## Supplemental Data for "Dysregulation of U12-Type Splicing in Lupus Neutrophils"

**Supplementary Material.**

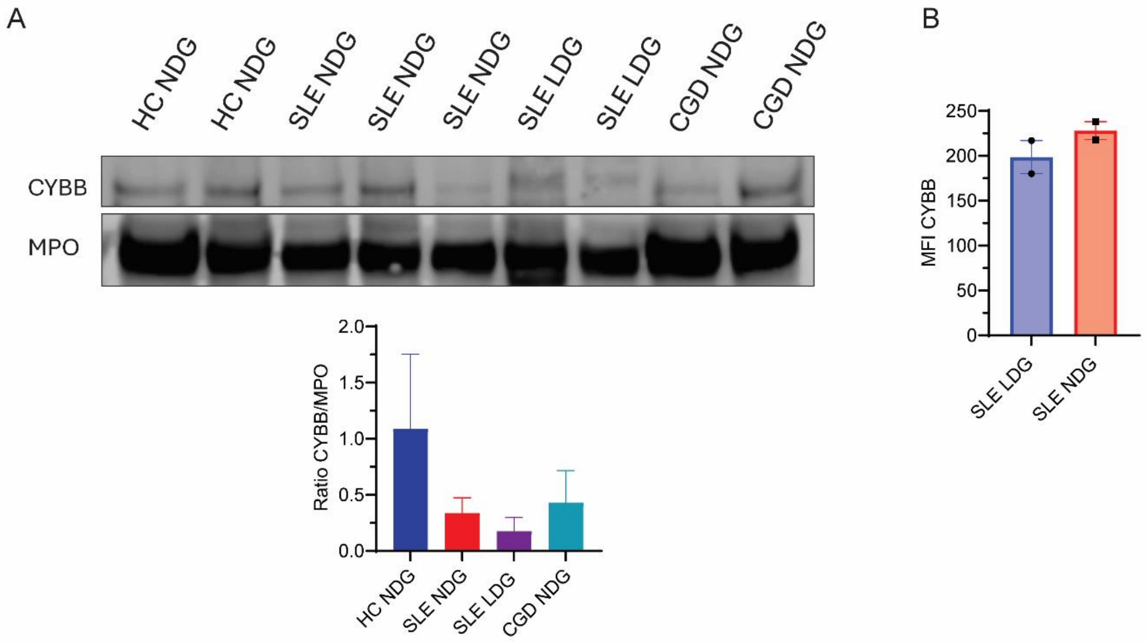

**Figure 1S: Reduced expression of the NADPH oxidase component CYBB (NOX2) in SLE neutrophils.** A) Western blot (top) and respective densitometry (Bottom) showing the expression of CYBB (NOX2) and MPO (as a housekeeping protein) in neutrophils from healthy controls or derived from SLE or CGD patients. B) Expression of NOX2 protein in neutrophils (NDGs and LDGs (from 2 SLE patients by flow cytometry. Shown average + SEM.

**Supplementary Data 1S**
**4_events_all**
***HC NDG versus SLE LDG***

| | | | | | | | | | | | | Inclusion counts | | | Skipping counts | | | Inclusion counts | | Skipping counts | | | | | | Inclusion_level(normalized) | | | | | |
|---|---|---|---|---|---|---|---|---|---|---|---|---|---|---|---|---|---|---|---|---|---|---|---|---|---|---|---|---|---|---|---|
| Splice_type | ID | Gene_ID | Gene_Symbol | Chromosome | Strand | LongExonStart_0base | LongExonEnd | ShortES | ShortEE | FlankingES | FlankingEE | control_s3 | control_s4 | control_s10 | control_s3 | control_s4 | control_s10 | ldg_14 | ldg_s16 | ldg_s14 | ldg_s16 | IncFormLen | SkipFormLen | p-value | adjust_p | IncLevel_s3 | IncLevel_s4 | IncLevel_s10 | IncLevel14 | IncLevel16 | Level_diff |
| SE | 8881 | CPEB2 | CPEB2 | chr4 | + | 15040463 | 15040487 | 15033160 | 15033211 | 15052413 | 15052584 | 14 | 15 | 2 | 1 | 0 | 0 | 0 | 1 | 5 | 1 | 73 | 49 | 1.71E-10 | 9.87E-07 | 0.904 | 1 | 1 | 0 | 0.402 | 0.767 |
| SE | 8677 | MAEA | MAEA | chr4 | + | 1322380 | 1322503 | 1315396 | 1315600 | 1332756 | 1332865 | 11 | 5 | 2 | 0 | 4 | 0 | 0 | 0 | 1 | 1 | 98 | 49 | 5.28E-08 | 0.000152595 | 1 | 0.385 | 1 | 0 | 0 | 0.795 |
| SE | 12478 | PQLC1 | PQLC1 | chr18 | - | 79933968 | 79934022 | 79919183 | 79919400 | 79943328 | 79943462 | 1 | 0 | 2 | 7 | 3 | 0 | 1 | 2 | 0 | 0 | 98 | 49 | 1.97E-07 | 0.000379884 | 0.067 | 0 | 1 | 1 | 1 | -0.644 |
| SE | 10324 | IRF7 | IRF7 | chr11 | - | 614173 | 614399 | 613784 | 613865 | 614475 | 614534 | 2 | 3 | 7 | 2 | 2 | 0 | 0 | 0 | 8 | 3 | 98 | 49 | 8.93E-07 | 0.00129026 | 0.333 | 0.429 | 1 | 0 | 0 | 0.587 |
| A5SS | 2439 | LILRA6 | LILRA6 | chr19 | - | 54241575 | 54241878 | 54241661 | 54241878 | 54240830 | 54241127 | 33 | 12 | 2 | 6 | 1 | 4 | 0 | 0 | 2 | 1 | 98 | 49 | 1.04E-06 | 0.000361711 | 0.733 | 0.857 | 0.2 | 0 | 0 | 0.597 |
| SE | 8882 | CPEB2 | CPEB2 | chr4 | + | 15059186 | 15059301 | 15058420 | 15058539 | 15062078 | 15062260 | 12 | 15 | 3 | 1 | 0 | 0 | 1 | 0 | 1 | 2 | 98 | 49 | 1.24E-06 | 0.001436638 | 0.857 | 1 | 1 | 0.333 | 0 | 0.786 |
| SE | 12726 | ADGRE5 | ADGRE5 | chr19 | + | 14397076 | 14397223 | 14396341 | 14396473 | 14397657 | 14397804 | 34 | 9 | 2 | 10 | 6 | 4 | 2 | 8 | 0 | 0 | 98 | 49 | 4.31E-06 | 0.004149582 | 0.63 | 0.429 | 0.2 | 1 | 1 | -0.58 |
| RI | 535 | TNFRSF10B | TNFRSF10B | chr8 | - | 23028330 | 23028602 | 23028330 | 23028441 | 23028528 | 23028602 | 22 | 4 | 2 | 4 | 7 | 1 | 1 | 2 | 0 | 0 | 98 | 49 | 4.88E-06 | 0.000804867 | 0.733 | 0.222 | 0.5 | 1 | 1 | -0.515 |
| SE | 17082 | FAM193B | FAM193B | chr5 | - | 177531281 | 177531521 | 177524184 | 177525205 | 177532442 | 177532641 | 21 | 5 | 2 | 10 | 4 | 0 | 3 | 7 | 0 | 0 | 98 | 49 | 1.30E-05 | 0.01058422 | 0.512 | 0.385 | 1 | 1 | 1 | -0.368 |
| SE | 3910 | AGTRAP | AGTRAP | chr1 | + | 11746126 | 11746223 | 11745802 | 11745837 | 11747439 | 11747545 | 5 | 4 | 2 | 5 | 2 | 0 | 0 | 0 | 1 | 2 | 98 | 49 | 1.47E-05 | 0.01058422 | 0.333 | 0.5 | 1 | 0 | 0 | 0.611 |
| SE | 5664 | WDFY3 | WDFY3 | chr4 | - | 84932269 | 84932363 | 84896910 | 84897010 | 84966208 | 84966391 | 36 | 18 | 3 | 4 | 11 | 1 | 2 | 13 | 0 | 0 | 98 | 49 | 2.47E-05 | 0.015846384 | 0.818 | 0.45 | 0.6 | 1 | 1 | -0.377 |
| A3SS | 1115 | PHF20L1 | PHF20L1 | chr8 | + | 132777791 | 132777911 | 132777795 | 132777911 | 132775357 | 132775645 | 32 | 9 | 5 | 10 | 5 | 3 | 0 | 1 | 3 | 3 | 53 | 49 | 2.54E-05 | 0.014675126 | 0.747 | 0.625 | 0.606 | 0 | 0.236 | 0.541 |
| RI | 948 | BCL2L11 | BCL2L11 | chr2 | + | 111123732 | 111124139 | 111123732 | 111123869 | 111124049 | 111124139 | 7 | 6 | 0 | 6 | 0 | 3 | 2 | 4 | 0 | 0 | 98 | 49 | 2.87E-05 | 0.002370232 | 0.368 | 1 | 0 | 1 | 1 | -0.544 |
| SE | 8257 | TNFSF13B | TNFSF13B | chr13 | + | 108286802 | 108286859 | 108270339 | 108270424 | 108303252 | 108303365 | 38 | 21 | 5 | 4 | 4 | 4 | 1 | 21 | 0 | 0 | 98 | 49 | 3.11E-05 | 0.017945737 | 0.826 | 0.724 | 0.385 | 1 | 1 | -0.355 |
| A3SS | 2900 | PGS1 | PGS1 | chr17 | + | 78414878 | 78415027 | 78414951 | 78415027 | 78403567 | 78404089 | 27 | 19 | 7 | 4 | 4 | 6 | 1 | 5 | 0 | 0 | 98 | 49 | 5.08E-05 | 0.014675126 | 0.771 | 0.704 | 0.368 | 1 | 1 | -0.386 |
| SE | 7275 | STK40 | STK40 | chr1 | - | 36367847 | 36368084 | 36361220 | 36361340 | 36385722 | 36385927 | 7 | 13 | 4 | 19 | 7 | 3 | 1 | 1 | 0 | 0 | 98 | 49 | 5.15E-05 | 0.027036581 | 0.156 | 0.481 | 0.4 | 1 | 1 | -0.654 |
| SE | 5316 | GATAD2A | GATAD2A | chr19 | + | 19492305 | 19492438 | 19465339 | 19465614 | 19492580 | 19492712 | 19 | 12 | 3 | 4 | 5 | 0 | 7 | 4 | 0 | 0 | 98 | 49 | 5.83E-05 | 0.028081837 | 0.704 | 0.545 | 1 | 1 | 1 | -0.25 |
| SE | 1880 | EEF1D | EEF1D | chr8 | - | 143586218 | 143586290 | 143581228 | 143581328 | 143586728 | 143586849 | 65 | 17 | 14 | 16 | 9 | 0 | 3 | 12 | 0 | 0 | 98 | 49 | 6.77E-05 | 0.029618192 | 0.67 | 0.486 | 1 | 1 | 1 | -0.281 |
| SE | 8102 | DEDD2 | DEDD2 | chr19 | - | 42216679 | 42217045 | 42215132 | 42215252 | 42217631 | 42217792 | 20 | 8 | 7 | 6 | 4 | 0 | 2 | 1 | 0 | 0 | 98 | 49 | 7.18E-05 | 0.029618192 | 0.625 | 0.5 | 1 | 1 | 1 | -0.292 |
| SE | 4618 | KIF13A | KIF13A | chr6 | - | 17789871 | 17789910 | 17787775 | 17787875 | 17794248 | 17794395 | 10 | 6 | 4 | 1 | 1 | 2 | 0 | 1 | 2 | 1 | 88 | 49 | 9.19E-05 | 0.035411696 | 0.848 | 0.77 | 0.527 | 0 | 0.358 | 0.536 |
| SE | 10413 | KLF6 | KLF6 | chr10 | - | 3780105 | 3780229 | 3775995 | 3779590 | 3781640 | 3782214 | 40 | 21 | 8 | 7 | 7 | 0 | 3 | 14 | 0 | 0 | 98 | 49 | 9.87E-05 | 0.035411696 | 0.741 | 0.6 | 1 | 1 | 1 | -0.22 |
| RI | 167 | WTAP | WTAP | chr6 | + | 159748190 | 159749421 | 159748190 | 159748369 | 159748589 | 159749421 | 39 | 9 | 9 | 12 | 2 | 0 | 1 | 13 | 0 | 0 | 98 | 49 | 0.000107296 | 0.005901273 | 0.619 | 0.692 | 1 | 1 | 1 | -0.23 |
| SE | 14148 | PSD4 | PSD4 | chr2 | + | 113198739 | 113198884 | 113197746 | 113197913 | 113199082 | 113199226 | 31 | 16 | 11 | 8 | 3 | 1 | 5 | 2 | 0 | 0 | 98 | 49 | 0.000108337 | 0.035411696 | 0.66 | 0.727 | 0.846 | 1 | 1 | -0.256 |
| SE | 9645 | MCTP1 | MCTP1 | chr5 | - | 94917895 | 94917973 | 94912805 | 94912976 | 94923961 | 94924021 | 17 | 27 | 15 | 13 | 3 | 1 | 2 | 3 | 0 | 0 | 98 | 49 | 0.000110298 | 0.035411696 | 0.395 | 0.818 | 0.882 | 1 | 1 | -0.302 |
| SE | 3227 | BANP | BANP | chr16 | + | 88004294 | 88004411 | 87984059 | 87984259 | 88006089 | 88006265 | 19 | 13 | 3 | 3 | 3 | 1 | 1 | 1 | 0 | 0 | 98 | 49 | 0.000123531 | 0.037572954 | 0.76 | 0.684 | 0.6 | 1 | 1 | -0.319 |
| A5SS | 1191 | PTPN22 | PTPN22 | chr1 | - | 113834308 | 113834439 | 113834392 | 113834439 | 113833110 | 113833138 | 16 | 24 | 53 | 8 | 1 | 10 | 1 | 10 | 0 | 0 | 98 | 49 | 0.000126122 | 0.020806987 | 0.5 | 0.923 | 0.726 | 1 | 1 | -0.284 |
| A3SS | 3574 | RALGAPB | RALGAPB | chr20 | + | 38535073 | 38535207 | 38535085 | 38535207 | 38532729 | 38532859 | 5 | 2 | 3 | 3 | 4 | 0 | 1 | 1 | 0 | 0 | 61 | 49 | 0.000130734 | 0.025188065 | 0.572 | 0.287 | 1 | 1 | 1 | -0.38 |
| SE | 14188 | ACOX1 | ACOX1 | chr17 | - | 75960214 | 75960375 | 75957458 | 75957566 | 75978533 | 75978693 | 35 | 8 | 22 | 9 | 4 | 0 | 1 | 11 | 0 | 0 | 98 | 49 | 0.000149043 | 0.043065865 | 0.66 | 0.5 | 1 | 1 | 1 | -0.28 |
| SE | 24 | G6PD | G6PD | chrX | - | 154532941 | 154533128 | 154532566 | 154532802 | 154533575 | 154533669 | 22 | 14 | 5 | 4 | 4 | 0 | 1 | 4 | 0 | 0 | 98 | 49 | 0.000172952 | 0.046713957 | 0.733 | 0.636 | 1 | 1 | 1 | -0.21 |
| A5SS | 237 | LAT2 | LAT2 | chr7 | + | 74214821 | 74215010 | 74214821 | 74214940 | 74215946 | 74216069 | 39 | 16 | 7 | 4 | 4 | 2 | 1 | 5 | 0 | 0 | 98 | 49 | 0.000179371 | 0.020806987 | 0.83 | 0.667 | 0.636 | 1 | 1 | -0.289 |
| SE | 5497 | SYNE1 | SYNE1 | chr6 | - | 152300640 | 152300781 | 152293959 | 152294127 | 152301868 | 152302063 | 6 | 3 | 0 | 0 | 1 | 1 | 0 | 1 | 1 | 1 | 98 | 49 | 0.000184481 | 0.046713957 | 1 | 0.6 | 0 | 0 | 0.333 | 0.367 |
| SE | 3520 | MTMR14 | MTMR14 | chr3 | + | 9689963 | 9690143 | 9688943 | 9689082 | 9701789 | 9702394 | 13 | 12 | 6 | 5 | 2 | 0 | 2 | 9 | 0 | 0 | 98 | 49 | 0.000189631 | 0.046713957 | 0.565 | 0.75 | 1 | 1 | 1 | -0.228 |
| SE | 1735 | VNN3 | VNN3 | chr6 | - | 132728740 | 132728979 | 132724605 | 132724979 | 132734252 | 132734383 | 44 | 14 | 6 | 7 | 6 | 0 | 1 | 2 | 0 | 0 | 98 | 49 | 0.000197256 | 0.046713957 | 0.759 | 0.538 | 1 | 1 | 1 | -0.234 |
| A3SS | 3415 | MAP4K4 | MAP4K4 | chr2 | + | 101866427 | 101866579 | 101866430 | 101866579 | 101864929 | 101865036 | 22 | 11 | 2 | 7 | 4 | 0 | 1 | 6 | 0 | 0 | 52 | 49 | 0.000199827 | 0.02887498 | 0.748 | 0.722 | 1 | 1 | 1 | -0.177 |
| SE | 1533 | WIPI2 | WIPI2 | chr7 | + | 5193117 | 5193171 | 5190203 | 5190493 | 5199575 | 5199658 | 3 | 1 | 0 | 10 | 7 | 2 | 1 | 3 | 0 | 1 | 98 | 49 | 0.000202085 | 0.046713957 | 0.13 | 0.067 | 0 | 1 | 0.6 | -0.734 |
| SE | 6524 | LYST | LYST | chr1 | - | 235728075 | 235728131 | 235724027 | 235724180 | 235729595 | 235729657 | 104 | 34 | 16 | 10 | 6 | 0 | 1 | 35 | 0 | 0 | 98 | 49 | 0.000210324 | 0.046748649 | 0.839 | 0.739 | 1 | 1 | 1 | -0.141 |
| SE | 3723 | ADGRG3 | ADGRG3 | chr16 | + | 57676199 | 57676338 | 57673320 | 57673468 | 57678169 | 57678316 | 109 | 54 | 15 | 13 | 11 | 0 | 2 | 14 | 0 | 0 | 98 | 49 | 0.000247814 | 0.051249962 | 0.807 | 0.711 | 1 | 1 | 1 | -0.161 |
| SE | 12145 | PGS1 | PGS1 | chr17 | + | 78414878 | 78415027 | 78403567 | 78404089 | 78419545 | 78419675 | 11 | 15 | 8 | 4 | 3 | 0 | 1 | 4 | 0 | 0 | 98 | 49 | 0.000253016 | 0.051249962 | 0.579 | 0.714 | 1 | 1 | 1 | -0.236 |
| SE | 1733 | VNN3 | VNN3 | chr6 | - | 132728740 | 132728933 | 132724605 | 132724979 | 132734252 | 132734383 | 51 | 21 | 4 | 7 | 6 | 0 | 1 | 5 | 0 | 0 | 98 | 49 | 0.000257181 | 0.051249962 | 0.785 | 0.636 | 1 | 1 | 1 | -0.193 |
| SE | 12146 | PGS1 | PGS1 | chr17 | + | 78414951 | 78415027 | 78403567 | 78404089 | 78419545 | 78419675 | 10 | 13 | 10 | 4 | 3 | 0 | 1 | 1 | 0 | 0 | 98 | 49 | 0.000268464 | 0.051715024 | 0.556 | 0.684 | 1 | 1 | 1 | -0.253 |
| SE | 3786 | RGS14 | RGS14 | chr5 | + | 177371474 | 177371589 | 177371349 | 177371396 | 177371872 | 177372598 | 22 | 11 | 8 | 4 | 3 | 0 | 1 | 10 | 0 | 0 | 98 | 49 | 0.000290269 | 0.052774163 | 0.733 | 0.647 | 1 | 1 | 1 | -0.207 |
| SE | 8838 | NR1H2 | NR1H2 | chr19 | + | 50378148 | 50378439 | 50377732 | 50377870 | 50378521 | 50378796 | 14 | 7 | 10 | 2 | 7 | 0 | 3 | 3 | 0 | 0 | 98 | 49 | 0.000293892 | 0.052774163 | 0.778 | 0.333 | 1 | 1 | 1 | -0.296 |
| SE | 8860 | GUK1 | GUK1 | chr1 | + | 228148370 | 228148456 | 228147614 | 228147699 | 228148664 | 228148954 | 32 | 21 | 5 | 0 | 1 | 0 | 0 | 8 | 1 | 3 | 98 | 49 | 0.000307457 | 0.052774163 | 1 | 0.913 | 1 | 0 | 0.571 | 0.686 |
| RI | 488 | STX10 | STX10 | chr19 | - | 13144057 | 13144671 | 13144057 | 13144486 | 13144576 | 13144671 | 15 | 16 | 0 | 9 | 10 | 13 | 1 | 6 | 0 | 1 | 98 | 49 | 0.000313634 | 0.0129374 | 0.455 | 0.444 | 0 | 1 | 0.75 | -0.575 |
| SE | 7426 | CD300A | CD300A | chr17 | + | 74473535 | 74473874 | 74466382 | 74466743 | 74474531 | 74474685 | 41 | 19 | 9 | 2 | 3 | 0 | 0 | 1 | 1 | 1 | 98 | 49 | 0.000315774 | 0.052774163 | 0.911 | 0.76 | 1 | 0 | 0.333 | 0.724 |
| SE | 9633 | ARIH1 | ARIH1 | chr15 | + | 72544819 | 72544964 | 72518066 | 72518134 | 72555270 | 72555363 | 17 | 7 | 5 | 2 | 7 | 0 | 1 | 2 | 0 | 0 | 98 | 49 | 0.000319622 | 0.052774163 | 0.81 | 0.333 | 1 | 1 | 1 | -0.286 |
| SE | 6469 | C1orf43 | C1orf43 | chr1 | - | 154219835 | 154219937 | 154214456 | 154214574 | 154220341 | 154220797 | 3 | 4 | 3 | 2 | 1 | 0 | 0 | 1 | 1 | 1 | 98 | 49 | 0.000389915 | 0.062553102 | 0.429 | 0.667 | 1 | 0 | 0.333 | 0.532 |
| SE | 12875 | ZFAS1 | ZFAS1 | chr20 | + | 49280902 | 49280964 | 49280484 | 49280570 | 49289044 | 49289258 | 45 | 37 | 17 | 12 | 14 | 13 | 1 | 12 | 0 | 1 | 98 | 49 | 0.000400496 | 0.062553102 | 0.652 | 0.569 | 0.395 | 1 | 0.857 | -0.39 |
| SE | 16934 | IRF1 | IRF1 | chr5 | - | 132486250 | 132486373 | 132484361 | 132484497 | 132486556 | 132486686 | 65 | 18 | 29 | 3 | 6 | 3 | 2 | 12 | 0 | 0 | 98 | 49 | 0.000423508 | 0.064406706 | 0.915 | 0.6 | 0.829 | 1 | 1 | -0.219 |
| A5SS | 2943 | MBD4 | MBD4 | chr3 | - | 129432502 | 129432606 | 129432555 | 129432606 | 129430943 | 129431578 | 33 | 10 | 3 | 6 | 2 | 0 | 0 | 1 | 1 | 1 | 98 | 49 | 0.0004824 | 0.041700018 | 0.733 | 0.714 | 1 | 0 | 0.333 | 0.649 |
| A3SS | 3084 | GMIP | GMIP | chr19 | - | 19636706 | 19636796 | 19636706 | 19636787 | 19636916 | 19637029 | 27 | 15 | 8 | 6 | 4 | 0 | 1 | 9 | 0 | 0 | 58 | 49 | 0.000488408 | 0.042850564 | 0.792 | 0.76 | 1 | 1 | 1 | -0.149 |
| A3SS | 3087 | CDV3 | CDV3 | chr3 | + | 133587158 | 133590274 | 133587895 | 133590274 | 133586562 | 133586722 | 17 | 15 | 11 | 11 | 1 | 0 | 1 | 2 | 0 | 0 | 98 | 49 | 0.000557132 | 0.042850564 | 0.436 | 0.882 | 1 | 1 | 1 | -0.227 |
| SE | 2174 | BAG6 | BAG6 | chr6 | - | 31647590 | 31647826 | 31646393 | 31646523 | 31648676 | 31648751 | 21 | 7 | 6 | 2 | 5 | 0 | 1 | 2 | 0 | 0 | 98 | 49 | 0.000561069 | 0.07959693 | 0.84 | 0.412 | 1 | 1 | 1 | -0.249 |
| SE | 12189 | HGS | HGS | chr17 | + | 81690181 | 81690234 | 81688703 | 81688827 | 81690673 | 81690742 | 32 | 18 | 10 | 2 | 9 | 0 | 1 | 1 | 0 | 0 | 98 | 49 | 0.000573304 | 0.07959693 | 0.889 | 0.5 | 1 | 1 | 1 | -0.204 |
| SE | 16397 | FAM13A-AS1 | FAM13A-AS1 | chr4 | + | 88722150 | 88722344 | 88711854 | 88712017 | 88723208 | 88723315 | 19 | 6 | 1 | 3 | 3 | 0 | 1 | 3 | 0 | 0 | 98 | 49 | 0.000577518 | 0.07959693 | 0.76 | 0.5 | 1 | 1 | 1 | -0.247 |
| SE | 16816 | AGTPBP1 | AGTPBP1 | chr9 | - | 85660935 | 85660973 | 85657434 | 85657643 | 85669484 | 85669578 | 44 | 26 | 26 | 8 | 5 | 0 | 1 | 9 | 0 | 0 | 87 | 49 | 0.000578486 | 0.07959693 | 0.756 | 0.745 | 1 | 1 | 1 | -0.166 |
| A3SS | 4077 | CNBP | CNBP | chr3 | - | 129171078 | 129171283 | 129171078 | 129171277 | 129171445 | 129171538 | 6 | 2 | 5 | 22 | 19 | 10 | 0 | 0 | 1 | 4 | 55 | 49 | 0.000580571 | 0.042850564 | 0.195 | 0.086 | 0.308 | 0 | 0 | 0.196 |
| A3SS | 971 | TPT1 | TPT1 | chr13 | - | 45333470 | 45337553 | 45333470 | 45337388 | 45338659 | 45338776 | 38 | 18 | 5 | 326 | 83 | 97 | 0 | 2 | 3 | 72 | 98 | 49 | 0.000593087 | 0.042850564 | 0.055 | 0.098 | 0.025 | 0 | 0.014 | 0.052 |
| A5SS | 1426 | TBXAS1 | TBXAS1 | chr7 | + | 139875584 | 139875666 | 139875584 | 139875637 | 139911224 | 139911321 | 20 | 11 | 2 | 53 | 44 | 9 | 0 | 1 | 1 | 15 | 78 | 49 | 0.000599138 | 0.041700018 | 0.192 | 0.136 | 0.122 | 0 | 0.04 | 0.13 |
| SE | 10242 | LST1 | LST1 | chr6 | + | 31587640 | 31587733 | 31587199 | 31587318 | 31588562 | 31588909 | 49 | 29 | 24 | 35 | 11 | 14 | 0 | 2 | 1 | 6 | 98 | 49 | 0.000674599 | 0.090662994 | 0.412 | 0.569 | 0.462 | 0 | 0.143 | 0.409 |
| RI | 408 | CPT1B | CPT1B | chr22 | - | 50568860 | 50569421 | 50568860 | 50569081 | 50569335 | 50569421 | 3 | 4 | 0 | 3 | 1 | 1 | 0 | 0 | 2 | 1 | 98 | 49 | 0.000688021 | 0.01974555 | 0.333 | 0.667 | 0 | 0 | 0 | 0.333 |
| RI | 529 | APH1A | APH1A | chr1 | - | 150265398 | 150266656 | 150265398 | 150266194 | 150266532 | 150266656 | 29 | 4 | 2 | 3 | 2 | 1 | 2 | 7 | 0 | 0 | 98 | 49 | 0.00071802 | 0.01974555 | 0.829 | 0.5 | 0.5 | 1 | 1 | -0.39 |
| SE | 9952 | SEC14L1 | SEC14L1 | chr17 | + | 77196201 | 77196311 | 77194676 | 77194911 | 77200483 | 77200673 | 180 | 90 | 52 | 9 | 7 | 2 | 11 | 34 | 0 | 0 | 98 | 49 | 0.000738806 | 0.095494005 | 0.909 | 0.865 | 0.929 | 1 | 1 | -0.099 |
| SE | 10018 | SBF2 | SBF2 | chr11 | - | 9998265 | 9998379 | 9993920 | 9993998 | 10000913 | 10001022 | 44 | 20 | 1 | 4 | 3 | 0 | 1 | 3 | 0 | 0 | 98 | 49 | 0.000771438 | 0.095494005 | 0.846 | 0.769 | 1 | 1 | 1 | -0.128 |
| SE | 6902 | CASP1 | CASP1 | chr11 | - | 105029123 | 105029267 | 105026841 | 105026951 | 105029664 | 105029899 | 48 | 13 | 23 | 6 | 3 | 0 | 1 | 17 | 0 | 0 | 98 | 49 | 0.000776107 | 0.095494005 | 0.8 | 0.684 | 1 | 1 | 1 | -0.172 |
| SE | 4285 | PLXNC1 | PLXNC1 | chr12 | + | 94298631 | 94298795 | 94297315 | 94297423 | 94300909 | 94301057 | 56 | 42 | 23 | 8 | 3 | 0 | 4 | 20 | 0 | 0 | 98 | 49 | 0.000802256 | 0.095494005 | 0.778 | 0.875 | 1 | 1 | 1 | -0.116 |
| SE | 5058 | ARAP1 | ARAP1 | chr11 | - | 72692752 | 72692785 | 72688454 | 72688537 | 72693324 | 72693470 | 13 | 7 | 12 | 26 | 12 | 17 | 0 | 2 | 1 | 16 | 82 | 49 | 0.000807769 | 0.095494005 | 0.23 | 0.258 | 0.297 | 0 | 0.07 | 0.227 |
| SE | 5663 | WDFY3 | WDFY3 | chr4 | - | 84836928 | 84837090 | 84831412 | 84831605 | 84841153 | 84841263 | 68 | 11 | 17 | 8 | 2 | 0 | 1 | 20 | 0 | 0 | 98 | 49 | 0.000809691 | 0.095494005 | 0.81 | 0.733 | 1 | 1 | 1 | -0.152 |
| A3SS | 3089 | CDV3 | CDV3 | chr3 | + | 133587161 | 133590274 | 133587895 | 133590274 | 133586562 | 133586722 | 10 | 15 | 10 | 11 | 1 | 0 | 1 | 2 | 0 | 0 | 98 | 49 | 0.000820412 | 0.052688694 | 0.312 | 0.882 | 1 | 1 | 1 | -0.269 |
| SE | 494 | ABCA1 | ABCA1 | chr9 | - | 104825682 | 104825887 | 104824464 | 104824578 | 104826947 | 104827169 | 31 | 8 | 7 | 9 | 0 | 0 | 4 | 6 | 0 | 0 | 98 | 49 | 0.001014686 | 0.117277454 | 0.633 | 1 | 1 | 1 | 1 | -0.122 |
| SE | 2588 | CDC42SE1 | CDC42SE1 | chr1 | - | 151056650 | 151056786 | 151055676 | 151055993 | 151059478 | 151059649 | 15 | 1 | 5 | 46 | 16 | 7 | 0 | 0 | 1 | 4 | 98 | 49 | 0.001070413 | 0.12129248 | 0.14 | 0.03 | 0.263 | 0 | 0 | 0.144 |
| SE | 1285 | SAMD9L | SAMD9L | chr7 | - | 93147087 | 93147146 | 93145384 | 93146040 | 93148193 | 93148388 | 6 | 7 | 3 | 7 | 1 | 3 | 1 | 7 | 0 | 1 | 98 | 49 | 0.001129194 | 0.125492512 | 0.3 | 0.778 | 0.333 | 1 | 0.778 | -0.419 |
| SE | 7935 | SUPT6H | SUPT6H | chr17 | + | 28682941 | 28683092 | 28682726 | 28682856 | 28683267 | 28683422 | 4 | 6 | 1 | 0 | 0 | 0 | 0 | 1 | 2 | 0 | 98 | 49 | 0.001189757 | 0.127567419 | 1 | 1 | 1 | 0 | 1 | 0.5 |
| SE | 6307 | CD82 | CD82 | chr11 | + | 44605354 | 44605429 | 44605057 | 44605182 | 44615271 | 44615373 | 44 | 22 | 10 | 0 | 0 | 0 | 0 | 1 | 2 | 0 | 98 | 49 | 0.001192013 | 0.127567419 | 1 | 1 | 1 | 0 | 1 | 0.5 |
| SE | 1284 | SAMD9L | SAMD9L | chr7 | - | 93146882 | 93147146 | 93145384 | 93146040 | 93148193 | 93148388 | 7 | 3 | 6 | 7 | 1 | 3 | 2 | 5 | 0 | 1 | 98 | 49 | 0.001254048 | 0.131766276 | 0.333 | 0.6 | 0.5 | 1 | 0.714 | -0.379 |
| SE | 14713 | PLEKHG3 | PLEKHG3 | chr14 | + | 64738741 | 64738855 | 64737355 | 64737375 | 64741035 | 64742455 | 48 | 3 | 10 | 5 | 3 | 0 | 2 | 14 | 0 | 0 | 98 | 49 | 0.001325462 | 0.136355346 | 0.828 | 0.333 | 1 | 1 | 1 | -0.28 |
| SE | 15669 | UBA7 | UBA7 | chr3 | - | 49812647 | 49812738 | 49812407 | 49812543 | 49813061 | 49813168 | 14 | 7 | 0 | 0 | 3 | 2 | 1 | 2 | 0 | 0 | 98 | 49 | 0.001349513 | 0.136355346 | 1 | 0.538 | 0 | 1 | 1 | -0.487 |
| SE | 16195 | UBE2R2 | UBE2R2 | chr9 | + | 33911963 | 33912098 | 33900173 | 33900271 | 33917017 | 33920403 | 21 | 10 | 8 | 1 | 0 | 0 | 0 | 6 | 2 | 1 | 98 | 49 | 0.001368508 | 0.136355346 | 0.913 | 1 | 1 | 0 | 0.75 | 0.596 |
| SE | 15849 | ARHGAP25 | ARHGAP25 | chr2 | + | 68813286 | 68813419 | 68807272 | 68807480 | 68816288 | 68816362 | 77 | 36 | 39 | 8 | 4 | 0 | 1 | 15 | 0 | 0 | 98 | 49 | 0.001401076 | 0.137234229 | 0.828 | 0.818 | 1 | 1 | 1 | -0.118 |
| SE | 8647 | SAMSN1 | SAMSN1 | chr21 | - | 14516891 | 14517041 | 14512443 | 14512573 | 14521149 | 14521221 | 26 | 9 | 10 | 0 | 1 | 0 | 1 | 12 | 1 | 3 | 98 | 49 | 0.001473673 | 0.141939264 | 1 | 0.818 | 1 | 0.333 | 0.667 | 0.439 |
| A5SS | 2947 | CSF2RA | CSF2RA | chrX | + | 1274754 | 1274842 | 1274754 | 1274818 | 1282677 | 1282779 | 43 | 28 | 3 | 6 | 3 | 0 | 1 | 3 | 0 | 0 | 73 | 49 | 0.001509767 | 0.07556475 | 0.828 | 0.862 | 1 | 1 | 1 | -0.103 |
| SE | 12923 | TYROBP | TYROBP | chr19 | - | 35907729 | 35907762 | 35907445 | 35907580 | 35908174 | 35908309 | 34 | 6 | 4 | 1 | 1 | 0 | 3 | 6 | 3 | 1 | 82 | 49 | 0.001510354 | 0.14308748 | 0.953 | 0.782 | 1 | 0.374 | 0.782 | 0.334 |
| SE | 6156 | LIMK2 | LIMK2 | chr22 | + | 31225719 | 31225819 | 31212263 | 31212424 | 31258290 | 31258426 | 118 | 52 | 19 | 7 | 4 | 0 | 14 | 29 | 0 | 0 | 98 | 49 | 0.001567896 | 0.146143067 | 0.894 | 0.867 | 1 | 1 | 1 | -0.08 |
| SE | 14889 | PREX1 | PREX1 | chr20 | - | 48659918 | 48660061 | 48658135 | 48658228 | 48666282 | 48666355 | 169 | 90 | 61 | 4 | 14 | 0 | 1 | 31 | 0 | 0 | 98 | 49 | 0.001683429 | 0.154421214 | 0.955 | 0.763 | 1 | 1 | 1 | -0.094 |
| A5SS | 1107 | SERPINA1 | SERPINA1 | chr14 | - | 94388800 | 94389010 | 94388818 | 94389010 | 94388559 | 94388660 | 38 | 15 | 12 | 18 | 3 | 11 | 1 | 6 | 0 | 1 | 67 | 49 | 0.001715222 | 0.07556475 | 0.607 | 0.785 | 0.444 | 1 | 0.814 | -0.295 |
| SE | 9038 | DENND5A | DENND5A | chr11 | - | 9170626 | 9170777 | 9169855 | 9169949 | 9178131 | 9178366 | 72 | 31 | 10 | 4 | 0 | 5 | 2 | 10 | 0 | 0 | 98 | 49 | 0.001799469 | 0.162486423 | 0.9 | 1 | 0.5 | 1 | 1 | -0.2 |
| SE | 13231 | RBM23 | RBM23 | chr14 | - | 22905605 | 22905659 | 22905335 | 22905453 | 22906194 | 22906368 | 38 | 23 | 7 | 1 | 3 | 3 | 0 | 2 | 2 | 1 | 98 | 49 | 0.001845941 | 0.1635565 | 0.95 | 0.793 | 0.538 | 0 | 0.5 | 0.51 |
| SE | 9349 | RAB18 | RAB18 | chr10 | + | 27531528 | 27531615 | 27526827 | 27526889 | 27532506 | 27532579 | 8 | 4 | 0 | 20 | 15 | 7 | 0 | 0 | 1 | 4 | 98 | 49 | 0.001867923 | 0.1635565 | 0.167 | 0.118 | 0 | 0 | 0 | 0.095 |
| A5SS | 2038 | ABTB1 | ABTB1 | chr3 | + | 127674453 | 127674593 | 127674390 | 127674418 | 127674544 | 127674600 | 57 | 37 | 13 | 15 | 18 | 7 | 2 | 6 | 0 | 1 | 84 | 49 | 0.001873218 | 0.07556475 | 0.689 | 0.545 | 0.52 | 1 | 0.778 | -0.304 |
| SE | 1093 | MGAM | MGAM | chr7 | + | 142027114 | 142027227 | 142025049 | 142025149 | 142027609 | 142027735 | 159 | 45 | 13 | 12 | 1 | 0 | 1 | 27 | 0 | 0 | 98 | 49 | 0.00193449 | 0.166856958 | 0.869 | 0.957 | 1 | 1 | 1 | -0.058 |
| A3SS | 3963 | CNN2 | CNN2 | chr19 | + | 1036066 | 1036246 | 1036129 | 1036246 | 1032558 | 1032696 | 13 | 10 | 0 | 83 | 43 | 21 | 2 | 9 | 1 | 10 | 98 | 49 | 0.001940249 | 0.109742062 | 0.073 | 0.104 | 0 | 0.5 | 0.31 | -0.346 |
| A5SS | 2352 | TYROBP | TYROBP | chr19 | - | 35908167 | 35908309 | 35908174 | 35908309 | 35907445 | 35907580 | 4 | 4 | 1 | 1 | 1 | 0 | 0 | 5 | 3 | 1 | 56 | 49 | 0.001954261 | 0.07556475 | 0.778 | 0.778 | 1 | 0 | 0.814 | 0.445 |
| SE | 11375 | LRRC75A-AS1 | LRRC75A-AS1 | chr17 | + | 16441073 | 16441130 | 16440110 | 16440253 | 16441367 | 16442026 | 23 | 8 | 9 | 5 | 1 | 0 | 1 | 8 | 0 | 0 | 98 | 49 | 0.001999476 | 0.169926028 | 0.697 | 0.8 | 1 | 1 | 1 | -0.168 |
| A3SS | 168 | TMEM71 | TMEM71 | chr8 | - | 132746941 | 132747114 | 132746941 | 132747057 | 132751784 | 132751997 | 107 | 66 | 50 | 4 | 3 | 5 | 5 | 22 | 0 | 0 | 98 | 49 | 0.002088517 | 0.109742062 | 0.93 | 0.917 | 0.833 | 1 | 1 | -0.107 |
| SE | 2482 | ARRB2 | ARRB2 | chr17 | + | 4716146 | 4716191 | 4715972 | 4716033 | 4716411 | 4716608 | 147 | 66 | 40 | 20 | 6 | 0 | 6 | 32 | 0 | 1 | 94 | 49 | 0.002174624 | 0.182132677 | 0.793 | 0.852 | 1 | 1 | 0.943 | -0.09 |
| SE | 9910 | PTP4A2 | PTP4A2 | chr1 | - | 31915894 | 31915987 | 31911695 | 31911826 | 31918969 | 31919658 | 21 | 15 | 12 | 4 | 1 | 1 | 4 | 13 | 0 | 0 | 98 | 49 | 0.002314755 | 0.189353778 | 0.724 | 0.882 | 0.857 | 1 | 1 | -0.179 |
| SE | 2943 | FCAR | FCAR | chr19 | + | 54888006 | 54888294 | 54885234 | 54885525 | 54889648 | 54891420 | 58 | 10 | 7 | 4 | 9 | 0 | 5 | 28 | 0 | 1 | 98 | 49 | 0.002326374 | 0.189353778 | 0.879 | 0.357 | 1 | 1 | 0.933 | -0.221 |
| A3SS | 700 | PCBP2 | PCBP2 | chr12 | + | 53462480 | 53462567 | 53462492 | 53462567 | 53461014 | 53461143 | 65 | 31 | 9 | 51 | 6 | 3 | 1 | 8 | 1 | 19 | 61 | 49 | 0.00238858 | 0.11504996 | 0.506 | 0.806 | 0.707 | 0.445 | 0.253 | 0.324 |
| A5SS | 760 | FAM160A2 | FAM160A2 | chr11 | - | 6218557 | 6218763 | 6218599 | 6218763 | 6217370 | 6218150 | 1 | 0 | 1 | 0 | 1 | 0 | 0 | 1 | 2 | 0 | 91 | 49 | 0.002461806 | 0.085670833 | 1 | 0 | 1 | 0 | 1 | 0.167 |
| SE | 2034 | TAP1 | TAP1 | chr6 | - | 32848989 | 32849118 | 32848651 | 32848840 | 32850319 | 32850517 | 61 | 38 | 37 | 6 | 2 | 0 | 2 | 34 | 0 | 0 | 98 | 49 | 0.002588194 | 0.207738482 | 0.836 | 0.905 | 1 | 1 | 1 | -0.086 |
| SE | 9654 | FAU | FAU | chr11 | - | 65120980 | 65121036 | 65120626 | 65120806 | 65121499 | 65121644 | 78 | 21 | 15 | 2 | 6 | 0 | 1 | 13 | 0 | 0 | 98 | 49 | 0.002676583 | 0.210269197 | 0.951 | 0.636 | 1 | 1 | 1 | -0.138 |
| SE | 11777 | FMNL1 | FMNL1 | chr17 | + | 45245631 | 45245733 | 45245252 | 45245416 | 45245877 | 45245973 | 108 | 41 | 29 | 8 | 3 | 1 | 5 | 21 | 3 | 6 | 98 | 49 | 0.002713341 | 0.210269197 | 0.871 | 0.872 | 0.935 | 0.455 | 0.636 | 0.347 |
| SE | 14886 | PREX1 | PREX1 | chr20 | - | 48644408 | 48644497 | 48642406 | 48642489 | 48645850 | 48646057 | 122 | 56 | 51 | 0 | 3 | 0 | 5 | 22 | 0 | 5 | 98 | 49 | 0.002756217 | 0.210269197 | 1 | 0.903 | 1 | 1 | 0.687 | 0.124 |
| SE | 9504 | HSH2D | HSH2D | chr19 | + | 16151446 | 16151578 | 16148723 | 16148875 | 16152551 | 16153208 | 12 | 4 | 0 | 44 | 18 | 3 | 0 | 1 | 4 | 15 | 98 | 49 | 0.002765264 | 0.210269197 | 0.12 | 0.1 | 0 | 0 | 0.032 | 0.057 |
| SE | 2825 | MEFV | MEFV | chr16 | - | 3249430 | 3249780 | 3247015 | 3247246 | 3254157 | 3254790 | 46 | 22 | 18 | 1 | 0 | 0 | 0 | 11 | 4 | 0 | 98 | 49 | 0.002963845 | 0.22054525 | 0.958 | 1 | 1 | 0 | 1 | 0.486 |
| SE | 2763 | PCBP2 | PCBP2 | chr12 | + | 53464759 | 53464852 | 53462480 | 53462567 | 53467220 | 53467293 | 54 | 17 | 19 | 0 | 1 | 0 | 0 | 13 | 1 | 3 | 98 | 49 | 0.002976731 | 0.22054525 | 1 | 0.895 | 1 | 0 | 0.684 | 0.623 |
| SE | 13280 | RBM5 | RBM5 | chr3 | + | 50092042 | 50092208 | 50090381 | 50090451 | 50093719 | 50093875 | 82 | 17 | 14 | 4 | 4 | 0 | 1 | 2 | 0 | 0 | 98 | 49 | 0.003118328 | 0.2281116 | 0.911 | 0.68 | 1 | 1 | 1 | -0.136 |
| SE | 16135 | IL1RAP | IL1RAP | chr3 | + | 190556129 | 190556216 | 190514050 | 190514219 | 190564288 | 190564353 | 22 | 20 | 1 | 5 | 1 | 0 | 3 | 7 | 0 | 1 | 98 | 49 | 0.003473051 | 0.250884516 | 0.687 | 0.909 | 1 | 1 | 0.778 | -0.024 |
| SE | 10388 | ZSWIM8 | ZSWIM8 | chr10 | + | 73788669 | 73788823 | 73785623 | 73786086 | 73789095 | 73789190 | 24 | 5 | 2 | 1 | 0 | 0 | 2 | 3 | 5 | 0 | 98 | 49 | 0.003538822 | 0.252479651 | 0.923 | 1 | 1 | 0.167 | 1 | 0.391 |
| RI | 1085 | UNC119 | UNC119 | chr17 | - | 28546706 | 28547849 | 28546706 | 28547409 | 28547676 | 28547849 | 36 | 22 | 3 | 3 | 3 | 6 | 1 | 10 | 0 | 1 | 98 | 49 | 0.003670791 | 0.086525783 | 0.857 | 0.786 | 0.2 | 1 | 0.833 | -0.302 |
| SE | 10573 | NFKBIA | NFKBIA | chr14 | - | 35402770 | 35402859 | 35402393 | 35402663 | 35403149 | 35403360 | 124 | 307 | 51 | 1 | 15 | 0 | 10 | 63 | 0 | 0 | 98 | 49 | 0.003690903 | 0.257474498 | 0.984 | 0.911 | 1 | 1 | 1 | -0.035 |
| SE | 7428 | CD300A | CD300A | chr17 | + | 74481725 | 74481833 | 74481288 | 74481326 | 74484000 | 74484798 | 38 | 29 | 35 | 5 | 6 | 3 | 4 | 15 | 0 | 1 | 98 | 49 | 0.003715696 | 0.257474498 | 0.792 | 0.707 | 0.854 | 1 | 0.882 | -0.157 |
| SE | 6933 | IL17RA | IL17RA | chr22 | + | 17105852 | 17105954 | 17105590 | 17105602 | 17107726 | 17107768 | 61 | 17 | 15 | 7 | 0 | 0 | 10 | 15 | 0 | 0 | 98 | 49 | 0.003761285 | 0.257474498 | 0.813 | 1 | 1 | 1 | 1 | -0.062 |
| SE | 10301 | NLRP6 | NLRP6 | chr11 | + | 280083 | 281839 | 279833 | 279872 | 282704 | 282797 | 10 | 11 | 4 | 1 | 1 | 3 | 1 | 2 | 0 | 0 | 98 | 49 | 0.003860898 | 0.257474498 | 0.833 | 0.846 | 0.4 | 1 | 1 | -0.307 |
| SE | 1138 | LINC-PINT | LINC-PINT | chr7 | - | 131002196 | 131002320 | 130984052 | 130984109 | 131052424 | 131052555 | 7 | 5 | 0 | 47 | 20 | 3 | 0 | 0 | 2 | 9 | 98 | 49 | 0.003875277 | 0.257474498 | 0.069 | 0.111 | 0 | 0 | 0 | 0.06 |
| SE | 10775 | PTK2B | PTK2B | chr8 | + | 27445793 | 27445919 | 27444205 | 27444271 | 27450748 | 27450895 | 8 | 1 | 0 | 22 | 5 | 8 | 0 | 0 | 1 | 2 | 98 | 49 | 0.003876152 | 0.257474498 | 0.154 | 0.091 | 0 | 0 | 0 | 0.082 |
| SE | 10245 | LST1 | LST1 | chr6 | + | 31587943 | 31587966 | 31587199 | 31587318 | 31588562 | 31588909 | 12 | 10 | 1 | 35 | 11 | 14 | 0 | 1 | 1 | 6 | 72 | 49 | 0.003941519 | 0.258841315 | 0.189 | 0.382 | 0.046 | 0 | 0.102 | 0.155 |
| A3SS | 805 | BANP | BANP | chr16 | + | 88035322 | 88035394 | 88035331 | 88035394 | 88033108 | 88033245 | 11 | 8 | 7 | 8 | 0 | 0 | 1 | 2 | 0 | 0 | 58 | 49 | 0.004108642 | 0.171485553 | 0.537 | 1 | 1 | 1 | 1 | -0.154 |
| A3SS | 2515 | ZSWIM8 | ZSWIM8 | chr10 | + | 73794540 | 73794639 | 73794555 | 73794639 | 73794146 | 73794330 | 19 | 5 | 0 | 9 | 3 | 1 | 1 | 7 | 1 | 0 | 64 | 49 | 0.004153629 | 0.171485553 | 0.618 | 0.561 | 0 | 0.434 | 1 | -0.324 |
| SE | 15293 | CSF2RB | CSF2RB | chr22 | + | 36936548 | 36936652 | 36935629 | 36935687 | 36937376 | 36940437 | 151 | 73 | 46 | 7 | 4 | 0 | 7 | 34 | 0 | 0 | 98 | 49 | 0.004519392 | 0.293455815 | 0.915 | 0.901 | 1 | 1 | 1 | -0.061 |
| SE | 1135 | LINC-PINT | LINC-PINT | chr7 | - | 130959868 | 130959970 | 130944159 | 130945835 | 130984052 | 130984109 | 21 | 1 | 0 | 14 | 1 | 2 | 0 | 1 | 1 | 2 | 98 | 49 | 0.004626486 | 0.297071834 | 0.429 | 0.333 | 0 | 0 | 0.2 | 0.154 |
| SE | 2941 | FCAR | FCAR | chr19 | + | 54888006 | 54888220 | 54885234 | 54885525 | 54889648 | 54891420 | 30 | 4 | 5 | 4 | 9 | 0 | 1 | 11 | 0 | 1 | 98 | 49 | 0.004730423 | 0.300407848 | 0.789 | 0.182 | 1 | 1 | 0.846 | -0.266 |
| A3SS | 1029 | OAS1 | OAS1 | chr12 | + | 112919388 | 112919907 | 112919486 | 112919907 | 112917546 | 112917700 | 1 | 1 | 9 | 0 | 0 | 0 | 3 | 5 | 0 | 4 | 98 | 49 | 0.004828658 | 0.186064281 | 1 | 1 | 1 | 1 | 0.385 | 0.307 |
| SE | 6536 | RBM39 | RBM39 | chr20 | - | 35740524 | 35740597 | 35738967 | 35739017 | 35740823 | 35740887 | 63 | 33 | 36 | 23 | 8 | 1 | 2 | 8 | 0 | 1 | 98 | 49 | 0.004971951 | 0.312314156 | 0.578 | 0.673 | 0.947 | 1 | 0.8 | -0.167 |
| A3SS | 4024 | RICTOR | RICTOR | chr5 | - | 38945490 | 38945724 | 38945490 | 38945610 | 38946467 | 38946552 | 100 | 32 | 12 | 12 | 8 | 0 | 1 | 15 | 0 | 1 | 98 | 49 | 0.005321448 | 0.192237298 | 0.806 | 0.667 | 1 | 1 | 0.882 | -0.117 |
| SE | 207 | PIM2 | PIM2 | chrX | - | 48914394 | 48914571 | 48913181 | 48914294 | 48915019 | 48915392 | 35 | 12 | 17 | 7 | 6 | 1 | 0 | 2 | 1 | 2 | 98 | 49 | 0.005370609 | 0.33372849 | 0.714 | 0.5 | 0.895 | 0 | 0.333 | 0.536 |
| SE | 7149 | ATP1A1 | ATP1A1 | chr1 | + | 116395109 | 116395285 | 116393530 | 116393723 | 116396597 | 116396734 | 17 | 4 | 8 | 6 | 0 | 0 | 1 | 11 | 0 | 0 | 98 | 49 | 0.005599832 | 0.344270522 | 0.586 | 1 | 1 | 1 | 1 | -0.138 |
| SE | 247 | DDX3X | DDX3X | chrX | + | 41347314 | 41347451 | 41346858 | 41347012 | 41347639 | 41350287 | 147 | 37 | 36 | 7 | 3 | 0 | 1 | 17 | 0 | 0 | 98 | 49 | 0.005759724 | 0.347607175 | 0.913 | 0.86 | 1 | 1 | 1 | -0.076 |
| SE | 3044 | STX16 | STX16 | chr20 | + | 58671153 | 58671297 | 58670511 | 58670603 | 58673630 | 58673711 | 37 | 27 | 7 | 4 | 7 | 0 | 1 | 16 | 0 | 1 | 98 | 49 | 0.005774405 | 0.347607175 | 0.822 | 0.659 | 1 | 1 | 0.889 | -0.118 |
| SE | 8799 | BROX | BROX | chr1 | + | 222731356 | 222731516 | 222730026 | 222730177 | 222732627 | 222735196 | 39 | 8 | 5 | 0 | 0 | 1 | 2 | 6 | 0 | 4 | 98 | 49 | 0.005905541 | 0.351836284 | 1 | 1 | 0.714 | 1 | 0.429 | 0.19 |
| SE | 11526 | CHFR | CHFR | chr12 | - | 132870723 | 132870783 | 132861466 | 132861634 | 132872284 | 132872394 | 19 | 3 | 6 | 0 | 4 | 1 | 3 | 5 | 0 | 0 | 98 | 49 | 0.006223406 | 0.366990437 | 1 | 0.273 | 0.75 | 1 | 1 | -0.326 |
| SE | 9240 | SVIL | SVIL | chr10 | - | 29526960 | 29527056 | 29524471 | 29524715 | 29529704 | 29529844 | 8 | 0 | 1 | 33 | 14 | 2 | 0 | 0 | 3 | 8 | 98 | 49 | 0.00637626 | 0.372206119 | 0.108 | 0 | 0.2 | 0 | 0 | 0.103 |
| SE | 9931 | MKNK1 | MKNK1 | chr1 | - | 46560233 | 46560277 | 46557406 | 46558800 | 46561477 | 46561642 | 16 | 9 | 2 | 5 | 2 | 0 | 0 | 2 | 1 | 2 | 93 | 49 | 0.006502959 | 0.375806004 | 0.628 | 0.703 | 1 | 0 | 0.345 | 0.605 |
| SE | 15577 | SUN2 | SUN2 | chr22 | - | 38750897 | 38751035 | 38750224 | 38750320 | 38751209 | 38751373 | 43 | 11 | 9 | 5 | 1 | 0 | 1 | 6 | 0 | 0 | 98 | 49 | 0.00671086 | 0.383255085 | 0.811 | 0.846 | 1 | 1 | 1 | -0.114 |
| SE | 4283 | PLXNC1 | PLXNC1 | chr12 | + | 94209589 | 94209704 | 94186372 | 94186473 | 94220015 | 94220163 | 77 | 36 | 26 | 7 | 5 | 0 | 4 | 27 | 0 | 1 | 98 | 49 | 0.006764495 | 0.383255085 | 0.846 | 0.783 | 1 | 1 | 0.931 | -0.089 |
| SE | 6938 | ARID4B | ARID4B | chr1 | - | 235213768 | 235214026 | 235196030 | 235196115 | 235219792 | 235219968 | 11 | 1 | 1 | 9 | 3 | 5 | 0 | 1 | 1 | 2 | 98 | 49 | 0.006951843 | 0.387975679 | 0.379 | 0.143 | 0.091 | 0 | 0.2 | 0.104 |
| SE | 236 | DDX3X | DDX3X | chrX | + | 41339035 | 41339083 | 41337407 | 41337465 | 41341483 | 41341616 | 120 | 66 | 28 | 5 | 4 | 0 | 2 | 15 | 0 | 0 | 97 | 49 | 0.006990144 | 0.387975679 | 0.924 | 0.893 | 1 | 1 | 1 | -0.061 |
| SE | 2824 | MEFV | MEFV | chr16 | - | 3248908 | 3249004 | 3247015 | 3247246 | 3254157 | 3254790 | 48 | 18 | 35 | 1 | 0 | 0 | 2 | 18 | 4 | 0 | 98 | 49 | 0.007049221 | 0.387975679 | 0.96 | 1 | 1 | 0.2 | 1 | 0.387 |
| SE | 7813 | PGD | PGD | chr1 | + | 10403070 | 10403136 | 10400392 | 10400572 | 10404160 | 10404279 | 93 | 46 | 31 | 1 | 0 | 0 | 7 | 13 | 4 | 0 | 98 | 49 | 0.007216115 | 0.39135749 | 0.979 | 1 | 1 | 0.467 | 1 | 0.259 |
| SE | 8990 | GLYR1 | GLYR1 | chr16 | - | 4821546 | 4821597 | 4821379 | 4821453 | 4822874 | 4822931 | 21 | 4 | 15 | 0 | 0 | 0 | 0 | 9 | 3 | 0 | 98 | 49 | 0.007246107 | 0.39135749 | 1 | 1 | 1 | 0 | 1 | 0.5 |
| SE | 13299 | TSEN34 | TSEN34 | chr19 | + | 54191720 | 54191964 | 54191213 | 54191607 | 54192115 | 54192373 | 62 | 31 | 46 | 3 | 7 | 3 | 5 | 17 | 0 | 1 | 98 | 49 | 0.007593268 | 0.406310123 | 0.912 | 0.689 | 0.885 | 1 | 0.895 | -0.119 |
| SE | 12576 | VAV1 | VAV1 | chr19 | + | 6847997 | 6848114 | 6843134 | 6843166 | 6850669 | 6850757 | 60 | 21 | 19 | 0 | 0 | 0 | 3 | 13 | 4 | 0 | 98 | 49 | 0.007817874 | 0.414490774 | 1 | 1 | 1 | 0.273 | 1 | 0.363 |
| SE | 245 | DDX3X | DDX3X | chrX | + | 41347311 | 41347451 | 41346858 | 41347012 | 41347639 | 41347915 | 173 | 52 | 46 | 7 | 3 | 0 | 1 | 19 | 0 | 0 | 98 | 49 | 0.008108954 | 0.426014944 | 0.925 | 0.897 | 1 | 1 | 1 | -0.059 |
| SE | 3570 | DIP2B | DIP2B | chr12 | + | 50692948 | 50693013 | 50691048 | 50691151 | 50695266 | 50695360 | 18 | 22 | 2 | 6 | 0 | 0 | 2 | 5 | 0 | 0 | 98 | 49 | 0.008244832 | 0.42925121 | 0.6 | 1 | 1 | 1 | 1 | -0.133 |
| SE | 9036 | DENND5A | DENND5A | chr11 | - | 9147029 | 9147151 | 9145669 | 9145815 | 9150080 | 9150209 | 66 | 31 | 24 | 7 | 0 | 0 | 3 | 13 | 0 | 0 | 98 | 49 | 0.008385906 | 0.432697767 | 0.825 | 1 | 1 | 1 | 1 | -0.058 |
| SE | 10695 | NPL | NPL | chr1 | + | 182822114 | 182822199 | 182818812 | 182818859 | 182825780 | 182825820 | 43 | 14 | 16 | 4 | 1 | 1 | 5 | 5 | 0 | 0 | 98 | 49 | 0.008492504 | 0.434320174 | 0.843 | 0.875 | 0.889 | 1 | 1 | -0.131 |
| SE | 9897 | ICAM3 | ICAM3 | chr19 | - | 10335065 | 10335353 | 10334527 | 10334782 | 10335670 | 10335976 | 209 | 102 | 117 | 19 | 7 | 4 | 1 | 23 | 0 | 1 | 98 | 49 | 0.008673586 | 0.439689953 | 0.846 | 0.879 | 0.936 | 1 | 0.92 | -0.073 |
| A5SS | 523 | BTNL8 | BTNL8 | chr5 | + | 180947511 | 180947755 | 180947511 | 180947625 | 180948354 | 180948375 | 13 | 4 | 0 | 4 | 0 | 3 | 2 | 2 | 0 | 1 | 98 | 49 | 0.009042031 | 0.277083706 | 0.619 | 1 | 0 | 1 | 0.5 | -0.21 |
| SE | 1187 | IFRD1 | IFRD1 | chr7 | + | 112462269 | 112462378 | 112462000 | 112462179 | 112467980 | 112468115 | 78 | 16 | 20 | 80 | 56 | 22 | 0 | 10 | 3 | 37 | 98 | 49 | 0.009088918 | 0.456737895 | 0.328 | 0.125 | 0.312 | 0 | 0.119 | 0.196 |
| SE | 6534 | RBM39 | RBM39 | chr20 | - | 35729461 | 35729527 | 35729311 | 35729365 | 35731940 | 35732135 | 186 | 68 | 19 | 4 | 0 | 5 | 3 | 27 | 0 | 0 | 98 | 49 | 0.009727315 | 0.484604766 | 0.959 | 1 | 0.655 | 1 | 1 | -0.129 |
| A5SS | 1150 | MYD88 | MYD88 | chr3 | + | 38140387 | 38140592 | 38140387 | 38140568 | 38140756 | 38140848 | 1 | 0 | 0 | 9 | 1 | 3 | 0 | 4 | 1 | 7 | 73 | 49 | 0.009750777 | 0.277083706 | 0.069 | 0 | 0 | 0 | 0.277 | -0.116 |

| | | | | | | | | | | | | | | | | | | | | | | | | | | | | | | | |
|---|---|---|---|---|---|---|---|---|---|---|---|---|---|---|---|---|---|---|---|---|---|---|---|---|---|---|---|---|---|---|---|
| RI | 1014 | OSCAR | OSCAR | chr19 | - | 54094667 | 54096153 | 54094667 | 54095357 | 54095871 | 54096153 | 5 | 1 | 1 | 0 | 2 | 3 | 2 | 1 | 0 | 1 | 98 | 49 | 0.009785372 | 0.201823302 | 1 | 0.2 | 0.143 | 1 | 0.333 | -0.219 |
| A3SS | 3474 | CFLAR | CFLAR | chr2 | + | 201162995 | 201172688 | 201163834 | 201172688 | 201160431 | 201160942 | 36 | 9 | 8 | 132 | 56 | 24 | 0 | 4 | 3 | 38 | 98 | 49 | 0.009823696 | 0.330007041 | 0.12 | 0.074 | 0.143 | 0 | 0.05 | 0.087 |
| SE | 10365 | AKNA | AKNA | chr9 | - | 114343707 | 114343803 | 114342008 | 114342125 | 114345862 | 114346009 | 63 | 18 | 18 | 2 | 4 | 0 | 0 | 10 | 2 | 2 | 98 | 49 | 0.009893908 | 0.48869139 | 0.94 | 0.692 | 1 | 0 | 0.714 | 0.52 |
| SE | 14778 | PLAUR | PLAUR | chr19 | - | 43655438 | 43655573 | 43652224 | 43652371 | 43656478 | 43656640 | 65 | 27 | 18 | 3 | 0 | 0 | 8 | 42 | 0 | 6 | 98 | 49 | 0.010169452 | 0.498044606 | 0.915 | 1 | 1 | 1 | 0.778 | 0.083 |
| A3SS | 1048 | DEF8 | DEF8 | chr16 | + | 89964169 | 89964310 | 89964220 | 89964310 | 89963362 | 89963443 | 19 | 8 | 3 | 2 | 2 | 3 | 3 | 3 | 0 | 1 | 98 | 49 | 0.010277036 | 0.330007041 | 0.826 | 0.667 | 0.333 | 1 | 0.6 | -0.191 |
| A5SS | 762 | PLEKHB2 | PLEKHB2 | chr2 | + | 131132901 | 131133015 | 131132901 | 131132991 | 131140166 | 131140275 | 1 | 0 | 0 | 18 | 5 | 2 | 0 | 4 | 1 | 3 | 73 | 49 | 0.010350828 | 0.277083706 | 0.036 | 0 | 0 | 0 | 0.472 | -0.224 |
| SE | 237 | DDX3X | DDX3X | chrX | + | 41343215 | 41343351 | 41342736 | 41342836 | 41343736 | 41343822 | 212 | 92 | 26 | 8 | 6 | 4 | 2 | 31 | 0 | 1 | 98 | 49 | 0.01082339 | 0.521780013 | 0.93 | 0.885 | 0.765 | 1 | 0.939 | -0.11 |
| SE | 12808 | ELL | ELL | chr19 | - | 18465796 | 18465918 | 18465411 | 18465575 | 18472834 | 18472882 | 69 | 32 | 3 | 2 | 0 | 0 | 2 | 13 | 4 | 0 | 98 | 49 | 0.010834678 | 0.521780013 | 0.945 | 1 | 1 | 0.2 | 1 | 0.382 |
| SE | 11623 | PSMB3 | PSMB3 | chr17 | + | 38760507 | 38760608 | 38755882 | 38755990 | 38762410 | 38762505 | 18 | 13 | 30 | 2 | 5 | 2 | 1 | 7 | 0 | 1 | 98 | 49 | 0.011365999 | 0.532195482 | 0.818 | 0.565 | 0.882 | 1 | 0.778 | -0.134 |
| SE | 6535 | RBM39 | RBM39 | chr20 | - | 35734185 | 35734257 | 35731940 | 35732135 | 35738967 | 35739017 | 2 | 2 | 8 | 56 | 49 | 9 | 0 | 0 | 2 | 9 | 98 | 49 | 0.011421734 | 0.532195482 | 0.018 | 0.02 | 0.308 | 0 | 0 | 0.115 |
| SE | 10891 | ADGRE2 | ADGRE2 | chr19 | - | 14755653 | 14755877 | 14752328 | 14752526 | 14756237 | 14756345 | 6 | 6 | 12 | 0 | 0 | 0 | 0 | 2 | 1 | 1 | 98 | 49 | 0.011489901 | 0.532195482 | 1 | 1 | 1 | 0 | 0.5 | 0.75 |
| SE | 7028 | TAOK3 | TAOK3 | chr12 | - | 118212913 | 118212995 | 118201295 | 118201463 | 118214016 | 118214110 | 28 | 5 | 5 | 3 | 3 | 0 | 2 | 4 | 0 | 1 | 98 | 49 | 0.011563355 | 0.532195482 | 0.824 | 0.455 | 1 | 1 | 0.667 | -0.074 |
| SE | 4824 | FRYL | FRYL | chr4 | - | 48512480 | 48512688 | 48510834 | 48510984 | 48515027 | 48515275 | 5 | 10 | 4 | 0 | 0 | 0 | 0 | 2 | 2 | 0 | 98 | 49 | 0.011681017 | 0.532195482 | 1 | 1 | 1 | 0 | 1 | 0.5 |
| SE | 2177 | PRRC2A | PRRC2A | chr6 | + | 31622726 | 31622901 | 31620672 | 31620858 | 31624260 | 31624360 | 16 | 3 | 2 | 0 | 0 | 0 | 0 | 2 | 2 | 0 | 98 | 49 | 0.011681454 | 0.532195482 | 1 | 1 | 1 | 0 | 1 | 0.5 |
| SE | 16224 | CIRBP | CIRBP | chr19 | + | 1271139 | 1271246 | 1270866 | 1271036 | 1271328 | 1271467 | 66 | 44 | 22 | 5 | 2 | 0 | 2 | 4 | 0 | 0 | 98 | 49 | 0.011717498 | 0.532195482 | 0.868 | 0.917 | 1 | 1 | 1 | -0.072 |
| SE | 9303 | PIK3CD | PIK3CD | chr1 | + | 9723968 | 9724092 | 9723124 | 9723292 | 9724275 | 9724421 | 67 | 52 | 19 | 2 | 5 | 0 | 3 | 12 | 0 | 0 | 98 | 49 | 0.011787683 | 0.532195482 | 0.944 | 0.839 | 1 | 1 | 1 | -0.072 |
| SE | 16189 | SH3BP2 | SH3BP2 | chr4 | + | 2831922 | 2831978 | 2831570 | 2831679 | 2832330 | 2832412 | 28 | 28 | 9 | 6 | 2 | 0 | 1 | 12 | 0 | 1 | 98 | 49 | 0.012193479 | 0.540043183 | 0.7 | 0.875 | 1 | 1 | 0.857 | -0.07 |
| SE | 4830 | CLEC7A | CLEC7A | chr12 | - | 10123244 | 10123363 | 10116780 | 10118590 | 10125086 | 10125173 | 96 | 47 | 30 | 2 | 0 | 1 | 0 | 38 | 2 | 1 | 98 | 49 | 0.012205539 | 0.540043183 | 0.96 | 1 | 0.937 | 0 | 0.95 | 0.491 |
| SE | 10339 | FIP1L1 | FIP1L1 | chr4 | + | 53414614 | 53414722 | 53399729 | 53399839 | 53425871 | 53425965 | 11 | 12 | 7 | 3 | 0 | 2 | 1 | 0 | 0 | 2 | 98 | 49 | 0.012241851 | 0.540043183 | 0.647 | 1 | 0.636 | 1 | 0 | 0.261 |
| SE | 15687 | MAPKAPK3 | MAPKAPK3 | chr3 | + | 50647122 | 50647203 | 50646739 | 50646825 | 50647893 | 50649297 | 43 | 30 | 7 | 5 | 1 | 0 | 2 | 11 | 0 | 0 | 98 | 49 | 0.012450865 | 0.545102635 | 0.811 | 0.937 | 1 | 1 | 1 | -0.084 |
| SE | 9203 | ACSL1 | ACSL1 | chr4 | - | 184773080 | 184773154 | 184770398 | 184770476 | 184773662 | 184773714 | 318 | 169 | 121 | 11 | 3 | 0 | 2 | 76 | 0 | 0 | 98 | 49 | 0.012737605 | 0.550988099 | 0.935 | 0.966 | 1 | 1 | 1 | -0.033 |
| SE | 14776 | PLAUR | PLAUR | chr19 | - | 43652224 | 43652371 | 43648570 | 43649143 | 43656478 | 43656640 | 23 | 4 | 2 | 1 | 4 | 0 | 2 | 23 | 0 | 0 | 98 | 49 | 0.012877679 | 0.550988099 | 0.92 | 0.333 | 1 | 1 | 1 | -0.249 |
| SE | 12252 | SMCHD1 | SMCHD1 | chr18 | + | 2778168 | 2778239 | 2777805 | 2777915 | 2784449 | 2784621 | 191 | 107 | 39 | 9 | 3 | 3 | 2 | 47 | 0 | 1 | 98 | 49 | 0.012958315 | 0.550988099 | 0.914 | 0.947 | 0.867 | 1 | 0.959 | -0.07 |
| SE | 13224 | FPR1 | FPR1 | chr19 | - | 51750202 | 51750253 | 51745769 | 51747005 | 51751813 | 51751897 | 47 | 21 | 16 | 13 | 7 | 3 | 1 | 2 | 0 | 1 | 98 | 49 | 0.012966669 | 0.550988099 | 0.644 | 0.6 | 0.727 | 1 | 0.5 | -0.093 |
| SE | 7858 | LRRK2 | LRRK2 | chr12 | + | 40305784 | 40305966 | 40303947 | 40304134 | 40308466 | 40308696 | 300 | 164 | 57 | 13 | 1 | 0 | 2 | 32 | 0 | 0 | 98 | 49 | 0.013129747 | 0.553845305 | 0.92 | 0.988 | 1 | 1 | 1 | -0.031 |
| SE | 2764 | PCBP2 | PCBP2 | chr12 | + | 53465931 | 53465973 | 53462480 | 53462567 | 53467220 | 53467293 | 72 | 31 | 6 | 0 | 1 | 0 | 2 | 30 | 1 | 3 | 91 | 49 | 0.013550016 | 0.567431451 | 1 | 0.943 | 1 | 0.519 | 0.843 | 0.3 |
| SE | 2605 | PXN | PXN | chr12 | - | 120215559 | 120215661 | 120215102 | 120215273 | 120221622 | 120221758 | 10 | 6 | 2 | 31 | 12 | 16 | 0 | 1 | 1 | 3 | 98 | 49 | 0.013824713 | 0.574769886 | 0.139 | 0.2 | 0.059 | 0 | 0.143 | 0.061 |
| SE | 6756 | SETD1B | SETD1B | chr12 | + | 121819403 | 121819895 | 121817798 | 121817904 | 121822489 | 121823749 | 8 | 9 | 3 | 0 | 0 | 0 | 3 | 1 | 0 | 3 | 98 | 49 | 0.014021703 | 0.578795883 | 1 | 1 | 1 | 1 | 0.143 | 0.428 |
| SE | 9544 | GMPR2 | GMPR2 | chr14 | + | 24238245 | 24238405 | 24237519 | 24237562 | 24238588 | 24239241 | 27 | 12 | 7 | 3 | 2 | 4 | 0 | 1 | 2 | 0 | 98 | 49 | 0.014211036 | 0.582450894 | 0.818 | 0.75 | 0.467 | 0 | 1 | 0.178 |
| SE | 7719 | CCDC88B | CCDC88B | chr11 | + | 64354003 | 64354170 | 64353714 | 64353813 | 64355193 | 64355400 | 36 | 17 | 6 | 0 | 2 | 0 | 4 | 4 | 1 | 2 | 98 | 49 | 0.014784184 | 0.600287427 | 1 | 0.81 | 1 | 0.667 | 0.5 | 0.353 |
| SE | 8321 | RNF10 | RNF10 | chr12 | + | 120571190 | 120571291 | 120566824 | 120566980 | 120575630 | 120575688 | 35 | 27 | 8 | 4 | 4 | 0 | 3 | 6 | 0 | 1 | 98 | 49 | 0.014853972 | 0.600287427 | 0.814 | 0.771 | 1 | 1 | 0.75 | -0.013 |
| SE | 12967 | DDX60L | DDX60L | chr4 | - | 168448637 | 168448779 | 168441336 | 168441492 | 168453123 | 168453282 | 92 | 48 | 7 | 10 | 0 | 0 | 2 | 53 | 0 | 1 | 98 | 49 | 0.015096044 | 0.605727518 | 0.821 | 1 | 1 | 1 | 0.964 | -0.042 |
| SE | 11622 | PSMB3 | PSMB3 | chr17 | + | 38760430 | 38760608 | 38755882 | 38755990 | 38762410 | 38762505 | 32 | 23 | 41 | 2 | 5 | 2 | 1 | 12 | 0 | 1 | 98 | 49 | 0.015198216 | 0.605727518 | 0.889 | 0.697 | 0.911 | 1 | 0.857 | -0.096 |
| SE | 4214 | ZBTB37 | ZBTB37 | chr1 | + | 173873466 | 173873566 | 173870196 | 173871148 | 173885635 | 173886636 | 15 | 3 | 2 | 5 | 0 | 0 | 1 | 6 | 0 | 0 | 98 | 49 | 0.015703593 | 0.621582645 | 0.6 | 1 | 1 | 1 | 1 | -0.133 |
| A3SS | 2442 | DNM2 | DNM2 | chr19 | + | 10830977 | 10831910 | 10830980 | 10831910 | 10830126 | 10830378 | 4 | 9 | 3 | 4 | 2 | 1 | 2 | 1 | 0 | 1 | 52 | 49 | 0.015869094 | 0.482754544 | 0.485 | 0.809 | 0.739 | 1 | 0.485 | -0.065 |
| RI | 113 | DDAH2 | DDAH2 | chr6 | - | 31728864 | 31730265 | 31728864 | 31729225 | 31730050 | 31730265 | 10 | 3 | 2 | 4 | 3 | 0 | 1 | 0 | 0 | 3 | 98 | 49 | 0.01604331 | 0.294127359 | 0.556 | 0.333 | 1 | 1 | 0 | 0.13 |
| SE | 11055 | THEMIS2 | THEMIS2 | chr1 | + | 27881970 | 27883043 | 27876587 | 27876728 | 27885294 | 27885451 | 25 | 14 | 1 | 3 | 2 | 6 | 0 | 2 | 3 | 1 | 98 | 49 | 0.016045206 | 0.627878418 | 0.806 | 0.778 | 0.077 | 0 | 0.5 | 0.304 |
| SE | 4833 | CLEC7A | CLEC7A | chr12 | - | 10125086 | 10125173 | 10123244 | 10123363 | 10125296 | 10125448 | 29 | 13 | 14 | 50 | 26 | 13 | 1 | 25 | 0 | 8 | 98 | 49 | 0.016142926 | 0.627878418 | 0.225 | 0.2 | 0.35 | 1 | 0.61 | -0.547 |
| SE | 8018 | RNF19B | RNF19B | chr1 | - | 32949568 | 32949774 | 32948221 | 32948363 | 32964050 | 32964813 | 56 | 29 | 8 | 1 | 1 | 0 | 0 | 7 | 2 | 0 | 98 | 49 | 0.016354916 | 0.627878418 | 0.966 | 0.935 | 1 | 0 | 1 | 0.467 |
| SE | 10036 | PRKCD | PRKCD | chr3 | + | 53185926 | 53186027 | 53185603 | 53185700 | 53186166 | 53186340 | 23 | 8 | 11 | 0 | 2 | 0 | 3 | 4 | 0 | 4 | 98 | 49 | 0.016356585 | 0.627878418 | 1 | 0.667 | 1 | 1 | 0.333 | 0.222 |
| SE | 5176 | SLC12A9 | SLC12A9 | chr7 | + | 100861137 | 100861262 | 100860149 | 100860232 | 100861391 | 100861584 | 62 | 24 | 16 | 1 | 2 | 3 | 4 | 10 | 0 | 0 | 98 | 49 | 0.01640589 | 0.627878418 | 0.969 | 0.857 | 0.727 | 1 | 1 | -0.149 |
| A5SS | 1792 | MLKL | MLKL | chr16 | - | 74675323 | 74675404 | 74675354 | 74675404 | 74674959 | 74675100 | 6 | 1 | 6 | 7 | 2 | 0 | 1 | 1 | 5 | 3 | 80 | 49 | 0.016544743 | 0.411255051 | 0.344 | 0.234 | 1 | 0.109 | 0.17 | 0.386 |
| A3SS | 2481 | LST1 | LST1 | chr6 | + | 31588496 | 31588909 | 31588562 | 31588909 | 31587943 | 31587966 | 67 | 45 | 41 | 5 | 2 | 0 | 4 | 10 | 0 | 0 | 98 | 49 | 0.016723378 | 0.482990798 | 0.87 | 0.918 | 1 | 1 | 1 | -0.071 |
| SE | 14578 | LRRFIP1 | LRRFIP1 | chr2 | + | 237751199 | 237751271 | 237749198 | 237749324 | 237753308 | 237753479 | 13 | 10 | 0 | 26 | 29 | 2 | 0 | 2 | 1 | 13 | 98 | 49 | 0.016726002 | 0.635918186 | 0.2 | 0.147 | 0 | 0 | 0.071 | 0.08 |
| SE | 10008 | NT5C2 | NT5C2 | chr10 | - | 103106588 | 103106706 | 103105705 | 103105801 | 103139405 | 103139479 | 62 | 22 | 12 | 3 | 3 | 0 | 1 | 7 | 0 | 0 | 98 | 49 | 0.017233157 | 0.650917757 | 0.912 | 0.786 | 1 | 1 | 1 | -0.101 |
| SE | 5342 | SF3B1 | SF3B1 | chr2 | - | 197392971 | 197393188 | 197391973 | 197392461 | 197396055 | 197396328 | 221 | 87 | 41 | 2 | 6 | 2 | 5 | 39 | 0 | 0 | 98 | 49 | 0.017359502 | 0.651432222 | 0.982 | 0.879 | 0.911 | 1 | 1 | -0.076 |
| A3SS | 2976 | MKNK2 | MKNK2 | chr19 | - | 2037470 | 2039856 | 2037470 | 2037829 | 2040133 | 2040177 | 127 | 48 | 42 | 2 | 2 | 4 | 6 | 15 | 0 | 0 | 98 | 49 | 0.017548109 | 0.482990798 | 0.969 | 0.923 | 0.84 | 1 | 1 | -0.089 |
| SE | 15943 | CSF2RA | CSF2RA | chrX | + | 1303922 | 1304019 | 1300490 | 1300626 | 1305445 | 1305527 | 20 | 7 | 17 | 8 | 0 | 0 | 1 | 11 | 0 | 1 | 98 | 49 | 0.017694655 | 0.659725219 | 0.556 | 1 | 1 | 1 | 0.846 | -0.071 |
| SE | 12824 | GMIP | GMIP | chr19 | - | 19635643 | 19635721 | 19635414 | 19635569 | 19636706 | 19636796 | 60 | 37 | 3 | 6 | 2 | 0 | 9 | 8 | 1 | 0 | 98 | 49 | 0.017993748 | 0.666576081 | 0.833 | 0.902 | 1 | 0.818 | 1 | 0.003 |
| SE | 14605 | HDLBP | HDLBP | chr2 | - | 241255373 | 241255580 | 241255050 | 241255158 | 241256183 | 241256399 | 29 | 5 | 1 | 3 | 2 | 0 | 1 | 1 | 0 | 0 | 98 | 49 | 0.018618823 | 0.685338728 | 0.829 | 0.556 | 1 | 1 | 1 | -0.205 |
| SE | 615 | TLN1 | TLN1 | chr9 | - | 35722860 | 35722921 | 35722118 | 35722223 | 35723951 | 35724079 | 103 | 87 | 32 | 4 | 4 | 0 | 3 | 26 | 0 | 0 | 98 | 49 | 0.019607496 | 0.717162791 | 0.928 | 0.916 | 1 | 1 | 1 | -0.052 |
| SE | 17072 | GRK6 | GRK6 | chr5 | + | 177431994 | 177432107 | 177430871 | 177430967 | 177432232 | 177432310 | 62 | 21 | 28 | 7 | 0 | 0 | 7 | 9 | 0 | 1 | 98 | 49 | 0.019813105 | 0.720125358 | 0.816 | 1 | 1 | 1 | 0.818 | 0.03 |
| SE | 9108 | MAN2C1 | MAN2C1 | chr15 | - | 75363998 | 75364188 | 75362641 | 75362748 | 75364487 | 75364665 | 10 | 0 | 3 | 0 | 1 | 0 | 2 | 0 | 0 | 2 | 98 | 49 | 0.020428857 | 0.737174147 | 1 | 0 | 1 | 1 | 0 | 0.167 |
| SE | 5207 | PTPRJ | PTPRJ | chr11 | + | 48121002 | 48121266 | 48112746 | 48112983 | 48123612 | 48123870 | 98 | 47 | 29 | 7 | 3 | 0 | 2 | 25 | 0 | 1 | 98 | 49 | 0.020572324 | 0.737174147 | 0.875 | 0.887 | 1 | 1 | 0.926 | -0.042 |
| SE | 7351 | FAM157A | FAM157A | chr3 | + | 198169784 | 198169872 | 198167675 | 198167871 | 198180754 | 198180857 | 63 | 20 | 3 | 5 | 2 | 9 | 4 | 48 | 0 | 3 | 98 | 49 | 0.020664858 | 0.737174147 | 0.863 | 0.833 | 0.143 | 1 | 0.889 | -0.332 |
| SE | 16115 | SIRPA | SIRPA | chr20 | + | 1921394 | 1921712 | 1915098 | 1915455 | 1924763 | 1924877 | 7 | 11 | 1 | 4 | 5 | 0 | 1 | 0 | 0 | 1 | 98 | 49 | 0.020848898 | 0.739176561 | 0.467 | 0.524 | 1 | 1 | 0 | 0.164 |
| A3SS | 1722 | MED11 | MED11 | chr17 | + | 4732938 | 4733610 | 4733049 | 4733610 | 4731775 | 4731906 | 12 | 8 | 4 | 5 | 0 | 0 | 1 | 4 | 0 | 0 | 98 | 49 | 0.020946769 | 0.550328744 | 0.545 | 1 | 1 | 1 | 1 | -0.152 |
| SE | 5052 | ST3GAL4 | ST3GAL4 | chr11 | + | 126409267 | 126409411 | 126408306 | 126408496 | 126413504 | 126413648 | 12 | 9 | 4 | 0 | 0 | 0 | 0 | 3 | 2 | 0 | 98 | 49 | 0.021635118 | 0.762374053 | 1 | 1 | 1 | 0 | 1 | 0.5 |
| SE | 11444 | USP22 | USP22 | chr17 | - | 21019083 | 21019185 | 21017941 | 21018111 | 21021112 | 21021226 | 20 | 8 | 3 | 0 | 0 | 6 | 1 | 14 | 0 | 0 | 98 | 49 | 0.022240574 | 0.777124869 | 1 | 1 | 0.2 | 1 | 1 | -0.267 |
| SE | 4936 | TNFRSF1A | TNFRSF1A | chr12 | - | 6334090 | 6334244 | 6333736 | 6333865 | 6341775 | 6342117 | 130 | 53 | 58 | 2 | 4 | 0 | 2 | 17 | 3 | 1 | 98 | 49 | 0.022373377 | 0.777124869 | 0.97 | 0.869 | 1 | 0.25 | 0.895 | 0.374 |
| SE | 16935 | IRF1 | IRF1 | chr5 | - | 132486250 | 132486373 | 132485666 | 132485716 | 132486556 | 132486686 | 113 | 50 | 55 | 8 | 2 | 0 | 2 | 36 | 0 | 1 | 98 | 49 | 0.022457147 | 0.777124869 | 0.876 | 0.926 | 1 | 1 | 0.947 | -0.039 |
| SE | 14480 | BAZ2A | BAZ2A | chr12 | - | 56605829 | 56606063 | 56605072 | 56605327 | 56606246 | 56606312 | 37 | 4 | 10 | 2 | 3 | 0 | 2 | 2 | 0 | 0 | 98 | 49 | 0.022681196 | 0.779565992 | 0.902 | 0.4 | 1 | 1 | 1 | -0.233 |
| SE | 11938 | MAP7D1 | MAP7D1 | chr1 | + | 36174897 | 36175008 | 36173363 | 36173478 | 36176198 | 36176321 | 24 | 17 | 3 | 8 | 1 | 1 | 0 | 4 | 2 | 2 | 98 | 49 | 0.022797483 | 0.779565992 | 0.6 | 0.895 | 0.6 | 0 | 0.5 | 0.448 |
| SE | 6610 | AKAP13 | AKAP13 | chr15 | + | 85658536 | 85658590 | 85655416 | 85655787 | 85664562 | 85664755 | 80 | 39 | 4 | 3 | 5 | 0 | 2 | 13 | 0 | 1 | 98 | 49 | 0.023450224 | 0.789226928 | 0.93 | 0.796 | 1 | 1 | 0.867 | -0.025 |
| SE | 4907 | GABARAPL1 | GABARAPL1 | chr12 | + | 10218062 | 10218141 | 10212889 | 10213219 | 10220439 | 10220558 | 45 | 31 | 26 | 2 | 0 | 0 | 0 | 8 | 2 | 0 | 98 | 49 | 0.023507475 | 0.789226928 | 0.918 | 1 | 1 | 0 | 1 | 0.473 |
| SE | 3726 | ADGRG3 | ADGRG3 | chr16 | + | 57684389 | 57684483 | 57680504 | 57680617 | 57685642 | 57685926 | 42 | 10 | 20 | 2 | 0 | 0 | 0 | 7 | 2 | 0 | 98 | 49 | 0.023598824 | 0.789226928 | 0.913 | 1 | 1 | 0 | 1 | 0.471 |
| SE | 1534 | WIPI2 | WIPI2 | chr7 | + | 5193117 | 5193345 | 5190203 | 5190493 | 5199575 | 5199658 | 3 | 0 | 0 | 10 | 7 | 2 | 1 | 1 | 0 | 1 | 98 | 49 | 0.023699285 | 0.789226928 | 0.13 | 0 | 0 | 1 | 0.333 | -0.623 |
| SE | 15851 | SYNE2 | SYNE2 | chr14 | + | 64029894 | 64030059 | 64027483 | 64027793 | 64031015 | 64031357 | 33 | 20 | 7 | 2 | 0 | 0 | 0 | 5 | 2 | 0 | 98 | 49 | 0.023762846 | 0.789226928 | 0.892 | 1 | 1 | 0 | 1 | 0.464 |
| A3SS | 4045 | SDCBP | SDCBP | chr8 | + | 58575899 | 58576061 | 58575902 | 58576061 | 58572204 | 58572314 | 53 | 62 | 38 | 10 | 10 | 8 | 3 | 20 | 0 | 2 | 52 | 49 | 0.024257968 | 0.609613283 | 0.833 | 0.854 | 0.817 | 1 | 0.904 | -0.117 |
| SE | 7851 | LRRK2 | LRRK2 | chr12 | + | 40298242 | 40298493 | 40295426 | 40295644 | 40299108 | 40299257 | 407 | 104 | 51 | 7 | 5 | 0 | 3 | 61 | 0 | 0 | 98 | 49 | 0.024357869 | 0.804366435 | 0.967 | 0.912 | 1 | 1 | 1 | -0.04 |
| A5SS | 2848 | TLE3 | TLE3 | chr15 | - | 70058158 | 70058291 | 70058185 | 70058291 | 70057458 | 70057658 | 31 | 10 | 15 | 5 | 2 | 1 | 1 | 4 | 0 | 1 | 76 | 49 | 0.024599774 | 0.570714761 | 0.8 | 0.763 | 0.906 | 1 | 0.721 | -0.037 |
| SE | 15838 | AOAH | AOAH | chr7 | - | 36686698 | 36686794 | 36673942 | 36674009 | 36724021 | 36724549 | 64 | 16 | 5 | 7 | 0 | 0 | 5 | 37 | 0 | 1 | 98 | 49 | 0.026174187 | 0.856138307 | 0.821 | 1 | 1 | 1 | 0.949 | -0.034 |
| SE | 7209 | NBEAL2 | NBEAL2 | chr3 | + | 47007065 | 47007155 | 47006332 | 47006449 | 47007240 | 47007350 | 85 | 26 | 16 | 1 | 1 | 4 | 2 | 16 | 0 | 0 | 98 | 49 | 0.026221921 | 0.856138307 | 0.977 | 0.929 | 0.667 | 1 | 1 | -0.142 |
| A3SS | 658 | PXN | PXN | chr12 | - | 120215102 | 120215417 | 120215102 | 120215273 | 120215559 | 120215661 | 20 | 9 | 4 | 3 | 1 | 1 | 1 | 7 | 0 | 1 | 98 | 49 | 0.026774513 | 0.644819526 | 0.769 | 0.818 | 0.667 | 1 | 0.778 | -0.138 |
| SE | 3034 | XPO6 | XPO6 | chr16 | - | 28180940 | 28181031 | 28177219 | 28177332 | 28211365 | 28211918 | 338 | 139 | 60 | 2 | 9 | 0 | 3 | 55 | 0 | 0 | 98 | 49 | 0.027362825 | 0.888369463 | 0.988 | 0.885 | 1 | 1 | 1 | -0.042 |
| A5SS | 1247 | CASS4 | CASS4 | chr20 | + | 56437163 | 56437586 | 56437163 | 56437424 | 56445899 | 56446001 | 44 | 9 | 8 | 1 | 1 | 3 | 2 | 10 | 0 | 0 | 98 | 49 | 0.027870677 | 0.606187224 | 0.957 | 0.818 | 0.571 | 1 | 1 | -0.218 |
| SE | 16895 | VASP | VASP | chr19 | + | 45521321 | 45521406 | 45517928 | 45518094 | 45522167 | 45522217 | 181 | 71 | 83 | 5 | 2 | 0 | 2 | 37 | 2 | 3 | 98 | 49 | 0.027956523 | 0.902539602 | 0.948 | 0.947 | 1 | 0.333 | 0.86 | 0.368 |
| SE | 12194 | SP110 | SP110 | chr2 | - | 230178156 | 230178255 | 230176472 | 230176724 | 230183571 | 230183640 | 34 | 16 | 10 | 0 | 1 | 4 | 1 | 8 | 0 | 0 | 98 | 49 | 0.028111633 | 0.902539602 | 1 | 0.889 | 0.556 | 1 | 1 | -0.185 |
| SE | 10325 | IRF7 | IRF7 | chr11 | - | 614173 | 614399 | 613950 | 614037 | 614475 | 614534 | 3 | 3 | 7 | 8 | 8 | 0 | 1 | 2 | 7 | 10 | 98 | 49 | 0.029017678 | 0.926481559 | 0.158 | 0.158 | 1 | 0.067 | 0.091 | 0.36 |
| SE | 11878 | SPAG9 | SPAG9 | chr17 | - | 50974770 | 50974947 | 50970706 | 50970856 | 50977107 | 50977221 | 35 | 17 | 6 | 3 | 2 | 0 | 1 | 3 | 0 | 0 | 98 | 49 | 0.030137537 | 0.956949587 | 0.854 | 0.81 | 1 | 1 | 1 | -0.112 |
| SE | 389 | RAPGEF1 | RAPGEF1 | chr9 | - | 131650128 | 131650242 | 131643238 | 131643417 | 131650809 | 131650949 | 18 | 3 | 1 | 4 | 2 | 0 | 2 | 1 | 0 | 1 | 98 | 49 | 0.030356618 | 0.958638776 | 0.692 | 0.429 | 1 | 1 | 0.333 | 0.04 |
| SE | 8171 | PECAM1 | PECAM1 | chr17 | - | 64354932 | 64355040 | 64353490 | 64353518 | 64356110 | 64356398 | 174 | 76 | 71 | 9 | 3 | 0 | 1 | 38 | 0 | 8 | 98 | 49 | 0.031092019 | 0.96633516 | 0.906 | 0.927 | 1 | 1 | 0.704 | 0.092 |
| SE | 3127 | CMIP | CMIP | chr16 | + | 81664268 | 81664349 | 81660883 | 81660946 | 81670141 | 81670245 | 27 | 11 | 4 | 0 | 0 | 0 | 0 | 5 | 2 | 0 | 98 | 49 | 0.031099492 | 0.96633516 | 1 | 1 | 1 | 0 | 1 | 0.5 |
| SE | 11704 | CASC3 | CASC3 | chr17 | + | 40167849 | 40167948 | 40167497 | 40167612 | 40168202 | 40168417 | 28 | 11 | 7 | 0 | 0 | 0 | 0 | 5 | 2 | 0 | 98 | 49 | 0.03110198 | 0.96633516 | 1 | 1 | 1 | 0 | 1 | 0.5 |
| A3SS | 3526 | SNRPB | SNRPB | chr20 | - | 2461634 | 2461939 | 2461634 | 2461793 | 2462635 | 2462761 | 22 | 10 | 1 | 7 | 1 | 0 | 0 | 3 | 1 | 2 | 98 | 49 | 0.031364389 | 0.725144665 | 0.611 | 0.833 | 1 | 0 | 0.429 | 0.6 |
| SE | 2485 | ARRB2 | ARRB2 | chr17 | + | 4720392 | 4720472 | 4720211 | 4720299 | 4720585 | 4720640 | 118 | 56 | 50 | 3 | 13 | 0 | 2 | 26 | 1 | 0 | 98 | 49 | 0.031561151 | 0.97535771 | 0.952 | 0.683 | 1 | 0.5 | 1 | 0.128 |
| SE | 4342 | ITPK1 | ITPK1 | chr14 | - | 93076594 | 93076619 | 93016675 | 93016801 | 93115068 | 93115184 | 55 | 27 | 12 | 4 | 2 | 0 | 4 | 6 | 0 | 0 | 74 | 49 | 0.032101744 | 0.984878326 | 0.901 | 0.899 | 1 | 1 | 1 | -0.067 |
| A5SS | 2353 | TYROBP | TYROBP | chr19 | - | 35908167 | 35908309 | 35908174 | 35908309 | 35907729 | 35907762 | 71 | 31 | 15 | 8 | 2 | 0 | 1 | 10 | 0 | 1 | 56 | 49 | 0.032118554 | 0.657485692 | 0.886 | 0.931 | 1 | 1 | 0.897 | -0.009 |
| SE | 10543 | ATF7IP | ATF7IP | chr12 | + | 14457206 | 14457295 | 14456560 | 14456634 | 14460494 | 14461133 | 27 | 8 | 3 | 0 | 0 | 5 | 1 | 3 | 0 | 0 | 98 | 49 | 0.032327505 | 0.984878326 | 1 | 1 | 0.231 | 1 | 1 | -0.256 |
| SE | 8341 | CASP8 | CASP8 | chr2 | + | 201272897 | 201272942 | 201272637 | 201272776 | 201274888 | 201274953 | 163 | 69 | 16 | 5 | 2 | 1 | 1 | 6 | 0 | 0 | 94 | 49 | 0.032380495 | 0.984878326 | 0.944 | 0.947 | 0.893 | 1 | 1 | -0.072 |
| A3SS | 2482 | LST1 | LST1 | chr6 | + | 31588517 | 31588909 | 31588562 | 31588909 | 31587943 | 31587966 | 78 | 54 | 55 | 5 | 2 | 0 | 3 | 16 | 0 | 0 | 94 | 49 | 0.032814021 | 0.729480928 | 0.89 | 0.934 | 1 | 1 | 1 | -0.059 |
| SE | 12922 | TYROBP | TYROBP | chr19 | - | 35907729 | 35907762 | 35907445 | 35907580 | 35908167 | 35908309 | 95 | 33 | 18 | 2 | 3 | 0 | 4 | 15 | 0 | 5 | 82 | 49 | 0.035068864 | 1 | 0.966 | 0.868 | 1 | 1 | 0.642 | 0.124 |
| SE | 5209 | PTPRJ | PTPRJ | chr11 | + | 48146875 | 48146963 | 48144999 | 48145124 | 48149446 | 48149488 | 36 | 5 | 11 | 0 | 5 | 0 | 6 | 5 | 0 | 0 | 98 | 49 | 0.035659347 | 1 | 1 | 0.333 | 1 | 1 | 1 | -0.222 |
| SE | 5658 | WDFY3 | WDFY3 | chr4 | - | 84741760 | 84741921 | 84740186 | 84740416 | 84743699 | 84743799 | 44 | 17 | 5 | 4 | 1 | 0 | 3 | 11 | 0 | 0 | 98 | 49 | 0.036395704 | 1 | 0.846 | 0.895 | 1 | 1 | 1 | -0.086 |
| A3SS | 710 | TMBIM4 | TMBIM4 | chr12 | - | 66153339 | 66153448 | 66153339 | 66153410 | 66169854 | 66170072 | 20 | 11 | 18 | 1 | 0 | 0 | 0 | 4 | 2 | 0 | 87 | 49 | 0.036512779 | 0.781643942 | 0.918 | 1 | 1 | 0 | 1 | 0.473 |
| A5SS | 1620 | RBM10 | RBM10 | chrX | + | 47145217 | 47145516 | 47145217 | 47145485 | 47147356 | 47147498 | 5 | 4 | 2 | 9 | 1 | 2 | 0 | 2 | 2 | 3 | 80 | 49 | 0.037869553 | 0.728051467 | 0.254 | 0.71 | 0.38 | 0 | 0.29 | 0.303 |
| SE | 8098 | DEDD2 | DEDD2 | chr19 | - | 42201988 | 42202072 | 42198592 | 42199829 | 42209699 | 42209840 | 2 | 0 | 1 | 16 | 25 | 11 | 1 | 2 | 3 | 5 | 98 | 49 | 0.037915884 | 1 | 0.059 | 0 | 0.043 | 0.143 | 0.167 | -0.121 |
| SE | 3160 | CTSA | CTSA | chr20 | + | 45893219 | 45893311 | 45892724 | 45892880 | 45893987 | 45894072 | 29 | 8 | 6 | 0 | 6 | 1 | 1 | 14 | 1 | 0 | 98 | 49 | 0.0382167 | 1 | 1 | 0.4 | 0.75 | 0.333 | 1 | 0.05 |
| SE | 15547 | ATG16L2 | ATG16L2 | chr11 | + | 72827187 | 72827293 | 72826702 | 72826823 | 72828358 | 72828508 | 132 | 85 | 43 | 7 | 2 | 1 | 1 | 41 | 0 | 1 | 98 | 49 | 0.038578466 | 1 | 0.904 | 0.955 | 0.956 | 1 | 0.953 | -0.038 |
| SE | 6790 | JMJD1C | JMJD1C | chr10 | - | 63264650 | 63264764 | 63217206 | 63217331 | 63268662 | 63269223 | 152 | 34 | 22 | 6 | 1 | 0 | 1 | 15 | 0 | 0 | 98 | 49 | 0.039711108 | 1 | 0.927 | 0.944 | 1 | 1 | 1 | -0.043 |
| SE | 11096 | ARHGAP15 | ARHGAP15 | chr2 | + | 143436912 | 143437042 | 143435600 | 143435699 | 143487372 | 143487495 | 197 | 49 | 30 | 0 | 0 | 0 | 0 | 19 | 2 | 0 | 98 | 49 | 0.039732904 | 1 | 1 | 1 | 1 | 0 | 1 | 0.5 |
| A5SS | 1739 | ADPGK | ADPGK | chr15 | - | 72783245 | 72783785 | 72783458 | 72783785 | 72774871 | 72775097 | 12 | 11 | 11 | 4 | 0 | 1 | 3 | 7 | 0 | 1 | 98 | 49 | 0.039749936 | 0.728051467 | 0.6 | 1 | 0.846 | 1 | 0.778 | -0.074 |
| SE | 11726 | BECN1 | BECN1 | chr17 | - | 42815907 | 42816054 | 42814523 | 42814673 | 42818220 | 42818415 | 14 | 14 | 6 | 0 | 0 | 0 | 2 | 7 | 0 | 3 | 98 | 49 | 0.040231039 | 1 | 1 | 1 | 1 | 1 | 0.538 | 0.231 |
| A3SS | 2480 | LST1 | LST1 | chr6 | + | 31588496 | 31588909 | 31588517 | 31588909 | 31587943 | 31587966 | 21 | 3 | 5 | 16 | 11 | 17 | 1 | 0 | 0 | 6 | 70 | 49 | 0.041875868 | 0.864437565 | 0.479 | 0.16 | 0.171 | 1 | 0 | -0.23 |
| SE | 5136 | MYL6 | MYL6 | chr12 | + | 56160625 | 56160670 | 56160242 | 56160320 | 56161386 | 56161582 | 10 | 3 | 2 | 45 | 57 | 61 | 0 | 1 | 3 | 15 | 94 | 49 | 0.041972875 | 1 | 0.104 | 0.027 | 0.017 | 0 | 0.034 | 0.032 |
| SE | 8284 | PLK3 | PLK3 | chr1 | + | 44801621 | 44801751 | 44801035 | 44801152 | 44801844 | 44801932 | 35 | 29 | 7 | 2 | 3 | 0 | 2 | 6 | 0 | 0 | 98 | 49 | 0.041980644 | 1 | 0.897 | 0.829 | 1 | 1 | 1 | -0.091 |
| SE | 14777 | PLAUR | PLAUR | chr19 | - | 43655438 | 43655573 | 43648566 | 43649143 | 43656478 | 43656640 | 35 | 17 | 4 | 1 | 4 | 0 | 8 | 20 | 0 | 0 | 98 | 49 | 0.04219046 | 1 | 0.946 | 0.68 | 1 | 1 | 1 | -0.125 |
| SE | 7861 | LRRK2 | LRRK2 | chr12 | + | 40313971 | 40314173 | 40310430 | 40310649 | 40315211 | 40315300 | 329 | 88 | 41 | 8 | 1 | 4 | 3 | 44 | 0 | 1 | 98 | 49 | 0.042582705 | 1 | 0.954 | 0.978 | 0.837 | 1 | 0.957 | -0.055 |
| SE | 16936 | IRF1 | IRF1 | chr5 | - | 132486794 | 132486844 | 132486556 | 132486686 | 132486953 | 132487130 | 173 | 106 | 45 | 11 | 1 | 0 | 2 | 21 | 0 | 1 | 98 | 49 | 0.042607206 | 1 | 0.887 | 0.981 | 1 | 1 | 0.913 | -0.001 |
| SE | 2762 | PCBP2 | PCBP2 | chr12 | + | 53464759 | 53464852 | 53462480 | 53462567 | 53465931 | 53465973 | 98 | 27 | 22 | 27 | 13 | 4 | 0 | 23 | 2 | 16 | 98 | 49 | 0.042684862 | 1 | 0.645 | 0.509 | 0.733 | 0 | 0.418 | 0.42 |
| SE | 9205 | ACSL1 | ACSL1 | chr4 | - | 184776483 | 184776662 | 184773842 | 184773875 | 184776883 | 184776983 | 213 | 52 | 116 | 7 | 5 | 0 | 3 | 33 | 0 | 1 | 98 | 49 | 0.043517605 | 1 | 0.938 | 0.839 | 1 | 1 | 0.943 | -0.046 |
| RI | 292 | DDIT3 | DDIT3 | chr12 | - | 57517268 | 57517753 | 57517268 | 57517438 | 57517705 | 57517753 | 91 | 57 | 8 | 5 | 4 | 0 | 2 | 7 | 0 | 1 | 98 | 49 | 0.044574092 | 0.735472513 | 0.901 | 0.877 | 1 | 1 | 0.778 | 0.037 |
| SE | 7558 | RFWD2 | RFWD2 | chr1 | - | 176175909 | 176176007 | 176163814 | 176163891 | 176184632 | 176184692 | 106 | 47 | 14 | 3 | 2 | 1 | 2 | 13 | 0 | 0 | 98 | 49 | 0.044718214 | 1 | 0.946 | 0.922 | 0.875 | 1 | 1 | -0.086 |
| SE | 1246 | LRCH4 | LRCH4 | chr7 | - | 100577085 | 100577154 | 100576901 | 100577005 | 100577272 | 100577389 | 34 | 27 | 13 | 2 | 3 | 0 | 4 | 5 | 0 | 0 | 98 | 49 | 0.045883646 | 1 | 0.895 | 0.818 | 1 | 1 | 1 | -0.096 |
| SE | 8398 | ATP2A3 | ATP2A3 | chr17 | - | 3953346 | 3953429 | 3951580 | 3951685 | 3953692 | 3953710 | 37 | 21 | 14 | 2 | 3 | 0 | 2 | 4 | 0 | 0 | 98 | 49 | 0.046992497 | 1 | 0.902 | 0.778 | 1 | 1 | 1 | -0.107 |
| SE | 12877 | LSM14A | LSM14A | chr19 | + | 34215123 | 34215300 | 34208928 | 34209051 | 34215595 | 34215661 | 31 | 12 | 8 | 5 | 0 | 0 | 1 | 7 | 0 | 0 | 98 | 49 | 0.047526789 | 1 | 0.756 | 1 | 1 | 1 | 1 | -0.081 |
| SE | 15922 | RPS6KA1 | RPS6KA1 | chr1 | + | 26555536 | 26555625 | 26555150 | 26555221 | 26556653 | 26556718 | 106 | 32 | 25 | 1 | 4 | 0 | 0 | 27 | 1 | 4 | 98 | 49 | 0.049990346 | 1 | 0.981 | 0.8 | 1 | 0 | 0.771 | 0.542 |
| SE | 5960 | SORL1 | SORL1 | chr11 | + | 121570156 | 121570270 | 121566939 | 121567113 | 121574240 | 121574363 | 515 | 176 | 96 | 23 | 8 | 4 | 0 | 56 | 1 | 12 | 98 | 49 | 0.050255826 | 1 | 0.918 | 0.917 | 0.923 | 0 | 0.7 | 0.569 |
| SE | 12750 | CYP4F3 | CYP4F3 | chr19 | + | 15645718 | 15645863 | 15641414 | 15641613 | 15647051 | 15647105 | 161 | 45 | 36 | 5 | 2 | 0 | 5 | 21 | 0 | 0 | 98 | 49 | 0.050941161 | 1 | 0.942 | 0.918 | 1 | 1 | 1 | -0.047 |
| SE | 11819 | PTPN6 | PTPN6 | chr12 | + | 6960805 | 6960945 | 6960343 | 6960435 | 6961125 | 6961316 | 199 | 83 | 40 | 8 | 3 | 0 | 1 | 28 | 0 | 1 | 98 | 49 | 0.05118402 | 1 | 0.926 | 0.933 | 1 | 1 | 0.933 | -0.014 |
| SE | 5411 | NCF2 | NCF2 | chr1 | - | 183567203 | 183567345 | 183565703 | 183565779 | 183569141 | 183569185 | 111 | 80 | 66 | 3 | 3 | 1 | 1 | 25 | 0 | 0 | 98 | 49 | 0.051634206 | 1 | 0.949 | 0.93 | 0.971 | 1 | 1 | -0.05 |
| SE | 5375 | MGEA5 | MGEA5 | chr10 | - | 101793912 | 101793998 | 101792838 | 101792943 | 101797979 | 101798154 | 19 | 7 | 7 | 8 | 3 | 0 | 0 | 3 | 2 | 1 | 98 | 49 | 0.052864731 | 1 | 0.543 | 0.538 | 1 | 0 | 0.6 | 0.394 |
| SE | 7210 | NBEAL2 | NBEAL2 | chr3 | + | 47007240 | 47007350 | 47006332 | 47006449 | 47007524 | 47007697 | 24 | 9 | 9 | 4 | 2 | 0 | 1 | 7 | 1 | 0 | 98 | 49 | 0.052977721 | 1 | 0.75 | 0.692 | 1 | 0.333 | 1 | 0.148 |
| SE | 1938 | CNPY3 | CNPY3 | chr6 | + | 42934474 | 42934598 | 42929121 | 42929721 | 42935573 | 42935670 | 37 | 18 | 18 | 1 | 4 | 0 | 3 | 10 | 0 | 0 | 98 | 49 | 0.053886737 | 1 | 0.949 | 0.692 | 1 | 1 | 1 | -0.12 |
| SE | 52 | IDS | IDS | chrX | - | 149503311 | 149503489 | 149500948 | 149501037 | 149504156 | 149504293 | 31 | 20 | 4 | 5 | 0 | 0 | 4 | 7 | 0 | 0 | 98 | 49 | 0.054213286 | 1 | 0.756 | 1 | 1 | 1 | 1 | -0.081 |
| SE | 1264 | ZNF394 | ZNF394 | chr7 | - | 99498715 | 99498842 | 99493230 | 99494631 | 99499637 | 99500254 | 14 | 22 | 1 | 2 | 0 | 2 | 2 | 5 | 0 | 0 | 98 | 49 | 0.054854194 | 1 | 0.778 | 1 | 0.2 | 1 | 1 | -0.341 |
| SE | 8832 | DOCK2 | DOCK2 | chr5 | + | 169759704 | 169759775 | 169747395 | 169747504 | 169761518 | 169761625 | 103 | 48 | 17 | 3 | 3 | 0 | 3 | 30 | 0 | 0 | 98 | 49 | 0.054985887 | 1 | 0.945 | 0.889 | 1 | 1 | 1 | -0.055 |
| SE | 15461 | RAF1 | RAF1 | chr3 | - | 12611949 | 12612062 | 12609232 | 12609335 | 12618514 | 12618747 | 159 | 54 | 30 | 3 | 0 | 4 | 3 | 15 | 0 | 0 | 98 | 49 | 0.055208514 | 1 | 0.964 | 1 | 0.789 | 1 | 1 | -0.082 |
| SE | 10771 | RILPL2 | RILPL2 | chr12 | - | 123430507 | 123430659 | 123423043 | 123423157 | 123436081 | 123436733 | 25 | 15 | 11 | 4 | 4 | 3 | 1 | 12 | 4 | 4 | 98 | 49 | 0.055941552 | 1 | 0.758 | 0.652 | 0.647 | 0.111 | 0.6 | 0.33 |
| SE | 51 | IDS | IDS | chrX | - | 149503311 | 149503445 | 149500948 | 149501037 | 149504156 | 149504293 | 34 | 20 | 4 | 5 | 0 | 0 | 4 | 7 | 0 | 0 | 98 | 49 | 0.056974615 | 1 | 0.773 | 1 | 1 | 1 | 1 | -0.076 |
| A3SS | 1540 | FLII | FLII | chr17 | - | 18251679 | 18251816 | 18251679 | 18251813 | 18251998 | 18252146 | 8 | 2 | 6 | 18 | 8 | 8 | 2 | 6 | 2 | 3 | 52 | 49 | 0.058311108 | 1 | 0.295 | 0.191 | 0.414 | 0.485 | 0.653 | -0.269 |
| SE | 5445 | BCL6 | BCL6 | chr3 | - | 187726730 | 187726898 | 187725498 | 187725629 | 187728359 | 187728544 | 72 | 29 | 20 | 15 | 5 | 0 | 1 | 25 | 0 | 2 | 98 | 49 | 0.060336665 | 1 | 0.706 | 0.744 | 1 | 1 | 0.862 | -0.114 |
| SE | 10383 | PIK3R5 | PIK3R5 | chr17 | - | 8889972 | 8890126 | 8889138 | 8889222 | 8890737 | 8890912 | 39 | 10 | 7 | 4 | 0 | 1 | 5 | 10 | 0 | 1 | 98 | 49 | 0.060499246 | 1 | 0.83 | 1 | 0.778 | 1 | 0.833 | -0.047 |
| SE | 9973 | DUSP1 | DUSP1 | chr5 | - | 172770160 | 172770306 | 172769574 | 172769794 | 172770585 | 172771200 | 974 | 462 | 420 | 12 | 6 | 3 | 105 | 424 | 1 | 1 | 98 | 49 | 0.061055612 | 1 | 0.976 | 0.975 | 0.986 | 0.981 | 0.995 | -0.009 |
| SE | 9605 | GLT1D1 | GLT1D1 | chr12 | + | 128875913 | 128876062 | 128853535 | 128853649 | 128888638 | 128888744 | 58 | 26 | 14 | 4 | 1 | 0 | 3 | 18 | 0 | 0 | 98 | 49 | 0.061072609 | 1 | 0.879 | 0.929 | 1 | 1 | 1 | -0.064 |
| SE | 9384 | AMICA1 | AMICA1 | chr11 | - | 118205881 | 118205991 | 118203427 | 118203665 | 118210486 | 118210712 | 265 | 109 | 117 | 7 | 3 | 0 | 1 | 9 | 0 | 0 | 98 | 49 | 0.061322729 | 1 | 0.95 | 0.948 | 1 | 1 | 1 | -0.034 |
| SE | 6836 | GAK | GAK | chr4 | - | 877614 | 877809 | 877089 | 877207 | 881906 | 882040 | 8 | 7 | 5 | 1 | 3 | 0 | 2 | 6 | 0 | 0 | 98 | 49 | 0.061544568 | 1 | 0.8 | 0.538 | 1 | 1 | 1 | -0.221 |
| RI | 637 | NXF1 | NXF1 | chr11 | - | 62798538 | 62800486 | 62798538 | 62798575 | 62800376 | 62800486 | 28 | 8 | 2 | 7 | 1 | 0 | 0 | 4 | 1 | 2 | 98 | 49 | 0.061822068 | 0.927331017 | 0.667 | 0.8 | 1 | 0 | 0.5 | 0.572 |
| A3SS | 2903 | TYMP | TYMP | chr22 | - | 50526000 | 50526156 | 50526000 | 50526141 | 50526245 | 50526476 | 4 | 5 | 0 | 13 | 14 | 13 | 2 | 0 | 5 | 13 | 64 | 49 | 0.062039036 | 1 | 0.191 | 0.215 | 0 | 0.234 | 0 | 0.018 |
| SE | 7830 | CD44 | CD44 | chr11 | + | 35180273 | 35180407 | 35176574 | 35176740 | 35186831 | 35186900 | 58 | 6 | 1 | 0 | 0 | 0 | 4 | 16 | 2 | 1 | 98 | 49 | 0.0624059 | 1 | 1 | 1 | 1 | 0.5 | 0.889 | 0.305 |
| SE | 10823 | HIF1A | HIF1A | chr14 | + | 61740754 | 61741188 | 61738086 | 61738373 | 61744704 | 61744813 | 35 | 38 | 6 | 0 | 0 | 0 | 1 | 13 | 0 | 3 | 98 | 49 | 0.063816629 | 1 | 1 | 1 | 1 | 1 | 0.684 | 0.158 |
| A3SS | 4076 | CNBP | CNBP | chr3 | - | 129171078 | 129171280 | 129171078 | 129171277 | 129171445 | 129171538 | 18 | 9 | 9 | 22 | 19 | 10 | 0 | 2 | 1 | 4 | 52 | 49 | 0.066637907 | 1 | 0.435 | 0.309 | 0.459 | 0 | 0.32 | 0.241 |
| A3SS | 1970 | FCGR3B | FCGR3B | chr1 | - | 161629777 | 161630035 | 161629777 | 161630032 | 161630367 | 161630388 | 57 | 9 | 5 | 6 | 0 | 0 | 3 | 1 | 0 | 0 | 52 | 49 | 0.067295016 | 1 | 0.9 | 1 | 1 | 1 | 1 | -0.033 |
| SE | 16938 | IRF1 | IRF1 | chr5 | - | 132487925 | 132488025 | 132486953 | 132487130 | 132490544 | 132490773 | 75 | 60 | 22 | 18 | 5 | 5 | 7 | 11 | 0 | 2 | 98 | 49 | 0.068165057 | 1 | 0.676 | 0.857 | 0.687 | 1 | 0.733 | -0.127 |
| SE | 5705 | SERPINA1 | SERPINA1 | chr14 | - | 94388800 | 94389010 | 94388559 | 94388663 | 94390456 | 94390692 | 66 | 13 | 9 | 28 | 18 | 5 | 1 | 17 | 0 | 4 | 98 | 49 | 0.068748003 | 1 | 0.541 | 0.265 | 0.474 | 1 | 0.68 | -0.413 |
| A3SS | 3007 | STXBP2 | STXBP2 | chr19 | + | 7640697 | 7640809 | 7640730 | 7640809 | 7639730 | 7639807 | 1 | 2 | 0 | 6 | 4 | 1 | 0 | 7 | 1 | 1 | 82 | 49 | 0.069429487 | 1 | 0.091 | 0.23 | 0 | 0 | 0.807 | -0.297 |
| SE | 4942 | KIAA1033 | KIAA1033 | chr12 | + | 105140293 | 105140401 | 105137885 | 105138011 | 105140898 | 105141045 | 34 | 14 | 3 | 3 | 0 | 2 | 3 | 12 | 0 | 1 | 98 | 49 | 0.069523012 | 1 | 0.85 | 1 | 0.429 | 1 | 0.857 | -0.169 |
| SE | 12748 | CYP4F3 | CYP4F3 | chr19 | + | 15643906 | 15644051 | 15641414 | 15641613 | 15645718 | 15645863 | 3 | 3 | 6 | 103 | 25 | 21 | 0 | 1 | 4 | 16 | 98 | 49 | 0.07067314 | 1 | 0.014 | 0.057 | 0.125 | 0 | 0.03 | 0.05 |
| SE | 495 | ABCA1 | ABCA1 | chr9 | - | 104884426 | 104884568 | 104883038 | 104883157 | 104889101 | 104889195 | 46 | 21 | 3 | 13 | 2 | 0 | 3 | 20 | 1 | 1 | 98 | 49 | 0.070809818 | 1 | 0.639 | 0.84 | 1 | 0.6 | 0.909 | 0.072 |
| SE | 7502 | SLC12A6 | SLC12A6 | chr15 | - | 34275344 | 34275389 | 34260925 | 34261020 | 34336409 | 34336601 | 155 | 85 | 27 | 2 | 2 | 3 | 6 | 36 | 0 | 0 | 94 | 49 | 0.072192319 | 1 | 0.976 | 0.957 | 0.824 | 1 | 1 | -0.081 |
| SE | 16969 | ETF1 | ETF1 | chr5 | - | 138517560 | 138517700 | 138513567 | 138513706 | 138518691 | 138518867 | 43 | 22 | 9 | 3 | 0 | 0 | 0 | 12 | 2 | 0 | 98 | 49 | 0.072739847 | 1 | 0.878 | 1 | 1 | 0 | 1 | 0.459 |
| SE | 5708 | SERPINA1 | SERPINA1 | chr14 | - | 94388818 | 94389010 | 94388559 | 94388663 | 94390456 | 94390692 | 59 | 15 | 11 | 28 | 18 | 5 | 1 | 17 | 0 | 4 | 98 | 49 | 0.073470529 | 1 | 0.513 | 0.294 | 0.524 | 1 | 0.68 | -0.396 |
| RI | 202 | ULK3 | ULK3 | chr15 | - | 74840233 | 74840641 | 74840233 | 74840316 | 74840497 | 74840641 | 4 | 2 | 0 | 1 | 0 | 1 | 2 | 1 | 0 | 3 | 98 | 49 | 0.074113907 | 1 | 0.667 | 1 | 0 | 1 | 0.143 | -0.016 |
| A5SS | 2605 | ACOX1 | ACOX1 | chr17 | - | 75978790 | 75979434 | 75978964 | 75979434 | 75978533 | 75978693 | 5 | 1 | 0 | 36 | 10 | 4 | 0 | 0 | 2 | 2 | 98 | 49 | 0.075943129 | 1 | 0.065 | 0.048 | 0 | 0 | 0 | 0.038 |
| SE | 4782 | CARD8 | CARD8 | chr19 | - | 48218870 | 48219012 | 48215339 | 48215384 | 48221729 | 48221855 | 62 | 26 | 16 | 3 | 2 | 0 | 2 | 6 | 0 | 0 | 98 | 49 | 0.076881264 | 1 | 0.912 | 0.867 | 1 | 1 | 1 | -0.074 |

| | | | | | | | | | | | | | | | | | | | | | | | | | | | | | | | |
|---|---|---|---|---|---|---|---|---|---|---|---|---|---|---|---|---|---|---|---|---|---|---|---|---|---|---|---|---|---|---|---|
| SE | 16211 | NSD1 | NSD1 | chr5 | + | 177238236 | 177238507 | 177235820 | 177235945 | 177239755 | 177239865 | 19 | 2 | 3 | 2 | 0 | 2 | 3 | 11 | 0 | 0 | 98 | 49 | 0.077838524 | 1 | 0.826 | 1 | 0.429 | 1 | 1 | -0.248 |
| SE | 12158 | CYTH1 | CYTH1 | chr17 | - | 78702537 | 78702604 | 78702121 | 78702240 | 78708196 | 78708261 | 32 | 12 | 16 | 6 | 6 | 0 | 0 | 6 | 1 | 3 | 98 | 49 | 0.079729274 | 1 | 0.727 | 0.5 | 1 | 0 | 0.5 | 0.492 |
| SE | 6674 | ARHGAP4 | ARHGAP4 | chrX | - | 153913777 | 153913879 | 153913408 | 153913600 | 153918831 | 153919053 | 42 | 16 | 19 | 5 | 0 | 0 | 1 | 11 | 0 | 0 | 98 | 49 | 0.081582192 | 1 | 0.808 | 1 | 1 | 1 | 1 | -0.064 |
| A5SS | 2845 | TLE3 | TLE3 | chr15 | - | 70057458 | 70057658 | 70057473 | 70057658 | 70056297 | 70056374 | 40 | 10 | 7 | 28 | 3 | 16 | 1 | 9 | 0 | 3 | 64 | 49 | 0.081597054 | 1 | 0.522 | 0.718 | 0.251 | 1 | 0.697 | -0.351 |
| SE | 1987 | SRPK1 | SRPK1 | chr6 | - | 35870280 | 35870494 | 35869481 | 35869901 | 35870933 | 35870959 | 19 | 4 | 7 | 8 | 10 | 0 | 0 | 4 | 1 | 5 | 98 | 49 | 0.082834818 | 1 | 0.543 | 0.167 | 1 | 0 | 0.286 | 0.427 |
| SE | 15984 | RNF13 | RNF13 | chr3 | + | 149813924 | 149814026 | 149812687 | 149813353 | 149846010 | 149846140 | 4 | 0 | 0 | 25 | 6 | 2 | 0 | 4 | 1 | 2 | 98 | 49 | 0.083683236 | 1 | 0.074 | 0 | 0 | 0 | 0.5 | -0.225 |
| SE | 3162 | PATL2 | PATL2 | chr15 | - | 44675485 | 44675691 | 44674149 | 44674230 | 44676474 | 44676565 | 1 | 1 | 0 | 0 | 0 | 5 | 1 | 3 | 0 | 1 | 98 | 49 | 0.084376737 | 1 | 1 | 1 | 0 | 1 | 0.6 | -0.133 |
| RI | 169 | STX4 | STX4 | chr16 | + | 31034225 | 31034536 | 31034225 | 31034325 | 31034461 | 31034536 | 18 | 13 | 0 | 1 | 0 | 1 | 1 | 5 | 0 | 3 | 98 | 49 | 0.088362273 | 1 | 0.9 | 1 | 0 | 1 | 0.455 | -0.094 |
| A5SS | 1475 | USF1 | USF1 | chr1 | - | 161042554 | 161042670 | 161042585 | 161042670 | 161042115 | 161042217 | 43 | 24 | 10 | 15 | 2 | 0 | 0 | 9 | 2 | 2 | 80 | 49 | 0.088876961 | 1 | 0.637 | 0.88 | 1 | 0 | 0.734 | 0.472 |
| SE | 8679 | MAEA | MAEA | chr4 | + | 1327626 | 1327703 | 1322380 | 1322503 | 1332756 | 1332865 | 13 | 8 | 8 | 0 | 3 | 1 | 3 | 2 | 0 | 0 | 98 | 49 | 0.089381693 | 1 | 1 | 0.571 | 0.8 | 1 | 1 | -0.21 |
| SE | 14389 | SRSF7 | SRSF7 | chr2 | - | 38746143 | 38746179 | 38743598 | 38745187 | 38746693 | 38746747 | 31 | 2 | 4 | 1 | 3 | 0 | 2 | 5 | 0 | 0 | 85 | 49 | 0.089682605 | 1 | 0.947 | 0.278 | 1 | 1 | 1 | -0.258 |
| SE | 9050 | STAT2 | STAT2 | chr12 | - | 56351291 | 56351450 | 56351097 | 56351190 | 56354465 | 56354614 | 10 | 6 | 1 | 2 | 2 | 0 | 1 | 7 | 0 | 0 | 98 | 49 | 0.09007975 | 1 | 0.714 | 0.6 | 1 | 1 | 1 | -0.229 |
| SE | 12179 | RNF213 | RNF213 | chr17 | + | 80389172 | 80389367 | 80388611 | 80388689 | 80389827 | 80389917 | 66 | 40 | 17 | 6 | 0 | 0 | 7 | 56 | 0 | 1 | 98 | 49 | 0.091311053 | 1 | 0.846 | 1 | 1 | 1 | 0.966 | -0.034 |
| SE | 12247 | SMCHD1 | SMCHD1 | chr18 | + | 2750042 | 2750122 | 2747521 | 2747647 | 2750349 | 2750507 | 354 | 198 | 64 | 10 | 13 | 2 | 2 | 54 | 0 | 8 | 98 | 49 | 0.091337788 | 1 | 0.947 | 0.884 | 0.941 | 1 | 0.771 | 0.038 |
| SE | 13736 | SMG5 | SMG5 | chr1 | - | 156266540 | 156266678 | 156265780 | 156266380 | 156267469 | 156267678 | 12 | 6 | 2 | 3 | 1 | 0 | 1 | 3 | 0 | 1 | 98 | 49 | 0.092398999 | 1 | 0.667 | 0.75 | 1 | 1 | 0.6 | 0.006 |
| SE | 12130 | SRSF2 | SRSF2 | chr17 | - | 76735771 | 76735875 | 76734114 | 76735158 | 76736153 | 76736464 | 26 | 9 | 1 | 16 | 3 | 2 | 1 | 2 | 2 | 3 | 98 | 49 | 0.092646108 | 1 | 0.448 | 0.6 | 0.2 | 0.2 | 0.25 | 0.191 |
| SE | 8463 | MAST3 | MAST3 | chr19 | + | 18124641 | 18124774 | 18124264 | 18124366 | 18128399 | 18128458 | 64 | 24 | 10 | 1 | 2 | 0 | 1 | 11 | 0 | 3 | 98 | 49 | 0.092711196 | 1 | 0.97 | 0.857 | 1 | 1 | 0.647 | 0.119 |
| SE | 14429 | CFLAR | CFLAR | chr2 | + | 201129728 | 201130146 | 201116103 | 201116481 | 201133028 | 201133134 | 356 | 120 | 62 | 4 | 2 | 3 | 1 | 63 | 0 | 0 | 98 | 49 | 0.093297333 | 1 | 0.978 | 0.968 | 0.912 | 1 | 1 | -0.047 |
| SE | 5253 | PTPRC | PTPRC | chr1 | + | 198699563 | 198699704 | 198692346 | 198692373 | 198703297 | 198703372 | 78 | 41 | 25 | 260 | 94 | 48 | 3 | 18 | 0 | 25 | 98 | 49 | 0.093994131 | 1 | 0.13 | 0.179 | 0.207 | 1 | 0.265 | -0.461 |
| A3SS | 709 | TMBIM4 | TMBIM4 | chr12 | - | 66138723 | 66138769 | 66138723 | 66138734 | 66145840 | 66145958 | 40 | 23 | 36 | 3 | 0 | 2 | 1 | 12 | 0 | 0 | 84 | 49 | 0.09406817 | 1 | 0.886 | 1 | 0.913 | 1 | 1 | -0.067 |
| SE | 2868 | NLRP12 | NLRP12 | chr19 | - | 53795858 | 53796029 | 53793600 | 53794136 | 53801226 | 53801397 | 19 | 6 | 5 | 1 | 3 | 0 | 2 | 2 | 0 | 0 | 98 | 49 | 0.095265544 | 1 | 0.905 | 0.5 | 1 | 1 | 1 | -0.198 |
| SE | 12833 | N4BP2L2 | N4BP2L2 | chr13 | - | 32527407 | 32527532 | 32522181 | 32522270 | 32535768 | 32537027 | 35 | 36 | 13 | 2 | 2 | 1 | 2 | 9 | 0 | 1 | 98 | 49 | 0.095937215 | 1 | 0.897 | 0.9 | 0.867 | 1 | 0.818 | -0.021 |
| SE | 4184 | GIT2 | GIT2 | chr12 | - | 109951211 | 109951316 | 109948061 | 109948846 | 109953091 | 109953234 | 19 | 5 | 5 | 1 | 1 | 2 | 1 | 2 | 0 | 0 | 98 | 49 | 0.096392161 | 1 | 0.905 | 0.714 | 0.556 | 1 | 1 | -0.275 |
| SE | 12725 | ADGRE5 | ADGRE5 | chr19 | + | 14396341 | 14396473 | 14390923 | 14391079 | 14397657 | 14397804 | 22 | 14 | 13 | 46 | 34 | 6 | 0 | 4 | 1 | 11 | 98 | 49 | 0.098212686 | 1 | 0.193 | 0.171 | 0.52 | 0 | 0.154 | 0.218 |
| SE | 12710 | CBX6 | CBX6 | chr22 | - | 38871901 | 38871945 | 38871691 | 38871757 | 38872121 | 38872334 | 2 | 6 | 1 | 0 | 0 | 0 | 0 | 1 | 1 | 0 | 93 | 49 | 0.098245981 | 1 | 1 | 1 | 1 | 0 | 1 | 0.5 |
| SE | 7842 | LRRK2 | LRRK2 | chr12 | + | 40243549 | 40243681 | 40240482 | 40240617 | 40249825 | 40249945 | 416 | 161 | 60 | 4 | 6 | 0 | 2 | 41 | 0 | 0 | 98 | 49 | 0.100960498 | 1 | 0.981 | 0.931 | 1 | 1 | 1 | -0.029 |
| SE | 2182 | PRRC2A | PRRC2A | chr6 | + | 31636732 | 31636945 | 31636509 | 31636608 | 31637041 | 31637136 | 7 | 5 | 1 | 0 | 0 | 0 | 0 | 2 | 1 | 0 | 98 | 49 | 0.101331953 | 1 | 1 | 1 | 1 | 0 | 1 | 0.5 |
| SE | 7319 | VPS11 | VPS11 | chr11 | + | 119078564 | 119078704 | 119078172 | 119078334 | 119078794 | 119078997 | 6 | 1 | 2 | 0 | 0 | 0 | 0 | 2 | 1 | 0 | 98 | 49 | 0.101331953 | 1 | 1 | 1 | 1 | 0 | 1 | 0.5 |
| SE | 10160 | BRMS1 | BRMS1 | chr11 | - | 66340120 | 66340213 | 66338720 | 66338785 | 66340773 | 66340870 | 21 | 4 | 9 | 0 | 0 | 0 | 0 | 2 | 1 | 0 | 98 | 49 | 0.101331953 | 1 | 1 | 1 | 1 | 0 | 1 | 0.5 |
| SE | 12459 | TTN-AS1 | TTN-AS1 | chr2 | + | 178571299 | 178571449 | 178542221 | 178542361 | 178597595 | 178597681 | 4 | 4 | 3 | 0 | 0 | 0 | 0 | 2 | 1 | 0 | 98 | 49 | 0.101331953 | 1 | 1 | 1 | 1 | 0 | 1 | 0.5 |
| SE | 17031 | PCGF3 | PCGF3 | chr4 | + | 764983 | 765064 | 761278 | 761416 | 766031 | 770639 | 30 | 12 | 2 | 0 | 0 | 0 | 0 | 2 | 1 | 0 | 98 | 49 | 0.101331953 | 1 | 1 | 1 | 1 | 0 | 1 | 0.5 |
| SE | 5503 | SYNE1 | SYNE1 | chr6 | - | 152455885 | 152456044 | 152455425 | 152455590 | 152458756 | 152458930 | 7 | 7 | 5 | 0 | 0 | 0 | 0 | 1 | 1 | 0 | 98 | 49 | 0.101440565 | 1 | 1 | 1 | 1 | 0 | 1 | 0.5 |
| SE | 7688 | TFIP11 | TFIP11 | chr22 | - | 26494138 | 26494304 | 26491928 | 26492368 | 26494796 | 26494939 | 15 | 12 | 4 | 0 | 0 | 0 | 1 | 0 | 0 | 1 | 98 | 49 | 0.101440565 | 1 | 1 | 1 | 1 | 1 | 0 | 0.5 |
| SE | 9457 | ZNF467 | ZNF467 | chr7 | - | 149770439 | 149770556 | 149769089 | 149769200 | 149770998 | 149771074 | 17 | 1 | 9 | 0 | 0 | 0 | 0 | 1 | 1 | 0 | 98 | 49 | 0.101440565 | 1 | 1 | 1 | 1 | 0 | 1 | 0.5 |
| SE | 3517 | MTMR14 | MTMR14 | chr3 | + | 9662266 | 9662375 | 9653620 | 9653769 | 9668718 | 9668794 | 34 | 19 | 7 | 0 | 0 | 0 | 0 | 1 | 1 | 0 | 98 | 49 | 0.101563741 | 1 | 1 | 1 | 1 | 0 | 1 | 0.5 |
| A5SS | 1110 | SERPINA1 | SERPINA1 | chr14 | - | 94390419 | 94390692 | 94390456 | 94390692 | 94388818 | 94389010 | 85 | 62 | 16 | 44 | 13 | 5 | 4 | 5 | 1 | 15 | 86 | 49 | 0.103524947 | 1 | 0.524 | 0.731 | 0.646 | 0.695 | 0.16 | 0.206 |
| RI | 626 | HSH2D | HSH2D | chr19 | + | 16152551 | 16153208 | 16152551 | 16152641 | 16153042 | 16153208 | 76 | 27 | 3 | 3 | 2 | 4 | 5 | 21 | 1 | 1 | 98 | 49 | 0.104386471 | 1 | 0.927 | 0.871 | 0.273 | 0.714 | 0.913 | -0.123 |
| A3SS | 3069 | LIME1 | LIME1 | chr20 | + | 63737972 | 63738060 | 63738007 | 63738060 | 63737820 | 63737902 | 2 | 0 | 3 | 2 | 1 | 0 | 0 | 1 | 1 | 0 | 84 | 49 | 0.10449659 | 1 | 0.368 | 0 | 1 | 0 | 1 | -0.044 |
| SE | 16882 | ALPL | ALPL | chr1 | + | 21561096 | 21561212 | 21560625 | 21560745 | 21563109 | 21563284 | 124 | 65 | 46 | 3 | 3 | 0 | 3 | 21 | 0 | 0 | 98 | 49 | 0.110341459 | 1 | 0.954 | 0.915 | 1 | 1 | 1 | -0.044 |
| SE | 4346 | OSBPL8 | OSBPL8 | chr12 | - | 76459858 | 76459895 | 76450850 | 76450988 | 76487509 | 76487618 | 1 | 4 | 4 | 15 | 11 | 1 | 1 | 0 | 0 | 2 | 86 | 49 | 0.110562647 | 1 | 0.037 | 0.172 | 0.695 | 1 | 0 | -0.199 |
| SE | 5410 | NCF2 | NCF2 | chr1 | - | 183566919 | 183566988 | 183565703 | 183565779 | 183569141 | 183569185 | 180 | 77 | 88 | 3 | 3 | 1 | 12 | 35 | 0 | 0 | 98 | 49 | 0.11452548 | 1 | 0.968 | 0.928 | 0.978 | 1 | 1 | -0.042 |
| SE | 1601 | TAGAP | TAGAP | chr6 | - | 159042077 | 159042244 | 159041353 | 159041515 | 159043588 | 159043655 | 143 | 66 | 40 | 6 | 1 | 0 | 4 | 19 | 0 | 1 | 98 | 49 | 0.116249145 | 1 | 0.923 | 0.971 | 1 | 1 | 0.905 | 0.012 |
| A3SS | 2499 | NLRP6 | NLRP6 | chr11 | + | 282701 | 282797 | 282704 | 282797 | 280083 | 281839 | 1 | 0 | 3 | 7 | 8 | 2 | 0 | 0 | 1 | 1 | 52 | 49 | 0.116389539 | 1 | 0.119 | 0 | 0.586 | 0 | 0 | 0.235 |
| A3SS | 415 | RXRB | RXRB | chr6 | - | 33195362 | 33195466 | 33195362 | 33195454 | 33195569 | 33195702 | 1 | 1 | 2 | 4 | 2 | 0 | 0 | 1 | 1 | 0 | 61 | 49 | 0.117574196 | 1 | 0.167 | 0.287 | 1 | 0 | 1 | -0.015 |
| A3SS | 639 | ARRB2 | ARRB2 | chr17 | + | 4720549 | 4720640 | 4720585 | 4720640 | 4720392 | 4720472 | 9 | 7 | 0 | 57 | 20 | 11 | 0 | 2 | 2 | 13 | 85 | 49 | 0.118308764 | 1 | 0.083 | 0.168 | 0 | 0 | 0.081 | 0.043 |
| SE | 3037 | MSRB1 | MSRB1 | chr16 | - | 1940777 | 1940892 | 1938232 | 1939143 | 1941256 | 1941405 | 52 | 44 | 33 | 3 | 2 | 0 | 2 | 10 | 0 | 0 | 98 | 49 | 0.11831167 | 1 | 0.897 | 0.917 | 1 | 1 | 1 | -0.062 |
| A5SS | 1477 | IFI6 | IFI6 | chr1 | - | 27669220 | 27669346 | 27669244 | 27669346 | 27668457 | 27668535 | 1 | 2 | 0 | 6 | 10 | 7 | 0 | 5 | 1 | 20 | 73 | 49 | 0.119011446 | 1 | 0.101 | 0.118 | 0 | 0 | 0.144 | 0.001 |
| A5SS | 1435 | U2AF2 | U2AF2 | chr19 | + | 55669082 | 55669181 | 55669082 | 55669169 | 55669443 | 55669692 | 3 | 0 | 1 | 5 | 4 | 2 | 0 | 0 | 2 | 1 | 61 | 49 | 0.119377499 | 1 | 0.325 | 0 | 0.287 | 0 | 0 | 0.204 |
| A3SS | 137 | FGD3 | FGD3 | chr9 | + | 93034540 | 93034681 | 93034543 | 93034681 | 93032768 | 93032873 | 47 | 25 | 14 | 4 | 0 | 2 | 6 | 5 | 0 | 1 | 52 | 49 | 0.119826628 | 1 | 0.917 | 1 | 0.868 | 1 | 0.825 | 0.016 |
| SE | 12237 | SMCHD1 | SMCHD1 | chr18 | + | 2694526 | 2694693 | 2688627 | 2688747 | 2697031 | 2697122 | 515 | 238 | 81 | 1 | 10 | 0 | 3 | 97 | 0 | 0 | 98 | 49 | 0.12013002 | 1 | 0.996 | 0.922 | 1 | 1 | 1 | -0.027 |
| SE | 9959 | HDAC7 | HDAC7 | chr12 | - | 47786584 | 47786703 | 47785751 | 47785885 | 47787711 | 47787809 | 20 | 7 | 4 | 1 | 0 | 0 | 0 | 4 | 1 | 1 | 98 | 49 | 0.121266089 | 1 | 0.909 | 1 | 1 | 0 | 0.667 | 0.636 |
| SE | 16445 | CNN2 | CNN2 | chr19 | + | 1036129 | 1036246 | 1032558 | 1032696 | 1036415 | 1036562 | 138 | 85 | 62 | 3 | 3 | 1 | 1 | 25 | 0 | 1 | 98 | 49 | 0.121520206 | 1 | 0.958 | 0.934 | 0.969 | 1 | 0.926 | -0.009 |
| SE | 3715 | TNIP1 | TNIP1 | chr5 | - | 151064959 | 151065131 | 151063612 | 151063747 | 151080879 | 151081084 | 17 | 4 | 13 | 6 | 2 | 0 | 0 | 5 | 1 | 2 | 98 | 49 | 0.123280066 | 1 | 0.586 | 0.5 | 1 | 0 | 0.556 | 0.417 |
| SE | 3455 | SLC15A3 | SLC15A3 | chr11 | - | 60941121 | 60941290 | 60939479 | 60939638 | 60942034 | 60942145 | 25 | 9 | 3 | 16 | 8 | 5 | 2 | 5 | 1 | 1 | 98 | 49 | 0.124092315 | 1 | 0.439 | 0.36 | 0.231 | 0.5 | 0.714 | -0.264 |
| SE | 12826 | GMIP | GMIP | chr19 | - | 19641809 | 19641867 | 19640445 | 19640571 | 19641975 | 19642051 | 98 | 64 | 17 | 2 | 0 | 0 | 1 | 27 | 0 | 3 | 98 | 49 | 0.1248425 | 1 | 0.961 | 1 | 1 | 1 | 0.818 | 0.078 |
| SE | 9165 | CELF2 | CELF2 | chr10 | + | 11217424 | 11217507 | 11165485 | 11165682 | 11249152 | 11249201 | 150 | 68 | 22 | 14 | 3 | 1 | 1 | 31 | 0 | 2 | 98 | 49 | 0.125461496 | 1 | 0.843 | 0.919 | 0.917 | 1 | 0.886 | -0.05 |
| SE | 9858 | NADK | NADK | chr1 | - | 1756502 | 1756608 | 1756257 | 1756343 | 1757180 | 1757310 | 75 | 38 | 35 | 0 | 0 | 0 | 1 | 24 | 0 | 3 | 98 | 49 | 0.126801704 | 1 | 1 | 1 | 1 | 1 | 0.8 | 0.1 |
| SE | 1502 | HNRNPA2B1 | HNRNPA2B1 | chr7 | - | 26197830 | 26197866 | 26197621 | 26197732 | 26200571 | 26200793 | 4 | 1 | 0 | 27 | 10 | 3 | 0 | 1 | 3 | 7 | 85 | 49 | 0.127514469 | 1 | 0.079 | 0.055 | 0 | 0 | 0.076 | 0.007 |
| A5SS | 1004 | SNX17 | SNX17 | chr2 | + | 27371268 | 27371468 | 27371268 | 27371343 | 27372622 | 27372740 | 2 | 0 | 2 | 4 | 1 | 1 | 0 | 0 | 1 | 3 | 98 | 49 | 0.127869113 | 1 | 0.2 | 0 | 0.5 | 0 | 0 | 0.233 |
| A3SS | 659 | PXN | PXN | chr12 | - | 120215102 | 120215417 | 120215102 | 120215273 | 120221622 | 120221758 | 21 | 9 | 5 | 31 | 12 | 16 | 1 | 7 | 1 | 3 | 98 | 49 | 0.127945795 | 1 | 0.253 | 0.273 | 0.135 | 0.333 | 0.538 | -0.215 |
| SE | 15052 | ABCG1 | ABCG1 | chr21 | + | 42276894 | 42276945 | 42273302 | 42273435 | 42282273 | 42282419 | 9 | 8 | 6 | 0 | 4 | 0 | 1 | 11 | 0 | 0 | 98 | 49 | 0.129179721 | 1 | 1 | 0.5 | 1 | 1 | 1 | -0.167 |
| SE | 4911 | ASAP1 | ASAP1 | chr8 | - | 130091972 | 130092143 | 130079901 | 130079971 | 130112093 | 130112322 | 12 | 4 | 3 | 6 | 0 | 0 | 1 | 0 | 0 | 1 | 98 | 49 | 0.129893977 | 1 | 0.5 | 1 | 1 | 1 | 0 | 0.333 |
| SE | 6639 | CSF3R | CSF3R | chr1 | - | 36473434 | 36473622 | 36472516 | 36472686 | 36473763 | 36473887 | 1105 | 330 | 219 | 31 | 13 | 3 | 19 | 197 | 1 | 3 | 98 | 49 | 0.134838662 | 1 | 0.947 | 0.927 | 0.973 | 0.905 | 0.97 | 0.011 |
| SE | 10004 | ADRBK1 | ADRBK1 | chr11 | + | 67283706 | 67283773 | 67283127 | 67283228 | 67283853 | 67283949 | 123 | 48 | 22 | 6 | 0 | 0 | 13 | 34 | 5 | 0 | 98 | 49 | 0.135600813 | 1 | 0.911 | 1 | 1 | 0.565 | 1 | 0.188 |
| A5SS | 102 | PTBP3 | PTBP3 | chr9 | - | 112297831 | 112297916 | 112297840 | 112297916 | 112275843 | 112276013 | 19 | 5 | 5 | 13 | 7 | 0 | 0 | 3 | 2 | 2 | 58 | 49 | 0.136047271 | 1 | 0.553 | 0.376 | 1 | 0 | 0.559 | 0.363 |
| SE | 14398 | STAT1 | STAT1 | chr2 | - | 190983641 | 190983740 | 190982382 | 190982518 | 190984309 | 190984393 | 88 | 37 | 23 | 4 | 1 | 0 | 2 | 25 | 0 | 0 | 98 | 49 | 0.136123209 | 1 | 0.917 | 0.949 | 1 | 1 | 1 | -0.045 |
| A3SS | 3122 | TYROBP | TYROBP | chr19 | - | 35907217 | 35907264 | 35907217 | 35907261 | 35907445 | 35907580 | 135 | 71 | 63 | 17 | 13 | 2 | 11 | 42 | 0 | 4 | 52 | 49 | 0.136563402 | 1 | 0.882 | 0.837 | 0.967 | 1 | 0.908 | -0.059 |
| SE | 5704 | SERPINA1 | SERPINA1 | chr14 | - | 94388800 | 94389010 | 94388559 | 94388660 | 94390456 | 94390692 | 46 | 13 | 6 | 33 | 5 | 10 | 1 | 17 | 1 | 3 | 98 | 49 | 0.137238291 | 1 | 0.411 | 0.565 | 0.231 | 0.333 | 0.739 | -0.134 |
| SE | 12236 | SMCHD1 | SMCHD1 | chr18 | + | 2674014 | 2674145 | 2673280 | 2673363 | 2688393 | 2688508 | 698 | 295 | 69 | 5 | 14 | 0 | 7 | 143 | 0 | 2 | 98 | 49 | 0.138416666 | 1 | 0.986 | 0.913 | 1 | 1 | 0.973 | -0.02 |
| SE | 4372 | SNRK | SNRK | chr3 | + | 43299753 | 43299815 | 43286511 | 43286675 | 43303097 | 43303792 | 81 | 42 | 6 | 1 | 3 | 1 | 1 | 7 | 0 | 0 | 98 | 49 | 0.141049021 | 1 | 0.976 | 0.875 | 0.75 | 1 | 1 | -0.133 |
| A3SS | 317 | GLIPR2 | GLIPR2 | chr9 | + | 36162231 | 36163913 | 36162361 | 36163913 | 36150871 | 36150949 | 17 | 3 | 2 | 51 | 13 | 20 | 1 | 6 | 0 | 5 | 98 | 49 | 0.14275805 | 1 | 0.143 | 0.103 | 0.048 | 1 | 0.375 | -0.59 |
| SE | 3986 | PSEN1 | PSEN1 | chr14 | + | 73198030 | 73198129 | 73192643 | 73192864 | 73206385 | 73206472 | 13 | 9 | 15 | 5 | 0 | 0 | 1 | 8 | 2 | 1 | 98 | 49 | 0.144842421 | 1 | 0.565 | 1 | 1 | 0.2 | 0.8 | 0.355 |
| SE | 14558 | CLK3 | CLK3 | chr15 | + | 74625801 | 74625968 | 74624901 | 74625018 | 74627351 | 74627446 | 25 | 8 | 3 | 2 | 1 | 1 | 3 | 4 | 0 | 0 | 98 | 49 | 0.146907852 | 1 | 0.862 | 0.8 | 0.6 | 1 | 1 | -0.246 |
| A5SS | 1914 | ADAR | ADAR | chr1 | - | 154590183 | 154590409 | 154590261 | 154590409 | 154589756 | 154589928 | 69 | 26 | 17 | 20 | 1 | 4 | 4 | 24 | 0 | 3 | 98 | 49 | 0.147328922 | 1 | 0.633 | 0.929 | 0.68 | 1 | 0.8 | -0.153 |
| SE | 3714 | TNIP1 | TNIP1 | chr5 | - | 151064959 | 151065127 | 151063612 | 151063747 | 151080879 | 151081088 | 12 | 2 | 8 | 6 | 2 | 0 | 0 | 4 | 1 | 2 | 98 | 49 | 0.150224689 | 1 | 0.5 | 0.333 | 1 | 0 | 0.5 | 0.361 |
| SE | 4600 | TUBA1A | TUBA1A | chr12 | - | 49186309 | 49186458 | 49184794 | 49185990 | 49186610 | 49186833 | 119 | 100 | 82 | 0 | 0 | 0 | 5 | 29 | 1 | 2 | 98 | 49 | 0.15233388 | 1 | 1 | 1 | 1 | 0.714 | 0.879 | 0.204 |
| A3SS | 2325 | ARPC5 | ARPC5 | chr1 | - | 183633081 | 183633163 | 183633081 | 183633154 | 183635516 | 183635941 | 7 | 1 | 0 | 103 | 77 | 27 | 0 | 0 | 4 | 20 | 58 | 49 | 0.153712068 | 1 | 0.054 | 0.011 | 0 | 0 | 0 | 0.022 |
| SE | 7711 | MDM4 | MDM4 | chr1 | + | 204537429 | 204537497 | 204532190 | 204532246 | 204538208 | 204538308 | 23 | 12 | 3 | 2 | 2 | 0 | 1 | 6 | 0 | 0 | 98 | 49 | 0.153839737 | 1 | 0.852 | 0.75 | 1 | 1 | 1 | -0.133 |
| SE | 11696 | OASL | OASL | chr12 | - | 121033460 | 121033743 | 121031441 | 121031617 | 121038773 | 121039242 | 15 | 18 | 2 | 2 | 5 | 0 | 0 | 16 | 2 | 2 | 98 | 49 | 0.154890071 | 1 | 0.789 | 0.643 | 1 | 0 | 0.8 | 0.411 |
| A3SS | 4117 | GRK6 | GRK6 | chr5 | + | 177441734 | 177442849 | 177441736 | 177442849 | 177440918 | 177441053 | 8 | 2 | 7 | 7 | 1 | 0 | 1 | 6 | 0 | 2 | 51 | 49 | 0.158599923 | 1 | 0.523 | 0.658 | 1 | 1 | 0.742 | -0.144 |
| A5SS | 2330 | N4BP2L2 | N4BP2L2 | chr13 | - | 32527407 | 32527532 | 32527457 | 32527532 | 32522181 | 32522270 | 70 | 73 | 47 | 6 | 4 | 10 | 2 | 17 | 0 | 2 | 98 | 49 | 0.1604111 | 1 | 0.854 | 0.901 | 0.701 | 1 | 0.81 | -0.086 |
| SE | 5959 | SORL1 | SORL1 | chr11 | + | 121557313 | 121557405 | 121555186 | 121555318 | 121558590 | 121558837 | 519 | 211 | 71 | 15 | 4 | 3 | 10 | 73 | 0 | 2 | 98 | 49 | 0.164593154 | 1 | 0.945 | 0.963 | 0.922 | 1 | 0.948 | -0.031 |
| SE | 3036 | XPO6 | XPO6 | chr16 | - | 28208986 | 28209122 | 28180940 | 28181031 | 28211365 | 28211918 | 8 | 10 | 3 | 48 | 22 | 8 | 1 | 1 | 0 | 17 | 98 | 49 | 0.166415202 | 1 | 0.077 | 0.185 | 0.158 | 1 | 0.029 | -0.374 |
| SE | 8074 | VPS29 | VPS29 | chr12 | - | 110492995 | 110493231 | 110491522 | 110492122 | 110496011 | 110496094 | 24 | 8 | 11 | 0 | 4 | 0 | 1 | 7 | 0 | 0 | 98 | 49 | 0.167858835 | 1 | 1 | 0.5 | 1 | 1 | 1 | -0.167 |
| SE | 11498 | SSH2 | SSH2 | chr17 | - | 29650653 | 29650800 | 29648143 | 29648344 | 29655560 | 29655607 | 128 | 82 | 14 | 0 | 0 | 0 | 1 | 32 | 0 | 3 | 98 | 49 | 0.168540954 | 1 | 1 | 1 | 1 | 1 | 0.842 | 0.079 |
| SE | 16957 | H2AFY | H2AFY | chr5 | - | 135352945 | 135353045 | 135345967 | 135346057 | 135360496 | 135360604 | 16 | 12 | 14 | 3 | 0 | 1 | 2 | 10 | 0 | 0 | 98 | 49 | 0.171430699 | 1 | 0.727 | 1 | 0.875 | 1 | 1 | -0.133 |
| A3SS | 2200 | STAT2 | STAT2 | chr12 | - | 56355707 | 56355803 | 56355707 | 56355791 | 56356131 | 56356285 | 14 | 3 | 1 | 2 | 0 | 0 | 4 | 17 | 0 | 3 | 61 | 49 | 0.174115106 | 1 | 0.849 | 1 | 1 | 1 | 0.82 | 0.04 |
| A5SS | 2171 | PGS1 | PGS1 | chr17 | + | 78419545 | 78419725 | 78419545 | 78419675 | 78424060 | 78424659 | 61 | 42 | 16 | 4 | 0 | 2 | 10 | 32 | 0 | 1 | 98 | 49 | 0.175839321 | 1 | 0.884 | 1 | 0.8 | 1 | 0.941 | -0.076 |
| SE | 9035 | DENND5A | DENND5A | chr11 | - | 9145669 | 9145815 | 9144994 | 9145113 | 9147029 | 9147151 | 81 | 32 | 23 | 5 | 4 | 1 | 0 | 13 | 1 | 0 | 98 | 49 | 0.176386755 | 1 | 0.89 | 0.8 | 0.92 | 0 | 1 | 0.37 |
| SE | 6835 | GAK | GAK | chr4 | - | 851749 | 851974 | 850935 | 851084 | 859605 | 859722 | 8 | 3 | 6 | 0 | 0 | 0 | 1 | 6 | 2 | 0 | 98 | 49 | 0.177931592 | 1 | 1 | 1 | 1 | 0.2 | 1 | 0.4 |
| SE | 3021 | XPO6 | XPO6 | chr16 | - | 28117317 | 28117462 | 28112903 | 28113050 | 28121669 | 28121762 | 267 | 159 | 66 | 4 | 3 | 1 | 1 | 55 | 0 | 1 | 98 | 49 | 0.179229462 | 1 | 0.971 | 0.964 | 0.971 | 1 | 0.965 | -0.014 |
| SE | 15911 | CEP63 | CEP63 | chr3 | + | 134558141 | 134558347 | 134551925 | 134552012 | 134561376 | 134565028 | 11 | 12 | 1 | 2 | 4 | 3 | 2 | 0 | 0 | 4 | 98 | 49 | 0.179730224 | 1 | 0.733 | 0.6 | 0.143 | 1 | 0 | -0.008 |
| SE | 15543 | ATG16L2 | ATG16L2 | chr11 | + | 72822847 | 72822961 | 72822477 | 72822543 | 72824059 | 72824122 | 161 | 82 | 52 | 3 | 3 | 0 | 9 | 41 | 0 | 1 | 98 | 49 | 0.182129729 | 1 | 0.964 | 0.932 | 1 | 1 | 0.953 | -0.011 |
| SE | 9063 | SIPA1 | SIPA1 | chr11 | + | 65644954 | 65645129 | 65642462 | 65642639 | 65646220 | 65646378 | 37 | 11 | 11 | 0 | 0 | 0 | 1 | 11 | 2 | 0 | 98 | 49 | 0.18440821 | 1 | 1 | 1 | 1 | 0.2 | 1 | 0.4 |
| SE | 10876 | PSMB4 | PSMB4 | chr1 | + | 151399980 | 151400187 | 151399564 | 151399727 | 151400441 | 151400588 | 24 | 9 | 14 | 2 | 2 | 0 | 1 | 3 | 0 | 0 | 98 | 49 | 0.185402396 | 1 | 0.857 | 0.692 | 1 | 1 | 1 | -0.15 |
| SE | 8604 | MAPKAPK2 | MAPKAPK2 | chr1 | + | 206731838 | 206731919 | 206731639 | 206731725 | 206732574 | 206734285 | 40 | 17 | 20 | 0 | 0 | 0 | 1 | 14 | 2 | 0 | 98 | 49 | 0.186541703 | 1 | 1 | 1 | 1 | 0.2 | 1 | 0.4 |
| SE | 11695 | OASL | OASL | chr12 | - | 121027575 | 121027817 | 121023989 | 121024137 | 121031441 | 121031617 | 19 | 18 | 4 | 0 | 0 | 0 | 0 | 26 | 1 | 2 | 98 | 49 | 0.186693868 | 1 | 1 | 1 | 1 | 0 | 0.867 | 0.567 |
| SE | 2951 | IQGAP1 | IQGAP1 | chr15 | + | 90439331 | 90439399 | 90433718 | 90433795 | 90440501 | 90440615 | 217 | 123 | 69 | 3 | 4 | 0 | 1 | 32 | 0 | 0 | 98 | 49 | 0.187913552 | 1 | 0.973 | 0.939 | 1 | 1 | 1 | -0.029 |
| SE | 2540 | MAP3K1 | MAP3K1 | chr5 | + | 56875031 | 56875310 | 56872824 | 56873005 | 56878979 | 56879101 | 66 | 10 | 4 | 8 | 0 | 0 | 0 | 5 | 1 | 1 | 98 | 49 | 0.188514258 | 1 | 0.805 | 1 | 1 | 0 | 0.714 | 0.578 |
| SE | 13009 | ARHGEF1 | ARHGEF1 | chr19 | + | 41905174 | 41905261 | 41904948 | 41905036 | 41905759 | 41905827 | 55 | 13 | 18 | 1 | 0 | 0 | 1 | 10 | 2 | 0 | 98 | 49 | 0.190810353 | 1 | 0.965 | 1 | 1 | 0.2 | 1 | 0.388 |
| A5SS | 1108 | SERPINA1 | SERPINA1 | chr14 | - | 94388800 | 94389010 | 94388818 | 94389010 | 94388559 | 94388663 | 58 | 15 | 15 | 15 | 2 | 6 | 1 | 6 | 0 | 2 | 67 | 49 | 0.193154337 | 1 | 0.739 | 0.846 | 0.646 | 1 | 0.687 | -0.1 |
| SE | 4705 | LCP1 | LCP1 | chr13 | - | 46158825 | 46158989 | 46158521 | 46158651 | 46159598 | 46159686 | 463 | 214 | 119 | 3 | 7 | 0 | 1 | 110 | 0 | 5 | 98 | 49 | 0.19318281 | 1 | 0.987 | 0.939 | 1 | 1 | 0.917 | 0.017 |
| SE | 13214 | FES | FES | chr15 | + | 90889516 | 90889636 | 90889305 | 90889443 | 90889839 | 90889962 | 14 | 10 | 6 | 0 | 4 | 0 | 1 | 7 | 0 | 0 | 98 | 49 | 0.193843347 | 1 | 1 | 0.556 | 1 | 1 | 1 | -0.148 |
| RI | 580 | ATG3 | ATG3 | chr3 | - | 112532506 | 112534337 | 112532506 | 112532780 | 112534268 | 112534337 | 10 | 6 | 4 | 28 | 11 | 3 | 0 | 3 | 3 | 5 | 98 | 49 | 0.199919419 | 1 | 0.152 | 0.214 | 0.4 | 0 | 0.231 | 0.14 |
| A3SS | 381 | MTCH1 | MTCH1 | chr6 | - | 36977633 | 36977784 | 36977633 | 36977691 | 36978077 | 36978155 | 18 | 5 | 1 | 18 | 5 | 4 | 1 | 1 | 0 | 3 | 98 | 49 | 0.205857876 | 1 | 0.333 | 0.333 | 0.111 | 1 | 0.143 | -0.312 |
| A3SS | 2637 | FERMT3 | FERMT3 | chr11 | + | 64219878 | 64220015 | 64219890 | 64220015 | 64219739 | 64219789 | 1 | 2 | 0 | 25 | 9 | 2 | 2 | 1 | 8 | 9 | 61 | 49 | 0.206700481 | 1 | 0.031 | 0.151 | 0 | 0.167 | 0.082 | -0.064 |
| SE | 5948 | SORL1 | SORL1 | chr11 | + | 121470006 | 121470123 | 121452202 | 121452616 | 121478117 | 121478243 | 354 | 200 | 103 | 3 | 5 | 1 | 9 | 67 | 0 | 1 | 98 | 49 | 0.20705801 | 1 | 0.983 | 0.952 | 0.981 | 1 | 0.971 | -0.014 |
| SE | 7874 | PAK1 | PAK1 | chr11 | - | 77325327 | 77325376 | 77322014 | 77323360 | 77332729 | 77332867 | 11 | 4 | 0 | 25 | 19 | 1 | 0 | 3 | 1 | 12 | 98 | 49 | 0.207915739 | 1 | 0.18 | 0.095 | 0 | 0 | 0.111 | 0.036 |
| A5SS | 579 | FCAR | FCAR | chr19 | + | 54888006 | 54888294 | 54888006 | 54888228 | 54889648 | 54891420 | 75 | 27 | 8 | 7 | 3 | 0 | 5 | 40 | 2 | 6 | 98 | 49 | 0.20797773 | 1 | 0.843 | 0.818 | 1 | 0.556 | 0.769 | 0.224 |
| SE | 8769 | ZCCHC2 | ZCCHC2 | chr18 | + | 62544801 | 62544873 | 62542497 | 62542574 | 62550347 | 62550460 | 26 | 30 | 4 | 0 | 0 | 0 | 0 | 29 | 1 | 2 | 98 | 49 | 0.211120851 | 1 | 1 | 1 | 1 | 0 | 0.879 | 0.56 |
| SE | 15206 | ADORA2A | ADORA2A | chr22 | + | 24433130 | 24433736 | 24427561 | 24427726 | 24440582 | 24442361 | 6 | 0 | 1 | 0 | 2 | 0 | 1 | 1 | 0 | 0 | 98 | 49 | 0.21534978 | 1 | 1 | 0 | 1 | 1 | 1 | -0.333 |
| SE | 8486 | SCYL3 | SCYL3 | chr1 | - | 169870254 | 169870357 | 169868927 | 169869039 | 169873695 | 169873752 | 7 | 3 | 0 | 0 | 0 | 2 | 2 | 1 | 0 | 0 | 98 | 49 | 0.215368466 | 1 | 1 | 1 | 0 | 1 | 1 | -0.333 |
| SE | 4182 | GIT2 | GIT2 | chr12 | - | 109951166 | 109951316 | 109948061 | 109948846 | 109953091 | 109953234 | 31 | 17 | 5 | 1 | 1 | 2 | 1 | 3 | 0 | 0 | 98 | 49 | 0.215861986 | 1 | 0.939 | 0.895 | 0.556 | 1 | 1 | -0.203 |
| SE | 11775 | FMNL1 | FMNL1 | chr17 | + | 45236135 | 45236244 | 45234071 | 45234200 | 45237280 | 45237357 | 115 | 36 | 19 | 1 | 0 | 0 | 8 | 42 | 2 | 1 | 98 | 49 | 0.216566122 | 1 | 0.983 | 1 | 1 | 0.667 | 0.955 | 0.183 |
| SE | 3020 | XPO6 | XPO6 | chr16 | - | 28111816 | 28112006 | 28107521 | 28107677 | 28112903 | 28113050 | 282 | 100 | 34 | 15 | 2 | 0 | 2 | 40 | 1 | 1 | 98 | 49 | 0.2195329 | 1 | 0.904 | 0.962 | 1 | 0.5 | 0.952 | 0.229 |
| RI | 603 | ATP6V0B | ATP6V0B | chr1 | + | 43977025 | 43978300 | 43977025 | 43977216 | 43977980 | 43978300 | 21 | 7 | 1 | 73 | 31 | 55 | 0 | 3 | 4 | 15 | 98 | 49 | 0.221518601 | 1 | 0.126 | 0.101 | 0.009 | 0 | 0.091 | 0.033 |
| A3SS | 3186 | STRN4 | STRN4 | chr19 | - | 46727451 | 46727567 | 46727451 | 46727546 | 46727893 | 46728007 | 7 | 3 | 1 | 6 | 5 | 0 | 1 | 1 | 0 | 4 | 70 | 49 | 0.22753096 | 1 | 0.45 | 0.296 | 1 | 1 | 0.149 | 0.007 |
| SE | 6722 | TNFRSF1B | TNFRSF1B | chr1 | + | 12188795 | 12188895 | 12167002 | 12167169 | 12190956 | 12191085 | 127 | 70 | 38 | 8 | 2 | 0 | 7 | 37 | 2 | 3 | 98 | 49 | 0.227588142 | 1 | 0.888 | 0.946 | 1 | 0.636 | 0.86 | 0.197 |
| A3SS | 3372 | GNAS | GNAS | chr20 | + | 58903527 | 58903585 | 58903530 | 58903585 | 58895611 | 58895684 | 37 | 18 | 9 | 10 | 3 | 1 | 3 | 4 | 0 | 2 | 52 | 49 | 0.228963552 | 1 | 0.777 | 0.85 | 0.895 | 1 | 0.653 | 0.014 |
| SE | 16897 | VASP | VASP | chr19 | + | 45523840 | 45523877 | 45523643 | 45523695 | 45524096 | 45524142 | 213 | 112 | 95 | 2 | 0 | 0 | 5 | 37 | 0 | 3 | 86 | 49 | 0.231255615 | 1 | 0.984 | 1 | 1 | 1 | 0.875 | 0.057 |
| A5SS | 640 | BANP | BANP | chr16 | + | 87975047 | 87975203 | 87975047 | 87975185 | 87981035 | 87981127 | 4 | 4 | 2 | 15 | 9 | 0 | 0 | 2 | 1 | 1 | 67 | 49 | 0.235724677 | 1 | 0.163 | 0.245 | 1 | 0 | 0.594 | 0.172 |
| SE | 701 | SLC45A4 | SLC45A4 | chr8 | - | 141221576 | 141221765 | 141219649 | 141219829 | 141253988 | 141254629 | 26 | 19 | 4 | 2 | 3 | 0 | 0 | 5 | 1 | 0 | 98 | 49 | 0.236450153 | 1 | 0.867 | 0.76 | 1 | 0 | 1 | 0.376 |
| SE | 13642 | EPSTI1 | EPSTI1 | chr13 | - | 42919299 | 42919332 | 42917540 | 42917624 | 42926335 | 42926429 | 0 | 1 | 1 | 5 | 3 | 0 | 0 | 4 | 4 | 14 | 82 | 49 | 0.238594202 | 1 | 0 | 0.166 | 1 | 0 | 0.146 | 0.316 |
| SE | 978 | DOCK5 | DOCK5 | chr8 | + | 25407982 | 25408154 | 25403557 | 25403724 | 25408801 | 25408940 | 105 | 24 | 4 | 5 | 5 | 1 | 1 | 23 | 1 | 1 | 98 | 49 | 0.239086319 | 1 | 0.913 | 0.706 | 0.667 | 0.333 | 0.92 | 0.135 |
| A3SS | 2222 | NFKB2 | NFKB2 | chr10 | + | 102402251 | 102402529 | 102402254 | 102402529 | 102402047 | 102402159 | 4 | 3 | 0 | 5 | 3 | 1 | 1 | 3 | 0 | 3 | 52 | 49 | 0.239433215 | 1 | 0.43 | 0.485 | 0 | 1 | 0.485 | -0.437 |
| SE | 13860 | FOXN2 | FOXN2 | chr2 | + | 48328560 | 48328702 | 48314655 | 48314814 | 48346200 | 48346751 | 4 | 2 | 0 | 2 | 0 | 1 | 1 | 2 | 0 | 1 | 98 | 49 | 0.243503547 | 1 | 0.5 | 1 | 0 | 1 | 0.5 | -0.25 |
| A5SS | 916 | TUBA1A | TUBA1A | chr12 | - | 49188416 | 49189324 | 49188976 | 49189324 | 49186610 | 49186833 | 18 | 4 | 1 | 41 | 18 | 9 | 1 | 1 | 1 | 3 | 98 | 49 | 0.24508394 | 1 | 0.18 | 0.1 | 0.053 | 0.333 | 0.143 | -0.127 |
| A5SS | 917 | TUBA1A | TUBA1A | chr12 | - | 49188888 | 49189324 | 49188976 | 49189324 | 49186610 | 49186833 | 18 | 4 | 1 | 41 | 18 | 9 | 1 | 1 | 1 | 3 | 98 | 49 | 0.24508394 | 1 | 0.18 | 0.1 | 0.053 | 0.333 | 0.143 | -0.127 |
| SE | 9048 | STAT2 | STAT2 | chr12 | - | 56349162 | 56349261 | 56348923 | 56349059 | 56349425 | 56349509 | 12 | 12 | 3 | 4 | 0 | 0 | 1 | 9 | 0 | 0 | 98 | 49 | 0.247044358 | 1 | 0.6 | 1 | 1 | 1 | 1 | -0.133 |
| SE | 17106 | NQO2 | NQO2 | chr6 | + | 3004494 | 3004651 | 2999815 | 3000085 | 3006467 | 3006559 | 15 | 4 | 1 | 17 | 7 | 0 | 0 | 3 | 5 | 2 | 98 | 49 | 0.248944006 | 1 | 0.306 | 0.222 | 1 | 0 | 0.429 | 0.295 |
| SE | 1581 | IGF2R | IGF2R | chr6 | + | 160045744 | 160045882 | 160044513 | 160044657 | 160046497 | 160046645 | 374 | 98 | 67 | 2 | 0 | 0 | 0 | 35 | 1 | 1 | 98 | 49 | 0.251838884 | 1 | 0.989 | 1 | 1 | 0 | 0.946 | 0.523 |
| A3SS | 2478 | LST1 | LST1 | chr6 | + | 31588496 | 31588909 | 31588562 | 31588909 | 31587640 | 31587733 | 66 | 44 | 40 | 10 | 5 | 2 | 3 | 10 | 0 | 2 | 98 | 49 | 0.255381106 | 1 | 0.767 | 0.815 | 0.909 | 1 | 0.714 | -0.027 |
| SE | 6853 | AMPD2 | AMPD2 | chr1 | + | 109628642 | 109628806 | 109628363 | 109628495 | 109629108 | 109629235 | 52 | 31 | 7 | 4 | 0 | 1 | 1 | 2 | 1 | 0 | 98 | 49 | 0.256881736 | 1 | 0.867 | 1 | 0.778 | 0.333 | 1 | 0.215 |
| SE | 13047 | CHD2 | CHD2 | chr15 | + | 92901166 | 92901299 | 92900320 | 92900824 | 92924320 | 92924552 | 40 | 24 | 8 | 9 | 0 | 0 | 4 | 13 | 0 | 2 | 98 | 49 | 0.257739962 | 1 | 0.69 | 1 | 1 | 1 | 0.765 | 0.014 |
| SE | 10856 | STAT6 | STAT6 | chr12 | - | 57107604 | 57107743 | 57107230 | 57107314 | 57108162 | 57108299 | 90 | 38 | 20 | 18 | 8 | 3 | 1 | 13 | 1 | 5 | 98 | 49 | 0.258500124 | 1 | 0.714 | 0.704 | 0.769 | 0.333 | 0.565 | 0.28 |
| A3SS | 2479 | LST1 | LST1 | chr6 | + | 31588517 | 31588909 | 31588562 | 31588909 | 31587640 | 31587733 | 65 | 47 | 42 | 10 | 5 | 2 | 3 | 10 | 0 | 2 | 94 | 49 | 0.25902872 | 1 | 0.772 | 0.831 | 0.916 | 1 | 0.723 | -0.022 |
| SE | 11921 | VMP1 | VMP1 | chr17 | + | 59737452 | 59737543 | 59735337 | 59735473 | 59738836 | 59738947 | 176 | 163 | 135 | 7 | 2 | 1 | 4 | 43 | 0 | 4 | 98 | 49 | 0.259710227 | 1 | 0.926 | 0.976 | 0.985 | 1 | 0.843 | 0.041 |
| SE | 5188 | SAP130 | SAP130 | chr2 | - | 127977984 | 127978089 | 127954985 | 127955344 | 127986784 | 127986962 | 1 | 2 | 4 | 0 | 0 | 3 | 2 | 2 | 0 | 0 | 98 | 49 | 0.260139245 | 1 | 1 | 1 | 0.4 | 1 | 1 | -0.2 |
| SE | 14545 | EIF4G2 | EIF4G2 | chr11 | - | 10801660 | 10801774 | 10800961 | 10801087 | 10802048 | 10802209 | 65 | 24 | 19 | 5 | 2 | 1 | 0 | 6 | 1 | 1 | 98 | 49 | 0.260458271 | 1 | 0.867 | 0.857 | 0.905 | 0 | 0.75 | 0.501 |
| SE | 12352 | DDX17 | DDX17 | chr22 | - | 38495795 | 38495937 | 38494885 | 38495046 | 38498084 | 38498150 | 182 | 92 | 23 | 2 | 5 | 0 | 1 | 31 | 0 | 1 | 98 | 49 | 0.262772824 | 1 | 0.978 | 0.902 | 1 | 1 | 0.939 | -0.01 |
| SE | 13235 | RBM23 | RBM23 | chr14 | - | 22909482 | 22909595 | 22908332 | 22908380 | 22911327 | 22911403 | 80 | 34 | 30 | 2 | 3 | 0 | 2 | 4 | 0 | 1 | 98 | 49 | 0.265544407 | 1 | 0.952 | 0.85 | 1 | 1 | 0.667 | 0.101 |
| SE | 9302 | PIK3CD | PIK3CD | chr1 | + | 9721443 | 9721587 | 9721126 | 9721248 | 9721760 | 9721860 | 72 | 39 | 29 | 1 | 3 | 0 | 1 | 19 | 0 | 3 | 98 | 49 | 0.265878817 | 1 | 0.973 | 0.867 | 1 | 1 | 0.76 | 0.067 |
| SE | 3576 | ACTN1 | ACTN1 | chr14 | - | 68909954 | 68910042 | 68909317 | 68909396 | 68912155 | 68912242 | 151 | 86 | 54 | 7 | 5 | 0 | 9 | 25 | 2 | 0 | 98 | 49 | 0.266375307 | 1 | 0.915 | 0.896 | 1 | 0.692 | 1 | 0.091 |
| SE | 5241 | CHI3L1 | CHI3L1 | chr1 | - | 203180469 | 203180652 | 203179760 | 203179877 | 203181161 | 203181285 | 82 | 30 | 43 | 1 | 1 | 2 | 4 | 12 | 0 | 0 | 98 | 49 | 0.270873952 | 1 | 0.976 | 0.937 | 0.915 | 1 | 1 | -0.057 |
| SE | 2677 | CEACAM3 | CEACAM3 | chr19 | + | 41809964 | 41810017 | 41808812 | 41808930 | 41810322 | 41810354 | 61 | 14 | 29 | 0 | 4 | 0 | 5 | 6 | 0 | 0 | 98 | 49 | 0.273594365 | 1 | 1 | 0.636 | 1 | 1 | 1 | -0.121 |
| A5SS | 2581 | HNRNPC | HNRNPC | chr14 | - | 21230957 | 21231072 | 21230996 | 21231072 | 21230318 | 21230366 | 65 | 22 | 16 | 60 | 15 | 22 | 6 | 6 | 1 | 5 | 88 | 49 | 0.276324142 | 1 | 0.376 | 0.45 | 0.288 | 0.77 | 0.401 | -0.214 |
| RI | 1035 | NCF4 | NCF4 | chr22 | + | 36875652 | 36876094 | 36875652 | 36875783 | 36876028 | 36876094 | 62 | 32 | 18 | 80 | 40 | 40 | 2 | 11 | 0 | 11 | 98 | 49 | 0.277848545 | 1 | 0.279 | 0.286 | 0.184 | 1 | 0.333 | -0.417 |
| SE | 7102 | ANKRD12 | ANKRD12 | chr18 | + | 9204475 | 9204544 | 9195550 | 9195698 | 9208656 | 9208803 | 28 | 7 | 0 | 20 | 8 | 3 | 1 | 3 | 0 | 2 | 98 | 49 | 0.279726536 | 1 | 0.412 | 0.304 | 0 | 1 | 0.429 | -0.476 |
| SE | 15047 | RAB11FIP1 | RAB11FIP1 | chr8 | - | 37871277 | 37873179 | 37870419 | 37870528 | 37874514 | 37875322 | 11 | 14 | 12 | 12 | 15 | 8 | 0 | 4 | 1 | 4 | 98 | 49 | 0.282239472 | 1 | 0.314 | 0.318 | 0.429 | 0 | 0.333 | 0.187 |
| A3SS | 3625 | RPS3 | RPS3 | chr11 | + | 75402303 | 75402446 | 75402351 | 75402446 | 75401639 | 75401733 | 9 | 3 | 1 | 76 | 13 | 8 | 0 | 2 | 2 | 7 | 97 | 49 | 0.287169766 | 1 | 0.056 | 0.104 | 0.059 | 0 | 0.126 | 0.01 |
| SE | 6114 | TSPAN2 | TSPAN2 | chr1 | - | 115060463 | 115060538 | 115058882 | 115058981 | 115062134 | 115062232 | 4 | 5 | 2 | 0 | 3 | 0 | 2 | 2 | 0 | 0 | 98 | 49 | 0.28728316 | 1 | 1 | 0.455 | 1 | 1 | 1 | -0.182 |
| SE | 7849 | LRRK2 | LRRK2 | chr12 | + | 40293544 | 40293663 | 40287350 | 40287539 | 40294844 | 40294914 | 513 | 192 | 66 | 3 | 6 | 0 | 2 | 74 | 0 | 0 | 98 | 49 | 0.292052716 | 1 | 0.988 | 0.941 | 1 | 1 | 1 | -0.024 |
| RI | 51 | GSTK1 | GSTK1 | chr7 | + | 143264991 | 143265296 | 143264991 | 143265092 | 143265260 | 143265296 | 26 | 2 | 5 | 15 | 4 | 4 | 2 | 3 | 1 | 6 | 98 | 49 | 0.297917554 | 1 | 0.464 | 0.2 | 0.385 | 0.5 | 0.2 | 0 |
| SE | 1947 | CCND3 | CCND3 | chr6 | - | 41940369 | 41940585 | 41937234 | 41937394 | 42048500 | 42048894 | 13 | 20 | 6 | 3 | 0 | 1 | 1 | 23 | 0 | 1 | 98 | 49 | 0.297942063 | 1 | 0.684 | 1 | 0.75 | 1 | 0.92 | -0.149 |
| SE | 13953 | DYSF | DYSF | chr2 | + | 71656161 | 71656290 | 71643964 | 71644063 | 71658877 | 71659033 | 52 | 21 | 7 | 0 | 0 | 0 | 2 | 17 | 2 | 0 | 98 | 49 | 0.298053201 | 1 | 1 | 1 | 1 | 0.333 | 1 | 0.334 |

| | | | | | | | | | | | | | | | | | | | | | | | | | | | | | | | |
|---|---|---|---|---|---|---|---|---|---|---|---|---|---|---|---|---|---|---|---|---|---|---|---|---|---|---|---|---|---|---|---|
| SE | 10845 | ITGAX | ITGAX | chr16 | + | 31371333 | 31371497 | 31371083 | 31371214 | 31371629 | 31371784 | 48 | 48 | 16 | 2 | 2 | 1 | 1 | 13 | 1 | 0 | 98 | 49 | 0.299662353 | 1 | 0.923 | 0.923 | 0.889 | 0.333 | 1 | 0.245 |
| RI | 401 | CDK5RAP3 | CDK5RAP3 | chr17 | + | 47975513 | 47976013 | 47975513 | 47975653 | 47975868 | 47976013 | 40 | 14 | 4 | 4 | 0 | 0 | 2 | 6 | 0 | 0 | 98 | 49 | 0.300766182 | 1 | 0.833 | 1 | 1 | 1 | 1 | -0.056 |
| SE | 7862 | LRRK2 | LRRK2 | chr12 | + | 40319987 | 40320175 | 40315211 | 40315300 | 40321033 | 40321188 | 398 | 102 | 74 | 35 | 7 | 2 | 2 | 64 | 4 | 2 | 98 | 49 | 0.301827803 | 1 | 0.85 | 0.879 | 0.949 | 0.2 | 0.941 | 0.322 |
| SE | 7112 | RGS18 | RGS18 | chr1 | + | 192159219 | 192159321 | 192158461 | 192158756 | 192160377 | 192160439 | 116 | 31 | 14 | 0 | 0 | 4 | 4 | 22 | 0 | 0 | 98 | 49 | 0.302925336 | 1 | 1 | 1 | 0.636 | 1 | 1 | -0.121 |
| SE | 13218 | SIGLEC5 | SIGLEC5 | chr19 | - | 51627461 | 51627746 | 51627148 | 51627248 | 51627833 | 51628091 | 15 | 19 | 18 | 0 | 0 | 0 | 2 | 8 | 2 | 0 | 98 | 49 | 0.30502781 | 1 | 1 | 1 | 1 | 0.333 | 1 | 0.334 |
| SE | 6675 | ARHGAP4 | ARHGAP4 | chrX | - | 153919566 | 153919686 | 153919154 | 153919283 | 153920625 | 153920808 | 4 | 4 | 2 | 21 | 15 | 2 | 2 | 0 | 2 | 2 | 98 | 49 | 0.305352645 | 1 | 0.087 | 0.118 | 0.333 | 0.333 | 0 | 0.013 |
| SE | 700 | SLC45A4 | SLC45A4 | chr8 | - | 141217089 | 141217189 | 141215758 | 141215970 | 141218010 | 141219029 | 34 | 25 | 4 | 4 | 0 | 0 | 4 | 2 | 0 | 0 | 98 | 49 | 0.305687075 | 1 | 0.81 | 1 | 1 | 1 | 1 | -0.063 |
| SE | 3766 | CD47 | CD47 | chr3 | - | 108051938 | 108051970 | 108043093 | 108047292 | 108057476 | 108057569 | 14 | 6 | 6 | 1 | 0 | 0 | 0 | 5 | 1 | 1 | 81 | 49 | 0.308504515 | 1 | 0.894 | 1 | 1 | 0 | 0.752 | 0.589 |
| A3SS | 2631 | HIF1A | HIF1A | chr14 | + | 61720378 | 61720572 | 61720381 | 61720572 | 61695400 | 61695839 | 25 | 8 | 3 | 40 | 22 | 5 | 1 | 4 | 1 | 14 | 52 | 49 | 0.310039241 | 1 | 0.371 | 0.255 | 0.361 | 0.485 | 0.212 | -0.019 |
| SE | 3738 | DDX58 | DDX58 | chr9 | - | 32500804 | 32500939 | 32493760 | 32493942 | 32526060 | 32526324 | 51 | 14 | 9 | 1 | 3 | 0 | 1 | 33 | 0 | 0 | 98 | 49 | 0.312908735 | 1 | 0.962 | 0.7 | 1 | 1 | 1 | -0.113 |
| SE | 15478 | SLC6A6 | SLC6A6 | chr3 | + | 14479084 | 14479185 | 14478465 | 14478568 | 14481670 | 14481841 | 129 | 53 | 23 | 4 | 1 | 0 | 3 | 12 | 0 | 0 | 98 | 49 | 0.313509545 | 1 | 0.942 | 0.964 | 1 | 1 | 1 | -0.031 |
| A3SS | 3149 | TMEM91 | TMEM91 | chr19 | + | 41383560 | 41384082 | 41383873 | 41384082 | 41382771 | 41382921 | 3 | 5 | 6 | 0 | 3 | 0 | 1 | 3 | 0 | 0 | 98 | 49 | 0.313679604 | 1 | 1 | 0.455 | 1 | 1 | 1 | -0.182 |
| SE | 4183 | GIT2 | GIT2 | chr12 | - | 109951211 | 109951316 | 109939164 | 109939247 | 109953091 | 109953234 | 17 | 5 | 5 | 0 | 0 | 3 | 1 | 2 | 0 | 0 | 98 | 49 | 0.315095602 | 1 | 1 | 1 | 0.455 | 1 | 1 | -0.182 |
| SE | 8978 | SAT1 | SAT1 | chrX | + | 23784293 | 23784403 | 23783799 | 23783883 | 23785327 | 23785429 | 114 | 53 | 25 | 115 | 88 | 54 | 5 | 44 | 10 | 78 | 98 | 49 | 0.315758853 | 1 | 0.331 | 0.231 | 0.188 | 0.2 | 0.22 | 0.04 |
| A5SS | 2846 | TLE3 | TLE3 | chr15 | - | 70058149 | 70058291 | 70058158 | 70058291 | 70057458 | 70057658 | 11 | 0 | 0 | 22 | 8 | 12 | 0 | 2 | 1 | 4 | 58 | 49 | 0.317935248 | 1 | 0.297 | 0 | 0 | 0 | 0.297 | -0.05 |
| A3SS | 2087 | MAPKAPK2 | MAPKAPK2 | chr1 | + | 206732037 | 206734285 | 206732574 | 206734285 | 206731838 | 206731919 | 19 | 8 | 0 | 21 | 9 | 11 | 0 | 4 | 1 | 9 | 98 | 49 | 0.319337215 | 1 | 0.311 | 0.308 | 0 | 0 | 0.182 | 0.115 |
| SE | 13061 | FOSB | FOSB | chr19 | + | 45471193 | 45471301 | 45470628 | 45470949 | 45472550 | 45475179 | 62 | 18 | 1 | 9 | 4 | 0 | 3 | 9 | 2 | 0 | 98 | 49 | 0.319578232 | 1 | 0.775 | 0.692 | 1 | 0.429 | 1 | 0.108 |
| SE | 5706 | SERPINA1 | SERPINA1 | chr14 | - | 94388818 | 94389010 | 94388559 | 94388660 | 94390456 | 94390692 | 62 | 16 | 16 | 33 | 5 | 10 | 1 | 16 | 1 | 3 | 98 | 49 | 0.319885457 | 1 | 0.484 | 0.615 | 0.444 | 0.333 | 0.727 | -0.016 |
| SE | 11222 | CASP4 | CASP4 | chr11 | - | 104950924 | 104951098 | 104949542 | 104949777 | 104954746 | 104955001 | 92 | 21 | 45 | 15 | 4 | 0 | 0 | 17 | 1 | 2 | 98 | 49 | 0.319912879 | 1 | 0.754 | 0.724 | 1 | 0 | 0.81 | 0.421 |
| RI | 295 | DDIT3 | DDIT3 | chr12 | - | 57517268 | 57517753 | 57517268 | 57517595 | 57517705 | 57517753 | 97 | 35 | 5 | 3 | 0 | 0 | 2 | 10 | 0 | 0 | 98 | 49 | 0.322152533 | 1 | 0.942 | 1 | 1 | 1 | 1 | -0.019 |
| SE | 4483 | USP15 | USP15 | chr12 | + | 62325871 | 62325933 | 62321463 | 62321609 | 62349220 | 62349307 | 283 | 86 | 43 | 5 | 0 | 1 | 2 | 32 | 0 | 1 | 98 | 49 | 0.323933701 | 1 | 0.966 | 1 | 0.956 | 1 | 0.941 | 0.003 |
| SE | 10726 | EIF1 | EIF1 | chr17 | + | 41689777 | 41689941 | 41688874 | 41689069 | 41690087 | 41690189 | 39 | 17 | 12 | 2 | 2 | 0 | 1 | 10 | 0 | 0 | 98 | 49 | 0.32396072 | 1 | 0.907 | 0.81 | 1 | 1 | 1 | -0.094 |
| SE | 11056 | THEMIS2 | THEMIS2 | chr1 | + | 27881970 | 27883043 | 27879643 | 27880054 | 27885294 | 27885451 | 71 | 18 | 19 | 2 | 0 | 1 | 1 | 19 | 0 | 1 | 98 | 49 | 0.326458678 | 1 | 0.947 | 1 | 0.905 | 1 | 0.905 | -0.002 |
| SE | 5702 | SERPINA1 | SERPINA1 | chr14 | - | 94388559 | 94388663 | 94382591 | 94383241 | 94390456 | 94390692 | 97 | 33 | 42 | 107 | 35 | 38 | 0 | 17 | 2 | 20 | 98 | 49 | 0.327279187 | 1 | 0.312 | 0.32 | 0.356 | 0 | 0.298 | 0.18 |
| SE | 12965 | DDX60L | DDX60L | chr4 | - | 168419301 | 168419397 | 168416681 | 168416797 | 168420260 | 168420380 | 26 | 27 | 8 | 3 | 0 | 0 | 1 | 32 | 0 | 1 | 98 | 49 | 0.327799215 | 1 | 0.812 | 1 | 1 | 1 | 0.941 | -0.033 |
| SE | 10741 | MYO1F | MYO1F | chr19 | - | 8548035 | 8548122 | 8545649 | 8545736 | 8548236 | 8548317 | 301 | 188 | 125 | 0 | 0 | 0 | 9 | 61 | 0 | 2 | 98 | 49 | 0.329790459 | 1 | 1 | 1 | 1 | 1 | 0.938 | 0.031 |
| SE | 2766 | PCBP2 | PCBP2 | chr12 | + | 53467804 | 53467843 | 53467220 | 53467293 | 53468776 | 53468832 | 60 | 29 | 10 | 14 | 0 | 2 | 1 | 9 | 0 | 7 | 88 | 49 | 0.331743206 | 1 | 0.705 | 1 | 0.736 | 1 | 0.417 | 0.105 |
| SE | 12248 | SMCHD1 | SMCHD1 | chr18 | + | 2751277 | 2751393 | 2750349 | 2750507 | 2752487 | 2752552 | 188 | 121 | 48 | 0 | 6 | 0 | 8 | 40 | 0 | 1 | 98 | 49 | 0.332012882 | 1 | 1 | 0.91 | 1 | 1 | 0.952 | -0.006 |
| SE | 12962 | DDX60L | DDX60L | chr4 | - | 168395958 | 168396124 | 168394460 | 168394617 | 168400825 | 168400978 | 83 | 27 | 6 | 3 | 0 | 0 | 6 | 32 | 0 | 0 | 98 | 49 | 0.332424323 | 1 | 0.933 | 1 | 1 | 1 | 1 | -0.022 |
| SE | 10367 | AKNA | AKNA | chr9 | - | 114381059 | 114381446 | 114376465 | 114377532 | 114394356 | 114394405 | 88 | 34 | 19 | 3 | 0 | 0 | 1 | 14 | 0 | 0 | 98 | 49 | 0.332785768 | 1 | 0.936 | 1 | 1 | 1 | 1 | -0.021 |
| RI | 773 | MAP7D1 | MAP7D1 | chr1 | + | 36176198 | 36176581 | 36176198 | 36176321 | 36176417 | 36176581 | 28 | 18 | 2 | 6 | 1 | 4 | 1 | 8 | 1 | 1 | 98 | 49 | 0.333527727 | 1 | 0.7 | 0.9 | 0.2 | 0.333 | 0.8 | 0.033 |
| A5SS | 744 | HAUS4 | HAUS4 | chr14 | - | 22956915 | 22957142 | 22956923 | 22957142 | 22955099 | 22955176 | 4 | 3 | 6 | 6 | 3 | 1 | 0 | 3 | 1 | 2 | 57 | 49 | 0.335219927 | 1 | 0.364 | 0.462 | 0.838 | 0 | 0.563 | 0.273 |
| SE | 10958 | CR1 | CR1 | chr1 | + | 207609289 | 207609688 | 207607250 | 207607336 | 207611676 | 207611853 | 63 | 11 | 18 | 1 | 3 | 0 | 1 | 20 | 0 | 0 | 98 | 49 | 0.335963878 | 1 | 0.969 | 0.647 | 1 | 1 | 1 | -0.128 |
| SE | 12631 | ALOX5 | ALOX5 | chr10 | + | 45382482 | 45382681 | 45374175 | 45374429 | 45395854 | 45395936 | 196 | 76 | 31 | 0 | 0 | 0 | 6 | 67 | 0 | 2 | 98 | 49 | 0.338181246 | 1 | 1 | 1 | 1 | 1 | 0.944 | 0.028 |
| SE | 15588 | CTNNB1 | CTNNB1 | chr3 | + | 41227207 | 41227352 | 41225659 | 41225861 | 41233340 | 41233444 | 34 | 12 | 26 | 4 | 0 | 0 | 1 | 9 | 0 | 0 | 98 | 49 | 0.339007481 | 1 | 0.81 | 1 | 1 | 1 | 1 | -0.063 |
| A5SS | 1106 | SERPINA1 | SERPINA1 | chr14 | - | 94390419 | 94390692 | 94390456 | 94390692 | 94382591 | 94383241 | 87 | 62 | 16 | 107 | 35 | 38 | 4 | 5 | 2 | 20 | 86 | 49 | 0.339487748 | 1 | 0.317 | 0.502 | 0.193 | 0.533 | 0.125 | 0.008 |
| SE | 7959 | TNRC6B | TNRC6B | chr22 | + | 40125771 | 40125862 | 40117054 | 40117128 | 40156114 | 40156182 | 38 | 24 | 8 | 4 | 0 | 0 | 1 | 5 | 0 | 0 | 98 | 49 | 0.340420177 | 1 | 0.826 | 1 | 1 | 1 | 1 | -0.058 |
| SE | 3287 | RNASET2 | RNASET2 | chr6 | - | 166948569 | 166948625 | 166946681 | 166946739 | 166952487 | 166952548 | 287 | 68 | 83 | 3 | 2 | 2 | 8 | 51 | 1 | 0 | 98 | 49 | 0.341060884 | 1 | 0.98 | 0.944 | 0.954 | 0.8 | 1 | 0.059 |
| A3SS | 988 | UBE2D3 | UBE2D3 | chr4 | - | 102826484 | 102826844 | 102826484 | 102826636 | 102827591 | 102827948 | 126 | 52 | 28 | 6 | 1 | 2 | 5 | 24 | 0 | 2 | 98 | 49 | 0.341867183 | 1 | 0.913 | 0.963 | 0.875 | 1 | 0.857 | -0.012 |
| SE | 3141 | EIF4A2 | EIF4A2 | chr3 | + | 186784961 | 186785101 | 186784563 | 186784696 | 186785882 | 186786051 | 32 | 7 | 7 | 25 | 4 | 0 | 1 | 6 | 2 | 3 | 98 | 49 | 0.342609716 | 1 | 0.39 | 0.467 | 1 | 0.2 | 0.5 | 0.269 |
| SE | 15542 | LTB | LTB | chr6 | - | 31581813 | 31581859 | 31581558 | 31581630 | 31582255 | 31582425 | 30 | 15 | 4 | 4 | 1 | 2 | 0 | 5 | 3 | 0 | 95 | 49 | 0.345211048 | 1 | 0.795 | 0.886 | 0.508 | 0 | 1 | 0.23 |
| SE | 516 | FGD3 | FGD3 | chr9 | + | 93002924 | 93003014 | 92976207 | 92976709 | 93004000 | 93004137 | 107 | 64 | 24 | 0 | 5 | 0 | 10 | 22 | 0 | 0 | 98 | 49 | 0.346206533 | 1 | 1 | 0.865 | 1 | 1 | 1 | -0.045 |
| SE | 10737 | MYO1F | MYO1F | chr19 | - | 8539946 | 8540028 | 8536948 | 8537055 | 8541905 | 8541991 | 210 | 144 | 78 | 0 | 0 | 0 | 6 | 55 | 0 | 2 | 98 | 49 | 0.346482776 | 1 | 1 | 1 | 1 | 1 | 0.932 | 0.034 |
| SE | 5964 | SORL1 | SORL1 | chr11 | + | 121577280 | 121577400 | 121574240 | 121574363 | 121583457 | 121583583 | 414 | 148 | 108 | 3 | 0 | 0 | 8 | 61 | 0 | 3 | 98 | 49 | 0.346526483 | 1 | 0.986 | 1 | 1 | 1 | 0.91 | 0.04 |
| SE | 976 | DOCK5 | DOCK5 | chr8 | + | 25368570 | 25368725 | 25366869 | 25366970 | 25369555 | 25369641 | 85 | 22 | 4 | 3 | 0 | 0 | 1 | 12 | 0 | 0 | 98 | 49 | 0.347461072 | 1 | 0.934 | 1 | 1 | 1 | 1 | -0.022 |
| SE | 11431 | MX1 | MX1 | chr21 | + | 41443787 | 41443866 | 41441715 | 41441914 | 41445447 | 41445570 | 50 | 61 | 8 | 0 | 0 | 0 | 4 | 62 | 0 | 2 | 98 | 49 | 0.348348103 | 1 | 1 | 1 | 1 | 1 | 0.939 | 0.03 |
| SE | 2179 | PRRC2A | PRRC2A | chr6 | + | 31625791 | 31625871 | 31625459 | 31625611 | 31626019 | 31626162 | 22 | 1 | 6 | 1 | 2 | 0 | 4 | 4 | 0 | 1 | 98 | 49 | 0.352306411 | 1 | 0.917 | 0.2 | 1 | 1 | 0.667 | -0.128 |
| SE | 3265 | DGAT2 | DGAT2 | chr11 | + | 75784617 | 75784746 | 75768732 | 75769112 | 75790187 | 75790295 | 33 | 26 | 25 | 0 | 4 | 0 | 7 | 3 | 0 | 0 | 98 | 49 | 0.352706146 | 1 | 1 | 0.765 | 1 | 1 | 1 | -0.078 |
| SE | 492 | ABCA1 | ABCA1 | chr9 | - | 104810799 | 104810924 | 104809465 | 104809564 | 104812573 | 104812722 | 18 | 1 | 4 | 6 | 5 | 2 | 0 | 3 | 1 | 1 | 98 | 49 | 0.353286236 | 1 | 0.6 | 0.091 | 0.5 | 0 | 0.6 | 0.097 |
| SE | 5920 | MYD88 | MYD88 | chr3 | + | 38140387 | 38140592 | 38139863 | 38139998 | 38140756 | 38140848 | 4 | 5 | 4 | 0 | 0 | 0 | 0 | 8 | 1 | 1 | 98 | 49 | 0.354202895 | 1 | 1 | 1 | 1 | 0 | 0.8 | 0.6 |
| SE | 4297 | PACSIN2 | PACSIN2 | chr22 | - | 42876887 | 42877010 | 42876136 | 42876333 | 42879047 | 42879169 | 37 | 20 | 13 | 2 | 7 | 1 | 0 | 5 | 3 | 0 | 98 | 49 | 0.354244445 | 1 | 0.902 | 0.588 | 0.867 | 0 | 1 | 0.286 |
| SE | 4305 | OAZ1 | OAZ1 | chr19 | + | 2271390 | 2271530 | 2269485 | 2269744 | 2271781 | 2271954 | 190 | 118 | 175 | 6 | 1 | 0 | 8 | 81 | 0 | 0 | 98 | 49 | 0.354607926 | 1 | 0.941 | 0.983 | 1 | 1 | 1 | -0.025 |
| SE | 3027 | XPO6 | XPO6 | chr16 | - | 28135215 | 28135324 | 28133840 | 28133933 | 28146093 | 28146203 | 317 | 187 | 67 | 1 | 0 | 0 | 5 | 50 | 0 | 2 | 98 | 49 | 0.357703086 | 1 | 0.994 | 1 | 1 | 1 | 0.926 | 0.035 |
| SE | 9881 | BAZ2B | BAZ2B | chr2 | - | 159438302 | 159438695 | 159432756 | 159433363 | 159439008 | 159439212 | 112 | 30 | 9 | 3 | 0 | 0 | 1 | 18 | 0 | 0 | 98 | 49 | 0.35775207 | 1 | 0.949 | 1 | 1 | 1 | 1 | -0.017 |
| SE | 33 | FLNA | FLNA | chrX | - | 154357250 | 154357274 | 154354824 | 154355072 | 154357433 | 154357623 | 18 | 6 | 0 | 23 | 5 | 3 | 1 | 4 | 7 | 5 | 73 | 49 | 0.359250106 | 1 | 0.344 | 0.446 | 0 | 0.087 | 0.349 | 0.045 |
| SE | 14571 | ADIPOR1 | ADIPOR1 | chr1 | - | 202954240 | 202954307 | 202950929 | 202951164 | 202958184 | 202958396 | 2 | 5 | 0 | 4 | 2 | 2 | 3 | 1 | 1 | 1 | 98 | 49 | 0.361687671 | 1 | 0.2 | 0.556 | 0 | 0.6 | 0.333 | -0.215 |
| SE | 7060 | CHD1 | CHD1 | chr5 | - | 98881974 | 98882123 | 98881278 | 98881375 | 98883087 | 98883237 | 48 | 28 | 8 | 0 | 4 | 0 | 2 | 14 | 0 | 0 | 98 | 49 | 0.361850765 | 1 | 1 | 0.778 | 1 | 1 | 1 | -0.074 |
| SE | 7781 | CYLD | CYLD | chr16 | + | 50781245 | 50781411 | 50779664 | 50780044 | 50782324 | 50782466 | 72 | 46 | 22 | 1 | 2 | 0 | 1 | 18 | 0 | 0 | 98 | 49 | 0.364325029 | 1 | 0.973 | 0.92 | 1 | 1 | 1 | -0.036 |
| A3SS | 3774 | SUN2 | SUN2 | chr22 | - | 38750224 | 38750383 | 38750224 | 38750320 | 38750897 | 38751035 | 6 | 3 | 1 | 20 | 9 | 6 | 0 | 2 | 1 | 1 | 98 | 49 | 0.365156177 | 1 | 0.13 | 0.143 | 0.077 | 0 | 0.5 | -0.133 |
| A3SS | 603 | BTN3A1 | BTN3A1 | chr6 | + | 26407670 | 26407952 | 26407826 | 26407952 | 26405908 | 26406256 | 33 | 15 | 3 | 0 | 4 | 0 | 2 | 3 | 0 | 0 | 98 | 49 | 0.365827954 | 1 | 1 | 0.652 | 1 | 1 | 1 | -0.116 |
| A3SS | 1073 | OSBPL8 | OSBPL8 | chr12 | - | 76450850 | 76450988 | 76450850 | 76450979 | 76487509 | 76487618 | 30 | 24 | 9 | 8 | 8 | 3 | 0 | 3 | 1 | 1 | 58 | 49 | 0.367474353 | 1 | 0.76 | 0.717 | 0.717 | 0 | 0.717 | 0.373 |
| SE | 6612 | AKAP13 | AKAP13 | chr15 | + | 85662387 | 85662453 | 85658536 | 85658590 | 85664562 | 85664755 | 4 | 1 | 1 | 22 | 14 | 3 | 0 | 2 | 2 | 11 | 98 | 49 | 0.367993412 | 1 | 0.083 | 0.034 | 0.143 | 0 | 0.083 | 0.045 |
| A3SS | 1953 | RNF19B | RNF19B | chr1 | - | 32936445 | 32937309 | 32936445 | 32937259 | 32938396 | 32938528 | 40 | 11 | 15 | 42 | 10 | 11 | 1 | 9 | 0 | 5 | 98 | 49 | 0.368999566 | 1 | 0.323 | 0.355 | 0.405 | 1 | 0.474 | -0.376 |
| SE | 316 | WWC3 | WWC3 | chrX | + | 10117123 | 10117689 | 10116417 | 10116507 | 10122618 | 10122739 | 40 | 23 | 14 | 4 | 0 | 0 | 1 | 4 | 0 | 0 | 98 | 49 | 0.373057457 | 1 | 0.833 | 1 | 1 | 1 | 1 | -0.056 |
| SE | 2394 | TMEM164 | TMEM164 | chrX | + | 110067346 | 110067396 | 110003500 | 110004164 | 110109079 | 110109146 | 65 | 27 | 1 | 3 | 0 | 0 | 2 | 14 | 0 | 0 | 98 | 49 | 0.376246788 | 1 | 0.915 | 1 | 1 | 1 | 1 | -0.028 |
| SE | 13855 | GNAS | GNAS | chr20 | + | 58898940 | 58898985 | 58895611 | 58895684 | 58903530 | 58903585 | 18 | 7 | 2 | 10 | 3 | 1 | 1 | 3 | 0 | 2 | 94 | 49 | 0.377418069 | 1 | 0.484 | 0.549 | 0.51 | 1 | 0.439 | -0.205 |
| SE | 7972 | LCOR | LCOR | chr10 | + | 96944112 | 96944245 | 96907667 | 96907747 | 96949007 | 96949295 | 17 | 8 | 4 | 4 | 0 | 0 | 2 | 0 | 0 | 1 | 98 | 49 | 0.37804421 | 1 | 0.68 | 1 | 1 | 1 | 0 | 0.393 |
| SE | 15173 | SHKBP1 | SHKBP1 | chr19 | + | 40589081 | 40589178 | 40588623 | 40588779 | 40590729 | 40590853 | 67 | 18 | 15 | 3 | 0 | 0 | 4 | 12 | 0 | 0 | 98 | 49 | 0.378060424 | 1 | 0.918 | 1 | 1 | 1 | 1 | -0.027 |
| SE | 3824 | PTPRE | PTPRE | chr10 | + | 128060938 | 128061015 | 128056122 | 128056213 | 128061678 | 128061715 | 118 | 64 | 26 | 2 | 2 | 0 | 3 | 11 | 0 | 0 | 98 | 49 | 0.378698267 | 1 | 0.967 | 0.941 | 1 | 1 | 1 | -0.031 |
| SE | 282 | SH3KBP1 | SH3KBP1 | chrX | - | 19683822 | 19684028 | 19645399 | 19645475 | 19695611 | 19695741 | 59 | 21 | 14 | 1 | 0 | 3 | 3 | 13 | 0 | 0 | 98 | 49 | 0.379406837 | 1 | 0.967 | 1 | 0.7 | 1 | 1 | -0.111 |
| SE | 12240 | SMCHD1 | SMCHD1 | chr18 | + | 2718157 | 2718235 | 2707806 | 2707920 | 2718314 | 2718434 | 320 | 153 | 43 | 3 | 0 | 4 | 2 | 71 | 0 | 0 | 98 | 49 | 0.37951052 | 1 | 0.982 | 1 | 0.843 | 1 | 1 | -0.058 |
| SE | 4221 | DEF8 | DEF8 | chr16 | + | 89962011 | 89962125 | 89961736 | 89961864 | 89963362 | 89963443 | 32 | 31 | 5 | 2 | 1 | 0 | 3 | 5 | 0 | 0 | 98 | 49 | 0.379976534 | 1 | 0.889 | 0.939 | 1 | 1 | 1 | -0.057 |
| SE | 2802 | IFI44L | IFI44L | chr1 | + | 78629719 | 78629915 | 78628950 | 78628999 | 78635336 | 78635489 | 9 | 3 | 1 | 0 | 1 | 0 | 3 | 61 | 0 | 2 | 98 | 49 | 0.382224179 | 1 | 1 | 0.6 | 1 | 1 | 0.938 | -0.102 |
| RI | 232 | ZFP36L1 | ZFP36L1 | chr14 | - | 68787654 | 68790492 | 68787654 | 68788809 | 68789526 | 68790492 | 173 | 46 | 11 | 5 | 5 | 2 | 4 | 17 | 0 | 2 | 98 | 49 | 0.382258712 | 1 | 0.945 | 0.821 | 0.733 | 1 | 0.81 | -0.072 |
| SE | 11429 | MX1 | MX1 | chr21 | + | 41430532 | 41430608 | 41427754 | 41427866 | 41432049 | 41432175 | 41 | 24 | 6 | 0 | 0 | 0 | 6 | 38 | 0 | 2 | 98 | 49 | 0.382268666 | 1 | 1 | 1 | 1 | 1 | 0.905 | 0.047 |
| SE | 7863 | LRRK2 | LRRK2 | chr12 | + | 40322034 | 40322181 | 40321033 | 40321188 | 40322318 | 40322510 | 353 | 137 | 46 | 0 | 0 | 0 | 1 | 45 | 0 | 2 | 98 | 49 | 0.388091869 | 1 | 1 | 1 | 1 | 1 | 0.918 | 0.041 |
| SE | 1579 | IGF2R | IGF2R | chr6 | + | 160029549 | 160029655 | 160027184 | 160027314 | 160032550 | 160032713 | 344 | 145 | 49 | 7 | 5 | 0 | 14 | 45 | 0 | 4 | 98 | 49 | 0.389795203 | 1 | 0.961 | 0.935 | 1 | 1 | 0.849 | 0.041 |
| A3SS | 2558 | SF1 | SF1 | chr11 | - | 64764603 | 64766155 | 64764603 | 64765518 | 64766899 | 64767079 | 16 | 4 | 1 | 4 | 0 | 0 | 3 | 7 | 0 | 0 | 98 | 49 | 0.390237161 | 1 | 0.667 | 1 | 1 | 1 | 1 | -0.111 |
| A3SS | 3621 | TPD52L2 | TPD52L2 | chr20 | + | 63882718 | 63882820 | 63882749 | 63882820 | 63875815 | 63875875 | 48 | 23 | 8 | 2 | 1 | 0 | 2 | 11 | 0 | 1 | 80 | 49 | 0.392016835 | 1 | 0.936 | 0.934 | 1 | 1 | 0.871 | 0.021 |
| A3SS | 3930 | FBXL5 | FBXL5 | chr4 | - | 15644492 | 15644708 | 15644492 | 15644657 | 15655203 | 15655412 | 81 | 58 | 25 | 1 | 2 | 0 | 1 | 40 | 0 | 0 | 98 | 49 | 0.393596853 | 1 | 0.976 | 0.935 | 1 | 1 | 1 | -0.03 |
| SE | 12020 | KLHL2 | KLHL2 | chr4 | + | 165238777 | 165238899 | 165228806 | 165228913 | 165263196 | 165263359 | 14 | 0 | 6 | 4 | 3 | 1 | 0 | 4 | 1 | 2 | 98 | 49 | 0.39720417 | 1 | 0.636 | 0 | 0.75 | 0 | 0.5 | 0.212 |
| SE | 11956 | CHD3 | CHD3 | chr17 | + | 7902936 | 7903061 | 7902609 | 7902727 | 7903271 | 7903503 | 9 | 7 | 2 | 0 | 0 | 0 | 1 | 3 | 0 | 2 | 98 | 49 | 0.39801607 | 1 | 1 | 1 | 1 | 1 | 0.429 | 0.285 |
| A3SS | 2329 | GMPR2 | GMPR2 | chr14 | + | 24238245 | 24239241 | 24238588 | 24239241 | 24237519 | 24237562 | 23 | 13 | 4 | 3 | 2 | 4 | 0 | 3 | 2 | 0 | 98 | 49 | 0.398116415 | 1 | 0.793 | 0.765 | 0.333 | 0 | 1 | 0.13 |
| A3SS | 752 | ISG20 | ISG20 | chr15 | + | 88650234 | 88652310 | 88652169 | 88652310 | 88639342 | 88639594 | 47 | 52 | 24 | 1 | 1 | 1 | 1 | 17 | 0 | 0 | 98 | 49 | 0.399153396 | 1 | 0.959 | 0.963 | 0.923 | 1 | 1 | -0.052 |
| SE | 14416 | ANKRD44 | ANKRD44 | chr2 | - | 197122649 | 197122792 | 197121331 | 197121544 | 197125380 | 197125468 | 151 | 81 | 24 | 2 | 2 | 0 | 4 | 11 | 0 | 0 | 98 | 49 | 0.399864693 | 1 | 0.974 | 0.953 | 1 | 1 | 1 | -0.024 |
| SE | 6529 | LYST | LYST | chr1 | - | 235791698 | 235792125 | 235788700 | 235788845 | 235793502 | 235793612 | 169 | 78 | 9 | 1 | 3 | 0 | 1 | 20 | 0 | 0 | 98 | 49 | 0.402422043 | 1 | 0.988 | 0.929 | 1 | 1 | 1 | -0.028 |
| SE | 10735 | MYO1F | MYO1F | chr19 | - | 8530458 | 8530573 | 8530195 | 8530365 | 8536251 | 8536396 | 77 | 70 | 51 | 1 | 3 | 0 | 3 | 27 | 0 | 0 | 98 | 49 | 0.403164003 | 1 | 0.975 | 0.921 | 1 | 1 | 1 | -0.035 |
| SE | 17074 | GRK6 | GRK6 | chr5 | + | 177433346 | 177433410 | 177433146 | 177433239 | 177433535 | 177433676 | 28 | 29 | 18 | 2 | 2 | 0 | 4 | 11 | 0 | 0 | 98 | 49 | 0.403207343 | 1 | 0.875 | 0.879 | 1 | 1 | 1 | -0.082 |
| SE | 6005 | NIN | NIN | chr14 | - | 50821873 | 50822077 | 50806736 | 50806818 | 50830463 | 50830517 | 61 | 31 | 6 | 3 | 1 | 0 | 3 | 8 | 0 | 1 | 98 | 49 | 0.403251294 | 1 | 0.91 | 0.939 | 1 | 1 | 0.8 | 0.05 |
| SE | 10818 | HIF1A | HIF1A | chr14 | + | 61720381 | 61720572 | 61695400 | 61695839 | 61721508 | 61721654 | 109 | 71 | 25 | 1 | 3 | 0 | 6 | 35 | 0 | 0 | 98 | 49 | 0.404683891 | 1 | 0.982 | 0.922 | 1 | 1 | 1 | -0.032 |
| SE | 15298 | TRIOBP | TRIOBP | chr22 | + | 37769261 | 37769375 | 37768073 | 37768176 | 37771649 | 37771736 | 12 | 2 | 1 | 0 | 0 | 0 | 2 | 3 | 0 | 2 | 98 | 49 | 0.404694173 | 1 | 1 | 1 | 1 | 1 | 0.429 | 0.285 |
| SE | 7631 | CSDE1 | CSDE1 | chr1 | - | 114739691 | 114739890 | 114737962 | 114738072 | 114749820 | 114750207 | 111 | 34 | 20 | 3 | 0 | 0 | 2 | 23 | 0 | 0 | 98 | 49 | 0.405048782 | 1 | 0.949 | 1 | 1 | 1 | 1 | -0.017 |
| SE | 10003 | ADRBK1 | ADRBK1 | chr11 | + | 67281092 | 67281184 | 67280731 | 67280783 | 67281458 | 67281558 | 152 | 66 | 46 | 2 | 2 | 0 | 2 | 32 | 0 | 0 | 98 | 49 | 0.405794829 | 1 | 0.974 | 0.943 | 1 | 1 | 1 | -0.028 |
| SE | 15609 | HIPK3 | HIPK3 | chr11 | + | 33348521 | 33348818 | 33347851 | 33348013 | 33349146 | 33349287 | 54 | 19 | 5 | 2 | 1 | 0 | 1 | 4 | 0 | 0 | 98 | 49 | 0.406461955 | 1 | 0.931 | 0.905 | 1 | 1 | 1 | -0.055 |
| SE | 8973 | FCGR2A | FCGR2A | chr1 | + | 161510833 | 161510956 | 161509819 | 161510074 | 161513894 | 161513932 | 349 | 205 | 165 | 1 | 3 | 0 | 1 | 66 | 0 | 2 | 98 | 49 | 0.40674608 | 1 | 0.994 | 0.972 | 1 | 1 | 0.943 | 0.017 |
| SE | 14733 | HCK | HCK | chr20 | + | 32054215 | 32054281 | 32052187 | 32052486 | 32071661 | 32071782 | 19 | 4 | 8 | 135 | 59 | 44 | 0 | 5 | 9 | 33 | 98 | 49 | 0.408086651 | 1 | 0.066 | 0.033 | 0.083 | 0 | 0.07 | 0.026 |
| SE | 16016 | CCNL1 | CCNL1 | chr3 | - | 157149286 | 157149385 | 157147555 | 157148589 | 157149484 | 157149596 | 172 | 75 | 20 | 2 | 2 | 0 | 1 | 41 | 0 | 0 | 98 | 49 | 0.408138584 | 1 | 0.977 | 0.949 | 1 | 1 | 1 | -0.025 |
| SE | 2024 | TAPBP | TAPBP | chr6 | - | 33303954 | 33303989 | 33299694 | 33301771 | 33304127 | 33304217 | 78 | 38 | 14 | 3 | 0 | 0 | 2 | 23 | 1 | 0 | 84 | 49 | 0.412371599 | 1 | 0.938 | 1 | 1 | 0.538 | 1 | 0.21 |
| SE | 15171 | SHKBP1 | SHKBP1 | chr19 | + | 40582350 | 40582466 | 40580745 | 40580936 | 40583397 | 40583485 | 110 | 30 | 27 | 3 | 0 | 0 | 1 | 14 | 0 | 0 | 98 | 49 | 0.414401449 | 1 | 0.948 | 1 | 1 | 1 | 1 | -0.017 |
| SE | 368 | RXRA | RXRA | chr9 | + | 134417157 | 134417327 | 134408939 | 134409119 | 134421675 | 134421805 | 69 | 31 | 17 | 3 | 0 | 0 | 3 | 11 | 0 | 0 | 98 | 49 | 0.41476143 | 1 | 0.92 | 1 | 1 | 1 | 1 | -0.027 |
| SE | 14779 | PLAUR | PLAUR | chr19 | - | 43665315 | 43665459 | 43656478 | 43656640 | 43667580 | 43667691 | 119 | 68 | 19 | 2 | 2 | 0 | 13 | 65 | 0 | 0 | 98 | 49 | 0.415108943 | 1 | 0.967 | 0.944 | 1 | 1 | 1 | -0.03 |
| SE | 12822 | MCL1 | MCL1 | chr1 | - | 150578243 | 150578491 | 150574550 | 150577491 | 150578842 | 150578963 | 544 | 221 | 161 | 66 | 59 | 10 | 5 | 123 | 4 | 15 | 98 | 49 | 0.417762806 | 1 | 0.805 | 0.652 | 0.89 | 0.385 | 0.804 | 0.188 |
| SE | 6782 | JMJD1C | JMJD1C | chr10 | - | 63186214 | 63186383 | 63185562 | 63185653 | 63189167 | 63189446 | 115 | 40 | 13 | 3 | 0 | 0 | 2 | 9 | 0 | 0 | 98 | 49 | 0.417962582 | 1 | 0.95 | 1 | 1 | 1 | 1 | -0.017 |
| SE | 611 | TLN1 | TLN1 | chr9 | - | 35704668 | 35704815 | 35704331 | 35704498 | 35705550 | 35705670 | 156 | 32 | 27 | 2 | 3 | 0 | 3 | 28 | 0 | 0 | 98 | 49 | 0.420055523 | 1 | 0.975 | 0.842 | 1 | 1 | 1 | -0.061 |
| SE | 7706 | TBXAS1 | TBXAS1 | chr7 | + | 139953367 | 139953456 | 139936190 | 139936307 | 139957633 | 139957764 | 90 | 67 | 13 | 3 | 0 | 0 | 1 | 32 | 0 | 0 | 98 | 49 | 0.420526793 | 1 | 0.938 | 1 | 1 | 1 | 1 | -0.021 |
| A3SS | 2476 | LST1 | LST1 | chr6 | + | 31588517 | 31588909 | 31588562 | 31588909 | 31587199 | 31587318 | 108 | 64 | 58 | 35 | 11 | 14 | 3 | 14 | 1 | 6 | 94 | 49 | 0.420817006 | 1 | 0.617 | 0.752 | 0.684 | 0.61 | 0.549 | 0.105 |
| SE | 8099 | DEDD2 | DEDD2 | chr19 | - | 42202050 | 42202072 | 42198592 | 42199829 | 42209699 | 42209840 | 3 | 0 | 1 | 16 | 25 | 11 | 0 | 3 | 3 | 5 | 71 | 49 | 0.421373914 | 1 | 0.115 | 0 | 0.059 | 0 | 0.293 | -0.088 |
| SE | 12176 | RNF213 | RNF213 | chr17 | + | 80369767 | 80369867 | 80369501 | 80369671 | 80371873 | 80371985 | 109 | 38 | 8 | 0 | 1 | 0 | 10 | 63 | 0 | 2 | 98 | 49 | 0.423196935 | 1 | 1 | 0.95 | 1 | 1 | 0.94 | 0.013 |
| SE | 12195 | SP110 | SP110 | chr2 | - | 230216780 | 230216928 | 230214949 | 230215118 | 230219873 | 230220112 | 27 | 29 | 6 | 1 | 2 | 0 | 5 | 11 | 0 | 0 | 98 | 49 | 0.423386143 | 1 | 0.931 | 0.879 | 1 | 1 | 1 | -0.063 |
| SE | 10846 | ITGAX | ITGAX | chr16 | + | 31372377 | 31372509 | 31371629 | 31371784 | 31372596 | 31372670 | 186 | 116 | 43 | 2 | 0 | 0 | 16 | 38 | 0 | 2 | 98 | 49 | 0.423798056 | 1 | 0.979 | 1 | 1 | 1 | 0.905 | 0.04 |
| A5SS | 2156 | HN1 | HN1 | chr17 | - | 75147519 | 75147653 | 75147555 | 75147653 | 75146665 | 75146684 | 0 | 1 | 4 | 34 | 19 | 8 | 0 | 1 | 1 | 2 | 85 | 49 | 0.424216982 | 1 | 0 | 0.029 | 0.224 | 0 | 0.224 | -0.028 |
| SE | 8525 | NFKBIZ | NFKBIZ | chr3 | + | 101855394 | 101855458 | 101855061 | 101855208 | 101855732 | 101855902 | 124 | 40 | 7 | 3 | 0 | 0 | 1 | 28 | 0 | 0 | 98 | 49 | 0.426137175 | 1 | 0.954 | 1 | 1 | 1 | 1 | -0.015 |
| SE | 3708 | IRF9 | IRF9 | chr14 | + | 24163877 | 24163959 | 24163377 | 24163508 | 24164062 | 24164134 | 46 | 27 | 7 | 4 | 6 | 0 | 3 | 25 | 1 | 5 | 98 | 49 | 0.426518129 | 1 | 0.852 | 0.692 | 1 | 0.6 | 0.714 | 0.191 |
| SE | 9065 | SIPA1 | SIPA1 | chr11 | + | 65649261 | 65649480 | 65647383 | 65647658 | 65649560 | 65649672 | 60 | 21 | 9 | 3 | 0 | 0 | 3 | 9 | 0 | 0 | 98 | 49 | 0.426604104 | 1 | 0.909 | 1 | 1 | 1 | 1 | -0.03 |
| A3SS | 605 | BTN3A1 | BTN3A1 | chr6 | + | 26412411 | 26415216 | 26413168 | 26415216 | 26411554 | 26411581 | 19 | 6 | 4 | 17 | 0 | 0 | 1 | 5 | 0 | 2 | 98 | 49 | 0.426625721 | 1 | 0.358 | 1 | 1 | 1 | 0.556 | 0.008 |
| SE | 13589 | FAM129A | FAM129A | chr1 | - | 184803584 | 184803692 | 184798078 | 184798190 | 184805945 | 184806056 | 214 | 78 | 64 | 2 | 4 | 0 | 1 | 38 | 0 | 0 | 98 | 49 | 0.428739516 | 1 | 0.982 | 0.907 | 1 | 1 | 1 | -0.037 |
| SE | 16420 | RHOA | RHOA | chr3 | - | 49368427 | 49368548 | 49362495 | 49362626 | 49375433 | 49375591 | 157 | 74 | 57 | 3 | 1 | 0 | 3 | 34 | 0 | 0 | 98 | 49 | 0.431166683 | 1 | 0.963 | 0.974 | 1 | 1 | 1 | -0.021 |
| SE | 5062 | ARAP1 | ARAP1 | chr11 | - | 72699004 | 72699107 | 72697910 | 72698106 | 72699416 | 72699552 | 163 | 116 | 39 | 0 | 1 | 0 | 8 | 34 | 0 | 2 | 98 | 49 | 0.434451175 | 1 | 1 | 0.983 | 1 | 1 | 0.895 | 0.047 |
| A3SS | 4087 | SPI1 | SPI1 | chr11 | - | 47375632 | 47375732 | 47375632 | 47375729 | 47378308 | 47378576 | 51 | 15 | 7 | 58 | 22 | 9 | 8 | 16 | 4 | 19 | 52 | 49 | 0.435975739 | 1 | 0.453 | 0.391 | 0.423 | 0.653 | 0.442 | -0.125 |
| SE | 2306 | HLA-A | HLA-A | chr6 | + | 29944499 | 29944616 | 29944121 | 29944397 | 29945058 | 29945091 | 734 | 514 | 291 | 1 | 4 | 0 | 41 | 187 | 1 | 2 | 98 | 49 | 0.436480895 | 1 | 0.997 | 0.985 | 1 | 0.953 | 0.979 | 0.028 |
| SE | 2689 | STX4 | STX4 | chr16 | + | 31034969 | 31035040 | 31034225 | 31034536 | 31037925 | 31038034 | 13 | 13 | 1 | 0 | 0 | 2 | 1 | 3 | 0 | 0 | 98 | 49 | 0.436872697 | 1 | 1 | 1 | 0.2 | 1 | 1 | -0.267 |
| SE | 14888 | PREX1 | PREX1 | chr20 | - | 48650893 | 48651055 | 48649995 | 48650206 | 48651395 | 48651583 | 129 | 77 | 48 | 0 | 0 | 0 | 3 | 32 | 0 | 2 | 98 | 49 | 0.43772807 | 1 | 1 | 1 | 1 | 1 | 0.889 | 0.055 |
| A3SS | 606 | BTN3A1 | BTN3A1 | chr6 | + | 26412704 | 26415216 | 26413168 | 26415216 | 26411554 | 26411581 | 21 | 8 | 4 | 17 | 0 | 0 | 1 | 5 | 0 | 2 | 98 | 49 | 0.439597639 | 1 | 0.382 | 1 | 1 | 1 | 0.556 | 0.016 |
| SE | 4482 | HBP1 | HBP1 | chr7 | + | 107189291 | 107189448 | 107186568 | 107186681 | 107190172 | 107190317 | 29 | 13 | 19 | 4 | 0 | 0 | 1 | 1 | 0 | 1 | 98 | 49 | 0.439686131 | 1 | 0.784 | 1 | 1 | 1 | 0.333 | 0.261 |
| A5SS | 1551 | OS9 | OS9 | chr12 | + | 57718145 | 57718421 | 57718145 | 57718376 | 57718992 | 57719182 | 99 | 50 | 22 | 1 | 2 | 0 | 6 | 6 | 0 | 0 | 94 | 49 | 0.443569573 | 1 | 0.981 | 0.929 | 1 | 1 | 1 | -0.03 |
| SE | 14047 | TMEM127 | TMEM127 | chr2 | - | 96265137 | 96265512 | 96254832 | 96254997 | 96265868 | 96266013 | 27 | 6 | 12 | 0 | 3 | 0 | 3 | 11 | 0 | 1 | 98 | 49 | 0.444110003 | 1 | 1 | 0.5 | 1 | 1 | 0.846 | -0.09 |
| SE | 13049 | CHD2 | CHD2 | chr15 | + | 92937517 | 92937625 | 92929029 | 92929091 | 92939577 | 92939718 | 65 | 24 | 3 | 4 | 0 | 0 | 1 | 7 | 0 | 0 | 98 | 49 | 0.446218154 | 1 | 0.89 | 1 | 1 | 1 | 1 | -0.037 |
| SE | 12793 | JAK3 | JAK3 | chr19 | - | 17835082 | 17835215 | 17834851 | 17835003 | 17835923 | 17836051 | 53 | 36 | 7 | 0 | 4 | 0 | 6 | 18 | 0 | 0 | 98 | 49 | 0.447951856 | 1 | 1 | 0.818 | 1 | 1 | 1 | -0.061 |
| RI | 787 | TYMP | TYMP | chr22 | - | 50529495 | 50530085 | 50529495 | 50529719 | 50529903 | 50530085 | 72 | 56 | 11 | 4 | 4 | 5 | 8 | 20 | 0 | 13 | 98 | 49 | 0.449170597 | 1 | 0.9 | 0.875 | 0.524 | 1 | 0.435 | 0.049 |
| A5SS | 313 | EEF1D | EEF1D | chr8 | - | 143581228 | 143581328 | 143581285 | 143581328 | 143581053 | 143581154 | 89 | 43 | 46 | 3 | 0 | 0 | 8 | 16 | 0 | 0 | 98 | 49 | 0.449604503 | 1 | 0.937 | 1 | 1 | 1 | 1 | -0.021 |
| SE | 7926 | EPB41 | EPB41 | chr1 | + | 29060421 | 29060484 | 29053103 | 29053312 | 29064981 | 29065158 | 3 | 0 | 0 | 21 | 11 | 11 | 0 | 0 | 2 | 10 | 98 | 49 | 0.449822548 | 1 | 0.067 | 0 | 0 | 0 | 0 | 0.022 |
| SE | 9197 | ACSL1 | ACSL1 | chr4 | - | 184764852 | 184764925 | 184763166 | 184763255 | 184766621 | 184766756 | 147 | 62 | 37 | 0 | 1 | 0 | 2 | 44 | 0 | 2 | 98 | 49 | 0.450670684 | 1 | 1 | 0.969 | 1 | 1 | 0.917 | 0.031 |
| SE | 15280 | TOM1 | TOM1 | chr22 | + | 35321958 | 35322037 | 35317876 | 35317961 | 35323027 | 35323177 | 24 | 34 | 9 | 1 | 2 | 0 | 4 | 11 | 0 | 1 | 98 | 49 | 0.45078102 | 1 | 0.923 | 0.895 | 1 | 1 | 0.846 | 0.016 |
| RI | 1042 | PLIN2 | PLIN2 | chr9 | - | 19126113 | 19126448 | 19126113 | 19126309 | 19126396 | 19126448 | 1 | 2 | 0 | 2 | 0 | 1 | 0 | 1 | 1 | 0 | 98 | 49 | 0.45174834 | 1 | 0.2 | 1 | 0 | 0 | 1 | -0.1 |
| SE | 1939 | UBR2 | UBR2 | chr6 | + | 42573733 | 42573993 | 42564021 | 42564397 | 42592150 | 42592229 | 156 | 72 | 10 | 0 | 4 | 1 | 5 | 9 | 0 | 0 | 98 | 49 | 0.452057563 | 1 | 1 | 0.9 | 0.833 | 1 | 1 | -0.089 |
| SE | 16878 | ALPL | ALPL | chr1 | + | 21553977 | 21554142 | 21509364 | 21509517 | 21560625 | 21560745 | 64 | 35 | 11 | 2 | 1 | 0 | 2 | 14 | 0 | 0 | 98 | 49 | 0.457733702 | 1 | 0.941 | 0.946 | 1 | 1 | 1 | -0.038 |
| SE | 3002 | R3HDM4 | R3HDM4 | chr19 | - | 901421 | 901546 | 900828 | 900952 | 901975 | 902130 | 128 | 109 | 50 | 5 | 0 | 0 | 6 | 19 | 0 | 1 | 98 | 49 | 0.458344066 | 1 | 0.928 | 1 | 1 | 1 | 0.905 | 0.023 |
| SE | 11816 | PTPN6 | PTPN6 | chr12 | + | 6957918 | 6958073 | 6957653 | 6957785 | 6959926 | 6959994 | 138 | 67 | 28 | 23 | 4 | 3 | 4 | 43 | 4 | 3 | 98 | 49 | 0.458449304 | 1 | 0.75 | 0.893 | 0.824 | 0.333 | 0.878 | 0.217 |
| SE | 8817 | ATXN2 | ATXN2 | chr12 | - | 111510384 | 111510582 | 111509890 | 111509998 | 111513356 | 111513539 | 7 | 5 | 0 | 2 | 0 | 1 | 0 | 2 | 1 | 0 | 98 | 49 | 0.460013403 | 1 | 0.636 | 1 | 0 | 0 | 1 | 0.045 |
| SE | 16308 | HOTAIRM1 | HOTAIRM1 | chr7 | + | 27099097 | 27099366 | 27096093 | 27096388 | 27099778 | 27100258 | 5 | 0 | 0 | 21 | 27 | 5 | 0 | 0 | 1 | 1 | 98 | 49 | 0.464250946 | 1 | 0.106 | 0 | 0 | 0 | 0 | 0.035 |
| SE | 1582 | IGF2R | IGF2R | chr6 | + | 160048374 | 160048543 | 160047791 | 160047907 | 160050472 | 160050652 | 233 | 104 | 38 | 0 | 2 | 0 | 3 | 42 | 0 | 2 | 98 | 49 | 0.464661849 | 1 | 1 | 0.963 | 1 | 1 | 0.913 | 0.031 |
| SE | 11898 | TRIM25 | TRIM25 | chr17 | - | 56895925 | 56895952 | 56895520 | 56895604 | 56899114 | 56899180 | 36 | 14 | 12 | 2 | 2 | 0 | 1 | 3 | 0 | 1 | 76 | 49 | 0.46806809 | 1 | 0.921 | 0.819 | 1 | 1 | 0.659 | 0.084 |
| A3SS | 3409 | TMEM127 | TMEM127 | chr2 | - | 96265137 | 96265512 | 96265137 | 96265489 | 96265868 | 96266013 | 21 | 3 | 3 | 10 | 7 | 1 | 4 | 2 | 3 | 4 | 72 | 49 | 0.469441269 | 1 | 0.588 | 0.226 | 0.671 | 0.476 | 0.254 | 0.13 |
| SE | 8049 | ADAM8 | ADAM8 | chr10 | - | 133263687 | 133263765 | 133262415 | 133263233 | 133267351 | 133267417 | 100 | 41 | 18 | 16 | 3 | 2 | 4 | 10 | 2 | 2 | 98 | 49 | 0.469863379 | 1 | 0.758 | 0.872 | 0.818 | 0.5 | 0.714 | 0.209 |
| A5SS | 740 | RNF31 | RNF31 | chr14 | + | 24148257 | 24148477 | 24148257 | 24148413 | 24148641 | 24148701 | 4 | 1 | 0 | 6 | 3 | 2 | 1 | 1 | 0 | 3 | 98 | 49 | 0.472635957 | 1 | 0.25 | 0.143 | 0 | 1 | 0.143 | -0.441 |
| SE | 5917 | MYD88 | MYD88 | chr3 | + | 38139863 | 38139998 | 38138477 | 38139028 | 38140756 | 38140848 | 50 | 12 | 5 | 3 | 0 | 0 | 6 | 12 | 0 | 0 | 98 | 49 | 0.473541925 | 1 | 0.893 | 1 | 1 | 1 | 1 | -0.036 |
| A5SS | 2716 | POLD4 | POLD4 | chr11 | - | 67353293 | 67353596 | 67353363 | 67353596 | 67352987 | 67353077 | 14 | 11 | 3 | 0 | 1 | 2 | 1 | 1 | 0 | 0 | 98 | 49 | 0.476503373 | 1 | 1 | 0.846 | 0.429 | 1 | 1 | -0.242 |
| SE | 1136 | LINC-PINT | LINC-PINT | chr7 | - | 130967388 | 130967517 | 130944159 | 130945835 | 130984052 | 130984109 | 36 | 7 | 0 | 14 | 1 | 2 | 0 | 5 | 1 | 2 | 98 | 49 | 0.476619732 | 1 | 0.562 | 0.778 | 0 | 0 | 0.556 | 0.169 |
| SE | 8831 | DOCK2 | DOCK2 | chr5 | + | 169714359 | 169714457 | 169714027 | 169714211 | 169716212 | 169716302 | 109 | 62 | 13 | 3 | 0 | 0 | 13 | 24 | 0 | 0 | 98 | 49 | 0.477632073 | 1 | 0.948 | 1 | 1 | 1 | 1 | -0.017 |
| SE | 5223 | MX2 | MX2 | chr21 | + | 41380016 | 41380151 | 41377788 | 41377981 | 41382409 | 41382564 | 32 | 26 | 37 | 5 | 1 | 0 | 4 | 25 | 0 | 2 | 98 | 49 | 0.478098223 | 1 | 0.762 | 0.929 | 1 | 1 | 0.862 | -0.034 |
| SE | 4593 | PARP8 | PARP8 | chr5 | + | 50821212 | 50821338 | 50815431 | 50815524 | 50822334 | 50822400 | 53 | 30 | 6 | 2 | 2 | 0 | 2 | 9 | 0 | 1 | 98 | 49 | 0.478298961 | 1 | 0.93 | 0.882 | 1 | 1 | 0.818 | 0.028 |
| SE | 12594 | STXBP2 | STXBP2 | chr19 | + | 7640730 | 7640809 | 7639730 | 7639807 | 7640899 | 7641003 | 28 | 16 | 11 | 2 | 1 | 0 | 1 | 7 | 0 | 0 | 98 | 49 | 0.478353872 | 1 | 0.875 | 0.889 | 1 | 1 | 1 | -0.079 |
| A3SS | 3788 | LPPR2 | LPPR2 | chr19 | + | 11359531 | 11359720 | 11359606 | 11359720 | 11357659 | 11357739 | 20 | 16 | 12 | 4 | 0 | 0 | 1 | 6 | 0 | 0 | 98 | 49 | 0.479378889 | 1 | 0.714 | 1 | 1 | 1 | 1 | -0.095 |
| SE | 4419 | ARHGAP9 | ARHGAP9 | chr12 | - | 57476870 | 57476963 | 57476589 | 57476651 | 57477155 | 57477269 | 192 | 72 | 60 | 7 | 5 | 0 | 2 | 25 | 0 | 2 | 98 | 49 | 0.480342678 | 1 | 0.932 | 0.878 | 1 | 1 | 0.862 | 0.006 |
| SE | 5177 | SLC12A9 | SLC12A9 | chr7 | + | 100861736 | 100861911 | 100861391 | 100861584 | 100862680 | 100862827 | 53 | 23 | 12 | 2 | 1 | 0 | 2 | 16 | 0 | 0 | 98 | 49 | 0.480487308 | 1 | 0.93 | 0.92 | 1 | 1 | 1 | -0.05 |

| | | | | | | | | | | | | | | | | | | | | | | | | | | | | | | |
|---|---|---|---|---|---|---|---|---|---|---|---|---|---|---|---|---|---|---|---|---|---|---|---|---|---|---|---|---|---|---|
| SE | 12573 VAV1 | VAV1 | chr19 | + | 6822418 | 6822514 | 6822220 | 6822329 | 6825052 | 6825121 | 91 | 45 | 14 | 3 | 7 | 0 | 0 | 15 | 2 | 0 | 98 | 49 | 0.481387676 | 1 | 0.938 | 0.763 | 1 | 0 | 1 | 0.4 |
| SE | 13803 PARP14 | PARP14 | chr3 | + | 122699389 | 122701635 | 122695425 | 122695662 | 122703741 | 122703978 | 78 | 24 | 1 | 0 | 0 | 0 | 8 | 30 | 0 | 2 | 98 | 49 | 0.481745882 | 1 | 1 | 1 | 1 | 1 | 0.882 | 0.059 |
| SE | 8919 ATP6V0B | ATP6V0B | chr1 | + | 43977025 | 43977216 | 43976772 | 43976824 | 43977980 | 43978300 | 91 | 62 | 65 | 1 | 3 | 0 | 4 | 22 | 0 | 0 | 98 | 49 | 0.483149345 | 1 | 0.978 | 0.912 | 1 | 1 | 1 | -0.037 |
| SE | 15965 DPEP2 | DPEP2 | chr16 | - | 67990046 | 67990131 | 67989322 | 67989398 | 67990820 | 67990997 | 105 | 42 | 37 | 3 | 0 | 0 | 1 | 15 | 0 | 0 | 98 | 49 | 0.483939387 | 1 | 0.946 | 1 | 1 | 1 | 1 | -0.018 |
| SE | 5221 BCL2A1 | BCL2A1 | chr15 | - | 79967636 | 79967692 | 79960889 | 79961174 | 79970699 | 79971301 | 10 | 4 | 6 | 104 | 148 | 64 | 0 | 2 | 1 | 19 | 98 | 49 | 0.485539689 | 1 | 0.046 | 0.013 | 0.045 | 0 | 0.05 | 0.01 |
| SE | 15339 TLE3 | TLE3 | chr15 | - | 70059409 | 70059460 | 70058662 | 70058815 | 70060529 | 70060649 | 157 | 61 | 34 | 0 | 0 | 0 | 4 | 31 | 2 | 0 | 98 | 49 | 0.486682137 | 1 | 1 | 1 | 1 | 0.5 | 1 | 0.25 |
| SE | 14401 STAT1 | STAT1 | chr2 | - | 191013524 | 191013678 | 191009875 | 191010004 | 191014017 | 191014250 | 89 | 53 | 9 | 3 | 4 | 0 | 5 | 34 | 0 | 2 | 98 | 49 | 0.491119546 | 1 | 0.937 | 0.869 | 1 | 1 | 0.895 | -0.012 |
| SE | 7748 ACAP1 | ACAP1 | chr17 | + | 7337311 | 7337369 | 7336528 | 7336787 | 7341947 | 7342067 | 24 | 18 | 5 | 2 | 1 | 0 | 1 | 3 | 0 | 0 | 98 | 49 | 0.491953713 | 1 | 0.857 | 0.9 | 1 | 1 | 1 | -0.081 |
| SE | 1248 PILRA | PILRA | chr7 | + | 100389887 | 100390106 | 100374043 | 100374433 | 100397878 | 100397912 | 91 | 45 | 54 | 17 | 6 | 18 | 4 | 21 | 0 | 5 | 98 | 49 | 0.492450455 | 1 | 0.728 | 0.789 | 0.6 | 1 | 0.677 | -0.133 |
| SE | 3064 DDX5 | DDX5 | chr17 | - | 64504676 | 64504842 | 64504221 | 64504318 | 64506075 | 64506366 | 293 | 110 | 22 | 1 | 4 | 0 | 2 | 38 | 0 | 0 | 98 | 49 | 0.493007314 | 1 | 0.993 | 0.932 | 1 | 1 | 1 | -0.025 |
| SE | 14933 OSBPL2 | OSBPL2 | chr20 | + | 62259980 | 62260125 | 62256056 | 62256221 | 62263615 | 62263691 | 30 | 14 | 10 | 2 | 1 | 0 | 2 | 3 | 0 | 0 | 98 | 49 | 0.494721497 | 1 | 0.882 | 0.875 | 1 | 1 | 1 | -0.081 |
| SE | 13839 PPM1B | PPM1B | chr2 | + | 44209209 | 44209327 | 44201185 | 44202045 | 44217966 | 44218078 | 27 | 9 | 12 | 0 | 3 | 0 | 3 | 7 | 0 | 0 | 98 | 49 | 0.494769315 | 1 | 1 | 0.6 | 1 | 1 | 1 | -0.133 |
| SE | 10608 MICAL2 | MICAL2 | chr11 | + | 12204249 | 12204457 | 12162078 | 12162419 | 12208022 | 12208139 | 74 | 30 | 7 | 3 | 0 | 0 | 1 | 17 | 0 | 1 | 98 | 49 | 0.49895827 | 1 | 0.925 | 1 | 1 | 1 | 0.895 | 0.027 |
| SE | 7206 NBEAL2 | NBEAL2 | chr3 | + | 47003173 | 47003309 | 47002956 | 47003081 | 47003815 | 47003976 | 120 | 36 | 9 | 0 | 0 | 0 | 14 | 31 | 0 | 2 | 98 | 49 | 0.499363766 | 1 | 1 | 1 | 1 | 1 | 0.886 | 0.057 |
| A3SS | 3712 TOM1 | TOM1 | chr22 | + | 35322998 | 35323177 | 35323027 | 35323177 | 35321958 | 35322037 | 7 | 5 | 0 | 14 | 19 | 3 | 1 | 5 | 4 | 6 | 78 | 49 | 0.499951338 | 1 | 0.239 | 0.142 | 0 | 0.136 | 0.344 | -0.113 |
| SE | 12634 ALOX5 | ALOX5 | chr10 | + | 45444115 | 45444286 | 45443727 | 45443828 | 45445507 | 45446119 | 96 | 37 | 30 | 0 | 1 | 0 | 2 | 41 | 0 | 2 | 98 | 49 | 0.500085279 | 1 | 1 | 0.949 | 1 | 1 | 0.911 | 0.027 |
| A5SS | 2045 RPL13A | RPL13A | chr19 | + | 49489849 | 49489922 | 49489849 | 49489912 | 49490231 | 49490297 | 99 | 13 | 1 | 3 | 1 | 0 | 3 | 11 | 0 | 1 | 59 | 49 | 0.501906011 | 1 | 0.965 | 0.915 | 1 | 1 | 0.901 | 0.009 |
| SE | 14397 STAT1 | STAT1 | chr2 | - | 190979771 | 190979866 | 190978855 | 190979001 | 190980619 | 190980669 | 41 | 25 | 5 | 2 | 4 | 0 | 1 | 20 | 1 | 2 | 98 | 49 | 0.502698128 | 1 | 0.911 | 0.758 | 1 | 0.333 | 0.833 | 0.307 |
| SE | 11487 FLOT2 | FLOT2 | chr17 | - | 28881191 | 28881375 | 28880712 | 28880862 | 28881813 | 28882028 | 144 | 64 | 62 | 0 | 0 | 0 | 3 | 27 | 0 | 2 | 98 | 49 | 0.502725055 | 1 | 1 | 1 | 1 | 1 | 0.871 | 0.065 |
| SE | 15289 NCF4 | NCF4 | chr22 | + | 36864044 | 36864129 | 36860987 | 36861203 | 36864918 | 36865072 | 197 | 128 | 96 | 5 | 0 | 0 | 8 | 28 | 0 | 0 | 98 | 49 | 0.502830419 | 1 | 0.952 | 1 | 1 | 1 | 1 | -0.016 |
| SE | 2484 ARRB2 | ARRB2 | chr17 | + | 4718260 | 4718345 | 4717887 | 4718023 | 4718611 | 4718684 | 106 | 27 | 85 | 30 | 9 | 10 | 0 | 26 | 1 | 3 | 98 | 49 | 0.504310317 | 1 | 0.639 | 0.6 | 0.81 | 0 | 0.812 | 0.277 |
| SE | 14887 PREX1 | PREX1 | chr20 | - | 48649995 | 48650206 | 48649299 | 48649576 | 48650893 | 48651055 | 184 | 95 | 70 | 2 | 3 | 0 | 8 | 40 | 0 | 0 | 98 | 49 | 0.505945346 | 1 | 0.979 | 0.941 | 1 | 1 | 1 | -0.027 |
| A3SS | 1764 GPBP1 | GPBP1 | chr5 | + | 57249408 | 57249576 | 57249453 | 57249576 | 57247074 | 57247215 | 42 | 25 | 25 | 2 | 0 | 1 | 1 | 5 | 0 | 0 | 94 | 49 | 0.506543634 | 1 | 0.916 | 1 | 0.929 | 1 | 1 | -0.052 |
| SE | 7869 LRRK2 | LRRK2 | chr12 | + | 40354298 | 40354492 | 40351538 | 40351733 | 40356114 | 40356187 | 206 | 77 | 31 | 1 | 0 | 0 | 2 | 26 | 0 | 2 | 98 | 49 | 0.510946613 | 1 | 0.99 | 1 | 1 | 1 | 0.867 | 0.063 |
| SE | 14024 SMAD2 | SMAD2 | chr18 | - | 47870474 | 47870564 | 47869242 | 47869436 | 47896520 | 47896809 | 55 | 18 | 19 | 3 | 0 | 0 | 1 | 1 | 0 | 0 | 98 | 49 | 0.516332408 | 1 | 0.902 | 1 | 1 | 1 | 1 | -0.033 |
| SE | 715 DENND3 | DENND3 | chr8 | + | 141190283 | 141190417 | 141188985 | 141189146 | 141192330 | 141192449 | 68 | 28 | 4 | 2 | 2 | 1 | 6 | 22 | 0 | 0 | 98 | 49 | 0.516511922 | 1 | 0.944 | 0.933 | 0.667 | 1 | 1 | -0.152 |
| SE | 5699 SERPINA1 | SERPINA1 | chr14 | - | 94388559 | 94388660 | 94382591 | 94383241 | 94390456 | 94390692 | 102 | 20 | 47 | 107 | 35 | 38 | 1 | 16 | 2 | 20 | 98 | 49 | 0.517955822 | 1 | 0.323 | 0.222 | 0.382 | 0.2 | 0.286 | 0.066 |
| SE | 3934 TPT1 | TPT1 | chr13 | - | 45339496 | 45339602 | 45338559 | 45338776 | 45339993 | 45340184 | 544 | 187 | 145 | 4 | 4 | 0 | 6 | 79 | 0 | 3 | 98 | 49 | 0.518168375 | 1 | 0.986 | 0.959 | 1 | 1 | 0.929 | 0.017 |
| SE | 5918 MYD88 | MYD88 | chr3 | + | 38140387 | 38140568 | 38138477 | 38139028 | 38140756 | 38140848 | 12 | 1 | 3 | 3 | 0 | 0 | 3 | 7 | 0 | 0 | 98 | 49 | 0.522673916 | 1 | 0.667 | 1 | 1 | 1 | 1 | -0.111 |
| SE | 5308 LDB1 | LDB1 | chr10 | - | 102110528 | 102110701 | 102109920 | 102110043 | 102110868 | 102110971 | 34 | 8 | 7 | 0 | 0 | 0 | 3 | 5 | 0 | 2 | 98 | 49 | 0.523770865 | 1 | 1 | 1 | 1 | 1 | 0.556 | 0.222 |
| SE | 2036 TAP1 | TAP1 | chr6 | - | 32850943 | 32851149 | 32850319 | 32850517 | 32852108 | 32852239 | 70 | 36 | 31 | 2 | 0 | 1 | 2 | 13 | 0 | 0 | 98 | 49 | 0.525221449 | 1 | 0.946 | 1 | 0.939 | 1 | 1 | -0.038 |
| SE | 6288 TCEB2 | TCEB2 | chr16 | - | 2775450 | 2775556 | 2771413 | 2772102 | 2776992 | 2777127 | 35 | 36 | 31 | 1 | 0 | 3 | 1 | 12 | 0 | 0 | 98 | 49 | 0.525837001 | 1 | 0.946 | 1 | 0.838 | 1 | 1 | -0.072 |
| SE | 11452 CARD16 | CARD16 | chr11 | - | 105043482 | 105043545 | 105041325 | 105041719 | 105044391 | 105044658 | 48 | 16 | 42 | 20 | 3 | 6 | 1 | 14 | 1 | 5 | 98 | 49 | 0.528217041 | 1 | 0.545 | 0.727 | 0.778 | 0.333 | 0.583 | 0.225 |
| SE | 4012 MTMR6 | MTMR6 | chr13 | - | 25251648 | 25251775 | 25246200 | 25249492 | 25251852 | 25251984 | 27 | 9 | 8 | 0 | 0 | 0 | 2 | 5 | 0 | 2 | 98 | 49 | 0.529097092 | 1 | 1 | 1 | 1 | 1 | 0.556 | 0.222 |
| SE | 16389 HERC5 | HERC5 | chr4 | + | 88462134 | 88462356 | 88460094 | 88460171 | 88463531 | 88463623 | 3 | 17 | 1 | 1 | 1 | 0 | 4 | 25 | 0 | 2 | 98 | 49 | 0.529188344 | 1 | 0.6 | 0.895 | 1 | 1 | 0.862 | -0.099 |
| SE | 7026 ELOVL5 | ELOVL5 | chr6 | - | 53287861 | 53287942 | 53276178 | 53276256 | 53291775 | 53291963 | 1 | 1 | 2 | 47 | 19 | 21 | 0 | 0 | 2 | 5 | 98 | 49 | 0.530437106 | 1 | 0.011 | 0.026 | 0.045 | 0 | 0 | 0.027 |
| A5SS | 598 STX16 | STX16 | chr20 | + | 58651252 | 58652138 | 58651252 | 58652087 | 58667489 | 58667597 | 13 | 8 | 3 | 3 | 2 | 2 | 4 | 7 | 0 | 3 | 98 | 49 | 0.531451593 | 1 | 0.684 | 0.667 | 0.429 | 1 | 0.538 | -0.176 |
| SE | 6530 LYST | LYST | chr1 | - | 235804503 | 235804665 | 235802907 | 235803064 | 235805742 | 235806772 | 186 | 55 | 10 | 0 | 0 | 0 | 4 | 25 | 0 | 2 | 98 | 49 | 0.533639671 | 1 | 1 | 1 | 1 | 1 | 0.862 | 0.069 |
| SE | 15466 STAT5A | STAT5A | chr17 | + | 42291971 | 42292061 | 42289865 | 42290022 | 42295618 | 42295793 | 9 | 5 | 6 | 0 | 2 | 1 | 1 | 2 | 2 | 0 | 98 | 49 | 0.535091642 | 1 | 1 | 0.556 | 0.75 | 0.2 | 1 | 0.169 |
| SE | 5626 CD74 | CD74 | chr5 | - | 150403120 | 150403312 | 150402562 | 150402625 | 150404679 | 150404767 | 21 | 2 | 0 | 114 | 6 | 7 | 2 | 7 | 2 | 32 | 98 | 49 | 0.537096648 | 1 | 0.084 | 0.143 | 0 | 0.333 | 0.099 | -0.14 |
| SE | 14925 ZBP1 | ZBP1 | chr20 | - | 57616243 | 57616468 | 57615511 | 57615580 | 57620261 | 57620576 | 13 | 7 | 4 | 3 | 0 | 0 | 1 | 9 | 0 | 0 | 98 | 49 | 0.538885454 | 1 | 0.684 | 1 | 1 | 1 | 1 | -0.105 |
| SE | 273 GK | GK | chrX | + | 30724100 | 30724181 | 30720851 | 30720995 | 30728731 | 30731460 | 69 | 41 | 22 | 2 | 2 | 0 | 4 | 5 | 0 | 0 | 98 | 49 | 0.541146332 | 1 | 0.945 | 0.911 | 1 | 1 | 1 | -0.048 |
| SE | 1731 VNN3 | VNN3 | chr6 | - | 132726721 | 132727012 | 132724605 | 132724979 | 132728740 | 132728933 | 79 | 48 | 5 | 22 | 9 | 1 | 0 | 11 | 1 | 1 | 98 | 49 | 0.54337128 | 1 | 0.642 | 0.727 | 0.714 | 0 | 0.846 | 0.271 |
| SE | 15760 LSP1 | LSP1 | chr11 | + | 1881431 | 1881596 | 1880086 | 1880224 | 1883418 | 1883560 | 223 | 169 | 106 | 1 | 1 | 3 | 7 | 58 | 0 | 0 | 98 | 49 | 0.544999045 | 1 | 0.991 | 0.988 | 0.946 | 1 | 1 | -0.025 |
| SE | 2829 STK38L | STK38L | chr12 | + | 27308338 | 27308461 | 27302136 | 27302188 | 27309113 | 27309197 | 42 | 12 | 7 | 2 | 1 | 0 | 3 | 8 | 0 | 0 | 98 | 49 | 0.545126619 | 1 | 0.913 | 0.857 | 1 | 1 | 1 | -0.077 |
| SE | 8154 CAP1 | CAP1 | chr1 | + | 40069869 | 40069874 | 40067539 | 40067717 | 40070158 | 40070282 | 72 | 34 | 18 | 0 | 4 | 0 | 2 | 10 | 0 | 0 | 98 | 49 | 0.545980056 | 1 | 1 | 0.81 | 1 | 1 | 1 | -0.063 |
| SE | 12089 ARRDC3 | ARRDC3 | chr5 | - | 91376620 | 91376768 | 91375510 | 91375613 | 91378693 | 91378775 | 106 | 33 | 48 | 18 | 10 | 1 | 1 | 13 | 0 | 6 | 98 | 49 | 0.54624919 | 1 | 0.746 | 0.623 | 0.96 | 1 | 0.52 | 0.016 |
| SE | 352 NOTCH1 | NOTCH1 | chr9 | - | 136515980 | 136516094 | 136515482 | 136515716 | 136517271 | 136517385 | 59 | 30 | 12 | 2 | 1 | 0 | 1 | 10 | 0 | 0 | 98 | 49 | 0.546699365 | 1 | 0.937 | 0.937 | 1 | 1 | 1 | -0.042 |
| SE | 10817 HIF1A | HIF1A | chr14 | + | 61720378 | 61720572 | 61695400 | 61695839 | 61721508 | 61721654 | 94 | 57 | 23 | 1 | 3 | 0 | 6 | 25 | 0 | 0 | 98 | 49 | 0.547705846 | 1 | 0.979 | 0.905 | 1 | 1 | 1 | -0.039 |
| SE | 10243 LST1 | LST1 | chr6 | + | 31587943 | 31587966 | 31587199 | 31587318 | 31588496 | 31588909 | 9 | 9 | 2 | 3 | 2 | 0 | 1 | 1 | 0 | 3 | 72 | 49 | 0.547785373 | 1 | 0.671 | 0.754 | 1 | 1 | 0.185 | 0.216 |
| SE | 953 SARAF | SARAF | chr8 | - | 30073876 | 30074055 | 30069641 | 30070059 | 30082846 | 30083208 | 72 | 48 | 39 | 20 | 10 | 11 | 3 | 13 | 1 | 5 | 98 | 49 | 0.549549436 | 1 | 0.643 | 0.706 | 0.639 | 0.6 | 0.565 | 0.08 |
| SE | 12701 GLUL | GLUL | chr1 | - | 182387130 | 182387292 | 182386255 | 182386402 | 182388571 | 182388750 | 1143 | 456 | 144 | 3 | 3 | 0 | 25 | 173 | 0 | 3 | 98 | 49 | 0.553318418 | 1 | 0.995 | 0.987 | 1 | 1 | 0.966 | 0.011 |
| SE | 9556 HNRNPU | HNRNPU | chr1 | - | 244858010 | 244858274 | 244857597 | 244857717 | 244858728 | 244858841 | 82 | 35 | 16 | 1 | 2 | 0 | 2 | 18 | 0 | 2 | 98 | 49 | 0.553341978 | 1 | 0.976 | 0.897 | 1 | 1 | 0.818 | 0.049 |
| SE | 4125 RPLP0 | RPLP0 | chr12 | - | 120198553 | 120198739 | 120197321 | 120197462 | 120198853 | 120199000 | 41 | 5 | 8 | 3 | 0 | 0 | 1 | 3 | 0 | 0 | 98 | 49 | 0.555263141 | 1 | 0.872 | 1 | 1 | 1 | 1 | -0.043 |
| SE | 8828 DOCK2 | DOCK2 | chr5 | + | 169684195 | 169684350 | 169681743 | 169681879 | 169689251 | 169689333 | 203 | 80 | 23 | 3 | 1 | 1 | 4 | 41 | 0 | 0 | 98 | 49 | 0.555294052 | 1 | 0.971 | 0.976 | 0.92 | 1 | 1 | -0.044 |
| SE | 10034 PRKCD | PRKCD | chr3 | + | 53165103 | 53165215 | 53161206 | 53161428 | 53178403 | 53178537 | 8 | 0 | 2 | 5 | 1 | 0 | 4 | 3 | 2 | 1 | 98 | 49 | 0.558514749 | 1 | 0.444 | 0 | 1 | 0.5 | 0.6 | -0.069 |
| SE | 517 FGD3 | FGD3 | chr9 | + | 93019830 | 93019861 | 93018135 | 93018215 | 93020316 | 93020424 | 85 | 35 | 25 | 2 | 2 | 1 | 3 | 10 | 0 | 1 | 80 | 49 | 0.559138062 | 1 | 0.963 | 0.955 | 1 | 1 | 0.86 | 0.043 |
| SE | 8751 NABP1 | NABP1 | chr2 | + | 191682491 | 191682595 | 191681945 | 191682017 | 191683728 | 191683804 | 42 | 11 | 10 | 48 | 10 | 4 | 2 | 4 | 0 | 5 | 98 | 49 | 0.559187208 | 1 | 0.304 | 0.355 | 0.556 | 1 | 0.286 | -0.238 |
| SE | 4151 PSTPIP1 | PSTPIP1 | chr15 | + | 77032297 | 77032394 | 77031179 | 77031278 | 77035507 | 77035563 | 11 | 5 | 4 | 3 | 0 | 0 | 1 | 1 | 0 | 0 | 98 | 49 | 0.562425178 | 1 | 0.647 | 1 | 1 | 1 | 1 | -0.118 |
| SE | 137 PGK1 | PGK1 | chrX | + | 78124873 | 78125051 | 78123194 | 78123374 | 78125326 | 78125425 | 78 | 40 | 29 | 1 | 2 | 0 | 2 | 30 | 0 | 0 | 98 | 49 | 0.565144298 | 1 | 0.975 | 0.909 | 1 | 1 | 1 | -0.039 |
| A3SS | 2905 TYMP | TYMP | chr22 | - | 50529495 | 50529754 | 50529495 | 50529719 | 50529903 | 50530085 | 38 | 29 | 7 | 4 | 4 | 5 | 3 | 9 | 0 | 13 | 84 | 49 | 0.565719428 | 1 | 0.847 | 0.809 | 0.45 | 1 | 0.288 | 0.058 |
| SE | 12963 DDX60L | DDX60L | chr4 | - | 168395958 | 168396127 | 168394460 | 168394617 | 168400825 | 168400978 | 62 | 58 | 6 | 3 | 0 | 0 | 4 | 29 | 0 | 0 | 98 | 49 | 0.566280009 | 1 | 0.912 | 1 | 1 | 1 | 1 | -0.029 |
| SE | 6242 APBB1IP | APBB1IP | chr10 | + | 26536073 | 26536217 | 26533438 | 26533525 | 26541581 | 26541692 | 99 | 17 | 22 | 1 | 3 | 0 | 1 | 12 | 0 | 0 | 98 | 49 | 0.566991124 | 1 | 0.98 | 0.739 | 1 | 1 | 1 | -0.094 |
| SE | 11052 THEMIS2 | THEMIS2 | chr1 | + | 27879643 | 27880054 | 27876587 | 27876728 | 27881970 | 27882125 | 155 | 32 | 32 | 0 | 5 | 0 | 7 | 42 | 0 | 1 | 98 | 49 | 0.56829042 | 1 | 1 | 0.762 | 1 | 1 | 0.955 | -0.057 |
| SE | 2382 FAM65B | FAM65B | chr6 | - | 24848024 | 24848154 | 24842861 | 24843554 | 24849801 | 24849950 | 314 | 131 | 97 | 3 | 3 | 0 | 3 | 57 | 0 | 0 | 98 | 49 | 0.569484819 | 1 | 0.981 | 0.956 | 1 | 1 | 1 | -0.021 |
| SE | 14618 USP32 | USP32 | chr17 | - | 60180544 | 60180637 | 60177329 | 60179428 | 60181323 | 60181748 | 67 | 31 | 5 | 3 | 0 | 0 | 0 | 12 | 1 | 0 | 98 | 49 | 0.569592115 | 1 | 0.918 | 1 | 1 | 0 | 1 | 0.473 |
| SE | 4932 TLE4 | TLE4 | chr9 | + | 79708592 | 79708786 | 79708117 | 79708250 | 79709622 | 79709699 | 41 | 4 | 11 | 3 | 0 | 0 | 3 | 7 | 0 | 0 | 98 | 49 | 0.571129463 | 1 | 0.872 | 1 | 1 | 1 | 1 | -0.043 |
| SE | 12509 ABCA7 | ABCA7 | chr19 | + | 1063759 | 1063863 | 1063543 | 1063678 | 1064160 | 1064253 | 2 | 5 | 1 | 2 | 0 | 0 | 1 | 3 | 0 | 0 | 98 | 49 | 0.572393395 | 1 | 0.333 | 1 | 1 | 1 | 1 | -0.222 |
| SE | 14792 PHF1 | PHF1 | chr6 | + | 33414954 | 33415144 | 33414724 | 33414829 | 33415234 | 33415329 | 2 | 5 | 1 | 2 | 0 | 0 | 1 | 1 | 0 | 0 | 98 | 49 | 0.572393395 | 1 | 0.333 | 1 | 1 | 1 | 1 | -0.222 |
| A3SS | 2653 IFITM3 | IFITM3 | chr11 | - | 319672 | 319990 | 319672 | 319985 | 320564 | 320914 | 22 | 6 | 8 | 0 | 0 | 3 | 1 | 33 | 0 | 0 | 54 | 49 | 0.573839549 | 1 | 1 | 1 | 0.708 | 1 | 1 | -0.097 |
| SE | 6426 RALGAPA2 | RALGAPA2 | chr20 | - | 20680690 | 20680801 | 20653529 | 20653587 | 20712374 | 20712623 | 23 | 10 | 2 | 2 | 1 | 0 | 2 | 7 | 0 | 0 | 98 | 49 | 0.574229202 | 1 | 0.852 | 0.833 | 1 | 1 | 1 | -0.105 |
| SE | 9970 DUSP1 | DUSP1 | chr5 | - | 172769574 | 172769794 | 172768089 | 172769132 | 172770160 | 172770306 | 842 | 360 | 330 | 3 | 7 | 4 | 17 | 269 | 0 | 3 | 98 | 49 | 0.576118237 | 1 | 0.993 | 0.963 | 0.976 | 1 | 0.978 | -0.012 |
| A5SS | 1592 TBC1D10C | TBC1D10C | chr11 | + | 67406626 | 67406738 | 67406626 | 67406692 | 67406826 | 67407016 | 38 | 9 | 8 | 8 | 5 | 0 | 1 | 2 | 0 | 4 | 95 | 49 | 0.577419673 | 1 | 0.71 | 0.481 | 1 | 1 | 0.205 | 0.128 |
| SE | 15458 RAF1 | RAF1 | chr3 | - | 12590797 | 12590974 | 12587590 | 12587637 | 12591707 | 12591792 | 180 | 64 | 32 | 2 | 0 | 0 | 1 | 24 | 0 | 2 | 98 | 49 | 0.579441471 | 1 | 0.978 | 1 | 1 | 1 | 0.857 | 0.064 |
| SE | 6704 LAPTM5 | LAPTM5 | chr1 | - | 30742455 | 30742549 | 30741639 | 30741716 | 30757658 | 30757836 | 90 | 35 | 46 | 1 | 0 | 0 | 3 | 22 | 0 | 2 | 98 | 49 | 0.580977501 | 1 | 0.978 | 1 | 1 | 1 | 0.846 | 0.07 |
| A3SS | 2839 PTPN6 | PTPN6 | chr12 | + | 6960662 | 6960945 | 6960805 | 6960945 | 6960343 | 6960435 | 26 | 31 | 0 | 95 | 46 | 27 | 1 | 8 | 1 | 17 | 98 | 49 | 0.581966841 | 1 | 0.12 | 0.252 | 0 | 0.333 | 0.19 | -0.138 |
| SE | 9269 NFKBID | NFKBID | chr19 | - | 35897650 | 35897856 | 35896912 | 35897058 | 35898471 | 35898532 | 18 | 4 | 1 | 0 | 1 | 0 | 7 | 10 | 2 | 0 | 98 | 49 | 0.582524393 | 1 | 1 | 0.667 | 1 | 0.636 | 1 | 0.071 |
| A3SS | 985 ELF1 | ELF1 | chr13 | - | 40949805 | 40949973 | 40949805 | 40949901 | 40951328 | 40951436 | 43 | 12 | 9 | 3 | 0 | 0 | 3 | 15 | 0 | 0 | 98 | 49 | 0.582990467 | 1 | 0.878 | 1 | 1 | 1 | 1 | -0.041 |
| SE | 14246 ITGAL | ITGAL | chr16 | + | 30489253 | 30489386 | 30489081 | 30489155 | 30494211 | 30494363 | 43 | 17 | 4 | 3 | 0 | 0 | 1 | 10 | 0 | 0 | 98 | 49 | 0.583106327 | 1 | 0.878 | 1 | 1 | 1 | 1 | -0.041 |
| SE | 10074 CD53 | CD53 | chr1 | + | 110897808 | 110897892 | 110891391 | 110891471 | 110899123 | 110899936 | 98 | 50 | 56 | 3 | 0 | 0 | 2 | 18 | 0 | 0 | 98 | 49 | 0.583622234 | 1 | 0.942 | 1 | 1 | 1 | 1 | -0.019 |
| SE | 15985 RNF13 | RNF13 | chr3 | + | 149911977 | 149912083 | 149902071 | 149902162 | 149921133 | 149921227 | 129 | 71 | 27 | 0 | 0 | 4 | 3 | 27 | 0 | 0 | 98 | 49 | 0.583653552 | 1 | 1 | 1 | 0.771 | 1 | 1 | -0.076 |
| SE | 10080 CAMK2G | CAMK2G | chr10 | - | 73839538 | 73839601 | 73837467 | 73837511 | 73842168 | 73842211 | 1 | 0 | 3 | 17 | 10 | 5 | 0 | 0 | 1 | 4 | 98 | 49 | 0.584918649 | 1 | 0.029 | 0 | 0.231 | 0 | 0 | 0.087 |
| SE | 12997 SASH3 | SASH3 | chrX | + | 129792327 | 129792476 | 129790939 | 129791081 | 129792626 | 129792836 | 51 | 31 | 9 | 3 | 0 | 0 | 3 | 17 | 0 | 0 | 98 | 49 | 0.586405367 | 1 | 0.895 | 1 | 1 | 1 | 1 | -0.035 |
| SE | 5115 N4BP1 | N4BP1 | chr16 | - | 48551385 | 48551482 | 48548006 | 48548114 | 48553538 | 48553669 | 47 | 10 | 7 | 10 | 5 | 1 | 1 | 10 | 0 | 3 | 98 | 49 | 0.588328632 | 1 | 0.701 | 0.5 | 0.778 | 1 | 0.625 | -0.153 |
| SE | 6964 HCLS1 | HCLS1 | chr3 | - | 121632331 | 121632563 | 121632100 | 121632184 | 121633066 | 121633171 | 174 | 114 | 99 | 2 | 2 | 0 | 10 | 51 | 0 | 2 | 98 | 49 | 0.591514184 | 1 | 0.978 | 0.966 | 1 | 1 | 0.927 | 0.018 |
| SE | 15923 RPS6KA1 | RPS6KA1 | chr1 | + | 26556653 | 26556718 | 26555150 | 26555221 | 26556997 | 26557100 | 45 | 24 | 23 | 3 | 1 | 0 | 4 | 8 | 0 | 0 | 98 | 49 | 0.593177157 | 1 | 0.882 | 0.923 | 1 | 1 | 1 | -0.065 |
| SE | 5916 MYD88 | MYD88 | chr3 | + | 38139863 | 38139998 | 38138477 | 38139028 | 38140387 | 38140568 | 54 | 17 | 9 | 3 | 0 | 0 | 5 | 19 | 2 | 0 | 98 | 49 | 0.594277976 | 1 | 0.9 | 1 | 1 | 0.556 | 1 | 0.189 |
| SE | 4181 GIT2 | GIT2 | chr12 | - | 109951166 | 109951316 | 109947255 | 109947504 | 109953091 | 109953234 | 21 | 13 | 10 | 0 | 3 | 0 | 1 | 5 | 0 | 1 | 98 | 49 | 0.595484855 | 1 | 1 | 0.684 | 1 | 1 | 0.714 | 0.038 |
| SE | 5444 BCL6 | BCL6 | chr3 | - | 187724940 | 187725078 | 187721376 | 187722601 | 187725498 | 187725629 | 143 | 71 | 32 | 1 | 4 | 0 | 8 | 53 | 0 | 0 | 98 | 49 | 0.596641777 | 1 | 0.986 | 0.899 | 1 | 1 | 1 | -0.038 |
| SE | 11633 FKBP8 | FKBP8 | chr19 | - | 18539550 | 18539720 | 18539370 | 18539459 | 18541678 | 18541995 | 119 | 51 | 36 | 3 | 0 | 0 | 2 | 16 | 0 | 0 | 98 | 49 | 0.59812016 | 1 | 0.952 | 1 | 1 | 1 | 1 | -0.016 |
| SE | 11189 CD300LF | CD300LF | chr17 | - | 74705203 | 74705377 | 74704477 | 74704816 | 74712823 | 74713000 | 3 | 2 | 0 | 7 | 4 | 1 | 0 | 0 | 2 | 0 | 98 | 49 | 0.600055751 | 1 | 0.176 | 0.2 | 0 | 0 | 1 | -0.375 |
| SE | 10847 ITGAX | ITGAX | chr16 | + | 31373248 | 31373390 | 31372596 | 31372670 | 31376798 | 31376915 | 156 | 142 | 31 | 10 | 6 | 0 | 12 | 40 | 0 | 5 | 98 | 49 | 0.600219064 | 1 | 0.886 | 0.922 | 1 | 1 | 0.8 | 0.036 |
| SE | 14417 ANKRD44 | ANKRD44 | chr2 | - | 197187022 | 197187106 | 197147026 | 197147105 | 197310577 | 197310797 | 189 | 78 | 20 | 2 | 3 | 0 | 3 | 25 | 0 | 0 | 98 | 49 | 0.600837139 | 1 | 0.979 | 0.929 | 1 | 1 | 1 | -0.031 |
| A3SS | 1461 SERPINA1 | SERPINA1 | chr14 | - | 94388559 | 94388663 | 94388559 | 94388660 | 94390456 | 94390692 | 31 | 18 | 5 | 33 | 5 | 10 | 0 | 5 | 1 | 3 | 52 | 49 | 0.601284271 | 1 | 0.47 | 0.772 | 0.32 | 0 | 0.611 | 0.215 |
| SE | 1108 MKRN1 | MKRN1 | chr7 | - | 140474380 | 140474489 | 140471882 | 140472011 | 140479159 | 140479569 | 0 | 0 | 3 | 5 | 3 | 3 | 0 | 0 | 1 | 1 | 98 | 49 | 0.601469268 | 1 | 0 | 0 | 0.333 | 0 | 0 | 0.111 |
| A3SS | 1888 MDM4 | MDM4 | chr1 | + | 204537429 | 204537497 | 204537458 | 204537497 | 204532190 | 204532246 | 31 | 9 | 2 | 3 | 0 | 0 | 1 | 10 | 0 | 0 | 78 | 49 | 0.60228108 | 1 | 0.867 | 1 | 1 | 1 | 1 | -0.044 |
| SE | 8583 RASSF3 | RASSF3 | chr12 | + | 64688215 | 64688453 | 64684786 | 64684894 | 64691469 | 64691579 | 91 | 31 | 62 | 1 | 0 | 0 | 1 | 21 | 0 | 2 | 98 | 49 | 0.603346308 | 1 | 0.978 | 1 | 1 | 1 | 0.84 | 0.073 |
| SE | 8824 AQP9 | AQP9 | chr15 | + | 58179127 | 58179345 | 58174917 | 58175036 | 58183960 | 58185911 | 229 | 142 | 134 | 4 | 2 | 0 | 1 | 35 | 0 | 0 | 98 | 49 | 0.604291852 | 1 | 0.966 | 0.979 | 1 | 1 | 1 | -0.02 |
| SE | 9289 MBNL1 | MBNL1 | chr3 | + | 152456266 | 152456361 | 152455541 | 152455577 | 152459270 | 152459345 | 51 | 22 | 22 | 3 | 0 | 0 | 1 | 10 | 0 | 0 | 98 | 49 | 0.608449796 | 1 | 0.895 | 1 | 1 | 1 | 1 | -0.035 |
| SE | 2787 ARHGAP27 | ARHGAP27 | chr17 | - | 45404268 | 45404334 | 45404028 | 45404096 | 45404444 | 45404528 | 39 | 9 | 5 | 18 | 5 | 1 | 2 | 4 | 0 | 3 | 98 | 49 | 0.608834548 | 1 | 0.52 | 0.474 | 0.714 | 1 | 0.4 | -0.131 |
| A5SS | 808 UBE2D3 | UBE2D3 | chr4 | - | 102827426 | 102827948 | 102827591 | 102827948 | 102826484 | 102826636 | 25 | 11 | 1 | 6 | 1 | 2 | 2 | 4 | 0 | 1 | 77 | 49 | 0.609205648 | 1 | 0.676 | 0.846 | 0.2 | 1 | 0.333 | -0.093 |
| SE | 8678 MAEA | MAEA | chr4 | + | 1327626 | 1327703 | 1315396 | 1315600 | 1332756 | 1332865 | 12 | 5 | 7 | 0 | 4 | 0 | 3 | 2 | 1 | 1 | 98 | 49 | 0.60937349 | 1 | 1 | 0.385 | 1 | 0.6 | 0.5 | 0.245 |
| SE | 4595 PARP8 | PARP8 | chr5 | + | 50834930 | 50835015 | 50833978 | 50834048 | 50841965 | 50846522 | 122 | 72 | 14 | 3 | 0 | 0 | 4 | 16 | 0 | 0 | 98 | 49 | 0.610101228 | 1 | 0.953 | 1 | 1 | 1 | 1 | -0.016 |
| SE | 17053 LCP2 | LCP2 | chr5 | - | 170270718 | 170270917 | 170268384 | 170268482 | 170274300 | 170274338 | 71 | 18 | 28 | 2 | 2 | 0 | 1 | 19 | 0 | 0 | 98 | 49 | 0.611178802 | 1 | 0.947 | 0.818 | 1 | 1 | 1 | -0.078 |
| SE | 12250 SMCHD1 | SMCHD1 | chr18 | + | 2762104 | 2762236 | 2760651 | 2760739 | 2763636 | 2763789 | 300 | 180 | 52 | 5 | 0 | 0 | 2 | 58 | 0 | 0 | 98 | 49 | 0.612583031 | 1 | 0.968 | 1 | 1 | 1 | 1 | -0.011 |
| A5SS | 1651 GAPVD1 | GAPVD1 | chr9 | + | 125321432 | 125321562 | 125321432 | 125321499 | 125323797 | 125323923 | 15 | 4 | 1 | 3 | 0 | 0 | 3 | 3 | 0 | 0 | 98 | 49 | 0.614143158 | 1 | 0.714 | 1 | 1 | 1 | 1 | -0.095 |
| SE | 15170 SHKBP1 | SHKBP1 | chr19 | + | 40580565 | 40580656 | 40580323 | 40580485 | 40580745 | 40580936 | 72 | 13 | 18 | 0 | 3 | 0 | 3 | 17 | 0 | 0 | 98 | 49 | 0.614265951 | 1 | 1 | 0.684 | 1 | 1 | 1 | -0.105 |
| SE | 1103 MGAM | MGAM | chr7 | + | 142064383 | 142064522 | 142063498 | 142063586 | 142065334 | 142065468 | 133 | 36 | 29 | 1 | 1 | 0 | 4 | 8 | 0 | 2 | 98 | 49 | 0.617391375 | 1 | 0.985 | 0.947 | 1 | 1 | 0.667 | 0.144 |
| SE | 6445 IRAK3 | IRAK3 | chr12 | + | 66226722 | 66226837 | 66217170 | 66217235 | 66228251 | 66228370 | 52 | 18 | 13 | 0 | 0 | 3 | 1 | 30 | 0 | 0 | 98 | 49 | 0.617973784 | 1 | 1 | 1 | 0.684 | 1 | 1 | -0.105 |
| A5SS | 2718 POLD4 | POLD4 | chr11 | - | 67353302 | 67353596 | 67353363 | 67353596 | 67352987 | 67353077 | 17 | 14 | 3 | 0 | 1 | 2 | 1 | 2 | 0 | 0 | 98 | 49 | 0.618730112 | 1 | 1 | 0.875 | 0.429 | 1 | 1 | -0.232 |
| A3SS | 2904 TYMP | TYMP | chr22 | - | 50529495 | 50529731 | 50529495 | 50529719 | 50529903 | 50530085 | 17 | 12 | 5 | 4 | 4 | 5 | 1 | 6 | 0 | 13 | 61 | 49 | 0.61890226 | 1 | 0.773 | 0.707 | 0.445 | 1 | 0.27 | 0.007 |
| SE | 923 KAT6A | KAT6A | chr8 | - | 41942792 | 41943000 | 41940841 | 41941044 | 41943747 | 41943979 | 50 | 26 | 16 | 3 | 0 | 0 | 3 | 9 | 0 | 1 | 98 | 49 | 0.61992813 | 1 | 0.893 | 1 | 1 | 1 | 0.818 | 0.055 |
| A3SS | 2190 FCGR2A | FCGR2A | chr1 | + | 161506333 | 161506591 | 161506336 | 161506591 | 161505986 | 161506007 | 99 | 41 | 17 | 88 | 27 | 14 | 1 | 20 | 0 | 17 | 52 | 49 | 0.62141272 | 1 | 0.515 | 0.589 | 0.534 | 1 | 0.526 | -0.217 |
| SE | 14085 MAP4K4 | MAP4K4 | chr2 | + | 101871493 | 101871685 | 101870294 | 101870415 | 101873646 | 101873764 | 23 | 5 | 4 | 12 | 6 | 6 | 0 | 7 | 1 | 4 | 98 | 49 | 0.622463836 | 1 | 0.489 | 0.294 | 0.25 | 0 | 0.467 | 0.111 |
| A5SS | 1478 IFI6 | IFI6 | chr1 | - | 27669232 | 27669346 | 27669244 | 27669346 | 27668457 | 27668535 | 0 | 2 | 0 | 6 | 10 | 7 | 0 | 2 | 1 | 20 | 61 | 49 | 0.62257771 | 1 | 0 | 0.138 | 0 | 0 | 0.074 | 0.009 |
| SE | 240 DDX3X | DDX3X | chrX | + | 41343736 | 41343822 | 41343215 | 41343351 | 41344029 | 41344128 | 233 | 80 | 33 | 10 | 0 | 0 | 4 | 36 | 2 | 0 | 98 | 49 | 0.623534899 | 1 | 0.921 | 1 | 1 | 0.5 | 1 | 0.224 |
| SE | 12825 GMIP | GMIP | chr19 | - | 19640295 | 19640360 | 19640084 | 19640192 | 19640445 | 19640571 | 161 | 65 | 30 | 4 | 1 | 0 | 9 | 25 | 0 | 2 | 98 | 49 | 0.624364766 | 1 | 0.953 | 0.97 | 1 | 1 | 0.862 | 0.043 |
| SE | 9199 ACSL1 | ACSL1 | chr4 | - | 184770254 | 184770332 | 184768315 | 184768450 | 184773080 | 184773154 | 18 | 3 | 6 | 1 | 2 | 0 | 2 | 3 | 0 | 1 | 98 | 49 | 0.625913977 | 1 | 0.9 | 0.429 | 1 | 1 | 0.6 | -0.024 |
| SE | 11778 FMNL1 | FMNL1 | chr17 | + | 45246504 | 45246604 | 45246209 | 45246330 | 45246866 | 45247316 | 2 | 2 | 0 | 58 | 31 | 6 | 1 | 0 | 1 | 15 | 98 | 49 | 0.626297292 | 1 | 0.017 | 0.031 | 0 | 0.333 | 0 | -0.151 |
| SE | 9386 AMICA1 | AMICA1 | chr11 | - | 118214823 | 118214886 | 118212406 | 118212561 | 118224940 | 118225094 | 229 | 73 | 31 | 30 | 15 | 6 | 7 | 17 | 1 | 5 | 98 | 49 | 0.626361724 | 1 | 0.792 | 0.709 | 0.721 | 0.778 | 0.63 | 0.037 |
| SE | 457 MEGF9 | MEGF9 | chr9 | - | 120607740 | 120608010 | 120600917 | 120605641 | 120612395 | 120612539 | 152 | 50 | 27 | 1 | 3 | 0 | 4 | 25 | 0 | 0 | 98 | 49 | 0.628685621 | 1 | 0.987 | 0.893 | 1 | 1 | 1 | -0.04 |
| SE | 7427 CD300A | CD300A | chr17 | + | 74481288 | 74481326 | 74477435 | 74477530 | 74484000 | 74484798 | 61 | 21 | 22 | 0 | 0 | 0 | 2 | 17 | 0 | 2 | 87 | 49 | 0.630034787 | 1 | 1 | 1 | 1 | 1 | 0.827 | 0.087 |
| SE | 15607 HIPK3 | HIPK3 | chr11 | + | 33348165 | 33348228 | 33347851 | 33348013 | 33348521 | 33348818 | 24 | 8 | 4 | 15 | 2 | 2 | 0 | 7 | 1 | 1 | 98 | 49 | 0.632723622 | 1 | 0.444 | 0.667 | 0.5 | 0 | 0.778 | 0.148 |
| RI | 441 SHISA5 | SHISA5 | chr3 | - | 48467797 | 48469573 | 48467797 | 48469186 | 48469360 | 48469573 | 4 | 1 | 2 | 31 | 7 | 10 | 3 | 1 | 5 | 13 | 98 | 49 | 0.634532702 | 1 | 0.061 | 0.067 | 0.091 | 0.231 | 0.037 | -0.061 |
| A3SS | 3511 INPP5D | INPP5D | chr2 | + | 233125744 | 233125919 | 233125747 | 233125919 | 233122106 | 233122257 | 8 | 2 | 8 | 15 | 8 | 0 | 0 | 7 | 1 | 5 | 52 | 49 | 0.634793545 | 1 | 0.334 | 0.191 | 1 | 0 | 0.569 | 0.224 |
| SE | 11506 CPD | CPD | chr17 | + | 30438974 | 30439077 | 30431771 | 30431881 | 30442307 | 30442450 | 106 | 39 | 36 | 2 | 2 | 0 | 1 | 10 | 0 | 2 | 98 | 49 | 0.636049511 | 1 | 0.964 | 0.907 | 1 | 1 | 0.714 | 0.1 |
| SE | 1596 SOD2 | SOD2 | chr6 | - | 159688125 | 159688242 | 159684853 | 159685033 | 159692660 | 159692863 | 781 | 510 | 412 | 10 | 9 | 2 | 7 | 82 | 2 | 0 | 98 | 49 | 0.639136682 | 1 | 0.975 | 0.966 | 0.99 | 0.636 | 1 | 0.159 |
| SE | 12976 GMFG | GMFG | chr19 | - | 39335260 | 39335310 | 39333076 | 39333126 | 39335434 | 39335531 | 157 | 104 | 81 | 4 | 0 | 0 | 1 | 18 | 0 | 1 | 98 | 49 | 0.639414084 | 1 | 0.952 | 1 | 1 | 1 | 0.9 | 0.034 |
| SE | 4304 OAZ1 | OAZ1 | chr19 | + | 2271384 | 2271530 | 2269485 | 2269744 | 2271781 | 2271954 | 353 | 218 | 283 | 6 | 1 | 0 | 18 | 139 | 0 | 0 | 98 | 49 | 0.644652121 | 1 | 0.967 | 0.991 | 1 | 1 | 1 | -0.014 |
| SE | 6125 NUMB | NUMB | chr14 | - | 73316389 | 73316422 | 73297210 | 73297285 | 73323129 | 73323204 | 4 | 0 | 0 | 35 | 28 | 6 | 0 | 2 | 3 | 8 | 82 | 49 | 0.644661198 | 1 | 0.064 | 0 | 0 | 0 | 0.13 | -0.044 |
| SE | 4325 GTPBP1 | GTPBP1 | chr22 | + | 38727212 | 38727348 | 38726257 | 38726440 | 38727982 | 38728161 | 44 | 22 | 10 | 3 | 0 | 0 | 1 | 10 | 0 | 0 | 98 | 49 | 0.646483805 | 1 | 0.88 | 1 | 1 | 1 | 1 | -0.04 |
| SE | 8390 FRY | FRY | chr13 | + | 32175545 | 32175632 | 32173366 | 32173549 | 32178176 | 32178436 | 62 | 14 | 4 | 0 | 0 | 0 | 2 | 7 | 0 | 2 | 98 | 49 | 0.646484019 | 1 | 1 | 1 | 1 | 1 | 0.636 | 0.182 |
| SE | 16511 HPCAL1 | HPCAL1 | chr2 | + | 10408561 | 10408734 | 10396834 | 10396920 | 10419733 | 10420135 | 0 | 0 | 4 | 3 | 3 | 5 | 0 | 2 | 2 | 7 | 98 | 49 | 0.647971128 | 1 | 0 | 0 | 0.286 | 0 | 0.125 | 0.033 |
| SE | 2734 RHOT1 | RHOT1 | chr17 | + | 32172936 | 32173060 | 32171042 | 32171101 | 32173830 | 32173912 | 16 | 2 | 18 | 0 | 0 | 3 | 5 | 3 | 0 | 0 | 98 | 49 | 0.648168093 | 1 | 1 | 1 | 0.75 | 1 | 1 | -0.083 |
| SE | 16237 BST1 | BST1 | chr4 | + | 15715284 | 15715361 | 15711806 | 15711889 | 15715706 | 15715799 | 51 | 25 | 24 | 3 | 0 | 0 | 2 | 33 | 0 | 0 | 98 | 49 | 0.65116662 | 1 | 0.895 | 1 | 1 | 1 | 1 | -0.035 |
| SE | 3274 USP4 | USP4 | chr3 | - | 49294406 | 49294598 | 49292509 | 49292598 | 49297869 | 49297964 | 19 | 7 | 13 | 0 | 0 | 0 | 1 | 7 | 0 | 2 | 98 | 49 | 0.652443727 | 1 | 1 | 1 | 1 | 1 | 0.636 | 0.182 |
| SE | 1206 NAMPT | NAMPT | chr7 | - | 106254363 | 106254504 | 106253016 | 106253151 | 106261587 | 106261707 | 1557 | 784 | 321 | 12 | 6 | 0 | 24 | 228 | 0 | 4 | 98 | 49 | 0.655738702 | 1 | 0.985 | 0.985 | 1 | 1 | 0.966 | 0.007 |
| SE | 5064 ARAP1 | ARAP1 | chr11 | - | 72711020 | 72711141 | 72710384 | 72710587 | 72711429 | 72711499 | 117 | 42 | 33 | 3 | 3 | 0 | 4 | 21 | 0 | 2 | 98 | 49 | 0.656734237 | 1 | 0.951 | 0.875 | 1 | 1 | 0.84 | 0.022 |
| SE | 6965 HCLS1 | HCLS1 | chr3 | - | 121637145 | 121637256 | 121636433 | 121636489 | 121642926 | 121642981 | 63 | 48 | 55 | 1 | 2 | 2 | 0 | 9 | 1 | 0 | 98 | 49 | 0.657047606 | 1 | 0.969 | 0.923 | 0.932 | 0 | 1 | 0.441 |
| SE | 12686 SLC38A2 | SLC38A2 | chr12 | - | 46370511 | 46370627 | 46366347 | 46367340 | 46370775 | 46370857 | 8 | 3 | 3 | 2 | 1 | 0 | 2 | 6 | 0 | 1 | 98 | 49 | 0.65813555 | 1 | 0.667 | 0.6 | 1 | 1 | 0.75 | -0.119 |
| SE | 14005 SRSF3 | SRSF3 | chr6 | + | 36599820 | 36600276 | 36598848 | 36598983 | 36601151 | 36601190 | 18 | 4 | 7 | 16 | 2 | 0 | 1 | 11 | 0 | 4 | 98 | 49 | 0.658575842 | 1 | 0.36 | 0.5 | 1 | 1 | 0.579 | -0.169 |
| SE | 3019 XPO6 | XPO6 | chr16 | - | 28104545 | 28104707 | 28101846 | 28101945 | 28106042 | 28106214 | 425 | 164 | 72 | 0 | 0 | 0 | 7 | 58 | 2 | 0 | 98 | 49 | 0.659208366 | 1 | 1 | 1 | 1 | 0.636 | 1 | 0.182 |
| A5SS | 327 MTCH1 | MTCH1 | chr6 | - | 36972651 | 36972796 | 36972702 | 36972796 | 36970646 | 36970694 | 33 | 34 | 13 | 24 | 16 | 7 | 2 | 9 | 0 | 8 | 98 | 49 | 0.661988709 | 1 | 0.407 | 0.515 | 0.481 | 1 | 0.36 | -0.212 |
| SE | 2554 RTFDC1 | RTFDC1 | chr20 | + | 56472325 | 56472415 | 56468584 | 56468766 | 56473300 | 56473395 | 4 | 0 | 0 | 21 | 14 | 4 | 0 | 0 | 2 | 1 | 98 | 49 | 0.662218871 | 1 | 0.087 | 0 | 0 | 0 | 0 | 0.029 |
| SE | 10104 PICALM | PICALM | chr11 | - | 86026291 | 86026367 | 86014869 | 86014963 | 86031468 | 86031611 | 144 | 66 | 56 | 4 | 0 | 1 | 1 | 14 | 0 | 0 | 98 | 49 | 0.664770716 | 1 | 0.947 | 1 | 0.966 | 1 | 1 | -0.029 |
| SE | 235 DDX3X | DDX3X | chrX | + | 41337407 | 41337465 | 41333307 | 41334297 | 41341483 | 41341616 | 84 | 33 | 3 | 5 | 0 | 0 | 2 | 10 | 0 | 2 | 98 | 49 | 0.666543847 | 1 | 0.894 | 1 | 1 | 1 | 0.714 | 0.108 |
| A3SS | 1957 ADAM8 | ADAM8 | chr10 | - | 133273253 | 133273443 | 133273253 | 133273430 | 133273761 | 133273838 | 43 | 23 | 4 | 4 | 0 | 1 | 6 | 11 | 0 | 2 | 62 | 49 | 0.666688927 | 1 | 0.895 | 1 | 0.76 | 1 | 0.813 | -0.021 |
| SE | 1587 IGF2R | IGF2R | chr6 | + | 160064803 | 160064901 | 160064400 | 160064531 | 160068248 | 160068385 | 112 | 38 | 21 | 6 | 5 | 0 | 2 | 17 | 0 | 3 | 98 | 49 | 0.671419462 | 1 | 0.903 | 0.792 | 1 | 1 | 0.739 | 0.029 |
| SE | 4832 CLEC7A | CLEC7A | chr12 | - | 10125086 | 10125173 | 10116780 | 10118590 | 10125296 | 10125448 | 26 | 2 | 10 | 12 | 2 | 1 | 3 | 18 | 0 | 8 | 98 | 49 | 0.675221418 | 1 | 0.52 | 0.333 | 0.833 | 1 | 0.529 | -0.203 |
| A5SS | 274 GLIPR2 | GLIPR2 | chr9 | + | 36147785 | 36147916 | 36147785 | 36147894 | 36148546 | 36148650 | 9 | 1 | 0 | 20 | 9 | 9 | 0 | 3 | 2 | 3 | 71 | 49 | 0.675450629 | 1 | 0.237 | 0.071 | 0 | 0 | 0.408 | -0.101 |
| SE | 12958 PSMD8 | PSMD8 | chr19 | + | 38379239 | 38379405 | 38376351 | 38376454 | 38380898 | 38380999 | 15 | 9 | 3 | 2 | 1 | 0 | 1 | 3 | 0 | 0 | 98 | 49 | 0.675480788 | 1 | 0.789 | 0.818 | 1 | 1 | 1 | -0.131 |
| SE | 5040 APLP2 | APLP2 | chr11 | + | 130137255 | 130137291 | 130135562 | 130135715 | 130140397 | 130140483 | 28 | 8 | 7 | 27 | 9 | 10 | 1 | 13 | 1 | 11 | 85 | 49 | 0.680652741 | 1 | 0.374 | 0.339 | 0.288 | 0.366 | 0.405 | -0.052 |
| SE | 14046 TMEM127 | TMEM127 | chr2 | - | 96265137 | 96265489 | 96254832 | 96254997 | 96265868 | 96266013 | 26 | 12 | 10 | 0 | 3 | 0 | 3 | 13 | 0 | 1 | 98 | 49 | 0.680667227 | 1 | 1 | 0.667 | 1 | 1 | 0.867 | -0.045 |
| SE | 3197 CCNDBP1 | CCNDBP1 | chr15 | + | 43194414 | 43194461 | 43192742 | 43192803 | 43194726 | 43197177 | 86 | 68 | 25 | 0 | 4 | 0 | 1 | 28 | 0 | 0 | 96 | 49 | 0.681207238 | 1 | 1 | 0.897 | 1 | 1 | 1 | -0.034 |
| SE | 14793 PHF1 | PHF1 | chr6 | + | 33414965 | 33415144 | 33414724 | 33414829 | 33415234 | 33415329 | 3 | 7 | 1 | 2 | 0 | 0 | 1 | 1 | 0 | 0 | 98 | 49 | 0.683136067 | 1 | 0.429 | 1 | 1 | 1 | 1 | -0.19 |

| | | | | | | | | | | | | | | | | | | | | | | | | | | | | | | |
|---|---|---|---|---|---|---|---|---|---|---|---|---|---|---|---|---|---|---|---|---|---|---|---|---|---|---|---|---|---|---|
| SE | 6430 LMBR1L | LMBR1L | chr12 | - | 49104447 | 49104551 | 49103686 | 49103813 | 49104745 | 49104885 | 9 | 3 | 2 | 0 | 2 | 0 | 1 | 2 | 0 | 0 | 98 | 49 | 0.683158228 | 1 | 1 | 0.429 | 1 | 1 | 1 | -0.19 |
| SE | 7519 MKL1 | MKL1 | chr22 | - | 40424205 | 40424381 | 40423535 | 40423685 | 40429605 | 40429767 | 12 | 3 | 1 | 0 | 2 | 0 | 2 | 2 | 0 | 0 | 98 | 49 | 0.683169487 | 1 | 1 | 0.429 | 1 | 1 | 1 | -0.19 |
| SE | 9962 HDAC7 | HDAC7 | chr12 | - | 47796206 | 47796298 | 47795586 | 47795767 | 47797016 | 47797142 | 15 | 3 | 2 | 0 | 2 | 0 | 1 | 1 | 0 | 0 | 98 | 49 | 0.683194702 | 1 | 1 | 0.429 | 1 | 1 | 1 | -0.19 |
| SE | 2629 NLRC5 | NLRC5 | chr16 | + | 57046551 | 57046641 | 57045447 | 57045492 | 57047544 | 57047628 | 23 | 19 | 1 | 3 | 0 | 0 | 1 | 10 | 0 | 0 | 98 | 49 | 0.687431362 | 1 | 0.793 | 1 | 1 | 1 | 1 | -0.069 |
| SE | 3026 XPO6 | XPO6 | chr16 | - | 28135215 | 28135324 | 28132333 | 28132403 | 28146093 | 28146203 | 125 | 73 | 21 | 2 | 1 | 0 | 1 | 25 | 0 | 0 | 98 | 49 | 0.688505845 | 1 | 0.969 | 0.973 | 1 | 1 | 1 | -0.019 |
| SE | 12874 ZFAS1 | ZFAS1 | chr20 | + | 49280484 | 49280570 | 49279103 | 49279208 | 49289044 | 49289258 | 54 | 41 | 23 | 3 | 1 | 0 | 1 | 10 | 0 | 0 | 98 | 49 | 0.688940427 | 1 | 0.9 | 0.953 | 1 | 1 | 1 | -0.049 |
| A3SS | 2605 LILRB2 | LILRB2 | chr19 | - | 54277549 | 54277600 | 54277549 | 54277597 | 54277888 | 54277939 | 24 | 5 | 2 | 11 | 3 | 0 | 1 | 2 | 0 | 6 | 52 | 49 | 0.69041571 | 1 | 0.673 | 0.611 | 1 | 1 | 0.239 | 0.142 |
| RI | 460 CSF3R | CSF3R | chr1 | - | 36466042 | 36466827 | 36466042 | 36466200 | 36466620 | 36466827 | 826 | 345 | 179 | 10 | 1 | 0 | 19 | 165 | 0 | 1 | 98 | 49 | 0.692461527 | 1 | 0.976 | 0.994 | 1 | 1 | 0.988 | -0.004 |
| SE | 3651 ERO1A | ERO1A | chr14 | - | 52683787 | 52683907 | 52682324 | 52682408 | 52695367 | 52695931 | 39 | 20 | 6 | 3 | 0 | 0 | 1 | 2 | 0 | 0 | 98 | 49 | 0.694855347 | 1 | 0.867 | 1 | 1 | 1 | 1 | -0.044 |
| A3SS | 241 LRCH4 | LRCH4 | chr7 | - | 100574010 | 100575304 | 100574010 | 100574356 | 100575704 | 100575782 | 12 | 5 | 3 | 19 | 3 | 7 | 1 | 2 | 0 | 4 | 98 | 49 | 0.69501919 | 1 | 0.24 | 0.455 | 0.176 | 1 | 0.2 | -0.31 |
| SE | 7089 MEGF6 | MEGF6 | chr1 | - | 3496987 | 3497119 | 3496654 | 3496783 | 3497232 | 3497361 | 6 | 3 | 1 | 8 | 4 | 0 | 3 | 0 | 0 | 2 | 98 | 49 | 0.696658222 | 1 | 0.273 | 0.273 | 1 | 1 | 0 | 0.015 |
| A5SS | 1109 SERPINA1 | SERPINA1 | chr14 | - | 94390419 | 94390692 | 94390456 | 94390692 | 94388559 | 94388663 | 84 | 62 | 16 | 28 | 18 | 5 | 4 | 5 | 0 | 4 | 86 | 49 | 0.696859738 | 1 | 0.631 | 0.662 | 0.646 | 1 | 0.416 | -0.062 |
| SE | 1945 CCND3 | CCND3 | chr6 | - | 41936558 | 41936695 | 41934932 | 41936107 | 41937234 | 41937394 | 26 | 8 | 17 | 1 | 2 | 0 | 3 | 19 | 2 | 0 | 98 | 49 | 0.696975016 | 1 | 0.929 | 0.667 | 1 | 0.429 | 1 | 0.151 |
| SE | 2180 PRRC2A | PRRC2A | chr6 | + | 31626771 | 31626862 | 31626019 | 31626162 | 31626981 | 31627198 | 33 | 12 | 2 | 3 | 0 | 0 | 1 | 5 | 0 | 0 | 98 | 49 | 0.697530193 | 1 | 0.846 | 1 | 1 | 1 | 1 | -0.051 |
| SE | 2603 CYTH4 | CYTH4 | chr22 | + | 37310987 | 37311064 | 37309211 | 37309323 | 37311455 | 37311527 | 172 | 85 | 41 | 0 | 0 | 4 | 2 | 30 | 0 | 0 | 98 | 49 | 0.698071195 | 1 | 1 | 1 | 0.837 | 1 | 1 | -0.054 |
| RI | 801 ARHGDIA | ARHGDIA | chr17 | - | 81869329 | 81869625 | 81869329 | 81869406 | 81869541 | 81869625 | 38 | 40 | 6 | 23 | 8 | 3 | 5 | 15 | 0 | 13 | 98 | 49 | 0.698940977 | 1 | 0.452 | 0.714 | 0.5 | 1 | 0.366 | -0.128 |
| SE | 6126 NUMB | NUMB | chr14 | - | 73366896 | 73366981 | 73355625 | 73355766 | 73409936 | 73410068 | 74 | 37 | 18 | 2 | 0 | 2 | 1 | 10 | 0 | 1 | 98 | 49 | 0.70020625 | 1 | 0.949 | 1 | 0.818 | 1 | 0.833 | 0.006 |
| SE | 13659 CLK1 | CLK1 | chr2 | - | 200854615 | 200854695 | 200853902 | 200853993 | 200855003 | 200855086 | 42 | 36 | 20 | 0 | 0 | 0 | 1 | 8 | 0 | 2 | 98 | 49 | 0.700781312 | 1 | 1 | 1 | 1 | 1 | 0.667 | 0.166 |
| SE | 2871 NLRP12 | NLRP12 | chr19 | - | 53807494 | 53807668 | 53805279 | 53805450 | 53809586 | 53811288 | 20 | 9 | 11 | 0 | 0 | 0 | 1 | 8 | 0 | 2 | 98 | 49 | 0.701325341 | 1 | 1 | 1 | 1 | 1 | 0.667 | 0.166 |
| SE | 12702 GLUL | GLUL | chr1 | - | 182390496 | 182390852 | 182388571 | 182388750 | 182391678 | 182392206 | 20 | 4 | 4 | 397 | 115 | 12 | 2 | 1 | 5 | 18 | 98 | 49 | 0.701769816 | 1 | 0.025 | 0.017 | 0.143 | 0.167 | 0.027 | -0.035 |
| SE | 5206 PTPRJ | PTPRJ | chr11 | + | 48112746 | 48112983 | 48110057 | 48110076 | 48121002 | 48121266 | 111 | 70 | 34 | 0 | 1 | 0 | 11 | 32 | 2 | 0 | 98 | 49 | 0.701904035 | 1 | 1 | 0.972 | 1 | 0.733 | 1 | 0.124 |
| SE | 2767 PCBP2 | PCBP2 | chr12 | + | 53467804 | 53467843 | 53467220 | 53467293 | 53468779 | 53468832 | 50 | 22 | 5 | 3 | 1 | 0 | 1 | 4 | 0 | 2 | 88 | 49 | 0.70273878 | 1 | 0.903 | 0.925 | 1 | 1 | 0.527 | 0.179 |
| SE | 9208 ACSL1 | ACSL1 | chr4 | - | 184803319 | 184803546 | 184788616 | 184788731 | 184825915 | 184826114 | 417 | 162 | 76 | 5 | 2 | 0 | 4 | 91 | 0 | 0 | 98 | 49 | 0.70655971 | 1 | 0.977 | 0.976 | 1 | 1 | 1 | -0.016 |
| A3SS | 2717 ABTB1 | ABTB1 | chr3 | + | 127674344 | 127674418 | 127674390 | 127674418 | 127672937 | 127673081 | 13 | 22 | 4 | 35 | 12 | 8 | 0 | 8 | 3 | 0 | 95 | 49 | 0.707794314 | 1 | 0.161 | 0.486 | 0.205 | 0 | 1 | -0.216 |
| SE | 12883 GRAMD1A | GRAMD1A | chr19 | + | 35013255 | 35013368 | 35011473 | 35011554 | 35013540 | 35013691 | 28 | 7 | 12 | 9 | 3 | 0 | 1 | 5 | 0 | 4 | 98 | 49 | 0.710171334 | 1 | 0.609 | 0.538 | 1 | 1 | 0.385 | 0.023 |
| SE | 4005 PYCARD | PYCARD | chr16 | - | 31202146 | 31202203 | 31201485 | 31201841 | 31202416 | 31202776 | 17 | 20 | 30 | 3 | 0 | 0 | 2 | 9 | 0 | 0 | 98 | 49 | 0.711092721 | 1 | 0.739 | 1 | 1 | 1 | 1 | -0.087 |
| SE | 16704 MICAL1 | MICAL1 | chr6 | - | 109447874 | 109447963 | 109447680 | 109447722 | 109448202 | 109448393 | 21 | 6 | 9 | 3 | 0 | 0 | 2 | 9 | 0 | 0 | 98 | 49 | 0.712273023 | 1 | 0.778 | 1 | 1 | 1 | 1 | -0.074 |
| A3SS | 3844 ARHGAP25 | ARHGAP25 | chr2 | + | 68807272 | 68807480 | 68807275 | 68807480 | 68787839 | 68787956 | 49 | 22 | 16 | 3 | 0 | 1 | 0 | 5 | 1 | 0 | 52 | 49 | 0.713208598 | 1 | 0.939 | 1 | 0.938 | 0 | 1 | 0.459 |
| SE | 12198 ARAP3 | ARAP3 | chr5 | - | 141655361 | 141655400 | 141654435 | 141654435 | 141655620 | 141655758 | 3 | 0 | 1 | 13 | 4 | 9 | 0 | 0 | 1 | 2 | 88 | 49 | 0.713267633 | 1 | 0.114 | 0 | 0.058 | 0 | 0 | 0.057 |
| SE | 8344 CASP8 | CASP8 | chr2 | + | 201274888 | 201274953 | 201272897 | 201272942 | 201276826 | 201276968 | 160 | 55 | 21 | 11 | 0 | 0 | 1 | 6 | 0 | 2 | 98 | 49 | 0.714017103 | 1 | 0.879 | 1 | 1 | 1 | 0.6 | 0.16 |
| RI | 506 RAB24 | RAB24 | chr5 | - | 177303171 | 177303344 | 177303171 | 177303320 | 177303406 | 177303744 | 23 | 19 | 8 | 7 | 1 | 2 | 6 | 10 | 2 | 1 | 98 | 49 | 0.714087862 | 1 | 0.622 | 0.905 | 0.667 | 0.6 | 0.833 | 0.015 |
| SE | 3730 ARID5A | ARID5A | chr2 | + | 96549320 | 96549459 | 96547401 | 96547517 | 96549752 | 96549805 | 12 | 3 | 4 | 8 | 8 | 0 | 2 | 2 | 0 | 3 | 98 | 49 | 0.715441443 | 1 | 0.429 | 0.158 | 1 | 1 | 0.25 | -0.096 |
| SE | 13149 FCGRT | FCGRT | chr19 | + | 49513386 | 49513473 | 49512278 | 49512750 | 49513881 | 49514133 | 99 | 37 | 27 | 0 | 1 | 3 | 9 | 14 | 0 | 0 | 98 | 49 | 0.717316717 | 1 | 1 | 0.949 | 0.818 | 1 | 1 | -0.078 |
| SE | 5532 NDE1 | NDE1 | chr16 | + | 15696708 | 15696860 | 15694164 | 15694256 | 15724190 | 15726351 | 21 | 28 | 5 | 3 | 0 | 0 | 1 | 4 | 0 | 0 | 98 | 49 | 0.717769788 | 1 | 0.778 | 1 | 1 | 1 | 1 | -0.074 |
| SE | 3285 RNASET2 | RNASET2 | chr6 | - | 166934090 | 166934136 | 166931043 | 166931118 | 166938894 | 166939008 | 231 | 74 | 117 | 2 | 4 | 0 | 3 | 45 | 0 | 0 | 95 | 49 | 0.719143808 | 1 | 0.983 | 0.905 | 1 | 1 | 1 | -0.037 |
| A3SS | 1695 CSF3R | CSF3R | chr1 | - | 36466042 | 36466908 | 36466042 | 36466827 | 36467229 | 36467311 | 99 | 28 | 16 | 479 | 165 | 105 | 3 | 22 | 17 | 96 | 98 | 49 | 0.71928138 | 1 | 0.094 | 0.078 | 0.071 | 0.081 | 0.103 | -0.011 |
| A3SS | 3626 RPS3 | RPS3 | chr11 | + | 75405582 | 75406913 | 75405613 | 75406913 | 75404671 | 75404868 | 39 | 8 | 8 | 61 | 21 | 21 | 0 | 4 | 1 | 4 | 80 | 49 | 0.719477917 | 1 | 0.281 | 0.189 | 0.189 | 0 | 0.38 | 0.03 |
| SE | 13854 GNAS | GNAS | chr20 | + | 58898940 | 58898985 | 58895611 | 58895684 | 58903527 | 58903585 | 10 | 4 | 3 | 28 | 12 | 5 | 1 | 2 | 3 | 2 | 94 | 49 | 0.72037482 | 1 | 0.157 | 0.148 | 0.238 | 0.148 | 0.343 | -0.065 |
| SE | 12234 SMCHD1 | SMCHD1 | chr18 | + | 2666156 | 2666232 | 2655886 | 2656261 | 2666869 | 2667031 | 415 | 229 | 66 | 4 | 0 | 3 | 20 | 98 | 0 | 0 | 98 | 49 | 0.720453135 | 1 | 0.981 | 1 | 0.917 | 1 | 1 | -0.034 |
| SE | 12238 SMCHD1 | SMCHD1 | chr18 | + | 2703691 | 2703886 | 2700734 | 2700918 | 2705693 | 2705807 | 514 | 273 | 65 | 0 | 0 | 5 | 4 | 139 | 0 | 0 | 98 | 49 | 0.720467692 | 1 | 1 | 1 | 0.867 | 1 | 1 | -0.044 |
| SE | 6636 CSF3R | CSF3R | chr1 | - | 36467821 | 36467962 | 36467557 | 36467651 | 36468074 | 36468221 | 634 | 178 | 131 | 4 | 3 | 0 | 15 | 149 | 0 | 0 | 98 | 49 | 0.721936982 | 1 | 0.988 | 0.967 | 1 | 1 | 1 | -0.015 |
| SE | 9241 SVIL | SVIL | chr10 | - | 29531253 | 29531288 | 29530606 | 29530668 | 29550596 | 29551263 | 30 | 14 | 2 | 3 | 0 | 0 | 2 | 4 | 0 | 0 | 84 | 49 | 0.722024346 | 1 | 0.854 | 1 | 1 | 1 | 1 | -0.049 |
| SE | 15940 CSF2RA | CSF2RA | chrX | + | 1285777 | 1285920 | 1282777 | 1282779 | 1288518 | 1288642 | 56 | 49 | 23 | 2 | 0 | 2 | 1 | 5 | 0 | 0 | 98 | 49 | 0.724133416 | 1 | 0.933 | 1 | 0.852 | 1 | 1 | -0.072 |
| SE | 8918 ATP6V0B | ATP6V0B | chr1 | + | 43975799 | 43975848 | 43974647 | 43975107 | 43976089 | 43976173 | 74 | 62 | 36 | 20 | 5 | 5 | 5 | 19 | 1 | 4 | 98 | 49 | 0.728043554 | 1 | 0.649 | 0.861 | 0.783 | 0.714 | 0.704 | 0.055 |
| SE | 5412 NCF2 | NCF2 | chr1 | - | 183567203 | 183567345 | 183566919 | 183566988 | 183569141 | 183569185 | 164 | 124 | 119 | 6 | 4 | 0 | 1 | 39 | 0 | 2 | 98 | 49 | 0.730433554 | 1 | 0.932 | 0.939 | 1 | 1 | 0.907 | 0.003 |
| SE | 970 DOCK5 | DOCK5 | chr8 | + | 25300577 | 25300657 | 25298943 | 25299101 | 25302324 | 25302454 | 135 | 60 | 13 | 0 | 4 | 0 | 3 | 24 | 0 | 0 | 98 | 49 | 0.730535785 | 1 | 1 | 0.882 | 1 | 1 | 1 | -0.039 |
| SE | 13949 DYSF | DYSF | chr2 | + | 71549349 | 71549391 | 71539156 | 71539239 | 71551040 | 71551156 | 10 | 11 | 1 | 38 | 16 | 10 | 2 | 2 | 1 | 11 | 91 | 49 | 0.731283912 | 1 | 0.124 | 0.27 | 0.051 | 0.519 | 0.089 | -0.156 |
| SE | 4341 ITPK1 | ITPK1 | chr14 | - | 92993879 | 92993997 | 92962750 | 92962849 | 93016675 | 93016801 | 54 | 46 | 22 | 1 | 0 | 0 | 2 | 10 | 0 | 2 | 98 | 49 | 0.731604336 | 1 | 0.964 | 1 | 1 | 1 | 0.714 | 0.131 |
| SE | 16958 H2AFY | H2AFY | chr5 | - | 135388921 | 135389126 | 135370035 | 135370142 | 135399061 | 135399238 | 38 | 12 | 15 | 3 | 0 | 0 | 2 | 8 | 0 | 0 | 98 | 49 | 0.732088065 | 1 | 0.864 | 1 | 1 | 1 | 1 | -0.045 |
| SE | 5243 PFKFB3 | PFKFB3 | chr10 | + | 6216124 | 6216191 | 6215220 | 6215317 | 6216705 | 6216780 | 157 | 110 | 11 | 3 | 2 | 0 | 19 | 68 | 1 | 1 | 98 | 49 | 0.733268371 | 1 | 0.963 | 0.965 | 1 | 0.905 | 0.971 | 0.038 |
| SE | 6533 RBM39 | RBM39 | chr20 | - | 35713018 | 35713096 | 35709223 | 35709256 | 35714184 | 35714389 | 71 | 48 | 21 | 1 | 3 | 0 | 4 | 24 | 0 | 0 | 98 | 49 | 0.733711812 | 1 | 0.973 | 0.889 | 1 | 1 | 1 | -0.046 |
| A3SS | 3083 RFXANK | RFXANK | chr19 | + | 19196962 | 19197046 | 19196965 | 19197046 | 19193938 | 19194133 | 2 | 1 | 0 | 6 | 4 | 2 | 0 | 1 | 1 | 1 | 52 | 49 | 0.734057817 | 1 | 0.239 | 0.191 | 0 | 0 | 0.485 | -0.099 |
| SE | 16459 TMC6 | TMC6 | chr17 | - | 78124888 | 78124985 | 78124523 | 78124781 | 78125157 | 78125263 | 14 | 10 | 6 | 2 | 1 | 0 | 4 | 2 | 1 | 0 | 98 | 49 | 0.734348789 | 1 | 0.778 | 0.833 | 1 | 0.667 | 1 | 0.037 |
| SE | 523 SEMA4D | SEMA4D | chr9 | - | 89388635 | 89388792 | 89387385 | 89387608 | 89388871 | 89389047 | 58 | 18 | 32 | 0 | 3 | 0 | 4 | 14 | 0 | 0 | 98 | 49 | 0.738125132 | 1 | 1 | 0.75 | 1 | 1 | 1 | -0.083 |
| SE | 3660 SOS2 | SOS2 | chr14 | - | 50158564 | 50158646 | 50156998 | 50157121 | 50159430 | 50160086 | 91 | 38 | 10 | 3 | 0 | 0 | 1 | 14 | 0 | 2 | 98 | 49 | 0.738538674 | 1 | 0.938 | 1 | 1 | 1 | 0.778 | 0.09 |
| SE | 16425 MANBA | MANBA | chr4 | - | 102714437 | 102714561 | 102690595 | 102690771 | 102722870 | 102723041 | 15 | 10 | 11 | 1 | 2 | 0 | 1 | 4 | 0 | 0 | 98 | 49 | 0.742914923 | 1 | 0.882 | 0.714 | 1 | 1 | 1 | -0.135 |
| A3SS | 693 PRKCB | PRKCB | chr16 | + | 24214657 | 24220611 | 24219960 | 24220611 | 24191089 | 24191230 | 37 | 23 | 9 | 1 | 0 | 2 | 2 | 14 | 0 | 0 | 98 | 49 | 0.747697883 | 1 | 0.949 | 1 | 0.692 | 1 | 1 | -0.12 |
| SE | 11424 ANKHD1 | ANKHD1 | chr5 | + | 140496519 | 140497278 | 140486957 | 140487060 | 140504820 | 140504966 | 20 | 7 | 2 | 2 | 3 | 0 | 2 | 6 | 0 | 3 | 98 | 49 | 0.747961443 | 1 | 0.833 | 0.538 | 1 | 1 | 0.5 | 0.04 |
| SE | 8606 TKT | TKT | chr3 | - | 53237796 | 53237820 | 53234982 | 53235174 | 53240250 | 53240348 | 5 | 0 | 11 | 75 | 47 | 49 | 2 | 1 | 1 | 33 | 73 | 49 | 0.748430679 | 1 | 0.043 | 0 | 0.131 | 0.573 | 0.02 | -0.238 |
| A3SS | 937 FCGR3A | FCGR3A | chr1 | - | 161548420 | 161548678 | 161548420 | 161548675 | 161549010 | 161549031 | 2 | 9 | 10 | 0 | 0 | 0 | 1 | 5 | 0 | 2 | 52 | 49 | 0.749944571 | 1 | 1 | 1 | 1 | 1 | 0.702 | 0.149 |
| SE | 15835 AOAH | AOAH | chr7 | - | 36621709 | 36621780 | 36620780 | 36620829 | 36623189 | 36623250 | 55 | 30 | 4 | 0 | 3 | 0 | 4 | 25 | 0 | 0 | 98 | 49 | 0.750059818 | 1 | 1 | 0.833 | 1 | 1 | 1 | -0.056 |
| SE | 4158 NLRP1 | NLRP1 | chr17 | - | 5520880 | 5521012 | 5517745 | 5517887 | 5521523 | 5521786 | 42 | 6 | 5 | 8 | 4 | 0 | 1 | 10 | 0 | 3 | 98 | 49 | 0.75216073 | 1 | 0.724 | 0.429 | 1 | 1 | 0.625 | -0.095 |
| SE | 1131 MKLN1 | MKLN1 | chr7 | + | 131445773 | 131445903 | 131443480 | 131443702 | 131463216 | 131463364 | 34 | 17 | 1 | 1 | 2 | 0 | 1 | 9 | 1 | 0 | 98 | 49 | 0.754100352 | 1 | 0.944 | 0.81 | 1 | 0.333 | 1 | 0.252 |
| A3SS | 1384 RASGRP4 | RASGRP4 | chr19 | - | 38419859 | 38420013 | 38419859 | 38419971 | 38420130 | 38420262 | 54 | 25 | 21 | 0 | 0 | 3 | 5 | 20 | 0 | 0 | 91 | 49 | 0.75562353 | 1 | 1 | 1 | 0.79 | 1 | 1 | -0.07 |
| A3SS | 2074 ITGAM | ITGAM | chr16 | + | 31297741 | 31297954 | 31297744 | 31297954 | 31297513 | 31297654 | 7 | 1 | 1 | 11 | 9 | 2 | 2 | 2 | 1 | 7 | 52 | 49 | 0.758089344 | 1 | 0.375 | 0.095 | 0.32 | 0.653 | 0.212 | -0.169 |
| SE | 684 DOCK8 | DOCK8 | chr9 | + | 386330 | 386426 | 382512 | 382685 | 390470 | 390566 | 93 | 23 | 16 | 2 | 2 | 0 | 4 | 13 | 0 | 0 | 98 | 49 | 0.759194701 | 1 | 0.959 | 0.852 | 1 | 1 | 1 | -0.063 |
| SE | 15578 SUN2 | SUN2 | chr22 | - | 38754641 | 38754706 | 38752506 | 38752665 | 38755762 | 38756019 | 13 | 5 | 3 | 14 | 1 | 3 | 0 | 7 | 1 | 2 | 98 | 49 | 0.760050741 | 1 | 0.317 | 0.714 | 0.333 | 0 | 0.636 | 0.137 |
| SE | 5254 PTPRC | PTPRC | chr1 | + | 198702386 | 198702530 | 198699563 | 198699704 | 198703297 | 198703372 | 16 | 0 | 8 | 35 | 24 | 9 | 0 | 4 | 3 | 6 | 98 | 49 | 0.762041288 | 1 | 0.186 | 0 | 0.308 | 0 | 0.25 | 0.04 |
| SE | 2792 USP25 | USP25 | chr21 | + | 15824038 | 15824166 | 15818697 | 15818846 | 15824965 | 15825061 | 22 | 16 | 1 | 3 | 0 | 0 | 1 | 3 | 0 | 2 | 98 | 49 | 0.762601432 | 1 | 0.786 | 1 | 1 | 1 | 0.429 | 0.214 |
| SE | 13744 NBN | NBN | chr8 | - | 89978219 | 89978323 | 89971172 | 89971290 | 89980733 | 89980893 | 34 | 16 | 5 | 3 | 0 | 0 | 2 | 17 | 0 | 0 | 98 | 49 | 0.763162175 | 1 | 0.85 | 1 | 1 | 1 | 1 | -0.05 |
| SE | 2804 UBN1 | UBN1 | chr16 | + | 4868832 | 4868903 | 4860663 | 4861102 | 4870211 | 4870341 | 30 | 33 | 10 | 3 | 0 | 0 | 0 | 4 | 1 | 0 | 98 | 49 | 0.764528525 | 1 | 0.833 | 1 | 1 | 0 | 1 | 0.444 |
| A3SS | 3648 ITGB2 | ITGB2 | chr21 | - | 44910283 | 44910415 | 44910283 | 44910372 | 44910724 | 44910785 | 10 | 2 | 3 | 92 | 29 | 19 | 1 | 3 | 1 | 16 | 92 | 49 | 0.765684671 | 1 | 0.055 | 0.035 | 0.078 | 0.348 | 0.091 | -0.163 |
| SE | 2761 CPPED1 | CPPED1 | chr16 | - | 12704623 | 12705049 | 12659798 | 12665115 | 12781184 | 12781403 | 87 | 37 | 15 | 1 | 1 | 4 | 1 | 12 | 0 | 3 | 98 | 49 | 0.766803703 | 1 | 0.978 | 0.949 | 0.652 | 1 | 0.667 | 0.026 |
| A5SS | 2776 ABCG1 | ABCG1 | chr21 | + | 42287888 | 42288073 | 42287888 | 42288037 | 42288210 | 42288312 | 21 | 10 | 4 | 2 | 2 | 0 | 2 | 3 | 0 | 2 | 85 | 49 | 0.767335084 | 1 | 0.858 | 0.742 | 1 | 1 | 0.464 | 0.135 |
| SE | 14190 ACOX1 | ACOX1 | chr17 | - | 75973624 | 75973785 | 75960214 | 75960375 | 75978533 | 75978693 | 10 | 3 | 1 | 16 | 3 | 10 | 0 | 3 | 1 | 2 | 98 | 49 | 0.767698723 | 1 | 0.238 | 0.333 | 0.048 | 0 | 0.429 | -0.008 |
| SE | 12161 TYMP | TYMP | chr22 | - | 50528511 | 50528610 | 50527587 | 50527717 | 50529135 | 50529338 | 73 | 24 | 32 | 0 | 0 | 3 | 1 | 41 | 0 | 0 | 98 | 49 | 0.768765459 | 1 | 1 | 1 | 0.842 | 1 | 1 | -0.053 |
| SE | 7501 SLC12A6 | SLC12A6 | chr15 | - | 34257641 | 34257788 | 34256228 | 34256283 | 34258812 | 34258944 | 64 | 22 | 9 | 10 | 11 | 3 | 1 | 11 | 0 | 4 | 98 | 49 | 0.771324392 | 1 | 0.762 | 0.5 | 0.6 | 1 | 0.579 | -0.169 |
| A3SS | 1471 LILRB3 | LILRB3 | chr19 | - | 54217318 | 54217477 | 54217318 | 54217474 | 54218360 | 54218413 | 15 | 12 | 4 | 44 | 15 | 2 | 2 | 8 | 3 | 11 | 52 | 49 | 0.772420962 | 1 | 0.243 | 0.43 | 0.653 | 0.386 | 0.407 | 0.046 |
| A3SS | 1670 RBM39 | RBM39 | chr20 | - | 35709223 | 35709274 | 35709223 | 35709256 | 35713018 | 35713096 | 38 | 12 | 8 | 24 | 4 | 4 | 0 | 7 | 1 | 1 | 67 | 49 | 0.773059786 | 1 | 0.537 | 0.687 | 0.594 | 0 | 0.837 | 0.188 |
| SE | 5059 ARAP1 | ARAP1 | chr11 | - | 72695717 | 72695865 | 72695541 | 72695628 | 72696548 | 72696654 | 95 | 77 | 31 | 0 | 3 | 0 | 1 | 24 | 0 | 0 | 98 | 49 | 0.778635347 | 1 | 1 | 0.928 | 1 | 1 | 1 | -0.024 |
| SE | 9620 GSK3B | GSK3B | chr3 | - | 119912703 | 119912810 | 119905754 | 119905852 | 119916043 | 119916174 | 40 | 5 | 10 | 1 | 2 | 0 | 2 | 8 | 0 | 2 | 98 | 49 | 0.780210798 | 1 | 0.952 | 0.556 | 1 | 1 | 0.667 | 0.002 |
| SE | 11293 MBD1 | MBD1 | chr18 | - | 50273333 | 50273471 | 50272955 | [illegible] | 50273563 | 50273860 | 4 | 4 | 2 | 2 | 0 | 0 | 1 | 1 | 0 | 1 | 98 | 49 | 0.780381124 | 1 | 0.5 | 1 | 1 | 1 | 0.333 | 0.167 |
| SE | 5415 NCF2 | NCF2 | chr1 | - | 183573184 | 183573292 | 183570779 | 183570839 | 183577598 | 183577707 | 304 | 147 | 80 | 3 | 2 | 0 | 5 | 53 | 0 | 1 | 98 | 49 | 0.783922926 | 1 | 0.981 | 0.974 | 1 | 1 | 0.964 | 0.003 |
| SE | 10100 PICALM | PICALM | chr11 | - | 85990249 | 85990399 | 85983865 | 85983973 | 85996825 | 85996929 | 104 | 59 | 9 | 71 | 25 | 20 | 1 | 25 | 0 | 21 | 98 | 49 | 0.784680554 | 1 | 0.423 | 0.541 | 0.184 | 1 | 0.373 | -0.304 |
| SE | 7840 LRRK2 | LRRK2 | chr12 | + | 40232273 | 40232383 | 40225554 | 40225640 | 40235625 | 40235714 | 338 | 169 | 65 | 0 | 0 | 4 | 6 | 38 | 0 | 0 | 98 | 49 | 0.785217176 | 1 | 1 | 1 | 0.89 | 1 | 1 | -0.037 |
| SE | 15545 ATG16L2 | ATG16L2 | chr11 | + | 72826517 | 72826589 | 72826172 | 72826243 | 72826702 | 72826823 | 222 | 145 | 48 | 1 | 3 | 0 | 12 | 63 | 0 | 0 | 98 | 49 | 0.785871628 | 1 | 0.991 | 0.96 | 1 | 1 | 1 | -0.016 |
| SE | 11053 THEMIS2 | THEMIS2 | chr1 | + | 27879643 | 27880054 | 27876587 | 27876728 | 27885294 | 27885451 | 111 | 23 | 15 | 3 | 2 | 6 | 6 | 25 | 3 | 1 | 98 | 49 | 0.787595271 | 1 | 0.949 | 0.852 | 0.556 | 0.5 | 0.926 | 0.073 |
| SE | 9009 HLA-B | HLA-B | chr6 | - | 31355106 | 31355223 | 31354632 | 31354665 | 31355316 | 31355592 | 2250 | 754 | 621 | 31 | 2 | 9 | 46 | 261 | 2 | 4 | 98 | 49 | 0.793423683 | 1 | 0.973 | 0.995 | 0.972 | 0.92 | 0.97 | 0.035 |
| A3SS | 2818 STAT3 | STAT3 | chr17 | - | 42316788 | 42316901 | 42316788 | 42316851 | 42317181 | 42317224 | 101 | 42 | 24 | 13 | 0 | 8 | 2 | 12 | 0 | 4 | 98 | 49 | 0.79351202 | 1 | 0.795 | 1 | 0.6 | 1 | 0.6 | -0.002 |
| SE | 9855 NADK | NADK | chr1 | - | 1754050 | 1754208 | 1753566 | 1753649 | 1754283 | 1754383 | 84 | 54 | 60 | 4 | 0 | 0 | 4 | 21 | 0 | 2 | 98 | 49 | 0.803552896 | 1 | 0.913 | 1 | 1 | 1 | 0.84 | 0.051 |
| SE | 14775 PLAUR | PLAUR | chr19 | - | 43652224 | 43652371 | 43648570 | 43649143 | 43655438 | 43655573 | 60 | 16 | 17 | 10 | 2 | 1 | 2 | 46 | 0 | 7 | 98 | 49 | 0.805341856 | 1 | 0.75 | 0.8 | 0.895 | 1 | 0.767 | -0.068 |
| SE | 2870 NLRP12 | NLRP12 | chr19 | - | 53807494 | 53807665 | 53805279 | 53805450 | 53809586 | 53811288 | 37 | 15 | 17 | 0 | 0 | 0 | 1 | 13 | 0 | 2 | 98 | 49 | 0.807194084 | 1 | 1 | 1 | 1 | 1 | 0.765 | 0.117 |
| A3SS | 753 ISG20 | ISG20 | chr15 | + | 88652109 | 88652310 | 88652169 | 88652310 | 88639342 | 88639594 | 60 | 67 | 37 | 1 | 1 | 1 | 1 | 28 | 0 | 0 | 98 | 49 | 0.807205421 | 1 | 0.968 | 0.971 | 0.949 | 1 | 1 | -0.037 |
| SE | 3709 IRF9 | IRF9 | chr14 | + | 24164613 | 24164955 | 24164062 | 24164134 | 24165846 | 24165962 | 41 | 31 | 26 | 3 | 0 | 0 | 2 | 13 | 0 | 0 | 98 | 49 | 0.810580629 | 1 | 0.872 | 1 | 1 | 1 | 1 | -0.043 |
| A3SS | 521 BAG6 | BAG6 | chr6 | - | 31647590 | 31647844 | 31647590 | 31647826 | 31648676 | 31648751 | 1 | 2 | 0 | 10 | 5 | 1 | 0 | 0 | 1 | 2 | 67 | 49 | 0.813743282 | 1 | 0.068 | 0.226 | 0 | 0 | 0 | 0.098 |
| SE | 9198 ACSL1 | ACSL1 | chr4 | - | 184765890 | 184765986 | 184764852 | 184764925 | 184766621 | 184766756 | 240 | 107 | 64 | 5 | 2 | 2 | 2 | 59 | 0 | 3 | 98 | 49 | 0.815627037 | 1 | 0.96 | 0.964 | 0.941 | 1 | 0.908 | 0.001 |
| A3SS | 2810 FKBP8 | FKBP8 | chr19 | - | 18538215 | 18538436 | 18538215 | 18538433 | 18539370 | 18539459 | 42 | 29 | 29 | 38 | 13 | 14 | 2 | 2 | 0 | 5 | 52 | 49 | 0.817161819 | 1 | 0.51 | 0.678 | 0.661 | 1 | 0.274 | -0.021 |
| A3SS | 2397 DBNL | DBNL | chr7 | + | 44058428 | 44058480 | 44058431 | 44058480 | 44058128 | 44058280 | 11 | 6 | 7 | 21 | 12 | 7 | 0 | 6 | 3 | 2 | 52 | 49 | 0.818390764 | 1 | 0.33 | 0.32 | 0.485 | 0 | 0.739 | 0.009 |
| SE | 266 GK | GK | chrX | + | 30665510 | 30665584 | 30653358 | 30653615 | 30677374 | 30677452 | 21 | 23 | 15 | 1 | 0 | 2 | 1 | 4 | 0 | 1 | 98 | 49 | 0.819518177 | 1 | 0.913 | 1 | 0.789 | 1 | 0.667 | 0.067 |
| SE | 13300 RPS9 | RPS9 | chr19 | + | 54206275 | 54206462 | 54201486 | 54201609 | 54207397 | 54207647 | 149 | 40 | 46 | 1 | 3 | 0 | 3 | 56 | 0 | 1 | 98 | 49 | 0.824162941 | 1 | 0.987 | 0.87 | 1 | 1 | 0.966 | -0.031 |
| SE | 9572 GNAI2 | GNAI2 | chr3 | + | 50253023 | 50253184 | 50252396 | 50252538 | 50256191 | 50256320 | 199 | 126 | 93 | 2 | 0 | 4 | 6 | 43 | 0 | 0 | 98 | 49 | 0.825744 | 1 | 0.98 | 1 | 0.921 | 1 | 1 | -0.033 |
| SE | 10099 PICALM | PICALM | chr11 | - | 85990249 | 85990378 | 85983865 | 85983973 | 85996825 | 85996929 | 78 | 43 | 4 | 71 | 25 | 20 | 1 | 18 | 0 | 21 | 98 | 49 | 0.825899785 | 1 | 0.355 | 0.462 | 0.091 | 1 | 0.3 | -0.347 |
| SE | 1589 IGF2R | IGF2R | chr6 | + | 160079579 | 160079787 | 160078200 | 160078362 | 160080128 | 160080275 | 264 | 84 | 44 | 4 | 0 | 1 | 4 | 40 | 0 | 1 | 98 | 49 | 0.826922372 | 1 | 0.971 | 1 | 0.957 | 1 | 0.952 | 0 |
| RI | 1017 POFUT2 | POFUT2 | chr21 | - | 45263927 | 45267713 | 45263927 | 45265635 | 45267589 | 45267713 | 6 | 7 | 0 | 5 | 3 | 1 | 0 | 3 | 1 | 0 | 98 | 49 | 0.827034304 | 1 | 0.375 | 0.538 | 0 | 0 | 1 | -0.196 |
| SE | 12784 MYO9B | MYO9B | chr19 | + | 17156908 | 17157038 | 17154314 | 17154415 | 17159394 | 17159484 | 95 | 46 | 13 | 0 | 3 | 0 | 6 | 18 | 0 | 0 | 98 | 49 | 0.827566797 | 1 | 1 | 0.885 | 1 | 1 | 1 | -0.038 |
| SE | 10738 MYO1F | MYO1F | chr19 | - | 8541905 | 8541991 | 8539946 | 8540028 | 8544296 | 8544464 | 163 | 108 | 78 | 2 | 9 | 0 | 11 | 31 | 0 | 3 | 98 | 49 | 0.83190545 | 1 | 0.976 | 0.857 | 1 | 1 | 0.838 | 0.025 |
| SE | 7211 NBEAL2 | NBEAL2 | chr3 | + | 47008069 | 47008186 | 47007815 | 47007910 | 47008282 | 47008441 | 97 | 42 | 14 | 0 | 3 | 0 | 9 | 40 | 0 | 0 | 98 | 49 | 0.832076213 | 1 | 1 | 0.875 | 1 | 1 | 1 | -0.042 |
| SE | 1410 IKZF1 | IKZF1 | chr7 | + | 50387344 | 50387470 | 50382539 | 50382707 | 50391728 | 50391833 | 29 | 5 | 9 | 3 | 0 | 0 | 2 | 10 | 0 | 1 | 98 | 49 | 0.832726723 | 1 | 0.829 | 1 | 1 | 1 | 0.833 | 0.026 |
| SE | 16143 KDM5C | KDM5C | chrX | - | 53218275 | 53218398 | 53217795 | 53217966 | 53220838 | 53220916 | 20 | 13 | 3 | 0 | 2 | 0 | 1 | 12 | 0 | 2 | 98 | 49 | 0.833157475 | 1 | 1 | 0.765 | 1 | 1 | 0.75 | 0.047 |
| A3SS | 3148 TMEM91 | TMEM91 | chr19 | + | 41383560 | 41384082 | 41383862 | 41384082 | 41382771 | 41382921 | 6 | 5 | 2 | 1 | 4 | 10 | 1 | 3 | 0 | 3 | 98 | 49 | 0.834081594 | 1 | 0.75 | 0.385 | 0.091 | 1 | 0.333 | -0.258 |
| SE | 239 DDX3X | DDX3X | chrX | + | 41343736 | 41343822 | 41342736 | 41342836 | 41344029 | 41344128 | 166 | 36 | 32 | 0 | 3 | 0 | 2 | 21 | 0 | 0 | 98 | 49 | 0.834518929 | 1 | 1 | 0.857 | 1 | 1 | 1 | -0.048 |
| SE | 16805 ZFAND5 | ZFAND5 | chr9 | - | 72363469 | 72363606 | 72360627 | 72360787 | 72364695 | 72365247 | 17 | 4 | 1 | 10 | 5 | 9 | 0 | 4 | 1 | 0 | 98 | 49 | 0.835172265 | 1 | 0.459 | 0.286 | 0.053 | 0 | 1 | -0.234 |
| SE | 12241 SMCHD1 | SMCHD1 | chr18 | + | 2722518 | 2722663 | 2718314 | 2718434 | 2724898 | 2724995 | 190 | 86 | 63 | 23 | 10 | 0 | 1 | 43 | 0 | 5 | 98 | 49 | 0.835663393 | 1 | 0.805 | 0.811 | 1 | 1 | 0.811 | -0.033 |
| A3SS | 2475 LST1 | LST1 | chr6 | + | 31588496 | 31588909 | 31588562 | 31588909 | 31587199 | 31587318 | 69 | 46 | 40 | 35 | 11 | 14 | 3 | 13 | 1 | 6 | 98 | 49 | 0.836902443 | 1 | 0.496 | 0.676 | 0.588 | 0.6 | 0.52 | 0.027 |
| SE | 3024 XPO6 | XPO6 | chr16 | - | 28133840 | 28133933 | 28132333 | 28132403 | 28135215 | 28135324 | 329 | 207 | 71 | 4 | 1 | 0 | 6 | 49 | 0 | 0 | 98 | 49 | 0.84163499 | 1 | 0.976 | 0.99 | 1 | 1 | 1 | -0.011 |
| SE | 16626 RTN4 | RTN4 | chr2 | - | 54974694 | 54974764 | 54973562 | 54973621 | 54982514 | 54982653 | 29 | 15 | 8 | 3 | 0 | 0 | 2 | 5 | 0 | 0 | 98 | 49 | 0.84261972 | 1 | 0.829 | 1 | 1 | 1 | 1 | -0.057 |
| SE | 6519 ACAP2 | ACAP2 | chr3 | - | 195392089 | 195392147 | 195381902 | 195382022 | 195442794 | 195443088 | 69 | 42 | 25 | 0 | 3 | 0 | 5 | 20 | 0 | 0 | 98 | 49 | 0.844595526 | 1 | 1 | 0.875 | 1 | 1 | 1 | -0.042 |
| A5SS | 234 SAMD9L | SAMD9L | chr7 | - | 93146882 | 93147146 | 93147087 | 93147146 | 93145384 | 93146040 | 18 | 1 | 3 | 3 | 4 | 0 | 4 | 10 | 0 | 6 | 98 | 49 | 0.844753468 | 1 | 0.75 | 0.111 | 1 | 1 | 0.455 | -0.107 |
| SE | 17083 FAM193B | FAM193B | chr5 | - | 177531952 | 177532200 | 177531281 | 177531521 | 177532442 | 177532641 | 19 | 5 | 2 | 7 | 4 | 0 | 2 | 0 | 0 | 1 | 98 | 49 | 0.844990272 | 1 | 0.576 | 0.385 | 1 | 1 | 0 | 0.154 |
| SE | 16046 ECE1 | ECE1 | chr1 | - | 21255946 | 21256138 | 21247220 | 21247363 | 21257524 | 21257590 | 47 | 15 | 9 | 1 | 2 | 0 | 1 | 5 | 0 | 0 | 98 | 49 | 0.849544904 | 1 | 0.959 | 0.789 | 1 | 1 | 1 | -0.084 |
| SE | 14665 CDS2 | CDS2 | chr20 | + | 5186686 | 5186839 | 5185757 | 5185826 | 5189066 | 5189186 | 34 | 16 | 10 | 3 | 0 | 0 | 3 | 8 | 0 | 0 | 98 | 49 | 0.850069193 | 1 | 0.85 | 1 | 1 | 1 | 1 | -0.05 |
| SE | 238 DDX3X | DDX3X | chrX | + | 41343215 | 41343351 | 41342736 | 41342836 | 41344029 | 41344128 | 147 | 42 | 21 | 0 | 3 | 0 | 2 | 15 | 0 | 0 | 98 | 49 | 0.854951073 | 1 | 1 | 0.875 | 1 | 1 | 1 | -0.042 |
| SE | 15493 NAIP | NAIP | chr5 | - | 71011274 | 71011374 | 71003775 | 71003857 | 71012347 | 71012918 | 63 | 16 | 5 | 0 | 1 | 2 | 1 | 9 | 0 | 0 | 98 | 49 | 0.855433741 | 1 | 1 | 0.889 | 0.556 | 1 | 1 | -0.185 |
| A5SS | 1405 RFWD2 | RFWD2 | chr1 | - | 176163814 | 176163891 | 176163826 | 176163891 | 176162868 | 176162988 | 95 | 45 | 5 | 33 | 11 | 1 | 1 | 17 | 1 | 5 | 61 | 49 | 0.856767086 | 1 | 0.698 | 0.767 | 0.801 | 0.445 | 0.732 | 0.167 |
| SE | 2881 SRRM2 | SRRM2 | chr16 | + | 2756333 | 2756606 | 2752328 | 2752846 | 2757471 | 2757579 | 30 | 25 | 2 | 3 | 0 | 0 | 0 | 7 | 1 | 0 | 98 | 49 | 0.857146712 | 1 | 0.833 | 1 | 1 | 0 | 1 | 0.444 |
| SE | 2723 PRKCB | PRKCB | chr16 | + | 24172269 | 24172361 | 24154683 | 24154857 | 24174517 | 24174580 | 105 | 58 | 23 | 1 | 3 | 0 | 9 | 27 | 0 | 1 | 98 | 49 | 0.85735006 | 1 | 0.981 | 0.906 | 1 | 1 | 0.931 | -0.003 |
| SE | 14489 ARPC2 | ARPC2 | chr2 | + | 218245419 | 218245546 | 218239390 | 218239484 | 218249263 | 218249464 | 357 | 133 | 74 | 1 | 1 | 0 | 3 | 57 | 0 | 0 | 98 | 49 | 0.858266198 | 1 | 0.994 | 1 | 0.902 | 1 | 1 | -0.035 |
| SE | 4306 DUSP6 | DUSP6 | chr12 | - | 89350587 | 89351025 | 89347824 | 89349561 | 89351639 | 89352859 | 41 | 7 | 7 | 4 | 0 | 2 | 3 | 21 | 2 | 1 | 98 | 49 | 0.860721438 | 1 | 0.837 | 1 | 0.636 | 0.429 | 0.913 | 0.153 |
| SE | 7308 MAX | MAX | chr14 | - | 65101545 | 65101572 | 65093707 | 65093815 | 65102303 | 65102544 | 17 | 5 | 3 | 2 | 3 | 3 | 1 | 2 | 0 | 3 | 76 | 49 | 0.861697781 | 1 | 0.846 | 0.518 | 0.392 | 1 | 0.301 | -0.065 |
| SE | 5224 MX2 | MX2 | chr21 | + | 41390564 | 41390703 | 41382409 | 41382564 | 41395586 | 41395785 | 59 | 36 | 25 | 23 | 16 | 2 | 4 | 23 | 0 | 13 | 98 | 49 | 0.863802513 | 1 | 0.562 | 0.529 | 0.862 | 1 | 0.469 | -0.083 |
| SE | 12966 DDX60L | DDX60L | chr4 | - | 168441336 | 168441492 | 168433009 | 168433115 | 168448637 | 168448779 | 96 | 56 | 4 | 0 | 3 | 0 | 3 | 44 | 0 | 0 | 98 | 49 | 0.864023371 | 1 | 1 | 0.903 | 1 | 1 | 1 | -0.032 |
| SE | 10313 JAK1 | JAK1 | chr1 | - | 64873369 | 64873523 | 64869310 | 64869474 | 64879024 | 64879148 | 84 | 27 | 32 | 4 | 0 | 0 | 1 | 13 | 0 | 0 | 98 | 49 | 0.867059698 | 1 | 0.913 | 1 | 1 | 1 | 1 | -0.029 |
| SE | 14738 HCK | HCK | chr20 | + | 32084390 | 32084543 | 32083893 | 32084043 | 32086627 | 32086807 | 173 | 75 | 64 | 19 | 6 | 2 | 3 | 55 | 1 | 5 | 98 | 49 | 0.868449242 | 1 | 0.82 | 0.862 | 0.941 | 0.6 | 0.846 | 0.151 |
| A3SS | 2398 DBNL | DBNL | chr7 | + | 44058877 | 44058983 | 44058901 | 44058983 | 44058428 | 44058480 | 3 | 2 | 1 | 42 | 19 | 9 | 0 | 2 | 1 | 3 | 73 | 49 | 0.869747429 | 1 | 0.046 | 0.066 | 0.069 | 0 | 0.309 | -0.094 |
| SE | 10048 ADD3 | ADD3 | chr10 | + | 110122109 | 110122292 | 110119465 | 110119564 | 110124016 | 110124274 | 39 | 17 | 5 | 0 | 0 | 2 | 2 | 5 | 0 | 0 | 98 | 49 | 0.874751865 | 1 | 1 | 1 | 0.556 | 1 | 1 | -0.148 |
| SE | 12305 FKBP1A | FKBP1A | chr20 | - | 1375490 | 1375603 | 1371711 | 1372240 | 1392833 | 1392881 | 12 | 6 | 5 | 0 | 0 | 2 | 1 | 1 | 0 | 0 | 98 | 49 | 0.874986718 | 1 | 1 | 1 | 0.556 | 1 | 1 | -0.148 |
| SE | 7551 RFWD2 | RFWD2 | chr1 | - | 176136487 | 176136547 | 176135009 | 176135086 | 176149005 | 176149074 | 156 | 44 | 20 | 29 | 6 | 0 | 1 | 16 | 0 | 4 | 98 | 49 | 0.876427787 | 1 | 0.729 | 0.786 | 1 | 1 | 0.667 | 0.005 |
| SE | 5085 SUCO | SUCO | chr1 | + | 172557279 | 172557417 | 172555868 | 172556023 | 172557643 | 172557794 | 46 | 22 | 16 | 1 | 2 | 0 | 4 | 9 | 0 | 0 | 98 | 49 | 0.881731767 | 1 | 0.958 | 0.846 | 1 | 1 | 1 | -0.065 |
| SE | 13947 DYSF | DYSF | chr2 | + | 71513239 | 71513332 | 71511806 | 71511921 | 71513715 | 71513921 | 53 | 16 | 16 | 1 | 2 | 0 | 1 | 10 | 0 | 1 | 98 | 49 | 0.881863743 | 1 | 0.964 | 0.8 | 1 | 1 | 0.833 | 0.005 |
| SE | 6855 AMPD2 | AMPD2 | chr1 | + | 109629795 | 109629916 | 109629326 | 109629490 | 109630232 | 109630406 | 51 | 15 | 17 | 4 | 14 | 0 | 1 | 7 | 0 | 3 | 98 | 49 | 0.883392835 | 1 | 0.864 | 0.349 | 1 | 1 | 0.538 | -0.031 |
| SE | 4794 HNRNPK | HNRNPK | chr9 | - | 83973901 | 83973973 | 83973285 | 83973399 | 83974516 | 83974589 | 185 | 110 | 61 | 5 | 3 | 4 | 1 | 53 | 0 | 3 | 98 | 49 | 0.884489462 | 1 | 0.949 | 0.948 | 0.884 | 1 | 0.898 | -0.022 |
| SE | 5373 RGS2 | RGS2 | chr1 | + | 192810369 | 192810431 | 192809038 | 192809181 | 192810980 | 192811147 | 620 | 414 | 275 | 2 | 5 | 1 | 6 | 89 | 0 | 0 | 98 | 49 | 0.885655317 | 1 | 0.994 | 0.976 | 0.993 | 1 | 1 | -0.012 |
| SE | 8848 CORO1A | CORO1A | chr16 | + | 30187941 | 30188087 | 30187724 | 30187829 | 30188191 | 30188249 | 327 | 150 | 138 | 1 | 4 | 0 | 18 | 59 | 0 | 0 | 98 | 49 | 0.888014184 | 1 | 0.994 | 0.949 | 1 | 1 | 1 | -0.019 |
| RI | 505 RAB24 | RAB24 | chr5 | - | 177302127 | 177302496 | 177302127 | 177302178 | 177302396 | 177302496 | 66 | 13 | 19 | 10 | 8 | 12 | 3 | 14 | 0 | 12 | 98 | 49 | 0.894120158 | 1 | 0.767 | 0.448 | 0.442 | 1 | 0.368 | -0.132 |
| A3SS | 2270 YY1AP1 | YY1AP1 | chr1 | - | 155668626 | 155668777 | 155668626 | 155668717 | 155670319 | 155670464 | 12 | 1 | 2 | 7 | 1 | 2 | 2 | 1 | 0 | 3 | 98 | 49 | 0.898227086 | 1 | 0.462 | 0.333 | 0.333 | 1 | 0.143 | -0.195 |
| SE | 12271 PPP4R1 | PPP4R1 | chr18 | - | 9549196 | 9549338 | 9546790 | 9547952 | 9550051 | 9550186 | 55 | 24 | 29 | 1 | 2 | 0 | 1 | 6 | 0 | 0 | 98 | 49 | 0.901736809 | 1 | 0.965 | 0.857 | 1 | 1 | 1 | -0.059 |
| SE | 5178 EIF4A1 | EIF4A1 | chr17 | + | 7575118 | 7575258 | 7574545 | 7574678 | 7576523 | 7576692 | 70 | 11 | 4 | 2 | 1 | 0 | 2 | 33 | 0 | 0 | 98 | 49 | 0.904424499 | 1 | 0.946 | 0.846 | 1 | 1 | 1 | -0.069 |
| A3SS | 2819 STAT3 | STAT3 | chr17 | - | 42348388 | 42348539 | 42348388 | 42348517 | 42388278 | 42388495 | 38 | 12 | 6 | 17 | 9 | 1 | 2 | 14 | 1 | 6 | 71 | 49 | 0.907213185 | 1 | 0.607 | 0.479 | 0.805 | 0.58 | 0.617 | 0.032 |
| SE | 16463 ZDHHC17 | ZDHHC17 | chr12 | + | 76797433 | 76797537 | 76764073 | 76764329 | 76805316 | 76805439 | 20 | 10 | 2 | 2 | 2 | 2 | 1 | 8 | 0 | 3 | 98 | 49 | 0.914990937 | 1 | 0.833 | 0.714 | 0.333 | 1 | 0.571 | -0.159 |
| A3SS | 1181 HNRNPK | HNRNPK | chr9 | - | 83968082 | 83969440 | 83968082 | 83969380 | 83970161 | 83970331 | 60 | 57 | 29 | 1 | 0 | 2 | 1 | 13 | 0 | 0 | 98 | 49 | 0.918678513 | 1 | 0.968 | 1 | 0.879 | 1 | 1 | -0.051 |
| SE | 2644 BRD2 | BRD2 | chr6 | + | 32975020 | 32975112 | 32974461 | 32974765 | 32975383 | 32975521 | 21 | 6 | 1 | 15 | 3 | 1 | 1 | 3 | 0 | 3 | 98 | 49 | 0.922814456 | 1 | 0.412 | 0.5 | 0.333 | 1 | 0.333 | -0.252 |
| SE | 246 DDX3X | DDX3X | chrX | + | 41347314 | 41347451 | 41346504 | 41346622 | 41347639 | 41350287 | 102 | 29 | 29 | 1 | 2 | 0 | 1 | 11 | 0 | 0 | 98 | 49 | 0.922903892 | 1 | 0.981 | 0.879 | 1 | 1 | 1 | -0.047 |
| SE | 13592 FAM129A | FAM129A | chr1 | - | 184823637 | 184823742 | 184823266 | 184823329 | 184831846 | 184831962 | 229 | 124 | 60 | 16 | 5 | 3 | 2 | 32 | 0 | 4 | 98 | 49 | 0.92335623 | 1 | 0.877 | 0.925 | 0.909 | 1 | 0.8 | 0.004 |
| SE | 15746 ERBB2IP | ERBB2IP | chr5 | + | 66075023 | 66075230 | 66053405 | 66054951 | 66076315 | 66076408 | 22 | 11 | 7 | 5 | 5 | 0 | 0 | 2 | 1 | 0 | 98 | 49 | 0.923482868 | 1 | 0.687 | 0.524 | 1 | 0 | 1 | 0.237 |
| SE | 5657 WDFY3 | WDFY3 | chr4 | - | 84726860 | 84726911 | 84724425 | 84724594 | 84733381 | 84733609 | 32 | 7 | 3 | 4 | 1 | 3 | 5 | 2 | 0 | 3 | 98 | 49 | 0.925440524 | 1 | 0.8 | 0.778 | 0.333 | 1 | 0.25 | 0.012 |

| | | | | | | | | | | | | | | | | | | | | | | | | | | | | | | | |
|---|---|---|---|---|---|---|---|---|---|---|---|---|---|---|---|---|---|---|---|---|---|---|---|---|---|---|---|---|---|---|---|
| A3SS | 600 | BTN2A1 | BTN2A1 | chr6 | + | 26467718 | 26469653 | 26467947 | 26469653 | 26466061 | 26466088 | 20 | 4 | 3 | 3 | 0 | 0 | 2 | 4 | 0 | 2 | 98 | 49 | 0.925814529 | 1 | 0.769 | 1 | 1 | 1 | 0.5 | 0.173 |
| SE | 4288 | PLXNC1 | PLXNC1 | chr12 | + | 94300909 | 94301057 | 94298631 | 94298795 | 94303755 | 94303896 | 85 | 44 | 32 | 0 | 1 | 2 | 4 | 17 | 0 | 0 | 98 | 49 | 0.929311539 | 1 | 1 | 0.957 | 0.889 | 1 | 1 | -0.051 |
| RI | 786 | SRSF2 | SRSF2 | chr17 | - | 76734114 | 76736464 | 76734114 | 76735158 | 76736153 | 76736464 | 55 | 21 | 6 | 16 | 3 | 2 | 7 | 12 | 2 | 3 | 98 | 49 | 0.930389012 | 1 | 0.632 | 0.778 | 0.6 | 0.636 | 0.667 | 0.019 |
| A3SS | 3977 | IGF1R | IGF1R | chr15 | + | 98929557 | 98929660 | 98929560 | 98929660 | 98924524 | 98924684 | 15 | 2 | 2 | 6 | 0 | 1 | 1 | 2 | 0 | 3 | 52 | 49 | 0.930997489 | 1 | 0.702 | 1 | 0.653 | 1 | 0.386 | 0.092 |
| SE | 10098 | PICALM | PICALM | chr11 | - | 85978069 | 85978093 | 85976622 | 85976682 | 85981128 | 85981228 | 28 | 12 | 12 | 58 | 39 | 13 | 0 | 12 | 2 | 14 | 73 | 49 | 0.931040519 | 1 | 0.245 | 0.171 | 0.383 | 0 | 0.365 | 0.084 |
| RI | 473 | CASP1 | CASP1 | chr11 | - | 105025509 | 105026356 | 105025509 | 105025600 | 105026225 | 105026356 | 20 | 12 | 13 | 44 | 9 | 13 | 0 | 15 | 2 | 6 | 98 | 49 | 0.931871799 | 1 | 0.185 | 0.4 | 0.333 | 0 | 0.556 | 0.028 |
| SE | 12122 | ST6GALNAC2 | ST6GALNAC2 | chr17 | - | 76578755 | 76578816 | 76574364 | 76574539 | 76585683 | 76586063 | 11 | 1 | 7 | 3 | 3 | 1 | 2 | 2 | 0 | 4 | 98 | 49 | 0.932313057 | 1 | 0.647 | 0.143 | 0.778 | 1 | 0.2 | -0.077 |
| A3SS | 4092 | NDUFA2 | NDUFA2 | chr5 | - | 140645362 | 140645725 | 140645362 | 140645678 | 140647255 | 140647362 | 5 | 0 | 5 | 1 | 2 | 18 | 0 | 3 | 1 | 2 | 96 | 49 | 0.938368028 | 1 | 0.718 | 0 | 0.124 | 0 | 0.434 | 0.064 |
| SE | 11300 | KDM6B | KDM6B | chr17 | + | 7845549 | 7845691 | 7845313 | 7845456 | 7845871 | 7845970 | 105 | 37 | 39 | 1 | 2 | 0 | 9 | 23 | 0 | 0 | 98 | 49 | 0.938610085 | 1 | 0.981 | 0.902 | 1 | 1 | 1 | -0.039 |
| SE | 1213 | NAMPT | NAMPT | chr7 | - | 106274945 | 106275049 | 106272529 | 106272658 | 106277022 | 106277179 | 2440 | 1189 | 341 | 16 | 7 | 0 | 15 | 313 | 0 | 2 | 98 | 49 | 0.94075831 | 1 | 0.987 | 0.988 | 1 | 1 | 0.987 | -0.002 |
| SE | 13786 | SPAST | SPAST | chr2 | + | 32126947 | 32127022 | 32116118 | 32116212 | 32128407 | 32128479 | 28 | 12 | 4 | 3 | 0 | 0 | 1 | 4 | 0 | 0 | 98 | 49 | 0.943048222 | 1 | 0.824 | 1 | 1 | 1 | 1 | -0.059 |
| SE | 12508 | ABCA7 | ABCA7 | chr19 | + | 1054569 | 1054694 | 1054192 | 1054341 | 1054779 | 1054878 | 13 | 12 | 1 | 7 | 4 | 0 | 1 | 4 | 0 | 3 | 98 | 49 | 0.949350037 | 1 | 0.481 | 0.6 | 1 | 1 | 0.4 | -0.006 |
| SE | 15924 | RPS6KA1 | RPS6KA1 | chr1 | + | 26560725 | 26560851 | 26558806 | 26558937 | 26561044 | 26561134 | 71 | 22 | 16 | 0 | 0 | 0 | 1 | 11 | 0 | 2 | 98 | 49 | 0.953350319 | 1 | 1 | 1 | 1 | 1 | 0.733 | 0.133 |
| SE | 11513 | SBNO2 | SBNO2 | chr19 | - | 1112188 | 1112301 | 1111995 | 1112067 | 1112401 | 1112537 | 61 | 21 | 22 | 10 | 4 | 0 | 8 | 15 | 4 | 0 | 98 | 49 | 0.956284839 | 1 | 0.753 | 0.724 | 1 | 0.5 | 1 | 0.076 |
| SE | 169 | IL2RG | IL2RG | chrX | - | 71108598 | 71108695 | 71108276 | 71108346 | 71109227 | 71109390 | 47 | 21 | 14 | 6 | 5 | 1 | 1 | 4 | 2 | 0 | 98 | 49 | 0.959685424 | 1 | 0.797 | 0.677 | 0.875 | 0.2 | 1 | 0.183 |
| SE | 8401 | C19orf38 | C19orf38 | chr19 | + | 10859914 | 10859958 | 10858315 | 10858343 | 10863169 | 10863207 | 26 | 4 | 3 | 0 | 0 | 0 | 2 | 11 | 0 | 2 | 93 | 49 | 0.969972169 | 1 | 1 | 1 | 1 | 1 | 0.743 | 0.129 |
| SE | 1585 | IGF2R | IGF2R | chr6 | + | 160061752 | 160061928 | 160061502 | 160061646 | 160062531 | 160062619 | 243 | 80 | 87 | 0 | 4 | 0 | 5 | 27 | 0 | 1 | 98 | 49 | 0.980810341 | 1 | 1 | 0.909 | 1 | 1 | 0.931 | 0.004 |
| SE | 5919 | MYD88 | MYD88 | chr3 | + | 38140387 | 38140568 | 38139863 | 38139998 | 38140756 | 38140848 | 13 | 6 | 7 | 0 | 0 | 0 | 1 | 15 | 1 | 1 | 98 | 49 | 0.983062405 | 1 | 1 | 1 | 1 | 0.333 | 0.882 | 0.392 |
| SE | 5962 | SORL1 | SORL1 | chr11 | + | 121574240 | 121574363 | 121566939 | 121567113 | 121583457 | 121583583 | 26 | 8 | 4 | 3 | 0 | 0 | 1 | 15 | 0 | 0 | 98 | 49 | 0.98862453 | 1 | 0.812 | 1 | 1 | 1 | 1 | -0.063 |
| A3SS | 1740 | AP1G2 | AP1G2 | chr14 | - | 23564045 | 23564221 | 23564045 | 23564159 | 23564332 | 23564388 | 9 | 4 | 0 | 4 | 0 | 1 | 3 | 1 | 0 | 3 | 98 | 49 | 0.98876747 | 1 | 0.529 | 1 | 0 | 1 | 0.143 | -0.062 |
| SE | 6589 | SETD2 | SETD2 | chr3 | - | 47101457 | 47101555 | 47097954 | 47098081 | 47103345 | 47103423 | 38 | 23 | 16 | 3 | 0 | 0 | 1 | 10 | 0 | 0 | 98 | 49 | 0.989533371 | 1 | 0.864 | 1 | 1 | 1 | 1 | -0.045 |
| A3SS | 5 | FAM122B | FAM122B | chrX | - | 134769565 | 134772283 | 134769565 | 134771354 | 134781820 | 134781917 | 2 | 0 | 0 | 0 | 2 | 0 | 0 | 3 | 0 | 0 | | | 1 | 1 | 1 | 0 | NA | NA | 1 | -0.5 |
| A3SS | 6 | FAM122B | FAM122B | chrX | - | 134785885 | 134785950 | 134785885 | 134785947 | 134787471 | 134787533 | 2 | 0 | 0 | 1 | 0 | 0 | 0 | 2 | 0 | 0 | 52 | 49 | 1 | 1 | 0.653 | NA | NA | NA | 1 | -0.347 |
| A3SS | 15 | MORF4L2 | MORF4L2 | chrX | - | 103678498 | 103678651 | 103678498 | 103678600 | 103684680 | 103684729 | 14 | 3 | 0 | 1 | 0 | 0 | 0 | 2 | 0 | 0 | 98 | 49 | 1 | 1 | 0.875 | 1 | NA | NA | 1 | -0.062 |
| A3SS | 25 | ARMCX3 | ARMCX3 | chrX | + | 101623785 | 101623893 | 101623813 | 101623893 | 101623129 | 101623276 | 0 | 0 | 0 | 1 | 1 | 0 | 0 | 1 | 0 | 0 | 77 | 49 | 1 | 1 | 0 | 0 | NA | NA | 1 | -1 |
| A3SS | 31 | ABCB7 | ABCB7 | chrX | - | 75114753 | 75114834 | 75114753 | 75114831 | 75156104 | 75156340 | 1 | 0 | 0 | 1 | 0 | 0 | 0 | 2 | 0 | 0 | 52 | 49 | 1 | 1 | 0.485 | NA | NA | NA | 1 | -0.515 |
| A3SS | 50 | TFE3 | TFE3 | chrX | - | 49039106 | 49039410 | 49039106 | 49039345 | 49040454 | 49040568 | 34 | 15 | 5 | 1 | 2 | 0 | 0 | 4 | 0 | 3 | 98 | 49 | 1 | 1 | 0.944 | 0.789 | 1 | NA | 0.4 | 0.511 |
| A3SS | 52 | FTSJ1 | FTSJ1 | chrX | + | 48482402 | 48482506 | 48482408 | 48482506 | 48481631 | 48481715 | 3 | 0 | 0 | 1 | 0 | 0 | 0 | 0 | 0 | 2 | 55 | 49 | 1 | 1 | 0.728 | NA | NA | NA | 0 | 0.728 |
| A3SS | 59 | DDX3X | DDX3X | chrX | + | 41347311 | 41347451 | 41347314 | 41347451 | 41346858 | 41347012 | 77 | 27 | 18 | 45 | 10 | 7 | 0 | 8 | 0 | 6 | 52 | 49 | 1 | 1 | 0.617 | 0.718 | 0.708 | NA | 0.557 | 0.124 |
| A3SS | 60 | BCOR | BCOR | chrX | - | 40063607 | 40063952 | 40063607 | 40063850 | 40064335 | 40064599 | 2 | 0 | 0 | 0 | 0 | 2 | 2 | 0 | 0 | 0 | 98 | 49 | 1 | 1 | 1 | NA | 0 | 1 | NA | -0.5 |
| A3SS | 66 | REPS2 | REPS2 | chrX | + | 17025058 | 17025185 | 17025061 | 17025185 | 17022122 | 17022271 | 3 | 0 | 3 | 9 | 2 | 6 | 0 | 1 | 0 | 0 | 52 | 49 | 1 | 1 | 0.239 | 0 | 0.32 | NA | 1 | -0.814 |
| A3SS | 69 | CD99 | CD99 | chrX | + | 2738199 | 2738256 | 2738202 | 2738256 | 2726259 | 2726373 | 20 | 3 | 2 | 1 | 0 | 0 | 0 | 10 | 0 | 1 | 52 | 49 | 1 | 1 | 0.95 | 1 | 1 | NA | 0.904 | 0.079 |
| A3SS | 84 | SEC16A | SEC16A | chr9 | - | 136446854 | 136447015 | 136446854 | 136446949 | 136447226 | 136447364 | 4 | 0 | 0 | 0 | 1 | 0 | 0 | 1 | 0 | 0 | 98 | 49 | 1 | 1 | 1 | 0 | NA | NA | 1 | -0.5 |
| A3SS | 126 | ZNF189 | ZNF189 | chr9 | + | 101399883 | 101400010 | 101399925 | 101400010 | 101398853 | 101399189 | 2 | 0 | 0 | 0 | 0 | 0 | 0 | 1 | 0 | 1 | 91 | 49 | 1 | 1 | 1 | NA | NA | NA | 0.35 | 0.65 |
| A3SS | 136 | FAM120A | FAM120A | chr9 | + | 93515666 | 93515767 | 93515669 | 93515767 | 93498789 | 93498886 | 9 | 4 | 1 | 2 | 0 | 0 | 0 | 1 | 0 | 2 | 52 | 49 | 1 | 1 | 0.809 | 1 | 1 | NA | 0.32 | 0.616 |
| A3SS | 139 | NOL8 | NOL8 | chr9 | - | 92299889 | 92300810 | 92299889 | 92300016 | 92301550 | 92301822 | 2 | 0 | 0 | 2 | 1 | 0 | 0 | 1 | 0 | 0 | 98 | 49 | 1 | 1 | 0.333 | 0 | NA | NA | 1 | -0.834 |
| A3SS | 140 | SECISBP2 | SECISBP2 | chr9 | + | 89325426 | 89325676 | 89325547 | 89325676 | 89319651 | 89319797 | 16 | 3 | 3 | 3 | 1 | 0 | 0 | 4 | 0 | 1 | 98 | 49 | 1 | 1 | 0.727 | 0.6 | 1 | NA | 0.667 | 0.109 |
| A3SS | 154 | FAM219A | FAM219A | chr9 | - | 34398183 | 34401122 | 34398183 | 34401119 | 34401665 | 34401720 | 5 | 0 | 2 | 1 | 0 | 3 | 0 | 0 | 1 | 0 | 52 | 49 | 1 | 1 | 0.825 | NA | 0.386 | 0 | NA | 0.605 |
| A3SS | 160 | KDM4C | KDM4C | chr9 | + | 6986324 | 6986666 | 6986343 | 6986666 | 6984165 | 6984404 | 2 | 0 | 0 | 0 | 2 | 0 | 0 | 0 | 1 | 2 | 68 | 49 | 1 | 1 | 1 | 0 | NA | 0 | 0 | 0.5 |
| A3SS | 182 | INTS8 | INTS8 | chr8 | + | 94838462 | 94838618 | 94838493 | 94838618 | 94836523 | 94836631 | 11 | 0 | 1 | 2 | 0 | 0 | 0 | 0 | 0 | 1 | 80 | 49 | 1 | 1 | 0.771 | NA | 1 | NA | 0 | 0.885 |
| A3SS | 194 | TM2D2 | TM2D2 | chr8 | - | 38995317 | 38995794 | 38995317 | 38995405 | 38996212 | 38996522 | 0 | 1 | 0 | 0 | 0 | 0 | 0 | 2 | 0 | 1 | 98 | 49 | 1 | 1 | NA | 1 | NA | NA | 0.5 | 0.5 |
| A3SS | 209 | PRKAG2 | PRKAG2 | chr7 | - | 151595344 | 151595454 | 151595344 | 151595451 | 151632068 | 151632138 | 6 | 1 | 6 | 0 | 8 | 0 | 0 | 1 | 0 | 0 | 52 | 49 | 1 | 1 | 1 | 0.105 | 1 | NA | 1 | -0.298 |
| A3SS | 228 | PNPLA8 | PNPLA8 | chr7 | - | 108514435 | 108515574 | 108514435 | 108515447 | 108521475 | 108521521 | 32 | 28 | 0 | 0 | 2 | 0 | 0 | 10 | 0 | 0 | 98 | 49 | 1 | 1 | 1 | 0.875 | NA | NA | 1 | -0.062 |
| A3SS | 239 | SRRT | SRRT | chr7 | + | 100885212 | 100885370 | 100885215 | 100885370 | 100884922 | 100885040 | 3 | 1 | 0 | 2 | 0 | 0 | 0 | 0 | 0 | 1 | 52 | 49 | 1 | 1 | 0.586 | 1 | NA | NA | 0 | 0.793 |
| A3SS | 240 | SRRT | SRRT | chr7 | + | 100888041 | 100888143 | 100888053 | 100888143 | 100887702 | 100887859 | 5 | 4 | 0 | 3 | 1 | 0 | 1 | 1 | 0 | 3 | 61 | 49 | 1 | 1 | 0.572 | 0.763 | NA | 1 | 0.211 | 0.062 |
| A3SS | 245 | STAG3L5P-PVRIG2P-PILRB | STAG3L5P-PVRIG2P-PILRB | chr7 | + | 100358689 | 100359079 | 100358898 | 100359079 | 100358219 | 100358366 | 12 | 2 | 3 | 0 | 0 | 1 | 0 | 1 | 0 | 0 | 98 | 49 | 1 | 1 | 1 | 1 | 0.6 | NA | 1 | -0.133 |
| A3SS | 251 | SMURF1 | SMURF1 | chr7 | - | 99033036 | 99033121 | 99033036 | 99033112 | 99035514 | 99035716 | 3 | 3 | 0 | 1 | 0 | 0 | 2 | 1 | 0 | 0 | 58 | 49 | 1 | 1 | 0.717 | 1 | NA | 1 | 1 | -0.141 |
| A3SS | 270 | GAB3 | GAB3 | chrX | - | 154713206 | 154713426 | 154713206 | 154713423 | 154716025 | 154716329 | 1 | 0 | 0 | 0 | 2 | 0 | 0 | 1 | 0 | 0 | 52 | 49 | 1 | 1 | 1 | 0 | NA | NA | 1 | -0.5 |
| A3SS | 280 | ZMIZ2 | ZMIZ2 | chr7 | + | 44756946 | 44757149 | 44757042 | 44757149 | 44756424 | 44756539 | 6 | 4 | 0 | 1 | 0 | 0 | 0 | 3 | 0 | 0 | 98 | 49 | 1 | 1 | 0.75 | 1 | NA | NA | 1 | -0.125 |
| A3SS | 282 | POLM | POLM | chr7 | - | 44072061 | 44073430 | 44072061 | 44073377 | 44073624 | 44073708 | 9 | 1 | 0 | 0 | 1 | 0 | 0 | 1 | 0 | 0 | 98 | 49 | 1 | 1 | 1 | 0.333 | NA | NA | 1 | -0.334 |
| A3SS | 286 | CDK13 | CDK13 | chr7 | + | 40092784 | 40093237 | 40092964 | 40093237 | 40088125 | 40088331 | 12 | 3 | 1 | 0 | 1 | 0 | 0 | 1 | 0 | 0 | 98 | 49 | 1 | 1 | 1 | 0.6 | 1 | NA | 1 | -0.133 |
| A3SS | 298 | RNF216 | RNF216 | chr7 | - | 5740972 | 5741815 | 5740972 | 5741644 | 5752845 | 5752979 | 21 | 9 | 0 | 3 | 1 | 0 | 0 | 4 | 0 | 0 | 98 | 49 | 1 | 1 | 0.778 | 0.818 | NA | NA | 1 | -0.202 |
| A3SS | 312 | IPCEF1 | IPCEF1 | chr6 | - | 154247448 | 154247488 | 154247448 | 154247485 | 154265911 | 154265964 | 2 | 3 | 0 | 6 | 0 | 0 | 0 | 0 | 0 | 1 | 52 | 49 | 1 | 1 | 0.239 | 1 | NA | NA | 0 | 0.619 |
| A3SS | 336 | VNN3 | VNN3 | chr6 | - | 132728740 | 132728979 | 132728740 | 132728933 | 132734252 | 132734383 | 109 | 54 | 13 | 29 | 12 | 3 | 0 | 9 | 0 | 4 | 95 | 49 | 1 | 1 | 0.66 | 0.699 | 0.691 | NA | 0.537 | 0.146 |
| A3SS | 337 | ARG1 | ARG1 | chr6 | + | 131579086 | 131579285 | 131579110 | 131579285 | 131576662 | 131576735 | 0 | 0 | 0 | 5 | 5 | 3 | 0 | 2 | 0 | 4 | 73 | 49 | 1 | 1 | 0 | 0 | 0 | NA | 0.251 | -0.251 |
| A3SS | 352 | FAM135A | FAM135A | chr6 | + | 70524342 | 70526698 | 70524930 | 70526698 | 70523966 | 70524121 | 0 | 0 | 0 | 1 | 1 | 0 | 0 | 1 | 0 | 0 | 98 | 49 | 1 | 1 | 0 | 0 | NA | NA | 1 | -1 |
| A3SS | 358 | HSP90AB1 | HSP90AB1 | chr6 | + | 44248616 | 44248776 | 44248629 | 44248776 | 44246957 | 44247195 | 0 | 0 | 0 | 3 | 0 | 0 | 0 | 2 | 0 | 0 | 62 | 49 | 1 | 1 | 0 | NA | NA | NA | 1 | -1 |
| A3SS | 369 | GBGT1 | GBGT1 | chr9 | - | 133155177 | 133155349 | 133155177 | 133155312 | 133155900 | 133155936 | 3 | 0 | 0 | 2 | 0 | 0 | 0 | 1 | 0 | 1 | 86 | 49 | 1 | 1 | 0.461 | NA | NA | NA | 0.363 | 0.098 |
| A3SS | 370 | GBGT1 | GBGT1 | chr9 | - | 133156014 | 133156102 | 133156014 | 133156065 | 133161466 | 133161532 | 2 | 3 | 0 | 1 | 4 | 0 | 0 | 0 | 0 | 2 | 86 | 49 | 1 | 1 | 0.533 | 0.299 | NA | NA | 0 | 0.416 |
| A3SS | 383 | STK38 | STK38 | chr6 | - | 36540071 | 36540207 | 36540071 | 36540204 | 36547189 | 36547516 | 4 | 1 | 2 | 8 | 3 | 6 | 0 | 0 | 0 | 1 | 52 | 49 | 1 | 1 | 0.32 | 0.239 | 0.239 | NA | 0 | 0.266 |
| A3SS | 384 | SRPK1 | SRPK1 | chr6 | - | 35890894 | 35891041 | 35890894 | 35891013 | 35920467 | 35920528 | 5 | 2 | 0 | 5 | 2 | 0 | 0 | 0 | 1 | 2 | 77 | 49 | 1 | 1 | 0.389 | 0.389 | NA | 0 | 0 | 0.389 |
| A3SS | 439 | PSMB8-AS1 | PSMB8-AS1 | chr6 | + | 32845329 | 32846500 | 32845417 | 32846500 | 32844085 | 32844274 | 67 | 37 | 39 | 7 | 0 | 0 | 0 | 17 | 0 | 0 | 98 | 49 | 1 | 1 | 0.827 | 1 | 1 | NA | 1 | -0.058 |
| A3SS | 500 | C6orf25 | C6orf25 | chr6 | + | 31724745 | 31724844 | 31724764 | 31724844 | 31724586 | 31724627 | 6 | 0 | 0 | 1 | 0 | 0 | 9 | 0 | 0 | 0 | 68 | 49 | 1 | 1 | 0.812 | NA | NA | 1 | NA | -0.188 |
| A3SS | 528 | PRRC2A | PRRC2A | chr6 | + | 31622689 | 31622901 | 31622726 | 31622901 | 31620672 | 31620858 | 8 | 0 | 0 | 16 | 2 | 2 | 0 | 2 | 0 | 2 | 86 | 49 | 1 | 1 | 0.222 | 0 | 0 | NA | 0.363 | -0.289 |
| A3SS | 543 | DDX39B | DDX39B | chr6 | - | 31540321 | 31540664 | 31540321 | 31540534 | 31541949 | 31542475 | 14 | 6 | 0 | 4 | 0 | 0 | 1 | 0 | 0 | 0 | 98 | 49 | 1 | 1 | 0.636 | 1 | NA | 1 | NA | -0.182 |
| A3SS | 582 | TRIM26 | TRIM26 | chr6 | - | 30198665 | 30199264 | 30198665 | 30199153 | 30201034 | 30201138 | 4 | 2 | 0 | 4 | 1 | 0 | 0 | 2 | 0 | 1 | 98 | 49 | 1 | 1 | 0.333 | 0.5 | NA | NA | 0.5 | -0.084 |
| A3SS | 604 | BTN3A1 | BTN3A1 | chr6 | + | 26412411 | 26415216 | 26412704 | 26415216 | 26411554 | 26411581 | 14 | 0 | 8 | 6 | 2 | 3 | 0 | 1 | 0 | 0 | 98 | 49 | 1 | 1 | 0.538 | 0 | 0.571 | NA | 1 | -0.63 |
| A3SS | 607 | BTN3A2 | BTN3A2 | chr6 | + | 26367974 | 26368050 | 26367989 | 26368050 | 26365158 | 26365352 | 0 | 0 | 0 | 11 | 4 | 0 | 0 | 1 | 0 | 2 | 64 | 49 | 1 | 1 | 0 | 0 | NA | NA | 0.277 | -0.277 |
| A3SS | 608 | BTN3A2 | BTN3A2 | chr6 | + | 26375514 | 26378320 | 26375796 | 26378320 | 26374770 | 26374798 | 7 | 2 | 0 | 3 | 1 | 2 | 0 | 0 | 0 | 2 | 98 | 49 | 1 | 1 | 0.538 | 0.5 | 0 | NA | 0 | 0.346 |
| A3SS | 611 | FAM65B | FAM65B | chr6 | - | 24839365 | 24840775 | 24839365 | 24839599 | 24842861 | 24843554 | 109 | 83 | 41 | 12 | 14 | 10 | 0 | 20 | 0 | 2 | 98 | 49 | 1 | 1 | 0.82 | 0.748 | 0.672 | NA | 0.833 | -0.086 |
| A3SS | 620 | PHF23 | PHF23 | chr17 | - | 7235929 | 7236767 | 7235929 | 7236566 | 7237384 | 7237477 | 10 | 7 | 3 | 1 | 0 | 0 | 1 | 1 | 0 | 0 | 98 | 49 | 1 | 1 | 0.833 | 1 | 1 | 1 | 1 | -0.056 |
| A3SS | 622 | C17orf49 | C17orf49 | chr17 | + | 7017256 | 7017524 | 7017260 | 7017524 | 7016909 | 7017014 | 4 | 0 | 0 | 6 | 0 | 0 | 0 | 0 | 0 | 2 | 53 | 49 | 1 | 1 | 0.381 | NA | NA | NA | 0 | 0.381 |
| A3SS | 632 | EML3 | EML3 | chr11 | - | 62611086 | 62611347 | 62611086 | 62611344 | 62611424 | 62611596 | 1 | 0 | 0 | 4 | 0 | 0 | 0 | 1 | 0 | 0 | 52 | 49 | 1 | 1 | 0.191 | NA | NA | NA | 1 | -0.809 |
| A3SS | 644 | BSDC1 | BSDC1 | chr1 | - | 32383829 | 32384048 | 32383829 | 32383997 | 32386778 | 32386895 | 3 | 1 | 0 | 8 | 5 | 3 | 0 | 0 | 0 | 1 | 98 | 49 | 1 | 1 | 0.158 | 0.091 | 0 | NA | 0 | 0.083 |
| A3SS | 649 | C12orf57 | C12orf57 | chr12 | + | 6944379 | 6944565 | 6944475 | 6944565 | 6943816 | 6944173 | 0 | 0 | 0 | 11 | 0 | 5 | 1 | 1 | 0 | 0 | 98 | 49 | 1 | 1 | 0 | NA | 0 | 1 | 1 | -1 |
| A3SS | 654 | PSME3 | PSME3 | chr17 | + | 42838691 | 42838799 | 42838730 | 42838799 | 42838092 | 42838205 | 1 | 0 | 0 | 17 | 9 | 5 | 0 | 1 | 0 | 4 | 88 | 49 | 1 | 1 | 0.032 | 0 | 0 | NA | 0.122 | -0.111 |
| A3SS | 669 | CIAPIN1 | CIAPIN1 | chr16 | - | 57439181 | 57439334 | 57439181 | 57439295 | 57440771 | 57440983 | 1 | 0 | 1 | 0 | 2 | 0 | 0 | 2 | 0 | 0 | 88 | 49 | 1 | 1 | 1 | 0 | 1 | NA | 1 | -0.333 |
| A3SS | 680 | STX4 | STX4 | chr16 | + | 31034203 | 31034325 | 31034225 | 31034325 | 31034012 | 31034114 | 2 | 0 | 0 | 4 | 5 | 0 | 0 | 3 | 1 | 2 | 71 | 49 | 1 | 1 | 0.257 | 0 | NA | 0 | 0.509 | -0.126 |
| A3SS | 685 | ATXN2L | ATXN2L | chr16 | + | 28832186 | 28832399 | 28832204 | 28832399 | 28830961 | 28831072 | 1 | 1 | 0 | 3 | 3 | 0 | 0 | 1 | 1 | 0 | 67 | 49 | 1 | 1 | 0.196 | 0.196 | NA | 0 | 1 | -0.304 |
| A3SS | 697 | WDTC1 | WDTC1 | chr1 | + | 27297047 | 27297156 | 27297050 | 27297156 | 27296325 | 27296401 | 19 | 6 | 10 | 1 | 0 | 0 | 0 | 4 | 0 | 1 | 52 | 49 | 1 | 1 | 0.947 | 1 | 1 | NA | 0.79 | 0.192 |
| A3SS | 701 | PCBP2 | PCBP2 | chr12 | + | 53468776 | 53468832 | 53468779 | 53468832 | 53467220 | 53467293 | 15 | 1 | 2 | 3 | 1 | 0 | 0 | 7 | 0 | 2 | 52 | 49 | 1 | 1 | 0.825 | 0.485 | 1 | NA | 0.767 | 0.003 |
| A3SS | 702 | PCBP2 | PCBP2 | chr12 | + | 53468776 | 53468832 | 53468779 | 53468832 | 53467804 | 53467843 | 28 | 9 | 7 | 14 | 1 | 2 | 0 | 5 | 0 | 0 | 52 | 49 | 1 | 1 | 0.653 | 0.895 | 0.767 | NA | 1 | -0.228 |
| A3SS | 711 | SBNO1 | SBNO1 | chr12 | - | 123345257 | 123345570 | 123345257 | 123345567 | 123348028 | 123348133 | 8 | 1 | 2 | 7 | 4 | 0 | 0 | 0 | 0 | 1 | 52 | 49 | 1 | 1 | 0.519 | 0.191 | 1 | NA | 0 | 0.57 |
| A3SS | 714 | MEFV | MEFV | chr16 | - | 3243859 | 3244067 | 3243859 | 3243892 | 3244253 | 3244286 | 5 | 3 | 3 | 10 | 8 | 13 | 0 | 8 | 0 | 3 | 98 | 49 | 1 | 1 | 0.2 | 0.158 | 0.103 | NA | 0.571 | -0.417 |
| A3SS | 719 | ZC3H14 | ZC3H14 | chr14 | + | 88609711 | 88609803 | 88609714 | 88609803 | 88609266 | 88609403 | 0 | 4 | 0 | 1 | 0 | 0 | 0 | 1 | 0 | 0 | 52 | 49 | 1 | 1 | 0 | 1 | NA | NA | 1 | -0.5 |
| A3SS | 727 | TSC2 | TSC2 | chr16 | + | 2079031 | 2079196 | 2079034 | 2079196 | 2076490 | 2076585 | 0 | 4 | 0 | 5 | 0 | 0 | 0 | 1 | 0 | 0 | 52 | 49 | 1 | 1 | 0 | 1 | NA | NA | 1 | -0.5 |
| A3SS | 732 | ATG5 | ATG5 | chr6 | - | 106248149 | 106248244 | 106248149 | 106248240 | 106279660 | 106279823 | 7 | 9 | 3 | 1 | 0 | 0 | 0 | 3 | 0 | 0 | 53 | 49 | 1 | 1 | 0.866 | 1 | 1 | NA | 1 | -0.045 |
| A3SS | 733 | NLRP12 | NLRP12 | chr19 | - | 53807494 | 53807668 | 53807494 | 53807665 | 53809586 | 53811288 | 4 | 1 | 0 | 21 | 7 | 6 | 0 | 1 | 0 | 6 | 52 | 49 | 1 | 1 | 0.152 | 0.119 | 0 | NA | 0.136 | -0.046 |
| A3SS | 736 | SERF2 | SERF2 | chr15 | + | 43793575 | 43796089 | 43793709 | 43796089 | 43792974 | 43793083 | 16 | 1 | 0 | 76 | 14 | 13 | 0 | 0 | 0 | 9 | 98 | 49 | 1 | 1 | 0.095 | 0.034 | 0 | NA | 0 | 0.043 |
| A3SS | 737 | SERF2 | SERF2 | chr15 | + | 43793686 | 43796089 | 43793709 | 43796089 | 43792974 | 43793083 | 12 | 1 | 0 | 76 | 14 | 13 | 0 | 0 | 0 | 9 | 72 | 49 | 1 | 1 | 0.097 | 0.046 | 0 | NA | 0 | 0.048 |
| A3SS | 748 | AP3S2 | AP3S2 | chr15 | - | 89889048 | 89889140 | 89889048 | 89889136 | 89893880 | 89894385 | 6 | 3 | 0 | 0 | 3 | 0 | 0 | 2 | 0 | 0 | 53 | 49 | 1 | 1 | 1 | 0.48 | NA | NA | 1 | -0.26 |
| A3SS | 751 | ISG20 | ISG20 | chr15 | + | 88650234 | 88652310 | 88652109 | 88652310 | 88639342 | 88639594 | 3 | 5 | 1 | 13 | 15 | 13 | 0 | 2 | 0 | 11 | 98 | 49 | 1 | 1 | 0.103 | 0.143 | 0.037 | NA | 0.083 | 0.011 |
| A3SS | 782 | ZNF143 | ZNF143 | chr11 | + | 9474549 | 9474633 | 9474552 | 9474633 | 9473940 | 9474024 | 9 | 1 | 0 | 4 | 0 | 0 | 0 | 1 | 0 | 3 | 52 | 49 | 1 | 1 | 0.68 | 1 | NA | NA | 0.239 | 0.601 |
| A3SS | 786 | TRPM7 | TRPM7 | chr15 | - | 50583088 | 50583159 | 50583088 | 50583156 | 50586391 | 50586488 | 1 | 1 | 6 | 1 | 0 | 1 | 0 | 1 | 0 | 0 | 52 | 49 | 1 | 1 | 0.485 | 1 | 0.85 | NA | 1 | -0.222 |
| A3SS | 790 | PLAGL1 | PLAGL1 | chr6 | - | 143960468 | 143960556 | 143960468 | 143960542 | 143964786 | 143964818 | 1 | 4 | 0 | 1 | 0 | 0 | 2 | 0 | 0 | 1 | 63 | 49 | 1 | 1 | 0.438 | 1 | NA | 1 | 0 | 0.219 |
| A3SS | 791 | PLAGL1 | PLAGL1 | chr6 | - | 143960468 | 143960580 | 143960468 | 143960542 | 143964786 | 143964818 | 1 | 3 | 0 | 1 | 0 | 0 | 2 | 0 | 0 | 1 | 87 | 49 | 1 | 1 | 0.36 | 1 | NA | 1 | 0 | 0.18 |
| A3SS | 831 | KIAA0430 | KIAA0430 | chr16 | - | 15622933 | 15623126 | 15622933 | 15623123 | 15624768 | 15624927 | 0 | 2 | 0 | 13 | 1 | 0 | 0 | 1 | 0 | 0 | 52 | 49 | 1 | 1 | 0 | 0.653 | NA | NA | 1 | -0.673 |
| A3SS | 832 | KIAA0430 | KIAA0430 | chr16 | - | 15625371 | 15625800 | 15625371 | 15625797 | 15630331 | 15630504 | 13 | 5 | 0 | 2 | 0 | 2 | 0 | 2 | 0 | 0 | 52 | 49 | 1 | 1 | 0.86 | 1 | 0 | NA | 1 | -0.38 |
| A3SS | 843 | CIITA | CIITA | chr16 | + | 10898921 | 10899002 | 10898924 | 10899002 | 10898669 | 10898732 | 4 | 0 | 0 | 0 | 0 | 0 | 0 | 2 | 0 | 2 | 52 | 49 | 1 | 1 | 1 | NA | NA | NA | 0.485 | 0.515 |
| A3SS | 844 | CIITA | CIITA | chr16 | + | 10908982 | 10909187 | 10909028 | 10909187 | 10906498 | 10908149 | 1 | 0 | 0 | 0 | 1 | 0 | 0 | 1 | 0 | 0 | 95 | 49 | 1 | 1 | 1 | 0 | NA | NA | 1 | -0.5 |
| A3SS | 850 | MMS19 | MMS19 | chr10 | - | 97466058 | 97466159 | 97466058 | 97466156 | 97466503 | 97466585 | 2 | 0 | 0 | 0 | 0 | 0 | 0 | 3 | 0 | 1 | 52 | 49 | 1 | 1 | 1 | NA | NA | NA | 0.739 | 0.261 |
| A3SS | 855 | DLST | DLST | chr14 | + | 74889274 | 74889349 | 74889276 | 74889349 | 74889094 | 74889147 | 2 | 5 | 1 | 1 | 0 | 0 | 0 | 1 | 0 | 0 | 51 | 49 | 1 | 1 | 0.658 | 1 | 1 | NA | 1 | -0.114 |
| A3SS | 858 | ZNF410 | ZNF410 | chr14 | + | 73896315 | 73896534 | 73896347 | 73896534 | 73893796 | 73893932 | 4 | 5 | 1 | 1 | 0 | 0 | 0 | 3 | 0 | 0 | 81 | 49 | 1 | 1 | 0.708 | 1 | 1 | NA | 1 | -0.097 |
| A3SS | 859 | ZNF410 | ZNF410 | chr14 | + | 73904901 | 73905083 | 73904930 | 73905083 | 73903959 | 73904110 | 9 | 2 | 3 | 1 | 0 | 0 | 0 | 2 | 0 | 0 | 78 | 49 | 1 | 1 | 0.85 | 1 | 1 | NA | 1 | -0.05 |
| A3SS | 862 | PCNX | PCNX | chr14 | + | 71012984 | 71013202 | 71012999 | 71013202 | 71011491 | 71011549 | 67 | 27 | 10 | 5 | 8 | 0 | 0 | 12 | 0 | 4 | 64 | 49 | 1 | 1 | 0.911 | 0.721 | 1 | NA | 0.697 | 0.18 |
| A3SS | 870 | NLRC3 | NLRC3 | chr16 | - | 3567242 | 3567385 | 3567242 | 3567324 | 3577148 | 3577404 | 1 | 0 | 0 | 2 | 0 | 0 | 0 | 1 | 0 | 0 | 98 | 49 | 1 | 1 | 0.2 | NA | NA | NA | 1 | -0.8 |
| A3SS | 878 | COPS2 | COPS2 | chr15 | - | 49137347 | 49137458 | 49137347 | 49137437 | 49139527 | 49139653 | 0 | 1 | 0 | 0 | 1 | 0 | 0 | 2 | 0 | 2 | 70 | 49 | 1 | 1 | NA | 0.412 | NA | NA | 0.412 | 0 |
| A3SS | 890 | LTB4R2 | LTB4R2 | chr14 | + | 24310598 | 24312053 | 24310654 | 24312053 | 24310150 | 24310255 | 4 | 1 | 0 | 1 | 0 | 0 | 0 | 1 | 0 | 0 | 98 | 49 | 1 | 1 | 0.667 | 1 | NA | NA | 1 | -0.166 |
| A3SS | 894 | TNIP1 | TNIP1 | chr5 | - | 151029942 | 151030747 | 151029942 | 151030673 | 151032286 | 151032383 | 21 | 13 | 13 | 1 | 1 | 0 | 0 | 6 | 0 | 0 | 98 | 49 | 1 | 1 | 0.913 | 0.867 | 1 | NA | 1 | -0.073 |
| A3SS | 899 | ARHGEF40 | ARHGEF40 | chr14 | + | 21076535 | 21076643 | 21076562 | 21076643 | 21076359 | 21076456 | 3 | 1 | 0 | 5 | 1 | 2 | 0 | 0 | 0 | 4 | 76 | 49 | 1 | 1 | 0.279 | 0.392 | 0 | NA | 0 | 0.224 |
| A3SS | 906 | NOP16 | NOP16 | chr5 | - | 176386839 | 176386943 | 176386839 | 176386909 | 176388234 | 176388343 | 3 | 0 | 0 | 0 | 0 | 0 | 0 | 0 | 0 | 1 | 83 | 49 | 1 | 1 | 1 | NA | NA | NA | 0 | 1 |
| A3SS | 918 | PLEKHB2 | PLEKHB2 | chr2 | + | 131140166 | 131140275 | 131140169 | 131140275 | 131132901 | 131132991 | 23 | 5 | 3 | 2 | 0 | 0 | 1 | 3 | 0 | 0 | 52 | 49 | 1 | 1 | 0.916 | 1 | 1 | 1 | 1 | -0.028 |
| A3SS | 919 | PLEKHB2 | PLEKHB2 | chr2 | + | 131140166 | 131140275 | 131140169 | 131140275 | 131132901 | 131133015 | 5 | 0 | 1 | 1 | 0 | 0 | 0 | 2 | 0 | 0 | 52 | 49 | 1 | 1 | 0.825 | NA | 1 | NA | 1 | -0.088 |
| A3SS | 921 | CARKD | CARKD | chr13 | + | 110638241 | 110639995 | 110638377 | 110639995 | 110637128 | 110637249 | 3 | 0 | 0 | 0 | 0 | 0 | 0 | 0 | 0 | 1 | 98 | 49 | 1 | 1 | 1 | NA | NA | NA | 0 | 1 |
| A3SS | 939 | TTC23 | TTC23 | chr15 | - | 99139316 | 99139467 | 99139316 | 99139399 | 99156147 | 99156297 | 0 | 1 | 0 | 0 | 0 | 0 | 0 | 0 | 0 | 1 | 98 | 49 | 1 | 1 | NA | 1 | NA | NA | 0 | 1 |
| A3SS | 942 | USP16 | USP16 | chr21 | + | 29037275 | 29037463 | 29037278 | 29037463 | 29036270 | 29036374 | 8 | 5 | 0 | 0 | 0 | 1 | 0 | 3 | 0 | 0 | 52 | 49 | 1 | 1 | 1 | 1 | 0 | NA | 1 | -0.333 |
| A3SS | 969 | ZNF18 | ZNF18 | chr17 | - | 11983296 | 11983407 | 11983296 | 11983404 | 11984112 | 11984197 | 0 | 0 | 2 | 0 | 0 | 0 | 0 | 0 | 0 | 1 | 52 | 49 | 1 | 1 | NA | NA | 1 | NA | 0 | 1 |
| A3SS | 974 | RBM26 | RBM26 | chr13 | - | 79365577 | 79365733 | 79365577 | 79365718 | 79366054 | 79366195 | 1 | 0 | 0 | 5 | 0 | 0 | 0 | 0 | 0 | 2 | 64 | 49 | 1 | 1 | 0.133 | NA | NA | NA | 0 | 0.133 |
| A3SS | 977 | SUPT20H | SUPT20H | chr13 | - | 37022010 | 37022317 | 37022010 | 37022080 | 37024034 | 37024193 | 5 | 0 | 0 | 2 | 0 | 0 | 0 | 2 | 0 | 1 | 98 | 49 | 1 | 1 | 0.556 | NA | NA | NA | 0.5 | 0.056 |
| A3SS | 987 | PPP4C | PPP4C | chr16 | + | 30081199 | 30081310 | 30081258 | 30081310 | 30076314 | 30076475 | 8 | 2 | 0 | 18 | 15 | 7 | 0 | 3 | 0 | 2 | 98 | 49 | 1 | 1 | 0.182 | 0.062 | 0 | NA | 0.429 | -0.348 |
| A3SS | 990 | PSPC1 | PSPC1 | chr13 | - | 19674751 | 19675653 | 19674751 | 19674984 | 19677723 | 19677823 | 2 | 1 | 1 | 2 | 1 | 1 | 0 | 0 | 0 | 1 | 98 | 49 | 1 | 1 | 0.333 | 0.333 | 0.333 | NA | 0 | 0.333 |
| A3SS | 1001 | TRAPPC13 | TRAPPC13 | chr5 | + | 65664233 | 65664384 | 65664236 | 65664384 | 65662049 | 65662150 | 1 | 4 | 0 | 0 | 2 | 0 | 0 | 0 | 0 | 1 | 52 | 49 | 1 | 1 | 1 | 0.653 | NA | NA | 0 | 0.827 |
| A3SS | 1003 | ARL6IP4 | ARL6IP4 | chr12 | + | 122981570 | 122981879 | 122981594 | 122981879 | 122981128 | 122981266 | 1 | 0 | 0 | 5 | 0 | 0 | 0 | 0 | 0 | 1 | 73 | 49 | 1 | 1 | 0.118 | NA | NA | NA | 0 | 0.118 |
| A3SS | 1004 | ARL6IP4 | ARL6IP4 | chr12 | + | 122981570 | 122981879 | 122981594 | 122981879 | 122981128 | 122981299 | 3 | 0 | 0 | 6 | 1 | 2 | 0 | 0 | 0 | 1 | 73 | 49 | 1 | 1 | 0.251 | 0 | 0 | NA | 0 | 0.084 |
| A3SS | 1005 | ARL6IP4 | ARL6IP4 | chr12 | + | 122981594 | 122981809 | 122981655 | 122981809 | 122981128 | 122981299 | 20 | 5 | 5 | 1 | 0 | 0 | 0 | 4 | 0 | 0 | 98 | 49 | 1 | 1 | 0.909 | 1 | 1 | NA | 1 | -0.03 |
| A3SS | 1018 | ATPIF1 | ATPIF1 | chr1 | + | 28237760 | 28238105 | 28237836 | 28238105 | 28236360 | 28236452 | 2 | 0 | 0 | 4 | 5 | 2 | 0 | 5 | 0 | 1 | 98 | 49 | 1 | 1 | 0.2 | 0 | 0 | NA | 0.714 | -0.647 |
| A3SS | 1031 | POLDIP3 | POLDIP3 | chr22 | - | 42601969 | 42602107 | 42601969 | 42602056 | 42602769 | 42603160 | 1 | 0 | 0 | 8 | 4 | 0 | 0 | 0 | 0 | 1 | 98 | 49 | 1 | 1 | 0.059 | 0 | NA | NA | 0 | 0.029 |
| A3SS | 1044 | CCDC53 | CCDC53 | chr12 | - | 102039867 | 102039978 | 102039867 | 102039975 | 102044104 | 102044212 | 9 | 2 | 0 | 2 | 0 | 0 | 0 | 1 | 0 | 0 | 52 | 49 | 1 | 1 | 0.809 | 1 | NA | NA | 1 | -0.095 |
| A3SS | 1049 | MARK3 | MARK3 | chr14 | + | 103468032 | 103468186 | 103468080 | 103468186 | 103467078 | 103467191 | 28 | 19 | 4 | 0 | 0 | 1 | 0 | 13 | 0 | 0 | 97 | 49 | 1 | 1 | 1 | 1 | 0.669 | NA | 1 | -0.11 |
| A3SS | 1064 | OAZ1 | OAZ1 | chr19 | + | 2271384 | 2271530 | 2271390 | 2271530 | 2269485 | 2269744 | 172 | 107 | 112 | 1 | 0 | 0 | 10 | 66 | 0 | 0 | 55 | 49 | 1 | 1 | 0.994 | 1 | 1 | 1 | 1 | -0.002 |
| A3SS | 1088 | TMCC1 | TMCC1 | chr3 | - | 129827802 | 129828508 | 129827802 | 129828077 | 129832773 | 129832826 | 35 | 29 | 11 | 0 | 0 | 1 | 0 | 6 | 0 | 0 | 98 | 49 | 1 | 1 | 1 | 1 | 0.846 | NA | 1 | -0.051 |
| A3SS | 1104 | DDIT3 | DDIT3 | chr12 | - | 57517268 | 57517519 | 57517268 | 57517438 | 57517705 | 57517753 | 37 | 34 | 5 | 5 | 4 | 0 | 0 | 5 | 0 | 1 | 98 | 49 | 1 | 1 | 0.787 | 0.81 | 1 | NA | 0.714 | 0.152 |
| A3SS | 1105 | DDIT3 | DDIT3 | chr12 | - | 57517268 | 57517542 | 57517268 | 57517438 | 57517705 | 57517753 | 37 | 34 | 5 | 5 | 4 | 0 | 0 | 5 | 0 | 1 | 98 | 49 | 1 | 1 | 0.787 | 0.81 | 1 | NA | 0.714 | 0.152 |
| A3SS | 1106 | DDIT3 | DDIT3 | chr12 | - | 57517268 | 57517595 | 57517268 | 57517438 | 57517705 | 57517753 | 40 | 34 | 5 | 5 | 4 | 0 | 0 | 5 | 0 | 1 | 98 | 49 | 1 | 1 | 0.8 | 0.81 | 1 | NA | 0.714 | 0.156 |
| A3SS | 1113 | STARD3 | STARD3 | chr17 | + | 39658708 | 39658820 | 39658721 | 39658820 | 39658404 | 39658522 | 0 | 1 | 0 | 4 | 5 | 0 | 0 | 0 | 1 | 1 | 62 | 49 | 1 | 1 | 0 | 0.136 | NA | 0 | 0 | 0.068 |
| A3SS | 1119 | ESYT1 | ESYT1 | chr12 | + | 56134079 | 56134181 | 56134109 | 56134181 | 56133780 | 56133873 | 2 | 0 | 0 | 2 | 1 | 0 | 0 | 1 | 0 | 1 | 79 | 49 | 1 | 1 | 0.383 | 0 | NA | NA | 0.383 | -0.192 |
| A3SS | 1126 | APPL2 | APPL2 | chr12 | - | 105207970 | 105208047 | 105207970 | 105208029 | 105208157 | 105208199 | 2 | 2 | 0 | 11 | 1 | 3 | 0 | 0 | 0 | 5 | 67 | 49 | 1 | 1 | 0.117 | 0.594 | 0 | NA | 0 | 0.237 |
| A3SS | 1130 | SURF4 | SURF4 | chr9 | - | 133367258 | 133367543 | 133367258 | 133367445 | 133375921 | 133376166 | 1 | 0 | 0 | 1 | 1 | 0 | 0 | 0 | 1 | 0 | 98 | 49 | 1 | 1 | 0.333 | 0 | NA | 0 | NA | 0.167 |
| A3SS | 1133 | MDM1 | MDM1 | chr12 | - | 68325004 | 68327510 | 68325004 | 68327021 | 68331106 | 68331221 | 2 | 3 | 1 | 3 | 0 | 0 | 0 | 0 | 0 | 1 | 98 | 49 | 1 | 1 | 0.25 | 1 | 1 | NA | 0 | 0.75 |
| A3SS | 1143 | TRIP12 | TRIP12 | chr2 | - | 229805729 | 229805883 | 229805729 | 229805802 | 229807707 | 229807864 | 25 | 6 | 4 | 1 | 1 | 0 | 0 | 8 | 0 | 0 | 98 | 49 | 1 | 1 | 0.926 | 0.75 | 1 | NA | 1 | -0.108 |
| A3SS | 1157 | FGD4 | FGD4 | chr12 | + | 32607937 | 32608095 | 32607956 | 32608095 | 32602160 | 32602317 | 6 | 3 | 0 | 24 | 7 | 15 | 0 | 0 | 0 | 4 | 68 | 49 | 1 | 1 | 0.153 | 0.236 | 0 | NA | 0 | 0.13 |
| A3SS | 1167 | DLG1 | DLG1 | chr3 | - | 197076585 | 197076685 | 197076585 | 197076682 | 197081050 | 197081117 | 1 | 0 | 0 | 1 | 0 | 0 | 0 | 4 | 0 | 0 | 52 | 49 | 1 | 1 | 0.485 | NA | NA | NA | 1 | -0.515 |
| A3SS | 1170 | LRMP | LRMP | chr12 | + | 25063719 | 25063816 | 25063722 | 25063816 | 25062819 | 25062900 | 50 | 29 | 6 | 24 | 6 | 0 | 0 | 3 | 0 | 2 | 52 | 49 | 1 | 1 | 0.663 | 0.82 | 1 | NA | 0.586 | 0.242 |
| A3SS | 1178 | CARD8 | CARD8 | chr19 | - | 48230437 | 48230700 | 48230437 | 48230697 | 48230776 | 48231006 | 25 | 9 | 7 | 5 | 2 | 0 | 0 | 4 | 0 | 0 | 52 | 49 | 1 | 1 | 0.825 | 0.809 | 1 | NA | 1 | -0.122 |
| A3SS | 1188 | QKI | QKI | chr6 | + | 163563419 | 163563719 | 163563443 | 163563719 | 163561981 | 163562069 | 39 | 22 | 18 | 1 | 2 | 0 | 0 | 7 | 0 | 0 | 73 | 49 | 1 | 1 | 0.963 | 0.881 | 1 | NA | 1 | -0.052 |
| A3SS | 1189 | QKI | QKI | chr6 | + | 163564666 | 163578596 | 163565945 | 163578596 | 163563419 | 163563719 | 14 | 4 | 4 | 8 | 0 | 0 | 0 | 3 | 0 | 0 | 98 | 49 | 1 | 1 | 0.467 | 1 | 1 | NA | 1 | -0.178 |
| A3SS | 1190 | QKI | QKI | chr6 | + | 163564666 | 163578596 | 163570693 | 163578596 | 163563419 | 163563719 | 8 | 0 | 3 | 1 | 0 | 0 | 0 | 7 | 0 | 0 | 98 | 49 | 1 | 1 | 0.8 | NA | 1 | NA | 1 | -0.1 |
| A3SS | 1191 | QKI | QKI | chr6 | + | 163565945 | 163578596 | 163570693 | 163578596 | 163563419 | 163563719 | 16 | 0 | 1 | 1 | 0 | 0 | 0 | 7 | 0 | 0 | 98 | 49 | 1 | 1 | 0.889 | NA | 1 | NA | 1 | -0.055 |
| A3SS | 1197 | CD163 | CD163 | chr12 | - | 7479859 | 7479996 | 7479859 | 7479913 | 7481160 | 7481256 | 0 | 0 | 0 | 2 | 0 | 0 | 0 | 5 | 0 | 3 | 98 | 49 | 1 | 1 | 0 | NA | NA | NA | 0.455 | -0.455 |
| A3SS | 1200 | C1RL | C1RL | chr12 | - | 7099685 | 7099807 | 7099685 | 7099760 | 7099900 | 7100026 | 3 | 1 | 0 | 12 | 3 | 2 | 0 | 2 | 0 | 2 | 96 | 49 | 1 | 1 | 0.113 | 0.145 | 0 | NA | 0.338 | -0.252 |
| A3SS | 1205 | TLE4 | TLE4 | chr9 | + | 79652589 | 79652794 | 79652592 | 79652794 | 79627373 | 79627448 | 3 | 0 | 3 | 6 | 1 | 0 | 0 | 0 | 0 | 2 | 52 | 49 | 1 | 1 | 0.32 | 0 | 1 | NA | 0 | 0.44 |
| A3SS | 1209 | SLTM | SLTM | chr15 | - | 58899468 | 58899937 | 58899468 | 58899883 | 58901259 | 58901287 | 10 | 5 | 1 | 1 | 0 | 0 | 0 | 6 | 0 | 0 | 98 | 49 | 1 | 1 | 0.833 | 1 | 1 | NA | 1 | -0.056 |
| A3SS | 1215 | WNK1 | WNK1 | chr12 | + | 878211 | 878361 | 878214 | 878361 | 862082 | 862270 | 6 | 0 | 0 | 1 | 1 | 0 | 0 | 1 | 0 | 2 | 52 | 49 | 1 | 1 | 0.85 | 0 | NA | NA | 0.32 | 0.105 |
| A3SS | 1216 | WNK1 | WNK1 | chr12 | + | 878211 | 878361 | 878214 | 878361 | 871264 | 871348 | 8 | 1 | 0 | 3 | 1 | 0 | 2 | 0 | 0 | 1 | 52 | 49 | 1 | 1 | 0.715 | 0.485 | NA | 1 | 0 | 0.1 |
| A3SS | 1221 | KANSL3 | KANSL3 | chr2 | - | 96631311 | 96631543 | 96631311 | 96631482 | 96636920 | 96637185 | 4 | 2 | 0 | 1 | 0 | 0 | 0 | 3 | 0 | 0 | 98 | 49 | 1 | 1 | 0.667 | 1 | NA | NA | 1 | -0.166 |

| | | | | | | | | | | | | | | | | | | | | | | | | | | | | | | | | |
|---|---|---|---|---|---|---|---|---|---|---|---|---|---|---|---|---|---|---|---|---|---|---|---|---|---|---|---|---|---|---|---|---|
| A3SS | 1225 | NFRKB | NFRKB | chr11 | - | 129884744 | 129884882 | 129884744 | 129884846 | 129885434 | 129885609 | 0 | 1 | 5 | 1 | 0 | 0 | 0 | 1 | 0 | 0 | 85 | 49 | 1 | 1 | 0 | 1 | 1 | NA | | 1 | -0.333 |
| A3SS | 1227 | MEF2C | MEF2C | chr5 | - | 88718240 | 88722925 | 88718240 | 88722829 | 88728492 | 88728628 | 5 | 0 | 0 | 2 | 0 | 0 | 0 | 1 | 0 | 1 | 98 | 49 | 1 | 1 | 0.556 | NA | NA | NA | | 0.333 | 0.223 |
| A3SS | 1228 | MEF2C | MEF2C | chr5 | - | 88823734 | 88823930 | 88823734 | 88823927 | 88882954 | 88883466 | 2 | 2 | 0 | 3 | 0 | 2 | 0 | 2 | 0 | 1 | 52 | 49 | 1 | 1 | 0.386 | 1 | 0 | NA | | 0.653 | -0.191 |
| A3SS | 1229 | ANAPC5 | ANAPC5 | chr12 | - | 121328315 | 121328497 | 121328315 | 121328458 | 121330582 | 121330672 | 9 | 1 | 0 | 0 | 0 | 1 | 0 | 3 | 0 | 0 | 88 | 49 | 1 | 1 | 1 | 1 | 0 | NA | | 1 | -0.333 |
| A3SS | 1235 | TMEM218 | TMEM218 | chr11 | - | 125101200 | 125101303 | 125101200 | 125101256 | 125102733 | 125102809 | 0 | 1 | 0 | 2 | 0 | 0 | 0 | 1 | 0 | 0 | 96 | 49 | 1 | 1 | 0 | 1 | NA | NA | | 1 | -0.5 |
| A3SS | 1242 | NRGN | NRGN | chr11 | + | 124745961 | 124745982 | 124745964 | 124745982 | 124745502 | 124745729 | 9 | 0 | 0 | 12 | 0 | 0 | 2 | 1 | 4 | 1 | 52 | 49 | 1 | 1 | 0.414 | NA | NA | | 0.32 | 0.485 | 0.012 |
| A3SS | 1257 | SLC25A45 | SLC25A45 | chr11 | - | 65380131 | 65380234 | 65380131 | 65380175 | 65380533 | 65380632 | 1 | 0 | 1 | 0 | 0 | 0 | 0 | 0 | 0 | 1 | 98 | 49 | 1 | 1 | 1 | NA | 1 | NA | | 0 | 1 |
| A3SS | 1263 | MPZL3 | MPZL3 | chr11 | - | 118240210 | 118240377 | 118240210 | 118240341 | 118252221 | 118252368 | 23 | 2 | 10 | 0 | 2 | 0 | 0 | 6 | 0 | 0 | 85 | 49 | 1 | 1 | 1 | 0.366 | 1 | NA | | 1 | -0.211 |
| A3SS | 1277 | MARK2 | MARK2 | chr11 | + | 63902600 | 63902782 | 63902603 | 63902782 | 63902197 | 63902330 | 1 | 0 | 0 | 25 | 4 | 8 | 0 | 0 | 0 | 6 | 52 | 49 | 1 | 1 | 0.036 | 0 | 0 | NA | | 0 | 0.012 |
| A3SS | 1363 | CCL4L2 | CCL4L2 | chr17 | + | 36212292 | 36212878 | 36212496 | 36212878 | 36211775 | 36211890 | 10 | 10 | 6 | 0 | 0 | 0 | 0 | 0 | 0 | 1 | 98 | 49 | 1 | 1 | 1 | 1 | 1 | NA | | 0 | 1 |
| A3SS | 1367 | CCL4L2 | CCL4L2 | chr17 | + | 36212302 | 36212878 | 36212496 | 36212878 | 36211775 | 36211890 | 10 | 10 | 6 | 0 | 0 | 0 | 0 | 0 | 0 | 1 | 98 | 49 | 1 | 1 | 1 | 1 | 1 | NA | | 0 | 1 |
| A3SS | 1370 | CCL4L2 | CCL4L2 | chr17 | + | 36212312 | 36212878 | 36212496 | 36212878 | 36211775 | 36211890 | 10 | 10 | 6 | 0 | 0 | 0 | 0 | 0 | 0 | 1 | 98 | 49 | 1 | 1 | 1 | 1 | 1 | NA | | 0 | 1 |
| A3SS | 1372 | CCL4L2 | CCL4L2 | chr17 | + | 36212333 | 36212878 | 36212496 | 36212878 | 36211775 | 36211890 | 10 | 10 | 6 | 0 | 0 | 0 | 0 | 0 | 0 | 1 | 98 | 49 | 1 | 1 | 1 | 1 | 1 | NA | | 0 | 1 |
| A3SS | 1373 | CCL4L2 | CCL4L2 | chr17 | + | 36212363 | 36212878 | 36212496 | 36212878 | 36211775 | 36211890 | 10 | 10 | 6 | 0 | 0 | 0 | 0 | 0 | 0 | 1 | 98 | 49 | 1 | 1 | 1 | 1 | 1 | NA | | 0 | 1 |
| A3SS | 1382 | RASGRP4 | RASGRP4 | chr19 | - | 38414847 | 38415123 | 38414847 | 38414916 | 38417051 | 38417168 | 55 | 18 | 9 | 0 | 1 | 1 | 2 | 16 | 0 | 0 | 98 | 49 | 1 | 1 | 1 | 0.9 | 0.818 | | 1 | 1 | -0.094 |
| A3SS | 1393 | RC3H1 | RC3H1 | chr1 | - | 173931083 | 173938874 | 173931083 | 173938871 | 173941264 | 173941380 | 6 | 6 | 2 | 8 | 2 | 4 | 0 | 0 | 0 | 4 | 52 | 49 | 1 | 1 | 0.414 | 0.739 | 0.32 | NA | | 0 | 0.491 |
| A3SS | 1394 | RC3H1 | RC3H1 | chr1 | - | 173943441 | 173943615 | 173943441 | 173943588 | 173946475 | 173946608 | 12 | 1 | 3 | 1 | 3 | 6 | 0 | 2 | 0 | 0 | 76 | 49 | 1 | 1 | 0.886 | 0.177 | 0.244 | NA | | 1 | -0.564 |
| A3SS | 1431 | COPA | COPA | chr1 | - | 160305432 | 160305598 | 160305432 | 160305571 | 160305687 | 160305773 | 0 | 0 | 0 | 9 | 10 | 0 | 0 | 4 | 0 | 3 | 76 | 49 | 1 | 1 | 0 | 0 | NA | NA | | 0.462 | -0.462 |
| A3SS | 1434 | KCNJ15 | KCNJ15 | chr21 | + | 38257045 | 38257185 | 38257103 | 38257185 | 38229914 | 38230023 | 44 | 6 | 10 | 2 | 0 | 0 | 0 | 1 | 0 | 0 | 98 | 49 | 1 | 1 | 0.917 | 1 | 1 | NA | | 1 | -0.028 |
| A3SS | 1440 | SDF4 | SDF4 | chr1 | - | 1216907 | 1217804 | 1216907 | 1217688 | 1218457 | 1218633 | 0 | 1 | 0 | 15 | 4 | 3 | 0 | 0 | 0 | 4 | 98 | 49 | 1 | 1 | 0 | 0.111 | 0 | NA | | 0 | 0.037 |
| A3SS | 1445 | METTL9 | METTL9 | chr16 | + | 21655226 | 21657473 | 21655229 | 21657473 | 21624930 | 21625115 | 2 | 6 | 1 | 1 | 0 | 1 | 0 | 2 | 0 | 2 | 52 | 49 | 1 | 1 | 0.653 | 1 | 0.485 | NA | | 0.485 | 0.228 |
| A3SS | 1459 | SERPINA1 | SERPINA1 | chr14 | - | 94388559 | 94388663 | 94388559 | 94388660 | 94388800 | 94389010 | 25 | 0 | 4 | 2 | 0 | 1 | 0 | 3 | 0 | 2 | 52 | 49 | 1 | 1 | 0.922 | NA | 0.79 | NA | | 0.586 | 0.27 |
| A3SS | 1460 | SERPINA1 | SERPINA1 | chr14 | - | 94388559 | 94388663 | 94388559 | 94388660 | 94388818 | 94389010 | 18 | 2 | 6 | 18 | 3 | 11 | 0 | 3 | 0 | 1 | 52 | 49 | 1 | 1 | 0.485 | 0.386 | 0.339 | NA | | 0.739 | -0.336 |
| A3SS | 1493 | SRSF10 | SRSF10 | chr1 | - | 23964346 | 23971439 | 23964346 | 23967723 | 23971572 | 23971623 | 4 | 1 | 0 | 4 | 0 | 2 | 0 | 1 | 0 | 0 | 98 | 49 | 1 | 1 | 0.333 | 1 | 0 | NA | | 1 | -0.556 |
| A3SS | 1494 | SRSF10 | SRSF10 | chr1 | - | 23964346 | 23971439 | 23964346 | 23967956 | 23971572 | 23971623 | 4 | 4 | 0 | 0 | 1 | 0 | 0 | 1 | 0 | 0 | 98 | 49 | 1 | 1 | 1 | 0.667 | NA | NA | | 1 | -0.166 |
| A3SS | 1498 | EDRF1 | EDRF1 | chr10 | + | 125730271 | 125730399 | 125730287 | 125730399 | 125729357 | 125729479 | 1 | 0 | 0 | 1 | 2 | 1 | 0 | 0 | 0 | 1 | 65 | 49 | 1 | 1 | 0.43 | 0 | 0 | NA | | 0 | 0.143 |
| A3SS | 1525 | FAM63A | FAM63A | chr1 | - | 151002164 | 151002706 | 151002164 | 151002323 | 151006311 | 151006909 | 24 | 4 | 3 | 2 | 0 | 0 | 3 | 4 | 0 | 1 | 98 | 49 | 1 | 1 | 0.857 | 1 | 1 | | 1 | 0.667 | 0.119 |
| A3SS | 1532 | CDIPT | CDIPT | chr16 | - | 29862585 | 29862720 | 29862585 | 29862635 | 29862814 | 29863288 | 23 | 8 | 11 | 1 | 0 | 0 | 0 | 1 | 0 | 0 | 98 | 49 | 1 | 1 | 0.92 | 1 | 1 | NA | | 1 | -0.027 |
| A3SS | 1554 | ALPK1 | ALPK1 | chr4 | + | 112377677 | 112377898 | 112377729 | 112377898 | 112315800 | 112315852 | 58 | 24 | 5 | 9 | 3 | 0 | 0 | 7 | 0 | 0 | 98 | 49 | 1 | 1 | 0.763 | 0.8 | 1 | NA | | 1 | -0.146 |
| A3SS | 1576 | TSC22D4 | TSC22D4 | chr7 | - | 100477276 | 100478204 | 100477276 | 100477459 | 100478793 | 100479279 | 27 | 8 | 9 | 1 | 1 | 2 | 0 | 9 | 0 | 0 | 98 | 49 | 1 | 1 | 0.931 | 0.8 | 0.692 | NA | | 1 | -0.192 |
| A3SS | 1577 | TSC22D4 | TSC22D4 | chr7 | - | 100477276 | 100478307 | 100477276 | 100477459 | 100478793 | 100479279 | 27 | 10 | 14 | 1 | 1 | 2 | 0 | 11 | 0 | 0 | 98 | 49 | 1 | 1 | 0.931 | 0.833 | 0.778 | NA | | 1 | -0.153 |
| A3SS | 1580 | TSC22D4 | TSC22D4 | chr7 | - | 100477276 | 100478307 | 100477276 | 100478204 | 100478793 | 100479279 | 11 | 6 | 7 | 2 | 0 | 1 | 0 | 2 | 0 | 0 | 98 | 49 | 1 | 1 | 0.733 | 1 | 0.778 | NA | | 1 | -0.163 |
| A3SS | 1592 | BRD7 | BRD7 | chr16 | - | 50320662 | 50320777 | 50320662 | 50320774 | 50321981 | 50322038 | 0 | 1 | 0 | 2 | 0 | 0 | 0 | 1 | 0 | 0 | 52 | 49 | 1 | 1 | 0 | 1 | NA | NA | | 1 | -0.5 |
| A3SS | 1604 | CEP170 | CEP170 | chr1 | - | 243156220 | 243156455 | 243156220 | 243156347 | 243164283 | 243166116 | 4 | 2 | 0 | 1 | 0 | 0 | 0 | 1 | 0 | 0 | 98 | 49 | 1 | 1 | 0.667 | 1 | NA | NA | | 1 | -0.166 |
| A3SS | 1607 | MAP2K7 | MAP2K7 | chr19 | + | 7911414 | 7911578 | 7911435 | 7911578 | 7911249 | 7911330 | 3 | 1 | 0 | 1 | 3 | 0 | 0 | 0 | 1 | 2 | 70 | 49 | 1 | 1 | 0.677 | 0.189 | NA | | 0 | 0 | 0.433 |
| A3SS | 1609 | ARHGEF2 | ARHGEF2 | chr1 | - | 155964987 | 155965131 | 155964987 | 155965128 | 155965302 | 155965412 | 16 | 4 | 0 | 15 | 5 | 0 | 0 | 2 | 0 | 4 | 52 | 49 | 1 | 1 | 0.501 | 0.43 | NA | NA | | 0.32 | 0.146 |
| A3SS | 1610 | TMEM59 | TMEM59 | chr1 | - | 54040755 | 54040840 | 54040755 | 54040837 | 54041723 | 54041805 | 11 | 10 | 5 | 30 | 16 | 18 | 0 | 0 | 0 | 2 | 52 | 49 | 1 | 1 | 0.257 | 0.371 | 0.207 | NA | | 0 | 0.278 |
| A3SS | 1613 | ADAT1 | ADAT1 | chr16 | - | 75618585 | 75618698 | 75618585 | 75618640 | 75620265 | 75620334 | 0 | 1 | 2 | 5 | 0 | 0 | 0 | 0 | 1 | 0 | 98 | 49 | 1 | 1 | 0 | 1 | 1 | | 0 | NA | 0.667 |
| A3SS | 1616 | SORT1 | SORT1 | chr1 | - | 109345750 | 109345881 | 109345750 | 109345878 | 109347482 | 109347532 | 5 | 0 | 2 | 3 | 0 | 0 | 0 | 5 | 0 | 2 | 52 | 49 | 1 | 1 | 0.611 | NA | 1 | NA | | 0.702 | 0.104 |
| A3SS | 1620 | DHDDS | DHDDS | chr1 | + | 26468891 | 26471304 | 26468894 | 26471304 | 26460036 | 26460144 | 0 | 0 | 0 | 2 | 0 | 1 | 0 | 1 | 0 | 0 | 52 | 49 | 1 | 1 | 0 | NA | 0 | NA | | 1 | -1 |
| A3SS | 1624 | SOAT1 | SOAT1 | chr1 | + | 179342009 | 179342192 | 179342113 | 179342192 | 179341027 | 179341310 | 2 | 0 | 0 | 4 | 1 | 2 | 0 | 0 | 0 | 3 | 98 | 49 | 1 | 1 | 0.2 | 0 | 0 | NA | | 0 | 0.067 |
| A3SS | 1639 | TGFBR3 | TGFBR3 | chr1 | - | 91719892 | 91720230 | 91719892 | 91720227 | 91721954 | 91722144 | 1 | 0 | 0 | 0 | 0 | 0 | 0 | 0 | 0 | 1 | 52 | 49 | 1 | 1 | 1 | NA | NA | NA | | 0 | 1 |
| A3SS | 1647 | HDDC3 | HDDC3 | chr15 | - | 90930917 | 90931556 | 90930917 | 90931405 | 90931703 | 90931944 | 0 | 4 | 0 | 0 | 0 | 0 | 4 | 1 | 0 | 2 | 98 | 49 | 1 | 1 | NA | 1 | NA | | 1 | 0.2 | 0.4 |
| A3SS | 1651 | LMBR1L | LMBR1L | chr12 | - | 49101249 | 49101323 | 49101249 | 49101263 | 49101471 | 49101549 | 8 | 3 | 4 | 1 | 1 | 0 | 1 | 4 | 0 | 0 | 98 | 49 | 1 | 1 | 0.8 | 0.6 | 1 | | 1 | 1 | -0.2 |
| A3SS | 1663 | SYTL1 | SYTL1 | chr1 | + | 27349079 | 27349152 | 27349115 | 27349152 | 27347966 | 27348012 | 6 | 4 | 5 | 1 | 1 | 0 | 0 | 3 | 0 | 0 | 85 | 49 | 1 | 1 | 0.776 | 0.698 | 1 | NA | | 1 | -0.175 |
| A3SS | 1666 | WDR47 | WDR47 | chr1 | - | 109004591 | 109004718 | 109004591 | 109004715 | 109010915 | 109011718 | 1 | 0 | 0 | 0 | 1 | 0 | 0 | 1 | 0 | 0 | 52 | 49 | 1 | 1 | 1 | 0 | NA | NA | | 1 | -0.5 |
| A3SS | 1678 | FDPS | FDPS | chr1 | + | 155310042 | 155310205 | 155310063 | 155310205 | 155308747 | 155308965 | 3 | 0 | 0 | 5 | 0 | 0 | 0 | 1 | 0 | 0 | 70 | 49 | 1 | 1 | 0.296 | NA | NA | NA | | 1 | -0.704 |
| A3SS | 1681 | AKAP13 | AKAP13 | chr15 | + | 85710578 | 85710645 | 85710581 | 85710645 | 85708018 | 85708086 | 66 | 11 | 12 | 4 | 0 | 0 | 0 | 13 | 0 | 0 | 52 | 49 | 1 | 1 | 0.94 | 1 | 1 | NA | | 1 | -0.02 |
| A3SS | 1698 | SRP19 | SRP19 | chr5 | + | 112867371 | 112868392 | 112867403 | 112868392 | 112864620 | 112864732 | 6 | 2 | 0 | 2 | 0 | 1 | 0 | 1 | 0 | 1 | 81 | 49 | 1 | 1 | 0.645 | 1 | 0 | NA | | 0.377 | 0.171 |
| A3SS | 1702 | TUG1 | TUG1 | chr22 | + | 30972854 | 30973601 | 30972923 | 30973601 | 30969210 | 30971779 | 13 | 3 | 5 | 2 | 1 | 2 | 0 | 1 | 0 | 0 | 98 | 49 | 1 | 1 | 0.765 | 0.6 | 0.556 | NA | | 1 | -0.36 |
| A3SS | 1704 | TRIM5 | TRIM5 | chr11 | - | 5663194 | 5665395 | 5663194 | 5664173 | 5665655 | 5665682 | 8 | 0 | 0 | 1 | 0 | 0 | 0 | 1 | 0 | 0 | 98 | 49 | 1 | 1 | 0.8 | NA | NA | NA | | 1 | -0.2 |
| A3SS | 1715 | DMXL2 | DMXL2 | chr15 | - | 51476588 | 51476719 | 51476588 | 51476716 | 51478270 | 51478347 | 16 | 1 | 0 | 14 | 2 | 0 | 0 | 8 | 0 | 2 | 52 | 49 | 1 | 1 | 0.519 | 0.32 | NA | NA | | 0.79 | -0.371 |
| A3SS | 1727 | CASP1 | CASP1 | chr11 | - | 105025509 | 105026356 | 105025509 | 105025600 | 105026841 | 105026951 | 62 | 20 | 21 | 1 | 2 | 0 | 0 | 29 | 0 | 0 | 98 | 49 | 1 | 1 | 0.969 | 0.833 | 1 | NA | | 1 | -0.066 |
| A3SS | 1736 | CD83 | CD83 | chr6 | + | 14135107 | 14136917 | 14135110 | 14136917 | 14133648 | 14133755 | 9 | 1 | 0 | 0 | 0 | 0 | 0 | 0 | 0 | 1 | 52 | 49 | 1 | 1 | 1 | 1 | NA | NA | | 0 | 1 |
| A3SS | 1742 | MVP | MVP | chr16 | + | 29830473 | 29830674 | 29830514 | 29830674 | 29820393 | 29820510 | 13 | 2 | 1 | 0 | 2 | 0 | 0 | 0 | 0 | 1 | 90 | 49 | 1 | 1 | 1 | 0.353 | 1 | NA | | 0 | 0.784 |
| A3SS | 1761 | SLC25A44 | SLC25A44 | chr1 | + | 156207861 | 156208013 | 156207885 | 156208013 | 156199834 | 156200472 | 1 | 0 | 0 | 8 | 9 | 2 | 0 | 0 | 0 | 1 | 73 | 49 | 1 | 1 | 0.077 | 0 | 0 | NA | | 0 | 0.026 |
| A3SS | 1765 | PAPD4 | PAPD4 | chr5 | + | 79642831 | 79642910 | 79642843 | 79642910 | 79641104 | 79641196 | 3 | 0 | 1 | 3 | 3 | 2 | 0 | 4 | 0 | 0 | 61 | 49 | 1 | 1 | 0.445 | 0 | 0.287 | NA | | 1 | -0.756 |
| A3SS | 1781 | RBMX | RBMX | chrX | - | 136879016 | 136879158 | 136879016 | 136879123 | 136879318 | 136879453 | 2 | 0 | 0 | 9 | 3 | 0 | 0 | 0 | 0 | 4 | 84 | 49 | 1 | 1 | 0.115 | 0 | NA | NA | | 0 | 0.058 |
| A3SS | 1790 | FAM217B | FAM217B | chr20 | + | 59943939 | 59948647 | 59944395 | 59948647 | 59942426 | 59942512 | 6 | 7 | 3 | 1 | 1 | 0 | 0 | 3 | 0 | 0 | 98 | 49 | 1 | 1 | 0.75 | 0.778 | 1 | NA | | 1 | -0.157 |
| A3SS | 1795 | SREK1 | SREK1 | chr5 | + | 66177513 | 66177658 | 66177516 | 66177658 | 66174945 | 66175041 | 6 | 7 | 0 | 2 | 2 | 0 | 0 | 1 | 0 | 0 | 52 | 49 | 1 | 1 | 0.739 | 0.767 | NA | NA | | 1 | -0.247 |
| A3SS | 1807 | STK40 | STK40 | chr1 | - | 36358736 | 36358837 | 36358736 | 36358822 | 36361220 | 36361340 | 2 | 0 | 0 | 11 | 6 | 12 | 0 | 1 | 0 | 1 | 64 | 49 | 1 | 1 | 0.122 | 0 | 0 | NA | | 0.434 | -0.393 |
| A3SS | 1810 | SHC1 | SHC1 | chr1 | - | 154965945 | 154966080 | 154965945 | 154966077 | 154966161 | 154966231 | 1 | 0 | 0 | 0 | 0 | 0 | 0 | 0 | 0 | 3 | 52 | 49 | 1 | 1 | 1 | NA | NA | NA | | 0 | 1 |
| A3SS | 1828 | ATP11A | ATP11A | chr13 | + | 112881815 | 112887168 | 112881875 | 112887168 | 112875775 | 112875941 | 11 | 1 | 0 | 0 | 3 | 0 | 3 | 1 | 0 | 1 | 98 | 49 | 1 | 1 | 1 | 0.143 | NA | | 1 | 0.333 | -0.095 |
| A3SS | 1842 | TPM3 | TPM3 | chr1 | - | 154155303 | 154159048 | 154155303 | 154157721 | 154170399 | 154170469 | 15 | 10 | 3 | 75 | 46 | 36 | 0 | 1 | 0 | 20 | 98 | 49 | 1 | 1 | 0.091 | 0.098 | 0.04 | NA | | 0.024 | 0.052 |
| A3SS | 1843 | TPM3 | TPM3 | chr1 | - | 154170399 | 154170469 | 154170399 | 154170465 | 154170648 | 154170711 | 114 | 42 | 46 | 0 | 2 | 0 | 0 | 19 | 0 | 1 | 53 | 49 | 1 | 1 | 1 | 0.951 | 1 | NA | | 0.946 | 0.038 |
| A3SS | 1845 | CDK12 | CDK12 | chr17 | + | 39530603 | 39534565 | 39530630 | 39534565 | 39525863 | 39526316 | 6 | 1 | 0 | 2 | 0 | 0 | 0 | 1 | 0 | 0 | 76 | 49 | 1 | 1 | 0.659 | 1 | NA | NA | | 1 | -0.17 |
| A3SS | 1862 | APTX | APTX | chr9 | - | 32988078 | 32988129 | 32988078 | 32988124 | 32989758 | 32989895 | 1 | 0 | 0 | 1 | 0 | 0 | 0 | 0 | 0 | 1 | 54 | 49 | 1 | 1 | 0.476 | NA | NA | NA | | 0 | 0.476 |
| A3SS | 1898 | SZRD1 | SZRD1 | chr1 | + | 16393227 | 16393482 | 16393230 | 16393482 | 16391374 | 16391424 | 0 | 0 | 1 | 1 | 0 | 0 | 1 | 0 | 0 | 0 | 52 | 49 | 1 | 1 | 0 | NA | 1 | | 1 | NA | -0.5 |
| A3SS | 1900 | FUS | FUS | chr16 | + | 31183857 | 31184002 | 31183860 | 31184002 | 31182512 | 31182664 | 6 | 6 | 5 | 3 | 6 | 0 | 0 | 5 | 0 | 3 | 52 | 49 | 1 | 1 | 0.653 | 0.485 | 1 | NA | | 0.611 | 0.102 |
| A3SS | 1901 | ZBTB17 | ZBTB17 | chr1 | - | 15942330 | 15942441 | 15942330 | 15942420 | 15942528 | 15942738 | 2 | 3 | 0 | 2 | 1 | 0 | 0 | 0 | 0 | 2 | 70 | 49 | 1 | 1 | 0.412 | 0.677 | NA | NA | | 0 | 0.544 |
| A3SS | 1902 | ZBTB17 | ZBTB17 | chr1 | - | 15946153 | 15946316 | 15946153 | 15946294 | 15946934 | 15947123 | 0 | 1 | 0 | 3 | 0 | 0 | 0 | 1 | 1 | 2 | 71 | 49 | 1 | 1 | 0 | 1 | NA | | 0 | 0.257 | 0.371 |
| A3SS | 1906 | APH1A | APH1A | chr1 | - | 150267956 | 150268127 | 150267956 | 150267991 | 150268697 | 150269209 | 19 | 12 | 1 | 2 | 3 | 1 | 0 | 6 | 0 | 0 | 98 | 49 | 1 | 1 | 0.826 | 0.667 | 0.333 | NA | | 1 | -0.391 |
| A3SS | 1913 | CD58 | CD58 | chr1 | - | 116519230 | 116519302 | 116519230 | 116519267 | 116521905 | 116521983 | 5 | 2 | 3 | 8 | 9 | 1 | 0 | 1 | 0 | 1 | 84 | 49 | 1 | 1 | 0.267 | 0.115 | 0.636 | NA | | 0.368 | -0.029 |
| A3SS | 1916 | RBBP5 | RBBP5 | chr1 | - | 205114788 | 205114961 | 205114788 | 205114905 | 205115857 | 205115883 | 7 | 8 | 0 | 0 | 1 | 0 | 0 | 5 | 0 | 0 | 98 | 49 | 1 | 1 | 1 | 0.8 | NA | NA | | 1 | -0.1 |
| A3SS | 1928 | CYB561D1 | CYB561D1 | chr1 | + | 109495678 | 109500441 | 109495755 | 109500441 | 109495142 | 109495180 | 2 | 2 | 0 | 1 | 0 | 0 | 0 | 0 | 0 | 1 | 98 | 49 | 1 | 1 | 0.5 | 1 | NA | NA | | 0 | 0.75 |
| A3SS | 1938 | RSRC2 | RSRC2 | chr12 | - | 122508217 | 122508450 | 122508217 | 122508447 | 122511108 | 122511188 | 0 | 1 | 0 | 2 | 1 | 0 | 0 | 0 | 0 | 1 | 52 | 49 | 1 | 1 | 0 | 0.485 | NA | NA | | 0 | 0.242 |
| A3SS | 1939 | RSRC2 | RSRC2 | chr12 | - | 122518838 | 122519051 | 122518838 | 122519029 | 122520503 | 122520581 | 0 | 1 | 0 | 1 | 0 | 0 | 0 | 0 | 1 | 1 | 71 | 49 | 1 | 1 | 0 | 1 | NA | | 0 | 0 | 0.5 |
| A3SS | 1940 | RSRC2 | RSRC2 | chr12 | - | 122518838 | 122519051 | 122518838 | 122519029 | 122521384 | 122521428 | 0 | 1 | 0 | 3 | 0 | 0 | 0 | 0 | 0 | 1 | 71 | 49 | 1 | 1 | 0 | 1 | NA | NA | | 0 | 0.5 |
| A3SS | 1954 | RNF19B | RNF19B | chr1 | - | 32948221 | 32948366 | 32948221 | 32948363 | 32949568 | 32949774 | 23 | 10 | 2 | 36 | 23 | 4 | 0 | 1 | 0 | 2 | 52 | 49 | 1 | 1 | 0.376 | 0.291 | 0.32 | NA | | 0.32 | 0.009 |
| A3SS | 1955 | PITPNM1 | PITPNM1 | chr11 | - | 67496177 | 67496348 | 67496177 | 67496345 | 67497230 | 67497436 | 3 | 0 | 0 | 2 | 0 | 0 | 0 | 2 | 0 | 0 | 52 | 49 | 1 | 1 | 0.586 | NA | NA | NA | | 1 | -0.414 |
| A3SS | 1997 | AES | AES | chr19 | - | 3052909 | 3054040 | 3052909 | 3054037 | 3054119 | 3054194 | 25 | 1 | 0 | 4 | 0 | 0 | 0 | 6 | 0 | 0 | 52 | 49 | 1 | 1 | 0.855 | 1 | NA | NA | | 1 | -0.073 |
| A3SS | 2001 | SYNRG | SYNRG | chr17 | - | 37541971 | 37542565 | 37541971 | 37542430 | 37553114 | 37554059 | 16 | 7 | 0 | 0 | 1 | 0 | 1 | 0 | 0 | 0 | 98 | 49 | 1 | 1 | 1 | 0.778 | NA | | 1 | NA | -0.111 |
| A3SS | 2005 | EWSR1 | EWSR1 | chr22 | + | 29291561 | 29291599 | 29291564 | 29291599 | 29288605 | 29288786 | 0 | 0 | 0 | 0 | 3 | 0 | 1 | 0 | 0 | 0 | 52 | 49 | 1 | 1 | NA | 0 | NA | | 1 | NA | -1 |
| A3SS | 2014 | NUDT13 | NUDT13 | chr10 | + | 73126634 | 73126827 | 73126672 | 73126827 | 73125397 | 73125509 | 3 | 1 | 0 | 0 | 0 | 0 | 0 | 0 | 0 | 1 | 87 | 49 | 1 | 1 | 1 | 1 | NA | NA | | 0 | 1 |
| A3SS | 2026 | IRAK1 | IRAK1 | chrX | - | 154013042 | 154013433 | 154013042 | 154013343 | 154014041 | 154014278 | 4 | 3 | 1 | 2 | 0 | 0 | 0 | 3 | 0 | 0 | 98 | 49 | 1 | 1 | 0.5 | 1 | 1 | NA | | 1 | -0.167 |
| A3SS | 2028 | C11orf68 | C11orf68 | chr11 | - | 65916811 | 65918218 | 65916811 | 65918215 | 65918909 | 65919060 | 1 | 0 | 0 | 0 | 0 | 0 | 0 | 0 | 0 | 1 | 52 | 49 | 1 | 1 | 1 | NA | NA | NA | | 0 | 1 |
| A3SS | 2032 | IL4R | IL4R | chr16 | + | 27352539 | 27352696 | 27352544 | 27352696 | 27346466 | 27346618 | 39 | 6 | 13 | 1 | 0 | 0 | 0 | 4 | 0 | 0 | 54 | 49 | 1 | 1 | 0.973 | 1 | 1 | NA | | 1 | -0.009 |
| A3SS | 2033 | ASPH | ASPH | chr8 | - | 61638321 | 61638363 | 61638321 | 61638360 | 61642887 | 61642920 | 4 | 2 | 1 | 4 | 0 | 0 | 0 | 3 | 0 | 0 | 52 | 49 | 1 | 1 | 0.485 | 1 | 1 | NA | | 1 | -0.172 |
| A3SS | 2034 | AKTIP | AKTIP | chr16 | - | 53491039 | 53492522 | 53491039 | 53492519 | 53492692 | 53492753 | 0 | 1 | 1 | 0 | 0 | 1 | 0 | 0 | 0 | 1 | 52 | 49 | 1 | 1 | NA | 1 | 0.485 | NA | | 0 | 0.742 |
| A3SS | 2046 | OS9 | OS9 | chr12 | + | 57716088 | 57716193 | 57716091 | 57716193 | 57715759 | 57715970 | 7 | 1 | 0 | 4 | 1 | 0 | 0 | 2 | 0 | 1 | 52 | 49 | 1 | 1 | 0.623 | 0.485 | NA | NA | | 0.653 | -0.099 |
| A3SS | 2062 | FOSL1 | FOSL1 | chr11 | - | 65894013 | 65894121 | 65894013 | 65894116 | 65896808 | 65897006 | 2 | 0 | 0 | 0 | 0 | 0 | 0 | 2 | 0 | 2 | 54 | 49 | 1 | 1 | 1 | NA | NA | NA | | 0.476 | 0.524 |
| A3SS | 2069 | ABI1 | ABI1 | chr10 | - | 26759061 | 26759238 | 26759061 | 26759235 | 26765217 | 26765318 | 4 | 2 | 1 | 1 | 3 | 0 | 0 | 1 | 0 | 2 | 52 | 49 | 1 | 1 | 0.79 | 0.386 | 1 | NA | | 0.32 | 0.405 |
| A3SS | 2082 | CNIH4 | CNIH4 | chr1 | + | 224371282 | 224371423 | 224371329 | 224371423 | 224365878 | 224365991 | 26 | 7 | 6 | 1 | 0 | 0 | 0 | 7 | 0 | 0 | 96 | 49 | 1 | 1 | 0.93 | 1 | 1 | NA | | 1 | -0.023 |
| A3SS | 2090 | CTDSP1 | CTDSP1 | chr2 | + | 218402110 | 218402215 | 218402113 | 218402215 | 218401563 | 218401712 | 5 | 2 | 3 | 13 | 5 | 6 | 0 | 0 | 0 | 7 | 52 | 49 | 1 | 1 | 0.266 | 0.274 | 0.32 | NA | | 0 | 0.287 |
| A3SS | 2100 | MEF2D | MEF2D | chr1 | - | 156483238 | 156483430 | 156483238 | 156483418 | 156500485 | 156500842 | 12 | 8 | 1 | 1 | 1 | 0 | 5 | 4 | 0 | 0 | 61 | 49 | 1 | 1 | 0.906 | 0.865 | 1 | | 1 | 1 | -0.076 |
| A3SS | 2116 | SNRNP70 | SNRNP70 | chr19 | + | 49107794 | 49108613 | 49107821 | 49108613 | 49107624 | 49107712 | 12 | 8 | 0 | 4 | 2 | 1 | 0 | 1 | 0 | 1 | 76 | 49 | 1 | 1 | 0.659 | 0.721 | 0 | NA | | 0.392 | 0.068 |
| A3SS | 2132 | TBC1D10C | TBC1D10C | chr11 | + | 67405902 | 67406017 | 67405906 | 67406017 | 67405594 | 67405701 | 29 | 7 | 9 | 1 | 6 | 0 | 0 | 7 | 0 | 2 | 53 | 49 | 1 | 1 | 0.964 | 0.519 | 1 | NA | | 0.764 | 0.064 |
| A3SS | 2140 | MRPS34 | MRPS34 | chr16 | - | 1771894 | 1772534 | 1771894 | 1772513 | 1772603 | 1772646 | 0 | 3 | 0 | 0 | 0 | 0 | 0 | 0 | 0 | 1 | 70 | 49 | 1 | 1 | NA | 1 | NA | NA | | 0 | 1 |
| A3SS | 2149 | ZBED5 | ZBED5 | chr11 | - | 10852703 | 10855118 | 10852703 | 10855086 | 10856143 | 10856257 | 1 | 2 | 0 | 2 | 0 | 0 | 0 | 1 | 0 | 0 | 81 | 49 | 1 | 1 | 0.232 | 1 | NA | NA | | 1 | -0.384 |
| A3SS | 2179 | ATP6V0B | ATP6V0B | chr1 | + | 43977025 | 43978300 | 43977980 | 43978300 | 43976772 | 43976824 | 22 | 37 | 10 | 1 | 3 | 0 | 0 | 10 | 0 | 0 | 98 | 49 | 1 | 1 | 0.917 | 0.86 | 1 | NA | | 1 | -0.074 |
| A3SS | 2191 | GLYR1 | GLYR1 | chr16 | - | 4814536 | 4814647 | 4814536 | 4814629 | 4817597 | 4817697 | 10 | 5 | 2 | 1 | 0 | 0 | 0 | 5 | 0 | 0 | 67 | 49 | 1 | 1 | 0.88 | 1 | 1 | NA | | 1 | -0.04 |
| A3SS | 2192 | PRELID1 | PRELID1 | chr5 | + | 177306421 | 177306959 | 177306454 | 177306959 | 177306097 | 177306176 | 32 | 11 | 29 | 1 | 0 | 0 | 1 | 7 | 0 | 0 | 82 | 49 | 1 | 1 | 0.95 | 1 | 1 | | 1 | 1 | -0.017 |
| A3SS | 2193 | ZDHHC20 | ZDHHC20 | chr13 | - | 21381433 | 21381549 | 21381433 | 21381546 | 21381900 | 21381945 | 1 | 0 | 0 | 1 | 0 | 0 | 0 | 2 | 0 | 0 | 52 | 49 | 1 | 1 | 0.485 | NA | NA | NA | | 1 | -0.515 |
| A3SS | 2194 | ZDHHC20 | ZDHHC20 | chr13 | - | 21381433 | 21381549 | 21381433 | 21381546 | 21382919 | 21383009 | 7 | 0 | 0 | 1 | 0 | 0 | 0 | 6 | 0 | 3 | 52 | 49 | 1 | 1 | 0.868 | NA | NA | NA | | 0.653 | 0.215 |
| A3SS | 2196 | C8orf58 | C8orf58 | chr8 | + | 22602536 | 22602643 | 22602560 | 22602643 | 22602199 | 22602312 | 2 | 0 | 0 | 1 | 0 | 0 | 1 | 0 | 0 | 0 | 73 | 49 | 1 | 1 | 0.573 | NA | NA | | 1 | NA | -0.427 |
| A3SS | 2225 | CHD4 | CHD4 | chr12 | - | 6601959 | 6602175 | 6601959 | 6602154 | 6602375 | 6602497 | 18 | 5 | 0 | 4 | 3 | 0 | 0 | 2 | 0 | 1 | 70 | 49 | 1 | 1 | 0.759 | 0.538 | NA | NA | | 0.583 | 0.066 |
| A3SS | 2226 | CELF2 | CELF2 | chr10 | + | 11321188 | 11321386 | 11321206 | 11321386 | 11314138 | 11314258 | 9 | 8 | 0 | 6 | 13 | 8 | 0 | 1 | 0 | 3 | 67 | 49 | 1 | 1 | 0.523 | 0.31 | 0 | NA | | 0.196 | 0.082 |
| A3SS | 2227 | TIAL1 | TIAL1 | chr10 | - | 119582458 | 119582608 | 119582458 | 119582557 | 119588151 | 119588248 | 4 | 10 | 0 | 4 | 4 | 0 | 0 | 0 | 0 | 1 | 98 | 49 | 1 | 1 | 0.333 | 0.556 | NA | NA | | 0 | 0.445 |
| A3SS | 2228 | BUB3 | BUB3 | chr10 | + | 123163819 | 123165370 | 123165046 | 123165370 | 123162611 | 123162828 | 11 | 2 | 4 | 1 | 0 | 0 | 0 | 5 | 0 | 0 | 98 | 49 | 1 | 1 | 0.846 | 1 | 1 | NA | | 1 | -0.051 |
| A3SS | 2259 | MGRN1 | MGRN1 | chr16 | + | 4686258 | 4690974 | 4688795 | 4690974 | 4683842 | 4683932 | 7 | 3 | 0 | 9 | 1 | 0 | 1 | 0 | 0 | 0 | 98 | 49 | 1 | 1 | 0.28 | 0.6 | NA | | 1 | NA | -0.56 |
| A3SS | 2279 | NFYC | NFYC | chr1 | + | 40770396 | 40771606 | 40770708 | 40771606 | 40769355 | 40769415 | 0 | 2 | 3 | 3 | 0 | 0 | 0 | 0 | 1 | 0 | 98 | 49 | 1 | 1 | 0 | 1 | 1 | | 0 | NA | 0.667 |
| A3SS | 2282 | TRMU | TRMU | chr22 | + | 46350290 | 46350463 | 46350345 | 46350463 | 46346421 | 46346544 | 3 | 1 | 0 | 0 | 1 | 0 | 0 | 1 | 0 | 0 | 98 | 49 | 1 | 1 | 1 | 0.333 | NA | NA | | 1 | -0.334 |
| A3SS | 2288 | RTN3 | RTN3 | chr11 | + | 63718701 | 63721032 | 63718980 | 63721032 | 63681449 | 63681778 | 14 | 8 | 1 | 0 | 0 | 1 | 1 | 0 | 0 | 0 | 98 | 49 | 1 | 1 | 1 | 1 | 0.333 | | 1 | NA | -0.222 |
| A3SS | 2293 | ZNF33A | ZNF33A | chr10 | + | 38034277 | 38034400 | 38034325 | 38034400 | 38017290 | 38017386 | 0 | 1 | 0 | 0 | 1 | 0 | 0 | 1 | 0 | 0 | 97 | 49 | 1 | 1 | NA | 0.336 | NA | NA | | 1 | -0.664 |
| A3SS | 2294 | ZNF33A | ZNF33A | chr10 | + | 38054374 | 38060067 | 38054377 | 38060067 | 38017290 | 38017386 | 9 | 5 | 3 | 3 | 4 | 0 | 0 | 1 | 0 | 1 | 52 | 49 | 1 | 1 | 0.739 | 0.541 | 1 | NA | | 0.485 | 0.275 |
| A3SS | 2296 | PRF1 | PRF1 | chr10 | - | 70600363 | 70600932 | 70600363 | 70600906 | 70602644 | 70602775 | 1 | 3 | 0 | 0 | 0 | 0 | 0 | 0 | 1 | 0 | 75 | 49 | 1 | 1 | 1 | 1 | NA | | 0 | NA | 1 |
| A3SS | 2306 | ELP4 | ELP4 | chr11 | + | 31603764 | 31603907 | 31603767 | 31603907 | 31594769 | 31594901 | 1 | 0 | 0 | 8 | 0 | 0 | 0 | 0 | 0 | 3 | 52 | 49 | 1 | 1 | 0.105 | NA | NA | NA | | 0 | 0.105 |
| A3SS | 2313 | C10orf128 | C10orf128 | chr10 | - | 49154723 | 49156259 | 49154723 | 49155909 | 49161598 | 49161622 | 2 | 6 | 0 | 0 | 0 | 0 | 0 | 0 | 0 | 1 | 98 | 49 | 1 | 1 | 1 | 1 | NA | NA | | 0 | 1 |
| A3SS | 2321 | XRN1 | XRN1 | chr3 | - | 142332374 | 142332534 | 142332374 | 142332531 | 142332966 | 142333089 | 9 | 0 | 1 | 1 | 0 | 0 | 0 | 1 | 0 | 0 | 52 | 49 | 1 | 1 | 0.895 | NA | 1 | NA | | 1 | -0.052 |
| A3SS | 2328 | GMPR2 | GMPR2 | chr14 | + | 24237041 | 24237152 | 24237070 | 24237152 | 24235966 | 24236140 | 19 | 9 | 0 | 13 | 5 | 8 | 0 | 0 | 0 | 1 | 78 | 49 | 1 | 1 | 0.479 | 0.531 | 0 | NA | | 0 | 0.337 |
| A3SS | 2331 | CIB1 | CIB1 | chr15 | - | 90232218 | 90232447 | 90232218 | 90232327 | 90233668 | 90233703 | 4 | 9 | 0 | 0 | 0 | 0 | 0 | 5 | 0 | 1 | 98 | 49 | 1 | 1 | 1 | 1 | NA | NA | | 0.714 | 0.286 |
| A3SS | 2340 | ADPGK | ADPGK | chr15 | - | 72751366 | 72752895 | 72751366 | 72752892 | 72755555 | 72755654 | 5 | 2 | 1 | 13 | 2 | 0 | 0 | 1 | 0 | 3 | 52 | 49 | 1 | 1 | 0.266 | 0.485 | 1 | NA | | 0.239 | 0.345 |
| A3SS | 2342 | TBC1D10A | TBC1D10A | chr22 | - | 30304530 | 30304651 | 30304530 | 30304630 | 30326672 | 30326966 | 0 | 0 | 1 | 3 | 1 | 0 | 0 | 1 | 0 | 0 | 70 | 49 | 1 | 1 | 0 | 0 | 1 | NA | | 1 | -0.667 |
| A3SS | 2348 | FUT11 | FUT11 | chr10 | + | 73775418 | 73776218 | 73775540 | 73776218 | 73773189 | 73773815 | 1 | 0 | 0 | 2 | 0 | 0 | 0 | 0 | 3 | 0 | 98 | 49 | 1 | 1 | 0.2 | NA | NA | | 0 | NA | 0.2 |
| A3SS | 2354 | TRPC4AP | TRPC4AP | chr20 | - | 35021189 | 35021356 | 35021189 | 35021332 | 35035122 | 35035308 | 12 | 5 | 0 | 2 | 5 | 1 | 0 | 3 | 0 | 1 | 73 | 49 | 1 | 1 | 0.801 | 0.402 | 0 | NA | | 0.668 | -0.267 |
| A3SS | 2356 | PNPLA6 | PNPLA6 | chr19 | + | 7553871 | 7554015 | 7553874 | 7554015 | 7551361 | 7551437 | 4 | 1 | 1 | 3 | 1 | 2 | 0 | 3 | 0 | 1 | 52 | 49 | 1 | 1 | 0.557 | 0.485 | 0.32 | NA | | 0.739 | -0.285 |
| A3SS | 2365 | IFIT1 | IFIT1 | chr10 | + | 89399889 | 89400003 | 89399894 | 89400003 | 89392545 | 89392717 | 2 | 0 | 0 | 0 | 0 | 0 | 0 | 0 | 1 | 1 | 54 | 49 | 1 | 1 | 1 | NA | NA | | 0 | 0 | 1 |
| A3SS | 2377 | RNH1 | RNH1 | chr11 | - | 502061 | 502249 | 502061 | 502181 | 504823 | 504996 | 2 | 0 | 0 | 1 | 2 | 0 | 0 | 5 | 0 | 2 | 98 | 49 | 1 | 1 | 0.5 | 0 | NA | NA | | 0.556 | -0.306 |
| A3SS | 2379 | RNH1 | RNH1 | chr11 | - | 502061 | 502249 | 502061 | 502181 | 507112 | 507283 | 2 | 0 | 0 | 0 | 1 | 0 | 0 | 5 | 0 | 0 | 98 | 49 | 1 | 1 | 1 | 0 | NA | NA | | 1 | -0.5 |
| A3SS | 2392 | BAZ2B | BAZ2B | chr2 | - | 159448241 | 159448409 | 159448241 | 159448403 | 159453612 | 159453801 | 43 | 9 | 0 | 6 | 1 | 0 | 0 | 5 | 0 | 0 | 55 | 49 | 1 | 1 | 0.865 | 0.889 | NA | NA | | 1 | -0.123 |
| A3SS | 2393 | BAZ2B | BAZ2B | chr2 | - | 159448241 | 159448409 | 159448241 | 159448403 | 159478574 | 159478721 | 8 | 7 | 0 | 2 | 0 | 0 | 0 | 5 | 0 | 0 | 55 | 49 | 1 | 1 | 0.781 | 1 | NA | NA | | 1 | -0.109 |
| A3SS | 2405 | SEC14L1 | SEC14L1 | chr17 | + | 77213917 | 77217099 | 77216548 | 77217099 | 77211949 | 77212201 | 164 | 43 | 8 | 0 | 2 | 0 | 8 | 10 | 0 | 0 | 98 | 49 | 1 | 1 | 1 | 0.915 | 1 | | 1 | 1 | -0.028 |
| A3SS | 2406 | CLEC16A | CLEC16A | chr16 | + | 10972931 | 10973061 | 10972937 | 10973061 | 10971124 | 10971230 | 0 | 0 | 0 | 5 | 3 | 0 | 0 | 2 | 0 | 0 | 55 | 49 | 1 | 1 | 0 | 0 | NA | NA | | 1 | -1 |
| A3SS | 2407 | GBF1 | GBF1 | chr10 | + | 102359266 | 102359435 | 102359269 | 102359435 | 102358505 | 102358729 | 1 | 0 | 0 | 4 | 0 | 0 | 0 | 0 | 0 | 1 | 52 | 49 | 1 | 1 | 0.191 | NA | NA | NA | | 0 | 0.191 |
| A3SS | 2413 | CALHM2 | CALHM2 | chr10 | - | 103451082 | 103451285 | 103451082 | 103451182 | 103452155 | 103452405 | 0 | 0 | 0 | 2 | 0 | 0 | 0 | 2 | 0 | 2 | 98 | 49 | 1 | 1 | 0 | NA | NA | NA | | 0.333 | -0.333 |
| A3SS | 2417 | ANKRD11 | ANKRD11 | chr16 | - | 89316932 | 89317078 | 89316932 | 89317075 | 89418283 | 89418368 | 0 | 2 | 0 | 0 | 5 | 0 | 1 | 2 | 0 | 0 | 52 | 49 | 1 | 1 | NA | 0.274 | NA | | 1 | 1 | -0.726 |
| A3SS | 2424 | PICALM | PICALM | chr11 | - | 85990249 | 85990399 | 85990249 | 85990378 | 85996825 | 85996929 | 125 | 58 | 10 | 4 | 3 | 0 | 0 | 18 | 0 | 1 | 70 | 49 | 1 | 1 | 0.956 | 0.931 | 1 | NA | | 0.926 | 0.036 |
| A3SS | 2426 | FAM89B | FAM89B | chr11 | + | 65573352 | 65574198 | 65573362 | 65574198 | 65572348 | 65572960 | 3 | 3 | 0 | 1 | 1 | 2 | 0 | 1 | 0 | 1 | 59 | 49 | 1 | 1 | 0.714 | 0.714 | 0 | NA | | 0.454 | 0.022 |
| A3SS | 2440 | TBCE | TBCE | chr1 | + | 235419416 | 235419561 | 235419472 | 235419561 | 235414432 | 235414618 | 1 | 0 | 0 | 0 | 1 | 0 | 1 | 0 | 0 | 0 | 98 | 49 | 1 | 1 | 1 | 0 | NA | | 1 | NA | -0.5 |
| A3SS | 2474 | LST1 | LST1 | chr6 | + | 31588496 | 31588909 | 31588517 | 31588909 | 31587199 | 31587318 | 23 | 4 | 4 | 46 | 21 | 20 | 0 | 3 | 0 | 4 | 70 | 49 | 1 | 1 | 0.259 | 0.118 | 0.123 | NA | | 0.344 | -0.177 |
| A3SS | 2495 | TUBGCP2 | TUBGCP2 | chr10 | - | 133283077 | 133283221 | 133283077 | 133283184 | 133283881 | 133284002 | 14 | 3 | 4 | 1 | 0 | 0 | 1 | 6 | 0 | 0 | 86 | 49 | 1 | 1 | 0.889 | 1 | 1 | | 1 | 1 | -0.037 |
| A3SS | 2514 | PIK3R5 | PIK3R5 | chr17 | - | 8888170 | 8888953 | 8888170 | 8888891 | 8889138 | 8889222 | 2 | 1 | 1 | 10 | 5 | 7 | 0 | 2 | 0 | 2 | 98 | 49 | 1 | 1 | 0.091 | 0.091 | 0.067 | NA | | 0.333 | -0.25 |
| A3SS | 2516 | ZSWIM8 | ZSWIM8 | chr10 | + | 73801016 | 73801195 | 73801084 | 73801195 | 73800639 | 73800759 | 14 | 6 | 1 | 2 | 0 | 0 | 0 | 2 | 0 | 0 | 98 | 49 | 1 | 1 | 0.778 | 1 | 1 | NA | | 1 | -0.074 |
| A3SS | 2517 | GCH1 | GCH1 | chr14 | - | 54842005 | 54844143 | 54842005 | 54843054 | 54845767 | 54845852 | 4 | 0 | 0 | 0 | 0 | 0 | 0 | 2 | 0 | 2 | 98 | 49 | 1 | 1 | 1 | NA | NA | NA | | 0.333 | 0.667 |
| A3SS | 2532 | LYPLAL1 | LYPLAL1 | chr1 | + | 219193081 | 219193251 | 219193129 | 219193251 | 219179146 | 219179246 | 7 | 2 | 3 | 0 | 2 | 0 | 0 | 1 | 0 | 0 | 97 | 49 | 1 | 1 | 1 | 0.336 | 1 | NA | | 1 | -0.221 |
| A3SS | 2560 | PPP1CA | PPP1CA | chr11 | - | 67401067 | 67401232 | 67401067 | 67401199 | 67401727 | 67401905 | 2 | 0 | 0 | 18 | 9 | 4 | 0 | 0 | 1 | 4 | 82 | 49 | 1 | 1 | 0.062 | 0 | 0 | | 0 | 0 | 0.021 |
| A3SS | 2564 | ATF7IP | ATF7IP | chr12 | + | 14434336 | 14434423 | 14434339 | 14434423 | 14423908 | 14425473 | 25 | 2 | 2 | 6 | 1 | 0 | 0 | 6 | 0 | 3 | 52 | 49 | 1 | 1 | 0.797 | 0.653 | 1 | NA | | 0.653 | 0.164 |
| A3SS | 2571 | SLC25A29 | SLC25A29 | chr14 | - | 100295625 | 100295993 | 100295625 | 100295697 | 100298841 | 100298885 | 1 | 1 | 0 | 2 | 0 | 0 | 3 | 0 | 0 | 0 | 98 | 49 | 1 | 1 | 0.2 | 1 | NA | | 1 | NA | -0.4 |
| A3SS | 2573 | KCNK7 | KCNK7 | chr11 | - | 65592854 | 65593383 | 65592854 | 65593210 | 65593475 | 65593874 | 8 | 4 | 0 | 3 | 0 | 0 | 0 | 5 | 0 | 0 | 98 | 49 | 1 | 1 | 0.571 | 1 | NA | NA | | 1 | -0.215 |
| A3SS | 2575 | SUPT7L | SUPT7L | chr2 | - | 27660983 | 27661601 | 27660983 | 27661388 | 27662178 | 27662281 | 0 | 1 | 0 | 0 | 1 | 0 | 0 | 0 | 0 | 1 | 98 | 49 | 1 | 1 | NA | 0.333 | NA | NA | | 0 | 0.333 |
| A3SS | 2583 | ADAR | ADAR | chr1 | - | 154596804 | 154596995 | 154596804 | 154596938 | 154597122 | 154597267 | 120 | 43 | 27 | 1 | 0 | 0 | 2 | 42 | 0 | 0 | 98 | 49 | 1 | 1 | 0.984 | 1 | 1 | | 1 | 1 | -0.005 |
| A3SS | 2594 | PTMA | PTMA | chr2 | + | 231711886 | 231711983 | 231711889 | 231711983 | 231711347 | 231711419 | 5 | 1 | 0 | 7 | 0 | 0 | 0 | 1 | 0 | 2 | 52 | 49 | 1 | 1 | 0.402 | 1 | NA | NA | | 0.32 | 0.381 |
| A3SS | 2604 | LILRB2 | LILRB2 | chr19 | - | 54276806 | 54276979 | 54276806 | 54276929 | 54277549 | 54277597 | 3 | 0 | 0 | 9 | 1 | 2 | 0 | 1 | 0 | 3 | 98 | 49 | 1 | 1 | 0.143 | 0 | 0 | NA | | 0.143 | -0.095 |
| A3SS | 2615 | ANKRD42 | ANKRD42 | chr11 | + | 83212652 | 83215200 | 83212978 | 83215200 | 83211294 | 83211430 | 3 | 0 | 0 | 0 | 0 | 0 | 0 | 0 | 0 | 1 | 98 | 49 | 1 | 1 | 1 | NA | NA | NA | | 0 | 1 |
| A3SS | 2623 | TNFSF13 | TNFSF13 | chr17 | + | 7559845 | 7559893 | 7559850 | 7559893 | 7558291 | 7559297 | 1 | 1 | 0 | 2 | 0 | 0 | 0 | 0 | 0 | 1 | 54 | 49 | 1 | 1 | 0.312 | 1 | NA | NA | | 0 | 0.656 |

| | | | | | | | | | | | | | | | | | | | | | | | | | | | | | | | | |
|---|---|---|---|---|---|---|---|---|---|---|---|---|---|---|---|---|---|---|---|---|---|---|---|---|---|---|---|---|---|---|---|---|
| A3SS | 2624 | TNFSF13 | TNFSF13 | chr17 | + | 7559847 | 7559893 | 7559850 | 7559893 | 7558291 | 7559297 | 0 | 1 | 0 | 2 | 0 | 0 | 0 | 0 | 0 | 1 | 52 | 49 | 1 | 1 | 0 | 1 | NA | | NA | 0 | 0.5 |
| A3SS | 2633 | SLC43A3 | SLC43A3 | chr11 | - | 57425988 | 57426389 | 57425988 | 57426352 | 57426587 | 57426652 | 0 | 0 | 0 | 0 | 1 | 0 | 0 | 2 | 0 | 0 | 86 | 49 | 1 | 1 | NA | 0 | NA | | NA | 1 | -1 |
| A3SS | 2638 | FAM111A | FAM111A | chr11 | + | 59148796 | 59148953 | 59148835 | 59148953 | 59145826 | 59145856 | 9 | 0 | 3 | 2 | 2 | 0 | 0 | 0 | 1 | 0 | 88 | 49 | 1 | 1 | 0.715 | 0 | 1 | | 0 | NA | 0.572 |
| A3SS | 2657 | PPP6R3 | PPP6R3 | chr11 | + | 68587926 | 68588024 | 68587944 | 68588024 | 68575957 | 68576043 | 2 | 3 | 3 | 0 | 0 | 0 | 0 | 0 | 0 | 1 | 67 | 49 | 1 | 1 | 1 | 1 | 1 | | NA | 0 | 1 |
| A3SS | 2658 | PPP6R3 | PPP6R3 | chr11 | + | 68587926 | 68588024 | 68587944 | 68588024 | 68583042 | 68583129 | 10 | 9 | 3 | 4 | 0 | 0 | 0 | 3 | 0 | 0 | 67 | 49 | 1 | 1 | 0.646 | 1 | 1 | | NA | 1 | -0.118 |
| A3SS | 2665 | IFIT3 | IFIT3 | chr10 | + | 89337420 | 89337542 | 89337424 | 89337542 | 89332478 | 89332627 | 2 | 4 | 0 | 0 | 1 | 0 | 0 | 3 | 0 | 0 | 53 | 49 | 1 | 1 | 1 | 0.787 | NA | | NA | 1 | -0.107 |
| A3SS | 2671 | KANSL1 | KANSL1 | chr17 | - | 46034160 | 46034285 | 46034160 | 46034282 | 46038537 | 46038686 | 13 | 1 | 0 | 4 | 0 | 0 | 1 | 1 | 0 | 0 | 52 | 49 | 1 | 1 | 0.754 | 1 | NA | | NA | 1 | -0.123 |
| A3SS | 2676 | THEMIS2 | THEMIS2 | chr1 | + | 27881970 | 27883043 | 27882512 | 27883043 | 27879643 | 27880054 | 191 | 107 | 53 | 1 | 0 | 0 | 17 | 67 | 0 | 0 | 98 | 49 | 1 | 1 | 0.99 | 1 | 1 | | 1 | 1 | -0.003 |
| A3SS | 2677 | PIGT | PIGT | chr20 | + | 45419472 | 45419590 | 45419503 | 45419590 | 45419294 | 45419395 | 1 | 0 | 0 | 1 | 2 | 0 | 0 | 0 | 0 | 2 | 80 | 49 | 1 | 1 | 0.38 | 0 | NA | | NA | 0 | 0.19 |
| A3SS | 2686 | ZYX | ZYX | chr7 | + | 143381556 | 143381779 | 143381559 | 143381779 | 143381266 | 143381409 | 1 | 0 | 4 | 1 | 0 | 0 | 1 | 1 | 1 | 0 | 52 | 49 | 1 | 1 | 0.485 | NA | 1 | | 0.485 | 1 | 0 |
| A3SS | 2701 | CD300LF | CD300LF | chr17 | - | 74703034 | 74703143 | 74703034 | 74703098 | 74704477 | 74704816 | 15 | 4 | 1 | 22 | 14 | 3 | 0 | 2 | 0 | 4 | 94 | 49 | 1 | 1 | 0.262 | 0.13 | 0.148 | | NA | 0.207 | -0.027 |
| A3SS | 2703 | ANAPC15 | ANAPC15 | chr11 | - | 72111156 | 72111286 | 72111156 | 72111281 | 72111411 | 72111496 | 1 | 0 | 0 | 1 | 0 | 0 | 0 | 1 | 0 | 0 | 54 | 49 | 1 | 1 | 0.476 | NA | NA | | NA | 1 | -0.524 |
| A3SS | 2714 | MAP4K2 | MAP4K2 | chr11 | - | 64797103 | 64797192 | 64797103 | 64797168 | 64797274 | 64797380 | 10 | 5 | 6 | 2 | 0 | 0 | 0 | 3 | 0 | 0 | 73 | 49 | 1 | 1 | 0.77 | 1 | 1 | | NA | 1 | -0.077 |
| A3SS | 2721 | ZNF276 | ZNF276 | chr16 | + | 89737720 | 89737905 | 89737805 | 89737905 | 89733920 | 89734038 | 2 | 1 | 0 | 7 | 2 | 0 | 1 | 1 | 0 | 5 | 98 | 49 | 1 | 1 | 0.125 | 0.2 | NA | NA | 1 | 0.091 | -0.383 |
| A3SS | 2729 | ST20 | ST20 | chr15 | - | 79907628 | 79907706 | 79907628 | 79907684 | 79914587 | 79915256 | 2 | 1 | 0 | 9 | 0 | 0 | 0 | 1 | 0 | 5 | 71 | 49 | 1 | 1 | 0.133 | 1 | NA | | NA | 0.121 | 0.446 |
| A3SS | 2735 | CTC1 | CTC1 | chr17 | - | 8237374 | 8237519 | 8237374 | 8237414 | 8238030 | 8238242 | 7 | 6 | 0 | 0 | 1 | 0 | 0 | 3 | 0 | 0 | 98 | 49 | 1 | 1 | 1 | 0.75 | NA | | NA | 1 | -0.125 |
| A3SS | 2738 | PAM | PAM | chr5 | + | 103028181 | 103028238 | 103028184 | 103028238 | 103025130 | 103025334 | 0 | 0 | 0 | 1 | 0 | 0 | 0 | 1 | 0 | 0 | 52 | 49 | 1 | 1 | 0 | NA | NA | | NA | 1 | -1 |
| A3SS | 2749 | LRRC75A-AS1 | LRRC75A-AS1 | chr17 | + | 16439527 | 16439703 | 16439659 | 16439703 | 16438986 | 16439414 | 28 | 10 | 8 | 2 | 0 | 0 | 1 | 4 | 0 | 0 | 98 | 49 | 1 | 1 | 0.875 | 1 | 1 | | 1 | 1 | -0.042 |
| A3SS | 2750 | LRRC75A-AS1 | LRRC75A-AS1 | chr17 | + | 16439580 | 16439703 | 16439659 | 16439703 | 16438986 | 16439414 | 28 | 10 | 8 | 2 | 0 | 0 | 1 | 4 | 0 | 0 | 98 | 49 | 1 | 1 | 0.875 | 1 | 1 | | 1 | 1 | -0.042 |
| A3SS | 2751 | LRRC75A-AS1 | LRRC75A-AS1 | chr17 | + | 16440110 | 16440253 | 16440184 | 16440253 | 16439326 | 16439414 | 5 | 0 | 0 | 1 | 1 | 0 | 0 | 2 | 0 | 1 | 98 | 49 | 1 | 1 | 0.714 | 0 | NA | | NA | 0.5 | -0.143 |
| A3SS | 2752 | LRRC75A-AS1 | LRRC75A-AS1 | chr17 | + | 16440110 | 16440253 | 16440184 | 16440253 | 16439659 | 16439703 | 5 | 0 | 0 | 2 | 2 | 0 | 0 | 2 | 0 | 0 | 98 | 49 | 1 | 1 | 0.556 | 0 | NA | | NA | 1 | -0.722 |
| A3SS | 2763 | MTA1 | MTA1 | chr14 | + | 105464415 | 105464567 | 105464462 | 105464567 | 105464031 | 105464147 | 2 | 0 | 0 | 0 | 1 | 0 | 0 | 1 | 0 | 0 | 96 | 49 | 1 | 1 | 1 | 0 | NA | | NA | 1 | -0.5 |
| A3SS | 2764 | LGALS9 | LGALS9 | chr17 | + | 27640571 | 27640773 | 27640642 | 27640773 | 27638262 | 27638354 | 21 | 4 | 6 | 1 | 0 | 0 | 2 | 25 | 0 | 0 | 98 | 49 | 1 | 1 | 0.913 | 1 | 1 | | 1 | 1 | -0.029 |
| A3SS | 2788 | LILRA1 | LILRA1 | chr19 | + | 54594196 | 54594476 | 54594440 | 54594476 | 54593575 | 54593781 | 0 | 3 | 1 | 11 | 6 | 4 | 0 | 0 | 0 | 3 | 98 | 49 | 1 | 1 | 0 | 0.2 | 0.111 | | NA | 0 | 0.104 |
| A3SS | 2813 | UBXN1 | UBXN1 | chr11 | - | 62676497 | 62677005 | 62676497 | 62676639 | 62677517 | 62677634 | 9 | 11 | 6 | 1 | 0 | 0 | 0 | 4 | 0 | 0 | 98 | 49 | 1 | 1 | 0.818 | 1 | 1 | | NA | 1 | -0.061 |
| A3SS | 2820 | ATP6V0A1 | ATP6V0A1 | chr17 | + | 42477638 | 42477742 | 42477659 | 42477742 | 42470089 | 42470218 | 1 | 3 | 3 | 5 | 1 | 0 | 0 | 0 | 0 | 1 | 70 | 49 | 1 | 1 | 0.123 | 0.677 | 1 | | NA | 0 | 0.6 |
| A3SS | 2826 | BRCA1 | BRCA1 | chr17 | - | 43076487 | 43076614 | 43076487 | 43076611 | 43082403 | 43082575 | 1 | 4 | 0 | 0 | 0 | 0 | 0 | 0 | 0 | 1 | 52 | 49 | 1 | 1 | 1 | 1 | NA | | NA | 0 | 1 |
| A3SS | 2843 | PHOSPHO1 | PHOSPHO1 | chr17 | - | 49223369 | 49225131 | 49223369 | 49225004 | 49226646 | 49226758 | 42 | 14 | 11 | 5 | 1 | 8 | 0 | 4 | 0 | 1 | 98 | 49 | 1 | 1 | 0.808 | 0.875 | 0.407 | | NA | 0.667 | 0.03 |
| A3SS | 2854 | MAP7D1 | MAP7D1 | chr1 | + | 36179511 | 36179557 | 36179514 | 36179557 | 36179261 | 36179315 | 2 | 4 | 0 | 26 | 18 | 14 | 0 | 0 | 0 | 3 | 52 | 49 | 1 | 1 | 0.068 | 0.173 | 0 | | NA | 0 | 0.08 |
| A3SS | 2881 | ZNF200 | ZNF200 | chr16 | - | 3222324 | 3224613 | 3222324 | 3224610 | 3232420 | 3232547 | 2 | 3 | 0 | 0 | 2 | 0 | 0 | 1 | 0 | 0 | 52 | 49 | 1 | 1 | 1 | 0.586 | NA | | NA | 1 | -0.207 |
| A3SS | 2895 | RHBDF2 | RHBDF2 | chr17 | - | 76479732 | 76479941 | 76479732 | 76479854 | 76481374 | 76481545 | 2 | 1 | 0 | 8 | 3 | 2 | 0 | 1 | 0 | 5 | 98 | 49 | 1 | 1 | 0.111 | 0.143 | 0 | | NA | 0.091 | -0.006 |
| A3SS | 2898 | PGS1 | PGS1 | chr17 | + | 78392475 | 78392665 | 78392513 | 78392665 | 78378617 | 78378808 | 34 | 19 | 3 | 2 | 0 | 0 | 0 | 15 | 0 | 0 | 87 | 49 | 1 | 1 | 0.905 | 1 | 1 | | NA | 1 | -0.032 |
| A3SS | 2906 | TYMP | TYMP | chr22 | - | 50529495 | 50529754 | 50529495 | 50529731 | 50529903 | 50530085 | 23 | 17 | 2 | 2 | 0 | 0 | 2 | 4 | 0 | 1 | 72 | 49 | 1 | 1 | 0.887 | 1 | 1 | | 1 | 0.731 | 0.097 |
| A3SS | 2921 | IL16 | IL16 | chr15 | + | 81306419 | 81306545 | 81306422 | 81306545 | 81305907 | 81306166 | 5 | 5 | 5 | 13 | 3 | 5 | 0 | 1 | 0 | 1 | 52 | 49 | 1 | 1 | 0.266 | 0.611 | 0.485 | | NA | 0.485 | -0.031 |
| A3SS | 2923 | HSDL1 | HSDL1 | chr16 | - | 84129985 | 84130431 | 84129985 | 84130171 | 84131101 | 84131327 | 1 | 1 | 0 | 0 | 1 | 0 | 0 | 3 | 0 | 0 | 98 | 49 | 1 | 1 | 1 | 0.333 | NA | | NA | 1 | -0.334 |
| A3SS | 2929 | HPS4 | HPS4 | chr22 | - | 26470718 | 26470922 | 26470718 | 26470813 | 26472301 | 26472418 | 1 | 0 | 0 | 1 | 1 | 0 | 1 | 0 | 0 | 0 | 98 | 49 | 1 | 1 | 0.333 | 0 | NA | | 1 | NA | -0.834 |
| A3SS | 2930 | PPP4R1 | PPP4R1 | chr18 | - | 9595017 | 9595153 | 9595017 | 9595102 | 9614225 | 9614270 | 10 | 4 | 3 | 9 | 0 | 2 | 0 | 4 | 0 | 0 | 98 | 49 | 1 | 1 | 0.357 | 1 | 0.429 | | NA | 1 | -0.405 |
| A3SS | 2940 | CARS | CARS | chr11 | - | 3040895 | 3041031 | 3040895 | 3040984 | 3042164 | 3042256 | 3 | 4 | 2 | 8 | 4 | 0 | 0 | 0 | 0 | 3 | 96 | 49 | 1 | 1 | 0.161 | 0.338 | 1 | | NA | 0 | 0.5 |
| A3SS | 2966 | DYRK1A | DYRK1A | chr21 | + | 37478180 | 37478300 | 37478207 | 37478300 | 37472683 | 37472880 | 6 | 4 | 0 | 24 | 4 | 2 | 0 | 0 | 0 | 3 | 76 | 49 | 1 | 1 | 0.139 | 0.392 | 0 | | NA | 0 | 0.177 |
| A3SS | 2969 | GPX4 | GPX4 | chr19 | + | 1106377 | 1106459 | 1106399 | 1106459 | 1106241 | 1106266 | 1 | 1 | 0 | 10 | 9 | 1 | 0 | 0 | 0 | 2 | 71 | 49 | 1 | 1 | 0.065 | 0.071 | 0 | | NA | 0 | 0.045 |
| A3SS | 2980 | RPL32 | RPL32 | chr3 | - | 12840141 | 12840341 | 12840141 | 12840242 | 12841493 | 12841582 | 7 | 2 | 0 | 1 | 0 | 1 | 0 | 1 | 0 | 1 | 98 | 49 | 1 | 1 | 0.778 | 1 | 0 | | NA | 0.333 | 0.26 |
| A3SS | 3006 | KIAA0226L | KIAA0226L | chr13 | - | 46371940 | 46372597 | 46371940 | 46372210 | 46387133 | 46387500 | 28 | 7 | 2 | 0 | 0 | 2 | 0 | 2 | 0 | 0 | 98 | 49 | 1 | 1 | 1 | 1 | 0.333 | | NA | 1 | -0.222 |
| A3SS | 3014 | IRF5 | IRF5 | chr7 | + | 128947229 | 128947535 | 128947277 | 128947535 | 128947022 | 128947056 | 1 | 0 | 0 | 1 | 1 | 0 | 0 | 1 | 0 | 1 | 97 | 49 | 1 | 1 | 0.336 | 0 | NA | | NA | 0.336 | -0.168 |
| A3SS | 3025 | ATG4D | ATG4D | chr19 | + | 10544708 | 10544866 | 10544782 | 10544866 | 10543893 | 10544325 | 3 | 2 | 0 | 1 | 0 | 0 | 2 | 1 | 0 | 0 | 98 | 49 | 1 | 1 | 0.6 | 1 | NA | | 1 | 1 | -0.2 |
| A3SS | 3031 | PRKCSH | PRKCSH | chr19 | + | 11447692 | 11447789 | 11447722 | 11447789 | 11447438 | 11447618 | 7 | 1 | 1 | 1 | 1 | 0 | 0 | 1 | 0 | 0 | 79 | 49 | 1 | 1 | 0.813 | 1 | 1 | | NA | 1 | -0.062 |
| A3SS | 3040 | TNPO2 | TNPO2 | chr19 | - | 12702071 | 12702177 | 12702071 | 12702147 | 12702822 | 12702918 | 0 | 0 | 0 | 2 | 0 | 0 | 1 | 0 | 0 | 0 | 79 | 49 | 1 | 1 | 0 | NA | NA | | 1 | NA | -1 |
| A3SS | 3066 | AUP1 | AUP1 | chr2 | - | 74526940 | 74527121 | 74526940 | 74527059 | 74527247 | 74527363 | 7 | 1 | 0 | 6 | 1 | 1 | 0 | 0 | 0 | 3 | 98 | 49 | 1 | 1 | 0.368 | 0.333 | 0 | | NA | 0 | 0.234 |
| A3SS | 3086 | CDV3 | CDV3 | chr3 | + | 133587158 | 133590274 | 133587161 | 133590274 | 133586562 | 133586722 | 11 | 3 | 2 | 3 | 3 | 1 | 0 | 4 | 0 | 1 | 52 | 49 | 1 | 1 | 0.776 | 0.485 | 0.653 | | NA | 0.79 | -0.152 |
| A3SS | 3088 | CDV3 | CDV3 | chr3 | + | 133587158 | 133590274 | 133588352 | 133590274 | 133586562 | 133586722 | 35 | 8 | 22 | 1 | 0 | 0 | 0 | 9 | 0 | 0 | 98 | 49 | 1 | 1 | 0.946 | 1 | 1 | | NA | 1 | -0.018 |
| A3SS | 3090 | CDV3 | CDV3 | chr3 | + | 133587161 | 133590274 | 133588352 | 133590274 | 133586562 | 133586722 | 28 | 8 | 21 | 1 | 0 | 0 | 0 | 9 | 0 | 0 | 98 | 49 | 1 | 1 | 0.933 | 1 | 1 | | NA | 1 | -0.022 |
| A3SS | 3091 | CDV3 | CDV3 | chr3 | + | 133587895 | 133590274 | 133588352 | 133590274 | 133586562 | 133586722 | 36 | 6 | 20 | 1 | 0 | 0 | 0 | 8 | 0 | 0 | 98 | 49 | 1 | 1 | 0.947 | 1 | 1 | | NA | 1 | -0.018 |
| A3SS | 3093 | ZNF493 | ZNF493 | chr19 | + | 21405757 | 21405856 | 21405760 | 21405856 | 21405128 | 21405255 | 0 | 2 | 0 | 0 | 3 | 1 | 0 | 1 | 0 | 0 | 52 | 49 | 1 | 1 | NA | 0.386 | 0 | | NA | 1 | -0.807 |
| A3SS | 3094 | ING4 | ING4 | chr12 | - | 6652661 | 6652770 | 6652661 | 6652767 | 6652935 | 6653050 | 2 | 0 | 0 | 2 | 2 | 0 | 0 | 2 | 0 | 2 | 52 | 49 | 1 | 1 | 0.485 | 0 | NA | | NA | 0.485 | -0.242 |
| A3SS | 3095 | ING4 | ING4 | chr12 | - | 6652661 | 6652770 | 6652661 | 6652767 | 6652944 | 6653050 | 0 | 0 | 0 | 1 | 0 | 2 | 0 | 1 | 0 | 0 | 52 | 49 | 1 | 1 | 0 | NA | 0 | | NA | 1 | -1 |
| A3SS | 3096 | TMEM150B | TMEM150B | chr19 | - | 55320968 | 55321093 | 55320968 | 55321044 | 55322647 | 55322743 | 1 | 0 | 0 | 0 | 0 | 0 | 0 | 0 | 0 | 1 | 98 | 49 | 1 | 1 | 1 | NA | NA | | NA | 0 | 1 |
| A3SS | 3105 | CUTA | CUTA | chr6 | - | 33417480 | 33417816 | 33417480 | 33417695 | 33417958 | 33418288 | 3 | 1 | 0 | 0 | 1 | 0 | 1 | 1 | 0 | 1 | 98 | 49 | 1 | 1 | 1 | 0.333 | NA | NA | 1 | 0.333 | 0 |
| A3SS | 3119 | AZI2 | AZI2 | chr3 | - | 28331688 | 28331954 | 28331688 | 28331950 | 28332368 | 28332427 | 1 | 0 | 0 | 2 | 0 | 0 | 0 | 2 | 0 | 0 | 53 | 49 | 1 | 1 | 0.316 | NA | NA | | NA | 1 | -0.684 |
| A3SS | 3121 | HCST | HCST | chr19 | + | 35904119 | 35904271 | 35904122 | 35904271 | 35903771 | 35903903 | 11 | 2 | 0 | 3 | 0 | 0 | 1 | 5 | 0 | 0 | 52 | 49 | 1 | 1 | 0.776 | 1 | NA | | 1 | 1 | -0.112 |
| A3SS | 3128 | INO80E | INO80E | chr16 | + | 30001211 | 30001530 | 30001413 | 30001530 | 30000928 | 30001040 | 4 | 1 | 1 | 5 | 0 | 0 | 0 | 1 | 0 | 0 | 98 | 49 | 1 | 1 | 0.286 | 1 | 1 | | NA | 1 | -0.238 |
| A3SS | 3140 | DDX60L | DDX60L | chr4 | - | 168395958 | 168396127 | 168395958 | 168396124 | 168400825 | 168400978 | 17 | 31 | 0 | 38 | 0 | 0 | 0 | 9 | 2 | 12 | 52 | 49 | 1 | 1 | 0.297 | 1 | NA | | 0 | 0.414 | 0.442 |
| A3SS | 3147 | TMEM91 | TMEM91 | chr19 | + | 41383560 | 41384082 | 41383714 | 41384082 | 41382771 | 41382921 | 1 | 5 | 1 | 0 | 1 | 0 | 1 | 1 | 0 | 0 | 98 | 49 | 1 | 1 | 1 | 0.714 | 1 | | 1 | 1 | -0.095 |
| A3SS | 3151 | TMEM91 | TMEM91 | chr19 | + | 41383714 | 41384082 | 41383862 | 41384082 | 41382771 | 41382921 | 5 | 1 | 1 | 1 | 4 | 10 | 0 | 2 | 0 | 3 | 98 | 49 | 1 | 1 | 0.714 | 0.111 | 0.048 | | NA | 0.25 | 0.041 |
| A3SS | 3152 | TMEM91 | TMEM91 | chr19 | + | 41383714 | 41384082 | 41383873 | 41384082 | 41382771 | 41382921 | 2 | 1 | 5 | 0 | 3 | 0 | 0 | 2 | 0 | 0 | 98 | 49 | 1 | 1 | 1 | 0.143 | 1 | | NA | 1 | -0.286 |
| A3SS | 3154 | TMEM91 | TMEM91 | chr19 | + | 41383862 | 41384082 | 41383873 | 41384082 | 41382771 | 41382921 | 1 | 4 | 14 | 0 | 3 | 0 | 0 | 3 | 0 | 0 | 60 | 49 | 1 | 1 | 1 | 0.521 | 1 | | NA | 1 | -0.16 |
| A3SS | 3162 | APBB3 | APBB3 | chr5 | - | 140561586 | 140561722 | 140561586 | 140561701 | 140562099 | 140562227 | 3 | 0 | 0 | 2 | 2 | 0 | 0 | 1 | 0 | 0 | 70 | 49 | 1 | 1 | 0.512 | 0 | NA | | NA | 1 | -0.744 |
| A3SS | 3163 | APBB3 | APBB3 | chr5 | - | 140561586 | 140561728 | 140561586 | 140561701 | 140562099 | 140562227 | 5 | 0 | 0 | 2 | 2 | 0 | 0 | 1 | 0 | 0 | 76 | 49 | 1 | 1 | 0.617 | 0 | NA | | NA | 1 | -0.692 |
| A3SS | 3175 | DGCR2 | DGCR2 | chr22 | - | 19064847 | 19065075 | 19064847 | 19065067 | 19068099 | 19068225 | 4 | 0 | 0 | 14 | 8 | 0 | 0 | 3 | 1 | 4 | 57 | 49 | 1 | 1 | 0.197 | 0 | NA | | 0 | 0.392 | -0.098 |
| A3SS | 3190 | AGFG1 | AGFG1 | chr2 | + | 227553703 | 227553795 | 227553709 | 227553795 | 227551958 | 227552117 | 7 | 7 | 4 | 0 | 0 | 0 | 0 | 2 | 0 | 4 | 55 | 49 | 1 | 1 | 1 | 1 | 1 | | NA | 0.308 | 0.692 |
| A3SS | 3194 | RPL18 | RPL18 | chr19 | - | 48617315 | 48617423 | 48617315 | 48617415 | 48617790 | 48617877 | 41 | 10 | 1 | 1 | 0 | 0 | 0 | 7 | 0 | 0 | 57 | 49 | 1 | 1 | 0.972 | 1 | 1 | | NA | 1 | -0.009 |
| A3SS | 3200 | PTOV1 | PTOV1 | chr19 | + | 49860565 | 49860744 | 49860613 | 49860744 | 49860267 | 49860336 | 9 | 4 | 0 | 6 | 1 | 2 | 0 | 3 | 0 | 3 | 97 | 49 | 1 | 1 | 0.431 | 0.669 | 0 | | NA | 0.336 | 0.031 |
| A3SS | 3210 | EIF2AK1 | EIF2AK1 | chr7 | - | 6042932 | 6042993 | 6042932 | 6042990 | 6044561 | 6044661 | 5 | 0 | 0 | 0 | 3 | 0 | 0 | 3 | 0 | 0 | 52 | 49 | 1 | 1 | 1 | 0 | NA | | NA | 1 | -0.5 |
| A3SS | 3213 | ALG13 | ALG13 | chrX | + | 111682131 | 111682294 | 111682235 | 111682294 | 111681169 | 111681424 | 2 | 5 | 0 | 0 | 0 | 0 | 0 | 0 | 2 | 0 | 98 | 49 | 1 | 1 | 1 | 1 | NA | NA | 0 | NA | 1 |
| A3SS | 3216 | LY6E | LY6E | chr8 | + | 143020882 | 143020991 | 143020887 | 143020991 | 143018484 | 143018586 | 1 | 2 | 0 | 0 | 0 | 0 | 0 | 2 | 0 | 1 | 54 | 49 | 1 | 1 | 1 | 1 | NA | | NA | 0.645 | 0.355 |
| A3SS | 3218 | ENTPD1 | ENTPD1 | chr10 | + | 95847445 | 95847706 | 95847568 | 95847706 | 95845356 | 95845596 | 82 | 29 | 11 | 0 | 0 | 0 | 4 | 26 | 0 | 1 | 98 | 49 | 1 | 1 | 1 | 1 | 1 | | 1 | 0.929 | 0.035 |
| A3SS | 3257 | LILRB1 | LILRB1 | chr19 | + | 54633970 | 54634021 | 54633973 | 54634021 | 54633637 | 54633688 | 4 | 1 | 0 | 1 | 0 | 0 | 0 | 0 | 0 | 3 | 52 | 49 | 1 | 1 | 0.79 | 1 | NA | | NA | 0 | 0.895 |
| A3SS | 3319 | EPN1 | EPN1 | chr19 | + | 55692950 | 55693037 | 55692953 | 55693037 | 55692685 | 55692796 | 5 | 2 | 0 | 2 | 0 | 0 | 0 | 1 | 0 | 0 | 52 | 49 | 1 | 1 | 0.702 | 1 | NA | | NA | 1 | -0.149 |
| A3SS | 3337 | ODC1 | ODC1 | chr2 | - | 10444473 | 10444647 | 10444473 | 10444547 | 10444930 | 10445049 | 11 | 0 | 4 | 1 | 0 | 0 | 0 | 2 | 0 | 1 | 98 | 49 | 1 | 1 | 0.846 | NA | 1 | | NA | 0.5 | 0.423 |
| A3SS | 3350 | EDF1 | EDF1 | chr9 | - | 136862113 | 136862496 | 136862113 | 136862345 | 136862905 | 136862999 | 7 | 5 | 0 | 16 | 2 | 3 | 0 | 3 | 0 | 1 | 98 | 49 | 1 | 1 | 0.179 | 0.556 | 0 | | NA | 0.6 | -0.355 |
| A3SS | 3351 | NCOA1 | NCOA1 | chr2 | + | 24768220 | 24770701 | 24768223 | 24770701 | 24762686 | 24762776 | 4 | 6 | 0 | 1 | 0 | 0 | 0 | 2 | 0 | 0 | 52 | 49 | 1 | 1 | 0.79 | 1 | NA | | NA | 1 | -0.105 |
| A3SS | 3352 | NCOA1 | NCOA1 | chr2 | + | 24768220 | 24770701 | 24768223 | 24770701 | 24768078 | 24768135 | 17 | 7 | 0 | 2 | 3 | 0 | 0 | 5 | 0 | 0 | 52 | 49 | 1 | 1 | 0.889 | 0.687 | NA | | NA | 1 | -0.212 |
| A3SS | 3355 | ACIN1 | ACIN1 | chr14 | - | 23064354 | 23064488 | 23064354 | 23064485 | 23065965 | 23066008 | 11 | 2 | 0 | 4 | 2 | 0 | 0 | 2 | 0 | 0 | 52 | 49 | 1 | 1 | 0.722 | 0.485 | NA | | NA | 1 | -0.397 |
| A3SS | 3358 | BRD9 | BRD9 | chr5 | - | 889586 | 889766 | 889586 | 889647 | 891154 | 891287 | 9 | 1 | 0 | 1 | 3 | 0 | 2 | 2 | 0 | 2 | 98 | 49 | 1 | 1 | 0.818 | 0.143 | NA | | 1 | 0.333 | -0.186 |
| A3SS | 3367 | PPM1B | PPM1B | chr2 | + | 44229992 | 44231473 | 44230412 | 44231473 | 44218479 | 44218537 | 5 | 2 | 0 | 3 | 1 | 0 | 0 | 3 | 0 | 2 | 98 | 49 | 1 | 1 | 0.455 | 0.5 | NA | | NA | 0.429 | 0.049 |
| A3SS | 3368 | FBXO7 | FBXO7 | chr22 | + | 32478980 | 32479275 | 32479132 | 32479275 | 32474719 | 32475124 | 8 | 6 | 0 | 0 | 1 | 0 | 1 | 1 | 0 | 0 | 98 | 49 | 1 | 1 | 1 | 0.75 | NA | | 1 | 1 | -0.125 |
| A3SS | 3370 | SH2D2A | SH2D2A | chr1 | - | 156814204 | 156814324 | 156814204 | 156814294 | 156815036 | 156815221 | 1 | 0 | 0 | 2 | 0 | 0 | 0 | 2 | 0 | 1 | 79 | 49 | 1 | 1 | 0.237 | NA | NA | | NA | 0.554 | -0.317 |
| A3SS | 3373 | GNAS | GNAS | chr20 | + | 58903527 | 58903585 | 58903530 | 58903585 | 58898940 | 58898985 | 9 | 8 | 5 | 8 | 5 | 0 | 0 | 2 | 0 | 1 | 52 | 49 | 1 | 1 | 0.515 | 0.601 | 1 | | NA | 0.653 | 0.052 |
| A3SS | 3379 | CCDC88A | CCDC88A | chr2 | - | 55328293 | 55328435 | 55328293 | 55328432 | 55332565 | 55332693 | 1 | 0 | 0 | 2 | 0 | 0 | 0 | 1 | 0 | 1 | 52 | 49 | 1 | 1 | 0.32 | NA | NA | | NA | 0.485 | -0.165 |
| A3SS | 3382 | VPS54 | VPS54 | chr2 | - | 63981645 | 63981887 | 63981645 | 63981851 | 63983863 | 63984019 | 0 | 0 | 0 | 3 | 0 | 0 | 2 | 1 | 0 | 0 | 85 | 49 | 1 | 1 | 0 | NA | NA | | 1 | 1 | -1 |
| A3SS | 3384 | AFTPH | AFTPH | chr2 | + | 64591884 | 64593004 | 64591887 | 64593004 | 64585421 | 64585545 | 5 | 1 | 1 | 2 | 2 | 4 | 0 | 1 | 0 | 0 | 52 | 49 | 1 | 1 | 0.702 | 0.191 | 1 | | NA | 1 | -0.369 |
| A3SS | 3395 | PSMA3-AS1 | PSMA3-AS1 | chr14 | - | 58273978 | 58274135 | 58273978 | 58274010 | 58291634 | 58291707 | 12 | 6 | 4 | 1 | 3 | 1 | 0 | 0 | 0 | 2 | 98 | 49 | 1 | 1 | 0.857 | 0.5 | 0.667 | | NA | 0 | 0.675 |
| A3SS | 3400 | TGOLN2 | TGOLN2 | chr2 | - | 85318017 | 85322758 | 85318017 | 85322741 | 85324914 | 85324998 | 5 | 0 | 0 | 17 | 6 | 0 | 0 | 0 | 0 | 2 | 66 | 49 | 1 | 1 | 0.179 | 0 | NA | | NA | 0 | 0.089 |
| A3SS | 3401 | TGOLN2 | TGOLN2 | chr2 | - | 85318017 | 85322799 | 85318017 | 85322741 | 85324914 | 85324998 | 10 | 3 | 0 | 17 | 6 | 0 | 0 | 0 | 0 | 2 | 98 | 49 | 1 | 1 | 0.227 | 0.2 | NA | | NA | 0 | 0.214 |
| A3SS | 3414 | MAP4K4 | MAP4K4 | chr2 | + | 101859642 | 101859864 | 101859735 | 101859864 | 101858995 | 101859082 | 24 | 14 | 10 | 14 | 4 | 4 | 0 | 4 | 0 | 1 | 98 | 49 | 1 | 1 | 0.462 | 0.636 | 0.556 | | NA | 0.667 | -0.116 |
| A3SS | 3422 | UXS1 | UXS1 | chr2 | - | 106164735 | 106164799 | 106164735 | 106164784 | 106166055 | 106166083 | 5 | 4 | 0 | 4 | 4 | 6 | 0 | 1 | 0 | 0 | 64 | 49 | 1 | 1 | 0.489 | 0.434 | 0 | | NA | 1 | -0.692 |
| A3SS | 3431 | MICU1 | MICU1 | chr10 | - | 72508154 | 72508275 | 72508154 | 72508269 | 72533745 | 72533789 | 4 | 0 | 0 | 7 | 5 | 5 | 0 | 0 | 0 | 2 | 55 | 49 | 1 | 1 | 0.337 | 0 | 0 | | NA | 0 | 0.112 |
| A3SS | 3441 | ITGAL | ITGAL | chr16 | + | 30505243 | 30505300 | 30505246 | 30505300 | 30504174 | 30504264 | 9 | 2 | 7 | 4 | 11 | 0 | 0 | 8 | 0 | 2 | 52 | 49 | 1 | 1 | 0.68 | 0.146 | 1 | | NA | 0.79 | -0.181 |
| A3SS | 3448 | SP3 | SP3 | chr2 | - | 173963760 | 173963883 | 173963760 | 173963874 | 173964404 | 173964553 | 10 | 5 | 1 | 0 | 0 | 0 | 0 | 2 | 0 | 1 | 58 | 49 | 1 | 1 | 1 | 1 | 1 | | NA | 0.628 | 0.372 |
| A3SS | 3450 | ATF2 | ATF2 | chr2 | - | 175072249 | 175074867 | 175072249 | 175074835 | 175080659 | 175080765 | 3 | 0 | 0 | 5 | 2 | 1 | 0 | 1 | 0 | 0 | 81 | 49 | 1 | 1 | 0.266 | 0 | 0 | | NA | 1 | -0.911 |
| A3SS | 3462 | WDR13 | WDR13 | chrX | + | 48598716 | 48598957 | 48598750 | 48598957 | 48597491 | 48598037 | 3 | 4 | 3 | 1 | 0 | 0 | 0 | 3 | 0 | 0 | 83 | 49 | 1 | 1 | 0.639 | 1 | 1 | | NA | 1 | -0.12 |
| A3SS | 3469 | DCTN3 | DCTN3 | chr9 | - | 34614041 | 34614132 | 34614041 | 34614101 | 34614709 | 34614768 | 0 | 5 | 0 | 3 | 4 | 1 | 1 | 0 | 0 | 0 | 80 | 49 | 1 | 1 | 0 | 0.434 | 0 | | 1 | NA | -0.855 |
| A3SS | 3470 | DCTN3 | DCTN3 | chr9 | - | 34614041 | 34614198 | 34614041 | 34614101 | 34614709 | 34614768 | 0 | 5 | 0 | 3 | 4 | 1 | 1 | 0 | 0 | 0 | 98 | 49 | 1 | 1 | 0 | 0.385 | 0 | | 1 | NA | -0.872 |
| A3SS | 3485 | SPATA20 | SPATA20 | chr17 | + | 50547664 | 50547767 | 50547719 | 50547767 | 50547088 | 50547285 | 0 | 0 | 0 | 2 | 0 | 0 | 0 | 1 | 0 | 0 | 98 | 49 | 1 | 1 | 0 | NA | NA | | NA | 1 | -1 |
| A3SS | 3513 | SERPINB1 | SERPINB1 | chr6 | - | 2835855 | 2836023 | 2835855 | 2836019 | 2836107 | 2836250 | 58 | 34 | 28 | 4 | 1 | 2 | 0 | 15 | 0 | 3 | 53 | 49 | 1 | 1 | 0.931 | 0.969 | 0.928 | | NA | 0.822 | 0.121 |
| A3SS | 3514 | USP19 | USP19 | chr3 | - | 49115723 | 49115944 | 49115723 | 49115938 | 49116046 | 49116162 | 3 | 1 | 0 | 1 | 3 | 0 | 0 | 1 | 0 | 0 | 55 | 49 | 1 | 1 | 0.728 | 0.229 | NA | | NA | 1 | -0.522 |
| A3SS | 3515 | RFC1 | RFC1 | chr4 | - | 39306591 | 39306704 | 39306591 | 39306701 | 39308635 | 39309032 | 1 | 0 | 0 | 1 | 2 | 0 | 0 | 0 | 0 | 2 | 52 | 49 | 1 | 1 | 0.485 | 0 | NA | | NA | 0 | 0.242 |
| A3SS | 3524 | NSFL1C | NSFL1C | chr20 | - | 1454966 | 1455132 | 1454966 | 1455037 | 1458199 | 1458274 | 19 | 10 | 12 | 2 | 0 | 0 | 0 | 6 | 0 | 1 | 98 | 49 | 1 | 1 | 0.826 | 1 | 1 | | NA | 0.75 | 0.192 |
| A3SS | 3566 | HCK | HCK | chr20 | + | 32073715 | 32073818 | 32073718 | 32073818 | 32073318 | 32073361 | 45 | 35 | 39 | 12 | 4 | 8 | 0 | 6 | 0 | 0 | 52 | 49 | 1 | 1 | 0.779 | 0.892 | 0.821 | | NA | 1 | -0.169 |
| A3SS | 3575 | RALGAPB | RALGAPB | chr20 | + | 38565355 | 38565478 | 38565358 | 38565478 | 38562531 | 38562697 | 1 | 2 | 1 | 0 | 0 | 0 | 0 | 0 | 0 | 2 | 52 | 49 | 1 | 1 | 1 | 1 | 1 | | NA | 0 | 1 |
| A3SS | 3586 | SH3BP5 | SH3BP5 | chr3 | - | 15304102 | 15304299 | 15304102 | 15304231 | 15330503 | 15330566 | 3 | 2 | 0 | 7 | 2 | 0 | 0 | 2 | 0 | 2 | 98 | 49 | 1 | 1 | 0.176 | 0.333 | NA | | NA | 0.333 | -0.079 |
| A3SS | 3587 | ELMO2 | ELMO2 | chr20 | - | 46394404 | 46394532 | 46394404 | 46394503 | 46398696 | 46398771 | 5 | 2 | 0 | 1 | 2 | 0 | 0 | 0 | 0 | 1 | 78 | 49 | 1 | 1 | 0.759 | 0.386 | NA | | NA | 0 | 0.573 |
| A3SS | 3591 | NCOA3 | NCOA3 | chr20 | + | 47634017 | 47634195 | 47634047 | 47634195 | 47633495 | 47633636 | 11 | 3 | 0 | 12 | 2 | 1 | 0 | 0 | 0 | 1 | 79 | 49 | 1 | 1 | 0.362 | 0.482 | 0 | | NA | 0 | 0.281 |
| A3SS | 3592 | NCOA3 | NCOA3 | chr20 | + | 47639576 | 47639822 | 47639621 | 47639822 | 47639007 | 47639202 | 15 | 14 | 0 | 4 | 1 | 0 | 0 | 11 | 0 | 0 | 94 | 49 | 1 | 1 | 0.662 | 0.879 | NA | | NA | 1 | -0.23 |
| A3SS | 3596 | STAU1 | STAU1 | chr20 | - | 49124374 | 49124605 | 49124374 | 49124587 | 49135832 | 49135931 | 4 | 1 | 0 | 8 | 1 | 1 | 0 | 3 | 0 | 0 | 67 | 49 | 1 | 1 | 0.268 | 0.422 | 0 | | NA | 1 | -0.77 |
| A3SS | 3603 | ZBP1 | ZBP1 | chr20 | - | 57613162 | 57613330 | 57613162 | 57613327 | 57614886 | 57615060 | 27 | 4 | 5 | 0 | 0 | 0 | 0 | 23 | 0 | 2 | 52 | 49 | 1 | 1 | 1 | 1 | 1 | | NA | 0.916 | 0.084 |
| A3SS | 3606 | OSBPL2 | OSBPL2 | chr20 | + | 62259980 | 62260125 | 62260016 | 62260125 | 62256056 | 62256221 | 43 | 24 | 12 | 1 | 0 | 0 | 2 | 13 | 0 | 0 | 85 | 49 | 1 | 1 | 0.961 | 1 | 1 | | 1 | 1 | -0.013 |
| A3SS | 3608 | PPP1R12C | PPP1R12C | chr19 | - | 55092782 | 55092868 | 55092782 | 55092865 | 55093015 | 55093076 | 2 | 2 | 0 | 8 | 5 | 0 | 0 | 0 | 0 | 1 | 52 | 49 | 1 | 1 | 0.191 | 0.274 | NA | | NA | 0 | 0.233 |
| A3SS | 3609 | PPP1R12C | PPP1R12C | chr19 | - | 55095444 | 55095603 | 55095444 | 55095600 | 55095866 | 55095940 | 7 | 0 | 0 | 6 | 2 | 2 | 0 | 1 | 0 | 0 | 52 | 49 | 1 | 1 | 0.524 | 0 | 0 | | NA | 1 | -0.825 |
| A3SS | 3623 | PCMTD2 | PCMTD2 | chr20 | + | 64267886 | 64268010 | 64267967 | 64268010 | 64265257 | 64265429 | 14 | 0 | 0 | 4 | 3 | 0 | 0 | 1 | 0 | 0 | 52 | 49 | 1 | 1 | 0.7 | NA | 1 | | NA | 1 | -0.15 |
| A3SS | 3649 | POFUT2 | POFUT2 | chr21 | - | 45263927 | 45267713 | 45263927 | 45265635 | 45269838 | 45270019 | 8 | 6 | 0 | 1 | 0 | 0 | 0 | 3 | 0 | 0 | 98 | 49 | 1 | 1 | 0.8 | 1 | NA | | NA | 1 | -0.1 |
| A3SS | 3684 | GUCD1 | GUCD1 | chr22 | - | 24543841 | 24544083 | 24543841 | 24543954 | 24546913 | 24547005 | 23 | 17 | 7 | 2 | 0 | 0 | 2 | 9 | 0 | 0 | 98 | 49 | 1 | 1 | 0.852 | 1 | 1 | | 1 | 1 | -0.049 |
| A3SS | 3706 | SIGLEC10 | SIGLEC10 | chr19 | - | 51416665 | 51416950 | 51416665 | 51416806 | 51417081 | 51417465 | 33 | 12 | 5 | 2 | 0 | 0 | 0 | 2 | 0 | 0 | 98 | 49 | 1 | 1 | 0.892 | 1 | 1 | | NA | 1 | -0.036 |
| A3SS | 3711 | TOM1 | TOM1 | chr22 | + | 35322998 | 35323177 | 35323027 | 35323177 | 35317876 | 35317961 | 7 | 4 | 0 | 1 | 2 | 0 | 1 | 5 | 0 | 1 | 78 | 49 | 1 | 1 | 0.815 | 0.557 | NA | | 1 | 0.759 | -0.194 |
| A3SS | 3717 | APOL3 | APOL3 | chr22 | - | 36149045 | 36149371 | 36149045 | 36149142 | 36160668 | 36160929 | 2 | 0 | 0 | 9 | 0 | 0 | 1 | 0 | 0 | 0 | 98 | 49 | 1 | 1 | 0.1 | NA | NA | | 1 | NA | -0.9 |
| A3SS | 3731 | PPP6R2 | PPP6R2 | chr22 | + | 50419345 | 50419462 | 50419348 | 50419462 | 50418866 | 50418979 | 0 | 0 | 0 | 3 | 2 | 0 | 1 | 0 | 0 | 1 | 52 | 49 | 1 | 1 | 0 | 0 | NA | | 1 | 0 | -0.5 |
| A3SS | 3732 | PPP6R2 | PPP6R2 | chr22 | + | 50439697 | 50439857 | 50439700 | 50439857 | 50438598 | 50438762 | 1 | 0 | 0 | 3 | 3 | 0 | 0 | 1 | 0 | 2 | 52 | 49 | 1 | 1 | 0.239 | 0 | NA | | NA | 0.32 | -0.201 |
| A3SS | 3756 | SLC6A6 | SLC6A6 | chr3 | + | 14443623 | 14443863 | 14443627 | 14443863 | 14416411 | 14416453 | 106 | 28 | 36 | 6 | 0 | 0 | 0 | 19 | 0 | 1 | 53 | 49 | 1 | 1 | 0.942 | 1 | 1 | | NA | 0.946 | 0.035 |
| A3SS | 3763 | PDCD6IP | PDCD6IP | chr3 | + | 33828837 | 33828969 | 33828852 | 33828969 | 33826479 | 33826580 | 3 | 1 | 0 | 1 | 0 | 0 | 0 | 2 | 0 | 0 | 64 | 49 | 1 | 1 | 0.697 | 1 | NA | | NA | 1 | -0.151 |
| A3SS | 3777 | CTNNB1 | CTNNB1 | chr3 | + | 41239659 | 41240448 | 41239818 | 41240448 | 41239133 | 41239354 | 18 | 10 | 6 | 1 | 0 | 1 | 0 | 11 | 0 | 0 | 98 | 49 | 1 | 1 | 0.9 | 1 | 0.75 | | NA | 1 | -0.117 |
| A3SS | 3797 | NCOR1 | NCOR1 | chr17 | - | 16101249 | 16101805 | 16101249 | 16101757 | 16108785 | 16108912 | 29 | 5 | 2 | 1 | 0 | 0 | 1 | 2 | 0 | 0 | 97 | 49 | 1 | 1 | 0.936 | 1 | 1 | | 1 | 1 | -0.021 |
| A3SS | 3809 | ENY2 | ENY2 | chr8 | + | 109340443 | 109340563 | 109340488 | 109340563 | 109339319 | 109339390 | 0 | 1 | 0 | 6 | 7 | 3 | 1 | 1 | 0 | 1 | 94 | 49 | 1 | 1 | 0 | 0.069 | 0 | | 1 | 0.343 | -0.648 |
| A3SS | 3810 | ENY2 | ENY2 | chr8 | + | 109340443 | 109340563 | 109340490 | 109340563 | 109339319 | 109339390 | 0 | 2 | 0 | 3 | 0 | 0 | 1 | 2 | 0 | 0 | 96 | 49 | 1 | 1 | 0 | 1 | NA | | 1 | 1 | -0.5 |
| A3SS | 3811 | ENY2 | ENY2 | chr8 | + | 109340488 | 109340563 | 109340490 | 109340563 | 109339319 | 109339390 | 6 | 8 | 3 | 3 | 0 | 0 | 0 | 2 | 0 | 0 | 51 | 49 | 1 | 1 | 0.658 | 1 | 1 | | NA | 1 | -0.114 |
| A3SS | 3815 | FAM208A | FAM208A | chr3 | - | 56633043 | 56633966 | 56633043 | 56633843 | 56638705 | 56638765 | 9 | 0 | 0 | 3 | 0 | 0 | 0 | 2 | 0 | 0 | 98 | 49 | 1 | 1 | 0.6 | NA | NA | | NA | 1 | -0.4 |
| A3SS | 3829 | CEP97 | CEP97 | chr3 | + | 101757633 | 101758423 | 101757810 | 101758423 | 101757062 | 101757196 | 1 | 1 | 0 | 0 | 1 | 0 | 0 | 1 | 0 | 0 | 98 | 49 | 1 | 1 | 1 | 0.333 | NA | | NA | 1 | -0.334 |
| A3SS | 3840 | NAA50 | NAA50 | chr3 | - | 113723958 | 113724095 | 113723958 | 113724092 | 113745941 | 113746299 | 14 | 12 | 1 | 4 | 3 | 3 | 0 | 2 | 0 | 3 | 52 | 49 | 1 | 1 | 0.767 | 0.79 | 0.239 | | NA | 0.386 | 0.213 |
| A3SS | 3845 | SYNE2 | SYNE2 | chr14 | + | 64221533 | 64221704 | 64221575 | 64221704 | 64220436 | 64220637 | 2 | 1 | 0 | 4 | 0 | 0 | 0 | 2 | 0 | 0 | 91 | 49 | 1 | 1 | 0.212 | 1 | NA | | NA | 1 | -0.394 |
| A3SS | 3853 | DGKZ | DGKZ | chr11 | + | 46379453 | 46379568 | 46379468 | 46379568 | 46379202 | 46379236 | 0 | 1 | 0 | 1 | 0 | 0 | 0 | 0 | 0 | 2 | 64 | 49 | 1 | 1 | 0 | 1 | NA | | NA | 0 | 0.5 |
| A3SS | 3854 | APRT | APRT | chr16 | - | 88809468 | 88809840 | 88809468 | 88809706 | 88810069 | 88810148 | 10 | 1 | 0 | 1 | 0 | 0 | 0 | 3 | 0 | 0 | 98 | 49 | 1 | 1 | 0.833 | 1 | NA | | NA | 1 | -0.084 |
| A3SS | 3857 | RYK | RYK | chr3 | - | 134191848 | 134191983 | 134191848 | 134191974 | 134195081 | 134195182 | 2 | 2 | 0 | 2 | 0 | 3 | 0 | 1 | 0 | 0 | 58 | 49 | 1 | 1 | 0.458 | 1 | 0 | | NA | 1 | -0.514 |
| A3SS | 3864 | CLINT1 | CLINT1 | chr5 | - | 157789362 | 157789567 | 157789362 | 157789513 | 157791702 | 157791995 | 1 | 0 | 0 | 3 | 0 | 0 | 0 | 2 | 0 | 3 | 98 | 49 | 1 | 1 | 0.143 | NA | NA | | NA | 0.25 | -0.107 |
| A3SS | 3865 | ARMC8 | ARMC8 | chr3 | + | 138188400 | 138188530 | 138188454 | 138188530 | 138187247 | 138187599 | 1 | 0 | 0 | 0 | 0 | 0 | 0 | 1 | 0 | 2 | 98 | 49 | 1 | 1 | 1 | NA | NA | | NA | 0.2 | 0.8 |
| A3SS | 3872 | MFSD1 | MFSD1 | chr3 | + | 158804318 | 158804371 | 158804322 | 158804371 | 158801925 | 158802315 | 15 | 1 | 0 | 1 | 1 | 0 | 0 | 5 | 0 | 0 | 53 | 49 | 1 | 1 | 0.933 | 0.48 | NA | | NA | 1 | -0.293 |
| A3SS | 3877 | ZNF124 | ZNF124 | chr1 | - | 247154988 | 247157403 | 247154988 | 247157217 | 247159005 | 247159066 | 3 | 3 | 0 | 2 | 0 | 0 | 0 | 0 | 0 | 2 | 98 | 49 | 1 | 1 | 0.429 | 1 | NA | | NA | 0 | 0.715 |
| A3SS | 3880 | TOR1AIP1 | TOR1AIP1 | chr1 | + | 179889309 | 179889369 | 179889312 | 179889369 | 179884691 | 179884769 | 1 | 1 | 0 | 5 | 1 | 0 | 0 | 0 | 0 | 3 | 52 | 49 | 1 | 1 | 0.159 | 0.485 | NA | | NA | 0 | 0.322 |
| A3SS | 3891 | EIF4G1 | EIF4G1 | chr3 | + | 184316071 | 184316218 | 184316131 | 184316218 | 184315762 | 184315856 | 6 | 0 | 0 | 1 | 0 | 0 | 0 | 2 | 0 | 0 | 98 | 49 | 1 | 1 | 0.75 | NA | NA | | NA | 1 | -0.25 |
| A3SS | 3892 | EIF4G1 | EIF4G1 | chr3 | + | 184323404 | 184323593 | 184323407 | 184323593 | 184323082 | 184323241 | 5 | 1 | 0 | 13 | 0 | 0 | 0 | 2 | 0 | 3 | 52 | 49 | 1 | 1 | 0.266 | 1 | NA | | NA | 0.386 | 0.247 |
| A3SS | 3906 | KDM5C | KDM5C | chrX | - | 53217142 | 53217277 | 53217142 | 53217274 | 53217795 | 53217966 | 13 | 6 | 3 | 4 | 0 | 0 | 0 | 5 | 0 | 0 | 52 | 49 | 1 | 1 | 0.754 | 1 | 1 | | NA | 1 | -0.082 |
| A3SS | 3914 | MFSD7 | MFSD7 | chr4 | - | 684482 | 684602 | 684482 | 684599 | 684718 | 684856 | 5 | 0 | 0 | 1 | 3 | 0 | 2 | 0 | 0 | 2 | 52 | 49 | 1 | 1 | 0.825 | 0 | NA | | 1 | 0 | -0.088 |
| A3SS | 3918 | TMEM175 | TMEM175 | chr4 | + | 951645 | 951717 | 951681 | 951717 | 951206 | 951258 | 2 | 2 | 0 | 10 | 0 | 3 | 0 | 1 | 0 | 1 | 85 | 49 | 1 | 1 | 0.103 | 1 | 0 | | NA | 0.366 | 0.002 |
| A3SS | 3929 | FBXL5 | FBXL5 | chr4 | - | 15636493 | 15636676 | 15636493 | 15636673 | 15638507 | 15638694 | 170 | 70 | 42 | 1 | 0 | 0 | 3 | 40 | 0 | 0 | 52 | 49 | 1 | 1 | 0.994 | 1 | 1 | | 1 | 1 | -0.002 |
| A3SS | 3951 | CXCL1 | CXCL1 | chr4 | + | 73869645 | 73869792 | 73869668 | 73869792 | 73869391 | 73869570 | 12 | 12 | 4 | 12 | 14 | 1 | 0 | 0 | 0 | 3 | 72 | 49 | 1 | 1 | 0.405 | 0.368 | 0.731 | | NA | 0 | 0.501 |
| A3SS | 3953 | OGT | OGT | chrX | + | 71536177 | 71536358 | 71536207 | 71536358 | 71533061 | 71533336 | 36 | 12 | 3 | 2 | 0 | 0 | 3 | 4 | 0 | 0 | 79 | 49 | 1 | 1 | 0.918 | 1 | 1 | | 1 | 1 | -0.027 |
| A3SS | 3962 | NFKB1 | NFKB1 | chr4 | + | 102533844 | 102533885 | 102533847 | 102533885 | 102529835 | 102529914 | 2 | 0 | 0 | 2 | 2 | 2 | 0 | 0 | 0 | 1 | 52 | 49 | 1 | 1 | 0.485 | 0 | 0 | | NA | 0 | 0.162 |
| A3SS | 3975 | ELF2 | ELF2 | chr4 | - | 139071865 | 139072039 | 139071865 | 139072003 | 139073453 | 139073567 | 16 | 8 | 6 | 34 | 11 | 7 | 0 | 4 | 0 | 0 | 85 | 49 | 1 | 1 | 0.213 | 0.295 | 0.331 | | NA | 1 | -0.72 |
| A3SS | 3980 | KIAA0922 | KIAA0922 | chr4 | + | 153584834 | 153584931 | 153584837 | 153584931 | 153583563 | 153583672 | 15 | 0 | 2 | 6 | 1 | 0 | 0 | 2 | 0 | 3 | 52 | 49 | 1 | 1 | 0.702 | 0 | 1 | | NA | 0.386 | 0.181 |

| | | | | | | | | | | | | | | | | | | | | | | | | | | | | | | | |
|---|---|---|---|---|---|---|---|---|---|---|---|---|---|---|---|---|---|---|---|---|---|---|---|---|---|---|---|---|---|---|---|
| A3SS | 3989 | RWDD4 | RWDD4 | chr4 | - | 183651217 | 183651327 | 183651217 | 183651323 | 183655880 | 183655961 | 0 | 0 | 0 | 0 | 2 | 0 | 0 | 1 | 0 | 0 | 53 | 49 | 1 | 1 | NA | 0 | NA | NA | 1 | -1 |
| A3SS | 3994 | CMTM3 | CMTM3 | chr16 | + | 66609758 | 66610003 | 66609882 | 66610003 | 66609434 | 66609530 | 0 | 0 | 0 | 6 | 0 | 2 | 0 | 1 | 0 | 1 | 98 | 49 | 1 | 1 | 0 | NA | 0 | NA | 0.333 | -0.333 |
| A3SS | 4007 | ATL2 | ATL2 | chr2 | - | 38293956 | 38296590 | 38293956 | 38296113 | 38298143 | 38298575 | 2 | 0 | 0 | 8 | 1 | 3 | 0 | 1 | 0 | 0 | 98 | 49 | 1 | 1 | 0.111 | 0 | 0 | NA | 1 | -0.963 |
| A3SS | 4025 | RICTOR | RICTOR | chr5 | - | 38945490 | 38945724 | 38945490 | 38945610 | 38947263 | 38947441 | 90 | 27 | 9 | 0 | 2 | 0 | 1 | 14 | 0 | 0 | 98 | 49 | 1 | 1 | 1 | 0.871 | 1 | 1 | 1 | -0.043 |
| A3SS | 4038 | SON | SON | chr21 | + | 33576364 | 33577514 | 33576417 | 33577514 | 33575777 | 33575893 | 12 | 4 | 1 | 7 | 1 | 1 | 0 | 5 | 0 | 0 | 98 | 49 | 1 | 1 | 0.462 | 0.667 | 0.333 | NA | 1 | -0.513 |
| A3SS | 4039 | GAPT | GAPT | chr5 | + | 58493618 | 58493838 | 58493710 | 58493838 | 58491434 | 58491541 | 3 | 0 | 0 | 2 | 0 | 0 | 0 | 1 | 0 | 0 | 98 | 49 | 1 | 1 | 0.429 | NA | NA | NA | 1 | -0.571 |
| A3SS | 4040 | GAPT | GAPT | chr5 | + | 58494147 | 58496358 | 58494246 | 58496358 | 58493710 | 58493838 | 8 | 5 | 2 | 7 | 1 | 0 | 0 | 1 | 0 | 2 | 98 | 49 | 1 | 1 | 0.364 | 0.714 | 1 | NA | 0.2 | 0.493 |
| A3SS | 4047 | ZFAND5 | ZFAND5 | chr9 | - | 72360627 | 72360787 | 72360627 | 72360784 | 72363469 | 72363606 | 13 | 4 | 1 | 1 | 0 | 0 | 0 | 3 | 0 | 0 | 52 | 49 | 1 | 1 | 0.925 | 1 | 1 | NA | 1 | -0.025 |
| A3SS | 4049 | AGTPBP1 | AGTPBP1 | chr9 | - | 85655142 | 85655320 | 85655142 | 85655200 | 85657434 | 85657643 | 47 | 28 | 17 | 25 | 13 | 0 | 0 | 20 | 0 | 4 | 98 | 49 | 1 | 1 | 0.485 | 0.519 | 1 | NA | 0.714 | -0.046 |
| A3SS | 4054 | GLRX | GLRX | chr5 | - | 95813848 | 95814807 | 95813848 | 95814389 | 95816506 | 95816626 | 1 | 0 | 0 | 10 | 3 | 1 | 0 | 0 | 0 | 3 | 98 | 49 | 1 | 1 | 0.048 | 0 | 0 | NA | 0 | 0.016 |
| A3SS | 4055 | GLRX | GLRX | chr5 | - | 95813848 | 95814957 | 95813848 | 95814389 | 95816506 | 95816626 | 3 | 0 | 0 | 10 | 3 | 1 | 0 | 0 | 0 | 3 | 98 | 49 | 1 | 1 | 0.13 | 0 | 0 | NA | 0 | 0.043 |
| A3SS | 4058 | CAST | CAST | chr5 | + | 96765220 | 96765325 | 96765223 | 96765325 | 96762273 | 96762372 | 19 | 3 | 1 | 2 | 0 | 0 | 0 | 8 | 0 | 3 | 52 | 49 | 1 | 1 | 0.9 | 1 | 1 | NA | 0.715 | 0.252 |
| A3SS | 4059 | CUX1 | CUX1 | chr7 | + | 102178468 | 102178657 | 102178474 | 102178657 | 102170445 | 102170550 | 9 | 7 | 2 | 2 | 0 | 0 | 0 | 5 | 0 | 0 | 55 | 49 | 1 | 1 | 0.8 | 1 | 1 | NA | 1 | -0.067 |
| A3SS | 4078 | CNBP | CNBP | chr3 | - | 129171078 | 129171283 | 129171078 | 129171280 | 129171445 | 129171538 | 0 | 0 | 2 | 12 | 7 | 6 | 0 | 0 | 0 | 2 | 52 | 49 | 1 | 1 | 0 | 0 | 0.239 | NA | 0 | 0.08 |
| A3SS | 4080 | H2AFY | H2AFY | chr5 | - | 135360496 | 135360607 | 135360496 | 135360604 | 135369405 | 135369603 | 18 | 4 | 7 | 14 | 5 | 10 | 0 | 3 | 0 | 5 | 52 | 49 | 1 | 1 | 0.548 | 0.43 | 0.397 | NA | 0.361 | 0.097 |
| A3SS | 4091 | SRA1 | SRA1 | chr5 | - | 140557146 | 140557332 | 140557146 | 140557272 | 140557427 | 140558093 | 9 | 2 | 0 | 0 | 0 | 0 | 0 | 1 | 0 | 1 | 98 | 49 | 1 | 1 | 1 | 1 | NA | NA | 0.333 | 0.667 |
| A3SS | 4093 | HARS2 | HARS2 | chr5 | + | 140693490 | 140693665 | 140693590 | 140693665 | 140691425 | 140691756 | 4 | 5 | 0 | 9 | 0 | 0 | 0 | 2 | 0 | 0 | 98 | 49 | 1 | 1 | 0.182 | 1 | NA | NA | 1 | -0.409 |
| A3SS | 4096 | SCARF1 | SCARF1 | chr17 | - | 1636708 | 1636855 | 1636708 | 1636818 | 1636940 | 1637062 | 13 | 19 | 3 | 2 | 0 | 0 | 0 | 2 | 0 | 0 | 86 | 49 | 1 | 1 | 0.787 | 1 | 1 | NA | 1 | -0.071 |
| A3SS | 4097 | SCARF1 | SCARF1 | chr17 | - | 1639911 | 1640040 | 1639911 | 1639970 | 1640447 | 1640666 | 4 | 10 | 0 | 0 | 1 | 0 | 0 | 2 | 0 | 0 | 98 | 49 | 1 | 1 | 1 | 0.833 | NA | NA | 1 | -0.084 |
| A3SS | 4104 | RNF111 | RNF111 | chr15 | + | 59092512 | 59092640 | 59092536 | 59092640 | 59091058 | 59091154 | 3 | 0 | 0 | 7 | 0 | 2 | 0 | 2 | 0 | 0 | 73 | 49 | 1 | 1 | 0.223 | NA | 0 | NA | 1 | -0.888 |
| A3SS | 4118 | PHYKPL | PHYKPL | chr5 | - | 178222354 | 178222580 | 178222354 | 178222492 | 178222851 | 178222934 | 9 | 0 | 0 | 3 | 0 | 0 | 0 | 5 | 0 | 0 | 98 | 49 | 1 | 1 | 0.6 | NA | NA | NA | 1 | -0.4 |
| A5SS | 5 | DNASE1L1 | DNASE1L1 | chrX | - | 154408557 | 154409278 | 154409111 | 154409278 | 154405433 | 154405655 | 0 | 0 | 0 | 3 | 0 | 0 | 0 | 1 | 0 | 0 | 98 | 49 | 1 | 1 | 0 | NA | NA | NA | 1 | -1 |
| A5SS | 24 | PSMD10 | PSMD10 | chrX | - | 108087685 | 108087852 | 108087764 | 108087852 | 108084204 | 108085127 | 4 | 0 | 0 | 0 | 0 | 0 | 0 | 0 | 0 | 2 | 98 | 49 | 1 | 1 | 1 | NA | NA | NA | 0 | 1 |
| A5SS | 40 | TAF1 | TAF1 | chrX | + | 71423116 | 71423264 | 71423116 | 71423239 | 71423973 | 71424066 | 0 | 1 | 0 | 0 | 0 | 0 | 0 | 0 | 0 | 2 | 74 | 49 | 1 | 1 | NA | 1 | NA | NA | 0 | 1 |
| A5SS | 64 | OTUD5 | OTUD5 | chrX | - | 48934695 | 48934867 | 48934710 | 48934867 | 48934463 | 48934612 | 1 | 0 | 0 | 11 | 7 | 8 | 0 | 3 | 0 | 2 | 64 | 49 | 1 | 1 | 0.065 | 0 | 0 | NA | 0.535 | -0.513 |
| A5SS | 66 | USP9X | USP9X | chrX | + | 41229566 | 41229827 | 41229566 | 41229779 | 41230500 | 41230596 | 13 | 1 | 1 | 17 | 2 | 0 | 0 | 0 | 0 | 1 | 97 | 49 | 1 | 1 | 0.279 | 0.202 | 1 | NA | 0 | 0.494 |
| A5SS | 73 | TRAPPC2 | TRAPPC2 | chrX | - | 13734043 | 13734185 | 13734049 | 13734185 | 13719870 | 13719982 | 2 | 3 | 1 | 0 | 2 | 0 | 0 | 2 | 0 | 0 | 55 | 49 | 1 | 1 | 1 | 0.572 | 1 | NA | 1 | -0.143 |
| A5SS | 95 | ODF2 | ODF2 | chr9 | + | 128460941 | 128461067 | 128460941 | 128461010 | 128469182 | 128469353 | 3 | 0 | 0 | 1 | 0 | 0 | 0 | 0 | 1 | 0 | 98 | 49 | 1 | 1 | 0.6 | NA | NA | 0 | NA | 0.6 |
| A5SS | 97 | SLC25A25 | SLC25A25 | chr9 | + | 128098481 | 128098768 | 128098481 | 128098725 | 128101095 | 128101222 | 0 | 0 | 0 | 2 | 0 | 0 | 0 | 1 | 0 | 0 | 92 | 49 | 1 | 1 | 0 | NA | NA | NA | 1 | -1 |
| A5SS | 115 | CTSL | CTSL | chr9 | + | 87726058 | 87726308 | 87726058 | 87726253 | 87727593 | 87727729 | 1 | 0 | 0 | 0 | 0 | 0 | 0 | 0 | 0 | 2 | 98 | 49 | 1 | 1 | 1 | NA | NA | NA | 0 | 1 |
| A5SS | 133 | ZCCHC6 | ZCCHC6 | chr9 | - | 86354270 | 86354487 | 86354380 | 86354487 | 86352679 | 86353230 | 90 | 62 | 15 | 0 | 2 | 0 | 3 | 11 | 0 | 0 | 98 | 49 | 1 | 1 | 1 | 0.939 | 1 | 1 | 1 | -0.02 |
| A5SS | 151 | GRINA | GRINA | chr8 | + | 143990057 | 143990311 | 143990057 | 143990199 | 143991199 | 143991602 | 1 | 1 | 7 | 3 | 0 | 0 | 1 | 0 | 0 | 0 | 98 | 49 | 1 | 1 | 0.143 | 1 | 1 | 1 | NA | -0.286 |
| A5SS | 156 | FAM49B | FAM49B | chr8 | - | 129896671 | 129896854 | 129896748 | 129896854 | 129879388 | 129879471 | 23 | 11 | 0 | 0 | 1 | 2 | 0 | 9 | 0 | 0 | 98 | 49 | 1 | 1 | 1 | 0.846 | 0 | NA | 1 | -0.385 |
| A5SS | 173 | CHCHD7 | CHCHD7 | chr8 | + | 56212761 | 56212909 | 56212761 | 56212889 | 56214597 | 56214667 | 2 | 0 | 0 | 0 | 0 | 0 | 0 | 0 | 0 | 1 | 69 | 49 | 1 | 1 | 1 | NA | NA | NA | 0 | 1 |
| A5SS | 191 | NUB1 | NUB1 | chr7 | + | 151374096 | 151374243 | 151374096 | 151374201 | 151375847 | 151375943 | 3 | 4 | 3 | 5 | 1 | 0 | 0 | 3 | 0 | 1 | 91 | 49 | 1 | 1 | 0.244 | 0.683 | 1 | NA | 0.618 | 0.024 |
| A5SS | 192 | FASTK | FASTK | chr7 | - | 151078841 | 151079021 | 151078922 | 151079021 | 151078561 | 151078701 | 3 | 0 | 1 | 1 | 0 | 0 | 0 | 1 | 0 | 0 | 98 | 49 | 1 | 1 | 0.6 | NA | 1 | NA | 1 | -0.2 |
| A5SS | 204 | TRIM24 | TRIM24 | chr7 | + | 138554697 | 138554966 | 138554697 | 138554864 | 138567480 | 138567654 | 9 | 1 | 1 | 1 | 0 | 0 | 0 | 2 | 0 | 0 | 98 | 49 | 1 | 1 | 0.818 | 1 | 1 | NA | 1 | -0.061 |
| A5SS | 207 | C7orf49 | C7orf49 | chr7 | - | 135168748 | 135169060 | 135168785 | 135169060 | 135167731 | 135167807 | 18 | 11 | 3 | 0 | 1 | 0 | 0 | 0 | 0 | 1 | 86 | 49 | 1 | 1 | 1 | 0.862 | 1 | NA | 0 | 0.954 |
| A5SS | 236 | AKAP9 | AKAP9 | chr7 | + | 92070029 | 92070206 | 92070029 | 92070182 | 92070904 | 92071009 | 4 | 0 | 0 | 2 | 0 | 0 | 2 | 0 | 0 | 0 | 73 | 49 | 1 | 1 | 0.573 | NA | NA | 1 | NA | -0.427 |
| A5SS | 247 | IKZF1 | IKZF1 | chr7 | + | 50391728 | 50391863 | 50391728 | 50391833 | 50399917 | 50405100 | 3 | 3 | 0 | 2 | 0 | 2 | 0 | 2 | 0 | 3 | 79 | 49 | 1 | 1 | 0.482 | 1 | 0 | NA | 0.293 | 0.201 |
| A5SS | 262 | WIPI2 | WIPI2 | chr7 | + | 5193117 | 5193345 | 5193117 | 5193171 | 5199575 | 5199658 | 7 | 3 | 0 | 0 | 1 | 0 | 0 | 0 | 0 | 2 | 98 | 49 | 1 | 1 | 1 | 0.6 | NA | NA | 0 | 0.8 |
| A5SS | 263 | WIPI2 | WIPI2 | chr7 | + | 5229607 | 5229738 | 5229607 | 5229705 | 5230834 | 5233855 | 12 | 1 | 2 | 3 | 2 | 3 | 0 | 1 | 0 | 0 | 82 | 49 | 1 | 1 | 0.705 | 0.23 | 0.285 | NA | 1 | -0.593 |
| A5SS | 272 | PCMT1 | PCMT1 | chr6 | + | 149802199 | 149802416 | 149802199 | 149802369 | 149810615 | 149811421 | 4 | 5 | 10 | 1 | 0 | 0 | 0 | 1 | 0 | 0 | 96 | 49 | 1 | 1 | 0.671 | 1 | 1 | NA | 1 | -0.11 |
| A5SS | 282 | TNFAIP3 | TNFAIP3 | chr6 | + | 137867187 | 137867518 | 137867187 | 137867494 | 137871212 | 137871522 | 10 | 13 | 0 | 2 | 2 | 0 | 3 | 1 | 2 | 0 | 73 | 49 | 1 | 1 | 0.77 | 0.814 | NA | 0.502 | 1 | 0.041 |
| A5SS | 283 | TNFAIP3 | TNFAIP3 | chr6 | + | 137867187 | 137867542 | 137867187 | 137867494 | 137871212 | 137871522 | 10 | 18 | 0 | 2 | 2 | 0 | 3 | 6 | 2 | 0 | 97 | 49 | 1 | 1 | 0.716 | 0.82 | NA | 0.431 | 1 | 0.052 |
| A5SS | 286 | VNN2 | VNN2 | chr6 | - | 132752460 | 132752749 | 132752672 | 132752749 | 132751144 | 132751518 | 402 | 191 | 203 | 1 | 1 | 0 | 5 | 36 | 0 | 0 | 98 | 49 | 1 | 1 | 0.995 | 0.99 | 1 | 1 | 1 | -0.005 |
| A5SS | 303 | SYTL3 | SYTL3 | chr6 | + | 158662749 | 158663378 | 158662749 | 158662812 | 158665394 | 158665613 | 13 | 7 | 1 | 1 | 1 | 0 | 1 | 2 | 0 | 1 | 98 | 49 | 1 | 1 | 0.867 | 0.778 | 1 | 1 | 0.5 | 0.132 |
| A5SS | 304 | HMGN3 | HMGN3 | chr6 | - | 79202275 | 79202389 | 79202316 | 79202389 | 79201244 | 79201726 | 5 | 0 | 0 | 1 | 0 | 0 | 0 | 1 | 0 | 0 | 90 | 49 | 1 | 1 | 0.731 | NA | NA | NA | 1 | -0.269 |
| A5SS | 305 | COX7A2 | COX7A2 | chr6 | - | 75241122 | 75241265 | 75241175 | 75241265 | 75240300 | 75240385 | 4 | 1 | 1 | 6 | 1 | 7 | 0 | 2 | 0 | 0 | 98 | 49 | 1 | 1 | 0.25 | 0.333 | 0.067 | NA | 1 | -0.783 |
| A5SS | 322 | TJAP1 | TJAP1 | chr6 | + | 43498977 | 43499136 | 43498977 | 43499100 | 43500743 | 43500772 | 2 | 0 | 0 | 2 | 0 | 0 | 0 | 0 | 1 | 0 | 85 | 49 | 1 | 1 | 0.366 | NA | NA | 0 | NA | 0.366 |
| A5SS | 324 | CCND3 | CCND3 | chr6 | - | 41940951 | 41941848 | 41941451 | 41941848 | 41940369 | 41940585 | 2 | 3 | 0 | 4 | 0 | 0 | 0 | 0 | 0 | 5 | 98 | 49 | 1 | 1 | 0.2 | 1 | NA | NA | 0 | 0.6 |
| A5SS | 330 | HMGA1 | HMGA1 | chr6 | + | 34240736 | 34240915 | 34240736 | 34240882 | 34242711 | 34242795 | 6 | 4 | 0 | 5 | 0 | 0 | 1 | 4 | 0 | 3 | 82 | 49 | 1 | 1 | 0.418 | 1 | NA | 1 | 0.443 | -0.013 |
| A5SS | 352 | VPS52 | VPS52 | chr6 | - | 33271136 | 33271965 | 33271585 | 33271965 | 33270198 | 33270283 | 2 | 0 | 2 | 3 | 1 | 0 | 0 | 0 | 0 | 1 | 98 | 49 | 1 | 1 | 0.25 | 0 | 1 | NA | 0 | 0.417 |
| A5SS | 423 | DDAH2 | DDAH2 | chr6 | - | 31728864 | 31730265 | 31730050 | 31730265 | 31728645 | 31728745 | 13 | 5 | 0 | 0 | 0 | 1 | 0 | 5 | 0 | 0 | 98 | 49 | 1 | 1 | 1 | 1 | 0 | NA | 1 | -0.333 |
| A5SS | 430 | APOM | APOM | chr6 | + | 31656471 | 31656633 | 31656471 | 31656626 | 31657224 | 31657298 | 0 | 1 | 0 | 0 | 0 | 0 | 0 | 0 | 0 | 1 | 56 | 49 | 1 | 1 | NA | 1 | NA | NA | 0 | 1 |
| A5SS | 437 | BAG6 | BAG6 | chr6 | - | 31652399 | 31652700 | 31652423 | 31652700 | 31651655 | 31651776 | 8 | 3 | 0 | 3 | 0 | 0 | 0 | 0 | 0 | 1 | 73 | 49 | 1 | 1 | 0.642 | 1 | NA | NA | 0 | 0.821 |
| A5SS | 459 | NRM | NRM | chr6 | - | 30690024 | 30690243 | 30690046 | 30690243 | 30689275 | 30689452 | 2 | 2 | 0 | 1 | 0 | 0 | 1 | 0 | 0 | 0 | 71 | 49 | 1 | 1 | 0.58 | 1 | NA | 1 | NA | -0.21 |
| A5SS | 491 | ARRB2 | ARRB2 | chr17 | + | 4716411 | 4716671 | 4716411 | 4716608 | 4717216 | 4717276 | 2 | 1 | 0 | 6 | 4 | 1 | 0 | 0 | 0 | 1 | 98 | 49 | 1 | 1 | 0.143 | 0.111 | 0 | NA | 0 | 0.085 |
| A5SS | 499 | CRK | CRK | chr17 | - | 1436619 | 1437155 | 1436789 | 1437155 | 1421352 | 1423650 | 4 | 5 | 4 | 3 | 4 | 1 | 0 | 6 | 0 | 1 | 98 | 49 | 1 | 1 | 0.4 | 0.385 | 0.667 | NA | 0.75 | -0.266 |
| A5SS | 503 | CHMP1A | CHMP1A | chr16 | - | 89647034 | 89647331 | 89647202 | 89647331 | 89646526 | 89646714 | 4 | 0 | 2 | 4 | 4 | 3 | 0 | 1 | 0 | 0 | 98 | 49 | 1 | 1 | 0.333 | 0 | 0.25 | NA | 1 | -0.806 |
| A5SS | 504 | C12orf57 | C12orf57 | chr12 | + | 6944379 | 6944652 | 6944379 | 6944565 | 6945770 | 6946003 | 12 | 0 | 1 | 2 | 0 | 0 | 2 | 1 | 0 | 1 | 98 | 49 | 1 | 1 | 0.75 | NA | 1 | 1 | 0.333 | 0.209 |
| A5SS | 508 | TSPAN17 | TSPAN17 | chr5 | + | 176656894 | 176656956 | 176656894 | 176656924 | 176657517 | 176659058 | 2 | 0 | 0 | 2 | 0 | 0 | 0 | 1 | 0 | 0 | 81 | 49 | 1 | 1 | 0.377 | NA | NA | NA | 1 | -0.623 |
| A5SS | 513 | ATRAID | ATRAID | chr2 | + | 27212350 | 27212607 | 27212350 | 27212467 | 27213176 | 27213298 | 1 | 0 | 0 | 1 | 1 | 0 | 1 | 3 | 0 | 0 | 98 | 49 | 1 | 1 | 0.333 | 0 | NA | 1 | 1 | -0.834 |
| A5SS | 518 | SH2B1 | SH2B1 | chr16 | + | 28872533 | 28872758 | 28872533 | 28872705 | 28873169 | 28873269 | 0 | 1 | 0 | 2 | 0 | 0 | 0 | 0 | 0 | 1 | 98 | 49 | 1 | 1 | 0 | 1 | NA | NA | 0 | 0.5 |
| A5SS | 521 | PRKD2 | PRKD2 | chr19 | - | 46716814 | 46717127 | 46716950 | 46717127 | 46716130 | 46716535 | 3 | 1 | 0 | 0 | 0 | 0 | 0 | 2 | 0 | 1 | 98 | 49 | 1 | 1 | 1 | 1 | NA | NA | 0.5 | 0.5 |
| A5SS | 525 | SLC35A2 | SLC35A2 | chrX | - | 48911506 | 48911958 | 48911545 | 48911958 | 48909813 | 48909996 | 3 | 3 | 0 | 0 | 0 | 0 | 0 | 0 | 0 | 3 | 88 | 49 | 1 | 1 | 1 | 1 | NA | NA | 0 | 1 |
| A5SS | 531 | NMRAL1 | NMRAL1 | chr16 | - | 4474409 | 4474650 | 4474422 | 4474650 | 4474092 | 4474166 | 0 | 1 | 0 | 1 | 0 | 0 | 0 | 1 | 0 | 0 | 62 | 49 | 1 | 1 | 0 | 1 | NA | NA | 1 | -0.5 |
| A5SS | 536 | TTC31 | TTC31 | chr2 | + | 74491480 | 74491672 | 74491480 | 74491666 | 74492003 | 74492055 | 1 | 0 | 0 | 2 | 2 | 0 | 0 | 1 | 3 | 0 | 55 | 49 | 1 | 1 | 0.308 | 0 | NA | 0 | 1 | -0.346 |
| A5SS | 539 | UNKL | UNKL | chr16 | - | 1369933 | 1370357 | 1370129 | 1370357 | 1367655 | 1367858 | 0 | 0 | 0 | 0 | 1 | 2 | 1 | 0 | 0 | 1 | 98 | 49 | 1 | 1 | NA | 0 | 0 | 1 | 0 | -0.5 |
| A5SS | 580 | TRAPPC5 | TRAPPC5 | chr19 | + | 7680820 | 7681026 | 7680820 | 7680878 | 7682241 | 7682862 | 2 | 0 | 0 | 0 | 0 | 0 | 0 | 0 | 0 | 1 | 98 | 49 | 1 | 1 | 1 | NA | NA | NA | 0 | 1 |
| A5SS | 581 | TRAPPC5 | TRAPPC5 | chr19 | + | 7680820 | 7681049 | 7680820 | 7680878 | 7682241 | 7682862 | 2 | 0 | 0 | 0 | 0 | 0 | 0 | 0 | 0 | 1 | 98 | 49 | 1 | 1 | 1 | NA | NA | NA | 0 | 1 |
| A5SS | 585 | ISG20 | ISG20 | chr15 | + | 88638742 | 88639178 | 88638742 | 88639076 | 88639342 | 88639594 | 0 | 2 | 0 | 12 | 6 | 1 | 0 | 1 | 0 | 1 | 98 | 49 | 1 | 1 | 0 | 0.143 | 0 | NA | 0.333 | -0.285 |
| A5SS | 597 | STX16 | STX16 | chr20 | + | 58651252 | 58652138 | 58651252 | 58652087 | 58659622 | 58659634 | 13 | 10 | 3 | 0 | 0 | 0 | 4 | 7 | 0 | 1 | 98 | 49 | 1 | 1 | 1 | 1 | 1 | 1 | 0.778 | 0.111 |
| A5SS | 608 | PML | PML | chr15 | + | 74033155 | 74033422 | 74033155 | 74033414 | 74034477 | 74034530 | 5 | 1 | 1 | 3 | 0 | 0 | 0 | 1 | 0 | 2 | 57 | 49 | 1 | 1 | 0.589 | 1 | 1 | NA | 0.301 | 0.562 |
| A5SS | 616 | USP8 | USP8 | chr15 | + | 50424376 | 50424654 | 50424376 | 50424514 | 50439008 | 50439177 | 6 | 3 | 0 | 3 | 0 | 0 | 0 | 1 | 0 | 0 | 98 | 49 | 1 | 1 | 0.5 | 1 | NA | NA | 1 | -0.25 |
| A5SS | 621 | PIGG | PIGG | chr4 | + | 499199 | 499725 | 499199 | 499489 | 500395 | 500601 | 5 | 0 | 0 | 7 | 0 | 0 | 0 | 1 | 0 | 3 | 98 | 49 | 1 | 1 | 0.263 | NA | NA | NA | 0.143 | 0.12 |
| A5SS | 622 | PIGG | PIGG | chr4 | + | 499199 | 500047 | 499199 | 499489 | 500395 | 500601 | 4 | 0 | 0 | 7 | 0 | 0 | 0 | 1 | 0 | 3 | 98 | 49 | 1 | 1 | 0.222 | NA | NA | NA | 0.143 | 0.079 |
| A5SS | 647 | ANAPC11 | ANAPC11 | chr17 | + | 81894466 | 81894624 | 81894466 | 81894586 | 81899919 | 81900533 | 0 | 2 | 0 | 10 | 1 | 3 | 0 | 0 | 1 | 0 | 87 | 49 | 1 | 1 | 0 | 0.53 | 0 | 0 | NA | 0.177 |
| A5SS | 672 | ZNF302 | ZNF302 | chr19 | + | 34677638 | 34677856 | 34677638 | 34677833 | 34678736 | 34678813 | 0 | 0 | 0 | 1 | 0 | 0 | 0 | 1 | 0 | 0 | 72 | 49 | 1 | 1 | 0 | NA | NA | NA | 1 | -1 |
| A5SS | 673 | ZNF302 | ZNF302 | chr19 | + | 34677638 | 34677862 | 34677638 | 34677833 | 34678736 | 34678813 | 0 | 0 | 0 | 1 | 0 | 0 | 0 | 1 | 0 | 0 | 78 | 49 | 1 | 1 | 0 | NA | NA | NA | 1 | -1 |
| A5SS | 674 | ZNF302 | ZNF302 | chr19 | + | 34677638 | 34678103 | 34677638 | 34677833 | 34678736 | 34678813 | 2 | 0 | 0 | 1 | 0 | 0 | 0 | 1 | 0 | 0 | 98 | 49 | 1 | 1 | 0.5 | NA | NA | NA | 1 | -0.5 |
| A5SS | 689 | NIPA2 | NIPA2 | chr15 | + | 22838640 | 22838947 | 22838640 | 22838921 | 22845145 | 22845267 | 0 | 0 | 1 | 0 | 1 | 0 | 2 | 1 | 0 | 0 | 75 | 49 | 1 | 1 | NA | 0 | 1 | 1 | 1 | -0.5 |
| A5SS | 725 | ARL5A | ARL5A | chr2 | - | 151828130 | 151828495 | 151828388 | 151828495 | 151815138 | 151815199 | 1 | 0 | 0 | 1 | 1 | 0 | 0 | 1 | 0 | 0 | 98 | 49 | 1 | 1 | 0.333 | 0 | NA | NA | 1 | -0.834 |
| A5SS | 731 | UQCR10 | UQCR10 | chr22 | + | 29767368 | 29767607 | 29767368 | 29767548 | 29769677 | 29770413 | 1 | 0 | 0 | 0 | 1 | 4 | 0 | 1 | 0 | 3 | 98 | 49 | 1 | 1 | 1 | 0 | 0 | NA | 0.143 | 0.19 |
| A5SS | 738 | TRRAP | TRRAP | chr7 | + | 98983263 | 98983492 | 98983263 | 98983459 | 98984092 | 98984358 | 1 | 0 | 0 | 1 | 5 | 0 | 0 | 3 | 0 | 1 | 82 | 49 | 1 | 1 | 0.374 | 0 | NA | NA | 0.642 | -0.455 |
| A5SS | 741 | PSME1 | PSME1 | chr14 | + | 24138473 | 24138568 | 24138473 | 24138560 | 24138735 | 24138967 | 3 | 5 | 2 | 37 | 24 | 15 | 0 | 1 | 0 | 11 | 57 | 49 | 1 | 1 | 0.065 | 0.152 | 0.103 | NA | 0.072 | 0.035 |
| A5SS | 746 | ARHGEF40 | ARHGEF40 | chr14 | + | 21073931 | 21075180 | 21073931 | 21074991 | 21075331 | 21075499 | 23 | 4 | 0 | 1 | 0 | 0 | 0 | 3 | 0 | 0 | 98 | 49 | 1 | 1 | 0.92 | 1 | NA | NA | 1 | -0.04 |
| A5SS | 748 | FANCA | FANCA | chr16 | - | 89739128 | 89739289 | 89739132 | 89739289 | 89738881 | 89738974 | 5 | 0 | 1 | 1 | 0 | 0 | 0 | 0 | 0 | 1 | 53 | 49 | 1 | 1 | 0.822 | NA | 1 | NA | 0 | 0.911 |
| A5SS | 763 | PLEKHB2 | PLEKHB2 | chr2 | + | 131132901 | 131133015 | 131132901 | 131132991 | 131140169 | 131140275 | 2 | 0 | 0 | 2 | 0 | 0 | 0 | 2 | 0 | 0 | 73 | 49 | 1 | 1 | 0.402 | NA | NA | NA | 1 | -0.598 |
| A5SS | 764 | AFF2 | AFF2 | chrX | + | 148651998 | 148652131 | 148651998 | 148652119 | 148661907 | 148662768 | 1 | 0 | 4 | 6 | 7 | 3 | 0 | 0 | 0 | 1 | 61 | 49 | 1 | 1 | 0.118 | 0 | 0.517 | NA | 0 | 0.212 |
| A5SS | 774 | PICK1 | PICK1 | chr22 | + | 38057254 | 38057587 | 38057254 | 38057489 | 38057752 | 38057850 | 0 | 0 | 0 | 1 | 0 | 0 | 1 | 0 | 0 | 0 | 98 | 49 | 1 | 1 | 0 | NA | NA | 1 | NA | -1 |
| A5SS | 793 | AGTRAP | AGTRAP | chr1 | + | 11748414 | 11748610 | 11748414 | 11748589 | 11750076 | 11750771 | 12 | 8 | 7 | 4 | 0 | 3 | 0 | 4 | 0 | 1 | 70 | 49 | 1 | 1 | 0.677 | 1 | 0.62 | NA | 0.737 | 0.029 |
| A5SS | 801 | PSEN1 | PSEN1 | chr14 | + | 73147966 | 73148106 | 73147966 | 73148094 | 73170796 | 73171047 | 10 | 6 | 2 | 25 | 3 | 0 | 0 | 2 | 0 | 2 | 61 | 49 | 1 | 1 | 0.243 | 0.616 | 1 | NA | 0.445 | 0.175 |
| A5SS | 827 | ARL6IP4 | ARL6IP4 | chr12 | + | 122981128 | 122981299 | 122981128 | 122981266 | 122981594 | 122981879 | 11 | 2 | 3 | 5 | 0 | 0 | 0 | 3 | 0 | 1 | 82 | 49 | 1 | 1 | 0.568 | 1 | 1 | NA | 0.642 | 0.214 |
| A5SS | 835 | PPP4R3A | PPP4R3A | chr14 | - | 91475810 | 91475966 | 91475849 | 91475966 | 91473238 | 91473370 | 15 | 12 | 4 | 1 | 0 | 0 | 0 | 3 | 0 | 2 | 88 | 49 | 1 | 1 | 0.893 | 1 | 1 | NA | 0.455 | 0.509 |
| A5SS | 838 | TRAPPC3 | TRAPPC3 | chr1 | - | 36149091 | 36149514 | 36149336 | 36149514 | 36140068 | 36140166 | 1 | 2 | 0 | 3 | 1 | 0 | 0 | 0 | 4 | 0 | 98 | 49 | 1 | 1 | 0.143 | 0.5 | NA | 0 | NA | 0.322 |
| A5SS | 844 | NLRP1 | NLRP1 | chr17 | - | 5533291 | 5533384 | 5533303 | 5533384 | 5532821 | 5532984 | 6 | 4 | 1 | 30 | 9 | 7 | 0 | 1 | 0 | 7 | 61 | 49 | 1 | 1 | 0.138 | 0.263 | 0.103 | NA | 0.103 | 0.065 |
| A5SS | 846 | MAPKAPK5 | MAPKAPK5 | chr12 | + | 111888884 | 111889006 | 111888884 | 111889000 | 111890039 | 111890144 | 0 | 1 | 1 | 2 | 0 | 0 | 0 | 2 | 0 | 2 | 55 | 49 | 1 | 1 | 0 | 1 | 1 | NA | 0.471 | 0.196 |
| A5SS | 847 | GIT2 | GIT2 | chr12 | - | 109951166 | 109951316 | 109951211 | 109951316 | 109948061 | 109948846 | 39 | 29 | 1 | 2 | 0 | 0 | 0 | 5 | 0 | 0 | 94 | 49 | 1 | 1 | 0.91 | 1 | 1 | NA | 1 | -0.03 |
| A5SS | 876 | ITFG1 | ITFG1 | chr16 | - | 47460837 | 47461274 | 47461114 | 47461274 | 47459102 | 47459175 | 3 | 4 | 0 | 0 | 2 | 0 | 0 | 2 | 0 | 0 | 98 | 49 | 1 | 1 | 1 | 0.5 | NA | NA | 1 | -0.25 |
| A5SS | 877 | IL32 | IL32 | chr16 | + | 3067376 | 3067613 | 3067376 | 3067415 | 3067983 | 3068010 | 16 | 3 | 0 | 3 | 1 | 0 | 0 | 1 | 0 | 0 | 98 | 49 | 1 | 1 | 0.727 | 0.6 | NA | NA | 1 | -0.337 |
| A5SS | 880 | KLF7 | KLF7 | chr2 | - | 207123773 | 207124404 | 207124343 | 207124404 | 207088457 | 207088581 | 26 | 13 | 1 | 2 | 0 | 0 | 4 | 5 | 0 | 0 | 98 | 49 | 1 | 1 | 0.867 | 1 | 1 | 1 | 1 | -0.044 |
| A5SS | 881 | KLF7 | KLF7 | chr2 | - | 207123975 | 207124404 | 207124343 | 207124404 | 207088457 | 207088581 | 24 | 12 | 0 | 2 | 0 | 0 | 4 | 4 | 0 | 0 | 98 | 49 | 1 | 1 | 0.857 | 1 | NA | 1 | 1 | -0.072 |
| A5SS | 889 | STARD3 | STARD3 | chr17 | + | 39657774 | 39657852 | 39657774 | 39657839 | 39657972 | 39658026 | 4 | 11 | 1 | 3 | 0 | 0 | 0 | 4 | 0 | 0 | 62 | 49 | 1 | 1 | 0.513 | 1 | 1 | NA | 1 | -0.162 |
| A5SS | 892 | PTGES3 | PTGES3 | chr12 | - | 56665631 | 56666266 | 56666203 | 56666266 | 56664775 | 56664800 | 0 | 2 | 0 | 6 | 13 | 1 | 0 | 0 | 0 | 5 | 98 | 49 | 1 | 1 | 0 | 0.071 | 0 | NA | 0 | 0.024 |
| A5SS | 952 | NDUFB11 | NDUFB11 | chrX | - | 47142583 | 47142744 | 47142613 | 47142744 | 47142215 | 47142440 | 1 | 0 | 0 | 7 | 1 | 0 | 0 | 0 | 0 | 3 | 79 | 49 | 1 | 1 | 0.081 | 0 | NA | NA | 0 | 0.041 |
| A5SS | 954 | CLEC2D | CLEC2D | chr12 | + | 9687901 | 9688161 | 9687901 | 9688086 | 9692827 | 9692931 | 11 | 0 | 0 | 18 | 3 | 0 | 0 | 1 | 0 | 0 | 98 | 49 | 1 | 1 | 0.234 | 0 | NA | NA | 1 | -0.883 |
| A5SS | 976 | KIAA1033 | KIAA1033 | chr12 | + | 105133769 | 105133899 | 105133769 | 105133896 | 105137885 | 105138011 | 1 | 2 | 0 | 29 | 17 | 4 | 0 | 0 | 0 | 6 | 52 | 49 | 1 | 1 | 0.031 | 0.1 | 0 | NA | 0 | 0.044 |
| A5SS | 982 | WNK1 | WNK1 | chr12 | + | 890452 | 890513 | 890452 | 890510 | 894561 | 894635 | 7 | 4 | 0 | 4 | 0 | 0 | 0 | 1 | 0 | 0 | 52 | 49 | 1 | 1 | 0.623 | 1 | NA | NA | 1 | -0.189 |
| A5SS | 1006 | NLRX1 | NLRX1 | chr11 | + | 119168712 | 119169302 | 119168712 | 119168894 | 119171355 | 119171473 | 0 | 0 | 0 | 1 | 0 | 0 | 1 | 0 | 0 | 0 | 98 | 49 | 1 | 1 | 0 | NA | NA | 1 | NA | -1 |
| A5SS | 1012 | SLC12A9 | SLC12A9 | chr7 | + | 100852718 | 100852839 | 100852718 | 100852835 | 100854155 | 100854378 | 4 | 2 | 1 | 3 | 2 | 1 | 0 | 0 | 0 | 2 | 53 | 49 | 1 | 1 | 0.552 | 0.48 | 0.48 | NA | 0 | 0.504 |
| A5SS | 1016 | POLG | POLG | chr15 | - | 89334672 | 89334795 | 89334684 | 89334795 | 89333095 | 89333913 | 3 | 0 | 0 | 1 | 0 | 0 | 0 | 0 | 0 | 3 | 61 | 49 | 1 | 1 | 0.707 | NA | NA | NA | 0 | 0.707 |
| A5SS | 1019 | SNX19 | SNX19 | chr11 | - | 130914265 | 130916487 | 130916279 | 130916487 | 130911632 | 130911771 | 7 | 8 | 0 | 1 | 0 | 0 | 0 | 3 | 0 | 1 | 98 | 49 | 1 | 1 | 0.778 | 1 | NA | NA | 0.6 | 0.289 |
| A5SS | 1051 | PTPN7 | PTPN7 | chr1 | - | 202160544 | 202161588 | 202161386 | 202161588 | 202159280 | 202159454 | 0 | 0 | 0 | 1 | 0 | 0 | 0 | 1 | 0 | 0 | 98 | 49 | 1 | 1 | 0 | NA | NA | NA | 1 | -1 |
| A5SS | 1067 | LOC374443 | LOC374443 | chr12 | + | 9648046 | 9648933 | 9648046 | 9648328 | 9653682 | 9653808 | 13 | 0 | 0 | 0 | 0 | 3 | 0 | 2 | 0 | 0 | 98 | 49 | 1 | 1 | 1 | NA | 0 | NA | 1 | -0.5 |
| A5SS | 1075 | IRF2BP2 | IRF2BP2 | chr1 | - | 234608446 | 234609525 | 234608494 | 234609525 | 234604268 | 234607852 | 16 | 2 | 0 | 9 | 1 | 0 | 0 | 5 | 1 | 5 | 97 | 49 | 1 | 1 | 0.473 | 0.503 | NA | 0 | 0.336 | 0.32 |
| A5SS | 1114 | TMEM55B | TMEM55B | chr14 | - | 20461162 | 20461612 | 20461183 | 20461612 | 20460654 | 20460845 | 0 | 2 | 0 | 0 | 1 | 0 | 0 | 2 | 0 | 0 | 70 | 49 | 1 | 1 | NA | 0.583 | NA | NA | 1 | -0.417 |
| A5SS | 1133 | WDR26 | WDR26 | chr1 | - | 224431476 | 224431581 | 224431524 | 224431581 | 224419517 | 224419615 | 33 | 17 | 3 | 1 | 1 | 0 | 0 | 10 | 0 | 0 | 97 | 49 | 1 | 1 | 0.943 | 0.896 | 1 | NA | 1 | -0.054 |
| A5SS | 1134 | WDR26 | WDR26 | chr1 | - | 224431476 | 224431581 | 224431524 | 224431581 | 224424517 | 224424654 | 40 | 25 | 3 | 10 | 11 | 1 | 0 | 11 | 0 | 4 | 97 | 49 | 1 | 1 | 0.669 | 0.534 | 0.602 | NA | 0.581 | 0.021 |
| A5SS | 1160 | PKM | PKM | chr15 | - | 72230895 | 72231386 | 72231115 | 72231386 | 72218943 | 72219110 | 1 | 0 | 0 | 6 | 0 | 0 | 0 | 0 | 0 | 1 | 98 | 49 | 1 | 1 | 0.077 | NA | NA | NA | 0 | 0.077 |
| A5SS | 1192 | CNOT1 | CNOT1 | chr16 | - | 58556846 | 58556993 | 58556861 | 58556993 | 58555783 | 58555908 | 9 | 1 | 0 | 3 | 0 | 0 | 0 | 1 | 0 | 1 | 64 | 49 | 1 | 1 | 0.697 | 1 | NA | NA | 0.434 | 0.415 |
| A5SS | 1203 | CANX | CANX | chr5 | + | 179698928 | 179699102 | 179698928 | 179699049 | 179705678 | 179705852 | 3 | 0 | 0 | 0 | 0 | 0 | 0 | 0 | 0 | 1 | 98 | 49 | 1 | 1 | 1 | NA | NA | NA | 0 | 1 |
| A5SS | 1239 | CHST11 | CHST11 | chr12 | + | 104456913 | 104457529 | 104456913 | 104457514 | 104601905 | 104601991 | 11 | 14 | 8 | 5 | 8 | 0 | 0 | 7 | 0 | 2 | 64 | 49 | 1 | 1 | 0.627 | 0.573 | 1 | NA | 0.728 | 0.005 |
| A5SS | 1248 | APAF1 | APAF1 | chr12 | + | 98648625 | 98648815 | 98648625 | 98648782 | 98649486 | 98649684 | 58 | 11 | 6 | 1 | 0 | 0 | 0 | 9 | 0 | 0 | 82 | 49 | 1 | 1 | 0.972 | 1 | 1 | NA | 1 | -0.009 |
| A5SS | 1249 | TRERF1 | TRERF1 | chr6 | - | 42263259 | 42263568 | 42263319 | 42263568 | 42259338 | 42259723 | 1 | 0 | 0 | 0 | 0 | 0 | 0 | 0 | 0 | 1 | 98 | 49 | 1 | 1 | 1 | NA | NA | NA | 0 | 1 |
| A5SS | 1255 | NBPF11 | NBPF11 | chr1 | - | 148110377 | 148110541 | 148110457 | 148110541 | 148109283 | 148109335 | 0 | 1 | 0 | 2 | 0 | 0 | 1 | 0 | 0 | 0 | 98 | 49 | 1 | 1 | 0 | 1 | NA | 1 | NA | -0.5 |
| A5SS | 1274 | TUG1 | TUG1 | chr22 | + | 30972854 | 30973601 | 30972854 | 30973587 | 30975169 | 30979394 | 9 | 4 | 0 | 4 | 2 | 0 | 1 | 4 | 0 | 0 | 63 | 49 | 1 | 1 | 0.636 | 0.609 | NA | 1 | 1 | -0.377 |
| A5SS | 1279 | TRIM5 | TRIM5 | chr11 | - | 5684867 | 5685109 | 5685015 | 5685109 | 5679760 | 5680238 | 3 | 2 | 0 | 0 | 2 | 0 | 0 | 1 | 0 | 0 | 98 | 49 | 1 | 1 | 1 | 0.333 | NA | NA | 1 | -0.334 |
| A5SS | 1303 | DAP3 | DAP3 | chr1 | + | 155689093 | 155689243 | 155689093 | 155689174 | 155709772 | 155709824 | 2 | 2 | 0 | 0 | 0 | 0 | 0 | 0 | 0 | 1 | 98 | 49 | 1 | 1 | 1 | 1 | NA | NA | 0 | 1 |
| A5SS | 1308 | AP1G2 | AP1G2 | chr14 | - | 23565815 | 23566160 | 23566063 | 23566160 | 23565605 | 23565701 | 1 | 0 | 0 | 0 | 0 | 0 | 0 | 3 | 0 | 1 | 98 | 49 | 1 | 1 | 1 | NA | NA | NA | 0.6 | 0.4 |
| A5SS | 1310 | SIRPB2 | SIRPB2 | chr20 | - | 1479699 | 1480065 | 1479993 | 1480065 | 1478265 | 1478607 | 21 | 3 | 5 | 5 | 2 | 2 | 0 | 8 | 0 | 2 | 98 | 49 | 1 | 1 | 0.677 | 0.429 | 0.556 | NA | 0.667 | -0.113 |
| A5SS | 1323 | THBS3 | THBS3 | chr1 | - | 155206192 | 155206406 | 155206199 | 155206406 | 155205059 | 155205316 | 0 | 0 | 0 | 2 | 0 | 0 | 0 | 1 | 0 | 0 | 56 | 49 | 1 | 1 | 0 | NA | NA | NA | 1 | -1 |
| A5SS | 1330 | AP3B1 | AP3B1 | chr5 | - | 78294439 | 78294755 | 78294451 | 78294755 | 78267519 | 78267595 | 2 | 1 | 0 | 13 | 5 | 3 | 0 | 0 | 0 | 2 | 61 | 49 | 1 | 1 | 0.11 | 0.138 | 0 | NA | 0 | 0.083 |
| A5SS | 1334 | PAPD4 | PAPD4 | chr5 | + | 79612419 | 79612769 | 79612419 | 79612735 | 79613010 | 79613075 | 7 | 8 | 0 | 0 | 0 | 0 | 2 | 1 | 0 | 1 | 83 | 49 | 1 | 1 | 1 | 1 | NA | 1 | 0.371 | 0.315 |
| A5SS | 1343 | TRAFD1 | TRAFD1 | chr12 | + | 112125544 | 112126149 | 112125544 | 112125618 | 112130510 | 112130569 | 0 | 0 | 0 | 2 | 0 | 0 | 0 | 1 | 0 | 0 | 98 | 49 | 1 | 1 | 0 | NA | NA | NA | 1 | -1 |
| A5SS | 1349 | STX10 | STX10 | chr19 | - | 13144763 | 13144870 | 13144767 | 13144870 | 13144576 | 13144671 | 38 | 11 | 6 | 0 | 0 | 1 | 4 | 7 | 0 | 0 | 53 | 49 | 1 | 1 | 1 | 1 | 0.847 | 1 | 1 | -0.051 |
| A5SS | 1359 | MIER1 | MIER1 | chr1 | + | 66984571 | 66984652 | 66984571 | 66984592 | 66986402 | 66988619 | 30 | 2 | 5 | 2 | 2 | 0 | 0 | 2 | 0 | 0 | 98 | 49 | 1 | 1 | 0.882 | 0.333 | 1 | NA | 1 | -0.262 |
| A5SS | 1366 | TSEN15 | TSEN15 | chr1 | + | 184072156 | 184072310 | 184072156 | 184072298 | 184072826 | 184074210 | 1 | 0 | 0 | 0 | 0 | 0 | 0 | 0 | 0 | 1 | 61 | 49 | 1 | 1 | 1 | NA | NA | NA | 0 | 1 |
| A5SS | 1375 | RAP1B | RAP1B | chr12 | + | 68610838 | 68611052 | 68610838 | 68611043 | 68648698 | 68648781 | 1 | 1 | 0 | 8 | 3 | 0 | 0 | 0 | 0 | 1 | 58 | 49 | 1 | 1 | 0.096 | 0.22 | NA | NA | 0 | 0.158 |
| A5SS | 1382 | ARID4A | ARID4A | chr14 | + | 58366023 | 58366230 | 58366023 | 58366068 | 58366882 | 58367029 | 22 | 7 | 2 | 0 | 0 | 1 | 0 | 4 | 0 | 0 | 98 | 49 | 1 | 1 | 1 | 1 | 0.5 | NA | 1 | -0.167 |
| A5SS | 1384 | MDM2 | MDM2 | chr12 | + | 68835828 | 68835984 | 68835828 | 68835843 | 68836671 | 68836749 | 57 | 22 | 17 | 3 | 0 | 0 | 0 | 9 | 0 | 0 | 98 | 49 | 1 | 1 | 0.905 | 1 | 1 | NA | 1 | -0.032 |
| A5SS | 1394 | MKL1 | MKL1 | chr22 | - | 40416975 | 40417046 | 40416985 | 40417046 | 40410280 | 40411907 | 1 | 0 | 0 | 3 | 5 | 0 | 0 | 0 | 4 | 0 | 59 | 49 | 1 | 1 | 0.217 | 0 | NA | 0 | NA | 0.108 |
| A5SS | 1397 | FLYWCH2 | FLYWCH2 | chr16 | + | 2883194 | 2883366 | 2883194 | 2883272 | 2895219 | 2895320 | 0 | 3 | 0 | 2 | 0 | 0 | 0 | 2 | 0 | 0 | 98 | 49 | 1 | 1 | 0 | 1 | NA | NA | 1 | -0.5 |
| A5SS | 1406 | UFM1 | UFM1 | chr13 | + | 38349998 | 38350227 | 38349998 | 38350055 | 38354238 | 38354296 | 2 | 0 | 0 | 1 | 1 | 0 | 0 | 0 | 0 | 2 | 98 | 49 | 1 | 1 | 0.5 | 0 | NA | NA | 0 | 0.25 |
| A5SS | 1418 | TMEM170A | TMEM170A | chr16 | - | 75451668 | 75451839 | 75451737 | 75451839 | 75443243 | 75447688 | 2 | 1 | 3 | 1 | 1 | 0 | 0 | 1 | 0 | 0 | 98 | 49 | 1 | 1 | 0.5 | 0.333 | 1 | NA | 1 | -0.389 |
| A5SS | 1424 | UBE2J2 | UBE2J2 | chr1 | - | 1267732 | 1267992 | 1267861 | 1267992 | 1263345 | 1263386 | 1 | 0 | 0 | 5 | 1 | 0 | 0 | 1 | 0 | 0 | 98 | 49 | 1 | 1 | 0.091 | 0 | NA | NA | 1 | -0.955 |
| A5SS | 1427 | RBMS1 | RBMS1 | chr2 | - | 160286968 | 160287084 | 160286977 | 160287084 | 160284994 | 160285044 | 15 | 4 | 2 | 13 | 6 | 1 | 0 | 2 | 0 | 2 | 58 | 49 | 1 | 1 | 0.494 | 0.36 | 0.628 | NA | 0.458 | 0.036 |
| A5SS | 1445 | RBBP5 | RBBP5 | chr1 | - | 205094872 | 205095064 | 205094986 | 205095064 | 205086141 | 205088815 | 5 | 5 | 0 | 0 | 7 | 0 | 0 | 1 | 0 | 0 | 98 | 49 | 1 | 1 | 1 | 0.263 | NA | NA | 1 | -0.369 |
| A5SS | 1453 | TAF12 | TAF12 | chr1 | - | 28642654 | 28643092 | 28642991 | 28643092 | 28621913 | 28622165 | 0 | 0 | 0 | 1 | 0 | 1 | 0 | 1 | 0 | 0 | 98 | 49 | 1 | 1 | 0 | NA | 0 | NA | 1 | -1 |
| A5SS | 1469 | KHDRBS1 | KHDRBS1 | chr1 | + | 32037834 | 32038119 | 32037834 | 32038036 | 32038551 | 32038619 | 2 | 1 | 0 | 4 | 2 | 1 | 0 | 0 | 0 | 2 | 98 | 49 | 1 | 1 | 0.2 | 0.2 | 0 | NA | 0 | 0.133 |
| A5SS | 1476 | IFI6 | IFI6 | chr1 | - | 27669220 | 27669346 | 27669232 | 27669346 | 27668457 | 27668535 | 1 | 0 | 0 | 0 | 0 | 0 | 0 | 4 | 0 | 1 | 61 | 49 | 1 | 1 | 1 | NA | NA | NA | 0.763 | 0.237 |
| A5SS | 1489 | NPEPL1 | NPEPL1 | chr20 | + | 58691723 | 58691830 | 58691723 | 58691780 | 58693736 | 58693922 | 13 | 4 | 0 | 1 | 0 | 0 | 0 | 5 | 0 | 1 | 98 | 49 | 1 | 1 | 0.867 | 1 | NA | NA | 0.714 | 0.22 |
| A5SS | 1490 | DEDD2 | DEDD2 | chr19 | - | 42201988 | 42202072 | 42202050 | 42202072 | 42198592 | 42199829 | 10 | 3 | 0 | 3 | 1 | 0 | 4 | 0 | 0 | 1 | 98 | 49 | 1 | 1 | 0.625 | 0.6 | NA | 1 | 0 | 0.113 |
| A5SS | 1495 | ADD1 | ADD1 | chr4 | + | 2904763 | 2905108 | 2904763 | 2905015 | 2907742 | 2907844 | 10 | 0 | 0 | 2 | 2 | 0 | 1 | 2 | 0 | 0 | 98 | 49 | 1 | 1 | 0.714 | 0 | NA | 1 | 1 | -0.643 |
| A5SS | 1500 | CAP1 | CAP1 | chr1 | + | 40040582 | 40040801 | 40040582 | 40040783 | 40059336 | 40059458 | 5 | 2 | 2 | 1 | 0 | 0 | 0 | 1 | 0 | 0 | 67 | 49 | 1 | 1 | 0.785 | 1 | 1 | NA | 1 | -0.072 |

| | | | | | | | | | | | | | | | | | | | | | | | | | | | | | | | | |
|---|---|---|---|---|---|---|---|---|---|---|---|---|---|---|---|---|---|---|---|---|---|---|---|---|---|---|---|---|---|---|---|---|
| A5SS | 1503 | ACAA1 | ACAA1 | chr3 | - | 38126161 | 38126700 | 38126509 | 38126700 | 38125825 | 38125881 | 7 | 5 | 5 | 2 | 0 | 0 | 0 | 3 | 0 | 0 | 98 | 49 | 1 | 1 | 0.636 | 1 | 1 | NA | | 1 | -0.121 |
| A5SS | 1520 | BTF3 | BTF3 | chr5 | + | 73498424 | 73498799 | 73498424 | 73498558 | 73499133 | 73499202 | 0 | 1 | 0 | 12 | 0 | 0 | 0 | 2 | 0 | 2 | 98 | 49 | 1 | 1 | 0 | 1 | NA | NA | | 0.333 | 0.167 |
| A5SS | 1534 | TTLL3 | TTLL3 | chr3 | + | 9826996 | 9827240 | 9826996 | 9827072 | 9828959 | 9829395 | 12 | 12 | 4 | 1 | 0 | 0 | 2 | 5 | 0 | 0 | 98 | 49 | 1 | 1 | 0.857 | 1 | 1 | | 1 | 1 | -0.048 |
| A5SS | 1536 | CASP8 | CASP8 | chr2 | + | 201276826 | 201277104 | 201276826 | 201276968 | 201284815 | 201285317 | 190 | 74 | 9 | 20 | 6 | 10 | 0 | 9 | 0 | 3 | 98 | 49 | 1 | 1 | 0.826 | 0.86 | 0.31 | NA | | 0.6 | 0.065 |
| A5SS | 1537 | EIF4B | EIF4B | chr12 | + | 53034611 | 53034724 | 53034611 | 53034709 | 53037408 | 53037622 | 8 | 2 | 0 | 51 | 13 | 14 | 0 | 0 | 0 | 3 | 64 | 49 | 1 | 1 | 0.107 | 0.105 | 0 | NA | | 0 | 0.071 |
| A5SS | 1546 | NR3C1 | NR3C1 | chr5 | - | 143313998 | 143314168 | 143314001 | 143314168 | 143310096 | 143310213 | 8 | 1 | 0 | 16 | 5 | 3 | 0 | 0 | 0 | 4 | 52 | 49 | 1 | 1 | 0.32 | 0.159 | 0 | NA | | 0 | 0.16 |
| A5SS | 1553 | NAT9 | NAT9 | chr17 | - | 74775598 | 74775707 | 74775621 | 74775707 | 74773575 | 74773685 | 1 | 0 | 0 | 2 | 0 | 0 | 0 | 1 | 0 | 0 | 72 | 49 | 1 | 1 | 0.254 | NA | NA | NA | | 1 | -0.746 |
| A5SS | 1576 | C12orf43 | C12orf43 | chr12 | - | 121011010 | 121011146 | 121011103 | 121011146 | 121010827 | 121010926 | 0 | 0 | 0 | 0 | 4 | 0 | 0 | 1 | 0 | 0 | 98 | 49 | 1 | 1 | NA | 0 | NA | NA | | 1 | -1 |
| A5SS | 1597 | RPA2 | RPA2 | chr1 | - | 27913911 | 27914169 | 27914062 | 27914169 | 27907180 | 27907282 | 7 | 5 | 0 | 4 | 0 | 0 | 0 | 0 | 0 | 2 | 98 | 49 | 1 | 1 | 0.467 | 1 | NA | NA | | 0 | 0.734 |
| A5SS | 1614 | CPEB2 | CPEB2 | chr4 | + | 15007304 | 15007595 | 15007304 | 15007586 | 15017187 | 15017278 | 2 | 3 | 0 | 3 | 1 | 0 | 0 | 1 | 0 | 2 | 58 | 49 | 1 | 1 | 0.36 | 0.717 | NA | NA | | 0.297 | 0.241 |
| A5SS | 1619 | PLCB2 | PLCB2 | chr15 | - | 40296745 | 40296908 | 40296757 | 40296908 | 40296521 | 40296634 | 16 | 7 | 1 | 3 | 6 | 0 | 0 | 7 | 0 | 3 | 61 | 49 | 1 | 1 | 0.811 | 0.484 | 1 | NA | | 0.652 | 0.113 |
| A5SS | 1621 | RBM10 | RBM10 | chrX | + | 47179879 | 47180040 | 47179879 | 47180037 | 47180211 | 47180309 | 0 | 0 | 0 | 3 | 3 | 0 | 0 | 3 | 0 | 0 | 52 | 49 | 1 | 1 | 0 | 0 | NA | NA | | 1 | -1 |
| A5SS | 1636 | DENND5A | DENND5A | chr11 | - | 9141939 | 9142108 | 9141973 | 9142108 | 9138825 | 9139854 | 41 | 25 | 22 | 0 | 1 | 1 | 1 | 6 | 0 | 0 | 83 | 49 | 1 | 1 | 1 | 0.937 | 0.929 | | 1 | 1 | -0.045 |
| A5SS | 1643 | CDC42EP3 | CDC42EP3 | chr2 | - | 37671425 | 37672199 | 37672084 | 37672199 | 37641881 | 37646822 | 22 | 3 | 4 | 1 | 0 | 0 | 4 | 6 | 0 | 0 | 98 | 49 | 1 | 1 | 0.917 | 1 | 1 | | 1 | 1 | -0.028 |
| A5SS | 1644 | CDC42EP3 | CDC42EP3 | chr2 | - | 37672060 | 37672199 | 37672084 | 37672199 | 37641881 | 37646822 | 3 | 0 | 3 | 1 | 0 | 0 | 0 | 1 | 0 | 0 | 73 | 49 | 1 | 1 | 0.668 | NA | 1 | NA | | 1 | -0.166 |
| A5SS | 1659 | CELF2 | CELF2 | chr10 | + | 11314138 | 11314270 | 11314138 | 11314258 | 11321206 | 11321386 | 9 | 11 | 3 | 6 | 13 | 8 | 0 | 1 | 0 | 3 | 61 | 49 | 1 | 1 | 0.546 | 0.405 | 0.231 | NA | | 0.211 | 0.183 |
| A5SS | 1677 | BPTF | BPTF | chr17 | + | 67945408 | 67946325 | 67945408 | 67945896 | 67947725 | 67947808 | 2 | 2 | 0 | 7 | 1 | 0 | 0 | 0 | 1 | 1 | 98 | 49 | 1 | 1 | 0.125 | 0.5 | NA | NA | 0 | 0 | 0.312 |
| A5SS | 1680 | CTNND1 | CTNND1 | chr11 | + | 57761761 | 57762119 | 57761761 | 57762046 | 57794009 | 57794081 | 0 | 2 | 0 | 0 | 0 | 0 | 0 | 1 | 0 | 1 | 98 | 49 | 1 | 1 | NA | 1 | NA | NA | | 0.333 | 0.667 |
| A5SS | 1697 | UBAP2L | UBAP2L | chr1 | + | 154234590 | 154234792 | 154234590 | 154234759 | 154235195 | 154235291 | 4 | 0 | 1 | 4 | 0 | 1 | 0 | 1 | 0 | 0 | 82 | 49 | 1 | 1 | 0.374 | NA | 0.374 | NA | | 1 | -0.626 |
| A5SS | 1705 | RCHY1 | RCHY1 | chr4 | - | 75491582 | 75491637 | 75491610 | 75491637 | 75490580 | 75490701 | 0 | 1 | 0 | 4 | 2 | 0 | 0 | 0 | 1 | 0 | 77 | 49 | 1 | 1 | 0 | 0.241 | NA | NA | 0 | NA | 0.12 |
| A5SS | 1706 | RCHY1 | RCHY1 | chr4 | - | 75508819 | 75508935 | 75508823 | 75508935 | 75494100 | 75494179 | 3 | 0 | 0 | 0 | 0 | 0 | 0 | 0 | 0 | 2 | 53 | 49 | 1 | 1 | 1 | NA | NA | NA | | 0 | 1 |
| A5SS | 1711 | AMICA1 | AMICA1 | chr11 | - | 118197967 | 118198091 | 118197997 | 118198091 | 118196734 | 118196821 | 130 | 46 | 31 | 45 | 39 | 38 | 0 | 14 | 0 | 6 | 79 | 49 | 1 | 1 | 0.642 | 0.422 | 0.336 | NA | | 0.591 | -0.124 |
| A5SS | 1720 | ZNF33A | ZNF33A | chr10 | + | 38016870 | 38017020 | 38016870 | 38017015 | 38017290 | 38017386 | 3 | 2 | 0 | 13 | 6 | 4 | 0 | 0 | 0 | 3 | 54 | 49 | 1 | 1 | 0.173 | 0.232 | 0 | NA | | 0 | 0.135 |
| A5SS | 1729 | GMPR2 | GMPR2 | chr14 | + | 24232421 | 24232847 | 24232421 | 24232788 | 24233218 | 24233340 | 21 | 11 | 1 | 9 | 4 | 3 | 0 | 2 | 0 | 4 | 98 | 49 | 1 | 1 | 0.538 | 0.579 | 0.143 | NA | | 0.2 | 0.22 |
| A5SS | 1730 | GMPR2 | GMPR2 | chr14 | + | 24233478 | 24233598 | 24233478 | 24233593 | 24235736 | 24235820 | 9 | 8 | 7 | 2 | 0 | 0 | 0 | 1 | 0 | 0 | 54 | 49 | 1 | 1 | 0.803 | 1 | 1 | NA | | 1 | -0.066 |
| A5SS | 1732 | UBE3B | UBE3B | chr12 | + | 109477622 | 109478109 | 109477622 | 109477764 | 109481636 | 109481742 | 5 | 0 | 0 | 1 | 1 | 0 | 0 | 1 | 0 | 0 | 98 | 49 | 1 | 1 | 0.714 | 0 | NA | NA | | 1 | -0.643 |
| A5SS | 1770 | CPEB3 | CPEB3 | chr10 | - | 92192476 | 92192636 | 92192545 | 92192636 | 92180962 | 92181019 | 2 | 0 | 0 | 0 | 0 | 0 | 0 | 0 | 0 | 1 | 98 | 49 | 1 | 1 | 1 | NA | NA | NA | | 0 | 1 |
| A5SS | 1771 | ARFIP1 | ARFIP1 | chr4 | + | 152779936 | 152780226 | 152779936 | 152780091 | 152790758 | 152790875 | 3 | 3 | 0 | 2 | 0 | 0 | 0 | 3 | 0 | 0 | 98 | 49 | 1 | 1 | 0.429 | 1 | NA | NA | | 1 | -0.285 |
| A5SS | 1772 | ARFIP1 | ARFIP1 | chr4 | + | 152779936 | 152780226 | 152779936 | 152780091 | 152829624 | 152829726 | 11 | 3 | 0 | 6 | 1 | 0 | 0 | 1 | 0 | 1 | 98 | 49 | 1 | 1 | 0.478 | 0.6 | NA | NA | | 0.333 | 0.206 |
| A5SS | 1778 | ZNF518A | ZNF518A | chr10 | + | 96155325 | 96155398 | 96155325 | 96155376 | 96155923 | 96156047 | 0 | 1 | 0 | 3 | 0 | 0 | 0 | 0 | 0 | 3 | 71 | 49 | 1 | 1 | 0 | 1 | NA | NA | | 0 | 0.5 |
| A5SS | 1783 | PIGBOS1 | PIGBOS1 | chr15 | - | 55318863 | 55319194 | 55318930 | 55319194 | 55317183 | 55317867 | 14 | 1 | 0 | 0 | 4 | 2 | 1 | 0 | 0 | 0 | 98 | 49 | 1 | 1 | 1 | 0.111 | 0 | | 1 | NA | -0.63 |
| A5SS | 1784 | PIGBOS1 | PIGBOS1 | chr15 | - | 55318906 | 55319194 | 55318930 | 55319194 | 55317183 | 55317867 | 13 | 0 | 0 | 0 | 4 | 2 | 1 | 0 | 0 | 0 | 73 | 49 | 1 | 1 | 1 | 0 | 0 | | 1 | NA | -0.667 |
| A5SS | 1788 | PDCD6 | PDCD6 | chr5 | + | 306601 | 306760 | 306601 | 306754 | 311292 | 311402 | 11 | 2 | 4 | 1 | 0 | 0 | 0 | 1 | 0 | 1 | 55 | 49 | 1 | 1 | 0.907 | 1 | 1 | NA | | 0.471 | 0.498 |
| A5SS | 1800 | TAZ | TAZ | chrX | + | 154411517 | 154412006 | 154411517 | 154411952 | 154412085 | 154412214 | 0 | 1 | 0 | 1 | 1 | 0 | 0 | 0 | 2 | 1 | 98 | 49 | 1 | 1 | 0 | 0.333 | NA | | 0 | 0 | 0.167 |
| A5SS | 1801 | TAZ | TAZ | chrX | + | 154413206 | 154413358 | 154413206 | 154413252 | 154413481 | 154413567 | 2 | 5 | 0 | 4 | 1 | 0 | 0 | 4 | 2 | 1 | 98 | 49 | 1 | 1 | 0.2 | 0.714 | NA | | 0 | 0.667 | 0.123 |
| A5SS | 1802 | GBF1 | GBF1 | chr10 | + | 102376934 | 102377140 | 102376934 | 102377128 | 102379283 | 102379434 | 1 | 0 | 0 | 2 | 0 | 0 | 0 | 2 | 0 | 2 | 61 | 49 | 1 | 1 | 0.287 | NA | NA | NA | | 0.445 | -0.158 |
| A5SS | 1810 | NAA60 | NAA60 | chr16 | + | 3484698 | 3485083 | 3484698 | 3485061 | 3485466 | 3486963 | 3 | 3 | 0 | 1 | 0 | 0 | 0 | 0 | 0 | 1 | 71 | 49 | 1 | 1 | 0.674 | 1 | NA | NA | | 0 | 0.837 |
| A5SS | 1813 | RNASEK | RNASEK | chr17 | + | 7012416 | 7012761 | 7012416 | 7012757 | 7013665 | 7013742 | 20 | 8 | 4 | 2 | 1 | 2 | 0 | 4 | 0 | 2 | 53 | 49 | 1 | 1 | 0.902 | 0.881 | 0.649 | NA | | 0.649 | 0.162 |
| A5SS | 1819 | CAMK2G | CAMK2G | chr10 | - | 73825278 | 73825347 | 73825320 | 73825347 | 73824039 | 73824084 | 26 | 12 | 10 | 0 | 1 | 0 | 0 | 2 | 0 | 1 | 91 | 49 | 1 | 1 | 1 | 0.866 | 1 | NA | | 0.519 | 0.436 |
| A5SS | 1823 | PABPC4 | PABPC4 | chr1 | - | 39564422 | 39564542 | 39564470 | 39564542 | 39563613 | 39563741 | 10 | 2 | 0 | 2 | 0 | 0 | 0 | 1 | 0 | 0 | 97 | 49 | 1 | 1 | 0.716 | 1 | NA | | | 1 | -0.142 |
| A5SS | 1832 | METTL10 | METTL10 | chr10 | - | 124789038 | 124789157 | 124789042 | 124789157 | 124774674 | 124774782 | 0 | 3 | 0 | 0 | 0 | 0 | 0 | 0 | 0 | 1 | 53 | 49 | 1 | 1 | NA | 1 | NA | | | 0 | 1 |
| A5SS | 1835 | UBE4A | UBE4A | chr11 | + | 118372506 | 118372687 | 118372506 | 118372666 | 118373085 | 118373288 | 2 | 1 | 0 | 14 | 1 | 0 | 0 | 0 | 0 | 1 | 70 | 49 | 1 | 1 | 0.091 | 0.412 | NA | | | 0 | 0.252 |
| A5SS | 1856 | NUP98 | NUP98 | chr11 | - | 3760487 | 3760626 | 3760538 | 3760626 | 3753315 | 3753408 | 40 | 18 | 0 | 24 | 18 | 4 | 0 | 4 | 0 | 2 | 98 | 49 | 1 | 1 | 0.455 | 0.333 | 0 | NA | | 0.5 | -0.237 |
| A5SS | 1859 | ZSWIM8 | ZSWIM8 | chr10 | + | 73800010 | 73800194 | 73800010 | 73800170 | 73800295 | 73800472 | 7 | 3 | 4 | 6 | 0 | 3 | 0 | 1 | 0 | 0 | 73 | 49 | 1 | 1 | 0.439 | 1 | 0.472 | NA | | 1 | -0.363 |
| A5SS | 1864 | KLF6 | KLF6 | chr10 | - | 3781640 | 3782214 | 3781766 | 3782214 | 3780105 | 3780229 | 132 | 44 | 18 | 2 | 0 | 0 | 3 | 35 | 0 | 0 | 98 | 49 | 1 | 1 | 0.971 | 1 | 1 | | 1 | 1 | -0.01 |
| A5SS | 1890 | SF1 | SF1 | chr11 | - | 64767566 | 64767844 | 64767570 | 64767844 | 64767191 | 64767251 | 0 | 3 | 0 | 8 | 0 | 1 | 0 | 0 | 0 | 3 | 53 | 49 | 1 | 1 | 0 | 1 | 0 | NA | | 0 | 0.333 |
| A5SS | 1901 | SUPT7L | SUPT7L | chr2 | - | 27663323 | 27663840 | 27663328 | 27663840 | 27662178 | 27662281 | 1 | 0 | 0 | 2 | 0 | 0 | 0 | 0 | 0 | 3 | 54 | 49 | 1 | 1 | 0.312 | NA | NA | NA | | 0 | 0.312 |
| A5SS | 1907 | P3H1 | P3H1 | chr1 | - | 42747252 | 42747412 | 42747271 | 42747412 | 42746334 | 42746852 | 0 | 0 | 0 | 1 | 0 | 0 | 0 | 1 | 0 | 0 | 68 | 49 | 1 | 1 | 0 | NA | NA | NA | | 1 | -1 |
| A5SS | 1909 | HDAC9 | HDAC9 | chr7 | + | 18591515 | 18591642 | 18591515 | 18591636 | 18593907 | 18594029 | 6 | 0 | 0 | 0 | 0 | 0 | 0 | 2 | 0 | 1 | 55 | 49 | 1 | 1 | 1 | NA | NA | NA | | 0.641 | 0.359 |
| A5SS | 1910 | MICAL2 | MICAL2 | chr11 | + | 12138389 | 12138464 | 12138389 | 12138460 | 12162078 | 12162419 | 7 | 2 | 0 | 2 | 5 | 0 | 0 | 1 | 0 | 0 | 53 | 49 | 1 | 1 | 0.764 | 0.27 | NA | NA | | 1 | -0.483 |
| A5SS | 1929 | NPL | NPL | chr1 | + | 182806144 | 182806531 | 182806144 | 182806232 | 182812155 | 182812213 | 12 | 8 | 0 | 37 | 9 | 6 | 0 | 10 | 0 | 0 | 98 | 49 | 1 | 1 | 0.14 | 0.308 | 0 | NA | | 1 | -0.851 |
| A5SS | 1936 | MADD | MADD | chr11 | + | 47284940 | 47285194 | 47284940 | 47285065 | 47285450 | 47285590 | 7 | 0 | 0 | 6 | 0 | 0 | 2 | 5 | 0 | 0 | 98 | 49 | 1 | 1 | 0.368 | NA | NA | | 1 | 1 | -0.632 |
| A5SS | 1950 | ABI3 | ABI3 | chr17 | + | 49216530 | 49216698 | 49216530 | 49216680 | 49217738 | 49217915 | 6 | 2 | 1 | 3 | 0 | 0 | 0 | 1 | 0 | 0 | 67 | 49 | 1 | 1 | 0.594 | 1 | 1 | NA | | 1 | -0.135 |
| A5SS | 1952 | JMJD4 | JMJD4 | chr1 | - | 227732880 | 227733027 | 227732928 | 227733027 | 227731188 | 227732676 | 1 | 0 | 0 | 1 | 0 | 0 | 0 | 1 | 0 | 0 | 97 | 49 | 1 | 1 | 0.336 | NA | NA | NA | | 1 | -0.664 |
| A5SS | 2000 | THTPA | THTPA | chr14 | + | 23555981 | 23557304 | 23555981 | 23557034 | 23558694 | 23559581 | 1 | 0 | 0 | 0 | 0 | 0 | 0 | 0 | 0 | 1 | 98 | 49 | 1 | 1 | 1 | NA | NA | NA | | 0 | 1 |
| A5SS | 2008 | ENTPD4 | ENTPD4 | chr8 | - | 23441568 | 23441723 | 23441592 | 23441723 | 23439748 | 23439915 | 5 | 2 | 0 | 4 | 0 | 1 | 0 | 2 | 0 | 0 | 73 | 49 | 1 | 1 | 0.456 | 1 | 0 | NA | | 1 | -0.515 |
| A5SS | 2014 | ANAPC15 | ANAPC15 | chr11 | - | 72111411 | 72111496 | 72111441 | 72111496 | 72111156 | 72111286 | 2 | 0 | 2 | 6 | 0 | 0 | 0 | 2 | 0 | 0 | 79 | 49 | 1 | 1 | 0.171 | NA | 1 | NA | | 1 | -0.414 |
| A5SS | 2035 | HPS5 | HPS5 | chr11 | - | 18321945 | 18322174 | 18322132 | 18322174 | 18317750 | 18317907 | 2 | 1 | 0 | 0 | 0 | 1 | 0 | 1 | 0 | 0 | 98 | 49 | 1 | 1 | 1 | 1 | 0 | NA | | 1 | -0.333 |
| A5SS | 2040 | ZNF276 | ZNF276 | chr16 | + | 89727278 | 89727357 | 89727278 | 89727325 | 89729234 | 89729318 | 11 | 3 | 1 | 4 | 2 | 0 | 0 | 8 | 0 | 2 | 81 | 49 | 1 | 1 | 0.625 | 0.476 | 1 | NA | | 0.708 | -0.008 |
| A5SS | 2055 | NDRG1 | NDRG1 | chr8 | - | 133296736 | 133297304 | 133297133 | 133297304 | 133284248 | 133284329 | 2 | 0 | 0 | 3 | 0 | 0 | 0 | 0 | 0 | 2 | 98 | 49 | 1 | 1 | 0.25 | NA | NA | NA | | 0 | 0.25 |
| A5SS | 2058 | TTC19 | TTC19 | chr17 | + | 16000117 | 16000382 | 16000117 | 16000245 | 16001914 | 16002025 | 0 | 3 | 0 | 0 | 0 | 0 | 0 | 0 | 0 | 1 | 98 | 49 | 1 | 1 | NA | 1 | NA | NA | | 0 | 1 |
| A5SS | 2073 | GOSR1 | GOSR1 | chr17 | + | 30481142 | 30481263 | 30481142 | 30481257 | 30484213 | 30484301 | 3 | 1 | 0 | 5 | 0 | 2 | 0 | 1 | 0 | 0 | 55 | 49 | 1 | 1 | 0.348 | 1 | 0 | NA | | 1 | -0.551 |
| A5SS | 2086 | STAT3 | STAT3 | chr17 | - | 42322281 | 42322494 | 42322284 | 42322494 | 42317181 | 42317224 | 61 | 32 | 30 | 16 | 10 | 10 | 0 | 19 | 0 | 4 | 52 | 49 | 1 | 1 | 0.782 | 0.751 | 0.739 | NA | | 0.817 | -0.06 |
| A5SS | 2095 | AOC2 | AOC2 | chr17 | + | 42849085 | 42849371 | 42849085 | 42849290 | 42849600 | 42849730 | 12 | 2 | 0 | 0 | 1 | 0 | 1 | 0 | 0 | 1 | 98 | 49 | 1 | 1 | 1 | 0.5 | NA | | 1 | 0 | 0.25 |
| A5SS | 2101 | TMEM230 | TMEM230 | chr20 | - | 5112729 | 5113087 | 5112960 | 5113087 | 5109331 | 5109445 | 0 | 2 | 0 | 1 | 0 | 0 | 0 | 0 | 0 | 2 | 98 | 49 | 1 | 1 | 0 | 1 | NA | NA | | 0 | 0.5 |
| A5SS | 2102 | TMEM230 | TMEM230 | chr20 | - | 5112742 | 5113087 | 5112960 | 5113087 | 5109331 | 5109445 | 2 | 2 | 0 | 1 | 0 | 0 | 0 | 0 | 0 | 2 | 98 | 49 | 1 | 1 | 0.5 | 1 | NA | NA | | 0 | 0.75 |
| A5SS | 2104 | KDM2A | KDM2A | chr11 | + | 67250085 | 67250798 | 67250085 | 67250346 | 67252693 | 67252857 | 30 | 10 | 3 | 1 | 0 | 0 | 2 | 7 | 0 | 0 | 98 | 49 | 1 | 1 | 0.937 | 1 | 1 | | 1 | 1 | -0.021 |
| A5SS | 2117 | BRPF1 | BRPF1 | chr3 | + | 9742024 | 9742171 | 9742024 | 9742153 | 9742943 | 9743253 | 6 | 0 | 1 | 1 | 3 | 0 | 0 | 1 | 0 | 2 | 67 | 49 | 1 | 1 | 0.814 | 0 | 1 | NA | | 0.268 | 0.337 |
| A5SS | 2120 | SUPT4H1 | SUPT4H1 | chr17 | - | 58352064 | 58352238 | 58352066 | 58352238 | 58351401 | 58351508 | 5 | 1 | 0 | 12 | 8 | 1 | 0 | 5 | 0 | 0 | 51 | 49 | 1 | 1 | 0.286 | 0.107 | 0 | NA | | 1 | -0.869 |
| A5SS | 2126 | CLTC | CLTC | chr17 | + | 59644275 | 59644495 | 59644275 | 59644483 | 59647397 | 59647666 | 2 | 0 | 0 | 12 | 3 | 3 | 0 | 0 | 0 | 1 | 61 | 49 | 1 | 1 | 0.118 | 0 | 0 | NA | | 0 | 0.039 |
| A5SS | 2135 | LYN | LYN | chr8 | + | 55941854 | 55941991 | 55941854 | 55941928 | 55946447 | 55946493 | 101 | 55 | 27 | 28 | 24 | 5 | 0 | 16 | 0 | 3 | 98 | 49 | 1 | 1 | 0.643 | 0.534 | 0.73 | NA | | 0.727 | -0.091 |
| A5SS | 2138 | SP140 | SP140 | chr2 | + | 230244987 | 230245080 | 230244987 | 230245071 | 230245862 | 230245940 | 3 | 6 | 0 | 3 | 0 | 0 | 2 | 3 | 0 | 0 | 58 | 49 | 1 | 1 | 0.458 | 1 | NA | | 1 | 1 | -0.271 |
| A5SS | 2147 | PRKAR1A | PRKAR1A | chr17 | + | 68512378 | 68512579 | 68512378 | 68512548 | 68515393 | 68515576 | 1 | 0 | 1 | 26 | 8 | 2 | 0 | 0 | 0 | 1 | 80 | 49 | 1 | 1 | 0.023 | 0 | 0.234 | NA | | 0 | 0.086 |
| A5SS | 2151 | CTAGE5 | CTAGE5 | chr14 | + | 39276933 | 39277065 | 39276933 | 39276955 | 39279336 | 39279358 | 43 | 21 | 4 | 2 | 0 | 1 | 1 | 6 | 0 | 0 | 98 | 49 | 1 | 1 | 0.915 | 1 | 0.667 | | 1 | 1 | -0.139 |
| A5SS | 2152 | CTAGE5 | CTAGE5 | chr14 | + | 39291018 | 39291111 | 39291018 | 39291096 | 39293270 | 39293381 | 1 | 1 | 0 | 2 | 5 | 0 | 0 | 0 | 0 | 3 | 64 | 49 | 1 | 1 | 0.277 | 0.133 | NA | NA | | 0 | 0.205 |
| A5SS | 2190 | ARHGDIA | ARHGDIA | chr17 | - | 81869329 | 81869625 | 81869541 | 81869625 | 81869172 | 81869236 | 21 | 15 | 3 | 1 | 0 | 0 | 3 | 16 | 0 | 1 | 98 | 49 | 1 | 1 | 0.913 | 1 | 1 | | 1 | 0.889 | 0.027 |
| A5SS | 2225 | DDX17 | DDX17 | chr22 | - | 38487872 | 38488115 | 38487878 | 38488115 | 38483437 | 38486440 | 8 | 1 | 2 | 27 | 5 | 0 | 0 | 0 | 0 | 2 | 55 | 49 | 1 | 1 | 0.209 | 0.151 | 1 | NA | | 1 | 0.453 |
| A5SS | 2247 | DYRK1A | DYRK1A | chr21 | + | 37506098 | 37506354 | 37506098 | 37506223 | 37511910 | 37515376 | 1 | 4 | 0 | 10 | 2 | 0 | 0 | 3 | 0 | 0 | 98 | 49 | 1 | 1 | 0.048 | 0.5 | NA | NA | | 1 | -0.726 |
| A5SS | 2270 | ADGRE1 | ADGRE1 | chr19 | + | 6928144 | 6928310 | 6928144 | 6928211 | 6934986 | 6935078 | 1 | 1 | 0 | 2 | 6 | 0 | 0 | 0 | 0 | 4 | 98 | 49 | 1 | 1 | 0.2 | 0.077 | NA | NA | | 0 | 0.139 |
| A5SS | 2278 | CREM | CREM | chr10 | + | 35195901 | 35196142 | 35195901 | 35196099 | 35206894 | 35207051 | 14 | 3 | 0 | 1 | 0 | 0 | 3 | 1 | 0 | 0 | 92 | 49 | 1 | 1 | 0.882 | 1 | NA | | 1 | 1 | -0.059 |
| A5SS | 2284 | TMEM205 | TMEM205 | chr19 | - | 11346007 | 11346305 | 11346126 | 11346305 | 11345519 | 11345628 | 1 | 0 | 2 | 3 | 0 | 0 | 0 | 2 | 0 | 1 | 98 | 49 | 1 | 1 | 0.143 | NA | 1 | NA | | 0.5 | 0.072 |
| A5SS | 2311 | CYP4F3 | CYP4F3 | chr19 | + | 15640896 | 15640945 | 15640896 | 15640929 | 15641414 | 15641613 | 1 | 1 | 1 | 0 | 4 | 0 | 4 | 0 | 0 | 0 | 65 | 49 | 1 | 1 | 1 | 0.159 | 1 | | 1 | NA | -0.28 |
| A5SS | 2312 | EPS15L1 | EPS15L1 | chr19 | - | 16440861 | 16440909 | 16440884 | 16440909 | 16437769 | 16437865 | 8 | 7 | 0 | 1 | 0 | 0 | 0 | 2 | 0 | 0 | 72 | 49 | 1 | 1 | 0.845 | 1 | NA | NA | | 1 | -0.078 |
| A5SS | 2314 | MYO9B | MYO9B | chr19 | + | 17211646 | 17211786 | 17211646 | 17211774 | 17211894 | 17213295 | 5 | 3 | 1 | 7 | 7 | 0 | 0 | 2 | 0 | 2 | 61 | 49 | 1 | 1 | 0.365 | 0.256 | 1 | NA | | 0.445 | 0.095 |
| A5SS | 2327 | CDV3 | CDV3 | chr3 | + | 133574433 | 133574756 | 133574433 | 133574592 | 133575038 | 133575115 | 10 | 0 | 0 | 2 | 0 | 0 | 0 | 1 | 0 | 0 | 98 | 49 | 1 | 1 | 0.714 | NA | NA | NA | | 1 | -0.286 |
| A5SS | 2334 | ING4 | ING4 | chr12 | - | 6652935 | 6653050 | 6652944 | 6653050 | 6652661 | 6652767 | 4 | 3 | 0 | 1 | 0 | 2 | 0 | 2 | 0 | 0 | 58 | 49 | 1 | 1 | 0.772 | 1 | 0 | NA | | 1 | -0.409 |
| A5SS | 2343 | CUTA | CUTA | chr6 | - | 33417868 | 33418288 | 33417958 | 33418288 | 33417480 | 33417695 | 1 | 1 | 0 | 0 | 1 | 0 | 0 | 0 | 0 | 1 | 98 | 49 | 1 | 1 | 1 | 0.333 | NA | NA | | 0 | 0.666 |
| A5SS | 2365 | C19orf47 | C19orf47 | chr19 | - | 40341785 | 40341890 | 40341838 | 40341890 | 40336319 | 40336407 | 0 | 1 | 0 | 0 | 0 | 0 | 0 | 0 | 0 | 1 | 98 | 49 | 1 | 1 | NA | 1 | NA | NA | | 0 | 1 |
| A5SS | 2383 | CLASRP | CLASRP | chr19 | + | 45068014 | 45068058 | 45068014 | 45068054 | 45068419 | 45068480 | 0 | 1 | 0 | 1 | 1 | 0 | 0 | 0 | 0 | 1 | 53 | 49 | 1 | 1 | 0 | 0.48 | NA | NA | | 0 | 0.24 |
| A5SS | 2389 | CNOT8 | CNOT8 | chr5 | + | 154858518 | 154858816 | 154858518 | 154858768 | 154863206 | 154863395 | 7 | 5 | 0 | 6 | 0 | 0 | 0 | 3 | 0 | 2 | 97 | 49 | 1 | 1 | 0.371 | 1 | NA | NA | | 0.431 | 0.255 |
| A5SS | 2403 | CD37 | CD37 | chr19 | + | 49335419 | 49335609 | 49335419 | 49335575 | 49335713 | 49335786 | 41 | 11 | 2 | 2 | 0 | 0 | 3 | 10 | 0 | 0 | 83 | 49 | 1 | 1 | 0.924 | 1 | 1 | | 1 | 1 | -0.025 |
| A5SS | 2509 | ZNF341 | ZNF341 | chr20 | + | 33757147 | 33757343 | 33757147 | 33757322 | 33758715 | 33758806 | 1 | 0 | 0 | 0 | 0 | 0 | 0 | 0 | 1 | 0 | 70 | 49 | 1 | 1 | 1 | NA | NA | | 0 | NA | 1 |
| A5SS | 2533 | ATAD2B | ATAD2B | chr2 | - | 23786026 | 23786223 | 23786041 | 23786223 | 23782868 | 23783028 | 7 | 2 | 1 | 0 | 3 | 0 | 0 | 0 | 0 | 1 | 64 | 49 | 1 | 1 | 1 | 0.338 | 1 | NA | | 0 | 0.779 |
| A5SS | 2545 | ZBTB8OS | ZBTB8OS | chr1 | - | 32650401 | 32650590 | 32650432 | 32650590 | 32634767 | 32634792 | 1 | 0 | 0 | 6 | 8 | 0 | 0 | 1 | 0 | 0 | 80 | 49 | 1 | 1 | 0.093 | 0 | NA | NA | | 1 | -0.954 |
| A5SS | 2548 | CLIP4 | CLIP4 | chr2 | + | 29157203 | 29157402 | 29157203 | 29157347 | 29160332 | 29160467 | 2 | 0 | 0 | 2 | 0 | 0 | 2 | 0 | 0 | 0 | 98 | 49 | 1 | 1 | 0.333 | NA | NA | | 1 | NA | -0.667 |
| A5SS | 2564 | SS18L1 | SS18L1 | chr20 | + | 62158671 | 62158942 | 62158671 | 62158748 | 62159876 | 62159961 | 3 | 0 | 0 | 0 | 3 | 2 | 0 | 0 | 0 | 1 | 98 | 49 | 1 | 1 | 1 | 0 | 0 | NA | | 0 | 0.333 |
| A5SS | 2565 | VRK2 | VRK2 | chr2 | + | 58046641 | 58046868 | 58046641 | 58046821 | 58048826 | 58048967 | 0 | 2 | 0 | 4 | 0 | 0 | 0 | 2 | 0 | 1 | 96 | 49 | 1 | 1 | 0 | 1 | NA | NA | | 0.505 | -0.005 |
| A5SS | 2585 | MAP4K4 | MAP4K4 | chr2 | + | 101887087 | 101887261 | 101887087 | 101887237 | 101887777 | 101887937 | 18 | 17 | 4 | 41 | 11 | 9 | 0 | 2 | 0 | 5 | 73 | 49 | 1 | 1 | 0.228 | 0.509 | 0.23 | NA | | 0.212 | 0.11 |
| A5SS | 2648 | GOLGB1 | GOLGB1 | chr3 | - | 121669196 | 121669355 | 121669211 | 121669355 | 121668060 | 121668158 | 1 | 0 | 0 | 5 | 0 | 0 | 0 | 1 | 0 | 1 | 64 | 49 | 1 | 1 | 0.133 | NA | NA | NA | | 0.434 | -0.301 |
| A5SS | 2649 | GOLGB1 | GOLGB1 | chr3 | - | 121730847 | 121730973 | 121730875 | 121730973 | 121729864 | 121730017 | 3 | 3 | 0 | 10 | 5 | 0 | 0 | 2 | 0 | 0 | 77 | 49 | 1 | 1 | 0.16 | 0.276 | NA | NA | | 1 | -0.782 |
| A5SS | 2650 | SP100 | SP100 | chr2 | + | 230467119 | 230467215 | 230467119 | 230467206 | 230469042 | 230469096 | 26 | 33 | 4 | 4 | 0 | 0 | 3 | 13 | 0 | 1 | 58 | 49 | 1 | 1 | 0.846 | 1 | 1 | | 1 | 0.917 | -0.01 |
| A5SS | 2658 | CCDC66 | CCDC66 | chr3 | + | 56566593 | 56566759 | 56566593 | 56566740 | 56566949 | 56567053 | 6 | 0 | 7 | 13 | 4 | 0 | 0 | 1 | 0 | 0 | 68 | 49 | 1 | 1 | 0.25 | 0 | 1 | NA | | 1 | -0.583 |
| A5SS | 2663 | ATG4B | ATG4B | chr2 | + | 241671311 | 241671425 | 241671311 | 241671405 | 241672190 | 241673856 | 2 | 3 | 0 | 0 | 2 | 3 | 0 | 2 | 0 | 0 | 69 | 49 | 1 | 1 | 1 | 0.516 | 0 | NA | | 1 | -0.495 |
| A5SS | 2710 | HM13 | HM13 | chr20 | + | 31568077 | 31568280 | 31568077 | 31568224 | 31569119 | 31569567 | 3 | 2 | 0 | 0 | 3 | 0 | 0 | 1 | 0 | 0 | 98 | 49 | 1 | 1 | 1 | 0.25 | NA | NA | | 1 | -0.375 |
| A5SS | 2715 | POLD4 | POLD4 | chr11 | - | 67353293 | 67353596 | 67353302 | 67353596 | 67352987 | 67353077 | 2 | 0 | 0 | 3 | 3 | 0 | 0 | 0 | 0 | 1 | 58 | 49 | 1 | 1 | 0.36 | 0 | NA | NA | | 0 | 0.18 |
| A5SS | 2722 | CPNE1 | CPNE1 | chr20 | - | 35664759 | 35664956 | 35664801 | 35664956 | 35632794 | 35632923 | 7 | 5 | 0 | 3 | 2 | 0 | 0 | 2 | 0 | 0 | 91 | 49 | 1 | 1 | 0.557 | 0.574 | NA | NA | | 1 | -0.434 |
| A5SS | 2725 | C20orf24 | C20orf24 | chr20 | + | 36609600 | 36609758 | 36609600 | 36609730 | 36612009 | 36612557 | 7 | 4 | 2 | 20 | 30 | 28 | 0 | 1 | 0 | 4 | 77 | 49 | 1 | 1 | 0.182 | 0.078 | 0.043 | NA | | 0.137 | -0.036 |
| A5SS | 2737 | NCOA3 | NCOA3 | chr20 | + | 47649004 | 47649109 | 47649004 | 47649097 | 47650981 | 47651276 | 6 | 3 | 0 | 0 | 2 | 0 | 0 | 2 | 0 | 0 | 61 | 49 | 1 | 1 | 1 | 0.546 | NA | NA | | 1 | -0.227 |
| A5SS | 2749 | ZGPAT | ZGPAT | chr20 | + | 63708027 | 63708188 | 63708027 | 63708118 | 63708552 | 63709164 | 1 | 0 | 0 | 2 | 1 | 0 | 0 | 2 | 0 | 0 | 98 | 49 | 1 | 1 | 0.2 | 0 | NA | NA | | 1 | -0.9 |
| A5SS | 2787 | SUMO3 | SUMO3 | chr21 | - | 44813861 | 44814104 | 44813975 | 44814104 | 44809046 | 44809118 | 1 | 0 | 0 | 3 | 0 | 1 | 0 | 0 | 0 | 1 | 98 | 49 | 1 | 1 | 0.143 | NA | 0 | NA | | 0 | 0.071 |
| A5SS | 2797 | TANGO2 | TANGO2 | chr22 | + | 20036759 | 20037094 | 20036759 | 20036854 | 20043354 | 20043443 | 3 | 5 | 0 | 4 | 2 | 3 | 0 | 2 | 0 | 7 | 98 | 49 | 1 | 1 | 0.273 | 0.556 | 0 | NA | | 0.125 | 0.151 |
| A5SS | 2798 | TANGO2 | TANGO2 | chr22 | + | 20061529 | 20061683 | 20061529 | 20061681 | 20063337 | 20063442 | 0 | 6 | 2 | 1 | 0 | 0 | 0 | 2 | 0 | 0 | 51 | 49 | 1 | 1 | 0 | 1 | 1 | NA | | 1 | -0.333 |
| A5SS | 2810 | SMARCB1 | SMARCB1 | chr22 | + | 23791755 | 23791894 | 23791755 | 23791867 | 23793558 | 23793688 | 1 | 0 | 0 | 1 | 1 | 0 | 0 | 1 | 0 | 1 | 76 | 49 | 1 | 1 | 0.392 | 0 | NA | NA | | 0.392 | -0.196 |
| A5SS | 2815 | ADORA2A | ADORA2A | chr22 | + | 24427561 | 24427746 | 24427561 | 24427726 | 24433130 | 24433736 | 3 | 1 | 0 | 2 | 0 | 0 | 3 | 0 | 0 | 0 | 69 | 49 | 1 | 1 | 0.516 | 1 | NA | | 1 | NA | -0.242 |
| A5SS | 2816 | ADORA2A | ADORA2A | chr22 | + | 24427561 | 24427746 | 24427561 | 24427726 | 24440582 | 24442361 | 2 | 2 | 0 | 0 | 2 | 0 | 3 | 0 | 0 | 0 | 69 | 49 | 1 | 1 | 1 | 0.415 | NA | | 1 | NA | -0.292 |
| A5SS | 2828 | FAM193A | FAM193A | chr4 | + | 2694945 | 2695129 | 2694945 | 2695082 | 2696362 | 2696593 | 7 | 3 | 2 | 1 | 0 | 0 | 2 | 2 | 0 | 0 | 96 | 49 | 1 | 1 | 0.781 | 1 | 1 | | 1 | 1 | -0.073 |
| A5SS | 2829 | FAM193A | FAM193A | chr4 | + | 2694945 | 2695142 | 2694945 | 2695082 | 2696362 | 2696593 | 5 | 2 | 2 | 1 | 0 | 0 | 0 | 2 | 0 | 0 | 98 | 49 | 1 | 1 | 0.714 | 1 | 1 | NA | | 1 | -0.095 |
| A5SS | 2831 | TOM1 | TOM1 | chr22 | + | 35346929 | 35346972 | 35346929 | 35346969 | 35347054 | 35347994 | 3 | 0 | 2 | 2 | 0 | 3 | 0 | 0 | 0 | 5 | 52 | 49 | 1 | 1 | 0.586 | NA | 0.386 | NA | | 0 | 0.486 |
| A5SS | 2835 | MAFF | MAFF | chr22 | + | 38201931 | 38202238 | 38201931 | 38202212 | 38213822 | 38213889 | 5 | 1 | 0 | 3 | 4 | 0 | 0 | 0 | 0 | 2 | 75 | 49 | 1 | 1 | 0.521 | 0.14 | NA | NA | | 0 | 0.331 |
| A5SS | 2847 | TLE3 | TLE3 | chr15 | - | 70058149 | 70058291 | 70058185 | 70058291 | 70057458 | 70057658 | 20 | 2 | 3 | 5 | 2 | 1 | 0 | 2 | 0 | 1 | 85 | 49 | 1 | 1 | 0.698 | 0.366 | 0.634 | NA | | 0.536 | 0.03 |
| A5SS | 2849 | TLE3 | TLE3 | chr15 | - | 70074502 | 70074607 | 70074532 | 70074607 | 70066013 | 70066218 | 21 | 12 | 3 | 27 | 10 | 5 | 0 | 7 | 0 | 4 | 79 | 49 | 1 | 1 | 0.325 | 0.427 | 0.271 | NA | | 0.52 | -0.179 |
| A5SS | 2855 | ARSA | ARSA | chr22 | - | 50627555 | 50628173 | 50627945 | 50628173 | 50627165 | 50627406 | 8 | 5 | 2 | 1 | 0 | 0 | 1 | 0 | 0 | 0 | 98 | 49 | 1 | 1 | 0.8 | 1 | 1 | | 1 | NA | -0.067 |
| A5SS | 2860 | ITPR1 | ITPR1 | chr3 | + | 4675067 | 4675248 | 4675067 | 4675221 | 4676613 | 4676801 | 1 | 3 | 0 | 4 | 1 | 0 | 0 | 1 | 0 | 3 | 76 | 49 | 1 | 1 | 0.139 | 0.659 | NA | NA | | 0.177 | 0.222 |
| A5SS | 2869 | CASP8AP2 | CASP8AP2 | chr6 | + | 89829899 | 89830106 | 89829899 | 89830067 | 89846561 | 89846644 | 1 | 2 | 0 | 1 | 0 | 0 | 3 | 0 | 0 | 0 | 88 | 49 | 1 | 1 | 0.358 | 1 | NA | | 1 | NA | -0.321 |
| A5SS | 2873 | FGD5-AS1 | FGD5-AS1 | chr3 | - | 14947086 | 14947505 | 14947113 | 14947505 | 14942778 | 14946154 | 3 | 2 | 1 | 1 | 1 | 0 | 0 | 1 | 0 | 0 | 76 | 49 | 1 | 1 | 0.659 | 0.563 | 1 | NA | | 1 | -0.259 |
| A5SS | 2877 | MS4A6A | MS4A6A | chr11 | - | 60175401 | 60175611 | 60175505 | 60175611 | 60173027 | 60173129 | 21 | 1 | 18 | 5 | 0 | 0 | 0 | 27 | 0 | 0 | 98 | 49 | 1 | 1 | 0.677 | 1 | 1 | NA | | 1 | -0.108 |
| A5SS | 2883 | GORASP1 | GORASP1 | chr3 | - | 39100746 | 39100877 | 39100787 | 39100877 | 39100304 | 39100503 | 2 | 0 | 1 | 2 | 0 | 0 | 1 | 0 | 0 | 0 | 90 | 49 | 1 | 1 | 0.353 | NA | | | 1 | NA | -0.324 |
| A5SS | 2889 | LPPR2 | LPPR2 | chr19 | + | 11363712 | 11363835 | 11363712 | 11363819 | 11364160 | 11364212 | 8 | 4 | 6 | 9 | 0 | 2 | 0 | 2 | 0 | 1 | 65 | 49 | 1 | 1 | 0.401 | 1 | 0.693 | NA | | 0.601 | 0.097 |
| A5SS | 2911 | ENY2 | ENY2 | chr8 | + | 109334322 | 109334474 | 109334322 | 109334402 | 109336127 | 109336204 | 1 | 1 | 3 | 0 | 0 | 0 | 0 | 1 | 0 | 2 | 98 | 49 | 1 | 1 | 1 | 1 | 1 | NA | | 0.2 | 0.8 |
| A5SS | 2921 | ERBB2IP | ERBB2IP | chr5 | + | 66044136 | 66044310 | 66044136 | 66044298 | 66046352 | 66046538 | 31 | 16 | 9 | 3 | 2 | 0 | 0 | 8 | 0 | 0 | 61 | 49 | 1 | 1 | 0.892 | 0.865 | 1 | NA | | 1 | -0.081 |
| A5SS | 2940 | PARP9 | PARP9 | chr3 | - | 122558328 | 122558467 | 122558433 | 122558467 | 122555285 | 122556121 | 8 | 7 | 0 | 20 | 9 | 2 | 0 | 10 | 0 | 6 | 98 | 49 | 1 | 1 | 0.167 | 0.28 | 0 | NA | | 0.455 | -0.306 |
| A5SS | 2941 | ZXDC | ZXDC | chr3 | - | 126461269 | 126462220 | 126461534 | 126462220 | 126459652 | 126459737 | 6 | 3 | 0 | 4 | 0 | 0 | 6 | 4 | 0 | 0 | 98 | 49 | 1 | 1 | 0.429 | 1 | NA | | 1 | 1 | -0.285 |
| A5SS | 2942 | ACAD9 | ACAD9 | chr3 | + | 128879489 | 128880017 | 128879489 | 128879841 | 128884652 | 128884746 | 1 | 2 | 0 | 1 | 1 | 0 | 0 | 4 | 0 | 2 | 98 | 49 | 1 | 1 | 0.333 | 0.5 | NA | NA | | 0.5 | -0.084 |
| A5SS | 2944 | MBD4 | MBD4 | chr3 | - | 129436442 | 129437308 | 129436460 | 129437308 | 129434061 | 129434136 | 4 | 0 | 0 | 24 | 4 | 2 | 0 | 0 | 0 | 1 | 67 | 49 | 1 | 1 | 0.109 | 0 | 0 | NA | | 0 | 0.036 |
| A5SS | 2945 | MBD4 | MBD4 | chr3 | - | 129437719 | 129437950 | 129437807 | 129437950 | 129436460 | 129437308 | 24 | 13 | 3 | 1 | 0 | 1 | 1 | 2 | 0 | 0 | 98 | 49 | 1 | 1 | 0.923 | 1 | 0.6 | | 1 | 1 | -0.159 |
| A5SS | 2963 | MYNN | MYNN | chr3 | + | 169774264 | 169774561 | 169774264 | 169774431 | 169778767 | 169779561 | 16 | 4 | 1 | 1 | 0 | 0 | 0 | 1 | 0 | 0 | 98 | 49 | 1 | 1 | 0.889 | 1 | 1 | NA | | 1 | -0.037 |
| A5SS | 2981 | SMARCA2 | SMARCA2 | chr9 | + | 2086828 | 2087071 | 2086828 | 2086897 | 2088499 | 2088613 | 14 | 3 | 2 | 2 | 0 | 0 | 0 | 1 | 0 | 0 | 98 | 49 | 1 | 1 | 0.778 | 1 | 1 | NA | | 1 | -0.074 |
| A5SS | 2984 | TMEM175 | TMEM175 | chr4 | + | 948115 | 948615 | 948115 | 948154 | 950420 | 950518 | 4 | 1 | 2 | 0 | 0 | 0 | 0 | 0 | 0 | 1 | 98 | 49 | 1 | 1 | 1 | 1 | 1 | NA | | 0 | 1 |
| A5SS | 2998 | EXOC1 | EXOC1 | chr4 | + | 55853649 | 55853953 | 55853649 | 55853797 | 55858313 | 55858447 | 5 | 2 | 2 | 4 | 4 | 0 | 0 | 1 | 0 | 0 | 98 | 49 | 1 | 1 | 0.385 | 0.2 | 1 | NA | | 1 | -0.472 |
| A5SS | 3003 | CNOT6L | CNOT6L | chr4 | - | 77819232 | 77819390 | 77819303 | 77819390 | 77776270 | 77776392 | 13 | 12 | 0 | 1 | 0 | 0 | 0 | 1 | 0 | 0 | 98 | 49 | 1 | 1 | 0.867 | 1 | NA | NA | | 1 | -0.067 |
| A5SS | 3006 | SNX10 | SNX10 | chr7 | + | 26291894 | 26292133 | 26291894 | 26292086 | 26346419 | 26346466 | 7 | 1 | 1 | 9 | 1 | 2 | 0 | 0 | 0 | 2 | 96 | 49 | 1 | 1 | 0.284 | 0.338 | 0.203 | NA | | 0 | 0.275 |
| A5SS | 3007 | ANTXR2 | ANTXR2 | chr4 | - | 80072408 | 80073323 | 80072708 | 80073323 | 80071582 | 80071654 | 46 | 15 | 16 | 1 | 0 | 0 | 10 | 10 | 0 | 0 | 98 | 49 | 1 | 1 | 0.958 | 1 | 1 | | 1 | 1 | -0.014 |
| A5SS | 3019 | CNN2 | CNN2 | chr19 | + | 1032558 | 1032696 | 1032558 | 1032663 | 1036129 | 1036246 | 117 | 73 | 37 | 0 | 3 | 0 | 8 | 20 | 0 | 0 | 82 | 49 | 1 | 1 | 1 | 0.936 | 1 | | 1 | 1 | -0.021 |
| A5SS | 3056 | PAIP1 | PAIP1 | chr5 | - | 43556581 | 43557093 | 43556818 | 43557093 | 43555829 | 43555999 | 2 | 0 | 0 | 1 | 1 | 0 | 1 | 1 | 0 | 0 | 98 | 49 | 1 | 1 | 0.5 | 0 | NA | | 1 | 1 | -0.75 |
| A5SS | 3059 | GAPT | GAPT | chr5 | + | 58493618 | 58493838 | 58493618 | 58493683 | 58494246 | 58496358 | 7 | 1 | 0 | 4 | 1 | 0 | 0 | 2 | 0 | 2 | 98 | 49 | 1 | 1 | 0.467 | 0.333 | NA | NA | | 0.333 | 0.067 |
| A5SS | 3065 | C14orf159 | C14orf159 | chr14 | + | 91173288 | 91173439 | 91173288 | 91173403 | 91175933 | 91176056 | 66 | 21 | 10 | 8 | 4 | 0 | 0 | 12 | 0 | 2 | 85 | 49 | 1 | 1 | 0.826 | 0.752 | 1 | NA | | 0.776 | 0.083 |
| A5SS | 3078 | GLRX | GLRX | chr5 | - | 95816495 | 95816626 | 95816506 | 95816626 | 95813848 | 95814389 | 0 | 3 | 0 | 10 | 3 | 1 | 0 | 0 | 0 | 3 | 60 | 49 | 1 | 1 | 0 | 0.45 | 0 | NA | | 0 | 0.15 |
| A5SS | 3089 | HINT1 | HINT1 | chr5 | - | 131165059 | 131165348 | 131165094 | 131165348 | 131162571 | 131162676 | 3 | 0 | 0 | 23 | 5 | 0 | 0 | 0 | 0 | 5 | 84 | 49 | 1 | 1 | 0.071 | 0 | NA | NA | | 0 | 0.035 |
| A5SS | 3090 | CNBP | CNBP | chr3 | - | 129171633 | 129171771 | 129171654 | 129171771 | 129171445 | 129171538 | 8 | 7 | 10 | 24 | 12 | 13 | 0 | 4 | 0 | 5 | 70 | 49 | 1 | 1 | 0.189 | 0.29 | 0.35 | NA | | 0.359 | -0.083 |
| A5SS | 3100 | ETF1 | ETF1 | chr5 | - | 138542633 | 138542936 | 138542832 | 138542936 | 138518691 | 138518867 | 36 | 7 | 0 | 20 | 18 | 0 | 0 | 2 | 2 | 4 | 98 | 49 | 1 | 1 | 0.474 | 0.163 | NA | | 0 | 0.2 | 0.218 |
| A5SS | 3106 | SCARF1 | SCARF1 | chr17 | - | 1645559 | 1645789 | 1645596 | 1645789 | 1645177 | 1645239 | 2 | 3 | 1 | 5 | 0 | 0 | 0 | 0 | 0 | 1 | 86 | 49 | 1 | 1 | 0.186 | 1 | 1 | NA | | 0 | 0.729 |
| A5SS | 3113 | ARHGAP26 | ARHGAP26 | chr5 | + | 143207197 | 143207473 | 143207197 | 143207308 | 143213996 | 143214088 | 8 | 0 | 2 | 6 | 4 | 0 | 0 | 0 | 0 | 1 | 98 | 49 | 1 | 1 | 0.4 | 0 | 1 | NA | | 0 | 0.467 |
| A5SS | 3114 | ZNF438 | ZNF438 | chr10 | - | 30848530 | 30850367 | 30850221 | 30850367 | 30844635 | 30845573 | 12 | 4 | 2 | 1 | 0 | 0 | 1 | 7 | 0 | 0 | 98 | 49 | 1 | 1 | 0.857 | 1 | 1 | | 1 | 1 | -0.048 |
| A5SS | 3115 | CSNK1A1 | CSNK1A1 | chr5 | - | 149505446 | 149505595 | 149505482 | 149505595 | 149495893 | 149496860 | 35 | 10 | 0 | 10 | 3 | 0 | 0 | 11 | 0 | 1 | 85 | 49 | 1 | 1 | 0.669 | 0.658 | NA | NA | | 0.864 | -0.201 |
| A5SS | 3122 | FAM193B | FAM193B | chr5 | - | 177553167 | 177554547 | 177554248 | 177554547 | 177538904 | 177539147 | 16 | 4 | 0 | 1 | 0 | 0 | 0 | 4 | 0 | 0 | 98 | 49 | 1 | 1 | 0.889 | 1 | NA | NA | | 1 | -0.055 |

| | | | | | | | | | | | | | | | | | | | | | | | | | | | | | | | |
|---|---|---|---|---|---|---|---|---|---|---|---|---|---|---|---|---|---|---|---|---|---|---|---|---|---|---|---|---|---|---|---|
| A5SS | 3123 | PHYKPL | PHYKPL | chr5 | - | 178231668 | 178232822 | 178232491 | 178232822 | 178231404 | 178231523 | 1 | 0 | 0 | 0 | 0 | 0 | 0 | 3 | 2 | 0 | 98 | 49 | 1 | 1 | 1 | NA | NA | 0 | 1 | 0.5 |
| A5SS | 3129 | MCMBP | MCMBP | chr10 | - | 119843247 | 119843426 | 119843253 | 119843426 | 119842471 | 119842595 | 0 | 2 | 7 | 2 | 1 | 2 | 0 | 0 | 0 | 3 | 55 | 49 | 1 | 1 | 0 | 0.641 | 0.757 | NA | 0 | 0.466 |
| SE | 20 | VAMP7 | VAMP7 | chrX | + | 155900496 | 155900587 | 155898111 | 155898249 | 155919812 | 155919880 | 12 | 7 | 3 | 1 | 0 | 0 | 0 | 2 | 0 | 0 | 98 | 49 | 1 | 1 | 0.857 | 1 | 1 | NA | 1 | -0.048 |
| SE | 30 | FLNA | FLNA | chrX | - | 154352552 | 154352675 | 154352180 | 154352447 | 154352771 | 154352924 | 85 | 28 | 9 | 1 | 0 | 0 | 9 | 44 | 0 | 1 | 98 | 49 | 1 | 1 | 0.977 | 1 | 1 | 1 | 0.957 | 0.014 |
| SE | 31 | FLNA | FLNA | chrX | - | 154353553 | 154353727 | 154353295 | 154353457 | 154353914 | 154354043 | 72 | 9 | 10 | 1 | 1 | 0 | 2 | 35 | 0 | 1 | 98 | 49 | 1 | 1 | 0.973 | 0.818 | 1 | 1 | 0.946 | -0.043 |
| SE | 32 | FLNA | FLNA | chrX | - | 154354615 | 154354711 | 154354380 | 154354483 | 154354824 | 154355072 | 79 | 21 | 7 | 0 | 0 | 0 | 2 | 25 | 0 | 1 | 98 | 49 | 1 | 1 | 1 | 1 | 1 | 1 | 0.926 | 0.037 |
| SE | 34 | FLNA | FLNA | chrX | - | 154359989 | 154360587 | 154359731 | 154359905 | 154361307 | 154361570 | 88 | 21 | 6 | 2 | 0 | 0 | 8 | 61 | 0 | 0 | 98 | 49 | 1 | 1 | 0.957 | 1 | 1 | 1 | 1 | -0.014 |
| SE | 35 | FLNA | FLNA | chrX | - | 154361669 | 154361787 | 154361307 | 154361570 | 154361978 | 154362148 | 49 | 27 | 7 | 2 | 0 | 0 | 0 | 22 | 0 | 0 | 98 | 49 | 1 | 1 | 0.925 | 1 | 1 | NA | 1 | -0.025 |
| SE | 38 | IDH3G | IDH3G | chrX | - | 153786800 | 153786947 | 153786354 | 153786449 | 153787050 | 153787153 | 8 | 0 | 3 | 1 | 0 | 3 | 2 | 2 | 0 | 0 | 98 | 49 | 1 | 1 | 0.8 | NA | 0.333 | 1 | 1 | -0.433 |
| SE | 49 | IDS | IDS | chrX | - | 149491537 | 149491667 | 149490313 | 149490440 | 149496345 | 149496516 | 0 | 0 | 1 | 60 | 26 | 19 | 0 | 1 | 0 | 7 | 98 | 49 | 1 | 1 | 0 | 0 | 0.026 | NA | 0.067 | -0.058 |
| SE | 50 | IDS | IDS | chrX | - | 149498106 | 149498307 | 149496345 | 149496516 | 149500948 | 149501037 | 102 | 48 | 33 | 0 | 1 | 2 | 0 | 9 | 0 | 0 | 98 | 49 | 1 | 1 | 1 | 0.96 | 0.892 | NA | 1 | -0.049 |
| SE | 53 | IDS | IDS | chrX | - | 149504156 | 149504293 | 149500948 | 149501037 | 149505034 | 149505354 | 42 | 12 | 17 | 1 | 1 | 0 | 1 | 14 | 0 | 1 | 98 | 49 | 1 | 1 | 0.955 | 0.857 | 1 | 1 | 0.875 | 0 |
| SE | 54 | ZNF75D | ZNF75D | chrX | - | 135293729 | 135294258 | 135291008 | 135291135 | 135295767 | 135296039 | 5 | 0 | 0 | 0 | 0 | 0 | 0 | 0 | 0 | 1 | 98 | 49 | 1 | 1 | 1 | NA | NA | NA | 0 | 1 |
| SE | 55 | MOSPD1 | MOSPD1 | chrX | - | 134891478 | 134891640 | 134887625 | 134889192 | 134896816 | 134897034 | 1 | 1 | 0 | 0 | 0 | 0 | 0 | 0 | 0 | 1 | 98 | 49 | 1 | 1 | 1 | 1 | NA | NA | 0 | 1 |
| SE | 62 | ELF4 | ELF4 | chrX | - | 130081255 | 130081539 | 130074580 | 130074752 | 130110324 | 130110500 | 9 | 9 | 2 | 2 | 0 | 0 | 2 | 5 | 1 | 0 | 98 | 49 | 1 | 1 | 0.692 | 1 | 1 | 0.5 | 1 | 0.147 |
| SE | 69 | STAG2 | STAG2 | chrX | + | 124021366 | 124021431 | 123960559 | 123960866 | 124022530 | 124022671 | 23 | 8 | 2 | 4 | 0 | 0 | 0 | 2 | 0 | 0 | 98 | 49 | 1 | 1 | 0.742 | 1 | 1 | NA | 1 | -0.086 |
| SE | 70 | STAG2 | STAG2 | chrX | + | 124021366 | 124021431 | 123961705 | 123961856 | 124022530 | 124022671 | 24 | 7 | 4 | 13 | 2 | 3 | 0 | 2 | 0 | 3 | 98 | 49 | 1 | 1 | 0.48 | 0.636 | 0.4 | NA | 0.25 | 0.255 |
| SE | 71 | STAG2 | STAG2 | chrX | + | 124090853 | 124090964 | 124090574 | 124090764 | 124094017 | 124094144 | 7 | 2 | 2 | 43 | 16 | 4 | 0 | 0 | 0 | 3 | 98 | 49 | 1 | 1 | 0.075 | 0.059 | 0.2 | NA | 0 | 0.111 |
| SE | 72 | XIAP | XIAP | chrX | + | 123885630 | 123886539 | 123860166 | 123860293 | 123888618 | 123888718 | 6 | 0 | 2 | 1 | 0 | 0 | 0 | 1 | 0 | 0 | 98 | 49 | 1 | 1 | 0.75 | NA | 1 | NA | 1 | -0.125 |
| SE | 83 | LONRF3 | LONRF3 | chrX | + | 119013038 | 119013201 | 119011814 | 119011973 | 119014206 | 119014356 | 0 | 0 | 0 | 1 | 0 | 0 | 0 | 1 | 0 | 0 | 98 | 49 | 1 | 1 | 0 | NA | NA | NA | 1 | -1 |
| SE | 84 | DOCK11 | DOCK11 | chrX | + | 118590209 | 118590302 | 118588412 | 118588478 | 118593213 | 118593337 | 77 | 26 | 27 | 7 | 3 | 0 | 0 | 15 | 0 | 1 | 98 | 49 | 1 | 1 | 0.846 | 0.812 | 1 | NA | 0.882 | 0.004 |
| SE | 85 | DOCK11 | DOCK11 | chrX | + | 118639434 | 118639577 | 118638079 | 118638127 | 118641189 | 118641305 | 46 | 22 | 1 | 2 | 0 | 0 | 0 | 4 | 0 | 0 | 98 | 49 | 1 | 1 | 0.92 | 1 | 1 | NA | 1 | -0.027 |
| SE | 86 | DOCK11 | DOCK11 | chrX | + | 118643456 | 118643594 | 118641189 | 118641305 | 118648944 | 118649127 | 42 | 22 | 10 | 0 | 1 | 0 | 0 | 10 | 0 | 0 | 98 | 49 | 1 | 1 | 1 | 0.917 | 1 | NA | 1 | -0.028 |
| SE | 87 | DOCK11 | DOCK11 | chrX | + | 118681693 | 118681794 | 118681057 | 118681248 | 118683078 | 118683217 | 40 | 19 | 8 | 1 | 0 | 0 | 1 | 6 | 0 | 0 | 98 | 49 | 1 | 1 | 0.952 | 1 | 1 | 1 | 1 | -0.016 |
| SE | 94 | ACSL4 | ACSL4 | chrX | - | 109661530 | 109661645 | 109659353 | 109659511 | 109663210 | 109663402 | 24 | 14 | 10 | 1 | 0 | 0 | 1 | 2 | 0 | 0 | 98 | 49 | 1 | 1 | 0.923 | 1 | 1 | 1 | 1 | -0.026 |
| SE | 123 | DIAPH2 | DIAPH2 | chrX | + | 96735757 | 96735790 | 96684662 | 96685190 | 96738585 | 96738762 | 2 | 1 | 2 | 1 | 1 | 0 | 0 | 1 | 0 | 0 | 82 | 49 | 1 | 1 | 0.544 | 0.374 | 1 | NA | 1 | -0.361 |
| SE | 136 | PGK1 | PGK1 | chrX | + | 78113743 | 78113899 | 78109866 | 78109917 | 78114015 | 78114160 | 33 | 16 | 18 | 1 | 0 | 0 | 0 | 6 | 0 | 0 | 98 | 49 | 1 | 1 | 0.943 | 1 | 1 | NA | 1 | -0.019 |
| SE | 139 | ATRX | ATRX | chrX | - | 77600433 | 77600564 | 77599731 | 77599820 | 77616612 | 77616730 | 13 | 7 | 0 | 0 | 0 | 0 | 0 | 2 | 0 | 2 | 98 | 49 | 1 | 1 | 1 | 1 | NA | NA | 0.333 | 0.667 |
| SE | 140 | ATRX | ATRX | chrX | - | 77693823 | 77693937 | 77688817 | 77688927 | 77696576 | 77696704 | 15 | 1 | 5 | 10 | 0 | 0 | 0 | 3 | 0 | 0 | 98 | 49 | 1 | 1 | 0.429 | 1 | 1 | NA | 1 | -0.19 |
| SE | 153 | ABCB7 | ABCB7 | chrX | - | 75114753 | 75114834 | 75112885 | 75112972 | 75156104 | 75156340 | 1 | 1 | 1 | 1 | 0 | 0 | 0 | 1 | 0 | 0 | 98 | 49 | 1 | 1 | 0.333 | 1 | 1 | NA | 1 | -0.222 |
| SE | 154 | RLIM | RLIM | chrX | - | 74612581 | 74612643 | 74595808 | 74596000 | 74614421 | 74614626 | 6 | 3 | 5 | 2 | 1 | 1 | 0 | 0 | 0 | 1 | 98 | 49 | 1 | 1 | 0.6 | 0.6 | 0.714 | NA | 0 | 0.638 |
| SE | 159 | NHSL2 | NHSL2 | chrX | + | 72137093 | 72137225 | 72134508 | 72134704 | 72138440 | 72140771 | 9 | 3 | 0 | 3 | 1 | 0 | 0 | 2 | 0 | 2 | 98 | 49 | 1 | 1 | 0.6 | 0.6 | NA | NA | 0.333 | 0.267 |
| SE | 160 | TAF1 | TAF1 | chrX | + | 71388236 | 71388378 | 71387260 | 71387461 | 71388737 | 71388868 | 9 | 0 | 2 | 0 | 0 | 0 | 0 | 3 | 0 | 1 | 98 | 49 | 1 | 1 | 1 | NA | 1 | NA | 0.6 | 0.4 |
| SE | 165 | NLGN3 | NLGN3 | chrX | + | 71153476 | 71153536 | 71147549 | 71148206 | 71155213 | 71155363 | 1 | 0 | 0 | 0 | 3 | 0 | 1 | 0 | 0 | 0 | 98 | 49 | 1 | 1 | 1 | 0 | NA | 1 | NA | -0.5 |
| SE | 167 | MED12 | MED12 | chrX | + | 71125350 | 71125495 | 71124975 | 71125146 | 71125662 | 71125713 | 11 | 4 | 7 | 2 | 0 | 0 | 0 | 3 | 0 | 0 | 98 | 49 | 1 | 1 | 0.733 | 1 | 1 | NA | 1 | -0.089 |
| SE | 168 | MED12 | MED12 | chrX | + | 71132844 | 71132956 | 71132376 | 71132538 | 71133122 | 71133212 | 5 | 9 | 1 | 1 | 0 | 0 | 0 | 6 | 0 | 0 | 98 | 49 | 1 | 1 | 0.714 | 1 | 1 | NA | 1 | -0.095 |
| SE | 170 | IL2RG | IL2RG | chrX | - | 71110155 | 71110295 | 71109227 | 71109390 | 71110503 | 71110688 | 52 | 18 | 18 | 5 | 2 | 0 | 0 | 7 | 0 | 0 | 98 | 49 | 1 | 1 | 0.839 | 0.818 | 1 | NA | 1 | -0.114 |
| SE | 171 | IL2RG | IL2RG | chrX | - | 71110896 | 71111050 | 71110503 | 71110688 | 71111424 | 71111631 | 58 | 14 | 18 | 0 | 1 | 0 | 1 | 12 | 0 | 0 | 98 | 49 | 1 | 1 | 1 | 0.875 | 1 | 1 | 1 | -0.042 |
| SE | 172 | SNX12 | SNX12 | chrX | - | 71061842 | 71061967 | 71059246 | 71061118 | 71062853 | 71062949 | 7 | 1 | 0 | 0 | 1 | 0 | 0 | 3 | 2 | 0 | 98 | 49 | 1 | 1 | 1 | 0.333 | NA | 0 | 1 | 0.166 |
| SE | 177 | IGBP1 | IGBP1 | chrX | + | 70146632 | 70146828 | 70134522 | 70134816 | 70148760 | 70148840 | 21 | 12 | 5 | 0 | 1 | 0 | 2 | 1 | 0 | 0 | 98 | 49 | 1 | 1 | 1 | 0.857 | 1 | 1 | 1 | -0.048 |
| SE | 189 | TSR2 | TSR2 | chrX | + | 54440689 | 54440780 | 54440419 | 54440502 | 54443399 | 54443491 | 0 | 0 | 0 | 0 | 2 | 0 | 0 | 1 | 0 | 0 | 98 | 49 | 1 | 1 | NA | 0 | NA | NA | 1 | -1 |
| SE | 190 | PHF8 | PHF8 | chrX | - | 53993600 | 53993903 | 53992735 | 53992839 | 53995692 | 53995782 | 8 | 2 | 1 | 1 | 0 | 0 | 0 | 0 | 0 | 1 | 98 | 49 | 1 | 1 | 0.8 | 1 | 1 | NA | 0 | 0.933 |
| SE | 191 | HUWE1 | HUWE1 | chrX | - | 53544559 | 53544762 | 53543840 | 53543968 | 53545028 | 53545161 | 5 | 2 | 0 | 0 | 2 | 0 | 1 | 5 | 0 | 0 | 98 | 49 | 1 | 1 | 1 | 0.333 | NA | 1 | 1 | -0.334 |
| SE | 192 | HUWE1 | HUWE1 | chrX | - | 53603367 | 53603501 | 53602563 | 53602658 | 53604588 | 53604834 | 12 | 2 | 0 | 0 | 0 | 0 | 0 | 1 | 0 | 2 | 98 | 49 | 1 | 1 | 1 | 1 | NA | NA | 0.2 | 0.8 |
| SE | 205 | WDR45 | WDR45 | chrX | - | 49075135 | 49075281 | 49074432 | 49074912 | 49075363 | 49075465 | 8 | 4 | 10 | 1 | 0 | 0 | 0 | 5 | 0 | 0 | 98 | 49 | 1 | 1 | 0.8 | 1 | 1 | NA | 1 | -0.067 |
| SE | 208 | PIM2 | PIM2 | chrX | - | 48918535 | 48918645 | 48917780 | 48917831 | 48918773 | 48919136 | 22 | 13 | 11 | 1 | 0 | 0 | 0 | 2 | 0 | 0 | 98 | 49 | 1 | 1 | 0.917 | 1 | 1 | NA | 1 | -0.028 |
| SE | 209 | TIMM17B | TIMM17B | chrX | - | 48895351 | 48895501 | 48895037 | 48895101 | 48896758 | 48896858 | 2 | 0 | 0 | 5 | 3 | 2 | 0 | 0 | 0 | 2 | 98 | 49 | 1 | 1 | 0.167 | 0 | 0 | NA | 0 | 0.056 |
| SE | 210 | TBC1D25 | TBC1D25 | chrX | + | 48544868 | 48545023 | 48541332 | 48541442 | 48558896 | 48559024 | 3 | 4 | 3 | 0 | 1 | 0 | 1 | 1 | 0 | 0 | 98 | 49 | 1 | 1 | 1 | 0.667 | 1 | 1 | 1 | -0.111 |
| SE | 225 | CDK16 | CDK16 | chrX | + | 47225771 | 47225866 | 47224987 | 47225102 | 47225964 | 47226027 | 15 | 8 | 1 | 2 | 0 | 0 | 1 | 4 | 0 | 0 | 98 | 49 | 1 | 1 | 0.789 | 1 | 1 | 1 | 1 | -0.07 |
| SE | 227 | UBA1 | UBA1 | chrX | + | 47199208 | 47199377 | 47199047 | 47199106 | 47199479 | 47199614 | 18 | 5 | 6 | 1 | 0 | 0 | 0 | 5 | 0 | 0 | 98 | 49 | 1 | 1 | 0.9 | 1 | 1 | NA | 1 | -0.033 |
| SE | 228 | UBA1 | UBA1 | chrX | + | 47211035 | 47211225 | 47210841 | 47210916 | 47212423 | 47212512 | 19 | 7 | 9 | 1 | 0 | 0 | 0 | 10 | 0 | 0 | 98 | 49 | 1 | 1 | 0.905 | 1 | 1 | NA | 1 | -0.032 |
| SE | 241 | DDX3X | DDX3X | chrX | + | 41346858 | 41347012 | 41346504 | 41346622 | 41347311 | 41347451 | 172 | 69 | 39 | 6 | 3 | 0 | 0 | 35 | 0 | 0 | 98 | 49 | 1 | 1 | 0.935 | 0.92 | 1 | NA | 1 | -0.048 |
| SE | 242 | DDX3X | DDX3X | chrX | + | 41346858 | 41347012 | 41346504 | 41346622 | 41347314 | 41347451 | 146 | 54 | 29 | 0 | 2 | 0 | 0 | 33 | 0 | 0 | 98 | 49 | 1 | 1 | 1 | 0.931 | 1 | NA | 1 | -0.023 |
| SE | 243 | DDX3X | DDX3X | chrX | + | 41346858 | 41347012 | 41346504 | 41346622 | 41347639 | 41347915 | 108 | 47 | 22 | 1 | 2 | 0 | 0 | 27 | 0 | 0 | 98 | 49 | 1 | 1 | 0.982 | 0.922 | 1 | NA | 1 | -0.032 |
| SE | 244 | DDX3X | DDX3X | chrX | + | 41347311 | 41347451 | 41346504 | 41346622 | 41347639 | 41347915 | 108 | 30 | 29 | 1 | 2 | 0 | 1 | 11 | 0 | 0 | 98 | 49 | 1 | 1 | 0.982 | 0.882 | 1 | 1 | 1 | -0.045 |
| SE | 248 | USP9X | USP9X | chrX | + | 41136803 | 41137022 | 41134724 | 41134837 | 41140655 | 41140771 | 53 | 19 | 6 | 0 | 1 | 0 | 1 | 4 | 0 | 0 | 98 | 49 | 1 | 1 | 1 | 0.905 | 1 | 1 | 1 | -0.032 |
| SE | 249 | USP9X | USP9X | chrX | + | 41140655 | 41140771 | 41136803 | 41137022 | 41140965 | 41141217 | 41 | 25 | 3 | 1 | 0 | 0 | 1 | 4 | 0 | 1 | 98 | 49 | 1 | 1 | 0.953 | 1 | 1 | 1 | 0.667 | 0.151 |
| SE | 250 | USP9X | USP9X | chrX | + | 41150920 | 41151057 | 41148368 | 41148575 | 41152947 | 41153081 | 29 | 11 | 2 | 4 | 0 | 0 | 0 | 10 | 0 | 0 | 98 | 49 | 1 | 1 | 0.784 | 1 | 1 | NA | 1 | -0.072 |
| SE | 251 | USP9X | USP9X | chrX | + | 41169994 | 41170235 | 41168006 | 41168218 | 41170469 | 41170619 | 37 | 15 | 1 | 0 | 1 | 0 | 0 | 5 | 0 | 0 | 98 | 49 | 1 | 1 | 1 | 0.882 | 1 | NA | 1 | -0.039 |
| SE | 252 | USP9X | USP9X | chrX | + | 41214567 | 41214709 | 41210508 | 41210682 | 41215898 | 41216652 | 53 | 12 | 2 | 1 | 0 | 0 | 0 | 2 | 0 | 0 | 98 | 49 | 1 | 1 | 0.964 | 1 | 1 | NA | 1 | -0.012 |
| SE | 253 | USP9X | USP9X | chrX | + | 41219101 | 41219231 | 41218371 | 41218597 | 41223216 | 41223402 | 32 | 6 | 7 | 1 | 0 | 0 | 0 | 1 | 0 | 0 | 98 | 49 | 1 | 1 | 0.941 | 1 | 1 | NA | 1 | -0.02 |
| SE | 254 | ATP6AP2 | ATP6AP2 | chrX | + | 40597248 | 40597344 | 40591233 | 40591365 | 40597526 | 40597664 | 35 | 4 | 3 | 1 | 0 | 0 | 4 | 11 | 0 | 0 | 98 | 49 | 1 | 1 | 0.946 | 1 | 1 | 1 | 1 | -0.018 |
| SE | 267 | GK | GK | chrX | + | 30668011 | 30668118 | 30665510 | 30665584 | 30677374 | 30677452 | 29 | 11 | 6 | 6 | 9 | 3 | 0 | 4 | 0 | 0 | 98 | 49 | 1 | 1 | 0.707 | 0.379 | 0.5 | NA | 1 | -0.471 |
| SE | 270 | GK | GK | chrX | + | 30694399 | 30694537 | 30677374 | 30677452 | 30696041 | 30696151 | 13 | 4 | 10 | 0 | 1 | 0 | 0 | 1 | 0 | 0 | 98 | 49 | 1 | 1 | 1 | 0.667 | 1 | NA | 1 | -0.111 |
| SE | 271 | GK | GK | chrX | + | 30694399 | 30694537 | 30691122 | 30691199 | 30696041 | 30696151 | 13 | 4 | 8 | 9 | 2 | 0 | 0 | 1 | 0 | 0 | 98 | 49 | 1 | 1 | 0.419 | 0.5 | 1 | NA | 1 | -0.36 |
| SE | 272 | GK | GK | chrX | + | 30697731 | 30697749 | 30696616 | 30696683 | 30700413 | 30700449 | 0 | 0 | 9 | 15 | 10 | 4 | 0 | 1 | 0 | 2 | 67 | 49 | 1 | 1 | 0 | 0 | 0.622 | NA | 0.268 | -0.061 |
| SE | 274 | GK | GK | chrX | + | 30727465 | 30727552 | 30724100 | 30724181 | 30728731 | 30731460 | 3 | 0 | 0 | 33 | 25 | 9 | 0 | 0 | 2 | 3 | 98 | 49 | 1 | 1 | 0.043 | 0 | 0 | 0 | 0 | 0.014 |
| SE | 275 | ACOT9 | ACOT9 | chrX | - | 23713134 | 23713208 | 23707876 | 23707944 | 23721880 | 23721984 | 18 | 11 | 2 | 6 | 0 | 0 | 0 | 9 | 0 | 0 | 98 | 49 | 1 | 1 | 0.6 | 1 | 1 | NA | 1 | -0.133 |
| SE | 281 | SH3KBP1 | SH3KBP1 | chrX | - | 19545921 | 19546050 | 19541924 | 19542193 | 19549973 | 19550083 | 21 | 6 | 4 | 0 | 1 | 0 | 0 | 2 | 0 | 0 | 98 | 49 | 1 | 1 | 1 | 0.75 | 1 | NA | 1 | -0.083 |
| SE | 283 | SH3KBP1 | SH3KBP1 | chrX | - | 19706880 | 19706984 | 19695611 | 19695741 | 19746317 | 19746441 | 54 | 27 | 15 | 1 | 0 | 0 | 0 | 3 | 0 | 0 | 98 | 49 | 1 | 1 | 0.964 | 1 | 1 | NA | 1 | -0.012 |
| SE | 293 | PHKA2 | PHKA2 | chrX | - | 18899172 | 18899226 | 18897162 | 18897333 | 18900669 | 18900699 | 3 | 1 | 0 | 4 | 0 | 3 | 0 | 1 | 0 | 0 | 98 | 49 | 1 | 1 | 0.273 | 1 | 0 | NA | 1 | -0.576 |
| SE | 294 | PHKA2 | PHKA2 | chrX | - | 18900669 | 18900699 | 18897162 | 18897333 | 18901484 | 18901603 | 8 | 0 | 3 | 3 | 0 | 0 | 1 | 2 | 0 | 0 | 79 | 49 | 1 | 1 | 0.623 | NA | 1 | 1 | 1 | -0.189 |
| SE | 295 | PHKA2 | PHKA2 | chrX | - | 18943709 | 18943808 | 18941528 | 18941675 | 18945077 | 18945158 | 6 | 1 | 0 | 1 | 0 | 0 | 2 | 4 | 0 | 0 | 98 | 49 | 1 | 1 | 0.75 | 1 | NA | 1 | 1 | -0.125 |
| SE | 300 | REPS2 | REPS2 | chrX | + | 17022122 | 17022271 | 17006220 | 17006344 | 17029525 | 17029623 | 12 | 4 | 5 | 1 | 0 | 0 | 0 | 0 | 0 | 1 | 98 | 49 | 1 | 1 | 0.857 | 1 | 1 | NA | 0 | 0.952 |
| SE | 301 | REPS2 | REPS2 | chrX | + | 17025058 | 17025185 | 17022122 | 17022271 | 17029525 | 17029623 | 12 | 5 | 3 | 1 | 1 | 4 | 0 | 2 | 0 | 0 | 98 | 49 | 1 | 1 | 0.857 | 0.714 | 0.273 | NA | 1 | -0.385 |
| SE | 302 | REPS2 | REPS2 | chrX | + | 17025061 | 17025185 | 17022122 | 17022271 | 17029525 | 17029623 | 20 | 7 | 6 | 1 | 1 | 4 | 0 | 1 | 0 | 0 | 98 | 49 | 1 | 1 | 0.909 | 0.778 | 0.429 | NA | 1 | -0.295 |
| SE | 303 | SYAP1 | SYAP1 | chrX | + | 16754944 | 16755093 | 16743700 | 16743840 | 16756662 | 16756721 | 13 | 9 | 0 | 2 | 0 | 0 | 1 | 19 | 0 | 0 | 98 | 49 | 1 | 1 | 0.765 | 1 | NA | 1 | 1 | -0.117 |
| SE | 305 | MOSPD2 | MOSPD2 | chrX | + | 14915667 | 14915764 | 14914502 | 14914599 | 14916196 | 14916326 | 12 | 3 | 2 | 0 | 2 | 0 | 0 | 1 | 0 | 0 | 98 | 49 | 1 | 1 | 1 | 0.429 | 1 | NA | 1 | -0.19 |
| SE | 317 | WWC3 | WWC3 | chrX | + | 10125023 | 10125110 | 10124264 | 10124339 | 10126113 | 10126306 | 26 | 7 | 2 | 0 | 1 | 0 | 0 | 3 | 0 | 0 | 98 | 49 | 1 | 1 | 1 | 0.778 | 1 | NA | 1 | -0.074 |
| SE | 318 | WWC3 | WWC3 | chrX | + | 10126113 | 10126306 | 10125023 | 10125110 | 10127987 | 10128161 | 20 | 4 | 8 | 1 | 0 | 0 | 0 | 3 | 0 | 0 | 98 | 49 | 1 | 1 | 0.909 | 1 | 1 | NA | 1 | -0.03 |
| SE | 319 | WWC3 | WWC3 | chrX | + | 10127987 | 10128161 | 10126113 | 10126306 | 10128556 | 10128708 | 23 | 8 | 16 | 0 | 1 | 0 | 0 | 2 | 0 | 0 | 98 | 49 | 1 | 1 | 1 | 0.8 | 1 | NA | 1 | -0.067 |
| SE | 321 | TBL1X | TBL1X | chrX | + | 9653544 | 9653689 | 9640272 | 9640360 | 9654214 | 9654322 | 6 | 6 | 9 | 4 | 3 | 1 | 0 | 1 | 0 | 3 | 98 | 49 | 1 | 1 | 0.429 | 0.5 | 0.818 | NA | 0.143 | 0.439 |
| SE | 322 | TBL1X | TBL1X | chrX | + | 9684042 | 9684188 | 9654214 | 9654322 | 9688016 | 9688275 | 39 | 9 | 7 | 0 | 2 | 0 | 0 | 4 | 0 | 0 | 98 | 49 | 1 | 1 | 1 | 0.692 | 1 | NA | 1 | -0.103 |
| SE | 330 | DHRSX | DHRSX | chrX | - | 2243022 | 2243230 | 2219513 | 2221229 | 2266739 | 2266947 | 4 | 0 | 2 | 0 | 1 | 0 | 0 | 2 | 0 | 0 | 98 | 49 | 1 | 1 | 1 | 0 | 1 | NA | 1 | -0.333 |
| SE | 347 | CCDC183-AS1 | CCDC183-AS1 | chr9 | - | 136806049 | 136806128 | 136803926 | 136805584 | 136806384 | 136806537 | 0 | 0 | 0 | 1 | 0 | 0 | 0 | 1 | 0 | 0 | 98 | 49 | 1 | 1 | 0 | NA | NA | NA | 1 | -1 |
| SE | 348 | SNHG7 | SNHG7 | chr9 | - | 136726310 | 136726416 | 136724593 | 136725110 | 136727045 | 136727086 | 1 | 0 | 0 | 0 | 1 | 0 | 0 | 2 | 0 | 0 | 98 | 49 | 1 | 1 | 1 | 0 | NA | NA | 1 | -0.5 |
| SE | 351 | NOTCH1 | NOTCH1 | chr9 | - | 136511151 | 136511271 | 136510652 | 136510805 | 136513020 | 136513134 | 138 | 61 | 22 | 0 | 1 | 0 | 9 | 22 | 0 | 0 | 98 | 49 | 1 | 1 | 1 | 0.968 | 1 | 1 | 1 | -0.011 |
| SE | 353 | NOTCH1 | NOTCH1 | chr9 | - | 136518136 | 136518292 | 136517751 | 136517937 | 136518590 | 136518824 | 89 | 54 | 24 | 1 | 0 | 0 | 7 | 26 | 0 | 0 | 98 | 49 | 1 | 1 | 0.978 | 1 | 1 | 1 | 1 | -0.007 |
| SE | 354 | NOTCH1 | NOTCH1 | chr9 | - | 136544023 | 136544102 | 136519442 | 136519565 | 136545725 | 136545786 | 60 | 27 | 10 | 0 | 1 | 0 | 6 | 16 | 0 | 0 | 98 | 49 | 1 | 1 | 1 | 0.931 | 1 | 1 | 1 | -0.023 |
| SE | 356 | SEC16A | SEC16A | chr9 | - | 136445644 | 136445719 | 136443822 | 136443900 | 136446854 | 136446949 | 6 | 3 | 2 | 0 | 1 | 0 | 0 | 2 | 0 | 0 | 98 | 49 | 1 | 1 | 1 | 0.6 | 1 | NA | 1 | -0.133 |
| SE | 357 | SEC16A | SEC16A | chr9 | - | 136468414 | 136468512 | 136466956 | 136467083 | 136471974 | 136472111 | 6 | 1 | 0 | 0 | 0 | 0 | 0 | 1 | 1 | 0 | 98 | 49 | 1 | 1 | 1 | 1 | NA | 0 | 1 | 0.5 |
| SE | 359 | PMPCA | PMPCA | chr9 | + | 136414552 | 136414647 | 136412809 | 136412892 | 136416290 | 136416391 | 3 | 0 | 0 | 0 | 0 | 0 | 0 | 0 | 0 | 1 | 98 | 49 | 1 | 1 | 1 | NA | NA | NA | 0 | 1 |
| SE | 360 | PMPCA | PMPCA | chr9 | + | 136414554 | 136414647 | 136412809 | 136412892 | 136416290 | 136416391 | 1 | 0 | 0 | 0 | 0 | 0 | 0 | 0 | 0 | 1 | 98 | 49 | 1 | 1 | 1 | NA | NA | NA | 0 | 1 |
| SE | 369 | BRD3 | BRD3 | chr9 | - | 134036031 | 134036324 | 134034700 | 134034829 | 134041759 | 134041951 | 2 | 1 | 0 | 0 | 1 | 0 | 0 | 1 | 0 | 0 | 98 | 49 | 1 | 1 | 1 | 0.333 | NA | NA | 1 | -0.334 |
| SE | 380 | SURF6 | SURF6 | chr9 | - | 133333717 | 133333806 | 133332547 | 133332760 | 133334391 | 133334601 | 0 | 0 | 2 | 0 | 0 | 0 | 0 | 1 | 0 | 1 | 98 | 49 | 1 | 1 | NA | NA | 1 | NA | 0.333 | 0.667 |
| SE | 383 | RALGDS | RALGDS | chr9 | - | 133107084 | 133107300 | 133106644 | 133106748 | 133107987 | 133108406 | 1 | 0 | 0 | 0 | 0 | 0 | 2 | 1 | 0 | 1 | 98 | 49 | 1 | 1 | 1 | NA | NA | 1 | 0.333 | 0.334 |
| SE | 388 | RAPGEF1 | RAPGEF1 | chr9 | - | 131630235 | 131630324 | 131629101 | 131629254 | 131638634 | 131638791 | 25 | 12 | 3 | 2 | 1 | 0 | 0 | 6 | 0 | 2 | 98 | 49 | 1 | 1 | 0.862 | 0.857 | 1 | NA | 0.6 | 0.306 |
| SE | 391 | PRRC2B | PRRC2B | chr9 | + | 131446400 | 131446642 | 131444184 | 131444328 | 131447084 | 131447206 | 7 | 3 | 0 | 1 | 0 | 0 | 0 | 2 | 0 | 1 | 98 | 49 | 1 | 1 | 0.778 | 1 | NA | NA | 0.5 | 0.389 |
| SE | 392 | PRRC2B | PRRC2B | chr9 | + | 131474453 | 131476535 | 131473507 | 131473724 | 131477743 | 131477949 | 5 | 0 | 2 | 1 | 0 | 0 | 0 | 3 | 0 | 0 | 98 | 49 | 1 | 1 | 0.714 | NA | 1 | NA | 1 | -0.143 |
| SE | 393 | PRRC2B | PRRC2B | chr9 | + | 131494404 | 131494486 | 131492168 | 131492260 | 131495739 | 131500188 | 0 | 2 | 0 | 2 | 0 | 0 | 0 | 1 | 0 | 0 | 98 | 49 | 1 | 1 | 0 | 1 | NA | NA | 1 | -0.5 |
| SE | 412 | GOLGA2 | GOLGA2 | chr9 | - | 128267193 | 128267274 | 128266286 | 128266325 | 128267457 | 128267517 | 6 | 2 | 3 | 0 | 2 | 0 | 0 | 3 | 0 | 0 | 98 | 49 | 1 | 1 | 1 | 0.333 | 1 | NA | 1 | -0.222 |
| SE | 422 | LRSAM1 | LRSAM1 | chr9 | + | 127497252 | 127497334 | 127495963 | 127496095 | 127501009 | 127501143 | 4 | 2 | 0 | 0 | 0 | 0 | 0 | 0 | 0 | 1 | 98 | 49 | 1 | 1 | 1 | 1 | NA | NA | 0 | 1 |
| SE | 425 | RALGPS1 | RALGPS1 | chr9 | + | 127050042 | 127050132 | 127034430 | 127034514 | 127052846 | 127052939 | 8 | 5 | 0 | 0 | 0 | 0 | 0 | 0 | 0 | 1 | 98 | 49 | 1 | 1 | 1 | 1 | NA | NA | 0 | 1 |
| SE | 430 | MAPKAP1 | MAPKAP1 | chr9 | - | 125672315 | 125672643 | 125669817 | 125669907 | 125706970 | 125707234 | 8 | 3 | 0 | 3 | 0 | 0 | 0 | 3 | 0 | 2 | 98 | 49 | 1 | 1 | 0.571 | 1 | NA | NA | 0.429 | 0.356 |
| SE | 435 | DENND1A | DENND1A | chr9 | - | 123450692 | 123450749 | 123440359 | 123440491 | 123452275 | 123452347 | 9 | 2 | 1 | 1 | 0 | 0 | 0 | 5 | 0 | 0 | 98 | 49 | 1 | 1 | 0.818 | 1 | 1 | NA | 1 | -0.061 |
| SE | 436 | DENND1A | DENND1A | chr9 | - | 123769513 | 123769563 | 123757702 | 123757822 | 123792586 | 123792630 | 12 | 8 | 1 | 1 | 0 | 0 | 0 | 3 | 0 | 0 | 98 | 49 | 1 | 1 | 0.857 | 1 | 1 | NA | 1 | -0.048 |
| SE | 442 | RBM18 | RBM18 | chr9 | - | 122254273 | 122254367 | 122251846 | 122251973 | 122261379 | 122261508 | 0 | 0 | 0 | 2 | 4 | 0 | 0 | 1 | 0 | 4 | 98 | 49 | 1 | 1 | 0 | 0 | NA | NA | 0.111 | -0.111 |
| SE | 448 | GSN | GSN | chr9 | + | 121282438 | 121282550 | 121281469 | 121281562 | 121301962 | 121302167 | 1 | 0 | 0 | 5 | 5 | 3 | 0 | 0 | 0 | 4 | 98 | 49 | 1 | 1 | 0.091 | 0 | 0 | NA | 0 | 0.03 |
| SE | 449 | GSN | GSN | chr9 | + | 121283318 | 121283392 | 121281469 | 121281562 | 121301962 | 121302167 | 0 | 0 | 1 | 5 | 5 | 3 | 0 | 0 | 0 | 4 | 98 | 49 | 1 | 1 | 0 | 0 | 0.143 | NA | 0 | 0.048 |
| SE | 450 | GSN | GSN | chr9 | + | 121300055 | 121300126 | 121281469 | 121281562 | 121301962 | 121302167 | 3 | 0 | 0 | 5 | 5 | 3 | 0 | 0 | 0 | 4 | 98 | 49 | 1 | 1 | 0.231 | 0 | 0 | NA | 0 | 0.077 |
| SE | 451 | GSN | GSN | chr9 | + | 121300064 | 121300126 | 121281469 | 121281562 | 121301962 | 121302167 | 3 | 0 | 0 | 5 | 5 | 3 | 0 | 0 | 0 | 4 | 98 | 49 | 1 | 1 | 0.231 | 0 | 0 | NA | 0 | 0.077 |
| SE | 452 | CNTRL | CNTRL | chr9 | + | 121140640 | 121140786 | 121138544 | 121138679 | 121142090 | 121142270 | 6 | 1 | 1 | 1 | 0 | 0 | 0 | 5 | 0 | 0 | 98 | 49 | 1 | 1 | 0.75 | 1 | 1 | NA | 1 | -0.083 |
| SE | 453 | CNTRL | CNTRL | chr9 | + | 121141380 | 121141588 | 121140640 | 121140786 | 121142090 | 121142270 | 3 | 6 | 4 | 4 | 1 | 0 | 0 | 2 | 0 | 0 | 98 | 49 | 1 | 1 | 0.273 | 0.75 | 1 | NA | 1 | -0.326 |
| SE | 462 | TLR4 | TLR4 | chr9 | + | 117708198 | 117708318 | 117704174 | 117704565 | 117708562 | 117708729 | 93 | 18 | 11 | 13 | 5 | 14 | 0 | 18 | 0 | 5 | 98 | 49 | 1 | 1 | 0.782 | 0.643 | 0.282 | NA | 0.643 | -0.074 |
| SE | 463 | TLR4 | TLR4 | chr9 | + | 117708198 | 117708318 | 117704174 | 117704565 | 117712388 | 117717491 | 80 | 13 | 10 | 66 | 14 | 7 | 0 | 14 | 0 | 10 | 98 | 49 | 1 | 1 | 0.377 | 0.317 | 0.417 | NA | 0.412 | -0.042 |
| SE | 464 | TLR4 | TLR4 | chr9 | + | 117708562 | 117708729 | 117704174 | 117704565 | 117712388 | 117717491 | 72 | 13 | 36 | 66 | 14 | 7 | 0 | 10 | 0 | 10 | 98 | 49 | 1 | 1 | 0.353 | 0.317 | 0.72 | NA | 0.333 | 0.13 |
| SE | 465 | TLR4 | TLR4 | chr9 | + | 117708562 | 117708729 | 117708198 | 117708318 | 117712388 | 117717491 | 74 | 13 | 23 | 1 | 0 | 0 | 0 | 9 | 0 | 0 | 98 | 49 | 1 | 1 | 0.974 | 1 | 1 | NA | 1 | -0.009 |
| SE | 470 | PTBP3 | PTBP3 | chr9 | - | 112224132 | 112224210 | 112217714 | 112223983 | 112227410 | 112227627 | 67 | 38 | 24 | 0 | 2 | 0 | 1 | 15 | 0 | 0 | 98 | 49 | 1 | 1 | 1 | 0.905 | 1 | 1 | 1 | -0.032 |
| SE | 471 | PTBP3 | PTBP3 | chr9 | - | 112262434 | 112262599 | 112252677 | 112252788 | 112268048 | 112268195 | 54 | 21 | 32 | 1 | 1 | 0 | 2 | 17 | 0 | 0 | 98 | 49 | 1 | 1 | 0.964 | 0.913 | 1 | 1 | 1 | -0.041 |
| SE | 473 | PTBP3 | PTBP3 | chr9 | - | 112330440 | 112330474 | 112297831 | 112297916 | 112333469 | 112333664 | 2 | 0 | 0 | 53 | 5 | 6 | 0 | 0 | 0 | 3 | 83 | 49 | 1 | 1 | 0.022 | 0 | 0 | NA | 0 | 0.007 |
| SE | 476 | UGCG | UGCG | chr9 | + | 111931270 | 111931357 | 111929499 | 111929678 | 111932169 | 111932359 | 9 | 10 | 2 | 1 | 0 | 0 | 0 | 3 | 0 | 0 | 98 | 49 | 1 | 1 | 0.818 | 1 | 1 | NA | 1 | -0.061 |
| SE | 478 | DNAJC25 | DNAJC25 | chr9 | + | 111642856 | 111643094 | 111631351 | 111631743 | 111647106 | 111647259 | 1 | 1 | 0 | 0 | 1 | 0 | 1 | 1 | 0 | 0 | 98 | 49 | 1 | 1 | 1 | 0.333 | NA | 1 | 1 | -0.334 |
| SE | 479 | KIAA0368 | KIAA0368 | chr9 | - | 111383210 | 111383332 | 111378579 | 111378730 | 111384521 | 111384569 | 6 | 2 | 2 | 1 | 0 | 0 | 0 | 4 | 0 | 0 | 98 | 49 | 1 | 1 | 0.75 | 1 | 1 | NA | 1 | -0.083 |
| SE | 488 | ABCA1 | ABCA1 | chr9 | - | 104785395 | 104785639 | 104781002 | 104784455 | 104786297 | 104786390 | 33 | 7 | 8 | 1 | 0 | 0 | 0 | 2 | 0 | 0 | 98 | 49 | 1 | 1 | 0.943 | 1 | 1 | NA | 1 | -0.019 |
| SE | 489 | ABCA1 | ABCA1 | chr9 | - | 104786872 | 104786976 | 104786297 | 104786390 | 104787919 | 104788054 | 9 | 2 | 0 | 8 | 1 | 0 | 0 | 1 | 0 | 1 | 98 | 49 | 1 | 1 | 0.36 | 0.5 | NA | NA | 0.333 | 0.097 |
| SE | 490 | ABCA1 | ABCA1 | chr9 | - | 104798420 | 104798598 | 104796308 | 104796424 | 104799818 | 104799988 | 11 | 10 | 6 | 3 | 0 | 0 | 0 | 2 | 0 | 0 | 98 | 49 | 1 | 1 | 0.647 | 1 | 1 | NA | 1 | -0.118 |
| SE | 493 | ABCA1 | ABCA1 | chr9 | - | 104819585 | 104819723 | 104818662 | 104818883 | 104819926 | 104820069 | 56 | 29 | 10 | 1 | 0 | 0 | 1 | 17 | 0 | 0 | 98 | 49 | 1 | 1 | 0.966 | 1 | 1 | 1 | 1 | -0.011 |
| SE | 498 | RNF20 | RNF20 | chr9 | + | 101544766 | 101544885 | 101540792 | 101540975 | 101546819 | 101546966 | 4 | 2 | 1 | 0 | 1 | 0 | 0 | 1 | 0 | 0 | 98 | 49 | 1 | 1 | 1 | 0.5 | 1 | NA | 1 | -0.167 |
| SE | 502 | ZNF189 | ZNF189 | chr9 | + | 101399394 | 101399530 | 101398853 | 101399189 | 101399925 | 101400010 | 1 | 0 | 0 | 0 | 0 | 0 | 0 | 0 | 0 | 1 | 98 | 49 | 1 | 1 | 1 | NA | NA | NA | 0 | 1 |
| SE | 511 | PHF2 | PHF2 | chr9 | + | 93645628 | 93645789 | 93636410 | 93636525 | 93649070 | 93649212 | 27 | 3 | 5 | 2 | 0 | 0 | 0 | 3 | 0 | 0 | 98 | 49 | 1 | 1 | 0.871 | 1 | 1 | NA | 1 | -0.043 |
| SE | 512 | FAM120A | FAM120A | chr9 | + | 93558580 | 93558718 | 93557826 | 93558010 | 93561108 | 93561250 | 34 | 12 | 14 | 0 | 0 | 0 | 0 | 12 | 0 | 1 | 98 | 49 | 1 | 1 | 1 | 1 | 1 | NA | 0.857 | 0.143 |
| SE | 524 | SEMA4D | SEMA4D | chr9 | - | 89418055 | 89418118 | 89405350 | 89405699 | 89455887 | 89455953 | 17 | 6 | 0 | 4 | 0 | 0 | 0 | 5 | 0 | 0 | 98 | 49 | 1 | 1 | 0.68 | 1 | NA | NA | 1 | -0.16 |
| SE | 525 | SEMA4D | SEMA4D | chr9 | - | 89418362 | 89418437 | 89418055 | 89418118 | 89455887 | 89455953 | 8 | 1 | 0 | 4 | 2 | 0 | 0 | 2 | 0 | 0 | 98 | 49 | 1 | 1 | 0.5 | 0.2 | NA | NA | 1 | -0.65 |
| SE | 526 | SECISBP2 | SECISBP2 | chr9 | + | 89357410 | 89357565 | 89350631 | 89350852 | 89357998 | 89358191 | 8 | 2 | 1 | 3 | 0 | 0 | 0 | 2 | 0 | 0 | 98 | 49 | 1 | 1 | 0.571 | 1 | 1 | NA | 1 | -0.143 |
| SE | 527 | SPIN1 | SPIN1 | chr9 | + | 88462495 | 88462749 | 88448940 | 88448989 | 88468371 | 88468605 | 0 | 0 | 0 | 2 | 0 | 0 | 0 | 2 | 0 | 0 | 98 | 49 | 1 | 1 | 0 | NA | NA | NA | 1 | -1 |
| SE | 530 | DAPK1 | DAPK1 | chr9 | + | 87686550 | 87686739 | 87681403 | 87681626 | 87697006 | 87697204 | 6 | 3 | 5 | 0 | 1 | 0 | 0 | 7 | 0 | 0 | 98 | 49 | 1 | 1 | 1 | 0.6 | 1 | NA | 1 | -0.133 |
| SE | 531 | DAPK1 | DAPK1 | chr9 | + | 87697006 | 87697204 | 87686550 | 87686739 | 87698655 | 87698794 | 8 | 4 | 0 | 1 | 1 | 0 | 0 | 3 | 0 | 0 | 98 | 49 | 1 | 1 | 0.8 | 0.667 | NA | NA | 1 | -0.266 |
| SE | 537 | UBQLN1 | UBQLN1 | chr9 | - | 83666349 | 83666433 | 83665029 | 83665145 | 83669184 | 83669327 | 17 | 3 | 10 | 3 | 0 | 0 | 0 | 4 | 0 | 1 | 98 | 49 | 1 | 1 | 0.739 | 1 | 1 | NA | 0.667 | 0.246 |
| SE | 539 | GNAQ | GNAQ | chr9 | - | 77794462 | 77794592 | 77728513 | 77728667 | 77797519 | 77797648 | 36 | 21 | 13 | 0 | 0 | 0 | 0 | 9 | 0 | 1 | 98 | 49 | 1 | 1 | 1 | 1 | 1 | NA | 0.818 | 0.182 |
| SE | 545 | TRPM6 | TRPM6 | chr9 | - | 74738406 | 74738612 | 74732684 | 74732736 | 74739366 | 74739449 | 5 | 2 | 6 | 1 | 0 | 0 | 0 | 2 | 0 | 0 | 98 | 49 | 1 | 1 | 0.714 | 1 | 1 | NA | 1 | -0.095 |
| SE | 546 | TRPM6 | TRPM6 | chr9 | - | 74761998 | 74763134 | 74761695 | 74761808 | 74771702 | 74771835 | 9 | 6 | 5 | 1 | 0 | 1 | 0 | 1 | 0 | 0 | 98 | 49 | 1 | 1 | 0.818 | 1 | 0.714 | NA | 1 | -0.156 |
| SE | 548 | TRPM6 | TRPM6 | chr9 | - | 74788613 | 74788742 | 74785873 | 74786125 | 74792623 | 74792770 | 2 | 6 | 3 | 4 | 2 | 0 | 0 | 0 | 1 | 0 | 98 | 49 | 1 | 1 | 0.2 | 0.6 | 1 | 0 | NA | 0.6 |
| SE | 550 | TRPM6 | TRPM6 | chr9 | - | 74821668 | 74821837 | 74820303 | 74820427 | 74827777 | 74827949 | 22 | 8 | 7 | 0 | 2 | 0 | 2 | 4 | 0 | 0 | 98 | 49 | 1 | 1 | 1 | 0.667 | 1 | 1 | 1 | -0.111 |
| SE | 551 | TRPM6 | TRPM6 | chr9 | - | 74842165 | 74842343 | 74840023 | 74840237 | 74855526 | 74855565 | 13 | 9 | 4 | 0 | 2 | 0 | 0 | 4 | 0 | 0 | 98 | 49 | 1 | 1 | 1 | 0.692 | 1 | NA | 1 | -0.103 |
| SE | 557 | PTAR1 | PTAR1 | chr9 | - | 69750780 | 69750950 | 69741791 | 69741858 | 69759852 | 69759960 | 4 | 4 | 0 | 0 | 1 | 0 | 0 | 1 | 0 | 0 | 98 | 49 | 1 | 1 | 1 | 0.667 | NA | NA | 1 | -0.166 |
| SE | 560 | LINC01506 | LINC01506 | chr9 | - | 68545716 | 68545836 | 68545500 | 68545624 | 68546558 | 68546602 | 9 | 8 | 17 | 0 | 0 | 1 | 0 | 1 | 0 | 0 | 98 | 49 | 1 | 1 | 1 | 1 | 0.895 | NA | 1 | -0.035 |
| SE | 565 | FRG1HP | FRG1HP | chr9 | + | 40993252 | 40993434 | 40992336 | 40992775 | 40994632 | 40994699 | 3 | 0 | 0 | 1 | 0 | 0 | 0 | 1 | 0 | 0 | 98 | 49 | 1 | 1 | 0.6 | NA | NA | NA | 1 | -0.4 |
| SE | 566 | SLC25A51 | SLC25A51 | chr9 | - | 37903530 | 37903557 | 37899828 | 37899950 | 37904067 | 37904353 | 2 | 0 | 2 | 1 | 0 | 0 | 0 | 0 | 0 | 1 | 76 | 49 | 1 | 1 | 0.563 | NA | 1 | NA | 0 | 0.781 |
| SE | 567 | DCAF10 | DCAF10 | chr9 | + | 37842088 | 37842286 | 37819287 | 37819401 | 37854779 | 37854982 | 1 | 2 | 0 | 2 | 0 | 0 | 0 | 1 | 0 | 0 | 98 | 49 | 1 | 1 | 0.2 | 1 | NA | NA | 1 | -0.4 |
| SE | 594 | RNF38 | RNF38 | chr9 | - | 36342324 | 36342424 | 36336400 | 36339814 | 36344831 | 36344953 | 10 | 4 | 0 | 0 | 0 | 0 | 0 | 1 | 0 | 2 | 98 | 49 | 1 | 1 | 1 | 1 | NA | NA | 0.2 | 0.8 |
| SE | 602 | ZCCHC6 | ZCCHC6 | chr9 | - | 86309462 | 86309576 | 86309211 | 86309289 | 86309927 | 86310017 | 48 | 40 | 12 | 2 | 0 | 0 | 0 | 12 | 0 | 0 | 98 | 49 | 1 | 1 | 0.923 | 1 | 1 | NA | 1 | -0.026 |
| SE | 603 | ZCCHC6 | ZCCHC6 | chr9 | - | 86318957 | 86319058 | 86317218 | 86317276 | 86319583 | 86319670 | 49 | 13 | 19 | 26 | 5 | 0 | 0 | 8 | 0 | 0 | 98 | 49 | 1 | 1 | 0.485 | 0.565 | 1 | NA | 1 | -0.317 |
| SE | 605 | ZCCHC6 | ZCCHC6 | chr9 | - | 86322324 | 86322475 | 86319583 | 86319670 | 86322872 | 86323960 | 66 | 41 | 27 | 1 | 0 | 0 | 0 | 6 | 0 | 0 | 98 | 49 | 1 | 1 | 0.971 | 1 | 1 | NA | 1 | -0.01 |
| SE | 606 | ZCCHC6 | ZCCHC6 | chr9 | - | 86346298 | 86346480 | 86345668 | 86345785 | 86352679 | 86353230 | 269 | 107 | 11 | 0 | 1 | 0 | 0 | 27 | 0 | 0 | 98 | 49 | 1 | 1 | 1 | 0.982 | 1 | NA | 1 | -0.006 |
| SE | 609 | TLN1 | TLN1 | chr9 | - | 35699355 | 35699461 | 35699031 | 35699156 | 35699973 | 35700081 | 195 | 53 | 21 | 0 | 3 | 0 | 9 | 45 | 0 | 0 | 98 | 49 | 1 | 1 | 1 | 0.898 | 1 | 1 | 1 | -0.034 |

| | | | | | | | | | | | | | | | | | | | | | | | | | | | | | | | |
|---|---|---|---|---|---|---|---|---|---|---|---|---|---|---|---|---|---|---|---|---|---|---|---|---|---|---|---|---|---|---|---|
| SE | 610 | TLN1 | TLN1 | chr9 | - | 35699973 | 35700081 | 35699355 | 35699461 | 35700190 | 35700376 | 207 | 50 | 28 | 1 | 0 | 0 | 11 | 39 | 0 | 0 | 98 | 49 | 1 | 1 | 0.99 | 1 | 1 | 1 | 1 | -0.003 |
| SE | 612 | TLN1 | TLN1 | chr9 | - | 35712834 | 35713043 | 35712004 | 35712124 | 35713195 | 35713298 | 75 | 34 | 30 | 0 | 1 | 0 | 11 | 20 | 0 | 0 | 98 | 49 | 1 | 1 | 1 | 0.944 | 1 | 1 | 1 | -0.019 |
| SE | 613 | TLN1 | TLN1 | chr9 | - | 35716389 | 35716556 | 35715058 | 35715187 | 35717145 | 35717440 | 178 | 63 | 47 | 2 | 0 | 0 | 14 | 44 | 0 | 0 | 98 | 49 | 1 | 1 | 0.978 | 1 | 1 | 1 | 1 | -0.007 |
| SE | 614 | TLN1 | TLN1 | chr9 | - | 35717145 | 35717440 | 35715058 | 35715187 | 35717618 | 35717786 | 40 | 14 | 13 | 1 | 0 | 0 | 0 | 10 | 0 | 0 | 98 | 49 | 1 | 1 | 0.952 | 1 | 1 | NA | 1 | -0.016 |
| SE | 616 | TLN1 | TLN1 | chr9 | - | 35725564 | 35725727 | 35725223 | 35725321 | 35732074 | 35732395 | 80 | 26 | 10 | 0 | 0 | 0 | 1 | 14 | 0 | 1 | 98 | 49 | 1 | 1 | 1 | 1 | 1 | 1 | 0.875 | 0.062 |
| SE | 619 | IL11RA | IL11RA | chr9 | + | 34657034 | 34657149 | 34656738 | 34656908 | 34657302 | 34657335 | 3 | 1 | 0 | 2 | 0 | 0 | 0 | 1 | 0 | 0 | 98 | 49 | 1 | 1 | 0.429 | 1 | NA | NA | 1 | -0.285 |
| SE | 627 | B4GALT1 | B4GALT1 | chr9 | - | 33115990 | 33116113 | 33113773 | 33113878 | 33120418 | 33120606 | 10 | 8 | 2 | 0 | 3 | 0 | 0 | 1 | 0 | 1 | 98 | 49 | 1 | 1 | 1 | 0.571 | 1 | NA | 0.333 | 0.524 |
| SE | 628 | DNAJA1 | DNAJA1 | chr9 | + | 33026812 | 33026990 | 33026474 | 33026616 | 33029884 | 33029989 | 11 | 14 | 3 | 2 | 0 | 0 | 0 | 6 | 0 | 1 | 98 | 49 | 1 | 1 | 0.733 | 1 | 1 | NA | 0.75 | 0.161 |
| SE | 664 | KDM4C | KDM4C | chr9 | + | 6849506 | 6849700 | 6814630 | 6814745 | 6880011 | 6880061 | 5 | 3 | 9 | 0 | 0 | 0 | 0 | 4 | 0 | 2 | 98 | 49 | 1 | 1 | 1 | 1 | 1 | NA | 0.5 | 0.5 |
| SE | 665 | KDM4C | KDM4C | chr9 | + | 6981841 | 6982765 | 6980924 | 6981118 | 6984165 | 6984404 | 1 | 0 | 0 | 2 | 2 | 0 | 0 | 0 | 0 | 3 | 98 | 49 | 1 | 1 | 0.2 | 0 | NA | NA | 0 | 0.1 |
| SE | 666 | UHRF2 | UHRF2 | chr9 | + | 6420911 | 6421142 | 6413150 | 6413643 | 6433913 | 6434173 | 14 | 1 | 0 | 0 | 0 | 1 | 0 | 4 | 0 | 0 | 98 | 49 | 1 | 1 | 1 | 1 | 0 | NA | 1 | -0.333 |
| SE | 667 | UHRF2 | UHRF2 | chr9 | + | 6433913 | 6434173 | 6420911 | 6421142 | 6460572 | 6460791 | 13 | 7 | 0 | 0 | 3 | 0 | 0 | 2 | 0 | 0 | 98 | 49 | 1 | 1 | 1 | 0.538 | NA | NA | 1 | -0.231 |
| SE | 670 | RIC1 | RIC1 | chr9 | + | 5689958 | 5690038 | 5656582 | 5656690 | 5713895 | 5714003 | 14 | 0 | 1 | 0 | 0 | 1 | 0 | 9 | 0 | 0 | 98 | 49 | 1 | 1 | 1 | NA | 0.333 | NA | 1 | -0.334 |
| SE | 671 | RIC1 | RIC1 | chr9 | + | 5745930 | 5746083 | 5743688 | 5743737 | 5747301 | 5747505 | 8 | 0 | 3 | 1 | 0 | 0 | 0 | 9 | 0 | 0 | 98 | 49 | 1 | 1 | 0.8 | NA | 1 | NA | 1 | -0.1 |
| SE | 672 | RIC1 | RIC1 | chr9 | + | 5753535 | 5753646 | 5753199 | 5753238 | 5754840 | 5754930 | 8 | 4 | 0 | 2 | 0 | 0 | 0 | 3 | 0 | 0 | 98 | 49 | 1 | 1 | 0.667 | 1 | NA | NA | 1 | -0.166 |
| SE | 677 | RFX3 | RFX3 | chr9 | - | 3270370 | 3270525 | 3266207 | 3266305 | 3271002 | 3271118 | 6 | 0 | 0 | 1 | 0 | 0 | 0 | 0 | 0 | 2 | 98 | 49 | 1 | 1 | 0.75 | NA | NA | NA | 0 | 0.75 |
| SE | 678 | RFX3 | RFX3 | chr9 | - | 3301545 | 3301620 | 3293076 | 3293258 | 3330258 | 3330517 | 0 | 0 | 0 | 3 | 0 | 0 | 0 | 4 | 0 | 0 | 98 | 49 | 1 | 1 | 0 | NA | NA | NA | 1 | -1 |
| SE | 682 | DOCK8 | DOCK8 | chr9 | + | 304580 | 304704 | 289509 | 289581 | 311953 | 312166 | 236 | 104 | 21 | 0 | 2 | 0 | 7 | 41 | 0 | 0 | 98 | 49 | 1 | 1 | 1 | 0.963 | 1 | 1 | 1 | -0.012 |
| SE | 683 | DOCK8 | DOCK8 | chr9 | + | 379770 | 379935 | 376976 | 377211 | 382512 | 382685 | 81 | 40 | 9 | 1 | 0 | 0 | 2 | 12 | 0 | 0 | 98 | 49 | 1 | 1 | 0.976 | 1 | 1 | 1 | 1 | -0.008 |
| SE | 685 | DOCK8 | DOCK8 | chr9 | + | 420948 | 421078 | 420400 | 420583 | 422047 | 422135 | 81 | 19 | 24 | 6 | 0 | 0 | 0 | 11 | 0 | 1 | 98 | 49 | 1 | 1 | 0.871 | 1 | 1 | NA | 0.846 | 0.111 |
| SE | 694 | GRINA | GRINA | chr8 | + | 143991877 | 143992078 | 143991691 | 143991804 | 143992244 | 143992373 | 9 | 2 | 9 | 2 | 0 | 0 | 0 | 5 | 0 | 0 | 98 | 49 | 1 | 1 | 0.692 | 1 | 1 | NA | 1 | -0.103 |
| SE | 695 | PARP10 | PARP10 | chr8 | - | 143983166 | 143983811 | 143982931 | 143983065 | 143984007 | 143984104 | 16 | 5 | 2 | 0 | 0 | 1 | 4 | 3 | 0 | 0 | 98 | 49 | 1 | 1 | 1 | 1 | 0.5 | 1 | 1 | -0.167 |
| SE | 696 | PARP10 | PARP10 | chr8 | - | 143985411 | 143985648 | 143984543 | 143985328 | 143985720 | 143985975 | 10 | 2 | 0 | 0 | 0 | 0 | 3 | 2 | 0 | 1 | 98 | 49 | 1 | 1 | 1 | 1 | NA | 1 | 0.5 | 0.25 |
| SE | 697 | PLEC | PLEC | chr8 | - | 143922503 | 143925884 | 143915146 | 143922395 | 143926783 | 143926882 | 4 | 2 | 0 | 0 | 0 | 2 | 0 | 5 | 0 | 0 | 98 | 49 | 1 | 1 | 1 | 1 | 0 | NA | 1 | -0.333 |
| SE | 698 | NAPRT | NAPRT | chr8 | - | 143577061 | 143577177 | 143576645 | 143576842 | 143577268 | 143577399 | 8 | 0 | 7 | 3 | 0 | 0 | 2 | 4 | 0 | 0 | 98 | 49 | 1 | 1 | 0.571 | NA | 1 | 1 | 1 | -0.215 |
| SE | 702 | DENND3 | DENND3 | chr8 | + | 141136557 | 141136791 | 141128620 | 141128809 | 141150833 | 141150953 | 4 | 2 | 0 | 0 | 3 | 0 | 0 | 1 | 0 | 0 | 98 | 49 | 1 | 1 | 1 | 0.25 | NA | NA | 1 | -0.375 |
| SE | 703 | DENND3 | DENND3 | chr8 | + | 141138021 | 141138137 | 141136557 | 141136791 | 141150833 | 141150953 | 64 | 64 | 11 | 1 | 0 | 0 | 3 | 16 | 0 | 0 | 98 | 49 | 1 | 1 | 0.97 | 1 | 1 | 1 | 1 | -0.01 |
| SE | 704 | DENND3 | DENND3 | chr8 | + | 141138021 | 141138137 | 141136557 | 141136791 | 141151618 | 141151837 | 53 | 54 | 11 | 1 | 0 | 0 | 3 | 15 | 0 | 0 | 98 | 49 | 1 | 1 | 0.964 | 1 | 1 | 1 | 1 | -0.012 |
| SE | 705 | DENND3 | DENND3 | chr8 | + | 141141202 | 141141324 | 141138021 | 141138137 | 141144147 | 141144259 | 87 | 36 | 18 | 0 | 0 | 0 | 0 | 17 | 0 | 1 | 98 | 49 | 1 | 1 | 1 | 1 | 1 | NA | 0.895 | 0.105 |
| SE | 706 | DENND3 | DENND3 | chr8 | + | 141144147 | 141144259 | 141138021 | 141138137 | 141150833 | 141150953 | 22 | 14 | 1 | 11 | 11 | 0 | 0 | 6 | 0 | 2 | 98 | 49 | 1 | 1 | 0.5 | 0.389 | 1 | NA | 0.6 | 0.03 |
| SE | 707 | DENND3 | DENND3 | chr8 | + | 141150833 | 141150953 | 141136557 | 141136791 | 141151618 | 141151837 | 84 | 39 | 10 | 1 | 0 | 0 | 0 | 20 | 0 | 0 | 98 | 49 | 1 | 1 | 0.977 | 1 | 1 | NA | 1 | -0.008 |
| SE | 708 | DENND3 | DENND3 | chr8 | + | 141150833 | 141150953 | 141138021 | 141138137 | 141151618 | 141151837 | 94 | 50 | 10 | 0 | 1 | 0 | 0 | 22 | 0 | 1 | 98 | 49 | 1 | 1 | 1 | 0.962 | 1 | NA | 0.917 | 0.07 |
| SE | 709 | DENND3 | DENND3 | chr8 | + | 141151618 | 141151837 | 141138021 | 141138137 | 141155848 | 141155970 | 73 | 51 | 17 | 1 | 0 | 0 | 1 | 31 | 0 | 0 | 98 | 49 | 1 | 1 | 0.973 | 1 | 1 | 1 | 1 | -0.009 |
| SE | 710 | DENND3 | DENND3 | chr8 | + | 141160631 | 141160787 | 141155848 | 141155970 | 141163332 | 141163429 | 75 | 32 | 15 | 5 | 0 | 0 | 0 | 19 | 0 | 0 | 98 | 49 | 1 | 1 | 0.882 | 1 | 1 | NA | 1 | -0.039 |
| SE | 711 | DENND3 | DENND3 | chr8 | + | 141160631 | 141160787 | 141155848 | 141155970 | 141165185 | 141165289 | 52 | 20 | 10 | 1 | 0 | 0 | 0 | 11 | 0 | 1 | 98 | 49 | 1 | 1 | 0.963 | 1 | 1 | NA | 0.846 | 0.142 |
| SE | 712 | DENND3 | DENND3 | chr8 | + | 141163332 | 141163429 | 141155848 | 141155970 | 141165185 | 141165289 | 60 | 17 | 13 | 1 | 0 | 0 | 0 | 8 | 0 | 1 | 98 | 49 | 1 | 1 | 0.968 | 1 | 1 | NA | 0.8 | 0.189 |
| SE | 713 | DENND3 | DENND3 | chr8 | + | 141163332 | 141163429 | 141160631 | 141160787 | 141165185 | 141165289 | 85 | 35 | 21 | 7 | 6 | 3 | 0 | 16 | 0 | 0 | 98 | 49 | 1 | 1 | 0.859 | 0.745 | 0.778 | NA | 1 | -0.206 |
| SE | 714 | DENND3 | DENND3 | chr8 | + | 141168003 | 141168525 | 141166189 | 141166389 | 141175199 | 141175459 | 151 | 71 | 16 | 2 | 0 | 0 | 0 | 36 | 0 | 0 | 98 | 49 | 1 | 1 | 0.974 | 1 | 1 | NA | 1 | -0.009 |
| SE | 716 | DENND3 | DENND3 | chr8 | + | 141192330 | 141192449 | 141190283 | 141190417 | 141192525 | 141192663 | 58 | 24 | 6 | 1 | 0 | 0 | 3 | 17 | 0 | 0 | 98 | 49 | 1 | 1 | 0.967 | 1 | 1 | 1 | 1 | -0.011 |
| SE | 719 | AGO2 | AGO2 | chr8 | - | 140558484 | 140558572 | 140557088 | 140557236 | 140559394 | 140559529 | 20 | 14 | 10 | 0 | 1 | 0 | 0 | 9 | 0 | 0 | 98 | 49 | 1 | 1 | 1 | 0.875 | 1 | NA | 1 | -0.042 |
| SE | 731 | SLA | SLA | chr8 | - | 133044983 | 133045115 | 133039997 | 133040130 | 133047829 | 133047933 | 51 | 33 | 34 | 6 | 2 | 1 | 0 | 35 | 0 | 6 | 98 | 49 | 1 | 1 | 0.81 | 0.892 | 0.944 | NA | 0.745 | 0.137 |
| SE | 734 | TMEM71 | TMEM71 | chr8 | - | 132722039 | 132722115 | 132714153 | 132714215 | 132727797 | 132727986 | 72 | 42 | 31 | 1 | 0 | 0 | 0 | 14 | 0 | 0 | 98 | 49 | 1 | 1 | 0.973 | 1 | 1 | NA | 1 | -0.009 |
| SE | 735 | TMEM71 | TMEM71 | chr8 | - | 132727797 | 132727986 | 132722039 | 132722115 | 132746941 | 132747057 | 47 | 25 | 29 | 2 | 2 | 5 | 0 | 9 | 0 | 0 | 98 | 49 | 1 | 1 | 0.922 | 0.862 | 0.744 | NA | 1 | -0.157 |
| SE | 736 | TMEM71 | TMEM71 | chr8 | - | 132757233 | 132757294 | 132751784 | 132751997 | 132758839 | 132758915 | 62 | 39 | 30 | 2 | 0 | 0 | 3 | 28 | 0 | 0 | 98 | 49 | 1 | 1 | 0.939 | 1 | 1 | 1 | 1 | -0.02 |
| SE | 737 | FAM49B | FAM49B | chr8 | - | 129862228 | 129862334 | 129855610 | 129855747 | 129871374 | 129871496 | 146 | 109 | 79 | 0 | 1 | 0 | 0 | 28 | 0 | 0 | 98 | 49 | 1 | 1 | 1 | 0.982 | 1 | NA | 1 | -0.006 |
| SE | 738 | FAM49B | FAM49B | chr8 | - | 129879388 | 129879471 | 129871374 | 129871496 | 129903311 | 129903350 | 118 | 67 | 33 | 8 | 10 | 2 | 0 | 14 | 0 | 1 | 98 | 49 | 1 | 1 | 0.881 | 0.77 | 0.892 | NA | 0.875 | -0.027 |
| SE | 740 | FAM49B | FAM49B | chr8 | - | 129896671 | 129896854 | 129879388 | 129879471 | 129903311 | 129903350 | 1 | 2 | 0 | 26 | 14 | 7 | 0 | 0 | 0 | 3 | 98 | 49 | 1 | 1 | 0.019 | 0.067 | 0 | NA | 0 | 0.029 |
| SE | 741 | FAM49B | FAM49B | chr8 | - | 129896748 | 129896854 | 129879388 | 129879471 | 129903311 | 129903350 | 0 | 3 | 2 | 26 | 14 | 7 | 0 | 0 | 0 | 3 | 98 | 49 | 1 | 1 | 0 | 0.097 | 0.125 | NA | 0 | 0.074 |
| SE | 742 | FAM49B | FAM49B | chr8 | - | 129903311 | 129903350 | 129879388 | 129879471 | 129970942 | 129970995 | 52 | 31 | 21 | 0 | 2 | 0 | 0 | 6 | 0 | 0 | 88 | 49 | 1 | 1 | 1 | 0.896 | 1 | NA | 1 | -0.035 |
| SE | 744 | FAM49B | FAM49B | chr8 | - | 129904498 | 129904585 | 129903311 | 129903350 | 129970942 | 129970995 | 45 | 17 | 5 | 27 | 22 | 15 | 0 | 3 | 0 | 3 | 98 | 49 | 1 | 1 | 0.455 | 0.279 | 0.143 | NA | 0.333 | -0.041 |
| SE | 747 | CNTNAP3 | CNTNAP3 | chr9 | - | 39140518 | 39140638 | 39132931 | 39133135 | 39144239 | 39144346 | 3 | 2 | 2 | 0 | 0 | 0 | 0 | 1 | 0 | 1 | 98 | 49 | 1 | 1 | 1 | 1 | 1 | NA | 0.333 | 0.667 |
| SE | 748 | KIAA0196 | KIAA0196 | chr8 | - | 125074997 | 125075111 | 125073152 | 125073324 | 125076347 | 125076500 | 6 | 1 | 0 | 0 | 0 | 0 | 0 | 2 | 0 | 1 | 98 | 49 | 1 | 1 | 1 | 1 | NA | NA | 0.5 | 0.5 |
| SE | 751 | TMEM65 | TMEM65 | chr8 | - | 124327353 | 124327421 | 124323320 | 124323375 | 124330747 | 124330792 | 3 | 1 | 4 | 2 | 0 | 0 | 0 | 2 | 0 | 0 | 98 | 49 | 1 | 1 | 0.429 | 1 | 1 | NA | 1 | -0.19 |
| SE | 752 | FAM91A1 | FAM91A1 | chr8 | + | 123774079 | 123774164 | 123768641 | 123768774 | 123775146 | 123775298 | 31 | 18 | 8 | 0 | 1 | 0 | 0 | 6 | 0 | 0 | 98 | 49 | 1 | 1 | 1 | 0.9 | 1 | NA | 1 | -0.033 |
| SE | 753 | FAM91A1 | FAM91A1 | chr8 | + | 123779984 | 123780075 | 123778658 | 123778772 | 123780479 | 123780542 | 28 | 15 | 2 | 1 | 0 | 0 | 0 | 6 | 0 | 0 | 98 | 49 | 1 | 1 | 0.933 | 1 | 1 | NA | 1 | -0.022 |
| SE | 754 | FAM91A1 | FAM91A1 | chr8 | + | 123786494 | 123786610 | 123785628 | 123785741 | 123787260 | 123787373 | 17 | 18 | 8 | 3 | 0 | 0 | 0 | 4 | 0 | 1 | 98 | 49 | 1 | 1 | 0.739 | 1 | 1 | NA | 0.667 | 0.246 |
| SE | 755 | FAM91A1 | FAM91A1 | chr8 | + | 123789612 | 123789745 | 123787663 | 123787750 | 123798089 | 123798238 | 13 | 3 | 3 | 0 | 2 | 0 | 0 | 2 | 0 | 0 | 98 | 49 | 1 | 1 | 1 | 0.429 | 1 | NA | 1 | -0.19 |
| SE | 756 | FAM91A1 | FAM91A1 | chr8 | + | 123806079 | 123806229 | 123805266 | 123805339 | 123808271 | 123808376 | 18 | 13 | 5 | 1 | 0 | 0 | 0 | 4 | 0 | 0 | 98 | 49 | 1 | 1 | 0.9 | 1 | 1 | NA | 1 | -0.033 |
| SE | 760 | ATAD2 | ATAD2 | chr8 | - | 123337624 | 123337821 | 123336372 | 123336532 | 123339310 | 123339446 | 1 | 0 | 0 | 0 | 0 | 0 | 0 | 0 | 0 | 1 | 98 | 49 | 1 | 1 | 1 | NA | NA | NA | 0 | 1 |
| SE | 767 | RAD21 | RAD21 | chr8 | - | 116854244 | 116854468 | 116852548 | 116852708 | 116856165 | 116856288 | 82 | 29 | 36 | 1 | 0 | 0 | 2 | 16 | 0 | 0 | 98 | 49 | 1 | 1 | 0.976 | 1 | 1 | 1 | 1 | -0.008 |
| SE | 768 | EIF3H | EIF3H | chr8 | - | 116726015 | 116726172 | 116658812 | 116658980 | 116755665 | 116755823 | 41 | 6 | 2 | 3 | 0 | 0 | 0 | 12 | 0 | 0 | 98 | 49 | 1 | 1 | 0.872 | 1 | 1 | NA | 1 | -0.043 |
| SE | 769 | TRPS1 | TRPS1 | chr8 | - | 115623600 | 115623758 | 115619131 | 115620060 | 115668544 | 115669028 | 5 | 1 | 0 | 3 | 0 | 0 | 0 | 2 | 0 | 0 | 98 | 49 | 1 | 1 | 0.455 | 1 | NA | NA | 1 | -0.272 |
| SE | 770 | TRPS1 | TRPS1 | chr8 | - | 115623614 | 115623758 | 115619131 | 115620060 | 115668544 | 115669028 | 0 | 0 | 0 | 3 | 0 | 0 | 0 | 1 | 0 | 0 | 98 | 49 | 1 | 1 | 0 | NA | NA | NA | 1 | -1 |
| SE | 779 | OXR1 | OXR1 | chr8 | + | 106745788 | 106745862 | 106742221 | 106742317 | 106750805 | 106752693 | 5 | 2 | 0 | 1 | 0 | 0 | 0 | 5 | 0 | 0 | 98 | 49 | 1 | 1 | 0.714 | 1 | NA | NA | 1 | -0.143 |
| SE | 783 | SLC25A32 | SLC25A32 | chr8 | - | 103404775 | 103404861 | 103403163 | 103403324 | 103407633 | 103407784 | 11 | 0 | 0 | 1 | 0 | 0 | 0 | 2 | 0 | 0 | 98 | 49 | 1 | 1 | 0.846 | NA | NA | NA | 1 | -0.154 |
| SE | 788 | UBR5 | UBR5 | chr8 | - | 102271133 | 102271245 | 102270038 | 102270183 | 102272528 | 102272772 | 20 | 4 | 1 | 0 | 2 | 0 | 0 | 2 | 0 | 0 | 98 | 49 | 1 | 1 | 1 | 0.5 | 1 | NA | 1 | -0.167 |
| SE | 789 | UBR5 | UBR5 | chr8 | - | 102278825 | 102278939 | 102276971 | 102277192 | 102279035 | 102279208 | 16 | 12 | 5 | 2 | 0 | 0 | 0 | 4 | 0 | 0 | 98 | 49 | 1 | 1 | 0.8 | 1 | 1 | NA | 1 | -0.067 |
| SE | 790 | UBR5 | UBR5 | chr8 | - | 102312341 | 102312485 | 102312133 | 102312245 | 102313803 | 102313916 | 17 | 15 | 3 | 0 | 1 | 0 | 1 | 4 | 0 | 0 | 98 | 49 | 1 | 1 | 1 | 0.882 | 1 | 1 | 1 | -0.039 |
| SE | 791 | YWHAZ | YWHAZ | chr8 | - | 100923954 | 100924050 | 100918575 | 100920752 | 100924134 | 100924298 | 25 | 16 | 6 | 1 | 0 | 0 | 0 | 5 | 0 | 0 | 98 | 49 | 1 | 1 | 0.926 | 1 | 1 | NA | 1 | -0.025 |
| SE | 792 | YWHAZ | YWHAZ | chr8 | - | 100924134 | 100924298 | 100918575 | 100920752 | 100924915 | 100925039 | 110 | 47 | 20 | 0 | 0 | 0 | 0 | 26 | 0 | 1 | 98 | 49 | 1 | 1 | 1 | 1 | 1 | NA | 0.929 | 0.071 |
| SE | 794 | RNF19A | RNF19A | chr8 | - | 100261541 | 100261755 | 100259853 | 100259997 | 100264033 | 100264195 | 16 | 2 | 5 | 1 | 0 | 0 | 0 | 1 | 0 | 0 | 98 | 49 | 1 | 1 | 0.889 | 1 | 1 | NA | 1 | -0.037 |
| SE | 795 | RNF19A | RNF19A | chr8 | - | 100264033 | 100264195 | 100261541 | 100261755 | 100264670 | 100264785 | 11 | 0 | 6 | 2 | 0 | 0 | 0 | 1 | 0 | 0 | 98 | 49 | 1 | 1 | 0.733 | NA | 1 | NA | 1 | -0.133 |
| SE | 797 | RNF19A | RNF19A | chr8 | - | 100287500 | 100288267 | 100269868 | 100270013 | 100309866 | 100310166 | 9 | 9 | 3 | 1 | 0 | 0 | 0 | 3 | 0 | 0 | 98 | 49 | 1 | 1 | 0.818 | 1 | 1 | NA | 1 | -0.061 |
| SE | 799 | SPAG1 | SPAG1 | chr8 | + | 100233410 | 100233537 | 100231155 | 100231288 | 100239239 | 100239404 | 0 | 8 | 3 | 0 | 2 | 0 | 0 | 1 | 0 | 0 | 98 | 49 | 1 | 1 | NA | 0.667 | 1 | NA | 1 | -0.166 |
| SE | 803 | CPQ | CPQ | chr8 | + | 96834972 | 96835180 | 96784863 | 96785330 | 96879797 | 96880005 | 64 | 40 | 16 | 1 | 0 | 0 | 1 | 16 | 0 | 0 | 98 | 49 | 1 | 1 | 0.97 | 1 | 1 | 1 | 1 | -0.01 |
| SE | 804 | CPQ | CPQ | chr8 | + | 96879797 | 96880005 | 96834972 | 96835180 | 96965934 | 96966046 | 57 | 39 | 14 | 3 | 0 | 0 | 0 | 7 | 0 | 0 | 98 | 49 | 1 | 1 | 0.905 | 1 | 1 | NA | 1 | -0.032 |
| SE | 817 | TP53INP1 | TP53INP1 | chr8 | - | 94932069 | 94932107 | 94925971 | 94930728 | 94939859 | 94940220 | 28 | 17 | 6 | 2 | 4 | 1 | 0 | 4 | 0 | 1 | 87 | 49 | 1 | 1 | 0.887 | 0.705 | 0.772 | NA | 0.693 | 0.095 |
| SE | 818 | INTS8 | INTS8 | chr8 | + | 94856776 | 94856978 | 94853804 | 94853915 | 94859510 | 94859632 | 7 | 1 | 1 | 1 | 0 | 0 | 0 | 2 | 0 | 0 | 98 | 49 | 1 | 1 | 0.778 | 1 | 1 | NA | 1 | -0.074 |
| SE | 822 | INTS8 | INTS8 | chr8 | + | 94868691 | 94868813 | 94867275 | 94867337 | 94871883 | 94872002 | 1 | 0 | 0 | 2 | 1 | 1 | 0 | 0 | 0 | 1 | 98 | 49 | 1 | 1 | 0.2 | 0 | 0 | NA | 0 | 0.067 |
| SE | 825 | PDP1 | PDP1 | chr8 | + | 93917910 | 93917945 | 93916933 | 93917079 | 93922015 | 93926068 | 1 | 0 | 0 | 3 | 0 | 0 | 0 | 0 | 0 | 2 | 84 | 49 | 1 | 1 | 0.163 | NA | NA | NA | 0 | 0.163 |
| SE | 849 | C8orf59 | C8orf59 | chr8 | - | 85214920 | 85215037 | 85214052 | 85214631 | 85217385 | 85217502 | 21 | 3 | 1 | 0 | 0 | 0 | 2 | 3 | 0 | 1 | 98 | 49 | 1 | 1 | 1 | 1 | 1 | 1 | 0.6 | 0.2 |
| SE | 853 | IMPA1 | IMPA1 | chr8 | - | 81670938 | 81671047 | 81660515 | 81660667 | 81673840 | 81673949 | 5 | 2 | 7 | 1 | 0 | 0 | 0 | 2 | 0 | 0 | 98 | 49 | 1 | 1 | 0.714 | 1 | 1 | NA | 1 | -0.095 |
| SE | 855 | PAG1 | PAG1 | chr8 | - | 80993102 | 80993307 | 80991478 | 80991530 | 81029995 | 81030089 | 9 | 8 | 1 | 0 | 0 | 0 | 0 | 2 | 0 | 1 | 98 | 49 | 1 | 1 | 1 | 1 | 1 | NA | 0.5 | 0.5 |
| SE | 856 | PAG1 | PAG1 | chr8 | - | 81070111 | 81070170 | 81029995 | 81030089 | 81111590 | 81112068 | 8 | 3 | 1 | 1 | 0 | 0 | 2 | 16 | 0 | 0 | 98 | 49 | 1 | 1 | 0.8 | 1 | 1 | 1 | 1 | -0.067 |
| SE | 869 | LY96 | LY96 | chr8 | + | 74004795 | 74004885 | 73991328 | 73991554 | 74010000 | 74010129 | 45 | 39 | 34 | 4 | 0 | 1 | 0 | 10 | 0 | 0 | 98 | 49 | 1 | 1 | 0.849 | 1 | 0.944 | NA | 1 | -0.069 |
| SE | 872 | UBE2W | UBE2W | chr8 | - | 73825146 | 73825249 | 73810473 | 73810629 | 73830380 | 73830472 | 32 | 23 | 8 | 1 | 0 | 0 | 0 | 4 | 0 | 0 | 98 | 49 | 1 | 1 | 0.941 | 1 | 1 | NA | 1 | -0.02 |
| SE | 878 | NCOA2 | NCOA2 | chr8 | - | 70144641 | 70144848 | 70141183 | 70141399 | 70148272 | 70148483 | 10 | 3 | 3 | 8 | 2 | 0 | 0 | 5 | 0 | 1 | 98 | 49 | 1 | 1 | 0.385 | 0.429 | 1 | NA | 0.714 | -0.109 |
| SE | 879 | NCOA2 | NCOA2 | chr8 | - | 70174755 | 70174859 | 70170201 | 70170379 | 70213902 | 70214075 | 55 | 23 | 11 | 0 | 2 | 1 | 0 | 9 | 0 | 0 | 98 | 49 | 1 | 1 | 1 | 0.852 | 0.846 | NA | 1 | -0.101 |
| SE | 880 | ARFGEF1 | ARFGEF1 | chr8 | - | 67259814 | 67259926 | 67258084 | 67258290 | 67266005 | 67266207 | 21 | 14 | 1 | 0 | 0 | 0 | 0 | 6 | 0 | 1 | 98 | 49 | 1 | 1 | 1 | 1 | 1 | NA | 0.75 | 0.25 |
| SE | 886 | PDE7A | PDE7A | chr8 | - | 65724254 | 65724351 | 65723540 | 65723621 | 65724776 | 65724921 | 11 | 10 | 3 | 2 | 0 | 0 | 0 | 2 | 0 | 0 | 98 | 49 | 1 | 1 | 0.733 | 1 | 1 | NA | 1 | -0.089 |
| SE | 888 | CHD7 | CHD7 | chr8 | + | 60801527 | 60801593 | 60800387 | 60800525 | 60808216 | 60808272 | 15 | 15 | 6 | 0 | 1 | 0 | 0 | 2 | 0 | 0 | 98 | 49 | 1 | 1 | 1 | 0.882 | 1 | NA | 1 | -0.039 |
| SE | 890 | GTF3C5 | GTF3C5 | chr9 | + | 133054943 | 133054964 | 133054711 | 133054809 | 133056011 | 133056094 | 0 | 0 | 0 | 2 | 0 | 0 | 1 | 0 | 0 | 0 | 70 | 49 | 1 | 1 | 0 | NA | NA | 1 | NA | -1 |
| SE | 892 | RAB2A | RAB2A | chr8 | + | 60572045 | 60572113 | 60558851 | 60558923 | 60584207 | 60584290 | 20 | 12 | 7 | 5 | 1 | 0 | 1 | 0 | 0 | 0 | 98 | 49 | 1 | 1 | 0.667 | 0.857 | 1 | 1 | NA | -0.159 |
| SE | 893 | CYSLTR1 | CYSLTR1 | chrX | - | 78283453 | 78283540 | 78271471 | 78273773 | 78327304 | 78327691 | 17 | 15 | 9 | 1 | 0 | 0 | 0 | 4 | 0 | 0 | 98 | 49 | 1 | 1 | 0.895 | 1 | 1 | NA | 1 | -0.035 |
| SE | 896 | CHCHD7 | CHCHD7 | chr8 | + | 56212761 | 56212889 | 56211755 | 56211837 | 56214597 | 56214667 | 0 | 0 | 0 | 0 | 3 | 0 | 0 | 1 | 0 | 0 | 98 | 49 | 1 | 1 | NA | 0 | NA | NA | 1 | -1 |
| SE | 899 | RB1CC1 | RB1CC1 | chr8 | - | 52628031 | 52628168 | 52624716 | 52624787 | 52630469 | 52630528 | 21 | 23 | 1 | 5 | 0 | 0 | 0 | 3 | 0 | 0 | 98 | 49 | 1 | 1 | 0.677 | 1 | 1 | NA | 1 | -0.108 |
| SE | 900 | RB1CC1 | RB1CC1 | chr8 | - | 52628040 | 52628168 | 52624716 | 52624787 | 52630469 | 52630528 | 17 | 19 | 0 | 5 | 0 | 0 | 0 | 3 | 0 | 0 | 98 | 49 | 1 | 1 | 0.63 | 1 | NA | NA | 1 | -0.185 |
| SE | 901 | RB1CC1 | RB1CC1 | chr8 | - | 52660595 | 52660657 | 52658872 | 52658976 | 52660925 | 52661007 | 34 | 13 | 8 | 4 | 9 | 0 | 0 | 3 | 0 | 0 | 98 | 49 | 1 | 1 | 0.81 | 0.419 | 1 | NA | 1 | -0.257 |
| SE | 902 | RB1CC1 | RB1CC1 | chr8 | - | 52686852 | 52686967 | 52685398 | 52685520 | 52714074 | 52714466 | 30 | 11 | 3 | 2 | 0 | 0 | 1 | 1 | 0 | 0 | 98 | 49 | 1 | 1 | 0.882 | 1 | 1 | 1 | 1 | -0.039 |
| SE | 905 | SPIDR | SPIDR | chr8 | + | 47396375 | 47396626 | 47293866 | 47294030 | 47407860 | 47407961 | 16 | 4 | 3 | 0 | 3 | 0 | 0 | 2 | 0 | 0 | 98 | 49 | 1 | 1 | 1 | 0.4 | 1 | NA | 1 | -0.2 |
| SE | 909 | POMK | POMK | chr8 | + | 43102504 | 43102600 | 43097523 | 43097615 | 43103527 | 43103830 | 8 | 1 | 0 | 0 | 0 | 0 | 0 | 0 | 0 | 2 | 98 | 49 | 1 | 1 | 1 | 1 | NA | NA | 0 | 1 |
| SE | 912 | RNF170 | RNF170 | chr8 | - | 42865415 | 42865489 | 42861744 | 42861855 | 42870003 | 42870112 | 0 | 0 | 0 | 0 | 0 | 3 | 0 | 6 | 0 | 0 | 98 | 49 | 1 | 1 | NA | NA | 0 | NA | 1 | -1 |
| SE | 914 | POLB | POLB | chr8 | + | 42339011 | 42339069 | 42338454 | 42338685 | 42344952 | 42345019 | 13 | 2 | 5 | 4 | 0 | 0 | 0 | 3 | 0 | 2 | 98 | 49 | 1 | 1 | 0.619 | 1 | 1 | NA | 0.429 | 0.444 |
| SE | 915 | POLB | POLB | chr8 | + | 42349015 | 42349090 | 42344952 | 42345019 | 42350006 | 42350065 | 16 | 20 | 1 | 1 | 0 | 0 | 0 | 1 | 0 | 0 | 98 | 49 | 1 | 1 | 0.889 | 1 | 1 | NA | 1 | -0.037 |
| SE | 916 | POLB | POLB | chr8 | + | 42350006 | 42350065 | 42344952 | 42345019 | 42352518 | 42352568 | 12 | 1 | 0 | 0 | 0 | 0 | 0 | 0 | 0 | 1 | 98 | 49 | 1 | 1 | 1 | 1 | NA | NA | 0 | 1 |
| SE | 917 | POLB | POLB | chr8 | + | 42362611 | 42362698 | 42361294 | 42361365 | 42369270 | 42369335 | 7 | 5 | 3 | 4 | 2 | 2 | 0 | 1 | 0 | 0 | 98 | 49 | 1 | 1 | 0.467 | 0.556 | 0.429 | NA | 1 | -0.516 |
| SE | 919 | IKBKB | IKBKB | chr8 | + | 42288633 | 42288728 | 42272082 | 42272205 | 42290155 | 42290273 | 4 | 1 | 0 | 0 | 1 | 0 | 0 | 3 | 0 | 0 | 98 | 49 | 1 | 1 | 1 | 0.333 | NA | NA | 1 | -0.334 |
| SE | 924 | KAT6A | KAT6A | chr8 | - | 41974703 | 41974822 | 41955295 | 41955411 | 41977007 | 41977327 | 47 | 39 | 14 | 1 | 0 | 0 | 2 | 10 | 0 | 0 | 98 | 49 | 1 | 1 | 0.959 | 1 | 1 | 1 | 1 | -0.014 |
| SE | 925 | KAT6A | KAT6A | chr8 | - | 41987454 | 41987563 | 41981838 | 41981954 | 42048377 | 42049302 | 94 | 32 | 16 | 1 | 0 | 0 | 0 | 17 | 0 | 0 | 98 | 49 | 1 | 1 | 0.979 | 1 | 1 | NA | 1 | -0.007 |
| SE | 938 | PLEKHA2 | PLEKHA2 | chr8 | + | 38943788 | 38943837 | 38935993 | 38936050 | 38946123 | 38946221 | 20 | 19 | 9 | 0 | 1 | 0 | 0 | 2 | 0 | 0 | 98 | 49 | 1 | 1 | 1 | 0.905 | 1 | NA | 1 | -0.032 |
| SE | 939 | PLEKHA2 | PLEKHA2 | chr8 | + | 38946123 | 38946221 | 38935993 | 38936050 | 38950849 | 38950990 | 5 | 8 | 4 | 0 | 0 | 1 | 0 | 4 | 0 | 0 | 98 | 49 | 1 | 1 | 1 | 1 | 0.667 | NA | 1 | -0.111 |
| SE | 940 | PLEKHA2 | PLEKHA2 | chr8 | + | 38952165 | 38952312 | 38950849 | 38950990 | 38952635 | 38952704 | 17 | 14 | 9 | 1 | 0 | 0 | 0 | 4 | 0 | 0 | 98 | 49 | 1 | 1 | 0.895 | 1 | 1 | NA | 1 | -0.035 |
| SE | 941 | PLEKHA2 | PLEKHA2 | chr8 | + | 38957322 | 38957386 | 38953296 | 38953367 | 38968591 | 38968669 | 9 | 13 | 3 | 0 | 0 | 0 | 0 | 5 | 0 | 1 | 98 | 49 | 1 | 1 | 1 | 1 | 1 | NA | 0.714 | 0.286 |
| SE | 943 | WHSC1L1 | WHSC1L1 | chr8 | - | 38299443 | 38299590 | 38295795 | 38295952 | 38304586 | 38304757 | 23 | 0 | 2 | 1 | 0 | 0 | 0 | 1 | 0 | 0 | 98 | 49 | 1 | 1 | 0.92 | NA | 1 | NA | 1 | -0.04 |
| SE | 944 | WHSC1L1 | WHSC1L1 | chr8 | - | 38299443 | 38299590 | 38295795 | 38295952 | 38314646 | 38314773 | 13 | 0 | 2 | 1 | 0 | 0 | 0 | 1 | 0 | 0 | 98 | 49 | 1 | 1 | 0.867 | NA | 1 | NA | 1 | -0.067 |
| SE | 945 | WHSC1L1 | WHSC1L1 | chr8 | - | 38326729 | 38326856 | 38321071 | 38321172 | 38329377 | 38329893 | 21 | 5 | 1 | 1 | 0 | 0 | 0 | 1 | 0 | 0 | 98 | 49 | 1 | 1 | 0.913 | 1 | 1 | NA | 1 | -0.029 |
| SE | 954 | INTS9 | INTS9 | chr8 | - | 28846746 | 28846809 | 28837636 | 28837776 | 28850212 | 28850273 | 0 | 1 | 0 | 1 | 0 | 0 | 0 | 1 | 0 | 0 | 98 | 49 | 1 | 1 | 0 | 1 | NA | NA | 1 | -0.5 |
| SE | 956 | EXTL3 | EXTL3 | chr8 | + | 28713456 | 28713550 | 28701472 | 28701659 | 28715584 | 28716066 | 4 | 3 | 4 | 1 | 0 | 0 | 0 | 1 | 0 | 0 | 98 | 49 | 1 | 1 | 0.667 | 1 | 1 | NA | 1 | -0.111 |
| SE | 966 | PPP2R2A | PPP2R2A | chr8 | + | 26363720 | 26363890 | 26362683 | 26362848 | 26366314 | 26366406 | 16 | 3 | 2 | 0 | 2 | 0 | 0 | 2 | 0 | 0 | 98 | 49 | 1 | 1 | 1 | 0.429 | 1 | NA | 1 | -0.19 |
| SE | 967 | DOCK5 | DOCK5 | chr8 | + | 25268844 | 25268885 | 25243673 | 25243757 | 25275385 | 25275441 | 148 | 62 | 18 | 1 | 0 | 0 | 1 | 18 | 0 | 0 | 90 | 49 | 1 | 1 | 0.988 | 1 | 1 | 1 | 1 | -0.004 |
| SE | 968 | DOCK5 | DOCK5 | chr8 | + | 25278568 | 25278665 | 25275385 | 25275441 | 25292023 | 25292172 | 229 | 94 | 12 | 0 | 3 | 0 | 5 | 32 | 0 | 0 | 98 | 49 | 1 | 1 | 1 | 0.94 | 1 | 1 | 1 | -0.02 |
| SE | 969 | DOCK5 | DOCK5 | chr8 | + | 25296512 | 25296648 | 25292023 | 25292172 | 25298943 | 25299101 | 181 | 82 | 14 | 2 | 1 | 0 | 3 | 29 | 0 | 0 | 98 | 49 | 1 | 1 | 0.978 | 0.976 | 1 | 1 | 1 | -0.015 |
| SE | 971 | DOCK5 | DOCK5 | chr8 | + | 25317006 | 25317131 | 25310406 | 25310532 | 25319577 | 25319676 | 132 | 75 | 18 | 1 | 0 | 0 | 2 | 30 | 0 | 0 | 98 | 49 | 1 | 1 | 0.985 | 1 | 1 | 1 | 1 | -0.005 |
| SE | 972 | DOCK5 | DOCK5 | chr8 | + | 25334095 | 25334196 | 25332602 | 25332692 | 25336238 | 25336373 | 175 | 73 | 14 | 1 | 0 | 0 | 1 | 36 | 0 | 0 | 98 | 49 | 1 | 1 | 0.989 | 1 | 1 | 1 | 1 | -0.004 |
| SE | 973 | DOCK5 | DOCK5 | chr8 | + | 25340876 | 25340988 | 25336238 | 25336373 | 25341738 | 25341809 | 115 | 49 | 6 | 8 | 7 | 0 | 0 | 19 | 0 | 0 | 98 | 49 | 1 | 1 | 0.878 | 0.778 | 1 | NA | 1 | -0.115 |
| SE | 974 | DOCK5 | DOCK5 | chr8 | + | 25342400 | 25342507 | 25341738 | 25341809 | 25345474 | 25345611 | 107 | 55 | 21 | 1 | 0 | 0 | 0 | 15 | 0 | 0 | 98 | 49 | 1 | 1 | 0.982 | 1 | 1 | NA | 1 | -0.006 |
| SE | 975 | DOCK5 | DOCK5 | chr8 | + | 25368191 | 25368250 | 25366869 | 25366970 | 25368570 | 25368725 | 86 | 32 | 10 | 3 | 0 | 0 | 0 | 11 | 0 | 5 | 98 | 49 | 1 | 1 | 0.935 | 1 | 1 | NA | 0.524 | 0.454 |
| SE | 977 | DOCK5 | DOCK5 | chr8 | + | 25373617 | 25373658 | 25372558 | 25372718 | 25374563 | 25374654 | 85 | 38 | 16 | 0 | 0 | 0 | 0 | 8 | 0 | 1 | 90 | 49 | 1 | 1 | 1 | 1 | 1 | NA | 0.813 | 0.187 |
| SE | 982 | TNFRSF10C | TNFRSF10C | chr8 | + | 23111719 | 23111825 | 23102813 | 23103181 | 23114656 | 23114770 | 340 | 217 | 190 | 33 | 15 | 0 | 0 | 51 | 0 | 4 | 98 | 49 | 1 | 1 | 0.837 | 0.879 | 1 | NA | 0.864 | 0.041 |
| SE | 983 | TNFRSF10C | TNFRSF10C | chr8 | + | 23115507 | 23115616 | 23114656 | 23114770 | 23116640 | 23117437 | 394 | 187 | 150 | 0 | 1 | 0 | 3 | 45 | 0 | 0 | 98 | 49 | 1 | 1 | 1 | 0.989 | 1 | 1 | 1 | -0.004 |
| SE | 984 | CCAR2 | CCAR2 | chr8 | + | 22618348 | 22618495 | 22617695 | 22617778 | 22618616 | 22618728 | 2 | 3 | 0 | 1 | 0 | 0 | 0 | 1 | 0 | 0 | 98 | 49 | 1 | 1 | 0.5 | 1 | NA | NA | 1 | -0.25 |
| SE | 985 | PDLIM2 | PDLIM2 | chr8 | + | 22589297 | 22589374 | 22585320 | 22585399 | 22589595 | 22589741 | 5 | 3 | 7 | 0 | 0 | 1 | 0 | 1 | 0 | 0 | 98 | 49 | 1 | 1 | 1 | 1 | 0.778 | NA | 1 | -0.074 |
| SE | 986 | PDLIM2 | PDLIM2 | chr8 | + | 22591550 | 22591668 | 22589595 | 22589741 | 22593732 | 22593832 | 20 | 11 | 13 | 2 | 0 | 0 | 0 | 6 | 0 | 0 | 98 | 49 | 1 | 1 | 0.833 | 1 | 1 | NA | 1 | -0.056 |
| SE | 988 | PPP3CC | PPP3CC | chr8 | + | 22539468 | 22539498 | 22532920 | 22533018 | 22540614 | 22541144 | 3 | 0 | 0 | 1 | 0 | 0 | 0 | 1 | 0 | 0 | 79 | 49 | 1 | 1 | 0.65 | NA | NA | NA | 1 | -0.35 |
| SE | 994 | XPO7 | XPO7 | chr8 | + | 21990810 | 21990919 | 21990343 | 21990407 | 21991867 | 21991974 | 3 | 0 | 3 | 1 | 0 | 0 | 0 | 2 | 0 | 0 | 98 | 49 | 1 | 1 | 0.6 | NA | 1 | NA | 1 | -0.2 |
| SE | 995 | DOK2 | DOK2 | chr8 | - | 21911900 | 21911988 | 21910672 | 21910857 | 21912228 | 21912510 | 10 | 2 | 1 | 1 | 0 | 0 | 0 | 7 | 0 | 2 | 98 | 49 | 1 | 1 | 0.833 | 1 | 1 | NA | 0.636 | 0.308 |
| SE | 998 | ATP6V1B2 | ATP6V1B2 | chr8 | + | 20210568 | 20210646 | 20210345 | 20210439 | 20211176 | 20211316 | 141 | 47 | 27 | 1 | 0 | 0 | 4 | 14 | 0 | 0 | 98 | 49 | 1 | 1 | 0.986 | 1 | 1 | 1 | 1 | -0.005 |
| SE | 999 | ATP6V1B2 | ATP6V1B2 | chr8 | + | 20214817 | 20214968 | 20212781 | 20212905 | 20216412 | 20216495 | 93 | 25 | 30 | 1 | 1 | 0 | 0 | 19 | 0 | 0 | 98 | 49 | 1 | 1 | 0.979 | 0.926 | 1 | NA | 1 | -0.032 |
| SE | 1000 | ATP6V1B2 | ATP6V1B2 | chr8 | + | 20216412 | 20216495 | 20212781 | 20212905 | 20217219 | 20217324 | 41 | 18 | 22 | 1 | 0 | 0 | 0 | 19 | 0 | 0 | 98 | 49 | 1 | 1 | 0.953 | 1 | 1 | NA | 1 | -0.016 |
| SE | 1015 | ASAH1 | ASAH1 | chr8 | - | 18062278 | 18062423 | 18061685 | 18061740 | 18063184 | 18063230 | 84 | 50 | 32 | 1 | 0 | 0 | 0 | 18 | 0 | 0 | 98 | 49 | 1 | 1 | 0.977 | 1 | 1 | NA | 1 | -0.008 |
| SE | 1016 | ASAH1 | ASAH1 | chr8 | - | 18063184 | 18063230 | 18062278 | 18062423 | 18064456 | 18064531 | 84 | 32 | 38 | 0 | 0 | 1 | 0 | 20 | 0 | 0 | 95 | 49 | 1 | 1 | 1 | 1 | 0.951 | NA | 1 | -0.016 |
| SE | 1018 | ASAH1 | ASAH1 | chr8 | - | 18073255 | 18073276 | 18071299 | 18071390 | 18075540 | 18075587 | 0 | 2 | 0 | 61 | 34 | 21 | 0 | 0 | 0 | 27 | 70 | 49 | 1 | 1 | 0 | 0.04 | 0 | NA | 0 | 0.013 |
| SE | 1019 | PCM1 | PCM1 | chr8 | + | 17924712 | 17924780 | 17922856 | 17923188 | 17935588 | 17935706 | 10 | 8 | 3 | 1 | 0 | 4 | 0 | 1 | 0 | 0 | 98 | 49 | 1 | 1 | 0.833 | 1 | 0.273 | NA | 1 | -0.298 |
| SE | 1020 | PCM1 | PCM1 | chr8 | + | 17963100 | 17963291 | 17962033 | 17962174 | 17964567 | 17964768 | 18 | 2 | 0 | 2 | 1 | 0 | 0 | 10 | 0 | 0 | 98 | 49 | 1 | 1 | 0.818 | 0.5 | NA | NA | 1 | -0.341 |
| SE | 1021 | PCM1 | PCM1 | chr8 | + | 17980590 | 17980755 | 17972328 | 17972687 | 17985446 | 17985619 | 4 | 1 | 1 | 2 | 0 | 0 | 0 | 1 | 0 | 0 | 98 | 49 | 1 | 1 | 0.5 | 1 | 1 | NA | 1 | -0.167 |
| SE | 1027 | ZDHHC2 | ZDHHC2 | chr8 | + | 17207959 | 17208092 | 17205654 | 17205775 | 17209931 | 17210058 | 8 | 3 | 9 | 0 | 0 | 2 | 0 | 2 | 0 | 0 | 98 | 49 | 1 | 1 | 1 | 1 | 0.692 | NA | 1 | -0.103 |
| SE | 1030 | CTSB | CTSB | chr8 | - | 11849045 | 11849164 | 11848066 | 11848152 | 11850865 | 11850980 | 21 | 9 | 3 | 0 | 0 | 0 | 0 | 32 | 0 | 2 | 98 | 49 | 1 | 1 | 1 | 1 | 1 | NA | 0.889 | 0.111 |
| SE | 1034 | CTSB | CTSB | chr8 | - | 11864375 | 11864463 | 11853328 | 11853479 | 11868000 | 11868137 | 2 | 0 | 0 | 0 | 1 | 0 | 0 | 1 | 2 | 1 | 98 | 49 | 1 | 1 | 1 | 0 | NA | 0 | 0.333 | 0.334 |
| SE | 1041 | TNKS | TNKS | chr8 | + | 9730889 | 9731035 | 9726640 | 9726720 | 9733278 | 9733444 | 4 | 2 | 0 | 0 | 1 | 0 | 0 | 1 | 0 | 1 | 98 | 49 | 1 | 1 | 1 | 1 | NA | NA | 0.333 | 0.667 |
| SE | 1051 | DNAJB6 | DNAJB6 | chr7 | + | 157363160 | 157363270 | 157358546 | 157358637 | 157366501 | 157366561 | 26 | 17 | 6 | 0 | 1 | 0 | 5 | 5 | 0 | 0 | 98 | 49 | 1 | 1 | 1 | 0.895 | 1 | 1 | 1 | -0.035 |
| SE | 1052 | RBM33 | RBM33 | chr7 | + | 155706859 | 155707068 | 155700772 | 155700944 | 155711202 | 155711455 | 31 | 9 | 7 | 1 | 0 | 0 | 0 | 4 | 0 | 0 | 98 | 49 | 1 | 1 | 0.939 | 1 | 1 | NA | 1 | -0.02 |
| SE | 1053 | RBM33 | RBM33 | chr7 | + | 155739714 | 155740026 | 155738059 | 155738403 | 155741818 | 155742106 | 8 | 1 | 0 | 2 | 0 | 0 | 0 | 2 | 0 | 0 | 98 | 49 | 1 | 1 | 0.667 | 1 | NA | NA | 1 | -0.166 |
| SE | 1054 | KMT2C | KMT2C | chr7 | - | 152146598 | 152146735 | 152145152 | 152145295 | 152148032 | 152149152 | 47 | 11 | 8 | 0 | 0 | 0 | 0 | 11 | 0 | 1 | 98 | 49 | 1 | 1 | 1 | 1 | 1 | NA | 0.846 | 0.154 |
| SE | 1055 | KMT2C | KMT2C | chr7 | - | 152182973 | 152183156 | 152180710 | 152182594 | 152185557 | 152185631 | 46 | 15 | 5 | 5 | 2 | 2 | 0 | 9 | 0 | 3 | 98 | 49 | 1 | 1 | 0.821 | 0.789 | 0.556 | NA | 0.6 | 0.122 |
| SE | 1056 | KMT2C | KMT2C | chr7 | - | 152195906 | 152196011 | 152194439 | 152194568 | 152199278 | 152199459 | 44 | 14 | 2 | 1 | 0 | 0 | 0 | 8 | 0 | 0 | 98 | 49 | 1 | 1 | 0.957 | 1 | 1 | NA | 1 | -0.014 |

| | | | | | | | | | | | | | | | | | | | | | | | | | | | | | | | | |
|---|---|---|---|---|---|---|---|---|---|---|---|---|---|---|---|---|---|---|---|---|---|---|---|---|---|---|---|---|---|---|---|---|
| SE | 1057 | KMT2C | KMT2C | chr7 | - | 152311797 | 152311946 | 152309965 | 152310075 | 152315137 | 152315338 | 99 | 35 | 7 | 1 | 0 | 0 | 0 | 12 | 0 | 0 | 98 | 49 | 1 | 1 | 0.98 | 1 | 1 | NA | | 1 | -0.007 |
| SE | 1061 | NUB1 | NUB1 | chr7 | + | 151355767 | 151355950 | 151352811 | 151352882 | 151360140 | 151360247 | 11 | 9 | 4 | 0 | 0 | 0 | 0 | 3 | 1 | 0 | 98 | 49 | 1 | 1 | 1 | 1 | 1 | | 0 | 1 | 0.5 |
| SE | 1062 | NUB1 | NUB1 | chr7 | + | 151356127 | 151356222 | 151355767 | 151355950 | 151360140 | 151360247 | 32 | 9 | 3 | 0 | 3 | 0 | 0 | 3 | 0 | 0 | 98 | 49 | 1 | 1 | 1 | 0.6 | 1 | NA | | 1 | -0.133 |
| SE | 1071 | ZNF862 | ZNF862 | chr7 | + | 149860382 | 149862494 | 149859421 | 149859526 | 149864108 | 149867479 | 1 | 0 | 0 | 2 | 0 | 0 | 0 | 1 | 0 | 0 | 98 | 49 | 1 | 1 | 0.2 | NA | NA | NA | | 1 | -0.8 |
| SE | 1078 | TPK1 | TPK1 | chr7 | - | 144835559 | 144835618 | 144765879 | 144765951 | 144835966 | 144836053 | 3 | 0 | 0 | 1 | 0 | 0 | 0 | 4 | 0 | 0 | 98 | 49 | 1 | 1 | 0.6 | NA | NA | NA | | 1 | -0.4 |
| SE | 1094 | MGAM | MGAM | chr7 | + | 142034261 | 142034379 | 142032824 | 142032909 | 142034669 | 142034841 | 154 | 87 | 31 | 2 | 0 | 0 | 2 | 32 | 0 | 0 | 98 | 49 | 1 | 1 | 0.975 | 1 | 1 | | 1 | 1 | -0.008 |
| SE | 1095 | MGAM | MGAM | chr7 | + | 142036168 | 142036285 | 142034669 | 142034841 | 142036822 | 142036977 | 158 | 41 | 27 | 0 | 2 | 0 | 2 | 23 | 0 | 0 | 98 | 49 | 1 | 1 | 1 | 0.911 | 1 | | 1 | 1 | -0.03 |
| SE | 1096 | MGAM | MGAM | chr7 | + | 142036168 | 142036285 | 142034669 | 142034841 | 142038530 | 142038615 | 110 | 29 | 10 | 1 | 0 | 0 | 2 | 18 | 0 | 0 | 98 | 49 | 1 | 1 | 0.982 | 1 | 1 | | 1 | 1 | -0.006 |
| SE | 1097 | MGAM | MGAM | chr7 | + | 142036822 | 142036977 | 142034669 | 142034841 | 142038530 | 142038615 | 97 | 19 | 10 | 1 | 0 | 0 | 0 | 14 | 0 | 0 | 98 | 49 | 1 | 1 | 0.98 | 1 | 1 | NA | | 1 | -0.007 |
| SE | 1098 | MGAM | MGAM | chr7 | + | 142036822 | 142036977 | 142036168 | 142036285 | 142038530 | 142038615 | 147 | 30 | 27 | 2 | 1 | 0 | 0 | 20 | 0 | 1 | 98 | 49 | 1 | 1 | 0.974 | 0.937 | 1 | NA | | 0.909 | 0.061 |
| SE | 1099 | MGAM | MGAM | chr7 | + | 142047784 | 142047873 | 142040721 | 142040846 | 142050234 | 142050284 | 176 | 46 | 39 | 2 | 0 | 0 | 2 | 16 | 0 | 0 | 98 | 49 | 1 | 1 | 0.978 | 1 | 1 | | 1 | 1 | -0.007 |
| SE | 1100 | MGAM | MGAM | chr7 | + | 142047784 | 142047873 | 142040721 | 142040846 | 142050696 | 142050864 | 85 | 20 | 17 | 1 | 2 | 0 | 2 | 7 | 0 | 0 | 98 | 49 | 1 | 1 | 0.977 | 0.833 | 1 | | 1 | 1 | -0.063 |
| SE | 1101 | MGAM | MGAM | chr7 | + | 142050234 | 142050284 | 142040721 | 142040846 | 142050696 | 142050864 | 124 | 20 | 11 | 1 | 2 | 0 | 0 | 9 | 0 | 0 | 98 | 49 | 1 | 1 | 0.984 | 0.833 | 1 | NA | | 1 | -0.061 |
| SE | 1102 | MGAM | MGAM | chr7 | + | 142050234 | 142050284 | 142047784 | 142047873 | 142050696 | 142050864 | 214 | 46 | 33 | 1 | 0 | 0 | 0 | 18 | 0 | 0 | 98 | 49 | 1 | 1 | 0.991 | 1 | 1 | NA | | 1 | -0.003 |
| SE | 1106 | BRWD3 | BRWD3 | chrX | - | 80681996 | 80682094 | 80681340 | 80681499 | 80682464 | 80682628 | 9 | 10 | 5 | 1 | 0 | 0 | 0 | 6 | 0 | 0 | 98 | 49 | 1 | 1 | 0.818 | 1 | 1 | NA | | 1 | -0.061 |
| SE | 1110 | SLC37A3 | SLC37A3 | chr7 | - | 140355667 | 140355764 | 140352061 | 140352146 | 140364407 | 140364491 | 12 | 7 | 4 | 1 | 0 | 0 | 0 | 2 | 0 | 0 | 98 | 49 | 1 | 1 | 0.857 | 1 | 1 | NA | | 1 | -0.048 |
| SE | 1111 | SLC37A3 | SLC37A3 | chr7 | - | 140358639 | 140358785 | 140355667 | 140355764 | 140364407 | 140364491 | 24 | 13 | 2 | 1 | 3 | 0 | 0 | 12 | 0 | 0 | 98 | 49 | 1 | 1 | 0.923 | 0.684 | 1 | NA | | 1 | -0.131 |
| SE | 1112 | SLC37A3 | SLC37A3 | chr7 | - | 140380281 | 140380390 | 140369589 | 140369682 | 140382437 | 140382596 | 14 | 6 | 0 | 3 | 0 | 0 | 1 | 1 | 0 | 0 | 98 | 49 | 1 | 1 | 0.7 | 1 | NA | | 1 | 1 | -0.15 |
| SE | 1113 | PARP12 | PARP12 | chr7 | - | 140027275 | 140027406 | 140026196 | 140026348 | 140028612 | 140028688 | 15 | 14 | 5 | 1 | 0 | 0 | 1 | 16 | 0 | 0 | 98 | 49 | 1 | 1 | 0.882 | 1 | 1 | | 1 | 1 | -0.039 |
| SE | 1115 | PARP12 | PARP12 | chr7 | - | 140057898 | 140058034 | 140056855 | 140057153 | 140062521 | 140063721 | 20 | 4 | 7 | 0 | 0 | 0 | 1 | 8 | 1 | 0 | 98 | 49 | 1 | 1 | 1 | 1 | 1 | | 0.333 | 1 | 0.334 |
| SE | 1117 | KIF13B | KIF13B | chr8 | - | 29186291 | 29186472 | 29181918 | 29182006 | 29188514 | 29188607 | 2 | 1 | 0 | 0 | 2 | 0 | 0 | 2 | 0 | 0 | 98 | 49 | 1 | 1 | 1 | 0.2 | NA | NA | | 1 | -0.4 |
| SE | 1132 | LINC-PINT | LINC-PINT | chr7 | - | 130912109 | 130912152 | 130884316 | 130884396 | 130945719 | 130945835 | 12 | 17 | 6 | 11 | 1 | 0 | 0 | 1 | 0 | 0 | 92 | 49 | 1 | 1 | 0.367 | 0.901 | 1 | NA | | 1 | -0.244 |
| SE | 1133 | LINC-PINT | LINC-PINT | chr7 | - | 130912109 | 130912152 | 130884316 | 130884396 | 130959868 | 130959970 | 7 | 10 | 0 | 1 | 0 | 0 | 0 | 1 | 0 | 0 | 92 | 49 | 1 | 1 | 0.789 | 1 | NA | NA | | 1 | -0.105 |
| SE | 1134 | LINC-PINT | LINC-PINT | chr7 | - | 130959868 | 130959970 | 130944159 | 130945835 | 130967388 | 130967517 | 22 | 3 | 0 | 18 | 3 | 0 | 0 | 1 | 0 | 3 | 98 | 49 | 1 | 1 | 0.379 | 0.333 | NA | NA | | 0.143 | 0.213 |
| SE | 1137 | LINC-PINT | LINC-PINT | chr7 | - | 130967388 | 130967517 | 130959868 | 130959970 | 130984052 | 130984109 | 22 | 7 | 0 | 3 | 1 | 0 | 0 | 2 | 0 | 0 | 98 | 49 | 1 | 1 | 0.786 | 0.778 | NA | NA | | 1 | -0.218 |
| SE | 1140 | LINC-PINT | LINC-PINT | chr7 | - | 131052424 | 131052555 | 130984052 | 130984109 | 131107719 | 131108803 | 56 | 24 | 8 | 1 | 0 | 0 | 2 | 11 | 0 | 0 | 98 | 49 | 1 | 1 | 0.966 | 1 | 1 | | 1 | 1 | -0.011 |
| SE | 1150 | NRF1 | NRF1 | chr7 | + | 129710373 | 129710571 | 129709074 | 129709233 | 129711474 | 129711576 | 4 | 4 | 5 | 1 | 2 | 0 | 0 | 1 | 0 | 0 | 98 | 49 | 1 | 1 | 0.667 | 0.5 | 1 | NA | | 1 | -0.278 |
| SE | 1152 | AHCYL2 | AHCYL2 | chr7 | + | 129400289 | 129400384 | 129397221 | 129397324 | 129403378 | 129403485 | 5 | 0 | 0 | 0 | 0 | 0 | 0 | 0 | 0 | 2 | 98 | 49 | 1 | 1 | 1 | NA | NA | NA | | 0 | 1 |
| SE | 1154 | TNPO3 | TNPO3 | chr7 | - | 128993806 | 128993914 | 128991998 | 128992090 | 128997388 | 128997535 | 12 | 5 | 10 | 0 | 0 | 0 | 0 | 0 | 0 | 2 | 98 | 49 | 1 | 1 | 1 | 1 | 1 | NA | | 0 | 1 |
| SE | 1157 | SND1 | SND1 | chr7 | + | 127707556 | 127707647 | 127704838 | 127704945 | 127721286 | 127721400 | 16 | 3 | 2 | 0 | 1 | 0 | 0 | 1 | 0 | 0 | 98 | 49 | 1 | 1 | 1 | 0.6 | 1 | NA | | 1 | -0.133 |
| SE | 1163 | POT1 | POT1 | chr7 | - | 124848539 | 124848659 | 124846941 | 124846998 | 124851871 | 124851951 | 1 | 0 | 0 | 2 | 0 | 0 | 0 | 0 | 1 | 0 | 98 | 49 | 1 | 1 | 0.2 | NA | NA | | 0 NA | | 0.2 |
| SE | 1170 | ING3 | ING3 | chr7 | + | 120964741 | 120964838 | 120955558 | 120955624 | 120966625 | 120966697 | 3 | 3 | 9 | 2 | 0 | 0 | 0 | 0 | 0 | 2 | 98 | 49 | 1 | 1 | 0.429 | 1 | 1 | NA | | 0 | 0.81 |
| SE | 1171 | ING3 | ING3 | chr7 | + | 120973204 | 120973243 | 120970687 | 120970880 | 120974727 | 120975657 | 8 | 4 | 5 | 0 | 0 | 0 | 0 | 3 | 0 | 1 | 88 | 49 | 1 | 1 | 1 | 1 | 1 | NA | | 0.626 | 0.374 |
| SE | 1174 | CAPZA2 | CAPZA2 | chr7 | + | 116906262 | 116906342 | 116904176 | 116904383 | 116910232 | 116910311 | 96 | 49 | 37 | 0 | 1 | 1 | 0 | 23 | 0 | 3 | 98 | 49 | 1 | 1 | 1 | 0.961 | 0.949 | NA | | 0.793 | 0.177 |
| SE | 1183 | IFRD1 | IFRD1 | chr7 | + | 112456913 | 112457038 | 112456001 | 112456086 | 112458860 | 112459018 | 274 | 84 | 26 | 3 | 0 | 0 | 6 | 52 | 0 | 0 | 98 | 49 | 1 | 1 | 0.979 | 1 | 1 | | 1 | 1 | -0.007 |
| SE | 1184 | IFRD1 | IFRD1 | chr7 | + | 112456913 | 112457038 | 112456001 | 112456086 | 112461865 | 112461916 | 216 | 63 | 15 | 0 | 2 | 0 | 6 | 38 | 0 | 0 | 98 | 49 | 1 | 1 | 1 | 0.94 | 1 | | 1 | 1 | -0.02 |
| SE | 1185 | IFRD1 | IFRD1 | chr7 | + | 112458860 | 112459018 | 112456001 | 112456086 | 112461865 | 112461916 | 61 | 44 | 24 | 0 | 2 | 0 | 0 | 28 | 0 | 0 | 98 | 49 | 1 | 1 | 1 | 0.917 | 1 | NA | | 1 | -0.028 |
| SE | 1186 | IFRD1 | IFRD1 | chr7 | + | 112458860 | 112459018 | 112456913 | 112457038 | 112461865 | 112461916 | 116 | 67 | 35 | 0 | 2 | 0 | 0 | 42 | 0 | 0 | 98 | 49 | 1 | 1 | 1 | 0.944 | 1 | NA | | 1 | -0.019 |
| SE | 1193 | PNPLA8 | PNPLA8 | chr7 | - | 108521475 | 108521521 | 108514435 | 108515574 | 108526028 | 108526318 | 17 | 15 | 0 | 1 | 0 | 0 | 0 | 3 | 0 | 0 | 95 | 49 | 1 | 1 | 0.898 | 1 | NA | NA | | 1 | -0.051 |
| SE | 1207 | NAMPT | NAMPT | chr7 | - | 106261587 | 106261707 | 106253016 | 106253151 | 106263391 | 106263617 | 12 | 6 | 0 | 0 | 1 | 0 | 0 | 4 | 0 | 0 | 98 | 49 | 1 | 1 | 1 | 0.75 | NA | NA | | 1 | -0.125 |
| SE | 1208 | NAMPT | NAMPT | chr7 | - | 106261587 | 106261707 | 106254363 | 106254504 | 106268463 | 106268600 | 3 | 3 | 0 | 1 | 6 | 0 | 0 | 0 | 0 | 1 | 98 | 49 | 1 | 1 | 0.6 | 0.2 | NA | NA | | 0 | 0.4 |
| SE | 1209 | NAMPT | NAMPT | chr7 | - | 106263391 | 106263617 | 106253016 | 106253151 | 106272529 | 106272658 | 0 | 1 | 0 | 0 | 0 | 1 | 0 | 1 | 0 | 0 | 98 | 49 | 1 | 1 | NA | 1 | 0 | NA | | 1 | -0.5 |
| SE | 1211 | NAMPT | NAMPT | chr7 | - | 106268463 | 106268600 | 106263391 | 106263617 | 106269153 | 106269312 | 678 | 328 | 111 | 0 | 1 | 0 | 9 | 124 | 0 | 0 | 98 | 49 | 1 | 1 | 1 | 0.994 | 1 | | 1 | 1 | -0.002 |
| SE | 1212 | NAMPT | NAMPT | chr7 | - | 106269153 | 106269312 | 106263391 | 106263617 | 106272529 | 106272658 | 0 | 1 | 0 | 0 | 0 | 0 | 0 | 0 | 0 | 1 | 98 | 49 | 1 | 1 | NA | 1 | NA | NA | | 0 | 1 |
| SE | 1214 | NAMPT | NAMPT | chr7 | - | 106277022 | 106277179 | 106272529 | 106272658 | 106284827 | 106285192 | 592 | 331 | 90 | 0 | 3 | 0 | 12 | 69 | 0 | 0 | 98 | 49 | 1 | 1 | 1 | 0.982 | 1 | | 1 | 1 | -0.006 |
| SE | 1216 | SRPK2 | SRPK2 | chr7 | - | 105133003 | 105133104 | 105132790 | 105132898 | 105142007 | 105142490 | 40 | 17 | 4 | 1 | 0 | 0 | 0 | 9 | 0 | 0 | 98 | 49 | 1 | 1 | 0.952 | 1 | 1 | NA | | 1 | -0.016 |
| SE | 1217 | SRPK2 | SRPK2 | chr7 | - | 105268805 | 105268869 | 105203627 | 105203785 | 105388647 | 105388702 | 4 | 4 | 1 | 10 | 2 | 5 | 0 | 0 | 0 | 1 | 98 | 49 | 1 | 1 | 0.167 | 0.5 | 0.091 | NA | | 0 | 0.253 |
| SE | 1223 | PMPCB | PMPCB | chr7 | + | 103307595 | 103307708 | 103304410 | 103304490 | 103308951 | 103309095 | 3 | 5 | 0 | 1 | 2 | 0 | 0 | 2 | 0 | 0 | 98 | 49 | 1 | 1 | 0.6 | 0.556 | NA | NA | | 1 | -0.422 |
| SE | 1236 | ORAI2 | ORAI2 | chr7 | + | 102436201 | 102436333 | 102433529 | 102433661 | 102438943 | 102439181 | 20 | 10 | 0 | 1 | 0 | 0 | 0 | 4 | 0 | 0 | 98 | 49 | 1 | 1 | 0.909 | 1 | NA | NA | | 1 | -0.045 |
| SE | 1237 | ORAI2 | ORAI2 | chr7 | + | 102436224 | 102436333 | 102433529 | 102433661 | 102438943 | 102439181 | 20 | 14 | 0 | 1 | 0 | 0 | 0 | 4 | 0 | 0 | 98 | 49 | 1 | 1 | 0.909 | 1 | NA | NA | | 1 | -0.045 |
| SE | 1238 | ORAI2 | ORAI2 | chr7 | + | 102438943 | 102439181 | 102436201 | 102436333 | 102446512 | 102456821 | 30 | 16 | 5 | 1 | 0 | 0 | 2 | 6 | 0 | 0 | 98 | 49 | 1 | 1 | 0.937 | 1 | 1 | | 1 | 1 | -0.021 |
| SE | 1247 | LRCH4 | LRCH4 | chr7 | - | 100577272 | 100577389 | 100576901 | 100577005 | 100577496 | 100577558 | 15 | 9 | 1 | 2 | 0 | 0 | 0 | 1 | 0 | 0 | 98 | 49 | 1 | 1 | 0.789 | 1 | 1 | NA | | 1 | -0.07 |
| SE | 1250 | STAG3L5P-PVRIG2P-PILRB | STAG3L5P-PVRIG2P-PILRB | chr7 | + | 100353372 | 100353483 | 100353209 | 100353270 | 100355142 | 100355240 | 4 | 1 | 0 | 0 | 2 | 0 | 1 | 1 | 0 | 0 | 98 | 49 | 1 | 1 | 1 | 0.2 | NA | | 1 | 1 | -0.4 |
| SE | 1251 | STAG3L5P-PVRIG2P-PILRB | STAG3L5P-PVRIG2P-PILRB | chr7 | + | 100353372 | 100354012 | 100353209 | 100353270 | 100355142 | 100355240 | 8 | 2 | 0 | 0 | 2 | 0 | 2 | 1 | 0 | 0 | 98 | 49 | 1 | 1 | 1 | 0.333 | NA | | 1 | 1 | -0.334 |
| SE | 1268 | TECPR1 | TECPR1 | chr7 | - | 98241069 | 98241244 | 98240850 | 98240951 | 98243466 | 98243592 | 10 | 3 | 0 | 0 | 0 | 0 | 0 | 2 | 0 | 1 | 98 | 49 | 1 | 1 | 1 | 1 | NA | NA | | 0.5 | 0.5 |
| SE | 1269 | TECPR1 | TECPR1 | chr7 | - | 98244884 | 98245067 | 98244570 | 98244693 | 98245921 | 98246165 | 11 | 12 | 0 | 2 | 0 | 0 | 0 | 1 | 0 | 0 | 98 | 49 | 1 | 1 | 0.733 | 1 | NA | NA | | 1 | -0.133 |
| SE | 1283 | SAMD9L | SAMD9L | chr7 | - | 93145908 | 93146040 | 93145384 | 93145532 | 93148193 | 93148388 | 8 | 1 | 3 | 1 | 1 | 0 | 0 | 1 | 0 | 0 | 98 | 49 | 1 | 1 | 0.8 | 0.333 | 1 | NA | | 1 | -0.289 |
| SE | 1292 | AKAP9 | AKAP9 | chr7 | + | 91992157 | 91992211 | 91980288 | 91980333 | 91992884 | 91993055 | 8 | 0 | 0 | 6 | 0 | 3 | 0 | 3 | 0 | 0 | 98 | 49 | 1 | 1 | 0.4 | NA | 0 | NA | | 1 | -0.8 |
| SE | 1293 | AKAP9 | AKAP9 | chr7 | + | 92000847 | 92003235 | 91995602 | 91995800 | 92012428 | 92012642 | 16 | 7 | 0 | 0 | 1 | 0 | 3 | 11 | 0 | 0 | 98 | 49 | 1 | 1 | 1 | 0.778 | NA | | 1 | 1 | -0.111 |
| SE | 1294 | AKAP9 | AKAP9 | chr7 | + | 92107292 | 92107422 | 92105677 | 92105763 | 92108493 | 92108633 | 3 | 2 | 3 | 1 | 0 | 0 | 1 | 1 | 0 | 0 | 98 | 49 | 1 | 1 | 0.6 | 1 | 1 | | 1 | 1 | -0.133 |
| SE | 1300 | STEAP4 | STEAP4 | chr7 | - | 88282640 | 88283168 | 88280914 | 88281079 | 88283813 | 88284271 | 130 | 54 | 39 | 0 | 2 | 0 | 2 | 17 | 0 | 0 | 98 | 49 | 1 | 1 | 1 | 0.931 | 1 | | 1 | 1 | -0.023 |
| SE | 1301 | STEAP4 | STEAP4 | chr7 | - | 88290915 | 88291014 | 88283813 | 88284271 | 88306791 | 88306913 | 12 | 7 | 2 | 5 | 8 | 0 | 0 | 4 | 0 | 0 | 98 | 49 | 1 | 1 | 0.545 | 0.304 | 1 | NA | | 1 | -0.384 |
| SE | 1305 | DMTF1 | DMTF1 | chr7 | + | 87164933 | 87165050 | 87163494 | 87163617 | 87166482 | 87166605 | 36 | 14 | 4 | 3 | 2 | 0 | 0 | 2 | 0 | 0 | 98 | 49 | 1 | 1 | 0.857 | 0.778 | 1 | NA | | 1 | -0.122 |
| SE | 1306 | DMTF1 | DMTF1 | chr7 | + | 87166482 | 87166605 | 87164933 | 87165050 | 87170994 | 87171089 | 42 | 16 | 0 | 3 | 0 | 0 | 0 | 1 | 0 | 0 | 98 | 49 | 1 | 1 | 0.875 | 1 | NA | NA | | 1 | -0.062 |
| SE | 1307 | DMTF1 | DMTF1 | chr7 | + | 87194683 | 87194828 | 87193724 | 87194102 | 87195030 | 87196332 | 13 | 3 | 0 | 4 | 1 | 0 | 0 | 1 | 0 | 2 | 98 | 49 | 1 | 1 | 0.619 | 0.6 | NA | NA | | 0.2 | 0.409 |
| SE | 1314 | PTPN12 | PTPN12 | chr7 | + | 77585542 | 77585581 | 77583554 | 77583650 | 77592184 | 77592256 | 38 | 19 | 22 | 0 | 0 | 0 | 1 | 9 | 0 | 1 | 88 | 49 | 1 | 1 | 1 | 1 | 1 | | 1 | 0.834 | 0.083 |
| SE | 1315 | GSAP | GSAP | chr7 | - | 77311840 | 77311940 | 77310750 | 77311449 | 77312100 | 77312202 | 6 | 14 | 0 | 0 | 0 | 0 | 1 | 2 | 0 | 1 | 98 | 49 | 1 | 1 | 1 | 1 | NA | | 1 | 0.5 | 0.25 |
| SE | 1316 | GSAP | GSAP | chr7 | - | 77349350 | 77349404 | 77330238 | 77330367 | 77352943 | 77353026 | 5 | 0 | 0 | 1 | 0 | 0 | 0 | 2 | 0 | 0 | 98 | 49 | 1 | 1 | 0.714 | NA | NA | NA | | 1 | -0.286 |
| SE | 1324 | DTX2 | DTX2 | chr7 | + | 76480420 | 76480777 | 76461654 | 76461836 | 76482507 | 76483147 | 4 | 1 | 0 | 1 | 0 | 0 | 0 | 6 | 0 | 0 | 98 | 49 | 1 | 1 | 0.667 | 1 | NA | NA | | 1 | -0.166 |
| SE | 1329 | TMEM120A | TMEM120A | chr7 | - | 75987537 | 75987602 | 75987359 | 75987428 | 75987717 | 75987810 | 2 | 0 | 1 | 2 | 1 | 0 | 0 | 1 | 0 | 0 | 98 | 49 | 1 | 1 | 0.333 | 0 | 1 | NA | | 1 | -0.556 |
| SE | 1330 | RHBDD2 | RHBDD2 | chr7 | + | 75881364 | 75881486 | 75878998 | 75879260 | 75881828 | 75882236 | 2 | 0 | 1 | 2 | 0 | 1 | 0 | 2 | 1 | 0 | 98 | 49 | 1 | 1 | 0.333 | NA | 0.333 | | 0 | 1 | -0.167 |
| SE | 1334 | GTF2I | GTF2I | chr7 | + | 74718878 | 74718941 | 74716893 | 74716950 | 74728785 | 74728896 | 1 | 1 | 0 | 3 | 0 | 0 | 0 | 1 | 0 | 1 | 98 | 49 | 1 | 1 | 0.143 | 1 | NA | NA | | 0.333 | 0.238 |
| SE | 1337 | LAT2 | LAT2 | chr7 | + | 74214821 | 74215010 | 74209756 | 74210088 | 74215946 | 74216069 | 22 | 12 | 5 | 0 | 2 | 0 | 1 | 2 | 0 | 0 | 98 | 49 | 1 | 1 | 1 | 0.75 | 1 | | 1 | 1 | -0.083 |
| SE | 1338 | LAT2 | LAT2 | chr7 | + | 74216824 | 74216864 | 74215946 | 74216069 | 74219743 | 74219787 | 42 | 29 | 18 | 1 | 0 | 0 | 0 | 13 | 0 | 0 | 89 | 49 | 1 | 1 | 0.959 | 1 | 1 | NA | | 1 | -0.014 |
| SE | 1339 | LAT2 | LAT2 | chr7 | + | 74221636 | 74221692 | 74220703 | 74220734 | 74223723 | 74223783 | 55 | 33 | 20 | 0 | 0 | 1 | 1 | 20 | 0 | 0 | 98 | 49 | 1 | 1 | 1 | 1 | 0.909 | | 1 | 1 | -0.03 |
| SE | 1340 | LAT2 | LAT2 | chr7 | + | 74224638 | 74224760 | 74224017 | 74224197 | 74228943 | 74229834 | 39 | 22 | 20 | 0 | 0 | 0 | 0 | 7 | 0 | 1 | 98 | 49 | 1 | 1 | 1 | 1 | 1 | NA | | 0.778 | 0.222 |
| SE | 1365 | RABGEF1 | RABGEF1 | chr7 | + | 66783674 | 66783841 | 66775226 | 66775393 | 66795510 | 66795592 | 9 | 3 | 5 | 0 | 2 | 1 | 0 | 9 | 0 | 0 | 98 | 49 | 1 | 1 | 1 | 0.429 | 0.714 | NA | | 1 | -0.286 |
| SE | 1366 | RABGEF1 | RABGEF1 | chr7 | + | 66795510 | 66795592 | 66775226 | 66775393 | 66797373 | 66797506 | 6 | 9 | 14 | 0 | 0 | 0 | 0 | 4 | 0 | 1 | 98 | 49 | 1 | 1 | 1 | 1 | 1 | NA | | 0.667 | 0.333 |
| SE | 1367 | TPST1 | TPST1 | chr7 | + | 66240324 | 66241270 | 66205271 | 66205522 | 66286510 | 66286709 | 19 | 5 | 4 | 0 | 0 | 1 | 0 | 7 | 0 | 0 | 98 | 49 | 1 | 1 | 1 | 1 | 0.667 | NA | | 1 | -0.111 |
| SE | 1368 | ASL | ASL | chr7 | + | 66086743 | 66086821 | 66086584 | 66086662 | 66087333 | 66087386 | 2 | 0 | 0 | 0 | 0 | 0 | 0 | 2 | 1 | 0 | 98 | 49 | 1 | 1 | 1 | NA | NA | | 0 | 1 | 0.5 |
| SE | 1385 | CHCHD2 | CHCHD2 | chr7 | - | 56104225 | 56104475 | 56102866 | 56103011 | 56106363 | 56106494 | 17 | 1 | 9 | 1 | 0 | 0 | 0 | 9 | 0 | 0 | 98 | 49 | 1 | 1 | 0.895 | 1 | 1 | NA | | 1 | -0.035 |
| SE | 1391 | GRB10 | GRB10 | chr7 | - | 50755886 | 50756056 | 50703820 | 50703908 | 50793223 | 50793462 | 0 | 0 | 2 | 1 | 0 | 0 | 0 | 1 | 0 | 0 | 98 | 49 | 1 | 1 | 0 | NA | 1 | NA | | 1 | -0.5 |
| SE | 1401 | IKZF1 | IKZF1 | chr7 | + | 50368005 | 50368065 | 50327637 | 50327757 | 50382539 | 50382707 | 10 | 0 | 0 | 12 | 1 | 3 | 0 | 1 | 0 | 2 | 98 | 49 | 1 | 1 | 0.294 | 0 | 0 | NA | | 0.2 | -0.102 |
| SE | 1402 | IKZF1 | IKZF1 | chr7 | + | 50376532 | 50376793 | 50327637 | 50327757 | 50382539 | 50382707 | 10 | 9 | 9 | 12 | 1 | 3 | 0 | 1 | 0 | 2 | 98 | 49 | 1 | 1 | 0.294 | 0.818 | 0.6 | NA | | 0.2 | 0.371 |
| SE | 1414 | IKZF1 | IKZF1 | chr7 | + | 50391728 | 50391833 | 50387344 | 50387470 | 50399917 | 50405100 | 7 | 2 | 7 | 1 | 0 | 0 | 0 | 6 | 0 | 0 | 98 | 49 | 1 | 1 | 0.778 | 1 | 1 | NA | | 1 | -0.074 |
| SE | 1417 | IKZF1 | IKZF1 | chr7 | + | 50391728 | 50391863 | 50387344 | 50387470 | 50399917 | 50405100 | 8 | 4 | 5 | 1 | 0 | 0 | 0 | 3 | 0 | 0 | 98 | 49 | 1 | 1 | 0.8 | 1 | 1 | NA | | 1 | -0.067 |
| SE | 1418 | NCBP1 | NCBP1 | chr9 | + | 97662041 | 97662144 | 97660945 | 97661068 | 97662953 | 97663047 | 6 | 3 | 0 | 0 | 0 | 0 | 1 | 1 | 0 | 2 | 98 | 49 | 1 | 1 | 1 | 1 | NA | | 1 | 0.2 | 0.4 |
| SE | 1419 | ABCA13 | ABCA13 | chr7 | + | 48372167 | 48372497 | 48367793 | 48367908 | 48374346 | 48374416 | 0 | 0 | 0 | 1 | 0 | 0 | 1 | 0 | 0 | 0 | 98 | 49 | 1 | 1 | NA | 0 | NA | | 1 NA | | -1 |
| SE | 1420 | UPP1 | UPP1 | chr7 | + | 48099669 | 48099787 | 48094762 | 48094827 | 48101823 | 48101982 | 2 | 3 | 1 | 1 | 0 | 0 | 0 | 1 | 0 | 0 | 98 | 49 | 1 | 1 | 0.5 | 1 | 1 | NA | | 1 | -0.167 |
| SE | 1421 | UPP1 | UPP1 | chr7 | + | 48101823 | 48101870 | 48094762 | 48094827 | 48106872 | 48107082 | 7 | 0 | 0 | 1 | 0 | 0 | 0 | 2 | 0 | 0 | 96 | 49 | 1 | 1 | 0.781 | NA | NA | NA | | 1 | -0.219 |
| SE | 1425 | LRRC6 | LRRC6 | chr8 | - | 132632739 | 132632963 | 132625271 | 132625454 | 132637934 | 132638107 | 7 | 2 | 0 | 0 | 1 | 0 | 0 | 1 | 0 | 1 | 98 | 49 | 1 | 1 | 1 | 0.5 | NA | NA | | 0.333 | 0.417 |
| SE | 1429 | FBXL13 | FBXL13 | chr7 | - | 102931880 | 102931933 | 102926273 | 102926374 | 102963532 | 102963665 | 42 | 16 | 3 | 2 | 0 | 0 | 0 | 8 | 0 | 0 | 98 | 49 | 1 | 1 | 0.913 | 1 | 1 | NA | | 1 | -0.029 |
| SE | 1430 | FBXL13 | FBXL13 | chr7 | - | 102963532 | 102963665 | 102931880 | 102931933 | 102968021 | 102968117 | 43 | 6 | 1 | 7 | 7 | 0 | 0 | 3 | 0 | 0 | 98 | 49 | 1 | 1 | 0.754 | 0.3 | 1 | NA | | 1 | -0.315 |
| SE | 1433 | TBRG4 | TBRG4 | chr7 | - | 45104098 | 45104256 | 45103332 | 45103443 | 45104537 | 45104709 | 3 | 0 | 0 | 0 | 0 | 0 | 0 | 0 | 0 | 1 | 98 | 49 | 1 | 1 | 1 | NA | NA | NA | | 0 | 1 |
| SE | 1435 | CCM2 | CCM2 | chr7 | + | 45038252 | 45038426 | 45000187 | 45000363 | 45063917 | 45064001 | 11 | 2 | 9 | 0 | 1 | 0 | 1 | 0 | 0 | 0 | 98 | 49 | 1 | 1 | 1 | 0.5 | 1 | | 1 NA | 1 | -0.167 |
| SE | 1436 | CCM2 | CCM2 | chr7 | + | 45038252 | 45038426 | 45027633 | 45027797 | 45063917 | 45064001 | 12 | 2 | 11 | 2 | 4 | 0 | 1 | 0 | 0 | 0 | 98 | 49 | 1 | 1 | 0.75 | 0.2 | 1 | | 1 NA | 1 | -0.35 |
| SE | 1437 | CCM2 | CCM2 | chr7 | + | 45068442 | 45068579 | 45064462 | 45064646 | 45069825 | 45069961 | 21 | 13 | 11 | 0 | 3 | 3 | 0 | 7 | 0 | 0 | 98 | 49 | 1 | 1 | 1 | 0.684 | 0.647 | NA | | 1 | -0.223 |
| SE | 1438 | CCM2 | CCM2 | chr7 | + | 45068442 | 45068579 | 45064462 | 45064646 | 45072725 | 45072783 | 9 | 8 | 6 | 0 | 0 | 2 | 0 | 4 | 0 | 0 | 98 | 49 | 1 | 1 | 1 | 1 | 0.6 | NA | | 1 | -0.133 |
| SE | 1439 | CCM2 | CCM2 | chr7 | + | 45069825 | 45069961 | 45064462 | 45064646 | 45072725 | 45072783 | 11 | 11 | 10 | 0 | 0 | 2 | 0 | 1 | 0 | 0 | 98 | 49 | 1 | 1 | 1 | 1 | 0.714 | NA | | 1 | -0.095 |
| SE | 1440 | CCM2 | CCM2 | chr7 | + | 45069825 | 45069961 | 45068442 | 45068579 | 45072725 | 45072783 | 24 | 13 | 12 | 1 | 0 | 0 | 0 | 4 | 0 | 0 | 98 | 49 | 1 | 1 | 0.923 | 1 | 1 | NA | | 1 | -0.026 |
| SE | 1441 | SNHG15 | SNHG15 | chr7 | - | 44986020 | 44986097 | 44984356 | 44984426 | 44986524 | 44986660 | 4 | 1 | 0 | 1 | 0 | 0 | 0 | 1 | 0 | 0 | 98 | 49 | 1 | 1 | 0.667 | 1 | NA | NA | | 1 | -0.166 |
| SE | 1445 | PPIA | PPIA | chr7 | + | 44798747 | 44798904 | 44796635 | 44796793 | 44799246 | 44799277 | 2 | 0 | 0 | 2 | 0 | 0 | 0 | 1 | 0 | 0 | 98 | 49 | 1 | 1 | 0.333 | NA | NA | NA | | 1 | -0.667 |
| SE | 1462 | VPS41 | VPS41 | chr7 | - | 38817816 | 38817882 | 38796744 | 38796864 | 38821202 | 38821265 | 22 | 6 | 2 | 1 | 0 | 0 | 0 | 4 | 0 | 0 | 98 | 49 | 1 | 1 | 0.917 | 1 | 1 | NA | | 1 | -0.028 |
| SE | 1470 | NT5C3A | NT5C3A | chr7 | - | 33024038 | 33024108 | 33022052 | 33022099 | 33026816 | 33026915 | 14 | 3 | 3 | 0 | 1 | 0 | 0 | 6 | 0 | 0 | 98 | 49 | 1 | 1 | 1 | 0.6 | 1 | NA | | 1 | -0.133 |
| SE | 1471 | NT5C3A | NT5C3A | chr7 | - | 33029628 | 33029714 | 33026816 | 33026915 | 33035933 | 33035988 | 2 | 0 | 0 | 12 | 3 | 0 | 0 | 0 | 0 | 1 | 98 | 49 | 1 | 1 | 0.077 | 0 | NA | NA | | 0 | 0.038 |
| SE | 1474 | KBTBD2 | KBTBD2 | chr7 | - | 32879434 | 32879942 | 32874991 | 32875157 | 32891535 | 32891856 | 28 | 11 | 2 | 2 | 2 | 0 | 0 | 6 | 0 | 0 | 98 | 49 | 1 | 1 | 0.875 | 0.733 | 1 | NA | | 1 | -0.131 |
| SE | 1479 | LOC401320 | LOC401320 | chr7 | - | 30552099 | 30552179 | 30548356 | 30550781 | 30561465 | 30562128 | 7 | 2 | 0 | 3 | 1 | 0 | 1 | 4 | 0 | 0 | 98 | 49 | 1 | 1 | 0.538 | 0.5 | NA | | 1 | 1 | -0.481 |
| SE | 1496 | CPVL | CPVL | chr7 | - | 29064060 | 29064234 | 29030576 | 29030759 | 29066022 | 29066121 | 29 | 0 | 0 | 1 | 0 | 0 | 0 | 9 | 0 | 0 | 98 | 49 | 1 | 1 | 0.935 | NA | NA | NA | | 1 | -0.065 |
| SE | 1500 | SKAP2 | SKAP2 | chr7 | - | 26684735 | 26684848 | 26670090 | 26670192 | 26690284 | 26690362 | 60 | 30 | 13 | 0 | 2 | 0 | 0 | 3 | 0 | 0 | 98 | 49 | 1 | 1 | 1 | 0.882 | 1 | NA | | 1 | -0.039 |
| SE | 1501 | HNRNPA2B1 | HNRNPA2B1 | chr7 | - | 26193574 | 26193694 | 26193250 | 26193373 | 26195846 | 26195909 | 30 | 7 | 1 | 5 | 1 | 0 | 0 | 1 | 0 | 0 | 98 | 49 | 1 | 1 | 0.75 | 0.778 | 1 | NA | | 1 | -0.157 |
| SE | 1509 | CCDC126 | CCDC126 | chr7 | + | 23597966 | 23598051 | 23597378 | 23597605 | 23611170 | 23611553 | 2 | 3 | 4 | 2 | 5 | 0 | 0 | 0 | 0 | 1 | 98 | 49 | 1 | 1 | 0.333 | 0.231 | 1 | NA | | 0 | 0.521 |
| SE | 1510 | FAM126A | FAM126A | chr7 | - | 22978268 | 22978448 | 22977340 | 22977421 | 22983943 | 22984045 | 9 | 5 | 6 | 1 | 0 | 0 | 0 | 2 | 0 | 0 | 98 | 49 | 1 | 1 | 0.818 | 1 | 1 | NA | | 1 | -0.061 |
| SE | 1514 | SNX13 | SNX13 | chr7 | - | 17845594 | 17845694 | 17839806 | 17840000 | 17868406 | 17868490 | 33 | 12 | 0 | 0 | 0 | 1 | 0 | 2 | 0 | 0 | 98 | 49 | 1 | 1 | 1 | 1 | 0 | NA | | 1 | -0.333 |
| SE | 1515 | SNX13 | SNX13 | chr7 | - | 17897333 | 17897446 | 17893331 | 17893434 | 17940283 | 17940508 | 13 | 2 | 3 | 1 | 0 | 0 | 0 | 2 | 0 | 0 | 98 | 49 | 1 | 1 | 0.867 | 1 | 1 | NA | | 1 | -0.044 |
| SE | 1521 | PHF14 | PHF14 | chr7 | + | 11013746 | 11013906 | 10990702 | 10990847 | 11022867 | 11022979 | 1 | 1 | 0 | 1 | 0 | 0 | 0 | 2 | 0 | 0 | 98 | 49 | 1 | 1 | 0.333 | 1 | NA | NA | | 1 | -0.334 |
| SE | 1524 | TDRD7 | TDRD7 | chr9 | + | 97470557 | 97470669 | 97464841 | 97465028 | 97472292 | 97472495 | 4 | 0 | 1 | 2 | 0 | 0 | 0 | 3 | 0 | 1 | 98 | 49 | 1 | 1 | 0.5 | NA | 1 | NA | | 0.6 | 0.15 |
| SE | 1525 | TDRD7 | TDRD7 | chr9 | + | 97475382 | 97475469 | 97473491 | 97473626 | 97478438 | 97478573 | 14 | 3 | 0 | 0 | 2 | 0 | 0 | 3 | 0 | 0 | 98 | 49 | 1 | 1 | 1 | 0.429 | NA | NA | | 1 | -0.285 |
| SE | 1527 | ZDHHC4 | ZDHHC4 | chr7 | + | 6580554 | 6580678 | 6578500 | 6578720 | 6581606 | 6581680 | 4 | 0 | 2 | 1 | 0 | 0 | 1 | 0 | 0 | 0 | 98 | 49 | 1 | 1 | 0.667 | NA | 1 | | 1 NA | | -0.166 |
| SE | 1532 | RNF216 | RNF216 | chr7 | - | 5721032 | 5721172 | 5716715 | 5716766 | 5725323 | 5725438 | 19 | 7 | 0 | 1 | 0 | 0 | 0 | 2 | 0 | 0 | 98 | 49 | 1 | 1 | 1 | 0.778 | NA | NA | | 1 | -0.111 |
| SE | 1544 | BRAT1 | BRAT1 | chr7 | - | 2544908 | 2545056 | 2543589 | 2543962 | 2547323 | 2547478 | 5 | 8 | 1 | 1 | 0 | 0 | 0 | 2 | 0 | 2 | 98 | 49 | 1 | 1 | 0.714 | 1 | 1 | NA | | 0.333 | 0.572 |
| SE | 1551 | CCDC22 | CCDC22 | chrX | + | 49249166 | 49249262 | 49248816 | 49248924 | 49249508 | 49249568 | 1 | 0 | 0 | 0 | 0 | 0 | 0 | 5 | 0 | 1 | 98 | 49 | 1 | 1 | 1 | NA | NA | NA | | 0.714 | 0.286 |
| SE | 1561 | ERMARD | ERMARD | chr6 | + | 169766637 | 169766667 | 169762428 | 169762531 | 169768102 | 169768171 | 5 | 1 | 1 | 0 | 0 | 0 | 0 | 1 | 0 | 1 | 79 | 49 | 1 | 1 | 1 | 1 | 1 | NA | | 0.383 | 0.617 |
| SE | 1578 | IGF2R | IGF2R | chr6 | + | 160027184 | 160027314 | 160024571 | 160024704 | 160029549 | 160029655 | 445 | 161 | 69 | 3 | 0 | 0 | 8 | 60 | 0 | 0 | 98 | 49 | 1 | 1 | 0.987 | 1 | 1 | | 1 | 1 | -0.004 |
| SE | 1580 | IGF2R | IGF2R | chr6 | + | 160034418 | 160034522 | 160032941 | 160033107 | 160040559 | 160040724 | 215 | 89 | 53 | 2 | 1 | 0 | 2 | 31 | 0 | 0 | 98 | 49 | 1 | 1 | 0.982 | 0.978 | 1 | | 1 | 1 | -0.013 |
| SE | 1583 | IGF2R | IGF2R | chr6 | + | 160060546 | 160060717 | 160058905 | 160059098 | 160061502 | 160061646 | 359 | 123 | 60 | 0 | 1 | 0 | 6 | 48 | 0 | 0 | 98 | 49 | 1 | 1 | 1 | 0.984 | 1 | | 1 | 1 | -0.005 |
| SE | 1584 | IGF2R | IGF2R | chr6 | + | 160061502 | 160061646 | 160060546 | 160060717 | 160062531 | 160062619 | 204 | 65 | 37 | 1 | 0 | 0 | 5 | 19 | 0 | 0 | 98 | 49 | 1 | 1 | 0.99 | 1 | 1 | | 1 | 1 | -0.003 |
| SE | 1586 | IGF2R | IGF2R | chr6 | + | 160063414 | 160063630 | 160062531 | 160062619 | 160064400 | 160064531 | 257 | 90 | 50 | 1 | 0 | 0 | 2 | 31 | 0 | 0 | 98 | 49 | 1 | 1 | 0.992 | 1 | 1 | | 1 | 1 | -0.003 |
| SE | 1588 | IGF2R | IGF2R | chr6 | + | 160072764 | 160072884 | 160071909 | 160072036 | 160073212 | 160073469 | 250 | 74 | 63 | 1 | 0 | 0 | 1 | 16 | 0 | 1 | 98 | 49 | 1 | 1 | 0.992 | 1 | 1 | | 1 | 0.889 | 0.053 |
| SE | 1590 | IGF2R | IGF2R | chr6 | + | 160084994 | 160085131 | 160083949 | 160084184 | 160089106 | 160089253 | 119 | 27 | 21 | 1 | 0 | 0 | 4 | 13 | 0 | 1 | 98 | 49 | 1 | 1 | 0.983 | 1 | 1 | | 1 | 0.867 | 0.061 |
| SE | 1591 | IGF2R | IGF2R | chr6 | + | 160088032 | 160088147 | 160084994 | 160085131 | 160089106 | 160089253 | 111 | 35 | 13 | 1 | 0 | 0 | 0 | 17 | 0 | 0 | 98 | 49 | 1 | 1 | 0.982 | 1 | 1 | NA | | 1 | -0.006 |
| SE | 1592 | TCP1 | TCP1 | chr6 | - | 159780934 | 159781110 | 159780442 | 159780566 | 159783940 | 159784067 | 13 | 7 | 3 | 3 | 0 | 0 | 0 | 1 | 0 | 0 | 98 | 49 | 1 | 1 | 0.684 | 1 | 1 | NA | | 1 | -0.105 |
| SE | 1594 | SOD2 | SOD2 | chr6 | - | 159682473 | 159682638 | 159679115 | 159679298 | 159684853 | 159685033 | 606 | 399 | 313 | 2 | 2 | 0 | 4 | 86 | 0 | 0 | 98 | 49 | 1 | 1 | 0.993 | 0.99 | 1 | | 1 | 1 | -0.006 |
| SE | 1595 | SOD2 | SOD2 | chr6 | - | 159684853 | 159685033 | 159681722 | 159682638 | 159688125 | 159688242 | 869 | 535 | 430 | 0 | 1 | 0 | 1 | 107 | 0 | 0 | 98 | 49 | 1 | 1 | 1 | 0.996 | 1 | | 1 | 1 | -0.001 |
| SE | 1602 | TAGAP | TAGAP | chr6 | - | 159043977 | 159044031 | 159043588 | 159043655 | 159044119 | 159044204 | 179 | 63 | 27 | 1 | 0 | 0 | 1 | 15 | 0 | 0 | 98 | 49 | 1 | 1 | 0.989 | 1 | 1 | | 1 | 1 | -0.004 |
| SE | 1606 | TMEM181 | TMEM181 | chr6 | + | 158583953 | 158584044 | 158580939 | 158580995 | 158585303 | 158585425 | 7 | 5 | 3 | 0 | 0 | 2 | 0 | 4 | 0 | 0 | 98 | 49 | 1 | 1 | 1 | 1 | 0.429 | NA | | 1 | -0.19 |
| SE | 1610 | ARID1B | ARID1B | chr6 | + | 156871613 | 156871652 | 156829226 | 156829421 | 156901375 | 156901525 | 0 | 1 | 0 | 12 | 0 | 0 | 0 | 0 | 1 | 1 | 88 | 49 | 1 | 1 | 0 | 1 | NA | | 0 | 0 | 0.5 |
| SE | 1611 | ARID1B | ARID1B | chr6 | + | 157110471 | 157110561 | 157084661 | 157084905 | 157133027 | 157133207 | 0 | 2 | 0 | 0 | 1 | 0 | 0 | 1 | 0 | 0 | 98 | 49 | 1 | 1 | NA | 0.5 | NA | NA | | 1 | -0.5 |
| SE | 1614 | ARMT1 | ARMT1 | chr6 | + | 151454547 | 151454653 | 151452257 | 151452586 | 151458327 | 151458572 | 2 | 1 | 0 | 0 | 3 | 0 | 0 | 2 | 0 | 0 | 98 | 49 | 1 | 1 | 1 | 0.143 | NA | NA | | 1 | -0.428 |
| SE | 1616 | CSGALNACT1 | CSGALNACT1 | chr8 | - | 19457529 | 19457816 | 19439829 | 19439931 | 19458425 | 19458642 | 1 | 0 | 0 | 1 | 1 | 0 | 0 | 3 | 1 | 5 | 98 | 49 | 1 | 1 | 0.333 | 0 | NA | | 0 | 0.231 | 0.051 |
| SE | 1617 | CSGALNACT1 | CSGALNACT1 | chr8 | - | 19585165 | 19585317 | 19505200 | 19506130 | 19601770 | 19601865 | 3 | 0 | 0 | 8 | 2 | 0 | 0 | 0 | 0 | 2 | 98 | 49 | 1 | 1 | 0.158 | 0 | NA | NA | | 0 | 0.079 |
| SE | 1618 | CSGALNACT1 | CSGALNACT1 | chr8 | - | 19591159 | 19591278 | 19505200 | 19506130 | 19601770 | 19601865 | 1 | 0 | 0 | 8 | 2 | 0 | 0 | 1 | 0 | 2 | 98 | 49 | 1 | 1 | 0.059 | 0 | NA | NA | | 0.2 | -0.171 |
| SE | 1623 | GLIPR2 | GLIPR2 | chr9 | + | 36148546 | 36148650 | 36147785 | 36147894 | 36150871 | 36150949 | 48 | 16 | 21 | 1 | 0 | 0 | 2 | 8 | 0 | 0 | 98 | 49 | 1 | 1 | 0.96 | 1 | 1 | | 1 | 1 | -0.013 |
| SE | 1629 | NUP43 | NUP43 | chr6 | - | 149743637 | 149743715 | 149742389 | 149742570 | 149745939 | 149746062 | 3 | 0 | 0 | 0 | 1 | 0 | 0 | 2 | 0 | 0 | 98 | 49 | 1 | 1 | 1 | 0 | NA | NA | | 1 | -0.5 |
| SE | 1632 | STXBP5 | STXBP5 | chr6 | + | 147314263 | 147314331 | 147313883 | 147314031 | 147314595 | 147314636 | 51 | 28 | 17 | 3 | 0 | 0 | 0 | 9 | 0 | 0 | 98 | 49 | 1 | 1 | 0.895 | 1 | 1 | NA | | 1 | -0.035 |
| SE | 1633 | STXBP5 | STXBP5 | chr6 | + | 147314595 | 147314636 | 147313883 | 147314031 | 147315514 | 147315735 | 26 | 3 | 1 | 0 | 1 | 0 | 0 | 4 | 0 | 0 | 90 | 49 | 1 | 1 | 1 | 0.62 | 1 | NA | | 1 | -0.127 |
| SE | 1634 | STXBP5 | STXBP5 | chr6 | + | 147339178 | 147339238 | 147334156 | 147334222 | 147353322 | 147353373 | 15 | 3 | 0 | 6 | 0 | 0 | 0 | 4 | 0 | 0 | 98 | 49 | 1 | 1 | 0.556 | 1 | NA | NA | | 1 | -0.222 |
| SE | 1635 | STXBP5 | STXBP5 | chr6 | + | 147339336 | 147339384 | 147339178 | 147339238 | 147353322 | 147353373 | 7 | 10 | 6 | 1 | 0 | 0 | 0 | 1 | 0 | 0 | 97 | 49 | 1 | 1 | 0.78 | 1 | 1 | NA | | 1 | -0.073 |
| SE | 1639 | UTRN | UTRN | chr6 | + | 144437564 | 144437746 | 144435934 | 144436138 | 144438744 | 144438895 | 57 | 20 | 10 | 3 | 0 | 0 | 0 | 19 | 0 | 1 | 98 | 49 | 1 | 1 | 0.905 | 1 | 1 | NA | | 0.905 | 0.063 |
| SE | 1640 | UTRN | UTRN | chr6 | + | 144827347 | 144827386 | 144820881 | 144821018 | 144827610 | 144827676 | 10 | 0 | 0 | 13 | 3 | 0 | 0 | 2 | 0 | 2 | 88 | 49 | 1 | 1 | 0.3 | 0 | NA | NA | | 0.358 | -0.208 |
| SE | 1641 | UTRN | UTRN | chr6 | + | 144839172 | 144839284 | 144836300 | 144836541 | 144840739 | 144840832 | 10 | 2 | 4 | 3 | 0 | 0 | 0 | 9 | 0 | 0 | 98 | 49 | 1 | 1 | 0.625 | 1 | 1 | NA | | 1 | -0.125 |
| SE | 1643 | PHACTR2 | PHACTR2 | chr6 | + | 143765260 | 143765798 | 143753753 | 143753912 | 143772257 | 143772457 | 10 | 0 | 0 | 0 | 0 | 0 | 1 | 4 | 0 | 1 | 98 | 49 | 1 | 1 | 1 | NA | NA | | 1 | 0.667 | 0.166 |
| SE | 1702 | TNFAIP3 | TNFAIP3 | chr6 | + | 137880070 | 137880252 | 137878431 | 137879351 | 137881034 | 137883314 | 26 | 16 | 1 | 1 | 0 | 0 | 0 | 12 | 0 | 0 | 98 | 49 | 1 | 1 | 0.929 | 1 | 1 | NA | | 1 | -0.024 |
| SE | 1705 | MAP3K5 | MAP3K5 | chr6 | - | 136601801 | 136601979 | 136601021 | 136601042 | 136605208 | 136605366 | 38 | 6 | 4 | 1 | 0 | 0 | 0 | 2 | 0 | 0 | 98 | 49 | 1 | 1 | 0.95 | 1 | 1 | NA | | 1 | -0.017 |
| SE | 1706 | MAP3K5 | MAP3K5 | chr6 | - | 136622847 | 136622981 | 136614158 | 136614286 | 136637324 | 136637406 | 53 | 13 | 17 | 2 | 2 | 0 | 0 | 8 | 0 | 0 | 98 | 49 | 1 | 1 | 0.93 | 0.765 | 1 | NA | | 1 | -0.102 |
| SE | 1707 | MAP3K5 | MAP3K5 | chr6 | - | 136650983 | 136651091 | 136642519 | 136642569 | 136656306 | 136656460 | 74 | 31 | 17 | 2 | 0 | 0 | 0 | 7 | 0 | 0 | 98 | 49 | 1 | 1 | 0.949 | 1 | 1 | NA | | 1 | -0.017 |

| | | | | | | | | | | | | | | | | | | | | | | | | | | | | | | | |
|---|---|---|---|---|---|---|---|---|---|---|---|---|---|---|---|---|---|---|---|---|---|---|---|---|---|---|---|---|---|---|---|
| SE | 1708 | MAP3K5 | MAP3K5 | chr6 | - | 136720449 | 136720589 | 136705109 | 136705133 | 136791709 | 136792518 | 43 | 16 | 20 | 1 | 0 | 0 | 1 | 12 | 0 | 1 | 98 | 49 | 1 | 1 | 0.956 | 1 | 1 | 1 | 0.857 | 0.057 |
| SE | 1715 | SGK1 | SGK1 | chr6 | - | 134171636 | 134171732 | 134171022 | 134171178 | 134172192 | 134172316 | 109 | 54 | 11 | 1 | 0 | 0 | 2 | 13 | 0 | 0 | 98 | 49 | 1 | 1 | 0.982 | 1 | 1 | 1 | 1 | -0.006 |
| SE | 1716 | SGK1 | SGK1 | chr6 | - | 134172192 | 134172316 | 134171636 | 134171732 | 134172661 | 134172774 | 100 | 69 | 18 | 0 | 1 | 1 | 0 | 15 | 0 | 0 | 98 | 49 | 1 | 1 | 1 | 0.972 | 0.9 | NA | 1 | -0.043 |
| SE | 1717 | SGK1 | SGK1 | chr6 | - | 134173022 | 134173154 | 134172661 | 134172774 | 134173273 | 134173357 | 177 | 96 | 37 | 0 | 1 | 0 | 3 | 25 | 0 | 0 | 98 | 49 | 1 | 1 | 1 | 0.98 | 1 | 1 | 1 | -0.007 |
| SE | 1720 | VNN2 | VNN2 | chr6 | - | 132751144 | 132751278 | 132749694 | 132749865 | 132755842 | 132756035 | 125 | 82 | 68 | 2 | 1 | 0 | 0 | 8 | 0 | 0 | 98 | 49 | 1 | 1 | 0.969 | 0.976 | 1 | NA | 1 | -0.018 |
| SE | 1721 | VNN2 | VNN2 | chr6 | - | 132751144 | 132751518 | 132749694 | 132749865 | 132755842 | 132756035 | 130 | 94 | 71 | 2 | 1 | 0 | 0 | 8 | 0 | 0 | 98 | 49 | 1 | 1 | 0.97 | 0.979 | 1 | NA | 1 | -0.017 |
| SE | 1723 | VNN2 | VNN2 | chr6 | - | 132752460 | 132752749 | 132751144 | 132751518 | 132755842 | 132756035 | 330 | 178 | 186 | 5 | 12 | 3 | 0 | 32 | 0 | 0 | 98 | 49 | 1 | 1 | 0.971 | 0.881 | 0.969 | NA | 1 | -0.06 |
| SE | 1724 | VNN2 | VNN2 | chr6 | - | 132752672 | 132752749 | 132751144 | 132751518 | 132753417 | 132753491 | 8 | 8 | 4 | 4 | 0 | 2 | 0 | 3 | 0 | 3 | 98 | 49 | 1 | 1 | 0.5 | 1 | 0.5 | NA | 0.333 | 0.334 |
| SE | 1725 | VNN2 | VNN2 | chr6 | - | 132752672 | 132752749 | 132751144 | 132751518 | 132755842 | 132756035 | 115 | 82 | 84 | 5 | 12 | 3 | 0 | 10 | 0 | 0 | 98 | 49 | 1 | 1 | 0.92 | 0.774 | 0.933 | NA | 1 | -0.124 |
| SE | 1726 | VNN2 | VNN2 | chr6 | - | 132753417 | 132753491 | 132751144 | 132751518 | 132755842 | 132756035 | 34 | 5 | 4 | 5 | 12 | 3 | 0 | 3 | 0 | 0 | 98 | 49 | 1 | 1 | 0.773 | 0.172 | 0.4 | NA | 1 | -0.552 |
| SE | 1727 | VNN2 | VNN2 | chr6 | - | 132753417 | 132753491 | 132752672 | 132752749 | 132755842 | 132756035 | 37 | 12 | 6 | 114 | 81 | 84 | 0 | 3 | 0 | 10 | 98 | 49 | 1 | 1 | 0.14 | 0.069 | 0.034 | NA | 0.13 | -0.049 |
| SE | 1729 | VNN2 | VNN2 | chr6 | - | 132755842 | 132756035 | 132751144 | 132751518 | 132757415 | 132757546 | 302 | 183 | 114 | 1 | 0 | 0 | 4 | 21 | 0 | 0 | 98 | 49 | 1 | 1 | 0.993 | 1 | 1 | 1 | 1 | -0.002 |
| SE | 1730 | VNN3 | VNN3 | chr6 | - | 132724605 | 132724979 | 132723733 | 132723907 | 132734252 | 132734383 | 38 | 17 | 5 | 1 | 0 | 0 | 0 | 5 | 0 | 0 | 98 | 49 | 1 | 1 | 0.95 | 1 | 1 | NA | 1 | -0.017 |
| SE | 1732 | VNN3 | VNN3 | chr6 | - | 132726721 | 132727012 | 132724605 | 132724979 | 132734252 | 132734383 | 39 | 26 | 1 | 7 | 6 | 0 | 0 | 9 | 0 | 0 | 98 | 49 | 1 | 1 | 0.736 | 0.684 | 1 | NA | 1 | -0.193 |
| SE | 1734 | VNN3 | VNN3 | chr6 | - | 132728740 | 132728933 | 132726721 | 132727012 | 132734252 | 132734383 | 70 | 34 | 7 | 1 | 0 | 0 | 0 | 6 | 0 | 0 | 98 | 49 | 1 | 1 | 0.972 | 1 | 1 | NA | 1 | -0.009 |
| SE | 1736 | VNN3 | VNN3 | chr6 | - | 132728740 | 132728979 | 132726721 | 132727012 | 132734252 | 132734383 | 63 | 27 | 9 | 1 | 0 | 0 | 0 | 3 | 0 | 0 | 98 | 49 | 1 | 1 | 0.969 | 1 | 1 | NA | 1 | -0.01 |
| SE | 1756 | ECHDC1 | ECHDC1 | chr6 | - | 127316449 | 127316502 | 127314815 | 127314896 | 127327001 | 127327144 | 8 | 5 | 0 | 1 | 0 | 0 | 0 | 1 | 0 | 0 | 98 | 49 | 1 | 1 | 0.8 | 1 | NA | NA | 1 | -0.1 |
| SE | 1761 | RNF146 | RNF146 | chr6 | + | 127286049 | 127286177 | 127280230 | 127280340 | 127286615 | 127288560 | 1 | 0 | 1 | 0 | 0 | 6 | 3 | 1 | 0 | 0 | 98 | 49 | 1 | 1 | 1 | NA | 0.077 | 1 | 1 | -0.462 |
| SE | 1762 | HDDC2 | HDDC2 | chr6 | - | 125298713 | 125298816 | 125292840 | 125292909 | 125300537 | 125300659 | 2 | 0 | 0 | 1 | 0 | 0 | 0 | 1 | 0 | 0 | 98 | 49 | 1 | 1 | 0.5 | NA | NA | NA | 1 | -0.5 |
| SE | 1773 | MAN1A1 | MAN1A1 | chr6 | - | 119204758 | 119204882 | 119201253 | 119201347 | 119290682 | 119290763 | 17 | 8 | 2 | 0 | 0 | 0 | 0 | 0 | 0 | 2 | 98 | 49 | 1 | 1 | 1 | 1 | 1 | NA | 0 | 1 |
| SE | 1774 | MAN1A1 | MAN1A1 | chr6 | - | 119248259 | 119248354 | 119204758 | 119204882 | 119290682 | 119290763 | 20 | 6 | 2 | 0 | 1 | 0 | 0 | 11 | 0 | 0 | 98 | 49 | 1 | 1 | 1 | 0.75 | 1 | NA | 1 | -0.083 |
| SE | 1777 | GOPC | GOPC | chr6 | - | 117570859 | 117570955 | 117569571 | 117569736 | 117573466 | 117573632 | 5 | 1 | 0 | 1 | 0 | 0 | 0 | 1 | 0 | 0 | 98 | 49 | 1 | 1 | 0.714 | 1 | NA | NA | 1 | -0.143 |
| SE | 1780 | KPNA5 | KPNA5 | chr6 | + | 116692292 | 116692392 | 116692054 | 116692156 | 116698703 | 116698798 | 5 | 6 | 0 | 3 | 0 | 0 | 0 | 2 | 0 | 0 | 98 | 49 | 1 | 1 | 0.455 | 1 | NA | NA | 1 | -0.272 |
| SE | 1786 | HDAC2 | HDAC2 | chr6 | - | 113953276 | 113953418 | 113949167 | 113949260 | 113956012 | 113956151 | 2 | 2 | 0 | 1 | 0 | 0 | 0 | 2 | 0 | 0 | 98 | 49 | 1 | 1 | 0.5 | 1 | NA | NA | 1 | -0.25 |
| SE | 1791 | FGFR1OP | FGFR1OP | chr6 | + | 167026546 | 167026615 | 167024781 | 167024894 | 167032602 | 167032642 | 14 | 4 | 0 | 0 | 1 | 0 | 0 | 1 | 0 | 0 | 98 | 49 | 1 | 1 | 1 | 0.667 | NA | NA | 1 | -0.166 |
| SE | 1794 | AMD1 | AMD1 | chr6 | + | 110887504 | 110887591 | 110874783 | 110875215 | 110888856 | 110888983 | 55 | 14 | 14 | 0 | 1 | 0 | 0 | 4 | 0 | 0 | 98 | 49 | 1 | 1 | 1 | 0.875 | 1 | NA | 1 | -0.042 |
| SE | 1802 | CD164 | CD164 | chr6 | - | 109370410 | 109370467 | 109366513 | 109369017 | 109376073 | 109376112 | 45 | 20 | 3 | 3 | 2 | 0 | 0 | 18 | 0 | 1 | 98 | 49 | 1 | 1 | 0.882 | 0.833 | 1 | NA | 0.9 | 0.005 |
| SE | 1808 | SNX3 | SNX3 | chr6 | - | 108214497 | 108214622 | 108211216 | 108212254 | 108222949 | 108223045 | 21 | 20 | 18 | 1 | 0 | 1 | 0 | 5 | 0 | 0 | 98 | 49 | 1 | 1 | 0.913 | 1 | 0.9 | NA | 1 | -0.062 |
| SE | 1810 | SEC63 | SEC63 | chr6 | - | 107897648 | 107897731 | 107893837 | 107893897 | 107901369 | 107901517 | 6 | 7 | 1 | 1 | 0 | 0 | 0 | 2 | 0 | 0 | 98 | 49 | 1 | 1 | 0.75 | 1 | 1 | NA | 1 | -0.083 |
| SE | 1818 | ASCC3 | ASCC3 | chr6 | - | 100509931 | 100510107 | 100508194 | 100509533 | 100512708 | 100512918 | 1 | 0 | 0 | 0 | 0 | 0 | 0 | 0 | 0 | 1 | 98 | 49 | 1 | 1 | 1 | NA | NA | NA | 0 | 1 |
| SE | 1825 | MAP3K7 | MAP3K7 | chr6 | - | 90544551 | 90544632 | 90536336 | 90536401 | 90547257 | 90547387 | 1 | 0 | 0 | 8 | 4 | 0 | 1 | 0 | 0 | 0 | 98 | 49 | 1 | 1 | 0.059 | 0 | NA | 1 | NA | -0.971 |
| SE | 1826 | UBE2J1 | UBE2J1 | chr6 | - | 89333085 | 89333205 | 89326624 | 89329957 | 89335301 | 89335431 | 58 | 17 | 8 | 1 | 0 | 0 | 2 | 6 | 0 | 0 | 98 | 49 | 1 | 1 | 0.967 | 1 | 1 | 1 | 1 | -0.011 |
| SE | 1828 | PM20D2 | PM20D2 | chr6 | + | 89154747 | 89154902 | 89153042 | 89153185 | 89158324 | 89158460 | 1 | 2 | 0 | 1 | 0 | 0 | 0 | 1 | 0 | 0 | 98 | 49 | 1 | 1 | 0.333 | 1 | NA | NA | 1 | -0.334 |
| SE | 1832 | RARS2 | RARS2 | chr6 | - | 87548590 | 87548646 | 87545615 | 87545699 | 87555407 | 87555505 | 4 | 10 | 2 | 1 | 0 | 0 | 0 | 2 | 0 | 0 | 98 | 49 | 1 | 1 | 0.667 | 1 | 1 | NA | 1 | -0.111 |
| SE | 1836 | SYNCRIP | SYNCRIP | chr6 | - | 85641291 | 85641451 | 85640445 | 85640564 | 85642796 | 85642890 | 3 | 2 | 0 | 2 | 0 | 0 | 0 | 1 | 0 | 0 | 98 | 49 | 1 | 1 | 0.429 | 1 | NA | NA | 1 | -0.285 |
| SE | 1837 | SNX14 | SNX14 | chr6 | - | 85528261 | 85528362 | 85526125 | 85526237 | 85530191 | 85530275 | 21 | 17 | 6 | 1 | 0 | 0 | 0 | 8 | 0 | 0 | 98 | 49 | 1 | 1 | 0.913 | 1 | 1 | NA | 1 | -0.029 |
| SE | 1838 | SNX14 | SNX14 | chr6 | - | 85538837 | 85538864 | 85536791 | 85536924 | 85541984 | 85542043 | 2 | 0 | 0 | 6 | 4 | 1 | 0 | 0 | 0 | 2 | 76 | 49 | 1 | 1 | 0.177 | 0 | 0 | NA | 0 | 0.059 |
| SE | 1839 | SNX14 | SNX14 | chr6 | - | 85557975 | 85558060 | 85549722 | 85549879 | 85565331 | 85565419 | 13 | 6 | 0 | 4 | 0 | 0 | 0 | 4 | 0 | 0 | 98 | 49 | 1 | 1 | 0.619 | 1 | NA | NA | 1 | -0.191 |
| SE | 1840 | SNX14 | SNX14 | chr6 | - | 85565331 | 85565419 | 85557975 | 85558060 | 85567533 | 85567577 | 16 | 5 | 3 | 2 | 0 | 0 | 0 | 5 | 0 | 0 | 98 | 49 | 1 | 1 | 0.8 | 1 | 1 | NA | 1 | -0.067 |
| SE | 1841 | SNX14 | SNX14 | chr6 | - | 85565331 | 85565419 | 85557975 | 85558060 | 85572136 | 85572215 | 13 | 6 | 0 | 2 | 0 | 0 | 0 | 2 | 0 | 0 | 98 | 49 | 1 | 1 | 0.765 | 1 | NA | NA | 1 | -0.117 |
| SE | 1843 | SNX14 | SNX14 | chr6 | - | 85567533 | 85567577 | 85565331 | 85565419 | 85572136 | 85572215 | 8 | 1 | 3 | 2 | 2 | 0 | 0 | 2 | 0 | 0 | 93 | 49 | 1 | 1 | 0.678 | 0.209 | 1 | NA | 1 | -0.371 |
| SE | 1851 | SYTL3 | SYTL3 | chr6 | + | 158663204 | 158663378 | 158662749 | 158662812 | 158665394 | 158665613 | 9 | 2 | 1 | 1 | 1 | 0 | 1 | 1 | 0 | 1 | 98 | 49 | 1 | 1 | 0.818 | 0.5 | 1 | 1 | 0.333 | 0.106 |
| SE | 1852 | HMGN3 | HMGN3 | chr6 | - | 79202062 | 79202155 | 79201244 | 79201726 | 79202275 | 79202389 | 1 | 0 | 0 | 0 | 0 | 0 | 0 | 0 | 0 | 1 | 98 | 49 | 1 | 1 | 1 | NA | NA | NA | 0 | 1 |
| SE | 1853 | PHIP | PHIP | chr6 | - | 78947622 | 78947775 | 78946710 | 78946874 | 78954813 | 78954963 | 28 | 16 | 0 | 1 | 1 | 0 | 0 | 6 | 0 | 0 | 98 | 49 | 1 | 1 | 0.933 | 0.889 | NA | NA | 1 | -0.089 |
| SE | 1854 | PHIP | PHIP | chr6 | - | 78965944 | 78966056 | 78965702 | 78965764 | 78969834 | 78969917 | 33 | 14 | 8 | 2 | 0 | 0 | 0 | 8 | 0 | 0 | 98 | 49 | 1 | 1 | 0.892 | 1 | 1 | NA | 1 | -0.036 |
| SE | 1855 | PHIP | PHIP | chr6 | - | 78970048 | 78970173 | 78969834 | 78969917 | 78970780 | 78970888 | 30 | 8 | 3 | 1 | 0 | 0 | 0 | 4 | 0 | 0 | 98 | 49 | 1 | 1 | 0.937 | 1 | 1 | NA | 1 | -0.021 |
| SE | 1856 | PHIP | PHIP | chr6 | - | 79042842 | 79043003 | 79025942 | 79026164 | 79060477 | 79060576 | 47 | 25 | 9 | 0 | 2 | 0 | 6 | 12 | 0 | 0 | 98 | 49 | 1 | 1 | 1 | 0.862 | 1 | 1 | 1 | -0.046 |
| SE | 1859 | SENP6 | SENP6 | chr6 | + | 75640683 | 75640704 | 75634706 | 75634811 | 75647730 | 75647801 | 6 | 4 | 0 | 2 | 0 | 0 | 0 | 1 | 0 | 1 | 70 | 49 | 1 | 1 | 0.677 | 1 | NA | NA | 0.412 | 0.427 |
| SE | 1860 | SENP6 | SENP6 | chr6 | + | 75647730 | 75647801 | 75634706 | 75634811 | 75659261 | 75659407 | 16 | 12 | 10 | 2 | 0 | 0 | 0 | 10 | 0 | 0 | 98 | 49 | 1 | 1 | 0.8 | 1 | 1 | NA | 1 | -0.067 |
| SE | 1861 | SENP6 | SENP6 | chr6 | + | 75702644 | 75703072 | 75697424 | 75697517 | 75709526 | 75709630 | 25 | 1 | 4 | 2 | 1 | 0 | 0 | 7 | 0 | 0 | 98 | 49 | 1 | 1 | 0.862 | 0.333 | 1 | NA | 1 | -0.268 |
| SE | 1865 | SLC17A5 | SLC17A5 | chr6 | - | 73638411 | 73638499 | 73636620 | 73636707 | 73641690 | 73641924 | 6 | 1 | 2 | 0 | 0 | 0 | 0 | 0 | 0 | 1 | 98 | 49 | 1 | 1 | 1 | 1 | 1 | NA | 0 | 1 |
| SE | 1869 | SMAP1 | SMAP1 | chr6 | + | 70791688 | 70791769 | 70773349 | 70773425 | 70798656 | 70798737 | 1 | 0 | 2 | 3 | 2 | 0 | 0 | 1 | 0 | 0 | 98 | 49 | 1 | 1 | 0.143 | 0 | 1 | NA | 1 | -0.619 |
| SE | 1875 | ZNF451 | ZNF451 | chr6 | + | 57090810 | 57090894 | 57090009 | 57090274 | 57099060 | 57099141 | 4 | 1 | 0 | 0 | 0 | 0 | 0 | 0 | 0 | 3 | 98 | 49 | 1 | 1 | 1 | 1 | NA | NA | 0 | 1 |
| SE | 1881 | EEF1D | EEF1D | chr8 | - | 143590607 | 143590738 | 143588990 | 143590081 | 143592646 | 143592660 | 2 | 0 | 0 | 1 | 2 | 0 | 2 | 0 | 0 | 0 | 98 | 49 | 1 | 1 | 0.5 | 0 | NA | 1 | NA | -0.75 |
| SE | 1900 | CDC5L | CDC5L | chr6 | + | 44426481 | 44426724 | 44426102 | 44426183 | 44429712 | 44429910 | 14 | 2 | 6 | 0 | 2 | 0 | 0 | 1 | 0 | 0 | 98 | 49 | 1 | 1 | 1 | 0.333 | 1 | NA | 1 | -0.222 |
| SE | 1916 | VEGFA | VEGFA | chr6 | + | 43781955 | 43782087 | 43778888 | 43778918 | 43784540 | 43786486 | 0 | 0 | 0 | 1 | 0 | 0 | 0 | 2 | 0 | 0 | 98 | 49 | 1 | 1 | 0 | NA | NA | NA | 1 | -1 |
| SE | 1921 | XPO5 | XPO5 | chr6 | - | 43533906 | 43534007 | 43531478 | 43531575 | 43546570 | 43546752 | 3 | 0 | 1 | 0 | 0 | 0 | 0 | 0 | 0 | 1 | 98 | 49 | 1 | 1 | 1 | NA | 1 | NA | 0 | 1 |
| SE | 1922 | YIPF3 | YIPF3 | chr6 | - | 43515594 | 43515701 | 43513587 | 43513633 | 43515888 | 43516095 | 33 | 23 | 14 | 0 | 0 | 0 | 0 | 5 | 0 | 1 | 98 | 49 | 1 | 1 | 1 | 1 | 1 | NA | 0.714 | 0.286 |
| SE | 1924 | TJAP1 | TJAP1 | chr6 | + | 43478658 | 43478718 | 43478086 | 43478232 | 43497880 | 43497977 | 1 | 0 | 0 | 1 | 0 | 0 | 0 | 1 | 0 | 0 | 98 | 49 | 1 | 1 | 0.333 | NA | NA | NA | 1 | -0.667 |
| SE | 1934 | KLHDC3 | KLHDC3 | chr6 | + | 43017518 | 43017695 | 43017133 | 43017346 | 43017852 | 43017968 | 13 | 8 | 4 | 2 | 0 | 0 | 0 | 4 | 0 | 0 | 98 | 49 | 1 | 1 | 0.765 | 1 | 1 | NA | 1 | -0.078 |
| SE | 1936 | KLHDC3 | KLHDC3 | chr6 | + | 43018144 | 43018216 | 43017852 | 43017968 | 43018342 | 43018556 | 3 | 6 | 6 | 1 | 0 | 0 | 0 | 2 | 0 | 0 | 98 | 49 | 1 | 1 | 0.6 | 1 | 1 | NA | 1 | -0.133 |
| SE | 1937 | PPP2R5D | PPP2R5D | chr6 | + | 43006910 | 43007110 | 43006445 | 43006679 | 43007195 | 43007306 | 4 | 4 | 0 | 0 | 1 | 0 | 0 | 1 | 0 | 0 | 98 | 49 | 1 | 1 | 1 | 0.667 | NA | NA | 1 | -0.166 |
| SE | 1940 | UBR2 | UBR2 | chr6 | + | 42605720 | 42605859 | 42603587 | 42603718 | 42606588 | 42606651 | 232 | 113 | 25 | 1 | 0 | 0 | 0 | 23 | 0 | 1 | 98 | 49 | 1 | 1 | 0.991 | 1 | 1 | NA | 0.92 | 0.077 |
| SE | 1941 | UBR2 | UBR2 | chr6 | + | 42665408 | 42665512 | 42663257 | 42663419 | 42666166 | 42666245 | 92 | 43 | 15 | 0 | 0 | 0 | 0 | 12 | 0 | 1 | 98 | 49 | 1 | 1 | 1 | 1 | 1 | NA | 0.857 | 0.143 |
| SE | 1942 | UBR2 | UBR2 | chr6 | + | 42673790 | 42673887 | 42670659 | 42670715 | 42674125 | 42674193 | 149 | 52 | 20 | 1 | 0 | 0 | 1 | 13 | 0 | 0 | 98 | 49 | 1 | 1 | 0.987 | 1 | 1 | 1 | 1 | -0.004 |
| SE | 1943 | UBR2 | UBR2 | chr6 | + | 42676055 | 42676191 | 42674125 | 42674193 | 42676782 | 42676873 | 67 | 40 | 8 | 1 | 0 | 0 | 0 | 20 | 0 | 1 | 98 | 49 | 1 | 1 | 0.971 | 1 | 1 | NA | 0.909 | 0.081 |
| SE | 1944 | UBR2 | UBR2 | chr6 | + | 42683054 | 42683111 | 42679723 | 42679832 | 42684793 | 42684871 | 88 | 57 | 29 | 2 | 0 | 0 | 0 | 7 | 0 | 0 | 98 | 49 | 1 | 1 | 0.957 | 1 | 1 | NA | 1 | -0.014 |
| SE | 1946 | CCND3 | CCND3 | chr6 | - | 41940369 | 41940585 | 41937234 | 41937394 | 41941451 | 41941848 | 17 | 14 | 5 | 2 | 0 | 0 | 0 | 22 | 0 | 0 | 98 | 49 | 1 | 1 | 0.81 | 1 | 1 | NA | 1 | -0.063 |
| SE | 1948 | CCND3 | CCND3 | chr6 | - | 42030008 | 42030190 | 41937234 | 41937394 | 42048500 | 42048894 | 0 | 0 | 0 | 3 | 0 | 1 | 0 | 1 | 0 | 1 | 98 | 49 | 1 | 1 | 0 | NA | 0 | NA | 0.333 | -0.333 |
| SE | 1955 | TREML1 | TREML1 | chr6 | - | 41150260 | 41150313 | 41149260 | 41149918 | 41150818 | 41150907 | 1 | 0 | 0 | 6 | 0 | 0 | 0 | 0 | 3 | 0 | 98 | 49 | 1 | 1 | 0.077 | NA | NA | 0 | NA | 0.077 |
| SE | 1957 | ADCY10P1 | ADCY10P1 | chr6 | + | 41118923 | 41119107 | 41118128 | 41118184 | 41120522 | 41120657 | 0 | 0 | 0 | 1 | 0 | 0 | 0 | 1 | 0 | 0 | 98 | 49 | 1 | 1 | 0 | NA | NA | NA | 1 | -1 |
| SE | 1960 | ADCY10P1 | ADCY10P1 | chr6 | + | 41127544 | 41127608 | 41124484 | 41124820 | 41132293 | 41132513 | 5 | 0 | 1 | 2 | 0 | 0 | 0 | 1 | 0 | 0 | 98 | 49 | 1 | 1 | 0.556 | NA | 1 | NA | 1 | -0.222 |
| SE | 1961 | NFYA | NFYA | chr6 | + | 41080810 | 41080897 | 41079028 | 41079164 | 41084045 | 41084192 | 0 | 0 | 0 | 2 | 2 | 0 | 0 | 1 | 0 | 1 | 98 | 49 | 1 | 1 | 0 | 0 | NA | NA | 0.333 | -0.333 |
| SE | 1962 | C9orf91 | C9orf91 | chr9 | + | 114624349 | 114624459 | 114617117 | 114617301 | 114626898 | 114627006 | 4 | 0 | 0 | 0 | 0 | 0 | 0 | 1 | 0 | 2 | 98 | 49 | 1 | 1 | 1 | NA | NA | NA | 0.2 | 0.8 |
| SE | 1966 | BTBD9 | BTBD9 | chr6 | - | 38592575 | 38592840 | 38580217 | 38580437 | 38593963 | 38594327 | 3 | 4 | 0 | 0 | 0 | 0 | 0 | 4 | 0 | 1 | 98 | 49 | 1 | 1 | 1 | 1 | NA | NA | 0.667 | 0.333 |
| SE | 1969 | MTCH1 | MTCH1 | chr6 | - | 36977633 | 36977691 | 36977198 | 36977250 | 36978077 | 36978155 | 65 | 27 | 13 | 1 | 0 | 0 | 2 | 12 | 0 | 0 | 98 | 49 | 1 | 1 | 0.97 | 1 | 1 | 1 | 1 | -0.01 |
| SE | 1970 | MTCH1 | MTCH1 | chr6 | - | 36977633 | 36977784 | 36977198 | 36977250 | 36978077 | 36978155 | 47 | 22 | 9 | 1 | 0 | 0 | 2 | 9 | 0 | 0 | 98 | 49 | 1 | 1 | 0.959 | 1 | 1 | 1 | 1 | -0.014 |
| SE | 1971 | MTCH1 | MTCH1 | chr6 | - | 36981587 | 36981672 | 36978504 | 36978611 | 36985852 | 36986551 | 33 | 15 | 9 | 0 | 1 | 0 | 2 | 15 | 0 | 0 | 98 | 49 | 1 | 1 | 1 | 0.882 | 1 | 1 | 1 | -0.039 |
| SE | 1974 | STK38 | STK38 | chr6 | - | 36498362 | 36498486 | 36497779 | 36497875 | 36506582 | 36506644 | 39 | 8 | 2 | 0 | 1 | 0 | 0 | 3 | 0 | 0 | 98 | 49 | 1 | 1 | 1 | 0.8 | 1 | NA | 1 | -0.067 |
| SE | 1975 | STK38 | STK38 | chr6 | - | 36499872 | 36499990 | 36498362 | 36498486 | 36506582 | 36506644 | 18 | 2 | 3 | 1 | 0 | 0 | 0 | 3 | 0 | 0 | 98 | 49 | 1 | 1 | 0.9 | 1 | 1 | NA | 1 | -0.033 |
| SE | 1977 | KCTD20 | KCTD20 | chr6 | + | 36470051 | 36470257 | 36442766 | 36443111 | 36479120 | 36479223 | 6 | 0 | 0 | 0 | 1 | 0 | 0 | 5 | 0 | 0 | 98 | 49 | 1 | 1 | 1 | 0 | NA | NA | 1 | -0.5 |
| SE | 1979 | KCTD20 | KCTD20 | chr6 | + | 36474788 | 36475062 | 36470051 | 36470257 | 36479120 | 36479223 | 12 | 2 | 1 | 1 | 0 | 0 | 0 | 2 | 0 | 4 | 98 | 49 | 1 | 1 | 0.857 | 1 | 1 | NA | 0.2 | 0.752 |
| SE | 1981 | KCTD20 | KCTD20 | chr6 | + | 36479120 | 36479223 | 36470051 | 36470257 | 36479590 | 36479711 | 5 | 0 | 0 | 0 | 3 | 0 | 0 | 6 | 0 | 0 | 98 | 49 | 1 | 1 | 1 | 0 | NA | NA | 1 | -0.5 |
| SE | 1985 | BRPF3 | BRPF3 | chr6 | + | 36222167 | 36222265 | 36217916 | 36218010 | 36225266 | 36225364 | 5 | 4 | 0 | 1 | 0 | 0 | 0 | 1 | 0 | 0 | 98 | 49 | 1 | 1 | 0.714 | 1 | NA | NA | 1 | -0.143 |
| SE | 1986 | SRPK1 | SRPK1 | chr6 | - | 35869009 | 35869110 | 35857260 | 35857368 | 35869481 | 35869901 | 49 | 15 | 8 | 2 | 1 | 0 | 0 | 5 | 0 | 0 | 98 | 49 | 1 | 1 | 0.925 | 0.882 | 1 | NA | 1 | -0.064 |
| SE | 1988 | FKBP5 | FKBP5 | chr6 | - | 35587033 | 35587117 | 35580035 | 35580221 | 35591129 | 35591220 | 9 | 4 | 2 | 0 | 0 | 0 | 0 | 12 | 0 | 1 | 98 | 49 | 1 | 1 | 1 | 1 | 1 | NA | 0.857 | 0.143 |
| SE | 1990 | FKBP5 | FKBP5 | chr6 | - | 35620131 | 35620274 | 35619095 | 35619210 | 35637013 | 35637158 | 11 | 2 | 5 | 2 | 0 | 0 | 0 | 13 | 0 | 0 | 98 | 49 | 1 | 1 | 0.733 | 1 | 1 | NA | 1 | -0.089 |
| SE | 1991 | FKBP5 | FKBP5 | chr6 | - | 35642719 | 35642843 | 35637013 | 35637158 | 35688803 | 35688942 | 7 | 5 | 4 | 0 | 0 | 2 | 0 | 3 | 0 | 0 | 98 | 49 | 1 | 1 | 1 | 1 | 0.5 | NA | 1 | -0.167 |
| SE | 2000 | TAF11 | TAF11 | chr6 | - | 34879966 | 34880063 | 34877777 | 34878720 | 34880288 | 34880376 | 2 | 0 | 0 | 1 | 0 | 0 | 0 | 0 | 0 | 2 | 98 | 49 | 1 | 1 | 0.5 | NA | NA | NA | 0 | 0.5 |
| SE | 2001 | YKT6 | YKT6 | chr7 | + | 44211022 | 44211124 | 44208132 | 44208198 | 44212246 | 44214294 | 7 | 4 | 4 | 0 | 1 | 0 | 0 | 1 | 0 | 0 | 98 | 49 | 1 | 1 | 1 | 0.667 | 1 | NA | 1 | -0.111 |
| SE | 2025 | TAPBP | TAPBP | chr6 | - | 33313216 | 33313477 | 33304988 | 33305387 | 33313693 | 33313864 | 50 | 15 | 8 | 1 | 0 | 0 | 9 | 3 | 0 | 0 | 98 | 49 | 1 | 1 | 0.962 | 1 | 1 | 1 | 1 | -0.013 |
| SE | 2026 | TAPBP | TAPBP | chr6 | - | 33313693 | 33313864 | 33313216 | 33313477 | 33314004 | 33314387 | 27 | 9 | 4 | 0 | 0 | 0 | 4 | 1 | 0 | 1 | 98 | 49 | 1 | 1 | 1 | 1 | 1 | 1 | 0.333 | 0.334 |
| SE | 2033 | TAP1 | TAP1 | chr6 | - | 32847918 | 32848092 | 32847512 | 32847675 | 32848651 | 32848840 | 69 | 36 | 37 | 1 | 0 | 0 | 4 | 21 | 0 | 0 | 98 | 49 | 1 | 1 | 0.972 | 1 | 1 | 1 | 1 | -0.009 |
| SE | 2062 | HLA-DQB1 | HLA-DQB1 | chr6 | - | 32661346 | 32661457 | 32659463 | 32660249 | 32661966 | 32662248 | 9 | 0 | 0 | 0 | 0 | 0 | 0 | 4 | 0 | 2 | 98 | 49 | 1 | 1 | 1 | NA | NA | NA | 0.5 | 0.5 |
| SE | 2063 | HLA-DQB1 | HLA-DQB1 | chr6 | - | 32664797 | 32665067 | 32661966 | 32662248 | 32666498 | 32666689 | 28 | 2 | 0 | 3 | 0 | 0 | 0 | 9 | 0 | 0 | 98 | 49 | 1 | 1 | 0.824 | 1 | NA | NA | 1 | -0.088 |
| SE | 2089 | PBX2 | PBX2 | chr6 | - | 32186812 | 32186901 | 32186603 | 32186690 | 32187241 | 32187395 | 88 | 43 | 26 | 1 | 1 | 0 | 1 | 11 | 0 | 0 | 98 | 49 | 1 | 1 | 0.978 | 0.956 | 1 | 1 | 1 | -0.022 |
| SE | 2133 | C6orf48 | C6orf48 | chr6 | + | 31836423 | 31836517 | 31835410 | 31835451 | 31837235 | 31837435 | 1 | 2 | 0 | 7 | 4 | 0 | 0 | 0 | 0 | 1 | 98 | 49 | 1 | 1 | 0.067 | 0.2 | NA | NA | 0 | 0.134 |
| SE | 2135 | DDAH2 | DDAH2 | chr6 | - | 31728864 | 31729225 | 31728645 | 31728745 | 31730050 | 31730265 | 13 | 6 | 0 | 0 | 0 | 1 | 0 | 8 | 0 | 0 | 98 | 49 | 1 | 1 | 1 | 1 | 0 | NA | 1 | -0.333 |
| SE | 2169 | BAG6 | BAG6 | chr6 | - | 31639499 | 31639646 | 31639027 | 31639226 | 31640198 | 31640306 | 12 | 1 | 3 | 11 | 2 | 3 | 0 | 4 | 0 | 2 | 98 | 49 | 1 | 1 | 0.353 | 0.2 | 0.333 | NA | 0.5 | -0.205 |
| SE | 2170 | BAG6 | BAG6 | chr6 | - | 31640384 | 31640528 | 31640198 | 31640306 | 31640644 | 31640704 | 37 | 7 | 9 | 0 | 0 | 2 | 1 | 16 | 0 | 0 | 98 | 49 | 1 | 1 | 1 | 1 | 0.692 | 1 | 1 | -0.103 |
| SE | 2171 | BAG6 | BAG6 | chr6 | - | 31640644 | 31640704 | 31640198 | 31640306 | 31640791 | 31640938 | 17 | 2 | 2 | 0 | 1 | 0 | 0 | 1 | 0 | 0 | 98 | 49 | 1 | 1 | 1 | 0.5 | 1 | NA | 1 | -0.167 |
| SE | 2172 | BAG6 | BAG6 | chr6 | - | 31643889 | 31643974 | 31643100 | 31643115 | 31644081 | 31644194 | 11 | 1 | 3 | 0 | 0 | 0 | 1 | 2 | 0 | 1 | 98 | 49 | 1 | 1 | 1 | 1 | 1 | 1 | 0.5 | 0.25 |
| SE | 2173 | BAG6 | BAG6 | chr6 | - | 31643889 | 31643977 | 31642828 | 31643115 | 31644081 | 31644194 | 12 | 1 | 4 | 0 | 0 | 0 | 1 | 3 | 0 | 1 | 98 | 49 | 1 | 1 | 1 | 1 | 1 | 1 | 0.6 | 0.2 |
| SE | 2176 | PRRC2A | PRRC2A | chr6 | + | 31622689 | 31622901 | 31620672 | 31620858 | 31624260 | 31624360 | 2 | 1 | 0 | 0 | 0 | 0 | 0 | 1 | 2 | 0 | 98 | 49 | 1 | 1 | 1 | 1 | NA | 0 | 1 | 0.5 |
| SE | 2178 | PRRC2A | PRRC2A | chr6 | + | 31623731 | 31623909 | 31622689 | 31622901 | 31624260 | 31624360 | 34 | 10 | 6 | 0 | 1 | 0 | 1 | 6 | 0 | 0 | 98 | 49 | 1 | 1 | 1 | 0.833 | 1 | 1 | 1 | -0.056 |
| SE | 2181 | PRRC2A | PRRC2A | chr6 | + | 31633378 | 31633647 | 31631138 | 31632992 | 31633858 | 31633989 | 9 | 5 | 1 | 1 | 0 | 0 | 0 | 4 | 0 | 0 | 98 | 49 | 1 | 1 | 0.818 | 1 | 1 | NA | 1 | -0.061 |
| SE | 2183 | AIF1 | AIF1 | chr6 | + | 31615669 | 31615736 | 31615520 | 31615582 | 31616343 | 31616506 | 105 | 37 | 22 | 1 | 0 | 0 | 2 | 21 | 0 | 0 | 98 | 49 | 1 | 1 | 0.981 | 1 | 1 | 1 | 1 | -0.006 |
| SE | 2184 | AIF1 | AIF1 | chr6 | + | 31616103 | 31616145 | 31615669 | 31615736 | 31616343 | 31616506 | 111 | 101 | 88 | 1 | 0 | 1 | 0 | 44 | 0 | 0 | 91 | 49 | 1 | 1 | 0.984 | 1 | 0.979 | NA | 1 | -0.012 |
| SE | 2185 | TNF | TNF | chr6 | + | 31576533 | 31576579 | 31575566 | 31575927 | 31576766 | 31576814 | 5 | 1 | 0 | 0 | 0 | 0 | 0 | 2 | 0 | 1 | 95 | 49 | 1 | 1 | 1 | 1 | NA | NA | 0.508 | 0.492 |
| SE | 2186 | DDX39B | DDX39B | chr6 | - | 31538762 | 31538855 | 31536499 | 31536683 | 31540321 | 31540534 | 15 | 10 | 0 | 2 | 0 | 0 | 0 | 1 | 0 | 0 | 98 | 49 | 1 | 1 | 0.789 | 1 | NA | NA | 1 | -0.105 |
| SE | 2187 | DDX39B | DDX39B | chr6 | - | 31539146 | 31539274 | 31538762 | 31538855 | 31540321 | 31540534 | 60 | 21 | 10 | 1 | 0 | 0 | 2 | 20 | 0 | 0 | 98 | 49 | 1 | 1 | 0.968 | 1 | 1 | 1 | 1 | -0.011 |
| SE | 2212 | FLOT1 | FLOT1 | chr6 | - | 30730021 | 30730186 | 30727733 | 30728145 | 30730427 | 30730565 | 37 | 19 | 26 | 0 | 5 | 0 | 0 | 6 | 0 | 0 | 98 | 49 | 1 | 1 | 1 | 0.655 | 1 | NA | 1 | -0.115 |
| SE | 2213 | FLOT1 | FLOT1 | chr6 | - | 30741189 | 30741333 | 30740678 | 30740798 | 30741613 | 30741704 | 50 | 24 | 20 | 1 | 0 | 0 | 0 | 12 | 0 | 0 | 98 | 49 | 1 | 1 | 0.962 | 1 | 1 | NA | 1 | -0.013 |
| SE | 2214 | FLOT1 | FLOT1 | chr6 | - | 30741189 | 30741333 | 30740678 | 30740798 | 30742146 | 30742203 | 39 | 12 | 17 | 0 | 1 | 1 | 0 | 12 | 0 | 0 | 98 | 49 | 1 | 1 | 1 | 0.857 | 0.895 | NA | 1 | -0.083 |
| SE | 2215 | FLOT1 | FLOT1 | chr6 | - | 30741613 | 30741704 | 30741189 | 30741333 | 30741791 | 30741867 | 34 | 26 | 16 | 0 | 2 | 0 | 1 | 6 | 0 | 1 | 98 | 49 | 1 | 1 | 1 | 0.867 | 1 | 1 | 0.75 | 0.081 |
| SE | 2216 | FLOT1 | FLOT1 | chr6 | - | 30741791 | 30741867 | 30741189 | 30741333 | 30742146 | 30742203 | 24 | 7 | 4 | 0 | 0 | 0 | 0 | 2 | 0 | 2 | 98 | 49 | 1 | 1 | 1 | 1 | 1 | NA | 0.333 | 0.667 |
| SE | 2294 | HLA-E | HLA-E | chr6 | + | 30491536 | 30491653 | 30491136 | 30491412 | 30492403 | 30492436 | 747 | 497 | 484 | 1 | 1 | 0 | 33 | 293 | 0 | 1 | 98 | 49 | 1 | 1 | 0.997 | 0.996 | 1 | 1 | 0.993 | 0.001 |
| SE | 2295 | HLA-E | HLA-E | chr6 | + | 30492403 | 30492436 | 30491536 | 30491653 | 30492540 | 30492583 | 584 | 384 | 406 | 0 | 4 | 0 | 26 | 172 | 0 | 1 | 82 | 49 | 1 | 1 | 1 | 0.983 | 1 | 1 | 0.99 | -0.001 |
| SE | 2296 | HLA-E | HLA-E | chr6 | + | 30492540 | 30492583 | 30491536 | 30491653 | 30492748 | 30494205 | 221 | 145 | 117 | 0 | 0 | 0 | 3 | 55 | 1 | 0 | 92 | 49 | 1 | 1 | 1 | 1 | 1 | 0.615 | 1 | 0.193 |
| SE | 2307 | HLA-H | HLA-H | chr6 | + | 29889706 | 29889823 | 29889327 | 29889604 | 29890263 | 29890296 | 613 | 40 | 312 | 0 | 0 | 1 | 18 | 52 | 0 | 0 | 98 | 49 | 1 | 1 | 1 | 1 | 0.994 | 1 | 1 | -0.002 |
| SE | 2314 | HLA-F | HLA-F | chr6 | + | 29725030 | 29725306 | 29724172 | 29724448 | 29725446 | 29725563 | 316 | 96 | 64 | 2 | 0 | 2 | 4 | 52 | 0 | 0 | 98 | 49 | 1 | 1 | 0.987 | 1 | 0.941 | 1 | 1 | -0.024 |
| SE | 2315 | HLA-F | HLA-F | chr6 | + | 29725446 | 29725563 | 29725030 | 29725306 | 29726402 | 29726526 | 197 | 46 | 33 | 1 | 1 | 0 | 4 | 28 | 0 | 0 | 98 | 49 | 1 | 1 | 0.99 | 0.958 | 1 | 1 | 1 | -0.017 |
| SE | 2316 | HLA-F | HLA-F | chr6 | + | 29726010 | 29726043 | 29725446 | 29725563 | 29726402 | 29726526 | 148 | 28 | 38 | 0 | 0 | 1 | 4 | 18 | 0 | 0 | 82 | 49 | 1 | 1 | 1 | 1 | 0.958 | 1 | 1 | -0.014 |
| SE | 2317 | HLA-F | HLA-F | chr6 | + | 29726010 | 29726043 | 29725446 | 29725563 | 29726882 | 29727296 | 74 | 16 | 14 | 0 | 0 | 1 | 3 | 16 | 0 | 0 | 82 | 49 | 1 | 1 | 1 | 1 | 0.893 | 1 | 1 | -0.036 |
| SE | 2365 | BTN2A1 | BTN2A1 | chr6 | + | 26458606 | 26458718 | 26457903 | 26458142 | 26459480 | 26459828 | 5 | 0 | 2 | 3 | 0 | 0 | 0 | 1 | 0 | 1 | 98 | 49 | 1 | 1 | 0.455 | NA | 1 | NA | 0.333 | 0.395 |
| SE | 2376 | FAM65B | FAM65B | chr6 | - | 24832255 | 24832391 | 24830508 | 24830670 | 24839090 | 24839272 | 18 | 16 | 12 | 0 | 0 | 0 | 1 | 4 | 0 | 1 | 98 | 49 | 1 | 1 | 1 | 1 | 1 | 1 | 0.667 | 0.166 |
| SE | 2377 | FAM65B | FAM65B | chr6 | - | 24835702 | 24835871 | 24832255 | 24832391 | 24839090 | 24839272 | 113 | 65 | 53 | 1 | 0 | 0 | 0 | 36 | 0 | 0 | 98 | 49 | 1 | 1 | 0.983 | 1 | 1 | NA | 1 | -0.006 |
| SE | 2378 | FAM65B | FAM65B | chr6 | - | 24835702 | 24835871 | 24832255 | 24832391 | 24842861 | 24843554 | 61 | 28 | 18 | 1 | 1 | 1 | 0 | 21 | 0 | 0 | 98 | 49 | 1 | 1 | 0.968 | 0.933 | 0.9 | NA | 1 | -0.066 |
| SE | 2379 | FAM65B | FAM65B | chr6 | - | 24839090 | 24839272 | 24832255 | 24832391 | 24842861 | 24843554 | 81 | 33 | 16 | 1 | 1 | 1 | 0 | 4 | 0 | 0 | 98 | 49 | 1 | 1 | 0.976 | 0.943 | 0.889 | NA | 1 | -0.064 |
| SE | 2380 | FAM65B | FAM65B | chr6 | - | 24839090 | 24839272 | 24835702 | 24835871 | 24842861 | 24843554 | 131 | 71 | 51 | 0 | 1 | 0 | 0 | 19 | 0 | 0 | 98 | 49 | 1 | 1 | 1 | 0.973 | 1 | NA | 1 | -0.009 |
| SE | 2383 | FAM65B | FAM65B | chr6 | - | 24870865 | 24870889 | 24869093 | 24869147 | 24872880 | 24872959 | 224 | 146 | 94 | 2 | 0 | 0 | 4 | 57 | 0 | 0 | 73 | 49 | 1 | 1 | 0.987 | 1 | 1 | 1 | 1 | -0.004 |
| SE | 2384 | KIAA0319 | KIAA0319 | chr6 | - | 24554540 | 24554631 | 24551433 | 24551525 | 24559012 | 24559155 | 2 | 1 | 0 | 1 | 0 | 0 | 0 | 1 | 0 | 0 | 98 | 49 | 1 | 1 | 0.5 | 1 | NA | NA | 1 | -0.25 |
| SE | 2386 | KIAA0319 | KIAA0319 | chr6 | - | 24572574 | 24572659 | 24569902 | 24570035 | 24576367 | 24576596 | 0 | 1 | 0 | 1 | 0 | 0 | 0 | 2 | 0 | 0 | 98 | 49 | 1 | 1 | 0 | 1 | NA | NA | 1 | -0.5 |
| SE | 2387 | KIAA0319 | KIAA0319 | chr6 | - | 24572574 | 24572698 | 24569902 | 24570035 | 24576367 | 24576596 | 1 | 1 | 2 | 1 | 0 | 0 | 0 | 2 | 0 | 0 | 98 | 49 | 1 | 1 | 0.333 | 1 | 1 | NA | 1 | -0.222 |
| SE | 2393 | TMEM164 | TMEM164 | chrX | + | 110003500 | 110004164 | 110002634 | 110002772 | 110067346 | 110067396 | 35 | 6 | 0 | 3 | 0 | 0 | 2 | 5 | 0 | 0 | 98 | 49 | 1 | 1 | 0.854 | 1 | NA | 1 | 1 | -0.073 |
| SE | 2395 | RNF144B | RNF144B | chr6 | + | 18427580 | 18427685 | 18399498 | 18399699 | 18439683 | 18439744 | 18 | 9 | 13 | 0 | 4 | 3 | 0 | 10 | 0 | 0 | 98 | 49 | 1 | 1 | 1 | 0.529 | 0.684 | NA | 1 | -0.262 |
| SE | 2396 | RNF144B | RNF144B | chr6 | + | 18457154 | 18457359 | 18439683 | 18439744 | 18459606 | 18459751 | 15 | 9 | 10 | 2 | 0 | 0 | 0 | 9 | 0 | 3 | 98 | 49 | 1 | 1 | 0.789 | 1 | 1 | NA | 0.6 | 0.33 |
| SE | 2398 | KDM1B | KDM1B | chr6 | + | 18162834 | 18162924 | 18161326 | 18161454 | 18166266 | 18166378 | 22 | 10 | 4 | 0 | 0 | 1 | 2 | 3 | 0 | 1 | 98 | 49 | 1 | 1 | 1 | 1 | 0.667 | 1 | 0.6 | 0.089 |
| SE | 2399 | KDM1B | KDM1B | chr6 | + | 18185771 | 18185810 | 18171362 | 18171479 | 18197056 | 18197233 | 3 | 6 | 3 | 1 | 0 | 0 | 0 | 5 | 0 | 0 | 88 | 49 | 1 | 1 | 0.626 | 1 | 1 | NA | 1 | -0.125 |
| SE | 2400 | NUP153 | NUP153 | chr6 | - | 17632649 | 17632844 | 17628654 | 17629539 | 17637152 | 17637770 | 23 | 10 | 12 | 1 | 0 | 0 | 0 | 6 | 0 | 0 | 98 | 49 | 1 | 1 | 0.92 | 1 | 1 | NA | 1 | -0.027 |
| SE | 2402 | NUP153 | NUP153 | chr6 | - | 17651833 | 17651926 | 17649162 | 17649300 | 17661652 | 17661779 | 0 | 2 | 3 | 10 | 1 | 1 | 0 | 0 | 0 | 6 | 98 | 49 | 1 | 1 | 0 | 0.5 | 0.6 | NA | 0 | 0.367 |
| SE | 2403 | NUP153 | NUP153 | chr6 | - | 17661652 | 17661779 | 17649162 | 17649300 | 17662017 | 17662070 | 27 | 3 | 3 | 1 | 0 | 0 | 0 | 7 | 0 | 0 | 98 | 49 | 1 | 1 | 0.931 | 1 | 1 | NA | 1 | -0.023 |
| SE | 2404 | NUP153 | NUP153 | chr6 | - | 17668974 | 17669028 | 17665238 | 17665385 | 17669292 | 17669337 | 48 | 20 | 4 | 0 | 0 | 0 | 3 | 7 | 0 | 1 | 98 | 49 | 1 | 1 | 1 | 1 | 1 | 1 | 0.778 | 0.111 |
| SE | 2405 | ATXN1 | ATXN1 | chr6 | - | 16753232 | 16753347 | 16657775 | 16657901 | 16761297 | 16761490 | 12 | 5 | 2 | 1 | 0 | 0 | 0 | 1 | 0 | 0 | 98 | 49 | 1 | 1 | 0.857 | 1 | 1 | NA | 1 | -0.048 |
| SE | 2414 | RANBP9 | RANBP9 | chr6 | - | 13642478 | 13642591 | 13641198 | 13641307 | 13644544 | 13644729 | 22 | 9 | 4 | 0 | 2 | 0 | 0 | 5 | 0 | 0 | 98 | 49 | 1 | 1 | 1 | 0.692 | 1 | NA | 1 | -0.103 |
| SE | 2415 | SIRT5 | SIRT5 | chr6 | + | 13588330 | 13588464 | 13584075 | 13584225 | 13591668 | 13591894 | 4 | 0 | 0 | 2 | 0 | 0 | 0 | 2 | 0 | 0 | 98 | 49 | 1 | 1 | 0.5 | NA | NA | NA | 1 | -0.5 |
| SE | 2418 | TBC1D7 | TBC1D7 | chr6 | - | 13325093 | 13325174 | 13320907 | 13321095 | 13326786 | 13326906 | 4 | 0 | 0 | 1 | 5 | 0 | 0 | 1 | 0 | 0 | 98 | 49 | 1 | 1 | 0.667 | 0 | NA | NA | 1 | -0.666 |
| SE | 2419 | HIVEP1 | HIVEP1 | chr6 | + | 12129758 | 12129892 | 12119889 | 12125870 | 12130766 | 12130942 | 21 | 1 | 2 | 0 | 1 | 0 | 0 | 3 | 0 | 1 | 98 | 49 | 1 | 1 | 1 | 0.333 | 1 | NA | 0.6 | 0.178 |
| SE | 2423 | NEDD9 | NEDD9 | chr6 | - | 11213280 | 11213727 | 11193590 | 11193692 | 11305991 | 11306155 | 24 | 11 | 15 | 1 | 0 | 0 | 3 | 2 | 0 | 0 | 98 | 49 | 1 | 1 | 0.923 | 1 | 1 | 1 | 1 | -0.026 |
| SE | 2424 | NEDD9 | NEDD9 | chr6 | - | 11305065 | 11305159 | 11213280 | 11213727 | 11305991 | 11306155 | 13 | 11 | 4 | 13 | 5 | 8 | 0 | 1 | 0 | 1 | 98 | 49 | 1 | 1 | 0.333 | 0.524 | 0.2 | NA | 0.333 | 0.019 |
| SE | 2425 | MAK | MAK | chr6 | - | 10770110 | 10770230 | 10762722 | 10764606 | 10775327 | 10775459 | 3 | 2 | 2 | 1 | 0 | 0 | 0 | 3 | 0 | 0 | 98 | 49 | 1 | 1 | 0.6 | 1 | 1 | NA | 1 | -0.133 |
| SE | 2426 | MAK | MAK | chr6 | - | 10773033 | 10773108 | 10770110 | 10770230 | 10775327 | 10775459 | 0 | 1 | 0 | 0 | 0 | 1 | 0 | 0 | 0 | 3 | 98 | 49 | 1 | 1 | NA | 1 | 0 | NA | 0 | 0.5 |
| SE | 2427 | MAK | MAK | chr6 | - | 10791674 | 10791847 | 10784423 | 10784572 | 10795997 | 10796309 | 2 | 4 | 1 | 3 | 0 | 0 | 0 | 4 | 0 | 0 | 98 | 49 | 1 | 1 | 0.25 | 1 | 1 | NA | 1 | -0.25 |
| SE | 2428 | MCU | MCU | chr10 | + | 72715110 | 72715226 | 72692130 | 72692301 | 72715853 | 72715902 | 15 | 7 | 3 | 0 | 2 | 0 | 0 | 1 | 0 | 0 | 98 | 49 | 1 | 1 | 1 | 0.636 | 1 | NA | 1 | -0.121 |
| SE | 2435 | TNFSF13 | TNFSF13 | chr17 | + | 7559845 | 7559893 | 7559623 | 7559702 | 7560048 | 7560167 | 18 | 3 | 1 | 0 | 0 | 0 | 0 | 11 | 0 | 1 | 97 | 49 | 1 | 1 | 1 | 1 | 1 | NA | 0.847 | 0.153 |

| | | | | | | | | | | | | | | | | | | | | | | | | | | | | | | |
|---|---|---|---|---|---|---|---|---|---|---|---|---|---|---|---|---|---|---|---|---|---|---|---|---|---|---|---|---|---|---|
| SE | 2446 XAF1 | XAF1 | chr17 | + | 6759661 | 6759718 | 6758088 | 6758224 | 6760405 | 6760601 | 14 | 20 | 1 | 0 | 2 | 0 | 0 | 19 | 0 | 0 | 98 | 49 | 1 | 1 | 1 | 0.833 | 1 NA | | 1 | -0.056 |
| SE | 2450 XAF1 | XAF1 | chr17 | + | 6760405 | 6760601 | 6759661 | 6759718 | 6762154 | 6762240 | 6 | 11 | 1 | 0 | 1 | 0 | 0 | 16 | 0 | 0 | 98 | 49 | 1 | 1 | 1 | 0.846 | 1 NA | | 1 | -0.051 |
| SE | 2451 XAF1 | XAF1 | chr17 | + | 6762154 | 6762240 | 6758088 | 6758224 | 6770642 | 6770984 | 5 | 5 | 0 | 1 | 0 | 0 | 0 | 11 | 0 | 0 | 98 | 49 | 1 | 1 | 0.714 | 1 NA | NA | | 1 | -0.143 |
| SE | 2459 DERL2 | DERL2 | chr17 | - | 5481295 | 5481389 | 5480386 | 5480582 | 5482808 | 5482882 | 4 | 1 | 4 | 0 | 0 | 5 | 0 | 1 | 0 | 0 | 98 | 49 | 1 | 1 | 1 | 1 | 0.286 NA | | 1 | -0.238 |
| SE | 2461 EML3 | EML3 | chr11 | - | 62605641 | 62605773 | 62605112 | 62605180 | 62605854 | 62605980 | 13 | 5 | 1 | 1 | 0 | 0 | 0 | 1 | 0 | 0 | 98 | 49 | 1 | 1 | 0.867 | 1 | 1 NA | | 1 | -0.044 |
| SE | 2465 CAMTA2 | CAMTA2 | chr17 | - | 4973584 | 4973769 | 4973174 | 4973253 | 4974384 | 4974500 | 6 | 8 | 7 | 2 | 0 | 0 | 0 | 3 | 0 | 0 | 98 | 49 | 1 | 1 | 0.6 | 1 | 1 NA | | 1 | -0.133 |
| SE | 2466 CAMTA2 | CAMTA2 | chr17 | - | 4982088 | 4982160 | 4981677 | 4981831 | 4982756 | 4982892 | 21 | 1 | 3 | 1 | 0 | 0 | 0 | 2 | 0 | 0 | 98 | 49 | 1 | 1 | 0.913 | 1 | 1 NA | | 1 | -0.029 |
| SE | 2481 ARRB2 | ARRB2 | chr17 | + | 4715012 | 4715043 | 4710493 | 4710744 | 4715972 | 4716033 | 32 | 34 | 5 | 7 | 1 | 0 | 0 | 7 | 0 | 0 | 80 | 49 | 1 | 1 | 0.737 | 0.954 | 1 NA | | 1 | -0.103 |
| SE | 2483 ARRB2 | ARRB2 | chr17 | + | 4716411 | 4716608 | 4715972 | 4716033 | 4717216 | 4717276 | 26 | 10 | 1 | 0 | 0 | 0 | 0 | 2 | 0 | 3 | 98 | 49 | 1 | 1 | 1 | 1 | 1 NA | | 0.25 | 0.75 |
| SE | 2486 DDX49 | DDX49 | chr19 | + | 18921662 | 18921748 | 18920579 | 18920703 | 18921842 | 18921964 | 5 | 3 | 0 | 0 | 3 | 0 | 0 | 7 | 0 | 0 | 98 | 49 | 1 | 1 | 1 | 0.333 NA | NA | | 1 | -0.334 |
| SE | 2490 PRPF8 | PRPF8 | chr17 | - | 1661610 | 1661790 | 1661270 | 1661406 | 1661905 | 1662153 | 19 | 10 | 9 | 1 | 0 | 0 | 0 | 5 | 0 | 0 | 98 | 49 | 1 | 1 | 0.905 | 1 | 1 NA | | 1 | -0.032 |
| SE | 2493 INPP5K | INPP5K | chr17 | - | 1509682 | 1509799 | 1509177 | 1509353 | 1513871 | 1513979 | 1 | 0 | 2 | 2 | 0 | 0 | 0 | 1 | 0 | 0 | 98 | 49 | 1 | 1 | 0.2 NA | | 1 NA | | 1 | -0.4 |
| SE | 2494 INPP5K | INPP5K | chr17 | - | 1513452 | 1513561 | 1509682 | 1509799 | 1513871 | 1513979 | 7 | 5 | 8 | 0 | 0 | 0 | 0 | 2 | 0 | 1 | 98 | 49 | 1 | 1 | 1 | 1 | 1 NA | | 0.5 | 0.5 |
| SE | 2496 INPP5K | INPP5K | chr17 | - | 1515426 | 1515585 | 1513871 | 1513979 | 1516455 | 1516888 | 0 | 1 | 0 | 1 | 1 | 3 | 1 | 0 | 0 | 1 | 98 | 49 | 1 | 1 | 0 | 0.333 | 0 | 1 | 0 | -0.389 |
| SE | 2500 YWHAE | YWHAE | chr17 | - | 1361901 | 1362008 | 1361091 | 1361298 | 1364858 | 1365058 | 27 | 3 | 5 | 1 | 0 | 0 | 0 | 5 | 0 | 0 | 98 | 49 | 1 | 1 | 0.931 | 1 | 1 NA | | 1 | -0.023 |
| SE | 2502 BIVM-ERCC5 | BIVM-ERCC5 | chr13 | + | 102865666 | 102865911 | 102862029 | 102863103 | 102866261 | 102866381 | 8 | 5 | 2 | 2 | 0 | 0 | 3 | 4 | 0 | 0 | 98 | 49 | 1 | 1 | 0.667 | 1 | 1 | 1 | 1 | -0.111 |
| SE | 2503 CCDC130 | CCDC130 | chr19 | + | 13754288 | 13754342 | 13751607 | 13751811 | 13756196 | 13756279 | 16 | 5 | 2 | 2 | 0 | 0 | 0 | 2 | 0 | 0 | 98 | 49 | 1 | 1 | 0.8 | 1 | 1 NA | | 1 | -0.067 |
| SE | 2504 GLOD4 | GLOD4 | chr17 | - | 775774 | 775919 | 771324 | 771461 | 776867 | 776988 | 7 | 0 | 0 | 0 | 0 | 0 | 0 | 0 | 0 | 2 | 98 | 49 | 1 | 1 | 1 NA | NA | NA | | 0 | 1 |
| SE | 2506 GNB4 | GNB4 | chr3 | - | 179419398 | 179419505 | 179416492 | 179416556 | 179420888 | 179420927 | 13 | 5 | 4 | 1 | 0 | 0 | 0 | 8 | 0 | 0 | 98 | 49 | 1 | 1 | 0.867 | 1 | 1 NA | | 1 | -0.044 |
| SE | 2508 CHMP1A | CHMP1A | chr16 | - | 89649350 | 89649497 | 89647034 | 89647331 | 89651568 | 89651646 | 27 | 11 | 13 | 0 | 0 | 1 | 0 | 4 | 0 | 0 | 98 | 49 | 1 | 1 | 1 | 1 | 0.867 NA | | 1 | -0.044 |
| SE | 2514 SLC4A7 | SLC4A7 | chr3 | - | 27421621 | 27421779 | 27420699 | 27420787 | 27424036 | 27424152 | 4 | 0 | 2 | 0 | 3 | 0 | 0 | 1 | 0 | 0 | 98 | 49 | 1 | 1 | 1 | 0 | 1 NA | | 1 | -0.333 |
| SE | 2517 BSDC1 | BSDC1 | chr1 | - | 32369229 | 32369338 | 32368446 | 32368550 | 32376261 | 32376741 | 0 | 0 | 3 | 2 | 3 | 0 | 0 | 0 | 0 | 1 | 98 | 49 | 1 | 1 | 0 | 0 | 1 NA | | 0 | 0.333 |
| SE | 2518 BSDC1 | BSDC1 | chr1 | - | 32383829 | 32383997 | 32381213 | 32381268 | 32386778 | 32386895 | 16 | 9 | 15 | 0 | 0 | 0 | 0 | 1 | 0 | 1 | 98 | 49 | 1 | 1 | 1 | 1 | 1 NA | | 0.333 | 0.667 |
| SE | 2519 BSDC1 | BSDC1 | chr1 | - | 32383829 | 32384048 | 32381213 | 32381268 | 32386778 | 32386895 | 8 | 5 | 12 | 0 | 0 | 0 | 0 | 0 | 0 | 1 | 98 | 49 | 1 | 1 | 1 | 1 | 1 NA | | 0 | 1 |
| SE | 2523 ZC3H18 | ZC3H18 | chr16 | + | 88587541 | 88587613 | 88586599 | 88586684 | 88598177 | 88598326 | 5 | 5 | 0 | 3 | 1 | 0 | 0 | 1 | 0 | 0 | 98 | 49 | 1 | 1 | 0.455 | 0.714 NA | NA | | 1 | -0.415 |
| SE | 2533 ANXA6 | ANXA6 | chr5 | - | 151110626 | 151110644 | 151109752 | 151109846 | 151117126 | 151117180 | 17 | 2 | 4 | 1 | 0 | 2 | 0 | 2 | 0 | 0 | 67 | 49 | 1 | 1 | 0.926 | 1 | 0.594 NA | | 1 | -0.16 |
| SE | 2534 FYN | FYN | chr6 | - | 111699514 | 111699670 | 111696276 | 111696456 | 111702884 | 111703034 | 5 | 3 | 0 | 3 | 0 | 0 | 0 | 7 | 0 | 0 | 98 | 49 | 1 | 1 | 0.455 | 1 NA | NA | | 1 | -0.272 |
| SE | 2536 EXOC6 | EXOC6 | chr10 | + | 92934143 | 92934190 | 92928338 | 92928422 | 92934309 | 92934430 | 23 | 12 | 4 | 1 | 0 | 0 | 0 | 5 | 0 | 0 | 96 | 49 | 1 | 1 | 0.922 | 1 | 1 NA | | 1 | -0.026 |
| SE | 2537 EXOC6 | EXOC6 | chr10 | + | 92934309 | 92934430 | 92934143 | 92934190 | 92935813 | 92935885 | 19 | 4 | 5 | 0 | 0 | 0 | 0 | 9 | 1 | 0 | 98 | 49 | 1 | 1 | 1 | 1 | 1 | 0 | 1 | 0.5 |
| SE | 2538 EXOC6 | EXOC6 | chr10 | + | 92935813 | 92935885 | 92934309 | 92934430 | 92940726 | 92940824 | 20 | 4 | 4 | 0 | 1 | 0 | 0 | 8 | 0 | 0 | 98 | 49 | 1 | 1 | 1 | 0.667 | 1 NA | | 1 | -0.111 |
| SE | 2539 EXOC6 | EXOC6 | chr10 | + | 92952272 | 92952382 | 92948273 | 92948379 | 92954629 | 92954741 | 20 | 5 | 3 | 0 | 2 | 0 | 0 | 2 | 0 | 0 | 98 | 49 | 1 | 1 | 1 | 0.556 | 1 NA | | 1 | -0.148 |
| SE | 2541 MAP3K1 | MAP3K1 | chr5 | + | 56887377 | 56887520 | 56885931 | 56886063 | 56888225 | 56888357 | 34 | 26 | 13 | 2 | 0 | 1 | 0 | 11 | 0 | 0 | 98 | 49 | 1 | 1 | 0.895 | 1 | 0.867 NA | | 1 | -0.079 |
| SE | 2545 PLCG2 | PLCG2 | chr16 | + | 81858262 | 81858356 | 81854443 | 81854587 | 81859115 | 81859163 | 19 | 21 | 6 | 1 | 0 | 0 | 0 | 9 | 0 | 0 | 98 | 49 | 1 | 1 | 0.905 | 1 | 1 NA | | 1 | -0.032 |
| SE | 2546 PLCG2 | PLCG2 | chr16 | + | 81880909 | 81880953 | 81870851 | 81870935 | 81889171 | 81889273 | 12 | 6 | 0 | 1 | 0 | 0 | 3 | 0 | 0 | 0 | 93 | 49 | 1 | 1 | 0.863 | 1 NA | | 1 NA | | -0.069 |
| SE | 2547 PLCG2 | PLCG2 | chr16 | + | 81883268 | 81883341 | 81880909 | 81880953 | 81889171 | 81889273 | 9 | 20 | 1 | 0 | 1 | 0 | 0 | 12 | 0 | 0 | 98 | 49 | 1 | 1 | 1 | 0.909 | 1 NA | | 1 | -0.03 |
| SE | 2548 PLCG2 | PLCG2 | chr16 | + | 81956694 | 81956879 | 81946174 | 81946263 | 81957955 | 81962693 | 42 | 15 | 6 | 2 | 0 | 0 | 0 | 7 | 0 | 0 | 98 | 49 | 1 | 1 | 0.913 | 1 | 1 NA | | 1 | -0.029 |
| SE | 2553 ANXA3 | ANXA3 | chr4 | + | 78586259 | 78586350 | 78582176 | 78582290 | 78591543 | 78591623 | 37 | 24 | 18 | 2 | 0 | 0 | 1 | 19 | 0 | 0 | 98 | 49 | 1 | 1 | 0.902 | 1 | 1 | 1 | 1 | -0.033 |
| SE | 2555 RTFDC1 | RTFDC1 | chr20 | + | 56516934 | 56516989 | 56513314 | 56513428 | 56517105 | 56517201 | 69 | 49 | 26 | 1 | 0 | 0 | 0 | 7 | 0 | 2 | 98 | 49 | 1 | 1 | 0.972 | 1 | 1 NA | | 0.636 | 0.355 |
| SE | 2562 ZMYM2 | ZMYM2 | chr13 | + | 19970024 | 19970071 | 19959957 | 19960026 | 19993062 | 19993919 | 6 | 4 | 0 | 2 | 0 | 0 | 0 | 1 | 0 | 0 | 96 | 49 | 1 | 1 | 0.605 | 1 NA | NA | | 1 | -0.198 |
| SE | 2563 PSME3 | PSME3 | chr17 | + | 42838092 | 42838205 | 42837648 | 42837697 | 42838730 | 42838799 | 34 | 18 | 12 | 1 | 0 | 0 | 0 | 5 | 0 | 0 | 98 | 49 | 1 | 1 | 0.944 | 1 | 1 NA | | 1 | -0.019 |
| SE | 2564 HNRNPR | HNRNPR | chr1 | - | 23318482 | 23318688 | 23313552 | 23313702 | 23321527 | 23321663 | 7 | 4 | 3 | 0 | 1 | 0 | 0 | 4 | 0 | 0 | 98 | 49 | 1 | 1 | 1 | 0.667 | 1 NA | | 1 | -0.111 |
| SE | 2565 HNRNPR | HNRNPR | chr1 | - | 23318482 | 23318697 | 23313552 | 23313702 | 23321527 | 23321663 | 3 | 1 | 3 | 0 | 1 | 0 | 0 | 4 | 0 | 0 | 98 | 49 | 1 | 1 | 1 | 0.333 | 1 NA | | 1 | -0.222 |
| SE | 2572 NEMF | NEMF | chr14 | - | 49829053 | 49829262 | 49828615 | 49828807 | 49829348 | 49829426 | 6 | 15 | 12 | 1 | 0 | 0 | 0 | 4 | 0 | 0 | 98 | 49 | 1 | 1 | 0.75 | 1 | 1 NA | | 1 | -0.083 |
| SE | 2573 NEMF | NEMF | chr14 | - | 49832050 | 49832126 | 49831298 | 49831361 | 49832206 | 49832277 | 6 | 6 | 1 | 1 | 0 | 0 | 0 | 3 | 0 | 0 | 98 | 49 | 1 | 1 | 0.75 | 1 | 1 NA | | 1 | -0.083 |
| SE | 2574 NEMF | NEMF | chr14 | - | 49833422 | 49833496 | 49832206 | 49832277 | 49834362 | 49834449 | 7 | 4 | 7 | 0 | 0 | 1 | 1 | 0 | 0 | 0 | 98 | 49 | 1 | 1 | 1 | 1 | 0.778 | 1 NA | | -0.074 |
| SE | 2575 NEMF | NEMF | chr14 | - | 49840717 | 49840866 | 49838138 | 49838206 | 49846139 | 49846265 | 7 | 11 | 0 | 0 | 2 | 0 | 0 | 2 | 0 | 0 | 98 | 49 | 1 | 1 | 1 | 0.733 NA | NA | | 1 | -0.133 |
| SE | 2583 RPL27 | RPL27 | chr17 | + | 42999932 | 43000102 | 42998748 | 42998831 | 43002672 | 43002783 | 97 | 25 | 13 | 1 | 0 | 0 | 3 | 13 | 0 | 0 | 98 | 49 | 1 | 1 | 0.98 | 1 | 1 | 1 | 1 | -0.007 |
| SE | 2585 CDC42SE1 | CDC42SE1 | chr1 | - | 151054230 | 151054321 | 151050970 | 151053327 | 151055676 | 151055993 | 126 | 56 | 40 | 1 | 0 | 0 | 0 | 20 | 0 | 0 | 98 | 49 | 1 | 1 | 0.984 | 1 | 1 NA | | 1 | -0.005 |
| SE | 2586 CDC42SE1 | CDC42SE1 | chr1 | - | 151055015 | 151055126 | 151054230 | 151054321 | 151055676 | 151055993 | 253 | 113 | 63 | 3 | 0 | 0 | 6 | 25 | 0 | 0 | 98 | 49 | 1 | 1 | 0.977 | 1 | 1 | 1 | 1 | -0.008 |
| SE | 2587 CDC42SE1 | CDC42SE1 | chr1 | - | 151055676 | 151055993 | 151055015 | 151055126 | 151059478 | 151059649 | 172 | 70 | 37 | 1 | 0 | 0 | 6 | 17 | 0 | 0 | 98 | 49 | 1 | 1 | 0.989 | 1 | 1 | 1 | 1 | -0.004 |
| SE | 2589 NFAT5 | NFAT5 | chr16 | + | 69568494 | 69568548 | 69565965 | 69566374 | 69626402 | 69626528 | 4 | 11 | 0 | 3 | 3 | 0 | 0 | 5 | 0 | 1 | 98 | 49 | 1 | 1 | 0.4 | 0.647 NA | NA | | 0.714 | -0.19 |
| SE | 2590 NFAT5 | NFAT5 | chr16 | + | 69646516 | 69646578 | 69626402 | 69626528 | 69647027 | 69647586 | 10 | 6 | 0 | 2 | 0 | 0 | 0 | 3 | 0 | 0 | 98 | 49 | 1 | 1 | 0.714 | 1 NA | NA | | 1 | -0.143 |
| SE | 2592 ERCC6-PGBD3 | ERCC6-PGBD3 | chr10 | - | 49530719 | 49530840 | 49528416 | 49528525 | 49532542 | 49532978 | 6 | 2 | 0 | 1 | 0 | 0 | 0 | 3 | 0 | 0 | 98 | 49 | 1 | 1 | 0.75 | 1 NA | NA | | 1 | -0.125 |
| SE | 2596 MS4A7 | MS4A7 | chr11 | + | 60385087 | 60385222 | 60383128 | 60383288 | 60386716 | 60386773 | 7 | 0 | 0 | 4 | 0 | 0 | 0 | 10 | 0 | 6 | 98 | 49 | 1 | 1 | 0.467 NA | NA | NA | | 0.455 | 0.012 |
| SE | 2597 MS4A7 | MS4A7 | chr11 | + | 60392684 | 60392786 | 60389389 | 60389596 | 60393786 | 60395953 | 7 | 0 | 2 | 1 | 0 | 0 | 0 | 13 | 0 | 5 | 98 | 49 | 1 | 1 | 0.778 NA | | 1 NA | | 0.565 | 0.324 |
| SE | 2598 EDC4 | EDC4 | chr16 | + | 67882678 | 67882865 | 67882428 | 67882594 | 67882957 | 67883177 | 11 | 2 | 1 | 1 | 0 | 0 | 0 | 1 | 0 | 0 | 98 | 49 | 1 | 1 | 0.846 | 1 | 1 NA | | 1 | -0.051 |
| SE | 2599 NAA20 | NAA20 | chr20 | + | 20032507 | 20032653 | 20026783 | 20026919 | 20033101 | 20033629 | 1 | 0 | 0 | 0 | 0 | 0 | 0 | 3 | 0 | 1 | 98 | 49 | 1 | 1 | 1 NA | NA | NA | | 0.6 | 0.4 |
| SE | 2601 SPG21 | SPG21 | chr15 | - | 64983506 | 64983593 | 64980863 | 64981025 | 64989664 | 64989946 | 22 | 7 | 15 | 0 | 0 | 0 | 1 | 6 | 0 | 1 | 98 | 49 | 1 | 1 | 1 | 1 | 1 | 1 | 0.75 | 0.125 |
| SE | 2602 CYTH4 | CYTH4 | chr22 | + | 37303253 | 37303402 | 37300906 | 37301019 | 37309211 | 37309323 | 107 | 76 | 29 | 0 | 2 | 0 | 2 | 12 | 0 | 0 | 98 | 49 | 1 | 1 | 1 | 0.95 | 1 | 1 | 1 | -0.017 |
| SE | 2604 CYTH4 | CYTH4 | chr22 | + | 37311455 | 37311527 | 37310987 | 37311064 | 37312019 | 37312174 | 146 | 103 | 62 | 1 | 0 | 0 | 2 | 23 | 0 | 0 | 98 | 49 | 1 | 1 | 0.986 | 1 | 1 | 1 | 1 | -0.005 |
| SE | 2607 ATP6V0D1 | ATP6V0D1 | chr16 | - | 67444527 | 67444706 | 67443098 | 67443178 | 67453543 | 67453715 | 40 | 22 | 21 | 2 | 1 | 1 | 0 | 13 | 0 | 0 | 98 | 49 | 1 | 1 | 0.909 | 0.917 | 0.913 NA | | 1 | -0.087 |
| SE | 2616 CKLF | CKLF | chr16 | + | 66558189 | 66558348 | 66552562 | 66552793 | 66563121 | 66563217 | 82 | 69 | 45 | 17 | 11 | 4 | 0 | 8 | 0 | 2 | 98 | 49 | 1 | 1 | 0.707 | 0.758 | 0.849 NA | | 0.667 | 0.104 |
| SE | 2620 CKLF | CKLF | chr16 | + | 66558189 | 66558348 | 66552562 | 66552793 | 66565885 | 66566287 | 71 | 53 | 18 | 1 | 0 | 1 | 0 | 9 | 0 | 0 | 98 | 49 | 1 | 1 | 0.973 | 1 | 0.9 NA | | 1 | -0.042 |
| SE | 2621 CKLF | CKLF | chr16 | + | 66563121 | 66563217 | 66552562 | 66552793 | 66565885 | 66566287 | 47 | 24 | 20 | 1 | 0 | 1 | 0 | 7 | 0 | 0 | 98 | 49 | 1 | 1 | 0.959 | 1 | 0.909 NA | | 1 | -0.044 |
| SE | 2622 CKLF | CKLF | chr16 | + | 66563121 | 66563217 | 66558189 | 66558348 | 66565885 | 66566287 | 43 | 33 | 43 | 2 | 4 | 0 | 0 | 6 | 0 | 2 | 98 | 49 | 1 | 1 | 0.915 | 0.805 | 1 NA | | 0.6 | 0.307 |
| SE | 2628 NLRC5 | NLRC5 | chr16 | + | 57028071 | 57028155 | 57025367 | 57027018 | 57028301 | 57028385 | 16 | 6 | 0 | 1 | 0 | 0 | 0 | 2 | 0 | 0 | 98 | 49 | 1 | 1 | 0.889 | 1 NA | NA | | 1 | -0.055 |
| SE | 2630 NLRC5 | NLRC5 | chr16 | + | 57054750 | 57054840 | 57051537 | 57051621 | 57055031 | 57055094 | 52 | 17 | 3 | 2 | 0 | 0 | 3 | 12 | 0 | 1 | 98 | 49 | 1 | 1 | 0.929 | 1 | 1 | 1 | 0.857 | 0.048 |
| SE | 2648 TLR1 | TLR1 | chr4 | - | 38800856 | 38800948 | 38796254 | 38798898 | 38804305 | 38804382 | 97 | 25 | 13 | 9 | 2 | 0 | 0 | 5 | 0 | 0 | 98 | 49 | 1 | 1 | 0.843 | 0.862 | 1 NA | | 1 | -0.098 |
| SE | 2653 ASB6 | ASB6 | chr9 | - | 129640540 | 129640722 | 129639401 | 129639508 | 129641886 | 129642169 | 6 | 1 | 0 | 0 | 0 | 0 | 0 | 1 | 0 | 1 | 98 | 49 | 1 | 1 | 1 | 1 NA | NA | | 0.333 | 0.667 |
| SE | 2661 ADCY7 | ADCY7 | chr16 | + | 50290456 | 50290660 | 50287911 | 50288350 | 50292675 | 50292825 | 14 | 5 | 2 | 0 | 0 | 0 | 0 | 6 | 0 | 1 | 98 | 49 | 1 | 1 | 1 | 1 | 1 NA | | 0.75 | 0.25 |
| SE | 2662 ADCY7 | ADCY7 | chr16 | + | 50291735 | 50291897 | 50290456 | 50290660 | 50292675 | 50292925 | 19 | 13 | 0 | 1 | 0 | 0 | 3 | 6 | 0 | 0 | 98 | 49 | 1 | 1 | 0.905 | 1 NA | | 1 | 1 | -0.047 |
| SE | 2663 ADCY7 | ADCY7 | chr16 | + | 50293353 | 50293502 | 50292675 | 50292825 | 50294639 | 50294751 | 9 | 2 | 5 | 1 | 0 | 0 | 0 | 3 | 0 | 0 | 98 | 49 | 1 | 1 | 0.818 | 1 | 1 NA | | 1 | -0.061 |
| SE | 2664 ADCY7 | ADCY7 | chr16 | + | 50310686 | 50310880 | 50309547 | 50309646 | 50311692 | 50311786 | 23 | 14 | 1 | 2 | 0 | 0 | 0 | 5 | 0 | 0 | 98 | 49 | 1 | 1 | 0.852 | 1 | 1 NA | | 1 | -0.049 |
| SE | 2665 ADCY7 | ADCY7 | chr16 | + | 50315013 | 50315138 | 50314291 | 50314406 | 50315358 | 50318134 | 26 | 12 | 0 | 1 | 0 | 0 | 1 | 9 | 0 | 0 | 98 | 49 | 1 | 1 | 1 | 0.857 NA | | 1 | 1 | -0.072 |
| SE | 2669 LONP2 | LONP2 | chr16 | + | 48256609 | 48256741 | 48252130 | 48252365 | 48258617 | 48258740 | 11 | 3 | 1 | 0 | 0 | 0 | 0 | 0 | 0 | 1 | 98 | 49 | 1 | 1 | 1 | 1 | 1 NA | | 0 | 1 |
| SE | 2670 LONP2 | LONP2 | chr16 | + | 48261423 | 48261587 | 48258617 | 48258740 | 48262777 | 48262872 | 5 | 0 | 3 | 0 | 0 | 0 | 0 | 4 | 0 | 1 | 98 | 49 | 1 | 1 | 1 NA | | 1 NA | | 0.667 | 0.333 |
| SE | 2673 PHKB | PHKB | chr16 | + | 47547432 | 47547548 | 47515520 | 47515601 | 47580294 | 47580358 | 3 | 0 | 0 | 0 | 0 | 0 | 0 | 2 | 0 | 1 | 98 | 49 | 1 | 1 | 1 NA | NA | 1 NA | | 0.333 | 0.667 |
| SE | 2675 PHKB | PHKB | chr16 | + | 47665941 | 47666032 | 47663676 | 47663734 | 47669214 | 47669417 | 2 | 0 | 0 | 1 | 0 | 0 | 0 | 2 | 0 | 0 | 98 | 49 | 1 | 1 | 0.5 NA | NA | 1 NA | | 1 | -0.5 |
| SE | 2676 CEACAM3 | CEACAM3 | chr19 | + | 41807110 | 41807617 | 41797588 | 41797948 | 41808812 | 41808930 | 4 | 1 | 0 | 15 | 4 | 6 | 0 | 2 | 0 | 2 | 98 | 49 | 1 | 1 | 0.118 | 0.111 | 0 NA | | 0.333 | -0.257 |
| SE | 2690 STX4 | STX4 | chr16 | + | 31039722 | 31039818 | 31039540 | 31039651 | 31039911 | 31040167 | 12 | 11 | 1 | 1 | 0 | 0 | 0 | 5 | 0 | 0 | 98 | 49 | 1 | 1 | 0.857 | 1 | 1 NA | | 1 | -0.048 |
| SE | 2695 TMEM189 | TMEM189 | chr20 | - | 50151211 | 50151315 | 50143501 | 50143621 | 50153516 | 50153798 | 3 | 0 | 0 | 2 | 0 | 0 | 0 | 0 | 0 | 2 | 98 | 49 | 1 | 1 | 0.429 NA | NA | NA | | 0 | 0.429 |
| SE | 2698 SLPI | SLPI | chr20 | - | 45253001 | 45253151 | 45252238 | 45252419 | 45253574 | 45253733 | 14 | 18 | 3 | 1 | 0 | 0 | 1 | 4 | 0 | 0 | 98 | 49 | 1 | 1 | 0.875 | 1 | 1 | 1 | 1 | -0.042 |
| SE | 2699 NR4A1 | NR4A1 | chr12 | + | 52052495 | 52052645 | 52051401 | 52051568 | 52054326 | 52055204 | 2 | 0 | 0 | 1 | 0 | 0 | 0 | 1 | 0 | 2 | 98 | 49 | 1 | 1 | 0.5 NA | NA | NA | | 0.2 | 0.3 |
| SE | 2704 MAZ | MAZ | chr16 | + | 29809539 | 29809764 | 29808569 | 29808741 | 29810076 | 29811183 | 9 | 0 | 9 | 12 | 1 | 0 | 0 | 1 | 0 | 0 | 98 | 49 | 1 | 1 | 0.273 | 0 | 1 NA | | 1 | -0.576 |
| SE | 2707 LY75-CD302 | LY75-CD302 | chr2 | - | 159882123 | 159882315 | 159881082 | 159881240 | 159885152 | 159885293 | 30 | 13 | 4 | 1 | 0 | 0 | 0 | 2 | 0 | 0 | 98 | 49 | 1 | 1 | 0.937 | 1 | 1 NA | | 1 | -0.021 |
| SE | 2708 LY75-CD302 | LY75-CD302 | chr2 | - | 159885152 | 159885293 | 159881082 | 159881240 | 159886419 | 159886530 | 23 | 4 | 6 | 0 | 2 | 0 | 0 | 1 | 0 | 0 | 98 | 49 | 1 | 1 | 1 | 0.5 | 1 NA | | 1 | -0.167 |
| SE | 2709 PIP4K2B | PIP4K2B | chr17 | - | 38784242 | 38784339 | 38780451 | 38780604 | 38786822 | 38786920 | 1 | 0 | 0 | 1 | 0 | 0 | 0 | 2 | 0 | 0 | 98 | 49 | 1 | 1 | 0.333 NA | NA | NA | | 1 | -0.667 |
| SE | 2711 SH2B1 | SH2B1 | chr16 | + | 28866011 | 28867033 | 28863667 | 28863754 | 28867330 | 28867432 | 2 | 0 | 0 | 0 | 0 | 0 | 0 | 0 | 0 | 1 | 98 | 49 | 1 | 1 | 1 NA | NA | NA | | 0 | 1 |
| SE | 2713 TUFM | TUFM | chr16 | - | 28845313 | 28845480 | 28844950 | 28845055 | 28845911 | 28846106 | 5 | 0 | 1 | 1 | 0 | 0 | 0 | 1 | 0 | 0 | 98 | 49 | 1 | 1 | 0.714 NA | | 1 NA | | 1 | -0.143 |
| SE | 2714 ATXN2L | ATXN2L | chr16 | + | 28826861 | 28826986 | 28826239 | 28826390 | 28829400 | 28829492 | 11 | 8 | 0 | 1 | 0 | 0 | 0 | 1 | 0 | 0 | 98 | 49 | 1 | 1 | 0.846 | 1 NA | NA | | 1 | -0.077 |
| SE | 2715 ATXN2L | ATXN2L | chr16 | + | 28832816 | 28832887 | 28832495 | 28832567 | 28833058 | 28833354 | 12 | 2 | 0 | 1 | 0 | 0 | 0 | 3 | 0 | 0 | 98 | 49 | 1 | 1 | 0.857 | 1 NA | NA | | 1 | -0.072 |
| SE | 2718 CD33 | CD33 | chr19 | + | 51226308 | 51226356 | 51225802 | 51226081 | 51235156 | 51235253 | 3 | 0 | 0 | 0 | 0 | 0 | 0 | 7 | 1 | 0 | 97 | 49 | 1 | 1 | 1 NA | NA | | 0 | 1 | 0.5 |
| SE | 2722 PRKCB | PRKCB | chr16 | + | 23837374 | 23837406 | 23835978 | 23836348 | 23988507 | 23988590 | 40 | 36 | 7 | 0 | 1 | 0 | 7 | 15 | 0 | 0 | 81 | 49 | 1 | 1 | 1 | 0.956 | 1 | 1 | 1 | -0.015 |
| SE | 2726 EIF2S2 | EIF2S2 | chr20 | - | 34093674 | 34093731 | 34090516 | 34090602 | 34096656 | 34096805 | 6 | 0 | 0 | 0 | 3 | 0 | 0 | 4 | 0 | 0 | 98 | 49 | 1 | 1 | 1 | 0 NA | NA | | 1 | -0.5 |
| SE | 2735 RHOT1 | RHOT1 | chr17 | + | 32193986 | 32194107 | 32193135 | 32193244 | 32198946 | 32199031 | 16 | 15 | 6 | 1 | 0 | 0 | 0 | 7 | 0 | 0 | 98 | 49 | 1 | 1 | 0.889 | 1 | 1 NA | | 1 | -0.037 |
| SE | 2736 RHOT1 | RHOT1 | chr17 | + | 32209349 | 32209445 | 32208106 | 32208309 | 32211115 | 32211238 | 4 | 5 | 1 | 1 | 2 | 0 | 0 | 2 | 0 | 0 | 98 | 49 | 1 | 1 | 0.5 | 1 | 1 NA | | 1 | -0.167 |
| SE | 2737 RHOT1 | RHOT1 | chr17 | + | 32209349 | 32209445 | 32208106 | 32208309 | 32224615 | 32225771 | 3 | 0 | 0 | 4 | 1 | 0 | 0 | 1 | 0 | 2 | 98 | 49 | 1 | 1 | 0.273 | 0 NA | NA | | 0.2 | -0.064 |
| SE | 2738 RHOT1 | RHOT1 | chr17 | + | 32211115 | 32211238 | 32208106 | 32208309 | 32224615 | 32225771 | 4 | 0 | 5 | 4 | 1 | 0 | 0 | 2 | 0 | 2 | 98 | 49 | 1 | 1 | 0.333 | 0 | 1 NA | | 0.333 | 0.111 |
| SE | 2740 RHOT1 | RHOT1 | chr17 | + | 32221048 | 32221186 | 32211115 | 32211238 | 32224615 | 32225771 | 1 | 0 | 0 | 2 | 0 | 5 | 0 | 2 | 0 | 2 | 98 | 49 | 1 | 1 | 0.2 NA | | 0 NA | | 0.333 | -0.233 |
| SE | 2746 MTHFD2 | MTHFD2 | chr2 | + | 74205704 | 74205889 | 74198562 | 74198742 | 74207703 | 74207826 | 7 | 1 | 0 | 1 | 0 | 0 | 0 | 2 | 0 | 0 | 98 | 49 | 1 | 1 | 0.778 | 1 NA | NA | | 1 | -0.111 |
| SE | 2747 MTHFD2 | MTHFD2 | chr2 | + | 74205704 | 74205889 | 74201381 | 74201479 | 74207703 | 74207826 | 7 | 1 | 0 | 0 | 2 | 0 | 0 | 2 | 0 | 0 | 98 | 49 | 1 | 1 | 1 | 0.2 NA | NA | | 1 | -0.4 |
| SE | 2753 WDTC1 | WDTC1 | chr1 | + | 27294021 | 27294116 | 27292214 | 27292397 | 27294513 | 27294629 | 19 | 9 | 7 | 1 | 2 | 0 | 0 | 7 | 0 | 0 | 98 | 49 | 1 | 1 | 0.905 | 0.692 | 1 NA | | 1 | -0.134 |
| SE | 2754 SMG1 | SMG1 | chr16 | - | 18838013 | 18838232 | 18837252 | 18837443 | 18838356 | 18838466 | 18 | 2 | 5 | 1 | 1 | 0 | 4 | 5 | 0 | 0 | 98 | 49 | 1 | 1 | 0.9 | 0.5 | 1 | 1 | 1 | -0.2 |
| SE | 2755 SMG1 | SMG1 | chr16 | - | 18879494 | 18879719 | 18877130 | 18877232 | 18882164 | 18882338 | 6 | 7 | 0 | 0 | 0 | 0 | 0 | 3 | 0 | 2 | 98 | 49 | 1 | 1 | 1 | 1 NA | NA | | 0.429 | 0.571 |
| SE | 2756 SMG1 | SMG1 | chr16 | - | 18889371 | 18889585 | 18885540 | 18885666 | 18890862 | 18890921 | 13 | 8 | 4 | 0 | 0 | 0 | 0 | 4 | 0 | 3 | 98 | 49 | 1 | 1 | 1 | 1 | 1 NA | | 0.4 | 0.6 |
| SE | 2758 ARL6IP1 | ARL6IP1 | chr16 | - | 18797924 | 18798044 | 18795463 | 18795581 | 18798700 | 18798834 | 29 | 5 | 22 | 0 | 1 | 0 | 1 | 3 | 0 | 0 | 98 | 49 | 1 | 1 | 1 | 0.714 | | 1 | 1 | -0.095 |
| SE | 2760 PMM2 | PMM2 | chr16 | + | 8811078 | 8811178 | 8806315 | 8806407 | 8811637 | 8811713 | 1 | 0 | 0 | 0 | 0 | 0 | 0 | 3 | 0 | 0 | 98 | 49 | 1 | 1 | 0.333 NA | NA | NA | | 1 | -0.667 |
| SE | 2765 PCBP2 | PCBP2 | chr12 | + | 53465931 | 53465973 | 53464759 | 53464852 | 53467220 | 53467293 | 92 | 28 | 6 | 3 | 0 | 1 | 0 | 24 | 0 | 0 | 91 | 49 | 1 | 1 | 0.943 | 1 | 0.764 NA | | 1 | -0.098 |
| SE | 2768 GSPT1 | GSPT1 | chr16 | - | 11887569 | 11887750 | 11886776 | 11886931 | 11891061 | 11891139 | 8 | 0 | 4 | 0 | 0 | 0 | 0 | 1 | 0 | 0 | 98 | 49 | 1 | 1 | 0.8 NA | | 1 NA | | 1 | -0.1 |
| SE | 2775 EBPL | EBPL | chr13 | - | 49663056 | 49663195 | 49660675 | 49661208 | 49691253 | 49691487 | 4 | 0 | 4 | 0 | 1 | 0 | 0 | 1 | 0 | 0 | 98 | 49 | 1 | 1 | 1 | 0 | 1 NA | | 1 | -0.333 |
| SE | 2784 ARHGAP27 | ARHGAP27 | chr17 | - | 45396486 | 45396585 | 45396206 | 45396284 | 45396667 | 45396790 | 56 | 30 | 10 | 0 | 0 | 0 | 4 | 14 | 0 | 1 | 98 | 49 | 1 | 1 | 1 | 1 | 1 NA | 1 | 0.875 | 0.062 |
| SE | 2785 ARHGAP27 | ARHGAP27 | chr17 | - | 45396667 | 45396790 | 45396206 | 45396284 | 45397948 | 45398047 | 0 | 1 | 1 | 0 | 2 | 0 | 0 | 1 | 0 | 0 | 98 | 49 | 1 | 1 NA | | 0.2 | 1 NA | | 1 | -0.4 |
| SE | 2786 ARHGAP27 | ARHGAP27 | chr17 | - | 45396915 | 45397024 | 45396667 | 45396790 | 45397948 | 45398047 | 38 | 16 | 15 | 0 | 1 | 1 | 1 | 12 | 0 | 0 | 98 | 49 | 1 | 1 | 1 | 0.889 | 0.882 | 1 | 1 | -0.076 |
| SE | 2788 ARHGAP27 | ARHGAP27 | chr17 | - | 45432215 | 45432349 | 45431620 | 45431752 | 45432751 | 45432916 | 12 | 6 | 3 | 0 | 1 | 0 | 0 | 4 | 0 | 0 | 98 | 49 | 1 | 1 | 1 | 0.75 | 1 NA | | 1 | -0.083 |
| SE | 2789 USP25 | USP25 | chr21 | + | 15777903 | 15778027 | 15765996 | 15766141 | 15791501 | 15791664 | 36 | 14 | 8 | 0 | 0 | 0 | 0 | 1 | 0 | 1 | 98 | 49 | 1 | 1 | 1 | 1 | 1 NA | | 0.333 | 0.667 |
| SE | 2790 USP25 | USP25 | chr21 | + | 15808808 | 15808885 | 15805120 | 15805258 | 15811136 | 15811210 | 34 | 9 | 1 | 0 | 7 | 0 | 0 | 6 | 0 | 6 | 98 | 49 | 1 | 1 | 1 | 0.391 | 1 NA | | 0.333 | 0.464 |
| SE | 2791 USP25 | USP25 | chr21 | + | 15811136 | 15811210 | 15805120 | 15805258 | 15818697 | 15818846 | 21 | 13 | 2 | 0 | 1 | 0 | 0 | 11 | 0 | 0 | 98 | 49 | 1 | 1 | 1 | 0.867 | 1 NA | | 1 | -0.044 |
| SE | 2793 USP25 | USP25 | chr21 | + | 15833347 | 15833548 | 15831400 | 15831629 | 15842397 | 15842540 | 39 | 4 | 6 | 1 | 0 | 0 | 0 | 11 | 0 | 0 | 98 | 49 | 1 | 1 | 0.951 | 1 | 1 NA | | 1 | -0.016 |
| SE | 2797 USP25 | USP25 | chr21 | + | 15864267 | 15864446 | 15830530 | 15830601 | 15866265 | 15866344 | 8 | 6 | 0 | 1 | 0 | 0 | 0 | 3 | 0 | 0 | 98 | 49 | 1 | 1 | 0.8 | 1 NA | NA | | 1 | -0.1 |
| SE | 2798 USP25 | USP25 | chr21 | + | 15864267 | 15864446 | 15842397 | 15842540 | 15866265 | 15866344 | 8 | 8 | 0 | 1 | 0 | 0 | 0 | 6 | 0 | 0 | 98 | 49 | 1 | 1 | 0.8 | 1 NA | NA | | 1 | -0.1 |
| SE | 2799 USP25 | USP25 | chr21 | + | 15864267 | 15864446 | 15849776 | 15849872 | 15866265 | 15866344 | 7 | 6 | 0 | 1 | 0 | 0 | 0 | 3 | 0 | 0 | 98 | 49 | 1 | 1 | 0.778 | 1 NA | NA | | 1 | -0.111 |
| SE | 2801 IFI44L | IFI44L | chr1 | + | 78627905 | 78628393 | 78620381 | 78620571 | 78628950 | 78628999 | 13 | 0 | 0 | 1 | 0 | 0 | 1 | 91 | 0 | 0 | 98 | 49 | 1 | 1 | 0.867 NA | NA | | 1 | 1 | -0.133 |
| SE | 2803 UBN1 | UBN1 | chr16 | + | 4857989 | 4858076 | 4852878 | 4853166 | 4858567 | 4858663 | 61 | 20 | 13 | 2 | 0 | 0 | 0 | 11 | 0 | 1 | 98 | 49 | 1 | 1 | 0.938 | 1 | 1 | 1 | 0.846 | 0.056 |
| SE | 2805 UBN1 | UBN1 | chr16 | + | 4870843 | 4870972 | 4870515 | 4870634 | 4872883 | 4872934 | 23 | 14 | 1 | 3 | 0 | 0 | 0 | 8 | 0 | 0 | 98 | 49 | 1 | 1 | 0.793 | 1 | 1 NA | | 1 | -0.069 |
| SE | 2807 UBN1 | UBN1 | chr16 | + | 4872883 | 4872934 | 4870515 | 4870634 | 4873030 | 4873073 | 49 | 24 | 12 | 0 | 0 | 0 | 1 | 8 | 0 | 1 | 98 | 49 | 1 | 1 | 1 | 1 | 1 | 1 | 0.8 | 0.1 |
| SE | 2808 UBN1 | UBN1 | chr16 | + | 4876870 | 4877111 | 4874210 | 4875434 | 4880082 | 4882401 | 5 | 6 | 0 | 0 | 1 | 0 | 0 | 3 | 0 | 1 | 98 | 49 | 1 | 1 | 1 | 0.75 NA | NA | | 0.6 | 0.275 |
| SE | 2809 UBN1 | UBN1 | chr16 | + | 4877384 | 4877474 | 4876870 | 4877111 | 4880082 | 4882401 | 13 | 16 | 5 | 0 | 0 | 0 | 0 | 1 | 0 | 1 | 98 | 49 | 1 | 1 | 1 | 1 | 1 NA | | 0.333 | 0.667 |
| SE | 2811 SLC15A4 | SLC15A4 | chr12 | - | 128808787 | 128808956 | 128800853 | 128801009 | 128809395 | 128809473 | 65 | 33 | 40 | 1 | 0 | 0 | 2 | 5 | 0 | 0 | 98 | 49 | 1 | 1 | 0.97 | 1 | 1 | 1 | 1 | -0.01 |
| SE | 2812 SLC15A4 | SLC15A4 | chr12 | - | 128809942 | 128810111 | 128809395 | 128809473 | 128814774 | 128815070 | 33 | 35 | 17 | 7 | 6 | 3 | 0 | 4 | 0 | 3 | 98 | 49 | 1 | 1 | 0.702 | 0.745 | 0.739 NA | | 0.4 | 0.329 |
| SE | 2813 CORO7 | CORO7 | chr16 | - | 4361984 | 4362160 | 4361360 | 4361469 | 4362611 | 4362738 | 19 | 8 | 2 | 1 | 0 | 0 | 1 | 6 | 0 | 0 | 98 | 49 | 1 | 1 | 0.905 | 1 | 1 | 1 | 1 | -0.032 |
| SE | 2816 USB1 | USB1 | chr16 | + | 58017333 | 58017439 | 58014272 | 58014326 | 58018971 | 58019055 | 12 | 3 | 3 | 0 | 1 | 0 | 0 | 5 | 0 | 0 | 98 | 49 | 1 | 1 | 1 | 0.6 | 1 NA | | 1 | -0.133 |
| SE | 2817 AGO4 | AGO4 | chr1 | + | 35822861 | 35822982 | 35816881 | 35817047 | 35825312 | 35825494 | 88 | 35 | 16 | 1 | 0 | 0 | 0 | 10 | 0 | 0 | 98 | 49 | 1 | 1 | 0.978 | 1 | 1 NA | | 1 | -0.007 |
| SE | 2818 AGO4 | AGO4 | chr1 | + | 35831426 | 35831574 | 35826747 | 35826835 | 35831811 | 35831931 | 84 | 26 | 20 | 0 | 1 | 2 | 0 | 12 | 0 | 0 | 98 | 49 | 1 | 1 | 1 | 0.929 | 0.833 NA | | 1 | -0.079 |
| SE | 2819 AGO4 | AGO4 | chr1 | + | 35832436 | 35832570 | 35832056 | 35832185 | 35833989 | 35834174 | 49 | 20 | 20 | 1 | 0 | 0 | 2 | 14 | 0 | 1 | 98 | 49 | 1 | 1 | 0.961 | 1 | 1 | 1 | 0.875 | 0.049 |
| SE | 2820 AGO4 | AGO4 | chr1 | + | 35833989 | 35834174 | 35832436 | 35832570 | 35835833 | 35835993 | 45 | 32 | 14 | 0 | 0 | 0 | 2 | 1 | 0 | 1 | 98 | 49 | 1 | 1 | 1 | 1 | 1 | 1 | 0.333 | 0.334 |
| SE | 2821 MEFV | MEFV | chr16 | - | 3244472 | 3244588 | 3244253 | 3244286 | 3246524 | 3246547 | 49 | 38 | 44 | 1 | 1 | 0 | 0 | 19 | 0 | 0 | 98 | 49 | 1 | 1 | 0.961 | 0.95 | 1 NA | | 1 | -0.03 |
| SE | 2822 MEFV | MEFV | chr16 | - | 3247015 | 3247246 | 3246524 | 3246547 | 3248908 | 3249004 | 131 | 74 | 74 | 0 | 0 | 2 | 3 | 43 | 0 | 0 | 98 | 49 | 1 | 1 | 1 | 1 | 0.949 | 1 | 1 | -0.017 |
| SE | 2823 MEFV | MEFV | chr16 | - | 3248908 | 3249004 | 3247015 | 3247246 | 3249430 | 3249780 | 70 | 28 | 45 | 0 | 2 | 0 | 3 | 23 | 0 | 0 | 98 | 49 | 1 | 1 | 1 | 0.875 | 1 | 1 | 1 | -0.042 |
| SE | 2826 MEFV | MEFV | chr16 | - | 3249430 | 3249780 | 3248908 | 3249004 | 3254157 | 3254790 | 69 | 30 | 28 | 1 | 0 | 0 | 1 | 16 | 0 | 0 | 98 | 49 | 1 | 1 | 0.972 | 1 | 1 | 1 | 1 | -0.009 |
| SE | 2827 MEFV | MEFV | chr16 | - | 3254157 | 3254790 | 3249430 | 3249780 | 3256310 | 3256627 | 126 | 46 | 48 | 0 | 0 | 1 | 0 | 28 | 0 | 0 | 98 | 49 | 1 | 1 | 1 | 1 | 0.96 NA | | 1 | -0.013 |
| SE | 2830 STK38L | STK38L | chr12 | + | 27312548 | 27312672 | 27309113 | 27309197 | 27314503 | 27314658 | 32 | 14 | 6 | 1 | 0 | 0 | 0 | 3 | 0 | 0 | 98 | 49 | 1 | 1 | 0.941 | 1 | 1 NA | | 1 | -0.02 |
| SE | 2831 STK38L | STK38L | chr12 | + | 27315014 | 27315117 | 27314503 | 27314658 | 27315288 | 27315350 | 7 | 4 | 1 | 3 | 0 | 0 | 0 | 2 | 0 | 3 | 98 | 49 | 1 | 1 | 0.538 | 1 | 1 NA | | 0.25 | 0.596 |
| SE | 2833 STK38L | STK38L | chr12 | + | 27322142 | 27322234 | 27319327 | 27319423 | 27322327 | 27325957 | 46 | 19 | 4 | 1 | 0 | 0 | 0 | 10 | 0 | 0 | 98 | 49 | 1 | 1 | 0.958 | 1 | 1 NA | | 1 | -0.014 |
| SE | 2836 MMP25 | MMP25 | chr16 | + | 3050253 | 3050546 | 3050008 | 3050144 | 3057032 | 3057209 | 153 | 63 | 44 | 1 | 0 | 0 | 7 | 19 | 0 | 1 | 98 | 49 | 1 | 1 | 0.987 | 1 | 1 | 1 | 0.905 | 0.043 |
| SE | 2837 MMP25 | MMP25 | chr16 | + | 3057309 | 3057394 | 3057032 | 3057209 | 3057530 | 3057613 | 162 | 58 | 58 | 0 | 0 | 0 | 1 | 33 | 0 | 1 | 98 | 49 | 1 | 1 | 1 | 1 | 1 | 1 | 0.943 | 0.028 |
| SE | 2838 MMP25 | MMP25 | chr16 | + | 3057530 | 3057613 | 3057032 | 3057209 | 3058180 | 3058333 | 41 | 12 | 11 | 1 | 0 | 0 | 0 | 8 | 0 | 0 | 98 | 49 | 1 | 1 | 0.953 | 1 | 1 NA | | 1 | -0.016 |
| SE | 2839 ZC3H14 | ZC3H14 | chr14 | + | 88563650 | 88563693 | 88562908 | 88563169 | 88568038 | 88568153 | 1 | 0 | 0 | 1 | 0 | 0 | 1 | 0 | 0 | 0 | 92 | 49 | 1 | 1 | 0.348 NA | NA | | 1 NA | | -0.652 |
| SE | 2849 NBAS | NBAS | chr2 | - | 15353552 | 15353710 | 15351991 | 15352081 | 15356302 | 15356416 | 1 | 0 | 0 | 1 | 0 | 0 | 0 | 0 | 0 | 1 | 98 | 49 | 1 | 1 | 0.333 NA | NA | NA | | 0 | 0.333 |
| SE | 2850 NBAS | NBAS | chr2 | - | 15488893 | 15489022 | 15478225 | 15478289 | 15504144 | 15504213 | 9 | 1 | 2 | 0 | 1 | 0 | 0 | 1 | 0 | 0 | 98 | 49 | 1 | 1 | 1 | 0.333 | 1 NA | | 1 | -0.222 |
| SE | 2856 DENND4C | DENND4C | chr9 | + | 19357964 | 19358160 | 19356971 | 19357154 | 19360243 | 19360489 | 5 | 0 | 0 | 0 | 0 | 0 | 0 | 3 | 0 | 2 | 98 | 49 | 1 | 1 | 1 NA | NA | NA | | 0.429 | 0.571 |
| SE | 2859 TTC31 | TTC31 | chr2 | + | 74490024 | 74490127 | 74483321 | 74483410 | 74490243 | 74490473 | 1 | 0 | 0 | 1 | 0 | 0 | 0 | 1 | 0 | 0 | 98 | 49 | 1 | 1 | 0.333 NA | NA | NA | | 1 | -0.667 |
| SE | 2869 NLRP12 | NLRP12 | chr19 | - | 53798242 | 53798413 | 53795858 | 53796029 | 53801226 | 53801397 | 14 | 9 | 7 | 1 | 1 | 0 | 0 | 2 | 0 | 0 | 98 | 49 | 1 | 1 | 0.875 | 0.818 | 1 NA | | 1 | -0.102 |
| SE | 2872 NLRP12 | NLRP12 | chr19 | - | 53809586 | 53811288 | 53807494 | 53807665 | 53814907 | 53814988 | 46 | 17 | 10 | 0 | 0 | 0 | 1 | 12 | 0 | 1 | 98 | 49 | 1 | 1 | 1 | 1 | 1 | 1 | 0.857 | 0.072 |
| SE | 2875 AXIN1 | AXIN1 | chr16 | - | 291189 | 291297 | 289439 | 289607 | 293487 | 293718 | 2 | 1 | 0 | 1 | 2 | 0 | 1 | 1 | 0 | 0 | 98 | 49 | 1 | 1 | 0.5 | 0.2 NA | | 1 | 1 | -0.65 |
| SE | 2882 WAC | WAC | chr10 | + | 28595732 | 28596041 | 28590719 | 28590832 | 28608185 | 28608431 | 38 | 18 | 6 | 5 | 0 | 0 | 0 | 6 | 0 | 0 | 98 | 49 | 1 | 1 | 0.792 | 1 | 1 NA | | 1 | -0.069 |

| | | | | | | | | | | | | | | | | | | | | | | | | | | | | | | | | |
|---|---|---|---|---|---|---|---|---|---|---|---|---|---|---|---|---|---|---|---|---|---|---|---|---|---|---|---|---|---|---|---|---|
| SE | 2883 | WAC | WAC | chr10 | + | 28608185 | 28608431 | 28595732 | 28596041 | 28610698 | 28610821 | 34 | 14 | 13 | 1 | 0 | 0 | 0 | 4 | 0 | 0 | 98 | 49 | 1 | 1 | 0.944 | 1 | 1 | NA | | 1 | -0.019 |
| SE | 2884 | TM2D3 | TM2D3 | chr15 | - | 101650003 | 101650161 | 101646724 | 101646899 | 101652270 | 101652391 | 10 | 2 | 0 | 1 | 0 | 0 | 4 | 4 | 0 | 0 | 98 | 49 | 1 | 1 | 0.833 | 1 | NA | | 1 | 1 | -0.084 |
| SE | 2890 | SLC7A6 | SLC7A6 | chr16 | + | 68273817 | 68273904 | 68266592 | 68266721 | 68274690 | 68275249 | 3 | 0 | 0 | 2 | 0 | 0 | 0 | 1 | 0 | 0 | 98 | 49 | 1 | 1 | 0.429 | NA | NA | NA | | 1 | -0.571 |
| SE | 2893 | MAGOH | MAGOH | chr1 | - | 53233541 | 53233652 | 53228871 | 53228954 | 53235576 | 53235635 | 9 | 11 | 1 | 2 | 2 | 0 | 0 | 1 | 0 | 0 | 98 | 49 | 1 | 1 | 0.692 | 0.733 | 1 | NA | | 1 | -0.192 |
| SE | 2939 | FCAR | FCAR | chr19 | + | 54875329 | 54875365 | 54874092 | 54874323 | 54885234 | 54885525 | 17 | 5 | 2 | 0 | 1 | 0 | 1 | 6 | 0 | 0 | 85 | 49 | 1 | 1 | 1 | 0.742 | 1 | | 1 | 1 | -0.086 |
| SE | 2942 | FCAR | FCAR | chr19 | + | 54888006 | 54888228 | 54885234 | 54885525 | 54889648 | 54891420 | 37 | 7 | 5 | 4 | 9 | 0 | 3 | 17 | 0 | 1 | 98 | 49 | 1 | 1 | 0.822 | 0.28 | 1 | | 1 | 0.895 | -0.247 |
| SE | 2946 | USP3 | USP3 | chr15 | + | 63529013 | 63529166 | 63504510 | 63504830 | 63532646 | 63532707 | 5 | 1 | 0 | 11 | 4 | 1 | 0 | 0 | 0 | 2 | 98 | 49 | 1 | 1 | 0.185 | 0.111 | 0 | NA | | 0 | 0.099 |
| SE | 2947 | USP3 | USP3 | chr15 | + | 63533802 | 63533918 | 63532646 | 63532707 | 63537024 | 63537156 | 26 | 13 | 13 | 12 | 9 | 8 | 0 | 6 | 0 | 6 | 98 | 49 | 1 | 1 | 0.52 | 0.419 | 0.448 | NA | | 0.333 | 0.129 |
| SE | 2949 | USP3 | USP3 | chr15 | + | 63558105 | 63558188 | 63556666 | 63556748 | 63559856 | 63559970 | 54 | 17 | 19 | 2 | 0 | 0 | 0 | 22 | 0 | 0 | 98 | 49 | 1 | 1 | 0.931 | 1 | 1 | NA | | 1 | -0.023 |
| SE | 2950 | MAT2B | MAT2B | chr5 | + | 163513554 | 163513669 | 163512001 | 163512196 | 163513841 | 163513994 | 34 | 22 | 17 | 0 | 2 | 0 | 0 | 6 | 0 | 0 | 98 | 49 | 1 | 1 | 1 | 0.846 | 1 | NA | | 1 | -0.051 |
| SE | 2952 | IQGAP1 | IQGAP1 | chr15 | + | 90441505 | 90441684 | 90440501 | 90440615 | 90443393 | 90443478 | 166 | 76 | 39 | 0 | 1 | 0 | 2 | 36 | 0 | 0 | 98 | 49 | 1 | 1 | 1 | 0.974 | 1 | | 1 | 1 | -0.009 |
| SE | 2953 | IQGAP1 | IQGAP1 | chr15 | + | 90466268 | 90466436 | 90466000 | 90466091 | 90467449 | 90467592 | 172 | 74 | 60 | 0 | 2 | 0 | 0 | 20 | 0 | 0 | 98 | 49 | 1 | 1 | 1 | 0.949 | 1 | NA | | 1 | -0.017 |
| SE | 2954 | IQGAP1 | IQGAP1 | chr15 | + | 90467449 | 90467592 | 90466000 | 90466091 | 90472839 | 90473010 | 71 | 42 | 17 | 1 | 0 | 0 | 0 | 16 | 0 | 0 | 98 | 49 | 1 | 1 | 0.973 | 1 | 1 | NA | | 1 | -0.009 |
| SE | 2957 | ANPEP | ANPEP | chr15 | - | 89790952 | 89791093 | 89790459 | 89790541 | 89792159 | 89792327 | 113 | 49 | 24 | 0 | 1 | 1 | 5 | 13 | 0 | 0 | 98 | 49 | 1 | 1 | 1 | 0.961 | 0.923 | | 1 | 1 | -0.039 |
| SE | 2958 | ANPEP | ANPEP | chr15 | - | 89793034 | 89793126 | 89792451 | 89792562 | 89799259 | 89799315 | 42 | 26 | 17 | 0 | 2 | 0 | 0 | 8 | 0 | 0 | 98 | 49 | 1 | 1 | 1 | 0.867 | 1 | NA | | 1 | -0.044 |
| SE | 2959 | ANPEP | ANPEP | chr15 | - | 89797574 | 89797722 | 89793034 | 89793126 | 89799259 | 89799315 | 131 | 65 | 23 | 1 | 0 | 0 | 0 | 50 | 0 | 0 | 98 | 49 | 1 | 1 | 0.985 | 1 | 1 | NA | | 1 | -0.005 |
| SE | 2960 | ANPEP | ANPEP | chr15 | - | 89805077 | 89805217 | 89804490 | 89804617 | 89805320 | 89805463 | 109 | 37 | 13 | 2 | 0 | 0 | 3 | 14 | 0 | 1 | 98 | 49 | 1 | 1 | 0.965 | 1 | 1 | | 1 | 0.875 | 0.051 |
| SE | 2962 | ABHD2 | ABHD2 | chr15 | + | 89104181 | 89104270 | 89103902 | 89104044 | 89113724 | 89113824 | 3 | 4 | 1 | 0 | 3 | 0 | 0 | 0 | 0 | 2 | 98 | 49 | 1 | 1 | 1 | 0.4 | 1 | NA | | 0 | 0.8 |
| SE | 2963 | ABHD2 | ABHD2 | chr15 | + | 89151676 | 89151852 | 89116321 | 89116521 | 89155366 | 89155534 | 77 | 56 | 17 | 0 | 1 | 0 | 4 | 15 | 0 | 0 | 98 | 49 | 1 | 1 | 1 | 0.966 | 1 | | 1 | 1 | -0.011 |
| SE | 2967 | RAB8B | RAB8B | chr15 | + | 63255506 | 63255584 | 63249644 | 63249705 | 63256504 | 63256594 | 37 | 14 | 24 | 2 | 0 | 0 | 0 | 9 | 0 | 0 | 98 | 49 | 1 | 1 | 0.902 | 1 | 1 | NA | | 1 | -0.033 |
| SE | 2978 | SEC11A | SEC11A | chr15 | - | 84691534 | 84691644 | 84687624 | 84687774 | 84716024 | 84716460 | 20 | 6 | 20 | 0 | 2 | 0 | 2 | 2 | 0 | 0 | 98 | 49 | 1 | 1 | 1 | 0.6 | 1 | | 1 | 1 | -0.133 |
| SE | 2984 | TM6SF1 | TM6SF1 | chr15 | + | 83115844 | 83115942 | 83112796 | 83112900 | 83119577 | 83119681 | 18 | 2 | 0 | 1 | 0 | 0 | 0 | 1 | 0 | 0 | 98 | 49 | 1 | 1 | 0.9 | 1 | NA | NA | | 1 | -0.05 |
| SE | 2985 | TM6SF1 | TM6SF1 | chr15 | + | 83124671 | 83124776 | 83119577 | 83119681 | 83126754 | 83126847 | 13 | 2 | 6 | 0 | 2 | 0 | 0 | 1 | 0 | 0 | 98 | 49 | 1 | 1 | 1 | 0.333 | 1 | NA | | 1 | -0.222 |
| SE | 2987 | BTBD1 | BTBD1 | chr15 | - | 83018706 | 83018853 | 83016428 | 83018225 | 83020674 | 83020762 | 6 | 2 | 1 | 2 | 0 | 0 | 0 | 6 | 0 | 0 | 98 | 49 | 1 | 1 | 0.6 | 1 | 1 | NA | | 1 | -0.133 |
| SE | 2994 | ZFAND6 | ZFAND6 | chr15 | + | 80098415 | 80098578 | 80059567 | 80059809 | 80120327 | 80120498 | 8 | 5 | 1 | 5 | 1 | 2 | 0 | 3 | 0 | 0 | 98 | 49 | 1 | 1 | 0.444 | 0.714 | 0.2 | NA | | 1 | -0.547 |
| SE | 3018 | ERAP2 | ERAP2 | chr5 | + | 96886654 | 96886789 | 96883791 | 96883930 | 96889184 | 96889305 | 9 | 0 | 0 | 0 | 0 | 0 | 0 | 2 | 2 | 0 | 98 | 49 | 1 | 1 | 1 | NA | NA | | 0 | 1 | 0.5 |
| SE | 3022 | XPO6 | XPO6 | chr16 | - | 28121669 | 28121762 | 28117317 | 28117462 | 28125688 | 28125848 | 334 | 215 | 81 | 0 | 4 | 0 | 6 | 79 | 0 | 0 | 98 | 49 | 1 | 1 | 1 | 0.964 | 1 | | 1 | 1 | -0.012 |
| SE | 3023 | XPO6 | XPO6 | chr16 | - | 28125688 | 28125848 | 28121669 | 28121762 | 28132333 | 28132403 | 311 | 194 | 61 | 1 | 0 | 0 | 9 | 69 | 0 | 0 | 98 | 49 | 1 | 1 | 0.994 | 1 | 1 | | 1 | 1 | -0.002 |
| SE | 3025 | XPO6 | XPO6 | chr16 | - | 28133840 | 28133933 | 28132333 | 28132403 | 28146093 | 28146203 | 134 | 92 | 25 | 2 | 1 | 0 | 2 | 26 | 0 | 0 | 98 | 49 | 1 | 1 | 0.971 | 0.979 | 1 | | 1 | 1 | -0.017 |
| SE | 3028 | XPO6 | XPO6 | chr16 | - | 28146093 | 28146203 | 28135215 | 28135324 | 28152658 | 28152785 | 247 | 115 | 41 | 2 | 1 | 0 | 2 | 47 | 0 | 0 | 98 | 49 | 1 | 1 | 0.984 | 0.983 | 1 | | 1 | 1 | -0.011 |
| SE | 3029 | XPO6 | XPO6 | chr16 | - | 28152658 | 28152785 | 28135215 | 28135324 | 28156073 | 28156527 | 51 | 33 | 19 | 1 | 1 | 0 | 1 | 13 | 0 | 0 | 98 | 49 | 1 | 1 | 0.962 | 0.943 | 1 | | 1 | 1 | -0.032 |
| SE | 3030 | XPO6 | XPO6 | chr16 | - | 28177219 | 28177332 | 28175897 | 28176095 | 28180940 | 28181031 | 306 | 142 | 58 | 2 | 2 | 1 | 3 | 53 | 0 | 1 | 98 | 49 | 1 | 1 | 0.987 | 0.973 | 0.967 | | 1 | 0.964 | -0.006 |
| SE | 3031 | XPO6 | XPO6 | chr16 | - | 28177219 | 28177332 | 28175897 | 28176095 | 28211365 | 28211918 | 18 | 34 | 6 | 0 | 0 | 2 | 0 | 15 | 0 | 0 | 98 | 49 | 1 | 1 | 1 | 1 | 0.6 | NA | | 1 | -0.133 |
| SE | 3032 | XPO6 | XPO6 | chr16 | - | 28180940 | 28181031 | 28175897 | 28176095 | 28211365 | 28211918 | 50 | 24 | 9 | 0 | 0 | 2 | 0 | 18 | 0 | 0 | 98 | 49 | 1 | 1 | 1 | 1 | 0.692 | NA | | 1 | -0.103 |
| SE | 3033 | XPO6 | XPO6 | chr16 | - | 28180940 | 28181031 | 28177219 | 28177332 | 28208986 | 28209122 | 292 | 121 | 54 | 2 | 0 | 0 | 3 | 39 | 0 | 0 | 98 | 49 | 1 | 1 | 0.986 | 1 | 1 | | 1 | 1 | -0.005 |
| SE | 3035 | XPO6 | XPO6 | chr16 | - | 28208986 | 28209122 | 28177219 | 28177332 | 28211365 | 28211918 | 8 | 6 | 1 | 2 | 9 | 0 | 1 | 0 | 0 | 0 | 98 | 49 | 1 | 1 | 0.667 | 0.25 | 1 | | 1 | NA | -0.361 |
| SE | 3041 | STX16 | STX16 | chr20 | + | 58659622 | 58659634 | 58651252 | 58652087 | 58667489 | 58667597 | 0 | 0 | 0 | 3 | 2 | 2 | 0 | 1 | 0 | 3 | 61 | 49 | 1 | 1 | 0 | 0 | 0 | NA | | 0.211 | -0.211 |
| SE | 3043 | STX16 | STX16 | chr20 | + | 58669290 | 58669453 | 58667986 | 58668127 | 58670511 | 58670603 | 45 | 7 | 4 | 26 | 14 | 3 | 0 | 9 | 0 | 3 | 98 | 49 | 1 | 1 | 0.464 | 0.2 | 0.4 | NA | | 0.6 | -0.245 |
| SE | 3045 | CSK | CSK | chr15 | + | 74798614 | 74798728 | 74798232 | 74798312 | 74799271 | 74799491 | 12 | 7 | 2 | 2 | 0 | 0 | 0 | 5 | 0 | 0 | 98 | 49 | 1 | 1 | 0.75 | 1 | 1 | NA | | 1 | -0.083 |
| SE | 3046 | CSK | CSK | chr15 | + | 74798825 | 74798938 | 74798614 | 74798728 | 74799271 | 74799491 | 38 | 13 | 16 | 0 | 1 | 0 | 2 | 10 | 0 | 0 | 98 | 49 | 1 | 1 | 1 | 0.867 | 1 | | 1 | 1 | -0.044 |
| SE | 3048 | EDC3 | EDC3 | chr15 | - | 74640465 | 74640619 | 74635408 | 74635626 | 74655732 | 74656068 | 2 | 2 | 0 | 1 | 0 | 0 | 3 | 1 | 0 | 0 | 98 | 49 | 1 | 1 | 0.5 | 1 | NA | | 1 | 1 | -0.25 |
| SE | 3062 | PML | PML | chr15 | + | 74033155 | 74033414 | 74024856 | 74024927 | 74034477 | 74034530 | 3 | 0 | 0 | 2 | 2 | 0 | 0 | 2 | 0 | 0 | 98 | 49 | 1 | 1 | 0.429 | 0 | NA | NA | | 1 | -0.785 |
| SE | 3063 | DDX5 | DDX5 | chr17 | - | 64500548 | 64500773 | 64498255 | 64500326 | 64502009 | 64502069 | 189 | 84 | 35 | 2 | 0 | 0 | 3 | 45 | 0 | 0 | 98 | 49 | 1 | 1 | 0.979 | 1 | 1 | | 1 | 1 | -0.007 |
| SE | 3069 | CELF6 | CELF6 | chr15 | - | 72315844 | 72315927 | 72304745 | 72304794 | 72319612 | 72319946 | 1 | 0 | 0 | 0 | 0 | 0 | 0 | 0 | 0 | 1 | 98 | 49 | 1 | 1 | 1 | NA | NA | NA | | 0 | 1 |
| SE | 3073 | XPC | XPC | chr3 | - | 14156334 | 14156495 | 14152334 | 14152416 | 14158010 | 14158892 | 8 | 6 | 3 | 2 | 0 | 0 | 0 | 3 | 0 | 0 | 98 | 49 | 1 | 1 | 0.667 | 1 | 1 | NA | | 1 | -0.111 |
| SE | 3084 | RNASEH1 | RNASEH1 | chr2 | - | 3556788 | 3556904 | 3552143 | 3552308 | 3558132 | 3558371 | 5 | 2 | 0 | 0 | 0 | 0 | 0 | 0 | 0 | 2 | 98 | 49 | 1 | 1 | 1 | 1 | NA | NA | | 0 | 1 |
| SE | 3085 | PAFAH1B1 | PAFAH1B1 | chr17 | + | 2665371 | 2665456 | 2638098 | 2638320 | 2666015 | 2666090 | 35 | 10 | 8 | 2 | 0 | 0 | 0 | 3 | 0 | 0 | 98 | 49 | 1 | 1 | 0.897 | 1 | 1 | NA | | 1 | -0.034 |
| SE | 3096 | HERC1 | HERC1 | chr15 | - | 63623724 | 63623890 | 63622814 | 63622891 | 63624157 | 63624327 | 16 | 0 | 2 | 5 | 0 | 3 | 0 | 2 | 0 | 2 | 98 | 49 | 1 | 1 | 0.615 | NA | 0.25 | NA | | 0.333 | 0.099 |
| SE | 3097 | HERC1 | HERC1 | chr15 | - | 63662983 | 63663204 | 63661752 | 63662021 | 63664469 | 63664594 | 17 | 3 | 0 | 0 | 0 | 0 | 0 | 1 | 0 | 1 | 98 | 49 | 1 | 1 | 1 | 1 | NA | NA | | 0.333 | 0.667 |
| SE | 3098 | HERC1 | HERC1 | chr15 | - | 63694773 | 63694894 | 63694311 | 63694549 | 63696123 | 63696339 | 30 | 8 | 5 | 0 | 4 | 0 | 0 | 2 | 0 | 0 | 98 | 49 | 1 | 1 | 1 | 0.5 | 1 | NA | | 1 | -0.167 |
| SE | 3100 | HERC1 | HERC1 | chr15 | - | 63758174 | 63758369 | 63756436 | 63756748 | 63764095 | 63764191 | 19 | 11 | 3 | 0 | 1 | 0 | 0 | 2 | 0 | 0 | 98 | 49 | 1 | 1 | 1 | 0.846 | 1 | NA | | 1 | -0.051 |
| SE | 3103 | 15-Sep | 15-Sep | chr1 | - | 86868052 | 86868102 | 86862444 | 86863605 | 86880661 | 86880725 | 58 | 21 | 16 | 0 | 0 | 0 | 0 | 12 | 1 | 0 | 98 | 49 | 1 | 1 | 1 | 1 | 1 | | 0 | 1 | 0.5 |
| SE | 3104 | ADAM10 | ADAM10 | chr15 | - | 58611807 | 58611991 | 58610998 | 58611107 | 58621470 | 58621621 | 52 | 25 | 8 | 0 | 0 | 0 | 0 | 11 | 0 | 1 | 98 | 49 | 1 | 1 | 1 | 1 | 1 | NA | | 0.846 | 0.154 |
| SE | 3105 | ADAM10 | ADAM10 | chr15 | - | 58643885 | 58643978 | 58640776 | 58640960 | 58646054 | 58646204 | 67 | 50 | 23 | 1 | 0 | 0 | 0 | 18 | 0 | 0 | 98 | 49 | 1 | 1 | 0.971 | 1 | 1 | NA | | 1 | -0.01 |
| SE | 3109 | COMMD9 | COMMD9 | chr11 | - | 36277088 | 36277123 | 36276136 | 36276240 | 36278476 | 36278616 | 4 | 3 | 0 | 1 | 0 | 0 | 0 | 1 | 0 | 0 | 84 | 49 | 1 | 1 | 0.7 | 1 | NA | NA | | 1 | -0.15 |
| SE | 3116 | SIN3B | SIN3B | chr19 | + | 16862883 | 16862979 | 16862351 | 16862559 | 16863679 | 16863796 | 2 | 0 | 0 | 1 | 2 | 0 | 0 | 4 | 0 | 0 | 98 | 49 | 1 | 1 | 0.5 | 0 | NA | NA | | 1 | -0.75 |
| SE | 3120 | MYO5A | MYO5A | chr15 | - | 52348817 | 52348826 | 52346360 | 52346461 | 52351253 | 52351481 | 1 | 2 | 1 | 0 | 0 | 0 | 0 | 0 | 0 | 1 | 58 | 49 | 1 | 1 | 1 | 1 | 1 | NA | | 0 | 1 |
| SE | 3121 | MYO5A | MYO5A | chr15 | - | 52407291 | 52407399 | 52405286 | 52405393 | 52408058 | 52408140 | 26 | 8 | 2 | 3 | 0 | 0 | 0 | 5 | 0 | 0 | 98 | 49 | 1 | 1 | 0.812 | 1 | 1 | NA | | 1 | -0.063 |
| SE | 3123 | MYO5A | MYO5A | chr15 | - | 52416144 | 52416301 | 52410332 | 52410476 | 52425829 | 52425974 | 54 | 15 | 6 | 1 | 0 | 0 | 0 | 7 | 0 | 0 | 98 | 49 | 1 | 1 | 0.964 | 1 | 1 | NA | | 1 | -0.012 |
| SE | 3124 | UFC1 | UFC1 | chr1 | + | 161158120 | 161158211 | 161157616 | 161157693 | 161158411 | 161158856 | 7 | 1 | 1 | 0 | 0 | 0 | 0 | 3 | 0 | 1 | 98 | 49 | 1 | 1 | 1 | 1 | 1 | NA | | 0.6 | 0.4 |
| SE | 3126 | ZNF207 | ZNF207 | chr17 | + | 32366664 | 32366757 | 32365329 | 32365487 | 32367771 | 32368014 | 23 | 9 | 1 | 4 | 2 | 1 | 0 | 0 | 0 | 2 | 98 | 49 | 1 | 1 | 0.742 | 0.692 | 0.333 | NA | | 0 | 0.589 |
| SE | 3128 | CMIP | CMIP | chr16 | + | 81699684 | 81699801 | 81696559 | 81696667 | 81701659 | 81701800 | 24 | 11 | 1 | 1 | 0 | 0 | 6 | 14 | 0 | 0 | 98 | 49 | 1 | 1 | 0.923 | 1 | 1 | | 1 | 1 | -0.026 |
| SE | 3131 | TRPM7 | TRPM7 | chr15 | - | 50628148 | 50628249 | 50624165 | 50624300 | 50631416 | 50631489 | 9 | 12 | 0 | 0 | 0 | 0 | 0 | 4 | 0 | 1 | 98 | 49 | 1 | 1 | 1 | 1 | NA | NA | | 0.667 | 0.333 |
| SE | 3133 | TRPM7 | TRPM7 | chr15 | - | 50662966 | 50663046 | 50657780 | 50657819 | 50686530 | 50686815 | 20 | 3 | 0 | 1 | 0 | 0 | 2 | 1 | 0 | 0 | 98 | 49 | 1 | 1 | 0.909 | 1 | NA | | 1 | 1 | -0.045 |
| SE | 3134 | USP8 | USP8 | chr15 | + | 50481480 | 50482065 | 50477275 | 50477499 | 50484274 | 50484361 | 10 | 8 | 0 | 1 | 0 | 0 | 0 | 5 | 0 | 0 | 98 | 49 | 1 | 1 | 0.833 | 1 | NA | NA | | 1 | -0.084 |
| SE | 3135 | USP8 | USP8 | chr15 | + | 50484274 | 50484361 | 50481480 | 50482065 | 50489800 | 50489881 | 8 | 5 | 2 | 8 | 1 | 1 | 0 | 2 | 0 | 0 | 98 | 49 | 1 | 1 | 0.333 | 0.714 | 0.5 | NA | | 1 | -0.484 |
| SE | 3136 | ATP8B4 | ATP8B4 | chr15 | - | 49901091 | 49901239 | 49898067 | 49898251 | 49916933 | 49917039 | 5 | 0 | 0 | 0 | 1 | 0 | 3 | 4 | 0 | 0 | 98 | 49 | 1 | 1 | 1 | 0 | NA | | 1 | 1 | -0.5 |
| SE | 3138 | ATP8B4 | ATP8B4 | chr15 | - | 49996676 | 49996759 | 49987390 | 49987549 | 50002152 | 50002223 | 5 | 9 | 0 | 0 | 0 | 0 | 1 | 2 | 0 | 1 | 98 | 49 | 1 | 1 | 1 | 1 | NA | | 1 | 0.5 | 0.25 |
| SE | 3139 | ATP8B4 | ATP8B4 | chr15 | - | 50002152 | 50002223 | 49987390 | 49987549 | 50010844 | 50010917 | 7 | 8 | 3 | 2 | 0 | 0 | 0 | 2 | 0 | 2 | 98 | 49 | 1 | 1 | 0.636 | 1 | 1 | NA | | 0.333 | 0.546 |
| SE | 3140 | ATP8B4 | ATP8B4 | chr15 | - | 50018888 | 50018976 | 50010844 | 50010917 | 50038767 | 50038829 | 4 | 8 | 1 | 9 | 5 | 8 | 0 | 0 | 0 | 5 | 98 | 49 | 1 | 1 | 0.182 | 0.444 | 0.059 | NA | | 0 | 0.228 |
| SE | 3142 | DERL1 | DERL1 | chr8 | - | 123030604 | 123030716 | 123024985 | 123025050 | 123041969 | 123042423 | 8 | 8 | 5 | 0 | 1 | 1 | 0 | 2 | 0 | 0 | 98 | 49 | 1 | 1 | 1 | 0.8 | 0.714 | NA | | 1 | -0.162 |
| SE | 3147 | PIGG | PIGG | chr4 | + | 521659 | 521941 | 521055 | 521273 | 523458 | 523913 | 1 | 2 | 0 | 2 | 0 | 0 | 0 | 1 | 0 | 0 | 98 | 49 | 1 | 1 | 0.2 | 1 | NA | NA | | 1 | -0.4 |
| SE | 3148 | SQRDL | SQRDL | chr15 | + | 45676100 | 45676310 | 45673606 | 45673801 | 45682477 | 45682661 | 37 | 20 | 14 | 0 | 2 | 0 | 0 | 14 | 0 | 0 | 98 | 49 | 1 | 1 | 1 | 0.833 | 1 | NA | | 1 | -0.056 |
| SE | 3149 | B4GALT5 | B4GALT5 | chr20 | - | 49639677 | 49639800 | 49637340 | 49637442 | 49640477 | 49640665 | 47 | 17 | 9 | 0 | 2 | 0 | 5 | 13 | 0 | 1 | 98 | 49 | 1 | 1 | 1 | 0.81 | 1 | | 1 | 0.867 | 0.003 |
| SE | 3158 | PDXDC1 | PDXDC1 | chr16 | + | 15026642 | 15026706 | 15022703 | 15022754 | 15028877 | 15028966 | 2 | 0 | 0 | 0 | 0 | 0 | 2 | 0 | 0 | 2 | 98 | 49 | 1 | 1 | 1 | NA | NA | | 1 | 0 | 0.5 |
| SE | 3161 | PATL2 | PATL2 | chr15 | - | 44667105 | 44667203 | 44666391 | 44666541 | 44668341 | 44668482 | 3 | 1 | 0 | 1 | 1 | 0 | 0 | 1 | 0 | 0 | 98 | 49 | 1 | 1 | 0.6 | 0.333 | NA | NA | | 1 | -0.533 |
| SE | 3183 | PLAGL1 | PLAGL1 | chr6 | - | 143985134 | 143985174 | 143968906 | 143968978 | 144064467 | 144064599 | 0 | 1 | 0 | 1 | 0 | 0 | 0 | 1 | 0 | 0 | 89 | 49 | 1 | 1 | 0 | 1 | NA | NA | | 1 | -0.5 |
| SE | 3194 | CCNDBP1 | CCNDBP1 | chr15 | + | 43185819 | 43185879 | 43185267 | 43185607 | 43189198 | 43189280 | 27 | 19 | 15 | 0 | 1 | 0 | 1 | 4 | 0 | 0 | 98 | 49 | 1 | 1 | 1 | 0.905 | 1 | | 1 | 1 | -0.032 |
| SE | 3195 | CCNDBP1 | CCNDBP1 | chr15 | + | 43186153 | 43186233 | 43185819 | 43185879 | 43189198 | 43189280 | 48 | 23 | 24 | 4 | 5 | 0 | 0 | 10 | 0 | 0 | 98 | 49 | 1 | 1 | 0.857 | 0.697 | 1 | NA | | 1 | -0.149 |
| SE | 3196 | CCNDBP1 | CCNDBP1 | chr15 | + | 43190054 | 43190151 | 43189198 | 43189280 | 43190324 | 43190398 | 34 | 10 | 15 | 20 | 2 | 3 | 0 | 7 | 0 | 5 | 98 | 49 | 1 | 1 | 0.459 | 0.714 | 0.714 | NA | | 0.412 | 0.217 |
| SE | 3199 | ABCD4 | ABCD4 | chr14 | - | 74288206 | 74288259 | 74287809 | 74287886 | 74288715 | 74288765 | 7 | 0 | 0 | 0 | 0 | 0 | 0 | 1 | 0 | 1 | 98 | 49 | 1 | 1 | 1 | NA | NA | NA | | 0.333 | 0.667 |
| SE | 3200 | ABCD4 | ABCD4 | chr14 | - | 74293153 | 74293248 | 74292747 | 74292869 | 74295147 | 74295198 | 3 | 0 | 0 | 1 | 0 | 0 | 0 | 1 | 0 | 0 | 98 | 49 | 1 | 1 | 0.6 | NA | NA | NA | | 1 | -0.4 |
| SE | 3207 | NFATC3 | NFATC3 | chr16 | + | 68214328 | 68214432 | 68190767 | 68191775 | 68226349 | 68229259 | 2 | 0 | 0 | 5 | 2 | 0 | 0 | 0 | 0 | 1 | 98 | 49 | 1 | 1 | 0.167 | 0 | NA | NA | | 0 | 0.084 |
| SE | 3208 | NFATC3 | NFATC3 | chr16 | + | 68221195 | 68221284 | 68190767 | 68191775 | 68226349 | 68229259 | 1 | 0 | 0 | 5 | 2 | 0 | 0 | 0 | 0 | 1 | 98 | 49 | 1 | 1 | 0.091 | 0 | NA | NA | | 0 | 0.045 |
| SE | 3211 | SEMA7A | SEMA7A | chr15 | - | 74414837 | 74414946 | 74414546 | 74414745 | 74415800 | 74415985 | 4 | 5 | 0 | 1 | 0 | 0 | 0 | 1 | 0 | 0 | 98 | 49 | 1 | 1 | 0.667 | 1 | NA | NA | | 1 | -0.166 |
| SE | 3246 | MTF2 | MTF2 | chr1 | + | 93129277 | 93129448 | 93127231 | 93127299 | 93133702 | 93133808 | 6 | 0 | 0 | 3 | 0 | 0 | 0 | 1 | 0 | 0 | 98 | 49 | 1 | 1 | 0.5 | NA | NA | NA | | 1 | -0.5 |
| SE | 3248 | YTHDF3 | YTHDF3 | chr8 | + | 63172652 | 63172798 | 63169386 | 63169411 | 63175330 | 63175416 | 1 | 0 | 0 | 3 | 6 | 0 | 0 | 0 | 0 | 3 | 98 | 49 | 1 | 1 | 0.143 | 0 | NA | NA | | 0 | 0.071 |
| SE | 3250 | YTHDF3 | YTHDF3 | chr8 | + | 63175330 | 63175416 | 63169386 | 63169411 | 63186146 | 63187745 | 9 | 12 | 3 | 3 | 0 | 0 | 0 | 5 | 0 | 0 | 98 | 49 | 1 | 1 | 0.6 | 1 | 1 | NA | | 1 | -0.133 |
| SE | 3252 | GANC | GANC | chr15 | + | 42297610 | 42297656 | 42292734 | 42292917 | 42306545 | 42306612 | 3 | 0 | 2 | 0 | 0 | 0 | 0 | 2 | 0 | 1 | 95 | 49 | 1 | 1 | 1 | NA | 1 | NA | | 0.508 | 0.492 |
| SE | 3258 | IRAK4 | IRAK4 | chr12 | + | 43771219 | 43771365 | 43768102 | 43768272 | 43772179 | 43772362 | 13 | 6 | 4 | 13 | 0 | 0 | 0 | 4 | 0 | 0 | 98 | 49 | 1 | 1 | 0.333 | 1 | 1 | NA | | 1 | -0.222 |
| SE | 3259 | IRAK4 | IRAK4 | chr12 | + | 43782306 | 43782490 | 43778192 | 43778302 | 43783661 | 43783724 | 17 | 5 | 2 | 1 | 0 | 0 | 0 | 6 | 0 | 0 | 98 | 49 | 1 | 1 | 0.895 | 1 | 1 | NA | | 1 | -0.035 |
| SE | 3260 | FCER1G | FCER1G | chr1 | + | 161217985 | 161218077 | 161215296 | 161215370 | 161218240 | 161218276 | 34 | 19 | 6 | 1 | 0 | 0 | 7 | 28 | 0 | 0 | 98 | 49 | 1 | 1 | 0.944 | 1 | 1 | | 1 | 1 | -0.019 |
| SE | 3261 | LPCAT4 | LPCAT4 | chr15 | - | 34361399 | 34361532 | 34360110 | 34360209 | 34362195 | 34362321 | 1 | 2 | 0 | 1 | 3 | 0 | 0 | 1 | 0 | 0 | 98 | 49 | 1 | 1 | 0.333 | 0.25 | NA | NA | | 1 | -0.709 |
| SE | 3269 | MTMR10 | MTMR10 | chr15 | - | 30942889 | 30943072 | 30938940 | 30942072 | 30947129 | 30947300 | 15 | 16 | 8 | 0 | 0 | 0 | 0 | 0 | 0 | 1 | 98 | 49 | 1 | 1 | 1 | 1 | 1 | NA | | 0 | 1 |
| SE | 3270 | MTMR10 | MTMR10 | chr15 | - | 30967919 | 30968010 | 30960880 | 30961073 | 30974313 | 30974456 | 43 | 18 | 12 | 1 | 2 | 0 | 0 | 8 | 0 | 0 | 98 | 49 | 1 | 1 | 0.956 | 0.818 | 1 | NA | | 1 | -0.075 |
| SE | 3271 | MTMR10 | MTMR10 | chr15 | - | 30976818 | 30976955 | 30974930 | 30975003 | 30990776 | 30990837 | 49 | 8 | 3 | 7 | 0 | 0 | 0 | 4 | 0 | 2 | 98 | 49 | 1 | 1 | 0.778 | 1 | 1 | NA | | 0.5 | 0.426 |
| SE | 3272 | USP4 | USP4 | chr3 | - | 49278813 | 49278902 | 49277143 | 49278451 | 49280743 | 49280847 | 27 | 17 | 9 | 3 | 3 | 0 | 0 | 9 | 0 | 0 | 98 | 49 | 1 | 1 | 0.818 | 0.739 | 1 | NA | | 1 | -0.148 |
| SE | 3273 | USP4 | USP4 | chr3 | - | 49283986 | 49284136 | 49280743 | 49280847 | 49284465 | 49284584 | 50 | 18 | 16 | 0 | 2 | 0 | 0 | 17 | 0 | 0 | 98 | 49 | 1 | 1 | 1 | 0.818 | 1 | NA | | 1 | -0.061 |
| SE | 3275 | USP4 | USP4 | chr3 | - | 49300466 | 49300691 | 49298551 | 49298635 | 49302383 | 49302542 | 22 | 9 | 2 | 2 | 4 | 0 | 1 | 0 | 0 | 0 | 98 | 49 | 1 | 1 | 0.846 | 0.529 | 1 | | 1 | NA | -0.208 |
| SE | 3276 | USP4 | USP4 | chr3 | - | 49310619 | 49310737 | 49305714 | 49305888 | 49324701 | 49324763 | 31 | 21 | 9 | 0 | 0 | 0 | 0 | 3 | 0 | 1 | 98 | 49 | 1 | 1 | 1 | 1 | 1 | NA | | 0.6 | 0.4 |
| SE | 3277 | USP4 | USP4 | chr3 | - | 49311513 | 49311654 | 49310619 | 49310737 | 49324701 | 49324763 | 5 | 12 | 2 | 8 | 10 | 1 | 0 | 1 | 0 | 1 | 98 | 49 | 1 | 1 | 0.238 | 0.375 | 0.5 | NA | | 0.333 | 0.038 |
| SE | 3278 | USP4 | USP4 | chr3 | - | 49324893 | 49325039 | 49324701 | 49324763 | 49327685 | 49327816 | 29 | 19 | 4 | 2 | 0 | 0 | 1 | 4 | 0 | 0 | 98 | 49 | 1 | 1 | 0.879 | 1 | 1 | | 1 | 1 | -0.04 |
| SE | 3279 | USP4 | USP4 | chr3 | - | 49325718 | 49325845 | 49324893 | 49325039 | 49327685 | 49327816 | 78 | 61 | 12 | 2 | 0 | 0 | 0 | 17 | 0 | 0 | 98 | 49 | 1 | 1 | 0.951 | 1 | 1 | NA | | 1 | -0.016 |
| SE | 3282 | PTTG1IP | PTTG1IP | chr21 | - | 44865394 | 44865447 | 44861162 | 44861271 | 44873501 | 44873903 | 13 | 3 | 9 | 1 | 0 | 0 | 2 | 4 | 0 | 0 | 98 | 49 | 1 | 1 | 0.867 | 1 | 1 | | 1 | 1 | -0.044 |
| SE | 3283 | CNOT2 | CNOT2 | chr12 | + | 70294070 | 70294294 | 70278131 | 70278274 | 70310894 | 70311017 | 2 | 1 | 1 | 4 | 2 | 0 | 0 | 0 | 0 | 1 | 98 | 49 | 1 | 1 | 0.2 | 0.2 | 1 | NA | | 0 | 0.467 |
| SE | 3286 | RNASET2 | RNASET2 | chr6 | - | 166938894 | 166939008 | 166934090 | 166934136 | 166943018 | 166943089 | 273 | 108 | 110 | 1 | 0 | 0 | 4 | 64 | 0 | 0 | 98 | 49 | 1 | 1 | 0.993 | 1 | 1 | | 1 | 1 | -0.002 |
| SE | 3290 | RAB37 | RAB37 | chr17 | + | 74745007 | 74745084 | 74744872 | 74744929 | 74745305 | 74747335 | 19 | 9 | 4 | 1 | 0 | 0 | 0 | 1 | 0 | 0 | 98 | 49 | 1 | 1 | 0.905 | 1 | 1 | NA | | 1 | -0.032 |
| SE | 3301 | GABPB1 | GABPB1 | chr15 | - | 50346371 | 50346395 | 50309690 | 50309798 | 50354984 | 50355408 | 1 | 2 | 4 | 2 | 1 | 0 | 0 | 0 | 0 | 1 | 73 | 49 | 1 | 1 | 0.251 | 0.573 | 1 | NA | | 0 | 0.608 |
| SE | 3306 | CDK17 | CDK17 | chr12 | - | 96311051 | 96311177 | 96300303 | 96300360 | 96313320 | 96313454 | 14 | 7 | 4 | 0 | 1 | 0 | 0 | 5 | 0 | 0 | 98 | 49 | 1 | 1 | 1 | 0.778 | 1 | NA | | 1 | -0.074 |
| SE | 3308 | GNL3L | GNL3L | chrX | + | 54554564 | 54554692 | 54552291 | 54552428 | 54558435 | 54558655 | 28 | 6 | 1 | 1 | 0 | 0 | 0 | 6 | 0 | 0 | 98 | 49 | 1 | 1 | 0.933 | 1 | 1 | NA | | 1 | -0.022 |
| SE | 3311 | VAPA | VAPA | chr18 | + | 9944923 | 9945058 | 9936985 | 9937066 | 9950394 | 9950568 | 4 | 0 | 0 | 16 | 0 | 4 | 0 | 1 | 0 | 3 | 98 | 49 | 1 | 1 | 0.111 | NA | 0 | NA | | 0.143 | -0.087 |
| SE | 3312 | VAPA | VAPA | chr18 | + | 9950394 | 9950568 | 9936985 | 9937066 | 9954052 | 9960021 | 47 | 19 | 25 | 1 | 0 | 0 | 0 | 10 | 0 | 0 | 98 | 49 | 1 | 1 | 0.959 | 1 | 1 | NA | | 1 | -0.014 |
| SE | 3316 | RAB1A | RAB1A | chr2 | - | 65090982 | 65091078 | 65088938 | 65089070 | 65097970 | 65098066 | 27 | 16 | 8 | 3 | 1 | 0 | 0 | 8 | 0 | 0 | 98 | 49 | 1 | 1 | 0.818 | 0.889 | 1 | NA | | 1 | -0.098 |
| SE | 3317 | PYGL | PYGL | chr14 | - | 50912155 | 50912303 | 50911977 | 50912036 | 50913028 | 50913130 | 125 | 85 | 57 | 1 | 0 | 0 | 1 | 14 | 0 | 1 | 98 | 49 | 1 | 1 | 0.984 | 1 | 1 | | 1 | 0.875 | 0.057 |
| SE | 3318 | PYGL | PYGL | chr14 | - | 50913028 | 50913130 | 50911977 | 50912036 | 50915335 | 50915499 | 1 | 0 | 1 | 1 | 0 | 0 | 0 | 1 | 0 | 0 | 98 | 49 | 1 | 1 | 0.333 | NA | 1 | NA | | 1 | -0.334 |
| SE | 3319 | PYGL | PYGL | chr14 | - | 50914700 | 50914815 | 50913028 | 50913130 | 50915335 | 50915499 | 119 | 57 | 34 | 0 | 0 | 1 | 1 | 20 | 0 | 0 | 98 | 49 | 1 | 1 | 1 | 1 | 0.944 | | 1 | 1 | -0.019 |
| SE | 3320 | PYGL | PYGL | chr14 | - | 50915335 | 50915499 | 50914700 | 50914815 | 50915824 | 50915971 | 97 | 79 | 29 | 0 | 1 | 0 | 1 | 25 | 0 | 0 | 98 | 49 | 1 | 1 | 1 | 0.975 | 1 | | 1 | 1 | -0.008 |
| SE | 3323 | NIPA2 | NIPA2 | chr15 | + | 22839654 | 22839790 | 22838640 | 22838921 | 22845145 | 22845267 | 0 | 0 | 0 | 0 | 1 | 0 | 0 | 1 | 0 | 0 | 98 | 49 | 1 | 1 | NA | 0 | NA | NA | | 1 | -1 |
| SE | 3324 | NIPA2 | NIPA2 | chr15 | + | 22839654 | 22839790 | 22838640 | 22838921 | 22851638 | 22851870 | 0 | 0 | 0 | 2 | 4 | 0 | 0 | 1 | 0 | 0 | 98 | 49 | 1 | 1 | 0 | 0 | NA | NA | | 1 | -1 |
| SE | 3330 | WDR11 | WDR11 | chr10 | + | 120860108 | 120860282 | 120858642 | 120858796 | 120862734 | 120862921 | 6 | 0 | 4 | 0 | 0 | 0 | 0 | 2 | 0 | 1 | 98 | 49 | 1 | 1 | 1 | NA | 1 | NA | | 0.5 | 0.5 |
| SE | 3331 | WDR11 | WDR11 | chr10 | + | 120867065 | 120867169 | 120866568 | 120866764 | 120871169 | 120871346 | 4 | 5 | 2 | 0 | 0 | 0 | 0 | 3 | 0 | 1 | 98 | 49 | 1 | 1 | 1 | 1 | 1 | NA | | 0.6 | 0.4 |
| SE | 3341 | UQCRH | UQCRH | chr1 | + | 46309896 | 46310044 | 46309100 | 46309127 | 46310154 | 46310316 | 8 | 0 | 0 | 4 | 2 | 0 | 0 | 1 | 2 | 0 | 98 | 49 | 1 | 1 | 0.5 | 0 | NA | | 0 | 1 | -0.25 |
| SE | 3343 | RABAC1 | RABAC1 | chr19 | - | 41957017 | 41957119 | 41956680 | 41956934 | 41958285 | 41958383 | 19 | 5 | 13 | 0 | 1 | 0 | 0 | 5 | 0 | 0 | 98 | 49 | 1 | 1 | 1 | 0.714 | 1 | NA | | 1 | -0.095 |
| SE | 3356 | CHST15 | CHST15 | chr10 | - | 124044579 | 124044919 | 124042300 | 124042447 | 124045666 | 124046724 | 136 | 60 | 56 | 1 | 0 | 0 | 0 | 34 | 0 | 0 | 98 | 49 | 1 | 1 | 0.986 | 1 | 1 | NA | | 1 | -0.005 |
| SE | 3357 | TNFAIP2 | TNFAIP2 | chr14 | + | 103127004 | 103127629 | 103126326 | 103126692 | 103129739 | 103129854 | 227 | 74 | 53 | 0 | 1 | 0 | 2 | 36 | 0 | 0 | 98 | 49 | 1 | 1 | 1 | 0.974 | 1 | | 1 | 1 | -0.009 |
| SE | 3358 | TNFAIP2 | TNFAIP2 | chr14 | + | 103129739 | 103129854 | 103126326 | 103126692 | 103130001 | 103130124 | 139 | 78 | 24 | 0 | 0 | 1 | 4 | 22 | 0 | 0 | 98 | 49 | 1 | 1 | 1 | 1 | 0.923 | | 1 | 1 | -0.026 |
| SE | 3359 | TNFAIP2 | TNFAIP2 | chr14 | + | 103130314 | 103130415 | 103130001 | 103130124 | 103131051 | 103131150 | 222 | 105 | 48 | 1 | 0 | 0 | 9 | 41 | 0 | 0 | 98 | 49 | 1 | 1 | 0.991 | 1 | 1 | | 1 | 1 | -0.003 |
| SE | 3360 | TNFAIP2 | TNFAIP2 | chr14 | + | 103133361 | 103133517 | 103132749 | 103132872 | 103133681 | 103133803 | 266 | 119 | 54 | 0 | 2 | 0 | 7 | 23 | 0 | 0 | 98 | 49 | 1 | 1 | 1 | 0.967 | 1 | | 1 | 1 | -0.011 |
| SE | 3389 | KIAA0430 | KIAA0430 | chr16 | - | 15600640 | 15600701 | 15600427 | 15600553 | 15601990 | 15602203 | 21 | 10 | 8 | 3 | 3 | 0 | 0 | 2 | 0 | 0 | 98 | 49 | 1 | 1 | 0.778 | 0.625 | 1 | NA | | 1 | -0.199 |
| SE | 3390 | KIAA0430 | KIAA0430 | chr16 | - | 15625371 | 15625797 | 15625015 | 15625173 | 15630331 | 15630504 | 22 | 11 | 7 | 1 | 0 | 0 | 0 | 5 | 0 | 0 | 98 | 49 | 1 | 1 | 0.917 | 1 | 1 | NA | | 1 | -0.028 |
| SE | 3391 | KIAA0430 | KIAA0430 | chr16 | - | 15625371 | 15625800 | 15625015 | 15625173 | 15630331 | 15630504 | 33 | 16 | 5 | 1 | 0 | 0 | 0 | 7 | 0 | 0 | 98 | 49 | 1 | 1 | 0.943 | 1 | 1 | NA | | 1 | -0.019 |
| SE | 3394 | SNHG6 | SNHG6 | chr8 | - | 66922323 | 66922392 | 66921929 | 66922114 | 66922613 | 66922725 | 22 | 6 | 1 | 0 | 0 | 0 | 0 | 2 | 0 | 1 | 98 | 49 | 1 | 1 | 1 | 1 | 1 | NA | | 0.5 | 0.5 |
| SE | 3395 | TECPR2 | TECPR2 | chr14 | + | 102424978 | 102425291 | 102414635 | 102414793 | 102434234 | 102435211 | 36 | 8 | 2 | 0 | 0 | 0 | 0 | 1 | 0 | 3 | 98 | 49 | 1 | 1 | 1 | 1 | 1 | NA | | 0.143 | 0.857 |
| SE | 3396 | TECPR2 | TECPR2 | chr14 | + | 102431795 | 102432128 | 102414635 | 102414793 | 102434234 | 102435211 | 19 | 5 | 3 | 0 | 0 | 0 | 0 | 1 | 0 | 3 | 98 | 49 | 1 | 1 | 1 | 1 | 1 | NA | | 0.143 | 0.857 |
| SE | 3397 | TECPR2 | TECPR2 | chr14 | + | 102452393 | 102452627 | 102450559 | 102450649 | 102465140 | 102465289 | 26 | 4 | 8 | 1 | 0 | 0 | 0 | 6 | 0 | 0 | 98 | 49 | 1 | 1 | 0.929 | 1 | 1 | NA | | 1 | -0.024 |
| SE | 3400 | ZZEF1 | ZZEF1 | chr17 | - | 4056215 | 4056345 | 4054056 | 4054195 | 4057993 | 4058155 | 18 | 7 | 5 | 0 | 0 | 0 | 0 | 2 | 0 | 1 | 98 | 49 | 1 | 1 | 1 | 1 | 1 | NA | | 0.5 | 0.5 |
| SE | 3401 | ZZEF1 | ZZEF1 | chr17 | - | 4072607 | 4072756 | 4070683 | 4070924 | 4074149 | 4074351 | 15 | 2 | 1 | 0 | 5 | 0 | 0 | 2 | 0 | 1 | 98 | 49 | 1 | 1 | 1 | 0.167 | 1 | NA | | 0.5 | 0.222 |
| SE | 3402 | ZZEF1 | ZZEF1 | chr17 | - | 4109652 | 4109863 | 4105692 | 4105809 | 4112608 | 4112808 | 30 | 11 | 3 | 1 | 3 | 0 | 0 | 2 | 0 | 2 | 98 | 49 | 1 | 1 | 0.937 | 0.647 | 1 | NA | | 0.333 | 0.528 |
| SE | 3405 | RSU1 | RSU1 | chr10 | - | 16816972 | 16817084 | 16782033 | 16782084 | 16817314 | 16817454 | 23 | 0 | 0 | 0 | 0 | 1 | 1 | 0 | 0 | 0 | 98 | 49 | 1 | 1 | 1 | NA | 0 | | 1 | NA | -0.5 |
| SE | 3410 | NME8 | NME8 | chr7 | + | 37850377 | 37850435 | 37850259 | 37850299 | 37850628 | 37850735 | 5 | 0 | 0 | 0 | 4 | 0 | 0 | 7 | 0 | 0 | 98 | 49 | 1 | 1 | 1 | 0 | NA | NA | | 1 | -0.5 |
| SE | 3453 | CIITA | CIITA | chr16 | + | 10922166 | 10922250 | 10918439 | 10918526 | 10922406 | 10922490 | 9 | 0 | 0 | 1 | 0 | 0 | 0 | 1 | 0 | 0 | 98 | 49 | 1 | 1 | 0.818 | NA | NA | NA | | 1 | -0.182 |
| SE | 3456 | SLC15A3 | SLC15A3 | chr11 | - | 60942034 | 60942145 | 60939479 | 60939638 | 60943688 | 60943836 | 32 | 13 | 10 | 1 | 0 | 0 | 2 | 6 | 0 | 0 | 98 | 49 | 1 | 1 | 0.941 | 1 | 1 | | 1 | 1 | -0.02 |
| SE | 3457 | PAPOLA | PAPOLA | chr14 | + | 96555846 | 96555947 | 96552479 | 96552622 | 96556174 | 96556413 | 34 | 8 | 15 | 2 | 0 | 0 | 0 | 7 | 0 | 0 | 98 | 49 | 1 | 1 | 0.895 | 1 | 1 | NA | | 1 | -0.035 |
| SE | 3458 | PAPOLA | PAPOLA | chr14 | + | 96555849 | 96555947 | 96552479 | 96552622 | 96556174 | 96556413 | 15 | 0 | 5 | 2 | 0 | 0 | 0 | 5 | 0 | 0 | 98 | 49 | 1 | 1 | 0.789 | NA | 1 | NA | | 1 | -0.105 |
| SE | 3459 | GPR107 | GPR107 | chr9 | + | 130104419 | 130104550 | 130101105 | 130101223 | 130107495 | 130107539 | 5 | 6 | 8 | 0 | 2 | 0 | 0 | 1 | 0 | 0 | 98 | 49 | 1 | 1 | 1 | 0.6 | 1 | NA | | 1 | -0.133 |
| SE | 3462 | CEP162 | CEP162 | chr6 | - | 84169327 | 84169433 | 84163143 | 84163270 | 84171605 | 84171718 | 2 | 1 | 0 | 2 | 0 | 0 | 0 | 1 | 0 | 0 | 98 | 49 | 1 | 1 | 0.333 | 1 | NA | NA | | 1 | -0.334 |
| SE | 3468 | HIATL2 | HIATL2 | chr9 | - | 96972852 | 96972948 | 96958834 | 96958897 | 97013258 | 97013580 | 0 | 0 | 0 | 1 | 0 | 0 | 0 | 1 | 0 | 0 | 98 | 49 | 1 | 1 | 0 | NA | NA | NA | | 1 | -1 |
| SE | 3475 | SUGP2 | SUGP2 | chr19 | - | 19001612 | 19001674 | 18995143 | 18995280 | 19004167 | 19004646 | 4 | 2 | 0 | 1 | 0 | 0 | 0 | 2 | 0 | 0 | 98 | 49 | 1 | 1 | 0.667 | 1 | NA | NA | | 1 | -0.166 |
| SE | 3477 | CTSS | CTSS | chr1 | - | 150750005 | 150750171 | 150747776 | 150747879 | 150751780 | 150752008 | 370 | 220 | 161 | 0 | 0 | 3 | 1 | 129 | 0 | 1 | 98 | 49 | 1 | 1 | 1 | 1 | 0.964 | | 1 | 0.985 | -0.004 |
| SE | 3478 | CTSS | CTSS | chr1 | - | 150750005 | 150750171 | 150747776 | 150747879 | 150755000 | 150755150 | 104 | 71 | 67 | 2 | 0 | 0 | 0 | 58 | 0 | 0 | 98 | 49 | 1 | 1 | 0.963 | 1 | 1 | NA | | 1 | -0.012 |
| SE | 3479 | CTSS | CTSS | chr1 | - | 150751780 | 150752008 | 150747776 | 150747879 | 150755000 | 150755150 | 261 | 144 | 134 | 2 | 0 | 0 | 2 | 110 | 0 | 0 | 98 | 49 | 1 | 1 | 0.985 | 1 | 1 | | 1 | 1 | -0.005 |
| SE | 3480 | CTSS | CTSS | chr1 | - | 150751780 | 150752008 | 150750005 | 150750171 | 150755000 | 150755150 | 527 | 293 | 226 | 0 | 0 | 1 | 3 | 181 | 0 | 1 | 98 | 49 | 1 | 1 | 1 | 1 | 0.991 | | 1 | 0.989 | 0.003 |
| SE | 3481 | CTSS | CTSS | chr1 | - | 150755000 | 150755150 | 150750005 | 150750171 | 150764637 | 150764764 | 1 | 0 | 1 | 0 | 0 | 2 | 0 | 1 | 0 | 0 | 98 | 49 | 1 | 1 | 1 | NA | 0.2 | NA | | 1 | -0.4 |
| SE | 3482 | CTSS | CTSS | chr1 | - | 150755000 | 150755150 | 150751780 | 150752008 | 150757857 | 150757980 | 673 | 276 | 253 | 2 | 0 | 2 | 4 | 234 | 0 | 1 | 98 | 49 | 1 | 1 | 0.994 | 1 | 0.984 | | 1 | 0.992 | -0.003 |
| SE | 3483 | CTSS | CTSS | chr1 | - | 150757857 | 150757980 | 150755000 | 150755150 | 150764637 | 150764764 | 820 | 292 | 228 | 1 | 0 | 0 | 3 | 247 | 0 | 0 | 98 | 49 | 1 | 1 | 0.998 | 1 | 1 | | 1 | 1 | -0.001 |

| | | | | | | | | | | | | | | | | | | | | | | | | | | | | | | | |
|---|---|---|---|---|---|---|---|---|---|---|---|---|---|---|---|---|---|---|---|---|---|---|---|---|---|---|---|---|---|---|---|
| SE | 3488 | PHC2 | PHC2 | chr1 | - | 33331347 | 33331462 | 33330070 | 33330212 | 33332274 | 33332404 | 50 | 47 | 51 | 0 | 3 | 0 | 0 | 12 | 0 | 0 | 98 | 49 | 1 | 1 | 1 | 0.887 | 1 | NA | 1 | -0.038 |
| SE | 3489 | HIPK1 | HIPK1 | chr1 | + | 113952765 | 113952889 | 113940381 | 113941459 | 113955562 | 113955649 | 22 | 12 | 8 | 0 | 2 | 0 | 0 | 5 | 0 | 0 | 98 | 49 | 1 | 1 | 1 | 0.75 | 1 | NA | 1 | -0.083 |
| SE | 3490 | HIPK1 | HIPK1 | chr1 | + | 113954650 | 113954770 | 113952765 | 113952889 | 113955562 | 113955649 | 18 | 24 | 5 | 2 | 4 | 0 | 0 | 6 | 0 | 0 | 98 | 49 | 1 | 1 | 0.818 | 0.75 | 1 | NA | 1 | -0.144 |
| SE | 3491 | HIPK1 | HIPK1 | chr1 | + | 113956626 | 113956811 | 113955562 | 113955649 | 113958065 | 113958291 | 25 | 9 | 1 | 0 | 1 | 0 | 0 | 8 | 0 | 0 | 98 | 49 | 1 | 1 | 1 | 0.818 | 1 | NA | 1 | -0.061 |
| SE | 3492 | HIPK1 | HIPK1 | chr1 | + | 113957123 | 113957286 | 113956626 | 113956811 | 113958065 | 113958291 | 23 | 9 | 4 | 1 | 0 | 0 | 0 | 7 | 0 | 0 | 98 | 49 | 1 | 1 | 0.92 | 1 | 1 | NA | 1 | -0.027 |
| SE | 3499 | MMS19 | MMS19 | chr10 | - | 97461495 | 97461622 | 97460906 | 97461007 | 97461827 | 97461896 | 4 | 1 | 0 | 1 | 0 | 0 | 1 | 3 | 0 | 0 | 98 | 49 | 1 | 1 | 0.667 | 1 | NA | 1 | 1 | -0.166 |
| SE | 3509 | ABCC1 | ABCC1 | chr16 | + | 16124788 | 16124915 | 16121974 | 16122174 | 16125809 | 16125911 | 3 | 6 | 5 | 1 | 0 | 0 | 0 | 1 | 0 | 0 | 98 | 49 | 1 | 1 | 0.6 | 1 | 1 | NA | 1 | -0.133 |
| SE | 3511 | VIPAS39 | VIPAS39 | chr14 | - | 77429680 | 77429767 | 77429005 | 77429095 | 77433841 | 77433931 | 3 | 2 | 0 | 0 | 0 | 1 | 0 | 1 | 0 | 0 | 98 | 49 | 1 | 1 | 1 | 1 | 0 | NA | 1 | -0.333 |
| SE | 3514 | LOC283922 | LOC283922 | chr16 | - | 74334891 | 74334986 | 74332405 | 74332656 | 74336593 | 74336706 | 4 | 0 | 0 | 1 | 0 | 0 | 0 | 1 | 0 | 0 | 98 | 49 | 1 | 1 | 0.667 | NA | NA | NA | 1 | -0.333 |
| SE | 3518 | MTMR14 | MTMR14 | chr3 | + | 9677983 | 9678058 | 9677316 | 9677387 | 9683177 | 9683244 | 32 | 19 | 18 | 10 | 2 | 0 | 0 | 2 | 0 | 1 | 98 | 49 | 1 | 1 | 0.615 | 0.826 | 1 | NA | 0.5 | 0.314 |
| SE | 3519 | MTMR14 | MTMR14 | chr3 | + | 9687820 | 9687891 | 9685210 | 9685247 | 9688695 | 9688754 | 15 | 17 | 6 | 0 | 1 | 0 | 1 | 10 | 0 | 0 | 98 | 49 | 1 | 1 | 1 | 0.895 | 1 | 1 | 1 | -0.035 |
| SE | 3521 | MTMR14 | MTMR14 | chr3 | + | 9697710 | 9697866 | 9689963 | 9690143 | 9701789 | 9702394 | 8 | 5 | 2 | 1 | 5 | 0 | 0 | 2 | 0 | 1 | 98 | 49 | 1 | 1 | 0.8 | 0.333 | 1 | NA | 0.5 | 0.211 |
| SE | 3527 | CFL1 | CFL1 | chr11 | - | 65855653 | 65855730 | 65854813 | 65855448 | 65855934 | 65856242 | 296 | 100 | 133 | 2 | 0 | 0 | 9 | 66 | 0 | 0 | 98 | 49 | 1 | 1 | 0.987 | 1 | 1 | 1 | 1 | -0.004 |
| SE | 3528 | NAA25 | NAA25 | chr12 | - | 112054387 | 112054568 | 112053557 | 112053657 | 112060269 | 112060359 | 1 | 0 | 0 | 0 | 0 | 0 | 0 | 0 | 0 | 1 | 98 | 49 | 1 | 1 | 1 | NA | NA | NA | 0 | 1 |
| SE | 3531 | DLST | DLST | chr14 | + | 74891055 | 74891167 | 74889896 | 74889952 | 74892833 | 74892986 | 2 | 2 | 0 | 2 | 0 | 0 | 0 | 1 | 0 | 0 | 98 | 49 | 1 | 1 | 0.333 | 1 | NA | NA | 1 | -0.334 |
| SE | 3532 | DLST | DLST | chr14 | + | 74894311 | 74894409 | 74893347 | 74893424 | 74898368 | 74898499 | 7 | 3 | 0 | 0 | 3 | 0 | 0 | 2 | 0 | 0 | 98 | 49 | 1 | 1 | 1 | 0.333 | NA | NA | 1 | -0.334 |
| SE | 3533 | NPC2 | NPC2 | chr14 | - | 74484414 | 74484587 | 74480701 | 74480779 | 74486328 | 74486436 | 58 | 22 | 23 | 0 | 0 | 1 | 2 | 26 | 0 | 0 | 98 | 49 | 1 | 1 | 1 | 1 | 0.92 | 1 | 1 | -0.027 |
| SE | 3534 | EIF2AK2 | EIF2AK2 | chr2 | - | 37122505 | 37122664 | 37119958 | 37120139 | 37126288 | 37126411 | 18 | 18 | 8 | 1 | 0 | 0 | 0 | 18 | 0 | 0 | 98 | 49 | 1 | 1 | 0.9 | 1 | 1 | NA | 1 | -0.033 |
| SE | 3539 | ZNF410 | ZNF410 | chr14 | + | 73894358 | 73894409 | 73893796 | 73893932 | 73896315 | 73896534 | 1 | 2 | 0 | 1 | 3 | 0 | 0 | 0 | 0 | 1 | 98 | 49 | 1 | 1 | 0.333 | 0.25 | NA | NA | 0 | 0.291 |
| SE | 3542 | ZNF410 | ZNF410 | chr14 | + | 73922065 | 73922206 | 73920979 | 73921105 | 73923394 | 73923522 | 2 | 0 | 7 | 3 | 3 | 0 | 0 | 2 | 0 | 4 | 98 | 49 | 1 | 1 | 0.25 | 0 | 1 | NA | 0.2 | 0.217 |
| SE | 3546 | PCNX | PCNX | chr14 | + | 70995740 | 70995925 | 70976941 | 70978648 | 71009633 | 71009724 | 5 | 4 | 3 | 5 | 2 | 0 | 0 | 2 | 0 | 0 | 98 | 49 | 1 | 1 | 0.333 | 0.5 | 1 | NA | 1 | -0.389 |
| SE | 3547 | PCNX | PCNX | chr14 | + | 71011491 | 71011549 | 71009633 | 71009724 | 71012999 | 71013202 | 38 | 22 | 2 | 0 | 0 | 0 | 0 | 10 | 0 | 1 | 98 | 49 | 1 | 1 | 1 | 1 | 1 | NA | 0.833 | 0.167 |
| SE | 3548 | PCNX | PCNX | chr14 | + | 71012984 | 71013202 | 71011491 | 71011549 | 71019008 | 71019162 | 107 | 31 | 11 | 0 | 1 | 0 | 2 | 12 | 0 | 0 | 98 | 49 | 1 | 1 | 1 | 0.939 | 1 | 1 | 1 | -0.02 |
| SE | 3549 | PCNX | PCNX | chr14 | + | 71012999 | 71013202 | 71011491 | 71011549 | 71019008 | 71019162 | 61 | 19 | 7 | 0 | 1 | 0 | 2 | 7 | 0 | 0 | 98 | 49 | 1 | 1 | 1 | 0.905 | 1 | 1 | 1 | -0.032 |
| SE | 3561 | SUSD6 | SUSD6 | chr14 | + | 69708676 | 69709104 | 69704603 | 69704742 | 69710953 | 69715144 | 54 | 23 | 15 | 2 | 0 | 0 | 0 | 7 | 0 | 0 | 98 | 49 | 1 | 1 | 0.931 | 1 | 1 | NA | 1 | -0.023 |
| SE | 3563 | SLC39A9 | SLC39A9 | chr14 | + | 69453240 | 69453309 | 69442068 | 69442266 | 69454811 | 69454897 | 3 | 0 | 4 | 0 | 0 | 0 | 0 | 0 | 0 | 1 | 98 | 49 | 1 | 1 | 1 | NA | 1 | NA | 0 | 1 |
| SE | 3568 | DIP2B | DIP2B | chr12 | + | 50671185 | 50671398 | 50660193 | 50660319 | 50674473 | 50674629 | 44 | 16 | 14 | 1 | 0 | 0 | 2 | 5 | 0 | 0 | 98 | 49 | 1 | 1 | 0.957 | 1 | 1 | 1 | 1 | -0.014 |
| SE | 3569 | DIP2B | DIP2B | chr12 | + | 50678678 | 50678876 | 50675328 | 50675448 | 50680671 | 50680763 | 68 | 34 | 8 | 2 | 0 | 0 | 1 | 8 | 0 | 0 | 98 | 49 | 1 | 1 | 0.944 | 1 | 1 | 1 | 1 | -0.019 |
| SE | 3571 | DIP2B | DIP2B | chr12 | + | 50723201 | 50723323 | 50721272 | 50721396 | 50724774 | 50724886 | 54 | 30 | 17 | 0 | 1 | 0 | 0 | 12 | 0 | 0 | 98 | 49 | 1 | 1 | 1 | 0.937 | 1 | NA | 1 | -0.021 |
| SE | 3572 | DIP2B | DIP2B | chr12 | + | 50731368 | 50731537 | 50728547 | 50728678 | 50732365 | 50732536 | 27 | 15 | 2 | 0 | 1 | 0 | 0 | 4 | 0 | 0 | 98 | 49 | 1 | 1 | 1 | 0.882 | 1 | NA | 1 | -0.039 |
| SE | 3573 | ACTN1 | ACTN1 | chr14 | - | 68878457 | 68878523 | 68877081 | 68877240 | 68878988 | 68879069 | 0 | 1 | 0 | 44 | 25 | 15 | 0 | 0 | 1 | 9 | 98 | 49 | 1 | 1 | 0 | 0.02 | 0 | 0 | 0 | 0.007 |
| SE | 3575 | ACTN1 | ACTN1 | chr14 | - | 68882872 | 68883055 | 68882457 | 68882592 | 68884167 | 68884308 | 68 | 39 | 22 | 2 | 0 | 0 | 0 | 14 | 0 | 0 | 98 | 49 | 1 | 1 | 0.944 | 1 | 1 | NA | 1 | -0.019 |
| SE | 3577 | MIDN | MIDN | chr19 | + | 1251561 | 1251649 | 1249889 | 1250529 | 1251838 | 1251901 | 60 | 24 | 6 | 0 | 0 | 0 | 6 | 6 | 0 | 1 | 98 | 49 | 1 | 1 | 1 | 1 | 1 | 1 | 0.75 | 0.125 |
| SE | 3581 | ERN1 | ERN1 | chr17 | - | 64066670 | 64066932 | 64065208 | 64065287 | 64068189 | 64068291 | 14 | 3 | 3 | 0 | 1 | 0 | 0 | 3 | 0 | 0 | 98 | 49 | 1 | 1 | 1 | 0.6 | 1 | NA | 1 | -0.133 |
| SE | 3582 | ERN1 | ERN1 | chr17 | - | 64068189 | 64068291 | 64065208 | 64065287 | 64071980 | 64072103 | 15 | 3 | 0 | 2 | 0 | 1 | 0 | 3 | 0 | 0 | 98 | 49 | 1 | 1 | 0.789 | 1 | 0 | NA | 1 | -0.404 |
| SE | 3583 | RDH11 | RDH11 | chr14 | - | 67685014 | 67685204 | 67676800 | 67678423 | 67690211 | 67690421 | 5 | 0 | 0 | 2 | 0 | 0 | 0 | 1 | 0 | 0 | 98 | 49 | 1 | 1 | 0.556 | NA | NA | NA | 1 | -0.444 |
| SE | 3585 | RDH11 | RDH11 | chr14 | - | 67692437 | 67692593 | 67691139 | 67691244 | 67692933 | 67693052 | 8 | 0 | 0 | 1 | 0 | 0 | 0 | 2 | 0 | 0 | 98 | 49 | 1 | 1 | 0.8 | NA | NA | NA | 1 | -0.2 |
| SE | 3593 | HDAC5 | HDAC5 | chr17 | - | 44085021 | 44085155 | 44084554 | 44084675 | 44086571 | 44086737 | 34 | 15 | 11 | 2 | 0 | 1 | 0 | 4 | 0 | 0 | 98 | 49 | 1 | 1 | 0.895 | 1 | 0.846 | NA | 1 | -0.086 |
| SE | 3595 | HDAC5 | HDAC5 | chr17 | - | 44117493 | 44117704 | 44110728 | 44110800 | 44123503 | 44123646 | 1 | 5 | 2 | 0 | 3 | 0 | 0 | 1 | 0 | 0 | 98 | 49 | 1 | 1 | 1 | 0.455 | 1 | NA | 1 | -0.182 |
| SE | 3596 | HDAC5 | HDAC5 | chr17 | - | 44117493 | 44117704 | 44110728 | 44110803 | 44123503 | 44123646 | 4 | 6 | 1 | 1 | 0 | 0 | 0 | 1 | 0 | 0 | 98 | 49 | 1 | 1 | 0.667 | 1 | 1 | NA | 1 | -0.111 |
| SE | 3603 | CHURC1 | CHURC1 | chr14 | + | 64926006 | 64926080 | 64923990 | 64924126 | 64932137 | 64935366 | 13 | 13 | 5 | 0 | 2 | 4 | 0 | 2 | 0 | 2 | 98 | 49 | 1 | 1 | 1 | 0.765 | 0.385 | NA | 0.333 | 0.384 |
| SE | 3604 | CHURC1 | CHURC1 | chr14 | + | 64926009 | 64926080 | 64923990 | 64924126 | 64932137 | 64935366 | 25 | 24 | 10 | 0 | 2 | 4 | 0 | 2 | 0 | 2 | 98 | 49 | 1 | 1 | 1 | 0.857 | 0.556 | NA | 0.333 | 0.471 |
| SE | 3620 | PPM1A | PPM1A | chr14 | + | 60250379 | 60250481 | 60249247 | 60249677 | 60282683 | 60283537 | 4 | 0 | 1 | 2 | 3 | 4 | 0 | 0 | 0 | 1 | 98 | 49 | 1 | 1 | 0.5 | 0 | 0.111 | NA | 0 | 0.204 |
| SE | 3621 | PPM1A | PPM1A | chr14 | + | 60282683 | 60283537 | 60249247 | 60249677 | 60285623 | 60285741 | 20 | 9 | 9 | 0 | 1 | 0 | 0 | 4 | 0 | 0 | 98 | 49 | 1 | 1 | 1 | 0.818 | 1 | NA | 1 | -0.061 |
| SE | 3623 | MIR22HG | MIR22HG | chr17 | - | 1714370 | 1714453 | 1713702 | 1714014 | 1716129 | 1716272 | 11 | 11 | 0 | 3 | 7 | 0 | 0 | 1 | 0 | 0 | 98 | 49 | 1 | 1 | 0.647 | 0.44 | NA | NA | 1 | -0.457 |
| SE | 3627 | AP5M1 | AP5M1 | chr14 | + | 57274243 | 57274889 | 57268887 | 57269388 | 57280194 | 57280422 | 9 | 5 | 1 | 0 | 0 | 0 | 0 | 3 | 0 | 1 | 98 | 49 | 1 | 1 | 1 | 1 | 1 | NA | 0.6 | 0.4 |
| SE | 3633 | SFT2D1 | SFT2D1 | chr6 | - | 166329506 | 166329589 | 166328275 | 166328357 | 166330160 | 166330247 | 15 | 11 | 3 | 1 | 0 | 0 | 1 | 4 | 0 | 0 | 98 | 49 | 1 | 1 | 0.882 | 1 | 1 | 1 | 1 | -0.039 |
| SE | 3635 | SFT2D1 | SFT2D1 | chr6 | - | 166331384 | 166331407 | 166330160 | 166330247 | 166342418 | 166342606 | 0 | 0 | 0 | 4 | 1 | 2 | 0 | 1 | 0 | 1 | 72 | 49 | 1 | 1 | 0 | 0 | 0 | NA | 0.405 | -0.405 |
| SE | 3659 | SOS2 | SOS2 | chr14 | - | 50145169 | 50145332 | 50139941 | 50140059 | 50145476 | 50145596 | 81 | 33 | 4 | 1 | 0 | 0 | 1 | 9 | 0 | 1 | 98 | 49 | 1 | 1 | 0.976 | 1 | 1 | 1 | 0.818 | 0.083 |
| SE | 3661 | SOS2 | SOS2 | chr14 | - | 50174453 | 50174552 | 50161481 | 50161609 | 50180571 | 50180682 | 91 | 37 | 15 | 2 | 0 | 0 | 1 | 8 | 0 | 0 | 98 | 49 | 1 | 1 | 0.958 | 1 | 1 | 1 | 1 | -0.014 |
| SE | 3662 | SOS2 | SOS2 | chr14 | - | 50180571 | 50180682 | 50174453 | 50174552 | 50182462 | 50182606 | 123 | 54 | 17 | 1 | 0 | 0 | 3 | 10 | 0 | 0 | 98 | 49 | 1 | 1 | 0.984 | 1 | 1 | 1 | 1 | -0.005 |
| SE | 3663 | SOS2 | SOS2 | chr14 | - | 50188496 | 50188700 | 50182462 | 50182606 | 50199690 | 50199855 | 72 | 25 | 16 | 0 | 1 | 0 | 1 | 13 | 0 | 0 | 98 | 49 | 1 | 1 | 1 | 0.926 | 1 | 1 | 1 | -0.025 |
| SE | 3664 | VCPKMT | VCPKMT | chr14 | - | 50112614 | 50112719 | 50108631 | 50109713 | 50114284 | 50114404 | 21 | 10 | 0 | 1 | 0 | 0 | 0 | 3 | 0 | 0 | 98 | 49 | 1 | 1 | 0.913 | 1 | NA | NA | 1 | -0.043 |
| SE | 3673 | MIS18BP1 | MIS18BP1 | chr14 | - | 45206082 | 45206170 | 45204398 | 45204453 | 45210379 | 45210528 | 6 | 1 | 0 | 0 | 0 | 0 | 0 | 5 | 0 | 2 | 98 | 49 | 1 | 1 | 1 | 1 | NA | NA | 0.556 | 0.444 |
| SE | 3686 | TPP2 | TPP2 | chr13 | + | 102635586 | 102635702 | 102633949 | 102634098 | 102636223 | 102636392 | 8 | 3 | 0 | 2 | 0 | 0 | 0 | 1 | 0 | 0 | 98 | 49 | 1 | 1 | 0.667 | 1 | NA | NA | 1 | -0.166 |
| SE | 3690 | G2E3 | G2E3 | chr14 | + | 30605504 | 30605812 | 30601998 | 30602131 | 30607887 | 30608069 | 1 | 2 | 7 | 1 | 0 | 0 | 0 | 2 | 0 | 0 | 98 | 49 | 1 | 1 | 0.333 | 1 | 1 | NA | 1 | -0.222 |
| SE | 3691 | TRRAP | TRRAP | chr7 | + | 98899678 | 98899767 | 98899421 | 98899499 | 98900623 | 98900720 | 1 | 1 | 0 | 1 | 0 | 0 | 0 | 3 | 0 | 0 | 98 | 49 | 1 | 1 | 0.333 | 1 | NA | NA | 1 | -0.334 |
| SE | 3692 | TRRAP | TRRAP | chr7 | + | 98945746 | 98945800 | 98942948 | 98943017 | 98948220 | 98948340 | 2 | 0 | 0 | 0 | 0 | 0 | 0 | 0 | 0 | 3 | 98 | 49 | 1 | 1 | 1 | NA | NA | NA | 0 | 1 |
| SE | 3693 | YY1 | YY1 | chr14 | + | 100276489 | 100276648 | 100274697 | 100274758 | 100277417 | 100279034 | 25 | 16 | 5 | 0 | 0 | 0 | 0 | 2 | 0 | 1 | 98 | 49 | 1 | 1 | 1 | 1 | 1 | NA | 0.5 | 0.5 |
| SE | 3701 | TCP11L2 | TCP11L2 | chr12 | + | 106335638 | 106335826 | 106323509 | 106323646 | 106336031 | 106336213 | 37 | 37 | 7 | 1 | 0 | 0 | 0 | 7 | 0 | 0 | 98 | 49 | 1 | 1 | 0.949 | 1 | 1 | NA | 1 | -0.017 |
| SE | 3704 | CDK9 | CDK9 | chr9 | + | 127787946 | 127788113 | 127787517 | 127787608 | 127788213 | 127788385 | 25 | 13 | 1 | 2 | 0 | 0 | 3 | 8 | 0 | 0 | 98 | 49 | 1 | 1 | 0.862 | 1 | 1 | 1 | 1 | -0.046 |
| SE | 3707 | IRF9 | IRF9 | chr14 | + | 24163377 | 24163508 | 24162965 | 24163149 | 24164062 | 24164134 | 55 | 65 | 7 | 1 | 1 | 0 | 3 | 26 | 0 | 0 | 98 | 49 | 1 | 1 | 0.965 | 0.97 | 1 | 1 | 1 | -0.022 |
| SE | 3710 | RNF31 | RNF31 | chr14 | + | 24159863 | 24159960 | 24158141 | 24158199 | 24160238 | 24160407 | 3 | 0 | 1 | 1 | 0 | 0 | 0 | 1 | 0 | 0 | 98 | 49 | 1 | 1 | 0.6 | NA | 1 | NA | 1 | -0.2 |
| SE | 3712 | TNIP1 | TNIP1 | chr5 | - | 151033607 | 151033799 | 151032286 | 151032383 | 151035001 | 151035067 | 18 | 14 | 8 | 0 | 0 | 1 | 1 | 5 | 0 | 0 | 98 | 49 | 1 | 1 | 1 | 1 | 0.8 | 1 | 1 | -0.067 |
| SE | 3713 | TNIP1 | TNIP1 | chr5 | - | 151045860 | 151045950 | 151042895 | 151042961 | 151049823 | 151049947 | 67 | 21 | 27 | 2 | 0 | 0 | 0 | 6 | 0 | 0 | 98 | 49 | 1 | 1 | 0.944 | 1 | 1 | NA | 1 | -0.019 |
| SE | 3716 | DCAF11 | DCAF11 | chr14 | + | 24116916 | 24117044 | 24115381 | 24115749 | 24117265 | 24117393 | 9 | 13 | 2 | 1 | 6 | 0 | 0 | 3 | 0 | 1 | 98 | 49 | 1 | 1 | 0.818 | 0.52 | 1 | NA | 0.6 | 0.179 |
| SE | 3717 | DCAF11 | DCAF11 | chr14 | + | 24116971 | 24117044 | 24115282 | 24115749 | 24117265 | 24117393 | 6 | 7 | 2 | 1 | 6 | 0 | 0 | 3 | 0 | 1 | 98 | 49 | 1 | 1 | 0.75 | 0.368 | 1 | NA | 0.6 | 0.106 |
| SE | 3718 | DCAF11 | DCAF11 | chr14 | + | 24117667 | 24117732 | 24117265 | 24117393 | 24118054 | 24118155 | 15 | 5 | 5 | 1 | 0 | 0 | 0 | 2 | 0 | 1 | 98 | 49 | 1 | 1 | 0.882 | 1 | 1 | NA | 0.5 | 0.461 |
| SE | 3719 | DCAF11 | DCAF11 | chr14 | + | 24120837 | 24120991 | 24119710 | 24119896 | 24121364 | 24121517 | 4 | 9 | 0 | 0 | 0 | 0 | 0 | 4 | 1 | 0 | 98 | 49 | 1 | 1 | 1 | 1 | NA | 0 | 1 | 0.5 |
| SE | 3724 | ADGRG3 | ADGRG3 | chr16 | + | 57679176 | 57679311 | 57678169 | 57678316 | 57679815 | 57679855 | 63 | 48 | 24 | 0 | 0 | 1 | 0 | 15 | 1 | 0 | 98 | 49 | 1 | 1 | 1 | 1 | 0.923 | 0 | 1 | 0.474 |
| SE | 3725 | ADGRG3 | ADGRG3 | chr16 | + | 57683931 | 57684212 | 57680504 | 57680617 | 57684389 | 57684483 | 41 | 4 | 9 | 6 | 0 | 3 | 0 | 1 | 0 | 1 | 98 | 49 | 1 | 1 | 0.774 | 1 | 0.6 | NA | 0.333 | 0.458 |
| SE | 3727 | ADGRG3 | ADGRG3 | chr16 | + | 57685642 | 57685926 | 57680504 | 57680617 | 57688351 | 57689378 | 41 | 13 | 13 | 1 | 0 | 0 | 4 | 8 | 0 | 0 | 98 | 49 | 1 | 1 | 0.953 | 1 | 1 | 1 | 1 | -0.016 |
| SE | 3731 | NDFIP1 | NDFIP1 | chr5 | + | 142137733 | 142137858 | 142135729 | 142135817 | 142140562 | 142140629 | 24 | 7 | 9 | 1 | 0 | 0 | 0 | 4 | 0 | 1 | 98 | 49 | 1 | 1 | 0.923 | 1 | 1 | NA | 0.667 | 0.307 |
| SE | 3732 | HAUS4 | HAUS4 | chr14 | - | 22951554 | 22951689 | 22950313 | 22950410 | 22952327 | 22952459 | 25 | 12 | 10 | 7 | 1 | 2 | 0 | 16 | 0 | 1 | 98 | 49 | 1 | 1 | 0.641 | 0.857 | 0.714 | NA | 0.889 | -0.152 |
| SE | 3733 | HAUS4 | HAUS4 | chr14 | - | 22952540 | 22952683 | 22952327 | 22952459 | 22955099 | 22955176 | 32 | 20 | 16 | 0 | 4 | 0 | 0 | 8 | 0 | 0 | 98 | 49 | 1 | 1 | 1 | 0.714 | 1 | NA | 1 | -0.095 |
| SE | 3734 | DDX58 | DDX58 | chr9 | - | 32466289 | 32466441 | 32459370 | 32459514 | 32467761 | 32467932 | 40 | 17 | 10 | 1 | 0 | 0 | 0 | 23 | 0 | 0 | 98 | 49 | 1 | 1 | 0.952 | 1 | 1 | NA | 1 | -0.016 |
| SE | 3735 | DDX58 | DDX58 | chr9 | - | 32467761 | 32467932 | 32459370 | 32459514 | 32472974 | 32473065 | 17 | 13 | 3 | 2 | 0 | 0 | 0 | 7 | 0 | 0 | 98 | 49 | 1 | 1 | 0.81 | 1 | 1 | NA | 1 | -0.063 |
| SE | 3736 | DDX58 | DDX58 | chr9 | - | 32488733 | 32488887 | 32487944 | 32488203 | 32489343 | 32489451 | 40 | 34 | 7 | 1 | 0 | 0 | 0 | 14 | 0 | 0 | 98 | 49 | 1 | 1 | 0.952 | 1 | 1 | NA | 1 | -0.016 |
| SE | 3737 | DDX58 | DDX58 | chr9 | - | 32492390 | 32492538 | 32491300 | 32491420 | 32493760 | 32493942 | 41 | 26 | 2 | 6 | 4 | 0 | 0 | 8 | 0 | 1 | 98 | 49 | 1 | 1 | 0.774 | 0.765 | 1 | NA | 0.8 | 0.046 |
| SE | 3739 | RPGRIP1 | RPGRIP1 | chr14 | + | 21343035 | 21343228 | 21334604 | 21334705 | 21345112 | 21345197 | 19 | 7 | 0 | 1 | 0 | 0 | 0 | 2 | 0 | 0 | 98 | 49 | 1 | 1 | 0.905 | 1 | NA | NA | 1 | -0.047 |
| SE | 3741 | ARHGEF40 | ARHGEF40 | chr14 | + | 21087319 | 21087463 | 21087000 | 21087105 | 21087967 | 21088098 | 13 | 8 | 5 | 0 | 0 | 1 | 0 | 4 | 0 | 1 | 98 | 49 | 1 | 1 | 1 | 1 | 0.714 | NA | 0.667 | 0.238 |
| SE | 3750 | FANCA | FANCA | chr16 | - | 89752137 | 89752222 | 89749729 | 89749902 | 89758576 | 89758705 | 0 | 2 | 1 | 0 | 0 | 1 | 0 | 1 | 0 | 0 | 98 | 49 | 1 | 1 | NA | 1 | 0.333 | NA | 1 | -0.334 |
| SE | 3751 | KDM5B | KDM5B | chr1 | - | 202746141 | 202746323 | 202745857 | 202745982 | 202748944 | 202749139 | 8 | 5 | 0 | 2 | 0 | 0 | 0 | 1 | 0 | 0 | 98 | 49 | 1 | 1 | 0.667 | 1 | NA | NA | 1 | -0.166 |
| SE | 3760 | TINF2 | TINF2 | chr14 | - | 24241889 | 24241994 | 24241674 | 24241776 | 24242140 | 24242674 | 10 | 13 | 0 | 7 | 4 | 0 | 1 | 0 | 0 | 0 | 98 | 49 | 1 | 1 | 0.417 | 0.619 | NA | 1 | NA | -0.482 |
| SE | 3765 | CD47 | CD47 | chr3 | - | 108050577 | 108050602 | 108043093 | 108047292 | 108051938 | 108051970 | 3 | 6 | 3 | 3 | 6 | 1 | 0 | 1 | 0 | 1 | 74 | 49 | 1 | 1 | 0.398 | 0.398 | 0.665 | NA | 0.398 | 0.089 |
| SE | 3769 | PSME2 | PSME2 | chr14 | - | 24143974 | 24144029 | 24143584 | 24143671 | 24144191 | 24144259 | 20 | 6 | 2 | 1 | 0 | 0 | 1 | 5 | 0 | 0 | 98 | 49 | 1 | 1 | 0.909 | 1 | 1 | 1 | 1 | -0.03 |
| SE | 3772 | PIEZO1 | PIEZO1 | chr16 | - | 88725008 | 88725080 | 88723870 | 88723971 | 88725415 | 88725519 | 3 | 0 | 0 | 1 | 0 | 0 | 0 | 1 | 0 | 0 | 98 | 49 | 1 | 1 | 0.6 | NA | NA | NA | 1 | -0.4 |
| SE | 3777 | CARS2 | CARS2 | chr13 | - | 110651033 | 110651100 | 110647100 | 110647239 | 110663450 | 110663518 | 21 | 15 | 14 | 2 | 0 | 0 | 0 | 5 | 0 | 0 | 98 | 49 | 1 | 1 | 0.84 | 1 | 1 | NA | 1 | -0.053 |
| SE | 3782 | PLEKHB2 | PLEKHB2 | chr2 | + | 131144407 | 131144449 | 131140166 | 131140275 | 131146636 | 131149852 | 3 | 1 | 0 | 10 | 1 | 5 | 0 | 0 | 0 | 3 | 91 | 49 | 1 | 1 | 0.139 | 0.35 | 0 | NA | 0 | 0.163 |
| SE | 3785 | RGS14 | RGS14 | chr5 | + | 177367713 | 177367825 | 177367413 | 177367557 | 177368156 | 177368266 | 23 | 14 | 6 | 0 | 1 | 0 | 1 | 12 | 0 | 0 | 98 | 49 | 1 | 1 | 1 | 0.875 | 1 | 1 | 1 | -0.042 |
| SE | 3801 | THBS1 | THBS1 | chr15 | + | 39588525 | 39588699 | 39588041 | 39588218 | 39588958 | 39589086 | 18 | 0 | 1 | 0 | 0 | 0 | 2 | 8 | 0 | 4 | 98 | 49 | 1 | 1 | 1 | NA | 1 | 1 | 0.5 | 0.25 |
| SE | 3804 | LINC00963 | LINC00963 | chr9 | + | 129493477 | 129493595 | 129488659 | 129489299 | 129503322 | 129503672 | 9 | 1 | 2 | 1 | 1 | 0 | 0 | 2 | 0 | 0 | 98 | 49 | 1 | 1 | 0.818 | 0.333 | 1 | NA | 1 | -0.283 |
| SE | 3805 | LINC00963 | LINC00963 | chr9 | + | 129502396 | 129503139 | 129493477 | 129493595 | 129503322 | 129503672 | 1 | 2 | 0 | 9 | 1 | 2 | 0 | 0 | 0 | 2 | 98 | 49 | 1 | 1 | 0.053 | 0.5 | 0 | NA | 0 | 0.184 |
| SE | 3807 | SLC25A1 | SLC25A1 | chr22 | - | 19177941 | 19178041 | 19177119 | 19177204 | 19178132 | 19178240 | 1 | 0 | 0 | 2 | 0 | 0 | 0 | 1 | 0 | 0 | 98 | 49 | 1 | 1 | 0.2 | NA | NA | NA | 1 | -0.8 |
| SE | 3808 | BIN2 | BIN2 | chr12 | - | 51295795 | 51295878 | 51291590 | 51292344 | 51297088 | 51297164 | 175 | 64 | 47 | 1 | 0 | 0 | 3 | 28 | 0 | 0 | 98 | 49 | 1 | 1 | 0.989 | 1 | 1 | 1 | 1 | -0.004 |
| SE | 3810 | BIN2 | BIN2 | chr12 | - | 51313822 | 51313903 | 51303086 | 51303141 | 51324021 | 51324198 | 60 | 24 | 22 | 1 | 0 | 0 | 0 | 5 | 0 | 0 | 98 | 49 | 1 | 1 | 0.968 | 1 | 1 | NA | 1 | -0.011 |
| SE | 3812 | MYO18A | MYO18A | chr17 | - | 29086437 | 29086577 | 29082315 | 29082438 | 29086935 | 29087121 | 0 | 2 | 0 | 0 | 2 | 0 | 0 | 3 | 0 | 0 | 98 | 49 | 1 | 1 | NA | 0.333 | NA | NA | 1 | -0.667 |
| SE | 3823 | ANP32A | ANP32A | chr15 | - | 68782955 | 68783053 | 68780409 | 68780473 | 68784396 | 68784595 | 42 | 11 | 11 | 0 | 1 | 0 | 2 | 5 | 0 | 0 | 98 | 49 | 1 | 1 | 1 | 0.846 | 1 | 1 | 1 | -0.051 |
| SE | 3825 | PTPRE | PTPRE | chr10 | + | 128066074 | 128066194 | 128063082 | 128063180 | 128068122 | 128068286 | 182 | 99 | 13 | 1 | 2 | 0 | 4 | 51 | 0 | 0 | 98 | 49 | 1 | 1 | 0.989 | 0.961 | 1 | 1 | 1 | -0.017 |
| SE | 3826 | PTPRE | PTPRE | chr10 | + | 128076602 | 128076728 | 128073336 | 128073471 | 128077616 | 128077783 | 147 | 62 | 23 | 0 | 1 | 0 | 3 | 39 | 0 | 0 | 98 | 49 | 1 | 1 | 1 | 0.969 | 1 | 1 | 1 | -0.01 |
| SE | 3827 | PTPRE | PTPRE | chr10 | + | 128079559 | 128079695 | 128077616 | 128077783 | 128082831 | 128085900 | 77 | 47 | 14 | 1 | 0 | 0 | 2 | 19 | 0 | 0 | 98 | 49 | 1 | 1 | 0.975 | 1 | 1 | 1 | 1 | -0.008 |
| SE | 3846 | C17orf85 | C17orf85 | chr17 | - | 3814321 | 3814483 | 3806750 | 3813279 | 3816115 | 3816270 | 9 | 3 | 1 | 0 | 0 | 0 | 1 | 1 | 0 | 1 | 98 | 49 | 1 | 1 | 1 | 1 | 1 | 1 | 0.333 | 0.334 |
| SE | 3852 | MYCBP2 | MYCBP2 | chr13 | - | 77095357 | 77095602 | 77093164 | 77093332 | 77096311 | 77096481 | 11 | 2 | 0 | 0 | 0 | 0 | 0 | 1 | 0 | 1 | 98 | 49 | 1 | 1 | 1 | 1 | NA | NA | 0.333 | 0.667 |
| SE | 3853 | MYCBP2 | MYCBP2 | chr13 | - | 77126317 | 77126542 | 77125335 | 77125468 | 77139195 | 77139336 | 5 | 1 | 0 | 2 | 2 | 0 | 0 | 2 | 1 | 0 | 98 | 49 | 1 | 1 | 0.556 | 0.2 | NA | 0 | 1 | -0.122 |
| SE | 3854 | MYCBP2 | MYCBP2 | chr13 | - | 77233155 | 77233263 | 77225434 | 77225554 | 77243058 | 77243160 | 20 | 6 | 3 | 1 | 0 | 0 | 0 | 2 | 0 | 0 | 98 | 49 | 1 | 1 | 0.909 | 1 | 1 | NA | 1 | -0.03 |
| SE | 3856 | ITCH | ITCH | chr20 | + | 34393790 | 34393881 | 34369393 | 34369470 | 34408650 | 34408792 | 31 | 16 | 4 | 1 | 0 | 0 | 0 | 6 | 0 | 0 | 98 | 49 | 1 | 1 | 0.939 | 1 | 1 | NA | 1 | -0.02 |
| SE | 3857 | ITCH | ITCH | chr20 | + | 34408650 | 34408792 | 34393790 | 34393881 | 34412514 | 34412639 | 49 | 38 | 20 | 0 | 0 | 0 | 0 | 5 | 0 | 1 | 98 | 49 | 1 | 1 | 1 | 1 | 1 | NA | 0.714 | 0.286 |
| SE | 3858 | ITCH | ITCH | chr20 | + | 34417116 | 34417239 | 34413741 | 34413879 | 34424479 | 34424525 | 1 | 0 | 0 | 30 | 36 | 4 | 0 | 3 | 0 | 3 | 98 | 49 | 1 | 1 | 0.016 | 0 | 0 | NA | 0.333 | -0.328 |
| SE | 3859 | ITCH | ITCH | chr20 | + | 34424479 | 34424525 | 34413741 | 34413879 | 34438473 | 34438631 | 67 | 44 | 15 | 4 | 0 | 0 | 0 | 5 | 0 | 0 | 95 | 49 | 1 | 1 | 0.896 | 1 | 1 | NA | 1 | -0.035 |
| SE | 3860 | ITCH | ITCH | chr20 | + | 34457389 | 34457474 | 34449410 | 34449480 | 34462092 | 34462221 | 33 | 28 | 9 | 0 | 0 | 0 | 0 | 3 | 0 | 1 | 98 | 49 | 1 | 1 | 1 | 1 | 1 | NA | 0.6 | 0.4 |
| SE | 3861 | COMMD6 | COMMD6 | chr13 | - | 75527812 | 75527851 | 75525213 | 75526639 | 75530113 | 75530266 | 0 | 2 | 0 | 13 | 2 | 2 | 0 | 0 | 1 | 2 | 88 | 49 | 1 | 1 | 0 | 0.358 | 0 | 0 | 0 | 0.119 |
| SE | 3865 | SYNJ1 | SYNJ1 | chr21 | - | 32699837 | 32700105 | 32695056 | 32695282 | 32701960 | 32702047 | 9 | 2 | 1 | 3 | 0 | 0 | 0 | 1 | 0 | 1 | 98 | 49 | 1 | 1 | 0.6 | 1 | 1 | NA | 0.333 | 0.534 |
| SE | 3866 | PSAP | PSAP | chr10 | - | 71819030 | 71819111 | 71818616 | 71818724 | 71819464 | 71819622 | 436 | 98 | 134 | 1 | 0 | 0 | 15 | 181 | 0 | 0 | 98 | 49 | 1 | 1 | 0.995 | 1 | 1 | 1 | 1 | -0.002 |
| SE | 3867 | PSAP | PSAP | chr10 | - | 71825836 | 71825893 | 71821875 | 71822007 | 71828013 | 71828157 | 520 | 148 | 225 | 0 | 0 | 1 | 10 | 255 | 0 | 0 | 98 | 49 | 1 | 1 | 1 | 1 | 0.991 | 1 | 1 | -0.003 |
| SE | 3876 | PPP1R16B | PPP1R16B | chr20 | + | 38900580 | 38900684 | 38895564 | 38895710 | 38902667 | 38902792 | 1 | 0 | 0 | 2 | 0 | 0 | 0 | 2 | 0 | 0 | 98 | 49 | 1 | 1 | 0.2 | NA | NA | NA | 1 | -0.8 |
| SE | 3878 | PSMB9 | PSMB9 | chr6 | + | 32857262 | 32857394 | 32856137 | 32856205 | 32858004 | 32858134 | 37 | 41 | 13 | 1 | 0 | 0 | 0 | 23 | 0 | 0 | 98 | 49 | 1 | 1 | 0.949 | 1 | 1 | NA | 1 | -0.017 |
| SE | 3879 | PSMB9 | PSMB9 | chr6 | + | 32858363 | 32858505 | 32858004 | 32858134 | 32859404 | 32859851 | 36 | 36 | 49 | 0 | 3 | 0 | 0 | 16 | 0 | 0 | 98 | 49 | 1 | 1 | 1 | 0.857 | 1 | NA | 1 | -0.048 |
| SE | 3889 | ST3GAL1 | ST3GAL1 | chr8 | - | 133475718 | 133476600 | 133465893 | 133466090 | 133499134 | 133499189 | 11 | 7 | 0 | 0 | 0 | 0 | 1 | 1 | 0 | 3 | 98 | 49 | 1 | 1 | 1 | 1 | NA | 1 | 0.143 | 0.428 |
| SE | 3893 | MCM9 | MCM9 | chr6 | - | 118931419 | 118931738 | 118923810 | 118924127 | 118934890 | 118935162 | 4 | 4 | 0 | 1 | 0 | 0 | 3 | 1 | 0 | 0 | 98 | 49 | 1 | 1 | 0.667 | 1 | NA | 1 | 1 | -0.166 |
| SE | 3895 | INTS6 | INTS6 | chr13 | - | 51421111 | 51421334 | 51395299 | 51395483 | 51423029 | 51423090 | 3 | 1 | 0 | 0 | 0 | 0 | 0 | 0 | 0 | 2 | 98 | 49 | 1 | 1 | 1 | 1 | NA | NA | 0 | 1 |
| SE | 3911 | ATF6 | ATF6 | chr1 | + | 161791407 | 161791537 | 161783989 | 161784096 | 161792123 | 161792327 | 97 | 30 | 3 | 2 | 0 | 0 | 0 | 16 | 0 | 0 | 98 | 49 | 1 | 1 | 0.96 | 1 | 1 | NA | 1 | -0.013 |
| SE | 3912 | ATF6 | ATF6 | chr1 | + | 161802051 | 161802272 | 161792123 | 161792327 | 161819632 | 161819818 | 47 | 7 | 7 | 2 | 0 | 0 | 0 | 8 | 0 | 0 | 98 | 49 | 1 | 1 | 0.922 | 1 | 1 | NA | 1 | -0.026 |
| SE | 3913 | ATF6 | ATF6 | chr1 | + | 161860206 | 161860277 | 161853223 | 161853323 | 161863197 | 161863312 | 53 | 15 | 4 | 2 | 0 | 0 | 0 | 17 | 0 | 0 | 98 | 49 | 1 | 1 | 0.93 | 1 | 1 | NA | 1 | -0.023 |
| SE | 3916 | EPG5 | EPG5 | chr18 | - | 45879012 | 45879214 | 45878375 | 45878448 | 45880074 | 45880223 | 5 | 1 | 8 | 0 | 1 | 0 | 0 | 3 | 0 | 0 | 98 | 49 | 1 | 1 | 1 | 0.333 | 1 | NA | 1 | -0.222 |
| SE | 3917 | EPG5 | EPG5 | chr18 | - | 45880074 | 45880223 | 45879012 | 45879214 | 45882273 | 45882487 | 2 | 4 | 3 | 0 | 0 | 1 | 0 | 4 | 0 | 0 | 98 | 49 | 1 | 1 | 1 | 1 | 0.6 | NA | 1 | -0.133 |
| SE | 3919 | EPG5 | EPG5 | chr18 | - | 45889797 | 45889940 | 45887750 | 45887907 | 45899403 | 45899566 | 10 | 3 | 0 | 1 | 0 | 0 | 0 | 1 | 0 | 0 | 98 | 49 | 1 | 1 | 0.833 | 1 | NA | NA | 1 | -0.084 |
| SE | 3920 | EPG5 | EPG5 | chr18 | - | 45916008 | 45916206 | 45915510 | 45915621 | 45916437 | 45916582 | 4 | 2 | 0 | 1 | 0 | 0 | 0 | 1 | 0 | 0 | 98 | 49 | 1 | 1 | 0.667 | 1 | NA | NA | 1 | -0.166 |
| SE | 3922 | SPG11 | SPG11 | chr15 | - | 44572682 | 44572820 | 44570524 | 44570658 | 44583813 | 44584558 | 16 | 0 | 2 | 0 | 1 | 0 | 0 | 10 | 0 | 0 | 98 | 49 | 1 | 1 | 1 | 0 | 1 | NA | 1 | -0.333 |
| SE | 3923 | SPG11 | SPG11 | chr15 | - | 44589251 | 44589414 | 44585635 | 44585850 | 44592330 | 44592438 | 17 | 4 | 7 | 1 | 0 | 0 | 0 | 2 | 0 | 0 | 98 | 49 | 1 | 1 | 0.895 | 1 | 1 | NA | 1 | -0.035 |
| SE | 3924 | SPG11 | SPG11 | chr15 | - | 44598630 | 44598836 | 44598264 | 44598373 | 44600466 | 44600632 | 16 | 3 | 0 | 0 | 1 | 0 | 0 | 1 | 0 | 0 | 98 | 49 | 1 | 1 | 1 | 0.6 | NA | NA | 1 | -0.2 |
| SE | 3925 | SPG11 | SPG11 | chr15 | - | 44606024 | 44606091 | 44600466 | 44600632 | 44608443 | 44608605 | 26 | 14 | 1 | 6 | 0 | 1 | 0 | 3 | 0 | 2 | 98 | 49 | 1 | 1 | 0.684 | 1 | 0.333 | NA | 0.429 | 0.243 |
| SE | 3926 | SPG11 | SPG11 | chr15 | - | 44613429 | 44613536 | 44610839 | 44610985 | 44615362 | 44615566 | 36 | 11 | 5 | 3 | 2 | 0 | 0 | 13 | 0 | 0 | 98 | 49 | 1 | 1 | 0.857 | 0.733 | 1 | NA | 1 | -0.137 |
| SE | 3932 | DAPP1 | DAPP1 | chr4 | + | 99863769 | 99863855 | 99863009 | 99863072 | 99866033 | 99866121 | 52 | 18 | 16 | 1 | 0 | 0 | 0 | 14 | 0 | 0 | 98 | 49 | 1 | 1 | 0.963 | 1 | 1 | NA | 1 | -0.012 |
| SE | 3933 | DAPP1 | DAPP1 | chr4 | + | 99866556 | 99866618 | 99866033 | 99866121 | 99868116 | 99870189 | 6 | 0 | 0 | 31 | 6 | 4 | 0 | 3 | 0 | 6 | 98 | 49 | 1 | 1 | 0.088 | 0 | 0 | NA | 0.2 | -0.171 |
| SE | 3935 | TPT1 | TPT1 | chr13 | - | 45340711 | 45340785 | 45339993 | 45340184 | 45341041 | 45341284 | 288 | 116 | 51 | 1 | 2 | 0 | 1 | 67 | 0 | 0 | 98 | 49 | 1 | 1 | 0.993 | 0.967 | 1 | 1 | 1 | -0.013 |
| SE | 3938 | DECR1 | DECR1 | chr8 | + | 90036840 | 90036940 | 90020908 | 90021056 | 90042727 | 90042800 | 36 | 16 | 9 | 1 | 0 | 0 | 0 | 6 | 0 | 0 | 98 | 49 | 1 | 1 | 0.947 | 1 | 1 | NA | 1 | -0.018 |
| SE | 3942 | SREBF1 | SREBF1 | chr17 | - | 17814614 | 17814747 | 17814244 | 17814410 | 17814834 | 17814944 | 3 | 1 | 0 | 2 | 0 | 0 | 0 | 0 | 2 | 0 | 98 | 49 | 1 | 1 | 0.429 | 1 | NA | 0 | NA | 0.715 |
| SE | 3943 | SREBF1 | SREBF1 | chr17 | - | 17817255 | 17817457 | 17816957 | 17817136 | 17817695 | 17817916 | 7 | 1 | 1 | 0 | 1 | 0 | 0 | 1 | 0 | 0 | 98 | 49 | 1 | 1 | 1 | 0.333 | 1 | NA | 1 | -0.222 |
| SE | 3944 | SREBF1 | SREBF1 | chr17 | - | 17819012 | 17819234 | 17818259 | 17818374 | 17819319 | 17819454 | 6 | 2 | 2 | 0 | 0 | 0 | 0 | 0 | 2 | 0 | 98 | 49 | 1 | 1 | 1 | 1 | 1 | 0 | NA | 1 |
| SE | 3965 | ARMC2 | ARMC2 | chr6 | + | 108868823 | 108868995 | 108858198 | 108858271 | 108876142 | 108876350 | 5 | 6 | 0 | 0 | 0 | 0 | 0 | 1 | 0 | 1 | 98 | 49 | 1 | 1 | 1 | 1 | NA | NA | 0.333 | 0.667 |
| SE | 3975 | SUPT20H | SUPT20H | chr13 | - | 37033448 | 37033588 | 37031738 | 37031895 | 37040404 | 37040458 | 20 | 3 | 13 | 1 | 0 | 0 | 0 | 4 | 0 | 0 | 98 | 49 | 1 | 1 | 0.909 | 1 | 1 | NA | 1 | -0.03 |
| SE | 3976 | SUPT20H | SUPT20H | chr13 | - | 37048563 | 37048599 | 37047877 | 37047936 | 37051487 | 37051583 | 19 | 2 | 10 | 4 | 0 | 1 | 0 | 0 | 0 | 1 | 85 | 49 | 1 | 1 | 0.732 | 1 | 0.852 | NA | 0 | 0.861 |
| SE | 3993 | PARVG | PARVG | chr22 | + | 44183317 | 44183408 | 44181741 | 44181917 | 44185807 | 44185872 | 97 | 32 | 18 | 0 | 2 | 0 | 7 | 24 | 0 | 0 | 98 | 49 | 1 | 1 | 1 | 0.889 | 1 | 1 | 1 | -0.037 |
| SE | 3995 | CIC | CIC | chr19 | + | 42289847 | 42289951 | 42289180 | 42289406 | 42290232 | 42291466 | 13 | 7 | 2 | 0 | 1 | 0 | 3 | 3 | 0 | 0 | 98 | 49 | 1 | 1 | 1 | 0.778 | 1 | 1 | 1 | -0.074 |
| SE | 3996 | CIC | CIC | chr19 | + | 42291557 | 42291745 | 42290232 | 42291466 | 42292085 | 42292207 | 4 | 0 | 1 | 1 | 0 | 0 | 0 | 1 | 0 | 2 | 98 | 49 | 1 | 1 | 0.667 | NA | 1 | NA | 0.2 | 0.633 |
| SE | 3997 | HMGB1 | HMGB1 | chr13 | - | 30463206 | 30463352 | 30462537 | 30462712 | 30463530 | 30463694 | 0 | 0 | 0 | 0 | 0 | 1 | 1 | 0 | 0 | 0 | 98 | 49 | 1 | 1 | NA | NA | 0 | 1 | NA | -1 |
| SE | 4000 | RPL24 | RPL24 | chr3 | - | 101682428 | 101682492 | 101681089 | 101681215 | 101682770 | 101682907 | 46 | 7 | 10 | 4 | 0 | 0 | 0 | 9 | 0 | 0 | 98 | 49 | 1 | 1 | 0.852 | 1 | 1 | NA | 1 | -0.049 |
| SE | 4001 | PAN3 | PAN3 | chr13 | + | 28177864 | 28177935 | 28176492 | 28176559 | 28220230 | 28220378 | 38 | 26 | 9 | 0 | 0 | 0 | 0 | 12 | 0 | 1 | 98 | 49 | 1 | 1 | 1 | 1 | 1 | NA | 0.857 | 0.143 |
| SE | 4002 | PAN3 | PAN3 | chr13 | + | 28197184 | 28197346 | 28177864 | 28177935 | 28220230 | 28220378 | 19 | 10 | 0 | 12 | 1 | 0 | 0 | 1 | 0 | 5 | 98 | 49 | 1 | 1 | 0.442 | 0.833 | NA | NA | 0.091 | 0.546 |
| SE | 4003 | PAN3 | PAN3 | chr13 | + | 28220230 | 28220378 | 28177864 | 28177935 | 28256291 | 28256539 | 26 | 1 | 1 | 1 | 0 | 0 | 0 | 6 | 0 | 0 | 98 | 49 | 1 | 1 | 0.929 | 1 | 1 | NA | 1 | -0.024 |
| SE | 4004 | PAN3 | PAN3 | chr13 | + | 28271980 | 28272071 | 28270700 | 28270866 | 28277236 | 28277376 | 12 | 5 | 8 | 0 | 0 | 0 | 0 | 0 | 0 | 1 | 98 | 49 | 1 | 1 | 1 | 1 | 1 | NA | 0 | 1 |
| SE | 4008 | NUPL1 | NUPL1 | chr13 | + | 25312882 | 25313032 | 25309246 | 25309282 | 25313613 | 25313751 | 19 | 8 | 4 | 0 | 0 | 0 | 0 | 3 | 0 | 1 | 98 | 49 | 1 | 1 | 1 | 1 | 1 | NA | 0.6 | 0.4 |
| SE | 4016 | SPATA13 | SPATA13 | chr13 | + | 24278613 | 24278821 | 24270686 | 24270912 | 24284134 | 24284271 | 3 | 0 | 0 | 4 | 4 | 0 | 0 | 1 | 1 | 3 | 98 | 49 | 1 | 1 | 0.273 | 0 | NA | 0 | 0.143 | 0.065 |
| SE | 4017 | SPATA13 | SPATA13 | chr13 | + | 24286764 | 24286950 | 24286213 | 24286393 | 24288998 | 24289178 | 9 | 5 | 6 | 1 | 0 | 1 | 0 | 5 | 0 | 2 | 98 | 49 | 1 | 1 | 0.818 | 1 | 0.75 | NA | 0.556 | 0.3 |
| SE | 4035 | CNOT7 | CNOT7 | chr8 | - | 17234715 | 17234860 | 17231512 | 17232537 | 17237211 | 17237373 | 17 | 19 | 3 | 1 | 0 | 0 | 2 | 1 | 0 | 0 | 98 | 49 | 1 | 1 | 0.895 | 1 | 1 | 1 | 1 | -0.035 |

| | | | | | | | | | | | | | | | | | | | | | | | | | | | | | | | |
|---|---|---|---|---|---|---|---|---|---|---|---|---|---|---|---|---|---|---|---|---|---|---|---|---|---|---|---|---|---|---|---|
| SE | 4053 | FBRSL1 | FBRSL1 | chr12 | + | 132581740 | 132581824 | 132581438 | 132581516 | 132582061 | 132582266 | 1 | 0 | 1 | 1 | 0 | 0 | 1 | 0 | 0 | 0 | 98 | 49 | 1 | 1 | 0.333 | NA | 1 | 1 | NA | -0.334 |
| SE | 4070 | RRP12 | RRP12 | chr10 | - | 97367040 | 97367132 | 97366741 | 97366909 | 97369424 | 97369582 | 14 | 6 | 4 | 0 | 3 | 0 | 0 | 1 | 0 | 0 | 98 | 49 | 1 | 1 | 1 | 0.5 | 1 | NA | 1 | -0.167 |
| SE | 4071 | RRP12 | RRP12 | chr10 | - | 97372735 | 97372803 | 97372072 | 97372166 | 97373045 | 97373200 | 15 | 11 | 2 | 1 | 1 | 0 | 0 | 4 | 0 | 2 | 98 | 49 | 1 | 1 | 0.882 | 0.846 | 1 | NA | 0.5 | 0.409 |
| SE | 4072 | RRP12 | RRP12 | chr10 | - | 97379627 | 97379770 | 97379292 | 97379414 | 97380798 | 97380913 | 37 | 8 | 1 | 1 | 0 | 0 | 6 | 8 | 0 | 0 | 98 | 49 | 1 | 1 | 0.949 | 1 | 1 | 1 | 1 | -0.017 |
| SE | 4073 | RRP12 | RRP12 | chr10 | - | 97381714 | 97381826 | 97381385 | 97381483 | 97385165 | 97385257 | 26 | 8 | 0 | 2 | 0 | 0 | 0 | 7 | 0 | 0 | 98 | 49 | 1 | 1 | 0.867 | 1 | NA | NA | 1 | -0.067 |
| SE | 4074 | RRP12 | RRP12 | chr10 | - | 97390422 | 97390539 | 97388488 | 97388624 | 97390738 | 97390844 | 53 | 20 | 5 | 0 | 0 | 0 | 7 | 7 | 0 | 1 | 98 | 49 | 1 | 1 | 1 | 1 | 1 | 1 | 0.778 | 0.111 |
| SE | 4076 | RRP12 | RRP12 | chr10 | - | 97390738 | 97390844 | 97390422 | 97390539 | 97393683 | 97393760 | 86 | 26 | 3 | 1 | 0 | 0 | 8 | 9 | 0 | 0 | 98 | 49 | 1 | 1 | 0.977 | 1 | 1 | 1 | 1 | -0.008 |
| SE | 4082 | NCOR2 | NCOR2 | chr12 | - | 124374412 | 124374463 | 124372021 | 124372610 | 124378236 | 124378384 | 3 | 0 | 0 | 1 | 0 | 1 | 0 | 2 | 0 | 0 | 98 | 49 | 1 | 1 | 0.6 | NA | 0 | NA | 1 | -0.7 |
| SE | 4083 | NCOR2 | NCOR2 | chr12 | - | 124385744 | 124385887 | 124378236 | 124378384 | 124398118 | 124398181 | 1 | 2 | 3 | 1 | 0 | 0 | 0 | 2 | 0 | 0 | 98 | 49 | 1 | 1 | 0.333 | 1 | 1 | NA | 1 | -0.222 |
| SE | 4084 | NCOR2 | NCOR2 | chr12 | - | 124457105 | 124457162 | 124449814 | 124449867 | 124466172 | 124466286 | 6 | 2 | 0 | 0 | 4 | 0 | 0 | 0 | 0 | 2 | 98 | 49 | 1 | 1 | 1 | 0.2 | NA | NA | 0 | 0.6 |
| SE | 4085 | NCOR2 | NCOR2 | chr12 | - | 124535564 | 124535611 | 124495146 | 124495368 | 124567307 | 124567464 | 3 | 2 | 0 | 1 | 0 | 0 | 0 | 3 | 0 | 0 | 96 | 49 | 1 | 1 | 0.605 | 1 | NA | NA | 1 | -0.198 |
| SE | 4094 | OGFOD2 | OGFOD2 | chr12 | + | 122976653 | 122976767 | 122975810 | 122975867 | 122976870 | 122976970 | 0 | 0 | 0 | 0 | 0 | 1 | 0 | 1 | 0 | 0 | 98 | 49 | 1 | 1 | NA | NA | 0 | NA | 1 | -1 |
| SE | 4101 | ZCCHC8 | ZCCHC8 | chr12 | - | 122482634 | 122482695 | 122481944 | 122482087 | 122489385 | 122489463 | 2 | 5 | 2 | 1 | 0 | 0 | 0 | 1 | 0 | 0 | 98 | 49 | 1 | 1 | 0.5 | 1 | 1 | NA | 1 | -0.167 |
| SE | 4108 | MLXIP | MLXIP | chr12 | + | 122130843 | 122130933 | 122129940 | 122130112 | 122132291 | 122132383 | 10 | 4 | 2 | 0 | 0 | 0 | 0 | 1 | 0 | 1 | 98 | 49 | 1 | 1 | 1 | 1 | 1 | NA | 0.333 | 0.667 |
| SE | 4112 | FAM219B | FAM219B | chr15 | - | 74906277 | 74906365 | 74905153 | 74905231 | 74906586 | 74907121 | 1 | 0 | 0 | 1 | 0 | 0 | 0 | 0 | 0 | 1 | 98 | 49 | 1 | 1 | 0.333 | NA | NA | NA | 0 | 0.333 |
| SE | 4116 | CAMKK2 | CAMKK2 | chr12 | - | 121248605 | 121248734 | 121245139 | 121245240 | 121249786 | 121249874 | 8 | 2 | 0 | 5 | 11 | 0 | 0 | 2 | 1 | 0 | 98 | 49 | 1 | 1 | 0.444 | 0.083 | NA | 0 | 1 | -0.236 |
| SE | 4117 | CAMKK2 | CAMKK2 | chr12 | - | 121249960 | 121250034 | 121249786 | 121249874 | 121252660 | 121252714 | 10 | 16 | 5 | 1 | 0 | 0 | 1 | 14 | 0 | 0 | 98 | 49 | 1 | 1 | 0.833 | 1 | 1 | 1 | 1 | -0.056 |
| SE | 4120 | P2RX4 | P2RX4 | chr12 | + | 121222729 | 121222767 | 121222093 | 121222166 | 121222946 | 121223043 | 1 | 0 | 0 | 3 | 0 | 0 | 0 | 0 | 0 | 5 | 87 | 49 | 1 | 1 | 0.158 | NA | NA | NA | 0 | 0.158 |
| SE | 4121 | CAMK2D | CAMK2D | chr4 | - | 113455725 | 113455821 | 113451031 | 113454515 | 113457334 | 113457563 | 2 | 0 | 0 | 2 | 0 | 0 | 0 | 1 | 0 | 0 | 98 | 49 | 1 | 1 | 0.333 | NA | NA | NA | 1 | -0.667 |
| SE | 4130 | HIRA | HIRA | chr22 | - | 19361726 | 19361931 | 19361236 | 19361341 | 19375630 | 19375792 | 14 | 8 | 0 | 0 | 1 | 0 | 0 | 4 | 0 | 0 | 98 | 49 | 1 | 1 | 1 | 0.8 | NA | NA | 1 | -0.1 |
| SE | 4131 | HIRA | HIRA | chr22 | - | 19387710 | 19387816 | 19385520 | 19385736 | 19388483 | 19388554 | 15 | 6 | 2 | 0 | 0 | 0 | 0 | 8 | 0 | 2 | 98 | 49 | 1 | 1 | 1 | 1 | 1 | NA | 0.667 | 0.333 |
| SE | 4132 | HIRA | HIRA | chr22 | - | 19388483 | 19388554 | 19385520 | 19385736 | 19392100 | 19392214 | 18 | 12 | 2 | 0 | 0 | 0 | 0 | 5 | 0 | 1 | 98 | 49 | 1 | 1 | 1 | 1 | 1 | NA | 0.714 | 0.286 |
| SE | 4138 | ZNRF2 | ZNRF2 | chr7 | + | 30323641 | 30323737 | 30284306 | 30285826 | 30355727 | 30355833 | 17 | 6 | 3 | 0 | 1 | 0 | 0 | 2 | 0 | 0 | 98 | 49 | 1 | 1 | 1 | 0.75 | 1 | NA | 1 | -0.083 |
| SE | 4139 | MED13L | MED13L | chr12 | - | 115968939 | 115969097 | 115966081 | 115966243 | 115970593 | 115970770 | 40 | 20 | 11 | 0 | 0 | 0 | 6 | 8 | 0 | 1 | 98 | 49 | 1 | 1 | 1 | 1 | 1 | 1 | 0.8 | 0.1 |
| SE | 4140 | MED13L | MED13L | chr12 | - | 115986265 | 115986489 | 115984179 | 115984372 | 115987108 | 115987288 | 20 | 15 | 8 | 0 | 3 | 0 | 0 | 4 | 0 | 0 | 98 | 49 | 1 | 1 | 1 | 0.714 | 1 | NA | 1 | -0.095 |
| SE | 4141 | MED13L | MED13L | chr12 | - | 116096668 | 116096752 | 116022455 | 116022601 | 116111427 | 116111512 | 100 | 56 | 21 | 0 | 1 | 0 | 0 | 23 | 0 | 0 | 98 | 49 | 1 | 1 | 1 | 0.966 | 1 | NA | 1 | -0.011 |
| SE | 4142 | MED13L | MED13L | chr12 | - | 116111427 | 116111512 | 116096668 | 116096752 | 116237467 | 116237705 | 130 | 65 | 32 | 2 | 1 | 0 | 0 | 17 | 0 | 0 | 98 | 49 | 1 | 1 | 0.97 | 0.97 | 1 | NA | 1 | -0.02 |
| SE | 4143 | MED13L | MED13L | chr12 | - | 116237467 | 116237705 | 116111427 | 116111512 | 116277059 | 116277186 | 138 | 53 | 19 | 1 | 0 | 0 | 0 | 18 | 0 | 0 | 98 | 49 | 1 | 1 | 0.986 | 1 | 1 | NA | 1 | -0.005 |
| SE | 4150 | PSTPIP1 | PSTPIP1 | chr15 | + | 77030501 | 77030581 | 77029528 | 77029574 | 77031179 | 77031278 | 41 | 18 | 9 | 0 | 3 | 0 | 0 | 3 | 0 | 0 | 98 | 49 | 1 | 1 | 1 | 0.75 | 1 | NA | 1 | -0.083 |
| SE | 4152 | PSTPIP1 | PSTPIP1 | chr15 | + | 77032861 | 77032952 | 77032297 | 77032394 | 77035507 | 77035563 | 26 | 15 | 9 | 0 | 1 | 0 | 4 | 6 | 0 | 0 | 98 | 49 | 1 | 1 | 1 | 0.882 | 1 | 1 | 1 | -0.039 |
| SE | 4153 | OAS2 | OAS2 | chr12 | + | 112987037 | 112987308 | 112978468 | 112978785 | 112995295 | 112995474 | 5 | 1 | 0 | 0 | 0 | 0 | 6 | 42 | 0 | 3 | 98 | 49 | 1 | 1 | 1 | 1 | NA | 1 | 0.875 | 0.062 |
| SE | 4154 | OAS2 | OAS2 | chr12 | + | 113004933 | 113005222 | 113002931 | 113003102 | 113006412 | 113006600 | 10 | 1 | 0 | 0 | 0 | 0 | 2 | 43 | 0 | 1 | 98 | 49 | 1 | 1 | 1 | 1 | NA | 1 | 0.956 | 0.022 |
| SE | 4155 | OAS3 | OAS3 | chr12 | + | 112941569 | 112941852 | 112938432 | 112938707 | 112944475 | 112944651 | 3 | 5 | 0 | 0 | 0 | 0 | 7 | 34 | 0 | 1 | 98 | 49 | 1 | 1 | 1 | 1 | NA | 1 | 0.944 | 0.028 |
| SE | 4156 | NLRP1 | NLRP1 | chr17 | - | 5515472 | 5515517 | 5501398 | 5501872 | 5517745 | 5517887 | 14 | 10 | 6 | 19 | 9 | 3 | 0 | 7 | 0 | 6 | 94 | 49 | 1 | 1 | 0.278 | 0.367 | 0.51 | NA | 0.378 | 0.007 |
| SE | 4157 | NLRP1 | NLRP1 | chr17 | - | 5517745 | 5517887 | 5515472 | 5515517 | 5530480 | 5530704 | 13 | 10 | 3 | 1 | 0 | 0 | 0 | 7 | 1 | 0 | 98 | 49 | 1 | 1 | 0.867 | 1 | 1 | 0 | 1 | 0.456 |
| SE | 4159 | NLRP1 | NLRP1 | chr17 | - | 5521523 | 5521786 | 5517745 | 5517887 | 5530480 | 5530704 | 22 | 22 | 0 | 1 | 0 | 0 | 0 | 9 | 0 | 0 | 98 | 49 | 1 | 1 | 0.917 | 1 | NA | NA | 1 | -0.041 |
| SE | 4161 | NLRP1 | NLRP1 | chr17 | - | 5533303 | 5533384 | 5532821 | 5532984 | 5536850 | 5536940 | 31 | 9 | 7 | 5 | 2 | 0 | 0 | 7 | 0 | 0 | 98 | 49 | 1 | 1 | 0.756 | 0.692 | 1 | NA | 1 | -0.184 |
| SE | 4162 | NLRP1 | NLRP1 | chr17 | - | 5533896 | 5533988 | 5533291 | 5533384 | 5536850 | 5536940 | 85 | 31 | 17 | 1 | 0 | 0 | 2 | 27 | 0 | 0 | 98 | 49 | 1 | 1 | 0.977 | 1 | 1 | 1 | 1 | -0.008 |
| SE | 4165 | OAS1 | OAS1 | chr12 | + | 112908535 | 112908824 | 112906933 | 112907219 | 112911050 | 112911235 | 7 | 5 | 0 | 0 | 1 | 0 | 1 | 25 | 0 | 0 | 98 | 49 | 1 | 1 | 1 | 0.714 | NA | 1 | 1 | -0.143 |
| SE | 4166 | OAS1 | OAS1 | chr12 | + | 112911050 | 112911235 | 112906933 | 112907219 | 112916508 | 112916738 | 7 | 5 | 0 | 0 | 0 | 0 | 0 | 15 | 0 | 2 | 98 | 49 | 1 | 1 | 1 | 1 | NA | NA | 0.789 | 0.211 |
| SE | 4167 | OAS1 | OAS1 | chr12 | + | 112917546 | 112917700 | 112916508 | 112916738 | 112919486 | 112919907 | 4 | 4 | 0 | 0 | 0 | 0 | 0 | 12 | 0 | 1 | 98 | 49 | 1 | 1 | 1 | 1 | NA | NA | 0.857 | 0.143 |
| SE | 4170 | POLDIP3 | POLDIP3 | chr22 | - | 42591954 | 42592084 | 42587505 | 42587572 | 42595536 | 42595603 | 9 | 6 | 7 | 0 | 0 | 0 | 0 | 0 | 0 | 2 | 98 | 49 | 1 | 1 | 1 | 1 | 1 | NA | 0 | 1 |
| SE | 4179 | GIT2 | GIT2 | chr12 | - | 109945259 | 109945349 | 109939164 | 109939247 | 109947255 | 109947504 | 2 | 1 | 0 | 1 | 0 | 1 | 0 | 0 | 0 | 0 | 98 | 49 | 1 | 1 | 0.5 | 1 | 0 | NA | 0 | 0.5 |
| SE | 4180 | GIT2 | GIT2 | chr12 | - | 109947255 | 109947504 | 109939164 | 109939247 | 109953091 | 109953234 | 1 | 3 | 1 | 0 | 0 | 3 | 0 | 2 | 0 | 0 | 98 | 49 | 1 | 1 | 1 | 1 | 0.143 | NA | 1 | -0.286 |
| SE | 4186 | GIT2 | GIT2 | chr12 | - | 109980951 | 109981046 | 109967457 | 109967503 | 109983372 | 109983503 | 81 | 22 | 28 | 0 | 1 | 0 | 2 | 20 | 0 | 0 | 98 | 49 | 1 | 1 | 1 | 0.917 | 1 | 1 | 1 | -0.028 |
| SE | 4187 | GIT2 | GIT2 | chr12 | - | 109991626 | 109991760 | 109989689 | 109989802 | 109996172 | 109996389 | 30 | 24 | 5 | 0 | 0 | 1 | 1 | 5 | 0 | 0 | 98 | 49 | 1 | 1 | 1 | 1 | 0.714 | 1 | 1 | -0.095 |
| SE | 4191 | DEPDC5 | DEPDC5 | chr22 | + | 31810520 | 31810641 | 31809610 | 31809647 | 31814991 | 31815212 | 4 | 0 | 0 | 0 | 0 | 0 | 0 | 1 | 0 | 0 | 98 | 49 | 1 | 1 | 1 | NA | NA | NA | 0.333 | 0.667 |
| SE | 4196 | YIPF1 | YIPF1 | chr1 | - | 53860055 | 53860153 | 53855000 | 53855051 | 53866199 | 53866382 | 11 | 5 | 6 | 0 | 0 | 0 | 0 | 1 | 0 | 1 | 98 | 49 | 1 | 1 | 1 | 1 | 1 | NA | 0.333 | 0.667 |
| SE | 4197 | YIPF1 | YIPF1 | chr1 | - | 53866199 | 53866382 | 53860055 | 53860153 | 53866757 | 53866924 | 19 | 4 | 8 | 1 | 0 | 0 | 0 | 1 | 0 | 0 | 98 | 49 | 1 | 1 | 0.905 | 1 | 1 | NA | 1 | -0.032 |
| SE | 4198 | YIPF1 | YIPF1 | chr1 | - | 53866199 | 53866382 | 53860055 | 53860153 | 53871371 | 53871488 | 9 | 2 | 3 | 2 | 3 | 0 | 0 | 2 | 0 | 0 | 98 | 49 | 1 | 1 | 0.692 | 0.25 | 1 | NA | 1 | -0.353 |
| SE | 4199 | YIPF1 | YIPF1 | chr1 | - | 53866757 | 53866924 | 53860055 | 53860153 | 53871371 | 53871488 | 13 | 2 | 7 | 2 | 3 | 0 | 0 | 4 | 0 | 0 | 98 | 49 | 1 | 1 | 0.765 | 0.25 | 1 | NA | 1 | -0.328 |
| SE | 4200 | YIPF1 | YIPF1 | chr1 | - | 53866757 | 53866924 | 53866199 | 53866382 | 53871371 | 53871488 | 22 | 4 | 12 | 0 | 0 | 0 | 0 | 4 | 0 | 0 | 98 | 49 | 1 | 1 | 1 | 1 | 1 | NA | 0.667 | 0.333 |
| SE | 4201 | YIPF1 | YIPF1 | chr1 | - | 53888906 | 53888986 | 53883112 | 53883276 | 53889712 | 53889814 | 1 | 6 | 0 | 1 | 0 | 0 | 1 | 1 | 0 | 0 | 98 | 49 | 1 | 1 | 0.333 | 1 | NA | 1 | 1 | -0.334 |
| SE | 4210 | SART3 | SART3 | chr12 | - | 108543027 | 108543152 | 108538933 | 108539089 | 108544426 | 108544478 | 5 | 0 | 8 | 2 | 0 | 0 | 0 | 1 | 0 | 0 | 98 | 49 | 1 | 1 | 0.556 | NA | 1 | NA | 1 | -0.222 |
| SE | 4211 | SART3 | SART3 | chr12 | - | 108544426 | 108544478 | 108543027 | 108543152 | 108545138 | 108545323 | 6 | 6 | 1 | 4 | 0 | 0 | 1 | 0 | 0 | 0 | 98 | 49 | 1 | 1 | 0.429 | 1 | 1 | 1 | NA | -0.19 |
| SE | 4212 | EIF3L | EIF3L | chr22 | + | 37858678 | 37858740 | 37855564 | 37855644 | 37862968 | 37863038 | 20 | 6 | 3 | 0 | 1 | 0 | 0 | 4 | 0 | 0 | 98 | 49 | 1 | 1 | 1 | 0.75 | 1 | NA | 1 | -0.083 |
| SE | 4213 | EIF3L | EIF3L | chr22 | + | 37886764 | 37886845 | 37877673 | 37878171 | 37888425 | 37888782 | 39 | 4 | 4 | 1 | 0 | 0 | 0 | 6 | 0 | 0 | 98 | 49 | 1 | 1 | 0.951 | 1 | 1 | NA | 1 | -0.016 |
| SE | 4220 | DEF8 | DEF8 | chr16 | + | 89961736 | 89961864 | 89960930 | 89961095 | 89963362 | 89963443 | 28 | 16 | 1 | 1 | 0 | 0 | 0 | 2 | 0 | 0 | 98 | 49 | 1 | 1 | 0.933 | 1 | 1 | NA | 1 | -0.022 |
| SE | 4226 | MARK3 | MARK3 | chr14 | + | 103448918 | 103448967 | 103428386 | 103428440 | 103451917 | 103451983 | 31 | 17 | 4 | 3 | 0 | 0 | 0 | 9 | 0 | 0 | 98 | 49 | 1 | 1 | 0.838 | 1 | 1 | NA | 1 | -0.054 |
| SE | 4227 | MARK3 | MARK3 | chr14 | + | 103465556 | 103465793 | 103462404 | 103462461 | 103465971 | 103466091 | 24 | 11 | 4 | 1 | 1 | 0 | 0 | 4 | 0 | 0 | 98 | 49 | 1 | 1 | 0.923 | 0.846 | 1 | NA | 1 | -0.077 |
| SE | 4228 | MARK3 | MARK3 | chr14 | + | 103465971 | 103466091 | 103465556 | 103465793 | 103466342 | 103466442 | 26 | 6 | 3 | 0 | 0 | 0 | 0 | 1 | 0 | 1 | 98 | 49 | 1 | 1 | 1 | 1 | 1 | NA | 0.333 | 0.667 |
| SE | 4229 | MARK3 | MARK3 | chr14 | + | 103474992 | 103475210 | 103468032 | 103468186 | 103480386 | 103480490 | 25 | 5 | 5 | 0 | 0 | 0 | 0 | 7 | 0 | 0 | 98 | 49 | 1 | 1 | 0.926 | 1 | 1 | NA | 1 | -0.025 |
| SE | 4230 | MARK3 | MARK3 | chr14 | + | 103491776 | 103492034 | 103480386 | 103480490 | 103502881 | 103503829 | 12 | 11 | 5 | 0 | 0 | 0 | 0 | 1 | 0 | 0 | 98 | 49 | 1 | 1 | 1 | 1 | 1 | NA | 0 | 1 |
| SE | 4235 | UHRF1BP1L | UHRF1BP1L | chr12 | - | 100058028 | 100059533 | 100057570 | 100057746 | 100059849 | 100060043 | 31 | 15 | 9 | 2 | 0 | 0 | 2 | 4 | 0 | 0 | 98 | 49 | 1 | 1 | 0.886 | 1 | 1 | 1 | 1 | -0.038 |
| SE | 4236 | UHRF1BP1L | UHRF1BP1L | chr12 | - | 100084475 | 100084658 | 100083015 | 100083134 | 100086274 | 100086358 | 37 | 13 | 1 | 6 | 1 | 0 | 0 | 4 | 0 | 0 | 98 | 49 | 1 | 1 | 0.755 | 0.867 | 1 | NA | 1 | -0.126 |
| SE | 4237 | UHRF1BP1L | UHRF1BP1L | chr12 | - | 100088914 | 100089095 | 100086274 | 100086358 | 100095666 | 100095837 | 48 | 20 | 9 | 0 | 0 | 0 | 0 | 11 | 0 | 3 | 98 | 49 | 1 | 1 | 1 | 1 | 1 | NA | 0.647 | 0.353 |
| SE | 4239 | C9orf89 | C9orf89 | chr9 | + | 93111878 | 93111938 | 93110567 | 93110721 | 93112217 | 93112289 | 8 | 5 | 5 | 3 | 1 | 2 | 0 | 6 | 0 | 0 | 98 | 49 | 1 | 1 | 0.571 | 0.714 | 0.556 | NA | 1 | -0.386 |
| SE | 4240 | TMEM86B | TMEM86B | chr19 | - | 55228190 | 55228437 | 55226633 | 55227563 | 55228690 | 55229264 | 0 | 0 | 2 | 0 | 0 | 0 | 0 | 1 | 1 | 0 | 98 | 49 | 1 | 1 | NA | NA | 1 | 0 | 1 | 0.5 |
| SE | 4261 | KATNBL1 | KATNBL1 | chr15 | - | 34146760 | 34146850 | 34145397 | 34145491 | 34147199 | 34147289 | 13 | 1 | 5 | 0 | 2 | 0 | 0 | 2 | 0 | 0 | 98 | 49 | 1 | 1 | 1 | 0.2 | 1 | NA | 1 | -0.267 |
| SE | 4264 | MTMR1 | MTMR1 | chrX | + | 150762564 | 150762737 | 150755688 | 150755865 | 150764511 | 150765103 | 4 | 4 | 0 | 0 | 0 | 0 | 0 | 1 | 1 | 0 | 98 | 49 | 1 | 1 | 1 | 1 | NA | 0 | 1 | 0.5 |
| SE | 4268 | LTA4H | LTA4H | chr12 | - | 96021084 | 96021137 | 96019167 | 96019240 | 96022146 | 96022251 | 60 | 15 | 8 | 1 | 0 | 0 | 1 | 16 | 0 | 0 | 98 | 49 | 1 | 1 | 0.968 | 1 | 1 | 1 | 1 | -0.011 |
| SE | 4271 | RSL24D1 | RSL24D1 | chr15 | - | 55190974 | 55191047 | 55185361 | 55185425 | 55192719 | 55192833 | 22 | 10 | 2 | 0 | 2 | 0 | 0 | 5 | 0 | 0 | 98 | 49 | 1 | 1 | 1 | 0.714 | 1 | NA | 1 | -0.095 |
| SE | 4273 | TOP3A | TOP3A | chr17 | - | 18290841 | 18291027 | 18290556 | 18290686 | 18294702 | 18294785 | 5 | 0 | 0 | 0 | 1 | 0 | 0 | 4 | 0 | 0 | 98 | 49 | 1 | 1 | 1 | 0 | NA | NA | 1 | -0.5 |
| SE | 4274 | TOP3A | TOP3A | chr17 | - | 18292644 | 18292852 | 18290841 | 18291027 | 18294702 | 18294785 | 19 | 6 | 2 | 1 | 0 | 0 | 0 | 1 | 0 | 1 | 98 | 49 | 1 | 1 | 0.905 | 1 | 1 | NA | 0.333 | 0.635 |
| SE | 4277 | IFI35 | IFI35 | chr17 | + | 43013775 | 43013882 | 43013475 | 43013662 | 43014107 | 43014459 | 10 | 6 | 0 | 0 | 1 | 0 | 0 | 5 | 0 | 0 | 98 | 49 | 1 | 1 | 1 | 0.75 | NA | NA | 1 | -0.125 |
| SE | 4282 | PLXNC1 | PLXNC1 | chr12 | + | 94181445 | 94181580 | 94169152 | 94169293 | 94186372 | 94186473 | 118 | 70 | 70 | 0 | 2 | 0 | 1 | 22 | 0 | 0 | 98 | 49 | 1 | 1 | 1 | 0.946 | 1 | 1 | 1 | -0.018 |
| SE | 4284 | PLXNC1 | PLXNC1 | chr12 | + | 94259336 | 94259375 | 94255192 | 94255296 | 94259609 | 94259734 | 103 | 64 | 33 | 2 | 2 | 0 | 0 | 26 | 0 | 3 | 88 | 49 | 1 | 1 | 0.966 | 0.947 | 1 | NA | 0.828 | 0.143 |
| SE | 4286 | PLXNC1 | PLXNC1 | chr12 | + | 94298631 | 94298795 | 94297315 | 94297423 | 94303755 | 94303896 | 31 | 26 | 12 | 2 | 0 | 0 | 0 | 14 | 0 | 0 | 98 | 49 | 1 | 1 | 0.886 | 1 | 1 | NA | 1 | -0.038 |
| SE | 4287 | PLXNC1 | PLXNC1 | chr12 | + | 94300909 | 94301057 | 94297315 | 94297423 | 94303755 | 94303896 | 68 | 30 | 19 | 2 | 0 | 0 | 0 | 11 | 0 | 0 | 98 | 49 | 1 | 1 | 0.944 | 1 | 1 | NA | 1 | -0.019 |
| SE | 4295 | CEP135 | CEP135 | chr4 | + | 56019352 | 56019555 | 56017647 | 56017857 | 56020675 | 56020780 | 6 | 0 | 0 | 0 | 0 | 0 | 0 | 2 | 0 | 1 | 98 | 49 | 1 | 1 | 1 | NA | NA | NA | 0.5 | 0.5 |
| SE | 4296 | PCID2 | PCID2 | chr13 | - | 113190871 | 113190975 | 113185484 | 113185560 | 113195070 | 113195125 | 5 | 1 | 0 | 1 | 0 | 0 | 0 | 2 | 0 | 0 | 98 | 49 | 1 | 1 | 0.714 | 1 | NA | NA | 1 | -0.143 |
| SE | 4298 | PACSIN2 | PACSIN2 | chr22 | - | 42882183 | 42882304 | 42879047 | 42879169 | 42884385 | 42884561 | 70 | 39 | 24 | 0 | 2 | 0 | 6 | 13 | 0 | 0 | 98 | 49 | 1 | 1 | 1 | 0.907 | 1 | 1 | 1 | -0.031 |
| SE | 4299 | PACSIN2 | PACSIN2 | chr22 | - | 42890946 | 42891182 | 42888642 | 42888798 | 42893456 | 42893613 | 59 | 47 | 13 | 1 | 0 | 0 | 1 | 8 | 0 | 0 | 98 | 49 | 1 | 1 | 0.967 | 1 | 1 | 1 | 1 | -0.011 |
| SE | 4303 | PFDN5 | PFDN5 | chr12 | + | 53296243 | 53296275 | 53295838 | 53295941 | 53297849 | 53297924 | 43 | 7 | 8 | 2 | 0 | 0 | 0 | 8 | 0 | 0 | 81 | 49 | 1 | 1 | 0.929 | 1 | 1 | NA | 1 | -0.024 |
| SE | 4313 | ZNF512 | ZNF512 | chr2 | + | 27600690 | 27600815 | 27599969 | 27600053 | 27601355 | 27601442 | 9 | 2 | 0 | 0 | 0 | 0 | 0 | 0 | 0 | 1 | 98 | 49 | 1 | 1 | 1 | 1 | NA | NA | 0 | 1 |
| SE | 4322 | GTPBP1 | GTPBP1 | chr22 | + | 38708844 | 38708956 | 38705801 | 38706147 | 38715906 | 38716087 | 31 | 8 | 3 | 4 | 7 | 1 | 0 | 8 | 0 | 1 | 98 | 49 | 1 | 1 | 0.795 | 0.364 | 0.6 | NA | 0.8 | -0.214 |
| SE | 4323 | GTPBP1 | GTPBP1 | chr22 | + | 38721741 | 38721865 | 38716651 | 38717000 | 38724296 | 38724411 | 32 | 15 | 1 | 0 | 1 | 0 | 0 | 4 | 0 | 0 | 98 | 49 | 1 | 1 | 1 | 0.882 | 1 | NA | 1 | -0.039 |
| SE | 4324 | GTPBP1 | GTPBP1 | chr22 | + | 38726257 | 38726440 | 38726005 | 38726150 | 38727982 | 38728161 | 22 | 19 | 5 | 0 | 0 | 0 | 0 | 12 | 0 | 1 | 98 | 49 | 1 | 1 | 1 | 1 | 1 | NA | 0.857 | 0.143 |
| SE | 4326 | GTPBP1 | GTPBP1 | chr22 | + | 38727982 | 38728161 | 38726005 | 38726150 | 38729461 | 38729662 | 24 | 12 | 0 | 1 | 0 | 0 | 0 | 8 | 0 | 0 | 98 | 49 | 1 | 1 | 0.923 | 1 | NA | NA | 1 | -0.038 |
| SE | 4327 | GTPBP1 | GTPBP1 | chr22 | + | 38727982 | 38728161 | 38727212 | 38727348 | 38729461 | 38729662 | 53 | 27 | 5 | 1 | 0 | 0 | 0 | 11 | 0 | 0 | 98 | 49 | 1 | 1 | 0.964 | 1 | 1 | NA | 1 | -0.012 |
| SE | 4352 | SLC8A1 | SLC8A1 | chr2 | - | 40170280 | 40170349 | 40164853 | 40164984 | 40174824 | 40174842 | 1 | 0 | 0 | 10 | 4 | 5 | 0 | 0 | 0 | 2 | 98 | 49 | 1 | 1 | 0.048 | 0 | 0 | NA | 0 | 0.016 |
| SE | 4354 | SLC8A1 | SLC8A1 | chr2 | - | 40177751 | 40177855 | 40174824 | 40174842 | 40428472 | 40430304 | 17 | 2 | 6 | 0 | 0 | 1 | 0 | 2 | 0 | 0 | 98 | 49 | 1 | 1 | 1 | 1 | 0.75 | NA | 1 | -0.083 |
| SE | 4357 | TBC1D15 | TBC1D15 | chr12 | + | 71880468 | 71880607 | 71872928 | 71873003 | 71884810 | 71885021 | 14 | 4 | 0 | 2 | 2 | 0 | 0 | 3 | 0 | 0 | 98 | 49 | 1 | 1 | 0.778 | 0.5 | NA | NA | 1 | -0.361 |
| SE | 4358 | TBC1D15 | TBC1D15 | chr12 | + | 71884810 | 71885021 | 71880468 | 71880607 | 71893221 | 71893324 | 10 | 7 | 0 | 1 | 0 | 0 | 0 | 3 | 0 | 0 | 98 | 49 | 1 | 1 | 0.833 | 1 | NA | NA | 1 | -0.084 |
| SE | 4359 | TBC1D15 | TBC1D15 | chr12 | + | 71894324 | 71894375 | 71893221 | 71893324 | 71894685 | 71894883 | 2 | 0 | 1 | 4 | 4 | 3 | 0 | 0 | 0 | 1 | 98 | 49 | 1 | 1 | 0.2 | 0 | 0.25 | NA | 0 | 0.15 |
| SE | 4364 | 2-Sep | 2-Sep | chr2 | + | 241335321 | 241335351 | 241335125 | 241335212 | 241335974 | 241336098 | 4 | 1 | 0 | 7 | 2 | 0 | 0 | 0 | 0 | 1 | 79 | 49 | 1 | 1 | 0.262 | 0.237 | NA | NA | 0 | 0.249 |
| SE | 4365 | 2-Sep | 2-Sep | chr2 | + | 241335974 | 241336098 | 241335125 | 241335212 | 241337381 | 241337516 | 17 | 3 | 0 | 0 | 1 | 0 | 0 | 4 | 0 | 0 | 98 | 49 | 1 | 1 | 1 | 0.6 | NA | NA | 1 | -0.2 |
| SE | 4373 | SNRK | SNRK | chr3 | + | 43332168 | 43332310 | 43303097 | 43303792 | 43340286 | 43340499 | 50 | 33 | 15 | 2 | 0 | 0 | 0 | 7 | 0 | 0 | 98 | 49 | 1 | 1 | 0.926 | 1 | 1 | NA | 1 | -0.025 |
| SE | 4376 | IL32 | IL32 | chr16 | + | 3067553 | 3067613 | 3067376 | 3067415 | 3067983 | 3068010 | 29 | 15 | 2 | 3 | 1 | 0 | 0 | 4 | 0 | 0 | 98 | 49 | 1 | 1 | 0.829 | 0.882 | 1 | NA | 1 | -0.096 |
| SE | 4379 | LYZ | LYZ | chr12 | + | 69352219 | 69352298 | 69350107 | 69350272 | 69353152 | 69354233 | 1515 | 182 | 52 | 5 | 0 | 0 | 40 | 456 | 0 | 1 | 98 | 49 | 1 | 1 | 0.993 | 1 | 1 | 1 | 0.996 | 0 |
| SE | 4385 | SIK3 | SIK3 | chr11 | - | 116867945 | 116868089 | 116863667 | 116863818 | 116870330 | 116870401 | 11 | 3 | 1 | 3 | 0 | 0 | 0 | 4 | 0 | 0 | 98 | 49 | 1 | 1 | 0.647 | 1 | 1 | NA | 1 | -0.118 |
| SE | 4386 | SIK3 | SIK3 | chr11 | - | 116875373 | 116875451 | 116875157 | 116875267 | 116876252 | 116876363 | 13 | 5 | 0 | 0 | 0 | 0 | 0 | 3 | 0 | 1 | 98 | 49 | 1 | 1 | 1 | 1 | NA | NA | 0.6 | 0.4 |
| SE | 4387 | SIK3 | SIK3 | chr11 | - | 116875865 | 116876009 | 116875373 | 116875451 | 116876252 | 116876363 | 7 | 1 | 5 | 1 | 4 | 0 | 0 | 1 | 0 | 2 | 98 | 49 | 1 | 1 | 0.778 | 0.111 | 1 | NA | 0.2 | 0.43 |
| SE | 4388 | SIK3 | SIK3 | chr11 | - | 116876923 | 116877042 | 116876252 | 116876363 | 116896252 | 116896376 | 20 | 10 | 10 | 1 | 1 | 0 | 0 | 8 | 0 | 0 | 98 | 49 | 1 | 1 | 0.909 | 0.833 | 1 | NA | 1 | -0.086 |
| SE | 4393 | ZNF692 | ZNF692 | chr1 | - | 248850681 | 248850781 | 248850003 | 248850516 | 248853936 | 248854051 | 12 | 3 | 0 | 0 | 0 | 0 | 0 | 0 | 0 | 3 | 98 | 49 | 1 | 1 | 1 | 1 | NA | NA | 0 | 1 |
| SE | 4407 | TOR2A | TOR2A | chr9 | - | 127732563 | 127732691 | 127731523 | 127732278 | 127733384 | 127733557 | 2 | 1 | 1 | 1 | 0 | 0 | 0 | 2 | 0 | 0 | 98 | 49 | 1 | 1 | 0.5 | 1 | 1 | NA | 1 | -0.167 |
| SE | 4410 | AP1G1 | AP1G1 | chr16 | - | 71748250 | 71748378 | 71746587 | 71746692 | 71749893 | 71749983 | 19 | 7 | 0 | 0 | 0 | 0 | 1 | 4 | 0 | 1 | 98 | 49 | 1 | 1 | 1 | 1 | NA | 1 | 0.667 | 0.166 |
| SE | 4411 | AP1G1 | AP1G1 | chr16 | - | 71756018 | 71756159 | 71753832 | 71753887 | 71758807 | 71758921 | 37 | 15 | 4 | 2 | 3 | 0 | 0 | 7 | 0 | 0 | 98 | 49 | 1 | 1 | 0.902 | 0.714 | 1 | NA | 1 | -0.128 |
| SE | 4412 | AP1G1 | AP1G1 | chr16 | - | 71771155 | 71771252 | 71769622 | 71769699 | 71773220 | 71773362 | 19 | 8 | 1 | 1 | 0 | 0 | 0 | 11 | 0 | 0 | 98 | 49 | 1 | 1 | 0.905 | 1 | 1 | NA | 1 | -0.032 |
| SE | 4413 | AP1G1 | AP1G1 | chr16 | - | 71789278 | 71789482 | 71774467 | 71774592 | 71808762 | 71809073 | 26 | 27 | 10 | 0 | 2 | 0 | 1 | 7 | 0 | 0 | 98 | 49 | 1 | 1 | 1 | 0.871 | 1 | 1 | 1 | -0.043 |
| SE | 4416 | ARHGAP9 | ARHGAP9 | chr12 | - | 57473602 | 57473708 | 57472254 | 57472688 | 57474041 | 57474176 | 289 | 109 | 70 | 30 | 11 | 8 | 0 | 32 | 0 | 3 | 98 | 49 | 1 | 1 | 0.828 | 0.832 | 0.814 | NA | 0.842 | -0.017 |
| SE | 4417 | ARHGAP9 | ARHGAP9 | chr12 | - | 57474041 | 57474176 | 57473602 | 57473708 | 57474422 | 57474476 | 362 | 134 | 91 | 5 | 0 | 0 | 0 | 40 | 0 | 0 | 98 | 49 | 1 | 1 | 0.973 | 1 | 1 | NA | 1 | -0.009 |
| SE | 4418 | ARHGAP9 | ARHGAP9 | chr12 | - | 57474874 | 57474973 | 57474625 | 57474703 | 57475290 | 57475398 | 204 | 55 | 68 | 1 | 1 | 0 | 3 | 37 | 0 | 0 | 98 | 49 | 1 | 1 | 0.99 | 0.965 | 1 | 1 | 1 | -0.015 |
| SE | 4423 | MPZL1 | MPZL1 | chr1 | + | 167772274 | 167772488 | 167765582 | 167765749 | 167773235 | 167773368 | 13 | 0 | 19 | 1 | 0 | 0 | 0 | 1 | 0 | 0 | 98 | 49 | 1 | 1 | 0.867 | NA | 1 | NA | 1 | -0.067 |
| SE | 4424 | MPZL1 | MPZL1 | chr1 | + | 167773235 | 167773368 | 167765582 | 167765749 | 167787819 | 167791919 | 10 | 5 | 10 | 0 | 0 | 3 | 0 | 1 | 0 | 0 | 98 | 49 | 1 | 1 | 1 | 1 | 0.625 | NA | 1 | -0.125 |
| SE | 4425 | MPZL1 | MPZL1 | chr1 | + | 167776063 | 167776166 | 167773235 | 167773368 | 167787819 | 167791919 | 5 | 0 | 0 | 9 | 5 | 10 | 0 | 1 | 0 | 0 | 98 | 49 | 1 | 1 | 0.217 | 0 | 0 | NA | 0.333 | -0.261 |
| SE | 4427 | STARD3 | STARD3 | chr17 | + | 39657972 | 39658026 | 39657774 | 39657852 | 39658404 | 39658522 | 8 | 15 | 2 | 0 | 2 | 0 | 0 | 4 | 0 | 0 | 98 | 49 | 1 | 1 | 1 | 0.789 | 1 | NA | 1 | -0.07 |
| SE | 4429 | LRP1 | LRP1 | chr12 | + | 57180666 | 57180807 | 57180329 | 57180479 | 57181156 | 57181291 | 7 | 0 | 0 | 2 | 0 | 0 | 0 | 9 | 0 | 0 | 98 | 49 | 1 | 1 | 0.636 | NA | NA | NA | 1 | -0.364 |
| SE | 4432 | MON2 | MON2 | chr12 | + | 62566013 | 62566031 | 62565236 | 62565380 | 62566321 | 62566450 | 5 | 1 | 0 | 1 | 0 | 0 | 0 | 0 | 0 | 1 | 67 | 49 | 1 | 1 | 0.785 | 1 | NA | NA | 0 | 0.893 |
| SE | 4438 | PTGES3 | PTGES3 | chr12 | - | 56665631 | 56666266 | 56664775 | 56664800 | 56670274 | 56670364 | 17 | 6 | 14 | 1 | 0 | 0 | 1 | 2 | 0 | 0 | 98 | 49 | 1 | 1 | 0.895 | 1 | 1 | 1 | 1 | -0.035 |
| SE | 4439 | PTGES3 | PTGES3 | chr12 | - | 56666203 | 56666266 | 56664775 | 56664800 | 56670274 | 56670364 | 23 | 19 | 15 | 1 | 0 | 0 | 1 | 7 | 0 | 0 | 98 | 49 | 1 | 1 | 0.92 | 1 | 1 | 1 | 1 | -0.027 |
| SE | 4445 | DYNC1H1 | DYNC1H1 | chr14 | + | 101991526 | 101991673 | 101988702 | 101988852 | 101994183 | 101994324 | 16 | 4 | 1 | 2 | 0 | 0 | 0 | 6 | 0 | 0 | 98 | 49 | 1 | 1 | 0.8 | 1 | 1 | 1 | 1 | -0.067 |
| SE | 4455 | ATP5B | ATP5B | chr12 | - | 56645170 | 56645353 | 56644780 | 56644955 | 56645836 | 56646068 | 35 | 9 | 6 | 2 | 0 | 1 | 0 | 21 | 0 | 0 | 98 | 49 | 1 | 1 | 0.897 | 1 | 0.75 | NA | 1 | -0.118 |
| SE | 4463 | ANKRD52 | ANKRD52 | chr12 | - | 56243784 | 56243876 | 56237806 | 56243392 | 56244050 | 56244133 | 6 | 1 | 0 | 0 | 0 | 0 | 0 | 0 | 0 | 1 | 98 | 49 | 1 | 1 | 1 | 1 | NA | NA | 0 | 1 |
| SE | 4473 | ANAPC4 | ANAPC4 | chr4 | + | 25413644 | 25413742 | 25409697 | 25409791 | 25414323 | 25414385 | 2 | 1 | 0 | 1 | 0 | 0 | 0 | 6 | 0 | 0 | 98 | 49 | 1 | 1 | 0.5 | 1 | NA | NA | 1 | -0.25 |
| SE | 4484 | USP15 | USP15 | chr12 | + | 62349220 | 62349307 | 62325871 | 62325933 | 62355330 | 62355475 | 170 | 68 | 40 | 5 | 5 | 1 | 0 | 15 | 0 | 0 | 98 | 49 | 1 | 1 | 0.944 | 0.872 | 0.952 | NA | 1 | -0.077 |
| SE | 4485 | USP15 | USP15 | chr12 | + | 62389604 | 62389699 | 62389430 | 62389514 | 62389796 | 62389988 | 194 | 66 | 36 | 1 | 0 | 0 | 4 | 22 | 0 | 0 | 98 | 49 | 1 | 1 | 0.99 | 1 | 1 | 1 | 1 | -0.003 |
| SE | 4486 | USP15 | USP15 | chr12 | + | 62396294 | 62396398 | 62393052 | 62393202 | 62401186 | 62401275 | 126 | 35 | 33 | 5 | 6 | 0 | 0 | 13 | 0 | 1 | 98 | 49 | 1 | 1 | 0.926 | 0.745 | 1 | NA | 0.867 | 0.023 |
| SE | 4488 | R3HDM2 | R3HDM2 | chr12 | - | 57288856 | 57288910 | 57283823 | 57284056 | 57289008 | 57289040 | 0 | 2 | 0 | 5 | 0 | 4 | 0 | 0 | 0 | 3 | 98 | 49 | 1 | 1 | 0 | 1 | 0 | NA | 0 | 0.333 |
| SE | 4489 | R3HDM2 | R3HDM2 | chr12 | - | 57289008 | 57289040 | 57283823 | 57284056 | 57295398 | 57295507 | 5 | 3 | 4 | 1 | 0 | 0 | 0 | 4 | 0 | 0 | 81 | 49 | 1 | 1 | 0.752 | 1 | 1 | NA | 1 | -0.083 |
| SE | 4490 | NCKAP1L | NCKAP1L | chr12 | + | 54523824 | 54523956 | 54523393 | 54523539 | 54526527 | 54526746 | 35 | 36 | 3 | 0 | 1 | 0 | 3 | 12 | 0 | 0 | 98 | 49 | 1 | 1 | 1 | 0.947 | 1 | 1 | 1 | -0.018 |
| SE | 4495 | NFE2 | NFE2 | chr12 | - | 54300146 | 54300276 | 54295134 | 54295304 | 54300800 | 54301037 | 8 | 1 | 7 | 9 | 9 | 1 | 0 | 0 | 0 | 2 | 98 | 49 | 1 | 1 | 0.308 | 0.053 | 0.778 | NA | 0 | 0.38 |
| SE | 4500 | ICAM1 | ICAM1 | chr19 | + | 10284782 | 10285028 | 10284402 | 10284657 | 10285114 | 10286615 | 10 | 13 | 6 | 0 | 1 | 0 | 0 | 7 | 0 | 0 | 98 | 49 | 1 | 1 | 1 | 0.867 | 1 | NA | 1 | -0.044 |
| SE | 4501 | MARS | MARS | chr12 | + | 57489895 | 57489971 | 57489423 | 57489558 | 57490206 | 57490379 | 15 | 4 | 0 | 5 | 1 | 0 | 0 | 5 | 0 | 0 | 98 | 49 | 1 | 1 | 0.6 | 0.667 | NA | NA | 1 | -0.367 |
| SE | 4502 | MARS | MARS | chr12 | + | 57498156 | 57498273 | 57490537 | 57490644 | 57498419 | 57498623 | 15 | 10 | 2 | 0 | 2 | 0 | 1 | 6 | 0 | 0 | 98 | 49 | 1 | 1 | 1 | 0.714 | 1 | 1 | 1 | -0.095 |
| SE | 4503 | MARS | MARS | chr12 | + | 57500320 | 57500522 | 57498419 | 57498623 | 57504224 | 57504299 | 34 | 20 | 8 | 1 | 0 | 0 | 1 | 6 | 0 | 0 | 98 | 49 | 1 | 1 | 0.944 | 1 | 1 | 1 | 1 | -0.019 |
| SE | 4508 | FIG4 | FIG4 | chr6 | + | 109727108 | 109727265 | 109716444 | 109716568 | 109732636 | 109732687 | 3 | 4 | 0 | 1 | 0 | 0 | 1 | 3 | 0 | 0 | 98 | 49 | 1 | 1 | 0.6 | 1 | NA | 1 | 1 | -0.2 |
| SE | 4510 | APPL2 | APPL2 | chr12 | - | 105203702 | 105203785 | 105199372 | 105199531 | 105207060 | 105207207 | 10 | 4 | 5 | 2 | 0 | 0 | 0 | 1 | 0 | 0 | 98 | 49 | 1 | 1 | 0.714 | 1 | 1 | NA | 1 | -0.095 |
| SE | 4511 | CALCOCO1 | CALCOCO1 | chr12 | - | 53715792 | 53716047 | 53715199 | 53715325 | 53719738 | 53719829 | 16 | 1 | 2 | 2 | 0 | 0 | 0 | 1 | 0 | 0 | 98 | 49 | 1 | 1 | 0.8 | 1 | 1 | NA | 1 | -0.067 |
| SE | 4516 | VTI1B | VTI1B | chr14 | - | 67662476 | 67662535 | 67659730 | 67659922 | 67674374 | 67674885 | 14 | 2 | 1 | 1 | 0 | 0 | 1 | 2 | 0 | 0 | 98 | 49 | 1 | 1 | 0.875 | 1 | 1 | 1 | 1 | -0.042 |
| SE | 4517 | TEP1 | TEP1 | chr14 | - | 20372732 | 20372857 | 20371488 | 20371632 | 20373010 | 20373147 | 12 | 0 | 0 | 4 | 3 | 0 | 0 | 3 | 0 | 0 | 98 | 49 | 1 | 1 | 0.6 | 0 | NA | NA | 1 | -0.7 |
| SE | 4518 | TEP1 | TEP1 | chr14 | - | 20378753 | 20378853 | 20378379 | 20378535 | 20378980 | 20379105 | 8 | 0 | 0 | 2 | 2 | 0 | 0 | 1 | 0 | 0 | 98 | 49 | 1 | 1 | 0.667 | 0 | NA | NA | 1 | -0.666 |
| SE | 4519 | TEP1 | TEP1 | chr14 | - | 20381552 | 20381686 | 20381312 | 20381401 | 20381912 | 20382063 | 2 | 1 | 0 | 1 | 0 | 0 | 0 | 1 | 0 | 0 | 98 | 49 | 1 | 1 | 0.5 | 1 | NA | NA | 1 | -0.25 |
| SE | 4533 | CTBP1 | CTBP1 | chr4 | - | 1241324 | 1241519 | 1238182 | 1238337 | 1248915 | 1249120 | 7 | 1 | 1 | 2 | 1 | 0 | 0 | 6 | 0 | 0 | 98 | 49 | 1 | 1 | 0.636 | 0.333 | 1 | NA | 1 | -0.344 |
| SE | 4537 | SPRYD3 | SPRYD3 | chr12 | - | 53074648 | 53074784 | 53073285 | 53073471 | 53075094 | 53075219 | 2 | 6 | 4 | 0 | 0 | 0 | 0 | 0 | 0 | 2 | 98 | 49 | 1 | 1 | 1 | 1 | 1 | NA | 0 | 1 |
| SE | 4539 | MDM1 | MDM1 | chr12 | - | 68316580 | 68316610 | 68316077 | 68316253 | 68321346 | 68321446 | 0 | 1 | 0 | 1 | 0 | 0 | 0 | 2 | 0 | 0 | 79 | 49 | 1 | 1 | 0 | 1 | NA | NA | 1 | -0.5 |
| SE | 4541 | C16orf93 | C16orf93 | chr16 | - | 30759011 | 30759101 | 30757422 | 30757654 | 30759177 | 30759247 | 0 | 0 | 0 | 1 | 1 | 0 | 1 | 0 | 0 | 0 | 98 | 49 | 1 | 1 | 0 | 0 | NA | 1 | NA | -1 |
| SE | 4542 | VCL | VCL | chr10 | + | 74111908 | 74112112 | 74108970 | 74109156 | 74114183 | 74114387 | 1 | 0 | 0 | 5 | 0 | 0 | 0 | 0 | 1 | 0 | 98 | 49 | 1 | 1 | 0.091 | NA | NA | 0 | NA | 0.091 |
| SE | 4549 | UBE2Q2 | UBE2Q2 | chr15 | + | 75859877 | 75859982 | 75854385 | 75854487 | 75868950 | 75869010 | 10 | 3 | 6 | 0 | 0 | 0 | 0 | 5 | 0 | 1 | 98 | 49 | 1 | 1 | 1 | 1 | 1 | NA | 0.714 | 0.286 |
| SE | 4550 | NDUFA11 | NDUFA11 | chr19 | - | 5901334 | 5901454 | 5896904 | 5896997 | 5903611 | 5904013 | 1 | 1 | 0 | 14 | 3 | 2 | 0 | 0 | 0 | 2 | 98 | 49 | 1 | 1 | 0.034 | 0.143 | 0 | NA | 0 | 0.059 |
| SE | 4553 | TFCP2 | TFCP2 | chr12 | - | 51098775 | 51098918 | 51095988 | 51096040 | 51099654 | 51099779 | 3 | 1 | 0 | 0 | 0 | 1 | 0 | 3 | 0 | 0 | 98 | 49 | 1 | 1 | 1 | 1 | 0 | NA | 1 | -0.333 |
| SE | 4554 | TFCP2 | TFCP2 | chr12 | - | 51109120 | 51109273 | 51107235 | 51107346 | 51110876 | 51110983 | 2 | 0 | 2 | 2 | 0 | 0 | 0 | 1 | 0 | 0 | 98 | 49 | 1 | 1 | 0.333 | NA | 1 | NA | 1 | -0.334 |
| SE | 4555 | RASSF2 | RASSF2 | chr20 | - | 4795814 | 4795966 | 4792538 | 4792627 | 4798009 | 4798085 | 136 | 98 | 66 | 0 | 0 | 0 | 4 | 29 | 0 | 1 | 98 | 49 | 1 | 1 | 1 | 1 | 1 | 1 | 0.935 | 0.032 |
| SE | 4556 | RASSF2 | RASSF2 | chr20 | - | 4822328 | 4822403 | 4800971 | 4801062 | 4823556 | 4823645 | 51 | 31 | 16 | 3 | 0 | 0 | 0 | 4 | 0 | 0 | 98 | 49 | 1 | 1 | 0.895 | 1 | 1 | NA | 1 | -0.035 |
| SE | 4557 | INPP5A | INPP5A | chr10 | + | 132645867 | 132645968 | 132607914 | 132607956 | 132650417 | 132650505 | 19 | 13 | 0 | 1 | 0 | 0 | 1 | 3 | 0 | 0 | 98 | 49 | 1 | 1 | 0.905 | 1 | NA | 1 | 1 | -0.047 |

| | | | | | | | | | | | | | | | | | | | | | | | | | | | | | | | |
|---|---|---|---|---|---|---|---|---|---|---|---|---|---|---|---|---|---|---|---|---|---|---|---|---|---|---|---|---|---|---|---|
| SE | 4558 | INPP5A | INPP5A | chr10 | + | 132690391 | 132690455 | 132650417 | 132650505 | 132697815 | 132697919 | 17 | 11 | 5 | 1 | 0 | 0 | 0 | 7 | 0 | 0 | 98 | 49 | 1 | 1 | 0.895 | 1 | 1 | NA | 1 | -0.035 |
| SE | 4563 | LRRC28 | LRRC28 | chr15 | + | 99333922 | 99334129 | 99287813 | 99287951 | 99352368 | 99352471 | 1 | 0 | 0 | 2 | 0 | 0 | 0 | 1 | 0 | 0 | 98 | 49 | 1 | 1 | 0.2 | NA | NA | NA | 1 | -0.8 |
| SE | 4564 | LRRC28 | LRRC28 | chr15 | + | 99363105 | 99363265 | 99361335 | 99361511 | 99386029 | 99386309 | 2 | 0 | 4 | 1 | 0 | 0 | 0 | 3 | 0 | 0 | 98 | 49 | 1 | 1 | 0.5 | NA | 1 | NA | 1 | -0.25 |
| SE | 4565 | HEXDC | HEXDC | chr17 | + | 82439630 | 82439713 | 82437167 | 82437363 | 82440996 | 82441266 | 2 | 3 | 0 | 0 | 0 | 0 | 0 | 1 | 0 | 1 | 98 | 49 | 1 | 1 | 1 | 1 | NA | NA | 0.333 | 0.667 |
| SE | 4577 | PI4KB | PI4KB | chr1 | - | 151324744 | 151324917 | 151315572 | 151316509 | 151326171 | 151326373 | 0 | 1 | 0 | 2 | 0 | 0 | 0 | 0 | 0 | 2 | 98 | 49 | 1 | 1 | 0 | 1 | NA | NA | 0 | 0.5 |
| SE | 4579 | PI4KB | PI4KB | chr1 | - | 151326171 | 151326373 | 151315572 | 151316509 | 151327270 | 151327657 | 3 | 1 | 0 | 1 | 3 | 0 | 0 | 4 | 0 | 0 | 98 | 49 | 1 | 1 | 0.6 | 0.143 | NA | NA | 1 | -0.629 |
| SE | 4580 | PI4KB | PI4KB | chr1 | - | 151326171 | 151326373 | 151324744 | 151324917 | 151327270 | 151327657 | 1 | 1 | 0 | 1 | 1 | 0 | 0 | 2 | 0 | 0 | 98 | 49 | 1 | 1 | 0.333 | 0.333 | NA | NA | 1 | -0.667 |
| SE | 4581 | TRIP12 | TRIP12 | chr2 | - | 229860405 | 229860531 | 229858771 | 229859574 | 229879981 | 229880128 | 14 | 5 | 0 | 9 | 3 | 0 | 0 | 3 | 0 | 2 | 98 | 49 | 1 | 1 | 0.437 | 0.455 | NA | NA | 0.429 | 0.017 |
| SE | 4583 | PSMA3 | PSMA3 | chr14 | + | 58252118 | 58252242 | 58247749 | 58247832 | 58257744 | 58257846 | 14 | 8 | 0 | 0 | 0 | 1 | 0 | 3 | 0 | 0 | 98 | 49 | 1 | 1 | 1 | 1 | 0 | NA | 1 | -0.333 |
| SE | 4584 | PSMA3 | PSMA3 | chr14 | + | 58270933 | 58270998 | 58270417 | 58270485 | 58271850 | 58272009 | 12 | 6 | 3 | 0 | 0 | 0 | 0 | 6 | 0 | 1 | 98 | 49 | 1 | 1 | 1 | 1 | 1 | NA | 0.75 | 0.25 |
| SE | 4589 | PARP8 | PARP8 | chr5 | + | 50759642 | 50759732 | 50750150 | 50750188 | 50761820 | 50761898 | 26 | 32 | 18 | 0 | 1 | 0 | 0 | 5 | 0 | 0 | 98 | 49 | 1 | 1 | 1 | 0.941 | 1 | NA | 1 | -0.02 |
| SE | 4590 | PARP8 | PARP8 | chr5 | + | 50760291 | 50760362 | 50759642 | 50759732 | 50761820 | 50761898 | 111 | 49 | 38 | 3 | 4 | 0 | 0 | 17 | 0 | 0 | 98 | 49 | 1 | 1 | 0.949 | 0.86 | 1 | NA | 1 | -0.064 |
| SE | 4591 | PARP8 | PARP8 | chr5 | + | 50763147 | 50763242 | 50761820 | 50761898 | 50778068 | 50778129 | 89 | 35 | 10 | 2 | 0 | 0 | 3 | 15 | 0 | 1 | 98 | 49 | 1 | 1 | 0.957 | 1 | 1 | 1 | 0.882 | 0.045 |
| SE | 4592 | PARP8 | PARP8 | chr5 | + | 50794852 | 50795417 | 50794206 | 50794332 | 50796981 | 50797032 | 152 | 56 | 11 | 6 | 2 | 0 | 0 | 30 | 0 | 0 | 98 | 49 | 1 | 1 | 0.927 | 0.933 | 1 | NA | 1 | -0.047 |
| SE | 4594 | PARP8 | PARP8 | chr5 | + | 50824907 | 50824975 | 50822334 | 50822400 | 50826754 | 50826803 | 109 | 45 | 14 | 0 | 0 | 0 | 0 | 14 | 0 | 1 | 98 | 49 | 1 | 1 | 1 | 1 | 1 | NA | 0.875 | 0.125 |
| SE | 4609 | YWHAB | YWHAB | chr20 | + | 44887647 | 44887742 | 44885598 | 44885886 | 44901530 | 44901833 | 5 | 0 | 0 | 9 | 5 | 4 | 0 | 0 | 0 | 1 | 98 | 49 | 1 | 1 | 0.217 | 0 | 0 | NA | 0 | 0.072 |
| SE | 4610 | P2RX7 | P2RX7 | chr12 | + | 121154784 | 121154953 | 121132827 | 121133095 | 121156078 | 121156147 | 5 | 0 | 0 | 2 | 0 | 0 | 0 | 1 | 0 | 0 | 98 | 49 | 1 | 1 | 0.556 | NA | NA | NA | 1 | -0.444 |
| SE | 4617 | KIF13A | KIF13A | chr6 | - | 17779591 | 17779684 | 17778946 | 17779099 | 17780729 | 17780906 | 13 | 3 | 4 | 2 | 0 | 0 | 1 | 2 | 0 | 0 | 98 | 49 | 1 | 1 | 0.765 | 1 | 1 | 1 | 1 | -0.078 |
| SE | 4619 | KIF13A | KIF13A | chr6 | - | 17794248 | 17794395 | 17787775 | 17787875 | 17794571 | 17794704 | 6 | 5 | 7 | 2 | 0 | 0 | 2 | 2 | 0 | 0 | 98 | 49 | 1 | 1 | 0.6 | 1 | 1 | 1 | 1 | -0.133 |
| SE | 4620 | KIF13A | KIF13A | chr6 | - | 17799951 | 17800113 | 17799265 | 17799439 | 17804360 | 17804510 | 36 | 15 | 2 | 1 | 0 | 0 | 0 | 2 | 0 | 0 | 98 | 49 | 1 | 1 | 0.947 | 1 | 1 | NA | 1 | -0.018 |
| SE | 4621 | KIF13A | KIF13A | chr6 | - | 17817019 | 17817233 | 17808767 | 17808930 | 17825767 | 17825934 | 21 | 7 | 4 | 0 | 0 | 0 | 0 | 5 | 0 | 1 | 98 | 49 | 1 | 1 | 1 | 1 | 1 | NA | 0.714 | 0.286 |
| SE | 4622 | KIF13A | KIF13A | chr6 | - | 17855436 | 17855617 | 17851954 | 17852042 | 17856029 | 17856122 | 30 | 20 | 3 | 0 | 1 | 0 | 0 | 9 | 0 | 1 | 98 | 49 | 1 | 1 | 1 | 0.909 | 1 | NA | 0.818 | 0.152 |
| SE | 4623 | KIF13A | KIF13A | chr6 | - | 17898167 | 17898180 | 17873376 | 17873437 | 17987053 | 17987144 | 16 | 14 | 2 | 0 | 0 | 0 | 0 | 6 | 0 | 1 | 62 | 49 | 1 | 1 | 1 | 1 | 1 | NA | 0.826 | 0.174 |
| SE | 4625 | AKAP11 | AKAP11 | chr13 | + | 42317527 | 42317688 | 42313893 | 42313940 | 42319087 | 42323267 | 7 | 0 | 0 | 0 | 0 | 0 | 0 | 1 | 0 | 1 | 98 | 49 | 1 | 1 | 1 | NA | NA | NA | 0.333 | 0.667 |
| SE | 4626 | INPP4A | INPP4A | chr2 | + | 98545968 | 98546073 | 98543876 | 98544007 | 98552785 | 98552969 | 18 | 2 | 0 | 0 | 0 | 0 | 3 | 0 | 0 | 1 | 98 | 49 | 1 | 1 | 1 | 1 | NA | 1 | 0 | 0.5 |
| SE | 4627 | INPP4A | INPP4A | chr2 | + | 98546585 | 98546694 | 98545968 | 98546073 | 98552785 | 98552969 | 14 | 11 | 1 | 1 | 0 | 0 | 0 | 1 | 0 | 0 | 98 | 49 | 1 | 1 | 0.875 | 1 | 1 | NA | 1 | -0.042 |
| SE | 4630 | HADHA | HADHA | chr2 | - | 26193576 | 26193772 | 26192309 | 26192424 | 26194569 | 26194638 | 25 | 3 | 4 | 1 | 0 | 0 | 1 | 11 | 0 | 1 | 98 | 49 | 1 | 1 | 0.926 | 1 | 1 | 1 | 0.846 | 0.052 |
| SE | 4631 | HADHA | HADHA | chr2 | - | 26204061 | 26204196 | 26201148 | 26201320 | 26209779 | 26209889 | 22 | 3 | 2 | 1 | 0 | 0 | 0 | 2 | 0 | 0 | 98 | 49 | 1 | 1 | 0.917 | 1 | 1 | NA | 1 | -0.028 |
| SE | 4633 | MS4A4A | MS4A4A | chr11 | + | 60301000 | 60301057 | 60297196 | 60297325 | 60306099 | 60306201 | 1 | 0 | 0 | 1 | 0 | 0 | 0 | 1 | 0 | 0 | 98 | 49 | 1 | 1 | 0.333 | NA | NA | NA | 1 | -0.667 |
| SE | 4637 | RPAP3 | RPAP3 | chr12 | - | 47679492 | 47679594 | 47670106 | 47670345 | 47679703 | 47679774 | 8 | 2 | 1 | 1 | 0 | 0 | 0 | 2 | 0 | 0 | 98 | 49 | 1 | 1 | 0.8 | 1 | 1 | NA | 1 | -0.067 |
| SE | 4646 | ANO6 | ANO6 | chr12 | + | 45416698 | 45416904 | 45409356 | 45409487 | 45421070 | 45421273 | 9 | 0 | 0 | 1 | 0 | 0 | 0 | 1 | 0 | 0 | 98 | 49 | 1 | 1 | 0.818 | NA | NA | NA | 1 | -0.182 |
| SE | 4647 | MACF1 | MACF1 | chr1 | + | 39378460 | 39378523 | 39372478 | 39372596 | 39379202 | 39379444 | 2 | 1 | 0 | 7 | 6 | 0 | 0 | 0 | 0 | 1 | 98 | 49 | 1 | 1 | 0.125 | 0.077 | NA | NA | 0 | 0.101 |
| SE | 4648 | MACF1 | MACF1 | chr1 | + | 39381952 | 39382152 | 39380243 | 39380373 | 39385433 | 39385929 | 11 | 3 | 0 | 1 | 0 | 0 | 0 | 1 | 0 | 0 | 98 | 49 | 1 | 1 | 0.846 | 1 | NA | NA | 1 | -0.077 |
| SE | 4651 | UBR1 | UBR1 | chr15 | - | 42970519 | 42970607 | 42966152 | 42966286 | 42976716 | 42976867 | 12 | 3 | 0 | 1 | 0 | 0 | 0 | 1 | 0 | 0 | 98 | 49 | 1 | 1 | 0.857 | 1 | NA | NA | 1 | -0.072 |
| SE | 4664 | ADGRE3 | ADGRE3 | chr19 | - | 14641418 | 14641616 | 14638104 | 14638340 | 14644107 | 14644275 | 87 | 42 | 15 | 0 | 2 | 0 | 3 | 9 | 0 | 0 | 98 | 49 | 1 | 1 | 1 | 0.913 | 1 | 1 | 1 | -0.029 |
| SE | 4665 | ADGRE3 | ADGRE3 | chr19 | - | 14654981 | 14655165 | 14651084 | 14651204 | 14658512 | 14658550 | 144 | 56 | 36 | 4 | 2 | 0 | 0 | 16 | 0 | 0 | 98 | 49 | 1 | 1 | 0.947 | 0.933 | 1 | NA | 1 | -0.04 |
| SE | 4667 | ADGRE3 | ADGRE3 | chr19 | - | 14658512 | 14658550 | 14654981 | 14655165 | 14661962 | 14662118 | 162 | 46 | 27 | 1 | 0 | 0 | 0 | 13 | 0 | 1 | 87 | 49 | 1 | 1 | 0.989 | 1 | 1 | NA | 0.88 | 0.116 |
| SE | 4668 | ADGRE3 | ADGRE3 | chr19 | - | 14661962 | 14662118 | 14658512 | 14658550 | 14663417 | 14663540 | 172 | 64 | 29 | 3 | 0 | 0 | 0 | 12 | 0 | 0 | 98 | 49 | 1 | 1 | 0.966 | 1 | 1 | NA | 1 | -0.011 |
| SE | 4669 | ADGRE3 | ADGRE3 | chr19 | - | 14663417 | 14663540 | 14661962 | 14662118 | 14668801 | 14668852 | 164 | 56 | 21 | 1 | 0 | 0 | 0 | 9 | 0 | 0 | 98 | 49 | 1 | 1 | 0.988 | 1 | 1 | NA | 1 | -0.004 |
| SE | 4670 | ADGRE3 | ADGRE3 | chr19 | - | 14668801 | 14668852 | 14663417 | 14663540 | 14674745 | 14674918 | 103 | 30 | 4 | 1 | 1 | 0 | 0 | 6 | 0 | 0 | 98 | 49 | 1 | 1 | 0.981 | 0.937 | 1 | NA | 1 | -0.027 |
| SE | 4675 | DNM1L | DNM1L | chr12 | + | 32705824 | 32705863 | 32701414 | 32701562 | 32707366 | 32707413 | 0 | 1 | 0 | 1 | 0 | 0 | 0 | 0 | 0 | 1 | 88 | 49 | 1 | 1 | 0 | 1 | NA | NA | 0 | 0.5 |
| SE | 4680 | FGD4 | FGD4 | chr12 | + | 32576265 | 32576449 | 32564136 | 32564289 | 32581959 | 32582467 | 6 | 1 | 0 | 0 | 1 | 0 | 0 | 5 | 0 | 0 | 98 | 49 | 1 | 1 | 1 | 0.333 | NA | NA | 1 | -0.334 |
| SE | 4681 | FGD4 | FGD4 | chr12 | + | 32581959 | 32582467 | 32564136 | 32564289 | 32598496 | 32598586 | 3 | 7 | 1 | 0 | 1 | 0 | 0 | 2 | 0 | 0 | 98 | 49 | 1 | 1 | 1 | 0.778 | 1 | NA | 1 | -0.074 |
| SE | 4682 | FGD4 | FGD4 | chr12 | + | 32600409 | 32600509 | 32598496 | 32598586 | 32601277 | 32601423 | 3 | 0 | 0 | 21 | 9 | 5 | 0 | 0 | 0 | 5 | 98 | 49 | 1 | 1 | 0.067 | 0 | 0 | NA | 0 | 0.022 |
| SE | 4683 | FGD4 | FGD4 | chr12 | + | 32607937 | 32608095 | 32602160 | 32602317 | 32610775 | 32610834 | 33 | 19 | 22 | 0 | 0 | 1 | 0 | 10 | 0 | 0 | 98 | 49 | 1 | 1 | 1 | 1 | 0.917 | NA | 1 | -0.028 |
| SE | 4684 | FGD4 | FGD4 | chr12 | + | 32607956 | 32608095 | 32602160 | 32602317 | 32610775 | 32610834 | 56 | 25 | 37 | 0 | 0 | 1 | 0 | 14 | 0 | 0 | 98 | 49 | 1 | 1 | 1 | 1 | 0.949 | NA | 1 | -0.017 |
| SE | 4685 | ZNF267 | ZNF267 | chr16 | + | 31875151 | 31875302 | 31873757 | 31873969 | 31884497 | 31884624 | 1 | 0 | 0 | 22 | 15 | 2 | 0 | 0 | 0 | 2 | 98 | 49 | 1 | 1 | 0.022 | 0 | 0 | NA | 0 | 0.007 |
| SE | 4686 | ZNF267 | ZNF267 | chr16 | + | 31885160 | 31885256 | 31884497 | 31884624 | 31914475 | 31917308 | 72 | 26 | 8 | 0 | 1 | 0 | 1 | 12 | 0 | 0 | 98 | 49 | 1 | 1 | 1 | 0.929 | 1 | 1 | 1 | -0.024 |
| SE | 4695 | SERPINB9 | SERPINB9 | chr6 | - | 2893410 | 2893553 | 2891832 | 2891988 | 2895390 | 2895508 | 7 | 1 | 0 | 0 | 0 | 0 | 0 | 3 | 0 | 1 | 98 | 49 | 1 | 1 | 1 | 1 | NA | NA | 0.6 | 0.4 |
| SE | 4696 | PHF3 | PHF3 | chr6 | + | 63646526 | 63646795 | 63635801 | 63636150 | 63679999 | 63680161 | 2 | 0 | 5 | 6 | 1 | 0 | 0 | 0 | 0 | 1 | 98 | 49 | 1 | 1 | 0.143 | 0 | 1 | NA | 0 | 0.381 |
| SE | 4698 | PHF3 | PHF3 | chr6 | + | 63679999 | 63680161 | 63635801 | 63636150 | 63684128 | 63685911 | 24 | 3 | 0 | 1 | 1 | 2 | 0 | 7 | 0 | 0 | 98 | 49 | 1 | 1 | 0.923 | 0.6 | 0 | NA | 1 | -0.492 |
| SE | 4700 | IPO8 | IPO8 | chr12 | - | 30662326 | 30662487 | 30661140 | 30661266 | 30663488 | 30663654 | 11 | 0 | 2 | 1 | 0 | 0 | 0 | 2 | 0 | 0 | 98 | 49 | 1 | 1 | 0.846 | NA | 1 | NA | 1 | -0.077 |
| SE | 4701 | LCP1 | LCP1 | chr13 | - | 46130813 | 46130938 | 46125922 | 46127723 | 46134126 | 46134250 | 450 | 237 | 209 | 0 | 0 | 1 | 5 | 131 | 0 | 0 | 98 | 49 | 1 | 1 | 1 | 1 | 0.991 | 1 | 1 | -0.003 |
| SE | 4702 | LCP1 | LCP1 | chr13 | - | 46146907 | 46147103 | 46144441 | 46144520 | 46148351 | 46148447 | 618 | 315 | 228 | 1 | 1 | 0 | 9 | 167 | 0 | 1 | 98 | 49 | 1 | 1 | 0.997 | 0.994 | 1 | 1 | 0.988 | 0.003 |
| SE | 4703 | LCP1 | LCP1 | chr13 | - | 46148351 | 46148447 | 46144441 | 46144520 | 46150935 | 46151078 | 82 | 45 | 35 | 0 | 0 | 0 | 0 | 37 | 0 | 3 | 98 | 49 | 1 | 1 | 1 | 1 | 1 | NA | 0.86 | 0.14 |
| SE | 4704 | LCP1 | LCP1 | chr13 | - | 46158521 | 46158651 | 46156437 | 46156570 | 46158825 | 46158989 | 663 | 268 | 190 | 5 | 0 | 0 | 1 | 128 | 0 | 0 | 98 | 49 | 1 | 1 | 0.985 | 1 | 1 | 1 | 1 | -0.005 |
| SE | 4706 | VPS13C | VPS13C | chr15 | - | 61920497 | 61920647 | 61920066 | 61920331 | 61921946 | 61922033 | 14 | 2 | 3 | 1 | 0 | 0 | 0 | 5 | 0 | 0 | 98 | 49 | 1 | 1 | 0.875 | 1 | 1 | NA | 1 | -0.042 |
| SE | 4709 | FGFR1OP2 | FGFR1OP2 | chr12 | + | 26960514 | 26960628 | 26957600 | 26957743 | 26963341 | 26963455 | 4 | 0 | 1 | 3 | 7 | 1 | 0 | 0 | 0 | 1 | 98 | 49 | 1 | 1 | 0.4 | 0 | 0.333 | NA | 0 | 0.244 |
| SE | 4710 | ASUN | ASUN | chr12 | - | 26914407 | 26914578 | 26913973 | 26914128 | 26916001 | 26916180 | 1 | 0 | 0 | 1 | 0 | 0 | 0 | 1 | 0 | 0 | 98 | 49 | 1 | 1 | 0.333 | NA | NA | NA | 1 | -0.667 |
| SE | 4716 | LPCAT2 | LPCAT2 | chr16 | + | 55534442 | 55534477 | 55532823 | 55532882 | 55537577 | 55537632 | 41 | 32 | 16 | 1 | 0 | 0 | 0 | 7 | 0 | 0 | 84 | 49 | 1 | 1 | 0.96 | 1 | 1 | NA | 1 | -0.013 |
| SE | 4717 | KRAS | KRAS | chr12 | - | 25215436 | 25215560 | 25204788 | 25209911 | 25225613 | 25225773 | 0 | 1 | 0 | 6 | 5 | 0 | 0 | 0 | 0 | 1 | 98 | 49 | 1 | 1 | 0 | 0.091 | NA | NA | 0 | 0.045 |
| SE | 4718 | KRAS | KRAS | chr12 | - | 25227233 | 25227412 | 25225613 | 25225773 | 25245273 | 25245395 | 10 | 4 | 1 | 0 | 1 | 0 | 0 | 1 | 0 | 0 | 98 | 49 | 1 | 1 | 1 | 0.667 | 1 | NA | 1 | -0.111 |
| SE | 4722 | LRMP | LRMP | chr12 | + | 25062819 | 25062900 | 25052246 | 25052956 | 25063719 | 25063816 | 52 | 35 | 13 | 8 | 0 | 2 | 0 | 10 | 0 | 0 | 98 | 49 | 1 | 1 | 0.765 | 1 | 0.765 | NA | 1 | -0.157 |
| SE | 4723 | LRMP | LRMP | chr12 | + | 25066364 | 25066512 | 25063719 | 25063816 | 25069349 | 25069431 | 121 | 46 | 10 | 5 | 1 | 0 | 0 | 17 | 0 | 0 | 98 | 49 | 1 | 1 | 0.924 | 0.958 | 1 | NA | 1 | -0.039 |
| SE | 4727 | HOOK3 | HOOK3 | chr8 | + | 42968012 | 42968214 | 42966472 | 42966613 | 42973288 | 42973399 | 5 | 1 | 0 | 0 | 0 | 0 | 0 | 1 | 0 | 3 | 98 | 49 | 1 | 1 | 1 | 1 | NA | NA | 0.143 | 0.857 |
| SE | 4735 | RECQL | RECQL | chr12 | - | 21477802 | 21477969 | 21476910 | 21476992 | 21483375 | 21483574 | 8 | 2 | 0 | 0 | 4 | 0 | 0 | 2 | 0 | 0 | 98 | 49 | 1 | 1 | 1 | 0.2 | NA | NA | 1 | -0.4 |
| SE | 4736 | CSAD | CSAD | chr12 | - | 53161126 | 53161191 | 53160762 | 53160844 | 53161272 | 53161389 | 8 | 3 | 6 | 4 | 5 | 0 | 0 | 2 | 0 | 4 | 98 | 49 | 1 | 1 | 0.5 | 0.231 | 1 | NA | 0.2 | 0.377 |
| SE | 4737 | PYROXD1 | PYROXD1 | chr12 | + | 21452080 | 21452154 | 21449562 | 21449691 | 21455131 | 21455292 | 1 | 0 | 3 | 1 | 0 | 0 | 0 | 1 | 0 | 0 | 98 | 49 | 1 | 1 | 0.333 | NA | 1 | NA | 1 | -0.334 |
| SE | 4744 | ARID2 | ARID2 | chr12 | + | 45836740 | 45836991 | 45836588 | 45836655 | 45837320 | 45837417 | 15 | 6 | 0 | 2 | 0 | 0 | 0 | 2 | 0 | 1 | 98 | 49 | 1 | 1 | 0.789 | 1 | NA | NA | 0.5 | 0.395 |
| SE | 4754 | SMARCC2 | SMARCC2 | chr12 | - | 56172936 | 56173029 | 56172584 | 56172704 | 56173695 | 56173849 | 1 | 3 | 0 | 3 | 7 | 0 | 0 | 0 | 0 | 1 | 98 | 49 | 1 | 1 | 0.143 | 0.176 | NA | NA | 0 | 0.159 |
| SE | 4755 | SMARCC2 | SMARCC2 | chr12 | - | 56178403 | 56178534 | 56178021 | 56178093 | 56178809 | 56178847 | 9 | 2 | 2 | 0 | 1 | 0 | 0 | 3 | 0 | 0 | 98 | 49 | 1 | 1 | 1 | 0.5 | 1 | NA | 1 | -0.167 |
| SE | 4756 | SMARCC2 | SMARCC2 | chr12 | - | 56183860 | 56183930 | 56182003 | 56182079 | 56184174 | 56184244 | 17 | 6 | 0 | 0 | 2 | 0 | 0 | 2 | 0 | 0 | 98 | 49 | 1 | 1 | 1 | 0.6 | NA | NA | 1 | -0.2 |
| SE | 4758 | SPG7 | SPG7 | chr16 | + | 89529476 | 89529579 | 89526328 | 89526468 | 89530682 | 89530808 | 7 | 6 | 2 | 1 | 2 | 0 | 0 | 2 | 0 | 0 | 98 | 49 | 1 | 1 | 0.778 | 0.6 | 1 | NA | 1 | -0.207 |
| SE | 4762 | ATP2C1 | ATP2C1 | chr3 | + | 130979248 | 130979419 | 130975331 | 130975488 | 130980581 | 130980679 | 1 | 0 | 0 | 2 | 0 | 0 | 1 | 2 | 0 | 0 | 98 | 49 | 1 | 1 | 0.2 | NA | NA | 1 | 1 | -0.8 |
| SE | 4773 | NDST2 | NDST2 | chr10 | - | 73805586 | 73805769 | 73804772 | 73804869 | 73805899 | 73806028 | 10 | 3 | 0 | 1 | 0 | 0 | 0 | 5 | 0 | 0 | 98 | 49 | 1 | 1 | 0.833 | 1 | NA | NA | 1 | -0.084 |
| SE | 4774 | NDST2 | NDST2 | chr10 | - | 73806288 | 73806474 | 73805899 | 73806028 | 73806656 | 73806811 | 8 | 3 | 6 | 1 | 0 | 0 | 0 | 8 | 0 | 0 | 98 | 49 | 1 | 1 | 0.8 | 1 | 1 | NA | 1 | -0.067 |
| SE | 4783 | CARD8 | CARD8 | chr19 | - | 48232452 | 48232493 | 48231659 | 48231810 | 48234402 | 48234543 | 23 | 9 | 0 | 6 | 5 | 1 | 0 | 3 | 0 | 0 | 90 | 49 | 1 | 1 | 0.676 | 0.495 | 0 | NA | 1 | -0.61 |
| SE | 4784 | CARD8 | CARD8 | chr19 | - | 48238382 | 48238532 | 48234402 | 48234543 | 48240961 | 48241063 | 57 | 18 | 20 | 18 | 6 | 7 | 0 | 7 | 0 | 1 | 98 | 49 | 1 | 1 | 0.613 | 0.6 | 0.588 | NA | 0.778 | -0.178 |
| SE | 4785 | CARD8 | CARD8 | chr19 | - | 48249522 | 48249633 | 48240961 | 48241063 | 48249750 | 48249847 | 30 | 12 | 5 | 3 | 0 | 0 | 0 | 1 | 0 | 1 | 98 | 49 | 1 | 1 | 0.833 | 1 | 1 | NA | 0.333 | 0.611 |
| SE | 4790 | HNRNPH3 | HNRNPH3 | chr10 | + | 68337857 | 68337996 | 68337198 | 68337333 | 68338502 | 68338642 | 8 | 5 | 0 | 7 | 0 | 0 | 0 | 1 | 0 | 0 | 98 | 49 | 1 | 1 | 0.364 | 1 | NA | NA | 1 | -0.318 |
| SE | 4795 | HNRNPK | HNRNPK | chr9 | - | 83975461 | 83975505 | 83974516 | 83974589 | 83976994 | 83977051 | 100 | 74 | 44 | 5 | 4 | 2 | 0 | 23 | 0 | 0 | 93 | 49 | 1 | 1 | 0.913 | 0.907 | 0.921 | NA | 1 | -0.086 |
| SE | 4796 | PADI2 | PADI2 | chr1 | - | 17071405 | 17071491 | 17070087 | 17070216 | 17074855 | 17074949 | 29 | 22 | 20 | 1 | 0 | 0 | 0 | 13 | 0 | 0 | 98 | 49 | 1 | 1 | 0.935 | 1 | 1 | NA | 1 | -0.022 |
| SE | 4797 | PADI2 | PADI2 | chr1 | - | 17079263 | 17079415 | 17075678 | 17075823 | 17082544 | 17082652 | 30 | 16 | 16 | 2 | 1 | 0 | 0 | 12 | 0 | 1 | 98 | 49 | 1 | 1 | 0.882 | 0.889 | 1 | NA | 0.857 | 0.067 |
| SE | 4798 | PADI2 | PADI2 | chr1 | - | 17086520 | 17086699 | 17084598 | 17084702 | 17092407 | 17092533 | 35 | 39 | 36 | 1 | 0 | 0 | 1 | 22 | 0 | 0 | 98 | 49 | 1 | 1 | 0.946 | 1 | 1 | 1 | 1 | -0.018 |
| SE | 4803 | YBX3 | YBX3 | chr12 | - | 10709907 | 10710114 | 10704050 | 10704148 | 10713210 | 10713333 | 8 | 4 | 1 | 3 | 1 | 0 | 0 | 5 | 0 | 0 | 98 | 49 | 1 | 1 | 0.571 | 0.667 | 1 | NA | 1 | -0.254 |
| SE | 4804 | KDM6A | KDM6A | chrX | + | 45020609 | 45020730 | 45010960 | 45011019 | 45034930 | 45034985 | 27 | 6 | 0 | 0 | 0 | 0 | 0 | 7 | 0 | 3 | 98 | 49 | 1 | 1 | 1 | 1 | NA | NA | 0.538 | 0.462 |
| SE | 4805 | KDM6A | KDM6A | chrX | + | 45059005 | 45059104 | 45053828 | 45053955 | 45059246 | 45059466 | 36 | 9 | 0 | 1 | 0 | 0 | 4 | 4 | 0 | 1 | 98 | 49 | 1 | 1 | 0.947 | 1 | NA | 1 | 0.667 | 0.14 |
| SE | 4806 | KDM6A | KDM6A | chrX | + | 45060021 | 45060156 | 45059246 | 45059466 | 45060608 | 45060764 | 6 | 0 | 0 | 7 | 0 | 0 | 0 | 0 | 0 | 3 | 98 | 49 | 1 | 1 | 0.3 | NA | NA | NA | 0 | 0.3 |
| SE | 4807 | KDM6A | KDM6A | chrX | + | 45060021 | 45060156 | 45059246 | 45059466 | 45061323 | 45061419 | 7 | 0 | 0 | 1 | 0 | 0 | 0 | 0 | 0 | 3 | 98 | 49 | 1 | 1 | 0.778 | NA | NA | NA | 0 | 0.778 |
| SE | 4808 | KDM6A | KDM6A | chrX | + | 45060608 | 45060764 | 45059246 | 45059466 | 45061323 | 45061419 | 11 | 8 | 2 | 1 | 0 | 0 | 0 | 4 | 0 | 3 | 98 | 49 | 1 | 1 | 0.846 | 1 | 1 | NA | 0.4 | 0.549 |
| SE | 4809 | KDM6A | KDM6A | chrX | + | 45060608 | 45060764 | 45060021 | 45060156 | 45061323 | 45061419 | 4 | 8 | 2 | 1 | 0 | 0 | 0 | 1 | 0 | 0 | 98 | 49 | 1 | 1 | 0.667 | 1 | 1 | NA | 1 | -0.111 |
| SE | 4810 | KDM6A | KDM6A | chrX | + | 45062646 | 45062748 | 45061323 | 45061419 | 45063421 | 45063817 | 20 | 2 | 1 | 1 | 1 | 1 | 0 | 6 | 0 | 0 | 98 | 49 | 1 | 1 | 0.909 | 0.5 | 0.333 | NA | 1 | -0.419 |
| SE | 4811 | KDM6A | KDM6A | chrX | + | 45110078 | 45110249 | 45107409 | 45107536 | 45111381 | 45112612 | 26 | 8 | 2 | 1 | 0 | 0 | 1 | 7 | 0 | 0 | 98 | 49 | 1 | 1 | 0.929 | 1 | 1 | 1 | 1 | -0.024 |
| SE | 4815 | FOXP1 | FOXP1 | chr3 | - | 71359149 | 71359244 | 71299819 | 71299880 | 71493425 | 71493555 | 16 | 5 | 2 | 0 | 0 | 0 | 0 | 4 | 0 | 1 | 98 | 49 | 1 | 1 | 1 | 1 | 1 | NA | 0.667 | 0.333 |
| SE | 4816 | QKI | QKI | chr6 | + | 163563419 | 163563719 | 163561981 | 163562069 | 163566720 | 163566795 | 39 | 13 | 15 | 0 | 2 | 0 | 0 | 4 | 0 | 0 | 98 | 49 | 1 | 1 | 1 | 0.765 | 1 | NA | 1 | -0.078 |
| SE | 4817 | QKI | QKI | chr6 | + | 163563443 | 163563719 | 163561981 | 163562069 | 163566720 | 163566795 | 13 | 4 | 5 | 0 | 2 | 0 | 0 | 1 | 0 | 0 | 98 | 49 | 1 | 1 | 1 | 0.5 | 1 | NA | 1 | -0.167 |
| SE | 4818 | QKI | QKI | chr6 | + | 163566720 | 163566795 | 163563419 | 163563719 | 163570693 | 163578596 | 22 | 4 | 10 | 1 | 0 | 0 | 0 | 4 | 0 | 0 | 98 | 49 | 1 | 1 | 0.917 | 1 | 1 | NA | 1 | -0.028 |
| SE | 4826 | FRYL | FRYL | chr4 | - | 48539970 | 48540068 | 48535656 | 48535827 | 48540352 | 48540960 | 11 | 0 | 0 | 0 | 1 | 0 | 0 | 2 | 0 | 0 | 98 | 49 | 1 | 1 | 1 | 0 | NA | NA | 1 | -0.5 |
| SE | 4827 | FRYL | FRYL | chr4 | - | 48609743 | 48609823 | 48608986 | 48609067 | 48619273 | 48619370 | 19 | 7 | 0 | 0 | 0 | 0 | 0 | 3 | 0 | 1 | 98 | 49 | 1 | 1 | 1 | 1 | NA | NA | 0.6 | 0.4 |
| SE | 4828 | FRYL | FRYL | chr4 | - | 48710518 | 48710698 | 48684672 | 48684795 | 48780077 | 48780299 | 5 | 9 | 0 | 3 | 0 | 1 | 0 | 1 | 0 | 0 | 98 | 49 | 1 | 1 | 0.455 | 1 | 0 | NA | 1 | -0.515 |
| SE | 4831 | CLEC7A | CLEC7A | chr12 | - | 10123244 | 10123363 | 10116780 | 10118590 | 10125296 | 10125448 | 141 | 62 | 38 | 12 | 2 | 1 | 0 | 38 | 0 | 8 | 98 | 49 | 1 | 1 | 0.855 | 0.939 | 0.95 | NA | 0.704 | 0.211 |
| SE | 4834 | CLEC7A | CLEC7A | chr12 | - | 10125296 | 10125448 | 10123244 | 10123363 | 10127746 | 10127845 | 86 | 43 | 27 | 1 | 0 | 0 | 0 | 19 | 0 | 0 | 98 | 49 | 1 | 1 | 0.977 | 1 | 1 | NA | 1 | -0.008 |
| SE | 4835 | CLEC7A | CLEC7A | chr12 | - | 10126570 | 10126708 | 10125296 | 10125448 | 10127746 | 10127845 | 63 | 57 | 12 | 36 | 17 | 14 | 0 | 18 | 0 | 11 | 98 | 49 | 1 | 1 | 0.467 | 0.626 | 0.3 | NA | 0.45 | 0.014 |
| SE | 4836 | CLEC7A | CLEC7A | chr12 | - | 10127746 | 10127845 | 10125296 | 10125448 | 10129979 | 10130269 | 234 | 91 | 50 | 2 | 0 | 0 | 6 | 55 | 0 | 0 | 98 | 49 | 1 | 1 | 0.983 | 1 | 1 | 1 | 1 | -0.006 |
| SE | 4837 | CLEC12A | CLEC12A | chr12 | + | 9978965 | 9979064 | 9971408 | 9971687 | 9979335 | 9979524 | 13 | 42 | 0 | 0 | 0 | 0 | 0 | 25 | 0 | 3 | 98 | 49 | 1 | 1 | 1 | 1 | NA | NA | 0.806 | 0.194 |
| SE | 4839 | COMMD2 | COMMD2 | chr3 | - | 149750677 | 149750851 | 149738469 | 149741718 | 149751402 | 149751485 | 14 | 0 | 0 | 0 | 1 | 0 | 0 | 3 | 0 | 0 | 98 | 49 | 1 | 1 | 1 | 0 | NA | NA | 1 | -0.5 |
| SE | 4843 | CLEC2D | CLEC2D | chr12 | + | 9693033 | 9693115 | 9692827 | 9692931 | 9694759 | 9699555 | 11 | 1 | 1 | 8 | 4 | 2 | 0 | 1 | 0 | 2 | 98 | 49 | 1 | 1 | 0.407 | 0.111 | 0.2 | NA | 0.2 | 0.039 |
| SE | 4845 | SH3BP1 | SH3BP1 | chr22 | + | 37641125 | 37641168 | 37639676 | 37639846 | 37641373 | 37641478 | 9 | 0 | 3 | 1 | 0 | 0 | 0 | 2 | 0 | 0 | 92 | 49 | 1 | 1 | 0.827 | NA | 1 | NA | 1 | -0.087 |
| SE | 4846 | SH3BP1 | SH3BP1 | chr22 | + | 37641373 | 37641478 | 37639676 | 37639846 | 37642538 | 37642615 | 11 | 6 | 0 | 0 | 1 | 0 | 0 | 4 | 0 | 0 | 98 | 49 | 1 | 1 | 1 | 0.75 | NA | NA | 1 | -0.125 |
| SE | 4847 | M6PR | M6PR | chr12 | - | 8943800 | 8943910 | 8943404 | 8943535 | 8945417 | 8945584 | 18 | 2 | 0 | 1 | 0 | 0 | 1 | 7 | 0 | 0 | 98 | 49 | 1 | 1 | 0.9 | 1 | NA | 1 | 1 | -0.05 |
| SE | 4852 | CLEC4E | CLEC4E | chr12 | - | 8536089 | 8536205 | 8533304 | 8534809 | 8539216 | 8539306 | 51 | 35 | 31 | 1 | 0 | 1 | 0 | 35 | 0 | 1 | 98 | 49 | 1 | 1 | 0.962 | 1 | 0.939 | NA | 0.946 | 0.021 |
| SE | 4853 | CLEC4E | CLEC4E | chr12 | - | 8537114 | 8537266 | 8536089 | 8536205 | 8539216 | 8539306 | 31 | 39 | 26 | 2 | 0 | 0 | 1 | 35 | 0 | 0 | 98 | 49 | 1 | 1 | 0.886 | 1 | 1 | 1 | 1 | -0.038 |
| SE | 4864 | NECAP1 | NECAP1 | chr12 | + | 8091768 | 8091850 | 8090194 | 8090299 | 8092675 | 8092784 | 8 | 3 | 3 | 5 | 0 | 0 | 0 | 2 | 0 | 0 | 98 | 49 | 1 | 1 | 0.444 | 1 | 1 | NA | 1 | -0.185 |
| SE | 4869 | SLC2A3 | SLC2A3 | chr12 | - | 7922820 | 7923024 | 7919227 | 7921631 | 7924409 | 7924511 | 143 | 75 | 22 | 0 | 1 | 0 | 2 | 34 | 0 | 0 | 98 | 49 | 1 | 1 | 1 | 0.974 | 1 | 1 | 1 | -0.009 |
| SE | 4870 | SLC2A3 | SLC2A3 | chr12 | - | 7931244 | 7931485 | 7930479 | 7930642 | 7932986 | 7933147 | 217 | 146 | 65 | 0 | 0 | 1 | 12 | 78 | 0 | 0 | 98 | 49 | 1 | 1 | 1 | 1 | 0.97 | 1 | 1 | -0.01 |
| SE | 4871 | SLC2A3 | SLC2A3 | chr12 | - | 7933809 | 7933902 | 7932986 | 7933147 | 7936019 | 7936296 | 419 | 206 | 58 | 2 | 0 | 0 | 21 | 107 | 0 | 0 | 98 | 49 | 1 | 1 | 0.991 | 1 | 1 | 1 | 1 | -0.003 |
| SE | 4876 | ANKFY1 | ANKFY1 | chr17 | - | 4173353 | 4173444 | 4172555 | 4172680 | 4173908 | 4174056 | 2 | 0 | 3 | 2 | 4 | 0 | 0 | 2 | 0 | 0 | 98 | 49 | 1 | 1 | 0.333 | 0 | 1 | NA | 1 | -0.556 |
| SE | 4877 | ANKFY1 | ANKFY1 | chr17 | - | 4189381 | 4189479 | 4184817 | 4185046 | 4194977 | 4195177 | 11 | 7 | 0 | 0 | 0 | 0 | 0 | 0 | 0 | 1 | 98 | 49 | 1 | 1 | 1 | 1 | NA | NA | 0 | 1 |
| SE | 4878 | ANKFY1 | ANKFY1 | chr17 | - | 4235771 | 4235890 | 4216982 | 4217118 | 4242255 | 4242448 | 13 | 6 | 6 | 2 | 3 | 0 | 0 | 6 | 0 | 0 | 98 | 49 | 1 | 1 | 0.765 | 0.5 | 1 | NA | 1 | -0.245 |
| SE | 4882 | FCHO2 | FCHO2 | chr5 | + | 73006444 | 73006549 | 72990711 | 72990864 | 73015625 | 73015724 | 19 | 7 | 1 | 1 | 0 | 0 | 0 | 6 | 0 | 0 | 98 | 49 | 1 | 1 | 0.905 | 1 | 1 | NA | 1 | -0.032 |
| SE | 4884 | FCHO2 | FCHO2 | chr5 | + | 73081782 | 73081982 | 73078179 | 73078312 | 73082760 | 73082825 | 15 | 9 | 11 | 0 | 1 | 0 | 2 | 0 | 0 | 0 | 98 | 49 | 1 | 1 | 1 | 0.818 | 1 | 1 | NA | -0.061 |
| SE | 4887 | C1RL-AS1 | C1RL-AS1 | chr12 | + | 7115385 | 7118677 | 7115129 | 7115219 | 7118898 | 7119026 | 2 | 5 | 0 | 3 | 0 | 0 | 0 | 1 | 0 | 0 | 98 | 49 | 1 | 1 | 0.25 | 1 | NA | NA | 1 | -0.375 |
| SE | 4889 | C1RL | C1RL | chr12 | - | 7108250 | 7108479 | 7101605 | 7102087 | 7109109 | 7109278 | 22 | 15 | 7 | 1 | 1 | 0 | 0 | 11 | 0 | 0 | 98 | 49 | 1 | 1 | 0.917 | 0.882 | 1 | NA | 1 | -0.067 |
| SE | 4908 | MLF2 | MLF2 | chr12 | - | 6751640 | 6751676 | 6750712 | 6750766 | 6751924 | 6752054 | 15 | 23 | 15 | 1 | 0 | 0 | 4 | 7 | 0 | 0 | 85 | 49 | 1 | 1 | 0.896 | 1 | 1 | 1 | 1 | -0.035 |
| SE | 4909 | BCR | BCR | chr22 | + | 23290338 | 23290413 | 23289516 | 23289621 | 23292540 | 23292638 | 7 | 0 | 3 | 0 | 1 | 0 | 0 | 3 | 0 | 0 | 98 | 49 | 1 | 1 | 1 | 0 | 1 | NA | 1 | -0.333 |
| SE | 4912 | ASAP1 | ASAP1 | chr8 | - | 130091972 | 130092143 | 130079901 | 130079971 | 130115627 | 130115735 | 5 | 5 | 2 | 1 | 0 | 0 | 1 | 0 | 0 | 0 | 98 | 49 | 1 | 1 | 0.714 | 1 | 1 | 1 | NA | -0.095 |
| SE | 4913 | ASAP1 | ASAP1 | chr8 | - | 130112093 | 130112322 | 130079901 | 130079971 | 130115627 | 130115735 | 37 | 7 | 0 | 1 | 0 | 0 | 0 | 12 | 0 | 0 | 98 | 49 | 1 | 1 | 0.949 | 1 | NA | NA | 1 | -0.026 |
| SE | 4914 | ASAP1 | ASAP1 | chr8 | - | 130112093 | 130112322 | 130091972 | 130092143 | 130115627 | 130115735 | 38 | 9 | 1 | 0 | 3 | 0 | 0 | 11 | 0 | 0 | 98 | 49 | 1 | 1 | 1 | 0.6 | 1 | NA | 1 | -0.133 |
| SE | 4921 | KTN1 | KTN1 | chr14 | + | 55663954 | 55664041 | 55661521 | 55661612 | 55667240 | 55667330 | 3 | 0 | 0 | 4 | 3 | 1 | 0 | 1 | 0 | 0 | 98 | 49 | 1 | 1 | 0.273 | 0 | 0 | NA | 1 | -0.909 |
| SE | 4923 | KTN1 | KTN1 | chr14 | + | 55680685 | 55680741 | 55679564 | 55679685 | 55684098 | 55684584 | 4 | 0 | 0 | 3 | 5 | 0 | 0 | 1 | 0 | 3 | 98 | 49 | 1 | 1 | 0.4 | 0 | NA | NA | 0.143 | 0.057 |
| SE | 4925 | TLE4 | TLE4 | chr9 | + | 79612655 | 79612718 | 79576132 | 79576177 | 79627373 | 79627448 | 31 | 9 | 5 | 1 | 0 | 0 | 0 | 4 | 0 | 0 | 98 | 49 | 1 | 1 | 0.939 | 1 | 1 | NA | 1 | -0.02 |
| SE | 4926 | TLE4 | TLE4 | chr9 | + | 79627373 | 79627448 | 79612655 | 79612718 | 79652589 | 79652794 | 12 | 5 | 2 | 1 | 0 | 0 | 0 | 4 | 0 | 0 | 98 | 49 | 1 | 1 | 0.857 | 1 | 1 | NA | 1 | -0.048 |
| SE | 4928 | TLE4 | TLE4 | chr9 | + | 79671208 | 79671338 | 79654058 | 79654075 | 79704782 | 79704902 | 1 | 1 | 0 | 14 | 9 | 4 | 0 | 0 | 0 | 7 | 98 | 49 | 1 | 1 | 0.034 | 0.053 | 0 | NA | 0 | 0.029 |
| SE | 4930 | TLE4 | TLE4 | chr9 | + | 79706746 | 79706899 | 79705888 | 79705942 | 79708117 | 79708250 | 41 | 9 | 2 | 3 | 0 | 0 | 0 | 9 | 0 | 0 | 98 | 49 | 1 | 1 | 0.872 | 1 | 1 | NA | 1 | -0.043 |
| SE | 4931 | TLE4 | TLE4 | chr9 | + | 79707122 | 79707218 | 79706746 | 79706899 | 79708117 | 79708250 | 0 | 0 | 2 | 18 | 5 | 1 | 0 | 0 | 0 | 2 | 98 | 49 | 1 | 1 | 0 | 0 | 0.5 | NA | 0 | 0.167 |
| SE | 4934 | TNFRSF1A | TNFRSF1A | chr12 | - | 6330852 | 6330926 | 6330597 | 6330711 | 6333068 | 6333147 | 104 | 35 | 58 | 0 | 0 | 0 | 0 | 21 | 0 | 1 | 98 | 49 | 1 | 1 | 1 | 1 | 1 | NA | 0.913 | 0.087 |
| SE | 4935 | TNFRSF1A | TNFRSF1A | chr12 | - | 6333736 | 6333865 | 6333366 | 6333516 | 6334090 | 6334244 | 131 | 58 | 49 | 1 | 0 | 0 | 0 | 16 | 0 | 0 | 98 | 49 | 1 | 1 | 0.985 | 1 | 1 | NA | 1 | -0.005 |
| SE | 4938 | UBE2D1 | UBE2D1 | chr10 | + | 58363608 | 58363686 | 58361494 | 58361526 | 58364770 | 58364876 | 42 | 8 | 17 | 1 | 0 | 0 | 1 | 16 | 0 | 0 | 98 | 49 | 1 | 1 | 0.955 | 1 | 1 | 1 | 1 | -0.015 |
| SE | 4940 | KIAA1033 | KIAA1033 | chr12 | + | 105126003 | 105126127 | 105122117 | 105122238 | 105127128 | 105127289 | 35 | 5 | 9 | 0 | 6 | 0 | 0 | 4 | 0 | 0 | 98 | 49 | 1 | 1 | 1 | 0.294 | 1 | NA | 1 | -0.235 |
| SE | 4941 | KIAA1033 | KIAA1033 | chr12 | + | 105126234 | 105126362 | 105126003 | 105126127 | 105127128 | 105127289 | 58 | 33 | 2 | 1 | 0 | 0 | 0 | 12 | 0 | 1 | 98 | 49 | 1 | 1 | 0.967 | 1 | 1 | NA | 0.857 | 0.132 |
| SE | 4943 | KIAA1033 | KIAA1033 | chr12 | + | 105142452 | 105142558 | 105141166 | 105141246 | 105143126 | 105143243 | 36 | 8 | 10 | 1 | 4 | 0 | 0 | 10 | 0 | 0 | 98 | 49 | 1 | 1 | 0.947 | 0.5 | 1 | NA | 1 | -0.184 |
| SE | 4945 | KIAA1033 | KIAA1033 | chr12 | + | 105144286 | 105144455 | 105143126 | 105143243 | 105144717 | 105144872 | 69 | 24 | 10 | 0 | 3 | 0 | 0 | 26 | 0 | 0 | 98 | 49 | 1 | 1 | 1 | 0.8 | 1 | NA | 1 | -0.067 |
| SE | 4946 | KIAA1033 | KIAA1033 | chr12 | + | 105156725 | 105156792 | 105152342 | 105152451 | 105157235 | 105157322 | 44 | 32 | 4 | 3 | 0 | 0 | 0 | 5 | 0 | 0 | 98 | 49 | 1 | 1 | 0.88 | 1 | 1 | NA | 1 | -0.04 |
| SE | 4947 | KIAA1033 | KIAA1033 | chr12 | + | 105160000 | 105160148 | 105157235 | 105157322 | 105162748 | 105162845 | 69 | 19 | 4 | 2 | 0 | 0 | 1 | 14 | 0 | 0 | 98 | 49 | 1 | 1 | 0.945 | 1 | 1 | 1 | 1 | -0.018 |
| SE | 4948 | KIAA1033 | KIAA1033 | chr12 | + | 105162748 | 105162845 | 105160000 | 105160148 | 105164110 | 105164307 | 31 | 20 | 5 | 0 | 0 | 0 | 1 | 7 | 1 | 0 | 98 | 49 | 1 | 1 | 1 | 1 | 1 | 0.333 | 1 | 0.334 |
| SE | 4949 | KIAA1033 | KIAA1033 | chr12 | + | 105164110 | 105164307 | 105162748 | 105162845 | 105164640 | 105164740 | 25 | 17 | 9 | 0 | 1 | 2 | 0 | 4 | 0 | 0 | 98 | 49 | 1 | 1 | 1 | 0.895 | 0.692 | NA | 1 | -0.138 |
| SE | 4952 | PARP11 | PARP11 | chr12 | - | 3863835 | 3863957 | 3829889 | 3830018 | 3873211 | 3873448 | 3 | 1 | 0 | 0 | 0 | 0 | 0 | 0 | 0 | 1 | 98 | 49 | 1 | 1 | 1 | 1 | NA | NA | 0 | 1 |
| SE | 4953 | HERPUD1 | HERPUD1 | chr16 | + | 56935400 | 56935475 | 56935234 | 56935312 | 56936686 | 56936817 | 12 | 5 | 2 | 1 | 0 | 0 | 1 | 5 | 0 | 0 | 98 | 49 | 1 | 1 | 0.857 | 1 | 1 | 1 | 1 | -0.048 |
| SE | 4954 | HERPUD1 | HERPUD1 | chr16 | + | 56935403 | 56935475 | 56935234 | 56935312 | 56936686 | 56936817 | 9 | 1 | 1 | 1 | 0 | 0 | 1 | 1 | 0 | 0 | 98 | 49 | 1 | 1 | 0.818 | 1 | 1 | 1 | 1 | -0.061 |
| SE | 4955 | HERPUD1 | HERPUD1 | chr16 | + | 56939236 | 56939359 | 56936686 | 56936817 | 56939894 | 56940088 | 19 | 12 | 8 | 0 | 1 | 0 | 0 | 3 | 0 | 0 | 98 | 49 | 1 | 1 | 1 | 0.857 | 1 | NA | 1 | -0.048 |
| SE | 4956 | SLTM | SLTM | chr15 | - | 58890280 | 58890461 | 58889429 | 58889554 | 58892896 | 58893060 | 7 | 0 | 0 | 1 | 2 | 0 | 0 | 2 | 0 | 1 | 98 | 49 | 1 | 1 | 0.778 | 0 | NA | NA | 0.5 | -0.111 |
| SE | 4957 | SLTM | SLTM | chr15 | - | 58899468 | 58899883 | 58898802 | 58898852 | 58901259 | 58901287 | 7 | 1 | 0 | 4 | 0 | 0 | 1 | 0 | 0 | 0 | 98 | 49 | 1 | 1 | 0.467 | 1 | NA | 1 | NA | -0.266 |
| SE | 4958 | SLTM | SLTM | chr15 | - | 58899468 | 58899937 | 58898802 | 58898852 | 58901259 | 58901287 | 7 | 2 | 0 | 4 | 0 | 0 | 1 | 0 | 0 | 0 | 98 | 49 | 1 | 1 | 0.467 | 1 | NA | 1 | NA | -0.266 |
| SE | 4959 | SLTM | SLTM | chr15 | - | 58913498 | 58913696 | 58912562 | 58912610 | 58916934 | 58916999 | 51 | 28 | 23 | 1 | 0 | 0 | 0 | 6 | 0 | 0 | 98 | 49 | 1 | 1 | 0.962 | 1 | 1 | NA | 1 | -0.013 |
| SE | 4972 | FNIP1 | FNIP1 | chr5 | - | 131651801 | 131651999 | 131647089 | 131647205 | 131670462 | 131670631 | 49 | 19 | 9 | 0 | 2 | 0 | 1 | 3 | 0 | 0 | 98 | 49 | 1 | 1 | 1 | 0.826 | 1 | 1 | 1 | -0.058 |

| | | | | | | | | | | | | | | | | | | | | | | | | | | | | | | | | |
|---|---|---|---|---|---|---|---|---|---|---|---|---|---|---|---|---|---|---|---|---|---|---|---|---|---|---|---|---|---|---|---|---|
| SE | 4973 | FNIP1 | FNIP1 | chr5 | - | 131677702 | 131677872 | 131671504 | 131672924 | 131679028 | 131679175 | 80 | 41 | 7 | 1 | 0 | 0 | 0 | 15 | 0 | 0 | 98 | 49 | 1 | 1 | 0.976 | 1 | 1 | NA | | 1 | -0.008 |
| SE | 4974 | FNIP1 | FNIP1 | chr5 | - | 131679028 | 131679175 | 131677702 | 131677872 | 131698916 | 131699002 | 62 | 28 | 7 | 1 | 0 | 0 | 0 | 10 | 0 | 0 | 98 | 49 | 1 | 1 | 0.969 | 1 | 1 | NA | | 1 | -0.01 |
| SE | 4979 | ERC1 | ERC1 | chr12 | + | 1371832 | 1371977 | 1289851 | 1290012 | 1408148 | 1408247 | 2 | 1 | 2 | 2 | 0 | 0 | 0 | 1 | 0 | 0 | 98 | 49 | 1 | 1 | 0.333 | 1 | 1 | NA | | 1 | -0.222 |
| SE | 4986 | WNK1 | WNK1 | chr12 | + | 871264 | 871348 | 862082 | 862270 | 878211 | 878361 | 17 | 4 | 0 | 6 | 0 | 0 | 2 | 4 | 0 | 1 | 98 | 49 | 1 | 1 | 0.586 | 1 | NA | | 1 | 0.667 | -0.041 |
| SE | 4987 | WNK1 | WNK1 | chr12 | + | 871264 | 871348 | 862082 | 862270 | 878214 | 878361 | 12 | 4 | 0 | 1 | 1 | 0 | 0 | 5 | 0 | 2 | 98 | 49 | 1 | 1 | 0.857 | 0.667 | NA | NA | | 0.556 | 0.206 |
| SE | 5004 | BAZ1A | BAZ1A | chr14 | - | 34785776 | 34786001 | 34783761 | 34783927 | 34786125 | 34786221 | 42 | 19 | 5 | 1 | 0 | 0 | 1 | 4 | 0 | 0 | 98 | 49 | 1 | 1 | 0.955 | 1 | 1 | | 1 | 1 | -0.015 |
| SE | 5005 | BAZ1A | BAZ1A | chr14 | - | 34786125 | 34786221 | 34785776 | 34786001 | 34792774 | 34792921 | 16 | 9 | 6 | 0 | 0 | 3 | 0 | 4 | 0 | 2 | 98 | 49 | 1 | 1 | 1 | 1 | 0.5 | NA | | 0.5 | 0.333 |
| SE | 5006 | BAZ1A | BAZ1A | chr14 | - | 34795669 | 34795765 | 34794748 | 34794887 | 34800223 | 34800390 | 60 | 25 | 14 | 1 | 0 | 0 | 0 | 9 | 0 | 0 | 98 | 49 | 1 | 1 | 0.968 | 1 | 1 | NA | | 1 | -0.011 |
| SE | 5007 | BAZ1A | BAZ1A | chr14 | - | 34826012 | 34826156 | 34810934 | 34811036 | 34874491 | 34874662 | 38 | 17 | 14 | 0 | 0 | 0 | 0 | 13 | 1 | 0 | 98 | 49 | 1 | 1 | 1 | 1 | 1 | | 0 | 1 | 0.5 |
| SE | 5008 | BAZ1A | BAZ1A | chr14 | - | 34862043 | 34862322 | 34826012 | 34826156 | 34874491 | 34874662 | 94 | 34 | 12 | 2 | 0 | 0 | 0 | 20 | 0 | 0 | 98 | 49 | 1 | 1 | 0.959 | 1 | 1 | NA | | 1 | -0.014 |
| SE | 5010 | ACAD8 | ACAD8 | chr11 | + | 134257087 | 134257257 | 134256547 | 134256648 | 134258514 | 134258624 | 1 | 1 | 2 | 0 | 1 | 0 | 0 | 1 | 0 | 0 | 98 | 49 | 1 | 1 | 1 | 0.333 | 1 | NA | | 1 | -0.222 |
| SE | 5011 | ACAD8 | ACAD8 | chr11 | + | 134261737 | 134261890 | 134261274 | 134261372 | 134262519 | 134262622 | 1 | 0 | 2 | 1 | 0 | 0 | 0 | 1 | 0 | 0 | 98 | 49 | 1 | 1 | 0.333 | NA | 1 | NA | | 1 | -0.334 |
| SE | 5015 | KANSL3 | KANSL3 | chr2 | - | 96611063 | 96611138 | 96610725 | 96610883 | 96612281 | 96612353 | 1 | 6 | 0 | 1 | 0 | 0 | 0 | 0 | 0 | 1 | 98 | 49 | 1 | 1 | 0.333 | 1 | NA | NA | | 0 | 0.666 |
| SE | 5018 | NCAPD3 | NCAPD3 | chr11 | - | 134209144 | 134209206 | 134208863 | 134208951 | 134209312 | 134209477 | 4 | 1 | 0 | 0 | 0 | 0 | 0 | 0 | 1 | 0 | 98 | 49 | 1 | 1 | 1 | 1 | NA | NA | 0 | NA | 1 |
| SE | 5023 | AHCTF1 | AHCTF1 | chr1 | - | 246843794 | 246843928 | 246842693 | 246842776 | 246849614 | 246851442 | 12 | 1 | 5 | 3 | 1 | 1 | 0 | 0 | 0 | 2 | 98 | 49 | 1 | 1 | 0.667 | 0.333 | 0.714 | NA | | 0 | 0.571 |
| SE | 5024 | AHCTF1 | AHCTF1 | chr1 | - | 246857690 | 246857814 | 246855729 | 246855827 | 246860898 | 246861295 | 27 | 2 | 5 | 1 | 1 | 0 | 0 | 6 | 0 | 0 | 98 | 49 | 1 | 1 | 0.931 | 0.5 | 1 | NA | | 1 | -0.19 |
| SE | 5025 | AHCTF1 | AHCTF1 | chr1 | - | 246863923 | 246864116 | 246861958 | 246862153 | 246867243 | 246867351 | 26 | 16 | 4 | 1 | 1 | 0 | 0 | 1 | 0 | 0 | 98 | 49 | 1 | 1 | 0.929 | 1 | 1 | NA | | 1 | -0.024 |
| SE | 5028 | AHCTF1 | AHCTF1 | chr1 | - | 246916141 | 246916395 | 246913231 | 246913412 | 246918249 | 246918377 | 12 | 12 | 2 | 5 | 1 | 1 | 0 | 4 | 0 | 1 | 98 | 49 | 1 | 1 | 0.545 | 0.857 | 0.5 | NA | | 0.667 | -0.033 |
| SE | 5029 | SPOP | SPOP | chr17 | - | 49636758 | 49636809 | 49622732 | 49622876 | 49668026 | 49668078 | 2 | 3 | 0 | 2 | 1 | 0 | 0 | 0 | 0 | 2 | 98 | 49 | 1 | 1 | 0.333 | 0.6 | NA | NA | | 0 | 0.467 |
| SE | 5032 | SPOP | SPOP | chr17 | - | 49668026 | 49668078 | 49622732 | 49622876 | 49677932 | 49678163 | 2 | 1 | 0 | 1 | 2 | 0 | 0 | 2 | 0 | 0 | 98 | 49 | 1 | 1 | 0.5 | 0.2 | NA | NA | | 1 | -0.65 |
| SE | 5039 | APLP2 | APLP2 | chr11 | + | 130123611 | 130123779 | 130122304 | 130122513 | 130126699 | 130126830 | 64 | 27 | 9 | 18 | 8 | 3 | 0 | 33 | 0 | 3 | 98 | 49 | 1 | 1 | 0.64 | 0.628 | 0.6 | NA | | 0.846 | -0.223 |
| SE | 5048 | ETS1 | ETS1 | chr11 | - | 128480190 | 128480451 | 128463508 | 128463627 | 128484822 | 128485071 | 27 | 8 | 5 | 0 | 0 | 1 | 2 | 2 | 0 | 0 | 98 | 49 | 1 | 1 | 1 | 1 | 0.714 | | 1 | 1 | -0.095 |
| SE | 5050 | UNC45A | UNC45A | chr15 | + | 90942436 | 90942605 | 90940305 | 90940473 | 90942911 | 90943082 | 3 | 3 | 5 | 0 | 0 | 0 | 0 | 0 | 0 | 2 | 98 | 49 | 1 | 1 | 1 | 1 | 1 | NA | | 0 | 1 |
| SE | 5054 | ARAP1 | ARAP1 | chr11 | - | 72686041 | 72686191 | 72685068 | 72685681 | 72687438 | 72687502 | 120 | 66 | 39 | 2 | 0 | 0 | 2 | 18 | 0 | 0 | 98 | 49 | 1 | 1 | 0.968 | 1 | 1 | | 1 | 1 | -0.011 |
| SE | 5055 | ARAP1 | ARAP1 | chr11 | - | 72686041 | 72686191 | 72685068 | 72685681 | 72687687 | 72687738 | 65 | 28 | 16 | 1 | 0 | 0 | 0 | 13 | 0 | 0 | 98 | 49 | 1 | 1 | 0.97 | 1 | 1 | NA | | 1 | -0.01 |
| SE | 5056 | ARAP1 | ARAP1 | chr11 | - | 72687438 | 72687502 | 72685068 | 72685681 | 72687687 | 72687738 | 29 | 18 | 25 | 1 | 0 | 0 | 1 | 5 | 0 | 0 | 98 | 49 | 1 | 1 | 0.935 | 1 | 1 | | 1 | 1 | -0.022 |
| SE | 5057 | ARAP1 | ARAP1 | chr11 | - | 72687438 | 72687502 | 72686041 | 72686191 | 72687687 | 72687738 | 83 | 56 | 48 | 2 | 0 | 0 | 3 | 10 | 0 | 0 | 98 | 49 | 1 | 1 | 0.954 | 1 | 1 | | 1 | 1 | -0.015 |
| SE | 5060 | ARAP1 | ARAP1 | chr11 | - | 72697322 | 72697486 | 72696982 | 72697195 | 72697597 | 72697649 | 141 | 72 | 50 | 1 | 2 | 0 | 2 | 24 | 1 | 0 | 98 | 49 | 1 | 1 | 0.986 | 0.947 | 1 | | 0.5 | 1 | 0.228 |
| SE | 5061 | ARAP1 | ARAP1 | chr11 | - | 72697910 | 72698106 | 72697597 | 72697649 | 72699000 | 72699107 | 92 | 69 | 31 | 0 | 1 | 0 | 7 | 25 | 0 | 0 | 98 | 49 | 1 | 1 | 1 | 0.972 | 1 | | 1 | 1 | -0.009 |
| SE | 5063 | ARAP1 | ARAP1 | chr11 | - | 72704151 | 72704334 | 72702904 | 72703079 | 72705804 | 72705890 | 79 | 69 | 42 | 6 | 2 | 3 | 0 | 25 | 0 | 0 | 98 | 49 | 1 | 1 | 0.868 | 0.945 | 0.875 | NA | | 1 | -0.104 |
| SE | 5065 | ARAP1 | ARAP1 | chr11 | - | 72732514 | 72732597 | 72726619 | 72727172 | 72752327 | 72752389 | 18 | 20 | 1 | 1 | 0 | 0 | 2 | 8 | 0 | 0 | 98 | 49 | 1 | 1 | 0.9 | 1 | 1 | | 1 | 1 | -0.033 |
| SE | 5071 | CYFIP1 | CYFIP1 | chr15 | - | 22873490 | 22873729 | 22872824 | 22872972 | 22874549 | 22874644 | 6 | 1 | 0 | 1 | 0 | 0 | 0 | 3 | 0 | 0 | 98 | 49 | 1 | 1 | 0.75 | 1 | NA | NA | | 1 | -0.125 |
| SE | 5074 | PPP4R3B | PPP4R3B | chr2 | - | 55573618 | 55573777 | 55568193 | 55568363 | 55577314 | 55577356 | 19 | 2 | 7 | 2 | 0 | 0 | 0 | 10 | 0 | 0 | 98 | 49 | 1 | 1 | 0.826 | 1 | 1 | NA | | 1 | -0.058 |
| SE | 5075 | PPP4R3B | PPP4R3B | chr2 | - | 55578246 | 55578342 | 55577314 | 55577356 | 55579678 | 55579781 | 3 | 0 | 1 | 6 | 0 | 0 | 0 | 0 | 1 | 1 | 98 | 49 | 1 | 1 | 0.2 | NA | 1 | | 0 | 0 | 0.6 |
| SE | 5076 | PPP4R3B | PPP4R3B | chr2 | - | 55586617 | 55586734 | 55585050 | 55585167 | 55588878 | 55588956 | 22 | 5 | 4 | 3 | 0 | 0 | 0 | 4 | 0 | 0 | 98 | 49 | 1 | 1 | 0.786 | 1 | 1 | NA | | 1 | -0.071 |
| SE | 5078 | LOC100288778 | LOC100288778 | chr12 | - | 18372 | 18484 | 18036 | 18183 | 26800 | 26954 | 0 | 2 | 0 | 0 | 0 | 0 | 0 | 0 | 0 | 1 | 98 | 49 | 1 | 1 | NA | 1 | NA | NA | | 0 | 1 |
| SE | 5089 | ITGB7 | ITGB7 | chr12 | - | 53192690 | 53192910 | 53192329 | 53192538 | 53193139 | 53193363 | 4 | 1 | 1 | 0 | 0 | 0 | 0 | 0 | 0 | 1 | 98 | 49 | 1 | 1 | 1 | 1 | 1 | NA | | 0 | 1 |
| SE | 5092 | TMEM218 | TMEM218 | chr11 | - | 125102131 | 125102302 | 125101200 | 125101303 | 125102733 | 125102809 | 1 | 0 | 0 | 0 | 1 | 0 | 0 | 1 | 0 | 0 | 98 | 49 | 1 | 1 | 1 | 0 | NA | NA | | 1 | -0.5 |
| SE | 5094 | TMEM218 | TMEM218 | chr11 | - | 125102131 | 125102317 | 125101200 | 125101303 | 125102733 | 125102809 | 1 | 0 | 0 | 0 | 1 | 0 | 0 | 1 | 0 | 0 | 98 | 49 | 1 | 1 | 1 | 0 | NA | NA | | 1 | -0.5 |
| SE | 5096 | TMEM218 | TMEM218 | chr11 | - | 125102131 | 125102351 | 125101200 | 125101303 | 125102733 | 125102809 | 1 | 0 | 0 | 0 | 1 | 0 | 0 | 1 | 0 | 0 | 98 | 49 | 1 | 1 | 1 | 0 | NA | NA | | 1 | -0.5 |
| SE | 5102 | GALC | GALC | chr14 | - | 87945552 | 87945733 | 87941394 | 87941558 | 87947727 | 87947878 | 7 | 2 | 1 | 0 | 0 | 0 | 0 | 3 | 0 | 1 | 98 | 49 | 1 | 1 | 1 | 1 | 1 | NA | | 0.6 | 0.4 |
| SE | 5103 | GALC | GALC | chr14 | - | 87988454 | 87988523 | 87988143 | 87988207 | 87992969 | 87993271 | 11 | 4 | 0 | 0 | 2 | 0 | 1 | 5 | 1 | 0 | 98 | 49 | 1 | 1 | 1 | 0.5 | NA | | 0.333 | 1 | 0.084 |
| SE | 5113 | FLI1 | FLI1 | chr11 | + | 128807179 | 128807239 | 128805365 | 128805431 | 128809156 | 128809204 | 38 | 15 | 23 | 0 | 0 | 0 | 1 | 11 | 0 | 1 | 98 | 49 | 1 | 1 | 1 | 1 | 1 | | 1 | 0.846 | 0.077 |
| SE | 5114 | N4BP1 | N4BP1 | chr16 | - | 48546146 | 48546254 | 48538725 | 48543261 | 48548006 | 48548114 | 24 | 3 | 5 | 0 | 0 | 0 | 0 | 2 | 0 | 1 | 98 | 49 | 1 | 1 | 1 | 1 | 1 | NA | | 0.5 | 0.5 |
| SE | 5116 | N4BP1 | N4BP1 | chr16 | - | 48560753 | 48562444 | 48553538 | 48553669 | 48609774 | 48610209 | 16 | 14 | 5 | 1 | 0 | 0 | 2 | 1 | 0 | 0 | 98 | 49 | 1 | 1 | 0.889 | 1 | 1 | | 1 | 1 | -0.037 |
| SE | 5122 | HSPA8 | HSPA8 | chr11 | - | 123060592 | 123060798 | 123060115 | 123060268 | 123061119 | 123061329 | 3 | 0 | 5 | 2 | 0 | 0 | 0 | 1 | 0 | 0 | 98 | 49 | 1 | 1 | 0.429 | NA | 1 | NA | | 1 | -0.285 |
| SE | 5133 | SNX17 | SNX17 | chr2 | + | 27373039 | 27373113 | 27372598 | 27372740 | 27373246 | 27373311 | 1 | 0 | 0 | 2 | 0 | 0 | 0 | 0 | 0 | 1 | 98 | 49 | 1 | 1 | 0.2 | NA | NA | NA | | 0 | 0.2 |
| SE | 5134 | SNX17 | SNX17 | chr2 | + | 27374688 | 27374758 | 27374345 | 27374433 | 27375060 | 27375153 | 23 | 4 | 3 | 0 | 3 | 0 | 0 | 6 | 0 | 0 | 98 | 49 | 1 | 1 | 1 | 0.4 | 1 | NA | | 1 | -0.2 |
| SE | 5135 | MYL6 | MYL6 | chr12 | + | 56159586 | 56159730 | 56158683 | 56158711 | 56159974 | 56160148 | 149 | 85 | 76 | 1 | 0 | 0 | 7 | 24 | 0 | 1 | 98 | 49 | 1 | 1 | 0.987 | 1 | 1 | | 1 | 0.923 | 0.034 |
| SE | 5138 | VPS35 | VPS35 | chr16 | - | 46668929 | 46669052 | 46662982 | 46663162 | 46671704 | 46671860 | 18 | 10 | 11 | 0 | 0 | 1 | 3 | 6 | 0 | 0 | 98 | 49 | 1 | 1 | 1 | 1 | 0.846 | | 1 | 1 | -0.051 |
| SE | 5157 | NLRX1 | NLRX1 | chr11 | + | 119172355 | 119172425 | 119171355 | 119171473 | 119172900 | 119172989 | 2 | 0 | 0 | 3 | 0 | 0 | 0 | 1 | 0 | 2 | 98 | 49 | 1 | 1 | 0.25 | NA | NA | NA | | 0.2 | 0.05 |
| SE | 5166 | RBM42 | RBM42 | chr19 | + | 35633686 | 35634019 | 35633076 | 35633252 | 35634255 | 35634373 | 5 | 2 | 2 | 0 | 0 | 0 | 2 | 0 | 1 | 0 | 98 | 49 | 1 | 1 | 1 | 1 | 1 | | 0.5 | NA | 0.5 |
| SE | 5167 | DDX6 | DDX6 | chr11 | - | 118758773 | 118758902 | 118757170 | 118757287 | 118759921 | 118760044 | 64 | 44 | 17 | 3 | 3 | 0 | 0 | 9 | 0 | 0 | 98 | 49 | 1 | 1 | 0.914 | 0.88 | 1 | NA | | 1 | -0.069 |
| SE | 5168 | DDX6 | DDX6 | chr11 | - | 118765208 | 118765355 | 118763211 | 118763306 | 118768222 | 118768352 | 40 | 25 | 7 | 2 | 0 | 0 | 0 | 12 | 0 | 0 | 98 | 49 | 1 | 1 | 0.909 | 1 | 1 | NA | | 1 | -0.03 |
| SE | 5169 | ARCN1 | ARCN1 | chr11 | + | 118581245 | 118581509 | 118572386 | 118572550 | 118583178 | 118583358 | 13 | 4 | 4 | 4 | 0 | 0 | 0 | 1 | 0 | 0 | 98 | 49 | 1 | 1 | 0.619 | 1 | 1 | NA | | 1 | -0.127 |
| SE | 5180 | TSPAN32 | TSPAN32 | chr11 | + | 2308735 | 2308810 | 2304106 | 2304204 | 2313653 | 2313755 | 11 | 4 | 0 | 2 | 0 | 0 | 2 | 3 | 0 | 0 | 98 | 49 | 1 | 1 | 0.733 | 1 | NA | | 1 | 1 | -0.133 |
| SE | 5181 | GPATCH8 | GPATCH8 | chr17 | - | 44464471 | 44464544 | 44436477 | 44436545 | 44474828 | 44474903 | 5 | 3 | 0 | 1 | 0 | 0 | 0 | 1 | 0 | 0 | 98 | 49 | 1 | 1 | 0.714 | 1 | NA | NA | | 1 | -0.143 |
| SE | 5184 | MPZL3 | MPZL3 | chr11 | - | 118235423 | 118235589 | 118233459 | 118233523 | 118240210 | 118240377 | 36 | 6 | 6 | 0 | 1 | 1 | 0 | 1 | 0 | 0 | 98 | 49 | 1 | 1 | 1 | 0.75 | 0.75 | NA | | 1 | -0.167 |
| SE | 5185 | MPZL3 | MPZL3 | chr11 | - | 118237049 | 118237260 | 118235423 | 118235589 | 118240210 | 118240341 | 13 | 9 | 14 | 4 | 1 | 1 | 0 | 2 | 0 | 0 | 98 | 49 | 1 | 1 | 0.619 | 0.818 | 0.875 | NA | | 1 | -0.229 |
| SE | 5189 | OSGEP | OSGEP | chr14 | - | 20447903 | 20447994 | 20447614 | 20447690 | 20448105 | 20448171 | 10 | 0 | 1 | 4 | 0 | 0 | 0 | 2 | 0 | 0 | 98 | 49 | 1 | 1 | 0.556 | NA | 1 | NA | | 1 | -0.222 |
| SE | 5200 | ARNTL | ARNTL | chr11 | + | 13349944 | 13350070 | 13309986 | 13310059 | 13354300 | 13354448 | 26 | 10 | 0 | 0 | 1 | 0 | 0 | 4 | 0 | 0 | 98 | 49 | 1 | 1 | 1 | 0.833 | NA | NA | | 1 | -0.084 |
| SE | 5201 | ARNTL | ARNTL | chr11 | + | 13354300 | 13354448 | 13349944 | 13350070 | 13356738 | 13356777 | 28 | 6 | 1 | 0 | 0 | 5 | 0 | 9 | 0 | 0 | 98 | 49 | 1 | 1 | 1 | 1 | 0.091 | NA | | 1 | -0.303 |
| SE | 5202 | ARNTL | ARNTL | chr11 | + | 13355225 | 13355296 | 13354300 | 13354448 | 13356738 | 13356777 | 1 | 0 | 0 | 10 | 1 | 1 | 0 | 1 | 0 | 5 | 98 | 49 | 1 | 1 | 0.048 | 0 | 0 | NA | | 0.091 | -0.075 |
| SE | 5203 | ARNTL | ARNTL | chr11 | + | 13355232 | 13355296 | 13354300 | 13354448 | 13356738 | 13356777 | 0 | 0 | 0 | 10 | 1 | 1 | 0 | 1 | 0 | 5 | 98 | 49 | 1 | 1 | 0 | 0 | 0 | NA | | 0.091 | -0.091 |
| SE | 5204 | ARNTL | ARNTL | chr11 | + | 13365499 | 13365592 | 13360344 | 13360424 | 13366664 | 13366782 | 16 | 12 | 3 | 0 | 1 | 0 | 0 | 1 | 0 | 2 | 98 | 49 | 1 | 1 | 1 | 0.857 | 1 | NA | | 0.2 | 0.752 |
| SE | 5208 | PTPRJ | PTPRJ | chr11 | + | 48144999 | 48145124 | 48144674 | 48144885 | 48146875 | 48146963 | 45 | 8 | 15 | 0 | 0 | 0 | 11 | 9 | 0 | 1 | 98 | 49 | 1 | 1 | 1 | 1 | 1 | | 1 | 0.818 | 0.091 |
| SE | 5210 | PTPRJ | PTPRJ | chr11 | + | 48164379 | 48164515 | 48163457 | 48163618 | 48167203 | 48170842 | 48 | 34 | 30 | 1 | 0 | 0 | 1 | 10 | 0 | 1 | 98 | 49 | 1 | 1 | 0.96 | 1 | 1 | | 1 | 0.833 | 0.07 |
| SE | 5211 | ETV6 | ETV6 | chr12 | + | 11752449 | 11752579 | 11649853 | 11650160 | 11839139 | 11839304 | 22 | 6 | 0 | 0 | 0 | 1 | 3 | 1 | 0 | 0 | 98 | 49 | 1 | 1 | 1 | 1 | 0 | | 1 | 1 | -0.333 |
| SE | 5216 | LPGAT1 | LPGAT1 | chr1 | - | 211787631 | 211787727 | 211783228 | 211783502 | 211793071 | 211793190 | 60 | 46 | 19 | 0 | 1 | 1 | 0 | 15 | 0 | 0 | 98 | 49 | 1 | 1 | 1 | 0.958 | 0.905 | NA | | 1 | -0.046 |
| SE | 5217 | PLEKHM2 | PLEKHM2 | chr1 | + | 15716706 | 15716816 | 15716236 | 15716343 | 15717892 | 15717992 | 5 | 4 | 3 | 2 | 0 | 0 | 0 | 3 | 0 | 0 | 98 | 49 | 1 | 1 | 0.556 | 1 | 1 | NA | | 1 | -0.148 |
| SE | 5218 | PLEKHM2 | PLEKHM2 | chr1 | + | 15719733 | 15719920 | 15718537 | 15718625 | 15721328 | 15721388 | 12 | 9 | 5 | 0 | 2 | 0 | 1 | 2 | 0 | 0 | 98 | 49 | 1 | 1 | 1 | 0.692 | 1 | | 1 | 1 | -0.103 |
| SE | 5228 | POLG | POLG | chr15 | - | 89322741 | 89322902 | 89321961 | 89322015 | 89323403 | 89323511 | 4 | 0 | 2 | 0 | 2 | 0 | 0 | 3 | 0 | 0 | 98 | 49 | 1 | 1 | 1 | 0 | 1 | NA | | 1 | -0.333 |
| SE | 5229 | POLG | POLG | chr15 | - | 89323403 | 89323511 | 89322741 | 89322902 | 89323814 | 89323901 | 10 | 1 | 2 | 1 | 0 | 0 | 0 | 4 | 0 | 0 | 98 | 49 | 1 | 1 | 0.833 | 1 | 1 | NA | | 1 | -0.056 |
| SE | 5230 | AKIP1 | AKIP1 | chr11 | + | 8912452 | 8912533 | 8911191 | 8911671 | 8914825 | 8914930 | 2 | 0 | 0 | 0 | 0 | 0 | 0 | 0 | 0 | 2 | 98 | 49 | 1 | 1 | 1 | NA | NA | NA | | 0 | 1 |
| SE | 5232 | SNX19 | SNX19 | chr11 | - | 130907952 | 130908083 | 130906624 | 130906721 | 130910017 | 130910137 | 4 | 4 | 1 | 1 | 0 | 0 | 0 | 1 | 0 | 0 | 98 | 49 | 1 | 1 | 0.667 | 1 | 1 | NA | | 1 | -0.111 |
| SE | 5240 | TRAF3IP3 | TRAF3IP3 | chr1 | + | 209780469 | 209780606 | 209779314 | 209779374 | 209781344 | 209781458 | 10 | 3 | 0 | 3 | 0 | 0 | 0 | 3 | 0 | 0 | 98 | 49 | 1 | 1 | 0.625 | 1 | NA | NA | | 1 | -0.188 |
| SE | 5242 | CHI3L1 | CHI3L1 | chr1 | - | 203183640 | 203183791 | 203182730 | 203182852 | 203184575 | 203184632 | 111 | 43 | 38 | 0 | 1 | 0 | 1 | 4 | 0 | 0 | 98 | 49 | 1 | 1 | 1 | 0.956 | 1 | | 1 | 1 | -0.015 |
| SE | 5244 | PFKFB3 | PFKFB3 | chr10 | + | 6217134 | 6217191 | 6216705 | 6216780 | 6219568 | 6219693 | 70 | 45 | 12 | 0 | 0 | 0 | 2 | 31 | 0 | 1 | 98 | 49 | 1 | 1 | 1 | 1 | 1 | | 1 | 0.939 | 0.03 |
| SE | 5245 | PFKFB3 | PFKFB3 | chr10 | + | 6221640 | 6221745 | 6221380 | 6221527 | 6222854 | 6222984 | 73 | 49 | 6 | 2 | 0 | 0 | 16 | 35 | 0 | 0 | 98 | 49 | 1 | 1 | 0.948 | 1 | 1 | | 1 | 1 | -0.017 |
| SE | 5246 | PFKFB3 | PFKFB3 | chr10 | + | 6221640 | 6221745 | 6221380 | 6221527 | 6223957 | 6224020 | 56 | 41 | 5 | 0 | 0 | 0 | 11 | 20 | 0 | 1 | 98 | 49 | 1 | 1 | 1 | 1 | 1 | | 1 | 0.909 | 0.045 |
| SE | 5247 | PFKFB3 | PFKFB3 | chr10 | + | 6222854 | 6222984 | 6221380 | 6221527 | 6223957 | 6224020 | 69 | 41 | 7 | 0 | 0 | 0 | 3 | 22 | 0 | 1 | 98 | 49 | 1 | 1 | 1 | 1 | 1 | | 1 | 0.917 | 0.041 |
| SE | 5248 | PFKFB3 | PFKFB3 | chr10 | + | 6222854 | 6222984 | 6221640 | 6221745 | 6223957 | 6224020 | 85 | 49 | 8 | 1 | 0 | 0 | 8 | 37 | 0 | 0 | 98 | 49 | 1 | 1 | 0.977 | 1 | 1 | | 1 | 1 | -0.008 |
| SE | 5249 | PFKFB3 | PFKFB3 | chr10 | + | 6224148 | 6224213 | 6223957 | 6224020 | 6226191 | 6226365 | 78 | 53 | 4 | 2 | 0 | 0 | 10 | 43 | 0 | 0 | 98 | 49 | 1 | 1 | 0.951 | 1 | 1 | | 1 | 1 | -0.016 |
| SE | 5252 | PTPRC | PTPRC | chr1 | + | 198696711 | 198696909 | 198692346 | 198692373 | 198699563 | 198699704 | 12 | 10 | 0 | 43 | 17 | 16 | 0 | 1 | 0 | 12 | 98 | 49 | 1 | 1 | 0.122 | 0.227 | 0 | NA | | 0.04 | 0.076 |
| SE | 5255 | PTPRC | PTPRC | chr1 | + | 198709686 | 198709824 | 198708132 | 198708261 | 198712952 | 198713072 | 276 | 128 | 85 | 1 | 0 | 0 | 0 | 36 | 0 | 0 | 98 | 49 | 1 | 1 | 0.993 | 1 | 1 | NA | | 1 | -0.002 |
| SE | 5256 | PTPRC | PTPRC | chr1 | + | 198741868 | 198742026 | 198735126 | 198735252 | 198742231 | 198742367 | 553 | 282 | 135 | 3 | 0 | 0 | 20 | 106 | 0 | 0 | 98 | 49 | 1 | 1 | 0.989 | 1 | 1 | | 1 | 1 | -0.004 |
| SE | 5257 | PTPRC | PTPRC | chr1 | + | 198742231 | 198742367 | 198741868 | 198742026 | 198744053 | 198744203 | 562 | 258 | 160 | 1 | 0 | 0 | 17 | 98 | 0 | 0 | 98 | 49 | 1 | 1 | 0.996 | 1 | 1 | | 1 | 1 | -0.001 |
| SE | 5258 | PTPRC | PTPRC | chr1 | + | 198749415 | 198749549 | 198748108 | 198748199 | 198750491 | 198750626 | 477 | 276 | 181 | 2 | 3 | 0 | 1 | 72 | 0 | 1 | 98 | 49 | 1 | 1 | 0.992 | 0.979 | 1 | | 1 | 0.973 | 0.004 |
| SE | 5266 | ARID3B | ARID3B | chr15 | + | 74573131 | 74573204 | 74572861 | 74572933 | 74589819 | 74590003 | 0 | 0 | 0 | 1 | 0 | 0 | 0 | 1 | 0 | 0 | 98 | 49 | 1 | 1 | 0 | NA | NA | NA | | 1 | -1 |
| SE | 5268 | TATDN3 | TATDN3 | chr1 | + | 212802700 | 212802763 | 212797111 | 212797196 | 212804319 | 212804429 | 4 | 2 | 5 | 0 | 1 | 0 | 0 | 2 | 0 | 1 | 98 | 49 | 1 | 1 | 1 | 0.5 | 1 | NA | | 0.5 | 0.333 |
| SE | 5269 | MARK2 | MARK2 | chr11 | + | 63902600 | 63902782 | 63902197 | 63902330 | 63903060 | 63903158 | 13 | 1 | 3 | 2 | 0 | 0 | 0 | 3 | 0 | 0 | 98 | 49 | 1 | 1 | 0.765 | 1 | 1 | NA | | 1 | -0.078 |
| SE | 5270 | MARK2 | MARK2 | chr11 | + | 63902603 | 63902782 | 63902197 | 63902330 | 63903060 | 63903158 | 37 | 5 | 11 | 2 | 0 | 0 | 0 | 9 | 0 | 0 | 98 | 49 | 1 | 1 | 0.902 | 1 | 1 | NA | | 1 | -0.033 |
| SE | 5271 | MARK2 | MARK2 | chr11 | + | 63903985 | 63904147 | 63903060 | 63903158 | 63904785 | 63905043 | 8 | 5 | 2 | 10 | 6 | 0 | 0 | 1 | 0 | 0 | 98 | 49 | 1 | 1 | 0.286 | 0.294 | 1 | NA | | 1 | -0.473 |
| SE | 5272 | MARK2 | MARK2 | chr11 | + | 63906087 | 63906114 | 63904785 | 63905043 | 63908259 | 63908304 | 2 | 0 | 0 | 18 | 2 | 1 | 0 | 0 | 0 | 1 | 76 | 49 | 1 | 1 | 0.067 | 0 | 0 | NA | | 0 | 0.022 |
| SE | 5284 | ZNRD1 | ZNRD1 | chr6 | + | 30061916 | 30062017 | 30061239 | 30061669 | 30062223 | 30062333 | 7 | 2 | 2 | 1 | 0 | 3 | 0 | 2 | 0 | 0 | 98 | 49 | 1 | 1 | 0.778 | 1 | 0.25 | NA | | 1 | -0.324 |
| SE | 5286 | CALML4 | CALML4 | chr15 | - | 68199540 | 68199681 | 68197439 | 68197628 | 68205120 | 68205151 | 3 | 0 | 0 | 2 | 0 | 0 | 0 | 1 | 0 | 0 | 98 | 49 | 1 | 1 | 0.429 | NA | NA | NA | | 1 | -0.571 |
| SE | 5290 | ZRANB1 | ZRANB1 | chr10 | + | 124974199 | 124974398 | 124973644 | 124973716 | 124981708 | 124981829 | 14 | 1 | 3 | 0 | 0 | 0 | 0 | 0 | 0 | 1 | 98 | 49 | 1 | 1 | 1 | 1 | 1 | NA | | 0 | 1 |
| SE | 5291 | ZRANB1 | ZRANB1 | chr10 | + | 124981708 | 124981829 | 124974199 | 124974398 | 124983174 | 124983304 | 18 | 11 | 2 | 1 | 0 | 0 | 1 | 1 | 0 | 1 | 98 | 49 | 1 | 1 | 0.9 | 1 | 1 | | 1 | 0.333 | 0.3 |
| SE | 5292 | ZRANB1 | ZRANB1 | chr10 | + | 124983174 | 124983304 | 124974199 | 124974398 | 124984773 | 124987436 | 3 | 0 | 0 | 0 | 1 | 0 | 0 | 2 | 0 | 0 | 98 | 49 | 1 | 1 | 1 | 0 | NA | NA | | 1 | -0.5 |
| SE | 5293 | ZRANB1 | ZRANB1 | chr10 | + | 124983458 | 124983688 | 124983174 | 124983304 | 124984773 | 124987436 | 10 | 12 | 6 | 2 | 0 | 0 | 0 | 0 | 0 | 1 | 98 | 49 | 1 | 1 | 0.714 | 1 | 1 | NA | | 0 | 0.905 |
| SE | 5294 | TGFB1 | TGFB1 | chr19 | - | 41341882 | 41342030 | 41332127 | 41332281 | 41342169 | 41342247 | 75 | 17 | 8 | 1 | 0 | 0 | 1 | 23 | 0 | 0 | 98 | 49 | 1 | 1 | 0.974 | 1 | 1 | | 1 | 1 | -0.009 |
| SE | 5304 | PTPN7 | PTPN7 | chr1 | - | 202153724 | 202153835 | 202152541 | 202152699 | 202155532 | 202155609 | 7 | 2 | 2 | 0 | 0 | 0 | 0 | 1 | 0 | 2 | 98 | 49 | 1 | 1 | 1 | 1 | 1 | NA | | 0.2 | 0.8 |
| SE | 5305 | PTPN7 | PTPN7 | chr1 | - | 202154185 | 202154323 | 202153724 | 202153835 | 202155532 | 202155609 | 9 | 2 | 6 | 1 | 0 | 0 | 1 | 0 | 0 | 0 | 98 | 49 | 1 | 1 | 0.818 | 1 | 1 | | 1 | NA | -0.061 |
| SE | 5310 | PPFIA1 | PPFIA1 | chr11 | + | 70348188 | 70348420 | 70343668 | 70343892 | 70354300 | 70354452 | 21 | 2 | 4 | 1 | 0 | 0 | 0 | 1 | 0 | 0 | 98 | 49 | 1 | 1 | 0.913 | 1 | 1 | NA | | 1 | -0.029 |
| SE | 5312 | PPFIA1 | PPFIA1 | chr11 | + | 70382087 | 70382158 | 70378029 | 70378195 | 70383002 | 70384501 | 18 | 2 | 1 | 3 | 6 | 1 | 0 | 4 | 0 | 0 | 98 | 49 | 1 | 1 | 0.75 | 0.143 | 0.333 | NA | | 1 | -0.591 |
| SE | 5313 | NDUFV1 | NDUFV1 | chr11 | + | 67611896 | 67611978 | 67611402 | 67611569 | 67612119 | 67612265 | 11 | 2 | 1 | 1 | 0 | 0 | 0 | 5 | 0 | 0 | 98 | 49 | 1 | 1 | 0.846 | 1 | 1 | NA | | 1 | -0.051 |
| SE | 5317 | GATAD2A | GATAD2A | chr19 | - | 19501968 | 19502043 | 19501117 | 19501416 | 19502330 | 19502526 | 14 | 3 | 4 | 0 | 4 | 0 | 0 | 3 | 0 | 0 | 98 | 49 | 1 | 1 | 1 | 0.273 | 1 | NA | | 1 | -0.242 |
| SE | 5325 | BAHD1 | BAHD1 | chr15 | + | 40461911 | 40462294 | 40458450 | 40459896 | 40463860 | 40464020 | 1 | 0 | 0 | 0 | 1 | 0 | 0 | 1 | 0 | 0 | 98 | 49 | 1 | 1 | 1 | 0 | NA | NA | | 1 | -0.5 |
| SE | 5343 | SF3B1 | SF3B1 | chr2 | - | 197409769 | 197410007 | 197408368 | 197408581 | 197416740 | 197416911 | 141 | 48 | 38 | 0 | 1 | 0 | 2 | 16 | 0 | 0 | 98 | 49 | 1 | 1 | 1 | 0.96 | 1 | | 1 | 1 | -0.013 |
| SE | 5353 | NBPF10 | NBPF10 | chr1 | - | 146133522 | 146133728 | 146132185 | 146132237 | 146134192 | 146134265 | 15 | 0 | 0 | 2 | 0 | 0 | 0 | 4 | 0 | 0 | 98 | 49 | 1 | 1 | 0.789 | NA | NA | NA | | 1 | -0.211 |
| SE | 5355 | TAB2 | TAB2 | chr6 | + | 149399103 | 149399184 | 149397968 | 149398062 | 149409576 | 149411613 | 59 | 22 | 4 | 1 | 0 | 0 | 1 | 9 | 0 | 0 | 98 | 49 | 1 | 1 | 0.967 | 1 | 1 | | 1 | 1 | -0.011 |
| SE | 5356 | CXCL8 | CXCL8 | chr4 | + | 73741541 | 73741677 | 73740505 | 73740722 | 73741948 | 73742032 | 1882 | 3505 | 450 | 5 | 7 | 0 | 26 | 133 | 1 | 0 | 98 | 49 | 1 | 1 | 0.995 | 0.996 | 1 | | 0.929 | 1 | 0.032 |
| SE | 5362 | TNRC18 | TNRC18 | chr7 | - | 5361593 | 5361722 | 5359397 | 5359569 | 5361896 | 5362033 | 24 | 5 | 1 | 0 | 3 | 3 | 0 | 4 | 0 | 2 | 98 | 49 | 1 | 1 | 1 | 0.455 | 0.143 | NA | | 0.5 | 0.033 |
| SE | 5364 | TROVE2 | TROVE2 | chr1 | + | 193069033 | 193069634 | 193059421 | 193059776 | 193075819 | 193076040 | 2 | 0 | 0 | 3 | 2 | 0 | 0 | 4 | 0 | 1 | 98 | 49 | 1 | 1 | 0.25 | 0 | NA | NA | | 0.667 | -0.542 |
| SE | 5372 | RGS2 | RGS2 | chr1 | + | 192810165 | 192810267 | 192809038 | 192809181 | 192810369 | 192810431 | 1491 | 624 | 334 | 1 | 0 | 0 | 11 | 186 | 0 | 0 | 98 | 49 | 1 | 1 | 0.999 | 1 | 1 | | 1 | 1 | 0 |
| SE | 5374 | RGS2 | RGS2 | chr1 | + | 192810980 | 192811147 | 192810369 | 192810431 | 192811401 | 192812277 | 1435 | 717 | 501 | 0 | 1 | 1 | 8 | 199 | 0 | 0 | 98 | 49 | 1 | 1 | 1 | 0.997 | 0.996 | | 1 | 1 | -0.002 |
| SE | 5377 | SLC9A8 | SLC9A8 | chr20 | + | 49839540 | 49839599 | 49823060 | 49823141 | 49845035 | 49845119 | 31 | 5 | 3 | 0 | 0 | 0 | 0 | 8 | 0 | 1 | 98 | 49 | 1 | 1 | 1 | 1 | 1 | NA | | 0.8 | 0.2 |
| SE | 5379 | SLC9A8 | SLC9A8 | chr20 | + | 49849578 | 49849680 | 49845035 | 49845119 | 49855437 | 49855581 | 32 | 13 | 0 | 3 | 0 | 0 | 0 | 10 | 0 | 2 | 98 | 49 | 1 | 1 | 0.842 | 1 | NA | NA | | 0.714 | 0.207 |
| SE | 5380 | SLC9A8 | SLC9A8 | chr20 | + | 49850761 | 49850844 | 49849578 | 49849680 | 49855437 | 49855581 | 7 | 4 | 0 | 14 | 9 | 0 | 0 | 4 | 0 | 4 | 98 | 49 | 1 | 1 | 0.2 | 0.182 | NA | NA | | 0.333 | -0.142 |
| SE | 5381 | SLC9A8 | SLC9A8 | chr20 | + | 49850809 | 49850844 | 49849578 | 49849680 | 49855437 | 49855581 | 7 | 6 | 0 | 14 | 9 | 0 | 0 | 4 | 0 | 4 | 84 | 49 | 1 | 1 | 0.226 | 0.28 | NA | NA | | 0.368 | -0.115 |
| SE | 5382 | SLC9A8 | SLC9A8 | chr20 | + | 49874704 | 49874821 | 49864727 | 49864844 | 49877980 | 49878063 | 23 | 10 | 8 | 2 | 0 | 0 | 0 | 4 | 0 | 0 | 98 | 49 | 1 | 1 | 0.852 | 1 | 1 | NA | | 1 | -0.049 |
| SE | 5388 | RRM2B | RRM2B | chr8 | - | 102224884 | 102225018 | 102224045 | 102224140 | 102225917 | 102226034 | 32 | 21 | 15 | 0 | 2 | 0 | 0 | 4 | 0 | 2 | 98 | 49 | 1 | 1 | 1 | 0.84 | 1 | NA | | 0.5 | 0.447 |
| SE | 5389 | RRM2B | RRM2B | chr8 | - | 102232148 | 102232304 | 102225917 | 102226034 | 102238826 | 102239118 | 24 | 7 | 1 | 0 | 0 | 2 | 0 | 1 | 0 | 0 | 98 | 49 | 1 | 1 | 1 | 1 | 0.2 | NA | | 1 | -0.267 |
| SE | 5392 | TMEM126B | TMEM126B | chr11 | + | 85631144 | 85631316 | 85628572 | 85628688 | 85631686 | 85631808 | 2 | 3 | 0 | 1 | 0 | 0 | 0 | 2 | 0 | 0 | 98 | 49 | 1 | 1 | 0.5 | 1 | NA | NA | | 1 | -0.25 |
| SE | 5405 | RASGRP1 | RASGRP1 | chr15 | - | 38501142 | 38501287 | 38500102 | 38500139 | 38502311 | 38502421 | 5 | 5 | 4 | 1 | 0 | 0 | 0 | 1 | 0 | 0 | 98 | 49 | 1 | 1 | 0.714 | 1 | 1 | NA | | 1 | -0.095 |
| SE | 5406 | RASGRP1 | RASGRP1 | chr15 | - | 38503271 | 38503376 | 38502311 | 38502421 | 38505839 | 38505920 | 9 | 1 | 3 | 0 | 1 | 0 | 0 | 2 | 0 | 0 | 98 | 49 | 1 | 1 | 1 | 0.333 | 1 | NA | | 1 | -0.222 |
| SE | 5408 | NCF2 | NCF2 | chr1 | - | 183565703 | 183565779 | 183564004 | 183564030 | 183566919 | 183566988 | 308 | 136 | 154 | 1 | 0 | 0 | 16 | 51 | 0 | 0 | 98 | 49 | 1 | 1 | 0.994 | 1 | 1 | | 1 | 1 | -0.002 |
| SE | 5409 | NCF2 | NCF2 | chr1 | - | 183566919 | 183566988 | 183565703 | 183565779 | 183567203 | 183567345 | 228 | 117 | 141 | 1 | 0 | 0 | 12 | 47 | 0 | 0 | 98 | 49 | 1 | 1 | 0.991 | 1 | 1 | | 1 | 1 | -0.003 |
| SE | 5413 | NCF2 | NCF2 | chr1 | - | 183569141 | 183569185 | 183565703 | 183565779 | 183570779 | 183570839 | 247 | 138 | 71 | 2 | 0 | 0 | 6 | 30 | 0 | 0 | 93 | 49 | 1 | 1 | 0.985 | 1 | 1 | | 1 | 1 | -0.005 |
| SE | 5414 | NCF2 | NCF2 | chr1 | - | 183573184 | 183573292 | 183570779 | 183570839 | 183574486 | 183574621 | 417 | 212 | 150 | 0 | 0 | 0 | 5 | 63 | 0 | 1 | 98 | 49 | 1 | 1 | 1 | 1 | 1 | | 1 | 0.969 | 0.016 |
| SE | 5416 | NCF2 | NCF2 | chr1 | - | 183574486 | 183574621 | 183570779 | 183570839 | 183577598 | 183577707 | 118 | 56 | 89 | 3 | 2 | 0 | 0 | 30 | 0 | 1 | 98 | 49 | 1 | 1 | 0.952 | 0.933 | 1 | | | 0.937 | 0.025 |
| SE | 5417 | NCF2 | NCF2 | chr1 | - | 183574486 | 183574621 | 183573184 | 183573292 | 183577598 | 183577707 | 255 | 137 | 159 | 24 | 16 | 0 | 0 | 51 | 0 | 12 | 98 | 49 | 1 | 1 | 0.842 | 0.811 | 1 | NA | | 0.68 | 0.204 |
| SE | 5418 | NCF2 | NCF2 | chr1 | - | 183577598 | 183577707 | 183573184 | 183573292 | 183586894 | 183586977 | 182 | 104 | 98 | 0 | 1 | 0 | 3 | 42 | 0 | 0 | 98 | 49 | 1 | 1 | 1 | 0.981 | 1 | | 1 | 1 | -0.006 |
| SE | 5419 | NCF2 | NCF2 | chr1 | - | 183586894 | 183586977 | 183577598 | 183577707 | 183590155 | 183590359 | 468 | 224 | 221 | 1 | 0 | 0 | 9 | 109 | 0 | 0 | 98 | 49 | 1 | 1 | 0.996 | 1 | 1 | | 1 | 1 | -0.001 |
| SE | 5426 | HBEGF | HBEGF | chr5 | - | 140345910 | 140346084 | 140342634 | 140342812 | 140346282 | 140346603 | 4 | 0 | 0 | 0 | 0 | 0 | 0 | 5 | 0 | 1 | 98 | 49 | 1 | 1 | 1 | NA | NA | NA | | 0.714 | 0.286 |
| SE | 5427 | PFKFB4 | PFKFB4 | chr3 | - | 48539253 | 48539310 | 48538497 | 48538619 | 48539696 | 48539771 | 20 | 7 | 2 | 1 | 1 | 0 | 0 | 7 | 0 | 1 | 98 | 49 | 1 | 1 | 0.909 | 0.778 | 1 | NA | | 0.778 | 0.118 |
| SE | 5431 | SMU1 | SMU1 | chr9 | - | 33053122 | 33053290 | 33048105 | 33048258 | 33056112 | 33056239 | 13 | 2 | 0 | 1 | 0 | 0 | 0 | 1 | 0 | 0 | 98 | 49 | 1 | 1 | 0.867 | 1 | NA | NA | | 1 | -0.067 |
| SE | 5432 | SMU1 | SMU1 | chr9 | - | 33071739 | 33071892 | 33068823 | 33068934 | 33073595 | 33073806 | 3 | 2 | 0 | 0 | 0 | 0 | 0 | 1 | 0 | 2 | 98 | 49 | 1 | 1 | 1 | 1 | NA | NA | | 0.2 | 0.8 |
| SE | 5433 | GLIPR1 | GLIPR1 | chr12 | + | 75490405 | 75490518 | 75481833 | 75482079 | 75495576 | 75495662 | 60 | 35 | 15 | 0 | 0 | 0 | 0 | 15 | 0 | 1 | 98 | 49 | 1 | 1 | 1 | 1 | 1 | NA | | 0.882 | 0.118 |
| SE | 5435 | FAR2 | FAR2 | chr12 | + | 29293299 | 29293475 | 29270411 | 29270638 | 29297020 | 29297200 | 20 | 6 | 11 | 1 | 0 | 0 | 0 | 1 | 0 | 0 | 98 | 49 | 1 | 1 | 0.909 | 1 | 1 | NA | | 1 | -0.03 |
| SE | 5436 | FAR2 | FAR2 | chr12 | + | 29297020 | 29297200 | 29293299 | 29293475 | 29307657 | 29307835 | 29 | 16 | 12 | 0 | 0 | 0 | 0 | 1 | 0 | 1 | 98 | 49 | 1 | 1 | 1 | 1 | 1 | NA | | 0.333 | 0.667 |
| SE | 5437 | FAR2 | FAR2 | chr12 | + | 29316840 | 29317012 | 29311882 | 29311950 | 29321794 | 29321924 | 9 | 0 | 1 | 0 | 0 | 0 | 0 | 1 | 0 | 1 | 98 | 49 | 1 | 1 | 1 | NA | 1 | NA | | 0.333 | 0.667 |
| SE | 5438 | TIMM8B | TIMM8B | chr11 | - | 112085977 | 112086310 | 112084810 | 112085462 | 112086639 | 112086798 | 0 | 0 | 0 | 0 | 0 | 3 | 0 | 1 | 0 | 0 | 98 | 49 | 1 | 1 | NA | NA | 0 | NA | | 1 | -1 |
| SE | 5439 | RASGRP4 | RASGRP4 | chr19 | - | 38414847 | 38414916 | 38413393 | 38413474 | 38417051 | 38417168 | 17 | 10 | 3 | 1 | 0 | 0 | 0 | 3 | 0 | 0 | 98 | 49 | 1 | 1 | 0.895 | 1 | 1 | NA | | 1 | -0.035 |
| SE | 5440 | RASGRP4 | RASGRP4 | chr19 | - | 38414847 | 38415123 | 38413393 | 38413474 | 38417051 | 38417168 | 56 | 20 | 10 | 1 | 0 | 0 | 2 | 16 | 0 | 0 | 98 | 49 | 1 | 1 | 0.966 | 1 | 1 | | 1 | 1 | -0.011 |
| SE | 5446 | BCL6 | BCL6 | chr3 | - | 187728359 | 187728544 | 187726730 | 187726898 | 187729049 | 187730021 | 81 | 30 | 25 | 0 | 0 | 0 | 2 | 24 | 0 | 1 | 98 | 49 | 1 | 1 | 1 | 1 | 1 | | 1 | 0.923 | 0.038 |
| SE | 5447 | RALGPS2 | RALGPS2 | chr1 | + | 178821611 | 178821704 | 178811314 | 178811404 | 178833423 | 178833550 | 8 | 1 | 5 | 1 | 0 | 0 | 0 | 1 | 0 | 0 | 98 | 49 | 1 | 1 | 0.8 | 1 | 1 | NA | | 1 | -0.067 |
| SE | 5448 | RALGPS2 | RALGPS2 | chr1 | + | 178892229 | 178892307 | 178889641 | 178889696 | 178893918 | 178894024 | 6 | 0 | 5 | 1 | 1 | 0 | 0 | 0 | 0 | 1 | 98 | 49 | 1 | 1 | 0.75 | 0 | 1 | NA | | 0 | 0.583 |
| SE | 5449 | RALGPS2 | RALGPS2 | chr1 | + | 178906775 | 178906867 | 178902105 | 178902211 | 178916329 | 178921842 | 8 | 6 | 0 | 1 | 0 | 0 | 0 | 1 | 0 | 0 | 98 | 49 | 1 | 1 | 0.8 | 1 | NA | NA | | 1 | -0.1 |
| SE | 5460 | SAMD8 | SAMD8 | chr10 | + | 75168540 | 75168658 | 75164644 | 75164740 | 75176065 | 75176216 | 11 | 10 | 2 | 1 | 0 | 0 | 0 | 2 | 0 | 1 | 98 | 49 | 1 | 1 | 0.846 | 1 | 1 | NA | | 0.5 | 0.449 |
| SE | 5461 | LOC374443 | LOC374443 | chr12 | + | 9653682 | 9653808 | 9648046 | 9648933 | 9655872 | 9655979 | 16 | 0 | 0 | 0 | 3 | 0 | 0 | 5 | 0 | 0 | 98 | 49 | 1 | 1 | 1 | 0 | NA | NA | | 1 | -0.5 |
| SE | 5464 | LOC374443 | LOC374443 | chr12 | + | 9656567 | 9656649 | 9655872 | 9655979 | 9656920 | 9658414 | 8 | 0 | 1 | 7 | 0 | 2 | 0 | 0 | 0 | 3 | 98 | 49 | 1 | 1 | 0.364 | NA | 0.2 | NA | | 0 | 0.282 |
| SE | 5465 | LOC374443 | LOC374443 | chr12 | + | 9656567 | 9656697 | 9655872 | 9655979 | 9656920 | 9658414 | 7 | 1 | 2 | 7 | 0 | 2 | 0 | 0 | 0 | 3 | 98 | 49 | 1 | 1 | 0.333 | 1 | 0.333 | NA | | 0 | 0.555 |
| SE | 5466 | RC3H1 | RC3H1 | chr1 | - | 173943441 | 173943588 | 173941264 | 173941380 | 173946475 | 173946608 | 6 | 6 | 6 | 2 | 3 | 0 | 0 | 2 | 0 | 3 | 98 | 49 | 1 | 1 | 0.6 | 0.5 | 1 | NA | | 0.25 | 0.45 |
| SE | 5467 | RC3H1 | RC3H1 | chr1 | - | 173943441 | 173943615 | 173941264 | 173941380 | 173946475 | 173946608 | 11 | 3 | 2 | 2 | 3 | 0 | 0 | 2 | 0 | 3 | 98 | 49 | 1 | 1 | 0.733 | 0.333 | 1 | NA | | 0.25 | 0.439 |

| | | | | | | | | | | | | | | | | | | | | | | | | | | | | | | | |
|---|---|---|---|---|---|---|---|---|---|---|---|---|---|---|---|---|---|---|---|---|---|---|---|---|---|---|---|---|---|---|---|
| SE | 5468 RC3H1 | RC3H1 | chr1 | - | 173951985 | 173952138 | 173947368 | 173947582 | 173961076 | 173961244 | 35 | 12 | 1 | 0 | 0 | 0 | 0 | 9 | 0 | 1 | 98 | 49 | 1 | 1 | 1 | 1 | 1 | NA | | 0.818 | 0.182 |
| SE | 5469 RC3H1 | RC3H1 | chr1 | - | 173964838 | 173965120 | 173963972 | 173964187 | 173970504 | 173970617 | 27 | 19 | 2 | 0 | 0 | 0 | 0 | 5 | 0 | 1 | 98 | 49 | 1 | 1 | 1 | 1 | 1 | NA | | 0.714 | 0.286 |
| SE | 5470 RC3H1 | RC3H1 | chr1 | - | 173972508 | 173972627 | 173970504 | 173970617 | 173978487 | 173978620 | 28 | 12 | 2 | 1 | 0 | 0 | 0 | 14 | 0 | 0 | 98 | 49 | 1 | 1 | 0.933 | 1 | 1 | NA | | 1 | -0.022 |
| SE | 5471 RC3H1 | RC3H1 | chr1 | - | 173983417 | 173983657 | 173982726 | 173982902 | 173984498 | 173984619 | 71 | 13 | 14 | 0 | 0 | 0 | 1 | 7 | 0 | 1 | 98 | 49 | 1 | 1 | 1 | 1 | 1 | | 1 | 0.778 | 0.111 |
| SE | 5480 DNAJB12 | DNAJB12 | chr10 | - | 72338201 | 72338311 | 72336523 | 72336696 | 72340788 | 72340868 | 11 | 7 | 3 | 0 | 0 | 0 | 0 | 2 | 0 | 1 | 98 | 49 | 1 | 1 | 1 | 1 | 1 | NA | | 0.5 | 0.5 |
| SE | 5485 TCIRG1 | TCIRG1 | chr11 | + | 68043370 | 68043497 | 68042945 | 68043031 | 68043570 | 68043653 | 76 | 29 | 21 | 1 | 0 | 0 | 2 | 17 | 0 | 0 | 98 | 49 | 1 | 1 | 0.974 | 1 | 1 | | 1 | 1 | -0.009 |
| SE | 5487 ZHX1-C8orf76 | ZHX1-C8orf76 | chr8 | - | 123231299 | 123231757 | 123226188 | 123226632 | 123237797 | 123237941 | 4 | 0 | 0 | 0 | 1 | 1 | 0 | 3 | 0 | 0 | 98 | 49 | 1 | 1 | 1 | 0 | 0 | NA | | 1 | -0.667 |
| SE | 5491 YWHAQ | YWHAQ | chr2 | - | 9588164 | 9588328 | 9587413 | 9587509 | 9591391 | 9591515 | 25 | 2 | 4 | 0 | 2 | 0 | 0 | 10 | 0 | 0 | 98 | 49 | 1 | 1 | 1 | 0.333 | 1 | NA | | 1 | -0.222 |
| SE | 5492 OGFRL1 | OGFRL1 | chr6 | + | 71296316 | 71296395 | 71293532 | 71293611 | 71296494 | 71296561 | 170 | 67 | 42 | 1 | 0 | 0 | 0 | 15 | 0 | 0 | 98 | 49 | 1 | 1 | 0.988 | 1 | 1 | NA | | 1 | -0.004 |
| SE | 5493 OGFRL1 | OGFRL1 | chr6 | + | 71296494 | 71296561 | 71293532 | 71293611 | 71296671 | 71296817 | 81 | 39 | 29 | 1 | 0 | 0 | 0 | 17 | 0 | 0 | 98 | 49 | 1 | 1 | 0.976 | 1 | 1 | NA | | 1 | -0.008 |
| SE | 5495 SYNE1 | SYNE1 | chr6 | - | 152145486 | 152145555 | 152143622 | 152143765 | 152148044 | 152148378 | 5 | 0 | 2 | 1 | 0 | 0 | 2 | 2 | 0 | 0 | 98 | 49 | 1 | 1 | 0.714 | NA | 1 | | 1 | 1 | -0.143 |
| SE | 5498 SYNE1 | SYNE1 | chr6 | - | 152325957 | 152326102 | 152325083 | 152325302 | 152326295 | 152326633 | 10 | 7 | 2 | 1 | 0 | 0 | 0 | 5 | 0 | 0 | 98 | 49 | 1 | 1 | 0.833 | 1 | 1 | NA | | 1 | -0.056 |
| SE | 5500 SYNE1 | SYNE1 | chr6 | - | 152352026 | 152352353 | 152350617 | 152350770 | 152353262 | 152353433 | 9 | 12 | 0 | 0 | 1 | 0 | 0 | 2 | 0 | 0 | 98 | 49 | 1 | 1 | 1 | 0.857 | NA | NA | | 1 | -0.072 |
| SE | 5502 SYNE1 | SYNE1 | chr6 | - | 152396774 | 152396980 | 152395515 | 152395671 | 152398618 | 152398731 | 12 | 5 | 1 | 1 | 0 | 0 | 0 | 2 | 0 | 0 | 98 | 49 | 1 | 1 | 0.857 | 1 | 1 | NA | | 1 | -0.048 |
| SE | 5512 RPAIN | RPAIN | chr17 | + | 5425970 | 5426082 | 5422768 | 5422829 | 5432541 | 5433020 | 0 | 2 | 0 | 0 | 1 | 0 | 0 | 2 | 0 | 1 | 98 | 49 | 1 | 1 | NA | 0.5 | NA | NA | | 0.5 | 0 |
| SE | 5517 RPAIN | RPAIN | chr17 | + | 5428070 | 5428211 | 5422768 | 5422829 | 5432541 | 5433020 | 2 | 0 | 0 | 0 | 1 | 0 | 0 | 0 | 0 | 1 | 98 | 49 | 1 | 1 | 1 | 0 | NA | NA | | 0 | 0.5 |
| SE | 5527 AMBRA1 | AMBRA1 | chr11 | - | 46512726 | 46512813 | 46508190 | 46508370 | 46541944 | 46542981 | 1 | 1 | 2 | 1 | 0 | 0 | 0 | 1 | 0 | 1 | 98 | 49 | 1 | 1 | 0.333 | 1 | 1 | NA | | 0.333 | 0.445 |
| SE | 5529 MICALL1 | MICALL1 | chr22 | + | 37927410 | 37927826 | 37925660 | 37926043 | 37931798 | 37931933 | 1 | 2 | 0 | 0 | 0 | 0 | 0 | 0 | 0 | 1 | 98 | 49 | 1 | 1 | 1 | 1 | NA | NA | | 0 | 1 |
| SE | 5537 PDE4B | PDE4B | chr1 | + | 66361614 | 66361793 | 66355526 | 66355620 | 66363167 | 66363266 | 55 | 43 | 14 | 1 | 0 | 0 | 0 | 8 | 0 | 0 | 98 | 49 | 1 | 1 | 0.965 | 1 | 1 | NA | | 1 | -0.012 |
| SE | 5538 PADI4 | PADI4 | chr1 | + | 17354532 | 17354687 | 17347940 | 17348048 | 17355982 | 17356127 | 50 | 18 | 47 | 1 | 0 | 0 | 4 | 8 | 0 | 0 | 98 | 49 | 1 | 1 | 0.962 | 1 | 1 | | 1 | 1 | -0.013 |
| SE | 5539 PADI4 | PADI4 | chr1 | + | 17355982 | 17356127 | 17347940 | 17348048 | 17356356 | 17356459 | 24 | 5 | 8 | 1 | 1 | 0 | 0 | 6 | 0 | 0 | 98 | 49 | 1 | 1 | 0.923 | 0.714 | 1 | NA | | 1 | -0.121 |
| SE | 5548 CRLF3 | CRLF3 | chr17 | - | 30793449 | 30793672 | 30792439 | 30792572 | 30796159 | 30796337 | 57 | 14 | 18 | 0 | 1 | 0 | 0 | 4 | 0 | 0 | 98 | 49 | 1 | 1 | 1 | 0.875 | 1 | NA | | 1 | -0.042 |
| SE | 5554 NAAA | NAAA | chr4 | - | 75918760 | 75918789 | 75914867 | 75914985 | 75919908 | 75919975 | 2 | 0 | 0 | 0 | 0 | 0 | 0 | 0 | 0 | 2 | 78 | 49 | 1 | 1 | 1 | NA | NA | NA | | 0 | 1 |
| SE | 5555 FCGR1B | FCGR1B | chr1 | + | 121094673 | 121094958 | 121092995 | 121093250 | 121095871 | 121097161 | 0 | 0 | 0 | 10 | 3 | 6 | 0 | 2 | 0 | 0 | 98 | 49 | 1 | 1 | 0 | 0 | 0 | NA | | 1 | -1 |
| SE | 5556 TAF2 | TAF2 | chr8 | - | 119793365 | 119793451 | 119791323 | 119791459 | 119795531 | 119795631 | 10 | 3 | 1 | 0 | 0 | 0 | 0 | 1 | 0 | 2 | 98 | 49 | 1 | 1 | 1 | 1 | 1 | NA | | 0.2 | 0.8 |
| SE | 5559 SDHAF2 | SDHAF2 | chr11 | + | 61437624 | 61437848 | 61430124 | 61430182 | 61438003 | 61438113 | 6 | 4 | 0 | 1 | 0 | 0 | 0 | 4 | 0 | 0 | 98 | 49 | 1 | 1 | 0.75 | 1 | NA | NA | | 1 | -0.125 |
| SE | 5560 SDHAF2 | SDHAF2 | chr11 | + | 61438003 | 61438113 | 61430124 | 61430182 | 61445940 | 61446767 | 3 | 1 | 6 | 1 | 1 | 1 | 0 | 2 | 0 | 0 | 98 | 49 | 1 | 1 | 0.6 | 0.333 | 0.75 | NA | | 1 | -0.439 |
| SE | 5561 KIAA0141 | KIAA0141 | chr5 | + | 141937197 | 141937357 | 141934493 | 141934586 | 141938520 | 141938760 | 10 | 4 | 0 | 1 | 0 | 0 | 0 | 2 | 0 | 0 | 98 | 49 | 1 | 1 | 0.833 | 1 | NA | NA | | 1 | -0.084 |
| SE | 5567 CERS2 | CERS2 | chr1 | - | 150968082 | 150968201 | 150967819 | 150967877 | 150968394 | 150968512 | 15 | 14 | 8 | 1 | 2 | 0 | 0 | 16 | 0 | 1 | 98 | 49 | 1 | 1 | 0.882 | 0.778 | 1 | NA | | 0.889 | -0.002 |
| SE | 5581 NSUN2 | NSUN2 | chr5 | - | 6622015 | 6622100 | 6620105 | 6620298 | 6623213 | 6623285 | 9 | 2 | 1 | 3 | 0 | 0 | 0 | 1 | 0 | 2 | 98 | 49 | 1 | 1 | 0.6 | 1 | 1 | NA | | 0.2 | 0.667 |
| SE | 5583 ROCK2 | ROCK2 | chr2 | - | 11222082 | 11222174 | 11221197 | 11221357 | 11224321 | 11224460 | 12 | 3 | 2 | 0 | 0 | 4 | 0 | 10 | 0 | 0 | 98 | 49 | 1 | 1 | 1 | 1 | 0.2 | NA | | 1 | -0.267 |
| SE | 5592 COPA | COPA | chr1 | - | 160294483 | 160294593 | 160293385 | 160293463 | 160294767 | 160294857 | 14 | 5 | 0 | 0 | 0 | 0 | 0 | 3 | 0 | 2 | 98 | 49 | 1 | 1 | 1 | 1 | NA | NA | | 0.429 | 0.571 |
| SE | 5593 COPA | COPA | chr1 | - | 160307162 | 160307245 | 160306353 | 160306493 | 160309100 | 160309176 | 16 | 5 | 7 | 2 | 0 | 0 | 0 | 6 | 0 | 0 | 98 | 49 | 1 | 1 | 0.8 | 1 | 1 | NA | | 1 | -0.067 |
| SE | 5594 COPA | COPA | chr1 | - | 160333602 | 160333679 | 160332447 | 160332557 | 160335241 | 160335322 | 17 | 6 | 3 | 0 | 2 | 0 | 0 | 4 | 0 | 0 | 98 | 49 | 1 | 1 | 1 | 0.6 | 1 | NA | | 1 | -0.133 |
| SE | 5600 DCAF8 | DCAF8 | chr1 | - | 160243959 | 160244034 | 160239221 | 160240370 | 160261284 | 160261325 | 8 | 2 | 0 | 1 | 0 | 0 | 0 | 3 | 0 | 0 | 98 | 49 | 1 | 1 | 0.8 | 1 | NA | NA | | 1 | -0.1 |
| SE | 5601 DCAF8 | DCAF8 | chr1 | - | 160243959 | 160244034 | 160239221 | 160240370 | 160262448 | 160262560 | 4 | 2 | 0 | 1 | 0 | 0 | 0 | 3 | 0 | 0 | 98 | 49 | 1 | 1 | 0.667 | 1 | NA | NA | | 1 | -0.166 |
| SE | 5611 KCNJ15 | KCNJ15 | chr21 | + | 38291605 | 38291747 | 38229914 | 38230023 | 38296925 | 38297023 | 35 | 2 | 4 | 8 | 0 | 0 | 0 | 2 | 0 | 0 | 98 | 49 | 1 | 1 | 0.686 | 1 | 1 | NA | | 1 | -0.105 |
| SE | 5612 KCNJ15 | KCNJ15 | chr21 | + | 38291605 | 38291747 | 38257045 | 38257185 | 38296925 | 38297023 | 40 | 2 | 3 | 8 | 2 | 2 | 0 | 2 | 0 | 0 | 98 | 49 | 1 | 1 | 0.714 | 0.333 | 0.429 | NA | | 1 | -0.508 |
| SE | 5613 KCNJ15 | KCNJ15 | chr21 | + | 38298143 | 38298282 | 38296925 | 38297023 | 38299243 | 38301824 | 15 | 1 | 0 | 39 | 7 | 14 | 0 | 1 | 0 | 7 | 98 | 49 | 1 | 1 | 0.161 | 0.067 | 0 | NA | | 0.067 | 0.009 |
| SE | 5619 VTI1A | VTI1A | chr10 | + | 112527086 | 112527164 | 112464546 | 112464657 | 112538245 | 112538330 | 21 | 7 | 1 | 4 | 0 | 0 | 0 | 13 | 0 | 0 | 98 | 49 | 1 | 1 | 0.724 | 1 | 1 | NA | | 1 | -0.092 |
| SE | 5622 AIM1 | AIM1 | chr6 | + | 106525132 | 106525180 | 106519130 | 106521453 | 106525267 | 106525386 | 23 | 8 | 4 | 0 | 1 | 0 | 0 | 11 | 0 | 0 | 97 | 49 | 1 | 1 | 1 | 0.802 | 1 | NA | | 1 | -0.066 |
| SE | 5627 CD74 | CD74 | chr5 | - | 150406880 | 150406960 | 150406258 | 150406321 | 150407151 | 150407324 | 145 | 8 | 14 | 1 | 1 | 0 | 7 | 68 | 0 | 1 | 98 | 49 | 1 | 1 | 0.986 | 0.8 | 1 | | 1 | 0.971 | -0.057 |
| SE | 5628 MORF4L1 | MORF4L1 | chr15 | + | 78884966 | 78885083 | 78880511 | 78880579 | 78886140 | 78886227 | 3 | 5 | 0 | 17 | 10 | 0 | 0 | 0 | 0 | 5 | 98 | 49 | 1 | 1 | 0.081 | 0.2 | NA | NA | | 0 | 0.141 |
| SE | 5629 MORF4L1 | MORF4L1 | chr15 | + | 78892211 | 78892313 | 78891483 | 78891572 | 78893538 | 78893627 | 18 | 8 | 2 | 1 | 0 | 0 | 1 | 4 | 0 | 0 | 98 | 49 | 1 | 1 | 0.9 | 1 | 1 | | 1 | 1 | -0.033 |
| SE | 5630 MORF4L1 | MORF4L1 | chr15 | + | 78894057 | 78894230 | 78893538 | 78893627 | 78894819 | 78894904 | 27 | 7 | 4 | 1 | 3 | 0 | 0 | 7 | 0 | 3 | 98 | 49 | 1 | 1 | 0.931 | 0.538 | 1 | NA | | 0.538 | 0.285 |
| SE | 5636 MNDA | MNDA | chr1 | + | 158843954 | 158844122 | 158843278 | 158843415 | 158845586 | 158846003 | 605 | 393 | 302 | 0 | 0 | 3 | 8 | 82 | 0 | 0 | 98 | 49 | 1 | 1 | 1 | 1 | 0.981 | | 1 | 1 | -0.006 |
| SE | 5637 MNDA | MNDA | chr1 | + | 158845586 | 158846003 | 158843954 | 158844122 | 158847727 | 158847916 | 679 | 552 | 400 | 4 | 1 | 4 | 6 | 105 | 0 | 0 | 98 | 49 | 1 | 1 | 0.988 | 0.996 | 0.98 | | 1 | 1 | -0.012 |
| SE | 5642 METTL9 | METTL9 | chr16 | + | 21612644 | 21612835 | 21599475 | 21599898 | 21617864 | 21618074 | 40 | 22 | 15 | 1 | 0 | 0 | 0 | 16 | 0 | 0 | 98 | 49 | 1 | 1 | 0.952 | 1 | 1 | NA | | 1 | -0.016 |
| SE | 5644 XPR1 | XPR1 | chr1 | + | 180803387 | 180803611 | 180787752 | 180787854 | 180806061 | 180806211 | 2 | 2 | 3 | 0 | 0 | 2 | 0 | 1 | 0 | 0 | 98 | 49 | 1 | 1 | 1 | 1 | 0.429 | NA | | 1 | -0.19 |
| SE | 5647 INO80 | INO80 | chr15 | - | 41074369 | 41074565 | 41073427 | 41073495 | 41079700 | 41079904 | 6 | 3 | 0 | 1 | 0 | 0 | 0 | 1 | 0 | 0 | 98 | 49 | 1 | 1 | 0.75 | 1 | NA | NA | | 1 | -0.125 |
| SE | 5655 WDFY3 | WDFY3 | chr4 | - | 84702352 | 84702506 | 84696731 | 84696823 | 84704337 | 84704444 | 35 | 12 | 9 | 1 | 0 | 0 | 0 | 21 | 0 | 0 | 98 | 49 | 1 | 1 | 0.946 | 1 | 1 | NA | | 1 | -0.018 |
| SE | 5656 WDFY3 | WDFY3 | chr4 | - | 84705393 | 84705511 | 84704337 | 84704444 | 84708908 | 84709028 | 53 | 17 | 2 | 2 | 1 | 0 | 0 | 11 | 0 | 1 | 98 | 49 | 1 | 1 | 0.93 | 0.895 | 1 | NA | | 0.846 | 0.096 |
| SE | 5659 WDFY3 | WDFY3 | chr4 | - | 84765809 | 84766027 | 84756925 | 84757161 | 84766251 | 84766372 | 51 | 16 | 10 | 0 | 1 | 0 | 0 | 8 | 0 | 0 | 98 | 49 | 1 | 1 | 1 | 0.889 | 1 | NA | | 1 | -0.037 |
| SE | 5660 WDFY3 | WDFY3 | chr4 | - | 84796520 | 84796752 | 84794878 | 84794979 | 84797995 | 84798108 | 75 | 23 | 16 | 1 | 0 | 0 | 1 | 13 | 0 | 0 | 98 | 49 | 1 | 1 | 0.974 | 1 | 1 | | 1 | 1 | -0.009 |
| SE | 5661 WDFY3 | WDFY3 | chr4 | - | 84808333 | 84808417 | 84803289 | 84803467 | 84809886 | 84810344 | 0 | 3 | 0 | 30 | 7 | 0 | 0 | 0 | 0 | 5 | 98 | 49 | 1 | 1 | 0 | 0.176 | NA | NA | | 0 | 0.088 |
| SE | 5662 WDFY3 | WDFY3 | chr4 | - | 84817391 | 84817585 | 84809886 | 84810344 | 84820084 | 84820186 | 100 | 28 | 9 | 2 | 0 | 0 | 1 | 19 | 1 | 0 | 98 | 49 | 1 | 1 | 0.962 | 1 | 1 | | 0.333 | 1 | 0.321 |
| SE | 5669 PPP2R5A | PPP2R5A | chr1 | + | 212329134 | 212329331 | 212285536 | 212286291 | 212333496 | 212333598 | 62 | 24 | 18 | 2 | 0 | 0 | 0 | 17 | 0 | 0 | 98 | 49 | 1 | 1 | 0.939 | 1 | 1 | NA | | 1 | -0.02 |
| SE | 5680 SELP | SELP | chr1 | - | 169593604 | 169593724 | 169591425 | 169591456 | 169594691 | 169594877 | 4 | 0 | 0 | 0 | 0 | 0 | 1 | 0 | 0 | 1 | 98 | 49 | 1 | 1 | 1 | NA | NA | | 1 | 0 | 0.5 |
| SE | 5683 PPP1R12A | PPP1R12A | chr12 | - | 79786373 | 79786478 | 79781814 | 79781862 | 79788647 | 79788783 | 15 | 8 | 11 | 0 | 1 | 0 | 0 | 2 | 0 | 0 | 98 | 49 | 1 | 1 | 1 | 0.8 | 1 | NA | | 1 | -0.067 |
| SE | 5684 PPP1R12A | PPP1R12A | chr12 | - | 79805591 | 79805768 | 79798493 | 79798584 | 79806165 | 79806333 | 14 | 2 | 0 | 17 | 6 | 2 | 0 | 2 | 0 | 4 | 98 | 49 | 1 | 1 | 0.292 | 0.143 | 0 | NA | | 0.2 | -0.055 |
| SE | 5685 PPP1R12A | PPP1R12A | chr12 | - | 79806165 | 79806333 | 79805591 | 79805768 | 79807225 | 79807330 | 12 | 6 | 1 | 1 | 0 | 0 | 0 | 4 | 0 | 0 | 98 | 49 | 1 | 1 | 0.857 | 1 | 1 | NA | | 1 | -0.048 |
| SE | 5686 PPP1R12A | PPP1R12A | chr12 | - | 79822115 | 79822190 | 79821077 | 79821166 | 79828319 | 79828464 | 40 | 19 | 8 | 2 | 0 | 0 | 0 | 13 | 0 | 0 | 98 | 49 | 1 | 1 | 0.909 | 1 | 1 | NA | | 1 | -0.03 |
| SE | 5696 SERPINA1 | SERPINA1 | chr14 | - | 94379463 | 94379611 | 94376746 | 94378640 | 94390456 | 94390692 | 142 | 65 | 104 | 2 | 0 | 0 | 6 | 51 | 0 | 0 | 98 | 49 | 1 | 1 | 0.973 | 1 | 1 | | 1 | 1 | -0.009 |
| SE | 5697 SERPINA1 | SERPINA1 | chr14 | - | 94382591 | 94383241 | 94376746 | 94378640 | 94390456 | 94390692 | 108 | 35 | 38 | 2 | 0 | 0 | 2 | 20 | 0 | 0 | 98 | 49 | 1 | 1 | 0.964 | 1 | 1 | | 1 | 1 | -0.012 |
| SE | 5698 SERPINA1 | SERPINA1 | chr14 | - | 94388559 | 94388660 | 94379463 | 94379611 | 94390456 | 94390692 | 34 | 6 | 10 | 1 | 0 | 0 | 1 | 3 | 0 | 0 | 98 | 49 | 1 | 1 | 0.944 | 1 | 1 | | 1 | 1 | -0.019 |
| SE | 5700 SERPINA1 | SERPINA1 | chr14 | - | 94388559 | 94388663 | 94379463 | 94379611 | 94390456 | 94390692 | 29 | 19 | 5 | 1 | 0 | 0 | 0 | 4 | 0 | 0 | 98 | 49 | 1 | 1 | 0.935 | 1 | 1 | NA | | 1 | -0.022 |
| SE | 5701 SERPINA1 | SERPINA1 | chr14 | - | 94388559 | 94388663 | 94382591 | 94383241 | 94390419 | 94390692 | 69 | 15 | 37 | 3 | 0 | 0 | 0 | 13 | 0 | 0 | 98 | 49 | 1 | 1 | 0.92 | 1 | 1 | NA | | 1 | -0.027 |
| SE | 5703 SERPINA1 | SERPINA1 | chr14 | - | 94388800 | 94389010 | 94376746 | 94378640 | 94390456 | 94390692 | 47 | 13 | 5 | 2 | 0 | 0 | 1 | 15 | 0 | 0 | 98 | 49 | 1 | 1 | 0.922 | 1 | 1 | | 1 | 1 | -0.026 |
| SE | 5717 ARPC3 | ARPC3 | chr12 | - | 110436556 | 110436683 | 110436109 | 110436204 | 110437083 | 110437152 | 164 | 84 | 70 | 0 | 1 | 0 | 3 | 30 | 0 | 0 | 98 | 49 | 1 | 1 | 1 | 0.977 | 1 | | 1 | 1 | -0.008 |
| SE | 5719 KIAA1324 | KIAA1324 | chr1 | + | 109173687 | 109173837 | 109173473 | 109173579 | 109185044 | 109185133 | 20 | 2 | 12 | 1 | 2 | 0 | 0 | 1 | 0 | 0 | 98 | 49 | 1 | 1 | 0.909 | 0.333 | 1 | NA | | 1 | -0.253 |
| SE | 5720 KIAA1324 | KIAA1324 | chr1 | + | 109185044 | 109185133 | 109173687 | 109173837 | 109188176 | 109188354 | 23 | 12 | 1 | 1 | 0 | 1 | 0 | 1 | 0 | 0 | 98 | 49 | 1 | 1 | 0.92 | 1 | 0.333 | NA | | 1 | -0.249 |
| SE | 5721 KIAA1324 | KIAA1324 | chr1 | + | 109197978 | 109198075 | 109197473 | 109197654 | 109198572 | 109198674 | 26 | 8 | 24 | 0 | 2 | 0 | 0 | 1 | 0 | 0 | 98 | 49 | 1 | 1 | 1 | 0.667 | 1 | NA | | 1 | -0.111 |
| SE | 5723 LILRB3 | LILRB3 | chr19 | - | 54222746 | 54222782 | 54222277 | 54222562 | 54222942 | 54223087 | 28 | 15 | 3 | 1 | 1 | 0 | 4 | 6 | 0 | 0 | 85 | 49 | 1 | 1 | 0.942 | 0.896 | 1 | | 1 | 1 | -0.054 |
| SE | 5726 WARS | WARS | chr14 | - | 100360553 | 100360662 | 100354446 | 100354566 | 100361707 | 100361921 | 16 | 5 | 3 | 0 | 1 | 0 | 0 | 7 | 0 | 0 | 98 | 49 | 1 | 1 | 1 | 0.714 | 1 | NA | | 1 | -0.095 |
| SE | 5739 WDR7 | WDR7 | chr18 | + | 56717963 | 56718159 | 56696241 | 56696462 | 56731382 | 56731597 | 25 | 5 | 1 | 2 | 0 | 0 | 0 | 2 | 0 | 0 | 98 | 49 | 1 | 1 | 0.862 | 1 | 1 | NA | | 1 | -0.046 |
| SE | 5741 EYA3 | EYA3 | chr1 | - | 27998283 | 27998351 | 27997319 | 27997378 | 27999959 | 28000049 | 5 | 3 | 0 | 0 | 0 | 0 | 0 | 0 | 0 | 1 | 98 | 49 | 1 | 1 | 1 | 1 | NA | NA | | 0 | 1 |
| SE | 5749 TMEM50A | TMEM50A | chr1 | + | 25342960 | 25343073 | 25340473 | 25340579 | 25351625 | 25351693 | 39 | 35 | 20 | 0 | 1 | 0 | 0 | 3 | 0 | 0 | 98 | 49 | 1 | 1 | 1 | 0.946 | 1 | NA | | 1 | -0.018 |
| SE | 5752 TPR | TPR | chr1 | - | 186317481 | 186317600 | 186314628 | 186314724 | 186318446 | 186318603 | 24 | 14 | 7 | 0 | 2 | 0 | 0 | 4 | 0 | 0 | 98 | 49 | 1 | 1 | 1 | 0.778 | 1 | NA | | 1 | -0.074 |
| SE | 5753 TPR | TPR | chr1 | - | 186335067 | 186335129 | 186334324 | 186334533 | 186335337 | 186335543 | 68 | 24 | 15 | 1 | 0 | 0 | 1 | 10 | 0 | 0 | 98 | 49 | 1 | 1 | 0.971 | 1 | 1 | | 1 | 1 | -0.01 |
| SE | 5754 TPR | TPR | chr1 | - | 186339641 | 186339772 | 186338032 | 186338243 | 186341027 | 186341159 | 58 | 25 | 7 | 1 | 0 | 0 | 0 | 1 | 0 | 0 | 98 | 49 | 1 | 1 | 0.967 | 1 | 1 | NA | | 1 | -0.011 |
| SE | 5755 TPR | TPR | chr1 | - | 186353687 | 186353850 | 186351975 | 186352110 | 186355409 | 186355558 | 72 | 36 | 3 | 0 | 2 | 0 | 0 | 5 | 0 | 0 | 98 | 49 | 1 | 1 | 1 | 0.9 | 1 | NA | | 1 | -0.033 |
| SE | 5756 TPR | TPR | chr1 | - | 186361621 | 186361709 | 186360764 | 186360905 | 186361788 | 186361869 | 28 | 15 | 2 | 0 | 2 | 0 | 0 | 5 | 0 | 0 | 98 | 49 | 1 | 1 | 1 | 0.789 | 1 | NA | | 1 | -0.07 |
| SE | 5757 TPR | TPR | chr1 | - | 186367885 | 186367982 | 186363341 | 186363445 | 186370969 | 186371043 | 53 | 11 | 7 | 1 | 0 | 0 | 0 | 3 | 0 | 0 | 98 | 49 | 1 | 1 | 0.964 | 1 | 1 | NA | | 1 | -0.012 |
| SE | 5767 SCAMP3 | SCAMP3 | chr1 | - | 155261656 | 155261734 | 155260536 | 155260659 | 155262085 | 155262385 | 0 | 1 | 3 | 2 | 0 | 0 | 0 | 1 | 0 | 0 | 98 | 49 | 1 | 1 | 0 | 1 | 1 | NA | | 1 | -0.333 |
| SE | 5778 TBC1D2B | TBC1D2B | chr15 | - | 78017846 | 78017957 | 78016545 | 78016739 | 78024155 | 78024539 | 5 | 3 | 4 | 1 | 0 | 0 | 0 | 2 | 0 | 0 | 98 | 49 | 1 | 1 | 0.714 | 1 | 1 | NA | | 1 | -0.095 |
| SE | 5786 SRSF10 | SRSF10 | chr1 | - | 23971337 | 23971439 | 23964346 | 23967723 | 23971572 | 23971626 | 4 | 1 | 0 | 4 | 0 | 2 | 0 | 1 | 0 | 0 | 98 | 49 | 1 | 1 | 0.333 | 1 | 0 | NA | | 1 | -0.556 |
| SE | 5793 RARA | RARA | chr17 | + | 40330856 | 40331396 | 40318220 | 40318448 | 40348315 | 40348464 | 34 | 14 | 3 | 1 | 0 | 0 | 1 | 16 | 0 | 0 | 98 | 49 | 1 | 1 | 0.944 | 1 | 1 | | 1 | 1 | -0.019 |
| SE | 5794 KIAA1109 | KIAA1109 | chr4 | + | 122173035 | 122173152 | 122170602 | 122170744 | 122174573 | 122174652 | 28 | 23 | 5 | 2 | 0 | 0 | 1 | 7 | 0 | 0 | 98 | 49 | 1 | 1 | 0.875 | 1 | 1 | | 1 | 1 | -0.042 |
| SE | 5795 KIAA1109 | KIAA1109 | chr4 | + | 122224497 | 122224677 | 122220332 | 122220470 | 122229120 | 122229240 | 13 | 13 | 4 | 0 | 1 | 0 | 0 | 2 | 0 | 0 | 98 | 49 | 1 | 1 | 1 | 0.867 | 1 | NA | | 1 | -0.044 |
| SE | 5796 KIAA1109 | KIAA1109 | chr4 | + | 122226706 | 122226835 | 122224497 | 122224677 | 122229120 | 122229240 | 32 | 18 | 2 | 7 | 3 | 0 | 0 | 1 | 0 | 1 | 98 | 49 | 1 | 1 | 0.696 | 0.75 | 1 | NA | | 0.333 | 0.482 |
| SE | 5797 KIAA1109 | KIAA1109 | chr4 | + | 122234772 | 122235023 | 122229930 | 122230211 | 122238091 | 122238350 | 41 | 11 | 2 | 3 | 1 | 0 | 0 | 9 | 0 | 0 | 98 | 49 | 1 | 1 | 0.872 | 0.846 | 1 | NA | | 1 | -0.094 |
| SE | 5798 KIAA1109 | KIAA1109 | chr4 | + | 122275971 | 122276022 | 122274342 | 122274436 | 122279774 | 122279981 | 2 | 7 | 0 | 6 | 3 | 3 | 0 | 4 | 0 | 0 | 98 | 49 | 1 | 1 | 0.143 | 0.538 | 0 | NA | | 1 | -0.773 |
| SE | 5799 KIAA1109 | KIAA1109 | chr4 | + | 122356620 | 122356755 | 122355795 | 122355990 | 122359556 | 122359722 | 43 | 19 | 10 | 0 | 0 | 0 | 0 | 14 | 0 | 0 | 98 | 49 | 1 | 1 | 1 | 1 | 1 | NA | | 0.875 | 0.125 |
| SE | 5800 WDR26 | WDR26 | chr1 | - | 224393827 | 224394013 | 224385142 | 224389860 | 224398096 | 224398226 | 44 | 22 | 8 | 3 | 2 | 0 | 0 | 9 | 0 | 0 | 98 | 49 | 1 | 1 | 0.88 | 0.846 | 1 | NA | | 1 | -0.091 |
| SE | 5801 WDR26 | WDR26 | chr1 | - | 224398096 | 224398226 | 224385142 | 224389860 | 224398514 | 224398593 | 13 | 14 | 3 | 1 | 0 | 0 | 0 | 4 | 0 | 0 | 98 | 49 | 1 | 1 | 0.867 | 1 | 1 | NA | | 1 | -0.044 |
| SE | 5802 WDR26 | WDR26 | chr1 | - | 224398888 | 224399034 | 224398514 | 224398593 | 224400949 | 224401069 | 49 | 24 | 15 | 1 | 0 | 0 | 0 | 5 | 0 | 1 | 98 | 49 | 1 | 1 | 0.961 | 1 | 1 | NA | | 0.714 | 0.273 |
| SE | 5803 WDR26 | WDR26 | chr1 | - | 224418259 | 224418416 | 224411426 | 224411565 | 224419517 | 224419615 | 50 | 26 | 15 | 1 | 6 | 0 | 0 | 16 | 0 | 0 | 98 | 49 | 1 | 1 | 0.962 | 0.684 | 1 | NA | | 1 | -0.118 |
| SE | 5804 WDR26 | WDR26 | chr1 | - | 224424517 | 224424654 | 224419517 | 224419615 | 224431476 | 224431581 | 23 | 25 | 1 | 0 | 1 | 0 | 2 | 12 | 0 | 0 | 98 | 49 | 1 | 1 | 1 | 0.926 | 1 | | 1 | 1 | -0.025 |
| SE | 5805 WDR26 | WDR26 | chr1 | - | 224424517 | 224424654 | 224419517 | 224419615 | 224431524 | 224431581 | 26 | 27 | 2 | 1 | 1 | 0 | 2 | 15 | 0 | 0 | 98 | 49 | 1 | 1 | 0.929 | 0.931 | 1 | | 1 | 1 | -0.047 |
| SE | 5806 WDR26 | WDR26 | chr1 | - | 224431681 | 224431781 | 224431476 | 224431581 | 224433683 | 224434299 | 44 | 51 | 5 | 2 | 0 | 0 | 1 | 6 | 0 | 1 | 98 | 49 | 1 | 1 | 0.917 | 1 | 1 | | 1 | 0.75 | 0.097 |
| SE | 5807 CCDC9 | CCDC9 | chr19 | + | 47270406 | 47270453 | 47266610 | 47266792 | 47270552 | 47270688 | 5 | 14 | 0 | 2 | 0 | 0 | 0 | 3 | 0 | 0 | 96 | 49 | 1 | 1 | 0.561 | 1 | NA | NA | | 1 | -0.22 |
| SE | 5813 CHTOP | CHTOP | chr1 | + | 153643226 | 153643364 | 153642242 | 153642429 | 153645063 | 153646306 | 2 | 3 | 2 | 0 | 0 | 1 | 0 | 1 | 0 | 0 | 98 | 49 | 1 | 1 | 1 | 1 | 0.5 | NA | | 1 | -0.167 |
| SE | 5814 EDRF1 | EDRF1 | chr10 | + | 125729002 | 125729104 | 125725681 | 125725838 | 125729357 | 125729479 | 2 | 0 | 0 | 0 | 0 | 2 | 0 | 0 | 0 | 1 | 98 | 49 | 1 | 1 | 1 | NA | 0 | NA | | 0 | 0.5 |
| SE | 5823 WAPL | WAPL | chr10 | - | 86472611 | 86472764 | 86472207 | 86472344 | 86473877 | 86473973 | 25 | 14 | 10 | 5 | 1 | 0 | 0 | 8 | 0 | 0 | 98 | 49 | 1 | 1 | 0.714 | 0.875 | 1 | NA | | 1 | -0.137 |
| SE | 5824 WAPL | WAPL | chr10 | - | 86497200 | 86497319 | 86473877 | 86473973 | 86499717 | 86500743 | 20 | 6 | 4 | 2 | 0 | 0 | 0 | 1 | 0 | 0 | 98 | 49 | 1 | 1 | 0.833 | 1 | 1 | NA | | 1 | -0.056 |
| SE | 5825 WAPL | WAPL | chr10 | - | 86517570 | 86518091 | 86499717 | 86500743 | 86521364 | 86521784 | 23 | 19 | 8 | 1 | 0 | 0 | 0 | 2 | 0 | 0 | 98 | 49 | 1 | 1 | 0.92 | 1 | 1 | NA | | 1 | -0.027 |
| SE | 5834 ITGB1 | ITGB1 | chr10 | - | 32932514 | 32932600 | 32929821 | 32930044 | 32935491 | 32935558 | 11 | 0 | 2 | 0 | 0 | 0 | 1 | 2 | 0 | 1 | 98 | 49 | 1 | 1 | 1 | NA | 1 | | 1 | 0.5 | 0.25 |
| SE | 5835 STX3 | STX3 | chr11 | + | 59787036 | 59787136 | 59773210 | 59773294 | 59788872 | 59788947 | 101 | 67 | 28 | 6 | 1 | 0 | 0 | 14 | 0 | 0 | 98 | 49 | 1 | 1 | 0.894 | 0.971 | 1 | NA | | 1 | -0.045 |
| SE | 5836 STX3 | STX3 | chr11 | + | 59787036 | 59787136 | 59773210 | 59773294 | 59790518 | 59790586 | 38 | 20 | 10 | 1 | 0 | 0 | 0 | 6 | 0 | 0 | 98 | 49 | 1 | 1 | 0.95 | 1 | 1 | NA | | 1 | -0.017 |
| SE | 5837 STX3 | STX3 | chr11 | + | 59788872 | 59788947 | 59773210 | 59773294 | 59790518 | 59790586 | 57 | 26 | 14 | 1 | 0 | 0 | 1 | 6 | 0 | 0 | 98 | 49 | 1 | 1 | 0.966 | 1 | 1 | | 1 | 1 | -0.011 |
| SE | 5838 STX3 | STX3 | chr11 | + | 59788872 | 59788947 | 59787036 | 59787136 | 59790518 | 59790586 | 114 | 73 | 32 | 0 | 1 | 0 | 1 | 14 | 0 | 0 | 98 | 49 | 1 | 1 | 1 | 0.973 | 1 | | 1 | 1 | -0.009 |
| SE | 5839 STX3 | STX3 | chr11 | + | 59795371 | 59795482 | 59793379 | 59793514 | 59797282 | 59797396 | 46 | 23 | 13 | 1 | 0 | 0 | 0 | 8 | 0 | 0 | 98 | 49 | 1 | 1 | 0.958 | 1 | 1 | NA | | 1 | -0.014 |
| SE | 5840 STX3 | STX3 | chr11 | + | 59797282 | 59797396 | 59795371 | 59795482 | 59800854 | 59805882 | 45 | 25 | 21 | 3 | 3 | 2 | 0 | 8 | 0 | 0 | 98 | 49 | 1 | 1 | 0.882 | 0.806 | 0.84 | NA | | 1 | -0.157 |
| SE | 5854 SNX27 | SNX27 | chr1 | + | 151662165 | 151662270 | 151660797 | 151660862 | 151665932 | 151666011 | 41 | 18 | 10 | 3 | 3 | 1 | 0 | 9 | 0 | 0 | 98 | 49 | 1 | 1 | 0.872 | 0.75 | 0.833 | NA | | 1 | -0.182 |
| SE | 5855 SNX27 | SNX27 | chr1 | + | 151665932 | 151666011 | 151662165 | 151662270 | 151668471 | 151668635 | 44 | 26 | 6 | 2 | 0 | 0 | 0 | 9 | 0 | 0 | 98 | 49 | 1 | 1 | 0.917 | 1 | 1 | NA | | 1 | -0.028 |
| SE | 5862 CTNNA1 | CTNNA1 | chr5 | + | 138783176 | 138783372 | 138781922 | 138782029 | 138810037 | 138810204 | 15 | 3 | 0 | 0 | 1 | 0 | 0 | 12 | 0 | 0 | 98 | 49 | 1 | 1 | 1 | 0.6 | NA | NA | | 1 | -0.2 |
| SE | 5866 PIP5K1A | PIP5K1A | chr1 | + | 151242122 | 151242269 | 151239954 | 151240039 | 151242437 | 151242567 | 1 | 0 | 0 | 0 | 0 | 0 | 0 | 0 | 0 | 1 | 98 | 49 | 1 | 1 | 1 | NA | NA | NA | | 0 | 1 |
| SE | 5869 PLEKHJ1 | PLEKHJ1 | chr19 | - | 2234149 | 2234240 | 2233997 | 2234061 | 2235761 | 2235828 | 6 | 6 | 2 | 0 | 0 | 0 | 0 | 0 | 0 | 1 | 98 | 49 | 1 | 1 | 1 | 1 | 1 | NA | | 0 | 1 |
| SE | 5870 ZDHHC13 | ZDHHC13 | chr11 | + | 19142977 | 19143123 | 19117144 | 19117276 | 19146180 | 19146303 | 2 | 0 | 0 | 1 | 0 | 0 | 0 | 2 | 0 | 0 | 98 | 49 | 1 | 1 | 0.5 | NA | NA | NA | | 1 | -0.5 |
| SE | 5871 ZDHHC13 | ZDHHC13 | chr11 | + | 19166301 | 19166385 | 19165051 | 19165145 | 19170410 | 19170568 | 2 | 2 | 0 | 2 | 0 | 0 | 0 | 2 | 0 | 0 | 98 | 49 | 1 | 1 | 0.333 | 1 | NA | NA | | 1 | -0.334 |
| SE | 5887 ATXN3 | ATXN3 | chr14 | - | 92093745 | 92093831 | 92093251 | 92093318 | 92096092 | 92096137 | 10 | 2 | 0 | 3 | 1 | 0 | 0 | 1 | 0 | 0 | 98 | 49 | 1 | 1 | 0.625 | 0.5 | NA | NA | | 1 | -0.438 |
| SE | 5901 FAM63A | FAM63A | chr1 | - | 150998081 | 150998273 | 150997623 | 150997779 | 150999368 | 150999511 | 27 | 10 | 8 | 2 | 0 | 0 | 0 | 1 | 0 | 0 | 98 | 49 | 1 | 1 | 0.871 | 1 | 1 | NA | | 1 | -0.043 |
| SE | 5902 FAM63A | FAM63A | chr1 | - | 151000456 | 151000615 | 150999861 | 150999964 | 151001249 | 151001314 | 10 | 8 | 14 | 2 | 0 | 0 | 0 | 4 | 0 | 0 | 98 | 49 | 1 | 1 | 0.714 | 1 | 1 | NA | | 1 | -0.095 |
| SE | 5904 RPN2 | RPN2 | chr20 | + | 37232295 | 37232391 | 37229972 | 37230059 | 37234019 | 37234095 | 19 | 0 | 5 | 0 | 0 | 0 | 0 | 2 | 0 | 1 | 98 | 49 | 1 | 1 | 1 | NA | 1 | NA | | 0.5 | 0.5 |
| SE | 5910 TP53BP2 | TP53BP2 | chr1 | - | 223818205 | 223818337 | 223814239 | 223814353 | 223821219 | 223821367 | 0 | 3 | 0 | 2 | 1 | 2 | 0 | 1 | 0 | 3 | 98 | 49 | 1 | 1 | 0 | 0.6 | 0 | NA | | 0.143 | 0.057 |
| SE | 5924 CDIPT | CDIPT | chr16 | - | 29859441 | 29859523 | 29858355 | 29859334 | 29860580 | 29860662 | 14 | 7 | 8 | 1 | 0 | 2 | 0 | 5 | 0 | 0 | 98 | 49 | 1 | 1 | 0.875 | 1 | 0.667 | NA | | 1 | -0.153 |
| SE | 5925 CDIPT | CDIPT | chr16 | - | 29862585 | 29862635 | 29861105 | 29861259 | 29862814 | 29863288 | 12 | 5 | 3 | 1 | 0 | 0 | 0 | 2 | 0 | 0 | 98 | 49 | 1 | 1 | 0.857 | 1 | 1 | NA | | 1 | -0.048 |
| SE | 5926 CDIPT | CDIPT | chr16 | - | 29862585 | 29862720 | 29861105 | 29861259 | 29862814 | 29863288 | 19 | 5 | 8 | 1 | 0 | 0 | 0 | 2 | 0 | 0 | 98 | 49 | 1 | 1 | 0.905 | 1 | 1 | NA | | 1 | -0.032 |
| SE | 5930 GDI2 | GDI2 | chr10 | - | 5794884 | 5795019 | 5785851 | 5786050 | 5796762 | 5796862 | 32 | 5 | 12 | 3 | 0 | 0 | 0 | 7 | 0 | 0 | 98 | 49 | 1 | 1 | 0.842 | 1 | 1 | NA | | 1 | -0.053 |
| SE | 5932 CREBBP | CREBBP | chr16 | - | 3738558 | 3738672 | 3736649 | 3736815 | 3739577 | 3739724 | 32 | 5 | 6 | 1 | 0 | 0 | 0 | 4 | 0 | 0 | 98 | 49 | 1 | 1 | 0.941 | 1 | 1 | NA | | 1 | -0.02 |
| SE | 5933 CREBBP | CREBBP | chr16 | - | 3739577 | 3739724 | 3736649 | 3736815 | 3740398 | 3740549 | 31 | 3 | 4 | 0 | 1 | 0 | 0 | 3 | 0 | 3 | 98 | 49 | 1 | 1 | 1 | 0.6 | 1 | NA | | 0.333 | 0.534 |
| SE | 5934 CREBBP | CREBBP | chr16 | - | 3777612 | 3777657 | 3774568 | 3774693 | 3778010 | 3778182 | 55 | 33 | 6 | 1 | 0 | 0 | 1 | 13 | 0 | 0 | 94 | 49 | 1 | 1 | 0.966 | 1 | 1 | | 1 | 1 | -0.011 |
| SE | 5935 CREBBP | CREBBP | chr16 | - | 3778699 | 3778817 | 3778010 | 3778182 | 3780731 | 3780878 | 61 | 17 | 8 | 0 | 1 | 0 | 1 | 10 | 0 | 0 | 98 | 49 | 1 | 1 | 1 | 0.895 | 1 | | 1 | 1 | -0.035 |
| SE | 5936 CREBBP | CREBBP | chr16 | - | 3780731 | 3780878 | 3778699 | 3778817 | 3781203 | 3781306 | 72 | 14 | 3 | 0 | 0 | 0 | 1 | 9 | 0 | 1 | 98 | 49 | 1 | 1 | 1 | 1 | 1 | | 1 | 0.818 | 0.091 |
| SE | 5937 CREBBP | CREBBP | chr16 | - | 3791980 | 3792094 | 3782683 | 3782926 | 3793385 | 3793626 | 57 | 18 | 12 | 1 | 0 | 0 | 1 | 9 | 0 | 0 | 98 | 49 | 1 | 1 | 0.966 | 1 | 1 | | 1 | 1 | -0.011 |
| SE | 5940 C11orf30 | C11orf30 | chr11 | + | 76459932 | 76460085 | 76458182 | 76458358 | 76463820 | 76464080 | 5 | 0 | 0 | 0 | 1 | 0 | 0 | 1 | 0 | 0 | 98 | 49 | 1 | 1 | 1 | 0 | NA | NA | | 1 | -0.5 |
| SE | 5941 C11orf30 | C11orf30 | chr11 | + | 76459935 | 76460085 | 76458182 | 76458358 | 76463820 | 76464080 | 4 | 0 | 0 | 0 | 1 | 0 | 0 | 1 | 0 | 0 | 98 | 49 | 1 | 1 | 1 | 0 | NA | NA | | 1 | -0.5 |
| SE | 5949 SORL1 | SORL1 | chr11 | + | 121478117 | 121478243 | 121470006 | 121470123 | 121488031 | 121488193 | 344 | 191 | 88 | 1 | 1 | 0 | 3 | 54 | 0 | 0 | 98 | 49 | 1 | 1 | 0.994 | 0.99 | 1 | | 1 | 1 | -0.005 |
| SE | 5950 SORL1 | SORL1 | chr11 | + | 121488031 | 121488193 | 121478117 | 121478243 | 121490042 | 121490110 | 458 | 227 | 186 | 3 | 2 | 0 | 0 | 79 | 0 | 0 | 98 | 49 | 1 | 1 | 0.987 | 0.983 | 1 | NA | | 1 | -0.01 |
| SE | 5951 SORL1 | SORL1 | chr11 | + | 121490042 | 121490110 | 121478117 | 121478243 | 121496868 | 121497049 | 317 | 113 | 79 | 1 | 0 | 0 | 5 | 41 | 0 | 0 | 98 | 49 | 1 | 1 | 0.994 | 1 | 1 | | 1 | 1 | -0.002 |
| SE | 5952 SORL1 | SORL1 | chr11 | + | 121513002 | 121513104 | 121496868 | 121497049 | 121514151 | 121514321 | 463 | 284 | 111 | 0 | 1 | 0 | 3 | 110 | 0 | 0 | 98 | 49 | 1 | 1 | 1 | 0.993 | 1 | | 1 | 1 | -0.002 |
| SE | 5953 SORL1 | SORL1 | chr11 | + | 121543547 | 121543726 | 121532463 | 121532552 | 121545242 | 121545429 | 373 | 144 | 88 | 1 | 0 | 0 | 4 | 58 | 0 | 0 | 98 | 49 | 1 | 1 | 0.995 | 1 | 1 | | 1 | 1 | -0.002 |
| SE | 5954 SORL1 | SORL1 | chr11 | + | 121545242 | 121545429 | 121543547 | 121543726 | 121549959 | 121550088 | 389 | 148 | 91 | 0 | 0 | 0 | 6 | 54 | 0 | 1 | 98 | 49 | 1 | 1 | 1 | 1 | 1 | | 1 | 0.964 | 0.018 |
| SE | 5955 SORL1 | SORL1 | chr11 | + | 121549959 | 121550088 | 121545242 | 121545429 | 121550584 | 121550670 | 332 | 120 | 72 | 1 | 3 | 0 | 2 | 33 | 0 | 0 | 98 | 49 | 1 | 1 | 0.994 | 0.952 | 1 | | 1 | 1 | -0.018 |
| SE | 5956 SORL1 | SORL1 | chr11 | + | 121550584 | 121550670 | 121545242 | 121545429 | 121553936 | 121554109 | 163 | 138 | 47 | 1 | 0 | 0 | 0 | 27 | 0 | 0 | 98 | 49 | 1 | 1 | 0.988 | 1 | 1 | NA | | 1 | -0.004 |
| SE | 5957 SORL1 | SORL1 | chr11 | + | 121553936 | 121554109 | 121545242 | 121545429 | 121555186 | 121555318 | 291 | 134 | 57 | 2 | 0 | 0 | 10 | 39 | 0 | 0 | 98 | 49 | 1 | 1 | 0.986 | 1 | 1 | | 1 | 1 | -0.005 |
| SE | 5958 SORL1 | SORL1 | chr11 | + | 121555186 | 121555318 | 121545242 | 121545429 | 121558590 | 121558837 | 17 | 4 | 3 | 1 | 0 | 0 | 0 | 2 | 0 | 0 | 98 | 49 | 1 | 1 | 0.895 | 1 | 1 | NA | | 1 | -0.035 |
| SE | 5961 SORL1 | SORL1 | chr11 | + | 121570156 | 121570270 | 121566939 | 121567113 | 121583457 | 121583583 | 170 | 45 | 28 | 3 | 0 | 0 | 0 | 32 | 0 | 0 | 98 | 49 | 1 | 1 | 0.966 | 1 | 1 | NA | | 1 | -0.011 |
| SE | 5963 SORL1 | SORL1 | chr11 | + | 121574240 | 121574363 | 121570156 | 121570270 | 121583457 | 121583583 | 349 | 131 | 68 | 1 | 0 | 0 | 0 | 27 | 0 | 0 | 98 | 49 | 1 | 1 | 0.994 | 1 | 1 | NA | | 1 | -0.002 |
| SE | 5965 SORL1 | SORL1 | chr11 | + | 121605401 | 121605571 | 121605112 | 121605239 | 121606844 | 121606957 | 524 | 184 | 145 | 2 | 4 | 0 | 11 | 57 | 0 | 1 | 98 | 49 | 1 | 1 | 0.992 | 0.958 | 1 | | 1 | 0.966 | 0 |
| SE | 5966 SORL1 | SORL1 | chr11 | + | 121608103 | 121608176 | 121607185 | 121607290 | 121611075 | 121611158 | 260 | 99 | 77 | 1 | 2 | 0 | 0 | 31 | 0 | 1 | 98 | 49 | 1 | 1 | 0.992 | 0.961 | 1 | NA | | 0.939 | 0.045 |
| SE | 5967 SORL1 | SORL1 | chr11 | + | 121611075 | 121611158 | 121608103 | 121608176 | 121612735 | 121612832 | 425 | 149 | 106 | 0 | 1 | 0 | 0 | 40 | 0 | 0 | 98 | 49 | 1 | 1 | 1 | 0.987 | 1 | NA | | 1 | -0.004 |
| SE | 5968 SORL1 | SORL1 | chr11 | + | 121625084 | 121625277 | 121622161 | 121622268 | 121627554 | 121627767 | 236 | 96 | 58 | 0 | 1 | 0 | 1 | 33 | 0 | 0 | 98 | 49 | 1 | 1 | 1 | 0.98 | 1 | | 1 | 1 | -0.007 |
| SE | 5974 CDK11B | CDK11B | chr1 | - | 1637080 | 1637202 | 1636896 | 1637004 | 1637407 | 1637513 | 12 | 0 | 4 | 0 | 0 | 0 | 1 | 0 | 2 | 0 | 98 | 49 | 1 | 1 | 1 | NA | 1 | | 0.2 | NA | 0.8 |
| SE | 5979 FLII | FLII | chr17 | - | 18246307 | 18246462 | 18246161 | 18246222 | 18246593 | 18246828 | 50 | 23 | 10 | 0 | 1 | 0 | 2 | 0 | 0 | 0 | 98 | 49 | 1 | 1 | 1 | 0.92 | 1 | | 1 | NA | -0.027 |

| | | | | | | | | | | | | | | | | | | | | | | | | | | | | | | | |
|---|---|---|---|---|---|---|---|---|---|---|---|---|---|---|---|---|---|---|---|---|---|---|---|---|---|---|---|---|---|---|---|
| SE | 5980 | FLII | FLII | chr17 | - | 18253543 | 18253719 | 18253300 | 18253458 | 18254078 | 18254182 | 9 | 3 | 0 | 3 | 0 | 0 | 3 | 1 | 0 | 0 | 98 | 49 | 1 | 1 | 0.6 | 1 | NA | 1 | 1 | -0.2 |
| SE | 5982 | FLII | FLII | chr17 | - | 18256525 | 18256597 | 18255182 | 18255263 | 18256908 | 18257019 | 44 | 20 | 6 | 2 | 0 | 0 | 7 | 15 | 0 | 0 | 98 | 49 | 1 | 1 | 0.917 | 1 | 1 | 1 | 1 | -0.028 |
| SE | 5990 | RPS6KB2 | RPS6KB2 | chr11 | + | 67431367 | 67431515 | 67429526 | 67429595 | 67432599 | 67432657 | 10 | 2 | 2 | 2 | 0 | 0 | 0 | 4 | 0 | 0 | 98 | 49 | 1 | 1 | 0.714 | 1 | 1 | NA | 1 | -0.095 |
| SE | 5992 | RPS6KB2 | RPS6KB2 | chr11 | + | 67433125 | 67433216 | 67432951 | 67433042 | 67433339 | 67433447 | 7 | 0 | 2 | 0 | 0 | 0 | 0 | 0 | 0 | 1 | 98 | 49 | 1 | 1 | 1 | NA | 1 | NA | 0 | 1 |
| SE | 5993 | DPP9 | DPP9 | chr19 | - | 4695377 | 4695555 | 4694660 | 4694823 | 4697550 | 4697651 | 4 | 0 | 0 | 1 | 0 | 0 | 0 | 4 | 0 | 0 | 98 | 49 | 1 | 1 | 0.667 | NA | NA | NA | | -0.333 |
| SE | 5994 | DPP9 | DPP9 | chr19 | - | 4702602 | 4702716 | 4702026 | 4702155 | 4703885 | 4704054 | 1 | 0 | 0 | 1 | 0 | 0 | 2 | 0 | 0 | 0 | 98 | 49 | 1 | 1 | 0.333 | NA | NA | 1 | NA | -0.667 |
| SE | 5999 | CHD1L | CHD1L | chr1 | + | 147270931 | 147271005 | 147268781 | 147268878 | 147272170 | 147272281 | 0 | 2 | 0 | 0 | 0 | 0 | 0 | 1 | 0 | 1 | 98 | 49 | 1 | 1 | NA | 1 | NA | NA | 0.333 | 0.667 |
| SE | 6001 | NIN | NIN | chr14 | - | 50756491 | 50758630 | 50754741 | 50754867 | 50759856 | 50760359 | 17 | 9 | 0 | 4 | 2 | 1 | 0 | 2 | 0 | 0 | 98 | 49 | 1 | 1 | 0.68 | 0.692 | 0 | NA | 1 | -0.543 |
| SE | 6002 | NIN | NIN | chr14 | - | 50759856 | 50760359 | 50754741 | 50754867 | 50761789 | 50761911 | 21 | 18 | 9 | 5 | 1 | 1 | 0 | 2 | 0 | 2 | 98 | 49 | 1 | 1 | 0.677 | 0.9 | 0.818 | NA | 0.333 | 0.465 |
| SE | 6003 | NIN | NIN | chr14 | - | 50766306 | 50766396 | 50763825 | 50763964 | 50766779 | 50766890 | 59 | 27 | 16 | 5 | 3 | 1 | 0 | 14 | 0 | 1 | 98 | 49 | 1 | 1 | 0.855 | 0.818 | 0.889 | NA | 0.875 | -0.021 |
| SE | 6004 | NIN | NIN | chr14 | - | 50772300 | 50772468 | 50771331 | 50771468 | 50772948 | 50773095 | 68 | 22 | 15 | 1 | 1 | 0 | 1 | 12 | 0 | 0 | 98 | 49 | 1 | 1 | 0.971 | 0.917 | 1 | 1 | 1 | -0.037 |
| SE | 6006 | NIN | NIN | chr14 | - | 50830463 | 50830517 | 50821873 | 50822077 | 50831005 | 50831121 | 29 | 13 | 2 | 0 | 0 | 0 | 3 | 3 | 0 | 1 | 98 | 49 | 1 | 1 | 1 | 1 | 1 | 1 | 0.6 | 0.2 |
| SE | 6008 | FOS | FOS | chr14 | + | 75279876 | 75280128 | 75278777 | 75279123 | 75280559 | 75280667 | 1366 | 319 | 192 | 0 | 0 | 0 | 18 | 153 | 0 | 1 | 98 | 49 | 1 | 1 | 1 | 1 | 1 | 1 | 0.987 | 0.006 |
| SE | 6009 | FOS | FOS | chr14 | + | 75280559 | 75280667 | 75279876 | 75280128 | 75280782 | 75282234 | 1634 | 373 | 175 | 5 | 1 | 2 | 25 | 189 | 1 | 0 | 98 | 49 | 1 | 1 | 0.994 | 0.995 | 0.978 | 0.926 | 1 | 0.026 |
| SE | 6019 | KSR1 | KSR1 | chr17 | + | 27601359 | 27601401 | 27597267 | 27597436 | 27603833 | 27603888 | 10 | 0 | 0 | 2 | 0 | 0 | 0 | 2 | 0 | 0 | 91 | 49 | 1 | 1 | 0.729 | NA | NA | NA | 1 | -0.271 |
| SE | 6026 | ZNF185 | ZNF185 | chrX | + | 152922719 | 152922809 | 152922172 | 152922256 | 152928574 | 152928661 | 5 | 1 | 2 | 4 | 1 | 0 | 1 | 0 | 0 | 0 | 98 | 49 | 1 | 1 | 0.385 | 0.333 | 1 | 1 | NA | -0.427 |
| SE | 6027 | ZNF185 | ZNF185 | chrX | + | 152922719 | 152922809 | 152922172 | 152922256 | 152931671 | 152931776 | 5 | 3 | 0 | 1 | 0 | 0 | 1 | 1 | 0 | 0 | 98 | 49 | 1 | 1 | 0.714 | 1 | NA | 1 | 1 | -0.143 |
| SE | 6029 | ZNF185 | ZNF185 | chrX | + | 152928574 | 152928661 | 152922719 | 152922809 | 152931671 | 152931776 | 7 | 2 | 2 | 0 | 3 | 0 | 0 | 0 | 0 | 1 | 98 | 49 | 1 | 1 | 1 | 0.25 | 1 | NA | 0 | 0.75 |
| SE | 6040 | USPL1 | USPL1 | chr13 | + | 30653147 | 30653305 | 30646931 | 30647057 | 30657473 | 30659549 | 5 | 5 | 2 | 1 | 0 | 0 | 0 | 1 | 0 | 0 | 98 | 49 | 1 | 1 | 0.714 | 1 | 1 | NA | 1 | -0.095 |
| SE | 6052 | NUDT5 | NUDT5 | chr10 | - | 12170899 | 12170908 | 12170716 | 12170770 | 12172764 | 12172866 | 0 | 1 | 2 | 0 | 1 | 0 | 0 | 1 | 0 | 0 | 58 | 49 | 1 | 1 | NA | 0.458 | 1 | NA | 1 | -0.271 |
| SE | 6054 | ALPK1 | ALPK1 | chr4 | + | 112423943 | 112424003 | 112411826 | 112412025 | 112425664 | 112425751 | 24 | 15 | 8 | 2 | 2 | 0 | 0 | 6 | 0 | 0 | 98 | 49 | 1 | 1 | 0.857 | 0.789 | 1 | NA | 1 | -0.118 |
| SE | 6055 | ALPK1 | ALPK1 | chr4 | + | 112430447 | 112432581 | 112429148 | 112429253 | 112435147 | 112435301 | 29 | 4 | 8 | 1 | 0 | 0 | 1 | 3 | 0 | 0 | 98 | 49 | 1 | 1 | 0.935 | 1 | 1 | 1 | 1 | -0.022 |
| SE | 6058 | TBC1D10B | TBC1D10B | chr16 | - | 30359171 | 30359361 | 30358662 | 30358817 | 30359537 | 30359603 | 10 | 2 | 2 | 4 | 2 | 0 | 0 | 2 | 0 | 0 | 98 | 49 | 1 | 1 | 0.556 | 0.333 | 1 | NA | 1 | -0.37 |
| SE | 6060 | THRAP3 | THRAP3 | chr1 | + | 36296582 | 36296770 | 36293850 | 36293935 | 36300885 | 36301084 | 26 | 18 | 7 | 0 | 1 | 0 | 0 | 6 | 0 | 0 | 98 | 49 | 1 | 1 | 1 | 0.9 | 1 | NA | 1 | -0.033 |
| SE | 6061 | THRAP3 | THRAP3 | chr1 | + | 36301552 | 36301696 | 36300885 | 36301084 | 36303795 | 36305356 | 27 | 24 | 11 | 0 | 0 | 0 | 0 | 3 | 0 | 2 | 98 | 49 | 1 | 1 | 1 | 1 | 1 | NA | 0.429 | 0.571 |
| SE | 6062 | HLA-DMB | HLA-DMB | chr6 | - | 32935535 | 32935652 | 32935341 | 32935377 | 32937171 | 32937456 | 9 | 0 | 0 | 0 | 0 | 0 | 0 | 4 | 0 | 1 | 98 | 49 | 1 | 1 | 1 | NA | NA | NA | 0.667 | 0.333 |
| SE | 6064 | NOTCH2 | NOTCH2 | chr1 | - | 119922635 | 119922778 | 119922235 | 119922446 | 119923636 | 119923984 | 68 | 17 | 10 | 0 | 0 | 0 | 2 | 5 | 0 | 1 | 98 | 49 | 1 | 1 | 1 | 1 | 1 | 1 | 0.714 | 0.143 |
| SE | 6065 | NOTCH2 | NOTCH2 | chr1 | - | 119923636 | 119923984 | 119922235 | 119922446 | 119925304 | 119925810 | 38 | 11 | 1 | 0 | 2 | 0 | 0 | 6 | 0 | 0 | 98 | 49 | 1 | 1 | 1 | 0.733 | 1 | NA | 1 | -0.089 |
| SE | 6066 | NOTCH2 | NOTCH2 | chr1 | - | 119925304 | 119925810 | 119923636 | 119923984 | 119926498 | 119926611 | 77 | 31 | 7 | 1 | 0 | 0 | 0 | 20 | 0 | 0 | 98 | 49 | 1 | 1 | 0.975 | 1 | 1 | NA | 1 | -0.008 |
| SE | 6067 | NOTCH2 | NOTCH2 | chr1 | - | 119959391 | 119959502 | 119955039 | 119955232 | 119963573 | 119963807 | 70 | 12 | 14 | 7 | 1 | 0 | 0 | 4 | 0 | 1 | 98 | 49 | 1 | 1 | 0.833 | 0.857 | 1 | NA | 0.667 | 0.23 |
| SE | 6068 | NOTCH2 | NOTCH2 | chr1 | - | 119968076 | 119968232 | 119967432 | 119967621 | 119969510 | 119969744 | 106 | 47 | 22 | 1 | 1 | 0 | 1 | 16 | 0 | 0 | 98 | 49 | 1 | 1 | 0.981 | 0.959 | 1 | 1 | 1 | -0.02 |
| SE | 6069 | NOTCH2 | NOTCH2 | chr1 | - | 120029905 | 120029987 | 120005328 | 120005588 | 120069333 | 120069703 | 50 | 22 | 2 | 0 | 1 | 0 | 0 | 9 | 0 | 0 | 98 | 49 | 1 | 1 | 1 | 0.917 | 1 | NA | 1 | -0.028 |
| SE | 6070 | FAM13A | FAM13A | chr4 | - | 88737471 | 88737555 | 88732001 | 88732198 | 88739029 | 88739125 | 2 | 0 | 0 | 0 | 0 | 0 | 0 | 0 | 0 | 2 | 98 | 49 | 1 | 1 | 1 | NA | NA | NA | 0 | 1 |
| SE | 6074 | CLIP1 | CLIP1 | chr12 | - | 122347374 | 122347479 | 122340752 | 122341697 | 122352725 | 122352786 | 2 | 2 | 0 | 4 | 3 | 0 | 0 | 1 | 0 | 0 | 98 | 49 | 1 | 1 | 0.2 | 0.25 | NA | NA | 1 | -0.775 |
| SE | 6076 | CLIP1 | CLIP1 | chr12 | - | 122355114 | 122355312 | 122354452 | 122354556 | 122360958 | 122361181 | 19 | 15 | 0 | 0 | 2 | 0 | 0 | 7 | 0 | 0 | 98 | 49 | 1 | 1 | 1 | 0.789 | NA | NA | 1 | -0.105 |
| SE | 6091 | C1orf162 | C1orf162 | chr1 | + | 111477650 | 111477775 | 111477333 | 111477428 | 111477982 | 111478512 | 3 | 1 | 0 | 0 | 0 | 0 | 1 | 9 | 0 | 1 | 98 | 49 | 1 | 1 | 1 | 1 | NA | 1 | 0.818 | 0.091 |
| SE | 6092 | C1orf162 | C1orf162 | chr1 | + | 111477725 | 111477775 | 111477333 | 111477428 | 111477982 | 111478512 | 12 | 1 | 0 | 0 | 0 | 0 | 1 | 9 | 0 | 1 | 98 | 49 | 1 | 1 | 1 | 1 | NA | 1 | 0.818 | 0.091 |
| SE | 6098 | UBE2K | UBE2K | chr4 | + | 39755656 | 39755739 | 39745751 | 39745810 | 39774833 | 39774933 | 12 | 8 | 4 | 0 | 0 | 0 | 0 | 2 | 0 | 1 | 98 | 49 | 1 | 1 | 1 | 1 | 1 | NA | 0.5 | 0.5 |
| SE | 6106 | ITPR2 | ITPR2 | chr12 | - | 26659112 | 26659285 | 26658010 | 26658130 | 26663684 | 26663846 | 16 | 6 | 1 | 0 | 2 | 0 | 0 | 6 | 0 | 0 | 98 | 49 | 1 | 1 | 1 | 0.6 | 1 | NA | 1 | -0.133 |
| SE | 6109 | ECT2 | ECT2 | chr3 | + | 172755482 | 172755575 | 172755294 | 172755374 | 172756982 | 172757165 | 1 | 0 | 0 | 0 | 0 | 0 | 0 | 0 | 1 | 0 | 98 | 49 | 1 | 1 | 1 | NA | NA | 0 | NA | 1 |
| SE | 6110 | GAB2 | GAB2 | chr11 | - | 78220318 | 78220444 | 78215289 | 78219415 | 78221676 | 78221779 | 23 | 15 | 18 | 2 | 0 | 0 | 1 | 4 | 0 | 0 | 98 | 49 | 1 | 1 | 0.852 | 1 | 1 | 1 | 1 | -0.049 |
| SE | 6111 | GAB2 | GAB2 | chr11 | - | 78250156 | 78250400 | 78226464 | 78227051 | 78280600 | 78280901 | 52 | 32 | 17 | 2 | 0 | 0 | 4 | 13 | 0 | 0 | 98 | 49 | 1 | 1 | 0.929 | 1 | 1 | 1 | 1 | -0.024 |
| SE | 6118 | CUTC | CUTC | chr10 | + | 99750368 | 99750396 | 99747256 | 99747390 | 99754528 | 99754634 | 10 | 6 | 0 | 1 | 0 | 0 | 0 | 0 | 0 | 3 | 77 | 49 | 1 | 1 | 0.864 | 1 | NA | NA | 0 | 0.932 |
| SE | 6119 | SENP1 | SENP1 | chr12 | - | 48074689 | 48074793 | 48074323 | 48074607 | 48083590 | 48083762 | 7 | 1 | 0 | 0 | 1 | 0 | 0 | 2 | 0 | 0 | 98 | 49 | 1 | 1 | 1 | 0.333 | NA | NA | 1 | -0.334 |
| SE | 6120 | SENP1 | SENP1 | chr12 | - | 48089162 | 48089293 | 48088800 | 48088960 | 48096342 | 48096427 | 0 | 1 | 0 | 2 | 0 | 0 | 0 | 0 | 0 | 1 | 98 | 49 | 1 | 1 | 0 | 1 | NA | NA | 0 | 0.5 |
| SE | 6121 | TIPRL | TIPRL | chr1 | + | 168191368 | 168191500 | 168184778 | 168184878 | 168196546 | 168196642 | 11 | 8 | 6 | 0 | 1 | 0 | 0 | 2 | 0 | 0 | 98 | 49 | 1 | 1 | 1 | 0.8 | 1 | NA | 1 | -0.067 |
| SE | 6123 | NUMB | NUMB | chr14 | - | 73279280 | 73279424 | 73275209 | 73277293 | 73282358 | 73282505 | 0 | 0 | 0 | 24 | 8 | 2 | 0 | 2 | 0 | 10 | 98 | 49 | 1 | 1 | 0 | 0 | 0 | NA | 0.091 | -0.091 |
| SE | 6124 | NUMB | NUMB | chr14 | - | 73284080 | 73284374 | 73282358 | 73282505 | 73287109 | 73287314 | 45 | 12 | 18 | 5 | 0 | 0 | 0 | 6 | 0 | 0 | 98 | 49 | 1 | 1 | 0.818 | 1 | 1 | NA | 1 | -0.061 |
| SE | 6128 | KDSR | KDSR | chr18 | - | 63335256 | 63335358 | 63327737 | 63331901 | 63338799 | 63338883 | 6 | 0 | 0 | 2 | 0 | 0 | 0 | 1 | 0 | 0 | 98 | 49 | 1 | 1 | 0.6 | NA | NA | NA | 1 | -0.4 |
| SE | 6134 | CD55 | CD55 | chr1 | + | 207324558 | 207324750 | 207322381 | 207322567 | 207325621 | 207325721 | 431 | 170 | 63 | 2 | 1 | 0 | 6 | 79 | 0 | 0 | 98 | 49 | 1 | 1 | 0.991 | 0.988 | 1 | 1 | 1 | -0.007 |
| SE | 6135 | CD55 | CD55 | chr1 | + | 207331107 | 207331296 | 207326751 | 207326837 | 207336692 | 207336818 | 181 | 53 | 39 | 0 | 3 | 0 | 2 | 28 | 0 | 0 | 98 | 49 | 1 | 1 | 1 | 0.898 | 1 | 1 | 1 | -0.034 |
| SE | 6136 | CD55 | CD55 | chr1 | + | 207340390 | 207340508 | 207339396 | 207339417 | 207359545 | 207360966 | 5 | 0 | 9 | 27 | 5 | 14 | 0 | 0 | 0 | 5 | 98 | 49 | 1 | 1 | 0.085 | 0 | 0.243 | NA | 0 | 0.109 |
| SE | 6137 | CD55 | CD55 | chr1 | + | 207347446 | 207347561 | 207339396 | 207339417 | 207359545 | 207360966 | 1 | 0 | 0 | 27 | 5 | 14 | 0 | 0 | 0 | 5 | 98 | 49 | 1 | 1 | 0.018 | 0 | 0 | NA | 0 | 0.006 |
| SE | 6139 | CD55 | CD55 | chr1 | + | 207354006 | 207354099 | 207339396 | 207339417 | 207359545 | 207360966 | 1 | 0 | 0 | 27 | 5 | 14 | 0 | 0 | 0 | 5 | 98 | 49 | 1 | 1 | 0.018 | 0 | 0 | NA | 0 | 0.006 |
| SE | 6143 | SRSF5 | SRSF5 | chr14 | + | 69768797 | 69768896 | 69768603 | 69768674 | 69769181 | 69769251 | 179 | 77 | 40 | 1 | 0 | 0 | 0 | 20 | 0 | 0 | 98 | 49 | 1 | 1 | 0.989 | 1 | 1 | NA | 1 | -0.004 |
| SE | 6146 | CNOT1 | CNOT1 | chr16 | - | 58547565 | 58547682 | 58547185 | 58547296 | 58549718 | 58549898 | 51 | 11 | 0 | 2 | 0 | 1 | 0 | 13 | 0 | 0 | 98 | 49 | 1 | 1 | 0.927 | 1 | 0 | NA | 1 | -0.358 |
| SE | 6147 | CNOT1 | CNOT1 | chr16 | - | 58555250 | 58555537 | 58553781 | 58553860 | 58555783 | 58555908 | 35 | 10 | 0 | 1 | 0 | 0 | 0 | 2 | 0 | 0 | 98 | 49 | 1 | 1 | 0.946 | 1 | NA | NA | 1 | -0.027 |
| SE | 6149 | CNOT1 | CNOT1 | chr16 | - | 58575006 | 58575129 | 58574608 | 58574760 | 58576462 | 58576582 | 23 | 11 | 7 | 1 | 0 | 0 | 2 | 6 | 0 | 0 | 98 | 49 | 1 | 1 | 0.92 | 1 | 1 | 1 | 1 | -0.027 |
| SE | 6150 | CNOT1 | CNOT1 | chr16 | - | 58583055 | 58583182 | 58582792 | 58582903 | 58585337 | 58585506 | 35 | 8 | 1 | 0 | 2 | 0 | 0 | 6 | 0 | 0 | 98 | 49 | 1 | 1 | 1 | 0.667 | 1 | NA | 1 | -0.111 |
| SE | 6151 | CNOT1 | CNOT1 | chr16 | - | 58599235 | 58599511 | 58588798 | 58588906 | 58629727 | 58629886 | 12 | 4 | 0 | 1 | 0 | 0 | 1 | 0 | 0 | 0 | 98 | 49 | 1 | 1 | 0.857 | 1 | NA | 1 | NA | -0.072 |
| SE | 6157 | LIMK2 | LIMK2 | chr22 | + | 31265945 | 31266132 | 31262594 | 31262791 | 31266983 | 31267070 | 68 | 29 | 16 | 0 | 1 | 0 | 2 | 21 | 0 | 0 | 98 | 49 | 1 | 1 | 1 | 0.935 | 1 | 1 | 1 | -0.022 |
| SE | 6158 | LIMK2 | LIMK2 | chr22 | + | 31266983 | 31267070 | 31265945 | 31266132 | 31267775 | 31267907 | 112 | 56 | 23 | 0 | 0 | 0 | 0 | 20 | 0 | 1 | 98 | 49 | 1 | 1 | 1 | 1 | 1 | NA | 0.909 | 0.091 |
| SE | 6159 | LIMK2 | LIMK2 | chr22 | + | 31268143 | 31268200 | 31267775 | 31267907 | 31271135 | 31271201 | 71 | 30 | 30 | 1 | 0 | 0 | 1 | 20 | 0 | 0 | 98 | 49 | 1 | 1 | 0.973 | 1 | 1 | 1 | 1 | -0.009 |
| SE | 6160 | LIMK2 | LIMK2 | chr22 | + | 31273451 | 31273507 | 31272529 | 31272704 | 31275150 | 31275308 | 201 | 114 | 21 | 1 | 0 | 0 | 13 | 40 | 0 | 0 | 98 | 49 | 1 | 1 | 0.99 | 1 | 1 | 1 | 1 | -0.003 |
| SE | 6165 | NBPF14 | NBPF14 | chr1 | - | 148593597 | 148593700 | 148592551 | 148592766 | 148595542 | 148595717 | 5 | 1 | 3 | 2 | 2 | 0 | 0 | 1 | 0 | 1 | 98 | 49 | 1 | 1 | 0.556 | 0.2 | 1 | NA | 0.333 | 0.252 |
| SE | 6166 | TCF25 | TCF25 | chr16 | + | 89893727 | 89893858 | 89892192 | 89892275 | 89895037 | 89895137 | 9 | 8 | 5 | 1 | 0 | 0 | 0 | 6 | 0 | 0 | 98 | 49 | 1 | 1 | 0.818 | 1 | 1 | NA | 1 | -0.061 |
| SE | 6174 | MAP2K7 | MAP2K7 | chr19 | + | 7905807 | 7905855 | 7903779 | 7904068 | 7909754 | 7909896 | 0 | 3 | 0 | 2 | 3 | 0 | 0 | 0 | 1 | 0 | 97 | 49 | 1 | 1 | 0 | 0.336 | NA | 0 | NA | 0.168 |
| SE | 6175 | PTGS2 | PTGS2 | chr1 | - | 186677648 | 186677830 | 186676832 | 186676916 | 186678260 | 186678404 | 526 | 106 | 24 | 2 | 0 | 0 | 1 | 45 | 0 | 0 | 98 | 49 | 1 | 1 | 0.992 | 1 | 1 | 1 | 1 | -0.003 |
| SE | 6185 | DDX60 | DDX60 | chr4 | - | 168272042 | 168272138 | 168268853 | 168268969 | 168273278 | 168273398 | 5 | 2 | 1 | 0 | 1 | 0 | 1 | 7 | 1 | 0 | 98 | 49 | 1 | 1 | 1 | 0.5 | 1 | 0.333 | 1 | 0.167 |
| SE | 6186 | ARHGEF2 | ARHGEF2 | chr1 | - | 155954901 | 155954969 | 155952627 | 155952828 | 155957712 | 155957882 | 53 | 26 | 10 | 2 | 0 | 0 | 0 | 12 | 0 | 0 | 98 | 49 | 1 | 1 | 0.93 | 1 | 1 | NA | 1 | -0.023 |
| SE | 6187 | ARHGEF2 | ARHGEF2 | chr1 | - | 155957712 | 155957882 | 155954901 | 155954969 | 155958319 | 155958396 | 35 | 29 | 20 | 0 | 2 | 0 | 0 | 10 | 0 | 0 | 98 | 49 | 1 | 1 | 1 | 0.879 | 1 | NA | 1 | -0.04 |
| SE | 6188 | ARHGEF2 | ARHGEF2 | chr1 | - | 155965630 | 155965760 | 155965302 | 155965412 | 155966415 | 155966479 | 37 | 4 | 9 | 6 | 2 | 0 | 0 | 8 | 0 | 0 | 98 | 49 | 1 | 1 | 0.755 | 0.5 | 1 | NA | 1 | -0.248 |
| SE | 6189 | ARHGEF2 | ARHGEF2 | chr1 | - | 155966819 | 155966887 | 155966415 | 155966479 | 155969155 | 155969300 | 45 | 23 | 7 | 1 | 1 | 0 | 0 | 4 | 0 | 0 | 98 | 49 | 1 | 1 | 0.957 | 0.92 | 1 | NA | 1 | -0.041 |
| SE | 6192 | TXNDC17 | TXNDC17 | chr17 | + | 6641752 | 6641834 | 6640901 | 6641227 | 6642248 | 6642324 | 1 | 1 | 5 | 0 | 0 | 0 | 0 | 1 | 0 | 2 | 98 | 49 | 1 | 1 | 1 | 1 | 1 | NA | 0.2 | 0.8 |
| SE | 6200 | CS | CS | chr12 | - | 56283791 | 56283857 | 56282859 | 56282991 | 56285915 | 56286023 | 9 | 6 | 0 | 0 | 1 | 0 | 0 | 7 | 0 | 0 | 98 | 49 | 1 | 1 | 0.818 | 1 | NA | NA | 1 | -0.091 |
| SE | 6208 | ADAT1 | ADAT1 | chr16 | - | 75618585 | 75618640 | 75617141 | 75617272 | 75620265 | 75620334 | 6 | 1 | 3 | 0 | 1 | 0 | 1 | 0 | 0 | 0 | 98 | 49 | 1 | 1 | 1 | 0.333 | 1 | 1 | NA | -0.222 |
| SE | 6210 | TRIM33 | TRIM33 | chr1 | - | 114397939 | 114397990 | 114392776 | 114397860 | 114399456 | 114399609 | 17 | 8 | 5 | 6 | 3 | 0 | 0 | 7 | 0 | 2 | 98 | 49 | 1 | 1 | 0.586 | 0.571 | 1 | NA | 0.636 | 0.083 |
| SE | 6217 | RIF1 | RIF1 | chr2 | + | 151462883 | 151466120 | 151462411 | 151462466 | 151467999 | 151468146 | 8 | 7 | 4 | 0 | 0 | 3 | 0 | 4 | 0 | 0 | 98 | 49 | 1 | 1 | 1 | 1 | 0.4 | NA | 1 | -0.2 |
| SE | 6238 | EHBP1L1 | EHBP1L1 | chr11 | + | 65589935 | 65589991 | 65589750 | 65589820 | 65590086 | 65590210 | 52 | 21 | 9 | 0 | 1 | 0 | 1 | 10 | 0 | 0 | 98 | 49 | 1 | 1 | 1 | 0.913 | 1 | 1 | 1 | -0.029 |
| SE | 6240 | APBB1IP | APBB1IP | chr10 | + | 26503196 | 26503274 | 26500818 | 26501111 | 26511746 | 26511906 | 165 | 44 | 23 | 3 | 0 | 0 | 0 | 28 | 0 | 0 | 98 | 49 | 1 | 1 | 0.965 | 1 | 1 | NA | 1 | -0.012 |
| SE | 6241 | APBB1IP | APBB1IP | chr10 | + | 26513538 | 26513660 | 26511746 | 26511906 | 26533438 | 26533525 | 80 | 38 | 24 | 1 | 1 | 0 | 0 | 20 | 0 | 0 | 98 | 49 | 1 | 1 | 0.976 | 0.95 | 1 | NA | 1 | -0.025 |
| SE | 6244 | PSMA5 | PSMA5 | chr1 | - | 109409927 | 109410014 | 109399030 | 109402090 | 109411010 | 109411113 | 11 | 8 | 5 | 1 | 0 | 0 | 0 | 4 | 0 | 0 | 98 | 49 | 1 | 1 | 0.846 | 1 | 1 | NA | 1 | -0.051 |
| SE | 6248 | STK24 | STK24 | chr13 | - | 98457167 | 98457304 | 98450198 | 98453209 | 98460371 | 98460440 | 30 | 9 | 4 | 0 | 0 | 0 | 0 | 5 | 0 | 1 | 98 | 49 | 1 | 1 | 1 | 1 | 1 | NA | 0.714 | 0.286 |
| SE | 6249 | STK24 | STK24 | chr13 | - | 98482264 | 98482321 | 98475249 | 98475358 | 98519242 | 98519473 | 42 | 8 | 6 | 0 | 2 | 0 | 0 | 2 | 0 | 0 | 98 | 49 | 1 | 1 | 1 | 0.667 | 1 | NA | 1 | -0.111 |
| SE | 6251 | SEC24B | SEC24B | chr4 | + | 109481676 | 109481781 | 109473003 | 109473186 | 109491326 | 109491407 | 4 | 1 | 0 | 5 | 0 | 0 | 0 | 2 | 0 | 0 | 98 | 49 | 1 | 1 | 0.286 | 1 | NA | NA | 1 | -0.357 |
| SE | 6252 | MAP2K1 | MAP2K1 | chr15 | + | 66484989 | 66485191 | 66481754 | 66481879 | 66487227 | 66487292 | 8 | 0 | 2 | 2 | 0 | 0 | 0 | 5 | 0 | 0 | 98 | 49 | 1 | 1 | 0.667 | NA | 1 | NA | 1 | -0.166 |
| SE | 6262 | CLASP1 | CLASP1 | chr2 | - | 121409023 | 121409047 | 121407470 | 121407715 | 121410865 | 121410969 | 2 | 0 | 1 | 1 | 1 | 0 | 0 | 0 | 0 | 2 | 73 | 49 | 1 | 1 | 0.573 | 0 | 1 | NA | 0 | 0.524 |
| SE | 6266 | CLASP1 | CLASP1 | chr2 | - | 121445448 | 121445496 | 121430072 | 121430177 | 121447336 | 121447507 | 5 | 0 | 0 | 6 | 2 | 0 | 0 | 0 | 0 | 1 | 97 | 49 | 1 | 1 | 0.296 | 0 | NA | NA | 0 | 0.148 |
| SE | 6267 | CLASP1 | CLASP1 | chr2 | - | 121459979 | 121460125 | 121458839 | 121458975 | 121461100 | 121461193 | 6 | 4 | 0 | 2 | 0 | 0 | 0 | 2 | 0 | 0 | 98 | 49 | 1 | 1 | 0.6 | 1 | NA | NA | 1 | -0.2 |
| SE | 6268 | CLASP1 | CLASP1 | chr2 | - | 121515664 | 121515762 | 121503166 | 121503234 | 121525844 | 121525920 | 20 | 11 | 2 | 0 | 0 | 0 | 0 | 1 | 0 | 2 | 98 | 49 | 1 | 1 | 1 | 1 | 1 | NA | 0.2 | 0.8 |
| SE | 6269 | CLASP1 | CLASP1 | chr2 | - | 121527798 | 121527890 | 121525844 | 121525920 | 121528676 | 121528780 | 41 | 15 | 5 | 1 | 0 | 0 | 0 | 5 | 0 | 0 | 98 | 49 | 1 | 1 | 0.953 | 1 | 1 | NA | 1 | -0.016 |
| SE | 6270 | CLASP1 | CLASP1 | chr2 | - | 121530246 | 121530325 | 121528676 | 121528780 | 121605700 | 121606180 | 34 | 9 | 1 | 3 | 0 | 0 | 0 | 2 | 0 | 0 | 98 | 49 | 1 | 1 | 0.85 | 1 | 1 | NA | 1 | -0.05 |
| SE | 6271 | CANX | CANX | chr5 | + | 179708977 | 179709059 | 179708238 | 179708380 | 179709872 | 179710065 | 32 | 5 | 2 | 1 | 0 | 0 | 3 | 6 | 0 | 0 | 98 | 49 | 1 | 1 | 0.941 | 1 | 1 | 1 | 1 | -0.02 |
| SE | 6272 | HIAT1 | HIAT1 | chr1 | + | 100068472 | 100068586 | 100067976 | 100068195 | 100069613 | 100069685 | 21 | 2 | 6 | 2 | 0 | 0 | 0 | 2 | 0 | 0 | 98 | 49 | 1 | 1 | 0.84 | 1 | 1 | NA | 1 | -0.053 |
| SE | 6287 | CBFB | CBFB | chr16 | + | 67036638 | 67036755 | 67029726 | 67029813 | 67066681 | 67066798 | 12 | 10 | 2 | 1 | 0 | 0 | 0 | 3 | 0 | 0 | 98 | 49 | 1 | 1 | 0.857 | 1 | 1 | NA | 1 | -0.048 |
| SE | 6293 | FXR1 | FXR1 | chr3 | + | 180970157 | 180970358 | 180968050 | 180968254 | 180976121 | 180982751 | 12 | 4 | 1 | 2 | 0 | 0 | 0 | 2 | 0 | 0 | 98 | 49 | 1 | 1 | 0.75 | 1 | 1 | NA | 1 | -0.083 |
| SE | 6296 | TMEM87A | TMEM87A | chr15 | - | 42272062 | 42272123 | 42267946 | 42268032 | 42273254 | 42273584 | 13 | 3 | 3 | 1 | 0 | 0 | 0 | 1 | 0 | 0 | 98 | 49 | 1 | 1 | 0.867 | 1 | 1 | NA | 1 | -0.044 |
| SE | 6297 | SOAT1 | SOAT1 | chr1 | + | 179302676 | 179302802 | 179293713 | 179293936 | 179323436 | 179323495 | 9 | 3 | 3 | 1 | 0 | 0 | 0 | 6 | 0 | 0 | 98 | 49 | 1 | 1 | 0.818 | 1 | 1 | NA | 1 | -0.061 |
| SE | 6301 | ADCK3 | ADCK3 | chr1 | + | 226965670 | 226965737 | 226964999 | 226965410 | 226977448 | 226977523 | 4 | 6 | 8 | 1 | 0 | 0 | 0 | 3 | 0 | 0 | 98 | 49 | 1 | 1 | 0.667 | 1 | 1 | NA | 1 | -0.111 |
| SE | 6318 | PTBP2 | PTBP2 | chr1 | + | 96813028 | 96813106 | 96812711 | 96812928 | 96813272 | 96815049 | 4 | 0 | 0 | 0 | 0 | 0 | 0 | 0 | 0 | 2 | 98 | 49 | 1 | 1 | 1 | NA | NA | NA | 0 | 1 |
| SE | 6326 | SHISA5 | SHISA5 | chr3 | - | 48503094 | 48503193 | 48501136 | 48501293 | 48504018 | 48504249 | 0 | 2 | 0 | 9 | 3 | 0 | 0 | 0 | 0 | 1 | 98 | 49 | 1 | 1 | 0 | 0.25 | NA | NA | 0 | 0.125 |
| SE | 6327 | SHISA5 | SHISA5 | chr3 | - | 48503094 | 48503193 | 48501136 | 48501293 | 48504422 | 48504814 | 0 | 0 | 0 | 1 | 5 | 0 | 0 | 2 | 0 | 0 | 98 | 49 | 1 | 1 | 0 | 0 | NA | NA | 1 | -1 |
| SE | 6335 | VWA8 | VWA8 | chr13 | - | 41949935 | 41950013 | 41912037 | 41912168 | 41960852 | 41961085 | 2 | 0 | 0 | 0 | 0 | 0 | 0 | 0 | 0 | 1 | 98 | 49 | 1 | 1 | 1 | NA | NA | NA | 0 | 1 |
| SE | 6336 | LINC01578 | LINC01578 | chr15 | + | 92883584 | 92883778 | 92882842 | 92883186 | 92885514 | 92885584 | 8 | 4 | 0 | 8 | 3 | 0 | 0 | 2 | 0 | 0 | 98 | 49 | 1 | 1 | 0.333 | 0.4 | NA | NA | 1 | -0.633 |
| SE | 6339 | ZFAND4 | ZFAND4 | chr10 | - | 45652983 | 45653059 | 45651965 | 45652033 | 45663541 | 45663842 | 4 | 0 | 0 | 0 | 0 | 0 | 0 | 0 | 0 | 2 | 98 | 49 | 1 | 1 | 1 | NA | NA | NA | 0 | 1 |
| SE | 6346 | CDC14A | CDC14A | chr1 | + | 100455404 | 100455492 | 100442933 | 100442996 | 100462650 | 100462881 | 12 | 10 | 2 | 2 | 0 | 0 | 0 | 1 | 0 | 1 | 98 | 49 | 1 | 1 | 0.75 | 1 | 1 | NA | 0.333 | 0.584 |
| SE | 6351 | EVI5 | EVI5 | chr1 | - | 92607580 | 92607727 | 92605306 | 92605402 | 92624175 | 92624334 | 5 | 0 | 0 | 0 | 1 | 0 | 0 | 6 | 0 | 0 | 98 | 49 | 1 | 1 | 1 | 0 | NA | NA | 1 | -0.5 |
| SE | 6353 | EVI5 | EVI5 | chr1 | - | 92693801 | 92693899 | 92677157 | 92677218 | 92694298 | 92694388 | 18 | 6 | 0 | 0 | 0 | 0 | 0 | 4 | 0 | 1 | 98 | 49 | 1 | 1 | 1 | 1 | NA | NA | 0.667 | 0.333 |
| SE | 6365 | DARS2 | DARS2 | chr1 | + | 173828332 | 173828399 | 173826686 | 173826786 | 173830659 | 173830761 | 0 | 0 | 0 | 2 | 0 | 0 | 0 | 1 | 0 | 0 | 98 | 49 | 1 | 1 | 0 | NA | NA | NA | 1 | -1 |
| SE | 6368 | ALDOA | ALDOA | chr16 | + | 30066884 | 30067038 | 30065672 | 30065927 | 30067233 | 30067366 | 0 | 0 | 0 | 44 | 14 | 9 | 0 | 1 | 0 | 7 | 98 | 49 | 1 | 1 | 0 | 0 | 0 | NA | 0.067 | -0.067 |
| SE | 6369 | ALDOA | ALDOA | chr16 | + | 30068645 | 30068700 | 30067449 | 30067661 | 30068817 | 30068978 | 149 | 91 | 77 | 0 | 0 | 0 | 3 | 41 | 0 | 1 | 98 | 49 | 1 | 1 | 1 | 1 | 1 | 1 | 0.953 | 0.024 |
| SE | 6384 | KMT2D | KMT2D | chr12 | - | 49028827 | 49028958 | 49028008 | 49028141 | 49029060 | 49029236 | 12 | 7 | 0 | 2 | 0 | 0 | 4 | 7 | 3 | 0 | 98 | 49 | 1 | 1 | 0.75 | 1 | NA | NA 0.4 | 1 | 0.175 |
| SE | 6385 | KMT2D | KMT2D | chr12 | - | 49030892 | 49031033 | 49030600 | 49030768 | 49031174 | 49033964 | 21 | 19 | 1 | 1 | 0 | 0 | 1 | 3 | 0 | 0 | 98 | 49 | 1 | 1 | 0.913 | 1 | 1 | 1 | 1 | -0.029 |
| SE | 6386 | KMT2D | KMT2D | chr12 | - | 49054899 | 49055026 | 49053977 | 49054140 | 49055275 | 49055324 | 11 | 21 | 0 | 0 | 1 | 0 | 0 | 3 | 0 | 0 | 98 | 49 | 1 | 1 | 1 | 0.913 | NA | NA | 1 | -0.043 |
| SE | 6387 | KMT2D | KMT2D | chr12 | - | 49054899 | 49055026 | 49054527 | 49054751 | 49055275 | 49055324 | 13 | 23 | 4 | 0 | 3 | 0 | 0 | 3 | 0 | 0 | 98 | 49 | 1 | 1 | 1 | 0.793 | 1 | NA | 1 | -0.069 |
| SE | 6400 | TGFBR3 | TGFBR3 | chr1 | - | 91800958 | 91801055 | 91797286 | 91797471 | 91861470 | 91861644 | 0 | 4 | 0 | 1 | 0 | 0 | 0 | 0 | 0 | 3 | 98 | 49 | 1 | 1 | 0 | 1 | NA | NA | 0 | 0.5 |
| SE | 6405 | ARID1A | ARID1A | chr1 | + | 26729650 | 26729863 | 26696030 | 26697540 | 26731151 | 26731604 | 12 | 5 | 0 | 1 | 0 | 0 | 0 | 3 | 0 | 0 | 98 | 49 | 1 | 1 | 0.857 | 1 | NA | NA | 1 | -0.072 |
| SE | 6408 | POU2F1 | POU2F1 | chr1 | + | 167396285 | 167396427 | 167389587 | 167389761 | 167397993 | 167398133 | 2 | 2 | 4 | 0 | 0 | 0 | 0 | 3 | 0 | 1 | 98 | 49 | 1 | 1 | 1 | 1 | 1 | NA | 0.6 | 0.4 |
| SE | 6412 | DGKA | DGKA | chr12 | + | 55941509 | 55941584 | 55941251 | 55941325 | 55941997 | 55942083 | 1 | 0 | 0 | 1 | 0 | 0 | 0 | 1 | 0 | 0 | 98 | 49 | 1 | 1 | 0.333 | NA | NA | NA | 1 | -0.667 |
| SE | 6416 | ECD | ECD | chr10 | - | 73154255 | 73154448 | 73152292 | 73152421 | 73156274 | 73156453 | 3 | 0 | 1 | 0 | 0 | 0 | 0 | 2 | 0 | 1 | 98 | 49 | 1 | 1 | 1 | NA | NA | NA | 0.5 | 0.5 |
| SE | 6417 | ECD | ECD | chr10 | - | 73163732 | 73163950 | 73160433 | 73160551 | 73167865 | 73168095 | 9 | 3 | 4 | 1 | 0 | 0 | 0 | 1 | 0 | 0 | 98 | 49 | 1 | 1 | 0.818 | 1 | 1 | NA | 1 | -0.061 |
| SE | 6418 | ZRANB2 | ZRANB2 | chr1 | - | 71065677 | 71065752 | 71063290 | 71065137 | 71066775 | 71066934 | 6 | 2 | 0 | 12 | 0 | 0 | 0 | 1 | 0 | 0 | 98 | 49 | 1 | 1 | 0.2 | 1 | NA | NA | 1 | -0.4 |
| SE | 6420 | RALGAPA2 | RALGAPA2 | chr20 | - | 20396694 | 20396734 | 20389627 | 20393253 | 20412026 | 20412148 | 11 | 2 | 6 | 0 | 0 | 3 | 0 | 3 | 0 | 0 | 89 | 49 | 1 | 1 | 1 | 1 | 0.524 | NA | 1 | -0.159 |
| SE | 6421 | RALGAPA2 | RALGAPA2 | chr20 | - | 20524405 | 20524543 | 20520916 | 20521100 | 20524829 | 20524898 | 12 | 9 | 2 | 0 | 3 | 0 | 0 | 2 | 0 | 1 | 98 | 49 | 1 | 1 | 1 | 0.6 | 1 | NA | 0.5 | 0.367 |
| SE | 6422 | RALGAPA2 | RALGAPA2 | chr20 | - | 20611314 | 20611426 | 20605174 | 20605412 | 20616042 | 20616191 | 33 | 4 | 0 | 0 | 0 | 0 | 0 | 6 | 0 | 1 | 98 | 49 | 1 | 1 | 1 | 1 | NA | NA | 0.75 | 0.25 |
| SE | 6423 | RALGAPA2 | RALGAPA2 | chr20 | - | 20629362 | 20629590 | 20620462 | 20620630 | 20635417 | 20635617 | 22 | 21 | 4 | 1 | 0 | 0 | 0 | 6 | 0 | 0 | 98 | 49 | 1 | 1 | 0.917 | 1 | 1 | NA | 1 | -0.028 |
| SE | 6424 | RALGAPA2 | RALGAPA2 | chr20 | - | 20637362 | 20637501 | 20635417 | 20635617 | 20639784 | 20639900 | 37 | 17 | 18 | 1 | 4 | 0 | 0 | 13 | 0 | 2 | 98 | 49 | 1 | 1 | 0.949 | 0.68 | 1 | NA | 0.765 | 0.111 |
| SE | 6425 | RALGAPA2 | RALGAPA2 | chr20 | - | 20676235 | 20676288 | 20653529 | 20653587 | 20680690 | 20680801 | 45 | 24 | 6 | 0 | 1 | 0 | 0 | 13 | 0 | 2 | 98 | 49 | 1 | 1 | 1 | 0.923 | 1 | NA | 0.765 | 0.209 |
| SE | 6427 | PRRG4 | PRRG4 | chr11 | + | 32840106 | 32840239 | 32838881 | 32838930 | 32853295 | 32858123 | 14 | 2 | 0 | 1 | 2 | 1 | 0 | 1 | 0 | 0 | 98 | 49 | 1 | 1 | 0.875 | 0.333 | 0 | NA | 1 | -0.597 |
| SE | 6428 | NPTN | NPTN | chr15 | - | 73597021 | 73597369 | 73591965 | 73592137 | 73633124 | 73633412 | 1 | 0 | 0 | 10 | 6 | 1 | 0 | 0 | 0 | 1 | 98 | 49 | 1 | 1 | 0.048 | 0 | 0 | NA | 0 | 0.016 |
| SE | 6429 | LMBR1L | LMBR1L | chr12 | - | 49102886 | 49102951 | 49102467 | 49102540 | 49103090 | 49103159 | 7 | 3 | 1 | 0 | 1 | 0 | 1 | 4 | 0 | 0 | 98 | 49 | 1 | 1 | 1 | 0.6 | 1 | 1 | 1 | -0.133 |
| SE | 6434 | SLK | SLK | chr10 | + | 104010815 | 104010908 | 104008176 | 104008356 | 104018159 | 104018289 | 0 | 1 | 0 | 7 | 14 | 4 | 0 | 0 | 0 | 4 | 98 | 49 | 1 | 1 | 0 | 0.034 | 0 | NA | 0 | 0.011 |
| SE | 6441 | NDUFB6 | NDUFB6 | chr9 | - | 32558909 | 32558954 | 32553525 | 32553944 | 32572880 | 32573184 | 6 | 15 | 12 | 0 | 0 | 0 | 0 | 5 | 0 | 3 | 94 | 49 | 1 | 1 | 1 | 1 | 1 | NA | 0.465 | 0.535 |
| SE | 6442 | NDUFB6 | NDUFB6 | chr9 | - | 32570959 | 32571052 | 32553525 | 32553944 | 32572880 | 32573184 | 2 | 2 | 2 | 0 | 0 | 0 | 0 | 5 | 0 | 3 | 98 | 49 | 1 | 1 | 1 | 1 | 1 | NA | 0 | 1 |
| SE | 6443 | NDUFB6 | NDUFB6 | chr9 | - | 32570959 | 32571052 | 32558909 | 32558954 | 32572880 | 32573184 | 7 | 11 | 4 | 0 | 4 | 2 | 1 | 0 | 0 | 3 | 98 | 49 | 1 | 1 | 1 | 0.579 | 0.5 | 1 | NA | -0.307 |
| SE | 6444 | IRAK3 | IRAK3 | chr12 | + | 66203710 | 66203893 | 66189197 | 66189432 | 66209455 | 66209520 | 24 | 8 | 5 | 1 | 1 | 0 | 0 | 13 | 0 | 0 | 98 | 49 | 1 | 1 | 0.923 | 0.8 | 1 | NA | 1 | -0.092 |
| SE | 6449 | ILF3 | ILF3 | chr19 | + | 10680345 | 10680443 | 10679815 | 10679927 | 10681018 | 10681130 | 5 | 1 | 1 | 0 | 0 | 2 | 0 | 7 | 0 | 0 | 98 | 49 | 1 | 1 | 1 | 1 | 0.2 | NA | 1 | -0.267 |
| SE | 6451 | ZMIZ1 | ZMIZ1 | chr10 | + | 79289774 | 79289889 | 79277180 | 79277325 | 79290958 | 79291176 | 10 | 3 | 1 | 1 | 0 | 0 | 0 | 5 | 0 | 0 | 98 | 49 | 1 | 1 | 0.833 | 1 | 1 | NA | 1 | -0.056 |
| SE | 6455 | NCF1 | NCF1 | chr7 | + | 74779081 | 74779157 | 74777266 | 74777347 | 74779256 | 74779422 | 17 | 13 | 5 | 2 | 1 | 0 | 0 | 4 | 0 | 0 | 98 | 49 | 1 | 1 | 0.81 | 0.867 | 1 | NA | 1 | -0.108 |
| SE | 6462 | DPH3 | DPH3 | chr3 | - | 16264154 | 16264229 | 16257060 | 16260829 | 16264768 | 16264989 | 8 | 2 | 1 | 3 | 1 | 7 | 0 | 1 | 0 | 0 | 98 | 49 | 1 | 1 | 0.571 | 0.5 | 0.067 | NA | 1 | -0.621 |
| SE | 6467 | C1orf43 | C1orf43 | chr1 | - | 154213892 | 154213946 | 154212457 | 154212624 | 154214456 | 154214574 | 12 | 6 | 10 | 2 | 1 | 0 | 0 | 3 | 0 | 1 | 98 | 49 | 1 | 1 | 0.75 | 0.75 | 1 | NA | 0.6 | 0.233 |
| SE | 6468 | C1orf43 | C1orf43 | chr1 | - | 154214456 | 154214574 | 154213892 | 154213946 | 154220341 | 154220797 | 8 | 1 | 3 | 1 | 0 | 0 | 1 | 3 | 0 | 0 | 98 | 49 | 1 | 1 | 0.8 | 1 | 1 | 1 | 1 | -0.067 |
| SE | 6474 | GBP5 | GBP5 | chr1 | - | 89262219 | 89262401 | 89258950 | 89260817 | 89263735 | 89263948 | 65 | 9 | 9 | 0 | 2 | 0 | 0 | 10 | 0 | 0 | 98 | 49 | 1 | 1 | 1 | 0.692 | 1 | NA | 1 | -0.103 |
| SE | 6475 | GBP5 | GBP5 | chr1 | - | 89262682 | 89262785 | 89262219 | 89262401 | 89263735 | 89263948 | 85 | 28 | 13 | 2 | 0 | 0 | 3 | 3 | 0 | 0 | 98 | 49 | 1 | 1 | 0.955 | 1 | 1 | 1 | 1 | -0.015 |
| SE | 6476 | GBP5 | GBP5 | chr1 | - | 89266956 | 89267153 | 89266345 | 89266588 | 89268728 | 89268856 | 62 | 19 | 18 | 1 | 0 | 0 | 0 | 7 | 0 | 0 | 98 | 49 | 1 | 1 | 0.969 | 1 | 1 | NA | 1 | -0.01 |
| SE | 6477 | GBP5 | GBP5 | chr1 | - | 89267416 | 89267526 | 89266956 | 89267153 | 89268728 | 89268856 | 136 | 36 | 23 | 1 | 0 | 0 | 2 | 14 | 0 | 0 | 98 | 49 | 1 | 1 | 0.986 | 1 | 1 | 1 | 1 | -0.005 |
| SE | 6478 | GBP5 | GBP5 | chr1 | - | 89268728 | 89268856 | 89267416 | 89267526 | 89269365 | 89269574 | 72 | 16 | 9 | 1 | 1 | 0 | 1 | 7 | 0 | 0 | 98 | 49 | 1 | 1 | 0.973 | 0.889 | 1 | 1 | 1 | -0.046 |
| SE | 6479 | GBP5 | GBP5 | chr1 | - | 89268728 | 89268856 | 89267416 | 89267526 | 89272451 | 89272861 | 48 | 11 | 5 | 1 | 1 | 0 | 1 | 3 | 0 | 0 | 98 | 49 | 1 | 1 | 0.96 | 0.846 | 1 | 1 | 1 | -0.065 |
| SE | 6481 | GBP5 | GBP5 | chr1 | - | 89269365 | 89269574 | 89268728 | 89268856 | 89272451 | 89272861 | 30 | 9 | 5 | 2 | 1 | 0 | 0 | 4 | 0 | 0 | 98 | 49 | 1 | 1 | 0.882 | 0.818 | 1 | NA | 1 | -0.1 |
| SE | 6482 | GBP5 | GBP5 | chr1 | - | 89270754 | 89270862 | 89269365 | 89269574 | 89272451 | 89272861 | 52 | 16 | 3 | 4 | 3 | 1 | 0 | 7 | 0 | 0 | 98 | 49 | 1 | 1 | 0.867 | 0.727 | 0.6 | NA | 1 | -0.269 |
| SE | 6501 | ARHGEF11 | ARHGEF11 | chr1 | - | 156947768 | 156947956 | 156947303 | 156947450 | 156948180 | 156948228 | 18 | 10 | 9 | 0 | 0 | 0 | 0 | 13 | 0 | 1 | 98 | 49 | 1 | 1 | 1 | 1 | 1 | NA | 0.867 | 0.133 |
| SE | 6508 | ADAM15 | ADAM15 | chr1 | + | 155061414 | 155061489 | 155060762 | 155060832 | 155062244 | 155062369 | 0 | 0 | 0 | 2 | 0 | 0 | 0 | 1 | 0 | 0 | 98 | 49 | 1 | 1 | 0 | NA | NA | NA | 1 | -1 |
| SE | 6517 | ACAP2 | ACAP2 | chr3 | - | 195289120 | 195289231 | 195285795 | 195285857 | 195291705 | 195291815 | 60 | 38 | 19 | 2 | 0 | 0 | 0 | 11 | 0 | 0 | 98 | 49 | 1 | 1 | 0.937 | 1 | 1 | NA | 1 | -0.021 |
| SE | 6518 | ACAP2 | ACAP2 | chr3 | - | 195291705 | 195291815 | 195289120 | 195289231 | 195292264 | 195292452 | 48 | 38 | 13 | 1 | 0 | 1 | 0 | 12 | 0 | 2 | 98 | 49 | 1 | 1 | 0.96 | 1 | 0.867 | NA | 0.75 | 0.192 |
| SE | 6520 | KLHL12 | KLHL12 | chr1 | - | 202893238 | 202893425 | 202891095 | 202892659 | 202894183 | 202894282 | 4 | 0 | 3 | 0 | 0 | 0 | 0 | 2 | 0 | 1 | 98 | 49 | 1 | 1 | 1 | NA | 1 | NA | 0.5 | 0.5 |

| | | | | | | | | | | | | | | | | | | | | | | | | | | | | | | | |
|---|---|---|---|---|---|---|---|---|---|---|---|---|---|---|---|---|---|---|---|---|---|---|---|---|---|---|---|---|---|---|---|
| SE | 6523 | LYST | LYST | chr1 | - | 235677479 | 235677619 | 235677090 | 235677188 | 235686948 | 235687047 | 90 | 32 | 16 | 1 | 0 | 0 | 0 | 11 | 0 | 0 | 98 | 49 | 1 | 1 | 0.978 | 1 | 1 | NA | 1 | -0.007 |
| SE | 6525 | LYST | LYST | chr1 | - | 235733502 | 235733691 | 235731031 | 235731177 | 235733829 | 235733906 | 121 | 59 | 17 | 0 | 3 | 0 | 0 | 3 | 0 | 0 | 98 | 49 | 1 | 1 | 1 | 0.908 | 1 | NA | 1 | -0.031 |
| SE | 6526 | LYST | LYST | chr1 | - | 235757280 | 235757458 | 235755477 | 235755647 | 235758971 | 235759599 | 59 | 61 | 8 | 4 | 0 | 0 | 0 | 13 | 0 | 0 | 98 | 49 | 1 | 1 | 0.881 | 1 | 1 | NA | 1 | -0.04 |
| SE | 6527 | LYST | LYST | chr1 | - | 235758971 | 235759599 | 235755477 | 235755647 | 235762719 | 235762851 | 61 | 30 | 8 | 0 | 1 | 0 | 0 | 6 | 0 | 0 | 98 | 49 | 1 | 1 | 1 | 0.937 | 1 | NA | 1 | -0.021 |
| SE | 6528 | LYST | LYST | chr1 | - | 235777062 | 235777308 | 235774912 | 235775086 | 235780864 | 235781055 | 124 | 53 | 18 | 1 | 0 | 0 | 2 | 15 | 0 | 0 | 98 | 49 | 1 | 1 | 0.984 | 1 | 1 | 1 | 1 | -0.005 |
| SE | 6531 | LYST | LYST | chr1 | - | 235812970 | 235813061 | 235808454 | 235810534 | 235830225 | 235830424 | 181 | 54 | 20 | 1 | 0 | 0 | 3 | 35 | 0 | 0 | 98 | 49 | 1 | 1 | 0.989 | 1 | 1 | 1 | 1 | -0.004 |
| SE | 6537 | RBM39 | RBM39 | chr20 | - | 35740823 | 35740887 | 35738967 | 35739017 | 35741940 | 35742336 | 73 | 28 | 8 | 1 | 0 | 0 | 2 | 10 | 0 | 0 | 98 | 49 | 1 | 1 | 0.973 | 1 | 1 | 1 | 1 | -0.009 |
| SE | 6538 | UBR4 | UBR4 | chr1 | - | 19077975 | 19078066 | 19076739 | 19076902 | 19081348 | 19081573 | 13 | 1 | 0 | 1 | 0 | 0 | 1 | 2 | 0 | 0 | 98 | 49 | 1 | 1 | 0.867 | 1 | NA | 1 | 1 | -0.067 |
| SE | 6539 | UBR4 | UBR4 | chr1 | - | 19092818 | 19092918 | 19088758 | 19088977 | 19093312 | 19093486 | 24 | 3 | 0 | 1 | 0 | 0 | 0 | 1 | 0 | 0 | 98 | 49 | 1 | 1 | 0.923 | 1 | NA | NA | 1 | -0.038 |
| SE | 6540 | UBR4 | UBR4 | chr1 | - | 19094905 | 19095025 | 19093948 | 19094139 | 19095544 | 19095652 | 4 | 1 | 1 | 0 | 0 | 0 | 0 | 3 | 0 | 1 | 98 | 49 | 1 | 1 | 1 | 1 | 1 | NA | 0.6 | 0.4 |
| SE | 6541 | UBR4 | UBR4 | chr1 | - | 19097192 | 19097280 | 19096522 | 19096650 | 19099596 | 19099677 | 8 | 3 | 0 | 0 | 1 | 0 | 0 | 2 | 0 | 0 | 98 | 49 | 1 | 1 | 1 | 0.6 | NA | NA | 1 | -0.2 |
| SE | 6542 | UBR4 | UBR4 | chr1 | - | 19099596 | 19099677 | 19096522 | 19096650 | 19100375 | 19100573 | 8 | 3 | 1 | 1 | 1 | 0 | 0 | 1 | 0 | 0 | 98 | 49 | 1 | 1 | 0.8 | 1 | 1 | NA | 1 | -0.067 |
| SE | 6543 | UBR4 | UBR4 | chr1 | - | 19148562 | 19148626 | 19147992 | 19148127 | 19150576 | 19150793 | 10 | 6 | 3 | 0 | 1 | 0 | 0 | 2 | 0 | 1 | 98 | 49 | 1 | 1 | 1 | 0.75 | 1 | NA | 0.5 | 0.417 |
| SE | 6544 | UBR4 | UBR4 | chr1 | - | 19150576 | 19150793 | 19148562 | 19148626 | 19151642 | 19151859 | 10 | 9 | 0 | 3 | 0 | 0 | 0 | 3 | 0 | 0 | 98 | 49 | 1 | 1 | 0.625 | 1 | NA | NA | 1 | -0.188 |
| SE | 6545 | UBR4 | UBR4 | chr1 | - | 19153767 | 19153939 | 19153300 | 19153502 | 19154917 | 19155075 | 20 | 0 | 0 | 0 | 0 | 0 | 0 | 1 | 0 | 1 | 98 | 49 | 1 | 1 | 1 | NA | NA | NA | 0.333 | 0.667 |
| SE | 6547 | UBR4 | UBR4 | chr1 | - | 19165655 | 19165757 | 19165248 | 19165349 | 19167021 | 19167231 | 20 | 9 | 0 | 1 | 0 | 0 | 0 | 4 | 0 | 1 | 98 | 49 | 1 | 1 | 0.909 | 1 | NA | NA | 0.667 | 0.287 |
| SE | 6548 | UBR4 | UBR4 | chr1 | - | 19168026 | 19168184 | 19167021 | 19167231 | 19169434 | 19169532 | 47 | 6 | 6 | 0 | 1 | 0 | 0 | 11 | 0 | 0 | 98 | 49 | 1 | 1 | 1 | 0.75 | 1 | NA | 1 | -0.083 |
| SE | 6549 | UBR4 | UBR4 | chr1 | - | 19174318 | 19174447 | 19173438 | 19173621 | 19174953 | 19175033 | 17 | 3 | 5 | 1 | 0 | 0 | 0 | 3 | 0 | 0 | 98 | 49 | 1 | 1 | 0.895 | 1 | 1 | NA | 1 | -0.035 |
| SE | 6550 | UBR4 | UBR4 | chr1 | - | 19185098 | 19185286 | 19184015 | 19184175 | 19186539 | 19186657 | 8 | 9 | 0 | 1 | 0 | 0 | 0 | 2 | 0 | 0 | 98 | 49 | 1 | 1 | 0.8 | 1 | NA | NA | 1 | -0.1 |
| SE | 6553 | APAF1 | APAF1 | chr12 | + | 98665552 | 98665791 | 98662674 | 98662806 | 98666189 | 98666357 | 17 | 0 | 10 | 7 | 0 | 0 | 0 | 2 | 0 | 1 | 98 | 49 | 1 | 1 | 0.548 | NA | 1 | NA | 0.5 | 0.274 |
| SE | 6554 | APAF1 | APAF1 | chr12 | + | 98677424 | 98677551 | 98671534 | 98671719 | 98680276 | 98680402 | 69 | 15 | 8 | 4 | 0 | 0 | 0 | 7 | 0 | 1 | 98 | 49 | 1 | 1 | 0.896 | 1 | 1 | NA | 0.778 | 0.187 |
| SE | 6555 | APAF1 | APAF1 | chr12 | + | 98703370 | 98703499 | 98699407 | 98699569 | 98706484 | 98706610 | 18 | 12 | 11 | 21 | 0 | 2 | 0 | 5 | 0 | 7 | 98 | 49 | 1 | 1 | 0.3 | 1 | 0.733 | NA | 0.263 | 0.415 |
| SE | 6556 | APAF1 | APAF1 | chr12 | + | 98706484 | 98706610 | 98699407 | 98699569 | 98708584 | 98708704 | 52 | 13 | 6 | 0 | 0 | 0 | 0 | 7 | 0 | 1 | 98 | 49 | 1 | 1 | 1 | 1 | 1 | NA | 0.778 | 0.222 |
| SE | 6557 | APAF1 | APAF1 | chr12 | + | 98725414 | 98725540 | 98723638 | 98723764 | 98727172 | 98727316 | 37 | 11 | 9 | 1 | 0 | 0 | 0 | 5 | 0 | 0 | 98 | 49 | 1 | 1 | 0.949 | 1 | 1 | NA | 1 | -0.017 |
| SE | 6558 | APAF1 | APAF1 | chr12 | + | 98727172 | 98727316 | 98725414 | 98725540 | 98732419 | 98735433 | 53 | 18 | 8 | 0 | 1 | 0 | 0 | 7 | 0 | 0 | 98 | 49 | 1 | 1 | 1 | 0.9 | 1 | NA | 1 | -0.033 |
| SE | 6561 | TRERF1 | TRERF1 | chr6 | - | 42263319 | 42263568 | 42259338 | 42259723 | 42264703 | 42264854 | 0 | 0 | 0 | 10 | 4 | 0 | 0 | 1 | 1 | 0 | 98 | 49 | 1 | 1 | 0 | 0 | NA | 0 | 1 | -0.5 |
| SE | 6570 | PSRC1 | PSRC1 | chr1 | - | 109280395 | 109280489 | 109279553 | 109280156 | 109280776 | 109280916 | 3 | 0 | 4 | 0 | 0 | 0 | 0 | 1 | 3 | 0 | 98 | 49 | 1 | 1 | 1 | NA | 1 | 0 | 1 | 0.5 |
| SE | 6577 | NOD2 | NOD2 | chr16 | + | 50719924 | 50720008 | 50716890 | 50716974 | 50722621 | 50722705 | 5 | 1 | 2 | 0 | 0 | 0 | 0 | 3 | 0 | 1 | 98 | 49 | 1 | 1 | 1 | 1 | 1 | NA | 0.6 | 0.4 |
| SE | 6586 | RGL4 | RGL4 | chr22 | + | 23697170 | 23697245 | 23696613 | 23696688 | 23697837 | 23697861 | 24 | 12 | 0 | 1 | 0 | 0 | 1 | 6 | 0 | 0 | 98 | 49 | 1 | 1 | 0.923 | 1 | NA | 1 | 1 | -0.038 |
| SE | 6587 | RGL4 | RGL4 | chr22 | + | 23697837 | 23697861 | 23696613 | 23696688 | 23698211 | 23698333 | 4 | 0 | 0 | 0 | 0 | 0 | 2 | 0 | 0 | 1 | 73 | 49 | 1 | 1 | 1 | NA | NA | 1 | 0 | 0.5 |
| SE | 6588 | SETD2 | SETD2 | chr3 | - | 47083719 | 47084382 | 47067069 | 47067118 | 47086194 | 47086314 | 15 | 5 | 0 | 0 | 1 | 0 | 1 | 1 | 0 | 0 | 98 | 49 | 1 | 1 | 1 | 0.714 | NA | 1 | 1 | -0.143 |
| SE | 6590 | PRKAA1 | PRKAA1 | chr5 | - | 40774927 | 40774972 | 40771718 | 40771863 | 40775409 | 40775503 | 4 | 3 | 0 | 6 | 0 | 2 | 0 | 0 | 0 | 1 | 94 | 49 | 1 | 1 | 0.258 | 1 | 0 | NA | 0 | 0.419 |
| SE | 6591 | CHPF2 | CHPF2 | chr7 | + | 151235047 | 151235612 | 151232488 | 151234274 | 151237373 | 151238827 | 4 | 3 | 4 | 0 | 1 | 0 | 0 | 2 | 0 | 0 | 98 | 49 | 1 | 1 | 1 | 0.6 | 1 | NA | 1 | -0.133 |
| SE | 6592 | CHPF2 | CHPF2 | chr7 | + | 151236407 | 151236590 | 151235047 | 151235612 | 151237373 | 151238827 | 10 | 2 | 3 | 2 | 1 | 0 | 0 | 4 | 0 | 0 | 98 | 49 | 1 | 1 | 0.714 | 0.5 | 1 | NA | 1 | -0.262 |
| SE | 6593 | KANSL2 | KANSL2 | chr12 | - | 48679035 | 48679150 | 48671798 | 48671962 | 48679654 | 48679833 | 5 | 1 | 0 | 3 | 3 | 1 | 0 | 6 | 0 | 2 | 98 | 49 | 1 | 1 | 0.455 | 0.143 | 0 | NA | 0.6 | -0.401 |
| SE | 6606 | NDUFB5 | NDUFB5 | chr3 | + | 179616982 | 179617044 | 179615982 | 179616049 | 179618414 | 179618521 | 11 | 2 | 3 | 0 | 0 | 0 | 0 | 7 | 0 | 1 | 98 | 49 | 1 | 1 | 1 | 1 | 1 | NA | 0.778 | 0.222 |
| SE | 6611 | AKAP13 | AKAP13 | chr15 | + | 85662387 | 85662453 | 85655416 | 85655787 | 85664562 | 85664755 | 3 | 1 | 0 | 3 | 5 | 0 | 0 | 0 | 0 | 1 | 98 | 49 | 1 | 1 | 0.333 | 0.091 | NA | NA | 0 | 0.212 |
| SE | 6613 | AKAP13 | AKAP13 | chr15 | + | 85715787 | 85715923 | 85710578 | 85710645 | 85717289 | 85717402 | 43 | 10 | 2 | 1 | 0 | 0 | 0 | 8 | 0 | 0 | 98 | 49 | 1 | 1 | 0.956 | 1 | 1 | NA | 1 | -0.015 |
| SE | 6614 | PPP1R10 | PPP1R10 | chr6 | - | 30604993 | 30605094 | 30604587 | 30604735 | 30605922 | 30606035 | 14 | 3 | 1 | 5 | 0 | 0 | 0 | 1 | 0 | 0 | 98 | 49 | 1 | 1 | 0.583 | 1 | 1 | NA | 1 | -0.139 |
| SE | 6619 | ILK | ILK | chr11 | + | 6608045 | 6608211 | 6603809 | 6604360 | 6608393 | 6608489 | 9 | 6 | 7 | 2 | 0 | 0 | 1 | 3 | 0 | 0 | 98 | 49 | 1 | 1 | 0.692 | 1 | 1 | 1 | 1 | -0.103 |
| SE | 6631 | PRUNE | PRUNE | chr1 | + | 151018466 | 151018669 | 151017811 | 151017904 | 151024610 | 151024795 | 2 | 0 | 6 | 4 | 0 | 0 | 0 | 1 | 0 | 0 | 98 | 49 | 1 | 1 | 0.2 | NA | 1 | NA | 1 | -0.4 |
| SE | 6637 | CSF3R | CSF3R | chr1 | - | 36469651 | 36469840 | 36469155 | 36469257 | 36471432 | 36471646 | 963 | 336 | 202 | 1 | 0 | 0 | 37 | 229 | 0 | 0 | 98 | 49 | 1 | 1 | 0.998 | 1 | 1 | 1 | 1 | -0.001 |
| SE | 6638 | CSF3R | CSF3R | chr1 | - | 36472065 | 36472139 | 36471432 | 36471646 | 36472237 | 36472391 | 930 | 461 | 214 | 3 | 0 | 0 | 94 | 256 | 0 | 1 | 98 | 49 | 1 | 1 | 0.994 | 1 | 1 | 1 | 0.992 | 0.002 |
| SE | 6640 | CSF3R | CSF3R | chr1 | - | 36473763 | 36473887 | 36472516 | 36472686 | 36479432 | 36479516 | 33 | 13 | 3 | 1 | 0 | 0 | 1 | 3 | 0 | 0 | 98 | 49 | 1 | 1 | 0.943 | 1 | 1 | 1 | 1 | -0.019 |
| SE | 6641 | CSF3R | CSF3R | chr1 | - | 36475376 | 36475673 | 36473763 | 36473887 | 36479432 | 36479516 | 826 | 284 | 151 | 2 | 0 | 0 | 19 | 114 | 0 | 0 | 98 | 49 | 1 | 1 | 0.995 | 1 | 1 | 1 | 1 | -0.002 |
| SE | 6642 | RPE | RPE | chr2 | + | 210003437 | 210003468 | 210002564 | 210002783 | 210009656 | 210009736 | 1 | 0 | 0 | 1 | 0 | 0 | 0 | 0 | 0 | 1 | 80 | 49 | 1 | 1 | 0.38 | NA | NA | NA | 0 | 0.38 |
| SE | 6664 | ERP29 | ERP29 | chr12 | + | 112019755 | 112019894 | 112013347 | 112013609 | 112022149 | 112023220 | 5 | 0 | 2 | 1 | 0 | 0 | 0 | 0 | 0 | 1 | 98 | 49 | 1 | 1 | 0.714 | NA | 1 | NA | 0 | 0.857 |
| SE | 6672 | ARHGAP4 | ARHGAP4 | chrX | - | 153910921 | 153910999 | 153910699 | 153910834 | 153911128 | 153911189 | 56 | 22 | 15 | 1 | 1 | 0 | 4 | 9 | 0 | 0 | 98 | 49 | 1 | 1 | 0.966 | 0.917 | 1 | 1 | 1 | -0.039 |
| SE | 6673 | ARHGAP4 | ARHGAP4 | chrX | - | 153913217 | 153913302 | 153913023 | 153913051 | 153913408 | 153913600 | 39 | 25 | 7 | 1 | 0 | 0 | 0 | 4 | 0 | 0 | 98 | 49 | 1 | 1 | 0.951 | 1 | 1 | NA | 1 | -0.016 |
| SE | 6676 | ARHGAP4 | ARHGAP4 | chrX | - | 153921096 | 153921159 | 153920625 | 153920808 | 153921364 | 153921527 | 34 | 22 | 6 | 2 | 0 | 0 | 3 | 6 | 0 | 0 | 98 | 49 | 1 | 1 | 0.895 | 1 | 1 | 1 | 1 | -0.035 |
| SE | 6679 | PKP4 | PKP4 | chr2 | + | 158676738 | 158676867 | 158673882 | 158674000 | 158678580 | 158678654 | 2 | 0 | 0 | 1 | 1 | 0 | 0 | 0 | 0 | 2 | 98 | 49 | 1 | 1 | 0.5 | 0 | NA | NA | 0 | 0.25 |
| SE | 6680 | NUDT22 | NUDT22 | chr11 | + | 64227567 | 64227666 | 64226634 | 64227132 | 64229246 | 64229344 | 3 | 1 | 0 | 0 | 0 | 2 | 0 | 2 | 0 | 0 | 98 | 49 | 1 | 1 | 1 | 1 | 0 | NA | 1 | -0.333 |
| SE | 6698 | GATAD2B | GATAD2B | chr1 | - | 153818039 | 153818171 | 153817371 | 153817542 | 153818790 | 153818922 | 16 | 7 | 3 | 0 | 0 | 0 | 0 | 2 | 0 | 1 | 98 | 49 | 1 | 1 | 1 | 1 | 1 | NA | 0.5 | 0.5 |
| SE | 6701 | LAPTM5 | LAPTM5 | chr1 | - | 30741639 | 30741716 | 30739808 | 30739937 | 30742455 | 30742549 | 209 | 62 | 86 | 0 | 0 | 0 | 4 | 37 | 0 | 1 | 98 | 49 | 1 | 1 | 1 | 1 | 1 | 1 | 0.949 | 0.026 |
| SE | 6702 | LAPTM5 | LAPTM5 | chr1 | - | 30741639 | 30741716 | 30739808 | 30739937 | 30757658 | 30757836 | 131 | 34 | 43 | 1 | 0 | 0 | 1 | 20 | 0 | 0 | 98 | 49 | 1 | 1 | 0.985 | 1 | 1 | 1 | 1 | -0.005 |
| SE | 6703 | LAPTM5 | LAPTM5 | chr1 | - | 30742455 | 30742549 | 30739808 | 30739937 | 30757658 | 30757836 | 11 | 7 | 3 | 1 | 0 | 0 | 0 | 4 | 0 | 0 | 98 | 49 | 1 | 1 | 0.846 | 1 | 1 | NA | 1 | -0.051 |
| SE | 6705 | STX12 | STX12 | chr1 | + | 27817850 | 27817923 | 27812162 | 27812268 | 27819649 | 27819732 | 5 | 6 | 1 | 0 | 0 | 0 | 0 | 5 | 0 | 1 | 98 | 49 | 1 | 1 | 1 | 1 | 1 | NA | 0.714 | 0.286 |
| SE | 6707 | MTMR3 | MTMR3 | chr22 | + | 29970975 | 29971062 | 29957035 | 29957088 | 29978441 | 29978531 | 31 | 37 | 4 | 3 | 3 | 0 | 0 | 9 | 0 | 1 | 98 | 49 | 1 | 1 | 0.838 | 0.86 | 1 | NA | 0.818 | 0.081 |
| SE | 6708 | MTMR3 | MTMR3 | chr22 | + | 30007113 | 30007319 | 30002879 | 30002993 | 30007900 | 30008032 | 48 | 15 | 5 | 0 | 1 | 0 | 0 | 11 | 0 | 0 | 98 | 49 | 1 | 1 | 1 | 0.882 | 1 | NA | 1 | -0.039 |
| SE | 6709 | MTMR3 | MTMR3 | chr22 | + | 30007900 | 30008032 | 30002879 | 30002993 | 30009017 | 30009129 | 33 | 31 | 13 | 1 | 1 | 0 | 1 | 14 | 0 | 0 | 98 | 49 | 1 | 1 | 0.943 | 0.939 | 1 | 1 | 1 | -0.039 |
| SE | 6710 | MTMR3 | MTMR3 | chr22 | + | 30012367 | 30012563 | 30009017 | 30009129 | 30013355 | 30013541 | 32 | 20 | 11 | 0 | 1 | 0 | 0 | 2 | 0 | 4 | 98 | 49 | 1 | 1 | 1 | 0.909 | 1 | NA | 0.2 | 0.77 |
| SE | 6711 | MTMR3 | MTMR3 | chr22 | + | 30013355 | 30013541 | 30009017 | 30009129 | 30016527 | 30016698 | 25 | 20 | 5 | 0 | 1 | 0 | 1 | 8 | 0 | 0 | 98 | 49 | 1 | 1 | 1 | 0.909 | 1 | 1 | 1 | -0.03 |
| SE | 6712 | MTMR3 | MTMR3 | chr22 | + | 30022028 | 30022139 | 30019479 | 30020884 | 30022608 | 30022697 | 31 | 7 | 12 | 3 | 5 | 0 | 0 | 10 | 0 | 0 | 98 | 49 | 1 | 1 | 0.838 | 0.412 | 1 | NA | 1 | -0.25 |
| SE | 6713 | MTMR3 | MTMR3 | chr22 | + | 30023456 | 30023483 | 30022608 | 30022697 | 30025629 | 30030868 | 10 | 6 | 3 | 12 | 4 | 4 | 0 | 5 | 0 | 2 | 76 | 49 | 1 | 1 | 0.35 | 0.492 | 0.326 | NA | 0.617 | -0.228 |
| SE | 6714 | TK2 | TK2 | chr16 | - | 66536963 | 66537017 | 66531379 | 66531469 | 66541878 | 66541953 | 30 | 8 | 0 | 1 | 0 | 0 | 0 | 3 | 0 | 0 | 98 | 49 | 1 | 1 | 0.937 | 1 | NA | NA | 1 | -0.031 |
| SE | 6716 | TK2 | TK2 | chr16 | - | 66547970 | 66548065 | 66541878 | 66541953 | 66548977 | 66549009 | 2 | 0 | 0 | 4 | 0 | 0 | 0 | 0 | 0 | 2 | 98 | 49 | 1 | 1 | 0.2 | NA | NA | NA | 0 | 0.2 |
| SE | 6719 | TRIM5 | TRIM5 | chr11 | - | 5665980 | 5666081 | 5663194 | 5665682 | 5667688 | 5667711 | 5 | 2 | 3 | 1 | 0 | 0 | 0 | 1 | 0 | 1 | 98 | 49 | 1 | 1 | 0.714 | 1 | 1 | NA | 0.333 | 0.572 |
| SE | 6720 | TRIM5 | TRIM5 | chr11 | - | 5679073 | 5679169 | 5678203 | 5678434 | 5679760 | 5680238 | 10 | 7 | 2 | 1 | 0 | 0 | 2 | 4 | 0 | 0 | 98 | 49 | 1 | 1 | 0.833 | 1 | 1 | 1 | 1 | -0.056 |
| SE | 6721 | C20orf24 | C20orf24 | chr20 | + | 36609600 | 36609730 | 36607714 | 36607818 | 36612009 | 36612557 | 29 | 41 | 54 | 1 | 0 | 1 | 0 | 8 | 0 | 0 | 98 | 49 | 1 | 1 | 0.935 | 1 | 0.964 | NA | 1 | -0.034 |
| SE | 6723 | TNFRSF1B | TNFRSF1B | chr1 | + | 12193954 | 12194032 | 12192862 | 12193098 | 12194583 | 12194618 | 112 | 87 | 55 | 1 | 0 | 0 | 8 | 48 | 0 | 0 | 98 | 49 | 1 | 1 | 0.982 | 1 | 1 | 1 | 1 | -0.006 |
| SE | 6724 | TNFRSF1B | TNFRSF1B | chr1 | + | 12201966 | 12202171 | 12194583 | 12194618 | 12206739 | 12209220 | 77 | 78 | 38 | 1 | 0 | 0 | 5 | 27 | 0 | 0 | 98 | 49 | 1 | 1 | 0.975 | 1 | 1 | 1 | 1 | -0.008 |
| SE | 6726 | DSCR3 | DSCR3 | chr21 | - | 37233361 | 37233442 | 37232376 | 37232451 | 37238459 | 37238609 | 7 | 12 | 2 | 1 | 0 | 0 | 0 | 3 | 0 | 0 | 98 | 49 | 1 | 1 | 0.778 | 1 | 1 | NA | 1 | -0.074 |
| SE | 6748 | DHRS12 | DHRS12 | chr13 | - | 51777059 | 51777121 | 51773929 | 51774034 | 51790010 | 51790092 | 3 | 7 | 3 | 1 | 0 | 7 | 0 | 1 | 0 | 0 | 98 | 49 | 1 | 1 | 0.6 | 1 | 0.176 | NA | 1 | -0.408 |
| SE | 6749 | DHRS12 | DHRS12 | chr13 | - | 51797825 | 51797908 | 51791164 | 51791257 | 51799533 | 51799667 | 1 | 0 | 0 | 6 | 1 | 0 | 0 | 2 | 0 | 6 | 98 | 49 | 1 | 1 | 0.077 | 0 | NA | NA | 0.143 | -0.104 |
| SE | 6754 | KMT2E | KMT2E | chr7 | + | 105038125 | 105038199 | 105014189 | 105014535 | 105040838 | 105041023 | 25 | 23 | 8 | 9 | 4 | 3 | 0 | 7 | 0 | 2 | 98 | 49 | 1 | 1 | 0.581 | 0.742 | 0.571 | NA | 0.636 | -0.005 |
| SE | 6755 | KMT2E | KMT2E | chr7 | + | 105062163 | 105062278 | 105040838 | 105041023 | 105063350 | 105063580 | 58 | 34 | 8 | 1 | 0 | 0 | 1 | 9 | 0 | 0 | 98 | 49 | 1 | 1 | 0.967 | 1 | 1 | 1 | 1 | -0.011 |
| SE | 6757 | SCCPDH | SCCPDH | chr1 | + | 246735974 | 246736055 | 246726891 | 246727004 | 246740171 | 246740301 | 0 | 3 | 2 | 1 | 0 | 0 | 0 | 5 | 0 | 0 | 98 | 49 | 1 | 1 | 0 | 1 | 1 | NA | 1 | -0.333 |
| SE | 6763 | OPA1 | OPA1 | chr3 | + | 193617177 | 193617285 | 193615673 | 193615770 | 193618868 | 193618936 | 11 | 0 | 1 | 4 | 0 | 0 | 0 | 2 | 0 | 0 | 98 | 49 | 1 | 1 | 0.579 | NA | 1 | NA | 1 | -0.211 |
| SE | 6765 | OPA1 | OPA1 | chr3 | + | 193617783 | 193617837 | 193617177 | 193617285 | 193618868 | 193618936 | 4 | 0 | 0 | 7 | 0 | 1 | 0 | 1 | 0 | 0 | 98 | 49 | 1 | 1 | 0.222 | NA | 0 | NA | 1 | -0.889 |
| SE | 6766 | OPA1 | OPA1 | chr3 | + | 193626091 | 193626202 | 193618868 | 193618936 | 193631611 | 193631665 | 2 | 0 | 0 | 3 | 0 | 0 | 0 | 1 | 0 | 0 | 98 | 49 | 1 | 1 | 0.25 | NA | NA | NA | 1 | -0.75 |
| SE | 6771 | PRPF19 | PRPF19 | chr11 | - | 60898861 | 60898931 | 60898540 | 60898626 | 60899148 | 60899304 | 1 | 3 | 0 | 2 | 0 | 0 | 0 | 1 | 0 | 0 | 98 | 49 | 1 | 1 | 0.2 | 1 | NA | NA | 1 | -0.4 |
| SE | 6775 | ZNF384 | ZNF384 | chr12 | - | 6672349 | 6672532 | 6670759 | 6670838 | 6673215 | 6673440 | 2 | 0 | 5 | 6 | 1 | 3 | 0 | 0 | 0 | 1 | 98 | 49 | 1 | 1 | 0.143 | 0 | 0.455 | NA | 0 | 0.199 |
| SE | 6776 | ZNF384 | ZNF384 | chr12 | - | 6673215 | 6673440 | 6670759 | 6670838 | 6678126 | 6678343 | 7 | 3 | 3 | 2 | 0 | 0 | 0 | 1 | 0 | 1 | 98 | 49 | 1 | 1 | 0.636 | 1 | 1 | NA | 0.333 | 0.546 |
| SE | 6777 | CSF1 | CSF1 | chr1 | + | 109914258 | 109914381 | 109910610 | 109911062 | 109915633 | 109915696 | 0 | 3 | 0 | 1 | 0 | 0 | 1 | 1 | 3 | 0 | 98 | 49 | 1 | 1 | 0 | 1 | NA | 0.143 | 1 | -0.072 |
| SE | 6778 | CSF1 | CSF1 | chr1 | + | 109915633 | 109915696 | 109910610 | 109911062 | 109917292 | 109917463 | 3 | 4 | 0 | 0 | 1 | 0 | 3 | 0 | 0 | 0 | 98 | 49 | 1 | 1 | 1 | 0.667 | NA | 1 | NA | -0.166 |
| SE | 6783 | JMJD1C | JMJD1C | chr10 | - | 63189167 | 63189446 | 63186214 | 63186383 | 63190893 | 63191108 | 189 | 51 | 23 | 0 | 1 | 0 | 3 | 24 | 0 | 0 | 98 | 49 | 1 | 1 | 1 | 0.962 | 1 | 1 | 1 | -0.013 |
| SE | 6784 | JMJD1C | JMJD1C | chr10 | - | 63198512 | 63198727 | 63197410 | 63197563 | 63206594 | 63208801 | 79 | 32 | 3 | 0 | 2 | 0 | 0 | 10 | 0 | 0 | 98 | 49 | 1 | 1 | 1 | 0.889 | 1 | NA | 1 | -0.037 |
| SE | 6785 | JMJD1C | JMJD1C | chr10 | - | 63200475 | 63200677 | 63198512 | 63198727 | 63206594 | 63208801 | 40 | 16 | 20 | 5 | 5 | 0 | 0 | 13 | 0 | 1 | 98 | 49 | 1 | 1 | 0.8 | 0.615 | 1 | NA | 0.867 | -0.062 |
| SE | 6786 | JMJD1C | JMJD1C | chr10 | - | 63215552 | 63215696 | 63215262 | 63215455 | 63217206 | 63217331 | 74 | 29 | 14 | 5 | 1 | 0 | 0 | 21 | 0 | 1 | 98 | 49 | 1 | 1 | 0.881 | 0.935 | 1 | NA | 0.913 | 0.026 |
| SE | 6787 | JMJD1C | JMJD1C | chr10 | - | 63217206 | 63217331 | 63215262 | 63215455 | 63264650 | 63264764 | 25 | 7 | 4 | 1 | 0 | 0 | 0 | 3 | 0 | 0 | 98 | 49 | 1 | 1 | 0.926 | 1 | 1 | NA | 1 | -0.025 |
| SE | 6788 | JMJD1C | JMJD1C | chr10 | - | 63219877 | 63219983 | 63217206 | 63217331 | 63264650 | 63264764 | 74 | 21 | 8 | 20 | 6 | 4 | 0 | 14 | 0 | 2 | 98 | 49 | 1 | 1 | 0.649 | 0.636 | 0.5 | NA | 0.778 | -0.183 |
| SE | 6789 | JMJD1C | JMJD1C | chr10 | - | 63219877 | 63219983 | 63217206 | 63217331 | 63268662 | 63269223 | 28 | 10 | 1 | 6 | 1 | 0 | 0 | 2 | 0 | 0 | 98 | 49 | 1 | 1 | 0.7 | 0.833 | 1 | NA | 1 | -0.156 |
| SE | 6791 | JMJD1C | JMJD1C | chr10 | - | 63264650 | 63264764 | 63219877 | 63219983 | 63268662 | 63269223 | 179 | 39 | 25 | 1 | 0 | 0 | 1 | 25 | 0 | 0 | 98 | 49 | 1 | 1 | 0.989 | 1 | 1 | 1 | 1 | -0.004 |
| SE | 6792 | JMJD1C | JMJD1C | chr10 | - | 63264650 | 63264764 | 63219877 | 63219983 | 63380317 | 63380482 | 48 | 11 | 8 | 2 | 0 | 0 | 1 | 12 | 0 | 0 | 98 | 49 | 1 | 1 | 0.923 | 1 | 1 | 1 | 1 | -0.026 |
| SE | 6793 | ZNF592 | ZNF592 | chr15 | + | 84778182 | 84778312 | 84764706 | 84764815 | 84782656 | 84784895 | 11 | 2 | 3 | 1 | 0 | 0 | 0 | 2 | 0 | 2 | 98 | 49 | 1 | 1 | 0.846 | 1 | 1 | NA | 0.333 | 0.616 |
| SE | 6794 | FAM96A | FAM96A | chr15 | - | 64081101 | 64081151 | 64075491 | 64075537 | 64088686 | 64088851 | 3 | 1 | 5 | 1 | 0 | 0 | 0 | 5 | 0 | 0 | 98 | 49 | 1 | 1 | 0.6 | 1 | 1 | NA | 1 | -0.133 |
| SE | 6796 | RRAGD | RRAGD | chr6 | - | 89380167 | 89380367 | 89379223 | 89379338 | 89387294 | 89387590 | 10 | 6 | 10 | 0 | 0 | 0 | 0 | 4 | 0 | 1 | 98 | 49 | 1 | 1 | 1 | 1 | 1 | NA | 0.667 | 0.333 |
| SE | 6799 | UBE2G1 | UBE2G1 | chr17 | - | 4282797 | 4282921 | 4269216 | 4272516 | 4289229 | 4289408 | 35 | 32 | 9 | 0 | 2 | 0 | 0 | 11 | 0 | 0 | 98 | 49 | 1 | 1 | 1 | 0.889 | 1 | NA | 1 | -0.037 |
| SE | 6800 | GAS5 | GAS5 | chr1 | - | 173864483 | 173864506 | 173864256 | 173864304 | 173864674 | 173864704 | 20 | 1 | 6 | 1 | 0 | 0 | 0 | 3 | 0 | 0 | 72 | 49 | 1 | 1 | 0.932 | 1 | 1 | NA | 1 | -0.023 |
| SE | 6807 | TMEM63A | TMEM63A | chr1 | - | 225860859 | 225860997 | 225859195 | 225859349 | 225862217 | 225862351 | 11 | 0 | 2 | 0 | 1 | 0 | 0 | 1 | 0 | 0 | 98 | 49 | 1 | 1 | 1 | 0 | 1 | NA | 1 | -0.333 |
| SE | 6809 | DMXL2 | DMXL2 | chr15 | - | 51451644 | 51451697 | 51450128 | 51450346 | 51453549 | 51453641 | 40 | 1 | 10 | 4 | 0 | 0 | 0 | 8 | 0 | 0 | 98 | 49 | 1 | 1 | 0.833 | 1 | 1 | NA | 1 | -0.056 |
| SE | 6810 | DMXL2 | DMXL2 | chr15 | - | 51457327 | 51457466 | 51456308 | 51456369 | 51458505 | 51458627 | 62 | 8 | 4 | 1 | 0 | 0 | 1 | 12 | 0 | 0 | 98 | 49 | 1 | 1 | 0.969 | 1 | 1 | 1 | 1 | -0.01 |
| SE | 6811 | DMXL2 | DMXL2 | chr15 | - | 51464674 | 51464876 | 51463378 | 51463496 | 51465565 | 51465651 | 46 | 7 | 3 | 4 | 3 | 0 | 0 | 15 | 0 | 1 | 98 | 49 | 1 | 1 | 0.852 | 0.538 | 1 | NA | 0.882 | -0.085 |
| SE | 6812 | DMXL2 | DMXL2 | chr15 | - | 51465565 | 51465651 | 51464674 | 51464876 | 51466183 | 51466311 | 52 | 3 | 3 | 2 | 0 | 0 | 0 | 15 | 0 | 0 | 98 | 49 | 1 | 1 | 0.929 | 1 | 1 | NA | 1 | -0.024 |
| SE | 6813 | DMXL2 | DMXL2 | chr15 | - | 51495023 | 51495134 | 51491577 | 51491747 | 51498551 | 51500231 | 43 | 7 | 9 | 1 | 0 | 0 | 1 | 12 | 0 | 0 | 98 | 49 | 1 | 1 | 0.956 | 1 | 1 | 1 | 1 | -0.015 |
| SE | 6814 | DMXL2 | DMXL2 | chr15 | - | 51565087 | 51565166 | 51564124 | 51564260 | 51568486 | 51568558 | 34 | 10 | 3 | 1 | 0 | 0 | 0 | 6 | 0 | 0 | 98 | 49 | 1 | 1 | 0.944 | 1 | 1 | NA | 1 | -0.019 |
| SE | 6819 | PRMT3 | PRMT3 | chr11 | + | 20397576 | 20397721 | 20395802 | 20395962 | 20402918 | 20402984 | 8 | 1 | 0 | 0 | 0 | 0 | 3 | 0 | 0 | 1 | 98 | 49 | 1 | 1 | 1 | 1 | NA | 1 | 0 | 0.5 |
| SE | 6820 | UBE3A | UBE3A | chr15 | - | 25354527 | 25354683 | 25354352 | 25354426 | 25355891 | 25356056 | 7 | 1 | 0 | 1 | 0 | 0 | 0 | 1 | 0 | 0 | 98 | 49 | 1 | 1 | 0.778 | 1 | NA | NA | 1 | -0.111 |
| SE | 6821 | UBE3A | UBE3A | chr15 | - | 25356690 | 25356896 | 25355891 | 25356056 | 25360382 | 25360527 | 3 | 0 | 0 | 1 | 0 | 0 | 0 | 1 | 0 | 0 | 98 | 49 | 1 | 1 | 0.6 | NA | NA | NA | 1 | -0.4 |
| SE | 6826 | GRIPAP1 | GRIPAP1 | chrX | - | 48990925 | 48991110 | 48990684 | 48990732 | 48993427 | 48993578 | 15 | 7 | 0 | 0 | 2 | 0 | 0 | 6 | 0 | 0 | 98 | 49 | 1 | 1 | 1 | 0.636 | NA | NA | 1 | -0.182 |
| SE | 6827 | LOC102724552 | LOC102724552 | chr1 | + | 8428481 | 8428588 | 8424644 | 8424701 | 8429826 | 8429972 | 1 | 0 | 0 | 0 | 0 | 0 | 0 | 0 | 0 | 1 | 98 | 49 | 1 | 1 | 1 | NA | NA | NA | 0 | 1 |
| SE | 6837 | GAK | GAK | chr4 | - | 913606 | 913668 | 904636 | 904779 | 932042 | 932386 | 9 | 2 | 0 | 1 | 1 | 0 | 3 | 2 | 0 | 0 | 98 | 49 | 1 | 1 | 0.818 | 0.5 | NA | 1 | 1 | -0.341 |
| SE | 6839 | BBIP1 | BBIP1 | chr10 | - | 110907679 | 110907836 | 110901537 | 110901612 | 110918120 | 110918213 | 1 | 0 | 0 | 6 | 2 | 0 | 0 | 1 | 0 | 0 | 98 | 49 | 1 | 1 | 0.077 | 0 | NA | NA | 1 | -0.962 |
| SE | 6843 | NAA16 | NAA16 | chr13 | + | 41331273 | 41331369 | 41328723 | 41328843 | 41355143 | 41355216 | 2 | 0 | 0 | 0 | 1 | 0 | 0 | 1 | 0 | 0 | 98 | 49 | 1 | 1 | 1 | 0 | NA | NA | 1 | -0.5 |
| SE | 6845 | TMCO3 | TMCO3 | chr13 | + | 113503495 | 113503588 | 113501751 | 113501880 | 113510237 | 113510424 | 11 | 1 | 4 | 0 | 0 | 0 | 0 | 1 | 0 | 2 | 98 | 49 | 1 | 1 | 1 | 1 | 1 | NA | 0.2 | 0.8 |
| SE | 6846 | TMCO3 | TMCO3 | chr13 | + | 113510237 | 113510424 | 113503495 | 113503588 | 113520615 | 113520733 | 7 | 2 | 5 | 0 | 2 | 0 | 0 | 1 | 0 | 0 | 98 | 49 | 1 | 1 | 1 | 0.333 | 1 | NA | 1 | -0.222 |
| SE | 6847 | TMCO3 | TMCO3 | chr13 | + | 113534044 | 113534240 | 113520615 | 113520733 | 113539356 | 113539507 | 2 | 2 | 1 | 1 | 0 | 0 | 0 | 2 | 0 | 0 | 98 | 49 | 1 | 1 | 0.5 | 1 | 1 | NA | 1 | -0.167 |
| SE | 6849 | AMPD2 | AMPD2 | chr1 | + | 109620913 | 109621266 | 109619812 | 109620278 | 109625661 | 109625792 | 4 | 3 | 0 | 11 | 6 | 2 | 0 | 0 | 0 | 3 | 98 | 49 | 1 | 1 | 0.154 | 0.2 | 0 | NA | 0 | 0.118 |
| SE | 6850 | AMPD2 | AMPD2 | chr1 | + | 109625302 | 109625433 | 109619812 | 109620278 | 109625661 | 109625792 | 26 | 6 | 5 | 11 | 6 | 2 | 0 | 2 | 0 | 3 | 98 | 49 | 1 | 1 | 0.542 | 0.333 | 0.556 | NA | 0.25 | 0.227 |
| SE | 6851 | AMPD2 | AMPD2 | chr1 | + | 109625302 | 109625433 | 109620580 | 109621266 | 109625661 | 109625792 | 24 | 5 | 2 | 4 | 3 | 0 | 0 | 2 | 0 | 0 | 98 | 49 | 1 | 1 | 0.75 | 0.455 | 1 | NA | 1 | -0.265 |
| SE | 6852 | AMPD2 | AMPD2 | chr1 | + | 109625661 | 109625792 | 109619812 | 109620278 | 109626159 | 109626228 | 33 | 38 | 5 | 0 | 0 | 0 | 2 | 10 | 0 | 1 | 98 | 49 | 1 | 1 | 1 | 1 | 1 | 1 | 0.833 | 0.084 |
| SE | 6854 | AMPD2 | AMPD2 | chr1 | + | 109629326 | 109629490 | 109629108 | 109629235 | 109629795 | 109629916 | 36 | 18 | 5 | 1 | 0 | 0 | 2 | 5 | 0 | 0 | 98 | 49 | 1 | 1 | 0.947 | 1 | 1 | 1 | 1 | -0.018 |
| SE | 6856 | AMPD2 | AMPD2 | chr1 | + | 109630682 | 109630793 | 109630232 | 109630406 | 109630942 | 109632055 | 93 | 61 | 25 | 3 | 3 | 1 | 0 | 9 | 0 | 0 | 98 | 49 | 1 | 1 | 0.939 | 0.91 | 0.926 | NA | 1 | -0.075 |
| SE | 6867 | TXNIP | TXNIP | chr1 | - | 145994543 | 145994800 | 145994280 | 145994437 | 145994928 | 145995031 | 796 | 229 | 218 | 0 | 0 | 0 | 17 | 121 | 0 | 1 | 98 | 49 | 1 | 1 | 1 | 1 | 1 | 1 | 0.984 | 0.008 |
| SE | 6869 | C11orf21 | C11orf21 | chr11 | - | 2300519 | 2300613 | 2299427 | 2299707 | 2300683 | 2300768 | 5 | 0 | 0 | 2 | 0 | 0 | 0 | 1 | 0 | 0 | 98 | 49 | 1 | 1 | 0.556 | NA | NA | NA | 1 | -0.444 |
| SE | 6871 | RNF4 | RNF4 | chr4 | + | 2470986 | 2471109 | 2469067 | 2469258 | 2490336 | 2490502 | 1 | 3 | 0 | 5 | 0 | 4 | 0 | 0 | 0 | 2 | 98 | 49 | 1 | 1 | 0.091 | 1 | 0 | NA | 0 | 0.364 |
| SE | 6872 | RNF4 | RNF4 | chr4 | + | 2511955 | 2511965 | 2500658 | 2500738 | 2512437 | 2512597 | 14 | 2 | 0 | 2 | 0 | 0 | 2 | 3 | 0 | 0 | 59 | 49 | 1 | 1 | 0.853 | 1 | NA | 1 | 1 | -0.074 |
| SE | 6882 | ZNF655 | ZNF655 | chr7 | + | 99571687 | 99571792 | 99560532 | 99560695 | 99572244 | 99576456 | 4 | 3 | 0 | 1 | 1 | 0 | 0 | 1 | 0 | 0 | 98 | 49 | 1 | 1 | 0.667 | 0.6 | NA | NA | 1 | -0.367 |
| SE | 6883 | ZNF655 | ZNF655 | chr7 | + | 99571687 | 99571792 | 99562367 | 99562494 | 99572244 | 99576456 | 5 | 4 | 0 | 1 | 0 | 2 | 0 | 4 | 0 | 0 | 98 | 49 | 1 | 1 | 0.714 | 1 | 0 | NA | 1 | -0.429 |
| SE | 6890 | MFSD10 | MFSD10 | chr4 | - | 2931554 | 2931663 | 2931374 | 2931465 | 2931824 | 2931936 | 2 | 2 | 0 | 2 | 0 | 0 | 0 | 2 | 0 | 0 | 98 | 49 | 1 | 1 | 0.333 | 1 | NA | NA | 1 | -0.334 |
| SE | 6896 | SLC43A2 | SLC43A2 | chr17 | - | 1591268 | 1591471 | 1590801 | 1590948 | 1591565 | 1591699 | 123 | 35 | 27 | 0 | 0 | 0 | 5 | 19 | 0 | 1 | 98 | 49 | 1 | 1 | 1 | 1 | 1 | 1 | 0.905 | 0.047 |
| SE | 6897 | SLC43A2 | SLC43A2 | chr17 | - | 1616561 | 1616769 | 1614978 | 1615034 | 1627714 | 1627920 | 74 | 25 | 23 | 7 | 1 | 5 | 0 | 9 | 0 | 1 | 98 | 49 | 1 | 1 | 0.841 | 0.926 | 0.697 | NA | 0.818 | 0.003 |
| SE | 6901 | CASP1 | CASP1 | chr11 | - | 105026225 | 105026356 | 105025509 | 105025600 | 105026841 | 105026951 | 97 | 20 | 29 | 1 | 2 | 0 | 2 | 29 | 0 | 0 | 98 | 49 | 1 | 1 | 0.98 | 0.833 | 1 | 1 | 1 | -0.062 |
| SE | 6903 | CASP1 | CASP1 | chr11 | - | 105033063 | 105033126 | 105031164 | 105031280 | 105034207 | 105034474 | 92 | 40 | 27 | 11 | 0 | 3 | 0 | 28 | 0 | 6 | 98 | 49 | 1 | 1 | 0.807 | 1 | 0.818 | NA | 0.7 | 0.175 |
| SE | 6904 | PPTC7 | PPTC7 | chr12 | - | 110538143 | 110538273 | 110534431 | 110537095 | 110539821 | 110539945 | 11 | 5 | 6 | 2 | 0 | 0 | 0 | 1 | 0 | 0 | 98 | 49 | 1 | 1 | 0.733 | 1 | 1 | NA | 1 | -0.089 |
| SE | 6917 | TMEM41B | TMEM41B | chr11 | - | 9286454 | 9286593 | 9280654 | 9283593 | 9287701 | 9287806 | 6 | 10 | 0 | 1 | 0 | 0 | 0 | 1 | 0 | 0 | 98 | 49 | 1 | 1 | 0.75 | 1 | NA | NA | 1 | -0.125 |
| SE | 6921 | DAP3 | DAP3 | chr1 | + | 155729041 | 155729122 | 155727607 | 155727738 | 155729207 | 155729366 | 11 | 3 | 1 | 0 | 2 | 0 | 0 | 4 | 0 | 0 | 98 | 49 | 1 | 1 | 1 | 0.429 | 1 | NA | 1 | -0.19 |
| SE | 6922 | KIF21B | KIF21B | chr1 | - | 200983055 | 200983094 | 200980959 | 200981096 | 200984858 | 200984972 | 2 | 0 | 0 | 3 | 0 | 0 | 0 | 1 | 0 | 0 | 88 | 49 | 1 | 1 | 0.271 | NA | NA | NA | 1 | -0.729 |
| SE | 6923 | KIF21B | KIF21B | chr1 | - | 201004765 | 201004933 | 201004339 | 201004455 | 201005307 | 201005442 | 25 | 8 | 7 | 1 | 0 | 1 | 2 | 3 | 0 | 0 | 98 | 49 | 1 | 1 | 0.926 | 1 | 0.778 | 1 | 1 | -0.099 |
| SE | 6927 | SNHG12 | SNHG12 | chr1 | - | 28581110 | 28581229 | 28580559 | 28580646 | 28581589 | 28581854 | 0 | 1 | 0 | 1 | 0 | 0 | 0 | 2 | 0 | 0 | 98 | 49 | 1 | 1 | 0 | 1 | NA | NA | 1 | -0.5 |
| SE | 6931 | IL17RA | IL17RA | chr22 | + | 17100354 | 17100481 | 17098774 | 17098887 | 17101995 | 17102043 | 64 | 25 | 25 | 8 | 1 | 0 | 0 | 15 | 0 | 0 | 98 | 49 | 1 | 1 | 0.8 | 0.926 | 1 | NA | 1 | -0.091 |
| SE | 6932 | IL17RA | IL17RA | chr22 | + | 17103493 | 17103577 | 17102138 | 17102302 | 17104725 | 17104810 | 73 | 13 | 14 | 1 | 0 | 0 | 0 | 8 | 0 | 0 | 98 | 49 | 1 | 1 | 0.973 | 1 | 1 | NA | 1 | -0.009 |
| SE | 6943 | CDA | CDA | chr1 | + | 20604927 | 20605039 | 20588950 | 20589283 | 20613841 | 20613899 | 26 | 20 | 25 | 0 | 1 | 0 | 0 | 15 | 0 | 0 | 98 | 49 | 1 | 1 | 1 | 0.909 | 1 | NA | 1 | -0.03 |
| SE | 6944 | CDA | CDA | chr1 | + | 20604927 | 20605039 | 20588950 | 20589283 | 20618451 | 20618907 | 15 | 12 | 15 | 0 | 0 | 0 | 0 | 14 | 0 | 1 | 98 | 49 | 1 | 1 | 1 | 1 | 1 | NA | 0.875 | 0.125 |
| SE | 6945 | CDA | CDA | chr1 | + | 20613841 | 20613899 | 20588950 | 20589283 | 20618451 | 20618907 | 26 | 7 | 10 | 0 | 0 | 0 | 0 | 3 | 0 | 1 | 98 | 49 | 1 | 1 | 1 | 1 | 1 | NA | 0.6 | 0.4 |
| SE | 6946 | CDA | CDA | chr1 | + | 20613841 | 20613899 | 20604927 | 20605039 | 20618451 | 20618907 | 37 | 14 | 20 | 0 | 0 | 0 | 0 | 6 | 0 | 2 | 98 | 49 | 1 | 1 | 1 | 1 | 1 | NA | 0.6 | 0.4 |
| SE | 6948 | SDHA | SDHA | chr5 | + | 230875 | 231000 | 228184 | 228333 | 233476 | 233645 | 4 | 3 | 2 | 1 | 0 | 0 | 1 | 1 | 0 | 0 | 98 | 49 | 1 | 1 | 0.667 | 1 | 1 | 1 | 1 | -0.111 |
| SE | 6950 | OSBPL9 | OSBPL9 | chr1 | + | 51669433 | 51669512 | 51651990 | 51652041 | 51714002 | 51714079 | 24 | 12 | 4 | 2 | 0 | 0 | 0 | 10 | 0 | 0 | 98 | 49 | 1 | 1 | 0.857 | 1 | 1 | NA | 1 | -0.048 |
| SE | 6951 | OSBPL9 | OSBPL9 | chr1 | + | 51740142 | 51740211 | 51714002 | 51714079 | 51745535 | 51745631 | 0 | 1 | 3 | 10 | 1 | 0 | 0 | 0 | 0 | 2 | 98 | 49 | 1 | 1 | 0 | 0.333 | 1 | NA | 0 | 0.444 |
| SE | 6966 | HCLS1 | HCLS1 | chr3 | - | 121647318 | 121647448 | 121644817 | 121644928 | 121657278 | 121657352 | 202 | 87 | 88 | 3 | 0 | 0 | 1 | 33 | 0 | 1 | 98 | 49 | 1 | 1 | 0.971 | 1 | 1 | 1 | 0.943 | 0.019 |
| SE | 6967 | AP1G2 | AP1G2 | chr14 | - | 23565815 | 23565892 | 23565605 | 23565701 | 23566063 | 23566160 | 0 | 3 | 0 | 0 | 0 | 0 | 0 | 2 | 0 | 1 | 98 | 49 | 1 | 1 | NA | 1 | NA | NA | 0.5 | 0.5 |
| SE | 6968 | AP1G2 | AP1G2 | chr14 | - | 23566561 | 23566686 | 23566277 | 23566419 | 23567110 | 23567319 | 10 | 4 | 5 | 3 | 1 | 2 | 0 | 4 | 0 | 0 | 98 | 49 | 1 | 1 | 0.625 | 0.667 | 0.556 | NA | 1 | -0.384 |
| SE | 6969 | EIF4G3 | EIF4G3 | chr1 | - | 20827616 | 20827698 | 20825099 | 20825198 | 20829146 | 20829272 | 6 | 1 | 2 | 1 | 0 | 0 | 0 | 1 | 0 | 0 | 98 | 49 | 1 | 1 | 0.75 | 1 | 1 | NA | 1 | -0.083 |
| SE | 6970 | EIF4G3 | EIF4G3 | chr1 | - | 20892618 | 20892729 | 20886200 | 20886371 | 20893516 | 20893636 | 6 | 1 | 0 | 2 | 2 | 0 | 0 | 0 | 0 | 1 | 98 | 49 | 1 | 1 | 0.6 | 0.2 | NA | NA | 0 | 0.4 |
| SE | 6976 | DSC2 | DSC2 | chr18 | - | 31068298 | 31068344 | 31065973 | 31068212 | 31068893 | 31069151 | 0 | 1 | 2 | 1 | 2 | 0 | 0 | 0 | 0 | 1 | 95 | 49 | 1 | 1 | 0 | 0.205 | 1 | NA | 0 | 0.402 |

| | | | | | | | | | | | | | | | | | | | | | | | | | | | | | | |
|---|---|---|---|---|---|---|---|---|---|---|---|---|---|---|---|---|---|---|---|---|---|---|---|---|---|---|---|---|---|---|
| SE | 6986 MVP | MVP | chr16 | + | 29841914 | 29842112 | 29841595 | 29841840 | 29844492 | 29844879 | 74 | 15 | 23 | 1 | 0 | 0 | 8 | 14 | 0 | 0 | 98 | 49 | 1 | 1 | 0.974 | 1 | 1 | 1 | 1 | -0.009 |
| SE | 6987 MVP | MVP | chr16 | + | 29844492 | 29844879 | 29841914 | 29842112 | 29845862 | 29845979 | 73 | 23 | 34 | 1 | 0 | 0 | 5 | 10 | 0 | 0 | 98 | 49 | 1 | 1 | 0.973 | 1 | 1 | 1 | 1 | -0.009 |
| SE | 6990 MPDU1 | MPDU1 | chr17 | + | 7586898 | 7587017 | 7586691 | 7586777 | 7587160 | 7587271 | 3 | 0 | 0 | 1 | 0 | 0 | 0 | 1 | 0 | 1 | 98 | 49 | 1 | 1 | 0.6 | NA | NA | NA | 0.333 | 0.267 |
| SE | 6996 SAMD9 | SAMD9 | chr7 | - | 93114794 | 93114895 | 93099512 | 93106105 | 93117862 | 93118023 | 48 | 33 | 1 | 3 | 0 | 0 | 0 | 9 | 0 | 0 | 98 | 49 | 1 | 1 | 0.889 | 1 | 1 | NA | 1 | -0.037 |
| SE | 6997 PUM2 | PUM2 | chr2 | - | 20263192 | 20263460 | 20260336 | 20260466 | 20282946 | 20283231 | 20 | 4 | 1 | 1 | 0 | 0 | 0 | 2 | 0 | 0 | 98 | 49 | 1 | 1 | 0.909 | 1 | 1 | NA | 1 | -0.03 |
| SE | 6999 PUM2 | PUM2 | chr2 | - | 20290651 | 20290790 | 20283342 | 20283486 | 20294375 | 20294518 | 24 | 12 | 5 | 1 | 1 | 0 | 0 | 1 | 0 | 0 | 98 | 49 | 1 | 1 | 0.923 | 0.857 | 1 | NA | 1 | -0.073 |
| SE | 7000 PUM2 | PUM2 | chr2 | - | 20294375 | 20294518 | 20283342 | 20283486 | 20297552 | 20297678 | 13 | 9 | 1 | 2 | 0 | 0 | 3 | 2 | 0 | 0 | 98 | 49 | 1 | 1 | 0.765 | 1 | 1 | 1 | 1 | -0.078 |
| SE | 7004 THBS3 | THBS3 | chr1 | - | 155200450 | 155200610 | 155199994 | 155200113 | 155200896 | 155201004 | 7 | 5 | 0 | 0 | 1 | 0 | 0 | 1 | 0 | 0 | 98 | 49 | 1 | 1 | 1 | 0.714 | NA | NA | 1 | -0.143 |
| SE | 7008 TACC1 | TACC1 | chr8 | + | 38827167 | 38827375 | 38825307 | 38825368 | 38831124 | 38831177 | 20 | 5 | 8 | 1 | 0 | 0 | 0 | 1 | 0 | 0 | 98 | 49 | 1 | 1 | 0.909 | 1 | 1 | NA | 1 | -0.03 |
| SE | 7009 SFI1 | SFI1 | chr22 | + | 31528689 | 31528863 | 31508254 | 31508376 | 31546860 | 31546971 | 6 | 13 | 3 | 1 | 3 | 0 | 0 | 0 | 0 | 1 | 98 | 49 | 1 | 1 | 0.75 | 0.684 | 1 | NA | 0 | 0.811 |
| SE | 7011 SFI1 | SFI1 | chr22 | + | 31546860 | 31546971 | 31528689 | 31528863 | 31550253 | 31550348 | 7 | 13 | 2 | 1 | 0 | 0 | 0 | 1 | 0 | 0 | 98 | 49 | 1 | 1 | 0.778 | 1 | 1 | NA | 1 | -0.074 |
| SE | 7012 SFI1 | SFI1 | chr22 | + | 31561289 | 31561392 | 31556941 | 31557059 | 31573057 | 31573214 | 22 | 16 | 3 | 1 | 0 | 0 | 0 | 4 | 0 | 0 | 98 | 49 | 1 | 1 | 0.917 | 1 | 1 | NA | 1 | -0.028 |
| SE | 7014 SFI1 | SFI1 | chr22 | + | 31613601 | 31613855 | 31613353 | 31613530 | 31614788 | 31614860 | 9 | 5 | 7 | 1 | 0 | 0 | 2 | 0 | 0 | 0 | 98 | 49 | 1 | 1 | 0.818 | 1 | 1 | 1 | NA | -0.061 |
| SE | 7024 ELOVL5 | ELOVL5 | chr6 | - | 53270592 | 53270727 | 53267397 | 53269270 | 53273219 | 53273344 | 79 | 34 | 58 | 0 | 0 | 0 | 0 | 15 | 0 | 2 | 98 | 49 | 1 | 1 | 1 | 1 | 1 | NA | 0.789 | 0.211 |
| SE | 7025 ELOVL5 | ELOVL5 | chr6 | - | 53273219 | 53273344 | 53270592 | 53270727 | 53275089 | 53275261 | 107 | 42 | 48 | 1 | 0 | 0 | 1 | 21 | 0 | 0 | 98 | 49 | 1 | 1 | 0.982 | 1 | 1 | 1 | 1 | -0.006 |
| SE | 7038 MPHOSPH9 | MPHOSPH9 | chr12 | - | 123203249 | 123203375 | 123202623 | 123203084 | 123210055 | 123210162 | 2 | 0 | 0 | 0 | 0 | 0 | 0 | 0 | 0 | 1 | 98 | 49 | 1 | 1 | 1 | NA | NA | NA | 0 | 1 |
| SE | 7039 MPHOSPH9 | MPHOSPH9 | chr12 | - | 123214743 | 123214834 | 123210055 | 123210162 | 123218375 | 123218499 | 3 | 0 | 0 | 0 | 0 | 0 | 0 | 0 | 0 | 3 | 98 | 49 | 1 | 1 | 1 | NA | NA | NA | 0 | 1 |
| SE | 7043 RHOT2 | RHOT2 | chr16 | + | 668487 | 668569 | 668352 | 668411 | 668655 | 668699 | 13 | 2 | 0 | 2 | 0 | 0 | 3 | 3 | 0 | 0 | 98 | 49 | 1 | 1 | 0.765 | 1 | NA | 1 | 1 | -0.117 |
| SE | 7044 RHOT2 | RHOT2 | chr16 | + | 669552 | 669606 | 668655 | 668699 | 670122 | 670175 | 5 | 2 | 0 | 0 | 0 | 0 | 0 | 1 | 2 | 0 | 98 | 49 | 1 | 1 | 1 | 1 | NA | 0 | 1 | 0.5 |
| SE | 7045 RHOT2 | RHOT2 | chr16 | + | 670674 | 670773 | 670455 | 670557 | 670891 | 671000 | 12 | 1 | 0 | 1 | 0 | 0 | 0 | 3 | 0 | 0 | 98 | 49 | 1 | 1 | 0.857 | 1 | NA | NA | 1 | -0.072 |
| SE | 7047 SLC25A44 | SLC25A44 | chr1 | + | 156196012 | 156196173 | 156193931 | 156194247 | 156199834 | 156200472 | 0 | 0 | 1 | 5 | 0 | 0 | 0 | 0 | 0 | 1 | 98 | 49 | 1 | 1 | 0 | NA | 1 | NA | 0 | 0.5 |
| SE | 7058 CHD1 | CHD1 | chr5 | - | 98858179 | 98858390 | 98855203 | 98856725 | 98858963 | 98859015 | 24 | 5 | 3 | 2 | 0 | 0 | 0 | 2 | 0 | 1 | 98 | 49 | 1 | 1 | 0.857 | 1 | 1 | NA | 0.5 | 0.452 |
| SE | 7059 CHD1 | CHD1 | chr5 | - | 98876397 | 98876558 | 98875071 | 98875113 | 98879551 | 98879728 | 17 | 7 | 0 | 4 | 5 | 0 | 1 | 2 | 0 | 0 | 98 | 49 | 1 | 1 | 0.68 | 0.412 | NA | 1 | 1 | -0.454 |
| SE | 7061 CHD1 | CHD1 | chr5 | - | 98896225 | 98896442 | 98894596 | 98894686 | 98897192 | 98897320 | 51 | 8 | 1 | 0 | 1 | 0 | 1 | 8 | 0 | 0 | 98 | 49 | 1 | 1 | 1 | 0.8 | 1 | 1 | 1 | -0.067 |
| SE | 7066 GPBP1 | GPBP1 | chr5 | + | 57221375 | 57221396 | 57214073 | 57214193 | 57230845 | 57230969 | 9 | 3 | 0 | 25 | 2 | 4 | 0 | 3 | 0 | 5 | 70 | 49 | 1 | 1 | 0.201 | 0.512 | 0 | NA | 0.296 | -0.058 |
| SE | 7067 GPBP1 | GPBP1 | chr5 | + | 57250953 | 57251141 | 57249408 | 57249576 | 57261179 | 57261282 | 52 | 20 | 21 | 0 | 1 | 0 | 0 | 10 | 0 | 0 | 98 | 49 | 1 | 1 | 1 | 0.909 | 1 | NA | 1 | -0.03 |
| SE | 7069 PAPD4 | PAPD4 | chr5 | + | 79641104 | 79641196 | 79640850 | 79640965 | 79642843 | 79642910 | 13 | 6 | 2 | 0 | 1 | 0 | 0 | 1 | 0 | 0 | 98 | 49 | 1 | 1 | 1 | 0.75 | 1 | NA | 1 | -0.083 |
| SE | 7070 PAPD4 | PAPD4 | chr5 | + | 79642831 | 79642910 | 79641104 | 79641196 | 79645122 | 79645192 | 8 | 4 | 5 | 0 | 0 | 0 | 0 | 4 | 0 | 1 | 98 | 49 | 1 | 1 | 1 | 1 | 1 | NA | 0.667 | 0.333 |
| SE | 7071 PAPD4 | PAPD4 | chr5 | + | 79642843 | 79642910 | 79641104 | 79641196 | 79645122 | 79645192 | 9 | 7 | 6 | 0 | 0 | 0 | 0 | 1 | 0 | 1 | 98 | 49 | 1 | 1 | 1 | 1 | 1 | NA | 0.333 | 0.667 |
| SE | 7072 PAPD4 | PAPD4 | chr5 | + | 79649061 | 79649190 | 79648616 | 79648693 | 79656957 | 79657001 | 24 | 5 | 2 | 4 | 0 | 0 | 0 | 0 | 0 | 1 | 98 | 49 | 1 | 1 | 0.75 | 1 | 1 | NA | 0 | 0.917 |
| SE | 7073 PAPD4 | PAPD4 | chr5 | + | 79656957 | 79657001 | 79649061 | 79649190 | 79668891 | 79669028 | 16 | 4 | 2 | 1 | 0 | 0 | 0 | 3 | 0 | 0 | 93 | 49 | 1 | 1 | 0.894 | 1 | 1 | NA | 1 | -0.035 |
| SE | 7083 PPP3CB | PPP3CB | chr10 | - | 73439871 | 73439901 | 73436427 | 73438420 | 73444724 | 73444822 | 11 | 1 | 0 | 2 | 0 | 0 | 0 | 1 | 0 | 2 | 79 | 49 | 1 | 1 | 0.773 | 1 | NA | NA | 0.237 | 0.65 |
| SE | 7087 GPR108 | GPR108 | chr19 | - | 6731890 | 6731934 | 6731472 | 6731522 | 6732024 | 6732155 | 17 | 0 | 4 | 1 | 0 | 0 | 0 | 5 | 0 | 0 | 93 | 49 | 1 | 1 | 0.9 | NA | 1 | NA | 1 | -0.05 |
| SE | 7088 GPR108 | GPR108 | chr19 | - | 6732986 | 6733062 | 6732475 | 6732549 | 6733167 | 6733301 | 17 | 11 | 15 | 2 | 0 | 0 | 0 | 7 | 0 | 0 | 98 | 49 | 1 | 1 | 0.81 | 1 | 1 | NA | 1 | -0.063 |
| SE | 7090 MEGF6 | MEGF6 | chr1 | - | 3498697 | 3498826 | 3498370 | 3498499 | 3499587 | 3499716 | 9 | 2 | 0 | 1 | 0 | 1 | 0 | 0 | 3 | 0 | 98 | 49 | 1 | 1 | 0.818 | 1 | 0 | 0 | NA | 0.606 |
| SE | 7091 MEGF6 | MEGF6 | chr1 | - | 3499137 | 3499266 | 3498697 | 3498826 | 3499587 | 3499716 | 15 | 6 | 2 | 1 | 0 | 0 | 1 | 3 | 0 | 0 | 98 | 49 | 1 | 1 | 0.882 | 1 | 1 | 1 | 1 | -0.039 |
| SE | 7092 MEGF6 | MEGF6 | chr1 | - | 3511549 | 3511687 | 3510782 | 3510902 | 3512005 | 3512128 | 14 | 13 | 3 | 1 | 0 | 0 | 0 | 1 | 0 | 0 | 98 | 49 | 1 | 1 | 0.875 | 1 | 1 | NA | 1 | -0.042 |
| SE | 7095 ABCC5 | ABCC5 | chr3 | - | 183987769 | 183987917 | 183985303 | 183985378 | 183988571 | 183988727 | 9 | 8 | 1 | 1 | 0 | 0 | 2 | 1 | 0 | 0 | 98 | 49 | 1 | 1 | 0.818 | 1 | 1 | 1 | 1 | -0.061 |
| SE | 7096 DPP8 | DPP8 | chr15 | - | 65450998 | 65451110 | 65445659 | 65447006 | 65451959 | 65452102 | 3 | 0 | 1 | 1 | 0 | 0 | 0 | 1 | 0 | 0 | 98 | 49 | 1 | 1 | 0.6 | NA | 1 | NA | 1 | -0.2 |
| SE | 7100 DPP8 | DPP8 | chr15 | - | 65500605 | 65500779 | 65497863 | 65498032 | 65507242 | 65507355 | 7 | 0 | 0 | 4 | 0 | 0 | 0 | 3 | 0 | 0 | 98 | 49 | 1 | 1 | 0.467 | NA | NA | NA | 1 | -0.533 |
| SE | 7103 ANKRD12 | ANKRD12 | chr18 | + | 9208656 | 9208803 | 9195550 | 9195698 | 9211583 | 9211784 | 52 | 32 | 9 | 1 | 0 | 0 | 1 | 5 | 0 | 0 | 98 | 49 | 1 | 1 | 0.963 | 1 | 1 | 1 | 1 | -0.012 |
| SE | 7108 USP47 | USP47 | chr11 | + | 11891967 | 11892106 | 11884466 | 11884580 | 11897596 | 11897693 | 11 | 2 | 0 | 0 | 0 | 0 | 0 | 1 | 0 | 2 | 98 | 49 | 1 | 1 | 1 | 1 | NA | NA | 0.2 | 0.8 |
| SE | 7109 USP47 | USP47 | chr11 | + | 11905398 | 11905548 | 11903262 | 11903342 | 11920155 | 11920251 | 11 | 7 | 1 | 0 | 0 | 0 | 0 | 0 | 0 | 1 | 98 | 49 | 1 | 1 | 1 | 1 | 1 | NA | 0 | 1 |
| SE | 7115 GNB1 | GNB1 | chr1 | - | 1804418 | 1804581 | 1793244 | 1793311 | 1806474 | 1806538 | 21 | 15 | 6 | 1 | 1 | 0 | 0 | 1 | 0 | 0 | 98 | 49 | 1 | 1 | 0.913 | 0.882 | 1 | NA | 1 | -0.068 |
| SE | 7116 GNB1 | GNB1 | chr1 | - | 1839189 | 1839238 | 1825396 | 1825499 | 1890819 | 1891117 | 27 | 19 | 12 | 1 | 0 | 0 | 2 | 5 | 1 | 0 | 98 | 49 | 1 | 1 | 0.931 | 1 | 1 | 0.5 | 1 | 0.227 |
| SE | 7119 PRDM2 | PRDM2 | chr1 | + | 13715540 | 13715614 | 13700239 | 13700300 | 13730999 | 13731117 | 2 | 4 | 1 | 0 | 0 | 0 | 0 | 0 | 0 | 2 | 98 | 49 | 1 | 1 | 1 | 1 | 1 | NA | 0 | 1 |
| SE | 7124 LAP3 | LAP3 | chr4 | + | 17597045 | 17597134 | 17595409 | 17595534 | 17598455 | 17598558 | 10 | 3 | 0 | 0 | 1 | 0 | 0 | 13 | 0 | 0 | 98 | 49 | 1 | 1 | 1 | 0.6 | NA | NA | 1 | -0.2 |
| SE | 7125 AMDHD2 | AMDHD2 | chr16 | + | 2527560 | 2527615 | 2520983 | 2521123 | 2527772 | 2527985 | 8 | 8 | 5 | 1 | 0 | 0 | 1 | 0 | 0 | 0 | 98 | 49 | 1 | 1 | 0.8 | 1 | 1 | 1 | NA | -0.067 |
| SE | 7129 MBD6 | MBD6 | chr12 | + | 57527847 | 57528017 | 57527506 | 57527660 | 57528146 | 57528560 | 22 | 7 | 1 | 1 | 0 | 0 | 1 | 4 | 0 | 0 | 98 | 49 | 1 | 1 | 0.917 | 1 | 1 | 1 | 1 | -0.028 |
| SE | 7131 SIVA1 | SIVA1 | chr14 | + | 104755629 | 104755824 | 104753132 | 104753319 | 104756603 | 104756760 | 7 | 1 | 0 | 1 | 0 | 0 | 0 | 1 | 0 | 0 | 98 | 49 | 1 | 1 | 0.778 | 1 | NA | NA | 1 | -0.111 |
| SE | 7135 PPM1M | PPM1M | chr3 | + | 52246973 | 52247228 | 52246694 | 52246812 | 52247681 | 52247794 | 18 | 3 | 5 | 2 | 0 | 0 | 2 | 4 | 0 | 0 | 98 | 49 | 1 | 1 | 0.818 | 1 | 1 | 1 | 1 | -0.061 |
| SE | 7136 PPM1M | PPM1M | chr3 | + | 52247681 | 52247794 | 52246694 | 52246812 | 52248152 | 52248237 | 13 | 2 | 9 | 1 | 1 | 0 | 0 | 0 | 0 | 1 | 98 | 49 | 1 | 1 | 0.867 | 0.5 | 1 | NA | 0 | 0.789 |
| SE | 7139 FANCM | FANCM | chr14 | + | 45140631 | 45140709 | 45137068 | 45137241 | 45148836 | 45148995 | 0 | 2 | 0 | 0 | 0 | 3 | 0 | 0 | 0 | 1 | 98 | 49 | 1 | 1 | NA | 1 | 0 | NA | 0 | 0.5 |
| SE | 7144 ISCU | ISCU | chr12 | + | 108567657 | 108567743 | 108567189 | 108567268 | 108568830 | 108569384 | 0 | 1 | 0 | 28 | 21 | 15 | 0 | 0 | 2 | 8 | 98 | 49 | 1 | 1 | 0 | 0.023 | 0 | 0 | 0 | 0.008 |
| SE | 7147 CDC73 | CDC73 | chr1 | + | 193236255 | 193236356 | 193232992 | 193233154 | 193249729 | 193249871 | 4 | 2 | 3 | 0 | 2 | 0 | 0 | 5 | 0 | 0 | 98 | 49 | 1 | 1 | 1 | 0.333 | 1 | NA | 1 | -0.222 |
| SE | 7152 FHL3 | FHL3 | chr1 | - | 37997962 | 37998132 | 37997683 | 37997870 | 37998973 | 37999148 | 7 | 4 | 4 | 1 | 0 | 0 | 0 | 7 | 0 | 0 | 98 | 49 | 1 | 1 | 0.778 | 1 | 1 | NA | 1 | -0.074 |
| SE | 7154 UROD | UROD | chr1 | + | 45013908 | 45014070 | 45013593 | 45013791 | 45014438 | 45014576 | 0 | 1 | 0 | 0 | 0 | 0 | 0 | 1 | 0 | 2 | 98 | 49 | 1 | 1 | NA | 1 | NA | NA | 0.2 | 0.8 |
| SE | 7155 FOXN3 | FOXN3 | chr14 | - | 89190383 | 89190449 | 89180700 | 89180806 | 89280949 | 89281014 | 3 | 6 | 0 | 3 | 1 | 0 | 1 | 1 | 0 | 1 | 98 | 49 | 1 | 1 | 0.333 | 0.75 | NA | 1 | 0.333 | -0.125 |
| SE | 7168 PSMA4 | PSMA4 | chr15 | + | 78542482 | 78542645 | 78542176 | 78542219 | 78544189 | 78544267 | 16 | 4 | 9 | 0 | 1 | 0 | 1 | 4 | 0 | 1 | 98 | 49 | 1 | 1 | 1 | 0.667 | 1 | 1 | 0.667 | 0.055 |
| SE | 7177 RGS12 | RGS12 | chr4 | + | 3428557 | 3428711 | 3428089 | 3428169 | 3430406 | 3430955 | 2 | 0 | 0 | 2 | 0 | 0 | 0 | 0 | 0 | 1 | 98 | 49 | 1 | 1 | 0.333 | NA | NA | NA | 0 | 0.333 |
| SE | 7178 CSNK1G1 | CSNK1G1 | chr15 | - | 64259200 | 64259241 | 64251511 | 64251581 | 64300318 | 64300723 | 3 | 6 | 0 | 2 | 0 | 0 | 0 | 2 | 0 | 1 | 90 | 49 | 1 | 1 | 0.45 | 1 | NA | NA | 0.521 | 0.204 |
| SE | 7181 TRPT1 | TRPT1 | chr11 | - | 64224099 | 64224210 | 64223798 | 64223967 | 64224284 | 64224341 | 2 | 1 | 0 | 0 | 0 | 0 | 0 | 0 | 0 | 1 | 98 | 49 | 1 | 1 | 1 | 1 | NA | NA | 0 | 1 |
| SE | 7182 TRPT1 | TRPT1 | chr11 | - | 64224099 | 64224213 | 64223798 | 64223967 | 64224284 | 64224341 | 2 | 0 | 0 | 0 | 0 | 0 | 0 | 0 | 0 | 1 | 98 | 49 | 1 | 1 | 1 | NA | NA | NA | 0 | 1 |
| SE | 7184 AFG3L2 | AFG3L2 | chr18 | - | 12367275 | 12367382 | 12366964 | 12367117 | 12370848 | 12370926 | 5 | 7 | 0 | 0 | 0 | 0 | 0 | 0 | 0 | 1 | 98 | 49 | 1 | 1 | 1 | 1 | NA | NA | 0 | 1 |
| SE | 7187 NEK4 | NEK4 | chr3 | - | 52749691 | 52749829 | 52746733 | 52746904 | 52751931 | 52752336 | 0 | 0 | 0 | 2 | 0 | 0 | 0 | 2 | 0 | 0 | 98 | 49 | 1 | 1 | 0 | NA | NA | NA | 1 | -1 |
| SE | 7189 SREK1 | SREK1 | chr5 | + | 66153462 | 66153596 | 66144217 | 66144537 | 66159218 | 66159334 | 6 | 2 | 0 | 0 | 1 | 0 | 0 | 1 | 0 | 0 | 98 | 49 | 1 | 1 | 1 | 0.5 | NA | NA | 1 | -0.25 |
| SE | 7190 SREK1 | SREK1 | chr5 | + | 66158808 | 66158932 | 66153462 | 66153596 | 66159218 | 66159334 | 1 | 0 | 1 | 4 | 0 | 0 | 0 | 1 | 0 | 0 | 98 | 49 | 1 | 1 | 0.111 | NA | 1 | NA | 1 | -0.445 |
| SE | 7191 SREK1 | SREK1 | chr5 | + | 66177513 | 66177658 | 66174945 | 66175041 | 66178718 | 66183616 | 18 | 8 | 0 | 0 | 3 | 0 | 0 | 2 | 0 | 0 | 98 | 49 | 1 | 1 | 1 | 0.571 | NA | NA | 1 | -0.215 |
| SE | 7192 SREK1 | SREK1 | chr5 | + | 66177516 | 66177658 | 66174945 | 66175041 | 66178718 | 66183616 | 14 | 3 | 0 | 0 | 3 | 0 | 0 | 1 | 0 | 0 | 98 | 49 | 1 | 1 | 1 | 0.333 | NA | NA | 1 | -0.334 |
| SE | 7194 RERE | RERE | chr1 | - | 8362682 | 8362844 | 8361762 | 8361876 | 8364055 | 8364255 | 36 | 14 | 6 | 0 | 0 | 0 | 2 | 6 | 0 | 1 | 98 | 49 | 1 | 1 | 1 | 1 | 1 | 1 | 0.75 | 0.125 |
| SE | 7201 NBEAL2 | NBEAL2 | chr3 | + | 46992474 | 46992555 | 46991839 | 46991946 | 46993936 | 46994020 | 32 | 12 | 3 | 29 | 2 | 9 | 0 | 10 | 0 | 9 | 98 | 49 | 1 | 1 | 0.356 | 0.75 | 0.143 | NA | 0.357 | 0.059 |
| SE | 7202 NBEAL2 | NBEAL2 | chr3 | + | 46996750 | 46996833 | 46996270 | 46996592 | 46996953 | 46997046 | 43 | 31 | 5 | 2 | 0 | 0 | 7 | 12 | 0 | 0 | 98 | 49 | 1 | 1 | 0.915 | 1 | 1 | 1 | 1 | -0.028 |
| SE | 7203 NBEAL2 | NBEAL2 | chr3 | + | 46998066 | 46998226 | 46997560 | 46997694 | 46998462 | 46998565 | 95 | 41 | 7 | 1 | 0 | 0 | 5 | 29 | 0 | 0 | 98 | 49 | 1 | 1 | 0.979 | 1 | 1 | 1 | 1 | -0.007 |
| SE | 7204 NBEAL2 | NBEAL2 | chr3 | + | 47001278 | 47001438 | 47001000 | 47001179 | 47001688 | 47001826 | 117 | 46 | 6 | 0 | 1 | 0 | 9 | 30 | 0 | 0 | 98 | 49 | 1 | 1 | 1 | 0.958 | 1 | 1 | 1 | -0.014 |
| SE | 7205 NBEAL2 | NBEAL2 | chr3 | + | 47001919 | 47002288 | 47001688 | 47001826 | 47002370 | 47002520 | 90 | 39 | 9 | 0 | 1 | 0 | 2 | 25 | 0 | 0 | 98 | 49 | 1 | 1 | 1 | 0.951 | 1 | 1 | 1 | -0.016 |
| SE | 7207 NBEAL2 | NBEAL2 | chr3 | + | 47003815 | 47003976 | 47003173 | 47003309 | 47004076 | 47004393 | 88 | 20 | 7 | 2 | 0 | 0 | 11 | 24 | 0 | 0 | 98 | 49 | 1 | 1 | 0.957 | 1 | 1 | 1 | 1 | -0.014 |
| SE | 7208 NBEAL2 | NBEAL2 | chr3 | + | 47005945 | 47006063 | 47005737 | 47005847 | 47006164 | 47006262 | 111 | 38 | 17 | 0 | 2 | 0 | 6 | 28 | 0 | 0 | 98 | 49 | 1 | 1 | 1 | 0.905 | 1 | 1 | 1 | -0.032 |
| SE | 7212 NBEAL2 | NBEAL2 | chr3 | + | 47008282 | 47008441 | 47007815 | 47007910 | 47008519 | 47008668 | 84 | 31 | 11 | 1 | 0 | 0 | 5 | 33 | 0 | 0 | 98 | 49 | 1 | 1 | 0.977 | 1 | 1 | 1 | 1 | -0.008 |
| SE | 7225 UPF2 | UPF2 | chr10 | - | 11979056 | 11979165 | 11967340 | 11967454 | 11997671 | 11997757 | 19 | 14 | 5 | 0 | 1 | 0 | 0 | 5 | 0 | 0 | 98 | 49 | 1 | 1 | 1 | 0.875 | 1 | NA | 1 | -0.042 |
| SE | 7227 UBE2Q1 | UBE2Q1 | chr1 | - | 154551396 | 154551492 | 154550937 | 154551004 | 154551770 | 154551819 | 9 | 6 | 6 | 1 | 0 | 0 | 0 | 1 | 0 | 0 | 98 | 49 | 1 | 1 | 0.818 | 1 | 1 | NA | 1 | -0.061 |
| SE | 7228 UBE2Q1 | UBE2Q1 | chr1 | - | 154552735 | 154552820 | 154552403 | 154552464 | 154553031 | 154553172 | 12 | 6 | 2 | 0 | 1 | 0 | 0 | 2 | 0 | 0 | 98 | 49 | 1 | 1 | 1 | 0.75 | 1 | NA | 1 | -0.083 |
| SE | 7234 PPP3CA | PPP3CA | chr4 | - | 101029165 | 101029195 | 101023429 | 101026061 | 101032266 | 101032364 | 3 | 0 | 0 | 22 | 10 | 3 | 0 | 0 | 0 | 2 | 79 | 49 | 1 | 1 | 0.078 | 0 | 0 | NA | 0 | 0.026 |
| SE | 7235 PPP3CA | PPP3CA | chr4 | - | 101063231 | 101063357 | 101061086 | 101061161 | 101080531 | 101080626 | 15 | 8 | 2 | 0 | 0 | 0 | 0 | 15 | 0 | 3 | 98 | 49 | 1 | 1 | 1 | 1 | 1 | NA | 0.714 | 0.286 |
| SE | 7240 MTR | MTR | chr1 | + | 236831965 | 236832078 | 236829188 | 236829268 | 236835546 | 236835687 | 6 | 4 | 0 | 5 | 0 | 0 | 0 | 2 | 0 | 0 | 98 | 49 | 1 | 1 | 0.375 | 1 | NA | NA | 1 | -0.312 |
| SE | 7241 MTR | MTR | chr1 | + | 236852520 | 236852637 | 236850343 | 236850523 | 236852947 | 236853088 | 4 | 0 | 0 | 1 | 0 | 0 | 0 | 3 | 0 | 0 | 98 | 49 | 1 | 1 | 0.667 | NA | NA | NA | 1 | -0.333 |
| SE | 7242 MTR | MTR | chr1 | + | 236861124 | 236861277 | 236859832 | 236859922 | 236862235 | 236862343 | 5 | 0 | 1 | 1 | 0 | 0 | 0 | 3 | 0 | 0 | 98 | 49 | 1 | 1 | 0.714 | NA | 1 | NA | 1 | -0.143 |
| SE | 7243 SSH1 | SSH1 | chr12 | - | 108798999 | 108799200 | 108791918 | 108792829 | 108800779 | 108800926 | 3 | 4 | 1 | 1 | 0 | 0 | 0 | 1 | 0 | 0 | 98 | 49 | 1 | 1 | 0.6 | 1 | 1 | NA | 1 | -0.133 |
| SE | 7244 SSH1 | SSH1 | chr12 | - | 108802321 | 108802368 | 108800779 | 108800926 | 108805055 | 108805184 | 7 | 4 | 0 | 1 | 2 | 1 | 0 | 1 | 0 | 0 | 96 | 49 | 1 | 1 | 0.781 | 0.505 | 0 | NA | 1 | -0.571 |
| SE | 7245 USP48 | USP48 | chr1 | - | 21721649 | 21721764 | 21721035 | 21721166 | 21723897 | 21724095 | 14 | 7 | 0 | 1 | 2 | 0 | 0 | 3 | 0 | 0 | 98 | 49 | 1 | 1 | 0.875 | 0.636 | NA | NA | 1 | -0.244 |
| SE | 7246 USP48 | USP48 | chr1 | - | 21736445 | 21736625 | 21729703 | 21729832 | 21747066 | 21747149 | 7 | 1 | 2 | 1 | 0 | 0 | 1 | 4 | 0 | 0 | 98 | 49 | 1 | 1 | 0.778 | 1 | 1 | 1 | 1 | -0.074 |
| SE | 7255 PANK4 | PANK4 | chr1 | - | 2512887 | 2513039 | 2511627 | 2511683 | 2514001 | 2514089 | 19 | 7 | 0 | 6 | 0 | 0 | 0 | 4 | 0 | 0 | 98 | 49 | 1 | 1 | 0.613 | 1 | NA | NA | 1 | -0.194 |
| SE | 7256 PANK4 | PANK4 | chr1 | - | 2515561 | 2515717 | 2514353 | 2514466 | 2518163 | 2518264 | 9 | 11 | 1 | 0 | 0 | 0 | 0 | 0 | 1 | 0 | 98 | 49 | 1 | 1 | 1 | 1 | 1 | 0 | NA | 1 |
| SE | 7268 HEXA | HEXA | chr15 | - | 72349078 | 72349259 | 72348047 | 72348134 | 72350517 | 72350650 | 10 | 3 | 0 | 0 | 0 | 0 | 0 | 3 | 0 | 2 | 98 | 49 | 1 | 1 | 1 | 1 | NA | NA | 0.429 | 0.571 |
| SE | 7270 SESN2 | SESN2 | chr1 | + | 28273357 | 28273508 | 28272580 | 28272793 | 28274039 | 28274158 | 0 | 2 | 2 | 0 | 4 | 0 | 0 | 1 | 0 | 0 | 98 | 49 | 1 | 1 | NA | 0.2 | 1 | NA | 1 | -0.4 |
| SE | 7271 STK40 | STK40 | chr1 | - | 36344119 | 36344264 | 36343859 | 36343979 | 36348699 | 36348815 | 32 | 23 | 29 | 0 | 2 | 0 | 3 | 6 | 0 | 0 | 98 | 49 | 1 | 1 | 1 | 0.852 | 1 | 1 | 1 | -0.049 |
| SE | 7272 STK40 | STK40 | chr1 | - | 36358736 | 36358822 | 36358238 | 36358382 | 36361220 | 36361340 | 40 | 24 | 21 | 0 | 1 | 0 | 1 | 4 | 0 | 0 | 98 | 49 | 1 | 1 | 1 | 0.923 | 1 | 1 | 1 | -0.026 |
| SE | 7273 STK40 | STK40 | chr1 | - | 36358736 | 36358837 | 36358238 | 36358382 | 36361220 | 36361340 | 29 | 18 | 9 | 0 | 1 | 0 | 1 | 3 | 0 | 0 | 98 | 49 | 1 | 1 | 1 | 0.9 | 1 | 1 | 1 | -0.033 |
| SE | 7274 STK40 | STK40 | chr1 | - | 36361220 | 36361340 | 36358736 | 36358822 | 36385722 | 36385927 | 30 | 13 | 15 | 1 | 0 | 0 | 0 | 1 | 0 | 0 | 98 | 49 | 1 | 1 | 0.937 | 1 | 1 | NA | 1 | -0.021 |
| SE | 7277 MFN2 | MFN2 | chr1 | + | 11989164 | 11989343 | 11981969 | 11982114 | 11992554 | 11992690 | 66 | 13 | 5 | 1 | 0 | 0 | 0 | 10 | 0 | 0 | 98 | 49 | 1 | 1 | 0.971 | 1 | 1 | NA | 1 | -0.01 |
| SE | 7278 MFN2 | MFN2 | chr1 | + | 12007052 | 12007249 | 12006537 | 12006693 | 12009591 | 12009726 | 25 | 16 | 10 | 2 | 0 | 0 | 0 | 5 | 1 | 0 | 98 | 49 | 1 | 1 | 0.862 | 1 | 1 | 0 | 1 | 0.454 |
| SE | 7279 SHC1 | SHC1 | chr1 | - | 154965542 | 154965781 | 154962297 | 154963931 | 154965945 | 154966077 | 6 | 1 | 0 | 0 | 2 | 0 | 0 | 1 | 0 | 0 | 98 | 49 | 1 | 1 | 1 | 0.2 | NA | NA | 1 | -0.4 |
| SE | 7280 SHC1 | SHC1 | chr1 | - | 154968203 | 154968257 | 154967979 | 154968031 | 154968494 | 154968614 | 7 | 1 | 2 | 0 | 2 | 0 | 0 | 3 | 0 | 0 | 98 | 49 | 1 | 1 | 1 | 0.2 | 1 | NA | 1 | -0.267 |
| SE | 7291 IL6R | IL6R | chr1 | + | 154435968 | 154436110 | 154434989 | 154435156 | 154448124 | 154448171 | 67 | 21 | 27 | 4 | 6 | 1 | 0 | 7 | 0 | 0 | 98 | 49 | 1 | 1 | 0.893 | 0.636 | 0.931 | NA | 1 | -0.18 |
| SE | 7292 IL6R | IL6R | chr1 | + | 154449910 | 154449980 | 154448124 | 154448171 | 154465133 | 154465450 | 25 | 9 | 9 | 0 | 0 | 6 | 0 | 5 | 0 | 0 | 98 | 49 | 1 | 1 | 1 | 1 | 0.429 | NA | 1 | -0.19 |
| SE | 7293 IL6R | IL6R | chr1 | + | 154454487 | 154454581 | 154449910 | 154449980 | 154465133 | 154469450 | 30 | 17 | 4 | 0 | 1 | 0 | 0 | 4 | 0 | 2 | 98 | 49 | 1 | 1 | 1 | 0.895 | 1 | NA | 0.5 | 0.465 |
| SE | 7296 IST1 | IST1 | chr16 | + | 71924768 | 71924817 | 71923287 | 71923380 | 71927613 | 71930637 | 4 | 10 | 5 | 1 | 0 | 0 | 0 | 1 | 0 | 0 | 98 | 49 | 1 | 1 | 0.667 | 1 | 1 | NA | 1 | -0.111 |
| SE | 7297 IST1 | IST1 | chr16 | + | 71924772 | 71924817 | 71923287 | 71923380 | 71927613 | 71930637 | 4 | 5 | 3 | 1 | 0 | 0 | 0 | 1 | 0 | 0 | 94 | 49 | 1 | 1 | 0.676 | 1 | 1 | NA | 1 | -0.108 |
| SE | 7298 KIAA0040 | KIAA0040 | chr1 | - | 175177610 | 175177684 | 175166561 | 175166737 | 175192639 | 175192716 | 46 | 29 | 2 | 2 | 0 | 0 | 2 | 7 | 0 | 0 | 98 | 49 | 1 | 1 | 0.92 | 1 | 1 | 1 | 1 | -0.027 |
| SE | 7307 MAX | MAX | chr14 | - | 65077327 | 65077428 | 65075123 | 65076663 | 65077912 | 65078036 | 16 | 3 | 9 | 15 | 4 | 2 | 0 | 0 | 0 | 1 | 98 | 49 | 1 | 1 | 0.348 | 0.273 | 0.692 | NA | 0 | 0.438 |
| SE | 7311 ANKDD1A | ANKDD1A | chr15 | + | 64942466 | 64942565 | 64934135 | 64934234 | 64943483 | 64943582 | 0 | 5 | 0 | 0 | 2 | 0 | 0 | 2 | 0 | 0 | 98 | 49 | 1 | 1 | NA | 0.556 | NA | NA | 1 | -0.444 |
| SE | 7316 MIER1 | MIER1 | chr1 | + | 66963087 | 66963160 | 66959678 | 66959743 | 66970807 | 66970959 | 38 | 31 | 20 | 0 | 3 | 0 | 0 | 11 | 0 | 0 | 98 | 49 | 1 | 1 | 1 | 0.838 | 1 | NA | 1 | -0.054 |
| SE | 7317 MSH5-SAPCD1 | MSH5-SAPCD1 | chr6 | + | 31758547 | 31758620 | 31758164 | 31758293 | 31758765 | 31758875 | 1 | 0 | 0 | 1 | 0 | 0 | 0 | 1 | 0 | 0 | 98 | 49 | 1 | 1 | 0.333 | NA | NA | NA | 1 | -0.667 |
| SE | 7318 VPS11 | VPS11 | chr11 | + | 119071595 | 119071843 | 119070233 | 119070397 | 119073197 | 119073399 | 10 | 0 | 0 | 0 | 0 | 0 | 0 | 0 | 0 | 2 | 98 | 49 | 1 | 1 | 1 | NA | NA | NA | 0 | 1 |
| SE | 7325 KIAA0513 | KIAA0513 | chr16 | + | 85078414 | 85078455 | 85077424 | 85077632 | 85078924 | 85079003 | 39 | 11 | 2 | 0 | 0 | 0 | 0 | 9 | 1 | 0 | 90 | 49 | 1 | 1 | 1 | 1 | 1 | 0 | 1 | 0.5 |
| SE | 7326 KIAA0513 | KIAA0513 | chr16 | + | 85082563 | 85082593 | 85081314 | 85081392 | 85086643 | 85086724 | 1 | 0 | 0 | 18 | 4 | 11 | 0 | 0 | 2 | 3 | 79 | 49 | 1 | 1 | 0.033 | 0 | 0 | 0 | 0 | 0.011 |
| SE | 7328 PI4KA | PI4KA | chr22 | - | 20727773 | 20727864 | 20727229 | 20727397 | 20729312 | 20729506 | 4 | 4 | 0 | 3 | 0 | 0 | 0 | 2 | 0 | 0 | 98 | 49 | 1 | 1 | 0.4 | 1 | NA | NA | 1 | -0.3 |
| SE | 7329 PI4KA | PI4KA | chr22 | - | 20744627 | 20744720 | 20742607 | 20742764 | 20747582 | 20747702 | 13 | 5 | 0 | 0 | 1 | 0 | 0 | 2 | 0 | 0 | 98 | 49 | 1 | 1 | 1 | 0.714 | NA | NA | 1 | -0.143 |
| SE | 7330 PI4KA | PI4KA | chr22 | - | 20764816 | 20764950 | 20761303 | 20761386 | 20765099 | 20765236 | 18 | 9 | 0 | 0 | 1 | 0 | 0 | 1 | 0 | 0 | 98 | 49 | 1 | 1 | 1 | 0.818 | NA | NA | 1 | -0.091 |
| SE | 7331 PI4KA | PI4KA | chr22 | - | 20796145 | 20796314 | 20793192 | 20793243 | 20798583 | 20798687 | 11 | 1 | 0 | 0 | 0 | 0 | 0 | 0 | 0 | 1 | 98 | 49 | 1 | 1 | 1 | 1 | NA | NA | 0 | 1 |
| SE | 7332 PI4KA | PI4KA | chr22 | - | 20818482 | 20818549 | 20813357 | 20813506 | 20819640 | 20819900 | 22 | 6 | 0 | 2 | 0 | 0 | 2 | 10 | 0 | 0 | 98 | 49 | 1 | 1 | 0.846 | 1 | NA | 1 | 1 | -0.077 |
| SE | 7336 TSEN15 | TSEN15 | chr1 | + | 184072156 | 184072298 | 184054727 | 184054863 | 184072826 | 184074210 | 0 | 0 | 0 | 1 | 1 | 0 | 0 | 1 | 0 | 0 | 98 | 49 | 1 | 1 | 0 | 0 | NA | NA | 1 | -1 |
| SE | 7342 ADA | ADA | chr20 | - | 44623006 | 44623078 | 44622828 | 44622930 | 44624201 | 44624329 | 1 | 0 | 0 | 3 | 0 | 0 | 0 | 4 | 0 | 0 | 98 | 49 | 1 | 1 | 0.143 | NA | NA | NA | 1 | -0.857 |
| SE | 7350 RASA3 | RASA3 | chr13 | - | 114040999 | 114041094 | 114029810 | 114029887 | 114052051 | 114052155 | 29 | 14 | 0 | 0 | 1 | 0 | 0 | 4 | 0 | 0 | 98 | 49 | 1 | 1 | 1 | 0.875 | NA | NA | 1 | -0.062 |
| SE | 7367 ZNF324 | ZNF324 | chr19 | + | 58469179 | 58469306 | 58467044 | 58467183 | 58469727 | 58469844 | 3 | 1 | 0 | 0 | 1 | 0 | 0 | 1 | 0 | 0 | 98 | 49 | 1 | 1 | 1 | 0.333 | NA | NA | 1 | -0.334 |
| SE | 7370 SMAP2 | SMAP2 | chr1 | + | 40406735 | 40406869 | 40373705 | 40374223 | 40408652 | 40408738 | 139 | 72 | 82 | 0 | 2 | 0 | 0 | 53 | 0 | 0 | 98 | 49 | 1 | 1 | 1 | 0.947 | 1 | NA | 1 | -0.018 |
| SE | 7377 ATP11A | ATP11A | chr13 | + | 112816082 | 112816211 | 112810618 | 112810726 | 112819303 | 112819407 | 43 | 25 | 0 | 1 | 0 | 0 | 0 | 8 | 0 | 0 | 98 | 49 | 1 | 1 | 0.956 | 1 | NA | NA | 1 | -0.022 |
| SE | 7378 ATP11A | ATP11A | chr13 | + | 112824343 | 112824425 | 112823344 | 112823409 | 112826693 | 112826891 | 26 | 9 | 1 | 0 | 1 | 0 | 0 | 3 | 0 | 0 | 98 | 49 | 1 | 1 | 1 | 0.818 | 1 | NA | 1 | -0.061 |
| SE | 7379 ATP11A | ATP11A | chr13 | + | 112825429 | 112825580 | 112824343 | 112824425 | 112826693 | 112826891 | 16 | 10 | 3 | 1 | 1 | 0 | 4 | 5 | 0 | 0 | 98 | 49 | 1 | 1 | 0.889 | 0.833 | 1 | 1 | 1 | -0.093 |
| SE | 7380 ATP11A | ATP11A | chr13 | + | 112851036 | 112851218 | 112842275 | 112842379 | 112854278 | 112854530 | 13 | 9 | 11 | 0 | 1 | 0 | 0 | 8 | 0 | 0 | 98 | 49 | 1 | 1 | 1 | 0.818 | 1 | NA | 1 | -0.061 |
| SE | 7381 ATP11A | ATP11A | chr13 | + | 112855910 | 112856085 | 112854278 | 112854530 | 112857817 | 112857920 | 7 | 8 | 3 | 2 | 0 | 0 | 0 | 4 | 0 | 0 | 98 | 49 | 1 | 1 | 0.636 | 1 | 1 | NA | 1 | -0.121 |
| SE | 7382 ATP11A | ATP11A | chr13 | + | 112858144 | 112858290 | 112857817 | 112857920 | 112859392 | 112859452 | 20 | 5 | 0 | 1 | 1 | 0 | 1 | 1 | 0 | 1 | 98 | 49 | 1 | 1 | 0.909 | 0.714 | NA | 1 | 0.333 | 0.145 |
| SE | 7383 ATP11A | ATP11A | chr13 | + | 112878216 | 112878303 | 112875775 | 112875941 | 112881875 | 112887168 | 2 | 2 | 3 | 0 | 3 | 0 | 0 | 0 | 0 | 1 | 98 | 49 | 1 | 1 | 1 | 0.25 | 1 | NA | 0 | 0.75 |
| SE | 7384 TRIM21 | TRIM21 | chr11 | - | 4388299 | 4388530 | 4386967 | 4386990 | 4389653 | 4389749 | 21 | 15 | 17 | 0 | 5 | 0 | 0 | 6 | 0 | 0 | 98 | 49 | 1 | 1 | 1 | 0.6 | 1 | NA | 1 | -0.133 |
| SE | 7387 CXorf38 | CXorf38 | chrX | - | 40636532 | 40636712 | 40630613 | 40630773 | 40637006 | 40637156 | 11 | 7 | 8 | 0 | 1 | 0 | 0 | 1 | 0 | 0 | 98 | 49 | 1 | 1 | 1 | 0.778 | 1 | NA | 1 | -0.074 |
| SE | 7388 CXorf38 | CXorf38 | chrX | - | 40647006 | 40647141 | 40639008 | 40639128 | 40647304 | 40647567 | 4 | 3 | 4 | 5 | 0 | 0 | 0 | 1 | 0 | 1 | 98 | 49 | 1 | 1 | 0.286 | 1 | 1 | NA | 0.333 | 0.429 |
| SE | 7391 DAZAP1 | DAZAP1 | chr19 | + | 1429966 | 1429996 | 1428841 | 1428995 | 1430221 | 1430362 | 5 | 1 | 0 | 0 | 1 | 0 | 0 | 4 | 0 | 0 | 79 | 49 | 1 | 1 | 1 | 1 | 0 | NA | 1 | -0.333 |
| SE | 7397 LOC100131564 | LOC100131564 | chr1 | - | 93326601 | 93326662 | 93325768 | 93325895 | 93337376 | 93339274 | 0 | 0 | 0 | 4 | 0 | 0 | 0 | 1 | 0 | 0 | 98 | 49 | 1 | 1 | 0 | NA | NA | NA | 1 | -1 |
| SE | 7404 LOC100289473 | LOC100289473 | chr20 | - | 1774510 | 1774665 | 1773364 | 1773542 | 1778519 | 1778848 | 5 | 2 | 2 | 1 | 0 | 0 | 0 | 1 | 0 | 0 | 98 | 49 | 1 | 1 | 0.714 | 1 | 1 | NA | 1 | -0.095 |
| SE | 7405 ZDHHC7 | ZDHHC7 | chr16 | - | 84988762 | 84988873 | 84981869 | 84981994 | 84990303 | 84990635 | 1 | 0 | 2 | 8 | 3 | 4 | 0 | 3 | 0 | 6 | 98 | 49 | 1 | 1 | 0.059 | 0 | 0.2 | NA | 0.2 | -0.114 |
| SE | 7406 KIN | KIN | chr10 | - | 7778837 | 7779019 | 7775750 | 7775799 | 7780055 | 7780178 | 21 | 4 | 0 | 1 | 0 | 0 | 0 | 2 | 0 | 0 | 98 | 49 | 1 | 1 | 0.913 | 1 | NA | NA | 1 | -0.043 |
| SE | 7408 HECTD1 | HECTD1 | chr14 | - | 31113876 | 31114059 | 31113276 | 31113484 | 31114285 | 31114340 | 6 | 1 | 1 | 0 | 1 | 0 | 0 | 0 | 0 | 1 | 98 | 49 | 1 | 1 | 1 | 0.333 | 1 | NA | 0 | 0.778 |
| SE | 7415 NARF | NARF | chr17 | + | 82483715 | 82483779 | 82481081 | 82481211 | 82484812 | 82484950 | 11 | 5 | 12 | 0 | 1 | 0 | 0 | 2 | 0 | 0 | 98 | 49 | 1 | 1 | 1 | 0.714 | 1 | NA | 1 | -0.095 |
| SE | 7421 TPM3 | TPM3 | chr1 | - | 154169304 | 154169383 | 154155303 | 154157721 | 154170399 | 154170469 | 9 | 2 | 0 | 75 | 46 | 36 | 0 | 4 | 0 | 20 | 98 | 49 | 1 | 1 | 0.057 | 0.021 | 0 | NA | 0.091 | -0.065 |
| SE | 7422 TPM3 | TPM3 | chr1 | - | 154169304 | 154169383 | 154161812 | 154167940 | 154170399 | 154170469 | 7 | 2 | 0 | 2 | 0 | 0 | 0 | 3 | 0 | 0 | 98 | 49 | 1 | 1 | 0.636 | 1 | NA | NA | 1 | -0.182 |
| SE | 7431 C9orf72 | C9orf72 | chr9 | - | 27560226 | 27560299 | 27558490 | 27558607 | 27561584 | 27561649 | 20 | 6 | 9 | 1 | 0 | 0 | 0 | 1 | 0 | 0 | 98 | 49 | 1 | 1 | 0.909 | 1 | 1 | NA | 1 | -0.03 |
| SE | 7435 ZNF100 | ZNF100 | chr19 | - | 21744940 | 21745067 | 21744016 | 21744115 | 21765693 | 21765786 | 4 | 2 | 0 | 0 | 2 | 0 | 0 | 1 | 0 | 0 | 98 | 49 | 1 | 1 | 0.5 | 1 | NA | NA | 1 | -0.25 |
| SE | 7439 TBC1D32 | TBC1D32 | chr6 | - | 121170404 | 121170527 | 121160947 | 121161056 | 121205074 | 121205163 | 0 | 0 | 0 | 1 | 0 | 0 | 0 | 1 | 0 | 0 | 98 | 49 | 1 | 1 | 0 | NA | NA | NA | 1 | -1 |
| SE | 7447 STX16-NPEPL1 | STX16-NPEPL1 | chr20 | + | 58673630 | 58673711 | 58671153 | 58671297 | 58691723 | 58691780 | 24 | 19 | 3 | 0 | 0 | 0 | 0 | 10 | 0 | 2 | 98 | 49 | 1 | 1 | 1 | 1 | 1 | NA | 0.714 | 0.286 |
| SE | 7448 STX16-NPEPL1 | STX16-NPEPL1 | chr20 | + | 58705497 | 58705569 | 58701015 | 58701158 | 58707122 | 58707200 | 0 | 6 | 1 | 11 | 1 | 0 | 0 | 0 | 0 | 1 | 98 | 49 | 1 | 1 | 0 | 0.75 | 1 | NA | 0 | 0.583 |
| SE | 7458 KIF1B | KIF1B | chr1 | + | 10337370 | 10337533 | 10337073 | 10337203 | 10339768 | 10339859 | 9 | 7 | 6 | 2 | 1 | 0 | 0 | 4 | 0 | 0 | 98 | 49 | 1 | 1 | 0.692 | 0.778 | 1 | NA | 1 | -0.177 |
| SE | 7459 KIF1B | KIF1B | chr1 | + | 10347760 | 10347827 | 10345844 | 10345953 | 10348648 | 10348733 | 8 | 9 | 5 | 0 | 1 | 0 | 0 | 1 | 0 | 0 | 98 | 49 | 1 | 1 | 1 | 0.818 | 1 | NA | 1 | -0.061 |
| SE | 7461 ARID4A | ARID4A | chr14 | + | 58301579 | 58301690 | 58299797 | 58299860 | 58304943 | 58305009 | 10 | 12 | 3 | 0 | 1 | 0 | 0 | 3 | 0 | 0 | 98 | 49 | 1 | 1 | 1 | 0.857 | 1 | NA | 1 | -0.048 |
| SE | 7463 ARID4A | ARID4A | chr14 | + | 58366023 | 58366230 | 58365517 | 58365622 | 58366882 | 58367029 | 13 | 5 | 2 | 3 | 0 | 0 | 0 | 1 | 0 | 0 | 98 | 49 | 1 | 1 | 0.684 | 1 | 1 | NA | 1 | -0.105 |
| SE | 7464 AATF | AATF | chr17 | + | 36986616 | 36986731 | 36953769 | 36953907 | 36988518 | 36988720 | 10 | 8 | 13 | 0 | 0 | 0 | 0 | 2 | 0 | 3 | 98 | 49 | 1 | 1 | 1 | 1 | 1 | NA | 0.25 | 0.75 |
| SE | 7467 VPS13D | VPS13D | chr1 | + | 12319496 | 12319630 | 12318071 | 12318337 | 12321808 | 12321964 | 8 | 2 | 1 | 0 | 0 | 1 | 0 | 2 | 0 | 0 | 98 | 49 | 1 | 1 | 1 | 1 | 0.333 | NA | 1 | -0.222 |

| | | | | | | | | | | | | | | | | | | | | | | | | | | | | | | | |
|---|---|---|---|---|---|---|---|---|---|---|---|---|---|---|---|---|---|---|---|---|---|---|---|---|---|---|---|---|---|---|---|
| SE | 7469 | VPS13D | VPS13D | chr1 | + | 12349163 | 12349374 | 12348822 | 12348973 | 12353973 | 12354221 | 3 | 2 | 0 | 0 | 0 | 0 | 0 | 3 | 0 | 1 | 98 | 49 | 1 | 1 | 1 | 1 | NA | NA | 0.6 | 0.4 |
| SE | 7475 | NRBP1 | NRBP1 | chr2 | + | 27434470 | 27434560 | 27433988 | 27434090 | 27434721 | 27434762 | 39 | 15 | 10 | 0 | 0 | 0 | 0 | 5 | 0 | 1 | 98 | 49 | 1 | 1 | 1 | 1 | 1 | NA | 0.714 | 0.286 |
| SE | 7476 | MDM2 | MDM2 | chr12 | + | 68816811 | 68816945 | 68813553 | 68813628 | 68820324 | 68820374 | 23 | 12 | 9 | 0 | 0 | 0 | 0 | 12 | 0 | 1 | 98 | 49 | 1 | 1 | 1 | 1 | 1 | NA | 0.857 | 0.143 |
| SE | 7477 | MDM2 | MDM2 | chr12 | + | 68828770 | 68828931 | 68824554 | 68824651 | 68835828 | 68835843 | 70 | 21 | 3 | 1 | 0 | 0 | 0 | 8 | 0 | 0 | 98 | 49 | 1 | 1 | 0.972 | 1 | 1 | NA | 1 | -0.009 |
| SE | 7493 | FAM126B | FAM126B | chr2 | - | 200992886 | 200993003 | 200992280 | 200992368 | 200997444 | 200997540 | 85 | 21 | 12 | 1 | 0 | 0 | 0 | 15 | 0 | 2 | 98 | 49 | 1 | 1 | 0.977 | 1 | 1 | NA | 0.789 | 0.203 |
| SE | 7494 | FAM126B | FAM126B | chr2 | - | 200992886 | 200993003 | 200992280 | 200992368 | 201008972 | 201009088 | 39 | 11 | 3 | 1 | 0 | 0 | 0 | 10 | 0 | 0 | 98 | 49 | 1 | 1 | 0.951 | 1 | 1 | NA | 1 | -0.016 |
| SE | 7495 | FAM126B | FAM126B | chr2 | - | 200997444 | 200997540 | 200992280 | 200992368 | 201008972 | 201009088 | 39 | 9 | 6 | 1 | 0 | 0 | 0 | 6 | 0 | 0 | 98 | 49 | 1 | 1 | 0.951 | 1 | 1 | NA | 1 | -0.016 |
| SE | 7496 | FAM126B | FAM126B | chr2 | - | 200997444 | 200997540 | 200992886 | 200993003 | 201008972 | 201009088 | 85 | 19 | 15 | 1 | 0 | 0 | 0 | 9 | 0 | 0 | 98 | 49 | 1 | 1 | 0.977 | 1 | 1 | NA | 1 | -0.008 |
| SE | 7497 | FAM126B | FAM126B | chr2 | - | 201016990 | 201017170 | 201011391 | 201011472 | 201022830 | 201022932 | 47 | 21 | 6 | 1 | 0 | 0 | 1 | 15 | 0 | 0 | 98 | 49 | 1 | 1 | 0.959 | 1 | 1 | 1 | 1 | -0.014 |
| SE | 7498 | SLC12A6 | SLC12A6 | chr15 | - | 34238964 | 34239160 | 34238231 | 34238401 | 34240660 | 34240829 | 71 | 12 | 3 | 0 | 1 | 0 | 1 | 11 | 0 | 0 | 98 | 49 | 1 | 1 | 1 | 0.857 | 1 | 1 | 1 | -0.048 |
| SE | 7499 | SLC12A6 | SLC12A6 | chr15 | - | 34242101 | 34242221 | 34241232 | 34241337 | 34243973 | 34244072 | 67 | 19 | 8 | 7 | 5 | 0 | 0 | 7 | 0 | 1 | 98 | 49 | 1 | 1 | 0.827 | 0.655 | 1 | NA | 0.778 | 0.049 |
| SE | 7500 | SLC12A6 | SLC12A6 | chr15 | - | 34245284 | 34245403 | 34243973 | 34244072 | 34245692 | 34245867 | 82 | 40 | 22 | 2 | 0 | 0 | 0 | 16 | 0 | 0 | 98 | 49 | 1 | 1 | 0.953 | 1 | 1 | NA | 1 | -0.016 |
| SE | 7506 | HAL | HAL | chr12 | - | 95990392 | 95990532 | 95988192 | 95988240 | 95992679 | 95992805 | 69 | 12 | 16 | 2 | 0 | 0 | 0 | 8 | 0 | 0 | 98 | 49 | 1 | 1 | 0.945 | 1 | 1 | NA | 1 | -0.018 |
| SE | 7507 | HAL | HAL | chr12 | - | 95993771 | 95993838 | 95993450 | 95993488 | 95993925 | 95993998 | 28 | 14 | 17 | 9 | 0 | 5 | 0 | 3 | 0 | 0 | 98 | 49 | 1 | 1 | 0.609 | 1 | 0.63 | NA | 1 | -0.254 |
| SE | 7508 | TMEM167B | TMEM167B | chr1 | + | 109092889 | 109093021 | 109090780 | 109090882 | 109094416 | 109096932 | 13 | 9 | 3 | 1 | 0 | 0 | 0 | 2 | 0 | 0 | 98 | 49 | 1 | 1 | 0.867 | 1 | 1 | NA | 1 | -0.044 |
| SE | 7512 | IFFO2 | IFFO2 | chr1 | - | 18918361 | 18918502 | 18916902 | 18917042 | 18919677 | 18919773 | 1 | 0 | 0 | 1 | 0 | 0 | 0 | 2 | 0 | 0 | 98 | 49 | 1 | 1 | 0.333 | NA | NA | NA | 1 | -0.667 |
| SE | 7513 | ZEB2 | ZEB2 | chr2 | - | 144398300 | 144400270 | 144396411 | 144396592 | 144401198 | 144401307 | 14 | 3 | 2 | 2 | 1 | 0 | 0 | 8 | 0 | 1 | 98 | 49 | 1 | 1 | 0.778 | 0.6 | 1 | NA | 0.8 | -0.007 |
| SE | 7514 | ZEB2 | ZEB2 | chr2 | - | 144404835 | 144405024 | 144403915 | 144404130 | 144429768 | 144430026 | 18 | 6 | 0 | 0 | 0 | 0 | 1 | 3 | 0 | 1 | 98 | 49 | 1 | 1 | 1 | 1 | NA | 1 | 0.6 | 0.2 |
| SE | 7515 | ZEB2 | ZEB2 | chr2 | - | 144424795 | 144424867 | 144404835 | 144405024 | 144429768 | 144430026 | 15 | 14 | 0 | 1 | 0 | 0 | 0 | 6 | 0 | 1 | 98 | 49 | 1 | 1 | 0.882 | 1 | NA | NA | 0.75 | 0.191 |
| SE | 7517 | MKL1 | MKL1 | chr22 | - | 40423535 | 40423685 | 40420846 | 40421100 | 40429605 | 40429767 | 5 | 6 | 0 | 0 | 1 | 0 | 0 | 1 | 0 | 0 | 98 | 49 | 1 | 1 | 1 | 0.75 | NA | NA | 1 | -0.125 |
| SE | 7518 | MKL1 | MKL1 | chr22 | - | 40424205 | 40424381 | 40420846 | 40421100 | 40429605 | 40429767 | 10 | 3 | 1 | 0 | 1 | 0 | 2 | 1 | 0 | 0 | 98 | 49 | 1 | 1 | 1 | 0.6 | 1 | 1 | 1 | -0.133 |
| SE | 7520 | MKL1 | MKL1 | chr22 | - | 40594673 | 40594735 | 40552105 | 40552367 | 40636477 | 40636719 | 7 | 4 | 0 | 0 | 0 | 0 | 0 | 4 | 0 | 2 | 98 | 49 | 1 | 1 | 1 | 1 | NA | NA | 0.5 | 0.5 |
| SE | 7521 | EDEM3 | EDEM3 | chr1 | - | 184723750 | 184723856 | 184721288 | 184721386 | 184726254 | 184726389 | 22 | 3 | 0 | 3 | 0 | 0 | 0 | 2 | 0 | 0 | 98 | 49 | 1 | 1 | 0.786 | 1 | NA | NA | 1 | -0.107 |
| SE | 7522 | FAM53B | FAM53B | chr10 | - | 124696157 | 124696212 | 124681606 | 124682379 | 124706635 | 124706887 | 17 | 17 | 5 | 2 | 0 | 0 | 1 | 12 | 0 | 0 | 98 | 49 | 1 | 1 | 0.81 | 1 | 1 | 1 | 1 | -0.063 |
| SE | 7524 | FAM53B | FAM53B | chr10 | - | 124706635 | 124706887 | 124681606 | 124682379 | 124744012 | 124744361 | 12 | 3 | 0 | 0 | 0 | 3 | 0 | 4 | 0 | 0 | 98 | 49 | 1 | 1 | 1 | 1 | 0 | NA | 1 | -0.333 |
| SE | 7525 | FAM53B | FAM53B | chr10 | - | 124706635 | 124706887 | 124696157 | 124696212 | 124744012 | 124744361 | 24 | 14 | 2 | 1 | 0 | 0 | 1 | 16 | 0 | 0 | 98 | 49 | 1 | 1 | 0.923 | 1 | 1 | 1 | 1 | -0.026 |
| SE | 7532 | 6-Mar | 6-Mar | chr5 | + | 10381799 | 10381943 | 10378758 | 10378832 | 10386993 | 10387066 | 13 | 6 | 0 | 1 | 0 | 0 | 0 | 5 | 0 | 0 | 98 | 49 | 1 | 1 | 0.867 | 1 | NA | NA | 1 | -0.067 |
| SE | 7533 | 6-Mar | 6-Mar | chr5 | + | 10386993 | 10387066 | 10378758 | 10378832 | 10390331 | 10390500 | 2 | 4 | 0 | 1 | 0 | 0 | 0 | 2 | 0 | 0 | 98 | 49 | 1 | 1 | 0.5 | 1 | NA | NA | 1 | -0.25 |
| SE | 7534 | 6-Mar | 6-Mar | chr5 | + | 10390331 | 10390500 | 10386993 | 10387066 | 10391541 | 10391731 | 8 | 5 | 0 | 0 | 0 | 0 | 0 | 6 | 0 | 2 | 98 | 49 | 1 | 1 | 1 | 1 | NA | NA | 0.6 | 0.4 |
| SE | 7535 | 6-Mar | 6-Mar | chr5 | + | 10407101 | 10407202 | 10405557 | 10405677 | 10410138 | 10410276 | 38 | 22 | 2 | 0 | 1 | 0 | 0 | 15 | 0 | 0 | 98 | 49 | 1 | 1 | 1 | 0.917 | 1 | NA | 1 | -0.028 |
| SE | 7536 | 6-Mar | 6-Mar | chr5 | + | 10426389 | 10426522 | 10423734 | 10423824 | 10429892 | 10430028 | 31 | 5 | 1 | 4 | 1 | 0 | 0 | 0 | 0 | 1 | 98 | 49 | 1 | 1 | 0.795 | 0.714 | 1 | NA | 0 | 0.836 |
| SE | 7537 | SSFA2 | SSFA2 | chr2 | + | 181896029 | 181896079 | 181892614 | 181892660 | 181896907 | 181896964 | 14 | 27 | 8 | 1 | 0 | 0 | 0 | 5 | 0 | 0 | 98 | 49 | 1 | 1 | 0.875 | 1 | 1 | NA | 1 | -0.042 |
| SE | 7538 | SSFA2 | SSFA2 | chr2 | + | 181920596 | 181920662 | 181919295 | 181919446 | 181921947 | 181922412 | 12 | 6 | 10 | 6 | 7 | 0 | 0 | 3 | 0 | 0 | 98 | 49 | 1 | 1 | 0.5 | 0.3 | 1 | NA | 1 | -0.4 |
| SE | 7542 | SFXN4 | SFXN4 | chr10 | - | 119155061 | 119155177 | 119147774 | 119147860 | 119156677 | 119156756 | 3 | 0 | 0 | 0 | 0 | 0 | 0 | 0 | 0 | 1 | 98 | 49 | 1 | 1 | 1 | NA | NA | NA | 0 | 1 |
| SE | 7545 | HEATR5A | HEATR5A | chr14 | - | 31304904 | 31305177 | 31302294 | 31302519 | 31306731 | 31306879 | 10 | 6 | 2 | 0 | 1 | 0 | 0 | 1 | 0 | 0 | 98 | 49 | 1 | 1 | 1 | 0.75 | 1 | NA | 1 | -0.083 |
| SE | 7547 | HEATR5A | HEATR5A | chr14 | - | 31364188 | 31364298 | 31358893 | 31359057 | 31371809 | 31371909 | 20 | 4 | 9 | 2 | 0 | 0 | 0 | 2 | 0 | 0 | 98 | 49 | 1 | 1 | 0.833 | 1 | 1 | NA | 1 | -0.056 |
| SE | 7549 | HEATR5A | HEATR5A | chr14 | - | 31400300 | 31400512 | 31398672 | 31398781 | 31402849 | 31403049 | 29 | 16 | 3 | 1 | 1 | 0 | 0 | 2 | 0 | 0 | 98 | 49 | 1 | 1 | 0.935 | 0.889 | 1 | NA | 1 | -0.059 |
| SE | 7550 | RFWD2 | RFWD2 | chr1 | - | 176135009 | 176135086 | 176116623 | 176116681 | 176149005 | 176149074 | 147 | 61 | 9 | 2 | 0 | 0 | 1 | 21 | 0 | 1 | 98 | 49 | 1 | 1 | 0.974 | 1 | 1 | 1 | 0.913 | 0.035 |
| SE | 7552 | RFWD2 | RFWD2 | chr1 | - | 176149005 | 176149074 | 176135009 | 176135086 | 176175909 | 176176007 | 30 | 7 | 0 | 0 | 2 | 0 | 0 | 4 | 0 | 0 | 98 | 49 | 1 | 1 | 1 | 0.636 | NA | NA | 1 | -0.182 |
| SE | 7553 | RFWD2 | RFWD2 | chr1 | - | 176149005 | 176149074 | 176136487 | 176136547 | 176184632 | 176184692 | 68 | 13 | 10 | 1 | 0 | 0 | 0 | 6 | 0 | 0 | 98 | 49 | 1 | 1 | 0.971 | 1 | 1 | NA | 1 | -0.01 |
| SE | 7554 | RFWD2 | RFWD2 | chr1 | - | 176162868 | 176162988 | 176149005 | 176149074 | 176163826 | 176163891 | 120 | 61 | 15 | 0 | 2 | 0 | 3 | 17 | 0 | 0 | 98 | 49 | 1 | 1 | 1 | 0.938 | 1 | 1 | 1 | -0.021 |
| SE | 7555 | RFWD2 | RFWD2 | chr1 | - | 176163826 | 176163891 | 176149005 | 176149074 | 176175909 | 176176007 | 74 | 35 | 11 | 1 | 1 | 0 | 2 | 13 | 0 | 0 | 98 | 49 | 1 | 1 | 0.974 | 0.946 | 1 | 1 | 1 | -0.027 |
| SE | 7559 | CDC16 | CDC16 | chr13 | + | 114257077 | 114257230 | 114250548 | 114250674 | 114259334 | 114259398 | 3 | 0 | 0 | 0 | 1 | 0 | 0 | 2 | 0 | 2 | 98 | 49 | 1 | 1 | 1 | 0 | NA | NA | 0.333 | 0.167 |
| SE | 7573 | KCNAB2 | KCNAB2 | chr1 | + | 6041830 | 6041872 | 6040516 | 6040645 | 6072754 | 6072798 | 5 | 5 | 0 | 3 | 1 | 0 | 0 | 1 | 0 | 0 | 91 | 49 | 1 | 1 | 0.473 | 0.729 | NA | NA | 1 | -0.399 |
| SE | 7584 | KIAA1551 | KIAA1551 | chr12 | + | 31960776 | 31960871 | 31959418 | 31959491 | 31970188 | 31970356 | 85 | 65 | 41 | 1 | 0 | 0 | 0 | 22 | 0 | 0 | 98 | 49 | 1 | 1 | 0.977 | 1 | 1 | NA | 1 | -0.008 |
| SE | 7585 | KIAA1551 | KIAA1551 | chr12 | + | 31960776 | 31960871 | 31959418 | 31959491 | 31980877 | 31985957 | 45 | 20 | 11 | 0 | 1 | 0 | 0 | 5 | 0 | 0 | 98 | 49 | 1 | 1 | 1 | 0.909 | 1 | NA | 1 | -0.03 |
| SE | 7586 | KIAA1551 | KIAA1551 | chr12 | + | 31970188 | 31970356 | 31959418 | 31959491 | 31980877 | 31985957 | 204 | 96 | 57 | 0 | 1 | 0 | 0 | 41 | 0 | 0 | 98 | 49 | 1 | 1 | 1 | 0.98 | 1 | NA | 1 | -0.007 |
| SE | 7587 | KIAA1551 | KIAA1551 | chr12 | + | 31970188 | 31970356 | 31960776 | 31960871 | 31980877 | 31985957 | 265 | 145 | 96 | 23 | 4 | 9 | 0 | 59 | 0 | 1 | 98 | 49 | 1 | 1 | 0.852 | 0.948 | 0.842 | NA | 0.967 | -0.086 |
| SE | 7588 | KIAA1551 | KIAA1551 | chr12 | + | 31980877 | 31985957 | 31970188 | 31970356 | 31987238 | 31987322 | 275 | 135 | 73 | 3 | 3 | 2 | 0 | 49 | 0 | 0 | 98 | 49 | 1 | 1 | 0.979 | 0.957 | 0.948 | NA | 1 | -0.039 |
| SE | 7589 | KIAA1551 | KIAA1551 | chr12 | + | 31980877 | 31985957 | 31970188 | 31970356 | 31992377 | 31993107 | 203 | 97 | 57 | 1 | 0 | 0 | 0 | 41 | 0 | 0 | 98 | 49 | 1 | 1 | 0.99 | 1 | 1 | NA | 1 | -0.003 |
| SE | 7590 | KIAA1551 | KIAA1551 | chr12 | + | 31987238 | 31987322 | 31970188 | 31970356 | 31992377 | 31993107 | 69 | 25 | 16 | 1 | 0 | 0 | 0 | 6 | 0 | 0 | 98 | 49 | 1 | 1 | 0.972 | 1 | 1 | NA | 1 | -0.009 |
| SE | 7591 | KIAA1551 | KIAA1551 | chr12 | + | 31987238 | 31987322 | 31980877 | 31985957 | 31992377 | 31993107 | 138 | 61 | 30 | 0 | 1 | 0 | 0 | 14 | 0 | 0 | 98 | 49 | 1 | 1 | 1 | 0.968 | 1 | NA | 1 | -0.011 |
| SE | 7592 | CSNK2A2 | CSNK2A2 | chr16 | - | 58167206 | 58167308 | 58166583 | 58166684 | 58167684 | 58167795 | 1 | 0 | 0 | 0 | 0 | 0 | 1 | 0 | 0 | 1 | 98 | 49 | 1 | 1 | 1 | NA | NA | 1 | 0 | 0.5 |
| SE | 7605 | CPSF2 | CPSF2 | chr14 | + | 92134010 | 92134170 | 92130950 | 92131133 | 92134249 | 92134355 | 12 | 1 | 1 | 0 | 0 | 0 | 0 | 2 | 0 | 3 | 98 | 49 | 1 | 1 | 1 | 1 | 1 | NA | 0.25 | 0.75 |
| SE | 7617 | NDUFA10 | NDUFA10 | chr2 | - | 240022171 | 240022340 | 240021196 | 240021412 | 240025226 | 240025402 | 4 | 3 | 3 | 0 | 1 | 0 | 0 | 3 | 0 | 0 | 98 | 49 | 1 | 1 | 1 | 0.6 | 1 | NA | 1 | -0.133 |
| SE | 7619 | DRAM2 | DRAM2 | chr1 | - | 111124741 | 111124881 | 111120515 | 111120693 | 111126226 | 111126294 | 5 | 1 | 2 | 1 | 0 | 0 | 0 | 6 | 0 | 0 | 98 | 49 | 1 | 1 | 0.714 | 1 | 1 | NA | 1 | -0.095 |
| SE | 7626 | CSDE1 | CSDE1 | chr1 | - | 114719578 | 114719742 | 114718612 | 114718745 | 114720538 | 114720717 | 55 | 29 | 5 | 0 | 1 | 0 | 0 | 2 | 0 | 0 | 98 | 49 | 1 | 1 | 1 | 0.935 | 1 | NA | 1 | -0.022 |
| SE | 7627 | CSDE1 | CSDE1 | chr1 | - | 114725220 | 114725333 | 114723882 | 114724002 | 114726210 | 114726386 | 83 | 40 | 13 | 3 | 1 | 0 | 0 | 22 | 0 | 0 | 98 | 49 | 1 | 1 | 0.933 | 0.952 | 1 | NA | 1 | -0.038 |
| SE | 7628 | CSDE1 | CSDE1 | chr1 | - | 114730257 | 114730422 | 114726982 | 114727090 | 114730507 | 114730648 | 73 | 35 | 18 | 0 | 0 | 0 | 0 | 21 | 0 | 1 | 98 | 49 | 1 | 1 | 1 | 1 | 1 | NA | 0.913 | 0.087 |
| SE | 7629 | CSDE1 | CSDE1 | chr1 | - | 114730507 | 114730648 | 114726982 | 114727090 | 114732603 | 114732816 | 43 | 32 | 14 | 2 | 0 | 0 | 0 | 6 | 0 | 0 | 98 | 49 | 1 | 1 | 0.915 | 1 | 1 | NA | 1 | -0.028 |
| SE | 7630 | CSDE1 | CSDE1 | chr1 | - | 114737470 | 114737563 | 114736757 | 114736855 | 114737962 | 114738072 | 9 | 2 | 0 | 60 | 36 | 2 | 0 | 0 | 0 | 18 | 98 | 49 | 1 | 1 | 0.07 | 0.027 | 0 | NA | 0 | 0.032 |
| SE | 7632 | CSDE1 | CSDE1 | chr1 | - | 114741526 | 114741673 | 114739691 | 114739890 | 114749820 | 114750207 | 2 | 0 | 0 | 71 | 15 | 15 | 0 | 0 | 2 | 17 | 98 | 49 | 1 | 1 | 0.014 | 0 | 0 | 0 | 0 | 0.005 |
| SE | 7633 | CSDE1 | CSDE1 | chr1 | - | 114749820 | 114750207 | 114739691 | 114739890 | 114757924 | 114758050 | 71 | 18 | 15 | 1 | 0 | 0 | 2 | 17 | 0 | 0 | 98 | 49 | 1 | 1 | 0.973 | 1 | 1 | 1 | 1 | -0.009 |
| SE | 7635 | CNNM3 | CNNM3 | chr2 | + | 96828098 | 96828195 | 96827730 | 96827900 | 96828566 | 96828700 | 3 | 0 | 0 | 1 | 0 | 0 | 0 | 1 | 0 | 0 | 98 | 49 | 1 | 1 | 0.6 | NA | NA | NA | 1 | -0.4 |
| SE | 7636 | PTEN | PTEN | chr10 | + | 87925512 | 87925557 | 87894024 | 87894109 | 87931045 | 87931089 | 77 | 45 | 51 | 1 | 0 | 0 | 0 | 17 | 0 | 0 | 94 | 49 | 1 | 1 | 0.976 | 1 | 1 | NA | 1 | -0.008 |
| SE | 7638 | PTEN | PTEN | chr10 | + | 87952117 | 87952259 | 87933012 | 87933251 | 87957852 | 87958019 | 16 | 9 | 10 | 3 | 0 | 2 | 0 | 4 | 0 | 1 | 98 | 49 | 1 | 1 | 0.727 | 1 | 0.714 | NA | 0.667 | 0.147 |
| SE | 7654 | CPSF3L | CPSF3L | chr1 | - | 1314300 | 1314365 | 1313731 | 1313921 | 1314823 | 1314962 | 7 | 2 | 1 | 0 | 1 | 0 | 0 | 1 | 0 | 0 | 98 | 49 | 1 | 1 | 1 | 0.5 | 1 | NA | 1 | -0.167 |
| SE | 7663 | PAPSS1 | PAPSS1 | chr4 | - | 107693770 | 107694006 | 107687038 | 107687177 | 107701170 | 107701285 | 33 | 4 | 10 | 1 | 0 | 0 | 0 | 3 | 0 | 0 | 98 | 49 | 1 | 1 | 0.943 | 1 | 1 | NA | 1 | -0.019 |
| SE | 7664 | TRAM2 | TRAM2 | chr6 | - | 52535782 | 52535846 | 52516627 | 52516737 | 52576795 | 52577064 | 1 | 0 | 0 | 1 | 0 | 0 | 0 | 1 | 0 | 0 | 98 | 49 | 1 | 1 | 0.333 | NA | NA | NA | 1 | -0.667 |
| SE | 7671 | PPP2R3C | PPP2R3C | chr14 | - | 35091069 | 35091207 | 35087950 | 35088010 | 35095047 | 35095184 | 13 | 10 | 4 | 1 | 1 | 0 | 0 | 5 | 0 | 0 | 98 | 49 | 1 | 1 | 0.867 | 0.833 | 1 | NA | 1 | -0.1 |
| SE | 7672 | PPP2R3C | PPP2R3C | chr14 | - | 35095047 | 35095184 | 35091069 | 35091207 | 35096557 | 35096633 | 9 | 5 | 1 | 3 | 0 | 0 | 0 | 6 | 0 | 0 | 98 | 49 | 1 | 1 | 0.6 | 1 | 1 | NA | 1 | -0.133 |
| SE | 7673 | PPP2R3C | PPP2R3C | chr14 | - | 35096708 | 35096764 | 35096557 | 35096633 | 35099251 | 35099384 | 24 | 17 | 6 | 1 | 0 | 0 | 0 | 6 | 0 | 0 | 98 | 49 | 1 | 1 | 0.923 | 1 | 1 | NA | 1 | -0.026 |
| SE | 7674 | PPP2R3C | PPP2R3C | chr14 | - | 35107303 | 35107374 | 35099251 | 35099384 | 35108138 | 35108236 | 33 | 15 | 8 | 1 | 0 | 0 | 0 | 6 | 0 | 0 | 98 | 49 | 1 | 1 | 0.943 | 1 | 1 | NA | 1 | -0.019 |
| SE | 7675 | PPP2R3C | PPP2R3C | chr14 | - | 35116609 | 35116737 | 35110524 | 35110629 | 35121901 | 35122542 | 15 | 6 | 2 | 0 | 0 | 0 | 1 | 4 | 1 | 0 | 98 | 49 | 1 | 1 | 1 | 1 | 1 | 0.333 | 1 | 0.334 |
| SE | 7678 | FNDC3B | FNDC3B | chr3 | + | 172247532 | 172247776 | 172226870 | 172226947 | 172251259 | 172251541 | 35 | 29 | 22 | 1 | 1 | 0 | 0 | 2 | 0 | 0 | 98 | 49 | 1 | 1 | 0.946 | 0.935 | 1 | NA | 1 | -0.04 |
| SE | 7679 | FNDC3B | FNDC3B | chr3 | + | 172247532 | 172247776 | 172226870 | 172226947 | 172285925 | 172285984 | 20 | 6 | 8 | 3 | 2 | 0 | 0 | 1 | 0 | 1 | 98 | 49 | 1 | 1 | 0.769 | 0.6 | 1 | NA | 0.333 | 0.457 |
| SE | 7680 | FNDC3B | FNDC3B | chr3 | + | 172251259 | 172251541 | 172226870 | 172226947 | 172285925 | 172285984 | 18 | 4 | 7 | 3 | 2 | 0 | 0 | 5 | 0 | 1 | 98 | 49 | 1 | 1 | 0.75 | 0.5 | 1 | NA | 0.714 | 0.036 |
| SE | 7681 | FNDC3B | FNDC3B | chr3 | + | 172251259 | 172251541 | 172247532 | 172247776 | 172285925 | 172285984 | 33 | 26 | 21 | 1 | 0 | 0 | 0 | 6 | 0 | 0 | 98 | 49 | 1 | 1 | 0.943 | 1 | 1 | NA | 1 | -0.019 |
| SE | 7682 | FNDC3B | FNDC3B | chr3 | + | 172328951 | 172329076 | 172310827 | 172310881 | 172330540 | 172330715 | 44 | 27 | 6 | 0 | 1 | 0 | 0 | 12 | 0 | 7 | 98 | 49 | 1 | 1 | 1 | 0.931 | 1 | NA | 0.462 | 0.515 |
| SE | 7683 | FNDC3B | FNDC3B | chr3 | + | 172333088 | 172333175 | 172330540 | 172330715 | 172334943 | 172335082 | 82 | 35 | 12 | 0 | 1 | 0 | 2 | 17 | 0 | 0 | 98 | 49 | 1 | 1 | 1 | 0.946 | 1 | 1 | 1 | -0.018 |
| SE | 7684 | FNDC3B | FNDC3B | chr3 | + | 172343010 | 172343116 | 172341112 | 172341231 | 172344085 | 172344258 | 16 | 5 | 6 | 1 | 1 | 0 | 0 | 10 | 0 | 0 | 98 | 49 | 1 | 1 | 0.889 | 0.714 | 1 | NA | 1 | -0.132 |
| SE | 7685 | FNDC3B | FNDC3B | chr3 | + | 172346326 | 172346440 | 172344085 | 172344258 | 172347211 | 172347361 | 36 | 16 | 5 | 0 | 1 | 0 | 1 | 6 | 0 | 0 | 98 | 49 | 1 | 1 | 1 | 0.889 | 1 | 1 | 1 | -0.037 |
| SE | 7686 | FNDC3B | FNDC3B | chr3 | + | 172347211 | 172347361 | 172344085 | 172344258 | 172352802 | 172353083 | 11 | 10 | 4 | 0 | 1 | 0 | 0 | 6 | 0 | 0 | 98 | 49 | 1 | 1 | 1 | 0.833 | 1 | NA | 1 | -0.056 |
| SE | 7687 | FNDC3B | FNDC3B | chr3 | + | 172352802 | 172353083 | 172347211 | 172347361 | 172362632 | 172362845 | 27 | 13 | 14 | 1 | 0 | 0 | 0 | 13 | 0 | 0 | 98 | 49 | 1 | 1 | 0.931 | 1 | 1 | NA | 1 | -0.023 |
| SE | 7696 | TREM1 | TREM1 | chr6 | - | 41280960 | 41281153 | 41275260 | 41276230 | 41282394 | 41282751 | 176 | 96 | 73 | 10 | 10 | 0 | 0 | 37 | 0 | 1 | 98 | 49 | 1 | 1 | 0.898 | 0.828 | 1 | NA | 0.949 | -0.04 |
| SE | 7700 | IKBIP | IKBIP | chr12 | - | 98634295 | 98634413 | 98613403 | 98614340 | 98644522 | 98645051 | 25 | 9 | 3 | 1 | 2 | 1 | 0 | 1 | 0 | 3 | 98 | 49 | 1 | 1 | 0.926 | 0.692 | 0.6 | NA | 0.143 | 0.596 |
| SE | 7704 | TBXAS1 | TBXAS1 | chr7 | + | 139911224 | 139911321 | 139875584 | 139875637 | 139936190 | 139936307 | 93 | 77 | 31 | 0 | 2 | 0 | 2 | 22 | 0 | 0 | 98 | 49 | 1 | 1 | 1 | 0.951 | 1 | 1 | 1 | -0.016 |
| SE | 7707 | TBXAS1 | TBXAS1 | chr7 | + | 139955458 | 139955607 | 139953367 | 139953456 | 139957633 | 139957764 | 65 | 38 | 25 | 1 | 0 | 0 | 1 | 15 | 0 | 0 | 98 | 49 | 1 | 1 | 0.97 | 1 | 1 | 1 | 1 | -0.01 |
| SE | 7708 | TBXAS1 | TBXAS1 | chr7 | + | 139957633 | 139957764 | 139955458 | 139955607 | 139961918 | 139962233 | 75 | 29 | 32 | 0 | 0 | 0 | 3 | 22 | 1 | 0 | 98 | 49 | 1 | 1 | 1 | 1 | 1 | 0.6 | 1 | 0.2 |
| SE | 7709 | TBXAS1 | TBXAS1 | chr7 | + | 140017670 | 140017833 | 140015722 | 140015860 | 140020024 | 140020325 | 18 | 34 | 7 | 0 | 2 | 0 | 0 | 14 | 0 | 1 | 98 | 49 | 1 | 1 | 1 | 0.895 | 1 | NA | 0.875 | 0.09 |
| SE | 7712 | MDM4 | MDM4 | chr1 | + | 204537458 | 204537497 | 204532190 | 204532246 | 204538208 | 204538308 | 11 | 9 | 1 | 2 | 2 | 0 | 0 | 1 | 0 | 0 | 88 | 49 | 1 | 1 | 0.754 | 0.715 | 1 | NA | 1 | -0.177 |
| SE | 7714 | MDM4 | MDM4 | chr1 | + | 204542783 | 204542944 | 204538208 | 204538308 | 204546796 | 204546877 | 13 | 11 | 1 | 2 | 0 | 0 | 0 | 2 | 0 | 0 | 98 | 49 | 1 | 1 | 0.765 | 1 | 1 | NA | 1 | -0.078 |
| SE | 7715 | MDM4 | MDM4 | chr1 | + | 204544534 | 204544684 | 204542783 | 204542944 | 204546796 | 204546877 | 35 | 27 | 1 | 0 | 0 | 0 | 0 | 3 | 0 | 1 | 98 | 49 | 1 | 1 | 1 | 1 | 1 | NA | 0.6 | 0.4 |
| SE | 7717 | CCDC88B | CCDC88B | chr11 | + | 64343996 | 64345157 | 64343777 | 64343914 | 64349330 | 64349458 | 35 | 12 | 6 | 1 | 0 | 0 | 4 | 4 | 0 | 0 | 98 | 49 | 1 | 1 | 0.946 | 1 | 1 | 1 | 1 | -0.018 |
| SE | 7718 | CCDC88B | CCDC88B | chr11 | + | 64351159 | 64351255 | 64349550 | 64349668 | 64351475 | 64351616 | 12 | 3 | 4 | 2 | 0 | 0 | 0 | 0 | 0 | 1 | 98 | 49 | 1 | 1 | 0.75 | 1 | 1 | NA | 0 | 0.917 |
| SE | 7724 | IFI16 | IFI16 | chr1 | + | 159053532 | 159053724 | 159051678 | 159052098 | 159054820 | 159055155 | 48 | 18 | 17 | 8 | 3 | 0 | 0 | 18 | 0 | 0 | 98 | 49 | 1 | 1 | 0.75 | 0.75 | 1 | NA | 1 | -0.167 |
| SE | 7749 | ACAP1 | ACAP1 | chr17 | + | 7349944 | 7350054 | 7348994 | 7349167 | 7350126 | 7350237 | 21 | 14 | 12 | 1 | 0 | 0 | 5 | 14 | 0 | 0 | 98 | 49 | 1 | 1 | 0.913 | 1 | 1 | 1 | 1 | -0.029 |
| SE | 7761 | PLCB3 | PLCB3 | chr11 | + | 64264020 | 64264112 | 64263690 | 64263795 | 64264950 | 64265104 | 0 | 0 | 0 | 1 | 0 | 0 | 0 | 2 | 0 | 0 | 98 | 49 | 1 | 1 | 0 | NA | NA | NA | 1 | -1 |
| SE | 7773 | APH1A | APH1A | chr1 | - | 150267956 | 150267991 | 150267715 | 150267789 | 150268697 | 150269209 | 7 | 6 | 1 | 2 | 0 | 0 | 8 | 4 | 0 | 0 | 84 | 49 | 1 | 1 | 0.671 | 1 | 1 | 1 | 1 | -0.11 |
| SE | 7775 | APH1A | APH1A | chr1 | - | 150267956 | 150268127 | 150267715 | 150267789 | 150268697 | 150269209 | 14 | 6 | 0 | 2 | 0 | 0 | 8 | 7 | 0 | 0 | 98 | 49 | 1 | 1 | 0.778 | 1 | NA | 1 | 1 | -0.111 |
| SE | 7779 | CYLD | CYLD | chr16 | + | 50744729 | 50744866 | 50742761 | 50742841 | 50749575 | 50750202 | 3 | 3 | 0 | 15 | 2 | 0 | 0 | 0 | 2 | 1 | 98 | 49 | 1 | 1 | 0.091 | 0.429 | NA | 0 | 0 | 0.26 |
| SE | 7780 | CYLD | CYLD | chr16 | + | 50776178 | 50776277 | 50754318 | 50754424 | 50777824 | 50777941 | 117 | 44 | 15 | 1 | 0 | 0 | 1 | 7 | 0 | 0 | 98 | 49 | 1 | 1 | 0.983 | 1 | 1 | 1 | 1 | -0.006 |
| SE | 7783 | RANBP2 | RANBP2 | chr2 | + | 108762100 | 108762195 | 108758412 | 108758548 | 108763236 | 108768388 | 2 | 0 | 0 | 0 | 2 | 0 | 0 | 1 | 0 | 0 | 98 | 49 | 1 | 1 | 1 | 0 | NA | NA | 1 | -0.5 |
| SE | 7784 | RANBP2 | RANBP2 | chr2 | + | 108775731 | 108775936 | 108772867 | 108773046 | 108777129 | 108777231 | 15 | 2 | 0 | 1 | 0 | 0 | 0 | 1 | 0 | 0 | 98 | 49 | 1 | 1 | 0.882 | 1 | NA | NA | 1 | -0.059 |
| SE | 7786 | WWP2 | WWP2 | chr16 | + | 69842023 | 69842120 | 69840125 | 69840263 | 69871803 | 69871931 | 58 | 20 | 9 | 0 | 2 | 0 | 1 | 4 | 0 | 0 | 98 | 49 | 1 | 1 | 1 | 0.833 | 1 | 1 | 1 | -0.056 |
| SE | 7787 | WWP2 | WWP2 | chr16 | + | 69917708 | 69917883 | 69908760 | 69908850 | 69925429 | 69925484 | 28 | 11 | 3 | 1 | 0 | 0 | 0 | 1 | 0 | 0 | 98 | 49 | 1 | 1 | 0.933 | 1 | 1 | NA | 1 | -0.022 |
| SE | 7789 | CC2D1B | CC2D1B | chr1 | - | 52359704 | 52359883 | 52359458 | 52359534 | 52360073 | 52360233 | 4 | 0 | 0 | 0 | 0 | 0 | 0 | 1 | 0 | 1 | 98 | 49 | 1 | 1 | 1 | NA | NA | NA | 0.333 | 0.667 |
| SE | 7794 | UBE2L3 | UBE2L3 | chr22 | + | 21614556 | 21614665 | 21610856 | 21611043 | 21621514 | 21624034 | 4 | 3 | 0 | 0 | 0 | 2 | 0 | 1 | 0 | 0 | 98 | 49 | 1 | 1 | 1 | 1 | 0 | NA | 1 | -0.333 |
| SE | 7799 | SLC22A15 | SLC22A15 | chr1 | + | 116026892 | 116027022 | 116020720 | 116020885 | 116031365 | 116031581 | 9 | 3 | 1 | 2 | 0 | 0 | 0 | 3 | 0 | 0 | 98 | 49 | 1 | 1 | 0.692 | 1 | 1 | NA | 1 | -0.103 |
| SE | 7800 | GS1-124K5.11 | GS1-124K5.11 | chr7 | - | 66560578 | 66560675 | 66553467 | 66554837 | 66573384 | 66573550 | 0 | 1 | 0 | 5 | 0 | 0 | 0 | 1 | 1 | 0 | 98 | 49 | 1 | 1 | 0 | 1 | NA | 0 | 1 | 0 |
| SE | 7806 | UBE4B | UBE4B | chr1 | + | 10106196 | 10106583 | 10105515 | 10105744 | 10117458 | 10117600 | 1 | 0 | 0 | 4 | 1 | 3 | 0 | 0 | 0 | 2 | 98 | 49 | 1 | 1 | 0.111 | 0 | 0 | NA | 0 | 0.037 |
| SE | 7807 | UBE4B | UBE4B | chr1 | + | 10121961 | 10122076 | 10119512 | 10119613 | 10126793 | 10126877 | 17 | 7 | 4 | 0 | 1 | 0 | 1 | 9 | 0 | 0 | 98 | 49 | 1 | 1 | 1 | 0.778 | 1 | 1 | 1 | -0.074 |
| SE | 7808 | UBE4B | UBE4B | chr1 | + | 10130499 | 10130616 | 10129391 | 10129448 | 10130714 | 10130813 | 24 | 10 | 4 | 1 | 0 | 0 | 0 | 2 | 0 | 0 | 98 | 49 | 1 | 1 | 0.923 | 1 | 1 | NA | 1 | -0.026 |
| SE | 7809 | UBE4B | UBE4B | chr1 | + | 10137066 | 10137205 | 10134987 | 10135186 | 10144939 | 10145039 | 20 | 7 | 2 | 1 | 0 | 0 | 0 | 7 | 0 | 0 | 98 | 49 | 1 | 1 | 0.909 | 1 | 1 | NA | 1 | -0.03 |
| SE | 7810 | UBE4B | UBE4B | chr1 | + | 10158355 | 10158482 | 10151325 | 10151561 | 10161141 | 10161286 | 24 | 3 | 3 | 0 | 0 | 0 | 0 | 0 | 0 | 1 | 98 | 49 | 1 | 1 | 1 | 1 | 1 | NA | 0 | 1 |
| SE | 7811 | NDUFS2 | NDUFS2 | chr1 | + | 161210110 | 161210188 | 161209856 | 161209931 | 161210303 | 161210389 | 22 | 1 | 4 | 0 | 1 | 0 | 3 | 9 | 0 | 0 | 98 | 49 | 1 | 1 | 1 | 0.333 | 1 | 1 | 1 | -0.222 |
| SE | 7818 | SELL | SELL | chr1 | - | 169696534 | 169696553 | 169690664 | 169691802 | 169701559 | 169701688 | 275 | 185 | 116 | 1 | 2 | 0 | 0 | 82 | 0 | 0 | 68 | 49 | 1 | 1 | 0.995 | 0.985 | 1 | NA | 1 | -0.007 |
| SE | 7819 | SELL | SELL | chr1 | - | 169696534 | 169696553 | 169690664 | 169691802 | 169703254 | 169703440 | 38 | 26 | 7 | 1 | 0 | 0 | 1 | 13 | 0 | 0 | 68 | 49 | 1 | 1 | 0.965 | 1 | 1 | 1 | 1 | -0.012 |
| SE | 7820 | SELL | SELL | chr1 | - | 169701559 | 169701688 | 169690664 | 169691802 | 169703254 | 169703440 | 124 | 55 | 47 | 1 | 0 | 0 | 2 | 26 | 0 | 0 | 98 | 49 | 1 | 1 | 0.984 | 1 | 1 | 1 | 1 | -0.005 |
| SE | 7821 | SELL | SELL | chr1 | - | 169701559 | 169701688 | 169696534 | 169696553 | 169703254 | 169703440 | 366 | 212 | 157 | 5 | 0 | 0 | 2 | 95 | 1 | 0 | 98 | 49 | 1 | 1 | 0.973 | 1 | 1 | 0.5 | 1 | 0.241 |
| SE | 7823 | CD58 | CD58 | chr1 | - | 116519230 | 116519267 | 116514533 | 116514822 | 116521905 | 116521983 | 20 | 16 | 3 | 2 | 0 | 0 | 0 | 1 | 0 | 0 | 86 | 49 | 1 | 1 | 0.851 | 1 | 1 | NA | 1 | -0.05 |
| SE | 7826 | ZNF687 | ZNF687 | chr1 | + | 151288527 | 151288706 | 151286274 | 151288406 | 151289094 | 151289271 | 7 | 0 | 4 | 0 | 0 | 2 | 0 | 1 | 0 | 0 | 98 | 49 | 1 | 1 | 1 | NA | 0.5 | NA | 1 | -0.25 |
| SE | 7828 | ZNF687 | ZNF687 | chr1 | + | 151289094 | 151289271 | 151286274 | 151288406 | 151289377 | 151289540 | 6 | 0 | 2 | 1 | 0 | 0 | 0 | 2 | 0 | 0 | 98 | 49 | 1 | 1 | 0.75 | NA | 1 | NA | 1 | -0.125 |
| SE | 7829 | ZNF687 | ZNF687 | chr1 | + | 151289094 | 151289271 | 151288527 | 151288706 | 151289377 | 151289540 | 9 | 0 | 0 | 0 | 1 | 1 | 0 | 3 | 0 | 0 | 98 | 49 | 1 | 1 | 1 | 0 | 0 | NA | 1 | -0.667 |
| SE | 7834 | CD44 | CD44 | chr11 | + | 35209964 | 35210054 | 35208104 | 35208206 | 35211245 | 35211449 | 4 | 1 | 0 | 1 | 0 | 0 | 0 | 4 | 0 | 0 | 98 | 49 | 1 | 1 | 0.667 | 1 | NA | NA | 1 | -0.166 |
| SE | 7835 | CD44 | CD44 | chr11 | + | 35211245 | 35211449 | 35189834 | 35190065 | 35214851 | 35214914 | 5 | 0 | 0 | 20 | 3 | 6 | 0 | 1 | 0 | 3 | 98 | 49 | 1 | 1 | 0.111 | 0 | 0 | NA | 0.143 | -0.106 |
| SE | 7836 | CD44 | CD44 | chr11 | + | 35211245 | 35211449 | 35209964 | 35210054 | 35214851 | 35214914 | 6 | 1 | 0 | 2 | 0 | 0 | 0 | 1 | 0 | 0 | 98 | 49 | 1 | 1 | 0.6 | 1 | NA | NA | 1 | -0.2 |
| SE | 7841 | LRRK2 | LRRK2 | chr12 | + | 40240482 | 40240617 | 40237968 | 40238103 | 40243549 | 40243681 | 732 | 253 | 82 | 0 | 1 | 0 | 5 | 60 | 0 | 0 | 98 | 49 | 1 | 1 | 1 | 0.992 | 1 | 1 | 1 | -0.003 |
| SE | 7843 | LRRK2 | LRRK2 | chr12 | + | 40257247 | 40257377 | 40252909 | 40253016 | 40259479 | 40259604 | 496 | 207 | 63 | 5 | 0 | 0 | 7 | 46 | 0 | 0 | 98 | 49 | 1 | 1 | 0.98 | 1 | 1 | 1 | 1 | -0.007 |
| SE | 7844 | LRRK2 | LRRK2 | chr12 | + | 40257247 | 40257377 | 40252909 | 40253016 | 40263788 | 40263901 | 175 | 69 | 34 | 2 | 0 | 0 | 2 | 19 | 0 | 1 | 98 | 49 | 1 | 1 | 0.978 | 1 | 1 | 1 | 0.905 | 0.04 |
| SE | 7845 | LRRK2 | LRRK2 | chr12 | + | 40259479 | 40259604 | 40252909 | 40253016 | 40263788 | 40263901 | 372 | 93 | 25 | 2 | 0 | 0 | 2 | 27 | 0 | 1 | 98 | 49 | 1 | 1 | 0.989 | 1 | 1 | 1 | 0.931 | 0.031 |
| SE | 7846 | LRRK2 | LRRK2 | chr12 | + | 40259479 | 40259604 | 40257247 | 40257377 | 40263788 | 40263901 | 689 | 231 | 54 | 1 | 0 | 0 | 7 | 54 | 0 | 0 | 98 | 49 | 1 | 1 | 0.997 | 1 | 1 | 1 | 1 | -0.001 |
| SE | 7847 | LRRK2 | LRRK2 | chr12 | + | 40278090 | 40278261 | 40277887 | 40278016 | 40283874 | 40284133 | 367 | 182 | 77 | 0 | 1 | 0 | 0 | 39 | 0 | 0 | 98 | 49 | 1 | 1 | 1 | 0.989 | 1 | NA | 1 | -0.004 |
| SE | 7848 | LRRK2 | LRRK2 | chr12 | + | 40283874 | 40284133 | 40278090 | 40278261 | 40287350 | 40287539 | 445 | 186 | 61 | 3 | 2 | 0 | 0 | 46 | 0 | 0 | 98 | 49 | 1 | 1 | 0.987 | 0.979 | 1 | NA | 1 | -0.011 |
| SE | 7850 | LRRK2 | LRRK2 | chr12 | + | 40295426 | 40295644 | 40294844 | 40294914 | 40298242 | 40298493 | 504 | 133 | 63 | 0 | 0 | 1 | 8 | 74 | 0 | 0 | 98 | 49 | 1 | 1 | 1 | 1 | 0.969 | 1 | 1 | -0.01 |
| SE | 7852 | LRRK2 | LRRK2 | chr12 | + | 40298242 | 40298493 | 40295426 | 40295644 | 40302788 | 40302882 | 192 | 59 | 33 | 2 | 2 | 0 | 0 | 32 | 0 | 1 | 98 | 49 | 1 | 1 | 0.98 | 0.937 | 1 | NA | 0.941 | 0.031 |
| SE | 7853 | LRRK2 | LRRK2 | chr12 | + | 40299108 | 40299257 | 40295426 | 40295644 | 40302788 | 40302882 | 390 | 124 | 39 | 2 | 2 | 0 | 2 | 64 | 0 | 1 | 98 | 49 | 1 | 1 | 0.99 | 0.969 | 1 | 1 | 0.97 | 0.001 |
| SE | 7854 | LRRK2 | LRRK2 | chr12 | + | 40299108 | 40299257 | 40298242 | 40298493 | 40302788 | 40302882 | 598 | 167 | 57 | 0 | 3 | 0 | 5 | 93 | 0 | 0 | 98 | 49 | 1 | 1 | 1 | 0.965 | 1 | 1 | 1 | -0.012 |
| SE | 7855 | LRRK2 | LRRK2 | chr12 | + | 40303947 | 40304134 | 40302788 | 40302882 | 40305784 | 40305966 | 405 | 135 | 63 | 1 | 0 | 1 | 2 | 47 | 0 | 0 | 98 | 49 | 1 | 1 | 0.995 | 1 | 0.969 | 1 | 1 | -0.012 |
| SE | 7856 | LRRK2 | LRRK2 | chr12 | + | 40303947 | 40304134 | 40302788 | 40302882 | 40308466 | 40308696 | 208 | 58 | 33 | 1 | 0 | 0 | 1 | 29 | 0 | 0 | 98 | 49 | 1 | 1 | 0.99 | 1 | 1 | 1 | 1 | -0.003 |
| SE | 7857 | LRRK2 | LRRK2 | chr12 | + | 40305784 | 40305966 | 40302788 | 40302882 | 40308466 | 40308696 | 91 | 86 | 28 | 1 | 0 | 0 | 1 | 14 | 0 | 0 | 98 | 49 | 1 | 1 | 0.978 | 1 | 1 | 1 | 1 | -0.007 |
| SE | 7859 | LRRK2 | LRRK2 | chr12 | + | 40308466 | 40308696 | 40305784 | 40305966 | 40309105 | 40309233 | 365 | 203 | 48 | 3 | 0 | 0 | 1 | 48 | 0 | 0 | 98 | 49 | 1 | 1 | 0.984 | 1 | 1 | 1 | 1 | -0.005 |
| SE | 7860 | LRRK2 | LRRK2 | chr12 | + | 40310430 | 40310649 | 40309105 | 40309233 | 40315211 | 40315300 | 161 | 70 | 29 | 3 | 0 | 0 | 0 | 24 | 0 | 0 | 98 | 49 | 1 | 1 | 0.964 | 1 | 1 | NA | 1 | -0.012 |
| SE | 7864 | LRRK2 | LRRK2 | chr12 | + | 40322318 | 40322510 | 40321033 | 40321188 | 40323159 | 40323306 | 237 | 64 | 26 | 0 | 0 | 1 | 6 | 37 | 0 | 0 | 98 | 49 | 1 | 1 | 1 | 1 | 0.929 | 1 | 1 | -0.024 |
| SE | 7865 | LRRK2 | LRRK2 | chr12 | + | 40340293 | 40340454 | 40334966 | 40335157 | 40346752 | 40346923 | 352 | 103 | 38 | 3 | 0 | 1 | 5 | 39 | 0 | 0 | 98 | 49 | 1 | 1 | 0.983 | 1 | 0.95 | 1 | 1 | -0.022 |
| SE | 7866 | LRRK2 | LRRK2 | chr12 | + | 40346752 | 40346923 | 40334966 | 40335157 | 40348408 | 40348509 | 146 | 58 | 53 | 1 | 0 | 0 | 1 | 17 | 0 | 0 | 98 | 49 | 1 | 1 | 0.986 | 1 | 1 | 1 | 1 | -0.005 |
| SE | 7867 | LRRK2 | LRRK2 | chr12 | + | 40348408 | 40348509 | 40346752 | 40346923 | 40351538 | 40351733 | 306 | 120 | 73 | 2 | 1 | 0 | 4 | 30 | 0 | 0 | 98 | 49 | 1 | 1 | 0.987 | 0.984 | 1 | 1 | 1 | -0.01 |
| SE | 7868 | LRRK2 | LRRK2 | chr12 | + | 40351538 | 40351733 | 40348408 | 40348509 | 40354298 | 40354492 | 244 | 99 | 31 | 1 | 0 | 0 | 3 | 25 | 0 | 0 | 98 | 49 | 1 | 1 | 0.992 | 1 | 1 | 1 | 1 | -0.003 |
| SE | 7870 | LRRK2 | LRRK2 | chr12 | + | 40367005 | 40367077 | 40364841 | 40365050 | 40367643 | 40369284 | 176 | 97 | 43 | 15 | 4 | 0 | 0 | 13 | 0 | 3 | 98 | 49 | 1 | 1 | 0.854 | 0.924 | 1 | NA | 0.684 | 0.242 |
| SE | 7875 | PAK1 | PAK1 | chr11 | - | 77332729 | 77332867 | 77322014 | 77323360 | 77336085 | 77336282 | 68 | 38 | 15 | 0 | 0 | 0 | 2 | 22 | 1 | 0 | 98 | 49 | 1 | 1 | 1 | 1 | 1 | 0.5 | 1 | 0.25 |
| SE | 7876 | PAK1 | PAK1 | chr11 | - | 77337323 | 77337423 | 77336085 | 77336282 | 77340645 | 77340763 | 47 | 32 | 22 | 1 | 1 | 0 | 0 | 12 | 0 | 0 | 98 | 49 | 1 | 1 | 0.959 | 0.941 | 1 | NA | 1 | -0.033 |
| SE | 7877 | PAK1 | PAK1 | chr11 | - | 77379240 | 77379388 | 77374327 | 77374365 | 77379893 | 77379994 | 55 | 23 | 22 | 0 | 1 | 0 | 0 | 15 | 0 | 0 | 98 | 49 | 1 | 1 | 1 | 0.92 | 1 | NA | 1 | -0.027 |
| SE | 7888 | BIRC2 | BIRC2 | chr11 | + | 102350843 | 102350943 | 102348597 | 102350749 | 102362895 | 102362974 | 17 | 8 | 4 | 1 | 0 | 0 | 0 | 4 | 0 | 0 | 98 | 49 | 1 | 1 | 0.895 | 1 | 1 | NA | 1 | -0.035 |
| SE | 7890 | CUBN | CUBN | chr10 | - | 16836234 | 16836382 | 16835013 | 16835195 | 16840329 | 16840535 | 0 | 0 | 1 | 0 | 0 | 0 | 0 | 0 | 0 | 1 | 98 | 49 | 1 | 1 | NA | NA | 1 | NA | 0 | 1 |
| SE | 7904 | FOXJ3 | FOXJ3 | chr1 | - | 42199101 | 42199230 | 42194889 | 42195064 | 42205761 | 42205863 | 9 | 0 | 0 | 0 | 0 | 0 | 0 | 0 | 0 | 1 | 98 | 49 | 1 | 1 | 1 | NA | NA | NA | 0 | 1 |

| | | | | | | | | | | | | | | | | | | | | | | | | | | | | | | | |
|---|---|---|---|---|---|---|---|---|---|---|---|---|---|---|---|---|---|---|---|---|---|---|---|---|---|---|---|---|---|---|---|
| SE | 7906 | GLG1 | GLG1 | chr16 | - | 74468945 | 74469063 | 74467755 | 74467848 | 74469984 | 74470073 | 20 | 1 | 4 | 1 | 0 | 0 | 0 | 5 | 0 | 0 | 98 | 49 | 1 | 1 | 0.909 | 1 | 1 | NA | 1 | -0.03 |
| SE | 7918 | EPB41 | EPB41 | chr1 | + | 28989355 | 28989435 | 28987430 | 28987905 | 28993329 | 28993542 | 3 | 3 | 0 | 45 | 17 | 5 | 0 | 0 | 0 | 2 | 98 | 49 | 1 | 1 | 0.032 | 0.081 | 0 | NA | 0 | 0.038 |
| SE | 7919 | EPB41 | EPB41 | chr1 | + | 28993329 | 28993542 | 28987430 | 28987905 | 28997214 | 28997319 | 90 | 37 | 10 | 0 | 1 | 0 | 0 | 3 | 0 | 0 | 98 | 49 | 1 | 1 | 1 | 0.949 | 1 | NA | 1 | -0.017 |
| SE | 7920 | EPB41 | EPB41 | chr1 | + | 28993329 | 28993542 | 28987430 | 28987905 | 29011864 | 29011907 | 46 | 19 | 5 | 6 | 2 | 0 | 0 | 2 | 0 | 0 | 98 | 49 | 1 | 1 | 0.793 | 0.826 | 1 | NA | 1 | -0.127 |
| SE | 7922 | EPB41 | EPB41 | chr1 | + | 28997214 | 28997319 | 28993329 | 28993542 | 29011864 | 29011907 | 72 | 26 | 12 | 1 | 2 | 0 | 0 | 1 | 0 | 0 | 98 | 49 | 1 | 1 | 0.973 | 0.867 | 1 | NA | 1 | -0.053 |
| SE | 7924 | EPB41 | EPB41 | chr1 | + | 29058588 | 29058645 | 29053103 | 29053312 | 29064981 | 29065158 | 0 | 2 | 0 | 21 | 11 | 11 | 0 | 1 | 2 | 10 | 98 | 49 | 1 | 1 | 0 | 0.083 | 0 | 0 | 0.048 | 0.004 |
| SE | 7928 | EPB41 | EPB41 | chr1 | + | 29097806 | 29097935 | 29064981 | 29065158 | 29109335 | 29109437 | 64 | 23 | 19 | 2 | 0 | 0 | 0 | 16 | 0 | 0 | 98 | 49 | 1 | 1 | 0.941 | 1 | 1 | NA | 1 | -0.02 |
| SE | 7944 | DHRS7 | DHRS7 | chr14 | - | 60150064 | 60150187 | 60149352 | 60149568 | 60152938 | 60153178 | 70 | 52 | 39 | 2 | 0 | 0 | 2 | 13 | 0 | 0 | 98 | 49 | 1 | 1 | 0.946 | 1 | 1 | 1 | 1 | -0.018 |
| SE | 7946 | CUL4A | CUL4A | chr13 | + | 113239432 | 113239551 | 113236822 | 113236890 | 113242967 | 113243160 | 7 | 2 | 0 | 0 | 0 | 0 | 0 | 5 | 0 | 1 | 98 | 49 | 1 | 1 | 1 | 1 | NA | NA | 0.714 | 0.286 |
| SE | 7951 | RSRC2 | RSRC2 | chr12 | - | 122520503 | 122520581 | 122518838 | 122519029 | 122521384 | 122521428 | 3 | 2 | 0 | 3 | 0 | 0 | 1 | 3 | 0 | 1 | 98 | 49 | 1 | 1 | 0.333 | 1 | NA | 1 | 0.6 | -0.134 |
| SE | 7961 | TNRC6B | TNRC6B | chr22 | + | 40270121 | 40270280 | 40264687 | 40267036 | 40273424 | 40273600 | 7 | 5 | 2 | 5 | 4 | 0 | 0 | 1 | 0 | 0 | 98 | 49 | 1 | 1 | 0.412 | 0.385 | 1 | NA | 1 | -0.401 |
| SE | 7971 | LCOR | LCOR | chr10 | + | 96833405 | 96833479 | 96832259 | 96832399 | 96907264 | 96907330 | 10 | 1 | 0 | 2 | 0 | 0 | 1 | 1 | 0 | 0 | 98 | 49 | 1 | 1 | 0.714 | 1 | NA | 1 | 1 | -0.143 |
| SE | 7975 | KHDRBS1 | KHDRBS1 | chr1 | + | 32031523 | 32031640 | 32030297 | 32030422 | 32033187 | 32033334 | 8 | 3 | 0 | 3 | 1 | 0 | 0 | 4 | 0 | 0 | 98 | 49 | 1 | 1 | 0.571 | 0.6 | NA | NA | 1 | -0.415 |
| SE | 7977 | TXLNA | TXLNA | chr1 | + | 32184524 | 32184616 | 32181241 | 32181577 | 32187953 | 32188124 | 2 | 0 | 0 | 0 | 0 | 0 | 0 | 1 | 0 | 1 | 98 | 49 | 1 | 1 | 1 | NA | NA | NA | 0.333 | 0.667 |
| SE | 7978 | MAN2A1 | MAN2A1 | chr5 | + | 109729341 | 109729513 | 109716119 | 109716264 | 109755328 | 109755456 | 17 | 2 | 0 | 1 | 1 | 0 | 0 | 3 | 0 | 0 | 98 | 49 | 1 | 1 | 0.895 | 0.5 | NA | NA | 1 | -0.302 |
| SE | 7979 | MAN2A1 | MAN2A1 | chr5 | + | 109817272 | 109817438 | 109789459 | 109789527 | 109819668 | 109819887 | 7 | 1 | 1 | 2 | 0 | 0 | 0 | 3 | 0 | 0 | 98 | 49 | 1 | 1 | 0.636 | 1 | 1 | NA | 1 | -0.121 |
| SE | 7980 | MAN2A1 | MAN2A1 | chr5 | + | 109820219 | 109820342 | 109819668 | 109819887 | 109823722 | 109823837 | 10 | 7 | 0 | 1 | 0 | 0 | 0 | 5 | 0 | 0 | 98 | 49 | 1 | 1 | 0.833 | 1 | NA | NA | 1 | -0.084 |
| SE | 7981 | MAN2A1 | MAN2A1 | chr5 | + | 109845864 | 109846006 | 109842327 | 109842461 | 109847656 | 109847790 | 3 | 2 | 0 | 0 | 3 | 0 | 0 | 4 | 0 | 0 | 98 | 49 | 1 | 1 | 1 | 0.25 | NA | NA | 1 | -0.375 |
| SE | 7984 | PTPN18 | PTPN18 | chr2 | + | 130372872 | 130372947 | 130372256 | 130372483 | 130373156 | 130375409 | 0 | 0 | 0 | 1 | 0 | 0 | 0 | 2 | 0 | 0 | 98 | 49 | 1 | 1 | 0 | NA | NA | NA | 1 | -1 |
| SE | 7985 | PAFAH1B2 | PAFAH1B2 | chr11 | + | 117152440 | 117152528 | 117144283 | 117144418 | 117159933 | 117160023 | 20 | 23 | 3 | 2 | 1 | 0 | 0 | 4 | 0 | 0 | 98 | 49 | 1 | 1 | 0.833 | 0.92 | 1 | NA | 1 | -0.082 |
| SE | 7987 | PAFAH1B2 | PAFAH1B2 | chr11 | + | 117161144 | 117161261 | 117159933 | 117160023 | 117163769 | 117163874 | 32 | 32 | 6 | 0 | 0 | 0 | 0 | 6 | 0 | 2 | 98 | 49 | 1 | 1 | 1 | 1 | 1 | NA | 0.6 | 0.4 |
| SE | 7998 | PMF1 | PMF1 | chr1 | + | 156225556 | 156225668 | 156212987 | 156213176 | 156232319 | 156232425 | 3 | 1 | 0 | 3 | 0 | 0 | 0 | 0 | 0 | 1 | 98 | 49 | 1 | 1 | 0.333 | 1 | NA | NA | 0 | 0.666 |
| SE | 8001 | USF1 | USF1 | chr1 | - | 161041650 | 161041846 | 161041323 | 161041411 | 161042115 | 161042217 | 24 | 9 | 4 | 2 | 1 | 0 | 0 | 5 | 0 | 0 | 98 | 49 | 1 | 1 | 0.857 | 0.818 | 1 | NA | 1 | -0.108 |
| SE | 8006 | ANO10 | ANO10 | chr3 | - | 43565652 | 43565727 | 43561219 | 43561402 | 43574808 | 43574864 | 12 | 6 | 2 | 1 | 0 | 0 | 2 | 1 | 0 | 0 | 98 | 49 | 1 | 1 | 0.857 | 1 | 1 | 1 | 1 | -0.048 |
| SE | 8009 | ANO10 | ANO10 | chr3 | - | 43600383 | 43600581 | 43598531 | 43598666 | 43605713 | 43605863 | 5 | 9 | 4 | 0 | 2 | 0 | 0 | 1 | 0 | 0 | 98 | 49 | 1 | 1 | 1 | 0.692 | 1 | NA | 1 | -0.103 |
| SE | 8014 | RNF19B | RNF19B | chr1 | - | 32938396 | 32938528 | 32936445 | 32937259 | 32942251 | 32942459 | 79 | 34 | 18 | 1 | 0 | 0 | 1 | 11 | 0 | 0 | 98 | 49 | 1 | 1 | 0.975 | 1 | 1 | 1 | 1 | -0.008 |
| SE | 8015 | RNF19B | RNF19B | chr1 | - | 32942251 | 32942459 | 32938396 | 32938528 | 32944018 | 32944159 | 91 | 49 | 45 | 1 | 1 | 0 | 2 | 25 | 0 | 0 | 98 | 49 | 1 | 1 | 0.978 | 0.961 | 1 | 1 | 1 | -0.02 |
| SE | 8016 | RNF19B | RNF19B | chr1 | - | 32945513 | 32945628 | 32944018 | 32944159 | 32946401 | 32946564 | 110 | 64 | 21 | 1 | 0 | 0 | 1 | 24 | 0 | 0 | 98 | 49 | 1 | 1 | 0.982 | 1 | 1 | 1 | 1 | -0.006 |
| SE | 8017 | RNF19B | RNF19B | chr1 | - | 32946401 | 32946564 | 32945513 | 32945628 | 32948221 | 32948363 | 124 | 54 | 35 | 1 | 0 | 0 | 1 | 19 | 0 | 0 | 98 | 49 | 1 | 1 | 0.984 | 1 | 1 | 1 | 1 | -0.005 |
| SE | 8019 | TBC1D9B | TBC1D9B | chr5 | - | 179867444 | 179867495 | 179865837 | 179865888 | 179867777 | 179867849 | 0 | 1 | 2 | 3 | 0 | 0 | 0 | 0 | 0 | 4 | 98 | 49 | 1 | 1 | 0 | 1 | 1 | NA | 0 | 0.667 |
| SE | 8020 | TBC1D9B | TBC1D9B | chr5 | - | 179875919 | 179876037 | 179874901 | 179875187 | 179878308 | 179878523 | 8 | 2 | 0 | 0 | 0 | 0 | 0 | 3 | 0 | 1 | 98 | 49 | 1 | 1 | 1 | 1 | NA | NA | 0.6 | 0.4 |
| SE | 8021 | SLC35D1 | SLC35D1 | chr1 | - | 67049781 | 67049850 | 67047264 | 67047367 | 67050432 | 67050504 | 1 | 2 | 0 | 0 | 0 | 0 | 0 | 2 | 0 | 1 | 98 | 49 | 1 | 1 | 1 | 1 | NA | NA | 0.5 | 0.5 |
| SE | 8037 | ARHGEF7 | ARHGEF7 | chr13 | + | 111205288 | 111205373 | 111153713 | 111153991 | 111209871 | 111210002 | 3 | 3 | 0 | 1 | 0 | 0 | 2 | 1 | 0 | 0 | 98 | 49 | 1 | 1 | 0.6 | 1 | NA | 1 | 1 | -0.2 |
| SE | 8038 | ARHGEF7 | ARHGEF7 | chr13 | + | 111209871 | 111210002 | 111205288 | 111205373 | 111217678 | 111217880 | 5 | 2 | 1 | 1 | 0 | 0 | 0 | 1 | 0 | 0 | 98 | 49 | 1 | 1 | 0.714 | 1 | 1 | NA | 1 | -0.095 |
| SE | 8041 | SRCAP | SRCAP | chr16 | + | 30712678 | 30712815 | 30712261 | 30712439 | 30713207 | 30713377 | 6 | 3 | 1 | 0 | 0 | 0 | 1 | 5 | 0 | 1 | 98 | 49 | 1 | 1 | 1 | 1 | 1 | 1 | 0.714 | 0.143 |
| SE | 8043 | AK2 | AK2 | chr1 | - | 33031543 | 33031661 | 33024441 | 33024567 | 33036735 | 33036911 | 1 | 0 | 0 | 4 | 1 | 0 | 0 | 1 | 0 | 0 | 98 | 49 | 1 | 1 | 0.111 | 0 | NA | NA | 1 | -0.945 |
| SE | 8050 | ADAM8 | ADAM8 | chr10 | - | 133268747 | 133268862 | 133267928 | 133268118 | 133269444 | 133269529 | 134 | 56 | 50 | 1 | 0 | 0 | 2 | 19 | 0 | 0 | 98 | 49 | 1 | 1 | 0.985 | 1 | 1 | 1 | 1 | -0.005 |
| SE | 8052 | ADAM8 | ADAM8 | chr10 | - | 133269896 | 133269974 | 133269444 | 133269529 | 133270359 | 133270510 | 208 | 103 | 61 | 2 | 0 | 0 | 7 | 30 | 0 | 0 | 98 | 49 | 1 | 1 | 0.981 | 1 | 1 | 1 | 1 | -0.006 |
| SE | 8053 | ADAM8 | ADAM8 | chr10 | - | 133270880 | 133271070 | 133270735 | 133270805 | 133271199 | 133271289 | 192 | 136 | 64 | 0 | 1 | 0 | 20 | 40 | 0 | 0 | 98 | 49 | 1 | 1 | 1 | 0.986 | 1 | 1 | 1 | -0.005 |
| SE | 8054 | ADAM8 | ADAM8 | chr10 | - | 133274158 | 133274235 | 133273950 | 133274029 | 133275483 | 133275587 | 116 | 70 | 41 | 1 | 0 | 0 | 14 | 29 | 0 | 0 | 98 | 49 | 1 | 1 | 0.983 | 1 | 1 | 1 | 1 | -0.006 |
| SE | 8066 | EIF5 | EIF5 | chr14 | + | 103336035 | 103336117 | 103335652 | 103335932 | 103336676 | 103336849 | 48 | 27 | 14 | 1 | 2 | 0 | 0 | 7 | 0 | 1 | 98 | 49 | 1 | 1 | 0.96 | 0.871 | 1 | NA | 0.778 | 0.166 |
| SE | 8067 | EIF5 | EIF5 | chr14 | + | 103339638 | 103339803 | 103339171 | 103339333 | 103340426 | 103340561 | 98 | 50 | 5 | 0 | 1 | 0 | 1 | 11 | 0 | 0 | 98 | 49 | 1 | 1 | 1 | 0.962 | 1 | 1 | 1 | -0.013 |
| SE | 8068 | TALDO1 | TALDO1 | chr11 | + | 758949 | 759057 | 755878 | 756002 | 760121 | 760253 | 161 | 136 | 132 | 0 | 1 | 0 | 1 | 25 | 0 | 0 | 98 | 49 | 1 | 1 | 1 | 0.986 | 1 | 1 | 1 | -0.005 |
| SE | 8069 | TALDO1 | TALDO1 | chr11 | + | 763343 | 763519 | 760121 | 760253 | 763746 | 763944 | 219 | 205 | 155 | 2 | 0 | 0 | 5 | 46 | 0 | 0 | 98 | 49 | 1 | 1 | 0.982 | 1 | 1 | 1 | 1 | -0.006 |
| SE | 8088 | HPS1 | HPS1 | chr10 | - | 98429790 | 98429889 | 98429145 | 98429642 | 98430570 | 98430670 | 14 | 1 | 2 | 0 | 0 | 3 | 0 | 1 | 0 | 1 | 98 | 49 | 1 | 1 | 1 | 1 | 0.25 | NA | 0.333 | 0.417 |
| SE | 8089 | HPS1 | HPS1 | chr10 | - | 98431130 | 98431291 | 98430570 | 98430670 | 98433982 | 98434091 | 15 | 0 | 3 | 2 | 0 | 0 | 0 | 6 | 0 | 0 | 98 | 49 | 1 | 1 | 0.789 | NA | 1 | NA | 1 | -0.105 |
| SE | 8090 | HPS1 | HPS1 | chr10 | - | 98435271 | 98435414 | 98433982 | 98434091 | 98435634 | 98435772 | 15 | 1 | 2 | 7 | 0 | 0 | 0 | 7 | 0 | 0 | 98 | 49 | 1 | 1 | 0.517 | 1 | 1 | NA | 1 | -0.161 |
| SE | 8091 | PANK2 | PANK2 | chr20 | + | 3912457 | 3912634 | 3910576 | 3910830 | 3916926 | 3917050 | 27 | 8 | 9 | 1 | 2 | 0 | 0 | 6 | 0 | 0 | 98 | 49 | 1 | 1 | 0.931 | 0.667 | 1 | NA | 1 | -0.134 |
| SE | 8092 | PANK2 | PANK2 | chr20 | + | 3918670 | 3918796 | 3916926 | 3917050 | 3923243 | 3923855 | 10 | 4 | 2 | 0 | 3 | 0 | 0 | 2 | 0 | 0 | 98 | 49 | 1 | 1 | 1 | 0.4 | 1 | NA | 1 | -0.2 |
| SE | 8096 | PARGP1 | PARGP1 | chr10 | - | 45939360 | 45939447 | 45916078 | 45916171 | 45946836 | 45946911 | 0 | 0 | 0 | 1 | 0 | 0 | 0 | 2 | 0 | 0 | 98 | 49 | 1 | 1 | 0 | NA | NA | NA | 1 | -1 |
| SE | 8097 | TMEM62 | TMEM62 | chr15 | + | 43178606 | 43178711 | 43169592 | 43169677 | 43181180 | 43181299 | 1 | 0 | 0 | 0 | 0 | 0 | 0 | 0 | 0 | 1 | 98 | 49 | 1 | 1 | 1 | NA | NA | NA | 0 | 1 |
| SE | 8100 | DEDD2 | DEDD2 | chr19 | - | 42204461 | 42204559 | 42198592 | 42199829 | 42209699 | 42209840 | 0 | 0 | 0 | 16 | 25 | 11 | 1 | 0 | 3 | 5 | 98 | 49 | 1 | 1 | 0 | 0 | 0 | 0.143 | 0 | -0.071 |
| SE | 8101 | DEDD2 | DEDD2 | chr19 | - | 42209699 | 42209840 | 42198592 | 42199829 | 42215132 | 42215252 | 32 | 36 | 19 | 2 | 0 | 0 | 3 | 17 | 0 | 0 | 98 | 49 | 1 | 1 | 0.889 | 1 | 1 | 1 | 1 | -0.037 |
| SE | 8105 | TAOK1 | TAOK1 | chr17 | + | 29498317 | 29498521 | 29495559 | 29495727 | 29502588 | 29502723 | 33 | 15 | 3 | 0 | 1 | 0 | 0 | 5 | 0 | 0 | 98 | 49 | 1 | 1 | 1 | 0.882 | 1 | NA | 1 | -0.039 |
| SE | 8114 | FCGR3B | FCGR3B | chr1 | - | 161626144 | 161626402 | 161623197 | 161624639 | 161629777 | 161630032 | 212 | 147 | 132 | 6 | 3 | 3 | 0 | 59 | 0 | 1 | 98 | 49 | 1 | 1 | 0.946 | 0.961 | 0.957 | NA | 0.967 | -0.012 |
| SE | 8115 | TMEM183A | TMEM183A | chr1 | + | 203007773 | 203007863 | 203007405 | 203007574 | 203008642 | 203008810 | 4 | 0 | 0 | 0 | 0 | 0 | 0 | 0 | 1 | 0 | 98 | 49 | 1 | 1 | 1 | NA | NA | 0 | NA | 1 |
| SE | 8121 | U2SURP | U2SURP | chr3 | + | 143023974 | 143024018 | 143022852 | 143023064 | 143027148 | 143027253 | 10 | 2 | 0 | 0 | 2 | 0 | 0 | 1 | 0 | 0 | 93 | 49 | 1 | 1 | 1 | 0.345 | NA | NA | 1 | -0.328 |
| SE | 8122 | U2SURP | U2SURP | chr3 | + | 143054942 | 143055119 | 143053675 | 143053794 | 143056311 | 143060725 | 12 | 3 | 1 | 1 | 0 | 0 | 0 | 1 | 0 | 0 | 98 | 49 | 1 | 1 | 0.857 | 1 | 1 | NA | 1 | -0.048 |
| SE | 8123 | ADD1 | ADD1 | chr4 | + | 2909338 | 2909431 | 2908514 | 2908604 | 2914883 | 2915040 | 2 | 0 | 0 | 10 | 2 | 5 | 0 | 0 | 0 | 2 | 98 | 49 | 1 | 1 | 0.091 | 0 | 0 | NA | 0 | 0.03 |
| SE | 8124 | ADD1 | ADD1 | chr4 | + | 2926641 | 2926675 | 2926013 | 2926112 | 2928170 | 2930075 | 1 | 0 | 0 | 21 | 8 | 3 | 0 | 0 | 2 | 3 | 83 | 49 | 1 | 1 | 0.027 | 0 | 0 | 0 | 0 | 0.009 |
| SE | 8125 | RCOR1 | RCOR1 | chr14 | + | 102714422 | 102714617 | 102710934 | 102711013 | 102721006 | 102721084 | 33 | 18 | 7 | 2 | 0 | 0 | 0 | 5 | 0 | 0 | 98 | 49 | 1 | 1 | 0.892 | 1 | 1 | NA | 1 | -0.036 |
| SE | 8126 | RCOR1 | RCOR1 | chr14 | + | 102721319 | 102721377 | 102721006 | 102721084 | 102722186 | 102722416 | 11 | 10 | 4 | 0 | 0 | 0 | 0 | 7 | 0 | 2 | 98 | 49 | 1 | 1 | 1 | 1 | 1 | NA | 0.636 | 0.364 |
| SE | 8130 | CNEP1R1 | CNEP1R1 | chr16 | + | 50029724 | 50029798 | 50026395 | 50026467 | 50033396 | 50033506 | 13 | 7 | 4 | 0 | 3 | 0 | 0 | 6 | 0 | 0 | 98 | 49 | 1 | 1 | 1 | 0.538 | 1 | NA | 1 | -0.154 |
| SE | 8133 | ATP2B4 | ATP2B4 | chr1 | + | 203707008 | 203707223 | 203703651 | 203703813 | 203707861 | 203708104 | 15 | 12 | 1 | 1 | 0 | 0 | 0 | 2 | 0 | 0 | 98 | 49 | 1 | 1 | 0.882 | 1 | 1 | NA | 1 | -0.039 |
| SE | 8134 | ATP2B4 | ATP2B4 | chr1 | + | 203721196 | 203721410 | 203720548 | 203720740 | 203722477 | 203722689 | 16 | 7 | 4 | 2 | 0 | 0 | 0 | 2 | 0 | 0 | 98 | 49 | 1 | 1 | 0.8 | 1 | 1 | NA | 1 | -0.067 |
| SE | 8136 | FAM160B1 | FAM160B1 | chr10 | + | 114848646 | 114848737 | 114847089 | 114847233 | 114855196 | 114855340 | 22 | 10 | 1 | 1 | 0 | 0 | 0 | 3 | 0 | 0 | 98 | 49 | 1 | 1 | 0.917 | 1 | 1 | NA | 1 | -0.028 |
| SE | 8137 | GNL2 | GNL2 | chr1 | - | 37574664 | 37574823 | 37574342 | 37574456 | 37575594 | 37575699 | 5 | 0 | 0 | 0 | 0 | 0 | 0 | 1 | 0 | 1 | 98 | 49 | 1 | 1 | 1 | NA | NA | NA | 0.333 | 0.667 |
| SE | 8143 | CD48 | CD48 | chr1 | - | 160684886 | 160685189 | 160680421 | 160681468 | 160711681 | 160711851 | 82 | 7 | 6 | 0 | 3 | 0 | 0 | 22 | 0 | 1 | 98 | 49 | 1 | 1 | 1 | 0.538 | 1 | NA | 0.917 | -0.071 |
| SE | 8149 | AKIRIN1 | AKIRIN1 | chr1 | + | 39000971 | 39001106 | 38998170 | 38998311 | 39003346 | 39003418 | 8 | 8 | 6 | 1 | 0 | 0 | 0 | 1 | 0 | 0 | 98 | 49 | 1 | 1 | 0.8 | 1 | 1 | NA | 1 | -0.067 |
| SE | 8150 | TFEC | TFEC | chr7 | - | 115974169 | 115974256 | 115956678 | 115956793 | 115984261 | 115984513 | 2 | 0 | 2 | 1 | 0 | 0 | 0 | 2 | 0 | 0 | 98 | 49 | 1 | 1 | 0.5 | NA | 1 | NA | 1 | -0.25 |
| SE | 8153 | CAP1 | CAP1 | chr1 | + | 40064226 | 40064370 | 40061734 | 40061812 | 40064473 | 40064559 | 175 | 135 | 62 | 1 | 0 | 0 | 2 | 29 | 0 | 0 | 98 | 49 | 1 | 1 | 0.989 | 1 | 1 | 1 | 1 | -0.004 |
| SE | 8157 | RBPJ | RBPJ | chr4 | + | 26420550 | 26420725 | 26415474 | 26415640 | 26424341 | 26424479 | 41 | 15 | 3 | 0 | 1 | 0 | 0 | 8 | 0 | 0 | 98 | 49 | 1 | 1 | 1 | 0.882 | 1 | NA | 1 | -0.039 |
| SE | 8158 | RBPJ | RBPJ | chr4 | + | 26429897 | 26430053 | 26428719 | 26428860 | 26430418 | 26430522 | 44 | 13 | 7 | 2 | 0 | 0 | 0 | 3 | 0 | 0 | 98 | 49 | 1 | 1 | 0.917 | 1 | 1 | NA | 1 | -0.028 |
| SE | 8160 | ACAA1 | ACAA1 | chr3 | - | 38126161 | 38126341 | 38125825 | 38125881 | 38126509 | 38126700 | 5 | 3 | 5 | 2 | 0 | 0 | 0 | 1 | 0 | 0 | 98 | 49 | 1 | 1 | 0.556 | 1 | 1 | NA | 1 | -0.148 |
| SE | 8161 | ACAA1 | ACAA1 | chr3 | - | 38129289 | 38129388 | 38127785 | 38127866 | 38131595 | 38131638 | 17 | 5 | 2 | 0 | 0 | 3 | 0 | 2 | 0 | 0 | 98 | 49 | 1 | 1 | 1 | 1 | 0.25 | NA | 1 | -0.25 |
| SE | 8163 | MAP3K11 | MAP3K11 | chr11 | - | 65607921 | 65608070 | 65607640 | 65607816 | 65608267 | 65608448 | 36 | 19 | 21 | 1 | 0 | 0 | 1 | 13 | 0 | 0 | 98 | 49 | 1 | 1 | 0.947 | 1 | 1 | 1 | 1 | -0.018 |
| SE | 8164 | MAP3K11 | MAP3K11 | chr11 | - | 65608267 | 65608448 | 65607921 | 65608070 | 65613017 | 65614249 | 44 | 27 | 21 | 1 | 0 | 0 | 1 | 10 | 0 | 0 | 98 | 49 | 1 | 1 | 0.957 | 1 | 1 | 1 | 1 | -0.014 |
| SE | 8166 | ATP5C1 | ATP5C1 | chr10 | + | 7802757 | 7802854 | 7802269 | 7802425 | 7806973 | 7807010 | 26 | 7 | 3 | 0 | 1 | 0 | 0 | 6 | 0 | 1 | 98 | 49 | 1 | 1 | 1 | 0.778 | 1 | NA | 0.75 | 0.176 |
| SE | 8168 | PECAM1 | PECAM1 | chr17 | - | 64329699 | 64329722 | 64319415 | 64323845 | 64341633 | 64341690 | 158 | 84 | 42 | 1 | 0 | 0 | 1 | 23 | 0 | 0 | 72 | 49 | 1 | 1 | 0.991 | 1 | 1 | 1 | 1 | -0.003 |
| SE | 8169 | PECAM1 | PECAM1 | chr17 | - | 64341633 | 64341690 | 64329699 | 64329722 | 64348259 | 64348322 | 124 | 89 | 59 | 0 | 0 | 1 | 0 | 17 | 0 | 0 | 98 | 49 | 1 | 1 | 1 | 1 | 0.967 | NA | 1 | -0.011 |
| SE | 8170 | PECAM1 | PECAM1 | chr17 | - | 64348259 | 64348322 | 64341633 | 64341690 | 64350379 | 64350433 | 62 | 64 | 67 | 0 | 5 | 0 | 0 | 15 | 0 | 0 | 98 | 49 | 1 | 1 | 1 | 0.865 | 1 | NA | 1 | -0.045 |
| SE | 8172 | PECAM1 | PECAM1 | chr17 | - | 64356110 | 64356398 | 64353490 | 64353518 | 64360139 | 64360415 | 127 | 62 | 42 | 0 | 2 | 0 | 1 | 25 | 0 | 0 | 98 | 49 | 1 | 1 | 1 | 0.939 | 1 | 1 | 1 | -0.02 |
| SE | 8173 | PECAM1 | PECAM1 | chr17 | - | 64360139 | 64360415 | 64356110 | 64356398 | 64363148 | 64363397 | 212 | 112 | 75 | 1 | 2 | 0 | 2 | 31 | 0 | 0 | 98 | 49 | 1 | 1 | 0.991 | 0.966 | 1 | 1 | 1 | -0.014 |
| SE | 8175 | NRIP1 | NRIP1 | chr21 | - | 15014343 | 15014466 | 14961234 | 14968526 | 15043494 | 15043574 | 3 | 1 | 0 | 1 | 0 | 0 | 0 | 1 | 0 | 0 | 98 | 49 | 1 | 1 | 0.6 | 1 | NA | NA | 1 | -0.2 |
| SE | 8187 | TMEM165 | TMEM165 | chr4 | + | 55403235 | 55403338 | 55395912 | 55396396 | 55411613 | 55411839 | 1 | 0 | 0 | 3 | 0 | 1 | 0 | 0 | 0 | 1 | 98 | 49 | 1 | 1 | 0.143 | NA | 0 | NA | 0 | 0.071 |
| SE | 8198 | DCLRE1C | DCLRE1C | chr10 | - | 14947524 | 14947609 | 14945104 | 14945189 | 14949035 | 14949087 | 3 | 0 | 0 | 0 | 2 | 2 | 0 | 1 | 0 | 3 | 98 | 49 | 1 | 1 | 1 | 0 | 0 | NA | 0.143 | 0.19 |
| SE | 8208 | SULF2 | SULF2 | chr20 | - | 47659398 | 47659452 | 47657405 | 47658392 | 47659696 | 47659721 | 48 | 27 | 15 | 1 | 0 | 4 | 0 | 9 | 0 | 0 | 98 | 49 | 1 | 1 | 0.96 | 1 | 0.652 | NA | 1 | -0.129 |
| SE | 8209 | SULF2 | SULF2 | chr20 | - | 47666259 | 47666488 | 47665856 | 47665953 | 47672197 | 47672393 | 28 | 16 | 11 | 0 | 0 | 0 | 1 | 5 | 0 | 1 | 98 | 49 | 1 | 1 | 1 | 1 | 1 | 1 | 0.714 | 0.143 |
| SE | 8210 | SULF2 | SULF2 | chr20 | - | 47702518 | 47702670 | 47690125 | 47690295 | 47736702 | 47736942 | 13 | 11 | 5 | 1 | 0 | 0 | 1 | 3 | 0 | 1 | 98 | 49 | 1 | 1 | 0.867 | 1 | 1 | 1 | 0.6 | 0.156 |
| SE | 8211 | SULF2 | SULF2 | chr20 | - | 47736702 | 47736942 | 47702518 | 47702670 | 47757188 | 47757463 | 38 | 13 | 6 | 1 | 0 | 0 | 7 | 14 | 0 | 0 | 98 | 49 | 1 | 1 | 0.95 | 1 | 1 | 1 | 1 | -0.017 |
| SE | 8213 | EFCAB14 | EFCAB14 | chr1 | - | 46688352 | 46688544 | 46686783 | 46686870 | 46689586 | 46689691 | 1 | 0 | 0 | 5 | 1 | 3 | 0 | 0 | 1 | 0 | 98 | 49 | 1 | 1 | 0.091 | 0 | 0 | 0 | NA | 0.03 |
| SE | 8214 | EFCAB14 | EFCAB14 | chr1 | - | 46716294 | 46716443 | 46707905 | 46708051 | 46717902 | 46719064 | 15 | 2 | 4 | 1 | 0 | 0 | 0 | 5 | 0 | 0 | 98 | 49 | 1 | 1 | 0.882 | 1 | 1 | NA | 1 | -0.039 |
| SE | 8216 | ACADVL | ACADVL | chr17 | + | 7220923 | 7221058 | 7220765 | 7220830 | 7221537 | 7221682 | 59 | 17 | 4 | 0 | 0 | 0 | 0 | 10 | 0 | 1 | 98 | 49 | 1 | 1 | 1 | 1 | 1 | NA | 0.833 | 0.167 |
| SE | 8217 | ACADVL | ACADVL | chr17 | + | 7224145 | 7224243 | 7223967 | 7224069 | 7224320 | 7224393 | 34 | 16 | 6 | 5 | 2 | 0 | 0 | 5 | 0 | 0 | 98 | 49 | 1 | 1 | 0.773 | 0.8 | 1 | NA | 1 | -0.142 |
| SE | 8220 | HELQ | HELQ | chr4 | - | 83446834 | 83447035 | 83446013 | 83446086 | 83448782 | 83448961 | 2 | 2 | 0 | 0 | 0 | 0 | 0 | 0 | 0 | 1 | 98 | 49 | 1 | 1 | 1 | 1 | NA | NA | 0 | 1 |
| SE | 8222 | EFHD2 | EFHD2 | chr1 | + | 15427149 | 15427284 | 15425870 | 15426018 | 15428592 | 15430343 | 150 | 117 | 81 | 1 | 0 | 0 | 8 | 39 | 0 | 0 | 98 | 49 | 1 | 1 | 0.987 | 1 | 1 | 1 | 1 | -0.004 |
| SE | 8228 | SYNRG | SYNRG | chr17 | - | 37529782 | 37529851 | 37520537 | 37520648 | 37535978 | 37536127 | 1 | 0 | 0 | 5 | 0 | 0 | 0 | 0 | 0 | 2 | 98 | 49 | 1 | 1 | 0.091 | NA | NA | NA | 0 | 0.091 |
| SE | 8234 | ELOVL1 | ELOVL1 | chr1 | - | 43365185 | 43365376 | 43364927 | 43365008 | 43365563 | 43365623 | 12 | 4 | 1 | 4 | 2 | 0 | 0 | 1 | 0 | 0 | 98 | 49 | 1 | 1 | 0.6 | 0.5 | 1 | NA | 1 | -0.3 |
| SE | 8235 | ZFC3H1 | ZFC3H1 | chr12 | - | 71619314 | 71619409 | 71615205 | 71615316 | 71619925 | 71620124 | 12 | 2 | 3 | 1 | 0 | 0 | 0 | 2 | 0 | 0 | 98 | 49 | 1 | 1 | 0.857 | 1 | 1 | NA | 1 | -0.048 |
| SE | 8236 | ZFC3H1 | ZFC3H1 | chr12 | - | 71624103 | 71624292 | 71623359 | 71623597 | 71626259 | 71626446 | 8 | 0 | 1 | 0 | 0 | 0 | 0 | 0 | 0 | 2 | 98 | 49 | 1 | 1 | 1 | NA | 1 | NA | 0 | 1 |
| SE | 8237 | ZFC3H1 | ZFC3H1 | chr12 | - | 71636849 | 71637059 | 71636489 | 71636654 | 71638417 | 71638515 | 27 | 17 | 1 | 0 | 0 | 0 | 0 | 3 | 0 | 1 | 98 | 49 | 1 | 1 | 1 | 1 | 1 | NA | 0.6 | 0.4 |
| SE | 8247 | SNX29P2 | SNX29P2 | chr16 | + | 29320927 | 29321095 | 29312349 | 29312589 | 29326346 | 29326483 | 1 | 2 | 0 | 0 | 4 | 0 | 1 | 0 | 0 | 0 | 98 | 49 | 1 | 1 | 1 | 0.2 | NA | 1 | NA | -0.4 |
| SE | 8249 | EWSR1 | EWSR1 | chr22 | + | 29282389 | 29282557 | 29278029 | 29278216 | 29286922 | 29287134 | 37 | 8 | 16 | 0 | 0 | 0 | 0 | 11 | 0 | 1 | 98 | 49 | 1 | 1 | 1 | 1 | 1 | NA | 0.846 | 0.154 |
| SE | 8250 | PPT1 | PPT1 | chr1 | - | 40076841 | 40076913 | 40072709 | 40074183 | 40078559 | 40078658 | 75 | 38 | 21 | 6 | 1 | 2 | 0 | 9 | 0 | 5 | 98 | 49 | 1 | 1 | 0.862 | 0.95 | 0.84 | NA | 0.474 | 0.41 |
| SE | 8254 | PKN2 | PKN2 | chr1 | + | 88771420 | 88771566 | 88770351 | 88770469 | 88771662 | 88771879 | 101 | 33 | 9 | 1 | 2 | 0 | 0 | 19 | 0 | 0 | 98 | 49 | 1 | 1 | 0.981 | 0.892 | 1 | NA | 1 | -0.042 |
| SE | 8255 | PKN2 | PKN2 | chr1 | + | 88804390 | 88804534 | 88786103 | 88786213 | 88804845 | 88804921 | 26 | 14 | 0 | 11 | 6 | 7 | 0 | 4 | 0 | 0 | 98 | 49 | 1 | 1 | 0.542 | 0.538 | 0 | NA | 1 | -0.64 |
| SE | 8262 | NLRP3 | NLRP3 | chr1 | + | 247429584 | 247429755 | 247423846 | 247425599 | 247434102 | 247434273 | 9 | 0 | 0 | 1 | 0 | 0 | 0 | 5 | 0 | 0 | 98 | 49 | 1 | 1 | 0.818 | NA | NA | NA | 1 | -0.182 |
| SE | 8263 | NLRP3 | NLRP3 | chr1 | + | 247434102 | 247434273 | 247429584 | 247429755 | 247443971 | 247444142 | 3 | 0 | 0 | 1 | 0 | 0 | 0 | 3 | 0 | 0 | 98 | 49 | 1 | 1 | 0.6 | NA | NA | NA | 1 | -0.4 |
| SE | 8264 | NLRP3 | NLRP3 | chr1 | + | 247435969 | 247436140 | 247434102 | 247434273 | 247443971 | 247444142 | 6 | 2 | 2 | 1 | 0 | 0 | 0 | 1 | 0 | 2 | 98 | 49 | 1 | 1 | 0.75 | 1 | 1 | NA | 0.2 | 0.717 |
| SE | 8266 | COL4A3BP | COL4A3BP | chr5 | - | 75459064 | 75459181 | 75426370 | 75426478 | 75505981 | 75506116 | 9 | 2 | 2 | 2 | 0 | 0 | 0 | 2 | 0 | 0 | 98 | 49 | 1 | 1 | 0.692 | 1 | 1 | NA | 1 | -0.103 |
| SE | 8282 | MLLT10 | MLLT10 | chr10 | + | 21617111 | 21617207 | 21614830 | 21614924 | 21651672 | 21651768 | 6 | 1 | 0 | 1 | 0 | 0 | 0 | 1 | 0 | 0 | 98 | 49 | 1 | 1 | 0.75 | 1 | NA | NA | 1 | -0.125 |
| SE | 8285 | PLK3 | PLK3 | chr1 | + | 44803930 | 44804024 | 44803599 | 44803691 | 44804338 | 44804501 | 10 | 0 | 2 | 2 | 0 | 0 | 0 | 2 | 0 | 0 | 98 | 49 | 1 | 1 | 0.714 | NA | 1 | NA | 1 | -0.143 |
| SE | 8286 | PLK3 | PLK3 | chr1 | + | 44804162 | 44804246 | 44803930 | 44804024 | 44804338 | 44804501 | 36 | 14 | 8 | 1 | 0 | 0 | 6 | 3 | 0 | 0 | 98 | 49 | 1 | 1 | 0.947 | 1 | 1 | 1 | 1 | -0.018 |
| SE | 8287 | PLK3 | PLK3 | chr1 | + | 44804649 | 44804779 | 44804338 | 44804501 | 44805265 | 44805379 | 9 | 6 | 1 | 20 | 9 | 2 | 0 | 2 | 0 | 3 | 98 | 49 | 1 | 1 | 0.184 | 0.25 | 0.2 | NA | 0.25 | -0.039 |
| SE | 8294 | TMED5 | TMED5 | chr1 | - | 93158814 | 93158885 | 93156299 | 93156483 | 93160128 | 93160226 | 3 | 0 | 0 | 8 | 0 | 0 | 0 | 1 | 0 | 0 | 98 | 49 | 1 | 1 | 0.158 | NA | NA | NA | 1 | -0.842 |
| SE | 8298 | ZNF821 | ZNF821 | chr16 | - | 71861775 | 71861942 | 71859679 | 71860672 | 71864137 | 71864242 | 1 | 0 | 0 | 0 | 0 | 0 | 0 | 0 | 0 | 1 | 98 | 49 | 1 | 1 | 1 | NA | NA | NA | 0 | 1 |
| SE | 8301 | IL10RA | IL10RA | chr11 | + | 117986683 | 117986813 | 117986390 | 117986534 | 117988381 | 117988502 | 4 | 2 | 2 | 1 | 2 | 1 | 0 | 0 | 0 | 2 | 98 | 49 | 1 | 1 | 0.667 | 0.333 | 0.5 | NA | 0 | 0.5 |
| SE | 8302 | IL10RA | IL10RA | chr11 | + | 117988381 | 117988502 | 117986390 | 117986534 | 117993240 | 117993410 | 1 | 2 | 1 | 0 | 0 | 0 | 0 | 3 | 0 | 1 | 98 | 49 | 1 | 1 | 1 | 1 | 1 | NA | 0.6 | 0.4 |
| SE | 8303 | IL10RA | IL10RA | chr11 | + | 117989441 | 117989620 | 117988381 | 117988502 | 117993240 | 117993410 | 18 | 0 | 0 | 0 | 0 | 0 | 2 | 14 | 0 | 1 | 98 | 49 | 1 | 1 | 1 | NA | NA | 1 | 0.875 | 0.062 |
| SE | 8310 | ZBTB7B | ZBTB7B | chr1 | + | 155014014 | 155014093 | 155010915 | 155011015 | 155014654 | 155015814 | 0 | 4 | 1 | 0 | 1 | 0 | 0 | 1 | 0 | 0 | 98 | 49 | 1 | 1 | NA | 0.667 | 1 | NA | 1 | -0.166 |
| SE | 8314 | RNF10 | RNF10 | chr12 | + | 120552498 | 120552698 | 120546404 | 120546601 | 120554717 | 120554808 | 32 | 18 | 10 | 0 | 0 | 0 | 1 | 5 | 0 | 1 | 98 | 49 | 1 | 1 | 1 | 1 | 1 | 1 | 0.714 | 0.143 |
| SE | 8315 | RNF10 | RNF10 | chr12 | + | 120560725 | 120560886 | 120557545 | 120557682 | 120562944 | 120563070 | 56 | 34 | 6 | 1 | 0 | 0 | 0 | 12 | 0 | 0 | 98 | 49 | 1 | 1 | 0.966 | 1 | 1 | NA | 1 | -0.011 |
| SE | 8316 | RNF10 | RNF10 | chr12 | + | 120563346 | 120563623 | 120562944 | 120563070 | 120563809 | 120563943 | 38 | 19 | 16 | 0 | 1 | 0 | 1 | 4 | 0 | 0 | 98 | 49 | 1 | 1 | 1 | 0.905 | 1 | 1 | 1 | -0.032 |
| SE | 8317 | RNF10 | RNF10 | chr12 | + | 120565071 | 120565189 | 120563809 | 120563943 | 120565427 | 120565529 | 68 | 29 | 6 | 1 | 0 | 0 | 1 | 10 | 0 | 0 | 98 | 49 | 1 | 1 | 0.971 | 1 | 1 | 1 | 1 | -0.01 |
| SE | 8318 | RNF10 | RNF10 | chr12 | + | 120566824 | 120566980 | 120565427 | 120565529 | 120571190 | 120571291 | 37 | 26 | 14 | 1 | 0 | 0 | 1 | 8 | 0 | 0 | 98 | 49 | 1 | 1 | 0.949 | 1 | 1 | 1 | 1 | -0.017 |
| SE | 8319 | RNF10 | RNF10 | chr12 | + | 120566824 | 120566980 | 120565427 | 120565529 | 120575630 | 120575688 | 19 | 19 | 11 | 1 | 0 | 0 | 0 | 6 | 0 | 0 | 98 | 49 | 1 | 1 | 0.905 | 1 | 1 | NA | 1 | -0.032 |
| SE | 8320 | RNF10 | RNF10 | chr12 | + | 120571190 | 120571291 | 120565427 | 120565529 | 120575630 | 120575688 | 14 | 16 | 5 | 1 | 0 | 0 | 2 | 3 | 0 | 0 | 98 | 49 | 1 | 1 | 0.875 | 1 | 1 | 1 | 1 | -0.042 |
| SE | 8325 | COPS7A | COPS7A | chr12 | + | 6727925 | 6728001 | 6724282 | 6724818 | 6728222 | 6728311 | 11 | 8 | 5 | 4 | 2 | 0 | 0 | 0 | 0 | 1 | 98 | 49 | 1 | 1 | 0.579 | 0.667 | 1 | NA | 0 | 0.749 |
| SE | 8332 | RAB3D | RAB3D | chr19 | - | 11335664 | 11335783 | 11335446 | 11335571 | 11337171 | 11337460 | 54 | 28 | 7 | 1 | 0 | 0 | 1 | 17 | 0 | 0 | 98 | 49 | 1 | 1 | 0.964 | 1 | 1 | 1 | 1 | -0.012 |
| SE | 8335 | DHTKD1 | DHTKD1 | chr10 | + | 12106245 | 12106396 | 12101041 | 12101181 | 12107908 | 12108015 | 10 | 5 | 8 | 0 | 1 | 0 | 0 | 1 | 0 | 0 | 98 | 49 | 1 | 1 | 1 | 0.714 | 1 | NA | 1 | -0.095 |
| SE | 8336 | TTLL3 | TTLL3 | chr3 | + | 9816073 | 9816202 | 9813247 | 9813345 | 9817644 | 9817759 | 6 | 4 | 0 | 4 | 2 | 1 | 0 | 1 | 0 | 0 | 98 | 49 | 1 | 1 | 0.429 | 0.5 | 0 | NA | 1 | -0.69 |
| SE | 8337 | TTLL3 | TTLL3 | chr3 | + | 9833103 | 9833245 | 9828959 | 9829395 | 9834680 | 9834907 | 4 | 3 | 4 | 1 | 3 | 3 | 0 | 2 | 0 | 0 | 98 | 49 | 1 | 1 | 0.667 | 0.333 | 0.4 | NA | 1 | -0.533 |
| SE | 8340 | CASP8 | CASP8 | chr2 | + | 201271515 | 201271621 | 201266460 | 201266791 | 201272637 | 201272776 | 142 | 47 | 33 | 7 | 10 | 0 | 0 | 9 | 0 | 1 | 98 | 49 | 1 | 1 | 0.91 | 0.701 | 1 | NA | 0.818 | 0.052 |
| SE | 8342 | CASP8 | CASP8 | chr2 | + | 201272897 | 201272942 | 201272637 | 201272776 | 201276826 | 201276968 | 106 | 49 | 12 | 4 | 2 | 0 | 0 | 6 | 0 | 1 | 94 | 49 | 1 | 1 | 0.932 | 0.927 | 1 | NA | 0.758 | 0.195 |
| SE | 8343 | CASP8 | CASP8 | chr2 | + | 201274888 | 201274953 | 201272637 | 201272776 | 201276826 | 201276968 | 89 | 36 | 17 | 4 | 2 | 0 | 0 | 3 | 0 | 1 | 98 | 49 | 1 | 1 | 0.918 | 0.9 | 1 | NA | 0.6 | 0.339 |
| SE | 8346 | EIF4B | EIF4B | chr12 | + | 53039237 | 53039343 | 53038355 | 53038411 | 53039629 | 53039702 | 16 | 4 | 0 | 1 | 0 | 0 | 0 | 1 | 0 | 0 | 98 | 49 | 1 | 1 | 0.889 | 1 | NA | NA | 1 | -0.055 |
| SE | 8350 | IL4R | IL4R | chr16 | + | 27330065 | 27330198 | 27313908 | 27314020 | 27340185 | 27340273 | 13 | 11 | 3 | 1 | 0 | 2 | 0 | 1 | 0 | 0 | 98 | 49 | 1 | 1 | 0.867 | 1 | 0.429 | NA | 1 | -0.235 |
| SE | 8351 | IL4R | IL4R | chr16 | + | 27340185 | 27340273 | 27330065 | 27330198 | 27342120 | 27342259 | 66 | 31 | 10 | 0 | 0 | 1 | 2 | 6 | 0 | 0 | 98 | 49 | 1 | 1 | 1 | 1 | 0.833 | 1 | 1 | -0.056 |
| SE | 8352 | IL4R | IL4R | chr16 | + | 27341069 | 27341313 | 27340185 | 27340273 | 27342120 | 27342259 | 2 | 0 | 1 | 53 | 21 | 10 | 0 | 0 | 2 | 5 | 98 | 49 | 1 | 1 | 0.019 | 0 | 0.048 | 0 | 0 | 0.022 |
| SE | 8353 | IL4R | IL4R | chr16 | + | 27346466 | 27346618 | 27344868 | 27345020 | 27352539 | 27352696 | 36 | 13 | 22 | 0 | 0 | 0 | 0 | 8 | 0 | 1 | 98 | 49 | 1 | 1 | 1 | 1 | 1 | NA | 0.8 | 0.2 |
| SE | 8354 | IL4R | IL4R | chr16 | + | 27355807 | 27352539 | 27352696 | 27358915 | 27358994 | 27358994 | 34 | 12 | 14 | 2 | 0 | 0 | 0 | 7 | 0 | 0 | 98 | 49 | 1 | 1 | 0.895 | 1 | 1 | NA | 1 | -0.035 |
| SE | 8355 | IL4R | IL4R | chr16 | + | 27360765 | 27360815 | 27358915 | 27358994 | 27362251 | 27364778 | 14 | 13 | 6 | 0 | 0 | 0 | 0 | 4 | 0 | 2 | 98 | 49 | 1 | 1 | 1 | 1 | 1 | NA | 0.5 | 0.5 |
| SE | 8358 | HNRNPDL | HNRNPDL | chr4 | - | 82425562 | 82425667 | 82422563 | 82424883 | 82426036 | 82426129 | 0 | 4 | 0 | 16 | 11 | 5 | 0 | 0 | 0 | 3 | 98 | 49 | 1 | 1 | 0 | 0.154 | 0 | NA | 0 | 0.051 |
| SE | 8359 | HNRNPDL | HNRNPDL | chr4 | - | 82426462 | 82426633 | 82426036 | 82426129 | 82427189 | 82427304 | 28 | 8 | 4 | 3 | 0 | 0 | 0 | 1 | 0 | 0 | 98 | 49 | 1 | 1 | 0.824 | 1 | 1 | NA | 1 | -0.059 |
| SE | 8360 | ASPH | ASPH | chr8 | - | 61553030 | 61553120 | 61548070 | 61548208 | 61555923 | 61556022 | 5 | 8 | 1 | 0 | 0 | 0 | 2 | 4 | 0 | 1 | 98 | 49 | 1 | 1 | 1 | 1 | 1 | 1 | 0.667 | 0.166 |
| SE | 8361 | ASPH | ASPH | chr8 | - | 61633682 | 61633727 | 61618977 | 61619019 | 61637946 | 61638003 | 9 | 10 | 6 | 1 | 0 | 0 | 0 | 1 | 0 | 0 | 94 | 49 | 1 | 1 | 0.824 | 1 | 1 | NA | 1 | -0.059 |
| SE | 8362 | ASPH | ASPH | chr8 | - | 61638321 | 61638360 | 61637946 | 61638003 | 61642887 | 61642920 | 11 | 12 | 3 | 1 | 1 | 0 | 1 | 6 | 0 | 0 | 88 | 49 | 1 | 1 | 0.86 | 0.87 | 1 | 1 | 1 | -0.09 |
| SE | 8363 | ASPH | ASPH | chr8 | - | 61638321 | 61638363 | 61637946 | 61638003 | 61642887 | 61642920 | 11 | 14 | 4 | 1 | 1 | 0 | 1 | 9 | 0 | 0 | 91 | 49 | 1 | 1 | 0.856 | 0.883 | 1 | 1 | 1 | -0.087 |
| SE | 8364 | ASPH | ASPH | chr8 | - | 61643944 | 61644001 | 61643385 | 61643433 | 61644599 | 61644632 | 12 | 19 | 2 | 1 | 0 | 0 | 0 | 2 | 0 | 0 | 98 | 49 | 1 | 1 | 0.857 | 1 | 1 | NA | 1 | -0.048 |
| SE | 8365 | ASPH | ASPH | chr8 | - | 61646749 | 61646878 | 61644599 | 61644632 | 61651049 | 61651124 | 21 | 6 | 0 | 0 | 1 | 1 | 1 | 8 | 0 | 0 | 98 | 49 | 1 | 1 | 1 | 0.75 | 0 | 1 | 1 | -0.417 |
| SE | 8366 | ASPH | ASPH | chr8 | - | 61682438 | 61682483 | 61680967 | 61681036 | 61684038 | 61684188 | 9 | 7 | 8 | 8 | 10 | 0 | 0 | 0 | 0 | 3 | 94 | 49 | 1 | 1 | 0.37 | 0.267 | 1 | NA | 0 | 0.546 |
| SE | 8378 | BCAS3 | BCAS3 | chr17 | + | 61026865 | 61026910 | 61015750 | 61015901 | 61034665 | 61034790 | 1 | 0 | 0 | 10 | 0 | 0 | 0 | 0 | 0 | 4 | 94 | 49 | 1 | 1 | 0.05 | NA | NA | NA | 0 | 0.05 |
| SE | 8388 | FRY | FRY | chr13 | + | 32131671 | 32131840 | 32124794 | 32124875 | 32134903 | 32134996 | 104 | 22 | 7 | 0 | 1 | 0 | 0 | 9 | 0 | 0 | 98 | 49 | 1 | 1 | 1 | 0.917 | 1 | NA | 1 | -0.028 |
| SE | 8389 | FRY | FRY | chr13 | + | 32147838 | 32147947 | 32147281 | 32147385 | 32149747 | 32149834 | 52 | 23 | 7 | 2 | 0 | 0 | 2 | 8 | 0 | 0 | 98 | 49 | 1 | 1 | 0.929 | 1 | 1 | 1 | 1 | -0.024 |
| SE | 8391 | FRY | FRY | chr13 | + | 32178176 | 32178436 | 32175545 | 32175632 | 32178843 | 32179033 | 65 | 17 | 5 | 1 | 0 | 0 | 3 | 7 | 0 | 0 | 98 | 49 | 1 | 1 | 0.97 | 1 | 1 | 1 | 1 | -0.01 |

| | | | | | | | | | | | | | | | | | | | | | | | | | | | | | | | |
|---|---|---|---|---|---|---|---|---|---|---|---|---|---|---|---|---|---|---|---|---|---|---|---|---|---|---|---|---|---|---|---|
| SE | 8392 | FRY | FRY | chr13 | + | 32194142 | 32194297 | 32187545 | 32187656 | 32201940 | 32202040 | 15 | 7 | 1 | 3 | 0 | 0 | 0 | 2 | 0 | 0 | 98 | 49 | 1 | 1 | 0.714 | 1 | 1 | NA | 1 | -0.095 |
| SE | 8397 | ATP2A3 | ATP2A3 | chr17 | - | 3951580 | 3951685 | 3951250 | 3951389 | 3953346 | 3953429 | 44 | 33 | 19 | 0 | 1 | 0 | 9 | 2 | 0 | 0 | 98 | 49 | 1 | 1 | 1 | 0.943 | 1 | 1 | 1 | -0.019 |
| SE | 8399 | ATP2A3 | ATP2A3 | chr17 | - | 3953692 | 3953710 | 3953346 | 3953429 | 3964173 | 3964464 | 22 | 10 | 13 | 0 | 1 | 0 | 2 | 5 | 0 | 0 | 67 | 49 | 1 | 1 | 1 | 0.88 | 1 | 1 | 1 | -0.04 |
| SE | 8411 | FCHO1 | FCHO1 | chr19 | + | 17755117 | 17755191 | 17754316 | 17754361 | 17762761 | 17762853 | 9 | 3 | 0 | 1 | 0 | 0 | 1 | 1 | 0 | 0 | 98 | 49 | 1 | 1 | 0.818 | 1 | NA | 1 | 1 | -0.091 |
| SE | 8412 | FCHO1 | FCHO1 | chr19 | + | 17781230 | 17781343 | 17778608 | 17778884 | 17781451 | 17781539 | 8 | 6 | 0 | 0 | 0 | 0 | 1 | 3 | 0 | 3 | 98 | 49 | 1 | 1 | 1 | 1 | NA | 1 | 0.333 | 0.334 |
| SE | 8413 | OS9 | OS9 | chr12 | + | 57694749 | 57694926 | 57693954 | 57694323 | 57695779 | 57695843 | 52 | 34 | 24 | 0 | 1 | 0 | 3 | 9 | 0 | 0 | 98 | 49 | 1 | 1 | 1 | 0.944 | 1 | 1 | 1 | -0.019 |
| SE | 8415 | OS9 | OS9 | chr12 | + | 57695961 | 57696038 | 57695779 | 57695843 | 57715759 | 57715970 | 22 | 16 | 12 | 0 | 1 | 0 | 0 | 1 | 0 | 0 | 98 | 49 | 1 | 1 | 1 | 0.889 | 1 | NA | 1 | -0.037 |
| SE | 8417 | OS9 | OS9 | chr12 | + | 57716088 | 57716193 | 57715759 | 57715970 | 57716411 | 57716512 | 27 | 17 | 4 | 1 | 0 | 0 | 0 | 12 | 0 | 0 | 98 | 49 | 1 | 1 | 0.931 | 1 | 1 | NA | 1 | -0.023 |
| SE | 8418 | OS9 | OS9 | chr12 | + | 57716091 | 57716193 | 57715759 | 57715970 | 57716411 | 57716512 | 26 | 17 | 4 | 1 | 0 | 0 | 0 | 11 | 0 | 0 | 98 | 49 | 1 | 1 | 0.929 | 1 | 1 | NA | 1 | -0.024 |
| SE | 8419 | OS9 | OS9 | chr12 | + | 57720098 | 57720263 | 57718992 | 57719182 | 57720405 | 57720518 | 24 | 14 | 22 | 25 | 13 | 21 | 0 | 2 | 0 | 3 | 98 | 49 | 1 | 1 | 0.324 | 0.35 | 0.344 | NA | 0.25 | 0.089 |
| SE | 8420 | NUCB2 | NUCB2 | chr11 | + | 17315385 | 17315475 | 17312027 | 17312120 | 17330126 | 17330297 | 1 | 0 | 0 | 0 | 0 | 0 | 0 | 0 | 1 | 0 | 98 | 49 | 1 | 1 | 1 | NA | NA | 0 | NA | 1 |
| SE | 8423 | MTDH | MTDH | chr8 | + | 97699753 | 97699852 | 97690951 | 97691188 | 97706625 | 97706750 | 10 | 3 | 1 | 2 | 0 | 0 | 0 | 8 | 0 | 0 | 98 | 49 | 1 | 1 | 0.714 | 1 | 1 | NA | 1 | -0.095 |
| SE | 8431 | AGO1 | AGO1 | chr1 | + | 35893673 | 35893810 | 35893096 | 35893278 | 35894036 | 35894171 | 12 | 0 | 4 | 1 | 1 | 0 | 0 | 4 | 0 | 0 | 98 | 49 | 1 | 1 | 0.857 | 0 | 1 | NA | 1 | -0.381 |
| SE | 8432 | AGO1 | AGO1 | chr1 | + | 35906934 | 35907119 | 35902203 | 35902337 | 35913841 | 35914001 | 1 | 2 | 1 | 0 | 0 | 0 | 0 | 2 | 0 | 1 | 98 | 49 | 1 | 1 | 1 | 1 | 1 | NA | 0.5 | 0.5 |
| SE | 8442 | CHD8 | CHD8 | chr14 | - | 21391462 | 21391642 | 21390946 | 21391063 | 21391832 | 21391946 | 18 | 7 | 4 | 2 | 1 | 0 | 0 | 1 | 0 | 0 | 98 | 49 | 1 | 1 | 0.818 | 0.778 | 1 | NA | 1 | -0.135 |
| SE | 8443 | CHD8 | CHD8 | chr14 | - | 21393105 | 21393254 | 21392506 | 21392809 | 21393475 | 21394195 | 25 | 1 | 2 | 0 | 0 | 0 | 2 | 2 | 0 | 1 | 98 | 49 | 1 | 1 | 1 | 1 | 1 | 1 | 0.5 | 0.25 |
| SE | 8444 | CHD8 | CHD8 | chr14 | - | 21408311 | 21408555 | 21406855 | 21407032 | 21408703 | 21408825 | 19 | 11 | 4 | 1 | 1 | 0 | 0 | 1 | 0 | 0 | 98 | 49 | 1 | 1 | 0.905 | 0.846 | 1 | NA | 1 | -0.083 |
| SE | 8445 | AEBP2 | AEBP2 | chr12 | + | 19514670 | 19514784 | 19512397 | 19512465 | 19518086 | 19518113 | 6 | 0 | 0 | 1 | 0 | 0 | 0 | 6 | 0 | 0 | 98 | 49 | 1 | 1 | 0.75 | NA | NA | NA | 1 | -0.25 |
| SE | 8446 | KDM3B | KDM3B | chr5 | + | 138379583 | 138379708 | 138377719 | 138377825 | 138381515 | 138381590 | 21 | 16 | 1 | 2 | 2 | 0 | 0 | 8 | 0 | 0 | 98 | 49 | 1 | 1 | 0.84 | 0.8 | 1 | NA | 1 | -0.12 |
| SE | 8447 | KDM3B | KDM3B | chr5 | + | 138386021 | 138386621 | 138381515 | 138381590 | 138391012 | 138392261 | 8 | 8 | 5 | 0 | 0 | 0 | 0 | 2 | 0 | 2 | 98 | 49 | 1 | 1 | 1 | 1 | 1 | NA | 0.333 | 0.667 |
| SE | 8448 | KDM3B | KDM3B | chr5 | + | 138391012 | 138392261 | 138381515 | 138381590 | 138393170 | 138393372 | 14 | 9 | 1 | 0 | 3 | 0 | 0 | 6 | 0 | 0 | 98 | 49 | 1 | 1 | 1 | 0.6 | 1 | NA | 1 | -0.133 |
| SE | 8453 | NRD1 | NRD1 | chr1 | - | 51790532 | 51790649 | 51789567 | 51789657 | 51790899 | 51790990 | 74 | 20 | 12 | 1 | 0 | 0 | 0 | 17 | 0 | 0 | 98 | 49 | 1 | 1 | 0.974 | 1 | 1 | NA | 1 | -0.009 |
| SE | 8454 | NRD1 | NRD1 | chr1 | - | 51791577 | 51791661 | 51790899 | 51790990 | 51792045 | 51792098 | 42 | 26 | 7 | 1 | 0 | 0 | 0 | 13 | 0 | 0 | 98 | 49 | 1 | 1 | 0.955 | 1 | 1 | NA | 1 | -0.015 |
| SE | 8455 | NRD1 | NRD1 | chr1 | - | 51800555 | 51800683 | 51798248 | 51798411 | 51803813 | 51803964 | 56 | 22 | 7 | 0 | 0 | 0 | 0 | 18 | 0 | 1 | 98 | 49 | 1 | 1 | 1 | 1 | 1 | NA | 0.9 | 0.1 |
| SE | 8456 | NRD1 | NRD1 | chr1 | - | 51819799 | 51819873 | 51818065 | 51818135 | 51821497 | 51821555 | 62 | 28 | 13 | 3 | 0 | 0 | 0 | 24 | 0 | 0 | 98 | 49 | 1 | 1 | 0.912 | 1 | 1 | NA | 1 | -0.029 |
| SE | 8464 | MAST3 | MAST3 | chr19 | + | 18128399 | 18128458 | 18124264 | 18124366 | 18128865 | 18128951 | 36 | 23 | 1 | 0 | 1 | 0 | 1 | 4 | 0 | 0 | 98 | 49 | 1 | 1 | 1 | 0.92 | 1 | 1 | 1 | -0.027 |
| SE | 8465 | MAST3 | MAST3 | chr19 | + | 18134816 | 18134982 | 18134578 | 18134711 | 18135739 | 18135841 | 57 | 17 | 9 | 2 | 3 | 0 | 0 | 12 | 0 | 0 | 98 | 49 | 1 | 1 | 0.934 | 0.739 | 1 | NA | 1 | -0.109 |
| SE | 8470 | ST8SIA4 | ST8SIA4 | chr5 | - | 100856102 | 100856396 | 100806934 | 100812129 | 100886342 | 100886600 | 82 | 44 | 51 | 5 | 2 | 1 | 0 | 12 | 0 | 0 | 98 | 49 | 1 | 1 | 0.891 | 0.917 | 0.962 | NA | 1 | -0.077 |
| SE | 8471 | ST8SIA4 | ST8SIA4 | chr5 | - | 100895653 | 100895785 | 100885310 | 100886600 | 100902842 | 100903285 | 87 | 58 | 27 | 0 | 2 | 0 | 4 | 14 | 0 | 0 | 98 | 49 | 1 | 1 | 1 | 0.935 | 1 | 1 | 1 | -0.022 |
| SE | 8473 | CCNH | CCNH | chr5 | - | 87404843 | 87405007 | 87401701 | 87401772 | 87407975 | 87408186 | 43 | 11 | 1 | 2 | 0 | 0 | 0 | 11 | 0 | 0 | 98 | 49 | 1 | 1 | 0.915 | 1 | 1 | NA | 1 | -0.028 |
| SE | 8481 | GABPB2 | GABPB2 | chr1 | + | 151103561 | 151103675 | 151097851 | 151098002 | 151107036 | 151107222 | 1 | 0 | 0 | 0 | 4 | 0 | 0 | 1 | 0 | 0 | 98 | 49 | 1 | 1 | 1 | 0 | NA | NA | 1 | -0.5 |
| SE | 8482 | GABPB2 | GABPB2 | chr1 | + | 151117391 | 151117516 | 151107036 | 151107222 | 151117956 | 151118531 | 0 | 0 | 0 | 1 | 0 | 0 | 0 | 2 | 0 | 0 | 98 | 49 | 1 | 1 | 0 | NA | NA | NA | 1 | -1 |
| SE | 8485 | SCYL3 | SCYL3 | chr1 | - | 169855795 | 169855957 | 169854269 | 169854964 | 169859040 | 169859212 | 1 | 0 | 0 | 0 | 0 | 1 | 0 | 0 | 0 | 1 | 98 | 49 | 1 | 1 | 1 | NA | 0 | NA | 0 | 0.5 |
| SE | 8500 | DPF2 | DPF2 | chr11 | + | 65343990 | 65344069 | 65343744 | 65343837 | 65345665 | 65345803 | 9 | 0 | 5 | 0 | 3 | 0 | 0 | 3 | 0 | 0 | 98 | 49 | 1 | 1 | 1 | 0 | 1 | NA | 1 | -0.333 |
| SE | 8504 | ABI1 | ABI1 | chr10 | - | 26755654 | 26755741 | 26751597 | 26751783 | 26759061 | 26759235 | 4 | 0 | 0 | 8 | 10 | 1 | 0 | 1 | 0 | 10 | 98 | 49 | 1 | 1 | 0.2 | 0 | 0 | NA | 0.048 | 0.019 |
| SE | 8505 | ABI1 | ABI1 | chr10 | - | 26755654 | 26755741 | 26751597 | 26751783 | 26765217 | 26765318 | 0 | 0 | 0 | 8 | 8 | 3 | 0 | 1 | 0 | 0 | 98 | 49 | 1 | 1 | 0 | 0 | 0 | NA | 1 | -1 |
| SE | 8506 | ABI1 | ABI1 | chr10 | - | 26759061 | 26759235 | 26751597 | 26751783 | 26765217 | 26765318 | 9 | 13 | 1 | 8 | 8 | 3 | 0 | 12 | 0 | 0 | 98 | 49 | 1 | 1 | 0.36 | 0.448 | 0.143 | NA | 1 | -0.683 |
| SE | 8508 | ABI1 | ABI1 | chr10 | - | 26759061 | 26759238 | 26751597 | 26751783 | 26765217 | 26765318 | 12 | 12 | 2 | 8 | 8 | 3 | 0 | 11 | 0 | 0 | 98 | 49 | 1 | 1 | 0.429 | 0.429 | 0.25 | NA | 1 | -0.631 |
| SE | 8510 | ABI1 | ABI1 | chr10 | - | 26763879 | 26763960 | 26759061 | 26759238 | 26765217 | 26765318 | 1 | 0 | 0 | 4 | 2 | 1 | 0 | 0 | 0 | 1 | 98 | 49 | 1 | 1 | 0.111 | 0 | 0 | NA | 0 | 0.037 |
| SE | 8511 | ABI1 | ABI1 | chr10 | - | 26771074 | 26771089 | 26770244 | 26770345 | 26777064 | 26777241 | 7 | 4 | 4 | 13 | 1 | 0 | 0 | 0 | 0 | 1 | 64 | 49 | 1 | 1 | 0.292 | 0.754 | 1 | NA | 0 | 0.682 |
| SE | 8519 | SELK | SELK | chr3 | - | 53885825 | 53885912 | 53885198 | 53885561 | 53886850 | 53886934 | 39 | 34 | 16 | 0 | 1 | 0 | 2 | 9 | 0 | 0 | 98 | 49 | 1 | 1 | 1 | 0.944 | 1 | 1 | 1 | -0.019 |
| SE | 8522 | RFC2 | RFC2 | chr7 | - | 74249011 | 74249118 | 74246661 | 74246763 | 74249738 | 74249780 | 5 | 0 | 4 | 3 | 0 | 0 | 3 | 0 | 0 | 0 | 98 | 49 | 1 | 1 | 0.455 | NA | 1 | 1 | NA | -0.272 |
| SE | 8523 | RFC2 | RFC2 | chr7 | - | 74249011 | 74249220 | 74246661 | 74246763 | 74249738 | 74249780 | 6 | 0 | 4 | 3 | 0 | 0 | 3 | 0 | 0 | 0 | 98 | 49 | 1 | 1 | 0.5 | NA | 1 | 1 | NA | -0.25 |
| SE | 8526 | NFKBIZ | NFKBIZ | chr3 | + | 101857072 | 101857183 | 101855732 | 101855902 | 101857291 | 101857459 | 56 | 36 | 6 | 2 | 2 | 5 | 0 | 19 | 0 | 3 | 98 | 49 | 1 | 1 | 0.933 | 0.9 | 0.375 | NA | 0.76 | -0.024 |
| SE | 8527 | KIAA0556 | KIAA0556 | chr16 | + | 27699533 | 27699599 | 27698327 | 27698500 | 27701588 | 27701695 | 14 | 3 | 0 | 2 | 0 | 0 | 2 | 3 | 0 | 0 | 98 | 49 | 1 | 1 | 0.778 | 1 | NA | 1 | 1 | -0.111 |
| SE | 8528 | KIAA0556 | KIAA0556 | chr16 | + | 27708704 | 27708920 | 27703895 | 27703998 | 27721557 | 27721695 | 11 | 1 | 0 | 1 | 0 | 0 | 0 | 1 | 0 | 0 | 98 | 49 | 1 | 1 | 0.846 | 1 | NA | NA | 1 | -0.077 |
| SE | 8529 | KIAA0556 | KIAA0556 | chr16 | + | 27754172 | 27754251 | 27751718 | 27751924 | 27761412 | 27761590 | 5 | 4 | 2 | 0 | 1 | 0 | 0 | 4 | 0 | 0 | 98 | 49 | 1 | 1 | 1 | 0.667 | 1 | NA | 1 | -0.111 |
| SE | 8530 | KIAA0556 | KIAA0556 | chr16 | + | 27777609 | 27777770 | 27776927 | 27777029 | 27777880 | 27777969 | 8 | 2 | 1 | 1 | 0 | 0 | 2 | 2 | 0 | 0 | 98 | 49 | 1 | 1 | 0.8 | 1 | 1 | 1 | 1 | -0.067 |
| SE | 8532 | ITGAM | ITGAM | chr16 | + | 31261691 | 31261797 | 31259966 | 31260092 | 31265394 | 31265498 | 29 | 18 | 4 | 0 | 2 | 0 | 0 | 9 | 0 | 0 | 98 | 49 | 1 | 1 | 1 | 0.818 | 1 | NA | 1 | -0.061 |
| SE | 8533 | ITGAM | ITGAM | chr16 | + | 31277966 | 31278109 | 31276919 | 31277049 | 31297513 | 31297654 | 53 | 36 | 11 | 1 | 1 | 0 | 0 | 11 | 0 | 0 | 98 | 49 | 1 | 1 | 0.964 | 0.947 | 1 | NA | 1 | -0.03 |
| SE | 8534 | ITGAM | ITGAM | chr16 | + | 31297741 | 31297954 | 31297513 | 31297654 | 31321240 | 31321371 | 30 | 7 | 3 | 2 | 0 | 0 | 3 | 14 | 0 | 0 | 98 | 49 | 1 | 1 | 0.882 | 1 | 1 | 1 | 1 | -0.039 |
| SE | 8535 | ITGAM | ITGAM | chr16 | + | 31297744 | 31297954 | 31297513 | 31297654 | 31321240 | 31321371 | 38 | 15 | 4 | 2 | 0 | 0 | 2 | 19 | 0 | 0 | 98 | 49 | 1 | 1 | 0.905 | 1 | 1 | 1 | 1 | -0.032 |
| SE | 8536 | ITGAM | ITGAM | chr16 | + | 31321463 | 31321627 | 31321240 | 31321371 | 31324398 | 31324553 | 52 | 25 | 23 | 0 | 0 | 0 | 0 | 12 | 0 | 1 | 98 | 49 | 1 | 1 | 1 | 1 | 1 | NA | 0.857 | 0.143 |
| SE | 8537 | ATG3 | ATG3 | chr3 | - | 112553279 | 112553329 | 112550191 | 112550262 | 112558375 | 112558417 | 31 | 9 | 9 | 1 | 3 | 0 | 0 | 4 | 0 | 3 | 98 | 49 | 1 | 1 | 0.939 | 0.6 | 1 | NA | 0.4 | 0.446 |
| SE | 8538 | CPSF6 | CPSF6 | chr12 | + | 69257731 | 69257905 | 69256696 | 69256842 | 69258589 | 69259094 | 7 | 5 | 0 | 0 | 0 | 1 | 0 | 2 | 0 | 0 | 98 | 49 | 1 | 1 | 1 | 1 | 0 | NA | 1 | -0.333 |
| SE | 8547 | TMEM180 | TMEM180 | chr10 | + | 102470664 | 102471117 | 102469992 | 102470075 | 102471272 | 102471396 | 0 | 0 | 0 | 1 | 2 | 0 | 0 | 1 | 0 | 0 | 98 | 49 | 1 | 1 | 0 | 0 | NA | NA | 1 | -1 |
| SE | 8551 | UBR3 | UBR3 | chr2 | + | 169877493 | 169877637 | 169875790 | 169875949 | 169878524 | 169878574 | 27 | 11 | 0 | 1 | 0 | 0 | 0 | 3 | 0 | 0 | 98 | 49 | 1 | 1 | 0.931 | 1 | NA | NA | 1 | -0.034 |
| SE | 8577 | MTHFR | MTHFR | chr1 | - | 11801160 | 11801399 | 11800211 | 11800322 | 11802880 | 11803129 | 7 | 2 | 0 | 1 | 0 | 0 | 0 | 1 | 0 | 0 | 98 | 49 | 1 | 1 | 0.778 | 1 | NA | NA | 1 | -0.111 |
| SE | 8582 | RASSF3 | RASSF3 | chr12 | + | 64684786 | 64684894 | 64610512 | 64610743 | 64691469 | 64691579 | 51 | 35 | 26 | 1 | 0 | 0 | 0 | 16 | 0 | 0 | 98 | 49 | 1 | 1 | 0.962 | 1 | 1 | NA | 1 | -0.013 |
| SE | 8595 | MINOS1 | MINOS1 | chr1 | + | 19625391 | 19625632 | 19623473 | 19623583 | 19626386 | 19629821 | 1 | 0 | 0 | 0 | 3 | 0 | 0 | 0 | 0 | 1 | 98 | 49 | 1 | 1 | 1 | 0 | NA | NA | 0 | 0.5 |
| SE | 8603 | MAPKAPK2 | MAPKAPK2 | chr1 | + | 206730687 | 206730763 | 206729971 | 206730098 | 206731137 | 206731262 | 32 | 21 | 5 | 0 | 1 | 0 | 1 | 6 | 0 | 0 | 98 | 49 | 1 | 1 | 1 | 0.913 | 1 | 1 | 1 | -0.029 |
| SE | 8607 | TKT | TKT | chr3 | - | 53240250 | 53240348 | 53234982 | 53235174 | 53241131 | 53241245 | 159 | 86 | 68 | 1 | 0 | 0 | 8 | 59 | 0 | 0 | 98 | 49 | 1 | 1 | 0.988 | 1 | 1 | 1 | 1 | -0.004 |
| SE | 8624 | EPS15 | EPS15 | chr1 | - | 51448045 | 51448135 | 51446959 | 51447105 | 51461090 | 51461150 | 16 | 6 | 5 | 1 | 0 | 0 | 0 | 8 | 0 | 0 | 98 | 49 | 1 | 1 | 0.889 | 1 | 1 | NA | 1 | -0.037 |
| SE | 8625 | CTR9 | CTR9 | chr11 | + | 10755677 | 10755795 | 10754957 | 10755197 | 10756748 | 10756838 | 2 | 3 | 4 | 0 | 0 | 0 | 0 | 2 | 0 | 1 | 98 | 49 | 1 | 1 | 1 | 1 | 1 | NA | 0.5 | 0.5 |
| SE | 8627 | ANKRD13C | ANKRD13C | chr1 | - | 70306223 | 70306290 | 70300763 | 70300908 | 70313744 | 70313790 | 8 | 0 | 0 | 0 | 0 | 1 | 0 | 1 | 0 | 0 | 98 | 49 | 1 | 1 | 1 | NA | 0 | NA | 1 | -0.5 |
| SE | 8633 | REC8 | REC8 | chr14 | + | 24175542 | 24175624 | 24173290 | 24173411 | 24176821 | 24176901 | 14 | 13 | 0 | 1 | 1 | 0 | 3 | 10 | 0 | 0 | 98 | 49 | 1 | 1 | 0.875 | 0.867 | NA | 1 | 1 | -0.129 |
| SE | 8634 | REC8 | REC8 | chr14 | + | 24179084 | 24179133 | 24178776 | 24178916 | 24179396 | 24179463 | 4 | 2 | 0 | 1 | 1 | 0 | 0 | 2 | 0 | 0 | 98 | 49 | 1 | 1 | 0.667 | 0.5 | NA | NA | 1 | -0.416 |
| SE | 8635 | OSBPL11 | OSBPL11 | chr3 | - | 125552180 | 125552679 | 125547405 | 125547592 | 125560378 | 125560519 | 11 | 4 | 4 | 2 | 0 | 2 | 0 | 5 | 0 | 1 | 98 | 49 | 1 | 1 | 0.733 | 1 | 0.5 | NA | 0.714 | 0.03 |
| SE | 8644 | ZFAND1 | ZFAND1 | chr8 | - | 81717248 | 81717288 | 81714986 | 81715114 | 81718181 | 81718224 | 2 | 0 | 0 | 2 | 0 | 0 | 0 | 2 | 0 | 1 | 89 | 49 | 1 | 1 | 0.355 | NA | NA | NA | 0.524 | -0.169 |
| SE | 8646 | SAMSN1 | SAMSN1 | chr21 | - | 14510309 | 14510461 | 14500528 | 14500735 | 14512443 | 14512573 | 34 | 13 | 0 | 4 | 0 | 0 | 0 | 29 | 0 | 0 | 98 | 49 | 1 | 1 | 0.81 | 1 | NA | NA | 1 | -0.095 |
| SE | 8649 | RABGGTB | RABGGTB | chr1 | + | 75792180 | 75792306 | 75791460 | 75791571 | 75794083 | 75794233 | 3 | 8 | 0 | 0 | 1 | 0 | 0 | 3 | 0 | 0 | 98 | 49 | 1 | 1 | 1 | 0.8 | NA | NA | 1 | -0.1 |
| SE | 8650 | DDX21 | DDX21 | chr10 | + | 68963290 | 68963469 | 68962081 | 68962157 | 68965376 | 68965494 | 36 | 20 | 1 | 1 | 0 | 0 | 1 | 5 | 0 | 0 | 98 | 49 | 1 | 1 | 0.947 | 1 | 1 | 1 | 1 | -0.018 |
| SE | 8651 | DDX21 | DDX21 | chr10 | + | 68970200 | 68970350 | 68968975 | 68969121 | 68973544 | 68973664 | 18 | 10 | 0 | 0 | 1 | 0 | 0 | 9 | 0 | 0 | 98 | 49 | 1 | 1 | 1 | 0.833 | NA | NA | 1 | -0.084 |
| SE | 8652 | DDX21 | DDX21 | chr10 | + | 68971890 | 68972052 | 68970200 | 68970350 | 68973544 | 68973664 | 6 | 4 | 0 | 2 | 0 | 0 | 0 | 3 | 0 | 0 | 98 | 49 | 1 | 1 | 0.6 | 1 | NA | NA | 1 | -0.2 |
| SE | 8665 | RNASE4 | RNASE4 | chr14 | + | 20693546 | 20693650 | 20684212 | 20684758 | 20699354 | 20700602 | 2 | 0 | 0 | 0 | 0 | 0 | 0 | 0 | 0 | 1 | 98 | 49 | 1 | 1 | 1 | NA | NA | NA | 0 | 1 |
| SE | 8680 | FGR | FGR | chr1 | - | 27614429 | 27614583 | 27613218 | 27613350 | 27614849 | 27614926 | 151 | 51 | 50 | 1 | 0 | 0 | 25 | 47 | 0 | 0 | 98 | 49 | 1 | 1 | 0.987 | 1 | 1 | 1 | 1 | -0.004 |
| SE | 8681 | FGR | FGR | chr1 | - | 27614849 | 27614926 | 27614429 | 27614583 | 27615433 | 27615613 | 131 | 57 | 34 | 1 | 0 | 0 | 9 | 55 | 0 | 0 | 98 | 49 | 1 | 1 | 0.985 | 1 | 1 | 1 | 1 | -0.005 |
| SE | 8682 | FGR | FGR | chr1 | - | 27617192 | 27617296 | 27616856 | 27617006 | 27621558 | 27621657 | 145 | 63 | 78 | 3 | 1 | 0 | 3 | 47 | 0 | 1 | 98 | 49 | 1 | 1 | 0.96 | 0.969 | 1 | 1 | 0.959 | -0.003 |
| SE | 8683 | KIAA0195 | KIAA0195 | chr17 | + | 75497112 | 75497198 | 75496729 | 75496807 | 75497780 | 75497862 | 0 | 0 | 2 | 0 | 2 | 0 | 0 | 1 | 0 | 0 | 98 | 49 | 1 | 1 | NA | 0 | 1 | NA | 1 | -0.5 |
| SE | 8692 | POGZ | POGZ | chr1 | - | 151424954 | 151425061 | 151423948 | 151424286 | 151430665 | 151430841 | 4 | 2 | 0 | 1 | 0 | 0 | 0 | 2 | 0 | 0 | 98 | 49 | 1 | 1 | 0.667 | 1 | NA | NA | 1 | -0.166 |
| SE | 8695 | POGZ | POGZ | chr1 | - | 151428122 | 151428413 | 151424954 | 151425061 | 151430665 | 151430841 | 2 | 0 | 0 | 0 | 0 | 0 | 0 | 0 | 0 | 2 | 98 | 49 | 1 | 1 | 1 | NA | NA | NA | 0 | 1 |
| SE | 8696 | POGZ | POGZ | chr1 | - | 151429602 | 151429711 | 151428122 | 151428413 | 151430665 | 151430841 | 1 | 1 | 0 | 1 | 0 | 0 | 0 | 3 | 0 | 0 | 98 | 49 | 1 | 1 | 0.333 | 1 | NA | NA | 1 | -0.334 |
| SE | 8699 | NAGK | NAGK | chr2 | + | 71071685 | 71071827 | 71070740 | 71070839 | 71072640 | 71072751 | 28 | 24 | 6 | 1 | 0 | 0 | 2 | 11 | 0 | 0 | 98 | 49 | 1 | 1 | 0.933 | 1 | 1 | 1 | 1 | -0.022 |
| SE | 8705 | PCSK7 | PCSK7 | chr11 | - | 117208896 | 117209053 | 117207969 | 117208064 | 117218465 | 117218568 | 8 | 8 | 0 | 2 | 0 | 0 | 0 | 0 | 0 | 3 | 98 | 49 | 1 | 1 | 0.667 | 1 | NA | NA | 0 | 0.834 |
| SE | 8708 | IFT88 | IFT88 | chr13 | + | 20615792 | 20615879 | 20605034 | 20605105 | 20625749 | 20625849 | 1 | 1 | 4 | 2 | 0 | 0 | 0 | 2 | 0 | 0 | 98 | 49 | 1 | 1 | 0.2 | 1 | 1 | NA | 1 | -0.267 |
| SE | 8718 | IFI44 | IFI44 | chr1 | + | 78660553 | 78660654 | 78659311 | 78659483 | 78662703 | 78662878 | 7 | 2 | 0 | 2 | 0 | 0 | 0 | 36 | 0 | 6 | 98 | 49 | 1 | 1 | 0.636 | 1 | NA | NA | 0.75 | 0.068 |
| SE | 8719 | IFI44 | IFI44 | chr1 | + | 78660553 | 78660654 | 78659311 | 78659483 | 78663764 | 78664078 | 5 | 1 | 0 | 1 | 0 | 0 | 0 | 31 | 1 | 1 | 98 | 49 | 1 | 1 | 0.714 | 1 | NA | 0 | 0.939 | 0.388 |
| SE | 8720 | IFI44 | IFI44 | chr1 | + | 78662703 | 78662878 | 78659311 | 78659483 | 78663764 | 78664078 | 12 | 6 | 0 | 1 | 0 | 0 | 6 | 51 | 1 | 1 | 98 | 49 | 1 | 1 | 0.857 | 1 | NA | 0.75 | 0.962 | 0.073 |
| SE | 8721 | IFI44 | IFI44 | chr1 | + | 78662703 | 78662878 | 78660553 | 78660654 | 78663764 | 78664078 | 12 | 7 | 0 | 0 | 0 | 0 | 6 | 52 | 0 | 2 | 98 | 49 | 1 | 1 | 1 | 1 | NA | 1 | 0.929 | 0.035 |
| SE | 8722 | CTSZ | CTSZ | chr20 | - | 58996638 | 58996801 | 58995186 | 58995759 | 58997602 | 58997753 | 62 | 27 | 18 | 0 | 0 | 1 | 0 | 22 | 0 | 0 | 98 | 49 | 1 | 1 | 1 | 1 | 0.9 | NA | 1 | -0.033 |
| SE | 8725 | RABGAP1L | RABGAP1L | chr1 | + | 174988640 | 174988838 | 174982833 | 174982905 | 174989848 | 174995308 | 21 | 9 | 10 | 0 | 5 | 0 | 0 | 4 | 0 | 0 | 98 | 49 | 1 | 1 | 1 | 0.474 | 1 | NA | 1 | -0.175 |
| SE | 8728 | TBC1D10C | TBC1D10C | chr11 | + | 67405398 | 67405506 | 67405084 | 67405184 | 67405594 | 67405701 | 48 | 38 | 14 | 0 | 0 | 1 | 0 | 7 | 0 | 0 | 98 | 49 | 1 | 1 | 1 | 1 | 0.875 | NA | 1 | -0.042 |
| SE | 8729 | TBC1D10C | TBC1D10C | chr11 | + | 67406626 | 67406692 | 67405902 | 67406017 | 67408978 | 67409133 | 12 | 14 | 9 | 0 | 0 | 1 | 0 | 5 | 0 | 0 | 98 | 49 | 1 | 1 | 1 | 1 | 0.818 | NA | 1 | -0.061 |
| SE | 8735 | PTBP1 | PTBP1 | chr19 | + | 805491 | 805569 | 805012 | 805187 | 806407 | 806556 | 7 | 0 | 0 | 2 | 0 | 0 | 0 | 1 | 0 | 0 | 98 | 49 | 1 | 1 | 0.636 | NA | NA | NA | 1 | -0.364 |
| SE | 8736 | PTBP1 | PTBP1 | chr19 | + | 805512 | 805569 | 805012 | 805187 | 806407 | 806556 | 1 | 0 | 0 | 2 | 0 | 0 | 0 | 1 | 0 | 0 | 98 | 49 | 1 | 1 | 0.2 | NA | NA | NA | 1 | -0.8 |
| SE | 8737 | PTBP1 | PTBP1 | chr19 | + | 807868 | 807902 | 806407 | 806556 | 808359 | 808452 | 4 | 0 | 0 | 0 | 0 | 0 | 0 | 3 | 0 | 1 | 83 | 49 | 1 | 1 | 1 | NA | NA | NA | 0.639 | 0.361 |
| SE | 8757 | ZNF106 | ZNF106 | chr15 | - | 42417804 | 42417951 | 42412436 | 42417360 | 42421060 | 42421132 | 50 | 5 | 5 | 4 | 4 | 0 | 0 | 5 | 0 | 0 | 98 | 49 | 1 | 1 | 0.862 | 0.385 | 1 | NA | 1 | -0.251 |
| SE | 8758 | ZNF106 | ZNF106 | chr15 | - | 42428017 | 42428134 | 42424833 | 42425025 | 42435383 | 42435518 | 53 | 15 | 17 | 0 | 0 | 0 | 0 | 7 | 0 | 1 | 98 | 49 | 1 | 1 | 1 | 1 | 1 | NA | 0.778 | 0.222 |
| SE | 8759 | ZNF106 | ZNF106 | chr15 | - | 42449770 | 42451954 | 42448071 | 42448705 | 42466052 | 42466114 | 19 | 7 | 4 | 2 | 0 | 0 | 0 | 3 | 0 | 0 | 98 | 49 | 1 | 1 | 0.826 | 1 | 1 | NA | 1 | -0.058 |
| SE | 8763 | RAB3GAP2 | RAB3GAP2 | chr1 | - | 220170891 | 220171120 | 220167501 | 220167675 | 220171888 | 220172049 | 5 | 0 | 0 | 1 | 0 | 0 | 0 | 0 | 1 | 0 | 98 | 49 | 1 | 1 | 0.714 | NA | NA | 0 | NA | 0.714 |
| SE | 8770 | ZCCHC2 | ZCCHC2 | chr18 | + | 62556202 | 62556297 | 62550347 | 62550460 | 62558686 | 62558770 | 28 | 33 | 4 | 0 | 0 | 0 | 0 | 20 | 0 | 2 | 98 | 49 | 1 | 1 | 1 | 1 | 1 | NA | 0.833 | 0.167 |
| SE | 8771 | ZCCHC2 | ZCCHC2 | chr18 | + | 62558686 | 62558770 | 62556202 | 62556297 | 62560586 | 62560644 | 17 | 12 | 4 | 0 | 3 | 0 | 0 | 13 | 0 | 2 | 98 | 49 | 1 | 1 | 1 | 0.667 | 1 | NA | 0.765 | 0.124 |
| SE | 8780 | SNW1 | SNW1 | chr14 | - | 77723180 | 77723277 | 77720710 | 77720828 | 77730987 | 77731129 | 12 | 8 | 5 | 1 | 1 | 0 | 2 | 0 | 0 | 0 | 98 | 49 | 1 | 1 | 0.857 | 0.8 | 1 | 1 | NA | -0.114 |
| SE | 8781 | SNW1 | SNW1 | chr14 | - | 77735936 | 77736006 | 77734946 | 77735012 | 77736970 | 77737075 | 16 | 16 | 11 | 1 | 0 | 0 | 0 | 5 | 0 | 0 | 98 | 49 | 1 | 1 | 0.889 | 1 | 1 | NA | 1 | -0.037 |
| SE | 8786 | ZKSCAN5 | ZKSCAN5 | chr7 | + | 99520168 | 99520304 | 99519826 | 99519909 | 99525812 | 99526418 | 0 | 5 | 0 | 0 | 3 | 0 | 0 | 0 | 0 | 1 | 98 | 49 | 1 | 1 | NA | 0.455 | NA | NA | 0 | 0.455 |
| SE | 8789 | CD46 | CD46 | chr1 | + | 207767606 | 207767651 | 207767012 | 207767195 | 207767778 | 207767823 | 16 | 7 | 1 | 68 | 16 | 11 | 0 | 7 | 0 | 0 | 94 | 49 | 1 | 1 | 0.109 | 0.186 | 0.045 | NA | 1 | -0.887 |
| SE | 8790 | CD46 | CD46 | chr1 | + | 207767778 | 207767823 | 207767012 | 207767195 | 207770320 | 207770362 | 140 | 36 | 38 | 11 | 2 | 9 | 0 | 12 | 0 | 0 | 94 | 49 | 1 | 1 | 0.869 | 0.904 | 0.688 | NA | 1 | -0.18 |
| SE | 8793 | CD46 | CD46 | chr1 | + | 207790252 | 207790345 | 207785618 | 207785682 | 207793518 | 207795516 | 88 | 36 | 23 | 89 | 37 | 22 | 0 | 12 | 0 | 6 | 98 | 49 | 1 | 1 | 0.331 | 0.327 | 0.343 | NA | 0.5 | -0.166 |
| SE | 8802 | CELF1 | CELF1 | chr11 | - | 47483452 | 47483532 | 47482694 | 47482856 | 47484388 | 47484520 | 31 | 1 | 5 | 0 | 1 | 0 | 0 | 4 | 0 | 0 | 98 | 49 | 1 | 1 | 1 | 0.333 | 1 | NA | 1 | -0.222 |
| SE | 8803 | CELF1 | CELF1 | chr11 | - | 47484388 | 47484520 | 47482694 | 47482856 | 47486749 | 47486798 | 0 | 1 | 0 | 0 | 0 | 0 | 0 | 0 | 0 | 1 | 98 | 49 | 1 | 1 | NA | 1 | NA | NA | 0 | 1 |
| SE | 8804 | CELF1 | CELF1 | chr11 | - | 47484388 | 47484523 | 47482694 | 47482856 | 47486749 | 47486798 | 6 | 3 | 0 | 0 | 0 | 0 | 0 | 2 | 0 | 1 | 98 | 49 | 1 | 1 | 1 | 1 | NA | NA | 0.5 | 0.5 |
| SE | 8805 | ASH1L | ASH1L | chr1 | - | 155415743 | 155415923 | 155395458 | 155395553 | 155438326 | 155439068 | 13 | 1 | 0 | 0 | 2 | 0 | 0 | 2 | 0 | 0 | 98 | 49 | 1 | 1 | 1 | 0.2 | NA | NA | 1 | -0.4 |
| SE | 8806 | TLR5 | TLR5 | chr1 | - | 223137177 | 223137263 | 223134681 | 223134864 | 223141647 | 223141763 | 4 | 0 | 0 | 2 | 0 | 1 | 0 | 7 | 0 | 0 | 98 | 49 | 1 | 1 | 0.5 | NA | 0 | NA | 1 | -0.75 |
| SE | 8807 | TLR5 | TLR5 | chr1 | - | 223141647 | 223141763 | 223137177 | 223137263 | 223143195 | 223143282 | 2 | 1 | 0 | 0 | 0 | 1 | 0 | 3 | 0 | 0 | 98 | 49 | 1 | 1 | 1 | 1 | 0 | NA | 1 | -0.333 |
| SE | 8808 | C1orf52 | C1orf52 | chr1 | - | 85258934 | 85259061 | 85258523 | 85258722 | 85259357 | 85259672 | 3 | 8 | 4 | 0 | 0 | 1 | 0 | 1 | 0 | 0 | 98 | 49 | 1 | 1 | 1 | 1 | 0.667 | NA | 1 | -0.111 |
| SE | 8820 | LBR | LBR | chr1 | - | 225410290 | 225410416 | 225406663 | 225406832 | 225411336 | 225411440 | 78 | 21 | 18 | 5 | 3 | 6 | 0 | 18 | 0 | 3 | 98 | 49 | 1 | 1 | 0.886 | 0.778 | 0.6 | NA | 0.75 | 0.005 |
| SE | 8821 | LBR | LBR | chr1 | - | 225411336 | 225411440 | 225406663 | 225406832 | 225412453 | 225412645 | 48 | 39 | 33 | 0 | 0 | 1 | 0 | 6 | 0 | 0 | 98 | 49 | 1 | 1 | 1 | 1 | 0.943 | NA | 1 | -0.019 |
| SE | 8822 | AQP9 | AQP9 | chr15 | + | 58173067 | 58173205 | 58166672 | 58166799 | 58174917 | 58175036 | 290 | 177 | 171 | 2 | 0 | 0 | 2 | 58 | 0 | 0 | 98 | 49 | 1 | 1 | 0.986 | 1 | 1 | 1 | 1 | -0.005 |
| SE | 8823 | AQP9 | AQP9 | chr15 | + | 58174917 | 58175036 | 58173067 | 58173205 | 58179127 | 58179345 | 289 | 151 | 147 | 0 | 0 | 0 | 1 | 43 | 0 | 1 | 98 | 49 | 1 | 1 | 1 | 1 | 1 | 1 | 0.956 | 0.022 |
| SE | 8826 | DOCK2 | DOCK2 | chr5 | + | 169670541 | 169670597 | 169669287 | 169669328 | 169671077 | 169671174 | 96 | 41 | 16 | 1 | 0 | 0 | 0 | 22 | 0 | 0 | 98 | 49 | 1 | 1 | 0.98 | 1 | 1 | NA | 1 | -0.007 |
| SE | 8827 | DOCK2 | DOCK2 | chr5 | + | 169671077 | 169671174 | 169670541 | 169670597 | 169674296 | 169674445 | 205 | 65 | 16 | 0 | 1 | 0 | 2 | 36 | 0 | 0 | 98 | 49 | 1 | 1 | 1 | 0.97 | 1 | 1 | 1 | -0.01 |
| SE | 8829 | DOCK2 | DOCK2 | chr5 | + | 169689251 | 169689333 | 169684195 | 169684350 | 169695802 | 169695938 | 176 | 92 | 10 | 2 | 0 | 0 | 5 | 49 | 0 | 0 | 98 | 49 | 1 | 1 | 0.978 | 1 | 1 | 1 | 1 | -0.007 |
| SE | 8830 | DOCK2 | DOCK2 | chr5 | + | 169712119 | 169712223 | 169711934 | 169712007 | 169714027 | 169714211 | 141 | 46 | 6 | 0 | 1 | 0 | 4 | 19 | 0 | 0 | 98 | 49 | 1 | 1 | 1 | 0.958 | 1 | 1 | 1 | -0.014 |
| SE | 8833 | DOCK2 | DOCK2 | chr5 | + | 169803057 | 169803206 | 169761518 | 169761625 | 169840756 | 169840852 | 143 | 72 | 20 | 0 | 2 | 0 | 2 | 21 | 0 | 0 | 98 | 49 | 1 | 1 | 1 | 0.947 | 1 | 1 | 1 | -0.018 |
| SE | 8834 | DOCK2 | DOCK2 | chr5 | + | 169983067 | 169983166 | 169840756 | 169840852 | 169985827 | 169985922 | 45 | 22 | 7 | 0 | 0 | 0 | 0 | 12 | 0 | 1 | 98 | 49 | 1 | 1 | 1 | 1 | 1 | NA | 0.857 | 0.143 |
| SE | 8835 | DOCK2 | DOCK2 | chr5 | + | 170008687 | 170008746 | 170008496 | 170008597 | 170018959 | 170019108 | 74 | 29 | 11 | 0 | 1 | 0 | 2 | 21 | 0 | 0 | 98 | 49 | 1 | 1 | 1 | 0.935 | 1 | 1 | 1 | -0.022 |
| SE | 8836 | DOCK2 | DOCK2 | chr5 | + | 170034398 | 170034555 | 170027862 | 170027948 | 170036514 | 170036555 | 61 | 8 | 18 | 1 | 0 | 0 | 0 | 16 | 0 | 0 | 98 | 49 | 1 | 1 | 0.968 | 1 | 1 | NA | 1 | -0.011 |
| SE | 8837 | DOCK2 | DOCK2 | chr5 | + | 170045815 | 170045905 | 170042012 | 170042132 | 170047509 | 170047614 | 49 | 19 | 5 | 0 | 1 | 0 | 1 | 14 | 0 | 0 | 98 | 49 | 1 | 1 | 1 | 0.905 | 1 | 1 | 1 | -0.032 |
| SE | 8841 | RPLP2 | RPLP2 | chr11 | + | 810233 | 810357 | 809935 | 810039 | 811596 | 811645 | 104 | 26 | 3 | 2 | 0 | 0 | 1 | 24 | 0 | 0 | 98 | 49 | 1 | 1 | 0.963 | 1 | 1 | 1 | 1 | -0.012 |
| SE | 8849 | CORO1A | CORO1A | chr16 | + | 30188360 | 30188576 | 30188191 | 30188249 | 30188859 | 30189076 | 248 | 133 | 94 | 0 | 1 | 0 | 2 | 46 | 0 | 0 | 98 | 49 | 1 | 1 | 1 | 0.985 | 1 | 1 | 1 | -0.005 |
| SE | 8852 | NCOA6 | NCOA6 | chr20 | - | 34727258 | 34727407 | 34714773 | 34715365 | 34732558 | 34732595 | 9 | 11 | 4 | 0 | 0 | 0 | 0 | 1 | 0 | 2 | 98 | 49 | 1 | 1 | 1 | 1 | 1 | NA | 0.2 | 0.8 |
| SE | 8857 | SEPSECS | SEPSECS | chr4 | - | 25145003 | 25145133 | 25144773 | 25144865 | 25151959 | 25152062 | 0 | 0 | 2 | 2 | 0 | 0 | 0 | 1 | 0 | 0 | 98 | 49 | 1 | 1 | 0 | NA | 1 | NA | 1 | -0.5 |
| SE | 8859 | GUK1 | GUK1 | chr1 | + | 228145510 | 228145624 | 228140227 | 228140363 | 228146025 | 228146067 | 26 | 8 | 4 | 2 | 0 | 0 | 0 | 3 | 0 | 0 | 98 | 49 | 1 | 1 | 0.867 | 1 | 1 | NA | 1 | -0.044 |
| SE | 8861 | IFNAR1 | IFNAR1 | chr21 | + | 33343267 | 33343422 | 33340998 | 33341174 | 33343534 | 33343676 | 51 | 46 | 6 | 0 | 0 | 0 | 0 | 6 | 0 | 1 | 98 | 49 | 1 | 1 | 1 | 1 | 1 | NA | 0.75 | 0.25 |
| SE | 8863 | ZC3HAV1 | ZC3HAV1 | chr7 | - | 139061035 | 139061138 | 139055204 | 139055295 | 139064878 | 139064999 | 22 | 5 | 2 | 0 | 1 | 0 | 1 | 2 | 0 | 0 | 98 | 49 | 1 | 1 | 1 | 0.714 | 1 | 1 | 1 | -0.095 |
| SE | 8864 | ZC3HAV1 | ZC3HAV1 | chr7 | - | 139076285 | 139076409 | 139073855 | 139074030 | 139078551 | 139078653 | 30 | 7 | 2 | 5 | 4 | 1 | 0 | 12 | 0 | 0 | 98 | 49 | 1 | 1 | 0.75 | 0.467 | 0.5 | NA | 1 | -0.428 |
| SE | 8866 | SYF2 | SYF2 | chr1 | - | 25228997 | 25229123 | 25228117 | 25228235 | 25232103 | 25232211 | 15 | 21 | 19 | 1 | 0 | 0 | 0 | 1 | 0 | 0 | 98 | 49 | 1 | 1 | 0.882 | 1 | 1 | NA | 1 | -0.039 |
| SE | 8869 | SRF | SRF | chr6 | + | 43175705 | 43175967 | 43173846 | 43174113 | 43176547 | 43176667 | 8 | 1 | 7 | 1 | 0 | 0 | 0 | 3 | 0 | 0 | 98 | 49 | 1 | 1 | 0.8 | 1 | 1 | NA | 1 | -0.067 |
| SE | 8872 | CLMN | CLMN | chr14 | - | 95196497 | 95196694 | 95194535 | 95194596 | 95202837 | 95204463 | 2 | 0 | 0 | 1 | 0 | 0 | 0 | 2 | 0 | 0 | 98 | 49 | 1 | 1 | 0.5 | NA | NA | NA | 1 | -0.5 |
| SE | 8874 | TRIM8 | TRIM8 | chr10 | + | 102654652 | 102654748 | 102644494 | 102645187 | 102655079 | 102655313 | 26 | 7 | 7 | 0 | 1 | 0 | 0 | 8 | 0 | 1 | 98 | 49 | 1 | 1 | 1 | 0.778 | 1 | NA | 0.8 | 0.126 |
| SE | 8878 | CPEB2 | CPEB2 | chr4 | + | 15007304 | 15007586 | 15002673 | 15004335 | 15017187 | 15017278 | 8 | 1 | 1 | 0 | 0 | 0 | 0 | 4 | 0 | 1 | 98 | 49 | 1 | 1 | 1 | 1 | 1 | NA | 0.667 | 0.333 |
| SE | 8879 | CPEB2 | CPEB2 | chr4 | + | 15007304 | 15007595 | 15002673 | 15004335 | 15017187 | 15017278 | 6 | 3 | 1 | 0 | 0 | 0 | 0 | 3 | 0 | 1 | 98 | 49 | 1 | 1 | 1 | 1 | 1 | NA | 0.6 | 0.4 |
| SE | 8880 | CPEB2 | CPEB2 | chr4 | + | 15008337 | 15008427 | 15007304 | 15007586 | 15017187 | 15017278 | 1 | 0 | 0 | 3 | 1 | 0 | 0 | 0 | 0 | 2 | 98 | 49 | 1 | 1 | 0.143 | 0 | NA | NA | 0 | 0.071 |
| SE | 8883 | DENND1B | DENND1B | chr1 | - | 197647054 | 197647114 | 197645689 | 197645743 | 197652234 | 197652315 | 8 | 1 | 2 | 7 | 0 | 4 | 1 | 0 | 0 | 0 | 98 | 49 | 1 | 1 | 0.364 | 1 | 0.2 | 1 | NA | -0.479 |
| SE | 8884 | DENND1B | DENND1B | chr1 | - | 197715030 | 197715074 | 197674119 | 197674169 | 197772867 | 197772932 | 4 | 0 | 2 | 1 | 0 | 0 | 2 | 1 | 0 | 0 | 93 | 49 | 1 | 1 | 0.678 | NA | 1 | 1 | 1 | -0.161 |
| SE | 8891 | PTPRA | PTPRA | chr20 | + | 2975214 | 2975241 | 2964860 | 2965202 | 2986764 | 2986849 | 1 | 0 | 0 | 5 | 2 | 0 | 0 | 0 | 0 | 3 | 76 | 49 | 1 | 1 | 0.114 | 0 | NA | NA | 0 | 0.057 |
| SE | 8892 | PTPRA | PTPRA | chr20 | + | 3022053 | 3022220 | 3021308 | 3021428 | 3022688 | 3022824 | 8 | 1 | 2 | 0 | 1 | 0 | 0 | 6 | 0 | 0 | 98 | 49 | 1 | 1 | 1 | 0.333 | 1 | NA | 1 | -0.222 |
| SE | 8896 | CASP5 | CASP5 | chr11 | - | 105007082 | 105007334 | 105003273 | 105003383 | 105008806 | 105008980 | 13 | 2 | 0 | 1 | 0 | 0 | 0 | 2 | 0 | 0 | 98 | 49 | 1 | 1 | 0.867 | 1 | NA | NA | 1 | -0.067 |
| SE | 8917 | PITHD1 | PITHD1 | chr1 | + | 23779437 | 23779481 | 23778385 | 23778713 | 23779863 | 23779941 | 0 | 1 | 2 | 1 | 0 | 0 | 1 | 0 | 0 | 0 | 93 | 49 | 1 | 1 | 0 | 1 | 1 | 1 | NA | -0.333 |
| SE | 8923 | DISC1 | DISC1 | chr1 | + | 231720847 | 231720924 | 231701954 | 231702024 | 231749925 | 231750619 | 1 | 0 | 0 | 4 | 7 | 3 | 0 | 0 | 0 | 3 | 98 | 49 | 1 | 1 | 0.111 | 0 | 0 | NA | 0 | 0.037 |
| SE | 8932 | PLCB2 | PLCB2 | chr15 | - | 40290024 | 40290082 | 40289271 | 40289358 | 40290576 | 40290672 | 63 | 24 | 12 | 0 | 3 | 0 | 0 | 8 | 0 | 0 | 98 | 49 | 1 | 1 | 1 | 0.8 | 1 | NA | 1 | -0.067 |
| SE | 8933 | PLCB2 | PLCB2 | chr15 | - | 40291605 | 40291650 | 40291264 | 40291487 | 40291848 | 40291924 | 10 | 5 | 0 | 10 | 3 | 1 | 0 | 1 | 0 | 2 | 94 | 49 | 1 | 1 | 0.343 | 0.465 | 0 | NA | 0.207 | 0.062 |
| SE | 8934 | PLCB2 | PLCB2 | chr15 | - | 40297876 | 40297959 | 40297520 | 40297605 | 40298222 | 40298380 | 59 | 15 | 5 | 1 | 0 | 0 | 1 | 6 | 0 | 0 | 98 | 49 | 1 | 1 | 0.967 | 1 | 1 | 1 | 1 | -0.011 |
| SE | 8940 | RBM10 | RBM10 | chrX | + | 47147356 | 47147498 | 47145217 | 47145485 | 47169314 | 47169498 | 14 | 2 | 2 | 0 | 1 | 0 | 2 | 3 | 0 | 0 | 98 | 49 | 1 | 1 | 1 | 0.5 | 1 | 1 | 1 | -0.167 |
| SE | 8942 | RBM10 | RBM10 | chrX | + | 47175018 | 47175092 | 47173127 | 47173197 | 47176499 | 47176586 | 2 | 4 | 0 | 3 | 0 | 0 | 0 | 1 | 0 | 0 | 98 | 49 | 1 | 1 | 0.25 | 1 | NA | NA | 1 | -0.375 |

| | | | | | | | | | | | | | | | | | | | | | | | | | | | | | | | |
|---|---|---|---|---|---|---|---|---|---|---|---|---|---|---|---|---|---|---|---|---|---|---|---|---|---|---|---|---|---|---|---|
| SE | 8967 | HEATR3 | HEATR3 | chr16 | + | 50072604 | 50072714 | 50070177 | 50070290 | 50075570 | 50075711 | 7 | 0 | 0 | 2 | 0 | 0 | 0 | 0 | 0 | 2 | 98 | 49 | 1 | 1 | 0.636 | NA | NA | NA | 0 | 0.636 |
| SE | 8971 | FCGR2A | FCGR2A | chr1 | + | 161506333 | 161506591 | 161505986 | 161506007 | 161509819 | 161510074 | 225 | 98 | 85 | 1 | 1 | 0 | 2 | 42 | 0 | 0 | 98 | 49 | 1 | 1 | 0.991 | 0.98 | 1 | 1 | 1 | -0.01 |
| SE | 8972 | FCGR2A | FCGR2A | chr1 | + | 161506336 | 161506591 | 161505986 | 161506007 | 161509819 | 161510074 | 241 | 93 | 87 | 1 | 1 | 0 | 1 | 45 | 0 | 0 | 98 | 49 | 1 | 1 | 0.992 | 0.979 | 1 | 1 | 1 | -0.01 |
| SE | 8974 | SGPL1 | SGPL1 | chr10 | + | 70854707 | 70854855 | 70851142 | 70851210 | 70857613 | 70857690 | 6 | 0 | 1 | 1 | 0 | 0 | 0 | 2 | 0 | 0 | 98 | 49 | 1 | 1 | 0.75 | NA | 1 | NA | 1 | -0.125 |
| SE | 8975 | SDCCAG8 | SDCCAG8 | chr1 | + | 243293090 | 243293219 | 243286271 | 243286397 | 243304712 | 243304777 | 5 | 1 | 0 | 0 | 0 | 0 | 0 | 0 | 0 | 1 | 98 | 49 | 1 | 1 | 1 | 1 | NA | NA | 0 | 1 |
| SE | 8977 | SAT1 | SAT1 | chrX | + | 23783799 | 23783883 | 23783657 | 23783709 | 23785327 | 23785429 | 590 | 359 | 141 | 2 | 3 | 0 | 20 | 265 | 1 | 1 | 98 | 49 | 1 | 1 | 0.993 | 0.984 | 1 | 0.909 | 0.993 | 0.041 |
| SE | 8982 | GTF2F1 | GTF2F1 | chr19 | - | 6383310 | 6383495 | 6381696 | 6381850 | 6387388 | 6387559 | 5 | 3 | 0 | 0 | 0 | 0 | 0 | 1 | 0 | 1 | 98 | 49 | 1 | 1 | 1 | 1 | NA | NA | 0.333 | 0.667 |
| SE | 8983 | RUNX2 | RUNX2 | chr6 | + | 45491940 | 45492114 | 45437946 | 45438051 | 45512245 | 45512407 | 5 | 1 | 5 | 1 | 0 | 0 | 0 | 1 | 0 | 2 | 98 | 49 | 1 | 1 | 0.714 | 1 | 1 | NA | 0.2 | 0.705 |
| SE | 8989 | GLYR1 | GLYR1 | chr16 | - | 4821379 | 4821453 | 4817597 | 4817697 | 4822874 | 4822931 | 13 | 4 | 2 | 1 | 0 | 0 | 3 | 6 | 0 | 0 | 98 | 49 | 1 | 1 | 0.867 | 1 | 1 | 1 | 1 | -0.044 |
| SE | 8991 | GLYR1 | GLYR1 | chr16 | - | 4831978 | 4832221 | 4823820 | 4823907 | 4832773 | 4832912 | 12 | 13 | 1 | 2 | 2 | 0 | 0 | 1 | 0 | 0 | 98 | 49 | 1 | 1 | 0.75 | 0.765 | 1 | NA | 1 | -0.162 |
| SE | 9001 | ZDHHC20 | ZDHHC20 | chr13 | - | 21378660 | 21378738 | 21372570 | 21376655 | 21381433 | 21381549 | 3 | 0 | 2 | 7 | 2 | 0 | 0 | 3 | 0 | 2 | 98 | 49 | 1 | 1 | 0.176 | 0 | 1 | NA | 0.429 | -0.037 |
| SE | 9002 | ZDHHC20 | ZDHHC20 | chr13 | - | 21381900 | 21381945 | 21381433 | 21381546 | 21382919 | 21383009 | 1 | 1 | 6 | 1 | 0 | 0 | 0 | 2 | 0 | 3 | 94 | 49 | 1 | 1 | 0.343 | 1 | 1 | NA | 0.258 | 0.523 |
| SE | 9003 | ZDHHC20 | ZDHHC20 | chr13 | - | 21381900 | 21381945 | 21381433 | 21381549 | 21382919 | 21383009 | 1 | 1 | 6 | 7 | 0 | 0 | 0 | 3 | 0 | 5 | 94 | 49 | 1 | 1 | 0.069 | 1 | 1 | NA | 0.238 | 0.452 |
| SE | 9004 | ZDHHC20 | ZDHHC20 | chr13 | - | 21400372 | 21400493 | 21391721 | 21391854 | 21401652 | 21401685 | 11 | 5 | 0 | 0 | 4 | 0 | 0 | 1 | 0 | 0 | 98 | 49 | 1 | 1 | 1 | 0.385 | NA | NA | 1 | -0.307 |
| SE | 9005 | ZDHHC20 | ZDHHC20 | chr13 | - | 21404224 | 21404342 | 21402796 | 21402866 | 21413651 | 21413772 | 0 | 0 | 0 | 6 | 0 | 0 | 0 | 1 | 0 | 3 | 98 | 49 | 1 | 1 | 0 | NA | NA | NA | 0.143 | -0.143 |
| SE | 9006 | ZDHHC20 | ZDHHC20 | chr13 | - | 21421060 | 21421164 | 21413651 | 21413772 | 21459053 | 21459370 | 7 | 4 | 2 | 0 | 1 | 0 | 0 | 3 | 0 | 0 | 98 | 49 | 1 | 1 | 1 | 0.667 | 1 | NA | 1 | -0.111 |
| SE | 9008 | HLA-B | HLA-B | chr6 | - | 31354632 | 31354665 | 31354478 | 31354526 | 31355106 | 31355223 | 1036 | 480 | 341 | 0 | 0 | 0 | 16 | 132 | 0 | 1 | 82 | 49 | 1 | 1 | 1 | 1 | 1 | 1 | 0.987 | 0.006 |
| SE | 9010 | HLA-B | HLA-B | chr6 | - | 31356166 | 31356442 | 31355316 | 31355592 | 31356687 | 31356957 | 1295 | 658 | 422 | 0 | 1 | 0 | 23 | 208 | 0 | 0 | 98 | 49 | 1 | 1 | 1 | 0.997 | 1 | 1 | 1 | -0.001 |
| SE | 9017 | HSD17B11 | HSD17B11 | chr4 | - | 87374698 | 87374830 | 87372708 | 87372815 | 87382254 | 87382362 | 170 | 78 | 84 | 1 | 0 | 0 | 3 | 26 | 0 | 1 | 98 | 49 | 1 | 1 | 0.988 | 1 | 1 | 1 | 0.929 | 0.031 |
| SE | 9023 | CDC123 | CDC123 | chr10 | + | 12249580 | 12249718 | 12246148 | 12246277 | 12250310 | 12250590 | 22 | 15 | 11 | 5 | 0 | 0 | 0 | 3 | 0 | 0 | 98 | 49 | 1 | 1 | 0.687 | 1 | 1 | NA | 1 | -0.104 |
| SE | 9031 | DENND5A | DENND5A | chr11 | - | 9142721 | 9142845 | 9141939 | 9142108 | 9143402 | 9143485 | 80 | 32 | 21 | 0 | 2 | 0 | 0 | 11 | 0 | 0 | 98 | 49 | 1 | 1 | 1 | 0.889 | 1 | NA | 1 | -0.037 |
| SE | 9032 | DENND5A | DENND5A | chr11 | - | 9142721 | 9142845 | 9141939 | 9142108 | 9144096 | 9144278 | 65 | 20 | 10 | 1 | 6 | 0 | 0 | 11 | 0 | 0 | 98 | 49 | 1 | 1 | 0.97 | 0.625 | 1 | NA | 1 | -0.135 |
| SE | 9033 | DENND5A | DENND5A | chr11 | - | 9143402 | 9143485 | 9141939 | 9142108 | 9144096 | 9144278 | 36 | 15 | 7 | 1 | 6 | 0 | 0 | 7 | 0 | 0 | 98 | 49 | 1 | 1 | 0.947 | 0.556 | 1 | NA | 1 | -0.166 |
| SE | 9034 | DENND5A | DENND5A | chr11 | - | 9143402 | 9143485 | 9142721 | 9142845 | 9144096 | 9144278 | 58 | 26 | 18 | 7 | 1 | 0 | 0 | 7 | 0 | 0 | 98 | 49 | 1 | 1 | 0.806 | 0.929 | 1 | NA | 1 | -0.088 |
| SE | 9037 | DENND5A | DENND5A | chr11 | - | 9150679 | 9150764 | 9150080 | 9150209 | 9152357 | 9152442 | 63 | 40 | 27 | 1 | 2 | 0 | 0 | 16 | 0 | 0 | 98 | 49 | 1 | 1 | 0.969 | 0.909 | 1 | NA | 1 | -0.041 |
| SE | 9043 | CCAR1 | CCAR1 | chr10 | + | 68756272 | 68756483 | 68755369 | 68755536 | 68757293 | 68757377 | 9 | 0 | 0 | 4 | 0 | 0 | 0 | 2 | 0 | 0 | 98 | 49 | 1 | 1 | 0.529 | NA | NA | NA | 1 | -0.471 |
| SE | 9045 | CEP57 | CEP57 | chr11 | + | 95795515 | 95795559 | 95790460 | 95790743 | 95799231 | 95799388 | 1 | 1 | 0 | 2 | 2 | 2 | 0 | 0 | 0 | 1 | 93 | 49 | 1 | 1 | 0.209 | 0.209 | 0 | NA | 0 | 0.139 |
| SE | 9046 | CEP57 | CEP57 | chr11 | + | 95822498 | 95822576 | 95821870 | 95821978 | 95827785 | 95828027 | 1 | 2 | 0 | 0 | 4 | 1 | 0 | 2 | 0 | 0 | 98 | 49 | 1 | 1 | 1 | 0.2 | 0 | NA | 1 | -0.6 |
| SE | 9047 | STAT2 | STAT2 | chr12 | - | 56346818 | 56346955 | 56346441 | 56346624 | 56348528 | 56348623 | 12 | 8 | 0 | 0 | 2 | 0 | 2 | 22 | 0 | 0 | 98 | 49 | 1 | 1 | 1 | 0.667 | NA | 1 | 1 | -0.166 |
| SE | 9049 | STAT2 | STAT2 | chr12 | - | 56350096 | 56350190 | 56349588 | 56349636 | 56350411 | 56350432 | 26 | 14 | 3 | 2 | 0 | 0 | 0 | 18 | 0 | 0 | 98 | 49 | 1 | 1 | 0.867 | 1 | 1 | NA | 1 | -0.044 |
| SE | 9054 | CPNE2 | CPNE2 | chr16 | + | 57137148 | 57137282 | 57134774 | 57134826 | 57146084 | 57146321 | 11 | 4 | 2 | 1 | 0 | 0 | 1 | 5 | 0 | 1 | 98 | 49 | 1 | 1 | 0.846 | 1 | 1 | 1 | 0.714 | 0.092 |
| SE | 9055 | AGAP2 | AGAP2 | chr12 | - | 57727680 | 57727771 | 57727359 | 57727582 | 57727936 | 57728085 | 15 | 2 | 1 | 4 | 0 | 0 | 0 | 3 | 0 | 0 | 98 | 49 | 1 | 1 | 0.652 | 1 | 1 | NA | 1 | -0.116 |
| SE | 9056 | AGAP2 | AGAP2 | chr12 | - | 57727936 | 57728085 | 57727359 | 57727582 | 57729638 | 57729767 | 7 | 1 | 0 | 0 | 0 | 0 | 0 | 0 | 1 | 0 | 98 | 49 | 1 | 1 | 1 | 1 | NA | 0 | NA | 1 |
| SE | 9058 | AGAP2 | AGAP2 | chr12 | - | 57729638 | 57729767 | 57727359 | 57727582 | 57730494 | 57730614 | 6 | 2 | 2 | 1 | 0 | 0 | 2 | 0 | 0 | 0 | 98 | 49 | 1 | 1 | 0.75 | 1 | 1 | 1 | NA | -0.083 |
| SE | 9059 | AGAP2 | AGAP2 | chr12 | - | 57730494 | 57730614 | 57729638 | 57729767 | 57730790 | 57730953 | 13 | 2 | 3 | 1 | 0 | 0 | 2 | 0 | 0 | 0 | 98 | 49 | 1 | 1 | 0.867 | 1 | 1 | 1 | NA | -0.044 |
| SE | 9060 | AGAP2 | AGAP2 | chr12 | - | 57731808 | 57731967 | 57731555 | 57731642 | 57732402 | 57732512 | 17 | 7 | 0 | 3 | 0 | 0 | 1 | 1 | 0 | 0 | 98 | 49 | 1 | 1 | 0.739 | 1 | NA | 1 | 1 | -0.131 |
| SE | 9061 | AGAP2 | AGAP2 | chr12 | - | 57732844 | 57732979 | 57732402 | 57732512 | 57734025 | 57734173 | 13 | 3 | 0 | 1 | 0 | 0 | 0 | 2 | 0 | 0 | 98 | 49 | 1 | 1 | 0.867 | 1 | NA | NA | 1 | -0.067 |
| SE | 9062 | SIPA1 | SIPA1 | chr11 | + | 65642249 | 65642377 | 65640800 | 65641600 | 65642462 | 65642639 | 40 | 21 | 14 | 4 | 1 | 0 | 0 | 14 | 0 | 0 | 98 | 49 | 1 | 1 | 0.833 | 0.913 | 1 | NA | 1 | -0.085 |
| SE | 9064 | SIPA1 | SIPA1 | chr11 | + | 65645853 | 65645957 | 65644954 | 65645129 | 65646220 | 65646378 | 49 | 27 | 20 | 0 | 0 | 0 | 0 | 19 | 0 | 2 | 98 | 49 | 1 | 1 | 1 | 1 | 1 | NA | 0.826 | 0.174 |
| SE | 9066 | SIPA1 | SIPA1 | chr11 | + | 65650400 | 65650479 | 65650137 | 65650192 | 65650568 | 65650920 | 54 | 31 | 18 | 0 | 0 | 0 | 0 | 9 | 1 | 0 | 98 | 49 | 1 | 1 | 1 | 1 | 1 | 0 | 1 | 0.5 |
| SE | 9071 | ESRRA | ESRRA | chr11 | + | 64307167 | 64307504 | 64305527 | 64305736 | 64313950 | 64314067 | 5 | 3 | 6 | 0 | 0 | 0 | 0 | 3 | 0 | 1 | 98 | 49 | 1 | 1 | 1 | 1 | 1 | NA | 0.6 | 0.4 |
| SE | 9074 | PRPF3 | PRPF3 | chr1 | + | 150334934 | 150335241 | 150332978 | 150333199 | 150338159 | 150338326 | 22 | 7 | 0 | 2 | 0 | 0 | 0 | 6 | 0 | 0 | 98 | 49 | 1 | 1 | 0.846 | 1 | NA | NA | 1 | -0.077 |
| SE | 9075 | PRPF3 | PRPF3 | chr1 | + | 150340397 | 150340477 | 150338159 | 150338326 | 150343308 | 150343452 | 9 | 8 | 1 | 1 | 0 | 0 | 0 | 4 | 0 | 0 | 98 | 49 | 1 | 1 | 0.818 | 1 | 1 | NA | 1 | -0.061 |
| SE | 9076 | PRPF3 | PRPF3 | chr1 | + | 150344433 | 150344547 | 150344161 | 150344261 | 150346017 | 150346136 | 11 | 1 | 5 | 2 | 0 | 0 | 0 | 3 | 0 | 0 | 98 | 49 | 1 | 1 | 0.733 | 1 | 1 | NA | 1 | -0.089 |
| SE | 9078 | BBX | BBX | chr3 | + | 107789786 | 107789876 | 107778370 | 107778519 | 107791239 | 107791299 | 6 | 0 | 2 | 1 | 0 | 0 | 0 | 0 | 0 | 2 | 98 | 49 | 1 | 1 | 0.75 | NA | 1 | NA | 0 | 0.875 |
| SE | 9087 | FBXL3 | FBXL3 | chr13 | - | 77021512 | 77021861 | 77018599 | 77018722 | 77026826 | 77027196 | 14 | 20 | 1 | 1 | 0 | 0 | 0 | 1 | 0 | 0 | 98 | 49 | 1 | 1 | 0.875 | 1 | 1 | NA | 1 | -0.042 |
| SE | 9103 | GAPVD1 | GAPVD1 | chr9 | + | 125295457 | 125295574 | 125268935 | 125268984 | 125298889 | 125299106 | 2 | 0 | 0 | 1 | 0 | 0 | 1 | 1 | 0 | 0 | 98 | 49 | 1 | 1 | 0.5 | NA | NA | 1 | 1 | -0.5 |
| SE | 9104 | GAPVD1 | GAPVD1 | chr9 | + | 125301982 | 125302826 | 125298889 | 125299106 | 125305062 | 125305149 | 28 | 8 | 3 | 1 | 0 | 0 | 0 | 11 | 0 | 0 | 98 | 49 | 1 | 1 | 0.933 | 1 | 1 | NA | 1 | -0.022 |
| SE | 9106 | GAPVD1 | GAPVD1 | chr9 | + | 125360527 | 125360725 | 125359419 | 125359492 | 125362605 | 125365011 | 13 | 4 | 1 | 0 | 2 | 0 | 0 | 5 | 0 | 2 | 98 | 49 | 1 | 1 | 1 | 0.5 | 1 | NA | 0.556 | 0.277 |
| SE | 9107 | MAN2C1 | MAN2C1 | chr15 | - | 75362641 | 75362748 | 75362342 | 75362453 | 75364487 | 75364665 | 4 | 2 | 0 | 0 | 0 | 0 | 0 | 4 | 0 | 1 | 98 | 49 | 1 | 1 | 1 | 1 | NA | NA | 0.667 | 0.333 |
| SE | 9110 | TPCN1 | TPCN1 | chr12 | + | 113284717 | 113284771 | 113284580 | 113284637 | 113285888 | 113285961 | 1 | 0 | 0 | 1 | 0 | 0 | 0 | 2 | 0 | 0 | 98 | 49 | 1 | 1 | 0.333 | NA | NA | NA | 1 | -0.667 |
| SE | 9114 | WDR37 | WDR37 | chr10 | + | 1077906 | 1078003 | 1072115 | 1072293 | 1080010 | 1080106 | 10 | 4 | 7 | 0 | 2 | 0 | 0 | 3 | 0 | 0 | 98 | 49 | 1 | 1 | 1 | 0.5 | 1 | NA | 1 | -0.167 |
| SE | 9115 | WDR37 | WDR37 | chr10 | + | 1103601 | 1103836 | 1096169 | 1096246 | 1124217 | 1124352 | 2 | 0 | 0 | 0 | 0 | 0 | 0 | 0 | 0 | 1 | 98 | 49 | 1 | 1 | 1 | NA | NA | NA | 0 | 1 |
| SE | 9116 | WDR37 | WDR37 | chr10 | + | 1105125 | 1105267 | 1103601 | 1103836 | 1124217 | 1124352 | 3 | 1 | 0 | 1 | 0 | 0 | 0 | 4 | 0 | 0 | 98 | 49 | 1 | 1 | 0.6 | 1 | NA | NA | 1 | -0.2 |
| SE | 9130 | DPM1 | DPM1 | chr20 | - | 50940864 | 50940933 | 50936147 | 50936262 | 50942030 | 50942126 | 18 | 4 | 3 | 0 | 0 | 1 | 0 | 3 | 0 | 0 | 98 | 49 | 1 | 1 | 1 | 1 | 0.6 | NA | 1 | -0.133 |
| SE | 9131 | DPM1 | DPM1 | chr20 | - | 50945736 | 50945762 | 50942030 | 50942126 | 50945846 | 50945923 | 3 | 7 | 5 | 0 | 1 | 0 | 0 | 1 | 0 | 0 | 75 | 49 | 1 | 1 | 1 | 0.821 | 1 | NA | 1 | -0.06 |
| SE | 9137 | HERC4 | HERC4 | chr10 | - | 67990210 | 67990400 | 67988662 | 67988835 | 67990903 | 67991015 | 8 | 4 | 3 | 1 | 0 | 0 | 0 | 1 | 0 | 2 | 98 | 49 | 1 | 1 | 0.8 | 1 | 1 | NA | 0.2 | 0.733 |
| SE | 9139 | HERC4 | HERC4 | chr10 | - | 68044403 | 68044563 | 68038092 | 68038169 | 68072882 | 68073186 | 20 | 10 | 0 | 1 | 0 | 0 | 0 | 1 | 0 | 0 | 98 | 49 | 1 | 1 | 0.909 | 1 | NA | NA | 1 | -0.045 |
| SE | 9141 | TLR10 | TLR10 | chr4 | - | 38775774 | 38775963 | 38772238 | 38775652 | 38776109 | 38776426 | 1 | 0 | 0 | 0 | 0 | 3 | 0 | 2 | 0 | 0 | 98 | 49 | 1 | 1 | 1 | NA | 0 | NA | 1 | -0.5 |
| SE | 9150 | FBXO18 | FBXO18 | chr10 | + | 5903019 | 5903175 | 5890202 | 5890346 | 5906036 | 5906632 | 2 | 3 | 0 | 0 | 1 | 0 | 0 | 3 | 0 | 0 | 98 | 49 | 1 | 1 | 1 | 0.6 | NA | NA | 1 | -0.2 |
| SE | 9151 | FBXO18 | FBXO18 | chr10 | + | 5918341 | 5918478 | 5917589 | 5917676 | 5921257 | 5921357 | 4 | 6 | 2 | 2 | 0 | 0 | 0 | 1 | 0 | 0 | 98 | 49 | 1 | 1 | 0.5 | 1 | 1 | NA | 1 | -0.167 |
| SE | 9161 | CHD4 | CHD4 | chr12 | - | 6582136 | 6582281 | 6581648 | 6581814 | 6582614 | 6582748 | 33 | 7 | 10 | 1 | 1 | 0 | 0 | 4 | 0 | 1 | 98 | 49 | 1 | 1 | 0.943 | 0.778 | 1 | NA | 0.667 | 0.24 |
| SE | 9162 | CHD4 | CHD4 | chr12 | - | 6583197 | 6583378 | 6583026 | 6583113 | 6587383 | 6587559 | 12 | 5 | 5 | 0 | 1 | 0 | 0 | 5 | 0 | 0 | 98 | 49 | 1 | 1 | 1 | 0.714 | 1 | NA | 1 | -0.095 |
| SE | 9163 | CHD4 | CHD4 | chr12 | - | 6600925 | 6601053 | 6600533 | 6600669 | 6601288 | 6601530 | 24 | 21 | 1 | 2 | 0 | 0 | 0 | 7 | 0 | 0 | 98 | 49 | 1 | 1 | 0.857 | 1 | 1 | NA | 1 | -0.048 |
| SE | 9166 | CELF2 | CELF2 | chr10 | + | 11266597 | 11266677 | 11257737 | 11257872 | 11270665 | 11270824 | 30 | 21 | 7 | 13 | 13 | 6 | 0 | 15 | 0 | 5 | 98 | 49 | 1 | 1 | 0.536 | 0.447 | 0.368 | NA | 0.6 | -0.15 |
| SE | 9167 | CELF2 | CELF2 | chr10 | + | 11266597 | 11266677 | 11257737 | 11257872 | 11275056 | 11275120 | 20 | 6 | 1 | 3 | 6 | 0 | 0 | 9 | 0 | 0 | 98 | 49 | 1 | 1 | 0.769 | 0.333 | 1 | NA | 1 | -0.299 |
| SE | 9168 | CELF2 | CELF2 | chr10 | + | 11270665 | 11270824 | 11257737 | 11257872 | 11275056 | 11275120 | 104 | 45 | 14 | 3 | 6 | 0 | 0 | 11 | 0 | 0 | 98 | 49 | 1 | 1 | 0.945 | 0.789 | 1 | NA | 1 | -0.089 |
| SE | 9169 | CELF2 | CELF2 | chr10 | + | 11270665 | 11270824 | 11266597 | 11266677 | 11275056 | 11275120 | 102 | 47 | 14 | 1 | 0 | 0 | 0 | 12 | 0 | 0 | 98 | 49 | 1 | 1 | 0.981 | 1 | 1 | NA | 1 | -0.006 |
| SE | 9170 | CELF2 | CELF2 | chr10 | + | 11288417 | 11288552 | 11275056 | 11275120 | 11314138 | 11314258 | 120 | 37 | 13 | 1 | 0 | 0 | 2 | 18 | 0 | 0 | 98 | 49 | 1 | 1 | 0.984 | 1 | 1 | 1 | 1 | -0.005 |
| SE | 9171 | CELF2 | CELF2 | chr10 | + | 11325835 | 11325979 | 11321188 | 11321386 | 11328925 | 11329097 | 46 | 30 | 19 | 0 | 1 | 0 | 0 | 9 | 0 | 0 | 98 | 49 | 1 | 1 | 1 | 0.937 | 1 | NA | 1 | -0.021 |
| SE | 9172 | TPP1 | TPP1 | chr11 | - | 6616004 | 6616074 | 6615441 | 6615562 | 6616314 | 6616503 | 16 | 3 | 4 | 0 | 0 | 0 | 0 | 13 | 0 | 1 | 98 | 49 | 1 | 1 | 1 | 1 | 1 | NA | 0.867 | 0.133 |
| SE | 9184 | TNNT1 | TNNT1 | chr19 | - | 55146433 | 55146466 | 55145543 | 55145565 | 55146680 | 55146707 | 1 | 1 | 0 | 0 | 0 | 0 | 0 | 0 | 0 | 1 | 82 | 49 | 1 | 1 | 1 | 1 | NA | NA | 0 | 1 |
| SE | 9187 | NCSTN | NCSTN | chr1 | + | 160351221 | 160351372 | 160350104 | 160350250 | 160351695 | 160351805 | 41 | 9 | 3 | 1 | 0 | 0 | 0 | 9 | 0 | 0 | 98 | 49 | 1 | 1 | 0.953 | 1 | 1 | NA | 1 | -0.016 |
| SE | 9188 | NCSTN | NCSTN | chr1 | + | 160355654 | 160355757 | 160354117 | 160354290 | 160355862 | 160355958 | 8 | 4 | 7 | 0 | 0 | 0 | 1 | 4 | 0 | 1 | 98 | 49 | 1 | 1 | 1 | 1 | 1 | 1 | 0.667 | 0.166 |
| SE | 9189 | NCSTN | NCSTN | chr1 | + | 160355862 | 160355958 | 160354117 | 160354290 | 160356259 | 160356347 | 17 | 2 | 2 | 1 | 0 | 0 | 0 | 9 | 0 | 1 | 98 | 49 | 1 | 1 | 0.895 | 1 | 1 | NA | 0.818 | 0.147 |
| SE | 9190 | NCSTN | NCSTN | chr1 | + | 160356259 | 160356347 | 160354117 | 160354290 | 160356599 | 160356754 | 13 | 3 | 5 | 0 | 0 | 0 | 0 | 4 | 0 | 1 | 98 | 49 | 1 | 1 | 1 | 1 | 1 | NA | 0.667 | 0.333 |
| SE | 9191 | C2CD3 | C2CD3 | chr11 | - | 74122987 | 74123135 | 74118227 | 74118382 | 74132843 | 74132972 | 7 | 0 | 0 | 1 | 0 | 0 | 0 | 5 | 0 | 0 | 98 | 49 | 1 | 1 | 0.778 | NA | NA | NA | 1 | -0.222 |
| SE | 9192 | C2CD3 | C2CD3 | chr11 | - | 74161398 | 74161556 | 74139604 | 74139828 | 74168343 | 74168613 | 8 | 2 | 0 | 0 | 0 | 0 | 0 | 2 | 0 | 1 | 98 | 49 | 1 | 1 | 1 | 1 | NA | NA | 0.5 | 0.5 |
| SE | 9200 | ACSL1 | ACSL1 | chr4 | - | 184770398 | 184770476 | 184768315 | 184768450 | 184773080 | 184773154 | 230 | 105 | 146 | 1 | 2 | 0 | 1 | 55 | 0 | 1 | 98 | 49 | 1 | 1 | 0.991 | 0.963 | 1 | 1 | 0.965 | 0.002 |
| SE | 9201 | ACSL1 | ACSL1 | chr4 | - | 184770398 | 184770476 | 184768315 | 184768450 | 184773662 | 184773714 | 85 | 32 | 78 | 1 | 0 | 0 | 1 | 16 | 0 | 1 | 98 | 49 | 1 | 1 | 0.977 | 1 | 1 | 1 | 0.889 | 0.048 |
| SE | 9202 | ACSL1 | ACSL1 | chr4 | - | 184773080 | 184773154 | 184768315 | 184768450 | 184773662 | 184773714 | 163 | 95 | 53 | 1 | 0 | 0 | 2 | 38 | 0 | 1 | 98 | 49 | 1 | 1 | 0.988 | 1 | 1 | 1 | 0.95 | 0.021 |
| SE | 9204 | ACSL1 | ACSL1 | chr4 | - | 184773842 | 184773875 | 184773662 | 184773714 | 184776483 | 184776662 | 231 | 70 | 108 | 1 | 0 | 0 | 2 | 51 | 0 | 0 | 82 | 49 | 1 | 1 | 0.993 | 1 | 1 | 1 | 1 | -0.002 |
| SE | 9206 | ACSL1 | ACSL1 | chr4 | - | 184780331 | 184780433 | 184776883 | 184776983 | 184783926 | 184783991 | 374 | 132 | 114 | 1 | 1 | 0 | 5 | 72 | 0 | 0 | 98 | 49 | 1 | 1 | 0.995 | 0.985 | 1 | 1 | 1 | -0.007 |
| SE | 9207 | ACSL1 | ACSL1 | chr4 | - | 184788378 | 184788513 | 184783926 | 184783991 | 184788616 | 184788731 | 1 | 3 | 1 | 158 | 76 | 30 | 0 | 0 | 1 | 55 | 98 | 49 | 1 | 1 | 0.003 | 0.019 | 0.016 | 0 | 0 | 0.013 |
| SE | 9212 | DIS3 | DIS3 | chr13 | - | 72781288 | 72781356 | 72780845 | 72781003 | 72781604 | 72782128 | 1 | 5 | 1 | 0 | 2 | 0 | 0 | 0 | 0 | 1 | 98 | 49 | 1 | 1 | 1 | 0.556 | 1 | NA | 0 | 0.852 |
| SE | 9214 | BPTF | BPTF | chr17 | + | 67875555 | 67875744 | 67874816 | 67875020 | 67886127 | 67886316 | 4 | 3 | 2 | 4 | 0 | 0 | 0 | 3 | 0 | 0 | 98 | 49 | 1 | 1 | 0.333 | 1 | 1 | NA | 1 | -0.222 |
| SE | 9215 | BPTF | BPTF | chr17 | + | 67875555 | 67875744 | 67874816 | 67875020 | 67891843 | 67892034 | 6 | 1 | 0 | 0 | 2 | 0 | 0 | 1 | 0 | 0 | 98 | 49 | 1 | 1 | 1 | 0.2 | NA | NA | 1 | -0.4 |
| SE | 9216 | BPTF | BPTF | chr17 | + | 67886127 | 67886316 | 67874816 | 67875020 | 67891843 | 67892034 | 10 | 0 | 0 | 0 | 2 | 0 | 0 | 3 | 0 | 0 | 98 | 49 | 1 | 1 | 1 | 0 | NA | NA | 1 | -0.5 |
| SE | 9217 | BPTF | BPTF | chr17 | + | 67886127 | 67886316 | 67875555 | 67875744 | 67891843 | 67892034 | 6 | 2 | 2 | 2 | 0 | 0 | 0 | 6 | 0 | 1 | 98 | 49 | 1 | 1 | 0.6 | 1 | 1 | NA | 0.75 | 0.117 |
| SE | 9218 | BPTF | BPTF | chr17 | + | 67966571 | 67966656 | 67964211 | 67964404 | 67975771 | 67975958 | 14 | 2 | 4 | 0 | 0 | 0 | 1 | 3 | 0 | 1 | 98 | 49 | 1 | 1 | 1 | 1 | 1 | 1 | 0.6 | 0.2 |
| SE | 9221 | CTNND1 | CTNND1 | chr11 | + | 57791384 | 57791673 | 57761761 | 57762046 | 57794009 | 57794081 | 1 | 0 | 0 | 0 | 0 | 0 | 0 | 0 | 0 | 1 | 98 | 49 | 1 | 1 | 1 | NA | NA | NA | 0 | 1 |
| SE | 9222 | CTNND1 | CTNND1 | chr11 | + | 57791493 | 57791673 | 57761761 | 57762046 | 57794009 | 57794081 | 1 | 0 | 0 | 0 | 0 | 0 | 0 | 0 | 0 | 1 | 98 | 49 | 1 | 1 | 1 | NA | NA | NA | 0 | 1 |
| SE | 9234 | FAM107B | FAM107B | chr10 | - | 14553321 | 14553387 | 14530331 | 14530515 | 14604239 | 14604418 | 1 | 4 | 3 | 1 | 0 | 0 | 1 | 0 | 0 | 0 | 98 | 49 | 1 | 1 | 0.333 | 1 | 1 | 1 | NA | -0.222 |
| SE | 9237 | SPEN | SPEN | chr1 | + | 15876201 | 15876678 | 15872815 | 15873136 | 15909320 | 15909481 | 17 | 2 | 1 | 0 | 0 | 2 | 0 | 3 | 0 | 0 | 98 | 49 | 1 | 1 | 1 | 1 | 0.2 | NA | 1 | -0.267 |
| SE | 9239 | SVIL | SVIL | chr10 | - | 29512734 | 29512861 | 29499115 | 29499263 | 29522409 | 29522635 | 34 | 9 | 16 | 2 | 2 | 0 | 0 | 3 | 0 | 0 | 98 | 49 | 1 | 1 | 0.895 | 0.692 | 1 | NA | 1 | -0.138 |
| SE | 9248 | CDC42SE2 | CDC42SE2 | chr5 | + | 131315975 | 131316144 | 131264008 | 131264166 | 131359208 | 131359547 | 17 | 3 | 6 | 0 | 1 | 0 | 0 | 1 | 0 | 0 | 98 | 49 | 1 | 1 | 1 | 0.6 | 1 | NA | 1 | -0.133 |
| SE | 9249 | CDC42SE2 | CDC42SE2 | chr5 | + | 131359208 | 131359547 | 131315975 | 131316144 | 131385542 | 131385644 | 59 | 14 | 16 | 1 | 0 | 0 | 1 | 10 | 0 | 0 | 98 | 49 | 1 | 1 | 0.967 | 1 | 1 | 1 | 1 | -0.011 |
| SE | 9252 | SEL1L | SEL1L | chr14 | - | 81492479 | 81492548 | 81490387 | 81490465 | 81495080 | 81495137 | 20 | 16 | 10 | 1 | 0 | 0 | 0 | 1 | 0 | 0 | 98 | 49 | 1 | 1 | 0.909 | 1 | 1 | NA | 1 | -0.03 |
| SE | 9262 | SLC38A9 | SLC38A9 | chr5 | - | 55649206 | 55649314 | 55645788 | 55645895 | 55652528 | 55652723 | 2 | 2 | 0 | 1 | 0 | 0 | 0 | 2 | 0 | 0 | 98 | 49 | 1 | 1 | 0.5 | 1 | NA | NA | 1 | -0.25 |
| SE | 9268 | OGDH | OGDH | chr7 | + | 44707224 | 44707388 | 44701542 | 44701615 | 44707581 | 44707736 | 6 | 1 | 4 | 0 | 4 | 0 | 0 | 1 | 0 | 0 | 98 | 49 | 1 | 1 | 1 | 0.111 | 1 | NA | 1 | -0.296 |
| SE | 9270 | PCF11 | PCF11 | chr11 | + | 83161326 | 83161452 | 83157094 | 83157631 | 83163678 | 83163867 | 14 | 20 | 0 | 0 | 0 | 1 | 0 | 3 | 0 | 0 | 98 | 49 | 1 | 1 | 1 | 1 | 0 | NA | 1 | -0.333 |
| SE | 9271 | PCF11 | PCF11 | chr11 | + | 83171817 | 83171914 | 83168427 | 83169995 | 83177084 | 83177204 | 48 | 9 | 5 | 0 | 0 | 1 | 0 | 9 | 0 | 0 | 98 | 49 | 1 | 1 | 1 | 1 | 0.714 | NA | 1 | -0.095 |
| SE | 9275 | VIM | VIM | chr10 | + | 17235845 | 17235889 | 17235168 | 17235389 | 17236293 | 17236379 | 301 | 80 | 44 | 4 | 0 | 0 | 12 | 171 | 0 | 0 | 93 | 49 | 1 | 1 | 0.975 | 1 | 1 | 1 | 1 | -0.008 |
| SE | 9276 | VIM | VIM | chr10 | + | 17235845 | 17235889 | 17235168 | 17235389 | 17237229 | 17237593 | 146 | 40 | 21 | 1 | 0 | 0 | 0 | 65 | 0 | 0 | 93 | 49 | 1 | 1 | 0.987 | 1 | 1 | NA | 1 | -0.004 |
| SE | 9277 | VIM | VIM | chr10 | + | 17236293 | 17236379 | 17235168 | 17235389 | 17237229 | 17237593 | 218 | 59 | 45 | 1 | 0 | 0 | 7 | 135 | 0 | 0 | 98 | 49 | 1 | 1 | 0.991 | 1 | 1 | 1 | 1 | -0.003 |
| SE | 9278 | VIM | VIM | chr10 | + | 17236293 | 17236379 | 17235845 | 17235889 | 17237229 | 17237593 | 384 | 100 | 70 | 3 | 0 | 0 | 19 | 241 | 0 | 0 | 98 | 49 | 1 | 1 | 0.985 | 1 | 1 | 1 | 1 | -0.005 |
| SE | 9280 | MGRN1 | MGRN1 | chr16 | + | 4677462 | 4677572 | 4673497 | 4673657 | 4681549 | 4681776 | 4 | 8 | 0 | 0 | 2 | 0 | 0 | 1 | 0 | 0 | 98 | 49 | 1 | 1 | 1 | 0.667 | NA | NA | 1 | -0.166 |
| SE | 9281 | MGRN1 | MGRN1 | chr16 | + | 4680031 | 4680097 | 4677462 | 4677572 | 4681549 | 4681776 | 18 | 8 | 1 | 1 | 3 | 0 | 0 | 3 | 0 | 0 | 98 | 49 | 1 | 1 | 0.9 | 0.571 | 1 | NA | 1 | -0.176 |
| SE | 9283 | RIPK2 | RIPK2 | chr8 | + | 89762828 | 89762982 | 89757746 | 89758233 | 89765340 | 89765496 | 3 | 1 | 0 | 6 | 2 | 0 | 0 | 1 | 0 | 0 | 98 | 49 | 1 | 1 | 0.2 | 0.2 | NA | NA | 1 | -0.8 |
| SE | 9284 | MBNL1 | MBNL1 | chr3 | + | 152299404 | 152300367 | 152268039 | 152269092 | 152414940 | 152415111 | 53 | 20 | 2 | 8 | 4 | 0 | 0 | 6 | 0 | 1 | 98 | 49 | 1 | 1 | 0.768 | 0.714 | 1 | NA | 0.75 | 0.077 |
| SE | 9285 | MBNL1 | MBNL1 | chr3 | + | 152432716 | 152432920 | 152414940 | 152415111 | 152445281 | 152445539 | 101 | 30 | 7 | 2 | 0 | 0 | 1 | 24 | 0 | 1 | 98 | 49 | 1 | 1 | 0.962 | 1 | 1 | 1 | 0.923 | 0.026 |
| SE | 9286 | MBNL1 | MBNL1 | chr3 | + | 152446703 | 152446757 | 152445281 | 152445539 | 152447619 | 152447773 | 3 | 0 | 0 | 51 | 23 | 13 | 0 | 0 | 0 | 10 | 98 | 49 | 1 | 1 | 0.029 | 0 | 0 | NA | 0 | 0.01 |
| SE | 9290 | MBNL1 | MBNL1 | chr3 | + | 152458129 | 152458193 | 152456266 | 152456361 | 152459270 | 152459345 | 1 | 0 | 1 | 35 | 7 | 9 | 0 | 0 | 0 | 5 | 98 | 49 | 1 | 1 | 0.014 | 0 | 0.053 | NA | 0 | 0.022 |
| SE | 9291 | SETD3 | SETD3 | chr14 | - | 99404224 | 99404310 | 99400098 | 99400259 | 99405204 | 99405371 | 2 | 3 | 0 | 1 | 0 | 2 | 0 | 2 | 0 | 0 | 98 | 49 | 1 | 1 | 0.5 | 1 | 0 | NA | 1 | -0.5 |
| SE | 9292 | SETD3 | SETD3 | chr14 | - | 99461191 | 99461340 | 99459112 | 99459185 | 99463485 | 99463578 | 21 | 8 | 0 | 2 | 0 | 0 | 0 | 4 | 0 | 1 | 98 | 49 | 1 | 1 | 0.84 | 1 | NA | NA | 0.667 | 0.253 |
| SE | 9293 | WLS | WLS | chr1 | - | 68153516 | 68153653 | 68150187 | 68150356 | 68155098 | 68155260 | 54 | 15 | 2 | 1 | 0 | 0 | 0 | 9 | 0 | 0 | 98 | 49 | 1 | 1 | 0.964 | 1 | 1 | NA | 1 | -0.012 |
| SE | 9299 | BEST1 | BEST1 | chr11 | + | 61959891 | 61960043 | 61959497 | 61959578 | 61962254 | 61962893 | 44 | 13 | 20 | 1 | 0 | 0 | 1 | 3 | 0 | 1 | 98 | 49 | 1 | 1 | 0.957 | 1 | 1 | 1 | 0.6 | 0.186 |
| SE | 9300 | PIK3CD | PIK3CD | chr1 | + | 9691466 | 9691571 | 9651731 | 9651802 | 9710423 | 9710596 | 11 | 6 | 1 | 0 | 3 | 2 | 0 | 2 | 0 | 0 | 98 | 49 | 1 | 1 | 1 | 0.5 | 0.2 | NA | 1 | -0.433 |
| SE | 9301 | PIK3CD | PIK3CD | chr1 | + | 9717536 | 9717626 | 9716958 | 9717108 | 9718693 | 9718915 | 93 | 40 | 40 | 1 | 0 | 0 | 1 | 8 | 0 | 0 | 98 | 49 | 1 | 1 | 0.979 | 1 | 1 | 1 | 1 | -0.007 |
| SE | 9304 | PIK3CD | PIK3CD | chr1 | + | 9723968 | 9724092 | 9723124 | 9723292 | 9724803 | 9724936 | 34 | 17 | 14 | 1 | 0 | 0 | 0 | 3 | 0 | 0 | 98 | 49 | 1 | 1 | 0.944 | 1 | 1 | NA | 1 | -0.019 |
| SE | 9305 | PIK3CD | PIK3CD | chr1 | + | 9724275 | 9724421 | 9723124 | 9723292 | 9724803 | 9724936 | 43 | 22 | 17 | 1 | 0 | 0 | 0 | 6 | 0 | 0 | 98 | 49 | 1 | 1 | 0.956 | 1 | 1 | NA | 1 | -0.015 |
| SE | 9306 | PIK3CD | PIK3CD | chr1 | + | 9724275 | 9724421 | 9723968 | 9724092 | 9724803 | 9724936 | 75 | 52 | 22 | 1 | 0 | 0 | 3 | 15 | 0 | 0 | 98 | 49 | 1 | 1 | 0.974 | 1 | 1 | 1 | 1 | -0.009 |
| SE | 9309 | PLXDC2 | PLXDC2 | chr10 | + | 20068169 | 20068239 | 20046868 | 20047015 | 20143294 | 20143417 | 73 | 48 | 13 | 0 | 0 | 0 | 0 | 21 | 0 | 1 | 98 | 49 | 1 | 1 | 1 | 1 | 1 | NA | 0.913 | 0.087 |
| SE | 9314 | UBAP2L | UBAP2L | chr1 | + | 154234590 | 154234738 | 154228614 | 154228725 | 154235195 | 154235291 | 3 | 3 | 0 | 4 | 4 | 0 | 0 | 1 | 0 | 0 | 98 | 49 | 1 | 1 | 0.273 | 0.273 | NA | NA | 1 | -0.727 |
| SE | 9315 | UBAP2L | UBAP2L | chr1 | + | 154234590 | 154234759 | 154228614 | 154228725 | 154235195 | 154235291 | 7 | 3 | 1 | 4 | 4 | 0 | 0 | 1 | 0 | 0 | 98 | 49 | 1 | 1 | 0.467 | 0.273 | 1 | NA | 1 | -0.42 |
| SE | 9316 | UBAP2L | UBAP2L | chr1 | + | 154234590 | 154234792 | 154228614 | 154228725 | 154235195 | 154235291 | 4 | 3 | 1 | 4 | 4 | 0 | 0 | 2 | 0 | 0 | 98 | 49 | 1 | 1 | 0.333 | 0.273 | 1 | NA | 1 | -0.465 |
| SE | 9318 | PIP4K2A | PIP4K2A | chr10 | - | 22591628 | 22591781 | 22573310 | 22573457 | 22607926 | 22608023 | 20 | 11 | 1 | 0 | 0 | 0 | 3 | 3 | 0 | 1 | 98 | 49 | 1 | 1 | 1 | 1 | 1 | 1 | 0.6 | 0.2 |
| SE | 9339 | ENKUR | ENKUR | chr10 | - | 24999400 | 24999546 | 24995645 | 24995869 | 25015859 | 25016156 | 0 | 0 | 0 | 1 | 0 | 0 | 0 | 1 | 0 | 0 | 98 | 49 | 1 | 1 | 0 | NA | NA | NA | 1 | -1 |
| SE | 9340 | FAM32A | FAM32A | chr19 | + | 16190519 | 16190573 | 16185623 | 16185765 | 16190886 | 16192046 | 13 | 4 | 6 | 5 | 0 | 1 | 0 | 2 | 0 | 0 | 98 | 49 | 1 | 1 | 0.565 | 1 | 0.75 | NA | 1 | -0.228 |
| SE | 9342 | RBM47 | RBM47 | chr4 | - | 40436440 | 40436647 | 40432650 | 40432862 | 40437770 | 40438924 | 31 | 16 | 8 | 3 | 2 | 0 | 0 | 6 | 0 | 0 | 98 | 49 | 1 | 1 | 0.838 | 0.8 | 1 | NA | 1 | -0.121 |
| SE | 9343 | RBM47 | RBM47 | chr4 | - | 40437770 | 40438924 | 40432650 | 40432862 | 40466576 | 40466699 | 29 | 10 | 8 | 0 | 0 | 0 | 2 | 9 | 0 | 1 | 98 | 49 | 1 | 1 | 1 | 1 | 1 | 1 | 0.818 | 0.091 |
| SE | 9347 | YME1L1 | YME1L1 | chr10 | - | 27147497 | 27147668 | 27145427 | 27145590 | 27148905 | 27149040 | 3 | 0 | 0 | 7 | 1 | 2 | 0 | 0 | 0 | 2 | 98 | 49 | 1 | 1 | 0.176 | 0 | 0 | NA | 0 | 0.059 |
| SE | 9351 | LGALS8 | LGALS8 | chr1 | + | 236542914 | 236543040 | 236542760 | 236542787 | 236543559 | 236543648 | 7 | 7 | 0 | 2 | 1 | 0 | 0 | 0 | 0 | 1 | 98 | 49 | 1 | 1 | 0.636 | 0.778 | NA | NA | 0 | 0.707 |
| SE | 9354 | LMTK2 | LMTK2 | chr7 | + | 98137314 | 98137442 | 98106884 | 98107280 | 98141396 | 98141541 | 18 | 29 | 15 | 1 | 0 | 0 | 1 | 0 | 0 | 0 | 98 | 49 | 1 | 1 | 0.9 | 1 | 1 | 1 | NA | -0.033 |
| SE | 9355 | LMTK2 | LMTK2 | chr7 | + | 98185050 | 98185135 | 98171540 | 98171674 | 98186876 | 98186998 | 8 | 4 | 4 | 0 | 2 | 0 | 0 | 5 | 0 | 0 | 98 | 49 | 1 | 1 | 1 | 0.5 | 1 | NA | 1 | -0.167 |
| SE | 9356 | LMTK2 | LMTK2 | chr7 | + | 98186876 | 98186998 | 98171540 | 98171674 | 98190727 | 98190877 | 5 | 4 | 0 | 0 | 2 | 0 | 0 | 2 | 0 | 0 | 98 | 49 | 1 | 1 | 1 | 0.5 | NA | NA | 1 | -0.25 |
| SE | 9357 | LMTK2 | LMTK2 | chr7 | + | 98203943 | 98204186 | 98203573 | 98203706 | 98205463 | 98209632 | 13 | 8 | 7 | 0 | 1 | 0 | 0 | 2 | 0 | 0 | 98 | 49 | 1 | 1 | 1 | 0.8 | 1 | NA | 1 | -0.067 |
| SE | 9360 | PPP4R1L | PPP4R1L | chr20 | - | 58248114 | 58248273 | 58247321 | 58247506 | 58250886 | 58250952 | 12 | 4 | 0 | 1 | 0 | 0 | 0 | 3 | 0 | 0 | 98 | 49 | 1 | 1 | 0.857 | 1 | NA | NA | 1 | -0.072 |
| SE | 9361 | PPP4R1L | PPP4R1L | chr20 | - | 58250886 | 58250952 | 58247321 | 58247506 | 58251756 | 58251864 | 11 | 2 | 0 | 1 | 0 | 0 | 0 | 4 | 0 | 0 | 98 | 49 | 1 | 1 | 0.846 | 1 | NA | NA | 1 | -0.077 |
| SE | 9372 | NFYC | NFYC | chr1 | + | 40766917 | 40766974 | 40766595 | 40766703 | 40769355 | 40769415 | 0 | 0 | 1 | 6 | 2 | 0 | 0 | 0 | 0 | 1 | 98 | 49 | 1 | 1 | 0 | 0 | 1 | NA | 0 | 0.333 |
| SE | 9377 | MAP3K8 | MAP3K8 | chr10 | + | 30437175 | 30437406 | 30434020 | 30434378 | 30438915 | 30439274 | 11 | 5 | 0 | 0 | 4 | 0 | 0 | 6 | 0 | 1 | 98 | 49 | 1 | 1 | 1 | 0.385 | NA | NA | 0.75 | -0.057 |
| SE | 9383 | EPC1 | EPC1 | chr10 | - | 32292495 | 32292644 | 32291162 | 32291322 | 32292987 | 32293194 | 23 | 11 | 6 | 2 | 0 | 0 | 0 | 3 | 0 | 0 | 98 | 49 | 1 | 1 | 0.852 | 1 | 1 | NA | 1 | -0.049 |
| SE | 9385 | AMICA1 | AMICA1 | chr11 | - | 118212406 | 118212561 | 118210486 | 118210712 | 118224940 | 118225094 | 168 | 114 | 70 | 1 | 0 | 0 | 4 | 27 | 0 | 0 | 98 | 49 | 1 | 1 | 0.988 | 1 | 1 | 1 | 1 | -0.004 |
| SE | 9395 | RTN3 | RTN3 | chr11 | + | 63718980 | 63721032 | 63681449 | 63681778 | 63749990 | 63750198 | 0 | 0 | 1 | 12 | 13 | 15 | 0 | 0 | 0 | 3 | 98 | 49 | 1 | 1 | 0 | 0 | 0.032 | NA | 0 | 0.011 |
| SE | 9401 | TXNRD1 | TXNRD1 | chr12 | + | 104321090 | 104321316 | 104319469 | 104319585 | 104325336 | 104325429 | 21 | 24 | 5 | 2 | 0 | 0 | 3 | 2 | 0 | 0 | 98 | 49 | 1 | 1 | 0.84 | 1 | 1 | 1 | 1 | -0.053 |
| SE | 9402 | TXNRD1 | TXNRD1 | chr12 | + | 104326346 | 104326423 | 104325336 | 104325429 | 104327514 | 104327671 | 37 | 22 | 11 | 2 | 0 | 0 | 0 | 5 | 0 | 0 | 98 | 49 | 1 | 1 | 0.902 | 1 | 1 | NA | 1 | -0.033 |
| SE | 9404 | KIAA0232 | KIAA0232 | chr4 | + | 6804522 | 6804606 | 6782731 | 6782841 | 6824184 | 6824684 | 1 | 0 | 0 | 8 | 4 | 0 | 0 | 0 | 0 | 1 | 98 | 49 | 1 | 1 | 0.059 | 0 | NA | NA | 0 | 0.029 |
| SE | 9407 | LDLR | LDLR | chr19 | + | 11105219 | 11105600 | 11102663 | 11102786 | 11106564 | 11106687 | 1 | 1 | 0 | 0 | 0 | 0 | 0 | 0 | 0 | 1 | 98 | 49 | 1 | 1 | 1 | 1 | NA | NA | 0 | 1 |
| SE | 9410 | CCNY | CCNY | chr10 | + | 35394815 | 35394878 | 35336873 | 35337207 | 35483403 | 35483478 | 1 | 0 | 0 | 13 | 1 | 8 | 0 | 0 | 2 | 1 | 98 | 49 | 1 | 1 | 0.037 | 0 | 0 | 0 | 0 | 0.012 |
| SE | 9412 | CCNY | CCNY | chr10 | + | 35529972 | 35530030 | 35525963 | 35525999 | 35530123 | 35530243 | 41 | 24 | 19 | 0 | 0 | 2 | 0 | 8 | 0 | 0 | 98 | 49 | 1 | 1 | 1 | 1 | 0.826 | NA | 1 | -0.058 |
| SE | 9416 | ZNF33A | ZNF33A | chr10 | + | 38025347 | 38025411 | 38017290 | 38017386 | 38054374 | 38060067 | 1 | 0 | 1 | 9 | 5 | 3 | 0 | 0 | 0 | 1 | 98 | 49 | 1 | 1 | 0.053 | 0 | 0.143 | NA | 0 | 0.065 |
| SE | 9417 | ZNF33A | ZNF33A | chr10 | + | 38034277 | 38034400 | 38017290 | 38017386 | 38054374 | 38060067 | 0 | 0 | 1 | 9 | 5 | 3 | 0 | 0 | 0 | 1 | 98 | 49 | 1 | 1 | 0 | 0 | 0.143 | NA | 0 | 0.048 |
| SE | 9419 | ZNF33A | ZNF33A | chr10 | + | 38034325 | 38034400 | 38017290 | 38017386 | 38054374 | 38060067 | 0 | 1 | 1 | 9 | 5 | 3 | 0 | 0 | 0 | 1 | 98 | 49 | 1 | 1 | 0 | 0.091 | 0.143 | NA | 0 | 0.078 |
| SE | 9420 | ZNF33A | ZNF33A | chr10 | + | 38039412 | 38039492 | 38017290 | 38017386 | 38054374 | 38060067 | 0 | 0 | 0 | 9 | 5 | 3 | 0 | 2 | 0 | 1 | 98 | 49 | 1 | 1 | 0 | 0 | 0 | NA | 0.5 | -0.5 |
| SE | 9430 | SCYL1 | SCYL1 | chr11 | + | 65526123 | 65526350 | 65525920 | 65526043 | 65526782 | 65526873 | 14 | 4 | 6 | 0 | 0 | 1 | 1 | 1 | 0 | 0 | 98 | 49 | 1 | 1 | 1 | 1 | 0.75 | 1 | 1 | -0.083 |

| | | | | | | | | | | | | | | | | | | | | | | | | | | | | | | | |
|---|---|---|---|---|---|---|---|---|---|---|---|---|---|---|---|---|---|---|---|---|---|---|---|---|---|---|---|---|---|---|---|
| SE | 9432 | TATDN2 | TATDN2 | chr3 | + | 10249194 | 10249614 | 10248492 | 10249067 | 10260136 | 10260670 | 8 | 2 | 2 | 0 | 0 | 0 | 3 | 6 | 0 | 1 | 98 | 49 | 1 | 1 | 1 | 1 | 1 | 1 | 0.75 | 0.125 |
| SE | 9433 | CUL5 | CUL5 | chr11 | + | 108049889 | 108050066 | 108046269 | 108046369 | 108052659 | 108052801 | 3 | 0 | 0 | 0 | 4 | 0 | 0 | 1 | 0 | 0 | 98 | 49 | 1 | 1 | 1 | 0 | NA | NA | 1 | -0.5 |
| SE | 9435 | ACBD5 | ACBD5 | chr10 | - | 27235091 | 27235212 | 27231747 | 27231820 | 27240318 | 27240484 | 15 | 3 | 4 | 1 | 0 | 0 | 0 | 2 | 0 | 0 | 98 | 49 | 1 | 1 | 0.882 | 1 | 1 | NA | 1 | -0.039 |
| SE | 9436 | TFRC | TFRC | chr3 | - | 196071395 | 196071498 | 196069454 | 196069568 | 196072002 | 196072152 | 9 | 4 | 0 | 0 | 2 | 0 | 0 | 3 | 0 | 0 | 98 | 49 | 1 | 1 | 1 | 0.5 | NA | NA | 1 | -0.25 |
| SE | 9446 | RAB1B | RAB1B | chr11 | + | 66272364 | 66272460 | 66272156 | 66272252 | 66275803 | 66275935 | 34 | 25 | 22 | 1 | 0 | 0 | 2 | 11 | 0 | 0 | 98 | 49 | 1 | 1 | 0.944 | 1 | 1 | 1 | 1 | -0.019 |
| SE | 9449 | ANKRD27 | ANKRD27 | chr19 | - | 32602014 | 32602126 | 32599971 | 32600050 | 32604262 | 32604424 | 2 | 0 | 0 | 0 | 0 | 2 | 0 | 1 | 0 | 0 | 98 | 49 | 1 | 1 | 1 | NA | 0 | NA | 1 | -0.5 |
| SE | 9450 | ANKRD27 | ANKRD27 | chr19 | - | 32646458 | 32646615 | 32644324 | 32644479 | 32649681 | 32649792 | 4 | 3 | 0 | 0 | 0 | 0 | 0 | 1 | 0 | 1 | 98 | 49 | 1 | 1 | 1 | 1 | NA | NA | 0.333 | 0.667 |
| SE | 9451 | ANKRD27 | ANKRD27 | chr19 | - | 32649681 | 32649792 | 32644324 | 32644479 | 32658913 | 32659045 | 3 | 1 | 0 | 0 | 0 | 0 | 0 | 2 | 0 | 2 | 98 | 49 | 1 | 1 | 1 | 1 | NA | NA | 0.333 | 0.667 |
| SE | 9454 | ZNF467 | ZNF467 | chr7 | - | 149769089 | 149769200 | 149764363 | 149766239 | 149770439 | 149770556 | 19 | 3 | 14 | 1 | 0 | 0 | 1 | 2 | 0 | 0 | 98 | 49 | 1 | 1 | 0.905 | 1 | 1 | 1 | 1 | -0.032 |
| SE | 9455 | ZNF467 | ZNF467 | chr7 | - | 149769089 | 149769200 | 149764363 | 149766239 | 149770998 | 149771074 | 9 | 3 | 5 | 1 | 0 | 0 | 2 | 1 | 0 | 0 | 98 | 49 | 1 | 1 | 0.818 | 1 | 1 | 1 | 1 | -0.061 |
| SE | 9459 | NCOA4 | NCOA4 | chr10 | - | 46009123 | 46009212 | 46005087 | 46006597 | 46009410 | 46009551 | 1 | 0 | 0 | 162 | 77 | 49 | 0 | 0 | 2 | 9 | 98 | 49 | 1 | 1 | 0.003 | 0 | 0 | 0 | 0 | 0.001 |
| SE | 9460 | NCOA4 | NCOA4 | chr10 | - | 46016539 | 46016694 | 46015125 | 46015266 | 46030525 | 46030714 | 102 | 33 | 29 | 2 | 1 | 0 | 0 | 5 | 0 | 0 | 98 | 49 | 1 | 1 | 0.962 | 0.943 | 1 | NA | 1 | -0.032 |
| SE | 9464 | LITAF | LITAF | chr16 | - | 11551706 | 11551804 | 11547721 | 11549745 | 11553532 | 11553689 | 3 | 0 | 2 | 92 | 51 | 66 | 0 | 0 | 2 | 12 | 98 | 49 | 1 | 1 | 0.016 | 0 | 0.015 | 0 | 0 | 0.01 |
| SE | 9467 | FAM131A | FAM131A | chr3 | + | 184337310 | 184337371 | 184335928 | 184336123 | 184341723 | 184341817 | 0 | 2 | 1 | 0 | 0 | 0 | 0 | 1 | 0 | 1 | 98 | 49 | 1 | 1 | NA | 1 | 1 | NA | 0.333 | 0.667 |
| SE | 9487 | PDCD4 | PDCD4 | chr10 | + | 110876669 | 110876753 | 110875965 | 110876070 | 110881232 | 110881535 | 5 | 3 | 1 | 2 | 0 | 1 | 0 | 0 | 0 | 1 | 98 | 49 | 1 | 1 | 0.556 | 1 | 0.333 | NA | 0 | 0.63 |
| SE | 9488 | PDCD4 | PDCD4 | chr10 | + | 110887664 | 110887886 | 110885252 | 110885366 | 110889532 | 110889630 | 48 | 22 | 9 | 0 | 1 | 0 | 1 | 4 | 0 | 0 | 98 | 49 | 1 | 1 | 1 | 0.917 | 1 | 1 | 1 | -0.028 |
| SE | 9489 | PDCD4 | PDCD4 | chr10 | + | 110890555 | 110890670 | 110889532 | 110889630 | 110894090 | 110894198 | 20 | 21 | 6 | 0 | 1 | 0 | 0 | 4 | 0 | 0 | 98 | 49 | 1 | 1 | 1 | 0.913 | 1 | NA | 1 | -0.029 |
| SE | 9490 | PDCD4 | PDCD4 | chr10 | + | 110894411 | 110894522 | 110894090 | 110894198 | 110895947 | 110896087 | 33 | 14 | 7 | 3 | 0 | 0 | 0 | 6 | 0 | 0 | 98 | 49 | 1 | 1 | 0.846 | 1 | 1 | NA | 1 | -0.051 |
| SE | 9494 | EMC4 | EMC4 | chr15 | + | 34225535 | 34225650 | 34224996 | 34225200 | 34227692 | 34227846 | 8 | 5 | 6 | 1 | 0 | 0 | 0 | 4 | 0 | 0 | 98 | 49 | 1 | 1 | 0.8 | 1 | 1 | NA | 1 | -0.067 |
| SE | 9496 | WDFY4 | WDFY4 | chr10 | + | 48727469 | 48727659 | 48725880 | 48726070 | 48729431 | 48729589 | 37 | 6 | 0 | 1 | 0 | 0 | 0 | 5 | 0 | 0 | 98 | 49 | 1 | 1 | 0.949 | 1 | NA | NA | 1 | -0.026 |
| SE | 9497 | WDFY4 | WDFY4 | chr10 | + | 48760346 | 48760440 | 48742967 | 48743548 | 48774457 | 48774672 | 28 | 13 | 13 | 0 | 0 | 0 | 0 | 11 | 0 | 1 | 98 | 49 | 1 | 1 | 1 | 1 | 1 | NA | 0.846 | 0.154 |
| SE | 9498 | WDFY4 | WDFY4 | chr10 | + | 48817244 | 48817409 | 48813956 | 48814082 | 48820233 | 48820437 | 21 | 1 | 5 | 1 | 0 | 0 | 1 | 7 | 0 | 0 | 98 | 49 | 1 | 1 | 0.913 | 1 | 1 | 1 | 1 | -0.029 |
| SE | 9499 | WDFY4 | WDFY4 | chr10 | + | 48826670 | 48826909 | 48822379 | 48822537 | 48828777 | 48828896 | 16 | 1 | 1 | 4 | 0 | 0 | 0 | 5 | 0 | 0 | 98 | 49 | 1 | 1 | 0.667 | 1 | 1 | NA | 1 | -0.111 |
| SE | 9503 | C10orf128 | C10orf128 | chr10 | - | 49165765 | 49165796 | 49161598 | 49161622 | 49166868 | 49167009 | 0 | 7 | 7 | 0 | 0 | 1 | 0 | 1 | 0 | 1 | 80 | 49 | 1 | 1 | NA | 1 | 0.811 | NA | 0.38 | 0.525 |
| SE | 9508 | BNIP2 | BNIP2 | chr15 | - | 59668891 | 59668990 | 59662862 | 59664120 | 59669275 | 59669362 | 72 | 49 | 20 | 1 | 0 | 0 | 1 | 13 | 0 | 0 | 98 | 49 | 1 | 1 | 0.973 | 1 | 1 | 1 | 1 | -0.009 |
| SE | 9509 | BNIP2 | BNIP2 | chr15 | - | 59669275 | 59669362 | 59668891 | 59668990 | 59671182 | 59671314 | 55 | 35 | 13 | 1 | 2 | 0 | 0 | 14 | 0 | 0 | 98 | 49 | 1 | 1 | 0.965 | 0.897 | 1 | NA | 1 | -0.046 |
| SE | 9510 | BNIP2 | BNIP2 | chr15 | - | 59679591 | 59679768 | 59677910 | 59678087 | 59680240 | 59680308 | 44 | 20 | 17 | 2 | 0 | 0 | 0 | 15 | 0 | 0 | 98 | 49 | 1 | 1 | 0.917 | 1 | 1 | NA | 1 | -0.028 |
| SE | 9515 | TBRG1 | TBRG1 | chr11 | + | 124626472 | 124626609 | 124625670 | 124625903 | 124626903 | 124627050 | 5 | 0 | 0 | 0 | 0 | 1 | 0 | 1 | 0 | 0 | 98 | 49 | 1 | 1 | 1 | NA | 0 | NA | 1 | -0.5 |
| SE | 9519 | XRN1 | XRN1 | chr3 | - | 142412543 | 142412663 | 142404906 | 142405076 | 142414134 | 142414291 | 51 | 17 | 2 | 1 | 0 | 0 | 0 | 14 | 0 | 0 | 98 | 49 | 1 | 1 | 0.962 | 1 | 1 | NA | 1 | -0.013 |
| SE | 9520 | XRN1 | XRN1 | chr3 | - | 142414134 | 142414291 | 142412543 | 142412663 | 142417139 | 142417229 | 55 | 17 | 3 | 0 | 3 | 0 | 0 | 11 | 0 | 0 | 98 | 49 | 1 | 1 | 1 | 0.739 | 1 | NA | 1 | -0.087 |
| SE | 9521 | XRN1 | XRN1 | chr3 | - | 142426743 | 142426841 | 142425428 | 142425538 | 142432660 | 142432893 | 36 | 11 | 1 | 0 | 2 | 0 | 0 | 8 | 0 | 1 | 98 | 49 | 1 | 1 | 1 | 0.733 | 1 | NA | 0.8 | 0.111 |
| SE | 9526 | LAMTOR2 | LAMTOR2 | chr1 | + | 156057977 | 156058067 | 156055262 | 156055425 | 156058314 | 156058510 | 6 | 1 | 5 | 0 | 0 | 1 | 0 | 3 | 0 | 0 | 98 | 49 | 1 | 1 | 1 | 1 | 0.714 | NA | 1 | -0.095 |
| SE | 9537 | GMPR2 | GMPR2 | chr14 | + | 24233218 | 24233340 | 24232421 | 24232847 | 24233478 | 24233593 | 22 | 9 | 4 | 0 | 5 | 0 | 0 | 4 | 0 | 0 | 98 | 49 | 1 | 1 | 1 | 0.474 | 1 | NA | 1 | -0.175 |
| SE | 9539 | GMPR2 | GMPR2 | chr14 | + | 24233478 | 24233598 | 24232421 | 24232847 | 24235736 | 24235820 | 8 | 9 | 2 | 0 | 1 | 0 | 0 | 1 | 0 | 0 | 98 | 49 | 1 | 1 | 1 | 0.818 | 1 | NA | 1 | -0.061 |
| SE | 9540 | GMPR2 | GMPR2 | chr14 | + | 24234106 | 24234241 | 24233478 | 24233598 | 24235736 | 24235820 | 1 | 0 | 0 | 8 | 4 | 2 | 0 | 0 | 0 | 1 | 98 | 49 | 1 | 1 | 0.059 | 0 | 0 | NA | 0 | 0.02 |
| SE | 9541 | GMPR2 | GMPR2 | chr14 | + | 24235736 | 24235820 | 24233478 | 24233598 | 24235966 | 24236140 | 13 | 11 | 5 | 1 | 0 | 0 | 0 | 1 | 0 | 0 | 98 | 49 | 1 | 1 | 0.867 | 1 | 1 | NA | 1 | -0.044 |
| SE | 9542 | GMPR2 | GMPR2 | chr14 | + | 24237041 | 24237152 | 24235966 | 24236140 | 24237244 | 24237351 | 6 | 3 | 3 | 0 | 2 | 0 | 0 | 3 | 0 | 0 | 98 | 49 | 1 | 1 | 1 | 0.429 | 1 | NA | 1 | -0.19 |
| SE | 9543 | GMPR2 | GMPR2 | chr14 | + | 24237070 | 24237152 | 24235966 | 24236140 | 24237244 | 24237351 | 19 | 8 | 11 | 0 | 2 | 0 | 0 | 4 | 0 | 0 | 98 | 49 | 1 | 1 | 1 | 0.667 | 1 | NA | 1 | -0.111 |
| SE | 9545 | FURIN | FURIN | chr15 | + | 90877526 | 90877615 | 90877134 | 90877211 | 90878131 | 90878304 | 29 | 17 | 18 | 0 | 0 | 0 | 1 | 6 | 0 | 1 | 98 | 49 | 1 | 1 | 1 | 1 | 1 | 1 | 0.75 | 0.125 |
| SE | 9546 | UBE3B | UBE3B | chr12 | + | 109530546 | 109530658 | 109529889 | 109530072 | 109533465 | 109533558 | 5 | 1 | 0 | 1 | 0 | 0 | 0 | 1 | 0 | 2 | 98 | 49 | 1 | 1 | 0.714 | 1 | NA | NA | 0.2 | 0.657 |
| SE | 9550 | ELP2 | ELP2 | chr18 | + | 36139418 | 36139613 | 36138794 | 36138872 | 36141136 | 36141201 | 0 | 6 | 0 | 3 | 0 | 0 | 0 | 0 | 0 | 2 | 98 | 49 | 1 | 1 | 0 | 1 | NA | NA | 0 | 0.5 |
| SE | 9553 | RBSN | RBSN | chr3 | - | 15082366 | 15082608 | 15080731 | 15080802 | 15084734 | 15084896 | 2 | 0 | 0 | 0 | 1 | 0 | 0 | 3 | 0 | 0 | 98 | 49 | 1 | 1 | 1 | 0 | NA | NA | 0 | -0.5 |
| SE | 9555 | HNRNPU | HNRNPU | chr1 | - | 244856727 | 244856856 | 244856456 | 244856625 | 244857597 | 244857717 | 116 | 47 | 31 | 1 | 3 | 0 | 0 | 23 | 0 | 0 | 98 | 49 | 1 | 1 | 0.983 | 0.887 | 1 | NA | 1 | -0.043 |
| SE | 9582 | SH3GLB1 | SH3GLB1 | chr1 | + | 86715723 | 86715865 | 86704569 | 86704971 | 86719506 | 86719635 | 55 | 24 | 4 | 1 | 1 | 0 | 0 | 15 | 0 | 0 | 98 | 49 | 1 | 1 | 0.965 | 0.923 | 1 | NA | 1 | -0.037 |
| SE | 9583 | SH3GLB1 | SH3GLB1 | chr1 | + | 86728402 | 86728489 | 86724312 | 86724405 | 86734601 | 86734691 | 1 | 1 | 4 | 14 | 9 | 9 | 0 | 1 | 0 | 7 | 98 | 49 | 1 | 1 | 0.034 | 0.053 | 0.182 | NA | 0.067 | 0.023 |
| SE | 9593 | ADPGK | ADPGK | chr15 | - | 72756250 | 72756447 | 72755555 | 72755654 | 72760406 | 72760527 | 17 | 4 | 10 | 5 | 0 | 0 | 0 | 5 | 0 | 0 | 98 | 49 | 1 | 1 | 0.63 | 1 | 1 | NA | 1 | -0.123 |
| SE | 9594 | ADPGK | ADPGK | chr15 | - | 72774871 | 72775097 | 72771782 | 72771845 | 72783245 | 72783785 | 37 | 7 | 6 | 2 | 2 | 1 | 0 | 5 | 0 | 2 | 98 | 49 | 1 | 1 | 0.902 | 0.636 | 0.75 | NA | 0.556 | 0.207 |
| SE | 9606 | GLT1D1 | GLT1D1 | chr12 | + | 128888638 | 128888744 | 128875913 | 128876062 | 128899235 | 128899287 | 76 | 42 | 18 | 2 | 0 | 0 | 6 | 18 | 0 | 1 | 98 | 49 | 1 | 1 | 0.95 | 1 | 1 | 1 | 0.9 | 0.033 |
| SE | 9607 | GLT1D1 | GLT1D1 | chr12 | + | 128899235 | 128899287 | 128888638 | 128888744 | 128945325 | 128945369 | 48 | 28 | 7 | 0 | 1 | 0 | 5 | 8 | 0 | 0 | 98 | 49 | 1 | 1 | 1 | 0.933 | 1 | 1 | 1 | -0.022 |
| SE | 9609 | SUPV3L1 | SUPV3L1 | chr10 | + | 69189266 | 69189435 | 69187641 | 69187756 | 69191654 | 69191766 | 3 | 1 | 0 | 0 | 0 | 0 | 0 | 0 | 1 | 0 | 98 | 49 | 1 | 1 | 1 | 1 | NA | 0 | NA | 1 |
| SE | 9611 | CTNNBL1 | CTNNBL1 | chr20 | + | 37842338 | 37842419 | 37840101 | 37840199 | 37859898 | 37860036 | 3 | 0 | 0 | 0 | 0 | 0 | 0 | 2 | 0 | 1 | 98 | 49 | 1 | 1 | 1 | NA | NA | NA | 0.5 | 0.5 |
| SE | 9614 | RMND5A | RMND5A | chr2 | + | 86771557 | 86771712 | 86770022 | 86770125 | 86773347 | 86778041 | 39 | 30 | 22 | 0 | 0 | 1 | 0 | 1 | 0 | 0 | 98 | 49 | 1 | 1 | 1 | 1 | 0.917 | NA | 1 | -0.028 |
| SE | 9617 | TMUB2 | TMUB2 | chr17 | + | 44189021 | 44189588 | 44186985 | 44187109 | 44190500 | 44191731 | 10 | 0 | 1 | 0 | 0 | 0 | 0 | 5 | 0 | 1 | 98 | 49 | 1 | 1 | 1 | NA | 1 | NA | 0.714 | 0.286 |
| SE | 9619 | GSK3B | GSK3B | chr3 | - | 119866589 | 119866628 | 119863418 | 119863605 | 119876412 | 119876508 | 0 | 5 | 0 | 10 | 3 | 1 | 0 | 0 | 0 | 2 | 88 | 49 | 1 | 1 | 0 | 0.481 | 0 | NA | 0 | 0.16 |
| SE | 9628 | CCBL2 | CCBL2 | chr1 | - | 88988251 | 88988351 | 88969408 | 88969467 | 88992584 | 88992960 | 7 | 5 | 1 | 5 | 1 | 0 | 0 | 1 | 0 | 0 | 98 | 49 | 1 | 1 | 0.412 | 0.714 | 1 | NA | 1 | -0.291 |
| SE | 9634 | ARIH1 | ARIH1 | chr15 | + | 72563393 | 72563500 | 72561482 | 72561549 | 72566562 | 72566605 | 19 | 11 | 1 | 2 | 2 | 0 | 0 | 1 | 0 | 0 | 98 | 49 | 1 | 1 | 0.826 | 0.733 | 1 | NA | 1 | -0.147 |
| SE | 9636 | ITGA5 | ITGA5 | chr12 | - | 54401584 | 54401665 | 54401372 | 54401478 | 54401775 | 54401855 | 28 | 26 | 11 | 1 | 0 | 0 | 0 | 14 | 0 | 0 | 98 | 49 | 1 | 1 | 0.933 | 1 | 1 | NA | 1 | -0.022 |
| SE | 9637 | ITGA5 | ITGA5 | chr12 | - | 54403624 | 54403779 | 54403186 | 54403324 | 54403910 | 54403966 | 30 | 11 | 14 | 1 | 0 | 0 | 0 | 12 | 0 | 0 | 98 | 49 | 1 | 1 | 0.937 | 1 | 1 | NA | 1 | -0.021 |
| SE | 9638 | ITGA5 | ITGA5 | chr12 | - | 54404144 | 54404246 | 54403910 | 54403966 | 54404429 | 54404475 | 51 | 22 | 7 | 0 | 1 | 0 | 7 | 15 | 0 | 0 | 98 | 49 | 1 | 1 | 1 | 0.917 | 1 | 1 | 1 | -0.028 |
| SE | 9639 | ITGA5 | ITGA5 | chr12 | - | 54409484 | 54409597 | 54409231 | 54409352 | 54411833 | 54411964 | 54 | 32 | 16 | 0 | 1 | 0 | 2 | 8 | 0 | 0 | 98 | 49 | 1 | 1 | 1 | 0.941 | 1 | 1 | 1 | -0.02 |
| SE | 9640 | ITGA5 | ITGA5 | chr12 | - | 54411833 | 54411964 | 54409484 | 54409597 | 54418980 | 54419266 | 40 | 22 | 10 | 0 | 1 | 0 | 3 | 6 | 0 | 0 | 98 | 49 | 1 | 1 | 1 | 0.917 | 1 | 1 | 1 | -0.028 |
| SE | 9641 | C10orf54 | C10orf54 | chr10 | - | 71755358 | 71755466 | 71752974 | 71753002 | 71760867 | 71760924 | 126 | 101 | 78 | 0 | 0 | 0 | 3 | 48 | 0 | 1 | 98 | 49 | 1 | 1 | 1 | 1 | 1 | 1 | 0.96 | 0.02 |
| SE | 9642 | C10orf54 | C10orf54 | chr10 | - | 71761597 | 71762026 | 71760867 | 71760924 | 71773357 | 71773580 | 131 | 70 | 76 | 1 | 0 | 0 | 4 | 72 | 0 | 0 | 98 | 49 | 1 | 1 | 0.985 | 1 | 1 | 1 | 1 | -0.005 |
| SE | 9643 | MCTP1 | MCTP1 | chr5 | - | 94799012 | 94799132 | 94779109 | 94779163 | 94868332 | 94868452 | 16 | 7 | 5 | 2 | 0 | 0 | 0 | 1 | 0 | 1 | 98 | 49 | 1 | 1 | 0.8 | 1 | 1 | NA | 0.333 | 0.6 |
| SE | 9644 | MCTP1 | MCTP1 | chr5 | - | 94870871 | 94870973 | 94870416 | 94870491 | 94871314 | 94871417 | 12 | 13 | 1 | 1 | 0 | 0 | 0 | 2 | 0 | 0 | 98 | 49 | 1 | 1 | 0.857 | 1 | 1 | NA | 1 | -0.048 |
| SE | 9646 | MCTP1 | MCTP1 | chr5 | - | 94917895 | 94917973 | 94912805 | 94912976 | 94940083 | 94940195 | 8 | 15 | 2 | 0 | 1 | 0 | 0 | 3 | 0 | 2 | 98 | 49 | 1 | 1 | 1 | 0.882 | 1 | NA | 0.429 | 0.532 |
| SE | 9647 | MCTP1 | MCTP1 | chr5 | - | 94923961 | 94924021 | 94912805 | 94912976 | 94931952 | 94931991 | 18 | 7 | 5 | 1 | 0 | 3 | 0 | 3 | 0 | 0 | 98 | 49 | 1 | 1 | 0.9 | 1 | 0.455 | NA | 1 | -0.215 |
| SE | 9648 | MCTP1 | MCTP1 | chr5 | - | 94923961 | 94924021 | 94912805 | 94912976 | 94940083 | 94940195 | 13 | 3 | 1 | 0 | 1 | 0 | 0 | 1 | 0 | 2 | 98 | 49 | 1 | 1 | 1 | 0.6 | 1 | NA | 0.2 | 0.667 |
| SE | 9649 | MCTP1 | MCTP1 | chr5 | - | 94923961 | 94924021 | 94917895 | 94917973 | 94940083 | 94940195 | 9 | 13 | 13 | 0 | 1 | 0 | 2 | 1 | 0 | 0 | 98 | 49 | 1 | 1 | 1 | 0.867 | 1 | 1 | 1 | -0.044 |
| SE | 9650 | MCTP1 | MCTP1 | chr5 | - | 94931952 | 94931991 | 94912805 | 94912976 | 94940083 | 94940195 | 25 | 9 | 6 | 0 | 1 | 0 | 0 | 6 | 0 | 2 | 88 | 49 | 1 | 1 | 1 | 0.834 | 1 | NA | 0.626 | 0.319 |
| SE | 9651 | MCTP1 | MCTP1 | chr5 | - | 94931952 | 94931991 | 94923961 | 94924021 | 94940083 | 94940195 | 29 | 13 | 7 | 0 | 0 | 0 | 0 | 9 | 0 | 1 | 88 | 49 | 1 | 1 | 1 | 1 | 1 | NA | 0.834 | 0.166 |
| SE | 9679 | SLC3A2 | SLC3A2 | chr11 | + | 62883988 | 62884112 | 62882907 | 62882999 | 62884456 | 62884525 | 2 | 0 | 0 | 11 | 4 | 6 | 0 | 1 | 0 | 4 | 98 | 49 | 1 | 1 | 0.083 | 0 | 0 | NA | 0.111 | -0.083 |
| SE | 9689 | MTHFS | MTHFS | chr15 | - | 79889092 | 79889354 | 79843546 | 79845442 | 79896871 | 79897049 | 13 | 8 | 2 | 1 | 0 | 0 | 0 | 3 | 0 | 0 | 98 | 49 | 1 | 1 | 0.867 | 1 | 1 | NA | 1 | -0.044 |
| SE | 9690 | MTHFS | MTHFS | chr15 | - | 79889092 | 79889354 | 79843546 | 79845442 | 79897160 | 79897285 | 9 | 7 | 0 | 1 | 0 | 0 | 0 | 2 | 0 | 0 | 98 | 49 | 1 | 1 | 0.818 | 1 | 1 | NA | 1 | -0.091 |
| SE | 9691 | PSMG4 | PSMG4 | chr6 | + | 3264208 | 3264325 | 3263683 | 3263759 | 3267590 | 3268066 | 0 | 2 | 0 | 1 | 0 | 0 | 0 | 0 | 0 | 1 | 98 | 49 | 1 | 1 | 0 | 1 | NA | NA | 0 | 0.5 |
| SE | 9703 | XPNPEP1 | XPNPEP1 | chr10 | - | 109893011 | 109893075 | 109891721 | 109891826 | 109907690 | 109907815 | 4 | 0 | 0 | 1 | 0 | 0 | 0 | 1 | 0 | 0 | 98 | 49 | 1 | 1 | 0.667 | NA | NA | NA | 1 | -0.333 |
| SE | 9705 | TSPAN14 | TSPAN14 | chr10 | + | 80509300 | 80509471 | 80507227 | 80507374 | 80512143 | 80512269 | 20 | 10 | 13 | 1 | 0 | 0 | 0 | 2 | 0 | 0 | 98 | 49 | 1 | 1 | 0.909 | 1 | 1 | NA | 1 | -0.03 |
| SE | 9713 | TRPC4AP | TRPC4AP | chr20 | - | 35013007 | 35013066 | 35010186 | 35010288 | 35016007 | 35016139 | 40 | 11 | 5 | 0 | 1 | 0 | 0 | 10 | 0 | 0 | 98 | 49 | 1 | 1 | 1 | 0.846 | 1 | NA | 1 | -0.051 |
| SE | 9714 | TRPC4AP | TRPC4AP | chr20 | - | 35021189 | 35021332 | 35016007 | 35016139 | 35035122 | 35035308 | 19 | 13 | 4 | 4 | 1 | 0 | 0 | 4 | 0 | 0 | 98 | 49 | 1 | 1 | 0.704 | 0.867 | 1 | NA | 1 | -0.143 |
| SE | 9715 | TRPC4AP | TRPC4AP | chr20 | - | 35021189 | 35021356 | 35016007 | 35016139 | 35035122 | 35035308 | 22 | 11 | 3 | 4 | 1 | 0 | 0 | 4 | 0 | 0 | 98 | 49 | 1 | 1 | 0.733 | 0.846 | 1 | NA | 1 | -0.14 |
| SE | 9716 | TRPC4AP | TRPC4AP | chr20 | - | 35069295 | 35069412 | 35057513 | 35057571 | 35078045 | 35078174 | 52 | 29 | 4 | 1 | 1 | 0 | 1 | 10 | 0 | 0 | 98 | 49 | 1 | 1 | 0.963 | 0.935 | 1 | 1 | 1 | -0.034 |
| SE | 9717 | NXF1 | NXF1 | chr11 | - | 62796286 | 62796345 | 62796065 | 62796181 | 62796459 | 62796567 | 20 | 6 | 6 | 0 | 2 | 0 | 0 | 1 | 0 | 0 | 98 | 49 | 1 | 1 | 1 | 0.6 | 1 | NA | 1 | -0.133 |
| SE | 9718 | NXF1 | NXF1 | chr11 | - | 62801093 | 62801201 | 62798538 | 62800486 | 62801328 | 62801417 | 48 | 17 | 8 | 1 | 0 | 0 | 4 | 11 | 0 | 0 | 98 | 49 | 1 | 1 | 0.96 | 1 | 1 | 1 | 1 | -0.013 |
| SE | 9719 | NXF1 | NXF1 | chr11 | - | 62803418 | 62803572 | 62802176 | 62802260 | 62803791 | 62803978 | 25 | 23 | 4 | 0 | 2 | 0 | 0 | 9 | 0 | 0 | 98 | 49 | 1 | 1 | 1 | 0.852 | 1 | NA | 1 | -0.049 |
| SE | 9722 | PNPLA6 | PNPLA6 | chr19 | + | 7536190 | 7536273 | 7535702 | 7536020 | 7536448 | 7536546 | 7 | 9 | 0 | 1 | 0 | 0 | 4 | 6 | 0 | 0 | 98 | 49 | 1 | 1 | 0.778 | 1 | NA | 1 | 1 | -0.111 |
| SE | 9724 | PNPLA6 | PNPLA6 | chr19 | + | 7555248 | 7555367 | 7554892 | 7555075 | 7555606 | 7555763 | 6 | 2 | 0 | 0 | 3 | 0 | 0 | 1 | 0 | 0 | 98 | 49 | 1 | 1 | 1 | 0.25 | NA | NA | 1 | -0.375 |
| SE | 9725 | PNPLA6 | PNPLA6 | chr19 | + | 7561013 | 7561110 | 7560647 | 7560764 | 7561207 | 7561317 | 14 | 6 | 6 | 0 | 1 | 0 | 0 | 3 | 0 | 0 | 98 | 49 | 1 | 1 | 1 | 0.75 | 1 | NA | 1 | -0.083 |
| SE | 9728 | DPYD | DPYD | chr1 | - | 97549559 | 97549744 | 97515725 | 97515941 | 97573759 | 97573970 | 166 | 35 | 16 | 3 | 0 | 0 | 4 | 25 | 0 | 1 | 98 | 49 | 1 | 1 | 0.965 | 1 | 1 | 1 | 0.926 | 0.025 |
| SE | 9729 | DPYD | DPYD | chr1 | - | 97593217 | 97593387 | 97573759 | 97573970 | 97595058 | 97595166 | 125 | 17 | 22 | 1 | 0 | 0 | 0 | 31 | 0 | 0 | 98 | 49 | 1 | 1 | 0.984 | 1 | 1 | NA | 1 | -0.005 |
| SE | 9730 | DPYD | DPYD | chr1 | - | 97828113 | 97828196 | 97740391 | 97740479 | 97883263 | 97883374 | 260 | 75 | 28 | 1 | 0 | 0 | 9 | 88 | 0 | 0 | 98 | 49 | 1 | 1 | 0.992 | 1 | 1 | 1 | 1 | -0.003 |
| SE | 9731 | CLEC12B | CLEC12B | chr12 | + | 10015251 | 10015406 | 10014522 | 10014741 | 10015611 | 10015727 | 0 | 0 | 0 | 0 | 2 | 0 | 0 | 2 | 0 | 0 | 98 | 49 | 1 | 1 | NA | 0 | NA | NA | 1 | -1 |
| SE | 9732 | AHCYL1 | AHCYL1 | chr1 | + | 110020730 | 110020851 | 110019547 | 110019626 | 110021673 | 110023742 | 1 | 0 | 1 | 1 | 1 | 0 | 0 | 4 | 0 | 0 | 98 | 49 | 1 | 1 | 0.333 | 0 | 1 | NA | 1 | -0.556 |
| SE | 9737 | CRBN | CRBN | chr3 | - | 3153959 | 3154075 | 3153423 | 3153488 | 3154746 | 3154831 | 9 | 5 | 5 | 1 | 0 | 0 | 0 | 7 | 0 | 0 | 98 | 49 | 1 | 1 | 0.818 | 1 | 1 | NA | 1 | -0.061 |
| SE | 9738 | CRBN | CRBN | chr3 | - | 3167633 | 3167793 | 3156218 | 3156281 | 3172775 | 3172925 | 15 | 3 | 7 | 0 | 1 | 0 | 0 | 1 | 0 | 0 | 98 | 49 | 1 | 1 | 1 | 0.6 | 1 | NA | 1 | -0.133 |
| SE | 9739 | VPS53 | VPS53 | chr17 | - | 586269 | 586364 | 562502 | 562745 | 601794 | 601896 | 9 | 1 | 0 | 2 | 0 | 0 | 0 | 1 | 0 | 0 | 98 | 49 | 1 | 1 | 0.692 | 1 | NA | NA | 1 | -0.154 |
| SE | 9743 | IGHMBP2 | IGHMBP2 | chr11 | + | 68935298 | 68935422 | 68934463 | 68934558 | 68936236 | 68937091 | 3 | 0 | 0 | 0 | 2 | 0 | 1 | 1 | 0 | 0 | 98 | 49 | 1 | 1 | 1 | 0 | NA | 1 | 1 | -0.5 |
| SE | 9745 | PRKACA | PRKACA | chr19 | - | 14093627 | 14093792 | 14091694 | 14093237 | 14097360 | 14097483 | 30 | 3 | 3 | 3 | 2 | 0 | 0 | 11 | 0 | 0 | 98 | 49 | 1 | 1 | 0.833 | 0.429 | 1 | NA | 1 | -0.246 |
| SE | 9751 | OSM | OSM | chr22 | - | 30265001 | 30265144 | 30262829 | 30264464 | 30266765 | 30266840 | 27 | 17 | 12 | 0 | 1 | 0 | 0 | 6 | 0 | 0 | 98 | 49 | 1 | 1 | 1 | 0.895 | 1 | NA | 1 | -0.035 |
| SE | 9753 | RELT | RELT | chr11 | + | 73391143 | 73391223 | 73390754 | 73390921 | 73392210 | 73392468 | 20 | 6 | 14 | 1 | 3 | 0 | 0 | 2 | 0 | 0 | 98 | 49 | 1 | 1 | 0.909 | 0.5 | 1 | NA | 1 | -0.197 |
| SE | 9754 | RELT | RELT | chr11 | + | 73391143 | 73391223 | 73390754 | 73390921 | 73393836 | 73393917 | 9 | 6 | 8 | 2 | 0 | 0 | 0 | 3 | 0 | 0 | 98 | 49 | 1 | 1 | 0.692 | 1 | 1 | NA | 1 | -0.103 |
| SE | 9755 | RELT | RELT | chr11 | + | 73392210 | 73392468 | 73390754 | 73390921 | 73393836 | 73393917 | 23 | 10 | 6 | 2 | 0 | 0 | 0 | 3 | 0 | 0 | 98 | 49 | 1 | 1 | 0.852 | 1 | 1 | NA | 1 | -0.049 |
| SE | 9756 | RELT | RELT | chr11 | + | 73392210 | 73392468 | 73391143 | 73391223 | 73393836 | 73393917 | 33 | 7 | 12 | 0 | 0 | 0 | 0 | 3 | 0 | 1 | 98 | 49 | 1 | 1 | 1 | 1 | 1 | NA | 0.6 | 0.4 |
| SE | 9757 | FAS | FAS | chr10 | + | 89007699 | 89007837 | 89003028 | 89003194 | 89010538 | 89010600 | 15 | 8 | 7 | 0 | 0 | 0 | 0 | 1 | 0 | 2 | 98 | 49 | 1 | 1 | 1 | 1 | 1 | NA | 0.2 | 0.8 |
| SE | 9758 | FAS | FAS | chr10 | + | 89008888 | 89008997 | 89007699 | 89007837 | 89010538 | 89010600 | 34 | 16 | 6 | 1 | 0 | 0 | 1 | 6 | 0 | 0 | 98 | 49 | 1 | 1 | 0.944 | 1 | 1 | 1 | 1 | -0.019 |
| SE | 9759 | FAS | FAS | chr10 | + | 89010752 | 89010815 | 89010538 | 89010600 | 89011998 | 89012081 | 20 | 13 | 3 | 8 | 15 | 1 | 0 | 2 | 0 | 2 | 98 | 49 | 1 | 1 | 0.556 | 0.302 | 0.6 | NA | 0.333 | 0.153 |
| SE | 9763 | IFIT1 | IFIT1 | chr10 | + | 89399889 | 89400003 | 89392545 | 89392717 | 89402280 | 89406487 | 5 | 2 | 0 | 1 | 6 | 0 | 0 | 11 | 0 | 5 | 98 | 49 | 1 | 1 | 0.714 | 0.143 | NA | NA | 0.524 | -0.096 |
| SE | 9764 | IFIT1 | IFIT1 | chr10 | + | 89399894 | 89400003 | 89392545 | 89392717 | 89402280 | 89406487 | 3 | 2 | 0 | 1 | 6 | 0 | 1 | 12 | 0 | 5 | 98 | 49 | 1 | 1 | 0.6 | 0.143 | NA | 1 | 0.545 | -0.401 |
| SE | 9773 | E2F4 | E2F4 | chr16 | + | 67194397 | 67194459 | 67192760 | 67192870 | 67194685 | 67194980 | 8 | 4 | 0 | 0 | 0 | 1 | 0 | 2 | 0 | 0 | 98 | 49 | 1 | 1 | 1 | 1 | 0 | NA | 1 | -0.333 |
| SE | 9775 | NADK2 | NADK2 | chr5 | - | 36219595 | 36219679 | 36217747 | 36217884 | 36225541 | 36225623 | 3 | 4 | 0 | 0 | 2 | 0 | 0 | 1 | 0 | 0 | 98 | 49 | 1 | 1 | 1 | 0.5 | NA | NA | 1 | -0.25 |
| SE | 9776 | PISD | PISD | chr22 | - | 31620552 | 31620713 | 31618490 | 31619836 | 31620995 | 31621142 | 26 | 34 | 9 | 1 | 0 | 0 | 0 | 8 | 0 | 0 | 98 | 49 | 1 | 1 | 0.929 | 1 | 1 | NA | 1 | -0.024 |
| SE | 9777 | PISD | PISD | chr22 | - | 31621333 | 31621472 | 31620995 | 31621142 | 31621648 | 31621885 | 50 | 25 | 20 | 0 | 2 | 0 | 0 | 12 | 0 | 0 | 98 | 49 | 1 | 1 | 1 | 0.862 | 1 | NA | 1 | -0.046 |
| SE | 9778 | PISD | PISD | chr22 | - | 31623683 | 31623849 | 31621648 | 31621885 | 31625762 | 31626016 | 23 | 7 | 12 | 1 | 0 | 0 | 0 | 6 | 0 | 0 | 98 | 49 | 1 | 1 | 0.92 | 1 | 1 | NA | 1 | -0.027 |
| SE | 9779 | ACSL5 | ACSL5 | chr10 | + | 112411455 | 112411529 | 112410583 | 112410635 | 112411901 | 112411979 | 5 | 1 | 0 | 1 | 0 | 0 | 0 | 1 | 0 | 0 | 98 | 49 | 1 | 1 | 0.714 | 1 | NA | NA | 1 | -0.143 |
| SE | 9781 | TNKS2 | TNKS2 | chr10 | + | 91827016 | 91827203 | 91822295 | 91822362 | 91828284 | 91828406 | 24 | 12 | 5 | 2 | 0 | 0 | 1 | 6 | 0 | 0 | 98 | 49 | 1 | 1 | 0.857 | 1 | 1 | 1 | 1 | -0.048 |
| SE | 9782 | BTAF1 | BTAF1 | chr10 | + | 91953736 | 91953873 | 91951402 | 91951566 | 91956527 | 91956657 | 30 | 3 | 0 | 1 | 0 | 0 | 1 | 5 | 0 | 1 | 98 | 49 | 1 | 1 | 0.937 | 1 | NA | 1 | 0.714 | 0.112 |
| SE | 9783 | BTAF1 | BTAF1 | chr10 | + | 91989153 | 91989580 | 91984200 | 91984404 | 91992118 | 91992309 | 19 | 2 | 2 | 0 | 1 | 0 | 0 | 2 | 0 | 0 | 98 | 49 | 1 | 1 | 1 | 0.5 | 1 | NA | 1 | -0.167 |
| SE | 9784 | BTAF1 | BTAF1 | chr10 | + | 92027129 | 92027300 | 92026591 | 92026751 | 92028789 | 92030323 | 19 | 1 | 0 | 0 | 0 | 0 | 0 | 4 | 0 | 3 | 98 | 49 | 1 | 1 | 1 | 1 | NA | NA | 0.4 | 0.6 |
| SE | 9787 | PIAS1 | PIAS1 | chr15 | + | 68145815 | 68145906 | 68142289 | 68142337 | 68146565 | 68146700 | 52 | 36 | 14 | 2 | 0 | 0 | 0 | 8 | 0 | 0 | 98 | 49 | 1 | 1 | 0.929 | 1 | 1 | NA | 1 | -0.024 |
| SE | 9788 | PIAS1 | PIAS1 | chr15 | + | 68173731 | 68173892 | 68164730 | 68164804 | 68175636 | 68175767 | 16 | 9 | 12 | 0 | 2 | 0 | 0 | 7 | 0 | 0 | 98 | 49 | 1 | 1 | 1 | 0.692 | 1 | NA | 1 | -0.103 |
| SE | 9789 | ARFIP1 | ARFIP1 | chr4 | + | 152790758 | 152790875 | 152779936 | 152780091 | 152829624 | 152829726 | 3 | 0 | 0 | 6 | 1 | 0 | 0 | 0 | 0 | 1 | 98 | 49 | 1 | 1 | 0.2 | 0 | NA | NA | 0 | 0.1 |
| SE | 9790 | ARFIP1 | ARFIP1 | chr4 | + | 152790758 | 152790875 | 152779936 | 152780226 | 152829624 | 152829726 | 3 | 0 | 0 | 10 | 0 | 0 | 0 | 2 | 0 | 0 | 98 | 49 | 1 | 1 | 0.13 | NA | NA | NA | 1 | -0.87 |
| SE | 9791 | ARFIP1 | ARFIP1 | chr4 | + | 152870752 | 152870848 | 152863605 | 152863714 | 152872451 | 152872564 | 22 | 7 | 2 | 9 | 4 | 1 | 0 | 3 | 0 | 0 | 98 | 49 | 1 | 1 | 0.55 | 0.467 | 0.5 | NA | 1 | -0.494 |
| SE | 9792 | ARFIP1 | ARFIP1 | chr4 | + | 152882722 | 152882880 | 152880962 | 152881184 | 152888132 | 152888307 | 10 | 4 | 5 | 1 | 0 | 0 | 0 | 1 | 0 | 0 | 98 | 49 | 1 | 1 | 0.833 | 1 | 1 | NA | 1 | -0.056 |
| SE | 9798 | RNH1 | RNH1 | chr11 | - | 504823 | 504996 | 502061 | 502181 | 507112 | 507283 | 1 | 2 | 0 | 0 | 1 | 0 | 0 | 2 | 0 | 0 | 98 | 49 | 1 | 1 | 1 | 0.5 | NA | NA | 1 | -0.25 |
| SE | 9830 | ARL8A | ARL8A | chr1 | - | 202135458 | 202135526 | 202135149 | 202135220 | 202135706 | 202135800 | 18 | 18 | 25 | 0 | 0 | 0 | 0 | 11 | 0 | 1 | 98 | 49 | 1 | 1 | 1 | 1 | 1 | NA | 0.846 | 0.154 |
| SE | 9836 | UBL3 | UBL3 | chr13 | - | 29777154 | 29777263 | 29772111 | 29772198 | 29849511 | 29850683 | 22 | 18 | 2 | 1 | 0 | 0 | 0 | 6 | 0 | 0 | 98 | 49 | 1 | 1 | 0.917 | 1 | 1 | NA | 1 | -0.028 |
| SE | 9837 | ZCCHC11 | ZCCHC11 | chr1 | - | 52458335 | 52458449 | 52446589 | 52446667 | 52461133 | 52461223 | 5 | 0 | 0 | 2 | 1 | 0 | 0 | 0 | 0 | 1 | 98 | 49 | 1 | 1 | 0.556 | 0 | NA | NA | 0 | 0.278 |
| SE | 9849 | MYH9 | MYH9 | chr22 | - | 36303994 | 36304155 | 36302567 | 36302676 | 36305032 | 36305102 | 184 | 98 | 47 | 0 | 1 | 0 | 6 | 33 | 0 | 0 | 98 | 49 | 1 | 1 | 1 | 0.98 | 1 | 1 | 1 | -0.007 |
| SE | 9854 | NADK | NADK | chr1 | - | 1753566 | 1753649 | 1751231 | 1753060 | 1754050 | 1754208 | 75 | 38 | 62 | 1 | 0 | 0 | 9 | 28 | 0 | 0 | 98 | 49 | 1 | 1 | 0.974 | 1 | 1 | 1 | 1 | -0.009 |
| SE | 9856 | NADK | NADK | chr1 | - | 1754543 | 1754698 | 1754283 | 1754383 | 1756257 | 1756343 | 98 | 57 | 38 | 0 | 2 | 0 | 4 | 24 | 0 | 0 | 98 | 49 | 1 | 1 | 1 | 0.934 | 1 | 1 | 1 | -0.022 |
| SE | 9857 | NADK | NADK | chr1 | - | 1755373 | 1755476 | 1754543 | 1754698 | 1756257 | 1756343 | 59 | 31 | 34 | 6 | 4 | 2 | 0 | 12 | 0 | 3 | 98 | 49 | 1 | 1 | 0.831 | 0.795 | 0.895 | NA | 0.667 | 0.173 |
| SE | 9859 | NADK | NADK | chr1 | - | 1765227 | 1765446 | 1761951 | 1762035 | 1779904 | 1780069 | 137 | 67 | 39 | 0 | 1 | 0 | 7 | 28 | 0 | 0 | 98 | 49 | 1 | 1 | 1 | 0.971 | 1 | 1 | 1 | -0.01 |
| SE | 9860 | STOM | STOM | chr9 | - | 121366115 | 121366245 | 121356052 | 121356156 | 121370126 | 121370304 | 0 | 0 | 0 | 6 | 2 | 0 | 2 | 0 | 6 | 0 | 98 | 49 | 1 | 1 | 0 | 0 | NA | 0.143 | NA | -0.143 |
| SE | 9862 | RNF40 | RNF40 | chr16 | + | 30766162 | 30766282 | 30765424 | 30765499 | 30766378 | 30766558 | 15 | 10 | 1 | 0 | 2 | 0 | 0 | 3 | 0 | 0 | 98 | 49 | 1 | 1 | 1 | 0.714 | 1 | NA | 1 | -0.095 |
| SE | 9866 | ZFYVE27 | ZFYVE27 | chr10 | + | 97743093 | 97743164 | 97738476 | 97738674 | 97744728 | 97744915 | 10 | 2 | 2 | 0 | 1 | 0 | 0 | 2 | 0 | 0 | 98 | 49 | 1 | 1 | 1 | 0.5 | 1 | NA | 1 | -0.167 |
| SE | 9873 | RCBTB2 | RCBTB2 | chr13 | - | 48522307 | 48522403 | 48521897 | 48521962 | 48524658 | 48524757 | 6 | 3 | 0 | 1 | 0 | 0 | 0 | 5 | 0 | 0 | 98 | 49 | 1 | 1 | 0.75 | 1 | NA | NA | 1 | -0.125 |
| SE | 9874 | VPS33A | VPS33A | chr12 | - | 122239877 | 122239945 | 122238586 | 122238724 | 122242381 | 122242508 | 1 | 3 | 3 | 2 | 0 | 0 | 0 | 2 | 0 | 0 | 98 | 49 | 1 | 1 | 0.2 | 1 | 1 | NA | 1 | -0.267 |
| SE | 9875 | VPS33A | VPS33A | chr12 | - | 122244568 | 122244762 | 122242381 | 122242508 | 122249870 | 122250045 | 12 | 2 | 0 | 0 | 0 | 0 | 0 | 1 | 2 | 0 | 98 | 49 | 1 | 1 | 1 | 1 | NA | 0 | 1 | 0.5 |
| SE | 9876 | BAZ2B | BAZ2B | chr2 | - | 159336941 | 159337077 | 159332539 | 159332686 | 159337566 | 159337772 | 10 | 6 | 1 | 6 | 2 | 3 | 0 | 4 | 0 | 0 | 98 | 49 | 1 | 1 | 0.455 | 0.6 | 0.143 | NA | 1 | -0.601 |
| SE | 9877 | BAZ2B | BAZ2B | chr2 | - | 159337566 | 159337772 | 159332539 | 159332686 | 159347485 | 159347646 | 44 | 21 | 6 | 0 | 0 | 0 | 1 | 9 | 0 | 1 | 98 | 49 | 1 | 1 | 1 | 1 | 1 | 1 | 0.818 | 0.091 |
| SE | 9878 | BAZ2B | BAZ2B | chr2 | - | 159389344 | 159389485 | 159386352 | 159386607 | 159395768 | 159395834 | 85 | 42 | 3 | 1 | 0 | 0 | 1 | 9 | 0 | 0 | 98 | 49 | 1 | 1 | 0.977 | 1 | 1 | 1 | 1 | -0.008 |
| SE | 9879 | BAZ2B | BAZ2B | chr2 | - | 159427940 | 159428042 | 159412334 | 159412545 | 159428310 | 159428419 | 57 | 12 | 2 | 19 | 1 | 0 | 0 | 5 | 0 | 0 | 98 | 49 | 1 | 1 | 0.6 | 0.857 | 1 | NA | 1 | -0.181 |
| SE | 9880 | BAZ2B | BAZ2B | chr2 | - | 159430862 | 159431156 | 159429199 | 159429260 | 159432756 | 159433363 | 82 | 34 | 2 | 5 | 8 | 0 | 0 | 18 | 0 | 3 | 98 | 49 | 1 | 1 | 0.891 | 0.68 | 1 | NA | 0.75 | 0.107 |
| SE | 9882 | BAZ2B | BAZ2B | chr2 | - | 159446781 | 159446975 | 159439008 | 159439212 | 159478574 | 159478721 | 77 | 22 | 2 | 1 | 0 | 0 | 0 | 16 | 0 | 0 | 98 | 49 | 1 | 1 | 0.975 | 1 | 1 | NA | 1 | -0.008 |
| SE | 9883 | BAZ2B | BAZ2B | chr2 | - | 159448241 | 159448403 | 159446781 | 159446975 | 159453612 | 159453801 | 41 | 23 | 5 | 1 | 0 | 0 | 0 | 9 | 0 | 0 | 98 | 49 | 1 | 1 | 0.953 | 1 | 1 | NA | 1 | -0.016 |
| SE | 9884 | BAZ2B | BAZ2B | chr2 | - | 159448241 | 159448403 | 159446781 | 159446975 | 159478574 | 159478721 | 37 | 22 | 5 | 5 | 3 | 0 | 0 | 9 | 0 | 0 | 98 | 49 | 1 | 1 | 0.787 | 0.786 | 1 | NA | 1 | -0.142 |
| SE | 9885 | BAZ2B | BAZ2B | chr2 | - | 159448241 | 159448409 | 159446781 | 159446975 | 159453612 | 159453801 | 78 | 30 | 5 | 1 | 0 | 0 | 0 | 14 | 0 | 0 | 98 | 49 | 1 | 1 | 0.975 | 1 | 1 | NA | 1 | -0.008 |
| SE | 9886 | BAZ2B | BAZ2B | chr2 | - | 159448241 | 159448409 | 159446781 | 159446975 | 159478574 | 159478721 | 43 | 28 | 5 | 5 | 3 | 0 | 0 | 14 | 0 | 0 | 98 | 49 | 1 | 1 | 0.811 | 0.824 | 1 | NA | 1 | -0.122 |
| SE | 9887 | BAZ2B | BAZ2B | chr2 | - | 159453612 | 159453801 | 159446781 | 159446975 | 159478574 | 159478721 | 36 | 15 | 2 | 5 | 3 | 0 | 0 | 6 | 0 | 0 | 98 | 49 | 1 | 1 | 0.783 | 0.714 | 1 | NA | 1 | -0.168 |
| SE | 9888 | BAZ2B | BAZ2B | chr2 | - | 159453612 | 159453801 | 159448241 | 159448403 | 159478574 | 159478721 | 41 | 16 | 2 | 2 | 0 | 0 | 0 | 6 | 0 | 0 | 98 | 49 | 1 | 1 | 0.911 | 1 | 1 | NA | 1 | -0.03 |
| SE | 9889 | BAZ2B | BAZ2B | chr2 | - | 159453612 | 159453801 | 159448241 | 159448409 | 159478574 | 159478721 | 78 | 23 | 2 | 8 | 6 | 0 | 0 | 11 | 0 | 5 | 98 | 49 | 1 | 1 | 0.83 | 0.657 | 1 | NA | 0.524 | 0.305 |
| SE | 9890 | MLKL | MLKL | chr16 | - | 74675354 | 74675404 | 74674959 | 74675100 | 74675612 | 74675764 | 19 | 5 | 2 | 1 | 0 | 0 | 5 | 6 | 0 | 0 | 98 | 49 | 1 | 1 | 0.905 | 1 | 1 | 1 | 1 | -0.032 |
| SE | 9891 | MLKL | MLKL | chr16 | - | 74678898 | 74678980 | 74675612 | 74675764 | 74682650 | 74682786 | 20 | 3 | 1 | 1 | 0 | 0 | 0 | 7 | 0 | 0 | 98 | 49 | 1 | 1 | 0.909 | 1 | 1 | NA | 1 | -0.03 |
| SE | 9892 | MLKL | MLKL | chr16 | - | 74678898 | 74678980 | 74675612 | 74675764 | 74685485 | 74685583 | 6 | 1 | 8 | 2 | 0 | 0 | 0 | 0 | 0 | 1 | 98 | 49 | 1 | 1 | 0.6 | 1 | 1 | NA | 0 | 0.867 |
| SE | 9893 | MLKL | MLKL | chr16 | - | 74682650 | 74682786 | 74675612 | 74675764 | 74685485 | 74685583 | 17 | 2 | 0 | 2 | 0 | 0 | 0 | 3 | 0 | 1 | 98 | 49 | 1 | 1 | 0.81 | 1 | NA | NA | 0.6 | 0.305 |

| | | | | | | | | | | | | | | | | | | | | | | | | | | | | | | | | |
|---|---|---|---|---|---|---|---|---|---|---|---|---|---|---|---|---|---|---|---|---|---|---|---|---|---|---|---|---|---|---|---|---|
| SE | 9894 | MLKL | MLKL | chr16 | - | 74682650 | 74682786 | 74678898 | 74678980 | 74685485 | 74685583 | 32 | 4 | 1 | 2 | 0 | 8 | 0 | 10 | 0 | 0 | 98 | 49 | 1 | 1 | 0.889 | 1 | 0.059 | NA | | 1 | -0.351 |
| SE | 9898 | ICAM3 | ICAM3 | chr19 | - | 10338681 | 10338948 | 10335670 | 10335976 | 10339538 | 10339669 | 126 | 79 | 79 | 1 | 0 | 1 | 0 | 15 | 0 | 0 | 98 | 49 | 1 | 1 | 0.984 | 1 | 0.975 | NA | | 1 | -0.014 |
| SE | 9900 | SLC30A6 | SLC30A6 | chr2 | + | 32175318 | 32175361 | 32174062 | 32174147 | 32184272 | 32184338 | 10 | 5 | 1 | 0 | 2 | 0 | 0 | 3 | 0 | 0 | 92 | 49 | 1 | 1 | 1 | 0.571 | 1 | NA | | 1 | -0.143 |
| SE | 9901 | SLC30A6 | SLC30A6 | chr2 | + | 32177476 | 32177596 | 32175318 | 32175361 | 32184272 | 32184338 | 1 | 0 | 0 | 6 | 1 | 1 | 0 | 0 | 0 | 1 | 98 | 49 | 1 | 1 | 0.077 | 0 | 0 | NA | | 0 | 0.026 |
| SE | 9909 | PTP4A2 | PTP4A2 | chr1 | - | 31910037 | 31910112 | 31906420 | 31908960 | 31911695 | 31911826 | 61 | 20 | 8 | 2 | 0 | 0 | 0 | 12 | 0 | 0 | 98 | 49 | 1 | 1 | 0.938 | 1 | 1 | NA | | 1 | -0.021 |
| SE | 9912 | PRKCQ | PRKCQ | chr10 | - | 6441892 | 6442081 | 6430809 | 6430938 | 6456673 | 6456812 | 5 | 0 | 0 | 2 | 0 | 0 | 0 | 2 | 0 | 0 | 98 | 49 | 1 | 1 | 0.556 | NA | NA | NA | | 1 | -0.444 |
| SE | 9914 | MFNG | MFNG | chr22 | - | 37472442 | 37472528 | 37469062 | 37470029 | 37474511 | 37474677 | 14 | 2 | 4 | 1 | 0 | 0 | 0 | 1 | 0 | 0 | 98 | 49 | 1 | 1 | 0.875 | 1 | 1 | NA | | 1 | -0.042 |
| SE | 9918 | DBNL | DBNL | chr7 | + | 44051829 | 44051942 | 44050224 | 44050280 | 44052866 | 44052941 | 88 | 25 | 26 | 1 | 0 | 0 | 5 | 6 | 0 | 0 | 98 | 49 | 1 | 1 | 0.978 | 1 | 1 | | 1 | 1 | -0.007 |
| SE | 9920 | DBNL | DBNL | chr7 | + | 44059353 | 44059449 | 44058877 | 44058983 | 44059542 | 44059658 | 79 | 28 | 18 | 0 | 1 | 0 | 5 | 13 | 0 | 0 | 98 | 49 | 1 | 1 | 1 | 0.933 | 1 | | 1 | 1 | -0.022 |
| SE | 9921 | ERLIN1 | ERLIN1 | chr10 | - | 100167347 | 100167406 | 100164003 | 100164095 | 100174207 | 100174281 | 2 | 3 | 0 | 1 | 0 | 0 | 0 | 3 | 0 | 0 | 98 | 49 | 1 | 1 | 0.5 | 1 | NA | NA | | 1 | -0.25 |
| SE | 9922 | CHUK | CHUK | chr10 | - | 100194424 | 100194521 | 100193983 | 100194131 | 100199970 | 100200020 | 5 | 2 | 9 | 0 | 0 | 0 | 0 | 2 | 0 | 2 | 98 | 49 | 1 | 1 | 1 | 1 | 1 | NA | | 0.333 | 0.667 |
| SE | 9923 | CHUK | CHUK | chr10 | - | 100219007 | 100219132 | 100218717 | 100218825 | 100219269 | 100219359 | 6 | 4 | 3 | 1 | 0 | 0 | 0 | 2 | 0 | 1 | 98 | 49 | 1 | 1 | 0.75 | 1 | 1 | NA | | 0.5 | 0.417 |
| SE | 9932 | MKNK1 | MKNK1 | chr1 | - | 46571416 | 46571539 | 46568442 | 46568498 | 46572062 | 46572167 | 1 | 0 | 0 | 28 | 5 | 0 | 0 | 0 | 2 | 4 | 98 | 49 | 1 | 1 | 0.018 | 0 | NA | | 0 | 0 | 0.009 |
| SE | 9934 | MKNK1 | MKNK1 | chr1 | - | 46580529 | 46580627 | 46576574 | 46576654 | 46583227 | 46583329 | 20 | 6 | 12 | 1 | 0 | 0 | 0 | 3 | 0 | 0 | 98 | 49 | 1 | 1 | 0.909 | 1 | 1 | NA | | 1 | -0.03 |
| SE | 9935 | MKNK1 | MKNK1 | chr1 | - | 46585873 | 46585974 | 46583227 | 46583329 | 46594112 | 46594280 | 1 | 1 | 1 | 2 | 0 | 2 | 0 | 1 | 0 | 0 | 98 | 49 | 1 | 1 | 0.2 | 1 | 0.2 | NA | | 1 | -0.533 |
| SE | 9937 | TRIM38 | TRIM38 | chr6 | + | 25973053 | 25973076 | 25971868 | 25972099 | 25973172 | 25973285 | 18 | 11 | 3 | 0 | 0 | 0 | 0 | 5 | 0 | 1 | 72 | 49 | 1 | 1 | 1 | 1 | 1 | NA | | 0.773 | 0.227 |
| SE | 9939 | SART1 | SART1 | chr11 | + | 65964073 | 65964131 | 65961688 | 65962093 | 65964514 | 65964570 | 3 | 10 | 3 | 0 | 1 | 0 | 0 | 2 | 0 | 0 | 98 | 49 | 1 | 1 | 1 | 0.833 | 1 | NA | | 1 | -0.056 |
| SE | 9942 | TMEM2 | TMEM2 | chr9 | - | 71697984 | 71698204 | 71694508 | 71694607 | 71700641 | 71700824 | 19 | 4 | 0 | 1 | 0 | 0 | 0 | 1 | 0 | 0 | 98 | 49 | 1 | 1 | 0.905 | 1 | NA | NA | | 1 | -0.047 |
| SE | 9943 | TMEM2 | TMEM2 | chr9 | - | 71734805 | 71734994 | 71732350 | 71732520 | 71740062 | 71740232 | 35 | 3 | 3 | 6 | 0 | 2 | 0 | 6 | 0 | 0 | 98 | 49 | 1 | 1 | 0.745 | 1 | 0.429 | NA | | 1 | -0.275 |
| SE | 9947 | MATR3 | MATR3 | chr5 | + | 139307238 | 139308327 | 139293643 | 139293805 | 139314674 | 139314736 | 12 | 7 | 5 | 0 | 0 | 0 | 0 | 0 | 1 | 0 | 98 | 49 | 1 | 1 | 1 | 1 | 1 | | 0 | NA | 1 |
| SE | 9949 | MATR3 | MATR3 | chr5 | + | 139318907 | 139319033 | 139317595 | 139317721 | 139319333 | 139319501 | 16 | 8 | 0 | 0 | 1 | 0 | 1 | 0 | 0 | 0 | 98 | 49 | 1 | 1 | 1 | 0.8 | NA | | 1 | NA | -0.1 |
| SE | 9951 | SEC14L1 | SEC14L1 | chr17 | + | 77193420 | 77193549 | 77191180 | 77191312 | 77194676 | 77194911 | 298 | 137 | 75 | 0 | 0 | 0 | 5 | 49 | 0 | 1 | 98 | 49 | 1 | 1 | 1 | 1 | 1 | | 1 | 0.961 | 0.02 |
| SE | 9953 | SEC14L1 | SEC14L1 | chr17 | + | 77213313 | 77213492 | 77211949 | 77212201 | 77213917 | 77214019 | 89 | 66 | 57 | 1 | 1 | 0 | 5 | 14 | 0 | 0 | 98 | 49 | 1 | 1 | 0.978 | 0.971 | 1 | | 1 | 1 | -0.017 |
| SE | 9956 | CLEC16A | CLEC16A | chr16 | + | 10977224 | 10977399 | 10972931 | 10973061 | 10979328 | 10979382 | 10 | 5 | 5 | 0 | 0 | 0 | 0 | 0 | 1 | 0 | 98 | 49 | 1 | 1 | 1 | 1 | 1 | | 0 | NA | 1 |
| SE | 9957 | ASCC1 | ASCC1 | chr10 | - | 72204421 | 72204505 | 72203426 | 72203524 | 72210731 | 72210831 | 0 | 1 | 0 | 0 | 0 | 0 | 0 | 0 | 0 | 1 | 98 | 49 | 1 | 1 | NA | 1 | NA | NA | | 0 | 1 |
| SE | 9958 | ASCC1 | ASCC1 | chr10 | - | 72210731 | 72210831 | 72203426 | 72203524 | 72213186 | 72213331 | 0 | 0 | 0 | 1 | 0 | 0 | 0 | 1 | 0 | 0 | 98 | 49 | 1 | 1 | 0 | NA | NA | NA | | 1 | -1 |
| SE | 9960 | HDAC7 | HDAC7 | chr12 | - | 47791258 | 47791308 | 47789812 | 47789920 | 47791585 | 47791706 | 9 | 17 | 3 | 2 | 0 | 0 | 0 | 2 | 0 | 0 | 98 | 49 | 1 | 1 | 0.692 | 1 | 1 | NA | | 1 | -0.103 |
| SE | 9961 | HDAC7 | HDAC7 | chr12 | - | 47795905 | 47796016 | 47795586 | 47795767 | 47796206 | 47796298 | 5 | 2 | 0 | 5 | 1 | 0 | 0 | 2 | 0 | 1 | 98 | 49 | 1 | 1 | 0.333 | 0.5 | NA | NA | | 0.5 | -0.084 |
| SE | 9965 | TAZ | TAZ | chrX | + | 154414100 | 154414190 | 154413481 | 154413567 | 154419542 | 154419623 | 0 | 2 | 0 | 1 | 1 | 0 | 0 | 0 | 0 | 1 | 98 | 49 | 1 | 1 | 0 | 0.5 | NA | NA | | 0 | 0.25 |
| SE | 9967 | TAZ | TAZ | chrX | + | 154420211 | 154420264 | 154420031 | 154420094 | 154420657 | 154420735 | 9 | 4 | 2 | 0 | 0 | 0 | 0 | 2 | 0 | 1 | 98 | 49 | 1 | 1 | 1 | 1 | 1 | NA | | 0.5 | 0.5 |
| SE | 9968 | DHX9 | DHX9 | chr1 | + | 182855576 | 182855713 | 182854029 | 182854178 | 182856531 | 182856578 | 19 | 5 | 0 | 13 | 2 | 0 | 2 | 1 | 0 | 5 | 98 | 49 | 1 | 1 | 0.422 | 0.556 | NA | | 1 | 0.091 | -0.056 |
| SE | 9969 | DHX9 | DHX9 | chr1 | + | 182856531 | 182856578 | 182855576 | 182855713 | 182858103 | 182858240 | 25 | 10 | 3 | 0 | 1 | 0 | 0 | 3 | 0 | 0 | 96 | 49 | 1 | 1 | 1 | 0.836 | 1 | NA | | 1 | -0.055 |
| SE | 9971 | DUSP1 | DUSP1 | chr5 | - | 172769574 | 172769794 | 172768089 | 172769132 | 172770585 | 172771200 | 438 | 200 | 178 | 0 | 2 | 0 | 6 | 110 | 1 | 0 | 98 | 49 | 1 | 1 | 1 | 0.98 | 1 | | 0.75 | 1 | 0.118 |
| SE | 9972 | DUSP1 | DUSP1 | chr5 | - | 172770160 | 172770306 | 172768089 | 172769132 | 172770585 | 172771200 | 561 | 303 | 269 | 0 | 2 | 0 | 93 | 267 | 1 | 0 | 98 | 49 | 1 | 1 | 1 | 0.987 | 1 | | 0.979 | 1 | 0.006 |
| SE | 9975 | GBF1 | GBF1 | chr10 | + | 102359266 | 102359435 | 102358505 | 102358729 | 102360183 | 102360395 | 2 | 0 | 1 | 1 | 1 | 0 | 0 | 1 | 0 | 0 | 98 | 49 | 1 | 1 | 0.5 | 0 | 1 | NA | | 1 | -0.5 |
| SE | 9976 | GBF1 | GBF1 | chr10 | + | 102359269 | 102359435 | 102358505 | 102358729 | 102360183 | 102360395 | 5 | 0 | 1 | 1 | 1 | 0 | 0 | 2 | 0 | 0 | 98 | 49 | 1 | 1 | 0.714 | 0 | 1 | NA | | 1 | -0.429 |
| SE | 9977 | GBF1 | GBF1 | chr10 | + | 102363255 | 102363396 | 102362474 | 102362664 | 102363709 | 102363798 | 8 | 2 | 0 | 0 | 2 | 0 | 1 | 0 | 0 | 0 | 98 | 49 | 1 | 1 | 1 | 0.333 | NA | | 1 | NA | -0.334 |
| SE | 9978 | GBF1 | GBF1 | chr10 | + | 102368217 | 102368454 | 102367477 | 102367560 | 102368738 | 102368832 | 12 | 3 | 4 | 3 | 0 | 0 | 0 | 4 | 0 | 0 | 98 | 49 | 1 | 1 | 0.667 | 1 | 1 | NA | | 1 | -0.111 |
| SE | 9979 | GBF1 | GBF1 | chr10 | + | 102368738 | 102368832 | 102368217 | 102368454 | 102369210 | 102369387 | 13 | 5 | 5 | 1 | 0 | 0 | 0 | 6 | 0 | 0 | 98 | 49 | 1 | 1 | 0.867 | 1 | 1 | NA | | 1 | -0.044 |
| SE | 9980 | GBF1 | GBF1 | chr10 | + | 102381126 | 102381255 | 102380505 | 102380686 | 102382055 | 102382899 | 6 | 1 | 6 | 1 | 0 | 0 | 0 | 1 | 0 | 0 | 98 | 49 | 1 | 1 | 0.75 | 1 | 1 | NA | | 1 | -0.083 |
| SE | 9984 | ACTR1A | ACTR1A | chr10 | - | 102484159 | 102484376 | 102483010 | 102483103 | 102485608 | 102485733 | 26 | 8 | 5 | 0 | 1 | 0 | 0 | 6 | 0 | 0 | 98 | 49 | 1 | 1 | 1 | 0.8 | 1 | NA | | 1 | -0.067 |
| SE | 10002 | TXNL1 | TXNL1 | chr18 | - | 56626161 | 56626266 | 56624287 | 56624461 | 56626360 | 56626457 | 3 | 0 | 0 | 6 | 0 | 3 | 0 | 3 | 0 | 1 | 98 | 49 | 1 | 1 | 0.2 | NA | 0 | NA | | 0.6 | -0.5 |
| SE | 10007 | NT5C2 | NT5C2 | chr10 | - | 103090610 | 103090787 | 103088016 | 103089908 | 103090935 | 103090996 | 71 | 22 | 11 | 8 | 4 | 2 | 0 | 14 | 0 | 5 | 98 | 49 | 1 | 1 | 0.816 | 0.733 | 0.733 | NA | | 0.583 | 0.178 |
| SE | 10009 | NT5C2 | NT5C2 | chr10 | - | 103139405 | 103139479 | 103106588 | 103106706 | 103174857 | 103174982 | 52 | 36 | 9 | 0 | 2 | 0 | 1 | 7 | 0 | 0 | 98 | 49 | 1 | 1 | 1 | 0.9 | 1 | | 1 | 1 | -0.033 |
| SE | 10013 | BSG | BSG | chr19 | + | 577773 | 578121 | 572453 | 572701 | 579499 | 579656 | 0 | 0 | 4 | 10 | 2 | 5 | 0 | 0 | 0 | 1 | 98 | 49 | 1 | 1 | 0 | 0 | 0.286 | NA | | 0 | 0.095 |
| SE | 10015 | SBF2 | SBF2 | chr11 | - | 9795830 | 9795957 | 9790555 | 9790683 | 9807999 | 9808185 | 5 | 1 | 1 | 0 | 3 | 0 | 0 | 1 | 0 | 1 | 98 | 49 | 1 | 1 | 1 | 0.143 | 1 | NA | | 0.333 | 0.381 |
| SE | 10016 | SBF2 | SBF2 | chr11 | - | 9853539 | 9853712 | 9852675 | 9852749 | 9856457 | 9856720 | 32 | 7 | 0 | 1 | 0 | 0 | 0 | 3 | 0 | 2 | 98 | 49 | 1 | 1 | 0.941 | 1 | NA | NA | | 0.429 | 0.541 |
| SE | 10017 | SBF2 | SBF2 | chr11 | - | 9992414 | 9992543 | 9989496 | 9989595 | 9992989 | 9993103 | 37 | 10 | 2 | 5 | 5 | 0 | 0 | 5 | 0 | 5 | 98 | 49 | 1 | 1 | 0.787 | 0.5 | 1 | NA | | 0.333 | 0.429 |
| SE | 10023 | DAPK2 | DAPK2 | chr15 | - | 63924815 | 63924861 | 63912107 | 63912197 | 63925940 | 63926093 | 24 | 18 | 8 | 1 | 0 | 0 | 0 | 6 | 0 | 0 | 95 | 49 | 1 | 1 | 0.925 | 1 | 1 | NA | | 1 | -0.025 |
| SE | 10024 | DAPK2 | DAPK2 | chr15 | - | 63939231 | 63939361 | 63930406 | 63930455 | 63971422 | 63971561 | 37 | 15 | 3 | 2 | 0 | 0 | 0 | 4 | 0 | 0 | 98 | 49 | 1 | 1 | 0.902 | 1 | 1 | NA | | 1 | -0.033 |
| SE | 10028 | ANKRD11 | ANKRD11 | chr16 | - | 89291680 | 89291777 | 89291012 | 89291183 | 89305205 | 89305344 | 2 | 3 | 0 | 9 | 4 | 2 | 0 | 0 | 0 | 4 | 98 | 49 | 1 | 1 | 0.1 | 0.273 | 0 | NA | | 0 | 0.124 |
| SE | 10030 | ANKRD11 | ANKRD11 | chr16 | - | 89431255 | 89431326 | 89418283 | 89418368 | 89490244 | 89490561 | 0 | 2 | 0 | 0 | 2 | 0 | 0 | 0 | 0 | 1 | 98 | 49 | 1 | 1 | NA | 0.333 | NA | NA | | 0 | 0.333 |
| SE | 10035 | PRKCD | PRKCD | chr3 | + | 53184873 | 53184974 | 53183451 | 53183581 | 53185603 | 53185700 | 74 | 31 | 6 | 1 | 1 | 0 | 1 | 23 | 0 | 0 | 98 | 49 | 1 | 1 | 0.974 | 0.939 | 1 | | 1 | 1 | -0.029 |
| SE | 10038 | TESK2 | TESK2 | chr1 | - | 45346978 | 45347062 | 45346692 | 45346779 | 45347608 | 45347693 | 0 | 1 | 0 | 0 | 0 | 0 | 3 | 3 | 1 | 0 | 98 | 49 | 1 | 1 | NA | 1 | NA | | 0.6 | 1 | 0.2 |
| SE | 10039 | TESK2 | TESK2 | chr1 | - | 45347917 | 45348000 | 45347608 | 45347693 | 45355302 | 45355449 | 7 | 3 | 0 | 0 | 1 | 0 | 0 | 1 | 0 | 0 | 98 | 49 | 1 | 1 | 1 | 0.6 | NA | NA | | 1 | -0.2 |
| SE | 10046 | SAFB2 | SAFB2 | chr19 | - | 5616421 | 5616486 | 5616131 | 5616335 | 5621308 | 5621396 | 21 | 15 | 1 | 2 | 0 | 0 | 0 | 1 | 0 | 0 | 98 | 49 | 1 | 1 | 0.84 | 1 | 1 | NA | | 1 | -0.053 |
| SE | 10047 | ADD3 | ADD3 | chr10 | + | 110116258 | 110116410 | 110112776 | 110112915 | 110117341 | 110117422 | 37 | 17 | 16 | 4 | 0 | 0 | 0 | 7 | 0 | 1 | 98 | 49 | 1 | 1 | 0.822 | 1 | 1 | NA | | 0.778 | 0.163 |
| SE | 10049 | ADD3 | ADD3 | chr10 | + | 110132304 | 110132400 | 110130362 | 110130486 | 110133325 | 110135565 | 22 | 3 | 8 | 0 | 0 | 0 | 0 | 9 | 0 | 2 | 98 | 49 | 1 | 1 | 1 | 1 | 1 | NA | | 0.692 | 0.308 |
| SE | 10059 | NRBF2 | NRBF2 | chr10 | + | 63146208 | 63146293 | 63133246 | 63133500 | 63152149 | 63152190 | 22 | 9 | 3 | 0 | 0 | 1 | 1 | 2 | 0 | 0 | 98 | 49 | 1 | 1 | 1 | 1 | 0.6 | | 1 | 1 | -0.133 |
| SE | 10069 | CD53 | CD53 | chr1 | + | 110892344 | 110892533 | 110891391 | 110891471 | 110894326 | 110894401 | 240 | 104 | 119 | 1 | 0 | 0 | 3 | 41 | 0 | 0 | 98 | 49 | 1 | 1 | 0.992 | 1 | 1 | | 1 | 1 | -0.003 |
| SE | 10070 | CD53 | CD53 | chr1 | + | 110892344 | 110892533 | 110891391 | 110891471 | 110899123 | 110899936 | 141 | 54 | 53 | 3 | 0 | 0 | 3 | 18 | 0 | 0 | 98 | 49 | 1 | 1 | 0.959 | 1 | 1 | | 1 | 1 | -0.014 |
| SE | 10071 | CD53 | CD53 | chr1 | + | 110896652 | 110896733 | 110891391 | 110891471 | 110897808 | 110897892 | 45 | 26 | 57 | 0 | 1 | 0 | 4 | 12 | 0 | 0 | 98 | 49 | 1 | 1 | 1 | 0.929 | 1 | | 1 | 1 | -0.024 |
| SE | 10072 | CD53 | CD53 | chr1 | + | 110896652 | 110896733 | 110891391 | 110891471 | 110899123 | 110899936 | 1 | 0 | 0 | 3 | 0 | 0 | 0 | 3 | 0 | 0 | 98 | 49 | 1 | 1 | 0.143 | NA | NA | NA | | 1 | -0.857 |
| SE | 10073 | CD53 | CD53 | chr1 | + | 110896652 | 110896733 | 110894959 | 110895055 | 110897808 | 110897892 | 77 | 55 | 92 | 1 | 0 | 0 | 4 | 21 | 0 | 0 | 98 | 49 | 1 | 1 | 0.975 | 1 | 1 | | 1 | 1 | -0.008 |
| SE | 10075 | CD53 | CD53 | chr1 | + | 110897808 | 110897892 | 110896652 | 110896733 | 110899123 | 110899936 | 143 | 75 | 113 | 1 | 0 | 0 | 6 | 27 | 0 | 0 | 98 | 49 | 1 | 1 | 0.986 | 1 | 1 | | 1 | 1 | -0.005 |
| SE | 10076 | CAMK2G | CAMK2G | chr10 | - | 73819531 | 73819645 | 73817478 | 73817554 | 73821681 | 73821730 | 6 | 3 | 2 | 12 | 5 | 4 | 0 | 2 | 0 | 0 | 98 | 49 | 1 | 1 | 0.2 | 0.231 | 0.2 | NA | | 1 | -0.79 |
| SE | 10077 | CAMK2G | CAMK2G | chr10 | - | 73825278 | 73825347 | 73824039 | 73824084 | 73837467 | 73837511 | 17 | 25 | 6 | 1 | 0 | 0 | 0 | 1 | 0 | 0 | 98 | 49 | 1 | 1 | 0.895 | 1 | 1 | NA | | 1 | -0.035 |
| SE | 10078 | CAMK2G | CAMK2G | chr10 | - | 73825320 | 73825347 | 73824039 | 73824084 | 73837467 | 73837511 | 7 | 15 | 2 | 1 | 0 | 0 | 0 | 1 | 0 | 0 | 76 | 49 | 1 | 1 | 0.819 | 1 | 1 | NA | | 1 | -0.06 |
| SE | 10081 | CAMK2G | CAMK2G | chr10 | - | 73847224 | 73847347 | 73842457 | 73842541 | 73847987 | 73848082 | 22 | 16 | 3 | 1 | 0 | 0 | 0 | 3 | 0 | 0 | 98 | 49 | 1 | 1 | 0.917 | 1 | 1 | NA | | 1 | -0.028 |
| SE | 10082 | CAMK2G | CAMK2G | chr10 | - | 73847224 | 73847347 | 73842457 | 73842541 | 73848525 | 73848609 | 15 | 1 | 2 | 1 | 0 | 0 | 0 | 2 | 0 | 0 | 98 | 49 | 1 | 1 | 0.882 | 1 | 1 | NA | | 1 | -0.039 |
| SE | 10084 | CAMK2G | CAMK2G | chr10 | - | 73847987 | 73848082 | 73847224 | 73847347 | 73848525 | 73848609 | 20 | 23 | 6 | 1 | 0 | 0 | 0 | 1 | 0 | 0 | 98 | 49 | 1 | 1 | 0.909 | 1 | 1 | NA | | 1 | -0.03 |
| SE | 10085 | CAMK2G | CAMK2G | chr10 | - | 73848525 | 73848609 | 73847987 | 73848082 | 73849012 | 73849115 | 34 | 24 | 22 | 5 | 0 | 0 | 0 | 2 | 0 | 2 | 98 | 49 | 1 | 1 | 0.773 | 1 | 1 | NA | | 0.333 | 0.591 |
| SE | 10086 | CAMK2G | CAMK2G | chr10 | - | 73853191 | 73853246 | 73852253 | 73852319 | 73860829 | 73860889 | 30 | 29 | 9 | 1 | 1 | 0 | 3 | 2 | 0 | 0 | 98 | 49 | 1 | 1 | 0.937 | 0.935 | 1 | | 1 | 1 | -0.043 |
| SE | 10090 | PHTF1 | PHTF1 | chr1 | - | 113706593 | 113706722 | 113705889 | 113706162 | 113710253 | 113710475 | 1 | 3 | 3 | 1 | 0 | 0 | 0 | 1 | 0 | 0 | 98 | 49 | 1 | 1 | 0.333 | 1 | 1 | NA | | 1 | -0.222 |
| SE | 10096 | PICALM | PICALM | chr11 | - | 85974707 | 85974812 | 85957170 | 85959060 | 85981128 | 85981228 | 54 | 19 | 8 | 0 | 0 | 1 | 0 | 13 | 0 | 0 | 98 | 49 | 1 | 1 | 1 | 1 | 0.8 | NA | | 1 | -0.067 |
| SE | 10097 | PICALM | PICALM | chr11 | - | 85976622 | 85976682 | 85974707 | 85974812 | 85981128 | 85981228 | 131 | 73 | 41 | 1 | 0 | 0 | 2 | 30 | 0 | 0 | 98 | 49 | 1 | 1 | 0.985 | 1 | 1 | | 1 | 1 | -0.005 |
| SE | 10101 | PICALM | PICALM | chr11 | - | 86001034 | 86001158 | 86000642 | 86000779 | 86003365 | 86003451 | 424 | 170 | 81 | 1 | 0 | 0 | 6 | 50 | 0 | 0 | 98 | 49 | 1 | 1 | 0.995 | 1 | 1 | | 1 | 1 | -0.002 |
| SE | 10102 | PICALM | PICALM | chr11 | - | 86022366 | 86022469 | 86014869 | 86014963 | 86026291 | 86026367 | 309 | 142 | 70 | 7 | 5 | 0 | 0 | 36 | 0 | 1 | 98 | 49 | 1 | 1 | 0.957 | 0.934 | 1 | NA | | 0.947 | 0.017 |
| SE | 10103 | PICALM | PICALM | chr11 | - | 86022366 | 86022469 | 86014869 | 86014963 | 86031468 | 86031611 | 188 | 89 | 37 | 4 | 0 | 1 | 0 | 16 | 0 | 0 | 98 | 49 | 1 | 1 | 0.959 | 1 | 0.949 | NA | | 1 | -0.031 |
| SE | 10105 | PICALM | PICALM | chr11 | - | 86026291 | 86026367 | 86022366 | 86022469 | 86031468 | 86031611 | 260 | 114 | 89 | 2 | 0 | 0 | 1 | 33 | 0 | 0 | 98 | 49 | 1 | 1 | 0.985 | 1 | 1 | | 1 | 1 | -0.005 |
| SE | 10106 | PICALM | PICALM | chr11 | - | 86031468 | 86031611 | 86014869 | 86014963 | 86068650 | 86069097 | 92 | 24 | 16 | 1 | 1 | 0 | 1 | 7 | 0 | 0 | 98 | 49 | 1 | 1 | 0.979 | 0.923 | 1 | | 1 | 1 | -0.033 |
| SE | 10109 | SUV420H1 | SUV420H1 | chr11 | - | 68179469 | 68179537 | 68175017 | 68175183 | 68180131 | 68180200 | 1 | 2 | 0 | 1 | 3 | 6 | 0 | 0 | 0 | 6 | 98 | 49 | 1 | 1 | 0.333 | 0.25 | 0 | NA | | 0 | 0.194 |
| SE | 10125 | PDZD8 | PDZD8 | chr10 | - | 117340979 | 117341102 | 117318871 | 117318974 | 117374355 | 117375467 | 8 | 8 | 1 | 2 | 1 | 0 | 0 | 4 | 0 | 0 | 98 | 49 | 1 | 1 | 0.667 | 0.8 | 1 | NA | | 1 | -0.178 |
| SE | 10127 | KMT2B | KMT2B | chr19 | + | 35726235 | 35726353 | 35725722 | 35725818 | 35727155 | 35727269 | 11 | 9 | 1 | 0 | 1 | 0 | 0 | 2 | 0 | 0 | 98 | 49 | 1 | 1 | 1 | 0.818 | 1 | NA | | 1 | -0.061 |
| SE | 10131 | PIK3C2A | PIK3C2A | chr11 | - | 17101277 | 17101434 | 17099859 | 17099969 | 17102661 | 17102831 | 7 | 2 | 1 | 2 | 0 | 0 | 0 | 1 | 0 | 0 | 98 | 49 | 1 | 1 | 0.636 | 1 | 1 | NA | | 1 | -0.121 |
| SE | 10132 | ABHD16A | ABHD16A | chr6 | - | 31701273 | 31701340 | 31700941 | 31701028 | 31703149 | 31703360 | 7 | 1 | 1 | 4 | 2 | 3 | 0 | 0 | 0 | 3 | 98 | 49 | 1 | 1 | 0.467 | 0.2 | 0.143 | NA | | 0 | 0.27 |
| SE | 10133 | ABHD16A | ABHD16A | chr6 | - | 31702073 | 31702130 | 31701273 | 31701340 | 31703149 | 31703360 | 11 | 1 | 9 | 1 | 0 | 0 | 0 | 3 | 0 | 0 | 98 | 49 | 1 | 1 | 0.846 | 1 | 1 | NA | | 1 | -0.051 |
| SE | 10135 | FNDC3A | FNDC3A | chr13 | + | 49191014 | 49191120 | 49188514 | 49188633 | 49191208 | 49191384 | 3 | 3 | 0 | 1 | 0 | 0 | 0 | 1 | 0 | 0 | 98 | 49 | 1 | 1 | 0.6 | 1 | NA | NA | | 1 | -0.2 |
| SE | 10136 | EIF3A | EIF3A | chr10 | - | 119061223 | 119061328 | 119060745 | 119060844 | 119065398 | 119065570 | 7 | 2 | 0 | 0 | 1 | 0 | 0 | 3 | 0 | 0 | 98 | 49 | 1 | 1 | 1 | 0.5 | NA | NA | | 1 | -0.25 |
| SE | 10146 | RGS10 | RGS10 | chr10 | - | 119515508 | 119515652 | 119499826 | 119500259 | 119526031 | 119526118 | 22 | 3 | 0 | 2 | 0 | 0 | 0 | 6 | 0 | 0 | 98 | 49 | 1 | 1 | 0.846 | 1 | NA | NA | | 1 | -0.077 |
| SE | 10149 | SEC23IP | SEC23IP | chr10 | + | 119929606 | 119929762 | 119926035 | 119926227 | 119930328 | 119930431 | 6 | 0 | 0 | 2 | 0 | 0 | 0 | 1 | 0 | 0 | 98 | 49 | 1 | 1 | 0.6 | NA | NA | NA | | 1 | -0.4 |
| SE | 10161 | DNM2 | DNM2 | chr19 | + | 10793719 | 10793855 | 10786563 | 10786706 | 10795371 | 10795439 | 55 | 19 | 14 | 2 | 0 | 0 | 1 | 15 | 0 | 0 | 98 | 49 | 1 | 1 | 0.932 | 1 | 1 | | 1 | 1 | -0.023 |
| SE | 10162 | DNM2 | DNM2 | chr19 | + | 10796060 | 10796199 | 10795371 | 10795439 | 10798485 | 10798572 | 12 | 5 | 1 | 2 | 0 | 0 | 0 | 2 | 0 | 0 | 98 | 49 | 1 | 1 | 0.75 | 1 | 1 | NA | | 1 | -0.083 |
| SE | 10163 | DNM2 | DNM2 | chr19 | + | 10797379 | 10797518 | 10795371 | 10795439 | 10798485 | 10798572 | 10 | 6 | 8 | 2 | 0 | 0 | 0 | 4 | 0 | 0 | 98 | 49 | 1 | 1 | 0.714 | 1 | 1 | NA | | 1 | -0.095 |
| SE | 10164 | DNM2 | DNM2 | chr19 | + | 10798485 | 10798572 | 10795371 | 10795439 | 10802287 | 10802358 | 38 | 14 | 3 | 2 | 0 | 0 | 1 | 10 | 0 | 0 | 98 | 49 | 1 | 1 | 0.905 | 1 | 1 | | 1 | 1 | -0.032 |
| SE | 10165 | DNM2 | DNM2 | chr19 | + | 10808568 | 10808580 | 10805915 | 10805967 | 10812263 | 10812377 | 23 | 13 | 2 | 0 | 0 | 0 | 0 | 4 | 0 | 1 | 61 | 49 | 1 | 1 | 1 | 1 | 1 | NA | | 0.763 | 0.237 |
| SE | 10168 | RIT1 | RIT1 | chr1 | - | 155910655 | 155910804 | 155910449 | 155910506 | 155911242 | 155911402 | 27 | 20 | 4 | 0 | 2 | 1 | 0 | 3 | 0 | 0 | 98 | 49 | 1 | 1 | 1 | 0.833 | 0.667 | NA | | 1 | -0.167 |
| SE | 10177 | CYFIP2 | CYFIP2 | chr5 | + | 157327972 | 157328049 | 157326170 | 157326267 | 157330741 | 157330850 | 21 | 7 | 2 | 1 | 0 | 0 | 0 | 3 | 0 | 0 | 98 | 49 | 1 | 1 | 0.913 | 1 | 1 | NA | | 1 | -0.029 |
| SE | 10182 | KIAA1468 | KIAA1468 | chr18 | + | 62252663 | 62252754 | 62244763 | 62244876 | 62255406 | 62255478 | 24 | 3 | 0 | 1 | 1 | 0 | 0 | 7 | 0 | 0 | 98 | 49 | 1 | 1 | 0.923 | 0.6 | NA | NA | | 1 | -0.238 |
| SE | 10183 | KIAA1468 | KIAA1468 | chr18 | + | 62280359 | 62280442 | 62275373 | 62275473 | 62280645 | 62280709 | 7 | 2 | 3 | 9 | 1 | 0 | 0 | 1 | 0 | 1 | 98 | 49 | 1 | 1 | 0.28 | 0.5 | 1 | NA | | 0.333 | 0.26 |
| SE | 10186 | CASP9 | CASP9 | chr1 | - | 15493891 | 15494001 | 15491400 | 15493035 | 15495272 | 15495452 | 1 | 1 | 1 | 0 | 0 | 3 | 0 | 3 | 0 | 0 | 98 | 49 | 1 | 1 | 1 | 1 | 0.143 | NA | | 1 | -0.286 |
| SE | 10188 | LGALS2 | LGALS2 | chr22 | - | 37571848 | 37571931 | 37570575 | 37570735 | 37579899 | 37580017 | 9 | 0 | 0 | 2 | 0 | 0 | 0 | 8 | 0 | 1 | 98 | 49 | 1 | 1 | 0.692 | NA | NA | NA | | 0.8 | -0.108 |
| SE | 10239 | LST1 | LST1 | chr6 | + | 31587640 | 31587733 | 31587199 | 31587318 | 31587943 | 31587966 | 63 | 29 | 32 | 11 | 10 | 2 | 0 | 1 | 0 | 2 | 98 | 49 | 1 | 1 | 0.741 | 0.592 | 0.889 | NA | | 0.2 | 0.541 |
| SE | 10240 | LST1 | LST1 | chr6 | + | 31587640 | 31587733 | 31587199 | 31587318 | 31588496 | 31588909 | 39 | 24 | 22 | 3 | 2 | 0 | 0 | 0 | 0 | 3 | 98 | 49 | 1 | 1 | 0.867 | 0.857 | 1 | NA | | 0 | 0.908 |
| SE | 10241 | LST1 | LST1 | chr6 | + | 31587640 | 31587733 | 31587199 | 31587318 | 31588517 | 31588909 | 42 | 28 | 26 | 46 | 21 | 20 | 0 | 0 | 0 | 4 | 98 | 49 | 1 | 1 | 0.313 | 0.4 | 0.394 | NA | | 0 | 0.369 |
| SE | 10244 | LST1 | LST1 | chr6 | + | 31587943 | 31587966 | 31587199 | 31587318 | 31588517 | 31588909 | 17 | 17 | 14 | 46 | 21 | 20 | 0 | 7 | 0 | 4 | 72 | 49 | 1 | 1 | 0.201 | 0.355 | 0.323 | NA | | 0.544 | -0.251 |
| SE | 10247 | LST1 | LST1 | chr6 | + | 31587943 | 31587966 | 31587640 | 31587733 | 31588517 | 31588909 | 31 | 14 | 19 | 3 | 4 | 4 | 0 | 6 | 0 | 0 | 72 | 49 | 1 | 1 | 0.876 | 0.704 | 0.764 | NA | | 1 | -0.219 |
| SE | 10248 | LST1 | LST1 | chr6 | + | 31587943 | 31587966 | 31587640 | 31587733 | 31588562 | 31588909 | 20 | 6 | 3 | 10 | 5 | 2 | 0 | 1 | 0 | 2 | 72 | 49 | 1 | 1 | 0.576 | 0.45 | 0.505 | NA | | 0.254 | 0.256 |
| SE | 10252 | OXA1L | OXA1L | chr14 | + | 22769790 | 22769934 | 22767957 | 22768171 | 22770192 | 22770278 | 12 | 7 | 1 | 0 | 1 | 0 | 0 | 5 | 0 | 0 | 98 | 49 | 1 | 1 | 1 | 0.778 | 1 | NA | | 1 | -0.074 |
| SE | 10259 | AP3D1 | AP3D1 | chr19 | - | 2108686 | 2108766 | 2100987 | 2102268 | 2109085 | 2109207 | 20 | 7 | 1 | 0 | 0 | 0 | 2 | 2 | 0 | 1 | 98 | 49 | 1 | 1 | 1 | 1 | 1 | | 1 | 0.5 | 0.25 |
| SE | 10260 | AP3D1 | AP3D1 | chr19 | - | 2114747 | 2114821 | 2114124 | 2114302 | 2115218 | 2115418 | 15 | 6 | 1 | 1 | 0 | 0 | 0 | 2 | 0 | 0 | 98 | 49 | 1 | 1 | 0.882 | 1 | 1 | NA | | 1 | -0.039 |
| SE | 10261 | AP3D1 | AP3D1 | chr19 | - | 2127151 | 2127201 | 2123829 | 2123879 | 2129089 | 2129163 | 13 | 5 | 0 | 1 | 0 | 0 | 0 | 7 | 0 | 0 | 98 | 49 | 1 | 1 | 0.867 | 1 | NA | NA | | 1 | -0.067 |
| SE | 10267 | NAP1L4 | NAP1L4 | chr11 | - | 2949226 | 2949264 | 2944429 | 2945646 | 2951258 | 2951315 | 3 | 4 | 0 | 2 | 0 | 0 | 0 | 0 | 0 | 1 | 87 | 49 | 1 | 1 | 0.458 | 1 | NA | NA | | 0 | 0.729 |
| SE | 10274 | TRAPPC6B | TRAPPC6B | chr14 | - | 39154210 | 39154294 | 39151745 | 39151839 | 39158284 | 39158402 | 18 | 6 | 1 | 1 | 0 | 0 | 0 | 2 | 0 | 0 | 98 | 49 | 1 | 1 | 0.9 | 1 | 1 | NA | | 1 | -0.033 |
| SE | 10280 | EIF3M | EIF3M | chr11 | + | 32587011 | 32587144 | 32583766 | 32583929 | 32588593 | 32588732 | 13 | 1 | 0 | 0 | 0 | 0 | 0 | 0 | 0 | 1 | 98 | 49 | 1 | 1 | 1 | 1 | NA | NA | | 0 | 1 |
| SE | 10285 | PPP2R2D | PPP2R2D | chr10 | + | 131911561 | 131912598 | 131901237 | 131901330 | 131934457 | 131934555 | 1 | 0 | 0 | 6 | 6 | 0 | 0 | 0 | 0 | 3 | 98 | 49 | 1 | 1 | 0.077 | 0 | NA | NA | | 0 | 0.038 |
| SE | 10286 | PPP2R2D | PPP2R2D | chr10 | + | 131934457 | 131934555 | 131901237 | 131901330 | 131940030 | 131940196 | 15 | 9 | 0 | 1 | 2 | 0 | 0 | 6 | 0 | 0 | 98 | 49 | 1 | 1 | 0.882 | 0.692 | NA | NA | | 1 | -0.213 |
| SE | 10287 | PPP2R2D | PPP2R2D | chr10 | + | 131934893 | 131935006 | 131934457 | 131934555 | 131940030 | 131940196 | 4 | 7 | 1 | 9 | 3 | 0 | 0 | 1 | 0 | 3 | 98 | 49 | 1 | 1 | 0.182 | 0.538 | 1 | NA | | 0.143 | 0.43 |
| SE | 10292 | DOK1 | DOK1 | chr2 | + | 74555574 | 74555668 | 74555153 | 74555453 | 74556307 | 74557551 | 7 | 0 | 5 | 0 | 0 | 0 | 1 | 4 | 0 | 1 | 98 | 49 | 1 | 1 | 1 | NA | 1 | | 1 | 0.667 | 0.166 |
| SE | 10293 | DOK1 | DOK1 | chr2 | + | 74555893 | 74556078 | 74555574 | 74555668 | 74556307 | 74557551 | 7 | 0 | 3 | 0 | 0 | 4 | 1 | 1 | 0 | 0 | 98 | 49 | 1 | 1 | 1 | NA | 0.273 | | 1 | 1 | -0.363 |
| SE | 10299 | NLRP6 | NLRP6 | chr11 | + | 279833 | 279872 | 279326 | 279607 | 282704 | 282797 | 5 | 7 | 4 | 0 | 1 | 2 | 1 | 0 | 0 | 0 | 88 | 49 | 1 | 1 | 1 | 0.796 | 0.527 | | 1 | NA | -0.226 |
| SE | 10300 | NLRP6 | NLRP6 | chr11 | + | 280083 | 281839 | 279833 | 279872 | 282701 | 282797 | 4 | 3 | 5 | 1 | 1 | 0 | 0 | 1 | 0 | 0 | 98 | 49 | 1 | 1 | 0.667 | 0.6 | 1 | NA | | 1 | -0.244 |
| SE | 10305 | MCTP2 | MCTP2 | chr15 | + | 94345128 | 94345164 | 94340812 | 94340924 | 94356136 | 94356301 | 41 | 15 | 10 | 1 | 2 | 0 | 0 | 2 | 0 | 0 | 85 | 49 | 1 | 1 | 0.959 | 0.812 | 1 | NA | | 1 | -0.076 |
| SE | 10306 | MCTP2 | MCTP2 | chr15 | + | 94398960 | 94399062 | 94385422 | 94385525 | 94399920 | 94399995 | 44 | 27 | 3 | 1 | 0 | 0 | 2 | 16 | 0 | 0 | 98 | 49 | 1 | 1 | 0.957 | 1 | 1 | | 1 | 1 | -0.014 |
| SE | 10307 | MCTP2 | MCTP2 | chr15 | + | 94440175 | 94440298 | 94401899 | 94402019 | 94442918 | 94442960 | 39 | 18 | 15 | 2 | 2 | 0 | 0 | 8 | 0 | 0 | 98 | 49 | 1 | 1 | 0.907 | 0.818 | 1 | NA | | 1 | -0.092 |
| SE | 10308 | MCTP2 | MCTP2 | chr15 | + | 94442918 | 94442960 | 94401899 | 94402019 | 94458136 | 94458246 | 27 | 16 | 5 | 2 | 0 | 0 | 0 | 4 | 0 | 1 | 91 | 49 | 1 | 1 | 0.879 | 1 | 1 | NA | | 0.683 | 0.277 |
| SE | 10311 | JAK1 | JAK1 | chr1 | - | 64839602 | 64839795 | 64838464 | 64838589 | 64841244 | 64841339 | 86 | 36 | 33 | 0 | 0 | 0 | 2 | 17 | 0 | 1 | 98 | 49 | 1 | 1 | 1 | 1 | 1 | | 1 | 0.895 | 0.052 |
| SE | 10312 | JAK1 | JAK1 | chr1 | - | 64860104 | 64860262 | 64857655 | 64857779 | 64864786 | 64864972 | 100 | 30 | 21 | 1 | 0 | 0 | 3 | 12 | 0 | 0 | 98 | 49 | 1 | 1 | 0.98 | 1 | 1 | | 1 | 1 | -0.007 |
| SE | 10315 | FRS2 | FRS2 | chr12 | + | 69480298 | 69480359 | 69470348 | 69470530 | 69530864 | 69530960 | 2 | 1 | 0 | 5 | 0 | 1 | 0 | 0 | 0 | 1 | 98 | 49 | 1 | 1 | 0.167 | 1 | 0 | NA | | 0 | 0.389 |
| SE | 10316 | FRS2 | FRS2 | chr12 | + | 69486201 | 69486297 | 69470348 | 69470530 | 69530864 | 69530960 | 1 | 0 | 2 | 5 | 0 | 1 | 0 | 0 | 0 | 1 | 98 | 49 | 1 | 1 | 0.091 | NA | 0.5 | NA | | 0 | 0.295 |
| SE | 10322 | IRF7 | IRF7 | chr11 | - | 613950 | 614037 | 613784 | 613865 | 614173 | 614399 | 6 | 4 | 0 | 0 | 1 | 0 | 2 | 6 | 0 | 0 | 98 | 49 | 1 | 1 | 1 | 0.667 | NA | | 1 | 1 | -0.166 |
| SE | 10323 | IRF7 | IRF7 | chr11 | - | 613950 | 614037 | 613784 | 613865 | 614475 | 614534 | 13 | 11 | 0 | 2 | 2 | 0 | 8 | 14 | 8 | 3 | 98 | 49 | 1 | 1 | 0.765 | 0.733 | NA | | 0.333 | 0.7 | 0.233 |
| SE | 10329 | RASSF5 | RASSF5 | chr1 | + | 206585179 | 206585295 | 206584386 | 206584684 | 206586825 | 206589284 | 52 | 36 | 23 | 1 | 4 | 3 | 0 | 10 | 0 | 2 | 98 | 49 | 1 | 1 | 0.963 | 0.818 | 0.793 | NA | | 0.714 | 0.144 |
| SE | 10330 | ANKRD13D | ANKRD13D | chr11 | + | 67289828 | 67289965 | 67289344 | 67289550 | 67290077 | 67290213 | 6 | 4 | 2 | 7 | 3 | 7 | 0 | 3 | 0 | 4 | 98 | 49 | 1 | 1 | 0.3 | 0.4 | 0.125 | NA | | 0.273 | 0.002 |
| SE | 10331 | ANKRD13D | ANKRD13D | chr11 | + | 67299057 | 67299124 | 67291990 | 67292180 | 67299529 | 67299611 | 21 | 14 | 19 | 0 | 0 | 1 | 0 | 3 | 0 | 1 | 98 | 49 | 1 | 1 | 1 | 1 | 0.905 | NA | | 0.6 | 0.368 |
| SE | 10332 | ANKRD13D | ANKRD13D | chr11 | + | 67301731 | 67301823 | 67301487 | 67301651 | 67302118 | 67302484 | 32 | 23 | 10 | 2 | 0 | 0 | 1 | 8 | 0 | 0 | 98 | 49 | 1 | 1 | 0.889 | 1 | 1 | | 1 | 1 | -0.037 |
| SE | 10337 | FIP1L1 | FIP1L1 | chr4 | + | 53379072 | 53379117 | 53377652 | 53377923 | 53379224 | 53379264 | 9 | 8 | 0 | 5 | 2 | 3 | 0 | 0 | 0 | 3 | 94 | 49 | 1 | 1 | 0.484 | 0.676 | 0 | NA | | 0 | 0.387 |
| SE | 10338 | FIP1L1 | FIP1L1 | chr4 | + | 53391429 | 53391498 | 53391008 | 53391139 | 53399729 | 53399839 | 4 | 5 | 3 | 2 | 0 | 0 | 0 | 1 | 0 | 0 | 98 | 49 | 1 | 1 | 0.5 | 1 | 1 | NA | | 1 | -0.167 |
| SE | 10350 | DYNC1LI2 | DYNC1LI2 | chr16 | - | 66749196 | 66749313 | 66742437 | 66742668 | 66751272 | 66751346 | 25 | 12 | 0 | 0 | 0 | 0 | 1 | 6 | 0 | 1 | 98 | 49 | 1 | 1 | 1 | 1 | NA | | 1 | 0.75 | 0.125 |
| SE | 10351 | TUBA4A | TUBA4A | chr2 | - | 219251564 | 219251713 | 219249710 | 219251323 | 219252007 | 219252230 | 48 | 21 | 26 | 0 | 0 | 1 | 6 | 4 | 0 | 0 | 98 | 49 | 1 | 1 | 1 | 1 | 0.929 | | 1 | 1 | -0.024 |
| SE | 10357 | GALNS | GALNS | chr16 | - | 88831997 | 88832101 | 88826701 | 88826838 | 88835212 | 88835352 | 16 | 5 | 6 | 1 | 0 | 0 | 1 | 3 | 0 | 0 | 98 | 49 | 1 | 1 | 0.889 | 1 | 1 | | 1 | 1 | -0.037 |
| SE | 10358 | GALNS | GALNS | chr16 | - | 88837621 | 88837765 | 88836200 | 88836267 | 88840991 | 88841094 | 4 | 3 | 0 | 1 | 0 | 0 | 0 | 1 | 0 | 2 | 98 | 49 | 1 | 1 | 0.667 | 1 | NA | NA | | 0.2 | 0.633 |
| SE | 10359 | GALNS | GALNS | chr16 | - | 88842705 | 88842829 | 88841896 | 88841971 | 88856757 | 88856966 | 2 | 2 | 3 | 4 | 1 | 1 | 0 | 0 | 0 | 1 | 98 | 49 | 1 | 1 | 0.2 | 0.5 | 0.6 | NA | | 0 | 0.433 |
| SE | 10364 | IL13RA1 | IL13RA1 | chrX | + | 118746953 | 118747092 | 118741016 | 118741156 | 118749657 | 118749778 | 58 | 17 | 13 | 0 | 1 | 0 | 0 | 11 | 0 | 0 | 98 | 49 | 1 | 1 | 1 | 0.895 | 1 | NA | | 1 | -0.035 |
| SE | 10366 | AKNA | AKNA | chr9 | - | 114376465 | 114377532 | 114374092 | 114374167 | 114381059 | 114381446 | 128 | 42 | 26 | 0 | 1 | 0 | 3 | 25 | 0 | 0 | 98 | 49 | 1 | 1 | 1 | 0.955 | 1 | | 1 | 1 | -0.015 |
| SE | 10370 | NUP98 | NUP98 | chr11 | - | 3731390 | 3731578 | 3725102 | 3725219 | 3735190 | 3735324 | 51 | 20 | 5 | 1 | 0 | 0 | 0 | 11 | 0 | 0 | 98 | 49 | 1 | 1 | 0.962 | 1 | 1 | NA | | 1 | -0.013 |
| SE | 10371 | NUP98 | NUP98 | chr11 | - | 3744508 | 3744649 | 3735190 | 3735324 | 3753315 | 3753408 | 28 | 28 | 13 | 3 | 2 | 0 | 0 | 13 | 0 | 0 | 98 | 49 | 1 | 1 | 0.824 | 0.875 | 1 | NA | | 1 | -0.1 |
| SE | 10372 | NUP98 | NUP98 | chr11 | - | 3782041 | 3782145 | 3779155 | 3779257 | 3797399 | 3797792 | 55 | 29 | 7 | 1 | 0 | 0 | 2 | 7 | 0 | 0 | 98 | 49 | 1 | 1 | 0.965 | 1 | 1 | | 1 | 1 | -0.012 |
| SE | 10373 | STIM1 | STIM1 | chr11 | + | 3967551 | 3967682 | 3855702 | 3856409 | 4023872 | 4023987 | 35 | 15 | 13 | 0 | 2 | 0 | 7 | 10 | 0 | 0 | 98 | 49 | 1 | 1 | 1 | 0.789 | 1 | | 1 | 1 | -0.07 |
| SE | 10384 | PIK3R5 | PIK3R5 | chr17 | - | 8890737 | 8890912 | 8889972 | 8890126 | 8893585 | 8893655 | 47 | 18 | 19 | 0 | 1 | 0 | 4 | 14 | 0 | 0 | 98 | 49 | 1 | 1 | 1 | 0.9 | 1 | | 1 | 1 | -0.033 |
| SE | 10389 | ZSWIM8 | ZSWIM8 | chr10 | + | 73791323 | 73791499 | 73790974 | 73791176 | 73791858 | 73792852 | 9 | 5 | 6 | 1 | 0 | 0 | 0 | 6 | 1 | 0 | 98 | 49 | 1 | 1 | 0.818 | 1 | 1 | | 0 | 1 | 0.439 |
| SE | 10390 | IP6K1 | IP6K1 | chr3 | - | 49738211 | 49738422 | 49732790 | 49732972 | 49786353 | 49786540 | 9 | 2 | 1 | 0 | 2 | 0 | 0 | 3 | 0 | 0 | 98 | 49 | 1 | 1 | 1 | 0.333 | 1 | NA | | 1 | -0.222 |
| SE | 10393 | C18orf25 | C18orf25 | chr18 | + | 46240004 | 46240187 | 46215810 | 46216595 | 46253696 | 46253823 | 2 | 0 | 0 | 1 | 0 | 0 | 0 | 2 | 0 | 0 | 98 | 49 | 1 | 1 | 0.5 | NA | NA | NA | | 1 | -0.5 |
| SE | 10396 | GCH1 | GCH1 | chr14 | - | 54844000 | 54844143 | 54842005 | 54843054 | 54845767 | 54845852 | 1 | 0 | 0 | 0 | 0 | 0 | 0 | 1 | 0 | 2 | 98 | 49 | 1 | 1 | 1 | NA | NA | NA | | 0.2 | 0.8 |
| SE | 10419 | ANKRD10 | ANKRD10 | chr13 | - | 110894108 | 110894202 | 110892691 | 110893263 | 110906032 | 110906124 | 3 | 0 | 0 | 1 | 1 | 1 | 0 | 0 | 0 | 4 | 98 | 49 | 1 | 1 | 0.6 | 0 | 0 | NA | | 0 | 0.2 |

| | | | | | | | | | | | | | | | | | | | | | | | | | | | | | |
|---|---|---|---|---|---|---|---|---|---|---|---|---|---|---|---|---|---|---|---|---|---|---|---|---|---|---|---|---|---|
| SE | 10526 TMEM33 | TMEM33 | chr4 | + | 41939195 | 41939383 | 41938601 | 41938696 | 41943746 | 41943814 | 13 | 6 | 3 | 6 | 0 | 0 | 0 | 4 | 0 | 0 | 98 | 49 | 1 | 1 | 0.52 | 1 | 1 | NA | 1 | -0.16 |
| SE | 10529 SF1 | SF1 | chr11 | - | 64765950 | 64766155 | 64764603 | 64765518 | 64766899 | 64767079 | 1 | 1 | 0 | 4 | 0 | 0 | 0 | 1 | 0 | 0 | 98 | 49 | 1 | 1 | 0.111 | 1 | NA | NA | 1 | -0.445 |
| SE | 10530 SF1 | SF1 | chr11 | - | 64773429 | 64773505 | 64770255 | 64770408 | 64776497 | 64776626 | 22 | 9 | 1 | 2 | 0 | 0 | 1 | 6 | 0 | 0 | 98 | 49 | 1 | 1 | 0.846 | 1 | 1 | 1 | 1 | -0.051 |
| SE | 10532 PPP1CA | PPP1CA | chr11 | - | 67401067 | 67401199 | 67400688 | 67400919 | 67401727 | 67401905 | 53 | 18 | 11 | 1 | 1 | 0 | 1 | 10 | 0 | 0 | 98 | 49 | 1 | 1 | 0.964 | 0.9 | 1 | 1 | 1 | -0.045 |
| SE | 10533 PPP1CA | PPP1CA | chr11 | - | 67401067 | 67401232 | 67400688 | 67400919 | 67401727 | 67401905 | 35 | 9 | 7 | 1 | 1 | 0 | 0 | 6 | 0 | 0 | 98 | 49 | 1 | 1 | 0.946 | 0.818 | 1 | NA | 1 | -0.079 |
| SE | 10542 ATF7IP | ATF7IP | chr12 | + | 14446987 | 14447053 | 14438129 | 14438267 | 14456560 | 14456634 | 43 | 7 | 2 | 1 | 0 | 0 | 4 | 17 | 0 | 0 | 98 | 49 | 1 | 1 | 0.956 | 1 | 1 | 1 | 1 | -0.015 |
| SE | 10544 RAD9A | RAD9A | chr11 | + | 67395715 | 67395825 | 67393690 | 67393790 | 67395911 | 67396021 | 1 | 0 | 2 | 0 | 0 | 0 | 0 | 0 | 0 | 2 | 98 | 49 | 1 | 1 | 1 | NA | 1 | NA | 0 | 1 |
| SE | 10551 FLAD1 | FLAD1 | chr1 | + | 154990338 | 154990528 | 154990158 | 154990257 | 154992712 | 154992786 | 2 | 0 | 0 | 0 | 2 | 0 | 1 | 7 | 0 | 0 | 98 | 49 | 1 | 1 | 1 | 0 | NA | 1 | 1 | -0.5 |
| SE | 10552 TMEM222 | TMEM222 | chr1 | + | 27330984 | 27331137 | 27330719 | 27330804 | 27332069 | 27332101 | 1 | 0 | 0 | 3 | 0 | 5 | 0 | 0 | 0 | 1 | 98 | 49 | 1 | 1 | 0.143 | NA | 0 | NA | 0 | 0.071 |
| SE | 10557 PBRM1 | PBRM1 | chr3 | - | 52615350 | 52615456 | 52609312 | 52609955 | 52617261 | 52617538 | 6 | 0 | 0 | 1 | 1 | 0 | 0 | 1 | 0 | 1 | 98 | 49 | 1 | 1 | 0.75 | NA | 1 | NA | 0.333 | 0.542 |
| SE | 10558 PBRM1 | PBRM1 | chr3 | - | 52643247 | 52643343 | 52641953 | 52642045 | 52644703 | 52644789 | 11 | 1 | 0 | 1 | 0 | 0 | 0 | 4 | 0 | 0 | 98 | 49 | 1 | 1 | 0.846 | 1 | NA | NA | 1 | -0.077 |
| SE | 10574 NFKBIA | NFKBIA | chr14 | - | 35403149 | 35403360 | 35402770 | 35402859 | 35403689 | 35403798 | 122 | 334 | 51 | 0 | 2 | 0 | 2 | 41 | 0 | 0 | 98 | 49 | 1 | 1 | 1 | 0.988 | 1 | 1 | 1 | -0.004 |
| SE | 10583 GZMH | GZMH | chr14 | - | 24607148 | 24607406 | 24606479 | 24606746 | 24608264 | 24608412 | 7 | 8 | 4 | 0 | 2 | 0 | 0 | 3 | 0 | 0 | 98 | 49 | 1 | 1 | 1 | 0.667 | 1 | NA | 1 | -0.111 |
| SE | 10585 GZMH | GZMH | chr14 | - | 24607340 | 24607406 | 24606479 | 24606746 | 24608264 | 24608412 | 0 | 0 | 0 | 0 | 2 | 0 | 0 | 3 | 0 | 0 | 98 | 49 | 1 | 1 | NA | 0 | NA | NA | 1 | -1 |
| SE | 10586 GZMH | GZMH | chr14 | - | 24607611 | 24607747 | 24606479 | 24606746 | 24608264 | 24608412 | 8 | 9 | 0 | 0 | 2 | 0 | 1 | 0 | 0 | 0 | 98 | 49 | 1 | 1 | 1 | 0.692 | NA | 1 | NA | -0.154 |
| SE | 10587 GZMH | GZMH | chr14 | - | 24607611 | 24607747 | 24607148 | 24607406 | 24608264 | 24608412 | 13 | 5 | 0 | 0 | 0 | 0 | 1 | 0 | 0 | 3 | 98 | 49 | 1 | 1 | 1 | 1 | NA | 1 | 0 | 0.5 |
| SE | 10604 HDAC9 | HDAC9 | chr7 | + | 18629349 | 18629481 | 18593907 | 18594029 | 18634626 | 18634742 | 2 | 0 | 2 | 2 | 0 | 0 | 0 | 1 | 0 | 1 | 98 | 49 | 1 | 1 | 0.333 | NA | 1 | NA | 0.333 | 0.333 |
| SE | 10606 MICAL2 | MICAL2 | chr11 | + | 12138389 | 12138460 | 12110575 | 12110726 | 12162078 | 12162419 | 14 | 10 | 1 | 2 | 0 | 0 | 0 | 2 | 0 | 1 | 98 | 49 | 1 | 1 | 0.778 | 1 | 1 | NA | 0.5 | 0.426 |
| SE | 10607 MICAL2 | MICAL2 | chr11 | + | 12138389 | 12138464 | 12110575 | 12110726 | 12162078 | 12162419 | 13 | 6 | 1 | 2 | 0 | 0 | 0 | 3 | 0 | 1 | 98 | 49 | 1 | 1 | 0.765 | 1 | 1 | NA | 0.6 | 0.322 |
| SE | 10609 MICAL2 | MICAL2 | chr11 | + | 12220200 | 12220458 | 12216218 | 12216319 | 12221643 | 12221759 | 9 | 6 | 0 | 0 | 0 | 0 | 0 | 3 | 0 | 1 | 98 | 49 | 1 | 1 | 1 | 1 | NA | NA | 0.6 | 0.4 |
| SE | 10610 MICAL2 | MICAL2 | chr11 | + | 12249183 | 12249246 | 12239435 | 12239585 | 12256784 | 12256971 | 10 | 6 | 2 | 7 | 13 | 0 | 0 | 1 | 0 | 0 | 98 | 49 | 1 | 1 | 0.417 | 0.187 | 1 | NA | 1 | -0.465 |
| SE | 10613 ZFAND3 | ZFAND3 | chr6 | + | 37929958 | 37929999 | 37819530 | 37820016 | 38061592 | 38061775 | 4 | 4 | 7 | 0 | 0 | 0 | 0 | 1 | 0 | 1 | 90 | 49 | 1 | 1 | 1 | 1 | 1 | NA | 0.353 | 0.647 |
| SE | 10614 CSNK1D | CSNK1D | chr17 | - | 82245975 | 82246039 | 82242660 | 82244831 | 82248874 | 82249014 | 2 | 0 | 0 | 1 | 2 | 0 | 0 | 2 | 0 | 1 | 98 | 49 | 1 | 1 | 0.5 | 0 | NA | NA | 0.5 | -0.25 |
| SE | 10615 CSNK1D | CSNK1D | chr17 | - | 82253015 | 82253244 | 82252433 | 82252604 | 82255428 | 82255577 | 24 | 9 | 7 | 0 | 0 | 0 | 2 | 3 | 1 | 0 | 98 | 49 | 1 | 1 | 1 | 1 | 1 | 0.5 | 1 | 0.25 |
| SE | 10616 CSNK1D | CSNK1D | chr17 | - | 82255428 | 82255577 | 82253015 | 82253244 | 82265685 | 82265796 | 20 | 10 | 7 | 0 | 0 | 0 | 0 | 3 | 0 | 2 | 98 | 49 | 1 | 1 | 1 | 1 | 1 | NA | 0.429 | 0.571 |
| SE | 10617 ATP5F1 | ATP5F1 | chr1 | + | 111454210 | 111454356 | 111449836 | 111449873 | 111456085 | 111456249 | 33 | 2 | 2 | 0 | 2 | 0 | 0 | 8 | 0 | 1 | 98 | 49 | 1 | 1 | 1 | 0.333 | 1 | NA | 0.8 | -0.022 |
| SE | 10625 ADAR | ADAR | chr1 | - | 154586180 | 154586363 | 154585752 | 154585865 | 154588124 | 154588258 | 119 | 57 | 37 | 8 | 3 | 1 | 0 | 38 | 0 | 2 | 98 | 49 | 1 | 1 | 0.881 | 0.905 | 0.949 | NA | 0.905 | 0.007 |
| SE | 10626 ADAR | ADAR | chr1 | - | 154589368 | 154589462 | 154588550 | 154588673 | 154589756 | 154589928 | 104 | 53 | 29 | 2 | 0 | 0 | 4 | 39 | 0 | 0 | 98 | 49 | 1 | 1 | 0.963 | 1 | 1 | 1 | 1 | -0.012 |
| SE | 10632 KATNB1 | KATNB1 | chr16 | + | 57750826 | 57750927 | 57744393 | 57744511 | 57751260 | 57751302 | 4 | 1 | 1 | 0 | 1 | 0 | 0 | 2 | 0 | 1 | 98 | 49 | 1 | 1 | 1 | 0.333 | 1 | NA | 0.5 | 0.278 |
| SE | 10638 HAGH | HAGH | chr16 | - | 1826506 | 1826602 | 1822864 | 1823037 | 1826711 | 1827194 | 1 | 0 | 0 | 1 | 1 | 0 | 0 | 0 | 1 | 0 | 98 | 49 | 1 | 1 | 0.333 | 0 | NA | 0 | NA | 0.167 |
| SE | 10639 EED | EED | chr11 | + | 86266082 | 86266216 | 86264171 | 86264263 | 86276979 | 86277138 | 4 | 1 | 0 | 0 | 1 | 0 | 0 | 2 | 0 | 0 | 98 | 49 | 1 | 1 | 1 | 0.333 | NA | NA | 1 | -0.334 |
| SE | 10648 TBC1D14 | TBC1D14 | chr4 | + | 6923372 | 6924111 | 6909767 | 6909951 | 6967303 | 6967424 | 52 | 35 | 23 | 1 | 0 | 0 | 2 | 5 | 0 | 0 | 98 | 49 | 1 | 1 | 0.963 | 1 | 1 | 1 | 1 | -0.012 |
| SE | 10649 TBC1D14 | TBC1D14 | chr4 | + | 6996324 | 6996407 | 6994183 | 6994302 | 6999084 | 6999202 | 83 | 50 | 52 | 1 | 0 | 0 | 0 | 7 | 0 | 0 | 98 | 49 | 1 | 1 | 0.976 | 1 | 1 | NA | 1 | -0.008 |
| SE | 10650 TBC1D14 | TBC1D14 | chr4 | + | 7004843 | 7004924 | 7001144 | 7001251 | 7006631 | 7006726 | 55 | 35 | 8 | 0 | 2 | 0 | 0 | 11 | 0 | 0 | 98 | 49 | 1 | 1 | 1 | 0.897 | 1 | NA | 1 | -0.034 |
| SE | 10651 TBC1D14 | TBC1D14 | chr4 | + | 7004843 | 7004924 | 7001144 | 7001251 | 7010652 | 7010781 | 33 | 21 | 4 | 0 | 0 | 0 | 0 | 3 | 0 | 3 | 98 | 49 | 1 | 1 | 1 | 1 | 1 | NA | 0.333 | 0.667 |
| SE | 10652 TBC1D14 | TBC1D14 | chr4 | + | 7006631 | 7006726 | 7001144 | 7001251 | 7010652 | 7010781 | 7 | 3 | 2 | 0 | 0 | 0 | 0 | 3 | 0 | 3 | 98 | 49 | 1 | 1 | 1 | 1 | 1 | NA | 0 | 1 |
| SE | 10653 TBC1D14 | TBC1D14 | chr4 | + | 7006631 | 7006726 | 7004843 | 7004924 | 7010652 | 7010781 | 29 | 15 | 6 | 0 | 0 | 0 | 0 | 9 | 0 | 1 | 98 | 49 | 1 | 1 | 1 | 1 | 1 | NA | 0.818 | 0.182 |
| SE | 10654 TBC1D14 | TBC1D14 | chr4 | + | 7009876 | 7009948 | 7006631 | 7006726 | 7010652 | 7010781 | 27 | 13 | 12 | 7 | 1 | 2 | 0 | 6 | 0 | 0 | 98 | 49 | 1 | 1 | 0.659 | 0.867 | 0.75 | NA | 1 | -0.241 |
| SE | 10656 MAPK3 | MAPK3 | chr16 | - | 30118046 | 30118163 | 30117669 | 30117784 | 30118348 | 30118538 | 10 | 2 | 9 | 0 | 0 | 0 | 0 | 0 | 0 | 2 | 98 | 49 | 1 | 1 | 1 | 1 | 1 | NA | 0 | 1 |
| SE | 10666 LILRB2 | LILRB2 | chr19 | - | 54276380 | 54276456 | 54276263 | 54276301 | 54276806 | 54276929 | 8 | 7 | 3 | 3 | 0 | 0 | 0 | 5 | 0 | 1 | 98 | 49 | 1 | 1 | 0.571 | 1 | 1 | NA | 0.714 | 0.143 |
| SE | 10667 LILRB2 | LILRB2 | chr19 | - | 54280263 | 54280299 | 54279790 | 54280075 | 54280462 | 54280544 | 41 | 17 | 10 | 1 | 0 | 0 | 0 | 8 | 0 | 0 | 85 | 49 | 1 | 1 | 0.959 | 1 | 1 | NA | 1 | -0.014 |
| SE | 10668 LILRB2 | LILRB2 | chr19 | - | 54280462 | 54280544 | 54279790 | 54280075 | 54280960 | 54281178 | 1 | 0 | 0 | 0 | 1 | 0 | 0 | 0 | 0 | 1 | 98 | 49 | 1 | 1 | 1 | 0 | NA | NA | 0 | 0.5 |
| SE | 10671 IFFO1 | IFFO1 | chr12 | - | 6549475 | 6549484 | 6548667 | 6548825 | 6549755 | 6549896 | 5 | 0 | 1 | 2 | 0 | 0 | 0 | 1 | 0 | 0 | 58 | 49 | 1 | 1 | 0.679 | NA | 1 | NA | 1 | -0.16 |
| SE | 10686 DICER1 | DICER1 | chr14 | - | 95129471 | 95129632 | 95126579 | 95126748 | 95130057 | 95130192 | 57 | 22 | 7 | 1 | 0 | 0 | 3 | 18 | 0 | 0 | 98 | 49 | 1 | 1 | 0.966 | 1 | 1 | 1 | 1 | -0.011 |
| SE | 10687 DICER1 | DICER1 | chr14 | - | 95131508 | 95131639 | 95130057 | 95130192 | 95132514 | 95132677 | 20 | 13 | 9 | 5 | 2 | 0 | 0 | 8 | 0 | 0 | 98 | 49 | 1 | 1 | 0.667 | 0.765 | 1 | NA | 1 | -0.189 |
| SE | 10688 THAP5 | THAP5 | chr7 | - | 108565829 | 108566022 | 108562143 | 108565105 | 108569489 | 108569768 | 2 | 0 | 0 | 0 | 0 | 0 | 0 | 0 | 0 | 2 | 98 | 49 | 1 | 1 | 1 | NA | NA | NA | 0 | 1 |
| SE | 10689 THAP5 | THAP5 | chr7 | - | 108565951 | 108566022 | 108562143 | 108565105 | 108569489 | 108569768 | 2 | 0 | 0 | 0 | 0 | 0 | 0 | 0 | 0 | 2 | 98 | 49 | 1 | 1 | 1 | NA | NA | NA | 0 | 1 |
| SE | 10691 NPL | NPL | chr1 | + | 182803697 | 182803771 | 182794355 | 182794439 | 182812155 | 182812213 | 6 | 2 | 0 | 0 | 3 | 0 | 0 | 1 | 0 | 0 | 98 | 49 | 1 | 1 | 1 | 0.25 | NA | NA | 1 | -0.375 |
| SE | 10692 NPL | NPL | chr1 | + | 182806144 | 182806232 | 182803697 | 182803771 | 182812155 | 182812213 | 58 | 17 | 23 | 0 | 2 | 0 | 0 | 2 | 0 | 0 | 98 | 49 | 1 | 1 | 1 | 0.81 | 1 | NA | 1 | -0.063 |
| SE | 10693 NPL | NPL | chr1 | + | 182806144 | 182806531 | 182803697 | 182803771 | 182812155 | 182812213 | 24 | 10 | 17 | 0 | 2 | 0 | 0 | 8 | 0 | 0 | 98 | 49 | 1 | 1 | 1 | 0.714 | 1 | NA | 1 | -0.095 |
| SE | 10699 PHF12 | PHF12 | chr17 | - | 28907589 | 28907672 | 28906855 | 28906994 | 28910225 | 28910369 | 13 | 8 | 0 | 0 | 1 | 0 | 0 | 4 | 0 | 0 | 98 | 49 | 1 | 1 | 1 | 0.8 | NA | NA | 1 | -0.1 |
| SE | 10700 PHF12 | PHF12 | chr17 | - | 28908782 | 28908881 | 28907589 | 28907672 | 28910225 | 28910369 | 26 | 11 | 3 | 2 | 1 | 0 | 0 | 5 | 0 | 1 | 98 | 49 | 1 | 1 | 0.867 | 0.846 | 1 | NA | 0.714 | 0.19 |
| SE | 10701 PHF12 | PHF12 | chr17 | - | 28919142 | 28919275 | 28917284 | 28917449 | 28921687 | 28921808 | 39 | 13 | 7 | 4 | 2 | 0 | 0 | 4 | 0 | 0 | 98 | 49 | 1 | 1 | 0.83 | 0.765 | 1 | NA | 1 | -0.135 |
| SE | 10702 PHF12 | PHF12 | chr17 | - | 28950064 | 28950246 | 28926990 | 28927063 | 28950894 | 28951490 | 10 | 3 | 6 | 0 | 3 | 0 | 0 | 3 | 0 | 0 | 98 | 49 | 1 | 1 | 1 | 0.333 | 1 | NA | 1 | -0.222 |
| SE | 10705 KLRD1 | KLRD1 | chr12 | + | 10309387 | 10309480 | 10307817 | 10308084 | 10309625 | 10309688 | 9 | 3 | 1 | 0 | 3 | 0 | 0 | 2 | 0 | 0 | 98 | 49 | 1 | 1 | 1 | 0.333 | 1 | NA | 1 | -0.222 |
| SE | 10706 PRR5L | PRR5L | chr11 | + | 36446299 | 36446440 | 36437384 | 36437476 | 36451208 | 36451335 | 8 | 5 | 1 | 1 | 0 | 0 | 0 | 1 | 0 | 0 | 98 | 49 | 1 | 1 | 0.8 | 1 | 1 | NA | 1 | -0.067 |
| SE | 10710 RASGRP2 | RASGRP2 | chr11 | - | 64727295 | 64727360 | 64726910 | 64727131 | 64728862 | 64729042 | 36 | 5 | 10 | 0 | 0 | 0 | 0 | 7 | 1 | 0 | 98 | 49 | 1 | 1 | 1 | 1 | 1 | 0 | 1 | 0.5 |
| SE | 10714 SRSF6 | SRSF6 | chr20 | + | 43459152 | 43459420 | 43458360 | 43458509 | 43459770 | 43459895 | 2 | 1 | 0 | 3 | 2 | 0 | 0 | 0 | 0 | 1 | 98 | 49 | 1 | 1 | 0.25 | 0.2 | NA | NA | 0 | 0.225 |
| SE | 10721 GAS7 | GAS7 | chr17 | - | 9940125 | 9940200 | 9934165 | 9934244 | 9943120 | 9943236 | 24 | 6 | 2 | 0 | 0 | 0 | 0 | 15 | 0 | 1 | 98 | 49 | 1 | 1 | 1 | 1 | 1 | NA | 0.882 | 0.118 |
| SE | 10722 GAS7 | GAS7 | chr17 | - | 9969676 | 9969762 | 9959201 | 9959255 | 9981803 | 9981884 | 18 | 4 | 2 | 1 | 0 | 0 | 1 | 9 | 1 | 0 | 98 | 49 | 1 | 1 | 0.9 | 1 | 1 | 0.333 | 1 | 0.3 |
| SE | 10730 ANXA5 | ANXA5 | chr4 | - | 121678414 | 121678494 | 121677893 | 121677950 | 121681670 | 121681761 | 35 | 25 | 3 | 0 | 0 | 0 | 0 | 22 | 0 | 1 | 98 | 49 | 1 | 1 | 1 | 1 | 1 | NA | 0.917 | 0.083 |
| SE | 10731 ANXA5 | ANXA5 | chr4 | - | 121684676 | 121684771 | 121683363 | 121683477 | 121686287 | 121686372 | 23 | 12 | 9 | 0 | 1 | 0 | 3 | 20 | 0 | 1 | 98 | 49 | 1 | 1 | 1 | 0.857 | 1 | 1 | 0.909 | -0.002 |
| SE | 10732 NUCB1 | NUCB1 | chr19 | + | 48921826 | 48921932 | 48921153 | 48921324 | 48922317 | 48923283 | 26 | 5 | 8 | 0 | 1 | 0 | 1 | 8 | 0 | 0 | 98 | 49 | 1 | 1 | 1 | 0.714 | 1 | 1 | 1 | -0.095 |
| SE | 10734 MYO1F | MYO1F | chr19 | - | 8527337 | 8527483 | 8526788 | 8526935 | 8530195 | 8530365 | 341 | 289 | 82 | 0 | 1 | 0 | 16 | 86 | 0 | 0 | 98 | 49 | 1 | 1 | 1 | 0.993 | 1 | 1 | 1 | -0.002 |
| SE | 10736 MYO1F | MYO1F | chr19 | - | 8536251 | 8536396 | 8530458 | 8530573 | 8536498 | 8536597 | 241 | 169 | 83 | 0 | 2 | 0 | 6 | 64 | 0 | 0 | 98 | 49 | 1 | 1 | 1 | 0.977 | 1 | 1 | 1 | -0.008 |
| SE | 10739 MYO1F | MYO1F | chr19 | - | 8544296 | 8544464 | 8539946 | 8540028 | 8548035 | 8548122 | 3 | 12 | 0 | 0 | 2 | 0 | 0 | 3 | 0 | 0 | 98 | 49 | 1 | 1 | 0.429 | 1 | NA | NA | 1 | -0.285 |
| SE | 10740 MYO1F | MYO1F | chr19 | - | 8545649 | 8545736 | 8544296 | 8544464 | 8548035 | 8548122 | 339 | 179 | 105 | 1 | 3 | 0 | 11 | 71 | 0 | 0 | 98 | 49 | 1 | 1 | 0.994 | 0.968 | 1 | 1 | 1 | -0.013 |
| SE | 10742 MYO1F | MYO1F | chr19 | - | 8550159 | 8550356 | 8548236 | 8548317 | 8550561 | 8550694 | 248 | 136 | 68 | 1 | 0 | 0 | 11 | 62 | 0 | 0 | 98 | 49 | 1 | 1 | 0.992 | 1 | 1 | 1 | 1 | -0.003 |
| SE | 10743 MYO1F | MYO1F | chr19 | - | 8552032 | 8552164 | 8551739 | 8551874 | 8553138 | 8553228 | 238 | 75 | 63 | 1 | 0 | 0 | 1 | 36 | 0 | 0 | 98 | 49 | 1 | 1 | 0.992 | 1 | 1 | 1 | 1 | -0.003 |
| SE | 10746 ATG13 | ATG13 | chr11 | + | 46618324 | 46618375 | 46617275 | 46617890 | 46630044 | 46630100 | 2 | 2 | 2 | 1 | 1 | 0 | 0 | 1 | 0 | 1 | 98 | 49 | 1 | 1 | 0.5 | 0.5 | 1 | NA | 0.333 | 0.334 |
| SE | 10747 ATG13 | ATG13 | chr11 | + | 46618324 | 46618375 | 46617275 | 46617890 | 46644278 | 46644360 | 0 | 0 | 0 | 1 | 0 | 0 | 0 | 2 | 0 | 0 | 98 | 49 | 1 | 1 | 0 | NA | NA | NA | 1 | -1 |
| SE | 10748 ATG13 | ATG13 | chr11 | + | 46630044 | 46630100 | 46617275 | 46617890 | 46644278 | 46644360 | 6 | 2 | 1 | 1 | 0 | 0 | 0 | 2 | 0 | 0 | 98 | 49 | 1 | 1 | 0.75 | 1 | 1 | NA | 1 | -0.083 |
| SE | 10749 ATG13 | ATG13 | chr11 | + | 46630044 | 46630100 | 46618324 | 46618375 | 46644278 | 46644360 | 7 | 3 | 3 | 0 | 0 | 0 | 0 | 1 | 0 | 1 | 98 | 49 | 1 | 1 | 1 | 1 | 1 | NA | 0.333 | 0.667 |
| SE | 10753 DOCK4 | DOCK4 | chr7 | - | 111735053 | 111735167 | 111732225 | 111732287 | 111736916 | 111736989 | 10 | 2 | 0 | 3 | 0 | 0 | 3 | 1 | 0 | 0 | 98 | 49 | 1 | 1 | 0.625 | 1 | NA | 1 | 1 | -0.188 |
| SE | 10754 DOCK4 | DOCK4 | chr7 | - | 111877029 | 111877186 | 111872466 | 111872564 | 111895611 | 111895718 | 20 | 5 | 1 | 1 | 0 | 0 | 0 | 10 | 0 | 0 | 98 | 49 | 1 | 1 | 0.909 | 1 | 1 | NA | 1 | -0.03 |
| SE | 10755 DOCK4 | DOCK4 | chr7 | - | 111945716 | 111945798 | 111944810 | 111944871 | 111977131 | 111977283 | 35 | 1 | 0 | 2 | 0 | 0 | 4 | 8 | 0 | 0 | 98 | 49 | 1 | 1 | 0.897 | 1 | NA | 1 | 1 | -0.051 |
| SE | 10759 TNFSF13 | TNFSF13 | chr17 | + | 7559623 | 7559702 | 7558291 | 7559297 | 7559845 | 7559893 | 13 | 3 | 0 | 1 | 0 | 0 | 0 | 6 | 0 | 0 | 98 | 49 | 1 | 1 | 0.867 | 1 | NA | NA | 1 | -0.067 |
| SE | 10769 NFXL1 | NFXL1 | chr4 | - | 47885497 | 47885657 | 47884345 | 47884437 | 47885878 | 47885999 | 30 | 4 | 2 | 1 | 0 | 0 | 0 | 2 | 0 | 0 | 98 | 49 | 1 | 1 | 0.937 | 1 | 1 | NA | 1 | -0.021 |
| SE | 10773 PTK2B | PTK2B | chr8 | + | 27436250 | 27436348 | 27435742 | 27435793 | 27437121 | 27437206 | 38 | 21 | 12 | 2 | 0 | 0 | 1 | 13 | 0 | 0 | 98 | 49 | 1 | 1 | 0.905 | 1 | 1 | 1 | 1 | -0.032 |
| SE | 10774 PTK2B | PTK2B | chr8 | + | 27437121 | 27437206 | 27435742 | 27435793 | 27437395 | 27437496 | 50 | 15 | 10 | 0 | 3 | 0 | 0 | 9 | 0 | 0 | 98 | 49 | 1 | 1 | 1 | 0.714 | 1 | NA | 1 | -0.095 |
| SE | 10776 PTK2B | PTK2B | chr8 | + | 27454530 | 27454611 | 27454153 | 27454291 | 27458293 | 27459391 | 87 | 62 | 19 | 1 | 0 | 0 | 7 | 21 | 0 | 0 | 98 | 49 | 1 | 1 | 0.978 | 1 | 1 | 1 | 1 | -0.007 |
| SE | 10779 TTC7A | TTC7A | chr2 | + | 47021861 | 47021979 | 47011330 | 47011435 | 47023407 | 47023465 | 9 | 0 | 0 | 2 | 0 | 0 | 0 | 8 | 0 | 0 | 98 | 49 | 1 | 1 | 0.692 | NA | NA | NA | 1 | -0.308 |
| SE | 10780 TTC7A | TTC7A | chr2 | + | 47047265 | 47047337 | 47046314 | 47046431 | 47049948 | 47050046 | 0 | 0 | 0 | 0 | 2 | 0 | 0 | 1 | 0 | 2 | 98 | 49 | 1 | 1 | NA | 0 | NA | NA | 0.2 | -0.2 |
| SE | 10781 FNBP4 | FNBP4 | chr11 | - | 47743952 | 47744163 | 47736615 | 47736740 | 47746055 | 47746394 | 17 | 4 | 0 | 1 | 0 | 0 | 0 | 1 | 0 | 0 | 98 | 49 | 1 | 1 | 0.895 | 1 | NA | NA | 1 | -0.052 |
| SE | 10784 BIN1 | BIN1 | chr2 | - | 127051153 | 127051243 | 127050801 | 127050912 | 127059010 | 127059155 | 1 | 0 | 0 | 0 | 3 | 0 | 0 | 2 | 0 | 0 | 98 | 49 | 1 | 1 | 1 | 0 | NA | NA | 1 | -0.5 |
| SE | 10792 PHF21A | PHF21A | chr11 | - | 45938156 | 45938312 | 45936493 | 45936569 | 45945839 | 45946003 | 27 | 2 | 1 | 0 | 1 | 0 | 0 | 6 | 0 | 3 | 98 | 49 | 1 | 1 | 1 | 0.5 | 1 | NA | 0.5 | 0.333 |
| SE | 10793 PHF21A | PHF21A | chr11 | - | 45945839 | 45946003 | 45938156 | 45938312 | 45948885 | 45948946 | 16 | 2 | 1 | 2 | 0 | 0 | 0 | 4 | 0 | 0 | 98 | 49 | 1 | 1 | 0.8 | 1 | 1 | NA | 1 | -0.067 |
| SE | 10794 PHF21A | PHF21A | chr11 | - | 45946075 | 45946098 | 45938156 | 45938312 | 45948885 | 45948946 | 7 | 6 | 0 | 2 | 0 | 0 | 1 | 0 | 0 | 0 | 72 | 49 | 1 | 1 | 0.704 | 1 | NA | 1 | NA | -0.148 |
| SE | 10796 PHF21A | PHF21A | chr11 | - | 45971115 | 45971367 | 45969814 | 45969904 | 45979759 | 45979966 | 18 | 4 | 7 | 1 | 0 | 0 | 0 | 2 | 0 | 0 | 98 | 49 | 1 | 1 | 0.9 | 1 | 1 | NA | 1 | -0.033 |
| SE | 10797 MAN2A2 | MAN2A2 | chr15 | + | 90906739 | 90906913 | 90906369 | 90906497 | 90907308 | 90907495 | 28 | 10 | 11 | 0 | 1 | 0 | 1 | 19 | 0 | 0 | 98 | 49 | 1 | 1 | 1 | 0.833 | 1 | 1 | 1 | -0.056 |
| SE | 10798 MAN2A2 | MAN2A2 | chr15 | + | 90910089 | 90910292 | 90909326 | 90909504 | 90910500 | 90910683 | 16 | 10 | 11 | 2 | 0 | 3 | 0 | 8 | 0 | 1 | 98 | 49 | 1 | 1 | 0.8 | 1 | 0.647 | NA | 0.8 | 0.016 |
| SE | 10799 MAN2A2 | MAN2A2 | chr15 | + | 90912541 | 90912664 | 90912042 | 90912279 | 90912876 | 90912991 | 21 | 12 | 4 | 0 | 0 | 0 | 0 | 2 | 0 | 1 | 98 | 49 | 1 | 1 | 1 | 1 | 1 | NA | 0.5 | 0.5 |
| SE | 10803 ERAP1 | ERAP1 | chr5 | - | 96788530 | 96788685 | 96786469 | 96786549 | 96790295 | 96790367 | 19 | 2 | 3 | 1 | 0 | 0 | 0 | 5 | 0 | 0 | 98 | 49 | 1 | 1 | 0.905 | 1 | 1 | NA | 1 | -0.032 |
| SE | 10811 LPCAT3 | LPCAT3 | chr12 | - | 6977597 | 6977745 | 6977366 | 6977525 | 6978340 | 6978507 | 8 | 3 | 2 | 1 | 0 | 0 | 0 | 2 | 0 | 0 | 98 | 49 | 1 | 1 | 0.8 | 1 | 1 | NA | 1 | -0.067 |
| SE | 10813 JMJD4 | JMJD4 | chr1 | - | 227734650 | 227734816 | 227733906 | 227734032 | 227735011 | 227735411 | 1 | 0 | 0 | 1 | 0 | 0 | 0 | 1 | 0 | 0 | 98 | 49 | 1 | 1 | 0.333 | NA | NA | NA | 1 | -0.667 |
| SE | 10819 HIF1A | HIF1A | chr14 | + | 61721738 | 61721823 | 61721508 | 61721654 | 61726705 | 61726818 | 142 | 125 | 35 | 2 | 0 | 0 | 1 | 43 | 0 | 0 | 98 | 49 | 1 | 1 | 0.973 | 1 | 1 | 1 | 1 | -0.009 |
| SE | 10820 HIF1A | HIF1A | chr14 | + | 61727452 | 61727655 | 61726705 | 61726818 | 61732417 | 61732524 | 264 | 166 | 27 | 0 | 1 | 0 | 5 | 67 | 0 | 0 | 98 | 49 | 1 | 1 | 1 | 0.988 | 1 | 1 | 1 | -0.004 |
| SE | 10821 HIF1A | HIF1A | chr14 | + | 61738086 | 61738373 | 61736888 | 61737109 | 61740504 | 61740627 | 62 | 29 | 14 | 0 | 1 | 0 | 1 | 10 | 0 | 1 | 98 | 49 | 1 | 1 | 1 | 0.935 | 1 | 1 | 0.833 | 0.062 |
| SE | 10822 HIF1A | HIF1A | chr14 | + | 61740504 | 61740627 | 61738086 | 61738373 | 61740754 | 61741188 | 64 | 37 | 8 | 0 | 0 | 0 | 0 | 14 | 1 | 0 | 98 | 49 | 1 | 1 | 1 | 1 | 1 | 0 | 1 | 0.5 |
| SE | 10824 HIF1A | HIF1A | chr14 | + | 61745690 | 61745817 | 61744704 | 61744813 | 61746933 | 61748259 | 124 | 77 | 15 | 0 | 1 | 1 | 4 | 49 | 0 | 0 | 98 | 49 | 1 | 1 | 1 | 0.975 | 0.882 | 1 | 1 | -0.048 |
| SE | 10825 SLC43A3 | SLC43A3 | chr11 | - | 57409934 | 57410121 | 57409174 | 57409298 | 57414614 | 57414731 | 3 | 0 | 0 | 1 | 0 | 0 | 0 | 1 | 0 | 0 | 98 | 49 | 1 | 1 | 0.6 | NA | NA | NA | 1 | -0.4 |
| SE | 10826 SLC43A3 | SLC43A3 | chr11 | - | 57425540 | 57425670 | 57423981 | 57424028 | 57425988 | 57426352 | 5 | 0 | 0 | 0 | 0 | 0 | 0 | 2 | 0 | 1 | 98 | 49 | 1 | 1 | 1 | NA | NA | NA | 0.5 | 0.5 |
| SE | 10827 SLC43A3 | SLC43A3 | chr11 | - | 57425540 | 57425709 | 57423981 | 57424028 | 57425988 | 57426389 | 1 | 0 | 0 | 0 | 0 | 0 | 0 | 0 | 0 | 1 | 98 | 49 | 1 | 1 | 1 | NA | NA | NA | 0 | 1 |
| SE | 10834 ZDHHC5 | ZDHHC5 | chr11 | + | 57695919 | 57696043 | 57693782 | 57693915 | 57696760 | 57696873 | 17 | 5 | 8 | 0 | 0 | 0 | 0 | 0 | 0 | 1 | 98 | 49 | 1 | 1 | 1 | 1 | 1 | NA | 0 | 1 |
| SE | 10836 PEX16 | PEX16 | chr11 | - | 45915467 | 45915568 | 45914603 | 45914684 | 45915702 | 45915836 | 19 | 14 | 1 | 1 | 0 | 0 | 0 | 1 | 0 | 0 | 98 | 49 | 1 | 1 | 0.905 | 1 | 1 | NA | 1 | -0.032 |
| SE | 10841 ITGAX | ITGAX | chr16 | + | 31355892 | 31355998 | 31355133 | 31355291 | 31356624 | 31356728 | 246 | 143 | 17 | 0 | 1 | 0 | 17 | 42 | 0 | 0 | 98 | 49 | 1 | 1 | 1 | 0.986 | 1 | 1 | 1 | -0.005 |
| SE | 10842 ITGAX | ITGAX | chr16 | + | 31356624 | 31356728 | 31355892 | 31355998 | 31357030 | 31357101 | 346 | 206 | 30 | 0 | 1 | 0 | 26 | 76 | 0 | 0 | 98 | 49 | 1 | 1 | 1 | 0.99 | 1 | 1 | 1 | -0.003 |
| SE | 10843 ITGAX | ITGAX | chr16 | + | 31357252 | 31357364 | 31357030 | 31357101 | 31359699 | 31359830 | 117 | 72 | 28 | 1 | 1 | 0 | 10 | 34 | 0 | 0 | 98 | 49 | 1 | 1 | 0.983 | 0.973 | 1 | 1 | 1 | -0.015 |
| SE | 10844 ITGAX | ITGAX | chr16 | + | 31371083 | 31371214 | 31363164 | 31363374 | 31371629 | 31371784 | 51 | 35 | 10 | 1 | 0 | 0 | 4 | 8 | 0 | 0 | 98 | 49 | 1 | 1 | 0.962 | 1 | 1 | 1 | 1 | -0.013 |
| SE | 10850 SRP54 | SRP54 | chr14 | + | 35000935 | 35001020 | 34999557 | 34999649 | 35007282 | 35007387 | 13 | 3 | 5 | 1 | 0 | 0 | 0 | 3 | 0 | 0 | 98 | 49 | 1 | 1 | 0.867 | 1 | 1 | NA | 1 | -0.044 |
| SE | 10851 SRP54 | SRP54 | chr14 | + | 35013345 | 35013494 | 35011508 | 35011659 | 35013801 | 35013902 | 8 | 2 | 4 | 1 | 0 | 0 | 0 | 2 | 0 | 1 | 98 | 49 | 1 | 1 | 0.8 | 1 | 1 | NA | 0.5 | 0.433 |
| SE | 10853 STAT6 | STAT6 | chr12 | - | 57099293 | 57099440 | 57099014 | 57099078 | 57099995 | 57100090 | 37 | 24 | 7 | 1 | 0 | 0 | 0 | 8 | 0 | 0 | 98 | 49 | 1 | 1 | 0.949 | 1 | 1 | NA | 1 | -0.017 |
| SE | 10854 STAT6 | STAT6 | chr12 | - | 57099766 | 57099903 | 57099293 | 57099440 | 57099995 | 57100090 | 74 | 45 | 23 | 3 | 0 | 0 | 0 | 13 | 0 | 0 | 98 | 49 | 1 | 1 | 0.925 | 1 | 1 | NA | 1 | -0.025 |
| SE | 10855 STAT6 | STAT6 | chr12 | - | 57104725 | 57104813 | 57104463 | 57104586 | 57105150 | 57105339 | 123 | 56 | 34 | 2 | 0 | 0 | 3 | 18 | 0 | 0 | 98 | 49 | 1 | 1 | 0.969 | 1 | 1 | 1 | 1 | -0.01 |
| SE | 10867 FAM111A | FAM111A | chr11 | + | 59145826 | 59145856 | 59143082 | 59143304 | 59148835 | 59148953 | 6 | 2 | 0 | 1 | 0 | 0 | 1 | 2 | 0 | 0 | 79 | 49 | 1 | 1 | 0.788 | 1 | NA | 1 | 1 | -0.106 |
| SE | 10873 PRDX5 | PRDX5 | chr11 | + | 64320660 | 64320792 | 64319733 | 64319868 | 64320864 | 64320903 | 14 | 31 | 17 | 1 | 0 | 0 | 0 | 4 | 0 | 1 | 98 | 49 | 1 | 1 | 0.875 | 1 | 1 | NA | 0.667 | 0.291 |
| SE | 10874 COPB2 | COPB2 | chr3 | - | 139373665 | 139373808 | 139373212 | 139373412 | 139374488 | 139374588 | 15 | 3 | 4 | 10 | 0 | 0 | 0 | 1 | 0 | 0 | 98 | 49 | 1 | 1 | 0.429 | 1 | 1 | NA | 1 | -0.19 |
| SE | 10875 INPP5B | INPP5B | chr1 | - | 37940687 | 37940798 | 37931912 | 37932053 | 37943639 | 37943669 | 3 | 0 | 0 | 0 | 0 | 2 | 0 | 2 | 0 | 0 | 98 | 49 | 1 | 1 | 1 | NA | 0 | NA | 1 | -0.5 |
| SE | 10877 PSMB4 | PSMB4 | chr1 | + | 151401238 | 151401355 | 151400763 | 151400845 | 151401541 | 151401630 | 54 | 27 | 17 | 1 | 0 | 0 | 0 | 11 | 0 | 0 | 98 | 49 | 1 | 1 | 0.964 | 1 | 1 | NA | 1 | -0.012 |
| SE | 10890 ADGRE2 | ADGRE2 | chr19 | - | 14754953 | 14755127 | 14752328 | 14752526 | 14756237 | 14756345 | 20 | 5 | 0 | 0 | 0 | 0 | 0 | 3 | 1 | 1 | 98 | 49 | 1 | 1 | 1 | 1 | NA | 0 | 0.6 | 0.7 |
| SE | 10892 ADGRE2 | ADGRE2 | chr19 | - | 14755653 | 14755877 | 14754953 | 14755127 | 14756237 | 14756345 | 24 | 6 | 12 | 2 | 0 | 0 | 0 | 4 | 0 | 0 | 98 | 49 | 1 | 1 | 0.857 | 1 | 1 | NA | 1 | -0.048 |
| SE | 10893 ADGRE2 | ADGRE2 | chr19 | - | 14764432 | 14764610 | 14756237 | 14756345 | 14765319 | 14765397 | 20 | 8 | 0 | 1 | 0 | 0 | 0 | 4 | 0 | 0 | 98 | 49 | 1 | 1 | 0.909 | 1 | NA | NA | 1 | -0.045 |
| SE | 10894 ADGRE2 | ADGRE2 | chr19 | - | 14765657 | 14765804 | 14765523 | 14765570 | 14772341 | 14772497 | 8 | 1 | 0 | 0 | 0 | 0 | 0 | 0 | 0 | 1 | 98 | 49 | 1 | 1 | 1 | 1 | NA | NA | 0 | 1 |
| SE | 10895 ADGRE2 | ADGRE2 | chr19 | - | 14766234 | 14766381 | 14765657 | 14765804 | 14766977 | 14767109 | 16 | 1 | 0 | 2 | 0 | 0 | 0 | 1 | 0 | 0 | 98 | 49 | 1 | 1 | 0.8 | 1 | NA | NA | 1 | -0.1 |
| SE | 10896 ADGRE2 | ADGRE2 | chr19 | - | 14766977 | 14767109 | 14765657 | 14765804 | 14772341 | 14772497 | 17 | 0 | 1 | 8 | 1 | 0 | 0 | 1 | 0 | 0 | 98 | 49 | 1 | 1 | 0.515 | 0 | 1 | NA | 1 | -0.495 |
| SE | 10897 PTS | PTS | chr11 | + | 112230207 | 112230230 | 112228593 | 112228673 | 112230625 | 112230682 | 1 | 0 | 0 | 10 | 6 | 0 | 0 | 0 | 0 | 4 | 72 | 49 | 1 | 1 | 0.064 | 0 | NA | NA | 0 | 0.032 |
| SE | 10898 PTS | PTS | chr11 | + | 112230625 | 112230682 | 112228593 | 112228673 | 112233162 | 112233233 | 23 | 8 | 0 | 0 | 0 | 0 | 0 | 5 | 0 | 1 | 98 | 49 | 1 | 1 | 1 | 1 | NA | NA | 0.714 | 0.286 |
| SE | 10899 KPNB1 | KPNB1 | chr17 | + | 47663088 | 47663178 | 47661118 | 47661178 | 47664158 | 47664269 | 58 | 19 | 19 | 2 | 0 | 0 | 0 | 10 | 0 | 0 | 98 | 49 | 1 | 1 | 0.935 | 1 | 1 | NA | 1 | -0.022 |
| SE | 10900 ZNF805 | ZNF805 | chr19 | + | 57243922 | 57244049 | 57240684 | 57240921 | 57248604 | 57248700 | 0 | 2 | 3 | 0 | 0 | 0 | 0 | 0 | 1 | 0 | 98 | 49 | 1 | 1 | NA | 1 | 1 | 0 | NA | 1 |
| SE | 10901 PACS1 | PACS1 | chr11 | + | 66221153 | 66221247 | 66220630 | 66220791 | 66227503 | 66227584 | 42 | 22 | 9 | 0 | 3 | 0 | 0 | 9 | 0 | 0 | 98 | 49 | 1 | 1 | 1 | 0.786 | 1 | NA | 1 | -0.071 |
| SE | 10902 PACS1 | PACS1 | chr11 | + | 66242911 | 66243031 | 66241426 | 66241653 | 66243164 | 66244747 | 33 | 5 | 4 | 1 | 2 | 1 | 0 | 8 | 0 | 0 | 98 | 49 | 1 | 1 | 0.943 | 0.556 | 0.667 | NA | 1 | -0.278 |
| SE | 10909 CTBS | CTBS | chr1 | - | 84569930 | 84570139 | 84565840 | 84566012 | 84570581 | 84570720 | 20 | 11 | 11 | 9 | 0 | 11 | 0 | 4 | 0 | 0 | 98 | 49 | 1 | 1 | 0.526 | 1 | 0.333 | NA | 1 | -0.38 |
| SE | 10912 RBL2 | RBL2 | chr16 | + | 53469915 | 53470185 | 53467057 | 53467169 | 53470382 | 53470663 | 38 | 16 | 11 | 0 | 2 | 0 | 0 | 3 | 0 | 0 | 98 | 49 | 1 | 1 | 1 | 0.8 | 1 | NA | 1 | -0.067 |
| SE | 10913 PLBD1 | PLBD1 | chr12 | - | 14553192 | 14553412 | 14542207 | 14542291 | 14567581 | 14567857 | 67 | 19 | 15 | 3 | 0 | 0 | 2 | 33 | 0 | 1 | 98 | 49 | 1 | 1 | 0.918 | 1 | 1 | 1 | 0.943 | 0.001 |
| SE | 10916 SPOPL | SPOPL | chr2 | + | 138550482 | 138550604 | 138550156 | 138550294 | 138550902 | 138551054 | 34 | 7 | 3 | 1 | 0 | 0 | 0 | 2 | 0 | 0 | 98 | 49 | 1 | 1 | 0.944 | 1 | 1 | NA | 1 | -0.019 |
| SE | 10921 GANAB | GANAB | chr11 | - | 62632564 | 62632745 | 62631029 | 62631183 | 62633004 | 62633101 | 14 | 1 | 0 | 0 | 0 | 0 | 1 | 3 | 0 | 1 | 98 | 49 | 1 | 1 | 1 | 1 | NA | 1 | 0.6 | 0.2 |
| SE | 10922 GANAB | GANAB | chr11 | - | 62634309 | 62634375 | 62633444 | 62633514 | 62634820 | 62635000 | 4 | 0 | 0 | 4 | 1 | 0 | 0 | 0 | 0 | 1 | 98 | 49 | 1 | 1 | 0.333 | 0 | NA | NA | 0 | 0.167 |
| SE | 10925 ZNF292 | ZNF292 | chr6 | + | 87218595 | 87218731 | 87216298 | 87216377 | 87233324 | 87233527 | 25 | 9 | 1 | 0 | 2 | 0 | 0 | 1 | 0 | 0 | 98 | 49 | 1 | 1 | 1 | 0.692 | 1 | NA | 1 | -0.103 |
| SE | 10926 ZNF292 | ZNF292 | chr6 | + | 87245502 | 87245644 | 87243474 | 87243611 | 87254649 | 87263688 | 13 | 13 | 3 | 0 | 1 | 0 | 0 | 7 | 0 | 0 | 98 | 49 | 1 | 1 | 1 | 0.867 | 1 | NA | 1 | -0.044 |
| SE | 10931 PRRC2C | PRRC2C | chr1 | + | 171512031 | 171512200 | 171485526 | 171485735 | 171513000 | 171513172 | 25 | 3 | 1 | 0 | 0 | 0 | 1 | 4 | 1 | 0 | 98 | 49 | 1 | 1 | 1 | 1 | 1 | 0.333 | 1 | 0.334 |
| SE | 10932 PRRC2C | PRRC2C | chr1 | + | 171545478 | 171545687 | 171539970 | 171542229 | 171550085 | 171550240 | 22 | 7 | 0 | 0 | 0 | 0 | 0 | 0 | 0 | 1 | 98 | 49 | 1 | 1 | 1 | 1 | NA | NA | 0 | 1 |
| SE | 10936 SIGIRR | SIGIRR | chr11 | - | 408694 | 408893 | 408072 | 408206 | 409867 | 410027 | 2 | 0 | 0 | 2 | 0 | 0 | 0 | 2 | 0 | 0 | 98 | 49 | 1 | 1 | 0.333 | NA | NA | NA | 1 | -0.667 |
| SE | 10937 TNPO1 | TNPO1 | chr5 | + | 72855773 | 72855923 | 72851243 | 72851319 | 72861807 | 72861914 | 18 | 6 | 6 | 1 | 0 | 0 | 0 | 1 | 0 | 0 | 98 | 49 | 1 | 1 | 0.9 | 1 | 1 | NA | 1 | -0.033 |
| SE | 10939 TNPO1 | TNPO1 | chr5 | + | 72900005 | 72900081 | 72897055 | 72897151 | 72900973 | 72901073 | 8 | 3 | 3 | 0 | 0 | 0 | 0 | 0 | 0 | 1 | 98 | 49 | 1 | 1 | 1 | 1 | 1 | NA | 0 | 1 |
| SE | 10944 TM9SF2 | TM9SF2 | chr13 | + | 99539445 | 99539557 | 99537738 | 99537863 | 99540713 | 99540793 | 64 | 22 | 27 | 0 | 0 | 0 | 0 | 11 | 0 | 1 | 98 | 49 | 1 | 1 | 1 | 1 | 1 | NA | 0.846 | 0.154 |
| SE | 10945 TM9SF2 | TM9SF2 | chr13 | + | 99546984 | 99547104 | 99543862 | 99543995 | 99549164 | 99549222 | 97 | 46 | 25 | 0 | 1 | 0 | 0 | 9 | 0 | 0 | 98 | 49 | 1 | 1 | 1 | 0.958 | 1 | NA | 1 | -0.014 |
| SE | 10946 TM9SF2 | TM9SF2 | chr13 | + | 99549164 | 99549222 | 99543862 | 99543995 | 99552166 | 99552326 | 34 | 17 | 13 | 1 | 0 | 0 | 0 | 19 | 0 | 0 | 98 | 49 | 1 | 1 | 0.944 | 1 | 1 | NA | 1 | -0.019 |
| SE | 10949 CERS5 | CERS5 | chr12 | - | 50144768 | 50144969 | 50143951 | 50144057 | 50167100 | 50167533 | 0 | 1 | 0 | 0 | 0 | 0 | 0 | 0 | 0 | 2 | 98 | 49 | 1 | 1 | NA | 1 | NA | NA | 0 | 1 |
| SE | 10953 ZC3H11A | ZC3H11A | chr1 | + | 203837965 | 203838064 | 203833790 | 203833853 | 203840305 | 203840374 | 35 | 25 | 8 | 1 | 0 | 0 | 2 | 2 | 0 | 0 | 98 | 49 | 1 | 1 | 0.946 | 1 | 1 | 1 | 1 | -0.018 |
| SE | 10957 CR1 | CR1 | chr1 | + | 207587385 | 207587565 | 207584648 | 207584676 | 207588674 | 207588774 | 158 | 63 | 21 | 1 | 0 | 0 | 2 | 47 | 0 | 0 | 98 | 49 | 1 | 1 | 0.987 | 1 | 1 | 1 | 1 | -0.004 |
| SE | 10959 CR1 | CR1 | chr1 | + | 207622992 | 207623068 | 207621972 | 207621996 | 207630516 | 207630621 | 49 | 24 | 31 | 0 | 2 | 0 | 0 | 11 | 0 | 0 | 98 | 49 | 1 | 1 | 1 | 0.857 | 1 | NA | 1 | -0.048 |
| SE | 10960 CR1 | CR1 | chr1 | + | 207630516 | 207630621 | 207622992 | 207623068 | 207639396 | 207641765 | 76 | 44 | 28 | 2 | 0 | 0 | 1 | 9 | 0 | 0 | 98 | 49 | 1 | 1 | 0.95 | 1 | 1 | 1 | 1 | -0.017 |
| SE | 10967 PPP6R3 | PPP6R3 | chr11 | + | 68551120 | 68551186 | 68548066 | 68548204 | 68554144 | 68554257 | 35 | 26 | 12 | 0 | 0 | 0 | 0 | 2 | 0 | 2 | 98 | 49 | 1 | 1 | 1 | 1 | 1 | NA | 0.333 | 0.667 |
| SE | 10968 PPP6R3 | PPP6R3 | chr11 | + | 68567013 | 68567166 | 68564302 | 68564432 | 68569747 | 68569897 | 17 | 8 | 6 | 0 | 3 | 0 | 0 | 6 | 0 | 0 | 98 | 49 | 1 | 1 | 1 | 0.571 | 1 | NA | 1 | -0.143 |
| SE | 10969 PPP6R3 | PPP6R3 | chr11 | + | 68583042 | 68583129 | 68575957 | 68576043 | 68587926 | 68588024 | 25 | 11 | 1 | 1 | 0 | 0 | 0 | 5 | 0 | 0 | 98 | 49 | 1 | 1 | 0.926 | 1 | 1 | NA | 1 | -0.025 |

| | | | | | | | | | | | | | | | | | | | | | | | | | | | | | | | |
|---|---|---|---|---|---|---|---|---|---|---|---|---|---|---|---|---|---|---|---|---|---|---|---|---|---|---|---|---|---|---|---|
| SE | 10970 | PPP6R3 | PPP6R3 | chr11 | + | 68583042 | 68583129 | 68575957 | 68576043 | 68587944 | 68588024 | 20 | 5 | 1 | 0 | 0 | 0 | 0 | 2 | 0 | 1 | 98 | 49 | 1 | 1 | 1 | 1 | 1 | NA | 0.5 | 0.5 |
| SE | 10971 | PPP6R3 | PPP6R3 | chr11 | + | 68587926 | 68588024 | 68575957 | 68576043 | 68590659 | 68590714 | 19 | 12 | 5 | 0 | 1 | 0 | 1 | 3 | 0 | 0 | 98 | 49 | 1 | 1 | 1 | 0.857 | 1 | 1 | 1 | -0.048 |
| SE | 10972 | PPP6R3 | PPP6R3 | chr11 | + | 68587944 | 68588024 | 68575957 | 68576043 | 68590659 | 68590714 | 18 | 12 | 5 | 0 | 1 | 0 | 1 | 4 | 0 | 0 | 98 | 49 | 1 | 1 | 1 | 0.857 | 1 | 1 | 1 | -0.048 |
| SE | 10976 | FRMD8 | FRMD8 | chr11 | + | 65393572 | 65393674 | 65389360 | 65389528 | 65394040 | 65394099 | 6 | 7 | 6 | 1 | 1 | 0 | 1 | 3 | 0 | 0 | 98 | 49 | 1 | 1 | 0.75 | 0.778 | 1 | 1 | 1 | -0.157 |
| SE | 10979 | PCNXL3 | PCNXL3 | chr11 | + | 65617469 | 65617509 | 65617249 | 65617349 | 65617939 | 65619067 | 4 | 3 | 0 | 0 | 2 | 0 | 0 | 1 | 0 | 0 | 89 | 49 | 1 | 1 | 1 | 0.452 | NA | NA | 1 | -0.274 |
| SE | 10980 | PCNXL3 | PCNXL3 | chr11 | + | 65617610 | 65617706 | 65617469 | 65617509 | 65617939 | 65619067 | 5 | 6 | 0 | 2 | 0 | 0 | 0 | 2 | 0 | 0 | 98 | 49 | 1 | 1 | 0.556 | 1 | NA | NA | 1 | -0.222 |
| SE | 10982 | PCNXL3 | PCNXL3 | chr11 | + | 65625651 | 65625744 | 65625404 | 65625510 | 65625903 | 65626054 | 4 | 1 | 1 | 1 | 0 | 0 | 0 | 1 | 0 | 0 | 98 | 49 | 1 | 1 | 0.667 | 1 | 1 | NA | 1 | -0.111 |
| SE | 11002 | TFAM | TFAM | chr10 | + | 58390764 | 58390860 | 58388669 | 58388819 | 58394357 | 58394414 | 4 | 7 | 6 | 0 | 4 | 0 | 0 | 1 | 0 | 0 | 98 | 49 | 1 | 1 | 1 | 0.467 | 1 | NA | 1 | -0.178 |
| SE | 11009 | MSH6 | MSH6 | chr2 | + | 47790926 | 47791123 | 47783081 | 47783493 | 47795893 | 47796063 | 5 | 0 | 0 | 1 | 0 | 0 | 0 | 1 | 0 | 0 | 98 | 49 | 1 | 1 | 0.714 | NA | NA | NA | 1 | -0.286 |
| SE | 11011 | ANKRD13A | ANKRD13A | chr12 | + | 110016387 | 110016433 | 110013124 | 110013249 | 110019138 | 110019328 | 36 | 20 | 9 | 1 | 0 | 0 | 0 | 9 | 0 | 0 | 95 | 49 | 1 | 1 | 0.949 | 1 | 1 | NA | 1 | -0.017 |
| SE | 11012 | ANKRD13A | ANKRD13A | chr12 | + | 110018344 | 110018488 | 110016387 | 110016433 | 110019138 | 110019328 | 55 | 14 | 25 | 2 | 2 | 0 | 0 | 18 | 0 | 1 | 98 | 49 | 1 | 1 | 0.932 | 0.778 | 1 | NA | 0.9 | 0.003 |
| SE | 11013 | ANKRD13A | ANKRD13A | chr12 | + | 110027704 | 110027766 | 110025741 | 110025823 | 110028511 | 110028642 | 71 | 33 | 35 | 1 | 0 | 0 | 0 | 8 | 0 | 0 | 98 | 49 | 1 | 1 | 0.973 | 1 | 1 | NA | 1 | -0.009 |
| SE | 11014 | ANKRD13A | ANKRD13A | chr12 | + | 110033796 | 110033957 | 110030644 | 110030758 | 110036260 | 110036328 | 38 | 19 | 14 | 1 | 0 | 0 | 2 | 4 | 0 | 1 | 98 | 49 | 1 | 1 | 0.95 | 1 | 1 | 1 | 0.667 | 0.15 |
| SE | 11015 | IFIT3 | IFIT3 | chr10 | + | 89337420 | 89337542 | 89332478 | 89332627 | 89338660 | 89340968 | 0 | 1 | 0 | 2 | 10 | 0 | 0 | 0 | 1 | 1 | 98 | 49 | 1 | 1 | 0 | 0.048 | NA | 0 | 0 | 0.024 |
| SE | 11016 | IFIT3 | IFIT3 | chr10 | + | 89337424 | 89337542 | 89327818 | 89328078 | 89338660 | 89340968 | 0 | 1 | 0 | 5 | 5 | 0 | 0 | 0 | 0 | 1 | 98 | 49 | 1 | 1 | 0 | 0.091 | NA | NA | 0 | 0.045 |
| SE | 11017 | IFIT3 | IFIT3 | chr10 | + | 89337424 | 89337542 | 89332478 | 89332627 | 89338660 | 89340968 | 0 | 2 | 0 | 2 | 10 | 0 | 0 | 0 | 1 | 1 | 98 | 49 | 1 | 1 | 0 | 0.091 | NA | 0 | 0 | 0.045 |
| SE | 11020 | RBM4 | RBM4 | chr11 | + | 66643449 | 66644140 | 66639699 | 66640123 | 66646026 | 66646473 | 13 | 5 | 2 | 1 | 0 | 1 | 0 | 2 | 0 | 0 | 98 | 49 | 1 | 1 | 0.867 | 1 | 0.5 | NA | 1 | -0.211 |
| SE | 11027 | KANSL1 | KANSL1 | chr17 | - | 46170854 | 46172232 | 46094559 | 46094701 | 46192822 | 46192906 | 38 | 12 | 0 | 0 | 1 | 0 | 0 | 7 | 0 | 0 | 98 | 49 | 1 | 1 | 1 | 0.857 | NA | NA | 1 | -0.072 |
| SE | 11032 | SIRT6 | SIRT6 | chr19 | - | 4180781 | 4180909 | 4179103 | 4179286 | 4182473 | 4182599 | 2 | 1 | 0 | 0 | 1 | 0 | 0 | 1 | 0 | 0 | 98 | 49 | 1 | 1 | 1 | 0.333 | NA | NA | 1 | -0.334 |
| SE | 11036 | DNAH1 | DNAH1 | chr3 | + | 52349988 | 52350108 | 52349194 | 52349420 | 52350507 | 52350590 | 3 | 2 | 0 | 0 | 1 | 0 | 0 | 3 | 0 | 0 | 98 | 49 | 1 | 1 | 1 | 0.5 | NA | NA | 1 | -0.25 |
| SE | 11038 | DNAH1 | DNAH1 | chr3 | + | 52398031 | 52398162 | 52397706 | 52397877 | 52398849 | 52399201 | 5 | 0 | 0 | 1 | 1 | 0 | 0 | 0 | 0 | 1 | 98 | 49 | 1 | 1 | 0.714 | 0 | NA | NA | 0 | 0.357 |
| SE | 11045 | PNISR | PNISR | chr6 | - | 99403828 | 99403882 | 99402539 | 99402710 | 99404602 | 99404702 | 23 | 7 | 3 | 0 | 4 | 1 | 0 | 4 | 0 | 0 | 98 | 49 | 1 | 1 | 1 | 0.467 | 0.6 | NA | 1 | -0.311 |
| SE | 11049 | THEMIS2 | THEMIS2 | chr1 | + | 27876587 | 27876728 | 27872542 | 27872665 | 27879643 | 27879963 | 181 | 71 | 24 | 1 | 0 | 0 | 13 | 31 | 0 | 1 | 98 | 49 | 1 | 1 | 0.989 | 1 | 1 | 1 | 0.939 | 0.027 |
| SE | 11050 | THEMIS2 | THEMIS2 | chr1 | + | 27876587 | 27876728 | 27872542 | 27872665 | 27885294 | 27885451 | 75 | 50 | 16 | 0 | 0 | 0 | 10 | 8 | 1 | 0 | 98 | 49 | 1 | 1 | 1 | 1 | 1 | 0.833 | 1 | 0.084 |
| SE | 11051 | THEMIS2 | THEMIS2 | chr1 | + | 27879643 | 27880054 | 27872542 | 27872665 | 27885294 | 27885451 | 3 | 0 | 1 | 0 | 0 | 0 | 0 | 2 | 1 | 0 | 98 | 49 | 1 | 1 | 1 | NA | 1 | 0 | 1 | 0.5 |
| SE | 11054 | THEMIS2 | THEMIS2 | chr1 | + | 27881970 | 27882125 | 27879643 | 27880054 | 27882512 | 27883043 | 46 | 9 | 18 | 1 | 0 | 0 | 1 | 18 | 0 | 0 | 98 | 49 | 1 | 1 | 0.958 | 1 | 1 | 1 | 1 | -0.014 |
| SE | 11057 | THEMIS2 | THEMIS2 | chr1 | + | 27882512 | 27883043 | 27879643 | 27880054 | 27885294 | 27885451 | 26 | 9 | 1 | 2 | 0 | 1 | 0 | 1 | 0 | 1 | 98 | 49 | 1 | 1 | 0.867 | 1 | 0.333 | NA | 0.333 | 0.4 |
| SE | 11059 | CNOT10 | CNOT10 | chr3 | + | 32713226 | 32713369 | 32708669 | 32708820 | 32716224 | 32716311 | 15 | 4 | 0 | 2 | 0 | 0 | 0 | 3 | 0 | 0 | 98 | 49 | 1 | 1 | 0.789 | 1 | NA | NA | 1 | -0.105 |
| SE | 11060 | CNOT10 | CNOT10 | chr3 | + | 32717153 | 32717237 | 32716224 | 32716311 | 32720113 | 32720231 | 16 | 2 | 5 | 0 | 1 | 0 | 0 | 4 | 0 | 0 | 98 | 49 | 1 | 1 | 1 | 0.5 | 1 | NA | 1 | -0.167 |
| SE | 11089 | FCHSD2 | FCHSD2 | chr11 | - | 73140030 | 73140128 | 73083694 | 73083740 | 73141856 | 73142098 | 9 | 9 | 2 | 1 | 1 | 0 | 0 | 1 | 0 | 0 | 98 | 49 | 1 | 1 | 0.818 | 0.818 | 1 | NA | 1 | -0.121 |
| SE | 11091 | PCMTD1 | PCMTD1 | chr8 | - | 51845660 | 51845763 | 51833517 | 51833689 | 51860844 | 51861246 | 18 | 13 | 22 | 0 | 0 | 1 | 0 | 8 | 0 | 0 | 98 | 49 | 1 | 1 | 1 | 1 | 0.917 | NA | 1 | -0.028 |
| SE | 11092 | PCMTD1 | PCMTD1 | chr8 | - | 51845660 | 51845763 | 51833517 | 51833689 | 51898929 | 51899186 | 9 | 7 | 13 | 0 | 0 | 0 | 0 | 4 | 0 | 1 | 98 | 49 | 1 | 1 | 1 | 1 | 1 | NA | 0.667 | 0.333 |
| SE | 11093 | PCMTD1 | PCMTD1 | chr8 | - | 51860844 | 51861246 | 51833517 | 51833689 | 51898929 | 51899186 | 12 | 12 | 2 | 0 | 0 | 0 | 1 | 1 | 0 | 1 | 98 | 49 | 1 | 1 | 1 | 1 | 1 | 1 | 0.333 | 0.334 |
| SE | 11095 | ARHGAP15 | ARHGAP15 | chr2 | + | 143202133 | 143202202 | 143155476 | 143155655 | 143216383 | 143216445 | 323 | 56 | 40 | 1 | 0 | 0 | 3 | 32 | 0 | 0 | 98 | 49 | 1 | 1 | 0.994 | 1 | 1 | 1 | 1 | -0.002 |
| SE | 11097 | ARHGAP15 | ARHGAP15 | chr2 | + | 143487372 | 143487495 | 143436912 | 143437042 | 143519265 | 143519364 | 193 | 53 | 16 | 3 | 0 | 0 | 0 | 19 | 0 | 0 | 98 | 49 | 1 | 1 | 0.97 | 1 | 1 | NA | 1 | -0.01 |
| SE | 11098 | ARHGAP15 | ARHGAP15 | chr2 | + | 143519265 | 143519364 | 143487372 | 143487495 | 143556407 | 143556485 | 160 | 33 | 14 | 0 | 2 | 0 | 1 | 15 | 0 | 0 | 98 | 49 | 1 | 1 | 1 | 0.892 | 1 | 1 | 1 | -0.036 |
| SE | 11105 | SF3A3 | SF3A3 | chr1 | - | 37978719 | 37978827 | 37976883 | 37976953 | 37978987 | 37979055 | 9 | 8 | 0 | 1 | 0 | 0 | 0 | 3 | 0 | 0 | 98 | 49 | 1 | 1 | 0.818 | 1 | NA | NA | 1 | -0.091 |
| SE | 11106 | CYB5R4 | CYB5R4 | chr6 | + | 83909008 | 83909090 | 83893521 | 83893622 | 83918004 | 83918065 | 12 | 15 | 18 | 0 | 2 | 0 | 0 | 3 | 0 | 0 | 98 | 49 | 1 | 1 | 1 | 0.789 | 1 | NA | 1 | -0.07 |
| SE | 11107 | CYB5R4 | CYB5R4 | chr6 | + | 83914415 | 83914448 | 83909008 | 83909090 | 83918004 | 83918065 | 113 | 38 | 42 | 0 | 0 | 1 | 0 | 17 | 0 | 0 | 82 | 49 | 1 | 1 | 1 | 1 | 0.962 | NA | 1 | -0.013 |
| SE | 11108 | CYB5R4 | CYB5R4 | chr6 | + | 83918004 | 83918065 | 83914415 | 83914448 | 83919396 | 83919454 | 125 | 40 | 36 | 1 | 0 | 0 | 0 | 16 | 0 | 0 | 98 | 49 | 1 | 1 | 0.984 | 1 | 1 | NA | 1 | -0.005 |
| SE | 11121 | RAVER1 | RAVER1 | chr19 | - | 10321047 | 10321259 | 10320654 | 10320951 | 10321530 | 10321618 | 7 | 2 | 0 | 0 | 0 | 1 | 0 | 1 | 0 | 0 | 98 | 49 | 1 | 1 | 1 | 1 | 0 | NA | 1 | -0.333 |
| SE | 11126 | CCT4 | CCT4 | chr2 | - | 61872457 | 61872588 | 61872081 | 61872316 | 61873001 | 61873112 | 6 | 0 | 1 | 0 | 0 | 0 | 0 | 0 | 0 | 1 | 98 | 49 | 1 | 1 | 1 | NA | 1 | NA | 0 | 1 |
| SE | 11139 | PER1 | PER1 | chr17 | - | 8145957 | 8146137 | 8144750 | 8144993 | 8146371 | 8146502 | 3 | 0 | 2 | 1 | 1 | 0 | 0 | 3 | 0 | 1 | 98 | 49 | 1 | 1 | 0.6 | 0 | 1 | NA | 0.6 | -0.067 |
| SE | 11140 | PER1 | PER1 | chr17 | - | 8146896 | 8147002 | 8146593 | 8146765 | 8147249 | 8147381 | 3 | 0 | 1 | 0 | 0 | 0 | 0 | 5 | 0 | 2 | 98 | 49 | 1 | 1 | 1 | NA | 1 | NA | 0.556 | 0.444 |
| SE | 11141 | ENTPD4 | ENTPD4 | chr8 | - | 23442006 | 23442066 | 23441568 | 23441723 | 23443849 | 23443953 | 19 | 10 | 4 | 3 | 4 | 0 | 0 | 2 | 0 | 1 | 98 | 49 | 1 | 1 | 0.76 | 0.556 | 1 | NA | 0.5 | 0.272 |
| SE | 11142 | SRGN | SRGN | chr10 | + | 69097083 | 69097231 | 69088071 | 69088236 | 69103870 | 69104811 | 1921 | 1250 | 628 | 5 | 6 | 1 | 28 | 397 | 0 | 0 | 98 | 49 | 1 | 1 | 0.995 | 0.99 | 0.997 | 1 | 1 | -0.006 |
| SE | 11143 | ZYX | ZYX | chr7 | + | 143382592 | 143382685 | 143382247 | 143382447 | 143382800 | 143383322 | 102 | 20 | 25 | 0 | 1 | 0 | 10 | 23 | 0 | 0 | 98 | 49 | 1 | 1 | 1 | 0.909 | 1 | 1 | 1 | -0.03 |
| SE | 11144 | ZYX | ZYX | chr7 | + | 143388218 | 143388339 | 143382800 | 143383322 | 143388488 | 143388658 | 89 | 33 | 35 | 1 | 0 | 0 | 5 | 34 | 0 | 0 | 98 | 49 | 1 | 1 | 0.978 | 1 | 1 | 1 | 1 | -0.007 |
| SE | 11145 | THADA | THADA | chr2 | - | 43556344 | 43556555 | 43552067 | 43552339 | 43560233 | 43560385 | 14 | 7 | 0 | 1 | 0 | 0 | 0 | 4 | 0 | 0 | 98 | 49 | 1 | 1 | 0.875 | 1 | NA | NA | 1 | -0.062 |
| SE | 11151 | FCRL6 | FCRL6 | chr1 | + | 159808960 | 159809245 | 159808177 | 159808444 | 159809401 | 159809683 | 2 | 0 | 0 | 3 | 0 | 0 | 0 | 0 | 0 | 1 | 98 | 49 | 1 | 1 | 0.25 | NA | NA | NA | 0 | 0.25 |
| SE | 11154 | RNF34 | RNF34 | chr12 | + | 121402741 | 121402807 | 121400117 | 121400218 | 121416158 | 121416377 | 1 | 0 | 0 | 1 | 0 | 1 | 0 | 0 | 0 | 1 | 98 | 49 | 1 | 1 | 0.333 | NA | 0 | NA | 0 | 0.167 |
| SE | 11155 | RNF34 | RNF34 | chr12 | + | 121420576 | 121420778 | 121420241 | 121420334 | 121423385 | 121424352 | 6 | 1 | 6 | 0 | 1 | 0 | 0 | 3 | 0 | 0 | 98 | 49 | 1 | 1 | 1 | 0.333 | 1 | NA | 1 | -0.222 |
| SE | 11161 | CBL | CBL | chr11 | + | 119232447 | 119232695 | 119206275 | 119206612 | 119271734 | 119271881 | 18 | 6 | 8 | 1 | 0 | 0 | 0 | 3 | 0 | 0 | 98 | 49 | 1 | 1 | 0.9 | 1 | 1 | NA | 1 | -0.033 |
| SE | 11162 | PHB2 | PHB2 | chr12 | - | 6967170 | 6967238 | 6966419 | 6966500 | 6967891 | 6968021 | 27 | 4 | 1 | 0 | 2 | 0 | 0 | 4 | 0 | 0 | 98 | 49 | 1 | 1 | 1 | 0.5 | 1 | NA | 1 | -0.167 |
| SE | 11167 | LARP4B | LARP4B | chr10 | - | 825064 | 825316 | 820799 | 820845 | 825763 | 825870 | 10 | 6 | 2 | 0 | 2 | 0 | 0 | 4 | 0 | 0 | 98 | 49 | 1 | 1 | 1 | 0.6 | 1 | NA | 1 | -0.133 |
| SE | 11177 | POLR2A | POLR2A | chr17 | + | 7496209 | 7496346 | 7495940 | 7496073 | 7496439 | 7496613 | 41 | 20 | 2 | 1 | 0 | 0 | 1 | 9 | 0 | 0 | 98 | 49 | 1 | 1 | 0.953 | 1 | 1 | 1 | 1 | -0.016 |
| SE | 11184 | CD300LF | CD300LF | chr17 | - | 74696194 | 74696217 | 74695724 | 74695859 | 74698368 | 74698481 | 12 | 2 | 2 | 2 | 1 | 0 | 0 | 2 | 0 | 0 | 72 | 49 | 1 | 1 | 0.803 | 0.576 | 1 | NA | 1 | -0.207 |
| SE | 11187 | CD300LF | CD300LF | chr17 | - | 74698368 | 74698481 | 74696194 | 74696217 | 74703034 | 74703098 | 24 | 8 | 10 | 0 | 1 | 0 | 0 | 1 | 0 | 0 | 98 | 49 | 1 | 1 | 1 | 0.8 | 1 | NA | 1 | -0.067 |
| SE | 11193 | MRE11A | MRE11A | chr11 | - | 94470470 | 94470642 | 94467812 | 94467893 | 94471573 | 94471759 | 6 | 0 | 0 | 1 | 0 | 0 | 0 | 2 | 0 | 0 | 98 | 49 | 1 | 1 | 0.75 | NA | NA | NA | 1 | -0.25 |
| SE | 11208 | PIP5K1C | PIP5K1C | chr19 | - | 3642906 | 3642939 | 3641704 | 3641809 | 3643242 | 3643381 | 2 | 0 | 0 | 0 | 0 | 0 | 0 | 2 | 0 | 2 | 82 | 49 | 1 | 1 | 1 | NA | NA | NA | 0.374 | 0.626 |
| SE | 11209 | SIPA1L1 | SIPA1L1 | chr14 | + | 71698980 | 71699127 | 71685361 | 71685631 | 71702380 | 71702505 | 42 | 17 | 0 | 1 | 0 | 0 | 0 | 6 | 0 | 0 | 98 | 49 | 1 | 1 | 0.955 | 1 | NA | NA | 1 | -0.022 |
| SE | 11211 | SIPA1L1 | SIPA1L1 | chr14 | + | 71730054 | 71730301 | 71724669 | 71724835 | 71733665 | 71733812 | 16 | 7 | 3 | 2 | 0 | 0 | 0 | 3 | 0 | 0 | 98 | 49 | 1 | 1 | 0.8 | 1 | 1 | NA | 1 | -0.067 |
| SE | 11212 | SIPA1L1 | SIPA1L1 | chr14 | + | 71735276 | 71735391 | 71733665 | 71733812 | 71738240 | 71738325 | 22 | 1 | 2 | 1 | 0 | 0 | 1 | 1 | 0 | 0 | 98 | 49 | 1 | 1 | 0.917 | 1 | 1 | 1 | 1 | -0.028 |
| SE | 11220 | HPS5 | HPS5 | chr11 | - | 18317750 | 18317907 | 18311913 | 18312024 | 18321945 | 18322174 | 2 | 1 | 0 | 0 | 1 | 0 | 0 | 1 | 0 | 0 | 98 | 49 | 1 | 1 | 1 | 0.333 | NA | NA | 1 | -0.334 |
| SE | 11223 | CASP4 | CASP4 | chr11 | - | 104951895 | 104952005 | 104950924 | 104951098 | 104954746 | 104955001 | 94 | 48 | 44 | 7 | 0 | 2 | 0 | 13 | 0 | 3 | 98 | 49 | 1 | 1 | 0.87 | 1 | 0.917 | NA | 0.684 | 0.245 |
| SE | 11232 | DPY19L3 | DPY19L3 | chr19 | + | 32437193 | 32437339 | 32436445 | 32436567 | 32439111 | 32439235 | 7 | 2 | 6 | 1 | 0 | 0 | 0 | 3 | 0 | 0 | 98 | 49 | 1 | 1 | 0.778 | 1 | 1 | NA | 1 | -0.074 |
| SE | 11237 | ATM | ATM | chr11 | + | 108235669 | 108235834 | 108229177 | 108229323 | 108243952 | 108244118 | 30 | 16 | 6 | 0 | 2 | 0 | 0 | 9 | 0 | 0 | 98 | 49 | 1 | 1 | 1 | 0.8 | 1 | NA | 1 | -0.067 |
| SE | 11238 | ATM | ATM | chr11 | + | 108244787 | 108245026 | 108243952 | 108244118 | 108246963 | 108247127 | 26 | 10 | 5 | 1 | 0 | 0 | 1 | 4 | 0 | 0 | 98 | 49 | 1 | 1 | 0.929 | 1 | 1 | 1 | 1 | -0.024 |
| SE | 11239 | ATM | ATM | chr11 | + | 108268409 | 108268609 | 108267170 | 108267342 | 108271063 | 108271146 | 31 | 6 | 10 | 0 | 0 | 0 | 0 | 9 | 0 | 1 | 98 | 49 | 1 | 1 | 1 | 1 | 1 | NA | 0.818 | 0.182 |
| SE | 11243 | MAP4K2 | MAP4K2 | chr11 | - | 64800899 | 64801031 | 64800763 | 64800826 | 64801110 | 64801183 | 7 | 7 | 0 | 1 | 0 | 0 | 1 | 0 | 0 | 0 | 98 | 49 | 1 | 1 | 0.778 | 1 | NA | 1 | NA | -0.111 |
| SE | 11244 | MTOR | MTOR | chr1 | - | 11233397 | 11233487 | 11232435 | 11232528 | 11234142 | 11234265 | 8 | 0 | 4 | 0 | 1 | 0 | 0 | 5 | 0 | 0 | 98 | 49 | 1 | 1 | 1 | 0 | 1 | NA | 1 | -0.333 |
| SE | 11245 | MTOR | MTOR | chr1 | - | 11256932 | 11257165 | 11255991 | 11256192 | 11258484 | 11258593 | 5 | 3 | 0 | 0 | 1 | 0 | 2 | 4 | 0 | 0 | 98 | 49 | 1 | 1 | 1 | 0.6 | NA | 1 | 1 | -0.2 |
| SE | 11246 | PROK2 | PROK2 | chr3 | - | 71774444 | 71774507 | 71771654 | 71772828 | 71781466 | 71781592 | 105 | 34 | 27 | 27 | 7 | 5 | 0 | 2 | 0 | 4 | 98 | 49 | 1 | 1 | 0.66 | 0.708 | 0.73 | NA | 0.2 | 0.499 |
| SE | 11253 | ABTB1 | ABTB1 | chr3 | + | 127674544 | 127674600 | 127674390 | 127674453 | 127675969 | 127676114 | 85 | 66 | 21 | 1 | 1 | 0 | 1 | 14 | 0 | 0 | 98 | 49 | 1 | 1 | 0.977 | 0.971 | 1 | 1 | 1 | -0.017 |
| SE | 11254 | ABTB1 | ABTB1 | chr3 | + | 127675969 | 127676114 | 127674544 | 127674600 | 127676535 | 127676581 | 61 | 43 | 14 | 1 | 0 | 0 | 1 | 12 | 0 | 0 | 98 | 49 | 1 | 1 | 0.968 | 1 | 1 | 1 | 1 | -0.011 |
| SE | 11255 | ABTB1 | ABTB1 | chr3 | + | 127676271 | 127676431 | 127675969 | 127676114 | 127676535 | 127676581 | 168 | 72 | 42 | 1 | 0 | 0 | 3 | 20 | 0 | 0 | 98 | 49 | 1 | 1 | 0.988 | 1 | 1 | 1 | 1 | -0.004 |
| SE | 11256 | ABTB1 | ABTB1 | chr3 | + | 127676966 | 127677083 | 127676535 | 127676581 | 127677167 | 127677286 | 96 | 44 | 25 | 0 | 0 | 2 | 1 | 20 | 0 | 0 | 98 | 49 | 1 | 1 | 1 | 1 | 0.862 | 1 | 1 | -0.046 |
| SE | 11260 | DLEU2 | DLEU2 | chr13 | - | 50029277 | 50029492 | 50027206 | 50027406 | 50044650 | 50044768 | 11 | 4 | 17 | 3 | 3 | 2 | 0 | 2 | 0 | 0 | 98 | 49 | 1 | 1 | 0.647 | 0.4 | 0.81 | NA | 1 | -0.381 |
| SE | 11261 | DLEU2 | DLEU2 | chr13 | - | 50044650 | 50044768 | 50027206 | 50027406 | 50049583 | 50049662 | 13 | 3 | 2 | 0 | 1 | 1 | 0 | 1 | 0 | 0 | 98 | 49 | 1 | 1 | 1 | 0.6 | 0.5 | NA | 1 | -0.3 |
| SE | 11263 | DLEU2 | DLEU2 | chr13 | - | 50075558 | 50075653 | 50068096 | 50068163 | 50125301 | 50125541 | 31 | 2 | 3 | 1 | 0 | 0 | 0 | 1 | 0 | 0 | 98 | 49 | 1 | 1 | 0.939 | 1 | 1 | NA | 1 | -0.02 |
| SE | 11264 | STXBP3 | STXBP3 | chr1 | + | 108800219 | 108800305 | 108798144 | 108798237 | 108807400 | 108807549 | 28 | 18 | 6 | 2 | 0 | 0 | 0 | 5 | 0 | 0 | 98 | 49 | 1 | 1 | 0.875 | 1 | 1 | NA | 1 | -0.042 |
| SE | 11270 | WBP11 | WBP11 | chr12 | - | 14794970 | 14795104 | 14794536 | 14794736 | 14796806 | 14797003 | 11 | 0 | 0 | 0 | 1 | 0 | 0 | 2 | 0 | 0 | 98 | 49 | 1 | 1 | 1 | 0 | NA | NA | 1 | -0.5 |
| SE | 11278 | FXR2 | FXR2 | chr17 | - | 7594237 | 7594347 | 7593917 | 7594004 | 7594678 | 7594757 | 4 | 0 | 5 | 1 | 1 | 0 | 0 | 1 | 0 | 0 | 98 | 49 | 1 | 1 | 0.667 | 0 | 1 | NA | 1 | -0.444 |
| SE | 11279 | SAT2 | SAT2 | chr17 | - | 7626942 | 7627044 | 7626752 | 7626793 | 7627142 | 7627226 | 0 | 0 | 0 | 4 | 0 | 0 | 0 | 2 | 0 | 1 | 98 | 49 | 1 | 1 | 0 | NA | NA | NA | 0.5 | -0.5 |
| SE | 11290 | TRIM39 | TRIM39 | chr6 | + | 30340271 | 30340361 | 30339907 | 30339930 | 30340504 | 30340620 | 0 | 0 | 1 | 6 | 4 | 4 | 0 | 0 | 1 | 0 | 98 | 49 | 1 | 1 | 0 | 0 | 0.111 | 0 | NA | 0.037 |
| SE | 11295 | MBD1 | MBD1 | chr18 | - | 50274185 | 50274353 | 50273563 | 50273863 | 50274976 | 50275045 | 0 | 7 | 1 | 1 | 0 | 0 | 0 | 1 | 0 | 0 | 98 | 49 | 1 | 1 | 0 | 1 | 1 | NA | 1 | -0.333 |
| SE | 11299 | KDM6B | KDM6B | chr17 | + | 7844900 | 7845020 | 7839916 | 7840024 | 7845313 | 7845456 | 88 | 56 | 22 | 1 | 0 | 0 | 3 | 14 | 0 | 1 | 98 | 49 | 1 | 1 | 0.978 | 1 | 1 | 1 | 0.875 | 0.055 |
| SE | 11301 | KDM6B | KDM6B | chr17 | + | 7851647 | 7851796 | 7851477 | 7851549 | 7851950 | 7852065 | 67 | 20 | 19 | 0 | 1 | 0 | 1 | 28 | 0 | 0 | 98 | 49 | 1 | 1 | 1 | 0.909 | 1 | 1 | 1 | -0.03 |
| SE | 11306 | YEATS2 | YEATS2 | chr3 | + | 183803255 | 183803335 | 183801454 | 183801528 | 183803986 | 183804188 | 2 | 0 | 0 | 0 | 0 | 0 | 0 | 0 | 0 | 1 | 98 | 49 | 1 | 1 | 1 | NA | NA | NA | 0 | 1 |
| SE | 11308 | CTC1 | CTC1 | chr17 | - | 8228726 | 8228892 | 8228502 | 8228629 | 8229301 | 8229446 | 0 | 0 | 0 | 4 | 0 | 0 | 0 | 1 | 0 | 0 | 98 | 49 | 1 | 1 | 0 | NA | NA | NA | 1 | -1 |
| SE | 11310 | CTC1 | CTC1 | chr17 | - | 8231275 | 8231469 | 8230562 | 8230651 | 8231725 | 8231815 | 5 | 7 | 0 | 2 | 0 | 0 | 0 | 2 | 0 | 0 | 98 | 49 | 1 | 1 | 0.556 | 1 | NA | NA | 1 | -0.222 |
| SE | 11311 | CTC1 | CTC1 | chr17 | - | 8238030 | 8238242 | 8237374 | 8237519 | 8238391 | 8238629 | 3 | 4 | 2 | 3 | 0 | 1 | 0 | 1 | 0 | 0 | 98 | 49 | 1 | 1 | 0.333 | 1 | 0.5 | NA | 1 | -0.389 |
| SE | 11315 | NDEL1 | NDEL1 | chr17 | + | 8445710 | 8445864 | 8444259 | 8444357 | 8446753 | 8446902 | 95 | 27 | 15 | 5 | 2 | 0 | 0 | 20 | 0 | 0 | 98 | 49 | 1 | 1 | 0.905 | 0.871 | 1 | NA | 1 | -0.075 |
| SE | 11316 | NDEL1 | NDEL1 | chr17 | + | 8446753 | 8446902 | 8445710 | 8445864 | 8448549 | 8448686 | 132 | 51 | 28 | 1 | 0 | 0 | 0 | 20 | 0 | 0 | 98 | 49 | 1 | 1 | 0.985 | 1 | 1 | NA | 1 | -0.005 |
| SE | 11317 | NDEL1 | NDEL1 | chr17 | + | 8454795 | 8454887 | 8450779 | 8450953 | 8460008 | 8460160 | 142 | 40 | 9 | 2 | 0 | 0 | 6 | 23 | 0 | 0 | 98 | 49 | 1 | 1 | 0.973 | 1 | 1 | 1 | 1 | -0.009 |
| SE | 11326 | PAM | PAM | chr5 | + | 103025130 | 103025334 | 103019789 | 103019843 | 103028886 | 103031104 | 2 | 3 | 0 | 0 | 0 | 1 | 0 | 1 | 0 | 0 | 98 | 49 | 1 | 1 | 1 | 1 | 0 | NA | 1 | -0.333 |
| SE | 11329 | PAM | PAM | chr5 | + | 103028181 | 103028238 | 103025130 | 103025334 | 103028886 | 103031104 | 2 | 2 | 0 | 1 | 0 | 0 | 0 | 1 | 0 | 0 | 98 | 49 | 1 | 1 | 0.5 | 1 | NA | NA | 1 | -0.25 |
| SE | 11335 | PABPC1 | PABPC1 | chr8 | - | 100718086 | 100718280 | 100717772 | 100717888 | 100721390 | 100722087 | 50 | 8 | 3 | 0 | 0 | 0 | 0 | 29 | 1 | 0 | 98 | 49 | 1 | 1 | 1 | 1 | 1 | 0 | 1 | 0.5 |
| SE | 11343 | AMPD3 | AMPD3 | chr11 | + | 10478525 | 10478730 | 10461514 | 10461740 | 10482062 | 10482225 | 9 | 2 | 0 | 3 | 0 | 0 | 0 | 1 | 0 | 0 | 98 | 49 | 1 | 1 | 0.6 | 1 | NA | NA | 1 | -0.2 |
| SE | 11344 | AMPD3 | AMPD3 | chr11 | + | 10482062 | 10482225 | 10478525 | 10478730 | 10484819 | 10485039 | 23 | 7 | 0 | 2 | 0 | 0 | 0 | 7 | 0 | 0 | 98 | 49 | 1 | 1 | 0.852 | 1 | NA | NA | 1 | -0.074 |
| SE | 11345 | AMPD3 | AMPD3 | chr11 | + | 10495569 | 10495733 | 10494898 | 10495030 | 10496811 | 10496938 | 18 | 10 | 1 | 0 | 3 | 0 | 0 | 4 | 0 | 0 | 98 | 49 | 1 | 1 | 1 | 0.625 | 1 | NA | 1 | -0.125 |
| SE | 11346 | AMPD3 | AMPD3 | chr11 | + | 10504548 | 10504659 | 10502720 | 10502894 | 10505707 | 10507579 | 27 | 7 | 4 | 1 | 0 | 0 | 0 | 3 | 0 | 0 | 98 | 49 | 1 | 1 | 0.931 | 1 | 1 | NA | 1 | -0.023 |
| SE | 11354 | NDRG1 | NDRG1 | chr8 | - | 133258365 | 133258426 | 133256776 | 133256863 | 133259167 | 133259230 | 9 | 2 | 5 | 0 | 0 | 0 | 0 | 6 | 0 | 1 | 98 | 49 | 1 | 1 | 1 | 1 | 1 | NA | 0.75 | 0.25 |
| SE | 11355 | NDRG1 | NDRG1 | chr8 | - | 133264546 | 133264652 | 133262046 | 133262167 | 133280231 | 133280267 | 26 | 7 | 9 | 1 | 0 | 0 | 0 | 9 | 0 | 0 | 98 | 49 | 1 | 1 | 0.929 | 1 | 1 | NA | 1 | -0.024 |
| SE | 11357 | ZNFX1 | ZNFX1 | chr20 | - | 49252719 | 49252830 | 49251526 | 49251622 | 49253665 | 49253811 | 61 | 18 | 9 | 1 | 0 | 0 | 0 | 12 | 0 | 0 | 98 | 49 | 1 | 1 | 0.968 | 1 | 1 | NA | 1 | -0.011 |
| SE | 11364 | SCAF11 | SCAF11 | chr12 | - | 45922935 | 45923154 | 45922462 | 45922582 | 45924727 | 45925074 | 12 | 6 | 2 | 2 | 0 | 0 | 0 | 5 | 0 | 0 | 98 | 49 | 1 | 1 | 0.75 | 1 | 1 | NA | 1 | -0.083 |
| SE | 11368 | TRPV2 | TRPV2 | chr17 | + | 16433573 | 16433698 | 16431965 | 16432300 | 16434889 | 16434969 | 5 | 1 | 0 | 0 | 0 | 4 | 0 | 1 | 0 | 0 | 98 | 49 | 1 | 1 | 1 | 1 | 0 | NA | 1 | -0.333 |
| SE | 11371 | LRRC75A-AS1 | LRRC75A-AS1 | chr17 | + | 16439527 | 16439703 | 16438986 | 16439414 | 16440184 | 16440253 | 2 | 2 | 0 | 1 | 1 | 0 | 0 | 0 | 0 | 1 | 98 | 49 | 1 | 1 | 0.5 | 0.5 | NA | NA | 0 | 0.5 |
| SE | 11372 | LRRC75A-AS1 | LRRC75A-AS1 | chr17 | + | 16439580 | 16439703 | 16438986 | 16439414 | 16440184 | 16440253 | 2 | 2 | 0 | 1 | 1 | 0 | 0 | 0 | 0 | 1 | 98 | 49 | 1 | 1 | 0.5 | 0.5 | NA | NA | 0 | 0.5 |
| SE | 11374 | LRRC75A-AS1 | LRRC75A-AS1 | chr17 | + | 16439659 | 16439703 | 16438986 | 16439414 | 16440184 | 16440253 | 3 | 2 | 0 | 1 | 1 | 0 | 0 | 0 | 0 | 1 | 93 | 49 | 1 | 1 | 0.613 | 0.513 | NA | NA | 0 | 0.563 |
| SE | 11393 | COPS3 | COPS3 | chr17 | - | 17247479 | 17247560 | 17246623 | 17247151 | 17248925 | 17249039 | 6 | 4 | 7 | 0 | 3 | 0 | 0 | 1 | 0 | 2 | 98 | 49 | 1 | 1 | 1 | 0.4 | 1 | NA | 0.2 | 0.6 |
| SE | 11398 | TOM1L2 | TOM1L2 | chr17 | - | 17882704 | 17882863 | 17879626 | 17879743 | 17893660 | 17893810 | 4 | 1 | 1 | 1 | 0 | 0 | 0 | 0 | 0 | 1 | 98 | 49 | 1 | 1 | 0.667 | 1 | 1 | NA | 0 | 0.889 |
| SE | 11399 | TOM1L2 | TOM1L2 | chr17 | - | 17884633 | 17884768 | 17879626 | 17879743 | 17893660 | 17893810 | 4 | 0 | 1 | 1 | 0 | 0 | 0 | 3 | 0 | 1 | 98 | 49 | 1 | 1 | 0.667 | NA | 1 | NA | 0.6 | 0.234 |
| SE | 11407 | RPS24 | RPS24 | chr10 | + | 78037961 | 78037982 | 78037193 | 78037304 | 78040203 | 78040225 | 0 | 1 | 0 | 99 | 14 | 7 | 0 | 0 | 0 | 10 | 70 | 49 | 1 | 1 | 0 | 0.048 | 0 | NA | 0 | 0.016 |
| SE | 11409 | RPS24 | RPS24 | chr10 | + | 78040200 | 78040225 | 78037193 | 78037304 | 78040614 | 78040715 | 62 | 12 | 6 | 72 | 17 | 11 | 0 | 1 | 0 | 26 | 74 | 49 | 1 | 1 | 0.363 | 0.319 | 0.265 | NA | 0.025 | 0.291 |
| SE | 11410 | RPS24 | RPS24 | chr10 | + | 78040203 | 78040225 | 78037193 | 78037304 | 78040614 | 78040715 | 161 | 26 | 13 | 72 | 17 | 11 | 0 | 11 | 0 | 26 | 71 | 49 | 1 | 1 | 0.607 | 0.514 | 0.449 | NA | 0.226 | 0.297 |
| SE | 11413 | LCMT1 | LCMT1 | chr16 | + | 25128474 | 25128566 | 25111725 | 25111996 | 25132401 | 25132523 | 1 | 0 | 0 | 0 | 0 | 0 | 0 | 0 | 0 | 1 | 98 | 49 | 1 | 1 | 1 | NA | NA | NA | 0 | 1 |
| SE | 11415 | PRPSAP2 | PRPSAP2 | chr17 | + | 18867281 | 18867334 | 18865801 | 18865952 | 18872582 | 18872649 | 7 | 0 | 1 | 3 | 0 | 0 | 0 | 1 | 0 | 0 | 98 | 49 | 1 | 1 | 0.538 | NA | 1 | NA | 1 | -0.231 |
| SE | 11423 | ANKHD1 | ANKHD1 | chr5 | + | 140485120 | 140485248 | 140482579 | 140482667 | 140485588 | 140485732 | 16 | 15 | 0 | 2 | 0 | 0 | 0 | 3 | 0 | 0 | 98 | 49 | 1 | 1 | 0.8 | 1 | NA | NA | 1 | -0.1 |
| SE | 11425 | ANKHD1 | ANKHD1 | chr5 | + | 140507784 | 140507998 | 140506834 | 140506977 | 140509636 | 140509812 | 44 | 9 | 8 | 1 | 1 | 0 | 1 | 11 | 0 | 0 | 98 | 49 | 1 | 1 | 0.957 | 0.818 | 1 | 1 | 1 | -0.075 |
| SE | 11426 | ANKHD1 | ANKHD1 | chr5 | + | 140538085 | 140538261 | 140537388 | 140537589 | 140538918 | 140539083 | 17 | 3 | 0 | 1 | 0 | 0 | 0 | 3 | 0 | 0 | 98 | 49 | 1 | 1 | 0.895 | 1 | NA | NA | 1 | -0.052 |
| SE | 11427 | QSOX1 | QSOX1 | chr1 | + | 180194212 | 180194392 | 180190432 | 180190580 | 180196261 | 180196599 | 6 | 3 | 8 | 0 | 0 | 0 | 1 | 3 | 0 | 1 | 98 | 49 | 1 | 1 | 1 | 1 | 1 | 1 | 0.6 | 0.2 |
| SE | 11430 | MX1 | MX1 | chr21 | + | 41439693 | 41439848 | 41437014 | 41437152 | 41440886 | 41441025 | 77 | 58 | 9 | 1 | 0 | 0 | 6 | 103 | 1 | 0 | 98 | 49 | 1 | 1 | 0.975 | 1 | 1 | 0.75 | 1 | 0.117 |
| SE | 11432 | MX1 | MX1 | chr21 | + | 41449136 | 41449295 | 41445999 | 41446141 | 41451166 | 41451243 | 24 | 19 | 4 | 1 | 3 | 0 | 0 | 30 | 0 | 0 | 98 | 49 | 1 | 1 | 0.923 | 0.76 | 1 | NA | 1 | -0.106 |
| SE | 11436 | AKAP10 | AKAP10 | chr17 | - | 19940886 | 19941010 | 19939712 | 19939849 | 19941825 | 19941910 | 12 | 4 | 0 | 2 | 6 | 0 | 0 | 4 | 0 | 4 | 98 | 49 | 1 | 1 | 0.75 | 0.25 | NA | NA | 0.333 | 0.167 |
| SE | 11437 | SPECC1 | SPECC1 | chr17 | + | 20204332 | 20205912 | 20110426 | 20110562 | 20227412 | 20227620 | 3 | 4 | 0 | 1 | 0 | 0 | 0 | 1 | 0 | 0 | 98 | 49 | 1 | 1 | 0.6 | 1 | NA | NA | 1 | -0.2 |
| SE | 11438 | DHX8 | DHX8 | chr17 | + | 43492680 | 43493040 | 43492182 | 43492292 | 43493444 | 43493589 | 25 | 12 | 0 | 0 | 2 | 0 | 0 | 4 | 0 | 0 | 98 | 49 | 1 | 1 | 1 | 0.75 | NA | NA | 1 | -0.125 |
| SE | 11439 | DHX8 | DHX8 | chr17 | + | 43493444 | 43493589 | 43492680 | 43493040 | 43493682 | 43493886 | 21 | 12 | 4 | 2 | 0 | 0 | 0 | 5 | 0 | 1 | 98 | 49 | 1 | 1 | 0.84 | 1 | 1 | NA | 0.714 | 0.233 |
| SE | 11440 | DHX8 | DHX8 | chr17 | + | 43508338 | 43508520 | 43507808 | 43508019 | 43513361 | 43513502 | 8 | 8 | 8 | 1 | 0 | 0 | 1 | 1 | 0 | 0 | 98 | 49 | 1 | 1 | 0.8 | 1 | 1 | 1 | 1 | -0.067 |
| SE | 11441 | DHX8 | DHX8 | chr17 | + | 43520750 | 43520879 | 43520129 | 43520267 | 43521368 | 43521565 | 15 | 5 | 16 | 0 | 1 | 0 | 0 | 1 | 0 | 0 | 98 | 49 | 1 | 1 | 1 | 0.714 | 1 | NA | 1 | -0.095 |
| SE | 11445 | MAP2K3 | MAP2K3 | chr17 | + | 21303182 | 21303234 | 21302142 | 21302259 | 21304425 | 21304553 | 57 | 30 | 11 | 0 | 2 | 0 | 0 | 14 | 0 | 0 | 98 | 49 | 1 | 1 | 1 | 0.882 | 1 | NA | 1 | -0.039 |
| SE | 11446 | MAP2K3 | MAP2K3 | chr17 | + | 21312141 | 21312281 | 21305050 | 21305128 | 21313491 | 21313537 | 31 | 9 | 2 | 1 | 0 | 0 | 1 | 10 | 0 | 0 | 98 | 49 | 1 | 1 | 0.939 | 1 | 1 | 1 | 1 | -0.02 |
| SE | 11453 | PIGS | PIGS | chr17 | - | 28561421 | 28561629 | 28560048 | 28560191 | 28563430 | 28563522 | 11 | 4 | 2 | 0 | 1 | 0 | 1 | 4 | 0 | 0 | 98 | 49 | 1 | 1 | 1 | 0.667 | 1 | 1 | 1 | -0.111 |
| SE | 11454 | NCK2 | NCK2 | chr2 | + | 105881327 | 105882049 | 105855047 | 105855289 | 105892981 | 105894274 | 11 | 4 | 2 | 1 | 1 | 0 | 0 | 1 | 0 | 0 | 98 | 49 | 1 | 1 | 0.846 | 0.667 | 1 | NA | 1 | -0.162 |
| SE | 11456 | KIAA0100 | KIAA0100 | chr17 | - | 28615663 | 28615798 | 28614439 | 28615246 | 28616095 | 28616197 | 13 | 1 | 4 | 1 | 0 | 0 | 0 | 5 | 0 | 0 | 98 | 49 | 1 | 1 | 0.867 | 1 | 1 | NA | 1 | -0.044 |
| SE | 11457 | KIAA0100 | KIAA0100 | chr17 | - | 28616888 | 28616999 | 28616617 | 28616753 | 28617231 | 28617318 | 23 | 7 | 4 | 0 | 0 | 0 | 0 | 7 | 0 | 1 | 98 | 49 | 1 | 1 | 1 | 1 | 1 | NA | 0.778 | 0.222 |
| SE | 11458 | KIAA0100 | KIAA0100 | chr17 | - | 28620508 | 28620630 | 28619859 | 28620015 | 28620987 | 28621170 | 3 | 7 | 1 | 1 | 0 | 0 | 0 | 3 | 0 | 0 | 98 | 49 | 1 | 1 | 0.6 | 1 | 1 | NA | 1 | -0.133 |
| SE | 11459 | KIAA0100 | KIAA0100 | chr17 | - | 28628264 | 28628559 | 28624232 | 28624390 | 28631478 | 28631681 | 20 | 5 | 0 | 0 | 1 | 1 | 0 | 7 | 0 | 1 | 98 | 49 | 1 | 1 | 1 | 0.714 | 0 | NA | 0.778 | -0.207 |
| SE | 11460 | KIAA0100 | KIAA0100 | chr17 | - | 28633282 | 28633401 | 28632941 | 28633162 | 28633548 | 28633751 | 5 | 1 | 0 | 2 | 0 | 0 | 0 | 4 | 0 | 0 | 98 | 49 | 1 | 1 | 0.556 | 1 | NA | NA | 1 | -0.222 |
| SE | 11461 | KIAA0100 | KIAA0100 | chr17 | - | 28641888 | 28642078 | 28640559 | 28640683 | 28642274 | 28642335 | 11 | 3 | 0 | 1 | 4 | 0 | 0 | 4 | 0 | 0 | 98 | 49 | 1 | 1 | 0.846 | 0.273 | NA | NA | 1 | -0.441 |
| SE | 11462 | KIAA0100 | KIAA0100 | chr17 | - | 28642274 | 28642335 | 28641888 | 28642078 | 28642868 | 28642987 | 14 | 3 | 0 | 1 | 0 | 0 | 0 | 5 | 0 | 0 | 98 | 49 | 1 | 1 | 0.875 | 1 | NA | NA | 1 | -0.062 |
| SE | 11479 | NFX1 | NFX1 | chr9 | + | 33351559 | 33351790 | 33347037 | 33347117 | 33352645 | 33352719 | 2 | 3 | 0 | 0 | 0 | 0 | 0 | 1 | 0 | 1 | 98 | 49 | 1 | 1 | 1 | 1 | NA | NA | 0.333 | 0.667 |
| SE | 11482 | BRD8 | BRD8 | chr5 | - | 138166517 | 138166610 | 138165827 | 138166108 | 138167933 | 138168078 | 2 | 0 | 2 | 1 | 0 | 0 | 0 | 1 | 0 | 0 | 98 | 49 | 1 | 1 | 0.5 | NA | 1 | NA | 1 | -0.25 |
| SE | 11483 | BRD8 | BRD8 | chr5 | - | 138166517 | 138166727 | 138165827 | 138166108 | 138167933 | 138168078 | 2 | 0 | 2 | 1 | 0 | 0 | 0 | 1 | 0 | 0 | 98 | 49 | 1 | 1 | 0.5 | NA | 1 | NA | 1 | -0.25 |
| SE | 11486 | EGLN1 | EGLN1 | chr1 | - | 231367568 | 231367636 | 231363750 | 231366475 | 231370561 | 231370698 | 57 | 42 | 33 | 6 | 4 | 0 | 0 | 7 | 0 | 1 | 98 | 49 | 1 | 1 | 0.826 | 0.84 | 1 | NA | 0.778 | 0.111 |
| SE | 11488 | FLOT2 | FLOT2 | chr17 | - | 28884224 | 28884315 | 28883107 | 28883231 | 28888944 | 28889026 | 57 | 43 | 33 | 6 | 6 | 2 | 0 | 13 | 0 | 2 | 98 | 49 | 1 | 1 | 0.826 | 0.782 | 0.892 | NA | 0.765 | 0.068 |
| SE | 11489 | FLOT2 | FLOT2 | chr17 | - | 28884224 | 28884315 | 28883107 | 28883231 | 28897525 | 28897697 | 33 | 28 | 26 | 1 | 0 | 0 | 0 | 10 | 0 | 0 | 98 | 49 | 1 | 1 | 0.943 | 1 | 1 | NA | 1 | -0.019 |
| SE | 11490 | FLOT2 | FLOT2 | chr17 | - | 28888944 | 28889026 | 28883107 | 28883231 | 28897525 | 28897697 | 26 | 7 | 14 | 1 | 0 | 0 | 1 | 4 | 0 | 0 | 98 | 49 | 1 | 1 | 0.929 | 1 | 1 | 1 | 1 | -0.024 |
| SE | 11491 | FLOT2 | FLOT2 | chr17 | - | 28888944 | 28889026 | 28884224 | 28884315 | 28897525 | 28897697 | 44 | 17 | 19 | 0 | 1 | 0 | 1 | 5 | 0 | 0 | 98 | 49 | 1 | 1 | 1 | 0.895 | 1 | 1 | 1 | -0.035 |
| SE | 11494 | DHRS13 | DHRS13 | chr17 | - | 28901492 | 28901616 | 28900989 | 28901301 | 28902582 | 28902701 | 8 | 9 | 17 | 1 | 4 | 0 | 0 | 5 | 0 | 3 | 98 | 49 | 1 | 1 | 0.8 | 0.529 | 1 | NA | 0.455 | 0.321 |
| SE | 11496 | SSH2 | SSH2 | chr17 | - | 29635967 | 29636802 | 29625938 | 29632931 | 29648143 | 29648344 | 140 | 48 | 23 | 2 | 0 | 0 | 2 | 21 | 0 | 0 | 98 | 49 | 1 | 1 | 0.972 | 1 | 1 | 1 | 1 | -0.009 |
| SE | 11497 | SSH2 | SSH2 | chr17 | - | 29648143 | 29648344 | 29625938 | 29632931 | 29650653 | 29650800 | 30 | 39 | 5 | 1 | 0 | 0 | 0 | 19 | 0 | 0 | 98 | 49 | 1 | 1 | 0.937 | 1 | 1 | NA | 1 | -0.021 |
| SE | 11499 | SSH2 | SSH2 | chr17 | - | 29695458 | 29695523 | 29684562 | 29684684 | 29702958 | 29703062 | 114 | 62 | 32 | 1 | 1 | 0 | 0 | 20 | 0 | 0 | 98 | 49 | 1 | 1 | 0.983 | 0.969 | 1 | NA | 1 | -0.016 |
| SE | 11500 | SSH2 | SSH2 | chr17 | - | 29702958 | 29703062 | 29695458 | 29695523 | 29761149 | 29761432 | 106 | 48 | 34 | 0 | 0 | 1 | 1 | 17 | 0 | 0 | 98 | 49 | 1 | 1 | 1 | 1 | 0.944 | 1 | 1 | -0.019 |
| SE | 11507 | CPD | CPD | chr17 | + | 30443801 | 30443971 | 30442307 | 30442450 | 30445690 | 30446020 | 63 | 33 | 11 | 0 | 0 | 1 | 0 | 1 | 0 | 0 | 98 | 49 | 1 | 1 | 1 | 1 | 0.846 | NA | 1 | -0.051 |

| | | | | | | | | | | | | | | | | | | | | | | | | | | | | | | | | |
|---|---|---|---|---|---|---|---|---|---|---|---|---|---|---|---|---|---|---|---|---|---|---|---|---|---|---|---|---|---|---|---|---|
| SE | 11508 | CPD | CPD | chr17 | + | 30449552 | 30449748 | 30445690 | 30446020 | 30451710 | 30451846 | 50 | 26 | 28 | 1 | 0 | 0 | 0 | 4 | 0 | 0 | 98 | 49 | 1 | 1 | 0.962 | 1 | 1 | NA | | 1 | -0.013 |
| SE | 11509 | CEP164 | CEP164 | chr11 | + | 117338565 | 117338668 | 117335604 | 117335680 | 117344165 | 117344277 | 4 | 3 | 1 | 1 | 0 | 0 | 0 | 3 | 0 | 0 | 98 | 49 | 1 | 1 | 0.667 | 1 | 1 | NA | | 1 | -0.111 |
| SE | 11512 | SBNO2 | SBNO2 | chr19 | - | 1111018 | 1111093 | 1110744 | 1110888 | 1111505 | 1111614 | 39 | 25 | 10 | 0 | 0 | 0 | 1 | 6 | 0 | 1 | 98 | 49 | 1 | 1 | 1 | 1 | 1 | | 1 | 0.75 | 0.125 |
| SE | 11514 | SBNO2 | SBNO2 | chr19 | - | 1113534 | 1113704 | 1112817 | 1112949 | 1114230 | 1114422 | 92 | 52 | 19 | 1 | 0 | 0 | 9 | 21 | 0 | 1 | 98 | 49 | 1 | 1 | 0.979 | 1 | 1 | | 1 | 0.913 | 0.036 |
| SE | 11516 | RNF135 | RNF135 | chr17 | + | 30987943 | 30988106 | 30984616 | 30984760 | 30997241 | 30997331 | 14 | 3 | 3 | 2 | 0 | 0 | 0 | 5 | 0 | 0 | 98 | 49 | 1 | 1 | 0.778 | 1 | 1 | NA | | 1 | -0.074 |
| SE | 11518 | NF1 | NF1 | chr17 | + | 31225094 | 31225250 | 31223443 | 31223567 | 31226434 | 31226684 | 14 | 2 | 0 | 1 | 0 | 0 | 0 | 1 | 0 | 0 | 98 | 49 | 1 | 1 | 0.875 | 1 | NA | NA | | 1 | -0.062 |
| SE | 11519 | NF1 | NF1 | chr17 | + | 31232699 | 31232881 | 31232072 | 31232189 | 31233001 | 31233213 | 5 | 4 | 1 | 1 | 0 | 0 | 0 | 2 | 0 | 0 | 98 | 49 | 1 | 1 | 0.714 | 1 | 1 | NA | | 1 | -0.095 |
| SE | 11520 | NF1 | NF1 | chr17 | + | 31252937 | 31253000 | 31248983 | 31249119 | 31258343 | 31258502 | 0 | 0 | 2 | 4 | 2 | 0 | 0 | 1 | 0 | 3 | 98 | 49 | 1 | 1 | 0 | 0 | 1 | NA | | 0.143 | 0.19 |
| SE | 11521 | CHFR | CHFR | chr12 | - | 132851617 | 132851734 | 132848640 | 132848724 | 132853430 | 132853573 | 9 | 3 | 4 | 1 | 0 | 0 | 0 | 3 | 0 | 0 | 98 | 49 | 1 | 1 | 0.818 | 1 | 1 | NA | | 1 | -0.061 |
| SE | 11522 | CHFR | CHFR | chr12 | - | 132851617 | 132851737 | 132848640 | 132848724 | 132853430 | 132853573 | 8 | 1 | 0 | 1 | 0 | 0 | 0 | 3 | 0 | 0 | 98 | 49 | 1 | 1 | 0.8 | 1 | NA | NA | | 1 | -0.1 |
| SE | 11524 | CHFR | CHFR | chr12 | - | 132869618 | 132869798 | 132861466 | 132861634 | 132870723 | 132870783 | 6 | 6 | 1 | 1 | 0 | 0 | 0 | 0 | 0 | 2 | 98 | 49 | 1 | 1 | 0.75 | 1 | 1 | NA | | 0 | 0.917 |
| SE | 11525 | CHFR | CHFR | chr12 | - | 132869618 | 132869834 | 132861466 | 132861634 | 132870723 | 132870783 | 4 | 4 | 1 | 1 | 0 | 0 | 0 | 0 | 0 | 2 | 98 | 49 | 1 | 1 | 0.667 | 1 | 1 | NA | | 0 | 0.889 |
| SE | 11527 | TCF7L2 | TCF7L2 | chr10 | + | 112964555 | 112964624 | 112951482 | 112951607 | 113040024 | 113040126 | 1 | 1 | 0 | 7 | 1 | 0 | 0 | 0 | 0 | 2 | 98 | 49 | 1 | 1 | 0.067 | 0.333 | NA | NA | | 0 | 0.2 |
| SE | 11538 | EVI2A | EVI2A | chr17 | - | 31320372 | 31320513 | 31316409 | 31319023 | 31321494 | 31321749 | 2 | 1 | 0 | 46 | 21 | 7 | 0 | 0 | 0 | 4 | 98 | 49 | 1 | 1 | 0.021 | 0.023 | 0 | NA | | 0 | 0.015 |
| SE | 11548 | TRNAU1AP | TRNAU1AP | chr1 | + | 28561112 | 28561260 | 28560632 | 28560732 | 28561345 | 28561398 | 2 | 1 | 0 | 1 | 2 | 0 | 0 | 1 | 0 | 1 | 98 | 49 | 1 | 1 | 0.5 | 0.2 | NA | NA | | 0.333 | 0.017 |
| SE | 11549 | NATD1 | NATD1 | chr17 | - | 21244105 | 21244224 | 21238870 | 21243429 | 21253158 | 21253266 | 7 | 2 | 0 | 2 | 0 | 0 | 0 | 1 | 0 | 0 | 98 | 49 | 1 | 1 | 0.636 | 1 | NA | NA | | 1 | -0.182 |
| SE | 11550 | CCDC69 | CCDC69 | chr5 | - | 151198996 | 151199084 | 151187385 | 151187459 | 151201581 | 151201688 | 27 | 17 | 1 | 0 | 0 | 1 | 0 | 5 | 0 | 0 | 98 | 49 | 1 | 1 | 1 | 1 | 0.333 | NA | | 1 | -0.222 |
| SE | 11551 | RFFL | RFFL | chr17 | - | 35017522 | 35017606 | 35016369 | 35016580 | 35021370 | 35021781 | 12 | 8 | 6 | 1 | 0 | 0 | 0 | 3 | 0 | 0 | 98 | 49 | 1 | 1 | 0.857 | 1 | 1 | NA | | 1 | -0.048 |
| SE | 11552 | RFFL | RFFL | chr17 | - | 35021370 | 35021781 | 35017522 | 35017606 | 35026373 | 35026561 | 10 | 5 | 2 | 5 | 0 | 1 | 0 | 6 | 0 | 1 | 98 | 49 | 1 | 1 | 0.5 | 1 | 0.5 | NA | | 0.75 | -0.083 |
| SE | 11554 | WDFY2 | WDFY2 | chr13 | + | 51660595 | 51660663 | 51584347 | 51584824 | 51675169 | 51675243 | 9 | 6 | 2 | 0 | 2 | 0 | 0 | 2 | 0 | 0 | 98 | 49 | 1 | 1 | 1 | 0.6 | 1 | NA | | 1 | -0.133 |
| SE | 11555 | WDFY2 | WDFY2 | chr13 | + | 51719197 | 51719348 | 51703595 | 51703650 | 51727677 | 51727790 | 20 | 3 | 3 | 0 | 0 | 0 | 0 | 3 | 0 | 1 | 98 | 49 | 1 | 1 | 1 | 1 | 1 | NA | | 0.6 | 0.4 |
| SE | 11556 | WDFY2 | WDFY2 | chr13 | + | 51739048 | 51739175 | 51727677 | 51727790 | 51751309 | 51751415 | 7 | 7 | 2 | 1 | 0 | 0 | 0 | 1 | 0 | 0 | 98 | 49 | 1 | 1 | 0.778 | 1 | 1 | NA | | 1 | -0.074 |
| SE | 11568 | PLB1 | PLB1 | chr2 | + | 28605852 | 28605948 | 28604654 | 28604759 | 28606495 | 28606567 | 13 | 2 | 2 | 2 | 0 | 0 | 0 | 1 | 0 | 0 | 98 | 49 | 1 | 1 | 0.765 | 1 | 1 | NA | | 1 | -0.078 |
| SE | 11569 | PLB1 | PLB1 | chr2 | + | 28620599 | 28620643 | 28620264 | 28620332 | 28620878 | 28620978 | 17 | 7 | 6 | 0 | 0 | 0 | 2 | 1 | 0 | 1 | 93 | 49 | 1 | 1 | 1 | 1 | 1 | | 1 | 0.345 | 0.328 |
| SE | 11570 | PLB1 | PLB1 | chr2 | + | 28640926 | 28641001 | 28632943 | 28633039 | 28642857 | 28643787 | 11 | 2 | 3 | 0 | 1 | 0 | 0 | 5 | 0 | 0 | 98 | 49 | 1 | 1 | 1 | 0.5 | 1 | NA | | 1 | -0.167 |
| SE | 11579 | LUCAT1 | LUCAT1 | chr5 | - | 91313077 | 91313229 | 91303872 | 91303971 | 91313637 | 91313777 | 6 | 1 | 5 | 1 | 0 | 0 | 2 | 3 | 0 | 1 | 98 | 49 | 1 | 1 | 0.75 | 1 | 1 | | 1 | 0.6 | 0.117 |
| SE | 11580 | LUCAT1 | LUCAT1 | chr5 | - | 91313077 | 91313229 | 91310913 | 91311032 | 91313637 | 91313777 | 7 | 1 | 5 | 0 | 1 | 0 | 2 | 4 | 0 | 0 | 98 | 49 | 1 | 1 | 1 | 0.333 | 1 | | 1 | 1 | -0.222 |
| SE | 11582 | PRDM8 | PRDM8 | chr4 | + | 80196962 | 80197216 | 80196351 | 80196601 | 80197624 | 80197763 | 2 | 0 | 1 | 0 | 2 | 0 | 0 | 2 | 0 | 0 | 98 | 49 | 1 | 1 | 1 | 0 | 1 | NA | | 1 | -0.333 |
| SE | 11601 | GGNBP2 | GGNBP2 | chr17 | + | 36545618 | 36545817 | 36544887 | 36545097 | 36554819 | 36554900 | 9 | 4 | 0 | 2 | 0 | 0 | 4 | 2 | 0 | 0 | 98 | 49 | 1 | 1 | 0.692 | 1 | NA | | 1 | 1 | -0.154 |
| SE | 11606 | P4HA1 | P4HA1 | chr10 | - | 73051052 | 73051249 | 73046924 | 73047101 | 73053350 | 73053590 | 6 | 2 | 0 | 0 | 1 | 0 | 0 | 1 | 0 | 0 | 98 | 49 | 1 | 1 | 1 | 0.5 | NA | NA | | 1 | -0.25 |
| SE | 11607 | P4HA1 | P4HA1 | chr10 | - | 73073730 | 73073827 | 73072028 | 73072180 | 73074807 | 73074915 | 5 | 0 | 4 | 1 | 0 | 0 | 0 | 1 | 0 | 0 | 98 | 49 | 1 | 1 | 0.714 | NA | 1 | NA | | 1 | -0.143 |
| SE | 11614 | ARHGAP12 | ARHGAP12 | chr10 | - | 31839304 | 31839319 | 31831738 | 31831800 | 31839636 | 31839711 | 1 | 0 | 0 | 0 | 2 | 0 | 0 | 2 | 0 | 0 | 64 | 49 | 1 | 1 | 1 | 0 | NA | NA | | 1 | -0.5 |
| SE | 11619 | MLLT6 | MLLT6 | chr17 | + | 38716366 | 38716981 | 38715611 | 38715828 | 38717431 | 38717613 | 16 | 1 | 6 | 1 | 0 | 0 | 2 | 4 | 0 | 0 | 98 | 49 | 1 | 1 | 0.889 | 1 | 1 | | 1 | 1 | -0.037 |
| SE | 11625 | CWC25 | CWC25 | chr17 | - | 38815624 | 38815741 | 38814860 | 38815097 | 38820900 | 38821073 | 0 | 3 | 0 | 18 | 20 | 0 | 0 | 0 | 1 | 0 | 98 | 49 | 1 | 1 | 0 | 0.07 | NA | | 0 | NA | 0.035 |
| SE | 11627 | LASP1 | LASP1 | chr17 | + | 38896901 | 38897071 | 38890419 | 38890504 | 38898411 | 38898519 | 3 | 0 | 0 | 67 | 19 | 16 | 0 | 0 | 0 | 7 | 98 | 49 | 1 | 1 | 0.022 | 0 | 0 | NA | | 0 | 0.007 |
| SE | 11628 | LASP1 | LASP1 | chr17 | + | 38915042 | 38915146 | 38914324 | 38914475 | 38918604 | 38921770 | 62 | 20 | 16 | 1 | 1 | 0 | 2 | 5 | 0 | 0 | 98 | 49 | 1 | 1 | 0.969 | 0.909 | 1 | | 1 | 1 | -0.041 |
| SE | 11629 | FKBP8 | FKBP8 | chr19 | - | 18533269 | 18533347 | 18532663 | 18532795 | 18537600 | 18537773 | 69 | 44 | 50 | 0 | 3 | 0 | 0 | 16 | 0 | 0 | 98 | 49 | 1 | 1 | 1 | 0.88 | 1 | NA | | 1 | -0.04 |
| SE | 11630 | FKBP8 | FKBP8 | chr19 | - | 18538215 | 18538433 | 18537600 | 18537773 | 18539370 | 18539459 | 110 | 48 | 32 | 2 | 0 | 0 | 4 | 20 | 0 | 0 | 98 | 49 | 1 | 1 | 0.965 | 1 | 1 | | 1 | 1 | -0.012 |
| SE | 11631 | FKBP8 | FKBP8 | chr19 | - | 18538215 | 18538436 | 18537600 | 18537773 | 18539370 | 18539459 | 111 | 64 | 47 | 2 | 0 | 0 | 6 | 17 | 0 | 0 | 98 | 49 | 1 | 1 | 0.965 | 1 | 1 | | 1 | 1 | -0.012 |
| SE | 11632 | FKBP8 | FKBP8 | chr19 | - | 18539370 | 18539459 | 18538215 | 18538436 | 18541678 | 18541995 | 42 | 29 | 29 | 0 | 1 | 0 | 2 | 2 | 0 | 0 | 98 | 49 | 1 | 1 | 1 | 0.935 | 1 | | 1 | 1 | -0.022 |
| SE | 11636 | FBXL20 | FBXL20 | chr17 | - | 39285477 | 39285573 | 39282728 | 39282855 | 39297126 | 39297195 | 20 | 3 | 3 | 0 | 1 | 0 | 0 | 1 | 0 | 0 | 98 | 49 | 1 | 1 | 1 | 0.6 | 1 | NA | | 1 | -0.133 |
| SE | 11637 | FBXL20 | FBXL20 | chr17 | - | 39301000 | 39301075 | 39298989 | 39299084 | 39303584 | 39303639 | 17 | 8 | 7 | 1 | 0 | 0 | 0 | 1 | 0 | 0 | 98 | 49 | 1 | 1 | 0.895 | 1 | 1 | NA | | 1 | -0.035 |
| SE | 11669 | PSMD3 | PSMD3 | chr17 | + | 39996182 | 39996338 | 39995423 | 39995527 | 39997329 | 39997380 | 10 | 1 | 2 | 1 | 0 | 0 | 0 | 5 | 0 | 0 | 98 | 49 | 1 | 1 | 0.833 | 1 | 1 | NA | | 1 | -0.056 |
| SE | 11670 | MSL1 | MSL1 | chr17 | + | 40131536 | 40131584 | 40129244 | 40129627 | 40132033 | 40132098 | 73 | 53 | 21 | 3 | 2 | 0 | 0 | 7 | 0 | 2 | 97 | 49 | 1 | 1 | 0.925 | 0.93 | 1 | NA | | 0.639 | 0.313 |
| SE | 11671 | UBXN1 | UBXN1 | chr11 | - | 62676812 | 62677005 | 62676497 | 62676639 | 62677517 | 62677634 | 8 | 6 | 10 | 1 | 0 | 0 | 0 | 11 | 0 | 0 | 98 | 49 | 1 | 1 | 0.8 | 1 | 1 | NA | | 1 | -0.067 |
| SE | 11672 | ARNT | ARNT | chr1 | - | 150842423 | 150842468 | 150839440 | 150839654 | 150846262 | 150846307 | 2 | 2 | 0 | 0 | 0 | 2 | 0 | 0 | 0 | 1 | 94 | 49 | 1 | 1 | 1 | 1 | 0 | NA | | 0 | 0.667 |
| SE | 11680 | TMEM70 | TMEM70 | chr8 | + | 73979078 | 73979171 | 73978755 | 73978861 | 73981154 | 73982783 | 0 | 4 | 0 | 2 | 1 | 2 | 0 | 1 | 0 | 1 | 98 | 49 | 1 | 1 | 0 | 0.667 | 0 | NA | | 0.333 | -0.111 |
| SE | 11687 | ZDHHC3 | ZDHHC3 | chr3 | - | 44929305 | 44929436 | 44915260 | 44926847 | 44932906 | 44932990 | 7 | 2 | 2 | 1 | 0 | 0 | 0 | 3 | 0 | 0 | 98 | 49 | 1 | 1 | 0.778 | 1 | 1 | NA | | 1 | -0.074 |
| SE | 11688 | ZDHHC3 | ZDHHC3 | chr3 | - | 44932906 | 44932990 | 44929305 | 44929436 | 44933117 | 44933199 | 2 | 1 | 1 | 10 | 1 | 0 | 0 | 2 | 0 | 0 | 98 | 49 | 1 | 1 | 0.091 | 0.333 | 1 | NA | | 1 | -0.525 |
| SE | 11690 | DNAL1 | DNAL1 | chr14 | + | 73661986 | 73662042 | 73658846 | 73658956 | 73671541 | 73671597 | 1 | 0 | 0 | 1 | 0 | 0 | 0 | 1 | 0 | 0 | 98 | 49 | 1 | 1 | 0.333 | NA | NA | NA | | 1 | -0.667 |
| SE | 11691 | SMARCA5 | SMARCA5 | chr4 | + | 143527867 | 143528023 | 143526280 | 143526460 | 143528582 | 143528714 | 29 | 8 | 4 | 1 | 0 | 0 | 0 | 4 | 0 | 0 | 98 | 49 | 1 | 1 | 0.935 | 1 | 1 | NA | | 1 | -0.022 |
| SE | 11692 | SMARCA5 | SMARCA5 | chr4 | + | 143540362 | 143540495 | 143538785 | 143538938 | 143543508 | 143543657 | 15 | 1 | 1 | 1 | 0 | 0 | 0 | 1 | 0 | 1 | 98 | 49 | 1 | 1 | 0.882 | 1 | 1 | NA | | 0.333 | 0.628 |
| SE | 11693 | SMARCA5 | SMARCA5 | chr4 | + | 143545924 | 143546047 | 143545469 | 143545583 | 143546775 | 143546908 | 9 | 3 | 2 | 0 | 1 | 0 | 0 | 3 | 0 | 0 | 98 | 49 | 1 | 1 | 1 | 0.6 | 1 | NA | | 1 | -0.133 |
| SE | 11698 | DNAJC7 | DNAJC7 | chr17 | - | 41982254 | 41982401 | 41981854 | 41982007 | 41983562 | 41983636 | 4 | 2 | 2 | 0 | 2 | 0 | 0 | 1 | 0 | 0 | 98 | 49 | 1 | 1 | 1 | 0.333 | 1 | NA | | 1 | -0.222 |
| SE | 11699 | DNAJC7 | DNAJC7 | chr17 | - | 41989403 | 41989557 | 41988731 | 41988896 | 41990263 | 41990382 | 16 | 7 | 0 | 0 | 1 | 0 | 0 | 1 | 0 | 0 | 98 | 49 | 1 | 1 | 1 | 0.778 | NA | NA | | 1 | -0.111 |
| SE | 11702 | STAT5B | STAT5B | chr17 | - | 42218722 | 42218878 | 42218150 | 42218330 | 42219311 | 42219463 | 45 | 21 | 22 | 1 | 0 | 0 | 1 | 4 | 0 | 0 | 98 | 49 | 1 | 1 | 0.957 | 1 | 1 | | 1 | 1 | -0.014 |
| SE | 11705 | STAT3 | STAT3 | chr17 | - | 42324710 | 42324846 | 42323572 | 42323625 | 42324962 | 42325061 | 97 | 43 | 40 | 1 | 0 | 0 | 1 | 28 | 0 | 0 | 98 | 49 | 1 | 1 | 0.98 | 1 | 1 | | 1 | 1 | -0.007 |
| SE | 11706 | STAT3 | STAT3 | chr17 | - | 42324710 | 42324846 | 42323572 | 42323625 | 42326115 | 42326199 | 49 | 25 | 19 | 0 | 0 | 0 | 1 | 17 | 0 | 1 | 98 | 49 | 1 | 1 | 1 | 1 | 1 | | 1 | 0.895 | 0.052 |
| SE | 11707 | STAT3 | STAT3 | chr17 | - | 42324962 | 42325061 | 42323572 | 42323625 | 42326115 | 42326199 | 27 | 18 | 26 | 0 | 0 | 0 | 0 | 11 | 0 | 1 | 98 | 49 | 1 | 1 | 1 | 1 | 1 | NA | | 0.846 | 0.154 |
| SE | 11708 | STAT3 | STAT3 | chr17 | - | 42324962 | 42325061 | 42324710 | 42324846 | 42326115 | 42326199 | 75 | 36 | 47 | 1 | 0 | 0 | 0 | 22 | 0 | 0 | 98 | 49 | 1 | 1 | 0.974 | 1 | 1 | NA | | 1 | -0.009 |
| SE | 11709 | STAT3 | STAT3 | chr17 | - | 42348388 | 42348517 | 42346568 | 42346713 | 42388278 | 42388495 | 68 | 44 | 19 | 0 | 0 | 2 | 1 | 28 | 0 | 0 | 98 | 49 | 1 | 1 | 1 | 1 | 0.826 | | 1 | 1 | -0.058 |
| SE | 11710 | STAT3 | STAT3 | chr17 | - | 42348388 | 42348539 | 42346568 | 42346713 | 42388278 | 42388387 | 62 | 40 | 21 | 0 | 0 | 2 | 2 | 29 | 0 | 0 | 98 | 49 | 1 | 1 | 1 | 1 | 0.84 | | 1 | 1 | -0.053 |
| SE | 11711 | ATP6V0A1 | ATP6V0A1 | chr17 | + | 42468009 | 42468107 | 42466428 | 42466507 | 42470089 | 42470218 | 19 | 13 | 2 | 1 | 0 | 0 | 0 | 12 | 0 | 0 | 98 | 49 | 1 | 1 | 0.905 | 1 | 1 | NA | | 1 | -0.032 |
| SE | 11715 | COASY | COASY | chr17 | + | 42563960 | 42564175 | 42562073 | 42563322 | 42564445 | 42564577 | 4 | 6 | 3 | 0 | 1 | 0 | 0 | 2 | 0 | 0 | 98 | 49 | 1 | 1 | 1 | 0.75 | 1 | NA | | 1 | -0.083 |
| SE | 11716 | MLX | MLX | chr17 | + | 42568469 | 42568559 | 42567618 | 42567655 | 42568836 | 42568943 | 2 | 1 | 0 | 2 | 2 | 0 | 1 | 0 | 0 | 0 | 98 | 49 | 1 | 1 | 0.333 | 0.2 | NA | | 1 | NA | -0.734 |
| SE | 11721 | EZH1 | EZH1 | chr17 | - | 42713208 | 42713389 | 42712288 | 42712485 | 42717975 | 42718067 | 27 | 4 | 1 | 0 | 0 | 1 | 0 | 4 | 0 | 0 | 98 | 49 | 1 | 1 | 1 | 1 | 0.333 | NA | | 1 | -0.222 |
| SE | 11722 | EZH1 | EZH1 | chr17 | - | 42724304 | 42724424 | 42722794 | 42722915 | 42727634 | 42727763 | 33 | 7 | 3 | 2 | 3 | 0 | 0 | 11 | 0 | 1 | 98 | 49 | 1 | 1 | 0.892 | 0.538 | 1 | NA | | 0.846 | -0.036 |
| SE | 11724 | BECN1 | BECN1 | chr17 | - | 42811654 | 42811797 | 42810131 | 42810928 | 42813947 | 42814008 | 13 | 3 | 6 | 0 | 0 | 2 | 0 | 5 | 0 | 0 | 98 | 49 | 1 | 1 | 1 | 1 | 0.6 | NA | | 1 | -0.133 |
| SE | 11725 | BECN1 | BECN1 | chr17 | - | 42813947 | 42814008 | 42810131 | 42810928 | 42814523 | 42814673 | 5 | 4 | 7 | 3 | 1 | 0 | 0 | 1 | 0 | 0 | 98 | 49 | 1 | 1 | 0.455 | 0.667 | 1 | NA | | 1 | -0.293 |
| SE | 11733 | NBR1 | NBR1 | chr17 | + | 43189587 | 43189802 | 43189041 | 43189119 | 43190608 | 43190776 | 24 | 7 | 14 | 0 | 2 | 0 | 0 | 9 | 0 | 0 | 98 | 49 | 1 | 1 | 1 | 0.636 | 1 | NA | | 1 | -0.121 |
| SE | 11734 | NBR1 | NBR1 | chr17 | + | 43191371 | 43191581 | 43190608 | 43190776 | 43193093 | 43193253 | 28 | 8 | 9 | 1 | 0 | 0 | 1 | 8 | 0 | 0 | 98 | 49 | 1 | 1 | 0.933 | 1 | 1 | | 1 | 1 | -0.022 |
| SE | 11735 | NBR1 | NBR1 | chr17 | + | 43196941 | 43197106 | 43196480 | 43196591 | 43200166 | 43200608 | 45 | 20 | 8 | 0 | 1 | 0 | 0 | 3 | 0 | 1 | 98 | 49 | 1 | 1 | 1 | 0.909 | 1 | NA | | 0.6 | 0.37 |
| SE | 11736 | NBR1 | NBR1 | chr17 | + | 43201685 | 43201780 | 43200166 | 43200608 | 43202654 | 43202712 | 28 | 13 | 10 | 0 | 1 | 0 | 0 | 10 | 0 | 0 | 98 | 49 | 1 | 1 | 1 | 0.867 | 1 | NA | | 1 | -0.044 |
| SE | 11747 | KDM2A | KDM2A | chr11 | + | 67121233 | 67121358 | 67119268 | 67120049 | 67180078 | 67180217 | 57 | 13 | 19 | 1 | 0 | 0 | 0 | 11 | 0 | 0 | 98 | 49 | 1 | 1 | 0.966 | 1 | 1 | NA | | 1 | -0.011 |
| SE | 11748 | KDM2A | KDM2A | chr11 | + | 67181319 | 67181398 | 67180078 | 67180217 | 67181845 | 67181892 | 39 | 17 | 7 | 3 | 3 | 0 | 0 | 7 | 0 | 1 | 98 | 49 | 1 | 1 | 0.867 | 0.739 | 1 | NA | | 0.778 | 0.091 |
| SE | 11749 | KDM2A | KDM2A | chr11 | + | 67215855 | 67215949 | 67215339 | 67215446 | 67217730 | 67217884 | 45 | 18 | 9 | 2 | 6 | 0 | 0 | 5 | 0 | 1 | 98 | 49 | 1 | 1 | 0.918 | 0.6 | 1 | NA | | 0.714 | 0.125 |
| SE | 11750 | KDM2A | KDM2A | chr11 | + | 67250085 | 67250346 | 67248280 | 67248370 | 67252693 | 67252857 | 27 | 12 | 2 | 0 | 1 | 0 | 1 | 5 | 0 | 0 | 98 | 49 | 1 | 1 | 1 | 0.857 | 1 | | 1 | 1 | -0.048 |
| SE | 11751 | KDM2A | KDM2A | chr11 | + | 67250085 | 67250798 | 67248280 | 67248370 | 67252693 | 67252857 | 41 | 18 | 4 | 0 | 1 | 0 | 1 | 11 | 0 | 0 | 98 | 49 | 1 | 1 | 1 | 0.9 | 1 | | 1 | 1 | -0.033 |
| SE | 11756 | GRN | GRN | chr17 | + | 44351549 | 44351795 | 44351362 | 44351460 | 44352014 | 44352248 | 119 | 44 | 41 | 1 | 0 | 0 | 7 | 58 | 0 | 0 | 98 | 49 | 1 | 1 | 0.983 | 1 | 1 | | 1 | 1 | -0.006 |
| SE | 11761 | EFTUD2 | EFTUD2 | chr17 | - | 44868286 | 44868350 | 44867806 | 44867897 | 44872445 | 44872570 | 10 | 2 | 0 | 1 | 0 | 0 | 1 | 4 | 0 | 0 | 98 | 49 | 1 | 1 | 0.833 | 1 | NA | | 1 | 1 | -0.084 |
| SE | 11776 | FMNL1 | FMNL1 | chr17 | + | 45245252 | 45245416 | 45244978 | 45245108 | 45245877 | 45245973 | 31 | 11 | 19 | 2 | 0 | 0 | 3 | 24 | 0 | 0 | 98 | 49 | 1 | 1 | 0.886 | 1 | 1 | | 1 | 1 | -0.038 |
| SE | 11786 | PPP6R1 | PPP6R1 | chr19 | - | 55232074 | 55232211 | 55231801 | 55231982 | 55236642 | 55236821 | 23 | 18 | 3 | 2 | 0 | 0 | 2 | 8 | 0 | 0 | 98 | 49 | 1 | 1 | 0.852 | 1 | 1 | | 1 | 1 | -0.049 |
| SE | 11811 | ANKRD17 | ANKRD17 | chr4 | - | 73147240 | 73147432 | 73146763 | 73146873 | 73148812 | 73149050 | 16 | 4 | 2 | 1 | 0 | 0 | 0 | 10 | 0 | 0 | 98 | 49 | 1 | 1 | 0.889 | 1 | 1 | NA | | 1 | -0.037 |
| SE | 11812 | PTPN6 | PTPN6 | chr12 | + | 6954804 | 6954994 | 6951982 | 6952177 | 6955371 | 6955485 | 49 | 28 | 32 | 1 | 3 | 0 | 0 | 15 | 0 | 0 | 98 | 49 | 1 | 1 | 0.961 | 0.824 | 1 | NA | | 1 | -0.072 |
| SE | 11813 | PTPN6 | PTPN6 | chr12 | + | 6955150 | 6955267 | 6954804 | 6954994 | 6955371 | 6955485 | 94 | 59 | 30 | 1 | 0 | 0 | 2 | 28 | 0 | 1 | 98 | 49 | 1 | 1 | 0.979 | 1 | 1 | | 1 | 0.933 | 0.026 |
| SE | 11814 | PTPN6 | PTPN6 | chr12 | + | 6956418 | 6956568 | 6956141 | 6956221 | 6959926 | 6959994 | 176 | 99 | 46 | 1 | 0 | 0 | 6 | 29 | 0 | 0 | 98 | 49 | 1 | 1 | 0.989 | 1 | 1 | | 1 | 1 | -0.004 |
| SE | 11815 | PTPN6 | PTPN6 | chr12 | + | 6957653 | 6957785 | 6956418 | 6956568 | 6959926 | 6959994 | 118 | 62 | 31 | 1 | 0 | 0 | 8 | 22 | 0 | 0 | 98 | 49 | 1 | 1 | 0.983 | 1 | 1 | | 1 | 1 | -0.006 |
| SE | 11817 | PTPN6 | PTPN6 | chr12 | + | 6960343 | 6960435 | 6960087 | 6960239 | 6960805 | 6960945 | 171 | 77 | 46 | 1 | 0 | 0 | 3 | 32 | 0 | 0 | 98 | 49 | 1 | 1 | 0.988 | 1 | 1 | | 1 | 1 | -0.004 |
| SE | 11818 | PTPN6 | PTPN6 | chr12 | + | 6960662 | 6960945 | 6960343 | 6960435 | 6961125 | 6961316 | 104 | 37 | 13 | 8 | 3 | 0 | 0 | 11 | 0 | 1 | 98 | 49 | 1 | 1 | 0.867 | 0.86 | 1 | NA | | 0.846 | 0.063 |
| SE | 11820 | NPEPPS | NPEPPS | chr17 | + | 47586147 | 47586244 | 47585499 | 47585700 | 47586364 | 47586398 | 16 | 13 | 1 | 1 | 0 | 0 | 0 | 6 | 0 | 1 | 98 | 49 | 1 | 1 | 0.889 | 1 | 1 | NA | | 0.75 | 0.213 |
| SE | 11828 | UTP15 | UTP15 | chr5 | + | 73572488 | 73572624 | 73570585 | 73570711 | 73576951 | 73577036 | 2 | 0 | 0 | 1 | 0 | 0 | 0 | 1 | 0 | 0 | 98 | 49 | 1 | 1 | 0.5 | NA | NA | NA | | 1 | -0.5 |
| SE | 11831 | HK3 | HK3 | chr5 | - | 176883769 | 176883869 | 176881943 | 176882127 | 176884038 | 176884134 | 26 | 20 | 11 | 2 | 0 | 0 | 1 | 12 | 0 | 0 | 98 | 49 | 1 | 1 | 0.867 | 1 | 1 | | 1 | 1 | -0.044 |
| SE | 11832 | HK3 | HK3 | chr5 | - | 176887200 | 176887337 | 176887001 | 176887121 | 176887450 | 176887746 | 34 | 31 | 3 | 0 | 0 | 0 | 10 | 17 | 1 | 0 | 98 | 49 | 1 | 1 | 1 | 1 | 1 | | 0.833 | 1 | 0.084 |
| SE | 11843 | ST3GAL6 | ST3GAL6 | chr3 | + | 98791840 | 98791993 | 98773919 | 98773983 | 98793674 | 98795845 | 10 | 0 | 0 | 3 | 0 | 0 | 0 | 2 | 0 | 0 | 98 | 49 | 1 | 1 | 0.625 | NA | NA | NA | | 1 | -0.375 |
| SE | 11846 | CALCOCO2 | CALCOCO2 | chr17 | + | 48846442 | 48846514 | 48841697 | 48841887 | 48848063 | 48848166 | 0 | 0 | 0 | 10 | 8 | 0 | 0 | 1 | 1 | 5 | 98 | 49 | 1 | 1 | 0 | 0 | NA | | 0 | 0.091 | -0.045 |
| SE | 11848 | CALCOCO2 | CALCOCO2 | chr17 | + | 48851088 | 48851177 | 48849251 | 48849377 | 48851558 | 48851628 | 33 | 14 | 3 | 0 | 3 | 0 | 0 | 2 | 0 | 0 | 98 | 49 | 1 | 1 | 1 | 0.7 | 1 | NA | | 1 | -0.1 |
| SE | 11849 | CALCOCO2 | CALCOCO2 | chr17 | + | 48852505 | 48852628 | 48851558 | 48851628 | 48852925 | 48853012 | 32 | 16 | 10 | 1 | 1 | 0 | 0 | 3 | 0 | 0 | 98 | 49 | 1 | 1 | 0.941 | 0.889 | 1 | NA | | 1 | -0.057 |
| SE | 11850 | HMBOX1 | HMBOX1 | chr8 | + | 28980070 | 28980156 | 28970042 | 28970519 | 29009071 | 29009182 | 11 | 3 | 0 | 2 | 0 | 0 | 0 | 2 | 0 | 0 | 98 | 49 | 1 | 1 | 0.733 | 1 | NA | NA | | 1 | -0.133 |
| SE | 11863 | KAT7 | KAT7 | chr17 | + | 49815802 | 49815913 | 49811475 | 49811574 | 49817819 | 49818011 | 17 | 11 | 1 | 1 | 0 | 0 | 0 | 1 | 0 | 0 | 98 | 49 | 1 | 1 | 0.895 | 1 | 1 | NA | | 1 | -0.035 |
| SE | 11864 | KAT7 | KAT7 | chr17 | + | 49826692 | 49826799 | 49825999 | 49826146 | 49827400 | 49829096 | 7 | 6 | 0 | 1 | 0 | 0 | 2 | 1 | 0 | 0 | 98 | 49 | 1 | 1 | 0.778 | 1 | NA | | 1 | 1 | -0.111 |
| SE | 11879 | SPAG9 | SPAG9 | chr17 | - | 50975862 | 50975901 | 50974770 | 50974947 | 50977107 | 50977221 | 4 | 0 | 0 | 8 | 1 | 1 | 0 | 0 | 0 | 2 | 88 | 49 | 1 | 1 | 0.218 | 0 | 0 | NA | | 0 | 0.073 |
| SE | 11880 | SPAG9 | SPAG9 | chr17 | - | 50984922 | 50984990 | 50982523 | 50982672 | 50985697 | 50985778 | 68 | 29 | 6 | 1 | 1 | 0 | 1 | 8 | 0 | 1 | 98 | 49 | 1 | 1 | 0.971 | 0.935 | 1 | | 1 | 0.8 | 0.069 |
| SE | 11881 | SPAG9 | SPAG9 | chr17 | - | 50984922 | 50984990 | 50982523 | 50982672 | 50987111 | 50987237 | 27 | 19 | 5 | 0 | 0 | 0 | 1 | 3 | 0 | 1 | 98 | 49 | 1 | 1 | 1 | 1 | 1 | | 1 | 0.6 | 0.2 |
| SE | 11882 | SPAG9 | SPAG9 | chr17 | - | 50985697 | 50985778 | 50982523 | 50982672 | 50987111 | 50987237 | 21 | 10 | 4 | 0 | 0 | 0 | 0 | 7 | 0 | 1 | 98 | 49 | 1 | 1 | 1 | 1 | 1 | NA | | 0.778 | 0.222 |
| SE | 11883 | SPAG9 | SPAG9 | chr17 | - | 50985697 | 50985778 | 50984922 | 50984990 | 50987111 | 50987237 | 62 | 19 | 5 | 1 | 0 | 0 | 0 | 11 | 0 | 0 | 98 | 49 | 1 | 1 | 0.969 | 1 | 1 | NA | | 1 | -0.01 |
| SE | 11884 | SPAG9 | SPAG9 | chr17 | - | 50996564 | 50996694 | 50995443 | 50995533 | 50998443 | 50998617 | 25 | 10 | 7 | 0 | 0 | 4 | 0 | 8 | 0 | 0 | 98 | 49 | 1 | 1 | 1 | 1 | 0.467 | NA | | 1 | -0.178 |
| SE | 11886 | SPAG9 | SPAG9 | chr17 | - | 51014231 | 51014353 | 51007268 | 51007326 | 51020158 | 51020258 | 111 | 61 | 27 | 0 | 0 | 0 | 0 | 8 | 1 | 0 | 98 | 49 | 1 | 1 | 1 | 1 | 1 | | 0 | 1 | 0.5 |
| SE | 11887 | SPAG9 | SPAG9 | chr17 | - | 51031680 | 51031722 | 51021157 | 51021365 | 51041500 | 51041651 | 17 | 6 | 3 | 17 | 15 | 7 | 0 | 0 | 0 | 8 | 91 | 49 | 1 | 1 | 0.35 | 0.177 | 0.188 | NA | | 0 | 0.238 |
| SE | 11888 | KIAA0319L | KIAA0319L | chr1 | - | 35442245 | 35442336 | 35441046 | 35441138 | 35442905 | 35443028 | 10 | 2 | 6 | 3 | 1 | 1 | 0 | 3 | 0 | 3 | 98 | 49 | 1 | 1 | 0.625 | 0.5 | 0.75 | NA | | 0.333 | 0.292 |
| SE | 11889 | KIAA0319L | KIAA0319L | chr1 | - | 35451627 | 35451776 | 35450357 | 35450509 | 35453556 | 35453689 | 7 | 1 | 3 | 0 | 0 | 0 | 0 | 0 | 0 | 1 | 98 | 49 | 1 | 1 | 1 | 1 | 1 | NA | | 0 | 1 |
| SE | 11890 | KIAA0319L | KIAA0319L | chr1 | - | 35456012 | 35456241 | 35454361 | 35454485 | 35460304 | 35460437 | 14 | 2 | 0 | 1 | 0 | 0 | 0 | 2 | 0 | 0 | 98 | 49 | 1 | 1 | 0.875 | 1 | NA | NA | | 1 | -0.062 |
| SE | 11891 | KIAA0319L | KIAA0319L | chr1 | - | 35506611 | 35507135 | 35478965 | 35479212 | 35554349 | 35554519 | 10 | 3 | 0 | 1 | 0 | 0 | 0 | 1 | 0 | 0 | 98 | 49 | 1 | 1 | 0.833 | 1 | NA | NA | | 1 | -0.084 |
| SE | 11895 | ME2 | ME2 | chr18 | + | 50916167 | 50916243 | 50912800 | 50912950 | 50917346 | 50917508 | 7 | 5 | 9 | 1 | 0 | 0 | 0 | 2 | 0 | 0 | 98 | 49 | 1 | 1 | 0.778 | 1 | 1 | NA | | 1 | -0.074 |
| SE | 11897 | TRIM25 | TRIM25 | chr17 | - | 56895520 | 56895604 | 56895342 | 56895441 | 56895925 | 56895952 | 56 | 7 | 12 | 0 | 0 | 0 | 0 | 13 | 0 | 1 | 98 | 49 | 1 | 1 | 1 | 1 | 1 | NA | | 0.867 | 0.133 |
| SE | 11899 | DYNLL2 | DYNLL2 | chr17 | + | 58087081 | 58087222 | 58083418 | 58083683 | 58089141 | 58090257 | 18 | 11 | 9 | 2 | 0 | 0 | 1 | 4 | 0 | 0 | 98 | 49 | 1 | 1 | 0.818 | 1 | 1 | | 1 | 1 | -0.061 |
| SE | 11909 | SUPT4H1 | SUPT4H1 | chr17 | - | 58347528 | 58347584 | 58347187 | 58347241 | 58351401 | 58351508 | 19 | 26 | 25 | 1 | 0 | 0 | 0 | 7 | 0 | 0 | 98 | 49 | 1 | 1 | 0.905 | 1 | 1 | NA | | 1 | -0.032 |
| SE | 11913 | GAA | GAA | chr17 | + | 80112875 | 80113027 | 80112577 | 80112711 | 80113217 | 80113366 | 16 | 0 | 3 | 0 | 1 | 0 | 0 | 11 | 0 | 0 | 98 | 49 | 1 | 1 | 1 | 0 | 1 | NA | | 1 | -0.333 |
| SE | 11916 | DHX40 | DHX40 | chr17 | + | 59566626 | 59566794 | 59565524 | 59565783 | 59570517 | 59570663 | 20 | 3 | 4 | 0 | 0 | 0 | 0 | 1 | 0 | 1 | 98 | 49 | 1 | 1 | 1 | 1 | 1 | NA | | 0.333 | 0.667 |
| SE | 11920 | VMP1 | VMP1 | chr17 | + | 59735337 | 59735473 | 59731420 | 59731522 | 59737452 | 59737543 | 112 | 95 | 120 | 1 | 0 | 0 | 2 | 24 | 0 | 0 | 98 | 49 | 1 | 1 | 0.982 | 1 | 1 | | 1 | 1 | -0.006 |
| SE | 11922 | VMP1 | VMP1 | chr17 | + | 59738836 | 59738947 | 59735337 | 59735473 | 59764970 | 59765138 | 142 | 54 | 63 | 0 | 2 | 2 | 0 | 29 | 0 | 1 | 98 | 49 | 1 | 1 | 1 | 0.931 | 0.94 | NA | | 0.935 | 0.022 |
| SE | 11923 | VMP1 | VMP1 | chr17 | + | 59764970 | 59765138 | 59738836 | 59738947 | 59773753 | 59773885 | 299 | 134 | 131 | 0 | 2 | 0 | 0 | 47 | 0 | 0 | 98 | 49 | 1 | 1 | 1 | 0.971 | 1 | NA | | 1 | -0.01 |
| SE | 11924 | VMP1 | VMP1 | chr17 | + | 59773753 | 59773885 | 59738836 | 59738947 | 59808795 | 59808876 | 237 | 109 | 110 | 0 | 1 | 0 | 2 | 43 | 0 | 0 | 98 | 49 | 1 | 1 | 1 | 0.982 | 1 | | 1 | 1 | -0.006 |
| SE | 11925 | VMP1 | VMP1 | chr17 | + | 59808795 | 59808876 | 59773753 | 59773885 | 59811669 | 59811786 | 367 | 190 | 180 | 2 | 0 | 0 | 2 | 76 | 0 | 0 | 98 | 49 | 1 | 1 | 0.989 | 1 | 1 | | 1 | 1 | -0.004 |
| SE | 11926 | VMP1 | VMP1 | chr17 | + | 59811669 | 59811786 | 59773753 | 59773885 | 59817711 | 59817773 | 90 | 72 | 55 | 0 | 0 | 0 | 1 | 26 | 0 | 1 | 98 | 49 | 1 | 1 | 1 | 1 | 1 | | 1 | 0.929 | 0.035 |
| SE | 11927 | VMP1 | VMP1 | chr17 | + | 59817711 | 59817773 | 59811669 | 59811786 | 59838294 | 59838397 | 187 | 135 | 114 | 0 | 2 | 0 | 1 | 36 | 0 | 0 | 98 | 49 | 1 | 1 | 1 | 0.971 | 1 | | 1 | 1 | -0.01 |
| SE | 11934 | PPM1D | PPM1D | chr17 | + | 60656598 | 60656841 | 60647891 | 60648082 | 60662994 | 60666279 | 0 | 1 | 1 | 0 | 3 | 0 | 0 | 1 | 0 | 0 | 98 | 49 | 1 | 1 | NA | 0.143 | 1 | NA | | 1 | -0.428 |
| SE | 11936 | KIAA1257 | KIAA1257 | chr3 | - | 128988727 | 128988918 | 128987615 | 128987829 | 128993042 | 128993336 | 4 | 0 | 0 | 0 | 1 | 0 | 0 | 1 | 0 | 0 | 98 | 49 | 1 | 1 | 1 | 0 | NA | NA | | 1 | -0.5 |
| SE | 11937 | MED13 | MED13 | chr17 | - | 61955679 | 61955838 | 61955381 | 61955567 | 61956338 | 61956481 | 9 | 5 | 1 | 1 | 0 | 0 | 0 | 2 | 0 | 1 | 98 | 49 | 1 | 1 | 0.818 | 1 | 1 | NA | | 0.5 | 0.439 |
| SE | 11939 | MAP7D1 | MAP7D1 | chr1 | + | 36176198 | 36176581 | 36173363 | 36173478 | 36176696 | 36176842 | 29 | 7 | 6 | 0 | 1 | 0 | 8 | 13 | 0 | 0 | 98 | 49 | 1 | 1 | 1 | 0.778 | 1 | | 1 | 1 | -0.074 |
| SE | 11951 | MAP3K3 | MAP3K3 | chr17 | + | 63634708 | 63634801 | 63632680 | 63632802 | 63646033 | 63646074 | 44 | 24 | 10 | 11 | 2 | 0 | 0 | 4 | 0 | 2 | 98 | 49 | 1 | 1 | 0.667 | 0.857 | 1 | NA | | 0.5 | 0.341 |
| SE | 11952 | MAP3K3 | MAP3K3 | chr17 | + | 63666939 | 63667060 | 63657793 | 63657907 | 63681765 | 63681899 | 39 | 30 | 19 | 1 | 0 | 0 | 0 | 5 | 0 | 0 | 98 | 49 | 1 | 1 | 0.951 | 1 | 1 | NA | | 1 | -0.016 |
| SE | 11953 | MAP3K3 | MAP3K3 | chr17 | + | 63688788 | 63688881 | 63688526 | 63688594 | 63689543 | 63689735 | 57 | 21 | 13 | 1 | 0 | 0 | 0 | 13 | 0 | 0 | 98 | 49 | 1 | 1 | 0.966 | 1 | 1 | NA | | 1 | -0.011 |
| SE | 11954 | MAP3K3 | MAP3K3 | chr17 | + | 63692241 | 63692419 | 63691732 | 63691862 | 63693548 | 63696310 | 53 | 28 | 15 | 1 | 0 | 0 | 0 | 8 | 0 | 0 | 98 | 49 | 1 | 1 | 0.964 | 1 | 1 | NA | | 1 | -0.012 |
| SE | 11957 | CHD3 | CHD3 | chr17 | + | 7907600 | 7907702 | 7907352 | 7907488 | 7907893 | 7908019 | 8 | 0 | 2 | 2 | 1 | 1 | 0 | 0 | 0 | 2 | 98 | 49 | 1 | 1 | 0.667 | 0 | 0.5 | NA | | 0 | 0.389 |
| SE | 11958 | STRADA | STRADA | chr17 | - | 63707246 | 63707418 | 63706634 | 63706739 | 63710490 | 63710614 | 1 | 3 | 1 | 3 | 0 | 0 | 0 | 2 | 0 | 0 | 98 | 49 | 1 | 1 | 0.143 | 1 | 1 | NA | | 1 | -0.286 |
| SE | 11962 | SP140 | SP140 | chr2 | + | 230238212 | 230238381 | 230237082 | 230237260 | 230241403 | 230241487 | 31 | 6 | 0 | 2 | 0 | 0 | 1 | 9 | 0 | 0 | 98 | 49 | 1 | 1 | 0.886 | 1 | NA | | 1 | 1 | -0.057 |
| SE | 11968 | SP140 | SP140 | chr2 | + | 230292645 | 230292788 | 230290459 | 230290564 | 230294270 | 230294318 | 4 | 0 | 4 | 3 | 1 | 0 | 0 | 2 | 0 | 0 | 98 | 49 | 1 | 1 | 0.4 | 0 | 1 | NA | | 1 | -0.533 |
| SE | 11969 | SMARCD2 | SMARCD2 | chr17 | - | 63833297 | 63833420 | 63833068 | 63833170 | 63833586 | 63833722 | 17 | 3 | 7 | 1 | 0 | 0 | 0 | 3 | 0 | 0 | 98 | 49 | 1 | 1 | 0.895 | 1 | 1 | NA | | 1 | -0.035 |
| SE | 11980 | FTH1 | FTH1 | chr11 | - | 61964986 | 61965112 | 61964284 | 61964891 | 61965368 | 61965515 | 399 | 143 | 55 | 0 | 1 | 0 | 8 | 114 | 0 | 1 | 98 | 49 | 1 | 1 | 1 | 0.986 | 1 | | 1 | 0.983 | 0.004 |
| SE | 11995 | CEP95 | CEP95 | chr17 | + | 64516722 | 64516828 | 64514247 | 64514358 | 64519320 | 64519436 | 0 | 0 | 1 | 0 | 0 | 0 | 0 | 0 | 0 | 1 | 98 | 49 | 1 | 1 | NA | NA | 1 | NA | | 0 | 1 |
| SE | 11996 | PLEKHM1P | PLEKHM1P | chr17 | - | 64804829 | 64805100 | 64799999 | 64800917 | 64821766 | 64822052 | 18 | 11 | 6 | 1 | 0 | 0 | 0 | 3 | 0 | 0 | 98 | 49 | 1 | 1 | 0.9 | 1 | 1 | NA | | 1 | -0.033 |
| SE | 11998 | LRRC37A3 | LRRC37A3 | chr17 | - | 64889680 | 64889752 | 64869094 | 64869166 | 64892528 | 64892600 | 1 | 0 | 0 | 0 | 0 | 0 | 0 | 0 | 0 | 1 | 98 | 49 | 1 | 1 | 1 | NA | NA | NA | | 0 | 1 |
| SE | 12002 | ITGB2-AS1 | ITGB2-AS1 | chr21 | + | 44922912 | 44922983 | 44921034 | 44921327 | 44926867 | 44927004 | 2 | 0 | 4 | 2 | 0 | 0 | 0 | 2 | 0 | 0 | 98 | 49 | 1 | 1 | 0.333 | NA | 1 | NA | | 1 | -0.334 |
| SE | 12003 | ITGB2-AS1 | ITGB2-AS1 | chr21 | + | 44927624 | 44927663 | 44926867 | 44927004 | 44927853 | 44929680 | 0 | 0 | 1 | 0 | 1 | 0 | 0 | 0 | 0 | 1 | 88 | 49 | 1 | 1 | NA | 0 | 1 | NA | | 0 | 0.5 |
| SE | 12004 | ITGB2-AS1 | ITGB2-AS1 | chr21 | + | 44927624 | 44927667 | 44926867 | 44927004 | 44927853 | 44929680 | 2 | 0 | 1 | 0 | 1 | 0 | 0 | 0 | 0 | 1 | 92 | 49 | 1 | 1 | 1 | 0 | 1 | NA | | 0 | 0.667 |
| SE | 12006 | UMAD1 | UMAD1 | chr7 | + | 7676101 | 7676208 | 7673308 | 7673453 | 7801669 | 7801743 | 1 | 1 | 1 | 0 | 0 | 0 | 0 | 0 | 0 | 1 | 98 | 49 | 1 | 1 | 1 | 1 | 1 | NA | | 0 | 1 |
| SE | 12007 | HELZ | HELZ | chr17 | - | 67135969 | 67136198 | 67128650 | 67128855 | 67137930 | 67138114 | 12 | 1 | 0 | 2 | 0 | 0 | 0 | 4 | 0 | 0 | 98 | 49 | 1 | 1 | 0.75 | 1 | NA | NA | | 1 | -0.125 |
| SE | 12008 | HELZ | HELZ | chr17 | - | 67149866 | 67149985 | 67148568 | 67148714 | 67151045 | 67151224 | 13 | 7 | 0 | 0 | 2 | 0 | 0 | 5 | 0 | 0 | 98 | 49 | 1 | 1 | 1 | 0.636 | NA | NA | | 1 | -0.182 |
| SE | 12009 | HELZ | HELZ | chr17 | - | 67167462 | 67167796 | 67166477 | 67166608 | 67178658 | 67178926 | 16 | 7 | 5 | 0 | 1 | 0 | 0 | 2 | 0 | 0 | 98 | 49 | 1 | 1 | 1 | 0.778 | 1 | NA | | 1 | -0.074 |
| SE | 12010 | SYS1 | SYS1 | chr20 | + | 45363528 | 45363693 | 45363099 | 45363315 | 45365618 | 45365686 | 1 | 0 | 0 | 1 | 0 | 0 | 1 | 0 | 0 | 0 | 98 | 49 | 1 | 1 | 0.333 | NA | NA | | 1 | NA | -0.667 |
| SE | 12019 | KLHL2 | KLHL2 | chr4 | + | 165228806 | 165228913 | 165219933 | 165220059 | 165238777 | 165238899 | 8 | 2 | 8 | 3 | 4 | 3 | 0 | 4 | 0 | 0 | 98 | 49 | 1 | 1 | 0.571 | 0.2 | 0.571 | NA | | 1 | -0.553 |
| SE | 12023 | PRKAR1A | PRKAR1A | chr17 | + | 68514986 | 68515119 | 68512378 | 68512548 | 68515393 | 68515576 | 2 | 0 | 0 | 26 | 8 | 2 | 0 | 3 | 0 | 1 | 98 | 49 | 1 | 1 | 0.037 | 0 | 0 | NA | | 0.6 | -0.588 |
| SE | 12024 | PRKAR1A | PRKAR1A | chr17 | + | 68525753 | 68525912 | 68524911 | 68524958 | 68527839 | 68527900 | 98 | 42 | 39 | 0 | 0 | 4 | 0 | 5 | 0 | 0 | 98 | 49 | 1 | 1 | 1 | 1 | 0.83 | NA | | 1 | -0.057 |
| SE | 12025 | WIPI1 | WIPI1 | chr17 | - | 68434555 | 68434626 | 68433467 | 68433575 | 68435619 | 68435712 | 15 | 3 | 0 | 0 | 2 | 0 | 0 | 4 | 0 | 0 | 98 | 49 | 1 | 1 | 1 | 0.429 | NA | NA | | 1 | -0.285 |
| SE | 12026 | WIPI1 | WIPI1 | chr17 | - | 68452909 | 68452992 | 68450727 | 68450897 | 68457341 | 68457512 | 18 | 7 | 3 | 0 | 0 | 1 | 0 | 1 | 0 | 0 | 98 | 49 | 1 | 1 | 1 | 1 | 0.6 | NA | | 1 | -0.133 |

| | | | | | | | | | | | | | | | | | | | | | | | | | | | | | | | |
|---|---|---|---|---|---|---|---|---|---|---|---|---|---|---|---|---|---|---|---|---|---|---|---|---|---|---|---|---|---|---|---|
| SE | 12047 | FAM104A | FAM104A | chr17 | - | 73227164 | 73227264 | 73207352 | 73209768 | 73232085 | 73232394 | 5 | 0 | 0 | 1 | 0 | 0 | 0 | 1 | 1 | 0 | 98 | 49 | 1 | 1 | 0.714 | NA | NA | 0 | 1 | 0.214 |
| SE | 12048 | ANXA7 | ANXA7 | chr10 | - | 73379878 | 73379954 | 73378910 | 73379023 | 73380030 | 73380201 | 32 | 7 | 16 | 1 | 0 | 0 | 0 | 9 | 0 | 0 | 98 | 49 | 1 | 1 | 0.941 | 1 | 1 | NA | 1 | -0.02 |
| SE | 12049 | ANXA7 | ANXA7 | chr10 | - | 73396043 | 73396109 | 73388311 | 73388414 | 73396518 | 73396583 | 0 | 4 | 0 | 5 | 3 | 3 | 0 | 0 | 0 | 1 | 98 | 49 | 1 | 1 | 0 | 0.4 | 0 | NA | 0 | 0.133 |
| SE | 12050 | CTAGE5 | CTAGE5 | chr14 | + | 39291018 | 39291096 | 39279448 | 39279537 | 39293270 | 39293381 | 6 | 8 | 2 | 0 | 0 | 1 | 0 | 5 | 0 | 0 | 98 | 49 | 1 | 1 | 1 | 1 | 0.5 | NA | 1 | -0.167 |
| SE | 12051 | CTAGE5 | CTAGE5 | chr14 | + | 39291018 | 39291111 | 39279448 | 39279537 | 39293270 | 39293381 | 5 | 3 | 2 | 0 | 0 | 1 | 0 | 2 | 0 | 0 | 98 | 49 | 1 | 1 | 1 | 1 | 0.5 | NA | 1 | -0.167 |
| SE | 12052 | CTAGE5 | CTAGE5 | chr14 | + | 39308448 | 39308587 | 39304290 | 39304381 | 39313339 | 39313441 | 14 | 4 | 3 | 1 | 0 | 0 | 0 | 4 | 0 | 0 | 98 | 49 | 1 | 1 | 0.875 | 1 | 1 | NA | 1 | -0.042 |
| SE | 12076 | ZNF200 | ZNF200 | chr16 | - | 3233505 | 3233836 | 3232832 | 3232921 | 3234986 | 3235456 | 4 | 3 | 0 | 2 | 0 | 0 | 0 | 1 | 0 | 0 | 98 | 49 | 1 | 1 | 0.5 | 1 | NA | NA | 1 | -0.25 |
| SE | 12093 | IQSEC1 | IQSEC1 | chr3 | - | 12902772 | 12902822 | 12897042 | 12901522 | 12908348 | 12908525 | 4 | 2 | 3 | 14 | 2 | 5 | 0 | 2 | 0 | 3 | 98 | 49 | 1 | 1 | 0.125 | 0.333 | 0.231 | NA | 0.25 | -0.02 |
| SE | 12094 | IQSEC1 | IQSEC1 | chr3 | - | 12920429 | 12920596 | 12915593 | 12915733 | 12922119 | 12922242 | 72 | 30 | 16 | 1 | 0 | 0 | 3 | 24 | 0 | 0 | 98 | 49 | 1 | 1 | 0.973 | 1 | 1 | 1 | 1 | -0.009 |
| SE | 12098 | SAP30BP | SAP30BP | chr17 | + | 75693439 | 75693482 | 75671815 | 75671863 | 75699782 | 75699871 | 32 | 13 | 6 | 0 | 2 | 0 | 1 | 5 | 0 | 0 | 92 | 49 | 1 | 1 | 1 | 0.776 | 1 | 1 | 1 | -0.075 |
| SE | 12101 | WBP2 | WBP2 | chr17 | - | 75851567 | 75851676 | 75849603 | 75849739 | 75855238 | 75855420 | 33 | 21 | 8 | 0 | 2 | 0 | 2 | 6 | 0 | 0 | 98 | 49 | 1 | 1 | 1 | 0.84 | 1 | 1 | 1 | -0.053 |
| SE | 12106 | HSPA4 | HSPA4 | chr5 | + | 133091192 | 133091374 | 133089561 | 133089695 | 133092699 | 133092789 | 12 | 1 | 0 | 1 | 0 | 0 | 0 | 3 | 0 | 0 | 98 | 49 | 1 | 1 | 0.857 | 1 | NA | NA | 1 | -0.072 |
| SE | 12110 | EXOC7 | EXOC7 | chr17 | - | 76090328 | 76090397 | 76089174 | 76089320 | 76094413 | 76094581 | 1 | 0 | 0 | 4 | 0 | 1 | 0 | 2 | 0 | 0 | 98 | 49 | 1 | 1 | 0.111 | NA | 0 | NA | 1 | -0.945 |
| SE | 12123 | PATL1 | PATL1 | chr11 | - | 59651543 | 59651641 | 59650753 | 59650813 | 59652463 | 59652587 | 23 | 5 | 4 | 1 | 0 | 0 | 0 | 7 | 0 | 0 | 98 | 49 | 1 | 1 | 0.92 | 1 | 1 | NA | 1 | -0.027 |
| SE | 12124 | PATL1 | PATL1 | chr11 | - | 59655955 | 59656045 | 59655522 | 59655740 | 59656498 | 59656600 | 5 | 1 | 2 | 0 | 3 | 2 | 0 | 3 | 0 | 0 | 98 | 49 | 1 | 1 | 1 | 0.143 | 0.333 | NA | 1 | -0.508 |
| SE | 12125 | JMJD6 | JMJD6 | chr17 | - | 76723771 | 76724058 | 76721797 | 76721933 | 76725466 | 76725855 | 25 | 9 | 1 | 0 | 1 | 0 | 0 | 4 | 0 | 1 | 98 | 49 | 1 | 1 | 1 | 0.818 | 1 | NA | 0.667 | 0.272 |
| SE | 12137 | EPB41L3 | EPB41L3 | chr18 | - | 5433902 | 5434121 | 5433468 | 5433556 | 5438034 | 5438110 | 3 | 0 | 0 | 0 | 0 | 0 | 0 | 2 | 0 | 1 | 98 | 49 | 1 | 1 | 1 | NA | NA | NA | 0.5 | 0.5 |
| SE | 12141 | PGS1 | PGS1 | chr17 | + | 78392475 | 78392665 | 78378617 | 78378808 | 78396307 | 78396385 | 31 | 24 | 7 | 1 | 0 | 0 | 0 | 4 | 0 | 0 | 98 | 49 | 1 | 1 | 0.939 | 1 | 1 | NA | 1 | -0.02 |
| SE | 12142 | PGS1 | PGS1 | chr17 | + | 78396307 | 78396385 | 78392475 | 78392665 | 78398251 | 78398351 | 31 | 34 | 8 | 2 | 5 | 1 | 0 | 5 | 0 | 1 | 98 | 49 | 1 | 1 | 0.886 | 0.773 | 0.8 | NA | 0.714 | 0.106 |
| SE | 12143 | PGS1 | PGS1 | chr17 | + | 78400676 | 78400855 | 78399347 | 78399537 | 78403567 | 78404089 | 48 | 30 | 19 | 1 | 0 | 0 | 0 | 13 | 0 | 0 | 98 | 49 | 1 | 1 | 0.96 | 1 | 1 | NA | 1 | -0.013 |
| SE | 12144 | PGS1 | PGS1 | chr17 | + | 78400734 | 78400855 | 78399347 | 78399537 | 78403567 | 78404089 | 30 | 17 | 5 | 1 | 0 | 0 | 0 | 9 | 0 | 0 | 98 | 49 | 1 | 1 | 0.937 | 1 | 1 | NA | 1 | -0.021 |
| SE | 12147 | DCP2 | DCP2 | chr5 | + | 113003941 | 113004077 | 113001566 | 113001674 | 113007937 | 113008042 | 22 | 10 | 10 | 5 | 1 | 1 | 0 | 10 | 0 | 0 | 98 | 49 | 1 | 1 | 0.687 | 0.833 | 0.833 | NA | 1 | -0.216 |
| SE | 12149 | ATG7 | ATG7 | chr3 | + | 11299361 | 11299416 | 11298685 | 11298855 | 11306942 | 11307060 | 67 | 30 | 7 | 0 | 0 | 0 | 0 | 7 | 1 | 0 | 98 | 49 | 1 | 1 | 1 | 1 | 1 | 0 | 1 | 0.5 |
| SE | 12151 | ATG7 | ATG7 | chr3 | + | 11332971 | 11333093 | 11331339 | 11331428 | 11340644 | 11340735 | 11 | 10 | 1 | 1 | 0 | 0 | 1 | 6 | 0 | 0 | 98 | 49 | 1 | 1 | 0.846 | 1 | 1 | 1 | 1 | -0.051 |
| SE | 12152 | ATG7 | ATG7 | chr3 | + | 11342134 | 11342279 | 11340644 | 11340735 | 11347876 | 11348035 | 13 | 12 | 7 | 0 | 1 | 0 | 0 | 16 | 0 | 1 | 98 | 49 | 1 | 1 | 1 | 0.857 | 1 | NA | 0.889 | 0.063 |
| SE | 12153 | ATG7 | ATG7 | chr3 | + | 11347876 | 11348035 | 11340644 | 11340735 | 11358417 | 11358612 | 8 | 8 | 1 | 0 | 0 | 1 | 0 | 5 | 0 | 0 | 98 | 49 | 1 | 1 | 1 | 1 | 0.333 | NA | 1 | -0.222 |
| SE | 12157 | CYTH1 | CYTH1 | chr17 | - | 78692416 | 78692493 | 78680970 | 78681042 | 78698268 | 78698380 | 28 | 5 | 3 | 0 | 1 | 0 | 0 | 7 | 0 | 0 | 98 | 49 | 1 | 1 | 1 | 0.714 | 1 | NA | 1 | -0.095 |
| SE | 12159 | TYMP | TYMP | chr22 | - | 50526000 | 50526141 | 50525752 | 50525918 | 50526245 | 50526476 | 33 | 36 | 25 | 0 | 1 | 0 | 8 | 30 | 1 | 0 | 98 | 49 | 1 | 1 | 1 | 0.947 | 1 | 0.8 | 1 | 0.082 |
| SE | 12160 | TYMP | TYMP | chr22 | - | 50526000 | 50526156 | 50525752 | 50525918 | 50526245 | 50526476 | 22 | 22 | 12 | 0 | 1 | 0 | 3 | 17 | 1 | 0 | 98 | 49 | 1 | 1 | 1 | 0.917 | 1 | 0.6 | 1 | 0.172 |
| SE | 12167 | EIF4A3 | EIF4A3 | chr17 | - | 80141318 | 80141381 | 80140007 | 80140140 | 80141781 | 80141848 | 19 | 2 | 1 | 0 | 2 | 0 | 0 | 3 | 0 | 0 | 98 | 49 | 1 | 1 | 1 | 0.333 | 1 | NA | 1 | -0.222 |
| SE | 12170 | RNF213 | RNF213 | chr17 | + | 80339200 | 80340356 | 80337832 | 80337997 | 80343131 | 80343325 | 67 | 45 | 9 | 0 | 1 | 0 | 4 | 50 | 0 | 0 | 98 | 49 | 1 | 1 | 1 | 0.957 | 1 | 1 | 1 | -0.014 |
| SE | 12171 | RNF213 | RNF213 | chr17 | + | 80349769 | 80349906 | 80344677 | 80348286 | 80350300 | 80350396 | 91 | 49 | 5 | 0 | 2 | 0 | 3 | 93 | 0 | 0 | 98 | 49 | 1 | 1 | 1 | 0.925 | 1 | 1 | 1 | -0.025 |
| SE | 12172 | RNF213 | RNF213 | chr17 | + | 80352939 | 80353059 | 80351684 | 80351803 | 80353511 | 80353666 | 37 | 34 | 4 | 1 | 0 | 0 | 1 | 35 | 0 | 0 | 98 | 49 | 1 | 1 | 0.949 | 1 | 1 | 1 | 1 | -0.017 |
| SE | 12173 | RNF213 | RNF213 | chr17 | + | 80360060 | 80360206 | 80358287 | 80358479 | 80361733 | 80361888 | 73 | 31 | 11 | 0 | 1 | 0 | 1 | 46 | 0 | 0 | 98 | 49 | 1 | 1 | 1 | 0.939 | 1 | 1 | 1 | -0.02 |
| SE | 12174 | RNF213 | RNF213 | chr17 | + | 80361733 | 80361888 | 80360060 | 80360206 | 80363101 | 80363314 | 74 | 53 | 13 | 0 | 1 | 0 | 2 | 52 | 0 | 0 | 98 | 49 | 1 | 1 | 1 | 0.964 | 1 | 1 | 1 | -0.012 |
| SE | 12175 | RNF213 | RNF213 | chr17 | + | 80363101 | 80363314 | 80361733 | 80361888 | 80363608 | 80363790 | 98 | 69 | 11 | 1 | 0 | 0 | 15 | 88 | 0 | 0 | 98 | 49 | 1 | 1 | 0.98 | 1 | 1 | 1 | 1 | -0.007 |
| SE | 12177 | RNF213 | RNF213 | chr17 | + | 80379619 | 80379714 | 80377761 | 80377796 | 80380830 | 80380987 | 78 | 29 | 10 | 0 | 0 | 0 | 3 | 53 | 0 | 1 | 98 | 49 | 1 | 1 | 1 | 1 | 1 | 1 | 0.964 | 0.018 |
| SE | 12178 | RNF213 | RNF213 | chr17 | + | 80385038 | 80385171 | 80383676 | 80383928 | 80385537 | 80385621 | 58 | 14 | 11 | 3 | 2 | 0 | 0 | 23 | 0 | 2 | 98 | 49 | 1 | 1 | 0.906 | 0.778 | 1 | NA | 0.852 | 0.043 |
| SE | 12180 | RNF213 | RNF213 | chr17 | + | 80390011 | 80390196 | 80389827 | 80389917 | 80393344 | 80398781 | 120 | 54 | 11 | 0 | 0 | 0 | 0 | 79 | 1 | 1 | 98 | 49 | 1 | 1 | 1 | 1 | 1 | 0 | 0.975 | 0.512 |
| SE | 12183 | AATK | AATK | chr17 | - | 81131060 | 81131205 | 81128469 | 81128549 | 81134367 | 81134501 | 20 | 20 | 6 | 2 | 0 | 0 | 0 | 4 | 0 | 0 | 98 | 49 | 1 | 1 | 0.833 | 1 | 1 | NA | 1 | -0.056 |
| SE | 12186 | CD36 | CD36 | chr7 | + | 80662989 | 80663169 | 80661062 | 80661210 | 80664405 | 80664497 | 28 | 0 | 0 | 2 | 0 | 0 | 1 | 28 | 0 | 2 | 98 | 49 | 1 | 1 | 0.875 | NA | NA | 1 | 0.875 | -0.062 |
| SE | 12188 | NPLOC4 | NPLOC4 | chr17 | - | 81610209 | 81610258 | 81608727 | 81608822 | 81613317 | 81613494 | 13 | 9 | 0 | 0 | 0 | 3 | 0 | 4 | 0 | 0 | 98 | 49 | 1 | 1 | 1 | 1 | 0 | NA | 1 | -0.333 |
| SE | 12191 | SP110 | SP110 | chr2 | - | 230170620 | 230170761 | 230168925 | 230169237 | 230172065 | 230172174 | 12 | 0 | 1 | 0 | 2 | 0 | 0 | 1 | 0 | 2 | 98 | 49 | 1 | 1 | 1 | 0 | 1 | NA | 0.2 | 0.467 |
| SE | 12193 | SP110 | SP110 | chr2 | - | 230177537 | 230177680 | 230176472 | 230176724 | 230178156 | 230178255 | 23 | 18 | 11 | 0 | 0 | 0 | 0 | 5 | 0 | 2 | 98 | 49 | 1 | 1 | 1 | 1 | 1 | NA | 0.556 | 0.444 |
| SE | 12199 | ARAP3 | ARAP3 | chr5 | - | 141656503 | 141656637 | 141656193 | 141656276 | 141656717 | 141656846 | 14 | 16 | 4 | 0 | 0 | 0 | 0 | 2 | 0 | 1 | 98 | 49 | 1 | 1 | 1 | 1 | 1 | NA | 0.5 | 0.5 |
| SE | 12201 | ARAP3 | ARAP3 | chr5 | - | 141669921 | 141670063 | 141669708 | 141669811 | 141670511 | 141670628 | 6 | 8 | 3 | 1 | 0 | 0 | 2 | 0 | 0 | 0 | 98 | 49 | 1 | 1 | 0.75 | 1 | 1 | 1 | NA | -0.083 |
| SE | 12202 | ARAP3 | ARAP3 | chr5 | - | 141671894 | 141671980 | 141671569 | 141671752 | 141672551 | 141672658 | 11 | 4 | 1 | 1 | 0 | 0 | 0 | 4 | 0 | 0 | 98 | 49 | 1 | 1 | 0.846 | 1 | 1 | NA | 1 | -0.051 |
| SE | 12203 | ARAP3 | ARAP3 | chr5 | - | 141672101 | 141672301 | 141671894 | 141671980 | 141672551 | 141672658 | 15 | 11 | 2 | 0 | 0 | 0 | 0 | 1 | 0 | 1 | 98 | 49 | 1 | 1 | 1 | 1 | 1 | NA | 0.333 | 0.667 |
| SE | 12205 | ARHGDIA | ARHGDIA | chr17 | - | 81869329 | 81869406 | 81869172 | 81869236 | 81869541 | 81869625 | 29 | 11 | 4 | 1 | 0 | 0 | 0 | 21 | 0 | 1 | 98 | 49 | 1 | 1 | 0.935 | 1 | 1 | NA | 0.913 | 0.065 |
| SE | 12208 | PCYT2 | PCYT2 | chr17 | - | 81908875 | 81909035 | 81908567 | 81908634 | 81909513 | 81909602 | 6 | 0 | 0 | 1 | 0 | 0 | 0 | 2 | 0 | 0 | 98 | 49 | 1 | 1 | 0.75 | NA | NA | NA | 1 | -0.25 |
| SE | 12209 | PCYT2 | PCYT2 | chr17 | - | 81908875 | 81909037 | 81908567 | 81908634 | 81909513 | 81909602 | 7 | 0 | 0 | 1 | 0 | 0 | 0 | 2 | 0 | 0 | 98 | 49 | 1 | 1 | 0.778 | NA | NA | NA | 1 | -0.222 |
| SE | 12210 | PCYT2 | PCYT2 | chr17 | - | 81908875 | 81909321 | 81908567 | 81908634 | 81909513 | 81909602 | 6 | 0 | 2 | 1 | 0 | 0 | 0 | 2 | 0 | 0 | 98 | 49 | 1 | 1 | 0.75 | NA | 1 | NA | 1 | -0.125 |
| SE | 12213 | GPR155 | GPR155 | chr2 | - | 174482872 | 174482918 | 174481496 | 174481987 | 174486872 | 174487088 | 3 | 2 | 3 | 2 | 0 | 0 | 0 | 0 | 0 | 3 | 95 | 49 | 1 | 1 | 0.436 | 1 | 1 | NA | 0 | 0.812 |
| SE | 12214 | GPR155 | GPR155 | chr2 | - | 174483009 | 174483158 | 174481496 | 174481987 | 174486872 | 174487088 | 3 | 0 | 0 | 2 | 0 | 0 | 0 | 0 | 0 | 3 | 98 | 49 | 1 | 1 | 0.429 | NA | NA | NA | 0 | 0.429 |
| SE | 12215 | IL1R1 | IL1R1 | chr2 | + | 102157718 | 102157785 | 102153940 | 102154017 | 102164773 | 102165008 | 5 | 2 | 2 | 2 | 0 | 0 | 0 | 1 | 0 | 0 | 98 | 49 | 1 | 1 | 0.556 | 1 | 1 | NA | 1 | -0.148 |
| SE | 12216 | IL1R1 | IL1R1 | chr2 | + | 102166112 | 102166281 | 102165114 | 102165304 | 102168597 | 102168663 | 9 | 4 | 0 | 1 | 3 | 0 | 0 | 2 | 0 | 0 | 98 | 49 | 1 | 1 | 0.818 | 0.4 | NA | NA | 1 | -0.391 |
| SE | 12219 | HSDL1 | HSDL1 | chr16 | - | 84130336 | 84130431 | 84129985 | 84130171 | 84131101 | 84131327 | 0 | 1 | 0 | 0 | 1 | 0 | 0 | 1 | 0 | 0 | 98 | 49 | 1 | 1 | NA | 0.333 | NA | NA | 1 | -0.667 |
| SE | 12233 | NR4A2 | NR4A2 | chr2 | - | 156328403 | 156328533 | 156327850 | 156328014 | 156329322 | 156330188 | 4 | 0 | 0 | 0 | 0 | 0 | 0 | 1 | 0 | 1 | 98 | 49 | 1 | 1 | 1 | NA | NA | NA | 0.333 | 0.667 |
| SE | 12235 | SMCHD1 | SMCHD1 | chr18 | + | 2673280 | 2673363 | 2666869 | 2667031 | 2688393 | 2688508 | 216 | 145 | 39 | 0 | 2 | 0 | 5 | 46 | 0 | 0 | 98 | 49 | 1 | 1 | 1 | 0.973 | 1 | 1 | 1 | -0.009 |
| SE | 12239 | SMCHD1 | SMCHD1 | chr18 | + | 2707562 | 2707645 | 2706363 | 2706470 | 2707806 | 2707920 | 430 | 242 | 69 | 0 | 1 | 0 | 2 | 82 | 0 | 0 | 98 | 49 | 1 | 1 | 1 | 0.992 | 1 | 1 | 1 | -0.003 |
| SE | 12242 | SMCHD1 | SMCHD1 | chr18 | + | 2722518 | 2722663 | 2718314 | 2718434 | 2726451 | 2726524 | 103 | 62 | 32 | 2 | 0 | 0 | 0 | 26 | 0 | 0 | 98 | 49 | 1 | 1 | 0.963 | 1 | 1 | NA | 1 | -0.012 |
| SE | 12243 | SMCHD1 | SMCHD1 | chr18 | + | 2724898 | 2724995 | 2718314 | 2718434 | 2726451 | 2726524 | 217 | 108 | 24 | 2 | 0 | 0 | 1 | 40 | 0 | 0 | 98 | 49 | 1 | 1 | 0.982 | 1 | 1 | 1 | 1 | -0.006 |
| SE | 12244 | SMCHD1 | SMCHD1 | chr18 | + | 2724898 | 2724995 | 2722518 | 2722663 | 2726451 | 2726524 | 282 | 123 | 55 | 1 | 1 | 0 | 2 | 53 | 0 | 1 | 98 | 49 | 1 | 1 | 0.993 | 0.984 | 1 | 1 | 0.964 | 0.01 |
| SE | 12245 | SMCHD1 | SMCHD1 | chr18 | + | 2732264 | 2732492 | 2729274 | 2729409 | 2738396 | 2738545 | 279 | 95 | 47 | 0 | 1 | 0 | 1 | 52 | 0 | 1 | 98 | 49 | 1 | 1 | 1 | 0.979 | 1 | 1 | 0.963 | 0.011 |
| SE | 12246 | SMCHD1 | SMCHD1 | chr18 | + | 2740702 | 2740821 | 2739431 | 2739520 | 2743760 | 2743928 | 427 | 196 | 66 | 1 | 0 | 0 | 3 | 91 | 0 | 0 | 98 | 49 | 1 | 1 | 0.995 | 1 | 1 | 1 | 1 | -0.002 |
| SE | 12249 | SMCHD1 | SMCHD1 | chr18 | + | 2760651 | 2760739 | 2752487 | 2752552 | 2762104 | 2762236 | 286 | 142 | 51 | 1 | 0 | 0 | 2 | 50 | 0 | 0 | 98 | 49 | 1 | 1 | 0.993 | 1 | 1 | 1 | 1 | -0.002 |
| SE | 12251 | SMCHD1 | SMCHD1 | chr18 | + | 2775733 | 2775924 | 2772249 | 2772372 | 2777805 | 2777915 | 335 | 159 | 66 | 3 | 0 | 0 | 1 | 70 | 0 | 0 | 98 | 49 | 1 | 1 | 0.982 | 1 | 1 | 1 | 1 | -0.006 |
| SE | 12253 | LPIN2 | LPIN2 | chr18 | - | 2923774 | 2923861 | 2922046 | 2922199 | 2924397 | 2924546 | 12 | 8 | 4 | 1 | 0 | 0 | 0 | 7 | 0 | 0 | 98 | 49 | 1 | 1 | 0.857 | 1 | 1 | NA | 1 | -0.048 |
| SE | 12254 | LPIN2 | LPIN2 | chr18 | - | 2925223 | 2925368 | 2924397 | 2924546 | 2926722 | 2926805 | 16 | 7 | 3 | 1 | 0 | 0 | 0 | 2 | 0 | 0 | 98 | 49 | 1 | 1 | 0.889 | 1 | 1 | NA | 1 | -0.037 |
| SE | 12255 | FBXO38 | FBXO38 | chr5 | + | 148427212 | 148427722 | 148425521 | 148425701 | 148433423 | 148433524 | 23 | 2 | 0 | 0 | 0 | 0 | 3 | 2 | 0 | 2 | 98 | 49 | 1 | 1 | 1 | 1 | NA | 1 | 0.333 | 0.334 |
| SE | 12256 | FBXO38 | FBXO38 | chr5 | + | 148439646 | 148439792 | 148438331 | 148438498 | 148440423 | 148440527 | 24 | 3 | 2 | 2 | 3 | 0 | 0 | 1 | 0 | 0 | 98 | 49 | 1 | 1 | 0.857 | 0.333 | 1 | NA | 1 | -0.27 |
| SE | 12264 | JARID2 | JARID2 | chr6 | + | 15452005 | 15452175 | 15410223 | 15410365 | 15468541 | 15468718 | 39 | 22 | 8 | 1 | 0 | 0 | 0 | 10 | 0 | 0 | 98 | 49 | 1 | 1 | 0.951 | 1 | 1 | NA | 1 | -0.016 |
| SE | 12265 | JARID2 | JARID2 | chr6 | + | 15496131 | 15497170 | 15487306 | 15487542 | 15500906 | 15501409 | 10 | 10 | 5 | 1 | 0 | 0 | 0 | 5 | 0 | 0 | 98 | 49 | 1 | 1 | 0.833 | 1 | 1 | NA | 1 | -0.056 |
| SE | 12272 | PPP4R1 | PPP4R1 | chr18 | - | 9553321 | 9553422 | 9550277 | 9550398 | 9557220 | 9557382 | 19 | 16 | 27 | 1 | 0 | 0 | 0 | 12 | 0 | 0 | 98 | 49 | 1 | 1 | 0.905 | 1 | 1 | NA | 1 | -0.032 |
| SE | 12273 | PPP4R1 | PPP4R1 | chr18 | - | 9562921 | 9563046 | 9561979 | 9562075 | 9563377 | 9563550 | 15 | 19 | 5 | 16 | 14 | 10 | 0 | 4 | 0 | 5 | 98 | 49 | 1 | 1 | 0.319 | 0.404 | 0.2 | NA | 0.286 | 0.022 |
| SE | 12274 | PPP4R1 | PPP4R1 | chr18 | - | 9593767 | 9593874 | 9588710 | 9588853 | 9595017 | 9595102 | 77 | 43 | 31 | 1 | 3 | 2 | 0 | 10 | 0 | 0 | 98 | 49 | 1 | 1 | 0.975 | 0.878 | 0.886 | NA | 1 | -0.087 |
| SE | 12279 | MPPE1 | MPPE1 | chr18 | - | 11884878 | 11885043 | 11883470 | 11884627 | 11885675 | 11885816 | 20 | 3 | 3 | 26 | 17 | 5 | 0 | 3 | 0 | 2 | 98 | 49 | 1 | 1 | 0.278 | 0.081 | 0.231 | NA | 0.429 | -0.232 |
| SE | 12280 | MPPE1 | MPPE1 | chr18 | - | 11885675 | 11885816 | 11883470 | 11884627 | 11886916 | 11887025 | 26 | 18 | 7 | 2 | 0 | 0 | 0 | 2 | 0 | 0 | 98 | 49 | 1 | 1 | 0.867 | 1 | 1 | NA | 1 | -0.044 |
| SE | 12281 | MPPE1 | MPPE1 | chr18 | - | 11886498 | 11886621 | 11885675 | 11885816 | 11886916 | 11887025 | 14 | 4 | 4 | 0 | 1 | 2 | 0 | 1 | 0 | 0 | 98 | 49 | 1 | 1 | 1 | 0.667 | 0.5 | NA | 1 | -0.278 |
| SE | 12282 | MPPE1 | MPPE1 | chr18 | - | 11886712 | 11886778 | 11886498 | 11886621 | 11886916 | 11887025 | 10 | 8 | 2 | 2 | 0 | 1 | 0 | 1 | 0 | 1 | 98 | 49 | 1 | 1 | 0.714 | 1 | 0.5 | NA | 0.333 | 0.405 |
| SE | 12283 | MPPE1 | MPPE1 | chr18 | - | 11886712 | 11886781 | 11886498 | 11886621 | 11886916 | 11887025 | 7 | 11 | 3 | 2 | 0 | 1 | 0 | 1 | 0 | 1 | 98 | 49 | 1 | 1 | 0.636 | 1 | 0.6 | NA | 0.333 | 0.412 |
| SE | 12285 | MPPE1 | MPPE1 | chr18 | - | 11906202 | 11906309 | 11896983 | 11897356 | 11908200 | 11908797 | 17 | 11 | 8 | 0 | 1 | 0 | 1 | 1 | 0 | 0 | 98 | 49 | 1 | 1 | 1 | 0.846 | 1 | 1 | 1 | -0.051 |
| SE | 12286 | SLC11A1 | SLC11A1 | chr2 | + | 218382959 | 218383102 | 218382028 | 218382375 | 218384242 | 218384365 | 55 | 28 | 4 | 9 | 3 | 3 | 0 | 13 | 0 | 0 | 98 | 49 | 1 | 1 | 0.753 | 0.824 | 0.4 | NA | 1 | -0.341 |
| SE | 12287 | SLC11A1 | SLC11A1 | chr2 | + | 218382959 | 218383102 | 218382028 | 218382375 | 218385146 | 218385266 | 27 | 17 | 0 | 1 | 0 | 0 | 0 | 2 | 0 | 0 | 98 | 49 | 1 | 1 | 0.931 | 1 | NA | NA | 1 | -0.034 |
| SE | 12288 | SLC11A1 | SLC11A1 | chr2 | + | 218384242 | 218384365 | 218382028 | 218382375 | 218385146 | 218385266 | 106 | 58 | 23 | 1 | 0 | 0 | 2 | 27 | 0 | 0 | 98 | 49 | 1 | 1 | 0.981 | 1 | 1 | 1 | 1 | -0.006 |
| SE | 12289 | SLC11A1 | SLC11A1 | chr2 | + | 218384242 | 218384365 | 218382959 | 218383102 | 218385146 | 218385266 | 125 | 67 | 24 | 0 | 1 | 0 | 2 | 38 | 0 | 0 | 98 | 49 | 1 | 1 | 1 | 0.971 | 1 | 1 | 1 | -0.01 |
| SE | 12294 | PSMG2 | PSMG2 | chr18 | + | 12706549 | 12706721 | 12702987 | 12703164 | 12712701 | 12712760 | 3 | 2 | 4 | 0 | 1 | 0 | 0 | 1 | 0 | 0 | 98 | 49 | 1 | 1 | 1 | 0.5 | 1 | NA | 1 | -0.167 |
| SE | 12304 | ESCO1 | ESCO1 | chr18 | - | 21567979 | 21568094 | 21566145 | 21566206 | 21573313 | 21575430 | 14 | 4 | 7 | 1 | 0 | 0 | 0 | 2 | 0 | 0 | 98 | 49 | 1 | 1 | 0.875 | 1 | 1 | NA | 1 | -0.042 |
| SE | 12306 | TRAPPC11 | TRAPPC11 | chr4 | + | 183682731 | 183682825 | 183680119 | 183680267 | 183683974 | 183684054 | 11 | 3 | 0 | 2 | 0 | 0 | 0 | 1 | 0 | 0 | 98 | 49 | 1 | 1 | 0.733 | 1 | NA | NA | 1 | -0.133 |
| SE | 12307 | TRAPPC11 | TRAPPC11 | chr4 | + | 183686617 | 183686748 | 183685270 | 183685403 | 183691315 | 183691471 | 12 | 2 | 3 | 1 | 0 | 0 | 0 | 3 | 0 | 0 | 98 | 49 | 1 | 1 | 0.857 | 1 | 1 | NA | 1 | -0.048 |
| SE | 12308 | ABHD3 | ABHD3 | chr18 | - | 21656860 | 21657026 | 21650896 | 21651763 | 21657103 | 21657152 | 57 | 33 | 17 | 1 | 0 | 0 | 0 | 13 | 0 | 0 | 98 | 49 | 1 | 1 | 0.966 | 1 | 1 | NA | 1 | -0.011 |
| SE | 12309 | ABHD3 | ABHD3 | chr18 | - | 21664117 | 21664230 | 21659169 | 21659343 | 21702315 | 21702498 | 13 | 3 | 2 | 2 | 1 | 0 | 0 | 2 | 0 | 1 | 98 | 49 | 1 | 1 | 0.765 | 0.6 | 1 | NA | 0.5 | 0.288 |
| SE | 12310 | ABHD3 | ABHD3 | chr18 | - | 21683919 | 21683965 | 21664117 | 21664230 | 21702315 | 21702498 | 38 | 13 | 8 | 4 | 0 | 1 | 0 | 6 | 0 | 0 | 95 | 49 | 1 | 1 | 0.831 | 1 | 0.805 | NA | 1 | -0.121 |
| SE | 12319 | TTC39C | TTC39C | chr18 | + | 24069156 | 24069271 | 24066011 | 24066140 | 24080584 | 24080939 | 20 | 6 | 2 | 1 | 0 | 0 | 1 | 2 | 0 | 0 | 98 | 49 | 1 | 1 | 0.909 | 1 | 1 | 1 | 1 | -0.03 |
| SE | 12321 | CARS | CARS | chr11 | - | 3017856 | 3017954 | 3017105 | 3017295 | 3018407 | 3018511 | 11 | 4 | 0 | 1 | 0 | 0 | 0 | 0 | 0 | 1 | 98 | 49 | 1 | 1 | 0.846 | 1 | NA | NA | 0 | 0.923 |
| SE | 12322 | CARS | CARS | chr11 | - | 3026675 | 3026797 | 3020219 | 3020332 | 3028995 | 3029084 | 20 | 13 | 9 | 2 | 0 | 0 | 0 | 7 | 0 | 0 | 98 | 49 | 1 | 1 | 0.833 | 1 | 1 | NA | 1 | -0.056 |
| SE | 12323 | CARS | CARS | chr11 | - | 3028995 | 3029084 | 3026675 | 3026797 | 3029302 | 3029443 | 35 | 15 | 9 | 2 | 0 | 0 | 0 | 10 | 0 | 0 | 98 | 49 | 1 | 1 | 0.897 | 1 | 1 | NA | 1 | -0.034 |
| SE | 12324 | CARS | CARS | chr11 | - | 3040895 | 3040984 | 3039834 | 3039931 | 3042164 | 3042256 | 11 | 6 | 0 | 0 | 1 | 0 | 0 | 3 | 0 | 0 | 98 | 49 | 1 | 1 | 1 | 0.75 | NA | NA | 1 | -0.125 |
| SE | 12338 | MAPK1 | MAPK1 | chr22 | - | 21788693 | 21788808 | 21788256 | 21788388 | 21799011 | 21799128 | 17 | 3 | 28 | 0 | 0 | 0 | 0 | 7 | 0 | 1 | 98 | 49 | 1 | 1 | 1 | 1 | 1 | NA | 0.778 | 0.222 |
| SE | 12342 | TRAPPC8 | TRAPPC8 | chr18 | - | 31913368 | 31913522 | 31909666 | 31909760 | 31916271 | 31916446 | 29 | 9 | 11 | 2 | 1 | 0 | 0 | 3 | 0 | 0 | 98 | 49 | 1 | 1 | 0.879 | 0.818 | 1 | NA | 1 | -0.101 |
| SE | 12347 | IQCB1 | IQCB1 | chr3 | - | 121788283 | 121788432 | 121781742 | 121781874 | 121790072 | 121790215 | 2 | 0 | 0 | 0 | 1 | 0 | 0 | 3 | 0 | 0 | 98 | 49 | 1 | 1 | 1 | 0 | NA | NA | 1 | -0.5 |
| SE | 12354 | RPRD1A | RPRD1A | chr18 | - | 36040806 | 36040865 | 36033707 | 36033837 | 36067253 | 36067594 | 4 | 4 | 0 | 1 | 4 | 2 | 0 | 2 | 0 | 0 | 98 | 49 | 1 | 1 | 0.667 | 0.333 | 0 | NA | 1 | -0.667 |
| SE | 12360 | PIK3C3 | PIK3C3 | chr18 | + | 41957569 | 41957758 | 41955197 | 41955359 | 41962488 | 41962632 | 34 | 11 | 1 | 0 | 0 | 4 | 0 | 2 | 0 | 0 | 98 | 49 | 1 | 1 | 1 | 1 | 0.111 | NA | 1 | -0.296 |
| SE | 12361 | PIK3C3 | PIK3C3 | chr18 | + | 41987811 | 41987898 | 41970326 | 41970456 | 41990458 | 41990554 | 26 | 17 | 0 | 1 | 0 | 0 | 0 | 2 | 0 | 0 | 98 | 49 | 1 | 1 | 0.929 | 1 | NA | NA | 1 | -0.035 |
| SE | 12367 | MAN2B2 | MAN2B2 | chr4 | + | 6598197 | 6598354 | 6597112 | 6597303 | 6600622 | 6600756 | 2 | 1 | 2 | 0 | 0 | 0 | 0 | 0 | 0 | 1 | 98 | 49 | 1 | 1 | 1 | 1 | 1 | NA | 0 | 1 |
| SE | 12372 | GPCPD1 | GPCPD1 | chr20 | - | 5559939 | 5560076 | 5558683 | 5558819 | 5561464 | 5561530 | 106 | 59 | 12 | 1 | 0 | 0 | 0 | 16 | 0 | 0 | 98 | 49 | 1 | 1 | 0.981 | 1 | 1 | NA | 1 | -0.006 |
| SE | 12373 | GPCPD1 | GPCPD1 | chr20 | - | 5570146 | 5570239 | 5567482 | 5567560 | 5573914 | 5573969 | 77 | 21 | 12 | 0 | 1 | 0 | 0 | 14 | 0 | 1 | 98 | 49 | 1 | 1 | 1 | 0.913 | 1 | NA | 0.875 | 0.096 |
| SE | 12374 | GPCPD1 | GPCPD1 | chr20 | - | 5580007 | 5580131 | 5578379 | 5578611 | 5584280 | 5584322 | 123 | 47 | 12 | 0 | 0 | 0 | 0 | 17 | 0 | 1 | 98 | 49 | 1 | 1 | 1 | 1 | 1 | NA | 0.895 | 0.105 |
| SE | 12375 | GPCPD1 | GPCPD1 | chr20 | - | 5586193 | 5586269 | 5584280 | 5584322 | 5593326 | 5593411 | 32 | 19 | 2 | 29 | 13 | 6 | 0 | 11 | 0 | 2 | 98 | 49 | 1 | 1 | 0.356 | 0.422 | 0.143 | NA | 0.733 | -0.426 |
| SE | 12376 | GPCPD1 | GPCPD1 | chr20 | - | 5593326 | 5593411 | 5584280 | 5584322 | 5598724 | 5598821 | 191 | 66 | 20 | 0 | 0 | 0 | 1 | 27 | 0 | 1 | 98 | 49 | 1 | 1 | 1 | 1 | 1 | 1 | 0.931 | 0.034 |
| SE | 12381 | PIAS2 | PIAS2 | chr18 | - | 46844733 | 46844839 | 46844053 | 46844127 | 46846706 | 46846841 | 0 | 0 | 0 | 1 | 0 | 0 | 0 | 2 | 0 | 0 | 98 | 49 | 1 | 1 | 0 | NA | NA | NA | 1 | -1 |
| SE | 12383 | ATG2A | ATG2A | chr11 | - | 64896461 | 64896616 | 64895289 | 64895442 | 64896747 | 64896869 | 23 | 19 | 1 | 1 | 0 | 0 | 0 | 5 | 0 | 0 | 98 | 49 | 1 | 1 | 0.92 | 1 | 1 | NA | 1 | -0.027 |
| SE | 12384 | ATG2A | ATG2A | chr11 | - | 64900883 | 64901092 | 64900493 | 64900629 | 64901961 | 64902176 | 16 | 9 | 0 | 0 | 0 | 0 | 0 | 0 | 0 | 1 | 98 | 49 | 1 | 1 | 1 | 1 | NA | NA | 0 | 1 |
| SE | 12385 | ATG2A | ATG2A | chr11 | - | 64902515 | 64902680 | 64902259 | 64902386 | 64903287 | 64903364 | 5 | 4 | 0 | 0 | 1 | 0 | 1 | 0 | 0 | 0 | 98 | 49 | 1 | 1 | 1 | 0.667 | NA | 1 | NA | -0.166 |
| SE | 12386 | ATG2A | ATG2A | chr11 | - | 64906112 | 64906193 | 64905741 | 64905848 | 64906333 | 64906533 | 23 | 20 | 4 | 0 | 1 | 0 | 1 | 7 | 0 | 0 | 98 | 49 | 1 | 1 | 1 | 0.909 | 1 | 1 | 1 | -0.03 |
| SE | 12387 | ATG2A | ATG2A | chr11 | - | 64906664 | 64906815 | 64906333 | 64906533 | 64907254 | 64907439 | 20 | 14 | 4 | 1 | 0 | 0 | 2 | 5 | 0 | 0 | 98 | 49 | 1 | 1 | 0.909 | 1 | 1 | 1 | 1 | -0.03 |
| SE | 12388 | ATG2A | ATG2A | chr11 | - | 64908990 | 64909150 | 64907747 | 64907890 | 64909270 | 64909367 | 24 | 18 | 9 | 0 | 5 | 0 | 0 | 2 | 0 | 0 | 98 | 49 | 1 | 1 | 1 | 0.643 | 1 | NA | 1 | -0.119 |
| SE | 12389 | ATG2A | ATG2A | chr11 | - | 64909270 | 64909367 | 64907747 | 64907890 | 64909680 | 64909924 | 11 | 10 | 3 | 0 | 1 | 0 | 0 | 6 | 0 | 0 | 98 | 49 | 1 | 1 | 1 | 0.833 | 1 | NA | 1 | -0.056 |
| SE | 12390 | ATG2A | ATG2A | chr11 | - | 64910615 | 64910708 | 64910039 | 64910195 | 64910806 | 64910954 | 43 | 46 | 13 | 0 | 1 | 0 | 8 | 7 | 0 | 0 | 98 | 49 | 1 | 1 | 1 | 0.958 | 1 | 1 | 1 | -0.014 |
| SE | 12391 | ATG2A | ATG2A | chr11 | - | 64912326 | 64912423 | 64912084 | 64912249 | 64913037 | 64913136 | 36 | 27 | 6 | 0 | 0 | 0 | 2 | 5 | 0 | 1 | 98 | 49 | 1 | 1 | 1 | 1 | 1 | 1 | 0.714 | 0.143 |
| SE | 12393 | DYM | DYM | chr18 | - | 49391592 | 49391645 | 49379664 | 49379758 | 49430254 | 49430447 | 16 | 11 | 1 | 1 | 0 | 0 | 3 | 1 | 0 | 0 | 98 | 49 | 1 | 1 | 0.889 | 1 | 1 | 1 | 1 | -0.037 |
| SE | 12404 | CLPTM1L | CLPTM1L | chr5 | - | 1330279 | 1330383 | 1325750 | 1325816 | 1331798 | 1331883 | 2 | 10 | 3 | 1 | 0 | 0 | 0 | 1 | 0 | 0 | 98 | 49 | 1 | 1 | 0.5 | 1 | 1 | NA | 1 | -0.167 |
| SE | 12405 | POLI | POLI | chr18 | + | 54280666 | 54280903 | 54277702 | 54277855 | 54282836 | 54283015 | 9 | 0 | 3 | 1 | 0 | 0 | 0 | 3 | 0 | 0 | 98 | 49 | 1 | 1 | 0.818 | NA | 1 | NA | 1 | -0.091 |
| SE | 12406 | RAB27B | RAB27B | chr18 | + | 54877566 | 54877738 | 54828476 | 54828700 | 54879368 | 54879454 | 10 | 0 | 0 | 1 | 0 | 0 | 1 | 0 | 0 | 0 | 98 | 49 | 1 | 1 | 0.833 | NA | NA | 1 | NA | -0.167 |
| SE | 12411 | ALAS1 | ALAS1 | chr3 | + | 52205838 | 52206023 | 52204692 | 52204915 | 52206571 | 52206751 | 2 | 3 | 9 | 1 | 0 | 0 | 0 | 2 | 0 | 0 | 98 | 49 | 1 | 1 | 0.5 | 1 | 1 | NA | 1 | -0.167 |
| SE | 12414 | WSB1 | WSB1 | chr17 | + | 27301787 | 27301956 | 27294079 | 27294435 | 27303366 | 27303635 | 47 | 23 | 0 | 7 | 7 | 0 | 0 | 12 | 0 | 1 | 98 | 49 | 1 | 1 | 0.77 | 0.622 | NA | NA | 0.857 | -0.161 |
| SE | 12416 | WSB1 | WSB1 | chr17 | + | 27310060 | 27310174 | 27309099 | 27309272 | 27311508 | 27311616 | 54 | 17 | 7 | 1 | 0 | 0 | 0 | 17 | 0 | 0 | 98 | 49 | 1 | 1 | 0.964 | 1 | 1 | NA | 1 | -0.012 |
| SE | 12434 | DYRK1A | DYRK1A | chr21 | + | 37420298 | 37420384 | 37367556 | 37367628 | 37472683 | 37472880 | 35 | 14 | 4 | 3 | 0 | 2 | 0 | 2 | 0 | 2 | 98 | 49 | 1 | 1 | 0.854 | 1 | 0.5 | NA | 0.333 | 0.452 |
| SE | 12448 | CFAP36 | CFAP36 | chr2 | + | 55522101 | 55522166 | 55519594 | 55519916 | 55523720 | 55523822 | 4 | 0 | 0 | 0 | 1 | 0 | 3 | 0 | 0 | 0 | 98 | 49 | 1 | 1 | 1 | 0 | NA | 1 | NA | -0.5 |
| SE | 12450 | NECAB2 | NECAB2 | chr16 | + | 83997215 | 83997269 | 83994608 | 83994688 | 83998204 | 83998317 | 2 | 2 | 4 | 0 | 1 | 0 | 0 | 3 | 0 | 0 | 98 | 49 | 1 | 1 | 1 | 0.5 | 1 | NA | 1 | -0.167 |
| SE | 12452 | RSBN1 | RSBN1 | chr1 | - | 113796185 | 113796303 | 113777670 | 113777808 | 113797362 | 113798036 | 1 | 0 | 0 | 4 | 4 | 0 | 0 | 0 | 0 | 1 | 98 | 49 | 1 | 1 | 0.111 | 0 | NA | NA | 0 | 0.056 |
| SE | 12458 | TTN-AS1 | TTN-AS1 | chr2 | + | 178538697 | 178539890 | 178537360 | 178537691 | 178540106 | 178540169 | 11 | 4 | 3 | 1 | 0 | 0 | 2 | 1 | 0 | 0 | 98 | 49 | 1 | 1 | 0.846 | 1 | 1 | 1 | 1 | -0.051 |
| SE | 12462 | ZNF516 | ZNF516 | chr18 | - | 76378854 | 76380303 | 76371466 | 76371571 | 76441244 | 76443211 | 20 | 9 | 2 | 1 | 0 | 0 | 3 | 4 | 0 | 0 | 98 | 49 | 1 | 1 | 0.909 | 1 | 1 | 1 | 1 | -0.03 |
| SE | 12463 | ZNF516 | ZNF516 | chr18 | - | 76463027 | 76463141 | 76441244 | 76443211 | 76495143 | 76495190 | 1 | 0 | 0 | 0 | 0 | 0 | 0 | 0 | 1 | 0 | 98 | 49 | 1 | 1 | 1 | NA | NA | 0 | NA | 1 |
| SE | 12466 | MBP | MBP | chr18 | - | 77066297 | 77066385 | 77012782 | 77017268 | 77105210 | 77105286 | 42 | 23 | 27 | 0 | 1 | 0 | 0 | 7 | 0 | 0 | 98 | 49 | 1 | 1 | 1 | 0.92 | 1 | NA | 1 | -0.027 |
| SE | 12471 | PANK3 | PANK3 | chr5 | - | 168563888 | 168564065 | 168561392 | 168561516 | 168566012 | 168566266 | 8 | 0 | 0 | 1 | 0 | 0 | 0 | 4 | 0 | 0 | 98 | 49 | 1 | 1 | 0.8 | NA | NA | NA | 1 | -0.2 |
| SE | 12475 | ATP9B | ATP9B | chr18 | + | 79348131 | 79348196 | 79347769 | 79347925 | 79359353 | 79359462 | 0 | 0 | 2 | 1 | 0 | 0 | 0 | 1 | 0 | 0 | 98 | 49 | 1 | 1 | 0 | NA | 1 | NA | 1 | -0.5 |
| SE | 12479 | TXNL4A | TXNL4A | chr18 | - | 79976717 | 79976787 | 79972866 | 79973856 | 79977597 | 79977701 | 0 | 0 | 0 | 8 | 0 | 0 | 0 | 1 | 0 | 0 | 98 | 49 | 1 | 1 | 0 | NA | NA | NA | 1 | -1 |
| SE | 12488 | RAB28 | RAB28 | chr4 | - | 13369870 | 13369965 | 13367722 | 13368650 | 13376544 | 13376622 | 0 | 0 | 0 | 1 | 2 | 0 | 0 | 2 | 0 | 0 | 98 | 49 | 1 | 1 | 0 | 0 | NA | NA | 1 | -1 |
| SE | 12491 | CLEC4A | CLEC4A | chr12 | + | 8125560 | 8125677 | 8123631 | 8123960 | 8129263 | 8129362 | 8 | 6 | 14 | 0 | 0 | 0 | 0 | 2 | 0 | 1 | 98 | 49 | 1 | 1 | 1 | 1 | 1 | NA | 0.5 | 0.5 |
| SE | 12495 | CLEC4A | CLEC4A | chr12 | + | 8135584 | 8135736 | 8129263 | 8129362 | 8136787 | 8136903 | 12 | 2 | 3 | 0 | 0 | 9 | 0 | 4 | 0 | 0 | 98 | 49 | 1 | 1 | 1 | 1 | 0.143 | NA | 1 | -0.286 |
| SE | 12497 | ARID3A | ARID3A | chr19 | + | 932417 | 932742 | 929261 | 929896 | 960091 | 960164 | 12 | 4 | 9 | 1 | 0 | 0 | 1 | 2 | 0 | 0 | 98 | 49 | 1 | 1 | 0.857 | 1 | 1 | 1 | 1 | -0.048 |
| SE | 12498 | ARID3A | ARID3A | chr19 | + | 964832 | 965080 | 964247 | 964431 | 966571 | 966868 | 12 | 6 | 3 | 2 | 0 | 0 | 2 | 4 | 0 | 0 | 98 | 49 | 1 | 1 | 0.75 | 1 | 1 | 1 | 1 | -0.083 |
| SE | 12499 | ARID3A | ARID3A | chr19 | + | 964832 | 965080 | 964247 | 964431 | 968404 | 968503 | 8 | 3 | 3 | 0 | 0 | 0 | 1 | 1 | 0 | 1 | 98 | 49 | 1 | 1 | 1 | 1 | 1 | 1 | 0.333 | 0.334 |
| SE | 12500 | ARID3A | ARID3A | chr19 | + | 966571 | 966868 | 964247 | 964431 | 968404 | 968503 | 17 | 4 | 3 | 0 | 0 | 0 | 0 | 0 | 0 | 1 | 98 | 49 | 1 | 1 | 1 | 1 | 1 | NA | 0 | 1 |
| SE | 12501 | ARID3A | ARID3A | chr19 | + | 966571 | 966868 | 964832 | 965080 | 968404 | 968503 | 19 | 7 | 3 | 0 | 0 | 0 | 1 | 4 | 0 | 1 | 98 | 49 | 1 | 1 | 1 | 1 | 1 | 1 | 0.667 | 0.166 |
| SE | 12513 | GPX4 | GPX4 | chr19 | + | 1106241 | 1106266 | 1105657 | 1105809 | 1106399 | 1106459 | 20 | 10 | 1 | 0 | 1 | 0 | 0 | 2 | 0 | 0 | 74 | 49 | 1 | 1 | 1 | 0.869 | 1 | NA | 1 | -0.044 |
| SE | 12526 | MKNK2 | MKNK2 | chr19 | - | 2043123 | 2043197 | 2042765 | 2042870 | 2043502 | 2043582 | 125 | 73 | 21 | 0 | 1 | 0 | 1 | 17 | 0 | 0 | 98 | 49 | 1 | 1 | 1 | 0.973 | 1 | 1 | 1 | -0.009 |
| SE | 12527 | MKNK2 | MKNK2 | chr19 | - | 2046185 | 2046283 | 2043502 | 2043582 | 2046366 | 2046468 | 123 | 61 | 23 | 0 | 2 | 0 | 4 | 17 | 0 | 0 | 98 | 49 | 1 | 1 | 1 | 0.938 | 1 | 1 | 1 | -0.021 |
| SE | 12528 | MOB3A | MOB3A | chr19 | - | 2085174 | 2085328 | 2078139 | 2078679 | 2096225 | 2096270 | 88 | 41 | 6 | 0 | 1 | 0 | 2 | 14 | 0 | 0 | 98 | 49 | 1 | 1 | 1 | 0.953 | 1 | 1 | 1 | -0.016 |
| SE | 12530 | GADD45B | GADD45B | chr19 | + | 2476528 | 2476630 | 2476124 | 2476402 | 2477028 | 2477251 | 32 | 45 | 18 | 0 | 2 | 0 | 7 | 19 | 0 | 0 | 98 | 49 | 1 | 1 | 1 | 0.918 | 1 | 1 | 1 | -0.027 |
| SE | 12531 | GADD45B | GADD45B | chr19 | + | 2477028 | 2477251 | 2476528 | 2476630 | 2477487 | 2478259 | 25 | 39 | 18 | 0 | 0 | 0 | 1 | 6 | 0 | 1 | 98 | 49 | 1 | 1 | 1 | 1 | 1 | 1 | 0.75 | 0.125 |
| SE | 12544 | FBXL4 | FBXL4 | chr6 | - | 98917373 | 98917719 | 98905425 | 98905670 | 98926476 | 98927060 | 1 | 0 | 0 | 0 | 0 | 0 | 0 | 0 | 0 | 1 | 98 | 49 | 1 | 1 | 1 | NA | NA | NA | 0 | 1 |
| SE | 12546 | PLIN3 | PLIN3 | chr19 | - | 4859825 | 4860024 | 4859589 | 4859672 | 4861328 | 4861411 | 18 | 17 | 13 | 0 | 1 | 0 | 0 | 10 | 0 | 0 | 98 | 49 | 1 | 1 | 1 | 0.895 | 1 | NA | 1 | -0.035 |
| SE | 12548 | KDM4B | KDM4B | chr19 | + | 5071009 | 5071059 | 5047475 | 5047669 | 5077366 | 5077470 | 32 | 20 | 10 | 0 | 2 | 0 | 0 | 3 | 0 | 0 | 98 | 49 | 1 | 1 | 1 | 0.833 | 1 | NA | 1 | -0.056 |

| | | | | | | | | | | | | | | | | | | | | | | | | | | | | | | | |
|---|---|---|---|---|---|---|---|---|---|---|---|---|---|---|---|---|---|---|---|---|---|---|---|---|---|---|---|---|---|---|---|
| SE | 12549 | KDM4B | KDM4B | chr19 | + | 5119652 | 5119852 | 5110621 | 5110818 | 5131075 | 5131545 | 17 | 23 | 1 | 0 | 1 | 0 | 0 | 3 | 0 | 0 | 98 | 49 | 1 | 1 | 1 | 0.92 | 1 | NA | 1 | -0.027 |
| SE | 12551 | SAFB | SAFB | chr19 | + | 5641739 | 5641946 | 5641593 | 5641658 | 5645336 | 5645399 | 24 | 17 | 4 | 1 | 0 | 0 | 0 | 3 | 0 | 0 | 98 | 49 | 1 | 1 | 0.923 | 1 | 1 | NA | 1 | -0.026 |
| SE | 12552 | SAFB | SAFB | chr19 | + | 5648988 | 5649499 | 5648015 | 5648043 | 5649925 | 5649975 | 13 | 2 | 0 | 0 | 0 | 0 | 0 | 1 | 0 | 3 | 98 | 49 | 1 | 1 | 1 | 1 | NA | NA | 0.143 | 0.857 |
| SE | 12559 | RNF24 | RNF24 | chr20 | - | 3974331 | 3974400 | 3963874 | 3964024 | 4015436 | 4015569 | 41 | 23 | 21 | 11 | 3 | 0 | 0 | 6 | 0 | 1 | 98 | 49 | 1 | 1 | 0.651 | 0.793 | 1 | NA | 0.75 | 0.065 |
| SE | 12567 | RFX2 | RFX2 | chr19 | - | 6008105 | 6008224 | 6007689 | 6007802 | 6010135 | 6010251 | 2 | 0 | 4 | 1 | 0 | 0 | 0 | 2 | 0 | 0 | 98 | 49 | 1 | 1 | 0.5 | NA | 1 | NA | 1 | -0.25 |
| SE | 12569 | OMA1 | OMA1 | chr1 | - | 58530600 | 58530729 | 58527260 | 58527335 | 58533952 | 58534060 | 13 | 5 | 8 | 3 | 0 | 0 | 0 | 1 | 0 | 0 | 98 | 49 | 1 | 1 | 0.684 | 1 | 1 | NA | 1 | -0.105 |
| SE | 12572 | VAV1 | VAV1 | chr19 | + | 6821790 | 6821859 | 6821621 | 6821680 | 6822220 | 6822329 | 84 | 32 | 11 | 0 | 1 | 0 | 4 | 4 | 0 | 0 | 98 | 49 | 1 | 1 | 1 | 0.941 | 1 | 1 | 1 | -0.02 |
| SE | 12574 | VAV1 | VAV1 | chr19 | + | 6826611 | 6826711 | 6825302 | 6825406 | 6828075 | 6828171 | 63 | 15 | 19 | 1 | 0 | 0 | 0 | 8 | 0 | 0 | 98 | 49 | 1 | 1 | 0.969 | 1 | 1 | NA | 1 | -0.01 |
| SE | 12577 | ADGRE1 | ADGRE1 | chr19 | + | 6897148 | 6897304 | 6896397 | 6896541 | 6897427 | 6897451 | 19 | 7 | 4 | 0 | 4 | 0 | 0 | 2 | 0 | 0 | 98 | 49 | 1 | 1 | 1 | 0.467 | 1 | NA | 1 | -0.178 |
| SE | 12578 | ADGRE1 | ADGRE1 | chr19 | + | 6924677 | 6924872 | 6921712 | 6921883 | 6926365 | 6926601 | 5 | 3 | 0 | 1 | 2 | 0 | 0 | 3 | 0 | 0 | 98 | 49 | 1 | 1 | 0.714 | 0.429 | NA | NA | 1 | -0.428 |
| SE | 12580 | KIAA0226L | KIAA0226L | chr13 | - | 46361440 | 46361573 | 46359485 | 46359631 | 46362537 | 46362598 | 99 | 18 | 15 | 2 | 0 | 0 | 0 | 6 | 0 | 0 | 98 | 49 | 1 | 1 | 0.961 | 1 | 1 | NA | 1 | -0.013 |
| SE | 12581 | KIAA0226L | KIAA0226L | chr13 | - | 46361440 | 46361573 | 46359485 | 46359631 | 46363114 | 46363213 | 64 | 9 | 9 | 2 | 1 | 0 | 0 | 6 | 0 | 1 | 98 | 49 | 1 | 1 | 0.941 | 0.818 | 1 | NA | 0.75 | 0.17 |
| SE | 12582 | KIAA0226L | KIAA0226L | chr13 | - | 46362537 | 46362598 | 46359485 | 46359631 | 46363114 | 46363213 | 42 | 11 | 10 | 2 | 1 | 0 | 0 | 3 | 0 | 1 | 98 | 49 | 1 | 1 | 0.913 | 0.846 | 1 | NA | 0.6 | 0.32 |
| SE | 12583 | KIAA0226L | KIAA0226L | chr13 | - | 46362537 | 46362598 | 46361440 | 46361573 | 46363114 | 46363213 | 83 | 20 | 16 | 8 | 0 | 0 | 0 | 3 | 0 | 0 | 98 | 49 | 1 | 1 | 0.838 | 1 | 1 | NA | 1 | -0.054 |
| SE | 12590 | KIAA0226L | KIAA0226L | chr13 | - | 46377889 | 46378005 | 46371940 | 46372597 | 46387133 | 46387500 | 10 | 3 | 4 | 5 | 3 | 0 | 0 | 1 | 0 | 0 | 98 | 49 | 1 | 1 | 0.5 | 0.333 | 1 | NA | 1 | -0.389 |
| SE | 12591 | PET100 | PET100 | chr19 | + | 7630572 | 7630659 | 7629784 | 7629860 | 7630822 | 7630846 | 34 | 9 | 13 | 0 | 1 | 0 | 0 | 9 | 0 | 0 | 98 | 49 | 1 | 1 | 1 | 0.818 | 1 | NA | 1 | -0.061 |
| SE | 12592 | PET100 | PET100 | chr19 | + | 7631231 | 7631321 | 7630822 | 7630846 | 7631472 | 7631624 | 2 | 0 | 1 | 32 | 10 | 5 | 0 | 0 | 0 | 8 | 98 | 49 | 1 | 1 | 0.03 | 0 | 0.091 | NA | 0 | 0.04 |
| SE | 12593 | STXBP2 | STXBP2 | chr19 | + | 7640697 | 7640809 | 7639730 | 7639807 | 7640899 | 7641003 | 22 | 12 | 10 | 2 | 1 | 0 | 0 | 6 | 0 | 0 | 98 | 49 | 1 | 1 | 0.846 | 0.857 | 1 | NA | 1 | -0.099 |
| SE | 12595 | STXBP2 | STXBP2 | chr19 | + | 7640739 | 7640809 | 7639730 | 7639807 | 7640899 | 7641003 | 22 | 12 | 10 | 2 | 1 | 0 | 0 | 6 | 0 | 0 | 98 | 49 | 1 | 1 | 0.846 | 0.857 | 1 | NA | 1 | -0.099 |
| SE | 12596 | STXBP2 | STXBP2 | chr19 | + | 7641704 | 7641853 | 7640899 | 7641003 | 7642033 | 7642118 | 55 | 38 | 17 | 0 | 2 | 0 | 2 | 13 | 0 | 0 | 98 | 49 | 1 | 1 | 1 | 0.905 | 1 | 1 | 1 | -0.032 |
| SE | 12598 | STXBP2 | STXBP2 | chr19 | + | 7644878 | 7644934 | 7644613 | 7644752 | 7645196 | 7645306 | 2 | 0 | 0 | 42 | 50 | 6 | 0 | 0 | 3 | 13 | 98 | 49 | 1 | 1 | 0.023 | 0 | 0 | 0 | 0 | 0.008 |
| SE | 12599 | STXBP2 | STXBP2 | chr19 | + | 7647353 | 7647511 | 7647161 | 7647247 | 7647724 | 7647874 | 16 | 11 | 10 | 1 | 1 | 0 | 0 | 14 | 0 | 0 | 98 | 49 | 1 | 1 | 0.889 | 0.846 | 1 | NA | 1 | -0.088 |
| SE | 12629 | DNMT1 | DNMT1 | chr19 | - | 10180186 | 10180234 | 10177291 | 10177367 | 10180349 | 10180569 | 4 | 0 | 0 | 2 | 0 | 0 | 0 | 0 | 1 | 0 | 97 | 49 | 1 | 1 | 0.503 | NA | NA | 0 | NA | 0.503 |
| SE | 12632 | ALOX5 | ALOX5 | chr10 | + | 45395854 | 45395936 | 45382482 | 45382681 | 45412190 | 45412313 | 225 | 79 | 37 | 0 | 2 | 0 | 5 | 67 | 0 | 1 | 98 | 49 | 1 | 1 | 1 | 0.952 | 1 | 1 | 0.971 | -0.002 |
| SE | 12633 | ALOX5 | ALOX5 | chr10 | + | 45428617 | 45428764 | 45424959 | 45425132 | 45440429 | 45440633 | 148 | 49 | 46 | 1 | 0 | 0 | 9 | 46 | 0 | 1 | 98 | 49 | 1 | 1 | 0.987 | 1 | 1 | 1 | 0.958 | 0.017 |
| SE | 12637 | TYK2 | TYK2 | chr19 | - | 10351051 | 10351162 | 10350527 | 10350968 | 10352433 | 10352551 | 15 | 9 | 7 | 0 | 2 | 0 | 2 | 4 | 0 | 0 | 98 | 49 | 1 | 1 | 1 | 0.692 | 1 | 1 | 1 | -0.103 |
| SE | 12638 | TYK2 | TYK2 | chr19 | - | 10362263 | 10362456 | 10362077 | 10362181 | 10362548 | 10362657 | 40 | 32 | 6 | 1 | 0 | 0 | 4 | 13 | 0 | 0 | 98 | 49 | 1 | 1 | 0.952 | 1 | 1 | 1 | 1 | -0.016 |
| SE | 12640 | COPG2 | COPG2 | chr7 | - | 130554480 | 130554724 | 130552354 | 130552430 | 130555036 | 130555132 | 5 | 0 | 0 | 0 | 4 | 0 | 0 | 4 | 0 | 0 | 98 | 49 | 1 | 1 | 1 | 0 | NA | NA | 1 | -0.5 |
| SE | 12648 | SLC44A2 | SLC44A2 | chr19 | + | 10631625 | 10631749 | 10631474 | 10631535 | 10631867 | 10631951 | 122 | 50 | 40 | 1 | 0 | 0 | 3 | 17 | 0 | 0 | 98 | 49 | 1 | 1 | 0.984 | 1 | 1 | 1 | 1 | -0.005 |
| SE | 12649 | SLC44A2 | SLC44A2 | chr19 | + | 10634973 | 10635073 | 10634755 | 10634887 | 10636322 | 10636585 | 30 | 16 | 14 | 2 | 0 | 0 | 3 | 8 | 0 | 1 | 98 | 49 | 1 | 1 | 0.882 | 1 | 1 | 1 | 0.8 | 0.061 |
| SE | 12650 | SLC44A2 | SLC44A2 | chr19 | + | 10636322 | 10636585 | 10634973 | 10635073 | 10636661 | 10636756 | 53 | 30 | 11 | 1 | 0 | 0 | 8 | 8 | 0 | 0 | 98 | 49 | 1 | 1 | 0.964 | 1 | 1 | 1 | 1 | -0.012 |
| SE | 12651 | SLC44A2 | SLC44A2 | chr19 | + | 10638022 | 10638093 | 10637855 | 10637929 | 10638226 | 10638315 | 157 | 45 | 30 | 1 | 0 | 0 | 4 | 16 | 0 | 0 | 98 | 49 | 1 | 1 | 0.987 | 1 | 1 | 1 | 1 | -0.004 |
| SE | 12653 | RNF123 | RNF123 | chr3 | + | 49712478 | 49712656 | 49706790 | 49706898 | 49713512 | 49713587 | 4 | 0 | 1 | 1 | 0 | 0 | 0 | 1 | 0 | 0 | 98 | 49 | 1 | 1 | 0.667 | NA | 1 | NA | 1 | -0.166 |
| SE | 12656 | SMARCA4 | SMARCA4 | chr19 | + | 11033766 | 11033865 | 11033289 | 11033517 | 11034122 | 11034200 | 0 | 1 | 0 | 4 | 0 | 0 | 0 | 0 | 0 | 1 | 98 | 49 | 1 | 1 | 0 | 1 | NA | NA | 0 | 0.5 |
| SE | 12658 | KDM7A | KDM7A | chr7 | - | 140113490 | 140113582 | 140111094 | 140111184 | 140119112 | 140119219 | 68 | 37 | 8 | 1 | 0 | 0 | 0 | 11 | 0 | 0 | 98 | 49 | 1 | 1 | 0.971 | 1 | 1 | NA | 1 | -0.01 |
| SE | 12666 | B3GNTL1 | B3GNTL1 | chr17 | - | 82965650 | 82965746 | 82961056 | 82961181 | 83005114 | 83005210 | 27 | 8 | 1 | 0 | 3 | 0 | 0 | 5 | 0 | 0 | 98 | 49 | 1 | 1 | 1 | 0.571 | 1 | NA | 1 | -0.143 |
| SE | 12677 | GABARAP | GABARAP | chr17 | - | 7241600 | 7241679 | 7241341 | 7241460 | 7242240 | 7242434 | 29 | 15 | 24 | 0 | 0 | 2 | 0 | 15 | 0 | 0 | 98 | 49 | 1 | 1 | 1 | 1 | 0.857 | NA | 1 | -0.048 |
| SE | 12681 | ZNF136 | ZNF136 | chr19 | + | 12185784 | 12185911 | 12163056 | 12163206 | 12186113 | 12186174 | 3 | 9 | 0 | 0 | 0 | 0 | 0 | 0 | 2 | 0 | 98 | 49 | 1 | 1 | 1 | 1 | NA | 0 | NA | 1 |
| SE | 12684 | SLC38A2 | SLC38A2 | chr12 | - | 46363020 | 46363145 | 46362493 | 46362638 | 46363725 | 46363826 | 11 | 14 | 5 | 1 | 2 | 0 | 0 | 6 | 0 | 0 | 98 | 49 | 1 | 1 | 0.846 | 0.778 | 1 | NA | 1 | -0.125 |
| SE | 12685 | SLC38A2 | SLC38A2 | chr12 | - | 46365106 | 46365189 | 46364643 | 46364702 | 46366863 | 46366945 | 19 | 10 | 2 | 1 | 0 | 0 | 0 | 5 | 0 | 0 | 98 | 49 | 1 | 1 | 0.905 | 1 | 1 | NA | 1 | -0.032 |
| SE | 12689 | ZNF564 | ZNF564 | chr19 | - | 12528303 | 12528364 | 12525369 | 12527916 | 12528569 | 12528696 | 14 | 4 | 1 | 0 | 1 | 0 | 0 | 1 | 0 | 0 | 98 | 49 | 1 | 1 | 1 | 0.667 | 1 | NA | 1 | -0.111 |
| SE | 12690 | KCNQ1 | KCNQ1 | chr11 | + | 2571324 | 2571403 | 2570627 | 2570754 | 2572012 | 2572109 | 6 | 3 | 2 | 0 | 2 | 0 | 0 | 3 | 0 | 0 | 98 | 49 | 1 | 1 | 1 | 0.429 | 1 | NA | 1 | -0.19 |
| SE | 12691 | KCNQ1 | KCNQ1 | chr11 | + | 2585211 | 2585307 | 2583434 | 2583545 | 2587569 | 2587692 | 11 | 3 | 1 | 0 | 0 | 0 | 0 | 4 | 1 | 0 | 98 | 49 | 1 | 1 | 1 | 1 | 1 | 0 | 1 | 0.5 |
| SE | 12692 | MAN2B1 | MAN2B1 | chr19 | - | 12658427 | 12658507 | 12658223 | 12658344 | 12661259 | 12661376 | 6 | 0 | 2 | 2 | 0 | 0 | 0 | 8 | 0 | 1 | 98 | 49 | 1 | 1 | 0.6 | NA | 1 | NA | 0.8 | 0 |
| SE | 12693 | MAN2B1 | MAN2B1 | chr19 | - | 12658427 | 12658510 | 12658223 | 12658344 | 12661259 | 12661376 | 20 | 5 | 2 | 2 | 0 | 0 | 0 | 10 | 0 | 1 | 98 | 49 | 1 | 1 | 0.833 | 1 | 1 | NA | 0.833 | 0.111 |
| SE | 12694 | MAN2B1 | MAN2B1 | chr19 | - | 12663316 | 12663462 | 12661259 | 12661376 | 12663702 | 12663835 | 32 | 8 | 2 | 0 | 2 | 0 | 2 | 11 | 0 | 0 | 98 | 49 | 1 | 1 | 1 | 0.667 | 1 | 1 | 1 | -0.111 |
| SE | 12696 | C19orf43 | C19orf43 | chr19 | - | 12731343 | 12731421 | 12730639 | 12731068 | 12734312 | 12734715 | 35 | 27 | 25 | 0 | 0 | 0 | 0 | 5 | 0 | 1 | 98 | 49 | 1 | 1 | 1 | 1 | 1 | NA | 0.714 | 0.286 |
| SE | 12698 | GLUL | GLUL | chr1 | - | 182385356 | 182385556 | 182378092 | 182384723 | 182385759 | 182385887 | 1476 | 554 | 212 | 6 | 0 | 0 | 32 | 197 | 0 | 0 | 98 | 49 | 1 | 1 | 0.992 | 1 | 1 | 1 | 1 | -0.003 |
| SE | 12699 | GLUL | GLUL | chr1 | - | 182385759 | 182385887 | 182385356 | 182385556 | 182386255 | 182386402 | 833 | 304 | 145 | 0 | 1 | 0 | 16 | 127 | 0 | 0 | 98 | 49 | 1 | 1 | 1 | 0.993 | 1 | 1 | 1 | -0.002 |
| SE | 12700 | GLUL | GLUL | chr1 | - | 182386255 | 182386402 | 182385759 | 182385887 | 182388571 | 182388750 | 659 | 195 | 81 | 0 | 2 | 0 | 10 | 80 | 0 | 0 | 98 | 49 | 1 | 1 | 1 | 0.98 | 1 | 1 | 1 | -0.007 |
| SE | 12706 | CLOCK | CLOCK | chr4 | - | 55478814 | 55478963 | 55475962 | 55476054 | 55479639 | 55479699 | 8 | 0 | 0 | 1 | 0 | 0 | 0 | 1 | 0 | 0 | 98 | 49 | 1 | 1 | 0.8 | NA | NA | NA | 1 | -0.2 |
| SE | 12711 | RNF145 | RNF145 | chr5 | - | 159181959 | 159182051 | 159176631 | 159176867 | 159194715 | 159194824 | 15 | 9 | 4 | 0 | 0 | 3 | 0 | 4 | 0 | 0 | 98 | 49 | 1 | 1 | 1 | 1 | 0.4 | NA | 1 | -0.2 |
| SE | 12728 | ADGRE5 | ADGRE5 | chr19 | + | 14402596 | 14402862 | 14401652 | 14401760 | 14404382 | 14404562 | 206 | 95 | 53 | 0 | 1 | 0 | 5 | 56 | 0 | 0 | 98 | 49 | 1 | 1 | 1 | 0.979 | 1 | 1 | 1 | -0.007 |
| SE | 12729 | ADGRE5 | ADGRE5 | chr19 | + | 14406330 | 14406557 | 14405747 | 14405939 | 14406699 | 14406766 | 132 | 72 | 50 | 1 | 0 | 0 | 4 | 63 | 0 | 0 | 98 | 49 | 1 | 1 | 0.985 | 1 | 1 | 1 | 1 | -0.005 |
| SE | 12730 | ADGRE5 | ADGRE5 | chr19 | + | 14407060 | 14407229 | 14406868 | 14406960 | 14407907 | 14408009 | 170 | 88 | 107 | 0 | 1 | 0 | 1 | 40 | 0 | 0 | 98 | 49 | 1 | 1 | 1 | 0.978 | 1 | 1 | 1 | -0.007 |
| SE | 12731 | ADGRE5 | ADGRE5 | chr19 | + | 14407907 | 14408009 | 14407060 | 14407229 | 14408091 | 14408725 | 184 | 68 | 100 | 0 | 0 | 2 | 0 | 44 | 0 | 0 | 98 | 49 | 1 | 1 | 1 | 1 | 0.962 | NA | 1 | -0.013 |
| SE | 12733 | PKN1 | PKN1 | chr19 | + | 14463827 | 14463969 | 14463666 | 14463742 | 14464059 | 14464153 | 22 | 7 | 6 | 0 | 1 | 0 | 0 | 5 | 0 | 0 | 98 | 49 | 1 | 1 | 1 | 0.778 | 1 | NA | 1 | -0.074 |
| SE | 12734 | PKN1 | PKN1 | chr19 | + | 14469751 | 14469814 | 14469340 | 14469517 | 14469911 | 14469988 | 24 | 28 | 4 | 2 | 0 | 0 | 1 | 5 | 0 | 1 | 98 | 49 | 1 | 1 | 0.857 | 1 | 1 | 1 | 0.714 | 0.095 |
| SE | 12742 | QARS | QARS | chr3 | - | 49101619 | 49101705 | 49101354 | 49101441 | 49101827 | 49101899 | 5 | 0 | 7 | 0 | 1 | 0 | 0 | 0 | 0 | 1 | 98 | 49 | 1 | 1 | 1 | 0 | 1 | NA | 0 | 0.667 |
| SE | 12743 | BRD4 | BRD4 | chr19 | - | 15267415 | 15267551 | 15265353 | 15265643 | 15268904 | 15269042 | 16 | 14 | 8 | 0 | 2 | 0 | 0 | 1 | 0 | 0 | 98 | 49 | 1 | 1 | 1 | 0.778 | 1 | NA | 1 | -0.074 |
| SE | 12744 | AKAP8L | AKAP8L | chr19 | - | 15398629 | 15398766 | 15397713 | 15397855 | 15399301 | 15399410 | 1 | 1 | 0 | 4 | 5 | 1 | 0 | 0 | 0 | 4 | 98 | 49 | 1 | 1 | 0.111 | 0.091 | 0 | NA | 0 | 0.067 |
| SE | 12746 | RASAL3 | RASAL3 | chr19 | - | 15452044 | 15452108 | 15451626 | 15451938 | 15452657 | 15452815 | 13 | 2 | 2 | 3 | 0 | 0 | 1 | 0 | 0 | 0 | 98 | 49 | 1 | 1 | 0.684 | 1 | 1 | 1 | NA | -0.105 |
| SE | 12747 | LRRK1 | LRRK1 | chr15 | + | 100973803 | 100973967 | 100924510 | 100924729 | 100983527 | 100983699 | 8 | 0 | 0 | 1 | 0 | 0 | 2 | 0 | 0 | 0 | 98 | 49 | 1 | 1 | 0.8 | NA | NA | 1 | NA | -0.2 |
| SE | 12749 | CYP4F3 | CYP4F3 | chr19 | + | 15643906 | 15644051 | 15641414 | 15641613 | 15647051 | 15647105 | 2 | 3 | 5 | 5 | 2 | 0 | 0 | 1 | 0 | 0 | 98 | 49 | 1 | 1 | 0.167 | 0.429 | 1 | NA | 1 | -0.468 |
| SE | 12760 | KLF2 | KLF2 | chr19 | + | 16325215 | 16326032 | 16324839 | 16324998 | 16326855 | 16327528 | 71 | 29 | 9 | 1 | 1 | 0 | 4 | 7 | 0 | 0 | 98 | 49 | 1 | 1 | 0.973 | 0.935 | 1 | 1 | 1 | -0.031 |
| SE | 12762 | EPS15L1 | EPS15L1 | chr19 | - | 16377121 | 16377254 | 16361577 | 16361984 | 16385128 | 16385211 | 6 | 4 | 0 | 0 | 0 | 3 | 0 | 1 | 0 | 0 | 98 | 49 | 1 | 1 | 1 | 1 | 0 | NA | 1 | -0.333 |
| SE | 12770 | CCDC125 | CCDC125 | chr5 | - | 69303846 | 69303929 | 69300011 | 69300127 | 69306816 | 69306902 | 10 | 3 | 3 | 0 | 0 | 3 | 0 | 4 | 0 | 0 | 98 | 49 | 1 | 1 | 1 | 1 | 0.333 | NA | 1 | -0.222 |
| SE | 12771 | CCDC125 | CCDC125 | chr5 | - | 69313984 | 69314043 | 69311117 | 69311204 | 69320236 | 69320345 | 8 | 9 | 6 | 1 | 0 | 0 | 0 | 3 | 0 | 0 | 98 | 49 | 1 | 1 | 0.8 | 1 | 1 | NA | 1 | -0.067 |
| SE | 12772 | CCDC125 | CCDC125 | chr5 | - | 69313984 | 69314046 | 69311117 | 69311204 | 69320236 | 69320580 | 11 | 6 | 5 | 1 | 0 | 0 | 0 | 3 | 0 | 0 | 98 | 49 | 1 | 1 | 0.846 | 1 | 1 | NA | 1 | -0.051 |
| SE | 12773 | CCDC125 | CCDC125 | chr5 | - | 69320236 | 69320345 | 69311117 | 69311204 | 69320464 | 69320580 | 6 | 1 | 1 | 1 | 0 | 0 | 0 | 1 | 0 | 0 | 98 | 49 | 1 | 1 | 0.75 | 1 | 1 | NA | 1 | -0.083 |
| SE | 12780 | CSNK1G3 | CSNK1G3 | chr5 | + | 123575728 | 123575963 | 123573392 | 123573541 | 123588067 | 123588153 | 9 | 8 | 0 | 1 | 0 | 0 | 0 | 3 | 0 | 0 | 98 | 49 | 1 | 1 | 0.818 | 1 | NA | NA | 1 | -0.091 |
| SE | 12785 | MYO9B | MYO9B | chr19 | + | 17159394 | 17159484 | 17156908 | 17157038 | 17162349 | 17162466 | 96 | 72 | 14 | 1 | 0 | 0 | 4 | 24 | 0 | 0 | 98 | 49 | 1 | 1 | 0.98 | 1 | 1 | 1 | 1 | -0.007 |
| SE | 12786 | MYO9B | MYO9B | chr19 | + | 17210332 | 17210380 | 17209585 | 17209709 | 17210714 | 17210848 | 1 | 1 | 0 | 22 | 7 | 2 | 0 | 0 | 0 | 3 | 97 | 49 | 1 | 1 | 0.022 | 0.067 | 0 | NA | 0 | 0.03 |
| SE | 12787 | PCGF5 | PCGF5 | chr10 | + | 91240483 | 91240580 | 91222688 | 91222983 | 91248504 | 91248560 | 12 | 6 | 1 | 1 | 0 | 0 | 0 | 2 | 0 | 0 | 98 | 49 | 1 | 1 | 0.857 | 1 | 1 | NA | 1 | -0.048 |
| SE | 12788 | PCGF5 | PCGF5 | chr10 | + | 91271637 | 91271697 | 91264430 | 91264520 | 91278268 | 91284331 | 9 | 0 | 2 | 1 | 0 | 0 | 0 | 7 | 0 | 0 | 98 | 49 | 1 | 1 | 0.818 | NA | 1 | NA | 1 | -0.091 |
| SE | 12794 | JAK3 | JAK3 | chr19 | - | 17843776 | 17843900 | 17843379 | 17843491 | 17844233 | 17844430 | 71 | 11 | 6 | 1 | 1 | 0 | 0 | 14 | 0 | 0 | 98 | 49 | 1 | 1 | 0.973 | 0.846 | 1 | NA | 1 | -0.06 |
| SE | 12798 | ARRDC2 | ARRDC2 | chr19 | + | 18009591 | 18009694 | 18008970 | 18009118 | 18009782 | 18010039 | 10 | 4 | 7 | 2 | 0 | 0 | 0 | 1 | 0 | 0 | 98 | 49 | 1 | 1 | 0.714 | 1 | 1 | NA | 1 | -0.095 |
| SE | 12799 | EDEM2 | EDEM2 | chr20 | - | 35118597 | 35118719 | 35115356 | 35115933 | 35123889 | 35124034 | 7 | 4 | 3 | 2 | 0 | 0 | 0 | 6 | 0 | 1 | 98 | 49 | 1 | 1 | 0.636 | 1 | 1 | NA | 0.75 | 0.129 |
| SE | 12800 | EDEM2 | EDEM2 | chr20 | - | 35131641 | 35131783 | 35126250 | 35126375 | 35134737 | 35134949 | 5 | 3 | 2 | 1 | 0 | 0 | 0 | 2 | 0 | 0 | 98 | 49 | 1 | 1 | 0.714 | 1 | 1 | NA | 1 | -0.095 |
| SE | 12801 | EDEM2 | EDEM2 | chr20 | - | 35144978 | 35145018 | 35142372 | 35142478 | 35146824 | 35146935 | 4 | 8 | 5 | 1 | 0 | 0 | 0 | 2 | 0 | 0 | 89 | 49 | 1 | 1 | 0.688 | 1 | 1 | NA | 1 | -0.104 |
| SE | 12804 | SSBP4 | SSBP4 | chr19 | + | 18431352 | 18431418 | 18430840 | 18430930 | 18431646 | 18431706 | 2 | 0 | 0 | 1 | 0 | 0 | 0 | 0 | 0 | 1 | 98 | 49 | 1 | 1 | 0.5 | NA | NA | NA | 0 | 0.5 |
| SE | 12811 | COPE | COPE | chr19 | - | 18906959 | 18907112 | 18905575 | 18905629 | 18910970 | 18911071 | 29 | 11 | 7 | 2 | 0 | 0 | 1 | 15 | 0 | 0 | 98 | 49 | 1 | 1 | 0.879 | 1 | 1 | 1 | 1 | -0.04 |
| SE | 12818 | ARMC6 | ARMC6 | chr19 | + | 19034130 | 19034238 | 19033577 | 19033930 | 19042710 | 19042877 | 0 | 0 | 0 | 1 | 1 | 0 | 0 | 2 | 0 | 0 | 98 | 49 | 1 | 1 | 0 | 0 | NA | NA | 1 | -1 |
| SE | 12823 | GMIP | GMIP | chr19 | - | 19635414 | 19635569 | 19635024 | 19635213 | 19636706 | 19636787 | 51 | 48 | 6 | 0 | 1 | 0 | 5 | 9 | 0 | 0 | 98 | 49 | 1 | 1 | 1 | 0.96 | 1 | 1 | 1 | -0.013 |
| SE | 12827 | ZNF101 | ZNF101 | chr19 | + | 19678725 | 19678786 | 19677863 | 19677990 | 19679180 | 19683506 | 8 | 1 | 0 | 0 | 0 | 0 | 0 | 1 | 0 | 1 | 98 | 49 | 1 | 1 | 1 | 1 | NA | NA | 0.333 | 0.667 |
| SE | 12839 | ING4 | ING4 | chr12 | - | 6653229 | 6653396 | 6652935 | 6653050 | 6656726 | 6656798 | 11 | 4 | 2 | 1 | 4 | 0 | 0 | 1 | 0 | 0 | 98 | 49 | 1 | 1 | 0.846 | 0.333 | 1 | NA | 1 | -0.274 |
| SE | 12848 | ANXA11 | ANXA11 | chr10 | - | 80168968 | 80169358 | 80167225 | 80167313 | 80170799 | 80170915 | 35 | 26 | 22 | 0 | 0 | 0 | 0 | 6 | 0 | 2 | 98 | 49 | 1 | 1 | 1 | 1 | 1 | NA | 0.6 | 0.4 |
| SE | 12850 | ANXA11 | ANXA11 | chr10 | - | 80176106 | 80176155 | 80172806 | 80172869 | 80205342 | 80205444 | 35 | 25 | 21 | 3 | 3 | 0 | 0 | 4 | 0 | 2 | 98 | 49 | 1 | 1 | 0.854 | 0.806 | 1 | NA | 0.5 | 0.387 |
| SE | 12858 | TLK2 | TLK2 | chr17 | + | 62524235 | 62524331 | 62523133 | 62523177 | 62536169 | 62536337 | 11 | 2 | 0 | 1 | 1 | 1 | 0 | 1 | 0 | 0 | 98 | 49 | 1 | 1 | 0.846 | 0.5 | 0 | NA | 1 | -0.551 |
| SE | 12861 | TLK2 | TLK2 | chr17 | + | 62574344 | 62574410 | 62573214 | 62573367 | 62576708 | 62576775 | 2 | 0 | 0 | 11 | 2 | 5 | 0 | 0 | 0 | 2 | 98 | 49 | 1 | 1 | 0.083 | 0 | 0 | NA | 0 | 0.028 |
| SE | 12862 | TLK2 | TLK2 | chr17 | + | 62602041 | 62602180 | 62600650 | 62600820 | 62606129 | 62606241 | 3 | 4 | 0 | 0 | 1 | 0 | 0 | 1 | 0 | 0 | 98 | 49 | 1 | 1 | 1 | 0.667 | NA | NA | 1 | -0.166 |
| SE | 12880 | GPI | GPI | chr19 | + | 34377502 | 34377586 | 34368582 | 34368702 | 34377734 | 34377881 | 13 | 7 | 1 | 0 | 0 | 0 | 0 | 3 | 0 | 1 | 98 | 49 | 1 | 1 | 1 | 1 | 1 | NA | 0.6 | 0.4 |
| SE | 12882 | GRAMD1A | GRAMD1A | chr19 | + | 35009422 | 35009443 | 35009118 | 35009329 | 35009887 | 35009972 | 24 | 7 | 9 | 1 | 0 | 0 | 1 | 5 | 0 | 0 | 70 | 49 | 1 | 1 | 0.944 | 1 | 1 | 1 | 1 | -0.019 |
| SE | 12885 | GRAMD1A | GRAMD1A | chr19 | + | 35023426 | 35023547 | 35023235 | 35023343 | 35026048 | 35026469 | 23 | 0 | 13 | 0 | 1 | 0 | 0 | 3 | 0 | 0 | 98 | 49 | 1 | 1 | 1 | 0 | 1 | NA | 1 | -0.333 |
| SE | 12909 | MMP8 | MMP8 | chr11 | - | 102714555 | 102714709 | 102713753 | 102713857 | 102715303 | 102715437 | 1 | 1 | 0 | 0 | 0 | 0 | 4 | 2 | 0 | 3 | 98 | 49 | 1 | 1 | 1 | 1 | NA | 1 | 0.25 | 0.375 |
| SE | 12910 | MMP8 | MMP8 | chr11 | - | 102722990 | 102723078 | 102722428 | 102722673 | 102724753 | 102724954 | 0 | 1 | 0 | 0 | 1 | 0 | 3 | 3 | 0 | 1 | 98 | 49 | 1 | 1 | NA | 0.333 | NA | 1 | 0.6 | -0.467 |
| SE | 12911 | MMP8 | MMP8 | chr11 | - | 102722990 | 102723081 | 102722428 | 102722673 | 102724753 | 102724954 | 0 | 1 | 0 | 0 | 1 | 0 | 3 | 3 | 0 | 1 | 98 | 49 | 1 | 1 | NA | 0.333 | NA | 1 | 0.6 | -0.467 |
| SE | 12912 | TREML3P | TREML3P | chr6 | - | 41217126 | 41217177 | 41215761 | 41215819 | 41217652 | 41217947 | 7 | 3 | 1 | 0 | 1 | 0 | 1 | 0 | 0 | 0 | 98 | 49 | 1 | 1 | 1 | 0.6 | 1 | 1 | NA | -0.133 |
| SE | 12917 | U2AF1L4 | U2AF1L4 | chr19 | - | 35744625 | 35744738 | 35744069 | 35744421 | 35745124 | 35745187 | 1 | 0 | 0 | 0 | 2 | 0 | 0 | 2 | 0 | 0 | 98 | 49 | 1 | 1 | 1 | 0 | NA | NA | 1 | -0.5 |
| SE | 12920 | TYROBP | TYROBP | chr19 | - | 35907217 | 35907261 | 35904400 | 35904634 | 35907445 | 35907580 | 126 | 73 | 76 | 0 | 0 | 0 | 7 | 39 | 0 | 1 | 93 | 49 | 1 | 1 | 1 | 1 | 1 | 1 | 0.954 | 0.023 |
| SE | 12921 | TYROBP | TYROBP | chr19 | - | 35907217 | 35907264 | 35904400 | 35904634 | 35907445 | 35907580 | 242 | 131 | 137 | 0 | 0 | 0 | 18 | 77 | 0 | 1 | 96 | 49 | 1 | 1 | 1 | 1 | 1 | 1 | 0.975 | 0.012 |
| SE | 12926 | INO80E | INO80E | chr16 | + | 30001211 | 30001530 | 30000928 | 30001040 | 30005220 | 30005790 | 4 | 1 | 5 | 0 | 1 | 0 | 0 | 2 | 0 | 0 | 98 | 49 | 1 | 1 | 1 | 0.333 | 1 | NA | 1 | -0.222 |
| SE | 12927 | INO80E | INO80E | chr16 | + | 30001413 | 30001530 | 30000928 | 30001040 | 30005220 | 30005790 | 5 | 0 | 5 | 0 | 1 | 0 | 0 | 2 | 0 | 0 | 98 | 49 | 1 | 1 | 1 | 0 | 1 | NA | 1 | -0.333 |
| SE | 12928 | INO80E | INO80E | chr16 | + | 30004567 | 30004657 | 30001211 | 30001530 | 30005220 | 30005790 | 1 | 0 | 0 | 0 | 0 | 5 | 0 | 0 | 0 | 2 | 98 | 49 | 1 | 1 | 1 | NA | 0 | NA | 0 | 0.5 |
| SE | 12934 | KDM1A | KDM1A | chr1 | + | 23044426 | 23044486 | 23030468 | 23030634 | 23050386 | 23050520 | 2 | 6 | 0 | 4 | 0 | 0 | 0 | 2 | 0 | 0 | 98 | 49 | 1 | 1 | 0.2 | 1 | NA | NA | 1 | -0.4 |
| SE | 12948 | PQBP1 | PQBP1 | chrX | + | 48902232 | 48902517 | 48901929 | 48902042 | 48902731 | 48902795 | 4 | 0 | 6 | 1 | 0 | 0 | 0 | 2 | 0 | 0 | 98 | 49 | 1 | 1 | 0.667 | NA | 1 | NA | 1 | -0.166 |
| SE | 12961 | DDX60L | DDX60L | chr4 | - | 168379725 | 168379830 | 168379362 | 168379504 | 168384611 | 168384812 | 28 | 10 | 8 | 0 | 0 | 0 | 1 | 18 | 0 | 1 | 98 | 49 | 1 | 1 | 1 | 1 | 1 | 1 | 0.9 | 0.05 |
| SE | 12964 | DDX60L | DDX60L | chr4 | - | 168415407 | 168415517 | 168406601 | 168406706 | 168415656 | 168415799 | 69 | 58 | 11 | 0 | 1 | 0 | 1 | 57 | 0 | 0 | 98 | 49 | 1 | 1 | 1 | 0.967 | 1 | 1 | 1 | -0.011 |
| SE | 12973 | GMFG | GMFG | chr19 | - | 39329543 | 39329626 | 39328999 | 39329073 | 39333076 | 39333126 | 288 | 205 | 223 | 2 | 1 | 0 | 0 | 53 | 0 | 0 | 98 | 49 | 1 | 1 | 0.986 | 0.99 | 1 | NA | 1 | -0.008 |
| SE | 12974 | GMFG | GMFG | chr19 | - | 39333076 | 39333126 | 39328999 | 39329073 | 39335434 | 39335531 | 6 | 1 | 0 | 1 | 0 | 0 | 0 | 1 | 0 | 0 | 98 | 49 | 1 | 1 | 0.75 | 1 | NA | NA | 1 | -0.125 |
| SE | 12975 | GMFG | GMFG | chr19 | - | 39333076 | 39333126 | 39329543 | 39329626 | 39335260 | 39335310 | 200 | 151 | 145 | 0 | 1 | 0 | 1 | 30 | 0 | 0 | 98 | 49 | 1 | 1 | 1 | 0.987 | 1 | 1 | 1 | -0.004 |
| SE | 12977 | GMFG | GMFG | chr19 | - | 39335260 | 39335310 | 39333076 | 39333126 | 39335973 | 39336086 | 97 | 63 | 46 | 1 | 0 | 0 | 1 | 12 | 0 | 0 | 98 | 49 | 1 | 1 | 0.98 | 1 | 1 | 1 | 1 | -0.007 |
| SE | 12978 | GMFG | GMFG | chr19 | - | 39335434 | 39335531 | 39333076 | 39333126 | 39335973 | 39336086 | 5 | 0 | 0 | 1 | 0 | 0 | 0 | 1 | 0 | 0 | 98 | 49 | 1 | 1 | 0.714 | NA | NA | NA | 1 | -0.286 |
| SE | 12981 | IFI30 | IFI30 | chr19 | + | 18177139 | 18177292 | 18175604 | 18175697 | 18177710 | 18177764 | 252 | 82 | 46 | 1 | 0 | 0 | 13 | 127 | 0 | 0 | 98 | 49 | 1 | 1 | 0.992 | 1 | 1 | 1 | 1 | -0.003 |
| SE | 12989 | CDK19 | CDK19 | chr6 | - | 110623289 | 110623362 | 110622814 | 110622912 | 110626775 | 110626845 | 23 | 2 | 0 | 3 | 0 | 0 | 3 | 1 | 0 | 0 | 98 | 49 | 1 | 1 | 0.793 | 1 | NA | 1 | 1 | -0.103 |
| SE | 12990 | CDK19 | CDK19 | chr6 | - | 110632029 | 110632161 | 110627001 | 110627145 | 110638648 | 110638706 | 11 | 4 | 4 | 1 | 0 | 0 | 0 | 4 | 0 | 0 | 98 | 49 | 1 | 1 | 0.846 | 1 | 1 | NA | 1 | -0.051 |
| SE | 12991 | CDK19 | CDK19 | chr6 | - | 110773875 | 110773937 | 110746125 | 110746201 | 110815008 | 110815316 | 5 | 2 | 0 | 4 | 1 | 1 | 0 | 0 | 0 | 1 | 98 | 49 | 1 | 1 | 0.385 | 0.5 | 0 | NA | 0 | 0.295 |
| SE | 12993 | ADCK4 | ADCK4 | chr19 | - | 40703714 | 40703855 | 40703540 | 40703622 | 40705095 | 40705181 | 2 | 1 | 0 | 0 | 1 | 0 | 0 | 3 | 0 | 0 | 98 | 49 | 1 | 1 | 1 | 0.333 | NA | NA | 1 | -0.334 |
| SE | 12995 | SASH3 | SASH3 | chrX | + | 129787974 | 129788070 | 129779915 | 129780154 | 129790939 | 129791081 | 41 | 14 | 8 | 2 | 0 | 0 | 0 | 8 | 0 | 0 | 98 | 49 | 1 | 1 | 0.911 | 1 | 1 | NA | 1 | -0.03 |
| SE | 12996 | SASH3 | SASH3 | chrX | + | 129788430 | 129788577 | 129787974 | 129788070 | 129790939 | 129791081 | 52 | 17 | 17 | 1 | 2 | 0 | 0 | 8 | 0 | 1 | 98 | 49 | 1 | 1 | 0.963 | 0.81 | 1 | NA | 0.8 | 0.124 |
| SE | 12998 | SASH3 | SASH3 | chrX | + | 129792626 | 129792836 | 129792327 | 129792476 | 129792988 | 129793139 | 41 | 25 | 8 | 2 | 0 | 0 | 0 | 20 | 0 | 0 | 98 | 49 | 1 | 1 | 0.911 | 1 | 1 | NA | 1 | -0.03 |
| SE | 13001 | CEACAM4 | CEACAM4 | chr19 | - | 41619669 | 41619711 | 41618970 | 41619395 | 41620210 | 41620242 | 29 | 10 | 30 | 1 | 0 | 0 | 1 | 10 | 0 | 0 | 91 | 49 | 1 | 1 | 0.94 | 1 | 1 | 1 | 1 | -0.02 |
| SE | 13002 | CEACAM4 | CEACAM4 | chr19 | - | 41621650 | 41621768 | 41620574 | 41620627 | 41625600 | 41625960 | 36 | 26 | 10 | 0 | 1 | 0 | 5 | 18 | 0 | 0 | 98 | 49 | 1 | 1 | 1 | 0.929 | 1 | 1 | 1 | -0.024 |
| SE | 13005 | RPS19 | RPS19 | chr19 | + | 41869030 | 41869214 | 41861111 | 41861212 | 41869698 | 41869753 | 205 | 40 | 27 | 0 | 0 | 0 | 6 | 58 | 0 | 1 | 98 | 49 | 1 | 1 | 1 | 1 | 1 | 1 | 0.967 | 0.016 |
| SE | 13006 | ARHGEF1 | ARHGEF1 | chr19 | + | 41888751 | 41888865 | 41888191 | 41888278 | 41892330 | 41892373 | 19 | 16 | 3 | 0 | 1 | 0 | 2 | 6 | 0 | 0 | 98 | 49 | 1 | 1 | 1 | 0.889 | 1 | 1 | 1 | -0.037 |
| SE | 13008 | ARHGEF1 | ARHGEF1 | chr19 | + | 41904948 | 41905036 | 41904215 | 41904383 | 41905174 | 41905261 | 38 | 13 | 4 | 0 | 1 | 0 | 3 | 4 | 0 | 0 | 98 | 49 | 1 | 1 | 1 | 0.867 | 1 | 1 | 1 | -0.044 |
| SE | 13010 | ARHGEF1 | ARHGEF1 | chr19 | + | 41906456 | 41906620 | 41905938 | 41906025 | 41906702 | 41906803 | 52 | 33 | 5 | 4 | 0 | 0 | 0 | 18 | 0 | 2 | 98 | 49 | 1 | 1 | 0.867 | 1 | 1 | NA | 0.818 | 0.138 |
| SE | 13028 | DGCR2 | DGCR2 | chr22 | - | 19063201 | 19063278 | 19056985 | 19057162 | 19064847 | 19065058 | 17 | 23 | 13 | 1 | 0 | 0 | 0 | 14 | 0 | 0 | 98 | 49 | 1 | 1 | 0.895 | 1 | 1 | NA | 1 | -0.035 |
| SE | 13043 | JTB | JTB | chr1 | - | 153976692 | 153976775 | 153975825 | 153975905 | 153976975 | 153977013 | 46 | 9 | 42 | 0 | 0 | 2 | 0 | 5 | 0 | 0 | 98 | 49 | 1 | 1 | 1 | 1 | 0.913 | NA | 1 | -0.029 |
| SE | 13046 | TRAPPC6A | TRAPPC6A | chr19 | - | 45165126 | 45165194 | 45164852 | 45164970 | 45178134 | 45178243 | 2 | 0 | 0 | 1 | 0 | 0 | 0 | 2 | 0 | 0 | 98 | 49 | 1 | 1 | 0.5 | NA | NA | NA | 1 | -0.5 |
| SE | 13048 | CHD2 | CHD2 | chr15 | + | 92924320 | 92924552 | 92901166 | 92901299 | 92927243 | 92927330 | 47 | 19 | 9 | 1 | 0 | 0 | 5 | 6 | 0 | 0 | 98 | 49 | 1 | 1 | 0.959 | 1 | 1 | 1 | 1 | -0.014 |
| SE | 13050 | CHD2 | CHD2 | chr15 | + | 92944414 | 92944515 | 92942842 | 92943068 | 92945820 | 92945865 | 45 | 12 | 4 | 0 | 2 | 0 | 0 | 5 | 0 | 0 | 98 | 49 | 1 | 1 | 1 | 0.75 | 1 | NA | 1 | -0.083 |
| SE | 13051 | CHD2 | CHD2 | chr15 | + | 92945820 | 92945865 | 92944414 | 92944515 | 92946037 | 92946216 | 67 | 20 | 12 | 1 | 0 | 0 | 2 | 13 | 0 | 0 | 94 | 49 | 1 | 1 | 0.972 | 1 | 1 | 1 | 1 | -0.009 |
| SE | 13052 | CHD2 | CHD2 | chr15 | + | 92956458 | 92956649 | 92955422 | 92955512 | 92967324 | 92967513 | 66 | 24 | 4 | 6 | 3 | 0 | 0 | 13 | 0 | 1 | 98 | 49 | 1 | 1 | 0.846 | 0.8 | 1 | NA | 0.867 | 0.015 |
| SE | 13053 | CHD2 | CHD2 | chr15 | + | 92978233 | 92978383 | 92974878 | 92974950 | 92979134 | 92979283 | 19 | 16 | 7 | 0 | 1 | 0 | 0 | 3 | 0 | 0 | 98 | 49 | 1 | 1 | 1 | 0.889 | 1 | NA | 1 | -0.037 |
| SE | 13054 | CHD2 | CHD2 | chr15 | + | 92992858 | 92992998 | 92991475 | 92991517 | 92996956 | 92997095 | 13 | 24 | 2 | 0 | 1 | 0 | 1 | 5 | 0 | 0 | 98 | 49 | 1 | 1 | 1 | 0.923 | 1 | 1 | 1 | -0.026 |
| SE | 13055 | CHD2 | CHD2 | chr15 | + | 93004616 | 93004751 | 93002176 | 93002317 | 93009144 | 93009323 | 21 | 13 | 0 | 1 | 0 | 0 | 0 | 6 | 0 | 0 | 98 | 49 | 1 | 1 | 0.913 | 1 | NA | NA | 1 | -0.043 |
| SE | 13056 | WAS | WAS | chrX | + | 48685733 | 48685836 | 48685546 | 48685633 | 48685945 | 48685987 | 112 | 59 | 30 | 0 | 0 | 2 | 5 | 20 | 0 | 0 | 98 | 49 | 1 | 1 | 1 | 1 | 0.882 | 1 | 1 | -0.039 |
| SE | 13057 | WAS | WAS | chrX | + | 48688659 | 48689066 | 48688299 | 48688453 | 48689319 | 48689434 | 64 | 13 | 22 | 2 | 0 | 0 | 1 | 15 | 0 | 0 | 98 | 49 | 1 | 1 | 0.941 | 1 | 1 | 1 | 1 | -0.02 |
| SE | 13059 | ERCC1 | ERCC1 | chr19 | - | 45413962 | 45414034 | 45413433 | 45413745 | 45414860 | 45414960 | 7 | 0 | 0 | 1 | 0 | 0 | 0 | 2 | 0 | 0 | 98 | 49 | 1 | 1 | 0.778 | NA | NA | NA | 1 | -0.222 |
| SE | 13060 | DOK3 | DOK3 | chr5 | - | 177508236 | 177508542 | 177505010 | 177505110 | 177509474 | 177509575 | 79 | 29 | 16 | 0 | 0 | 0 | 7 | 32 | 0 | 1 | 98 | 49 | 1 | 1 | 1 | 1 | 1 | 1 | 0.941 | 0.03 |
| SE | 13070 | CMTR1 | CMTR1 | chr6 | + | 37461969 | 37462102 | 37461548 | 37461645 | 37462828 | 37463008 | 6 | 2 | 0 | 0 | 0 | 0 | 0 | 3 | 0 | 1 | 98 | 49 | 1 | 1 | 1 | 1 | NA | NA | 0.6 | 0.4 |
| SE | 13071 | CMTR1 | CMTR1 | chr6 | + | 37462828 | 37463008 | 37461548 | 37461645 | 37471020 | 37471077 | 8 | 0 | 0 | 0 | 0 | 0 | 0 | 3 | 1 | 0 | 98 | 49 | 1 | 1 | 1 | NA | NA | 0 | 1 | 0.5 |
| SE | 13073 | CNOT8 | CNOT8 | chr5 | + | 154863206 | 154863395 | 154858518 | 154858768 | 154865191 | 154865385 | 12 | 3 | 4 | 2 | 0 | 1 | 0 | 2 | 0 | 0 | 98 | 49 | 1 | 1 | 0.75 | 1 | 0.667 | NA | 1 | -0.194 |
| SE | 13074 | CNOT8 | CNOT8 | chr5 | + | 154863206 | 154863395 | 154858518 | 154858768 | 154870660 | 154870822 | 6 | 0 | 0 | 0 | 0 | 0 | 0 | 2 | 0 | 1 | 98 | 49 | 1 | 1 | 1 | NA | NA | NA | 0.5 | 0.5 |
| SE | 13075 | CNOT8 | CNOT8 | chr5 | + | 154865191 | 154865385 | 154858518 | 154858768 | 154870660 | 154870822 | 6 | 2 | 6 | 0 | 0 | 0 | 0 | 0 | 0 | 1 | 98 | 49 | 1 | 1 | 1 | 1 | 1 | NA | 0 | 1 |
| SE | 13080 | CNOT8 | CNOT8 | chr5 | + | 154870660 | 154870822 | 154867691 | 154867792 | 154871729 | 154871874 | 10 | 3 | 2 | 0 | 0 | 1 | 0 | 3 | 0 | 0 | 98 | 49 | 1 | 1 | 1 | 1 | 0.5 | NA | 1 | -0.167 |
| SE | 13085 | STT3B | STT3B | chr3 | + | 31624913 | 31625085 | 31623673 | 31623861 | 31625953 | 31626127 | 15 | 7 | 1 | 2 | 0 | 0 | 0 | 3 | 0 | 0 | 98 | 49 | 1 | 1 | 0.789 | 1 | 1 | NA | 1 | -0.07 |
| SE | 13087 | TSC1 | TSC1 | chr9 | - | 132903650 | 132903817 | 132902604 | 132902787 | 132904410 | 132904454 | 1 | 0 | 3 | 1 | 0 | 0 | 0 | 1 | 0 | 0 | 98 | 49 | 1 | 1 | 0.333 | NA | 1 | NA | 1 | -0.334 |
| SE | 13093 | USP33 | USP33 | chr1 | - | 77701368 | 77701471 | 77697862 | 77697931 | 77711746 | 77711855 | 9 | 2 | 2 | 1 | 2 | 0 | 0 | 5 | 0 | 2 | 98 | 49 | 1 | 1 | 0.818 | 0.333 | 1 | NA | 0.556 | 0.161 |
| SE | 13094 | USP33 | USP33 | chr1 | - | 77745420 | 77745599 | 77741616 | 77741748 | 77759642 | 77759879 | 2 | 2 | 0 | 8 | 3 | 0 | 0 | 1 | 0 | 0 | 98 | 49 | 1 | 1 | 0.111 | 0.25 | NA | NA | 1 | -0.82 |
| SE | 13098 | DHX34 | DHX34 | chr19 | + | 47357865 | 47358120 | 47355038 | 47355350 | 47359967 | 47360070 | 58 | 21 | 2 | 0 | 0 | 0 | 2 | 24 | 0 | 1 | 98 | 49 | 1 | 1 | 1 | 1 | 1 | 1 | 0.923 | 0.038 |
| SE | 13099 | DHX34 | DHX34 | chr19 | + | 47373598 | 47373700 | 47372729 | 47372923 | 47375465 | 47375708 | 21 | 21 | 7 | 1 | 0 | 0 | 1 | 14 | 0 | 0 | 98 | 49 | 1 | 1 | 0.913 | 1 | 1 | 1 | 1 | -0.029 |
| SE | 13107 | KIDINS220 | KIDINS220 | chr2 | - | 8793809 | 8793987 | 8791059 | 8791224 | 8796770 | 8796869 | 18 | 7 | 4 | 0 | 1 | 0 | 0 | 2 | 0 | 0 | 98 | 49 | 1 | 1 | 1 | 0.778 | 1 | NA | 1 | -0.074 |
| SE | 13108 | KIDINS220 | KIDINS220 | chr2 | - | 8826985 | 8827129 | 8818694 | 8818793 | 8837479 | 8837625 | 14 | 8 | 3 | 0 | 1 | 0 | 0 | 3 | 0 | 0 | 98 | 49 | 1 | 1 | 1 | 0.8 | 1 | NA | 1 | -0.067 |

| | | | | | | | | | | | | | | | | | | | | | | | | | | | | | | | |
|---|---|---|---|---|---|---|---|---|---|---|---|---|---|---|---|---|---|---|---|---|---|---|---|---|---|---|---|---|---|---|---|
| SE | 13109 | GBP3 | GBP3 | chr1 | - | 89020531 | 89020743 | 89015286 | 89015414 | 89022683 | 89022866 | 8 | 1 | 0 | 1 | 0 | 0 | 1 | 0 | 0 | 0 | 98 | 49 | 1 | 1 | 0.8 | 1 | NA | 1 | NA | -0.1 |
| SE | 13113 | LRRC42 | LRRC42 | chr1 | + | 53962033 | 53962122 | 53960355 | 53960474 | 53962295 | 53962409 | 0 | 0 | 0 | 0 | 1 | 0 | 0 | 1 | 0 | 0 | 98 | 49 | 1 | 1 | NA | 0 | NA | NA | 1 | -1 |
| SE | 13120 | PRPF18 | PRPF18 | chr10 | + | 13613740 | 13613881 | 13611614 | 13611683 | 13614014 | 13614086 | 9 | 7 | 1 | 3 | 0 | 0 | 0 | 3 | 0 | 0 | 98 | 49 | 1 | 1 | 0.6 | 1 | 1 | NA | 1 | -0.133 |
| SE | 13123 | EHMT1 | EHMT1 | chr9 | + | 137752330 | 137752408 | 137743901 | 137744090 | 137754170 | 137754291 | 9 | 0 | 0 | 2 | 0 | 0 | 0 | 4 | 0 | 0 | 98 | 49 | 1 | 1 | 0.692 | NA | NA | NA | 1 | -0.308 |
| SE | 13126 | CEP350 | CEP350 | chr1 | + | 179990506 | 179990621 | 179987239 | 179987286 | 179992061 | 179992221 | 17 | 4 | 0 | 1 | 0 | 2 | 0 | 5 | 0 | 0 | 98 | 49 | 1 | 1 | 0.895 | 1 | 0 | NA | 1 | -0.368 |
| SE | 13127 | CEP350 | CEP350 | chr1 | + | 180036925 | 180037089 | 180033861 | 180034082 | 180041137 | 180041248 | 5 | 7 | 4 | 0 | 0 | 0 | 0 | 1 | 0 | 1 | 98 | 49 | 1 | 1 | 1 | 1 | 1 | NA | 0.333 | 0.667 |
| SE | 13128 | CEP350 | CEP350 | chr1 | + | 180043055 | 180043192 | 180041661 | 180041802 | 180044050 | 180044173 | 13 | 3 | 0 | 2 | 0 | 0 | 2 | 5 | 0 | 0 | 98 | 49 | 1 | 1 | 0.765 | 1 | NA | 1 | 1 | -0.117 |
| SE | 13137 | RPS6KA3 | RPS6KA3 | chrX | - | 20167588 | 20167747 | 20164898 | 20165060 | 20169401 | 20169491 | 26 | 19 | 12 | 0 | 4 | 0 | 0 | 11 | 0 | 0 | 98 | 49 | 1 | 1 | 1 | 0.704 | 1 | NA | 1 | -0.099 |
| SE | 13141 | CD37 | CD37 | chr19 | + | 49339329 | 49339413 | 49338699 | 49338936 | 49340250 | 49340604 | 29 | 17 | 24 | 2 | 0 | 0 | 1 | 21 | 0 | 0 | 98 | 49 | 1 | 1 | 0.879 | 1 | 1 | 1 | 1 | -0.04 |
| SE | 13146 | IL1RN | IL1RN | chr2 | + | 113131044 | 113131157 | 113129575 | 113129664 | 113132655 | 113134016 | 37 | 37 | 14 | 0 | 1 | 0 | 0 | 12 | 0 | 0 | 98 | 49 | 1 | 1 | 1 | 0.949 | 1 | NA | 1 | -0.017 |
| SE | 13150 | FCGRT | FCGRT | chr19 | + | 49524506 | 49524776 | 49514210 | 49514486 | 49526009 | 49526428 | 73 | 30 | 27 | 1 | 0 | 0 | 3 | 16 | 0 | 0 | 98 | 49 | 1 | 1 | 0.973 | 1 | 1 | 1 | 1 | -0.009 |
| SE | 13151 | FCGRT | FCGRT | chr19 | + | 49525456 | 49525573 | 49524506 | 49524776 | 49526009 | 49526428 | 66 | 64 | 45 | 2 | 0 | 0 | 6 | 18 | 0 | 0 | 98 | 49 | 1 | 1 | 0.943 | 1 | 1 | 1 | 1 | -0.019 |
| SE | 13160 | AP2A1 | AP2A1 | chr19 | + | 49800960 | 49801058 | 49799967 | 49800150 | 49801389 | 49801621 | 20 | 9 | 5 | 0 | 0 | 0 | 1 | 3 | 1 | 0 | 98 | 49 | 1 | 1 | 1 | 1 | 1 | 0.333 | 1 | 0.334 |
| SE | 13163 | SMG7 | SMG7 | chr1 | + | 183512836 | 183512868 | 183472370 | 183472649 | 183515873 | 183515991 | 3 | 2 | 1 | 0 | 1 | 0 | 0 | 2 | 0 | 0 | 81 | 49 | 1 | 1 | 1 | 0.547 | 1 | NA | 1 | -0.151 |
| SE | 13166 | GPATCH2L | GPATCH2L | chr14 | + | 76178207 | 76178322 | 76177987 | 76178042 | 76180763 | 76180849 | 0 | 3 | 0 | 6 | 6 | 0 | 0 | 0 | 0 | 2 | 98 | 49 | 1 | 1 | 0 | 0.2 | NA | NA | 0 | 0.1 |
| SE | 13167 | GPATCH2L | GPATCH2L | chr14 | + | 76180763 | 76180849 | 76177987 | 76178042 | 76195877 | 76195972 | 16 | 7 | 0 | 2 | 0 | 0 | 1 | 4 | 0 | 0 | 98 | 49 | 1 | 1 | 0.8 | 1 | NA | 1 | 1 | -0.1 |
| SE | 13168 | PNKP | PNKP | chr19 | - | 49861607 | 49861695 | 49861448 | 49861510 | 49861771 | 49861881 | 3 | 0 | 6 | 0 | 1 | 0 | 1 | 2 | 0 | 0 | 98 | 49 | 1 | 1 | 1 | 0 | 1 | 1 | 1 | -0.333 |
| SE | 13172 | VRK3 | VRK3 | chr19 | - | 49997503 | 49997570 | 49995190 | 49995275 | 50000789 | 50000854 | 20 | 14 | 8 | 1 | 1 | 0 | 0 | 5 | 0 | 0 | 98 | 49 | 1 | 1 | 0.909 | 0.875 | 1 | NA | 1 | -0.072 |
| SE | 13173 | VRK3 | VRK3 | chr19 | - | 50009235 | 50009385 | 50007568 | 50007826 | 50016023 | 50016163 | 25 | 3 | 2 | 2 | 3 | 1 | 0 | 4 | 0 | 0 | 98 | 49 | 1 | 1 | 0.862 | 0.333 | 0.5 | NA | 1 | -0.435 |
| SE | 13183 | EMC10 | EMC10 | chr19 | + | 50481860 | 50481947 | 50480883 | 50480977 | 50482148 | 50483526 | 2 | 0 | 0 | 3 | 0 | 1 | 0 | 1 | 0 | 2 | 98 | 49 | 1 | 1 | 0.25 | NA | 0 | NA | 0.2 | -0.075 |
| SE | 13194 | LONRF1 | LONRF1 | chr8 | - | 12725726 | 12725879 | 12721896 | 12723254 | 12728900 | 12729063 | 2 | 1 | 0 | 0 | 1 | 0 | 0 | 1 | 0 | 0 | 98 | 49 | 1 | 1 | 1 | 0.333 | NA | NA | 1 | -0.334 |
| SE | 13204 | ARHGAP30 | ARHGAP30 | chr1 | - | 161051313 | 161051715 | 161049423 | 161049689 | 161052285 | 161052363 | 62 | 18 | 14 | 1 | 0 | 0 | 0 | 6 | 0 | 0 | 98 | 49 | 1 | 1 | 0.969 | 1 | 1 | NA | 1 | -0.01 |
| SE | 13205 | ARHGAP30 | ARHGAP30 | chr1 | - | 161056387 | 161056532 | 161054622 | 161054705 | 161059613 | 161059716 | 92 | 62 | 21 | 5 | 3 | 2 | 0 | 26 | 0 | 3 | 98 | 49 | 1 | 1 | 0.902 | 0.912 | 0.84 | NA | 0.812 | 0.073 |
| SE | 13206 | ARHGAP30 | ARHGAP30 | chr1 | - | 161056387 | 161056532 | 161054622 | 161054705 | 161069527 | 161069970 | 54 | 33 | 7 | 1 | 1 | 0 | 0 | 18 | 0 | 2 | 98 | 49 | 1 | 1 | 0.964 | 0.943 | 1 | NA | 0.818 | 0.151 |
| SE | 13207 | ARHGAP30 | ARHGAP30 | chr1 | - | 161059613 | 161059716 | 161054622 | 161054705 | 161069527 | 161069970 | 40 | 18 | 10 | 1 | 1 | 0 | 1 | 18 | 0 | 2 | 98 | 49 | 1 | 1 | 0.952 | 0.9 | 1 | 1 | 0.818 | 0.042 |
| SE | 13208 | ARHGAP30 | ARHGAP30 | chr1 | - | 161059613 | 161059716 | 161056387 | 161056532 | 161069527 | 161069970 | 75 | 44 | 22 | 2 | 0 | 0 | 1 | 23 | 0 | 0 | 98 | 49 | 1 | 1 | 0.949 | 1 | 1 | 1 | 1 | -0.017 |
| SE | 13213 | FES | FES | chr15 | + | 90885411 | 90885585 | 90885036 | 90885258 | 90886960 | 90887057 | 23 | 13 | 6 | 1 | 0 | 0 | 1 | 3 | 0 | 1 | 98 | 49 | 1 | 1 | 0.92 | 1 | 1 | 1 | 0.6 | 0.173 |
| SE | 13215 | FES | FES | chr15 | + | 90890981 | 90891191 | 90890400 | 90890484 | 90891553 | 90891676 | 10 | 9 | 5 | 1 | 0 | 0 | 0 | 7 | 0 | 0 | 98 | 49 | 1 | 1 | 0.833 | 1 | 1 | NA | 1 | -0.056 |
| SE | 13216 | FES | FES | chr15 | + | 90891553 | 90891676 | 90890400 | 90890484 | 90892057 | 90892111 | 13 | 6 | 4 | 1 | 0 | 0 | 0 | 8 | 0 | 0 | 98 | 49 | 1 | 1 | 0.867 | 1 | 1 | NA | 1 | -0.044 |
| SE | 13217 | SIGLEC5 | SIGLEC5 | chr19 | - | 51626031 | 51626113 | 51611502 | 51612422 | 51627148 | 51627248 | 23 | 18 | 26 | 1 | 0 | 1 | 0 | 13 | 0 | 1 | 98 | 49 | 1 | 1 | 0.92 | 1 | 0.929 | NA | 0.867 | 0.083 |
| SE | 13219 | ALG13 | ALG13 | chrX | + | 111686136 | 111686174 | 111684964 | 111685103 | 111687886 | 111690395 | 4 | 0 | 0 | 4 | 0 | 0 | 0 | 2 | 0 | 0 | 87 | 49 | 1 | 1 | 0.36 | NA | NA | NA | 1 | -0.64 |
| SE | 13225 | AFF4 | AFF4 | chr5 | - | 132893029 | 132893118 | 132892163 | 132892404 | 132896322 | 132897240 | 7 | 10 | 0 | 0 | 1 | 0 | 0 | 2 | 0 | 0 | 98 | 49 | 1 | 1 | 1 | 0.833 | NA | NA | 1 | -0.084 |
| SE | 13226 | AFF4 | AFF4 | chr5 | - | 132898229 | 132898392 | 132896322 | 132897240 | 132899103 | 132899141 | 15 | 7 | 7 | 1 | 0 | 0 | 0 | 2 | 0 | 0 | 98 | 49 | 1 | 1 | 0.882 | 1 | 1 | NA | 1 | -0.039 |
| SE | 13230 | PPP2R1A | PPP2R1A | chr19 | + | 52220188 | 52220249 | 52219690 | 52219864 | 52220978 | 52221133 | 5 | 0 | 4 | 0 | 0 | 0 | 0 | 6 | 0 | 1 | 98 | 49 | 1 | 1 | 1 | NA | 1 | NA | 0.75 | 0.25 |
| SE | 13232 | RBM23 | RBM23 | chr14 | - | 22908332 | 22908380 | 22906194 | 22906368 | 22909482 | 22909595 | 25 | 12 | 2 | 7 | 6 | 8 | 0 | 2 | 0 | 0 | 97 | 49 | 1 | 1 | 0.643 | 0.503 | 0.112 | NA | 1 | -0.581 |
| SE | 13236 | RBM23 | RBM23 | chr14 | - | 22911327 | 22911403 | 22908332 | 22908380 | 22918998 | 22919187 | 9 | 16 | 3 | 0 | 1 | 0 | 0 | 2 | 0 | 0 | 98 | 49 | 1 | 1 | 1 | 0.889 | 1 | NA | 1 | -0.037 |
| SE | 13244 | RELL1 | RELL1 | chr4 | - | 37638446 | 37638504 | 37634886 | 37635123 | 37647367 | 37647439 | 56 | 55 | 36 | 2 | 0 | 0 | 0 | 7 | 0 | 0 | 98 | 49 | 1 | 1 | 0.933 | 1 | 1 | NA | 1 | -0.022 |
| SE | 13245 | RELL1 | RELL1 | chr4 | - | 37649275 | 37649500 | 37647367 | 37647439 | 37686199 | 37686377 | 26 | 15 | 12 | 0 | 0 | 2 | 1 | 7 | 0 | 0 | 98 | 49 | 1 | 1 | 1 | 1 | 0.75 | 1 | 1 | -0.083 |
| SE | 13249 | TSR3 | TSR3 | chr16 | - | 1349888 | 1349952 | 1349239 | 1349608 | 1350057 | 1350234 | 12 | 9 | 5 | 0 | 0 | 0 | 0 | 0 | 0 | 1 | 98 | 49 | 1 | 1 | 1 | 1 | 1 | NA | 0 | 1 |
| SE | 13252 | ENTPD1 | ENTPD1 | chr10 | + | 95842343 | 95842494 | 95839690 | 95839808 | 95844475 | 95844635 | 38 | 26 | 11 | 3 | 4 | 1 | 0 | 10 | 0 | 0 | 98 | 49 | 1 | 1 | 0.864 | 0.765 | 0.846 | NA | 1 | -0.175 |
| SE | 13253 | ENTPD1 | ENTPD1 | chr10 | + | 95845356 | 95845596 | 95844475 | 95844635 | 95847445 | 95847706 | 49 | 20 | 10 | 1 | 1 | 0 | 1 | 12 | 0 | 0 | 98 | 49 | 1 | 1 | 0.961 | 0.909 | 1 | 1 | 1 | -0.043 |
| SE | 13266 | PSMF1 | PSMF1 | chr20 | + | 1127425 | 1127508 | 1125497 | 1125650 | 1135120 | 1135306 | 16 | 6 | 5 | 0 | 2 | 0 | 1 | 1 | 0 | 0 | 98 | 49 | 1 | 1 | 1 | 0.6 | 1 | 1 | 1 | -0.133 |
| SE | 13268 | CYP20A1 | CYP20A1 | chr2 | + | 203246754 | 203246921 | 203245845 | 203245895 | 203251966 | 203252109 | 1 | 0 | 0 | 0 | 0 | 0 | 0 | 0 | 1 | 0 | 98 | 49 | 1 | 1 | 1 | NA | NA | 0 | NA | 1 |
| SE | 13269 | CYP20A1 | CYP20A1 | chr2 | + | 203272669 | 203272748 | 203266513 | 203266681 | 203278572 | 203278688 | 2 | 0 | 0 | 3 | 0 | 0 | 0 | 1 | 0 | 0 | 98 | 49 | 1 | 1 | 0.25 | NA | NA | NA | 1 | -0.75 |
| SE | 13270 | CYP20A1 | CYP20A1 | chr2 | + | 203278572 | 203278688 | 203266513 | 203266681 | 203280058 | 203280113 | 4 | 1 | 0 | 0 | 1 | 0 | 0 | 1 | 0 | 0 | 98 | 49 | 1 | 1 | 1 | 0.333 | NA | NA | 1 | -0.334 |
| SE | 13271 | BCKDK | BCKDK | chr16 | + | 31110687 | 31110761 | 31110400 | 31110499 | 31111090 | 31111219 | 11 | 3 | 7 | 1 | 2 | 0 | 0 | 1 | 0 | 1 | 98 | 49 | 1 | 1 | 0.846 | 0.429 | 1 | NA | 0.333 | 0.425 |
| SE | 13281 | RBM5 | RBM5 | chr3 | + | 50100531 | 50100605 | 50099981 | 50100051 | 50103082 | 50103166 | 8 | 1 | 5 | 14 | 6 | 1 | 0 | 0 | 0 | 2 | 98 | 49 | 1 | 1 | 0.222 | 0.077 | 0.714 | NA | 0 | 0.338 |
| SE | 13282 | RBM5 | RBM5 | chr3 | + | 50104247 | 50104308 | 50103082 | 50103166 | 50105076 | 50105142 | 49 | 24 | 14 | 1 | 0 | 0 | 2 | 6 | 0 | 1 | 98 | 49 | 1 | 1 | 0.961 | 1 | 1 | 1 | 0.75 | 0.112 |
| SE | 13283 | RBM5 | RBM5 | chr3 | + | 50105548 | 50105709 | 50105076 | 50105142 | 50106766 | 50106864 | 70 | 35 | 32 | 0 | 1 | 0 | 1 | 9 | 0 | 0 | 98 | 49 | 1 | 1 | 1 | 0.946 | 1 | 1 | 1 | -0.018 |
| SE | 13284 | RBM5 | RBM5 | chr3 | + | 50110378 | 50110463 | 50109602 | 50109688 | 50110678 | 50110770 | 37 | 6 | 17 | 25 | 5 | 2 | 0 | 3 | 0 | 0 | 98 | 49 | 1 | 1 | 0.425 | 0.375 | 0.81 | NA | 1 | -0.463 |
| SE | 13285 | RBM5 | RBM5 | chr3 | + | 50115427 | 50115607 | 50114179 | 50114251 | 50115905 | 50115980 | 93 | 20 | 12 | 0 | 1 | 0 | 6 | 16 | 0 | 0 | 98 | 49 | 1 | 1 | 1 | 0.909 | 1 | 1 | 1 | -0.03 |
| SE | 13286 | RBM5 | RBM5 | chr3 | + | 50117249 | 50117379 | 50117073 | 50117171 | 50118330 | 50118964 | 49 | 8 | 21 | 0 | 1 | 0 | 2 | 10 | 0 | 0 | 98 | 49 | 1 | 1 | 1 | 0.8 | 1 | 1 | 1 | -0.067 |
| SE | 13287 | TARM1 | TARM1 | chr19 | - | 54075882 | 54075918 | 54074823 | 54075114 | 54081306 | 54081365 | 0 | 0 | 0 | 2 | 0 | 0 | 0 | 1 | 0 | 0 | 85 | 49 | 1 | 1 | 0 | NA | NA | NA | 1 | -1 |
| SE | 13294 | MBOAT7 | MBOAT7 | chr19 | - | 54180772 | 54181133 | 54178274 | 54178941 | 54183520 | 54183680 | 69 | 32 | 18 | 1 | 1 | 0 | 0 | 9 | 0 | 0 | 98 | 49 | 1 | 1 | 0.972 | 0.941 | 1 | NA | 1 | -0.029 |
| SE | 13295 | MBOAT7 | MBOAT7 | chr19 | - | 54187160 | 54187287 | 54183520 | 54183680 | 54188216 | 54188346 | 115 | 46 | 59 | 1 | 0 | 0 | 10 | 24 | 0 | 1 | 98 | 49 | 1 | 1 | 0.983 | 1 | 1 | 1 | 0.923 | 0.033 |
| SE | 13296 | MBOAT7 | MBOAT7 | chr19 | - | 54188216 | 54188346 | 54187160 | 54187287 | 54188432 | 54188511 | 68 | 38 | 34 | 1 | 0 | 0 | 3 | 20 | 0 | 0 | 98 | 49 | 1 | 1 | 0.971 | 1 | 1 | 1 | 1 | -0.01 |
| SE | 13297 | MBOAT7 | MBOAT7 | chr19 | - | 54188432 | 54188511 | 54188216 | 54188346 | 54189337 | 54189882 | 118 | 64 | 47 | 4 | 4 | 0 | 9 | 20 | 0 | 1 | 98 | 49 | 1 | 1 | 0.937 | 0.889 | 1 | 1 | 0.909 | -0.012 |
| SE | 13298 | TSEN34 | TSEN34 | chr19 | + | 54191360 | 54191607 | 54190267 | 54190450 | 54191720 | 54191964 | 18 | 11 | 29 | 0 | 1 | 0 | 2 | 5 | 0 | 0 | 98 | 49 | 1 | 1 | 1 | 0.846 | 1 | 1 | 1 | -0.051 |
| SE | 13301 | LILRA6 | LILRA6 | chr19 | - | 54239900 | 54239951 | 54238821 | 54239089 | 54240273 | 54240576 | 3 | 5 | 0 | 0 | 2 | 0 | 1 | 4 | 0 | 0 | 98 | 49 | 1 | 1 | 1 | 0.556 | NA | 1 | 1 | -0.222 |
| SE | 13316 | LILRA5 | LILRA5 | chr19 | - | 54311863 | 54312148 | 54311164 | 54311716 | 54312536 | 54312621 | 57 | 36 | 42 | 0 | 0 | 0 | 1 | 18 | 0 | 1 | 98 | 49 | 1 | 1 | 1 | 1 | 1 | 1 | 0.9 | 0.05 |
| SE | 13317 | LILRA5 | LILRA5 | chr19 | - | 54312334 | 54312370 | 54311863 | 54312148 | 54312536 | 54312621 | 39 | 27 | 49 | 6 | 7 | 2 | 0 | 15 | 0 | 2 | 85 | 49 | 1 | 1 | 0.789 | 0.69 | 0.934 | NA | 0.812 | -0.008 |
| SE | 13318 | LILRA5 | LILRA5 | chr19 | - | 54312536 | 54312621 | 54312334 | 54312370 | 54313016 | 54313139 | 33 | 20 | 36 | 2 | 0 | 0 | 0 | 14 | 0 | 0 | 98 | 49 | 1 | 1 | 0.892 | 1 | 1 | NA | 1 | -0.036 |
| SE | 13319 | LENG8 | LENG8 | chr19 | + | 54451289 | 54451382 | 54449161 | 54449310 | 54452092 | 54452267 | 17 | 15 | 2 | 0 | 1 | 0 | 1 | 4 | 0 | 0 | 98 | 49 | 1 | 1 | 1 | 0.882 | 1 | 1 | 1 | -0.039 |
| SE | 13320 | LENG8 | LENG8 | chr19 | + | 54453545 | 54453656 | 54452650 | 54452752 | 54454429 | 54454682 | 20 | 5 | 1 | 1 | 0 | 0 | 0 | 5 | 0 | 0 | 98 | 49 | 1 | 1 | 0.909 | 1 | 1 | NA | 0.714 | 0.256 |
| SE | 13331 | LILRB1 | LILRB1 | chr19 | + | 54633637 | 54633688 | 54633015 | 54633318 | 54633970 | 54634021 | 5 | 1 | 0 | 0 | 0 | 0 | 0 | 2 | 0 | 1 | 98 | 49 | 1 | 1 | 1 | 1 | NA | NA | 0.5 | 0.5 |
| SE | 13571 | CEP192 | CEP192 | chr18 | + | 13038369 | 13038579 | 13037236 | 13037301 | 13040829 | 13040956 | 1 | 1 | 0 | 1 | 0 | 0 | 0 | 2 | 0 | 0 | 98 | 49 | 1 | 1 | 0.333 | 1 | NA | NA | 1 | -0.334 |
| SE | 13573 | CEP192 | CEP192 | chr18 | + | 13059081 | 13059312 | 13055779 | 13056698 | 13067830 | 13067956 | 2 | 5 | 0 | 2 | 0 | 0 | 1 | 1 | 0 | 0 | 98 | 49 | 1 | 1 | 0.333 | 1 | NA | 1 | 1 | -0.334 |
| SE | 13583 | PIK3AP1 | PIK3AP1 | chr10 | - | 96602279 | 96602398 | 96593311 | 96595634 | 96603978 | 96604049 | 36 | 19 | 18 | 0 | 1 | 0 | 2 | 27 | 0 | 0 | 98 | 49 | 1 | 1 | 1 | 0.905 | 1 | 1 | 1 | -0.032 |
| SE | 13584 | PIK3AP1 | PIK3AP1 | chr10 | - | 96603978 | 96604049 | 96593311 | 96595634 | 96609711 | 96609867 | 18 | 5 | 6 | 1 | 0 | 0 | 0 | 9 | 0 | 0 | 98 | 49 | 1 | 1 | 0.9 | 1 | 1 | NA | 1 | -0.033 |
| SE | 13585 | PIK3AP1 | PIK3AP1 | chr10 | - | 96609711 | 96609867 | 96603978 | 96604049 | 96616638 | 96616711 | 32 | 26 | 11 | 0 | 1 | 0 | 1 | 11 | 0 | 0 | 98 | 49 | 1 | 1 | 1 | 0.929 | 1 | 1 | 1 | -0.024 |
| SE | 13586 | PIK3AP1 | PIK3AP1 | chr10 | - | 96652697 | 96652842 | 96651508 | 96651651 | 96656797 | 96656934 | 27 | 16 | 3 | 1 | 0 | 0 | 1 | 15 | 0 | 0 | 98 | 49 | 1 | 1 | 0.931 | 1 | 1 | 1 | 1 | -0.023 |
| SE | 13590 | FAM129A | FAM129A | chr1 | - | 184805945 | 184806056 | 184803584 | 184803692 | 184808073 | 184808235 | 167 | 67 | 46 | 0 | 1 | 0 | 1 | 25 | 0 | 0 | 98 | 49 | 1 | 1 | 1 | 0.971 | 1 | 1 | 1 | -0.01 |
| SE | 13591 | FAM129A | FAM129A | chr1 | - | 184818637 | 184818825 | 184808073 | 184808235 | 184823166 | 184823329 | 163 | 107 | 53 | 0 | 0 | 1 | 0 | 16 | 0 | 0 | 98 | 49 | 1 | 1 | 1 | 1 | 0.964 | NA | 1 | -0.012 |
| SE | 13593 | FAM129A | FAM129A | chr1 | - | 184831846 | 184831962 | 184823637 | 184823742 | 184884632 | 184884800 | 227 | 89 | 67 | 0 | 3 | 0 | 0 | 32 | 0 | 0 | 98 | 49 | 1 | 1 | 1 | 0.937 | 1 | NA | 1 | -0.021 |
| SE | 13594 | HIATL1 | HIATL1 | chr9 | + | 94456239 | 94456337 | 94453957 | 94454070 | 94458321 | 94458464 | 23 | 5 | 5 | 0 | 0 | 0 | 0 | 10 | 0 | 1 | 98 | 49 | 1 | 1 | 1 | 1 | 1 | NA | 0.833 | 0.167 |
| SE | 13612 | GTF3C2 | GTF3C2 | chr2 | - | 27335597 | 27335706 | 27333973 | 27333999 | 27335916 | 27336028 | 6 | 0 | 0 | 0 | 1 | 0 | 0 | 5 | 0 | 0 | 98 | 49 | 1 | 1 | 1 | 0 | NA | NA | 1 | -0.5 |
| SE | 13615 | ZNF274 | ZNF274 | chr19 | + | 58209960 | 58210073 | 58206719 | 58207202 | 58211559 | 58211686 | 1 | 1 | 0 | 0 | 0 | 0 | 1 | 0 | 3 | 0 | 98 | 49 | 1 | 1 | 1 | 1 | NA | 0.143 | NA | 0.857 |
| SE | 13623 | VPS8 | VPS8 | chr3 | + | 184866875 | 184866950 | 184862896 | 184863067 | 184868023 | 184868059 | 94 | 40 | 4 | 0 | 1 | 0 | 1 | 11 | 0 | 0 | 98 | 49 | 1 | 1 | 1 | 0.952 | 1 | 1 | 1 | -0.016 |
| SE | 13624 | VPS8 | VPS8 | chr3 | + | 184868945 | 184869036 | 184868023 | 184868059 | 184869481 | 184869528 | 93 | 22 | 7 | 2 | 0 | 0 | 0 | 12 | 0 | 0 | 98 | 49 | 1 | 1 | 0.959 | 1 | 1 | NA | 1 | -0.014 |
| SE | 13625 | VPS8 | VPS8 | chr3 | + | 184914980 | 184915053 | 184913518 | 184913561 | 184915354 | 184915474 | 51 | 32 | 11 | 2 | 0 | 0 | 2 | 12 | 0 | 0 | 98 | 49 | 1 | 1 | 0.927 | 1 | 1 | 1 | 1 | -0.024 |
| SE | 13628 | SH3YL1 | SH3YL1 | chr2 | - | 224863 | 224920 | 218135 | 219001 | 229965 | 230044 | 0 | 0 | 0 | 1 | 0 | 0 | 0 | 3 | 0 | 0 | 98 | 49 | 1 | 1 | 0 | NA | NA | NA | 1 | -1 |
| SE | 13636 | CAMLG | CAMLG | chr5 | + | 134741062 | 134741523 | 134738479 | 134738792 | 134743986 | 134744052 | 8 | 2 | 4 | 0 | 0 | 0 | 0 | 0 | 0 | 1 | 98 | 49 | 1 | 1 | 1 | 1 | 1 | NA | 0 | 1 |
| SE | 13638 | CAMLG | CAMLG | chr5 | + | 134743986 | 134744052 | 134738479 | 134738792 | 134750758 | 134752160 | 4 | 4 | 1 | 2 | 2 | 0 | 0 | 1 | 0 | 0 | 98 | 49 | 1 | 1 | 0.5 | 0.5 | 1 | NA | 1 | -0.333 |
| SE | 13640 | EPSTI1 | EPSTI1 | chr13 | - | 42889210 | 42889242 | 42886387 | 42888502 | 42895008 | 42895108 | 0 | 1 | 0 | 0 | 0 | 0 | 0 | 8 | 0 | 4 | 81 | 49 | 1 | 1 | NA | 1 | NA | NA | 0.547 | 0.453 |
| SE | 13641 | EPSTI1 | EPSTI1 | chr13 | - | 42895008 | 42895108 | 42886387 | 42888502 | 42900309 | 42900383 | 2 | 5 | 0 | 0 | 0 | 0 | 1 | 9 | 0 | 2 | 98 | 49 | 1 | 1 | 1 | 1 | NA | 1 | 0.692 | 0.154 |
| SE | 13643 | EPSTI1 | EPSTI1 | chr13 | - | 42970611 | 42970670 | 42969093 | 42969177 | 42991977 | 42992271 | 5 | 5 | 0 | 0 | 0 | 0 | 0 | 25 | 1 | 0 | 98 | 49 | 1 | 1 | 1 | 1 | NA | 0 | 1 | 0.5 |
| SE | 13644 | DOPEY2 | DOPEY2 | chr21 | + | 36211552 | 36211651 | 36208714 | 36208904 | 36211973 | 36212097 | 3 | 1 | 2 | 0 | 0 | 0 | 2 | 1 | 0 | 1 | 98 | 49 | 1 | 1 | 1 | 1 | 1 | 1 | 0.333 | 0.334 |
| SE | 13645 | DOPEY2 | DOPEY2 | chr21 | + | 36214080 | 36214190 | 36211973 | 36212097 | 36214441 | 36214556 | 3 | 1 | 0 | 0 | 0 | 0 | 2 | 6 | 0 | 1 | 98 | 49 | 1 | 1 | 1 | 1 | NA | 1 | 0.75 | 0.125 |
| SE | 13660 | CLK1 | CLK1 | chr2 | - | 200857717 | 200857884 | 200856890 | 200856985 | 200857972 | 200858089 | 45 | 41 | 28 | 3 | 2 | 1 | 0 | 12 | 0 | 0 | 98 | 49 | 1 | 1 | 0.882 | 0.911 | 0.933 | NA | 1 | -0.091 |
| SE | 13661 | CLK1 | CLK1 | chr2 | - | 200857972 | 200858089 | 200856890 | 200856985 | 200859679 | 200859746 | 53 | 18 | 16 | 1 | 0 | 0 | 1 | 1 | 0 | 0 | 98 | 49 | 1 | 1 | 0.964 | 1 | 1 | 1 | 1 | -0.012 |
| SE | 13662 | CLK1 | CLK1 | chr2 | - | 200860124 | 200860215 | 200859679 | 200859746 | 200861237 | 200861466 | 7 | 1 | 3 | 18 | 9 | 6 | 0 | 0 | 0 | 2 | 98 | 49 | 1 | 1 | 0.163 | 0.053 | 0.2 | NA | 0 | 0.139 |
| SE | 13669 | STYXL1 | STYXL1 | chr7 | - | 76013741 | 76013887 | 76005258 | 76005404 | 76021850 | 76021992 | 2 | 2 | 0 | 0 | 0 | 3 | 0 | 1 | 0 | 0 | 98 | 49 | 1 | 1 | 1 | 1 | 0 | NA | 1 | -0.333 |
| SE | 13672 | LOC101929567 | LOC101929567 | chr2 | - | 8579910 | 8580047 | 8578569 | 8578711 | 8583716 | 8583792 | 0 | 1 | 0 | 0 | 0 | 0 | 0 | 0 | 0 | 2 | 98 | 49 | 1 | 1 | NA | 1 | NA | NA | 0 | 1 |
| SE | 13674 | ADAM17 | ADAM17 | chr2 | - | 9492897 | 9492986 | 9491100 | 9491151 | 9493746 | 9493825 | 26 | 3 | 3 | 1 | 0 | 0 | 0 | 7 | 0 | 0 | 98 | 49 | 1 | 1 | 0.929 | 1 | 1 | NA | 1 | -0.024 |
| SE | 13675 | ADAM17 | ADAM17 | chr2 | - | 9518102 | 9518247 | 9517900 | 9517989 | 9521202 | 9521316 | 18 | 10 | 3 | 0 | 0 | 0 | 0 | 10 | 0 | 0 | 98 | 49 | 1 | 1 | 0.75 | 1 | 1 | NA | 1 | -0.083 |
| SE | 13676 | ADAM17 | ADAM17 | chr2 | - | 9535833 | 9535922 | 9527785 | 9527954 | 9536697 | 9536828 | 17 | 2 | 3 | 0 | 0 | 0 | 0 | 3 | 0 | 1 | 98 | 49 | 1 | 1 | 1 | 1 | 1 | NA | 0.6 | 0.4 |
| SE | 13677 | ADAM17 | ADAM17 | chr2 | - | 9536697 | 9536828 | 9527785 | 9527954 | 9543152 | 9543285 | 5 | 2 | 4 | 2 | 0 | 0 | 0 | 2 | 0 | 0 | 98 | 49 | 1 | 1 | 0.556 | 1 | 1 | NA | 1 | -0.148 |
| SE | 13683 | XPO1 | XPO1 | chr2 | - | 61485767 | 61485962 | 61483936 | 61484105 | 61488164 | 61488271 | 8 | 0 | 2 | 1 | 0 | 0 | 0 | 4 | 0 | 0 | 98 | 49 | 1 | 1 | 0.8 | NA | 1 | NA | 1 | -0.1 |
| SE | 13684 | XPO1 | XPO1 | chr2 | - | 61488587 | 61488771 | 61488164 | 61488271 | 61490641 | 61490776 | 24 | 3 | 6 | 3 | 0 | 0 | 0 | 8 | 0 | 0 | 98 | 49 | 1 | 1 | 0.8 | 1 | 1 | NA | 1 | -0.067 |
| SE | 13685 | XPO1 | XPO1 | chr2 | - | 61492914 | 61493053 | 61492566 | 61492748 | 61493893 | 61494091 | 18 | 16 | 0 | 0 | 0 | 0 | 0 | 4 | 0 | 2 | 98 | 49 | 1 | 1 | 1 | 1 | NA | NA | 0.5 | 0.5 |
| SE | 13686 | PQLC3 | PQLC3 | chr2 | + | 11171924 | 11172045 | 11164203 | 11164261 | 11174967 | 11175009 | 3 | 2 | 3 | 0 | 1 | 0 | 0 | 1 | 0 | 1 | 98 | 49 | 1 | 1 | 1 | 0.5 | 1 | NA | 0.333 | 0.5 |
| SE | 13693 | AAED1 | AAED1 | chr9 | - | 96645892 | 96646024 | 96641250 | 96641886 | 96651389 | 96651495 | 2 | 0 | 0 | 0 | 1 | 0 | 0 | 1 | 0 | 0 | 98 | 49 | 1 | 1 | 1 | 0 | NA | NA | 0.333 | 0.167 |
| SE | 13694 | FAM49A | FAM49A | chr2 | - | 16562004 | 16562141 | 16561455 | 16561533 | 16563988 | 16564094 | 24 | 8 | 2 | 0 | 2 | 0 | 3 | 10 | 0 | 0 | 98 | 49 | 1 | 1 | 1 | 0.667 | 1 | 1 | 1 | -0.111 |
| SE | 13695 | FAM49A | FAM49A | chr2 | - | 16563988 | 16564094 | 16561455 | 16561533 | 16565645 | 16565767 | 9 | 10 | 2 | 0 | 0 | 0 | 0 | 11 | 0 | 2 | 98 | 49 | 1 | 1 | 1 | 1 | 1 | NA | 0.733 | 0.267 |
| SE | 13697 | FAM49A | FAM49A | chr2 | - | 16565645 | 16565767 | 16561455 | 16561533 | 16588049 | 16588129 | 16 | 6 | 5 | 1 | 0 | 0 | 0 | 4 | 0 | 0 | 98 | 49 | 1 | 1 | 0.889 | 1 | 1 | NA | 1 | -0.037 |
| SE | 13698 | FAM49A | FAM49A | chr2 | - | 16565645 | 16565767 | 16563988 | 16564094 | 16588049 | 16588129 | 25 | 14 | 7 | 1 | 0 | 0 | 0 | 13 | 0 | 0 | 98 | 49 | 1 | 1 | 0.926 | 1 | 1 | NA | 1 | -0.025 |
| SE | 13700 | ARPP19 | ARPP19 | chr15 | - | 52564189 | 52564246 | 52557099 | 52557222 | 52568847 | 52568902 | 6 | 0 | 0 | 20 | 1 | 0 | 0 | 2 | 0 | 0 | 98 | 49 | 1 | 1 | 0.13 | 0 | NA | NA | 1 | -0.935 |
| SE | 13701 | ROCK1 | ROCK1 | chr18 | - | 21042564 | 21042709 | 21042096 | 21042235 | 21044101 | 21044186 | 40 | 18 | 2 | 0 | 1 | 0 | 0 | 4 | 0 | 0 | 98 | 49 | 1 | 1 | 1 | 0.9 | 1 | NA | 1 | -0.033 |
| SE | 13706 | LAPTM4A | LAPTM4A | chr2 | - | 20037537 | 20037614 | 20037315 | 20037438 | 20040890 | 20041011 | 30 | 14 | 3 | 0 | 1 | 0 | 0 | 11 | 0 | 0 | 98 | 49 | 1 | 1 | 1 | 0.875 | 1 | NA | 1 | -0.042 |
| SE | 13713 | RBM38 | RBM38 | chr20 | + | 57393278 | 57393333 | 57392653 | 57392777 | 57407542 | 57409330 | 4 | 1 | 1 | 1 | 0 | 0 | 0 | 1 | 0 | 0 | 98 | 49 | 1 | 1 | 0.667 | 1 | 1 | NA | 1 | -0.111 |
| SE | 13715 | FAM228B | FAM228B | chr2 | + | 24137908 | 24138100 | 24135118 | 24135187 | 24139369 | 24139450 | 6 | 1 | 5 | 1 | 0 | 1 | 0 | 1 | 0 | 0 | 98 | 49 | 1 | 1 | 0.75 | 1 | 0.714 | NA | 1 | -0.179 |
| SE | 13718 | TP53I3 | TP53I3 | chr2 | - | 24079443 | 24079640 | 24077432 | 24077761 | 24080818 | 24081031 | 0 | 2 | 0 | 1 | 0 | 0 | 0 | 1 | 0 | 0 | 98 | 49 | 1 | 1 | 0 | 1 | NA | NA | 1 | -0.5 |
| SE | 13719 | ITSN2 | ITSN2 | chr2 | - | 24284762 | 24284843 | 24275712 | 24275849 | 24286211 | 24286351 | 2 | 1 | 0 | 29 | 15 | 2 | 0 | 0 | 0 | 3 | 98 | 49 | 1 | 1 | 0.033 | 0.032 | 0 | NA | 0 | 0.022 |
| SE | 13720 | NCOA1 | NCOA1 | chr2 | + | 24705085 | 24705233 | 24697657 | 24697798 | 24706567 | 24707888 | 77 | 48 | 17 | 1 | 0 | 0 | 1 | 11 | 0 | 1 | 98 | 49 | 1 | 1 | 0.975 | 1 | 1 | 1 | 0.846 | 0.069 |
| SE | 13721 | NCOA1 | NCOA1 | chr2 | + | 24762686 | 24762776 | 24757972 | 24758156 | 24768078 | 24768135 | 47 | 14 | 7 | 1 | 0 | 0 | 0 | 6 | 0 | 0 | 98 | 49 | 1 | 1 | 0.959 | 1 | 1 | NA | 1 | -0.014 |
| SE | 13722 | NCOA1 | NCOA1 | chr2 | + | 24768078 | 24768135 | 24762686 | 24762776 | 24768220 | 24770701 | 31 | 14 | 2 | 4 | 4 | 0 | 0 | 8 | 0 | 2 | 98 | 49 | 1 | 1 | 0.795 | 0.636 | 1 | NA | 0.667 | 0.143 |
| SE | 13723 | NCOA1 | NCOA1 | chr2 | + | 24768078 | 24768135 | 24762686 | 24762776 | 24768223 | 24770701 | 16 | 12 | 2 | 1 | 0 | 0 | 0 | 3 | 0 | 0 | 98 | 49 | 1 | 1 | 0.889 | 1 | 1 | NA | 1 | -0.037 |
| SE | 13725 | RAPGEF2 | RAPGEF2 | chr4 | + | 159355852 | 159356158 | 159353486 | 159354046 | 159358113 | 159360149 | 20 | 14 | 4 | 0 | 1 | 0 | 0 | 2 | 0 | 0 | 98 | 49 | 1 | 1 | 1 | 0.875 | 1 | NA | 1 | -0.042 |
| SE | 13729 | DNMT3A | DNMT3A | chr2 | - | 25241561 | 25241707 | 25240639 | 25240730 | 25243897 | 25243982 | 5 | 6 | 0 | 0 | 0 | 0 | 0 | 0 | 0 | 1 | 98 | 49 | 1 | 1 | 1 | 1 | NA | NA | 0 | 1 |
| SE | 13735 | SMG5 | SMG5 | chr1 | - | 156250857 | 156250996 | 156249223 | 156250670 | 156251402 | 156251477 | 6 | 4 | 0 | 0 | 0 | 2 | 0 | 1 | 0 | 0 | 98 | 49 | 1 | 1 | 1 | 1 | 0 | NA | 1 | -0.333 |
| SE | 13737 | SMG5 | SMG5 | chr1 | - | 156273360 | 156273450 | 156272319 | 156272398 | 156274596 | 156274686 | 11 | 3 | 2 | 0 | 0 | 0 | 0 | 3 | 0 | 1 | 98 | 49 | 1 | 1 | 1 | 1 | 1 | NA | 0.6 | 0.4 |
| SE | 13740 | BIRC6 | BIRC6 | chr2 | + | 32435495 | 32435585 | 32433643 | 32433804 | 32436052 | 32436184 | 5 | 2 | 0 | 4 | 0 | 0 | 0 | 1 | 0 | 0 | 98 | 49 | 1 | 1 | 0.385 | 1 | NA | NA | 1 | -0.307 |
| SE | 13741 | BIRC6 | BIRC6 | chr2 | + | 32464508 | 32464823 | 32463193 | 32463381 | 32465064 | 32465164 | 22 | 23 | 3 | 0 | 0 | 0 | 0 | 1 | 0 | 1 | 98 | 49 | 1 | 1 | 1 | 1 | 1 | NA | 0.333 | 0.667 |
| SE | 13742 | BIRC6 | BIRC6 | chr2 | + | 32499546 | 32500109 | 32493539 | 32493667 | 32501712 | 32501888 | 20 | 14 | 0 | 1 | 0 | 0 | 0 | 13 | 0 | 0 | 98 | 49 | 1 | 1 | 0.909 | 1 | NA | NA | 1 | -0.045 |
| SE | 13743 | NBN | NBN | chr8 | - | 89946139 | 89946295 | 89943252 | 89943366 | 89947823 | 89947892 | 30 | 16 | 13 | 1 | 0 | 0 | 0 | 6 | 0 | 0 | 98 | 49 | 1 | 1 | 0.937 | 1 | 1 | NA | 1 | -0.021 |
| SE | 13745 | NBN | NBN | chr8 | - | 89980733 | 89980893 | 89971172 | 89971290 | 89981374 | 89981523 | 10 | 1 | 1 | 1 | 0 | 0 | 0 | 1 | 0 | 0 | 98 | 49 | 1 | 1 | 0.833 | 1 | 1 | NA | 1 | -0.056 |
| SE | 13746 | NBN | NBN | chr8 | - | 89981952 | 89982002 | 89981374 | 89981523 | 89982721 | 89982855 | 3 | 2 | 2 | 3 | 0 | 1 | 0 | 2 | 0 | 2 | 98 | 49 | 1 | 1 | 0.333 | 1 | 0.5 | NA | 0.333 | 0.278 |
| SE | 13751 | ZBTB8OS | ZBTB8OS | chr1 | - | 32631826 | 32631879 | 32627507 | 32627544 | 32633644 | 32633727 | 6 | 7 | 8 | 0 | 2 | 0 | 0 | 1 | 0 | 0 | 98 | 49 | 1 | 1 | 1 | 0.636 | 1 | NA | 1 | -0.121 |
| SE | 13756 | ACIN1 | ACIN1 | chr14 | - | 23089981 | 23090101 | 23081747 | 23081836 | 23090521 | 23090633 | 17 | 4 | 13 | 2 | 0 | 0 | 1 | 5 | 0 | 0 | 98 | 49 | 1 | 1 | 0.81 | 1 | 1 | 1 | 1 | -0.063 |
| SE | 13758 | ERGIC1 | ERGIC1 | chr5 | + | 172909666 | 172909761 | 172897001 | 172897074 | 172914713 | 172914838 | 92 | 46 | 44 | 0 | 0 | 0 | 1 | 13 | 0 | 1 | 98 | 49 | 1 | 1 | 1 | 1 | 1 | 1 | 0.867 | 0.067 |
| SE | 13759 | ERGIC1 | ERGIC1 | chr5 | + | 172924004 | 172924109 | 172914713 | 172914838 | 172926508 | 172926569 | 58 | 26 | 32 | 0 | 0 | 4 | 0 | 24 | 0 | 0 | 98 | 49 | 1 | 1 | 1 | 1 | 0.8 | NA | 1 | -0.067 |
| SE | 13760 | MRPL33 | MRPL33 | chr2 | + | 27774423 | 27774530 | 27772673 | 27772692 | 27779432 | 27779741 | 4 | 1 | 0 | 1 | 0 | 0 | 0 | 0 | 0 | 1 | 98 | 49 | 1 | 1 | 0.667 | 1 | NA | NA | 0 | 0.834 |
| SE | 13762 | GFER | GFER | chr16 | + | 1984746 | 1984943 | 1984148 | 1984476 | 1985865 | 1987749 | 1 | 0 | 0 | 2 | 0 | 0 | 0 | 1 | 0 | 0 | 98 | 49 | 1 | 1 | 0.2 | NA | NA | NA | 1 | -0.8 |
| SE | 13763 | BRE | BRE | chr2 | + | 27892382 | 27892448 | 27890614 | 27890842 | 27894532 | 27894684 | 3 | 0 | 0 | 0 | 0 | 0 | 0 | 1 | 0 | 3 | 98 | 49 | 1 | 1 | 1 | NA | NA | NA | 0.143 | 0.857 |
| SE | 13765 | BRE | BRE | chr2 | + | 28327273 | 28327447 | 28298337 | 28298491 | 28338449 | 28338900 | 1 | 0 | 0 | 1 | 0 | 0 | 0 | 1 | 0 | 0 | 98 | 49 | 1 | 1 | 0.333 | NA | NA | NA | 1 | -0.667 |
| SE | 13767 | ZSWIM6 | ZSWIM6 | chr5 | + | 61490785 | 61490934 | 61472680 | 61473037 | 61494259 | 61494410 | 17 | 6 | 1 | 1 | 0 | 0 | 0 | 4 | 0 | 0 | 98 | 49 | 1 | 1 | 0.895 | 1 | 1 | NA | 1 | -0.035 |
| SE | 13778 | MTRR | MTRR | chr5 | + | 7886614 | 7886703 | 7885700 | 7885854 | 7889094 | 7889275 | 3 | 1 | 0 | 0 | 0 | 0 | 0 | 0 | 0 | 1 | 98 | 49 | 1 | 1 | 1 | 1 | NA | NA | 0 | 1 |
| SE | 13779 | GALNT14 | GALNT14 | chr2 | - | 30944857 | 30944942 | 30942200 | 30942304 | 30945782 | 30945870 | 1 | 1 | 1 | 0 | 1 | 0 | 0 | 1 | 0 | 0 | 98 | 49 | 1 | 1 | 1 | 0.333 | 1 | NA | 1 | -0.222 |
| SE | 13781 | GALNT14 | GALNT14 | chr2 | - | 30992837 | 30993007 | 30966203 | 30966302 | 31137957 | 31138726 | 3 | 2 | 0 | 0 | 1 | 0 | 1 | 3 | 0 | 0 | 98 | 49 | 1 | 1 | 1 | 0.5 | NA | 1 | 1 | -0.25 |
| SE | 13785 | SPAST | SPAST | chr2 | + | 32098795 | 32098891 | 32089521 | 32089605 | 32114637 | 32114825 | 25 | 10 | 5 | 2 | 0 | 0 | 0 | 2 | 0 | 0 | 98 | 49 | 1 | 1 | 0.862 | 1 | 1 | NA | 1 | -0.046 |
| SE | 13804 | PARP14 | PARP14 | chr3 | + | 122703741 | 122703978 | 122699389 | 122701635 | 122704526 | 122704748 | 66 | 23 | 3 | 0 | 0 | 0 | 6 | 21 | 1 | 0 | 98 | 49 | 1 | 1 | 1 | 1 | 1 | 0.75 | 1 | 0.125 |
| SE | 13805 | PARP14 | PARP14 | chr3 | + | 122704526 | 122704748 | 122699389 | 122701635 | 122708189 | 122708268 | 18 | 2 | 0 | 1 | 1 | 0 | 2 | 3 | 0 | 1 | 98 | 49 | 1 | 1 | 0.9 | 0.5 | NA | 1 | 0.6 | -0.1 |
| SE | 13809 | HNRNPLL | HNRNPLL | chr2 | - | 38569803 | 38569925 | 38569132 | 38569334 | 38573209 | 38573427 | 4 | 6 | 1 | 1 | 0 | 0 | 0 | 2 | 0 | 0 | 98 | 49 | 1 | 1 | 0.667 | 1 | 1 | NA | 1 | -0.111 |
| SE | 13811 | APEH | APEH | chr3 | + | 49676776 | 49676817 | 49676608 | 49676700 | 49676902 | 49677024 | 5 | 10 | 4 | 0 | 0 | 0 | 0 | 4 | 0 | 1 | 98 | 49 | 1 | 1 | 1 | 1 | 1 | NA | 0.685 | 0.315 |
| SE | 13812 | APEH | APEH | chr3 | + | 49679592 | 49679644 | 49678851 | 49678949 | 49680540 | 49680629 | 10 | 0 | 6 | 0 | 0 | 1 | 0 | 4 | 0 | 0 | 90 | 49 | 1 | 1 | 1 | NA | 0.75 | NA | 1 | -0.125 |
| SE | 13813 | APEH | APEH | chr3 | + | 49681721 | 49681805 | 49681100 | 49681239 | 49681886 | 49681967 | 6 | 1 | 2 | 2 | 0 | 0 | 0 | 2 | 0 | 0 | 98 | 49 | 1 | 1 | 0.6 | 1 | 1 | NA | 1 | -0.133 |
| SE | 13814 | MAP4K3 | MAP4K3 | chr2 | - | 39307942 | 39308005 | 39299742 | 39299801 | 39309460 | 39309519 | 1 | 0 | 0 | 0 | 0 | 1 | 0 | 3 | 0 | 0 | 98 | 49 | 1 | 1 | 1 | NA | 0 | NA | 1 | -0.5 |
| SE | 13815 | MEF2A | MEF2A | chr15 | + | 99598429 | 99598511 | 99565927 | 99566104 | 99632977 | 99633173 | 4 | 0 | 0 | 1 | 0 | 0 | 0 | 2 | 0 | 0 | 98 | 49 | 1 | 1 | 0.667 | NA | NA | NA | 1 | -0.333 |
| SE | 13818 | MEF2A | MEF2A | chr15 | + | 99690240 | 99690428 | 99675398 | 99675458 | 99710633 | 99710760 | 13 | 1 | 7 | 0 | 0 | 0 | 0 | 7 | 0 | 1 | 98 | 49 | 1 | 1 | 1 | 1 | 1 | NA | 0.778 | 0.222 |
| SE | 13820 | MEF2A | MEF2A | chr15 | + | 99706728 | 99706855 | 99690240 | 99690428 | 99710633 | 99710760 | 9 | 0 | 0 | 0 | 0 | 0 | 0 | 6 | 0 | 1 | 98 | 49 | 1 | 1 | 1 | NA | NA | NA | 0.75 | 0.25 |
| SE | 13829 | EML4 | EML4 | chr2 | + | 42301240 | 42301392 | 42295380 | 42295516 | 42303103 | 42303229 | 17 | 11 | 2 | 1 | 0 | 0 | 0 | 6 | 0 | 0 | 98 | 49 | 1 | 1 | 0.895 | 1 | 1 | NA | 1 | -0.035 |
| SE | 13830 | SYK | SYK | chr9 | + | 90867130 | 90867199 | 90865047 | 90865097 | 90874203 | 90874291 | 31 | 22 | 12 | 1 | 0 | 0 | 0 | 3 | 0 | 0 | 98 | 49 | 1 | 1 | 0.939 | 1 | 1 | NA | 1 | -0.02 |
| SE | 13844 | SRBD1 | SRBD1 | chr2 | - | 45476992 | 45477075 | 45419787 | 45419894 | 45488239 | 45488331 | 11 | 4 | 1 | 0 | 1 | 0 | 0 | 2 | 0 | 0 | 98 | 49 | 1 | 1 | 1 | 0.667 | 1 | NA | 1 | -0.111 |
| SE | 13845 | SRBD1 | SRBD1 | chr2 | - | 45547521 | 45547612 | 45546731 | 45546839 | 45551124 | 45551282 | 15 | 9 | 8 | 0 | 1 | 0 | 0 | 3 | 0 | 0 | 98 | 49 | 1 | 1 | 1 | 0.818 | 1 | NA | 1 | -0.061 |
| SE | 13846 | RHOQ | RHOQ | chr2 | + | 46543753 | 46543812 | 46542727 | 46543188 | 46576086 | 46576251 | 13 | 7 | 2 | 0 | 0 | 0 | 0 | 0 | 0 | 1 | 98 | 49 | 1 | 1 | 1 | 1 | 1 | NA | 0 | 1 |

| | | | | | | | | | | | | | | | | | | | | | | | | | | | | | | |
|---|---|---|---|---|---|---|---|---|---|---|---|---|---|---|---|---|---|---|---|---|---|---|---|---|---|---|---|---|---|---|
| SE | 13847 MYLIP | MYLIP | chr6 | + | 16130556 | 16130747 | 16129085 | 16129409 | 16141624 | 16141810 | 15 | 10 | 9 | 0 | 2 | 0 | 0 | 3 | 0 | 0 | 98 | 49 | 1 | 1 | 1 | 0.714 | 1 NA | | 1 | -0.095 |
| SE | 13856 GNAS | GNAS | chr20 | + | 58909683 | 58909804 | 58909520 | 58909579 | 58909950 | 58910081 | 83 | 39 | 33 | 1 | 0 | 0 | 8 | 35 | 0 | 0 | 98 | 49 | 1 | 1 | 0.976 | 1 | 1 | 1 | 1 | -0.008 |
| SE | 13857 CALM2 | CALM2 | chr2 | - | 47172413 | 47172459 | 47170733 | 47170764 | 47176440 | 47176623 | 1 | 0 | 0 | 15 | 11 | 3 | 0 | 0 | 0 | 1 | 95 | 49 | 1 | 1 | 0.033 | 0 | 0 NA | | 0 | 0.011 |
| SE | 13861 TBCK | TBCK | chr4 | - | 106254946 | 106255047 | 106251865 | 106252007 | 106260436 | 106260510 | 0 | 0 | 0 | 1 | 0 | 0 | 0 | 3 | 0 | 0 | 98 | 49 | 1 | 1 | 0 NA | NA | NA | | 1 | -1 |
| SE | 13862 PPP1R21 | PPP1R21 | chr2 | + | 48471086 | 48471188 | 48465492 | 48465642 | 48479923 | 48480016 | 8 | 4 | 3 | 1 | 0 | 0 | 0 | 2 | 0 | 0 | 98 | 49 | 1 | 1 | 0.8 | 1 | 1 NA | | 1 | -0.067 |
| SE | 13874 EIF4E3 | EIF4E3 | chr3 | - | 71696459 | 71696520 | 71693874 | 71693941 | 71699613 | 71699708 | 60 | 32 | 41 | 1 | 0 | 0 | 0 | 13 | 0 | 1 | 98 | 49 | 1 | 1 | 0.968 | 1 | 1 NA | | 0.867 | 0.122 |
| SE | 13875 EIF4E3 | EIF4E3 | chr3 | - | 71696459 | 71696520 | 71693874 | 71693941 | 71710411 | 71710484 | 29 | 15 | 15 | 2 | 2 | 0 | 0 | 3 | 0 | 0 | 98 | 49 | 1 | 1 | 0.879 | 0.789 | 1 NA | | 1 | -0.111 |
| SE | 13876 EIF4E3 | EIF4E3 | chr3 | - | 71699613 | 71699708 | 71693874 | 71693941 | 71710411 | 71710484 | 11 | 12 | 8 | 2 | 2 | 0 | 0 | 2 | 0 | 0 | 98 | 49 | 1 | 1 | 0.733 | 0.75 | 1 NA | | 1 | -0.172 |
| SE | 13877 EIF4E3 | EIF4E3 | chr3 | - | 71699613 | 71699708 | 71696459 | 71696520 | 71710411 | 71710484 | 44 | 29 | 34 | 3 | 0 | 0 | 0 | 11 | 0 | 0 | 98 | 49 | 1 | 1 | 0.88 | 1 | 1 NA | | 1 | -0.04 |
| SE | 13878 EIF4E3 | EIF4E3 | chr3 | - | 71728475 | 71728623 | 71710411 | 71710484 | 71754214 | 71754383 | 12 | 7 | 2 | 0 | 0 | 0 | 0 | 10 | 0 | 1 | 98 | 49 | 1 | 1 | 1 | 1 | 1 NA | | 0.833 | 0.167 |
| SE | 13879 PSME4 | PSME4 | chr2 | - | 53899880 | 53900017 | 53898300 | 53898354 | 53901349 | 53901559 | 8 | 1 | 2 | 1 | 2 | 0 | 0 | 2 | 0 | 0 | 98 | 49 | 1 | 1 | 0.8 | 0.2 | 1 NA | | 1 | -0.333 |
| SE | 13880 PPIP5K2 | PPIP5K2 | chr5 | + | 103176865 | 103176910 | 103173857 | 103173972 | 103177666 | 103177774 | 4 | 1 | 0 | 4 | 1 | 0 | 0 | 2 | 0 | 1 | 94 | 49 | 1 | 1 | 0.343 | 0.343 NA | NA | | 0.51 | -0.167 |
| SE | 13881 PPIP5K2 | PPIP5K2 | chr5 | + | 103183233 | 103183407 | 103180020 | 103180188 | 103184671 | 103184744 | 2 | 0 | 0 | 6 | 4 | 0 | 0 | 0 | 1 | 0 | 98 | 49 | 1 | 1 | 0.143 | 0 NA | NA | 0 NA | | 0.071 |
| SE | 13882 PPIP5K2 | PPIP5K2 | chr5 | + | 103186319 | 103186439 | 103184671 | 103184744 | 103190841 | 103190982 | 1 | 0 | 0 | 3 | 1 | 0 | 0 | 1 | 0 | 0 | 98 | 49 | 1 | 1 | 0.143 | 0 NA | NA | | 1 | -0.928 |
| SE | 13883 PPIP5K2 | PPIP5K2 | chr5 | + | 103187313 | 103187376 | 103186319 | 103186439 | 103190841 | 103190982 | 0 | 6 | 1 | 0 | 0 | 0 | 0 | 0 | 0 | 1 | 98 | 49 | 1 | 1 NA | | 1 | 1 NA | | 0 | 1 |
| SE | 13886 CCDC88A | CCDC88A | chr2 | - | 55312433 | 55312579 | 55309161 | 55309254 | 55315927 | 55316114 | 0 | 0 | 0 | 2 | 0 | 0 | 0 | 6 | 0 | 0 | 98 | 49 | 1 | 1 | 0 NA | NA | NA | | 1 | -1 |
| SE | 13891 VRK2 | VRK2 | chr2 | + | 58154753 | 58154802 | 58146315 | 58146474 | 58159348 | 58159920 | 0 | 1 | 0 | 1 | 1 | 0 | 0 | 2 | 0 | 1 | 98 | 49 | 1 | 1 | 0 | 0.333 NA | NA | | 0.5 | -0.334 |
| SE | 13892 GPBP1L1 | GPBP1L1 | chr1 | - | 45634095 | 45634236 | 45633488 | 45633647 | 45640209 | 45640403 | 28 | 8 | 1 | 0 | 0 | 2 | 0 | 13 | 0 | 0 | 98 | 49 | 1 | 1 | 1 | 1 | 0.2 NA | | 1 | -0.267 |
| SE | 13893 GPBP1L1 | GPBP1L1 | chr1 | - | 45685575 | 45685620 | 45660183 | 45661225 | 45686411 | 45686630 | 17 | 14 | 0 | 1 | 1 | 0 | 0 | 4 | 0 | 0 | 94 | 49 | 1 | 1 | 0.899 | 0.879 NA | NA | | 1 | -0.111 |
| SE | 13896 USP34 | USP34 | chr2 | - | 61192900 | 61192980 | 61190517 | 61190658 | 61203139 | 61203263 | 32 | 4 | 3 | 0 | 0 | 0 | 0 | 3 | 0 | 1 | 98 | 49 | 1 | 1 | 1 | 1 | 1 NA | | 0.6 | 0.4 |
| SE | 13897 USP34 | USP34 | chr2 | - | 61204255 | 61204380 | 61203139 | 61203263 | 61204496 | 61204601 | 12 | 7 | 9 | 1 | 0 | 0 | 0 | 4 | 0 | 0 | 98 | 49 | 1 | 1 | 0.857 | 1 | 1 NA | | 1 | -0.048 |
| SE | 13898 USP34 | USP34 | chr2 | - | 61214059 | 61214694 | 61211771 | 61211929 | 61220309 | 61220457 | 16 | 2 | 3 | 3 | 0 | 0 | 0 | 2 | 0 | 0 | 98 | 49 | 1 | 1 | 0.727 | 1 | 1 NA | | 1 | -0.091 |
| SE | 13900 USP34 | USP34 | chr2 | - | 61265396 | 61265557 | 61259710 | 61259776 | 61265983 | 61266167 | 16 | 19 | 4 | 1 | 0 | 0 | 0 | 3 | 0 | 0 | 98 | 49 | 1 | 1 | 0.889 | 1 | 1 NA | | 1 | -0.037 |
| SE | 13909 ATP8A1 | ATP8A1 | chr4 | - | 42485495 | 42485668 | 42464892 | 42465076 | 42503449 | 42503514 | 4 | 6 | 1 | 1 | 0 | 0 | 0 | 1 | 0 | 1 | 98 | 49 | 1 | 1 | 0.667 | 1 | 1 NA | | 0.333 | 0.556 |
| SE | 13912 ATP8A1 | ATP8A1 | chr4 | - | 42594288 | 42594362 | 42588259 | 42588329 | 42600477 | 42600518 | 8 | 5 | 0 | 2 | 1 | 1 | 0 | 1 | 0 | 0 | 98 | 49 | 1 | 1 | 0.667 | 0.714 | 0 NA | | 1 | -0.54 |
| SE | 13913 AFTPH | AFTPH | chr2 | + | 64569091 | 64569218 | 64567561 | 64567713 | 64569622 | 64569679 | 32 | 6 | 14 | 2 | 0 | 0 | 0 | 3 | 0 | 0 | 98 | 49 | 1 | 1 | 0.889 | 1 | 1 NA | | 1 | -0.037 |
| SE | 13921 NEURL4 | NEURL4 | chr17 | - | 7327164 | 7327230 | 7326710 | 7327009 | 7327439 | 7327884 | 1 | 0 | 3 | 1 | 1 | 0 | 0 | 1 | 0 | 0 | 98 | 49 | 1 | 1 | 0.333 | 0 | 1 NA | | 1 | -0.556 |
| SE | 13929 G3BP2 | G3BP2 | chr4 | - | 75653982 | 75654081 | 75648638 | 75648741 | 75655065 | 75655246 | 3 | 0 | 0 | 17 | 7 | 2 | 0 | 1 | 0 | 3 | 98 | 49 | 1 | 1 | 0.081 | 0 | 0 NA | | 0.143 | -0.116 |
| SE | 13930 G3BP2 | G3BP2 | chr4 | - | 75656923 | 75657014 | 75655767 | 75655870 | 75657556 | 75657730 | 20 | 5 | 0 | 1 | 0 | 0 | 0 | 6 | 0 | 0 | 98 | 49 | 1 | 1 | 0.909 | 1 NA | NA | | 1 | -0.045 |
| SE | 13933 TRAPPC9 | TRAPPC9 | chr8 | - | 140426614 | 140426641 | 140405576 | 140405698 | 140435111 | 140435240 | 5 | 4 | 0 | 2 | 0 | 0 | 0 | 4 | 0 | 0 | 76 | 49 | 1 | 1 | 0.617 | 1 NA | NA | | 1 | -0.192 |
| SE | 13935 AAK1 | AAK1 | chr2 | - | 69532040 | 69532162 | 69530624 | 69530706 | 69542522 | 69542665 | 10 | 1 | 2 | 1 | 0 | 0 | 0 | 1 | 0 | 0 | 98 | 49 | 1 | 1 | 0.833 | 1 | 1 NA | | 1 | -0.056 |
| SE | 13939 ATP2B1 | ATP2B1 | chr12 | - | 89603711 | 89603925 | 89603042 | 89603254 | 89604154 | 89604346 | 11 | 11 | 2 | 1 | 0 | 0 | 0 | 6 | 0 | 0 | 98 | 49 | 1 | 1 | 0.846 | 1 | 1 NA | | 1 | -0.051 |
| SE | 13940 ATP2B1 | ATP2B1 | chr12 | - | 89616801 | 89617039 | 89611192 | 89611372 | 89619998 | 89620240 | 39 | 13 | 12 | 3 | 0 | 0 | 0 | 11 | 0 | 0 | 98 | 49 | 1 | 1 | 0.867 | 1 | 1 NA | | 1 | -0.044 |
| SE | 13945 ZNF638 | ZNF638 | chr2 | + | 71408121 | 71408247 | 71406127 | 71406262 | 71418601 | 71418639 | 12 | 3 | 4 | 1 | 0 | 0 | 0 | 3 | 0 | 0 | 98 | 49 | 1 | 1 | 0.857 | 1 | 1 NA | | 1 | -0.048 |
| SE | 13946 DYSF | DYSF | chr2 | + | 71511806 | 71511921 | 71503213 | 71503319 | 71513715 | 71513921 | 43 | 15 | 11 | 0 | 0 | 0 | 0 | 5 | 0 | 1 | 98 | 49 | 1 | 1 | 1 | 1 | 1 NA | | 0.714 | 0.286 |
| SE | 13948 DYSF | DYSF | chr2 | + | 71520177 | 71520208 | 71516988 | 71517039 | 71520788 | 71520904 | 73 | 27 | 16 | 0 | 0 | 0 | 1 | 12 | 1 | 0 | 80 | 49 | 1 | 1 | 1 | 1 | 1 | 0.38 | 1 | 0.31 |
| SE | 13950 DYSF | DYSF | chr2 | + | 71570228 | 71570334 | 71569819 | 71569934 | 71570598 | 71570741 | 107 | 69 | 18 | 0 | 1 | 0 | 5 | 39 | 0 | 0 | 98 | 49 | 1 | 1 | 1 | 0.972 | 1 | 1 | 1 | -0.009 |
| SE | 13951 DYSF | DYSF | chr2 | + | 71574197 | 71574371 | 71570598 | 71570741 | 71589592 | 71589686 | 85 | 27 | 20 | 0 | 2 | 0 | 5 | 17 | 0 | 0 | 98 | 49 | 1 | 1 | 1 | 0.871 | 1 | 1 | 1 | -0.043 |
| SE | 13952 DYSF | DYSF | chr2 | + | 71620546 | 71620609 | 71613333 | 71613410 | 71643964 | 71644063 | 19 | 13 | 4 | 9 | 13 | 8 | 0 | 13 | 0 | 3 | 98 | 49 | 1 | 1 | 0.514 | 0.333 | 0.2 NA | | 0.684 | -0.335 |
| SE | 13954 DYSF | DYSF | chr2 | + | 71668753 | 71668842 | 71667375 | 71667515 | 71669604 | 71669746 | 9 | 3 | 1 | 1 | 0 | 0 | 0 | 8 | 0 | 0 | 98 | 49 | 1 | 1 | 0.818 | 1 | 1 NA | | 1 | -0.061 |
| SE | 13955 DYSF | DYSF | chr2 | + | 71669111 | 71669207 | 71668753 | 71668842 | 71669604 | 71669746 | 54 | 16 | 13 | 0 | 0 | 0 | 0 | 17 | 0 | 3 | 98 | 49 | 1 | 1 | 1 | 1 | 1 NA | | 0.739 | 0.261 |
| SE | 13956 DYSF | DYSF | chr2 | + | 71674196 | 71674296 | 71669604 | 71669746 | 71679056 | 71679235 | 94 | 46 | 36 | 0 | 1 | 0 | 3 | 26 | 0 | 0 | 98 | 49 | 1 | 1 | 1 | 0.958 | 1 | 1 | 1 | -0.014 |
| SE | 13964 CCT7 | CCT7 | chr2 | + | 73243996 | 73244049 | 73243003 | 73243129 | 73244543 | 73244715 | 6 | 2 | 12 | 0 | 0 | 0 | 0 | 0 | 1 | 0 | 98 | 49 | 1 | 1 | 1 | 1 | 1 | 0 NA | | 1 |
| SE | 13965 CCT7 | CCT7 | chr2 | + | 73249818 | 73249916 | 73248990 | 73249179 | 73250305 | 73250438 | 5 | 2 | 3 | 2 | 0 | 0 | 1 | 0 | 0 | 0 | 98 | 49 | 1 | 1 | 0.556 | 1 | 1 | 1 NA | | -0.148 |
| SE | 13966 CCT7 | CCT7 | chr2 | + | 73251225 | 73251432 | 73250305 | 73250438 | 73252639 | 73253022 | 10 | 1 | 1 | 0 | 1 | 0 | 0 | 1 | 0 | 0 | 98 | 49 | 1 | 1 | 1 | 0.333 | 1 NA | | 1 | -0.222 |
| SE | 13969 MSN | MSN | chrX | + | 65731837 | 65731984 | 65731106 | 65731190 | 65733183 | 65733280 | 186 | 98 | 81 | 0 | 1 | 0 | 1 | 29 | 0 | 0 | 98 | 49 | 1 | 1 | 1 | 0.98 | 1 | 1 | 1 | -0.007 |
| SE | 13970 MSN | MSN | chrX | + | 65738524 | 65738617 | 65737177 | 65737338 | 65738969 | 65739194 | 138 | 48 | 47 | 0 | 0 | 0 | 3 | 22 | 0 | 1 | 98 | 49 | 1 | 1 | 1 | 1 | 1 | 1 | 0.917 | 0.041 |
| SE | 13973 PSMA3-AS1 | PSMA3-AS1 | chr14 | - | 58285590 | 58285756 | 58273978 | 58274010 | 58291634 | 58291707 | 6 | 3 | 0 | 1 | 3 | 1 | 0 | 1 | 0 | 2 | 98 | 49 | 1 | 1 | 0.75 | 0.333 | 0 NA | | 0.2 | 0.161 |
| SE | 13974 PSMA3-AS1 | PSMA3-AS1 | chr14 | - | 58285590 | 58285756 | 58273978 | 58274135 | 58291634 | 58291707 | 13 | 5 | 4 | 6 | 1 | 4 | 0 | 1 | 0 | 0 | 98 | 49 | 1 | 1 | 0.52 | 0.714 | 0.333 NA | | 1 | -0.478 |
| SE | 13975 INTS3 | INTS3 | chr1 | + | 153763232 | 153763362 | 153762727 | 153762847 | 153763831 | 153763886 | 29 | 4 | 1 | 1 | 0 | 0 | 0 | 4 | 0 | 0 | 98 | 49 | 1 | 1 | 0.935 | 1 | 1 NA | | 1 | -0.022 |
| SE | 13976 STAMBP | STAMBP | chr2 | + | 73847386 | 73847753 | 73845166 | 73845262 | 73849362 | 73849487 | 8 | 0 | 7 | 0 | 2 | 0 | 0 | 2 | 0 | 0 | 98 | 49 | 1 | 1 | 1 | 0 | 1 NA | | 1 | -0.333 |
| SE | 13981 TET3 | TET3 | chr2 | + | 74046277 | 74048411 | 74003109 | 74003166 | 74073548 | 74073639 | 17 | 4 | 6 | 0 | 1 | 0 | 0 | 4 | 0 | 0 | 98 | 49 | 1 | 1 | 1 | 0.667 | 1 NA | | 1 | -0.111 |
| SE | 13989 APMAP | APMAP | chr20 | - | 24968891 | 24969084 | 24962943 | 24964022 | 24969525 | 24969660 | 47 | 18 | 23 | 0 | 0 | 1 | 2 | 2 | 0 | 0 | 98 | 49 | 1 | 1 | 1 | 1 | 0.92 | 1 | 1 | -0.027 |
| SE | 13990 APMAP | APMAP | chr20 | - | 24978766 | 24978882 | 24973644 | 24973737 | 24983902 | 24984019 | 52 | 17 | 38 | 0 | 0 | 0 | 1 | 5 | 0 | 1 | 98 | 49 | 1 | 1 | 1 | 1 | 1 | 1 | 0.714 | 0.143 |
| SE | 13991 APMAP | APMAP | chr20 | - | 24983902 | 24984019 | 24973644 | 24973737 | 24992593 | 24992789 | 13 | 9 | 2 | 1 | 0 | 0 | 0 | 3 | 0 | 0 | 98 | 49 | 1 | 1 | 0.867 | 1 | 1 NA | | 1 | -0.044 |
| SE | 13993 VPS13B | VPS13B | chr8 | + | 99134631 | 99134727 | 99121176 | 99121445 | 99135014 | 99135137 | 94 | 29 | 11 | 0 | 2 | 0 | 2 | 12 | 0 | 0 | 98 | 49 | 1 | 1 | 1 | 0.879 | 1 | 1 | 1 | -0.04 |
| SE | 13994 VPS13B | VPS13B | chr8 | + | 99156548 | 99156743 | 99147840 | 99148010 | 99170038 | 99170163 | 90 | 46 | 2 | 0 | 1 | 0 | 0 | 24 | 0 | 0 | 98 | 49 | 1 | 1 | 1 | 0.958 | 1 NA | | 1 | -0.014 |
| SE | 13995 VPS13B | VPS13B | chr8 | + | 99507136 | 99507203 | 99502835 | 99502950 | 99507769 | 99507911 | 26 | 5 | 0 | 1 | 0 | 0 | 0 | 3 | 0 | 0 | 98 | 49 | 1 | 1 | 0.929 | 1 NA | NA | | 1 | -0.035 |
| SE | 13996 VPS13B | VPS13B | chr8 | + | 99507136 | 99507203 | 99502835 | 99502950 | 99511103 | 99511512 | 38 | 11 | 0 | 1 | 0 | 0 | 0 | 5 | 0 | 0 | 98 | 49 | 1 | 1 | 0.95 | 1 NA | NA | | 1 | -0.025 |
| SE | 13997 VPS13B | VPS13B | chr8 | + | 99507769 | 99507911 | 99502835 | 99502950 | 99511103 | 99511512 | 4 | 1 | 0 | 1 | 0 | 0 | 0 | 2 | 0 | 0 | 98 | 49 | 1 | 1 | 0.667 | 1 NA | NA | | 1 | -0.166 |
| SE | 13998 VPS13B | VPS13B | chr8 | + | 99507769 | 99507911 | 99507136 | 99507203 | 99511103 | 99511512 | 4 | 1 | 0 | 13 | 6 | 0 | 0 | 2 | 0 | 2 | 98 | 49 | 1 | 1 | 0.133 | 0.077 NA | NA | | 0.333 | -0.228 |
| SE | 14004 HK2 | HK2 | chr2 | + | 74882119 | 74882239 | 74881710 | 74881859 | 74885493 | 74885589 | 4 | 10 | 1 | 1 | 0 | 0 | 0 | 3 | 0 | 0 | 98 | 49 | 1 | 1 | 0.667 | 1 | 1 NA | | 1 | -0.111 |
| SE | 14011 ELMOD3 | ELMOD3 | chr2 | + | 85355554 | 85355598 | 85355075 | 85355170 | 85356966 | 85357252 | 1 | 0 | 0 | 0 | 0 | 0 | 0 | 1 | 0 | 1 | 93 | 49 | 1 | 1 | 1 NA | NA | | | 0.345 | 0.655 |
| SE | 14015 RNF181 | RNF181 | chr2 | + | 85596821 | 85596931 | 85596518 | 85596649 | 85597103 | 85597178 | 12 | 22 | 10 | 0 | 0 | 5 | 0 | 1 | 0 | 0 | 98 | 49 | 1 | 1 | 1 | 1 | 0.5 NA | | 1 | -0.167 |
| SE | 14016 RNF181 | RNF181 | chr2 | + | 85597103 | 85597178 | 85596518 | 85596649 | 85597444 | 85597708 | 21 | 4 | 11 | 0 | 1 | 0 | 1 | 4 | 0 | 0 | 98 | 49 | 1 | 1 | 1 | 0.667 | 1 | 1 | | -0.111 |
| SE | 14021 GADD45A | GADD45A | chr1 | + | 67686024 | 67686126 | 67685176 | 67685538 | 67687660 | 67688338 | 4 | 0 | 2 | 0 | 0 | 0 | 0 | 0 | 0 | 1 | 98 | 49 | 1 | 1 | 1 NA | | 1 NA | | 0 | 1 |
| SE | 14022 GADD45A | GADD45A | chr1 | + | 67686349 | 67686587 | 67685176 | 67685538 | 67687660 | 67688338 | 3 | 4 | 1 | 0 | 0 | 0 | 0 | 0 | 0 | 1 | 98 | 49 | 1 | 1 | 1 | 1 | 1 NA | | 0 | 1 |
| SE | 14026 PTCD3 | PTCD3 | chr2 | + | 86125454 | 86125515 | 86124994 | 86125082 | 86125794 | 86125880 | 11 | 5 | 1 | 1 | 0 | 0 | 0 | 2 | 0 | 0 | 98 | 49 | 1 | 1 | 0.846 | 1 | 1 NA | | 1 | -0.051 |
| SE | 14028 TPCN2 | TPCN2 | chr11 | + | 69054032 | 69054097 | 69048881 | 69049106 | 69062883 | 69062990 | 6 | 12 | 1 | 1 | 0 | 0 | 2 | 2 | 0 | 1 | 98 | 49 | 1 | 1 | 0.75 | 1 | 1 | 1 | 0.5 | 0.167 |
| SE | 14029 TPCN2 | TPCN2 | chr11 | + | 69054720 | 69054797 | 69054032 | 69054097 | 69055174 | 69055352 | 0 | 3 | 3 | 0 | 0 | 0 | 0 | 2 | 0 | 1 | 98 | 49 | 1 | 1 NA | | 1 | 1 NA | | 0.5 | 0.5 |
| SE | 14030 TPCN2 | TPCN2 | chr11 | + | 69055174 | 69055352 | 69054032 | 69054097 | 69057577 | 69057694 | 3 | 0 | 0 | 0 | 1 | 0 | 0 | 3 | 0 | 1 | 98 | 49 | 1 | 1 | 1 | 0 NA | NA | | 0.6 | -0.1 |
| SE | 14031 TPCN2 | TPCN2 | chr11 | + | 69057577 | 69057694 | 69054032 | 69054097 | 69062883 | 69062990 | 0 | 3 | 1 | 0 | 4 | 0 | 0 | 2 | 0 | 0 | 98 | 49 | 1 | 1 NA | | 0.273 | 1 NA | | 1 | -0.363 |
| SE | 14032 TPCN2 | TPCN2 | chr11 | + | 69072626 | 69072708 | 69071922 | 69072023 | 69072914 | 69073001 | 7 | 5 | 0 | 0 | 0 | 0 | 2 | 0 | 0 | 1 | 98 | 49 | 1 | 1 | 1 | 1 NA | | 1 | 0 | 0.5 |
| SE | 14033 TPCN2 | TPCN2 | chr11 | + | 69072914 | 69073001 | 69071922 | 69072023 | 69078481 | 69078601 | 5 | 2 | 0 | 0 | 1 | 0 | 0 | 2 | 0 | 2 | 98 | 49 | 1 | 1 | 1 | 0.5 NA | NA | | 0.333 | 0.417 |
| SE | 14034 TPCN2 | TPCN2 | chr11 | + | 69079833 | 69079883 | 69078891 | 69079020 | 69081399 | 69081499 | 7 | 2 | 2 | 0 | 1 | 0 | 0 | 2 | 0 | 1 | 98 | 49 | 1 | 1 | 1 | 0.5 | 1 NA | | 1 | -0.167 |
| SE | 14037 CHMP3 | CHMP3 | chr2 | - | 86542251 | 86542312 | 86529217 | 86529397 | 86563303 | 86563497 | 18 | 5 | 8 | 2 | 0 | 0 | 0 | 1 | 0 | 0 | 98 | 49 | 1 | 1 | 0.818 | 1 | 1 NA | | 1 | -0.061 |
| SE | 14042 HNRNPC | HNRNPC | chr14 | - | 21211405 | 21211566 | 21209135 | 21211306 | 21211809 | 21211923 | 69 | 43 | 28 | 0 | 0 | 0 | 0 | 6 | 0 | 2 | 98 | 49 | 1 | 1 | 1 | 1 | 1 NA | | 0.6 | 0.4 |
| SE | 14045 IRAK2 | IRAK2 | chr3 | + | 10238747 | 10239039 | 10234458 | 10234659 | 10242115 | 10243743 | 1 | 0 | 0 | 1 | 0 | 0 | 0 | 2 | 0 | 0 | 98 | 49 | 1 | 1 | 0.333 NA | NA | NA | | 1 | -0.667 |
| SE | 14054 TMEM131 | TMEM131 | chr2 | - | 97759651 | 97759749 | 97758892 | 97759053 | 97760592 | 97760689 | 5 | 0 | 0 | 1 | 0 | 0 | 0 | 2 | 0 | 0 | 98 | 49 | 1 | 1 | 0.714 NA | NA | NA | | 1 | -0.286 |
| SE | 14055 TMEM131 | TMEM131 | chr2 | - | 97794929 | 97795115 | 97793394 | 97793553 | 97796217 | 97796404 | 5 | 0 | 0 | 0 | 0 | 0 | 0 | 2 | 0 | 2 | 98 | 49 | 1 | 1 | 1 NA | NA | NA | | 0.333 | 0.667 |
| SE | 14056 TMEM131 | TMEM131 | chr2 | - | 97796217 | 97796404 | 97793394 | 97793553 | 97796843 | 97796986 | 4 | 3 | 0 | 0 | 0 | 0 | 0 | 3 | 0 | 2 | 98 | 49 | 1 | 1 | 1 | 1 NA | NA | | 0.429 | 0.571 |
| SE | 14057 TMEM131 | TMEM131 | chr2 | - | 97797364 | 97797516 | 97796843 | 97796986 | 97801894 | 97801961 | 16 | 5 | 0 | 1 | 0 | 0 | 0 | 1 | 0 | 0 | 98 | 49 | 1 | 1 | 0.889 | 1 NA | NA | | 1 | -0.055 |
| SE | 14058 TMEM131 | TMEM131 | chr2 | - | 97844144 | 97844261 | 97841814 | 97841937 | 97859303 | 97859427 | 5 | 2 | 1 | 1 | 0 | 0 | 0 | 1 | 0 | 0 | 98 | 49 | 1 | 1 | 0.714 | 1 | 1 NA | | 1 | -0.095 |
| SE | 14059 PAXBP1 | PAXBP1 | chr21 | - | 32750916 | 32751032 | 32748498 | 32748698 | 32751118 | 32751218 | 6 | 0 | 0 | 1 | 0 | 0 | 0 | 3 | 0 | 0 | 98 | 49 | 1 | 1 | 0.75 NA | NA | NA | | 1 | -0.25 |
| SE | 14070 TBC1D8 | TBC1D8 | chr2 | - | 101036017 | 101036168 | 101033543 | 101033758 | 101037531 | 101037708 | 7 | 0 | 3 | 1 | 0 | 0 | 0 | 2 | 0 | 0 | 98 | 49 | 1 | 1 | 0.778 NA | | 1 NA | | 1 | -0.111 |
| SE | 14072 RNF149 | RNF149 | chr2 | - | 101286080 | 101286177 | 101281858 | 101282057 | 101288972 | 101289055 | 112 | 84 | 53 | 8 | 6 | 0 | 0 | 25 | 0 | 0 | 98 | 49 | 1 | 1 | 0.875 | 0.875 | 1 NA | | 1 | -0.083 |
| SE | 14078 MAP4K4 | MAP4K4 | chr2 | + | 101831720 | 101831851 | 101829503 | 101829594 | 101834408 | 101834463 | 54 | 39 | 9 | 1 | 0 | 0 | 2 | 6 | 0 | 0 | 98 | 49 | 1 | 1 | 0.964 | 1 | 1 | 1 | 1 | -0.012 |
| SE | 14079 MAP4K4 | MAP4K4 | chr2 | + | 101855976 | 101856138 | 101844100 | 101844311 | 101858995 | 101859082 | 28 | 13 | 4 | 3 | 0 | 0 | 0 | 1 | 0 | 0 | 98 | 49 | 1 | 1 | 0.824 | 1 | 1 NA | | 1 | -0.059 |
| SE | 14081 MAP4K4 | MAP4K4 | chr2 | + | 101860824 | 101860986 | 101859642 | 101859864 | 101864929 | 101865036 | 8 | 3 | 2 | 10 | 3 | 1 | 0 | 1 | 0 | 2 | 98 | 49 | 1 | 1 | 0.286 | 0.333 | 0.5 NA | | 0.2 | 0.173 |
| SE | 14083 MAP4K4 | MAP4K4 | chr2 | + | 101863820 | 101864051 | 101860824 | 101860986 | 101864929 | 101865036 | 1 | 0 | 0 | 4 | 0 | 2 | 0 | 0 | 0 | 0 | 98 | 49 | 1 | 1 | 0.111 NA | | 0 NA | | 0 | 0.056 |
| SE | 14084 MAP4K4 | MAP4K4 | chr2 | + | 101868028 | 101868037 | 101867211 | 101867309 | 101869621 | 101869797 | 4 | 1 | 0 | 17 | 16 | 9 | 0 | 0 | 0 | 5 | 58 | 49 | 1 | 1 | 0.166 | 0.05 | 0 NA | | 0 | 0.072 |
| SE | 14086 MAP4K4 | MAP4K4 | chr2 | + | 101874081 | 101874252 | 101873646 | 101873764 | 101877002 | 101877146 | 28 | 24 | 5 | 1 | 0 | 0 | 0 | 14 | 0 | 0 | 98 | 49 | 1 | 1 | 0.933 | 1 | 1 NA | | 1 | -0.022 |
| SE | 14087 MAP4K4 | MAP4K4 | chr2 | + | 101887087 | 101887237 | 101885186 | 101885287 | 101887777 | 101887937 | 63 | 23 | 22 | 2 | 0 | 0 | 0 | 10 | 0 | 0 | 98 | 49 | 1 | 1 | 0.94 | 1 | 1 NA | | 1 | -0.02 |
| SE | 14088 MAP4K4 | MAP4K4 | chr2 | + | 101887087 | 101887261 | 101885186 | 101885287 | 101887777 | 101887937 | 30 | 19 | 17 | 2 | 0 | 0 | 0 | 6 | 0 | 0 | 98 | 49 | 1 | 1 | 0.882 | 1 | 1 NA | | 1 | -0.039 |
| SE | 14089 MAP4K4 | MAP4K4 | chr2 | + | 101888935 | 101888995 | 101887777 | 101887937 | 101891165 | 101894690 | 60 | 19 | 26 | 1 | 0 | 0 | 0 | 8 | 0 | 0 | 98 | 49 | 1 | 1 | 0.968 | 1 | 1 NA | | 1 | -0.011 |
| SE | 14092 IL18R1 | IL18R1 | chr2 | + | 102373936 | 102373993 | 102371952 | 102372118 | 102375906 | 102376063 | 2 | 0 | 0 | 1 | 0 | 0 | 0 | 2 | 0 | 4 | 98 | 49 | 1 | 1 | 0.5 NA | NA | NA | | 0.2 | 0.3 |
| SE | 14093 IL18RAP | IL18RAP | chr2 | + | 102423810 | 102424135 | 102423177 | 102423347 | 102424230 | 102424414 | 9 | 9 | 2 | 2 | 0 | 8 | 1 | 0 | 0 | 0 | 98 | 49 | 1 | 1 | 0.692 | 1 | 0.111 | 1 NA | | -0.399 |
| SE | 14094 IL18RAP | IL18RAP | chr2 | + | 102443199 | 102443323 | 102441311 | 102441377 | 102445188 | 102445340 | 5 | 7 | 14 | 1 | 0 | 0 | 0 | 1 | 0 | 0 | 98 | 49 | 1 | 1 | 0.714 | 1 | 1 NA | | 1 | -0.095 |
| SE | 14095 IL18RAP | IL18RAP | chr2 | + | 102450847 | 102451021 | 102447069 | 102447207 | 102451765 | 102452564 | 20 | 12 | 22 | 0 | 4 | 0 | 0 | 1 | 0 | 0 | 98 | 49 | 1 | 1 | 1 | 0.6 | 1 NA | | 1 | -0.133 |
| SE | 14104 HADHB | HADHB | chr2 | + | 26260980 | 26261063 | 26254429 | 26254474 | 26263379 | 26263479 | 1 | 1 | 0 | 10 | 1 | 3 | 0 | 1 | 0 | 2 | 98 | 49 | 1 | 1 | 0.048 | 0.333 | 0 NA | | 0.2 | -0.073 |
| SE | 14106 HADHB | HADHB | chr2 | + | 26285406 | 26285571 | 26284882 | 26284957 | 26289917 | 26290465 | 10 | 7 | 1 | 1 | 1 | 0 | 2 | 11 | 0 | 0 | 98 | 49 | 1 | 1 | 0.833 | 0.778 | 1 | 1 | 1 | -0.13 |
| SE | 14109 TPM4 | TPM4 | chr19 | + | 16093535 | 16093598 | 16089044 | 16089120 | 16093683 | 16093753 | 80 | 7 | 14 | 1 | 0 | 0 | 1 | 10 | 0 | 0 | 98 | 49 | 1 | 1 | 0.976 | 1 | 1 | 1 | 1 | -0.008 |
| SE | 14122 BCL2L11 | BCL2L11 | chr2 | + | 111123732 | 111123869 | 111120913 | 111121188 | 111150043 | 111150147 | 3 | 1 | 0 | 2 | 0 | 0 | 0 | 1 | 0 | 1 | 98 | 49 | 1 | 1 | 0.429 | 1 NA | NA | | 0.333 | 0.382 |
| SE | 14123 BCL2L11 | BCL2L11 | chr2 | + | 111123732 | 111124139 | 111120913 | 111121188 | 111150043 | 111150147 | 9 | 0 | 1 | 2 | 0 | 0 | 0 | 1 | 0 | 1 | 98 | 49 | 1 | 1 | 0.692 NA | | 1 NA | | 0.333 | 0.513 |
| SE | 14136 TMEM87B | TMEM87B | chr2 | + | 112077191 | 112077282 | 112074911 | 112074962 | 112081056 | 112081118 | 9 | 0 | 0 | 1 | 0 | 0 | 0 | 1 | 0 | 0 | 98 | 49 | 1 | 1 | 0.818 NA | NA | NA | | 1 | -0.182 |
| SE | 14137 TMEM87B | TMEM87B | chr2 | + | 112081334 | 112081518 | 112081056 | 112081118 | 112086004 | 112086104 | 3 | 0 | 0 | 1 | 0 | 0 | 0 | 1 | 0 | 0 | 98 | 49 | 1 | 1 | 0.6 NA | NA | NA | | 1 | -0.4 |
| SE | 14142 CHCHD5 | CHCHD5 | chr2 | + | 112585973 | 112586114 | 112584436 | 112584649 | 112586199 | 112586365 | 2 | 1 | 0 | 1 | 0 | 0 | 0 | 1 | 0 | 0 | 98 | 49 | 1 | 1 | 0.5 | 1 NA | NA | | 1 | -0.25 |
| SE | 14143 SLC20A1 | SLC20A1 | chr2 | + | 112647323 | 112647464 | 112646562 | 112647162 | 112647652 | 112647738 | 3 | 3 | 2 | 0 | 2 | 0 | 2 | 0 | 0 | 0 | 98 | 49 | 1 | 1 | 1 | 0.429 | 1 | 1 NA | | -0.19 |
| SE | 14144 SLC20A1 | SLC20A1 | chr2 | + | 112660386 | 112660572 | 112659203 | 112659762 | 112661141 | 112661226 | 27 | 12 | 8 | 0 | 1 | 0 | 1 | 2 | 0 | 0 | 98 | 49 | 1 | 1 | 1 | 0.857 | 1 | 1 | | -0.048 |
| SE | 14145 IL1B | IL1B | chr2 | - | 112835565 | 112835617 | 112833373 | 112833575 | 112836182 | 112836244 | 72 | 61 | 3 | 2 | 0 | 0 | 0 | 3 | 0 | 0 | 98 | 49 | 1 | 1 | 0.947 | 1 | 1 NA | | 1 | -0.018 |
| SE | 14146 PSD4 | PSD4 | chr2 | + | 113192379 | 113192589 | 113185876 | 113186255 | 113193047 | 113193128 | 31 | 16 | 13 | 0 | 3 | 0 | 0 | 11 | 0 | 0 | 98 | 49 | 1 | 1 | 1 | 0.727 | 1 NA | | 1 | -0.091 |
| SE | 14147 PSD4 | PSD4 | chr2 | + | 113195726 | 113195770 | 113193858 | 113193948 | 113196146 | 113196307 | 24 | 29 | 14 | 1 | 1 | 0 | 2 | 12 | 0 | 0 | 93 | 49 | 1 | 1 | 0.927 | 0.939 | 1 | 1 | 1 | -0.045 |
| SE | 14152 SLC35F5 | SLC35F5 | chr2 | - | 113717752 | 113717850 | 113712791 | 113715195 | 113719153 | 113719308 | 8 | 1 | 3 | 0 | 2 | 0 | 0 | 1 | 0 | 0 | 98 | 49 | 1 | 1 | 1 | 0.2 | 1 NA | | 1 | -0.267 |
| SE | 14154 EMB | EMB | chr5 | - | 50410876 | 50410965 | 50405724 | 50405852 | 50411196 | 50411383 | 84 | 43 | 30 | 2 | 0 | 0 | 1 | 22 | 0 | 1 | 98 | 49 | 1 | 1 | 0.955 | 1 | 1 | 1 | 0.917 | 0.026 |
| SE | 14155 F5 | F5 | chr1 | - | 169523800 | 169523904 | 169523196 | 169523352 | 169524836 | 169524908 | 7 | 9 | 0 | 1 | 0 | 0 | 0 | 2 | 0 | 0 | 98 | 49 | 1 | 1 | 0.778 | 1 NA | NA | | 1 | -0.111 |
| SE | 14165 ACSF3 | ACSF3 | chr16 | + | 89100661 | 89101347 | 89093808 | 89093996 | 89102603 | 89102759 | 4 | 0 | 0 | 0 | 2 | 0 | 0 | 1 | 0 | 0 | 98 | 49 | 1 | 1 | 1 | 0 NA | NA | | 1 | -0.5 |
| SE | 14167 FAM188A | FAM188A | chr10 | - | 15838227 | 15838279 | 15837203 | 15837318 | 15841425 | 15841599 | 1 | 0 | 1 | 3 | 3 | 7 | 0 | 0 | 0 | 1 | 98 | 49 | 1 | 1 | 0.143 | 0 | 0.067 NA | | 0 | 0.07 |
| SE | 14177 MAP3K2 | MAP3K2 | chr2 | - | 127325727 | 127325807 | 127324173 | 127324241 | 127326686 | 127326817 | 78 | 26 | 17 | 4 | 5 | 0 | 0 | 17 | 0 | 0 | 98 | 49 | 1 | 1 | 0.907 | 0.722 | 1 NA | | 1 | -0.124 |
| SE | 14178 MAP3K2 | MAP3K2 | chr2 | - | 127330391 | 127330505 | 127329920 | 127330008 | 127335869 | 127335969 | 56 | 19 | 17 | 1 | 0 | 0 | 0 | 7 | 0 | 0 | 98 | 49 | 1 | 1 | 0.966 | 1 | 1 NA | | 1 | -0.011 |
| SE | 14186 UGGT1 | UGGT1 | chr2 | + | 128180889 | 128181072 | 128179785 | 128179870 | 128182129 | 128182290 | 13 | 4 | 6 | 2 | 0 | 0 | 0 | 3 | 0 | 0 | 98 | 49 | 1 | 1 | 0.765 | 1 | 1 NA | | 1 | -0.078 |
| SE | 14189 ACOX1 | ACOX1 | chr17 | - | 75973624 | 75973785 | 75957458 | 75957566 | 75978533 | 75978693 | 8 | 3 | 1 | 9 | 4 | 0 | 0 | 3 | 0 | 0 | 98 | 49 | 1 | 1 | 0.308 | 0.273 | 1 NA | | 1 | -0.473 |
| SE | 14193 CNST | CNST | chr1 | + | 246591511 | 246591941 | 246566336 | 246566663 | 246621428 | 246621634 | 7 | 2 | 0 | 1 | 0 | 0 | 0 | 1 | 0 | 0 | 98 | 49 | 1 | 1 | 0.778 | 1 NA | NA | | 1 | -0.111 |
| SE | 14201 INSR | INSR | chr19 | - | 7166153 | 7166404 | 7163031 | 7163199 | 7167967 | 7168094 | 2 | 2 | 0 | 1 | 0 | 0 | 0 | 1 | 0 | 0 | 98 | 49 | 1 | 1 | 0.5 | 1 NA | NA | | 1 | -0.25 |
| SE | 14215 MGAT5 | MGAT5 | chr2 | + | 134341589 | 134341759 | 134338258 | 134338420 | 134344929 | 134345064 | 20 | 1 | 2 | 1 | 0 | 0 | 1 | 2 | 0 | 0 | 98 | 49 | 1 | 1 | 0.909 | 1 | 1 | 1 | | -0.03 |
| SE | 14217 RRBP1 | RRBP1 | chr20 | - | 17616731 | 17616839 | 17615925 | 17616009 | 17618595 | 17618679 | 13 | 3 | 0 | 0 | 0 | 0 | 1 | 0 | 0 | 1 | 98 | 49 | 1 | 1 | 1 | 1 NA | | 1 | 0 | 0.5 |
| SE | 14218 RRBP1 | RRBP1 | chr20 | - | 17679998 | 17680075 | 17659979 | 17660528 | 17682027 | 17682283 | 7 | 2 | 0 | 2 | 1 | 0 | 0 | 0 | 1 | 0 | 98 | 49 | 1 | 1 | 0.636 | 0.5 NA | | 0 NA | | 0.568 |
| SE | 14222 RAB3GAP1 | RAB3GAP1 | chr2 | + | 135164593 | 135164696 | 135162985 | 135163101 | 135168544 | 135170709 | 15 | 9 | 5 | 0 | 0 | 1 | 0 | 2 | 0 | 0 | 98 | 49 | 1 | 1 | 1 | 1 | 0.714 NA | | 1 | -0.095 |
| SE | 14237 ACVR2A | ACVR2A | chr2 | + | 147922972 | 147923111 | 147920229 | 147920344 | 147926030 | 147926161 | 1 | 0 | 0 | 1 | 0 | 0 | 0 | 2 | 0 | 0 | 98 | 49 | 1 | 1 | 0.333 NA | NA | NA | | 1 | -0.667 |
| SE | 14239 GALK2 | GALK2 | chr15 | + | 49281986 | 49282085 | 49239220 | 49239367 | 49283565 | 49283718 | 4 | 1 | 0 | 0 | 1 | 0 | 0 | 2 | 0 | 0 | 98 | 49 | 1 | 1 | 1 | 0.333 NA | NA | | 1 | -0.334 |
| SE | 14241 ORC4 | ORC4 | chr2 | - | 147981927 | 147982081 | 147975901 | 147975975 | 148020632 | 148020747 | 3 | 0 | 0 | 2 | 2 | 0 | 0 | 2 | 0 | 0 | 98 | 49 | 1 | 1 | 0.429 | 0 NA | NA | | 1 | -0.785 |
| SE | 14242 ITGAL | ITGAL | chr16 | + | 30475305 | 30475400 | 30474195 | 30474298 | 30479090 | 30479208 | 40 | 17 | 2 | 0 | 1 | 0 | 0 | 12 | 0 | 0 | 98 | 49 | 1 | 1 | 1 | 0.895 | 1 NA | | 1 | -0.035 |
| SE | 14243 ITGAL | ITGAL | chr16 | + | 30475512 | 30475580 | 30475305 | 30475400 | 30479090 | 30479208 | 44 | 17 | 15 | 1 | 0 | 0 | 0 | 17 | 0 | 0 | 98 | 49 | 1 | 1 | 0.957 | 1 | 1 NA | | 1 | -0.014 |
| SE | 14244 ITGAL | ITGAL | chr16 | + | 30479090 | 30479208 | 30475512 | 30475580 | 30481438 | 30481584 | 42 | 10 | 11 | 4 | 0 | 0 | 0 | 18 | 0 | 1 | 98 | 49 | 1 | 1 | 0.84 | 1 | 1 NA | | 0.9 | 0.047 |
| SE | 14245 ITGAL | ITGAL | chr16 | + | 30479330 | 30479461 | 30479090 | 30479208 | 30481438 | 30481584 | 64 | 37 | 10 | 11 | 1 | 0 | 0 | 11 | 0 | 4 | 98 | 49 | 1 | 1 | 0.744 | 0.949 | 1 NA | | 0.579 | 0.319 |
| SE | 14247 ITGAL | ITGAL | chr16 | + | 30510880 | 30510960 | 30510360 | 30510471 | 30511049 | 30511136 | 65 | 20 | 9 | 1 | 0 | 0 | 2 | 13 | 0 | 1 | 98 | 49 | 1 | 1 | 0.97 | 1 | 1 | 1 | 0.867 | 0.056 |
| SE | 14251 SLC2A9 | SLC2A9 | chr4 | - | 9941912 | 9942045 | 9920384 | 9920572 | 9980591 | 9980737 | 1 | 0 | 0 | 0 | 0 | 0 | 0 | 0 | 0 | 2 | 98 | 49 | 1 | 1 | 1 NA | NA | NA | | 0 | 1 |
| SE | 14252 SLC2A9 | SLC2A9 | chr4 | - | 9985668 | 9985793 | 9980591 | 9980737 | 9996780 | 9996941 | 1 | 0 | 0 | 1 | 0 | 0 | 0 | 2 | 0 | 0 | 98 | 49 | 1 | 1 | 0.333 NA | NA | NA | | 1 | -0.667 |
| SE | 14259 SH2D3C | SH2D3C | chr9 | - | 127741787 | 127741959 | 127740257 | 127740369 | 127742848 | 127742964 | 31 | 24 | 3 | 0 | 0 | 0 | 2 | 7 | 1 | 0 | 98 | 49 | 1 | 1 | 1 | 1 | 1 | 0.5 | 1 | 0.25 |
| SE | 14260 SH2D3C | SH2D3C | chr9 | - | 127761610 | 127761650 | 127751171 | 127751300 | 127773989 | 127774467 | 12 | 14 | 8 | 1 | 1 | 1 | 0 | 1 | 0 | 1 | 89 | 49 | 1 | 1 | 0.869 | 0.885 | 0.815 NA | | 0.355 | 0.501 |
| SE | 14262 SH2D3C | SH2D3C | chr9 | - | 127766900 | 127767271 | 127761610 | 127761650 | 127773989 | 127774467 | 2 | 3 | 0 | 3 | 13 | 6 | 0 | 0 | 0 | 1 | 98 | 49 | 1 | 1 | 0.25 | 0.103 | 0 NA | | 0 | 0.118 |
| SE | 14263 CYTIP | CYTIP | chr2 | - | 157418522 | 157418589 | 157414618 | 157416143 | 157427350 | 157427420 | 156 | 108 | 46 | 0 | 0 | 0 | 5 | 36 | 0 | 1 | 98 | 49 | 1 | 1 | 1 | 1 | 1 | 1 | 0.947 | 0.026 |
| SE | 14264 CYTIP | CYTIP | chr2 | - | 157430859 | 157430962 | 157430558 | 157430652 | 157434369 | 157434424 | 437 | 195 | 67 | 0 | 2 | 0 | 11 | 97 | 0 | 0 | 98 | 49 | 1 | 1 | 1 | 0.98 | 1 | 1 | 1 | -0.007 |
| SE | 14265 CYTIP | CYTIP | chr2 | - | 157434369 | 157434424 | 157430859 | 157430962 | 157434697 | 157434747 | 436 | 196 | 66 | 2 | 0 | 0 | 9 | 93 | 0 | 0 | 98 | 49 | 1 | 1 | 0.991 | 1 | 1 | 1 | 1 | -0.003 |
| SE | 14281 HDAC10 | HDAC10 | chr22 | - | 50247691 | 50247776 | 50246874 | 50246966 | 50247889 | 50248145 | 5 | 2 | 1 | 0 | 0 | 0 | 0 | 0 | 0 | 1 | 98 | 49 | 1 | 1 | 1 | 1 | 1 NA | | 0 | 1 |
| SE | 14287 MAP2K4 | MAP2K4 | chr17 | + | 12032265 | 12032298 | 12020817 | 12021001 | 12054888 | 12054991 | 0 | 1 | 0 | 2 | 0 | 0 | 1 | 1 | 0 | 0 | 82 | 49 | 1 | 1 | 0 | 1 NA | | 1 | 1 | -0.5 |
| SE | 14289 MAP2K4 | MAP2K4 | chr17 | + | 12081355 | 12081530 | 12054888 | 12054991 | 12095574 | 12095694 | 10 | 5 | 5 | 0 | 5 | 0 | 0 | 6 | 0 | 0 | 98 | 49 | 1 | 1 | 1 | 0.333 | 1 NA | | 1 | -0.222 |
| SE | 14291 MAP2K4 | MAP2K4 | chr17 | + | 12129138 | 12129287 | 12125293 | 12125371 | 12139838 | 12139884 | 13 | 8 | 7 | 1 | 0 | 0 | 0 | 1 | 0 | 0 | 98 | 49 | 1 | 1 | 0.867 | 1 | 1 NA | | 1 | -0.044 |
| SE | 14298 ARHGDIB | ARHGDIB | chr12 | - | 14949801 | 14949885 | 14947872 | 14947949 | 14950531 | 14950724 | 464 | 197 | 202 | 0 | 0 | 0 | 13 | 90 | 0 | 1 | 98 | 49 | 1 | 1 | 1 | 1 | 1 | 1 | 0.978 | 0.011 |
| SE | 14311 UBE2F | UBE2F | chr2 | + | 237973648 | 237973747 | 237973091 | 237973225 | 237987962 | 237987992 | 1 | 0 | 0 | 4 | 0 | 0 | 0 | 0 | 0 | 1 | 98 | 49 | 1 | 1 | 0.111 NA | NA | NA | | 0 | 0.111 |
| SE | 14312 UBE2F | UBE2F | chr2 | + | 238016565 | 238016633 | 237994743 | 237994809 | 238025341 | 238025412 | 6 | 2 | 0 | 2 | 0 | 0 | 5 | 1 | 0 | 0 | 98 | 49 | 1 | 1 | 0.6 | 1 NA | | 1 | 1 | -0.2 |
| SE | 14313 UBE2F | UBE2F | chr2 | + | 238035877 | 238035940 | 238032221 | 238032254 | 238041287 | 238042782 | 9 | 9 | 0 | 1 | 0 | 0 | 0 | 3 | 0 | 0 | 98 | 49 | 1 | 1 | 0.818 | 1 NA | NA | | 1 | -0.091 |
| SE | 14317 EIF6 | EIF6 | chr20 | - | 35280653 | 35280829 | 35279941 | 35280118 | 35284175 | 35284261 | 5 | 2 | 0 | 1 | 1 | 0 | 0 | 5 | 0 | 0 | 98 | 49 | 1 | 1 | 0.714 | 0.5 NA | NA | | 1 | -0.393 |
| SE | 14319 PDK1 | PDK1 | chr2 | + | 172562740 | 172562800 | 172562219 | 172562291 | 172564502 | 172564687 | 4 | 3 | 1 | 4 | 0 | 0 | 0 | 1 | 0 | 0 | 98 | 49 | 1 | 1 | 0.333 | 1 | 1 NA | | 1 | -0.222 |
| SE | 14320 PDK1 | PDK1 | chr2 | + | 172562740 | 172562800 | 172562219 | 172562291 | 172564977 | 172565073 | 3 | 3 | 0 | 1 | 2 | 0 | 1 | 0 | 0 | 1 | 98 | 49 | 1 | 1 | 0.6 | 0.429 NA | | 1 | 0 | 0.014 |
| SE | 14321 PDK1 | PDK1 | chr2 | + | 172564502 | 172564687 | 172562219 | 172562291 | 172564977 | 172565073 | 8 | 0 | 2 | 1 | 2 | 0 | 0 | 0 | 0 | 1 | 98 | 49 | 1 | 1 | 0.8 | 0 | 1 NA | | 0 | 0.6 |

| | | | | | | | | | | | | | | | | | | | | | | | | | | | | | | | | |
|---|---|---|---|---|---|---|---|---|---|---|---|---|---|---|---|---|---|---|---|---|---|---|---|---|---|---|---|---|---|---|---|---|
| SE | 14322 | PDK1 | PDK1 | chr2 | + | 172564502 | 172564687 | 172562740 | 172562800 | 172564977 | 172565073 | 6 | 0 | 3 | 1 | 0 | 0 | 0 | 1 | 1 | 0 | 98 | 49 | 1 | 1 | 0.75 | NA | 1 | 0 | | 1 | 0.375 |
| SE | 14323 | PDK1 | PDK1 | chr2 | + | 172568740 | 172568817 | 172566855 | 172566933 | 172570725 | 172570824 | 9 | 5 | 1 | 1 | 1 | 0 | 0 | 2 | 0 | 0 | 98 | 49 | 1 | 1 | 0.818 | 0.714 | 1 | NA | | 1 | -0.156 |
| SE | 14326 | SP3 | SP3 | chr2 | - | 173918592 | 173918785 | 173913069 | 173913266 | 173954872 | 173956232 | 72 | 44 | 13 | 0 | 0 | 0 | 0 | 19 | 0 | 1 | 98 | 49 | 1 | 1 | 1 | 1 | 1 | NA | | 0.905 | 0.095 |
| SE | 14334 | ATF2 | ATF2 | chr2 | - | 175121443 | 175121540 | 175118250 | 175118369 | 175130137 | 175130207 | 16 | 6 | 4 | 2 | 3 | 0 | 0 | 4 | 0 | 1 | 98 | 49 | 1 | 1 | 0.8 | 0.5 | 1 | NA | | 0.667 | 0.1 |
| SE | 14336 | ATF2 | ATF2 | chr2 | - | 175147945 | 175147980 | 175136411 | 175136486 | 175151059 | 175151158 | 2 | 0 | 0 | 8 | 2 | 0 | 0 | 0 | 0 | 2 | 84 | 49 | 1 | 1 | 0.127 | 0 | NA | NA | | 0 | 0.064 |
| SE | 14345 | SETX | SETX | chr9 | - | 132288235 | 132288351 | 132286422 | 132286494 | 132288549 | 132288651 | 76 | 23 | 10 | 0 | 1 | 0 | 0 | 7 | 0 | 0 | 98 | 49 | 1 | 1 | 1 | 0.92 | 1 | NA | | 1 | -0.027 |
| SE | 14346 | SETX | SETX | chr9 | - | 132298079 | 132298312 | 132296886 | 132297054 | 132300629 | 132300803 | 103 | 13 | 3 | 0 | 0 | 0 | 0 | 16 | 0 | 1 | 98 | 49 | 1 | 1 | 1 | 1 | 1 | NA | | 0.889 | 0.111 |
| SE | 14347 | SETX | SETX | chr9 | - | 132326323 | 132330499 | 132311756 | 132311856 | 132331051 | 132331139 | 110 | 34 | 10 | 1 | 0 | 0 | 0 | 18 | 0 | 0 | 98 | 49 | 1 | 1 | 0.982 | 1 | 1 | NA | | 1 | -0.006 |
| SE | 14348 | SETX | SETX | chr9 | - | 132334607 | 132334727 | 132331276 | 132331448 | 132336295 | 132336515 | 171 | 39 | 6 | 0 | 3 | 0 | 0 | 17 | 0 | 0 | 98 | 49 | 1 | 1 | 1 | 0.867 | 1 | NA | | 1 | -0.044 |
| SE | 14349 | SETX | SETX | chr9 | - | 132334607 | 132334727 | 132331276 | 132331448 | 132342689 | 132342799 | 80 | 20 | 2 | 1 | 0 | 0 | 0 | 8 | 0 | 0 | 98 | 49 | 1 | 1 | 0.976 | 1 | 1 | NA | | 1 | -0.008 |
| SE | 14350 | SETX | SETX | chr9 | - | 132336295 | 132336515 | 132331276 | 132331448 | 132342689 | 132342799 | 116 | 29 | 8 | 1 | 0 | 0 | 0 | 9 | 0 | 0 | 98 | 49 | 1 | 1 | 0.983 | 1 | 1 | NA | | 1 | -0.006 |
| SE | 14351 | SETX | SETX | chr9 | - | 132336295 | 132336515 | 132334607 | 132334727 | 132342689 | 132342799 | 207 | 45 | 12 | 1 | 0 | 0 | 0 | 18 | 0 | 0 | 98 | 49 | 1 | 1 | 0.99 | 1 | 1 | NA | | 1 | -0.003 |
| SE | 14352 | SETX | SETX | chr9 | - | 132342689 | 132342799 | 132336295 | 132336515 | 132346260 | 132346471 | 187 | 43 | 10 | 0 | 1 | 0 | 1 | 22 | 0 | 0 | 98 | 49 | 1 | 1 | 1 | 0.956 | 1 | 1 | | 1 | -0.015 |
| SE | 14355 | SESTD1 | SESTD1 | chr2 | - | 179116667 | 179116790 | 179115064 | 179115256 | 179117531 | 179117613 | 3 | 0 | 0 | 0 | 0 | 0 | 0 | 5 | 0 | 1 | 98 | 49 | 1 | 1 | 1 | NA | NA | NA | | 0.714 | 0.286 |
| SE | 14365 | ITGA4 | ITGA4 | chr2 | + | 181475158 | 181475288 | 181474959 | 181475066 | 181478756 | 181478824 | 13 | 8 | 7 | 1 | 1 | 0 | 0 | 2 | 0 | 0 | 98 | 49 | 1 | 1 | 0.867 | 0.8 | 1 | NA | | 1 | -0.111 |
| SE | 14366 | ITGA4 | ITGA4 | chr2 | + | 181530523 | 181530649 | 181529540 | 181529648 | 181531656 | 181531776 | 26 | 11 | 0 | 0 | 0 | 0 | 0 | 7 | 0 | 2 | 98 | 49 | 1 | 1 | 1 | 1 | NA | NA | | 0.636 | 0.364 |
| SE | 14375 | PUM1 | PUM1 | chr1 | - | 30941997 | 30942123 | 30941150 | 30941272 | 30945345 | 30945483 | 7 | 0 | 5 | 1 | 0 | 0 | 0 | 2 | 0 | 2 | 98 | 49 | 1 | 1 | 0.778 | NA | 1 | NA | | 0.333 | 0.556 |
| SE | 14376 | PUM1 | PUM1 | chr1 | - | 30964673 | 30964910 | 30953713 | 30953981 | 30965981 | 30966278 | 5 | 0 | 2 | 0 | 0 | 0 | 0 | 0 | 0 | 1 | 98 | 49 | 1 | 1 | 1 | NA | 1 | NA | | 0 | 1 |
| SE | 14377 | PUM1 | PUM1 | chr1 | - | 30980061 | 30980163 | 30974650 | 30974802 | 30981311 | 30981405 | 6 | 0 | 2 | 2 | 1 | 0 | 0 | 2 | 0 | 0 | 98 | 49 | 1 | 1 | 0.6 | 0 | 1 | NA | | 1 | -0.467 |
| SE | 14383 | DNAJC10 | DNAJC10 | chr2 | + | 182752548 | 182752623 | 182752071 | 182752188 | 182755002 | 182755104 | 1 | 0 | 0 | 7 | 2 | 0 | 0 | 0 | 0 | 3 | 98 | 49 | 1 | 1 | 0.067 | 0 | NA | NA | | 0 | 0.034 |
| SE | 14385 | SMPDL3A | SMPDL3A | chr6 | + | 122795676 | 122795890 | 122789048 | 122789458 | 122796823 | 122796968 | 2 | 0 | 0 | 0 | 0 | 0 | 0 | 1 | 0 | 1 | 98 | 49 | 1 | 1 | 1 | NA | NA | NA | | 0.333 | 0.667 |
| SE | 14386 | ABHD14B | ABHD14B | chr3 | - | 51971459 | 51971698 | 51969942 | 51970184 | 51973964 | 51974630 | 2 | 0 | 0 | 0 | 0 | 0 | 0 | 0 | 0 | 1 | 98 | 49 | 1 | 1 | 1 | NA | NA | NA | | 0 | 1 |
| SE | 14387 | ABHD14B | ABHD14B | chr3 | - | 51971459 | 51971892 | 51969942 | 51970184 | 51973964 | 51974630 | 2 | 0 | 0 | 0 | 0 | 0 | 0 | 0 | 0 | 1 | 98 | 49 | 1 | 1 | 1 | NA | NA | NA | | 0 | 1 |
| SE | 14391 | HIBCH | HIBCH | chr2 | - | 190294545 | 190294630 | 190290404 | 190290485 | 190296812 | 190296953 | 3 | 0 | 0 | 1 | 0 | 0 | 0 | 1 | 0 | 0 | 98 | 49 | 1 | 1 | 0.6 | NA | NA | NA | | 1 | -0.4 |
| SE | 14399 | STAT1 | STAT1 | chr2 | - | 190998216 | 190998308 | 190997855 | 190998007 | 190999625 | 190999704 | 112 | 56 | 20 | 3 | 5 | 0 | 0 | 33 | 0 | 2 | 98 | 49 | 1 | 1 | 0.949 | 0.848 | 1 | NA | | 0.892 | 0.04 |
| SE | 14400 | STAT1 | STAT1 | chr2 | - | 191001073 | 191001163 | 190999625 | 190999704 | 191007562 | 191007661 | 162 | 84 | 22 | 1 | 6 | 0 | 0 | 49 | 0 | 1 | 98 | 49 | 1 | 1 | 0.988 | 0.875 | 1 | NA | | 0.961 | -0.007 |
| SE | 14407 | HSDL2 | HSDL2 | chr9 | + | 112416840 | 112416944 | 112408906 | 112409021 | 112418859 | 112418958 | 7 | 10 | 5 | 1 | 0 | 0 | 0 | 1 | 0 | 0 | 98 | 49 | 1 | 1 | 0.778 | 1 | 1 | NA | | 1 | -0.074 |
| SE | 14410 | RPL32P3 | RPL32P3 | chr3 | - | 129397039 | 129397140 | 129393634 | 129393795 | 129399007 | 129399439 | 0 | 0 | 2 | 0 | 0 | 1 | 0 | 0 | 0 | 1 | 98 | 49 | 1 | 1 | NA | NA | 0.5 | NA | | 0 | 0.5 |
| SE | 14411 | ANKRD44 | ANKRD44 | chr2 | - | 196998906 | 196999052 | 196998336 | 196998419 | 197000418 | 197000502 | 57 | 21 | 15 | 2 | 2 | 0 | 0 | 9 | 0 | 5 | 98 | 49 | 1 | 1 | 0.934 | 0.84 | 1 | NA | | 0.474 | 0.451 |
| SE | 14412 | ANKRD44 | ANKRD44 | chr2 | - | 197000418 | 197000502 | 196998906 | 196999052 | 197001752 | 197001840 | 77 | 25 | 8 | 0 | 2 | 0 | 0 | 8 | 0 | 0 | 98 | 49 | 1 | 1 | 1 | 0.862 | 1 | NA | | 1 | -0.046 |
| SE | 14413 | ANKRD44 | ANKRD44 | chr2 | - | 197013510 | 197013712 | 197008943 | 197009031 | 197025195 | 197025267 | 83 | 36 | 15 | 1 | 3 | 0 | 0 | 9 | 0 | 0 | 98 | 49 | 1 | 1 | 0.976 | 0.857 | 1 | NA | | 1 | -0.056 |
| SE | 14414 | ANKRD44 | ANKRD44 | chr2 | - | 197078702 | 197078814 | 197025195 | 197025267 | 197081644 | 197081725 | 114 | 32 | 21 | 2 | 0 | 0 | 0 | 14 | 0 | 0 | 98 | 49 | 1 | 1 | 0.966 | 1 | 1 | NA | | 1 | -0.011 |
| SE | 14415 | ANKRD44 | ANKRD44 | chr2 | - | 197086679 | 197086748 | 197083368 | 197083509 | 197088710 | 197088774 | 58 | 38 | 25 | 1 | 1 | 0 | 2 | 9 | 0 | 0 | 98 | 49 | 1 | 1 | 0.967 | 0.95 | 1 | 1 | | 1 | -0.028 |
| SE | 14423 | SLC10A7 | SLC10A7 | chr4 | - | 146256825 | 146256936 | 146253980 | 146256520 | 146258691 | 146258837 | 0 | 0 | 0 | 2 | 0 | 0 | 0 | 2 | 0 | 0 | 98 | 49 | 1 | 1 | 0 | NA | NA | NA | | 1 | -1 |
| SE | 14427 | ORC2 | ORC2 | chr2 | - | 200925835 | 200925932 | 200920992 | 200921139 | 200931338 | 200931448 | 4 | 0 | 0 | 0 | 0 | 0 | 0 | 0 | 0 | 1 | 98 | 49 | 1 | 1 | 1 | NA | NA | NA | | 0 | 1 |
| SE | 14430 | CFLAR | CFLAR | chr2 | + | 201145377 | 201145432 | 201140274 | 201140439 | 201149002 | 201149052 | 277 | 153 | 108 | 2 | 0 | 0 | 0 | 64 | 0 | 0 | 98 | 49 | 1 | 1 | 0.986 | 1 | 1 | NA | | 1 | -0.005 |
| SE | 14441 | INO80D | INO80D | chr2 | - | 206063150 | 206063244 | 206062798 | 206063045 | 206085900 | 206086182 | 6 | 0 | 0 | 1 | 0 | 0 | 0 | 2 | 0 | 0 | 98 | 49 | 1 | 1 | 0.75 | NA | NA | NA | | 1 | -0.25 |
| SE | 14445 | DCTN4 | DCTN4 | chr5 | - | 150742113 | 150742157 | 150733367 | 150733475 | 150753478 | 150753657 | 12 | 4 | 5 | 2 | 4 | 0 | 0 | 6 | 0 | 0 | 93 | 49 | 1 | 1 | 0.76 | 0.345 | 1 | NA | | 1 | -0.298 |
| SE | 14446 | CCNYL1 | CCNYL1 | chr2 | + | 207733946 | 207734047 | 207726841 | 207726876 | 207737410 | 207737446 | 8 | 2 | 2 | 3 | 1 | 0 | 0 | 1 | 0 | 0 | 98 | 49 | 1 | 1 | 0.571 | 0.5 | 1 | NA | | 1 | -0.31 |
| SE | 14448 | PIKFYVE | PIKFYVE | chr2 | + | 208273583 | 208273733 | 208271510 | 208271691 | 208285725 | 208285933 | 11 | 0 | 3 | 1 | 0 | 0 | 0 | 2 | 0 | 0 | 98 | 49 | 1 | 1 | 0.846 | NA | 1 | NA | | 1 | -0.077 |
| SE | 14450 | PIKFYVE | PIKFYVE | chr2 | + | 208277536 | 208277708 | 208276711 | 208276830 | 208285725 | 208285933 | 10 | 4 | 3 | 2 | 0 | 0 | 0 | 5 | 0 | 0 | 98 | 49 | 1 | 1 | 0.714 | 1 | 1 | NA | | 1 | -0.095 |
| SE | 14452 | PIKFYVE | PIKFYVE | chr2 | + | 208285725 | 208285933 | 208276711 | 208276830 | 208288728 | 208288818 | 13 | 9 | 0 | 1 | 0 | 0 | 0 | 1 | 0 | 0 | 98 | 49 | 1 | 1 | 0.867 | 1 | NA | NA | | 1 | -0.067 |
| SE | 14453 | PIKFYVE | PIKFYVE | chr2 | + | 208285725 | 208285933 | 208277536 | 208277708 | 208288728 | 208288818 | 18 | 11 | 0 | 5 | 0 | 0 | 0 | 6 | 0 | 0 | 98 | 49 | 1 | 1 | 0.643 | 1 | NA | NA | | 1 | -0.178 |
| SE | 14454 | PIKFYVE | PIKFYVE | chr2 | + | 208304845 | 208305013 | 208304170 | 208304318 | 208312235 | 208312295 | 4 | 1 | 2 | 2 | 7 | 0 | 0 | 1 | 0 | 4 | 98 | 49 | 1 | 1 | 0.5 | 0.067 | 1 | NA | | 0.111 | 0.411 |
| SE | 14455 | PIKFYVE | PIKFYVE | chr2 | + | 208333314 | 208333493 | 208330522 | 208330694 | 208335305 | 208335419 | 3 | 0 | 2 | 0 | 2 | 0 | 0 | 1 | 0 | 0 | 98 | 49 | 1 | 1 | 1 | 0 | 1 | NA | | 1 | -0.333 |
| SE | 14456 | PIKFYVE | PIKFYVE | chr2 | + | 208347858 | 208348023 | 208346049 | 208346147 | 208350023 | 208350083 | 9 | 2 | 0 | 0 | 0 | 0 | 0 | 2 | 0 | 2 | 98 | 49 | 1 | 1 | 1 | 1 | NA | NA | | 0.333 | 0.667 |
| SE | 14457 | PIKFYVE | PIKFYVE | chr2 | + | 208351351 | 208351455 | 208350770 | 208350947 | 208352653 | 208352782 | 10 | 6 | 0 | 0 | 0 | 0 | 0 | 3 | 0 | 1 | 98 | 49 | 1 | 1 | 1 | 1 | NA | NA | | 0.6 | 0.4 |
| SE | 14464 | BARD1 | BARD1 | chr2 | - | 214792296 | 214792445 | 214780559 | 214781509 | 214809411 | 214809711 | 0 | 0 | 0 | 0 | 1 | 0 | 0 | 2 | 0 | 0 | 98 | 49 | 1 | 1 | NA | 0 | NA | NA | | 1 | -1 |
| SE | 14466 | BARD1 | BARD1 | chr2 | - | 214797060 | 214797117 | 214780559 | 214781509 | 214809411 | 214809711 | 0 | 0 | 0 | 0 | 1 | 0 | 0 | 1 | 0 | 0 | 98 | 49 | 1 | 1 | NA | 0 | NA | NA | | 1 | -1 |
| SE | 14472 | UBXN7 | UBXN7 | chr3 | - | 196369420 | 196369511 | 196368027 | 196368155 | 196371895 | 196372042 | 9 | 14 | 3 | 0 | 3 | 0 | 0 | 8 | 0 | 0 | 98 | 49 | 1 | 1 | 1 | 0.7 | 1 | NA | | 1 | -0.1 |
| SE | 14473 | RELA | RELA | chr11 | - | 65658286 | 65658499 | 65655854 | 65655935 | 65658717 | 65658822 | 12 | 7 | 1 | 0 | 0 | 1 | 0 | 4 | 0 | 0 | 98 | 49 | 1 | 1 | 1 | 1 | 0.333 | NA | | 1 | -0.222 |
| SE | 14474 | RELA | RELA | chr11 | - | 65658717 | 65658822 | 65655854 | 65655935 | 65659665 | 65659788 | 28 | 2 | 4 | 8 | 1 | 0 | 0 | 2 | 0 | 0 | 98 | 49 | 1 | 1 | 0.636 | 0.5 | 1 | NA | | 1 | -0.288 |
| SE | 14479 | BAZ2A | BAZ2A | chr12 | - | 56604584 | 56604799 | 56604216 | 56604291 | 56605072 | 56605327 | 31 | 14 | 0 | 1 | 2 | 0 | 0 | 3 | 0 | 0 | 98 | 49 | 1 | 1 | 0.939 | 0.778 | NA | NA | | 1 | -0.141 |
| SE | 14481 | BAZ2A | BAZ2A | chr12 | - | 56610408 | 56610517 | 56610113 | 56610215 | 56611572 | 56611633 | 34 | 13 | 4 | 0 | 2 | 0 | 1 | 2 | 0 | 0 | 98 | 49 | 1 | 1 | 1 | 0.765 | 1 | 1 | | 1 | -0.078 |
| SE | 14482 | ATP5L | ATP5L | chr11 | + | 118405656 | 118405709 | 118401388 | 118401717 | 118406936 | 118407097 | 0 | 0 | 0 | 9 | 4 | 0 | 0 | 2 | 0 | 0 | 98 | 49 | 1 | 1 | 0 | 0 | NA | NA | | 1 | -1 |
| SE | 14486 | ARPC2 | ARPC2 | chr2 | + | 218234351 | 218234397 | 218228737 | 218228850 | 218238663 | 218238850 | 168 | 112 | 91 | 1 | 0 | 0 | 4 | 32 | 0 | 0 | 95 | 49 | 1 | 1 | 0.989 | 1 | 1 | 1 | | 1 | -0.004 |
| SE | 14487 | ARPC2 | ARPC2 | chr2 | + | 218238663 | 218238850 | 218228737 | 218228850 | 218245419 | 218245546 | 2 | 0 | 0 | 1 | 1 | 0 | 0 | 1 | 0 | 1 | 98 | 49 | 1 | 1 | 0.5 | 0 | NA | NA | | 0.333 | -0.083 |
| SE | 14488 | ARPC2 | ARPC2 | chr2 | + | 218239390 | 218239484 | 218238663 | 218238850 | 218245419 | 218245546 | 338 | 115 | 89 | 1 | 0 | 0 | 0 | 56 | 0 | 1 | 98 | 49 | 1 | 1 | 0.994 | 1 | 1 | NA | | 0.966 | 0.032 |
| SE | 14492 | FRMD4B | FRMD4B | chr3 | - | 69249225 | 69249248 | 69224606 | 69224690 | 69250042 | 69250099 | 1 | 0 | 0 | 1 | 0 | 0 | 0 | 1 | 0 | 0 | 72 | 49 | 1 | 1 | 0.405 | NA | NA | NA | | 1 | -0.595 |
| SE | 14500 | ZFAND2B | ZFAND2B | chr2 | + | 219207647 | 219207779 | 219207326 | 219207421 | 219207886 | 219208038 | 11 | 2 | 3 | 0 | 2 | 0 | 0 | 2 | 0 | 0 | 98 | 49 | 1 | 1 | 1 | 0.333 | 1 | NA | | 1 | -0.222 |
| SE | 14501 | ZFAND2B | ZFAND2B | chr2 | + | 219208976 | 219209049 | 219208571 | 219208639 | 219209261 | 219209651 | 9 | 5 | 4 | 2 | 0 | 0 | 0 | 5 | 0 | 0 | 98 | 49 | 1 | 1 | 0.692 | 1 | 1 | NA | | 1 | -0.103 |
| SE | 14502 | ATG9A | ATG9A | chr2 | - | 219227921 | 219228053 | 219227769 | 219227813 | 219228433 | 219228485 | 8 | 3 | 0 | 1 | 0 | 0 | 0 | 1 | 0 | 0 | 98 | 49 | 1 | 1 | 0.8 | 1 | NA | NA | | 1 | -0.1 |
| SE | 14506 | GOLGB1 | GOLGB1 | chr3 | - | 121699811 | 121699885 | 121693740 | 121698929 | 121702480 | 121702595 | 12 | 2 | 0 | 1 | 0 | 0 | 0 | 2 | 0 | 1 | 98 | 49 | 1 | 1 | 0.857 | 1 | NA | NA | | 0.5 | 0.428 |
| SE | 14507 | GOLGB1 | GOLGB1 | chr3 | - | 121702480 | 121702595 | 121693740 | 121698929 | 121714860 | 121714976 | 5 | 8 | 1 | 1 | 0 | 0 | 0 | 5 | 0 | 0 | 98 | 49 | 1 | 1 | 0.714 | 1 | 1 | NA | | 1 | -0.095 |
| SE | 14508 | GOLGB1 | GOLGB1 | chr3 | - | 121719645 | 121719753 | 121718387 | 121718501 | 121722261 | 121722378 | 1 | 0 | 0 | 3 | 0 | 2 | 0 | 0 | 0 | 2 | 98 | 49 | 1 | 1 | 0.143 | NA | 0 | NA | | 0 | 0.071 |
| SE | 14509 | GOLGB1 | GOLGB1 | chr3 | - | 121719645 | 121719768 | 121718387 | 121718501 | 121722261 | 121722378 | 1 | 0 | 0 | 3 | 0 | 2 | 0 | 0 | 0 | 2 | 98 | 49 | 1 | 1 | 0.143 | NA | 0 | NA | | 0 | 0.071 |
| SE | 14512 | ACSL3 | ACSL3 | chr2 | + | 222900673 | 222900780 | 222887829 | 222887888 | 222908732 | 222909150 | 3 | 7 | 0 | 1 | 0 | 0 | 0 | 6 | 0 | 0 | 98 | 49 | 1 | 1 | 0.6 | 1 | NA | NA | | 1 | -0.2 |
| SE | 14513 | ACSL3 | ACSL3 | chr2 | + | 222921279 | 222921430 | 222919063 | 222919202 | 222922707 | 222922831 | 4 | 4 | 0 | 0 | 2 | 0 | 0 | 9 | 0 | 0 | 98 | 49 | 1 | 1 | 1 | 0.5 | NA | NA | | 1 | -0.25 |
| SE | 14517 | CCNJL | CCNJL | chr5 | - | 160254128 | 160254335 | 160251651 | 160253798 | 160255548 | 160255708 | 1 | 0 | 0 | 14 | 7 | 4 | 0 | 0 | 0 | 4 | 98 | 49 | 1 | 1 | 0.034 | 0 | 0 | NA | | 0 | 0.011 |
| SE | 14521 | MFF | MFF | chr2 | + | 227328677 | 227328789 | 227325150 | 227325427 | 227330625 | 227330846 | 1 | 0 | 0 | 1 | 0 | 0 | 1 | 0 | 0 | 0 | 98 | 49 | 1 | 1 | 0.333 | NA | NA | 1 | NA | | -0.667 |
| SE | 14523 | MFF | MFF | chr2 | + | 227330625 | 227330846 | 227325150 | 227325427 | 227332418 | 227332588 | 6 | 0 | 0 | 1 | 1 | 0 | 0 | 0 | 0 | 1 | 98 | 49 | 1 | 1 | 0.75 | 0 | NA | NA | | 0 | 0.375 |
| SE | 14535 | SP100 | SP100 | chr2 | + | 230440475 | 230440557 | 230417590 | 230417665 | 230442936 | 230443099 | 0 | 1 | 0 | 26 | 36 | 1 | 0 | 1 | 1 | 3 | 98 | 49 | 1 | 1 | 0 | 0.014 | 0 | 0 | | 0.143 | -0.067 |
| SE | 14536 | SP100 | SP100 | chr2 | + | 230449560 | 230449710 | 230449087 | 230449150 | 230450171 | 230450255 | 86 | 64 | 10 | 1 | 0 | 0 | 7 | 33 | 0 | 0 | 98 | 49 | 1 | 1 | 0.977 | 1 | 1 | 1 | | 1 | -0.008 |
| SE | 14537 | SP100 | SP100 | chr2 | + | 230450171 | 230450255 | 230449087 | 230449150 | 230461261 | 230461414 | 52 | 32 | 10 | 1 | 1 | 0 | 2 | 23 | 1 | 0 | 98 | 49 | 1 | 1 | 0.963 | 0.941 | 1 | 0.5 | | 1 | 0.218 |
| SE | 14538 | SP100 | SP100 | chr2 | + | 230464066 | 230464150 | 230462434 | 230462518 | 230466300 | 230466354 | 71 | 39 | 23 | 5 | 8 | 0 | 0 | 25 | 0 | 2 | 98 | 49 | 1 | 1 | 0.877 | 0.709 | 1 | NA | | 0.862 | 0 |
| SE | 14539 | SP100 | SP100 | chr2 | + | 230498460 | 230498535 | 230494415 | 230494460 | 230503065 | 230503110 | 42 | 29 | 5 | 1 | 0 | 0 | 0 | 11 | 0 | 0 | 98 | 49 | 1 | 1 | 0.955 | 1 | 1 | NA | | 1 | -0.015 |
| SE | 14540 | SP100 | SP100 | chr2 | + | 230507992 | 230508031 | 230506302 | 230506445 | 230511124 | 230511166 | 25 | 15 | 4 | 1 | 0 | 0 | 0 | 4 | 0 | 0 | 88 | 49 | 1 | 1 | 0.933 | 1 | 1 | NA | | 1 | -0.022 |
| SE | 14546 | EIF4G2 | EIF4G2 | chr11 | - | 10803479 | 10803590 | 10803210 | 10803294 | 10803898 | 10804050 | 152 | 46 | 33 | 0 | 1 | 0 | 1 | 22 | 0 | 0 | 98 | 49 | 1 | 1 | 1 | 0.958 | 1 | 1 | | 1 | -0.014 |
| SE | 14547 | EIF4G2 | EIF4G2 | chr11 | - | 10804912 | 10805015 | 10804286 | 10804418 | 10805906 | 10806047 | 114 | 18 | 23 | 3 | 0 | 0 | 0 | 14 | 0 | 0 | 98 | 49 | 1 | 1 | 0.95 | 1 | 1 | NA | | 1 | -0.017 |
| SE | 14553 | GIGYF2 | GIGYF2 | chr2 | + | 232734416 | 232734486 | 232703423 | 232703489 | 232735154 | 232735238 | 0 | 2 | 3 | 1 | 0 | 0 | 0 | 3 | 0 | 0 | 98 | 49 | 1 | 1 | 0 | 1 | 1 | NA | | 1 | -0.333 |
| SE | 14559 | CLK3 | CLK3 | chr15 | + | 74627351 | 74627446 | 74624901 | 74625018 | 74627538 | 74627668 | 18 | 5 | 4 | 1 | 0 | 0 | 0 | 5 | 0 | 0 | 98 | 49 | 1 | 1 | 0.9 | 1 | 1 | NA | | 1 | -0.033 |
| SE | 14560 | CLK3 | CLK3 | chr15 | + | 74627969 | 74628052 | 74627538 | 74627668 | 74628603 | 74628683 | 24 | 3 | 13 | 0 | 0 | 0 | 0 | 3 | 0 | 1 | 98 | 49 | 1 | 1 | 1 | 1 | 1 | NA | | 0.6 | 0.4 |
| SE | 14562 | SEC23A | SEC23A | chr14 | - | 39074414 | 39074530 | 39067172 | 39067296 | 39075934 | 39076093 | 18 | 3 | 2 | 1 | 0 | 0 | 2 | 0 | 0 | 0 | 98 | 49 | 1 | 1 | 0.9 | 1 | 1 | 1 | NA | | -0.033 |
| SE | 14563 | DGKD | DGKD | chr2 | + | 233434768 | 233434901 | 233434379 | 233434484 | 233435817 | 233435924 | 21 | 5 | 8 | 1 | 0 | 0 | 0 | 13 | 0 | 0 | 98 | 49 | 1 | 1 | 0.913 | 1 | 1 | NA | | 1 | -0.029 |
| SE | 14564 | DGKD | DGKD | chr2 | + | 233435817 | 233435924 | 233434768 | 233434901 | 233436315 | 233436441 | 12 | 9 | 12 | 0 | 0 | 0 | 0 | 12 | 0 | 1 | 98 | 49 | 1 | 1 | 1 | 1 | 1 | NA | | 0.857 | 0.143 |
| SE | 14572 | SERPINB1 | SERPINB1 | chr6 | - | 2839334 | 2839466 | 2838548 | 2838686 | 2840418 | 2840594 | 0 | 0 | 0 | 11 | 11 | 9 | 0 | 2 | 0 | 7 | 98 | 49 | 1 | 1 | 0 | 0 | 0 | NA | | 0.125 | -0.125 |
| SE | 14574 | LRRFIP1 | LRRFIP1 | chr2 | + | 237708543 | 237708630 | 237692163 | 237692521 | 237748363 | 237748399 | 15 | 14 | 1 | 0 | 0 | 0 | 0 | 2 | 0 | 2 | 98 | 49 | 1 | 1 | 1 | 1 | 1 | NA | | 0.333 | 0.667 |
| SE | 14576 | LRRFIP1 | LRRFIP1 | chr2 | + | 237717761 | 237717809 | 237708543 | 237708630 | 237719522 | 237719567 | 2 | 2 | 2 | 1 | 0 | 0 | 0 | 1 | 0 | 0 | 97 | 49 | 1 | 1 | 0.503 | 1 | 1 | NA | | 1 | -0.166 |
| SE | 14579 | UNC13D | UNC13D | chr17 | - | 75835646 | 75835777 | 75835408 | 75835529 | 75835854 | 75835906 | 47 | 14 | 10 | 2 | 0 | 0 | 4 | 11 | 0 | 0 | 98 | 49 | 1 | 1 | 0.922 | 1 | 1 | 1 | | 1 | -0.026 |
| SE | 14580 | UNC13D | UNC13D | chr17 | - | 75840506 | 75840576 | 75840224 | 75840329 | 75840761 | 75840830 | 39 | 19 | 9 | 0 | 2 | 0 | 3 | 7 | 0 | 0 | 98 | 49 | 1 | 1 | 1 | 0.826 | 1 | 1 | | 1 | -0.058 |
| SE | 14584 | HDAC4 | HDAC4 | chr2 | - | 239053459 | 239053601 | 239048167 | 239053136 | 239054748 | 239054833 | 6 | 1 | 1 | 1 | 0 | 0 | 0 | 6 | 0 | 0 | 98 | 49 | 1 | 1 | 0.75 | 1 | 1 | NA | | 1 | -0.083 |
| SE | 14586 | HDAC4 | HDAC4 | chr2 | - | 239144582 | 239144714 | 239139683 | 239139796 | 239156651 | 239156773 | 12 | 3 | 3 | 0 | 0 | 0 | 0 | 3 | 0 | 2 | 98 | 49 | 1 | 1 | 1 | 1 | 1 | NA | | 0.429 | 0.571 |
| SE | 14587 | CAPN10 | CAPN10 | chr2 | + | 240596318 | 240596521 | 240595023 | 240595304 | 240597887 | 240598087 | 1 | 2 | 1 | 0 | 0 | 0 | 0 | 2 | 0 | 1 | 98 | 49 | 1 | 1 | 1 | 1 | 1 | NA | | 0.5 | 0.5 |
| SE | 14591 | RFC1 | RFC1 | chr4 | - | 39306591 | 39306701 | 39304813 | 39304928 | 39308635 | 39309032 | 8 | 2 | 0 | 1 | 0 | 0 | 0 | 2 | 0 | 0 | 98 | 49 | 1 | 1 | 0.8 | 1 | NA | NA | | 1 | -0.1 |
| SE | 14594 | GOLPH3 | GOLPH3 | chr5 | - | 32143748 | 32143880 | 32135571 | 32135686 | 32173809 | 32174319 | 17 | 14 | 9 | 0 | 0 | 2 | 0 | 3 | 0 | 0 | 98 | 49 | 1 | 1 | 1 | 1 | 0.692 | NA | | 1 | -0.103 |
| SE | 14595 | CCDC66 | CCDC66 | chr3 | + | 56559568 | 56559594 | 56558845 | 56558910 | 56563683 | 56564125 | 2 | 0 | 0 | 1 | 0 | 0 | 0 | 1 | 0 | 0 | 75 | 49 | 1 | 1 | 0.566 | NA | NA | NA | | 1 | -0.434 |
| SE | 14612 | ATG4B | ATG4B | chr2 | + | 241655270 | 241655343 | 241654545 | 241654647 | 241659107 | 241659187 | 3 | 5 | 1 | 3 | 0 | 0 | 0 | 1 | 0 | 0 | 98 | 49 | 1 | 1 | 0.333 | 1 | 1 | NA | | 1 | -0.222 |
| SE | 14619 | USP32 | USP32 | chr17 | - | 60288522 | 60288682 | 60271349 | 60271481 | 60294682 | 60294801 | 75 | 33 | 12 | 1 | 0 | 0 | 0 | 12 | 0 | 0 | 98 | 49 | 1 | 1 | 0.974 | 1 | 1 | NA | | 1 | -0.009 |
| SE | 14622 | RBCK1 | RBCK1 | chr20 | + | 409880 | 410025 | 408064 | 408779 | 417525 | 417619 | 25 | 15 | 16 | 1 | 0 | 0 | 0 | 5 | 0 | 0 | 98 | 49 | 1 | 1 | 0.926 | 1 | 1 | NA | | 1 | -0.025 |
| SE | 14623 | RBCK1 | RBCK1 | chr20 | + | 420870 | 421031 | 419557 | 419731 | 422126 | 422238 | 7 | 8 | 4 | 0 | 1 | 0 | 2 | 3 | 0 | 1 | 98 | 49 | 1 | 1 | 1 | 0.8 | 1 | 1 | | 0.6 | 0.133 |
| SE | 14627 | TBC1D20 | TBC1D20 | chr20 | - | 447888 | 448074 | 445049 | 445130 | 462335 | 462553 | 8 | 4 | 0 | 2 | 0 | 0 | 1 | 6 | 0 | 0 | 98 | 49 | 1 | 1 | 0.667 | 1 | NA | 1 | | 1 | -0.166 |
| SE | 14637 | SNRPB | SNRPB | chr20 | - | 2465707 | 2465819 | 2463746 | 2463899 | 2467606 | 2467758 | 25 | 14 | 8 | 0 | 0 | 0 | 1 | 10 | 0 | 1 | 98 | 49 | 1 | 1 | 1 | 1 | 1 | 1 | | 0.833 | 0.084 |
| SE | 14656 | NUMA1 | NUMA1 | chr11 | - | 72012400 | 72012442 | 72010785 | 72010854 | 72012894 | 72016260 | 10 | 0 | 0 | 4 | 2 | 1 | 0 | 0 | 0 | 1 | 91 | 49 | 1 | 1 | 0.574 | 0 | 0 | NA | | 0 | 0.191 |
| SE | 14668 | CSF1R | CSF1R | chr5 | - | 150059699 | 150059862 | 150057503 | 150057592 | 150060861 | 150060972 | 20 | 1 | 3 | 1 | 0 | 0 | 0 | 7 | 0 | 0 | 98 | 49 | 1 | 1 | 0.909 | 1 | 1 | NA | | 1 | -0.03 |
| SE | 14673 | LRRFIP2 | LRRFIP2 | chr3 | - | 37091466 | 37091538 | 37083635 | 37083806 | 37094791 | 37094908 | 4 | 1 | 1 | 5 | 0 | 0 | 0 | 2 | 0 | 0 | 98 | 49 | 1 | 1 | 0.286 | 1 | 1 | NA | | 1 | -0.238 |
| SE | 14678 | PCIF1 | PCIF1 | chr20 | + | 45943339 | 45943423 | 45943096 | 45943244 | 45943665 | 45943765 | 6 | 4 | 4 | 0 | 0 | 0 | 0 | 6 | 0 | 1 | 98 | 49 | 1 | 1 | 1 | 1 | 1 | NA | | 0.75 | 0.25 |
| SE | 14679 | PCIF1 | PCIF1 | chr20 | + | 45943665 | 45943765 | 45943096 | 45943244 | 45944867 | 45945030 | 5 | 0 | 4 | 0 | 3 | 0 | 0 | 2 | 0 | 0 | 98 | 49 | 1 | 1 | 1 | 0 | 1 | NA | | 1 | -0.333 |
| SE | 14680 | PCIF1 | PCIF1 | chr20 | + | 45945710 | 45945883 | 45944867 | 45945030 | 45946028 | 45946115 | 3 | 1 | 0 | 1 | 0 | 3 | 0 | 1 | 0 | 1 | 98 | 49 | 1 | 1 | 0.6 | 1 | 0 | NA | | 0.333 | 0.2 |
| SE | 14681 | TIAM1 | TIAM1 | chr21 | - | 31202907 | 31203012 | 31195223 | 31195305 | 31210044 | 31210215 | 2 | 0 | 0 | 1 | 0 | 0 | 0 | 4 | 0 | 0 | 98 | 49 | 1 | 1 | 0.5 | NA | NA | NA | | 1 | -0.5 |
| SE | 14696 | NAPB | NAPB | chr20 | - | 23395138 | 23395185 | 23394921 | 23394999 | 23397071 | 23397188 | 0 | 0 | 0 | 1 | 0 | 0 | 0 | 2 | 0 | 0 | 96 | 49 | 1 | 1 | 0 | NA | NA | NA | | 1 | -1 |
| SE | 14701 | STK10 | STK10 | chr5 | - | 172093411 | 172093960 | 172090231 | 172090362 | 172105655 | 172105737 | 42 | 3 | 6 | 2 | 0 | 0 | 1 | 8 | 0 | 0 | 98 | 49 | 1 | 1 | 0.913 | 1 | 1 | 1 | | 1 | -0.029 |
| SE | 14702 | STK10 | STK10 | chr5 | - | 172096425 | 172096560 | 172093411 | 172093960 | 172105655 | 172105737 | 59 | 25 | 13 | 1 | 0 | 0 | 2 | 8 | 0 | 0 | 98 | 49 | 1 | 1 | 0.967 | 1 | 1 | 1 | | 1 | -0.011 |
| SE | 14703 | STK10 | STK10 | chr5 | - | 172107779 | 172107852 | 172106619 | 172106814 | 172117480 | 172117630 | 107 | 56 | 14 | 0 | 1 | 0 | 6 | 14 | 0 | 0 | 98 | 49 | 1 | 1 | 1 | 0.966 | 1 | 1 | | 1 | -0.011 |
| SE | 14710 | PLEKHG3 | PLEKHG3 | chr14 | + | 64728995 | 64729093 | 64727592 | 64727982 | 64730641 | 64730688 | 21 | 11 | 5 | 2 | 0 | 0 | 1 | 2 | 0 | 0 | 98 | 49 | 1 | 1 | 0.84 | 1 | 1 | 1 | | 1 | -0.053 |
| SE | 14711 | PLEKHG3 | PLEKHG3 | chr14 | + | 64730242 | 64730312 | 64728995 | 64729093 | 64730641 | 64730688 | 37 | 27 | 18 | 5 | 6 | 0 | 0 | 4 | 0 | 0 | 98 | 49 | 1 | 1 | 0.787 | 0.692 | 1 | NA | | 1 | -0.174 |
| SE | 14712 | PLEKHG3 | PLEKHG3 | chr14 | + | 64731360 | 64731543 | 64731037 | 64731169 | 64731713 | 64731806 | 35 | 20 | 25 | 2 | 0 | 0 | 0 | 6 | 0 | 0 | 98 | 49 | 1 | 1 | 0.897 | 1 | 1 | NA | | 1 | -0.034 |
| SE | 14727 | HM13 | HM13 | chr20 | + | 31568077 | 31568224 | 31566209 | 31566295 | 31569119 | 31569567 | 0 | 3 | 0 | 4 | 0 | 0 | 0 | 0 | 0 | 1 | 98 | 49 | 1 | 1 | 0 | 1 | NA | NA | | 0 | 0.5 |
| SE | 14734 | HCK | HCK | chr20 | + | 32069687 | 32069751 | 32052187 | 32052486 | 32071661 | 32071782 | 2 | 0 | 0 | 135 | 59 | 44 | 0 | 0 | 9 | 33 | 98 | 49 | 1 | 1 | 0.007 | 0 | 0 | 0 | | 0 | 0.002 |
| SE | 14735 | HCK | HCK | chr20 | + | 32073318 | 32073361 | 32071661 | 32071782 | 32073715 | 32073818 | 146 | 80 | 82 | 3 | 3 | 0 | 0 | 24 | 0 | 0 | 92 | 49 | 1 | 1 | 0.963 | 0.934 | 1 | NA | | 1 | -0.034 |
| SE | 14736 | HCK | HCK | chr20 | + | 32073318 | 32073361 | 32071661 | 32071782 | 32073718 | 32073818 | 114 | 49 | 55 | 2 | 0 | 0 | 0 | 19 | 0 | 0 | 92 | 49 | 1 | 1 | 0.968 | 1 | 1 | NA | | 1 | -0.011 |
| SE | 14737 | HCK | HCK | chr20 | + | 32079773 | 32079877 | 32074622 | 32074721 | 32083893 | 32084043 | 186 | 83 | 65 | 1 | 0 | 0 | 0 | 38 | 0 | 0 | 98 | 49 | 1 | 1 | 0.989 | 1 | 1 | NA | | 1 | -0.004 |
| SE | 14739 | TM9SF4 | TM9SF4 | chr20 | + | 32157793 | 32157969 | 32155102 | 32155186 | 32158450 | 32158514 | 6 | 0 | 0 | 1 | 0 | 0 | 1 | 3 | 0 | 0 | 98 | 49 | 1 | 1 | 0.75 | NA | NA | 1 | | 1 | -0.25 |
| SE | 14740 | TM9SF4 | TM9SF4 | chr20 | + | 32159991 | 32160111 | 32158450 | 32158514 | 32161275 | 32161365 | 9 | 1 | 0 | 0 | 0 | 0 | 2 | 4 | 1 | 0 | 98 | 49 | 1 | 1 | 1 | 1 | NA | 0.5 | | 1 | 0.25 |
| SE | 14741 | GPR132 | GPR132 | chr14 | - | 105055386 | 105056166 | 105049388 | 105052102 | 105057166 | 105057280 | 3 | 0 | 0 | 5 | 2 | 0 | 0 | 0 | 0 | 2 | 98 | 49 | 1 | 1 | 0.231 | 0 | NA | NA | | 0 | 0.116 |
| SE | 14744 | IDUA | IDUA | chr4 | + | 1000881 | 1000989 | 1000611 | 1000697 | 1001467 | 1001563 | 1 | 0 | 0 | 0 | 0 | 0 | 0 | 0 | 0 | 2 | 98 | 49 | 1 | 1 | 1 | NA | NA | NA | | 0 | 1 |
| SE | 14750 | ASXL1 | ASXL1 | chr20 | + | 32429900 | 32430053 | 32429337 | 32429431 | 32431320 | 32431484 | 6 | 2 | 0 | 1 | 3 | 0 | 0 | 1 | 0 | 0 | 98 | 49 | 1 | 1 | 0.75 | 0.25 | NA | NA | | 1 | -0.5 |
| SE | 14753 | RALY | RALY | chr20 | + | 34031521 | 34031604 | 33993651 | 33994131 | 34072065 | 34072330 | 9 | 10 | 6 | 1 | 0 | 0 | 3 | 10 | 0 | 0 | 98 | 49 | 1 | 1 | 0.818 | 1 | 1 | 1 | | 1 | -0.061 |
| SE | 14754 | RALY | RALY | chr20 | + | 34073818 | 34073866 | 34073562 | 34073635 | 34075873 | 34076040 | 11 | 3 | 7 | 1 | 0 | 0 | 0 | 8 | 0 | 0 | 97 | 49 | 1 | 1 | 0.847 | 1 | 1 | NA | | 1 | -0.051 |
| SE | 14761 | MAP1LC3A | MAP1LC3A | chr20 | + | 34559207 | 34559263 | 34558696 | 34558908 | 34559346 | 34559453 | 4 | 2 | 2 | 1 | 0 | 0 | 0 | 1 | 0 | 0 | 98 | 49 | 1 | 1 | 0.667 | 1 | 1 | NA | | 1 | -0.111 |
| SE | 14770 | CPNE1 | CPNE1 | chr20 | - | 35630740 | 35630795 | 35630438 | 35630490 | 35630900 | 35631034 | 31 | 7 | 24 | 1 | 0 | 2 | 0 | 4 | 0 | 0 | 98 | 49 | 1 | 1 | 0.939 | 1 | 0.857 | NA | | 1 | -0.068 |
| SE | 14771 | CPNE1 | CPNE1 | chr20 | - | 35632162 | 35632234 | 35631944 | 35632025 | 35632310 | 35632385 | 16 | 11 | 4 | 5 | 1 | 2 | 4 | 0 | 0 | 0 | 98 | 49 | 1 | 1 | 0.615 | 0.846 | 0.5 | 1 | NA | | -0.346 |
| SE | 14772 | CPNE1 | CPNE1 | chr20 | - | 35655201 | 35655344 | 35632794 | 35632923 | 35658929 | 35658982 | 12 | 5 | 0 | 0 | 3 | 0 | 0 | 0 | 0 | 2 | 98 | 49 | 1 | 1 | 1 | 0.455 | NA | NA | | 0 | 0.728 |
| SE | 14773 | CPNE1 | CPNE1 | chr20 | - | 35658929 | 35658982 | 35632794 | 35632923 | 35664759 | 35664937 | 3 | 4 | 0 | 3 | 1 | 0 | 0 | 2 | 0 | 0 | 98 | 49 | 1 | 1 | 0.333 | 0.667 | NA | NA | | 1 | -0.5 |
| SE | 14774 | CPNE1 | CPNE1 | chr20 | - | 35658929 | 35659014 | 35632794 | 35632923 | 35664759 | 35664956 | 3 | 5 | 1 | 3 | 1 | 0 | 0 | 2 | 0 | 0 | 98 | 49 | 1 | 1 | 0.333 | 0.714 | 1 | NA | | 1 | -0.318 |
| SE | 14782 | PHF20 | PHF20 | chr20 | + | 35863012 | 35863400 | 35858301 | 35858381 | 35869437 | 35869551 | 19 | 8 | 4 | 3 | 0 | 0 | 0 | 2 | 0 | 0 | 98 | 49 | 1 | 1 | 0.76 | 1 | 1 | NA | | 1 | -0.08 |
| SE | 14783 | PHF20 | PHF20 | chr20 | + | 35870954 | 35871134 | 35869437 | 35869551 | 35871649 | 35871829 | 14 | 2 | 4 | 1 | 0 | 0 | 0 | 1 | 0 | 0 | 98 | 49 | 1 | 1 | 0.875 | 1 | 1 | NA | | 1 | -0.042 |
| SE | 14784 | PHF20 | PHF20 | chr20 | + | 35927779 | 35927879 | 35917483 | 35917662 | 35931248 | 35931444 | 14 | 4 | 3 | 0 | 1 | 0 | 0 | 1 | 0 | 0 | 98 | 49 | 1 | 1 | 1 | 0.667 | 1 | NA | | 1 | -0.111 |
| SE | 14785 | DHX36 | DHX36 | chr3 | - | 154280762 | 154280862 | 154280578 | 154280669 | 154283187 | 154283271 | 7 | 4 | 0 | 2 | 0 | 0 | 0 | 3 | 0 | 0 | 98 | 49 | 1 | 1 | 0.636 | 1 | NA | NA | | 1 | -0.182 |
| SE | 14786 | DHX36 | DHX36 | chr3 | - | 154306215 | 154306295 | 154305093 | 154305168 | 154309652 | 154309823 | 5 | 2 | 0 | 0 | 0 | 0 | 0 | 1 | 0 | 1 | 98 | 49 | 1 | 1 | 1 | 1 | NA | NA | | 0.333 | 0.667 |
| SE | 14787 | MYL9 | MYL9 | chr20 | + | 36548031 | 36548193 | 36544858 | 36545068 | 36549076 | 36549823 | 7 | 0 | 2 | 1 | 0 | 0 | 4 | 0 | 0 | 0 | 98 | 49 | 1 | 1 | 0.778 | NA | 1 | 1 | NA | | -0.111 |
| SE | 14789 | PHF1 | PHF1 | chr6 | + | 33412507 | 33412589 | 33412247 | 33412422 | 33413195 | 33413296 | 2 | 1 | 0 | 2 | 0 | 0 | 2 | 2 | 0 | 0 | 98 | 49 | 1 | 1 | 0.333 | 1 | NA | 1 | | 1 | -0.334 |
| SE | 14790 | PHF1 | PHF1 | chr6 | + | 33412697 | 33412793 | 33412507 | 33412589 | 33413195 | 33413296 | 10 | 8 | 1 | 1 | 0 | 0 | 0 | 4 | 0 | 0 | 98 | 49 | 1 | 1 | 0.833 | 1 | 1 | NA | | 1 | -0.056 |
| SE | 14795 | C20orf24 | C20orf24 | chr20 | + | 36607889 | 36608000 | 36607714 | 36607818 | 36609600 | 36609730 | 14 | 8 | 3 | 9 | 11 | 26 | 0 | 3 | 0 | 4 | 98 | 49 | 1 | 1 | 0.437 | 0.267 | 0.055 | NA | | 0.273 | -0.02 |
| SE | 14796 | C20orf24 | C20orf24 | chr20 | + | 36607889 | 36608000 | 36607714 | 36607818 | 36612009 | 36612557 | 3 | 3 | 1 | 1 | 0 | 1 | 0 | 1 | 0 | 0 | 98 | 49 | 1 | 1 | 0.6 | 1 | 0.333 | NA | | 1 | -0.356 |
| SE | 14797 | C20orf24 | C20orf24 | chr20 | + | 36609600 | 36609730 | 36607889 | 36608000 | 36612009 | 36612557 | 32 | 36 | 30 | 1 | 1 | 0 | 0 | 6 | 0 | 0 | 98 | 49 | 1 | 1 | 0.941 | 0.947 | 1 | NA | | 1 | -0.037 |
| SE | 14798 | C20orf24 | C20orf24 | chr20 | + | 36609600 | 36609758 | 36607714 | 36607818 | 36612009 | 36612557 | 9 | 11 | 26 | 1 | 0 | 1 | 0 | 4 | 0 | 0 | 98 | 49 | 1 | 1 | 0.818 | 1 | 0.929 | NA | | 1 | -0.084 |
| SE | 14806 | SAMHD1 | SAMHD1 | chr20 | - | 36898439 | 36898544 | 36897821 | 36897959 | 36904156 | 36904249 | 35 | 12 | 6 | 0 | 1 | 0 | 0 | 25 | 0 | 1 | 98 | 49 | 1 | 1 | 1 | 0.857 | 1 | NA | | 0.926 | 0.026 |
| SE | 14807 | SAMHD1 | SAMHD1 | chr20 | - | 36935028 | 36935189 | 36930759 | 36930875 | 36941038 | 36941111 | 27 | 1 | 2 | 0 | 0 | 0 | 0 | 21 | 0 | 1 | 98 | 49 | 1 | 1 | 1 | 1 | 1 | NA | | 0.913 | 0.087 |
| SE | 14809 | HNRNPH1 | HNRNPH1 | chr5 | - | 179615545 | 179615595 | 179614177 | 179614959 | 179616125 | 179616218 | 83 | 9 | 12 | 4 | 2 | 0 | 0 | 4 | 0 | 0 | 98 | 49 | 1 | 1 | 0.912 | 0.692 | 1 | NA | | 1 | -0.132 |
| SE | 14810 | HNRNPH1 | HNRNPH1 | chr5 | - | 179619268 | 179619407 | 179618144 | 179618323 | 179620891 | 179621035 | 23 | 5 | 0 | 12 | 4 | 0 | 0 | 3 | 0 | 0 | 98 | 49 | 1 | 1 | 0.489 | 0.385 | NA | NA | | 1 | -0.563 |
| SE | 14811 | HNRNPH1 | HNRNPH1 | chr5 | - | 179623036 | 179623164 | 179621241 | 179621397 | 179623594 | 179623710 | 43 | 5 | 1 | 0 | 2 | 0 | 0 | 3 | 0 | 0 | 98 | 49 | 1 | 1 | 1 | 0.556 | 1 | NA | | 1 | -0.148 |
| SE | 14819 | RALGAPB | RALGAPB | chr20 | + | 38548688 | 38548795 | 38546242 | 38546430 | 38551070 | 38551223 | 18 | 11 | 3 | 0 | 1 | 0 | 0 | 4 | 0 | 3 | 98 | 49 | 1 | 1 | 1 | 0.846 | 1 | NA | | 0.4 | 0.549 |
| SE | 14820 | RALGAPB | RALGAPB | chr20 | + | 38565355 | 38565478 | 38562531 | 38562697 | 38567095 | 38567232 | 8 | 4 | 6 | 0 | 0 | 0 | 0 | 0 | 0 | 4 | 98 | 49 | 1 | 1 | 1 | 1 | 1 | NA | | 0 | 1 |
| SE | 14821 | RALGAPB | RALGAPB | chr20 | + | 38565358 | 38565478 | 38562531 | 38562697 | 38567095 | 38567232 | 7 | 2 | 5 | 0 | 0 | 0 | 0 | 2 | 0 | 4 | 98 | 49 | 1 | 1 | 1 | 1 | 1 | NA | | 0.2 | 0.8 |
| SE | 14827 | TOP1 | TOP1 | chr20 | + | 41076170 | 41076294 | 41061393 | 41061490 | 41077581 | 41077637 | 9 | 6 | 0 | 0 | 1 | 0 | 1 | 3 | 0 | 0 | 98 | 49 | 1 | 1 | 1 | 0.75 | NA | 1 | | 1 | -0.125 |

| | | | | | | | | | | | | | | | | | | | | | | | | | | | | | | | |
|---|---|---|---|---|---|---|---|---|---|---|---|---|---|---|---|---|---|---|---|---|---|---|---|---|---|---|---|---|---|---|---|
| SE | 14828 | MBD5 | MBD5 | chr2 | + | 148233244 | 148233395 | 148178699 | 148178793 | 148342213 | 148342336 | 13 | 1 | 0 | 1 | 0 | 0 | 0 | 1 | 0 | 0 | 98 | 49 | 1 | 1 | 0.867 | 1 | NA | NA | 1 | -0.067 |
| SE | 14838 | SERINC3 | SERINC3 | chr20 | - | 44511288 | 44511368 | 44509890 | 44510028 | 44512800 | 44512994 | 46 | 6 | 28 | 2 | 0 | 0 | 0 | 13 | 0 | 0 | 98 | 49 | 1 | 1 | 0.92 | 1 | 1 | NA | 1 | -0.027 |
| SE | 14842 | STK4 | STK4 | chr20 | + | 45001166 | 45001353 | 45000391 | 45000520 | 45024972 | 45025130 | 116 | 59 | 21 | 0 | 1 | 0 | 0 | 11 | 0 | 0 | 98 | 49 | 1 | 1 | 1 | 0.967 | 1 | NA | 1 | -0.011 |
| SE | 14874 | ZMYND8 | ZMYND8 | chr20 | - | 47212641 | 47212725 | 47209213 | 47210897 | 47220257 | 47220324 | 2 | 0 | 0 | 3 | 0 | 0 | 0 | 0 | 0 | 1 | 98 | 49 | 1 | 1 | 0.25 | NA | NA | NA | 0 | 0.25 |
| SE | 14876 | ZMYND8 | ZMYND8 | chr20 | - | 47236325 | 47236516 | 47229725 | 47229806 | 47246007 | 47246517 | 10 | 3 | 4 | 1 | 0 | 0 | 0 | 3 | 0 | 0 | 98 | 49 | 1 | 1 | 0.833 | 1 | 1 | NA | 1 | -0.056 |
| SE | 14880 | ZMYND8 | ZMYND8 | chr20 | - | 47276313 | 47276795 | 47262287 | 47262428 | 47282101 | 47282217 | 18 | 11 | 1 | 0 | 1 | 0 | 0 | 1 | 0 | 0 | 98 | 49 | 1 | 1 | 1 | 0.846 | 1 | NA | 1 | -0.051 |
| SE | 14881 | ZMYND8 | ZMYND8 | chr20 | - | 47282101 | 47282217 | 47262287 | 47262428 | 47283570 | 47283648 | 4 | 5 | 2 | 0 | 1 | 0 | 0 | 3 | 0 | 0 | 98 | 49 | 1 | 1 | 1 | 0.714 | 1 | NA | 1 | -0.095 |
| SE | 14885 | PREX1 | PREX1 | chr20 | - | 48627547 | 48627615 | 48624254 | 48625927 | 48627860 | 48627963 | 156 | 84 | 50 | 2 | 0 | 0 | 6 | 30 | 0 | 0 | 98 | 49 | 1 | 1 | 0.975 | 1 | 1 | 1 | 1 | -0.008 |
| SE | 14890 | PREX1 | PREX1 | chr20 | - | 48676192 | 48676268 | 48666282 | 48666355 | 48679359 | 48679409 | 252 | 177 | 111 | 1 | 0 | 0 | 11 | 96 | 0 | 0 | 98 | 49 | 1 | 1 | 0.992 | 1 | 1 | 1 | 1 | -0.003 |
| SE | 14891 | PREX1 | PREX1 | chr20 | - | 48679359 | 48679409 | 48676192 | 48676268 | 48679650 | 48679754 | 183 | 119 | 92 | 1 | 0 | 0 | 9 | 74 | 0 | 0 | 98 | 49 | 1 | 1 | 0.989 | 1 | 1 | 1 | 1 | -0.004 |
| SE | 14892 | PREX1 | PREX1 | chr20 | - | 48690946 | 48691096 | 48688656 | 48688804 | 48692671 | 48692790 | 182 | 122 | 60 | 0 | 1 | 0 | 5 | 38 | 0 | 0 | 98 | 49 | 1 | 1 | 1 | 0.984 | 1 | 1 | 1 | -0.005 |
| SE | 14893 | PREX1 | PREX1 | chr20 | - | 48726289 | 48726391 | 48708259 | 48708421 | 48734545 | 48734650 | 330 | 204 | 94 | 0 | 1 | 0 | 23 | 82 | 0 | 0 | 98 | 49 | 1 | 1 | 1 | 0.99 | 1 | 1 | 1 | -0.003 |
| SE | 14894 | PREX1 | PREX1 | chr20 | - | 48734545 | 48734650 | 48708259 | 48708421 | 48745024 | 48745147 | 112 | 69 | 46 | 0 | 1 | 0 | 7 | 26 | 0 | 0 | 98 | 49 | 1 | 1 | 1 | 0.972 | 1 | 1 | 1 | -0.009 |
| SE | 14895 | PREX1 | PREX1 | chr20 | - | 48747808 | 48747880 | 48745024 | 48745147 | 48827641 | 48827883 | 97 | 107 | 30 | 1 | 0 | 1 | 20 | 45 | 0 | 1 | 98 | 49 | 1 | 1 | 0.98 | 1 | 0.937 | 1 | 0.957 | -0.006 |
| SE | 14897 | DOPEY1 | DOPEY1 | chr6 | + | 83155950 | 83156103 | 83154179 | 83154241 | 83157181 | 83157318 | 1 | 0 | 0 | 1 | 0 | 0 | 0 | 1 | 0 | 0 | 98 | 49 | 1 | 1 | 0.333 | NA | NA | NA | 1 | -0.667 |
| SE | 14901 | CSE1L | CSE1L | chr20 | + | 49085282 | 49085386 | 49084025 | 49084162 | 49088008 | 49088106 | 5 | 2 | 0 | 2 | 0 | 0 | 0 | 2 | 0 | 0 | 98 | 49 | 1 | 1 | 0.556 | 1 | NA | NA | 1 | -0.222 |
| SE | 14905 | TMEM8A | TMEM8A | chr16 | - | 375132 | 375256 | 374755 | 374892 | 375344 | 375435 | 6 | 9 | 8 | 0 | 0 | 1 | 1 | 5 | 0 | 0 | 98 | 49 | 1 | 1 | 1 | 1 | 0.8 | 1 | 1 | -0.067 |
| SE | 14906 | TMEM8A | TMEM8A | chr16 | - | 376543 | 376812 | 376135 | 376455 | 377036 | 377164 | 16 | 9 | 5 | 0 | 1 | 0 | 2 | 7 | 0 | 0 | 98 | 49 | 1 | 1 | 1 | 0.818 | 1 | 1 | 1 | -0.061 |
| SE | 14907 | DCP1A | DCP1A | chr3 | - | 53304176 | 53304290 | 53292068 | 53292827 | 53312240 | 53312379 | 2 | 1 | 0 | 1 | 0 | 0 | 0 | 2 | 0 | 0 | 98 | 49 | 1 | 1 | 0.5 | 1 | NA | NA | 1 | -0.25 |
| SE | 14919 | ZNF217 | ZNF217 | chr20 | - | 53571720 | 53571853 | 53567070 | 53569264 | 53575726 | 53577280 | 19 | 15 | 4 | 3 | 0 | 1 | 0 | 4 | 0 | 1 | 98 | 49 | 1 | 1 | 0.76 | 1 | 0.667 | NA | 0.667 | 0.142 |
| SE | 14921 | UBA3 | UBA3 | chr3 | - | 69057255 | 69057309 | 69056778 | 69056815 | 69061813 | 69061927 | 9 | 2 | 8 | 2 | 0 | 0 | 0 | 7 | 0 | 0 | 98 | 49 | 1 | 1 | 0.692 | 1 | 1 | NA | 1 | -0.103 |
| SE | 14922 | UBA3 | UBA3 | chr3 | - | 69071534 | 69071617 | 69067927 | 69068008 | 69075429 | 69075510 | 16 | 6 | 14 | 4 | 0 | 0 | 0 | 3 | 0 | 0 | 98 | 49 | 1 | 1 | 0.667 | 1 | 1 | NA | 1 | -0.111 |
| SE | 14926 | ATP6V1D | ATP6V1D | chr14 | - | 67349036 | 67349104 | 67347408 | 67347453 | 67350610 | 67350690 | 21 | 18 | 1 | 2 | 1 | 0 | 0 | 2 | 0 | 0 | 98 | 49 | 1 | 1 | 0.84 | 0.9 | 1 | NA | 1 | -0.087 |
| SE | 14930 | SYCP2 | SYCP2 | chr20 | - | 59877384 | 59877555 | 59875270 | 59875469 | 59878007 | 59878045 | 2 | 0 | 0 | 1 | 0 | 0 | 0 | 1 | 0 | 0 | 98 | 49 | 1 | 1 | 0.5 | NA | NA | NA | 1 | -0.5 |
| SE | 14934 | OSBPL2 | OSBPL2 | chr20 | + | 62260016 | 62260125 | 62256056 | 62256221 | 62263615 | 62263691 | 27 | 6 | 5 | 2 | 1 | 0 | 0 | 3 | 0 | 0 | 98 | 49 | 1 | 1 | 0.871 | 0.75 | 1 | NA | 1 | -0.126 |
| SE | 14935 | OSBPL2 | OSBPL2 | chr20 | + | 62264628 | 62264718 | 62263615 | 62263691 | 62272124 | 62272259 | 13 | 5 | 0 | 17 | 28 | 8 | 0 | 2 | 0 | 3 | 98 | 49 | 1 | 1 | 0.277 | 0.082 | 0 | NA | 0.25 | -0.13 |
| SE | 14936 | OSBPL2 | OSBPL2 | chr20 | + | 62273308 | 62273406 | 62272124 | 62272259 | 62279156 | 62279339 | 13 | 12 | 6 | 0 | 2 | 0 | 0 | 1 | 0 | 0 | 98 | 49 | 1 | 1 | 1 | 0.75 | 1 | NA | 1 | -0.083 |
| SE | 14937 | OSBPL2 | OSBPL2 | chr20 | + | 62289206 | 62289330 | 62286582 | 62286711 | 62291702 | 62291793 | 24 | 19 | 7 | 1 | 0 | 2 | 0 | 7 | 0 | 0 | 98 | 49 | 1 | 1 | 0.923 | 1 | 0.636 | NA | 1 | -0.147 |
| SE | 14938 | ADRM1 | ADRM1 | chr20 | + | 62304460 | 62304577 | 62303567 | 62303781 | 62306196 | 62306320 | 6 | 3 | 0 | 3 | 0 | 0 | 0 | 5 | 0 | 0 | 98 | 49 | 1 | 1 | 0.5 | 1 | NA | NA | 1 | -0.25 |
| SE | 14939 | ADRM1 | ADRM1 | chr20 | + | 62308367 | 62308470 | 62308020 | 62308178 | 62308654 | 62308862 | 2 | 2 | 2 | 0 | 2 | 1 | 0 | 4 | 0 | 1 | 98 | 49 | 1 | 1 | 1 | 0.333 | 0.5 | NA | 0.667 | -0.056 |
| SE | 14940 | ERCC6L2 | ERCC6L2 | chr9 | + | 95907077 | 95907271 | 95897848 | 95897971 | 95915667 | 95915829 | 8 | 1 | 0 | 4 | 0 | 0 | 0 | 0 | 0 | 1 | 98 | 49 | 1 | 1 | 0.5 | 1 | NA | NA | 0 | 0.75 |
| SE | 14942 | EIF3K | EIF3K | chr19 | + | 38620336 | 38620435 | 38619073 | 38619327 | 38624076 | 38624197 | 31 | 7 | 4 | 0 | 0 | 0 | 0 | 8 | 0 | 1 | 98 | 49 | 1 | 1 | 1 | 1 | 1 | NA | 0.8 | 0.2 |
| SE | 14943 | EIF3K | EIF3K | chr19 | + | 38632600 | 38632678 | 38632429 | 38632496 | 38634992 | 38635118 | 30 | 9 | 10 | 0 | 0 | 0 | 0 | 5 | 0 | 1 | 98 | 49 | 1 | 1 | 1 | 1 | 1 | NA | 0.714 | 0.286 |
| SE | 14951 | OGFR | OGFR | chr20 | + | 62809584 | 62809663 | 62808246 | 62808325 | 62810498 | 62810565 | 31 | 15 | 8 | 0 | 2 | 0 | 2 | 5 | 0 | 0 | 98 | 49 | 1 | 1 | 1 | 0.789 | 1 | 1 | 1 | -0.07 |
| SE | 14952 | CGGBP1 | CGGBP1 | chr3 | - | 88057190 | 88057292 | 88051949 | 88055999 | 88058018 | 88058223 | 2 | 6 | 0 | 15 | 1 | 4 | 0 | 1 | 0 | 1 | 98 | 49 | 1 | 1 | 0.062 | 0.75 | 0 | NA | 0.333 | -0.062 |
| SE | 14954 | ARFGAP1 | ARFGAP1 | chr20 | + | 63277204 | 63277305 | 63276479 | 63276651 | 63278116 | 63278203 | 5 | 3 | 4 | 1 | 0 | 0 | 0 | 1 | 0 | 0 | 98 | 49 | 1 | 1 | 0.714 | 1 | 1 | NA | 1 | -0.095 |
| SE | 14955 | ARFGAP1 | ARFGAP1 | chr20 | + | 63282818 | 63282851 | 63281290 | 63281347 | 63284865 | 63284922 | 3 | 1 | 1 | 2 | 0 | 0 | 0 | 1 | 0 | 0 | 82 | 49 | 1 | 1 | 0.473 | 1 | 1 | NA | 1 | -0.176 |
| SE | 14964 | HELZ2 | HELZ2 | chr20 | - | 63561600 | 63561745 | 63561349 | 63561466 | 63561822 | 63561984 | 22 | 11 | 8 | 1 | 0 | 0 | 2 | 25 | 0 | 0 | 98 | 49 | 1 | 1 | 0.917 | 1 | 1 | 1 | 1 | -0.028 |
| SE | 14965 | HELZ2 | HELZ2 | chr20 | - | 63562071 | 63562203 | 63561822 | 63561984 | 63562287 | 63562382 | 8 | 12 | 1 | 0 | 0 | 3 | 0 | 11 | 0 | 0 | 98 | 49 | 1 | 1 | 1 | 1 | 0.143 | NA | 1 | -0.286 |
| SE | 14971 | TPD52L2 | TPD52L2 | chr20 | + | 63875815 | 63875875 | 63873667 | 63873816 | 63882718 | 63882820 | 39 | 21 | 14 | 2 | 0 | 1 | 0 | 17 | 0 | 0 | 98 | 49 | 1 | 1 | 0.907 | 1 | 0.875 | NA | 1 | -0.073 |
| SE | 14974 | TPD52L2 | TPD52L2 | chr20 | + | 63887563 | 63887605 | 63882718 | 63882820 | 63889189 | 63889238 | 0 | 0 | 0 | 13 | 7 | 4 | 0 | 1 | 0 | 8 | 91 | 49 | 1 | 1 | 0 | 0 | 0 | NA | 0.063 | -0.063 |
| SE | 14977 | RGS19 | RGS19 | chr20 | - | 64076856 | 64076954 | 64076524 | 64076646 | 64079293 | 64079492 | 18 | 4 | 3 | 5 | 1 | 0 | 0 | 7 | 0 | 2 | 98 | 49 | 1 | 1 | 0.643 | 0.667 | 1 | NA | 0.636 | 0.134 |
| SE | 14979 | OPRL1 | OPRL1 | chr20 | + | 64092687 | 64092953 | 64080117 | 64080352 | 64097801 | 64098157 | 0 | 1 | 0 | 1 | 0 | 0 | 0 | 1 | 0 | 0 | 98 | 49 | 1 | 1 | 0 | 1 | NA | NA | 1 | -0.5 |
| SE | 14980 | PCMTD2 | PCMTD2 | chr20 | + | 64265257 | 64265429 | 64264428 | 64264531 | 64267886 | 64268010 | 10 | 1 | 6 | 0 | 1 | 0 | 0 | 2 | 0 | 0 | 98 | 49 | 1 | 1 | 1 | 0.333 | 1 | NA | 1 | -0.222 |
| SE | 14988 | RPS3 | RPS3 | chr11 | + | 75404019 | 75404207 | 75402303 | 75402446 | 75404671 | 75404868 | 77 | 16 | 16 | 1 | 0 | 0 | 1 | 7 | 0 | 0 | 98 | 49 | 1 | 1 | 0.975 | 1 | 1 | 1 | 1 | -0.008 |
| SE | 14997 | MGAM2 | MGAM2 | chr7 | + | 142154728 | 142154845 | 142154017 | 142154189 | 142157936 | 142158091 | 1 | 2 | 1 | 1 | 0 | 0 | 0 | 1 | 0 | 0 | 98 | 49 | 1 | 1 | 0.333 | 1 | 1 | NA | 1 | -0.222 |
| SE | 14998 | MGAM2 | MGAM2 | chr7 | + | 142172097 | 142172194 | 142171271 | 142171440 | 142172651 | 142172764 | 3 | 2 | 0 | 1 | 0 | 0 | 0 | 2 | 0 | 0 | 98 | 49 | 1 | 1 | 0.6 | 1 | NA | NA | 1 | -0.2 |
| SE | 15008 | UBP1 | UBP1 | chr3 | - | 33408689 | 33408797 | 33402800 | 33402904 | 33409235 | 33409346 | 6 | 0 | 0 | 3 | 1 | 1 | 0 | 2 | 0 | 0 | 98 | 49 | 1 | 1 | 0.5 | 0 | 0 | NA | 1 | -0.833 |
| SE | 15011 | APP | APP | chr21 | - | 25905023 | 25905077 | 25897572 | 25897673 | 25911740 | 25911962 | 11 | 0 | 1 | 2 | 1 | 0 | 0 | 6 | 0 | 1 | 98 | 49 | 1 | 1 | 0.733 | 0 | 1 | NA | 0.75 | -0.172 |
| SE | 15012 | APP | APP | chr21 | - | 25997359 | 25997416 | 25982343 | 25982477 | 26000014 | 26000182 | 1 | 0 | 0 | 2 | 0 | 0 | 0 | 1 | 0 | 3 | 98 | 49 | 1 | 1 | 0.2 | NA | NA | NA | 0.143 | 0.057 |
| SE | 15015 | N6AMT1 | N6AMT1 | chr21 | - | 28879869 | 28879953 | 28878191 | 28878333 | 28882159 | 28882250 | 0 | 0 | 0 | 0 | 2 | 0 | 1 | 0 | 0 | 0 | 98 | 49 | 1 | 1 | NA | 0 | NA | 1 | NA | -1 |
| SE | 15016 | FBRS | FBRS | chr16 | + | 30661179 | 30661215 | 30660262 | 30660442 | 30662419 | 30662468 | 6 | 5 | 0 | 0 | 1 | 0 | 0 | 4 | 0 | 0 | 85 | 49 | 1 | 1 | 1 | 0.742 | NA | NA | 1 | -0.129 |
| SE | 15017 | FBRS | FBRS | chr16 | + | 30661303 | 30661333 | 30661179 | 30661215 | 30662419 | 30662468 | 10 | 7 | 0 | 0 | 1 | 0 | 0 | 1 | 0 | 1 | 79 | 49 | 1 | 1 | 1 | 0.813 | NA | NA | 0.383 | 0.523 |
| SE | 15018 | FBRS | FBRS | chr16 | + | 30666511 | 30666541 | 30665637 | 30665706 | 30666918 | 30666990 | 14 | 10 | 0 | 0 | 1 | 0 | 4 | 0 | 0 | 0 | 79 | 49 | 1 | 1 | 1 | 0.861 | NA | 1 | NA | -0.07 |
| SE | 15019 | FBRS | FBRS | chr16 | + | 30667319 | 30667437 | 30666918 | 30666990 | 30667541 | 30667622 | 30 | 14 | 11 | 1 | 0 | 0 | 5 | 5 | 0 | 0 | 98 | 49 | 1 | 1 | 0.937 | 1 | 1 | 1 | 1 | -0.021 |
| SE | 15026 | BACH1 | BACH1 | chr21 | + | 29305203 | 29305322 | 29299415 | 29299598 | 29321220 | 29321514 | 14 | 3 | 0 | 6 | 5 | 0 | 0 | 3 | 0 | 2 | 98 | 49 | 1 | 1 | 0.538 | 0.231 | NA | NA | 0.429 | -0.044 |
| SE | 15027 | BACH1 | BACH1 | chr21 | + | 29351633 | 29351813 | 29329486 | 29329693 | 29361551 | 29361896 | 14 | 3 | 0 | 1 | 0 | 0 | 0 | 3 | 0 | 0 | 98 | 49 | 1 | 1 | 0.875 | 1 | NA | NA | 1 | -0.062 |
| SE | 15029 | BACH1 | BACH1 | chr21 | + | 29361030 | 29361243 | 29352578 | 29352663 | 29361551 | 29361896 | 0 | 1 | 0 | 2 | 0 | 0 | 0 | 2 | 0 | 0 | 98 | 49 | 1 | 1 | 0 | 1 | NA | NA | 1 | -0.5 |
| SE | 15030 | ATP5A1 | ATP5A1 | chr18 | - | 46087340 | 46087492 | 46087007 | 46087232 | 46088108 | 46088257 | 14 | 3 | 3 | 1 | 0 | 0 | 0 | 16 | 0 | 0 | 98 | 49 | 1 | 1 | 0.875 | 1 | 1 | NA | 1 | -0.042 |
| SE | 15038 | HERC3 | HERC3 | chr4 | + | 88605794 | 88606049 | 88595556 | 88595614 | 88649839 | 88649999 | 76 | 39 | 3 | 4 | 0 | 0 | 0 | 14 | 0 | 0 | 98 | 49 | 1 | 1 | 0.905 | 1 | 1 | NA | 1 | -0.032 |
| SE | 15039 | HERC3 | HERC3 | chr4 | + | 88667891 | 88668081 | 88667376 | 88667488 | 88669859 | 88670023 | 91 | 34 | 3 | 3 | 3 | 0 | 0 | 20 | 0 | 0 | 98 | 49 | 1 | 1 | 0.938 | 0.85 | 1 | NA | 1 | -0.071 |
| SE | 15040 | HERC3 | HERC3 | chr4 | + | 88676217 | 88676241 | 88670138 | 88670252 | 88676333 | 88676423 | 26 | 21 | 4 | 2 | 2 | 0 | 0 | 6 | 0 | 3 | 73 | 49 | 1 | 1 | 0.897 | 0.876 | 1 | NA | 0.573 | 0.351 |
| SE | 15041 | HERC3 | HERC3 | chr4 | + | 88677963 | 88678134 | 88676333 | 88676423 | 88680092 | 88680236 | 35 | 13 | 2 | 2 | 0 | 0 | 0 | 10 | 0 | 0 | 98 | 49 | 1 | 1 | 0.897 | 1 | 1 | NA | 1 | -0.034 |
| SE | 15042 | HERC3 | HERC3 | chr4 | + | 88681158 | 88681325 | 88680092 | 88680236 | 88686735 | 88686802 | 75 | 16 | 16 | 4 | 9 | 0 | 0 | 12 | 0 | 2 | 98 | 49 | 1 | 1 | 0.904 | 0.471 | 1 | NA | 0.75 | 0.042 |
| SE | 15048 | IFNAR2 | IFNAR2 | chr21 | + | 33248708 | 33248854 | 33246717 | 33246890 | 33252661 | 33252830 | 56 | 12 | 3 | 0 | 0 | 0 | 1 | 7 | 0 | 1 | 98 | 49 | 1 | 1 | 1 | 1 | 1 | 1 | 0.778 | 0.111 |
| SE | 15049 | IFNAR2 | IFNAR2 | chr21 | + | 33260596 | 33260727 | 33252661 | 33252830 | 33262560 | 33262757 | 10 | 12 | 0 | 0 | 5 | 0 | 0 | 7 | 0 | 0 | 98 | 49 | 1 | 1 | 1 | 0.545 | NA | NA | 1 | -0.228 |
| SE | 15050 | IFNAR2 | IFNAR2 | chr21 | + | 33260596 | 33260727 | 33252661 | 33252830 | 33262792 | 33264513 | 14 | 13 | 1 | 1 | 0 | 0 | 0 | 7 | 0 | 0 | 98 | 49 | 1 | 1 | 0.875 | 1 | 1 | NA | 1 | -0.042 |
| SE | 15053 | IFNGR2 | IFNGR2 | chr21 | + | 33432176 | 33432336 | 33426883 | 33427032 | 33432713 | 33432871 | 80 | 42 | 45 | 0 | 0 | 0 | 0 | 16 | 0 | 1 | 98 | 49 | 1 | 1 | 1 | 1 | 1 | NA | 0.889 | 0.111 |
| SE | 15059 | RER1 | RER1 | chr1 | + | 2399414 | 2399514 | 2397115 | 2397220 | 2400856 | 2400935 | 25 | 16 | 13 | 1 | 0 | 0 | 0 | 3 | 0 | 3 | 98 | 49 | 1 | 1 | 0.926 | 1 | 1 | NA | 0.333 | 0.642 |
| SE | 15062 | MORC3 | MORC3 | chr21 | + | 36356619 | 36356724 | 36349310 | 36349408 | 36359954 | 36360077 | 18 | 6 | 11 | 1 | 0 | 0 | 0 | 2 | 0 | 0 | 98 | 49 | 1 | 1 | 0.9 | 1 | 1 | NA | 1 | -0.033 |
| SE | 15070 | ETS2 | ETS2 | chr21 | + | 38818424 | 38818646 | 38817007 | 38817091 | 38819502 | 38819766 | 21 | 8 | 2 | 1 | 0 | 0 | 3 | 18 | 0 | 0 | 98 | 49 | 1 | 1 | 0.913 | 1 | 1 | 1 | 1 | -0.029 |
| SE | 15071 | ETS2 | ETS2 | chr21 | + | 38821585 | 38821704 | 38819502 | 38819766 | 38822673 | 38824954 | 37 | 4 | 16 | 0 | 0 | 0 | 0 | 8 | 0 | 3 | 98 | 49 | 1 | 1 | 1 | 1 | 1 | NA | 0.571 | 0.429 |
| SE | 15072 | HMHA1 | HMHA1 | chr19 | + | 1068413 | 1068744 | 1067165 | 1067495 | 1073148 | 1073292 | 65 | 43 | 18 | 1 | 0 | 0 | 8 | 14 | 0 | 1 | 98 | 49 | 1 | 1 | 0.97 | 1 | 1 | 1 | 0.875 | 0.052 |
| SE | 15073 | HMHA1 | HMHA1 | chr19 | + | 1073673 | 1073746 | 1073505 | 1073590 | 1073947 | 1074014 | 99 | 51 | 38 | 1 | 0 | 3 | 0 | 23 | 0 | 1 | 98 | 49 | 1 | 1 | 0.98 | 1 | 0.864 | NA | 0.92 | 0.028 |
| SE | 15074 | HMHA1 | HMHA1 | chr19 | + | 1083142 | 1083353 | 1082839 | 1083066 | 1084237 | 1084346 | 152 | 62 | 42 | 0 | 1 | 0 | 10 | 40 | 0 | 0 | 98 | 49 | 1 | 1 | 1 | 0.969 | 1 | 1 | 1 | -0.01 |
| SE | 15075 | BRWD1 | BRWD1 | chr21 | - | 39218151 | 39218272 | 39215236 | 39215362 | 39218504 | 39218660 | 17 | 3 | 2 | 0 | 1 | 0 | 0 | 2 | 0 | 0 | 98 | 49 | 1 | 1 | 1 | 0.6 | 1 | NA | 1 | -0.133 |
| SE | 15090 | C2CD2 | C2CD2 | chr21 | - | 41921971 | 41922085 | 41918855 | 41918960 | 41942146 | 41942245 | 0 | 1 | 0 | 0 | 0 | 0 | 0 | 0 | 0 | 3 | 98 | 49 | 1 | 1 | NA | 1 | NA | NA | 0 | 1 |
| SE | 15094 | TRAPPC10 | TRAPPC10 | chr21 | + | 44037791 | 44037927 | 44032090 | 44032172 | 44052279 | 44052476 | 14 | 13 | 12 | 0 | 3 | 0 | 0 | 5 | 0 | 0 | 98 | 49 | 1 | 1 | 1 | 0.684 | 1 | NA | 1 | -0.105 |
| SE | 15095 | TRAPPC10 | TRAPPC10 | chr21 | + | 44074323 | 44074470 | 44063537 | 44063785 | 44075038 | 44075153 | 16 | 12 | 2 | 0 | 2 | 0 | 0 | 4 | 0 | 0 | 98 | 49 | 1 | 1 | 1 | 0.75 | 1 | NA | 1 | -0.083 |
| SE | 15118 | ITGB2 | ITGB2 | chr21 | - | 44889275 | 44889495 | 44888692 | 44888895 | 44889977 | 44890222 | 214 | 174 | 89 | 0 | 3 | 0 | 4 | 78 | 0 | 0 | 98 | 49 | 1 | 1 | 1 | 0.967 | 1 | 1 | 1 | -0.011 |
| SE | 15123 | POFUT2 | POFUT2 | chr21 | - | 45266021 | 45266227 | 45263927 | 45265635 | 45267589 | 45267713 | 0 | 0 | 1 | 5 | 3 | 1 | 0 | 0 | 1 | 0 | 98 | 49 | 1 | 1 | 0 | 0 | 0.333 | 0 | NA | 0.111 |
| SE | 15124 | POFUT2 | POFUT2 | chr21 | - | 45267589 | 45267713 | 45263927 | 45265635 | 45269838 | 45270019 | 10 | 4 | 1 | 1 | 0 | 0 | 1 | 0 | 0 | 0 | 98 | 49 | 1 | 1 | 0.833 | 1 | 1 | 1 | NA | -0.056 |
| SE | 15125 | POFUT2 | POFUT2 | chr21 | - | 45283382 | 45283527 | 45282348 | 45282459 | 45285677 | 45285928 | 2 | 5 | 0 | 1 | 0 | 0 | 0 | 1 | 0 | 0 | 98 | 49 | 1 | 1 | 0.5 | 1 | NA | NA | 1 | -0.25 |
| SE | 15132 | DIP2A | DIP2A | chr21 | + | 46532126 | 46532237 | 46529091 | 46529183 | 46533523 | 46533647 | 5 | 9 | 0 | 1 | 0 | 0 | 0 | 5 | 0 | 0 | 98 | 49 | 1 | 1 | 0.714 | 1 | NA | NA | 1 | -0.143 |
| SE | 15133 | PRMT2 | PRMT2 | chr21 | + | 46636438 | 46636547 | 46635594 | 46635763 | 46636895 | 46636990 | 4 | 3 | 0 | 0 | 2 | 1 | 0 | 1 | 0 | 0 | 98 | 49 | 1 | 1 | 1 | 0.429 | 0 | NA | 1 | -0.524 |
| SE | 15134 | PRMT2 | PRMT2 | chr21 | + | 46658744 | 46658920 | 46649574 | 46649739 | 46664294 | 46665124 | 9 | 2 | 5 | 0 | 0 | 0 | 0 | 2 | 0 | 1 | 98 | 49 | 1 | 1 | 1 | 1 | 1 | NA | 0.5 | 0.5 |
| SE | 15135 | PRMT2 | PRMT2 | chr21 | + | 46661799 | 46661936 | 46649574 | 46649739 | 46663382 | 46663554 | 9 | 4 | 3 | 1 | 5 | 0 | 0 | 3 | 0 | 0 | 98 | 49 | 1 | 1 | 0.818 | 0.286 | 1 | NA | 1 | -0.299 |
| SE | 15136 | PRMT2 | PRMT2 | chr21 | + | 46663382 | 46663554 | 46649574 | 46649739 | 46664294 | 46665124 | 4 | 9 | 2 | 0 | 0 | 0 | 0 | 1 | 0 | 1 | 98 | 49 | 1 | 1 | 1 | 1 | 1 | NA | 0.333 | 0.667 |
| SE | 15137 | BID | BID | chr22 | - | 17743802 | 17744013 | 17739348 | 17739488 | 17750104 | 17750174 | 28 | 26 | 19 | 1 | 0 | 0 | 4 | 10 | 0 | 0 | 98 | 49 | 1 | 1 | 0.933 | 1 | 1 | 1 | 1 | -0.022 |
| SE | 15145 | PARP4 | PARP4 | chr13 | - | 24459971 | 24460136 | 24459263 | 24459310 | 24469023 | 24469110 | 27 | 18 | 9 | 0 | 1 | 0 | 1 | 15 | 0 | 0 | 98 | 49 | 1 | 1 | 1 | 0.9 | 1 | 1 | 1 | -0.033 |
| SE | 15154 | TANGO2 | TANGO2 | chr22 | + | 20053436 | 20053551 | 20052464 | 20052584 | 20055942 | 20056013 | 5 | 10 | 8 | 2 | 3 | 1 | 0 | 4 | 0 | 0 | 98 | 49 | 1 | 1 | 0.556 | 0.625 | 0.8 | NA | 1 | -0.34 |
| SE | 15157 | TANGO2 | TANGO2 | chr22 | + | 20055942 | 20056013 | 20052464 | 20052584 | 20063337 | 20063442 | 2 | 3 | 1 | 2 | 0 | 0 | 0 | 1 | 0 | 0 | 98 | 49 | 1 | 1 | 0.333 | 1 | 1 | NA | 1 | -0.222 |
| SE | 15158 | TANGO2 | TANGO2 | chr22 | + | 20055942 | 20056013 | 20053436 | 20053551 | 20063337 | 20063442 | 4 | 6 | 1 | 0 | 2 | 0 | 0 | 5 | 0 | 0 | 98 | 49 | 1 | 1 | 1 | 0.6 | 1 | NA | 1 | -0.133 |
| SE | 15159 | TANGO2 | TANGO2 | chr22 | + | 20061529 | 20061681 | 20055942 | 20056013 | 20063337 | 20063442 | 2 | 1 | 2 | 0 | 0 | 0 | 0 | 0 | 0 | 1 | 98 | 49 | 1 | 1 | 1 | 1 | 1 | NA | 0 | 1 |
| SE | 15160 | TANGO2 | TANGO2 | chr22 | + | 20061529 | 20061683 | 20052464 | 20052584 | 20063337 | 20063442 | 0 | 3 | 2 | 2 | 0 | 0 | 0 | 2 | 0 | 0 | 98 | 49 | 1 | 1 | 0 | 1 | 1 | NA | 1 | -0.333 |
| SE | 15161 | TANGO2 | TANGO2 | chr22 | + | 20061529 | 20061683 | 20055942 | 20056013 | 20063337 | 20063442 | 1 | 4 | 4 | 0 | 0 | 0 | 0 | 2 | 0 | 1 | 98 | 49 | 1 | 1 | 1 | 1 | 1 | NA | 0.5 | 0.5 |
| SE | 15162 | TANGO2 | TANGO2 | chr22 | + | 20063337 | 20063442 | 20055942 | 20056013 | 20064541 | 20065926 | 5 | 7 | 5 | 0 | 1 | 0 | 0 | 4 | 0 | 0 | 98 | 49 | 1 | 1 | 1 | 0.778 | 1 | NA | 1 | -0.074 |
| SE | 15165 | LZTR1 | LZTR1 | chr22 | + | 20989624 | 20989682 | 20988788 | 20988872 | 20990385 | 20990525 | 3 | 0 | 0 | 0 | 3 | 0 | 0 | 4 | 0 | 0 | 98 | 49 | 1 | 1 | 1 | 0 | NA | NA | 1 | -0.5 |
| SE | 15172 | SHKBP1 | SHKBP1 | chr19 | + | 40583397 | 40583485 | 40582350 | 40582466 | 40583600 | 40583717 | 151 | 50 | 36 | 2 | 1 | 0 | 1 | 9 | 0 | 0 | 98 | 49 | 1 | 1 | 0.974 | 0.962 | 1 | 1 | 1 | -0.021 |
| SE | 15174 | SHKBP1 | SHKBP1 | chr19 | + | 40590243 | 40590422 | 40589081 | 40589178 | 40590729 | 40590853 | 51 | 5 | 15 | 2 | 0 | 0 | 0 | 6 | 0 | 0 | 98 | 49 | 1 | 1 | 0.927 | 1 | 1 | NA | 1 | -0.024 |
| SE | 15181 | NME6 | NME6 | chr3 | - | 48296118 | 48296158 | 48295074 | 48295235 | 48296726 | 48296829 | 2 | 1 | 0 | 1 | 0 | 0 | 0 | 2 | 0 | 0 | 89 | 49 | 1 | 1 | 0.524 | 1 | NA | NA | 1 | -0.238 |
| SE | 15188 | GUSBP11 | GUSBP11 | chr22 | - | 23700725 | 23700845 | 23694118 | 23695517 | 23705428 | 23705541 | 5 | 6 | 2 | 5 | 3 | 0 | 0 | 6 | 0 | 0 | 98 | 49 | 1 | 1 | 0.333 | 0.5 | 1 | NA | 1 | -0.389 |
| SE | 15189 | GUSBP11 | GUSBP11 | chr22 | - | 23714186 | 23714342 | 23709785 | 23709867 | 23715008 | 23715194 | 46 | 24 | 3 | 0 | 2 | 0 | 0 | 4 | 0 | 0 | 98 | 49 | 1 | 1 | 1 | 0.857 | 1 | NA | 1 | -0.048 |
| SE | 15199 | CABIN1 | CABIN1 | chr22 | + | 24168446 | 24168521 | 24166638 | 24167313 | 24171712 | 24171995 | 14 | 15 | 1 | 1 | 0 | 0 | 0 | 4 | 0 | 0 | 98 | 49 | 1 | 1 | 0.875 | 1 | 1 | NA | 1 | -0.042 |
| SE | 15207 | ADORA2A | ADORA2A | chr22 | + | 24433130 | 24433736 | 24427561 | 24427746 | 24440582 | 24442361 | 5 | 0 | 1 | 0 | 1 | 0 | 1 | 1 | 0 | 0 | 98 | 49 | 1 | 1 | 1 | 0 | 1 | 1 | 1 | -0.333 |
| SE | 15212 | GUCD1 | GUCD1 | chr22 | - | 24547907 | 24548073 | 24546913 | 24547005 | 24548916 | 24549001 | 13 | 22 | 4 | 1 | 0 | 0 | 0 | 1 | 0 | 0 | 98 | 49 | 1 | 1 | 0.867 | 1 | 1 | NA | 1 | -0.044 |
| SE | 15213 | GUCD1 | GUCD1 | chr22 | - | 24547907 | 24548073 | 24546913 | 24547005 | 24554948 | 24555307 | 13 | 7 | 0 | 0 | 0 | 0 | 0 | 1 | 0 | 1 | 98 | 49 | 1 | 1 | 1 | 1 | NA | NA | 0.333 | 0.667 |
| SE | 15214 | GUCD1 | GUCD1 | chr22 | - | 24548916 | 24549001 | 24546913 | 24547005 | 24554948 | 24555307 | 5 | 4 | 0 | 0 | 0 | 0 | 2 | 0 | 0 | 1 | 98 | 49 | 1 | 1 | 1 | 1 | NA | 1 | 0 | 0.5 |
| SE | 15215 | GUCD1 | GUCD1 | chr22 | - | 24548916 | 24549001 | 24547907 | 24548073 | 24554948 | 24555307 | 7 | 19 | 4 | 3 | 0 | 0 | 2 | 0 | 0 | 0 | 98 | 49 | 1 | 1 | 0.538 | 1 | 1 | 1 | NA | -0.154 |
| SE | 15224 | CRYBB2P1 | CRYBB2P1 | chr22 | + | 25453315 | 25453434 | 25448086 | 25448284 | 25455711 | 25455840 | 3 | 0 | 3 | 1 | 0 | 0 | 0 | 1 | 0 | 0 | 98 | 49 | 1 | 1 | 0.6 | NA | 1 | NA | 1 | -0.2 |
| SE | 15225 | ADAM19 | ADAM19 | chr5 | - | 157502802 | 157502980 | 157499572 | 157499662 | 157505668 | 157505808 | 11 | 0 | 2 | 0 | 1 | 3 | 0 | 1 | 0 | 0 | 98 | 49 | 1 | 1 | 1 | 0 | 0.25 | NA | 1 | -0.583 |
| SE | 15226 | DIAPH1 | DIAPH1 | chr5 | - | 141529602 | 141529697 | 141529171 | 141529273 | 141534334 | 141534433 | 20 | 17 | 0 | 2 | 2 | 0 | 0 | 8 | 0 | 0 | 98 | 49 | 1 | 1 | 0.833 | 0.81 | NA | NA | 1 | -0.178 |
| SE | 15227 | DIAPH1 | DIAPH1 | chr5 | - | 141583484 | 141583615 | 141583205 | 141583292 | 141584123 | 141584225 | 41 | 10 | 5 | 2 | 0 | 0 | 3 | 26 | 0 | 0 | 98 | 49 | 1 | 1 | 0.911 | 1 | 1 | 1 | 1 | -0.03 |
| SE | 15228 | DIAPH1 | DIAPH1 | chr5 | - | 141588223 | 141588250 | 141587041 | 141587197 | 141618797 | 141619055 | 0 | 1 | 0 | 16 | 10 | 3 | 0 | 0 | 0 | 7 | 76 | 49 | 1 | 1 | 0 | 0.061 | 0 | NA | 0 | 0.02 |
| SE | 15229 | XBP1 | XBP1 | chr22 | - | 28796046 | 28796192 | 28794559 | 28795732 | 28797076 | 28797205 | 15 | 0 | 2 | 0 | 0 | 0 | 0 | 6 | 1 | 0 | 98 | 49 | 1 | 1 | 1 | NA | 1 | 0 | 1 | 0.5 |
| SE | 15230 | AP1B1 | AP1B1 | chr22 | - | 29329711 | 29329720 | 29327679 | 29328895 | 29330377 | 29330532 | 1 | 0 | 0 | 2 | 0 | 0 | 0 | 1 | 0 | 0 | 58 | 49 | 1 | 1 | 0.297 | NA | NA | NA | 1 | -0.703 |
| SE | 15231 | AP1B1 | AP1B1 | chr22 | - | 29339753 | 29339774 | 29338989 | 29339133 | 29340655 | 29340857 | 0 | 0 | 0 | 3 | 0 | 0 | 0 | 3 | 0 | 3 | 70 | 49 | 1 | 1 | 0 | NA | NA | NA | 0.412 | -0.412 |
| SE | 15232 | AP1B1 | AP1B1 | chr22 | - | 29356425 | 29356616 | 29354649 | 29354871 | 29358725 | 29358971 | 2 | 1 | 0 | 1 | 0 | 0 | 0 | 3 | 0 | 0 | 98 | 49 | 1 | 1 | 0.5 | 1 | NA | NA | 1 | -0.25 |
| SE | 15238 | THOC5 | THOC5 | chr22 | - | 29550727 | 29550843 | 29549051 | 29549158 | 29553670 | 29553747 | 5 | 1 | 2 | 2 | 0 | 0 | 0 | 1 | 0 | 0 | 98 | 49 | 1 | 1 | 0.556 | 1 | 1 | NA | 1 | -0.148 |
| SE | 15239 | GIGYF1 | GIGYF1 | chr7 | - | 100683549 | 100683632 | 100683303 | 100683444 | 100683817 | 100683918 | 21 | 5 | 1 | 0 | 0 | 0 | 0 | 0 | 1 | 0 | 98 | 49 | 1 | 1 | 1 | 1 | 1 | 0 | NA | 1 |
| SE | 15241 | ZMAT5 | ZMAT5 | chr22 | - | 29754748 | 29754863 | 29748417 | 29748571 | 29766871 | 29766980 | 2 | 0 | 0 | 0 | 0 | 1 | 0 | 1 | 0 | 0 | 98 | 49 | 1 | 1 | 1 | NA | 0 | NA | 1 | -0.5 |
| SE | 15246 | DUSP18 | DUSP18 | chr22 | - | 30664954 | 30665121 | 30661457 | 30664080 | 30667461 | 30667890 | 0 | 0 | 0 | 4 | 0 | 0 | 0 | 1 | 0 | 2 | 98 | 49 | 1 | 1 | 0 | NA | NA | NA | 0.2 | -0.2 |
| SE | 15266 | SIGLEC10 | SIGLEC10 | chr19 | - | 51414823 | 51415108 | 51414421 | 51414515 | 51415180 | 51415438 | 13 | 19 | 1 | 10 | 7 | 0 | 0 | 4 | 0 | 0 | 98 | 49 | 1 | 1 | 0.394 | 0.576 | 1 | NA | 1 | -0.343 |
| SE | 15267 | SIGLEC10 | SIGLEC10 | chr19 | - | 51415897 | 51416167 | 51415567 | 51415615 | 51416309 | 51416357 | 8 | 5 | 6 | 2 | 0 | 1 | 0 | 1 | 0 | 0 | 98 | 49 | 1 | 1 | 0.667 | 1 | 0.75 | NA | 1 | -0.194 |
| SE | 15269 | PRR14L | PRR14L | chr22 | - | 31738386 | 31738911 | 31725537 | 31725610 | 31749992 | 31750134 | 4 | 3 | 0 | 2 | 0 | 0 | 0 | 1 | 0 | 0 | 98 | 49 | 1 | 1 | 0.5 | 1 | NA | NA | 1 | -0.25 |
| SE | 15270 | FAM193A | FAM193A | chr4 | + | 2655057 | 2655129 | 2646684 | 2646832 | 2657802 | 2657880 | 3 | 2 | 0 | 4 | 0 | 0 | 0 | 1 | 0 | 2 | 98 | 49 | 1 | 1 | 0.273 | 1 | NA | NA | 0.2 | 0.437 |
| SE | 15276 | RTCB | RTCB | chr22 | - | 32397940 | 32398100 | 32396073 | 32396249 | 32399602 | 32399759 | 3 | 1 | 0 | 0 | 0 | 0 | 0 | 4 | 0 | 1 | 98 | 49 | 1 | 1 | 1 | 1 | NA | NA | 0.667 | 0.333 |
| SE | 15290 | CSF2RB | CSF2RB | chr22 | + | 36925986 | 36926177 | 36923243 | 36923367 | 36929401 | 36929559 | 203 | 99 | 92 | 3 | 0 | 0 | 1 | 39 | 0 | 1 | 98 | 49 | 1 | 1 | 0.971 | 1 | 1 | 1 | 0.951 | 0.015 |
| SE | 15291 | CSF2RB | CSF2RB | chr22 | + | 36929401 | 36929559 | 36925986 | 36926177 | 36929638 | 36929807 | 249 | 107 | 99 | 1 | 0 | 0 | 6 | 42 | 0 | 0 | 98 | 49 | 1 | 1 | 0.992 | 1 | 1 | 1 | 1 | -0.003 |
| SE | 15292 | CSF2RB | CSF2RB | chr22 | + | 36930374 | 36930510 | 36929638 | 36929807 | 36930672 | 36930830 | 68 | 27 | 46 | 1 | 1 | 0 | 2 | 8 | 0 | 0 | 98 | 49 | 1 | 1 | 0.971 | 0.931 | 1 | 1 | 1 | -0.033 |
| SE | 15297 | TRIOBP | TRIOBP | chr22 | + | 37769027 | 37769187 | 37768073 | 37768176 | 37769261 | 37769375 | 25 | 20 | 20 | 1 | 0 | 0 | 0 | 6 | 0 | 0 | 98 | 49 | 1 | 1 | 0.926 | 1 | 1 | NA | 1 | -0.025 |
| SE | 15308 | TSTD1 | TSTD1 | chr1 | - | 161038550 | 161038673 | 161037631 | 161038075 | 161038879 | 161038984 | 1 | 0 | 0 | 1 | 0 | 0 | 0 | 1 | 0 | 0 | 98 | 49 | 1 | 1 | 0.333 | NA | NA | NA | 1 | -0.667 |
| SE | 15314 | EP300 | EP300 | chr22 | + | 41135812 | 41135906 | 41131387 | 41131633 | 41137652 | 41137790 | 46 | 21 | 15 | 0 | 0 | 0 | 2 | 14 | 0 | 1 | 98 | 49 | 1 | 1 | 1 | 1 | 1 | 1 | 0.875 | 0.062 |
| SE | 15315 | EP300 | EP300 | chr22 | + | 41146738 | 41146816 | 41141047 | 41141222 | 41147836 | 41147946 | 56 | 5 | 2 | 0 | 0 | 1 | 0 | 8 | 0 | 0 | 98 | 49 | 1 | 1 | 1 | 1 | 0.5 | NA | 1 | -0.167 |
| SE | 15320 | ADAMTSL4 | ADAMTSL4 | chr1 | + | 150556620 | 150556793 | 150556161 | 150556366 | 150556938 | 150556976 | 5 | 0 | 0 | 4 | 0 | 0 | 0 | 5 | 0 | 0 | 98 | 49 | 1 | 1 | 0.385 | NA | NA | NA | 1 | -0.615 |
| SE | 15321 | UTP11L | UTP11L | chr1 | + | 38016358 | 38016420 | 38012711 | 38012865 | 38017667 | 38017770 | 4 | 0 | 0 | 1 | 0 | 0 | 0 | 1 | 0 | 0 | 98 | 49 | 1 | 1 | 0.667 | NA | NA | NA | 1 | -0.333 |
| SE | 15323 | SCAF8 | SCAF8 | chr6 | + | 154787860 | 154788022 | 154778000 | 154778045 | 154792822 | 154792976 | 15 | 10 | 4 | 0 | 0 | 3 | 0 | 3 | 0 | 0 | 98 | 49 | 1 | 1 | 1 | 1 | 0.4 | NA | 1 | -0.2 |
| SE | 15324 | SCAF8 | SCAF8 | chr6 | + | 154822275 | 154822409 | 154820176 | 154820333 | 154824233 | 154824378 | 47 | 8 | 9 | 1 | 0 | 0 | 0 | 3 | 0 | 0 | 98 | 49 | 1 | 1 | 0.959 | 1 | 1 | NA | 1 | -0.014 |
| SE | 15325 | SCAF8 | SCAF8 | chr6 | + | 154824233 | 154824378 | 154822275 | 154822409 | 154827171 | 154827240 | 44 | 4 | 1 | 1 | 0 | 0 | 0 | 2 | 0 | 0 | 98 | 49 | 1 | 1 | 0.957 | 1 | 1 | NA | 1 | -0.014 |
| SE | 15327 | C16orf72 | C16orf72 | chr16 | + | 9092920 | 9093001 | 9091679 | 9092270 | 9102983 | 9103259 | 102 | 57 | 22 | 0 | 1 | 0 | 1 | 22 | 1 | 1 | 98 | 49 | 1 | 1 | 1 | 0.966 | 1 | 0.333 | 0.917 | 0.364 |
| SE | 15334 | ARFGAP3 | ARFGAP3 | chr22 | - | 42810812 | 42810944 | 42808766 | 42808890 | 42817141 | 42817264 | 14 | 4 | 3 | 0 | 1 | 0 | 0 | 3 | 0 | 1 | 98 | 49 | 1 | 1 | 1 | 0.667 | 1 | NA | 0.6 | 0.289 |
| SE | 15335 | ARFGAP3 | ARFGAP3 | chr22 | - | 42834241 | 42834325 | 42831548 | 42831636 | 42835361 | 42835493 | 28 | 19 | 6 | 0 | 0 | 0 | 0 | 8 | 1 | 0 | 98 | 49 | 1 | 1 | 1 | 1 | 1 | 0 | 1 | 0.5 |
| SE | 15337 | TLE3 | TLE3 | chr15 | - | 70054437 | 70054685 | 70053226 | 70053374 | 70055048 | 70055298 | 92 | 29 | 26 | 1 | 1 | 0 | 4 | 15 | 0 | 4 | 98 | 49 | 1 | 1 | 0.979 | 0.935 | 1 | 1 | 0.652 | 0.145 |
| SE | 15338 | TLE3 | TLE3 | chr15 | - | 70056297 | 70056374 | 70055048 | 70055298 | 70057473 | 70057658 | 43 | 8 | 21 | 2 | 0 | 0 | 0 | 3 | 0 | 0 | 98 | 49 | 1 | 1 | 0.915 | 1 | 1 | NA | 1 | -0.028 |
| SE | 15340 | TLE3 | TLE3 | chr15 | - | 70064453 | 70064470 | 70060529 | 70060649 | 70066013 | 70066218 | 72 | 39 | 17 | 0 | 0 | 0 | 3 | 13 | 0 | 1 | 66 | 49 | 1 | 1 | 1 | 1 | 1 | 1 | 0.906 | 0.047 |
| SE | 15341 | TLE3 | TLE3 | chr15 | - | 70066013 | 70066218 | 70060529 | 70060649 | 70074532 | 70074607 | 27 | 10 | 5 | 3 | 0 | 0 | 0 | 5 | 0 | 0 | 98 | 49 | 1 | 1 | 0.818 | 1 | 1 | NA | 1 | -0.061 |
| SE | 15342 | TLE3 | TLE3 | chr15 | - | 70074502 | 70074607 | 70066013 | 70066218 | 70076095 | 70076158 | 71 | 30 | 16 | 1 | 1 | 0 | 0 | 15 | 0 | 0 | 98 | 49 | 1 | 1 | 0.973 | 0.937 | 1 | NA | 1 | -0.03 |
| SE | 15343 | TLE3 | TLE3 | chr15 | - | 70074532 | 70074607 | 70066013 | 70066218 | 70076095 | 70076158 | 92 | 36 | 18 | 1 | 1 | 0 | 0 | 15 | 0 | 0 | 98 | 49 | 1 | 1 | 0.979 | 0.947 | 1 | NA | 1 | -0.025 |
| SE | 15344 | TLE3 | TLE3 | chr15 | - | 70095577 | 70095641 | 70094531 | 70094576 | 70096160 | 70096261 | 106 | 18 | 28 | 2 | 0 | 0 | 0 | 2 | 0 | 1 | 98 | 49 | 1 | 1 | 0.964 | 1 | 1 | NA | 0.5 | 0.488 |
| SE | 15345 | SZT2 | SZT2 | chr1 | + | 43415959 | 43416101 | 43415081 | 43415213 | 43416534 | 43416641 | 6 | 4 | 0 | 2 | 0 | 0 | 1 | 0 | 0 | 0 | 98 | 49 | 1 | 1 | 0.6 | 1 | NA | 1 | NA | -0.2 |
| SE | 15346 | SZT2 | SZT2 | chr1 | + | 43443596 | 43443796 | 43443351 | 43443477 | 43445893 | 43445984 | 4 | 0 | 0 | 3 | 0 | 0 | 0 | 1 | 0 | 0 | 98 | 49 | 1 | 1 | 0.4 | NA | NA | NA | 1 | -0.6 |
| SE | 15351 | TAX1BP1 | TAX1BP1 | chr7 | + | 27765833 | 27766021 | 27758030 | 27758133 | 27769675 | 27769834 | 60 | 42 | 24 | 1 | 0 | 0 | 0 | 11 | 0 | 0 | 98 | 49 | 1 | 1 | 0.968 | 1 | 1 | NA | 1 | -0.011 |
| SE | 15352 | NUP50 | NUP50 | chr22 | + | 45170987 | 45171177 | 45168167 | 45168246 | 45171599 | 45171683 | 7 | 3 | 2 | 6 | 9 | 1 | 0 | 2 | 0 | 0 | 98 | 49 | 1 | 1 | 0.368 | 0.143 | 0.5 | NA | 1 | -0.663 |
| SE | 15356 | NUP50 | NUP50 | chr22 | + | 45178237 | 45178900 | 45171599 | 45171683 | 45181285 | 45181367 | 12 | 12 | 3 | 1 | 1 | 0 | 0 | 1 | 0 | 0 | 98 | 49 | 1 | 1 | 0.857 | 0.857 | 1 | NA | 1 | -0.095 |
| SE | 15357 | NUP50 | NUP50 | chr22 | + | 45178237 | 45178900 | 45175893 | 45176080 | 45181285 | 45181367 | 20 | 16 | 3 | 0 | 0 | 1 | 0 | 1 | 0 | 0 | 98 | 49 | 1 | 1 | 1 | 1 | 0.6 | NA | 1 | -0.133 |

| | | | | | | | | | | | | | | | | | | | | | | | | | | | | | | | |
|---|---|---|---|---|---|---|---|---|---|---|---|---|---|---|---|---|---|---|---|---|---|---|---|---|---|---|---|---|---|---|---|
| SE | 15362 | CERK | CERK | chr22 | - | 46689991 | 46690200 | 46684409 | 46687206 | 46691571 | 46691777 | 26 | 10 | 0 | 2 | 0 | 0 | 0 | 2 | 0 | 0 | 98 | 49 | 1 | 1 | 0.867 | 1 | NA | NA | 1 | -0.067 |
| SE | 15366 | TBC1D22A | TBC1D22A | chr22 | + | 46878652 | 46878723 | 46797443 | 46797620 | 46891265 | 46891394 | 25 | 15 | 0 | 1 | 0 | 0 | 3 | 5 | 0 | 0 | 98 | 49 | 1 | 1 | 0.926 | 1 | NA | 1 | 1 | -0.037 |
| SE | 15368 | TBC1D22A | TBC1D22A | chr22 | + | 46974289 | 46974399 | 46912073 | 46912188 | 46997633 | 46997709 | 16 | 5 | 10 | 1 | 0 | 0 | 0 | 3 | 0 | 0 | 98 | 49 | 1 | 1 | 0.889 | 1 | 1 | NA | 1 | -0.037 |
| SE | 15369 | TBC1D22A | TBC1D22A | chr22 | + | 46997633 | 46997709 | 46974289 | 46974399 | 47037070 | 47037198 | 18 | 11 | 14 | 1 | 0 | 0 | 0 | 3 | 0 | 0 | 98 | 49 | 1 | 1 | 0.9 | 1 | 1 | NA | 1 | -0.033 |
| SE | 15377 | PIM3 | PIM3 | chr22 | + | 49961124 | 49961234 | 49960494 | 49961032 | 49961317 | 49961368 | 39 | 20 | 17 | 1 | 0 | 0 | 8 | 15 | 1 | 0 | 98 | 49 | 1 | 1 | 0.951 | 1 | 1 | 0.8 | 1 | 0.084 |
| SE | 15383 | PLXNB2 | PLXNB2 | chr22 | - | 50294718 | 50294778 | 50289516 | 50290597 | 50307552 | 50307572 | 0 | 0 | 0 | 1 | 0 | 0 | 0 | 1 | 0 | 0 | 98 | 49 | 1 | 1 | 0 | NA | NA | NA | 1 | -1 |
| SE | 15384 | FBXW11 | FBXW11 | chr5 | - | 171872871 | 171872990 | 171870747 | 171870858 | 171876284 | 171876534 | 13 | 4 | 1 | 0 | 2 | 0 | 0 | 2 | 0 | 0 | 98 | 49 | 1 | 1 | 1 | 0.5 | 1 | NA | 1 | -0.167 |
| SE | 15387 | PPP6R2 | PPP6R2 | chr22 | + | 50436987 | 50437068 | 50436366 | 50436452 | 50437505 | 50437603 | 2 | 4 | 1 | 2 | 1 | 0 | 0 | 2 | 0 | 0 | 98 | 49 | 1 | 1 | 0.333 | 0.667 | 1 | NA | 1 | -0.333 |
| SE | 15388 | SBF1 | SBF1 | chr22 | - | 50457033 | 50457111 | 50456491 | 50456673 | 50459254 | 50459392 | 4 | 1 | 0 | 2 | 0 | 0 | 0 | 4 | 0 | 0 | 98 | 49 | 1 | 1 | 0.5 | 1 | NA | NA | 1 | -0.25 |
| SE | 15389 | LMF2 | LMF2 | chr22 | - | 50505056 | 50505153 | 50504801 | 50504984 | 50505228 | 50505334 | 10 | 4 | 8 | 1 | 0 | 0 | 1 | 12 | 0 | 0 | 98 | 49 | 1 | 1 | 0.833 | 1 | 1 | 1 | 1 | -0.056 |
| SE | 15391 | JOSD2 | JOSD2 | chr19 | - | 50507573 | 50507699 | 50506377 | 50506572 | 50510285 | 50510445 | 1 | 0 | 0 | 0 | 0 | 0 | 0 | 2 | 0 | 1 | 98 | 49 | 1 | 1 | 1 | NA | NA | NA | 0.5 | 0.5 |
| SE | 15393 | ZNF346 | ZNF346 | chr5 | + | 177023159 | 177023234 | 177022679 | 177022913 | 177041125 | 177041229 | 0 | 0 | 0 | 4 | 0 | 0 | 1 | 0 | 0 | 0 | 98 | 49 | 1 | 1 | 0 | NA | NA | 1 | NA | -1 |
| SE | 15418 | BHLHE40 | BHLHE40 | chr3 | + | 4981391 | 4981515 | 4980300 | 4980408 | 4982835 | 4985180 | 11 | 10 | 12 | 0 | 0 | 0 | 3 | 3 | 0 | 1 | 98 | 49 | 1 | 1 | 1 | 1 | 1 | 1 | 0.6 | 0.2 |
| SE | 15423 | SETD5 | SETD5 | chr3 | + | 9436849 | 9436888 | 9435727 | 9435906 | 9440455 | 9440698 | 2 | 5 | 0 | 0 | 0 | 0 | 0 | 0 | 0 | 3 | 88 | 49 | 1 | 1 | 1 | 1 | NA | NA | 0 | 1 |
| SE | 15428 | ARPC4 | ARPC4 | chr3 | + | 9801660 | 9801756 | 9800184 | 9800296 | 9803842 | 9804013 | 132 | 94 | 52 | 0 | 0 | 0 | 3 | 21 | 0 | 1 | 98 | 49 | 1 | 1 | 1 | 1 | 1 | 1 | 0.913 | 0.043 |
| SE | 15437 | RPRD2 | RPRD2 | chr1 | + | 150441880 | 150441958 | 150440922 | 150441023 | 150443230 | 150443283 | 5 | 4 | 0 | 1 | 0 | 0 | 2 | 0 | 0 | 0 | 98 | 49 | 1 | 1 | 0.714 | 1 | NA | 1 | NA | -0.143 |
| SE | 15448 | SEC13 | SEC13 | chr3 | - | 10312578 | 10312730 | 10311576 | 10312098 | 10315320 | 10315436 | 18 | 2 | 6 | 1 | 0 | 0 | 0 | 4 | 0 | 0 | 98 | 49 | 1 | 1 | 0.9 | 1 | 1 | NA | 1 | -0.033 |
| SE | 15459 | RAF1 | RAF1 | chr3 | - | 12600151 | 12600279 | 12599690 | 12599808 | 12600387 | 12600415 | 162 | 48 | 33 | 2 | 1 | 0 | 2 | 19 | 0 | 0 | 98 | 49 | 1 | 1 | 0.976 | 0.96 | 1 | 1 | 1 | -0.021 |
| SE | 15460 | RAF1 | RAF1 | chr3 | - | 12606200 | 12606299 | 12604135 | 12604289 | 12608765 | 12608923 | 154 | 59 | 46 | 39 | 3 | 3 | 0 | 20 | 0 | 5 | 98 | 49 | 1 | 1 | 0.664 | 0.908 | 0.885 | NA | 0.667 | 0.152 |
| SE | 15470 | SLC6A6 | SLC6A6 | chr3 | + | 14416411 | 14416453 | 14402575 | 14402847 | 14443623 | 14443863 | 175 | 62 | 37 | 1 | 0 | 0 | 3 | 30 | 0 | 0 | 91 | 49 | 1 | 1 | 0.989 | 1 | 1 | 1 | 1 | -0.004 |
| SE | 15471 | SLC6A6 | SLC6A6 | chr3 | + | 14443623 | 14443863 | 14416411 | 14416453 | 14445716 | 14445851 | 155 | 60 | 57 | 0 | 0 | 1 | 5 | 30 | 0 | 1 | 98 | 49 | 1 | 1 | 1 | 1 | 0.966 | 1 | 0.937 | 0.02 |
| SE | 15472 | SLC6A6 | SLC6A6 | chr3 | + | 14443627 | 14443863 | 14416411 | 14416453 | 14445716 | 14445851 | 80 | 39 | 28 | 0 | 0 | 1 | 5 | 14 | 0 | 1 | 98 | 49 | 1 | 1 | 1 | 1 | 0.933 | 1 | 0.875 | 0.04 |
| SE | 15473 | SLC6A6 | SLC6A6 | chr3 | + | 14457949 | 14458082 | 14447581 | 14447816 | 14466515 | 14466650 | 191 | 46 | 45 | 1 | 1 | 0 | 3 | 36 | 0 | 0 | 98 | 49 | 1 | 1 | 0.99 | 0.958 | 1 | 1 | 1 | -0.017 |
| SE | 15474 | SLC6A6 | SLC6A6 | chr3 | + | 14468087 | 14468212 | 14467852 | 14467956 | 14472204 | 14472317 | 40 | 15 | 13 | 5 | 4 | 2 | 0 | 3 | 0 | 1 | 98 | 49 | 1 | 1 | 0.8 | 0.652 | 0.765 | NA | 0.6 | 0.139 |
| SE | 15475 | SLC6A6 | SLC6A6 | chr3 | + | 14472204 | 14472317 | 14467852 | 14467956 | 14477204 | 14477342 | 41 | 22 | 15 | 2 | 0 | 0 | 0 | 5 | 0 | 0 | 98 | 49 | 1 | 1 | 0.911 | 1 | 1 | NA | 1 | -0.03 |
| SE | 15476 | SLC6A6 | SLC6A6 | chr3 | + | 14472204 | 14472317 | 14467852 | 14467956 | 14481670 | 14481841 | 5 | 5 | 2 | 1 | 0 | 0 | 0 | 1 | 0 | 0 | 98 | 49 | 1 | 1 | 0.714 | 1 | 1 | NA | 1 | -0.095 |
| SE | 15477 | SLC6A6 | SLC6A6 | chr3 | + | 14478465 | 14478568 | 14477204 | 14477342 | 14479084 | 14479185 | 88 | 33 | 20 | 1 | 0 | 0 | 2 | 14 | 0 | 0 | 98 | 49 | 1 | 1 | 0.978 | 1 | 1 | 1 | 1 | -0.007 |
| SE | 15479 | TRANK1 | TRANK1 | chr3 | - | 36838371 | 36838504 | 36834761 | 36834907 | 36838612 | 36838716 | 31 | 9 | 2 | 2 | 7 | 0 | 0 | 3 | 0 | 0 | 98 | 49 | 1 | 1 | 0.886 | 0.391 | 1 | NA | 1 | -0.241 |
| SE | 15480 | TRANK1 | TRANK1 | chr3 | - | 36842621 | 36842710 | 36838612 | 36838716 | 36846247 | 36846404 | 47 | 26 | 9 | 1 | 0 | 0 | 0 | 3 | 0 | 0 | 98 | 49 | 1 | 1 | 0.959 | 1 | 1 | NA | 1 | -0.014 |
| SE | 15481 | TRANK1 | TRANK1 | chr3 | - | 36847199 | 36847346 | 36846247 | 36846404 | 36851718 | 36851856 | 66 | 11 | 15 | 1 | 0 | 0 | 0 | 12 | 0 | 0 | 98 | 49 | 1 | 1 | 0.971 | 1 | 1 | NA | 1 | -0.01 |
| SE | 15482 | TRANK1 | TRANK1 | chr3 | - | 36855172 | 36858049 | 36852145 | 36852345 | 36858717 | 36858894 | 27 | 12 | 5 | 0 | 0 | 0 | 0 | 15 | 0 | 2 | 98 | 49 | 1 | 1 | 1 | 1 | 1 | NA | 0.789 | 0.211 |
| SE | 15483 | TRANK1 | TRANK1 | chr3 | - | 36874125 | 36874296 | 36864318 | 36864480 | 36889828 | 36889960 | 63 | 26 | 5 | 1 | 0 | 0 | 3 | 13 | 0 | 1 | 98 | 49 | 1 | 1 | 0.969 | 1 | 1 | 1 | 0.867 | 0.056 |
| SE | 15484 | TRANK1 | TRANK1 | chr3 | - | 36892201 | 36892340 | 36889828 | 36889960 | 36892900 | 36892984 | 42 | 16 | 6 | 0 | 2 | 0 | 0 | 20 | 0 | 0 | 98 | 49 | 1 | 1 | 1 | 0.8 | 1 | NA | 1 | -0.067 |
| SE | 15485 | TRANK1 | TRANK1 | chr3 | - | 36892900 | 36892984 | 36892201 | 36892340 | 36899108 | 36899259 | 27 | 10 | 5 | 0 | 0 | 0 | 0 | 15 | 0 | 2 | 98 | 49 | 1 | 1 | 1 | 1 | 1 | NA | 0.789 | 0.211 |
| SE | 15486 | TRANK1 | TRANK1 | chr3 | - | 36895639 | 36895758 | 36892900 | 36892984 | 36899108 | 36899259 | 58 | 12 | 1 | 1 | 0 | 0 | 0 | 8 | 0 | 3 | 98 | 49 | 1 | 1 | 0.967 | 1 | 1 | NA | 0.571 | 0.418 |
| SE | 15496 | NR2C2 | NR2C2 | chr3 | + | 15004556 | 15004613 | 15003875 | 15003986 | 15013588 | 15013789 | 1 | 0 | 0 | 1 | 0 | 0 | 0 | 2 | 0 | 0 | 98 | 49 | 1 | 1 | 0.333 | NA | NA | NA | 1 | -0.667 |
| SE | 15497 | EAF1 | EAF1 | chr3 | + | 15432086 | 15432223 | 15429912 | 15430007 | 15434347 | 15434538 | 13 | 4 | 1 | 2 | 0 | 2 | 0 | 3 | 0 | 0 | 98 | 49 | 1 | 1 | 0.765 | 1 | 0.2 | NA | 1 | -0.345 |
| SE | 15501 | HACL1 | HACL1 | chr3 | - | 15585247 | 15585342 | 15582876 | 15582989 | 15586524 | 15586602 | 1 | 0 | 0 | 1 | 0 | 0 | 0 | 1 | 0 | 0 | 98 | 49 | 1 | 1 | 0.333 | NA | NA | NA | 1 | -0.667 |
| SE | 15508 | STAB1 | STAB1 | chr3 | + | 52502631 | 52502727 | 52502158 | 52502228 | 52502998 | 52503109 | 8 | 1 | 0 | 1 | 0 | 0 | 0 | 4 | 0 | 0 | 98 | 49 | 1 | 1 | 0.8 | 1 | NA | NA | 1 | -0.1 |
| SE | 15509 | STAB1 | STAB1 | chr3 | + | 52512340 | 52512436 | 52511649 | 52511745 | 52512595 | 52512643 | 9 | 0 | 0 | 0 | 0 | 0 | 0 | 13 | 0 | 2 | 98 | 49 | 1 | 1 | 1 | NA | NA | NA | 0.765 | 0.235 |
| SE | 15514 | KAT2B | KAT2B | chr3 | + | 20125904 | 20126113 | 20122667 | 20122804 | 20136941 | 20137052 | 12 | 3 | 0 | 0 | 0 | 0 | 0 | 2 | 0 | 1 | 98 | 49 | 1 | 1 | 1 | 1 | NA | NA | 0.5 | 0.5 |
| SE | 15515 | KAT2B | KAT2B | chr3 | + | 20127422 | 20127549 | 20125904 | 20126113 | 20136941 | 20137052 | 27 | 11 | 8 | 0 | 2 | 0 | 0 | 12 | 0 | 0 | 98 | 49 | 1 | 1 | 1 | 0.733 | 1 | NA | 1 | -0.089 |
| SE | 15519 | TOP2B | TOP2B | chr3 | - | 25604759 | 25604870 | 25601099 | 25601225 | 25606042 | 25606122 | 24 | 5 | 5 | 2 | 1 | 0 | 0 | 10 | 0 | 1 | 98 | 49 | 1 | 1 | 0.857 | 0.714 | 1 | NA | 0.833 | 0.024 |
| SE | 15520 | TOP2B | TOP2B | chr3 | - | 25618417 | 25618509 | 25615430 | 25615586 | 25618653 | 25618849 | 17 | 5 | 0 | 1 | 0 | 0 | 0 | 1 | 0 | 0 | 98 | 49 | 1 | 1 | 0.895 | 1 | NA | NA | 1 | -0.052 |
| SE | 15521 | TOP2B | TOP2B | chr3 | - | 25618653 | 25618849 | 25615430 | 25615586 | 25619861 | 25620062 | 14 | 2 | 0 | 0 | 0 | 0 | 0 | 2 | 0 | 2 | 98 | 49 | 1 | 1 | 1 | 1 | NA | NA | 0.333 | 0.667 |
| SE | 15524 | NGLY1 | NGLY1 | chr3 | - | 25732318 | 25732483 | 25729132 | 25729318 | 25733871 | 25733982 | 6 | 0 | 2 | 1 | 0 | 4 | 0 | 1 | 0 | 0 | 98 | 49 | 1 | 1 | 0.75 | NA | 0.2 | NA | 1 | -0.525 |
| SE | 15525 | NGLY1 | NGLY1 | chr3 | - | 25737333 | 25737455 | 25736308 | 25736400 | 25739576 | 25739799 | 2 | 0 | 0 | 0 | 0 | 0 | 0 | 0 | 0 | 1 | 98 | 49 | 1 | 1 | 1 | NA | NA | NA | 0 | 1 |
| SE | 15527 | TGFBR2 | TGFBR2 | chr3 | + | 30623198 | 30623273 | 30606501 | 30606977 | 30644746 | 30644915 | 24 | 5 | 3 | 9 | 3 | 4 | 0 | 3 | 0 | 2 | 98 | 49 | 1 | 1 | 0.571 | 0.455 | 0.273 | NA | 0.429 | 0.004 |
| SE | 15528 | MS4A6A | MS4A6A | chr11 | - | 60179830 | 60179965 | 60178259 | 60178316 | 60181580 | 60181741 | 27 | 2 | 6 | 3 | 1 | 0 | 0 | 20 | 0 | 4 | 98 | 49 | 1 | 1 | 0.818 | 0.5 | 1 | NA | 0.714 | 0.059 |
| SE | 15529 | CMTM7 | CMTM7 | chr3 | + | 32449453 | 32449552 | 32441839 | 32442013 | 32452391 | 32452473 | 16 | 0 | 0 | 0 | 0 | 2 | 0 | 2 | 0 | 0 | 98 | 49 | 1 | 1 | 1 | NA | 0 | NA | 1 | -0.5 |
| SE | 15530 | DYNC1LI1 | DYNC1LI1 | chr3 | - | 32530283 | 32530328 | 32529539 | 32529660 | 32530460 | 32530520 | 49 | 39 | 10 | 1 | 0 | 0 | 1 | 7 | 0 | 0 | 94 | 49 | 1 | 1 | 0.962 | 1 | 1 | 1 | 1 | -0.013 |
| SE | 15531 | DYNC1LI1 | DYNC1LI1 | chr3 | - | 32544875 | 32545106 | 32541036 | 32541206 | 32545848 | 32545965 | 22 | 4 | 10 | 2 | 0 | 0 | 0 | 4 | 0 | 0 | 98 | 49 | 1 | 1 | 0.846 | 1 | 1 | NA | 1 | -0.051 |
| SE | 15532 | DYNC1LI1 | DYNC1LI1 | chr3 | - | 32545848 | 32545965 | 32541036 | 32541206 | 32570345 | 32570419 | 15 | 9 | 6 | 2 | 0 | 0 | 0 | 2 | 0 | 0 | 98 | 49 | 1 | 1 | 0.789 | 1 | 1 | NA | 1 | -0.07 |
| SE | 15534 | GLB1 | GLB1 | chr3 | - | 33065462 | 33065557 | 33058088 | 33058269 | 33068229 | 33068290 | 19 | 15 | 11 | 0 | 0 | 0 | 0 | 3 | 0 | 1 | 98 | 49 | 1 | 1 | 1 | 1 | 1 | NA | 0.6 | 0.4 |
| SE | 15544 | ATG16L2 | ATG16L2 | chr11 | + | 72824733 | 72824842 | 72824059 | 72824122 | 72825301 | 72825407 | 355 | 162 | 46 | 0 | 1 | 0 | 13 | 68 | 0 | 0 | 98 | 49 | 1 | 1 | 1 | 0.988 | 1 | 1 | 1 | -0.004 |
| SE | 15546 | ATG16L2 | ATG16L2 | chr11 | + | 72826702 | 72826823 | 72826172 | 72826243 | 72827187 | 72827293 | 55 | 44 | 20 | 1 | 0 | 0 | 1 | 24 | 0 | 0 | 98 | 49 | 1 | 1 | 0.965 | 1 | 1 | 1 | 1 | -0.012 |
| SE | 15548 | PDCD6IP | PDCD6IP | chr3 | + | 33812071 | 33812126 | 33798570 | 33798937 | 33813558 | 33813628 | 20 | 10 | 4 | 1 | 0 | 0 | 1 | 6 | 0 | 0 | 98 | 49 | 1 | 1 | 0.909 | 1 | 1 | 1 | 1 | -0.03 |
| SE | 15549 | PDCD6IP | PDCD6IP | chr3 | + | 33825186 | 33825340 | 33821954 | 33822082 | 33826479 | 33826580 | 21 | 4 | 10 | 1 | 0 | 0 | 0 | 8 | 0 | 0 | 98 | 49 | 1 | 1 | 0.913 | 1 | 1 | NA | 1 | -0.029 |
| SE | 15550 | PDCD6IP | PDCD6IP | chr3 | + | 33828837 | 33828969 | 33826479 | 33826580 | 33836043 | 33836266 | 15 | 6 | 3 | 0 | 0 | 0 | 0 | 1 | 0 | 1 | 98 | 49 | 1 | 1 | 1 | 1 | 1 | NA | 0.333 | 0.667 |
| SE | 15551 | PDCD6IP | PDCD6IP | chr3 | + | 33828852 | 33828969 | 33826479 | 33826580 | 33836043 | 33836266 | 16 | 5 | 3 | 0 | 0 | 0 | 0 | 1 | 0 | 1 | 98 | 49 | 1 | 1 | 1 | 1 | 1 | NA | 0.333 | 0.667 |
| SE | 15559 | MLH1 | MLH1 | chr3 | + | 37017505 | 37017599 | 37014431 | 37014544 | 37020309 | 37020463 | 2 | 3 | 0 | 2 | 0 | 0 | 0 | 1 | 0 | 0 | 98 | 49 | 1 | 1 | 0.333 | 1 | NA | NA | 1 | -0.334 |
| SE | 15564 | KLHL6 | KLHL6 | chr3 | - | 183499589 | 183499827 | 183494078 | 183494281 | 183508058 | 183508508 | 2 | 4 | 0 | 0 | 1 | 0 | 0 | 1 | 0 | 0 | 98 | 49 | 1 | 1 | 1 | 0.667 | NA | NA | 1 | -0.166 |
| SE | 15566 | GOLGA4 | GOLGA4 | chr3 | + | 37337142 | 37337163 | 37335052 | 37335166 | 37337665 | 37337734 | 4 | 3 | 0 | 2 | 0 | 0 | 0 | 3 | 0 | 0 | 70 | 49 | 1 | 1 | 0.583 | 1 | NA | NA | 1 | -0.209 |
| SE | 15567 | GOLGA4 | GOLGA4 | chr3 | + | 37347192 | 37347296 | 37340123 | 37340199 | 37355100 | 37355187 | 2 | 3 | 0 | 1 | 0 | 0 | 0 | 1 | 0 | 0 | 98 | 49 | 1 | 1 | 0.5 | 1 | NA | NA | 1 | -0.25 |
| SE | 15575 | MTHFSD | MTHFSD | chr16 | - | 86548034 | 86548111 | 86546558 | 86546649 | 86548463 | 86548577 | 0 | 0 | 1 | 1 | 0 | 0 | 0 | 1 | 0 | 0 | 98 | 49 | 1 | 1 | 0 | NA | 1 | NA | 1 | -0.5 |
| SE | 15576 | SUN2 | SUN2 | chr22 | - | 38738872 | 38738988 | 38738586 | 38738754 | 38739341 | 38739426 | 45 | 18 | 11 | 0 | 3 | 0 | 0 | 6 | 0 | 4 | 98 | 49 | 1 | 1 | 1 | 0.75 | 1 | NA | 0.429 | 0.488 |
| SE | 15589 | TRAK1 | TRAK1 | chr3 | + | 42176813 | 42176890 | 42125419 | 42125614 | 42184684 | 42184801 | 9 | 4 | 6 | 1 | 0 | 0 | 0 | 3 | 0 | 0 | 98 | 49 | 1 | 1 | 0.818 | 1 | 1 | NA | 1 | -0.061 |
| SE | 15595 | ABHD5 | ABHD5 | chr3 | + | 43702214 | 43702587 | 43699275 | 43699361 | 43711708 | 43711863 | 85 | 16 | 18 | 1 | 0 | 0 | 1 | 13 | 0 | 0 | 98 | 49 | 1 | 1 | 0.977 | 1 | 1 | 1 | 1 | -0.008 |
| SE | 15596 | ABHD5 | ABHD5 | chr3 | + | 43714946 | 43715058 | 43711708 | 43711863 | 43717670 | 43717857 | 89 | 26 | 26 | 1 | 0 | 0 | 0 | 18 | 0 | 0 | 98 | 49 | 1 | 1 | 0.978 | 1 | 1 | NA | 1 | -0.007 |
| SE | 15597 | ABHD5 | ABHD5 | chr3 | + | 43717670 | 43717857 | 43714946 | 43715058 | 43718442 | 43722725 | 74 | 17 | 10 | 7 | 1 | 0 | 0 | 17 | 0 | 3 | 98 | 49 | 1 | 1 | 0.841 | 0.895 | 1 | NA | 0.739 | 0.173 |
| SE | 15598 | LARS2 | LARS2 | chr3 | + | 45491516 | 45491800 | 45488696 | 45488812 | 45496274 | 45496373 | 2 | 0 | 3 | 1 | 0 | 0 | 0 | 1 | 0 | 0 | 98 | 49 | 1 | 1 | 0.5 | NA | 1 | NA | 1 | -0.25 |
| SE | 15599 | SACM1L | SACM1L | chr3 | + | 45735234 | 45735373 | 45732052 | 45732151 | 45737582 | 45737652 | 16 | 5 | 0 | 1 | 0 | 0 | 0 | 2 | 0 | 0 | 98 | 49 | 1 | 1 | 0.889 | 1 | NA | NA | 1 | -0.055 |
| SE | 15605 | YDJC | YDJC | chr22 | - | 21629009 | 21629187 | 21628089 | 21628787 | 21629307 | 21629407 | 3 | 0 | 0 | 1 | 5 | 0 | 0 | 0 | 0 | 2 | 98 | 49 | 1 | 1 | 0.6 | 0 | NA | NA | 0 | 0.3 |
| SE | 15606 | HIPK3 | HIPK3 | chr11 | + | 33337074 | 33337194 | 33328509 | 33328633 | 33338756 | 33338843 | 93 | 20 | 18 | 2 | 0 | 0 | 1 | 27 | 0 | 0 | 98 | 49 | 1 | 1 | 0.959 | 1 | 1 | 1 | 1 | -0.014 |
| SE | 15608 | HIPK3 | HIPK3 | chr11 | + | 33348165 | 33348228 | 33347851 | 33348013 | 33349146 | 33349287 | 14 | 4 | 2 | 2 | 1 | 0 | 0 | 6 | 0 | 0 | 98 | 49 | 1 | 1 | 0.778 | 0.667 | 1 | NA | 1 | -0.185 |
| SE | 15610 | HIPK3 | HIPK3 | chr11 | + | 33348521 | 33348818 | 33348165 | 33348228 | 33349146 | 33349287 | 52 | 23 | 5 | 3 | 2 | 0 | 0 | 4 | 0 | 0 | 98 | 49 | 1 | 1 | 0.897 | 0.852 | 1 | NA | 1 | -0.084 |
| SE | 15611 | HIPK3 | HIPK3 | chr11 | + | 33349146 | 33349287 | 33348521 | 33348818 | 33351607 | 33351843 | 68 | 32 | 7 | 1 | 0 | 0 | 0 | 9 | 0 | 0 | 98 | 49 | 1 | 1 | 0.971 | 1 | 1 | NA | 1 | -0.01 |
| SE | 15612 | SCAP | SCAP | chr3 | - | 47417303 | 47417826 | 47417121 | 47417207 | 47418133 | 47418249 | 5 | 2 | 4 | 1 | 0 | 0 | 1 | 0 | 0 | 0 | 98 | 49 | 1 | 1 | 0.714 | 1 | 1 | 1 | NA | -0.095 |
| SE | 15617 | SMARCC1 | SMARCC1 | chr3 | - | 47662333 | 47662592 | 47661293 | 47661455 | 47670657 | 47670717 | 22 | 3 | 1 | 1 | 0 | 0 | 1 | 1 | 0 | 0 | 98 | 49 | 1 | 1 | 0.917 | 1 | 1 | 1 | 1 | -0.028 |
| SE | 15620 | SMARCC1 | SMARCC1 | chr3 | - | 47729024 | 47729094 | 47720665 | 47720735 | 47736033 | 47736126 | 24 | 11 | 4 | 0 | 2 | 0 | 0 | 2 | 0 | 0 | 98 | 49 | 1 | 1 | 1 | 0.733 | 1 | NA | 1 | -0.089 |
| SE | 15621 | MAP4 | MAP4 | chr3 | - | 47918718 | 47918841 | 47915950 | 47917174 | 47921764 | 47921878 | 0 | 1 | 2 | 0 | 0 | 0 | 0 | 0 | 0 | 1 | 98 | 49 | 1 | 1 | NA | 1 | 1 | NA | 0 | 1 |
| SE | 15630 | UQCRC1 | UQCRC1 | chr3 | - | 48604232 | 48604431 | 48603563 | 48603643 | 48604650 | 48604780 | 15 | 2 | 9 | 0 | 1 | 0 | 0 | 2 | 0 | 0 | 98 | 49 | 1 | 1 | 1 | 0.5 | 1 | NA | 1 | -0.167 |
| SE | 15631 | UQCRC1 | UQCRC1 | chr3 | - | 48605769 | 48605856 | 48604650 | 48604780 | 48609161 | 48609302 | 7 | 0 | 1 | 2 | 0 | 0 | 0 | 6 | 0 | 0 | 98 | 49 | 1 | 1 | 0.636 | NA | 1 | NA | 1 | -0.182 |
| SE | 15632 | NCOR1 | NCOR1 | chr17 | - | 16039432 | 16039654 | 16034764 | 16034944 | 16040440 | 16040494 | 32 | 0 | 2 | 1 | 0 | 0 | 0 | 3 | 0 | 0 | 98 | 49 | 1 | 1 | 0.941 | NA | 1 | NA | 1 | -0.03 |
| SE | 15633 | NCOR1 | NCOR1 | chr17 | - | 16048844 | 16048988 | 16046950 | 16047093 | 16057513 | 16057737 | 12 | 4 | 3 | 0 | 0 | 0 | 0 | 3 | 0 | 2 | 98 | 49 | 1 | 1 | 1 | 1 | 1 | NA | 0.429 | 0.571 |
| SE | 15634 | NCOR1 | NCOR1 | chr17 | - | 16067893 | 16068121 | 16065484 | 16065694 | 16070164 | 16070525 | 15 | 6 | 4 | 1 | 1 | 0 | 1 | 2 | 0 | 1 | 98 | 49 | 1 | 1 | 0.882 | 0.75 | 1 | 1 | 0.5 | 0.127 |
| SE | 15635 | NCOR1 | NCOR1 | chr17 | - | 16072144 | 16072228 | 16071408 | 16071665 | 16073428 | 16073569 | 14 | 9 | 4 | 1 | 0 | 0 | 0 | 4 | 0 | 0 | 98 | 49 | 1 | 1 | 0.875 | 1 | 1 | NA | 1 | -0.042 |
| SE | 15636 | NCOR1 | NCOR1 | chr17 | - | 16080407 | 16080509 | 16079963 | 16080064 | 16080606 | 16080727 | 29 | 9 | 1 | 2 | 0 | 0 | 0 | 8 | 0 | 0 | 98 | 49 | 1 | 1 | 0.879 | 1 | 1 | NA | 1 | -0.04 |
| SE | 15637 | NCOR1 | NCOR1 | chr17 | - | 16126081 | 16126206 | 16121051 | 16121269 | 16137310 | 16137412 | 14 | 9 | 2 | 0 | 2 | 0 | 0 | 1 | 0 | 0 | 98 | 49 | 1 | 1 | 1 | 0.692 | 1 | NA | 1 | -0.103 |
| SE | 15643 | ARIH2 | ARIH2 | chr3 | + | 48922747 | 48922811 | 48918847 | 48918998 | 48927461 | 48927813 | 4 | 4 | 0 | 1 | 0 | 0 | 0 | 1 | 0 | 0 | 98 | 49 | 1 | 1 | 0.667 | 1 | NA | NA | 1 | -0.166 |
| SE | 15645 | NMT2 | NMT2 | chr10 | - | 15130141 | 15130312 | 15128349 | 15128458 | 15132816 | 15132933 | 4 | 0 | 2 | 1 | 0 | 0 | 0 | 1 | 0 | 0 | 98 | 49 | 1 | 1 | 0.667 | NA | 1 | NA | 1 | -0.166 |
| SE | 15647 | QRICH1 | QRICH1 | chr3 | - | 49044389 | 49044504 | 49033119 | 49033228 | 49046424 | 49046579 | 8 | 0 | 0 | 0 | 0 | 1 | 0 | 5 | 0 | 0 | 98 | 49 | 1 | 1 | 1 | NA | 0 | NA | 1 | -0.5 |
| SE | 15648 | QRICH1 | QRICH1 | chr3 | - | 49092396 | 49092456 | 49076708 | 49077038 | 49093911 | 49094071 | 1 | 2 | 0 | 4 | 5 | 0 | 0 | 0 | 1 | 0 | 98 | 49 | 1 | 1 | 0.111 | 0.167 | NA | 0 | NA | 0.139 |
| SE | 15667 | UBA7 | UBA7 | chr3 | - | 49809389 | 49809464 | 49808975 | 49809159 | 49809541 | 49809725 | 7 | 3 | 0 | 1 | 0 | 0 | 0 | 2 | 0 | 1 | 98 | 49 | 1 | 1 | 0.778 | 1 | NA | NA | 0.5 | 0.389 |
| SE | 15671 | RBM6 | RBM6 | chr3 | + | 49967469 | 49968748 | 49962575 | 49962685 | 49972058 | 49972148 | 14 | 1 | 6 | 2 | 0 | 0 | 0 | 4 | 0 | 0 | 98 | 49 | 1 | 1 | 0.778 | 1 | 1 | NA | 1 | -0.074 |
| SE | 15672 | RBM6 | RBM6 | chr3 | + | 49999439 | 49999513 | 49975322 | 49975392 | 50048244 | 50048319 | 13 | 7 | 0 | 3 | 0 | 0 | 0 | 1 | 0 | 0 | 98 | 49 | 1 | 1 | 0.684 | 1 | NA | NA | 1 | -0.158 |
| SE | 15673 | RBM6 | RBM6 | chr3 | + | 50059648 | 50059746 | 50058401 | 50058562 | 50060955 | 50060998 | 10 | 9 | 1 | 0 | 0 | 0 | 0 | 2 | 0 | 2 | 98 | 49 | 1 | 1 | 1 | 1 | 1 | NA | 0.333 | 0.667 |
| SE | 15677 | HEXB | HEXB | chr5 | + | 74713505 | 74713635 | 74705218 | 74705320 | 74715509 | 74715690 | 12 | 10 | 5 | 0 | 0 | 0 | 0 | 12 | 0 | 2 | 98 | 49 | 1 | 1 | 1 | 1 | 1 | NA | 0.75 | 0.25 |
| SE | 15685 | MAPKAPK3 | MAPKAPK3 | chr3 | + | 50644408 | 50644532 | 50642252 | 50642332 | 50645709 | 50645785 | 34 | 23 | 12 | 2 | 5 | 0 | 0 | 11 | 0 | 5 | 98 | 49 | 1 | 1 | 0.895 | 0.697 | 1 | NA | 0.524 | 0.34 |
| SE | 15686 | MAPKAPK3 | MAPKAPK3 | chr3 | + | 50645709 | 50645785 | 50644408 | 50644532 | 50646139 | 50646264 | 18 | 16 | 7 | 0 | 1 | 0 | 1 | 3 | 0 | 0 | 98 | 49 | 1 | 1 | 1 | 0.889 | 1 | 1 | 1 | -0.037 |
| SE | 15691 | DSE | DSE | chr6 | + | 116433342 | 116433550 | 116430953 | 116431193 | 116435586 | 116438281 | 8 | 1 | 3 | 2 | 0 | 0 | 0 | 1 | 0 | 0 | 98 | 49 | 1 | 1 | 0.667 | 1 | 1 | NA | 1 | -0.111 |
| SE | 15693 | DENND6A | DENND6A | chr3 | - | 57630420 | 57630523 | 57628810 | 57628885 | 57630714 | 57630824 | 9 | 18 | 5 | 2 | 0 | 0 | 0 | 8 | 0 | 0 | 98 | 49 | 1 | 1 | 0.692 | 1 | 1 | NA | 1 | -0.103 |
| SE | 15695 | WDR82 | WDR82 | chr3 | - | 52270711 | 52270809 | 52266951 | 52267018 | 52278200 | 52278643 | 20 | 13 | 5 | 0 | 0 | 1 | 0 | 5 | 0 | 1 | 98 | 49 | 1 | 1 | 1 | 1 | 0.714 | NA | 0.714 | 0.191 |
| SE | 15696 | BAP1 | BAP1 | chr3 | - | 52405109 | 52405294 | 52404452 | 52404586 | 52405764 | 52405912 | 5 | 0 | 2 | 0 | 2 | 0 | 0 | 3 | 0 | 0 | 98 | 49 | 1 | 1 | 1 | 0 | 1 | NA | 1 | -0.333 |
| SE | 15698 | BAP1 | BAP1 | chr3 | - | 52408473 | 52408606 | 52407957 | 52408077 | 52409553 | 52409608 | 7 | 3 | 1 | 0 | 0 | 0 | 1 | 1 | 0 | 0 | 98 | 49 | 1 | 1 | 1 | 0.6 | 1 | 1 | 1 | -0.133 |
| SE | 15703 | C4orf29 | C4orf29 | chr4 | + | 127989720 | 127989821 | 127984338 | 127984423 | 128008919 | 128008998 | 4 | 0 | 2 | 1 | 1 | 0 | 0 | 3 | 0 | 0 | 98 | 49 | 1 | 1 | 0.667 | 0 | 1 | NA | 1 | -0.444 |
| SE | 15709 | FAM208A | FAM208A | chr3 | - | 56663540 | 56663572 | 56662384 | 56662490 | 56668396 | 56668558 | 2 | 0 | 0 | 0 | 0 | 0 | 0 | 1 | 0 | 0 | 81 | 49 | 1 | 1 | 1 | NA | NA | NA | 0 | 1 |
| SE | 15710 | FAM208A | FAM208A | chr3 | - | 56666259 | 56666384 | 56663540 | 56663572 | 56668396 | 56668558 | 12 | 0 | 2 | 1 | 0 | 0 | 0 | 1 | 0 | 0 | 98 | 49 | 1 | 1 | 0.857 | NA | 1 | NA | 1 | -0.072 |
| SE | 15729 | ATXN7 | ATXN7 | chr3 | + | 63952378 | 63952483 | 63913156 | 63913225 | 63979914 | 63980167 | 35 | 5 | 5 | 1 | 0 | 0 | 0 | 5 | 0 | 0 | 98 | 49 | 1 | 1 | 0.946 | 1 | 1 | NA | 1 | -0.018 |
| SE | 15730 | ATXN7 | ATXN7 | chr3 | + | 63982185 | 63982445 | 63979914 | 63980167 | 63982938 | 63983021 | 21 | 12 | 0 | 1 | 0 | 0 | 0 | 9 | 0 | 0 | 98 | 49 | 1 | 1 | 0.913 | 1 | NA | NA | 1 | -0.043 |
| SE | 15732 | ATXN7 | ATXN7 | chr3 | + | 63990175 | 63990374 | 63988058 | 63988324 | 63990737 | 63990859 | 13 | 5 | 0 | 3 | 0 | 0 | 0 | 2 | 0 | 0 | 98 | 49 | 1 | 1 | 0.684 | 1 | NA | NA | 1 | -0.158 |
| SE | 15733 | ATXN7 | ATXN7 | chr3 | + | 63997620 | 63997687 | 63995504 | 63996483 | 63999449 | 64003460 | 6 | 6 | 0 | 1 | 0 | 1 | 0 | 2 | 0 | 2 | 98 | 49 | 1 | 1 | 0.75 | 1 | 0 | NA | 0.333 | 0.25 |
| SE | 15740 | ARL6IP5 | ARL6IP5 | chr3 | + | 69101838 | 69102056 | 69084938 | 69085223 | 69104463 | 69106088 | 25 | 12 | 7 | 0 | 1 | 0 | 0 | 12 | 0 | 0 | 98 | 49 | 1 | 1 | 1 | 0.857 | 1 | NA | 1 | -0.048 |
| SE | 15741 | ERBB2IP | ERBB2IP | chr5 | + | 66028273 | 66028343 | 66026301 | 66026417 | 66038382 | 66038482 | 60 | 18 | 13 | 1 | 0 | 0 | 0 | 7 | 0 | 0 | 98 | 49 | 1 | 1 | 0.968 | 1 | 1 | NA | 1 | -0.011 |
| SE | 15742 | ERBB2IP | ERBB2IP | chr5 | + | 66068876 | 66069020 | 66053405 | 66054951 | 66072168 | 66072291 | 2 | 2 | 0 | 5 | 2 | 0 | 0 | 0 | 0 | 2 | 98 | 49 | 1 | 1 | 0.167 | 0.333 | NA | NA | 0 | 0.25 |
| SE | 15743 | ERBB2IP | ERBB2IP | chr5 | + | 66068876 | 66069020 | 66053405 | 66054951 | 66075023 | 66075230 | 2 | 0 | 0 | 9 | 5 | 5 | 0 | 0 | 0 | 1 | 98 | 49 | 1 | 1 | 0.1 | 0 | 0 | NA | 0 | 0.033 |
| SE | 15744 | ERBB2IP | ERBB2IP | chr5 | + | 66072168 | 66072291 | 66053405 | 66054951 | 66075023 | 66075230 | 6 | 9 | 0 | 9 | 5 | 5 | 0 | 2 | 0 | 1 | 98 | 49 | 1 | 1 | 0.25 | 0.474 | 0 | NA | 0.5 | -0.259 |
| SE | 15748 | COQ2 | COQ2 | chr4 | - | 83269859 | 83269993 | 83267585 | 83267774 | 83272086 | 83272172 | 8 | 2 | 0 | 1 | 0 | 0 | 0 | 4 | 0 | 0 | 98 | 49 | 1 | 1 | 0.8 | 1 | NA | NA | 1 | -0.1 |
| SE | 15749 | PPP4R2 | PPP4R2 | chr3 | + | 73047185 | 73047356 | 72998076 | 72998158 | 73059036 | 73059130 | 24 | 10 | 12 | 1 | 0 | 0 | 0 | 13 | 0 | 0 | 98 | 49 | 1 | 1 | 0.923 | 1 | 1 | NA | 1 | -0.026 |
| SE | 15751 | GBE1 | GBE1 | chr3 | - | 81646391 | 81646482 | 81642780 | 81642990 | 81648855 | 81648991 | 8 | 2 | 3 | 1 | 0 | 0 | 0 | 0 | 0 | 1 | 98 | 49 | 1 | 1 | 0.8 | 1 | 1 | NA | 0 | 0.933 |
| SE | 15752 | GBE1 | GBE1 | chr3 | - | 81705443 | 81705613 | 81670837 | 81670953 | 81761374 | 81761799 | 5 | 6 | 1 | 0 | 1 | 0 | 1 | 1 | 0 | 0 | 98 | 49 | 1 | 1 | 1 | 0.75 | 1 | 1 | 1 | -0.083 |
| SE | 15761 | LSP1 | LSP1 | chr11 | + | 1883418 | 1883560 | 1880086 | 1880224 | 1883931 | 1884024 | 182 | 100 | 71 | 0 | 1 | 1 | 4 | 21 | 0 | 0 | 98 | 49 | 1 | 1 | 1 | 0.98 | 0.973 | 1 | 1 | -0.016 |
| SE | 15762 | LSP1 | LSP1 | chr11 | + | 1884279 | 1884323 | 1883931 | 1884024 | 1884499 | 1884581 | 322 | 196 | 157 | 2 | 0 | 0 | 5 | 71 | 0 | 1 | 93 | 49 | 1 | 1 | 0.988 | 1 | 1 | 1 | 0.974 | 0.009 |
| SE | 15763 | LSP1 | LSP1 | chr11 | + | 1887236 | 1887314 | 1886731 | 1886866 | 1887473 | 1887576 | 130 | 85 | 44 | 1 | 0 | 0 | 6 | 18 | 0 | 1 | 98 | 49 | 1 | 1 | 0.985 | 1 | 1 | 1 | 0.9 | 0.045 |
| SE | 15769 | CRYBG3 | CRYBG3 | chr3 | + | 97892859 | 97892993 | 97889354 | 97889390 | 97895958 | 97896085 | 1 | 0 | 0 | 0 | 0 | 0 | 0 | 3 | 0 | 0 | 98 | 49 | 1 | 1 | 1 | NA | 0 | NA | 1 | -0.5 |
| SE | 15771 | UBTF | UBTF | chr17 | - | 44212343 | 44212454 | 44211872 | 44212006 | 44212818 | 44212939 | 4 | 2 | 1 | 9 | 1 | 1 | 0 | 1 | 0 | 0 | 98 | 49 | 1 | 1 | 0.182 | 0.5 | 0.333 | NA | 0.333 | 0.005 |
| SE | 15773 | CWF19L1 | CWF19L1 | chr10 | - | 100256261 | 100256476 | 100253420 | 100253539 | 100260217 | 100260319 | 7 | 9 | 1 | 1 | 1 | 0 | 0 | 4 | 0 | 0 | 98 | 49 | 1 | 1 | 0.778 | 0.818 | 1 | NA | 1 | -0.135 |
| SE | 15774 | CWF19L1 | CWF19L1 | chr10 | - | 100260217 | 100260319 | 100256261 | 100256476 | 100260965 | 100261028 | 14 | 8 | 1 | 1 | 1 | 0 | 0 | 3 | 0 | 0 | 98 | 49 | 1 | 1 | 0.875 | 0.8 | 1 | NA | 1 | -0.108 |
| SE | 15778 | TBC1D23 | TBC1D23 | chr3 | + | 100310402 | 100310542 | 100306436 | 100306543 | 100316098 | 100316187 | 6 | 2 | 2 | 0 | 0 | 0 | 0 | 1 | 0 | 1 | 98 | 49 | 1 | 1 | 1 | 1 | 1 | NA | 0 | 1 |
| SE | 15783 | SENP7 | SENP7 | chr3 | - | 101417592 | 101417790 | 101398860 | 101399055 | 101458954 | 101459052 | 11 | 14 | 2 | 1 | 0 | 0 | 0 | 0 | 0 | 0 | 98 | 49 | 1 | 1 | 0.846 | 1 | 1 | NA | 0 | 0.949 |
| SE | 15785 | HSP90AA1 | HSP90AA1 | chr14 | - | 102082110 | 102082444 | 102080737 | 102081821 | 102083033 | 102083302 | 29 | 17 | 4 | 0 | 0 | 0 | 0 | 5 | 0 | 1 | 98 | 49 | 1 | 1 | 1 | 1 | 1 | NA | 0.714 | 0.286 |
| SE | 15786 | HSP90AA1 | HSP90AA1 | chr14 | - | 102085757 | 102086124 | 102085297 | 102085431 | 102086216 | 102086378 | 20 | 14 | 7 | 0 | 2 | 0 | 0 | 10 | 0 | 0 | 98 | 49 | 1 | 1 | 1 | 0.778 | 1 | NA | 1 | -0.074 |
| SE | 15787 | TRA2B | TRA2B | chr3 | - | 185923795 | 185923984 | 185922010 | 185922126 | 185925463 | 185925626 | 19 | 6 | 8 | 1 | 0 | 0 | 0 | 5 | 0 | 0 | 98 | 49 | 1 | 1 | 0.905 | 1 | 1 | NA | 1 | -0.032 |
| SE | 15799 | FKBP15 | FKBP15 | chr9 | - | 113196931 | 113197078 | 113194026 | 113194169 | 113198854 | 113198923 | 16 | 8 | 6 | 1 | 0 | 0 | 0 | 4 | 0 | 0 | 98 | 49 | 1 | 1 | 0.889 | 1 | 1 | NA | 1 | -0.037 |
| SE | 15813 | MRVI1 | MRVI1 | chr11 | - | 10580454 | 10580589 | 10573090 | 10576575 | 10581866 | 10581986 | 13 | 12 | 6 | 1 | 0 | 0 | 0 | 6 | 0 | 0 | 98 | 49 | 1 | 1 | 0.867 | 1 | 1 | NA | 1 | -0.044 |
| SE | 15815 | MRVI1 | MRVI1 | chr11 | - | 10600917 | 10601059 | 10594145 | 10594195 | 10603119 | 10603251 | 47 | 14 | 16 | 1 | 2 | 0 | 0 | 4 | 0 | 1 | 98 | 49 | 1 | 1 | 0.959 | 0.778 | 1 | NA | 0.667 | 0.245 |
| SE | 15833 | AOAH | AOAH | chr7 | - | 36514471 | 36514590 | 36512940 | 36513380 | 36522038 | 36522115 | 5 | 0 | 0 | 13 | 2 | 0 | 0 | 1 | 0 | 6 | 98 | 49 | 1 | 1 | 0.161 | 0 | NA | NA | 0.077 | 0.004 |
| SE | 15834 | AOAH | AOAH | chr7 | - | 36548611 | 36548686 | 36540318 | 36540491 | 36549438 | 36549475 | 87 | 24 | 15 | 10 | 0 | 2 | 0 | 16 | 0 | 1 | 98 | 49 | 1 | 1 | 0.813 | 1 | 0.789 | NA | 0.889 | -0.022 |
| SE | 15836 | AOAH | AOAH | chr7 | - | 36637850 | 36637910 | 36632035 | 36632106 | 36659165 | 36659265 | 60 | 21 | 5 | 0 | 2 | 0 | 6 | 12 | 0 | 0 | 98 | 49 | 1 | 1 | 1 | 0.84 | 1 | 1 | 1 | -0.053 |
| SE | 15837 | AOAH | AOAH | chr7 | - | 36659165 | 36659265 | 36637850 | 36637910 | 36673942 | 36674009 | 48 | 12 | 1 | 2 | 2 | 0 | 0 | 10 | 0 | 1 | 98 | 49 | 1 | 1 | 0.923 | 0.75 | 1 | NA | 0.833 | 0.058 |
| SE | 15842 | GON4L | GON4L | chr1 | - | 155777621 | 155777820 | 155776394 | 155776481 | 155783985 | 155784089 | 8 | 2 | 1 | 0 | 0 | 0 | 1 | 5 | 0 | 1 | 98 | 49 | 1 | 1 | 1 | 1 | 1 | 1 | 0.714 | 0.143 |
| SE | 15846 | ATP6V0E1 | ATP6V0E1 | chr5 | + | 172994774 | 172994822 | 172983759 | 172983964 | 173020237 | 173020367 | 33 | 29 | 33 | 0 | 1 | 0 | 4 | 18 | 0 | 0 | 97 | 49 | 1 | 1 | 1 | 0.936 | 1 | 1 | 1 | -0.021 |
| SE | 15847 | ARHGAP25 | ARHGAP25 | chr2 | + | 68787839 | 68787956 | 68782232 | 68782320 | 68807272 | 68807480 | 59 | 31 | 30 | 3 | 1 | 2 | 0 | 6 | 0 | 0 | 98 | 49 | 1 | 1 | 0.908 | 0.939 | 0.882 | NA | 1 | -0.09 |
| SE | 15850 | SYNE2 | SYNE2 | chr14 | + | 64009965 | 64010116 | 64007042 | 64007222 | 64016472 | 64016631 | 28 | 22 | 16 | 0 | 0 | 0 | 0 | 3 | 0 | 2 | 98 | 49 | 1 | 1 | 1 | 1 | 1 | NA | 0.429 | 0.571 |
| SE | 15852 | SYNE2 | SYNE2 | chr14 | + | 64075944 | 64076100 | 64073967 | 64074136 | 64078465 | 64078606 | 39 | 10 | 5 | 0 | 2 | 0 | 0 | 3 | 0 | 0 | 98 | 49 | 1 | 1 | 1 | 0.714 | 1 | NA | 1 | -0.095 |
| SE | 15853 | SYNE2 | SYNE2 | chr14 | + | 64163401 | 64163581 | 64162071 | 64162276 | 64165284 | 64165410 | 20 | 14 | 8 | 0 | 0 | 0 | 0 | 1 | 0 | 1 | 98 | 49 | 1 | 1 | 1 | 1 | 1 | NA | 0.333 | 0.667 |
| SE | 15854 | SYNE2 | SYNE2 | chr14 | + | 64215285 | 64215354 | 64214140 | 64214470 | 64216247 | 64216387 | 3 | 0 | 0 | 1 | 1 | 0 | 0 | 0 | 0 | 4 | 98 | 49 | 1 | 1 | 0.6 | 0 | NA | NA | 0 | 0.3 |
| SE | 15855 | SYNE2 | SYNE2 | chr14 | + | 64221533 | 64221704 | 64220436 | 64220637 | 64223188 | 64223380 | 10 | 3 | 0 | 1 | 0 | 0 | 0 | 2 | 0 | 0 | 98 | 49 | 1 | 1 | 0.833 | 1 | NA | NA | 1 | -0.084 |
| SE | 15856 | SYNE2 | SYNE2 | chr14 | + | 64221575 | 64221704 | 64220436 | 64220637 | 64223188 | 64223380 | 14 | 3 | 0 | 1 | 0 | 0 | 0 | 2 | 0 | 0 | 98 | 49 | 1 | 1 | 0.875 | 1 | NA | NA | 1 | -0.062 |
| SE | 15857 | EAF2 | EAF2 | chr3 | + | 121854686 | 121854823 | 121844452 | 121844547 | 121857010 | 121857156 | 3 | 1 | 2 | 1 | 0 | 0 | 0 | 1 | 0 | 0 | 98 | 49 | 1 | 1 | 0.6 | 1 | 1 | NA | 1 | -0.133 |
| SE | 15872 | SMURF2 | SMURF2 | chr17 | - | 64591083 | 64591149 | 64586085 | 64586170 | 64593439 | 64593573 | 17 | 3 | 0 | 1 | 0 | 0 | 0 | 1 | 0 | 0 | 98 | 49 | 1 | 1 | 0.895 | 1 | NA | NA | 1 | -0.052 |
| SE | 15879 | MR1 | MR1 | chr1 | + | 181053572 | 181053677 | 181050010 | 181050286 | 181055224 | 181061938 | 6 | 0 | 3 | 1 | 0 | 0 | 0 | 1 | 0 | 0 | 98 | 49 | 1 | 1 | 0.75 | NA | 1 | NA | 1 | -0.125 |
| SE | 15883 | ZXDC | ZXDC | chr3 | - | 126466154 | 126466325 | 126458900 | 126462220 | 126470894 | 126471025 | 20 | 5 | 1 | 2 | 0 | 0 | 0 | 5 | 0 | 0 | 98 | 49 | 1 | 1 | 0.833 | 1 | 1 | NA | 1 | -0.056 |
| SE | 15884 | HDAC1 | HDAC1 | chr1 | + | 32330577 | 32330686 | 32329067 | 32329160 | 32330767 | 32330908 | 14 | 6 | 0 | 1 | 0 | 0 | 0 | 3 | 0 | 0 | 98 | 49 | 1 | 1 | 0.875 | 1 | NA | NA | 1 | -0.062 |
| SE | 15885 | HDAC1 | HDAC1 | chr1 | + | 32332700 | 32332749 | 32332089 | 32332242 | 32333016 | 32333623 | 9 | 3 | 0 | 0 | 1 | 0 | 0 | 4 | 0 | 0 | 98 | 49 | 1 | 1 | 1 | 0.6 | NA | NA | 1 | -0.2 |

| | | | | | | | | | | | | | | | | | | | | | | | | | | | | | | | |
|---|---|---|---|---|---|---|---|---|---|---|---|---|---|---|---|---|---|---|---|---|---|---|---|---|---|---|---|---|---|---|---|
| SE | 15889 | ULK4 | ULK4 | chr3 | - | 41715446 | 41715568 | 41715236 | 41715293 | 41717727 | 41717861 | 0 | 0 | 2 | 1 | 0 | 0 | 0 | 1 | 0 | 0 | 98 | 49 | 1 | 1 | 0 | NA | 1 | NA | 1 | -0.5 |
| SE | 15890 | ULK4 | ULK4 | chr3 | - | 41915976 | 41916052 | 41912806 | 41912899 | 41918456 | 41918540 | 1 | 0 | 0 | 4 | 5 | 0 | 0 | 0 | 0 | 1 | 98 | 49 | 1 | 1 | 0.111 | 0 | NA | NA | 0 | 0.056 |
| SE | 15891 | DGKZ | DGKZ | chr11 | + | 46369940 | 46370009 | 46369493 | 46369550 | 46371312 | 46371384 | 17 | 5 | 2 | 4 | 2 | 0 | 0 | 5 | 0 | 0 | 98 | 49 | 1 | 1 | 0.68 | 0.556 | 1 | NA | 1 | -0.255 |
| SE | 15906 | TOPBP1 | TOPBP1 | chr3 | - | 133624051 | 133624175 | 133623310 | 133623457 | 133628361 | 133628471 | 5 | 4 | 2 | 0 | 0 | 0 | 0 | 2 | 0 | 2 | 98 | 49 | 1 | 1 | 1 | 1 | 1 | NA | 0.333 | 0.667 |
| SE | 15908 | CEP63 | CEP63 | chr3 | + | 134532777 | 134532900 | 134531844 | 134531940 | 134537154 | 134537268 | 27 | 15 | 1 | 2 | 0 | 0 | 1 | 1 | 0 | 0 | 98 | 49 | 1 | 1 | 0.871 | 1 | 1 | 1 | 1 | -0.043 |
| SE | 15909 | CEP63 | CEP63 | chr3 | + | 134547334 | 134547472 | 134546148 | 134546288 | 134549061 | 134549176 | 13 | 10 | 1 | 13 | 4 | 6 | 0 | 4 | 0 | 6 | 98 | 49 | 1 | 1 | 0.333 | 0.556 | 0.077 | NA | 0.25 | 0.072 |
| SE | 15910 | CEP63 | CEP63 | chr3 | + | 134558141 | 134558347 | 134551925 | 134552012 | 134559149 | 134559429 | 20 | 16 | 2 | 0 | 2 | 0 | 2 | 0 | 0 | 0 | 98 | 49 | 1 | 1 | 1 | 0.8 | 1 | 1 | NA | -0.067 |
| SE | 15912 | CEP63 | CEP63 | chr3 | + | 134559149 | 134559429 | 134551925 | 134552012 | 134561376 | 134565028 | 5 | 2 | 3 | 2 | 4 | 3 | 0 | 0 | 0 | 4 | 98 | 49 | 1 | 1 | 0.556 | 0.2 | 0.333 | NA | 0 | 0.363 |
| SE | 15914 | FMR1 | FMR1 | chrX | + | 147938098 | 147938161 | 147937465 | 147937600 | 147940575 | 147940662 | 8 | 1 | 0 | 2 | 2 | 0 | 0 | 0 | 0 | 1 | 98 | 49 | 1 | 1 | 0.667 | 0.2 | NA | NA | 0 | 0.433 |
| SE | 15919 | STAG1 | STAG1 | chr3 | - | 136568764 | 136568861 | 136542118 | 136542195 | 136604308 | 136604473 | 15 | 7 | 3 | 2 | 0 | 0 | 0 | 4 | 0 | 0 | 98 | 49 | 1 | 1 | 0.789 | 1 | 1 | NA | 1 | -0.07 |
| SE | 15920 | RPS6KA1 | RPS6KA1 | chr1 | + | 26546866 | 26546983 | 26545851 | 26546035 | 26547188 | 26547270 | 60 | 36 | 24 | 1 | 5 | 1 | 3 | 19 | 0 | 1 | 98 | 49 | 1 | 1 | 0.968 | 0.783 | 0.923 | 1 | 0.905 | -0.061 |
| SE | 15921 | RPS6KA1 | RPS6KA1 | chr1 | + | 26551396 | 26551477 | 26547188 | 26547270 | 26551643 | 26551723 | 88 | 46 | 31 | 1 | 0 | 0 | 1 | 19 | 0 | 0 | 98 | 49 | 1 | 1 | 0.978 | 1 | 1 | 1 | 1 | -0.007 |
| SE | 15925 | RPS6KA1 | RPS6KA1 | chr1 | + | 26561504 | 26561663 | 26561044 | 26561134 | 26561448 | 26571610 | 41 | 38 | 38 | 0 | 0 | 0 | 0 | 12 | 0 | 1 | 98 | 49 | 1 | 1 | 1 | 1 | 1 | NA | 0.857 | 0.143 |
| SE | 15926 | RPS6KA1 | RPS6KA1 | chr1 | + | 26572175 | 26572293 | 26571848 | 26571925 | 26573223 | 26573361 | 49 | 24 | 31 | 1 | 0 | 0 | 0 | 13 | 0 | 0 | 98 | 49 | 1 | 1 | 0.961 | 1 | 1 | NA | 1 | -0.013 |
| SE | 15927 | ARMC8 | ARMC8 | chr3 | + | 138188400 | 138188530 | 138187247 | 138187599 | 138209816 | 138209893 | 2 | 0 | 0 | 0 | 2 | 0 | 0 | 2 | 0 | 0 | 98 | 49 | 1 | 1 | 1 | 0 | NA | NA | 1 | -0.5 |
| SE | 15929 | ARMC8 | ARMC8 | chr3 | + | 138188454 | 138188530 | 138187247 | 138187599 | 138209816 | 138209893 | 2 | 0 | 0 | 0 | 2 | 0 | 0 | 4 | 0 | 0 | 98 | 49 | 1 | 1 | 1 | 0 | NA | NA | 1 | -0.5 |
| SE | 15932 | ARMC8 | ARMC8 | chr3 | + | 138209816 | 138209893 | 138188400 | 138188530 | 138221925 | 138221997 | 12 | 0 | 2 | 0 | 3 | 0 | 0 | 4 | 0 | 0 | 98 | 49 | 1 | 1 | 1 | 0 | 1 | NA | 1 | -0.333 |
| SE | 15933 | ARMC8 | ARMC8 | chr3 | + | 138223388 | 138223531 | 138221925 | 138221997 | 138223635 | 138223733 | 13 | 5 | 0 | 0 | 1 | 0 | 0 | 2 | 0 | 0 | 98 | 49 | 1 | 1 | 1 | 0.714 | NA | NA | 1 | -0.143 |
| SE | 15935 | ARMC8 | ARMC8 | chr3 | + | 138237308 | 138237384 | 138235033 | 138235114 | 138237481 | 138237572 | 20 | 3 | 1 | 0 | 0 | 0 | 0 | 1 | 0 | 2 | 98 | 49 | 1 | 1 | 1 | 1 | 1 | NA | 0.2 | 0.8 |
| SE | 15941 | CSF2RA | CSF2RA | chrX | + | 1300490 | 1300626 | 1295426 | 1295456 | 1303922 | 1304019 | 19 | 15 | 15 | 0 | 0 | 0 | 0 | 19 | 0 | 4 | 98 | 49 | 1 | 1 | 1 | 1 | 1 | NA | 0.704 | 0.296 |
| SE | 15942 | CSF2RA | CSF2RA | chrX | + | 1303260 | 1303362 | 1300490 | 1300626 | 1303922 | 1304019 | 4 | 1 | 0 | 12 | 4 | 7 | 0 | 1 | 0 | 5 | 98 | 49 | 1 | 1 | 0.143 | 0.111 | 0 | NA | 0.091 | -0.006 |
| SE | 15944 | CSF2RA | CSF2RA | chrX | + | 1305445 | 1305527 | 1300490 | 1300626 | 1309401 | 1309935 | 25 | 8 | 4 | 0 | 1 | 0 | 0 | 1 | 0 | 0 | 98 | 49 | 1 | 1 | 1 | 0.8 | 1 | NA | 1 | -0.067 |
| SE | 15950 | RASA2 | RASA2 | chr3 | + | 141573943 | 141574067 | 141573146 | 141573221 | 141576999 | 141577106 | 11 | 4 | 1 | 1 | 0 | 0 | 0 | 1 | 0 | 0 | 98 | 49 | 1 | 1 | 0.846 | 1 | 1 | NA | 1 | -0.051 |
| SE | 15952 | RASA2 | RASA2 | chr3 | + | 141609876 | 141610066 | 141609419 | 141609523 | 141612282 | 141615363 | 14 | 8 | 9 | 0 | 3 | 0 | 0 | 2 | 0 | 0 | 98 | 49 | 1 | 1 | 1 | 0.571 | 1 | NA | 1 | -0.143 |
| SE | 15956 | ATR | ATR | chr3 | - | 142542664 | 142542757 | 142540903 | 142541034 | 142547724 | 142547910 | 4 | 3 | 0 | 1 | 0 | 0 | 0 | 2 | 0 | 0 | 98 | 49 | 1 | 1 | 0.667 | 1 | NA | NA | 1 | -0.166 |
| SE | 15958 | NEK9 | NEK9 | chr14 | - | 75121118 | 75121174 | 75120509 | 75120580 | 75124045 | 75124223 | 6 | 2 | 0 | 0 | 1 | 0 | 0 | 2 | 0 | 0 | 98 | 49 | 1 | 1 | 1 | 0.5 | NA | NA | 1 | -0.25 |
| SE | 15959 | AREL1 | AREL1 | chr14 | - | 74667464 | 74667594 | 74667318 | 74667377 | 74669648 | 74669774 | 23 | 6 | 1 | 1 | 0 | 0 | 0 | 3 | 0 | 0 | 98 | 49 | 1 | 1 | 0.92 | 1 | 1 | NA | 1 | -0.027 |
| SE | 15960 | AREL1 | AREL1 | chr14 | - | 74669946 | 74670126 | 74669648 | 74669774 | 74670761 | 74670871 | 21 | 12 | 4 | 1 | 1 | 0 | 2 | 2 | 0 | 3 | 98 | 49 | 1 | 1 | 0.913 | 0.857 | 1 | 1 | 0.25 | 0.298 |
| SE | 15961 | AREL1 | AREL1 | chr14 | - | 74674033 | 74674111 | 74673076 | 74673218 | 74675698 | 74675946 | 23 | 18 | 4 | 8 | 3 | 0 | 0 | 2 | 0 | 0 | 98 | 49 | 1 | 1 | 0.59 | 0.75 | 1 | NA | 1 | -0.22 |
| SE | 15962 | AREL1 | AREL1 | chr14 | - | 74684453 | 74684680 | 74683295 | 74683533 | 74685599 | 74685660 | 18 | 14 | 2 | 2 | 0 | 2 | 0 | 4 | 0 | 0 | 98 | 49 | 1 | 1 | 0.818 | 1 | 0.333 | NA | 1 | -0.283 |
| SE | 15964 | DPEP2 | DPEP2 | chr16 | - | 67987851 | 67987987 | 67987389 | 67987760 | 67989322 | 67989398 | 61 | 31 | 21 | 6 | 0 | 2 | 0 | 18 | 0 | 0 | 98 | 49 | 1 | 1 | 0.836 | 1 | 0.84 | NA | 1 | -0.108 |
| SE | 15966 | DPEP2 | DPEP2 | chr16 | - | 67991837 | 67991979 | 67991114 | 67991184 | 67992063 | 67992193 | 21 | 13 | 7 | 7 | 0 | 0 | 0 | 4 | 0 | 0 | 98 | 49 | 1 | 1 | 0.6 | 1 | 1 | NA | 1 | -0.133 |
| SE | 15968 | PLSCR1 | PLSCR1 | chr3 | - | 146528613 | 146528831 | 146525604 | 146525647 | 146533469 | 146533550 | 8 | 4 | 7 | 0 | 2 | 0 | 0 | 8 | 0 | 2 | 98 | 49 | 1 | 1 | 1 | 0.5 | 1 | NA | 0.667 | 0.166 |
| SE | 15976 | 7-Mar | 7-Mar | chr2 | + | 159745769 | 159745937 | 159743060 | 159743253 | 159747804 | 159748903 | 63 | 39 | 21 | 0 | 2 | 0 | 0 | 4 | 0 | 0 | 98 | 49 | 1 | 1 | 1 | 0.907 | 1 | NA | 1 | -0.031 |
| SE | 15977 | 7-Mar | 7-Mar | chr2 | + | 159762879 | 159762993 | 159759225 | 159759335 | 159764625 | 159764674 | 56 | 31 | 17 | 1 | 3 | 0 | 0 | 8 | 0 | 0 | 98 | 49 | 1 | 1 | 0.966 | 0.838 | 1 | NA | 1 | -0.065 |
| SE | 15982 | FUBP3 | FUBP3 | chr9 | + | 130634666 | 130634738 | 130632201 | 130632278 | 130635998 | 130636126 | 3 | 0 | 0 | 0 | 1 | 0 | 0 | 1 | 0 | 0 | 98 | 49 | 1 | 1 | 1 | 0 | NA | NA | 1 | -0.5 |
| SE | 15988 | NSUN4 | NSUN4 | chr1 | + | 46352867 | 46353028 | 46344800 | 46345144 | 46360703 | 46360828 | 1 | 0 | 0 | 1 | 0 | 0 | 0 | 1 | 0 | 0 | 98 | 49 | 1 | 1 | 0.333 | NA | NA | NA | 1 | -0.667 |
| SE | 15989 | TSC22D2 | TSC22D2 | chr3 | + | 150423048 | 150423120 | 150408334 | 150411308 | 150457075 | 150457127 | 1 | 0 | 0 | 0 | 3 | 0 | 0 | 4 | 0 | 1 | 98 | 49 | 1 | 1 | 1 | 0 | NA | NA | 0.667 | -0.167 |
| SE | 15991 | SERP1 | SERP1 | chr3 | - | 150545702 | 150545778 | 150541992 | 150544498 | 150546051 | 150546641 | 58 | 20 | 6 | 1 | 1 | 0 | 1 | 14 | 0 | 0 | 98 | 49 | 1 | 1 | 0.967 | 0.909 | 1 | 1 | 1 | -0.041 |
| SE | 15992 | EIF2A | EIF2A | chr3 | + | 150571957 | 150572529 | 150568175 | 150568292 | 150575648 | 150575762 | 14 | 0 | 2 | 1 | 0 | 0 | 0 | 6 | 0 | 0 | 98 | 49 | 1 | 1 | 0.875 | NA | 1 | NA | 1 | -0.062 |
| SE | 15993 | EIF2A | EIF2A | chr3 | + | 150575648 | 150575762 | 150571957 | 150572529 | 150581617 | 150581746 | 14 | 1 | 0 | 1 | 0 | 0 | 0 | 8 | 0 | 0 | 98 | 49 | 1 | 1 | 0.875 | 1 | NA | NA | 1 | -0.062 |
| SE | 15998 | MME | MME | chr3 | + | 155114993 | 155115155 | 155085058 | 155085094 | 155116478 | 155116559 | 236 | 111 | 57 | 2 | 0 | 0 | 0 | 30 | 0 | 0 | 98 | 49 | 1 | 1 | 0.983 | 1 | 1 | NA | 1 | -0.006 |
| SE | 15999 | MME | MME | chr3 | + | 155141990 | 155142127 | 155140190 | 155140292 | 155142236 | 155142330 | 84 | 27 | 32 | 0 | 1 | 0 | 0 | 12 | 0 | 0 | 98 | 49 | 1 | 1 | 1 | 0.931 | 1 | NA | 1 | -0.023 |
| SE | 16000 | MME | MME | chr3 | + | 155141990 | 155142127 | 155140190 | 155140292 | 155143442 | 155143571 | 67 | 18 | 23 | 3 | 1 | 2 | 0 | 9 | 0 | 0 | 98 | 49 | 1 | 1 | 0.918 | 0.9 | 0.852 | NA | 1 | -0.11 |
| SE | 16001 | MME | MME | chr3 | + | 155142236 | 155142330 | 155140190 | 155140292 | 155143442 | 155143571 | 95 | 36 | 23 | 3 | 1 | 2 | 0 | 7 | 0 | 0 | 98 | 49 | 1 | 1 | 0.941 | 0.947 | 0.852 | NA | 1 | -0.087 |
| SE | 16002 | MME | MME | chr3 | + | 155142236 | 155142330 | 155141990 | 155142127 | 155143442 | 155143571 | 117 | 46 | 32 | 5 | 2 | 0 | 0 | 10 | 0 | 0 | 98 | 49 | 1 | 1 | 0.921 | 0.92 | 1 | NA | 1 | -0.053 |
| SE | 16003 | MME | MME | chr3 | + | 155168491 | 155168625 | 155166901 | 155167021 | 155168731 | 155168797 | 177 | 57 | 55 | 10 | 0 | 0 | 0 | 10 | 0 | 3 | 98 | 49 | 1 | 1 | 0.898 | 1 | 1 | NA | 0.625 | 0.341 |
| SE | 16004 | MME | MME | chr3 | + | 155172116 | 155172212 | 155168731 | 155168797 | 155172535 | 155172612 | 117 | 74 | 37 | 1 | 0 | 1 | 0 | 17 | 0 | 0 | 98 | 49 | 1 | 1 | 0.983 | 1 | 0.949 | NA | 1 | -0.023 |
| SE | 16010 | SLC33A1 | SLC33A1 | chr3 | - | 155833856 | 155834041 | 155833467 | 155833585 | 155842431 | 155842619 | 3 | 1 | 0 | 1 | 0 | 0 | 0 | 1 | 0 | 0 | 98 | 49 | 1 | 1 | 0.6 | 1 | NA | NA | 1 | -0.2 |
| SE | 16017 | MFSD1 | MFSD1 | chr3 | + | 158805361 | 158805474 | 158804318 | 158804371 | 158807039 | 158807082 | 41 | 13 | 11 | 3 | 1 | 2 | 0 | 10 | 0 | 0 | 98 | 49 | 1 | 1 | 0.872 | 0.867 | 0.733 | NA | 1 | -0.176 |
| SE | 16018 | MFSD1 | MFSD1 | chr3 | + | 158821983 | 158822140 | 158821596 | 158821653 | 158823427 | 158823525 | 38 | 4 | 7 | 1 | 0 | 0 | 2 | 16 | 0 | 0 | 98 | 49 | 1 | 1 | 0.95 | 1 | 1 | 1 | 1 | -0.017 |
| SE | 16024 | NMD3 | NMD3 | chr3 | + | 161246335 | 161246448 | 161242507 | 161242653 | 161247257 | 161247330 | 7 | 0 | 0 | 1 | 3 | 0 | 3 | 5 | 0 | 0 | 98 | 49 | 1 | 1 | 0.778 | 0 | NA | 1 | 1 | -0.611 |
| SE | 16027 | ATP11C | ATP11C | chrX | - | 139731650 | 139731755 | 139726345 | 139728962 | 139737915 | 139738069 | 0 | 0 | 0 | 2 | 0 | 0 | 0 | 2 | 0 | 0 | 98 | 49 | 1 | 1 | 0 | NA | NA | NA | 1 | -1 |
| SE | 16028 | RNPEPL1 | RNPEPL1 | chr2 | + | 240574112 | 240574348 | 240573774 | 240573891 | 240574514 | 240574628 | 18 | 9 | 3 | 3 | 0 | 0 | 0 | 6 | 0 | 1 | 98 | 49 | 1 | 1 | 0.75 | 1 | 1 | NA | 0.75 | 0.167 |
| SE | 16035 | PDCD10 | PDCD10 | chr3 | - | 167725400 | 167725473 | 167720061 | 167720273 | 167734805 | 167734863 | 5 | 1 | 0 | 4 | 0 | 0 | 0 | 5 | 0 | 1 | 98 | 49 | 1 | 1 | 0.385 | 1 | NA | NA | 0.714 | -0.021 |
| SE | 16036 | YTHDC2 | YTHDC2 | chr5 | + | 113540967 | 113541116 | 113539073 | 113539181 | 113542367 | 113542503 | 9 | 6 | 0 | 0 | 1 | 0 | 0 | 1 | 0 | 0 | 98 | 49 | 1 | 1 | 1 | 0.75 | NA | NA | 1 | -0.125 |
| SE | 16038 | RASGEF1B | RASGEF1B | chr4 | - | 81447503 | 81447578 | 81445742 | 81445838 | 81448068 | 81448281 | 5 | 0 | 0 | 1 | 0 | 0 | 0 | 5 | 0 | 0 | 98 | 49 | 1 | 1 | 0.714 | NA | NA | NA | 1 | -0.286 |
| SE | 16039 | RASGEF1B | RASGEF1B | chr4 | - | 81457498 | 81457621 | 81456650 | 81456788 | 81459331 | 81459514 | 16 | 0 | 0 | 4 | 0 | 0 | 0 | 27 | 0 | 0 | 98 | 49 | 1 | 1 | 0.667 | NA | NA | NA | 1 | -0.333 |
| SE | 16041 | MYNN | MYNN | chr3 | + | 169782464 | 169782643 | 169780589 | 169780749 | 169783476 | 169783560 | 3 | 1 | 2 | 0 | 4 | 0 | 0 | 4 | 0 | 0 | 98 | 49 | 1 | 1 | 1 | 0.111 | 1 | NA | 1 | -0.296 |
| SE | 16044 | ECE1 | ECE1 | chr1 | - | 21221746 | 21221842 | 21217246 | 21220131 | 21227158 | 21227226 | 29 | 9 | 10 | 1 | 0 | 0 | 3 | 9 | 0 | 0 | 98 | 49 | 1 | 1 | 0.935 | 1 | 1 | 1 | 1 | -0.022 |
| SE | 16045 | ECE1 | ECE1 | chr1 | - | 21225249 | 21225440 | 21221746 | 21221842 | 21227158 | 21227226 | 41 | 24 | 13 | 0 | 3 | 0 | 0 | 8 | 0 | 1 | 98 | 49 | 1 | 1 | 1 | 0.8 | 1 | NA | 0.8 | 0.133 |
| SE | 16047 | ECE1 | ECE1 | chr1 | - | 21279190 | 21279332 | 21272698 | 21272911 | 21290069 | 21290156 | 46 | 22 | 11 | 2 | 0 | 0 | 2 | 12 | 0 | 0 | 98 | 49 | 1 | 1 | 0.92 | 1 | 1 | 1 | 1 | -0.027 |
| SE | 16048 | PHC3 | PHC3 | chr3 | - | 170145422 | 170145521 | 170136418 | 170136665 | 170171372 | 170171450 | 11 | 8 | 14 | 1 | 1 | 0 | 0 | 1 | 0 | 0 | 98 | 49 | 1 | 1 | 0.846 | 0.8 | 1 | NA | 1 | -0.118 |
| SE | 16049 | PHC3 | PHC3 | chr3 | - | 170149085 | 170149244 | 170145422 | 170145521 | 170171372 | 170171450 | 14 | 12 | 13 | 1 | 0 | 0 | 0 | 3 | 0 | 0 | 98 | 49 | 1 | 1 | 0.875 | 1 | 1 | NA | 1 | -0.042 |
| SE | 16051 | ZMYM5 | ZMYM5 | chr13 | - | 19851354 | 19851448 | 19838699 | 19838985 | 19851688 | 19852190 | 9 | 2 | 3 | 1 | 0 | 0 | 0 | 1 | 0 | 0 | 98 | 49 | 1 | 1 | 0.818 | 1 | 1 | NA | 1 | -0.061 |
| SE | 16052 | ZMYM5 | ZMYM5 | chr13 | - | 19862398 | 19862466 | 19851688 | 19852190 | 19863449 | 19863636 | 19 | 4 | 2 | 1 | 0 | 0 | 0 | 2 | 0 | 0 | 98 | 49 | 1 | 1 | 0.905 | 1 | 1 | NA | 1 | -0.032 |
| SE | 16056 | PIK3CA | PIK3CA | chr3 | + | 179220985 | 179221157 | 179219948 | 179220052 | 179224080 | 179224187 | 20 | 17 | 6 | 0 | 0 | 0 | 0 | 4 | 0 | 1 | 98 | 49 | 1 | 1 | 1 | 1 | 1 | NA | 0.667 | 0.333 |
| SE | 16068 | ATP11B | ATP11B | chr3 | + | 182885950 | 182886010 | 182884752 | 182884898 | 182887585 | 182887713 | 43 | 22 | 24 | 1 | 0 | 0 | 0 | 3 | 0 | 0 | 98 | 49 | 1 | 1 | 0.956 | 1 | 1 | NA | 1 | -0.015 |
| SE | 16074 | SIPA1L2 | SIPA1L2 | chr1 | - | 232404124 | 232404178 | 232403447 | 232403571 | 232415493 | 232415625 | 1 | 0 | 0 | 11 | 0 | 0 | 0 | 0 | 0 | 2 | 98 | 49 | 1 | 1 | 0.043 | NA | NA | NA | 0 | 0.043 |
| SE | 16075 | SIPA1L2 | SIPA1L2 | chr1 | - | 232441767 | 232441868 | 232441290 | 232441394 | 232443601 | 232443685 | 9 | 3 | 2 | 3 | 0 | 0 | 0 | 2 | 0 | 0 | 98 | 49 | 1 | 1 | 0.6 | 1 | 1 | NA | 1 | -0.133 |
| SE | 16076 | SIPA1L2 | SIPA1L2 | chr1 | - | 232471370 | 232471528 | 232464839 | 232465416 | 232479626 | 232479730 | 29 | 1 | 4 | 1 | 0 | 0 | 0 | 3 | 0 | 0 | 98 | 49 | 1 | 1 | 0.935 | 1 | 1 | NA | 1 | -0.022 |
| SE | 16081 | UNC119 | UNC119 | chr17 | - | 28547998 | 28548101 | 28546706 | 28547849 | 28548591 | 28548705 | 38 | 16 | 3 | 1 | 0 | 0 | 0 | 8 | 0 | 0 | 98 | 49 | 1 | 1 | 0.95 | 1 | 1 | NA | 1 | -0.017 |
| SE | 16085 | EIF4G1 | EIF4G1 | chr3 | + | 184315488 | 184315545 | 184314494 | 184314674 | 184315762 | 184315856 | 4 | 0 | 0 | 1 | 1 | 0 | 0 | 2 | 0 | 1 | 98 | 49 | 1 | 1 | 0.667 | 0 | NA | NA | 0.5 | -0.166 |
| SE | 16101 | XRN2 | XRN2 | chr20 | + | 21328558 | 21328670 | 21326489 | 21326601 | 21330480 | 21330539 | 60 | 20 | 9 | 2 | 0 | 0 | 0 | 13 | 0 | 0 | 98 | 49 | 1 | 1 | 0.937 | 1 | 1 | NA | 1 | -0.021 |
| SE | 16102 | XRN2 | XRN2 | chr20 | + | 21356585 | 21356665 | 21356079 | 21356177 | 21357735 | 21357792 | 16 | 9 | 10 | 0 | 0 | 0 | 0 | 12 | 0 | 2 | 98 | 49 | 1 | 1 | 1 | 1 | 1 | NA | 0.75 | 0.25 |
| SE | 16103 | XRN2 | XRN2 | chr20 | + | 21357735 | 21357792 | 21356585 | 21356665 | 21365420 | 21365489 | 30 | 9 | 7 | 0 | 0 | 0 | 0 | 9 | 0 | 1 | 98 | 49 | 1 | 1 | 1 | 1 | 1 | NA | 0.818 | 0.182 |
| SE | 16106 | STK17B | STK17B | chr2 | - | 196139619 | 196139799 | 196133582 | 196137729 | 196141248 | 196141297 | 171 | 95 | 23 | 0 | 2 | 0 | 1 | 70 | 0 | 0 | 98 | 49 | 1 | 1 | 1 | 0.96 | 1 | 1 | 1 | -0.013 |
| SE | 16107 | STK17B | STK17B | chr2 | - | 196163261 | 196163427 | 196156438 | 196156651 | 196171332 | 196171612 | 408 | 196 | 43 | 0 | 1 | 0 | 21 | 136 | 0 | 0 | 98 | 49 | 1 | 1 | 1 | 0.99 | 1 | 1 | 1 | -0.003 |
| SE | 16109 | HSPA9 | HSPA9 | chr5 | - | 138566625 | 138566718 | 138561579 | 138561789 | 138567000 | 138567163 | 7 | 6 | 0 | 2 | 0 | 0 | 0 | 2 | 0 | 0 | 98 | 49 | 1 | 1 | 0.636 | 1 | NA | NA | 1 | -0.182 |
| SE | 16113 | GGA3 | GGA3 | chr17 | - | 75244618 | 75244717 | 75243446 | 75243570 | 75246508 | 75246584 | 9 | 0 | 10 | 4 | 0 | 1 | 0 | 0 | 0 | 1 | 98 | 49 | 1 | 1 | 0.529 | NA | 0.833 | NA | 0 | 0.681 |
| SE | 16116 | SIRPA | SIRPA | chr20 | + | 1922312 | 1922645 | 1921394 | 1921712 | 1924763 | 1924877 | 48 | 17 | 9 | 2 | 0 | 0 | 3 | 7 | 0 | 0 | 98 | 49 | 1 | 1 | 0.923 | 1 | 1 | 1 | 1 | -0.026 |
| SE | 16118 | LPP | LPP | chr3 | + | 188341662 | 188341719 | 188225404 | 188225527 | 188406111 | 188406313 | 21 | 4 | 0 | 0 | 0 | 0 | 0 | 0 | 0 | 1 | 98 | 49 | 1 | 1 | 1 | 1 | NA | NA | 0 | 1 |
| SE | 16133 | IL1RAP | IL1RAP | chr3 | + | 190550341 | 190550505 | 190514050 | 190514219 | 190556129 | 190556216 | 3 | 0 | 0 | 11 | 8 | 0 | 0 | 0 | 3 | 3 | 98 | 49 | 1 | 1 | 0.12 | 0 | NA | 0 | 0 | 0.06 |
| SE | 16134 | IL1RAP | IL1RAP | chr3 | + | 190550341 | 190550505 | 190514050 | 190514219 | 190564288 | 190564353 | 0 | 1 | 0 | 5 | 1 | 0 | 0 | 0 | 0 | 1 | 98 | 49 | 1 | 1 | 0 | 0.333 | NA | NA | 0 | 0.167 |
| SE | 16136 | IL1RAP | IL1RAP | chr3 | + | 190556129 | 190556216 | 190550341 | 190550505 | 190564288 | 190564353 | 14 | 12 | 1 | 0 | 1 | 0 | 0 | 4 | 0 | 0 | 98 | 49 | 1 | 1 | 1 | 0.857 | 1 | NA | 1 | -0.048 |
| SE | 16137 | IL1RAP | IL1RAP | chr3 | + | 190564288 | 190564353 | 190514050 | 190514219 | 190604127 | 190604413 | 39 | 17 | 10 | 1 | 0 | 0 | 2 | 14 | 0 | 0 | 98 | 49 | 1 | 1 | 0.951 | 1 | 1 | 1 | 1 | -0.016 |
| SE | 16138 | IL1RAP | IL1RAP | chr3 | + | 190608994 | 190609181 | 190604127 | 190604413 | 190620274 | 190620440 | 40 | 14 | 22 | 0 | 0 | 2 | 0 | 8 | 0 | 0 | 98 | 49 | 1 | 1 | 1 | 1 | 0.846 | NA | 1 | -0.051 |
| SE | 16139 | IL1RAP | IL1RAP | chr3 | + | 190620274 | 190620440 | 190604127 | 190604413 | 190623343 | 190623415 | 16 | 23 | 21 | 1 | 0 | 0 | 0 | 4 | 0 | 0 | 98 | 49 | 1 | 1 | 0.889 | 1 | 1 | NA | 1 | -0.037 |
| SE | 16141 | KDM5C | KDM5C | chrX | - | 53198489 | 53198637 | 53197770 | 53197876 | 53198763 | 53198888 | 15 | 7 | 5 | 1 | 0 | 0 | 1 | 3 | 0 | 0 | 98 | 49 | 1 | 1 | 0.882 | 1 | 1 | 1 | 1 | -0.039 |
| SE | 16142 | KDM5C | KDM5C | chrX | - | 53210675 | 53210857 | 53210413 | 53210576 | 53211496 | 53211655 | 34 | 10 | 2 | 1 | 1 | 0 | 0 | 10 | 0 | 0 | 98 | 49 | 1 | 1 | 0.944 | 0.833 | 1 | NA | 1 | -0.074 |
| SE | 16149 | TNK2 | TNK2 | chr3 | - | 195882050 | 195882328 | 195879048 | 195879175 | 195883156 | 195883309 | 12 | 9 | 2 | 2 | 0 | 0 | 0 | 3 | 0 | 1 | 98 | 49 | 1 | 1 | 0.75 | 1 | 1 | NA | 0.6 | 0.317 |
| SE | 16152 | PCYT1A | PCYT1A | chr3 | - | 196283474 | 196283538 | 196270414 | 196270541 | 196287614 | 196287713 | 1 | 0 | 0 | 1 | 0 | 0 | 0 | 1 | 0 | 1 | 98 | 49 | 1 | 1 | 0.333 | NA | NA | NA | 0.333 | 0 |
| SE | 16154 | CEP19 | CEP19 | chr3 | - | 196708527 | 196708727 | 196706276 | 196707912 | 196711928 | 196712294 | 7 | 2 | 8 | 1 | 0 | 0 | 0 | 2 | 0 | 0 | 98 | 49 | 1 | 1 | 0.778 | 1 | 1 | NA | 1 | -0.074 |
| SE | 16155 | SMARCA2 | SMARCA2 | chr9 | + | 2115821 | 2116049 | 2110253 | 2110417 | 2119457 | 2119535 | 26 | 4 | 4 | 0 | 1 | 0 | 0 | 3 | 0 | 0 | 98 | 49 | 1 | 1 | 1 | 0.667 | 1 | NA | 1 | -0.111 |
| SE | 16156 | SMARCA2 | SMARCA2 | chr9 | + | 2170418 | 2170472 | 2161685 | 2161903 | 2181570 | 2181676 | 6 | 2 | 1 | 6 | 1 | 4 | 0 | 2 | 0 | 0 | 98 | 49 | 1 | 1 | 0.333 | 0.5 | 0.111 | NA | 1 | -0.685 |
| SE | 16165 | KIAA0226 | KIAA0226 | chr3 | - | 197699186 | 197699231 | 197696953 | 197697049 | 197700612 | 197701146 | 1 | 0 | 0 | 2 | 0 | 0 | 0 | 1 | 0 | 2 | 94 | 49 | 1 | 1 | 0.207 | NA | NA | NA | 0.207 | 0 |
| SE | 16166 | KIAA0226 | KIAA0226 | chr3 | - | 197717976 | 197718130 | 197705091 | 197705175 | 197736654 | 197736926 | 3 | 0 | 3 | 1 | 0 | 0 | 0 | 2 | 1 | 0 | 98 | 49 | 1 | 1 | 0.6 | NA | 1 | 0 | 1 | 0.3 |
| SE | 16167 | LRCH3 | LRCH3 | chr3 | + | 197858833 | 197858905 | 197854391 | 197854445 | 197865422 | 197865471 | 3 | 1 | 0 | 1 | 0 | 0 | 0 | 3 | 0 | 1 | 98 | 49 | 1 | 1 | 0.6 | 1 | NA | NA | 0.6 | 0.2 |
| SE | 16168 | AFF1 | AFF1 | chr4 | + | 87046165 | 87046286 | 87007000 | 87007424 | 87046694 | 87047594 | 28 | 8 | 7 | 1 | 1 | 0 | 2 | 9 | 0 | 0 | 98 | 49 | 1 | 1 | 0.933 | 0.8 | 1 | 1 | 1 | -0.089 |
| SE | 16169 | AFF1 | AFF1 | chr4 | + | 87089983 | 87090070 | 87084119 | 87084164 | 87091792 | 87091829 | 23 | 7 | 2 | 0 | 0 | 0 | 1 | 12 | 0 | 1 | 98 | 49 | 1 | 1 | 1 | 1 | 1 | 1 | 0.857 | 0.072 |
| SE | 16170 | AFF1 | AFF1 | chr4 | + | 87132270 | 87132408 | 87131792 | 87131864 | 87134470 | 87134694 | 5 | 9 | 1 | 0 | 5 | 0 | 0 | 3 | 0 | 0 | 98 | 49 | 1 | 1 | 1 | 0.474 | 1 | NA | 1 | -0.175 |
| SE | 16174 | HLA-C | HLA-C | chr6 | - | 31269965 | 31270085 | 31269492 | 31269525 | 31270209 | 31270485 | 734 | 697 | 301 | 0 | 1 | 0 | 48 | 112 | 0 | 0 | 98 | 49 | 1 | 1 | 1 | 0.997 | 1 | 1 | 1 | -0.001 |
| SE | 16177 | TMEM175 | TMEM175 | chr4 | + | 948410 | 948615 | 948115 | 948154 | 950420 | 950518 | 4 | 0 | 2 | 0 | 0 | 0 | 0 | 0 | 0 | 1 | 98 | 49 | 1 | 1 | 1 | NA | 1 | NA | 0 | 1 |
| SE | 16181 | DUSP22 | DUSP22 | chr6 | + | 335113 | 335163 | 311879 | 311962 | 345853 | 345928 | 11 | 7 | 1 | 0 | 0 | 0 | 0 | 2 | 0 | 1 | 98 | 49 | 1 | 1 | 1 | 1 | 1 | NA | 0.5 | 0.5 |
| SE | 16185 | SLC25A13 | SLC25A13 | chr7 | - | 96189580 | 96189674 | 96189293 | 96189378 | 96191108 | 96191247 | 6 | 2 | 0 | 0 | 0 | 0 | 0 | 1 | 0 | 1 | 98 | 49 | 1 | 1 | 1 | 1 | NA | NA | 0.333 | 0.667 |
| SE | 16190 | HTT | HTT | chr4 | + | 3140509 | 3140656 | 3136225 | 3136326 | 3142765 | 3142886 | 8 | 0 | 0 | 1 | 0 | 0 | 0 | 4 | 0 | 0 | 98 | 49 | 1 | 1 | 0.8 | NA | NA | NA | 1 | -0.2 |
| SE | 16200 | SH3TC1 | SH3TC1 | chr4 | + | 8216956 | 8217167 | 8216110 | 8216257 | 8218270 | 8218347 | 3 | 0 | 0 | 0 | 0 | 0 | 0 | 1 | 0 | 5 | 98 | 49 | 1 | 1 | 1 | NA | NA | NA | 0.091 | 0.909 |
| SE | 16201 | SH3TC1 | SH3TC1 | chr4 | + | 8236277 | 8236428 | 8235432 | 8235555 | 8237473 | 8237670 | 2 | 0 | 0 | 3 | 0 | 0 | 0 | 1 | 0 | 0 | 98 | 49 | 1 | 1 | 0.25 | NA | NA | NA | 1 | -0.75 |
| SE | 16203 | DMXL1 | DMXL1 | chr5 | + | 119110150 | 119110283 | 119105179 | 119105258 | 119114474 | 119114541 | 10 | 1 | 2 | 2 | 0 | 0 | 0 | 1 | 0 | 0 | 98 | 49 | 1 | 1 | 0.714 | 1 | 1 | NA | 1 | -0.095 |
| SE | 16204 | DMXL1 | DMXL1 | chr5 | + | 119110154 | 119110283 | 119105179 | 119105258 | 119114474 | 119114541 | 6 | 1 | 2 | 2 | 0 | 0 | 0 | 1 | 0 | 0 | 98 | 49 | 1 | 1 | 0.6 | 1 | 1 | NA | 1 | -0.133 |
| SE | 16206 | DMXL1 | DMXL1 | chr5 | + | 119238988 | 119239080 | 119237321 | 119237414 | 119240418 | 119240471 | 3 | 0 | 0 | 0 | 0 | 2 | 0 | 1 | 0 | 0 | 98 | 49 | 1 | 1 | 1 | NA | 0 | NA | 1 | -0.5 |
| SE | 16207 | NIPBL | NIPBL | chr5 | + | 37000372 | 37000570 | 36995621 | 36995804 | 37000816 | 37000888 | 26 | 13 | 1 | 1 | 0 | 0 | 0 | 1 | 0 | 1 | 98 | 49 | 1 | 1 | 0.929 | 1 | 1 | NA | 0.333 | 0.643 |
| SE | 16209 | NIPBL | NIPBL | chr5 | + | 37060843 | 37061018 | 37052365 | 37052566 | 37063789 | 37063978 | 4 | 2 | 1 | 0 | 1 | 0 | 0 | 1 | 0 | 0 | 98 | 49 | 1 | 1 | 1 | 0.5 | 1 | NA | 1 | -0.167 |
| SE | 16210 | ATG2B | ATG2B | chr14 | - | 96322111 | 96322254 | 96317697 | 96317855 | 96322539 | 96322735 | 3 | 5 | 2 | 0 | 1 | 0 | 0 | 0 | 1 | 0 | 98 | 49 | 1 | 1 | 1 | 0.714 | 1 | 0 | NA | 0.905 |
| SE | 16212 | NSD1 | NSD1 | chr5 | + | 177269601 | 177269807 | 177267561 | 177267718 | 177273671 | 177273784 | 14 | 4 | 1 | 1 | 0 | 0 | 0 | 1 | 0 | 0 | 98 | 49 | 1 | 1 | 0.875 | 1 | 1 | NA | 1 | -0.042 |
| SE | 16213 | NSD1 | NSD1 | chr5 | + | 177282464 | 177282581 | 177280564 | 177280834 | 177283786 | 177283928 | 9 | 0 | 5 | 1 | 0 | 0 | 0 | 1 | 0 | 0 | 98 | 49 | 1 | 1 | 0.818 | NA | 1 | NA | 1 | -0.091 |
| SE | 16215 | WDR1 | WDR1 | chr4 | - | 10077752 | 10077926 | 10077303 | 10077448 | 10078890 | 10079001 | 83 | 27 | 26 | 2 | 0 | 0 | 12 | 27 | 0 | 0 | 98 | 49 | 1 | 1 | 0.954 | 1 | 1 | 1 | 1 | -0.015 |
| SE | 16217 | WDR1 | WDR1 | chr4 | - | 10088663 | 10088741 | 10088292 | 10088373 | 10097710 | 10097891 | 55 | 15 | 17 | 2 | 0 | 0 | 3 | 4 | 0 | 0 | 98 | 49 | 1 | 1 | 0.932 | 1 | 1 | 1 | 1 | -0.023 |
| SE | 16218 | WDR1 | WDR1 | chr4 | - | 10097710 | 10097891 | 10088663 | 10088741 | 10116112 | 10116234 | 37 | 7 | 7 | 2 | 0 | 0 | 2 | 3 | 0 | 0 | 98 | 49 | 1 | 1 | 0.902 | 1 | 1 | 1 | 1 | -0.033 |
| SE | 16219 | WDR1 | WDR1 | chr4 | - | 10098991 | 10099139 | 10097710 | 10097891 | 10116112 | 10116234 | 35 | 23 | 17 | 2 | 0 | 0 | 1 | 6 | 0 | 0 | 98 | 49 | 1 | 1 | 0.897 | 1 | 1 | 1 | 1 | -0.034 |
| SE | 16220 | WDR1 | WDR1 | chr4 | - | 10103895 | 10103986 | 10088663 | 10088741 | 10116112 | 10116234 | 36 | 16 | 13 | 2 | 0 | 0 | 4 | 6 | 0 | 0 | 98 | 49 | 1 | 1 | 0.9 | 1 | 1 | 1 | 1 | -0.033 |
| SE | 16221 | WDR1 | WDR1 | chr4 | - | 10103895 | 10103986 | 10098991 | 10099139 | 10116112 | 10116234 | 64 | 30 | 25 | 0 | 0 | 0 | 5 | 16 | 0 | 1 | 98 | 49 | 1 | 1 | 1 | 1 | 1 | 1 | 0.889 | 0.055 |
| SE | 16222 | SHTN1 | SHTN1 | chr10 | - | 116929848 | 116930002 | 116927791 | 116927891 | 116940465 | 116940612 | 6 | 0 | 0 | 1 | 0 | 0 | 0 | 4 | 0 | 0 | 98 | 49 | 1 | 1 | 0.75 | NA | NA | NA | 1 | -0.25 |
| SE | 16228 | BOD1L1 | BOD1L1 | chr4 | - | 13595859 | 13595944 | 13591922 | 13591966 | 13597103 | 13597168 | 43 | 7 | 2 | 0 | 2 | 0 | 1 | 12 | 0 | 0 | 98 | 49 | 1 | 1 | 1 | 0.636 | 1 | 1 | 1 | -0.121 |
| SE | 16229 | BOD1L1 | BOD1L1 | chr4 | - | 13610933 | 13611100 | 13609294 | 13609406 | 13613511 | 13613661 | 86 | 25 | 8 | 1 | 0 | 0 | 2 | 9 | 0 | 0 | 98 | 49 | 1 | 1 | 0.977 | 1 | 1 | 1 | 1 | -0.008 |
| SE | 16232 | FBXL5 | FBXL5 | chr4 | - | 15638507 | 15638694 | 15636493 | 15636676 | 15640787 | 15640883 | 366 | 142 | 81 | 3 | 0 | 0 | 3 | 71 | 0 | 0 | 98 | 49 | 1 | 1 | 0.984 | 1 | 1 | 1 | 1 | -0.005 |
| SE | 16233 | FBXL5 | FBXL5 | chr4 | - | 15641523 | 15641627 | 15640787 | 15640883 | 15644492 | 15644708 | 5 | 5 | 2 | 96 | 46 | 54 | 0 | 1 | 0 | 31 | 98 | 49 | 1 | 1 | 0.025 | 0.052 | 0.018 | NA | 0.016 | 0.016 |
| SE | 16234 | FBXL5 | FBXL5 | chr4 | - | 15644492 | 15644657 | 15640787 | 15640883 | 15655203 | 15655412 | 97 | 48 | 54 | 0 | 2 | 0 | 0 | 31 | 0 | 0 | 98 | 49 | 1 | 1 | 1 | 0.923 | 1 | NA | 1 | -0.026 |
| SE | 16235 | FBXL5 | FBXL5 | chr4 | - | 15644492 | 15644708 | 15640787 | 15640883 | 15655203 | 15655412 | 108 | 49 | 63 | 0 | 2 | 0 | 1 | 42 | 0 | 0 | 98 | 49 | 1 | 1 | 1 | 0.925 | 1 | 1 | 1 | -0.025 |
| SE | 16236 | HLA-DRB1 | HLA-DRB1 | chr6 | - | 32581556 | 32581838 | 32580745 | 32580856 | 32584108 | 32584170 | 88 | 0 | 4 | 0 | 2 | 0 | 0 | 14 | 0 | 0 | 98 | 49 | 1 | 1 | 1 | 0 | 1 | NA | 1 | -0.333 |
| SE | 16238 | BST1 | BST1 | chr4 | + | 15715706 | 15715799 | 15715284 | 15715361 | 15718906 | 15718993 | 55 | 35 | 29 | 0 | 2 | 0 | 2 | 21 | 0 | 0 | 98 | 49 | 1 | 1 | 1 | 0.897 | 1 | 1 | 1 | -0.034 |
| SE | 16244 | TAPT1 | TAPT1 | chr4 | - | 16213767 | 16213898 | 16191360 | 16191523 | 16226258 | 16226538 | 2 | 2 | 0 | 1 | 0 | 0 | 0 | 1 | 0 | 0 | 98 | 49 | 1 | 1 | 0.5 | 1 | NA | NA | 1 | -0.25 |
| SE | 16249 | DHX15 | DHX15 | chr4 | - | 24576242 | 24576678 | 24570653 | 24570847 | 24584322 | 24584561 | 10 | 5 | 8 | 0 | 1 | 0 | 0 | 1 | 0 | 0 | 98 | 49 | 1 | 1 | 1 | 0.714 | 1 | NA | 1 | -0.095 |
| SE | 16251 | PI4K2B | PI4K2B | chr4 | + | 25252320 | 25252475 | 25234030 | 25234431 | 25255064 | 25255265 | 2 | 0 | 0 | 1 | 0 | 0 | 0 | 1 | 0 | 0 | 98 | 49 | 1 | 1 | 0.5 | NA | NA | NA | 1 | -0.5 |
| SE | 16260 | ARAP2 | ARAP2 | chr4 | - | 36177826 | 36178005 | 36166931 | 36167047 | 36187450 | 36187571 | 10 | 3 | 1 | 1 | 0 | 0 | 1 | 2 | 0 | 0 | 98 | 49 | 1 | 1 | 0.833 | 1 | 1 | 1 | 1 | -0.056 |
| SE | 16261 | ARAP2 | ARAP2 | chr4 | - | 36212395 | 36212487 | 36210389 | 36210743 | 36213242 | 36213319 | 16 | 6 | 15 | 0 | 0 | 0 | 0 | 3 | 0 | 2 | 98 | 49 | 1 | 1 | 1 | 1 | 1 | NA | 0.429 | 0.571 |
| SE | 16262 | PGM2 | PGM2 | chr4 | + | 37844363 | 37844553 | 37840065 | 37840259 | 37845632 | 37845730 | 14 | 3 | 1 | 1 | 4 | 0 | 0 | 6 | 0 | 0 | 98 | 49 | 1 | 1 | 0.875 | 0.273 | 1 | NA | 1 | -0.284 |
| SE | 16263 | PGM2 | PGM2 | chr4 | + | 37848521 | 37848651 | 37847201 | 37847295 | 37850183 | 37850373 | 12 | 21 | 7 | 1 | 0 | 0 | 2 | 3 | 0 | 0 | 98 | 49 | 1 | 1 | 0.857 | 1 | 1 | 1 | 1 | -0.048 |
| SE | 16264 | PGM2 | PGM2 | chr4 | + | 37850183 | 37850373 | 37848521 | 37848651 | 37855607 | 37855741 | 28 | 13 | 6 | 1 | 0 | 0 | 2 | 6 | 0 | 0 | 98 | 49 | 1 | 1 | 0.933 | 1 | 1 | 1 | 1 | -0.022 |
| SE | 16265 | TBC1D1 | TBC1D1 | chr4 | + | 38118032 | 38118192 | 38115709 | 38115954 | 38124961 | 38125131 | 18 | 6 | 4 | 0 | 0 | 1 | 4 | 10 | 0 | 0 | 98 | 49 | 1 | 1 | 1 | 1 | 0.667 | 1 | 1 | -0.111 |
| SE | 16275 | RIPK1 | RIPK1 | chr6 | + | 3085258 | 3085408 | 3083084 | 3083313 | 3089580 | 3089657 | 15 | 5 | 9 | 0 | 0 | 0 | 0 | 0 | 0 | 1 | 98 | 49 | 1 | 1 | 1 | 1 | 1 | NA | 0 | 1 |
| SE | 16283 | SPTAN1 | SPTAN1 | chr9 | + | 128609650 | 128609665 | 128609121 | 128609284 | 128611713 | 128611845 | 2 | 2 | 0 | 4 | 1 | 0 | 0 | 0 | 0 | 2 | 64 | 49 | 1 | 1 | 0.277 | 0.605 | NA | NA | 0 | 0.441 |
| SE | 16292 | DDX46 | DDX46 | chr5 | + | 134782944 | 134783065 | 134781920 | 134782086 | 134784365 | 134784541 | 11 | 3 | 3 | 1 | 0 | 0 | 2 | 3 | 0 | 0 | 98 | 49 | 1 | 1 | 0.846 | 1 | 1 | 1 | 1 | -0.051 |
| SE | 16298 | EXOC1 | EXOC1 | chr4 | + | 55871848 | 55871958 | 55871100 | 55871233 | 55877916 | 55878066 | 8 | 3 | 4 | 0 | 0 | 0 | 0 | 2 | 0 | 2 | 98 | 49 | 1 | 1 | 1 | 1 | 1 | NA | 0.333 | 0.667 |
| SE | 16300 | EXOC1 | EXOC1 | chr4 | + | 55891314 | 55891422 | 55890222 | 55890386 | 55892634 | 55892711 | 16 | 5 | 2 | 1 | 0 | 0 | 0 | 3 | 0 | 0 | 98 | 49 | 1 | 1 | 0.889 | 1 | 1 | NA | 1 | -0.037 |
| SE | 16301 | EXOC1 | EXOC1 | chr4 | + | 55893551 | 55893780 | 55892634 | 55892711 | 55896716 | 55896900 | 9 | 4 | 3 | 0 | 0 | 0 | 0 | 7 | 0 | 1 | 98 | 49 | 1 | 1 | 1 | 1 | 1 | NA | 0.778 | 0.222 |
| SE | 16311 | YTHDC1 | YTHDC1 | chr4 | - | 68330201 | 68330310 | 68330001 | 68330119 | 68332102 | 68332197 | 27 | 18 | 4 | 0 | 1 | 0 | 0 | 1 | 0 | 0 | 98 | 49 | 1 | 1 | 1 | 0.9 | 1 | NA | 1 | -0.033 |
| SE | 16312 | YTHDC1 | YTHDC1 | chr4 | - | 68332793 | 68332847 | 68332102 | 68332197 | 68333307 | 68333397 | 39 | 26 | 3 | 3 | 0 | 0 | 0 | 2 | 0 | 0 | 98 | 49 | 1 | 1 | 0.867 | 1 | 1 | NA | 1 | -0.044 |
| SE | 16314 | PARN | PARN | chr16 | - | 14627105 | 14627187 | 14617589 | 14617650 | 14628171 | 14628251 | 8 | 4 | 0 | 3 | 0 | 0 | 0 | 1 | 0 | 0 | 98 | 49 | 1 | 1 | 0.571 | 1 | NA | NA | 1 | -0.215 |
| SE | 16315 | PARN | PARN | chr16 | - | 14627268 | 14627336 | 14627105 | 14627187 | 14628171 | 14628251 | 8 | 9 | 2 | 0 | 0 | 0 | 0 | 4 | 0 | 1 | 98 | 49 | 1 | 1 | 1 | 1 | 1 | NA | 0.667 | 0.333 |
| SE | 16333 | SCARB2 | SCARB2 | chr4 | - | 76163224 | 76163383 | 76158738 | 76161751 | 76166249 | 76166301 | 3 | 0 | 0 | 1 | 0 | 0 | 0 | 2 | 0 | 0 | 98 | 49 | 1 | 1 | 0.6 | NA | NA | NA | 1 | -0.4 |
| SE | 16334 | SCARB2 | SCARB2 | chr4 | - | 76179516 | 76179705 | 76176436 | 76176528 | 76180953 | 76181101 | 1 | 0 | 0 | 0 | 1 | 0 | 0 | 8 | 0 | 0 | 98 | 49 | 1 | 1 | 1 | 0 | NA | NA | 1 | -0.5 |
| SE | 16341 | BCAP31 | BCAP31 | chrX | - | 153702934 | 153703058 | 153702006 | 153702107 | 153703958 | 153704094 | 37 | 21 | 11 | 0 | 0 | 0 | 0 | 11 | 0 | 1 | 98 | 49 | 1 | 1 | 1 | 1 | 1 | NA | 0.846 | 0.154 |
| SE | 16343 | BMP2K | BMP2K | chr4 | + | 78847187 | 78847269 | 78844927 | 78845049 | 78850923 | 78851056 | 15 | 5 | 5 | 1 | 0 | 0 | 0 | 8 | 0 | 0 | 98 | 49 | 1 | 1 | 0.882 | 1 | 1 | NA | 1 | -0.039 |
| SE | 16344 | BMP2K | BMP2K | chr4 | + | 78872613 | 78872798 | 78871849 | 78871948 | 78878733 | 78878891 | 19 | 3 | 4 | 1 | 0 | 0 | 0 | 5 | 0 | 0 | 98 | 49 | 1 | 1 | 0.905 | 1 | 1 | NA | 1 | -0.032 |

| | | | | | | | | | | | | | | | | | | | | | | | | | | | | | | |
|---|---|---|---|---|---|---|---|---|---|---|---|---|---|---|---|---|---|---|---|---|---|---|---|---|---|---|---|---|---|---|
| SE | 16345 | BMP2K | BMP2K | chr4 | + | 78887173 | 78887284 | 78878733 | 78878891 | 78910609 | 78912187 | 8 | 0 | 1 | 1 | 0 | 0 | 0 | 1 | 0 | 0 | 98 | 49 | 1 | 1 | 0.8 | NA | 1 | NA | 1 | -0.1 |
| SE | 16346 | SNX10 | SNX10 | chr7 | + | 26357006 | 26357127 | 26346419 | 26346466 | 26360974 | 26361061 | 2 | 0 | 0 | 9 | 3 | 5 | 0 | 3 | 0 | 5 | 98 | 49 | 1 | 1 | 0.1 | 0 | 0 | NA | 0.231 | -0.198 |
| SE | 16347 | TRABD | TRABD | chr22 | + | 50197240 | 50197351 | 50194899 | 50195040 | 50197448 | 50197588 | 39 | 12 | 10 | 0 | 1 | 0 | 0 | 3 | 0 | 0 | 98 | 49 | 1 | 1 | 1 | 0.857 | 1 | NA | 1 | -0.048 |
| SE | 16348 | ANTXR2 | ANTXR2 | chr4 | - | 80033471 | 80033570 | 80031622 | 80031692 | 80054271 | 80054352 | 46 | 13 | 12 | 0 | 0 | 0 | 0 | 4 | 0 | 1 | 98 | 49 | 1 | 1 | 1 | 1 | 1 | NA | 0.667 | 0.333 |
| SE | 16349 | ANTXR2 | ANTXR2 | chr4 | - | 80035971 | 80036032 | 80033471 | 80033570 | 80054271 | 80054352 | 65 | 9 | 11 | 1 | 0 | 0 | 0 | 6 | 0 | 0 | 98 | 49 | 1 | 1 | 0.97 | 1 | 1 | NA | 1 | -0.01 |
| SE | 16351 | HNRNPD | HNRNPD | chr4 | - | 82356536 | 82356683 | 82355303 | 82355401 | 82356795 | 82356895 | 10 | 11 | 0 | 6 | 6 | 2 | 0 | 1 | 0 | 0 | 98 | 49 | 1 | 1 | 0.455 | 0.478 | 0 | NA | 1 | -0.689 |
| SE | 16356 | SEC31A | SEC31A | chr4 | - | 82848803 | 82848977 | 82844385 | 82844509 | 82851430 | 82851604 | 14 | 3 | 2 | 0 | 0 | 0 | 0 | 3 | 0 | 1 | 98 | 49 | 1 | 1 | 1 | 1 | 1 | NA | 0.6 | 0.4 |
| SE | 16361 | THAP9-AS1 | THAP9-AS1 | chr4 | - | 82899712 | 82899737 | 82897988 | 82898062 | 82900076 | 82900223 | 11 | 7 | 0 | 3 | 0 | 0 | 0 | 3 | 0 | 0 | 74 | 49 | 1 | 1 | 0.708 | 1 | NA | NA | 1 | -0.146 |
| SE | 16362 | EDEM1 | EDEM1 | chr3 | + | 5205066 | 5205241 | 5202965 | 5203149 | 5207152 | 5207273 | 11 | 11 | 0 | 1 | 0 | 0 | 0 | 2 | 0 | 0 | 98 | 49 | 1 | 1 | 0.846 | 1 | NA | NA | 1 | -0.077 |
| SE | 16363 | EDEM1 | EDEM1 | chr3 | + | 5210174 | 5210248 | 5208092 | 5208263 | 5211119 | 5211216 | 10 | 14 | 0 | 4 | 0 | 0 | 0 | 3 | 0 | 0 | 98 | 49 | 1 | 1 | 0.556 | 1 | NA | NA | 1 | -0.222 |
| SE | 16364 | OGT | OGT | chrX | + | 71544566 | 71544635 | 71537828 | 71538072 | 71547906 | 71548023 | 8 | 4 | 3 | 0 | 0 | 4 | 0 | 3 | 0 | 0 | 98 | 49 | 1 | 1 | 1 | 1 | 0.273 | NA | 1 | -0.242 |
| SE | 16365 | OGT | OGT | chrX | + | 71555189 | 71555385 | 71554512 | 71554592 | 71555953 | 71556094 | 32 | 7 | 4 | 8 | 7 | 0 | 0 | 2 | 0 | 0 | 98 | 49 | 1 | 1 | 0.667 | 0.333 | 1 | NA | 1 | -0.333 |
| SE | 16374 | PDLIM7 | PDLIM7 | chr5 | - | 177489392 | 177489627 | 177488067 | 177488248 | 177489770 | 177489832 | 37 | 17 | 12 | 0 | 0 | 0 | 3 | 11 | 0 | 1 | 98 | 49 | 1 | 1 | 1 | 1 | 1 | 1 | 0.846 | 0.077 |
| SE | 16375 | PDLIM7 | PDLIM7 | chr5 | - | 177491403 | 177491420 | 177491009 | 177491146 | 177491806 | 177491925 | 3 | 0 | 0 | 8 | 5 | 6 | 0 | 0 | 0 | 4 | 66 | 49 | 1 | 1 | 0.218 | 0 | 0 | NA | 0 | 0.073 |
| SE | 16390 | HERC5 | HERC5 | chr4 | + | 88463854 | 88463985 | 88463531 | 88463623 | 88467058 | 88467204 | 18 | 20 | 0 | 1 | 0 | 0 | 2 | 27 | 0 | 1 | 98 | 49 | 1 | 1 | 0.9 | 1 | NA | 1 | 0.931 | -0.016 |
| SE | 16391 | HERC5 | HERC5 | chr4 | + | 88470613 | 88470673 | 88469156 | 88469260 | 88472408 | 88472502 | 13 | 26 | 2 | 0 | 2 | 0 | 2 | 23 | 0 | 1 | 98 | 49 | 1 | 1 | 1 | 0.867 | 1 | 1 | 0.92 | -0.004 |
| SE | 16392 | HERC5 | HERC5 | chr4 | + | 88472408 | 88472502 | 88470613 | 88470673 | 88475840 | 88476030 | 11 | 17 | 2 | 1 | 0 | 0 | 2 | 29 | 0 | 0 | 98 | 49 | 1 | 1 | 0.846 | 1 | 1 | 1 | 1 | -0.051 |
| SE | 16393 | HERC5 | HERC5 | chr4 | + | 88486114 | 88486228 | 88479352 | 88479507 | 88489165 | 88489336 | 6 | 21 | 0 | 1 | 0 | 0 | 0 | 8 | 0 | 1 | 98 | 49 | 1 | 1 | 0.75 | 1 | NA | NA | 0.8 | 0.075 |
| SE | 16394 | HERC5 | HERC5 | chr4 | + | 88487068 | 88487179 | 88486114 | 88486228 | 88489165 | 88489336 | 3 | 17 | 1 | 0 | 0 | 0 | 1 | 19 | 0 | 1 | 98 | 49 | 1 | 1 | 1 | 1 | 1 | 1 | 0.905 | 0.047 |
| SE | 16395 | HERC5 | HERC5 | chr4 | + | 88489165 | 88489336 | 88479352 | 88479507 | 88493011 | 88493155 | 14 | 15 | 1 | 0 | 1 | 0 | 1 | 15 | 0 | 0 | 98 | 49 | 1 | 1 | 1 | 0.882 | 1 | 1 | 1 | -0.039 |
| SE | 16396 | HERC5 | HERC5 | chr4 | + | 88499925 | 88499992 | 88494164 | 88494331 | 88500914 | 88500985 | 15 | 27 | 0 | 0 | 0 | 0 | 3 | 48 | 1 | 1 | 98 | 49 | 1 | 1 | 1 | 1 | NA | 0.6 | 0.96 | 0.22 |
| SE | 16422 | NFKB1 | NFKB1 | chr4 | + | 102578880 | 102579039 | 102576875 | 102577039 | 102580534 | 102580639 | 15 | 2 | 5 | 1 | 0 | 0 | 0 | 4 | 0 | 0 | 98 | 49 | 1 | 1 | 0.882 | 1 | 1 | NA | 1 | -0.039 |
| SE | 16423 | MANBA | MANBA | chr4 | - | 102635864 | 102636007 | 102634787 | 102635045 | 102639712 | 102639857 | 12 | 7 | 1 | 1 | 0 | 0 | 1 | 2 | 0 | 0 | 98 | 49 | 1 | 1 | 0.857 | 1 | 1 | 1 | 1 | -0.048 |
| SE | 16424 | MANBA | MANBA | chr4 | - | 102650536 | 102650701 | 102639712 | 102639857 | 102657681 | 102657900 | 17 | 2 | 6 | 0 | 0 | 0 | 0 | 4 | 0 | 1 | 98 | 49 | 1 | 1 | 1 | 1 | 1 | NA | 0.667 | 0.333 |
| SE | 16427 | PELI1 | PELI1 | chr2 | - | 64100397 | 64100499 | 64096412 | 64096610 | 64108239 | 64108379 | 129 | 27 | 17 | 0 | 1 | 0 | 2 | 12 | 0 | 0 | 98 | 49 | 1 | 1 | 1 | 0.931 | 1 | 1 | 1 | -0.023 |
| SE | 16428 | PELI1 | PELI1 | chr2 | - | 64104700 | 64104830 | 64100397 | 64100499 | 64108239 | 64108379 | 77 | 46 | 44 | 0 | 2 | 0 | 0 | 13 | 0 | 0 | 98 | 49 | 1 | 1 | 1 | 0.92 | 1 | NA | 1 | -0.027 |
| SE | 16433 | TET2 | TET2 | chr4 | + | 105190359 | 105190505 | 105146684 | 105146979 | 105233896 | 105237351 | 65 | 21 | 15 | 5 | 2 | 0 | 0 | 6 | 0 | 0 | 98 | 49 | 1 | 1 | 0.867 | 0.84 | 1 | NA | 1 | -0.098 |
| SE | 16434 | TET2 | TET2 | chr4 | + | 105241338 | 105241429 | 105233896 | 105237351 | 105242833 | 105242927 | 32 | 8 | 0 | 0 | 0 | 0 | 0 | 6 | 0 | 2 | 98 | 49 | 1 | 1 | 1 | 1 | NA | NA | 0.6 | 0.4 |
| SE | 16435 | TET2 | TET2 | chr4 | + | 105243569 | 105243778 | 105242833 | 105242927 | 105259618 | 105259769 | 26 | 17 | 12 | 7 | 0 | 0 | 0 | 3 | 0 | 0 | 98 | 49 | 1 | 1 | 0.65 | 1 | 1 | NA | 1 | -0.117 |
| SE | 16436 | RAP1GAP2 | RAP1GAP2 | chr17 | + | 2962669 | 2962714 | 2957758 | 2957794 | 2963429 | 2963462 | 8 | 8 | 1 | 10 | 11 | 3 | 0 | 3 | 0 | 6 | 94 | 49 | 1 | 1 | 0.294 | 0.275 | 0.148 | NA | 0.207 | 0.032 |
| SE | 16437 | ZFR | ZFR | chr5 | - | 32380074 | 32380172 | 32379114 | 32379210 | 32385507 | 32385649 | 17 | 2 | 1 | 0 | 0 | 0 | 0 | 4 | 0 | 2 | 98 | 49 | 1 | 1 | 1 | 1 | 1 | NA | 0.5 | 0.5 |
| SE | 16443 | CNN2 | CNN2 | chr19 | + | 1032558 | 1032696 | 1032391 | 1032458 | 1036415 | 1036562 | 72 | 35 | 36 | 2 | 0 | 0 | 1 | 17 | 0 | 0 | 98 | 49 | 1 | 1 | 0.947 | 1 | 1 | 1 | 1 | -0.018 |
| SE | 16444 | CNN2 | CNN2 | chr19 | + | 1036066 | 1036246 | 1032558 | 1032696 | 1036415 | 1036562 | 55 | 42 | 41 | 3 | 3 | 1 | 0 | 15 | 0 | 1 | 98 | 49 | 1 | 1 | 0.902 | 0.875 | 0.953 | NA | 0.882 | 0.028 |
| SE | 16454 | KDM5A | KDM5A | chr12 | - | 297040 | 297200 | 295572 | 295793 | 306945 | 307089 | 40 | 16 | 9 | 1 | 0 | 0 | 0 | 4 | 0 | 0 | 98 | 49 | 1 | 1 | 0.952 | 1 | 1 | NA | 1 | -0.016 |
| SE | 16455 | KDM5A | KDM5A | chr12 | - | 306945 | 307089 | 297040 | 297200 | 307453 | 308005 | 43 | 15 | 10 | 0 | 0 | 0 | 0 | 1 | 0 | 1 | 98 | 49 | 1 | 1 | 1 | 1 | 1 | NA | 0.333 | 0.667 |
| SE | 16456 | KDM5A | KDM5A | chr12 | - | 333486 | 333649 | 331818 | 331938 | 334240 | 334422 | 46 | 23 | 6 | 0 | 1 | 0 | 0 | 8 | 0 | 0 | 98 | 49 | 1 | 1 | 1 | 0.92 | 1 | NA | 1 | -0.027 |
| SE | 16457 | KDM5A | KDM5A | chr12 | - | 354075 | 354234 | 352204 | 352324 | 355157 | 355249 | 41 | 12 | 13 | 1 | 0 | 0 | 3 | 7 | 0 | 0 | 98 | 49 | 1 | 1 | 0.953 | 1 | 1 | 1 | 1 | -0.016 |
| SE | 16462 | LOC645513 | LOC645513 | chr4 | + | 119497810 | 119497903 | 119494432 | 119494523 | 119499560 | 119499592 | 9 | 0 | 6 | 2 | 0 | 0 | 0 | 1 | 0 | 0 | 98 | 49 | 1 | 1 | 0.692 | NA | 1 | NA | 1 | -0.154 |
| SE | 16470 | JADE1 | JADE1 | chr4 | + | 128867855 | 128867973 | 128861703 | 128862225 | 128871354 | 128875224 | 5 | 0 | 8 | 1 | 0 | 0 | 0 | 2 | 0 | 0 | 98 | 49 | 1 | 1 | 0.714 | NA | 1 | NA | 1 | -0.143 |
| SE | 16471 | SCLT1 | SCLT1 | chr4 | - | 128942995 | 128943188 | 128936654 | 128936851 | 128946006 | 128946152 | 11 | 9 | 3 | 4 | 1 | 0 | 0 | 0 | 0 | 2 | 98 | 49 | 1 | 1 | 0.579 | 0.818 | 1 | NA | 0 | 0.799 |
| SE | 16472 | SCLT1 | SCLT1 | chr4 | - | 128965226 | 128965318 | 128959599 | 128959777 | 128970377 | 128970468 | 20 | 15 | 6 | 3 | 3 | 3 | 0 | 3 | 0 | 1 | 98 | 49 | 1 | 1 | 0.769 | 0.714 | 0.5 | NA | 0.6 | 0.061 |
| SE | 16476 | ELF2 | ELF2 | chr4 | - | 139061864 | 139062057 | 139060323 | 139060674 | 139067683 | 139067770 | 52 | 21 | 3 | 2 | 0 | 0 | 0 | 7 | 0 | 0 | 98 | 49 | 1 | 1 | 0.929 | 1 | 1 | NA | 1 | -0.024 |
| SE | 16478 | ELF2 | ELF2 | chr4 | - | 139137629 | 139137867 | 139125163 | 139125329 | 139139412 | 139139476 | 63 | 35 | 7 | 3 | 0 | 0 | 0 | 10 | 0 | 0 | 98 | 49 | 1 | 1 | 0.913 | 1 | 1 | NA | 1 | -0.029 |
| SE | 16482 | IGF1R | IGF1R | chr15 | + | 98891324 | 98891637 | 98707561 | 98708107 | 98896756 | 98896905 | 32 | 7 | 1 | 1 | 0 | 0 | 0 | 1 | 0 | 0 | 98 | 49 | 1 | 1 | 0.941 | 1 | 1 | NA | 1 | -0.02 |
| SE | 16484 | IGF1R | IGF1R | chr15 | + | 98934823 | 98935053 | 98930234 | 98930305 | 98935315 | 98935426 | 25 | 11 | 3 | 1 | 1 | 0 | 0 | 2 | 0 | 0 | 98 | 49 | 1 | 1 | 0.926 | 0.846 | 1 | NA | 1 | -0.076 |
| SE | 16492 | GAB1 | GAB1 | chr4 | + | 143457679 | 143457769 | 143440078 | 143440382 | 143459384 | 143459478 | 1 | 0 | 0 | 2 | 0 | 0 | 0 | 0 | 0 | 3 | 98 | 49 | 1 | 1 | 0.2 | NA | NA | NA | 0 | 0.2 |
| SE | 16509 | FBXW7 | FBXW7 | chr4 | - | 152412479 | 152412529 | 152411159 | 152411872 | 152534940 | 152535033 | 17 | 8 | 0 | 3 | 0 | 0 | 0 | 0 | 2 | 0 | 98 | 49 | 1 | 1 | 0.739 | 1 | NA | 0 | NA | 0.869 |
| SE | 16510 | CIDEB | CIDEB | chr14 | - | 24305946 | 24306137 | 24305186 | 24305765 | 24306373 | 24306523 | 4 | 0 | 2 | 0 | 0 | 0 | 0 | 3 | 0 | 2 | 98 | 49 | 1 | 1 | 1 | NA | 1 | NA | 0.429 | 0.571 |
| SE | 16515 | KIAA0922 | KIAA0922 | chr4 | + | 153587741 | 153587811 | 153586208 | 153586379 | 153588889 | 153589007 | 22 | 4 | 3 | 0 | 2 | 0 | 0 | 3 | 0 | 2 | 98 | 49 | 1 | 1 | 1 | 0.5 | 1 | NA | 0.429 | 0.404 |
| SE | 16516 | KIAA0922 | KIAA0922 | chr4 | + | 153588889 | 153589007 | 153586208 | 153586379 | 153593798 | 153593871 | 2 | 2 | 0 | 1 | 0 | 0 | 0 | 2 | 0 | 0 | 98 | 49 | 1 | 1 | 0.5 | 1 | NA | NA | 1 | -0.25 |
| SE | 16517 | KIAA0922 | KIAA0922 | chr4 | + | 153591052 | 153591194 | 153588889 | 153589007 | 153592474 | 153592584 | 30 | 15 | 2 | 1 | 2 | 0 | 0 | 6 | 0 | 0 | 98 | 49 | 1 | 1 | 0.937 | 0.789 | 1 | NA | 1 | -0.091 |
| SE | 16518 | KIAA0922 | KIAA0922 | chr4 | + | 153591052 | 153591194 | 153588889 | 153589007 | 153593798 | 153593871 | 19 | 14 | 1 | 2 | 0 | 0 | 0 | 3 | 0 | 0 | 98 | 49 | 1 | 1 | 0.826 | 1 | 1 | NA | 1 | -0.058 |
| SE | 16519 | KIAA0922 | KIAA0922 | chr4 | + | 153592474 | 153592584 | 153588889 | 153589007 | 153593798 | 153593871 | 9 | 13 | 0 | 2 | 0 | 0 | 0 | 4 | 0 | 0 | 98 | 49 | 1 | 1 | 0.692 | 1 | NA | NA | 1 | -0.154 |
| SE | 16520 | KIAA0922 | KIAA0922 | chr4 | + | 153592474 | 153592584 | 153591052 | 153591194 | 153593798 | 153593871 | 23 | 12 | 1 | 4 | 0 | 0 | 0 | 7 | 0 | 0 | 98 | 49 | 1 | 1 | 0.742 | 1 | 1 | NA | 1 | -0.086 |
| SE | 16521 | KIAA0922 | KIAA0922 | chr4 | + | 153632717 | 153632838 | 153627604 | 153627687 | 153634191 | 153634280 | 11 | 3 | 8 | 1 | 0 | 0 | 0 | 1 | 0 | 0 | 98 | 49 | 1 | 1 | 0.846 | 1 | 1 | NA | 1 | -0.051 |
| SE | 16522 | PLRG1 | PLRG1 | chr4 | - | 154537285 | 154537479 | 154534996 | 154536744 | 154537968 | 154538108 | 5 | 4 | 0 | 1 | 0 | 0 | 0 | 2 | 0 | 0 | 98 | 49 | 1 | 1 | 0.714 | 1 | NA | NA | 1 | -0.143 |
| SE | 16574 | LILRA2 | LILRA2 | chr19 | + | 54574075 | 54574111 | 54573697 | 54573912 | 54574300 | 54574582 | 29 | 5 | 8 | 6 | 0 | 0 | 0 | 5 | 0 | 1 | 85 | 49 | 1 | 1 | 0.736 | 1 | 1 | NA | 0.742 | 0.17 |
| SE | 16575 | LILRA2 | LILRA2 | chr19 | + | 54574300 | 54574582 | 54573830 | 54573912 | 54574730 | 54575033 | 54 | 25 | 20 | 1 | 0 | 0 | 1 | 22 | 0 | 0 | 98 | 49 | 1 | 1 | 0.964 | 1 | 1 | 1 | 1 | -0.012 |
| SE | 16576 | LILRA2 | LILRA2 | chr19 | + | 54575255 | 54575552 | 54574730 | 54575033 | 54575806 | 54576109 | 33 | 16 | 16 | 1 | 0 | 0 | 0 | 9 | 0 | 0 | 98 | 49 | 1 | 1 | 0.943 | 1 | 1 | NA | 1 | -0.019 |
| SE | 16577 | LILRA2 | LILRA2 | chr19 | + | 54577464 | 54577723 | 54575806 | 54576109 | 54587009 | 54587060 | 1 | 1 | 3 | 5 | 0 | 2 | 0 | 4 | 0 | 0 | 98 | 49 | 1 | 1 | 0.091 | 1 | 0.429 | NA | 1 | -0.493 |
| SE | 16578 | LILRA2 | LILRA2 | chr19 | + | 54577464 | 54577723 | 54575806 | 54576109 | 54587200 | 54587561 | 5 | 4 | 4 | 12 | 9 | 9 | 0 | 3 | 0 | 3 | 98 | 49 | 1 | 1 | 0.172 | 0.182 | 0.182 | NA | 0.333 | -0.154 |
| SE | 16579 | LILRA2 | LILRA2 | chr19 | + | 54587009 | 54587060 | 54575806 | 54576109 | 54587200 | 54587561 | 11 | 0 | 6 | 12 | 9 | 9 | 0 | 1 | 0 | 3 | 98 | 49 | 1 | 1 | 0.314 | 0 | 0.25 | NA | 0.143 | 0.045 |
| SE | 16580 | LILRA2 | LILRA2 | chr19 | + | 54587009 | 54587060 | 54577464 | 54577723 | 54587200 | 54587561 | 6 | 0 | 4 | 4 | 3 | 1 | 0 | 2 | 0 | 0 | 98 | 49 | 1 | 1 | 0.429 | 0 | 0.667 | NA | 1 | -0.635 |
| SE | 16583 | IRF2 | IRF2 | chr4 | - | 184408157 | 184408275 | 184398914 | 184399079 | 184418166 | 184418213 | 47 | 15 | 19 | 1 | 0 | 0 | 0 | 8 | 0 | 0 | 98 | 49 | 1 | 1 | 0.959 | 1 | 1 | NA | 1 | -0.014 |
| SE | 16584 | IRF2 | IRF2 | chr4 | - | 184418166 | 184418213 | 184398914 | 184399079 | 184418531 | 184418708 | 71 | 9 | 27 | 1 | 0 | 0 | 2 | 7 | 0 | 0 | 96 | 49 | 1 | 1 | 0.973 | 1 | 1 | 1 | 1 | -0.009 |
| SE | 16585 | IRF2 | IRF2 | chr4 | - | 184419468 | 184419568 | 184418531 | 184418708 | 184428977 | 184429070 | 48 | 21 | 42 | 1 | 0 | 0 | 0 | 12 | 0 | 0 | 98 | 49 | 1 | 1 | 0.96 | 1 | 1 | NA | 1 | -0.013 |
| SE | 16586 | GPSM3 | GPSM3 | chr6 | - | 32191708 | 32191908 | 32190765 | 32191503 | 32192147 | 32192250 | 95 | 38 | 58 | 1 | 0 | 0 | 6 | 21 | 0 | 0 | 98 | 49 | 1 | 1 | 0.979 | 1 | 1 | 1 | 1 | -0.007 |
| SE | 16587 | GPSM3 | GPSM3 | chr6 | - | 32192147 | 32192250 | 32191708 | 32191908 | 32192471 | 32192530 | 64 | 29 | 41 | 0 | 1 | 0 | 4 | 8 | 0 | 0 | 98 | 49 | 1 | 1 | 1 | 0.935 | 1 | 1 | 1 | -0.022 |
| SE | 16608 | EXOC3 | EXOC3 | chr5 | + | 456888 | 457006 | 453369 | 454051 | 457899 | 458025 | 12 | 5 | 4 | 1 | 0 | 0 | 0 | 7 | 0 | 0 | 98 | 49 | 1 | 1 | 0.857 | 1 | 1 | NA | 1 | -0.048 |
| SE | 16612 | CPEB4 | CPEB4 | chr5 | + | 173932449 | 173932500 | 173910522 | 173910604 | 173944966 | 173945140 | 11 | 2 | 9 | 2 | 0 | 1 | 0 | 5 | 0 | 0 | 98 | 49 | 1 | 1 | 0.733 | 1 | 0.818 | NA | 1 | -0.15 |
| SE | 16615 | CPEB4 | CPEB4 | chr5 | + | 173944966 | 173945140 | 173932449 | 173932500 | 173949507 | 173949597 | 29 | 12 | 11 | 0 | 1 | 0 | 1 | 18 | 0 | 0 | 98 | 49 | 1 | 1 | 1 | 0.857 | 1 | 1 | 1 | -0.048 |
| SE | 16619 | IMPDH1 | IMPDH1 | chr7 | - | 128405766 | 128405865 | 128403705 | 128403754 | 128409755 | 128409982 | 18 | 10 | 5 | 2 | 0 | 0 | 0 | 5 | 0 | 0 | 98 | 49 | 1 | 1 | 0.818 | 1 | 1 | NA | 1 | -0.061 |
| SE | 16620 | IMPDH1 | IMPDH1 | chr7 | - | 128409288 | 128409366 | 128405766 | 128405865 | 128409755 | 128409982 | 2 | 3 | 0 | 12 | 6 | 0 | 0 | 0 | 0 | 4 | 98 | 49 | 1 | 1 | 0.077 | 0.2 | NA | NA | 0 | 0.139 |
| SE | 16625 | RTN4 | RTN4 | chr2 | - | 54973820 | 54973867 | 54973562 | 54973621 | 54974694 | 54974764 | 38 | 8 | 8 | 0 | 3 | 0 | 0 | 9 | 0 | 0 | 96 | 49 | 1 | 1 | 1 | 0.576 | 1 | NA | 1 | -0.141 |
| SE | 16629 | MPP1 | MPP1 | chrX | - | 154784027 | 154784108 | 154783426 | 154783507 | 154785050 | 154785157 | 32 | 10 | 10 | 1 | 0 | 0 | 0 | 10 | 0 | 0 | 98 | 49 | 1 | 1 | 0.941 | 1 | 1 | NA | 1 | -0.02 |
| SE | 16630 | MPP1 | MPP1 | chrX | - | 154799744 | 154799893 | 154792141 | 154792285 | 154805271 | 154805527 | 0 | 0 | 0 | 10 | 5 | 5 | 0 | 1 | 0 | 6 | 98 | 49 | 1 | 1 | 0 | 0 | 0 | NA | 0.077 | -0.077 |
| SE | 16632 | HNRNPM | HNRNPM | chr19 | + | 8483157 | 8483211 | 8474166 | 8474244 | 8485602 | 8486405 | 9 | 3 | 0 | 2 | 0 | 0 | 0 | 2 | 0 | 0 | 98 | 49 | 1 | 1 | 0.692 | 1 | NA | NA | 1 | -0.154 |
| SE | 16644 | ITM2A | ITM2A | chrX | - | 79363422 | 79363554 | 79362941 | 79363139 | 79367104 | 79367552 | 6 | 0 | 0 | 0 | 1 | 0 | 0 | 2 | 0 | 0 | 98 | 49 | 1 | 1 | 1 | 0 | NA | NA | 1 | -0.5 |
| SE | 16650 | FAM105A | FAM105A | chr5 | + | 14601350 | 14601442 | 14601212 | 14601244 | 14602182 | 14602332 | 18 | 4 | 0 | 1 | 0 | 0 | 2 | 3 | 0 | 2 | 98 | 49 | 1 | 1 | 0.9 | 1 | NA | 1 | 0.429 | 0.235 |
| SE | 16651 | OTULIN | OTULIN | chr5 | + | 14673641 | 14673718 | 14664673 | 14664977 | 14678680 | 14678775 | 16 | 5 | 1 | 1 | 0 | 0 | 1 | 8 | 0 | 0 | 98 | 49 | 1 | 1 | 0.889 | 1 | 1 | 1 | 1 | -0.037 |
| SE | 16665 | PCBP1-AS1 | PCBP1-AS1 | chr2 | - | 70083537 | 70083687 | 70059642 | 70059680 | 70085546 | 70085624 | 6 | 0 | 2 | 2 | 3 | 0 | 0 | 1 | 0 | 0 | 98 | 49 | 1 | 1 | 0.6 | 0 | 1 | NA | 1 | -0.467 |
| SE | 16669 | ZNF335 | ZNF335 | chr20 | - | 45969450 | 45969691 | 45968284 | 45968362 | 45971209 | 45971460 | 9 | 2 | 1 | 0 | 0 | 0 | 0 | 0 | 0 | 1 | 98 | 49 | 1 | 1 | 1 | 1 | 1 | NA | 0 | 1 |
| SE | 16685 | NUP155 | NUP155 | chr5 | - | 37307296 | 37307432 | 37305056 | 37305210 | 37309128 | 37309267 | 4 | 0 | 0 | 0 | 1 | 0 | 0 | 1 | 0 | 1 | 98 | 49 | 1 | 1 | 1 | 0 | NA | NA | 0.333 | 0.167 |
| SE | 16687 | CFAP54 | CFAP54 | chr12 | + | 96535514 | 96535600 | 96534061 | 96534227 | 96538383 | 96538518 | 2 | 1 | 0 | 1 | 0 | 0 | 0 | 4 | 0 | 0 | 98 | 49 | 1 | 1 | 0.5 | 1 | NA | NA | 1 | -0.25 |
| SE | 16689 | RICTOR | RICTOR | chr5 | - | 38944445 | 38944569 | 38942832 | 38942971 | 38944912 | 38945068 | 105 | 34 | 10 | 1 | 1 | 0 | 0 | 15 | 0 | 0 | 98 | 49 | 1 | 1 | 0.981 | 0.944 | 1 | NA | 1 | -0.025 |
| SE | 16690 | RICTOR | RICTOR | chr5 | - | 38946467 | 38946552 | 38945490 | 38945610 | 38947263 | 38947441 | 70 | 29 | 10 | 0 | 2 | 0 | 0 | 9 | 0 | 0 | 98 | 49 | 1 | 1 | 1 | 0.879 | 1 | NA | 1 | -0.04 |
| SE | 16691 | RICTOR | RICTOR | chr5 | - | 38946467 | 38946552 | 38945490 | 38945724 | 38947263 | 38947441 | 71 | 26 | 13 | 3 | 0 | 0 | 0 | 9 | 0 | 0 | 98 | 49 | 1 | 1 | 0.922 | 1 | 1 | NA | 1 | -0.026 |
| SE | 16692 | RICTOR | RICTOR | chr5 | - | 38949355 | 38949427 | 38947263 | 38947441 | 38949711 | 38950720 | 18 | 20 | 2 | 37 | 23 | 1 | 0 | 0 | 0 | 11 | 98 | 49 | 1 | 1 | 0.196 | 0.303 | 0.5 | NA | 0 | 0.333 |
| SE | 16693 | RICTOR | RICTOR | chr5 | - | 38975536 | 38975604 | 38971876 | 38971959 | 38978582 | 38978648 | 133 | 60 | 7 | 0 | 0 | 0 | 0 | 12 | 0 | 1 | 98 | 49 | 1 | 1 | 1 | 1 | 1 | NA | 0.857 | 0.143 |
| SE | 16694 | RICTOR | RICTOR | chr5 | - | 38978582 | 38978648 | 38975536 | 38975604 | 38981866 | 38982036 | 100 | 32 | 6 | 0 | 1 | 0 | 0 | 7 | 0 | 0 | 98 | 49 | 1 | 1 | 1 | 0.941 | 1 | NA | 1 | -0.02 |
| SE | 16695 | RICTOR | RICTOR | chr5 | - | 38978582 | 38978650 | 38975536 | 38975604 | 38981866 | 38982036 | 155 | 57 | 25 | 0 | 1 | 0 | 0 | 18 | 0 | 0 | 98 | 49 | 1 | 1 | 1 | 0.966 | 1 | NA | 1 | -0.011 |
| SE | 16696 | RICTOR | RICTOR | chr5 | - | 39074110 | 39074158 | 39021038 | 39021136 | 39074328 | 39074408 | 21 | 14 | 0 | 1 | 0 | 0 | 1 | 0 | 0 | 0 | 97 | 49 | 1 | 1 | 0.914 | 1 | NA | 1 | NA | -0.043 |
| SE | 16697 | FYB | FYB | chr5 | - | 39125997 | 39126135 | 39124252 | 39124278 | 39127740 | 39127807 | 145 | 61 | 23 | 11 | 12 | 8 | 0 | 17 | 0 | 9 | 98 | 49 | 1 | 1 | 0.868 | 0.718 | 0.59 | NA | 0.486 | 0.239 |
| SE | 16698 | FYB | FYB | chr5 | - | 39127740 | 39127807 | 39125997 | 39126135 | 39130589 | 39130612 | 167 | 59 | 24 | 1 | 0 | 0 | 0 | 25 | 0 | 0 | 98 | 49 | 1 | 1 | 0.988 | 1 | 1 | NA | 1 | -0.004 |
| SE | 16699 | FYB | FYB | chr5 | - | 39137599 | 39137720 | 39134854 | 39135014 | 39138656 | 39138691 | 105 | 57 | 24 | 2 | 0 | 0 | 0 | 20 | 0 | 0 | 98 | 49 | 1 | 1 | 0.963 | 1 | 1 | NA | 1 | -0.012 |
| SE | 16700 | FYB | FYB | chr5 | - | 39153447 | 39153604 | 39141094 | 39141141 | 39201825 | 39202987 | 189 | 65 | 25 | 1 | 0 | 0 | 7 | 35 | 0 | 0 | 98 | 49 | 1 | 1 | 0.99 | 1 | 1 | 1 | 1 | -0.003 |
| SE | 16701 | FYB | FYB | chr5 | - | 39201825 | 39202987 | 39153447 | 39153604 | 39219442 | 39219578 | 151 | 41 | 15 | 1 | 2 | 0 | 8 | 22 | 0 | 0 | 98 | 49 | 1 | 1 | 0.987 | 0.911 | 1 | 1 | 1 | -0.034 |
| SE | 16702 | MICAL1 | MICAL1 | chr6 | - | 109445196 | 109445290 | 109444895 | 109444995 | 109445415 | 109445529 | 30 | 16 | 9 | 2 | 0 | 0 | 1 | 9 | 0 | 0 | 98 | 49 | 1 | 1 | 0.882 | 1 | 1 | 1 | 1 | -0.039 |
| SE | 16703 | MICAL1 | MICAL1 | chr6 | - | 109447680 | 109447722 | 109447356 | 109447440 | 109448202 | 109448393 | 5 | 4 | 3 | 1 | 0 | 0 | 2 | 2 | 0 | 0 | 91 | 49 | 1 | 1 | 0.729 | 1 | 1 | 1 | 1 | -0.09 |
| SE | 16705 | MICAL1 | MICAL1 | chr6 | - | 109450299 | 109450557 | 109449969 | 109450085 | 109451599 | 109451700 | 6 | 9 | 1 | 1 | 0 | 0 | 2 | 4 | 0 | 1 | 98 | 49 | 1 | 1 | 0.75 | 1 | 1 | 1 | 0.667 | 0.083 |
| SE | 16716 | HMGCS1 | HMGCS1 | chr5 | - | 43298008 | 43298134 | 43297001 | 43297166 | 43298517 | 43298975 | 3 | 0 | 2 | 0 | 1 | 0 | 0 | 2 | 0 | 0 | 98 | 49 | 1 | 1 | 1 | 0 | 1 | NA | 1 | -0.333 |
| SE | 16725 | SKIV2L | SKIV2L | chr6 | + | 31960431 | 31960546 | 31960219 | 31960337 | 31960665 | 31960740 | 17 | 9 | 2 | 1 | 0 | 2 | 0 | 1 | 0 | 0 | 98 | 49 | 1 | 1 | 0.895 | 1 | 0.333 | NA | 1 | -0.257 |
| SE | 16726 | SKIV2L | SKIV2L | chr6 | + | 31961195 | 31961386 | 31961040 | 31961094 | 31961546 | 31961675 | 15 | 7 | 5 | 0 | 3 | 0 | 0 | 1 | 0 | 1 | 98 | 49 | 1 | 1 | 1 | 0.538 | 1 | NA | 0.333 | 0.513 |
| SE | 16727 | SKIV2L | SKIV2L | chr6 | + | 31961908 | 31962054 | 31961546 | 31961675 | 31962438 | 31962585 | 16 | 9 | 0 | 0 | 1 | 0 | 1 | 3 | 0 | 0 | 98 | 49 | 1 | 1 | 1 | 0.818 | NA | 1 | 1 | -0.091 |
| SE | 16728 | SKIV2L | SKIV2L | chr6 | + | 31967003 | 31967141 | 31966708 | 31966846 | 31967272 | 31967377 | 22 | 4 | 1 | 1 | 0 | 0 | 0 | 4 | 0 | 0 | 98 | 49 | 1 | 1 | 0.917 | 1 | 1 | NA | 1 | -0.028 |
| SE | 16730 | FNBP1 | FNBP1 | chr9 | - | 129923843 | 129924026 | 129908889 | 129908999 | 129924959 | 129925157 | 1 | 1 | 0 | 18 | 1 | 0 | 0 | 0 | 0 | 1 | 98 | 49 | 1 | 1 | 0.027 | 0.333 | NA | NA | 0 | 0.18 |
| SE | 16731 | FNBP1 | FNBP1 | chr9 | - | 129958490 | 129958553 | 129957359 | 129957464 | 129978464 | 129978612 | 73 | 11 | 13 | 2 | 0 | 0 | 1 | 8 | 0 | 0 | 98 | 49 | 1 | 1 | 0.948 | 1 | 1 | 1 | 1 | -0.017 |
| SE | 16734 | IL6ST | IL6ST | chr5 | - | 55959625 | 55959671 | 55957208 | 55957291 | 55960401 | 55960561 | 2 | 2 | 0 | 0 | 0 | 1 | 0 | 0 | 0 | 5 | 95 | 49 | 1 | 1 | 1 | 1 | 0 | NA | 0 | 0.667 |
| SE | 16736 | SON | SON | chr21 | + | 33572550 | 33572630 | 33568970 | 33569087 | 33573307 | 33573455 | 5 | 0 | 0 | 8 | 0 | 5 | 0 | 0 | 0 | 1 | 98 | 49 | 1 | 1 | 0.238 | NA | 0 | NA | 0 | 0.119 |
| SE | 16740 | C14orf159 | C14orf159 | chr14 | + | 91145220 | 91145313 | 91114012 | 91114283 | 91147780 | 91147858 | 26 | 4 | 0 | 1 | 0 | 0 | 0 | 1 | 0 | 0 | 98 | 49 | 1 | 1 | 0.929 | 1 | NA | NA | 1 | -0.035 |
| SE | 16741 | C14orf159 | C14orf159 | chr14 | + | 91145220 | 91145313 | 91114012 | 91114283 | 91157638 | 91157690 | 16 | 1 | 2 | 1 | 1 | 0 | 0 | 2 | 0 | 1 | 98 | 49 | 1 | 1 | 0.889 | 0.333 | 1 | NA | 0.5 | 0.241 |
| SE | 16743 | C14orf159 | C14orf159 | chr14 | + | 91145220 | 91145313 | 91114634 | 91114994 | 91157638 | 91157690 | 35 | 8 | 5 | 22 | 12 | 0 | 0 | 4 | 0 | 5 | 98 | 49 | 1 | 1 | 0.443 | 0.25 | 1 | NA | 0.286 | 0.278 |
| SE | 16744 | C14orf159 | C14orf159 | chr14 | + | 91147780 | 91147858 | 91114012 | 91114283 | 91157638 | 91157690 | 29 | 9 | 0 | 1 | 1 | 0 | 0 | 6 | 0 | 1 | 98 | 49 | 1 | 1 | 0.935 | 0.818 | NA | NA | 0.75 | 0.127 |
| SE | 16745 | C14orf159 | C14orf159 | chr14 | + | 91147780 | 91147858 | 91145220 | 91145313 | 91157638 | 91157690 | 54 | 13 | 0 | 16 | 1 | 2 | 0 | 6 | 0 | 1 | 98 | 49 | 1 | 1 | 0.628 | 0.867 | 0 | NA | 0.75 | -0.252 |
| SE | 16747 | C14orf159 | C14orf159 | chr14 | + | 91170002 | 91170186 | 91167224 | 91167378 | 91173288 | 91173403 | 13 | 7 | 6 | 3 | 6 | 1 | 0 | 4 | 0 | 1 | 98 | 49 | 1 | 1 | 0.684 | 0.368 | 0.75 | NA | 0.667 | -0.066 |
| SE | 16748 | C14orf159 | C14orf159 | chr14 | + | 91170002 | 91170201 | 91167224 | 91167378 | 91173288 | 91173439 | 31 | 17 | 9 | 3 | 6 | 1 | 0 | 5 | 0 | 1 | 98 | 49 | 1 | 1 | 0.838 | 0.586 | 0.818 | NA | 0.714 | 0.033 |
| SE | 16749 | C14orf159 | C14orf159 | chr14 | + | 91204705 | 91204825 | 91199756 | 91199905 | 91215404 | 91215556 | 94 | 17 | 19 | 8 | 11 | 0 | 0 | 17 | 0 | 0 | 98 | 49 | 1 | 1 | 0.855 | 0.436 | 1 | NA | 1 | -0.236 |
| SE | 16761 | SLC30A5 | SLC30A5 | chr5 | + | 69128003 | 69128132 | 69123198 | 69123425 | 69129446 | 69131072 | 6 | 0 | 2 | 1 | 0 | 0 | 0 | 1 | 0 | 0 | 98 | 49 | 1 | 1 | 0.75 | NA | 1 | NA | 1 | -0.125 |
| SE | 16778 | PYGM | PYGM | chr11 | - | 64751596 | 64751655 | 64751324 | 64751466 | 64751923 | 64752071 | 3 | 1 | 6 | 1 | 1 | 0 | 0 | 2 | 0 | 0 | 98 | 49 | 1 | 1 | 0.6 | 0.333 | 1 | NA | 1 | -0.356 |
| SE | 16779 | PYGM | PYGM | chr11 | - | 64758245 | 64758349 | 64757778 | 64757910 | 64758436 | 64758515 | 2 | 0 | 0 | 0 | 0 | 0 | 0 | 1 | 0 | 2 | 98 | 49 | 1 | 1 | 1 | NA | NA | NA | 0.2 | 0.8 |
| SE | 16787 | MCCC2 | MCCC2 | chr5 | + | 71649096 | 71649253 | 71646210 | 71646277 | 71650068 | 71650183 | 0 | 2 | 0 | 0 | 1 | 0 | 0 | 3 | 0 | 0 | 98 | 49 | 1 | 1 | NA | 0.5 | NA | NA | 1 | -0.5 |
| SE | 16793 | ANKRA2 | ANKRA2 | chr5 | - | 73554860 | 73554986 | 73554321 | 73554388 | 73555487 | 73555585 | 5 | 3 | 0 | 0 | 1 | 0 | 0 | 0 | 0 | 3 | 98 | 49 | 1 | 1 | 1 | 0.6 | NA | NA | 0 | 0.8 |
| SE | 16795 | GFM2 | GFM2 | chr5 | - | 74730259 | 74730398 | 74725940 | 74726126 | 74733021 | 74733098 | 0 | 0 | 0 | 4 | 0 | 0 | 0 | 1 | 0 | 0 | 98 | 49 | 1 | 1 | 0 | NA | NA | NA | 1 | -1 |
| SE | 16797 | GFM2 | GFM2 | chr5 | - | 74750578 | 74750667 | 74747691 | 74747780 | 74751367 | 74751493 | 9 | 0 | 2 | 1 | 0 | 0 | 0 | 1 | 0 | 0 | 98 | 49 | 1 | 1 | 0.818 | NA | 1 | NA | 1 | -0.091 |
| SE | 16801 | ZFAND5 | ZFAND5 | chr9 | - | 72359417 | 72359521 | 72356930 | 72357056 | 72360109 | 72360221 | 61 | 15 | 4 | 2 | 0 | 0 | 0 | 17 | 0 | 0 | 98 | 49 | 1 | 1 | 0.938 | 1 | 1 | NA | 1 | -0.021 |
| SE | 16802 | ZFAND5 | ZFAND5 | chr9 | - | 72360109 | 72360221 | 72359417 | 72359521 | 72360627 | 72360784 | 20 | 10 | 0 | 1 | 1 | 0 | 0 | 9 | 0 | 0 | 98 | 49 | 1 | 1 | 0.909 | 0.833 | NA | NA | 1 | -0.129 |
| SE | 16806 | POLK | POLK | chr5 | + | 75593877 | 75594049 | 75590343 | 75590440 | 75596221 | 75597178 | 5 | 0 | 0 | 9 | 0 | 0 | 0 | 2 | 0 | 0 | 98 | 49 | 1 | 1 | 0.217 | NA | NA | NA | 1 | -0.783 |
| SE | 16809 | IQGAP2 | IQGAP2 | chr5 | + | 76683775 | 76683917 | 76683114 | 76683217 | 76693354 | 76693442 | 40 | 9 | 8 | 1 | 2 | 0 | 0 | 6 | 0 | 0 | 98 | 49 | 1 | 1 | 0.952 | 0.692 | 1 | NA | 1 | -0.119 |
| SE | 16812 | ZFYVE16 | ZFYVE16 | chr5 | + | 80427491 | 80427545 | 80408012 | 80408153 | 80434108 | 80434217 | 3 | 5 | 0 | 4 | 0 | 0 | 0 | 2 | 0 | 0 | 98 | 49 | 1 | 1 | 0.273 | 1 | NA | NA | 1 | -0.363 |
| SE | 16813 | ZFYVE16 | ZFYVE16 | chr5 | + | 80448025 | 80448404 | 80445262 | 80445405 | 80449590 | 80449713 | 23 | 13 | 1 | 12 | 9 | 5 | 0 | 8 | 0 | 0 | 98 | 49 | 1 | 1 | 0.489 | 0.419 | 0.091 | NA | 1 | -0.667 |
| SE | 16814 | ZFYVE16 | ZFYVE16 | chr5 | + | 80472760 | 80472923 | 80459413 | 80459494 | 80473753 | 80473859 | 29 | 9 | 12 | 4 | 1 | 0 | 0 | 2 | 0 | 1 | 98 | 49 | 1 | 1 | 0.784 | 0.818 | 1 | NA | 0.5 | 0.367 |
| SE | 16815 | AGTPBP1 | AGTPBP1 | chr9 | - | 85588297 | 85588478 | 85586830 | 85586960 | 85589527 | 85589681 | 55 | 32 | 22 | 4 | 2 | 0 | 0 | 12 | 0 | 1 | 98 | 49 | 1 | 1 | 0.873 | 0.889 | 1 | NA | 0.857 | 0.064 |
| SE | 16817 | AGTPBP1 | AGTPBP1 | chr9 | - | 85692688 | 85692813 | 85681267 | 85681335 | 85741774 | 85742029 | 88 | 35 | 10 | 0 | 0 | 0 | 3 | 9 | 0 | 1 | 98 | 49 | 1 | 1 | 1 | 1 | 1 | 1 | 0.818 | 0.091 |
| SE | 16818 | AGTPBP1 | AGTPBP1 | chr9 | - | 85712501 | 85712566 | 85692688 | 85692813 | 85741774 | 85742029 | 56 | 29 | 17 | 2 | 0 | 0 | 2 | 7 | 0 | 0 | 98 | 49 | 1 | 1 | 0.933 | 1 | 1 | 1 | 1 | -0.022 |
| SE | 16824 | SSBP2 | SSBP2 | chr5 | - | 81489249 | 81489309 | 81474495 | 81474562 | 81513627 | 81513717 | 9 | 2 | 3 | 3 | 0 | 0 | 0 | 0 | 0 | 1 | 98 | 49 | 1 | 1 | 0.6 | 1 | 1 | NA | 0 | 0.867 |
| SE | 16825 | SSBP2 | SSBP2 | chr5 | - | 81513627 | 81513717 | 81489249 | 81489309 | 81615472 | 81615557 | 10 | 7 | 0 | 0 | 0 | 0 | 0 | 4 | 0 | 3 | 98 | 49 | 1 | 1 | 1 | 1 | NA | NA | 0.4 | 0.6 |
| SE | 16826 | TMEM167A | TMEM167A | chr5 | - | 83061876 | 83061911 | 83052845 | 83057154 | 83077320 | 83077453 | 5 | 5 | 1 | 0 | 0 | 0 | 0 | 3 | 0 | 1 | 84 | 49 | 1 | 1 | 1 | 1 | 1 | NA | 0.636 | 0.364 |
| SE | 16827 | TMEM167A | TMEM167A | chr5 | - | 83065007 | 83065117 | 83061876 | 83061911 | 83077320 | 83077453 | 7 | 5 | 0 | 1 | 0 | 0 | 0 | 3 | 0 | 0 | 98 | 49 | 1 | 1 | 0.778 | 1 | NA | NA | 1 | -0.111 |
| SE | 16828 | VCAN | VCAN | chr5 | + | 83519348 | 83522309 | 83512102 | 83512396 | 83537006 | 83542268 | 1 | 0 | 0 | 14 | 0 | 0 | 0 | 1 | 0 | 19 | 98 | 49 | 1 | 1 | 0.034 | NA | NA | NA | 0.026 | 0.008 |
| SE | 16829 | VCAN | VCAN | chr5 | + | 83519348 | 83522309 | 83512102 | 83512396 | 83545536 | 83545650 | 1 | 0 | 0 | 1 | 0 | 0 | 0 | 1 | 0 | 1 | 98 | 49 | 1 | 1 | 0.333 | NA | NA | NA | 0.333 | 0 |
| SE | 16830 | VCAN | VCAN | chr5 | + | 83537006 | 83542268 | 83512102 | 83512396 | 83545536 | 83545650 | 56 | 0 | 0 | 1 | 0 | 0 | 2 | 51 | 0 | 1 | 98 | 49 | 1 | 1 | 0.966 | NA | NA | 1 | 0.962 | -0.015 |
| SE | 16832 | DNAJC13 | DNAJC13 | chr3 | + | 132461049 | 132461205 | 132460249 | 132460357 | 132462466 | 132462523 | 9 | 2 | 0 | 2 | 0 | 0 | 1 | 2 | 0 | 0 | 98 | 49 | 1 | 1 | 0.692 | 1 | NA | 1 | 1 | -0.154 |
| SE | 16841 | FAM172A | FAM172A | chr5 | - | 93958776 | 93958968 | 93881469 | 93881688 | 93964457 | 93964523 | 17 | 8 | 0 | 0 | 0 | 0 | 0 | 5 | 0 | 2 | 98 | 49 | 1 | 1 | 1 | 1 | NA | NA | 0.556 | 0.444 |
| SE | 16842 | FAM172A | FAM172A | chr5 | - | 94050730 | 94050831 | 93964457 | 93964523 | 94053125 | 94053227 | 11 | 6 | 1 | 2 | 0 | 0 | 0 | 3 | 0 | 1 | 98 | 49 | 1 | 1 | 0.733 | 1 | 1 | NA | 0.6 | 0.311 |
| SE | 16843 | FAM172A | FAM172A | chr5 | - | 94074645 | 94074769 | 94053125 | 94053227 | 94111510 | 94111699 | 2 | 0 | 0 | 4 | 1 | 0 | 0 | 2 | 0 | 1 | 98 | 49 | 1 | 1 | 0.2 | 0 | NA | NA | 0.5 | -0.4 |
| SE | 16844 | KIAA0825 | KIAA0825 | chr5 | - | 94582432 | 94582583 | 94536995 | 94537127 | 94618499 | 94618604 | 9 | 3 | 2 | 1 | 0 | 0 | 0 | 1 | 0 | 0 | 98 | 49 | 1 | 1 | 0.818 | 1 | 1 | NA | 1 | -0.061 |
| SE | 16845 | SLF1 | SLF1 | chr5 | + | 94643272 | 94643435 | 94630502 | 94630743 | 94649453 | 94649597 | 0 | 2 | 1 | 0 | 2 | 0 | 0 | 1 | 0 | 0 | 98 | 49 | 1 | 1 | NA | 0.333 | 1 | NA | 1 | -0.334 |
| SE | 16846 | SLF1 | SLF1 | chr5 | + | 94653271 | 94653421 | 94651701 | 94651845 | 94654629 | 94654752 | 8 | 0 | 6 | 0 | 0 | 0 | 0 | 5 | 0 | 1 | 98 | 49 | 1 | 1 | 1 | NA | 1 | NA | 0.714 | 0.286 |
| SE | 16855 | CAST | CAST | chr5 | + | 96730779 | 96730860 | 96729611 | 96729725 | 96736171 | 96736240 | 16 | 2 | 3 | 2 | 0 | 0 | 0 | 8 | 0 | 1 | 98 | 49 | 1 | 1 | 0.8 | 1 | 1 | NA | 0.8 | 0.133 |
| SE | 16857 | CAST | CAST | chr5 | + | 96740744 | 96740783 | 96740037 | 96740118 | 96741265 | 96741358 | 14 | 3 | 0 | 10 | 3 | 3 | 0 | 8 | 0 | 0 | 88 | 49 | 1 | 1 | 0.438 | 0.358 | 0 | NA | 1 | -0.735 |
| SE | 16858 | CAST | CAST | chr5 | + | 96746341 | 96746425 | 96742654 | 96742756 | 96747344 | 96747392 | 25 | 2 | 4 | 2 | 0 | 0 | 0 | 11 | 0 | 0 | 98 | 49 | 1 | 1 | 0.862 | 1 | 1 | NA | 1 | -0.046 |
| SE | 16868 | ANKZF1 | ANKZF1 | chr2 | + | 219231927 | 219232040 | 219230227 | 219230405 | 219232259 | 219232362 | 10 | 5 | 5 | 1 | 0 | 2 | 1 | 0 | 0 | 0 | 98 | 49 | 1 | 1 | 0.833 | 1 | 0.556 | 1 | NA | -0.204 |
| SE | 16872 | APC | APC | chr5 | + | 112818965 | 112819344 | 112815494 | 112815593 | 112821895 | 112821991 | 14 | 14 | 7 | 0 | 2 | 0 | 0 | 4 | 0 | 0 | 98 | 49 | 1 | 1 | 1 | 0.778 | 1 | NA | 1 | -0.074 |
| SE | 16873 | APC | APC | chr5 | + | 112834950 | 112835165 | 112828855 | 112828972 | 112837552 | 112846239 | 17 | 13 | 4 | 1 | 0 | 2 | 0 | 1 | 0 | 0 | 98 | 49 | 1 | 1 | 0.895 | 1 | 0.5 | NA | 1 | -0.202 |
| SE | 16879 | ALPL | ALPL | chr1 | + | 21560625 | 21560745 | 21553977 | 21554142 | 21561096 | 21561212 | 95 | 52 | 20 | 1 | 0 | 0 | 3 | 11 | 0 | 0 | 98 | 49 | 1 | 1 | 0.979 | 1 | 1 | 1 | 1 | -0.007 |

| | | | | | | | | | | | | | | | | | | | | | | | | | | | | | | | | | |
|---|---|---|---|---|---|---|---|---|---|---|---|---|---|---|---|---|---|---|---|---|---|---|---|---|---|---|---|---|---|---|---|---|---|
| SE | 16880 | ALPL | ALPL | chr1 | + | 21560625 | 21560745 | 21553977 | 21554142 | 21563109 | 21563284 | 43 | 31 | 5 | 1 | 0 | 0 | 0 | 8 | 0 | 0 | 98 | 49 | 1 | 1 | 0.956 | 1 | 1 | NA | | | 1 | -0.015 |
| SE | 16881 | ALPL | ALPL | chr1 | + | 21561096 | 21561212 | 21553977 | 21554142 | 21563109 | 21563284 | 70 | 41 | 31 | 1 | 0 | 0 | 0 | 18 | 0 | 0 | 98 | 49 | 1 | 1 | 0.972 | 1 | 1 | NA | | | 1 | -0.009 |
| SE | 16883 | ALPL | ALPL | chr1 | + | 21573664 | 21573799 | 21570304 | 21570374 | 21575732 | 21575924 | 155 | 52 | 67 | 1 | 0 | 0 | 0 | 18 | 0 | 1 | 98 | 49 | 1 | 1 | 0.987 | 1 | 1 | NA | | | 0.9 | 0.096 |
| SE | 16890 | HSD17B4 | HSD17B4 | chr5 | + | 119531265 | 119531404 | 119529893 | 119529980 | 119536422 | 119536550 | 2 | 0 | 0 | 1 | 0 | 0 | 0 | 3 | 0 | 0 | 98 | 49 | 1 | 1 | 0.5 | NA | NA | NA | | | 1 | -0.5 |
| SE | 16891 | ITPKC | ITPKC | chr19 | + | 40733159 | 40733364 | 40729201 | 40729415 | 40736985 | 40737087 | 1 | 0 | 0 | 2 | 0 | 0 | 0 | 1 | 0 | 0 | 98 | 49 | 1 | 1 | 0.2 | NA | NA | NA | | | 1 | -0.8 |
| SE | 16894 | VASP | VASP | chr19 | + | 45517928 | 45518094 | 45517662 | 45517834 | 45521321 | 45521406 | 200 | 80 | 69 | 1 | 0 | 0 | 7 | 36 | 0 | 0 | 98 | 49 | 1 | 1 | 0.99 | 1 | 1 | | 1 | | 1 | -0.003 |
| SE | 16896 | VASP | VASP | chr19 | + | 45522339 | 45522581 | 45517662 | 45517834 | 45522717 | 45522818 | 17 | 6 | 7 | 1 | 0 | 0 | 2 | 6 | 0 | 0 | 98 | 49 | 1 | 1 | 0.895 | 1 | 1 | | 1 | | 1 | -0.035 |
| SE | 16906 | LMNB1 | LMNB1 | chr5 | + | 126822780 | 126822885 | 126820909 | 126821135 | 126825987 | 126826107 | 126 | 35 | 30 | 0 | 2 | 0 | 2 | 18 | 0 | 0 | 98 | 49 | 1 | 1 | 1 | 0.897 | 1 | | 1 | | 1 | -0.034 |
| SE | 16918 | RAPGEF6 | RAPGEF6 | chr5 | - | 131561977 | 131562047 | 131548046 | 131548190 | 131592382 | 131592466 | 11 | 8 | 1 | 2 | 2 | 0 | 0 | 1 | 0 | 0 | 98 | 49 | 1 | 1 | 0.733 | 0.667 | 1 | NA | | | 1 | -0.2 |
| SE | 16921 | SLC22A4 | SLC22A4 | chr5 | + | 132312160 | 132312264 | 132294451 | 132295009 | 132313613 | 132313768 | 15 | 5 | 2 | 0 | 0 | 0 | 0 | 4 | 1 | 0 | 98 | 49 | 1 | 1 | 1 | 1 | 1 | | 0 | | 1 | 0.5 |
| SE | 16922 | SLC22A4 | SLC22A4 | chr5 | + | 132313613 | 132313768 | 132294451 | 132295009 | 132327276 | 132327403 | 2 | 1 | 2 | 0 | 1 | 0 | 1 | 0 | 0 | 0 | 98 | 49 | 1 | 1 | 1 | 0.333 | 1 | | 1 | NA | 1 | -0.222 |
| SE | 16923 | SLC22A4 | SLC22A4 | chr5 | + | 132322183 | 132322355 | 132313613 | 132313768 | 132327276 | 132327403 | 9 | 5 | 4 | 2 | 1 | 2 | 0 | 2 | 0 | 0 | 98 | 49 | 1 | 1 | 0.692 | 0.714 | 0.5 | NA | | | 1 | -0.365 |
| SE | 16926 | IRF1 | IRF1 | chr5 | - | 132484361 | 132484497 | 132481608 | 132484075 | 132485666 | 132485716 | 105 | 52 | 26 | 3 | 2 | 0 | 0 | 18 | 0 | 0 | 98 | 49 | 1 | 1 | 0.946 | 0.929 | 1 | NA | | | 1 | -0.042 |
| SE | 16927 | IRF1 | IRF1 | chr5 | - | 132484361 | 132484497 | 132481608 | 132484075 | 132486250 | 132486373 | 43 | 18 | 10 | 2 | 3 | 0 | 0 | 8 | 0 | 0 | 98 | 49 | 1 | 1 | 0.915 | 0.75 | 1 | NA | | | 1 | -0.112 |
| SE | 16928 | IRF1 | IRF1 | chr5 | - | 132484361 | 132484497 | 132481608 | 132484075 | 132486556 | 132486686 | 44 | 24 | 13 | 0 | 0 | 0 | 0 | 8 | 0 | 1 | 98 | 49 | 1 | 1 | 1 | 1 | 1 | NA | | | 0.8 | 0.2 |
| SE | 16929 | IRF1 | IRF1 | chr5 | - | 132485666 | 132485716 | 132481608 | 132484075 | 132486250 | 132486373 | 53 | 34 | 26 | 2 | 3 | 0 | 0 | 24 | 0 | 0 | 98 | 49 | 1 | 1 | 0.93 | 0.85 | 1 | NA | | | 1 | -0.073 |
| SE | 16930 | IRF1 | IRF1 | chr5 | - | 132485666 | 132485716 | 132481608 | 132484075 | 132486556 | 132486686 | 11 | 4 | 0 | 0 | 0 | 0 | 0 | 1 | 0 | 1 | 98 | 49 | 1 | 1 | 1 | 1 | NA | NA | | | 0.333 | 0.667 |
| SE | 16931 | IRF1 | IRF1 | chr5 | - | 132485666 | 132485716 | 132484361 | 132484497 | 132486250 | 132486373 | 114 | 66 | 42 | 2 | 0 | 0 | 0 | 34 | 0 | 0 | 98 | 49 | 1 | 1 | 0.966 | 1 | 1 | NA | | | 1 | -0.011 |
| SE | 16932 | IRF1 | IRF1 | chr5 | - | 132485666 | 132485716 | 132484361 | 132484497 | 132486556 | 132486686 | 72 | 36 | 16 | 3 | 6 | 3 | 0 | 11 | 0 | 0 | 98 | 49 | 1 | 1 | 0.923 | 0.75 | 0.727 | NA | | | 1 | -0.2 |
| SE | 16933 | IRF1 | IRF1 | chr5 | - | 132486250 | 132486373 | 132481608 | 132484075 | 132486556 | 132486686 | 65 | 21 | 29 | 0 | 0 | 0 | 2 | 12 | 0 | 1 | 98 | 49 | 1 | 1 | 1 | 1 | 1 | | 1 | | 0.857 | 0.072 |
| SE | 16937 | IRF1 | IRF1 | chr5 | - | 132487925 | 132488025 | 132486953 | 132487130 | 132489391 | 132489483 | 85 | 62 | 26 | 0 | 3 | 0 | 7 | 14 | 0 | 1 | 98 | 49 | 1 | 1 | 1 | 0.912 | 1 | | 1 | | 0.875 | 0.033 |
| SE | 16939 | IRF1 | IRF1 | chr5 | - | 132489391 | 132489483 | 132486953 | 132487130 | 132490544 | 132490773 | 43 | 25 | 13 | 18 | 5 | 5 | 0 | 6 | 0 | 2 | 98 | 49 | 1 | 1 | 0.544 | 0.714 | 0.565 | NA | | | 0.6 | 0.008 |
| SE | 16940 | IRF1 | IRF1 | chr5 | - | 132489391 | 132489483 | 132487925 | 132488025 | 132490544 | 132490773 | 55 | 28 | 18 | 2 | 4 | 1 | 0 | 8 | 0 | 0 | 98 | 49 | 1 | 1 | 0.932 | 0.778 | 0.9 | NA | | | 1 | -0.13 |
| SE | 16949 | PPP2CA | PPP2CA | chr5 | - | 134200334 | 134200496 | 134199085 | 134199204 | 134200984 | 134201074 | 18 | 11 | 8 | 1 | 2 | 0 | 0 | 4 | 0 | 3 | 98 | 49 | 1 | 1 | 0.9 | 0.733 | 1 | NA | | | 0.4 | 0.478 |
| SE | 16956 | H2AFY | H2AFY | chr5 | - | 135350822 | 135350913 | 135345967 | 135346057 | 135360496 | 135360607 | 5 | 3 | 3 | 3 | 0 | 1 | 0 | 3 | 0 | 0 | 98 | 49 | 1 | 1 | 0.455 | 1 | 0.6 | NA | | | 1 | -0.315 |
| SE | 16963 | FAM13B | FAM13B | chr5 | - | 137946227 | 137946311 | 137945901 | 137945997 | 137948954 | 137949184 | 9 | 7 | 4 | 1 | 4 | 0 | 0 | 5 | 0 | 0 | 98 | 49 | 1 | 1 | 0.818 | 0.467 | 1 | NA | | | 1 | -0.238 |
| SE | 16964 | FAM13B | FAM13B | chr5 | - | 137956476 | 137956542 | 137954165 | 137954376 | 137959615 | 137959763 | 0 | 0 | 0 | 6 | 0 | 0 | 0 | 1 | 0 | 0 | 98 | 49 | 1 | 1 | 0 | NA | NA | NA | | | 1 | -1 |
| SE | 16965 | FAM13B | FAM13B | chr5 | - | 138018954 | 138019146 | 138018301 | 138018514 | 138021030 | 138021197 | 4 | 5 | 0 | 2 | 1 | 0 | 0 | 1 | 0 | 0 | 98 | 49 | 1 | 1 | 0.5 | 0.714 | NA | NA | | | 1 | -0.393 |
| SE | 16971 | ETF1 | ETF1 | chr5 | - | 138529554 | 138529678 | 138518691 | 138518867 | 138542832 | 138542936 | 0 | 1 | 0 | 20 | 18 | 0 | 0 | 0 | 2 | 4 | 98 | 49 | 1 | 1 | 0 | 0.027 | NA | NA | 0 | | 0 | 0.013 |
| SE | 16972 | SPI1 | SPI1 | chr11 | - | 47359852 | 47360040 | 47358843 | 47359006 | 47375632 | 47375729 | 283 | 126 | 83 | 0 | 0 | 2 | 14 | 70 | 0 | 0 | 98 | 49 | 1 | 1 | 1 | 1 | 0.954 | | 1 | | 1 | -0.015 |
| SE | 16976 | IK | IK | chr5 | + | 140655828 | 140655992 | 140654680 | 140654727 | 140657553 | 140657662 | 30 | 7 | 6 | 3 | 0 | 0 | 0 | 2 | 0 | 2 | 98 | 49 | 1 | 1 | 0.833 | 1 | 1 | NA | | | 0.333 | 0.611 |
| SE | 16979 | CASC4 | CASC4 | chr15 | + | 44402886 | 44403054 | 44380805 | 44380976 | 44413335 | 44415761 | 23 | 5 | 14 | 8 | 1 | 2 | 0 | 1 | 0 | 0 | 98 | 49 | 1 | 1 | 0.59 | 0.714 | 0.778 | NA | | | 1 | -0.306 |
| SE | 16982 | HARS2 | HARS2 | chr5 | + | 140696102 | 140696201 | 140695737 | 140695845 | 140696520 | 140696614 | 7 | 2 | 1 | 1 | 0 | 0 | 0 | 1 | 0 | 0 | 98 | 49 | 1 | 1 | 0.778 | 1 | 1 | NA | | | 1 | -0.074 |
| SE | 16986 | HDAC3 | HDAC3 | chr5 | - | 141626012 | 141626071 | 141625684 | 141625764 | 141626193 | 141626283 | 16 | 1 | 2 | 0 | 0 | 0 | 0 | 6 | 0 | 2 | 98 | 49 | 1 | 1 | 1 | 1 | 1 | NA | | | 0.6 | 0.4 |
| SE | 16987 | FCHSD1 | FCHSD1 | chr5 | - | 141643000 | 141643088 | 141641701 | 141641757 | 141644217 | 141644438 | 7 | 7 | 5 | 2 | 0 | 0 | 3 | 5 | 0 | 0 | 98 | 49 | 1 | 1 | 0.636 | 1 | 1 | | 1 | | 1 | -0.121 |
| SE | 16988 | FCHSD1 | FCHSD1 | chr5 | - | 141646602 | 141646722 | 141646086 | 141646191 | 141647134 | 141647230 | 6 | 4 | 3 | 0 | 0 | 0 | 0 | 1 | 0 | 1 | 98 | 49 | 1 | 1 | 1 | 1 | 1 | NA | | | 0.333 | 0.667 |
| SE | 16989 | FCHSD1 | FCHSD1 | chr5 | - | 141649394 | 141649536 | 141649171 | 141649308 | 141649886 | 141649954 | 4 | 1 | 2 | 1 | 0 | 0 | 0 | 2 | 0 | 0 | 98 | 49 | 1 | 1 | 0.667 | 1 | 1 | NA | | | 1 | -0.111 |
| SE | 16993 | ARHGAP26 | ARHGAP26 | chr5 | + | 142873399 | 142873495 | 142770726 | 142770915 | 142879373 | 142879445 | 51 | 28 | 8 | 0 | 0 | 0 | 0 | 11 | 0 | 1 | 98 | 49 | 1 | 1 | 1 | 1 | 1 | NA | | | 0.846 | 0.154 |
| SE | 16994 | ARHGAP26 | ARHGAP26 | chr5 | + | 142875109 | 142875171 | 142873399 | 142873495 | 142879373 | 142879445 | 232 | 139 | 55 | 2 | 0 | 0 | 5 | 59 | 0 | 0 | 98 | 49 | 1 | 1 | 0.983 | 1 | 1 | | 1 | | 1 | -0.006 |
| SE | 16995 | ARHGAP26 | ARHGAP26 | chr5 | + | 142894237 | 142894348 | 142885297 | 142885399 | 142903539 | 142903669 | 104 | 70 | 17 | 0 | 1 | 0 | 0 | 26 | 0 | 0 | 98 | 49 | 1 | 1 | 1 | 0.972 | 1 | NA | | | 1 | -0.009 |
| SE | 16996 | ARHGAP26 | ARHGAP26 | chr5 | + | 142901934 | 142902039 | 142894237 | 142894348 | 142903539 | 142903669 | 282 | 160 | 34 | 3 | 1 | 0 | 2 | 101 | 0 | 0 | 98 | 49 | 1 | 1 | 0.979 | 0.988 | 1 | | 1 | | 1 | -0.011 |
| SE | 16997 | ARHGAP26 | ARHGAP26 | chr5 | + | 143014079 | 143014116 | 142932046 | 142932125 | 143037195 | 143037261 | 231 | 77 | 21 | 1 | 0 | 0 | 1 | 47 | 0 | 0 | 86 | 49 | 1 | 1 | 0.992 | 1 | 1 | | 1 | | 1 | -0.003 |
| SE | 16998 | ARHGAP26 | ARHGAP26 | chr5 | + | 143037195 | 143037261 | 143014079 | 143014116 | 143041815 | 143041890 | 150 | 75 | 13 | 3 | 1 | 0 | 0 | 32 | 0 | 0 | 98 | 49 | 1 | 1 | 0.962 | 0.974 | 1 | NA | | | 1 | -0.021 |
| SE | 16999 | ARHGAP26 | ARHGAP26 | chr5 | + | 143207197 | 143207308 | 143147230 | 143147381 | 143213996 | 143214088 | 23 | 14 | 0 | 1 | 1 | 1 | 0 | 4 | 0 | 0 | 98 | 49 | 1 | 1 | 0.92 | 0.875 | 0 | NA | | | 1 | -0.402 |
| SE | 17000 | ARHGAP26 | ARHGAP26 | chr5 | + | 143207197 | 143207473 | 143147230 | 143147381 | 143213996 | 143214088 | 17 | 10 | 0 | 1 | 1 | 1 | 0 | 3 | 0 | 0 | 98 | 49 | 1 | 1 | 0.895 | 0.833 | 0 | NA | | | 1 | -0.424 |
| SE | 17003 | WDSUB1 | WDSUB1 | chr2 | - | 159248371 | 159248512 | 159235792 | 159236190 | 159256195 | 159256375 | 1 | 0 | 0 | 1 | 0 | 0 | 0 | 0 | 0 | 2 | 98 | 49 | 1 | 1 | 0.333 | NA | NA | NA | | | 0 | 0.333 |
| SE | 17008 | ZNF438 | ZNF438 | chr10 | - | 30857679 | 30857778 | 30848530 | 30850367 | 30876997 | 30877065 | 2 | 5 | 0 | 6 | 3 | 0 | 0 | 0 | 0 | 2 | 98 | 49 | 1 | 1 | 0.143 | 0.455 | NA | NA | | | 0 | 0.299 |
| SE | 17011 | ZNF438 | ZNF438 | chr10 | - | 30910359 | 30910440 | 30908932 | 30909015 | 30941574 | 30941651 | 1 | 1 | 0 | 4 | 4 | 0 | 0 | 0 | 0 | 1 | 98 | 49 | 1 | 1 | 0.111 | 0.111 | NA | NA | | | 0 | 0.111 |
| SE | 17012 | ZNF438 | ZNF438 | chr10 | - | 30923286 | 30923415 | 30908932 | 30909015 | 30941574 | 30941651 | 5 | 4 | 0 | 4 | 4 | 0 | 0 | 1 | 0 | 1 | 98 | 49 | 1 | 1 | 0.385 | 0.333 | NA | NA | | | 0.333 | 0.026 |
| SE | 17013 | ZNF438 | ZNF438 | chr10 | - | 30923286 | 30923415 | 30910359 | 30910440 | 30941574 | 30941651 | 1 | 4 | 0 | 1 | 0 | 0 | 0 | 1 | 0 | 0 | 98 | 49 | 1 | 1 | 0.333 | 1 | NA | NA | | | 1 | -0.334 |
| SE | 17014 | ZNF438 | ZNF438 | chr10 | - | 30941574 | 30941651 | 30923286 | 30923415 | 30984368 | 30984426 | 8 | 9 | 1 | 1 | 0 | 0 | 0 | 3 | 0 | 0 | 98 | 49 | 1 | 1 | 0.8 | 1 | 1 | NA | | | 1 | -0.067 |
| SE | 17015 | ZNF438 | ZNF438 | chr10 | - | 30984368 | 30984426 | 30941574 | 30941651 | 31031832 | 31031937 | 7 | 6 | 1 | 0 | 0 | 1 | 0 | 2 | 0 | 0 | 98 | 49 | 1 | 1 | 1 | 1 | 0.333 | NA | | | 1 | -0.222 |
| SE | 17017 | CSNK1A1 | CSNK1A1 | chr5 | - | 149517793 | 149517877 | 149513069 | 149513209 | 149520289 | 149520388 | 3 | 0 | 0 | 14 | 6 | 2 | 0 | 0 | 0 | 5 | 98 | 49 | 1 | 1 | 0.097 | 0 | 0 | NA | | | 0 | 0.032 |
| SE | 17023 | ARRB1 | ARRB1 | chr11 | - | 75276838 | 75276911 | 75274073 | 75274211 | 75277363 | 75277448 | 10 | 2 | 0 | 1 | 0 | 0 | 0 | 2 | 0 | 0 | 98 | 49 | 1 | 1 | 0.833 | 1 | NA | NA | | | 1 | -0.084 |
| SE | 17029 | FAM114A2 | FAM114A2 | chr5 | - | 154029488 | 154029580 | 154028148 | 154028283 | 154033790 | 154033883 | 2 | 0 | 1 | 0 | 0 | 0 | 1 | 0 | 0 | 2 | 98 | 49 | 1 | 1 | 1 | NA | 1 | | 1 | | 0 | 0.5 |
| SE | 17030 | GALNT10 | GALNT10 | chr5 | + | 154329571 | 154329738 | 154297940 | 154298079 | 154376276 | 154376462 | 8 | 3 | 2 | 2 | 0 | 0 | 0 | 1 | 0 | 0 | 98 | 49 | 1 | 1 | 0.667 | 1 | 1 | NA | | | 1 | -0.111 |
| SE | 17033 | SAP30L | SAP30L | chr5 | + | 154452455 | 154452499 | 154451090 | 154451213 | 154453401 | 154453500 | 10 | 1 | 0 | 2 | 0 | 0 | 0 | 1 | 0 | 0 | 93 | 49 | 1 | 1 | 0.725 | 1 | NA | NA | | | 1 | -0.137 |
| SE | 17039 | RENBP | RENBP | chrX | - | 153941949 | 153942031 | 153941477 | 153941653 | 153942854 | 153943079 | 8 | 5 | 2 | 0 | 1 | 0 | 0 | 3 | 0 | 0 | 98 | 49 | 1 | 1 | 1 | 0.714 | 1 | NA | | | 1 | -0.095 |
| SE | 17040 | ITK | ITK | chr5 | + | 157214190 | 157214319 | 157211286 | 157211368 | 157217866 | 157217907 | 10 | 0 | 0 | 0 | 0 | 0 | 0 | 1 | 0 | 0 | 98 | 49 | 1 | 1 | 0.833 | NA | NA | NA | | | 1 | -0.167 |
| SE | 17054 | LCP2 | LCP2 | chr5 | - | 170275319 | 170275351 | 170274300 | 170274338 | 170275794 | 170275860 | 63 | 36 | 5 | 3 | 0 | 0 | 0 | 23 | 0 | 0 | 81 | 49 | 1 | 1 | 0.927 | 1 | 1 | NA | | | 1 | -0.024 |
| SE | 17058 | CIR1 | CIR1 | chr2 | - | 174378922 | 174379040 | 174351618 | 174351725 | 174380201 | 174380250 | 45 | 30 | 9 | 0 | 0 | 1 | 0 | 8 | 0 | 0 | 98 | 49 | 1 | 1 | 1 | 1 | 0.818 | NA | | | 1 | -0.061 |
| SE | 17073 | GRK6 | GRK6 | chr5 | + | 177432705 | 177432806 | 177432232 | 177432310 | 177433146 | 177433239 | 81 | 54 | 35 | 1 | 0 | 0 | 9 | 23 | 0 | 0 | 98 | 49 | 1 | 1 | 0.976 | 1 | 1 | | 1 | | 1 | -0.008 |
| SE | 17078 | PRKDC | PRKDC | chr8 | - | 47782161 | 47782254 | 47779003 | 47779093 | 47782377 | 47782598 | 11 | 3 | 6 | 2 | 2 | 2 | 0 | 4 | 0 | 0 | 98 | 49 | 1 | 1 | 0.733 | 0.429 | 0.6 | NA | | | 1 | -0.413 |
| SE | 17079 | PRKDC | PRKDC | chr8 | - | 47826655 | 47826861 | 47823857 | 47823996 | 47828167 | 47828347 | 13 | 4 | 7 | 2 | 0 | 0 | 0 | 7 | 0 | 0 | 98 | 49 | 1 | 1 | 0.765 | 1 | 1 | NA | | | 1 | -0.078 |
| SE | 17080 | PRKDC | PRKDC | chr8 | - | 47836337 | 47836527 | 47834195 | 47834396 | 47837211 | 47837419 | 15 | 6 | 12 | 0 | 0 | 0 | 0 | 11 | 0 | 2 | 98 | 49 | 1 | 1 | 1 | 1 | 1 | NA | | | 0.733 | 0.267 |
| SE | 17081 | PRKDC | PRKDC | chr8 | - | 47839147 | 47839246 | 47837211 | 47837419 | 47840015 | 47840189 | 16 | 5 | 3 | 1 | 0 | 0 | 3 | 15 | 0 | 0 | 98 | 49 | 1 | 1 | 0.889 | 1 | 1 | | 1 | | 1 | -0.037 |
| SE | 17087 | PHYKPL | PHYKPL | chr5 | - | 178212972 | 178213103 | 178211889 | 178211970 | 178214795 | 178214885 | 11 | 5 | 1 | 3 | 0 | 0 | 0 | 1 | 0 | 1 | 98 | 49 | 1 | 1 | 0.647 | 1 | 1 | NA | | | 0 | 0.882 |
| SE | 17092 | CLK4 | CLK4 | chr5 | - | 178617343 | 178617434 | 178616881 | 178616948 | 178618555 | 178618778 | 5 | 0 | 2 | 12 | 3 | 9 | 0 | 0 | 0 | 1 | 98 | 49 | 1 | 1 | 0.172 | 0 | 0.1 | NA | | | 0 | 0.091 |
| SE | 17096 | CASP10 | CASP10 | chr2 | + | 201203729 | 201203766 | 201195841 | 201195948 | 201208074 | 201208183 | 2 | 0 | 0 | 0 | 0 | 0 | 0 | 1 | 0 | 1 | 86 | 49 | 1 | 1 | 1 | NA | NA | NA | | | 0.363 | 0.637 |
| SE | 17099 | NEK7 | NEK7 | chr1 | + | 198277960 | 198278069 | 198264124 | 198264235 | 198278953 | 198279061 | 40 | 12 | 15 | 0 | 2 | 0 | 0 | 4 | 0 | 0 | 98 | 49 | 1 | 1 | 1 | 0.75 | 1 | NA | | | 1 | -0.083 |
| SE | 17102 | NQO2 | NQO2 | chr6 | + | 3002049 | 3002286 | 2999815 | 3000085 | 3003668 | 3003789 | 7 | 0 | 0 | 1 | 1 | 0 | 0 | 3 | 0 | 0 | 98 | 49 | 1 | 1 | 0.778 | 0 | NA | NA | | | 1 | -0.611 |
| SE | 17105 | NQO2 | NQO2 | chr6 | + | 3003668 | 3003789 | 3002049 | 3002286 | 3004494 | 3004651 | 11 | 0 | 0 | 1 | 1 | 0 | 0 | 3 | 0 | 0 | 98 | 49 | 1 | 1 | 0.846 | 0 | NA | NA | | | 1 | -0.577 |
| SE | 17107 | NQO2 | NQO2 | chr6 | + | 3015529 | 3015643 | 3012543 | 3012674 | 3016883 | 3016985 | 31 | 12 | 6 | 4 | 0 | 0 | 0 | 4 | 0 | 1 | 98 | 49 | 1 | 1 | 0.795 | 1 | 1 | NA | | | 0.667 | 0.265 |
| SE | 17108 | ECI2 | ECI2 | chr6 | - | 4117307 | 4117451 | 4115692 | 4116029 | 4119185 | 4119275 | 2 | 0 | 0 | 0 | 0 | 0 | 0 | 2 | 0 | 1 | 98 | 49 | 1 | 1 | 1 | NA | NA | NA | | | 0.5 | 0.5 |
| SE | 17117 | MCMBP | MCMBP | chr10 | - | 119836895 | 119837029 | 119835539 | 119835704 | 119838534 | 119838700 | 17 | 9 | 27 | 1 | 0 | 0 | 0 | 12 | 0 | 0 | 98 | 49 | 1 | 1 | 0.895 | 1 | 1 | NA | | | 1 | -0.035 |
| SE | 17118 | MCMBP | MCMBP | chr10 | - | 119842471 | 119842595 | 119840842 | 119840960 | 119843247 | 119843426 | 2 | 5 | 14 | 0 | 0 | 0 | 0 | 1 | 0 | 1 | 98 | 49 | 1 | 1 | 1 | 1 | 1 | | 0 | NA | | 1 |
| RI | 0 | FAM3A | FAM3A | chrX | - | 154515759 | 154516242 | 154515759 | 154515812 | 154515909 | 154516242 | 0 | 1 | 0 | 0 | 0 | 0 | 0 | 0 | 0 | 1 | 98 | 49 | 1 | 1 | NA | 1 | NA | NA | 0 | NA | 0 | 1 |
| RI | 2 | IDH3G | IDH3G | chrX | - | 153785765 | 153786272 | 153785765 | 153785973 | 153786211 | 153786272 | 3 | 0 | 0 | 3 | 0 | 0 | 1 | 0 | 0 | 0 | 98 | 49 | 1 | 1 | 0.333 | NA | NA | | 1 | NA | 1 | -0.667 |
| RI | 56 | MCM7 | MCM7 | chr7 | - | 100099022 | 100099403 | 100099022 | 100099203 | 100099278 | 100099403 | 2 | 2 | 0 | 0 | 2 | 0 | 0 | 1 | 0 | 0 | 98 | 49 | 1 | 1 | 1 | 0.333 | NA | NA | | | 1 | -0.334 |
| RI | 58 | SAMD9L | SAMD9L | chr7 | - | 93145384 | 93146040 | 93145384 | 93145532 | 93145908 | 93146040 | 15 | 3 | 5 | 1 | 0 | 0 | 3 | 10 | 0 | 0 | 98 | 49 | 1 | 1 | 0.882 | 1 | 1 | | 1 | | 1 | -0.039 |
| RI | 70 | TMED4 | TMED4 | chr7 | - | 44581722 | 44582295 | 44581722 | 44581823 | 44582046 | 44582295 | 10 | 1 | 0 | 7 | 0 | 2 | 0 | 1 | 0 | 0 | 98 | 49 | 1 | 1 | 0.417 | 1 | 0 | NA | | | 1 | -0.528 |
| RI | 79 | CD164 | CD164 | chr6 | - | 109366513 | 109369017 | 109366513 | 109368325 | 109368921 | 109369017 | 36 | 34 | 6 | 0 | 1 | 0 | 1 | 31 | 0 | 1 | 98 | 49 | 1 | 1 | 1 | 0.944 | 1 | | 1 | | 0.939 | 0.012 |
| RI | 81 | SYTL3 | SYTL3 | chr6 | + | 158662749 | 158663378 | 158662749 | 158662812 | 158663204 | 158663378 | 17 | 13 | 3 | 2 | 0 | 0 | 2 | 1 | 0 | 0 | 98 | 49 | 1 | 1 | 0.81 | 1 | 1 | | 1 | | 1 | -0.063 |
| RI | 82 | HMGN3 | HMGN3 | chr6 | - | 79201244 | 79202389 | 79201244 | 79201726 | 79202275 | 79202389 | 7 | 4 | 0 | 0 | 0 | 0 | 0 | 2 | 0 | 1 | 98 | 49 | 1 | 1 | 1 | 1 | NA | NA | | | 0.5 | 0.5 |
| RI | 83 | HMGN3 | HMGN3 | chr6 | - | 79201244 | 79202389 | 79201244 | 79201726 | 79202316 | 79202389 | 9 | 4 | 0 | 1 | 0 | 0 | 0 | 2 | 0 | 0 | 98 | 49 | 1 | 1 | 0.818 | 1 | NA | NA | | | 1 | -0.091 |
| RI | 93 | SLC39A7 | SLC39A7 | chr6 | + | 33200825 | 33201656 | 33200825 | 33200938 | 33201241 | 33201656 | 1 | 2 | 0 | 2 | 0 | 0 | 0 | 1 | 0 | 0 | 98 | 49 | 1 | 1 | 0.2 | 1 | NA | NA | | | 1 | -0.4 |
| RI | 133 | POU5F1 | POU5F1 | chr6 | - | 31165570 | 31167170 | 31165570 | 31165701 | 31165926 | 31167170 | 0 | 0 | 0 | 2 | 0 | 0 | 0 | 1 | 0 | 0 | 98 | 49 | 1 | 1 | 0 | NA | NA | NA | | | 1 | -1 |
| RI | 164 | BSDC1 | BSDC1 | chr1 | - | 32364632 | 32368550 | 32364632 | 32366654 | 32368446 | 32368550 | 1 | 4 | 1 | 5 | 1 | 3 | 3 | 0 | 0 | 0 | 98 | 49 | 1 | 1 | 0.091 | 0.667 | 0.143 | | 1 | NA | | -0.7 |
| RI | 170 | TAOK2 | TAOK2 | chr16 | + | 29986264 | 29988405 | 29986264 | 29986504 | 29986843 | 29988405 | 2 | 1 | 0 | 1 | 0 | 0 | 0 | 0 | 0 | 0 | 98 | 49 | 1 | 1 | 0.5 | 1 | NA | NA | | | 1 | -0.25 |
| RI | 180 | PRKD2 | PRKD2 | chr19 | - | 46716130 | 46717127 | 46716130 | 46716535 | 46716814 | 46717127 | 11 | 3 | 0 | 2 | 0 | 0 | 0 | 3 | 0 | 0 | 98 | 49 | 1 | 1 | 0.733 | 1 | NA | NA | | | 1 | -0.133 |
| RI | 181 | PRKD2 | PRKD2 | chr19 | - | 46716130 | 46717127 | 46716130 | 46716535 | 46716950 | 46717127 | 7 | 4 | 0 | 0 | 0 | 0 | 0 | 4 | 0 | 1 | 98 | 49 | 1 | 1 | 1 | 1 | NA | NA | | | 0.667 | 0.333 |
| RI | 189 | SERF2 | SERF2 | chr15 | + | 43793709 | 43796089 | 43793709 | 43793936 | 43794373 | 43796089 | 52 | 25 | 17 | 0 | 1 | 0 | 1 | 27 | 0 | 0 | 98 | 49 | 1 | 1 | 1 | 0.926 | 1 | | 1 | | 1 | -0.025 |
| RI | 199 | KAT8 | KAT8 | chr16 | + | 31130745 | 31131393 | 31130745 | 31130900 | 31131194 | 31131393 | 6 | 5 | 4 | 6 | 4 | 0 | 0 | 0 | 1 | 0 | 98 | 49 | 1 | 1 | 0.333 | 0.385 | 1 | | 0 | NA | | 0.573 |
| RI | 205 | PML | PML | chr15 | + | 74034477 | 74036394 | 74034477 | 74034530 | 74035171 | 74036394 | 14 | 7 | 2 | 1 | 0 | 0 | 1 | 7 | 0 | 0 | 98 | 49 | 1 | 1 | 0.875 | 1 | 1 | | 1 | | 1 | -0.042 |
| RI | 206 | CCPG1 | CCPG1 | chr15 | - | 55352222 | 55360944 | 55352222 | 55356409 | 55359538 | 55360944 | 15 | 17 | 7 | 9 | 4 | 1 | 0 | 1 | 0 | 0 | 98 | 49 | 1 | 1 | 0.455 | 0.68 | 0.778 | NA | | | 0.333 | 0.305 |
| RI | 219 | CDK17 | CDK17 | chr12 | - | 96278260 | 96280885 | 96278260 | 96280279 | 96280807 | 96280885 | 9 | 1 | 6 | 2 | 2 | 0 | 0 | 13 | 0 | 0 | 98 | 49 | 1 | 1 | 0.692 | 0.2 | 1 | NA | | | 1 | -0.369 |
| RI | 244 | PSME1 | PSME1 | chr14 | + | 24138473 | 24138967 | 24138473 | 24138560 | 24138735 | 24138967 | 9 | 7 | 5 | 37 | 24 | 15 | 0 | 2 | 0 | 11 | 98 | 49 | 1 | 1 | 0.108 | 0.127 | 0.143 | NA | | | 0.083 | 0.043 |
| RI | 254 | BIN2 | BIN2 | chr12 | - | 51302019 | 51302780 | 51302019 | 51302115 | 51302685 | 51302780 | 26 | 11 | 2 | 23 | 19 | 17 | 0 | 1 | 4 | 5 | 98 | 49 | 1 | 1 | 0.361 | 0.224 | 0.056 | | 0 | | 0.091 | 0.168 |
| RI | 255 | LUC7L3 | LUC7L3 | chr17 | + | 50750500 | 50752711 | 50750500 | 50750694 | 50752199 | 50752711 | 23 | 2 | 1 | 2 | 0 | 0 | 0 | 2 | 0 | 2 | 98 | 49 | 1 | 1 | 0.852 | 1 | 1 | NA | | | 0.333 | 0.618 |
| RI | 273 | ARL6IP4 | ARL6IP4 | chr12 | + | 122982468 | 122982913 | 122982468 | 122982538 | 122982619 | 122982913 | 2 | 0 | 2 | 5 | 1 | 0 | 0 | 0 | 0 | 1 | 98 | 49 | 1 | 1 | 0.167 | 0 | 1 | NA | | | 0 | 0.389 |
| RI | 275 | ATPIF1 | ATPIF1 | chr1 | - | 28236360 | 28238105 | 28236360 | 28236452 | 28237836 | 28238105 | 2 | 0 | 0 | 4 | 5 | 2 | 0 | 5 | 0 | 1 | 98 | 49 | 1 | 1 | 0.2 | 0 | 0 | NA | | | 0.714 | -0.647 |
| RI | 277 | PPP1CC | PPP1CC | chr12 | - | 110719807 | 110721165 | 110719807 | 110720204 | 110721104 | 110721165 | 28 | 7 | 24 | 0 | 0 | 2 | 1 | 4 | 0 | 0 | 98 | 49 | 1 | 1 | 1 | 1 | 0.857 | | 1 | | 1 | -0.048 |
| RI | 288 | IL32 | IL32 | chr16 | + | 3067376 | 3067613 | 3067376 | 3067415 | 3067553 | 3067613 | 3 | 4 | 0 | 14 | 13 | 2 | 0 | 1 | 0 | 3 | 98 | 49 | 1 | 1 | 0.097 | 0.133 | 0 | NA | | | 0.143 | -0.066 |
| RI | 306 | P2RX7 | P2RX7 | chr12 | + | 121177146 | 121177446 | 121177146 | 121177212 | 121177296 | 121177446 | 0 | 1 | 0 | 3 | 0 | 0 | 0 | 2 | 0 | 0 | 98 | 49 | 1 | 1 | 0 | 1 | NA | NA | | | 1 | -0.5 |
| RI | 311 | C2CD5 | C2CD5 | chr12 | - | 22544060 | 22544546 | 22544060 | 22544179 | 22544319 | 22544546 | 1 | 0 | 0 | 9 | 2 | 0 | 0 | 2 | 0 | 0 | 98 | 49 | 1 | 1 | 0.053 | 0 | NA | NA | | | 1 | -0.974 |
| RI | 312 | RECQL | RECQL | chr12 | - | 21501169 | 21501669 | 21501169 | 21501215 | 21501412 | 21501669 | 4 | 1 | 6 | 2 | 0 | 0 | 0 | 0 | 0 | 1 | 98 | 49 | 1 | 1 | 0.5 | 1 | 1 | NA | | | 0 | 0.833 |
| RI | 314 | RPL28 | RPL28 | chr19 | + | 55387929 | 55392083 | 55387929 | 55388048 | 55388242 | 55392083 | 6 | 1 | 0 | 17 | 9 | 2 | 0 | 1 | 0 | 13 | 98 | 49 | 1 | 1 | 0.15 | 0.053 | 0 | NA | | | 0.037 | 0.031 |
| RI | 320 | QKI | QKI | chr6 | + | 163563419 | 163578596 | 163563419 | 163563719 | 163564666 | 163578596 | 15 | 4 | 5 | 0 | 0 | 2 | 0 | 2 | 0 | 0 | 98 | 49 | 1 | 1 | 1 | 1 | 0.556 | NA | | | 1 | -0.148 |
| RI | 321 | QKI | QKI | chr6 | + | 163563419 | 163578596 | 163563419 | 163563719 | 163565945 | 163578596 | 23 | 7 | 4 | 8 | 0 | 0 | 0 | 4 | 0 | 0 | 98 | 49 | 1 | 1 | 0.59 | 1 | 1 | NA | | | 1 | -0.137 |
| RI | 322 | QKI | QKI | chr6 | + | 163563419 | 163578596 | 163563419 | 163563719 | 163570693 | 163578596 | 17 | 3 | 3 | 1 | 0 | 0 | 0 | 8 | 0 | 0 | 98 | 49 | 1 | 1 | 0.895 | 1 | 1 | NA | | | 1 | -0.035 |
| RI | 354 | ZNRD1 | ZNRD1 | chr6 | + | 30061239 | 30061669 | 30061239 | 30061389 | 30061514 | 30061669 | 2 | 0 | 0 | 0 | 0 | 2 | 3 | 0 | 0 | 0 | 98 | 49 | 1 | 1 | 1 | NA | 0 | | 1 | NA | | -0.5 |
| RI | 391 | CCL4L2 | CCL4L2 | chr17 | + | 36211775 | 36212878 | 36211775 | 36211890 | 36212496 | 36212878 | 11 | 14 | 6 | 0 | 0 | 0 | 0 | 0 | 0 | 1 | 98 | 49 | 1 | 1 | 1 | 1 | 1 | NA | | | 0 | 1 |
| RI | 394 | SAMD8 | SAMD8 | chr10 | + | 75176065 | 75182123 | 75176065 | 75176216 | 75176387 | 75182123 | 18 | 6 | 0 | 10 | 2 | 0 | 0 | 0 | 0 | 4 | 98 | 49 | 1 | 1 | 0.474 | 0.6 | NA | NA | | | 0 | 0.537 |
| RI | 395 | DNAJB12 | DNAJB12 | chr10 | - | 72332829 | 72335931 | 72332829 | 72334617 | 72335779 | 72335931 | 10 | 7 | 1 | 4 | 0 | 4 | 0 | 1 | 0 | 3 | 98 | 49 | 1 | 1 | 0.556 | 1 | 0.111 | NA | | | 0.143 | 0.413 |
| RI | 400 | CDK5RAP3 | CDK5RAP3 | chr17 | + | 47974399 | 47975337 | 47974399 | 47974448 | 47975158 | 47975337 | 66 | 15 | 4 | 0 | 1 | 0 | 7 | 10 | 0 | 1 | 98 | 49 | 1 | 1 | 1 | 0.882 | 1 | | 1 | | 0.833 | 0.044 |
| RI | 411 | DNAJB2 | DNAJB2 | chr2 | + | 219284631 | 219286900 | 219284631 | 219284835 | 219285986 | 219286900 | 3 | 0 | 0 | 0 | 0 | 3 | 1 | 0 | 0 | 0 | 98 | 49 | 1 | 1 | 1 | NA | 0 | | 1 | NA | | -0.5 |
| RI | 413 | CDKN1C | CDKN1C | chr11 | - | 2884669 | 2885765 | 2884669 | 2885466 | 2885633 | 2885765 | 1 | 3 | 0 | 0 | 0 | 0 | 1 | 2 | 0 | 2 | 98 | 49 | 1 | 1 | 1 | 1 | NA | | 1 | | 0.333 | 0.334 |
| RI | 416 | OTUB1 | OTUB1 | chr11 | + | 63996529 | 63996941 | 63996529 | 63996648 | 63996856 | 63996941 | 7 | 4 | 2 | 5 | 0 | 1 | 0 | 0 | 0 | 1 | 98 | 49 | 1 | 1 | 0.412 | 1 | 0.5 | NA | | | 0 | 0.637 |
| RI | 417 | SPRTN | SPRTN | chr1 | + | 231351303 | 231355027 | 231351303 | 231351571 | 231352609 | 231355027 | 8 | 3 | 4 | 0 | 0 | 0 | 0 | 1 | 0 | 2 | 98 | 49 | 1 | 1 | 1 | 1 | 1 | NA | | | 0.2 | 0.8 |
| RI | 419 | SLC46A3 | SLC46A3 | chr13 | - | 28700080 | 28701581 | 28700080 | 28700956 | 28701500 | 28701581 | 2 | 0 | 0 | 3 | 0 | 0 | 0 | 1 | 0 | 0 | 98 | 49 | 1 | 1 | 0.25 | NA | NA | NA | | | 1 | -0.75 |
| RI | 433 | VAMP4 | VAMP4 | chr1 | - | 171700155 | 171704534 | 171700155 | 171702084 | 171704386 | 171704534 | 5 | 0 | 0 | 1 | 0 | 0 | 2 | 0 | 0 | 0 | 98 | 49 | 1 | 1 | 0.714 | NA | NA | | 1 | NA | | -0.286 |
| RI | 442 | SNRNP27 | SNRNP27 | chr2 | + | 69904253 | 69905236 | 69904253 | 69904425 | 69904511 | 69905236 | 2 | 0 | 1 | 0 | 0 | 2 | 0 | 3 | 0 | 0 | 98 | 49 | 1 | 1 | 1 | NA | 0.2 | NA | | | 1 | -0.4 |
| RI | 452 | ATP5D | ATP5D | chr19 | + | 1244314 | 1244825 | 1244314 | 1244440 | 1244736 | 1244825 | 1 | 0 | 0 | 5 | 0 | 1 | 0 | 3 | 0 | 0 | 98 | 49 | 1 | 1 | 0.091 | NA | 0 | NA | | | 1 | -0.955 |
| RI | 462 | TK2 | TK2 | chr16 | - | 66548977 | 66550412 | 66548977 | 66549009 | 66549937 | 66550412 | 2 | 0 | 0 | 5 | 0 | 0 | 0 | 0 | 0 | 1 | 98 | 49 | 1 | 1 | 0.167 | NA | NA | NA | | | 0 | 0.167 |
| RI | 463 | TRIM5 | TRIM5 | chr11 | - | 5663194 | 5665682 | 5663194 | 5664173 | 5665655 | 5665682 | 3 | 0 | 0 | 1 | 0 | 0 | 0 | 1 | 0 | 0 | 98 | 49 | 1 | 1 | 0.6 | NA | NA | NA | | | 1 | -0.4 |
| RI | 464 | TRIM5 | TRIM5 | chr11 | - | 5663194 | 5665682 | 5663194 | 5665395 | 5665655 | 5665682 | 2 | 0 | 0 | 6 | 0 | 0 | 0 | 2 | 0 | 1 | 98 | 49 | 1 | 1 | 0.143 | NA | NA | NA | | | 0.5 | -0.357 |
| RI | 466 | DHRS12 | DHRS12 | chr13 | - | 51767992 | 51769293 | 51767992 | 51768296 | 51769155 | 51769293 | 4 | 8 | 3 | 1 | 3 | 0 | 0 | 6 | 0 | 0 | 98 | 49 | 1 | 1 | 0.667 | 0.571 | 1 | NA | | | 1 | -0.254 |
| RI | 474 | TMEM41B | TMEM41B | chr11 | - | 9280654 | 9283593 | 9280654 | 9281041 | 9283370 | 9283593 | 8 | 2 | 0 | 1 | 1 | 0 | 0 | 2 | 0 | 0 | 98 | 49 | 1 | 1 | 0.8 | 0.5 | NA | NA | | | 1 | -0.35 |
| RI | 475 | AP1G2 | AP1G2 | chr14 | - | 23565815 | 23566160 | 23565815 | 23565892 | 23566063 | 23566160 | 6 | 0 | 0 | 0 | 3 | 0 | 0 | 4 | 0 | 2 | 98 | 49 | 1 | 1 | 1 | 0 | NA | NA | | | 0.5 | 0 |
| RI | 483 | PAPD4 | PAPD4 | chr5 | + | 79612419 | 79613075 | 79612419 | 79612735 | 79613010 | 79613075 | 43 | 22 | 7 | 0 | 0 | 0 | 3 | 7 | 0 | 1 | 98 | 49 | 1 | 1 | 1 | 1 | 1 | | 1 | | 0.778 | 0.111 |
| RI | 490 | MIER1 | MIER1 | chr1 | + | 66984571 | 66988619 | 66984571 | 66984592 | 66986402 | 66988619 | 48 | 10 | 7 | 2 | 2 | 0 | 0 | 6 | 0 | 0 | 98 | 49 | 1 | 1 | 0.923 | 0.714 | 1 | NA | | | 1 | -0.121 |
| RI | 496 | THUMPD1 | THUMPD1 | chr16 | - | 20733665 | 20737286 | 20733665 | 20736358 | 20736859 | 20737286 | 12 | 3 | 1 | 2 | 0 | 0 | 0 | 2 | 0 | 0 | 98 | 49 | 1 | 1 | 0.75 | 1 | 1 | NA | | | 1 | -0.083 |
| RI | 498 | CSTF3 | CSTF3 | chr11 | - | 33141430 | 33141986 | 33141430 | 33141762 | 33141884 | 33141986 | 3 | 0 | 0 | 0 | 3 | 0 | 0 | 0 | 0 | 1 | 98 | 49 | 1 | 1 | 1 | 0 | NA | NA | | | 0 | 0.5 |
| RI | 503 | BICD2 | BICD2 | chr9 | - | 92711362 | 92715463 | 92711362 | 92713461 | 92715252 | 92715463 | 9 | 4 | 2 | 11 | 0 | 0 | 0 | 0 | 0 | 2 | 98 | 49 | 1 | 1 | 0.29 | 1 | 1 | NA | | | 0 | 0.763 |
| RI | 507 | CDC16 | CDC16 | chr13 | + | 114272183 | 114272675 | 114272183 | 114272481 | 114272563 | 114272675 | 3 | 2 | 2 | 0 | 0 | 0 | 0 | 7 | 0 | 2 | 98 | 49 | 1 | 1 | 1 | 1 | 1 | NA | | | 0.636 | 0.364 |
| RI | 531 | MYCL | MYCL | chr1 | - | 39900938 | 39902015 | 39900938 | 39901443 | 39901807 | 39902015 | 1 | 0 | 0 | 1 | 0 | 0 | 0 | 1 | 0 | 0 | 98 | 49 | 1 | 1 | 0.333 | NA | NA | NA | | | 1 | -0.667 |
| RI | 533 | EGR2 | EGR2 | chr10 | - | 62815860 | 62816366 | 62815860 | 62815969 | 62816069 | 62816366 | 85 | 8 | 1 | 1 | 0 | 0 | 0 | 10 | 0 | 0 | 98 | 49 | 1 | 1 | 0.977 | 1 | 1 | NA | | | 1 | -0.008 |
| RI | 537 | NDUFS2 | NDUFS2 | chr1 | + | 161213863 | 161214394 | 161213863 | 161213921 | 161214155 | 161214394 | 2 | 0 | 0 | 1 | 1 | 0 | 0 | 2 | 0 | 0 | 98 | 49 | 1 | 1 | 0.5 | 0 | NA | NA | | | 1 | -0.75 |
| RI | 552 | ACAA1 | ACAA1 | chr3 | - | 38126161 | 38126700 | 38126161 | 38126341 | 38126509 | 38126700 | 9 | 6 | 0 | 2 | 0 | 0 | 0 | 9 | 0 | 1 | 98 | 49 | 1 | 1 | 0.692 | 1 | NA | NA | | | 0.818 | 0.028 |
| RI | 554 | ACADVL | ACADVL | chr17 | + | 7219830 | 7220197 | 7219830 | 7220046 | 7220121 | 7220197 | 6 | 5 | 0 | 5 | 12 | 0 | 2 | 1 | 2 | 3 | 98 | 49 | 1 | 1 | 0.375 | 0.172 | NA | | 0.333 | | 0.143 | 0.035 |
| RI | 555 | VPS28 | VPS28 | chr8 | - | 144423600 | 144424132 | 144423600 | 144423922 | 144424040 | 144424132 | 7 | 5 | 0 | 2 | 7 | 7 | 0 | 0 | 0 | 3 | 98 | 49 | 1 | 1 | 0.636 | 0.263 | 0 | NA | | | 0 | 0.3 |
| RI | 574 | HCFC1R1 | HCFC1R1 | chr16 | - | 3023232 | 3023530 | 3023232 | 3023361 | 3023473 | 3023530 | 1 | 0 | 0 | 1 | 0 | 0 | 0 | 1 | 0 | 0 | 98 | 49 | 1 | 1 | 0.333 | NA | NA | NA | | | 1 | -0.667 |
| RI | 578 | NAP1L1 | NAP1L1 | chr12 | - | 76045382 | 76048464 | 76045382 | 76047718 | 76048392 | 76048464 | 3 | 0 | 0 | 27 | 1 | 5 | 0 | 0 | 0 | 1 | 98 | 49 | 1 | 1 | 0.053 | 0 | 0 | NA | | | 0 | 0.018 |
| RI | 583 | REC8 | REC8 | chr14 | + | 24172024 | 24172608 | 24172024 | 24172365 | 24172466 | 24172608 | 2 | 8 | 0 | 0 | 0 | 1 | 1 | 7 | 0 | 0 | 98 | 49 | 1 | 1 | 1 | 1 | 0 | | 1 | | 1 | -0.333 |
| RI | 611 | NFKB2 | NFKB2 | chr10 | + | 102395595 | 102395980 | 102395595 | 102395796 | 102395887 | 102395980 | 11 | 1 | 3 | 0 | 4 | 0 | 0 | 0 | 0 | 1 | 98 | 49 | 1 | 1 | 1 | 0.111 | 1 | NA | | | 0 | 0.704 |
| RI | 614 | CELF2 | CELF2 | chr10 | + | 11328925 | 11336673 | 11328925 | 11329097 | 11330455 | 11336673 | 106 | 44 | 20 | 9 | 1 | 3 | 0 | 29 | 0 | 1 | 98 | 49 | 1 | 1 | 0.855 | 0.957 | 0.769 | NA | | | 0.935 | -0.075 |
| RI | 618 | TRIM52-AS1 | TRIM52-AS1 | chr5 | + | 181263627 | 181264566 | 181263627 | 181263766 | 181264051 | 181264566 | 1 | 2 | 3 | 2 | 0 | 0 | 0 | 2 | 0 | 0 | 98 | 49 | 1 | 1 | 0.2 | 1 | 1 | NA | | | 1 | -0.267 |
| RI | 630 | GMPR2 | GMPR2 | chr14 | + | 24238245 | 24239241 | 24238245 | 24238405 | 24238588 | 24239241 | 23 | 11 | 3 | 13 | 4 | 5 | 0 | 7 | 0 | 1 | 98 | 49 | 1 | 1 | 0.469 | 0.579 | 0.231 | NA | | | 0.778 | -0.352 |
| RI | 648 | SEC14L1 | SEC14L1 | chr17 | + | 77213917 | 77217099 | 77213917 | 77214019 | 77216548 | 77217099 | 197 | 72 | 25 | 1 | 0 | 1 | 13 | 22 | 0 | 0 | 98 | 49 | 1 | 1 | 0.99 | 1 | 0.926 | | 1 | | 1 | -0.028 |
| RI | 663 | LRTOMT | LRTOMT | chr11 | + | 72094947 | 72096892 | 72094947 | 72095096 | 72095378 | 72096892 | 2 | 0 | 0 | 0 | 1 | 0 | 0 | 1 | 0 | 0 | 98 | 49 | 1 | 1 | 1 | 0 | NA | NA | | | 1 | -0.5 |
| RI | 673 | KCNK7 | KCNK7 | chr11 | - | 65592854 | 65593874 | 65592854 | 65593210 | 65593475 | 65593874 | 12 | 10 | 4 | 3 | 0 | 0 | 0 | 5 | 0 | 0 | 98 | 49 | 1 | 1 | 0.667 | 1 | 1 | NA | | | 1 | -0.111 |
| RI | 677 | P3H1 | P3H1 | chr1 | - | 42746334 | 42747412 | 42746334 | 42746852 | 42747271 | 42747412 | 0 | 0 | 0 | 1 | 0 | 0 | 0 | 1 | 0 | 0 | 98 | 49 | 1 | 1 | 0 | NA | NA | NA | | | 1 | -1 |
| RI | 712 | CREBZF | CREBZF | chr11 | - | 85657563 | 85665138 | 85657563 | 85662457 | 85663523 | 85665138 | 9 | 12 | 0 | 4 | 1 | 1 | 0 | 4 | 0 | 2 | 98 | 49 | 1 | 1 | 0.529 | 0.857 | 0 | NA | | | 0.5 | -0.038 |
| RI | 713 | CREBZF | CREBZF | chr11 | - | 85663523 | 85665138 | 85663523 | 85663601 | 85663814 | 85665138 | 7 | 3 | 0 | 3 | 0 | 0 | 0 | 2 | 0 | 0 | 98 | 49 | 1 | 1 | 0.538 | 1 | NA | NA | | | 1 | -0.231 |
| RI | 716 | BET1L | BET1L | chr11 | - | 202923 | 205667 | 202923 | 205469 | 205610 | 205667 | 17 | 1 | 0 | 1 | 2 | 0 | 0 | 7 | 0 | 0 | 98 | 49 | 1 | 1 | 0.895 | 0.2 | NA | NA | | | 1 | -0.453 |

| | | | | | | | | | | | | | | | | | | | | | | | | | | | | | | | |
|---|---|---|---|---|---|---|---|---|---|---|---|---|---|---|---|---|---|---|---|---|---|---|---|---|---|---|---|---|---|---|---|
| RI | 728 | PTPMT1 | PTPMT1 | chr11 | + | 47565429 | 47565986 | 47565429 | 47565796 | 47565905 | 47565986 | 2 | 1 | 0 | 1 | 0 | 0 | 0 | 2 | 1 | 0 | 98 | 49 | 1 | 1 | 0.5 | 1 | NA | | 0 | 1 | 0.25 |
| RI | 730 | WRAP53 | WRAP53 | chr17 | + | 7688348 | 7689079 | 7688348 | 7688561 | 7688647 | 7689079 | 3 | 0 | 0 | 0 | 0 | 0 | 0 | 0 | 0 | 2 | 98 | 49 | 1 | 1 | 1 | NA | NA | NA | 0 | 1 |
| RI | 731 | RANGRF | RANGRF | chr17 | + | 8289257 | 8290091 | 8289257 | 8289414 | 8289502 | 8290091 | 1 | 0 | 0 | 2 | 0 | 0 | 0 | 1 | 0 | 0 | 98 | 49 | 1 | 1 | 0.2 | NA | NA | NA | 1 | -0.8 |
| RI | 741 | QSOX1 | QSOX1 | chr1 | + | 180196261 | 180198034 | 180196261 | 180196599 | 180197332 | 180198034 | 12 | 5 | 11 | 0 | 0 | 0 | 0 | 11 | 0 | 3 | 98 | 49 | 1 | 1 | 1 | 1 | 1 | NA | 0.647 | 0.353 |
| RI | 743 | ZNF248 | ZNF248 | chr10 | - | 37828764 | 37833116 | 37828764 | 37831312 | 37833024 | 37833116 | 0 | 0 | 0 | 1 | 0 | 0 | 0 | 1 | 0 | 0 | 98 | 49 | 1 | 1 | 0 | NA | NA | NA | 1 | -1 |
| RI | 754 | UBXN1 | UBXN1 | chr11 | - | 62676497 | 62677005 | 62676497 | 62676639 | 62676812 | 62677005 | 7 | 7 | 0 | 1 | 2 | 4 | 0 | 0 | 0 | 7 | 98 | 49 | 1 | 1 | 0.778 | 0.636 | 0 | NA | 0 | 0.471 |
| RI | 765 | SRSF1 | SRSF1 | chr17 | - | 58000918 | 58005973 | 58000918 | 58004001 | 58004922 | 58005973 | 31 | 20 | 14 | 3 | 1 | 0 | 0 | 6 | 0 | 0 | 98 | 49 | 1 | 1 | 0.838 | 0.909 | 1 | NA | 1 | -0.084 |
| RI | 766 | SRSF1 | SRSF1 | chr17 | - | 58000918 | 58005973 | 58000918 | 58005600 | 58005800 | 58005973 | 10 | 3 | 5 | 12 | 2 | 1 | 0 | 1 | 0 | 5 | 98 | 49 | 1 | 1 | 0.294 | 0.429 | 0.714 | NA | 0.091 | 0.388 |
| RI | 767 | GLYCTK | GLYCTK | chr3 | + | 52292259 | 52293192 | 52292259 | 52292569 | 52293045 | 52293192 | 4 | 0 | 2 | 0 | 0 | 1 | 2 | 1 | 0 | 0 | 98 | 49 | 1 | 1 | 1 | NA | 0.5 | 1 | 1 | -0.25 |
| RI | 775 | DCAF7 | DCAF7 | chr17 | + | 63588999 | 63594281 | 63588999 | 63589237 | 63594202 | 63594281 | 3 | 11 | 1 | 1 | 1 | 0 | 0 | 5 | 0 | 2 | 98 | 49 | 1 | 1 | 0.6 | 0.846 | 1 | NA | 0.556 | 0.259 |
| RI | 788 | TYMP | TYMP | chr22 | - | 50529495 | 50530085 | 50529495 | 50529731 | 50529903 | 50530085 | 70 | 48 | 6 | 2 | 0 | 0 | 7 | 17 | 0 | 1 | 98 | 49 | 1 | 1 | 0.946 | 1 | 1 | 1 | 0.895 | 0.034 |
| RI | 790 | CANT1 | CANT1 | chr17 | - | 78996991 | 78997963 | 78996991 | 78997644 | 78997839 | 78997963 | 24 | 16 | 4 | 6 | 1 | 0 | 0 | 8 | 0 | 0 | 98 | 49 | 1 | 1 | 0.667 | 0.889 | 1 | NA | 1 | -0.148 |
| RI | 794 | OXLD1 | OXLD1 | chr17 | - | 81665035 | 81666637 | 81665035 | 81665584 | 81666154 | 81666637 | 2 | 0 | 0 | 0 | 0 | 0 | 0 | 1 | 0 | 2 | 98 | 49 | 1 | 1 | 1 | NA | NA | NA | 0.2 | 0.8 |
| RI | 797 | OXLD1 | OXLD1 | chr17 | - | 81665035 | 81666637 | 81665035 | 81665584 | 81666517 | 81666637 | 1 | 0 | 0 | 0 | 0 | 0 | 0 | 1 | 0 | 1 | 98 | 49 | 1 | 1 | 1 | NA | NA | NA | 0.333 | 0.667 |
| RI | 798 | FAM195B | FAM195B | chr17 | - | 81822360 | 81823311 | 81822360 | 81822907 | 81823208 | 81823311 | 3 | 2 | 0 | 2 | 0 | 0 | 0 | 3 | 0 | 0 | 98 | 49 | 1 | 1 | 0.429 | 1 | NA | NA | 1 | -0.285 |
| RI | 833 | TMEM205 | TMEM205 | chr19 | - | 11345519 | 11346305 | 11345519 | 11345628 | 11346126 | 11346305 | 1 | 1 | 2 | 3 | 0 | 0 | 0 | 4 | 0 | 1 | 98 | 49 | 1 | 1 | 0.143 | 1 | 1 | NA | 0.667 | 0.047 |
| RI | 840 | AKAP8L | AKAP8L | chr19 | - | 15403474 | 15403715 | 15403474 | 15403555 | 15403657 | 15403715 | 31 | 5 | 6 | 0 | 2 | 5 | 0 | 4 | 0 | 1 | 98 | 49 | 1 | 1 | 1 | 0.556 | 0.375 | NA | 0.667 | -0.023 |
| RI | 843 | AUP1 | AUP1 | chr2 | - | 74528750 | 74529282 | 74528750 | 74528935 | 74529131 | 74529282 | 6 | 0 | 0 | 0 | 0 | 1 | 1 | 0 | 0 | 2 | 98 | 49 | 1 | 1 | 1 | NA | 0 | 1 | 0 | 0 |
| RI | 845 | CCDC125 | CCDC125 | chr5 | - | 69320236 | 69320580 | 69320236 | 69320345 | 69320464 | 69320580 | 33 | 7 | 0 | 5 | 1 | 1 | 0 | 0 | 0 | 1 | 98 | 49 | 1 | 1 | 0.767 | 0.778 | 0 | NA | 0 | 0.515 |
| RI | 847 | EDEM2 | EDEM2 | chr20 | - | 35146824 | 35147358 | 35146824 | 35146935 | 35147151 | 35147358 | 7 | 0 | 0 | 4 | 6 | 0 | 0 | 0 | 0 | 1 | 98 | 49 | 1 | 1 | 0.467 | 0 | NA | NA | 0 | 0.234 |
| RI | 849 | MCL1 | MCL1 | chr1 | - | 150578842 | 150579738 | 150578842 | 150578963 | 150579422 | 150579738 | 1364 | 513 | 216 | 3 | 0 | 0 | 41 | 264 | 0 | 2 | 98 | 49 | 1 | 1 | 0.996 | 1 | 1 | 1 | 0.985 | 0.006 |
| RI | 856 | CUTA | CUTA | chr6 | - | 33417480 | 33418288 | 33417480 | 33417695 | 33417958 | 33418288 | 4 | 2 | 0 | 0 | 1 | 0 | 1 | 1 | 0 | 1 | 98 | 49 | 1 | 1 | 1 | 0.5 | NA | 1 | 0.333 | 0.084 |
| RI | 865 | U2AF1L4 | U2AF1L4 | chr19 | - | 35744069 | 35744421 | 35744069 | 35744145 | 35744322 | 35744421 | 5 | 13 | 0 | 2 | 1 | 4 | 0 | 1 | 0 | 1 | 98 | 49 | 1 | 1 | 0.556 | 0.867 | 0 | NA | 0.333 | 0.141 |
| RI | 871 | NFKBIB | NFKBIB | chr19 | + | 38907398 | 38908894 | 38907398 | 38907659 | 38908730 | 38908894 | 9 | 3 | 1 | 2 | 0 | 0 | 0 | 0 | 0 | 1 | 98 | 49 | 1 | 1 | 0.692 | 1 | 1 | NA | 0 | 0.897 |
| RI | 881 | NAPA | NAPA | chr19 | - | 47487633 | 47488389 | 47487633 | 47487890 | 47488234 | 47488389 | 16 | 17 | 11 | 1 | 0 | 0 | 0 | 17 | 0 | 0 | 98 | 49 | 1 | 1 | 0.889 | 1 | 1 | NA | 1 | -0.037 |
| RI | 919 | PTPRO | PTPRO | chr12 | + | 15594936 | 15598331 | 15594936 | 15595057 | 15596089 | 15598331 | 1 | 0 | 0 | 0 | 0 | 0 | 0 | 0 | 0 | 1 | 98 | 49 | 1 | 1 | 1 | NA | NA | NA | 0 | 1 |
| RI | 925 | CLK1 | CLK1 | chr2 | - | 200859679 | 200861466 | 200859679 | 200859746 | 200861237 | 200861466 | 27 | 16 | 2 | 18 | 9 | 6 | 0 | 5 | 0 | 2 | 98 | 49 | 1 | 1 | 0.429 | 0.471 | 0.143 | NA | 0.556 | -0.208 |
| RI | 944 | SUMO1 | SUMO1 | chr2 | - | 202206179 | 202207321 | 202206179 | 202206803 | 202207252 | 202207321 | 23 | 13 | 9 | 0 | 0 | 0 | 1 | 10 | 0 | 1 | 98 | 49 | 1 | 1 | 1 | 1 | 1 | 1 | 0.833 | 0.084 |
| RI | 945 | RPL31 | RPL31 | chr2 | + | 101005958 | 101006423 | 101005958 | 101006071 | 101006349 | 101006423 | 6 | 1 | 3 | 53 | 13 | 6 | 0 | 4 | 0 | 5 | 98 | 49 | 1 | 1 | 0.054 | 0.037 | 0.2 | NA | 0.286 | -0.189 |
| RI | 947 | C2orf49 | C2orf49 | chr2 | + | 105342847 | 105343223 | 105342847 | 105342974 | 105343100 | 105343223 | 5 | 3 | 6 | 1 | 0 | 0 | 0 | 2 | 0 | 0 | 98 | 49 | 1 | 1 | 0.714 | 1 | 1 | NA | 1 | -0.095 |
| RI | 953 | EIF6 | EIF6 | chr20 | - | 35284175 | 35284492 | 35284175 | 35284261 | 35284380 | 35284492 | 4 | 0 | 0 | 7 | 1 | 0 | 0 | 1 | 0 | 0 | 98 | 49 | 1 | 1 | 0.222 | 0 | NA | NA | 1 | -0.889 |
| RI | 956 | ATP5G3 | ATP5G3 | chr2 | - | 175176257 | 175179250 | 175176257 | 175178402 | 175179056 | 175179250 | 1 | 2 | 0 | 2 | 0 | 0 | 0 | 0 | 0 | 1 | 98 | 49 | 1 | 1 | 0.2 | 1 | NA | NA | 0 | 0.6 |
| RI | 974 | SMOX | SMOX | chr20 | + | 4182088 | 4182848 | 4182088 | 4182323 | 4182482 | 4182848 | 3 | 0 | 0 | 0 | 1 | 0 | 1 | 0 | 1 | 0 | 98 | 49 | 1 | 1 | 1 | 0 | NA | 0.333 | NA | 0.167 |
| RI | 991 | DCTN1 | DCTN1 | chr2 | - | 74366243 | 74366620 | 74366243 | 74366375 | 74366458 | 74366620 | 4 | 0 | 0 | 8 | 0 | 1 | 0 | 1 | 1 | 1 | 98 | 49 | 1 | 1 | 0.2 | NA | 0 | 0 | 0.333 | -0.067 |
| RI | 996 | PAQR6 | PAQR6 | chr1 | - | 156245781 | 156246250 | 156245781 | 156245987 | 156246122 | 156246250 | 0 | 0 | 1 | 0 | 0 | 6 | 2 | 0 | 0 | 0 | 98 | 49 | 1 | 1 | NA | NA | 0.077 | 1 | NA | -0.923 |
| RI | 1007 | DKC1 | DKC1 | chrX | + | 154774601 | 154775273 | 154774601 | 154774705 | 154775194 | 154775273 | 5 | 4 | 0 | 2 | 3 | 0 | 0 | 1 | 0 | 0 | 98 | 49 | 1 | 1 | 0.556 | 0.4 | NA | NA | 1 | -0.522 |
| RI | 1010 | TAF1C | TAF1C | chr16 | - | 84181327 | 84181663 | 84181327 | 84181463 | 84181591 | 84181663 | 8 | 1 | 0 | 0 | 0 | 0 | 0 | 3 | 0 | 1 | 98 | 49 | 1 | 1 | 1 | 1 | NA | NA | 0.6 | 0.4 |
| RI | 1030 | XBP1 | XBP1 | chr22 | - | 28796046 | 28796192 | 28796046 | 28796123 | 28796149 | 28796192 | 11 | 3 | 0 | 2 | 0 | 0 | 0 | 4 | 0 | 0 | 75 | 49 | 1 | 1 | 0.782 | 1 | NA | NA | 1 | -0.109 |
| RI | 1036 | TSTD1 | TSTD1 | chr1 | - | 161037631 | 161038075 | 161037631 | 161037826 | 161037912 | 161038075 | 2 | 0 | 0 | 4 | 7 | 2 | 0 | 0 | 0 | 1 | 98 | 49 | 1 | 1 | 0.2 | 0 | 0 | NA | 0 | 0.067 |
| RI | 1052 | IL17RC | IL17RC | chr3 | + | 9923880 | 9924291 | 9923880 | 9924020 | 9924231 | 9924291 | 0 | 0 | 0 | 2 | 0 | 0 | 1 | 0 | 0 | 0 | 98 | 49 | 1 | 1 | 0 | NA | NA | 1 | 0 | -0.5 |
| RI | 1053 | RPRD2 | RPRD2 | chr1 | + | 150470560 | 150476565 | 150470560 | 150471742 | 150471882 | 150476565 | 0 | 0 | 0 | 1 | 0 | 0 | 1 | 0 | 0 | 0 | 98 | 49 | 1 | 1 | 0 | NA | NA | 1 | NA | -1 |
| RI | 1068 | CTNNB1 | CTNNB1 | chr3 | + | 41239133 | 41240448 | 41239133 | 41239354 | 41239659 | 41240448 | 19 | 13 | 0 | 7 | 0 | 6 | 0 | 6 | 0 | 3 | 98 | 49 | 1 | 1 | 0.576 | 1 | 0 | NA | 0.5 | 0.025 |
| RI | 1069 | CTNNB1 | CTNNB1 | chr3 | + | 41239133 | 41240448 | 41239133 | 41239354 | 41239818 | 41240448 | 19 | 13 | 0 | 1 | 0 | 1 | 0 | 13 | 0 | 0 | 98 | 49 | 1 | 1 | 0.905 | 1 | 0 | NA | 1 | -0.365 |
| RI | 1080 | NDUFB4 | NDUFB4 | chr3 | + | 120601110 | 120602411 | 120601110 | 120601257 | 120602207 | 120602411 | 1 | 1 | 0 | 1 | 0 | 3 | 0 | 0 | 0 | 1 | 98 | 49 | 1 | 1 | 0.333 | 1 | 0 | NA | 0 | 0.444 |
| RI | 1089 | ROPN1L | ROPN1L | chr5 | + | 10441861 | 10442298 | 10441861 | 10442003 | 10442126 | 10442298 | 5 | 3 | 1 | 2 | 0 | 0 | 0 | 4 | 0 | 0 | 98 | 49 | 1 | 1 | 0.556 | 1 | 1 | NA | 1 | -0.148 |
| RI | 1090 | NIPBL | NIPBL | chr5 | + | 37063789 | 37065819 | 37063789 | 37063978 | 37064526 | 37065819 | 15 | 5 | 0 | 6 | 1 | 0 | 0 | 4 | 0 | 0 | 98 | 49 | 1 | 1 | 0.556 | 0.714 | NA | NA | 1 | -0.365 |
| RI | 1091 | NSD1 | NSD1 | chr5 | + | 177135086 | 177136030 | 177135086 | 177135200 | 177135940 | 177136030 | 11 | 5 | 1 | 17 | 1 | 0 | 0 | 1 | 0 | 0 | 98 | 49 | 1 | 1 | 0.244 | 0.714 | 1 | NA | 1 | -0.347 |
| RI | 1096 | CIRBP | CIRBP | chr19 | - | 1271980 | 1273172 | 1271980 | 1272051 | 1272426 | 1273172 | 32 | 2 | 2 | 22 | 6 | 3 | 0 | 3 | 0 | 1 | 98 | 49 | 1 | 1 | 0.421 | 0.143 | 0.25 | NA | 0.6 | -0.329 |
| RI | 1110 | HNRNPM | HNRNPM | chr19 | + | 8465323 | 8465515 | 8465323 | 8465362 | 8465479 | 8465515 | 13 | 6 | 1 | 5 | 2 | 3 | 0 | 8 | 0 | 1 | 98 | 49 | 1 | 1 | 0.565 | 0.6 | 0.143 | NA | 0.8 | -0.364 |
| RI | 1123 | SKP1 | SKP1 | chr5 | - | 134156390 | 134158595 | 134156390 | 134157768 | 134158454 | 134158595 | 18 | 3 | 3 | 2 | 0 | 0 | 0 | 1 | 0 | 0 | 98 | 49 | 1 | 1 | 0.818 | 1 | 1 | NA | 1 | -0.061 |
| RI | 1131 | NDUFA2 | NDUFA2 | chr5 | - | 140647255 | 140647785 | 140647255 | 140647362 | 140647482 | 140647785 | 8 | 2 | 5 | 8 | 5 | 1 | 0 | 0 | 0 | 2 | 98 | 49 | 1 | 1 | 0.333 | 0.167 | 0.714 | NA | 0 | 0.405 |
| RI | 1133 | RELL2 | RELL2 | chr5 | + | 141636949 | 141638409 | 141636949 | 141637741 | 141638169 | 141638409 | 6 | 2 | 0 | 2 | 0 | 0 | 2 | 0 | 0 | 0 | 98 | 49 | 1 | 1 | 0.6 | 1 | NA | 1 | NA | -0.2 |
| RI | 1134 | ABLIM3 | ABLIM3 | chr5 | + | 149141482 | 149142108 | 149141482 | 149141654 | 149142008 | 149142108 | 1 | 0 | 0 | 0 | 0 | 0 | 0 | 1 | 1 | 0 | 98 | 49 | 1 | 1 | 1 | NA | NA | 0 | 1 | 0.5 |
| RI | 1135 | ZNF438 | ZNF438 | chr10 | - | 30848530 | 30850367 | 30848530 | 30849443 | 30850221 | 30850367 | 10 | 8 | 4 | 1 | 0 | 0 | 1 | 12 | 0 | 0 | 98 | 49 | 1 | 1 | 0.833 | 1 | 1 | 1 | 1 | -0.056 |

**U12_4_filtered**

| | | | | | | | | | | | | | Inclusion counts | | | Skipping counts | | | Inclusion counts | | Skipping counts | | | | | | | Inclusion_level(normalized) | | | |
|---|---|---|---|---|---|---|---|---|---|---|---|---|---|---|---|---|---|---|---|---|---|---|---|---|---|---|---|---|---|---|---|
| Splice_type | ID | Gene_ID | Gene_Symbol | Chromosome | Strand | LongExonStart_0base | LongExonEnd | ShortES | ShortEE | FlankingES | FlankingEE | control_s3 | control_s4 | control_s10 | control_s3 | control_s4 | control_s10 | ldg_14 | ldg_s16 | ldg_s14 | ldg_s16 | IncFormLen | SkipFormLen | p-value | adjust_p | IncLevel_s3 | IncLevel_s4 | IncLevel_s10 | IncLevel14 | IncLevel16 | Level_diff |
| SE | 8677 | MAEA | MAEA | chr4 | + | 1322380 | 1322503 | 1315396 | 1315600 | 1332756 | 1332865 | 11 | 5 | 2 | 0 | 4 | 0 | 0 | 0 | 1 | 1 | 98 | 49 | 5.28102E-08 | 0.000152595 | 1 | 0.385 | 1 | 0 | 0 | 0.795 |
| RI | 488 | STX10 | STX10 | chr19 | - | 13144057 | 13144671 | 13144057 | 13144486 | 13144576 | 13144671 | 15 | 16 | 0 | 9 | 10 | 13 | 1 | 6 | 0 | 1 | 98 | 49 | 0.000313634 | 0.0129374 | 0.455 | 0.444 | 0 | 1 | 0.75 | -0.575 |
| SE | 6756 | SETD1B | SETD1B | chr12 | + | 121819403 | 121819895 | 121817798 | 121817904 | 121822489 | 121823749 | 8 | 9 | 3 | 0 | 0 | 0 | 3 | 1 | 0 | 3 | 98 | 49 | 0.014021703 | 0.578795883 | 1 | 1 | 1 | 1 | 0.143 | 0.428 |
| SE | 5176 | SLC12A9 | SLC12A9 | chr7 | + | 100861137 | 100861262 | 100860149 | 100860232 | 100861391 | 100861584 | 62 | 24 | 16 | 1 | 2 | 3 | 4 | 10 | 0 | 3 | 98 | 49 | 0.01640589 | 0.627878418 | 0.969 | 0.857 | 0.727 | 1 | 1 | -0.149 |
| SE | 389 | RAPGEF1 | RAPGEF1 | chr9 | - | 131650128 | 131650242 | 131643238 | 131643417 | 131650809 | 131650949 | 18 | 3 | 1 | 4 | 2 | 0 | 2 | 1 | 0 | 1 | 98 | 49 | 0.030356618 | 0.958638776 | 0.692 | 0.429 | 1 | 1 | 0.333 | 0.04 |
| SE | 8832 | DOCK2 | DOCK2 | chr5 | + | 169759704 | 169759775 | 169747395 | 169747504 | 169761518 | 169761625 | 103 | 48 | 17 | 3 | 3 | 0 | 3 | 30 | 0 | 0 | 98 | 49 | 0.054985887 | 1 | 0.945 | 0.889 | 1 | 1 | 1 | -0.055 |
| SE | 15461 | RAF1 | RAF1 | chr3 | - | 12611949 | 12612062 | 12609232 | 12609335 | 12618514 | 12618747 | 159 | 54 | 30 | 3 | 0 | 4 | 3 | 15 | 0 | 0 | 98 | 49 | 0.055208514 | 1 | 0.964 | 1 | 0.789 | 1 | 1 | -0.082 |
| SE | 6836 | GAK | GAK | chr4 | - | 877614 | 877809 | 877089 | 877207 | 881906 | 882040 | 8 | 7 | 5 | 1 | 3 | 0 | 2 | 6 | 0 | 0 | 98 | 49 | 0.061544568 | 1 | 0.8 | 0.538 | 1 | 1 | 1 | -0.221 |
| SE | 7502 | SLC12A6 | SLC12A6 | chr15 | - | 34275344 | 34275389 | 34260925 | 34261020 | 34336409 | 34336601 | 155 | 85 | 27 | 2 | 2 | 3 | 6 | 36 | 0 | 0 | 94 | 49 | 0.072192319 | 1 | 0.976 | 0.957 | 0.824 | 1 | 1 | -0.081 |
| SE | 1987 | SRPK1 | SRPK1 | chr6 | - | 35870280 | 35870494 | 35869481 | 35869901 | 35870933 | 35870959 | 19 | 4 | 7 | 8 | 10 | 0 | 0 | 4 | 1 | 5 | 98 | 49 | 0.082834818 | 1 | 0.543 | 0.167 | 1 | 0 | 0.286 | 0.427 |
| SE | 8679 | MAEA | MAEA | chr4 | + | 1327626 | 1327703 | 1322380 | 1322503 | 1332756 | 1332865 | 13 | 8 | 8 | 0 | 3 | 1 | 3 | 2 | 0 | 0 | 98 | 49 | 0.089381693 | 1 | 1 | 0.571 | 0.8 | 1 | 1 | -0.21 |
| SE | 7319 | VPS11 | VPS11 | chr11 | + | 119078564 | 119078704 | 119078172 | 119078334 | 119078794 | 119078997 | 6 | 1 | 2 | 0 | 0 | 0 | 0 | 2 | 1 | 0 | 98 | 49 | 0.101331953 | 1 | 1 | 1 | 1 | 0 | 1 | 0.5 |
| SE | 10160 | BRMS1 | BRMS1 | chr11 | - | 66340120 | 66340213 | 66338720 | 66338785 | 66340773 | 66340870 | 21 | 4 | 9 | 0 | 0 | 0 | 0 | 2 | 1 | 0 | 98 | 49 | 0.101331953 | 1 | 1 | 1 | 1 | 0 | 1 | 0.5 |
| A3SS | 2325 | ARPC5 | ARPC5 | chr1 | - | 183633081 | 183633163 | 183633081 | 183633154 | 183635516 | 183635941 | 7 | 1 | 0 | 103 | 77 | 27 | 0 | 0 | 4 | 20 | 58 | 49 | 0.153712068 | 1 | 0.054 | 0.011 | 0 | 0 | 0 | 0.022 |
| SE | 6835 | GAK | GAK | chr4 | - | 851749 | 851974 | 850935 | 851084 | 859605 | 859722 | 8 | 3 | 6 | 0 | 0 | 0 | 1 | 6 | 2 | 0 | 98 | 49 | 0.177931592 | 1 | 1 | 1 | 1 | 0.2 | 1 | 0.4 |
| RI | 580 | ATG3 | ATG3 | chr3 | - | 112532506 | 112534337 | 112532506 | 112532780 | 112534268 | 112534337 | 10 | 6 | 4 | 28 | 11 | 3 | 0 | 3 | 3 | 5 | 98 | 49 | 0.199919419 | 1 | 0.152 | 0.214 | 0.4 | 0 | 0.231 | 0.14 |
| SE | 978 | DOCK5 | DOCK5 | chr8 | + | 25407982 | 25408154 | 25403557 | 25403724 | 25408801 | 25408940 | 105 | 24 | 4 | 5 | 5 | 1 | 1 | 23 | 1 | 1 | 98 | 49 | 0.239086319 | 1 | 0.913 | 0.706 | 0.667 | 0.333 | 0.92 | 0.135 |
| SE | 976 | DOCK5 | DOCK5 | chr8 | + | 25368570 | 25368725 | 25366869 | 25366970 | 25369555 | 25369641 | 85 | 22 | 4 | 3 | 0 | 0 | 1 | 12 | 0 | 0 | 98 | 49 | 0.347461072 | 1 | 0.934 | 1 | 1 | 1 | 1 | -0.022 |
| SE | 11956 | CHD3 | CHD3 | chr17 | + | 7902936 | 7903061 | 7902609 | 7902727 | 7903271 | 7903503 | 9 | 7 | 2 | 0 | 0 | 0 | 1 | 3 | 0 | 2 | 98 | 49 | 0.39801607 | 1 | 1 | 1 | 1 | 1 | 0.429 | 0.285 |
| SE | 8831 | DOCK2 | DOCK2 | chr5 | + | 169714359 | 169714457 | 169714027 | 169714211 | 169716212 | 169716302 | 109 | 62 | 13 | 3 | 0 | 0 | 13 | 24 | 0 | 0 | 98 | 49 | 0.477632073 | 1 | 0.948 | 1 | 1 | 1 | 1 | -0.017 |
| SE | 5177 | SLC12A9 | SLC12A9 | chr7 | + | 100861736 | 100861911 | 100861391 | 100861584 | 100862680 | 100862827 | 53 | 23 | 12 | 2 | 1 | 0 | 2 | 16 | 0 | 0 | 98 | 49 | 0.480487308 | 1 | 0.93 | 0.92 | 1 | 1 | 1 | -0.05 |
| SE | 8828 | DOCK2 | DOCK2 | chr5 | + | 169684195 | 169684350 | 169681743 | 169681879 | 169689251 | 169689333 | 203 | 80 | 23 | 3 | 1 | 1 | 4 | 41 | 0 | 0 | 98 | 49 | 0.555294052 | 1 | 0.971 | 0.976 | 0.92 | 1 | 1 | -0.044 |
| SE | 6426 | RALGAPA2 | RALGAPA2 | chr20 | - | 20680690 | 20680801 | 20653529 | 20653587 | 20712374 | 20712623 | 23 | 10 | 2 | 2 | 1 | 0 | 2 | 7 | 0 | 0 | 98 | 49 | 0.574229202 | 1 | 0.852 | 0.833 | 1 | 1 | 1 | -0.105 |
| SE | 15458 | RAF1 | RAF1 | chr3 | - | 12590797 | 12590974 | 12587590 | 12587637 | 12591707 | 12591792 | 180 | 64 | 32 | 2 | 0 | 0 | 1 | 24 | 0 | 2 | 98 | 49 | 0.579441471 | 1 | 0.978 | 1 | 1 | 1 | 0.857 | 0.064 |
| SE | 8678 | MAEA | MAEA | chr4 | + | 1327626 | 1327703 | 1315396 | 1315600 | 1332756 | 1332865 | 12 | 5 | 7 | 0 | 4 | 0 | 3 | 2 | 1 | 1 | 98 | 49 | 0.60937349 | 1 | 1 | 0.385 | 1 | 0.6 | 0.5 | 0.245 |
| SE | 6430 | LMBR1L | LMBR1L | chr12 | - | 49104447 | 49104551 | 49103686 | 49103813 | 49104745 | 49104885 | 9 | 3 | 2 | 0 | 2 | 0 | 1 | 2 | 0 | 0 | 98 | 49 | 0.683158228 | 1 | 1 | 0.429 | 1 | 1 | 1 | -0.19 |
| SE | 970 | DOCK5 | DOCK5 | chr8 | + | 25300575 | 25300657 | 25298943 | 25299101 | 25302324 | 25302454 | 135 | 60 | 13 | 0 | 4 | 0 | 3 | 24 | 0 | 0 | 98 | 49 | 0.730535785 | 1 | 1 | 0.882 | 1 | 1 | 1 | -0.039 |
| A3SS | 1384 | RASGRP4 | RASGRP4 | chr19 | - | 38419859 | 38420013 | 38419859 | 38419971 | 38420130 | 38420262 | 54 | 25 | 21 | 0 | 0 | 3 | 5 | 20 | 0 | 0 | 91 | 49 | 0.75562353 | 1 | 1 | 1 | 0.79 | 1 | 1 | -0.07 |
| SE | 7501 | SLC12A6 | SLC12A6 | chr15 | - | 34257641 | 34257788 | 34256228 | 34256283 | 34258812 | 34258944 | 64 | 22 | 9 | 10 | 11 | 3 | 1 | 11 | 0 | 4 | 98 | 49 | 0.771324392 | 1 | 0.762 | 0.5 | 0.6 | 1 | 0.579 | -0.169 |
| SE | 16463 | ZDHHC17 | ZDHHC17 | chr12 | + | 76797433 | 76797537 | 76764073 | 76764329 | 76805316 | 76805439 | 20 | 10 | 2 | 2 | 2 | 2 | 1 | 8 | 0 | 3 | 98 | 49 | 0.914990937 | 1 | 0.833 | 0.714 | 0.333 | 1 | 0.571 | -0.159 |
| SE | 11513 | SBNO2 | SBNO2 | chr19 | - | 1112188 | 1112301 | 1111995 | 1112067 | 1112401 | 1112537 | 61 | 21 | 22 | 10 | 4 | 0 | 8 | 15 | 4 | 0 | 98 | 49 | 0.956284839 | 1 | 0.753 | 0.724 | 1 | 0.5 | 1 | 0.076 |
| A3SS | 139 | NOL8 | NOL8 | chr9 | - | 92299889 | 92300810 | 92299889 | 92300016 | 92301550 | 92301822 | 2 | 0 | 0 | 2 | 1 | 0 | 0 | 1 | 0 | 0 | 98 | 49 | 1 | 1 | 0.333 | 0 | NA | NA | 1 | -0.834 |
| A3SS | 182 | INTS8 | INTS8 | chr8 | + | 94838462 | 94838618 | 94838493 | 94838618 | 94836523 | 94836631 | 11 | 0 | 1 | 2 | 1 | 0 | 0 | 0 | 0 | 1 | 80 | 49 | 1 | 1 | 0.771 | NA | 1 | NA | 0 | 0.885 |
| A3SS | 384 | SRPK1 | SRPK1 | chr6 | - | 35890894 | 35891041 | 35890894 | 35891013 | 35920467 | 35920528 | 5 | 2 | 0 | 5 | 2 | 0 | 0 | 0 | 1 | 2 | 77 | 49 | 1 | 1 | 0.389 | 0.389 | NA | 0 | 0 | 0.389 |
| A3SS | 632 | EML3 | EML3 | chr11 | - | 62611086 | 62611347 | 62611086 | 62611344 | 62611424 | 62611596 | 1 | 0 | 0 | 4 | 0 | 0 | 0 | 1 | 0 | 0 | 52 | 49 | 1 | 1 | 0.191 | NA | NA | NA | 1 | -0.809 |
| A3SS | 711 | SBNO1 | SBNO1 | chr12 | - | 123345257 | 123345570 | 123345257 | 123345567 | 123348028 | 123348133 | 8 | 1 | 2 | 7 | 4 | 0 | 0 | 0 | 0 | 1 | 52 | 49 | 1 | 1 | 0.519 | 0.191 | 1 | NA | 0 | 0.57 |
| A3SS | 939 | TTC23 | TTC23 | chr15 | - | 99139316 | 99139467 | 99139316 | 99139399 | 99156147 | 99156297 | 0 | 1 | 0 | 0 | 0 | 0 | 0 | 0 | 0 | 1 | 98 | 49 | 1 | 1 | NA | 1 | NA | NA | 0 | 1 |
| A3SS | 1382 | RASGRP4 | RASGRP4 | chr19 | - | 38414847 | 38415123 | 38414847 | 38414916 | 38417051 | 38417168 | 55 | 18 | 9 | 0 | 1 | 1 | 2 | 16 | 0 | 0 | 98 | 49 | 1 | 1 | 1 | 0.9 | 0.818 | 1 | 1 | -0.094 |
| A3SS | 1493 | SRSF10 | SRSF10 | chr1 | - | 23964346 | 23971439 | 23964346 | 23967723 | 23971572 | 23971623 | 4 | 1 | 0 | 4 | 0 | 2 | 0 | 1 | 0 | 0 | 98 | 49 | 1 | 1 | 0.333 | 1 | 0 | NA | 1 | -0.556 |
| A3SS | 1494 | SRSF10 | SRSF10 | chr1 | - | 23964346 | 23971439 | 23964346 | 23967956 | 23971572 | 23971623 | 4 | 4 | 0 | 0 | 1 | 0 | 0 | 1 | 0 | 0 | 98 | 49 | 1 | 1 | 1 | 0.667 | NA | NA | 1 | -0.166 |
| A3SS | 1604 | CEP170 | CEP170 | chr1 | - | 243156220 | 243156455 | 243156220 | 243156347 | 243164283 | 243166116 | 4 | 2 | 0 | 1 | 0 | 0 | 0 | 1 | 0 | 0 | 98 | 49 | 1 | 1 | 0.667 | 1 | NA | NA | 1 | -0.166 |
| A3SS | 1651 | LMBR1L | LMBR1L | chr12 | - | 49101249 | 49101323 | 49101249 | 49101263 | 49101471 | 49101549 | 8 | 3 | 4 | 1 | 1 | 0 | 1 | 4 | 0 | 0 | 98 | 49 | 1 | 1 | 0.8 | 0.6 | 1 | 1 | 1 | -0.2 |
| A3SS | 2082 | CNIH4 | CNIH4 | chr1 | + | 224371282 | 224371423 | 224371329 | 224371423 | 224365878 | 224365991 | 26 | 7 | 6 | 1 | 0 | 0 | 0 | 7 | 0 | 0 | 96 | 49 | 1 | 1 | 0.93 | 1 | 1 | NA | 1 | -0.023 |
| A3SS | 2225 | CHD4 | CHD4 | chr12 | - | 6601959 | 6602175 | 6601959 | 6602154 | 6602375 | 6602497 | 18 | 5 | 0 | 4 | 3 | 0 | 0 | 2 | 0 | 1 | 70 | 49 | 1 | 1 | 0.759 | 0.538 | NA | NA | 0.583 | 0.066 |
| A3SS | 3040 | TNPO2 | TNPO2 | chr19 | - | 12702071 | 12702177 | 12702071 | 12702147 | 12702822 | 12702918 | 0 | 0 | 0 | 2 | 0 | 0 | 1 | 0 | 0 | 0 | 79 | 49 | 1 | 1 | 0 | NA | NA | 1 | NA | -1 |
| A3SS | 3096 | TMEM150B | TMEM150B | chr19 | - | 55320968 | 55321093 | 55320968 | 55321044 | 55322647 | 55322743 | 1 | 0 | 0 | 0 | 0 | 0 | 0 | 0 | 0 | 1 | 98 | 49 | 1 | 1 | 1 | NA | NA | NA | 0 | 1 |
| A3SS | 3469 | DCTN3 | DCTN3 | chr9 | - | 34614041 | 34614132 | 34614041 | 34614101 | 34614709 | 34614768 | 0 | 5 | 0 | 3 | 4 | 1 | 1 | 0 | 0 | 0 | 80 | 49 | 1 | 1 | 0 | 0.434 | 0 | 1 | NA | -0.855 |
| A3SS | 3470 | DCTN3 | DCTN3 | chr9 | - | 34614041 | 34614198 | 34614041 | 34614101 | 34614709 | 34614768 | 0 | 5 | 0 | 3 | 4 | 1 | 1 | 0 | 0 | 0 | 98 | 49 | 1 | 1 | 0 | 0.385 | 0 | 1 | NA | -0.872 |
| A3SS | 4093 | HARS2 | HARS2 | chr5 | + | 140693490 | 140693665 | 140693590 | 140693665 | 140691425 | 140691756 | 4 | 5 | 0 | 9 | 0 | 0 | 0 | 2 | 0 | 0 | 98 | 49 | 1 | 1 | 0.182 | 1 | NA | NA | 1 | -0.409 |
| A5SS | 73 | TRAPPC2 | TRAPPC2 | chrX | - | 13734043 | 13734185 | 13734049 | 13734185 | 13719870 | 13719982 | 2 | 3 | 1 | 0 | 2 | 0 | 0 | 2 | 0 | 0 | 55 | 49 | 1 | 1 | 1 | 0.572 | 1 | NA | 1 | -0.143 |
| A5SS | 352 | VPS52 | VPS52 | chr6 | - | 33271136 | 33271965 | 33271585 | 33271965 | 33270198 | 33270283 | 2 | 0 | 2 | 3 | 1 | 0 | 0 | 0 | 0 | 1 | 98 | 49 | 1 | 1 | 0.25 | 0 | 1 | NA | 0 | 0.417 |
| A5SS | 1012 | SLC12A9 | SLC12A9 | chr7 | + | 100852718 | 100852839 | 100852718 | 100852835 | 100854155 | 100854378 | 4 | 2 | 1 | 3 | 2 | 1 | 0 | 0 | 0 | 2 | 53 | 49 | 1 | 1 | 0.552 | 0.48 | 0.48 | NA | 0 | 0.504 |
| A5SS | 1303 | DAP3 | DAP3 | chr1 | + | 155689093 | 155689243 | 155689093 | 155689174 | 155709772 | 155709824 | 2 | 2 | 0 | 0 | 0 | 0 | 0 | 0 | 0 | 1 | 98 | 49 | 1 | 1 | 1 | 1 | NA | NA | 0 | 1 |
| A5SS | 1349 | STX10 | STX10 | chr19 | - | 13144763 | 13144870 | 13144767 | 13144870 | 13144576 | 13144671 | 38 | 11 | 6 | 0 | 0 | 1 | 4 | 7 | 0 | 0 | 53 | 49 | 1 | 1 | 1 | 1 | 0.847 | 1 | 1 | -0.051 |
| A5SS | 1619 | PLCB2 | PLCB2 | chr15 | - | 40296745 | 40296908 | 40296757 | 40296908 | 40296521 | 40296634 | 16 | 7 | 1 | 3 | 6 | 0 | 0 | 7 | 0 | 3 | 61 | 49 | 1 | 1 | 0.811 | 0.484 | 1 | NA | 0.652 | 0.113 |
| A5SS | 1810 | NAA60 | NAA60 | chr16 | + | 3484698 | 3485083 | 3484698 | 3485061 | 3485466 | 3486963 | 3 | 3 | 0 | 1 | 0 | 0 | 0 | 0 | 0 | 1 | 71 | 49 | 1 | 1 | 0.674 | 1 | NA | NA | 0 | 0.837 |
| A5SS | 2073 | GOSR1 | GOSR1 | chr17 | + | 30481142 | 30481263 | 30481142 | 30481257 | 30484213 | 30484301 | 3 | 1 | 0 | 5 | 0 | 2 | 0 | 1 | 0 | 0 | 55 | 49 | 1 | 1 | 0.348 | 1 | 0 | NA | 1 | -0.551 |
| A5SS | 3129 | MCMBP | MCMBP | chr10 | - | 119843247 | 119843426 | 119843253 | 119843426 | 119842471 | 119842595 | 0 | 2 | 7 | 2 | 1 | 2 | 0 | 0 | 0 | 3 | 55 | 49 | 1 | 1 | 0 | 0.641 | 0.757 | NA | 0 | 0.466 |
| SE | 123 | DIAPH2 | DIAPH2 | chrX | + | 96735757 | 96735790 | 96684662 | 96685190 | 96738585 | 96738762 | 2 | 1 | 2 | 1 | 1 | 0 | 0 | 1 | 0 | 0 | 82 | 49 | 1 | 1 | 0.544 | 0.374 | 1 | NA | 1 | -0.361 |
| SE | 388 | RAPGEF1 | RAPGEF1 | chr9 | - | 131630235 | 131630324 | 131629101 | 131629254 | 131638634 | 131638791 | 25 | 12 | 3 | 2 | 1 | 0 | 0 | 6 | 0 | 2 | 98 | 49 | 1 | 1 | 0.862 | 0.857 | 1 | NA | 0.6 | 0.306 |
| SE | 818 | INTS8 | INTS8 | chr8 | + | 94856776 | 94856978 | 94853804 | 94853915 | 94859510 | 94859632 | 7 | 1 | 1 | 1 | 0 | 0 | 0 | 2 | 0 | 0 | 98 | 49 | 1 | 1 | 0.778 | 1 | 1 | NA | 1 | -0.074 |
| SE | 822 | INTS8 | INTS8 | chr8 | + | 94868691 | 94868813 | 94867275 | 94867337 | 94871883 | 94872002 | 1 | 0 | 0 | 2 | 1 | 1 | 0 | 0 | 0 | 1 | 98 | 49 | 1 | 1 | 0.2 | 0 | 0 | NA | 0 | 0.067 |
| SE | 966 | PPP2R2A | PPP2R2A | chr8 | + | 26363720 | 26363890 | 26362683 | 26362848 | 26366314 | 26366406 | 16 | 3 | 2 | 0 | 2 | 0 | 0 | 2 | 0 | 0 | 98 | 49 | 1 | 1 | 1 | 0.429 | 1 | NA | 1 | -0.19 |
| SE | 967 | DOCK5 | DOCK5 | chr8 | + | 25268844 | 25268885 | 25243673 | 25243757 | 25275385 | 25275441 | 148 | 62 | 18 | 1 | 0 | 0 | 1 | 18 | 0 | 0 | 90 | 49 | 1 | 1 | 0.988 | 1 | 1 | 1 | 1 | -0.004 |
| SE | 968 | DOCK5 | DOCK5 | chr8 | + | 25278568 | 25278665 | 25275385 | 25275441 | 25292023 | 25292172 | 229 | 94 | 12 | 0 | 3 | 0 | 5 | 32 | 0 | 0 | 98 | 49 | 1 | 1 | 1 | 0.94 | 1 | 1 | 1 | -0.02 |
| SE | 969 | DOCK5 | DOCK5 | chr8 | + | 25296512 | 25296648 | 25292023 | 25292172 | 25298943 | 25299101 | 181 | 82 | 14 | 2 | 1 | 0 | 3 | 29 | 0 | 0 | 98 | 49 | 1 | 1 | 0.978 | 0.976 | 1 | 1 | 1 | -0.015 |
| SE | 971 | DOCK5 | DOCK5 | chr8 | + | 25317006 | 25317131 | 25310406 | 25310532 | 25319577 | 25319676 | 132 | 75 | 18 | 1 | 0 | 0 | 2 | 30 | 0 | 0 | 98 | 49 | 1 | 1 | 0.985 | 1 | 1 | 1 | 1 | -0.005 |
| SE | 972 | DOCK5 | DOCK5 | chr8 | + | 25334095 | 25334196 | 25332602 | 25332692 | 25336238 | 25336373 | 175 | 73 | 14 | 1 | 0 | 0 | 1 | 36 | 0 | 0 | 98 | 49 | 1 | 1 | 0.989 | 1 | 1 | 1 | 1 | -0.004 |
| SE | 973 | DOCK5 | DOCK5 | chr8 | + | 25340876 | 25340988 | 25336238 | 25336373 | 25341738 | 25341809 | 115 | 49 | 6 | 8 | 7 | 0 | 0 | 19 | 0 | 0 | 98 | 49 | 1 | 1 | 0.878 | 0.778 | 1 | NA | 1 | -0.115 |
| SE | 974 | DOCK5 | DOCK5 | chr8 | + | 25342400 | 25342507 | 25341738 | 25341809 | 25345474 | 25345611 | 107 | 55 | 21 | 1 | 0 | 0 | 0 | 15 | 0 | 0 | 98 | 49 | 1 | 1 | 0.982 | 1 | 1 | NA | 1 | -0.006 |
| SE | 975 | DOCK5 | DOCK5 | chr8 | + | 25368191 | 25368250 | 25366869 | 25366970 | 25368570 | 25368725 | 86 | 32 | 10 | 3 | 0 | 0 | 0 | 11 | 0 | 5 | 98 | 49 | 1 | 1 | 0.935 | 1 | 1 | NA | 0.524 | 0.454 |
| SE | 977 | DOCK5 | DOCK5 | chr8 | + | 25373617 | 25373658 | 25372558 | 25372718 | 25374563 | 25374654 | 85 | 38 | 16 | 0 | 0 | 0 | 0 | 8 | 0 | 1 | 90 | 49 | 1 | 1 | 1 | 1 | 1 | NA | 0.813 | 0.187 |
| SE | 994 | XPO7 | XPO7 | chr8 | + | 21990810 | 21990919 | 21990343 | 21990407 | 21991867 | 21991974 | 3 | 0 | 3 | 1 | 0 | 0 | 0 | 2 | 0 | 0 | 98 | 49 | 1 | 1 | 0.6 | NA | 1 | NA | 1 | -0.2 |
| SE | 1216 | SRPK2 | SRPK2 | chr7 | - | 105133003 | 105133104 | 105132790 | 105132898 | 105142007 | 105142490 | 40 | 17 | 4 | 1 | 0 | 0 | 0 | 9 | 0 | 0 | 98 | 49 | 1 | 1 | 0.952 | 1 | 1 | NA | 1 | -0.016 |
| SE | 1217 | SRPK2 | SRPK2 | chr7 | - | 105268805 | 105268869 | 105203627 | 105203785 | 105388647 | 105388702 | 4 | 4 | 1 | 10 | 2 | 5 | 0 | 0 | 0 | 1 | 98 | 49 | 1 | 1 | 0.167 | 0.5 | 0.091 | NA | 0 | 0.253 |
| SE | 1330 | RHBDD2 | RHBDD2 | chr7 | + | 75881364 | 75881486 | 75878998 | 75879260 | 75881828 | 75882236 | 2 | 0 | 1 | 2 | 0 | 1 | 0 | 2 | 1 | 0 | 98 | 49 | 1 | 1 | 0.333 | NA | 0.333 | 0 | 1 | -0.167 |
| SE | 1418 | NCBP1 | NCBP1 | chr9 | + | 97662041 | 97662144 | 97660945 | 97661068 | 97662953 | 97663047 | 6 | 3 | 0 | 0 | 0 | 0 | 1 | 1 | 0 | 2 | 98 | 49 | 1 | 1 | 1 | 1 | NA | 1 | 0.2 | 0.4 |
| SE | 1592 | TCP1 | TCP1 | chr6 | - | 159780934 | 159781110 | 159780442 | 159780566 | 159783940 | 159784067 | 13 | 7 | 3 | 3 | 0 | 0 | 0 | 1 | 0 | 0 | 98 | 49 | 1 | 1 | 0.684 | 1 | 1 | NA | 1 | -0.105 |
| SE | 1606 | TMEM181 | TMEM181 | chr6 | + | 158583953 | 158584044 | 158580939 | 158580995 | 158585303 | 158585425 | 7 | 5 | 3 | 0 | 0 | 2 | 0 | 4 | 0 | 0 | 98 | 49 | 1 | 1 | 1 | 1 | 0.429 | NA | 1 | -0.19 |
| SE | 1632 | STXBP5 | STXBP5 | chr6 | + | 147314263 | 147314331 | 147313883 | 147314031 | 147314595 | 147314636 | 51 | 28 | 17 | 3 | 0 | 0 | 0 | 9 | 0 | 0 | 98 | 49 | 1 | 1 | 0.895 | 1 | 1 | NA | 1 | -0.035 |
| SE | 1633 | STXBP5 | STXBP5 | chr6 | + | 147314595 | 147314636 | 147313883 | 147314031 | 147315514 | 147315735 | 26 | 3 | 1 | 0 | 0 | 1 | 0 | 4 | 0 | 0 | 90 | 49 | 1 | 1 | 1 | 0.62 | 1 | NA | 1 | -0.127 |
| SE | 1634 | STXBP5 | STXBP5 | chr6 | + | 147339178 | 147339238 | 147334156 | 147334222 | 147353322 | 147353373 | 15 | 3 | 0 | 6 | 0 | 0 | 0 | 4 | 0 | 0 | 98 | 49 | 1 | 1 | 0.556 | 1 | NA | NA | 1 | -0.222 |
| SE | 1635 | STXBP5 | STXBP5 | chr6 | + | 147339336 | 147339384 | 147339178 | 147339238 | 147353322 | 147353373 | 7 | 10 | 6 | 1 | 0 | 0 | 0 | 1 | 0 | 0 | 97 | 49 | 1 | 1 | 0.78 | 1 | 1 | NA | 1 | -0.073 |
| SE | 1869 | SMAP1 | SMAP1 | chr6 | + | 70791688 | 70791769 | 70773349 | 70773425 | 70798656 | 70798737 | 1 | 0 | 2 | 3 | 2 | 0 | 0 | 1 | 0 | 0 | 98 | 49 | 1 | 1 | 0.143 | 0 | 1 | NA | 1 | -0.619 |
| SE | 1921 | XPO5 | XPO5 | chr6 | - | 43533906 | 43534007 | 43531478 | 43531575 | 43546570 | 43546752 | 3 | 0 | 1 | 0 | 0 | 0 | 0 | 0 | 0 | 1 | 98 | 49 | 1 | 1 | 1 | NA | 1 | NA | 0 | 1 |
| SE | 1986 | SRPK1 | SRPK1 | chr6 | - | 35869009 | 35869110 | 35857260 | 35857368 | 35869481 | 35869901 | 49 | 15 | 8 | 2 | 1 | 0 | 0 | 5 | 0 | 0 | 98 | 49 | 1 | 1 | 0.925 | 0.882 | 1 | NA | 1 | -0.064 |
| SE | 2459 | DERL2 | DERL2 | chr17 | - | 5481295 | 5481389 | 5480386 | 5480582 | 5482808 | 5482882 | 4 | 1 | 4 | 0 | 0 | 5 | 0 | 1 | 0 | 0 | 98 | 49 | 1 | 1 | 1 | 1 | 0.286 | NA | 1 | -0.238 |
| SE | 2461 | EML3 | EML3 | chr11 | - | 62605641 | 62605773 | 62605112 | 62605180 | 62605854 | 62605980 | 13 | 5 | 1 | 1 | 0 | 0 | 0 | 1 | 0 | 0 | 98 | 49 | 1 | 1 | 0.867 | 1 | 1 | NA | 1 | -0.044 |
| SE | 2598 | EDC4 | EDC4 | chr16 | + | 67882678 | 67882865 | 67882428 | 67882594 | 67882957 | 67883177 | 11 | 2 | 1 | 1 | 0 | 0 | 0 | 1 | 0 | 0 | 98 | 49 | 1 | 1 | 0.846 | 1 | 1 | NA | 1 | -0.051 |
| SE | 2599 | NAA20 | NAA20 | chr20 | + | 20032507 | 20032653 | 20026783 | 20026919 | 20033101 | 20033629 | 1 | 0 | 0 | 0 | 0 | 0 | 0 | 3 | 0 | 1 | 98 | 49 | 1 | 1 | 1 | NA | NA | NA | 0.6 | 0.4 |
| SE | 3126 | ZNF207 | ZNF207 | chr17 | + | 32366664 | 32366757 | 32365329 | 32365487 | 32367771 | 32368014 | 23 | 9 | 1 | 4 | 2 | 1 | 0 | 0 | 0 | 2 | 98 | 49 | 1 | 1 | 0.742 | 0.692 | 0.333 | NA | 0 | 0.589 |
| SE | 3142 | DERL1 | DERL1 | chr8 | - | 123030604 | 123030716 | 123024985 | 123025050 | 123041969 | 123042423 | 8 | 8 | 5 | 0 | 1 | 1 | 0 | 2 | 0 | 0 | 98 | 49 | 1 | 1 | 1 | 0.8 | 0.714 | NA | 1 | -0.162 |
| SE | 3330 | WDR11 | WDR11 | chr10 | + | 120860108 | 120860282 | 120858642 | 120858796 | 120862734 | 120862921 | 6 | 0 | 4 | 0 | 0 | 0 | 0 | 2 | 0 | 1 | 98 | 49 | 1 | 1 | 1 | NA | 1 | NA | 0.5 | 0.5 |
| SE | 3331 | WDR11 | WDR11 | chr10 | + | 120867065 | 120867169 | 120866568 | 120866764 | 120871169 | 120871346 | 4 | 5 | 2 | 0 | 0 | 0 | 0 | 3 | 0 | 1 | 98 | 49 | 1 | 1 | 1 | 1 | 1 | NA | 0.6 | 0.4 |
| SE | 3922 | SPG11 | SPG11 | chr15 | - | 44572682 | 44572820 | 44570524 | 44570658 | 44583813 | 44584558 | 16 | 0 | 2 | 0 | 1 | 0 | 0 | 10 | 0 | 0 | 98 | 49 | 1 | 1 | 1 | 0 | 1 | NA | 1 | -0.333 |
| SE | 3923 | SPG11 | SPG11 | chr15 | - | 44589251 | 44589414 | 44585635 | 44585850 | 44592330 | 44592438 | 17 | 4 | 7 | 1 | 0 | 0 | 0 | 2 | 0 | 0 | 98 | 49 | 1 | 1 | 0.895 | 1 | 1 | NA | 1 | -0.035 |
| SE | 3924 | SPG11 | SPG11 | chr15 | - | 44598630 | 44598836 | 44598264 | 44598373 | 44600466 | 44600632 | 16 | 3 | 0 | 0 | 1 | 0 | 0 | 1 | 0 | 0 | 98 | 49 | 1 | 1 | 1 | 0.6 | NA | NA | 1 | -0.2 |

| | | | | | | | | | | | | | | | | | | | | | | | | | | | | | | | | | |
|---|---|---|---|---|---|---|---|---|---|---|---|---|---|---|---|---|---|---|---|---|---|---|---|---|---|---|---|---|---|---|---|---|---|
| SE | 3925 | SPG11 | SPG11 | chr15 | - | 44606024 | 44606091 | 44600466 | 44600632 | 44608443 | 44608605 | 26 | 14 | 1 | 6 | 0 | 1 | 0 | 3 | 0 | 2 | 98 | 49 | 1 | 1 | 0.684 | 1 | 0.333 | NA | | | 0.429 | 0.243 |
| SE | 3926 | SPG11 | SPG11 | chr15 | - | 44613429 | 44613536 | 44610839 | 44610985 | 44615362 | 44615566 | 36 | 11 | 5 | 3 | 2 | 0 | 0 | 13 | 0 | 0 | 98 | 49 | 1 | 1 | 0.857 | 0.733 | 1 | NA | | | 1 | -0.137 |
| SE | 4101 | ZCCHC8 | ZCCHC8 | chr12 | - | 122482634 | 122482695 | 122481944 | 122482087 | 122489385 | 122489463 | 2 | 5 | 2 | 1 | 0 | 0 | 0 | 1 | 0 | 0 | 98 | 49 | 1 | 1 | 0.5 | 1 | 1 | NA | | | 1 | -0.167 |
| SE | 4108 | MLXIP | MLXIP | chr12 | + | 122130843 | 122130933 | 122129940 | 122130112 | 122132291 | 122132383 | 10 | 4 | 2 | 0 | 0 | 0 | 0 | 1 | 0 | 1 | 98 | 49 | 1 | 1 | 1 | 1 | 1 | NA | | | 0.333 | 0.667 |
| SE | 4303 | PFDN5 | PFDN5 | chr12 | + | 53296243 | 53296275 | 53295838 | 53295941 | 53297849 | 53297924 | 43 | 7 | 8 | 2 | 0 | 0 | 0 | 8 | 0 | 0 | 81 | 49 | 1 | 1 | 0.929 | 1 | 1 | NA | | | 1 | -0.024 |
| SE | 4508 | FIG4 | FIG4 | chr6 | + | 109727108 | 109727265 | 109716444 | 109716568 | 109732636 | 109732687 | 3 | 4 | 0 | 1 | 0 | 0 | 1 | 3 | 0 | 0 | 98 | 49 | 1 | 1 | 0.6 | 1 | NA | | 1 | | 1 | -0.2 |
| SE | 4882 | FCHO2 | FCHO2 | chr5 | + | 73006444 | 73006549 | 72990711 | 72990864 | 73015625 | 73015724 | 19 | 7 | 1 | 1 | 0 | 0 | 0 | 6 | 0 | 0 | 98 | 49 | 1 | 1 | 0.905 | | 1 | NA | | | 1 | -0.032 |
| SE | 4884 | FCHO2 | FCHO2 | chr5 | + | 73081782 | 73081982 | 73078179 | 73078312 | 73082760 | 73082825 | 15 | 9 | 11 | 0 | 1 | 0 | 2 | 0 | 0 | 0 | 98 | 49 | 1 | 1 | 1 | 0.818 | 1 | | 1 | NA | 1 | -0.061 |
| SE | 5023 | AHCTF1 | AHCTF1 | chr1 | - | 246843794 | 246843928 | 246842693 | 246842776 | 246849614 | 246851442 | 12 | 1 | 5 | 3 | 1 | 1 | 0 | 0 | 0 | 2 | 98 | 49 | 1 | 1 | 0.667 | 0.333 | 0.714 | NA | | | 0 | 0.571 |
| SE | 5024 | AHCTF1 | AHCTF1 | chr1 | - | 246857690 | 246857814 | 246855729 | 246855827 | 246860898 | 246861295 | 27 | 2 | 5 | 1 | 1 | 0 | 0 | 6 | 0 | 0 | 98 | 49 | 1 | 1 | 0.931 | 0.5 | 1 | NA | | | 1 | -0.19 |
| SE | 5025 | AHCTF1 | AHCTF1 | chr1 | - | 246863923 | 246864116 | 246861958 | 246862153 | 246867243 | 246867351 | 26 | 16 | 4 | 1 | 0 | 0 | 0 | 1 | 0 | 0 | 98 | 49 | 1 | 1 | 0.929 | 1 | 1 | NA | | | 1 | -0.024 |
| SE | 5028 | AHCTF1 | AHCTF1 | chr1 | - | 246916141 | 246916395 | 246913231 | 246913412 | 246918249 | 246918377 | 12 | 12 | 2 | 5 | 1 | 1 | 0 | 4 | 0 | 1 | 98 | 49 | 1 | 1 | 0.545 | 0.857 | 0.5 | NA | | | 0.667 | -0.033 |
| SE | 5138 | VPS35 | VPS35 | chr16 | - | 46668929 | 46669052 | 46662982 | 46663162 | 46671704 | 46671860 | 18 | 10 | 11 | 0 | 0 | 1 | 3 | 6 | 0 | 0 | 98 | 49 | 1 | 1 | 1 | 1 | | 0.846 | 1 | | 1 | -0.051 |
| SE | 5377 | SLC9A8 | SLC9A8 | chr20 | + | 49839540 | 49839599 | 49823060 | 49823141 | 49845035 | 49845119 | 31 | 5 | 3 | 0 | 0 | 0 | 0 | 8 | 0 | 1 | 98 | 49 | 1 | 1 | 1 | 1 | 1 | NA | | | 0.8 | 0.2 |
| SE | 5379 | SLC9A8 | SLC9A8 | chr20 | + | 49849578 | 49849680 | 49845035 | 49845119 | 49855437 | 49855581 | 32 | 13 | 0 | 3 | 0 | 0 | 0 | 10 | 0 | 2 | 98 | 49 | 1 | 1 | 0.842 | 1 | NA | NA | | | 0.714 | 0.207 |
| SE | 5380 | SLC9A8 | SLC9A8 | chr20 | + | 49850761 | 49850844 | 49849578 | 49849680 | 49855437 | 49855581 | 7 | 4 | 0 | 14 | 9 | 0 | 0 | 4 | 0 | 4 | 98 | 49 | 1 | 1 | 0.2 | 0.182 | NA | NA | | | 0.333 | -0.142 |
| SE | 5381 | SLC9A8 | SLC9A8 | chr20 | + | 49850809 | 49850844 | 49849578 | 49849680 | 49855437 | 49855581 | 7 | 6 | 0 | 14 | 9 | 0 | 0 | 4 | 0 | 4 | 84 | 49 | 1 | 1 | 0.226 | 0.28 | NA | NA | | | 0.368 | -0.115 |
| SE | 5382 | SLC9A8 | SLC9A8 | chr20 | + | 49874704 | 49874821 | 49864727 | 49864844 | 49877980 | 49878063 | 23 | 10 | 8 | 2 | 0 | 0 | 0 | 4 | 0 | 0 | 98 | 49 | 1 | 1 | 0.852 | 1 | 1 | NA | | | 1 | -0.049 |
| SE | 5405 | RASGRP1 | RASGRP1 | chr15 | - | 38501142 | 38501287 | 38500102 | 38500139 | 38502311 | 38502421 | 5 | 5 | 4 | 1 | 0 | 0 | 0 | 1 | 0 | 0 | 98 | 49 | 1 | 1 | 0.714 | 1 | 1 | NA | | | 1 | -0.095 |
| SE | 5406 | RASGRP1 | RASGRP1 | chr15 | - | 38503271 | 38503376 | 38502311 | 38502421 | 38505839 | 38505920 | 9 | 1 | 3 | 0 | 1 | 0 | 0 | 2 | 0 | 0 | 98 | 49 | 1 | 1 | 1 | 0.333 | 1 | NA | | | 1 | -0.222 |
| SE | 5439 | RASGRP4 | RASGRP4 | chr19 | - | 38414847 | 38414916 | 38413393 | 38413474 | 38417051 | 38417168 | 17 | 10 | 3 | 1 | 0 | 0 | 0 | 3 | 0 | 0 | 98 | 49 | 1 | 1 | 0.895 | 1 | 1 | NA | | | 1 | -0.035 |
| SE | 5440 | RASGRP4 | RASGRP4 | chr19 | - | 38414847 | 38415123 | 38413393 | 38413474 | 38417051 | 38417168 | 56 | 20 | 10 | 1 | 0 | 0 | 2 | 16 | 0 | 0 | 98 | 49 | 1 | 1 | 0.966 | 1 | 1 | | 1 | | 1 | -0.011 |
| SE | 5556 | TAF2 | TAF2 | chr8 | - | 119793365 | 119793451 | 119791323 | 119791459 | 119795531 | 119795631 | 10 | 3 | 1 | 0 | 0 | 0 | 0 | 1 | 0 | 2 | 98 | 49 | 1 | 1 | 1 | 1 | 1 | NA | | | 0.2 | 0.8 |
| SE | 5786 | SRSF10 | SRSF10 | chr1 | - | 23971337 | 23971439 | 23964346 | 23967723 | 23971572 | 23971626 | 4 | 1 | 0 | 0 | 0 | 2 | 0 | 1 | 0 | 0 | 98 | 49 | 1 | 1 | 0.333 | 1 | 0 | NA | | | 1 | -0.556 |
| SE | 5870 | ZDHHC13 | ZDHHC13 | chr11 | + | 19142977 | 19143123 | 19117144 | 19117276 | 19146180 | 19146303 | 2 | 0 | 0 | 1 | 0 | 0 | 0 | 2 | 0 | 0 | 98 | 49 | 1 | 1 | 0.5 | NA | NA | NA | | | 1 | -0.5 |
| SE | 5871 | ZDHHC13 | ZDHHC13 | chr11 | + | 19166301 | 19166385 | 19165051 | 19165145 | 19170410 | 19170568 | 2 | 2 | 0 | 2 | 0 | 0 | 0 | 2 | 0 | 0 | 98 | 49 | 1 | 1 | 0.333 | 1 | NA | NA | | | 1 | -0.334 |
| SE | 5999 | CHD1L | CHD1L | chr1 | + | 147270931 | 147271005 | 147268781 | 147268878 | 147272170 | 147272281 | 0 | 2 | 0 | 0 | 0 | 0 | 0 | 1 | 0 | 1 | 98 | 49 | 1 | 1 | NA | 1 | NA | NA | | | 0.333 | 0.667 |
| SE | 6296 | TMEM87A | TMEM87A | chr15 | - | 42272062 | 42272123 | 42267946 | 42268032 | 42273254 | 42273584 | 13 | 3 | 3 | 1 | 0 | 0 | 0 | 1 | 0 | 0 | 98 | 49 | 1 | 1 | 0.867 | | 1 | NA | | | 1 | -0.044 |
| SE | 6420 | RALGAPA2 | RALGAPA2 | chr20 | - | 20396694 | 20396734 | 20389627 | 20393253 | 20412026 | 20412148 | 11 | 2 | 6 | 0 | 0 | 3 | 0 | 3 | 0 | 0 | 89 | 49 | 1 | 1 | 1 | 1 | 0.524 | NA | | | 1 | -0.159 |
| SE | 6421 | RALGAPA2 | RALGAPA2 | chr20 | - | 20524405 | 20524543 | 20520916 | 20521100 | 20524829 | 20524898 | 12 | 9 | 2 | 0 | 3 | 0 | 0 | 2 | 0 | 1 | 98 | 49 | 1 | 1 | 1 | 0.6 | 1 | NA | | | 0.5 | 0.367 |
| SE | 6422 | RALGAPA2 | RALGAPA2 | chr20 | - | 20611314 | 20611426 | 20605174 | 20605412 | 20616042 | 20616191 | 33 | 4 | 0 | 0 | 0 | 0 | 0 | 6 | 0 | 1 | 98 | 49 | 1 | 1 | 1 | 1 | NA | NA | | | 0.75 | 0.25 |
| SE | 6423 | RALGAPA2 | RALGAPA2 | chr20 | - | 20629362 | 20629590 | 20620462 | 20620630 | 20635417 | 20635617 | 22 | 21 | 4 | 1 | 0 | 0 | 0 | 6 | 0 | 0 | 98 | 49 | 1 | 1 | 0.917 | | 1 | NA | | | 1 | -0.028 |
| SE | 6424 | RALGAPA2 | RALGAPA2 | chr20 | - | 20637362 | 20637501 | 20635417 | 20635617 | 20639784 | 20639900 | 37 | 17 | 18 | 1 | 4 | 0 | 0 | 13 | 0 | 2 | 98 | 49 | 1 | 1 | 0.949 | 0.68 | 1 | NA | | | 0.765 | 0.111 |
| SE | 6425 | RALGAPA2 | RALGAPA2 | chr20 | - | 20676235 | 20676288 | 20653529 | 20653587 | 20680690 | 20680801 | 45 | 24 | 6 | 0 | 1 | 0 | 0 | 13 | 0 | 2 | 98 | 49 | 1 | 1 | 1 | 0.923 | 1 | NA | | | 0.765 | 0.209 |
| SE | 6429 | LMBR1L | LMBR1L | chr12 | - | 49102886 | 49102951 | 49102467 | 49102540 | 49103090 | 49103159 | 7 | 3 | 1 | 0 | 1 | 0 | 1 | 4 | 0 | 0 | 98 | 49 | 1 | 1 | 1 | 0.6 | 1 | | 1 | | 1 | -0.133 |
| SE | 6474 | GBP5 | GBP5 | chr1 | - | 89262219 | 89262401 | 89258950 | 89260817 | 89263735 | 89263948 | 65 | 9 | 9 | 0 | 2 | 0 | 0 | 10 | 0 | 0 | 98 | 49 | 1 | 1 | 1 | 0.692 | 1 | NA | | | 1 | -0.103 |
| SE | 6475 | GBP5 | GBP5 | chr1 | - | 89262682 | 89262785 | 89262219 | 89262401 | 89263735 | 89263948 | 85 | 28 | 13 | 2 | 0 | 0 | 3 | 3 | 0 | 0 | 98 | 49 | 1 | 1 | 0.955 | 1 | 1 | | 1 | | 1 | -0.015 |
| SE | 6476 | GBP5 | GBP5 | chr1 | - | 89266956 | 89267153 | 89266345 | 89266588 | 89268728 | 89268856 | 62 | 19 | 18 | 1 | 0 | 0 | 0 | 7 | 0 | 0 | 98 | 49 | 1 | 1 | 0.969 | 1 | 1 | NA | | | 1 | -0.01 |
| SE | 6477 | GBP5 | GBP5 | chr1 | - | 89267416 | 89267526 | 89266956 | 89267153 | 89268728 | 89268856 | 136 | 36 | 23 | 1 | 0 | 0 | 2 | 14 | 0 | 0 | 98 | 49 | 1 | 1 | 0.986 | 1 | 1 | | 1 | | 1 | -0.005 |
| SE | 6478 | GBP5 | GBP5 | chr1 | - | 89268728 | 89268856 | 89267416 | 89267526 | 89269365 | 89269574 | 72 | 16 | 9 | 1 | 1 | 0 | 1 | 7 | 0 | 0 | 98 | 49 | 1 | 1 | 0.973 | 0.889 | 1 | | 1 | | 1 | -0.046 |
| SE | 6479 | GBP5 | GBP5 | chr1 | - | 89268728 | 89268856 | 89267416 | 89267526 | 89272451 | 89272861 | 48 | 11 | 5 | 1 | 1 | 0 | 1 | 3 | 0 | 0 | 98 | 49 | 1 | 1 | 0.96 | 0.846 | 1 | | 1 | | 1 | -0.065 |
| SE | 6481 | GBP5 | GBP5 | chr1 | - | 89269365 | 89269574 | 89268728 | 89268856 | 89272451 | 89272861 | 30 | 9 | 5 | 2 | 1 | 0 | 0 | 4 | 0 | 0 | 98 | 49 | 1 | 1 | 0.882 | 0.818 | 1 | NA | | | 1 | -0.1 |
| SE | 6482 | GBP5 | GBP5 | chr1 | - | 89270754 | 89270862 | 89269365 | 89269574 | 89272451 | 89272861 | 52 | 16 | 3 | 4 | 3 | 1 | 0 | 7 | 0 | 0 | 98 | 49 | 1 | 1 | 0.867 | 0.727 | 0.6 | NA | | | 1 | -0.269 |
| SE | 6593 | KANSL2 | KANSL2 | chr12 | - | 48679035 | 48679150 | 48671798 | 48671962 | 48679654 | 48679833 | 5 | 1 | 0 | 3 | 3 | 1 | 0 | 6 | 0 | 2 | 98 | 49 | 1 | 1 | 0.455 | 0.143 | 0 | NA | | | 0.6 | -0.401 |
| SE | 6837 | GAK | GAK | chr4 | - | 913606 | 913668 | 904636 | 904779 | 932042 | 932386 | 9 | 2 | 0 | 1 | 1 | 0 | 3 | 2 | 0 | 0 | 98 | 49 | 1 | 1 | 0.818 | 0.5 | NA | | 1 | | 1 | -0.341 |
| SE | 6843 | NAA16 | NAA16 | chr13 | + | 41331273 | 41331369 | 41328723 | 41328843 | 41355143 | 41355216 | 2 | 0 | 0 | 1 | 0 | 0 | 0 | 1 | 0 | 0 | 98 | 49 | 1 | 1 | 1 | 0 | NA | NA | | | 1 | -0.5 |
| SE | 6917 | TMEM41B | TMEM41B | chr11 | - | 9286454 | 9286593 | 9280654 | 9283593 | 9287701 | 9287806 | 6 | 10 | 0 | 1 | 0 | 0 | 0 | 1 | 0 | 0 | 98 | 49 | 1 | 1 | 0.75 | 1 | NA | NA | | | 1 | -0.125 |
| SE | 6921 | DAP3 | DAP3 | chr1 | + | 155729041 | 155729122 | 155727607 | 155727738 | 155729207 | 155729366 | 11 | 3 | 1 | 0 | 2 | 0 | 0 | 4 | 0 | 0 | 98 | 49 | 1 | 1 | 1 | 0.429 | 1 | NA | | | 1 | -0.19 |
| SE | 7009 | SFI1 | SFI1 | chr22 | + | 31528689 | 31528863 | 31508254 | 31508376 | 31546860 | 31546971 | 6 | 13 | 3 | 1 | 3 | 0 | 0 | 0 | 0 | 1 | 98 | 49 | 1 | 1 | 0.75 | 0.684 | 1 | NA | | | 0 | 0.811 |
| SE | 7011 | SFI1 | SFI1 | chr22 | + | 31546860 | 31546971 | 31528689 | 31528863 | 31550253 | 31550348 | 7 | 13 | 2 | 1 | 0 | 0 | 0 | 1 | 0 | 0 | 98 | 49 | 1 | 1 | 0.778 | 1 | 1 | NA | | | 1 | -0.074 |
| SE | 7012 | SFI1 | SFI1 | chr22 | + | 31561289 | 31561392 | 31556941 | 31557059 | 31573057 | 31573214 | 22 | 16 | 3 | 1 | 0 | 0 | 0 | 4 | 0 | 0 | 98 | 49 | 1 | 1 | 0.917 | 1 | 1 | NA | | | 1 | -0.028 |
| SE | 7014 | SFI1 | SFI1 | chr22 | + | 31613601 | 31613855 | 31613353 | 31613530 | 31614788 | 31614860 | 9 | 5 | 7 | 1 | 0 | 0 | 2 | 0 | 0 | 0 | 98 | 49 | 1 | 1 | 0.818 | 1 | 1 | | 1 | NA | | -0.061 |
| SE | 7318 | VPS11 | VPS11 | chr11 | + | 119071595 | 119071843 | 119070233 | 119070397 | 119073197 | 119073399 | 10 | 0 | 0 | 0 | 0 | 0 | 0 | 0 | 0 | 2 | 98 | 49 | 1 | 1 | 1 | NA | NA | NA | | | 0 | 1 |
| SE | 7370 | SMAP2 | SMAP2 | chr1 | + | 40406735 | 40406869 | 40373705 | 40374223 | 40408652 | 40408738 | 139 | 72 | 82 | 0 | 2 | 0 | 0 | 53 | 0 | 0 | 98 | 49 | 1 | 1 | 1 | 0.947 | 1 | NA | | | 1 | -0.018 |
| SE | 7439 | TBC1D32 | TBC1D32 | chr6 | - | 121170404 | 121170527 | 121160947 | 121161056 | 121205074 | 121205163 | 0 | 0 | 0 | 1 | 0 | 0 | 0 | 1 | 0 | 0 | 98 | 49 | 1 | 1 | 0 | NA | NA | NA | | | 1 | -1 |
| SE | 7498 | SLC12A6 | SLC12A6 | chr15 | - | 34238964 | 34239160 | 34238231 | 34238401 | 34240660 | 34240829 | 71 | 12 | 3 | 0 | 1 | 0 | 1 | 11 | 0 | 0 | 98 | 49 | 1 | 1 | 1 | 0.857 | 1 | | 1 | | 1 | -0.048 |
| SE | 7499 | SLC12A6 | SLC12A6 | chr15 | - | 34242101 | 34242221 | 34241232 | 34241337 | 34243973 | 34244072 | 67 | 19 | 8 | 7 | 5 | 0 | 0 | 7 | 0 | 1 | 98 | 49 | 1 | 1 | 0.827 | 0.655 | 1 | NA | | | 0.778 | 0.049 |
| SE | 7500 | SLC12A6 | SLC12A6 | chr15 | - | 34245284 | 34245403 | 34243973 | 34244072 | 34245692 | 34245867 | 82 | 40 | 22 | 2 | 0 | 0 | 0 | 16 | 0 | 0 | 98 | 49 | 1 | 1 | 0.953 | 1 | 1 | NA | | | 1 | -0.016 |
| SE | 7506 | HAL | HAL | chr12 | - | 95990392 | 95990532 | 95988192 | 95988240 | 95992679 | 95992805 | 69 | 12 | 16 | 2 | 0 | 0 | 0 | 8 | 0 | 0 | 98 | 49 | 1 | 1 | 0.945 | 1 | 1 | NA | | | 1 | -0.018 |
| SE | 7507 | HAL | HAL | chr12 | - | 95993771 | 95993838 | 95993450 | 95993488 | 95993925 | 95993998 | 28 | 14 | 17 | 9 | 0 | 5 | 0 | 3 | 0 | 0 | 98 | 49 | 1 | 1 | 0.609 | 1 | 0.63 | NA | | | 1 | -0.254 |
| SE | 7619 | DRAM2 | DRAM2 | chr1 | - | 111124741 | 111124881 | 111120515 | 111120693 | 111126226 | 111126294 | 5 | 1 | 2 | 1 | 0 | 0 | 0 | 6 | 0 | 0 | 98 | 49 | 1 | 1 | 0.714 | 1 | 1 | NA | | | 1 | -0.095 |
| SE | 7636 | PTEN | PTEN | chr10 | + | 87925512 | 87925557 | 87894024 | 87894109 | 87931045 | 87931089 | 77 | 45 | 51 | 1 | 0 | 0 | 0 | 17 | 0 | 0 | 94 | 49 | 1 | 1 | 0.976 | 1 | 1 | NA | | | 1 | -0.008 |
| SE | 7638 | PTEN | PTEN | chr10 | + | 87952117 | 87952259 | 87933012 | 87933251 | 87957852 | 87958019 | 16 | 9 | 10 | 3 | 0 | 2 | 0 | 4 | 0 | 1 | 98 | 49 | 1 | 1 | 0.727 | 1 | 0.714 | NA | | | 0.667 | 0.147 |
| SE | 7761 | PLCB3 | PLCB3 | chr11 | + | 64264020 | 64264112 | 64263690 | 64263795 | 64264950 | 64265104 | 0 | 0 | 0 | 1 | 0 | 0 | 0 | 2 | 0 | 0 | 98 | 49 | 1 | 1 | | 0 | NA | NA | | | 1 | -1 |
| SE | 7946 | CUL4A | CUL4A | chr13 | + | 113239432 | 113239551 | 113236822 | 113236890 | 113242967 | 113243160 | 7 | 2 | 0 | 1 | 0 | 0 | 0 | 5 | 0 | 0 | 98 | 49 | 1 | 1 | 1 | 1 | NA | NA | | | 0.714 | 0.286 |
| SE | 8088 | HPS1 | HPS1 | chr10 | - | 98429790 | 98429889 | 98429145 | 98429642 | 98430570 | 98430670 | 14 | 1 | 2 | 0 | 0 | 3 | 0 | 1 | 0 | 1 | 98 | 49 | 1 | 1 | 1 | 1 | 0.25 | NA | | | 0.333 | 0.417 |
| SE | 8089 | HPS1 | HPS1 | chr10 | - | 98431130 | 98431291 | 98430570 | 98430670 | 98433982 | 98434091 | 15 | 0 | 3 | 2 | 0 | 0 | 0 | 6 | 0 | 0 | 98 | 49 | 1 | 1 | 0.789 | NA | 1 | NA | | | 1 | -0.105 |
| SE | 8090 | HPS1 | HPS1 | chr10 | - | 98435271 | 98435414 | 98433982 | 98434091 | 98435634 | 98435772 | 15 | 1 | 2 | 7 | 0 | 0 | 0 | 7 | 0 | 0 | 98 | 49 | 1 | 1 | 0.517 | | 1 | NA | | | 1 | -0.161 |
| SE | 8130 | CNEP1R1 | CNEP1R1 | chr16 | + | 50029724 | 50029798 | 50026395 | 50026467 | 50033396 | 50033506 | 13 | 7 | 4 | 0 | 3 | 0 | 0 | 6 | 0 | 0 | 98 | 49 | 1 | 1 | 1 | 0.538 | 1 | NA | | | 1 | -0.154 |
| SE | 8235 | ZFC3H1 | ZFC3H1 | chr12 | - | 71619314 | 71619409 | 71615205 | 71615316 | 71619925 | 71620124 | 12 | 2 | 3 | 1 | 0 | 0 | 0 | 2 | 0 | 0 | 98 | 49 | 1 | 1 | 0.857 | 1 | 1 | NA | | | 1 | -0.048 |
| SE | 8236 | ZFC3H1 | ZFC3H1 | chr12 | - | 71624103 | 71624292 | 71623359 | 71623597 | 71626259 | 71626446 | 8 | 0 | 1 | 0 | 0 | 0 | 0 | 0 | 0 | 2 | 98 | 49 | 1 | 1 | 1 | NA | 1 | NA | | | 0 | 1 |
| SE | 8237 | ZFC3H1 | ZFC3H1 | chr12 | - | 71636849 | 71637059 | 71636489 | 71636654 | 71638417 | 71638515 | 27 | 17 | 1 | 0 | 0 | 0 | 0 | 3 | 0 | 1 | 98 | 49 | 1 | 1 | 1 | 1 | 1 | NA | | | 0.6 | 0.4 |
| SE | 8411 | FCHO1 | FCHO1 | chr19 | + | 17755117 | 17755191 | 17754316 | 17754361 | 17762761 | 17762853 | 9 | 3 | 0 | 1 | 0 | 0 | 1 | 1 | 0 | 0 | 98 | 49 | 1 | 1 | 0.818 | 1 | NA | | 1 | | 1 | -0.091 |
| SE | 8412 | FCHO1 | FCHO1 | chr19 | + | 17781230 | 17781343 | 17778608 | 17778884 | 17781451 | 17781539 | 8 | 6 | 0 | 0 | 0 | 0 | 1 | 3 | 0 | 3 | 98 | 49 | 1 | 1 | 1 | 1 | NA | | 1 | | 0.333 | 0.334 |
| SE | 8527 | KIAA0556 | KIAA0556 | chr16 | + | 27699533 | 27699599 | 27698327 | 27698500 | 27701588 | 27701695 | 14 | 3 | 0 | 2 | 0 | 0 | 2 | 3 | 0 | 0 | 98 | 49 | 1 | 1 | 0.778 | 1 | NA | | 1 | | 1 | -0.111 |
| SE | 8528 | KIAA0556 | KIAA0556 | chr16 | + | 27708704 | 27708920 | 27703895 | 27703998 | 27721557 | 27721695 | 11 | 1 | 0 | 1 | 0 | 0 | 0 | 1 | 0 | 0 | 98 | 49 | 1 | 1 | 0.846 | 1 | NA | NA | | | 1 | -0.077 |
| SE | 8529 | KIAA0556 | KIAA0556 | chr16 | + | 27754172 | 27754251 | 27751718 | 27751924 | 27761412 | 27761590 | 5 | 4 | 2 | 0 | 1 | 0 | 0 | 4 | 0 | 0 | 98 | 49 | 1 | 1 | 1 | 0.667 | 1 | NA | | | 1 | -0.111 |
| SE | 8530 | KIAA0556 | KIAA0556 | chr16 | + | 27777609 | 27777770 | 27776927 | 27777029 | 27777880 | 27777969 | 8 | 2 | 1 | 1 | 0 | 0 | 2 | 2 | 0 | 0 | 98 | 49 | 1 | 1 | 0.8 | 1 | 1 | | 1 | | 1 | -0.067 |
| SE | 8537 | ATG3 | ATG3 | chr3 | - | 112553279 | 112553329 | 112550191 | 112550262 | 112558375 | 112558417 | 31 | 9 | 9 | 1 | 3 | 0 | 0 | 4 | 0 | 3 | 98 | 49 | 1 | 1 | 0.939 | 0.6 | 1 | NA | | | 0.4 | 0.446 |
| SE | 8551 | UBR3 | UBR3 | chr2 | + | 169877493 | 169877637 | 169875790 | 169875949 | 169878524 | 169878574 | 27 | 11 | 0 | 1 | 0 | 0 | 0 | 3 | 0 | 0 | 98 | 49 | 1 | 1 | 0.931 | 1 | NA | NA | | | 1 | -0.034 |
| SE | 8708 | IFT88 | IFT88 | chr13 | + | 20615792 | 20615879 | 20605034 | 20605105 | 20625749 | 20625849 | 1 | 1 | 4 | 2 | 0 | 0 | 0 | 2 | 0 | 0 | 98 | 49 | 1 | 1 | 0.2 | 1 | 1 | NA | | | 1 | -0.267 |
| SE | 8826 | DOCK2 | DOCK2 | chr5 | + | 169670541 | 169670597 | 169669287 | 169669328 | 169671077 | 169671174 | 96 | 41 | 16 | 1 | 0 | 0 | 0 | 22 | 0 | 0 | 98 | 49 | 1 | 1 | 0.98 | 1 | 1 | NA | | | 1 | -0.007 |
| SE | 8827 | DOCK2 | DOCK2 | chr5 | + | 169671077 | 169671174 | 169670541 | 169670597 | 169674296 | 169674445 | 205 | 65 | 16 | 0 | 1 | 0 | 2 | 36 | 0 | 0 | 98 | 49 | 1 | 1 | 1 | 0.97 | 1 | | 1 | | 1 | -0.01 |
| SE | 8829 | DOCK2 | DOCK2 | chr5 | + | 169689251 | 169689333 | 169684195 | 169684350 | 169695802 | 169695938 | 176 | 92 | 10 | 2 | 0 | 0 | 5 | 49 | 0 | 0 | 98 | 49 | 1 | 1 | 0.978 | 1 | 1 | | 1 | | 1 | -0.007 |
| SE | 8830 | DOCK2 | DOCK2 | chr5 | + | 169712119 | 169712223 | 169711934 | 169712007 | 169714027 | 169714211 | 141 | 46 | 6 | 0 | 1 | 0 | 4 | 19 | 0 | 0 | 98 | 49 | 1 | 1 | 1 | 0.958 | 1 | | 1 | | 1 | -0.014 |
| SE | 8833 | DOCK2 | DOCK2 | chr5 | + | 169803057 | 169803206 | 169761518 | 169761625 | 169840756 | 169840852 | 143 | 72 | 20 | 0 | 2 | 0 | 2 | 21 | 0 | 0 | 98 | 49 | 1 | 1 | 1 | 0.947 | 1 | | 1 | | 1 | -0.018 |
| SE | 8834 | DOCK2 | DOCK2 | chr5 | + | 169983067 | 169983166 | 169840756 | 169840852 | 169985827 | 169985922 | 45 | 22 | 7 | 0 | 0 | 0 | 0 | 12 | 0 | 1 | 98 | 49 | 1 | 1 | 1 | 1 | 1 | NA | | | 0.857 | 0.143 |
| SE | 8835 | DOCK2 | DOCK2 | chr5 | + | 170008687 | 170008746 | 170008496 | 170008597 | 170018959 | 170019108 | 74 | 29 | 11 | 0 | 0 | 0 | 2 | 21 | 0 | 0 | 98 | 49 | 1 | 1 | 1 | 0.935 | 1 | | 1 | | 1 | -0.022 |
| SE | 8836 | DOCK2 | DOCK2 | chr5 | + | 170034398 | 170034555 | 170027862 | 170027948 | 170036514 | 170036555 | 61 | 8 | 18 | 1 | 0 | 0 | 0 | 16 | 0 | 0 | 98 | 49 | 1 | 1 | 0.968 | 1 | 1 | NA | | | 1 | -0.011 |
| SE | 8837 | DOCK2 | DOCK2 | chr5 | + | 170045815 | 170045905 | 170042012 | 170042132 | 170047509 | 170047614 | 49 | 19 | 5 | 0 | 1 | 0 | 1 | 14 | 0 | 0 | 98 | 49 | 1 | 1 | 1 | 0.905 | 1 | | 1 | | 1 | -0.032 |
| SE | 8923 | DISC1 | DISC1 | chr1 | + | 231720847 | 231720924 | 231701954 | 231702024 | 231749925 | 231750619 | 1 | 0 | 0 | 4 | 7 | 3 | 0 | 0 | 0 | 3 | 98 | 49 | 1 | 1 | 0.111 | 0 | 0 | NA | | | 0 | 0.037 |
| SE | 8932 | PLCB2 | PLCB2 | chr15 | - | 40290024 | 40290082 | 40289271 | 40289358 | 40290576 | 40290672 | 63 | 24 | 12 | 0 | 3 | 0 | 0 | 8 | 0 | 0 | 98 | 49 | 1 | 1 | 1 | 0.8 | 1 | NA | | | 1 | -0.067 |
| SE | 8933 | PLCB2 | PLCB2 | chr15 | - | 40291605 | 40291650 | 40291264 | 40291487 | 40291848 | 40291924 | 10 | 5 | 0 | 10 | 3 | 1 | 0 | 1 | 0 | 2 | 94 | 49 | 1 | 1 | 0.343 | 0.465 | 0 | NA | | | 0.207 | 0.062 |
| SE | 8934 | PLCB2 | PLCB2 | chr15 | - | 40297876 | 40297959 | 40297520 | 40297605 | 40298222 | 40298380 | 59 | 15 | 5 | 1 | 0 | 0 | 1 | 6 | 0 | 0 | 98 | 49 | 1 | 1 | 0.967 | 1 | 1 | | 1 | | 1 | -0.011 |
| SE | 9161 | CHD4 | CHD4 | chr12 | - | 6582136 | 6582281 | 6581648 | 6581814 | 6582614 | 6582748 | 33 | 7 | 10 | 1 | 1 | 0 | 0 | 4 | 0 | 1 | 98 | 49 | 1 | 1 | 0.943 | 0.778 | 1 | NA | | | 0.667 | 0.24 |
| SE | 9162 | CHD4 | CHD4 | chr12 | - | 6583197 | 6583378 | 6583026 | 6583113 | 6587383 | 6587559 | 12 | 5 | 5 | 0 | 1 | 0 | 0 | 5 | 0 | 0 | 98 | 49 | 1 | 1 | 1 | 0.714 | 1 | NA | | | 1 | -0.095 |
| SE | 9163 | CHD4 | CHD4 | chr12 | - | 6600925 | 6601053 | 6600533 | 6600669 | 6601288 | 6601530 | 24 | 21 | 1 | 2 | 0 | 0 | 0 | 7 | 0 | 0 | 98 | 49 | 1 | 1 | 0.857 | 1 | 1 | NA | | | 1 | -0.048 |
| SE | 9191 | C2CD3 | C2CD3 | chr11 | - | 74122987 | 74123135 | 74118227 | 74118382 | 74132843 | 74132972 | 7 | 0 | 0 | 1 | 0 | 0 | 0 | 5 | 0 | 0 | 98 | 49 | 1 | 1 | 0.778 | NA | NA | NA | | | 1 | -0.222 |
| SE | 9192 | C2CD3 | C2CD3 | chr11 | - | 74161398 | 74161556 | 74139604 | 74139828 | 74168343 | 74168613 | 8 | 2 | 0 | 0 | 0 | 0 | 0 | 2 | 0 | 1 | 98 | 49 | 1 | 1 | 1 | 1 | NA | NA | | | 0.5 | 0.5 |
| SE | 9293 | WLS | WLS | chr1 | - | 68153516 | 68153653 | 68150187 | 68150356 | 68155098 | 68155260 | 54 | 15 | 2 | 1 | 0 | 0 | 0 | 9 | 0 | 0 | 98 | 49 | 1 | 1 | 0.964 | 1 | 1 | NA | | | 1 | -0.012 |
| SE | 9611 | CTNNBL1 | CTNNBL1 | chr20 | + | 37842338 | 37842419 | 37840101 | 37840199 | 37859898 | 37860036 | 3 | 0 | 0 | 0 | 0 | 0 | 0 | 2 | 0 | 1 | 98 | 49 | 1 | 1 | 1 | NA | NA | NA | | | 0.5 | 0.5 |
| SE | 9773 | E2F4 | E2F4 | chr16 | + | 67194397 | 67194459 | 67192760 | 67192870 | 67194685 | 67194980 | 8 | 4 | 0 | 0 | 0 | 0 | 1 | 2 | 0 | 0 | 98 | 49 | 1 | 1 | 1 | 1 | 0 | NA | | | 1 | -0.333 |
| SE | 9782 | BTAF1 | BTAF1 | chr10 | + | 91953736 | 91953873 | 91951402 | 91951566 | 91956527 | 91956657 | 30 | 3 | 0 | 1 | 0 | 0 | 1 | 5 | 0 | 1 | 98 | 49 | 1 | 1 | 0.937 | 1 | NA | | 1 | | 0.714 | 0.112 |
| SE | 9783 | BTAF1 | BTAF1 | chr10 | + | 91989153 | 91989580 | 91984200 | 91984404 | 91992118 | 91992309 | 19 | 2 | 2 | 0 | 1 | 0 | 0 | 2 | 0 | 0 | 98 | 49 | 1 | 1 | 1 | 0.5 | 1 | NA | | | 1 | -0.167 |
| SE | 9784 | BTAF1 | BTAF1 | chr10 | + | 92027129 | 92027300 | 92026591 | 92026751 | 92028789 | 92030323 | 19 | 1 | 0 | 0 | 0 | 0 | 0 | 4 | 0 | 3 | 98 | 49 | 1 | 1 | 1 | 1 | NA | NA | | | 0.4 | 0.6 |
| SE | 9849 | MYH9 | MYH9 | chr22 | - | 36303994 | 36304155 | 36302567 | 36302676 | 36305032 | 36305102 | 184 | 98 | 47 | 0 | 1 | 0 | 6 | 33 | 0 | 0 | 98 | 49 | 1 | 1 | 1 | 0.98 | 1 | | 1 | | 1 | -0.007 |
| SE | 10285 | PPP2R2D | PPP2R2D | chr10 | + | 131911561 | 131912598 | 131901237 | 131901330 | 131934457 | 131934555 | 1 | 0 | 0 | 6 | 6 | 0 | 0 | 0 | 0 | 3 | 98 | 49 | 1 | 1 | 0.077 | 0 | NA | NA | | | 0 | 0.038 |
| SE | 10286 | PPP2R2D | PPP2R2D | chr10 | + | 131934457 | 131934555 | 131901237 | 131901330 | 131940030 | 131940196 | 15 | 9 | 0 | 1 | 2 | 0 | 0 | 6 | 0 | 0 | 98 | 49 | 1 | 1 | 0.882 | 0.692 | NA | NA | | | 1 | -0.213 |
| SE | 10287 | PPP2R2D | PPP2R2D | chr10 | + | 131934893 | 131935006 | 131934457 | 131934555 | 131940030 | 131940196 | 4 | 7 | 1 | 9 | 3 | 0 | 0 | 1 | 0 | 3 | 98 | 49 | 1 | 1 | 0.182 | 0.538 | 1 | NA | | | 0.143 | 0.43 |
| SE | 10350 | DYNC1LI2 | DYNC1LI2 | chr16 | - | 66749196 | 66749313 | 66742437 | 66742668 | 66751272 | 66751346 | 25 | 12 | 0 | 0 | 0 | 0 | 1 | 6 | 0 | 1 | 98 | 49 | 1 | 1 | 1 | 1 | NA | | 1 | | 0.75 | 0.125 |
| SE | 10390 | IP6K1 | IP6K1 | chr3 | - | 49738211 | 49738422 | 49732790 | 49732972 | 49786353 | 49786540 | 9 | 2 | 1 | 0 | 2 | 0 | 0 | 3 | 0 | 0 | 98 | 49 | 1 | 1 | 1 | 0.333 | 1 | NA | | | 1 | -0.222 |
| SE | 10526 | TMEM33 | TMEM33 | chr4 | + | 41939195 | 41939601 | 41938601 | 41938696 | 41943746 | 41943814 | 13 | 6 | 3 | 6 | 0 | 0 | 0 | 4 | 0 | 0 | 98 | 49 | 1 | 1 | 0.52 | | 1 | NA | | | 1 | -0.16 |
| SE | 10639 | EED | EED | chr11 | + | 86266082 | 86266216 | 86264171 | 86264263 | 86276979 | 86277138 | 4 | 1 | 0 | 0 | 1 | 0 | 0 | 2 | 0 | 0 | 98 | 49 | 1 | 1 | 1 | 0.333 | NA | NA | | | 1 | -0.334 |
| SE | 10710 | RASGRP2 | RASGRP2 | chr11 | - | 64727295 | 64727360 | 64726910 | 64727131 | 64728862 | 64729042 | 36 | 5 | 10 | 0 | 0 | 0 | 0 | 7 | 1 | 0 | 98 | 49 | 1 | 1 | 1 | 1 | 1 | | 0 | | 1 | 0.5 |
| SE | 10834 | ZDHHC5 | ZDHHC5 | chr11 | + | 57695919 | 57696043 | 57693782 | 57693915 | 57696760 | 57696873 | 17 | 5 | 8 | 0 | 0 | 0 | 0 | 0 | 0 | 1 | 98 | 49 | 1 | 1 | 1 | 1 | 1 | NA | | | 0 | 1 |
| SE | 10836 | PEX16 | PEX16 | chr11 | - | 45915467 | 45915568 | 45914603 | 45914684 | 45915702 | 45915836 | 19 | 14 | 1 | 1 | 0 | 0 | 0 | 1 | 0 | 0 | 98 | 49 | 1 | 1 | 0.905 | 1 | 1 | NA | | | 1 | -0.032 |
| SE | 10875 | INPP5B | INPP5B | chr1 | - | 37940687 | 37940798 | 37931912 | 37932053 | 37943639 | 37943669 | 3 | 0 | 0 | 0 | 0 | 2 | 0 | 2 | 0 | 0 | 98 | 49 | 1 | 1 | 1 | NA | 0 | NA | | | 1 | -0.5 |
| SE | 10937 | TNPO1 | TNPO1 | chr5 | + | 72855773 | 72855923 | 72851243 | 72851319 | 72861807 | 72861914 | 18 | 6 | 6 | 1 | 0 | 0 | 0 | 1 | 0 | 0 | 98 | 49 | 1 | 1 | 0.9 | 1 | 1 | NA | | | 1 | -0.033 |
| SE | 10939 | TNPO1 | TNPO1 | chr5 | + | 72900005 | 72900081 | 72897055 | 72897151 | 72900973 | 72901073 | 8 | 3 | 3 | 0 | 0 | 0 | 0 | 0 | 0 | 1 | 98 | 49 | 1 | 1 | 1 | 1 | 1 | NA | | | 0 | 1 |
| SE | 11162 | PHB2 | PHB2 | chr12 | - | 6967170 | 6967238 | 6966419 | 6966500 | 6967891 | 6968021 | 27 | 4 | 1 | 0 | 2 | 0 | 0 | 4 | 0 | 0 | 98 | 49 | 1 | 1 | 1 | 0.5 | 1 | NA | | | 1 | -0.167 |
| SE | 11509 | CEP164 | CEP164 | chr11 | + | 117338565 | 117338668 | 117335604 | 117335680 | 117344165 | 117344277 | 4 | 3 | 1 | 1 | 0 | 0 | 0 | 3 | 0 | 0 | 98 | 49 | 1 | 1 | 0.667 | 1 | 1 | NA | | | 1 | -0.111 |
| SE | 11512 | SBNO2 | SBNO2 | chr19 | - | 1111018 | 1111093 | 1110744 | 1110888 | 1111505 | 1111614 | 39 | 25 | 10 | 0 | 0 | 0 | 1 | 6 | 0 | 1 | 98 | 49 | 1 | 1 | 1 | 1 | 1 | | 1 | | 0.75 | 0.125 |
| SE | 11514 | SBNO2 | SBNO2 | chr19 | - | 1113534 | 1113704 | 1112817 | 1112949 | 1114230 | 1114422 | 92 | 52 | 19 | 1 | 0 | 0 | 9 | 21 | 0 | 1 | 98 | 49 | 1 | 1 | 0.979 | 1 | 1 | | 1 | | 0.913 | 0.036 |
| SE | 11554 | WDFY2 | WDFY2 | chr13 | + | 51660595 | 51660663 | 51584347 | 51584824 | 51675169 | 51675243 | 9 | 6 | 2 | 0 | 2 | 0 | 0 | 2 | 0 | 0 | 98 | 49 | 1 | 1 | 1 | 0.6 | 1 | NA | | | 1 | -0.133 |
| SE | 11555 | WDFY2 | WDFY2 | chr13 | + | 51719197 | 51719348 | 51703595 | 51703650 | 51727677 | 51727790 | 20 | 3 | 3 | 0 | 0 | 0 | 0 | 3 | 0 | 1 | 98 | 49 | 1 | 1 | 1 | 1 | 1 | NA | | | 0.6 | 0.4 |
| SE | 11556 | WDFY2 | WDFY2 | chr13 | + | 51739048 | 51739175 | 51727677 | 51727790 | 51751309 | 51751415 | 7 | 7 | 2 | 1 | 0 | 0 | 0 | 1 | 0 | 0 | 98 | 49 | 1 | 1 | 0.778 | 1 | 1 | NA | | | 1 | -0.074 |
| SE | 11957 | CHD3 | CHD3 | chr17 | + | 7907600 | 7907702 | 7907352 | 7907488 | 7907893 | 7908019 | 8 | 0 | 2 | 2 | 1 | 1 | 0 | 0 | 0 | 2 | 98 | 49 | 1 | 1 | 0.667 | 0 | 0.5 | NA | | | 0 | 0.389 |
| SE | 12147 | DCP2 | DCP2 | chr5 | + | 113003941 | 113004077 | 113001566 | 113001674 | 113007937 | 113008042 | 22 | 10 | 10 | 5 | 1 | 1 | 0 | 10 | 0 | 0 | 98 | 49 | 1 | 1 | 0.687 | 0.833 | 0.833 | NA | | | 1 | -0.216 |
| SE | 12208 | PCYT2 | PCYT2 | chr17 | - | 81908875 | 81909035 | 81908567 | 81908634 | 81909513 | 81909602 | 6 | 0 | 0 | 1 | 0 | 0 | 0 | 2 | 0 | 0 | 98 | 49 | 1 | 1 | 0.75 | NA | NA | NA | | | 1 | -0.25 |
| SE | 12209 | PCYT2 | PCYT2 | chr17 | - | 81908875 | 81909037 | 81908567 | 81908634 | 81909513 | 81909602 | 7 | 0 | 0 | 1 | 0 | 0 | 0 | 2 | 0 | 0 | 98 | 49 | 1 | 1 | 0.778 | NA | NA | NA | | | 1 | -0.222 |
| SE | 12210 | PCYT2 | PCYT2 | chr17 | - | 81908875 | 81909321 | 81908567 | 81908634 | 81909513 | 81909602 | 6 | 0 | 2 | 1 | 0 | 0 | 0 | 2 | 0 | 0 | 98 | 49 | 1 | 1 | 0.75 | NA | 1 | NA | | | 1 | -0.125 |
| SE | 12338 | MAPK1 | MAPK1 | chr22 | - | 21788693 | 21788808 | 21788256 | 21788388 | 21799011 | 21799128 | 17 | 3 | 28 | 0 | 0 | 0 | 0 | 7 | 0 | 1 | 98 | 49 | 1 | 1 | 1 | 1 | 1 | NA | | | 0.778 | 0.222 |
| SE | 12342 | TRAPPC8 | TRAPPC8 | chr18 | - | 31913368 | 31913522 | 31909666 | 31909760 | 31916271 | 31916446 | 29 | 9 | 11 | 2 | 1 | 0 | 0 | 3 | 0 | 0 | 98 | 49 | 1 | 1 | 0.879 | 0.818 | 1 | NA | | | 1 | -0.101 |
| SE | 12653 | RNF123 | RNF123 | chr3 | + | 49712478 | 49712656 | 49706790 | 49706898 | 49713512 | 49713587 | 4 | 0 | 1 | 1 | 0 | 0 | 0 | 1 | 0 | 0 | 98 | 49 | 1 | 1 | 0.667 | NA | 1 | NA | | | 1 | -0.166 |
| SE | 13109 | GBP3 | GBP3 | chr1 | - | 89020531 | 89020743 | 89015286 | 89015414 | 89022683 | 89022866 | 8 | 1 | 0 | 1 | 0 | 0 | 1 | 0 | 0 | 0 | 98 | 49 | 1 | 1 | 0.8 | 1 | NA | | 1 | NA | | -0.1 |
| SE | 13160 | AP2A1 | AP2A1 | chr19 | + | 49800960 | 49801058 | 49799967 | 49800150 | 49801389 | 49801621 | 20 | 9 | 5 | 0 | 0 | 0 | 1 | 3 | 1 | 0 | 98 | 49 | 1 | 1 | 1 | 1 | 1 | | 0.333 | | 1 | 0.334 |
| SE | 13686 | PQLC3 | PQLC3 | chr2 | + | 11171924 | 11172045 | 11164203 | 11164261 | 11174967 | 11175009 | 3 | 2 | 3 | 0 | 1 | 0 | 0 | 1 | 0 | 1 | 98 | 49 | 1 | 1 | 1 | 0.5 | 1 | NA | | | 0.333 | 0.5 |
| SE | 13809 | HNRNPLL | HNRNPLL | chr2 | - | 38569803 | 38569925 | 38569132 | 38569334 | 38573209 | 38573427 | 4 | 6 | 1 | 1 | 0 | 0 | 0 | 2 | 0 | 0 | 98 | 49 | 1 | 1 | 0.667 | 1 | 1 | NA | | | 1 | -0.111 |
| SE | 13829 | EML4 | EML4 | chr2 | + | 42301240 | 42301392 | 42295380 | 42295516 | 42303103 | 42303229 | 17 | 11 | 2 | 1 | 0 | 0 | 0 | 6 | 0 | 0 | 98 | 49 | 1 | 1 | 0.895 | 1 | 1 | NA | | | 1 | -0.035 |
| SE | 13861 | TBCK | TBCK | chr4 | - | 106254946 | 106255047 | 106251865 | 106252007 | 106260436 | 106260510 | 0 | 2 | 0 | 1 | 0 | 0 | 0 | 3 | 0 | 0 | 98 | 49 | 1 | 1 | 0 | NA | NA | NA | | | 1 | -1 |
| SE | 13933 | TRAPPC9 | TRAPPC9 | chr8 | - | 140426614 | 140426641 | 140405576 | 140405698 | 140435111 | 140435240 | 5 | 4 | 0 | 2 | 0 | 0 | 0 | 4 | 0 | 0 | 76 | 49 | 1 | 1 | 0.617 | 1 | NA | NA | | | 1 | -0.192 |
| SE | 14136 | TMEM87B | TMEM87B | chr2 | + | 112077191 | 112077282 | 112074911 | 112074962 | 112081056 | 112081118 | 9 | 0 | 0 | 1 | 0 | 0 | 0 | 2 | 0 | 0 | 98 | 49 | 1 | 1 | 0.818 | NA | NA | NA | | | 1 | -0.182 |
| SE | 14137 | TMEM87B | TMEM87B | chr2 | + | 112081334 | 112081518 | 112081056 | 112081118 | 112086004 | 112086104 | 3 | 0 | 0 | 1 | 0 | 0 | 0 | 2 | 0 | 0 | 98 | 49 | 1 | 1 | 0.6 | NA | NA | NA | | | 1 | -0.4 |
| SE | 14193 | CNST | CNST | chr1 | + | 246591511 | 246591941 | 246566336 | 246566663 | 246621428 | 246621634 | 7 | 2 | 0 | 1 | 0 | 0 | 0 | 1 | 0 | 0 | 98 | 49 | 1 | 1 | 0.778 | 1 | NA | NA | | | 1 | -0.111 |
| SE | 14441 | INO80D | INO80D | chr2 | - | 206063150 | 206063244 | 206062798 | 206063045 | 206085900 | 206086182 | 6 | 0 | 0 | 1 | 0 | 0 | 0 | 2 | 0 | 0 | 98 | 49 | 1 | 1 | 0.75 | NA | NA | NA | | | 1 | -0.25 |
| SE | 14502 | ATG9A | ATG9A | chr2 | - | 219227921 | 219228053 | 219227769 | 219227813 | 219228433 | 219228485 | 8 | 3 | 0 | 1 | 0 | 0 | 0 | 1 | 0 | 0 | 98 | 49 | 1 | 1 | 0.8 | 1 | NA | NA | | | 1 | -0.1 |
| SE | 14921 | UBA3 | UBA3 | chr3 | - | 69057255 | 69057309 | 69056778 | 69056815 | 69061813 | 69061927 | 9 | 2 | 8 | 2 | 0 | 0 | 0 | 7 | 0 | 0 | 98 | 49 | 1 | 1 | 0.692 | 1 | 1 | NA | | | 1 | -0.103 |
| SE | 14922 | UBA3 | UBA3 | chr3 | - | 69071534 | 69071617 | 69067927 | 69068008 | 69075429 | 69075510 | 16 | 6 | 14 | 4 | 0 | 0 | 0 | 3 | 0 | 0 | 98 | 49 | 1 | 1 | 0.667 | 1 | 1 | NA | | | 1 | -0.111 |
| SE | 14942 | EIF3K | EIF3K | chr19 | + | 38620336 | 38620435 | 38619073 | 38619327 | 38624076 | 38624197 | 31 | 7 | 4 | 0 | 0 | 0 | 0 | 8 | 0 | 1 | 98 | 49 | 1 | 1 | 1 | 1 | 1 | NA | | | 0.8 | 0.2 |
| SE | 14943 | EIF3K | EIF3K | chr19 | + | 38632600 | 38632678 | 38632429 | 38632496 | 38634992 | 38635118 | 30 | 9 | 10 | 0 | 0 | 0 | 0 | 5 | 0 | 1 | 98 | 49 | 1 | 1 | 1 | 1 | 1 | NA | | | 0.714 | 0.286 |
| SE | 15062 | MORC3 | MORC3 | chr21 | + | 36356619 | 36356724 | 36349310 | 36349408 | 36359954 | 36360077 | 18 | 6 | 11 | 1 | 0 | 0 | 0 | 2 | 0 | 0 | 98 | 49 | 1 | 1 | 0.9 | 1 | 1 | NA | | | 1 | -0.033 |
| SE | 15094 | TRAPPC10 | TRAPPC10 | chr21 | + | 44037791 | 44037927 | 44032090 | 44032172 | 44052279 | 44052476 | 14 | 13 | 12 | 0 | 3 | 0 | 0 | 5 | 0 | 0 | 98 | 49 | 1 | 1 | 1 | 0.684 | 1 | NA | | | 1 | -0.105 |
| SE | 15095 | TRAPPC10 | TRAPPC10 | chr21 | + | 44074323 | 44074470 | 44063537 | 44063785 | 44075038 | 44075153 | 16 | 12 | 2 | 0 | 2 | 0 | 0 | 4 | 0 | 0 | 98 | 49 | 1 | 1 | 1 | 0.75 | 1 | NA | | | 1 | -0.083 |
| SE | 15165 | LZTR1 | LZTR1 | chr22 | + | 20989624 | 20989682 | 20988788 | 20988872 | 20990385 | 20990525 | 3 | 0 | 0 | 0 | 3 | 0 | 0 | 4 | 0 | 0 | 98 | 49 | 1 | 1 | 1 | 0 | NA | NA | | | 1 | -0.5 |
| SE | 15226 | DIAPH1 | DIAPH1 | chr5 | - | 141529602 | 141529697 | 141529171 | 141529273 | 141534334 | 141534433 | 20 | 17 | 0 | 2 | 2 | 0 | 0 | 8 | 0 | 0 | 98 | 49 | 1 | 1 | 0.833 | 0.81 | NA | NA | | | 1 | -0.178 |

| | | | | | | | | | | | | | | | | | | | | | | | | | | | | | | | |
|---|---|---|---|---|---|---|---|---|---|---|---|---|---|---|---|---|---|---|---|---|---|---|---|---|---|---|---|---|---|---|---|
| SE | 15227 | DIAPH1 | DIAPH1 | chr5 | - | 141583484 | 141583615 | 141583205 | 141583292 | 141584123 | 141584225 | 41 | 10 | 5 | 2 | 0 | 0 | 3 | 26 | 0 | 0 | 98 | 49 | 1 | 1 | 0.911 | 1 | 1 | 1 | 1 | -0.03 |
| SE | 15228 | DIAPH1 | DIAPH1 | chr5 | - | 141588223 | 141588250 | 141587041 | 141587197 | 141618797 | 141619055 | 0 | 1 | 0 | 16 | 10 | 3 | 0 | 0 | 0 | 7 | 76 | 49 | 1 | 1 | 0 | 0.061 | 0 | NA | 0 | 0.02 |
| SE | 15459 | RAF1 | RAF1 | chr3 | - | 12600151 | 12600279 | 12599690 | 12599808 | 12600387 | 12600415 | 162 | 48 | 33 | 2 | 1 | 0 | 2 | 19 | 0 | 0 | 98 | 49 | 1 | 1 | 0.976 | 0.96 | 1 | 1 | 1 | -0.021 |
| SE | 15460 | RAF1 | RAF1 | chr3 | - | 12606200 | 12606299 | 12604135 | 12604289 | 12608765 | 12608923 | 154 | 59 | 46 | 39 | 3 | 3 | 0 | 20 | 0 | 5 | 98 | 49 | 1 | 1 | 0.664 | 0.908 | 0.885 | NA | 0.667 | 0.152 |
| SE | 15530 | DYNC1LI1 | DYNC1LI1 | chr3 | - | 32530283 | 32530328 | 32529539 | 32529660 | 32530460 | 32530520 | 49 | 39 | 10 | 1 | 0 | 0 | 1 | 7 | 0 | 0 | 94 | 49 | 1 | 1 | 0.962 | 1 | 1 | 1 | 1 | -0.013 |
| SE | 15531 | DYNC1LI1 | DYNC1LI1 | chr3 | - | 32544875 | 32545106 | 32541036 | 32541206 | 32545848 | 32545965 | 22 | 4 | 10 | 2 | 0 | 0 | 0 | 4 | 0 | 0 | 98 | 49 | 1 | 1 | 0.846 | 1 | 1 | NA | 1 | -0.051 |
| SE | 15532 | DYNC1LI1 | DYNC1LI1 | chr3 | - | 32545848 | 32545965 | 32541036 | 32541206 | 32570345 | 32570419 | 15 | 9 | 6 | 2 | 0 | 0 | 0 | 2 | 0 | 0 | 98 | 49 | 1 | 1 | 0.789 | 1 | 1 | NA | 1 | -0.07 |
| SE | 15599 | SACM1L | SACM1L | chr3 | + | 45735234 | 45735373 | 45732052 | 45732151 | 45737582 | 45737652 | 16 | 5 | 0 | 1 | 0 | 0 | 0 | 2 | 0 | 0 | 98 | 49 | 1 | 1 | 0.889 | 1 | NA | NA | 1 | -0.055 |
| SE | 15751 | GBE1 | GBE1 | chr3 | - | 81646391 | 81646482 | 81642780 | 81642990 | 81648855 | 81648991 | 8 | 2 | 3 | 1 | 0 | 0 | 0 | 0 | 0 | 1 | 98 | 49 | 1 | 1 | 0.8 | 1 | 1 | NA | 0 | 0.933 |
| SE | 15752 | GBE1 | GBE1 | chr3 | - | 81705443 | 81705613 | 81670837 | 81670953 | 81761374 | 81761799 | 5 | 6 | 1 | 0 | 1 | 0 | 1 | 1 | 0 | 0 | 98 | 49 | 1 | 1 | 1 | 0.75 | 1 | 1 | 1 | -0.083 |
| SE | 16190 | HTT | HTT | chr4 | + | 3140509 | 3140656 | 3136225 | 3136326 | 3142765 | 3142886 | 8 | 0 | 0 | 1 | 0 | 0 | 0 | 4 | 0 | 0 | 98 | 49 | 1 | 1 | 0.8 | NA | NA | NA | 1 | -0.2 |
| SE | 16215 | WDR1 | WDR1 | chr4 | - | 10077752 | 10077926 | 10077303 | 10077448 | 10078890 | 10079001 | 83 | 27 | 26 | 2 | 0 | 0 | 12 | 27 | 0 | 0 | 98 | 49 | 1 | 1 | 0.954 | 1 | 1 | 1 | 1 | -0.015 |
| SE | 16217 | WDR1 | WDR1 | chr4 | - | 10088663 | 10088741 | 10088292 | 10088373 | 10097710 | 10097891 | 55 | 15 | 17 | 2 | 0 | 0 | 3 | 4 | 0 | 0 | 98 | 49 | 1 | 1 | 0.932 | 1 | 1 | 1 | 1 | -0.023 |
| SE | 16218 | WDR1 | WDR1 | chr4 | - | 10097710 | 10097891 | 10088663 | 10088741 | 10116112 | 10116234 | 37 | 7 | 7 | 2 | 0 | 0 | 2 | 3 | 0 | 0 | 98 | 49 | 1 | 1 | 0.902 | 1 | 1 | 1 | 1 | -0.033 |
| SE | 16219 | WDR1 | WDR1 | chr4 | - | 10098991 | 10099139 | 10097710 | 10097891 | 10116112 | 10116234 | 35 | 23 | 17 | 2 | 0 | 0 | 1 | 6 | 0 | 0 | 98 | 49 | 1 | 1 | 0.897 | 1 | 1 | 1 | 1 | -0.034 |
| SE | 16220 | WDR1 | WDR1 | chr4 | - | 10103895 | 10103986 | 10088663 | 10088741 | 10116112 | 10116234 | 36 | 16 | 13 | 2 | 0 | 0 | 4 | 6 | 0 | 0 | 98 | 49 | 1 | 1 | 0.9 | 1 | 1 | 1 | 1 | -0.033 |
| SE | 16221 | WDR1 | WDR1 | chr4 | - | 10103895 | 10103986 | 10098991 | 10099139 | 10116112 | 10116234 | 64 | 30 | 25 | 0 | 0 | 0 | 5 | 16 | 0 | 1 | 98 | 49 | 1 | 1 | 1 | 1 | 1 | 1 | 0.889 | 0.055 |
| SE | 16244 | TAPT1 | TAPT1 | chr4 | - | 16213767 | 16213898 | 16191360 | 16191523 | 16226258 | 16226538 | 2 | 2 | 0 | 1 | 0 | 0 | 0 | 1 | 0 | 0 | 98 | 49 | 1 | 1 | 0.5 | 1 | NA | NA | 1 | -0.25 |
| SE | 16632 | HNRNPM | HNRNPM | chr19 | + | 8483157 | 8483211 | 8474166 | 8474244 | 8485602 | 8486405 | 9 | 3 | 0 | 2 | 0 | 0 | 0 | 2 | 0 | 0 | 98 | 49 | 1 | 1 | 0.692 | 1 | NA | NA | 1 | -0.154 |
| SE | 16761 | SLC30A5 | SLC30A5 | chr5 | + | 69128003 | 69128132 | 69123198 | 69123425 | 69129446 | 69131072 | 6 | 0 | 2 | 1 | 0 | 0 | 0 | 1 | 0 | 0 | 98 | 49 | 1 | 1 | 0.75 | NA | 1 | NA | 1 | -0.125 |
| SE | 16976 | IK | IK | chr5 | + | 140655828 | 140655992 | 140654680 | 140654727 | 140657553 | 140657662 | 30 | 7 | 6 | 3 | 0 | 0 | 0 | 2 | 0 | 2 | 98 | 49 | 1 | 1 | 0.833 | 1 | 1 | NA | 0.333 | 0.611 |
| SE | 16982 | HARS2 | HARS2 | chr5 | + | 140696102 | 140696201 | 140695737 | 140695845 | 140696520 | 140696614 | 7 | 2 | 1 | 1 | 0 | 0 | 0 | 1 | 0 | 0 | 98 | 49 | 1 | 1 | 0.778 | 1 | 1 | NA | 1 | -0.074 |
| SE | 17117 | MCMBP | MCMBP | chr10 | - | 119836895 | 119837029 | 119835539 | 119835704 | 119838534 | 119838700 | 17 | 9 | 27 | 1 | 0 | 0 | 0 | 12 | 0 | 0 | 98 | 49 | 1 | 1 | 0.895 | 1 | 1 | NA | 1 | -0.035 |
| SE | 17118 | MCMBP | MCMBP | chr10 | - | 119842471 | 119842595 | 119840842 | 119840960 | 119843247 | 119843426 | 2 | 5 | 14 | 0 | 0 | 0 | 0 | 0 | 1 | 0 | 98 | 49 | 1 | 1 | 1 | 1 | 1 | 0 | NA | 1 |
| RI | 474 | TMEM41B | TMEM41B | chr11 | - | 9280654 | 9283593 | 9280654 | 9281041 | 9283370 | 9283593 | 8 | 2 | 0 | 1 | 1 | 0 | 0 | 2 | 0 | 0 | 98 | 49 | 1 | 1 | 0.8 | 0.5 | NA | NA | 1 | -0.35 |
| RI | 1010 | TAF1C | TAF1C | chr16 | - | 84181327 | 84181663 | 84181327 | 84181463 | 84181591 | 84181663 | 8 | 1 | 0 | 0 | 0 | 0 | 0 | 3 | 0 | 1 | 98 | 49 | 1 | 1 | 1 | 1 | NA | NA | 0.6 | 0.4 |
| RI | 1110 | HNRNPM | HNRNPM | chr19 | + | 8465323 | 8465515 | 8465323 | 8465362 | 8465479 | 8465515 | 13 | 6 | 1 | 5 | 2 | 3 | 0 | 8 | 0 | 1 | 98 | 49 | 1 | 1 | 0.565 | 0.6 | 0.143 | NA | 0.8 | -0.364 |

**MXE_all**

| | | | | | | | | | | | | | Inclusion counts | | | Skipping counts | | | Inclusion counts | | Skipping counts | | | | | | Inclusion_level(normalized) | | | | | |
|---|---|---|---|---|---|---|---|---|---|---|---|---|---|---|---|---|---|---|---|---|---|---|---|---|---|---|---|---|---|---|---|---|
| ID | Gene_ID | Gene_Symbol | Chromosome | Strand | 1stExonStart_0base | 1stExonEnd | 2ndExonStart_0base | 2ndExonEnd | UpstreamES | UpstreamEE | DownstreamES | DownstreamEE | control_s3 | control_s4 | control_s10 | control_s3 | control_s4 | control_s10 | ldg_14 | lldg_s16 | ldg_s14 | ldg_s16 | IncFormLen | SkipFormLen | p-value | adjust_p | IncLevel_s3 | IncLevel_s4 | IncLevel_s10 | IncLevel14 | IncLevel16 | Level_diff |
| 767 | ATP5C1 | ATP5C1 | chr10 | + | 7802757 | 7802854 | 7806973 | 7807010 | 7802269 | 7802425 | 7807658 | 7807799 | 20 | 7 | 2 | 3 | 1 | 3 | 0 | 0 | 1 | 8 | 98 | 86 | 9.92E-09 | 6.58E-06 | 0.854 | 0.86 | 0.369 | 0 | 0 | 0.694 |
| 69 | CPQ | CPQ | chr8 | + | 96834972 | 96835180 | 96879797 | 96880005 | 96784863 | 96785330 | 96965934 | 96966046 | 33 | 16 | 7 | 24 | 15 | 5 | 1 | 9 | 0 | 0 | 98 | 98 | 1.45E-06 | 0.000481929 | 0.579 | 0.516 | 0.583 | 1 | 1 | -0.441 |
| 973 | LST1 | LST1 | chr6 | + | 31587640 | 31587733 | 31587943 | 31587966 | 31587199 | 31587318 | 31588496 | 31588909 | 39 | 24 | 22 | 9 | 9 | 2 | 0 | 0 | 1 | 1 | 98 | 72 | 1.71E-05 | 0.00253914 | 0.761 | 0.662 | 0.89 | 0 | 0 | 0.771 |
| 700 | RFWD2 | RFWD2 | chr1 | - | 176136487 | 176136547 | 176175909 | 176176007 | 176135009 | 176135086 | 176184632 | 176184692 | 32 | 16 | 3 | 89 | 32 | 10 | 0 | 0 | 1 | 10 | 98 | 98 | 1.93E-05 | 0.00253914 | 0.264 | 0.333 | 0.231 | 0 | 0 | 0.276 |
| 1169 | ARID3A | ARID3A | chr19 | + | 964832 | 965080 | 966571 | 966868 | 964247 | 964431 | 968404 | 968503 | 8 | 3 | 3 | 17 | 4 | 3 | 1 | 1 | 0 | 0 | 98 | 98 | 2.60E-05 | 0.00253914 | 0.32 | 0.429 | 0.5 | 1 | 1 | -0.584 |
| 983 | IRF7 | IRF7 | chr11 | - | 613950 | 614037 | 614173 | 614399 | 613784 | 613865 | 614475 | 614534 | 2 | 3 | 7 | 13 | 11 | 0 | 0 | 0 | 8 | 14 | 98 | 98 | 2.64E-05 | 0.00253914 | 0.133 | 0.214 | 1 | 0 | 0 | 0.449 |
| 814 | MAEA | MAEA | chr4 | + | 1322380 | 1322503 | 1327626 | 1327703 | 1315396 | 1315600 | 1332756 | 1332865 | 11 | 5 | 2 | 12 | 5 | 7 | 0 | 0 | 3 | 2 | 98 | 98 | 2.68E-05 | 0.00253914 | 0.478 | 0.5 | 0.222 | 0 | 0 | 0.4 |
| 672 | SLBP | SLBP | chr4 | - | 1703595 | 1703700 | 1711873 | 1711990 | 1700010 | 1700070 | 1712129 | 1712741 | 1 | 0 | 1 | 10 | 6 | 0 | 1 | 2 | 0 | 1 | 98 | 98 | 4.66E-05 | 0.003870084 | 0.091 | 0 | 1 | 1 | 0.667 | -0.47 |
| 933 | DUSP1 | DUSP1 | chr5 | - | 172769574 | 172769794 | 172770160 | 172770306 | 172768089 | 172769132 | 172770585 | 172771200 | 561 | 303 | 269 | 438 | 200 | 178 | 93 | 267 | 6 | 110 | 98 | 98 | 7.32E-05 | 0.005400629 | 0.562 | 0.602 | 0.602 | 0.939 | 0.708 | -0.235 |
| 677 | STK40 | STK40 | chr1 | - | 36358736 | 36358822 | 36361220 | 36361340 | 36358238 | 36358382 | 36385722 | 36385927 | 19 | 8 | 3 | 30 | 18 | 9 | 0 | 0 | 1 | 3 | 98 | 98 | 9.01E-05 | 0.005983417 | 0.388 | 0.308 | 0.25 | 0 | 0 | 0.315 |
| 1210 | CHD2 | CHD2 | chr15 | + | 92944414 | 92944515 | 92945820 | 92945865 | 92942842 | 92943068 | 92946037 | 92946216 | 23 | 3 | 0 | 44 | 13 | 8 | 0 | 0 | 2 | 8 | 98 | 94 | 0.000158716 | 0.009580676 | 0.334 | 0.181 | 0 | 0 | 0 | 0.172 |
| 887 | ZNF467 | ZNF467 | chr7 | - | 149769089 | 149769200 | 149770439 | 149770556 | 149764363 | 149766239 | 149770998 | 149771074 | 8 | 1 | 0 | 9 | 3 | 5 | 0 | 0 | 2 | 1 | 98 | 98 | 0.000367755 | 0.020349121 | 0.471 | 0.25 | 0 | 0 | 0 | 0.24 |
| 603 | C1orf43 | C1orf43 | chr1 | - | 154213892 | 154213946 | 154214456 | 154214574 | 154212457 | 154212624 | 154220341 | 154220797 | 4 | 2 | 0 | 7 | 6 | 7 | 1 | 2 | 0 | 1 | 98 | 98 | 0.000476501 | 0.024338219 | 0.364 | 0.25 | 0 | 1 | 0.667 | -0.629 |
| 275 | SEC11A | SEC11A | chr15 | - | 84680712 | 84680832 | 84687624 | 84687774 | 84670724 | 84670782 | 84691534 | 84691644 | 15 | 4 | 20 | 3 | 2 | 4 | 2 | 2 | 0 | 0 | 98 | 98 | 0.000719589 | 0.030060115 | 0.833 | 0.667 | 0.833 | 1 | 1 | -0.222 |
| 1520 | ALPL | ALPL | chr1 | + | 21553977 | 21554142 | 21560625 | 21560745 | 21509364 | 21509517 | 21563109 | 21563284 | 25 | 7 | 6 | 5 | 4 | 0 | 2 | 6 | 0 | 0 | 98 | 98 | 0.000721258 | 0.030060115 | 0.833 | 0.636 | 1 | 1 | 1 | -0.177 |
| 720 | APH1A | APH1A | chr1 | - | 150267715 | 150267789 | 150267956 | 150267991 | 150267355 | 150267478 | 150268697 | 150269209 | 2 | 3 | 1 | 11 | 2 | 6 | 0 | 0 | 1 | 1 | 84 | 98 | 0.00072434 | 0.030060115 | 0.175 | 0.636 | 0.163 | 0 | 0 | 0.325 |
| 1140 | SAP30BP | SAP30BP | chr17 | + | 75671815 | 75671863 | 75693439 | 75693482 | 75668515 | 75668625 | 75699782 | 75699871 | 25 | 6 | 5 | 15 | 5 | 5 | 1 | 0 | 1 | 4 | 97 | 92 | 0.000838883 | 0.03276579 | 0.613 | 0.532 | 0.487 | 0.487 | 0 | 0.301 |
| 439 | KIAA1033 | KIAA1033 | chr12 | + | 105160000 | 105160148 | 105162748 | 105162845 | 105157235 | 105157322 | 105164110 | 105164307 | 44 | 7 | 1 | 8 | 8 | 2 | 1 | 8 | 0 | 1 | 98 | 98 | 0.001281181 | 0.047261349 | 0.846 | 0.467 | 0.333 | 1 | 0.889 | -0.396 |
| 347 | PLEKHB2 | PLEKHB2 | chr2 | + | 131126683 | 131126786 | 131132901 | 131132991 | 131125752 | 131125905 | 131140166 | 131140275 | 24 | 9 | 2 | 18 | 5 | 2 | 1 | 0 | 1 | 3 | 98 | 98 | 0.001418793 | 0.048685792 | 0.571 | 0.643 | 0.5 | 0.5 | 0 | 0.321 |
| 272 | FCAR | FCAR | chr19 | + | 54885234 | 54885525 | 54888006 | 54888294 | 54874092 | 54874323 | 54889648 | 54891420 | 4 | 10 | 0 | 28 | 6 | 2 | 0 | 1 | 4 | 17 | 98 | 98 | 0.001508121 | 0.048685792 | 0.125 | 0.625 | 0 | 0 | 0.056 | 0.222 |
| 1172 | GADD45B | GADD45B | chr19 | + | 2476528 | 2476630 | 2477028 | 2477251 | 2476124 | 2476402 | 2477487 | 2478259 | 29 | 30 | 14 | 22 | 26 | 14 | 7 | 19 | 1 | 5 | 98 | 98 | 0.001539761 | 0.048685792 | 0.569 | 0.536 | 0.5 | 0.875 | 0.792 | -0.298 |
| 866 | CDC42SE2 | CDC42SE2 | chr5 | + | 131315975 | 131316144 | 131359208 | 131359547 | 131264008 | 131264166 | 131385542 | 131385644 | 9 | 2 | 0 | 50 | 14 | 10 | 0 | 0 | 1 | 9 | 98 | 98 | 0.001681804 | 0.050759907 | 0.153 | 0.125 | 0 | 0 | 0 | 0.093 |
| 605 | GBP5 | GBP5 | chr1 | - | 89268728 | 89268856 | 89269365 | 89269574 | 89267416 | 89267526 | 89272451 | 89272861 | 5 | 4 | 1 | 48 | 11 | 5 | 0 | 0 | 1 | 3 | 98 | 98 | 0.001832725 | 0.05290996 | 0.094 | 0.267 | 0.167 | 0 | 0 | 0.176 |
| 1043 | THEMIS2 | THEMIS2 | chr1 | + | 27879643 | 27880054 | 27881970 | 27883043 | 27876587 | 27876728 | 27885294 | 27885451 | 111 | 23 | 15 | 25 | 14 | 1 | 6 | 25 | 0 | 2 | 98 | 98 | 0.002730415 | 0.075541495 | 0.816 | 0.622 | 0.937 | 1 | 0.926 | -0.171 |
| 752 | ARHGEF7 | ARHGEF7 | chr13 | + | 111205288 | 111205373 | 111209871 | 111210002 | 111153713 | 111153991 | 111217678 | 111217880 | 4 | 2 | 0 | 6 | 1 | 1 | 2 | 1 | 0 | 1 | 98 | 98 | 0.002961552 | 0.078658825 | 0.4 | 0.667 | 0 | 1 | 0.5 | -0.394 |
| 924 | NADK | NADK | chr1 | - | 1753566 | 1753649 | 1754050 | 1754208 | 1751231 | 1753060 | 1754283 | 1754383 | 60 | 35 | 36 | 54 | 19 | 38 | 2 | 6 | 7 | 15 | 98 | 98 | 0.00311261 | 0.079491271 | 0.526 | 0.648 | 0.486 | 0.222 | 0.286 | 0.299 |
| 1492 | GPSM3 | GPSM3 | chr6 | - | 32191708 | 32191908 | 32192147 | 32192250 | 32190765 | 32191503 | 32192471 | 32192530 | 20 | 13 | 8 | 50 | 23 | 25 | 1 | 1 | 3 | 14 | 98 | 98 | 0.004345138 | 0.106858211 | 0.286 | 0.361 | 0.242 | 0.25 | 0.067 | 0.138 |
| 812 | MAEA | MAEA | chr4 | + | 1315396 | 1315600 | 1322380 | 1322503 | 1311978 | 1312161 | 1327626 | 1327703 | 15 | 13 | 2 | 1 | 4 | 1 | 2 | 1 | 0 | 0 | 98 | 98 | 0.005546739 | 0.13153696 | 0.937 | 0.765 | 0.667 | 1 | 1 | -0.21 |
| 1232 | TSEN34 | TSEN34 | chr19 | + | 54191360 | 54191607 | 54191720 | 54191964 | 54190267 | 54190450 | 54192115 | 54192373 | 3 | 7 | 3 | 44 | 21 | 17 | 0 | 1 | 3 | 12 | 98 | 98 | 0.007192046 | 0.164673055 | 0.064 | 0.25 | 0.15 | 0 | 0.077 | 0.116 |
| 1156 | SLC11A1 | SLC11A1 | chr2 | + | 218382959 | 218383102 | 218384242 | 218384365 | 218382028 | 218382375 | 218385146 | 218385266 | 27 | 17 | 0 | 106 | 58 | 23 | 0 | 2 | 2 | 27 | 98 | 98 | 0.007791047 | 0.172441837 | 0.203 | 0.227 | 0 | 0 | 0.069 | 0.109 |
| 694 | MKL1 | MKL1 | chr22 | - | 40423535 | 40423685 | 40424205 | 40424381 | 40420846 | 40421100 | 40429605 | 40429767 | 10 | 3 | 1 | 5 | 6 | 0 | 2 | 1 | 0 | 1 | 98 | 98 | 0.008293126 | 0.177633408 | 0.667 | 0.333 | 1 | 1 | 0.5 | -0.083 |
| 995 | NFKBIA | NFKBIA | chr14 | - | 35402770 | 35402859 | 35403149 | 35403360 | 35402393 | 35402663 | 35403689 | 35403798 | 77 | 218 | 23 | 78 | 178 | 23 | 0 | 24 | 8 | 46 | 98 | 98 | 0.008601596 | 0.178483112 | 0.497 | 0.551 | 0.5 | 0 | 0.343 | 0.345 |
| 580 | LIMK2 | LIMK2 | chr22 | + | 31265945 | 31266132 | 31266983 | 31267070 | 31262594 | 31262791 | 31267775 | 31267907 | 24 | 9 | 2 | 68 | 37 | 9 | 2 | 8 | 0 | 6 | 98 | 98 | 0.008893756 | 0.178953152 | 0.261 | 0.196 | 0.182 | 1 | 0.571 | -0.573 |
| 469 | PLEKHM2 | PLEKHM2 | chr1 | + | 15719733 | 15719920 | 15721328 | 15721388 | 15718537 | 15718625 | 15725316 | 15725545 | 10 | 6 | 5 | 2 | 2 | 1 | 1 | 2 | 0 | 0 | 98 | 98 | 0.00958895 | 0.187266563 | 0.833 | 0.75 | 0.833 | 1 | 1 | -0.195 |
| 478 | ZRANB1 | ZRANB1 | chr10 | + | 124974199 | 124974398 | 124981708 | 124981829 | 124973644 | 124973716 | 124983174 | 124983304 | 9 | 0 | 3 | 12 | 10 | 2 | 0 | 1 | 1 | 2 | 98 | 98 | 0.010790876 | 0.204718338 | 0.429 | 0 | 0.6 | 0 | 0.333 | 0.176 |
| 128 | IKZF1 | IKZF1 | chr7 | + | 50387344 | 50387470 | 50391728 | 50391863 | 50382539 | 50382707 | 50399917 | 50405100 | 25 | 3 | 4 | 6 | 2 | 0 | 2 | 7 | 0 | 1 | 98 | 98 | 0.013093288 | 0.241498429 | 0.806 | 0.6 | 1 | 1 | 0.875 | -0.135 |
| 428 | CLEC7A | CLEC7A | chr12 | - | 10123244 | 10123363 | 10125086 | 10125173 | 10116780 | 10118590 | 10125296 | 10125448 | 26 | 2 | 10 | 141 | 62 | 38 | 3 | 18 | 0 | 38 | 98 | 98 | 0.014111252 | 0.25323976 | 0.156 | 0.031 | 0.208 | 1 | 0.321 | -0.529 |
| 1058 | C18orf8 | C18orf8 | chr18 | + | 23504370 | 23504447 | 23509192 | 23509279 | 23503469 | 23503720 | 23515855 | 23515996 | 1 | 7 | 0 | 15 | 5 | 2 | 4 | 3 | 0 | 5 | 98 | 98 | 0.026036619 | 0.454955651 | 0.062 | 0.583 | 0 | 1 | 0.375 | -0.473 |
| 654 | MVP | MVP | chr16 | + | 29841914 | 29842112 | 29844492 | 29844879 | 29841595 | 29841840 | 29845862 | 29845979 | 31 | 8 | 7 | 30 | 16 | 18 | 4 | 10 | 1 | 6 | 98 | 98 | 0.027787914 | 0.473107055 | 0.508 | 0.333 | 0.28 | 0.8 | 0.625 | -0.339 |
| 986 | AKNA | AKNA | chr9 | - | 114376465 | 114377532 | 114381059 | 114381446 | 114374092 | 114374167 | 114394356 | 114394405 | 13 | 9 | 1 | 56 | 16 | 8 | 0 | 2 | 2 | 13 | 98 | 98 | 0.030058921 | 0.498978088 | 0.188 | 0.36 | 0.111 | 0 | 0.133 | 0.153 |
| 111 | ORAI2 | ORAI2 | chr7 | + | 102436224 | 102436333 | 102438943 | 102439181 | 102433529 | 102433661 | 102446512 | 102456821 | 1 | 4 | 0 | 11 | 6 | 5 | 0 | 0 | 2 | 2 | 98 | 98 | 0.032538655 | 0.526967491 | 0.083 | 0.4 | 0 | 0 | 0 | 0.161 |
| 1371 | SHKBP1 | SHKBP1 | chr19 | + | 40582350 | 40582466 | 40583397 | 40583485 | 40580745 | 40580936 | 40583600 | 40583717 | 65 | 23 | 14 | 107 | 42 | 23 | 1 | 10 | 1 | 5 | 98 | 98 | 0.038106412 | 0.600128823 | 0.378 | 0.354 | 0.378 | 0.5 | 0.667 | -0.214 |
| 743 | TNRC6B | TNRC6B | chr22 | + | 40264687 | 40267036 | 40270121 | 40270280 | 40261831 | 40262173 | 40273424 | 40273600 | 27 | 14 | 0 | 6 | 2 | 1 | 1 | 4 | 0 | 1 | 98 | 98 | 0.038863764 | 0.600128823 | 0.818 | 0.875 | 0 | 1 | 0.8 | -0.336 |
| 433 | ASAP1 | ASAP1 | chr8 | - | 130091972 | 130092143 | 130112093 | 130112322 | 130079901 | 130079971 | 130115627 | 130115735 | 37 | 7 | 0 | 5 | 5 | 2 | 0 | 12 | 1 | 0 | 98 | 98 | 0.040307474 | 0.608276428 | 0.881 | 0.583 | 0 | 0 | 1 | -0.012 |
| 774 | SULF2 | SULF2 | chr20 | - | 47702518 | 47702670 | 47736702 | 47736942 | 47690125 | 47690295 | 47757188 | 47757463 | 32 | 7 | 4 | 7 | 5 | 3 | 6 | 15 | 0 | 3 | 98 | 98 | 0.042284294 | 0.623928249 | 0.821 | 0.583 | 0.571 | 1 | 0.833 | -0.258 |
| 744 | LCOR | LCOR | chr10 | + | 96944112 | 96944245 | 96949007 | 96949295 | 96907667 | 96907747 | 96952102 | 96952196 | 7 | 2 | 3 | 11 | 4 | 1 | 2 | 0 | 0 | 1 | 98 | 98 | 0.043446552 | 0.627141534 | 0.389 | 0.333 | 0.75 | 1 | 0 | -0.009 |
| 996 | GZMH | GZMH | chr14 | - | 24607148 | 24607406 | 24607611 | 24607747 | 24606479 | 24606746 | 24608264 | 24608412 | 8 | 9 | 0 | 7 | 8 | 4 | 1 | 0 | 0 | 3 | 98 | 98 | 0.046773203 | 0.657258859 | 0.533 | 0.529 | 0 | 1 | 0 | -0.146 |
| 438 | UBE2D1 | UBE2D1 | chr10 | + | 58361494 | 58361526 | 58363608 | 58363686 | 58361337 | 58361401 | 58364770 | 58364876 | 12 | 0 | 5 | 33 | 4 | 13 | 0 | 2 | 1 | 13 | 81 | 98 | 0.047512689 | 0.657258859 | 0.306 | 0 | 0.318 | 0 | 0.157 | 0.13 |
| 382 | SART3 | SART3 | chr12 | - | 108543027 | 108543152 | 108544426 | 108544478 | 108538933 | 108539089 | 108545138 | 108545323 | 5 | 6 | 1 | 6 | 0 | 8 | 1 | 0 | 0 | 1 | 98 | 98 | 0.048879289 | 0.662364246 | 0.455 | 1 | 0.111 | 1 | 0 | 0.022 |
| 788 | ATP2A3 | ATP2A3 | chr17 | - | 3953346 | 3953429 | 3953692 | 3953710 | 3951580 | 3951685 | 3964173 | 3964464 | 6 | 4 | 5 | 19 | 13 | 6 | 1 | 2 | 1 | 1 | 67 | 98 | 0.052742755 | 0.70042378 | 0.316 | 0.31 | 0.549 | 0.594 | 0.745 | -0.278 |
| 403 | USP15 | USP15 | chr12 | + | 62325871 | 62325933 | 62349220 | 62349307 | 62321463 | 62321609 | 62355330 | 62355475 | 193 | 56 | 26 | 80 | 33 | 23 | 2 | 22 | 0 | 6 | 98 | 98 | 0.055556301 | 0.713311355 | 0.707 | 0.629 | 0.531 | 1 | 0.786 | -0.271 |
| 430 | CLEC7A | CLEC7A | chr12 | - | 10125296 | 10125448 | 10127746 | 10127845 | 10123244 | 10123363 | 10129979 | 10130269 | 199 | 74 | 36 | 52 | 26 | 13 | 6 | 44 | 0 | 8 | 98 | 98 | 0.055861733 | 0.713311355 | 0.793 | 0.74 | 0.735 | 1 | 0.846 | -0.167 |
| 60 | DENND3 | DENND3 | chr8 | + | 141190283 | 141190417 | 141192330 | 141192449 | 141188985 | 141189146 | 141192525 | 141192663 | 38 | 11 | 3 | 29 | 8 | 6 | 3 | 9 | 0 | 4 | 98 | 98 | 0.056978495 | 0.71384378 | 0.567 | 0.579 | 0.333 | 1 | 0.692 | -0.353 |
| 1017 | PHF21A | PHF21A | chr11 | - | 45945839 | 45946003 | 45946075 | 45946098 | 45938156 | 45938312 | 45948885 | 45948946 | 7 | 6 | 0 | 16 | 2 | 1 | 1 | 0 | 0 | 4 | 72 | 98 | 0.058600631 | 0.720570725 | 0.373 | 0.803 | 0 | 1 | 0 | -0.108 |
| 1519 | ALPL | ALPL | chr1 | + | 21553977 | 21554142 | 21560625 | 21560745 | 21509364 | 21509517 | 21561096 | 21561212 | 25 | 7 | 6 | 57 | 25 | 15 | 2 | 6 | 3 | 3 | 98 | 98 | 0.067066433 | 0.79747995 | 0.305 | 0.219 | 0.286 | 0.4 | 0.667 | -0.264 |
| 638 | JMJD1C | JMJD1C | chr10 | - | 63219877 | 63219983 | 63264650 | 63264764 | 63217206 | 63217331 | 63268662 | 63269223 | 152 | 34 | 22 | 28 | 10 | 1 | 1 | 15 | 0 | 2 | 98 | 98 | 0.067819904 | 0.79747995 | 0.844 | 0.773 | 0.957 | 1 | 0.882 | -0.083 |
| 1024 | ITGAX | ITGAX | chr16 | + | 31355892 | 31355998 | 31356624 | 31356728 | 31355133 | 31355291 | 31357030 | 31357101 | 12 | 18 | 3 | 112 | 81 | 16 | 0 | 4 | 9 | 38 | 98 | 98 | 0.068458369 | 0.79747995 | 0.097 | 0.182 | 0.158 | 0 | 0.095 | 0.098 |
| 251 | PCBP2 | PCBP2 | chr12 | + | 53464759 | 53464852 | 53465931 | 53465973 | 53462480 | 53462567 | 53467220 | 53467293 | 54 | 17 | 19 | 72 | 31 | 6 | 0 | 13 | 2 | 30 | 98 | 91 | 0.073459606 | 0.821205033 | 0.411 | 0.337 | 0.746 | 0 | 0.287 | 0.355 |
| 1149 | RNF213 | RNF213 | chr17 | + | 80361733 | 80361888 | 80363101 | 80363314 | 80360060 | 80360206 | 80363608 | 80363790 | 43 | 13 | 9 | 66 | 30 | 7 | 1 | 27 | 14 | 63 | 98 | 98 | 0.074169176 | 0.821205033 | 0.394 | 0.302 | 0.562 | 0.067 | 0.3 | 0.236 |
| 527 | SERPINA1 | SERPINA1 | chr14 | - | 94382591 | 94383241 | 94388800 | 94389010 | 94376746 | 94378640 | 94390456 | 94390692 | 47 | 13 | 5 | 108 | 35 | 38 | 1 | 15 | 2 | 20 | 98 | 98 | 0.074205274 | 0.821205033 | 0.303 | 0.271 | 0.116 | 0.333 | 0.429 | -0.151 |
| 241 | CYTH4 | CYTH4 | chr22 | + | 37310987 | 37311064 | 37311455 | 37311527 | 37309211 | 37309323 | 37312019 | 37312174 | 82 | 23 | 19 | 55 | 41 | 44 | 2 | 12 | 2 | 5 | 98 | 98 | 0.076746722 | 0.835406946 | 0.599 | 0.359 | 0.302 | 0.5 | 0.706 | -0.183 |
| 930 | DBNL | DBNL | chr7 | + | 44050224 | 44050280 | 44052866 | 44052941 | 44044639 | 44044820 | 44056756 | 44056903 | 11 | 3 | 5 | 24 | 6 | 9 | 1 | 1 | 2 | 5 | 98 | 98 | 0.083414571 | 0.893343144 | 0.314 | 0.333 | 0.357 | 0.333 | 0.167 | 0.085 |
| 550 | SORL1 | SORL1 | chr11 | + | 121470006 | 121470123 | 121478117 | 121478243 | 121452202 | 121452616 | 121488031 | 121488193 | 174 | 85 | 67 | 166 | 80 | 53 | 6 | 35 | 0 | 23 | 98 | 98 | 0.086250776 | 0.896644508 | 0.512 | 0.515 | 0.558 | 1 | 0.603 | -0.273 |
| 832 | DOCK2 | DOCK2 | chr5 | + | 169684195 | 169684350 | 169689251 | 169689333 | 169681743 | 169681879 | 169695802 | 169695938 | 109 | 35 | 17 | 83 | 48 | 5 | 2 | 13 | 3 | 21 | 98 | 98 | 0.086423567 | 0.896644508 | 0.568 | 0.422 | 0.773 | 0.4 | 0.382 | 0.197 |
| 330 | CTSS | CTSS | chr1 | - | 150751780 | 150752008 | 150755000 | 150755150 | 150750005 | 150750171 | 150757857 | 150757980 | 412 | 132 | 123 | 268 | 149 | 97 | 2 | 126 | 1 | 73 | 98 | 98 | 0.091023357 | 0.929838596 | 0.606 | 0.47 | 0.559 | 0.667 | 0.633 | -0.105 |
| 1114 | CALCOCO2 | CALCOCO2 | chr17 | + | 48849251 | 48849377 | 48851088 | 48851177 | 48848321 | 48848455 | 48851558 | 48851628 | 15 | 15 | 10 | 11 | 5 | 3 | 2 | 4 | 0 | 2 | 98 | 98 | 0.095466674 | 0.960452598 | 0.577 | 0.75 | 0.769 | 1 | 0.667 | -0.135 |
| 747 | RNF19B | RNF19B | chr1 | - | 32938396 | 32938528 | 32942251 | 32942459 | 32936445 | 32937259 | 32944018 | 32944159 | 55 | 25 | 38 | 43 | 11 | 11 | 1 | 19 | 0 | 5 | 98 | 98 | 0.098734743 | 0.978505508 | 0.561 | 0.694 | 0.776 | 1 | 0.792 | -0.219 |
| 1187 | CYP4F3 | CYP4F3 | chr19 | + | 15643906 | 15644051 | 15645718 | 15645863 | 15641414 | 15641613 | 15647051 | 15647105 | 2 | 3 | 5 | 161 | 45 | 36 | 0 | 1 | 5 | 21 | 98 | 98 | 0.10473881 | 1 | 0.012 | 0.062 | 0.122 | 0 | 0.045 | 0.043 |
| 213 | HLA-E | HLA-E | chr6 | + | 30491536 | 30491653 | 30492403 | 30492436 | 30491136 | 30491412 | 30492540 | 30492583 | 302 | 247 | 202 | 143 | 135 | 125 | 12 | 185 | 7 | 64 | 98 | 82 | 0.10716204 | 1 | 0.639 | 0.605 | 0.575 | 0.589 | 0.707 | -0.042 |
| 524 | SERPINA1 | SERPINA1 | chr14 | - | 94379463 | 94379611 | 94382591 | 94383241 | 94376746 | 94378640 | 94388559 | 94388660 | 70 | 15 | 37 | 142 | 66 | 104 | 0 | 13 | 6 | 51 | 98 | 98 | 0.113078522 | 1 | 0.33 | 0.185 | 0.262 | 0 | 0.203 | 0.158 |
| 548 | MYD88 | MYD88 | chr3 | + | 38139863 | 38139998 | 38140387 | 38140568 | 38138477 | 38139028 | 38140756 | 38140848 | 50 | 12 | 5 | 12 | 1 | 3 | 6 | 12 | 3 | 7 | 98 | 98 | 0.117799117 | 1 | 0.806 | 0.923 | 0.625 | 0.667 | 0.632 | 0.135 |
| 1008 | MYO1F | MYO1F | chr19 | - | 8545649 | 8545736 | 8548035 | 8548122 | 8544296 | 8544464 | 8548236 | 8548317 | 127 | 112 | 51 | 164 | 100 | 31 | 3 | 27 | 5 | 39 | 98 | 98 | 0.118730396 | 1 | 0.436 | 0.528 | 0.622 | 0.375 | 0.409 | 0.137 |
| 556 | FOS | FOS | chr14 | + | 75279876 | 75280128 | 75280559 | 75280667 | 75278777 | 75279123 | 75280782 | 75282234 | 619 | 124 | 110 | 881 | 177 | 91 | 6 | 55 | 12 | 92 | 98 | 98 | 0.121335262 | 1 | 0.413 | 0.412 | 0.547 | 0.333 | 0.374 | 0.104 |
| 637 | JMJD1C | JMJD1C | chr10 | - | 63186214 | 63186383 | 63189167 | 63189446 | 63185562 | 63185653 | 63190893 | 63191108 | 140 | 33 | 16 | 63 | 23 | 6 | 3 | 18 | 2 | 3 | 98 | 98 | 0.122157015 | 1 | 0.69 | 0.589 | 0.727 | 0.6 | 0.857 | -0.06 |
| 768 | PECAM1 | PECAM1 | chr17 | - | 64329699 | 64329722 | 64341633 | 64341690 | 64319415 | 64323845 | 64348259 | 64348322 | 21 | 26 | 28 | 56 | 21 | 12 | 0 | 6 | 1 | 12 | 98 | 72 | 0.123750931 | 1 | 0.216 | 0.476 | 0.632 | 0 | 0.269 | 0.307 |
| 847 | DENND5A | DENND5A | chr11 | - | 9145669 | 9145815 | 9147029 | 9147151 | 9144994 | 9145113 | 9150080 | 9150209 | 26 | 15 | 10 | 43 | 12 | 8 | 4 | 5 | 0 | 5 | 98 | 98 | 0.125778306 | 1 | 0.377 | 0.556 | 0.556 | 1 | 0.5 | -0.254 |
| 869 | VIM | VIM | chr10 | + | 17235845 | 17235889 | 17236293 | 17236379 | 17235168 | 17235389 | 17237229 | 17237593 | 146 | 40 | 21 | 218 | 59 | 45 | 0 | 65 | 7 | 135 | 93 | 98 | 0.129220765 | 1 | 0.414 | 0.417 | 0.33 | 0 | 0.337 | 0.218 |
| 639 | JMJD1C | JMJD1C | chr10 | - | 63219877 | 63219983 | 63264650 | 63264764 | 63217206 | 63217331 | 63380317 | 63380482 | 21 | 6 | 5 | 29 | 10 | 1 | 1 | 2 | 0 | 2 | 98 | 98 | 0.132304353 | 1 | 0.42 | 0.375 | 0.833 | 1 | 0.5 | -0.207 |
| 303 | RNASET2 | RNASET2 | chr6 | - | 166934090 | 166934136 | 166938894 | 166939008 | 166931043 | 166931118 | 166943018 | 166943089 | 162 | 68 | 52 | 119 | 30 | 59 | 1 | 31 | 0 | 12 | 98 | 95 | 0.132843371 | 1 | 0.569 | 0.687 | 0.461 | 1 | 0.715 | -0.285 |
| 1522 | VASP | VASP | chr19 | + | 45517928 | 45518094 | 45521321 | 45521406 | 45517662 | 45517834 | 45522167 | 45522217 | 121 | 49 | 30 | 98 | 38 | 44 | 7 | 22 | 0 | 20 | 98 | 98 | 0.139102038 | 1 | 0.553 | 0.563 | 0.405 | 1 | 0.524 | -0.255 |
| 490 | NCF2 | NCF2 | chr1 | - | 183566919 | 183566988 | 183567203 | 183567345 | 183565703 | 183565779 | 183569141 | 183569185 | 111 | 80 | 66 | 180 | 77 | 88 | 1 | 25 | 12 | 35 | 98 | 98 | 0.143527964 | 1 | 0.381 | 0.51 | 0.429 | 0.077 | 0.417 | 0.193 |
| 553 | SORL1 | SORL1 | chr11 | + | 121545242 | 121545429 | 121549959 | 121550088 | 121543547 | 121543726 | 121550584 | 121550670 | 254 | 80 | 55 | 196 | 49 | 36 | 4 | 41 | 0 | 21 | 98 | 98 | 0.154456522 | 1 | 0.564 | 0.62 | 0.604 | 1 | 0.661 | -0.235 |
| 289 | XPO6 | XPO6 | chr16 | - | 28180940 | 28181031 | 28208986 | 28209122 | 28177219 | 28177332 | 28211365 | 28211918 | 8 | 6 | 1 | 338 | 139 | 60 | 1 | 0 | 3 | 55 | 98 | 98 | 0.159407357 | 1 | 0.023 | 0.041 | 0.016 | 0.25 | 0 | -0.098 |
| 1521 | VASP | VASP | chr19 | + | 45517662 | 45517834 | 45522167 | 45522217 | 45507429 | 45507776 | 45522339 | 45522581 | 125 | 46 | 31 | 50 | 19 | 25 | 6 | 30 | 2 | 9 | 98 | 98 | 0.160587804 | 1 | 0.714 | 0.708 | 0.554 | 0.75 | 0.769 | -0.101 |
| 144 | SOD2 | SOD2 | chr6 | - | 159682473 | 159682638 | 159684853 | 159685033 | 159679115 | 159679298 | 159688125 | 159688242 | 413 | 247 | 217 | 148 | 110 | 100 | 1 | 49 | 4 | 28 | 98 | 98 | 0.165128506 | 1 | 0.736 | 0.692 | 0.685 | 0.2 | 0.636 | 0.286 |
| 148 | GLIPR2 | GLIPR2 | chr9 | + | 36148546 | 36148650 | 36150871 | 36150949 | 36147785 | 36147894 | 36162361 | 36163913 | 20 | 9 | 9 | 52 | 13 | 20 | 2 | 3 | 0 | 5 | 98 | 98 | 0.167785562 | 1 | 0.278 | 0.409 | 0.31 | 1 | 0.375 | -0.355 |
| 1528 | IRF1 | IRF1 | chr5 | - | 132487925 | 132488025 | 132489391 | 132489483 | 132486953 | 132487130 | 132490544 | 132490773 | 43 | 25 | 13 | 75 | 60 | 22 | 0 | 6 | 7 | 11 | 98 | 98 | 0.172303606 | 1 | 0.364 | 0.294 | 0.371 | 0 | 0.353 | 0.166 |
| 718 | TBXAS1 | TBXAS1 | chr7 | + | 139955458 | 139955607 | 139957633 | 139957764 | 139953367 | 139953456 | 139961918 | 139962233 | 27 | 25 | 5 | 38 | 16 | 12 | 1 | 5 | 2 | 12 | 98 | 98 | 0.172540394 | 1 | 0.415 | 0.61 | 0.294 | 0.333 | 0.294 | 0.126 |
| 167 | VNN2 | VNN2 | chr6 | - | 132751144 | 132751518 | 132755842 | 132756035 | 132749694 | 132749865 | 132757415 | 132757546 | 299 | 172 | 111 | 126 | 82 | 68 | 4 | 21 | 0 | 8 | 98 | 98 | 0.174346114 | 1 | 0.704 | 0.677 | 0.62 | 1 | 0.724 | -0.195 |
| 1110 | PTPN6 | PTPN6 | chr12 | + | 6960343 | 6960435 | 6960805 | 6960945 | 6960087 | 6960239 | 6961125 | 6961316 | 84 | 34 | 19 | 105 | 37 | 13 | 2 | 16 | 0 | 11 | 98 | 98 | 0.17819376 | 1 | 0.444 | 0.479 | 0.594 | 1 | 0.593 | -0.291 |
| 657 | ELOVL5 | ELOVL5 | chr6 | - | 53270592 | 53270727 | 53273219 | 53273344 | 53267397 | 53269270 | 53275089 | 53275261 | 79 | 38 | 24 | 52 | 30 | 34 | 1 | 16 | 0 | 8 | 98 | 98 | 0.187130875 | 1 | 0.603 | 0.559 | 0.414 | 1 | 0.667 | -0.308 |
| 551 | SORL1 | SORL1 | chr11 | + | 121478117 | 121478243 | 121488031 | 121488193 | 121470006 | 121470123 | 121490042 | 121490110 | 184 | 118 | 36 | 296 | 153 | 134 | 3 | 32 | 0 | 57 | 98 | 98 | 0.193262821 | 1 | 0.383 | 0.435 | 0.212 | 1 | 0.36 | -0.337 |
| 1131 | KLHL2 | KLHL2 | chr4 | + | 165228806 | 165228913 | 165238777 | 165238899 | 165219933 | 165220059 | 165263196 | 165263359 | 12 | 5 | 9 | 17 | 4 | 9 | 1 | 6 | 0 | 4 | 98 | 98 | 0.193339356 | 1 | 0.414 | 0.556 | 0.5 | 1 | 0.6 | -0.31 |
| 525 | SERPINA1 | SERPINA1 | chr14 | - | 94379463 | 94379611 | 94382591 | 94383241 | 94376746 | 94378640 | 94390456 | 94390692 | 108 | 35 | 38 | 142 | 65 | 104 | 2 | 20 | 6 | 51 | 98 | 98 | 0.194500005 | 1 | 0.432 | 0.35 | 0.268 | 0.25 | 0.282 | 0.084 |
| 1299 | SETX | SETX | chr9 | - | 132336295 | 132336515 | 132342689 | 132342799 | 132334607 | 132334727 | 132346260 | 132346471 | 72 | 17 | 2 | 91 | 20 | 4 | 1 | 13 | 0 | 9 | 98 | 98 | 0.195265172 | 1 | 0.442 | 0.459 | 0.333 | 1 | 0.591 | -0.384 |
| 283 | XPO6 | XPO6 | chr16 | - | 28117317 | 28117462 | 28121669 | 28121762 | 28112903 | 28113050 | 28125688 | 28125848 | 186 | 130 | 42 | 115 | 75 | 26 | 5 | 50 | 0 | 25 | 98 | 98 | 0.203788993 | 1 | 0.618 | 0.634 | 0.618 | 1 | 0.667 | -0.21 |
| 237 | EXOC6 | EXOC6 | chr10 | + | 92934309 | 92934430 | 92935813 | 92935885 | 92934143 | 92934190 | 92940726 | 92940824 | 9 | 4 | 2 | 10 | 3 | 1 | 0 | 2 | 1 | 1 | 98 | 98 | 0.213519627 | 1 | 0.474 | 0.571 | 0.667 | 0 | 0.667 | 0.237 |
| 493 | NCF2 | NCF2 | chr1 | - | 183577598 | 183577707 | 183586894 | 183586977 | 183573184 | 183573292 | 183590155 | 183590604 | 310 | 137 | 123 | 25 | 16 | 0 | 6 | 79 | 0 | 12 | 98 | 98 | 0.214252004 | 1 | 0.925 | 0.895 | 1 | 1 | 0.868 | 0.006 |
| 643 | AMPD2 | AMPD2 | chr1 | + | 109629326 | 109629490 | 109629795 | 109629916 | 109629108 | 109629235 | 109630232 | 109630406 | 19 | 25 | 0 | 31 | 8 | 12 | 2 | 5 | 1 | 4 | 98 | 98 | 0.215680739 | 1 | 0.38 | 0.758 | 0 | 0.667 | 0.556 | -0.232 |
| 474 | PTPRC | PTPRC | chr1 | + | 198741868 | 198742026 | 198742231 | 198742367 | 198735126 | 198735252 | 198744053 | 198744203 | 196 | 110 | 52 | 207 | 86 | 77 | 5 | 34 | 2 | 26 | 98 | 98 | 0.216034204 | 1 | 0.486 | 0.561 | 0.403 | 0.714 | 0.567 | -0.157 |
| 458 | ARAP1 | ARAP1 | chr11 | - | 72697910 | 72698106 | 72699004 | 72699107 | 72697597 | 72697649 | 72699416 | 72699552 | 97 | 76 | 23 | 26 | 29 | 15 | 4 | 19 | 3 | 12 | 98 | 98 | 0.216993421 | 1 | 0.789 | 0.724 | 0.605 | 0.571 | 0.613 | 0.114 |
| 1047 | ARHGAP15 | ARHGAP15 | chr2 | + | 143436912 | 143437042 | 143487372 | 143487495 | 143435600 | 143435699 | 143519265 | 143519364 | 72 | 17 | 18 | 65 | 21 | 4 | 0 | 10 | 2 | 10 | 98 | 98 | 0.223268285 | 1 | 0.526 | 0.447 | 0.818 | 0 | 0.5 | 0.347 |
| 782 | IL4R | IL4R | chr16 | + | 27330065 | 27330198 | 27340185 | 27340273 | 27313908 | 27314020 | 27342120 | 27342259 | 0 | 1 | 4 | 54 | 21 | 12 | 0 | 0 | 2 | 5 | 98 | 98 | 0.225979246 | 1 | 0 | 0.045 | 0.25 | 0 | 0 | 0.098 |
| 473 | PFKFB3 | PFKFB3 | chr10 | + | 6221640 | 6221745 | 6222854 | 6222984 | 6221380 | 6221527 | 6223957 | 6224020 | 56 | 41 | 5 | 69 | 41 | 7 | 11 | 20 | 3 | 22 | 98 | 98 | 0.231324144 | 1 | 0.448 | 0.5 | 0.417 | 0.786 | 0.476 | -0.176 |
| 223 | FAM65B | FAM65B | chr6 | - | 24832255 | 24832391 | 24839090 | 24839272 | 24830508 | 24830670 | 24842861 | 24843554 | 80 | 33 | 16 | 18 | 17 | 13 | 0 | 5 | 1 | 4 | 98 | 98 | 0.243139221 | 1 | 0.816 | 0.66 | 0.552 | 0 | 0.556 | 0.398 |
| 1435 | AOAH | AOAH | chr7 | - | 36637850 | 36637910 | 36659165 | 36659265 | 36632035 | 36632106 | 36673942 | 36674009 | 18 | 4 | 0 | 32 | 13 | 4 | 0 | 3 | 6 | 6 | 98 | 98 | 0.246598151 | 1 | 0.36 | 0.235 | 0 | 0 | 0.333 | 0.032 |
| 257 | MEFV | MEFV | chr16 | - | 3248908 | 3249004 | 3249430 | 3249780 | 3247015 | 3247246 | 3254157 | 3254790 | 46 | 22 | 18 | 48 | 18 | 35 | 0 | 11 | 2 | 18 | 98 | 98 | 0.248207938 | 1 | 0.489 | 0.55 | 0.34 | 0 | 0.379 | 0.27 |
| 1256 | PARP14 | PARP14 | chr3 | + | 122699389 | 122701635 | 122703741 | 122703978 | 122695425 | 122695662 | 122704526 | 122704748 | 46 | 12 | 0 | 34 | 11 | 2 | 7 | 16 | 4 | 9 | 98 | 98 | 0.250710902 | 1 | 0.575 | 0.522 | 0 | 0.636 | 0.64 | -0.272 |
| 724 | SELL | SELL | chr1 | - | 169696534 | 169696553 | 169701559 | 169701688 | 169690664 | 169691802 | 169703254 | 169703440 | 124 | 55 | 47 | 38 | 26 | 7 | 2 | 26 | 1 | 13 | 98 | 68 | 0.257332021 | 1 | 0.694 | 0.595 | 0.823 | 0.581 | 0.581 | 0.123 |
| 143 | IGF2R | IGF2R | chr6 | + | 160027184 | 160027314 | 160029549 | 160029655 | 160024571 | 160024704 | 160032550 | 160032713 | 252 | 84 | 46 | 147 | 63 | 26 | 1 | 40 | 7 | 21 | 98 | 98 | 0.25857467 | 1 | 0.632 | 0.571 | 0.639 | 0.125 | 0.656 | 0.223 |
| 998 | CSNK1D | CSNK1D | chr17 | - | 82253015 | 82253244 | 82255428 | 82255577 | 82252433 | 82252604 | 82265685 | 82265796 | 14 | 8 | 5 | 18 | 7 | 5 | 1 | 2 | 2 | 4 | 98 | 98 | 0.26440277 | 1 | 0.437 | 0.533 | 0.5 | 0.333 | 0.333 | 0.157 |
| 732 | LRRK2 | LRRK2 | chr12 | + | 40303947 | 40304134 | 40305784 | 40305966 | 40302788 | 40302882 | 40308466 | 40308696 | 208 | 58 | 33 | 91 | 86 | 28 | 1 | 29 | 1 | 14 | 98 | 98 | 0.271535111 | 1 | 0.696 | 0.403 | 0.541 | 0.5 | 0.674 | -0.04 |
| 858 | ACSL1 | ACSL1 | chr4 | - | 184770398 | 184770476 | 184773080 | 184773154 | 184768315 | 184768450 | 184773662 | 184773714 | 163 | 95 | 53 | 85 | 32 | 78 | 2 | 38 | 1 | 16 | 98 | 98 | 0.274471942 | 1 | 0.657 | 0.748 | 0.405 | 0.667 | 0.704 | -0.082 |
| 1206 | ARHGEF1 | ARHGEF1 | chr19 | + | 41904948 | 41905036 | 41905174 | 41905261 | 41904215 | 41904383 | 41905759 | 41905827 | 19 | 9 | 0 | 35 | 10 | 14 | 4 | 1 | 0 | 7 | 98 | 98 | 0.276065057 | 1 | 0.352 | 0.474 | 0 | 1 | 0.125 | -0.287 |
| 815 | FGR | FGR | chr1 | - | 27614429 | 27614583 | 27614849 | 27614926 | 27613218 | 27613350 | 27615433 | 27615613 | 67 | 28 | 22 | 87 | 22 | 38 | 1 | 35 | 17 | 27 | 98 | 98 | 0.278358891 | 1 | 0.435 | 0.56 | 0.367 | 0.056 | 0.565 | 0.144 |
| 650 | OSBPL9 | OSBPL9 | chr1 | + | 51651990 | 51652041 | 51669433 | 51669512 | 51616873 | 51617221 | 51714002 | 51714079 | 4 | 7 | 0 | 11 | 8 | 3 | 1 | 1 | 0 | 8 | 98 | 98 | 0.282183756 | 1 | 0.267 | 0.467 | 0 | 1 | 0.111 | -0.311 |
| 813 | MAEA | MAEA | chr4 | + | 1315396 | 1315600 | 1322380 | 1322503 | 1311978 | 1312161 | 1332756 | 1332865 | 15 | 16 | 2 | 0 | 3 | 1 | 3 | 2 | 0 | 0 | 98 | 98 | 0.297185281 | 1 | 1 | 0.842 | 0.667 | 1 | 1 | -0.164 |
| 1527 | IRF1 | IRF1 | chr5 | - | 132485666 | 132485716 | 132486250 | 132486373 | 132484361 | 132484497 | 132486556 | 132486686 | 65 | 18 | 29 | 72 | 36 | 16 | 2 | 12 | 0 | 11 | 98 | 98 | 0.306476344 | 1 | 0.474 | 0.333 | 0.644 | 1 | 0.522 | -0.277 |
| 871 | BEST1 | BEST1 | chr11 | + | 61959497 | 61959578 | 61959891 | 61960043 | 61958145 | 61958298 | 61962254 | 61962893 | 7 | 1 | 5 | 26 | 7 | 17 | 0 | 2 | 1 | 3 | 98 | 98 | 0.306596855 | 1 | 0.212 | 0.125 | 0.227 | 0 | 0.4 | -0.012 |
| 1309 | ARPC2 | ARPC2 | chr2 | + | 218239390 | 218239484 | 218245419 | 218245546 | 218238663 | 218238850 | 218249363 | 218249464 | 144 | 58 | 65 | 163 | 76 | 46 | 0 | 29 | 3 | 31 | 98 | 98 | 0.307941549 | 1 | 0.469 | 0.433 | 0.586 | 0 | 0.483 | 0.255 |
| 754 | ADAM8 | ADAM8 | chr10 | - | 133274158 | 133274235 | 133275483 | 133275587 | 133273950 | 133274029 | 133276771 | 133276903 | 18 | 15 | 1 | 84 | 46 | 27 | 1 | 3 | 8 | 19 | 98 | 98 | 0.314113239 | 1 | 0.176 | 0.246 | 0.036 | 0.111 | 0.136 | 0.029 |
| 829 | AQP9 | AQP9 | chr15 | + | 58173067 | 58173205 | 58174917 | 58175036 | 58166672 | 58166799 | 58179127 | 58179345 | 144 | 111 | 100 | 145 | 85 | 76 | 1 | 40 | 0 | 24 | 98 | 98 | 0.323281869 | 1 | 0.498 | 0.566 | 0.568 | 1 | 0.625 | -0.268 |
| 400 | PTGES3 | PTGES3 | chr12 | - | 56670274 | 56670364 | 56671748 | 56671847 | 56666203 | 56666266 | 56672739 | 56672809 | 13 | 2 | 0 | 17 | 6 | 14 | 0 | 2 | 1 | 2 | 98 | 98 | 0.333454073 | 1 | 0.433 | 0.25 | 0 | 0 | 0.5 | -0.022 |
| 1151 | C17orf62 | C17orf62 | chr17 | - | 82446622 | 82446696 | 82447579 | 82447621 | 82445863 | 82445960 | 82449169 | 82449292 | 28 | 15 | 1 | 47 | 20 | 15 | 2 | 4 | 2 | 5 | 91 | 98 | 0.342520084 | 1 | 0.391 | 0.447 | 0.067 | 0.519 | 0.463 | -0.189 |

| | | | | | | | | | | | | | | | | | | | | | | | | | | | | | | | | |
|---|---|---|---|---|---|---|---|---|---|---|---|---|---|---|---|---|---|---|---|---|---|---|---|---|---|---|---|---|---|---|---|---|
| 1397 | SLC6A6 | SLC6A6 | chr3 | + | 14478465 | 14478568 | 14479084 | 14479185 | 14477204 | 14477342 | 14481670 | 14481841 | 59 | 20 | 15 | 97 | 39 | 18 | 1 | 11 | 2 | 9 | 98 | 98 | 0.346752966 | 1 | 0.378 | 0.339 | 0.455 | 0.333 | 0.55 | -0.051 |
| 557 | PKM | PKM | chr15 | - | 72202453 | 72202620 | 72203021 | 72203188 | 72200473 | 72200655 | 72206727 | 72206880 | 3 | 0 | 0 | 36 | 6 | 7 | 0 | 0 | 3 | 15 | 98 | 98 | 0.350414649 | 1 | 0.077 | 0 | 0 | 0 | 0 | 0.026 |
| 165 | SGK1 | SGK1 | chr6 | - | 134171636 | 134171732 | 134172192 | 134172316 | 134171022 | 134171178 | 134172661 | 134172774 | 82 | 52 | 12 | 90 | 38 | 6 | 0 | 13 | 2 | 11 | 98 | 98 | 0.354258391 | 1 | 0.477 | 0.578 | 0.667 | 0 | 0.542 | 0.303 |
| 1497 | CPEB4 | CPEB4 | chr5 | + | 173932449 | 173932500 | 173944966 | 173945140 | 173910522 | 173910604 | 173949507 | 173949597 | 8 | 3 | 7 | 28 | 12 | 10 | 0 | 5 | 1 | 18 | 98 | 98 | 0.363102749 | 1 | 0.222 | 0.2 | 0.412 | 0 | 0.217 | 0.17 |
| 500 | BCL6 | BCL6 | chr3 | - | 187726730 | 187726898 | 187728359 | 187728544 | 187725498 | 187725629 | 187729049 | 187730021 | 47 | 22 | 12 | 23 | 16 | 7 | 2 | 13 | 1 | 13 | 98 | 98 | 0.363508365 | 1 | 0.671 | 0.579 | 0.632 | 0.667 | 0.5 | 0.044 |
| 779 | RNF10 | RNF10 | chr12 | + | 120566824 | 120566980 | 120571190 | 120571291 | 120565427 | 120565529 | 120575630 | 120575688 | 19 | 19 | 11 | 14 | 16 | 5 | 0 | 6 | 2 | 3 | 98 | 98 | 0.367190598 | 1 | 0.576 | 0.543 | 0.687 | 0 | 0.667 | 0.268 |
| 106 | IFRD1 | IFRD1 | chr7 | + | 112456913 | 112457038 | 112458860 | 112459018 | 112456001 | 112456086 | 112461865 | 112461916 | 216 | 63 | 15 | 61 | 44 | 24 | 6 | 38 | 0 | 28 | 98 | 98 | 0.368297315 | 1 | 0.78 | 0.589 | 0.385 | 1 | 0.576 | -0.203 |
| 57 | DENND3 | DENND3 | chr8 | + | 141138021 | 141138137 | 141150833 | 141150953 | 141136557 | 141136791 | 141151618 | 141151837 | 53 | 54 | 11 | 84 | 39 | 10 | 3 | 15 | 0 | 20 | 98 | 98 | 0.372123371 | 1 | 0.387 | 0.581 | 0.524 | 1 | 0.429 | -0.217 |
| 1178 | ALOX5 | ALOX5 | chr10 | + | 45382482 | 45382681 | 45395854 | 45395936 | 45374175 | 45374429 | 45412190 | 45412313 | 45 | 36 | 5 | 74 | 37 | 11 | 1 | 30 | 0 | 31 | 98 | 98 | 0.380866482 | 1 | 0.378 | 0.493 | 0.312 | 1 | 0.492 | -0.352 |
| 1526 | IRF1 | IRF1 | chr5 | - | 132484361 | 132484497 | 132486250 | 132486373 | 132481608 | 132484075 | 132486556 | 132486686 | 65 | 21 | 29 | 44 | 24 | 13 | 2 | 12 | 0 | 8 | 98 | 98 | 0.381480785 | 1 | 0.596 | 0.467 | 0.69 | 1 | 0.6 | -0.216 |
| 526 | SERPINA1 | SERPINA1 | chr14 | - | 94379463 | 94379611 | 94388800 | 94389010 | 94376746 | 94378640 | 94390456 | 94390692 | 47 | 13 | 5 | 142 | 65 | 104 | 1 | 15 | 6 | 51 | 98 | 98 | 0.381777559 | 1 | 0.249 | 0.167 | 0.046 | 0.143 | 0.227 | -0.031 |
| 101 | MGAM | MGAM | chr7 | + | 142036168 | 142036285 | 142036822 | 142036977 | 142034669 | 142034841 | 142038530 | 142038615 | 110 | 29 | 10 | 97 | 19 | 10 | 2 | 18 | 0 | 14 | 98 | 98 | 0.385226423 | 1 | 0.531 | 0.604 | 0.5 | 1 | 0.562 | -0.236 |
| 491 | NCF2 | NCF2 | chr1 | - | 183573184 | 183573292 | 183574486 | 183574621 | 183570779 | 183570839 | 183577598 | 183577707 | 118 | 56 | 89 | 304 | 147 | 80 | 0 | 30 | 5 | 53 | 98 | 98 | 0.390483373 | 1 | 0.28 | 0.276 | 0.527 | 0 | 0.361 | 0.181 |
| 623 | AKAP13 | AKAP13 | chr15 | + | 85658536 | 85658590 | 85662387 | 85662453 | 85655416 | 85655787 | 85664562 | 85664755 | 80 | 39 | 4 | 3 | 1 | 0 | 2 | 13 | 0 | 0 | 98 | 98 | 0.394292383 | 1 | 0.964 | 0.975 | 1 | 1 | 1 | -0.02 |
| 873 | PLXDC2 | PLXDC2 | chr10 | + | 20046868 | 20047015 | 20068169 | 20068239 | 20001774 | 20001986 | 20143294 | 20143417 | 36 | 9 | 0 | 45 | 27 | 6 | 1 | 6 | 0 | 11 | 98 | 98 | 0.394870873 | 1 | 0.444 | 0.25 | 0 | 1 | 0.353 | -0.445 |
| 1223 | ARHGAP30 | ARHGAP30 | chr1 | - | 161056387 | 161056532 | 161059613 | 161059716 | 161054622 | 161054705 | 161069527 | 161069970 | 40 | 18 | 10 | 54 | 33 | 7 | 1 | 18 | 0 | 18 | 98 | 98 | 0.399292684 | 1 | 0.426 | 0.353 | 0.588 | 1 | 0.5 | -0.294 |
| 102 | MGAM | MGAM | chr7 | + | 142047784 | 142047873 | 142050234 | 142050284 | 142040721 | 142040846 | 142050696 | 142050864 | 85 | 20 | 17 | 124 | 20 | 11 | 2 | 7 | 0 | 9 | 98 | 98 | 0.399781628 | 1 | 0.407 | 0.5 | 0.607 | 1 | 0.437 | -0.214 |
| 1464 | WDR1 | WDR1 | chr4 | - | 10097710 | 10097891 | 10103895 | 10103986 | 10088663 | 10088741 | 10116112 | 10116234 | 36 | 16 | 13 | 37 | 7 | 7 | 4 | 6 | 2 | 3 | 98 | 98 | 0.401096861 | 1 | 0.493 | 0.696 | 0.65 | 0.667 | 0.667 | -0.054 |
| 197 | TAPBP | TAPBP | chr6 | - | 33313216 | 33313477 | 33313693 | 33313864 | 33304988 | 33305387 | 33314004 | 33314387 | 4 | 0 | 0 | 26 | 6 | 4 | 0 | 1 | 5 | 4 | 98 | 98 | 0.40341675 | 1 | 0.133 | 0 | 0 | 0 | 0.2 | -0.056 |
| 1007 | MYO1F | MYO1F | chr19 | - | 8539946 | 8540028 | 8541905 | 8541991 | 8536948 | 8537055 | 8544296 | 8544464 | 66 | 33 | 35 | 115 | 78 | 35 | 6 | 17 | 1 | 42 | 98 | 98 | 0.414774608 | 1 | 0.365 | 0.297 | 0.5 | 0.857 | 0.288 | -0.185 |
| 786 | FRY | FRY | chr13 | + | 32175545 | 32175632 | 32178176 | 32178436 | 32173366 | 32173549 | 32178843 | 32179033 | 49 | 12 | 4 | 51 | 15 | 5 | 2 | 3 | 3 | 5 | 98 | 98 | 0.42175388 | 1 | 0.49 | 0.444 | 0.444 | 0.4 | 0.375 | 0.072 |
| 1226 | RBM23 | RBM23 | chr14 | - | 22908332 | 22908380 | 22909482 | 22909595 | 22906194 | 22906368 | 22911327 | 22911403 | 67 | 35 | 36 | 7 | 10 | 0 | 2 | 3 | 0 | 2 | 98 | 97 | 0.428534074 | 1 | 0.905 | 0.776 | 1 | 1 | 0.598 | 0.095 |
| 1102 | STAT3 | STAT3 | chr17 | - | 42324710 | 42324846 | 42324962 | 42325061 | 42323572 | 42323625 | 42326115 | 42326199 | 27 | 18 | 26 | 49 | 25 | 19 | 0 | 11 | 1 | 17 | 98 | 98 | 0.432753276 | 1 | 0.355 | 0.419 | 0.578 | 0 | 0.393 | 0.254 |
| 1041 | THEMIS2 | THEMIS2 | chr1 | + | 27876587 | 27876728 | 27879643 | 27880054 | 27872542 | 27872665 | 27885294 | 27885451 | 75 | 50 | 16 | 3 | 0 | 1 | 10 | 8 | 0 | 2 | 98 | 98 | 0.443747177 | 1 | 0.962 | 1 | 0.941 | 1 | 0.8 | 0.068 |
| 1420 | MAPKAPK3 | MAPKAPK3 | chr3 | + | 50644408 | 50644532 | 50645709 | 50645785 | 50642252 | 50642332 | 50646139 | 50646264 | 29 | 17 | 9 | 15 | 14 | 4 | 0 | 10 | 1 | 7 | 98 | 98 | 0.445444298 | 1 | 0.659 | 0.548 | 0.692 | 0 | 0.588 | 0.339 |
| 857 | ACSL1 | ACSL1 | chr4 | - | 184770254 | 184770332 | 184770398 | 184770476 | 184768315 | 184768450 | 184773080 | 184773154 | 230 | 105 | 146 | 18 | 3 | 6 | 1 | 55 | 2 | 3 | 98 | 98 | 0.452276841 | 1 | 0.927 | 0.972 | 0.961 | 0.333 | 0.948 | 0.313 |
| 1122 | VMP1 | VMP1 | chr17 | + | 59735337 | 59735473 | 59737452 | 59737543 | 59731420 | 59731522 | 59738836 | 59738947 | 65 | 35 | 48 | 124 | 101 | 62 | 0 | 14 | 2 | 29 | 98 | 98 | 0.455253454 | 1 | 0.344 | 0.257 | 0.436 | 0 | 0.326 | 0.183 |
| 987 | PIK3R5 | PIK3R5 | chr17 | - | 8889972 | 8890126 | 8890737 | 8890912 | 8889138 | 8889222 | 8893585 | 8893655 | 16 | 11 | 14 | 4 | 4 | 1 | 0 | 6 | 1 | 1 | 98 | 98 | 0.460982421 | 1 | 0.8 | 0.733 | 0.933 | 0 | 0.857 | 0.394 |
| 440 | KIAA1033 | KIAA1033 | chr12 | + | 105162748 | 105162845 | 105164110 | 105164307 | 105160000 | 105160148 | 105164640 | 105164740 | 25 | 13 | 5 | 19 | 9 | 7 | 1 | 6 | 1 | 3 | 98 | 98 | 0.46252304 | 1 | 0.568 | 0.591 | 0.417 | 0.5 | 0.667 | -0.058 |
| 489 | NCF2 | NCF2 | chr1 | - | 183565703 | 183565779 | 183566919 | 183566988 | 183564004 | 183564030 | 183567203 | 183567345 | 55 | 44 | 53 | 135 | 63 | 66 | 0 | 14 | 4 | 18 | 98 | 98 | 0.46267558 | 1 | 0.289 | 0.411 | 0.445 | 0 | 0.437 | 0.163 |
| 731 | LRRK2 | LRRK2 | chr12 | + | 40298242 | 40298493 | 40299108 | 40299257 | 40295426 | 40295644 | 40302788 | 40302882 | 192 | 59 | 33 | 390 | 124 | 39 | 0 | 32 | 2 | 64 | 98 | 98 | 0.472031432 | 1 | 0.33 | 0.322 | 0.458 | 0 | 0.333 | 0.204 |
| 1529 | H2AFY | H2AFY | chr5 | - | 135350822 | 135350913 | 135352945 | 135353045 | 135345967 | 135346057 | 135360496 | 135360607 | 16 | 12 | 14 | 5 | 3 | 3 | 2 | 10 | 0 | 3 | 98 | 98 | 0.472653985 | 1 | 0.762 | 0.8 | 0.824 | 1 | 0.769 | -0.089 |
| 898 | GLT1D1 | GLT1D1 | chr12 | + | 128875913 | 128876062 | 128888638 | 128888744 | 128853535 | 128853649 | 128899235 | 128899287 | 31 | 12 | 3 | 51 | 29 | 7 | 2 | 9 | 5 | 8 | 98 | 98 | 0.478814744 | 1 | 0.378 | 0.293 | 0.3 | 0.286 | 0.529 | -0.084 |
| 1440 | CSF2RA | CSF2RA | chrX | + | 1300490 | 1300626 | 1303922 | 1304019 | 1295426 | 1295456 | 1305445 | 1305527 | 15 | 11 | 8 | 8 | 3 | 10 | 0 | 15 | 1 | 10 | 98 | 98 | 0.482127935 | 1 | 0.652 | 0.786 | 0.444 | 0 | 0.6 | 0.327 |
| 1502 | FYB | FYB | chr5 | - | 39153447 | 39153604 | 39201825 | 39202987 | 39141094 | 39141141 | 39219442 | 39219578 | 52 | 17 | 0 | 90 | 43 | 10 | 4 | 8 | 3 | 21 | 98 | 98 | 0.490580949 | 1 | 0.366 | 0.283 | 0 | 0.571 | 0.276 | -0.207 |
| 286 | XPO6 | XPO6 | chr16 | - | 28135215 | 28135324 | 28146093 | 28146203 | 28133840 | 28133933 | 28152658 | 28152785 | 127 | 43 | 20 | 198 | 116 | 46 | 1 | 24 | 4 | 25 | 98 | 98 | 0.502162687 | 1 | 0.391 | 0.27 | 0.303 | 0.2 | 0.49 | -0.024 |
| 859 | ACSL1 | ACSL1 | chr4 | - | 184773842 | 184773875 | 184776483 | 184776662 | 184773662 | 184773714 | 184776883 | 184776983 | 82 | 24 | 41 | 106 | 47 | 33 | 2 | 14 | 1 | 33 | 98 | 82 | 0.513367948 | 1 | 0.393 | 0.299 | 0.51 | 0.626 | 0.262 | -0.043 |
| 904 | ITGA5 | ITGA5 | chr12 | - | 54409484 | 54409597 | 54411833 | 54411964 | 54409231 | 54409352 | 54418980 | 54419266 | 6 | 5 | 2 | 20 | 15 | 8 | 1 | 1 | 0 | 3 | 98 | 98 | 0.514001261 | 1 | 0.231 | 0.25 | 0.2 | 1 | 0.25 | -0.398 |
| 284 | XPO6 | XPO6 | chr16 | - | 28121669 | 28121762 | 28125688 | 28125848 | 28117317 | 28117462 | 28132333 | 28132403 | 129 | 71 | 20 | 153 | 88 | 40 | 4 | 20 | 1 | 30 | 98 | 98 | 0.524444894 | 1 | 0.457 | 0.447 | 0.333 | 0.8 | 0.4 | -0.188 |
| 1380 | CSF2RB | CSF2RB | chr22 | + | 36925986 | 36926177 | 36929401 | 36929559 | 36923243 | 36923367 | 36929638 | 36929807 | 46 | 34 | 37 | 94 | 42 | 44 | 1 | 13 | 6 | 17 | 98 | 98 | 0.526871726 | 1 | 0.329 | 0.447 | 0.457 | 0.143 | 0.433 | 0.123 |
| 302 | PTTG1IP | PTTG1IP | chr21 | - | 44861162 | 44861271 | 44865394 | 44865447 | 44856192 | 44856364 | 44873501 | 44873903 | 10 | 3 | 6 | 27 | 15 | 4 | 1 | 3 | 0 | 7 | 98 | 98 | 0.534318244 | 1 | 0.27 | 0.167 | 0.6 | 1 | 0.3 | -0.304 |
| 1405 | HIPK3 | HIPK3 | chr11 | + | 33348165 | 33348228 | 33348521 | 33348818 | 33347851 | 33348013 | 33349146 | 33349287 | 14 | 4 | 2 | 54 | 19 | 5 | 0 | 6 | 1 | 4 | 98 | 98 | 0.539792793 | 1 | 0.206 | 0.174 | 0.286 | 0 | 0.6 | -0.078 |
| 1292 | CYTIP | CYTIP | chr2 | - | 157430859 | 157430962 | 157434369 | 157434424 | 157430558 | 157430652 | 157434697 | 157434747 | 184 | 88 | 26 | 187 | 85 | 27 | 3 | 30 | 5 | 34 | 98 | 98 | 0.540075756 | 1 | 0.496 | 0.509 | 0.491 | 0.375 | 0.469 | 0.077 |
| 1244 | FAM129A | FAM129A | chr1 | - | 184823637 | 184823742 | 184831846 | 184831962 | 184823166 | 184823329 | 184884632 | 184884800 | 136 | 33 | 28 | 122 | 66 | 18 | 0 | 23 | 2 | 19 | 98 | 98 | 0.547893713 | 1 | 0.527 | 0.333 | 0.609 | 0 | 0.548 | 0.216 |
| 1042 | THEMIS2 | THEMIS2 | chr1 | + | 27879643 | 27880054 | 27881970 | 27882125 | 27876587 | 27876728 | 27882512 | 27883043 | 110 | 23 | 14 | 0 | 5 | 0 | 6 | 24 | 0 | 1 | 98 | 98 | 0.548720612 | 1 | 1 | 0.821 | 1 | 1 | 0.96 | -0.04 |
| 1530 | ARHGAP26 | ARHGAP26 | chr5 | + | 143014079 | 143014116 | 143037261 | 143037261 | 142932046 | 142932125 | 143041815 | 143041890 | 134 | 37 | 12 | 50 | 33 | 4 | 1 | 28 | 0 | 13 | 86 | 98 | 0.550512855 | 1 | 0.753 | 0.561 | 0.774 | 1 | 0.711 | -0.159 |
| 844 | HLA-B | HLA-B | chr6 | - | 31354632 | 31354665 | 31355106 | 31355223 | 31354478 | 31354526 | 31355316 | 31355592 | 1587 | 500 | 504 | 407 | 232 | 231 | 40 | 238 | 12 | 113 | 98 | 82 | 0.551571777 | 1 | 0.765 | 0.643 | 0.646 | 0.736 | 0.638 | -0.002 |
| 727 | LRRK2 | LRRK2 | chr12 | + | 40240482 | 40240617 | 40243549 | 40243681 | 40237968 | 40238103 | 40249825 | 40249945 | 491 | 161 | 45 | 171 | 64 | 23 | 4 | 41 | 1 | 22 | 98 | 98 | 0.572355298 | 1 | 0.742 | 0.716 | 0.662 | 0.8 | 0.651 | -0.019 |
| 519 | MNDA | MNDA | chr1 | + | 158843954 | 158844122 | 158845586 | 158846003 | 158843278 | 158843415 | 158847727 | 158847916 | 358 | 171 | 135 | 428 | 329 | 232 | 4 | 45 | 2 | 68 | 98 | 98 | 0.576278334 | 1 | 0.455 | 0.342 | 0.368 | 0.667 | 0.398 | -0.144 |
| 1021 | HIF1A | HIF1A | chr14 | + | 61738086 | 61738373 | 61740504 | 61740627 | 61736888 | 61737109 | 61740754 | 61741188 | 45 | 18 | 13 | 47 | 27 | 7 | 2 | 6 | 0 | 11 | 98 | 98 | 0.577069499 | 1 | 0.489 | 0.4 | 0.65 | 1 | 0.353 | -0.163 |
| 1048 | ARHGAP15 | ARHGAP15 | chr2 | + | 143487372 | 143487495 | 143519265 | 143519364 | 143436912 | 143437042 | 143556407 | 143556485 | 128 | 34 | 12 | 98 | 12 | 10 | 0 | 9 | 1 | 5 | 98 | 98 | 0.577192634 | 1 | 0.566 | 0.739 | 0.545 | 0 | 0.643 | 0.295 |
| 259 | MEFV | MEFV | chr16 | - | 3249430 | 3249780 | 3254157 | 3254790 | 3248908 | 3249004 | 3256310 | 3256627 | 81 | 26 | 30 | 23 | 10 | 11 | 0 | 16 | 1 | 5 | 98 | 98 | 0.579281899 | 1 | 0.779 | 0.722 | 0.732 | 0 | 0.762 | 0.363 |
| 1347 | PREX1 | PREX1 | chr20 | - | 48676192 | 48676268 | 48679359 | 48679409 | 48666282 | 48666355 | 48679650 | 48679754 | 40 | 25 | 28 | 109 | 83 | 47 | 1 | 12 | 3 | 34 | 98 | 98 | 0.59053247 | 1 | 0.268 | 0.231 | 0.373 | 0.25 | 0.261 | 0.035 |
| 1026 | STAT6 | STAT6 | chr12 | - | 57107604 | 57107743 | 57108162 | 57108299 | 57107230 | 57107314 | 57111128 | 57111413 | 28 | 11 | 3 | 36 | 20 | 11 | 1 | 5 | 1 | 6 | 98 | 98 | 0.590630237 | 1 | 0.437 | 0.355 | 0.214 | 0.5 | 0.455 | -0.142 |
| 948 | PICALM | PICALM | chr11 | - | 86022366 | 86022469 | 86026291 | 86026367 | 86014869 | 86014963 | 86031468 | 86031611 | 144 | 66 | 56 | 188 | 89 | 37 | 1 | 14 | 0 | 16 | 98 | 98 | 0.593652134 | 1 | 0.434 | 0.426 | 0.602 | 1 | 0.467 | -0.246 |
| 552 | SORL1 | SORL1 | chr11 | + | 121543547 | 121543726 | 121545242 | 121545429 | 121532463 | 121532552 | 121549959 | 121550088 | 120 | 67 | 33 | 137 | 71 | 36 | 0 | 18 | 2 | 13 | 98 | 98 | 0.596956742 | 1 | 0.467 | 0.486 | 0.478 | 0 | 0.581 | 0.187 |
| 1179 | SLC44A2 | SLC44A2 | chr19 | + | 10634973 | 10635073 | 10636322 | 10636585 | 10634755 | 10634887 | 10636661 | 10636756 | 29 | 14 | 12 | 53 | 28 | 9 | 1 | 8 | 6 | 9 | 98 | 98 | 0.603913668 | 1 | 0.354 | 0.333 | 0.571 | 0.143 | 0.471 | 0.112 |
| 377 | GIT2 | GIT2 | chr12 | - | 109947255 | 109947504 | 109951211 | 109951316 | 109939164 | 109939247 | 109953091 | 109953234 | 17 | 5 | 5 | 1 | 3 | 1 | 1 | 2 | 0 | 2 | 98 | 98 | 0.607057526 | 1 | 0.944 | 0.625 | 0.833 | 1 | 0.5 | 0.051 |
| 1153 | SMCHD1 | SMCHD1 | chr18 | + | 2722518 | 2722663 | 2724898 | 2724995 | 2718314 | 2718434 | 2726451 | 2726524 | 103 | 62 | 32 | 217 | 108 | 24 | 0 | 26 | 1 | 40 | 98 | 98 | 0.620617635 | 1 | 0.322 | 0.365 | 0.571 | 0 | 0.394 | 0.222 |
| 467 | PTPRJ | PTPRJ | chr11 | + | 48112746 | 48112983 | 48121002 | 48121266 | 48110057 | 48110076 | 48123612 | 48123870 | 63 | 43 | 22 | 43 | 18 | 17 | 11 | 21 | 4 | 13 | 98 | 98 | 0.623407942 | 1 | 0.594 | 0.705 | 0.564 | 0.733 | 0.618 | -0.054 |
| 1205 | SASH3 | SASH3 | chrX | + | 129792327 | 129792476 | 129792626 | 129792836 | 129790939 | 129791081 | 129792988 | 129793139 | 33 | 18 | 3 | 24 | 12 | 2 | 3 | 5 | 0 | 8 | 98 | 98 | 0.625998837 | 1 | 0.579 | 0.6 | 0.6 | 1 | 0.385 | -0.1 |
| 19 | DDX3X | DDX3X | chrX | + | 41343215 | 41343351 | 41343736 | 41343822 | 41342736 | 41342836 | 41344029 | 41344128 | 147 | 42 | 21 | 166 | 36 | 32 | 2 | 15 | 2 | 21 | 98 | 98 | 0.627654747 | 1 | 0.47 | 0.538 | 0.396 | 0.5 | 0.417 | 0.009 |
| 1075 | FLOT2 | FLOT2 | chr17 | - | 28884224 | 28884315 | 28888944 | 28889026 | 28883107 | 28883231 | 28897525 | 28897697 | 26 | 7 | 14 | 33 | 28 | 26 | 1 | 4 | 0 | 10 | 98 | 98 | 0.633526331 | 1 | 0.441 | 0.2 | 0.35 | 1 | 0.286 | -0.313 |
| 1385 | TBC1D22A | TBC1D22A | chr22 | + | 46797443 | 46797620 | 46878652 | 46878723 | 46793500 | 46793841 | 46891265 | 46891394 | 24 | 2 | 2 | 15 | 6 | 0 | 1 | 5 | 3 | 3 | 98 | 98 | 0.641897603 | 1 | 0.615 | 0.25 | 1 | 0.25 | 0.625 | 0.184 |
| 256 | MEFV | MEFV | chr16 | - | 3247015 | 3247246 | 3248908 | 3249004 | 3246524 | 3246547 | 3249430 | 3249780 | 23 | 10 | 12 | 84 | 58 | 39 | 1 | 5 | 1 | 25 | 98 | 98 | 0.643337978 | 1 | 0.215 | 0.147 | 0.235 | 0.5 | 0.167 | -0.135 |
| 1076 | SSH2 | SSH2 | chr17 | - | 29695458 | 29695523 | 29702958 | 29703062 | 29684562 | 29684684 | 29761149 | 29761432 | 41 | 20 | 13 | 48 | 33 | 12 | 1 | 4 | 0 | 7 | 98 | 98 | 0.656275231 | 1 | 0.461 | 0.377 | 0.52 | 1 | 0.364 | -0.229 |
| 604 | GBP5 | GBP5 | chr1 | - | 89267416 | 89267526 | 89268728 | 89268856 | 89266956 | 89267153 | 89269365 | 89269574 | 27 | 6 | 4 | 91 | 27 | 18 | 0 | 4 | 1 | 11 | 98 | 98 | 0.659255602 | 1 | 0.229 | 0.182 | 0.182 | 0 | 0.267 | 0.064 |
| 52 | TLN1 | TLN1 | chr9 | - | 35699355 | 35699461 | 35699973 | 35700081 | 35699031 | 35699156 | 35700190 | 35700376 | 85 | 21 | 17 | 74 | 21 | 10 | 2 | 17 | 0 | 23 | 98 | 98 | 0.659351734 | 1 | 0.535 | 0.5 | 0.63 | 1 | 0.425 | -0.157 |
| 127 | IKZF1 | IKZF1 | chr7 | + | 50387344 | 50387470 | 50391728 | 50391833 | 50382539 | 50382707 | 50399917 | 50405100 | 25 | 3 | 4 | 5 | 0 | 2 | 2 | 7 | 0 | 4 | 98 | 98 | 0.669024756 | 1 | 0.833 | 1 | 0.667 | 1 | 0.636 | 0.015 |
| 831 | DOCK2 | DOCK2 | chr5 | + | 169670541 | 169670597 | 169671077 | 169671174 | 169669287 | 169669328 | 169674296 | 169674445 | 32 | 11 | 5 | 142 | 34 | 5 | 0 | 8 | 2 | 22 | 98 | 98 | 0.672793032 | 1 | 0.184 | 0.244 | 0.5 | 0 | 0.267 | 0.176 |
| 899 | GLT1D1 | GLT1D1 | chr12 | + | 128888638 | 128888744 | 128899235 | 128899287 | 128875913 | 128876062 | 128945325 | 128945369 | 29 | 15 | 11 | 3 | 0 | 0 | 1 | 10 | 0 | 1 | 98 | 98 | 0.676978034 | 1 | 0.906 | 1 | 1 | 1 | 0.909 | 0.014 |
| 1288 | ACOX1 | ACOX1 | chr17 | - | 75960214 | 75960375 | 75973624 | 75973785 | 75957458 | 75957566 | 75978533 | 75978693 | 8 | 3 | 1 | 35 | 8 | 22 | 0 | 3 | 1 | 11 | 98 | 98 | 0.677706498 | 1 | 0.186 | 0.273 | 0.043 | 0 | 0.214 | 0.06 |
| 334 | PCNX | PCNX | chr14 | + | 71011491 | 71011549 | 71012999 | 71013202 | 71009633 | 71009724 | 71019008 | 71019162 | 33 | 15 | 2 | 56 | 11 | 7 | 0 | 6 | 2 | 4 | 98 | 98 | 0.689326839 | 1 | 0.371 | 0.577 | 0.222 | 0 | 0.6 | 0.09 |
| 631 | LAPTM5 | LAPTM5 | chr1 | - | 30741639 | 30741716 | 30742455 | 30742549 | 30739808 | 30739937 | 30757658 | 30757836 | 11 | 7 | 3 | 131 | 34 | 43 | 0 | 4 | 1 | 20 | 98 | 98 | 0.695609396 | 1 | 0.077 | 0.171 | 0.065 | 0 | 0.167 | 0.021 |
| 535 | STX3 | STX3 | chr11 | + | 59787036 | 59787136 | 59788872 | 59788947 | 59773210 | 59773294 | 59790518 | 59790586 | 38 | 20 | 10 | 57 | 26 | 14 | 0 | 6 | 1 | 6 | 98 | 98 | 0.713148432 | 1 | 0.4 | 0.435 | 0.417 | 0 | 0.5 | 0.167 |
| 1201 | GMFG | GMFG | chr19 | - | 39329543 | 39329626 | 39333076 | 39333126 | 39328999 | 39329073 | 39335260 | 39335310 | 99 | 64 | 46 | 185 | 118 | 124 | 1 | 11 | 0 | 34 | 98 | 98 | 0.716396186 | 1 | 0.349 | 0.352 | 0.271 | 1 | 0.244 | -0.298 |
| 1406 | HIPK3 | HIPK3 | chr11 | + | 33348521 | 33348818 | 33349146 | 33349287 | 33347851 | 33348013 | 33351607 | 33351843 | 16 | 2 | 2 | 31 | 16 | 4 | 1 | 1 | 0 | 6 | 98 | 98 | 0.724062162 | 1 | 0.34 | 0.111 | 0.333 | 1 | 0.143 | -0.31 |
| 1148 | RNF213 | RNF213 | chr17 | + | 80360060 | 80360206 | 80361733 | 80361888 | 80358287 | 80358479 | 80363101 | 80363314 | 31 | 19 | 2 | 32 | 41 | 4 | 0 | 19 | 1 | 25 | 98 | 98 | 0.753935402 | 1 | 0.492 | 0.317 | 0.333 | 0 | 0.432 | 0.165 |
| 943 | CD53 | CD53 | chr1 | + | 110896652 | 110896733 | 110897808 | 110897892 | 110894959 | 110895055 | 110899123 | 110899936 | 33 | 29 | 35 | 99 | 49 | 56 | 0 | 12 | 2 | 18 | 98 | 98 | 0.754948443 | 1 | 0.25 | 0.372 | 0.385 | 0 | 0.4 | 0.136 |
| 392 | GTPBP1 | GTPBP1 | chr22 | + | 38727212 | 38727348 | 38727982 | 38728161 | 38726257 | 38726440 | 38729461 | 38729662 | 16 | 7 | 5 | 27 | 12 | 0 | 1 | 6 | 0 | 7 | 98 | 98 | 0.764160975 | 1 | 0.372 | 0.368 | 1 | 1 | 0.462 | -0.151 |
| 721 | APH1A | APH1A | chr1 | - | 150267715 | 150267789 | 150267956 | 150268127 | 150267355 | 150267478 | 150268697 | 150269209 | 9 | 0 | 0 | 11 | 2 | 6 | 0 | 3 | 1 | 1 | 98 | 98 | 0.766857327 | 1 | 0.45 | 0 | 0 | 0 | 0.75 | -0.225 |
| 328 | CTSS | CTSS | chr1 | - | 150750005 | 150750171 | 150751780 | 150752008 | 150747776 | 150747879 | 150755000 | 150755150 | 261 | 144 | 134 | 104 | 71 | 67 | 2 | 110 | 0 | 58 | 98 | 98 | 0.767195433 | 1 | 0.715 | 0.67 | 0.667 | 1 | 0.655 | -0.143 |
| 669 | NBEAL2 | NBEAL2 | chr3 | + | 47003173 | 47003309 | 47003815 | 47003976 | 47002956 | 47003081 | 47004076 | 47004393 | 47 | 20 | 6 | 13 | 4 | 4 | 3 | 16 | 0 | 11 | 98 | 98 | 0.768824062 | 1 | 0.783 | 0.833 | 0.6 | 1 | 0.593 | -0.058 |
| 735 | LRRK2 | LRRK2 | chr12 | + | 40351538 | 40351733 | 40354298 | 40354492 | 40348408 | 40348509 | 40356114 | 40356187 | 164 | 62 | 21 | 126 | 40 | 21 | 3 | 15 | 2 | 14 | 98 | 98 | 0.771638748 | 1 | 0.566 | 0.608 | 0.5 | 0.6 | 0.517 | -0.001 |
| 1189 | MYO9B | MYO9B | chr19 | + | 17156908 | 17157038 | 17159394 | 17159484 | 17154314 | 17154415 | 17162349 | 17162466 | 57 | 25 | 4 | 57 | 54 | 5 | 6 | 9 | 4 | 15 | 98 | 98 | 0.775648212 | 1 | 0.5 | 0.316 | 0.444 | 0.6 | 0.375 | -0.068 |
| 932 | DHX9 | DHX9 | chr1 | + | 182855576 | 182855713 | 182856531 | 182856578 | 182854029 | 182854178 | 182858103 | 182858240 | 9 | 4 | 0 | 28 | 10 | 3 | 2 | 0 | 0 | 7 | 98 | 96 | 0.777970856 | 1 | 0.239 | 0.282 | 0 | 1 | 0 | -0.326 |
| 1040 | THEMIS2 | THEMIS2 | chr1 | + | 27876587 | 27876728 | 27879643 | 27880054 | 27872542 | 27872665 | 27881970 | 27882125 | 72 | 53 | 10 | 47 | 9 | 18 | 7 | 8 | 1 | 19 | 98 | 98 | 0.786188136 | 1 | 0.605 | 0.855 | 0.357 | 0.875 | 0.296 | 0.02 |
| 733 | LRRK2 | LRRK2 | chr12 | + | 40305784 | 40305966 | 40308466 | 40308696 | 40303947 | 40304134 | 40309105 | 40309233 | 213 | 78 | 30 | 288 | 118 | 21 | 1 | 18 | 0 | 34 | 98 | 98 | 0.79075928 | 1 | 0.425 | 0.398 | 0.588 | 1 | 0.346 | -0.203 |
| 368 | RRP12 | RRP12 | chr10 | - | 97390422 | 97390539 | 97390738 | 97390844 | 97388488 | 97388624 | 97393683 | 97393760 | 59 | 13 | 2 | 27 | 7 | 4 | 2 | 7 | 1 | 4 | 98 | 98 | 0.791114293 | 1 | 0.686 | 0.65 | 0.333 | 0.667 | 0.636 | -0.095 |
| 830 | AQP9 | AQP9 | chr15 | + | 58174917 | 58175036 | 58179127 | 58179345 | 58173067 | 58173205 | 58183960 | 58185911 | 150 | 68 | 71 | 86 | 57 | 58 | 1 | 19 | 1 | 12 | 98 | 98 | 0.802867631 | 1 | 0.636 | 0.544 | 0.55 | 0.5 | 0.613 | 0.02 |
| 198 | FLOT1 | FLOT1 | chr6 | - | 30741189 | 30741333 | 30741613 | 30741704 | 30740678 | 30740798 | 30741791 | 30741867 | 24 | 14 | 13 | 39 | 14 | 17 | 1 | 4 | 0 | 11 | 98 | 98 | 0.810040084 | 1 | 0.381 | 0.5 | 0.433 | 1 | 0.267 | -0.195 |
| 145 | SOD2 | SOD2 | chr6 | - | 159684853 | 159685033 | 159688125 | 159688242 | 159682473 | 159682638 | 159692660 | 159692863 | 370 | 266 | 195 | 468 | 299 | 215 | 6 | 33 | 2 | 58 | 98 | 98 | 0.8101779 | 1 | 0.442 | 0.471 | 0.476 | 0.75 | 0.363 | -0.093 |
| 787 | ATP2A3 | ATP2A3 | chr17 | - | 3951580 | 3951685 | 3953346 | 3953429 | 3951250 | 3951389 | 3953692 | 3953710 | 18 | 10 | 8 | 27 | 24 | 13 | 1 | 3 | 8 | 1 | 98 | 98 | 0.810828389 | 1 | 0.4 | 0.294 | 0.381 | 0.111 | 0.75 | -0.072 |
| 457 | ARAP1 | ARAP1 | chr11 | - | 72686041 | 72686191 | 72687438 | 72687502 | 72685068 | 72685681 | 72687687 | 72687738 | 29 | 18 | 25 | 65 | 28 | 16 | 1 | 5 | 0 | 13 | 98 | 98 | 0.813010615 | 1 | 0.309 | 0.391 | 0.61 | 1 | 0.278 | -0.202 |
| 1346 | PREX1 | PREX1 | chr20 | - | 48649995 | 48650206 | 48650893 | 48651055 | 48649299 | 48649576 | 48651395 | 48651583 | 31 | 22 | 13 | 84 | 37 | 35 | 1 | 11 | 6 | 21 | 98 | 98 | 0.814486416 | 1 | 0.27 | 0.373 | 0.271 | 0.143 | 0.344 | 0.061 |
| 1185 | ADGRE5 | ADGRE5 | chr19 | + | 14407060 | 14407229 | 14407907 | 14408009 | 14406868 | 14406960 | 14408091 | 14408725 | 45 | 51 | 44 | 59 | 32 | 34 | 1 | 15 | 0 | 19 | 98 | 98 | 0.816695615 | 1 | 0.433 | 0.614 | 0.564 | 1 | 0.441 | -0.184 |
| 374 | NLRP1 | NLRP1 | chr17 | - | 5532821 | 5532984 | 5533896 | 5533988 | 5530480 | 5530704 | 5536850 | 5536940 | 55 | 19 | 11 | 47 | 39 | 5 | 2 | 20 | 4 | 20 | 98 | 98 | 0.823687728 | 1 | 0.539 | 0.328 | 0.687 | 0.333 | 0.5 | 0.102 |
| 730 | LRRK2 | LRRK2 | chr12 | + | 40295426 | 40295644 | 40298242 | 40298493 | 40294844 | 40294914 | 40299108 | 40299257 | 319 | 82 | 30 | 215 | 48 | 19 | 8 | 42 | 3 | 29 | 98 | 98 | 0.825500335 | 1 | 0.597 | 0.631 | 0.612 | 0.727 | 0.592 | -0.046 |
| 728 | LRRK2 | LRRK2 | chr12 | + | 40257247 | 40257377 | 40259479 | 40259604 | 40252909 | 40253016 | 40263788 | 40263901 | 175 | 69 | 34 | 372 | 93 | 25 | 2 | 19 | 2 | 27 | 98 | 98 | 0.828099365 | 1 | 0.32 | 0.426 | 0.576 | 0.5 | 0.413 | -0.016 |
| 331 | CTSS | CTSS | chr1 | - | 150755000 | 150755150 | 150757857 | 150757980 | 150751780 | 150752008 | 150764637 | 150764764 | 410 | 160 | 108 | 262 | 144 | 131 | 1 | 123 | 2 | 109 | 98 | 98 | 0.829639519 | 1 | 0.61 | 0.526 | 0.452 | 0.333 | 0.53 | 0.098 |
| 236 | EXOC6 | EXOC6 | chr10 | + | 92934143 | 92934190 | 92934309 | 92934430 | 92928338 | 92928422 | 92935813 | 92935885 | 14 | 9 | 2 | 11 | 1 | 3 | 1 | 3 | 0 | 7 | 96 | 98 | 0.832162999 | 1 | 0.565 | 0.902 | 0.405 | 1 | 0.304 | -0.028 |
| 343 | SOS2 | SOS2 | chr14 | - | 50174453 | 50174552 | 50180571 | 50180682 | 50161481 | 50161609 | 50182462 | 50182606 | 70 | 30 | 7 | 37 | 13 | 5 | 3 | 9 | 1 | 7 | 98 | 98 | 0.834511645 | 1 | 0.654 | 0.698 | 0.583 | 0.75 | 0.562 | -0.011 |
| 449 | BAZ1A | BAZ1A | chr14 | - | 34785776 | 34786001 | 34786125 | 34786221 | 34783761 | 34783927 | 34792774 | 34792921 | 13 | 6 | 4 | 38 | 16 | 6 | 0 | 4 | 1 | 6 | 98 | 98 | 0.836937733 | 1 | 0.255 | 0.273 | 0.4 | 0 | 0.4 | 0.109 |
| 1396 | SLC6A6 | SLC6A6 | chr3 | + | 14416411 | 14416453 | 14443623 | 14443863 | 14402575 | 14402847 | 14445716 | 14445851 | 95 | 41 | 9 | 75 | 39 | 28 | 3 | 14 | 5 | 13 | 98 | 91 | 0.85311396 | 1 | 0.577 | 0.531 | 0.257 | 0.393 | 0.537 | -0.01 |
| 240 | CDC42SE1 | CDC42SE1 | chr1 | - | 151055015 | 151055126 | 151055676 | 151055993 | 151054230 | 151054321 | 151059478 | 151059649 | 49 | 16 | 7 | 128 | 59 | 33 | 1 | 4 | 1 | 12 | 98 | 98 | 0.858739689 | 1 | 0.277 | 0.213 | 0.175 | 0.5 | 0.25 | -0.153 |
| 942 | CD53 | CD53 | chr1 | + | 110892344 | 110892533 | 110897808 | 110897892 | 110891391 | 110891471 | 110899123 | 110899936 | 141 | 54 | 53 | 98 | 50 | 56 | 3 | 18 | 2 | 18 | 98 | 98 | 0.865568364 | 1 | 0.59 | 0.519 | 0.486 | 0.6 | 0.5 | -0.018 |
| 437 | TNFRSF1A | TNFRSF1A | chr12 | - | 6333736 | 6333865 | 6334090 | 6334244 | 6333366 | 6333516 | 6341775 | 6342117 | 62 | 26 | 20 | 64 | 36 | 11 | 2 | 11 | 3 | 11 | 98 | 98 | 0.873353503 | 1 | 0.492 | 0.419 | 0.645 | 0.4 | 0.5 | 0.069 |
| 498 | RASGRP4 | RASGRP4 | chr19 | - | 38417051 | 38417168 | 38419859 | 38420013 | 38414847 | 38415123 | 38420130 | 38420262 | 20 | 18 | 4 | 39 | 11 | 9 | 4 | 7 | 2 | 13 | 98 | 98 | 0.882368191 | 1 | 0.339 | 0.621 | 0.308 | 0.667 | 0.35 | -0.086 |
| 1117 | SPAG9 | SPAG9 | chr17 | - | 50984922 | 50984990 | 50985697 | 50985778 | 50982523 | 50982676 | 50987111 | 50987237 | 21 | 10 | 4 | 27 | 19 | 5 | 0 | 7 | 1 | 3 | 98 | 98 | 0.895684561 | 1 | 0.438 | 0.345 | 0.444 | 0 | 0.7 | 0.059 |
| 1199 | DDX60L | DDX60L | chr4 | - | 168441336 | 168441492 | 168448637 | 168448779 | 168433009 | 168433115 | 168453123 | 168453282 | 30 | 22 | 3 | 44 | 27 | 0 | 1 | 22 | 2 | 14 | 98 | 98 | 0.895713753 | 1 | 0.405 | 0.449 | 1 | 0.333 | 0.611 | 0.146 |
| 1006 | MYO1F | MYO1F | chr19 | - | 8530458 | 8530573 | 8536251 | 8536396 | 8530195 | 8530365 | 8536498 | 8536597 | 221 | 148 | 54 | 56 | 48 | 22 | 6 | 53 | 3 | 16 | 98 | 98 | 0.899193342 | 1 | 0.798 | 0.755 | 0.711 | 0.667 | 0.768 | 0.037 |
| 58 | DENND3 | DENND3 | chr8 | + | 141138021 | 141138137 | 141151618 | 141151837 | 141136557 | 141136791 | 141155848 | 141155970 | 54 | 53 | 11 | 74 | 50 | 17 | 3 | 14 | 1 | 30 | 98 | 98 | 0.901910799 | 1 | 0.422 | 0.515 | 0.393 | 0.75 | 0.318 | -0.091 |
| 177 | CD164 | CD164 | chr6 | - | 109370410 | 109370467 | 109376073 | 109376112 | 109366513 | 109369017 | 109377899 | 109377971 | 34 | 17 | 3 | 20 | 10 | 0 | 1 | 6 | 0 | 5 | 88 | 98 | 0.903877373 | 1 | 0.654 | 0.654 | 1 | 1 | 0.572 | -0.017 |
| 468 | PTPRJ | PTPRJ | chr11 | + | 48144999 | 48145124 | 48146875 | 48146963 | 48144674 | 48144885 | 48149446 | 48149488 | 35 | 12 | 5 | 26 | 4 | 1 | 8 | 8 | 3 | 5 | 98 | 98 | 0.904773974 | 1 | 0.574 | 0.75 | 0.833 | 0.727 | 0.615 | 0.048 |
| 699 | RFWD2 | RFWD2 | chr1 | - | 176136487 | 176136547 | 176149005 | 176149074 | 176135009 | 176135086 | 176184632 | 176184692 | 29 | 7 | 0 | 89 | 32 | 10 | 0 | 4 | 1 | 10 | 98 | 98 | 0.907396354 | 1 | 0.246 | 0.179 | 0 | 0 | 0.286 | -0.001 |
| 1182 | GLUL | GLUL | chr1 | - | 182385356 | 182385556 | 182385759 | 182385887 | 182378092 | 182384723 | 182386255 | 182386402 | 662 | 192 | 81 | 1299 | 443 | 148 | 10 | 77 | 26 | 147 | 98 | 98 | 0.913655916 | 1 | 0.338 | 0.302 | 0.354 | 0.278 | 0.344 | 0.02 |
| 734 | LRRK2 | LRRK2 | chr12 | + | 40348408 | 40348509 | 40351538 | 40351733 | 40346752 | 40346923 | 40354298 | 40354492 | 144 | 58 | 52 | 83 | 38 | 10 | 1 | 17 | 0 | 12 | 98 | 98 | 0.915533828 | 1 | 0.634 | 0.604 | 0.839 | 1 | 0.586 | -0.101 |
| 1118 | TRIM25 | TRIM25 | chr17 | - | 56895520 | 56895604 | 56895925 | 56895952 | 56895342 | 56895441 | 56899114 | 56899180 | 21 | 14 | 11 | 43 | 9 | 11 | 1 | 4 | 0 | 14 | 98 | 76 | 0.930125935 | 1 | 0.386 | 0.667 | 0.563 | 1 | 0.269 | -0.096 |
| 1209 | CHD2 | CHD2 | chr15 | + | 92901166 | 92901299 | 92924320 | 92924552 | 92900320 | 92900824 | 92927243 | 92927330 | 30 | 20 | 8 | 45 | 15 | 9 | 0 | 9 | 1 | 4 | 98 | 98 | 0.932025488 | 1 | 0.4 | 0.571 | 0.471 | 0 | 0.692 | 0.135 |
| 285 | XPO6 | XPO6 | chr16 | - | 28133840 | 28133933 | 28135215 | 28135324 | 28132333 | 28132403 | 28146093 | 28146203 | 125 | 73 | 21 | 134 | 92 | 25 | 1 | 25 | 2 | 26 | 98 | 98 | 0.937455275 | 1 | 0.483 | 0.442 | 0.457 | 0.333 | 0.49 | 0.049 |
| 1033 | CR1 | CR1 | chr1 | + | 207622992 | 207623068 | 207630516 | 207630621 | 207621972 | 207621996 | 207639396 | 207641765 | 16 | 5 | 8 | 41 | 25 | 5 | 0 | 6 | 1 | 4 | 98 | 98 | 0.939008769 | 1 | 0.281 | 0.167 | 0.615 | 0 | 0.6 | 0.054 |
| 116 | PTPN12 | PTPN12 | chr7 | + | 77583554 | 77583650 | 77585542 | 77585581 | 77581426 | 77581503 | 77592184 | 77592256 | 45 | 22 | 13 | 20 | 16 | 11 | 0 | 9 | 1 | 1 | 98 | 88 | 0.945708507 | 1 | 0.669 | 0.553 | 0.515 | 0 | 0.89 | 0.134 |
| 170 | VNN3 | VNN3 | chr6 | - | 132726721 | 132727012 | 132728740 | 132728933 | 132724605 | 132724979 | 132734252 | 132734383 | 51 | 21 | 4 | 39 | 26 | 1 | 1 | 5 | 0 | 9 | 98 | 98 | 0.94976096 | 1 | 0.567 | 0.447 | 0.8 | 1 | 0.357 | -0.074 |
| 21 | DDX3X | DDX3X | chrX | + | 41346858 | 41347012 | 41347314 | 41347451 | 41346504 | 41346622 | 41347639 | 41350287 | 108 | 47 | 22 | 102 | 29 | 29 | 0 | 27 | 1 | 11 | 98 | 98 | 0.95277589 | 1 | 0.514 | 0.618 | 0.431 | 0 | 0.711 | 0.166 |
| 889 | BNIP2 | BNIP2 | chr15 | - | 59668891 | 59668990 | 59669275 | 59669362 | 59662862 | 59664120 | 59671182 | 59671314 | 27 | 15 | 3 | 44 | 31 | 10 | 0 | 8 | 1 | 7 | 98 | 98 | 0.963645053 | 1 | 0.38 | 0.326 | 0.231 | 0 | 0.533 | 0.046 |
| 1231 | MBOAT7 | MBOAT7 | chr19 | - | 54187160 | 54187287 | 54188216 | 54188346 | 54183520 | 54183680 | 54188432 | 54188511 | 15 | 16 | 8 | 62 | 24 | 33 | 1 | 6 | 8 | 9 | 98 | 98 | 0.967774437 | 1 | 0.195 | 0.4 | 0.195 | 0.111 | 0.4 | 0.008 |
| 1257 | MEF2A | MEF2A | chr15 | + | 99671322 | 99671454 | 99671519 | 99671657 | 99645560 | 99645764 | 99674392 | 99674612 | 20 | 3 | 2 | 6 | 0 | 0 | 1 | 5 | 0 | 3 | 98 | 98 | 0.969117258 | 1 | 0.769 | 1 | 1 | 1 | 0.625 | 0.111 |
| 20 | DDX3X | DDX3X | chrX | + | 41346858 | 41347012 | 41347311 | 41347451 | 41346504 | 41346622 | 41347639 | 41347915 | 108 | 47 | 22 | 108 | 30 | 29 | 0 | 27 | 1 | 11 | 98 | 98 | 0.96964034 | 1 | 0.5 | 0.61 | 0.431 | 0 | 0.711 | 0.158 |
| 171 | VNN3 | VNN3 | chr6 | - | 132726721 | 132727012 | 132728740 | 132728979 | 132724605 | 132724979 | 132734252 | 132734383 | 44 | 14 | 6 | 39 | 26 | 1 | 1 | 2 | 0 | 9 | 98 | 98 | 0.975627308 | 1 | 0.53 | 0.35 | 0.857 | 1 | 0.182 | -0.012 |
| 492 | NCF2 | NCF2 | chr1 | - | 183573184 | 183573292 | 183577598 | 183577707 | 183570779 | 183570839 | 183586894 | 183586977 | 161 | 90 | 98 | 280 | 132 | 80 | 3 | 31 | 5 | 41 | 98 | 98 | 0.981091398 | 1 | 0.365 | 0.405 | 0.551 | 0.375 | 0.431 | 0.037 |
| 261 | NLRP12 | NLRP12 | chr19 | - | 53807494 | 53807665 | 53809586 | 53811288 | 53805279 | 53805450 | 53814907 | 53814988 | 25 | 10 | 4 | 16 | 8 | 11 | 1 | 8 | 1 | 8 | 98 | 98 | 0.993526134 | 1 | 0.61 | 0.556 | 0.267 | 0.5 | 0.5 | -0.022 |
| 1383 | ARFGAP3 | ARFGAP3 | chr22 | - | 42834241 | 42834325 | 42835361 | 42835493 | 42831548 | 42831636 | 42840943 | 42841016 | 8 | 9 | 6 | 11 | 7 | 5 | 1 | 0 | 0 | 1 | 98 | 98 | 0.995820004 | 1 | 0.421 | 0.562 | 0.545 | 1 | 0 | 0.009 |
| 2 | VAMP7 | VAMP7 | chrX | + | 155900496 | 155900587 | 155919812 | 155919880 | 155898111 | 155898249 | 155939700 | 155939793 | 9 | 2 | 0 | 1 | 3 | 0 | 0 | 2 | 0 | 0 | 98 | 98 | 1 | 1 | 0.9 | 0.4 | NA | NA | 1 | -0.35 |
| 6 | MORF4L2 | MORF4L2 | chrX | - | 103678498 | 103678600 | 103684680 | 103684729 | 103675497 | 103677051 | 103685170 | 103685260 | 1 | 2 | 0 | 10 | 2 | 3 | 0 | 0 | 0 | 5 | 98 | 98 | 1 | 1 | 0.091 | 0.5 | 0 | NA | 0 | 0.197 |
| 7 | MORF4L2 | MORF4L2 | chrX | - | 103678498 | 103678620 | 103684680 | 103684729 | 103675497 | 103677051 | 103685170 | 103685260 | 1 | 2 | 0 | 10 | 2 | 3 | 0 | 0 | 0 | 5 | 98 | 98 | 1 | 1 | 0.091 | 0.5 | 0 | NA | 0 | 0.197 |
| 8 | MORF4L2 | MORF4L2 | chrX | - | 103678498 | 103678651 | 103684680 | 103684729 | 103675497 | 103677051 | 103685170 | 103685260 | 1 | 2 | 0 | 12 | 2 | 3 | 0 | 0 | 0 | 5 | 98 | 98 | 1 | 1 | 0.077 | 0.5 | 0 | NA | 0 | 0.192 |
| 22 | USP9X | USP9X | chrX | + | 41136803 | 41137022 | 41140655 | 41140771 | 41134724 | 41134837 | 41140965 | 41141217 | 19 | 2 | 4 | 6 | 9 | 1 | 0 | 4 | 0 | 3 | 98 | 98 | 1 | 1 | 0.76 | 0.182 | 0.8 | NA | 0.571 | 0.01 |
| 23 | GK | GK | chrX | + | 30691122 | 30691199 | 30694399 | 30694537 | 30677374 | 30677452 | 30696041 | 30696151 | 9 | 2 | 0 | 13 | 4 | 10 | 0 | 0 | 0 | 1 | 98 | 98 | 1 | 1 | 0.409 | 0.333 | 0 | NA | 0 | 0.247 |
| 25 | WWC3 | WWC3 | chrX | + | 10125023 | 10125110 | 10126113 | 10126306 | 10124264 | 10124339 | 10127987 | 10128161 | 16 | 3 | 0 | 9 | 1 | 6 | 0 | 2 | 0 | 2 | 98 | 98 | 1 | 1 | 0.64 | 0.75 | 0 | NA | 0.5 | -0.037 |
| 26 | WWC3 | WWC3 | chrX | + | 10126113 | 10126306 | 10127987 | 10128161 | 10125023 | 10125110 | 10128556 | 10128708 | 11 | 5 | 2 | 15 | 8 | 10 | 0 | 1 | 0 | 0 | 98 | 98 | 1 | 1 | 0.423 | 0.385 | 0.167 | NA | 1 | -0.675 |
| 43 | TLR4 | TLR4 | chr9 | + | 117708198 | 117708318 | 117708562 | 117708729 | 117704174 | 117704565 | 117712388 | 117717491 | 80 | 13 | 10 | 72 | 13 | 36 | 0 | 14 | 0 | 10 | 98 | 98 | 1 | 1 | 0.526 | 0.5 | 0.217 | NA | 0.583 | -0.169 |
| 44 | DAPK1 | DAPK1 | chr9 | + | 87686550 | 87686739 | 87697006 | 87697204 | 87681403 | 87681626 | 87698655 | 87698794 | 4 | 4 | 5 | 5 | 5 | 0 | 0 | 4 | 0 | 0 | 98 | 98 | 1 | 1 | 0.444 | 0.444 | 1 | NA | 1 | -0.371 |
| 55 | UHRF2 | UHRF2 | chr9 | + | 6420911 | 6421142 | 6433913 | 6434173 | 6413150 | 6413643 | 6460572 | 6460791 | 4 | 3 | 0 | 3 | 6 | 1 | 0 | 2 | 0 | 0 | 98 | 98 | 1 | 1 | 0.571 | 0.333 | 0 | NA | 1 | -0.699 |
| 56 | DENND3 | DENND3 | chr8 | + | 141136557 | 141136791 | 141150833 | 141150953 | 141128620 | 141128809 | 141151618 | 141151837 | 4 | 2 | 0 | 83 | 42 | 10 | 0 | 1 | 0 | 20 | 98 | 98 | 1 | 1 | 0.046 | 0.045 | 0 | NA | 0.048 | -0.018 |
| 59 | DENND3 | DENND3 | chr8 | + | 141160631 | 141160787 | 141163332 | 141163429 | 141155848 | 141155970 | 141165185 | 141165289 | 52 | 20 | 10 | 60 | 17 | 13 | 0 | 11 | 0 | 8 | 98 | 98 | 1 | 1 | 0.464 | 0.541 | 0.435 | NA | 0.579 | -0.099 |
| 64 | FAM49B | FAM49B | chr8 | - | 129879388 | 129879471 | 129903311 | 129903350 | 129871374 | 129871496 | 129970942 | 129970995 | 33 | 27 | 14 | 92 | 55 | 26 | 0 | 4 | 0 | 11 | 88 | 98 | 1 | 1 | 0.285 | 0.353 | 0.375 | NA | 0.288 | 0.05 |
| 82 | RAB2A | RAB2A | chr8 | + | 60558851 | 60558923 | 60572045 | 60572113 | 60516909 | 60517253 | 60584207 | 60584290 | 14 | 7 | 0 | 9 | 6 | 4 | 0 | 1 | 0 | 0 | 98 | 98 | 1 | 1 | 0.609 | 0.538 | 0 | NA | 1 | -0.618 |
| 84 | IKBKB | IKBKB | chr8 | + | 42272082 | 42272205 | 42288633 | 42288728 | 42271301 | 42271469 | 42290155 | 42290273 | 0 | 1 | 0 | 1 | 0 | 0 | 0 | 1 | 0 | 2 | 98 | 98 | 1 | 1 | 0 | 1 | NA | NA | 0.333 | 0.167 |
| 85 | ADAM9 | ADAM9 | chr8 | + | 39082967 | 39083073 | 39090046 | 39090188 | 39082640 | 39082721 | 39091258 | 39091346 | 2 | 3 | 0 | 11 | 0 | 0 | 0 | 1 | 1 | 3 | 98 | 98 | 1 | 1 | 0.154 | 1 | NA | 0 | 0.25 | 0.452 |
| 93 | ASAH1 | ASAH1 | chr8 | - | 18062278 | 18062423 | 18063184 | 18063230 | 18061685 | 18061740 | 18064456 | 18064531 | 40 | 11 | 20 | 39 | 29 | 15 | 0 | 4 | 0 | 1 | 95 | 98 | 1 | 1 | 0.514 | 0.281 | 0.579 | NA | 0.805 | -0.347 |
| 104 | LINC-PINT | LINC-PINT | chr7 | - | 130959868 | 130959970 | 130967388 | 130967517 | 130944159 | 130945835 | 130984052 | 130984109 | 36 | 7 | 0 | 21 | 1 | 0 | 0 | 5 | 0 | 1 | 98 | 98 | 1 | 1 | 0.632 | 0.875 | NA | NA | 0.833 | -0.079 |
| 107 | NAMPT | NAMPT | chr7 | - | 106254363 | 106254504 | 106263391 | 106263617 | 106253016 | 106253151 | 106268463 | 106268600 | 0 | 1 | 0 | 1558 | 790 | 321 | 0 | 0 | 24 | 229 | 98 | 98 | 1 | 1 | 0 | 0.001 | 0 | 0 | 0 | 0 |
| 108 | NAMPT | NAMPT | chr7 | - | 106263391 | 106263617 | 106268463 | 106268600 | 106261587 | 106261707 | 106269153 | 106269312 | 680 | 331 | 111 | 1 | 1 | 1 | 9 | 124 | 0 | 0 | 98 | 98 | 1 | 1 | 0.999 | 0.997 | 0.991 | 1 | 1 | -0.004 |
| 115 | DMTF1 | DMTF1 | chr7 | + | 87164933 | 87165050 | 87166482 | 87166605 | 87163494 | 87163617 | 87170994 | 87171089 | 9 | 7 | 4 | 15 | 11 | 0 | 0 | 1 | 0 | 0 | 98 | 98 | 1 | 1 | 0.375 | 0.389 | 1 | NA | 1 | -0.412 |
| 122 | IKZF1 | IKZF1 | chr7 | + | 50376532 | 50376793 | 50391728 | 50391833 | 50327637 | 50327757 | 50399917 | 50405100 | 3 | 1 | 5 | 2 | 0 | 2 | 0 | 0 | 0 | 3 | 98 | 98 | 1 | 1 | 0.6 | 1 | 0.714 | NA | 0 | 0.771 |

| | | | | | | | | | | | | | | | | | | | | | | | | | | | | | | | | | |
|---|---|---|---|---|---|---|---|---|---|---|---|---|---|---|---|---|---|---|---|---|---|---|---|---|---|---|---|---|---|---|---|---|---|
| 124 | IKZF1 | IKZF1 | chr7 | + | 50382539 | 50382707 | 50387344 | 50387470 | 50327637 | 50327757 | 50391728 | 50391833 | 15 | 1 | 3 | 5 | 2 | 5 | 0 | 3 | 0 | 3 | 98 | 98 | 1 | 1 | 0.75 | 0.333 | 0.375 | NA | | 0.5 | -0.014 |
| 125 | IKZF1 | IKZF1 | chr7 | + | 50387344 | 50387470 | 50391728 | 50391833 | 50327637 | 50327757 | 50399917 | 50405100 | 1 | 0 | 0 | 2 | 0 | 2 | 0 | 0 | 0 | 3 | 98 | 98 | 1 | 1 | 0.333 | NA | 0 | NA | | 0 | 0.167 |
| 131 | FBXL13 | FBXL13 | chr7 | - | 102931880 | 102931933 | 102963532 | 102963665 | 102926273 | 102926374 | 102968021 | 102968117 | 26 | 1 | 0 | 30 | 18 | 2 | 0 | 0 | 0 | 5 | 98 | 98 | 1 | 1 | 0.464 | 0.053 | 0 | NA | | 0 | 0.172 |
| 132 | CCM2 | CCM2 | chr7 | + | 45068442 | 45068579 | 45069825 | 45069961 | 45064462 | 45064646 | 45072725 | 45072783 | 9 | 8 | 6 | 11 | 11 | 10 | 0 | 4 | 0 | 1 | 98 | 98 | 1 | 1 | 0.45 | 0.421 | 0.375 | NA | | 0.8 | -0.385 |
| 134 | H2AFV | H2AFV | chr7 | - | 44840898 | 44841012 | 44843276 | 44843354 | 44835528 | 44835658 | 44847968 | 44848126 | 4 | 8 | 1 | 7 | 1 | 10 | 0 | 0 | 0 | 3 | 98 | 98 | 1 | 1 | 0.364 | 0.889 | 0.091 | NA | | 0 | 0.448 |
| 136 | GGCT | GGCT | chr7 | - | 30498802 | 30498938 | 30500535 | 30500681 | 30496620 | 30497235 | 30504568 | 30504841 | 2 | 0 | 0 | 2 | 0 | 0 | 0 | 2 | 0 | 0 | 98 | 98 | 1 | 1 | 0.5 | NA | NA | NA | | 1 | -0.5 |
| 147 | CSGALNACT1 | CSGALNACT1 | chr8 | - | 19585165 | 19585317 | 19591159 | 19591278 | 19505200 | 19506130 | 19601770 | 19601865 | 1 | 0 | 0 | 3 | 0 | 0 | 0 | 1 | 0 | 0 | 98 | 98 | 1 | 1 | 0.25 | NA | NA | NA | | 1 | -0.75 |
| 149 | NUP43 | NUP43 | chr6 | - | 149742389 | 149742570 | 149743637 | 149743715 | 149738642 | 149738778 | 149745939 | 149746062 | 2 | 1 | 0 | 0 | 1 | 0 | 0 | 2 | 0 | 0 | 98 | 98 | 1 | 1 | 1 | 0.5 | NA | NA | | 1 | -0.25 |
| 168 | VNN2 | VNN2 | chr6 | - | 132752460 | 132752749 | 132753417 | 132753491 | 132751144 | 132751518 | 132755842 | 132756035 | 34 | 5 | 4 | 330 | 178 | 186 | 0 | 3 | 0 | 32 | 98 | 98 | 1 | 1 | 0.093 | 0.027 | 0.021 | NA | | 0.086 | -0.039 |
| 169 | VNN2 | VNN2 | chr6 | - | 132752672 | 132752749 | 132753417 | 132753491 | 132751144 | 132751518 | 132755842 | 132756035 | 34 | 5 | 4 | 115 | 82 | 84 | 0 | 3 | 0 | 10 | 98 | 98 | 1 | 1 | 0.228 | 0.057 | 0.045 | NA | | 0.231 | -0.121 |
| 179 | SNX3 | SNX3 | chr6 | - | 108214497 | 108214622 | 108222949 | 108223045 | 108211216 | 108212254 | 108260759 | 108261260 | 6 | 5 | 3 | 13 | 14 | 5 | 0 | 0 | 0 | 1 | 98 | 98 | 1 | 1 | 0.316 | 0.263 | 0.375 | NA | | 0 | 0.318 |
| 180 | SNX14 | SNX14 | chr6 | - | 85557975 | 85558060 | 85565331 | 85565419 | 85549722 | 85549879 | 85567533 | 85567577 | 9 | 1 | 3 | 4 | 2 | 0 | 0 | 3 | 0 | 2 | 98 | 98 | 1 | 1 | 0.692 | 0.333 | 1 | NA | | 0.6 | 0.075 |
| 181 | SNX14 | SNX14 | chr6 | - | 85565331 | 85565419 | 85567533 | 85567577 | 85557975 | 85558060 | 85572136 | 85572215 | 5 | 0 | 0 | 13 | 6 | 0 | 0 | 0 | 0 | 2 | 93 | 98 | 1 | 1 | 0.288 | 0 | NA | NA | | 0 | 0.144 |
| 189 | KLHDC3 | KLHDC3 | chr6 | + | 43017852 | 43017968 | 43018144 | 43018216 | 43017518 | 43017695 | 43018342 | 43018556 | 7 | 7 | 0 | 3 | 2 | 0 | 0 | 4 | 0 | 2 | 98 | 98 | 1 | 1 | 0.7 | 0.778 | NA | NA | | 0.667 | 0.072 |
| 190 | CCND3 | CCND3 | chr6 | - | 41940369 | 41940585 | 42030008 | 42030190 | 41937234 | 41937394 | 42048500 | 42048894 | 0 | 0 | 0 | 13 | 20 | 6 | 0 | 1 | 1 | 23 | 98 | 98 | 1 | 1 | 0 | 0 | 0 | | 0 | 0.042 | -0.021 |
| 192 | KCTD20 | KCTD20 | chr6 | + | 36470051 | 36470257 | 36479120 | 36479223 | 36442766 | 36443111 | 36479590 | 36479711 | 5 | 3 | 0 | 4 | 1 | 0 | 0 | 1 | 0 | 2 | 98 | 98 | 1 | 1 | 0.556 | 0.75 | NA | NA | | 0.333 | 0.32 |
| 193 | KCTD20 | KCTD20 | chr6 | + | 36470051 | 36470257 | 36479120 | 36479223 | 36442766 | 36443111 | 36481561 | 36481759 | 5 | 0 | 0 | 0 | 1 | 0 | 0 | 1 | 0 | 0 | 98 | 98 | 1 | 1 | 1 | 0 | NA | NA | | 1 | -0.5 |
| 194 | KCTD20 | KCTD20 | chr6 | + | 36479120 | 36479223 | 36479590 | 36479711 | 36470051 | 36470257 | 36481561 | 36481759 | 1 | 0 | 0 | 8 | 5 | 2 | 0 | 4 | 0 | 1 | 98 | 98 | 1 | 1 | 0.111 | 0 | 0 | NA | | 0.8 | -0.763 |
| 214 | HLA-F | HLA-F | chr6 | + | 29725030 | 29725306 | 29725446 | 29725563 | 29724172 | 29724448 | 29726402 | 29726526 | 120 | 51 | 32 | 2 | 0 | 3 | 0 | 24 | 0 | 0 | 98 | 98 | 1 | 1 | 0.984 | 1 | 0.914 | NA | | 1 | -0.034 |
| 224 | FAM65B | FAM65B | chr6 | - | 24835702 | 24835871 | 24839090 | 24839272 | 24832255 | 24832391 | 24842861 | 24843554 | 81 | 33 | 16 | 61 | 28 | 18 | 0 | 4 | 0 | 21 | 98 | 98 | 1 | 1 | 0.57 | 0.541 | 0.471 | NA | | 0.16 | 0.367 |
| 228 | XAF1 | XAF1 | chr17 | + | 6759661 | 6759718 | 6760405 | 6760601 | 6758088 | 6758224 | 6762154 | 6762240 | 11 | 14 | 1 | 3 | 6 | 1 | 0 | 13 | 0 | 10 | 98 | 98 | 1 | 1 | 0.786 | 0.7 | 0.5 | NA | | 0.565 | 0.097 |
| 229 | XAF1 | XAF1 | chr17 | + | 6760405 | 6760601 | 6762154 | 6762240 | 6758088 | 6758224 | 6770642 | 6770984 | 0 | 2 | 0 | 5 | 5 | 0 | 0 | 0 | 0 | 11 | 98 | 98 | 1 | 1 | 0 | 0.286 | NA | NA | | 0 | 0.143 |
| 230 | XAF1 | XAF1 | chr17 | + | 6760405 | 6760601 | 6762154 | 6762240 | 6759661 | 6759718 | 6770642 | 6770984 | 3 | 7 | 0 | 5 | 6 | 0 | 0 | 6 | 0 | 11 | 98 | 98 | 1 | 1 | 0.375 | 0.538 | NA | NA | | 0.353 | 0.104 |
| 235 | FYN | FYN | chr6 | - | 111699514 | 111699670 | 111700103 | 111700268 | 111696276 | 111696456 | 111702884 | 111703034 | 0 | 0 | 1 | 5 | 3 | 0 | 0 | 0 | 0 | 7 | 98 | 98 | 1 | 1 | 0 | 0 | 1 | NA | | 0 | 0.333 |
| 238 | HNRNPR | HNRNPR | chr1 | - | 23323555 | 23323732 | 23333517 | 23333631 | 23321527 | 23321663 | 23337753 | 23337861 | 7 | 0 | 0 | 6 | 3 | 0 | 0 | 0 | 0 | 1 | 98 | 98 | 1 | 1 | 0.538 | 0 | NA | NA | | 0 | 0.269 |
| 243 | CKLF-CMTM1 | CKLF-CMTM1 | chr16 | + | 66558189 | 66558348 | 66563121 | 66563217 | 66552562 | 66552793 | 66577103 | 66577202 | 69 | 49 | 18 | 17 | 11 | 4 | 0 | 7 | 0 | 2 | 98 | 98 | 1 | 1 | 0.802 | 0.817 | 0.818 | NA | | 0.778 | 0.034 |
| 244 | CKLF | CKLF | chr16 | + | 66558189 | 66558348 | 66563121 | 66563217 | 66552562 | 66552793 | 66565885 | 66566287 | 71 | 53 | 18 | 47 | 24 | 20 | 0 | 9 | 0 | 7 | 98 | 98 | 1 | 1 | 0.602 | 0.688 | 0.474 | NA | | 0.562 | 0.026 |
| 245 | ASB6 | ASB6 | chr9 | - | 129638572 | 129638659 | 129639201 | 129639310 | 129634603 | 129638457 | 129639401 | 129639508 | 2 | 0 | 4 | 4 | 2 | 0 | 0 | 1 | 0 | 1 | 98 | 98 | 1 | 1 | 0.333 | 0 | 1 | NA | | 0.5 | -0.056 |
| 246 | GUSB | GUSB | chr7 | - | 65976014 | 65976202 | 65979833 | 65979911 | 65974918 | 65975071 | 65980223 | 65980409 | 1 | 0 | 0 | 0 | 0 | 1 | 1 | 0 | 0 | 0 | 98 | 98 | 1 | 1 | 1 | NA | 0 | | 1 | NA | -0.5 |
| 248 | RHOT1 | RHOT1 | chr17 | + | 32209349 | 32209445 | 32211115 | 32211238 | 32208106 | 32208309 | 32224615 | 32225771 | 3 | 0 | 0 | 4 | 0 | 5 | 0 | 1 | 0 | 2 | 98 | 98 | 1 | 1 | 0.429 | NA | 0 | NA | | 0.333 | -0.119 |
| 249 | MTHFD2 | MTHFD2 | chr2 | + | 74201381 | 74201479 | 74205704 | 74205889 | 74198562 | 74198742 | 74207703 | 74207826 | 0 | 2 | 0 | 7 | 1 | 0 | 0 | 0 | 0 | 2 | 98 | 98 | 1 | 1 | 0 | 0.667 | NA | NA | | 0 | 0.334 |
| 253 | USP25 | USP25 | chr21 | + | 15849776 | 15849872 | 15864267 | 15864446 | 15842397 | 15842540 | 15866265 | 15866344 | 1 | 0 | 0 | 8 | 8 | 0 | 0 | 0 | 0 | 6 | 98 | 98 | 1 | 1 | 0.111 | 0 | NA | NA | | 0 | 0.056 |
| 254 | USB1 | USB1 | chr16 | + | 58014272 | 58014326 | 58017333 | 58017439 | 58009928 | 58010112 | 58018971 | 58019055 | 14 | 4 | 1 | 6 | 1 | 0 | 0 | 3 | 0 | 4 | 98 | 98 | 1 | 1 | 0.7 | 0.8 | 1 | NA | | 0.429 | 0.404 |
| 255 | AGO4 | AGO4 | chr1 | + | 35832436 | 35832570 | 35833989 | 35834174 | 35832056 | 35832185 | 35835833 | 35835993 | 21 | 12 | 18 | 18 | 24 | 12 | 0 | 15 | 0 | 2 | 98 | 98 | 1 | 1 | 0.538 | 0.333 | 0.6 | NA | | 0.882 | -0.392 |
| 258 | MEFV | MEFV | chr16 | - | 3249430 | 3249780 | 3254157 | 3254790 | 3247015 | 3247246 | 3256310 | 3256627 | 81 | 26 | 30 | 0 | 2 | 1 | 4 | 16 | 0 | 0 | 98 | 98 | 1 | 1 | 1 | 0.929 | 0.968 | | 1 | 1 | -0.034 |
| 260 | STK38L | STK38L | chr12 | + | 27315014 | 27315117 | 27315288 | 27315350 | 27314503 | 27314658 | 27317335 | 27317453 | 2 | 0 | 1 | 16 | 5 | 0 | 0 | 2 | 0 | 3 | 98 | 98 | 1 | 1 | 0.111 | 0 | 1 | NA | | 0.4 | -0.03 |
| 262 | WAC | WAC | chr10 | + | 28595732 | 28596041 | 28608185 | 28608431 | 28590719 | 28590832 | 28610698 | 28610821 | 26 | 13 | 2 | 26 | 9 | 9 | 0 | 3 | 0 | 1 | 98 | 98 | 1 | 1 | 0.5 | 0.591 | 0.182 | NA | | 0.75 | -0.326 |
| 273 | SEC11A | SEC11A | chr15 | - | 84670724 | 84670782 | 84680712 | 84680832 | 84669536 | 84670069 | 84687624 | 84687774 | 0 | 0 | 0 | 6 | 4 | 6 | 0 | 4 | 0 | 5 | 98 | 98 | 1 | 1 | 0 | 0 | 0 | NA | | 0.444 | -0.444 |
| 276 | SEC11A | SEC11A | chr15 | - | 84687624 | 84687774 | 84691534 | 84691644 | 84680712 | 84680832 | 84716024 | 84716460 | 5 | 2 | 0 | 0 | 2 | 0 | 0 | 0 | 0 | 4 | 98 | 98 | 1 | 1 | 1 | 0.5 | NA | NA | | 0 | 0.75 |
| 277 | TM6SF1 | TM6SF1 | chr15 | + | 83124671 | 83124776 | 83126754 | 83126847 | 83119577 | 83119681 | 83127357 | 83127477 | 3 | 0 | 0 | 11 | 10 | 8 | 0 | 0 | 0 | 1 | 98 | 98 | 1 | 1 | 0.214 | 0 | 0 | NA | | 0 | 0.071 |
| 278 | BTBD1 | BTBD1 | chr15 | - | 83018706 | 83018853 | 83020674 | 83020762 | 83016428 | 83018225 | 83030135 | 83030328 | 3 | 4 | 0 | 5 | 2 | 0 | 0 | 0 | 0 | 3 | 98 | 98 | 1 | 1 | 0.375 | 0.667 | NA | NA | | 0 | 0.521 |
| 287 | XPO6 | XPO6 | chr16 | - | 28177219 | 28177332 | 28180940 | 28181031 | 28175897 | 28176095 | 28208986 | 28209122 | 4 | 6 | 3 | 18 | 25 | 6 | 0 | 2 | 0 | 15 | 98 | 98 | 1 | 1 | 0.182 | 0.194 | 0.333 | NA | | 0.118 | 0.118 |
| 288 | XPO6 | XPO6 | chr16 | - | 28177219 | 28177332 | 28180940 | 28181031 | 28175897 | 28176095 | 28211365 | 28211918 | 50 | 24 | 9 | 18 | 34 | 6 | 0 | 18 | 0 | 15 | 98 | 98 | 1 | 1 | 0.735 | 0.414 | 0.6 | NA | | 0.545 | 0.038 |
| 290 | EDC3 | EDC3 | chr15 | - | 74635408 | 74635626 | 74640465 | 74640619 | 74630557 | 74632946 | 74655732 | 74656068 | 1 | 2 | 0 | 1 | 1 | 0 | 3 | 1 | 0 | 0 | 98 | 98 | 1 | 1 | 0.5 | 0.667 | NA | NA | 1 | 1 | -0.416 |
| 293 | USP8 | USP8 | chr15 | + | 50481480 | 50482065 | 50484274 | 50484361 | 50477275 | 50477499 | 50489800 | 50489881 | 13 | 6 | 1 | 4 | 2 | 2 | 0 | 5 | 0 | 2 | 98 | 98 | 1 | 1 | 0.765 | 0.75 | 0.333 | NA | | 0.714 | -0.098 |
| 294 | PLAGL1 | PLAGL1 | chr6 | - | 143947984 | 143948460 | 143960468 | 143960542 | 143940299 | 143942663 | 143964786 | 143964818 | 5 | 3 | 0 | 2 | 0 | 0 | 0 | 1 | 0 | 0 | 98 | 98 | 1 | 1 | 0.714 | 1 | NA | NA | | 1 | -0.143 |
| 301 | YTHDF3 | YTHDF3 | chr8 | + | 63175330 | 63175416 | 63186146 | 63187745 | 63169386 | 63169411 | 63209682 | 63212788 | 3 | 6 | 0 | 19 | 3 | 0 | 0 | 3 | 0 | 5 | 98 | 98 | 1 | 1 | 0.136 | 0.667 | NA | NA | | 0.375 | 0.027 |
| 306 | PYGL | PYGL | chr14 | - | 50914700 | 50914815 | 50915335 | 50915499 | 50913028 | 50913130 | 50915824 | 50915971 | 46 | 46 | 19 | 68 | 25 | 23 | 0 | 19 | 0 | 14 | 98 | 98 | 1 | 1 | 0.404 | 0.648 | 0.452 | NA | | 0.576 | -0.075 |
| 307 | NIPA2 | NIPA2 | chr15 | + | 22839654 | 22839790 | 22845145 | 22845267 | 22838640 | 22838921 | 22851638 | 22851870 | 0 | 0 | 0 | 0 | 2 | 0 | 0 | 1 | 0 | 0 | 98 | 98 | 1 | 1 | NA | 0 | NA | NA | | 1 | -1 |
| 311 | TECPR2 | TECPR2 | chr14 | + | 102424978 | 102425291 | 102431795 | 102432128 | 102414635 | 102414793 | 102434234 | 102435211 | 36 | 8 | 2 | 19 | 5 | 3 | 0 | 1 | 0 | 1 | 98 | 98 | 1 | 1 | 0.655 | 0.615 | 0.4 | NA | | 0.5 | 0.057 |
| 314 | NME8 | NME8 | chr7 | + | 37864347 | 37864421 | 37867701 | 37867898 | 37863395 | 37863462 | 37876831 | 37877007 | 8 | 9 | 4 | 1 | 1 | 1 | 0 | 4 | 0 | 1 | 98 | 98 | 1 | 1 | 0.889 | 0.9 | 0.8 | NA | | 0.8 | 0.063 |
| 318 | FOPNL | FOPNL | chr16 | - | 15873490 | 15873627 | 15879803 | 15879888 | 15865718 | 15867516 | 15884007 | 15884205 | 1 | 1 | 0 | 0 | 1 | 0 | 0 | 0 | 0 | 1 | 98 | 98 | 1 | 1 | 1 | 0.5 | NA | NA | | 0 | 0.75 |
| 329 | CTSS | CTSS | chr1 | - | 150751780 | 150752008 | 150755000 | 150755150 | 150747776 | 150747879 | 150757857 | 150757980 | 414 | 132 | 122 | 2 | 0 | 5 | 2 | 125 | 0 | 2 | 98 | 98 | 1 | 1 | 0.995 | 1 | 0.961 | | 1 | 0.984 | -0.007 |
| 332 | EIF2AK2 | EIF2AK2 | chr2 | - | 37122505 | 37122664 | 37126288 | 37126411 | 37119958 | 37120139 | 37135483 | 37135546 | 17 | 7 | 0 | 12 | 13 | 2 | 0 | 9 | 0 | 6 | 98 | 98 | 1 | 1 | 0.586 | 0.35 | 0 | NA | | 0.6 | -0.288 |
| 336 | ACTN1 | ACTN1 | chr14 | - | 68878457 | 68878523 | 68878988 | 68879069 | 68877081 | 68877240 | 68879961 | 68880108 | 85 | 49 | 27 | 0 | 1 | 0 | 1 | 15 | 0 | 0 | 98 | 98 | 1 | 1 | 1 | 0.98 | 1 | | 1 | 1 | -0.007 |
| 337 | RDH11 | RDH11 | chr14 | - | 67685014 | 67685204 | 67690211 | 67690421 | 67676800 | 67678423 | 67691139 | 67691244 | 9 | 0 | 0 | 1 | 0 | 0 | 0 | 0 | 0 | 1 | 98 | 98 | 1 | 1 | 0.9 | NA | NA | | | 0 | 0.9 |
| 349 | ITCH | ITCH | chr20 | + | 34393790 | 34393881 | 34408650 | 34408792 | 34369393 | 34369470 | 34412514 | 34412639 | 9 | 6 | 0 | 28 | 28 | 16 | 0 | 4 | 0 | 2 | 98 | 98 | 1 | 1 | 0.243 | 0.176 | 0 | NA | | 0.667 | -0.527 |
| 356 | EPG5 | EPG5 | chr18 | - | 45879012 | 45879214 | 45880074 | 45880223 | 45878375 | 45878448 | 45882273 | 45882487 | 1 | 5 | 0 | 4 | 1 | 6 | 0 | 1 | 0 | 0 | 98 | 98 | 1 | 1 | 0.2 | 0.833 | 0 | NA | | 1 | -0.656 |
| 357 | EPG5 | EPG5 | chr18 | - | 45887750 | 45887907 | 45889797 | 45889940 | 45884616 | 45884811 | 45899403 | 45899566 | 6 | 3 | 0 | 3 | 0 | 6 | 0 | 1 | 0 | 0 | 98 | 98 | 1 | 1 | 0.667 | 1 | 0 | NA | | 1 | -0.444 |
| 359 | ARMC2 | ARMC2 | chr6 | + | 108858198 | 108858271 | 108868823 | 108868995 | 108854224 | 108854485 | 108876142 | 108876350 | 3 | 0 | 1 | 5 | 3 | 0 | 0 | 3 | 0 | 1 | 98 | 98 | 1 | 1 | 0.375 | 0 | 1 | NA | | 0.75 | -0.292 |
| 362 | PAN3 | PAN3 | chr13 | + | 28177864 | 28177935 | 28220230 | 28220378 | 28176492 | 28176559 | 28256291 | 28256539 | 27 | 25 | 9 | 14 | 0 | 1 | 0 | 7 | 0 | 2 | 98 | 98 | 1 | 1 | 0.659 | 1 | 0.9 | NA | | 0.778 | 0.075 |
| 363 | NUPL1 | NUPL1 | chr13 | + | 25309246 | 25309282 | 25312882 | 25313032 | 25307805 | 25307948 | 25313613 | 25313751 | 10 | 3 | 3 | 15 | 2 | 2 | 0 | 5 | 0 | 2 | 85 | 98 | 1 | 1 | 0.435 | 0.634 | 0.634 | NA | | 0.742 | -0.174 |
| 365 | ZNF268 | ZNF268 | chr12 | + | 133187871 | 133188072 | 133191488 | 133191615 | 133181945 | 133182030 | 133191907 | 133192003 | 0 | 0 | 1 | 0 | 2 | 0 | 0 | 0 | 0 | 2 | 98 | 98 | 1 | 1 | NA | 0 | 1 | NA | | 0 | 0.5 |
| 369 | OGFOD2 | OGFOD2 | chr12 | + | 122975810 | 122975867 | 122976653 | 122976767 | 122974702 | 122975121 | 122976870 | 122976970 | 0 | 0 | 1 | 0 | 0 | 0 | 0 | 0 | 0 | 1 | 98 | 98 | 1 | 1 | NA | NA | 1 | NA | | 0 | 1 |
| 372 | MED13L | MED13L | chr12 | - | 116096668 | 116096752 | 116111427 | 116111512 | 116022455 | 116022601 | 116237467 | 116237705 | 84 | 36 | 16 | 56 | 27 | 5 | 0 | 10 | 0 | 16 | 98 | 98 | 1 | 1 | 0.6 | 0.571 | 0.762 | NA | | 0.385 | 0.259 |
| 373 | MED13L | MED13L | chr12 | - | 116111427 | 116111512 | 116237467 | 116237705 | 116096668 | 116096752 | 116277059 | 116277186 | 56 | 19 | 3 | 47 | 30 | 16 | 0 | 8 | 0 | 7 | 98 | 98 | 1 | 1 | 0.544 | 0.388 | 0.158 | NA | | 0.533 | -0.17 |
| 375 | NLRP1 | NLRP1 | chr17 | - | 5533896 | 5533988 | 5536850 | 5536940 | 5533303 | 5533384 | 5539414 | 5539585 | 36 | 25 | 5 | 31 | 12 | 6 | 0 | 10 | 0 | 7 | 98 | 98 | 1 | 1 | 0.537 | 0.676 | 0.455 | NA | | 0.588 | -0.032 |
| 378 | YIPF1 | YIPF1 | chr1 | - | 53855000 | 53855051 | 53860055 | 53860153 | 53851718 | 53852270 | 53866199 | 53866382 | 9 | 2 | 3 | 4 | 0 | 0 | 0 | 1 | 0 | 2 | 98 | 98 | 1 | 1 | 0.692 | 1 | 1 | NA | | 0.333 | 0.564 |
| 379 | YIPF1 | YIPF1 | chr1 | - | 53860055 | 53860153 | 53866199 | 53866382 | 53855000 | 53855051 | 53866757 | 53866924 | 10 | 2 | 5 | 3 | 3 | 3 | 0 | 1 | 0 | 0 | 98 | 98 | 1 | 1 | 0.769 | 0.4 | 0.625 | NA | | 1 | -0.402 |
| 380 | YIPF1 | YIPF1 | chr1 | - | 53866199 | 53866382 | 53866757 | 53866924 | 53860055 | 53860153 | 53871371 | 53871488 | 13 | 2 | 7 | 9 | 2 | 3 | 0 | 4 | 0 | 2 | 98 | 98 | 1 | 1 | 0.591 | 0.5 | 0.7 | NA | | 0.667 | -0.07 |
| 384 | MARK3 | MARK3 | chr14 | + | 103465556 | 103465793 | 103465971 | 103466091 | 103462404 | 103462461 | 103466342 | 103466442 | 10 | 5 | 4 | 15 | 1 | 3 | 0 | 5 | 0 | 1 | 98 | 98 | 1 | 1 | 0.4 | 0.833 | 0.571 | NA | | 0.833 | -0.232 |
| 388 | NF2 | NF2 | chr22 | + | 29636750 | 29636876 | 29639089 | 29639212 | 29603555 | 29604112 | 29642201 | 29642285 | 2 | 0 | 0 | 1 | 0 | 0 | 0 | 1 | 0 | 0 | 98 | 98 | 1 | 1 | 0.667 | NA | NA | NA | | 1 | -0.333 |
| 390 | PLXNC1 | PLXNC1 | chr12 | + | 94298631 | 94298795 | 94300909 | 94301057 | 94297315 | 94297423 | 94303755 | 94303896 | 31 | 26 | 12 | 68 | 30 | 19 | 0 | 14 | 0 | 11 | 98 | 98 | 1 | 1 | 0.313 | 0.464 | 0.387 | NA | | 0.56 | -0.172 |
| 391 | ZNF512 | ZNF512 | chr2 | + | 27599969 | 27600053 | 27600690 | 27600815 | 27599582 | 27599678 | 27601355 | 27601442 | 6 | 3 | 0 | 4 | 2 | 0 | 0 | 1 | 0 | 0 | 98 | 98 | 1 | 1 | 0.6 | 0.6 | NA | NA | | 1 | -0.4 |
| 395 | IL32 | IL32 | chr16 | + | 3067553 | 3067613 | 3067983 | 3068010 | 3067376 | 3067415 | 3068179 | 3068239 | 14 | 13 | 2 | 11 | 2 | 0 | 0 | 3 | 0 | 1 | 98 | 76 | 1 | 1 | 0.497 | 0.834 | 1 | NA | | 0.699 | 0.078 |
| 397 | ARHGAP9 | ARHGAP9 | chr12 | - | 57473602 | 57473708 | 57474041 | 57474176 | 57472254 | 57472688 | 57474422 | 57474476 | 214 | 71 | 70 | 116 | 35 | 41 | 0 | 27 | 0 | 16 | 98 | 98 | 1 | 1 | 0.648 | 0.67 | 0.631 | NA | | 0.628 | 0.022 |
| 399 | PTGES3 | PTGES3 | chr12 | - | 56666203 | 56666266 | 56670274 | 56670364 | 56664775 | 56664800 | 56671748 | 56671847 | 6 | 3 | 0 | 6 | 13 | 1 | 0 | 0 | 0 | 5 | 98 | 98 | 1 | 1 | 0.5 | 0.187 | 0 | NA | | 0 | 0.229 |
| 408 | PI4KB | PI4KB | chr1 | - | 151324744 | 151324917 | 151326171 | 151326373 | 151315572 | 151316509 | 151327270 | 151327657 | 3 | 1 | 0 | 1 | 2 | 0 | 0 | 4 | 0 | 0 | 98 | 98 | 1 | 1 | 0.75 | 0.333 | NA | NA | | 1 | -0.459 |
| 409 | PSMA3 | PSMA3 | chr14 | + | 58247749 | 58247832 | 58252118 | 58252242 | 58244804 | 58244941 | 58257744 | 58257846 | 0 | 1 | 1 | 11 | 4 | 0 | 0 | 2 | 0 | 2 | 98 | 98 | 1 | 1 | 0 | 0.2 | 1 | NA | | 0.5 | -0.1 |
| 411 | P2RX7 | P2RX7 | chr12 | + | 121160901 | 121160974 | 121162423 | 121162520 | 121156078 | 121156147 | 121165356 | 121165437 | 4 | 0 | 0 | 0 | 0 | 0 | 0 | 5 | 0 | 1 | 98 | 98 | 1 | 1 | 1 | NA | NA | NA | | 0.833 | 0.167 |
| 414 | ADGRE3 | ADGRE3 | chr19 | - | 14654981 | 14655165 | 14658512 | 14658550 | 14651084 | 14651204 | 14661962 | 14662118 | 98 | 26 | 12 | 77 | 34 | 21 | 0 | 7 | 0 | 11 | 87 | 98 | 1 | 1 | 0.589 | 0.463 | 0.392 | NA | | 0.418 | 0.063 |
| 415 | ADGRE3 | ADGRE3 | chr19 | - | 14658512 | 14658550 | 14661962 | 14662118 | 14654981 | 14655165 | 14663417 | 14663540 | 72 | 31 | 17 | 66 | 13 | 15 | 0 | 5 | 0 | 5 | 98 | 87 | 1 | 1 | 0.492 | 0.679 | 0.502 | NA | | 0.47 | 0.088 |
| 416 | ADGRE3 | ADGRE3 | chr19 | - | 14661962 | 14662118 | 14663417 | 14663540 | 14658512 | 14658550 | 14668801 | 14668852 | 96 | 25 | 4 | 102 | 33 | 12 | 0 | 5 | 0 | 8 | 98 | 98 | 1 | 1 | 0.485 | 0.431 | 0.25 | NA | | 0.385 | 0.004 |
| 417 | ADGRE3 | ADGRE3 | chr19 | - | 14663417 | 14663540 | 14668801 | 14668852 | 14661962 | 14662118 | 14674745 | 14674918 | 11 | 5 | 0 | 72 | 32 | 17 | 0 | 1 | 0 | 4 | 98 | 98 | 1 | 1 | 0.133 | 0.135 | 0 | NA | | 0.2 | -0.111 |
| 419 | LCP1 | LCP1 | chr13 | - | 46158521 | 46158651 | 46158825 | 46158989 | 46156437 | 46156570 | 46159598 | 46159686 | 50 | 39 | 15 | 250 | 100 | 86 | 0 | 23 | 0 | 46 | 98 | 98 | 1 | 1 | 0.167 | 0.281 | 0.149 | NA | | 0.333 | -0.134 |
| 421 | DLG1 | DLG1 | chr3 | - | 197075836 | 197075870 | 197076585 | 197076685 | 197069218 | 197069260 | 197081050 | 197081117 | 1 | 0 | 0 | 2 | 0 | 0 | 0 | 4 | 0 | 0 | 98 | 83 | 1 | 1 | 0.297 | NA | NA | NA | | 1 | -0.703 |
| 423 | KDM6A | KDM6A | chrX | + | 45060021 | 45060156 | 45060608 | 45060764 | 45059246 | 45059466 | 45061323 | 45061419 | 7 | 0 | 0 | 11 | 8 | 2 | 0 | 0 | 0 | 4 | 98 | 98 | 1 | 1 | 0.389 | 0 | 0 | NA | | 0 | 0.13 |
| 426 | QKI | QKI | chr6 | + | 163563419 | 163563719 | 163566720 | 163566795 | 163561981 | 163562069 | 163570693 | 163578596 | 28 | 11 | 10 | 10 | 4 | 5 | 0 | 3 | 0 | 3 | 98 | 98 | 1 | 1 | 0.737 | 0.733 | 0.667 | NA | | 0.5 | 0.212 |
| 427 | QKI | QKI | chr6 | + | 163563443 | 163563719 | 163566720 | 163566795 | 163561981 | 163562069 | 163570693 | 163578596 | 2 | 2 | 0 | 10 | 4 | 5 | 0 | 0 | 0 | 3 | 98 | 98 | 1 | 1 | 0.167 | 0.333 | 0 | NA | | 0 | 0.167 |
| 429 | CLEC7A | CLEC7A | chr12 | - | 10123244 | 10123363 | 10125246 | 10125448 | 10116780 | 10118590 | 10127746 | 10127845 | 48 | 19 | 15 | 92 | 36 | 25 | 0 | 19 | 0 | 30 | 98 | 98 | 1 | 1 | 0.343 | 0.345 | 0.375 | NA | | 0.388 | -0.034 |
| 432 | FCHO2 | FCHO2 | chr5 | + | 73006444 | 73006549 | 73015625 | 73015724 | 72990711 | 72990864 | 73017211 | 73017308 | 15 | 3 | 0 | 22 | 9 | 1 | 0 | 6 | 0 | 0 | 98 | 98 | 1 | 1 | 0.405 | 0.25 | 0 | NA | | 0.75 | -0.532 |
| 434 | SYNJ2BP-COX16 | SYNJ2BP-COX16 | chr14 | - | 70342657 | 70342729 | 70373032 | 70373131 | 70329173 | 70329236 | 70375675 | 70375771 | 8 | 0 | 0 | 0 | 0 | 0 | 0 | 0 | 0 | 2 | 98 | 98 | 1 | 1 | 1 | NA | NA | NA | | 0 | 1 |
| 436 | TLE4 | TLE4 | chr9 | + | 79612655 | 79612718 | 79627373 | 79627448 | 79576132 | 79576177 | 79652592 | 79652794 | 22 | 4 | 3 | 7 | 1 | 0 | 0 | 0 | 0 | 2 | 98 | 98 | 1 | 1 | 0.759 | 0.8 | 1 | NA | | 0 | 0.853 |
| 446 | FNIP1 | FNIP1 | chr5 | - | 131677702 | 131677872 | 131679028 | 131679175 | 131671504 | 131672924 | 131698916 | 131699002 | 16 | 2 | 2 | 34 | 15 | 2 | 0 | 1 | 0 | 6 | 98 | 98 | 1 | 1 | 0.32 | 0.118 | 0.5 | NA | | 0.143 | 0.17 |
| 450 | KANSL3 | KANSL3 | chr2 | - | 96602115 | 96602338 | 96602752 | 96602862 | 96601642 | 96601776 | 96604249 | 96604380 | 3 | 1 | 0 | 1 | 0 | 0 | 0 | 0 | 1 | 2 | 98 | 98 | 1 | 1 | 0.75 | 1 | NA | | 0 | 0 | 0.875 |
| 451 | AHCTF1 | AHCTF1 | chr1 | - | 246863923 | 246864116 | 246867243 | 246867351 | 246861958 | 246862153 | 246867660 | 246867811 | 1 | 0 | 0 | 26 | 16 | 4 | 0 | 0 | 0 | 3 | 98 | 98 | 1 | 1 | 0.037 | 0 | 0 | NA | | 0 | 0.012 |
| 452 | SPOP | SPOP | chr17 | - | 49636758 | 49636809 | 49668026 | 49668078 | 49622732 | 49622876 | 49677932 | 49678163 | 2 | 1 | 0 | 2 | 5 | 0 | 0 | 2 | 0 | 0 | 98 | 98 | 1 | 1 | 0.5 | 0.167 | NA | NA | | 1 | -0.666 |
| 455 | APLP2 | APLP2 | chr11 | + | 130109428 | 130109505 | 130120705 | 130120818 | 130069820 | 130070082 | 130121613 | 130121810 | 21 | 12 | 5 | 53 | 34 | 10 | 0 | 9 | 0 | 20 | 98 | 98 | 1 | 1 | 0.284 | 0.261 | 0.333 | NA | | 0.31 | -0.017 |
| 456 | MEF2C | MEF2C | chr5 | - | 88760987 | 88761125 | 88761184 | 88761328 | 88751856 | 88752043 | 88804597 | 88804801 | 5 | 2 | 0 | 1 | 0 | 0 | 0 | 5 | 0 | 0 | 98 | 98 | 1 | 1 | 0.833 | 1 | NA | NA | | 1 | -0.084 |
| 460 | SNX17 | SNX17 | chr2 | + | 27371268 | 27371468 | 27372622 | 27372740 | 27370495 | 27370806 | 27373246 | 27373311 | 0 | 1 | 0 | 2 | 0 | 0 | 0 | 0 | 0 | 1 | 98 | 98 | 1 | 1 | 0 | 1 | NA | NA | | 0 | 0.5 |
| 461 | SNX17 | SNX17 | chr2 | + | 27372622 | 27372740 | 27373039 | 27373113 | 27371268 | 27371343 | 27373246 | 27373311 | 6 | 1 | 1 | 1 | 0 | 0 | 1 | 4 | 0 | 0 | 98 | 98 | 1 | 1 | 0.857 | 1 | 1 | | 1 | 1 | -0.048 |
| 464 | GPATCH8 | GPATCH8 | chr17 | - | 44464471 | 44464544 | 44474828 | 44474903 | 44436477 | 44436545 | 44503325 | 44503602 | 3 | 0 | 0 | 3 | 0 | 0 | 0 | 0 | 0 | 1 | 98 | 98 | 1 | 1 | 0.5 | NA | NA | NA | | 0 | 0.5 |
| 466 | ARNTL | ARNTL | chr11 | + | 13349944 | 13350070 | 13354300 | 13354448 | 13309986 | 13310059 | 13356738 | 13356777 | 8 | 5 | 5 | 10 | 2 | 1 | 0 | 0 | 0 | 5 | 98 | 98 | 1 | 1 | 0.444 | 0.714 | 0.833 | NA | | 0 | 0.664 |
| 470 | POLG | POLG | chr15 | - | 89322741 | 89322902 | 89323403 | 89323511 | 89321961 | 89322015 | 89323814 | 89323901 | 7 | 3 | 0 | 2 | 0 | 0 | 0 | 3 | 0 | 2 | 98 | 98 | 1 | 1 | 0.778 | 1 | NA | NA | | 0.6 | 0.289 |
| 472 | TRAF3IP3 | TRAF3IP3 | chr1 | + | 209779314 | 209779374 | 209780469 | 209780606 | 209778110 | 209778173 | 209781344 | 209781458 | 23 | 6 | 0 | 8 | 2 | 0 | 0 | 0 | 0 | 3 | 98 | 98 | 1 | 1 | 0.742 | 0.75 | NA | NA | | 0 | 0.746 |
| 485 | SLC9A8 | SLC9A8 | chr20 | + | 49839540 | 49839599 | 49845035 | 49845119 | 49823060 | 49823141 | 49849578 | 49849680 | 21 | 3 | 3 | 18 | 4 | 0 | 0 | 6 | 0 | 7 | 98 | 98 | 1 | 1 | 0.538 | 0.429 | 1 | NA | | 0.462 | 0.194 |
| 486 | SLC9A8 | SLC9A8 | chr20 | + | 49845035 | 49845119 | 49849578 | 49849680 | 49839540 | 49839599 | 49855437 | 49855581 | 13 | 2 | 2 | 14 | 9 | 0 | 0 | 4 | 0 | 4 | 98 | 98 | 1 | 1 | 0.481 | 0.182 | NA | NA | | 0.5 | -0.168 |
| 494 | NDUFS1 | NDUFS1 | chr2 | - | 206149817 | 206149925 | 206152418 | 206152510 | 206149019 | 206149096 | 206153617 | 206153682 | 3 | 4 | 2 | 10 | 6 | 0 | 0 | 1 | 0 | 0 | 98 | 98 | 1 | 1 | 0.231 | 0.4 | 1 | NA | | 1 | -0.456 |
| 496 | FAR2 | FAR2 | chr12 | + | 29293299 | 29293475 | 29297020 | 29297200 | 29270411 | 29270638 | 29307657 | 29307835 | 2 | 3 | 2 | 12 | 13 | 3 | 0 | 2 | 0 | 1 | 98 | 98 | 1 | 1 | 0.143 | 0.187 | 0.4 | NA | | 0.667 | -0.424 |
| 497 | RASGRP4 | RASGRP4 | chr19 | - | 38417051 | 38417168 | 38419859 | 38420013 | 38413393 | 38413474 | 38420130 | 38420262 | 20 | 18 | 4 | 1 | 0 | 1 | 4 | 7 | 0 | 0 | 98 | 98 | 1 | 1 | 0.952 | 1 | 0.8 | | 1 | 0 | -0.083 |
| 501 | ELP3 | ELP3 | chr8 | + | 28097218 | 28097318 | 28099827 | 28099966 | 28093066 | 28093233 | 28106712 | 28106783 | 1 | 0 | 0 | 4 | 0 | 0 | 0 | 0 | 0 | 2 | 98 | 98 | 1 | 1 | 0.2 | NA | NA | NA | | 0 | 0.2 |
| 502 | RPAIN | RPAIN | chr17 | + | 5425970 | 5426082 | 5426235 | 5426299 | 5422768 | 5422829 | 5428070 | 5428211 | 0 | 2 | 0 | 2 | 0 | 2 | 0 | 2 | 0 | 1 | 98 | 98 | 1 | 1 | 0 | 1 | 0 | NA | | 0.667 | -0.334 |
| 504 | RPAIN | RPAIN | chr17 | + | 5425970 | 5426082 | 5428070 | 5428211 | 5422768 | 5422829 | 5432541 | 5433020 | 0 | 2 | 0 | 2 | 0 | 0 | 0 | 2 | 0 | 0 | 98 | 98 | 1 | 1 | 0 | 1 | NA | NA | | 1 | -0.5 |
| 507 | SCP2 | SCP2 | chr1 | + | 52941795 | 52941853 | 52948008 | 52948080 | 52927228 | 52927465 | 52950754 | 52950886 | 0 | 0 | 0 | 1 | 1 | 0 | 0 | 1 | 0 | 0 | 98 | 98 | 1 | 1 | 0 | 0 | NA | NA | | 1 | -1 |
| 511 | NAAA | NAAA | chr4 | - | 75914867 | 75914985 | 75918760 | 75918789 | 75913655 | 75914338 | 75919908 | 75919975 | 1 | 0 | 0 | 3 | 0 | 1 | 0 | 0 | 0 | 2 | 78 | 98 | 1 | 1 | 0.295 | NA | 0 | NA | | 0 | 0.147 |
| 514 | DCAF8 | DCAF8 | chr1 | - | 160243959 | 160244034 | 160261284 | 160261358 | 160239696 | 160240370 | 160262116 | 160262528 | 1 | 0 | 0 | 3 | 2 | 0 | 0 | 0 | 0 | 3 | 98 | 98 | 1 | 1 | 0.25 | 0 | NA | NA | | 0.25 | -0.125 |
| 515 | DCAF8 | DCAF8 | chr1 | - | 160243959 | 160244034 | 160261284 | 160261358 | 160239696 | 160240370 | 160262448 | 160262560 | 1 | 2 | 0 | 4 | 2 | 0 | 0 | 0 | 0 | 3 | 98 | 98 | 1 | 1 | 0.2 | 0.5 | NA | NA | | 0 | 0.35 |
| 523 | PPP1R12A | PPP1R12A | chr12 | - | 79805591 | 79805768 | 79806165 | 79806333 | 79798493 | 79798584 | 79807225 | 79807330 | 22 | 12 | 3 | 8 | 2 | 0 | 0 | 6 | 0 | 0 | 98 | 98 | 1 | 1 | 0.733 | 0.857 | 1 | NA | | 1 | -0.137 |
| 533 | SLC50A1 | SLC50A1 | chr1 | + | 155136827 | 155136951 | 155137560 | 155137722 | 155136298 | 155136376 | 155137978 | 155138098 | 2 | 0 | 0 | 0 | 0 | 0 | 0 | 0 | 0 | 1 | 98 | 98 | 1 | 1 | 1 | NA | NA | NA | | 0 | 1 |
| 536 | STX3 | STX3 | chr11 | + | 59795371 | 59795482 | 59797282 | 59797396 | 59793379 | 59793514 | 59800854 | 59805882 | 24 | 12 | 8 | 21 | 11 | 14 | 0 | 5 | 0 | 5 | 98 | 98 | 1 | 1 | 0.533 | 0.522 | 0.364 | NA | | 0.5 | -0.027 |
| 538 | SNX27 | SNX27 | chr1 | + | 151662165 | 151662270 | 151665932 | 151666011 | 151660797 | 151660862 | 151668471 | 151668635 | 17 | 0 | 6 | 21 | 11 | 3 | 0 | 5 | 0 | 5 | 98 | 98 | 1 | 1 | 0.447 | 0 | 0.667 | NA | | 0.5 | -0.129 |
| 541 | ATXN3 | ATXN3 | chr14 | - | 92093251 | 92093318 | 92096092 | 92096137 | 92088729 | 92088817 | 92096673 | 92096838 | 7 | 0 | 2 | 9 | 1 | 0 | 0 | 1 | 0 | 1 | 94 | 98 | 1 | 1 | 0.448 | 0 | 1 | NA | | 0.51 | -0.027 |
| 542 | ATXN3 | ATXN3 | chr14 | - | 92093745 | 92093831 | 92096092 | 92096137 | 92093251 | 92093318 | 92096673 | 92096838 | 10 | 1 | 2 | 4 | 2 | 0 | 0 | 1 | 0 | 1 | 94 | 98 | 1 | 1 | 0.723 | 0.343 | 1 | NA | | 0.51 | 0.179 |
| 543 | ATXN3 | ATXN3 | chr14 | - | 92093745 | 92093831 | 92096092 | 92096137 | 92093251 | 92093318 | 92106528 | 92106621 | 3 | 1 | 0 | 4 | 2 | 0 | 0 | 0 | 0 | 1 | 94 | 98 | 1 | 1 | 0.439 | 0.343 | NA | NA | | 0 | 0.391 |
| 549 | CREBBP | CREBBP | chr16 | - | 3778699 | 3778817 | 3780731 | 3780878 | 3778010 | 3778182 | 3781203 | 3781306 | 37 | 7 | 1 | 26 | 9 | 6 | 0 | 5 | 0 | 7 | 98 | 98 | 1 | 1 | 0.587 | 0.437 | 0.143 | NA | | 0.417 | -0.028 |
| 554 | SORL1 | SORL1 | chr11 | + | 121608103 | 121608176 | 121611075 | 121611158 | 121607185 | 121607290 | 121612735 | 121612832 | 73 | 47 | 30 | 239 | 98 | 59 | 0 | 12 | 0 | 22 | 98 | 98 | 1 | 1 | 0.234 | 0.324 | 0.337 | NA | | 0.353 | -0.055 |
| 555 | NIN | NIN | chr14 | - | 50821873 | 50822077 | 50830463 | 50830517 | 50806736 | 50806818 | 50831005 | 50831121 | 5 | 7 | 0 | 34 | 24 | 4 | 0 | 1 | 0 | 6 | 98 | 98 | 1 | 1 | 0.128 | 0.226 | 0 | NA | | 0.143 | -0.025 |
| 558 | ZNF185 | ZNF185 | chrX | + | 152922719 | 152922809 | 152928574 | 152928661 | 152922172 | 152922256 | 152931671 | 152931776 | 5 | 3 | 0 | 11 | 2 | 0 | 1 | 1 | 0 | 0 | 98 | 98 | 1 | 1 | 0.312 | 0.6 | NA | | 1 | 1 | -0.544 |
| 559 | R3HCC1L | R3HCC1L | chr10 | + | 98208100 | 98209899 | 98231511 | 98231687 | 98163292 | 98163397 | 98234445 | 98234516 | 4 | 2 | 0 | 2 | 3 | 0 | 0 | 2 | 0 | 0 | 98 | 98 | 1 | 1 | 0.667 | 0.4 | NA | NA | | 1 | -0.466 |
| 568 | C1orf162 | C1orf162 | chr1 | + | 111477333 | 111477428 | 111477650 | 111477775 | 111476797 | 111476867 | 111477982 | 111478512 | 7 | 2 | 0 | 3 | 1 | 0 | 0 | 5 | 1 | 9 | 98 | 98 | 1 | 1 | 0.7 | 0.667 | NA | | 0 | 0.357 | 0.505 |
| 584 | ARHGEF2 | ARHGEF2 | chr1 | - | 155954901 | 155954969 | 155957712 | 155957882 | 155952627 | 155952828 | 155958319 | 155958396 | 9 | 15 | 12 | 25 | 14 | 2 | 0 | 5 | 0 | 7 | 98 | 98 | 1 | 1 | 0.265 | 0.517 | 0.857 | NA | | 0.417 | 0.129 |
| 590 | SOAT1 | SOAT1 | chr1 | + | 179302676 | 179302802 | 179323436 | 179323495 | 179293713 | 179293936 | 179335505 | 179335657 | 1 | 1 | 0 | 8 | 6 | 0 | 0 | 3 | 0 | 0 | 98 | 98 | 1 | 1 | 0.111 | 0.143 | NA | NA | | 1 | -0.873 |
| 598 | BLOC1S6 | BLOC1S6 | chr15 | + | 45603099 | 45603187 | 45605427 | 45605514 | 45592134 | 45592276 | 45606394 | 45609716 | 1 | 0 | 1 | 2 | 3 | 0 | 0 | 0 | 0 | 4 | 98 | 98 | 1 | 1 | 0.333 | 0 | 1 | NA | | 0 | 0.444 |
| 601 | ECD | ECD | chr10 | - | 73152292 | 73152421 | 73154255 | 73154448 | 73148275 | 73148404 | 73156274 | 73156453 | 3 | 0 | 0 | 4 | 0 | 0 | 0 | 1 | 0 | 1 | 98 | 98 | 1 | 1 | 0.429 | NA | NA | NA | | 0.5 | -0.071 |
| 602 | NDUFB6 | NDUFB6 | chr9 | - | 32558909 | 32558954 | 32570959 | 32571052 | 32553525 | 32553944 | 32572880 | 32573184 | 2 | 2 | 2 | 6 | 15 | 12 | 0 | 0 | 0 | 5 | 98 | 94 | 1 | 1 | 0.242 | 0.113 | 0.138 | NA | | 0 | 0.164 |
| 616 | ACAP2 | ACAP2 | chr3 | - | 195289120 | 195289231 | 195291705 | 195291815 | 195285795 | 195285857 | 195292264 | 195292452 | 20 | 13 | 2 | 31 | 13 | 9 | 0 | 6 | 0 | 7 | 98 | 98 | 1 | 1 | 0.392 | 0.5 | 0.182 | NA | | 0.462 | -0.104 |
| 618 | UBR4 | UBR4 | chr1 | - | 19148562 | 19148626 | 19150576 | 19150793 | 19147992 | 19148127 | 19151642 | 19151859 | 5 | 8 | 0 | 8 | 4 | 3 | 0 | 3 | 0 | 1 | 98 | 98 | 1 | 1 | 0.385 | 0.667 | 0 | NA | | 0.75 | -0.399 |
| 619 | APAF1 | APAF1 | chr12 | + | 98725414 | 98725540 | 98727172 | 98727316 | 98723638 | 98723764 | 98732419 | 98735433 | 15 | 4 | 5 | 32 | 10 | 4 | 0 | 2 | 0 | 4 | 98 | 98 | 1 | 1 | 0.319 | 0.286 | 0.556 | NA | | 0.333 | 0.054 |
| 622 | NDUFB5 | NDUFB5 | chr3 | + | 179616982 | 179617044 | 179618414 | 179618521 | 179615982 | 179616049 | 179623919 | 179624500 | 8 | 0 | 3 | 5 | 0 | 0 | 0 | 5 | 1 | 5 | 98 | 98 | 1 | 1 | 0.615 | NA | 1 | | 0 | 0.5 | 0.557 |
| 626 | PRUNE | PRUNE | chr1 | + | 151017811 | 151017904 | 151025514 | 151025673 | 151008390 | 151008671 | 151027232 | 151027327 | 2 | 0 | 0 | 5 | 3 | 0 | 0 | 0 | 0 | 1 | 98 | 98 | 1 | 1 | 0.286 | 0 | NA | NA | | 0 | 0.143 |
| 627 | PRUNE | PRUNE | chr1 | + | 151018466 | 151018669 | 151025514 | 151025673 | 151017811 | 151017904 | 151027232 | 151027327 | 0 | 0 | 3 | 5 | 3 | 0 | 0 | 1 | 0 | 1 | 98 | 98 | 1 | 1 | 0 | 0 | 1 | NA | | 0.5 | -0.167 |
| 630 | SRP19 | SRP19 | chr5 | + | 112862507 | 112862583 | 112864456 | 112864528 | 112861187 | 112861417 | 112864620 | 112864732 | 1 | 0 | 0 | 2 | 2 | 0 | 0 | 1 | 0 | 0 | 98 | 98 | 1 | 1 | 0.333 | 0 | NA | NA | | 1 | -0.834 |
| 632 | TK2 | TK2 | chr16 | - | 66536963 | 66537017 | 66541878 | 66541953 | 66531379 | 66531469 | 66548977 | 66549009 | 5 | 0 | 0 | 23 | 4 | 0 | 0 | 2 | 0 | 2 | 98 | 98 | 1 | 1 | 0.179 | 0 | NA | NA | | 0.5 | -0.41 |
| 634 | OPA1 | OPA1 | chr3 | + | 193617177 | 193617285 | 193617783 | 193617837 | 193615673 | 193615770 | 193618868 | 193618936 | 11 | 0 | 1 | 3 | 0 | 0 | 0 | 2 | 0 | 0 | 98 | 98 | 1 | 1 | 0.786 | NA | 1 | NA | | 1 | -0.107 |
| 640 | DMXL2 | DMXL2 | chr15 | - | 51464674 | 51464876 | 51465565 | 51465651 | 51463378 | 51463496 | 51466183 | 51466311 | 33 | 5 | 3 | 25 | 6 | 3 | 0 | 9 | 0 | 8 | 98 | 98 | 1 | 1 | 0.569 | 0.455 | 0.5 | NA | | 0.529 | -0.021 |
| 642 | TMCO3 | TMCO3 | chr13 | + | 113503495 | 113503588 | 113510237 | 113510424 | 113501751 | 113501880 | 113520615 | 113520733 | 5 | 3 | 3 | 1 | 2 | 4 | 0 | 1 | 0 | 3 | 98 | 98 | 1 | 1 | 0.833 | 0.6 | 0.429 | NA | | 0.25 | 0.371 |
| 644 | C11orf21 | C11orf21 | chr11 | - | 2300519 | 2300613 | 2300683 | 2300768 | 2299427 | 2299707 | 2301755 | 2301913 | 2 | 0 | 0 | 6 | 0 | 0 | 0 | 0 | 0 | 1 | 98 | 98 | 1 | 1 | 0.25 | NA | NA | NA | | 0 | 0.25 |
| 648 | DAP3 | DAP3 | chr1 | + | 155717005 | 155717128 | 155721516 | 155721618 | 155709772 | 155709824 | 155725381 | 155725490 | 6 | 1 | 2 | 5 | 7 | 1 | 0 | 0 | 0 | 3 | 98 | 98 | 1 | 1 | 0.545 | 0.125 | 0.667 | NA | | 0 | 0.446 |
| 649 | CDA | CDA | chr1 | + | 20604927 | 20605039 | 20613841 | 20613899 | 20588950 | 20589283 | 20618451 | 20618907 | 15 | 12 | 15 | 26 | 7 | 10 | 0 | 14 | 0 | 3 | 98 | 98 | 1 | 1 | 0.366 | 0.632 | 0.6 | NA | | 0.824 | -0.291 |
| 656 | SFI1 | SFI1 | chr22 | + | 31528689 | 31528863 | 31546860 | 31546971 | 31508254 | 31508376 | 31550253 | 31550348 | 3 | 5 | 1 | 4 | 8 | 0 | 0 | 0 | 0 | 2 | 98 | 98 | 1 | 1 | 0.429 | 0.385 | 1 | NA | | 0 | 0.605 |
| 658 | MPHOSPH9 | MPHOSPH9 | chr12 | - | 123202623 | 123203084 | 123203113 | 123203375 | 123202163 | 123202319 | 123210055 | 123210162 | 1 | 0 | 0 | 4 | 0 | 0 | 0 | 0 | 0 | 1 | 98 | 98 | 1 | 1 | 0.2 | NA | NA | NA | | 0 | 0.2 |
| 660 | PAPD4 | PAPD4 | chr5 | + | 79641104 | 79641196 | 79642843 | 79642910 | 79640850 | 79640965 | 79645122 | 79645192 | 10 | 3 | 0 | 6 | 5 | 4 | 0 | 2 | 0 | 1 | 98 | 98 | 1 | 1 | 0.625 | 0.375 | 0 | NA | | 0.667 | -0.334 |
| 661 | PAPD4 | PAPD4 | chr5 | + | 79649061 | 79649190 | 79656957 | 79657001 | 79648616 | 79648693 | 79668891 | 79669028 | 18 | 4 | 0 | 13 | 3 | 0 | 0 | 0 | 0 | 4 | 98 | 93 | 1 | 1 | 0.568 | 0.559 | NA | NA | | 0 | 0.564 |
| 662 | ABCC5 | ABCC5 | chr3 | - | 183985303 | 183985378 | 183987769 | 183987917 | 183983752 | 183984894 | 183988571 | 183988727 | 8 | 9 | 0 | 1 | 0 | 0 | 2 | 1 | 0 | 0 | 98 | 98 | 1 | 1 | 0.889 | 1 | NA | | 1 | 1 | -0.055 |
| 663 | DPP8 | DPP8 | chr15 | - | 65450998 | 65451110 | 65451959 | 65452102 | 65445659 | 65447006 | 65456224 | 65456371 | 1 | 1 | 0 | 1 | 0 | 1 | 0 | 0 | 0 | 1 | 98 | 98 | 1 | 1 | 0.5 | 1 | 0 | NA | | 0 | 0.5 |

| | | | | | | | | | | | | | | | | | | | | | | | | | | | | | | | | |
|---|---|---|---|---|---|---|---|---|---|---|---|---|---|---|---|---|---|---|---|---|---|---|---|---|---|---|---|---|---|---|---|---|
| 678 | MFN2 | MFN2 | chr1 | + | 11981969 | 11982114 | 11989164 | 11989343 | 11980180 | 11980484 | 11992554 | 11992690 | 7 | 3 | 0 | 29 | 10 | 0 | 0 | 1 | 0 | 7 | 98 | 98 | 1 | 1 | 0.194 | 0.231 | NA | NA | 0.125 | 0.088 |
| 679 | IST1 | IST1 | chr16 | + | 71923287 | 71923380 | 71924768 | 71924817 | 71922473 | 71922680 | 71927613 | 71930637 | 20 | 8 | 6 | 4 | 5 | 3 | 0 | 1 | 0 | 1 | 98 | 98 | 1 | 1 | 0.833 | 0.615 | 0.667 | NA | 0.5 | 0.205 |
| 690 | TPM3 | TPM3 | chr1 | - | 154171412 | 154171488 | 154172028 | 154172104 | 154170648 | 154170711 | 154172907 | 154172978 | 159 | 96 | 73 | 3 | 0 | 0 | 2 | 34 | 0 | 0 | 98 | 98 | 1 | 1 | 0.981 | 1 | 1 | 1 | 1 | -0.006 |
| 693 | FAM126B | FAM126B | chr2 | - | 200992886 | 200993003 | 200997444 | 200997540 | 200992280 | 200992368 | 201008972 | 201009088 | 39 | 9 | 6 | 39 | 11 | 3 | 0 | 6 | 0 | 10 | 98 | 98 | 1 | 1 | 0.5 | 0.45 | 0.667 | NA | 0.375 | 0.164 |
| 695 | FAM53B | FAM53B | chr10 | - | 124696157 | 124696212 | 124706635 | 124706887 | 124681606 | 124682379 | 124744012 | 124744361 | 12 | 3 | 0 | 4 | 6 | 3 | 0 | 4 | 0 | 0 | 98 | 98 | 1 | 1 | 0.75 | 0.333 | 0 | NA | 1 | -0.639 |
| 696 | 6-Mar | 6-Mar | chr5 | + | 10386993 | 10387066 | 10390331 | 10390500 | 10378758 | 10378832 | 10391541 | 10391731 | 1 | 0 | 0 | 8 | 1 | 0 | 0 | 2 | 0 | 4 | 98 | 98 | 1 | 1 | 0.111 | 0 | NA | NA | 0.333 | -0.278 |
| 698 | RFWD2 | RFWD2 | chr1 | - | 176135009 | 176135086 | 176149005 | 176149074 | 176116623 | 176116681 | 176175909 | 176176007 | 3 | 1 | 0 | 118 | 57 | 9 | 0 | 1 | 1 | 17 | 98 | 98 | 1 | 1 | 0.025 | 0.017 | 0 | 0 | 0.056 | -0.014 |
| 701 | RFWD2 | RFWD2 | chr1 | - | 176149005 | 176149074 | 176175909 | 176176007 | 176135009 | 176135086 | 176184632 | 176184692 | 32 | 16 | 3 | 29 | 7 | 0 | 0 | 0 | 0 | 4 | 98 | 98 | 1 | 1 | 0.525 | 0.696 | 1 | NA | 0 | 0.74 |
| 702 | KIAA1551 | KIAA1551 | chr12 | + | 31960776 | 31960871 | 31970188 | 31970356 | 31959418 | 31959491 | 31980877 | 31985957 | 45 | 20 | 11 | 204 | 96 | 57 | 0 | 5 | 0 | 41 | 98 | 98 | 1 | 1 | 0.181 | 0.172 | 0.162 | NA | 0.109 | 0.063 |
| 703 | KIAA1551 | KIAA1551 | chr12 | + | 31970188 | 31970356 | 31980877 | 31985957 | 31960776 | 31960871 | 31987238 | 31987322 | 65 | 52 | 41 | 95 | 43 | 25 | 0 | 18 | 0 | 9 | 98 | 98 | 1 | 1 | 0.406 | 0.547 | 0.621 | NA | 0.667 | -0.142 |
| 704 | KIAA1551 | KIAA1551 | chr12 | + | 31980877 | 31985957 | 31987238 | 31987322 | 31970188 | 31970356 | 31992377 | 31993107 | 203 | 97 | 57 | 69 | 25 | 16 | 0 | 41 | 0 | 6 | 98 | 98 | 1 | 1 | 0.746 | 0.795 | 0.781 | NA | 0.872 | -0.098 |
| 711 | CSDE1 | CSDE1 | chr1 | - | 114739691 | 114739890 | 114749820 | 114750207 | 114737962 | 114738072 | 114757924 | 114758050 | 3 | 3 | 0 | 41 | 19 | 5 | 0 | 0 | 0 | 6 | 98 | 98 | 1 | 1 | 0.068 | 0.136 | 0 | NA | 0 | 0.068 |
| 715 | PPP2R3C | PPP2R3C | chr14 | - | 35091069 | 35091207 | 35095047 | 35095184 | 35087950 | 35088010 | 35096557 | 35096633 | 3 | 1 | 1 | 9 | 5 | 4 | 0 | 3 | 0 | 2 | 98 | 98 | 1 | 1 | 0.25 | 0.167 | 0.2 | NA | 0.6 | -0.394 |
| 716 | FNDC3B | FNDC3B | chr3 | + | 172247532 | 172247776 | 172251259 | 172251541 | 172226870 | 172226947 | 172285925 | 172285984 | 20 | 6 | 8 | 18 | 4 | 7 | 0 | 1 | 0 | 5 | 98 | 98 | 1 | 1 | 0.526 | 0.6 | 0.533 | NA | 0.167 | 0.386 |
| 725 | ZNF687 | ZNF687 | chr1 | + | 151288527 | 151288706 | 151289094 | 151289271 | 151286274 | 151288406 | 151289377 | 151289540 | 4 | 1 | 5 | 6 | 0 | 2 | 0 | 0 | 0 | 2 | 98 | 98 | 1 | 1 | 0.4 | 1 | 0.714 | NA | 0 | 0.705 |
| 726 | CD44 | CD44 | chr11 | + | 35209964 | 35210054 | 35211245 | 35211449 | 35208104 | 35208206 | 35214851 | 35214914 | 5 | 0 | 0 | 6 | 0 | 0 | 0 | 4 | 0 | 1 | 98 | 98 | 1 | 1 | 0.455 | NA | NA | NA | 0.8 | -0.345 |
| 729 | LRRK2 | LRRK2 | chr12 | + | 40278090 | 40278261 | 40283874 | 40284133 | 40277887 | 40278016 | 40287350 | 40287539 | 102 | 91 | 48 | 177 | 94 | 32 | 0 | 22 | 0 | 29 | 98 | 98 | 1 | 1 | 0.366 | 0.492 | 0.6 | NA | 0.431 | 0.055 |
| 737 | FOXJ3 | FOXJ3 | chr1 | - | 42199101 | 42199230 | 42205761 | 42205863 | 42194889 | 42195064 | 42227882 | 42227966 | 2 | 1 | 0 | 8 | 0 | 0 | 0 | 1 | 0 | 0 | 98 | 98 | 1 | 1 | 0.2 | 1 | NA | NA | 1 | -0.4 |
| 738 | CYB561D1 | CYB561D1 | chr1 | + | 109494519 | 109494623 | 109495142 | 109495180 | 109494078 | 109494287 | 109495755 | 109500441 | 1 | 0 | 0 | 1 | 0 | 0 | 0 | 0 | 0 | 1 | 98 | 87 | 1 | 1 | 0.47 | NA | NA | NA | 0 | 0.47 |
| 739 | EPB41 | EPB41 | chr1 | + | 28993329 | 28993542 | 28997214 | 28997319 | 28987430 | 28987905 | 29011864 | 29011907 | 46 | 19 | 5 | 27 | 7 | 7 | 0 | 2 | 0 | 0 | 98 | 98 | 1 | 1 | 0.63 | 0.731 | 0.417 | NA | 1 | -0.407 |
| 740 | EPB41 | EPB41 | chr1 | + | 29058588 | 29058645 | 29060421 | 29060484 | 29053103 | 29053312 | 29064981 | 29065158 | 0 | 2 | 0 | 3 | 0 | 0 | 0 | 1 | 0 | 0 | 98 | 98 | 1 | 1 | 0 | 1 | NA | NA | 1 | -0.5 |
| 741 | EPB41 | EPB41 | chr1 | + | 29097806 | 29097935 | 29109335 | 29109437 | 29064981 | 29065158 | 29112367 | 29112448 | 31 | 14 | 5 | 13 | 12 | 5 | 0 | 8 | 0 | 0 | 98 | 98 | 1 | 1 | 0.705 | 0.538 | 0.5 | NA | 1 | -0.419 |
| 748 | RNF19B | RNF19B | chr1 | - | 32945513 | 32945628 | 32946401 | 32946564 | 32944018 | 32944159 | 32948221 | 32948363 | 67 | 25 | 18 | 53 | 35 | 4 | 0 | 7 | 0 | 12 | 98 | 98 | 1 | 1 | 0.558 | 0.417 | 0.818 | NA | 0.368 | 0.23 |
| 759 | DEDD2 | DEDD2 | chr19 | - | 42201988 | 42202072 | 42204461 | 42204559 | 42198592 | 42199829 | 42209699 | 42209840 | 0 | 0 | 0 | 2 | 0 | 1 | 1 | 0 | 1 | 2 | 98 | 98 | 1 | 1 | 0 | NA | 0 | 0.5 | 0 | -0.25 |
| 760 | DEDD2 | DEDD2 | chr19 | - | 42202050 | 42202072 | 42204461 | 42204559 | 42198592 | 42199829 | 42209699 | 42209840 | 0 | 0 | 0 | 3 | 0 | 1 | 1 | 0 | 0 | 3 | 98 | 71 | 1 | 1 | 0 | NA | 0 | 1 | 0 | -0.5 |
| 763 | RAB6A | RAB6A | chr11 | - | 73718612 | 73718718 | 73718784 | 73718890 | 73716250 | 73716362 | 73720845 | 73720899 | 0 | 0 | 1 | 11 | 3 | 4 | 0 | 0 | 0 | 2 | 98 | 98 | 1 | 1 | 0 | 0 | 0.2 | NA | 0 | 0.067 |
| 765 | ACAA1 | ACAA1 | chr3 | - | 38129289 | 38129388 | 38131595 | 38131638 | 38127785 | 38127866 | 38131925 | 38132005 | 3 | 0 | 3 | 9 | 8 | 2 | 0 | 0 | 0 | 2 | 92 | 98 | 1 | 1 | 0.262 | 0 | 0.615 | NA | 0 | 0.292 |
| 766 | MAP3K11 | MAP3K11 | chr11 | - | 65607921 | 65608070 | 65608267 | 65608448 | 65607640 | 65607816 | 65613017 | 65614249 | 25 | 18 | 11 | 17 | 10 | 11 | 0 | 4 | 0 | 7 | 98 | 98 | 1 | 1 | 0.595 | 0.643 | 0.5 | NA | 0.364 | 0.215 |
| 769 | PECAM1 | PECAM1 | chr17 | - | 64341633 | 64341690 | 64348259 | 64348322 | 64329699 | 64329722 | 64350379 | 64350433 | 42 | 38 | 40 | 104 | 68 | 31 | 0 | 9 | 0 | 11 | 98 | 98 | 1 | 1 | 0.288 | 0.358 | 0.563 | NA | 0.45 | -0.047 |
| 770 | CD86 | CD86 | chr3 | + | 122091600 | 122091650 | 122103511 | 122103847 | 122055361 | 122055503 | 122106197 | 122106500 | 3 | 0 | 0 | 2 | 0 | 0 | 0 | 1 | 0 | 0 | 98 | 98 | 1 | 1 | 0.6 | NA | NA | NA | 1 | -0.4 |
| 772 | DCLRE1C | DCLRE1C | chr10 | - | 14945104 | 14945189 | 14949035 | 14949087 | 14939809 | 14939869 | 14953901 | 14954432 | 4 | 1 | 0 | 6 | 1 | 0 | 0 | 2 | 0 | 0 | 98 | 98 | 1 | 1 | 0.4 | 0.5 | NA | NA | 1 | -0.55 |
| 775 | ELOVL1 | ELOVL1 | chr1 | - | 43364927 | 43365008 | 43365185 | 43365376 | 43364737 | 43364794 | 43365563 | 43365623 | 0 | 3 | 0 | 6 | 2 | 0 | 0 | 0 | 0 | 5 | 98 | 98 | 1 | 1 | 0 | 0.6 | NA | NA | 0 | 0.3 |
| 776 | NLRP3 | NLRP3 | chr1 | + | 247429584 | 247429755 | 247434102 | 247434273 | 247423846 | 247425599 | 247443971 | 247444142 | 8 | 0 | 0 | 2 | 0 | 0 | 0 | 4 | 0 | 2 | 98 | 98 | 1 | 1 | 0.8 | NA | NA | NA | 0.667 | 0.133 |
| 780 | CASP8 | CASP8 | chr2 | + | 201272897 | 201272942 | 201274888 | 201274953 | 201272637 | 201272776 | 201276826 | 201276968 | 106 | 49 | 12 | 89 | 36 | 17 | 0 | 6 | 0 | 3 | 94 | 98 | 1 | 1 | 0.554 | 0.587 | 0.424 | NA | 0.676 | -0.154 |
| 796 | ABI1 | ABI1 | chr10 | - | 26755654 | 26755741 | 26759061 | 26759235 | 26751597 | 26751783 | 26765217 | 26765318 | 9 | 13 | 1 | 0 | 0 | 0 | 0 | 12 | 0 | 1 | 98 | 98 | 1 | 1 | 1 | 1 | 1 | NA | 0.923 | 0.077 |
| 797 | ABI1 | ABI1 | chr10 | - | 26755654 | 26755741 | 26759061 | 26759238 | 26751597 | 26751783 | 26765217 | 26765318 | 12 | 12 | 2 | 0 | 0 | 0 | 0 | 11 | 0 | 1 | 98 | 98 | 1 | 1 | 1 | 1 | 1 | NA | 0.917 | 0.083 |
| 799 | RFC2 | RFC2 | chr7 | - | 74246661 | 74246763 | 74249011 | 74249220 | 74243145 | 74243246 | 74249738 | 74249780 | 1 | 0 | 0 | 5 | 3 | 2 | 0 | 0 | 0 | 1 | 98 | 98 | 1 | 1 | 0.167 | 0 | 0 | NA | 0 | 0.056 |
| 805 | CNIH4 | CNIH4 | chr1 | + | 224360494 | 224360563 | 224364108 | 224364455 | 224356810 | 224356993 | 224365878 | 224365991 | 3 | 2 | 7 | 2 | 1 | 0 | 0 | 2 | 0 | 0 | 98 | 98 | 1 | 1 | 0.6 | 0.667 | 1 | NA | 1 | -0.244 |
| 816 | POGZ | POGZ | chr1 | - | 151428122 | 151428413 | 151430665 | 151430841 | 151424954 | 151425061 | 151440927 | 151441086 | 0 | 0 | 0 | 1 | 0 | 0 | 0 | 3 | 0 | 0 | 98 | 98 | 1 | 1 | 0 | NA | NA | NA | 1 | -1 |
| 817 | POGZ | POGZ | chr1 | - | 151430665 | 151430841 | 151440927 | 151441086 | 151428122 | 151428413 | 151442080 | 151442205 | 4 | 0 | 0 | 1 | 0 | 0 | 0 | 2 | 0 | 0 | 98 | 98 | 1 | 1 | 0.8 | NA | NA | NA | 1 | -0.2 |
| 822 | IFI44 | IFI44 | chr1 | + | 78660553 | 78660654 | 78662703 | 78662878 | 78659311 | 78659483 | 78663764 | 78664078 | 5 | 1 | 0 | 12 | 6 | 0 | 0 | 31 | 6 | 51 | 98 | 98 | 1 | 1 | 0.294 | 0.143 | NA | 0 | 0.378 | 0.029 |
| 825 | ZCCHC2 | ZCCHC2 | chr18 | + | 62556202 | 62556297 | 62558686 | 62558770 | 62550347 | 62550460 | 62560586 | 62560644 | 23 | 29 | 2 | 12 | 5 | 2 | 0 | 16 | 0 | 9 | 98 | 98 | 1 | 1 | 0.657 | 0.853 | 0.5 | NA | 0.64 | 0.03 |
| 828 | TLR5 | TLR5 | chr1 | - | 223137177 | 223137263 | 223141647 | 223141763 | 223134681 | 223134864 | 223143195 | 223143282 | 3 | 1 | 1 | 3 | 0 | 1 | 0 | 0 | 0 | 4 | 98 | 98 | 1 | 1 | 0.5 | 1 | 0.5 | NA | 0 | 0.667 |
| 835 | CASP5 | CASP5 | chr11 | - | 105007082 | 105007334 | 105008806 | 105008980 | 105003273 | 105003383 | 105023129 | 105023168 | 1 | 0 | 0 | 10 | 0 | 0 | 0 | 0 | 0 | 2 | 98 | 98 | 1 | 1 | 0.091 | NA | NA | NA | 0 | 0.091 |
| 836 | CASP5 | CASP5 | chr11 | - | 105007082 | 105007334 | 105008806 | 105009019 | 105003273 | 105003383 | 105023129 | 105023168 | 1 | 0 | 0 | 10 | 0 | 0 | 0 | 0 | 0 | 2 | 98 | 98 | 1 | 1 | 0.091 | NA | NA | NA | 0 | 0.091 |
| 846 | DENND5A | DENND5A | chr11 | - | 9142721 | 9142845 | 9143402 | 9143485 | 9141939 | 9142108 | 9144096 | 9144278 | 36 | 15 | 7 | 65 | 20 | 10 | 0 | 7 | 0 | 11 | 98 | 98 | 1 | 1 | 0.356 | 0.429 | 0.412 | NA | 0.389 | 0.01 |
| 848 | WDR37 | WDR37 | chr10 | + | 1072115 | 1072293 | 1077906 | 1078003 | 1056835 | 1056968 | 1080010 | 1080106 | 2 | 4 | 2 | 9 | 2 | 6 | 0 | 0 | 0 | 3 | 98 | 98 | 1 | 1 | 0.182 | 0.667 | 0.25 | NA | 0 | 0.366 |
| 849 | CLTA | CLTA | chr9 | + | 36197550 | 36197588 | 36198978 | 36199096 | 36190855 | 36191273 | 36204067 | 36204179 | 11 | 0 | 2 | 3 | 1 | 4 | 0 | 0 | 0 | 4 | 87 | 98 | 1 | 1 | 0.805 | 0 | 0.36 | NA | 0 | 0.388 |
| 852 | FBXO18 | FBXO18 | chr10 | + | 5918341 | 5918478 | 5921257 | 5921357 | 5917589 | 5917676 | 5921447 | 5921569 | 4 | 4 | 1 | 6 | 5 | 0 | 0 | 1 | 0 | 0 | 98 | 98 | 1 | 1 | 0.4 | 0.444 | 1 | NA | 1 | -0.385 |
| 853 | CELF2 | CELF2 | chr10 | + | 11266597 | 11266677 | 11270665 | 11270824 | 11257737 | 11257872 | 11275056 | 11275120 | 20 | 6 | 1 | 104 | 45 | 14 | 0 | 9 | 0 | 11 | 98 | 98 | 1 | 1 | 0.161 | 0.118 | 0.067 | NA | 0.45 | -0.335 |
| 860 | DIS3 | DIS3 | chr13 | - | 72780845 | 72781003 | 72781288 | 72781356 | 72778186 | 72778380 | 72781604 | 72782128 | 1 | 2 | 1 | 2 | 2 | 0 | 0 | 0 | 0 | 1 | 98 | 98 | 1 | 1 | 0.333 | 0.5 | 1 | NA | 0 | 0.611 |
| 861 | BPTF | BPTF | chr17 | + | 67875555 | 67875744 | 67886127 | 67886316 | 67874816 | 67875020 | 67891843 | 67892034 | 6 | 1 | 0 | 10 | 0 | 0 | 0 | 1 | 0 | 3 | 98 | 98 | 1 | 1 | 0.375 | 1 | NA | NA | 0.25 | 0.438 |
| 868 | OGDH | OGDH | chr7 | + | 44647443 | 44647534 | 44647656 | 44647759 | 44645326 | 44645518 | 44666735 | 44666851 | 1 | 5 | 0 | 11 | 7 | 4 | 0 | 2 | 0 | 0 | 98 | 98 | 1 | 1 | 0.083 | 0.417 | 0 | NA | 1 | -0.833 |
| 870 | MBNL1 | MBNL1 | chr3 | + | 152455541 | 152455577 | 152456266 | 152456361 | 152447619 | 152447773 | 152459270 | 152459345 | 17 | 1 | 8 | 35 | 7 | 9 | 0 | 10 | 0 | 5 | 85 | 98 | 1 | 1 | 0.359 | 0.141 | 0.506 | NA | 0.698 | -0.363 |
| 872 | PIK3CD | PIK3CD | chr1 | + | 9723968 | 9724092 | 9724275 | 9724421 | 9723124 | 9723292 | 9724803 | 9724936 | 34 | 17 | 14 | 43 | 22 | 17 | 0 | 3 | 0 | 6 | 98 | 98 | 1 | 1 | 0.442 | 0.436 | 0.452 | NA | 0.333 | 0.11 |
| 875 | FBXO28 | FBXO28 | chr1 | + | 224134073 | 224134212 | 224153141 | 224153337 | 224130471 | 224130581 | 224157351 | 224162047 | 13 | 3 | 5 | 1 | 1 | 0 | 0 | 3 | 0 | 0 | 98 | 98 | 1 | 1 | 0.929 | 0.75 | 1 | NA | 1 | -0.107 |
| 878 | TXNRD1 | TXNRD1 | chr12 | + | 104288930 | 104289040 | 104311289 | 104311412 | 104286681 | 104287109 | 104313244 | 104313317 | 0 | 1 | 0 | 27 | 20 | 2 | 0 | 0 | 0 | 2 | 98 | 98 | 1 | 1 | 0 | 0.048 | 0 | NA | 0 | 0.016 |
| 879 | LDLR | LDLR | chr19 | + | 11102663 | 11102786 | 11105219 | 11105600 | 11100222 | 11100345 | 11106564 | 11106687 | 2 | 0 | 0 | 1 | 0 | 0 | 0 | 2 | 0 | 0 | 98 | 98 | 1 | 1 | 0.667 | NA | NA | NA | 1 | -0.333 |
| 882 | ZNF33A | ZNF33A | chr10 | + | 38025347 | 38025411 | 38039412 | 38039492 | 38017290 | 38017386 | 38054374 | 38060067 | 1 | 0 | 1 | 0 | 0 | 0 | 0 | 0 | 0 | 2 | 98 | 98 | 1 | 1 | 1 | NA | 1 | NA | 0 | 1 |
| 883 | ZNF33A | ZNF33A | chr10 | + | 38034277 | 38034400 | 38039412 | 38039492 | 38017290 | 38017386 | 38054374 | 38060067 | 0 | 0 | 1 | 0 | 0 | 0 | 0 | 0 | 0 | 2 | 98 | 98 | 1 | 1 | NA | NA | 1 | NA | 0 | 1 |
| 884 | ZNF33A | ZNF33A | chr10 | + | 38034325 | 38034400 | 38039412 | 38039492 | 38017290 | 38017386 | 38054374 | 38060067 | 0 | 1 | 1 | 0 | 0 | 0 | 0 | 0 | 0 | 2 | 98 | 98 | 1 | 1 | NA | 1 | 1 | NA | 0 | 1 |
| 890 | CAPRIN2 | CAPRIN2 | chr12 | - | 30719078 | 30719225 | 30720810 | 30720915 | 30716507 | 30716676 | 30723258 | 30723314 | 1 | 0 | 0 | 0 | 3 | 0 | 0 | 1 | 0 | 0 | 98 | 98 | 1 | 1 | 1 | 0 | NA | NA | 1 | -0.5 |
| 891 | XRN1 | XRN1 | chr3 | - | 142412543 | 142412663 | 142414134 | 142414291 | 142404906 | 142405076 | 142417139 | 142417229 | 21 | 11 | 2 | 16 | 14 | 1 | 0 | 4 | 0 | 7 | 98 | 98 | 1 | 1 | 0.568 | 0.44 | 0.667 | NA | 0.364 | 0.194 |
| 892 | ELP2 | ELP2 | chr18 | + | 36138794 | 36138872 | 36139418 | 36139613 | 36138269 | 36138426 | 36141136 | 36141201 | 7 | 2 | 0 | 0 | 5 | 0 | 0 | 4 | 0 | 0 | 98 | 98 | 1 | 1 | 1 | 0.286 | NA | NA | 1 | -0.357 |
| 905 | MCTP1 | MCTP1 | chr5 | - | 94917895 | 94917973 | 94931952 | 94931991 | 94912805 | 94912976 | 94940083 | 94940195 | 25 | 9 | 6 | 8 | 15 | 2 | 0 | 6 | 0 | 3 | 88 | 98 | 1 | 1 | 0.777 | 0.401 | 0.77 | NA | 0.69 | -0.041 |
| 912 | RELT | RELT | chr11 | + | 73391143 | 73391223 | 73392210 | 73392468 | 73390754 | 73390921 | 73393836 | 73393917 | 9 | 6 | 8 | 23 | 10 | 6 | 0 | 3 | 0 | 3 | 98 | 98 | 1 | 1 | 0.281 | 0.375 | 0.571 | NA | 0.5 | -0.091 |
| 926 | RCBTB2 | RCBTB2 | chr13 | - | 48521897 | 48521962 | 48522307 | 48522403 | 48515585 | 48515741 | 48524658 | 48524757 | 2 | 1 | 0 | 6 | 3 | 0 | 0 | 0 | 0 | 2 | 98 | 98 | 1 | 1 | 0.25 | 0.25 | NA | NA | 0 | 0.25 |
| 927 | BAZ2B | BAZ2B | chr2 | - | 159448241 | 159448403 | 159453612 | 159453801 | 159446781 | 159446975 | 159478574 | 159478721 | 36 | 15 | 2 | 37 | 22 | 5 | 0 | 6 | 0 | 9 | 98 | 98 | 1 | 1 | 0.493 | 0.405 | 0.286 | NA | 0.4 | -0.005 |
| 928 | BAZ2B | BAZ2B | chr2 | - | 159448241 | 159448409 | 159453612 | 159453801 | 159446781 | 159446975 | 159478574 | 159478721 | 36 | 15 | 2 | 43 | 28 | 5 | 0 | 6 | 0 | 14 | 98 | 98 | 1 | 1 | 0.456 | 0.349 | 0.286 | NA | 0.3 | 0.064 |
| 929 | MLKL | MLKL | chr16 | - | 74678898 | 74678980 | 74682650 | 74682786 | 74675612 | 74675764 | 74685485 | 74685583 | 17 | 2 | 0 | 6 | 1 | 8 | 0 | 3 | 0 | 0 | 98 | 98 | 1 | 1 | 0.739 | 0.667 | 0 | NA | 1 | -0.531 |
| 934 | GBF1 | GBF1 | chr10 | + | 102368217 | 102368454 | 102368738 | 102368832 | 102367477 | 102367560 | 102369210 | 102369387 | 7 | 1 | 0 | 10 | 3 | 1 | 0 | 2 | 0 | 4 | 98 | 98 | 1 | 1 | 0.412 | 0.25 | 0 | NA | 0.333 | -0.112 |
| 939 | NT5C2 | NT5C2 | chr10 | - | 103106588 | 103106706 | 103139405 | 103139479 | 103105705 | 103105801 | 103174857 | 103174982 | 36 | 30 | 8 | 43 | 15 | 11 | 0 | 7 | 0 | 7 | 98 | 98 | 1 | 1 | 0.456 | 0.667 | 0.421 | NA | 0.5 | 0.015 |
| 940 | USMG5 | USMG5 | chr10 | - | 103394198 | 103394394 | 103395745 | 103396032 | 103392370 | 103392466 | 103396408 | 103396513 | 1 | 0 | 0 | 0 | 0 | 0 | 0 | 0 | 0 | 1 | 98 | 98 | 1 | 1 | 1 | NA | NA | NA | 0 | 1 |
| 946 | CAMK2G | CAMK2G | chr10 | - | 73847224 | 73847347 | 73847987 | 73848082 | 73842457 | 73842541 | 73848525 | 73848609 | 13 | 8 | 5 | 15 | 1 | 2 | 0 | 0 | 0 | 2 | 98 | 98 | 1 | 1 | 0.464 | 0.889 | 0.714 | NA | 0 | 0.689 |
| 947 | CAMK2G | CAMK2G | chr10 | - | 73847987 | 73848082 | 73848525 | 73848609 | 73847224 | 73847347 | 73849012 | 73849115 | 23 | 16 | 17 | 13 | 15 | 1 | 0 | 2 | 0 | 3 | 98 | 98 | 1 | 1 | 0.639 | 0.516 | 0.944 | NA | 0.4 | 0.3 |
| 952 | ATE1 | ATE1 | chr10 | - | 121898840 | 121898969 | 121899865 | 121899994 | 121870005 | 121870038 | 121902390 | 121902620 | 4 | 0 | 0 | 3 | 0 | 0 | 0 | 1 | 0 | 0 | 98 | 98 | 1 | 1 | 0.571 | NA | NA | NA | 1 | -0.429 |
| 953 | ATE1 | ATE1 | chr10 | - | 121913789 | 121913893 | 121922348 | 121922411 | 121910905 | 121911151 | 121924265 | 121924329 | 2 | 0 | 0 | 15 | 6 | 0 | 0 | 1 | 0 | 0 | 98 | 98 | 1 | 1 | 0.118 | 0 | NA | NA | 1 | -0.941 |
| 954 | DNM2 | DNM2 | chr19 | + | 10796060 | 10796199 | 10797379 | 10797518 | 10795371 | 10795439 | 10798485 | 10798572 | 12 | 5 | 1 | 10 | 6 | 8 | 0 | 2 | 0 | 4 | 98 | 98 | 1 | 1 | 0.545 | 0.455 | 0.111 | NA | 0.333 | 0.037 |
| 955 | FAM134C | FAM134C | chr17 | - | 42586764 | 42586891 | 42587833 | 42587864 | 42586052 | 42586137 | 42592055 | 42592162 | 11 | 2 | 0 | 3 | 0 | 2 | 0 | 1 | 0 | 0 | 80 | 98 | 1 | 1 | 0.818 | 1 | 0 | NA | 1 | -0.394 |
| 958 | GSR | GSR | chr8 | - | 30689160 | 30689319 | 30692968 | 30693055 | 30684087 | 30684199 | 30696379 | 30696479 | 11 | 1 | 1 | 6 | 5 | 7 | 0 | 2 | 0 | 3 | 98 | 98 | 1 | 1 | 0.647 | 0.167 | 0.125 | NA | 0.4 | -0.087 |
| 974 | LST1 | LST1 | chr6 | + | 31587640 | 31587733 | 31587943 | 31587966 | 31587199 | 31587318 | 31588517 | 31588909 | 42 | 28 | 26 | 17 | 17 | 14 | 0 | 0 | 0 | 7 | 98 | 72 | 1 | 1 | 0.645 | 0.548 | 0.577 | NA | 0 | 0.59 |
| 979 | MAPK14 | MAPK14 | chr6 | + | 36095986 | 36096066 | 36100187 | 36100267 | 36076536 | 36076608 | 36102570 | 36102649 | 31 | 21 | 14 | 18 | 11 | 4 | 0 | 8 | 0 | 1 | 98 | 98 | 1 | 1 | 0.633 | 0.656 | 0.778 | NA | 0.889 | -0.2 |
| 985 | DYNC1LI2 | DYNC1LI2 | chr16 | - | 66742437 | 66742668 | 66749196 | 66749313 | 66736074 | 66736244 | 66751272 | 66751346 | 19 | 2 | 0 | 2 | 5 | 1 | 0 | 2 | 0 | 1 | 98 | 98 | 1 | 1 | 0.905 | 0.286 | 0 | NA | 0.667 | -0.27 |
| 997 | GZMH | GZMH | chr14 | - | 24607340 | 24607406 | 24607611 | 24607747 | 24606479 | 24606746 | 24608264 | 24608412 | 8 | 9 | 0 | 0 | 0 | 0 | 1 | 0 | 0 | 3 | 98 | 98 | 1 | 1 | 1 | 1 | NA | 1 | 0 | 0.5 |
| 1000 | SDHD | SDHD | chr11 | + | 112087856 | 112087973 | 112088866 | 112089011 | 112086823 | 112086959 | 112094804 | 112095801 | 1 | 1 | 0 | 2 | 0 | 5 | 0 | 1 | 0 | 0 | 98 | 98 | 1 | 1 | 0.333 | 1 | 0 | NA | 1 | -0.556 |
| 1003 | PRR5L | PRR5L | chr11 | + | 36446299 | 36446440 | 36451208 | 36451335 | 36437384 | 36437476 | 36462341 | 36465204 | 4 | 5 | 1 | 1 | 0 | 0 | 0 | 1 | 0 | 0 | 98 | 98 | 1 | 1 | 0.8 | 1 | 1 | NA | 1 | -0.067 |
| 1010 | ATG13 | ATG13 | chr11 | + | 46618324 | 46618375 | 46630044 | 46630100 | 46617275 | 46617890 | 46644278 | 46644360 | 0 | 0 | 0 | 6 | 2 | 1 | 0 | 2 | 0 | 2 | 98 | 98 | 1 | 1 | 0 | 0 | 0 | NA | 0.5 | -0.5 |
| 1012 | TNFSF13 | TNFSF13 | chr17 | + | 7559623 | 7559702 | 7559845 | 7559893 | 7558291 | 7559297 | 7560048 | 7560167 | 6 | 3 | 0 | 12 | 3 | 1 | 0 | 4 | 0 | 8 | 98 | 97 | 1 | 1 | 0.331 | 0.497 | 0 | NA | 0.331 | -0.055 |
| 1013 | TNFSF13 | TNFSF13 | chr17 | + | 7559623 | 7559702 | 7559847 | 7559893 | 7558291 | 7559297 | 7560048 | 7560167 | 6 | 3 | 0 | 11 | 3 | 1 | 0 | 4 | 0 | 8 | 98 | 95 | 1 | 1 | 0.346 | 0.492 | 0 | NA | 0.326 | -0.047 |
| 1014 | TNFSF13 | TNFSF13 | chr17 | + | 7559623 | 7559702 | 7559850 | 7559893 | 7558291 | 7559297 | 7560048 | 7560167 | 6 | 3 | 0 | 13 | 2 | 1 | 0 | 4 | 0 | 9 | 98 | 92 | 1 | 1 | 0.302 | 0.585 | 0 | NA | 0.294 | 0.002 |
| 1016 | PHF21A | PHF21A | chr11 | - | 45938156 | 45938312 | 45945839 | 45946003 | 45936493 | 45936569 | 45948885 | 45948946 | 8 | 3 | 0 | 21 | 2 | 0 | 0 | 6 | 0 | 5 | 98 | 98 | 1 | 1 | 0.276 | 0.6 | NA | NA | 0.545 | -0.107 |
| 1025 | SRP54 | SRP54 | chr14 | + | 34999557 | 34999649 | 35000935 | 35001020 | 34996676 | 34996787 | 35007282 | 35007387 | 6 | 2 | 0 | 10 | 3 | 0 | 2 | 0 | 0 | 2 | 98 | 98 | 1 | 1 | 0.375 | 0.4 | NA | 1 | 0 | -0.112 |
| 1027 | ZEB1 | ZEB1 | chr10 | + | 31461036 | 31461237 | 31495778 | 31495838 | 31319171 | 31319292 | 31502347 | 31502509 | 2 | 0 | 0 | 8 | 2 | 0 | 0 | 0 | 0 | 2 | 98 | 98 | 1 | 1 | 0.2 | 0 | NA | NA | 0 | 0.1 |
| 1029 | ADGRE2 | ADGRE2 | chr19 | - | 14754953 | 14755127 | 14755653 | 14755877 | 14752328 | 14752526 | 14756237 | 14756345 | 6 | 6 | 12 | 20 | 5 | 0 | 0 | 2 | 0 | 3 | 98 | 98 | 1 | 1 | 0.231 | 0.545 | 1 | NA | 0.4 | 0.192 |
| 1034 | FRMD8 | FRMD8 | chr11 | + | 65389360 | 65389528 | 65393572 | 65393674 | 65387036 | 65387121 | 65394040 | 65394099 | 4 | 3 | 0 | 4 | 3 | 6 | 0 | 0 | 0 | 1 | 98 | 98 | 1 | 1 | 0.5 | 0.5 | 0 | NA | 0 | 0.333 |
| 1038 | TMEM135 | TMEM135 | chr11 | + | 87091361 | 87091395 | 87157340 | 87157406 | 87071522 | 87071615 | 87236637 | 87236684 | 1 | 0 | 0 | 0 | 0 | 0 | 0 | 0 | 1 | 0 | 83 | 98 | 1 | 1 | 1 | NA | NA | 0 | NA | 1 |
| 1039 | CLN3 | CLN3 | chr16 | - | 28487661 | 28487741 | 28488590 | 28488662 | 28487455 | 28487541 | 28489289 | 28489386 | 4 | 2 | 0 | 8 | 6 | 2 | 0 | 4 | 0 | 1 | 98 | 98 | 1 | 1 | 0.333 | 0.25 | 0 | NA | 0.8 | -0.606 |
| 1049 | CYB5R4 | CYB5R4 | chr6 | + | 83914415 | 83914448 | 83918004 | 83918065 | 83909008 | 83909090 | 83919396 | 83919454 | 65 | 27 | 25 | 68 | 29 | 20 | 0 | 10 | 0 | 6 | 82 | 98 | 1 | 1 | 0.533 | 0.527 | 0.599 | NA | 0.666 | -0.113 |
| 1052 | C11orf54 | C11orf54 | chr11 | + | 93755209 | 93755386 | 93757315 | 93757465 | 93753935 | 93754037 | 93759741 | 93759858 | 7 | 1 | 0 | 11 | 3 | 0 | 0 | 5 | 0 | 0 | 98 | 98 | 1 | 1 | 0.389 | 0.25 | NA | NA | 1 | -0.68 |
| 1053 | CD300LF | CD300LF | chr17 | - | 74696194 | 74696217 | 74698368 | 74698481 | 74695724 | 74695859 | 74703034 | 74703098 | 15 | 7 | 10 | 1 | 1 | 2 | 0 | 0 | 0 | 1 | 98 | 72 | 1 | 1 | 0.917 | 0.837 | 0.786 | NA | 0 | 0.847 |
| 1059 | MBD1 | MBD1 | chr18 | - | 50274185 | 50274353 | 50274976 | 50275045 | 50273563 | 50273863 | 50275128 | 50275245 | 7 | 0 | 1 | 0 | 4 | 0 | 0 | 1 | 0 | 0 | 98 | 98 | 1 | 1 | 1 | 0 | 1 | NA | 1 | -0.333 |
| 1060 | NDEL1 | NDEL1 | chr17 | + | 8445710 | 8445864 | 8446753 | 8446902 | 8444259 | 8444357 | 8448549 | 8448686 | 28 | 12 | 1 | 69 | 38 | 14 | 0 | 8 | 0 | 8 | 98 | 98 | 1 | 1 | 0.289 | 0.24 | 0.067 | NA | 0.5 | -0.301 |
| 1062 | PAM | PAM | chr5 | + | 103025130 | 103025334 | 103028184 | 103028238 | 103019789 | 103019843 | 103028886 | 103031104 | 2 | 3 | 0 | 2 | 2 | 0 | 0 | 1 | 0 | 0 | 98 | 98 | 1 | 1 | 0.5 | 0.6 | NA | NA | 1 | -0.45 |
| 1064 | AMPD3 | AMPD3 | chr11 | + | 10478525 | 10478730 | 10482062 | 10482225 | 10461514 | 10461740 | 10484819 | 10485039 | 3 | 0 | 0 | 18 | 5 | 0 | 0 | 0 | 0 | 6 | 98 | 98 | 1 | 1 | 0.143 | 0 | NA | NA | 0 | 0.071 |
| 1065 | LRRC75A-AS1 | LRRC75A-AS1 | chr17 | + | 16441073 | 16441130 | 16460892 | 16461245 | 16440184 | 16440253 | 16462832 | 16462941 | 1 | 0 | 3 | 0 | 0 | 0 | 0 | 0 | 0 | 1 | 98 | 98 | 1 | 1 | 1 | NA | 1 | NA | 0 | 1 |
| 1066 | TOM1L2 | TOM1L2 | chr17 | - | 17882704 | 17882863 | 17884633 | 17884768 | 17879626 | 17879743 | 17893660 | 17893810 | 4 | 0 | 1 | 4 | 1 | 1 | 0 | 3 | 0 | 0 | 98 | 98 | 1 | 1 | 0.5 | 0 | 0.5 | NA | 1 | -0.667 |
| 1067 | TOM1L2 | TOM1L2 | chr17 | - | 17884633 | 17884768 | 17893660 | 17893810 | 17882704 | 17882863 | 17898595 | 17898674 | 4 | 0 | 0 | 0 | 0 | 0 | 0 | 1 | 1 | 0 | 98 | 98 | 1 | 1 | 1 | NA | NA | 0 | 1 | 0.5 |
| 1068 | GYS1 | GYS1 | chr19 | - | 48987193 | 48987385 | 48991301 | 48991483 | 48985849 | 48986035 | 48992994 | 48993353 | 0 | 1 | 0 | 2 | 0 | 3 | 0 | 0 | 3 | 2 | 98 | 98 | 1 | 1 | 0 | 1 | 0 | 0 | 0 | 0.333 |
| 1069 | DHX8 | DHX8 | chr17 | + | 43492680 | 43493040 | 43493444 | 43493589 | 43492182 | 43492292 | 43493682 | 43493886 | 21 | 9 | 0 | 15 | 11 | 4 | 0 | 2 | 0 | 2 | 98 | 98 | 1 | 1 | 0.583 | 0.45 | 0 | NA | 0.5 | -0.156 |
| 1070 | KIAA0100 | KIAA0100 | chr17 | - | 28641888 | 28642078 | 28642274 | 28642335 | 28640559 | 28640683 | 28642868 | 28642987 | 9 | 7 | 0 | 6 | 3 | 0 | 0 | 3 | 0 | 2 | 98 | 98 | 1 | 1 | 0.6 | 0.7 | NA | NA | 0.6 | 0.05 |
| 1077 | ANP32E | ANP32E | chr1 | - | 150229071 | 150229237 | 150230570 | 150230693 | 150226607 | 150226795 | 150231776 | 150231926 | 3 | 3 | 2 | 2 | 0 | 1 | 0 | 1 | 0 | 0 | 98 | 98 | 1 | 1 | 0.6 | 1 | 0.667 | NA | 1 | -0.244 |
| 1085 | RFFL | RFFL | chr17 | - | 35017522 | 35017606 | 35021370 | 35021781 | 35016369 | 35016580 | 35026373 | 35026561 | 9 | 1 | 0 | 15 | 4 | 5 | 0 | 3 | 0 | 1 | 98 | 98 | 1 | 1 | 0.375 | 0.2 | 0 | NA | 0.75 | -0.558 |
| 1101 | ZDHHC3 | ZDHHC3 | chr3 | - | 44929305 | 44929436 | 44932906 | 44932990 | 44915260 | 44926847 | 44933117 | 44933199 | 1 | 1 | 1 | 15 | 3 | 2 | 0 | 2 | 0 | 3 | 98 | 98 | 1 | 1 | 0.062 | 0.25 | 0.333 | NA | 0.4 | -0.185 |
| 1120 | DHX40 | DHX40 | chr17 | + | 59566626 | 59566709 | 59570517 | 59570663 | 59565524 | 59565783 | 59573115 | 59573235 | 7 | 3 | 0 | 13 | 6 | 4 | 0 | 1 | 0 | 1 | 98 | 98 | 1 | 1 | 0.35 | 0.333 | 0 | NA | 0.5 | -0.272 |
| 1121 | DHX40 | DHX40 | chr17 | + | 59566626 | 59566794 | 59570517 | 59570663 | 59565524 | 59565783 | 59573115 | 59573235 | 8 | 3 | 0 | 13 | 6 | 4 | 0 | 1 | 0 | 1 | 98 | 98 | 1 | 1 | 0.381 | 0.333 | 0 | NA | 0.5 | -0.262 |
| 1128 | MILR1 | MILR1 | chr17 | + | 64452596 | 64452866 | 64457399 | 64457684 | 64449313 | 64449355 | 64460821 | 64460932 | 4 | 3 | 0 | 7 | 0 | 0 | 0 | 1 | 0 | 0 | 98 | 98 | 1 | 1 | 0.364 | 1 | NA | NA | 1 | -0.318 |
| 1129 | MILR1 | MILR1 | chr17 | + | 64457399 | 64457684 | 64460821 | 64460932 | 64452596 | 64452866 | 64465451 | 64465541 | 6 | 2 | 0 | 6 | 0 | 0 | 0 | 6 | 0 | 0 | 98 | 98 | 1 | 1 | 0.5 | 1 | NA | NA | 1 | -0.25 |
| 1141 | EXOC7 | EXOC7 | chr17 | - | 76090328 | 76090397 | 76091142 | 76091235 | 76089174 | 76089320 | 76094413 | 76094581 | 1 | 0 | 0 | 1 | 0 | 0 | 0 | 0 | 0 | 2 | 98 | 98 | 1 | 1 | 0.5 | NA | NA | NA | 0 | 0.5 |
| 1144 | PGS1 | PGS1 | chr17 | + | 78392475 | 78392665 | 78396307 | 78396385 | 78378617 | 78378808 | 78398251 | 78398351 | 12 | 14 | 1 | 11 | 19 | 1 | 0 | 4 | 0 | 4 | 98 | 98 | 1 | 1 | 0.522 | 0.424 | 0.5 | NA | 0.5 | -0.018 |
| 1145 | PGS1 | PGS1 | chr17 | + | 78392513 | 78392665 | 78396307 | 78396385 | 78378617 | 78378808 | 78398251 | 78398351 | 4 | 5 | 1 | 11 | 19 | 1 | 0 | 1 | 0 | 4 | 98 | 98 | 1 | 1 | 0.267 | 0.208 | 0.5 | NA | 0.2 | 0.125 |
| 1146 | DCP2 | DCP2 | chr5 | + | 113003941 | 113004077 | 113007937 | 113008042 | 113001566 | 113001674 | 113010755 | 113010807 | 12 | 8 | 7 | 26 | 7 | 2 | 0 | 5 | 0 | 1 | 98 | 98 | 1 | 1 | 0.316 | 0.533 | 0.778 | NA | 0.833 | -0.291 |
| 1147 | ATG7 | ATG7 | chr3 | + | 11379971 | 11380052 | 11426803 | 11426926 | 11364658 | 11364734 | 11554810 | 11557665 | 16 | 8 | 2 | 3 | 5 | 0 | 0 | 6 | 0 | 2 | 98 | 98 | 1 | 1 | 0.842 | 0.615 | 1 | NA | 0.75 | 0.069 |
| 1154 | SMCHD1 | SMCHD1 | chr18 | + | 2760651 | 2760739 | 2762104 | 2762236 | 2752487 | 2752552 | 2763636 | 2763789 | 110 | 56 | 32 | 120 | 94 | 33 | 0 | 22 | 0 | 30 | 98 | 98 | 1 | 1 | 0.478 | 0.373 | 0.492 | NA | 0.423 | 0.025 |
| 1158 | CARS | CARS | chr11 | - | 3026675 | 3026797 | 3028995 | 3029084 | 3020219 | 3020332 | 3029302 | 3029443 | 27 | 10 | 3 | 12 | 8 | 3 | 0 | 6 | 0 | 3 | 98 | 98 | 1 | 1 | 0.692 | 0.556 | 0.5 | NA | 0.667 | -0.084 |
| 1162 | PPCDC | PPCDC | chr15 | + | 75043440 | 75043536 | 75044385 | 75044514 | 75028246 | 75028453 | 75048552 | 75048721 | 0 | 2 | 0 | 16 | 0 | 4 | 0 | 0 | 0 | 2 | 98 | 98 | 1 | 1 | 0 | 1 | 0 | NA | 0 | 0.333 |
| 1166 | CLEC4A | CLEC4A | chr12 | + | 8125560 | 8125677 | 8129263 | 8129362 | 8123631 | 8123960 | 8135584 | 8135736 | 3 | 4 | 1 | 8 | 1 | 0 | 0 | 0 | 0 | 2 | 98 | 98 | 1 | 1 | 0.273 | 0.8 | 1 | NA | 0 | 0.691 |
| 1167 | CLEC4A | CLEC4A | chr12 | + | 8129263 | 8129362 | 8135584 | 8135736 | 8123631 | 8123960 | 8136787 | 8136903 | 0 | 0 | 9 | 4 | 1 | 3 | 0 | 1 | 0 | 3 | 98 | 98 | 1 | 1 | 0 | 0 | 0.75 | NA | 0.25 | 0 |
| 1168 | CLEC4A | CLEC4A | chr12 | + | 8129263 | 8129362 | 8135584 | 8135736 | 8125560 | 8125677 | 8136787 | 8136903 | 5 | 2 | 22 | 4 | 1 | 3 | 0 | 2 | 0 | 3 | 98 | 98 | 1 | 1 | 0.556 | 0.667 | 0.88 | NA | 0.4 | 0.301 |
| 1173 | KIAA0226L | KIAA0226L | chr13 | - | 46361440 | 46361573 | 46362537 | 46362598 | 46359485 | 46359631 | 46363114 | 46363213 | 42 | 11 | 10 | 64 | 9 | 9 | 0 | 3 | 0 | 6 | 98 | 98 | 1 | 1 | 0.396 | 0.55 | 0.526 | NA | 0.333 | 0.158 |
| 1190 | EDEM2 | EDEM2 | chr20 | - | 35144978 | 35145018 | 35146824 | 35146935 | 35142372 | 35142478 | 35147151 | 35147358 | 5 | 6 | 0 | 0 | 5 | 5 | 0 | 1 | 0 | 0 | 98 | 89 | 1 | 1 | 1 | 0.521 | 0 | NA | 1 | -0.493 |
| 1200 | GMFG | GMFG | chr19 | - | 39328999 | 39329073 | 39335260 | 39335310 | 39328356 | 39328548 | 39335434 | 39335531 | 60 | 41 | 35 | 85 | 105 | 133 | 0 | 7 | 0 | 24 | 98 | 98 | 1 | 1 | 0.414 | 0.281 | 0.208 | NA | 0.226 | 0.075 |
| 1202 | GMFG | GMFG | chr19 | - | 39333076 | 39333126 | 39335260 | 39335310 | 39329543 | 39329626 | 39335434 | 39335531 | 60 | 42 | 35 | 107 | 88 | 99 | 0 | 6 | 0 | 20 | 98 | 98 | 1 | 1 | 0.359 | 0.323 | 0.261 | NA | 0.231 | 0.083 |
| 1203 | GMFG | GMFG | chr19 | - | 39333076 | 39333126 | 39335260 | 39335310 | 39329543 | 39329626 | 39335973 | 39336086 | 0 | 1 | 0 | 104 | 88 | 99 | 0 | 0 | 0 | 19 | 98 | 98 | 1 | 1 | 0 | 0.011 | 0 | NA | 0 | 0.004 |
| 1204 | GMFG | GMFG | chr19 | - | 39335260 | 39335310 | 39335434 | 39335531 | 39333076 | 39333126 | 39335973 | 39336086 | 5 | 0 | 0 | 97 | 63 | 46 | 0 | 1 | 1 | 12 | 98 | 98 | 1 | 1 | 0.049 | 0 | 0 | 0 | 0.077 | -0.022 |
| 1211 | CNOT8 | CNOT8 | chr5 | + | 154865191 | 154865385 | 154870660 | 154870822 | 154863206 | 154863395 | 154871729 | 154871874 | 6 | 3 | 4 | 9 | 2 | 2 | 0 | 0 | 0 | 3 | 98 | 98 | 1 | 1 | 0.4 | 0.6 | 0.667 | NA | 0 | 0.556 |
| 1212 | CNOT8 | CNOT8 | chr5 | + | 154867691 | 154867792 | 154870660 | 154870822 | 154865191 | 154865385 | 154871729 | 154871874 | 0 | 0 | 1 | 13 | 4 | 7 | 0 | 0 | 0 | 3 | 98 | 98 | 1 | 1 | 0 | 0 | 0.125 | NA | 0 | 0.042 |
| 1221 | GPATCH2L | GPATCH2L | chr14 | + | 76177891 | 76177931 | 76177987 | 76178042 | 76176622 | 76176690 | 76180763 | 76180849 | 2 | 0 | 0 | 9 | 13 | 0 | 0 | 0 | 0 | 2 | 89 | 98 | 1 | 1 | 0.197 | 0 | NA | NA | 0 | 0.099 |
| 1225 | ALG13 | ALG13 | chrX | + | 111682131 | 111682235 | 111682235 | 111682294 | 111681169 | 111681424 | 111684964 | 111685103 | 1 | 0 | 0 | 14 | 0 | 2 | 0 | 0 | 3 | 0 | 98 | 98 | 1 | 1 | 0.067 | NA | 0 | 0 | NA | 0.034 |
| 1227 | RBM23 | RBM23 | chr14 | - | 22908332 | 22908380 | 22911327 | 22911403 | 22906194 | 22906368 | 22918998 | 22919187 | 7 | 13 | 3 | 5 | 8 | 0 | 0 | 1 | 0 | 1 | 98 | 97 | 1 | 1 | 0.581 | 0.617 | 1 | NA | 0.497 | 0.236 |
| 1233 | LILRA5 | LILRA5 | chr19 | - | 54312334 | 54312370 | 54312536 | 54312621 | 54311863 | 54312148 | 54313016 | 54313139 | 21 | 14 | 12 | 25 | 14 | 23 | 0 | 6 | 0 | 5 | 98 | 85 | 1 | 1 | 0.421 | 0.464 | 0.312 | NA | 0.51 | -0.111 |
| 1238 | C2 | C2 | chr6 | + | 31933866 | 31933965 | 31935922 | 31936061 | 31933609 | 31933783 | 31937318 | 31937459 | 0 | 0 | 0 | 1 | 0 | 0 | 0 | 2 | 0 | 0 | 98 | 98 | 1 | 1 | 0 | NA | NA | NA | 1 | -1 |
| 1241 | VPS33B | VPS33B | chr15 | - | 91016962 | 91017024 | 91017804 | 91017885 | 91014383 | 91014433 | 91022153 | 91022621 | 2 | 0 | 0 | 1 | 0 | 0 | 0 | 2 | 0 | 1 | 98 | 98 | 1 | 1 | 0.667 | NA | NA | NA | 0.667 | 0 |
| 1243 | FAM129A | FAM129A | chr1 | - | 184803584 | 184803692 | 184805945 | 184806056 | 184798078 | 184798190 | 184808073 | 184808235 | 59 | 34 | 14 | 104 | 42 | 32 | 0 | 4 | 0 | 17 | 98 | 98 | 1 | 1 | 0.362 | 0.447 | 0.304 | NA | 0.19 | 0.181 |
| 1246 | CAMLG | CAMLG | chr5 | + | 134741062 | 134741523 | 134743986 | 134744052 | 134738479 | 134738792 | 134750758 | 134752160 | 4 | 0 | 4 | 4 | 4 | 1 | 0 | 0 | 0 | 1 | 98 | 98 | 1 | 1 | 0.5 | 0 | 0.8 | NA | 0 | 0.433 |
| 1250 | PQLC3 | PQLC3 | chr2 | + | 11171924 | 11172045 | 11174967 | 11175009 | 11164203 | 11164261 | 11177736 | 11178872 | 2 | 0 | 3 | 0 | 1 | 0 | 0 | 1 | 0 | 1 | 98 | 91 | 1 | 1 | 1 | 0 | 1 | NA | 0.481 | 0.186 |
| 1251 | NCOA1 | NCOA1 | chr2 | + | 24762686 | 24762776 | 24768078 | 24768135 | 24757972 | 24758156 | 24768220 | 24770701 | 37 | 9 | 5 | 18 | 5 | 0 | 0 | 5 | 0 | 5 | 98 | 98 | 1 | 1 | 0.673 | 0.643 | 1 | NA | 0.5 | 0.272 |
| 1253 | BRE | BRE | chr2 | + | 28310081 | 28310153 | 28327273 | 28327447 | 28298337 | 28298491 | 28338449 | 28338900 | 1 | 0 | 0 | 1 | 0 | 0 | 0 | 0 | 0 | 1 | 98 | 98 | 1 | 1 | 0.5 | NA | NA | NA | 0 | 0.5 |
| 1259 | PPM1B | PPM1B | chr2 | + | 44201185 | 44202045 | 44209209 | 44209327 | 44168802 | 44169274 | 44217966 | 44218078 | 4 | 7 | 3 | 19 | 7 | 7 | 0 | 2 | 0 | 3 | 98 | 98 | 1 | 1 | 0.174 | 0.5 | 0.3 | NA | 0.4 | -0.075 |
| 1260 | PREPL | PREPL | chr2 | - | 44332458 | 44332656 | 44338350 | 44338536 | 44328936 | 44329112 | 44339146 | 44339363 | 2 | 0 | 1 | 1 | 0 | 0 | 0 | 1 | 0 | 0 | 98 | 98 | 1 | 1 | 0.667 | NA | 1 | NA | 1 | -0.166 |
| 1261 | EIF4E3 | EIF4E3 | chr3 | - | 71696459 | 71696520 | 71699613 | 71699708 | 71693874 | 71693941 | 71710411 | 71710484 | 11 | 12 | 8 | 29 | 15 | 15 | 0 | 2 | 0 | 3 | 98 | 98 | 1 | 1 | 0.275 | 0.444 | 0.348 | NA | 0.4 | -0.044 |
| 1266 | PSMA3-AS1 | PSMA3-AS1 | chr14 | - | 58273978 | 58274010 | 58285590 | 58285756 | 58265364 | 58267393 | 58291634 | 58291707 | 5 | 3 | 1 | 3 | 6 | 1 | 0 | 1 | 0 | 2 | 98 | 81 | 1 | 1 | 0.579 | 0.292 | 0.453 | NA | 0.292 | 0.149 |
| 1267 | PSMA3-AS1 | PSMA3-AS1 | chr14 | - | 58273978 | 58274135 | 58285590 | 58285756 | 58265364 | 58267393 | 58291634 | 58291707 | 5 | 3 | 1 | 10 | 5 | 4 | 0 | 1 | 0 | 0 | 98 | 98 | 1 | 1 | 0.333 | 0.375 | 0.2 | NA | 1 | -0.697 |
| 1268 | VPS13B | VPS13B | chr8 | + | 99507136 | 99507203 | 99507769 | 99507911 | 99502835 | 99502950 | 99511103 | 99511512 | 38 | 11 | 0 | 4 | 1 | 0 | 0 | 5 | 0 | 2 | 98 | 98 | 1 | 1 | 0.905 | 0.917 | NA | NA | 0.714 | 0.197 |
| 1271 | PTCD3 | PTCD3 | chr2 | + | 86125454 | 86125515 | 86125794 | 86125880 | 86124994 | 86125082 | 86127160 | 86127305 | 11 | 3 | 0 | 3 | 0 | 0 | 0 | 2 | 0 | 0 | 98 | 98 | 1 | 1 | 0.786 | 1 | NA | NA | 1 | -0.107 |
| 1277 | BCL2L11 | BCL2L11 | chr2 | + | 111123732 | 111123869 | 111150043 | 111150147 | 111120913 | 111121188 | 111164132 | 111168445 | 3 | 0 | 0 | 3 | 2 | 0 | 0 | 1 | 0 | 1 | 98 | 98 | 1 | 1 | 0.5 | 0 | NA | NA | 0.5 | -0.25 |

| | | | | | | | | | | | | | | | | | | | | | | | | | | | | | | | | |
|---|---|---|---|---|---|---|---|---|---|---|---|---|---|---|---|---|---|---|---|---|---|---|---|---|---|---|---|---|---|---|---|---|
| 1278 | BCL2L11 | BCL2L11 | chr2 | + | 111123732 | 111124139 | 111150043 | 111150147 | 111120913 | 111121188 | 111153779 | 111153904 | 3 | 0 | 0 | 5 | 0 | 0 | 0 | 1 | 0 | 1 | 98 | 98 | 1 | 1 | 0.375 | NA | NA | NA | 0.5 | -0.125 |
| 1279 | BCL2L11 | BCL2L11 | chr2 | + | 111123732 | 111124139 | 111150043 | 111150147 | 111120913 | 111121188 | 111164132 | 111168445 | 3 | 0 | 0 | 3 | 2 | 0 | 0 | 1 | 0 | 1 | 98 | 98 | 1 | 1 | 0.5 | 0 | NA | NA | 0.5 | -0.25 |
| 1289 | SMPD4 | SMPD4 | chr2 | - | 130173278 | 130173354 | 130173513 | 130173656 | 130172786 | 130172895 | 130174913 | 130175000 | 0 | 0 | 0 | 0 | 5 | 0 | 0 | 2 | 0 | 0 | 98 | 98 | 1 | 1 | NA | 0 | NA | NA | 1 | -1 |
| 1295 | PDK1 | PDK1 | chr2 | + | 172562740 | 172562800 | 172564502 | 172564687 | 172562219 | 172562291 | 172564977 | 172565073 | 3 | 3 | 0 | 8 | 0 | 2 | 1 | 0 | 0 | 0 | 98 | 98 | 1 | 1 | 0.273 | 1 | 0 | 1 | NA | -0.576 |
| 1298 | SETX | SETX | chr9 | - | 132334607 | 132334727 | 132336295 | 132336515 | 132331276 | 132331448 | 132342689 | 132342799 | 116 | 29 | 8 | 80 | 20 | 2 | 0 | 9 | 0 | 8 | 98 | 98 | 1 | 1 | 0.592 | 0.592 | 0.8 | NA | 0.529 | 0.132 |
| 1303 | ANKRD44 | ANKRD44 | chr2 | - | 196998906 | 196999052 | 197000418 | 197000502 | 196998336 | 196998419 | 197001752 | 197001840 | 47 | 11 | 6 | 25 | 7 | 13 | 0 | 9 | 0 | 5 | 98 | 98 | 1 | 1 | 0.653 | 0.611 | 0.316 | NA | 0.643 | -0.116 |
| 1304 | PPIL3 | PPIL3 | chr2 | - | 200882341 | 200882435 | 200885261 | 200885367 | 200881420 | 200881488 | 200885697 | 200885772 | 0 | 0 | 0 | 1 | 2 | 0 | 0 | 1 | 0 | 0 | 98 | 98 | 1 | 1 | 0 | 0 | NA | NA | 1 | -1 |
| 1305 | PIKFYVE | PIKFYVE | chr2 | + | 208276711 | 208276830 | 208285725 | 208285933 | 208273583 | 208273733 | 208288728 | 208288818 | 10 | 1 | 0 | 12 | 9 | 0 | 0 | 0 | 0 | 1 | 98 | 98 | 1 | 1 | 0.455 | 0.1 | NA | NA | 0 | 0.278 |
| 1306 | PIKFYVE | PIKFYVE | chr2 | + | 208277536 | 208277708 | 208285725 | 208285933 | 208276711 | 208276830 | 208288728 | 208288818 | 8 | 2 | 3 | 13 | 9 | 0 | 0 | 0 | 0 | 1 | 98 | 98 | 1 | 1 | 0.381 | 0.182 | 1 | NA | 0 | 0.521 |
| 1307 | BARD1 | BARD1 | chr2 | - | 214780559 | 214781509 | 214792296 | 214792445 | 214769231 | 214769312 | 214797060 | 214797117 | 1 | 0 | 0 | 0 | 0 | 0 | 0 | 0 | 0 | 1 | 98 | 98 | 1 | 1 | 1 | NA | NA | NA | 0 | 1 |
| 1317 | DGKD | DGKD | chr2 | + | 233434768 | 233434901 | 233435817 | 233435924 | 233434379 | 233434484 | 233436315 | 233436441 | 11 | 3 | 0 | 3 | 7 | 4 | 0 | 5 | 0 | 3 | 98 | 98 | 1 | 1 | 0.786 | 0.3 | 0 | NA | 0.625 | -0.263 |
| 1319 | STK25 | STK25 | chr2 | - | 241501477 | 241501708 | 241508005 | 241508135 | 241500739 | 241500796 | 241508442 | 241508695 | 0 | 0 | 0 | 1 | 1 | 0 | 0 | 1 | 0 | 1 | 98 | 98 | 1 | 1 | 0 | 0 | NA | NA | 0.5 | -0.5 |
| 1324 | NAPB | NAPB | chr20 | - | 23395138 | 23395185 | 23397071 | 23397182 | 23394921 | 23394999 | 23402992 | 23403072 | 1 | 0 | 0 | 0 | 0 | 0 | 0 | 0 | 0 | 2 | 98 | 96 | 1 | 1 | 1 | NA | NA | NA | 0 | 1 |
| 1325 | NAPB | NAPB | chr20 | - | 23395138 | 23395185 | 23397071 | 23397188 | 23394921 | 23394999 | 23402992 | 23403072 | 1 | 0 | 0 | 0 | 0 | 0 | 0 | 0 | 0 | 2 | 98 | 96 | 1 | 1 | 1 | NA | NA | NA | 0 | 1 |
| 1329 | HCK | HCK | chr20 | + | 32054215 | 32054281 | 32069687 | 32069751 | 32052187 | 32052486 | 32071661 | 32071782 | 19 | 4 | 8 | 2 | 0 | 0 | 0 | 5 | 0 | 0 | 98 | 98 | 1 | 1 | 0.905 | 1 | 1 | NA | 1 | -0.032 |
| 1332 | C20orf24 | C20orf24 | chr20 | + | 36607889 | 36608000 | 36609600 | 36609730 | 36607714 | 36607818 | 36612009 | 36612557 | 3 | 3 | 1 | 29 | 41 | 54 | 0 | 1 | 0 | 8 | 98 | 98 | 1 | 1 | 0.094 | 0.068 | 0.018 | NA | 0.111 | -0.051 |
| 1333 | C20orf24 | C20orf24 | chr20 | + | 36607889 | 36608000 | 36609600 | 36609758 | 36607714 | 36607818 | 36612009 | 36612557 | 3 | 3 | 1 | 9 | 11 | 26 | 0 | 1 | 0 | 4 | 98 | 98 | 1 | 1 | 0.25 | 0.214 | 0.037 | NA | 0.2 | -0.033 |
| 1345 | ZMYND8 | ZMYND8 | chr20 | - | 47262287 | 47262428 | 47276313 | 47276795 | 47249286 | 47249439 | 47282101 | 47282217 | 5 | 7 | 1 | 1 | 3 | 0 | 0 | 1 | 0 | 1 | 98 | 98 | 1 | 1 | 0.833 | 0.7 | 1 | NA | 0.5 | 0.344 |
| 1365 | PRMT2 | PRMT2 | chr21 | + | 46658744 | 46658920 | 46663382 | 46663554 | 46649574 | 46649739 | 46664294 | 46665124 | 9 | 2 | 5 | 4 | 9 | 2 | 0 | 2 | 0 | 1 | 98 | 98 | 1 | 1 | 0.692 | 0.182 | 0.714 | NA | 0.667 | -0.138 |
| 1366 | BID | BID | chr22 | - | 17743802 | 17744013 | 17750104 | 17750174 | 17739348 | 17739488 | 17774380 | 17774495 | 7 | 0 | 0 | 11 | 15 | 12 | 0 | 0 | 0 | 9 | 98 | 98 | 1 | 1 | 0.389 | 0 | 0 | NA | 0 | 0.13 |
| 1367 | TANGO2 | TANGO2 | chr22 | + | 20053436 | 20053551 | 20055942 | 20056013 | 20052464 | 20052584 | 20063337 | 20063442 | 1 | 6 | 7 | 2 | 3 | 1 | 0 | 0 | 0 | 1 | 98 | 98 | 1 | 1 | 0.333 | 0.667 | 0.875 | NA | 0 | 0.625 |
| 1368 | TANGO2 | TANGO2 | chr22 | + | 20053436 | 20053551 | 20061529 | 20061683 | 20052464 | 20052584 | 20063337 | 20063442 | 1 | 6 | 7 | 0 | 3 | 2 | 0 | 0 | 0 | 2 | 98 | 98 | 1 | 1 | 1 | 0.667 | 0.778 | NA | 0 | 0.815 |
| 1369 | TANGO2 | TANGO2 | chr22 | + | 20055942 | 20056013 | 20061529 | 20061683 | 20052464 | 20052584 | 20063337 | 20063442 | 2 | 3 | 1 | 0 | 3 | 2 | 0 | 1 | 0 | 2 | 98 | 98 | 1 | 1 | 1 | 0.5 | 0.333 | NA | 0.333 | 0.278 |
| 1370 | TANGO2 | TANGO2 | chr22 | + | 20055942 | 20056013 | 20063337 | 20063442 | 20053436 | 20053551 | 20064541 | 20065926 | 4 | 7 | 1 | 5 | 9 | 5 | 0 | 4 | 0 | 3 | 98 | 98 | 1 | 1 | 0.444 | 0.437 | 0.167 | NA | 0.571 | -0.222 |
| 1372 | GUCD1 | GUCD1 | chr22 | - | 24547907 | 24548073 | 24548916 | 24549001 | 24546913 | 24547005 | 24554948 | 24555307 | 5 | 4 | 0 | 13 | 7 | 0 | 2 | 0 | 0 | 1 | 98 | 98 | 1 | 1 | 0.278 | 0.364 | NA | 1 | 0 | -0.179 |
| 1373 | GUCD1 | GUCD1 | chr22 | - | 24547907 | 24548073 | 24548916 | 24549001 | 24546913 | 24547005 | 24555614 | 24555935 | 2 | 1 | 0 | 11 | 7 | 0 | 0 | 0 | 0 | 1 | 98 | 98 | 1 | 1 | 0.154 | 0.125 | NA | NA | 0 | 0.14 |
| 1377 | UBAP1 | UBAP1 | chr9 | + | 34220907 | 34220948 | 34234215 | 34234340 | 34179004 | 34179240 | 34241184 | 34242108 | 7 | 3 | 4 | 4 | 0 | 4 | 0 | 1 | 0 | 0 | 90 | 98 | 1 | 1 | 0.656 | 1 | 0.521 | NA | 1 | -0.274 |
| 1382 | SCAF8 | SCAF8 | chr6 | + | 154822275 | 154822409 | 154824233 | 154824378 | 154820176 | 154820333 | 154827171 | 154827240 | 16 | 4 | 8 | 13 | 0 | 0 | 0 | 1 | 0 | 0 | 98 | 98 | 1 | 1 | 0.552 | 1 | 1 | NA | 1 | -0.149 |
| 1384 | NUP50 | NUP50 | chr22 | + | 45175893 | 45176080 | 45178237 | 45178900 | 45171599 | 45171683 | 45181285 | 45181367 | 19 | 3 | 3 | 12 | 12 | 3 | 0 | 0 | 0 | 1 | 98 | 98 | 1 | 1 | 0.613 | 0.2 | 0.5 | NA | 0 | 0.438 |
| 1386 | TBC1D22A | TBC1D22A | chr22 | + | 46913333 | 46913446 | 46974289 | 46974399 | 46912073 | 46912188 | 46997633 | 46997709 | 0 | 0 | 2 | 16 | 5 | 10 | 0 | 0 | 0 | 3 | 98 | 98 | 1 | 1 | 0 | 0 | 0.167 | NA | 0 | 0.056 |
| 1387 | TBC1D22A | TBC1D22A | chr22 | + | 46974289 | 46974399 | 46997633 | 46997709 | 46912073 | 46912188 | 47037070 | 47037198 | 11 | 2 | 0 | 13 | 8 | 4 | 0 | 3 | 0 | 3 | 98 | 98 | 1 | 1 | 0.458 | 0.2 | 0 | NA | 0.5 | -0.281 |
| 1391 | ZNF346 | ZNF346 | chr5 | + | 177041777 | 177041870 | 177044391 | 177044533 | 177041125 | 177041229 | 177050750 | 177050936 | 1 | 1 | 0 | 1 | 0 | 0 | 0 | 1 | 0 | 0 | 98 | 98 | 1 | 1 | 0.5 | 1 | NA | NA | 1 | -0.25 |
| 1398 | HACL1 | HACL1 | chr3 | - | 15585247 | 15585342 | 15586524 | 15586602 | 15582876 | 15582989 | 15589539 | 15589612 | 1 | 0 | 1 | 0 | 0 | 0 | 0 | 0 | 0 | 1 | 98 | 98 | 1 | 1 | 1 | NA | 1 | NA | 0 | 1 |
| 1404 | ABHD5 | ABHD5 | chr3 | + | 43714946 | 43715058 | 43717670 | 43717857 | 43711708 | 43711863 | 43718442 | 43722725 | 61 | 17 | 23 | 40 | 7 | 7 | 0 | 13 | 0 | 9 | 98 | 98 | 1 | 1 | 0.604 | 0.708 | 0.767 | NA | 0.591 | 0.102 |
| 1407 | HIPK3 | HIPK3 | chr11 | + | 33348521 | 33348818 | 33349146 | 33349287 | 33348165 | 33348228 | 33351607 | 33351843 | 14 | 6 | 2 | 32 | 17 | 4 | 0 | 1 | 0 | 6 | 98 | 98 | 1 | 1 | 0.304 | 0.261 | 0.333 | NA | 0.143 | 0.156 |
| 1425 | ERBB2IP | ERBB2IP | chr5 | + | 66068876 | 66069020 | 66072168 | 66072291 | 66053405 | 66054951 | 66075023 | 66075230 | 2 | 0 | 0 | 6 | 9 | 0 | 0 | 0 | 0 | 2 | 98 | 98 | 1 | 1 | 0.25 | 0 | NA | NA | 0 | 0.125 |
| 1427 | SENP7 | SENP7 | chr3 | - | 101398860 | 101399055 | 101417592 | 101417790 | 101372007 | 101372126 | 101458954 | 101459052 | 11 | 8 | 2 | 13 | 4 | 0 | 0 | 0 | 0 | 1 | 98 | 98 | 1 | 1 | 0.458 | 0.667 | 1 | NA | 0 | 0.708 |
| 1432 | MRVI1 | MRVI1 | chr11 | - | 10594145 | 10594195 | 10600917 | 10601059 | 10593491 | 10593599 | 10603119 | 10603251 | 27 | 9 | 9 | 10 | 2 | 4 | 0 | 4 | 0 | 1 | 98 | 98 | 1 | 1 | 0.73 | 0.818 | 0.692 | NA | 0.8 | -0.053 |
| 1437 | CEP63 | CEP63 | chr3 | + | 134558141 | 134558347 | 134559149 | 134559429 | 134551925 | 134552012 | 134561376 | 134565028 | 11 | 12 | 1 | 5 | 2 | 3 | 2 | 0 | 0 | 0 | 98 | 98 | 1 | 1 | 0.687 | 0.857 | 0.25 | 1 | NA | -0.402 |
| 1438 | ARMC8 | ARMC8 | chr3 | + | 138188400 | 138188530 | 138209816 | 138209893 | 138187247 | 138187599 | 138221925 | 138221997 | 0 | 3 | 0 | 10 | 2 | 2 | 0 | 2 | 0 | 2 | 98 | 98 | 1 | 1 | 0 | 0.6 | 0 | NA | 0 | 0.2 |
| 1439 | ARMC8 | ARMC8 | chr3 | + | 138188454 | 138188530 | 138209816 | 138209893 | 138187247 | 138187599 | 138221925 | 138221997 | 0 | 3 | 0 | 10 | 2 | 2 | 0 | 2 | 0 | 2 | 98 | 98 | 1 | 1 | 0 | 0.6 | 0 | NA | 0.5 | -0.3 |
| 1442 | 7-Mar | 7-Mar | chr2 | + | 159762879 | 159762993 | 159764625 | 159764674 | 159759225 | 159759335 | 159767283 | 159768583 | 16 | 10 | 0 | 26 | 16 | 15 | 0 | 2 | 0 | 5 | 98 | 98 | 1 | 1 | 0.381 | 0.385 | 0 | NA | 0.286 | -0.031 |
| 1443 | EIF2A | EIF2A | chr3 | + | 150571957 | 150572529 | 150575648 | 150575762 | 150568175 | 150568292 | 150581617 | 150581746 | 6 | 0 | 2 | 6 | 1 | 0 | 0 | 0 | 0 | 1 | 98 | 98 | 1 | 1 | 0.5 | 0 | 1 | NA | 0 | 0.5 |
| 1445 | MME | MME | chr3 | + | 155141990 | 155142127 | 155142236 | 155142330 | 155140190 | 155140292 | 155143442 | 155143571 | 67 | 18 | 23 | 95 | 36 | 23 | 0 | 9 | 0 | 7 | 98 | 98 | 1 | 1 | 0.414 | 0.333 | 0.5 | NA | 0.562 | -0.146 |
| 1447 | ZMYM5 | ZMYM5 | chr13 | - | 19838699 | 19838985 | 19851354 | 19851448 | 19837452 | 19837821 | 19851688 | 19852190 | 6 | 1 | 0 | 2 | 2 | 0 | 0 | 0 | 0 | 1 | 98 | 98 | 1 | 1 | 0.75 | 0.333 | NA | NA | 0 | 0.541 |
| 1450 | XRN2 | XRN2 | chr20 | + | 21356585 | 21356665 | 21357735 | 21357792 | 21356079 | 21356177 | 21365420 | 21365489 | 8 | 4 | 4 | 22 | 4 | 1 | 0 | 8 | 0 | 6 | 98 | 98 | 1 | 1 | 0.267 | 0.5 | 0.8 | NA | 0.571 | -0.049 |
| 1457 | IL1RAP | IL1RAP | chr3 | + | 190550341 | 190550505 | 190556129 | 190556216 | 190514050 | 190514219 | 190564288 | 190564353 | 0 | 1 | 0 | 22 | 20 | 1 | 0 | 0 | 3 | 7 | 98 | 98 | 1 | 1 | 0 | 0.048 | 0 | 0 | 0 | 0.016 |
| 1460 | DUSP22 | DUSP22 | chr6 | + | 311879 | 311962 | 335113 | 335163 | 304627 | 304661 | 345853 | 345928 | 5 | 4 | 3 | 7 | 7 | 0 | 0 | 1 | 0 | 1 | 98 | 98 | 1 | 1 | 0.417 | 0.364 | 1 | NA | 0.5 | 0.094 |
| 1463 | NIPBL | NIPBL | chr5 | + | 37057185 | 37057332 | 37060843 | 37061018 | 37052365 | 37052566 | 37063789 | 37063978 | 16 | 6 | 0 | 4 | 2 | 1 | 0 | 0 | 0 | 1 | 98 | 98 | 1 | 1 | 0.8 | 0.75 | 0 | NA | 0 | 0.517 |
| 1465 | BST1 | BST1 | chr4 | + | 15715284 | 15715361 | 15715706 | 15715799 | 15711806 | 15711889 | 15718906 | 15718993 | 30 | 11 | 11 | 37 | 19 | 16 | 0 | 15 | 0 | 3 | 98 | 98 | 1 | 1 | 0.448 | 0.367 | 0.407 | NA | 0.833 | -0.426 |
| 1468 | PGM2 | PGM2 | chr4 | + | 37848521 | 37848651 | 37850183 | 37850373 | 37847201 | 37847295 | 37855607 | 37855741 | 7 | 18 | 3 | 23 | 10 | 2 | 0 | 0 | 0 | 3 | 98 | 98 | 1 | 1 | 0.233 | 0.643 | 0.6 | NA | 0 | 0.492 |
| 1476 | HERC5 | HERC5 | chr4 | + | 88470613 | 88470673 | 88472408 | 88472502 | 88469156 | 88469260 | 88475840 | 88476030 | 7 | 18 | 1 | 4 | 11 | 1 | 0 | 9 | 0 | 16 | 98 | 98 | 1 | 1 | 0.636 | 0.621 | 0.5 | NA | 0.36 | 0.226 |
| 1480 | KDM5A | KDM5A | chr12 | - | 297040 | 297200 | 306945 | 307089 | 295572 | 295793 | 307453 | 308005 | 17 | 2 | 3 | 13 | 3 | 2 | 0 | 0 | 0 | 4 | 98 | 98 | 1 | 1 | 0.567 | 0.4 | 0.6 | NA | 0 | 0.522 |
| 1481 | ELF2 | ELF2 | chr4 | - | 139061864 | 139062057 | 139067683 | 139067770 | 139060323 | 139060674 | 139071865 | 139072003 | 33 | 3 | 2 | 49 | 12 | 2 | 0 | 8 | 0 | 5 | 98 | 98 | 1 | 1 | 0.402 | 0.2 | 0.5 | NA | 0.615 | -0.248 |
| 1484 | KIAA0922 | KIAA0922 | chr4 | + | 153591052 | 153591194 | 153592474 | 153592584 | 153588889 | 153589007 | 153593798 | 153593871 | 19 | 14 | 1 | 9 | 13 | 0 | 0 | 3 | 0 | 4 | 98 | 98 | 1 | 1 | 0.679 | 0.519 | 1 | NA | 0.429 | 0.304 |
| 1491 | LILRA2 | LILRA2 | chr19 | + | 54577464 | 54577723 | 54587009 | 54587060 | 54575806 | 54576109 | 54587200 | 54587561 | 5 | 4 | 4 | 11 | 0 | 6 | 0 | 3 | 0 | 1 | 98 | 98 | 1 | 1 | 0.312 | 1 | 0.4 | NA | 0.75 | -0.179 |
| 1496 | CPEB4 | CPEB4 | chr5 | + | 173932449 | 173932500 | 173943025 | 173943049 | 173910522 | 173910604 | 173944966 | 173945140 | 11 | 2 | 9 | 1 | 0 | 0 | 0 | 5 | 0 | 0 | 98 | 73 | 1 | 1 | 0.891 | 1 | 1 | NA | 1 | -0.036 |
| 1498 | PCBP1-AS1 | PCBP1-AS1 | chr2 | - | 70059642 | 70059680 | 70083537 | 70083687 | 70051202 | 70051305 | 70085546 | 70085624 | 8 | 0 | 0 | 5 | 3 | 0 | 0 | 1 | 0 | 0 | 98 | 87 | 1 | 1 | 0.587 | 0 | NA | NA | 1 | -0.707 |
| 1499 | RICTOR | RICTOR | chr5 | - | 38975536 | 38975604 | 38978582 | 38978648 | 38971876 | 38971959 | 38981866 | 38982036 | 0 | 0 | 0 | 33 | 29 | 1 | 0 | 1 | 0 | 5 | 98 | 98 | 1 | 1 | 0 | 0 | 0 | NA | 0.167 | -0.167 |
| 1500 | RICTOR | RICTOR | chr5 | - | 38975536 | 38975604 | 38978582 | 38978650 | 38971876 | 38971959 | 38981866 | 38982036 | 55 | 25 | 19 | 33 | 29 | 1 | 0 | 12 | 0 | 5 | 98 | 98 | 1 | 1 | 0.625 | 0.463 | 0.95 | NA | 0.706 | -0.027 |
| 1501 | FYB | FYB | chr5 | - | 39125997 | 39126135 | 39127740 | 39127807 | 39124252 | 39124278 | 39130589 | 39130612 | 91 | 29 | 17 | 59 | 19 | 8 | 0 | 25 | 0 | 8 | 98 | 98 | 1 | 1 | 0.607 | 0.604 | 0.68 | NA | 0.758 | -0.128 |
| 1505 | GAPT | GAPT | chr5 | + | 58493618 | 58493683 | 58493710 | 58493838 | 58491434 | 58491541 | 58494246 | 58496358 | 4 | 1 | 0 | 9 | 1 | 0 | 0 | 2 | 0 | 2 | 98 | 98 | 1 | 1 | 0.308 | 0.5 | NA | NA | 0.5 | -0.096 |
| 1507 | RAD17 | RAD17 | chr5 | + | 69372033 | 69372217 | 69373829 | 69374087 | 69371034 | 69371171 | 69374627 | 69374711 | 4 | 0 | 0 | 1 | 0 | 0 | 0 | 0 | 0 | 1 | 98 | 98 | 1 | 1 | 0.8 | NA | NA | NA | 0 | 0.8 |
| 1508 | MRPS18C | MRPS18C | chr4 | + | 83458345 | 83458429 | 83459739 | 83459797 | 83456908 | 83456958 | 83460972 | 83461032 | 3 | 0 | 1 | 2 | 2 | 5 | 0 | 3 | 0 | 5 | 98 | 98 | 1 | 1 | 0.6 | 0 | 0.167 | NA | 0.375 | -0.119 |
| 1510 | ZFAND5 | ZFAND5 | chr9 | - | 72359417 | 72359521 | 72360109 | 72360221 | 72356930 | 72357056 | 72360627 | 72360784 | 2 | 0 | 0 | 42 | 6 | 4 | 0 | 0 | 0 | 8 | 98 | 98 | 1 | 1 | 0.045 | 0 | 0 | NA | 0 | 0.015 |
| 1513 | SSBP2 | SSBP2 | chr5 | - | 81489249 | 81489309 | 81513627 | 81513717 | 81474495 | 81474562 | 81615472 | 81615557 | 8 | 6 | 0 | 4 | 1 | 3 | 0 | 5 | 0 | 3 | 98 | 98 | 1 | 1 | 0.667 | 0.857 | 0 | NA | 0.625 | -0.117 |
| 1514 | VCAN | VCAN | chr5 | + | 83519348 | 83522309 | 83537006 | 83542268 | 83512102 | 83512396 | 83545536 | 83545650 | 1 | 0 | 0 | 56 | 0 | 0 | 0 | 1 | 2 | 51 | 98 | 98 | 1 | 1 | 0.018 | NA | NA | 0 | 0.019 | 0.008 |
| 1523 | PPP2R4 | PPP2R4 | chr9 | + | 129123051 | 129123138 | 129128984 | 129129110 | 129120512 | 129120610 | 129131521 | 129131639 | 1 | 0 | 0 | 2 | 0 | 0 | 0 | 1 | 2 | 1 | 98 | 98 | 1 | 1 | 0.333 | NA | NA | 0 | 0.5 | 0.083 |
| 1524 | RAPGEF6 | RAPGEF6 | chr5 | - | 131561977 | 131562047 | 131592382 | 131592466 | 131548046 | 131548190 | 131603270 | 131603327 | 4 | 3 | 0 | 8 | 6 | 1 | 0 | 0 | 0 | 1 | 98 | 98 | 1 | 1 | 0.333 | 0.333 | 0 | NA | 0 | 0.222 |
| 1525 | IRF1 | IRF1 | chr5 | - | 132484361 | 132484497 | 132485666 | 132485716 | 132481608 | 132484075 | 132486250 | 132486373 | 53 | 34 | 26 | 43 | 18 | 10 | 0 | 24 | 0 | 8 | 98 | 98 | 1 | 1 | 0.552 | 0.654 | 0.722 | NA | 0.75 | -0.107 |
| 1533 | ZNF438 | ZNF438 | chr10 | - | 30910359 | 30910440 | 30923286 | 30923415 | 30908932 | 30909015 | 30941574 | 30941651 | 5 | 4 | 0 | 1 | 1 | 0 | 0 | 1 | 0 | 0 | 98 | 98 | 1 | 1 | 0.833 | 0.8 | NA | NA | 1 | -0.183 |
| 1534 | ZNF438 | ZNF438 | chr10 | - | 30923286 | 30923415 | 30941574 | 30941651 | 30908932 | 30909015 | 30984368 | 30984426 | 11 | 10 | 1 | 5 | 1 | 0 | 0 | 3 | 0 | 0 | 98 | 98 | 1 | 1 | 0.687 | 0.909 | 1 | NA | 1 | -0.135 |
| 1545 | MAPK9 | MAPK9 | chr5 | - | 180247438 | 180247510 | 180247854 | 180247926 | 180242572 | 180242755 | 180248972 | 180249138 | 0 | 1 | 0 | 3 | 0 | 0 | 0 | 0 | 0 | 1 | 98 | 98 | 1 | 1 | 0 | 1 | NA | NA | 0 | 0.5 |
| 1546 | CASP10 | CASP10 | chr2 | + | 201203729 | 201203766 | 201208074 | 201208183 | 201195841 | 201195948 | 201209069 | 201209562 | 2 | 0 | 0 | 2 | 0 | 0 | 0 | 1 | 0 | 1 | 86 | 98 | 1 | 1 | 0.533 | NA | NA | NA | 0.533 | 0 |
| 1548 | RREB1 | RREB1 | chr6 | + | 7240437 | 7240602 | 7246423 | 7247221 | 7228996 | 7231907 | 7248510 | 7251980 | 2 | 0 | 0 | 1 | 0 | 0 | 0 | 0 | 1 | 0 | 98 | 98 | 1 | 1 | 0.667 | NA | NA | 0 | NA | 0.667 |

**U12_MXE_filtered**

| | | | | | | | | | | | | | Inclusion counts | | | Skipping counts | | | Inclusion counts | | Skipping counts | | | | | | | | Inclusion_level(normalized) | | | |
|---|---|---|---|---|---|---|---|---|---|---|---|---|---|---|---|---|---|---|---|---|---|---|---|---|---|---|---|---|---|---|---|---|
| ID | Gene_ID | Gene_Symbol | Chromosome | Strand | 1stExonStart_0base | 1stExonEnd | 2ndExonStart_0base | 2ndExonEnd | UpstreamES | UpstreamEE | DownstreamES | DownstreamEE | control_s3 | control_s4 | control_s10 | control_s3 | control_s4 | control_s10 | Idg_14 | Ildg_s16 | Idg_s14 | Idg_s16 | IncFormLen | SkipFormLen | p-value | adjust_p | IncLevel_s3 | IncLevel_s4 | IncLevel_s10 | IncLevel14 | IncLevel16 | Level_diff |
| 814 | MAEA | MAEA | chr4 | + | 1322380 | 1322503 | 1327626 | 1327703 | 1315396 | 1315600 | 1332756 | 1332865 | 11 | 5 | 2 | 12 | 5 | 7 | 0 | 0 | 3 | 2 | 98 | 98 | 2.6768E-05 | 0.00253914 | 0.478 | 0.5 | 0.222 | 0 | 0 | 0.4 |
| 605 | GBP5 | GBP5 | chr1 | - | 89268728 | 89268856 | 89269365 | 89269574 | 89267416 | 89267526 | 89272451 | 89272861 | 5 | 4 | 1 | 48 | 11 | 5 | 0 | 0 | 1 | 3 | 98 | 98 | 0.001832725 | 0.05290996 | 0.094 | 0.267 | 0.167 | 0 | 0 | 0.176 |
| 812 | MAEA | MAEA | chr4 | + | 1315396 | 1315600 | 1322380 | 1322503 | 1311978 | 1312161 | 1327626 | 1327703 | 15 | 13 | 2 | 1 | 4 | 1 | 2 | 1 | 0 | 0 | 98 | 98 | 0.005546739 | 0.13153696 | 0.937 | 0.765 | 0.667 | 1 | 1 | -0.21 |
| 832 | DOCK2 | DOCK2 | chr5 | + | 169684195 | 169684350 | 169689251 | 169689333 | 169681743 | 169681879 | 169695802 | 169695938 | 109 | 35 | 17 | 83 | 48 | 5 | 2 | 13 | 3 | 21 | 98 | 98 | 0.086423567 | 0.896644508 | 0.568 | 0.422 | 0.773 | 0.4 | 0.382 | 0.197 |
| 813 | MAEA | MAEA | chr4 | + | 1315396 | 1315600 | 1322380 | 1322503 | 1311978 | 1312161 | 1332756 | 1332865 | 15 | 16 | 2 | 0 | 3 | 1 | 3 | 2 | 0 | 0 | 98 | 98 | 0.297185281 | 1 | 1 | 0.842 | 0.667 | 1 | 1 | -0.164 |
| 1464 | WDR1 | WDR1 | chr4 | - | 10097710 | 10097891 | 10103895 | 10103986 | 10088663 | 10088741 | 10116112 | 10116234 | 36 | 16 | 13 | 37 | 7 | 7 | 4 | 6 | 2 | 3 | 98 | 98 | 0.401096861 | 1 | 0.493 | 0.696 | 0.65 | 0.667 | 0.667 | -0.054 |
| 604 | GBP5 | GBP5 | chr1 | - | 89267416 | 89267526 | 89268728 | 89268856 | 89266956 | 89267153 | 89269365 | 89269574 | 27 | 6 | 4 | 91 | 27 | 18 | 0 | 4 | 1 | 11 | 98 | 98 | 0.659255602 | 1 | 0.229 | 0.182 | 0.182 | 0 | 0.267 | 0.064 |
| 831 | DOCK2 | DOCK2 | chr5 | + | 169670541 | 169670597 | 169671077 | 169671174 | 169669287 | 169669328 | 169674296 | 169674445 | 32 | 11 | 5 | 142 | 34 | 5 | 0 | 8 | 2 | 22 | 98 | 98 | 0.672793032 | 1 | 0.184 | 0.244 | 0.5 | 0 | 0.267 | 0.176 |
| 498 | RASGRP4 | RASGRP4 | chr19 | - | 38417051 | 38417168 | 38419859 | 38420013 | 38414847 | 38415123 | 38420130 | 38420262 | 20 | 18 | 4 | 39 | 11 | 9 | 4 | 7 | 2 | 13 | 98 | 98 | 0.882368191 | 1 | 0.339 | 0.621 | 0.308 | 0.667 | 0.35 | -0.086 |
| 432 | FCHO2 | FCHO2 | chr5 | + | 73006444 | 73006549 | 73015625 | 73015724 | 72990711 | 72990864 | 73017211 | 73017308 | 15 | 3 | 0 | 22 | 9 | 1 | 0 | 6 | 0 | 2 | 98 | 98 | 1 | 1 | 0.405 | 0.25 | 0 | NA | 0.75 | -0.532 |
| 451 | AHCTF1 | AHCTF1 | chr1 | - | 246863923 | 246864116 | 246867243 | 246867351 | 246861958 | 246862153 | 246867660 | 246867811 | 1 | 0 | 0 | 26 | 16 | 4 | 0 | 0 | 0 | 3 | 98 | 98 | 1 | 1 | 0.037 | 0 | 0 | NA | 0 | 0.012 |
| 485 | SLC9A8 | SLC9A8 | chr20 | + | 49839540 | 49839599 | 49845035 | 49845119 | 49823060 | 49823141 | 49849578 | 49849680 | 21 | 3 | 3 | 18 | 4 | 0 | 0 | 6 | 0 | 7 | 98 | 98 | 1 | 1 | 0.538 | 0.429 | 1 | NA | 0.462 | 0.194 |
| 486 | SLC9A8 | SLC9A8 | chr20 | + | 49845035 | 49845119 | 49849578 | 49849680 | 49839540 | 49839599 | 49855437 | 49855581 | 13 | 2 | 0 | 14 | 9 | 0 | 0 | 4 | 0 | 4 | 98 | 98 | 1 | 1 | 0.481 | 0.182 | NA | NA | 0.5 | -0.168 |
| 497 | RASGRP4 | RASGRP4 | chr19 | - | 38417051 | 38417168 | 38419859 | 38420013 | 38413393 | 38413474 | 38420130 | 38420262 | 20 | 18 | 4 | 1 | 0 | 1 | 4 | 7 | 0 | 0 | 98 | 98 | 1 | 1 | 0.952 | 1 | 0.8 | 1 | 1 | -0.083 |
| 648 | DAP3 | DAP3 | chr1 | + | 155717005 | 155717128 | 155721516 | 155721618 | 155709772 | 155709824 | 155725381 | 155725490 | 6 | 1 | 2 | 5 | 7 | 1 | 0 | 0 | 0 | 3 | 98 | 98 | 1 | 1 | 0.545 | 0.125 | 0.667 | NA | 0 | 0.446 |
| 656 | SFI1 | SFI1 | chr22 | + | 31528689 | 31528863 | 31546860 | 31546971 | 31508254 | 31508376 | 31550253 | 31550348 | 3 | 5 | 1 | 4 | 8 | 0 | 0 | 0 | 0 | 2 | 98 | 98 | 1 | 1 | 0.429 | 0.385 | 1 | NA | 0 | 0.605 |
| 763 | RAB6A | RAB6A | chr11 | - | 73718612 | 73718718 | 73718784 | 73718890 | 73716250 | 73716362 | 73720845 | 73720899 | 0 | 0 | 1 | 11 | 3 | 4 | 0 | 0 | 0 | 2 | 98 | 98 | 1 | 1 | 0 | 0 | 0.2 | NA | 0 | 0.067 |
| 805 | CNIH4 | CNIH4 | chr1 | + | 224360494 | 224360563 | 224364108 | 224364455 | 224356810 | 224356993 | 224365878 | 224365991 | 3 | 2 | 7 | 2 | 1 | 0 | 0 | 2 | 0 | 0 | 98 | 98 | 1 | 1 | 0.6 | 0.667 | 1 | NA | 1 | -0.244 |
| 979 | MAPK14 | MAPK14 | chr6 | + | 36095986 | 36096066 | 36100187 | 36100267 | 36076536 | 36076608 | 36102570 | 36102649 | 31 | 21 | 14 | 18 | 11 | 4 | 0 | 8 | 0 | 1 | 98 | 98 | 1 | 1 | 0.633 | 0.656 | 0.778 | NA | 0.889 | -0.2 |
| 985 | DYNC1LI2 | DYNC1LI2 | chr16 | - | 66742437 | 66742668 | 66749196 | 66749313 | 66736074 | 66736244 | 66751272 | 66751346 | 19 | 2 | 0 | 2 | 5 | 1 | 0 | 2 | 0 | 1 | 98 | 98 | 1 | 1 | 0.905 | 0.286 | 0 | NA | 0.667 | -0.27 |
| 1146 | DCP2 | DCP2 | chr5 | + | 113003941 | 113004077 | 113007937 | 113008042 | 113001566 | 113001674 | 113010755 | 113010807 | 12 | 8 | 7 | 26 | 7 | 2 | 0 | 5 | 0 | 1 | 98 | 98 | 1 | 1 | 0.316 | 0.533 | 0.778 | NA | 0.833 | -0.291 |
| 1250 | PQLC3 | PQLC3 | chr2 | + | 11171924 | 11172045 | 11174967 | 11175009 | 11164203 | 11164261 | 11177736 | 11178872 | 2 | 0 | 3 | 0 | 1 | 0 | 0 | 1 | 0 | 1 | 98 | 91 | 1 | 1 | 1 | 0 | 1 | NA | 0.481 | 0.186 |
| 1289 | SMPD4 | SMPD4 | chr2 | - | 130173278 | 130173354 | 130173513 | 130173656 | 130172786 | 130172895 | 130174913 | 130175000 | 0 | 0 | 0 | 0 | 5 | 0 | 0 | 2 | 0 | 0 | 98 | 98 | 1 | 1 | NA | 0 | NA | NA | 1 | -1 |
| 1545 | MAPK9 | MAPK9 | chr5 | - | 180247438 | 180247510 | 180247854 | 180247926 | 180242572 | 180242755 | 180248972 | 180249138 | 0 | 1 | 0 | 3 | 0 | 0 | 0 | 0 | 0 | 1 | 98 | 98 | 1 | 1 | 0 | 1 | NA | NA | 0 | 0.5 |

**Supplementary Data 2S**
**4_events_all**
**SLE LDG versus SLE NDG**

| | | | | | | | | | | | IJC_SAMPLE_1 | | | SJC_SAMPLE_1 | | | IJC_SAMPLE_2 | | | SJC_SAMPLE_2 | | | | | | | IncLevel1 | | | IncLevel2 | | | |
|---|---|---|---|---|---|---|---|---|---|---|---|---|---|---|---|---|---|---|---|---|---|---|---|---|---|---|---|---|---|---|---|---|---|
| **Event** | **ID** | **Gene_ID** | **Chromosome** | ***Strand*** | ***LongExonStart_0base*** | ***LongExonEnd*** | ***ShortES*** | ***ShortEE*** | ***FlankingES*** | ***FlankingEE*** | **ldg_12** | **ldg_14** | **ldg_16** | **ldg_12** | **ldg_14** | **ldg_16** | **pmn_13** | **pmn_15** | **pmn_17** | **pmn_13** | **pmn_15** | **pmn_17** | **IncFormLen** | **SkipFormLen** | **p value** | **FDR** | **ldg_12** | **ldg_14** | **ldg_16** | **pmn_13** | **pmn_15** | **pmn_17** | **IncLevelDifference** |
| SE | 13845 | C10orf54 | chr10 | - | 71761597 | 71762026 | 71760867 | 71760924 | 71773357 | 71773580 | 0 | 4 | 72 | 0 | 0 | 0 | 1 | 16 | 24 | 0 | 1 | 3 | 98 | 49 | 0.001516497 | 0.170760496 | NA | 1 | 1 | 1 | 0.889 | 0.8 | 0.104 |
| SE | 1661 | FAM193B | chr5 | - | 177531281 | 177531521 | 177524184 | 177525205 | 177532442 | 177532641 | 0 | 3 | 7 | 0 | 0 | 0 | 0 | 1 | 2 | 0 | 1 | 2 | 98 | 49 | 0.001603576 | 0.170760496 | NA | 1 | 1 | NA | 0.333 | 0.333 | 0.667 |
| SE | 7639 | IRF7 | chr11 | - | 614173 | 614399 | 613784 | 613865 | 614475 | 614534 | 0 | 0 | 0 | 0 | 8 | 3 | 0 | 4 | 1 | 0 | 3 | 2 | 98 | 49 | 0.001756837 | 0.170760496 | NA | 0 | 0 | NA | 0.4 | 0.2 | -0.3 |
| SE | 6866 | ATG2A | chr11 | - | 64906664 | 64906815 | 64906333 | 64906533 | 64907254 | 64907439 | 0 | 2 | 5 | 0 | 0 | 0 | 0 | 6 | 5 | 0 | 2 | 1 | 98 | 49 | 0.001882581 | 0.170760496 | NA | 1 | 1 | NA | 0.6 | 0.714 | 0.343 |
| SE | 9706 | REC8 | chr14 | + | 24175542 | 24175624 | 24173290 | 24173411 | 24176821 | 24176901 | 0 | 3 | 10 | 0 | 0 | 0 | 1 | 3 | 1 | 0 | 2 | 4 | 98 | 49 | 0.002515847 | 0.170760496 | NA | 1 | 1 | 1 | 0.429 | 0.111 | 0.487 |
| SE | 13860 | ACSL1 | chr4 | - | 184764852 | 184764925 | 184763166 | 184763255 | 184765890 | 184765986 | 0 | 2 | 50 | 0 | 0 | 0 | 7 | 51 | 14 | 0 | 2 | 2 | 98 | 49 | 0.003078802 | 0.170760496 | NA | 1 | 1 | 1 | 0.927 | 0.778 | 0.098 |
| SE | 8555 | PIK3AP1 | chr10 | - | 96652697 | 96652842 | 96651508 | 96651651 | 96656797 | 96656934 | 0 | 1 | 15 | 0 | 0 | 0 | 0 | 4 | 7 | 0 | 2 | 1 | 98 | 49 | 0.003471069 | 0.170760496 | NA | 1 | 1 | NA | 0.5 | 0.778 | 0.361 |
| SE | 11710 | SENP3-EIF4A1 | chr17 | + | 7575118 | 7575258 | 7574545 | 7574678 | 7576523 | 7576692 | 0 | 2 | 33 | 0 | 0 | 0 | 3 | 11 | 2 | 0 | 1 | 1 | 98 | 49 | 0.004779846 | 0.170760496 | NA | 1 | 1 | 1 | 0.846 | 0.5 | 0.218 |
| SE | 12939 | SMCHD1 | chr18 | + | 2666156 | 2666232 | 2655886 | 2656261 | 2666869 | 2667031 | 0 | 20 | 98 | 0 | 0 | 0 | 4 | 61 | 8 | 0 | 1 | 1 | 98 | 49 | 0.00491362 | 0.170760496 | NA | 1 | 1 | 1 | 0.968 | 0.8 | 0.077 |
| A3SS | 1863 | SRPK1 | chr6 | - | 35890894 | 35891041 | 35890894 | 35891013 | 35920467 | 35920528 | 0 | 0 | 0 | 0 | 1 | 2 | 0 | 1 | 1 | 0 | 4 | 1 | 77 | 49 | 0.005762586 | 0.197717869 | NA | 0 | 0 | NA | 0.137 | 0.389 | -0.263 |
| SE | 12824 | SEC14L1 | chr17 | + | 77196201 | 77196311 | 77194676 | 77194911 | 77200483 | 77200673 | 0 | 11 | 34 | 0 | 0 | 0 | 0 | 20 | 0 | 0 | 11 | 2 | 98 | 49 | 0.005971369 | 0.170760496 | NA | 1 | 1 | NA | 0.476 | 0 | 0.762 |
| A5SS | 72 | CSF2RA | chrX | + | 1274754 | 1274842 | 1274754 | 1274818 | 1282677 | 1282779 | 0 | 1 | 3 | 0 | 0 | 0 | 0 | 2 | 0 | 0 | 2 | 1 | 73 | 49 | 0.006411688 | 0.170414363 | NA | 1 | 1 | NA | 0.402 | 0 | 0.799 |
| SE | 4425 | PREX1 | chr20 | - | 48659918 | 48660061 | 48658135 | 48658228 | 48666282 | 48666355 | 0 | 1 | 31 | 0 | 0 | 0 | 0 | 16 | 2 | 0 | 1 | 1 | 98 | 49 | 0.008019007 | 0.170760496 | NA | 1 | 1 | NA | 0.889 | 0.5 | 0.305 |
| SE | 11393 | ADGRG3 | chr16 | + | 57676199 | 57676338 | 57673320 | 57673468 | 57678169 | 57678316 | 0 | 2 | 14 | 0 | 0 | 0 | 0 | 25 | 2 | 0 | 3 | 3 | 98 | 49 | 0.008245186 | 0.170760496 | NA | 1 | 1 | NA | 0.806 | 0.25 | 0.472 |
| SE | 9930 | PTPRE | chr10 | + | 128060938 | 128061015 | 128056122 | 128056213 | 128061678 | 128061715 | 0 | 3 | 11 | 0 | 0 | 0 | 0 | 22 | 0 | 0 | 7 | 0 | 98 | 49 | 0.009650972 | 0.170760496 | NA | 1 | 1 | NA | 0.611 | NA | 0.389 |
| SE | 5497 | ADAM8 | chr10 | - | 133270880 | 133271070 | 133270735 | 133270805 | 133271199 | 133271289 | 1 | 20 | 40 | 0 | 0 | 0 | 4 | 55 | 21 | 0 | 1 | 2 | 98 | 49 | 0.010036062 | 0.170760496 | 1 | 1 | 1 | 1 | 0.965 | 0.84 | 0.065 |
| SE | 11923 | RBM39 | chr20 | - | 35740524 | 35740597 | 35738967 | 35739017 | 35740823 | 35740887 | 0 | 2 | 8 | 0 | 0 | 1 | 0 | 8 | 0 | 0 | 8 | 0 | 98 | 49 | 0.010118399 | 0.170760496 | NA | 1 | 0.8 | NA | 0.333 | NA | 0.567 |
| SE | 14093 | NLRP6 | chr11 | + | 280083 | 281839 | 279833 | 279872 | 282704 | 282797 | 0 | 1 | 2 | 0 | 0 | 0 | 0 | 1 | 0 | 0 | 1 | 2 | 98 | 49 | 0.010649553 | 0.170760496 | NA | 1 | 1 | NA | 0.333 | 0 | 0.834 |
| SE | 10848 | IRF1 | chr5 | - | 132486250 | 132486373 | 132485666 | 132485716 | 132486556 | 132486686 | 0 | 2 | 36 | 0 | 0 | 1 | 7 | 13 | 0 | 0 | 6 | 0 | 98 | 49 | 0.011205299 | 0.170760496 | NA | 1 | 0.947 | 1 | 0.52 | NA | 0.214 |
| SE | 2075 | HERC5 | chr4 | + | 88462134 | 88462356 | 88460094 | 88460171 | 88463531 | 88463623 | 0 | 4 | 25 | 0 | 0 | 2 | 0 | 11 | 0 | 0 | 5 | 0 | 98 | 49 | 0.011241877 | 0.170760496 | NA | 1 | 0.862 | NA | 0.524 | NA | 0.407 |
| SE | 8188 | EPSTI1 | chr13 | - | 42919299 | 42919332 | 42917540 | 42917624 | 42926335 | 42926429 | 0 | 0 | 4 | 0 | 4 | 14 | 0 | 6 | 0 | 0 | 10 | 0 | 82 | 49 | 0.011354222 | 0.170760496 | NA | 0 | 0.146 | NA | 0.264 | NA | -0.191 |
| SE | 4426 | PREX1 | chr20 | - | 48726289 | 48726391 | 48708259 | 48708421 | 48734545 | 48734650 | 1 | 23 | 82 | 0 | 0 | 0 | 8 | 65 | 15 | 0 | 2 | 3 | 98 | 49 | 0.01202775 | 0.170760496 | 1 | 1 | 1 | 1 | 0.942 | 0.714 | 0.115 |
| SE | 3738 | BRD2 | chr6 | + | 32975020 | 32975112 | 32974461 | 32974765 | 32975383 | 32975521 | 0 | 1 | 3 | 0 | 0 | 3 | 0 | 4 | 0 | 0 | 8 | 0 | 98 | 49 | 0.012603077 | 0.170760496 | NA | 1 | 0.333 | NA | 0.2 | NA | 0.466 |
| SE | 3103 | MX2 | chr21 | + | 41380016 | 41380151 | 41377788 | 41377981 | 41382409 | 41382564 | 0 | 4 | 25 | 0 | 0 | 2 | 0 | 16 | 5 | 0 | 11 | 2 | 98 | 49 | 0.013190421 | 0.170760496 | NA | 1 | 0.862 | NA | 0.421 | 0.556 | 0.443 |
| SE | 10602 | DMXL2 | chr15 | - | 51495023 | 51495134 | 51491577 | 51491747 | 51498551 | 51500231 | 0 | 1 | 12 | 0 | 0 | 0 | 0 | 7 | 0 | 0 | 5 | 0 | 98 | 49 | 0.014099979 | 0.170760496 | NA | 1 | 1 | NA | 0.412 | NA | 0.588 |
| SE | 3545 | OSBPL2 | chr20 | + | 62259980 | 62260125 | 62256056 | 62256221 | 62263615 | 62263691 | 0 | 2 | 3 | 0 | 0 | 0 | 0 | 5 | 0 | 0 | 4 | 0 | 98 | 49 | 0.014587636 | 0.170760496 | NA | 1 | 1 | NA | 0.385 | NA | 0.615 |
| SE | 12803 | CYP4F3 | chr19 | + | 15645718 | 15645863 | 15641414 | 15641613 | 15647051 | 15647105 | 0 | 5 | 21 | 0 | 0 | 0 | 7 | 13 | 0 | 0 | 3 | 0 | 98 | 49 | 0.014732939 | 0.170760496 | NA | 1 | 1 | 1 | 0.684 | NA | 0.158 |
| SE | 1826 | MCTP1 | chr5 | - | 94917895 | 94917973 | 94912805 | 94912976 | 94923961 | 94924021 | 0 | 2 | 3 | 0 | 0 | 0 | 0 | 4 | 0 | 0 | 4 | 0 | 98 | 49 | 0.014963682 | 0.170760496 | NA | 1 | 1 | NA | 0.333 | NA | 0.667 |
| SE | 6336 | FCAR | chr19 | + | 54888006 | 54888294 | 54885234 | 54885525 | 54889648 | 54891420 | 0 | 5 | 28 | 0 | 0 | 1 | 0 | 19 | 0 | 0 | 3 | 0 | 98 | 49 | 0.01511042 | 0.170760496 | NA | 1 | 0.933 | NA | 0.76 | NA | 0.207 |
| SE | 2203 | CPEB2 | chr4 | + | 15040463 | 15040487 | 15033160 | 15033211 | 15052413 | 15052584 | 0 | 0 | 1 | 0 | 5 | 1 | 0 | 5 | 0 | 0 | 2 | 0 | 73 | 49 | 0.015421435 | 0.170760496 | NA | 0 | 0.402 | NA | 0.627 | NA | -0.426 |
| SE | 12815 | BST2 | chr19 | - | 17404370 | 17404437 | 17404128 | 17404189 | 17405290 | 17405648 | 0 | 4 | 23 | 0 | 0 | 0 | 0 | 4 | 0 | 0 | 3 | 0 | 98 | 49 | 0.015554946 | 0.170760496 | NA | 1 | 1 | NA | 0.4 | NA | 0.6 |
| SE | 11842 | RHOT1 | chr17 | + | 32172936 | 32173060 | 32171042 | 32171101 | 32173830 | 32173912 | 0 | 5 | 3 | 0 | 0 | 0 | 0 | 3 | 0 | 0 | 3 | 0 | 98 | 49 | 0.015643271 | 0.170760496 | NA | 1 | 1 | NA | 0.333 | NA | 0.667 |
| SE | 6335 | FCAR | chr19 | + | 54888006 | 54888228 | 54885234 | 54885525 | 54889648 | 54891420 | 0 | 3 | 17 | 0 | 0 | 1 | 0 | 12 | 0 | 0 | 3 | 0 | 98 | 49 | 0.015872783 | 0.170760496 | NA | 1 | 0.895 | NA | 0.667 | NA | 0.28 |
| SE | 5621 | ANKRD12 | chr18 | + | 9204475 | 9204544 | 9195550 | 9195698 | 9208656 | 9208803 | 0 | 1 | 3 | 0 | 0 | 2 | 0 | 5 | 0 | 0 | 4 | 0 | 98 | 49 | 0.016160337 | 0.170760496 | NA | 1 | 0.429 | NA | 0.385 | NA | 0.33 |
| SE | 10536 | SGK1 | chr6 | - | 134170825 | 134170915 | 134169245 | 134170435 | 134171022 | 134171178 | 0 | 1 | 11 | 0 | 0 | 0 | 1 | 16 | 0 | 0 | 4 | 0 | 98 | 49 | 0.016182238 | 0.170760496 | NA | 1 | 1 | 1 | 0.667 | NA | 0.166 |
| SE | 6177 | ABCA1 | chr9 | - | 104825682 | 104825887 | 104824464 | 104824578 | 104826947 | 104827169 | 0 | 4 | 6 | 0 | 0 | 0 | 0 | 21 | 1 | 0 | 5 | 0 | 98 | 49 | 0.016495032 | 0.170760496 | NA | 1 | 1 | NA | 0.677 | 1 | 0.161 |
| SE | 799 | TBXAS1 | chr7 | + | 139911224 | 139911321 | 139875584 | 139875637 | 139936190 | 139936307 | 0 | 2 | 22 | 0 | 0 | 0 | 0 | 7 | 0 | 0 | 3 | 0 | 98 | 49 | 0.016691194 | 0.170760496 | NA | 1 | 1 | NA | 0.538 | NA | 0.462 |
| A5SS | 1875 | N4BP2L2 | chr13 | - | 32527407 | 32527532 | 32527457 | 32527532 | 32522181 | 32522270 | 0 | 2 | 17 | 0 | 0 | 2 | 0 | 9 | 0 | 0 | 3 | 0 | 98 | 49 | 0.016742666 | 0.170414363 | NA | 1 | 0.81 | NA | 0.6 | NA | 0.305 |
| SE | 7593 | ATG16L2 | chr11 | + | 72822847 | 72822961 | 72822477 | 72822543 | 72824059 | 72824122 | 0 | 9 | 41 | 0 | 0 | 1 | 4 | 19 | 16 | 0 | 4 | 2 | 98 | 49 | 0.017045614 | 0.170760496 | NA | 1 | 0.953 | 1 | 0.704 | 0.8 | 0.142 |
| A3SS | 2805 | TPT1 | chr13 | - | 45333470 | 45337553 | 45333470 | 45337388 | 45338659 | 45338776 | 0 | 0 | 2 | 1 | 3 | 72 | 0 | 4 | 0 | 0 | 47 | 0 | 98 | 49 | 0.017197054 | 0.197717869 | 0 | 0 | 0.014 | NA | 0.041 | NA | -0.036 |
| SE | 883 | SH2B2 | chr7 | + | 102300521 | 102301279 | 102285061 | 102285258 | 102306720 | 102306822 | 0 | 4 | 0 | 0 | 0 | 0 | 0 | 2 | 1 | 0 | 3 | 0 | 98 | 49 | 0.017202771 | 0.170760496 | NA | 1 | NA | NA | 0.25 | 1 | 0.375 |
| SE | 1452 | PRRC2A | chr6 | + | 31625791 | 31625871 | 31625459 | 31625611 | 31626019 | 31626162 | 0 | 4 | 4 | 0 | 0 | 1 | 0 | 1 | 0 | 0 | 5 | 0 | 98 | 49 | 0.017880599 | 0.170760496 | NA | 1 | 0.667 | NA | 0.091 | NA | 0.743 |
| SE | 6626 | PCBP2 | chr12 | + | 53464759 | 53464852 | 53462480 | 53462567 | 53465931 | 53465973 | 0 | 0 | 23 | 0 | 2 | 16 | 0 | 11 | 0 | 0 | 1 | 0 | 98 | 49 | 0.017987559 | 0.170760496 | NA | 0 | 0.418 | NA | 0.846 | NA | -0.637 |
| SE | 6174 | ABCA1 | chr9 | - | 104810799 | 104810924 | 104809465 | 104809564 | 104812573 | 104812722 | 0 | 0 | 3 | 0 | 1 | 1 | 0 | 4 | 0 | 2 | 4 | 0 | 98 | 49 | 0.018518561 | 0.170760496 | NA | 0 | 0.6 | 0 | 0.333 | NA | 0.133 |
| SE | 11731 | TAOK3 | chr12 | - | 118212913 | 118212995 | 118201295 | 118201463 | 118214016 | 118214110 | 1 | 2 | 4 | 0 | 0 | 1 | 0 | 2 | 0 | 0 | 3 | 0 | 98 | 49 | 0.018766874 | 0.170760496 | 1 | 1 | 0.667 | NA | 0.25 | NA | 0.639 |
| SE | 3208 | STX16-NPEPL1 | chr20 | + | 58671153 | 58671297 | 58670511 | 58670603 | 58673630 | 58673711 | 0 | 1 | 16 | 0 | 0 | 1 | 0 | 5 | 0 | 0 | 3 | 0 | 98 | 49 | 0.018766876 | 0.170760496 | NA | 1 | 0.889 | NA | 0.455 | NA | 0.489 |
| SE | 1982 | DDX60L | chr4 | - | 168419301 | 168419397 | 168416681 | 168416797 | 168420260 | 168420380 | 0 | 1 | 32 | 0 | 0 | 1 | 0 | 8 | 0 | 0 | 3 | 0 | 98 | 49 | 0.018767562 | 0.170760496 | NA | 1 | 0.941 | NA | 0.571 | NA | 0.399 |
| SE | 6334 | FCAR | chr19 | + | 54888006 | 54888220 | 54885234 | 54885525 | 54889648 | 54891420 | 0 | 1 | 11 | 0 | 0 | 1 | 0 | 10 | 0 | 0 | 3 | 0 | 98 | 49 | 0.018937654 | 0.170760496 | NA | 1 | 0.846 | NA | 0.625 | NA | 0.298 |
| SE | 4057 | APBB1IP | chr10 | + | 26536073 | 26536217 | 26533438 | 26533525 | 26541581 | 26541692 | 0 | 1 | 12 | 0 | 0 | 0 | 3 | 4 | 0 | 0 | 3 | 0 | 98 | 49 | 0.018984944 | 0.170760496 | NA | 1 | 1 | 1 | 0.4 | NA | 0.3 |
| SE | 3889 | MAP7D1 | chr1 | + | 36174897 | 36175008 | 36173363 | 36173478 | 36176198 | 36176321 | 0 | 0 | 4 | 0 | 2 | 2 | 0 | 8 | 0 | 0 | 1 | 0 | 98 | 49 | 0.019005986 | 0.170760496 | NA | 0 | 0.5 | NA | 0.8 | NA | -0.55 |
| A3SS | 1378 | OSBPL8 | chr12 | - | 76450850 | 76450988 | 76450850 | 76450979 | 76487509 | 76487618 | 0 | 0 | 3 | 0 | 1 | 1 | 0 | 5 | 0 | 0 | 2 | 0 | 58 | 49 | 0.019031697 | 0.197717869 | NA | 0 | 0.717 | NA | 0.679 | NA | -0.321 |
| SE | 5821 | MGEA5 | chr10 | - | 101793912 | 101793998 | 101792838 | 101792943 | 101797979 | 101798154 | 0 | 0 | 3 | 0 | 2 | 1 | 0 | 3 | 0 | 0 | 4 | 0 | 98 | 49 | 0.019064109 | 0.170760496 | NA | 0 | 0.6 | NA | 0.273 | NA | 0.027 |
| SE | 5241 | PLXNC1 | chr12 | + | 94209589 | 94209704 | 94186372 | 94186473 | 94220015 | 94220163 | 0 | 4 | 27 | 0 | 0 | 1 | 0 | 5 | 0 | 0 | 2 | 0 | 98 | 49 | 0.019110193 | 0.170760496 | NA | 1 | 0.931 | NA | 0.556 | NA | 0.409 |
| A3SS | 3389 | RBM39 | chr20 | - | 35709223 | 35709274 | 35709223 | 35709256 | 35713018 | 35713096 | 0 | 0 | 7 | 0 | 1 | 1 | 0 | 4 | 0 | 0 | 3 | 0 | 67 | 49 | 0.019156561 | 0.197717869 | NA | 0 | 0.837 | NA | 0.494 | NA | -0.076 |
| SE | 10591 | RNF213 | chr17 | + | 80383676 | 80383928 | 80382978 | 80383070 | 80385537 | 80385621 | 0 | 1 | 30 | 0 | 0 | 0 | 1 | 12 | 0 | 0 | 3 | 0 | 98 | 49 | 0.019182317 | 0.170760496 | NA | 1 | 1 | 1 | 0.667 | NA | 0.166 |
| SE | 11022 | TAP1 | chr6 | - | 32847918 | 32848092 | 32847512 | 32847675 | 32848651 | 32848840 | 0 | 4 | 21 | 0 | 0 | 0 | 0 | 7 | 0 | 0 | 2 | 0 | 98 | 49 | 0.019207741 | 0.170760496 | NA | 1 | 1 | NA | 0.636 | NA | 0.364 |
| A5SS | 2325 | DEDD2 | chr19 | - | 42201988 | 42202072 | 42202050 | 42202072 | 42198592 | 42199829 | 0 | 4 | 0 | 0 | 0 | 1 | 2 | 3 | 0 | 0 | 2 | 0 | 98 | 49 | 0.019484918 | 0.170414363 | NA | 1 | 0 | 1 | 0.429 | NA | -0.215 |
| SE | 5270 | BID | chr22 | - | 17750104 | 17750174 | 17743802 | 17744013 | 17774380 | 17774495 | 0 | 4 | 1 | 0 | 0 | 0 | 0 | 3 | 0 | 0 | 2 | 0 | 98 | 49 | 0.019484918 | 0.170760496 | NA | 1 | 1 | NA | 0.429 | NA | 0.571 |
| SE | 7455 | FPR1 | chr19 | - | 51750202 | 51750253 | 51745769 | 51747005 | 51751813 | 51751897 | 0 | 1 | 2 | 0 | 0 | 1 | 0 | 3 | 0 | 0 | 3 | 0 | 98 | 49 | 0.019485123 | 0.170760496 | NA | 1 | 0.5 | NA | 0.333 | NA | 0.417 |
| SE | 7415 | VAV1 | chr19 | + | 6821790 | 6821859 | 6821621 | 6821680 | 6822220 | 6822329 | 0 | 4 | 4 | 0 | 0 | 0 | 3 | 11 | 0 | 0 | 2 | 0 | 98 | 49 | 0.01972669 | 0.170760496 | NA | 1 | 1 | 1 | 0.733 | NA | 0.133 |
| SE | 9526 | EIF4G2 | chr11 | - | 10801660 | 10801774 | 10800961 | 10801087 | 10802048 | 10802209 | 0 | 0 | 6 | 0 | 1 | 1 | 2 | 10 | 0 | 0 | 1 | 0 | 98 | 49 | 0.01979002 | 0.170760496 | NA | 0 | 0.75 | 1 | 0.833 | NA | -0.541 |
| SE | 9353 | GBP5 | chr1 | - | 89262682 | 89262785 | 89262219 | 89262401 | 89263735 | 89263948 | 0 | 3 | 3 | 0 | 0 | 0 | 1 | 6 | 0 | 0 | 2 | 0 | 98 | 49 | 0.019996576 | 0.170760496 | NA | 1 | 1 | 1 | 0.6 | NA | 0.2 |
| SE | 12946 | SMCHD1 | chr18 | + | 2750042 | 2750122 | 2747521 | 2747647 | 2750349 | 2750507 | 0 | 2 | 54 | 0 | 0 | 8 | 0 | 46 | 0 | 0 | 3 | 0 | 98 | 49 | 0.020053524 | 0.170760496 | NA | 1 | 0.771 | NA | 0.885 | NA | 0 |
| SE | 496 | EEF1D | chr8 | - | 143586218 | 143586290 | 143581228 | 143581328 | 143586728 | 143586849 | 0 | 3 | 12 | 0 | 0 | 0 | 0 | 8 | 0 | 0 | 2 | 0 | 98 | 49 | 0.020218801 | 0.170760496 | NA | 1 | 1 | NA | 0.667 | NA | 0.333 |
| SE | 4780 | TXNRD1 | chr12 | + | 104321090 | 104321316 | 104319469 | 104319585 | 104325336 | 104325429 | 0 | 3 | 2 | 0 | 0 | 0 | 0 | 8 | 0 | 0 | 2 | 0 | 98 | 49 | 0.020218801 | 0.170760496 | NA | 1 | 1 | NA | 0.667 | NA | 0.333 |
| SE | 4967 | DDX3X | chrX | + | 41347314 | 41347451 | 41346858 | 41347012 | 41347639 | 41350287 | 0 | 1 | 17 | 0 | 0 | 0 | 0 | 20 | 0 | 0 | 3 | 0 | 98 | 49 | 0.020293915 | 0.170760496 | NA | 1 | 1 | NA | 0.769 | NA | 0.231 |
| SE | 1577 | USP9X | chrX | + | 41123470 | 41123724 | 41085634 | 41086109 | 41128999 | 41129145 | 0 | 0 | 4 | 0 | 0 | 0 | 0 | 1 | 2 | 0 | 0 | 2 | 98 | 49 | 0.020294338 | 0.170760496 | NA | NA | 1 | NA | 1 | 0.333 | 0.334 |
| A5SS | 976 | CTAGE5 | chr14 | + | 39276933 | 39277065 | 39276933 | 39276955 | 39279336 | 39279358 | 0 | 1 | 6 | 0 | 0 | 0 | 0 | 2 | 0 | 0 | 3 | 0 | 98 | 49 | 0.020627966 | 0.170414363 | NA | 1 | 1 | NA | 0.25 | NA | 0.75 |
| SE | 9058 | FAM107B | chr10 | - | 14553321 | 14553387 | 14530331 | 14530515 | 14604239 | 14604418 | 0 | 1 | 0 | 0 | 0 | 0 | 0 | 2 | 0 | 0 | 3 | 0 | 98 | 49 | 0.020627966 | 0.170760496 | NA | 1 | NA | NA | 0.25 | NA | 0.75 |
| SE | 11109 | USP25 | chr21 | + | 15824038 | 15824166 | 15818697 | 15818846 | 15824965 | 15825061 | 0 | 1 | 3 | 0 | 0 | 2 | 0 | 2 | 0 | 0 | 3 | 0 | 98 | 49 | 0.020627966 | 0.170760496 | NA | 1 | 0.429 | NA | 0.25 | NA | 0.465 |
| SE | 7818 | DPEP2 | chr16 | - | 67990046 | 67990131 | 67989322 | 67989398 | 67990820 | 67990997 | 0 | 1 | 15 | 0 | 0 | 0 | 0 | 5 | 0 | 0 | 1 | 1 | 98 | 49 | 0.020811238 | 0.170760496 | NA | 1 | 1 | NA | 0.714 | 0 | 0.643 |
| SE | 3457 | CDS2 | chr20 | + | 5186686 | 5186839 | 5185757 | 5185826 | 5189066 | 5189186 | 0 | 3 | 8 | 0 | 0 | 0 | 0 | 1 | 0 | 0 | 3 | 0 | 98 | 49 | 0.020882905 | 0.170760496 | NA | 1 | 1 | NA | 0.143 | NA | 0.857 |
| A5SS | 2345 | HN1 | chr17 | - | 75147519 | 75147653 | 75147555 | 75147653 | 75146665 | 75146684 | 2 | 0 | 1 | 0 | 1 | 2 | 0 | 0 | 0 | 4 | 5 | 0 | 85 | 49 | 0.020913098 | 0.170414363 | 1 | 0 | 0.224 | 0 | 0 | NA | 0.408 |
| SE | 6630 | PCBP2 | chr12 | + | 53467804 | 53467843 | 53467220 | 53467293 | 53468776 | 53468832 | 0 | 1 | 9 | 0 | 0 | 7 | 8 | 2 | 0 | 1 | 3 | 0 | 88 | 49 | 0.020946855 | 0.170760496 | NA | 1 | 0.417 | 0.817 | 0.271 | NA | 0.164 |
| SE | 1457 | ATP6V1G2-DDX39B | chr6 | - | 31539146 | 31539274 | 31538762 | 31538855 | 31540321 | 31540664 | 0 | 2 | 20 | 0 | 0 | 0 | 0 | 4 | 0 | 0 | 2 | 0 | 98 | 49 | 0.021248483 | 0.170760496 | NA | 1 | 1 | NA | 0.5 | NA | 0.5 |
| SE | 6867 | ATG2A | chr11 | - | 64908990 | 64909150 | 64907747 | 64907890 | 64909270 | 64909367 | 0 | 0 | 2 | 0 | 0 | 0 | 0 | 10 | 4 | 0 | 1 | 2 | 98 | 49 | 0.021248483 | 0.170760496 | NA | NA | 1 | NA | 0.833 | 0.5 | 0.334 |
| A5SS | 2264 | MTCH1 | chr6 | - | 36972651 | 36972796 | 36972702 | 36972796 | 36970646 | 36970694 | 0 | 2 | 9 | 0 | 0 | 8 | 0 | 6 | 0 | 0 | 2 | 0 | 98 | 49 | 0.021315983 | 0.170414363 | NA | 1 | 0.36 | NA | 0.6 | NA | 0.08 |
| SE | 12526 | GUCD1 | chr22 | - | 24548916 | 24549001 | 24547907 | 24548073 | 24554948 | 24555307 | 0 | 2 | 0 | 0 | 0 | 0 | 0 | 6 | 2 | 0 | 2 | 0 | 98 | 49 | 0.021315983 | 0.170760496 | NA | 1 | NA | NA | 0.6 | 1 | 0.2 |
| SE | 5012 | ADRBK1 | chr11 | + | 67281092 | 67281184 | 67280731 | 67280783 | 67281458 | 67281558 | 0 | 2 | 32 | 0 | 0 | 0 | 0 | 7 | 0 | 0 | 2 | 0 | 98 | 49 | 0.021459804 | 0.170760496 | NA | 1 | 1 | NA | 0.636 | NA | 0.364 |
| SE | 2371 | MFSD1 | chr3 | + | 158821983 | 158822140 | 158821596 | 158821653 | 158823427 | 158823525 | 0 | 2 | 16 | 0 | 0 | 0 | 0 | 3 | 0 | 0 | 2 | 0 | 98 | 49 | 0.021489596 | 0.170760496 | NA | 1 | 1 | NA | 0.429 | NA | 0.571 |
| SE | 10746 | MAN2C1 | chr15 | - | 75363998 | 75364188 | 75362641 | 75362748 | 75364487 | 75364665 | 0 | 2 | 0 | 0 | 0 | 2 | 0 | 3 | 0 | 0 | 2 | 0 | 98 | 49 | 0.021489596 | 0.170760496 | NA | 1 | 0 | NA | 0.429 | NA | 0.071 |
| SE | 13957 | DIP2B | chr12 | + | 50671185 | 50671398 | 50660193 | 50660319 | 50674473 | 50674629 | 0 | 2 | 5 | 0 | 0 | 0 | 0 | 3 | 0 | 0 | 2 | 0 | 98 | 49 | 0.021489596 | 0.170760496 | NA | 1 | 1 | NA | 0.429 | NA | 0.571 |
| SE | 12161 | HSH2D | chr19 | + | 16151446 | 16151578 | 16148723 | 16148875 | 16152551 | 16153208 | 0 | 0 | 1 | 0 | 4 | 15 | 0 | 2 | 0 | 0 | 9 | 0 | 98 | 49 | 0.021888757 | 0.170760496 | NA | 0 | 0.032 | NA | 0.1 | NA | -0.084 |
| SE | 6970 | MFN2 | chr1 | + | 12007052 | 12007249 | 12006537 | 12006693 | 12009591 | 12009726 | 0 | 0 | 5 | 0 | 1 | 0 | 0 | 6 | 0 | 0 | 1 | 0 | 98 | 49 | 0.021892715 | 0.170760496 | NA | 0 | 1 | NA | 0.75 | NA | -0.25 |
| A3SS | 689 | CNBP | chr3 | - | 129171078 | 129171280 | 129171078 | 129171277 | 129171445 | 129171538 | 0 | 0 | 2 | 0 | 1 | 4 | 0 | 3 | 0 | 0 | 11 | 0 | 52 | 49 | 0.021942316 | 0.197717869 | NA | 0 | 0.32 | NA | 0.204 | NA | -0.044 |
| A3SS | 490 | GRK6 | chr5 | + | 177441734 | 177442849 | 177441736 | 177442849 | 177440918 | 177441053 | 0 | 1 | 6 | 0 | 0 | 2 | 0 | 0 | 0 | 0 | 2 | 2 | 51 | 49 | 0.021987328 | 0.197717869 | NA | 1 | 0.742 | NA | 0 | 0 | 0.871 |
| SE | 4999 | WIPI2 | chr7 | + | 5193117 | 5193171 | 5190203 | 5190493 | 5199575 | 5199658 | 0 | 1 | 3 | 0 | 0 | 1 | 0 | 0 | 0 | 0 | 2 | 2 | 98 | 49 | 0.021999349 | 0.170760496 | NA | 1 | 0.6 | NA | 0 | 0 | 0.8 |
| SE | 2204 | CPEB2 | chr4 | + | 15059186 | 15059301 | 15058420 | 15058539 | 15062078 | 15062260 | 0 | 1 | 0 | 0 | 1 | 2 | 0 | 3 | 3 | 0 | 0 | 0 | 98 | 49 | 0.022049079 | 0.170760496 | NA | 0.333 | 0 | NA | 1 | 1 | -0.834 |
| A5SS | 2821 | LILRA6 | chr19 | - | 54241575 | 54241878 | 54241661 | 54241878 | 54240830 | 54241127 | 0 | 0 | 0 | 0 | 2 | 1 | 0 | 4 | 0 | 0 | 1 | 0 | 98 | 49 | 0.022093415 | 0.170414363 | NA | 0 | 0 | NA | 0.667 | NA | -0.667 |
| SE | 9285 | VIM | chr10 | + | 17235845 | 17235889 | 17235168 | 17235389 | 17236293 | 17236379 | 0 | 12 | 171 | 0 | 0 | 0 | 3 | 36 | 3 | 0 | 3 | 0 | 93 | 49 | 0.022225644 | 0.170760496 | NA | 1 | 1 | 1 | 0.863 | 1 | 0.046 |
| SE | 13688 | BAZ2A | chr12 | - | 56605829 | 56606063 | 56605072 | 56605327 | 56606246 | 56606312 | 0 | 2 | 2 | 0 | 0 | 0 | 0 | 2 | 0 | 0 | 2 | 0 | 98 | 49 | 0.022262485 | 0.170760496 | NA | 1 | 1 | NA | 0.333 | NA | 0.667 |
| SE | 14090 | AREL1 | chr14 | - | 74669946 | 74670126 | 74669648 | 74669774 | 74670761 | 74670871 | 0 | 2 | 2 | 0 | 0 | 3 | 0 | 2 | 0 | 0 | 2 | 0 | 98 | 49 | 0.022262485 | 0.170760496 | NA | 1 | 0.25 | NA | 0.333 | NA | 0.292 |
| SE | 6878 | UBN1 | chr16 | + | 4868832 | 4868903 | 4860663 | 4861102 | 4870211 | 4870341 | 0 | 0 | 4 | 0 | 1 | 0 | 0 | 5 | 0 | 0 | 1 | 0 | 98 | 49 | 0.022832961 | 0.170760496 | NA | 0 | 1 | NA | 0.714 | NA | -0.214 |
| SE | 11349 | ADGRE2 | chr19 | - | 14754953 | 14755127 | 14752328 | 14752526 | 14756237 | 14756345 | 0 | 0 | 3 | 0 | 1 | 1 | 0 | 5 | 0 | 0 | 1 | 0 | 98 | 49 | 0.022832961 | 0.170760496 | NA | 0 | 0.6 | NA | 0.714 | NA | -0.414 |
| SE | 3680 | USP15 | chr12 | + | 62325871 | 62325933 | 62321463 | 62321609 | 62349220 | 62349307 | 0 | 2 | 32 | 0 | 0 | 1 | 0 | 15 | 0 | 0 | 2 | 0 | 98 | 49 | 0.023010064 | 0.170760496 | NA | 1 | 0.941 | NA | 0.789 | NA | 0.181 |
| SE | 10210 | RFWD2 | chr1 | - | 176175909 | 176176007 | 176163814 | 176163891 | 176184632 | 176184692 | 0 | 2 | 13 | 0 | 0 | 0 | 0 | 17 | 0 | 0 | 2 | 0 | 98 | 49 | 0.023408389 | 0.170760496 | NA | 1 | 1 | NA | 0.81 | NA | 0.19 |
| SE | 558 | PABPC1 | chr8 | - | 100713086 | 100713181 | 100712651 | 100712789 | 100715461 | 100715601 | 0 | 6 | 63 | 0 | 0 | 0 | 5 | 32 | 2 | 0 | 3 | 0 | 98 | 49 | 0.0234244 | 0.170760496 | NA | 1 | 1 | 1 | 0.842 | 1 | 0.053 |
| SE | 17 | FLNA | chrX | - | 154353553 | 154353727 | 154353295 | 154353457 | 154353914 | 154354043 | 0 | 2 | 35 | 0 | 0 | 1 | 0 | 18 | 0 | 2 | 2 | 0 | 98 | 49 | 0.023592394 | 0.170760496 | NA | 1 | 0.946 | 0 | 0.818 | NA | 0.564 |
| A5SS | 93 | GAPVD1 | chr9 | + | 125321432 | 125321562 | 125321432 | 125321499 | 125323797 | 125323923 | 0 | 3 | 3 | 0 | 0 | 0 | 0 | 1 | 0 | 0 | 2 | 0 | 98 | 49 | 0.023673881 | 0.170414363 | NA | 1 | 1 | NA | 0.2 | NA | 0.8 |
| SE | 2652 | SETD2 | chr3 | - | 47101457 | 47101555 | 47097954 | 47098081 | 47103345 | 47103423 | 0 | 1 | 10 | 0 | 0 | 0 | 0 | 4 | 0 | 0 | 2 | 0 | 98 | 49 | 0.023775553 | 0.170760496 | NA | 1 | 1 | NA | 0.5 | NA | 0.5 |
| SE | 4064 | FBXL5 | chr4 | - | 15638507 | 15638694 | 15636493 | 15636676 | 15640787 | 15640883 | 0 | 3 | 71 | 0 | 0 | 0 | 6 | 32 | 0 | 0 | 2 | 0 | 98 | 49 | 0.0238404 | 0.170760496 | NA | 1 | 1 | 1 | 0.889 | NA | 0.055 |
| A3SS | 681 | CDV3 | chr3 | + | 133587158 | 133590274 | 133587895 | 133590274 | 133586562 | 133586722 | 0 | 1 | 2 | 0 | 0 | 0 | 0 | 5 | 0 | 0 | 2 | 0 | 98 | 49 | 0.023845162 | 0.197717869 | NA | 1 | 1 | NA | 0.556 | NA | 0.444 |
| SE | 3215 | PDCD6IP | chr3 | + | 33812071 | 33812126 | 33798570 | 33798937 | 33813558 | 33813628 | 0 | 1 | 6 | 0 | 0 | 0 | 0 | 1 | 0 | 0 | 3 | 0 | 98 | 49 | 0.023847213 | 0.170760496 | NA | 1 | 1 | NA | 0.143 | NA | 0.857 |
| SE | 4847 | TINF2 | chr14 | - | 24241889 | 24241994 | 24241674 | 24241776 | 24242140 | 24242674 | 0 | 1 | 0 | 0 | 0 | 0 | 0 | 1 | 0 | 0 | 3 | 0 | 98 | 49 | 0.023847213 | 0.170760496 | NA | 1 | NA | NA | 0.143 | NA | 0.857 |
| SE | 2082 | HERC6 | chr4 | + | 88393487 | 88393582 | 88390651 | 88390879 | 88396014 | 88396142 | 0 | 1 | 12 | 0 | 0 | 0 | 0 | 3 | 0 | 0 | 2 | 0 | 98 | 49 | 0.023922591 | 0.170760496 | NA | 1 | 1 | NA | 0.429 | NA | 0.571 |
| SE | 7201 | MDM4 | chr1 | + | 204537429 | 204537497 | 204532190 | 204532246 | 204538208 | 204538308 | 0 | 1 | 6 | 0 | 0 | 0 | 0 | 3 | 0 | 0 | 2 | 0 | 98 | 49 | 0.023922591 | 0.170760496 | NA | 1 | 1 | NA | 0.429 | NA | 0.571 |
| SE | 10725 | PTMA | chr2 | + | 231711889 | 231711983 | 231711347 | 231711419 | 231712442 | 231712516 | 0 | 1 | 3 | 0 | 0 | 0 | 0 | 3 | 0 | 0 | 2 | 0 | 98 | 49 | 0.023922591 | 0.170760496 | NA | 1 | 1 | NA | 0.429 | NA | 0.571 |
| SE | 6590 | UQCRH | chr1 | + | 46309896 | 46310044 | 46309100 | 46309127 | 46310154 | 46310316 | 0 | 0 | 1 | 0 | 2 | 0 | 0 | 2 | 0 | 3 | 3 | 0 | 98 | 49 | 0.024004538 | 0.170760496 | NA | 0 | 1 | 0 | 0.25 | NA | 0.375 |
| SE | 4111 | ATP2B1 | chr12 | - | 89601325 | 89601433 | 89599116 | 89599299 | 89603042 | 89603254 | 0 | 1 | 2 | 0 | 0 | 0 | 2 | 6 | 0 | 0 | 2 | 0 | 98 | 49 | 0.024023827 | 0.170760496 | NA | 1 | 1 | 1 | 0.6 | NA | 0.2 |
| SE | 5016 | ARHGAP25 | chr2 | + | 68813286 | 68813419 | 68807272 | 68807480 | 68816288 | 68816362 | 0 | 1 | 15 | 0 | 0 | 0 | 0 | 6 | 0 | 0 | 2 | 0 | 98 | 49 | 0.024023827 | 0.170760496 | NA | 1 | 1 | NA | 0.6 | NA | 0.4 |
| SE | 7011 | HIPK3 | chr11 | + | 33348521 | 33348818 | 33347851 | 33348013 | 33349146 | 33349287 | 0 | 1 | 4 | 0 | 0 | 0 | 0 | 6 | 0 | 0 | 2 | 0 | 98 | 49 | 0.024023827 | 0.170760496 | NA | 1 | 1 | NA | 0.6 | NA | 0.4 |
| SE | 10363 | SOS2 | chr14 | - | 50145169 | 50145332 | 50139941 | 50140059 | 50145476 | 50145596 | 0 | 1 | 9 | 0 | 0 | 1 | 0 | 6 | 0 | 0 | 2 | 0 | 98 | 49 | 0.024023827 | 0.170760496 | NA | 1 | 0.818 | NA | 0.6 | NA | 0.309 |
| SE | 10724 | PTMA | chr2 | + | 231711886 | 231711983 | 231711347 | 231711419 | 231712442 | 231712516 | 0 | 1 | 2 | 0 | 0 | 0 | 0 | 6 | 0 | 0 | 2 | 0 | 98 | 49 | 0.024023827 | 0.170760496 | NA | 1 | 1 | NA | 0.6 | NA | 0.4 |
| SE | 11386 | NLRC5 | chr16 | + | 57046551 | 57046641 | 57045447 | 57045492 | 57047544 | 57047628 | 0 | 1 | 10 | 0 | 0 | 0 | 0 | 6 | 0 | 0 | 2 | 0 | 98 | 49 | 0.024023827 | 0.170760496 | NA | 1 | 1 | NA | 0.6 | NA | 0.4 |
| SE | 690 | CDK16 | chrX | + | 47225771 | 47225866 | 47224987 | 47225102 | 47225964 | 47226027 | 0 | 1 | 4 | 0 | 0 | 0 | 0 | 7 | 0 | 0 | 2 | 0 | 98 | 49 | 0.02426018 | 0.170760496 | NA | 1 | 1 | NA | 0.636 | NA | 0.364 |
| SE | 7907 | IFIT1 | chr10 | + | 89399894 | 89400003 | 89392545 | 89392717 | 89402280 | 89406487 | 0 | 1 | 12 | 0 | 0 | 5 | 0 | 7 | 0 | 4 | 2 | 0 | 98 | 49 | 0.02426018 | 0.170760496 | NA | 1 | 0.545 | 0 | 0.636 | NA | 0.454 |
| SE | 8359 | ARHGAP30 | chr1 | - | 161059613 | 161059716 | 161054622 | 161054705 | 161069527 | 161069970 | 0 | 1 | 18 | 0 | 0 | 2 | 0 | 7 | 0 | 0 | 2 | 0 | 98 | 49 | 0.02426018 | 0.170760496 | NA | 1 | 0.818 | NA | 0.636 | NA | 0.273 |
| SE | 9209 | CELF2 | chr10 | + | 11217424 | 11217507 | 11165485 | 11165682 | 11249152 | 11249201 | 0 | 1 | 31 | 0 | 0 | 2 | 5 | 7 | 0 | 0 | 2 | 0 | 98 | 49 | 0.02426018 | 0.170760496 | NA | 1 | 0.886 | 1 | 0.636 | NA | 0.125 |
| SE | 8140 | ITGAX | chr16 | + | 31373248 | 31373390 | 31372596 | 31372670 | 31376798 | 31376915 | 0 | 12 | 40 | 0 | 0 | 5 | 6 | 46 | 13 | 0 | 0 | 0 | 98 | 49 | 0.024390083 | 0.170760496 | NA | 1 | 0.8 | 1 | 1 | 1 | -0.1 |

| | | | | | | | | | | | | | | | | | | | | | | | | | | | | | | | | | |
|---|---|---|---|---|---|---|---|---|---|---|---|---|---|---|---|---|---|---|---|---|---|---|---|---|---|---|---|---|---|---|---|---|---|
| SE | 12583 | SBNO2 | chr19 | - | 1109134 | 1109211 | 1108778 | 1108969 | 1109291 | 1109423 | 0 | 4 | 9 | 0 | 0 | 0 | 0 | 25 | 2 | 0 | 3 | 0 | 98 | 49 | 0.024422917 | 0.170760496 | NA | | 1 | 1 | NA | 0.806 | 1 | 0.097 |
| SE | 13765 | ABCG1 | chr21 | + | 42276894 | 42276945 | 42273302 | 42273435 | 42282273 | 42282419 | 0 | 1 | 11 | 0 | 0 | 0 | 0 | 8 | 0 | 0 | 2 | 0 | 98 | 49 | 0.024517355 | 0.170760496 | NA | | 1 | 1 | NA | 0.667 | NA | 0.333 |
| A3SS | 683 | CDV3 | chr3 | + | 133587161 | 133590274 | 133587895 | 133590274 | 133586562 | 133586722 | 0 | 1 | 2 | 0 | 0 | 0 | 0 | 2 | 0 | 0 | 2 | 0 | 98 | 49 | 0.024627853 | 0.197717869 | NA | | 1 | 1 | NA | 0.333 | NA | 0.667 |
| SE | 1480 | FLOT1 | chr6 | - | 30741613 | 30741704 | 30741189 | 30741333 | 30741791 | 30741867 | 0 | 1 | 6 | 0 | 0 | 1 | 0 | 2 | 0 | 0 | 2 | 0 | 98 | 49 | 0.024627853 | 0.170760496 | NA | | 1 | 0.75 | NA | 0.333 | NA | 0.542 |
| SE | 3632 | USP4 | chr3 | - | 49300466 | 49300691 | 49298551 | 49298635 | 49302383 | 49302542 | 0 | 1 | 0 | 0 | 0 | 0 | 0 | 2 | 0 | 0 | 2 | 0 | 98 | 49 | 0.024627853 | 0.170760496 | NA | | 1 | NA | NA | 0.333 | NA | 0.667 |
| SE | 4078 | PHF1 | chr6 | + | 33414954 | 33415144 | 33414724 | 33414829 | 33415234 | 33415329 | 0 | 1 | 1 | 0 | 0 | 0 | 0 | 2 | 0 | 0 | 2 | 0 | 98 | 49 | 0.024627853 | 0.170760496 | NA | | 1 | 1 | NA | 0.333 | NA | 0.667 |
| SE | 4079 | PHF1 | chr6 | + | 33414965 | 33415144 | 33414724 | 33414829 | 33415234 | 33415329 | 0 | 1 | 1 | 0 | 0 | 0 | 0 | 2 | 0 | 0 | 2 | 0 | 98 | 49 | 0.024627853 | 0.170760496 | NA | | 1 | 1 | NA | 0.333 | NA | 0.667 |
| SE | 6133 | STXBP3 | chr1 | + | 108779285 | 108779410 | 108776332 | 108776423 | 108782421 | 108782517 | 0 | 2 | 7 | 0 | 0 | 0 | 0 | 1 | 0 | 0 | 2 | 0 | 98 | 49 | 0.02492004 | 0.170760496 | NA | | 1 | 1 | NA | 0.2 | NA | 0.8 |
| SE | 7313 | WNK1 | chr12 | + | 871264 | 871348 | 862082 | 862270 | 878211 | 878361 | 0 | 2 | 4 | 0 | 0 | 1 | 3 | 1 | 0 | 0 | 2 | 0 | 98 | 49 | 0.02492004 | 0.170760496 | NA | | 1 | 0.667 | 1 | 0.2 | NA | 0.234 |
| SE | 8790 | VPS13D | chr1 | + | 12311453 | 12311625 | 12308430 | 12308641 | 12311812 | 12311925 | 0 | 2 | 6 | 0 | 0 | 0 | 0 | 1 | 0 | 0 | 2 | 0 | 98 | 49 | 0.02492004 | 0.170760496 | NA | | 1 | 1 | NA | 0.2 | NA | 0.8 |
| A5SS | 1139 | TBC1D10C | chr11 | + | 67406626 | 67406738 | 67406626 | 67406692 | 67406826 | 67407016 | 0 | 1 | 2 | 0 | 0 | 4 | 0 | 3 | 3 | 0 | 0 | 0 | 95 | 49 | 0.024974929 | 0.170414363 | NA | | 1 | 0.205 | NA | 1 | 1 | -0.397 |
| SE | 5184 | MYO9B | chr19 | + | 17145396 | 17145491 | 17101659 | 17102557 | 17152643 | 17152706 | 0 | 3 | 23 | 0 | 0 | 0 | 0 | 19 | 2 | 0 | 4 | 0 | 98 | 49 | 0.024989397 | 0.170760496 | NA | | 1 | 1 | NA | 0.704 | 1 | 0.148 |
| SE | 13164 | PPP1CB | chr2 | + | 28788657 | 28788809 | 28783906 | 28783978 | 28793862 | 28793997 | 0 | 1 | 6 | 0 | 0 | 0 | 0 | 10 | 0 | 0 | 2 | 0 | 98 | 49 | 0.025074234 | 0.170760496 | NA | | 1 | 1 | NA | 0.714 | NA | 0.286 |
| SE | 2167 | FYB | chr5 | - | 39201825 | 39202987 | 39153447 | 39153604 | 39219442 | 39219578 | 0 | 8 | 22 | 0 | 0 | 0 | 1 | 10 | 1 | 0 | 2 | 0 | 98 | 49 | 0.025260336 | 0.170760496 | NA | | 1 | 1 | 1 | 0.714 | 1 | 0.095 |
| SE | 2076 | HERC5 | chr4 | + | 88463854 | 88463985 | 88463531 | 88463623 | 88467058 | 88467204 | 0 | 2 | 27 | 0 | 0 | 1 | 0 | 28 | 0 | 0 | 2 | 0 | 98 | 49 | 0.02534026 | 0.170760496 | NA | | 1 | 0.931 | NA | 0.875 | NA | 0.091 |
| SE | 4015 | RPLP0 | chr12 | - | 120198553 | 120198739 | 120197321 | 120197462 | 120198853 | 120199000 | 0 | 1 | 3 | 0 | 0 | 0 | 0 | 12 | 0 | 0 | 2 | 0 | 98 | 49 | 0.025646206 | 0.170760496 | NA | | 1 | 1 | NA | 0.75 | NA | 0.25 |
| SE | 5192 | PICALM | chr11 | - | 85976622 | 85976682 | 85974707 | 85974812 | 85981128 | 85981228 | 0 | 2 | 30 | 0 | 0 | 0 | 3 | 30 | 0 | 0 | 2 | 0 | 98 | 49 | 0.025657506 | 0.170760496 | NA | | 1 | 1 | 1 | 0.882 | NA | 0.059 |
| A3SS | 932 | CXCL1 | chr4 | + | 73869645 | 73869792 | 73869668 | 73869792 | 73869391 | 73869570 | 0 | 0 | 0 | 0 | 0 | 3 | 3 | 1 | 2 | 0 | 1 | 1 | 72 | 49 | 0.025891313 | 0.197717869 | NA | NA | | 0 | 1 | 0.405 | 0.576 | -0.66 |
| SE | 7720 | THEMIS2 | chr1 | + | 27881970 | 27883043 | 27876587 | 27876728 | 27885294 | 27885451 | 0 | 0 | 2 | 0 | 3 | 1 | 0 | 2 | 0 | 0 | 1 | 0 | 98 | 49 | 0.025897097 | 0.170760496 | NA | | 0 | 0.5 | NA | 0.5 | NA | -0.25 |
| SE | 13194 | MBOAT7 | chr19 | - | 54188432 | 54188511 | 54188216 | 54188346 | 54189337 | 54189882 | 0 | 9 | 20 | 0 | 0 | 1 | 1 | 12 | 31 | 0 | 2 | 6 | 98 | 49 | 0.025921935 | 0.170760496 | NA | | 1 | 0.909 | 1 | 0.75 | 0.721 | 0.131 |
| SE | 6180 | ABCA1 | chr9 | - | 104884426 | 104884568 | 104883038 | 104883157 | 104889101 | 104889195 | 0 | 3 | 20 | 0 | 1 | 1 | 0 | 16 | 2 | 0 | 7 | 3 | 98 | 49 | 0.026099399 | 0.170760496 | NA | | 0.6 | 0.909 | NA | 0.533 | 0.25 | 0.363 |
| SE | 2583 | LCP2 | chr5 | - | 170270718 | 170270917 | 170268384 | 170268482 | 170274300 | 170274338 | 1 | 1 | 19 | 0 | 0 | 0 | 0 | 14 | 0 | 0 | 2 | 0 | 98 | 49 | 0.026208434 | 0.170760496 | 1 | | 1 | 1 | NA | 0.778 | NA | 0.222 |
| A5SS | 2038 | TRAPPC5 | chr19 | + | 7680820 | 7681026 | 7680820 | 7680878 | 7682241 | 7682862 | 0 | 0 | 0 | 0 | 0 | 1 | 0 | 4 | 3 | 0 | 0 | 1 | 98 | 49 | 0.026257417 | 0.170414363 | NA | NA | | 0 | NA | 1 | 0.6 | -0.8 |
| A5SS | 2039 | TRAPPC5 | chr19 | + | 7680820 | 7681049 | 7680820 | 7680878 | 7682241 | 7682862 | 0 | 0 | 0 | 0 | 0 | 1 | 0 | 3 | 3 | 0 | 0 | 1 | 98 | 49 | 0.026257417 | 0.170414363 | NA | NA | | 0 | NA | 1 | 0.6 | -0.8 |
| SE | 6627 | PCBP2 | chr12 | + | 53464759 | 53464852 | 53462480 | 53462567 | 53467220 | 53467293 | 0 | 0 | 13 | 0 | 1 | 3 | 0 | 3 | 0 | 0 | 1 | 0 | 98 | 49 | 0.026257417 | 0.170760496 | NA | | 0 | 0.684 | NA | 0.6 | NA | -0.258 |
| SE | 7010 | HIPK3 | chr11 | + | 33348165 | 33348228 | 33347851 | 33348013 | 33348521 | 33348818 | 0 | 0 | 7 | 0 | 1 | 1 | 0 | 3 | 0 | 0 | 1 | 0 | 98 | 49 | 0.026257417 | 0.170760496 | NA | | 0 | 0.778 | NA | 0.6 | NA | -0.211 |
| SE | 12300 | CYTH1 | chr17 | - | 78702537 | 78702604 | 78702121 | 78702240 | 78708196 | 78708261 | 0 | 0 | 6 | 0 | 1 | 3 | 0 | 3 | 0 | 0 | 1 | 0 | 98 | 49 | 0.026257417 | 0.170760496 | NA | | 0 | 0.5 | NA | 0.6 | NA | -0.35 |
| SE | 6175 | ABCA1 | chr9 | - | 104812573 | 104812722 | 104809465 | 104809564 | 104814117 | 104814231 | 0 | 1 | 3 | 0 | 0 | 0 | 2 | 15 | 0 | 0 | 2 | 0 | 98 | 49 | 0.026481025 | 0.170760496 | NA | | 1 | 1 | 1 | 0.789 | NA | 0.105 |
| SE | 5194 | PICALM | chr11 | - | 85990249 | 85990378 | 85983865 | 85983973 | 85996825 | 85996929 | 0 | 1 | 18 | 0 | 0 | 21 | 0 | 16 | 0 | 0 | 2 | 0 | 98 | 49 | 0.026734839 | 0.170760496 | NA | | 1 | 0.3 | NA | 0.8 | NA | -0.15 |
| SE | 5975 | TAP2 | chr6 | - | 32835153 | 32835359 | 32832626 | 32832824 | 32835642 | 32835773 | 0 | 0 | 11 | 0 | 0 | 0 | 0 | 11 | 1 | 0 | 0 | 1 | 98 | 49 | 0.026878049 | 0.170760496 | NA | NA | | 1 | NA | 1 | 0.333 | 0.334 |
| SE | 9687 | IFRD1 | chr7 | + | 112456913 | 112457038 | 112456001 | 112456086 | 112458860 | 112459018 | 0 | 6 | 52 | 0 | 0 | 0 | 0 | 84 | 4 | 0 | 3 | 0 | 98 | 49 | 0.026897321 | 0.170760496 | NA | | 1 | 1 | NA | 0.933 | 1 | 0.033 |
| SE | 6887 | SRPK1 | chr6 | - | 35870280 | 35870494 | 35869481 | 35869901 | 35870933 | 35870959 | 0 | 0 | 4 | 0 | 1 | 5 | 0 | 2 | 0 | 0 | 3 | 0 | 98 | 49 | 0.027065308 | 0.170760496 | NA | | 0 | 0.286 | NA | 0.25 | NA | -0.107 |
| A3SS | 3741 | TMEM91 | chr19 | + | 41383560 | 41384082 | 41383862 | 41384082 | 41382771 | 41382921 | 0 | 1 | 3 | 0 | 0 | 3 | 0 | 1 | 0 | 0 | 2 | 0 | 98 | 49 | 0.027259421 | 0.197717869 | NA | | 1 | 0.333 | NA | 0.2 | NA | 0.466 |
| SE | 2726 | SATB1 | chr3 | - | 18351991 | 18352195 | 18347640 | 18349682 | 18378169 | 18378325 | 0 | 1 | 3 | 0 | 0 | 0 | 0 | 1 | 0 | 0 | 2 | 0 | 98 | 49 | 0.027259421 | 0.170760496 | NA | | 1 | 1 | NA | 0.2 | NA | 0.8 |
| SE | 9126 | HGS | chr17 | + | 81690181 | 81690234 | 81688703 | 81688827 | 81690673 | 81690742 | 0 | 1 | 1 | 0 | 0 | 0 | 0 | 1 | 0 | 0 | 2 | 0 | 98 | 49 | 0.027259421 | 0.170760496 | NA | | 1 | 1 | NA | 0.2 | NA | 0.8 |
| SE | 14102 | SLC12A4 | chr16 | - | 67951822 | 67952037 | 67951139 | 67951304 | 67952183 | 67952425 | 0 | 1 | 0 | 0 | 0 | 0 | 0 | 1 | 0 | 0 | 2 | 0 | 98 | 49 | 0.027259421 | 0.170760496 | NA | | 1 | NA | NA | 0.2 | NA | 0.8 |
| A5SS | 1672 | PTBP3 | chr9 | - | 112297831 | 112297916 | 112297840 | 112297916 | 112275843 | 112276013 | 0 | 0 | 3 | 0 | 2 | 2 | 0 | 2 | 0 | 0 | 1 | 0 | 58 | 49 | 0.027334905 | 0.170414363 | NA | | 0 | 0.559 | NA | 0.628 | NA | -0.348 |
| SE | 8906 | FBXW7 | chr4 | - | 152412479 | 152412529 | 152411159 | 152411872 | 152534940 | 152535033 | 0 | 0 | 0 | 0 | 2 | 0 | 0 | 2 | 0 | 0 | 1 | 1 | 98 | 49 | 0.02734801 | 0.170760496 | NA | | 0 | NA | NA | 0.5 | 0 | -0.25 |
| SE | 3622 | ITGAM | chr16 | + | 31266029 | 31266147 | 31265810 | 31265881 | 31270953 | 31271084 | 0 | 6 | 27 | 0 | 0 | 0 | 0 | 3 | 0 | 0 | 1 | 0 | 98 | 49 | 0.027600753 | 0.170760496 | NA | | 1 | 1 | NA | 0.6 | NA | 0.4 |
| A3SS | 405 | BAG6 | chr6 | - | 31647590 | 31647844 | 31647590 | 31647826 | 31648676 | 31648751 | 0 | 0 | 0 | 0 | 1 | 2 | 0 | 2 | 0 | 0 | 2 | 0 | 67 | 49 | 0.027623127 | 0.197717869 | NA | | 0 | 0 | NA | 0.422 | NA | -0.422 |
| SE | 3281 | RAB11FIP1 | chr8 | - | 37871277 | 37873179 | 37870419 | 37870528 | 37874514 | 37875322 | 0 | 0 | 4 | 0 | 1 | 4 | 0 | 2 | 0 | 0 | 2 | 0 | 98 | 49 | 0.027623276 | 0.170760496 | NA | | 0 | 0.333 | NA | 0.333 | NA | -0.167 |
| SE | 5922 | MKNK1 | chr1 | - | 46560233 | 46560277 | 46557406 | 46558800 | 46561477 | 46561642 | 0 | 0 | 2 | 0 | 1 | 2 | 0 | 2 | 0 | 0 | 2 | 0 | 93 | 49 | 0.027624169 | 0.170760496 | NA | | 0 | 0.345 | NA | 0.345 | NA | -0.172 |
| SE | 472 | DOCK8 | chr9 | + | 372184 | 372286 | 371427 | 371566 | 376209 | 376305 | 0 | 10 | 33 | 0 | 0 | 0 | 0 | 9 | 0 | 0 | 1 | 0 | 98 | 49 | 0.027770234 | 0.170760496 | NA | | 1 | 1 | NA | 0.818 | NA | 0.182 |
| SE | 20 | FLNA | chrX | - | 154362417 | 154362578 | 154362241 | 154362332 | 154362660 | 154362784 | 0 | 5 | 37 | 0 | 0 | 0 | 2 | 18 | 1 | 0 | 2 | 0 | 98 | 49 | 0.027864407 | 0.170760496 | NA | | 1 | 1 | 1 | 0.818 | 1 | 0.061 |
| SE | 5181 | ANKRD13A | chr12 | + | 110024045 | 110024112 | 110019138 | 110019328 | 110025741 | 110025823 | 0 | 6 | 10 | 0 | 0 | 0 | 0 | 4 | 0 | 0 | 1 | 0 | 98 | 49 | 0.027908407 | 0.170760496 | NA | | 1 | 1 | NA | 0.667 | NA | 0.333 |
| SE | 6868 | ATG2A | chr11 | - | 64909270 | 64909367 | 64907747 | 64907890 | 64909680 | 64909924 | 0 | 0 | 6 | 0 | 0 | 0 | 0 | 7 | 4 | 0 | 0 | 1 | 98 | 49 | 0.027908407 | 0.170760496 | NA | NA | | 1 | NA | 1 | 0.667 | 0.166 |
| SE | 4661 | AMICA1 | chr11 | - | 118205881 | 118205991 | 118203427 | 118203665 | 118210486 | 118210712 | 0 | 1 | 9 | 0 | 0 | 0 | 0 | 21 | 0 | 0 | 2 | 0 | 98 | 49 | 0.028047628 | 0.170760496 | NA | | 1 | 1 | NA | 0.84 | NA | 0.16 |
| SE | 5195 | PICALM | chr11 | - | 85990249 | 85990399 | 85983865 | 85983973 | 85996825 | 85996929 | 0 | 1 | 25 | 0 | 0 | 21 | 0 | 21 | 0 | 0 | 2 | 0 | 98 | 49 | 0.028047628 | 0.170760496 | NA | | 1 | 0.373 | NA | 0.84 | NA | -0.153 |
| SE | 13787 | PSD4 | chr2 | + | 113198739 | 113198884 | 113197746 | 113197913 | 113199082 | 113199226 | 0 | 5 | 2 | 0 | 0 | 0 | 0 | 7 | 1 | 0 | 3 | 0 | 98 | 49 | 0.028063223 | 0.170760496 | NA | | 1 | 1 | NA | 0.538 | 1 | 0.231 |
| SE | 16 | FLNA | chrX | - | 154352552 | 154352675 | 154352180 | 154352447 | 154352771 | 154352924 | 0 | 9 | 44 | 0 | 0 | 1 | 0 | 9 | 0 | 0 | 1 | 0 | 98 | 49 | 0.028070044 | 0.170760496 | NA | | 1 | 0.957 | NA | 0.818 | NA | 0.16 |
| SE | 688 | PTK2B | chr8 | + | 27397547 | 27397788 | 27325478 | 27325681 | 27419894 | 27420073 | 0 | 7 | 27 | 0 | 0 | 0 | 0 | 6 | 0 | 0 | 1 | 0 | 98 | 49 | 0.028077309 | 0.170760496 | NA | | 1 | 1 | NA | 0.75 | NA | 0.25 |
| SE | 8356 | ARHGAP30 | chr1 | - | 161052625 | 161052797 | 161052439 | 161052543 | 161053257 | 161053385 | 0 | 7 | 26 | 0 | 0 | 0 | 0 | 6 | 0 | 0 | 1 | 0 | 98 | 49 | 0.028077309 | 0.170760496 | NA | | 1 | 1 | NA | 0.75 | NA | 0.25 |
| SE | 12701 | ARFGEF1 | chr8 | - | 67275975 | 67276109 | 67271701 | 67271936 | 67277281 | 67277457 | 0 | 5 | 2 | 0 | 0 | 0 | 7 | 2 | 0 | 0 | 1 | 0 | 98 | 49 | 0.028306385 | 0.170760496 | NA | | 1 | 1 | 1 | 0.5 | NA | 0.25 |
| SE | 4456 | CLEC7A | chr12 | - | 10123244 | 10123363 | 10116780 | 10118590 | 10125086 | 10125173 | 0 | 0 | 38 | 0 | 2 | 1 | 0 | 15 | 3 | 0 | 0 | 0 | 98 | 49 | 0.028599911 | 0.170760496 | NA | | 0 | 0.95 | NA | 1 | 1 | -0.525 |
| A5SS | 672 | STX16 | chr20 | + | 58651252 | 58652138 | 58651252 | 58652087 | 58667489 | 58667597 | 0 | 4 | 7 | 0 | 0 | 3 | 1 | 2 | 0 | 0 | 1 | 0 | 98 | 49 | 0.029221508 | 0.170414363 | NA | | 1 | 0.538 | 1 | 0.5 | NA | 0.019 |
| SE | 5228 | NFKBIA | chr14 | - | 35402770 | 35402859 | 35402393 | 35402663 | 35403149 | 35403360 | 0 | 10 | 63 | 0 | 0 | 0 | 7 | 167 | 19 | 0 | 4 | 0 | 98 | 49 | 0.029442582 | 0.170760496 | NA | | 1 | 1 | 1 | 0.954 | 1 | 0.015 |
| SE | 5000 | WIPI2 | chr7 | + | 5193117 | 5193345 | 5190203 | 5190493 | 5199575 | 5199658 | 0 | 1 | 1 | 0 | 0 | 1 | 0 | 0 | 0 | 0 | 2 | 2 | 98 | 49 | 0.029568334 | 0.170760496 | NA | | 1 | 0.333 | NA | 0 | 0 | 0.666 |
| SE | 4834 | ULK3 | chr15 | - | 74838265 | 74838344 | 74838151 | 74838192 | 74838439 | 74838504 | 0 | 5 | 0 | 0 | 0 | 0 | 0 | 1 | 0 | 0 | 1 | 0 | 98 | 49 | 0.029576123 | 0.170760496 | NA | | 1 | NA | NA | 0.333 | NA | 0.667 |
| SE | 4835 | ULK3 | chr15 | - | 74838269 | 74838344 | 74838151 | 74838192 | 74838439 | 74838504 | 0 | 5 | 0 | 0 | 0 | 0 | 0 | 1 | 0 | 0 | 1 | 0 | 98 | 49 | 0.029576123 | 0.170760496 | NA | | 1 | NA | NA | 0.333 | NA | 0.667 |
| SE | 7841 | B4GALT5 | chr20 | - | 49639677 | 49639800 | 49637340 | 49637442 | 49640477 | 49640665 | 0 | 5 | 13 | 0 | 0 | 1 | 0 | 6 | 0 | 0 | 1 | 0 | 98 | 49 | 0.029628726 | 0.170760496 | NA | | 1 | 0.867 | NA | 0.75 | NA | 0.183 |
| SE | 11675 | ULK1 | chr12 | + | 131915334 | 131915421 | 131915082 | 131915231 | 131915890 | 131916159 | 0 | 0 | 5 | 0 | 0 | 0 | 0 | 3 | 6 | 0 | 0 | 1 | 98 | 49 | 0.029628726 | 0.170760496 | NA | NA | | 1 | NA | 1 | 0.75 | 0.125 |
| SE | 6193 | FCER1G | chr1 | + | 161217985 | 161218077 | 161215296 | 161215370 | 161218240 | 161218276 | 0 | 7 | 28 | 0 | 0 | 0 | 4 | 10 | 0 | 0 | 1 | 0 | 98 | 49 | 0.029643012 | 0.170760496 | NA | | 1 | 1 | 1 | 0.833 | NA | 0.084 |
| A5SS | 1841 | OS9 | chr12 | + | 57718145 | 57718421 | 57718145 | 57718376 | 57718992 | 57719182 | 0 | 6 | 6 | 0 | 0 | 0 | 3 | 8 | 0 | 0 | 1 | 0 | 94 | 49 | 0.029835799 | 0.170414363 | NA | | 1 | 1 | 1 | 0.807 | NA | 0.097 |
| SE | 11769 | FLII | chr17 | - | 18255182 | 18255263 | 18254768 | 18254854 | 18256525 | 18256597 | 0 | 6 | 12 | 0 | 0 | 0 | 0 | 9 | 0 | 0 | 1 | 0 | 98 | 49 | 0.03004548 | 0.170760496 | NA | | 1 | 1 | NA | 0.818 | NA | 0.182 |
| A3SS | 3631 | SURF4 | chr9 | - | 133367258 | 133367543 | 133367258 | 133367445 | 133375921 | 133376166 | 0 | 0 | 0 | 0 | 1 | 0 | 1 | 2 | 0 | 0 | 1 | 0 | 98 | 49 | 0.030221083 | 0.197717869 | NA | | 0 | NA | 1 | 0.5 | NA | -0.75 |
| SE | 388 | ZFAND5 | chr9 | - | 72363469 | 72363606 | 72360627 | 72360787 | 72364695 | 72365247 | 0 | 0 | 4 | 0 | 1 | 0 | 0 | 2 | 0 | 0 | 1 | 0 | 98 | 49 | 0.030221083 | 0.170760496 | NA | | 0 | 1 | NA | 0.5 | NA | 0 |
| SE | 11350 | ADGRE2 | chr19 | - | 14755653 | 14755877 | 14752328 | 14752526 | 14756237 | 14756345 | 0 | 0 | 2 | 0 | 1 | 1 | 0 | 2 | 0 | 0 | 1 | 0 | 98 | 49 | 0.030221083 | 0.170760496 | NA | | 0 | 0.5 | NA | 0.5 | NA | -0.25 |
| SE | 5198 | PICALM | chr11 | - | 86026291 | 86026367 | 86022366 | 86022469 | 86031468 | 86031611 | 0 | 1 | 33 | 0 | 0 | 0 | 3 | 24 | 1 | 0 | 3 | 0 | 98 | 49 | 0.03042033 | 0.170760496 | NA | | 1 | 1 | 1 | 0.8 | 1 | 0.067 |
| SE | 1451 | BAG6 | chr6 | - | 31648910 | 31648964 | 31648676 | 31648751 | 31649198 | 31649395 | 0 | 3 | 2 | 0 | 0 | 0 | 0 | 2 | 0 | 0 | 1 | 0 | 98 | 49 | 0.030496725 | 0.170760496 | NA | | 1 | 1 | NA | 0.5 | NA | 0.5 |
| SE | 8360 | ARHGAP30 | chr1 | - | 161059613 | 161059716 | 161056387 | 161056532 | 161069527 | 161069970 | 0 | 1 | 23 | 0 | 0 | 0 | 0 | 8 | 0 | 0 | 1 | 1 | 98 | 49 | 0.030607676 | 0.170760496 | NA | | 1 | 1 | NA | 0.8 | 0 | 0.6 |
| SE | 143 | NONO | chrX | + | 71297835 | 71297938 | 71297376 | 71297461 | 71298468 | 71298508 | 0 | 3 | 2 | 0 | 0 | 0 | 0 | 3 | 0 | 0 | 1 | 0 | 98 | 49 | 0.030693475 | 0.170760496 | NA | | 1 | 1 | NA | 0.6 | NA | 0.4 |
| SE | 9279 | PIK3CD | chr1 | + | 9723968 | 9724092 | 9723124 | 9723292 | 9724275 | 9724421 | 0 | 3 | 12 | 0 | 0 | 0 | 0 | 3 | 0 | 0 | 1 | 0 | 98 | 49 | 0.030693475 | 0.170760496 | NA | | 1 | 1 | NA | 0.6 | NA | 0.4 |
| SE | 9304 | SNRNP200 | chr2 | - | 96298283 | 96298420 | 96297636 | 96297720 | 96298602 | 96298702 | 0 | 3 | 3 | 0 | 0 | 0 | 0 | 3 | 0 | 0 | 1 | 0 | 98 | 49 | 0.030693475 | 0.170760496 | NA | | 1 | 1 | NA | 0.6 | NA | 0.4 |
| SE | 6670 | FGR | chr1 | - | 27616856 | 27617006 | 27615688 | 27615844 | 27621558 | 27621657 | 0 | 2 | 21 | 0 | 0 | 0 | 0 | 9 | 0 | 0 | 1 | 1 | 98 | 49 | 0.030846049 | 0.170760496 | NA | | 1 | 1 | NA | 0.818 | 0 | 0.591 |
| A3SS | 2290 | SF1 | chr11 | - | 64764603 | 64766155 | 64764603 | 64765518 | 64766899 | 64767079 | 0 | 3 | 7 | 0 | 0 | 0 | 0 | 4 | 0 | 0 | 1 | 0 | 98 | 49 | 0.03121249 | 0.197717869 | NA | | 1 | 1 | NA | 0.667 | NA | 0.333 |
| SE | 6445 | RALGDS | chr9 | - | 133102475 | 133102571 | 133101937 | 133102139 | 133102778 | 133102900 | 0 | 3 | 0 | 0 | 0 | 0 | 0 | 4 | 0 | 0 | 1 | 0 | 98 | 49 | 0.03121249 | 0.170760496 | NA | | 1 | NA | NA | 0.667 | NA | 0.333 |
| SE | 54 | ELF4 | chrX | - | 130071039 | 130071232 | 130069299 | 130069677 | 130071335 | 130071419 | 0 | 3 | 7 | 0 | 0 | 0 | 0 | 1 | 0 | 0 | 1 | 0 | 98 | 49 | 0.031636388 | 0.170760496 | NA | | 1 | 1 | NA | 0.333 | NA | 0.667 |
| SE | 8271 | ADAM9 | chr8 | + | 39025802 | 39025884 | 39023155 | 39023325 | 39026676 | 39026810 | 0 | 3 | 4 | 0 | 0 | 0 | 0 | 5 | 0 | 0 | 1 | 0 | 98 | 49 | 0.031777706 | 0.170760496 | NA | | 1 | 1 | NA | 0.714 | NA | 0.286 |
| SE | 11767 | KDM5A | chr12 | - | 354075 | 354234 | 352204 | 352324 | 355157 | 355249 | 0 | 3 | 7 | 0 | 0 | 0 | 1 | 5 | 0 | 0 | 1 | 0 | 98 | 49 | 0.031777706 | 0.170760496 | NA | | 1 | 1 | 1 | 0.714 | NA | 0.143 |
| SE | 12649 | TLN1 | chr9 | - | 35722860 | 35722921 | 35722118 | 35722223 | 35723951 | 35724079 | 0 | 3 | 26 | 0 | 0 | 0 | 2 | 5 | 0 | 0 | 1 | 0 | 98 | 49 | 0.031777706 | 0.170760496 | NA | | 1 | 1 | 1 | 0.714 | NA | 0.143 |
| SE | 13848 | GNLY | chr2 | + | 85697505 | 85697677 | 85695957 | 85696056 | 85698563 | 85698851 | 0 | 3 | 2 | 0 | 0 | 0 | 0 | 5 | 0 | 0 | 1 | 0 | 98 | 49 | 0.031777706 | 0.170760496 | NA | | 1 | 1 | NA | 0.714 | NA | 0.286 |
| SE | 8338 | CHD2 | chr15 | + | 92901166 | 92901299 | 92900320 | 92900824 | 92924320 | 92924552 | 0 | 4 | 13 | 0 | 0 | 2 | 0 | 8 | 0 | 0 | 1 | 0 | 98 | 49 | 0.031803979 | 0.170760496 | NA | | 1 | 0.765 | NA | 0.8 | NA | 0.083 |
| SE | 11875 | MGAM | chr7 | + | 142036168 | 142036285 | 142034669 | 142034841 | 142038530 | 142038615 | 0 | 2 | 18 | 0 | 0 | 0 | 2 | 5 | 1 | 0 | 4 | 0 | 98 | 49 | 0.032046501 | 0.170760496 | NA | | 1 | 1 | 1 | 0.385 | 1 | 0.205 |
| A5SS | 2853 | CD37 | chr19 | + | 49335419 | 49335609 | 49335419 | 49335575 | 49335713 | 49335786 | 0 | 3 | 10 | 0 | 0 | 0 | 0 | 4 | 0 | 0 | 1 | 0 | 83 | 49 | 0.032094074 | 0.170414363 | NA | | 1 | 1 | NA | 0.703 | NA | 0.297 |
| SE | 2478 | GSK3B | chr3 | - | 119912703 | 119912810 | 119905754 | 119905852 | 119916043 | 119916174 | 0 | 2 | 8 | 0 | 0 | 2 | 0 | 2 | 0 | 0 | 1 | 0 | 98 | 49 | 0.032484896 | 0.170760496 | NA | | 1 | 0.667 | NA | 0.5 | NA | 0.334 |
| SE | 6900 | PSME3 | chr17 | + | 42838691 | 42838799 | 42838092 | 42838205 | 42838944 | 42839012 | 0 | 2 | 1 | 0 | 0 | 0 | 0 | 2 | 0 | 0 | 1 | 0 | 98 | 49 | 0.032484896 | 0.170760496 | NA | | 1 | 1 | NA | 0.5 | NA | 0.5 |
| SE | 9190 | GIT2 | chr12 | - | 109980951 | 109981046 | 109967457 | 109967503 | 109983372 | 109983503 | 0 | 2 | 20 | 0 | 0 | 0 | 0 | 2 | 0 | 0 | 1 | 0 | 98 | 49 | 0.032484896 | 0.170760496 | NA | | 1 | 1 | NA | 0.5 | NA | 0.5 |
| SE | 13501 | HADHB | chr2 | + | 26285406 | 26285571 | 26284882 | 26284957 | 26289917 | 26290465 | 0 | 2 | 11 | 0 | 0 | 0 | 0 | 2 | 0 | 0 | 1 | 0 | 98 | 49 | 0.032484896 | 0.170760496 | NA | | 1 | 1 | NA | 0.5 | NA | 0.5 |
| SE | 13782 | SLC20A1 | chr2 | + | 112647323 | 112647464 | 112646562 | 112647162 | 112647652 | 112647738 | 0 | 2 | 0 | 0 | 0 | 0 | 0 | 2 | 0 | 0 | 1 | 0 | 98 | 49 | 0.032484896 | 0.170760496 | NA | | 1 | NA | NA | 0.5 | NA | 0.5 |
| SE | 4966 | DDX3X | chrX | + | 41347311 | 41347451 | 41346858 | 41347012 | 41347639 | 41347915 | 0 | 1 | 19 | 0 | 0 | 0 | 0 | 34 | 1 | 0 | 3 | 0 | 98 | 49 | 0.032593127 | 0.170760496 | NA | | 1 | 1 | NA | 0.85 | 1 | 0.075 |
| SE | 5243 | PLXNC1 | chr12 | + | 94298631 | 94298795 | 94297315 | 94297423 | 94300909 | 94301057 | 0 | 4 | 20 | 0 | 0 | 0 | 1 | 10 | 0 | 0 | 1 | 0 | 98 | 49 | 0.03278556 | 0.170760496 | NA | | 1 | 1 | 1 | 0.833 | NA | 0.084 |
| A5SS | 1502 | MADD | chr11 | + | 47284940 | 47285194 | 47284940 | 47285065 | 47285450 | 47285590 | 0 | 2 | 5 | 0 | 0 | 0 | 3 | 3 | 0 | 0 | 1 | 0 | 98 | 49 | 0.032835937 | 0.170414363 | NA | | 1 | 1 | 1 | 0.6 | NA | 0.2 |
| SE | 1141 | IGF2R | chr6 | + | 160064803 | 160064901 | 160064400 | 160064531 | 160068248 | 160068385 | 0 | 2 | 17 | 0 | 0 | 3 | 0 | 3 | 0 | 0 | 1 | 0 | 98 | 49 | 0.032835937 | 0.170760496 | NA | | 1 | 0.739 | NA | 0.6 | NA | 0.269 |
| SE | 2318 | FAM105A | chr5 | + | 14601350 | 14601442 | 14601212 | 14601244 | 14602182 | 14602332 | 0 | 2 | 3 | 0 | 0 | 2 | 0 | 3 | 0 | 0 | 1 | 0 | 98 | 49 | 0.032835937 | 0.170760496 | NA | | 1 | 0.429 | NA | 0.6 | NA | 0.115 |
| SE | 2543 | GAK | chr4 | - | 877614 | 877809 | 877089 | 877207 | 881906 | 882040 | 0 | 2 | 6 | 0 | 0 | 0 | 0 | 3 | 0 | 0 | 1 | 0 | 98 | 49 | 0.032835937 | 0.170760496 | NA | | 1 | 1 | NA | 0.6 | NA | 0.4 |
| SE | 11659 | NLRP1 | chr17 | - | 5536850 | 5536940 | 5533896 | 5533988 | 5539414 | 5539585 | 0 | 2 | 30 | 0 | 0 | 0 | 0 | 3 | 0 | 0 | 1 | 0 | 98 | 49 | 0.032835937 | 0.170760496 | NA | | 1 | 1 | NA | 0.6 | NA | 0.4 |
| SE | 652 | KAT6A | chr8 | - | 41942792 | 41943000 | 41940841 | 41941444 | 41943747 | 41943979 | 0 | 3 | 9 | 0 | 0 | 1 | 0 | 7 | 0 | 0 | 1 | 0 | 98 | 49 | 0.032969828 | 0.170760496 | NA | | 1 | 0.818 | NA | 0.778 | NA | 0.131 |
| SE | 1651 | CANX | chr5 | + | 179708977 | 179709059 | 179708238 | 179708380 | 179709872 | 179710065 | 1 | 3 | 6 | 0 | 0 | 0 | 0 | 7 | 0 | 0 | 1 | 0 | 98 | 49 | 0.032969828 | 0.170760496 | 1 | | 1 | 1 | NA | 0.778 | NA | 0.222 |
| A3SS | 3558 | TMEM127 | chr2 | - | 96265137 | 96265512 | 96265137 | 96265489 | 96265868 | 96266013 | 0 | 4 | 2 | 0 | 3 | 4 | 0 | 6 | 1 | 0 | 0 | 0 | 72 | 49 | 0.033183051 | 0.197717869 | NA | | 0.476 | 0.254 | NA | 1 | 1 | -0.635 |
| SE | 13035 | VASP | chr19 | + | 45522339 | 45522581 | 45522167 | 45522217 | 45522717 | 45522818 | 0 | 4 | 13 | 0 | 0 | 0 | 0 | 11 | 0 | 0 | 1 | 0 | 98 | 49 | 0.033237301 | 0.170760496 | NA | | 1 | 1 | NA | 0.846 | NA | 0.154 |
| SE | 10971 | ARHGAP15 | chr2 | + | 143436912 | 143437042 | 143435600 | 143435699 | 143487372 | 143487495 | 0 | 0 | 19 | 0 | 2 | 0 | 0 | 21 | 2 | 0 | 0 | 0 | 98 | 49 | 0.033416613 | 0.170760496 | NA | | 0 | 1 | NA | 1 | 1 | -0.5 |
| SE | 2607 | RBM5 | chr3 | + | 50104247 | 50104308 | 50103082 | 50103166 | 50105076 | 50105142 | 0 | 2 | 6 | 0 | 0 | 1 | 0 | 1 | 0 | 3 | 1 | 0 | 98 | 49 | 0.033473796 | 0.170760496 | NA | | 1 | 0.75 | 0 | 0.333 | NA | 0.709 |
| SE | 7140 | KIF21B | chr1 | - | 201004765 | 201004933 | 201004339 | 201004455 | 201005307 | 201005442 | 0 | 2 | 3 | 0 | 0 | 0 | 0 | 1 | 0 | 0 | 1 | 0 | 98 | 49 | 0.033473796 | 0.170760496 | NA | | 1 | 1 | NA | 0.333 | NA | 0.667 |
| SE | 7425 | H2AFY | chr5 | - | 135352945 | 135353045 | 135345967 | 135346057 | 135360496 | 135360604 | 0 | 2 | 10 | 0 | 0 | 0 | 0 | 1 | 0 | 0 | 1 | 0 | 98 | 49 | 0.033473796 | 0.170760496 | NA | | 1 | 1 | NA | 0.333 | NA | 0.667 |
| SE | 7587 | DHX9 | chr1 | + | 182855576 | 182855713 | 182854029 | 182854178 | 182856531 | 182856578 | 0 | 2 | 1 | 0 | 0 | 5 | 0 | 1 | 0 | 0 | 1 | 0 | 98 | 49 | 0.033473796 | 0.170760496 | NA | | 1 | 0.091 | NA | 0.333 | NA | 0.212 |
| SE | 9970 | SCYL3 | chr1 | - | 169870254 | 169870357 | 169868927 | 169869039 | 169873695 | 169873752 | 0 | 2 | 1 | 0 | 0 | 0 | 0 | 1 | 0 | 0 | 1 | 0 | 98 | 49 | 0.033473796 | 0.170760496 | NA | | 1 | 1 | NA | 0.333 | NA | 0.667 |
| SE | 12042 | TMC8 | chr17 | + | 78137234 | 78137358 | 78134869 | 78135009 | 78137716 | 78137814 | 0 | 2 | 4 | 0 | 0 | 0 | 0 | 1 | 0 | 0 | 1 | 0 | 98 | 49 | 0.033473796 | 0.170760496 | NA | | 1 | 1 | NA | 0.333 | NA | 0.667 |
| SE | 12085 | DCAF8 | chr1 | - | 160225061 | 160225119 | 160224441 | 160224549 | 160225590 | 160225663 | 0 | 2 | 9 | 0 | 0 | 0 | 2 | 1 | 0 | 0 | 1 | 0 | 98 | 49 | 0.033473796 | 0.170760496 | NA | | 1 | 1 | 1 | 0.333 | NA | 0.334 |
| SE | 12285 | ST6GALNAC2 | chr17 | - | 76578755 | 76578816 | 76574364 | 76574539 | 76585683 | 76586063 | 0 | 2 | 2 | 0 | 0 | 4 | 0 | 1 | 0 | 0 | 1 | 0 | 98 | 49 | 0.033473796 | 0.170760496 | NA | | 1 | 0.2 | NA | 0.333 | NA | 0.267 |
| SE | 12756 | ADGRE5 | chr19 | + | 14397076 | 14397223 | 14396341 | 14396473 | 14397657 | 14397804 | 0 | 2 | 8 | 0 | 0 | 0 | 1 | 1 | 0 | 5 | 1 | 0 | 98 | 49 | 0.033473796 | 0.170760496 | NA | | 1 | 1 | 0.091 | 0.333 | NA | 0.788 |
| SE | 13958 | DIP2B | chr12 | + | 50692948 | 50693013 | 50691048 | 50691151 | 50695266 | 50695360 | 0 | 2 | 5 | 0 | 0 | 0 | 0 | 1 | 0 | 0 | 1 | 0 | 98 | 49 | 0.033473796 | 0.170760496 | NA | | 1 | 1 | NA | 0.333 | NA | 0.667 |
| SE | 10454 | THBS1 | chr15 | + | 39588525 | 39588699 | 39588041 | 39588218 | 39588958 | 39589086 | 0 | 2 | 8 | 0 | 0 | 4 | 0 | 4 | 0 | 0 | 1 | 0 | 98 | 49 | 0.033475981 | 0.170760496 | NA | | 1 | 0.5 | NA | 0.667 | NA | 0.083 |
| SE | 13548 | STRN | chr2 | - | 36855211 | 36855352 | 36850999 | 36851107 | 36857855 | 36858023 | 0 | 2 | 0 | 0 | 0 | 0 | 0 | 4 | 0 | 0 | 1 | 0 | 98 | 49 | 0.033475981 | 0.170760496 | NA | | 1 | NA | NA | 0.667 | NA | 0.333 |
| RI | 452 | BIN2 | chr12 | - | 51302019 | 51302780 | 51302019 | 51302115 | 51302685 | 51302780 | 0 | 0 | 1 | 0 | 4 | 5 | 1 | 1 | 0 | 0 | 4 | 0 | 98 | 49 | 0.033604201 | 0.297702651 | NA | | 0 | 0.091 | 1 | 0.111 | NA | -0.51 |
| SE | 8340 | CHD2 | chr15 | + | 92945820 | 92945865 | 92942842 | 92943068 | 92946037 | 92946216 | 0 | 2 | 8 | 0 | 0 | 0 | 0 | 4 | 0 | 0 | 1 | 0 | 94 | 49 | 0.033727313 | 0.170760496 | NA | | 1 | 1 | NA | 0.676 | NA | 0.324 |
| SE | 4559 | SLC6A6 | chr3 | + | 14457949 | 14458082 | 14447581 | 14447816 | 14466515 | 14466650 | 0 | 3 | 36 | 0 | 0 | 0 | 0 | 9 | 2 | 0 | 3 | 0 | 98 | 49 | 0.033971646 | 0.170760496 | NA | | 1 | 1 | NA | 0.6 | 1 | 0.2 |
| SE | 2911 | PPP6C | chr9 | - | 125153532 | 125153742 | 125146572 | 125149921 | 125153905 | 125153985 | 0 | 3 | 7 | 0 | 0 | 0 | 0 | 9 | 0 | 0 | 1 | 0 | 98 | 49 | 0.034107533 | 0.170760496 | NA | | 1 | 1 | NA | 0.818 | NA | 0.182 |
| SE | 8654 | SVIL | chr10 | - | 29555050 | 29555108 | 29554782 | 29554934 | 29563200 | 29563292 | 0 | 3 | 13 | 0 | 0 | 0 | 0 | 9 | 0 | 0 | 1 | 0 | 98 | 49 | 0.034107533 | 0.170760496 | NA | | 1 | 1 | NA | 0.818 | NA | 0.182 |
| SE | 5187 | MYO9B | chr19 | + | 17192745 | 17193062 | 17191096 | 17191219 | 17194555 | 17195473 | 0 | 2 | 5 | 0 | 0 | 0 | 0 | 5 | 0 | 0 | 1 | 0 | 98 | 49 | 0.03419934 | 0.170760496 | NA | | 1 | 1 | NA | 0.714 | NA | 0.286 |
| SE | 1137 | IGF2R | chr6 | + | 160058022 | 160058124 | 160056423 | 160056525 | 160058905 | 160059098 | 0 | 2 | 35 | 0 | 0 | 0 | 4 | 21 | 1 | 0 | 2 | 0 | 98 | 49 | 0.034257364 | 0.170760496 | NA | | 1 | 1 | 1 | 0.84 | 1 | 0.053 |
| SE | 6669 | FGR | chr1 | - | 27614429 | 27614583 | 27613218 | 27613350 | 27614849 | 27614926 | 0 | 25 | 47 | 0 | 0 | 0 | 2 | 22 | 2 | 0 | 1 | 0 | 98 | 49 | 0.034275909 | 0.170760496 | NA | | 1 | 1 | 1 | 0.917 | 1 | 0.028 |
| SE | 8631 | SMG7 | chr1 | + | 183512836 | 183512868 | 183472370 | 183472649 | 183515873 | 183515991 | 0 | 0 | 2 | 0 | 0 | 0 | 0 | 1 | 1 | 0 | 0 | 1 | 81 | 49 | 0.034410483 | 0.170760496 | NA | NA | | 1 | NA | 1 | 0.377 | 0.311 |
| SE | 4362 | LRRK2 | chr12 | + | 40305784 | 40305966 | 40303947 | 40304134 | 40308466 | 40308696 | 0 | 2 | 32 | 0 | 0 | 0 | 6 | 23 | 1 | 0 | 2 | 0 | 98 | 49 | 0.034647673 | 0.170760496 | NA | | 1 | 1 | 1 | 0.852 | 1 | 0.049 |

| | | | | | | | | | | | | | | | | | | | | | | | | | | | | | | | | | |
|---|---|---|---|---|---|---|---|---|---|---|---|---|---|---|---|---|---|---|---|---|---|---|---|---|---|---|---|---|---|---|---|---|---|
| SE | 691 | DOCK5 | chr8 | + | 25300575 | 25300657 | 25298943 | 25299101 | 25302324 | 25302454 | 0 | 3 | 24 | 0 | 0 | 0 | 0 | 10 | 0 | 0 | 1 | 0 | 98 | 49 | 0.03465568 | 0.170760496 | NA | 1 | 1 | NA | 0.833 | NA | 0.167 |
| A3SS | 806 | TYMP | chr22 | - | 50529495 | 50529754 | 50529495 | 50529731 | 50529903 | 50530085 | 0 | 2 | 4 | 0 | 0 | 1 | 3 | 3 | 0 | 0 | 1 | 0 | 72 | 49 | 0.034683351 | 0.197717869 | NA | 1 | 0.731 | 1 | 0.671 | NA | 0.03 |
| SE | 4962 | DDX3X | chrX | + | 41339035 | 41339083 | 41337407 | 41337465 | 41341483 | 41341616 | 0 | 2 | 15 | 0 | 0 | 0 | 0 | 19 | 3 | 0 | 3 | 0 | 97 | 49 | 0.034732632 | 0.170760496 | NA | 1 | 1 | NA | 0.762 | 1 | 0.119 |
| SE | 9028 | MYD88 | chr3 | + | 38139863 | 38139998 | 38138477 | 38139028 | 38140756 | 38140848 | 0 | 6 | 12 | 0 | 0 | 0 | 0 | 11 | 3 | 0 | 2 | 0 | 98 | 49 | 0.03475926 | 0.170760496 | NA | 1 | 1 | NA | 0.733 | 1 | 0.133 |
| A3SS | 2212 | GNAS | chr20 | + | 58903527 | 58903585 | 58903530 | 58903585 | 58895611 | 58895684 | 0 | 3 | 4 | 0 | 0 | 2 | 0 | 4 | 0 | 0 | 1 | 0 | 52 | 49 | 0.034903266 | 0.197717869 | NA | 1 | 0.653 | NA | 0.79 | NA | 0.036 |
| SE | 3591 | TRIP12 | chr2 | - | 229778875 | 229778990 | 229778432 | 229778587 | 229785756 | 229785855 | 0 | 2 | 11 | 0 | 0 | 0 | 0 | 6 | 0 | 0 | 1 | 0 | 98 | 49 | 0.034934183 | 0.170760496 | NA | 1 | 1 | NA | 0.75 | NA | 0.25 |
| SE | 863 | NAMPT | chr7 | - | 106268463 | 106268600 | 106263391 | 106263617 | 106269153 | 106269312 | 0 | 9 | 124 | 0 | 0 | 0 | 0 | 88 | 7 | 0 | 2 | 0 | 98 | 49 | 0.035105524 | 0.170760496 | NA | 1 | 1 | NA | 0.957 | 1 | 0.022 |
| SE | 7287 | HDAC5 | chr17 | - | 44078794 | 44078879 | 44078499 | 44078665 | 44079143 | 44079277 | 0 | 1 | 2 | 0 | 0 | 0 | 0 | 5 | 1 | 0 | 3 | 0 | 98 | 49 | 0.035115165 | 0.170760496 | NA | 1 | 1 | NA | 0.455 | 1 | 0.272 |
| SE | 9359 | FERMT3 | chr11 | + | 64219523 | 64219658 | 64219250 | 64219358 | 64219739 | 64219789 | 0 | 3 | 13 | 0 | 0 | 0 | 0 | 11 | 0 | 0 | 1 | 0 | 98 | 49 | 0.03520334 | 0.170760496 | NA | 1 | 1 | NA | 0.846 | NA | 0.154 |
| SE | 7828 | SRGN | chr10 | + | 69097083 | 69097231 | 69088071 | 69088236 | 69103870 | 69104811 | 0 | 28 | 397 | 0 | 0 | 0 | 31 | 320 | 22 | 0 | 2 | 0 | 98 | 49 | 0.035427964 | 0.170760496 | NA | 1 | 1 | 1 | 0.988 | 1 | 0.004 |
| SE | 8959 | N4BP2L2 | chr13 | - | 32527407 | 32527532 | 32522181 | 32522270 | 32535768 | 32537027 | 0 | 2 | 9 | 0 | 0 | 1 | 0 | 7 | 0 | 0 | 1 | 0 | 98 | 49 | 0.0356372 | 0.170760496 | NA | 1 | 0.818 | NA | 0.778 | NA | 0.131 |
| A3SS | 1851 | OAS1 | chr12 | + | 112919388 | 112919907 | 112919486 | 112919907 | 112917546 | 112917700 | 0 | 3 | 5 | 0 | 0 | 4 | 0 | 15 | 2 | 0 | 0 | 0 | 98 | 49 | 0.035922391 | 0.197717869 | NA | 1 | 0.385 | NA | 1 | 1 | -0.307 |
| A5SS | 478 | UBE2D3 | chr4 | - | 102827426 | 102827948 | 102827591 | 102827948 | 102826484 | 102826636 | 0 | 2 | 2 | 0 | 0 | 2 | 1 | 9 | 3 | 0 | 0 | 0 | 98 | 49 | 0.036267739 | 0.170414363 | NA | 1 | 0.333 | 1 | 1 | 1 | -0.334 |
| SE | 14036 | CFLAR | chr2 | + | 201135971 | 201136107 | 201132964 | 201133134 | 201140356 | 201140439 | 1 | 6 | 52 | 0 | 0 | 0 | 0 | 29 | 0 | 0 | 1 | 0 | 98 | 49 | 0.03628875 | 0.170760496 | 1 | 1 | 1 | NA | 0.935 | NA | 0.065 |
| SE | 1368 | RGL2 | chr6 | - | 33294687 | 33294762 | 33294033 | 33294066 | 33294976 | 33295045 | 0 | 2 | 17 | 0 | 0 | 0 | 2 | 8 | 0 | 0 | 1 | 0 | 98 | 49 | 0.036328243 | 0.170760496 | NA | 1 | 1 | 1 | 0.8 | NA | 0.1 |
| SE | 4793 | CASP1 | chr11 | - | 105026225 | 105026356 | 105025509 | 105025600 | 105026841 | 105026951 | 0 | 2 | 29 | 0 | 0 | 0 | 0 | 8 | 0 | 1 | 1 | 0 | 98 | 49 | 0.036328243 | 0.170760496 | NA | 1 | 1 | 0 | 0.8 | NA | 0.6 |
| SE | 6901 | PSME3 | chr17 | + | 42838730 | 42838799 | 42838092 | 42838205 | 42838944 | 42839012 | 0 | 2 | 5 | 0 | 0 | 0 | 0 | 8 | 0 | 0 | 1 | 0 | 98 | 49 | 0.036328243 | 0.170760496 | NA | 1 | 1 | NA | 0.8 | NA | 0.2 |
| SE | 338 | DNAJC25 | chr9 | + | 111642856 | 111643094 | 111631351 | 111631743 | 111647106 | 111647259 | 0 | 1 | 1 | 0 | 0 | 0 | 0 | 2 | 0 | 0 | 1 | 0 | 98 | 49 | 0.036515846 | 0.170760496 | NA | 1 | 1 | NA | 0.5 | NA | 0.5 |
| SE | 2653 | CCDC12 | chr3 | - | 46923606 | 46923668 | 46923328 | 46923363 | 46925455 | 46925535 | 0 | 1 | 1 | 0 | 0 | 0 | 0 | 2 | 0 | 0 | 1 | 0 | 98 | 49 | 0.036515846 | 0.170760496 | NA | 1 | 1 | NA | 0.5 | NA | 0.5 |
| SE | 3010 | MED15 | chr22 | + | 20584854 | 20585015 | 20584358 | 20584425 | 20585100 | 20585267 | 0 | 1 | 1 | 0 | 0 | 0 | 0 | 2 | 0 | 0 | 1 | 0 | 98 | 49 | 0.036515846 | 0.170760496 | NA | 1 | 1 | NA | 0.5 | NA | 0.5 |
| SE | 6389 | SUPT6H | chr17 | + | 28690920 | 28691063 | 28690081 | 28690229 | 28693695 | 28693836 | 0 | 1 | 2 | 0 | 0 | 0 | 0 | 2 | 0 | 0 | 1 | 0 | 98 | 49 | 0.036515846 | 0.170760496 | NA | 1 | 1 | NA | 0.5 | NA | 0.5 |
| SE | 6793 | PLEKHM2 | chr1 | + | 15719733 | 15719920 | 15718537 | 15718625 | 15721328 | 15721388 | 0 | 1 | 2 | 0 | 0 | 0 | 0 | 2 | 0 | 0 | 1 | 0 | 98 | 49 | 0.036515846 | 0.170760496 | NA | 1 | 1 | NA | 0.5 | NA | 0.5 |
| SE | 7794 | G6PD | chrX | - | 154532941 | 154533128 | 154532566 | 154532802 | 154533575 | 154533669 | 0 | 1 | 4 | 0 | 0 | 0 | 0 | 2 | 0 | 0 | 1 | 0 | 98 | 49 | 0.036515846 | 0.170760496 | NA | 1 | 1 | NA | 0.5 | NA | 0.5 |
| SE | 8149 | TTC39C | chr18 | + | 24069156 | 24069271 | 24066011 | 24066140 | 24080584 | 24080939 | 0 | 1 | 2 | 0 | 0 | 0 | 0 | 2 | 0 | 0 | 1 | 0 | 98 | 49 | 0.036515846 | 0.170760496 | NA | 1 | 1 | NA | 0.5 | NA | 0.5 |
| SE | 8182 | LSM14A | chr19 | + | 34215123 | 34215300 | 34208928 | 34209051 | 34215595 | 34215661 | 0 | 1 | 7 | 0 | 0 | 0 | 0 | 2 | 0 | 0 | 1 | 0 | 98 | 49 | 0.036515846 | 0.170760496 | NA | 1 | 1 | NA | 0.5 | NA | 0.5 |
| SE | 9120 | CR1 | chr1 | + | 207609289 | 207609688 | 207607250 | 207607336 | 207611676 | 207611853 | 0 | 1 | 20 | 0 | 0 | 0 | 0 | 2 | 0 | 0 | 1 | 0 | 98 | 49 | 0.036515846 | 0.170760496 | NA | 1 | 1 | NA | 0.5 | NA | 0.5 |
| SE | 9391 | ZNF346 | chr5 | + | 177023159 | 177023234 | 177022679 | 177022913 | 177041125 | 177041229 | 0 | 1 | 0 | 0 | 0 | 0 | 0 | 2 | 0 | 0 | 1 | 0 | 98 | 49 | 0.036515846 | 0.170760496 | NA | 1 | NA | NA | 0.5 | NA | 0.5 |
| SE | 10262 | SNX17 | chr2 | + | 27372622 | 27372740 | 27371268 | 27371343 | 27373039 | 27373113 | 0 | 1 | 3 | 0 | 0 | 0 | 0 | 2 | 0 | 0 | 1 | 0 | 98 | 49 | 0.036515846 | 0.170760496 | NA | 1 | 1 | NA | 0.5 | NA | 0.5 |
| SE | 10380 | NFKBIE | chr6 | - | 44262559 | 44262662 | 44261625 | 44261848 | 44264981 | 44265788 | 0 | 1 | 2 | 0 | 0 | 0 | 0 | 2 | 0 | 0 | 1 | 0 | 98 | 49 | 0.036515846 | 0.170760496 | NA | 1 | 1 | NA | 0.5 | NA | 0.5 |
| SE | 12595 | GBF1 | chr10 | + | 102363255 | 102363396 | 102362474 | 102362664 | 102363709 | 102363798 | 0 | 1 | 0 | 0 | 0 | 0 | 0 | 2 | 0 | 0 | 1 | 0 | 98 | 49 | 0.036515846 | 0.170760496 | NA | 1 | NA | NA | 0.5 | NA | 0.5 |
| SE | 12839 | DENND1B | chr1 | - | 197647054 | 197647114 | 197645689 | 197645743 | 197652234 | 197652315 | 0 | 1 | 0 | 0 | 0 | 0 | 0 | 2 | 0 | 0 | 1 | 0 | 98 | 49 | 0.036515846 | 0.170760496 | NA | 1 | NA | NA | 0.5 | NA | 0.5 |
| SE | 13738 | SERPINA1 | chr14 | - | 94388818 | 94389010 | 94388559 | 94388663 | 94390456 | 94390692 | 0 | 1 | 17 | 0 | 0 | 4 | 0 | 2 | 0 | 0 | 1 | 0 | 98 | 49 | 0.036515846 | 0.170760496 | NA | 1 | 0.68 | NA | 0.5 | NA | 0.34 |
| SE | 347 | OGT | chrX | + | 71561774 | 71561900 | 71559587 | 71559677 | 71562846 | 71563017 | 0 | 3 | 24 | 0 | 0 | 0 | 5 | 14 | 0 | 0 | 1 | 0 | 98 | 49 | 0.036639257 | 0.170760496 | NA | 1 | 1 | 1 | 0.875 | NA | 0.062 |
| SE | 7594 | ATG16L2 | chr11 | + | 72826517 | 72826589 | 72826172 | 72826243 | 72826702 | 72826823 | 0 | 12 | 63 | 0 | 0 | 0 | 11 | 29 | 8 | 0 | 2 | 0 | 98 | 49 | 0.036924565 | 0.170760496 | NA | 1 | 1 | 1 | 0.879 | 1 | 0.04 |
| SE | 3590 | TRIP12 | chr2 | - | 229771518 | 229771632 | 229769230 | 229769325 | 229774096 | 229774261 | 0 | 2 | 2 | 0 | 0 | 0 | 0 | 9 | 0 | 0 | 1 | 0 | 98 | 49 | 0.037002078 | 0.170760496 | NA | 1 | 1 | NA | 0.818 | NA | 0.182 |
| SE | 8756 | ZC3H11A | chr1 | + | 203848330 | 203848407 | 203847183 | 203847687 | 203849710 | 203850026 | 0 | 2 | 10 | 0 | 0 | 0 | 0 | 9 | 0 | 0 | 1 | 0 | 98 | 49 | 0.037002078 | 0.170760496 | NA | 1 | 1 | NA | 0.818 | NA | 0.182 |
| SE | 279 | EXOSC2 | chr9 | + | 130698161 | 130698251 | 130697581 | 130697627 | 130699328 | 130699394 | 0 | 1 | 0 | 0 | 0 | 0 | 0 | 1 | 0 | 0 | 1 | 0 | 98 | 49 | 0.037136978 | 0.170760496 | NA | 1 | NA | NA | 0.333 | NA | 0.667 |
| SE | 3510 | HDLBP | chr2 | - | 241255373 | 241255580 | 241255050 | 241255158 | 241256183 | 241256399 | 0 | 1 | 1 | 0 | 0 | 0 | 0 | 1 | 0 | 0 | 1 | 0 | 98 | 49 | 0.037136978 | 0.170760496 | NA | 1 | 1 | NA | 0.333 | NA | 0.667 |
| SE | 4017 | AGAP2 | chr12 | - | 57731808 | 57731967 | 57731555 | 57731642 | 57732402 | 57732512 | 0 | 1 | 1 | 0 | 0 | 0 | 4 | 1 | 0 | 0 | 1 | 0 | 98 | 49 | 0.037136978 | 0.170760496 | NA | 1 | 1 | 1 | 0.333 | NA | 0.334 |
| SE | 4033 | TRAFD1 | chr12 | + | 112151800 | 112152140 | 112149750 | 112149871 | 112152426 | 112152499 | 0 | 1 | 5 | 0 | 0 | 0 | 0 | 1 | 0 | 0 | 1 | 0 | 98 | 49 | 0.037136978 | 0.170760496 | NA | 1 | 1 | NA | 0.333 | NA | 0.667 |
| SE | 5121 | CCM2 | chr7 | + | 45038252 | 45038426 | 45027633 | 45027797 | 45063917 | 45064001 | 0 | 1 | 0 | 0 | 0 | 0 | 0 | 1 | 0 | 0 | 1 | 0 | 98 | 49 | 0.037136978 | 0.170760496 | NA | 1 | NA | NA | 0.333 | NA | 0.667 |
| SE | 6443 | ATP11A | chr13 | + | 112858144 | 112858290 | 112857817 | 112857920 | 112859392 | 112859452 | 0 | 1 | 1 | 0 | 0 | 1 | 0 | 1 | 0 | 0 | 1 | 0 | 98 | 49 | 0.037136978 | 0.170760496 | NA | 1 | 0.333 | NA | 0.333 | NA | 0.333 |
| SE | 6888 | SMC3 | chr10 | + | 110590990 | 110591132 | 110590411 | 110590572 | 110593072 | 110593223 | 0 | 1 | 3 | 0 | 0 | 0 | 0 | 1 | 0 | 0 | 1 | 0 | 98 | 49 | 0.037136978 | 0.170760496 | NA | 1 | 1 | NA | 0.333 | NA | 0.667 |
| SE | 10311 | KLC1 | chr14 | + | 103657545 | 103657776 | 103654563 | 103654825 | 103662115 | 103662194 | 0 | 1 | 8 | 0 | 0 | 0 | 0 | 1 | 0 | 0 | 1 | 0 | 98 | 49 | 0.037136978 | 0.170760496 | NA | 1 | 1 | NA | 0.333 | NA | 0.667 |
| SE | 11210 | GYPC | chr2 | + | 126690254 | 126690311 | 126655934 | 126656312 | 126693863 | 126693947 | 0 | 1 | 0 | 0 | 0 | 0 | 0 | 1 | 0 | 0 | 1 | 0 | 98 | 49 | 0.037136978 | 0.170760496 | NA | 1 | NA | NA | 0.333 | NA | 0.667 |
| SE | 13735 | SERPINA1 | chr14 | - | 94388800 | 94389010 | 94388559 | 94388663 | 94390456 | 94390692 | 0 | 1 | 17 | 0 | 0 | 4 | 0 | 1 | 0 | 0 | 1 | 0 | 98 | 49 | 0.037136978 | 0.170760496 | NA | 1 | 0.68 | NA | 0.333 | NA | 0.507 |
| SE | 13855 | FBXW11 | chr5 | - | 171914342 | 171914405 | 171910571 | 171910797 | 172006457 | 172006873 | 0 | 0 | 1 | 0 | 0 | 0 | 0 | 1 | 1 | 0 | 0 | 1 | 98 | 49 | 0.037136978 | 0.170760496 | NA | NA | 1 | NA | 1 | 0.333 | 0.334 |
| SE | 1807 | ASAP1 | chr8 | - | 130091972 | 130092143 | 130079901 | 130079971 | 130112093 | 130112322 | 0 | 1 | 0 | 0 | 0 | 1 | 0 | 3 | 0 | 0 | 1 | 0 | 98 | 49 | 0.03722674 | 0.170760496 | NA | 1 | 0 | NA | 0.6 | NA | -0.1 |
| SE | 3240 | ZFAS1 | chr20 | + | 49280902 | 49280964 | 49280484 | 49280570 | 49289044 | 49289258 | 0 | 1 | 12 | 0 | 0 | 1 | 0 | 3 | 0 | 0 | 1 | 0 | 98 | 49 | 0.03722674 | 0.170760496 | NA | 1 | 0.857 | NA | 0.6 | NA | 0.329 |
| SE | 5144 | KIAA0226L | chr13 | - | 46371940 | 46372210 | 46368732 | 46368815 | 46387133 | 46387500 | 0 | 1 | 1 | 0 | 0 | 0 | 0 | 3 | 0 | 0 | 1 | 0 | 98 | 49 | 0.03722674 | 0.170760496 | NA | 1 | 1 | NA | 0.6 | NA | 0.4 |
| SE | 5505 | SBF2 | chr11 | - | 10031047 | 10031170 | 10029764 | 10029875 | 10042843 | 10042981 | 0 | 1 | 12 | 0 | 0 | 0 | 0 | 3 | 0 | 0 | 1 | 0 | 98 | 49 | 0.03722674 | 0.170760496 | NA | 1 | 1 | NA | 0.6 | NA | 0.4 |
| SE | 6391 | KMT2B | chr19 | + | 35736689 | 35736827 | 35733762 | 35733872 | 35736911 | 35736986 | 0 | 1 | 1 | 0 | 0 | 0 | 0 | 3 | 0 | 0 | 1 | 0 | 98 | 49 | 0.03722674 | 0.170760496 | NA | 1 | 1 | NA | 0.6 | NA | 0.4 |
| SE | 8166 | ARL5B | chr10 | + | 18672621 | 18672705 | 18668529 | 18668677 | 18673983 | 18674135 | 0 | 1 | 5 | 0 | 0 | 0 | 0 | 3 | 0 | 0 | 1 | 0 | 98 | 49 | 0.03722674 | 0.170760496 | NA | 1 | 1 | NA | 0.6 | NA | 0.4 |
| SE | 9178 | SH3BP1 | chr22 | + | 37644869 | 37644960 | 37644636 | 37644705 | 37645364 | 37645510 | 0 | 1 | 4 | 0 | 0 | 0 | 2 | 3 | 0 | 0 | 1 | 0 | 98 | 49 | 0.03722674 | 0.170760496 | NA | 1 | 1 | 1 | 0.6 | NA | 0.2 |
| SE | 10631 | VPS13C | chr15 | - | 61951823 | 61951980 | 61950944 | 61951024 | 61954420 | 61954554 | 0 | 1 | 3 | 0 | 0 | 0 | 0 | 3 | 0 | 0 | 1 | 0 | 98 | 49 | 0.03722674 | 0.170760496 | NA | 1 | 1 | NA | 0.6 | NA | 0.4 |
| SE | 10953 | FES | chr15 | + | 90889516 | 90889636 | 90889305 | 90889443 | 90889839 | 90889962 | 0 | 1 | 7 | 0 | 0 | 0 | 0 | 3 | 0 | 0 | 1 | 0 | 98 | 49 | 0.03722674 | 0.170760496 | NA | 1 | 1 | NA | 0.6 | NA | 0.4 |
| SE | 11935 | STK40 | chr1 | - | 36367847 | 36368084 | 36361220 | 36361340 | 36385722 | 36385927 | 0 | 1 | 1 | 0 | 0 | 0 | 0 | 3 | 0 | 0 | 1 | 0 | 98 | 49 | 0.03722674 | 0.170760496 | NA | 1 | 1 | NA | 0.6 | NA | 0.4 |
| SE | 13687 | SIRPA | chr20 | + | 1924763 | 1924877 | 1915098 | 1915455 | 1927874 | 1927899 | 0 | 1 | 10 | 0 | 0 | 0 | 0 | 3 | 0 | 0 | 1 | 0 | 98 | 49 | 0.03722674 | 0.170760496 | NA | 1 | 1 | NA | 0.6 | NA | 0.4 |
| SE | 331 | AKNA | chr9 | - | 114343707 | 114343803 | 114342008 | 114342125 | 114345862 | 114346009 | 0 | 0 | 10 | 0 | 2 | 2 | 0 | 1 | 0 | 0 | 1 | 0 | 98 | 49 | 0.037261148 | 0.170760496 | NA | 0 | 0.714 | NA | 0.333 | NA | 0.024 |
| SE | 6388 | SUPT6H | chr17 | + | 28682941 | 28683092 | 28682726 | 28682856 | 28683267 | 28683422 | 0 | 0 | 1 | 0 | 2 | 0 | 0 | 1 | 0 | 0 | 1 | 0 | 98 | 49 | 0.037261148 | 0.170760496 | NA | 0 | 1 | NA | 0.333 | NA | 0.167 |
| SE | 7325 | SREBF1 | chr17 | - | 17814614 | 17814747 | 17814244 | 17814410 | 17814834 | 17814944 | 0 | 0 | 0 | 0 | 2 | 0 | 0 | 1 | 0 | 0 | 1 | 0 | 98 | 49 | 0.037261148 | 0.170760496 | NA | 0 | NA | NA | 0.333 | NA | -0.333 |
| SE | 559 | PABPC1 | chr8 | - | 100718086 | 100718280 | 100717772 | 100717888 | 100721390 | 100722087 | 0 | 0 | 29 | 0 | 1 | 0 | 1 | 19 | 6 | 0 | 0 | 0 | 98 | 49 | 0.037496585 | 0.170760496 | NA | 0 | 1 | 1 | 1 | 1 | -0.5 |
| SE | 371 | AGTPBP1 | chr9 | - | 85678334 | 85678398 | 85677435 | 85677582 | 85681267 | 85681335 | 0 | 2 | 31 | 0 | 0 | 0 | 0 | 10 | 0 | 0 | 1 | 0 | 98 | 49 | 0.037649757 | 0.170760496 | NA | 1 | 1 | NA | 0.833 | NA | 0.167 |
| SE | 5702 | PPP4C | chr16 | + | 30083654 | 30083781 | 30083393 | 30083567 | 30084665 | 30084855 | 0 | 2 | 14 | 0 | 0 | 0 | 0 | 10 | 0 | 0 | 1 | 0 | 98 | 49 | 0.037649757 | 0.170760496 | NA | 1 | 1 | NA | 0.833 | NA | 0.167 |
| SE | 11023 | TAP1 | chr6 | - | 32848989 | 32849118 | 32848651 | 32848840 | 32850319 | 32850517 | 1 | 2 | 34 | 0 | 0 | 0 | 0 | 10 | 0 | 0 | 1 | 0 | 98 | 49 | 0.037649757 | 0.170760496 | 1 | 1 | 1 | NA | 0.833 | NA | 0.167 |
| A3SS | 1947 | SNRPB | chr20 | - | 2461634 | 2461939 | 2461634 | 2461793 | 2462635 | 2462761 | 0 | 0 | 3 | 0 | 1 | 2 | 7 | 5 | 1 | 0 | 0 | 0 | 98 | 49 | 0.037878306 | 0.197717869 | NA | 0 | 0.429 | 1 | 1 | 1 | -0.785 |
| SE | 370 | AGTPBP1 | chr9 | - | 85660935 | 85660973 | 85657434 | 85657643 | 85669484 | 85669578 | 0 | 1 | 9 | 0 | 0 | 0 | 0 | 3 | 0 | 0 | 1 | 0 | 87 | 49 | 0.038097905 | 0.170760496 | NA | 1 | 1 | NA | 0.628 | NA | 0.372 |
| SE | 5887 | SIPA1L1 | chr14 | + | 71650334 | 71650509 | 71624047 | 71624236 | 71658332 | 71658436 | 0 | 1 | 14 | 0 | 0 | 0 | 0 | 4 | 0 | 0 | 1 | 0 | 98 | 49 | 0.038206571 | 0.170760496 | NA | 1 | 1 | NA | 0.667 | NA | 0.333 |
| SE | 8614 | TM9SF3 | chr10 | - | 96533050 | 96533190 | 96530539 | 96530608 | 96544075 | 96544206 | 0 | 1 | 11 | 0 | 0 | 0 | 0 | 4 | 0 | 0 | 1 | 0 | 98 | 49 | 0.038206571 | 0.170760496 | NA | 1 | 1 | NA | 0.667 | NA | 0.333 |
| SE | 11155 | NDE1 | chr16 | + | 15694164 | 15694256 | 15691143 | 15691323 | 15696708 | 15696860 | 0 | 1 | 16 | 0 | 0 | 0 | 0 | 4 | 0 | 0 | 1 | 0 | 98 | 49 | 0.038206571 | 0.170760496 | NA | 1 | 1 | NA | 0.667 | NA | 0.333 |
| SE | 11295 | CREBBP | chr16 | - | 3782683 | 3782926 | 3781203 | 3781306 | 3791980 | 3792094 | 0 | 1 | 6 | 0 | 0 | 0 | 0 | 4 | 0 | 0 | 1 | 0 | 98 | 49 | 0.038206571 | 0.170760496 | NA | 1 | 1 | NA | 0.667 | NA | 0.333 |
| SE | 12432 | SPAG9 | chr17 | - | 50974770 | 50974947 | 50970706 | 50970856 | 50977107 | 50977221 | 0 | 1 | 3 | 0 | 0 | 0 | 0 | 4 | 0 | 0 | 1 | 0 | 98 | 49 | 0.038206571 | 0.170760496 | NA | 1 | 1 | NA | 0.667 | NA | 0.333 |
| SE | 3199 | TNIP1 | chr5 | - | 151060317 | 151060395 | 151056765 | 151056957 | 151062126 | 151062212 | 0 | 2 | 8 | 0 | 0 | 0 | 0 | 11 | 0 | 0 | 1 | 0 | 98 | 49 | 0.038274566 | 0.170760496 | NA | 1 | 1 | NA | 0.846 | NA | 0.154 |
| SE | 349 | NR4A3 | chr9 | + | 99832688 | 99832818 | 99828040 | 99828993 | 99833281 | 99833454 | 0 | 1 | 3 | 0 | 0 | 0 | 0 | 7 | 1 | 0 | 2 | 0 | 98 | 49 | 0.038321227 | 0.170760496 | NA | 1 | 1 | NA | 0.636 | 1 | 0.182 |
| SE | 11639 | ARRB2 | chr17 | + | 4716146 | 4716191 | 4715972 | 4716033 | 4716411 | 4716608 | 0 | 6 | 32 | 0 | 0 | 1 | 0 | 13 | 1 | 0 | 5 | 2 | 94 | 49 | 0.038650672 | 0.170760496 | NA | 1 | 0.943 | NA | 0.575 | 0.207 | 0.581 |
| SE | 9931 | PTPRE | chr10 | + | 128066074 | 128066194 | 128063082 | 128063180 | 128068122 | 128068286 | 0 | 4 | 51 | 0 | 0 | 0 | 0 | 47 | 9 | 0 | 3 | 0 | 98 | 49 | 0.038824113 | 0.170760496 | NA | 1 | 1 | NA | 0.887 | 1 | 0.056 |
| SE | 11065 | CORO1A | chr16 | + | 30187941 | 30188087 | 30187724 | 30187829 | 30188191 | 30188249 | 0 | 18 | 59 | 0 | 0 | 0 | 5 | 24 | 3 | 0 | 1 | 0 | 98 | 49 | 0.038917622 | 0.170760496 | NA | 1 | 1 | 1 | 0.923 | 1 | 0.026 |
| A5SS | 3018 | SERPINA1 | chr14 | - | 94388800 | 94389010 | 94388818 | 94389010 | 94388559 | 94388663 | 0 | 1 | 6 | 0 | 0 | 2 | 0 | 2 | 0 | 0 | 1 | 0 | 67 | 49 | 0.039150683 | 0.170414363 | NA | 1 | 0.687 | NA | 0.594 | NA | 0.25 |
| A3SS | 3771 | STRN4 | chr19 | - | 46727451 | 46727567 | 46727451 | 46727546 | 46727893 | 46728007 | 0 | 1 | 1 | 0 | 0 | 4 | 0 | 1 | 0 | 0 | 1 | 0 | 70 | 49 | 0.039208323 | 0.197717869 | NA | 1 | 0.149 | NA | 0.412 | NA | 0.163 |
| SE | 2205 | BOD1L1 | chr4 | - | 13595859 | 13595944 | 13591922 | 13591966 | 13597103 | 13597168 | 0 | 1 | 12 | 0 | 0 | 0 | 0 | 5 | 0 | 0 | 1 | 0 | 98 | 49 | 0.039248064 | 0.170760496 | NA | 1 | 1 | NA | 0.714 | NA | 0.286 |
| SE | 2551 | ATXN7 | chr3 | + | 63913156 | 63913225 | 63912587 | 63912923 | 63952378 | 63952483 | 0 | 1 | 7 | 0 | 0 | 0 | 0 | 5 | 0 | 0 | 1 | 0 | 98 | 49 | 0.039248064 | 0.170760496 | NA | 1 | 1 | NA | 0.714 | NA | 0.286 |
| SE | 4158 | SH3BP2 | chr4 | + | 2831922 | 2831978 | 2831570 | 2831679 | 2832330 | 2832412 | 0 | 1 | 12 | 0 | 0 | 1 | 1 | 5 | 0 | 0 | 1 | 0 | 98 | 49 | 0.039248064 | 0.170760496 | NA | 1 | 0.857 | 1 | 0.714 | NA | 0.072 |
| SE | 4319 | BIRC6 | chr2 | + | 32441328 | 32441462 | 32439507 | 32439686 | 32442064 | 32442226 | 0 | 1 | 5 | 0 | 0 | 0 | 0 | 5 | 0 | 0 | 1 | 0 | 98 | 49 | 0.039248064 | 0.170760496 | NA | 1 | 1 | NA | 0.714 | NA | 0.286 |
| SE | 5146 | KIAA0226L | chr13 | - | 46371940 | 46372597 | 46368732 | 46368815 | 46387133 | 46387500 | 0 | 1 | 1 | 0 | 0 | 0 | 0 | 5 | 0 | 0 | 1 | 0 | 98 | 49 | 0.039248064 | 0.170760496 | NA | 1 | 1 | NA | 0.714 | NA | 0.286 |
| SE | 5222 | CLEC4E | chr12 | - | 8537114 | 8537266 | 8536089 | 8536205 | 8539216 | 8539306 | 0 | 1 | 35 | 0 | 0 | 0 | 0 | 5 | 0 | 0 | 1 | 0 | 98 | 49 | 0.039248064 | 0.170760496 | NA | 1 | 1 | NA | 0.714 | NA | 0.286 |
| SE | 5861 | DISC1 | chr1 | + | 231795241 | 231795296 | 231770834 | 231771070 | 231818328 | 231819244 | 0 | 1 | 7 | 0 | 0 | 0 | 0 | 5 | 0 | 0 | 1 | 0 | 98 | 49 | 0.039248064 | 0.170760496 | NA | 1 | 1 | NA | 0.714 | NA | 0.286 |
| SE | 6191 | FOXN2 | chr2 | + | 48328560 | 48328702 | 48314655 | 48314814 | 48346200 | 48346751 | 0 | 1 | 2 | 0 | 0 | 1 | 0 | 5 | 0 | 0 | 1 | 0 | 98 | 49 | 0.039248064 | 0.170760496 | NA | 1 | 0.5 | NA | 0.714 | NA | 0.036 |
| SE | 8721 | NOD2 | chr16 | + | 50707854 | 50707960 | 50699349 | 50699954 | 50710557 | 50712373 | 0 | 1 | 3 | 0 | 0 | 0 | 0 | 5 | 0 | 0 | 1 | 0 | 98 | 49 | 0.039248064 | 0.170760496 | NA | 1 | 1 | NA | 0.714 | NA | 0.286 |
| SE | 8817 | KDM2A | chr11 | + | 67243008 | 67243092 | 67231565 | 67231960 | 67245188 | 67245458 | 0 | 1 | 11 | 0 | 0 | 0 | 2 | 5 | 0 | 0 | 1 | 0 | 98 | 49 | 0.039248064 | 0.170760496 | NA | 1 | 1 | 1 | 0.714 | NA | 0.143 |
| SE | 9383 | PECAM1 | chr17 | - | 64354932 | 64355040 | 64353490 | 64353518 | 64356110 | 64356398 | 0 | 1 | 38 | 0 | 0 | 8 | 0 | 5 | 0 | 0 | 1 | 0 | 98 | 49 | 0.039248064 | 0.170760496 | NA | 1 | 0.704 | NA | 0.714 | NA | 0.138 |
| SE | 11702 | POLR2A | chr17 | + | 7496209 | 7496346 | 7495940 | 7496073 | 7496439 | 7496613 | 0 | 1 | 9 | 0 | 0 | 0 | 0 | 5 | 0 | 0 | 1 | 0 | 98 | 49 | 0.039248064 | 0.170760496 | NA | 1 | 1 | NA | 0.714 | NA | 0.286 |
| SE | 11979 | TSEN34 | chr19 | + | 54191720 | 54191964 | 54191213 | 54191607 | 54192115 | 54192373 | 0 | 5 | 17 | 0 | 0 | 1 | 0 | 8 | 4 | 0 | 3 | 1 | 98 | 49 | 0.039311924 | 0.170760496 | NA | 1 | 0.895 | NA | 0.571 | 0.667 | 0.329 |
| SE | 7521 | AQP9 | chr15 | + | 58173067 | 58173205 | 58166672 | 58166799 | 58179127 | 58179345 | 0 | 1 | 40 | 0 | 0 | 0 | 0 | 21 | 1 | 0 | 2 | 0 | 98 | 49 | 0.039675176 | 0.170760496 | NA | 1 | 1 | NA | 0.84 | 1 | 0.08 |
| A3SS | 462 | BTN3A1 | chr6 | + | 26412704 | 26415216 | 26413168 | 26415216 | 26411554 | 26411581 | 0 | 1 | 5 | 0 | 0 | 2 | 0 | 6 | 0 | 0 | 1 | 0 | 98 | 49 | 0.040279671 | 0.197717869 | NA | 1 | 0.556 | NA | 0.75 | NA | 0.028 |
| SE | 1761 | FAM53C | chr5 | + | 138341808 | 138341866 | 138341183 | 138341413 | 138344824 | 138345609 | 0 | 1 | 7 | 0 | 0 | 0 | 0 | 6 | 0 | 0 | 1 | 0 | 98 | 49 | 0.040279671 | 0.170760496 | NA | 1 | 1 | NA | 0.75 | NA | 0.25 |
| SE | 3245 | SULF2 | chr20 | - | 47665198 | 47665293 | 47664129 | 47664189 | 47665856 | 47665953 | 0 | 1 | 14 | 0 | 0 | 0 | 0 | 6 | 0 | 0 | 1 | 0 | 98 | 49 | 0.040279671 | 0.170760496 | NA | 1 | 1 | NA | 0.75 | NA | 0.25 |
| SE | 8677 | MANBA | chr4 | - | 102714437 | 102714561 | 102690595 | 102690771 | 102722870 | 102723041 | 0 | 1 | 4 | 0 | 0 | 0 | 0 | 6 | 0 | 0 | 1 | 0 | 98 | 49 | 0.040279671 | 0.170760496 | NA | 1 | 1 | NA | 0.75 | NA | 0.25 |
| SE | 12664 | STXBP2 | chr19 | + | 7643164 | 7643245 | 7642982 | 7643048 | 7644613 | 7644752 | 0 | 1 | 4 | 0 | 0 | 0 | 1 | 6 | 0 | 0 | 1 | 0 | 98 | 49 | 0.040279671 | 0.170760496 | NA | 1 | 1 | 1 | 0.75 | NA | 0.125 |
| SE | 19 | FLNA | chrX | - | 154357250 | 154357274 | 154354824 | 154355072 | 154357433 | 154357623 | 0 | 1 | 4 | 0 | 7 | 5 | 2 | 10 | 0 | 0 | 4 | 0 | 73 | 49 | 0.040614468 | 0.170760496 | NA | 0.087 | 0.349 | 1 | 0.627 | NA | -0.596 |
| SE | 12838 | PTTG1IP | chr21 | - | 44865394 | 44865447 | 44861162 | 44861271 | 44873501 | 44873903 | 0 | 2 | 4 | 0 | 0 | 0 | 0 | 3 | 1 | 0 | 2 | 0 | 98 | 49 | 0.040702408 | 0.170760496 | NA | 1 | 1 | NA | 0.429 | 1 | 0.285 |
| SE | 4624 | SORL1 | chr11 | + | 121543547 | 121543726 | 121532463 | 121532552 | 121545242 | 121545429 | 0 | 4 | 58 | 0 | 0 | 0 | 0 | 34 | 0 | 0 | 1 | 0 | 98 | 49 | 0.04109126 | 0.170760496 | NA | 1 | 1 | NA | 0.944 | NA | 0.056 |
| SE | 1140 | IGF2R | chr6 | + | 160063414 | 160063630 | 160062531 | 160062619 | 160064400 | 160064531 | 0 | 2 | 31 | 0 | 0 | 0 | 0 | 16 | 0 | 0 | 1 | 0 | 98 | 49 | 0.041121367 | 0.170760496 | NA | 1 | 1 | NA | 0.889 | NA | 0.111 |
| SE | 5495 | ADAM8 | chr10 | - | 133269444 | 133269529 | 133267928 | 133268118 | 133269896 | 133269974 | 0 | 2 | 12 | 0 | 0 | 0 | 0 | 16 | 0 | 0 | 1 | 0 | 98 | 49 | 0.041121367 | 0.170760496 | NA | 1 | 1 | NA | 0.889 | NA | 0.111 |
| SE | 2907 | RAF1 | chr3 | - | 12600151 | 12600279 | 12599690 | 12599808 | 12600387 | 12600415 | 0 | 2 | 19 | 0 | 0 | 0 | 0 | 22 | 3 | 0 | 2 | 0 | 98 | 49 | 0.041304338 | 0.170760496 | NA | 1 | 1 | NA | 0.846 | 1 | 0.077 |
| SE | 5124 | DNM2 | chr19 | + | 10759737 | 10759811 | 10718052 | 10718403 | 10772478 | 10772628 | 0 | 11 | 23 | 0 | 0 | 0 | 0 | 12 | 3 | 0 | 1 | 0 | 98 | 49 | 0.041331042 | 0.170760496 | NA | 1 | 1 | NA | 0.857 | 1 | 0.072 |
| A3SS | 3207 | ANKRD11 | chr16 | - | 89316932 | 89317078 | 89316932 | 89317075 | 89418283 | 89418368 | 0 | 1 | 2 | 0 | 0 | 0 | 0 | 1 | 0 | 0 | 1 | 0 | 52 | 49 | 0.041336327 | 0.197717869 | NA | 1 | 1 | NA | 0.485 | NA | 0.515 |
| SE | 8601 | PYGL | chr14 | - | 50912155 | 50912303 | 50911977 | 50912036 | 50913028 | 50913130 | 0 | 1 | 14 | 0 | 0 | 1 | 0 | 7 | 0 | 0 | 1 | 0 | 98 | 49 | 0.041336891 | 0.170760496 | NA | 1 | 0.875 | NA | 0.778 | NA | 0.159 |
| SE | 9493 | SAP30BP | chr17 | + | 75668515 | 75668625 | 75667115 | 75667478 | 75671815 | 75671863 | 0 | 1 | 1 | 0 | 0 | 0 | 0 | 7 | 0 | 0 | 1 | 0 | 98 | 49 | 0.041336891 | 0.170760496 | NA | 1 | 1 | NA | 0.778 | NA | 0.222 |
| SE | 10364 | SOS2 | chr14 | - | 50158564 | 50158646 | 50156998 | 50157121 | 50159430 | 50160086 | 0 | 1 | 14 | 0 | 0 | 2 | 0 | 7 | 0 | 0 | 1 | 0 | 98 | 49 | 0.041336891 | 0.170760496 | NA | 1 | 0.778 | NA | 0.778 | NA | 0.111 |
| SE | 11078 | TCEB2 | chr16 | - | 2775450 | 2775556 | 2771413 | 2772102 | 2776992 | 2777127 | 0 | 1 | 12 | 0 | 0 | 0 | 1 | 7 | 0 | 0 | 1 | 0 | 98 | 49 | 0.041336891 | 0.170760496 | NA | 1 | 1 | 1 | 0.778 | NA | 0.111 |
| SE | 10594 | RNF213 | chr17 | + | 80390011 | 80390196 | 80389827 | 80389917 | 80393344 | 80398781 | 0 | 0 | 79 | 0 | 1 | 1 | 0 | 30 | 3 | 0 | 0 | 0 | 98 | 49 | 0.041591024 | 0.170760496 | NA | 0 | 0.975 | NA | 1 | 1 | -0.512 |
| SE | 1662 | FAM193B | chr5 | - | 177531952 | 177532200 | 177531281 | 177531521 | 177532442 | 177532641 | 0 | 2 | 0 | 0 | 0 | 1 | 0 | 0 | 1 | 0 | 0 | 2 | 98 | 49 | 0.041785228 | 0.170760496 | NA | 1 | 0 | NA | NA | 0.2 | 0.3 |
| SE | 9538 | CAMK2G | chr10 | - | 73839538 | 73839601 | 73837467 | 73837511 | 73842168 | 73842211 | 0 | 0 | 0 | 0 | 1 | 4 | 0 | 1 | 0 | 3 | 2 | 0 | 98 | 49 | 0.041785228 | 0.170760496 | NA | 0 | 0 | 0 | 0.2 | NA | -0.1 |
| A5SS | 2638 | BPTF | chr17 | + | 67945408 | 67946325 | 67945408 | 67945896 | 67947725 | 67947808 | 0 | 0 | 0 | 0 | 1 | 1 | 0 | 1 | 0 | 0 | 1 | 0 | 98 | 49 | 0.041803869 | 0.170414363 | NA | 0 | 0 | NA | 0.333 | NA | -0.333 |
| RI | 287 | EDEM2 | chr20 | - | 35146824 | 35147358 | 35146824 | 35146935 | 35147151 | 35147358 | 0 | 0 | 0 | 0 | 0 | 1 | 0 | 0 | 1 | 0 | 0 | 1 | 98 | 49 | 0.041803869 | 0.297702651 | NA | NA | 0 | NA | NA | 0.333 | -0.333 |
| SE | 888 | GIGYF1 | chr7 | - | 100683549 | 100683632 | 100683303 | 100683444 | 100683817 | 100683918 | 0 | 0 | 0 | 0 | 1 | 0 | 0 | 1 | 0 | 0 | 1 | 0 | 98 | 49 | 0.041803869 | 0.170760496 | NA | 0 | NA | NA | 0.333 | NA | -0.333 |
| SE | 6999 | AGTRAP | chr1 | + | 11746126 | 11746223 | 11745802 | 11745837 | 11747439 | 11747545 | 0 | 0 | 0 | 0 | 1 | 2 | 0 | 1 | 0 | 0 | 1 | 0 | 98 | 49 | 0.041803869 | 0.170760496 | NA | 0 | 0 | NA | 0.333 | NA | -0.333 |
| SE | 8430 | GS1-124K5.11 | chr7 | - | 66560578 | 66560675 | 66553467 | 66554837 | 66573384 | 66573550 | 0 | 0 | 1 | 0 | 1 | 0 | 0 | 1 | 0 | 0 | 1 | 0 | 98 | 49 | 0.041803869 | 0.170760496 | NA | 0 | 1 | NA | 0.333 | NA | 0.167 |
| SE | 9926 | PPP4R3B | chr2 | - | 55578246 | 55578342 | 55577314 | 55577356 | 55579678 | 55579781 | 0 | 0 | 0 | 0 | 1 | 1 | 0 | 1 | 0 | 0 | 1 | 0 | 98 | 49 | 0.041803869 | 0.170760496 | NA | 0 | 0 | NA | 0.333 | NA | -0.333 |
| SE | 7544 | TKT | chr3 | - | 53237796 | 53237820 | 53234982 | 53235174 | 53240250 | 53240348 | 0 | 2 | 1 | 0 | 1 | 33 | 0 | 0 | 0 | 0 | 10 | 6 | 73 | 49 | 0.041838974 | 0.170760496 | NA | 0.573 | 0.02 | NA | 0 | 0 | 0.296 |
| SE | 2150 | YTHDC1 | chr4 | - | 68322748 | 68322915 | 68320122 | 68320205 | 68324138 | 68324223 | 0 | 1 | 3 | 0 | 0 | 0 | 0 | 8 | 0 | 0 | 1 | 0 | 98 | 49 | 0.04224465 | 0.170760496 | NA | 1 | 1 | NA | 0.8 | NA | 0.2 |
| SE | 2915 | LPCAT1 | chr5 | - | 1483427 | 1483486 | 1480941 | 1480976 | 1488390 | 1488451 | 0 | 1 | 2 | 0 | 0 | 0 | 0 | 8 | 0 | 0 | 1 | 0 | 98 | 49 | 0.04224465 | 0.170760496 | NA | 1 | 1 | NA | 0.8 | NA | 0.2 |
| SE | 3672 | CALCOCO2 | chr17 | + | 48848063 | 48848166 | 48841697 | 48841887 | 48848321 | 48848455 | 0 | 1 | 8 | 0 | 0 | 0 | 0 | 8 | 0 | 0 | 1 | 0 | 98 | 49 | 0.04224465 | 0.170760496 | NA | 1 | 1 | NA | 0.8 | NA | 0.2 |

| | | | | | | | | | | | | | | | | | | | | | | | | | | | | | | | | | |
|---|---|---|---|---|---|---|---|---|---|---|---|---|---|---|---|---|---|---|---|---|---|---|---|---|---|---|---|---|---|---|---|---|---|
| SE | 1138 | IGF2R | chr6 | + | 160061502 | 160061646 | 160060546 | 160060717 | 160062531 | 160062619 | 0 | 5 | 19 | 0 | 0 | 0 | 2 | 9 | 3 | 0 | 4 | 0 | 98 | 49 | 0.042663026 | 0.170760496 | NA | 1 | 1 | 1 | 0.529 | 1 | 0.157 |
| SE | 5043 | CFL1 | chr11 | - | 65855653 | 65855730 | 65854813 | 65855448 | 65855934 | 65856242 | 0 | 9 | 66 | 0 | 0 | 0 | 10 | 26 | 1 | 0 | 1 | 0 | 98 | 49 | 0.042685592 | 0.170760496 | NA | 1 | 1 | 1 | 0.929 | 1 | 0.024 |
| SE | 7672 | SF3B1 | chr2 | - | 197392971 | 197393188 | 197391973 | 197392461 | 197396055 | 197396328 | 0 | 5 | 39 | 0 | 0 | 0 | 0 | 9 | 1 | 0 | 1 | 0 | 98 | 49 | 0.042920223 | 0.170760496 | NA | 1 | 1 | NA | 0.818 | 1 | 0.091 |
| SE | 9384 | PECAM1 | chr17 | - | 64375050 | 64375356 | 64369749 | 64370025 | 64377823 | 64378117 | 0 | 2 | 51 | 0 | 0 | 0 | 8 | 20 | 0 | 0 | 1 | 0 | 98 | 49 | 0.043140749 | 0.170760496 | NA | 1 | 1 | 1 | 0.909 | NA | 0.045 |
| SE | 8137 | ITGAX | chr16 | + | 31356624 | 31356728 | 31355892 | 31355998 | 31357030 | 31357101 | 0 | 26 | 76 | 0 | 0 | 0 | 8 | 82 | 30 | 0 | 2 | 0 | 98 | 49 | 0.043231591 | 0.170760496 | NA | 1 | 1 | 1 | 0.953 | 1 | 0.016 |
| SE | 22 | ARHGAP4 | chrX | - | 153919566 | 153919686 | 153919154 | 153919283 | 153920625 | 153920808 | 0 | 2 | 0 | 2 | 2 | 2 | 0 | 0 | 0 | 0 | 6 | 3 | 98 | 49 | 0.043694816 | 0.170760496 | 0 | 0.333 | 0 | NA | 0 | 0 | 0.111 |
| A3SS | 4015 | PLEKHB2 | chr2 | + | 131140166 | 131140275 | 131140169 | 131140275 | 131132901 | 131132991 | 0 | 1 | 3 | 0 | 0 | 0 | 0 | 4 | 0 | 0 | 1 | 0 | 52 | 49 | 0.043807749 | 0.197717869 | NA | 1 | 1 | NA | 0.79 | NA | 0.21 |
| A3SS | 461 | BTN3A1 | chr6 | + | 26412411 | 26415216 | 26413168 | 26415216 | 26411554 | 26411581 | 0 | 1 | 5 | 0 | 0 | 2 | 0 | 10 | 0 | 0 | 1 | 0 | 98 | 49 | 0.044092288 | 0.197717869 | NA | 1 | 0.556 | NA | 0.833 | NA | -0.055 |
| SE | 7268 | IVNS1ABP | chr1 | - | 185309002 | 185309172 | 185308799 | 185308875 | 185309382 | 185309511 | 0 | 1 | 10 | 0 | 0 | 0 | 0 | 10 | 0 | 0 | 1 | 0 | 98 | 49 | 0.044092288 | 0.170760496 | NA | 1 | 1 | NA | 0.833 | NA | 0.167 |
| SE | 5494 | ADAM8 | chr10 | - | 133268747 | 133268862 | 133267928 | 133268118 | 133269444 | 133269529 | 0 | 2 | 19 | 0 | 0 | 0 | 0 | 25 | 4 | 0 | 2 | 0 | 98 | 49 | 0.044237698 | 0.170760496 | NA | 1 | 1 | NA | 0.862 | 1 | 0.069 |
| SE | 1818 | CHD1 | chr5 | - | 98881974 | 98882123 | 98881278 | 98881375 | 98883087 | 98883237 | 0 | 2 | 14 | 0 | 0 | 0 | 0 | 23 | 0 | 0 | 1 | 0 | 98 | 49 | 0.044532178 | 0.170760496 | NA | 1 | 1 | NA | 0.92 | NA | 0.08 |
| SE | 814 | LINC-PINT | chr7 | - | 130959868 | 130959970 | 130944159 | 130945835 | 130984052 | 130984109 | 0 | 0 | 1 | 0 | 1 | 2 | 0 | 1 | 0 | 0 | 4 | 0 | 98 | 49 | 0.044611886 | 0.170760496 | NA | 0 | 0.2 | NA | 0.111 | NA | -0.011 |
| SE | 815 | LINC-PINT | chr7 | - | 130967388 | 130967517 | 130944159 | 130945835 | 130984052 | 130984109 | 0 | 0 | 5 | 0 | 1 | 2 | 0 | 1 | 0 | 0 | 4 | 0 | 98 | 49 | 0.044611886 | 0.170760496 | NA | 0 | 0.556 | NA | 0.111 | NA | 0.167 |
| A5SS | 3046 | PLEKHB2 | chr2 | + | 131132901 | 131133015 | 131132901 | 131132991 | 131140166 | 131140275 | 0 | 0 | 4 | 0 | 1 | 3 | 0 | 1 | 0 | 0 | 4 | 0 | 73 | 49 | 0.044612593 | 0.170414363 | NA | 0 | 0.472 | NA | 0.144 | NA | 0.092 |
| SE | 11394 | ADGRG3 | chr16 | + | 57679176 | 57679311 | 57678169 | 57678316 | 57679815 | 57679855 | 0 | 0 | 15 | 0 | 1 | 0 | 0 | 11 | 3 | 0 | 0 | 0 | 98 | 49 | 0.044729667 | 0.170760496 | NA | 0 | 1 | NA | 1 | 1 | -0.5 |
| SE | 2103 | WDFY3 | chr4 | - | 84836928 | 84837090 | 84831412 | 84831605 | 84841153 | 84841263 | 0 | 1 | 20 | 0 | 0 | 0 | 0 | 11 | 0 | 0 | 1 | 0 | 98 | 49 | 0.044972537 | 0.170760496 | NA | 1 | 1 | NA | 0.846 | NA | 0.154 |
| SE | 7942 | IFI44L | chr1 | + | 78629719 | 78629915 | 78628950 | 78628999 | 78635336 | 78635489 | 0 | 3 | 61 | 0 | 0 | 2 | 4 | 38 | 0 | 0 | 1 | 0 | 98 | 49 | 0.045281167 | 0.170760496 | NA | 1 | 0.938 | 1 | 0.95 | NA | -0.006 |
| SE | 7117 | MED13L | chr12 | - | 116007410 | 116007636 | 116006305 | 116006411 | 116008400 | 116009132 | 0 | 2 | 11 | 0 | 0 | 0 | 0 | 25 | 0 | 0 | 1 | 0 | 98 | 49 | 0.045416917 | 0.170760496 | NA | 1 | 1 | NA | 0.926 | NA | 0.074 |
| SE | 4382 | STK38L | chr12 | + | 27308338 | 27308461 | 27302136 | 27302188 | 27309113 | 27309197 | 0 | 3 | 8 | 0 | 0 | 0 | 0 | 5 | 1 | 0 | 1 | 0 | 98 | 49 | 0.045453113 | 0.170760496 | NA | 1 | 1 | NA | 0.714 | 1 | 0.143 |
| SE | 1691 | DOCK2 | chr5 | + | 169761518 | 169761625 | 169759704 | 169759775 | 169803057 | 169803206 | 0 | 4 | 19 | 0 | 0 | 0 | 0 | 12 | 1 | 0 | 1 | 0 | 98 | 49 | 0.045620082 | 0.170760496 | NA | 1 | 1 | NA | 0.857 | 1 | 0.072 |
| SE | 6818 | OAS1 | chr12 | + | 112917546 | 112917700 | 112916508 | 112916738 | 112919388 | 112919907 | 0 | 3 | 8 | 0 | 0 | 0 | 0 | 8 | 1 | 0 | 1 | 0 | 98 | 49 | 0.045786327 | 0.170760496 | NA | 1 | 1 | NA | 0.8 | 1 | 0.1 |
| SE | 7899 | IFIT3 | chr10 | + | 89337420 | 89337542 | 89332478 | 89332627 | 89338660 | 89340968 | 0 | 0 | 0 | 0 | 1 | 1 | 0 | 1 | 0 | 0 | 5 | 0 | 98 | 49 | 0.046158728 | 0.170760496 | NA | 0 | 0 | NA | 0.091 | NA | -0.091 |
| SE | 9005 | LYST | chr1 | - | 235709090 | 235709308 | 235702746 | 235702977 | 235712056 | 235712197 | 0 | 3 | 10 | 0 | 0 | 0 | 0 | 4 | 1 | 0 | 1 | 0 | 98 | 49 | 0.046507952 | 0.170760496 | NA | 1 | 1 | NA | 0.667 | 1 | 0.166 |
| A5SS | 2803 | TYROBP | chr19 | - | 35908167 | 35908309 | 35908174 | 35908309 | 35907445 | 35907580 | 0 | 0 | 5 | 0 | 3 | 1 | 0 | 1 | 2 | 0 | 0 | 0 | 56 | 49 | 0.046582587 | 0.170414363 | NA | 0 | 0.814 | NA | 1 | 1 | -0.593 |
| SE | 5273 | NIPBL | chr5 | + | 37036378 | 37036487 | 37027358 | 37027412 | 37038601 | 37038738 | 0 | 1 | 11 | 0 | 0 | 0 | 2 | 13 | 0 | 0 | 1 | 0 | 98 | 49 | 0.046638092 | 0.170760496 | NA | 1 | 1 | 1 | 0.867 | NA | 0.067 |
| SE | 9385 | CLTC | chr17 | + | 59682270 | 59682428 | 59681646 | 59681839 | 59682628 | 59682793 | 0 | 1 | 12 | 0 | 0 | 0 | 0 | 13 | 0 | 0 | 1 | 0 | 98 | 49 | 0.046638092 | 0.170760496 | NA | 1 | 1 | NA | 0.867 | NA | 0.133 |
| SE | 4948 | ARAP1 | chr11 | - | 72712195 | 72712339 | 72711429 | 72711499 | 72712437 | 72712568 | 0 | 7 | 9 | 0 | 0 | 0 | 0 | 6 | 3 | 0 | 1 | 0 | 98 | 49 | 0.047039397 | 0.170760496 | NA | 1 | 1 | NA | 0.75 | 1 | 0.125 |
| SE | 9916 | CASP8 | chr2 | + | 201274888 | 201274953 | 201272897 | 201272942 | 201276826 | 201276968 | 0 | 1 | 6 | 0 | 0 | 2 | 0 | 14 | 0 | 0 | 1 | 0 | 98 | 49 | 0.047441935 | 0.170760496 | NA | 1 | 0.6 | NA | 0.875 | NA | -0.075 |
| SE | 11995 | STAT3 | chr17 | - | 42348388 | 42348517 | 42346568 | 42346713 | 42388278 | 42388495 | 0 | 1 | 28 | 0 | 0 | 0 | 0 | 14 | 0 | 0 | 1 | 0 | 98 | 49 | 0.047441935 | 0.170760496 | NA | 1 | 1 | NA | 0.875 | NA | 0.125 |
| SE | 9286 | VIM | chr10 | + | 17236293 | 17236379 | 17235168 | 17235389 | 17237229 | 17237593 | 0 | 7 | 135 | 0 | 0 | 0 | 15 | 37 | 1 | 0 | 1 | 0 | 98 | 49 | 0.0475366 | 0.170760496 | NA | 1 | 1 | 1 | 0.949 | 1 | 0.017 |
| SE | 510 | ST3GAL1 | chr8 | - | 133475718 | 133476600 | 133465893 | 133466090 | 133499134 | 133499189 | 0 | 1 | 1 | 0 | 0 | 3 | 0 | 3 | 1 | 0 | 0 | 0 | 98 | 49 | 0.047617 | 0.170760496 | NA | 1 | 0.143 | NA | 1 | 1 | -0.428 |
| SE | 12755 | ADGRE5 | chr19 | + | 14396341 | 14396473 | 14390923 | 14391079 | 14397657 | 14397804 | 0 | 0 | 4 | 0 | 1 | 11 | 5 | 1 | 0 | 0 | 6 | 0 | 98 | 49 | 0.047680582 | 0.170760496 | NA | 0 | 0.154 | 1 | 0.077 | NA | -0.461 |
| SE | 5193 | PICALM | chr11 | - | 85978069 | 85978093 | 85976622 | 85976682 | 85981128 | 85981228 | 0 | 0 | 12 | 0 | 2 | 14 | 0 | 1 | 0 | 0 | 14 | 0 | 73 | 49 | 0.047713059 | 0.170760496 | NA | 0 | 0.365 | NA | 0.046 | NA | 0.137 |
| SE | 12573 | R3HDM4 | chr19 | - | 899600 | 899686 | 899439 | 899495 | 900060 | 900146 | 0 | 6 | 13 | 0 | 0 | 0 | 0 | 32 | 2 | 0 | 1 | 0 | 98 | 49 | 0.047912713 | 0.170760496 | NA | 1 | 1 | NA | 0.941 | 1 | 0.03 |
| SE | 6173 | ABCA1 | chr9 | - | 104804625 | 104804720 | 104803283 | 104803316 | 104806240 | 104806430 | 0 | 4 | 7 | 0 | 0 | 0 | 0 | 25 | 1 | 0 | 1 | 0 | 98 | 49 | 0.048012831 | 0.170760496 | NA | 1 | 1 | NA | 0.926 | 1 | 0.037 |
| SE | 11306 | SP100 | chr2 | + | 230461261 | 230461414 | 230450171 | 230450255 | 230462434 | 230462518 | 0 | 3 | 35 | 0 | 0 | 0 | 8 | 15 | 5 | 0 | 2 | 0 | 98 | 49 | 0.048106877 | 0.170760496 | NA | 1 | 1 | 1 | 0.789 | 1 | 0.07 |
| A3SS | 1145 | FBXL5 | chr4 | - | 15644498 | 15644708 | 15644492 | 15644657 | 15655203 | 15655412 | 0 | 1 | 40 | 0 | 0 | 0 | 0 | 15 | 0 | 0 | 1 | 0 | 98 | 49 | 0.048230843 | 0.197717869 | NA | 1 | 1 | NA | 0.882 | NA | 0.118 |
| SE | 8157 | KMT2E | chr7 | + | 105078845 | 105078963 | 105077302 | 105077433 | 105081687 | 105081797 | 0 | 1 | 12 | 0 | 0 | 0 | 0 | 15 | 0 | 0 | 1 | 0 | 98 | 49 | 0.048230843 | 0.170760496 | NA | 1 | 1 | NA | 0.882 | NA | 0.118 |
| SE | 10588 | RNF213 | chr17 | + | 80363608 | 80363790 | 80363101 | 80363314 | 80364432 | 80364553 | 0 | 15 | 83 | 0 | 0 | 0 | 0 | 31 | 8 | 0 | 1 | 0 | 98 | 49 | 0.048276428 | 0.170760496 | NA | 1 | 1 | NA | 0.939 | 1 | 0.03 |
| SE | 12401 | CSF2RB | chr22 | + | 36936548 | 36936652 | 36935629 | 36935687 | 36937376 | 36940437 | 0 | 7 | 34 | 0 | 0 | 0 | 4 | 32 | 2 | 0 | 1 | 0 | 98 | 49 | 0.048373702 | 0.170760496 | NA | 1 | 1 | 1 | 0.941 | 1 | 0.02 |
| SE | 7883 | HIF1A | chr14 | + | 61740504 | 61740627 | 61738086 | 61738373 | 61740754 | 61741188 | 0 | 0 | 14 | 0 | 1 | 0 | 0 | 18 | 1 | 0 | 0 | 0 | 98 | 49 | 0.048559241 | 0.170760496 | NA | 0 | 1 | NA | 1 | 1 | -0.5 |
| SE | 4361 | LRRK2 | chr12 | + | 40293544 | 40293663 | 40287350 | 40287539 | 40294844 | 40294914 | 0 | 2 | 74 | 0 | 0 | 0 | 0 | 33 | 0 | 0 | 1 | 0 | 98 | 49 | 0.048704034 | 0.170760496 | NA | 1 | 1 | NA | 0.943 | NA | 0.057 |
| SE | 11375 | ARHGEF1 | chr19 | + | 41894625 | 41894661 | 41894450 | 41894547 | 41895348 | 41895486 | 0 | 5 | 22 | 0 | 0 | 0 | 0 | 16 | 2 | 0 | 1 | 0 | 85 | 49 | 0.048992317 | 0.170760496 | NA | 1 | 1 | NA | 0.902 | 1 | 0.049 |
| A3SS | 555 | RICTOR | chr5 | - | 38945490 | 38945724 | 38945490 | 38945610 | 38946467 | 38946552 | 0 | 1 | 15 | 0 | 0 | 1 | 0 | 16 | 0 | 0 | 1 | 0 | 98 | 49 | 0.048997106 | 0.197717869 | NA | 1 | 0.882 | NA | 0.889 | NA | 0.052 |
| SE | 9006 | LYST | chr1 | - | 235728075 | 235728131 | 235724027 | 235724180 | 235729595 | 235729657 | 0 | 1 | 35 | 0 | 0 | 0 | 2 | 16 | 0 | 0 | 1 | 0 | 98 | 49 | 0.048997106 | 0.170760496 | NA | 1 | 1 | 1 | 0.889 | NA | 0.055 |
| SE | 6618 | AMPD2 | chr1 | + | 109629326 | 109629490 | 109629108 | 109629235 | 109629795 | 109629916 | 0 | 2 | 5 | 0 | 0 | 0 | 0 | 6 | 1 | 0 | 1 | 0 | 98 | 49 | 0.049093709 | 0.170760496 | NA | 1 | 1 | NA | 0.75 | 1 | 0.125 |
| SE | 6896 | ITGB2 | chr21 | - | 44889275 | 44889495 | 44888692 | 44888895 | 44889977 | 44890222 | 0 | 4 | 78 | 0 | 0 | 0 | 3 | 11 | 2 | 0 | 1 | 0 | 98 | 49 | 0.049219655 | 0.170760496 | NA | 1 | 1 | 1 | 0.846 | 1 | 0.051 |
| SE | 11046 | HLA-B | chr6 | - | 31355106 | 31355223 | 31354478 | 31354526 | 31355316 | 31355592 | 0 | 40 | 238 | 0 | 0 | 0 | 10 | 193 | 14 | 0 | 1 | 0 | 98 | 49 | 0.049293349 | 0.170760496 | NA | 1 | 1 | 1 | 0.99 | 1 | 0.003 |
| SE | 9374 | FMNL1 | chr17 | + | 45233647 | 45233731 | 45233223 | 45233297 | 45234071 | 45234200 | 0 | 4 | 23 | 0 | 0 | 0 | 0 | 21 | 1 | 0 | 1 | 0 | 98 | 49 | 0.049821364 | 0.170760496 | NA | 1 | 1 | NA | 0.913 | 1 | 0.043 |
| SE | 7884 | HIF1A | chr14 | + | 61740754 | 61741188 | 61738086 | 61738373 | 61744704 | 61744813 | 0 | 1 | 13 | 0 | 0 | 3 | 0 | 34 | 1 | 0 | 0 | 0 | 98 | 49 | 0.049963886 | 0.170760496 | NA | 1 | 0.684 | NA | 1 | 1 | -0.158 |
| SE | 9513 | SIPA1 | chr11 | + | 65644954 | 65645129 | 65642462 | 65642639 | 65646220 | 65646378 | 0 | 1 | 11 | 0 | 2 | 0 | 0 | 7 | 7 | 0 | 0 | 0 | 98 | 49 | 0.050052195 | 0.170760496 | NA | 0.2 | 1 | NA | 1 | 1 | -0.4 |
| SE | 6178 | ABCA1 | chr9 | - | 104828915 | 104829138 | 104826947 | 104827169 | 104830924 | 104831101 | 0 | 4 | 14 | 0 | 0 | 0 | 0 | 25 | 1 | 0 | 1 | 0 | 98 | 49 | 0.050181472 | 0.170760496 | NA | 1 | 1 | NA | 0.926 | 1 | 0.037 |
| SE | 1146 | IGF2R | chr6 | + | 160102518 | 160102671 | 160096438 | 160096625 | 160103745 | 160103815 | 0 | 6 | 13 | 0 | 0 | 0 | 0 | 14 | 7 | 0 | 2 | 0 | 98 | 49 | 0.050333626 | 0.170760496 | NA | 1 | 1 | NA | 0.778 | 1 | 0.111 |
| SE | 10067 | ACTN1 | chr14 | - | 68884774 | 68884883 | 68884167 | 68884308 | 68885424 | 68885575 | 0 | 1 | 15 | 0 | 0 | 0 | 0 | 18 | 0 | 0 | 1 | 0 | 98 | 49 | 0.050460196 | 0.170760496 | NA | 1 | 1 | NA | 0.9 | NA | 0.1 |
| SE | 4427 | PREX1 | chr20 | - | 48734545 | 48734650 | 48726289 | 48726391 | 48745024 | 48745147 | 0 | 12 | 55 | 0 | 0 | 0 | 3 | 75 | 3 | 0 | 1 | 0 | 98 | 49 | 0.050469046 | 0.170760496 | NA | 1 | 1 | 1 | 0.974 | 1 | 0.009 |
| SE | 12942 | SMCHD1 | chr18 | + | 2703691 | 2703886 | 2700734 | 2700918 | 2705693 | 2705807 | 1 | 4 | 139 | 0 | 0 | 0 | 0 | 77 | 0 | 0 | 1 | 0 | 98 | 49 | 0.05052881 | 0.170760496 | 1 | 1 | 1 | NA | 0.975 | NA | 0.025 |
| SE | 7030 | ARPC2 | chr2 | + | 218217462 | 218217544 | 218217093 | 218217254 | 218225919 | 218225954 | 0 | 4 | 9 | 0 | 0 | 0 | 0 | 24 | 2 | 0 | 1 | 0 | 98 | 49 | 0.050622824 | 0.170760496 | NA | 1 | 1 | NA | 0.923 | 1 | 0.038 |
| SE | 12325 | MAPKAPK2 | chr1 | + | 206731838 | 206731919 | 206731639 | 206731725 | 206732574 | 206734285 | 0 | 1 | 14 | 0 | 2 | 0 | 5 | 16 | 3 | 0 | 0 | 0 | 98 | 49 | 0.050687516 | 0.170760496 | NA | 0.2 | 1 | 1 | 1 | 1 | -0.4 |
| SE | 11256 | SRSF7 | chr2 | - | 38746143 | 38746179 | 38743598 | 38745187 | 38746693 | 38746747 | 0 | 2 | 5 | 0 | 0 | 0 | 0 | 7 | 1 | 0 | 1 | 0 | 85 | 49 | 0.05078876 | 0.170760496 | NA | 1 | 1 | NA | 0.801 | 1 | 0.099 |
| SE | 5864 | NFKBID | chr19 | - | 35896912 | 35897058 | 35896725 | 35896831 | 35897650 | 35897856 | 0 | 6 | 7 | 0 | 0 | 0 | 0 | 4 | 7 | 0 | 0 | 1 | 98 | 49 | 0.050792748 | 0.170760496 | NA | 1 | 1 | NA | 1 | 0.778 | 0.111 |
| SE | 9455 | PFKFB3 | chr10 | + | 6216124 | 6216191 | 6215220 | 6215317 | 6216705 | 6216780 | 0 | 19 | 68 | 0 | 1 | 1 | 1 | 72 | 24 | 0 | 0 | 0 | 98 | 49 | 0.050850888 | 0.170760496 | NA | 0.905 | 0.971 | 1 | 1 | 1 | -0.062 |
| SE | 8552 | EHBP1L1 | chr11 | + | 65581538 | 65583765 | 65581210 | 65581373 | 65584240 | 65584398 | 0 | 2 | 12 | 0 | 0 | 0 | 0 | 5 | 1 | 0 | 1 | 0 | 98 | 49 | 0.051092621 | 0.170760496 | NA | 1 | 1 | NA | 0.714 | 1 | 0.143 |
| SE | 5410 | TAGAP | chr6 | - | 159043588 | 159043655 | 159042077 | 159042244 | 159043977 | 159044031 | 0 | 1 | 14 | 0 | 0 | 0 | 0 | 19 | 0 | 0 | 1 | 0 | 98 | 49 | 0.051175801 | 0.170760496 | NA | 1 | 1 | NA | 0.905 | NA | 0.095 |
| RI | 1060 | SEC14L1 | chr17 | + | 77213917 | 77217099 | 77213917 | 77214019 | 77216548 | 77217099 | 0 | 13 | 22 | 0 | 0 | 0 | 6 | 16 | 9 | 0 | 1 | 0 | 98 | 49 | 0.051358786 | 0.297702651 | NA | 1 | 1 | 1 | 0.889 | 1 | 0.037 |
| SE | 4670 | TNFRSF1B | chr1 | + | 12201966 | 12202171 | 12194583 | 12194618 | 12206739 | 12209220 | 0 | 5 | 27 | 0 | 0 | 0 | 0 | 27 | 2 | 0 | 1 | 0 | 98 | 49 | 0.051369383 | 0.170760496 | NA | 1 | 1 | NA | 0.931 | 1 | 0.034 |
| SE | 4501 | PTPN6 | chr12 | + | 6951608 | 6951731 | 6951270 | 6951520 | 6951982 | 6952177 | 0 | 5 | 25 | 0 | 0 | 0 | 0 | 8 | 3 | 0 | 1 | 0 | 98 | 49 | 0.051433279 | 0.170760496 | NA | 1 | 1 | NA | 0.8 | 1 | 0.1 |
| SE | 7417 | VAV1 | chr19 | + | 6836984 | 6837050 | 6836431 | 6836568 | 6843134 | 6843166 | 0 | 1 | 12 | 0 | 0 | 0 | 3 | 20 | 0 | 0 | 1 | 0 | 98 | 49 | 0.051859277 | 0.170760496 | NA | 1 | 1 | 1 | 0.909 | NA | 0.045 |
| SE | 12684 | HNRNPK | chr9 | - | 83973901 | 83973973 | 83973285 | 83973399 | 83974516 | 83974589 | 0 | 1 | 53 | 0 | 0 | 3 | 1 | 20 | 0 | 0 | 1 | 0 | 98 | 49 | 0.051859277 | 0.170760496 | NA | 1 | 0.898 | 1 | 0.909 | NA | -0.005 |
| SE | 3134 | IFNGR2 | chr21 | + | 33414887 | 33415020 | 33402895 | 33403616 | 33421479 | 33421685 | 9 | 14 | 31 | 0 | 0 | 0 | 0 | 15 | 47 | 0 | 0 | 1 | 98 | 49 | 0.052010728 | 0.170760496 | 1 | 1 | 1 | NA | 1 | 0.959 | 0.02 |
| SE | 7031 | ARPC2 | chr2 | + | 218239390 | 218239484 | 218238663 | 218238850 | 218245419 | 218245546 | 0 | 0 | 56 | 0 | 0 | 1 | 1 | 33 | 2 | 0 | 1 | 2 | 98 | 49 | 0.052035219 | 0.170760496 | NA | NA | 0.966 | 1 | 0.943 | 0.333 | 0.207 |
| SE | 1737 | ARHGAP26 | chr5 | + | 143014079 | 143014116 | 142932046 | 142932125 | 143037195 | 143037261 | 0 | 1 | 47 | 0 | 0 | 0 | 0 | 23 | 4 | 0 | 2 | 0 | 86 | 49 | 0.052066191 | 0.170760496 | NA | 1 | 1 | NA | 0.868 | 1 | 0.066 |
| SE | 13867 | ACSL1 | chr4 | - | 184776483 | 184776662 | 184773842 | 184773875 | 184776883 | 184776983 | 0 | 3 | 33 | 0 | 0 | 1 | 1 | 36 | 1 | 0 | 6 | 0 | 98 | 49 | 0.052282739 | 0.170760496 | NA | 1 | 0.943 | 1 | 0.75 | 1 | 0.055 |
| SE | 4669 | TNFRSF1B | chr1 | + | 12188795 | 12188895 | 12167002 | 12167169 | 12190956 | 12191085 | 0 | 7 | 37 | 0 | 2 | 3 | 5 | 25 | 32 | 0 | 0 | 1 | 98 | 49 | 0.052384975 | 0.170760496 | NA | 0.636 | 0.86 | 1 | 1 | 0.941 | -0.232 |
| SE | 9376 | FMNL1 | chr17 | + | 45245252 | 45245416 | 45244978 | 45245108 | 45245877 | 45245973 | 0 | 3 | 24 | 0 | 0 | 0 | 0 | 14 | 0 | 0 | 1 | 2 | 98 | 49 | 0.052499618 | 0.170760496 | NA | 1 | 1 | NA | 0.875 | 0 | 0.562 |
| SE | 10927 | DEDD2 | chr19 | - | 42201988 | 42202072 | 42198592 | 42199829 | 42209699 | 42209840 | 0 | 1 | 2 | 0 | 3 | 5 | 0 | 1 | 0 | 0 | 16 | 3 | 98 | 49 | 0.052585138 | 0.170760496 | NA | 0.143 | 0.167 | NA | 0.03 | 0 | 0.14 |
| SE | 1854 | DBNL | chr7 | + | 44058431 | 44058480 | 44058128 | 44058280 | 44058901 | 44058983 | 0 | 4 | 5 | 0 | 0 | 0 | 0 | 3 | 2 | 0 | 1 | 0 | 98 | 49 | 0.052858149 | 0.170760496 | NA | 1 | 1 | NA | 0.6 | 1 | 0.2 |
| RI | 576 | SRSF2 | chr17 | - | 76734114 | 76736464 | 76734114 | 76735158 | 76736153 | 76736464 | 0 | 7 | 12 | 0 | 2 | 3 | 0 | 16 | 1 | 0 | 12 | 1 | 98 | 49 | 0.052909456 | 0.297702651 | NA | 0.636 | 0.667 | NA | 0.4 | 0.333 | 0.285 |
| A3SS | 459 | BTN3A1 | chr6 | + | 26407670 | 26407952 | 26407826 | 26407952 | 26405908 | 26406256 | 0 | 2 | 3 | 0 | 0 | 0 | 2 | 2 | 1 | 0 | 1 | 0 | 98 | 49 | 0.05291102 | 0.197717869 | NA | 1 | 1 | 1 | 0.5 | 1 | 0.167 |
| SE | 8417 | GRAMD1A | chr19 | + | 35021950 | 35022038 | 35021690 | 35021864 | 35023235 | 35023343 | 0 | 4 | 17 | 0 | 0 | 0 | 0 | 4 | 12 | 0 | 0 | 1 | 98 | 49 | 0.052978196 | 0.170760496 | NA | 1 | 1 | NA | 1 | 0.857 | 0.072 |
| SE | 7910 | ATP6V0B | chr1 | + | 43977025 | 43977216 | 43976772 | 43976824 | 43977980 | 43978300 | 0 | 4 | 22 | 0 | 0 | 0 | 1 | 11 | 3 | 0 | 1 | 0 | 98 | 49 | 0.053042143 | 0.170760496 | NA | 1 | 1 | 1 | 0.846 | 1 | 0.051 |
| SE | 10586 | RNF213 | chr17 | + | 80360060 | 80360206 | 80358287 | 80358479 | 80361733 | 80361888 | 0 | 1 | 46 | 0 | 0 | 0 | 2 | 41 | 4 | 0 | 2 | 0 | 98 | 49 | 0.053139356 | 0.170760496 | NA | 1 | 1 | 1 | 0.911 | 1 | 0.03 |
| SE | 6938 | CD55 | chr1 | + | 207331107 | 207331296 | 207326751 | 207326837 | 207336692 | 207336818 | 0 | 2 | 28 | 0 | 0 | 0 | 0 | 46 | 0 | 0 | 1 | 0 | 98 | 49 | 0.053445903 | 0.170760496 | NA | 1 | 1 | NA | 0.958 | NA | 0.042 |
| SE | 7392 | IRAK3 | chr12 | + | 66211445 | 66211597 | 66210146 | 66210201 | 66217170 | 66217235 | 0 | 2 | 30 | 0 | 0 | 0 | 2 | 10 | 1 | 0 | 1 | 0 | 98 | 49 | 0.053461329 | 0.170760496 | NA | 1 | 1 | 1 | 0.833 | 1 | 0.056 |
| SE | 9740 | PXN | chr12 | - | 120222548 | 120222750 | 120221622 | 120221758 | 120222862 | 120222999 | 0 | 3 | 23 | 0 | 0 | 0 | 0 | 6 | 2 | 0 | 1 | 0 | 98 | 49 | 0.053502796 | 0.170760496 | NA | 1 | 1 | NA | 0.75 | 1 | 0.125 |
| SE | 4088 | TNFRSF1A | chr12 | - | 6334090 | 6334244 | 6333736 | 6333865 | 6341775 | 6342117 | 0 | 2 | 17 | 0 | 3 | 1 | 0 | 5 | 3 | 0 | 0 | 0 | 98 | 49 | 0.053505644 | 0.170760496 | NA | 0.25 | 0.895 | NA | 1 | 1 | -0.427 |
| SE | 13863 | ACSL1 | chr4 | - | 184765890 | 184765986 | 184764852 | 184764925 | 184766621 | 184766756 | 0 | 2 | 59 | 0 | 0 | 3 | 4 | 48 | 12 | 0 | 0 | 0 | 98 | 49 | 0.053534738 | 0.170760496 | NA | 1 | 0.908 | 1 | 1 | 1 | -0.046 |
| SE | 1144 | IGF2R | chr6 | + | 160084994 | 160085131 | 160083949 | 160084184 | 160088032 | 160088147 | 0 | 4 | 17 | 0 | 0 | 0 | 1 | 20 | 3 | 0 | 1 | 0 | 98 | 49 | 0.05376828 | 0.170760496 | NA | 1 | 1 | 1 | 0.909 | 1 | 0.03 |
| SE | 8590 | LTB | chr6 | - | 31581813 | 31581859 | 31581558 | 31581630 | 31582255 | 31582425 | 0 | 0 | 5 | 0 | 3 | 0 | 0 | 11 | 0 | 0 | 0 | 0 | 95 | 49 | 0.054362012 | 0.170760496 | NA | 0 | 1 | NA | 1 | NA | -0.5 |
| SE | 4241 | ITGA5 | chr12 | - | 54403624 | 54403779 | 54403186 | 54403324 | 54403910 | 54403966 | 0 | 0 | 12 | 0 | 0 | 0 | 0 | 7 | 0 | 0 | 0 | 2 | 98 | 49 | 0.054536123 | 0.170760496 | NA | NA | 1 | NA | 1 | 0 | 0.5 |
| A3SS | 690 | CNBP | chr3 | - | 129171078 | 129171283 | 129171078 | 129171277 | 129171445 | 129171538 | 0 | 0 | 0 | 0 | 1 | 4 | 0 | 1 | 0 | 0 | 11 | 0 | 55 | 49 | 0.054675117 | 0.197717869 | NA | 0 | 0 | NA | 0.075 | NA | -0.075 |
| SE | 5196 | PICALM | chr11 | - | 86001034 | 86001158 | 86000642 | 86000779 | 86003365 | 86003451 | 0 | 6 | 50 | 0 | 0 | 0 | 2 | 55 | 1 | 0 | 1 | 0 | 98 | 49 | 0.054703796 | 0.170760496 | NA | 1 | 1 | 1 | 0.965 | 1 | 0.012 |
| SE | 11996 | STAT3 | chr17 | - | 42348388 | 42348539 | 42346568 | 42346713 | 42388278 | 42388387 | 0 | 2 | 29 | 0 | 0 | 0 | 0 | 12 | 1 | 0 | 1 | 0 | 98 | 49 | 0.054806328 | 0.170760496 | NA | 1 | 1 | NA | 0.857 | 1 | 0.072 |
| SE | 7497 | IFI44 | chr1 | + | 78660553 | 78660654 | 78659311 | 78659483 | 78663764 | 78664078 | 0 | 0 | 31 | 0 | 1 | 1 | 0 | 20 | 1 | 0 | 0 | 0 | 98 | 49 | 0.054840236 | 0.170760496 | NA | 0 | 0.939 | NA | 1 | 1 | -0.53 |
| SE | 7726 | FGD3 | chr9 | + | 93002924 | 93003014 | 92976207 | 92976709 | 93004000 | 93004137 | 0 | 10 | 22 | 0 | 0 | 0 | 0 | 16 | 9 | 0 | 1 | 0 | 98 | 49 | 0.054857277 | 0.170760496 | NA | 1 | 1 | NA | 0.889 | 1 | 0.055 |
| SE | 4940 | ARAP1 | chr11 | - | 72687438 | 72687502 | 72686041 | 72686191 | 72687687 | 72687738 | 0 | 3 | 10 | 0 | 0 | 0 | 0 | 4 | 2 | 0 | 1 | 0 | 98 | 49 | 0.055117945 | 0.170760496 | NA | 1 | 1 | NA | 0.667 | 1 | 0.166 |
| SE | 12992 | PLAUR | chr19 | - | 43656478 | 43656640 | 43655438 | 43655573 | 43665315 | 43665459 | 0 | 14 | 45 | 0 | 0 | 0 | 0 | 84 | 14 | 0 | 1 | 0 | 98 | 49 | 0.055245993 | 0.170760496 | NA | 1 | 1 | NA | 0.977 | 1 | 0.012 |
| SE | 11304 | SP100 | chr2 | + | 230449560 | 230449710 | 230449087 | 230449150 | 230450171 | 230450255 | 0 | 7 | 33 | 0 | 0 | 0 | 0 | 28 | 7 | 0 | 1 | 0 | 98 | 49 | 0.055357429 | 0.170760496 | NA | 1 | 1 | NA | 0.933 | 1 | 0.033 |
| SE | 3739 | BRD2 | chr6 | + | 32976561 | 32976936 | 32976249 | 32976464 | 32977441 | 32977570 | 0 | 0 | 8 | 0 | 0 | 0 | 0 | 8 | 0 | 0 | 0 | 2 | 98 | 49 | 0.055397548 | 0.170760496 | NA | NA | 1 | NA | 1 | 0 | 0.5 |
| A3SS | 1302 | GMPR2 | chr14 | + | 24238245 | 24239241 | 24238588 | 24239241 | 24237519 | 24237562 | 1 | 0 | 3 | 0 | 2 | 0 | 0 | 10 | 0 | 0 | 0 | 0 | 98 | 49 | 0.055620739 | 0.197717869 | 1 | 0 | 1 | NA | 1 | NA | -0.333 |
| SE | 2096 | WDFY3 | chr4 | - | 84678918 | 84679242 | 84678167 | 84678279 | 84682373 | 84682470 | 0 | 2 | 8 | 0 | 0 | 0 | 0 | 2 | 1 | 0 | 1 | 0 | 98 | 49 | 0.055836208 | 0.170760496 | NA | 1 | 1 | NA | 0.5 | 1 | 0.25 |
| SE | 6103 | GLUL | chr1 | - | 182385759 | 182385887 | 182385356 | 182385556 | 182386255 | 182386402 | 0 | 16 | 127 | 0 | 0 | 0 | 6 | 106 | 11 | 0 | 1 | 0 | 98 | 49 | 0.055843452 | 0.170760496 | NA | 1 | 1 | 1 | 0.981 | 1 | 0.006 |
| SE | 10624 | RRP12 | chr10 | - | 97366107 | 97366233 | 97363853 | 97363903 | 97366445 | 97366621 | 0 | 0 | 2 | 0 | 0 | 0 | 0 | 1 | 0 | 0 | 0 | 5 | 98 | 49 | 0.055847068 | 0.170760496 | NA | NA | 1 | NA | 1 | 0 | 0.5 |
| SE | 2782 | ATG7 | chr3 | + | 11299361 | 11299416 | 11298665 | 11298855 | 11306942 | 11307060 | 0 | 0 | 7 | 0 | 1 | 0 | 0 | 9 | 1 | 0 | 0 | 0 | 98 | 49 | 0.055867249 | 0.170760496 | NA | 0 | 1 | NA | 1 | 1 | -0.5 |
| SE | 11397 | ADGRG3 | chr16 | + | 57684389 | 57684483 | 57680504 | 57680617 | 57685642 | 57685926 | 0 | 0 | 7 | 0 | 2 | 0 | 0 | 9 | 0 | 0 | 0 | 0 | 98 | 49 | 0.055954714 | 0.170760496 | NA | 0 | 1 | NA | 1 | NA | -0.5 |
| SE | 2283 | IL1RAP | chr3 | + | 190550341 | 190550505 | 190514050 | 190514219 | 190556129 | 190556216 | 0 | 0 | 0 | 0 | 3 | 3 | 0 | 3 | 0 | 0 | 4 | 2 | 98 | 49 | 0.05597701 | 0.170760496 | NA | 0 | 0 | NA | 0.273 | 0 | -0.137 |
| SE | 1506 | HLA-E | chr6 | + | 30491536 | 30491653 | 30491136 | 30491412 | 30492540 | 30492583 | 0 | 12 | 185 | 0 | 0 | 0 | 11 | 70 | 10 | 0 | 1 | 0 | 98 | 49 | 0.056110759 | 0.170760496 | NA | 1 | 1 | 1 | 0.972 | 1 | 0.009 |
| SE | 3382 | HCK | chr20 | + | 32084390 | 32084543 | 32083893 | 32084043 | 32086627 | 32086807 | 0 | 3 | 55 | 0 | 1 | 5 | 5 | 17 | 4 | 0 | 0 | 0 | 98 | 49 | 0.056127965 | 0.170760496 | NA | 0.6 | 0.846 | 1 | 1 | 1 | -0.277 |
| SE | 3057 | COL18A1 | chr21 | + | 45505133 | 45505278 | 45504521 | 45504556 | 45505357 | 45505431 | 0 | 0 | 6 | 0 | 0 | 0 | 0 | 2 | 0 | 0 | 0 | 2 | 98 | 49 | 0.056129395 | 0.170760496 | NA | NA | 1 | NA | 1 | 0 | 0.5 |
| SE | 10626 | RRP12 | chr10 | - | 97379292 | 97379414 | 97373829 | 97373894 | 97379627 | 97379770 | 0 | 2 | 7 | 0 | 0 | 0 | 0 | 9 | 2 | 0 | 1 | 0 | 98 | 49 | 0.056251461 | 0.170760496 | NA | 1 | 1 | NA | 0.818 | 1 | 0.091 |
| SE | 1507 | HLA-E | chr6 | + | 30492403 | 30492436 | 30491136 | 30491412 | 30492540 | 30492583 | 0 | 7 | 64 | 0 | 0 | 0 | 0 | 48 | 3 | 0 | 1 | 0 | 82 | 49 | 0.05627258 | 0.170760496 | NA | 1 | 1 | NA | 0.966 | 1 | 0.017 |
| SE | 9587 | ZNF467 | chr7 | - | 149770439 | 149770556 | 149769089 | 149769200 | 149770998 | 149771074 | 0 | 0 | 1 | 0 | 1 | 0 | 0 | 2 | 3 | 0 | 0 | 0 | 98 | 49 | 0.056299876 | 0.170760496 | NA | 0 | 1 | NA | 1 | 1 | -0.5 |
| SE | 3981 | NCOR2 | chr12 | - | 124374412 | 124374463 | 124372021 | 124372610 | 124378236 | 124378384 | 0 | 0 | 2 | 0 | 0 | 0 | 0 | 0 | 0 | 0 | 0 | 4 | 98 | 49 | 0.056348041 | 0.170760496 | NA | NA | 1 | NA | NA | 0 | 1 |
| SE | 6199 | RPRD2 | chr1 | + | 150441880 | 150441958 | 150440922 | 150441023 | 150443230 | 150443283 | 0 | 2 | 0 | 0 | 0 | 0 | 0 | 0 | 0 | 0 | 4 | 0 | 98 | 49 | 0.056348041 | 0.170760496 | NA | 1 | NA | NA | 0 | NA | 1 |
| SE | 636 | RB1CC1 | chr8 | - | 52660595 | 52660657 | 52658872 | 52658976 | 52660925 | 52661007 | 0 | 0 | 3 | 0 | 0 | 0 | 0 | 4 | 0 | 0 | 0 | 3 | 98 | 49 | 0.056460428 | 0.170760496 | NA | NA | 1 | NA | 1 | 0 | 0.5 |
| SE | 12437 | SPAG9 | chr17 | - | 51014231 | 51014353 | 51007268 | 51007326 | 51020158 | 51020258 | 0 | 0 | 8 | 0 | 1 | 0 | 0 | 14 | 0 | 0 | 0 | 0 | 98 | 49 | 0.056479947 | 0.170760496 | NA | 0 | 1 | NA | 1 | NA | -0.5 |
| SE | 6672 | FGR | chr1 | - | 27621558 | 27621657 | 27615688 | 27615844 | 27623041 | 27623144 | 0 | 3 | 15 | 0 | 0 | 0 | 0 | 10 | 3 | 0 | 1 | 0 | 98 | 49 | 0.0565412 | 0.170760496 | NA | 1 | 1 | NA | 0.833 | 1 | 0.084 |
| SE | 3944 | ZDHHC20 | chr13 | - | 21421060 | 21421164 | 21413651 | 21413772 | 21425651 | 21425678 | 0 | 0 | 5 | 0 | 0 | 0 | 0 | 0 | 0 | 0 | 1 | 2 | 98 | 49 | 0.056609827 | 0.170760496 | NA | NA | 1 | NA | 0 | 0 | 1 |
| SE | 10040 | SYNE2 | chr14 | + | 64029894 | 64030059 | 64027483 | 64027793 | 64031015 | 64031357 | 0 | 0 | 5 | 0 | 2 | 0 | 0 | 7 | 0 | 0 | 0 | 0 | 98 | 49 | 0.056662448 | 0.170760496 | NA | 0 | 1 | NA | 1 | NA | -0.5 |
| SE | 3801 | ZCCHC2 | chr18 | + | 62544801 | 62544873 | 62542497 | 62542574 | 62550347 | 62550460 | 0 | 0 | 29 | 0 | 1 | 2 | 3 | 13 | 0 | 0 | 0 | 0 | 98 | 49 | 0.056686045 | 0.170760496 | NA | 0 | 0.879 | 1 | 1 | NA | -0.56 |
| SE | 1817 | CHD1 | chr5 | - | 98876397 | 98876558 | 98875071 | 98875113 | 98879551 | 98879728 | 0 | 1 | 2 | 0 | 0 | 0 | 0 | 8 | 2 | 0 | 9 | 0 | 98 | 49 | 0.056740038 | 0.170760496 | NA | 1 | 1 | NA | 0.308 | 1 | 0.346 |
| SE | 2275 | VNN3 | chr6 | - | 132728740 | 132728979 | 132724605 | 132724979 | 132734252 | 132734383 | 0 | 1 | 2 | 0 | 0 | 0 | 0 | 7 | 1 | 0 | 1 | 0 | 98 | 49 | 0.056900121 | 0.170760496 | NA | 1 | 1 | NA | 0.778 | 1 | 0.111 |
| A5SS | 2964 | USF1 | chr1 | - | 161042554 | 161042670 | 161042585 | 161042670 | 161042115 | 161042217 | 0 | 0 | 9 | 0 | 2 | 2 | 0 | 6 | 0 | 0 | 0 | 0 | 80 | 49 | 0.057135855 | 0.170414363 | NA | 0 | 0.734 | NA | 1 | NA | -0.633 |
| SE | 3101 | MX1 | chr21 | + | 41451166 | 41451243 | 41445999 | 41446141 | 41452620 | 41452869 | 0 | 2 | 24 | 0 | 0 | 0 | 1 | 15 | 1 | 0 | 1 | 0 | 98 | 49 | 0.057325035 | 0.170760496 | NA | 1 | 1 | 1 | 0.882 | 1 | 0.039 |

| | | | | | | | | | | | | | | | | | | | | | | | | | | | | | | | | |
|---|---|---|---|---|---|---|---|---|---|---|---|---|---|---|---|---|---|---|---|---|---|---|---|---|---|---|---|---|---|---|---|---|
| SE | 12164 | USP32 | chr17 | - | 60180544 | 60180637 | 60177329 | 60179428 | 60181323 | 60181748 | 0 | 0 | 12 | 0 | 1 | 0 | 0 | 10 | 0 | 0 | 0 | 0 | 98 | 49 | 0.057462192 | 0.170760496 | NA | 0 | 1 | NA | 1 | NA | -0.5 |
| SE | 6865 | ATG2A | chr11 | - | 64901961 | 64902176 | 64900493 | 64900629 | 64902259 | 64902386 | 0 | 0 | 2 | 0 | 0 | 0 | 0 | 2 | 0 | 0 | 1 | 3 | 98 | 49 | 0.057547789 | 0.170760496 | NA | NA | 1 | NA | 0.5 | 0 | 0.75 |
| SE | 8248 | PRRC2C | chr1 | + | 171513000 | 171513172 | 171485526 | 171485735 | 171514535 | 171514645 | 0 | 1 | 1 | 0 | 0 | 0 | 2 | 2 | 1 | 0 | 1 | 0 | 98 | 49 | 0.057579548 | 0.170760496 | NA | 1 | 1 | 1 | 0.5 | 1 | 0.167 |
| SE | 8660 | DUSP1 | chr5 | - | 172770160 | 172770306 | 172769574 | 172769794 | 172770585 | 172771200 | 4 | 105 | 424 | 0 | 1 | 1 | 27 | 197 | 285 | 0 | 2 | 6 | 98 | 49 | 0.057728535 | 0.170760496 | 1 | 0.981 | 0.995 | 1 | 0.98 | 0.96 | 0.012 |
| SE | 5304 | CD82 | chr11 | + | 44605354 | 44605429 | 44605057 | 44605182 | 44615271 | 44615373 | 0 | 0 | 1 | 0 | 2 | 0 | 0 | 5 | 0 | 0 | 0 | 0 | 98 | 49 | 0.057729 | 0.170760496 | NA | 0 | 1 | NA | 1 | NA | -0.5 |
| SE | 4803 | CASP4 | chr11 | - | 104950924 | 104951098 | 104949542 | 104949777 | 104954746 | 104955001 | 0 | 0 | 17 | 0 | 1 | 2 | 4 | 9 | 0 | 0 | 0 | 0 | 98 | 49 | 0.057774487 | 0.170760496 | NA | 0 | 0.81 | 1 | 1 | NA | -0.595 |
| SE | 5518 | DENND5A | chr11 | - | 9145669 | 9145815 | 9144994 | 9145113 | 9147029 | 9147151 | 0 | 0 | 13 | 0 | 1 | 0 | 2 | 9 | 0 | 0 | 0 | 0 | 98 | 49 | 0.057774487 | 0.170760496 | NA | 0 | 1 | 1 | 1 | NA | -0.5 |
| SE | 5966 | 15-Sep | chr1 | - | 86868052 | 86868102 | 86862444 | 86863605 | 86880661 | 86880725 | 0 | 0 | 12 | 0 | 1 | 0 | 0 | 9 | 0 | 0 | 0 | 0 | 98 | 49 | 0.057774487 | 0.170760496 | NA | 0 | 1 | NA | 1 | NA | -0.5 |
| SE | 6091 | SRRM2 | chr16 | + | 2756333 | 2756606 | 2752328 | 2752846 | 2757471 | 2757579 | 0 | 0 | 7 | 0 | 1 | 0 | 0 | 9 | 0 | 0 | 0 | 0 | 98 | 49 | 0.057774487 | 0.170760496 | NA | 0 | 1 | NA | 1 | NA | -0.5 |
| SE | 11055 | ARFGAP3 | chr22 | - | 42834241 | 42834325 | 42831548 | 42831636 | 42835361 | 42835493 | 0 | 0 | 8 | 0 | 1 | 0 | 0 | 9 | 0 | 0 | 0 | 0 | 98 | 49 | 0.057774487 | 0.170760496 | NA | 0 | 1 | NA | 1 | NA | -0.5 |
| SE | 15 | FLNA | chrX | - | 154352180 | 154352447 | 154351883 | 154352021 | 154352771 | 154352924 | 0 | 1 | 15 | 0 | 0 | 0 | 5 | 8 | 3 | 0 | 2 | 0 | 98 | 49 | 0.058033181 | 0.170760496 | NA | 1 | 1 | 1 | 0.667 | 1 | 0.111 |
| SE | 9987 | TRANK1 | chr3 | - | 36838371 | 36838504 | 36834761 | 36834907 | 36838612 | 36838716 | 0 | 0 | 3 | 0 | 0 | 0 | 0 | 3 | 0 | 0 | 0 | 2 | 98 | 49 | 0.058049942 | 0.170760496 | NA | NA | 1 | NA | 1 | 0 | 0.5 |
| SE | 8136 | ITGAX | chr16 | + | 31355892 | 31355998 | 31355133 | 31355291 | 31356624 | 31356728 | 0 | 17 | 42 | 0 | 0 | 0 | 8 | 52 | 24 | 0 | 1 | 0 | 98 | 49 | 0.058165611 | 0.170760496 | NA | 1 | 1 | 1 | 0.963 | 1 | 0.012 |
| SE | 4886 | DCAF11 | chr14 | + | 24120837 | 24120991 | 24119710 | 24119896 | 24121364 | 24121517 | 0 | 0 | 4 | 0 | 1 | 0 | 0 | 8 | 0 | 0 | 0 | 0 | 98 | 49 | 0.058181256 | 0.170760496 | NA | 0 | 1 | NA | 1 | NA | -0.5 |
| RI | 523 | DDAH2 | chr6 | - | 31728864 | 31730265 | 31728864 | 31729225 | 31730050 | 31730265 | 0 | 1 | 0 | 0 | 0 | 3 | 0 | 0 | 3 | 0 | 0 | 0 | 98 | 49 | 0.058431081 | 0.297702651 | NA | 1 | 0 | NA | NA | 1 | -0.5 |
| SE | 4634 | GMPR2 | chr14 | + | 24238245 | 24238405 | 24237519 | 24237562 | 24238588 | 24239241 | 0 | 0 | 1 | 0 | 2 | 0 | 0 | 4 | 0 | 0 | 0 | 0 | 98 | 49 | 0.058521801 | 0.170760496 | NA | 0 | 1 | NA | 1 | NA | -0.5 |
| SE | 11517 | CMIP | chr16 | + | 81664268 | 81664349 | 81660883 | 81660946 | 81670141 | 81670245 | 0 | 0 | 5 | 0 | 2 | 0 | 0 | 4 | 0 | 0 | 0 | 0 | 98 | 49 | 0.058521801 | 0.170760496 | NA | 0 | 1 | NA | 1 | NA | -0.5 |
| SE | 3630 | USP4 | chr3 | - | 49286097 | 49286325 | 49284848 | 49284919 | 49292509 | 49292598 | 0 | 2 | 22 | 0 | 0 | 0 | 6 | 10 | 2 | 0 | 1 | 0 | 98 | 49 | 0.058749844 | 0.170760496 | NA | 1 | 1 | 1 | 0.833 | 1 | 0.056 |
| SE | 14092 | NLRP6 | chr11 | + | 279833 | 279872 | 279326 | 279607 | 280083 | 281839 | 0 | 1 | 1 | 0 | 0 | 0 | 0 | 0 | 0 | 0 | 0 | 4 | 88 | 49 | 0.058768543 | 0.170760496 | NA | 1 | 1 | NA | NA | 0 | 1 |
| SE | 13371 | CCT7 | chr2 | + | 73243996 | 73244049 | 73243003 | 73243129 | 73244543 | 73244715 | 1 | 0 | 0 | 0 | 1 | 0 | 6 | 1 | 0 | 0 | 0 | 0 | 98 | 49 | 0.058830341 | 0.170760496 | 1 | 0 | NA | 1 | 1 | NA | -0.5 |
| SE | 1853 | DBNL | chr7 | + | 44058428 | 44058480 | 44058128 | 44058280 | 44058901 | 44058983 | 0 | 1 | 3 | 0 | 0 | 0 | 0 | 3 | 1 | 0 | 1 | 0 | 98 | 49 | 0.058903155 | 0.170760496 | NA | 1 | 1 | NA | 0.6 | 1 | 0.2 |
| SE | 3097 | MX1 | chr21 | + | 41437014 | 41437152 | 41435836 | 41436029 | 41439693 | 41439848 | 0 | 4 | 56 | 0 | 0 | 0 | 7 | 31 | 3 | 0 | 1 | 0 | 98 | 49 | 0.05891182 | 0.170760496 | NA | 1 | 1 | 1 | 0.939 | 1 | 0.02 |
| SE | 2611 | KIAA0922 | chr4 | + | 153592474 | 153592584 | 153591052 | 153591194 | 153593798 | 153593871 | 0 | 0 | 7 | 0 | 0 | 0 | 0 | 0 | 0 | 0 | 0 | 1 | 98 | 49 | 0.05894622 | 0.170760496 | NA | NA | 1 | NA | NA | 0 | 1 |
| SE | 6096 | CLN3 | chr16 | - | 28491481 | 28491560 | 28489289 | 28489386 | 28491713 | 28491835 | 0 | 0 | 7 | 0 | 0 | 0 | 0 | 0 | 0 | 0 | 0 | 1 | 98 | 49 | 0.05894622 | 0.170760496 | NA | NA | 1 | NA | NA | 0 | 1 |
| SE | 9258 | MOV10 | chr1 | + | 112694748 | 112694896 | 112694452 | 112694629 | 112695415 | 112695574 | 0 | 5 | 7 | 0 | 0 | 0 | 0 | 0 | 0 | 0 | 0 | 1 | 98 | 49 | 0.05894622 | 0.170760496 | NA | 1 | 1 | NA | NA | 0 | 1 |
| SE | 12991 | PLAUR | chr19 | - | 43655438 | 43655573 | 43652224 | 43652371 | 43656478 | 43656640 | 0 | 8 | 42 | 0 | 0 | 6 | 0 | 48 | 10 | 0 | 1 | 0 | 98 | 49 | 0.059088669 | 0.170760496 | NA | 1 | 0.778 | NA | 0.96 | 1 | -0.091 |
| SE | 13915 | CD300A | chr17 | + | 74473535 | 74473874 | 74466382 | 74466743 | 74474531 | 74474685 | 0 | 0 | 1 | 0 | 1 | 1 | 0 | 6 | 0 | 0 | 0 | 0 | 98 | 49 | 0.05910855 | 0.170760496 | NA | 0 | 0.333 | NA | 1 | NA | -0.834 |
| SE | 4341 | SLC38A2 | chr12 | - | 46370511 | 46370627 | 46367266 | 46367340 | 46370775 | 46370857 | 0 | 2 | 6 | 0 | 0 | 1 | 0 | 0 | 0 | 0 | 2 | 0 | 98 | 49 | 0.05919998 | 0.170760496 | NA | 1 | 0.75 | NA | 0 | NA | 0.875 |
| SE | 5039 | RELT | chr11 | + | 73391143 | 73391223 | 73390754 | 73390921 | 73392210 | 73392468 | 0 | 0 | 2 | 0 | 0 | 0 | 0 | 6 | 0 | 0 | 1 | 2 | 98 | 49 | 0.05919998 | 0.170760496 | NA | NA | 1 | NA | 0.75 | 0 | 0.625 |
| SE | 7302 | PPP6R2 | chr22 | + | 50436987 | 50437068 | 50436366 | 50436452 | 50437505 | 50437603 | 0 | 0 | 2 | 0 | 0 | 0 | 0 | 0 | 0 | 0 | 0 | 2 | 98 | 49 | 0.05919998 | 0.170760496 | NA | NA | 1 | NA | NA | 0 | 1 |
| SE | 12511 | WDR7 | chr18 | + | 56717963 | 56718159 | 56696241 | 56696462 | 56731382 | 56731597 | 0 | 0 | 2 | 0 | 0 | 0 | 0 | 3 | 0 | 0 | 0 | 2 | 98 | 49 | 0.05919998 | 0.170760496 | NA | NA | 1 | NA | 1 | 0 | 0.5 |
| SE | 8184 | EVI2A | chr17 | - | 31320372 | 31320513 | 31316409 | 31319023 | 31321494 | 31321749 | 0 | 0 | 0 | 0 | 0 | 4 | 0 | 0 | 2 | 0 | 0 | 0 | 98 | 49 | 0.059276699 | 0.170760496 | NA | NA | 0 | NA | NA | 1 | -1 |
| SE | 9171 | KLF6 | chr10 | - | 3780105 | 3780229 | 3775995 | 3779590 | 3781640 | 3782214 | 0 | 3 | 14 | 1 | 0 | 0 | 0 | 6 | 3 | 0 | 1 | 0 | 98 | 49 | 0.059302042 | 0.170760496 | 0 | 1 | 1 | NA | 0.75 | 1 | -0.208 |
| SE | 7343 | FAM129A | chr1 | - | 184803584 | 184803692 | 184798078 | 184798190 | 184805945 | 184806056 | 0 | 1 | 38 | 0 | 0 | 0 | 0 | 7 | 1 | 0 | 1 | 0 | 98 | 49 | 0.059353638 | 0.170760496 | NA | 1 | 1 | NA | 0.778 | 1 | 0.111 |
| A5SS | 553 | HMGA1 | chr6 | + | 34240736 | 34240915 | 34240736 | 34240882 | 34242711 | 34242795 | 0 | 1 | 4 | 0 | 0 | 3 | 0 | 5 | 1 | 0 | 0 | 0 | 82 | 49 | 0.059398863 | 0.170414363 | NA | 1 | 0.443 | NA | 1 | 1 | -0.278 |
| A3SS | 414 | DCTN3 | chr9 | - | 34614041 | 34614198 | 34614041 | 34614101 | 34614709 | 34614768 | 1 | 1 | 0 | 0 | 0 | 0 | 0 | 0 | 0 | 0 | 3 | 0 | 98 | 49 | 0.059555994 | 0.197717869 | 1 | 1 | NA | NA | 0 | NA | 1 |
| A5SS | 2673 | RNF31 | chr14 | + | 24148257 | 24148477 | 24148257 | 24148413 | 24148641 | 24148701 | 0 | 1 | 1 | 0 | 0 | 3 | 0 | 0 | 0 | 0 | 3 | 0 | 98 | 49 | 0.059555994 | 0.170414363 | NA | 1 | 0.143 | NA | 0 | NA | 0.572 |
| SE | 6361 | ZSWIM8 | chr10 | + | 73789366 | 73789539 | 73789095 | 73789190 | 73789716 | 73789824 | 0 | 1 | 4 | 0 | 0 | 0 | 0 | 0 | 0 | 0 | 3 | 0 | 98 | 49 | 0.059555994 | 0.170760496 | NA | 1 | 1 | NA | 0 | NA | 1 |
| SE | 7469 | SLTM | chr15 | - | 58899468 | 58899883 | 58898802 | 58898852 | 58901259 | 58901287 | 0 | 1 | 0 | 0 | 0 | 0 | 0 | 0 | 0 | 0 | 3 | 0 | 98 | 49 | 0.059555994 | 0.170760496 | NA | 1 | NA | NA | 0 | NA | 1 |
| SE | 7470 | SLTM | chr15 | - | 58899468 | 58899937 | 58898802 | 58898852 | 58901259 | 58901287 | 0 | 1 | 0 | 0 | 0 | 0 | 0 | 0 | 0 | 0 | 3 | 0 | 98 | 49 | 0.059555994 | 0.170760496 | NA | 1 | NA | NA | 0 | NA | 1 |
| SE | 9472 | USP33 | chr1 | - | 77745420 | 77745599 | 77741616 | 77741748 | 77759642 | 77759879 | 0 | 0 | 1 | 0 | 0 | 0 | 0 | 0 | 0 | 0 | 0 | 3 | 98 | 49 | 0.059555994 | 0.170760496 | NA | NA | 1 | NA | NA | 0 | 1 |
| SE | 13888 | RIF1 | chr2 | + | 151410413 | 151410527 | 151409882 | 151410033 | 151411259 | 151411338 | 0 | 0 | 1 | 0 | 0 | 0 | 0 | 2 | 0 | 0 | 0 | 3 | 98 | 49 | 0.059555994 | 0.170760496 | NA | NA | 1 | NA | 1 | 0 | 0.5 |
| SE | 4181 | DDIT3 | chr12 | - | 57517268 | 57517595 | 57516587 | 57517180 | 57517705 | 57517753 | 0 | 1 | 5 | 0 | 0 | 0 | 0 | 4 | 1 | 0 | 1 | 0 | 98 | 49 | 0.059568296 | 0.170760496 | NA | 1 | 1 | NA | 0.667 | 1 | 0.166 |
| SE | 12744 | DYSF | chr2 | + | 71561751 | 71561944 | 71555964 | 71556071 | 71564057 | 71564213 | 0 | 5 | 41 | 0 | 0 | 0 | 2 | 10 | 26 | 0 | 0 | 1 | 98 | 49 | 0.059610573 | 0.170760496 | NA | 1 | 1 | 1 | 1 | 0.929 | 0.024 |
| SE | 2865 | POLDIP3 | chr22 | - | 42591954 | 42592084 | 42587505 | 42587572 | 42595536 | 42595603 | 0 | 0 | 0 | 0 | 0 | 2 | 0 | 3 | 3 | 0 | 0 | 0 | 98 | 49 | 0.059642855 | 0.170760496 | NA | NA | 0 | NA | 1 | 1 | -1 |
| SE | 7379 | RNF19B | chr1 | - | 32949568 | 32949774 | 32948221 | 32948363 | 32964050 | 32964813 | 0 | 0 | 7 | 0 | 2 | 0 | 0 | 3 | 0 | 0 | 0 | 0 | 98 | 49 | 0.059642855 | 0.170760496 | NA | 0 | 1 | NA | 1 | NA | -0.5 |
| SE | 7408 | RBM23 | chr14 | - | 22905605 | 22905659 | 22905335 | 22905453 | 22906194 | 22906368 | 0 | 0 | 2 | 0 | 2 | 1 | 0 | 3 | 0 | 0 | 0 | 0 | 98 | 49 | 0.059642855 | 0.170760496 | NA | 0 | 0.5 | NA | 1 | NA | -0.75 |
| SE | 9772 | FRYL | chr4 | - | 48512480 | 48512688 | 48510834 | 48510984 | 48515027 | 48515275 | 0 | 0 | 2 | 0 | 2 | 0 | 7 | 3 | 0 | 0 | 0 | 0 | 98 | 49 | 0.059642855 | 0.170760496 | NA | 0 | 1 | 1 | 1 | NA | -0.5 |
| SE | 11703 | POLR2A | chr17 | + | 7511810 | 7512014 | 7511405 | 7511588 | 7512157 | 7512344 | 0 | 2 | 19 | 0 | 0 | 0 | 0 | 5 | 2 | 0 | 1 | 0 | 98 | 49 | 0.059650539 | 0.170760496 | NA | 1 | 1 | NA | 0.714 | 1 | 0.143 |
| SE | 6864 | ATG2A | chr11 | - | 64900883 | 64901092 | 64900493 | 64900629 | 64901961 | 64902176 | 0 | 0 | 0 | 0 | 0 | 1 | 0 | 5 | 5 | 0 | 0 | 0 | 98 | 49 | 0.059786145 | 0.170760496 | NA | NA | 0 | NA | 1 | 1 | -1 |
| A5SS | 2457 | SCARF1 | chr17 | - | 1645559 | 1645789 | 1645596 | 1645789 | 1645177 | 1645239 | 0 | 0 | 0 | 0 | 0 | 1 | 0 | 3 | 5 | 0 | 3 | 0 | 86 | 49 | 0.059797532 | 0.170414363 | NA | NA | 0 | NA | 0.363 | 1 | -0.681 |
| A3SS | 1375 | ARHGAP25 | chr2 | + | 68807272 | 68807480 | 68807275 | 68807480 | 68787839 | 68787956 | 0 | 0 | 5 | 0 | 1 | 0 | 0 | 5 | 0 | 0 | 0 | 0 | 52 | 49 | 0.059800864 | 0.197717869 | NA | 0 | 1 | NA | 1 | NA | -0.5 |
| SE | 12948 | SMCHD1 | chr18 | + | 2778168 | 2778239 | 2777805 | 2777915 | 2784449 | 2784621 | 0 | 2 | 47 | 0 | 0 | 1 | 2 | 28 | 1 | 0 | 4 | 0 | 98 | 49 | 0.059897198 | 0.170760496 | NA | 1 | 0.959 | 1 | 0.778 | 1 | 0.053 |
| SE | 5027 | OSBPL8 | chr12 | - | 76459858 | 76459895 | 76450850 | 76450988 | 76487509 | 76487618 | 0 | 1 | 0 | 0 | 0 | 2 | 0 | 0 | 0 | 0 | 3 | 0 | 86 | 49 | 0.059993631 | 0.170760496 | NA | 1 | 0 | NA | 0 | NA | 0.5 |
| SE | 9705 | ABTB1 | chr3 | + | 127676966 | 127677083 | 127676535 | 127676581 | 127677167 | 127677286 | 0 | 1 | 20 | 0 | 0 | 0 | 5 | 4 | 1 | 0 | 1 | 0 | 98 | 49 | 0.060016261 | 0.170760496 | NA | 1 | 1 | 1 | 0.667 | 1 | 0.111 |
| SE | 4178 | DDIT3 | chr12 | - | 57517268 | 57517438 | 57516587 | 57517180 | 57517705 | 57517753 | 0 | 1 | 6 | 0 | 0 | 0 | 0 | 5 | 1 | 0 | 1 | 0 | 98 | 49 | 0.060034631 | 0.170760496 | NA | 1 | 1 | NA | 0.714 | 1 | 0.143 |
| SE | 364 | SEMA4D | chr9 | - | 89418055 | 89418118 | 89405350 | 89405699 | 89455887 | 89455953 | 0 | 0 | 5 | 0 | 0 | 0 | 0 | 7 | 0 | 1 | 0 | 1 | 98 | 49 | 0.060038159 | 0.170760496 | NA | NA | 1 | 0 | 1 | 0 | 0.667 |
| A5SS | 1404 | RBMS1 | chr2 | - | 160286968 | 160287084 | 160286977 | 160287084 | 160284994 | 160285044 | 1 | 0 | 2 | 0 | 0 | 2 | 0 | 1 | 0 | 4 | 0 | 0 | 58 | 49 | 0.060055861 | 0.170414363 | 1 | NA | 0.458 | 0 | 1 | NA | 0.229 |
| A3SS | 413 | DCTN3 | chr9 | - | 34614041 | 34614132 | 34614041 | 34614101 | 34614709 | 34614768 | 1 | 1 | 0 | 0 | 0 | 0 | 0 | 0 | 0 | 0 | 3 | 0 | 80 | 49 | 0.060211792 | 0.197717869 | 1 | 1 | NA | NA | 0 | NA | 1 |
| SE | 12010 | GPR108 | chr19 | - | 6731890 | 6731934 | 6731472 | 6731522 | 6732024 | 6732155 | 0 | 0 | 5 | 0 | 0 | 0 | 0 | 5 | 0 | 0 | 0 | 1 | 93 | 49 | 0.060224876 | 0.170760496 | NA | NA | 1 | NA | 1 | 0 | 0.5 |
| SE | 8079 | NADK | chr1 | - | 1757180 | 1757310 | 1756257 | 1756343 | 1761951 | 1762035 | 0 | 1 | 24 | 0 | 0 | 0 | 0 | 6 | 1 | 0 | 1 | 0 | 98 | 49 | 0.060230863 | 0.170760496 | NA | 1 | 1 | NA | 0.75 | 1 | 0.125 |
| SE | 2349 | FNDC3B | chr3 | + | 172346326 | 172346440 | 172344085 | 172344258 | 172347211 | 172347361 | 0 | 1 | 6 | 0 | 0 | 0 | 0 | 7 | 3 | 0 | 2 | 0 | 98 | 49 | 0.060484453 | 0.170760496 | NA | 1 | 1 | NA | 0.636 | 1 | 0.182 |
| SE | 4177 | DDIT3 | chr12 | - | 57517268 | 57517424 | 57516587 | 57517180 | 57517705 | 57517753 | 0 | 1 | 5 | 0 | 0 | 0 | 0 | 3 | 1 | 0 | 1 | 0 | 98 | 49 | 0.060546566 | 0.170760496 | NA | 1 | 1 | NA | 0.6 | 1 | 0.2 |
| SE | 4179 | DDIT3 | chr12 | - | 57517268 | 57517519 | 57516587 | 57517180 | 57517705 | 57517753 | 0 | 1 | 5 | 0 | 0 | 0 | 0 | 3 | 1 | 0 | 1 | 0 | 98 | 49 | 0.060546566 | 0.170760496 | NA | 1 | 1 | NA | 0.6 | 1 | 0.2 |
| SE | 4180 | DDIT3 | chr12 | - | 57517268 | 57517542 | 57516587 | 57517180 | 57517705 | 57517753 | 0 | 1 | 5 | 0 | 0 | 0 | 0 | 3 | 1 | 0 | 1 | 0 | 98 | 49 | 0.060546566 | 0.170760496 | NA | 1 | 1 | NA | 0.6 | 1 | 0.2 |
| SE | 13577 | STK17B | chr2 | - | 196163261 | 196163427 | 196156438 | 196156651 | 196171332 | 196171612 | 0 | 21 | 136 | 0 | 0 | 0 | 6 | 68 | 29 | 0 | 1 | 0 | 98 | 49 | 0.060570323 | 0.170760496 | NA | 1 | 1 | 1 | 0.971 | 1 | 0.01 |
| SE | 1540 | HLA-F | chr6 | + | 29725030 | 29725306 | 29724172 | 29724448 | 29725446 | 29725563 | 0 | 4 | 52 | 0 | 0 | 0 | 0 | 23 | 5 | 0 | 1 | 0 | 98 | 49 | 0.06060798 | 0.170760496 | NA | 1 | 1 | NA | 0.92 | 1 | 0.04 |
| SE | 11556 | KIAA0513 | chr16 | + | 85078414 | 85078455 | 85077424 | 85077632 | 85078924 | 85079003 | 0 | 0 | 9 | 0 | 1 | 0 | 0 | 4 | 0 | 0 | 0 | 0 | 90 | 49 | 0.060658984 | 0.170760496 | NA | 0 | 1 | NA | 1 | NA | -0.5 |
| A3SS | 3134 | STX4 | chr16 | + | 31034203 | 31034325 | 31034225 | 31034325 | 31034012 | 31034114 | 0 | 0 | 3 | 0 | 1 | 2 | 1 | 4 | 0 | 0 | 0 | 0 | 71 | 49 | 0.06066392 | 0.197717869 | NA | 0 | 0.509 | 1 | 1 | NA | -0.746 |
| SE | 2842 | HDAC10 | chr22 | - | 50247691 | 50247776 | 50246874 | 50246966 | 50247889 | 50248145 | 0 | 0 | 0 | 0 | 0 | 1 | 0 | 0 | 4 | 0 | 2 | 0 | 98 | 49 | 0.060672385 | 0.170760496 | NA | NA | 0 | NA | 0 | 1 | -0.5 |
| SE | 6993 | MATR3 | chr5 | + | 139307238 | 139308327 | 139293643 | 139293805 | 139314674 | 139314736 | 0 | 0 | 0 | 0 | 1 | 0 | 0 | 4 | 0 | 0 | 0 | 0 | 98 | 49 | 0.060672385 | 0.170760496 | NA | 0 | NA | NA | 1 | NA | -1 |
| SE | 7078 | ERBB2IP | chr5 | + | 66075023 | 66075230 | 66053405 | 66054951 | 66076315 | 66076408 | 0 | 0 | 2 | 0 | 1 | 0 | 0 | 4 | 0 | 0 | 0 | 0 | 98 | 49 | 0.060672385 | 0.170760496 | NA | 0 | 1 | NA | 1 | NA | -0.5 |
| SE | 1858 | AP3B1 | chr5 | - | 78228143 | 78228239 | 78227371 | 78227532 | 78240861 | 78240936 | 0 | 1 | 13 | 0 | 0 | 0 | 0 | 5 | 1 | 0 | 1 | 0 | 98 | 49 | 0.060716183 | 0.170760496 | NA | 1 | 1 | NA | 0.714 | 1 | 0.143 |
| A3SS | 2183 | PPP4R1 | chr18 | - | 9595017 | 9595153 | 9595017 | 9595102 | 9614225 | 9614270 | 0 | 0 | 4 | 0 | 0 | 0 | 0 | 2 | 0 | 0 | 2 | 1 | 98 | 49 | 0.060734355 | 0.197717869 | NA | NA | 1 | NA | 0.333 | 0 | 0.834 |
| SE | 788 | PARP10 | chr8 | - | 143984007 | 143984104 | 143983166 | 143983811 | 143984209 | 143984431 | 0 | 0 | 4 | 0 | 0 | 0 | 0 | 4 | 0 | 0 | 0 | 1 | 98 | 49 | 0.060734355 | 0.170760496 | NA | NA | 1 | NA | 1 | 0 | 0.5 |
| SE | 5889 | SIPA1L1 | chr14 | + | 71723646 | 71723886 | 71709221 | 71709664 | 71724669 | 71724835 | 0 | 0 | 4 | 0 | 0 | 0 | 0 | 0 | 0 | 0 | 0 | 1 | 98 | 49 | 0.060734355 | 0.170760496 | NA | NA | 1 | NA | NA | 0 | 1 |
| A5SS | 3026 | ABCG1 | chr21 | + | 42287888 | 42288073 | 42287888 | 42288037 | 42288210 | 42288312 | 0 | 2 | 3 | 0 | 0 | 2 | 0 | 8 | 2 | 0 | 0 | 0 | 85 | 49 | 0.060789913 | 0.170414363 | NA | 1 | 0.464 | NA | 1 | 1 | -0.268 |
| SE | 4502 | PTPN6 | chr12 | + | 6954804 | 6954994 | 6951982 | 6952177 | 6955150 | 6955267 | 0 | 2 | 31 | 0 | 0 | 0 | 0 | 21 | 1 | 0 | 1 | 0 | 98 | 49 | 0.061131927 | 0.170760496 | NA | 1 | 1 | NA | 0.913 | 1 | 0.043 |
| SE | 10395 | HES4 | chr1 | - | 999691 | 999787 | 999525 | 999613 | 999865 | 1000172 | 0 | 0 | 0 | 0 | 1 | 1 | 0 | 1 | 0 | 0 | 4 | 1 | 98 | 49 | 0.061306069 | 0.170760496 | NA | 0 | 0 | NA | 0.111 | 0 | -0.056 |
| RI | 672 | PDE4D | chr5 | - | 59038858 | 59039512 | 59038858 | 59038971 | 59039057 | 59039512 | 0 | 0 | 1 | 0 | 0 | 0 | 3 | 3 | 0 | 0 | 0 | 2 | 98 | 49 | 0.061401334 | 0.297702651 | NA | NA | 1 | 1 | 1 | 0 | 0.333 |
| SE | 7868 | VRK3 | chr19 | - | 49989638 | 49989771 | 49988371 | 49988492 | 49992859 | 49992952 | 0 | 1 | 2 | 0 | 0 | 0 | 0 | 0 | 0 | 0 | 2 | 0 | 98 | 49 | 0.061401334 | 0.170760496 | NA | 1 | 1 | NA | 0 | NA | 1 |
| SE | 8054 | CUL4B | chrX | - | 120544480 | 120544643 | 120544113 | 120544203 | 120545443 | 120545517 | 0 | 0 | 1 | 0 | 0 | 0 | 0 | 1 | 0 | 0 | 0 | 2 | 98 | 49 | 0.061401334 | 0.170760496 | NA | NA | 1 | NA | 1 | 0 | 0.5 |
| SE | 11362 | RSRC2 | chr12 | - | 122520503 | 122520581 | 122518838 | 122519029 | 122521384 | 122521428 | 0 | 1 | 3 | 0 | 0 | 1 | 0 | 0 | 0 | 0 | 2 | 0 | 98 | 49 | 0.061401334 | 0.170760496 | NA | 1 | 0.6 | NA | 0 | NA | 0.8 |
| SE | 12007 | DHX8 | chr17 | + | 43508338 | 43508520 | 43507808 | 43508019 | 43513361 | 43513502 | 0 | 1 | 1 | 0 | 0 | 0 | 0 | 0 | 0 | 0 | 2 | 0 | 98 | 49 | 0.061401334 | 0.170760496 | NA | 1 | 1 | NA | 0 | NA | 1 |
| SE | 9457 | PFKFB3 | chr10 | + | 6221640 | 6221745 | 6221380 | 6221527 | 6222854 | 6222984 | 0 | 16 | 35 | 0 | 0 | 0 | 0 | 44 | 26 | 0 | 1 | 0 | 98 | 49 | 0.0614161 | 0.170760496 | NA | 1 | 1 | NA | 0.957 | 1 | 0.022 |
| A3SS | 3638 | MAP2K7 | chr19 | + | 7911414 | 7911578 | 7911435 | 7911578 | 7911249 | 7911330 | 0 | 0 | 0 | 0 | 1 | 2 | 0 | 0 | 2 | 0 | 0 | 0 | 70 | 49 | 0.061460457 | 0.197717869 | NA | 0 | 0 | NA | NA | 1 | -1 |
| A5SS | 725 | FAM193A | chr4 | + | 2694945 | 2695142 | 2694945 | 2695129 | 2696362 | 2696593 | 0 | 0 | 0 | 0 | 2 | 0 | 0 | 2 | 0 | 0 | 0 | 0 | 62 | 49 | 0.061467502 | 0.170414363 | NA | 0 | NA | NA | 1 | NA | -1 |
| A3SS | 16 | ALG13 | chrX | + | 111682131 | 111682294 | 111682235 | 111682294 | 111681169 | 111681424 | 0 | 0 | 0 | 0 | 2 | 0 | 0 | 2 | 0 | 0 | 0 | 0 | 98 | 49 | 0.061489123 | 0.197717869 | NA | 0 | NA | NA | 1 | NA | -1 |
| A5SS | 1081 | WIPI2 | chr7 | + | 5193117 | 5193345 | 5193117 | 5193171 | 5199575 | 5199658 | 0 | 0 | 0 | 0 | 0 | 2 | 0 | 0 | 2 | 0 | 0 | 0 | 98 | 49 | 0.061489123 | 0.170414363 | NA | NA | 0 | NA | NA | 1 | -1 |
| RI | 843 | FAM96A | chr15 | - | 64072558 | 64075537 | 64072558 | 64073028 | 64075491 | 64075537 | 0 | 0 | 0 | 0 | 0 | 2 | 1 | 0 | 2 | 0 | 1 | 0 | 98 | 49 | 0.061489123 | 0.297702651 | NA | NA | 0 | 1 | 0 | 1 | -0.667 |
| SE | 4053 | CASC3 | chr17 | + | 40167849 | 40167948 | 40167497 | 40167612 | 40168202 | 40168417 | 0 | 0 | 5 | 0 | 2 | 0 | 0 | 2 | 0 | 0 | 0 | 0 | 98 | 49 | 0.061489123 | 0.170760496 | NA | 0 | 1 | NA | 1 | NA | -0.5 |
| SE | 10886 | AKAP13 | chr15 | + | 85662387 | 85662453 | 85658536 | 85658590 | 85664562 | 85664755 | 0 | 0 | 2 | 0 | 2 | 11 | 0 | 2 | 0 | 0 | 0 | 0 | 98 | 49 | 0.061489123 | 0.170760496 | NA | 0 | 0.083 | NA | 1 | NA | -0.959 |
| SE | 11013 | RHOT2 | chr16 | + | 669552 | 669606 | 668655 | 668699 | 670122 | 670175 | 0 | 0 | 1 | 0 | 2 | 0 | 0 | 2 | 0 | 0 | 0 | 0 | 98 | 49 | 0.061489123 | 0.170760496 | NA | 0 | 1 | NA | 1 | NA | -0.5 |
| SE | 11296 | CREBBP | chr16 | - | 3791980 | 3792094 | 3782683 | 3782926 | 3793385 | 3793626 | 0 | 1 | 9 | 0 | 0 | 0 | 0 | 3 | 1 | 0 | 1 | 0 | 98 | 49 | 0.061515403 | 0.170760496 | NA | 1 | 1 | NA | 0.6 | 1 | 0.2 |
| A3SS | 3506 | DYRK1B | chr19 | - | 39826671 | 39826987 | 39826671 | 39826867 | 39827284 | 39827425 | 0 | 0 | 3 | 0 | 0 | 0 | 0 | 0 | 0 | 0 | 2 | 1 | 98 | 49 | 0.06170155 | 0.197717869 | NA | NA | 1 | NA | 0 | 0 | 1 |
| A5SS | 1533 | BTF3 | chr5 | + | 73498424 | 73498799 | 73498424 | 73498558 | 73499133 | 73499202 | 3 | 0 | 2 | 0 | 0 | 2 | 0 | 2 | 0 | 1 | 3 | 0 | 98 | 49 | 0.06170155 | 0.170414363 | 1 | NA | 0.333 | 0 | 0.25 | NA | 0.541 |
| SE | 7580 | MMP8 | chr11 | - | 102722990 | 102723078 | 102722428 | 102722673 | 102724753 | 102724954 | 0 | 3 | 3 | 0 | 0 | 1 | 0 | 0 | 0 | 0 | 1 | 0 | 98 | 49 | 0.06170155 | 0.170760496 | NA | 1 | 0.6 | NA | 0 | NA | 0.8 |
| SE | 7581 | MMP8 | chr11 | - | 102722990 | 102723081 | 102722428 | 102722673 | 102724753 | 102724954 | 0 | 3 | 3 | 0 | 0 | 1 | 0 | 0 | 0 | 0 | 1 | 0 | 98 | 49 | 0.06170155 | 0.170760496 | NA | 1 | 0.6 | NA | 0 | NA | 0.8 |
| SE | 8154 | CD302 | chr2 | - | 159783358 | 159783469 | 159780881 | 159780998 | 159798131 | 159798255 | 0 | 0 | 3 | 0 | 0 | 0 | 0 | 3 | 0 | 0 | 0 | 1 | 98 | 49 | 0.06170155 | 0.170760496 | NA | NA | 1 | NA | 1 | 0 | 0.5 |
| SE | 9121 | KDM4B | chr19 | + | 5071009 | 5071059 | 5047475 | 5047669 | 5077366 | 5077470 | 0 | 0 | 3 | 0 | 0 | 0 | 0 | 4 | 0 | 0 | 0 | 1 | 98 | 49 | 0.06170155 | 0.170760496 | NA | NA | 1 | NA | 1 | 0 | 0.5 |
| SE | 10516 | ZNF316 | chr7 | + | 6642381 | 6642764 | 6641824 | 6641962 | 6642963 | 6643073 | 0 | 3 | 0 | 0 | 0 | 0 | 0 | 0 | 0 | 0 | 1 | 0 | 98 | 49 | 0.06170155 | 0.170760496 | NA | 1 | NA | NA | 0 | NA | 1 |
| SE | 11106 | MGRN1 | chr16 | + | 4680031 | 4680097 | 4677462 | 4677572 | 4681549 | 4681776 | 0 | 0 | 3 | 0 | 0 | 0 | 0 | 1 | 0 | 0 | 0 | 1 | 98 | 49 | 0.06170155 | 0.170760496 | NA | NA | 1 | NA | 1 | 0 | 0.5 |
| A3SS | 910 | LIME1 | chr20 | + | 63737972 | 63738060 | 63738007 | 63738060 | 63737820 | 63737902 | 0 | 0 | 1 | 0 | 1 | 0 | 0 | 3 | 0 | 0 | 0 | 0 | 84 | 49 | 0.061899751 | 0.197717869 | NA | 0 | 1 | NA | 1 | NA | -0.5 |
| A3SS | 2475 | ZBTB17 | chr1 | - | 15946153 | 15946316 | 15946153 | 15946294 | 15946934 | 15947123 | 0 | 0 | 1 | 0 | 1 | 2 | 0 | 3 | 0 | 0 | 0 | 0 | 71 | 49 | 0.061903 | 0.197717869 | NA | 0 | 0.257 | NA | 1 | NA | -0.871 |
| A3SS | 122 | FAM219A | chr9 | - | 34398183 | 34401122 | 34398183 | 34401119 | 34401665 | 34401720 | 0 | 0 | 0 | 0 | 1 | 0 | 0 | 3 | 0 | 0 | 0 | 0 | 52 | 49 | 0.061903774 | 0.197717869 | NA | 0 | NA | NA | 1 | NA | -1 |
| A3SS | 3899 | PPP1R12C | chr19 | - | 55092782 | 55092868 | 55092782 | 55092865 | 55093015 | 55093076 | 0 | 0 | 0 | 0 | 0 | 1 | 0 | 0 | 3 | 0 | 1 | 0 | 52 | 49 | 0.061903774 | 0.197717869 | NA | NA | 0 | NA | 0 | 1 | -0.5 |
| RI | 528 | NFKB2 | chr10 | + | 102395595 | 102395980 | 102395595 | 102395796 | 102395887 | 102395980 | 0 | 0 | 0 | 0 | 0 | 1 | 0 | 5 | 3 | 0 | 0 | 0 | 98 | 49 | 0.061911422 | 0.297702651 | NA | NA | 0 | NA | 1 | 1 | -1 |
| SE | 4260 | SLC22A4 | chr5 | + | 132312160 | 132312264 | 132294451 | 132295009 | 132313613 | 132313768 | 0 | 0 | 4 | 0 | 1 | 0 | 0 | 3 | 0 | 0 | 0 | 0 | 98 | 49 | 0.061911422 | 0.170760496 | NA | 0 | 1 | NA | 1 | NA | -0.5 |
| SE | 5368 | BAZ1A | chr14 | - | 34826012 | 34826156 | 34810934 | 34811036 | 34874491 | 34874662 | 0 | 0 | 13 | 0 | 1 | 0 | 2 | 3 | 0 | 0 | 0 | 0 | 98 | 49 | 0.061911422 | 0.170760496 | NA | 0 | 1 | 1 | 1 | NA | -0.5 |
| SE | 6045 | GADD45A | chr1 | + | 67686349 | 67686587 | 67685176 | 67685538 | 67687660 | 67688338 | 0 | 0 | 0 | 0 | 0 | 1 | 0 | 1 | 3 | 0 | 0 | 0 | 98 | 49 | 0.061911422 | 0.170760496 | NA | NA | 0 | NA | 1 | 1 | -1 |
| SE | 12169 | HDAC7 | chr12 | - | 47786584 | 47786703 | 47785751 | 47785885 | 47787711 | 47787809 | 0 | 0 | 4 | 0 | 1 | 1 | 0 | 3 | 0 | 0 | 0 | 0 | 98 | 49 | 0.061911422 | 0.170760496 | NA | 0 | 0.667 | NA | 1 | NA | -0.666 |
| SE | 12235 | FAM104A | chr17 | - | 73227164 | 73227264 | 73207352 | 73209768 | 73232085 | 73232394 | 0 | 0 | 1 | 0 | 1 | 0 | 0 | 3 | 0 | 0 | 0 | 0 | 98 | 49 | 0.061911422 | 0.170760496 | NA | 0 | 1 | NA | 1 | NA | -0.5 |
| A5SS | 2118 | CSNK1A1 | chr5 | - | 149505446 | 149505595 | 149505482 | 149505595 | 149495893 | 149496860 | 3 | 0 | 11 | 0 | 0 | 1 | 0 | 5 | 0 | 1 | 1 | 0 | 85 | 49 | 0.062188913 | 0.170414363 | 1 | NA | 0.864 | 0 | 0.742 | NA | 0.561 |
| SE | 12245 | DYRK1B | chr19 | - | 39826951 | 39826987 | 39826671 | 39826867 | 39827284 | 39827425 | 0 | 0 | 3 | 0 | 0 | 0 | 0 | 0 | 0 | 0 | 2 | 1 | 85 | 49 | 0.062188913 | 0.170760496 | NA | NA | 1 | NA | 0 | 0 | 1 |
| A3SS | 2080 | AP2S1 | chr19 | - | 46839464 | 46839620 | 46839464 | 46839578 | 46845992 | 46846142 | 0 | 0 | 0 | 0 | 0 | 5 | 0 | 0 | 1 | 5 | 2 | 0 | 91 | 49 | 0.062394347 | 0.197717869 | NA | NA | 0 | 0 | 0 | 1 | -0.333 |
| SE | 4424 | PREX1 | chr20 | - | 48658135 | 48658228 | 48657039 | 48657188 | 48666282 | 48666355 | 0 | 2 | 10 | 0 | 0 | 0 | 0 | 20 | 2 | 0 | 1 | 0 | 98 | 49 | 0.062524268 | 0.170760496 | NA | 1 | 1 | NA | 0.909 | 1 | 0.045 |
| SE | 2274 | VNN3 | chr6 | - | 132728740 | 132728933 | 132724605 | 132724979 | 132734252 | 132734383 | 0 | 1 | 5 | 0 | 0 | 0 | 0 | 9 | 1 | 0 | 1 | 0 | 98 | 49 | 0.062727245 | 0.170760496 | NA | 1 | 1 | NA | 0.818 | 1 | 0.091 |
| SE | 13163 | PPP1CB | chr2 | + | 28752037 | 28752176 | 28751747 | 28751820 | 28776850 | 28776982 | 0 | 2 | 2 | 0 | 0 | 0 | 1 | 4 | 3 | 0 | 1 | 0 | 98 | 49 | 0.06302451 | 0.170760496 | NA | 1 | 1 | 1 | 0.667 | 1 | 0.111 |
| A3SS | 2776 | MYADM | chr19 | + | 53869689 | 53869838 | 53869753 | 53869838 | 53867864 | 53867943 | 0 | 0 | 2 | 0 | 0 | 0 | 3 | 0 | 0 | 0 | 0 | 1 | 98 | 49 | 0.063117334 | 0.197717869 | NA | NA | 1 | 1 | NA | 0 | 0.5 |
| A5SS | 1568 | POLD4 | chr11 | - | 67353302 | 67353596 | 67353363 | 67353596 | 67352987 | 67353077 | 0 | 1 | 2 | 0 | 0 | 0 | 0 | 0 | 0 | 0 | 0 | 1 | 98 | 49 | 0.063117334 | 0.170414363 | NA | 1 | 1 | NA | NA | 0 | 1 |
| SE | 498 | NAPRT | chr8 | - | 143577061 | 143577177 | 143576645 | 143576842 | 143577268 | 143577399 | 0 | 2 | 4 | 0 | 0 | 0 | 0 | 0 | 0 | 0 | 1 | 0 | 98 | 49 | 0.063117334 | 0.170760496 | NA | 1 | 1 | NA | 0 | NA | 1 |
| SE | 3230 | TMEM189 | chr20 | - | 50130855 | 50130947 | 50129545 | 50129690 | 50143501 | 50143621 | 0 | 2 | 1 | 0 | 0 | 0 | 0 | 0 | 0 | 0 | 1 | 0 | 98 | 49 | 0.063117334 | 0.170760496 | NA | 1 | 1 | NA | 0 | NA | 1 |
| SE | 3231 | TMEM189 | chr20 | - | 50130864 | 50130947 | 50129545 | 50129690 | 50143501 | 50143621 | 0 | 2 | 0 | 0 | 0 | 0 | 0 | 0 | 0 | 0 | 1 | 0 | 98 | 49 | 0.063117334 | 0.170760496 | NA | 1 | NA | NA | 0 | NA | 1 |

| | | | | | | | | | | | | | | | | | | | | | | | | | | | | | | | | | |
|---|---|---|---|---|---|---|---|---|---|---|---|---|---|---|---|---|---|---|---|---|---|---|---|---|---|---|---|---|---|---|---|---|---|
| SE | 3810 | CPNE1 | chr20 | - | 35658929 | 35658982 | 35632794 | 35632923 | 35664759 | 35664937 | 0 | 0 | 2 | 0 | 0 | 0 | 0 | 0 | 0 | 0 | 0 | 1 | 98 | 49 | 0.063117334 | 0.170760496 | NA | NA | 1 | NA | NA | 0 | 1 |
| SE | 3811 | CPNE1 | chr20 | - | 35658929 | 35659014 | 35632794 | 35632923 | 35664759 | 35664956 | 0 | 0 | 2 | 0 | 0 | 0 | 0 | 0 | 0 | 0 | 0 | 1 | 98 | 49 | 0.063117334 | 0.170760496 | NA | NA | 1 | NA | NA | 0 | 1 |
| SE | 6736 | MTOR | chr1 | - | 11256932 | 11257165 | 11255991 | 11256192 | 11258484 | 11258593 | 0 | 2 | 4 | 0 | 0 | 0 | 0 | 0 | 0 | 0 | 1 | 0 | 98 | 49 | 0.063117334 | 0.170760496 | NA | 1 | 1 | NA | 0 | NA | 1 |
| SE | 6925 | DMTN | chr8 | + | 22067084 | 22067159 | 22066704 | 22066893 | 22067526 | 22067682 | 0 | 2 | 1 | 0 | 0 | 0 | 0 | 0 | 0 | 0 | 1 | 0 | 98 | 49 | 0.063117334 | 0.170760496 | NA | 1 | 1 | NA | 0 | NA | 1 |
| SE | 9827 | SHISA5 | chr3 | - | 48503094 | 48503193 | 48501136 | 48501293 | 48504422 | 48504814 | 0 | 0 | 2 | 0 | 0 | 0 | 0 | 0 | 0 | 0 | 0 | 1 | 98 | 49 | 0.063117334 | 0.170760496 | NA | NA | 1 | NA | NA | 0 | 1 |
| SE | 10041 | SYNE2 | chr14 | + | 64031015 | 64031357 | 64027483 | 64027793 | 64047999 | 64048155 | 0 | 2 | 0 | 0 | 0 | 0 | 0 | 0 | 0 | 0 | 1 | 0 | 98 | 49 | 0.063117334 | 0.170760496 | NA | 1 | NA | NA | 0 | NA | 1 |
| SE | 10233 | SYNE3 | chr14 | - | 95467794 | 95467967 | 95465930 | 95466240 | 95475677 | 95475836 | 0 | 2 | 0 | 0 | 0 | 0 | 0 | 0 | 0 | 0 | 1 | 0 | 98 | 49 | 0.063117334 | 0.170760496 | NA | 1 | NA | NA | 0 | NA | 1 |
| SE | 12575 | ARID3A | chr19 | + | 964832 | 965080 | 964247 | 964431 | 966571 | 966868 | 0 | 2 | 4 | 0 | 0 | 0 | 0 | 0 | 0 | 0 | 1 | 0 | 98 | 49 | 0.063117334 | 0.170760496 | NA | 1 | 1 | NA | 0 | NA | 1 |
| SE | 13045 | CNOT7 | chr8 | - | 17232426 | 17232537 | 17229230 | 17230848 | 17234715 | 17234860 | 0 | 2 | 2 | 0 | 0 | 0 | 0 | 0 | 0 | 0 | 1 | 0 | 98 | 49 | 0.063117334 | 0.170760496 | NA | 1 | 1 | NA | 0 | NA | 1 |
| A3SS | 1438 | FGD4 | chr12 | + | 32607937 | 32608095 | 32607956 | 32608095 | 32602160 | 32602317 | 1 | 0 | 0 | 0 | 0 | 4 | 0 | 2 | 1 | 0 | 0 | 0 | 68 | 49 | 0.063176479 | 0.197717869 | 1 | NA | 0 | NA | 1 | 1 | -0.5 |
| A5SS | 1600 | TRAPPC3 | chr1 | - | 36149091 | 36149514 | 36149336 | 36149514 | 36140068 | 36140166 | 0 | 0 | 0 | 0 | 4 | 0 | 8 | 1 | 0 | 0 | 0 | 0 | 98 | 49 | 0.063182848 | 0.170414363 | NA | 0 | NA | 1 | 1 | NA | -1 |
| SE | 5519 | DENND5A | chr11 | - | 9203659 | 9204317 | 9193493 | 9193681 | 9206672 | 9206782 | 0 | 2 | 20 | 0 | 0 | 0 | 0 | 18 | 2 | 0 | 1 | 0 | 98 | 49 | 0.063221519 | 0.170760496 | NA | 1 | 1 | NA | 0.9 | 1 | 0.05 |
| SE | 7498 | IFI44 | chr1 | + | 78662703 | 78662878 | 78659311 | 78659483 | 78663764 | 78664078 | 0 | 6 | 51 | 0 | 1 | 1 | 0 | 87 | 6 | 0 | 0 | 0 | 98 | 49 | 0.063259916 | 0.170760496 | NA | 0.75 | 0.962 | NA | 1 | 1 | -0.144 |
| SE | 12385 | JAK3 | chr19 | - | 17831227 | 17831400 | 17830502 | 17830620 | 17831673 | 17831798 | 0 | 2 | 10 | 0 | 0 | 0 | 0 | 7 | 3 | 0 | 1 | 0 | 98 | 49 | 0.063527186 | 0.170760496 | NA | 1 | 1 | NA | 0.778 | 1 | 0.111 |
| A3SS | 3618 | CYB561D1 | chr1 | + | 109495678 | 109500441 | 109495755 | 109500441 | 109495142 | 109495180 | 0 | 0 | 0 | 0 | 0 | 1 | 0 | 0 | 2 | 0 | 0 | 0 | 98 | 49 | 0.063770311 | 0.197717869 | NA | NA | 0 | NA | NA | 1 | -1 |
| A5SS | 1161 | GAPDH | chr12 | + | 6534404 | 6534661 | 6534404 | 6534569 | 6534809 | 6534861 | 0 | 0 | 0 | 0 | 0 | 1 | 0 | 0 | 2 | 0 | 1 | 0 | 98 | 49 | 0.063770311 | 0.170414363 | NA | NA | 0 | NA | 0 | 1 | -0.5 |
| A5SS | 2679 | OXLD1 | chr17 | - | 81666145 | 81666637 | 81666517 | 81666637 | 81665035 | 81665584 | 0 | 0 | 0 | 0 | 0 | 1 | 0 | 0 | 2 | 0 | 0 | 0 | 98 | 49 | 0.063770311 | 0.170414363 | NA | NA | 0 | NA | NA | 1 | -1 |
| A5SS | 2683 | OXLD1 | chr17 | - | 81666149 | 81666637 | 81666517 | 81666637 | 81665035 | 81665584 | 0 | 0 | 0 | 0 | 0 | 1 | 0 | 0 | 2 | 0 | 0 | 0 | 98 | 49 | 0.063770311 | 0.170414363 | NA | NA | 0 | NA | NA | 1 | -1 |
| A5SS | 2688 | OXLD1 | chr17 | - | 81666163 | 81666637 | 81666517 | 81666637 | 81665035 | 81665584 | 0 | 0 | 0 | 0 | 0 | 1 | 0 | 0 | 2 | 0 | 0 | 0 | 98 | 49 | 0.063770311 | 0.170414363 | NA | NA | 0 | NA | NA | 1 | -1 |
| A5SS | 2689 | OXLD1 | chr17 | - | 81666276 | 81666637 | 81666517 | 81666637 | 81665035 | 81665584 | 0 | 0 | 0 | 0 | 0 | 1 | 0 | 0 | 2 | 0 | 0 | 0 | 98 | 49 | 0.063770311 | 0.170414363 | NA | NA | 0 | NA | NA | 1 | -1 |
| A5SS | 2944 | UNKL | chr16 | - | 1369933 | 1370357 | 1370129 | 1370357 | 1367655 | 1367858 | 0 | 1 | 0 | 0 | 0 | 1 | 0 | 0 | 2 | 0 | 0 | 0 | 98 | 49 | 0.063770311 | 0.170414363 | NA | 1 | 0 | NA | NA | 1 | -0.5 |
| RI | 746 | PTPMT1 | chr11 | + | 47565429 | 47565986 | 47565429 | 47565796 | 47565905 | 47565986 | 1 | 0 | 2 | 0 | 1 | 0 | 0 | 2 | 0 | 0 | 0 | 0 | 98 | 49 | 0.063770311 | 0.297702651 | 1 | 0 | 1 | NA | 1 | NA | -0.333 |
| RI | 790 | CCDC125 | chr5 | - | 69320236 | 69320580 | 69320236 | 69320345 | 69320464 | 69320580 | 0 | 0 | 0 | 0 | 0 | 1 | 1 | 1 | 2 | 0 | 0 | 0 | 98 | 49 | 0.063770311 | 0.297702651 | NA | NA | 0 | 1 | 1 | 1 | -1 |
| SE | 255 | SEC16A | chr9 | - | 136468414 | 136468512 | 136466956 | 136467083 | 136471974 | 136472111 | 0 | 0 | 1 | 0 | 1 | 0 | 0 | 2 | 0 | 0 | 0 | 0 | 98 | 49 | 0.063770311 | 0.170760496 | NA | 0 | 1 | NA | 1 | NA | -0.5 |
| SE | 506 | SLC45A4 | chr8 | - | 141221576 | 141221765 | 141219649 | 141219829 | 141253988 | 141254629 | 0 | 0 | 5 | 0 | 1 | 0 | 2 | 2 | 0 | 0 | 0 | 0 | 98 | 49 | 0.063770311 | 0.170760496 | NA | 0 | 1 | 1 | 1 | NA | -0.5 |
| SE | 3157 | XBP1 | chr22 | - | 28796046 | 28796192 | 28794559 | 28795732 | 28797076 | 28797205 | 0 | 0 | 6 | 0 | 1 | 0 | 0 | 2 | 0 | 0 | 0 | 0 | 98 | 49 | 0.063770311 | 0.170760496 | NA | 0 | 1 | NA | 1 | NA | -0.5 |
| SE | 3781 | MYCBP2 | chr13 | - | 77126317 | 77126542 | 77125335 | 77125468 | 77139195 | 77139336 | 0 | 0 | 2 | 0 | 1 | 0 | 0 | 2 | 0 | 0 | 0 | 0 | 98 | 49 | 0.063770311 | 0.170760496 | NA | 0 | 1 | NA | 1 | NA | -0.5 |
| SE | 5029 | BRMS1 | chr11 | - | 66340120 | 66340213 | 66338720 | 66338785 | 66340773 | 66340870 | 0 | 0 | 2 | 0 | 1 | 0 | 0 | 2 | 0 | 0 | 0 | 0 | 98 | 49 | 0.063770311 | 0.170760496 | NA | 0 | 1 | NA | 1 | NA | -0.5 |
| SE | 10606 | ANKFY1 | chr17 | - | 4189381 | 4189479 | 4184817 | 4185046 | 4194977 | 4195177 | 0 | 0 | 0 | 0 | 0 | 1 | 0 | 0 | 2 | 0 | 0 | 0 | 98 | 49 | 0.063770311 | 0.170760496 | NA | NA | 0 | NA | NA | 1 | -1 |
| SE | 13383 | ZNF805 | chr19 | + | 57243922 | 57244049 | 57240684 | 57240921 | 57248604 | 57248700 | 0 | 0 | 0 | 0 | 1 | 0 | 0 | 2 | 0 | 0 | 0 | 0 | 98 | 49 | 0.063770311 | 0.170760496 | NA | 0 | NA | NA | 1 | NA | -1 |
| SE | 12368 | NBEAL2 | chr3 | + | 47007065 | 47007155 | 47006332 | 47006449 | 47007240 | 47007350 | 0 | 2 | 16 | 0 | 0 | 0 | 0 | 8 | 3 | 0 | 1 | 0 | 98 | 49 | 0.063873063 | 0.170760496 | NA | 1 | 1 | NA | 0.8 | 1 | 0.1 |
| SE | 8540 | ARAP3 | chr5 | - | 141655361 | 141655400 | 141653400 | 141654435 | 141655620 | 141655758 | 0 | 0 | 0 | 0 | 1 | 2 | 0 | 2 | 0 | 0 | 0 | 0 | 88 | 49 | 0.063945674 | 0.170760496 | NA | 0 | 0 | NA | 1 | NA | -1 |
| A3SS | 530 | GLRX | chr5 | - | 95813848 | 95814807 | 95813848 | 95814389 | 95816506 | 95816626 | 0 | 0 | 0 | 0 | 0 | 3 | 0 | 0 | 1 | 0 | 1 | 0 | 98 | 49 | 0.064268453 | 0.197717869 | NA | NA | 0 | NA | 0 | 1 | -0.5 |
| A3SS | 531 | GLRX | chr5 | - | 95813848 | 95814957 | 95813848 | 95814389 | 95816506 | 95816626 | 0 | 0 | 0 | 0 | 0 | 3 | 0 | 0 | 1 | 0 | 1 | 0 | 98 | 49 | 0.064268453 | 0.197717869 | NA | NA | 0 | NA | 0 | 1 | -0.5 |
| SE | 5261 | PCSK7 | chr11 | - | 117208896 | 117209053 | 117207969 | 117208064 | 117218465 | 117218568 | 0 | 0 | 0 | 0 | 0 | 3 | 0 | 0 | 1 | 0 | 0 | 0 | 98 | 49 | 0.064268453 | 0.170760496 | NA | NA | 0 | NA | NA | 1 | -1 |
| SE | 6387 | PSRC1 | chr1 | - | 109280395 | 109280489 | 109279553 | 109280156 | 109280776 | 109280916 | 0 | 0 | 1 | 0 | 3 | 0 | 0 | 1 | 0 | 0 | 0 | 0 | 98 | 49 | 0.064268453 | 0.170760496 | NA | 0 | 1 | NA | 1 | NA | -0.5 |
| A5SS | 1969 | SLC35A2 | chrX | - | 48911506 | 48911958 | 48911545 | 48911958 | 48909813 | 48909996 | 0 | 0 | 0 | 0 | 0 | 3 | 0 | 0 | 1 | 0 | 0 | 0 | 88 | 49 | 0.064272896 | 0.170414363 | NA | NA | 0 | NA | NA | 1 | -1 |
| A5SS | 1774 | SF1 | chr11 | - | 64767566 | 64767844 | 64767570 | 64767844 | 64767191 | 64767251 | 0 | 0 | 0 | 0 | 0 | 3 | 6 | 0 | 1 | 0 | 2 | 0 | 53 | 49 | 0.064284969 | 0.170414363 | NA | NA | 0 | 1 | 0 | 1 | -0.667 |
| SE | 10332 | CYTIP | chr2 | - | 157434369 | 157434424 | 157430859 | 157430962 | 157434697 | 157434747 | 0 | 9 | 93 | 0 | 0 | 0 | 18 | 103 | 8 | 0 | 1 | 0 | 98 | 49 | 0.064528457 | 0.170760496 | NA | 1 | 1 | 1 | 0.981 | 1 | 0.006 |
| SE | 11921 | RBM39 | chr20 | - | 35729461 | 35729527 | 35729311 | 35729365 | 35731940 | 35732135 | 0 | 3 | 27 | 0 | 0 | 0 | 2 | 37 | 2 | 0 | 1 | 0 | 98 | 49 | 0.064578009 | 0.170760496 | NA | 1 | 1 | 1 | 0.949 | 1 | 0.017 |
| SE | 8709 | CD74 | chr5 | - | 150403120 | 150403312 | 150402562 | 150402625 | 150404679 | 150404767 | 0 | 2 | 7 | 0 | 2 | 32 | 1 | 1 | 0 | 0 | 20 | 0 | 98 | 49 | 0.064910289 | 0.170760496 | NA | 0.333 | 0.099 | 1 | 0.024 | NA | -0.296 |
| SE | 250 | NOTCH1 | chr9 | - | 136515980 | 136516094 | 136515482 | 136515716 | 136517271 | 136517385 | 1 | 1 | 10 | 0 | 0 | 0 | 0 | 2 | 1 | 0 | 1 | 0 | 98 | 49 | 0.065113604 | 0.170760496 | 1 | 1 | 1 | NA | 0.5 | 1 | 0.25 |
| SE | 6697 | FCGR2A | chr1 | + | 161506333 | 161506591 | 161505986 | 161506007 | 161509819 | 161510074 | 0 | 2 | 42 | 0 | 0 | 0 | 1 | 17 | 3 | 0 | 1 | 0 | 98 | 49 | 0.065339874 | 0.170760496 | NA | 1 | 1 | 1 | 0.895 | 1 | 0.035 |
| SE | 12941 | SMCHD1 | chr18 | + | 2694526 | 2694693 | 2688627 | 2688747 | 2697031 | 2697122 | 0 | 3 | 97 | 0 | 0 | 0 | 0 | 55 | 1 | 0 | 1 | 0 | 98 | 49 | 0.065369701 | 0.170760496 | NA | 1 | 1 | NA | 0.965 | 1 | 0.018 |
| A3SS | 2057 | FBXO7 | chr22 | + | 32478980 | 32479275 | 32479132 | 32479275 | 32474719 | 32475124 | 0 | 1 | 1 | 0 | 0 | 0 | 0 | 0 | 0 | 0 | 1 | 0 | 98 | 49 | 0.065790241 | 0.197717869 | NA | 1 | 1 | NA | 0 | NA | 1 |
| A3SS | 2058 | FBXO7 | chr22 | + | 32478980 | 32479275 | 32479132 | 32479275 | 32475236 | 32475398 | 0 | 1 | 1 | 0 | 0 | 0 | 0 | 0 | 0 | 0 | 1 | 0 | 98 | 49 | 0.065790241 | 0.197717869 | NA | 1 | 1 | NA | 0 | NA | 1 |
| A3SS | 2827 | APOL3 | chr22 | - | 36149045 | 36149371 | 36149045 | 36149142 | 36160668 | 36160929 | 1 | 1 | 0 | 0 | 0 | 0 | 0 | 0 | 0 | 0 | 1 | 0 | 98 | 49 | 0.065790241 | 0.197717869 | 1 | 1 | NA | NA | 0 | NA | 1 |
| A3SS | 3029 | POLM | chr7 | - | 44072061 | 44073430 | 44072061 | 44073377 | 44073624 | 44073708 | 1 | 0 | 1 | 0 | 0 | 0 | 0 | 3 | 0 | 0 | 0 | 1 | 98 | 49 | 0.065790241 | 0.197717869 | 1 | NA | 1 | NA | 1 | 0 | 0.5 |
| A5SS | 1566 | POLD4 | chr11 | - | 67353293 | 67353596 | 67353363 | 67353596 | 67352987 | 67353077 | 0 | 1 | 1 | 0 | 0 | 0 | 0 | 0 | 0 | 0 | 0 | 1 | 98 | 49 | 0.065790241 | 0.170414363 | NA | 1 | 1 | NA | NA | 0 | 1 |
| A5SS | 2228 | USP8 | chr15 | + | 50424376 | 50424654 | 50424376 | 50424514 | 50439008 | 50439177 | 0 | 0 | 1 | 0 | 0 | 0 | 0 | 0 | 0 | 0 | 3 | 1 | 98 | 49 | 0.065790241 | 0.170414363 | NA | NA | 1 | NA | 0 | 0 | 1 |
| A5SS | 2719 | NBPF11 | chr1 | - | 148110377 | 148110541 | 148110457 | 148110541 | 148109283 | 148109335 | 0 | 1 | 0 | 0 | 0 | 0 | 0 | 0 | 0 | 0 | 1 | 1 | 98 | 49 | 0.065790241 | 0.170414363 | NA | 1 | NA | NA | 0 | 0 | 1 |
| SE | 1021 | PPIA | chr7 | + | 44798747 | 44798904 | 44796635 | 44796793 | 44799246 | 44799277 | 0 | 0 | 1 | 0 | 0 | 0 | 0 | 0 | 0 | 0 | 1 | 1 | 98 | 49 | 0.065790241 | 0.170760496 | NA | NA | 1 | NA | 0 | 0 | 1 |
| SE | 2879 | ACO2 | chr22 | + | 41522829 | 41522987 | 41520170 | 41520276 | 41523204 | 41523278 | 0 | 1 | 2 | 0 | 0 | 0 | 0 | 0 | 0 | 0 | 1 | 0 | 98 | 49 | 0.065790241 | 0.170760496 | NA | 1 | 1 | NA | 0 | NA | 1 |
| SE | 4101 | FAM13B | chr5 | - | 138018954 | 138019146 | 138018301 | 138018514 | 138021030 | 138021197 | 0 | 0 | 1 | 0 | 0 | 0 | 0 | 1 | 0 | 3 | 0 | 1 | 98 | 49 | 0.065790241 | 0.170760496 | NA | NA | 1 | 0 | 1 | 0 | 0.667 |
| SE | 4610 | HSPA8 | chr11 | - | 123060592 | 123060798 | 123060115 | 123060268 | 123061119 | 123061329 | 0 | 0 | 1 | 0 | 0 | 0 | 0 | 0 | 0 | 0 | 2 | 1 | 98 | 49 | 0.065790241 | 0.170760496 | NA | NA | 1 | NA | 0 | 0 | 1 |
| SE | 6531 | FRMD8 | chr11 | + | 65393572 | 65393674 | 65389360 | 65389528 | 65394040 | 65394099 | 0 | 1 | 3 | 0 | 0 | 0 | 0 | 0 | 0 | 0 | 1 | 0 | 98 | 49 | 0.065790241 | 0.170760496 | NA | 1 | 1 | NA | 0 | NA | 1 |
| SE | 7131 | GGA3 | chr17 | - | 75243062 | 75243166 | 75242830 | 75242911 | 75243446 | 75243570 | 0 | 1 | 5 | 0 | 0 | 0 | 0 | 0 | 0 | 0 | 1 | 0 | 98 | 49 | 0.065790241 | 0.170760496 | NA | 1 | 1 | NA | 0 | NA | 1 |
| SE | 7595 | ATG16L2 | chr11 | + | 72826702 | 72826823 | 72826172 | 72826243 | 72828358 | 72828508 | 0 | 0 | 1 | 0 | 0 | 0 | 0 | 2 | 0 | 0 | 1 | 1 | 98 | 49 | 0.065790241 | 0.170760496 | NA | NA | 1 | NA | 0.5 | 0 | 0.75 |
| SE | 8118 | POLE | chr12 | - | 132648928 | 132649072 | 132643836 | 132643977 | 132649305 | 132649515 | 0 | 1 | 1 | 0 | 0 | 0 | 0 | 0 | 0 | 0 | 1 | 0 | 98 | 49 | 0.065790241 | 0.170760496 | NA | 1 | 1 | NA | 0 | NA | 1 |
| SE | 8179 | MLXIP | chr12 | + | 122137468 | 122137590 | 122135488 | 122135666 | 122138193 | 122138295 | 0 | 0 | 1 | 0 | 0 | 0 | 0 | 0 | 0 | 0 | 0 | 1 | 98 | 49 | 0.065790241 | 0.170760496 | NA | NA | 1 | NA | NA | 0 | 1 |
| SE | 8364 | STRAP | chr12 | + | 15883540 | 15883676 | 15882353 | 15882819 | 15889927 | 15890009 | 0 | 0 | 1 | 0 | 0 | 0 | 0 | 2 | 0 | 0 | 1 | 1 | 98 | 49 | 0.065790241 | 0.170760496 | NA | NA | 1 | NA | 0.5 | 0 | 0.75 |
| SE | 10100 | GGA1 | chr22 | + | 37625796 | 37625949 | 37624968 | 37625076 | 37629461 | 37629526 | 0 | 1 | 1 | 0 | 0 | 0 | 0 | 0 | 0 | 0 | 1 | 0 | 98 | 49 | 0.065790241 | 0.170760496 | NA | 1 | 1 | NA | 0 | NA | 1 |
| SE | 10460 | OSBPL11 | chr3 | - | 125547405 | 125547592 | 125538450 | 125538633 | 125560378 | 125560519 | 0 | 1 | 2 | 0 | 0 | 0 | 0 | 0 | 0 | 0 | 1 | 0 | 98 | 49 | 0.065790241 | 0.170760496 | NA | 1 | 1 | NA | 0 | NA | 1 |
| SE | 10539 | P4HA1 | chr10 | - | 73051052 | 73051249 | 73046924 | 73047101 | 73053350 | 73053590 | 0 | 0 | 1 | 0 | 0 | 0 | 0 | 0 | 0 | 0 | 0 | 1 | 98 | 49 | 0.065790241 | 0.170760496 | NA | NA | 1 | NA | NA | 0 | 1 |
| SE | 11584 | FANCA | chr16 | - | 89742799 | 89742938 | 89740803 | 89740866 | 89744958 | 89745071 | 0 | 1 | 1 | 0 | 0 | 0 | 0 | 0 | 0 | 0 | 1 | 0 | 98 | 49 | 0.065790241 | 0.170760496 | NA | 1 | 1 | NA | 0 | NA | 1 |
| SE | 13943 | PDK1 | chr2 | + | 172562740 | 172562800 | 172562219 | 172562291 | 172564977 | 172565073 | 0 | 1 | 0 | 0 | 0 | 1 | 0 | 0 | 0 | 0 | 1 | 0 | 98 | 49 | 0.065790241 | 0.170760496 | NA | 1 | 0 | NA | 0 | NA | 0.5 |
| A3SS | 2853 | TGOLN2 | chr2 | - | 85318017 | 85322799 | 85318017 | 85322741 | 85324914 | 85324998 | 0 | 0 | 0 | 0 | 0 | 2 | 0 | 2 | 1 | 2 | 4 | 0 | 98 | 49 | 0.0658956 | 0.197717869 | NA | NA | 0 | 0 | 0.2 | 1 | -0.4 |
| A5SS | 1409 | PHYKPL | chr5 | - | 178231668 | 178232822 | 178232491 | 178232822 | 178231404 | 178231523 | 0 | 0 | 3 | 0 | 2 | 0 | 0 | 1 | 0 | 0 | 0 | 0 | 98 | 49 | 0.0658956 | 0.170414363 | NA | 0 | 1 | NA | 1 | NA | -0.5 |
| SE | 145 | SNX12 | chrX | - | 71061842 | 71061967 | 71059246 | 71061118 | 71062853 | 71062949 | 0 | 0 | 3 | 0 | 2 | 0 | 0 | 1 | 0 | 0 | 0 | 0 | 98 | 49 | 0.0658956 | 0.170760496 | NA | 0 | 1 | NA | 1 | NA | -0.5 |
| SE | 239 | NDOR1 | chr9 | + | 137215406 | 137215521 | 137215094 | 137215202 | 137215658 | 137215805 | 0 | 0 | 0 | 0 | 2 | 0 | 0 | 1 | 0 | 0 | 0 | 0 | 98 | 49 | 0.0658956 | 0.170760496 | NA | 0 | NA | NA | 1 | NA | -1 |
| SE | 436 | UBE2R2 | chr9 | + | 33911963 | 33912098 | 33900173 | 33900271 | 33917017 | 33920403 | 0 | 0 | 6 | 0 | 2 | 1 | 0 | 1 | 0 | 0 | 0 | 0 | 98 | 49 | 0.0658956 | 0.170760496 | NA | 0 | 0.75 | NA | 1 | NA | -0.625 |
| SE | 4582 | ST3GAL4 | chr11 | + | 126409267 | 126409411 | 126408306 | 126408496 | 126413504 | 126413648 | 0 | 0 | 3 | 0 | 2 | 0 | 0 | 1 | 0 | 0 | 0 | 0 | 98 | 49 | 0.0658956 | 0.170760496 | NA | 0 | 1 | NA | 1 | NA | -0.5 |
| SE | 7326 | SREBF1 | chr17 | - | 17819012 | 17819234 | 17818259 | 17818374 | 17819319 | 17819454 | 0 | 0 | 0 | 0 | 2 | 0 | 0 | 1 | 0 | 0 | 0 | 0 | 98 | 49 | 0.0658956 | 0.170760496 | NA | 0 | NA | NA | 1 | NA | -1 |
| SE | 8223 | MAPK3 | chr16 | - | 30118046 | 30118163 | 30117669 | 30117784 | 30118348 | 30118538 | 0 | 0 | 0 | 0 | 0 | 2 | 0 | 0 | 1 | 0 | 0 | 0 | 98 | 49 | 0.0658956 | 0.170760496 | NA | NA | 0 | NA | NA | 1 | -1 |
| SE | 11719 | CHD3 | chr17 | + | 7907600 | 7907702 | 7907352 | 7907488 | 7907893 | 7908019 | 0 | 0 | 0 | 0 | 0 | 2 | 0 | 0 | 1 | 0 | 0 | 0 | 98 | 49 | 0.0658956 | 0.170760496 | NA | NA | 0 | NA | NA | 1 | -1 |
| A3SS | 81 | GBGT1 | chr9 | - | 133156014 | 133156102 | 133156014 | 133156065 | 133161466 | 133161532 | 0 | 0 | 0 | 0 | 0 | 2 | 0 | 0 | 1 | 0 | 1 | 0 | 86 | 49 | 0.065901144 | 0.197717869 | NA | NA | 0 | NA | 0 | 1 | -0.5 |
| SE | 4442 | CORO7 | chr16 | - | 4357967 | 4358103 | 4357167 | 4357259 | 4358366 | 4358483 | 0 | 1 | 4 | 0 | 0 | 0 | 5 | 8 | 2 | 0 | 1 | 0 | 98 | 49 | 0.067252723 | 0.171142152 | NA | 1 | 1 | 1 | 0.8 | 1 | 0.067 |
| SE | 8028 | GNAS | chr20 | + | 58898940 | 58898985 | 58895611 | 58895684 | 58903530 | 58903585 | 0 | 1 | 3 | 0 | 0 | 2 | 0 | 0 | 0 | 0 | 1 | 0 | 94 | 49 | 0.067458351 | 0.171142152 | NA | 1 | 0.439 | NA | 0 | NA | 0.72 |
| SE | 911 | LMTK2 | chr7 | + | 98171540 | 98171674 | 98159337 | 98159425 | 98185050 | 98185135 | 0 | 1 | 2 | 0 | 0 | 0 | 0 | 3 | 2 | 0 | 2 | 0 | 98 | 49 | 0.068013122 | 0.171142152 | NA | 1 | 1 | NA | 0.429 | 1 | 0.285 |
| SE | 2940 | LIMK2 | chr22 | + | 31273451 | 31273507 | 31272529 | 31272704 | 31275150 | 31275308 | 0 | 13 | 40 | 0 | 0 | 0 | 3 | 93 | 36 | 0 | 1 | 0 | 98 | 49 | 0.068060835 | 0.171142152 | NA | 1 | 1 | 1 | 0.979 | 1 | 0.007 |
| SE | 2968 | AP1B1 | chr22 | - | 29329711 | 29329720 | 29327679 | 29328895 | 29330377 | 29330532 | 0 | 0 | 1 | 0 | 0 | 0 | 0 | 0 | 0 | 0 | 0 | 1 | 58 | 49 | 0.068063814 | 0.171142152 | NA | NA | 1 | NA | NA | 0 | 1 |
| SE | 12329 | ARHGDIA | chr17 | - | 81869329 | 81869406 | 81869172 | 81869236 | 81869541 | 81869625 | 0 | 0 | 21 | 0 | 0 | 1 | 0 | 11 | 1 | 0 | 0 | 2 | 98 | 49 | 0.068249658 | 0.171142152 | NA | NA | 0.913 | NA | 1 | 0.2 | 0.313 |
| SE | 5783 | DENND3 | chr8 | + | 141138021 | 141138137 | 141136557 | 141136791 | 141151618 | 141151837 | 0 | 3 | 15 | 0 | 0 | 0 | 0 | 19 | 7 | 0 | 1 | 0 | 98 | 49 | 0.068459154 | 0.171142152 | NA | 1 | 1 | NA | 0.905 | 1 | 0.047 |
| A3SS | 1319 | P2RX7 | chr12 | + | 121162270 | 121162520 | 121162423 | 121162520 | 121160901 | 121160974 | 0 | 0 | 0 | 0 | 0 | 1 | 0 | 0 | 1 | 0 | 0 | 0 | 98 | 49 | 0.068700612 | 0.197717869 | NA | NA | 0 | NA | NA | 1 | -1 |
| A3SS | 3364 | CCL4L2 | chr17 | + | 36212292 | 36212878 | 36212496 | 36212878 | 36211775 | 36211890 | 0 | 0 | 0 | 0 | 0 | 1 | 2 | 4 | 1 | 0 | 0 | 0 | 98 | 49 | 0.068700612 | 0.197717869 | NA | NA | 0 | 1 | 1 | 1 | -1 |
| A3SS | 3368 | CCL4L2 | chr17 | + | 36212302 | 36212878 | 36212496 | 36212878 | 36211775 | 36211890 | 0 | 0 | 0 | 0 | 0 | 1 | 2 | 4 | 1 | 0 | 0 | 0 | 98 | 49 | 0.068700612 | 0.197717869 | NA | NA | 0 | 1 | 1 | 1 | -1 |
| A3SS | 3371 | CCL4L2 | chr17 | + | 36212312 | 36212878 | 36212496 | 36212878 | 36211775 | 36211890 | 0 | 0 | 0 | 0 | 0 | 1 | 2 | 4 | 1 | 0 | 0 | 0 | 98 | 49 | 0.068700612 | 0.197717869 | NA | NA | 0 | 1 | 1 | 1 | -1 |
| A3SS | 3373 | CCL4L2 | chr17 | + | 36212333 | 36212878 | 36212496 | 36212878 | 36211775 | 36211890 | 0 | 0 | 0 | 0 | 0 | 1 | 2 | 4 | 1 | 0 | 0 | 0 | 98 | 49 | 0.068700612 | 0.197717869 | NA | NA | 0 | 1 | 1 | 1 | -1 |
| A3SS | 3374 | CCL4L2 | chr17 | + | 36212363 | 36212878 | 36212496 | 36212878 | 36211775 | 36211890 | 0 | 0 | 0 | 0 | 0 | 1 | 2 | 4 | 1 | 0 | 0 | 0 | 98 | 49 | 0.068700612 | 0.197717869 | NA | NA | 0 | 1 | 1 | 1 | -1 |
| A3SS | 3842 | CARKD | chr13 | + | 110638241 | 110639995 | 110638377 | 110639995 | 110637128 | 110637249 | 0 | 0 | 0 | 0 | 0 | 1 | 0 | 0 | 1 | 0 | 0 | 0 | 98 | 49 | 0.068700612 | 0.197717869 | NA | NA | 0 | NA | NA | 1 | -1 |
| A3SS | 3925 | C10orf128 | chr10 | - | 49154723 | 49156259 | 49154723 | 49155909 | 49161598 | 49161622 | 0 | 0 | 0 | 0 | 0 | 1 | 7 | 0 | 1 | 0 | 0 | 0 | 98 | 49 | 0.068700612 | 0.197717869 | NA | NA | 0 | 1 | NA | 1 | -1 |
| A5SS | 407 | ARHGAP26 | chr5 | + | 143207197 | 143207473 | 143207197 | 143207308 | 143213996 | 143214088 | 0 | 0 | 0 | 0 | 0 | 1 | 0 | 0 | 1 | 0 | 1 | 0 | 98 | 49 | 0.068700612 | 0.170414363 | NA | NA | 0 | NA | 0 | 1 | -0.5 |
| A5SS | 1221 | C6orf203 | chr6 | + | 107028171 | 107028797 | 107028171 | 107028271 | 107039748 | 107040230 | 0 | 0 | 0 | 0 | 0 | 1 | 0 | 0 | 1 | 0 | 0 | 0 | 98 | 49 | 0.068700612 | 0.170414363 | NA | NA | 0 | NA | NA | 1 | -1 |
| A5SS | 1816 | POLR2F | chr22 | + | 37956772 | 37956901 | 37956772 | 37956842 | 37959345 | 37959476 | 0 | 0 | 0 | 0 | 0 | 1 | 0 | 0 | 1 | 2 | 0 | 0 | 98 | 49 | 0.068700612 | 0.170414363 | NA | NA | 0 | 0 | NA | 1 | -0.5 |
| A5SS | 2218 | TRIM69 | chr15 | + | 44758612 | 44758854 | 44758612 | 44758803 | 44759639 | 44759662 | 0 | 0 | 0 | 0 | 0 | 1 | 0 | 2 | 1 | 0 | 0 | 0 | 98 | 49 | 0.068700612 | 0.170414363 | NA | NA | 0 | NA | 1 | 1 | -1 |
| RI | 334 | ARL6IP4 | chr12 | + | 122982468 | 122982913 | 122982468 | 122982538 | 122982619 | 122982913 | 0 | 0 | 0 | 0 | 0 | 1 | 0 | 0 | 1 | 0 | 1 | 0 | 98 | 49 | 0.068700612 | 0.297702651 | NA | NA | 0 | NA | 0 | 1 | -0.5 |
| RI | 369 | RECQL | chr12 | - | 21501169 | 21501669 | 21501169 | 21501215 | 21501412 | 21501669 | 0 | 0 | 0 | 0 | 0 | 1 | 2 | 1 | 1 | 0 | 1 | 0 | 98 | 49 | 0.068700612 | 0.297702651 | NA | NA | 0 | 1 | 0.333 | 1 | -0.778 |
| RI | 581 | TK2 | chr16 | - | 66548977 | 66550412 | 66548977 | 66549009 | 66549937 | 66550412 | 0 | 0 | 0 | 0 | 0 | 1 | 0 | 3 | 1 | 0 | 0 | 0 | 98 | 49 | 0.068700612 | 0.297702651 | NA | NA | 0 | NA | 1 | 1 | -1 |
| RI | 682 | NAP1L1 | chr12 | - | 76045382 | 76048464 | 76045382 | 76047718 | 76048392 | 76048464 | 0 | 0 | 0 | 0 | 0 | 1 | 0 | 2 | 1 | 0 | 2 | 0 | 98 | 49 | 0.068700612 | 0.297702651 | NA | NA | 0 | NA | 0.333 | 1 | -0.666 |
| RI | 968 | CCL4L2 | chr17 | + | 36211775 | 36212878 | 36211775 | 36211890 | 36212496 | 36212878 | 0 | 0 | 0 | 0 | 0 | 1 | 2 | 4 | 1 | 0 | 0 | 0 | 98 | 49 | 0.068700612 | 0.297702651 | NA | NA | 0 | 1 | 1 | 1 | -1 |
| RI | 1065 | CPT1B | chr22 | - | 50568860 | 50569421 | 50568860 | 50569081 | 50569335 | 50569421 | 0 | 0 | 0 | 0 | 2 | 1 | 0 | 0 | 1 | 2 | 0 | 0 | 98 | 49 | 0.068700612 | 0.297702651 | NA | 0 | 0 | 0 | NA | 1 | -0.5 |
| RI | 1071 | NFKBIB | chr19 | + | 38907398 | 38908894 | 38907398 | 38907659 | 38908730 | 38908894 | 0 | 0 | 0 | 0 | 0 | 1 | 0 | 2 | 1 | 0 | 0 | 0 | 98 | 49 | 0.068700612 | 0.297702651 | NA | NA | 0 | NA | 1 | 1 | -1 |
| RI | 1115 | DCTN1 | chr2 | - | 74366243 | 74366620 | 74366243 | 74366375 | 74366458 | 74366620 | 0 | 0 | 1 | 0 | 1 | 1 | 1 | 1 | 0 | 0 | 0 | 0 | 98 | 49 | 0.068700612 | 0.297702651 | NA | 0 | 0.333 | 1 | 1 | NA | -0.834 |
| SE | 689 | PTK2B | chr8 | + | 27445793 | 27445919 | 27444205 | 27444271 | 27450748 | 27450895 | 0 | 0 | 0 | 0 | 1 | 2 | 0 | 1 | 0 | 0 | 0 | 0 | 98 | 49 | 0.068700612 | 0.171142152 | NA | 0 | 0 | NA | 1 | NA | -1 |
| SE | 969 | RHBDD2 | chr7 | + | 75881364 | 75881486 | 75878998 | 75879260 | 75881828 | 75882236 | 0 | 0 | 2 | 0 | 1 | 0 | 2 | 1 | 0 | 0 | 0 | 0 | 98 | 49 | 0.068700612 | 0.171142152 | NA | 0 | 1 | 1 | 1 | NA | -0.5 |
| SE | 3923 | HAGH | chr16 | - | 1826506 | 1826602 | 1822864 | 1823037 | 1826711 | 1827194 | 0 | 0 | 0 | 0 | 1 | 0 | 0 | 1 | 0 | 0 | 0 | 0 | 98 | 49 | 0.068700612 | 0.171142152 | NA | 0 | NA | NA | 1 | NA | -1 |
| SE | 3999 | ASL | chr7 | + | 66086743 | 66086821 | 66086584 | 66086662 | 66087333 | 66087386 | 0 | 0 | 2 | 0 | 1 | 0 | 0 | 1 | 0 | 0 | 0 | 0 | 98 | 49 | 0.068700612 | 0.171142152 | NA | 0 | 1 | NA | 1 | NA | -0.5 |
| SE | 4928 | FCRL6 | chr1 | + | 159808960 | 159809245 | 159808177 | 159808444 | 159809401 | 159809683 | 0 | 0 | 0 | 0 | 0 | 1 | 0 | 0 | 1 | 0 | 0 | 0 | 98 | 49 | 0.068700612 | 0.171142152 | NA | NA | 0 | NA | NA | 1 | -1 |
| SE | 5379 | SYNE1 | chr6 | - | 152455885 | 152456044 | 152455425 | 152455590 | 152458756 | 152458930 | 0 | 0 | 1 | 0 | 1 | 0 | 0 | 1 | 0 | 0 | 0 | 0 | 98 | 49 | 0.068700612 | 0.171142152 | NA | 0 | 1 | NA | 1 | NA | -0.5 |
| SE | 6602 | CCDC88B | chr11 | + | 64351159 | 64351255 | 64349550 | 64349668 | 64351475 | 64351616 | 0 | 0 | 0 | 0 | 0 | 1 | 0 | 5 | 1 | 0 | 0 | 0 | 98 | 49 | 0.068700612 | 0.171142152 | NA | NA | 0 | NA | 1 | 1 | -1 |
| SE | 6680 | TTN-AS1 | chr2 | + | 178571299 | 178571449 | 178542221 | 178542361 | 178597595 | 178597681 | 0 | 0 | 2 | 0 | 1 | 0 | 0 | 1 | 0 | 0 | 0 | 0 | 98 | 49 | 0.068700612 | 0.171142152 | NA | 0 | 1 | NA | 1 | NA | -0.5 |
| SE | 7590 | UBE4B | chr1 | + | 10158355 | 10158482 | 10151325 | 10151561 | 10161141 | 10161286 | 0 | 0 | 0 | 0 | 0 | 1 | 0 | 0 | 1 | 0 | 1 | 0 | 98 | 49 | 0.068700612 | 0.171142152 | NA | NA | 0 | NA | 0 | 1 | -0.5 |
| SE | 10301 | MARK3 | chr14 | + | 103491776 | 103492034 | 103480386 | 103480490 | 103502881 | 103503829 | 0 | 0 | 0 | 0 | 0 | 1 | 0 | 1 | 1 | 0 | 0 | 0 | 98 | 49 | 0.068700612 | 0.171142152 | NA | NA | 0 | NA | 1 | 1 | -1 |
| SE | 11137 | CLEC16A | chr16 | + | 10977224 | 10977399 | 10972931 | 10973061 | 10979328 | 10979382 | 0 | 0 | 0 | 0 | 1 | 0 | 0 | 1 | 0 | 0 | 0 | 0 | 98 | 49 | 0.068700612 | 0.171142152 | NA | 0 | NA | NA | 1 | NA | -1 |
| SE | 11367 | LONP2 | chr16 | + | 48256609 | 48256741 | 48252130 | 48252365 | 48258617 | 48258740 | 0 | 0 | 0 | 0 | 0 | 1 | 0 | 0 | 1 | 0 | 1 | 0 | 98 | 49 | 0.068700612 | 0.171142152 | NA | NA | 0 | NA | 0 | 1 | -0.5 |
| SE | 12842 | SMARCA4 | chr19 | + | 11033766 | 11033865 | 11033289 | 11033517 | 11034122 | 11034200 | 0 | 0 | 0 | 0 | 0 | 1 | 0 | 0 | 1 | 0 | 0 | 0 | 98 | 49 | 0.068700612 | 0.171142152 | NA | NA | 0 | NA | NA | 1 | -1 |
| SE | 13356 | TMEM86B | chr19 | - | 55228190 | 55228437 | 55226633 | 55227563 | 55228690 | 55229264 | 0 | 0 | 1 | 0 | 1 | 0 | 0 | 1 | 0 | 0 | 0 | 0 | 98 | 49 | 0.068700612 | 0.171142152 | NA | 0 | 1 | NA | 1 | NA | -0.5 |
| SE | 13996 | QRICH1 | chr3 | - | 49092396 | 49092456 | 49076708 | 49077038 | 49093911 | 49094071 | 0 | 0 | 0 | 0 | 1 | 0 | 0 | 1 | 0 | 0 | 0 | 0 | 98 | 49 | 0.068700612 | 0.171142152 | NA | 0 | NA | NA | 1 | NA | -1 |
| SE | 14058 | ATG2B | chr14 | - | 96322111 | 96322254 | 96317697 | 96317855 | 96322539 | 96322735 | 0 | 0 | 0 | 0 | 1 | 0 | 0 | 1 | 0 | 0 | 0 | 0 | 98 | 49 | 0.068700612 | 0.171142152 | NA | 0 | NA | NA | 1 | NA | -1 |
| A5SS | 1424 | C7orf49 | chr7 | - | 135168748 | 135169060 | 135168785 | 135169060 | 135167731 | 135167807 | 0 | 0 | 0 | 0 | 0 | 1 | 0 | 6 | 1 | 0 | 0 | 0 | 86 | 49 | 0.068703243 | 0.170414363 | NA | NA | 0 | NA | 1 | 1 | -1 |
| A5SS | 3076 | NCEH1 | chr3 | - | 172647861 | 172648114 | 172647885 | 172648114 | 172645622 | 172645692 | 0 | 0 | 0 | 0 | 0 | 1 | 0 | 0 | 1 | 0 | 0 | 0 | 73 | 49 | 0.06870403 | 0.170414363 | NA | NA | 0 | NA | NA | 1 | -1 |
| A5SS | 903 | ATF7 | chr12 | - | 53543296 | 53543448 | 53543329 | 53543448 | 53537414 | 53537552 | 1 | 0 | 0 | 0 | 0 | 1 | 0 | 0 | 1 | 0 | 0 | 0 | 82 | 49 | 0.068705526 | 0.170414363 | 1 | NA | 0 | NA | NA | 1 | -0.5 |
| A3SS | 2808 | SRRT | chr7 | + | 100885212 | 100885370 | 100885215 | 100885370 | 100884922 | 100885040 | 0 | 0 | 0 | 0 | 0 | 1 | 0 | 4 | 1 | 0 | 2 | 0 | 52 | 49 | 0.068726673 | 0.197717869 | NA | NA | 0 | NA | 0.653 | 1 | -0.827 |

| | | | | | | | | | | | | | | | | | | | | | | | | | | | | | | | | | |
|---|---|---|---|---|---|---|---|---|---|---|---|---|---|---|---|---|---|---|---|---|---|---|---|---|---|---|---|---|---|---|---|---|---|
| A3SS | 3999 | SERPINA1 | chr14 | - | 94388559 | 94388663 | 94388559 | 94388660 | 94390456 | 94390692 | 0 | 0 | 5 | 0 | 1 | 3 | 0 | 1 | 0 | 4 | 0 | 0 | 52 | 49 | 0.068726673 | 0.197717869 | NA | 0 | 0.611 | 0 | 1 | NA | -0.195 |
| SE | 9686 | IFRD1 | chr7 | + | 112450507 | 112450782 | 112423143 | 112423432 | 112455762 | 112455867 | 0 | 3 | 58 | 0 | 0 | 0 | 0 | 33 | 5 | 0 | 1 | 0 | 98 | 49 | 0.068775688 | 0.171142152 | NA | 1 | 1 | NA | 0.943 | 1 | 0.028 |
| A5SS | 2089 | CLASRP | chr19 | + | 45068014 | 45068058 | 45068014 | 45068054 | 45068419 | 45068480 | 0 | 0 | 0 | 0 | 0 | 1 | 0 | 1 | 1 | 0 | 0 | 0 | 53 | 49 | 0.068946269 | 0.170414363 | NA | NA | 0 | NA | 1 | 1 | -1 |
| SE | 11969 | RARA | chr17 | + | 40348315 | 40348464 | 40342018 | 40342867 | 40349783 | 40349925 | 0 | 2 | 18 | 0 | 0 | 0 | 0 | 9 | 4 | 0 | 1 | 0 | 98 | 49 | 0.069039705 | 0.17126162 | NA | 1 | 1 | NA | 0.818 | 1 | 0.091 |
| SE | 3617 | NIN | chr14 | - | 50772300 | 50772468 | 50771331 | 50771468 | 50772948 | 50773095 | 0 | 1 | 12 | 0 | 0 | 0 | 0 | 13 | 1 | 0 | 1 | 0 | 98 | 49 | 0.069097631 | 0.17126162 | NA | 1 | 1 | NA | 0.867 | 1 | 0.067 |
| SE | 784 | ZYX | chr7 | + | 143388218 | 143388339 | 143382800 | 143383322 | 143388488 | 143388658 | 0 | 5 | 34 | 0 | 0 | 0 | 0 | 9 | 7 | 0 | 1 | 0 | 98 | 49 | 0.069165538 | 0.17126162 | NA | 1 | 1 | NA | 0.818 | 1 | 0.091 |
| A3SS | 2597 | FERMT3 | chr11 | + | 64219878 | 64220015 | 64219890 | 64220015 | 64219739 | 64219789 | 0 | 2 | 1 | 2 | 8 | 9 | 0 | 0 | 0 | 3 | 6 | 6 | 61 | 49 | 0.069201254 | 0.197717869 | 0 | 0.167 | 0.082 | 0 | 0 | 0 | 0.083 |
| SE | 2329 | ATP11B | chr3 | + | 182842074 | 182842122 | 182837070 | 182837174 | 182845457 | 182845522 | 0 | 1 | 22 | 0 | 0 | 0 | 0 | 6 | 2 | 0 | 1 | 0 | 97 | 49 | 0.069320212 | 0.171362302 | NA | 1 | 1 | NA | 0.752 | 1 | 0.124 |
| SE | 5624 | ETF1 | chr5 | - | 138517560 | 138517700 | 138513567 | 138513706 | 138518691 | 138518867 | 0 | 0 | 12 | 0 | 2 | 0 | 0 | 10 | 1 | 0 | 1 | 0 | 98 | 49 | 0.069479149 | 0.17144651 | NA | 0 | 1 | NA | 0.833 | 1 | -0.416 |
| SE | 1355 | MAPK14 | chr6 | + | 36102570 | 36102649 | 36095986 | 36096066 | 36107454 | 36107628 | 0 | 1 | 19 | 0 | 0 | 0 | 0 | 14 | 1 | 0 | 1 | 0 | 98 | 49 | 0.069582416 | 0.17144651 | NA | 1 | 1 | NA | 0.875 | 1 | 0.062 |
| SE | 4942 | ARAP1 | chr11 | - | 72695717 | 72695865 | 72695541 | 72695628 | 72696548 | 72696654 | 0 | 1 | 24 | 0 | 0 | 0 | 0 | 15 | 1 | 0 | 1 | 0 | 98 | 49 | 0.06991611 | 0.171507349 | NA | 1 | 1 | NA | 0.882 | 1 | 0.059 |
| SE | 10851 | IRF1 | chr5 | - | 132489391 | 132489483 | 132486953 | 132487130 | 132490544 | 132490773 | 0 | 0 | 6 | 0 | 0 | 2 | 0 | 8 | 2 | 0 | 2 | 7 | 98 | 49 | 0.069995497 | 0.171507349 | NA | NA | 0.6 | NA | 0.667 | 0.125 | 0.204 |
| SE | 4877 | TPP1 | chr11 | - | 6618775 | 6618915 | 6617625 | 6617776 | 6619195 | 6619267 | 1 | 2 | 13 | 0 | 0 | 0 | 0 | 1 | 1 | 0 | 1 | 0 | 98 | 49 | 0.07001504 | 0.171507349 | 1 | 1 | 1 | NA | 0.333 | 1 | 0.334 |
| SE | 6551 | SESN2 | chr1 | + | 28271673 | 28271871 | 28269182 | 28269248 | 28272283 | 28272466 | 0 | 1 | 1 | 0 | 0 | 0 | 0 | 7 | 4 | 0 | 1 | 0 | 98 | 49 | 0.070207347 | 0.171507349 | NA | 1 | 1 | NA | 0.778 | 1 | 0.111 |
| SE | 8748 | ADIPOR1 | chr1 | - | 202954240 | 202954307 | 202950929 | 202951164 | 202958184 | 202958396 | 0 | 3 | 1 | 0 | 1 | 1 | 0 | 2 | 2 | 0 | 0 | 0 | 98 | 49 | 0.070257706 | 0.171507349 | NA | 0.6 | 0.333 | NA | 1 | 1 | -0.533 |
| SE | 13036 | VASP | chr19 | + | 45523840 | 45523877 | 45523643 | 45523695 | 45524096 | 45524142 | 0 | 5 | 37 | 0 | 0 | 3 | 0 | 33 | 3 | 0 | 0 | 0 | 86 | 49 | 0.070291768 | 0.171507349 | NA | 1 | 0.875 | NA | 1 | 1 | -0.062 |
| SE | 7499 | IFI44 | chr1 | + | 78662703 | 78662878 | 78660553 | 78660654 | 78663764 | 78664078 | 0 | 6 | 52 | 0 | 0 | 2 | 0 | 85 | 6 | 0 | 0 | 0 | 98 | 49 | 0.070496338 | 0.171727708 | NA | 1 | 0.929 | NA | 1 | 1 | -0.035 |
| SE | 12903 | TYROBP | chr19 | - | 35907729 | 35907762 | 35907445 | 35907580 | 35908174 | 35908309 | 0 | 3 | 6 | 0 | 3 | 1 | 0 | 5 | 1 | 0 | 0 | 0 | 82 | 49 | 0.070693174 | 0.171928544 | NA | 0.374 | 0.782 | NA | 1 | 1 | -0.422 |
| A3SS | 581 | LRCH4 | chr7 | - | 100574010 | 100575304 | 100574010 | 100574356 | 100575704 | 100575782 | 0 | 1 | 2 | 0 | 0 | 4 | 0 | 3 | 9 | 0 | 1 | 1 | 98 | 49 | 0.070727262 | 0.198669639 | NA | 1 | 0.2 | NA | 0.6 | 0.818 | -0.109 |
| SE | 12640 | CYTH4 | chr22 | + | 37310987 | 37311064 | 37309211 | 37309323 | 37311455 | 37311527 | 0 | 2 | 30 | 0 | 0 | 0 | 0 | 38 | 1 | 0 | 1 | 0 | 98 | 49 | 0.07087784 | 0.172099182 | NA | 1 | 1 | NA | 0.95 | 1 | 0.025 |
| SE | 12746 | DYSF | chr2 | + | 71656161 | 71656290 | 71643964 | 71644063 | 71658877 | 71659033 | 0 | 2 | 17 | 0 | 2 | 0 | 0 | 7 | 6 | 0 | 0 | 0 | 98 | 49 | 0.071270681 | 0.172773926 | NA | 0.333 | 1 | NA | 1 | 1 | -0.334 |
| SE | 333 | AKNA | chr9 | - | 114356862 | 114356969 | 114355924 | 114356136 | 114357920 | 114358167 | 0 | 1 | 3 | 0 | 0 | 0 | 6 | 8 | 3 | 0 | 1 | 0 | 98 | 49 | 0.071722008 | 0.173368597 | NA | 1 | 1 | 1 | 0.8 | 1 | 0.067 |
| A3SS | 3730 | RASGRP4 | chr19 | - | 38414847 | 38415123 | 38414847 | 38414916 | 38417051 | 38417168 | 0 | 2 | 16 | 0 | 0 | 0 | 1 | 4 | 3 | 0 | 1 | 0 | 98 | 49 | 0.071741814 | 0.198669639 | NA | 1 | 1 | 1 | 0.667 | 1 | 0.111 |
| SE | 4363 | LRRK2 | chr12 | + | 40313971 | 40314173 | 40310430 | 40310649 | 40315211 | 40315300 | 0 | 3 | 44 | 0 | 0 | 1 | 1 | 19 | 1 | 0 | 3 | 0 | 98 | 49 | 0.071746685 | 0.173368597 | NA | 1 | 0.957 | 1 | 0.76 | 1 | 0.058 |
| A3SS | 590 | UBE2D3 | chr4 | - | 102826484 | 102826844 | 102826484 | 102826636 | 102827591 | 102827948 | 0 | 5 | 24 | 0 | 0 | 2 | 2 | 32 | 7 | 0 | 0 | 0 | 98 | 49 | 0.073395949 | 0.200170771 | NA | 1 | 0.857 | 1 | 1 | 1 | -0.072 |
| SE | 4359 | LRRK2 | chr12 | + | 40274582 | 40274727 | 40263788 | 40263901 | 40274853 | 40274993 | 0 | 1 | 45 | 0 | 0 | 0 | 0 | 7 | 3 | 0 | 1 | 0 | 98 | 49 | 0.073501365 | 0.177323519 | NA | 1 | 1 | NA | 0.778 | 1 | 0.111 |
| SE | 6869 | ATG2A | chr11 | - | 64910615 | 64910708 | 64910039 | 64910195 | 64910806 | 64910954 | 0 | 8 | 7 | 0 | 0 | 0 | 8 | 26 | 23 | 0 | 1 | 0 | 98 | 49 | 0.07395786 | 0.178138883 | NA | 1 | 1 | 1 | 0.929 | 1 | 0.024 |
| SE | 1736 | ARHGAP26 | chr5 | + | 142901934 | 142902039 | 142894237 | 142894348 | 142903539 | 142903669 | 0 | 2 | 101 | 0 | 0 | 0 | 0 | 61 | 1 | 0 | 1 | 0 | 98 | 49 | 0.074241382 | 0.178535676 | NA | 1 | 1 | NA | 0.968 | 1 | 0.016 |
| SE | 8074 | BCL2A1 | chr15 | - | 79967636 | 79967692 | 79960889 | 79961174 | 79970699 | 79971301 | 0 | 0 | 2 | 0 | 1 | 19 | 0 | 0 | 0 | 1 | 40 | 1 | 98 | 49 | 0.074407684 | 0.178649759 | NA | 0 | 0.05 | 0 | 0 | 0 | 0.025 |
| SE | 13869 | ACSL1 | chr4 | - | 184788378 | 184788513 | 184783926 | 184783991 | 184788616 | 184788731 | 0 | 0 | 0 | 0 | 1 | 55 | 0 | 2 | 0 | 1 | 37 | 9 | 98 | 49 | 0.074750662 | 0.178951805 | NA | 0 | 0 | 0 | 0.026 | 0 | -0.009 |
| SE | 11135 | ECE1 | chr1 | - | 21279190 | 21279332 | 21272698 | 21272911 | 21290069 | 21290156 | 0 | 2 | 12 | 0 | 0 | 0 | 0 | 8 | 5 | 0 | 1 | 0 | 98 | 49 | 0.074771613 | 0.178951805 | NA | 1 | 1 | NA | 0.8 | 1 | 0.1 |
| SE | 2078 | HERC5 | chr4 | + | 88472408 | 88472502 | 88470613 | 88470673 | 88475840 | 88476030 | 0 | 2 | 29 | 0 | 0 | 0 | 0 | 34 | 3 | 0 | 1 | 0 | 98 | 49 | 0.075329785 | 0.180001061 | NA | 1 | 1 | NA | 0.944 | 1 | 0.028 |
| A5SS | 2702 | IFI6 | chr1 | - | 27669220 | 27669346 | 27669244 | 27669346 | 27668457 | 27668535 | 0 | 0 | 5 | 0 | 1 | 20 | 0 | 0 | 0 | 0 | 9 | 1 | 73 | 49 | 0.075687381 | 0.183611979 | NA | 0 | 0.144 | NA | 0 | 0 | 0.072 |
| SE | 801 | TBXAS1 | chr7 | + | 139955458 | 139955607 | 139953367 | 139953456 | 139957633 | 139957764 | 0 | 1 | 15 | 0 | 0 | 0 | 0 | 3 | 2 | 0 | 1 | 0 | 98 | 49 | 0.07582917 | 0.180642255 | NA | 1 | 1 | NA | 0.6 | 1 | 0.2 |
| SE | 12943 | SMCHD1 | chr18 | + | 2722518 | 2722663 | 2718314 | 2718434 | 2724898 | 2724995 | 0 | 1 | 43 | 0 | 0 | 5 | 1 | 37 | 1 | 0 | 12 | 0 | 98 | 49 | 0.075958015 | 0.180642255 | NA | 1 | 0.811 | 1 | 0.607 | 1 | 0.036 |
| SE | 6058 | HCLS1 | chr3 | - | 121637145 | 121637256 | 121636433 | 121636489 | 121642926 | 121642981 | 0 | 0 | 9 | 0 | 1 | 0 | 1 | 4 | 1 | 0 | 0 | 0 | 98 | 49 | 0.075958686 | 0.180642255 | NA | 0 | 1 | 1 | 1 | 1 | -0.5 |
| RI | 461 | WTAP | chr6 | + | 159748190 | 159749421 | 159748190 | 159748369 | 159748589 | 159749421 | 0 | 1 | 13 | 0 | 0 | 0 | 3 | 9 | 5 | 0 | 2 | 0 | 98 | 49 | 0.076502744 | 0.314063896 | NA | 1 | 1 | 1 | 0.692 | 1 | 0.103 |
| SE | 11482 | AP1G1 | chr16 | - | 71789278 | 71789482 | 71774467 | 71774592 | 71808762 | 71809073 | 0 | 1 | 7 | 0 | 0 | 0 | 0 | 7 | 3 | 0 | 1 | 0 | 98 | 49 | 0.076633988 | 0.181960321 | NA | 1 | 1 | NA | 0.778 | 1 | 0.111 |
| SE | 4413 | MEGF6 | chr1 | - | 3496987 | 3497119 | 3496654 | 3496783 | 3497232 | 3497361 | 0 | 3 | 0 | 0 | 0 | 2 | 0 | 1 | 1 | 0 | 0 | 0 | 98 | 49 | 0.076772626 | 0.182001983 | NA | 1 | 0 | NA | 1 | 1 | -0.5 |
| SE | 5396 | MTMR14 | chr3 | + | 9662266 | 9662375 | 9653620 | 9653769 | 9668718 | 9668794 | 0 | 0 | 1 | 0 | 1 | 0 | 0 | 1 | 3 | 0 | 0 | 0 | 98 | 49 | 0.077074472 | 0.182348728 | NA | 0 | 1 | NA | 1 | 1 | -0.5 |
| SE | 5680 | GMIP | chr19 | - | 19641809 | 19641867 | 19640445 | 19640571 | 19641975 | 19642051 | 0 | 1 | 27 | 0 | 0 | 3 | 0 | 15 | 6 | 0 | 0 | 0 | 98 | 49 | 0.077161537 | 0.182348728 | NA | 1 | 0.818 | NA | 1 | 1 | -0.091 |
| SE | 6871 | ATG2A | chr11 | - | 64914080 | 64914233 | 64913820 | 64913923 | 64914337 | 64914500 | 0 | 2 | 8 | 0 | 0 | 0 | 4 | 20 | 8 | 0 | 1 | 0 | 98 | 49 | 0.077718556 | 0.183327985 | NA | 1 | 1 | 1 | 0.909 | 1 | 0.03 |
| SE | 10716 | GADD45B | chr19 | + | 2477028 | 2477251 | 2476528 | 2476630 | 2477487 | 2478259 | 0 | 1 | 6 | 0 | 0 | 1 | 0 | 22 | 4 | 0 | 0 | 0 | 98 | 49 | 0.077819863 | 0.183327985 | NA | 1 | 0.75 | NA | 1 | 1 | -0.125 |
| SE | 10342 | PDCD4 | chr10 | + | 110887664 | 110887886 | 110885252 | 110885366 | 110889532 | 110889630 | 0 | 1 | 4 | 0 | 0 | 0 | 0 | 21 | 2 | 0 | 1 | 0 | 98 | 49 | 0.078019098 | 0.183509709 | NA | 1 | 1 | NA | 0.913 | 1 | 0.043 |
| A3SS | 1541 | GMIP | chr19 | - | 19636706 | 19636796 | 19636706 | 19636787 | 19636916 | 19637029 | 0 | 1 | 9 | 0 | 0 | 0 | 0 | 5 | 2 | 0 | 1 | 0 | 58 | 49 | 0.078029018 | 0.208872272 | NA | 1 | 1 | NA | 0.809 | 1 | 0.095 |
| SE | 11436 | PLK3 | chr1 | + | 44803267 | 44803391 | 44802954 | 44803153 | 44803599 | 44803691 | 0 | 2 | 8 | 0 | 0 | 0 | 0 | 6 | 5 | 0 | 1 | 0 | 98 | 49 | 0.078722929 | 0.184875878 | NA | 1 | 1 | NA | 0.75 | 1 | 0.125 |
| A3SS | 1062 | ATXN2L | chr16 | + | 28832186 | 28832399 | 28832204 | 28832399 | 28830961 | 28831072 | 0 | 0 | 1 | 0 | 1 | 0 | 0 | 0 | 0 | 0 | 1 | 2 | 67 | 49 | 0.078907303 | 0.208872272 | NA | 0 | 1 | NA | 0 | 0 | 0.5 |
| SE | 9029 | MYD88 | chr3 | + | 38140387 | 38140568 | 38138477 | 38139028 | 38140756 | 38140848 | 0 | 3 | 7 | 0 | 0 | 0 | 0 | 2 | 2 | 0 | 2 | 0 | 98 | 49 | 0.079080192 | 0.185334725 | NA | 1 | 1 | NA | 0.333 | 1 | 0.334 |
| SE | 9375 | FMNL1 | chr17 | + | 45236135 | 45236244 | 45234071 | 45234200 | 45237280 | 45237357 | 0 | 8 | 42 | 0 | 2 | 1 | 3 | 17 | 7 | 0 | 0 | 0 | 98 | 49 | 0.079164932 | 0.185334725 | NA | 0.667 | 0.955 | 1 | 1 | 1 | -0.189 |
| SE | 4098 | SETD1B | chr12 | + | 121819403 | 121819895 | 121817798 | 121817904 | 121822489 | 121823749 | 0 | 3 | 1 | 0 | 0 | 3 | 1 | 6 | 1 | 0 | 0 | 0 | 98 | 49 | 0.079530423 | 0.185900818 | NA | 1 | 0.143 | 1 | 1 | 1 | -0.428 |
| SE | 5185 | MYO9B | chr19 | + | 17159394 | 17159484 | 17156908 | 17157038 | 17162349 | 17162466 | 0 | 4 | 24 | 0 | 0 | 0 | 1 | 16 | 12 | 0 | 1 | 0 | 98 | 49 | 0.079811629 | 0.186229709 | NA | 1 | 1 | 1 | 0.889 | 1 | 0.037 |
| SE | 6698 | FCGR2A | chr1 | + | 161506336 | 161506591 | 161505986 | 161506007 | 161509819 | 161510074 | 0 | 1 | 45 | 0 | 0 | 0 | 1 | 23 | 2 | 0 | 1 | 0 | 98 | 49 | 0.079918937 | 0.186229709 | NA | 1 | 1 | 1 | 0.92 | 1 | 0.027 |
| SE | 7345 | FAM129A | chr1 | - | 184823637 | 184823742 | 184823166 | 184823329 | 184831846 | 184831962 | 0 | 2 | 32 | 0 | 0 | 4 | 1 | 19 | 1 | 0 | 0 | 0 | 98 | 49 | 0.080074419 | 0.186303176 | NA | 1 | 0.8 | 1 | 1 | 1 | -0.1 |
| SE | 7416 | VAV1 | chr19 | + | 6822418 | 6822514 | 6822220 | 6822329 | 6825052 | 6825121 | 0 | 0 | 15 | 0 | 2 | 0 | 0 | 11 | 4 | 0 | 2 | 0 | 98 | 49 | 0.080433731 | 0.18684992 | NA | 0 | 1 | NA | 0.733 | 1 | -0.367 |
| SE | 12584 | SBNO2 | chr19 | - | 1111018 | 1111093 | 1110744 | 1110888 | 1111505 | 1111614 | 0 | 1 | 6 | 0 | 0 | 1 | 1 | 26 | 2 | 0 | 0 | 0 | 98 | 49 | 0.080803043 | 0.187418168 | NA | 1 | 0.75 | 1 | 1 | 1 | -0.125 |
| SE | 10297 | TNFAIP2 | chr14 | + | 103127004 | 103127629 | 103126326 | 103126692 | 103129739 | 103129854 | 0 | 2 | 36 | 0 | 0 | 0 | 0 | 35 | 5 | 0 | 1 | 0 | 98 | 49 | 0.081018716 | 0.187601066 | NA | 1 | 1 | NA | 0.946 | 1 | 0.027 |
| SE | 4657 | PRKD2 | chr19 | - | 46681649 | 46681748 | 46678395 | 46678663 | 46689536 | 46689698 | 0 | 1 | 4 | 0 | 0 | 0 | 1 | 17 | 4 | 0 | 1 | 0 | 98 | 49 | 0.081231413 | 0.187601066 | NA | 1 | 1 | 1 | 0.895 | 1 | 0.035 |
| SE | 11395 | ADGRG3 | chr16 | + | 57680504 | 57680617 | 57680264 | 57680365 | 57684389 | 57684483 | 0 | 3 | 11 | 0 | 0 | 0 | 0 | 12 | 14 | 0 | 0 | 1 | 98 | 49 | 0.081296035 | 0.187601066 | NA | 1 | 1 | NA | 1 | 0.875 | 0.062 |
| SE | 5698 | RPLP2 | chr11 | + | 812534 | 812633 | 811596 | 811645 | 812759 | 812876 | 0 | 2 | 43 | 0 | 0 | 0 | 0 | 41 | 4 | 0 | 1 | 0 | 98 | 49 | 0.081381168 | 0.187601066 | NA | 1 | 1 | NA | 0.953 | 1 | 0.024 |
| SE | 10951 | IQGAP1 | chr15 | + | 90482196 | 90482281 | 90481959 | 90482100 | 90483360 | 90483593 | 0 | 1 | 25 | 0 | 0 | 0 | 0 | 12 | 4 | 0 | 1 | 0 | 98 | 49 | 0.083074196 | 0.191210592 | NA | 1 | 1 | NA | 0.857 | 1 | 0.072 |
| SE | 3399 | ELOVL5 | chr6 | - | 53273219 | 53273344 | 53270592 | 53270727 | 53275089 | 53275261 | 0 | 1 | 21 | 0 | 0 | 0 | 3 | 9 | 4 | 0 | 1 | 0 | 98 | 49 | 0.083349562 | 0.191551057 | NA | 1 | 1 | 1 | 0.818 | 1 | 0.061 |
| SE | 5156 | PSAP | chr10 | - | 71819030 | 71819111 | 71818616 | 71818724 | 71819464 | 71819622 | 1 | 15 | 181 | 0 | 0 | 0 | 10 | 44 | 9 | 0 | 1 | 0 | 98 | 49 | 0.084300652 | 0.193441038 | 1 | 1 | 1 | 1 | 0.957 | 1 | 0.014 |
| SE | 9929 | PTPRE | chr10 | + | 128056122 | 128056213 | 128049529 | 128049666 | 128060938 | 128061015 | 1 | 1 | 31 | 0 | 0 | 0 | 0 | 31 | 1 | 0 | 1 | 0 | 98 | 49 | 0.084566773 | 0.193502494 | 1 | 1 | 1 | NA | 0.939 | 1 | 0.03 |
| SE | 6870 | ATG2A | chr11 | - | 64912326 | 64912423 | 64912084 | 64912249 | 64913037 | 64913136 | 0 | 2 | 5 | 0 | 0 | 1 | 4 | 16 | 7 | 0 | 0 | 0 | 98 | 49 | 0.084584923 | 0.193502494 | NA | 1 | 0.714 | 1 | 1 | 1 | -0.143 |
| A5SS | 886 | DDIT3 | chr12 | - | 57517268 | 57517753 | 57517705 | 57517753 | 57516587 | 57517180 | 1 | 3 | 7 | 0 | 0 | 0 | 2 | 20 | 3 | 0 | 1 | 0 | 98 | 49 | 0.084700702 | 0.201741673 | 1 | 1 | 1 | 1 | 0.909 | 1 | 0.03 |
| SE | 10384 | SIGLEC5 | chr19 | - | 51627461 | 51627746 | 51627148 | 51627248 | 51627833 | 51628091 | 0 | 2 | 8 | 0 | 2 | 0 | 0 | 4 | 5 | 0 | 0 | 0 | 98 | 49 | 0.084994123 | 0.19414311 | NA | 0.333 | 1 | NA | 1 | 1 | -0.334 |
| SE | 12730 | PILRA | chr7 | + | 100397878 | 100397912 | 100374043 | 100374433 | 100399290 | 100399340 | 0 | 1 | 14 | 0 | 0 | 0 | 0 | 3 | 2 | 0 | 3 | 0 | 83 | 49 | 0.085434257 | 0.194790215 | NA | 1 | 1 | NA | 0.371 | 1 | 0.315 |
| SE | 4324 | KMT2D | chr12 | - | 49026181 | 49027322 | 49024809 | 49024946 | 49027802 | 49027930 | 0 | 1 | 7 | 0 | 0 | 0 | 0 | 2 | 2 | 0 | 1 | 0 | 98 | 49 | 0.085536621 | 0.194790215 | NA | 1 | 1 | NA | 0.5 | 1 | 0.25 |
| SE | 11837 | RAB11FIP4 | chr17 | + | 31517650 | 31517877 | 31488000 | 31488272 | 31521165 | 31521360 | 0 | 1 | 2 | 0 | 0 | 0 | 0 | 6 | 5 | 0 | 1 | 0 | 98 | 49 | 0.086321731 | 0.196280728 | NA | 1 | 1 | NA | 0.75 | 1 | 0.125 |
| RI | 359 | DDIT3 | chr12 | - | 57517268 | 57517753 | 57517268 | 57517595 | 57517705 | 57517753 | 4 | 2 | 10 | 0 | 0 | 0 | 8 | 27 | 2 | 0 | 1 | 0 | 98 | 49 | 0.086408407 | 0.336992786 | 1 | 1 | 1 | 1 | 0.931 | 1 | 0.023 |
| SE | 10863 | ARHGAP9 | chr12 | - | 57473602 | 57473708 | 57472254 | 57472688 | 57474041 | 57474176 | 0 | 0 | 32 | 0 | 0 | 3 | 0 | 6 | 1 | 0 | 2 | 2 | 98 | 49 | 0.086690204 | 0.196820811 | NA | NA | 0.842 | NA | 0.6 | 0.2 | 0.442 |
| SE | 9932 | PTPRE | chr10 | + | 128072137 | 128072214 | 128070807 | 128070901 | 128073336 | 128073471 | 0 | 1 | 29 | 0 | 0 | 0 | 0 | 27 | 2 | 0 | 1 | 0 | 98 | 49 | 0.087167679 | 0.197606366 | NA | 1 | 1 | NA | 0.931 | 1 | 0.034 |
| SE | 10447 | ELL | chr19 | - | 18465796 | 18465918 | 18465411 | 18465575 | 18472834 | 18472882 | 0 | 2 | 13 | 0 | 4 | 0 | 0 | 10 | 0 | 0 | 0 | 0 | 98 | 49 | 0.088641831 | 0.200598284 | NA | 0.2 | 1 | NA | 1 | NA | -0.4 |
| SE | 10822 | C16orf72 | chr16 | + | 9092920 | 9093001 | 9091679 | 9092270 | 9102983 | 9103259 | 0 | 1 | 22 | 0 | 1 | 1 | 0 | 20 | 0 | 0 | 0 | 0 | 98 | 49 | 0.088754397 | 0.200598284 | NA | 0.333 | 0.917 | NA | 1 | NA | -0.375 |
| SE | 13628 | SUN2 | chr22 | - | 38754641 | 38754706 | 38752506 | 38752665 | 38755762 | 38756019 | 0 | 0 | 7 | 0 | 1 | 2 | 0 | 0 | 0 | 0 | 0 | 6 | 98 | 49 | 0.088928004 | 0.200688873 | NA | 0 | 0.636 | NA | NA | 0 | 0.318 |
| SE | 1903 | MAP3K1 | chr5 | + | 56875031 | 56875310 | 56872824 | 56873005 | 56878979 | 56879101 | 0 | 0 | 5 | 0 | 1 | 1 | 1 | 3 | 2 | 0 | 0 | 0 | 98 | 49 | 0.089518138 | 0.201717783 | NA | 0 | 0.714 | 1 | 1 | 1 | -0.643 |
| SE | 7640 | IRF7 | chr11 | - | 614173 | 614399 | 613950 | 614037 | 614475 | 614534 | 0 | 1 | 2 | 0 | 7 | 10 | 0 | 5 | 1 | 0 | 3 | 3 | 98 | 49 | 0.089929548 | 0.202173556 | NA | 0.067 | 0.091 | NA | 0.455 | 0.143 | -0.22 |
| SE | 5219 | TPM4 | chr19 | + | 16081912 | 16082046 | 16076324 | 16076697 | 16086422 | 16086540 | 0 | 4 | 6 | 0 | 0 | 0 | 0 | 10 | 7 | 0 | 0 | 1 | 98 | 49 | 0.09016239 | 0.202173556 | NA | 1 | 1 | NA | 1 | 0.778 | 0.111 |
| SE | 6816 | OAS1 | chr12 | + | 112908535 | 112908824 | 112906933 | 112907219 | 112911050 | 112911235 | 0 | 1 | 25 | 0 | 0 | 0 | 5 | 3 | 3 | 0 | 1 | 0 | 98 | 49 | 0.090232466 | 0.202173556 | NA | 1 | 1 | 1 | 0.6 | 1 | 0.133 |
| SE | 2081 | HERC5 | chr4 | + | 88499925 | 88499992 | 88494164 | 88494331 | 88500914 | 88500985 | 0 | 3 | 48 | 0 | 1 | 1 | 0 | 33 | 0 | 0 | 2 | 0 | 98 | 49 | 0.090258454 | 0.202173556 | NA | 0.6 | 0.96 | NA | 0.892 | NA | -0.112 |
| A3SS | 872 | DGCR2 | chr22 | - | 19064847 | 19065075 | 19064847 | 19065067 | 19068099 | 19068225 | 0 | 0 | 3 | 0 | 1 | 4 | 0 | 0 | 0 | 0 | 2 | 2 | 57 | 49 | 0.092162455 | 0.240423795 | NA | 0 | 0.392 | NA | 0 | 0 | 0.196 |
| RI | 832 | HES4 | chr1 | - | 999691 | 1000172 | 999691 | 999787 | 999865 | 1000172 | 1 | 2 | 1 | 0 | 0 | 0 | 3 | 13 | 4 | 0 | 1 | 0 | 98 | 49 | 0.092178476 | 0.342377196 | 1 | 1 | 1 | 1 | 0.867 | 1 | 0.044 |
| SE | 1266 | MICAL1 | chr6 | - | 109447356 | 109447440 | 109447072 | 109447229 | 109447680 | 109447722 | 0 | 3 | 5 | 0 | 0 | 0 | 0 | 2 | 1 | 0 | 0 | 1 | 98 | 49 | 0.094107379 | 0.210481236 | NA | 1 | 1 | NA | 1 | 0.333 | 0.334 |
| SE | 4124 | XPO6 | chr16 | - | 28135215 | 28135324 | 28133840 | 28133933 | 28146093 | 28146203 | 0 | 5 | 50 | 0 | 0 | 2 | 3 | 60 | 4 | 0 | 0 | 0 | 98 | 49 | 0.094351496 | 0.210563613 | NA | 1 | 0.926 | 1 | 1 | 1 | -0.037 |
| SE | 3589 | SP110 | chr2 | - | 230202578 | 230202728 | 230200884 | 230200965 | 230207990 | 230208059 | 0 | 1 | 8 | 0 | 0 | 0 | 0 | 10 | 6 | 0 | 1 | 0 | 98 | 49 | 0.094424401 | 0.210563613 | NA | 1 | 1 | NA | 0.833 | 1 | 0.084 |
| SE | 10651 | SLC43A2 | chr17 | - | 1616561 | 1616769 | 1614978 | 1615034 | 1627714 | 1627920 | 0 | 0 | 9 | 0 | 0 | 1 | 0 | 9 | 0 | 0 | 0 | 2 | 98 | 49 | 0.094824344 | 0.211028906 | NA | NA | 0.818 | NA | 1 | 0 | 0.318 |
| SE | 7418 | VAV1 | chr19 | + | 6847997 | 6848114 | 6843134 | 6843166 | 6850669 | 6850757 | 0 | 3 | 13 | 0 | 4 | 0 | 3 | 10 | 0 | 0 | 0 | 0 | 98 | 49 | 0.095128464 | 0.211028906 | NA | 0.273 | 1 | 1 | 1 | NA | -0.363 |
| SE | 10849 | IRF1 | chr5 | - | 132486794 | 132486844 | 132486556 | 132486686 | 132486953 | 132487130 | 0 | 2 | 21 | 0 | 0 | 1 | 0 | 36 | 5 | 0 | 8 | 0 | 98 | 49 | 0.095196923 | 0.211028906 | NA | 1 | 0.913 | NA | 0.692 | 1 | 0.111 |
| SE | 4963 | DDX3X | chrX | + | 41343215 | 41343351 | 41342736 | 41342836 | 41343736 | 41343822 | 0 | 2 | 31 | 0 | 0 | 1 | 0 | 24 | 1 | 0 | 3 | 0 | 98 | 49 | 0.095320071 | 0.211028906 | NA | 1 | 0.939 | NA | 0.8 | 1 | 0.07 |
| SE | 1143 | IGF2R | chr6 | + | 160079579 | 160079787 | 160078200 | 160078362 | 160080128 | 160080275 | 0 | 4 | 40 | 0 | 0 | 1 | 0 | 6 | 2 | 0 | 3 | 0 | 98 | 49 | 0.095335081 | 0.211028906 | NA | 1 | 0.952 | NA | 0.5 | 1 | 0.226 |
| SE | 10701 | TLE3 | chr15 | - | 70059409 | 70059460 | 70058662 | 70058815 | 70060529 | 70060649 | 0 | 4 | 31 | 0 | 2 | 0 | 0 | 21 | 0 | 0 | 0 | 0 | 98 | 49 | 0.095684167 | 0.211266553 | NA | 0.5 | 1 | NA | 1 | NA | -0.25 |
| SE | 6176 | ABCA1 | chr9 | - | 104819585 | 104819723 | 104818662 | 104818883 | 104819926 | 104820069 | 0 | 1 | 17 | 0 | 0 | 0 | 0 | 29 | 3 | 0 | 1 | 0 | 98 | 49 | 0.095723568 | 0.211266553 | NA | 1 | 1 | NA | 0.935 | 1 | 0.032 |
| SE | 8187 | EPSTI1 | chr13 | - | 42895008 | 42895108 | 42886387 | 42888502 | 42900309 | 42900383 | 0 | 1 | 9 | 0 | 0 | 2 | 4 | 11 | 1 | 0 | 0 | 0 | 98 | 49 | 0.096411782 | 0.212473472 | NA | 1 | 0.692 | 1 | 1 | 1 | -0.154 |
| SE | 12365 | NBEAL2 | chr3 | + | 46995931 | 46996051 | 46995713 | 46995846 | 46996270 | 46996592 | 0 | 8 | 19 | 0 | 0 | 0 | 0 | 5 | 4 | 0 | 0 | 2 | 98 | 49 | 0.09699006 | 0.213434934 | NA | 1 | 1 | NA | 1 | 0.5 | 0.25 |
| SE | 8710 | CD74 | chr5 | - | 150406880 | 150406960 | 150406258 | 150406321 | 150407151 | 150407324 | 0 | 7 | 68 | 0 | 0 | 1 | 2 | 12 | 4 | 0 | 3 | 0 | 98 | 49 | 0.097385214 | 0.213991193 | NA | 1 | 0.971 | 1 | 0.667 | 1 | 0.097 |
| SE | 11126 | PSEN1 | chr14 | + | 73198030 | 73198129 | 73192643 | 73192864 | 73206385 | 73206472 | 0 | 1 | 8 | 0 | 2 | 1 | 0 | 5 | 0 | 0 | 0 | 0 | 98 | 49 | 0.098181291 | 0.215425518 | NA | 0.2 | 0.8 | NA | 1 | NA | -0.5 |
| A3SS | 1986 | PGS1 | chr17 | + | 78414878 | 78415027 | 78414951 | 78415027 | 78403567 | 78404089 | 0 | 1 | 5 | 0 | 0 | 0 | 3 | 5 | 5 | 0 | 1 | 0 | 98 | 49 | 0.098812002 | 0.253724157 | NA | 1 | 1 | 1 | 0.714 | 1 | 0.095 |
| SE | 13029 | FOSB | chr19 | + | 45471193 | 45471301 | 45470628 | 45470949 | 45472550 | 45475179 | 0 | 3 | 9 | 0 | 2 | 0 | 0 | 13 | 0 | 0 | 0 | 0 | 98 | 49 | 0.099118539 | 0.217164962 | NA | 0.429 | 1 | NA | 1 | NA | -0.285 |
| SE | 13040 | MAP3K8 | chr10 | + | 30437175 | 30437406 | 30434020 | 30434378 | 30438915 | 30439274 | 0 | 0 | 6 | 0 | 0 | 1 | 0 | 2 | 0 | 0 | 2 | 2 | 98 | 49 | 0.099502086 | 0.217687969 | NA | NA | 0.75 | NA | 0.333 | 0 | 0.584 |
| SE | 12128 | LRRC59 | chr17 | - | 50392738 | 50392897 | 50392397 | 50392502 | 50394928 | 50394988 | 1 | 1 | 3 | 0 | 0 | 0 | 0 | 1 | 1 | 0 | 3 | 0 | 98 | 49 | 0.099753465 | 0.217812352 | 1 | 1 | 1 | NA | 0.143 | 1 | 0.428 |
| SE | 55 | ELF4 | chrX | - | 130081255 | 130081539 | 130074580 | 130074752 | 130110324 | 130110500 | 0 | 2 | 5 | 0 | 1 | 0 | 0 | 0 | 1 | 0 | 1 | 4 | 98 | 49 | 0.100072297 | 0.217812352 | NA | 0.5 | 1 | NA | 0 | 0.111 | 0.695 |
| A3SS | 2860 | TPM3 | chr1 | - | 154155303 | 154159048 | 154155303 | 154157721 | 154170399 | 154170469 | 1 | 0 | 1 | 0 | 0 | 20 | 0 | 2 | 1 | 0 | 14 | 0 | 98 | 49 | 0.100080084 | 0.253724157 | 1 | NA | 0.024 | NA | 0.067 | 1 | -0.021 |
| SE | 9378 | FMNL1 | chr17 | + | 45246504 | 45246604 | 45246209 | 45246330 | 45246866 | 45247316 | 0 | 1 | 0 | 0 | 1 | 15 | 5 | 0 | 0 | 0 | 13 | 2 | 98 | 49 | 0.100134734 | 0.217812352 | NA | 0.333 | 0 | 1 | 0 | 0 | -0.167 |
| SE | 6719 | KIF1B | chr1 | + | 10345844 | 10345953 | 10343231 | 10343287 | 10348648 | 10348733 | 0 | 4 | 2 | 0 | 0 | 0 | 0 | 1 | 2 | 0 | 1 | 0 | 98 | 49 | 0.100138613 | 0.217812352 | NA | 1 | 1 | NA | 0.333 | 1 | 0.334 |
| SE | 13866 | ACSL1 | chr4 | - | 184773080 | 184773154 | 184768315 | 184768450 | 184773662 | 184773714 | 0 | 2 | 38 | 0 | 0 | 1 | 3 | 28 | 2 | 0 | 3 | 0 | 98 | 49 | 0.100289399 | 0.217825096 | NA | 1 | 0.95 | 1 | 0.824 | 1 | 0.034 |
| SE | 13870 | ACSL1 | chr4 | - | 184803319 | 184803546 | 184788616 | 184788731 | 184825915 | 184826114 | 0 | 4 | 91 | 0 | 0 | 0 | 8 | 83 | 44 | 0 | 3 | 0 | 98 | 49 | 0.100452402 | 0.217864301 | NA | 1 | 1 | 1 | 0.933 | 1 | 0.022 |
| SE | 13051 | NAPA | chr19 | - | 47500632 | 47500749 | 47495549 | 47495596 | 47514842 | 47515258 | 0 | 1 | 13 | 0 | 0 | 0 | 0 | 11 | 7 | 0 | 1 | 0 | 98 | 49 | 0.100911104 | 0.218543788 | NA | 1 | 1 | NA | 0.846 | 1 | 0.077 |
| SE | 7577 | SPATS2L | chr2 | + | 200439121 | 200439328 | 200419249 | 200419496 | 200440648 | 200440784 | 0 | 0 | 3 | 0 | 0 | 1 | 0 | 1 | 0 | 0 | 0 | 3 | 98 | 49 | 0.101162648 | 0.218773323 | NA | NA | 0.6 | NA | 1 | 0 | 0.1 |
| SE | 12742 | DYSF | chr2 | + | 71520177 | 71520208 | 71516988 | 71517039 | 71520788 | 71520904 | 0 | 1 | 12 | 0 | 1 | 0 | 2 | 7 | 0 | 0 | 0 | 0 | 80 | 49 | 0.10199177 | 0.220129306 | NA | 0.38 | 1 | 1 | 1 | NA | -0.31 |
| SE | 10929 | DEDD2 | chr19 | - | 42204461 | 42204559 | 42198592 | 42199829 | 42209699 | 42209840 | 0 | 1 | 0 | 0 | 3 | 5 | 0 | 0 | 0 | 0 | 16 | 3 | 98 | 49 | 0.102192668 | 0.220129306 | NA | 0.143 | 0 | NA | 0 | 0 | 0.071 |
| SE | 3914 | ALOX5AP | chr13 | + | 30755943 | 30756025 | 30752051 | 30752122 | 30763943 | 30764428 | 0 | 2 | 74 | 0 | 0 | 0 | 0 | 36 | 17 | 0 | 1 | 0 | 98 | 49 | 0.102229046 | 0.220129306 | NA | 1 | 1 | NA | 0.947 | 1 | 0.026 |
| SE | 3886 | LCP1 | chr13 | - | 46146907 | 46147103 | 46144441 | 46144520 | 46148351 | 46148447 | 0 | 9 | 167 | 0 | 0 | 1 | 8 | 67 | 3 | 0 | 2 | 0 | 98 | 49 | 0.102410835 | 0.220205272 | NA | 1 | 0.988 | 1 | 0.944 | 1 | 0.013 |
| SE | 3619 | NIN | chr14 | - | 50830463 | 50830517 | 50821873 | 50822077 | 50831005 | 50831121 | 0 | 3 | 3 | 0 | 0 | 1 | 2 | 0 | 18 | 0 | 0 | 0 | 98 | 49 | 0.10387195 | 0.223027916 | NA | 1 | 0.6 | 1 | NA | 1 | -0.2 |
| SE | 803 | TBXAS1 | chr7 | + | 140017670 | 140017833 | 140015722 | 140015860 | 140020024 | 140020325 | 0 | 0 | 14 | 0 | 0 | 1 | 0 | 2 | 0 | 0 | 1 | 1 | 98 | 49 | 0.104215706 | 0.223185153 | NA | NA | 0.875 | NA | 0.5 | 0 | 0.625 |
| SE | 10226 | ACAP1 | chr17 | + | 7337311 | 7337369 | 7336528 | 7336787 | 7341947 | 7342067 | 2 | 1 | 3 | 0 | 0 | 0 | 0 | 2 | 3 | 0 | 1 | 0 | 98 | 49 | 0.104242167 | 0.223185153 | 1 | 1 | 1 | NA | 0.5 | 1 | 0.25 |
| SE | 6858 | SELL | chr1 | - | 169701559 | 169701688 | 169696534 | 169696553 | 169703254 | 169703440 | 0 | 2 | 95 | 0 | 1 | 0 | 12 | 29 | 3 | 0 | 0 | 0 | 98 | 49 | 0.105494926 | 0.225138011 | NA | 0.5 | 1 | 1 | 1 | 1 | -0.25 |
| SE | 796 | PARP12 | chr7 | - | 140057898 | 140058034 | 140056855 | 140057153 | 140062521 | 140063721 | 0 | 1 | 8 | 0 | 1 | 0 | 0 | 6 | 0 | 0 | 0 | 0 | 98 | 49 | 0.105498898 | 0.225138011 | NA | 0.333 | 1 | NA | 1 | NA | -0.334 |
| SE | 4946 | ARAP1 | chr11 | - | 72705804 | 72705890 | 72702904 | 72703079 | 72707174 | 72707374 | 0 | 2 | 13 | 0 | 0 | 0 | 0 | 5 | 7 | 0 | 1 | 0 | 98 | 49 | 0.105603658 | 0.225138011 | NA | 1 | 1 | NA | 0.714 | 1 | 0.143 |
| SE | 7968 | OAS3 | chr12 | + | 112967417 | 112967593 | 112964234 | 112964408 | 112967935 | 112968174 | 0 | 0 | 27 | 0 | 0 | 1 | 0 | 7 | 1 | 0 | 2 | 1 | 98 | 49 | 0.106158162 | 0.225817599 | NA | NA | 0.931 | NA | 0.636 | 0.333 | 0.447 |
| SE | 4056 | FOS | chr14 | + | 75280559 | 75280667 | 75279876 | 75280128 | 75280782 | 75282234 | 0 | 25 | 189 | 0 | 1 | 0 | 3 | 248 | 16 | 0 | 0 | 0 | 98 | 49 | 0.106325146 | 0.225817599 | NA | 0.926 | 1 | 1 | 1 | 1 | -0.037 |
| SE | 1509 | HLA-E | chr6 | + | 30492540 | 30492583 | 30491536 | 30491653 | 30492748 | 30494205 | 0 | 3 | 55 | 0 | 1 | 0 | 0 | 33 | 4 | 0 | 0 | 0 | 92 | 49 | 0.106659671 | 0.225817599 | NA | 0.615 | 1 | NA | 1 | 1 | -0.193 |
| SE | 11045 | HLA-B | chr6 | - | 31354632 | 31354665 | 31354478 | 31354526 | 31355316 | 31355592 | 1 | 12 | 113 | 0 | 0 | 0 | 8 | 112 | 7 | 0 | 1 | 0 | 82 | 49 | 0.106709745 | 0.225817599 | 1 | 1 | 1 | 1 | 0.985 | 1 | 0.005 |
| SE | 7902 | PACSIN2 | chr22 | - | 42890946 | 42891182 | 42888642 | 42888798 | 42893456 | 42893613 | 0 | 1 | 8 | 0 | 0 | 0 | 0 | 10 | 8 | 0 | 1 | 0 | 98 | 49 | 0.106828817 | 0.225817599 | NA | 1 | 1 | NA | 0.833 | 1 | 0.084 |

| | | | | | | | | | | | | | | | | | | | | | | | | | | | | | | | | | |
|---|---|---|---|---|---|---|---|---|---|---|---|---|---|---|---|---|---|---|---|---|---|---|---|---|---|---|---|---|---|---|---|---|---|
| SE | 7882 | HIF1A | chr14 | + | 61738086 | 61738373 | 61736888 | 61737109 | 61740504 | 61740627 | 0 | 1 | 10 | 0 | 0 | 1 | 0 | 34 | 1 | 0 | 0 | 0 | 98 | 49 | 0.106940257 | 0.225817599 | NA | 1 | 0.833 | NA | 1 | 1 | -0.084 |
| SE | 7143 | CPD | chr17 | + | 30438974 | 30439077 | 30431771 | 30431881 | 30442307 | 30442450 | 0 | 1 | 10 | 0 | 0 | 2 | 0 | 11 | 1 | 0 | 0 | 0 | 98 | 49 | 0.107022882 | 0.225817599 | NA | 1 | 0.714 | NA | 1 | 1 | -0.143 |
| SE | 5742 | ENTPD1 | chr10 | + | 95860468 | 95860582 | 95847445 | 95847706 | 95864723 | 95864861 | 0 | 1 | 5 | 0 | 0 | 0 | 0 | 2 | 3 | 0 | 1 | 0 | 98 | 49 | 0.107124383 | 0.225817599 | NA | 1 | 1 | NA | 0.5 | 1 | 0.25 |
| A3SS | 166 | SDCBP | chr8 | + | 58575899 | 58576061 | 58575902 | 58576061 | 58572204 | 58572314 | 0 | 3 | 20 | 0 | 0 | 2 | 0 | 9 | 1 | 0 | 4 | 0 | 52 | 49 | 0.10775843 | 0.269396074 | NA | 1 | 0.904 | NA | 0.68 | 1 | 0.112 |
| SE | 4628 | SORL1 | chr11 | + | 121577280 | 121577400 | 121574240 | 121574363 | 121583457 | 121583583 | 0 | 8 | 61 | 0 | 0 | 3 | 0 | 33 | 7 | 0 | 0 | 0 | 98 | 49 | 0.107936364 | 0.227210581 | NA | 1 | 0.91 | NA | 1 | 1 | -0.045 |
| SE | 9584 | ANXA11 | chr10 | - | 80176106 | 80176155 | 80172806 | 80172869 | 80205342 | 80205444 | 0 | 0 | 4 | 0 | 0 | 2 | 0 | 4 | 0 | 0 | 0 | 3 | 98 | 49 | 0.108478756 | 0.228032965 | NA | NA | 0.5 | NA | 1 | 0 | 0 |
| SE | 10587 | RNF213 | chr17 | + | 80361733 | 80361888 | 80358287 | 80358479 | 80363101 | 80363314 | 0 | 1 | 25 | 0 | 0 | 0 | 0 | 32 | 6 | 0 | 1 | 0 | 98 | 49 | 0.108649764 | 0.228073457 | NA | 1 | 1 | NA | 0.941 | 1 | 0.03 |
| SE | 9787 | HERC4 | chr10 | - | 67990210 | 67990400 | 67988662 | 67988835 | 67990903 | 67991015 | 0 | 0 | 1 | 0 | 0 | 2 | 0 | 2 | 3 | 0 | 0 | 0 | 98 | 49 | 0.108971078 | 0.228428912 | NA | NA | 0.2 | NA | 1 | 1 | -0.8 |
| A5SS | 1408 | HNRNPC | chr14 | - | 21230957 | 21231072 | 21230996 | 21231072 | 21230318 | 21230366 | 0 | 6 | 6 | 0 | 1 | 5 | 4 | 15 | 0 | 1 | 7 | 0 | 88 | 49 | 0.108976627 | 0.254927467 | NA | 0.77 | 0.401 | 0.69 | 0.544 | NA | -0.031 |
| SE | 11640 | ARRB2 | chr17 | + | 4716411 | 4716608 | 4715972 | 4716033 | 4717216 | 4717276 | 0 | 0 | 2 | 0 | 0 | 3 | 0 | 5 | 4 | 0 | 0 | 0 | 98 | 49 | 0.1093759 | 0.228660806 | NA | NA | 0.25 | NA | 1 | 1 | -0.75 |
| SE | 7570 | PGD | chr1 | + | 10403070 | 10403136 | 10400392 | 10400572 | 10404160 | 10404279 | 0 | 7 | 13 | 0 | 4 | 0 | 0 | 8 | 0 | 0 | 0 | 0 | 98 | 49 | 0.109641955 | 0.228660806 | NA | 0.467 | 1 | NA | 1 | NA | -0.266 |
| SE | 5385 | CD44 | chr11 | + | 35180273 | 35180407 | 35176574 | 35176740 | 35186831 | 35186900 | 1 | 4 | 16 | 0 | 2 | 1 | 0 | 9 | 0 | 0 | 0 | 0 | 98 | 49 | 0.109674831 | 0.228660806 | 1 | 0.5 | 0.889 | NA | 1 | NA | -0.204 |
| SE | 1986 | DDX60 | chr4 | - | 168272042 | 168272138 | 168268853 | 168268969 | 168273278 | 168273398 | 0 | 1 | 7 | 0 | 1 | 0 | 0 | 5 | 0 | 0 | 0 | 0 | 98 | 49 | 0.109690247 | 0.228660806 | NA | 0.333 | 1 | NA | 1 | NA | -0.334 |
| SE | 1038 | AOAH | chr7 | - | 36659165 | 36659265 | 36637850 | 36637910 | 36673942 | 36674009 | 0 | 0 | 10 | 0 | 0 | 1 | 1 | 8 | 0 | 0 | 1 | 1 | 98 | 49 | 0.110142389 | 0.229285332 | NA | NA | 0.833 | 1 | 0.8 | 0 | 0.233 |
| SE | 5785 | DENND3 | chr8 | + | 141144147 | 141144259 | 141138021 | 141138137 | 141150833 | 141150953 | 0 | 0 | 6 | 0 | 0 | 2 | 1 | 0 | 0 | 0 | 4 | 2 | 98 | 49 | 0.111042654 | 0.230839709 | NA | NA | 0.6 | 1 | 0 | 0 | 0.267 |
| SE | 5585 | WTAP | chr6 | + | 159736257 | 159736295 | 159726997 | 159727333 | 159738989 | 159739045 | 0 | 3 | 10 | 0 | 0 | 0 | 0 | 6 | 3 | 0 | 0 | 1 | 87 | 49 | 0.111205631 | 0.230859204 | NA | 1 | 1 | NA | 1 | 0.628 | 0.186 |
| SE | 13176 | ZNF274 | chr19 | + | 58209960 | 58210073 | 58206719 | 58207202 | 58211559 | 58211686 | 0 | 1 | 0 | 0 | 3 | 0 | 0 | 2 | 0 | 0 | 0 | 0 | 98 | 49 | 0.111677163 | 0.231518312 | NA | 0.143 | NA | NA | 1 | NA | -0.857 |
| SE | 422 | FAM214B | chr9 | - | 35106537 | 35106643 | 35106251 | 35106410 | 35106832 | 35106891 | 0 | 1 | 10 | 0 | 0 | 0 | 0 | 4 | 4 | 0 | 2 | 0 | 98 | 49 | 0.112112283 | 0.232100222 | NA | 1 | 1 | NA | 0.5 | 1 | 0.25 |
| SE | 13719 | HMHA1 | chr19 | + | 1073673 | 1073746 | 1073505 | 1073590 | 1073947 | 1074014 | 0 | 0 | 23 | 0 | 0 | 1 | 0 | 6 | 1 | 0 | 3 | 1 | 98 | 49 | 0.112614145 | 0.232818515 | NA | NA | 0.92 | NA | 0.5 | 0.333 | 0.504 |
| SE | 8977 | RPS9 | chr19 | + | 54206275 | 54206462 | 54201486 | 54201609 | 54207397 | 54207647 | 0 | 3 | 56 | 0 | 0 | 1 | 0 | 22 | 2 | 0 | 2 | 0 | 98 | 49 | 0.113459253 | 0.234243486 | NA | 1 | 0.966 | NA | 0.846 | 1 | 0.06 |
| SE | 12370 | NBEAL2 | chr3 | + | 47007524 | 47007697 | 47006332 | 47006449 | 47007815 | 47007910 | 0 | 14 | 33 | 0 | 0 | 0 | 2 | 7 | 17 | 0 | 1 | 0 | 98 | 49 | 0.114232158 | 0.235336193 | NA | 1 | 1 | 1 | 0.778 | 1 | 0.074 |
| SE | 4364 | LRRK2 | chr12 | + | 40319987 | 40320175 | 40315211 | 40315300 | 40321033 | 40321188 | 0 | 2 | 64 | 0 | 4 | 2 | 0 | 16 | 1 | 0 | 3 | 0 | 98 | 49 | 0.114457785 | 0.235336193 | NA | 0.2 | 0.941 | NA | 0.727 | 1 | -0.293 |
| SE | 11194 | PARP14 | chr3 | + | 122703741 | 122703978 | 122699389 | 122701635 | 122704526 | 122704748 | 0 | 6 | 21 | 0 | 1 | 0 | 0 | 22 | 0 | 0 | 0 | 0 | 98 | 49 | 0.114458255 | 0.235336193 | NA | 0.75 | 1 | NA | 1 | NA | -0.125 |
| A5SS | 1123 | MYO9B | chr19 | + | 17211646 | 17211786 | 17211646 | 17211774 | 17211894 | 17213295 | 0 | 0 | 2 | 0 | 0 | 2 | 0 | 0 | 0 | 0 | 0 | 4 | 61 | 49 | 0.11468766 | 0.263580411 | NA | NA | 0.445 | NA | NA | 0 | 0.445 |
| SE | 10702 | TLE3 | chr15 | - | 70064453 | 70064470 | 70060529 | 70060649 | 70066013 | 70066218 | 0 | 3 | 13 | 0 | 0 | 1 | 0 | 11 | 1 | 0 | 3 | 0 | 66 | 49 | 0.114960195 | 0.235829003 | NA | 1 | 0.906 | NA | 0.731 | 1 | 0.088 |
| SE | 2899 | GTPBP1 | chr22 | + | 38708844 | 38708956 | 38705801 | 38706147 | 38715906 | 38716087 | 0 | 0 | 8 | 0 | 0 | 1 | 0 | 9 | 0 | 0 | 6 | 1 | 98 | 49 | 0.115011749 | 0.235829003 | NA | NA | 0.8 | NA | 0.429 | 0 | 0.586 |
| SE | 7731 | KIAA1033 | chr12 | + | 105162748 | 105162845 | 105160000 | 105160148 | 105164110 | 105164307 | 0 | 1 | 7 | 0 | 1 | 0 | 0 | 4 | 0 | 0 | 0 | 0 | 98 | 49 | 0.115352023 | 0.235883116 | NA | 0.333 | 1 | NA | 1 | NA | -0.334 |
| SE | 13111 | AP2A1 | chr19 | + | 49800960 | 49801058 | 49799967 | 49800150 | 49801389 | 49801621 | 0 | 1 | 3 | 0 | 1 | 0 | 0 | 4 | 0 | 0 | 0 | 0 | 98 | 49 | 0.115352023 | 0.235883116 | NA | 0.333 | 1 | NA | 1 | NA | -0.334 |
| SE | 3524 | DYRK1A | chr21 | + | 37420298 | 37420384 | 37367556 | 37367628 | 37472683 | 37472880 | 0 | 0 | 2 | 0 | 0 | 2 | 0 | 2 | 4 | 0 | 0 | 0 | 98 | 49 | 0.115941495 | 0.236766395 | NA | NA | 0.333 | NA | 1 | 1 | -0.667 |
| SE | 7378 | RNF19B | chr1 | - | 32945513 | 32945628 | 32944018 | 32944159 | 32946401 | 32946564 | 0 | 1 | 24 | 0 | 0 | 0 | 1 | 30 | 9 | 0 | 1 | 0 | 98 | 49 | 0.116142758 | 0.236855585 | NA | 1 | 1 | 1 | 0.937 | 1 | 0.021 |
| SE | 6360 | ZSWIM8 | chr10 | + | 73788669 | 73788823 | 73785623 | 73786086 | 73789095 | 73789190 | 0 | 2 | 3 | 0 | 5 | 0 | 0 | 2 | 0 | 0 | 0 | 0 | 98 | 49 | 0.116399553 | 0.237028277 | NA | 0.167 | 1 | NA | 1 | NA | -0.416 |
| SE | 12686 | MYO1F | chr19 | - | 8541905 | 8541991 | 8539946 | 8540028 | 8544296 | 8544464 | 0 | 11 | 31 | 0 | 0 | 3 | 0 | 21 | 4 | 0 | 0 | 0 | 98 | 49 | 0.116542846 | 0.237028277 | NA | 1 | 0.838 | NA | 1 | 1 | -0.081 |
| SE | 6861 | CARD8 | chr19 | - | 48238382 | 48238532 | 48234402 | 48234543 | 48240961 | 48241063 | 0 | 0 | 7 | 0 | 0 | 1 | 0 | 0 | 0 | 0 | 0 | 1 | 98 | 49 | 0.11681427 | 0.237259253 | NA | NA | 0.778 | NA | NA | 0 | 0.778 |
| A3SS | 412 | PRRC2A | chr6 | + | 31622689 | 31622901 | 31622726 | 31622901 | 31620672 | 31620858 | 0 | 0 | 2 | 0 | 0 | 2 | 0 | 0 | 0 | 0 | 3 | 4 | 86 | 49 | 0.117982531 | 0.29091583 | NA | NA | 0.363 | NA | 0 | 0 | 0.363 |
| SE | 11993 | STAT5A | chr17 | + | 42291971 | 42292061 | 42289865 | 42290022 | 42295618 | 42295793 | 0 | 1 | 2 | 0 | 2 | 0 | 0 | 4 | 0 | 0 | 1 | 0 | 98 | 49 | 0.118118135 | 0.239583747 | NA | 0.2 | 1 | NA | 0.667 | NA | -0.067 |
| SE | 12348 | CSNK1D | chr17 | - | 82255428 | 82255577 | 82253015 | 82253244 | 82265685 | 82265796 | 0 | 0 | 3 | 0 | 0 | 2 | 0 | 2 | 5 | 0 | 0 | 0 | 98 | 49 | 0.118962034 | 0.240970265 | NA | NA | 0.429 | NA | 1 | 1 | -0.571 |
| SE | 4455 | GABARAPL1 | chr12 | + | 10218062 | 10218141 | 10212889 | 10213219 | 10220439 | 10220558 | 0 | 0 | 8 | 0 | 2 | 0 | 0 | 6 | 3 | 0 | 2 | 0 | 98 | 49 | 0.119908107 | 0.242559736 | NA | 0 | 1 | NA | 0.6 | 1 | -0.3 |
| SE | 11089 | IQSEC1 | chr3 | - | 12902772 | 12902822 | 12897042 | 12901522 | 12908348 | 12908525 | 0 | 0 | 2 | 0 | 0 | 3 | 0 | 0 | 0 | 0 | 5 | 5 | 98 | 49 | 0.120556389 | 0.24354335 | NA | NA | 0.25 | NA | 0 | 0 | 0.25 |
| SE | 7230 | NFKBIZ | chr3 | + | 101855394 | 101855458 | 101855061 | 101855208 | 101855732 | 101855902 | 0 | 1 | 28 | 0 | 0 | 0 | 5 | 65 | 2 | 0 | 1 | 0 | 98 | 49 | 0.120755712 | 0.243618571 | NA | 1 | 1 | 1 | 0.97 | 1 | 0.01 |
| SE | 6272 | BAZ2B | chr2 | - | 159430862 | 159431156 | 159429199 | 159429260 | 159432756 | 159433363 | 0 | 0 | 18 | 0 | 0 | 3 | 0 | 5 | 0 | 0 | 0 | 1 | 98 | 49 | 0.12105318 | 0.243891326 | NA | NA | 0.75 | NA | 1 | 0 | 0.25 |
| SE | 4944 | ARAP1 | chr11 | - | 72699004 | 72699107 | 72697910 | 72698106 | 72699416 | 72699552 | 2 | 8 | 34 | 0 | 0 | 2 | 0 | 30 | 7 | 0 | 0 | 0 | 98 | 49 | 0.121497957 | 0.24431631 | 1 | 1 | 0.895 | NA | 1 | 1 | -0.035 |
| SE | 11629 | ATP2A3 | chr17 | - | 3953346 | 3953429 | 3951580 | 3951685 | 3953692 | 3953710 | 0 | 2 | 4 | 0 | 0 | 0 | 0 | 8 | 4 | 0 | 0 | 1 | 98 | 49 | 0.121589221 | 0.24431631 | NA | 1 | 1 | NA | 1 | 0.667 | 0.166 |
| SE | 3974 | GLT1D1 | chr12 | + | 128875913 | 128876062 | 128853535 | 128853649 | 128888638 | 128888744 | 0 | 3 | 18 | 0 | 0 | 0 | 0 | 7 | 3 | 0 | 0 | 1 | 98 | 49 | 0.122907287 | 0.246635051 | NA | 1 | 1 | NA | 1 | 0.6 | 0.2 |
| SE | 3455 | GPCPD1 | chr20 | - | 5586193 | 5586269 | 5584280 | 5584322 | 5593326 | 5593411 | 0 | 0 | 11 | 0 | 0 | 2 | 0 | 7 | 0 | 0 | 6 | 1 | 98 | 49 | 0.123901232 | 0.24829807 | NA | NA | 0.733 | NA | 0.368 | 0 | 0.549 |
| SE | 812 | MKLN1 | chr7 | + | 131445773 | 131445903 | 131443480 | 131443702 | 131463216 | 131463364 | 0 | 1 | 9 | 0 | 1 | 0 | 2 | 3 | 0 | 0 | 0 | 0 | 98 | 49 | 0.124958526 | 0.24977771 | NA | 0.333 | 1 | 1 | 1 | NA | -0.334 |
| SE | 8076 | NADK | chr1 | - | 1754050 | 1754208 | 1753566 | 1753649 | 1754283 | 1754383 | 0 | 4 | 21 | 0 | 0 | 2 | 4 | 17 | 5 | 0 | 0 | 0 | 98 | 49 | 0.125009625 | 0.24977771 | NA | 1 | 0.84 | 1 | 1 | 1 | -0.08 |
| SE | 6057 | HCLS1 | chr3 | - | 121632331 | 121632563 | 121632100 | 121632184 | 121633066 | 121633171 | 0 | 10 | 51 | 0 | 0 | 2 | 0 | 41 | 11 | 0 | 0 | 0 | 98 | 49 | 0.125175468 | 0.24977771 | NA | 1 | 0.927 | NA | 1 | 1 | -0.036 |
| SE | 11932 | FBRS | chr16 | + | 30661303 | 30661333 | 30661179 | 30661215 | 30662419 | 30662468 | 0 | 0 | 1 | 0 | 0 | 1 | 0 | 0 | 3 | 0 | 0 | 0 | 79 | 49 | 0.12530432 | 0.24977771 | NA | NA | 0.383 | NA | NA | 1 | -0.617 |
| A3SS | 1733 | ZSWIM8 | chr10 | + | 73794540 | 73794639 | 73794555 | 73794639 | 73794146 | 73794330 | 0 | 1 | 7 | 0 | 1 | 0 | 0 | 3 | 0 | 5 | 0 | 0 | 64 | 49 | 0.125310145 | 0.30480846 | NA | 0.434 | 1 | 0 | 1 | NA | 0.217 |
| SE | 7879 | ALPL | chr1 | + | 21573664 | 21573799 | 21570304 | 21570374 | 21575732 | 21575924 | 0 | 0 | 18 | 0 | 0 | 1 | 0 | 14 | 1 | 0 | 1 | 1 | 98 | 49 | 0.125632388 | 0.250099972 | NA | NA | 0.9 | NA | 0.875 | 0.333 | 0.296 |
| SE | 8247 | PRRC2C | chr1 | + | 171512031 | 171512200 | 171485526 | 171485735 | 171513000 | 171513172 | 0 | 1 | 4 | 0 | 1 | 0 | 0 | 8 | 2 | 0 | 0 | 0 | 98 | 49 | 0.127922639 | 0.254322389 | NA | 0.333 | 1 | NA | 1 | 1 | -0.334 |
| SE | 10699 | TLE3 | chr15 | - | 70054437 | 70054685 | 70053226 | 70053374 | 70055048 | 70055298 | 0 | 4 | 15 | 0 | 0 | 4 | 0 | 21 | 6 | 0 | 1 | 0 | 98 | 49 | 0.128234493 | 0.254605606 | NA | 1 | 0.652 | NA | 0.913 | 1 | -0.13 |
| SE | 3888 | LCP1 | chr13 | - | 46158825 | 46158989 | 46158521 | 46158651 | 46159598 | 46159686 | 0 | 1 | 110 | 0 | 0 | 5 | 0 | 59 | 2 | 0 | 6 | 0 | 98 | 49 | 0.129117546 | 0.256020675 | NA | 1 | 0.917 | NA | 0.831 | 1 | 0.043 |
| SE | 6628 | PCBP2 | chr12 | + | 53465931 | 53465973 | 53462480 | 53462567 | 53467220 | 53467293 | 0 | 2 | 30 | 0 | 1 | 3 | 0 | 11 | 0 | 0 | 1 | 0 | 91 | 49 | 0.129961836 | 0.257034707 | NA | 0.519 | 0.843 | NA | 0.856 | NA | -0.175 |
| SE | 7727 | FGD3 | chr9 | + | 93019830 | 93019861 | 93018135 | 93018215 | 93020316 | 93020424 | 0 | 3 | 10 | 0 | 0 | 1 | 0 | 10 | 1 | 0 | 3 | 0 | 80 | 49 | 0.129970976 | 0.257034707 | NA | 1 | 0.86 | NA | 0.671 | 1 | 0.094 |
| SE | 9562 | OASL | chr12 | - | 121027575 | 121027817 | 121023989 | 121024137 | 121031441 | 121031617 | 0 | 0 | 26 | 0 | 1 | 2 | 1 | 10 | 1 | 0 | 0 | 0 | 98 | 49 | 0.130217062 | 0.257182975 | NA | 0 | 0.867 | 1 | 1 | 1 | -0.567 |
| SE | 8139 | ITGAX | chr16 | + | 31372377 | 31372509 | 31371629 | 31371784 | 31372596 | 31372670 | 0 | 16 | 38 | 0 | 0 | 2 | 0 | 24 | 19 | 0 | 0 | 0 | 98 | 49 | 0.132864223 | 0.262066834 | NA | 1 | 0.905 | NA | 1 | 1 | -0.047 |
| SE | 6225 | AHCTF1 | chr1 | - | 246916141 | 246916395 | 246913231 | 246913412 | 246918249 | 246918377 | 0 | 0 | 4 | 0 | 0 | 1 | 0 | 0 | 0 | 0 | 0 | 1 | 98 | 49 | 0.133877166 | 0.263718717 | NA | NA | 0.667 | NA | NA | 0 | 0.667 |
| SE | 5499 | ADAM8 | chr10 | - | 133275483 | 133275587 | 133274158 | 133274235 | 133276771 | 133276903 | 0 | 7 | 13 | 0 | 0 | 0 | 1 | 20 | 14 | 0 | 0 | 2 | 98 | 49 | 0.134548885 | 0.264504128 | NA | 1 | 1 | 1 | 1 | 0.778 | 0.074 |
| SE | 12648 | TLN1 | chr9 | - | 35704668 | 35704815 | 35704331 | 35704498 | 35705550 | 35705670 | 0 | 3 | 28 | 0 | 0 | 0 | 1 | 6 | 8 | 0 | 2 | 0 | 98 | 49 | 0.134879669 | 0.264504128 | NA | 1 | 1 | 1 | 0.6 | 1 | 0.133 |
| SE | 5934 | RASGRP2 | chr11 | - | 64727295 | 64727360 | 64726910 | 64727131 | 64728862 | 64729042 | 0 | 0 | 7 | 0 | 1 | 0 | 0 | 1 | 1 | 0 | 0 | 0 | 98 | 49 | 0.134979818 | 0.264504128 | NA | 0 | 1 | NA | 1 | 1 | -0.5 |
| SE | 9259 | MOV10 | chr1 | + | 112695415 | 112695574 | 112694452 | 112694629 | 112696147 | 112696251 | 0 | 0 | 7 | 0 | 1 | 0 | 0 | 1 | 1 | 0 | 0 | 0 | 98 | 49 | 0.134979818 | 0.264504128 | NA | 0 | 1 | NA | 1 | 1 | -0.5 |
| SE | 7950 | PPP4R1 | chr18 | - | 9562921 | 9563046 | 9561979 | 9562075 | 9563377 | 9563550 | 0 | 0 | 4 | 0 | 0 | 5 | 0 | 2 | 2 | 0 | 1 | 0 | 98 | 49 | 0.135910558 | 0.265981209 | NA | NA | 0.286 | NA | 0.5 | 1 | -0.464 |
| A3SS | 2930 | LRRC75A-AS1 | chr17 | + | 16440110 | 16440253 | 16440184 | 16440253 | 16439326 | 16439414 | 0 | 0 | 2 | 0 | 0 | 1 | 0 | 1 | 4 | 0 | 1 | 0 | 98 | 49 | 0.136565787 | 0.324235571 | NA | NA | 0.5 | NA | 0.333 | 1 | -0.166 |
| SE | 865 | NAMPT | chr7 | - | 106274945 | 106275049 | 106272529 | 106272658 | 106277022 | 106277179 | 0 | 15 | 313 | 0 | 0 | 2 | 3 | 467 | 4 | 0 | 6 | 0 | 98 | 49 | 0.136632841 | 0.267047022 | NA | 1 | 0.987 | 1 | 0.975 | 1 | 0.002 |
| A3SS | 1281 | CHD4 | chr12 | - | 6601959 | 6602175 | 6601959 | 6602154 | 6602375 | 6602497 | 0 | 0 | 2 | 0 | 0 | 1 | 0 | 17 | 4 | 4 | 4 | 0 | 70 | 49 | 0.136899463 | 0.324235571 | NA | NA | 0.583 | 0 | 0.748 | 1 | 0 |
| A5SS | 1981 | C12orf57 | chr12 | + | 6944379 | 6944652 | 6944379 | 6944565 | 6945770 | 6946003 | 0 | 2 | 1 | 0 | 0 | 1 | 0 | 4 | 3 | 0 | 0 | 0 | 98 | 49 | 0.137146559 | 0.309762055 | NA | 1 | 0.333 | NA | 1 | 1 | -0.334 |
| SE | 8155 | KMT2E | chr7 | + | 105038125 | 105038199 | 105014189 | 105014535 | 105040838 | 105041023 | 0 | 0 | 7 | 0 | 0 | 2 | 0 | 12 | 0 | 0 | 2 | 1 | 98 | 49 | 0.137637656 | 0.268661555 | NA | NA | 0.636 | NA | 0.75 | 0 | 0.261 |
| SE | 11184 | MEF2D | chr1 | - | 156480833 | 156480971 | 156479585 | 156479796 | 156482436 | 156482640 | 1 | 4 | 3 | 0 | 0 | 0 | 0 | 3 | 5 | 0 | 2 | 0 | 98 | 49 | 0.139889993 | 0.272539469 | 1 | 1 | 1 | NA | 0.429 | 1 | 0.285 |
| SE | 1447 | BAG6 | chr6 | - | 31643889 | 31643974 | 31643100 | 31643115 | 31644081 | 31644194 | 0 | 1 | 2 | 0 | 0 | 1 | 0 | 6 | 1 | 0 | 0 | 0 | 98 | 49 | 0.139987006 | 0.272539469 | NA | 1 | 0.5 | NA | 1 | 1 | -0.25 |
| SE | 1213 | VNN2 | chr6 | - | 132753417 | 132753491 | 132752672 | 132752749 | 132755842 | 132756035 | 0 | 0 | 3 | 0 | 0 | 10 | 0 | 2 | 1 | 0 | 3 | 0 | 98 | 49 | 0.140487467 | 0.273115403 | NA | NA | 0.13 | NA | 0.25 | 1 | -0.495 |
| SE | 4623 | SORL1 | chr11 | + | 121470006 | 121470123 | 121452202 | 121452616 | 121478117 | 121478243 | 0 | 9 | 67 | 0 | 0 | 1 | 1 | 40 | 3 | 0 | 2 | 0 | 98 | 49 | 0.140646255 | 0.273115403 | NA | 1 | 0.971 | 1 | 0.909 | 1 | 0.016 |
| SE | 2848 | PIM3 | chr22 | + | 49961124 | 49961234 | 49960494 | 49961032 | 49961317 | 49961368 | 0 | 8 | 15 | 0 | 1 | 0 | 0 | 32 | 7 | 0 | 0 | 0 | 98 | 49 | 0.14293986 | 0.277211109 | NA | 0.8 | 1 | NA | 1 | 1 | -0.1 |
| RI | 341 | CDKN1C | chr11 | - | 2884669 | 2885765 | 2884669 | 2885466 | 2885633 | 2885765 | 1 | 1 | 2 | 0 | 0 | 2 | 2 | 0 | 1 | 0 | 0 | 0 | 98 | 49 | 0.143188949 | 0.496482453 | 1 | 1 | 0.333 | 1 | NA | 1 | -0.222 |
| SE | 2268 | TNK2 | chr3 | - | 195882050 | 195882328 | 195879048 | 195879175 | 195883156 | 195883309 | 0 | 0 | 3 | 0 | 0 | 1 | 2 | 2 | 0 | 0 | 2 | 1 | 98 | 49 | 0.143961591 | 0.278473967 | NA | NA | 0.6 | 1 | 0.333 | 0 | 0.156 |
| SE | 8090 | SCLT1 | chr4 | - | 128965226 | 128965318 | 128959599 | 128959777 | 128970377 | 128970468 | 0 | 0 | 3 | 0 | 0 | 1 | 0 | 6 | 0 | 0 | 0 | 1 | 98 | 49 | 0.143961591 | 0.278473967 | NA | NA | 0.6 | NA | 1 | 0 | 0.1 |
| SE | 9704 | ABTB1 | chr3 | + | 127676271 | 127676431 | 127675969 | 127676114 | 127676535 | 127676581 | 0 | 3 | 20 | 0 | 0 | 0 | 1 | 11 | 4 | 0 | 0 | 1 | 98 | 49 | 0.145196396 | 0.280100163 | NA | 1 | 1 | 1 | 1 | 0.667 | 0.111 |
| A5SS | 1928 | PIGG | chr4 | + | 499199 | 499725 | 499199 | 499489 | 500395 | 500601 | 0 | 0 | 1 | 0 | 0 | 3 | 0 | 0 | 1 | 0 | 0 | 0 | 98 | 49 | 0.145772881 | 0.312928566 | NA | NA | 0.143 | NA | NA | 1 | -0.857 |
| A5SS | 1929 | PIGG | chr4 | + | 499199 | 500047 | 499199 | 499489 | 500395 | 500601 | 0 | 0 | 1 | 0 | 0 | 3 | 0 | 0 | 1 | 0 | 0 | 0 | 98 | 49 | 0.145772881 | 0.312928566 | NA | NA | 0.143 | NA | NA | 1 | -0.857 |
| SE | 7876 | TLR4 | chr9 | + | 117708562 | 117708729 | 117704174 | 117704565 | 117712388 | 117717491 | 0 | 0 | 10 | 0 | 0 | 10 | 0 | 2 | 0 | 0 | 6 | 2 | 98 | 49 | 0.146224332 | 0.280100163 | NA | NA | 0.333 | NA | 0.143 | 0 | 0.262 |
| SE | 324 | STOM | chr9 | - | 121366115 | 121366245 | 121356052 | 121356156 | 121370126 | 121370304 | 0 | 2 | 0 | 0 | 6 | 0 | 0 | 1 | 0 | 0 | 0 | 0 | 98 | 49 | 0.146368947 | 0.280100163 | NA | 0.143 | NA | NA | 1 | NA | -0.857 |
| A5SS | 468 | ARFIP1 | chr4 | + | 152779936 | 152780226 | 152779936 | 152780091 | 152829624 | 152829726 | 0 | 0 | 1 | 0 | 0 | 1 | 0 | 2 | 2 | 0 | 0 | 0 | 98 | 49 | 0.146398672 | 0.312928566 | NA | NA | 0.333 | NA | 1 | 1 | -0.667 |
| RI | 1023 | OXLD1 | chr17 | - | 81665035 | 81666637 | 81665035 | 81665584 | 81666517 | 81666637 | 0 | 0 | 1 | 0 | 0 | 1 | 0 | 0 | 2 | 0 | 0 | 0 | 98 | 49 | 0.146398672 | 0.496482453 | NA | NA | 0.333 | NA | NA | 1 | -0.667 |
| SE | 3770 | KIAA1109 | chr4 | + | 122226706 | 122226835 | 122224497 | 122224677 | 122229120 | 122229240 | 0 | 0 | 1 | 0 | 0 | 1 | 0 | 9 | 2 | 0 | 1 | 0 | 98 | 49 | 0.146398672 | 0.280100163 | NA | NA | 0.333 | NA | 0.818 | 1 | -0.576 |
| SE | 6502 | GAS7 | chr17 | - | 9969676 | 9969762 | 9959201 | 9959255 | 9981803 | 9981884 | 0 | 1 | 9 | 0 | 1 | 0 | 0 | 2 | 0 | 0 | 0 | 0 | 98 | 49 | 0.146398672 | 0.280100163 | NA | 0.333 | 1 | NA | 1 | NA | -0.334 |
| SE | 11396 | ADGRG3 | chr16 | + | 57683931 | 57684212 | 57680504 | 57680617 | 57684389 | 57684483 | 0 | 0 | 1 | 0 | 0 | 1 | 0 | 3 | 2 | 0 | 3 | 0 | 98 | 49 | 0.146398672 | 0.280100163 | NA | NA | 0.333 | NA | 0.333 | 1 | -0.333 |
| SE | 13361 | SMARCA5 | chr4 | + | 143540362 | 143540495 | 143538785 | 143538938 | 143543508 | 143543657 | 0 | 0 | 1 | 0 | 0 | 1 | 0 | 3 | 2 | 0 | 0 | 0 | 98 | 49 | 0.146398672 | 0.280100163 | NA | NA | 0.333 | NA | 1 | 1 | -0.667 |
| SE | 13734 | SERPINA1 | chr14 | - | 94388800 | 94389010 | 94388559 | 94388660 | 94390456 | 94390692 | 0 | 1 | 17 | 0 | 1 | 3 | 0 | 2 | 0 | 4 | 0 | 0 | 98 | 49 | 0.146398672 | 0.280100163 | NA | 0.333 | 0.739 | 0 | 1 | NA | 0.036 |
| SE | 4332 | NCF2 | chr1 | - | 183567203 | 183567345 | 183566919 | 183566988 | 183569141 | 183569185 | 0 | 1 | 39 | 0 | 0 | 2 | 2 | 30 | 2 | 0 | 0 | 0 | 98 | 49 | 0.146479526 | 0.280100163 | NA | 1 | 0.907 | 1 | 1 | 1 | -0.046 |
| SE | 9574 | FAS | chr10 | + | 89010752 | 89010815 | 89010538 | 89010600 | 89011998 | 89012081 | 0 | 0 | 2 | 0 | 0 | 2 | 0 | 6 | 2 | 0 | 0 | 0 | 98 | 49 | 0.147583451 | 0.281852512 | NA | NA | 0.333 | NA | 1 | 1 | -0.667 |
| A5SS | 2262 | TLE3 | chr15 | - | 70074502 | 70074607 | 70074532 | 70074607 | 70066013 | 70066218 | 0 | 0 | 7 | 0 | 0 | 4 | 0 | 2 | 3 | 0 | 2 | 0 | 79 | 49 | 0.148103596 | 0.312928566 | NA | NA | 0.52 | NA | 0.383 | 1 | -0.171 |
| SE | 2581 | PRKCD | chr3 | + | 53165103 | 53165215 | 53161206 | 53161428 | 53178403 | 53178537 | 0 | 4 | 3 | 0 | 2 | 1 | 0 | 1 | 2 | 0 | 1 | 3 | 98 | 49 | 0.148640738 | 0.283511458 | NA | 0.5 | 0.6 | NA | 0.333 | 0.25 | 0.259 |
| SE | 12586 | SBNO2 | chr19 | - | 1113534 | 1113704 | 1112817 | 1112949 | 1114230 | 1114422 | 0 | 9 | 21 | 0 | 0 | 1 | 0 | 40 | 18 | 0 | 0 | 0 | 98 | 49 | 0.149179961 | 0.284179318 | NA | 1 | 0.913 | NA | 1 | 1 | -0.043 |
| SE | 8708 | CTSS | chr1 | - | 150764637 | 150764764 | 150757857 | 150757980 | 150765697 | 150765957 | 0 | 1 | 129 | 0 | 0 | 0 | 20 | 18 | 6 | 0 | 0 | 1 | 98 | 49 | 0.150561932 | 0.28644884 | NA | 1 | 1 | 1 | 1 | 0.75 | 0.083 |
| SE | 759 | RBM33 | chr7 | + | 155665174 | 155665253 | 155644508 | 155644919 | 155672866 | 155672915 | 0 | 4 | 8 | 0 | 0 | 0 | 0 | 7 | 2 | 0 | 0 | 1 | 98 | 49 | 0.15316134 | 0.291025909 | NA | 1 | 1 | NA | 1 | 0.5 | 0.25 |
| SE | 6532 | CARD16 | chr11 | - | 105043482 | 105043545 | 105041325 | 105041719 | 105044391 | 105044658 | 0 | 1 | 14 | 0 | 1 | 5 | 4 | 1 | 0 | 2 | 5 | 0 | 98 | 49 | 0.15394085 | 0.292137749 | NA | 0.333 | 0.583 | 0.5 | 0.091 | NA | 0.162 |
| SE | 7042 | SRSF2 | chr17 | - | 76735771 | 76735875 | 76734114 | 76735158 | 76736153 | 76736464 | 0 | 1 | 2 | 0 | 2 | 3 | 0 | 3 | 0 | 0 | 12 | 1 | 98 | 49 | 0.154541762 | 0.292908283 | NA | 0.2 | 0.25 | NA | 0.111 | 0 | 0.17 |
| SE | 9278 | PIK3CD | chr1 | + | 9721443 | 9721587 | 9721126 | 9721248 | 9721760 | 9721860 | 0 | 1 | 19 | 0 | 0 | 3 | 0 | 8 | 1 | 0 | 0 | 0 | 98 | 49 | 0.156377277 | 0.296013914 | NA | 1 | 0.76 | NA | 1 | 1 | -0.12 |
| SE | 6507 | TRIM33 | chr1 | - | 114397939 | 114397990 | 114392776 | 114397860 | 114399456 | 114399609 | 0 | 0 | 7 | 0 | 0 | 2 | 0 | 4 | 5 | 0 | 0 | 0 | 98 | 49 | 0.156576702 | 0.296018595 | NA | NA | 0.636 | NA | 1 | 1 | -0.364 |
| A3SS | 3032 | NME4 | chr16 | + | 400108 | 400754 | 400218 | 400754 | 399626 | 399739 | 2 | 0 | 0 | 1 | 0 | 0 | 3 | 0 | 0 | 0 | 0 | 0 | 98 | 49 | 0.156977715 | 0.366960892 | 0.5 | NA | NA | 1 | NA | NA | -0.5 |
| SE | 5476 | MICAL2 | chr11 | + | 12138389 | 12138460 | 12110575 | 12110726 | 12162078 | 12162419 | 0 | 0 | 2 | 0 | 0 | 1 | 1 | 2 | 3 | 0 | 2 | 0 | 98 | 49 | 0.156977715 | 0.296403901 | NA | NA | 0.5 | 1 | 0.333 | 1 | -0.278 |
| SE | 4367 | LRRK2 | chr12 | + | 40367005 | 40367077 | 40364841 | 40365050 | 40367643 | 40369284 | 0 | 0 | 13 | 0 | 0 | 3 | 0 | 13 | 6 | 0 | 3 | 0 | 98 | 49 | 0.157632777 | 0.297267333 | NA | NA | 0.684 | NA | 0.684 | 1 | -0.158 |
| SE | 9609 | TAPBP | chr6 | - | 33303954 | 33303989 | 33299694 | 33301771 | 33304127 | 33304217 | 0 | 2 | 23 | 0 | 1 | 0 | 4 | 10 | 1 | 0 | 0 | 0 | 84 | 49 | 0.158787547 | 0.299069779 | NA | 0.538 | 1 | 1 | 1 | 1 | -0.231 |
| A5SS | 3020 | SERPINA1 | chr14 | - | 94390419 | 94390692 | 94390456 | 94390692 | 94388818 | 94389010 | 0 | 4 | 5 | 0 | 1 | 15 | 3 | 3 | 2 | 0 | 1 | 0 | 86 | 49 | 0.159752672 | 0.332184127 | NA | 0.695 | 0.16 | 1 | 0.631 | 1 | -0.45 |
| SE | 8027 | GNAS | chr20 | + | 58898940 | 58898985 | 58895611 | 58895684 | 58903527 | 58903585 | 0 | 1 | 2 | 0 | 3 | 2 | 0 | 0 | 0 | 0 | 4 | 0 | 94 | 49 | 0.16039283 | 0.301490206 | NA | 0.148 | 0.343 | NA | 0 | NA | 0.245 |
| SE | 1448 | BAG6 | chr6 | - | 31643889 | 31643977 | 31642828 | 31643115 | 31644081 | 31644194 | 0 | 1 | 3 | 0 | 0 | 1 | 0 | 7 | 1 | 0 | 0 | 0 | 98 | 49 | 0.160473829 | 0.301490206 | NA | 1 | 0.6 | NA | 1 | 1 | -0.2 |
| SE | 4626 | SORL1 | chr11 | + | 121557313 | 121557405 | 121555186 | 121555318 | 121558590 | 121558837 | 0 | 10 | 73 | 0 | 0 | 2 | 0 | 39 | 5 | 0 | 3 | 0 | 98 | 49 | 0.162347432 | 0.304629451 | NA | 1 | 0.948 | NA | 0.867 | 1 | 0.04 |
| A3SS | 1989 | PPP6R2 | chr22 | + | 50419345 | 50419462 | 50419348 | 50419462 | 50418866 | 50418979 | 0 | 1 | 0 | 0 | 0 | 1 | 0 | 2 | 1 | 0 | 0 | 0 | 52 | 49 | 0.163314358 | 0.376879289 | NA | 1 | 0 | NA | 1 | 1 | -0.5 |
| SE | 8529 | CKLF | chr16 | + | 66558189 | 66558348 | 66552562 | 66552793 | 66563121 | 66563217 | 0 | 0 | 8 | 0 | 0 | 2 | 0 | 3 | 5 | 0 | 0 | 0 | 98 | 49 | 0.164123685 | 0.307578427 | NA | NA | 0.667 | NA | 1 | 1 | -0.333 |
| SE | 695 | DOCK5 | chr8 | + | 25368191 | 25368250 | 25366869 | 25366970 | 25368570 | 25368725 | 0 | 0 | 11 | 0 | 0 | 5 | 0 | 5 | 3 | 0 | 1 | 0 | 98 | 49 | 0.166703165 | 0.311886731 | NA | NA | 0.524 | NA | 0.714 | 1 | -0.333 |
| SE | 7636 | IRF7 | chr11 | - | 612998 | 613117 | 612554 | 612800 | 613205 | 613595 | 0 | 9 | 20 | 0 | 0 | 0 | 0 | 20 | 5 | 0 | 0 | 1 | 98 | 49 | 0.16703421 | 0.311886731 | NA | 1 | 1 | NA | 1 | 0.714 | 0.143 |
| SE | 2487 | ATG3 | chr3 | - | 112553279 | 112553329 | 112550191 | 112550262 | 112558375 | 112558417 | 0 | 0 | 4 | 0 | 0 | 3 | 1 | 4 | 2 | 0 | 0 | 0 | 98 | 49 | 0.167045122 | 0.311886731 | NA | NA | 0.4 | 1 | 1 | 1 | -0.6 |
| SE | 471 | DOCK8 | chr9 | + | 317042 | 317128 | 311953 | 312166 | 325670 | 325737 | 0 | 10 | 33 | 0 | 0 | 0 | 2 | 15 | 7 | 0 | 0 | 2 | 98 | 49 | 0.167661893 | 0.311930048 | NA | 1 | 1 | 1 | 1 | 0.636 | 0.121 |
| SE | 4637 | ARID5A | chr2 | + | 96549320 | 96549459 | 96547401 | 96547517 | 96549752 | 96549805 | 0 | 2 | 2 | 0 | 0 | 3 | 0 | 1 | 0 | 0 | 6 | 4 | 98 | 49 | 0.167831186 | 0.311930048 | NA | 1 | 0.25 | NA | 0.077 | 0 | 0.587 |
| RI | 566 | SPRTN | chr1 | + | 231351303 | 231355027 | 231351303 | 231351571 | 231352609 | 231355027 | 0 | 0 | 1 | 0 | 0 | 2 | 2 | 2 | 1 | 0 | 0 | 0 | 98 | 49 | 0.167898476 | 0.545670046 | NA | NA | 0.2 | 1 | 1 | 1 | -0.8 |
| SE | 5473 | ARNTL | chr11 | + | 13365499 | 13365592 | 13360344 | 13360424 | 13366664 | 13366782 | 0 | 0 | 1 | 0 | 0 | 2 | 2 | 4 | 1 | 0 | 3 | 0 | 98 | 49 | 0.167898476 | 0.311930048 | NA | NA | 0.2 | 1 | 0.4 | 1 | -0.6 |
| SE | 11688 | CDK11B | chr1 | - | 1637080 | 1637202 | 1636896 | 1637004 | 1637407 | 1637513 | 0 | 1 | 0 | 0 | 2 | 0 | 0 | 1 | 1 | 0 | 0 | 0 | 98 | 49 | 0.167898476 | 0.311930048 | NA | 0.2 | NA | NA | 1 | 1 | -0.8 |
| A3SS | 3932 | OTOF | chr2 | - | 26472518 | 26472649 | 26472518 | 26472589 | 26473131 | 26473294 | 0 | 0 | 1 | 0 | 1 | 1 | 0 | 0 | 0 | 0 | 2 | 2 | 98 | 49 | 0.167980116 | 0.379975466 | NA | 0 | 0.333 | NA | 0 | 0 | 0.167 |
| A3SS | 2018 | RNF19B | chr1 | - | 32948221 | 32948366 | 32948221 | 32948363 | 32949568 | 32949774 | 0 | 0 | 1 | 0 | 0 | 2 | 0 | 1 | 1 | 0 | 1 | 0 | 52 | 49 | 0.168877985 | 0.379975466 | NA | NA | 0.32 | NA | 0.485 | 1 | -0.422 |
| SE | 7215 | LSP1 | chr11 | + | 1880086 | 1880224 | 1852969 | 1853197 | 1881431 | 1881596 | 0 | 7 | 39 | 0 | 0 | 0 | 2 | 37 | 11 | 0 | 0 | 1 | 98 | 49 | 0.17099036 | 0.31689431 | NA | 1 | 1 | 1 | 1 | 0.846 | 0.051 |
| SE | 1069 | HNRNPA2B1 | chr7 | - | 26197830 | 26197866 | 26197621 | 26197732 | 26200571 | 26200793 | 0 | 0 | 1 | 0 | 3 | 7 | 0 | 2 | 0 | 0 | 2 | 1 | 85 | 49 | 0.170992206 | 0.31689431 | NA | 0 | 0.076 | NA | 0.366 | 0 | -0.145 |
| SE | 4615 | PTPRJ | chr11 | + | 48112746 | 48112983 | 48110057 | 48110076 | 48121002 | 48121266 | 2 | 11 | 32 | 0 | 2 | 0 | 0 | 9 | 5 | 0 | 0 | 0 | 98 | 49 | 0.171767102 | 0.317938368 | 1 | 0.733 | 1 | NA | 1 | 1 | -0.089 |
| A3SS | 502 | ARG1 | chr6 | + | 131579086 | 131579285 | 131579110 | 131579285 | 131576662 | 131576735 | 0 | 0 | 2 | 0 | 0 | 4 | 0 | 0 | 1 | 0 | 0 | 0 | 73 | 49 | 0.173440158 | 0.385422574 | NA | NA | 0.251 | NA | NA | 1 | -0.749 |

| | | | | | | | | | | | | | | | | | | | | | | | | | | | | | | | | | |
|---|---|---|---|---|---|---|---|---|---|---|---|---|---|---|---|---|---|---|---|---|---|---|---|---|---|---|---|---|---|---|---|---|---|
| SE | 11874 | MGAM | chr7 | + | 142036168 | 142036285 | 142034669 | 142034841 | 142036822 | 142036977 | 0 | 2 | 23 | 0 | 0 | 0 | 6 | 6 | 3 | 0 | 0 | 2 | 98 | 49 | 0.173570524 | 0.3208813 | NA | 1 | 1 | 1 | 1 | 0.429 | 0.19 |
| A5SS | 1004 | GMPR2 | chr14 | + | 24232421 | 24232847 | 24232421 | 24232788 | 24233218 | 24233340 | 3 | 0 | 2 | 0 | 0 | 4 | 0 | 4 | 1 | 0 | 1 | 0 | 98 | 49 | 0.174787188 | 0.357767525 | 1 | NA | 0.2 | NA | 0.667 | 1 | -0.234 |
| SE | 7320 | IL18R1 | chr2 | + | 102373936 | 102373993 | 102371952 | 102372118 | 102375906 | 102376063 | 0 | 0 | 2 | 0 | 0 | 4 | 0 | 0 | 1 | 0 | 0 | 0 | 98 | 49 | 0.174787188 | 0.322733591 | NA | NA | 0.2 | NA | NA | 1 | -0.8 |
| SE | 4120 | XPO6 | chr16 | - | 28104545 | 28104707 | 28101846 | 28101945 | 28106042 | 28106214 | 0 | 7 | 58 | 0 | 2 | 0 | 1 | 60 | 4 | 0 | 2 | 0 | 98 | 49 | 0.176059763 | 0.324652462 | NA | 0.636 | 1 | 1 | 0.937 | 1 | -0.161 |
| SE | 235 | CSF2RA | chrX | + | 1303922 | 1304019 | 1300490 | 1300626 | 1305445 | 1305527 | 0 | 1 | 11 | 0 | 0 | 1 | 0 | 2 | 0 | 0 | 5 | 1 | 98 | 49 | 0.176258423 | 0.324652462 | NA | 1 | 0.846 | NA | 0.167 | 0 | 0.84 |
| SE | 12687 | MYO1F | chr19 | - | 8548035 | 8548122 | 8545649 | 8545736 | 8548236 | 8548317 | 0 | 9 | 61 | 0 | 0 | 2 | 0 | 38 | 5 | 0 | 0 | 0 | 98 | 49 | 0.177681081 | 0.326872294 | NA | 1 | 0.938 | NA | 1 | 1 | -0.031 |
| SE | 10039 | SYNE2 | chr14 | + | 64009965 | 64010116 | 64007042 | 64007222 | 64016472 | 64016631 | 0 | 0 | 3 | 0 | 0 | 2 | 0 | 7 | 2 | 0 | 0 | 0 | 98 | 49 | 0.179363678 | 0.329564313 | NA | NA | 0.429 | NA | 1 | 1 | -0.571 |
| SE | 5557 | SPATA13 | chr13 | + | 24278613 | 24278821 | 24270686 | 24270912 | 24284134 | 24284271 | 0 | 0 | 1 | 0 | 1 | 3 | 0 | 0 | 0 | 0 | 3 | 4 | 98 | 49 | 0.18048442 | 0.331070043 | NA | 0 | 0.143 | NA | 0 | 0 | 0.071 |
| SE | 10589 | RNF213 | chr17 | + | 80369767 | 80369867 | 80369501 | 80369671 | 80371873 | 80371985 | 0 | 10 | 63 | 0 | 0 | 2 | 3 | 33 | 13 | 0 | 0 | 0 | 98 | 49 | 0.180942085 | 0.331070043 | NA | 1 | 0.94 | 1 | 1 | 1 | -0.03 |
| SE | 10850 | IRF1 | chr5 | - | 132487925 | 132488025 | 132486953 | 132487130 | 132489391 | 132489483 | 0 | 7 | 14 | 0 | 0 | 1 | 0 | 12 | 1 | 0 | 2 | 0 | 98 | 49 | 0.181005184 | 0.331070043 | NA | 1 | 0.875 | NA | 0.75 | 1 | 0.062 |
| SE | 4941 | ARAP1 | chr11 | - | 72692752 | 72692785 | 72688454 | 72688537 | 72693324 | 72693470 | 0 | 0 | 2 | 0 | 1 | 16 | 0 | 2 | 0 | 0 | 2 | 1 | 82 | 49 | 0.181064255 | 0.331070043 | NA | 0 | 0.07 | NA | 0.374 | 0 | -0.152 |
| SE | 6699 | FCGR2A | chr1 | + | 161510833 | 161510956 | 161509819 | 161510074 | 161513894 | 161513932 | 0 | 1 | 66 | 0 | 0 | 2 | 0 | 40 | 2 | 0 | 0 | 0 | 98 | 49 | 0.181721355 | 0.331867797 | NA | 1 | 0.943 | NA | 1 | 1 | -0.028 |
| SE | 7908 | ATP6V0B | chr1 | + | 43975799 | 43975848 | 43974647 | 43975107 | 43976089 | 43976173 | 0 | 5 | 19 | 0 | 1 | 4 | 1 | 10 | 5 | 0 | 3 | 3 | 98 | 49 | 0.183348939 | 0.334433805 | NA | 0.714 | 0.704 | 1 | 0.625 | 0.455 | 0.016 |
| A3SS | 457 | BTN2A1 | chr6 | + | 26467718 | 26469653 | 26467947 | 26469653 | 26466061 | 26466088 | 0 | 2 | 4 | 0 | 0 | 2 | 0 | 3 | 1 | 0 | 0 | 0 | 98 | 49 | 0.184424674 | 0.403832852 | NA | 1 | 0.5 | NA | 1 | 1 | -0.25 |
| SE | 2077 | HERC5 | chr4 | + | 88470613 | 88470673 | 88469156 | 88469260 | 88472408 | 88472502 | 0 | 2 | 23 | 0 | 0 | 1 | 0 | 41 | 1 | 0 | 3 | 0 | 98 | 49 | 0.18526689 | 0.337522588 | NA | 1 | 0.92 | NA | 0.872 | 1 | 0.024 |
| SE | 8553 | EHBP1L1 | chr11 | + | 65589935 | 65589991 | 65589750 | 65589820 | 65590086 | 65590210 | 0 | 1 | 10 | 0 | 0 | 0 | 3 | 7 | 2 | 0 | 0 | 1 | 98 | 49 | 0.185747044 | 0.337684951 | NA | 1 | 1 | 1 | 1 | 0.5 | 0.167 |
| SE | 5679 | GMIP | chr19 | - | 19640295 | 19640360 | 19640084 | 19640192 | 19640445 | 19640571 | 0 | 9 | 25 | 0 | 0 | 2 | 2 | 12 | 3 | 0 | 3 | 0 | 98 | 49 | 0.185805359 | 0.337684951 | NA | 1 | 0.862 | 1 | 0.667 | 1 | 0.042 |
| SE | 9611 | TAPBP | chr6 | - | 33313693 | 33313864 | 33313216 | 33313477 | 33314004 | 33314387 | 0 | 4 | 1 | 0 | 0 | 1 | 0 | 9 | 3 | 0 | 0 | 0 | 98 | 49 | 0.186099817 | 0.337811625 | NA | 1 | 0.333 | NA | 1 | 1 | -0.334 |
| SE | 6577 | RILPL2 | chr12 | - | 123430507 | 123430659 | 123423043 | 123423157 | 123436081 | 123436733 | 0 | 1 | 12 | 0 | 4 | 4 | 0 | 13 | 6 | 0 | 3 | 3 | 98 | 49 | 0.189327441 | 0.343255903 | NA | 0.111 | 0.6 | NA | 0.684 | 0.5 | -0.237 |
| SE | 6105 | GLUL | chr1 | - | 182390496 | 182390852 | 182388571 | 182388750 | 182391678 | 182392206 | 0 | 2 | 1 | 0 | 5 | 18 | 0 | 1 | 1 | 3 | 24 | 22 | 98 | 49 | 0.189877016 | 0.343454188 | NA | 0.167 | 0.027 | 0 | 0.02 | 0.022 | 0.083 |
| SE | 2138 | CXCL8 | chr4 | + | 73741541 | 73741677 | 73740505 | 73740722 | 73741948 | 73742032 | 1 | 26 | 133 | 0 | 1 | 0 | 6 | 985 | 18 | 0 | 1 | 0 | 98 | 49 | 0.189893833 | 0.343454188 | 1 | 0.929 | 1 | 1 | 0.998 | 1 | -0.023 |
| RI | 624 | MCL1 | chr1 | - | 150578842 | 150579738 | 150578842 | 150578963 | 150579422 | 150579738 | 0 | 41 | 264 | 0 | 0 | 2 | 13 | 148 | 29 | 0 | 0 | 0 | 98 | 49 | 0.190371362 | 0.586344893 | NA | 1 | 0.985 | 1 | 1 | 1 | -0.008 |
| A3SS | 504 | NDUFA2 | chr5 | - | 140645362 | 140645725 | 140645362 | 140645678 | 140647255 | 140647362 | 0 | 0 | 3 | 0 | 1 | 2 | 0 | 0 | 0 | 0 | 0 | 1 | 96 | 49 | 0.190583999 | 0.403832852 | NA | 0 | 0.434 | NA | NA | 0 | 0.217 |
| A5SS | 1894 | MYD88 | chr3 | + | 38140387 | 38140592 | 38140387 | 38140568 | 38140756 | 38140848 | 0 | 0 | 4 | 0 | 1 | 7 | 0 | 0 | 0 | 0 | 0 | 2 | 73 | 49 | 0.190693809 | 0.381630589 | NA | 0 | 0.277 | NA | NA | 0 | 0.139 |
| SE | 12366 | NBEAL2 | chr3 | + | 46996750 | 46996833 | 46996270 | 46996592 | 46996953 | 46997046 | 0 | 7 | 12 | 0 | 0 | 0 | 0 | 4 | 6 | 0 | 4 | 0 | 98 | 49 | 0.19095347 | 0.344955607 | NA | 1 | 1 | NA | 0.333 | 1 | 0.334 |
| A3SS | 2130 | FGD3 | chr9 | + | 93034540 | 93034681 | 93034543 | 93034681 | 93032768 | 93032873 | 0 | 6 | 5 | 0 | 0 | 1 | 0 | 11 | 5 | 0 | 0 | 0 | 52 | 49 | 0.19141247 | 0.403832852 | NA | 1 | 0.825 | NA | 1 | 1 | -0.088 |
| SE | 5865 | NFKBID | chr19 | - | 35897650 | 35897856 | 35896912 | 35897058 | 35898471 | 35898532 | 0 | 7 | 10 | 0 | 2 | 0 | 0 | 4 | 6 | 0 | 0 | 0 | 98 | 49 | 0.192464506 | 0.34726789 | NA | 0.636 | 1 | NA | 1 | 1 | -0.182 |
| A3SS | 4000 | MAP4K4 | chr2 | + | 101859642 | 101859864 | 101859735 | 101859864 | 101858995 | 101859082 | 0 | 0 | 4 | 0 | 0 | 1 | 0 | 1 | 4 | 0 | 0 | 0 | 98 | 49 | 0.193038976 | 0.403832852 | NA | NA | 0.667 | NA | 1 | 1 | -0.333 |
| SE | 3456 | GPCPD1 | chr20 | - | 5593326 | 5593411 | 5584280 | 5584322 | 5598724 | 5598821 | 0 | 1 | 27 | 0 | 0 | 1 | 0 | 38 | 4 | 0 | 0 | 0 | 98 | 49 | 0.193523387 | 0.348759773 | NA | 1 | 0.931 | NA | 1 | 1 | -0.034 |
| A3SS | 1806 | PCBP2 | chr12 | + | 53462480 | 53462567 | 53462492 | 53462567 | 53461014 | 53461143 | 1 | 1 | 8 | 0 | 1 | 19 | 0 | 9 | 0 | 4 | 7 | 0 | 61 | 49 | 0.194077531 | 0.403832852 | 1 | 0.445 | 0.253 | 0 | 0.508 | NA | 0.312 |
| SE | 8686 | PTPRC | chr1 | + | 198702386 | 198702530 | 198699563 | 198699704 | 198703297 | 198703372 | 1 | 0 | 4 | 0 | 3 | 6 | 4 | 3 | 2 | 1 | 11 | 0 | 98 | 49 | 0.195041125 | 0.351074025 | 1 | 0 | 0.25 | 0.667 | 0.12 | 1 | -0.179 |
| A5SS | 1245 | TUBA1A | chr12 | - | 49188416 | 49189324 | 49188976 | 49189324 | 49186610 | 49186833 | 0 | 1 | 1 | 0 | 1 | 3 | 0 | 1 | 0 | 0 | 4 | 0 | 98 | 49 | 0.195185111 | 0.381630589 | NA | 0.333 | 0.143 | NA | 0.111 | NA | 0.127 |
| A5SS | 1246 | TUBA1A | chr12 | - | 49188888 | 49189324 | 49188976 | 49189324 | 49186610 | 49186833 | 0 | 1 | 1 | 0 | 1 | 3 | 0 | 1 | 0 | 0 | 4 | 0 | 98 | 49 | 0.195185111 | 0.381630589 | NA | 0.333 | 0.143 | NA | 0.111 | NA | 0.127 |
| A3SS | 570 | DDX60L | chr4 | - | 168395958 | 168396127 | 168395958 | 168396124 | 168400825 | 168400978 | 0 | 0 | 9 | 0 | 2 | 12 | 0 | 16 | 0 | 0 | 5 | 1 | 52 | 49 | 0.195185878 | 0.403832852 | NA | 0 | 0.414 | NA | 0.751 | 0 | -0.169 |
| RI | 52 | GSTK1 | chr7 | + | 143264991 | 143265296 | 143264991 | 143265092 | 143265260 | 143265296 | 0 | 2 | 3 | 0 | 1 | 6 | 2 | 4 | 1 | 9 | 2 | 0 | 98 | 49 | 0.195448298 | 0.586344893 | NA | 0.5 | 0.2 | 0.1 | 0.5 | 1 | -0.183 |
| SE | 5839 | TUBA1A | chr12 | - | 49186309 | 49186458 | 49184794 | 49185990 | 49186610 | 49186833 | 0 | 5 | 29 | 0 | 1 | 2 | 6 | 17 | 0 | 0 | 1 | 0 | 98 | 49 | 0.195483437 | 0.35144929 | NA | 0.714 | 0.879 | 1 | 0.895 | NA | -0.151 |
| SE | 12944 | SMCHD1 | chr18 | + | 2724898 | 2724995 | 2722518 | 2722663 | 2726451 | 2726524 | 0 | 2 | 53 | 0 | 0 | 1 | 0 | 50 | 1 | 0 | 2 | 0 | 98 | 49 | 0.196703531 | 0.353220319 | NA | 1 | 0.964 | NA | 0.926 | 1 | 0.019 |
| SE | 2289 | BCL6 | chr3 | - | 187726730 | 187726898 | 187725498 | 187725629 | 187728359 | 187728544 | 0 | 1 | 25 | 0 | 0 | 2 | 3 | 7 | 3 | 0 | 3 | 0 | 98 | 49 | 0.197315011 | 0.353895539 | NA | 1 | 0.862 | 1 | 0.538 | 1 | 0.085 |
| SE | 12685 | MYO1F | chr19 | - | 8539946 | 8540028 | 8536948 | 8537055 | 8541905 | 8541991 | 0 | 6 | 55 | 0 | 0 | 2 | 1 | 30 | 5 | 0 | 0 | 0 | 98 | 49 | 0.197961525 | 0.354631909 | NA | 1 | 0.932 | 1 | 1 | 1 | -0.034 |
| SE | 5477 | MICAL2 | chr11 | + | 12138389 | 12138464 | 12110575 | 12110726 | 12162078 | 12162419 | 0 | 0 | 3 | 0 | 0 | 1 | 1 | 3 | 3 | 0 | 2 | 0 | 98 | 49 | 0.199033101 | 0.354859729 | NA | NA | 0.6 | 1 | 0.429 | 1 | -0.21 |
| SE | 5479 | MICAL2 | chr11 | + | 12220200 | 12220458 | 12216218 | 12216319 | 12221643 | 12221759 | 0 | 0 | 3 | 0 | 0 | 1 | 0 | 1 | 3 | 0 | 0 | 0 | 98 | 49 | 0.199033101 | 0.354859729 | NA | NA | 0.6 | NA | 1 | 1 | -0.4 |
| SE | 7788 | CLK3 | chr15 | + | 74627969 | 74628052 | 74627538 | 74627668 | 74628603 | 74628683 | 0 | 0 | 3 | 0 | 0 | 1 | 0 | 0 | 3 | 0 | 0 | 0 | 98 | 49 | 0.199033101 | 0.354859729 | NA | NA | 0.6 | NA | NA | 1 | -0.4 |
| SE | 8840 | C1orf43 | chr1 | - | 154213892 | 154213946 | 154212457 | 154212624 | 154214456 | 154214574 | 0 | 0 | 3 | 0 | 0 | 1 | 0 | 5 | 3 | 0 | 0 | 0 | 98 | 49 | 0.199033101 | 0.354859729 | NA | NA | 0.6 | NA | 1 | 1 | -0.4 |
| SE | 9582 | ANXA11 | chr10 | - | 80168968 | 80169358 | 80167225 | 80167313 | 80170799 | 80170915 | 0 | 0 | 6 | 0 | 0 | 2 | 0 | 2 | 3 | 0 | 0 | 0 | 98 | 49 | 0.200611555 | 0.357250198 | NA | NA | 0.6 | NA | 1 | 1 | -0.4 |
| SE | 10668 | LAPTM5 | chr1 | - | 30741639 | 30741716 | 30739808 | 30739937 | 30742455 | 30742549 | 0 | 4 | 37 | 0 | 0 | 1 | 6 | 52 | 6 | 0 | 0 | 0 | 98 | 49 | 0.201003059 | 0.357523784 | NA | 1 | 0.949 | 1 | 1 | 1 | -0.026 |
| SE | 1136 | IGF2R | chr6 | + | 160048374 | 160048543 | 160047791 | 160047907 | 160050472 | 160050652 | 0 | 3 | 42 | 0 | 0 | 2 | 0 | 21 | 4 | 0 | 0 | 0 | 98 | 49 | 0.201533872 | 0.357884259 | NA | 1 | 0.913 | NA | 1 | 1 | -0.043 |
| SE | 5119 | SLC3A2 | chr11 | + | 62883988 | 62884112 | 62882907 | 62882999 | 62884456 | 62884525 | 0 | 0 | 1 | 0 | 0 | 4 | 0 | 0 | 0 | 0 | 5 | 4 | 98 | 49 | 0.201681948 | 0.357884259 | NA | NA | 0.111 | NA | 0 | 0 | 0.111 |
| A3SS | 3948 | ALOX5 | chr10 | + | 45443037 | 45443216 | 45443133 | 45443216 | 45441343 | 45441430 | 2 | 7 | 33 | 0 | 0 | 0 | 1 | 6 | 7 | 0 | 3 | 0 | 98 | 49 | 0.202413488 | 0.410636839 | 1 | 1 | 1 | 1 | 0.5 | 1 | 0.167 |
| SE | 13916 | CD300A | chr17 | + | 74481288 | 74481326 | 74477435 | 74477530 | 74484000 | 74484798 | 0 | 2 | 17 | 0 | 0 | 2 | 0 | 5 | 6 | 0 | 0 | 0 | 87 | 49 | 0.203130618 | 0.359943165 | NA | 1 | 0.827 | NA | 1 | 1 | -0.087 |
| SE | 11717 | KDM6B | chr17 | + | 7844900 | 7845020 | 7839916 | 7840024 | 7845313 | 7845456 | 0 | 3 | 14 | 0 | 0 | 1 | 0 | 15 | 9 | 0 | 0 | 0 | 98 | 49 | 0.203321189 | 0.359943165 | NA | 1 | 0.875 | NA | 1 | 1 | -0.062 |
| A5SS | 519 | TNFAIP3 | chr6 | + | 137867187 | 137867518 | 137867187 | 137867494 | 137871212 | 137871522 | 0 | 3 | 1 | 0 | 2 | 0 | 0 | 8 | 4 | 0 | 1 | 0 | 73 | 49 | 0.204508982 | 0.388363884 | NA | 0.502 | 1 | NA | 0.843 | 1 | -0.17 |
| A3SS | 1274 | PTPN6 | chr12 | + | 6960662 | 6960945 | 6960805 | 6960945 | 6960343 | 6960435 | 0 | 1 | 8 | 0 | 1 | 17 | 0 | 2 | 0 | 3 | 6 | 3 | 98 | 49 | 0.204565043 | 0.410636839 | NA | 0.333 | 0.19 | 0 | 0.143 | 0 | 0.214 |
| SE | 13219 | LILRA5 | chr19 | - | 54312334 | 54312370 | 54311863 | 54312148 | 54312536 | 54312621 | 0 | 0 | 15 | 0 | 0 | 2 | 1 | 2 | 1 | 0 | 3 | 1 | 85 | 49 | 0.205560222 | 0.36347884 | NA | NA | 0.812 | 1 | 0.278 | 0.366 | 0.264 |
| SE | 5498 | ADAM8 | chr10 | - | 133274158 | 133274235 | 133273950 | 133274029 | 133276771 | 133276903 | 0 | 8 | 19 | 0 | 0 | 0 | 1 | 26 | 10 | 0 | 0 | 2 | 98 | 49 | 0.206600072 | 0.364888259 | NA | 1 | 1 | 1 | 1 | 0.714 | 0.095 |
| A3SS | 3540 | MAPKAPK2 | chr1 | + | 206732037 | 206734285 | 206732574 | 206734285 | 206731838 | 206731919 | 0 | 0 | 4 | 0 | 1 | 9 | 0 | 8 | 9 | 5 | 10 | 2 | 98 | 49 | 0.208792326 | 0.410636839 | NA | 0 | 0.182 | 0 | 0.286 | 0.692 | -0.235 |
| A3SS | 1990 | PPP6R2 | chr22 | + | 50439697 | 50439857 | 50439700 | 50439857 | 50438598 | 50438762 | 0 | 0 | 1 | 0 | 0 | 2 | 0 | 0 | 0 | 0 | 3 | 2 | 52 | 49 | 0.209323809 | 0.410636839 | NA | NA | 0.32 | NA | 0 | 0 | 0.32 |
| A5SS | 1572 | ACAD9 | chr3 | + | 128879489 | 128880017 | 128879489 | 128879841 | 128884652 | 128884746 | 0 | 0 | 4 | 0 | 0 | 2 | 0 | 0 | 2 | 0 | 0 | 0 | 98 | 49 | 0.209418448 | 0.388363884 | NA | NA | 0.5 | NA | NA | 1 | -0.5 |
| SE | 3047 | FAM157A | chr3 | + | 198169784 | 198169872 | 198167675 | 198167871 | 198180754 | 198180857 | 0 | 4 | 48 | 0 | 0 | 3 | 0 | 7 | 4 | 0 | 1 | 4 | 98 | 49 | 0.209675253 | 0.368087771 | NA | 1 | 0.889 | NA | 0.778 | 0.333 | 0.389 |
| A3SS | 2544 | TRIM26 | chr6 | - | 30198665 | 30199264 | 30198665 | 30199153 | 30201034 | 30201138 | 0 | 0 | 2 | 0 | 0 | 1 | 1 | 2 | 2 | 0 | 2 | 0 | 98 | 49 | 0.209881051 | 0.410636839 | NA | NA | 0.5 | 1 | 0.333 | 1 | -0.278 |
| A5SS | 2686 | OXLD1 | chr17 | - | 81666154 | 81666637 | 81666517 | 81666637 | 81665035 | 81665584 | 0 | 0 | 2 | 0 | 0 | 1 | 0 | 0 | 2 | 0 | 0 | 0 | 98 | 49 | 0.209881051 | 0.388363884 | NA | NA | 0.5 | NA | NA | 1 | -0.5 |
| SE | 533 | PRRC2B | chr9 | + | 131446400 | 131446642 | 131444184 | 131444328 | 131447084 | 131447206 | 0 | 0 | 2 | 0 | 0 | 1 | 0 | 1 | 2 | 0 | 0 | 0 | 98 | 49 | 0.209881051 | 0.368087771 | NA | NA | 0.5 | NA | 1 | 1 | -0.5 |
| SE | 3379 | TM9SF4 | chr20 | + | 32159991 | 32160111 | 32158450 | 32158514 | 32161275 | 32161365 | 0 | 2 | 4 | 0 | 1 | 0 | 0 | 2 | 0 | 0 | 0 | 0 | 98 | 49 | 0.209881051 | 0.368087771 | NA | 0.5 | 1 | NA | 1 | NA | -0.25 |
| SE | 3413 | RALGAPA2 | chr20 | - | 20524405 | 20524543 | 20520916 | 20521100 | 20524829 | 20524898 | 0 | 0 | 2 | 0 | 0 | 1 | 0 | 0 | 2 | 0 | 0 | 0 | 98 | 49 | 0.209881051 | 0.368087771 | NA | NA | 0.5 | NA | NA | 1 | -0.5 |
| SE | 10410 | RBM42 | chr19 | + | 35633686 | 35634019 | 35633076 | 35633252 | 35634255 | 35634373 | 0 | 2 | 0 | 0 | 1 | 0 | 0 | 2 | 2 | 0 | 0 | 0 | 98 | 49 | 0.209881051 | 0.368087771 | NA | 0.5 | NA | NA | 1 | 1 | -0.5 |
| SE | 12955 | MAN2A2 | chr15 | + | 90912541 | 90912664 | 90912042 | 90912279 | 90912876 | 90912991 | 0 | 0 | 2 | 0 | 0 | 1 | 0 | 2 | 2 | 0 | 0 | 0 | 98 | 49 | 0.209881051 | 0.368087771 | NA | NA | 0.5 | NA | 1 | 1 | -0.5 |
| A5SS | 2442 | RNASEK | chr17 | + | 7012416 | 7012761 | 7012416 | 7012757 | 7013665 | 7013742 | 0 | 0 | 4 | 0 | 0 | 2 | 1 | 2 | 2 | 0 | 1 | 0 | 53 | 49 | 0.210487296 | 0.388363884 | NA | NA | 0.649 | 1 | 0.649 | 1 | -0.234 |
| SE | 5409 | TAGAP | chr6 | - | 159042077 | 159042244 | 159041353 | 159041515 | 159043588 | 159043655 | 0 | 4 | 19 | 0 | 0 | 1 | 0 | 28 | 2 | 0 | 0 | 0 | 98 | 49 | 0.21465984 | 0.37603 | NA | 1 | 0.905 | NA | 1 | 1 | -0.047 |
| SE | 4197 | CPPED1 | chr16 | - | 12704623 | 12705049 | 12659798 | 12665115 | 12781184 | 12781403 | 0 | 1 | 12 | 0 | 0 | 3 | 0 | 2 | 1 | 0 | 3 | 0 | 98 | 49 | 0.215298398 | 0.376709536 | NA | 1 | 0.667 | NA | 0.25 | 1 | 0.209 |
| SE | 3187 | RGS19 | chr20 | - | 64076856 | 64076954 | 64076524 | 64076646 | 64079293 | 64079492 | 0 | 0 | 7 | 0 | 0 | 2 | 0 | 1 | 3 | 0 | 0 | 0 | 98 | 49 | 0.220448488 | 0.384909961 | NA | NA | 0.636 | NA | 1 | 1 | -0.364 |
| SE | 8747 | ADIPOR1 | chr1 | - | 202943757 | 202943945 | 202942024 | 202942218 | 202944982 | 202945169 | 0 | 1 | 9 | 0 | 0 | 0 | 0 | 5 | 1 | 0 | 0 | 1 | 98 | 49 | 0.220497323 | 0.384909961 | NA | 1 | 1 | NA | 1 | 0.333 | 0.334 |
| SE | 6274 | BAZ2B | chr2 | - | 159453612 | 159453801 | 159448241 | 159448409 | 159478574 | 159478721 | 0 | 0 | 11 | 0 | 0 | 5 | 0 | 3 | 1 | 0 | 0 | 2 | 98 | 49 | 0.220936482 | 0.385229157 | NA | NA | 0.524 | NA | 1 | 0.2 | -0.076 |
| SE | 838 | CAPZA2 | chr7 | + | 116906262 | 116906342 | 116904176 | 116904383 | 116910232 | 116910311 | 0 | 0 | 23 | 0 | 0 | 3 | 0 | 10 | 6 | 0 | 1 | 0 | 98 | 49 | 0.221735441 | 0.386174239 | NA | NA | 0.793 | NA | 0.833 | 1 | -0.123 |
| SE | 276 | RAPGEF1 | chr9 | - | 131650128 | 131650242 | 131643238 | 131643417 | 131650809 | 131650949 | 0 | 2 | 1 | 0 | 0 | 1 | 3 | 10 | 1 | 0 | 7 | 0 | 98 | 49 | 0.223239446 | 0.388343619 | NA | 1 | 0.333 | 1 | 0.417 | 1 | -0.139 |
| SE | 7716 | THEMIS2 | chr1 | + | 27876587 | 27876728 | 27872542 | 27872665 | 27885294 | 27885451 | 0 | 10 | 8 | 0 | 1 | 0 | 6 | 6 | 15 | 3 | 0 | 0 | 98 | 49 | 0.223535788 | 0.388409582 | NA | 0.833 | 1 | 0.5 | 1 | 1 | 0.083 |
| SE | 7875 | TLR4 | chr9 | + | 117708198 | 117708318 | 117704174 | 117704565 | 117708562 | 117708729 | 0 | 0 | 18 | 0 | 0 | 5 | 0 | 9 | 3 | 0 | 0 | 0 | 98 | 49 | 0.224282921 | 0.389257772 | NA | NA | 0.643 | NA | 1 | 1 | -0.357 |
| SE | 4169 | RNF10 | chr12 | + | 120552498 | 120552698 | 120546404 | 120546601 | 120554717 | 120554808 | 0 | 1 | 5 | 0 | 0 | 1 | 0 | 6 | 2 | 0 | 0 | 0 | 98 | 49 | 0.225232695 | 0.390455294 | NA | 1 | 0.714 | NA | 1 | 1 | -0.143 |
| RI | 991 | HSH2D | chr19 | + | 16152551 | 16153208 | 16152551 | 16152641 | 16153042 | 16153208 | 0 | 5 | 21 | 0 | 1 | 1 | 4 | 8 | 1 | 0 | 0 | 0 | 98 | 49 | 0.226524618 | 0.654404451 | NA | 0.714 | 0.913 | 1 | 1 | 1 | -0.186 |
| A5SS | 1015 | AMICA1 | chr11 | - | 118197967 | 118198091 | 118197997 | 118198091 | 118196734 | 118196821 | 0 | 0 | 14 | 0 | 0 | 6 | 0 | 9 | 2 | 0 | 1 | 0 | 79 | 49 | 0.228281084 | 0.409690921 | NA | NA | 0.591 | NA | 0.848 | 1 | -0.333 |
| SE | 12396 | TMEM127 | chr2 | - | 96265137 | 96265512 | 96254832 | 96254997 | 96265868 | 96266013 | 0 | 3 | 11 | 0 | 0 | 1 | 0 | 13 | 4 | 0 | 0 | 0 | 98 | 49 | 0.229343062 | 0.397122837 | NA | 1 | 0.846 | NA | 1 | 1 | -0.077 |
| SE | 10331 | CYTIP | chr2 | - | 157418522 | 157418589 | 157414618 | 157416143 | 157427350 | 157427420 | 0 | 5 | 36 | 0 | 0 | 1 | 4 | 50 | 1 | 0 | 0 | 0 | 98 | 49 | 0.231211633 | 0.398828525 | NA | 1 | 0.947 | 1 | 1 | 1 | -0.026 |
| SE | 6671 | FGR | chr1 | - | 27617192 | 27617296 | 27616856 | 27617006 | 27621558 | 27621657 | 0 | 3 | 47 | 0 | 0 | 1 | 5 | 12 | 2 | 0 | 1 | 0 | 98 | 49 | 0.231633345 | 0.398828525 | NA | 1 | 0.959 | 1 | 0.857 | 1 | 0.027 |
| A3SS | 2782 | GCH1 | chr14 | - | 54842005 | 54844143 | 54842005 | 54843054 | 54845767 | 54845852 | 0 | 0 | 2 | 0 | 0 | 2 | 0 | 2 | 0 | 1 | 0 | 1 | 98 | 49 | 0.232250281 | 0.43959786 | NA | NA | 0.333 | 0 | 1 | 0 | 0 |
| SE | 3236 | SLC9A8 | chr20 | + | 49850761 | 49850844 | 49849578 | 49849680 | 49855437 | 49855581 | 0 | 0 | 4 | 0 | 0 | 4 | 0 | 0 | 0 | 0 | 4 | 1 | 98 | 49 | 0.23228173 | 0.398828525 | NA | NA | 0.333 | NA | 0 | 0 | 0.333 |
| SE | 10641 | USP3 | chr15 | + | 63533802 | 63533918 | 63532646 | 63532707 | 63537024 | 63537156 | 0 | 0 | 6 | 0 | 0 | 6 | 0 | 3 | 0 | 0 | 2 | 1 | 98 | 49 | 0.232454602 | 0.398828525 | NA | NA | 0.333 | NA | 0.429 | 0 | 0.119 |
| SE | 4196 | FAM49B | chr8 | - | 129904498 | 129904585 | 129903311 | 129903350 | 129970942 | 129970995 | 0 | 0 | 3 | 0 | 0 | 3 | 0 | 1 | 0 | 0 | 3 | 1 | 98 | 49 | 0.232595516 | 0.398828525 | NA | NA | 0.333 | NA | 0.143 | 0 | 0.262 |
| SE | 3237 | SLC9A8 | chr20 | + | 49850809 | 49850844 | 49849578 | 49849680 | 49855437 | 49855581 | 0 | 0 | 4 | 0 | 0 | 4 | 0 | 0 | 0 | 0 | 4 | 1 | 84 | 49 | 0.232676511 | 0.398828525 | NA | NA | 0.368 | NA | 0 | 0 | 0.368 |
| SE | 4286 | RASSF3 | chr12 | + | 64688215 | 64688453 | 64684786 | 64684894 | 64691469 | 64691579 | 0 | 1 | 21 | 0 | 0 | 2 | 1 | 10 | 1 | 0 | 0 | 0 | 98 | 49 | 0.233640332 | 0.398828525 | NA | 1 | 0.84 | 1 | 1 | 1 | -0.08 |
| SE | 10884 | AKAP13 | chr15 | + | 85658536 | 85658590 | 85655416 | 85655787 | 85664562 | 85664755 | 0 | 2 | 13 | 0 | 0 | 1 | 0 | 4 | 3 | 0 | 2 | 0 | 98 | 49 | 0.233666262 | 0.398828525 | NA | 1 | 0.867 | NA | 0.5 | 1 | 0.183 |
| A3SS | 650 | TMEM175 | chr4 | + | 951645 | 951717 | 951681 | 951717 | 951206 | 951258 | 0 | 0 | 1 | 0 | 0 | 1 | 0 | 0 | 1 | 0 | 0 | 0 | 85 | 49 | 0.234374295 | 0.43959786 | NA | NA | 0.366 | NA | NA | 1 | -0.634 |
| A3SS | 1341 | FAM89B | chr11 | + | 65573352 | 65574198 | 65573362 | 65574198 | 65572348 | 65572960 | 0 | 0 | 1 | 0 | 0 | 1 | 0 | 0 | 1 | 0 | 1 | 0 | 59 | 49 | 0.234434278 | 0.43959786 | NA | NA | 0.454 | NA | 0 | 1 | -0.046 |
| A3SS | 193 | ZYX | chr7 | + | 143381556 | 143381779 | 143381559 | 143381779 | 143381266 | 143381409 | 2 | 1 | 1 | 0 | 1 | 0 | 0 | 1 | 0 | 0 | 0 | 0 | 52 | 49 | 0.234452192 | 0.43959786 | 1 | 0.485 | 1 | NA | 1 | NA | -0.172 |
| A5SS | 696 | PDCD6 | chr5 | + | 306601 | 306760 | 306601 | 306754 | 311292 | 311402 | 0 | 0 | 1 | 0 | 0 | 1 | 0 | 0 | 1 | 0 | 0 | 0 | 55 | 49 | 0.234460292 | 0.409690921 | NA | NA | 0.471 | NA | NA | 1 | -0.529 |
| A5SS | 2321 | ISG20 | chr15 | + | 88638742 | 88639178 | 88638742 | 88639076 | 88639342 | 88639594 | 0 | 0 | 1 | 0 | 0 | 1 | 0 | 1 | 1 | 0 | 2 | 0 | 98 | 49 | 0.234545893 | 0.409690921 | NA | NA | 0.333 | NA | 0.2 | 1 | -0.267 |
| SE | 739 | CTSB | chr8 | - | 11864375 | 11864463 | 11853328 | 11853479 | 11868000 | 11868137 | 0 | 0 | 1 | 0 | 2 | 1 | 0 | 0 | 1 | 0 | 0 | 0 | 98 | 49 | 0.234545893 | 0.398828525 | NA | 0 | 0.333 | NA | NA | 1 | -0.834 |
| SE | 5272 | NIPBL | chr5 | + | 37000372 | 37000570 | 36995621 | 36995804 | 37000816 | 37000888 | 0 | 0 | 1 | 0 | 0 | 1 | 0 | 2 | 1 | 0 | 0 | 0 | 98 | 49 | 0.234545893 | 0.398828525 | NA | NA | 0.333 | NA | 1 | 1 | -0.667 |
| SE | 7330 | NEDD9 | chr6 | - | 11305065 | 11305159 | 11213280 | 11213727 | 11305991 | 11306155 | 0 | 0 | 1 | 0 | 0 | 1 | 0 | 2 | 1 | 0 | 0 | 0 | 98 | 49 | 0.234545893 | 0.398828525 | NA | NA | 0.333 | NA | 1 | 1 | -0.667 |
| SE | 9136 | TRIM5 | chr11 | - | 5665980 | 5666081 | 5663194 | 5665682 | 5667688 | 5667711 | 0 | 0 | 1 | 0 | 0 | 1 | 0 | 0 | 1 | 0 | 2 | 0 | 98 | 49 | 0.234545893 | 0.398828525 | NA | NA | 0.333 | NA | 0 | 1 | -0.167 |
| SE | 10646 | HERC1 | chr15 | - | 63662983 | 63663204 | 63661752 | 63662021 | 63664469 | 63664594 | 0 | 0 | 1 | 0 | 0 | 1 | 0 | 2 | 1 | 0 | 1 | 0 | 98 | 49 | 0.234545893 | 0.398828525 | NA | NA | 0.333 | NA | 0.5 | 1 | -0.417 |
| SE | 11342 | BCKDK | chr16 | + | 31110687 | 31110761 | 31110400 | 31110499 | 31111090 | 31111219 | 0 | 0 | 1 | 0 | 0 | 1 | 0 | 0 | 1 | 0 | 0 | 0 | 98 | 49 | 0.234545893 | 0.398828525 | NA | NA | 0.333 | NA | NA | 1 | -0.667 |
| SE | 13736 | SERPINA1 | chr14 | - | 94388818 | 94389010 | 94388559 | 94388660 | 94390456 | 94390692 | 0 | 1 | 16 | 0 | 1 | 3 | 0 | 1 | 0 | 4 | 0 | 0 | 98 | 49 | 0.234545893 | 0.398828525 | NA | 0.333 | 0.727 | 0 | 1 | NA | 0.03 |
| A5SS | 2440 | SMARCB1 | chr22 | + | 23791755 | 23791894 | 23791755 | 23791867 | 23793558 | 23793688 | 2 | 0 | 1 | 0 | 0 | 1 | 0 | 2 | 1 | 0 | 0 | 0 | 76 | 49 | 0.234555871 | 0.409690921 | 1 | NA | 0.392 | NA | 1 | 1 | -0.304 |
| SE | 11000 | TBC1D14 | chr4 | + | 7004843 | 7004924 | 7001144 | 7001251 | 7010652 | 7010781 | 0 | 0 | 3 | 0 | 0 | 3 | 0 | 1 | 1 | 0 | 0 | 0 | 98 | 49 | 0.234573797 | 0.398828525 | NA | NA | 0.333 | NA | 1 | 1 | -0.667 |
| SE | 2249 | PCGF3 | chr4 | + | 764983 | 765064 | 761278 | 761416 | 766031 | 770639 | 0 | 0 | 2 | 0 | 1 | 0 | 0 | 1 | 1 | 0 | 0 | 0 | 98 | 49 | 0.236755697 | 0.402083405 | NA | 0 | 1 | NA | 1 | 1 | -0.5 |
| SE | 12650 | TLN1 | chr9 | - | 35725564 | 35725727 | 35725223 | 35725321 | 35732074 | 35732395 | 0 | 1 | 14 | 0 | 0 | 1 | 0 | 15 | 3 | 0 | 0 | 0 | 98 | 49 | 0.239010624 | 0.405454817 | NA | 1 | 0.875 | NA | 1 | 1 | -0.062 |
| SE | 9492 | SLC15A3 | chr11 | - | 60941121 | 60941290 | 60939479 | 60939638 | 60942034 | 60942145 | 0 | 2 | 5 | 0 | 1 | 1 | 0 | 2 | 3 | 0 | 2 | 2 | 98 | 49 | 0.239519623 | 0.405735216 | NA | 0.5 | 0.714 | NA | 0.333 | 0.429 | 0.226 |
| SE | 13178 | NLRP12 | chr19 | - | 53807494 | 53807665 | 53805279 | 53805450 | 53809586 | 53811288 | 0 | 1 | 13 | 0 | 0 | 2 | 0 | 3 | 4 | 0 | 0 | 0 | 98 | 49 | 0.23984512 | 0.405735216 | NA | 1 | 0.765 | NA | 1 | 1 | -0.117 |
| SE | 6632 | MRVI1 | chr11 | - | 10600917 | 10601059 | 10594145 | 10594195 | 10603119 | 10603251 | 0 | 0 | 4 | 0 | 0 | 1 | 0 | 6 | 3 | 0 | 0 | 0 | 98 | 49 | 0.239985767 | 0.405735216 | NA | NA | 0.667 | NA | 1 | 1 | -0.333 |
| SE | 13627 | SUN2 | chr22 | - | 38750897 | 38751035 | 38750224 | 38750320 | 38751209 | 38751373 | 0 | 1 | 6 | 0 | 0 | 0 | 0 | 1 | 3 | 0 | 2 | 0 | 98 | 49 | 0.241094036 | 0.407150939 | NA | 1 | 1 | NA | 0.2 | 1 | 0.4 |
| SE | 3099 | MX1 | chr21 | + | 41443787 | 41443866 | 41441715 | 41441914 | 41445447 | 41445570 | 0 | 4 | 62 | 0 | 0 | 2 | 18 | 50 | 6 | 0 | 3 | 0 | 98 | 49 | 0.244079828 | 0.411554843 | NA | 1 | 0.939 | 1 | 0.893 | 1 | 0.005 |
| SE | 11814 | FLOT2 | chr17 | - | 28881191 | 28881375 | 28880712 | 28880862 | 28881813 | 28882028 | 0 | 3 | 27 | 0 | 0 | 2 | 0 | 12 | 1 | 0 | 2 | 0 | 98 | 49 | 0.244249447 | 0.411554843 | NA | 1 | 0.871 | NA | 0.75 | 1 | 0.06 |
| A5SS | 1373 | FCAR | chr19 | + | 54888006 | 54888294 | 54888006 | 54888228 | 54889648 | 54891420 | 1 | 5 | 40 | 0 | 2 | 6 | 9 | 17 | 4 | 0 | 2 | 0 | 98 | 49 | 0.244859012 | 0.422059613 | 1 | 0.556 | 0.769 | 1 | 0.81 | 1 | -0.162 |
| SE | 1039 | AOAH | chr7 | - | 36686698 | 36686794 | 36673942 | 36674009 | 36724021 | 36724549 | 0 | 5 | 37 | 0 | 0 | 1 | 0 | 10 | 2 | 0 | 1 | 0 | 98 | 49 | 0.245974583 | 0.413770577 | NA | 1 | 0.949 | NA | 0.833 | 1 | 0.058 |
| SE | 12364 | NBEAL2 | chr3 | + | 46992474 | 46992555 | 46991839 | 46991946 | 46993936 | 46994020 | 0 | 0 | 10 | 0 | 0 | 9 | 0 | 2 | 1 | 0 | 1 | 0 | 98 | 49 | 0.246115034 | 0.413770577 | NA | NA | 0.357 | NA | 0.5 | 1 | -0.393 |
| SE | 12574 | R3HDM4 | chr19 | - | 901421 | 901546 | 900828 | 900952 | 901975 | 902130 | 0 | 6 | 19 | 0 | 0 | 1 | 5 | 27 | 1 | 0 | 0 | 0 | 98 | 49 | 0.247240838 | 0.41489777 | NA | 1 | 0.905 | 1 | 1 | 1 | -0.047 |
| SE | 516 | SLA | chr8 | - | 133044983 | 133045115 | 133039997 | 133040130 | 133047829 | 133047933 | 0 | 0 | 35 | 0 | 0 | 6 | 0 | 8 | 4 | 0 | 1 | 0 | 98 | 49 | 0.247337593 | 0.41489777 | NA | NA | 0.745 | NA | 0.8 | 1 | -0.155 |
| SE | 4618 | PTPRJ | chr11 | + | 48164379 | 48164515 | 48163457 | 48163618 | 48167203 | 48170842 | 0 | 1 | 10 | 0 | 0 | 1 | 0 | 10 | 3 | 0 | 0 | 0 | 98 | 49 | 0.248230356 | 0.415931131 | NA | 1 | 0.833 | NA | 1 | 1 | -0.084 |
| SE | 4055 | FOS | chr14 | + | 75279876 | 75280128 | 75278777 | 75279123 | 75280559 | 75280667 | 0 | 18 | 153 | 0 | 0 | 1 | 11 | 176 | 10 | 0 | 0 | 0 | 98 | 49 | 0.249706448 | 0.417938521 | NA | 1 | 0.987 | 1 | 1 | 1 | -0.006 |
| A3SS | 3814 | NLRP12 | chr19 | - | 53807494 | 53807668 | 53807494 | 53807665 | 53809586 | 53811288 | 0 | 0 | 1 | 0 | 0 | 6 | 0 | 0 | 0 | 0 | 2 | 4 | 52 | 49 | 0.250107194 | 0.464116443 | NA | NA | 0.136 | NA | 0 | 0 | 0.136 |
| SE | 2700 | MBNL1 | chr3 | + | 152432716 | 152432920 | 152414940 | 152415111 | 152445281 | 152445539 | 0 | 1 | 24 | 0 | 0 | 1 | 7 | 7 | 1 | 0 | 1 | 0 | 98 | 49 | 0.252076189 | 0.421435498 | NA | 1 | 0.923 | 1 | 0.778 | 1 | 0.035 |
| SE | 12940 | SMCHD1 | chr18 | + | 2674014 | 2674145 | 2673280 | 2673363 | 2688393 | 2688508 | 0 | 7 | 143 | 0 | 0 | 2 | 5 | 100 | 1 | 0 | 3 | 0 | 98 | 49 | 0.254061338 | 0.424282435 | NA | 1 | 0.973 | 1 | 0.943 | 1 | 0.005 |
| A3SS | 3504 | FCGR3A | chr1 | - | 161548420 | 161548678 | 161548420 | 161548675 | 161549010 | 161549031 | 0 | 1 | 5 | 0 | 0 | 2 | 0 | 2 | 1 | 0 | 0 | 0 | 52 | 49 | 0.25427504 | 0.467035788 | NA | 1 | 0.702 | NA | 1 | 1 | -0.149 |
| SE | 4164 | NT5C2 | chr10 | - | 103090610 | 103090787 | 103088016 | 103089908 | 103090935 | 103090996 | 0 | 0 | 14 | 0 | 0 | 5 | 0 | 2 | 2 | 0 | 0 | 0 | 98 | 49 | 0.254640733 | 0.424778049 | NA | NA | 0.583 | NA | 1 | 1 | -0.417 |
| A5SS | 2258 | TLE3 | chr15 | - | 70057458 | 70057658 | 70057473 | 70057658 | 70056297 | 70056374 | 0 | 1 | 9 | 0 | 0 | 3 | 0 | 3 | 1 | 0 | 3 | 0 | 64 | 49 | 0.254834903 | 0.430953855 | NA | 1 | 0.697 | NA | 0.434 | 1 | 0.132 |

| | | | | | | | | | | | | | | | | | | | | | | | | | | | | | | | | | |
|---|---|---|---|---|---|---|---|---|---|---|---|---|---|---|---|---|---|---|---|---|---|---|---|---|---|---|---|---|---|---|---|---|---|
| SE | 13858 | MMP25 | chr16 | + | 3050253 | 3050546 | 3050008 | 3050144 | 3057032 | 3057209 | 0 | 7 | 19 | 0 | 0 | 1 | 0 | 17 | 10 | 0 | 0 | 0 | 98 | 49 | 0.255901705 | 0.426408273 | NA | 1 | 0.905 | NA | 1 | 1 | -0.047 |
| A5SS | 403 | CNOT8 | chr5 | + | 154858518 | 154858816 | 154858518 | 154858768 | 154863206 | 154863395 | 1 | 0 | 3 | 0 | 0 | 2 | 1 | 0 | 1 | 0 | 0 | 0 | 97 | 49 | 0.256598479 | 0.430953855 | 1 | NA | 0.431 | 1 | NA | 1 | -0.284 |
| SE | 12369 | NBEAL2 | chr3 | + | 47007240 | 47007350 | 47006332 | 47006449 | 47007524 | 47007697 | 0 | 1 | 7 | 0 | 1 | 0 | 0 | 5 | 0 | 0 | 2 | 0 | 98 | 49 | 0.256741576 | 0.427007803 | NA | 0.333 | 1 | NA | 0.556 | NA | 0.11 |
| SE | 12947 | SMCHD1 | chr18 | + | 2751277 | 2751393 | 2750349 | 2750507 | 2752487 | 2752552 | 1 | 8 | 40 | 0 | 0 | 1 | 0 | 46 | 4 | 0 | 0 | 0 | 98 | 49 | 0.25682971 | 0.427007803 | 1 | 1 | 0.952 | NA | 1 | 1 | -0.016 |
| SE | 2166 | FYB | chr5 | - | 39125997 | 39126135 | 39124252 | 39124278 | 39127740 | 39127807 | 0 | 0 | 17 | 0 | 0 | 9 | 6 | 25 | 1 | 0 | 6 | 2 | 98 | 49 | 0.257579272 | 0.427780824 | NA | NA | 0.486 | 1 | 0.676 | 0.2 | -0.139 |
| SE | 7217 | LSP1 | chr11 | + | 1887236 | 1887314 | 1886731 | 1886866 | 1887473 | 1887576 | 0 | 6 | 18 | 0 | 0 | 1 | 0 | 22 | 2 | 0 | 0 | 0 | 98 | 49 | 0.260907854 | 0.432830579 | NA | 1 | 0.9 | NA | 1 | 1 | -0.05 |
| SE | 7231 | NFKBIZ | chr3 | + | 101857072 | 101857183 | 101855732 | 101855902 | 101857291 | 101857459 | 0 | 0 | 19 | 0 | 0 | 3 | 5 | 34 | 4 | 0 | 7 | 0 | 98 | 49 | 0.261478977 | 0.432929688 | NA | NA | 0.76 | 1 | 0.708 | 1 | -0.143 |
| SE | 5891 | CSF1 | chr1 | + | 109914258 | 109914381 | 109910610 | 109911062 | 109915633 | 109915696 | 0 | 1 | 1 | 0 | 3 | 0 | 0 | 0 | 0 | 0 | 2 | 0 | 98 | 49 | 0.261543684 | 0.432929688 | NA | 0.143 | 1 | NA | 0 | NA | 0.572 |
| SE | 6056 | HCLS1 | chr3 | - | 121632100 | 121632184 | 121631397 | 121631982 | 121632331 | 121632563 | 0 | 3 | 39 | 0 | 0 | 0 | 1 | 28 | 4 | 0 | 0 | 1 | 98 | 49 | 0.261910932 | 0.433060651 | NA | 1 | 1 | 1 | 1 | 0.667 | 0.111 |
| SE | 1667 | GRK6 | chr5 | + | 177431994 | 177432107 | 177430871 | 177430967 | 177432232 | 177432310 | 0 | 7 | 9 | 0 | 0 | 1 | 0 | 4 | 3 | 0 | 3 | 0 | 98 | 49 | 0.26379167 | 0.435653879 | NA | 1 | 0.818 | NA | 0.4 | 1 | 0.209 |
| SE | 10928 | DEDD2 | chr19 | - | 42202050 | 42202072 | 42198592 | 42199829 | 42209699 | 42209840 | 0 | 0 | 3 | 0 | 3 | 5 | 0 | 2 | 0 | 0 | 16 | 3 | 71 | 49 | 0.264511909 | 0.435653879 | NA | 0 | 0.293 | NA | 0.079 | 0 | 0.107 |
| RI | 607 | TAF1C | chr16 | - | 84181327 | 84181663 | 84181327 | 84181463 | 84181591 | 84181663 | 0 | 0 | 3 | 0 | 0 | 1 | 0 | 1 | 2 | 0 | 0 | 0 | 98 | 49 | 0.264638717 | 0.737207854 | NA | NA | 0.6 | NA | 1 | 1 | -0.4 |
| SE | 3872 | WDFY2 | chr13 | + | 51719197 | 51719348 | 51703595 | 51703650 | 51727677 | 51727790 | 0 | 0 | 3 | 0 | 0 | 1 | 3 | 2 | 2 | 0 | 0 | 0 | 98 | 49 | 0.264638717 | 0.435653879 | NA | NA | 0.6 | 1 | 1 | 1 | -0.4 |
| SE | 7725 | ZFC3H1 | chr12 | - | 71636849 | 71637059 | 71636489 | 71636654 | 71638417 | 71638515 | 0 | 0 | 3 | 0 | 0 | 1 | 0 | 9 | 2 | 0 | 0 | 0 | 98 | 49 | 0.264638717 | 0.435653879 | NA | NA | 0.6 | NA | 1 | 1 | -0.4 |
| A5SS | 1432 | ADPGK | chr15 | - | 72783245 | 72783785 | 72783458 | 72783785 | 72774871 | 72775097 | 0 | 3 | 7 | 0 | 0 | 1 | 2 | 2 | 8 | 0 | 0 | 0 | 98 | 49 | 0.265624158 | 0.440465375 | NA | 1 | 0.778 | 1 | 1 | 1 | -0.111 |
| SE | 5576 | SAT1 | chrX | + | 23784293 | 23784403 | 23783799 | 23783883 | 23785327 | 23785429 | 0 | 5 | 44 | 0 | 10 | 78 | 0 | 46 | 0 | 0 | 89 | 9 | 98 | 49 | 0.267669304 | 0.440160792 | NA | 0.2 | 0.22 | NA | 0.205 | 0 | 0.108 |
| A3SS | 1360 | DDX3X | chrX | + | 41347311 | 41347451 | 41347314 | 41347451 | 41346858 | 41347012 | 0 | 0 | 8 | 0 | 0 | 6 | 0 | 16 | 1 | 0 | 2 | 0 | 52 | 49 | 0.267844114 | 0.486989299 | NA | NA | 0.557 | NA | 0.883 | 1 | -0.384 |
| SE | 7628 | RALGAPB | chr20 | + | 38548688 | 38548795 | 38546242 | 38546430 | 38551070 | 38551223 | 0 | 0 | 4 | 0 | 0 | 3 | 0 | 0 | 1 | 0 | 1 | 0 | 98 | 49 | 0.268677998 | 0.441336646 | NA | NA | 0.4 | NA | 0 | 1 | -0.1 |
| SE | 6197 | TPCN2 | chr11 | + | 69072626 | 69072708 | 69071922 | 69072023 | 69072914 | 69073001 | 0 | 2 | 0 | 0 | 0 | 1 | 0 | 3 | 1 | 0 | 0 | 0 | 98 | 49 | 0.27131636 | 0.444793103 | NA | 1 | 0 | NA | 1 | 1 | -0.5 |
| A5SS | 435 | PAPD4 | chr5 | + | 79612419 | 79612769 | 79612419 | 79612735 | 79613010 | 79613075 | 0 | 2 | 1 | 0 | 0 | 1 | 0 | 2 | 1 | 0 | 0 | 0 | 83 | 49 | 0.271326485 | 0.444297119 | NA | 1 | 0.371 | NA | 1 | 1 | -0.315 |
| SE | 1339 | CMTR1 | chr6 | + | 37462828 | 37463008 | 37461548 | 37461645 | 37471020 | 37471077 | 0 | 0 | 3 | 0 | 1 | 0 | 0 | 2 | 5 | 0 | 0 | 0 | 98 | 49 | 0.271849638 | 0.444793103 | NA | 0 | 1 | NA | 1 | 1 | -0.5 |
| SE | 9456 | PFKFB3 | chr10 | + | 6217134 | 6217191 | 6216705 | 6216780 | 6219568 | 6219693 | 0 | 2 | 31 | 0 | 0 | 1 | 0 | 32 | 1 | 0 | 0 | 0 | 98 | 49 | 0.271902353 | 0.444793103 | NA | 1 | 0.939 | NA | 1 | 1 | -0.03 |
| SE | 9007 | LYST | chr1 | - | 235804503 | 235804665 | 235802907 | 235803064 | 235805742 | 235806772 | 0 | 4 | 25 | 0 | 0 | 2 | 0 | 11 | 1 | 0 | 0 | 0 | 98 | 49 | 0.271988796 | 0.444793103 | NA | 1 | 0.862 | NA | 1 | 1 | -0.069 |
| SE | 12740 | DYSF | chr2 | + | 71511806 | 71511921 | 71503213 | 71503319 | 71513715 | 71513921 | 0 | 0 | 5 | 0 | 0 | 1 | 0 | 4 | 3 | 0 | 0 | 0 | 98 | 49 | 0.272557849 | 0.444793103 | NA | NA | 0.714 | NA | 1 | 1 | -0.286 |
| SE | 13209 | LILRB2 | chr19 | - | 54276380 | 54276456 | 54276263 | 54276301 | 54276806 | 54276929 | 0 | 0 | 5 | 0 | 0 | 1 | 0 | 2 | 3 | 0 | 0 | 0 | 98 | 49 | 0.272557849 | 0.444793103 | NA | NA | 0.714 | NA | 1 | 1 | -0.286 |
| A3SS | 831 | TOM1 | chr22 | + | 35322998 | 35323177 | 35323027 | 35323177 | 35317876 | 35317961 | 0 | 1 | 5 | 0 | 0 | 1 | 0 | 0 | 3 | 0 | 0 | 0 | 78 | 49 | 0.272864991 | 0.491156984 | NA | 1 | 0.759 | NA | NA | 1 | -0.121 |
| SE | 12823 | SEC14L1 | chr17 | + | 77193420 | 77193549 | 77191180 | 77191312 | 77194676 | 77194911 | 0 | 5 | 49 | 0 | 0 | 1 | 2 | 50 | 3 | 0 | 0 | 0 | 98 | 49 | 0.274578151 | 0.44760408 | NA | 1 | 0.961 | 1 | 1 | 1 | -0.02 |
| SE | 4882 | DCAF11 | chr14 | + | 24116916 | 24117044 | 24115381 | 24115749 | 24117265 | 24117393 | 0 | 0 | 3 | 0 | 0 | 1 | 0 | 2 | 2 | 0 | 2 | 2 | 98 | 49 | 0.277220408 | 0.450933196 | NA | NA | 0.6 | NA | 0.333 | 0.333 | 0.267 |
| SE | 4883 | DCAF11 | chr14 | + | 24116971 | 24117044 | 24115282 | 24115749 | 24117265 | 24117393 | 0 | 0 | 3 | 0 | 0 | 1 | 0 | 0 | 2 | 0 | 2 | 2 | 98 | 49 | 0.277220408 | 0.450933196 | NA | NA | 0.6 | NA | 0 | 0.333 | 0.433 |
| A3SS | 3597 | EIF4G1 | chr3 | + | 184323404 | 184323593 | 184323407 | 184323593 | 184323082 | 184323241 | 0 | 0 | 2 | 0 | 0 | 3 | 0 | 0 | 0 | 0 | 1 | 1 | 52 | 49 | 0.278939638 | 0.491656631 | NA | NA | 0.386 | NA | 0 | 0 | 0.386 |
| SE | 8081 | CSF3R | chr1 | - | 36472065 | 36472139 | 36471432 | 36471646 | 36472237 | 36472391 | 0 | 94 | 256 | 0 | 0 | 1 | 16 | 183 | 131 | 0 | 0 | 0 | 98 | 49 | 0.278963874 | 0.453278598 | NA | 1 | 0.992 | 1 | 1 | 1 | -0.004 |
| A3SS | 1548 | PPP4C | chr16 | + | 30081199 | 30081310 | 30081258 | 30081310 | 30076314 | 30076475 | 0 | 0 | 3 | 0 | 0 | 2 | 0 | 2 | 1 | 0 | 5 | 0 | 98 | 49 | 0.280935793 | 0.491656631 | NA | NA | 0.429 | NA | 0.167 | 1 | -0.155 |
| SE | 3522 | HDAC4 | chr2 | - | 239144582 | 239144714 | 239139683 | 239139796 | 239156651 | 239156773 | 0 | 0 | 3 | 0 | 0 | 2 | 0 | 0 | 1 | 0 | 0 | 0 | 98 | 49 | 0.280935793 | 0.454548838 | NA | NA | 0.429 | NA | NA | 1 | -0.571 |
| SE | 4254 | MAPKAP1 | chr9 | - | 125672315 | 125672643 | 125669817 | 125669907 | 125706970 | 125707234 | 0 | 0 | 3 | 0 | 0 | 2 | 8 | 0 | 1 | 0 | 0 | 0 | 98 | 49 | 0.280935793 | 0.454548838 | NA | NA | 0.429 | 1 | NA | 1 | -0.571 |
| SE | 4531 | DUSP6 | chr12 | - | 89350587 | 89351025 | 89347824 | 89349561 | 89351639 | 89352859 | 0 | 3 | 21 | 0 | 2 | 1 | 0 | 1 | 0 | 0 | 0 | 0 | 98 | 49 | 0.280935793 | 0.454548838 | NA | 0.429 | 0.913 | NA | 1 | NA | -0.329 |
| SE | 10627 | RRP12 | chr10 | - | 97390422 | 97390539 | 97388488 | 97388624 | 97390738 | 97390844 | 0 | 7 | 7 | 0 | 0 | 1 | 5 | 9 | 3 | 0 | 0 | 0 | 98 | 49 | 0.281057395 | 0.454548838 | NA | 1 | 0.778 | 1 | 1 | 1 | -0.111 |
| SE | 13626 | SUN2 | chr22 | - | 38738872 | 38738988 | 38738586 | 38738754 | 38739341 | 38739426 | 0 | 0 | 6 | 0 | 0 | 4 | 0 | 6 | 1 | 0 | 0 | 0 | 98 | 49 | 0.281257764 | 0.454548838 | NA | NA | 0.429 | NA | 1 | 1 | -0.571 |
| A3SS | 1827 | FCGR2A | chr1 | + | 161506333 | 161506591 | 161506336 | 161506591 | 161505986 | 161506007 | 0 | 1 | 20 | 0 | 0 | 17 | 0 | 6 | 2 | 0 | 10 | 1 | 52 | 49 | 0.28133685 | 0.491656631 | NA | 1 | 0.526 | NA | 0.361 | 0.653 | 0.256 |
| SE | 12951 | GMFG | chr19 | - | 39335260 | 39335310 | 39333076 | 39333126 | 39335434 | 39335531 | 0 | 1 | 18 | 0 | 0 | 1 | 13 | 16 | 2 | 0 | 0 | 0 | 98 | 49 | 0.283872459 | 0.458281747 | NA | 1 | 0.9 | 1 | 1 | 1 | -0.05 |
| SE | 3214 | RTFDC1 | chr20 | + | 56516934 | 56516989 | 56513314 | 56513428 | 56517105 | 56517201 | 0 | 0 | 7 | 0 | 0 | 2 | 0 | 13 | 2 | 0 | 0 | 0 | 98 | 49 | 0.286074243 | 0.461340759 | NA | NA | 0.636 | NA | 1 | 1 | -0.364 |
| SE | 9377 | FMNL1 | chr17 | + | 45245631 | 45245733 | 45245252 | 45245416 | 45245877 | 45245973 | 0 | 5 | 21 | 0 | 3 | 6 | 0 | 24 | 7 | 0 | 7 | 0 | 98 | 49 | 0.287117418 | 0.462526773 | NA | 0.455 | 0.636 | NA | 0.632 | 1 | -0.271 |
| SE | 9458 | PFKFB3 | chr10 | + | 6222854 | 6222984 | 6221380 | 6221527 | 6223957 | 6224020 | 0 | 3 | 22 | 0 | 0 | 1 | 0 | 21 | 7 | 0 | 2 | 0 | 98 | 49 | 0.28811154 | 0.463631312 | NA | 1 | 0.917 | NA | 0.84 | 1 | 0.039 |
| SE | 10611 | PIK3R5 | chr17 | - | 8889972 | 8890126 | 8889138 | 8889222 | 8890737 | 8890912 | 0 | 5 | 10 | 0 | 0 | 1 | 0 | 3 | 10 | 0 | 0 | 0 | 98 | 49 | 0.288581572 | 0.463891019 | NA | 1 | 0.833 | NA | 1 | 1 | -0.084 |
| SE | 816 | LINC-PINT | chr7 | - | 131002196 | 131002320 | 130984052 | 130984109 | 131052424 | 131052555 | 0 | 0 | 0 | 0 | 2 | 9 | 0 | 0 | 1 | 0 | 8 | 1 | 98 | 49 | 0.290996266 | 0.46727285 | NA | 0 | 0 | NA | 0 | 0.333 | -0.167 |
| A3SS | 2039 | H2AFY | chr5 | - | 135360496 | 135360607 | 135360496 | 135360604 | 135369405 | 135369603 | 0 | 0 | 3 | 0 | 0 | 5 | 6 | 1 | 0 | 0 | 0 | 1 | 52 | 49 | 0.292512865 | 0.506272265 | NA | NA | 0.361 | 1 | 1 | 0 | -0.306 |
| SE | 7721 | THEMIS2 | chr1 | + | 27881970 | 27883043 | 27879643 | 27880054 | 27885294 | 27885451 | 0 | 1 | 19 | 0 | 0 | 1 | 0 | 15 | 3 | 0 | 0 | 0 | 98 | 49 | 0.292531456 | 0.469236689 | NA | 1 | 0.905 | NA | 1 | 1 | -0.047 |
| SE | 11044 | HLA-B | chr6 | - | 31354632 | 31354665 | 31354478 | 31354526 | 31355106 | 31355223 | 1 | 16 | 132 | 0 | 0 | 1 | 8 | 135 | 6 | 0 | 0 | 0 | 82 | 49 | 0.294858598 | 0.472465323 | 1 | 1 | 0.987 | 1 | 1 | 1 | -0.004 |
| SE | 2178 | WAS | chrX | + | 48685733 | 48685836 | 48685546 | 48685633 | 48685945 | 48685987 | 0 | 5 | 20 | 0 | 0 | 0 | 0 | 20 | 2 | 0 | 0 | 1 | 98 | 49 | 0.297924203 | 0.476869091 | NA | 1 | 1 | NA | 1 | 0.5 | 0.25 |
| RI | 186 | PAPD4 | chr5 | + | 79612419 | 79613075 | 79612419 | 79612735 | 79613010 | 79613075 | 2 | 3 | 7 | 0 | 0 | 1 | 2 | 7 | 2 | 0 | 0 | 0 | 98 | 49 | 0.299346388 | 0.80513856 | 1 | 1 | 0.778 | 1 | 1 | 1 | -0.074 |
| A3SS | 2683 | PXN | chr12 | - | 120215102 | 120215417 | 120215102 | 120215273 | 120221622 | 120221758 | 0 | 1 | 7 | 0 | 1 | 3 | 0 | 5 | 3 | 0 | 2 | 0 | 98 | 49 | 0.29950383 | 0.513435137 | NA | 0.333 | 0.538 | NA | 0.556 | 1 | -0.343 |
| SE | 8361 | TNFRSF10C | chr8 | + | 23111719 | 23111825 | 23102813 | 23103181 | 23114656 | 23114770 | 0 | 0 | 51 | 0 | 0 | 4 | 0 | 12 | 6 | 0 | 1 | 0 | 98 | 49 | 0.302014034 | 0.482901162 | NA | NA | 0.864 | NA | 0.857 | 1 | -0.065 |
| SE | 9507 | FOXP1 | chr3 | - | 71359149 | 71359244 | 71299819 | 71299880 | 71493425 | 71493555 | 0 | 0 | 4 | 0 | 0 | 1 | 0 | 5 | 2 | 0 | 0 | 0 | 98 | 49 | 0.303158096 | 0.484215322 | NA | NA | 0.667 | NA | 1 | 1 | -0.333 |
| A5SS | 2161 | SIRPB2 | chr20 | - | 1479699 | 1480065 | 1479993 | 1480065 | 1478265 | 1478607 | 0 | 0 | 8 | 0 | 0 | 2 | 0 | 1 | 2 | 0 | 0 | 0 | 98 | 49 | 0.304701638 | 0.492789069 | NA | NA | 0.667 | NA | 1 | 1 | -0.333 |
| SE | 7847 | RPS6KA1 | chr1 | + | 26555536 | 26555625 | 26555150 | 26555221 | 26556653 | 26556718 | 0 | 0 | 27 | 0 | 1 | 4 | 4 | 8 | 0 | 0 | 3 | 1 | 98 | 49 | 0.305107208 | 0.48651819 | NA | 0 | 0.771 | 1 | 0.571 | 0 | -0.138 |
| SE | 4904 | PAK1 | chr11 | - | 77325327 | 77325376 | 77322014 | 77323360 | 77332729 | 77332867 | 0 | 0 | 3 | 0 | 1 | 12 | 0 | 0 | 0 | 0 | 2 | 1 | 98 | 49 | 0.305247274 | 0.48651819 | NA | 0 | 0.111 | NA | 0 | 0 | 0.056 |
| A3SS | 3600 | CNN2 | chr19 | + | 1036066 | 1036246 | 1036129 | 1036246 | 1032558 | 1032696 | 0 | 2 | 9 | 0 | 1 | 10 | 11 | 5 | 10 | 0 | 5 | 1 | 98 | 49 | 0.309557191 | 0.525663154 | NA | 0.5 | 0.31 | 1 | 0.333 | 0.833 | -0.317 |
| SE | 8217 | FURIN | chr15 | + | 90877526 | 90877615 | 90877134 | 90877211 | 90878131 | 90878304 | 1 | 1 | 6 | 0 | 0 | 1 | 0 | 2 | 4 | 0 | 0 | 0 | 98 | 49 | 0.310862931 | 0.494727732 | 1 | 1 | 0.75 | NA | 1 | 1 | -0.083 |
| SE | 12945 | SMCHD1 | chr18 | + | 2732264 | 2732492 | 2729274 | 2729409 | 2738396 | 2738545 | 0 | 1 | 52 | 0 | 0 | 1 | 1 | 43 | 2 | 0 | 0 | 0 | 98 | 49 | 0.311165715 | 0.494727732 | NA | 1 | 0.963 | 1 | 1 | 1 | -0.018 |
| SE | 5935 | RASGRP2 | chr11 | - | 64735902 | 64735980 | 64735541 | 64735664 | 64736752 | 64737034 | 0 | 1 | 5 | 0 | 0 | 0 | 1 | 3 | 1 | 0 | 0 | 3 | 98 | 49 | 0.311385518 | 0.494727732 | NA | 1 | 1 | 1 | 1 | 0.143 | 0.286 |
| SE | 13180 | NLRP12 | chr19 | - | 53809586 | 53811288 | 53807494 | 53807665 | 53814907 | 53814988 | 0 | 1 | 12 | 0 | 0 | 1 | 0 | 5 | 6 | 0 | 0 | 0 | 98 | 49 | 0.311762966 | 0.49480437 | NA | 1 | 0.857 | NA | 1 | 1 | -0.072 |
| SE | 3534 | LRRFIP1 | chr2 | + | 237751199 | 237751271 | 237749198 | 237749324 | 237753308 | 237753479 | 0 | 0 | 2 | 0 | 1 | 13 | 0 | 7 | 0 | 0 | 4 | 2 | 98 | 49 | 0.314453037 | 0.498547379 | NA | 0 | 0.071 | NA | 0.467 | 0 | -0.198 |
| SE | 6694 | RBCK1 | chr20 | + | 420870 | 421031 | 419557 | 419731 | 422126 | 422238 | 0 | 2 | 3 | 0 | 0 | 1 | 0 | 1 | 3 | 0 | 0 | 0 | 98 | 49 | 0.315518625 | 0.499709688 | NA | 1 | 0.6 | NA | 1 | 1 | -0.2 |
| SE | 1664 | PDLIM7 | chr5 | - | 177489392 | 177489627 | 177488067 | 177488248 | 177489770 | 177489832 | 0 | 3 | 11 | 0 | 0 | 1 | 0 | 6 | 5 | 0 | 0 | 0 | 98 | 49 | 0.31710227 | 0.50058381 | NA | 1 | 0.846 | NA | 1 | 1 | -0.077 |
| SE | 3246 | SULF2 | chr20 | - | 47666259 | 47666488 | 47665856 | 47665953 | 47672197 | 47672393 | 0 | 1 | 5 | 0 | 0 | 1 | 0 | 3 | 2 | 0 | 0 | 0 | 98 | 49 | 0.317618048 | 0.50058381 | NA | 1 | 0.714 | NA | 1 | 1 | -0.143 |
| A5SS | 1014 | PRKD2 | chr19 | - | 46716814 | 46717127 | 46716950 | 46717127 | 46716130 | 46716535 | 0 | 0 | 2 | 0 | 0 | 1 | 0 | 0 | 1 | 0 | 0 | 0 | 98 | 49 | 0.318735 | 0.509198598 | NA | NA | 0.5 | NA | NA | 1 | -0.5 |
| RI | 727 | TRIM5 | chr11 | - | 5663194 | 5665682 | 5663194 | 5665395 | 5665655 | 5665682 | 0 | 0 | 2 | 0 | 0 | 1 | 4 | 0 | 1 | 0 | 0 | 0 | 98 | 49 | 0.318735 | 0.805848647 | NA | NA | 0.5 | 1 | NA | 1 | -0.5 |
| SE | 702 | ENTPD4 | chr8 | - | 23442006 | 23442066 | 23441568 | 23441723 | 23443849 | 23443953 | 0 | 0 | 2 | 0 | 0 | 1 | 0 | 2 | 1 | 0 | 1 | 0 | 98 | 49 | 0.318735 | 0.50058381 | NA | NA | 0.5 | NA | 0.5 | 1 | -0.25 |
| SE | 2612 | UBA7 | chr3 | - | 49809389 | 49809464 | 49808975 | 49809159 | 49809541 | 49809725 | 0 | 0 | 2 | 0 | 0 | 1 | 0 | 1 | 1 | 0 | 0 | 0 | 98 | 49 | 0.318735 | 0.50058381 | NA | NA | 0.5 | NA | 1 | 1 | -0.5 |
| SE | 2722 | KAT2B | chr3 | + | 20125904 | 20126113 | 20122667 | 20122804 | 20136941 | 20137052 | 0 | 0 | 2 | 0 | 0 | 1 | 2 | 1 | 1 | 0 | 0 | 0 | 98 | 49 | 0.318735 | 0.50058381 | NA | NA | 0.5 | 1 | 1 | 1 | -0.5 |
| SE | 4806 | GNB1 | chr1 | - | 1839189 | 1839238 | 1825396 | 1825499 | 1890819 | 1891117 | 0 | 2 | 5 | 0 | 1 | 0 | 0 | 1 | 0 | 0 | 0 | 0 | 98 | 49 | 0.318735 | 0.50058381 | NA | 0.5 | 1 | NA | 1 | NA | -0.25 |
| SE | 6222 | DNAJB12 | chr10 | - | 72338201 | 72338311 | 72336523 | 72336696 | 72340788 | 72340868 | 0 | 0 | 2 | 0 | 0 | 1 | 0 | 0 | 1 | 0 | 0 | 0 | 98 | 49 | 0.318735 | 0.50058381 | NA | NA | 0.5 | NA | NA | 1 | -0.5 |
| SE | 12813 | SRP54 | chr14 | + | 35013345 | 35013494 | 35011508 | 35011659 | 35013801 | 35013902 | 0 | 0 | 2 | 0 | 0 | 1 | 0 | 1 | 1 | 0 | 0 | 0 | 98 | 49 | 0.318735 | 0.50058381 | NA | NA | 0.5 | NA | 1 | 1 | -0.5 |
| SE | 10593 | RNF213 | chr17 | + | 80389172 | 80389367 | 80388611 | 80388689 | 80389827 | 80389917 | 0 | 7 | 56 | 0 | 0 | 1 | 2 | 23 | 1 | 0 | 1 | 0 | 98 | 49 | 0.320121262 | 0.502236176 | NA | 1 | 0.966 | 1 | 0.92 | 1 | 0.01 |
| RI | 389 | GMPR2 | chr14 | + | 24238245 | 24239241 | 24238245 | 24238405 | 24238588 | 24239241 | 1 | 0 | 7 | 0 | 0 | 1 | 0 | 8 | 3 | 0 | 0 | 0 | 98 | 49 | 0.32027318 | 0.805848647 | 1 | NA | 0.778 | NA | 1 | 1 | -0.111 |
| A3SS | 3968 | SUN2 | chr22 | - | 38750224 | 38750383 | 38750224 | 38750320 | 38750897 | 38751035 | 0 | 0 | 2 | 0 | 1 | 1 | 0 | 2 | 1 | 0 | 0 | 1 | 98 | 49 | 0.320307546 | 0.538835124 | NA | 0 | 0.5 | NA | 1 | 0.333 | -0.416 |
| SE | 7718 | THEMIS2 | chr1 | + | 27879643 | 27880054 | 27876587 | 27876728 | 27881970 | 27882125 | 0 | 7 | 42 | 0 | 0 | 1 | 1 | 25 | 11 | 0 | 0 | 0 | 98 | 49 | 0.322009305 | 0.504671517 | NA | 1 | 0.955 | 1 | 1 | 1 | -0.022 |
| SE | 372 | AGTPBP1 | chr9 | - | 85692688 | 85692813 | 85681267 | 85681335 | 85741774 | 85742029 | 0 | 3 | 9 | 0 | 0 | 1 | 3 | 8 | 1 | 0 | 0 | 0 | 98 | 49 | 0.326777752 | 0.511611417 | NA | 1 | 0.818 | 1 | 1 | 1 | -0.091 |
| SE | 13193 | MBOAT7 | chr19 | - | 54187160 | 54187287 | 54183520 | 54183680 | 54188216 | 54188346 | 0 | 10 | 24 | 0 | 0 | 1 | 2 | 13 | 11 | 0 | 0 | 0 | 98 | 49 | 0.327855035 | 0.51276391 | NA | 1 | 0.923 | 1 | 1 | 1 | -0.038 |
| SE | 2938 | LIMK2 | chr22 | + | 31225719 | 31225819 | 31212263 | 31212424 | 31258290 | 31258426 | 0 | 14 | 29 | 0 | 0 | 0 | 3 | 19 | 8 | 0 | 0 | 5 | 98 | 49 | 0.328408932 | 0.512945188 | NA | 1 | 1 | 1 | 1 | 0.444 | 0.185 |
| SE | 3975 | GLT1D1 | chr12 | + | 128888638 | 128888744 | 128875913 | 128876062 | 128899235 | 128899287 | 0 | 6 | 18 | 0 | 0 | 1 | 0 | 7 | 2 | 0 | 1 | 0 | 98 | 49 | 0.328653504 | 0.512945188 | NA | 1 | 0.9 | NA | 0.778 | 1 | 0.061 |
| SE | 2767 | LST1 | chr6 | + | 31587943 | 31587966 | 31587199 | 31587318 | 31588517 | 31588909 | 0 | 0 | 7 | 0 | 0 | 4 | 0 | 0 | 1 | 0 | 4 | 0 | 72 | 49 | 0.329153667 | 0.513192906 | NA | NA | 0.544 | NA | 0 | 1 | 0.044 |
| SE | 4457 | CLEC7A | chr12 | - | 10123244 | 10123363 | 10116780 | 10118590 | 10125296 | 10125448 | 0 | 0 | 38 | 0 | 0 | 8 | 0 | 16 | 2 | 0 | 1 | 0 | 98 | 49 | 0.329686827 | 0.513491504 | NA | NA | 0.704 | NA | 0.889 | 1 | -0.241 |
| SE | 10068 | ACTN1 | chr14 | - | 68909954 | 68910042 | 68909317 | 68909396 | 68912155 | 68912242 | 0 | 9 | 25 | 0 | 2 | 0 | 0 | 18 | 1 | 0 | 2 | 1 | 98 | 49 | 0.331082125 | 0.515130884 | NA | 0.692 | 1 | NA | 0.818 | 0.333 | 0.27 |
| A3SS | 1213 | CIB1 | chr15 | - | 90232218 | 90232447 | 90232218 | 90232327 | 90233668 | 90233703 | 0 | 0 | 5 | 0 | 0 | 1 | 2 | 0 | 2 | 0 | 0 | 0 | 98 | 49 | 0.332444824 | 0.545043142 | NA | NA | 0.714 | 1 | NA | 1 | -0.286 |
| A3SS | 1850 | OAS1 | chr12 | + | 112919388 | 112919907 | 112919486 | 112919907 | 112916508 | 112916738 | 0 | 0 | 5 | 0 | 0 | 1 | 0 | 12 | 2 | 0 | 1 | 0 | 98 | 49 | 0.332444824 | 0.545043142 | NA | NA | 0.714 | NA | 0.857 | 1 | -0.215 |
| RI | 641 | ATPIF1 | chr1 | + | 28236360 | 28238105 | 28236360 | 28236452 | 28237836 | 28238105 | 0 | 0 | 5 | 0 | 0 | 1 | 0 | 0 | 2 | 0 | 0 | 0 | 98 | 49 | 0.332444824 | 0.810334259 | NA | NA | 0.714 | NA | NA | 1 | -0.286 |
| SE | 7846 | RPS6KA1 | chr1 | + | 26546866 | 26546983 | 26545851 | 26546035 | 26547188 | 26547270 | 0 | 3 | 19 | 0 | 0 | 1 | 0 | 10 | 6 | 0 | 0 | 0 | 98 | 49 | 0.332491859 | 0.516037235 | NA | 1 | 0.905 | NA | 1 | 1 | -0.047 |
| SE | 9991 | TRANK1 | chr3 | - | 36892900 | 36892984 | 36892201 | 36892340 | 36899108 | 36899259 | 0 | 0 | 15 | 0 | 0 | 2 | 1 | 3 | 3 | 0 | 0 | 0 | 98 | 49 | 0.332635761 | 0.516037235 | NA | NA | 0.789 | 1 | 1 | 1 | -0.211 |
| SE | 10092 | NUMB | chr14 | - | 73279280 | 73279424 | 73275209 | 73277293 | 73282358 | 73282505 | 0 | 0 | 2 | 0 | 0 | 10 | 0 | 0 | 0 | 0 | 0 | 2 | 98 | 49 | 0.332694665 | 0.516037235 | NA | NA | 0.091 | NA | NA | 0 | 0.091 |
| A3SS | 653 | VNN3 | chr6 | - | 132728740 | 132728979 | 132728740 | 132728933 | 132734252 | 132734383 | 0 | 0 | 9 | 0 | 0 | 4 | 0 | 23 | 1 | 0 | 2 | 0 | 95 | 49 | 0.33308192 | 0.545043142 | NA | NA | 0.537 | NA | 0.856 | 1 | -0.391 |
| SE | 13633 | CCND3 | chr6 | - | 41936558 | 41936695 | 41934932 | 41936107 | 41937234 | 41937394 | 0 | 3 | 19 | 0 | 2 | 0 | 0 | 2 | 2 | 0 | 0 | 2 | 98 | 49 | 0.333586462 | 0.516887065 | NA | 0.429 | 1 | NA | 1 | 0.333 | 0.048 |
| SE | 9234 | RPS24 | chr10 | + | 78040203 | 78040225 | 78037193 | 78037304 | 78040614 | 78040715 | 0 | 0 | 11 | 0 | 0 | 26 | 5 | 22 | 0 | 0 | 6 | 1 | 71 | 49 | 0.336551286 | 0.520943957 | NA | NA | 0.226 | 1 | 0.717 | 0 | -0.346 |
| SE | 2900 | DDX17 | chr22 | - | 38495795 | 38495937 | 38494885 | 38495046 | 38498084 | 38498150 | 0 | 1 | 31 | 0 | 0 | 1 | 3 | 22 | 2 | 0 | 0 | 0 | 98 | 49 | 0.337825803 | 0.522378788 | NA | 1 | 0.939 | 1 | 1 | 1 | -0.03 |
| SE | 12582 | CNN2 | chr19 | + | 1036129 | 1036246 | 1032558 | 1032696 | 1036415 | 1036562 | 0 | 1 | 25 | 0 | 0 | 1 | 0 | 17 | 2 | 0 | 0 | 0 | 98 | 49 | 0.338322411 | 0.522609027 | NA | 1 | 0.926 | NA | 1 | 1 | -0.037 |
| SE | 7966 | OAS3 | chr12 | + | 112941569 | 112941852 | 112938432 | 112938707 | 112944475 | 112944651 | 0 | 7 | 34 | 0 | 0 | 1 | 0 | 17 | 11 | 0 | 0 | 0 | 98 | 49 | 0.338676564 | 0.52261897 | NA | 1 | 0.944 | NA | 1 | 1 | -0.028 |
| SE | 12901 | TYROBP | chr19 | - | 35907217 | 35907264 | 35904400 | 35904634 | 35907445 | 35907580 | 0 | 18 | 77 | 0 | 0 | 1 | 0 | 30 | 10 | 0 | 1 | 0 | 96 | 49 | 0.342984435 | 0.527529386 | NA | 1 | 0.975 | NA | 0.939 | 1 | 0.018 |
| SE | 4627 | SORL1 | chr11 | + | 121570156 | 121570270 | 121566939 | 121567113 | 121574240 | 121574363 | 0 | 0 | 56 | 0 | 1 | 12 | 0 | 27 | 2 | 4 | 1 | 1 | 98 | 49 | 0.343393791 | 0.527529386 | NA | 0 | 0.7 | 0 | 0.931 | 0.5 | -0.127 |
| SE | 5558 | SPATA13 | chr13 | + | 24286764 | 24286950 | 24286213 | 24286393 | 24288998 | 24289178 | 0 | 0 | 5 | 0 | 0 | 2 | 0 | 1 | 1 | 0 | 1 | 0 | 98 | 49 | 0.34342743 | 0.527529386 | NA | NA | 0.556 | NA | 0.333 | 1 | -0.11 |
| SE | 11729 | PER1 | chr17 | - | 8146896 | 8147002 | 8146593 | 8146765 | 8147249 | 8147381 | 0 | 0 | 5 | 0 | 0 | 2 | 0 | 1 | 1 | 0 | 2 | 0 | 98 | 49 | 0.34342743 | 0.527529386 | NA | NA | 0.556 | NA | 0.2 | 1 | -0.044 |
| SE | 12395 | TMEM127 | chr2 | - | 96265137 | 96265489 | 96254832 | 96254997 | 96265868 | 96266013 | 0 | 3 | 13 | 0 | 0 | 1 | 0 | 7 | 4 | 0 | 0 | 0 | 98 | 49 | 0.343613619 | 0.527529386 | NA | 1 | 0.867 | NA | 1 | 1 | -0.067 |
| SE | 13844 | C10orf54 | chr10 | - | 71755358 | 71755466 | 71752974 | 71753002 | 71760867 | 71760924 | 0 | 3 | 48 | 0 | 0 | 1 | 2 | 32 | 4 | 0 | 0 | 0 | 98 | 49 | 0.344174583 | 0.527680329 | NA | 1 | 0.96 | 1 | 1 | 1 | -0.02 |
| SE | 13536 | ALOX5 | chr10 | + | 45382482 | 45382681 | 45374175 | 45374429 | 45395854 | 45395936 | 1 | 6 | 67 | 0 | 0 | 2 | 1 | 16 | 6 | 0 | 0 | 0 | 98 | 49 | 0.344414107 | 0.527680329 | 1 | 1 | 0.944 | 1 | 1 | 1 | -0.019 |
| SE | 12400 | CSF2RB | chr22 | + | 36925986 | 36926177 | 36923243 | 36923367 | 36929401 | 36929559 | 0 | 1 | 39 | 0 | 0 | 1 | 1 | 24 | 4 | 0 | 0 | 0 | 98 | 49 | 0.345199125 | 0.528344486 | NA | 1 | 0.951 | 1 | 1 | 1 | -0.024 |
| SE | 4506 | PTPN6 | chr12 | + | 6960805 | 6960945 | 6960343 | 6960435 | 6961125 | 6961316 | 0 | 1 | 28 | 0 | 0 | 1 | 7 | 18 | 3 | 0 | 0 | 0 | 98 | 49 | 0.346027303 | 0.528422573 | NA | 1 | 0.933 | 1 | 1 | 1 | -0.033 |
| SE | 9933 | PTPRE | chr10 | + | 128076602 | 128076728 | 128073336 | 128073471 | 128077616 | 128077783 | 0 | 3 | 39 | 0 | 0 | 0 | 0 | 24 | 2 | 0 | 0 | 1 | 98 | 49 | 0.346119905 | 0.528422573 | NA | 1 | 1 | NA | 1 | 0.5 | 0.25 |
| SE | 7719 | THEMIS2 | chr1 | + | 27879643 | 27880054 | 27876587 | 27876728 | 27885294 | 27885451 | 0 | 6 | 25 | 0 | 3 | 1 | 1 | 12 | 8 | 0 | 1 | 0 | 98 | 49 | 0.346439311 | 0.528422573 | NA | 0.5 | 0.926 | 1 | 0.857 | 1 | -0.239 |
| SE | 12745 | DYSF | chr2 | + | 71620546 | 71620609 | 71613333 | 71613410 | 71643964 | 71644063 | 0 | 0 | 13 | 0 | 0 | 3 | 0 | 3 | 2 | 0 | 2 | 0 | 98 | 49 | 0.346656459 | 0.528422573 | NA | NA | 0.684 | NA | 0.429 | 1 | -0.03 |
| SE | 13437 | HBP1 | chr7 | + | 107189291 | 107189448 | 107186568 | 107186681 | 107190172 | 107190317 | 0 | 1 | 1 | 0 | 0 | 1 | 0 | 1 | 1 | 0 | 3 | 0 | 98 | 49 | 0.348964996 | 0.531353191 | NA | 1 | 0.333 | NA | 0.143 | 1 | 0.095 |
| SE | 9997 | GCH1 | chr14 | - | 54844000 | 54844143 | 54842005 | 54843054 | 54845767 | 54845852 | 0 | 0 | 1 | 0 | 0 | 2 | 0 | 0 | 0 | 1 | 0 | 1 | 98 | 49 | 0.349286063 | 0.531353191 | NA | NA | 0.2 | 0 | NA | 0 | 0.2 |
| SE | 3661 | ETS2 | chr21 | + | 38821585 | 38821704 | 38819502 | 38819766 | 38822673 | 38824954 | 0 | 0 | 8 | 0 | 0 | 3 | 3 | 13 | 1 | 0 | 0 | 0 | 98 | 49 | 0.35298525 | 0.535895789 | NA | NA | 0.571 | 1 | 1 | 1 | -0.429 |
| SE | 9992 | TRANK1 | chr3 | - | 36895639 | 36895758 | 36892900 | 36892984 | 36899108 | 36899259 | 0 | 0 | 8 | 0 | 0 | 3 | 0 | 4 | 1 | 0 | 1 | 0 | 98 | 49 | 0.35298525 | 0.535895789 | NA | NA | 0.571 | NA | 0.667 | 1 | -0.263 |
| SE | 11324 | ITGAL | chr16 | + | 30479330 | 30479461 | 30479090 | 30479208 | 30481438 | 30481584 | 0 | 0 | 11 | 0 | 0 | 4 | 0 | 6 | 1 | 0 | 0 | 0 | 98 | 49 | 0.355559836 | 0.539081642 | NA | NA | 0.579 | NA | 1 | 1 | -0.421 |
| SE | 2458 | MPP1 | chrX | - | 154799744 | 154799893 | 154792141 | 154792285 | 154805271 | 154805527 | 0 | 0 | 1 | 0 | 0 | 6 | 0 | 0 | 0 | 0 | 0 | 2 | 98 | 49 | 0.355801057 | 0.539081642 | NA | NA | 0.077 | NA | NA | 0 | 0.077 |
| A3SS | 564 | PHF20L1 | chr8 | + | 132777791 | 132777911 | 132777795 | 132777911 | 132775357 | 132775645 | 0 | 0 | 1 | 0 | 3 | 3 | 0 | 3 | 0 | 2 | 5 | 3 | 53 | 49 | 0.358798866 | 0.579927566 | NA | 0 | 0.236 | 0 | 0.357 | 0 | -0.001 |
| SE | 2884 | MKL1 | chr22 | - | 40594673 | 40594735 | 40552105 | 40552367 | 40636477 | 40636719 | 0 | 0 | 4 | 0 | 0 | 2 | 0 | 0 | 1 | 0 | 0 | 0 | 98 | 49 | 0.359545146 | 0.544205795 | NA | NA | 0.5 | NA | NA | 1 | -0.5 |
| SE | 3751 | ARHGAP27 | chr17 | - | 45396486 | 45396585 | 45396206 | 45396284 | 45396667 | 45396790 | 0 | 4 | 14 | 0 | 0 | 1 | 0 | 8 | 3 | 0 | 0 | 0 | 98 | 49 | 0.360380165 | 0.544920914 | NA | 1 | 0.875 | NA | 1 | 1 | -0.062 |
| A3SS | 140 | LY6E | chr8 | + | 143020882 | 143020991 | 143020887 | 143020991 | 143018484 | 143018586 | 0 | 0 | 2 | 0 | 0 | 1 | 2 | 1 | 1 | 0 | 1 | 0 | 54 | 49 | 0.360843819 | 0.579927566 | NA | NA | 0.645 | 1 | 0.476 | 1 | -0.18 |
| SE | 18 | FLNA | chrX | - | 154354615 | 154354711 | 154354380 | 154354483 | 154354824 | 154355072 | 0 | 2 | 25 | 0 | 0 | 1 | 0 | 15 | 2 | 0 | 0 | 0 | 98 | 49 | 0.361294614 | 0.545754578 | NA | 1 | 0.926 | NA | 1 | 1 | -0.037 |
| SE | 12741 | DYSF | chr2 | + | 71513239 | 71513332 | 71511806 | 71511921 | 71513715 | 71513921 | 0 | 1 | 10 | 0 | 0 | 1 | 0 | 3 | 2 | 0 | 1 | 0 | 98 | 49 | 0.362088102 | 0.546301307 | NA | 1 | 0.833 | NA | 0.6 | 1 | 0.116 |
| SE | 12142 | TRIM25 | chr17 | - | 56895520 | 56895604 | 56895342 | 56895441 | 56895925 | 56895952 | 0 | 0 | 13 | 0 | 0 | 1 | 0 | 15 | 4 | 0 | 0 | 0 | 98 | 49 | 0.362383502 | 0.546301307 | NA | NA | 0.867 | NA | 1 | 1 | -0.133 |
| SE | 275 | RAPGEF1 | chr9 | - | 131630235 | 131630324 | 131629101 | 131629254 | 131638634 | 131638791 | 0 | 0 | 6 | 0 | 0 | 2 | 0 | 8 | 1 | 0 | 1 | 0 | 98 | 49 | 0.365054683 | 0.547059405 | NA | NA | 0.6 | NA | 0.8 | 1 | -0.3 |
| SE | 5253 | SEC31A | chr4 | - | 82848803 | 82848977 | 82844385 | 82844509 | 82851430 | 82851604 | 0 | 0 | 3 | 0 | 0 | 1 | 0 | 4 | 1 | 0 | 0 | 0 | 98 | 49 | 0.365118555 | 0.547059405 | NA | NA | 0.6 | NA | 1 | 1 | -0.4 |
| SE | 6132 | ANKRD13D | chr11 | + | 67299057 | 67299124 | 67291990 | 67292180 | 67299529 | 67299611 | 0 | 0 | 3 | 0 | 0 | 1 | 0 | 8 | 1 | 0 | 0 | 0 | 98 | 49 | 0.365118555 | 0.547059405 | NA | NA | 0.6 | NA | 1 | 1 | -0.4 |

| | | | | | | | | | | | | | | | | | | | | | | | | | | | | | | | | | |
|---|---|---|---|---|---|---|---|---|---|---|---|---|---|---|---|---|---|---|---|---|---|---|---|---|---|---|---|---|---|---|---|---|---|
| SE | 7202 | MDM4 | chr1 | + | 204544534 | 204544684 | 204542783 | 204542944 | 204546796 | 204546877 | 0 | 0 | 3 | 0 | 0 | 1 | 0 | 7 | 1 | 0 | 0 | 0 | 98 | 49 | 0.365118555 | 0.547059405 | NA | NA | 0.6 | NA | 1 | 1 | -0.4 |
| SE | 13919 | LUCAT1 | chr5 | - | 91313077 | 91313229 | 91303872 | 91303971 | 91313637 | 91313777 | 0 | 2 | 3 | 0 | 0 | 1 | 0 | 0 | 1 | 0 | 0 | 0 | 98 | 49 | 0.365118555 | 0.547059405 | NA | 1 | 0.6 | NA | NA | 1 | -0.2 |
| SE | 1533 | HLA-A | chr6 | + | 29944499 | 29944616 | 29944121 | 29944397 | 29945058 | 29945091 | 1 | 41 | 187 | 0 | 1 | 2 | 5 | 230 | 19 | 0 | 6 | 0 | 98 | 49 | 0.3653902 | 0.547059405 | 1 | 0.953 | 0.979 | 1 | 0.95 | 1 | -0.006 |
| SE | 3397 | APMAP | chr20 | - | 24978766 | 24978882 | 24973644 | 24973737 | 24983902 | 24984019 | 0 | 1 | 5 | 0 | 0 | 1 | 0 | 2 | 2 | 0 | 0 | 0 | 98 | 49 | 0.365434227 | 0.547059405 | NA | 1 | 0.714 | NA | 1 | 1 | -0.143 |
| SE | 11877 | MGAM | chr7 | + | 142036822 | 142036977 | 142036168 | 142036285 | 142038530 | 142038615 | 0 | 0 | 20 | 0 | 0 | 1 | 4 | 3 | 6 | 0 | 0 | 0 | 98 | 49 | 0.366623914 | 0.548294271 | NA | NA | 0.909 | 1 | 1 | 1 | -0.091 |
| SE | 2284 | IL1RAP | chr3 | + | 190556129 | 190556216 | 190514050 | 190514219 | 190564288 | 190564353 | 0 | 3 | 7 | 0 | 0 | 1 | 0 | 4 | 2 | 0 | 0 | 0 | 98 | 49 | 0.367396868 | 0.548904067 | NA | 1 | 0.778 | NA | 1 | 1 | -0.111 |
| SE | 12976 | RPS19 | chr19 | + | 41869030 | 41869214 | 41861111 | 41861212 | 41869698 | 41869753 | 1 | 6 | 58 | 0 | 0 | 1 | 3 | 38 | 4 | 0 | 0 | 0 | 98 | 49 | 0.368911011 | 0.550618918 | 1 | 1 | 0.967 | 1 | 1 | 1 | -0.011 |
| SE | 8375 | OAS2 | chr12 | + | 112987037 | 112987308 | 112978468 | 112978785 | 112995295 | 112995474 | 0 | 6 | 42 | 0 | 0 | 3 | 3 | 7 | 1 | 0 | 0 | 0 | 98 | 49 | 0.370767489 | 0.552840809 | NA | 1 | 0.875 | 1 | 1 | 1 | -0.062 |
| SE | 6279 | IDS | chrX | - | 149491537 | 149491667 | 149490313 | 149490440 | 149496345 | 149496516 | 0 | 0 | 1 | 0 | 0 | 7 | 0 | 0 | 0 | 0 | 2 | 2 | 98 | 49 | 0.37479931 | 0.558298674 | NA | NA | 0.067 | NA | 0 | 0 | 0.067 |
| SE | 6619 | AMPD2 | chr1 | + | 109629795 | 109629916 | 109629326 | 109629490 | 109630232 | 109630406 | 0 | 1 | 7 | 0 | 0 | 3 | 0 | 7 | 1 | 0 | 4 | 0 | 98 | 49 | 0.376779701 | 0.560692961 | NA | 1 | 0.538 | NA | 0.467 | 1 | 0.035 |
| SE | 12898 | TXNIP | chr1 | - | 145994543 | 145994800 | 145994280 | 145994437 | 145994928 | 145995031 | 0 | 17 | 121 | 0 | 0 | 1 | 4 | 61 | 19 | 0 | 0 | 0 | 98 | 49 | 0.378921937 | 0.563323118 | NA | 1 | 0.984 | 1 | 1 | 1 | -0.008 |
| A3SS | 3632 | STXBP2 | chr19 | + | 7640697 | 7640809 | 7640730 | 7640809 | 7639730 | 7639807 | 0 | 0 | 7 | 0 | 1 | 1 | 2 | 0 | 2 | 0 | 0 | 0 | 82 | 49 | 0.379879433 | 0.605117681 | NA | 0 | 0.807 | 1 | NA | 1 | -0.597 |
| RI | 545 | CDK5RAP3 | chr17 | + | 47974399 | 47975337 | 47974399 | 47974448 | 47975158 | 47975337 | 0 | 7 | 10 | 0 | 0 | 1 | 0 | 4 | 5 | 0 | 0 | 0 | 98 | 49 | 0.38169658 | 0.878475733 | NA | 1 | 0.833 | NA | 1 | 1 | -0.084 |
| SE | 8578 | RC3H1 | chr1 | - | 173983417 | 173983657 | 173982726 | 173982902 | 173984498 | 173984619 | 0 | 1 | 7 | 0 | 0 | 1 | 0 | 3 | 1 | 0 | 1 | 0 | 98 | 49 | 0.382084955 | 0.567464117 | NA | 1 | 0.778 | NA | 0.6 | 1 | 0.089 |
| SE | 11360 | TATDN2 | chr3 | + | 10249194 | 10249614 | 10248492 | 10249067 | 10260136 | 10260670 | 0 | 3 | 6 | 0 | 0 | 1 | 2 | 4 | 1 | 0 | 0 | 0 | 98 | 49 | 0.382895649 | 0.568106772 | NA | 1 | 0.75 | 1 | 1 | 1 | -0.125 |
| RI | 660 | CSF3R | chr1 | - | 36466042 | 36466827 | 36466042 | 36466200 | 36466620 | 36466827 | 1 | 19 | 165 | 0 | 0 | 1 | 7 | 83 | 24 | 0 | 0 | 0 | 98 | 49 | 0.382925319 | 0.878475733 | 1 | 1 | 0.988 | 1 | 1 | 1 | -0.004 |
| SE | 7522 | AQP9 | chr15 | + | 58174917 | 58175036 | 58173067 | 58173205 | 58179127 | 58179345 | 0 | 1 | 43 | 0 | 0 | 1 | 3 | 20 | 6 | 0 | 0 | 0 | 98 | 49 | 0.383614897 | 0.568612613 | NA | 1 | 0.956 | 1 | 1 | 1 | -0.022 |
| SE | 802 | TBXAS1 | chr7 | + | 139957633 | 139957764 | 139955458 | 139955607 | 139961918 | 139962233 | 0 | 3 | 22 | 0 | 1 | 0 | 2 | 4 | 2 | 0 | 0 | 0 | 98 | 49 | 0.385851284 | 0.571364019 | NA | 0.6 | 1 | 1 | 1 | 1 | -0.2 |
| SE | 8267 | DPYD | chr1 | - | 97549559 | 97549744 | 97515725 | 97515941 | 97573759 | 97573970 | 0 | 4 | 25 | 0 | 0 | 1 | 0 | 11 | 4 | 0 | 1 | 0 | 98 | 49 | 0.389051763 | 0.57553622 | NA | 1 | 0.926 | NA | 0.846 | 1 | 0.04 |
| A5SS | 1166 | IRF2BP2 | chr1 | - | 234608446 | 234609525 | 234608494 | 234609525 | 234604268 | 234607852 | 0 | 0 | 5 | 0 | 1 | 5 | 0 | 4 | 0 | 0 | 10 | 1 | 97 | 49 | 0.389300363 | 0.614437923 | NA | 0 | 0.336 | NA | 0.168 | 0 | 0.084 |
| SE | 306 | SH2D3C | chr9 | - | 127741787 | 127741959 | 127740257 | 127740369 | 127742848 | 127742964 | 0 | 2 | 7 | 0 | 1 | 0 | 0 | 2 | 2 | 0 | 0 | 0 | 98 | 49 | 0.390828967 | 0.577596791 | NA | 0.5 | 1 | NA | 1 | 1 | -0.25 |
| SE | 4458 | CLEC7A | chr12 | - | 10125086 | 10125173 | 10116780 | 10118590 | 10125296 | 10125448 | 0 | 3 | 18 | 0 | 0 | 8 | 0 | 4 | 3 | 0 | 1 | 0 | 98 | 49 | 0.391525682 | 0.578058056 | NA | 1 | 0.529 | NA | 0.667 | 1 | -0.069 |
| SE | 6890 | EP300 | chr22 | + | 41135812 | 41135906 | 41131387 | 41131633 | 41137652 | 41137790 | 0 | 2 | 14 | 0 | 0 | 1 | 0 | 8 | 1 | 0 | 0 | 0 | 98 | 49 | 0.392944473 | 0.579315862 | NA | 1 | 0.875 | NA | 1 | 1 | -0.062 |
| SE | 3233 | SLC9A8 | chr20 | + | 49839540 | 49839599 | 49823060 | 49823141 | 49845035 | 49845119 | 0 | 0 | 8 | 0 | 0 | 1 | 0 | 0 | 2 | 0 | 0 | 0 | 98 | 49 | 0.393533929 | 0.579315862 | NA | NA | 0.8 | NA | NA | 1 | -0.2 |
| SE | 11134 | ECE1 | chr1 | - | 21225249 | 21225440 | 21221746 | 21221842 | 21227158 | 21227226 | 1 | 0 | 8 | 0 | 0 | 1 | 0 | 12 | 2 | 0 | 0 | 0 | 98 | 49 | 0.393533929 | 0.579315862 | 1 | NA | 0.8 | NA | 1 | 1 | -0.1 |
| SE | 11656 | NLRP1 | chr17 | - | 5517745 | 5517887 | 5515472 | 5515517 | 5530480 | 5530704 | 0 | 0 | 7 | 0 | 1 | 0 | 0 | 2 | 3 | 0 | 0 | 0 | 98 | 49 | 0.39950276 | 0.587426728 | NA | 0 | 1 | NA | 1 | 1 | -0.5 |
| SE | 1505 | HLA-E | chr6 | + | 30491536 | 30491653 | 30491136 | 30491412 | 30492403 | 30492436 | 0 | 33 | 293 | 0 | 0 | 1 | 21 | 137 | 29 | 0 | 0 | 0 | 98 | 49 | 0.399825378 | 0.587426728 | NA | 1 | 0.993 | 1 | 1 | 1 | -0.004 |
| SE | 5153 | FTH1 | chr11 | - | 61964986 | 61965112 | 61964284 | 61964891 | 61965368 | 61965515 | 0 | 8 | 114 | 0 | 0 | 1 | 7 | 66 | 2 | 0 | 0 | 0 | 98 | 49 | 0.400407723 | 0.587707819 | NA | 1 | 0.983 | 1 | 1 | 1 | -0.008 |
| SE | 11835 | NCOA1 | chr2 | + | 24768078 | 24768135 | 24762686 | 24762776 | 24768220 | 24770701 | 0 | 0 | 8 | 0 | 0 | 2 | 0 | 2 | 1 | 0 | 0 | 0 | 98 | 49 | 0.401159139 | 0.58823628 | NA | NA | 0.667 | NA | 1 | 1 | -0.333 |
| A3SS | 909 | TPD52L2 | chr20 | + | 63882718 | 63882820 | 63882749 | 63882820 | 63875815 | 63875875 | 0 | 2 | 11 | 0 | 0 | 1 | 3 | 5 | 2 | 0 | 0 | 0 | 80 | 49 | 0.401186798 | 0.63162252 | NA | 1 | 0.871 | 1 | 1 | 1 | -0.065 |
| SE | 9689 | IFRD1 | chr7 | + | 112462269 | 112462378 | 112462000 | 112462179 | 112467980 | 112468115 | 0 | 0 | 10 | 0 | 3 | 37 | 4 | 3 | 0 | 2 | 22 | 2 | 98 | 49 | 0.40162165 | 0.588340487 | NA | 0 | 0.119 | 0.5 | 0.064 | 0 | -0.129 |
| A3SS | 4073 | CFLAR | chr2 | + | 201162995 | 201172688 | 201163834 | 201172688 | 201160431 | 201160942 | 0 | 0 | 4 | 0 | 3 | 38 | 2 | 3 | 2 | 7 | 14 | 3 | 98 | 49 | 0.40353661 | 0.63162252 | NA | 0 | 0.05 | 0.125 | 0.097 | 0.25 | -0.132 |
| SE | 13946 | AP3D1 | chr19 | - | 2108686 | 2108766 | 2100987 | 2102268 | 2109085 | 2109207 | 0 | 2 | 2 | 0 | 0 | 1 | 0 | 1 | 1 | 0 | 0 | 0 | 98 | 49 | 0.409859466 | 0.599823542 | NA | 1 | 0.5 | NA | 1 | 1 | -0.25 |
| SE | 13218 | LILRA5 | chr19 | - | 54311863 | 54312148 | 54311164 | 54311716 | 54312536 | 54312621 | 0 | 1 | 18 | 0 | 0 | 1 | 0 | 9 | 1 | 0 | 0 | 0 | 98 | 49 | 0.410993926 | 0.600898707 | NA | 1 | 0.9 | NA | 1 | 1 | -0.05 |
| SE | 6789 | RAB4B-EGLN2 | chr19 | + | 40806554 | 40806674 | 40800338 | 40801415 | 40807137 | 40807274 | 0 | 2 | 3 | 0 | 0 | 0 | 0 | 1 | 0 | 0 | 0 | 1 | 98 | 49 | 0.412150794 | 0.601420042 | NA | 1 | 1 | NA | 1 | 0 | 0.5 |
| SE | 7891 | FHL3 | chr1 | - | 37999256 | 37999432 | 37998973 | 37999148 | 38005356 | 38005515 | 0 | 2 | 3 | 0 | 0 | 0 | 0 | 1 | 0 | 0 | 0 | 1 | 98 | 49 | 0.412150794 | 0.601420042 | NA | 1 | 1 | NA | 1 | 0 | 0.5 |
| SE | 12902 | TYROBP | chr19 | - | 35907729 | 35907762 | 35907445 | 35907580 | 35908167 | 35908309 | 0 | 4 | 15 | 0 | 0 | 5 | 0 | 7 | 3 | 0 | 0 | 0 | 82 | 49 | 0.416321308 | 0.606916515 | NA | 1 | 0.642 | NA | 0.807 | 1 | -0.083 |
| RI | 981 | ACAA1 | chr3 | - | 38126161 | 38126700 | 38126161 | 38126341 | 38126509 | 38126700 | 0 | 0 | 9 | 0 | 0 | 1 | 0 | 1 | 3 | 0 | 1 | 0 | 98 | 49 | 0.421251047 | 0.934008656 | NA | NA | 0.818 | NA | 0.333 | 1 | 0.151 |
| SE | 5079 | WDR13 | chrX | + | 48599586 | 48599717 | 48599352 | 48599462 | 48600318 | 48600626 | 0 | 1 | 4 | 0 | 0 | 0 | 0 | 1 | 0 | 0 | 0 | 1 | 98 | 49 | 0.421888846 | 0.613929698 | NA | 1 | 1 | NA | 1 | 0 | 0.5 |
| SE | 11377 | ARHGEF1 | chr19 | + | 41906456 | 41906620 | 41905938 | 41906025 | 41906702 | 41906803 | 0 | 0 | 18 | 0 | 0 | 2 | 0 | 5 | 3 | 0 | 0 | 0 | 98 | 49 | 0.422105527 | 0.613929698 | NA | NA | 0.818 | NA | 1 | 1 | -0.182 |
| SE | 3692 | DOK3 | chr5 | - | 177508236 | 177508542 | 177505010 | 177505110 | 177509474 | 177509575 | 0 | 7 | 32 | 0 | 0 | 1 | 0 | 13 | 5 | 0 | 0 | 0 | 98 | 49 | 0.42235749 | 0.613929698 | NA | 1 | 0.941 | NA | 1 | 1 | -0.03 |
| SE | 13638 | RNF114 | chr20 | + | 49941560 | 49941711 | 49936376 | 49936552 | 49945381 | 49945488 | 0 | 1 | 2 | 0 | 0 | 0 | 0 | 0 | 1 | 0 | 1 | 0 | 98 | 49 | 0.423875417 | 0.615287988 | NA | 1 | 1 | NA | 0 | 1 | 0.5 |
| SE | 5214 | SUCO | chr1 | + | 172551511 | 172551626 | 172533117 | 172533497 | 172553259 | 172553370 | 0 | 1 | 1 | 0 | 0 | 0 | 1 | 1 | 0 | 0 | 0 | 1 | 98 | 49 | 0.424110682 | 0.615287988 | NA | 1 | 1 | 1 | 1 | 0 | 0.333 |
| SE | 5013 | ADRBK1 | chr11 | + | 67283706 | 67283773 | 67283127 | 67283228 | 67283853 | 67283949 | 0 | 13 | 34 | 0 | 5 | 0 | 4 | 7 | 17 | 0 | 1 | 0 | 98 | 49 | 0.425222253 | 0.6162202 | NA | 0.565 | 1 | 1 | 0.778 | 1 | -0.144 |
| SE | 2290 | BCL6 | chr3 | - | 187728359 | 187728544 | 187726730 | 187726898 | 187729049 | 187730021 | 0 | 2 | 24 | 0 | 0 | 1 | 3 | 8 | 4 | 0 | 0 | 0 | 98 | 49 | 0.425573232 | 0.6162202 | NA | 1 | 0.923 | 1 | 1 | 1 | -0.038 |
| A5SS | 1299 | WDR26 | chr1 | - | 224431476 | 224431581 | 224431524 | 224431581 | 224424517 | 224424654 | 0 | 0 | 11 | 0 | 0 | 4 | 4 | 5 | 1 | 0 | 2 | 0 | 97 | 49 | 0.426432576 | 0.665031756 | NA | NA | 0.581 | 1 | 0.558 | 1 | -0.272 |
| SE | 6186 | HERC3 | chr4 | + | 88681158 | 88681325 | 88680092 | 88680236 | 88686735 | 88686802 | 0 | 0 | 12 | 0 | 0 | 2 | 0 | 8 | 2 | 0 | 0 | 0 | 98 | 49 | 0.429038403 | 0.620451665 | NA | NA | 0.75 | NA | 1 | 1 | -0.25 |
| SE | 2247 | MAEA | chr4 | + | 1327626 | 1327703 | 1315396 | 1315600 | 1332756 | 1332865 | 0 | 3 | 2 | 0 | 1 | 1 | 0 | 1 | 0 | 0 | 1 | 0 | 98 | 49 | 0.429432607 | 0.620451665 | NA | 0.6 | 0.5 | NA | 0.333 | NA | 0.217 |
| SE | 12900 | TYROBP | chr19 | - | 35907217 | 35907261 | 35904400 | 35904634 | 35907445 | 35907580 | 0 | 7 | 39 | 0 | 0 | 1 | 0 | 22 | 1 | 0 | 1 | 0 | 93 | 49 | 0.429733987 | 0.620451665 | NA | 1 | 0.954 | NA | 0.921 | 1 | 0.016 |
| SE | 679 | SARAF | chr8 | - | 30073876 | 30074055 | 30069641 | 30070059 | 30082846 | 30083208 | 0 | 3 | 13 | 0 | 1 | 5 | 0 | 17 | 4 | 0 | 5 | 6 | 98 | 49 | 0.430454253 | 0.620561858 | NA | 0.6 | 0.565 | NA | 0.63 | 0.25 | 0.143 |
| SE | 5660 | KCNQ1 | chr11 | + | 2585211 | 2585307 | 2583434 | 2583545 | 2587569 | 2587692 | 0 | 0 | 4 | 0 | 1 | 0 | 0 | 1 | 1 | 0 | 0 | 0 | 98 | 49 | 0.430636073 | 0.620561858 | NA | 0 | 1 | NA | 1 | 1 | -0.5 |
| RI | 694 | UNC119 | chr17 | - | 28546706 | 28547849 | 28546706 | 28547409 | 28547676 | 28547849 | 0 | 1 | 10 | 0 | 0 | 1 | 2 | 3 | 2 | 1 | 0 | 0 | 98 | 49 | 0.431080918 | 0.934008656 | NA | 1 | 0.833 | 0.5 | 1 | 1 | 0.083 |
| A3SS | 3011 | ISG20 | chr15 | + | 88650234 | 88652310 | 88652109 | 88652310 | 88639342 | 88639594 | 0 | 0 | 2 | 0 | 0 | 11 | 0 | 3 | 1 | 0 | 1 | 2 | 98 | 49 | 0.432135137 | 0.670554523 | NA | NA | 0.083 | NA | 0.6 | 0.2 | -0.317 |
| SE | 13731 | SERPINA1 | chr14 | - | 94388559 | 94388660 | 94382591 | 94383241 | 94390456 | 94390692 | 0 | 1 | 16 | 0 | 2 | 20 | 4 | 5 | 1 | 0 | 5 | 0 | 98 | 49 | 0.432495196 | 0.621784006 | NA | 0.2 | 0.286 | 1 | 0.333 | 1 | -0.535 |
| SE | 1921 | PARP8 | chr5 | + | 50821212 | 50821338 | 50815431 | 50815524 | 50822334 | 50822400 | 0 | 2 | 9 | 0 | 0 | 1 | 0 | 4 | 1 | 0 | 0 | 0 | 98 | 49 | 0.432725263 | 0.621784006 | NA | 1 | 0.818 | NA | 1 | 1 | -0.091 |
| SE | 2181 | RBM47 | chr4 | - | 40437770 | 40438924 | 40432650 | 40432862 | 40466576 | 40466699 | 0 | 2 | 9 | 0 | 0 | 1 | 0 | 4 | 1 | 0 | 0 | 0 | 98 | 49 | 0.432725263 | 0.621784006 | NA | 1 | 0.818 | NA | 1 | 1 | -0.091 |
| A5SS | 1440 | MLKL | chr16 | - | 74675323 | 74675404 | 74675354 | 74675404 | 74674959 | 74675100 | 1 | 1 | 1 | 0 | 5 | 3 | 0 | 3 | 0 | 1 | 1 | 0 | 80 | 49 | 0.432994858 | 0.667321487 | 1 | 0.109 | 0.17 | 0 | 0.648 | NA | 0.102 |
| SE | 4557 | SLC6A6 | chr3 | + | 14443623 | 14443863 | 14416411 | 14416453 | 14445716 | 14445851 | 0 | 5 | 30 | 0 | 0 | 1 | 0 | 16 | 5 | 0 | 1 | 0 | 98 | 49 | 0.434955535 | 0.624391756 | NA | 1 | 0.937 | NA | 0.889 | 1 | 0.024 |
| SE | 5786 | DENND3 | chr8 | + | 141150833 | 141150953 | 141138021 | 141138137 | 141151618 | 141151837 | 0 | 0 | 22 | 0 | 0 | 1 | 0 | 10 | 4 | 0 | 0 | 0 | 98 | 49 | 0.435694354 | 0.624855547 | NA | NA | 0.917 | NA | 1 | 1 | -0.083 |
| SE | 3704 | LYZ | chr12 | + | 69352219 | 69352298 | 69350107 | 69350272 | 69353152 | 69354233 | 0 | 40 | 456 | 0 | 0 | 1 | 43 | 267 | 1 | 0 | 0 | 0 | 98 | 49 | 0.441484717 | 0.632034884 | NA | 1 | 0.996 | 1 | 1 | 1 | -0.002 |
| SE | 2699 | MBNL1 | chr3 | + | 152299404 | 152300367 | 152268039 | 152269092 | 152414940 | 152415111 | 0 | 0 | 6 | 0 | 0 | 1 | 5 | 3 | 1 | 0 | 1 | 0 | 98 | 49 | 0.441541336 | 0.632034884 | NA | NA | 0.75 | 1 | 0.6 | 1 | -0.117 |
| SE | 13538 | ALOX5 | chr10 | + | 45428617 | 45428764 | 45424959 | 45425132 | 45440429 | 45440633 | 0 | 9 | 46 | 0 | 0 | 1 | 0 | 19 | 4 | 0 | 0 | 0 | 98 | 49 | 0.442803979 | 0.633239183 | NA | 1 | 0.958 | NA | 1 | 1 | -0.021 |
| SE | 3207 | STX16 | chr20 | + | 58669290 | 58669453 | 58667986 | 58668127 | 58670511 | 58670603 | 0 | 0 | 9 | 0 | 0 | 3 | 0 | 9 | 1 | 0 | 4 | 0 | 98 | 49 | 0.44356605 | 0.633726019 | NA | NA | 0.6 | NA | 0.529 | 1 | -0.164 |
| SE | 10590 | RNF213 | chr17 | + | 80379619 | 80379714 | 80377761 | 80377796 | 80380830 | 80380987 | 0 | 3 | 53 | 0 | 0 | 1 | 0 | 22 | 3 | 0 | 0 | 0 | 98 | 49 | 0.44486502 | 0.634978277 | NA | 1 | 0.964 | NA | 1 | 1 | -0.018 |
| A5SS | 2190 | HES4 | chr1 | - | 999691 | 1000172 | 999865 | 1000172 | 999525 | 999613 | 1 | 1 | 1 | 0 | 1 | 1 | 3 | 11 | 4 | 0 | 4 | 1 | 98 | 49 | 0.445770867 | 0.679023064 | 1 | 0.333 | 0.333 | 1 | 0.579 | 0.667 | -0.193 |
| SE | 1983 | DDX60L | chr4 | - | 168448637 | 168448779 | 168441336 | 168441492 | 168453123 | 168453282 | 0 | 2 | 53 | 0 | 0 | 1 | 0 | 35 | 2 | 0 | 1 | 0 | 98 | 49 | 0.445818676 | 0.63573574 | NA | 1 | 0.964 | NA | 0.946 | 1 | 0.009 |
| SE | 4191 | FAM49B | chr8 | - | 129879388 | 129879471 | 129871374 | 129871496 | 129903311 | 129903350 | 0 | 0 | 14 | 0 | 0 | 1 | 4 | 13 | 4 | 0 | 4 | 0 | 98 | 49 | 0.449034561 | 0.639399819 | NA | NA | 0.875 | 1 | 0.619 | 1 | 0.002 |
| SE | 13835 | SPG21 | chr15 | - | 64983506 | 64983593 | 64980863 | 64981025 | 64989664 | 64989946 | 0 | 1 | 6 | 0 | 0 | 1 | 0 | 2 | 1 | 0 | 0 | 0 | 98 | 49 | 0.449609977 | 0.639399819 | NA | 1 | 0.75 | NA | 1 | 1 | -0.125 |
| SE | 11826 | ARHGDIB | chr12 | - | 14949801 | 14949885 | 14947872 | 14947949 | 14950531 | 14950724 | 0 | 13 | 90 | 0 | 0 | 1 | 12 | 34 | 7 | 0 | 0 | 0 | 98 | 49 | 0.44966441 | 0.639399819 | NA | 1 | 0.978 | 1 | 1 | 1 | -0.011 |
| RI | 780 | SHISA5 | chr3 | - | 48467797 | 48469573 | 48467797 | 48469186 | 48469360 | 48469573 | 0 | 3 | 1 | 0 | 5 | 13 | 5 | 0 | 1 | 0 | 10 | 2 | 98 | 49 | 0.455153121 | 0.94773154 | NA | 0.231 | 0.037 | 1 | 0 | 0.2 | -0.266 |
| SE | 2102 | WDFY3 | chr4 | - | 84817391 | 84817585 | 84809886 | 84810344 | 84820084 | 84820186 | 0 | 1 | 19 | 0 | 1 | 0 | 1 | 6 | 3 | 0 | 0 | 0 | 98 | 49 | 0.458220849 | 0.650950791 | NA | 0.333 | 1 | 1 | 1 | 1 | -0.334 |
| SE | 13718 | HMHA1 | chr19 | + | 1068413 | 1068744 | 1067165 | 1067495 | 1073148 | 1073292 | 0 | 8 | 14 | 0 | 0 | 1 | 0 | 2 | 6 | 0 | 0 | 0 | 98 | 49 | 0.460039403 | 0.652917113 | NA | 1 | 0.875 | NA | 1 | 1 | -0.062 |
| SE | 8333 | NCSTN | chr1 | + | 160355654 | 160355757 | 160354117 | 160354290 | 160355862 | 160355958 | 0 | 1 | 4 | 0 | 0 | 1 | 0 | 1 | 1 | 0 | 0 | 0 | 98 | 49 | 0.46501066 | 0.659350021 | NA | 1 | 0.667 | NA | 1 | 1 | -0.166 |
| A3SS | 2675 | ABTB1 | chr3 | + | 127674344 | 127674418 | 127674390 | 127674418 | 127672937 | 127673081 | 0 | 0 | 8 | 0 | 3 | 0 | 0 | 2 | 0 | 0 | 3 | 2 | 95 | 49 | 0.468058768 | 0.720090412 | NA | 0 | 1 | NA | 0.256 | 0 | 0.372 |
| SE | 1195 | MAP3K5 | chr6 | - | 136720449 | 136720589 | 136705109 | 136705133 | 136791709 | 136792518 | 0 | 1 | 12 | 0 | 0 | 1 | 0 | 4 | 1 | 0 | 0 | 0 | 98 | 49 | 0.46879248 | 0.66408586 | NA | 1 | 0.857 | NA | 1 | 1 | -0.072 |
| A5SS | 3016 | SERPINA1 | chr14 | - | 94390419 | 94390692 | 94390456 | 94390692 | 94382591 | 94383241 | 0 | 4 | 5 | 0 | 2 | 20 | 3 | 3 | 2 | 0 | 5 | 0 | 86 | 49 | 0.47090828 | 0.709068789 | NA | 0.533 | 0.125 | 1 | 0.255 | 1 | -0.423 |
| SE | 9545 | PRKCB | chr16 | + | 24172269 | 24172361 | 24154683 | 24154857 | 24174517 | 24174580 | 0 | 9 | 27 | 0 | 0 | 1 | 0 | 11 | 2 | 0 | 0 | 0 | 98 | 49 | 0.472851524 | 0.669205122 | NA | 1 | 0.931 | NA | 1 | 1 | -0.034 |
| A3SS | 1443 | TBC1D10C | chr11 | + | 67405902 | 67406017 | 67405906 | 67406017 | 67405594 | 67405701 | 0 | 0 | 7 | 0 | 0 | 2 | 0 | 0 | 1 | 0 | 0 | 0 | 53 | 49 | 0.472854846 | 0.721304002 | NA | NA | 0.764 | NA | NA | 1 | -0.236 |
| RI | 556 | CDC16 | chr13 | + | 114272183 | 114272675 | 114272183 | 114272481 | 114272563 | 114272675 | 0 | 0 | 7 | 0 | 0 | 2 | 0 | 1 | 1 | 0 | 0 | 0 | 98 | 49 | 0.47352014 | 0.94773154 | NA | NA | 0.636 | NA | 1 | 1 | -0.364 |
| SE | 11966 | MSL1 | chr17 | - | 40131536 | 40131584 | 40129244 | 40129627 | 40132033 | 40132098 | 0 | 0 | 7 | 0 | 0 | 2 | 0 | 4 | 1 | 0 | 1 | 0 | 97 | 49 | 0.474814393 | 0.671350925 | NA | NA | 0.639 | NA | 0.669 | 1 | -0.196 |
| SE | 2939 | LIMK2 | chr22 | + | 31266983 | 31267070 | 31265945 | 31266132 | 31267775 | 31267907 | 0 | 0 | 20 | 0 | 0 | 1 | 0 | 34 | 5 | 0 | 0 | 0 | 98 | 49 | 0.475908434 | 0.671934183 | NA | NA | 0.909 | NA | 1 | 1 | -0.091 |
| SE | 1139 | IGF2R | chr6 | + | 160061752 | 160061928 | 160061502 | 160061646 | 160062531 | 160062619 | 0 | 5 | 27 | 0 | 0 | 1 | 0 | 13 | 8 | 0 | 1 | 0 | 98 | 49 | 0.476121028 | 0.671934183 | NA | 1 | 0.931 | NA | 0.867 | 1 | 0.032 |
| SE | 3270 | CTSA | chr20 | + | 45893219 | 45893311 | 45892724 | 45892880 | 45893987 | 45894072 | 0 | 1 | 14 | 0 | 1 | 0 | 0 | 4 | 2 | 2 | 0 | 0 | 98 | 49 | 0.476656296 | 0.672058549 | NA | 0.333 | 1 | 0 | 1 | 1 | 0 |
| SE | 5812 | HAUS4 | chr14 | - | 22951554 | 22951689 | 22950313 | 22950410 | 22952327 | 22952459 | 0 | 0 | 16 | 0 | 0 | 1 | 0 | 3 | 2 | 0 | 0 | 0 | 98 | 49 | 0.480086309 | 0.676260283 | NA | NA | 0.889 | NA | 1 | 1 | -0.111 |
| RI | 1055 | STX10 | chr19 | - | 13144057 | 13144671 | 13144057 | 13144486 | 13144576 | 13144671 | 0 | 1 | 6 | 0 | 0 | 1 | 0 | 0 | 1 | 0 | 1 | 0 | 98 | 49 | 0.482241355 | 0.94773154 | NA | 1 | 0.75 | NA | 0 | 1 | 0.375 |
| A5SS | 2804 | TYROBP | chr19 | - | 35908167 | 35908309 | 35908174 | 35908309 | 35907729 | 35907762 | 0 | 1 | 10 | 0 | 0 | 1 | 0 | 3 | 4 | 0 | 1 | 0 | 56 | 49 | 0.482508271 | 0.718279358 | NA | 1 | 0.897 | NA | 0.724 | 1 | 0.087 |
| SE | 8138 | ITGAX | chr16 | + | 31371333 | 31371497 | 31371083 | 31371214 | 31371629 | 31371784 | 0 | 1 | 13 | 0 | 1 | 0 | 0 | 10 | 6 | 0 | 3 | 1 | 98 | 49 | 0.482836455 | 0.67949737 | NA | 0.333 | 1 | NA | 0.625 | 0.75 | -0.021 |
| SE | 12743 | DYSF | chr2 | + | 71549349 | 71549391 | 71539156 | 71539239 | 71551040 | 71551156 | 0 | 2 | 2 | 0 | 1 | 11 | 0 | 2 | 1 | 0 | 4 | 0 | 91 | 49 | 0.489990933 | 0.688920834 | NA | 0.519 | 0.089 | NA | 0.212 | 1 | -0.302 |
| A3SS | 1013 | INPP5D | chr2 | + | 233125744 | 233125919 | 233125747 | 233125919 | 233122106 | 233122257 | 1 | 0 | 7 | 0 | 1 | 5 | 0 | 5 | 0 | 1 | 2 | 0 | 52 | 49 | 0.490380444 | 0.74107425 | 1 | 0 | 0.569 | 0 | 0.702 | NA | 0.172 |
| SE | 10208 | RFWD2 | chr1 | - | 176135009 | 176135086 | 176116623 | 176116681 | 176149005 | 176149074 | 0 | 1 | 21 | 0 | 0 | 1 | 0 | 7 | 1 | 0 | 0 | 0 | 98 | 49 | 0.493056216 | 0.692582704 | NA | 1 | 0.913 | NA | 1 | 1 | -0.043 |
| RI | 322 | MAP7D1 | chr1 | + | 36176198 | 36176581 | 36176198 | 36176321 | 36176417 | 36176581 | 0 | 1 | 8 | 0 | 1 | 1 | 3 | 5 | 6 | 0 | 1 | 0 | 98 | 49 | 0.496928876 | 0.94773154 | NA | 0.333 | 0.8 | 1 | 0.714 | 1 | -0.338 |
| SE | 4467 | CLEC12A | chr12 | + | 9978965 | 9979064 | 9971408 | 9971687 | 9979335 | 9979524 | 0 | 0 | 25 | 0 | 0 | 3 | 0 | 0 | 2 | 0 | 0 | 0 | 98 | 49 | 0.497638055 | 0.696506389 | NA | NA | 0.806 | NA | NA | 1 | -0.194 |
| SE | 4625 | SORL1 | chr11 | + | 121545242 | 121545429 | 121543547 | 121543726 | 121549959 | 121550088 | 0 | 6 | 54 | 0 | 0 | 1 | 0 | 38 | 2 | 0 | 1 | 0 | 98 | 49 | 0.497995567 | 0.696506389 | NA | 1 | 0.964 | NA | 0.95 | 1 | 0.007 |
| A3SS | 1731 | SECISBP2 | chr9 | + | 89325426 | 89325676 | 89325547 | 89325676 | 89319651 | 89319797 | 0 | 0 | 4 | 0 | 0 | 1 | 0 | 4 | 1 | 0 | 0 | 0 | 98 | 49 | 0.498166579 | 0.74107425 | NA | NA | 0.667 | NA | 1 | 1 | -0.333 |
| A3SS | 3445 | PHOSPHO1 | chr17 | - | 49223369 | 49225131 | 49223369 | 49225004 | 49226646 | 49226758 | 0 | 0 | 4 | 0 | 0 | 1 | 0 | 2 | 1 | 0 | 0 | 0 | 98 | 49 | 0.498166579 | 0.74107425 | NA | NA | 0.667 | NA | 1 | 1 | -0.333 |
| RI | 392 | PRKD2 | chr19 | - | 46716130 | 46717127 | 46716130 | 46716535 | 46716950 | 46717127 | 0 | 0 | 4 | 0 | 0 | 1 | 0 | 0 | 1 | 0 | 0 | 0 | 98 | 49 | 0.498166579 | 0.94773154 | NA | NA | 0.667 | NA | NA | 1 | -0.333 |
| SE | 3691 | ARHGEF40 | chr14 | + | 21087319 | 21087463 | 21087000 | 21087105 | 21087967 | 21088098 | 0 | 0 | 4 | 0 | 0 | 1 | 0 | 0 | 1 | 0 | 0 | 0 | 98 | 49 | 0.498166579 | 0.696506389 | NA | NA | 0.667 | NA | NA | 1 | -0.333 |
| SE | 6752 | UBR4 | chr1 | - | 19165655 | 19165757 | 19165248 | 19165349 | 19167021 | 19167231 | 0 | 0 | 4 | 0 | 0 | 1 | 0 | 4 | 1 | 0 | 1 | 0 | 98 | 49 | 0.498166579 | 0.696506389 | NA | NA | 0.667 | NA | 0.667 | 1 | -0.166 |
| SE | 7087 | PRDX5 | chr11 | + | 64320660 | 64320792 | 64319733 | 64319868 | 64320864 | 64320903 | 0 | 0 | 4 | 0 | 0 | 1 | 0 | 5 | 1 | 0 | 0 | 0 | 98 | 49 | 0.498166579 | 0.696506389 | NA | NA | 0.667 | NA | 1 | 1 | -0.333 |
| SE | 4125 | XPO6 | chr16 | - | 28177219 | 28177332 | 28175897 | 28176095 | 28180940 | 28181031 | 0 | 3 | 53 | 0 | 0 | 1 | 0 | 35 | 8 | 0 | 1 | 0 | 98 | 49 | 0.499162452 | 0.69697095 | NA | 1 | 0.964 | NA | 0.946 | 1 | 0.009 |
| SE | 13537 | ALOX5 | chr10 | + | 45395854 | 45395936 | 45382482 | 45382681 | 45412190 | 45412313 | 0 | 5 | 67 | 0 | 0 | 1 | 1 | 17 | 7 | 0 | 0 | 0 | 98 | 49 | 0.49942629 | 0.69697095 | NA | 1 | 0.971 | 1 | 1 | 1 | -0.014 |
| A5SS | 2491 | ZNF276 | chr16 | + | 89727278 | 89727357 | 89727278 | 89727325 | 89729234 | 89729318 | 0 | 0 | 8 | 0 | 0 | 2 | 0 | 3 | 1 | 0 | 4 | 0 | 81 | 49 | 0.50024699 | 0.736318604 | NA | NA | 0.708 | NA | 0.312 | 1 | 0.052 |
| SE | 10650 | SLC43A2 | chr17 | - | 1591268 | 1591471 | 1590801 | 1590948 | 1591565 | 1591699 | 0 | 5 | 19 | 0 | 0 | 1 | 0 | 5 | 8 | 0 | 1 | 0 | 98 | 49 | 0.500994407 | 0.698510756 | NA | 1 | 0.905 | NA | 0.714 | 1 | 0.096 |
| A3SS | 1434 | PICALM | chr11 | - | 85990249 | 85990399 | 85990249 | 85990378 | 85996825 | 85996929 | 0 | 0 | 18 | 0 | 0 | 1 | 0 | 21 | 4 | 0 | 1 | 0 | 70 | 49 | 0.504274672 | 0.744011811 | NA | NA | 0.926 | NA | 0.936 | 1 | -0.042 |
| SE | 12757 | PKN1 | chr19 | + | 14469751 | 14469814 | 14469340 | 14469517 | 14469911 | 14469988 | 0 | 1 | 5 | 0 | 0 | 1 | 2 | 1 | 1 | 0 | 0 | 0 | 98 | 49 | 0.507272183 | 0.706608055 | NA | 1 | 0.714 | 1 | 1 | 1 | -0.143 |
| SE | 6543 | NRD1 | chr1 | - | 51800555 | 51800683 | 51798248 | 51798411 | 51803813 | 51803964 | 0 | 0 | 18 | 0 | 0 | 1 | 0 | 10 | 4 | 0 | 0 | 0 | 98 | 49 | 0.507742839 | 0.706608784 | NA | NA | 0.9 | NA | 1 | 1 | -0.1 |
| SE | 1150 | SOD2 | chr6 | - | 159688125 | 159688242 | 159684853 | 159685033 | 159692660 | 159692863 | 3 | 7 | 82 | 0 | 2 | 0 | 18 | 158 | 13 | 0 | 13 | 1 | 98 | 49 | 0.509005729 | 0.707237006 | 1 | 0.636 | 1 | 1 | 0.859 | 0.867 | -0.03 |
| SE | 13859 | MMP25 | chr16 | + | 3057309 | 3057394 | 3057032 | 3057209 | 3057530 | 3057613 | 0 | 1 | 33 | 0 | 0 | 1 | 3 | 8 | 3 | 0 | 0 | 0 | 98 | 49 | 0.509135356 | 0.707237006 | NA | 1 | 0.943 | 1 | 1 | 1 | -0.028 |
| SE | 8685 | PTPRC | chr1 | + | 198699563 | 198699704 | 198692346 | 198692373 | 198703297 | 198703372 | 0 | 3 | 18 | 0 | 0 | 25 | 2 | 15 | 1 | 0 | 11 | 6 | 98 | 49 | 0.509721028 | 0.707396773 | NA | 1 | 0.265 | 1 | 0.405 | 0.077 | 0.139 |
| SE | 8376 | OAS2 | chr12 | + | 113004933 | 113005222 | 113002931 | 113003102 | 113006412 | 113006600 | 0 | 2 | 43 | 0 | 0 | 1 | 0 | 11 | 3 | 0 | 0 | 0 | 98 | 49 | 0.512035658 | 0.709568442 | NA | 1 | 0.956 | NA | 1 | 1 | -0.022 |
| SE | 4905 | PAK1 | chr11 | - | 77332729 | 77332867 | 77322014 | 77323360 | 77336085 | 77336282 | 0 | 2 | 22 | 0 | 1 | 0 | 0 | 7 | 1 | 0 | 0 | 0 | 98 | 49 | 0.512585144 | 0.709568442 | NA | 0.5 | 1 | NA | 1 | 1 | -0.25 |
| SE | 5056 | SRSF3 | chr6 | + | 36599820 | 36600276 | 36598848 | 36598983 | 36601151 | 36601190 | 0 | 1 | 11 | 0 | 0 | 4 | 0 | 7 | 1 | 0 | 1 | 0 | 98 | 49 | 0.512702147 | 0.709568442 | NA | 1 | 0.579 | NA | 0.778 | 1 | -0.1 |
| A5SS | 3021 | MAP4K4 | chr2 | + | 101887087 | 101887261 | 101887087 | 101887237 | 101887777 | 101887937 | 0 | 0 | 2 | 0 | 0 | 5 | 0 | 5 | 1 | 0 | 2 | 1 | 73 | 49 | 0.51637545 | 0.751613155 | NA | NA | 0.212 | NA | 0.627 | 0.402 | -0.302 |
| SE | 2564 | HK3 | chr5 | - | 176887200 | 176887337 | 176887001 | 176887121 | 176887450 | 176887746 | 0 | 10 | 17 | 0 | 1 | 0 | 2 | 5 | 2 | 0 | 0 | 0 | 98 | 49 | 0.517806019 | 0.715972812 | NA | 0.833 | 1 | 1 | 1 | 1 | -0.084 |
| SE | 12585 | SBNO2 | chr19 | - | 1112188 | 1112301 | 1111995 | 1112067 | 1112401 | 1112537 | 0 | 8 | 15 | 0 | 4 | 0 | 0 | 8 | 3 | 0 | 2 | 1 | 98 | 49 | 0.521310981 | 0.720156622 | NA | 0.5 | 1 | NA | 0.667 | 0.6 | 0.117 |
| SE | 4558 | SLC6A6 | chr3 | + | 14443627 | 14443863 | 14416411 | 14416453 | 14445716 | 14445851 | 0 | 5 | 14 | 0 | 0 | 1 | 0 | 9 | 4 | 0 | 1 | 0 | 98 | 49 | 0.523615544 | 0.722675999 | NA | 1 | 0.875 | NA | 0.818 | 1 | 0.028 |
| SE | 5960 | LILRA2 | chr19 | + | 54574075 | 54574111 | 54573697 | 54573912 | 54574300 | 54574582 | 0 | 0 | 5 | 0 | 0 | 1 | 1 | 1 | 1 | 0 | 0 | 0 | 85 | 49 | 0.5248053 | 0.723653547 | NA | NA | 0.742 | 1 | 1 | 1 | -0.258 |
| SE | 7023 | HPCAL1 | chr2 | + | 10408561 | 10408734 | 10396834 | 10396920 | 10419733 | 10420135 | 0 | 0 | 2 | 0 | 2 | 7 | 0 | 2 | 0 | 0 | 1 | 2 | 98 | 49 | 0.527686141 | 0.726959 | NA | 0 | 0.125 | NA | 0.5 | 0 | -0.188 |
| SE | 9067 | ERGIC1 | chr5 | + | 172909666 | 172909761 | 172897001 | 172897074 | 172914713 | 172914838 | 0 | 1 | 13 | 0 | 0 | 1 | 0 | 1 | 3 | 0 | 0 | 0 | 98 | 49 | 0.528398477 | 0.727273728 | NA | 1 | 0.867 | NA | 1 | 1 | -0.067 |
| SE | 11047 | HLA-B | chr6 | - | 31355106 | 31355223 | 31354632 | 31354665 | 31355316 | 31355592 | 0 | 46 | 261 | 0 | 2 | 4 | 10 | 220 | 15 | 0 | 4 | 2 | 98 | 49 | 0.529614267 | 0.728280186 | NA | 0.92 | 0.97 | 1 | 0.965 | 0.789 | 0.027 |
| SE | 7715 | THEMIS2 | chr1 | + | 27876587 | 27876728 | 27872542 | 27872665 | 27879643 | 27879963 | 0 | 13 | 31 | 0 | 0 | 1 | 7 | 17 | 23 | 0 | 0 | 2 | 98 | 49 | 0.530434112 | 0.728740832 | NA | 1 | 0.939 | 1 | 1 | 0.852 | 0.019 |
| A5SS | 2259 | TLE3 | chr15 | - | 70058149 | 70058291 | 70058158 | 70058291 | 70057458 | 70057658 | 0 | 0 | 2 | 0 | 1 | 4 | 0 | 2 | 0 | 0 | 3 | 1 | 58 | 49 | 0.531185703 | 0.764673924 | NA | 0 | 0.297 | NA | 0.36 | 0 | -0.032 |
| A3SS | 1208 | SDF4 | chr1 | - | 1216907 | 1217804 | 1216907 | 1217688 | 1218457 | 1218633 | 1 | 0 | 0 | 0 | 0 | 4 | 1 | 0 | 1 | 2 | 0 | 0 | 98 | 49 | 0.53213915 | 0.762573929 | 1 | NA | 0 | 0.2 | NA | 1 | -0.1 |

| | | | | | | | | | | | | | | | | | | | | | | | | | | | | | | | | | |
|---|---|---|---|---|---|---|---|---|---|---|---|---|---|---|---|---|---|---|---|---|---|---|---|---|---|---|---|---|---|---|---|---|---|
| SE | 4662 | AMICA1 | chr11 | - | 118214823 | 118214886 | 118212406 | 118212561 | 118224940 | 118225094 | 0 | 7 | 17 | 0 | 1 | 5 | 0 | 13 | 2 | 0 | 2 | 0 | 98 | 49 | 0.537451897 | 0.737707946 | NA | 0.778 | 0.63 | NA | 0.765 | 1 | -0.179 |
| A3SS | 1184 | DDIT3 | chr12 | - | 57517268 | 57517519 | 57517268 | 57517438 | 57517705 | 57517753 | 0 | 0 | 5 | 0 | 0 | 1 | 0 | 11 | 1 | 0 | 2 | 0 | 98 | 49 | 0.538906714 | 0.762573929 | NA | NA | 0.714 | NA | 0.733 | 1 | -0.153 |
| A3SS | 1185 | DDIT3 | chr12 | - | 57517268 | 57517542 | 57517268 | 57517438 | 57517705 | 57517753 | 0 | 0 | 5 | 0 | 0 | 1 | 0 | 11 | 1 | 0 | 2 | 0 | 98 | 49 | 0.538906714 | 0.762573929 | NA | NA | 0.714 | NA | 0.733 | 1 | -0.153 |
| A3SS | 1186 | DDIT3 | chr12 | - | 57517268 | 57517595 | 57517268 | 57517438 | 57517705 | 57517753 | 0 | 0 | 5 | 0 | 0 | 1 | 0 | 12 | 1 | 0 | 2 | 0 | 98 | 49 | 0.538906714 | 0.762573929 | NA | NA | 0.714 | NA | 0.75 | 1 | -0.161 |
| SE | 8577 | RC3H1 | chr1 | - | 173964838 | 173965120 | 173963972 | 173964187 | 173970504 | 173970617 | 0 | 0 | 5 | 0 | 0 | 1 | 0 | 5 | 1 | 0 | 0 | 0 | 98 | 49 | 0.538906714 | 0.737743862 | NA | NA | 0.714 | NA | 1 | 1 | -0.286 |
| SE | 1576 | TRIM38 | chr6 | + | 25973053 | 25973076 | 25971868 | 25972099 | 25973172 | 25973285 | 0 | 0 | 5 | 0 | 0 | 1 | 0 | 9 | 1 | 0 | 1 | 0 | 72 | 49 | 0.539026433 | 0.737743862 | NA | NA | 0.773 | NA | 0.86 | 1 | -0.157 |
| SE | 3235 | SLC9A8 | chr20 | + | 49849578 | 49849680 | 49845035 | 49845119 | 49855437 | 49855581 | 0 | 0 | 10 | 0 | 0 | 2 | 0 | 6 | 1 | 0 | 2 | 0 | 98 | 49 | 0.539441453 | 0.737743862 | NA | NA | 0.714 | NA | 0.6 | 1 | -0.086 |
| SE | 5886 | RASSF5 | chr1 | + | 206585179 | 206585295 | 206584386 | 206584684 | 206586825 | 206589284 | 0 | 0 | 10 | 0 | 0 | 2 | 0 | 10 | 1 | 0 | 6 | 0 | 98 | 49 | 0.539441453 | 0.737743862 | NA | NA | 0.714 | NA | 0.455 | 1 | -0.014 |
| A3SS | 3256 | FLII | chr17 | - | 18251679 | 18251816 | 18251679 | 18251813 | 18251998 | 18252146 | 0 | 2 | 6 | 0 | 2 | 3 | 0 | 1 | 0 | 0 | 2 | 0 | 52 | 49 | 0.542191316 | 0.762573929 | NA | 0.485 | 0.653 | NA | 0.32 | NA | 0.249 |
| A3SS | 116 | AGTPBP1 | chr9 | - | 85655142 | 85655320 | 85655142 | 85655200 | 85657434 | 85657643 | 0 | 0 | 20 | 0 | 0 | 4 | 0 | 8 | 1 | 0 | 2 | 0 | 98 | 49 | 0.542274794 | 0.762573929 | NA | NA | 0.714 | NA | 0.667 | 1 | -0.12 |
| SE | 4629 | SORL1 | chr11 | + | 121605401 | 121605571 | 121605112 | 121605239 | 121606844 | 121606957 | 0 | 11 | 57 | 0 | 0 | 1 | 0 | 45 | 4 | 0 | 1 | 0 | 98 | 49 | 0.544440604 | 0.743903843 | NA | 1 | 0.966 | NA | 0.957 | 1 | 0.005 |
| SE | 2099 | WDFY3 | chr4 | - | 84726860 | 84726911 | 84724425 | 84724594 | 84733381 | 84733609 | 0 | 5 | 2 | 0 | 0 | 3 | 0 | 5 | 3 | 0 | 1 | 0 | 98 | 49 | 0.545895829 | 0.745214742 | NA | 1 | 0.25 | NA | 0.714 | 1 | -0.232 |
| SE | 6349 | FLI1 | chr11 | + | 128807179 | 128807239 | 128805365 | 128805431 | 128809156 | 128809204 | 0 | 1 | 11 | 0 | 0 | 1 | 1 | 2 | 1 | 0 | 0 | 0 | 98 | 49 | 0.547491166 | 0.746714358 | NA | 1 | 0.846 | 1 | 1 | 1 | -0.077 |
| A5SS | 1194 | ETF1 | chr5 | - | 138542633 | 138542936 | 138542832 | 138542936 | 138518691 | 138518867 | 0 | 0 | 2 | 0 | 2 | 4 | 0 | 4 | 3 | 0 | 3 | 0 | 98 | 49 | 0.547780994 | 0.779992502 | NA | 0 | 0.2 | NA | 0.4 | 1 | -0.6 |
| SE | 1134 | IGF2R | chr6 | + | 160029549 | 160029655 | 160027184 | 160027314 | 160032550 | 160032713 | 0 | 14 | 45 | 0 | 0 | 4 | 0 | 27 | 6 | 0 | 1 | 0 | 98 | 49 | 0.549943569 | 0.749379133 | NA | 1 | 0.849 | NA | 0.931 | 1 | -0.041 |
| SE | 11237 | APLP2 | chr11 | + | 130123611 | 130123779 | 130122304 | 130122513 | 130126699 | 130126830 | 0 | 0 | 33 | 0 | 0 | 3 | 0 | 5 | 2 | 0 | 0 | 0 | 98 | 49 | 0.550569813 | 0.749552925 | NA | NA | 0.846 | NA | 1 | 1 | -0.154 |
| SE | 8917 | IRF9 | chr14 | + | 24163877 | 24163959 | 24163377 | 24163508 | 24164062 | 24164134 | 0 | 3 | 25 | 0 | 1 | 5 | 0 | 7 | 3 | 0 | 8 | 0 | 98 | 49 | 0.551132735 | 0.749640272 | NA | 0.6 | 0.714 | NA | 0.304 | 1 | 0.005 |
| SE | 5788 | DENND3 | chr8 | + | 141160631 | 141160787 | 141155848 | 141155970 | 141165185 | 141165289 | 0 | 0 | 11 | 0 | 0 | 1 | 0 | 5 | 2 | 0 | 0 | 0 | 98 | 49 | 0.551645974 | 0.749659945 | NA | NA | 0.846 | NA | 1 | 1 | -0.154 |
| RI | 675 | PSME1 | chr14 | + | 24138473 | 24138967 | 24138473 | 24138560 | 24138735 | 24138967 | 0 | 0 | 2 | 0 | 0 | 11 | 3 | 0 | 0 | 0 | 6 | 1 | 98 | 49 | 0.557006469 | 0.999202116 | NA | NA | 0.083 | 1 | 0 | 0 | -0.25 |
| A3SS | 2017 | RNF19B | chr1 | - | 32936445 | 32937309 | 32936445 | 32937259 | 32938396 | 32938528 | 0 | 1 | 9 | 0 | 0 | 5 | 0 | 3 | 2 | 0 | 5 | 0 | 98 | 49 | 0.558523697 | 0.779335391 | NA | 1 | 0.474 | NA | 0.231 | 1 | 0.121 |
| SE | 13539 | ALOX5 | chr10 | + | 45444115 | 45444286 | 45443727 | 45443828 | 45445507 | 45446119 | 0 | 2 | 41 | 0 | 0 | 2 | 1 | 4 | 7 | 0 | 0 | 1 | 98 | 49 | 0.559152751 | 0.759174873 | NA | 1 | 0.911 | 1 | 1 | 0.778 | 0.029 |
| SE | 4964 | DDX3X | chrX | + | 41343736 | 41343822 | 41343215 | 41343351 | 41344029 | 41344128 | 0 | 4 | 36 | 0 | 2 | 0 | 0 | 34 | 1 | 0 | 4 | 1 | 98 | 49 | 0.565342278 | 0.76688578 | NA | 0.5 | 1 | NA | 0.81 | 0.333 | 0.178 |
| A5SS | 520 | TNFAIP3 | chr6 | + | 137867187 | 137867542 | 137867187 | 137867494 | 137871212 | 137871522 | 0 | 3 | 6 | 0 | 2 | 0 | 0 | 9 | 4 | 0 | 1 | 0 | 97 | 49 | 0.570399439 | 0.803465876 | NA | 0.431 | 1 | NA | 0.82 | 1 | -0.194 |
| A3SS | 1102 | FKBP8 | chr19 | - | 18538215 | 18538436 | 18538215 | 18538433 | 18539370 | 18539459 | 0 | 2 | 2 | 0 | 0 | 5 | 0 | 2 | 1 | 2 | 5 | 0 | 52 | 49 | 0.570761829 | 0.79028561 | NA | 1 | 0.274 | 0 | 0.274 | 1 | 0.212 |
| SE | 7496 | IFI44 | chr1 | + | 78660553 | 78660654 | 78659311 | 78659483 | 78662703 | 78662878 | 0 | 0 | 36 | 0 | 0 | 6 | 0 | 23 | 1 | 0 | 5 | 0 | 98 | 49 | 0.575224839 | 0.778766917 | NA | NA | 0.75 | NA | 0.697 | 1 | -0.099 |
| SE | 8238 | LBR | chr1 | - | 225410290 | 225410416 | 225406663 | 225406832 | 225411336 | 225411440 | 0 | 0 | 18 | 0 | 0 | 3 | 1 | 10 | 1 | 0 | 1 | 0 | 98 | 49 | 0.575643233 | 0.778766917 | NA | NA | 0.75 | 1 | 0.833 | 1 | -0.194 |
| SE | 11307 | SP100 | chr2 | + | 230464066 | 230464150 | 230462434 | 230462518 | 230466300 | 230466354 | 0 | 0 | 25 | 0 | 0 | 2 | 0 | 12 | 2 | 0 | 2 | 0 | 98 | 49 | 0.575655386 | 0.778766917 | NA | NA | 0.862 | NA | 0.75 | 1 | -0.013 |
| A5SS | 3019 | SERPINA1 | chr14 | - | 94390419 | 94390692 | 94390456 | 94390692 | 94388559 | 94388663 | 0 | 4 | 5 | 0 | 0 | 4 | 3 | 3 | 2 | 0 | 1 | 0 | 86 | 49 | 0.578375981 | 0.804333654 | NA | 1 | 0.416 | 1 | 0.631 | 1 | -0.169 |
| A5SS | 1552 | PGS1 | chr17 | + | 78419545 | 78419725 | 78419545 | 78419675 | 78424060 | 78424659 | 0 | 10 | 32 | 0 | 0 | 1 | 0 | 27 | 6 | 0 | 1 | 0 | 98 | 49 | 0.583295398 | 0.804333654 | NA | 1 | 0.941 | NA | 0.931 | 1 | 0.005 |
| SE | 6362 | ZSWIM8 | chr10 | + | 73791323 | 73791499 | 73790974 | 73791176 | 73791858 | 73792852 | 0 | 0 | 6 | 0 | 1 | 0 | 0 | 1 | 0 | 0 | 0 | 1 | 98 | 49 | 0.584480703 | 0.789995051 | NA | 0 | 1 | NA | 1 | 0 | 0 |
| SE | 3417 | RALGAPA2 | chr20 | - | 20676235 | 20676288 | 20653529 | 20653587 | 20680690 | 20680801 | 0 | 0 | 13 | 0 | 0 | 2 | 0 | 5 | 1 | 0 | 0 | 0 | 98 | 49 | 0.588967448 | 0.794630228 | NA | NA | 0.765 | NA | 1 | 1 | -0.235 |
| SE | 3803 | ZCCHC2 | chr18 | + | 62558686 | 62558770 | 62556202 | 62556297 | 62560586 | 62560644 | 0 | 0 | 13 | 0 | 0 | 2 | 5 | 11 | 1 | 0 | 0 | 0 | 98 | 49 | 0.588967448 | 0.794630228 | NA | NA | 0.765 | 1 | 1 | 1 | -0.235 |
| SE | 1922 | PARP8 | chr5 | + | 50824907 | 50824975 | 50822334 | 50822400 | 50826754 | 50826803 | 0 | 0 | 14 | 0 | 0 | 1 | 0 | 5 | 2 | 0 | 0 | 0 | 98 | 49 | 0.592731575 | 0.797560929 | NA | NA | 0.875 | NA | 1 | 1 | -0.125 |
| SE | 3771 | KIAA1109 | chr4 | + | 122356620 | 122356755 | 122355795 | 122355990 | 122359556 | 122359722 | 0 | 0 | 14 | 0 | 0 | 1 | 0 | 7 | 2 | 0 | 0 | 0 | 98 | 49 | 0.592731575 | 0.797560929 | NA | NA | 0.875 | NA | 1 | 1 | -0.125 |
| SE | 4732 | CDA | chr1 | + | 20604927 | 20605039 | 20588950 | 20589283 | 20618451 | 20618907 | 0 | 0 | 14 | 0 | 0 | 1 | 0 | 5 | 2 | 0 | 0 | 0 | 98 | 49 | 0.592731575 | 0.797560929 | NA | NA | 0.875 | NA | 1 | 1 | -0.125 |
| SE | 7937 | TMC6 | chr17 | - | 78124888 | 78124985 | 78124523 | 78124781 | 78125157 | 78125263 | 1 | 4 | 2 | 0 | 1 | 0 | 0 | 3 | 3 | 0 | 2 | 0 | 98 | 49 | 0.596766039 | 0.802271339 | 1 | 0.667 | 1 | NA | 0.429 | 1 | 0.174 |
| A5SS | 2516 | NLRP1 | chr17 | - | 5533291 | 5533384 | 5533303 | 5533384 | 5532821 | 5532984 | 0 | 0 | 1 | 0 | 0 | 7 | 0 | 5 | 0 | 0 | 4 | 1 | 61 | 49 | 0.597621998 | 0.815505018 | NA | NA | 0.103 | NA | 0.501 | 0 | -0.148 |
| SE | 11312 | ALDOA | chr16 | + | 30066884 | 30067038 | 30065672 | 30065927 | 30067233 | 30067366 | 0 | 0 | 1 | 0 | 0 | 7 | 0 | 0 | 0 | 0 | 5 | 1 | 98 | 49 | 0.59822268 | 0.803510892 | NA | NA | 0.067 | NA | 0 | 0 | 0.067 |
| A3SS | 1568 | ENTPD1 | chr10 | + | 95847445 | 95847706 | 95847568 | 95847706 | 95845356 | 95845596 | 0 | 4 | 26 | 0 | 0 | 1 | 2 | 4 | 1 | 0 | 0 | 0 | 98 | 49 | 0.600161541 | 0.824649445 | NA | 1 | 0.929 | 1 | 1 | 1 | -0.035 |
| SE | 4503 | PTPN6 | chr12 | + | 6955150 | 6955267 | 6954804 | 6954994 | 6955371 | 6955485 | 0 | 2 | 28 | 0 | 0 | 1 | 0 | 24 | 1 | 0 | 1 | 0 | 98 | 49 | 0.600780878 | 0.805924887 | NA | 1 | 0.933 | NA | 0.923 | 1 | 0.005 |
| SE | 2258 | KIAA0226 | chr3 | - | 197717976 | 197718130 | 197705091 | 197705175 | 197736654 | 197736926 | 0 | 0 | 2 | 0 | 1 | 0 | 0 | 3 | 1 | 0 | 0 | 1 | 98 | 49 | 0.601092347 | 0.805924887 | NA | 0 | 1 | NA | 1 | 0.333 | -0.166 |
| SE | 2636 | PFKFB4 | chr3 | - | 48539253 | 48539310 | 48538497 | 48538619 | 48539696 | 48539771 | 0 | 0 | 7 | 0 | 0 | 1 | 0 | 1 | 1 | 0 | 0 | 0 | 98 | 49 | 0.602141516 | 0.806612031 | NA | NA | 0.778 | NA | 1 | 1 | -0.222 |
| SE | 9030 | MYD88 | chr3 | + | 38140387 | 38140568 | 38139863 | 38139998 | 38140756 | 38140848 | 0 | 1 | 15 | 0 | 1 | 1 | 0 | 0 | 2 | 0 | 0 | 0 | 98 | 49 | 0.608707381 | 0.814681384 | NA | 0.333 | 0.882 | NA | NA | 1 | -0.392 |
| SE | 4423 | PREX1 | chr20 | - | 48650893 | 48651055 | 48649995 | 48650206 | 48651395 | 48651583 | 0 | 3 | 32 | 0 | 0 | 2 | 0 | 37 | 5 | 0 | 1 | 0 | 98 | 49 | 0.611249391 | 0.817355724 | NA | 1 | 0.889 | NA | 0.949 | 1 | -0.03 |
| SE | 8705 | CTSS | chr1 | - | 150750005 | 150750171 | 150747776 | 150747879 | 150751780 | 150752008 | 0 | 1 | 129 | 0 | 0 | 1 | 5 | 24 | 1 | 0 | 0 | 0 | 98 | 49 | 0.612568689 | 0.818391768 | NA | 1 | 0.985 | 1 | 1 | 1 | -0.008 |
| SE | 8707 | CTSS | chr1 | - | 150755000 | 150755150 | 150751780 | 150752008 | 150757857 | 150757980 | 0 | 4 | 234 | 0 | 0 | 1 | 6 | 39 | 7 | 0 | 0 | 0 | 98 | 49 | 0.614855148 | 0.820716951 | NA | 1 | 0.992 | 1 | 1 | 1 | -0.004 |
| SE | 56 | SASH3 | chrX | + | 129788430 | 129788577 | 129787974 | 129788070 | 129790939 | 129791081 | 0 | 0 | 8 | 0 | 0 | 1 | 0 | 12 | 1 | 0 | 1 | 0 | 98 | 49 | 0.622026252 | 0.829552314 | NA | NA | 0.8 | NA | 0.857 | 1 | -0.128 |
| SE | 11031 | STAT1 | chr2 | - | 190998216 | 190998308 | 190997855 | 190998007 | 190999625 | 190999704 | 0 | 0 | 33 | 0 | 0 | 2 | 0 | 14 | 2 | 0 | 4 | 0 | 98 | 49 | 0.624060418 | 0.831527312 | NA | NA | 0.892 | NA | 0.636 | 1 | 0.074 |
| RI | 30 | PLIN2 | chr9 | - | 19126113 | 19126448 | 19126113 | 19126309 | 19126396 | 19126448 | 2 | 0 | 1 | 0 | 1 | 0 | 4 | 1 | 0 | 0 | 1 | 0 | 98 | 49 | 0.627465042 | 0.999202116 | 1 | 0 | 1 | 1 | 0.333 | NA | 0 |
| SE | 12347 | CSNK1D | chr17 | - | 82253015 | 82253244 | 82252433 | 82252604 | 82255428 | 82255577 | 0 | 2 | 3 | 0 | 1 | 0 | 0 | 4 | 7 | 0 | 1 | 0 | 98 | 49 | 0.630476403 | 0.839332182 | NA | 0.5 | 1 | NA | 0.667 | 1 | -0.084 |
| SE | 11030 | STAT1 | chr2 | - | 190979771 | 190979866 | 190978855 | 190979001 | 190980619 | 190980669 | 0 | 1 | 20 | 0 | 1 | 2 | 0 | 2 | 0 | 0 | 3 | 0 | 98 | 49 | 0.633284155 | 0.842323969 | NA | 0.333 | 0.833 | NA | 0.25 | NA | 0.333 |
| SE | 381 | GNAQ | chr9 | - | 77794462 | 77794592 | 77728513 | 77728667 | 77797519 | 77797648 | 0 | 0 | 9 | 0 | 0 | 1 | 0 | 2 | 1 | 0 | 0 | 0 | 98 | 49 | 0.642008309 | 0.852419159 | NA | NA | 0.818 | NA | 1 | 1 | -0.182 |
| SE | 8334 | NCSTN | chr1 | + | 160355862 | 160355958 | 160354117 | 160354290 | 160356259 | 160356347 | 0 | 0 | 9 | 0 | 0 | 1 | 0 | 2 | 1 | 0 | 2 | 0 | 98 | 49 | 0.642008309 | 0.852419159 | NA | NA | 0.818 | NA | 0.333 | 1 | 0.151 |
| SE | 5678 | GMIP | chr19 | - | 19635643 | 19635721 | 19635414 | 19635569 | 19636706 | 19636796 | 0 | 9 | 8 | 1 | 1 | 0 | 0 | 5 | 1 | 0 | 0 | 1 | 98 | 49 | 0.642808803 | 0.852728712 | 0 | 0.818 | 1 | NA | 1 | 0.333 | -0.06 |
| SE | 8358 | ARHGAP30 | chr1 | - | 161056387 | 161056532 | 161054622 | 161054705 | 161069527 | 161069970 | 0 | 0 | 18 | 0 | 0 | 2 | 0 | 11 | 1 | 0 | 2 | 0 | 98 | 49 | 0.643507131 | 0.852902309 | NA | NA | 0.818 | NA | 0.733 | 1 | -0.049 |
| SE | 4961 | DDX3X | chrX | + | 41337407 | 41337465 | 41333307 | 41334297 | 41341483 | 41341616 | 0 | 2 | 10 | 0 | 0 | 2 | 0 | 23 | 2 | 0 | 2 | 0 | 98 | 49 | 0.648092299 | 0.858222665 | NA | 1 | 0.714 | NA | 0.852 | 1 | -0.069 |
| SE | 11305 | SP100 | chr2 | + | 230450171 | 230450255 | 230449087 | 230449150 | 230461261 | 230461414 | 0 | 2 | 23 | 0 | 1 | 0 | 8 | 15 | 5 | 0 | 1 | 0 | 98 | 49 | 0.652122993 | 0.862079722 | NA | 0.5 | 1 | 1 | 0.882 | 1 | -0.211 |
| SE | 4428 | PREX1 | chr20 | - | 48747808 | 48747880 | 48745024 | 48745147 | 48827641 | 48827883 | 0 | 20 | 45 | 0 | 0 | 1 | 2 | 45 | 9 | 0 | 1 | 0 | 98 | 49 | 0.652532646 | 0.862079722 | NA | 1 | 0.957 | 1 | 0.957 | 1 | -0.007 |
| SE | 9514 | SIPA1 | chr11 | + | 65645853 | 65645957 | 65644954 | 65645129 | 65646220 | 65646378 | 0 | 0 | 19 | 0 | 0 | 2 | 0 | 5 | 1 | 0 | 0 | 0 | 98 | 49 | 0.652725698 | 0.862079722 | NA | NA | 0.826 | NA | 1 | 1 | -0.174 |
| RI | 230 | ATG3 | chr3 | - | 112532506 | 112534337 | 112532506 | 112532780 | 112534268 | 112534337 | 0 | 0 | 3 | 0 | 3 | 5 | 0 | 0 | 1 | 0 | 4 | 0 | 98 | 49 | 0.654603078 | 0.999202116 | NA | 0 | 0.231 | NA | 0 | 1 | -0.385 |
| SE | 6617 | AMPD2 | chr1 | + | 109628642 | 109628806 | 109628363 | 109628495 | 109629108 | 109629235 | 0 | 1 | 2 | 0 | 1 | 0 | 0 | 7 | 2 | 0 | 5 | 1 | 98 | 49 | 0.656411303 | 0.866186294 | NA | 0.333 | 1 | NA | 0.412 | 0.5 | 0.211 |
| RI | 644 | ATP6V0B | chr1 | + | 43977025 | 43978300 | 43977025 | 43977216 | 43977980 | 43978300 | 3 | 0 | 3 | 0 | 4 | 15 | 0 | 0 | 4 | 1 | 7 | 3 | 98 | 49 | 0.657605162 | 0.999202116 | 1 | 0 | 0.091 | 0 | 0 | 0.4 | 0.23 |
| SE | 2839 | TYMP | chr22 | - | 50526000 | 50526141 | 50525752 | 50525918 | 50526245 | 50526476 | 1 | 8 | 30 | 0 | 1 | 0 | 2 | 8 | 14 | 0 | 0 | 1 | 98 | 49 | 0.661103457 | 0.871612716 | 1 | 0.8 | 1 | 1 | 1 | 0.875 | -0.025 |
| SE | 3332 | SAMHD1 | chr20 | - | 36935028 | 36935189 | 36930759 | 36930875 | 36941038 | 36941111 | 0 | 0 | 21 | 0 | 0 | 1 | 0 | 3 | 2 | 0 | 1 | 0 | 98 | 49 | 0.661813092 | 0.871783591 | NA | NA | 0.913 | NA | 0.6 | 1 | 0.113 |
| SE | 8706 | CTSS | chr1 | - | 150751780 | 150752008 | 150750005 | 150750171 | 150755000 | 150755150 | 0 | 3 | 181 | 0 | 0 | 1 | 7 | 24 | 1 | 0 | 0 | 0 | 98 | 49 | 0.66301558 | 0.872602817 | NA | 1 | 0.989 | 1 | 1 | 1 | -0.006 |
| SE | 11325 | ITGAL | chr16 | + | 30510880 | 30510960 | 30510360 | 30510471 | 30511049 | 30511136 | 0 | 2 | 13 | 0 | 0 | 1 | 0 | 1 | 1 | 0 | 0 | 0 | 98 | 49 | 0.665452763 | 0.875044185 | NA | 1 | 0.867 | NA | 1 | 1 | -0.067 |
| SE | 7596 | ATG16L2 | chr11 | + | 72827187 | 72827293 | 72826702 | 72826823 | 72828358 | 72828508 | 0 | 1 | 41 | 0 | 0 | 1 | 1 | 4 | 1 | 0 | 0 | 0 | 98 | 49 | 0.668079787 | 0.8777307 | NA | 1 | 0.953 | 1 | 1 | 1 | -0.024 |
| SE | 9411 | WDFY4 | chr10 | + | 48760346 | 48760440 | 48742967 | 48743548 | 48774457 | 48774672 | 0 | 0 | 11 | 0 | 0 | 1 | 0 | 1 | 1 | 0 | 0 | 0 | 98 | 49 | 0.668749011 | 0.877842588 | NA | NA | 0.846 | NA | 1 | 1 | -0.154 |
| A5SS | 1756 | PSME1 | chr14 | + | 24138473 | 24138568 | 24138473 | 24138560 | 24138735 | 24138967 | 0 | 0 | 1 | 0 | 0 | 11 | 0 | 0 | 0 | 0 | 6 | 1 | 57 | 49 | 0.670811132 | 0.905940807 | NA | NA | 0.072 | NA | 0 | 0 | 0.072 |
| SE | 4459 | CLEC7A | chr12 | - | 10125086 | 10125173 | 10123244 | 10123363 | 10125296 | 10125448 | 0 | 1 | 25 | 0 | 0 | 8 | 0 | 7 | 4 | 0 | 4 | 0 | 98 | 49 | 0.674199264 | 0.884224689 | NA | 1 | 0.61 | NA | 0.467 | 1 | 0.071 |
| SE | 7535 | MCL1 | chr1 | - | 150578243 | 150578491 | 150574550 | 150577491 | 150578842 | 150578963 | 0 | 5 | 123 | 1 | 4 | 15 | 1 | 70 | 5 | 0 | 30 | 0 | 98 | 49 | 0.678459838 | 0.889036737 | 0 | 0.385 | 0.804 | 1 | 0.538 | 1 | -0.45 |
| SE | 8082 | CSF3R | chr1 | - | 36473434 | 36473622 | 36472516 | 36472686 | 36473763 | 36473887 | 0 | 19 | 197 | 0 | 1 | 3 | 20 | 148 | 32 | 0 | 2 | 1 | 98 | 49 | 0.679743226 | 0.889942569 | NA | 0.905 | 0.97 | 1 | 0.974 | 0.941 | -0.034 |
| A5SS | 2261 | TLE3 | chr15 | - | 70058158 | 70058291 | 70058185 | 70058291 | 70057458 | 70057658 | 1 | 1 | 4 | 0 | 0 | 1 | 0 | 4 | 1 | 0 | 1 | 0 | 76 | 49 | 0.682465604 | 0.91227545 | 1 | 1 | 0.721 | NA | 0.721 | 1 | 0.046 |
| SE | 6482 | PADI2 | chr1 | - | 17079263 | 17079415 | 17075678 | 17075823 | 17082544 | 17082652 | 0 | 0 | 12 | 0 | 0 | 1 | 0 | 0 | 1 | 0 | 0 | 0 | 98 | 49 | 0.685696462 | 0.895643369 | NA | NA | 0.857 | NA | NA | 1 | -0.143 |
| SE | 6819 | OAS1 | chr12 | + | 112917546 | 112917700 | 112916508 | 112916738 | 112919486 | 112919907 | 0 | 0 | 12 | 0 | 0 | 1 | 0 | 4 | 1 | 0 | 1 | 0 | 98 | 49 | 0.685696462 | 0.895643369 | NA | NA | 0.857 | NA | 0.667 | 1 | 0.023 |
| SE | 8684 | PTPRC | chr1 | + | 198696711 | 198696909 | 198692346 | 198692373 | 198699563 | 198699704 | 0 | 0 | 1 | 0 | 0 | 12 | 0 | 3 | 0 | 1 | 4 | 1 | 98 | 49 | 0.685885241 | 0.895643369 | NA | NA | 0.04 | 0 | 0.273 | 0 | -0.051 |
| SE | 3887 | LCP1 | chr13 | - | 46148351 | 46148447 | 46144441 | 46144520 | 46150935 | 46151078 | 0 | 0 | 37 | 0 | 0 | 3 | 0 | 8 | 1 | 0 | 0 | 0 | 98 | 49 | 0.687307489 | 0.89672149 | NA | NA | 0.86 | NA | 1 | 1 | -0.14 |
| A3SS | 803 | TYMP | chr22 | - | 50526000 | 50526156 | 50526000 | 50526141 | 50526245 | 50526476 | 0 | 2 | 0 | 0 | 5 | 13 | 1 | 0 | 6 | 2 | 5 | 5 | 64 | 49 | 0.687426061 | 0.937399174 | NA | 0.234 | 0 | 0.277 | 0 | 0.479 | -0.135 |
| SE | 13917 | CD300A | chr17 | + | 74481725 | 74481833 | 74481288 | 74481326 | 74484000 | 74484798 | 0 | 4 | 15 | 0 | 0 | 1 | 0 | 6 | 0 | 0 | 0 | 1 | 98 | 49 | 0.688861707 | 0.89796977 | NA | 1 | 0.882 | NA | 1 | 0 | 0.441 |
| SE | 2765 | LST1 | chr6 | + | 31587640 | 31587733 | 31587199 | 31587318 | 31588562 | 31588909 | 0 | 0 | 2 | 0 | 1 | 6 | 0 | 1 | 0 | 0 | 2 | 1 | 98 | 49 | 0.690552858 | 0.899394233 | NA | 0 | 0.143 | NA | 0.2 | 0 | -0.029 |
| SE | 1145 | IGF2R | chr6 | + | 160088032 | 160088147 | 160083949 | 160084184 | 160089106 | 160089253 | 0 | 0 | 13 | 0 | 0 | 1 | 0 | 2 | 1 | 0 | 0 | 0 | 98 | 49 | 0.699215366 | 0.909888048 | NA | NA | 0.867 | NA | 1 | 1 | -0.133 |
| SE | 13865 | ACSL1 | chr4 | - | 184770398 | 184770476 | 184768315 | 184768450 | 184773080 | 184773154 | 0 | 1 | 55 | 0 | 0 | 1 | 4 | 29 | 0 | 0 | 0 | 0 | 98 | 49 | 0.700770989 | 0.911123526 | NA | 1 | 0.965 | 1 | 1 | NA | -0.018 |
| SE | 12581 | CNN2 | chr19 | + | 1036066 | 1036246 | 1032558 | 1032696 | 1036415 | 1036562 | 0 | 0 | 15 | 0 | 0 | 1 | 0 | 12 | 1 | 0 | 0 | 0 | 98 | 49 | 0.713986072 | 0.927503083 | NA | NA | 0.882 | NA | 1 | 1 | -0.118 |
| SE | 8658 | DUSP1 | chr5 | - | 172769574 | 172769794 | 172768089 | 172769132 | 172770585 | 172771200 | 0 | 6 | 110 | 0 | 1 | 0 | 3 | 47 | 10 | 0 | 2 | 0 | 98 | 49 | 0.7163398 | 0.929757097 | NA | 0.75 | 1 | 1 | 0.922 | 1 | -0.099 |
| A3SS | 775 | LST1 | chr6 | + | 31588496 | 31588909 | 31588562 | 31588909 | 31587199 | 31587318 | 0 | 3 | 13 | 0 | 1 | 6 | 4 | 11 | 0 | 0 | 2 | 1 | 98 | 49 | 0.718836501 | 0.97286143 | NA | 0.6 | 0.52 | 1 | 0.733 | 0 | -0.018 |
| SE | 11238 | APLP2 | chr11 | + | 130137255 | 130137291 | 130135562 | 130135715 | 130140397 | 130140483 | 0 | 1 | 13 | 0 | 1 | 11 | 0 | 3 | 3 | 0 | 10 | 0 | 85 | 49 | 0.719023331 | 0.932434915 | NA | 0.366 | 0.405 | NA | 0.147 | 1 | -0.188 |
| SE | 1142 | IGF2R | chr6 | + | 160072764 | 160072884 | 160071909 | 160072036 | 160073212 | 160073469 | 0 | 1 | 16 | 0 | 0 | 1 | 0 | 18 | 2 | 0 | 1 | 0 | 98 | 49 | 0.724649909 | 0.938921391 | NA | 1 | 0.889 | NA | 0.9 | 1 | -0.005 |
| SE | 1037 | AOAH | chr7 | - | 36548611 | 36548686 | 36540318 | 36540491 | 36549438 | 36549475 | 0 | 0 | 16 | 0 | 0 | 1 | 0 | 3 | 1 | 0 | 0 | 0 | 98 | 49 | 0.727128809 | 0.941321792 | NA | NA | 0.889 | NA | 1 | 1 | -0.111 |
| RI | 251 | TYMP | chr22 | - | 50529495 | 50530085 | 50529495 | 50529731 | 50529903 | 50530085 | 1 | 7 | 17 | 0 | 0 | 1 | 7 | 16 | 4 | 0 | 1 | 0 | 98 | 49 | 0.727141068 | 0.999202116 | 1 | 1 | 0.895 | 1 | 0.889 | 1 | 0.002 |
| SE | 5784 | DENND3 | chr8 | + | 141141202 | 141141324 | 141138021 | 141138137 | 141144147 | 141144259 | 0 | 0 | 17 | 0 | 0 | 1 | 2 | 10 | 1 | 0 | 0 | 0 | 98 | 49 | 0.73240269 | 0.947333256 | NA | NA | 0.895 | 1 | 1 | 1 | -0.105 |
| SE | 5221 | CLEC4E | chr12 | - | 8536089 | 8536205 | 8533304 | 8534809 | 8539216 | 8539306 | 0 | 0 | 35 | 0 | 0 | 1 | 0 | 8 | 2 | 0 | 0 | 0 | 98 | 49 | 0.736969319 | 0.952420366 | NA | NA | 0.946 | NA | 1 | 1 | -0.054 |
| A5SS | 2690 | ARHGDIA | chr17 | - | 81869329 | 81869625 | 81869541 | 81869625 | 81869172 | 81869236 | 1 | 3 | 16 | 0 | 0 | 1 | 0 | 7 | 3 | 0 | 0 | 2 | 98 | 49 | 0.739861649 | 0.968908208 | 1 | 1 | 0.889 | NA | 1 | 0.429 | 0.249 |
| A5SS | 2132 | RFWD2 | chr1 | - | 176163814 | 176163891 | 176163826 | 176163891 | 176162868 | 176162988 | 0 | 1 | 17 | 0 | 1 | 5 | 0 | 5 | 0 | 0 | 6 | 0 | 61 | 49 | 0.745584037 | 0.968908208 | NA | 0.445 | 0.732 | NA | 0.401 | NA | 0.188 |
| A3SS | 1291 | SLC6A6 | chr3 | + | 14443623 | 14443863 | 14443627 | 14443863 | 14416411 | 14416453 | 0 | 0 | 19 | 0 | 0 | 1 | 0 | 9 | 1 | 0 | 0 | 0 | 53 | 49 | 0.746035627 | 0.999202116 | NA | NA | 0.946 | NA | 1 | 1 | -0.054 |
| A5SS | 2703 | IFI6 | chr1 | - | 27669232 | 27669346 | 27669244 | 27669346 | 27668457 | 27668535 | 0 | 0 | 2 | 0 | 1 | 20 | 0 | 1 | 0 | 0 | 9 | 1 | 61 | 49 | 0.747020832 | 0.968908208 | NA | 0 | 0.074 | NA | 0.082 | 0 | -0.004 |
| SE | 862 | NAMPT | chr7 | - | 106254363 | 106254504 | 106253016 | 106253151 | 106261587 | 106261707 | 0 | 24 | 228 | 0 | 0 | 4 | 14 | 336 | 15 | 0 | 3 | 0 | 98 | 49 | 0.748257683 | 0.966178091 | NA | 1 | 0.966 | 1 | 0.982 | 1 | -0.011 |
| SE | 7082 | UBR2 | chr6 | + | 42676055 | 42676191 | 42674125 | 42674193 | 42676782 | 42676873 | 0 | 0 | 20 | 0 | 0 | 1 | 0 | 6 | 1 | 0 | 0 | 0 | 98 | 49 | 0.751735186 | 0.96983518 | NA | NA | 0.909 | NA | 1 | 1 | -0.091 |
| SE | 8416 | GRAMD1A | chr19 | + | 35014188 | 35014387 | 35013540 | 35013691 | 35015823 | 35015967 | 0 | 3 | 13 | 0 | 0 | 0 | 0 | 1 | 0 | 0 | 0 | 2 | 98 | 49 | 0.752841225 | 0.970277462 | NA | 1 | 1 | NA | 1 | 0 | 0.5 |
| SE | 4184 | STAT6 | chr12 | - | 57107604 | 57107743 | 57107230 | 57107314 | 57108162 | 57108299 | 0 | 1 | 13 | 0 | 1 | 5 | 0 | 3 | 1 | 0 | 3 | 0 | 98 | 49 | 0.753369127 | 0.970277462 | NA | 0.333 | 0.565 | NA | 0.333 | 1 | -0.218 |
| SE | 3752 | ARHGAP27 | chr17 | - | 45404268 | 45404334 | 45404028 | 45404096 | 45404444 | 45404528 | 0 | 2 | 4 | 0 | 0 | 3 | 0 | 0 | 1 | 0 | 5 | 0 | 98 | 49 | 0.759645396 | 0.976832296 | NA | 1 | 0.4 | NA | 0 | 1 | 0.2 |
| SE | 9027 | MYD88 | chr3 | + | 38139863 | 38139998 | 38138477 | 38139028 | 38140387 | 38140568 | 0 | 5 | 19 | 0 | 2 | 0 | 0 | 11 | 3 | 0 | 2 | 0 | 98 | 49 | 0.759758452 | 0.976832296 | NA | 0.556 | 1 | NA | 0.733 | 1 | -0.089 |
| SE | 2840 | TYMP | chr22 | - | 50526000 | 50526156 | 50525752 | 50525918 | 50526245 | 50526476 | 1 | 3 | 17 | 0 | 1 | 0 | 0 | 3 | 9 | 0 | 0 | 1 | 98 | 49 | 0.762114536 | 0.979024058 | 1 | 0.6 | 1 | NA | 1 | 0.818 | -0.042 |
| A3SS | 3416 | STAT3 | chr17 | - | 42348388 | 42348539 | 42348388 | 42348517 | 42388278 | 42388495 | 0 | 2 | 14 | 0 | 1 | 6 | 0 | 2 | 1 | 0 | 2 | 0 | 71 | 49 | 0.764799772 | 0.999202116 | NA | 0.58 | 0.617 | NA | 0.408 | 1 | -0.105 |
| RI | 356 | DDIT3 | chr12 | - | 57517268 | 57517753 | 57517268 | 57517438 | 57517705 | 57517753 | 1 | 2 | 7 | 0 | 0 | 1 | 2 | 28 | 3 | 0 | 2 | 0 | 98 | 49 | 0.777903504 | 0.999202116 | 1 | 1 | 0.778 | 1 | 0.875 | 1 | -0.032 |
| SE | 9233 | RPS24 | chr10 | + | 78040200 | 78040225 | 78037193 | 78037304 | 78040614 | 78040715 | 0 | 0 | 1 | 0 | 0 | 26 | 0 | 8 | 0 | 0 | 6 | 1 | 74 | 49 | 0.781756708 | 0.999202116 | NA | NA | 0.025 | NA | 0.469 | 0 | -0.209 |
| RI | 733 | CELF2 | chr10 | + | 11328925 | 11336673 | 11328925 | 11329097 | 11330455 | 11336673 | 0 | 0 | 29 | 0 | 0 | 1 | 1 | 10 | 1 | 0 | 1 | 0 | 98 | 49 | 0.794160768 | 0.999202116 | NA | NA | 0.935 | 1 | 0.833 | 1 | -0.009 |
| SE | 8415 | GRAMD1A | chr19 | + | 35013255 | 35013368 | 35011473 | 35011554 | 35013540 | 35013691 | 0 | 1 | 5 | 0 | 0 | 4 | 0 | 0 | 1 | 0 | 1 | 0 | 98 | 49 | 0.795544674 | 0.999202116 | NA | 1 | 0.385 | NA | 0 | 1 | 0.193 |
| RI | 1027 | ARHGDIA | chr17 | - | 81869329 | 81869625 | 81869329 | 81869406 | 81869541 | 81869625 | 1 | 5 | 15 | 0 | 0 | 13 | 5 | 7 | 6 | 0 | 8 | 1 | 98 | 49 | 0.797337051 | 0.999202116 | 1 | 1 | 0.366 | 1 | 0.304 | 0.75 | 0.104 |
| SE | 11387 | NLRC5 | chr16 | + | 57054750 | 57054840 | 57051537 | 57051621 | 57055031 | 57055094 | 0 | 3 | 12 | 0 | 0 | 1 | 0 | 9 | 11 | 0 | 1 | 0 | 98 | 49 | 0.799295763 | 0.999202116 | NA | 1 | 0.857 | NA | 0.818 | 1 | 0.019 |
| A5SS | 2652 | ACOX1 | chr17 | - | 75978790 | 75979434 | 75978964 | 75979434 | 75978533 | 75978693 | 0 | 0 | 0 | 0 | 2 | 2 | 1 | 2 | 0 | 0 | 0 | 1 | 98 | 49 | 0.808907745 | 0.999208248 | NA | 0 | 0 | 1 | 1 | 0 | -0.667 |
| A3SS | 3230 | ARRB2 | chr17 | + | 4720549 | 4720640 | 4720585 | 4720640 | 4720392 | 4720472 | 0 | 0 | 2 | 0 | 2 | 13 | 0 | 1 | 0 | 2 | 5 | 8 | 85 | 49 | 0.814988864 | 0.999202116 | NA | 0 | 0.081 | 0 | 0.103 | 0 | 0.006 |
| SE | 1331 | TREM1 | chr6 | - | 41280960 | 41281153 | 41275260 | 41276230 | 41282394 | 41282751 | 0 | 0 | 37 | 0 | 0 | 1 | 0 | 38 | 1 | 0 | 3 | 0 | 98 | 49 | 0.817902743 | 0.999202116 | NA | NA | 0.949 | NA | 0.864 | 1 | 0.017 |
| SE | 8687 | PTPRC | chr1 | + | 198749415 | 198749549 | 198748108 | 198748199 | 198750491 | 198750626 | 0 | 1 | 72 | 0 | 0 | 1 | 9 | 51 | 0 | 0 | 0 | 0 | 98 | 49 | 0.818606032 | 0.999202116 | NA | 1 | 0.973 | 1 | 1 | NA | -0.014 |
| SE | 6603 | CCDC88B | chr11 | + | 64354003 | 64354170 | 64353714 | 64353813 | 64355193 | 64355400 | 0 | 4 | 4 | 0 | 1 | 2 | 0 | 2 | 2 | 0 | 0 | 1 | 98 | 49 | 0.823294092 | 0.999202116 | NA | 0.667 | 0.5 | NA | 1 | 0.5 | -0.166 |
| A5SS | 1623 | ADAR | chr1 | - | 154590183 | 154590409 | 154590261 | 154590409 | 154589756 | 154589928 | 0 | 4 | 24 | 0 | 0 | 3 | 0 | 17 | 2 | 0 | 1 | 0 | 98 | 49 | 0.823441962 | 0.999208248 | NA | 1 | 0.8 | NA | 0.895 | 1 | -0.047 |
| SE | 5377 | SYNE1 | chr6 | - | 152300640 | 152300781 | 152293959 | 152294127 | 152301868 | 152302063 | 0 | 0 | 1 | 0 | 1 | 1 | 0 | 5 | 0 | 0 | 0 | 1 | 98 | 49 | 0.827502283 | 0.999202116 | NA | 0 | 0.333 | NA | 1 | 0 | -0.334 |
| SE | 4122 | XPO6 | chr16 | - | 28117317 | 28117462 | 28112903 | 28113050 | 28121669 | 28121762 | 0 | 1 | 55 | 0 | 0 | 1 | 8 | 46 | 0 | 0 | 0 | 0 | 98 | 49 | 0.827647446 | 0.999202116 | NA | 1 | 0.965 | 1 | 1 | NA | -0.018 |
| SE | 12729 | PILRA | chr7 | + | 100389887 | 100390106 | 100374043 | 100374433 | 100397878 | 100397912 | 0 | 4 | 21 | 0 | 0 | 5 | 7 | 11 | 3 | 0 | 2 | 0 | 98 | 49 | 0.837221542 | 0.999202116 | NA | 1 | 0.677 | 1 | 0.733 | 1 | -0.073 |
| SE | 11878 | MGAM | chr7 | + | 142064383 | 142064522 | 142063498 | 142063586 | 142065334 | 142065468 | 0 | 4 | 8 | 0 | 0 | 2 | 3 | 8 | 2 | 0 | 1 | 0 | 98 | 49 | 0.840992744 | 0.999202116 | NA | 1 | 0.667 | 1 | 0.8 | 1 | -0.1 |
| A3SS | 1500 | ADAM8 | chr10 | - | 133273253 | 133273443 | 133273253 | 133273430 | 133273761 | 133273838 | 0 | 6 | 11 | 0 | 0 | 2 | 2 | 5 | 3 | 0 | 1 | 0 | 62 | 49 | 0.846278337 | 0.999202116 | NA | 1 | 0.813 | 1 | 0.798 | 1 | -0.026 |
| A3SS | 537 | DBNL | chr7 | + | 44058428 | 44058480 | 44058431 | 44058480 | 44058128 | 44058280 | 0 | 0 | 6 | 0 | 3 | 2 | 0 | 1 | 0 | 0 | 1 | 1 | 52 | 49 | 0.847104365 | 0.999202116 | NA | 0 | 0.739 | NA | 0.485 | 0 | 0.127 |
| SE | 9515 | SIPA1 | chr11 | + | 65650400 | 65650479 | 65650137 | 65650192 | 65650568 | 65650920 | 0 | 0 | 9 | 0 | 1 | 0 | 0 | 4 | 2 | 0 | 0 | 2 | 98 | 49 | 0.852675535 | 0.999202116 | NA | 0 | 1 | NA | 1 | 0.333 | -0.166 |
| SE | 6906 | FCGR3B | chr1 | - | 161626144 | 161626402 | 161623197 | 161624639 | 161629777 | 161630032 | 0 | 0 | 59 | 0 | 0 | 1 | 2 | 19 | 1 | 0 | 2 | 0 | 98 | 49 | 0.85537265 | 0.999202116 | NA | NA | 0.967 | 1 | 0.826 | 1 | 0.025 |

| | | | | | | | | | | | | | | | | | | | | | | | | | | | | | | | | | | |
|---|---|---|---|---|---|---|---|---|---|---|---|---|---|---|---|---|---|---|---|---|---|---|---|---|---|---|---|---|---|---|---|---|---|---|
| A3SS | 3415 | STAT3 | chr17 | - | 42316788 | 42316901 | 42316788 | 42316851 | 42317181 | 42317224 | 3 | 2 | 12 | 0 | 0 | 4 | 0 | 8 | 2 | 0 | 2 | 0 | 98 | 49 | 0.866784393 | 0.999202116 | 1 | 1 | 0.6 | NA | | 0.667 | 1 | 0.033 |
| SE | 2906 | RAF1 | chr3 | - | 12590797 | 12590974 | 12587590 | 12587637 | 12591707 | 12591792 | 1 | 1 | 24 | 0 | 0 | 2 | 4 | 25 | 0 | 0 | 0 | 0 | 98 | 49 | 0.867186844 | 0.999202116 | 1 | 1 | 0.857 | | 1 | 1 | NA | -0.048 |
| SE | 144 | IL2RG | chrX | - | 71108598 | 71108695 | 71108276 | 71108346 | 71109227 | 71109390 | 0 | 1 | 4 | 0 | 2 | 0 | 0 | 12 | 1 | 0 | 10 | 0 | 98 | 49 | 0.876241607 | 0.999202116 | NA | 0.2 | 1 | NA | | 0.375 | 1 | -0.088 |
| SE | 4365 | LRRK2 | chr12 | + | 40322034 | 40322181 | 40321033 | 40321188 | 40322318 | 40322510 | 0 | 1 | 45 | 0 | 0 | 2 | 0 | 21 | 0 | 0 | 0 | 0 | 98 | 49 | 0.881723484 | 0.999202116 | NA | 1 | 0.918 | NA | | 1 | NA | -0.041 |
| RI | 809 | ZFP36L1 | chr14 | - | 68787654 | 68790492 | 68787654 | 68788809 | 68789526 | 68790492 | 0 | 4 | 17 | 0 | 0 | 2 | 8 | 27 | 5 | 0 | 2 | 0 | 98 | 49 | 0.883440896 | 0.999202116 | NA | 1 | 0.81 | | 1 | 0.871 | 1 | -0.052 |
| SE | 9293 | ABCA7 | chr19 | + | 1054569 | 1054694 | 1054192 | 1054341 | 1054779 | 1054878 | 0 | 1 | 4 | 0 | 0 | 3 | 0 | 4 | 1 | 0 | 2 | 0 | 98 | 49 | 0.891129146 | 0.999202116 | NA | 1 | 0.4 | NA | | 0.5 | 1 | -0.05 |
| SE | 6059 | HCLS1 | chr3 | - | 121647318 | 121647448 | 121644817 | 121644928 | 121657278 | 121657352 | 0 | 1 | 33 | 0 | 0 | 1 | 3 | 18 | 0 | 0 | 0 | 0 | 98 | 49 | 0.89175776 | 0.999202116 | NA | 1 | 0.943 | | 1 | 1 | NA | -0.028 |
| SE | 9137 | CLK1 | chr2 | - | 200854615 | 200854695 | 200853902 | 200853993 | 200855003 | 200855086 | 0 | 1 | 8 | 0 | 0 | 2 | 0 | 18 | 0 | 0 | 0 | 0 | 98 | 49 | 0.89175776 | 0.999202116 | NA | 1 | 0.667 | NA | | 1 | NA | -0.166 |
| A3SS | 1625 | RPS3 | chr11 | + | 75402303 | 75402446 | 75402351 | 75402446 | 75401639 | 75401733 | 0 | 0 | 2 | 0 | 2 | 7 | 3 | 0 | 0 | 0 | 15 | 0 | 97 | 49 | 0.906255181 | 0.999202116 | NA | 0 | 0.126 | | 1 | 0 | NA | -0.437 |
| SE | 1126 | RNASET2 | chr6 | - | 166948569 | 166948625 | 166946681 | 166946739 | 166952487 | 166952548 | 0 | 8 | 51 | 0 | 1 | 0 | 0 | 18 | 2 | 0 | 1 | 0 | 98 | 49 | 0.909429153 | 0.999202116 | NA | 0.8 | 1 | NA | | 0.9 | 1 | -0.05 |
| SE | 11303 | SP100 | chr2 | + | 230440475 | 230440557 | 230417590 | 230417665 | 230442936 | 230443099 | 0 | 0 | 1 | 0 | 1 | 3 | 0 | 0 | 0 | 0 | 7 | 0 | 98 | 49 | 0.909814497 | 0.999202116 | NA | 0 | 0.143 | NA | | 0 | NA | 0.071 |
| SE | 1834 | ARRDC3 | chr5 | - | 91376620 | 91376768 | 91375510 | 91375613 | 91378693 | 91378775 | 0 | 1 | 13 | 0 | 0 | 6 | 0 | 13 | 0 | 0 | 0 | 0 | 98 | 49 | 0.910129019 | 0.999202116 | NA | 1 | 0.52 | NA | | 1 | NA | -0.24 |
| SE | 11821 | SSH2 | chr17 | - | 29650653 | 29650800 | 29648143 | 29648344 | 29655560 | 29655607 | 0 | 1 | 32 | 0 | 0 | 3 | 0 | 13 | 0 | 0 | 0 | 0 | 98 | 49 | 0.910129019 | 0.999202116 | NA | 1 | 0.842 | NA | | 1 | NA | -0.079 |
| A3SS | 3715 | TYROBP | chr19 | - | 35907217 | 35907264 | 35907217 | 35907261 | 35907445 | 35907580 | 0 | 11 | 42 | 0 | 0 | 4 | 0 | 9 | 9 | 0 | 1 | 0 | 52 | 49 | 0.912303287 | 0.999202116 | NA | 1 | 0.908 | NA | | 0.895 | 1 | 0.006 |
| SE | 3136 | IFNAR2 | chr21 | + | 33248708 | 33248854 | 33246717 | 33246890 | 33252661 | 33252830 | 0 | 1 | 7 | 0 | 0 | 1 | 0 | 12 | 0 | 0 | 0 | 0 | 98 | 49 | 0.91423345 | 0.999202116 | NA | 1 | 0.778 | NA | | 1 | NA | -0.111 |
| SE | 6130 | WDR26 | chr1 | - | 224431681 | 224431781 | 224431476 | 224431581 | 224433683 | 224434299 | 0 | 1 | 6 | 0 | 0 | 1 | 0 | 12 | 0 | 0 | 0 | 0 | 98 | 49 | 0.91423345 | 0.999202116 | NA | 1 | 0.75 | NA | | 1 | NA | -0.125 |
| SE | 12143 | TRIM25 | chr17 | - | 56895925 | 56895952 | 56895520 | 56895604 | 56899114 | 56899180 | 0 | 1 | 3 | 0 | 0 | 1 | 0 | 9 | 0 | 0 | 0 | 0 | 76 | 49 | 0.917397557 | 0.999202116 | NA | 1 | 0.659 | NA | | 1 | NA | -0.17 |
| A3SS | 1626 | RPS3 | chr11 | + | 75405582 | 75406913 | 75405613 | 75406913 | 75404671 | 75404868 | 1 | 0 | 4 | 0 | 1 | 4 | 0 | 0 | 0 | 0 | 6 | 0 | 80 | 49 | 0.917875723 | 0.999202116 | 1 | 0 | 0.38 | NA | | 0 | NA | 0.46 |
| SE | 3618 | NIN | chr14 | - | 50821873 | 50822077 | 50806736 | 50806818 | 50830463 | 50830517 | 0 | 3 | 8 | 0 | 0 | 1 | 0 | 7 | 6 | 0 | 0 | 1 | 98 | 49 | 0.918267371 | 0.999202116 | NA | 1 | 0.8 | NA | | 1 | 0.75 | 0.025 |
| SE | 8078 | NADK | chr1 | - | 1756502 | 1756608 | 1756257 | 1756343 | 1757180 | 1757310 | 0 | 1 | 24 | 0 | 0 | 3 | 0 | 11 | 0 | 0 | 0 | 0 | 98 | 49 | 0.918460091 | 0.999202116 | NA | 1 | 0.8 | NA | | 1 | NA | -0.1 |
| A3SS | 2809 | SRRT | chr7 | + | 100888041 | 100888143 | 100888053 | 100888143 | 100887702 | 100887859 | 0 | 1 | 1 | 0 | 0 | 3 | 0 | 7 | 0 | 0 | 0 | 0 | 61 | 49 | 0.919108663 | 0.999202116 | NA | 1 | 0.211 | NA | | 1 | NA | -0.394 |
| SE | 10061 | TPT1 | chr13 | - | 45339496 | 45339602 | 45338659 | 45338776 | 45339993 | 45340184 | 0 | 6 | 79 | 0 | 0 | 3 | 10 | 61 | 0 | 0 | 0 | 0 | 98 | 49 | 0.922183161 | 0.999202116 | NA | 1 | 0.929 | | 1 | 1 | NA | -0.035 |
| RI | 894 | STX4 | chr16 | + | 31034225 | 31034536 | 31034225 | 31034325 | 31034461 | 31034536 | 0 | 1 | 5 | 0 | 0 | 3 | 6 | 10 | 0 | 0 | 0 | 0 | 98 | 49 | 0.922793683 | 0.999202116 | NA | 1 | 0.455 | | 1 | 1 | NA | -0.272 |
| SE | 5493 | ADAM8 | chr10 | - | 133263687 | 133263765 | 133262415 | 133263233 | 133267351 | 133267417 | 0 | 4 | 10 | 0 | 2 | 2 | 3 | 19 | 13 | 0 | 5 | 4 | 98 | 49 | 0.923886696 | 0.999202116 | NA | 0.5 | 0.714 | | 1 | 0.655 | 0.619 | -0.151 |
| SE | 1583 | SYTL3 | chr6 | + | 158663204 | 158663378 | 158662749 | 158662812 | 158665394 | 158665613 | 0 | 1 | 1 | 0 | 0 | 1 | 0 | 2 | 0 | 0 | 0 | 1 | 98 | 49 | 0.925071736 | 0.999202116 | NA | 1 | 0.333 | NA | | 1 | 0 | 0.166 |
| SE | 4333 | NCF2 | chr1 | - | 183573184 | 183573292 | 183570779 | 183570839 | 183574486 | 183574621 | 2 | 5 | 63 | 0 | 0 | 1 | 0 | 47 | 0 | 0 | 0 | 0 | 98 | 49 | 0.925576519 | 0.999202116 | 1 | 1 | 0.969 | NA | | 1 | NA | -0.01 |
| SE | 8261 | ARHGEF11 | chr1 | - | 156957791 | 156957815 | 156956419 | 156956564 | 156958741 | 156958864 | 0 | 2 | 7 | 0 | 0 | 0 | 0 | 2 | 0 | 0 | 0 | 1 | 73 | 49 | 0.92640979 | 0.999202116 | NA | 1 | 1 | NA | | 1 | 0 | 0.5 |
| A5SS | 2405 | SP100 | chr2 | + | 230467119 | 230467215 | 230467119 | 230467206 | 230469042 | 230469096 | 0 | 3 | 13 | 0 | 0 | 1 | 0 | 15 | 0 | 0 | 0 | 0 | 58 | 49 | 0.926498131 | 0.999208248 | NA | 1 | 0.917 | NA | | 1 | NA | -0.041 |
| RI | 1099 | RPL28 | chr19 | + | 55387929 | 55392083 | 55387929 | 55388048 | 55388242 | 55392083 | 1 | 0 | 1 | 0 | 0 | 13 | 0 | 0 | 1 | 4 | 0 | 0 | 98 | 49 | 0.927341297 | 0.999202116 | 1 | NA | 0.037 | | 0 | NA | 1 | 0.018 |
| SE | 11228 | PTPN12 | chr7 | + | 77585542 | 77585581 | 77583554 | 77583650 | 77592184 | 77592256 | 0 | 1 | 9 | 0 | 0 | 1 | 0 | 8 | 0 | 0 | 0 | 0 | 88 | 49 | 0.9284209 | 0.999202116 | NA | 1 | 0.834 | NA | | 1 | NA | -0.083 |
| SE | 10669 | LAPTM5 | chr1 | - | 30742455 | 30742549 | 30741639 | 30741716 | 30757658 | 30757836 | 0 | 3 | 22 | 0 | 0 | 2 | 1 | 26 | 0 | 0 | 0 | 0 | 98 | 49 | 0.928473035 | 0.999202116 | NA | 1 | 0.846 | | 1 | 1 | NA | -0.077 |
| SE | 4334 | NCF2 | chr1 | - | 183573184 | 183573292 | 183570779 | 183570839 | 183577598 | 183577707 | 2 | 5 | 53 | 0 | 0 | 1 | 0 | 44 | 0 | 0 | 0 | 0 | 98 | 49 | 0.928484533 | 0.999202116 | 1 | 1 | 0.964 | NA | | 1 | NA | -0.012 |
| SE | 11373 | CYLD | chr16 | + | 50744729 | 50744866 | 50742761 | 50742841 | 50749575 | 50750202 | 0 | 0 | 0 | 0 | 2 | 1 | 2 | 0 | 0 | 0 | 7 | 2 | 98 | 49 | 0.928492447 | 0.999202116 | NA | 0 | 0 | | 1 | 0 | 0 | -0.333 |
| A5SS | 2858 | RPL13A | chr19 | + | 49489849 | 49489922 | 49489849 | 49489912 | 49490231 | 49490297 | 0 | 3 | 11 | 0 | 0 | 1 | 4 | 17 | 0 | 0 | 0 | 0 | 59 | 49 | 0.928715497 | 0.999208248 | NA | 1 | 0.901 | | 1 | 1 | NA | -0.049 |
| SE | 1923 | EMB | chr5 | - | 50410876 | 50410965 | 50405724 | 50405852 | 50411196 | 50411383 | 1 | 1 | 22 | 0 | 0 | 1 | 1 | 9 | 0 | 0 | 0 | 0 | 98 | 49 | 0.928729465 | 0.999202116 | 1 | 1 | 0.917 | | 1 | 1 | NA | -0.028 |
| SE | 3671 | CALCOCO2 | chr17 | + | 48846442 | 48846514 | 48841697 | 48841887 | 48848063 | 48848166 | 0 | 0 | 1 | 0 | 1 | 5 | 0 | 0 | 0 | 0 | 4 | 0 | 98 | 49 | 0.930162674 | 0.999202116 | NA | 0 | 0.091 | NA | | 0 | NA | 0.045 |
| SE | 10146 | GALC | chr14 | - | 87988454 | 87988523 | 87988143 | 87988207 | 87992969 | 87993271 | 0 | 1 | 5 | 0 | 1 | 0 | 0 | 5 | 12 | 0 | 0 | 2 | 98 | 49 | 0.93135784 | 0.999202116 | NA | 0.333 | 1 | NA | | 1 | 0.75 | -0.209 |
| A3SS | 538 | DBNL | chr7 | + | 44058877 | 44058983 | 44058901 | 44058983 | 44058428 | 44058480 | 0 | 0 | 2 | 0 | 1 | 3 | 3 | 1 | 0 | 0 | 2 | 1 | 73 | 49 | 0.931399982 | 0.999202116 | NA | 0 | 0.309 | | 1 | 0.251 | 0 | -0.262 |
| SE | 10209 | RFWD2 | chr1 | - | 176136487 | 176136547 | 176135009 | 176135086 | 176149005 | 176149074 | 0 | 1 | 16 | 0 | 0 | 4 | 3 | 15 | 3 | 0 | 3 | 0 | 98 | 49 | 0.931793753 | 0.999202116 | NA | 1 | 0.667 | | 1 | 0.714 | 1 | -0.071 |
| SE | 5478 | MICAL2 | chr11 | + | 12204249 | 12204457 | 12162078 | 12162419 | 12208022 | 12208139 | 0 | 1 | 17 | 0 | 0 | 1 | 0 | 8 | 0 | 0 | 0 | 0 | 98 | 49 | 0.932280727 | 0.999202116 | NA | 1 | 0.895 | NA | | 1 | NA | -0.052 |
| SE | 11480 | AP1G1 | chr16 | - | 71748250 | 71748378 | 71746587 | 71746692 | 71749893 | 71749983 | 0 | 1 | 4 | 0 | 0 | 1 | 0 | 8 | 0 | 0 | 0 | 0 | 98 | 49 | 0.932280727 | 0.999202116 | NA | 1 | 0.667 | NA | | 1 | NA | -0.166 |
| SE | 13003 | HNRNPU | chr1 | - | 244858010 | 244858274 | 244857597 | 244857717 | 244858728 | 244858841 | 0 | 2 | 18 | 0 | 0 | 2 | 2 | 13 | 0 | 0 | 0 | 0 | 98 | 49 | 0.932958395 | 0.999202116 | NA | 1 | 0.818 | | 1 | 1 | NA | -0.091 |
| SE | 7216 | LSP1 | chr11 | + | 1884279 | 1884323 | 1883931 | 1884024 | 1884499 | 1884581 | 0 | 5 | 71 | 0 | 0 | 1 | 0 | 36 | 0 | 0 | 0 | 0 | 93 | 49 | 0.933472812 | 0.999202116 | NA | 1 | 0.974 | NA | | 1 | NA | -0.013 |
| SE | 4366 | LRRK2 | chr12 | + | 40354298 | 40354492 | 40351538 | 40351733 | 40356114 | 40356187 | 0 | 2 | 26 | 0 | 0 | 2 | 0 | 15 | 0 | 0 | 0 | 0 | 98 | 49 | 0.934689043 | 0.999202116 | NA | 1 | 0.867 | NA | | 1 | NA | -0.067 |
| SE | 13049 | DHX34 | chr19 | + | 47357865 | 47358120 | 47355038 | 47355350 | 47359967 | 47360070 | 0 | 2 | 24 | 0 | 0 | 1 | 0 | 15 | 0 | 0 | 0 | 0 | 98 | 49 | 0.934689043 | 0.999202116 | NA | 1 | 0.923 | NA | | 1 | NA | -0.038 |
| SE | 3908 | MYL6 | chr12 | + | 56160625 | 56160670 | 56160242 | 56160320 | 56161386 | 56161582 | 0 | 0 | 1 | 0 | 3 | 15 | 0 | 0 | 0 | 0 | 11 | 0 | 94 | 49 | 0.935467898 | 0.999202116 | NA | 0 | 0.034 | NA | | 0 | NA | 0.017 |
| A3SS | 2959 | MTCH1 | chr6 | - | 36977633 | 36977784 | 36977633 | 36977691 | 36978077 | 36978155 | 0 | 1 | 1 | 0 | 0 | 3 | 2 | 6 | 0 | 0 | 0 | 0 | 98 | 49 | 0.936414148 | 0.999202116 | NA | 1 | 0.143 | | 1 | 1 | NA | -0.428 |
| RI | 864 | SUMO1 | chr2 | - | 202206179 | 202207321 | 202206179 | 202206803 | 202207252 | 202207321 | 0 | 1 | 10 | 0 | 0 | 1 | 0 | 6 | 0 | 0 | 0 | 0 | 98 | 49 | 0.936414148 | 0.999202116 | NA | 1 | 0.833 | NA | | 1 | NA | -0.084 |
| SE | 11834 | NCOA1 | chr2 | + | 24705085 | 24705233 | 24697657 | 24697798 | 24706567 | 24707888 | 0 | 1 | 11 | 0 | 0 | 1 | 0 | 6 | 0 | 0 | 0 | 0 | 98 | 49 | 0.936414148 | 0.999202116 | NA | 1 | 0.846 | NA | | 1 | NA | -0.077 |
| SE | 12434 | SPAG9 | chr17 | - | 50984922 | 50984990 | 50982523 | 50982672 | 50985697 | 50985778 | 0 | 1 | 8 | 0 | 0 | 1 | 0 | 6 | 0 | 0 | 0 | 0 | 98 | 49 | 0.936414148 | 0.999202116 | NA | 1 | 0.8 | NA | | 1 | NA | -0.1 |
| SE | 13879 | ZEB2 | chr2 | - | 144404835 | 144405024 | 144403915 | 144404130 | 144429768 | 144430026 | 0 | 1 | 3 | 0 | 0 | 1 | 0 | 6 | 0 | 0 | 0 | 0 | 98 | 49 | 0.936414148 | 0.999202116 | NA | 1 | 0.6 | NA | | 1 | NA | -0.2 |
| A3SS | 3220 | LILRB3 | chr19 | - | 54217318 | 54217477 | 54217318 | 54217474 | 54218360 | 54218413 | 0 | 2 | 8 | 0 | 3 | 11 | 0 | 1 | 3 | 4 | 6 | 1 | 52 | 49 | 0.937019423 | 0.999202116 | NA | 0.386 | 0.407 | | 0 | 0.136 | 0.739 | 0.105 |
| RI | 712 | CD164 | chr6 | - | 109366513 | 109369017 | 109366513 | 109368325 | 109368921 | 109369017 | 0 | 1 | 31 | 0 | 0 | 1 | 4 | 7 | 0 | 0 | 0 | 0 | 98 | 49 | 0.937374675 | 0.999202116 | NA | 1 | 0.939 | | 1 | 1 | NA | -0.03 |
| SE | 2080 | HERC5 | chr4 | + | 88487068 | 88487179 | 88486114 | 88486228 | 88489165 | 88489336 | 0 | 1 | 19 | 0 | 0 | 1 | 0 | 7 | 0 | 0 | 0 | 0 | 98 | 49 | 0.937374675 | 0.999202116 | NA | 1 | 0.905 | NA | | 1 | NA | -0.047 |
| SE | 2091 | HSD17B11 | chr4 | - | 87374698 | 87374830 | 87372708 | 87372815 | 87382254 | 87382362 | 0 | 3 | 26 | 0 | 0 | 1 | 2 | 20 | 0 | 0 | 0 | 0 | 98 | 49 | 0.939243248 | 0.999202116 | NA | 1 | 0.929 | | 1 | 1 | NA | -0.035 |
| RI | 861 | JUP | chr17 | - | 41754606 | 41755895 | 41754606 | 41755432 | 41755729 | 41755895 | 1 | 0 | 0 | 0 | 0 | 0 | 5 | 1 | 0 | 0 | 1 | 0 | 98 | 49 | 0.94318207 | 0.999202116 | 1 | NA | NA | | 1 | 0.333 | NA | 0.334 |
| SE | 8746 | BEST1 | chr11 | + | 61959891 | 61960043 | 61959497 | 61959578 | 61962254 | 61962893 | 0 | 1 | 3 | 0 | 0 | 1 | 0 | 5 | 0 | 0 | 0 | 0 | 98 | 49 | 0.94318207 | 0.999202116 | NA | 1 | 0.6 | NA | | 1 | NA | -0.2 |
| SE | 9188 | GIT2 | chr12 | - | 109951166 | 109951316 | 109947255 | 109947504 | 109953091 | 109953234 | 0 | 1 | 5 | 0 | 0 | 1 | 0 | 5 | 0 | 0 | 0 | 0 | 98 | 49 | 0.94318207 | 0.999202116 | NA | 1 | 0.714 | NA | | 1 | NA | -0.143 |
| SE | 1843 | VCAN | chr5 | + | 83537006 | 83542268 | 83512102 | 83512396 | 83545536 | 83545650 | 0 | 2 | 51 | 0 | 0 | 1 | 0 | 10 | 0 | 0 | 0 | 0 | 98 | 49 | 0.944027198 | 0.999202116 | NA | 1 | 0.962 | NA | | 1 | NA | -0.019 |
| SE | 3459 | RASSF2 | chr20 | - | 4795814 | 4795966 | 4792538 | 4792627 | 4798009 | 4798085 | 0 | 4 | 29 | 0 | 0 | 1 | 0 | 20 | 0 | 0 | 0 | 0 | 98 | 49 | 0.944343057 | 0.999202116 | NA | 1 | 0.935 | NA | | 1 | NA | -0.032 |
| SE | 3950 | ARID4B | chr1 | - | 235213768 | 235214026 | 235196030 | 235196115 | 235219792 | 235219968 | 0 | 0 | 1 | 0 | 1 | 2 | 0 | 0 | 0 | 0 | 3 | 0 | 98 | 49 | 0.94451324 | 0.999202116 | NA | 0 | 0.2 | NA | | 0 | NA | 0.1 |
| SE | 8841 | C1orf43 | chr1 | - | 154219835 | 154219937 | 154214456 | 154214574 | 154220341 | 154220797 | 0 | 0 | 1 | 0 | 1 | 1 | 0 | 0 | 0 | 0 | 3 | 0 | 98 | 49 | 0.94451324 | 0.999202116 | NA | 0 | 0.333 | NA | | 0 | NA | 0.167 |
| SE | 12024 | KLHL2 | chr4 | + | 165238777 | 165238899 | 165228806 | 165228913 | 165263196 | 165263359 | 0 | 0 | 4 | 0 | 1 | 2 | 0 | 0 | 0 | 0 | 3 | 0 | 98 | 49 | 0.94451324 | 0.999202116 | NA | 0 | 0.5 | NA | | 0 | NA | 0.25 |
| A5SS | 192 | TBXAS1 | chr7 | + | 139875584 | 139875666 | 139875584 | 139875637 | 139911224 | 139911321 | 0 | 0 | 1 | 0 | 1 | 15 | 5 | 0 | 0 | 0 | 3 | 0 | 78 | 49 | 0.945027512 | 0.999208248 | NA | 0 | 0.04 | | 1 | 0 | NA | -0.48 |
| RI | 396 | CASP1 | chr11 | - | 105025509 | 105026356 | 105025509 | 105025600 | 105026225 | 105026356 | 0 | 0 | 15 | 0 | 2 | 6 | 0 | 0 | 0 | 0 | 6 | 0 | 98 | 49 | 0.945187435 | 0.999202116 | NA | 0 | 0.556 | NA | | 0 | NA | 0.278 |
| SE | 11313 | ALDOA | chr16 | + | 30068645 | 30068700 | 30067449 | 30067661 | 30068817 | 30068978 | 0 | 3 | 41 | 0 | 0 | 1 | 4 | 14 | 0 | 0 | 0 | 0 | 98 | 49 | 0.946228808 | 0.999202116 | NA | 1 | 0.953 | | 1 | 1 | NA | -0.024 |
| SE | 10440 | SLC12A6 | chr15 | - | 34257641 | 34257788 | 34256228 | 34256283 | 34258812 | 34258944 | 0 | 1 | 11 | 0 | 0 | 4 | 0 | 2 | 4 | 0 | 2 | 0 | 98 | 49 | 0.949596688 | 0.999202116 | NA | 1 | 0.579 | NA | | 0.333 | 1 | 0.123 |
| SE | 10864 | ARHGAP9 | chr12 | - | 57476870 | 57476963 | 57476589 | 57476651 | 57477155 | 57477269 | 0 | 2 | 25 | 0 | 0 | 2 | 0 | 9 | 0 | 0 | 0 | 0 | 98 | 49 | 0.950138287 | 0.999202116 | NA | 1 | 0.862 | NA | | 1 | NA | -0.069 |
| SE | 7901 | PACSIN2 | chr22 | - | 42876887 | 42877010 | 42876136 | 42876333 | 42879047 | 42879169 | 0 | 0 | 5 | 0 | 3 | 0 | 0 | 0 | 0 | 0 | 8 | 0 | 98 | 49 | 0.950176987 | 0.999202116 | NA | 0 | 1 | NA | | 0 | NA | 0.5 |
| SE | 7638 | IRF7 | chr11 | - | 613950 | 614037 | 613784 | 613865 | 614475 | 614534 | 0 | 8 | 14 | 0 | 8 | 3 | 0 | 7 | 3 | 0 | 3 | 2 | 98 | 49 | 0.950918257 | 0.999202116 | NA | 0.333 | 0.7 | NA | | 0.538 | 0.429 | 0.033 |
| A3SS | 779 | LST1 | chr6 | + | 31588517 | 31588909 | 31588562 | 31588909 | 31587640 | 31587733 | 0 | 3 | 10 | 0 | 0 | 2 | 4 | 12 | 0 | 0 | 0 | 0 | 94 | 49 | 0.951788859 | 0.999202116 | NA | 1 | 0.723 | | 1 | 1 | NA | -0.139 |
| A5SS | 1230 | STRADA | chr17 | - | 63704340 | 63704582 | 63704431 | 63704582 | 63704004 | 63704047 | 1 | 0 | 0 | 0 | 0 | 0 | 4 | 1 | 0 | 0 | 1 | 0 | 98 | 49 | 0.952968863 | 0.999208248 | 1 | NA | NA | | 1 | 0.333 | NA | 0.334 |
| RI | 174 | NDUFA2 | chr5 | - | 140647255 | 140647785 | 140647255 | 140647362 | 140647482 | 140647785 | 1 | 0 | 0 | 0 | 0 | 2 | 4 | 3 | 0 | 0 | 1 | 0 | 98 | 49 | 0.952968863 | 0.999202116 | 1 | NA | 0 | | 1 | 0.6 | NA | -0.3 |
| SE | 1981 | DDX60L | chr4 | - | 168379725 | 168379830 | 168379362 | 168379504 | 168384611 | 168384812 | 0 | 1 | 18 | 0 | 0 | 1 | 0 | 4 | 0 | 0 | 0 | 0 | 98 | 49 | 0.952968863 | 0.999202116 | NA | 1 | 0.9 | NA | | 1 | NA | -0.05 |
| SE | 2114 | KDM5C | chrX | - | 53218275 | 53218398 | 53217795 | 53217966 | 53220838 | 53220916 | 0 | 1 | 12 | 0 | 0 | 2 | 0 | 4 | 0 | 0 | 0 | 0 | 98 | 49 | 0.952968863 | 0.999202116 | NA | 1 | 0.75 | NA | | 1 | NA | -0.125 |
| SE | 7631 | BTAF1 | chr10 | + | 91953736 | 91953873 | 91951402 | 91951566 | 91956527 | 91956657 | 0 | 1 | 5 | 0 | 0 | 1 | 0 | 4 | 0 | 0 | 0 | 0 | 98 | 49 | 0.952968863 | 0.999202116 | NA | 1 | 0.714 | NA | | 1 | NA | -0.143 |
| SE | 10952 | FES | chr15 | + | 90885411 | 90885585 | 90885036 | 90885258 | 90886960 | 90887057 | 0 | 1 | 3 | 0 | 0 | 1 | 0 | 4 | 0 | 0 | 0 | 0 | 98 | 49 | 0.952968863 | 0.999202116 | NA | 1 | 0.6 | NA | | 1 | NA | -0.2 |
| SE | 11294 | CREBBP | chr16 | - | 3780731 | 3780878 | 3778699 | 3778817 | 3781203 | 3781306 | 0 | 1 | 9 | 0 | 0 | 1 | 2 | 4 | 0 | 0 | 0 | 0 | 98 | 49 | 0.952968863 | 0.999202116 | NA | 1 | 0.818 | | 1 | 1 | NA | -0.091 |
| SE | 11440 | PSMB3 | chr17 | + | 38760430 | 38760608 | 38755882 | 38755990 | 38762410 | 38762505 | 0 | 1 | 12 | 0 | 0 | 1 | 0 | 4 | 0 | 0 | 0 | 0 | 98 | 49 | 0.952968863 | 0.999202116 | NA | 1 | 0.857 | NA | | 1 | NA | -0.072 |
| A5SS | 3017 | SERPINA1 | chr14 | - | 94388800 | 94389010 | 94388818 | 94389010 | 94388559 | 94388660 | 0 | 1 | 6 | 0 | 0 | 1 | 0 | 3 | 0 | 0 | 0 | 0 | 67 | 49 | 0.954109788 | 0.999208248 | NA | 1 | 0.814 | NA | | 1 | NA | -0.093 |
| A5SS | 399 | BTNL8 | chr5 | + | 180947511 | 180947755 | 180947511 | 180947625 | 180948354 | 180948375 | 0 | 2 | 2 | 0 | 0 | 1 | 0 | 7 | 0 | 0 | 0 | 0 | 98 | 49 | 0.957490202 | 0.999208248 | NA | 1 | 0.5 | NA | | 1 | NA | -0.25 |
| SE | 5182 | ANKRD13A | chr12 | + | 110033796 | 110033957 | 110030644 | 110030758 | 110036260 | 110036328 | 0 | 2 | 4 | 0 | 0 | 1 | 0 | 7 | 0 | 0 | 0 | 0 | 98 | 49 | 0.957490202 | 0.999202116 | NA | 1 | 0.667 | NA | | 1 | NA | -0.166 |
| SE | 6877 | UBN1 | chr16 | + | 4857989 | 4858076 | 4852878 | 4853166 | 4858567 | 4858663 | 0 | 2 | 11 | 0 | 0 | 1 | 1 | 7 | 0 | 0 | 0 | 0 | 98 | 49 | 0.957490202 | 0.999202116 | NA | 1 | 0.846 | | 1 | 1 | NA | -0.077 |
| SE | 9478 | JAK1 | chr1 | - | 64839602 | 64839795 | 64838464 | 64838589 | 64841244 | 64841339 | 0 | 2 | 17 | 0 | 0 | 1 | 0 | 7 | 0 | 0 | 0 | 0 | 98 | 49 | 0.957490202 | 0.999202116 | NA | 1 | 0.895 | NA | | 1 | NA | -0.052 |
| SE | 11195 | PARP14 | chr3 | + | 122704526 | 122704748 | 122699389 | 122701635 | 122708189 | 122708268 | 0 | 2 | 3 | 0 | 0 | 1 | 0 | 7 | 0 | 0 | 0 | 0 | 98 | 49 | 0.957490202 | 0.999202116 | NA | 1 | 0.6 | NA | | 1 | NA | -0.2 |
| A3SS | 778 | LST1 | chr6 | + | 31588496 | 31588909 | 31588562 | 31588909 | 31587640 | 31587733 | 0 | 3 | 10 | 0 | 0 | 2 | 4 | 11 | 0 | 0 | 0 | 0 | 98 | 49 | 0.957822245 | 0.999202116 | NA | 1 | 0.714 | | 1 | 1 | NA | -0.143 |
| SE | 11576 | PPIF | chr10 | + | 79349075 | 79349106 | 79347463 | 79347743 | 79349664 | 79349753 | 0 | 0 | 2 | 0 | 0 | 0 | 0 | 13 | 6 | 0 | 2 | 0 | 80 | 49 | 0.958077329 | 0.999202116 | NA | NA | 1 | NA | | 0.799 | 1 | 0.101 |
| SE | 6631 | PCBP2 | chr12 | + | 53467804 | 53467843 | 53467220 | 53467293 | 53468779 | 53468832 | 0 | 1 | 4 | 0 | 0 | 2 | 8 | 3 | 0 | 0 | 0 | 0 | 88 | 49 | 0.960459131 | 0.999202116 | NA | 1 | 0.527 | | 1 | 1 | NA | -0.236 |
| RI | 557 | QSOX1 | chr1 | + | 180196261 | 180198034 | 180196261 | 180196599 | 180197332 | 180198034 | 1 | 0 | 11 | 0 | 0 | 3 | 3 | 8 | 0 | 0 | 1 | 0 | 98 | 49 | 0.961336908 | 0.999202116 | 1 | NA | 0.647 | | 1 | 0.8 | NA | -0.077 |
| RI | 1064 | FXYD5 | chr19 | + | 35154768 | 35155611 | 35154768 | 35154869 | 35155550 | 35155611 | 1 | 0 | 0 | 0 | 0 | 0 | 3 | 1 | 0 | 0 | 1 | 0 | 98 | 49 | 0.961336908 | 0.999202116 | 1 | NA | NA | | 1 | 0.333 | NA | 0.334 |
| SE | 1449 | BAG6 | chr6 | - | 31647590 | 31647826 | 31646393 | 31646523 | 31648676 | 31648751 | 0 | 1 | 2 | 1 | 0 | 0 | 0 | 3 | 0 | 0 | 0 | 0 | 98 | 49 | 0.961336908 | 0.999202116 | 0 | 1 | 1 | NA | | 1 | NA | -0.333 |
| SE | 3433 | RRBP1 | chr20 | - | 17616731 | 17616839 | 17615925 | 17616009 | 17618595 | 17618679 | 0 | 1 | 0 | 0 | 0 | 1 | 0 | 3 | 0 | 0 | 0 | 0 | 98 | 49 | 0.961336908 | 0.999202116 | NA | 1 | 0 | NA | | 1 | NA | -0.5 |
| SE | 4548 | GZMH | chr14 | - | 24607611 | 24607747 | 24607148 | 24607406 | 24608264 | 24608412 | 0 | 1 | 0 | 0 | 0 | 3 | 0 | 3 | 0 | 0 | 0 | 0 | 98 | 49 | 0.961336908 | 0.999202116 | NA | 1 | 0 | NA | | 1 | NA | -0.5 |
| SE | 5312 | AFF1 | chr4 | + | 87089983 | 87090070 | 87084119 | 87084164 | 87091792 | 87091829 | 0 | 1 | 12 | 0 | 0 | 1 | 0 | 3 | 0 | 0 | 0 | 0 | 98 | 49 | 0.961336908 | 0.999202116 | NA | 1 | 0.857 | NA | | 1 | NA | -0.072 |
| SE | 6569 | HADHA | chr2 | - | 26193576 | 26193772 | 26192309 | 26192424 | 26194569 | 26194638 | 0 | 1 | 11 | 0 | 0 | 1 | 0 | 3 | 0 | 0 | 0 | 0 | 98 | 49 | 0.961336908 | 0.999202116 | NA | 1 | 0.846 | NA | | 1 | NA | -0.077 |
| SE | 7389 | ZRANB1 | chr10 | + | 124981708 | 124981829 | 124974199 | 124974398 | 124983174 | 124983304 | 0 | 1 | 1 | 0 | 0 | 1 | 0 | 3 | 0 | 0 | 0 | 0 | 98 | 49 | 0.961336908 | 0.999202116 | NA | 1 | 0.333 | NA | | 1 | NA | -0.334 |
| SE | 7848 | RPS6KA1 | chr1 | + | 26560725 | 26560851 | 26558806 | 26558937 | 26561044 | 26561134 | 0 | 1 | 11 | 0 | 0 | 2 | 0 | 3 | 0 | 0 | 0 | 0 | 98 | 49 | 0.961336908 | 0.999202116 | NA | 1 | 0.733 | NA | | 1 | NA | -0.133 |
| SE | 7862 | C1orf162 | chr1 | + | 111477725 | 111477775 | 111477333 | 111477428 | 111477982 | 111478512 | 0 | 1 | 9 | 0 | 0 | 1 | 0 | 3 | 0 | 0 | 0 | 0 | 98 | 49 | 0.961336908 | 0.999202116 | NA | 1 | 0.818 | NA | | 1 | NA | -0.091 |
| SE | 11425 | DYNC1LI2 | chr16 | - | 66749196 | 66749313 | 66742437 | 66742668 | 66751272 | 66751346 | 0 | 1 | 6 | 0 | 0 | 1 | 0 | 3 | 0 | 0 | 0 | 0 | 98 | 49 | 0.961336908 | 0.999202116 | NA | 1 | 0.75 | NA | | 1 | NA | -0.125 |
| SE | 12705 | GANAB | chr11 | - | 62632564 | 62632745 | 62631029 | 62631183 | 62633004 | 62633101 | 0 | 1 | 3 | 0 | 0 | 1 | 0 | 3 | 0 | 0 | 0 | 0 | 98 | 49 | 0.961336908 | 0.999202116 | NA | 1 | 0.6 | NA | | 1 | NA | -0.2 |
| SE | 10094 | NUMB | chr14 | - | 73316389 | 73316422 | 73297210 | 73297285 | 73323129 | 73323204 | 0 | 0 | 2 | 0 | 3 | 8 | 0 | 0 | 0 | 0 | 6 | 0 | 82 | 49 | 0.961535291 | 0.999202116 | NA | 0 | 0.13 | NA | | 0 | NA | 0.065 |
| A3SS | 3132 | ITGAL | chr16 | + | 30505243 | 30505300 | 30505246 | 30505300 | 30504174 | 30504264 | 1 | 0 | 8 | 0 | 0 | 2 | 2 | 3 | 0 | 0 | 0 | 0 | 52 | 49 | 0.961575457 | 0.999202116 | 1 | NA | 0.79 | | 1 | 1 | NA | -0.105 |
| A5SS | 2154 | SNX17 | chr2 | + | 27371268 | 27371468 | 27371268 | 27371343 | 27372622 | 27372740 | 1 | 0 | 0 | 0 | 1 | 3 | 0 | 0 | 0 | 0 | 2 | 0 | 98 | 49 | 0.962033383 | 0.999208248 | 1 | 0 | 0 | NA | | 0 | NA | 0.333 |
| SE | 9739 | PXN | chr12 | - | 120215559 | 120215661 | 120215102 | 120215273 | 120221622 | 120221758 | 0 | 0 | 1 | 0 | 1 | 3 | 0 | 0 | 0 | 0 | 2 | 0 | 98 | 49 | 0.962033383 | 0.999202116 | NA | 0 | 0.143 | NA | | 0 | NA | 0.071 |
| SE | 11604 | INPP5K | chr17 | - | 1515426 | 1515585 | 1513871 | 1513979 | 1516455 | 1516888 | 0 | 1 | 0 | 0 | 0 | 1 | 0 | 0 | 0 | 0 | 0 | 2 | 98 | 49 | 0.962033383 | 0.999202116 | NA | 1 | 0 | NA | | NA | 0 | 0.5 |
| SE | 13744 | MAP4K4 | chr2 | + | 101871493 | 101871685 | 101870294 | 101870415 | 101873646 | 101873764 | 0 | 0 | 7 | 0 | 1 | 4 | 0 | 0 | 0 | 0 | 2 | 0 | 98 | 49 | 0.962033383 | 0.999202116 | NA | 0 | 0.467 | NA | | 0 | NA | 0.234 |
| SE | 6797 | BHLHE40 | chr3 | + | 4981391 | 4981515 | 4980300 | 4980408 | 4982835 | 4985180 | 0 | 3 | 3 | 0 | 0 | 1 | 0 | 9 | 0 | 0 | 0 | 0 | 98 | 49 | 0.962139286 | 0.999202116 | NA | 1 | 0.6 | NA | | 1 | NA | -0.2 |
| SE | 2917 | TOM1 | chr22 | + | 35321958 | 35322037 | 35317876 | 35317961 | 35323027 | 35323177 | 0 | 4 | 11 | 0 | 0 | 1 | 0 | 12 | 0 | 0 | 0 | 0 | 98 | 49 | 0.962709327 | 0.999202116 | NA | 1 | 0.846 | NA | | 1 | NA | -0.077 |
| SE | 1267 | MICAL1 | chr6 | - | 109450299 | 109450557 | 109449969 | 109450085 | 109451599 | 109451700 | 0 | 2 | 4 | 0 | 0 | 1 | 0 | 6 | 0 | 0 | 0 | 0 | 98 | 49 | 0.963061436 | 0.999202116 | NA | 1 | 0.667 | NA | | 1 | NA | -0.166 |
| SE | 4007 | FRY | chr13 | + | 32175545 | 32175632 | 32173366 | 32173549 | 32178176 | 32178436 | 0 | 2 | 7 | 0 | 0 | 2 | 0 | 6 | 0 | 0 | 0 | 0 | 98 | 49 | 0.963061436 | 0.999202116 | NA | 1 | 0.636 | NA | | 1 | NA | -0.182 |
| SE | 6402 | AGO4 | chr1 | + | 35832436 | 35832570 | 35832056 | 35832185 | 35833989 | 35834174 | 0 | 2 | 14 | 0 | 0 | 1 | 0 | 6 | 0 | 0 | 0 | 0 | 98 | 49 | 0.963061436 | 0.999202116 | NA | 1 | 0.875 | NA | | 1 | NA | -0.062 |
| SE | 7412 | RBM23 | chr14 | - | 22909482 | 22909595 | 22908332 | 22908380 | 22911327 | 22911403 | 0 | 2 | 4 | 0 | 0 | 1 | 0 | 6 | 0 | 0 | 0 | 0 | 98 | 49 | 0.963061436 | 0.999202116 | NA | 1 | 0.667 | NA | | 1 | NA | -0.166 |
| SE | 1919 | PARP8 | chr5 | + | 50763147 | 50763242 | 50761820 | 50761898 | 50778068 | 50778129 | 0 | 3 | 15 | 0 | 0 | 1 | 0 | 8 | 0 | 0 | 0 | 0 | 98 | 49 | 0.964425677 | 0.999202116 | NA | 1 | 0.882 | NA | | 1 | NA | -0.059 |
| SE | 12367 | NBEAL2 | chr3 | + | 47003173 | 47003309 | 47002956 | 47003081 | 47003815 | 47003976 | 0 | 14 | 31 | 0 | 0 | 2 | 2 | 12 | 14 | 0 | 1 | 0 | 98 | 49 | 0.965210998 | 0.999202116 | NA | 1 | 0.886 | | 1 | 0.857 | 1 | -0.009 |
| SE | 12157 | VMP1 | chr17 | + | 59737452 | 59737543 | 59735337 | 59735473 | 59738836 | 59738947 | 0 | 4 | 43 | 0 | 0 | 4 | 0 | 10 | 0 | 0 | 0 | 0 | 98 | 49 | 0.967766702 | 0.999202116 | NA | 1 | 0.843 | NA | | 1 | NA | -0.079 |
| SE | 1590 | KDM1B | chr6 | + | 18162834 | 18162924 | 18161326 | 18161454 | 18166266 | 18166378 | 0 | 2 | 3 | 0 | 0 | 1 | 0 | 5 | 0 | 0 | 0 | 0 | 98 | 49 | 0.967767969 | 0.999202116 | NA | 1 | 0.6 | NA | | 1 | NA | -0.2 |
| SE | 4616 | PTPRJ | chr11 | + | 48121002 | 48121266 | 48112746 | 48112983 | 48123612 | 48123870 | 0 | 2 | 25 | 0 | 0 | 1 | 0 | 5 | 0 | 0 | 0 | 0 | 98 | 49 | 0.967767969 | 0.999202116 | NA | 1 | 0.926 | NA | | 1 | NA | -0.037 |
| SE | 6403 | AGO4 | chr1 | + | 35833989 | 35834174 | 35832436 | 35832570 | 35835833 | 35835993 | 0 | 2 | 1 | 0 | 0 | 1 | 0 | 5 | 0 | 0 | 0 | 0 | 98 | 49 | 0.967767969 | 0.999202116 | NA | 1 | 0.333 | NA | | 1 | NA | -0.334 |
| SE | 6446 | RALGDS | chr9 | - | 133107084 | 133107300 | 133106644 | 133106748 | 133107987 | 133108406 | 0 | 2 | 1 | 0 | 0 | 1 | 0 | 5 | 0 | 0 | 0 | 0 | 98 | 49 | 0.967767969 | 0.999202116 | NA | 1 | 0.333 | NA | | 1 | NA | -0.334 |
| SE | 7363 | NOTCH2 | chr1 | - | 119922635 | 119922778 | 119922235 | 119922446 | 119923636 | 119923984 | 0 | 2 | 5 | 0 | 0 | 1 | 0 | 5 | 0 | 0 | 0 | 0 | 98 | 49 | 0.967767969 | 0.999202116 | NA | 1 | 0.714 | NA | | 1 | NA | -0.143 |
| SE | 13360 | BROX | chr1 | + | 222731356 | 222731516 | 222730026 | 222730177 | 222732627 | 222735196 | 0 | 2 | 6 | 0 | 0 | 4 | 0 | 5 | 0 | 0 | 0 | 0 | 98 | 49 | 0.967767969 | 0.999202116 | NA | 1 | 0.429 | NA | | 1 | NA | -0.285 |
| A5SS | 2323 | POLG | chr15 | - | 89334672 | 89334795 | 89334684 | 89334795 | 89333095 | 89333913 | 0 | 0 | 0 | 0 | 0 | 3 | 0 | 1 | 0 | 0 | 0 | 4 | 61 | 49 | 0.969523928 | 0.999208248 | NA | NA | 0 | NA | | 1 | 0 | -0.5 |
| SE | 3907 | MYL6 | chr12 | + | 56159586 | 56159730 | 56158683 | 56158711 | 56159974 | 56160148 | 0 | 7 | 24 | 0 | 0 | 1 | 0 | 11 | 0 | 0 | 0 | 0 | 98 | 49 | 0.969652412 | 0.999202116 | NA | 1 | 0.923 | NA | | 1 | NA | -0.038 |

| | | | | | | | | | | | | | | | | | | | | | | | | | | | | | | | | | |
|---|---|---|---|---|---|---|---|---|---|---|---|---|---|---|---|---|---|---|---|---|---|---|---|---|---|---|---|---|---|---|---|---|---|
| SE | 9995 | UPP1 | chr7 | + | 48101823 | 48101870 | 48094762 | 48094827 | 48106872 | 48107082 | 0 | 0 | 2 | 0 | 0 | 0 | 0 | 3 | 3 | 0 | 3 | 0 | 96 | 49 | 0.970230495 | 0.999202116 | NA | NA | 1 | NA | 0.338 | 1 | 0.331 |
| RI | 401 | CTNNB1 | chr3 | + | 41239133 | 41240448 | 41239133 | 41239354 | 41239659 | 41240448 | 4 | 0 | 6 | 0 | 0 | 3 | 6 | 8 | 0 | 0 | 2 | 0 | 98 | 49 | 0.970780408 | 0.999202116 | 1 | NA | 0.5 | 1 | 0.667 | NA | -0.084 |
| SE | 813 | LINC-PINT | chr7 | - | 130912109 | 130912152 | 130884316 | 130884396 | 130945719 | 130945835 | 0 | 0 | 1 | 0 | 0 | 0 | 0 | 4 | 2 | 0 | 1 | 0 | 92 | 49 | 0.970840435 | 0.999202116 | NA | NA | 1 | NA | 0.681 | 1 | 0.159 |
| SE | 3380 | HCK | chr20 | + | 32054215 | 32054281 | 32052187 | 32052486 | 32071661 | 32071782 | 0 | 0 | 5 | 0 | 9 | 33 | 0 | 0 | 0 | 2 | 14 | 0 | 98 | 49 | 0.971752688 | 0.999202116 | NA | 0 | 0.07 | 0 | 0 | NA | 0.035 |
| A3SS | 80 | GBGT1 | chr9 | - | 133155177 | 133155349 | 133155177 | 133155312 | 133155900 | 133155936 | 1 | 0 | 1 | 0 | 0 | 1 | 2 | 0 | 0 | 0 | 0 | 0 | 86 | 49 | 0.972021626 | 0.999202116 | 1 | NA | 0.363 | 1 | NA | NA | -0.319 |
| A3SS | 2931 | LRRC75A-AS1 | chr17 | + | 16440110 | 16440253 | 16440184 | 16440253 | 16439659 | 16439703 | 0 | 0 | 2 | 0 | 0 | 0 | 0 | 1 | 4 | 0 | 3 | 0 | 98 | 49 | 0.97202508 | 0.999202116 | NA | NA | 1 | NA | 0.143 | 1 | 0.428 |
| RI | 848 | ARSA | chr22 | - | 50627555 | 50628173 | 50627555 | 50627792 | 50627897 | 50628173 | 0 | 0 | 2 | 0 | 0 | 0 | 0 | 0 | 4 | 0 | 2 | 0 | 98 | 49 | 0.97202508 | 0.999202116 | NA | NA | 1 | NA | 0 | 1 | 0.5 |
| SE | 3390 | ANKHD1-EIF4EBP3 | chr5 | + | 140496519 | 140497278 | 140486957 | 140487060 | 140504820 | 140504966 | 0 | 2 | 6 | 0 | 0 | 3 | 0 | 4 | 0 | 0 | 0 | 0 | 98 | 49 | 0.97202508 | 0.999202116 | NA | 1 | 0.5 | NA | 1 | NA | -0.25 |
| SE | 4583 | BECN1 | chr17 | - | 42815907 | 42816054 | 42814523 | 42814673 | 42818220 | 42818415 | 0 | 2 | 7 | 0 | 0 | 3 | 0 | 4 | 0 | 0 | 0 | 0 | 98 | 49 | 0.97202508 | 0.999202116 | NA | 1 | 0.538 | NA | 1 | NA | -0.231 |
| A5SS | 2532 | TTC19 | chr17 | + | 16000117 | 16000382 | 16000117 | 16000245 | 16001914 | 16002025 | 1 | 0 | 0 | 0 | 0 | 1 | 2 | 0 | 0 | 0 | 0 | 0 | 98 | 49 | 0.9723413 | 0.999208248 | 1 | NA | 0 | 1 | NA | NA | -0.5 |
| SE | 1578 | USP9X | chrX | + | 41140655 | 41140771 | 41136803 | 41137022 | 41140965 | 41141217 | 0 | 1 | 4 | 0 | 0 | 1 | 0 | 2 | 0 | 0 | 0 | 0 | 98 | 49 | 0.9723413 | 0.999202116 | NA | 1 | 0.667 | NA | 1 | NA | -0.166 |
| SE | 3631 | USP4 | chr3 | - | 49294406 | 49294598 | 49292509 | 49292598 | 49297869 | 49297964 | 0 | 1 | 7 | 0 | 0 | 2 | 4 | 2 | 0 | 0 | 0 | 0 | 98 | 49 | 0.9723413 | 0.999202116 | NA | 1 | 0.636 | 1 | 1 | NA | -0.182 |
| SE | 3782 | MYCBP2 | chr13 | - | 77278757 | 77278911 | 77273471 | 77273668 | 77288160 | 77288376 | 0 | 1 | 2 | 0 | 0 | 0 | 0 | 2 | 0 | 5 | 0 | 0 | 98 | 49 | 0.9723413 | 0.999202116 | NA | 1 | 1 | 0 | 1 | NA | 0.5 |
| SE | 6270 | BAZ2B | chr2 | - | 159337566 | 159337772 | 159332539 | 159332686 | 159347485 | 159347646 | 0 | 1 | 9 | 0 | 0 | 1 | 0 | 2 | 0 | 0 | 0 | 0 | 98 | 49 | 0.9723413 | 0.999202116 | NA | 1 | 0.818 | NA | 1 | NA | -0.091 |
| SE | 9517 | SMG5 | chr1 | - | 156266540 | 156266678 | 156265780 | 156266380 | 156267469 | 156267678 | 0 | 1 | 3 | 0 | 0 | 1 | 0 | 2 | 0 | 0 | 0 | 0 | 98 | 49 | 0.9723413 | 0.999202116 | NA | 1 | 0.6 | NA | 1 | NA | -0.2 |
| SE | 9988 | TRANK1 | chr3 | - | 36838612 | 36838716 | 36834761 | 36834907 | 36842621 | 36842710 | 0 | 0 | 1 | 0 | 0 | 0 | 0 | 1 | 2 | 0 | 1 | 0 | 98 | 49 | 0.9723413 | 0.999202116 | NA | NA | 1 | NA | 0.333 | 1 | 0.334 |
| SE | 10095 | NUMB | chr14 | - | 73366896 | 73366981 | 73355625 | 73355766 | 73409936 | 73410068 | 0 | 1 | 10 | 0 | 0 | 1 | 2 | 2 | 0 | 0 | 0 | 0 | 98 | 49 | 0.9723413 | 0.999202116 | NA | 1 | 0.833 | 1 | 1 | NA | -0.084 |
| SE | 11441 | PSMB3 | chr17 | + | 38760507 | 38760608 | 38755882 | 38755990 | 38762410 | 38762505 | 0 | 1 | 7 | 0 | 0 | 1 | 0 | 2 | 0 | 0 | 0 | 0 | 98 | 49 | 0.9723413 | 0.999202116 | NA | 1 | 0.778 | NA | 1 | NA | -0.111 |
| SE | 13354 | TFIP11 | chr22 | - | 26494138 | 26494304 | 26491928 | 26492368 | 26494796 | 26494939 | 0 | 1 | 0 | 0 | 0 | 1 | 0 | 2 | 0 | 0 | 0 | 0 | 98 | 49 | 0.9723413 | 0.999202116 | NA | 1 | 0 | NA | 1 | NA | -0.5 |
| SE | 1601 | NUP153 | chr6 | - | 17668974 | 17669028 | 17665238 | 17665385 | 17669292 | 17669337 | 0 | 3 | 7 | 0 | 0 | 1 | 0 | 6 | 0 | 0 | 0 | 0 | 98 | 49 | 0.973770068 | 0.999202116 | NA | 1 | 0.778 | NA | 1 | NA | -0.111 |
| SE | 332 | AKNA | chr9 | - | 114355924 | 114356136 | 114350858 | 114351021 | 114357920 | 114358167 | 0 | 2 | 12 | 0 | 0 | 1 | 0 | 3 | 0 | 0 | 0 | 0 | 98 | 49 | 0.974386813 | 0.999202116 | NA | 1 | 0.857 | NA | 1 | NA | -0.072 |
| SE | 3924 | MTMR6 | chr13 | - | 25251648 | 25251775 | 25246200 | 25249492 | 25251852 | 25251984 | 0 | 2 | 5 | 0 | 0 | 2 | 0 | 3 | 0 | 0 | 0 | 0 | 98 | 49 | 0.974386813 | 0.999202116 | NA | 1 | 0.556 | NA | 1 | NA | -0.222 |
| SE | 11516 | PDXDC1 | chr16 | + | 15026642 | 15026706 | 15022703 | 15022754 | 15028877 | 15028966 | 0 | 2 | 0 | 0 | 0 | 2 | 0 | 3 | 0 | 0 | 0 | 0 | 98 | 49 | 0.974386813 | 0.999202116 | NA | 1 | 0 | NA | 1 | NA | -0.5 |
| SE | 13864 | ACSL1 | chr4 | - | 184770254 | 184770332 | 184768315 | 184768450 | 184773080 | 184773154 | 0 | 2 | 3 | 0 | 0 | 1 | 0 | 3 | 0 | 0 | 0 | 0 | 98 | 49 | 0.974386813 | 0.999202116 | NA | 1 | 0.6 | NA | 1 | NA | -0.2 |
| A3SS | 1029 | ITGAM | chr16 | + | 31297741 | 31297954 | 31297744 | 31297954 | 31297513 | 31297654 | 0 | 2 | 2 | 0 | 1 | 7 | 0 | 3 | 0 | 6 | 1 | 2 | 52 | 49 | 0.975810876 | 0.999202116 | NA | 0.653 | 0.212 | 0 | 0.739 | 0 | 0.186 |
| SE | 7116 | MED13L | chr12 | - | 115968939 | 115969097 | 115966081 | 115966243 | 115970593 | 115970770 | 0 | 6 | 8 | 0 | 0 | 1 | 0 | 10 | 0 | 0 | 0 | 0 | 98 | 49 | 0.976427731 | 0.999202116 | NA | 1 | 0.8 | NA | 1 | NA | -0.1 |
| SE | 12588 | MIDN | chr19 | + | 1251561 | 1251649 | 1249889 | 1250529 | 1251838 | 1251901 | 0 | 6 | 6 | 0 | 0 | 1 | 0 | 10 | 0 | 0 | 0 | 0 | 98 | 49 | 0.976427731 | 0.999202116 | NA | 1 | 0.75 | NA | 1 | NA | -0.125 |
| SE | 9816 | LDB1 | chr10 | - | 102110528 | 102110701 | 102109920 | 102110043 | 102110868 | 102110971 | 0 | 3 | 5 | 0 | 0 | 2 | 3 | 5 | 0 | 0 | 0 | 0 | 98 | 49 | 0.976610097 | 0.999202116 | NA | 1 | 0.556 | 1 | 1 | NA | -0.222 |
| SE | 8808 | WDR1 | chr4 | - | 10103895 | 10103986 | 10098991 | 10099139 | 10116112 | 10116234 | 0 | 5 | 16 | 0 | 0 | 1 | 1 | 9 | 0 | 0 | 0 | 0 | 98 | 49 | 0.976663141 | 0.999202116 | NA | 1 | 0.889 | 1 | 1 | NA | -0.055 |
| SE | 3259 | ELMO2 | chr20 | - | 46387337 | 46387437 | 46386123 | 46386275 | 46389038 | 46389220 | 0 | 0 | 3 | 0 | 0 | 0 | 0 | 0 | 4 | 0 | 2 | 0 | 98 | 49 | 0.977411915 | 0.999202116 | NA | NA | 1 | NA | 0 | 1 | 0.5 |
| SE | 186 | KDM6A | chrX | + | 45059005 | 45059104 | 45053828 | 45053955 | 45059246 | 45059466 | 0 | 4 | 4 | 0 | 0 | 1 | 0 | 3 | 0 | 0 | 0 | 0 | 98 | 49 | 0.99285348 | 0.999202116 | NA | 1 | 0.667 | NA | 1 | NA | -0.166 |
| SE | 1798 | FBXO38 | chr5 | + | 148427212 | 148427722 | 148425521 | 148425701 | 148433423 | 148433524 | 0 | 3 | 2 | 0 | 0 | 2 | 0 | 2 | 0 | 0 | 0 | 0 | 98 | 49 | 0.993205848 | 0.999202116 | NA | 1 | 0.333 | NA | 1 | NA | -0.334 |
| SE | 5041 | RELT | chr11 | + | 73392210 | 73392468 | 73390754 | 73390921 | 73393836 | 73393917 | 0 | 0 | 3 | 0 | 0 | 0 | 0 | 6 | 2 | 2 | 3 | 0 | 98 | 49 | 0.993205848 | 0.999202116 | NA | NA | 1 | 0 | 0.5 | 1 | 0.5 |
| SE | 5295 | PEX16 | chr11 | - | 45916226 | 45916303 | 45915702 | 45915836 | 45917457 | 45917493 | 0 | 0 | 3 | 0 | 0 | 0 | 0 | 5 | 2 | 0 | 1 | 0 | 98 | 49 | 0.993205848 | 0.999202116 | NA | NA | 1 | NA | 0.714 | 1 | 0.143 |
| A3SS | 107 | ZNF189 | chr9 | + | 101399883 | 101400010 | 101399925 | 101400010 | 101398853 | 101399189 | 0 | 0 | 1 | 0 | 0 | 1 | 3 | 0 | 1 | 0 | 0 | 1 | 91 | 49 | 0.997361002 | 0.999202116 | NA | NA | 0.35 | 1 | NA | 0.35 | -0.325 |
| A5SS | 2698 | ANAPC11 | chr17 | + | 81894466 | 81894624 | 81894466 | 81894586 | 81899919 | 81900533 | 1 | 0 | 0 | 0 | 1 | 0 | 0 | 0 | 0 | 0 | 1 | 0 | 87 | 49 | 0.997625179 | 0.999208248 | 1 | 0 | NA | NA | 0 | NA | 0.5 |
| A3SS | 70 | MAN1B1 | chr9 | + | 137107249 | 137107447 | 137107274 | 137107447 | 137106688 | 137106809 | 0 | 0 | 3 | 0 | 0 | 0 | 0 | 0 | 2 | 0 | 1 | 0 | 74 | 49 | 0.999202116 | 0.999202116 | NA | NA | 1 | NA | 0 | 1 | 0.5 |
| A3SS | 338 | CUTA | chr6 | - | 33417480 | 33417816 | 33417480 | 33417695 | 33417958 | 33418288 | 0 | 1 | 1 | 0 | 0 | 1 | 0 | 1 | 0 | 0 | 0 | 0 | 98 | 49 | 0.999202116 | 0.999202116 | NA | 1 | 0.333 | NA | 1 | NA | -0.334 |
| A3SS | 776 | LST1 | chr6 | + | 31588517 | 31588909 | 31588562 | 31588909 | 31587199 | 31587318 | 0 | 3 | 14 | 0 | 1 | 6 | 4 | 15 | 0 | 0 | 2 | 1 | 94 | 49 | 0.999202116 | 0.999202116 | NA | 0.61 | 0.549 | 1 | 0.796 | 0 | -0.019 |
| A3SS | 780 | LST1 | chr6 | + | 31588496 | 31588909 | 31588517 | 31588909 | 31587943 | 31587966 | 0 | 1 | 0 | 0 | 0 | 6 | 0 | 0 | 0 | 0 | 0 | 1 | 70 | 49 | 0.999202116 | 0.999202116 | NA | 1 | 0 | NA | NA | 0 | 0.5 |
| A3SS | 804 | TYMP | chr22 | - | 50529495 | 50529731 | 50529495 | 50529719 | 50529903 | 50530085 | 0 | 1 | 6 | 0 | 0 | 13 | 0 | 20 | 1 | 0 | 3 | 2 | 61 | 49 | 0.999202116 | 0.999202116 | NA | 1 | 0.27 | NA | 0.843 | 0.287 | 0.07 |
| A3SS | 805 | TYMP | chr22 | - | 50529495 | 50529754 | 50529495 | 50529719 | 50529903 | 50530085 | 0 | 3 | 9 | 0 | 0 | 13 | 3 | 22 | 1 | 0 | 3 | 2 | 84 | 49 | 0.999202116 | 0.999202116 | NA | 1 | 0.288 | 1 | 0.811 | 0.226 | -0.035 |
| A3SS | 832 | TOM1 | chr22 | + | 35322998 | 35323177 | 35323027 | 35323177 | 35321958 | 35322037 | 0 | 1 | 5 | 0 | 4 | 6 | 0 | 0 | 3 | 0 | 4 | 0 | 78 | 49 | 0.999202116 | 0.999202116 | NA | 0.136 | 0.344 | NA | 0 | 1 | -0.26 |
| A3SS | 837 | TUG1 | chr22 | + | 30972854 | 30973601 | 30972923 | 30973601 | 30969210 | 30971779 | 0 | 0 | 1 | 0 | 0 | 0 | 3 | 2 | 1 | 0 | 2 | 0 | 98 | 49 | 0.999202116 | 0.999202116 | NA | NA | 1 | 1 | 0.333 | 1 | 0.222 |
| A3SS | 845 | GAS2L1 | chr22 | + | 29308058 | 29308738 | 29308086 | 29308738 | 29306994 | 29307153 | 2 | 0 | 0 | 0 | 1 | 0 | 2 | 0 | 0 | 0 | 0 | 0 | 77 | 49 | 0.999202116 | 0.999202116 | 1 | 0 | NA | 1 | NA | NA | -0.5 |
| A3SS | 1313 | BRD9 | chr5 | - | 889586 | 889766 | 889586 | 889647 | 891154 | 891287 | 0 | 2 | 2 | 0 | 0 | 2 | 0 | 2 | 0 | 0 | 0 | 0 | 98 | 49 | 0.999202116 | 0.999202116 | NA | 1 | 0.333 | NA | 1 | NA | -0.334 |
| A3SS | 1412 | GLIPR2 | chr9 | + | 36162231 | 36163913 | 36162361 | 36163913 | 36150871 | 36150949 | 0 | 1 | 6 | 0 | 0 | 5 | 0 | 2 | 0 | 0 | 5 | 2 | 98 | 49 | 0.999202116 | 0.999202116 | NA | 1 | 0.375 | NA | 0.167 | 0 | 0.604 |
| A3SS | 1450 | SPI1 | chr11 | - | 47375632 | 47375732 | 47375632 | 47375729 | 47378308 | 47378576 | 0 | 8 | 16 | 0 | 4 | 19 | 0 | 9 | 1 | 0 | 9 | 2 | 52 | 49 | 0.999202116 | 0.999202116 | NA | 0.653 | 0.442 | NA | 0.485 | 0.32 | 0.145 |
| A3SS | 1515 | KRCC1 | chr2 | - | 88034133 | 88034292 | 88034133 | 88034231 | 88036942 | 88037051 | 0 | 0 | 1 | 0 | 0 | 0 | 0 | 0 | 1 | 0 | 1 | 0 | 98 | 49 | 0.999202116 | 0.999202116 | NA | NA | 1 | NA | 0 | 1 | 0.5 |
| A3SS | 1711 | NFKB2 | chr10 | + | 102402251 | 102402529 | 102402254 | 102402529 | 102402047 | 102402159 | 0 | 1 | 3 | 0 | 0 | 3 | 0 | 6 | 0 | 0 | 0 | 1 | 52 | 49 | 0.999202116 | 0.999202116 | NA | 1 | 0.485 | NA | 1 | 0 | 0.242 |
| A3SS | 1716 | YY1AP1 | chr1 | - | 155668626 | 155668777 | 155668626 | 155668717 | 155670319 | 155670464 | 0 | 2 | 1 | 0 | 0 | 3 | 0 | 2 | 0 | 0 | 0 | 0 | 98 | 49 | 0.999202116 | 0.999202116 | NA | 1 | 0.143 | NA | 1 | NA | -0.428 |
| A3SS | 1761 | ATP11A | chr13 | + | 112881815 | 112887168 | 112881875 | 112887168 | 112875775 | 112875941 | 1 | 3 | 1 | 0 | 0 | 1 | 0 | 1 | 0 | 0 | 0 | 0 | 98 | 49 | 0.999202116 | 0.999202116 | 1 | 1 | 0.333 | NA | 1 | NA | -0.222 |
| A3SS | 1775 | PHYKPL | chr5 | - | 178222354 | 178222580 | 178222354 | 178222492 | 178222851 | 178222934 | 0 | 0 | 5 | 0 | 0 | 0 | 0 | 5 | 1 | 0 | 1 | 0 | 98 | 49 | 0.999202116 | 0.999202116 | NA | NA | 1 | NA | 0.714 | 1 | 0.143 |
| A3SS | 1875 | ITGB2 | chr21 | - | 44910283 | 44910415 | 44910283 | 44910372 | 44910724 | 44910785 | 0 | 1 | 3 | 0 | 1 | 16 | 0 | 1 | 1 | 2 | 6 | 1 | 92 | 49 | 0.999202116 | 0.999202116 | NA | 0.348 | 0.091 | 0 | 0.082 | 0.348 | 0.076 |
| A3SS | 1896 | ADAT1 | chr16 | - | 75618585 | 75618698 | 75618585 | 75618640 | 75620265 | 75620334 | 0 | 0 | 0 | 0 | 1 | 0 | 0 | 0 | 1 | 0 | 1 | 0 | 98 | 49 | 0.999202116 | 0.999202116 | NA | 0 | NA | NA | 0 | 1 | -0.5 |
| A3SS | 2090 | STAT2 | chr12 | - | 56355707 | 56355803 | 56355707 | 56355791 | 56356131 | 56356285 | 0 | 4 | 17 | 0 | 0 | 3 | 0 | 6 | 2 | 0 | 1 | 0 | 61 | 49 | 0.999202116 | 0.999202116 | NA | 1 | 0.82 | NA | 0.828 | 1 | -0.004 |
| A3SS | 2169 | HIF1A | chr14 | + | 61720378 | 61720572 | 61720381 | 61720572 | 61695400 | 61695839 | 0 | 1 | 4 | 0 | 1 | 14 | 0 | 11 | 0 | 1 | 25 | 3 | 52 | 49 | 0.999202116 | 0.999202116 | NA | 0.485 | 0.212 | 0 | 0.293 | 0 | 0.251 |
| A3SS | 2223 | CSF3R | chr1 | - | 36466042 | 36466908 | 36466042 | 36466827 | 36467229 | 36467311 | 1 | 3 | 22 | 0 | 17 | 96 | 3 | 23 | 5 | 11 | 73 | 20 | 98 | 49 | 0.999202116 | 0.999202116 | 1 | 0.081 | 0.103 | 0.12 | 0.136 | 0.111 | 0.272 |
| A3SS | 2815 | DEF8 | chr16 | + | 89964169 | 89964310 | 89964220 | 89964310 | 89963362 | 89963443 | 0 | 3 | 3 | 0 | 0 | 1 | 0 | 1 | 0 | 0 | 0 | 0 | 98 | 49 | 0.999202116 | 0.999202116 | NA | 1 | 0.6 | NA | 1 | NA | -0.2 |
| A3SS | 3095 | MEF2D | chr1 | - | 156483238 | 156483430 | 156483238 | 156483418 | 156500485 | 156500842 | 0 | 5 | 4 | 0 | 0 | 0 | 0 | 10 | 0 | 0 | 0 | 1 | 61 | 49 | 0.999202116 | 0.999202116 | NA | 1 | 1 | NA | 1 | 0 | 0.5 |
| A3SS | 3209 | ZNF276 | chr16 | + | 89737720 | 89737905 | 89737805 | 89737905 | 89733920 | 89734038 | 0 | 1 | 1 | 0 | 0 | 5 | 0 | 1 | 0 | 0 | 0 | 0 | 98 | 49 | 0.999202116 | 0.999202116 | NA | 1 | 0.091 | NA | 1 | NA | -0.455 |
| A3SS | 3689 | DDX49 | chr19 | + | 18920579 | 18920703 | 18920650 | 18920703 | 18919674 | 18919856 | 0 | 0 | 1 | 0 | 0 | 0 | 0 | 1 | 1 | 0 | 1 | 0 | 98 | 49 | 0.999202116 | 0.999202116 | NA | NA | 1 | NA | 0.333 | 1 | 0.334 |
| A3SS | 3839 | LILRB2 | chr19 | - | 54277549 | 54277600 | 54277549 | 54277597 | 54277888 | 54277939 | 0 | 1 | 2 | 0 | 0 | 6 | 1 | 1 | 0 | 2 | 1 | 1 | 52 | 49 | 0.999202116 | 0.999202116 | NA | 1 | 0.239 | 0.32 | 0.485 | 0 | 0.351 |
| A5SS | 59 | RBM10 | chrX | + | 47145217 | 47145516 | 47145217 | 47145485 | 47147356 | 47147498 | 0 | 0 | 2 | 0 | 2 | 3 | 0 | 0 | 0 | 0 | 2 | 0 | 80 | 49 | 0.999202116 | 0.999208248 | NA | 0 | 0.29 | NA | 0 | NA | 0.145 |
| A5SS | 140 | GRINA | chr8 | + | 143990057 | 143990311 | 143990057 | 143990199 | 143991199 | 143991602 | 0 | 1 | 0 | 0 | 0 | 0 | 0 | 1 | 8 | 0 | 0 | 1 | 98 | 49 | 0.999202116 | 0.999208248 | NA | 1 | NA | NA | 1 | 0.8 | 0.1 |
| A5SS | 211 | SAMD9L | chr7 | - | 93146882 | 93147146 | 93147087 | 93147146 | 93145384 | 93146040 | 0 | 4 | 10 | 0 | 0 | 6 | 2 | 1 | 0 | 0 | 0 | 0 | 98 | 49 | 0.999202116 | 0.999208248 | NA | 1 | 0.455 | 1 | 1 | NA | -0.272 |
| A5SS | 388 | SYTL3 | chr6 | + | 158662749 | 158663378 | 158662749 | 158662812 | 158665394 | 158665613 | 0 | 1 | 2 | 0 | 0 | 1 | 2 | 2 | 1 | 0 | 0 | 1 | 98 | 49 | 0.999202116 | 0.999208248 | NA | 1 | 0.5 | 1 | 1 | 0.333 | -0.028 |
| A5SS | 615 | TOM1 | chr22 | + | 35346929 | 35346972 | 35346929 | 35346969 | 35347054 | 35347994 | 0 | 0 | 0 | 0 | 0 | 5 | 0 | 2 | 0 | 0 | 4 | 1 | 52 | 49 | 0.999202116 | 0.999208248 | NA | NA | 0 | NA | 0.32 | 0 | -0.16 |
| A5SS | 671 | STX16 | chr20 | + | 58651252 | 58652138 | 58651252 | 58652087 | 58659622 | 58659634 | 0 | 4 | 7 | 0 | 0 | 1 | 1 | 1 | 0 | 0 | 0 | 0 | 98 | 49 | 0.999202116 | 0.999208248 | NA | 1 | 0.778 | 1 | 1 | NA | -0.111 |
| A5SS | 745 | TAZ | chrX | + | 154413206 | 154413358 | 154413206 | 154413252 | 154413481 | 154413567 | 0 | 0 | 4 | 0 | 2 | 1 | 0 | 0 | 0 | 0 | 1 | 0 | 98 | 49 | 0.999202116 | 0.999208248 | NA | 0 | 0.667 | NA | 0 | NA | 0.334 |
| A5SS | 791 | SLAIN1 | chr13 | + | 77744282 | 77744516 | 77744282 | 77744432 | 77746513 | 77746855 | 1 | 0 | 0 | 0 | 0 | 1 | 1 | 0 | 0 | 0 | 0 | 0 | 98 | 49 | 0.999202116 | 0.999208248 | 1 | NA | 0 | 1 | NA | NA | -0.5 |
| A5SS | 1115 | GLIPR2 | chr9 | + | 36147785 | 36147916 | 36147785 | 36147894 | 36148546 | 36148650 | 0 | 0 | 3 | 0 | 2 | 3 | 0 | 0 | 0 | 0 | 1 | 0 | 71 | 49 | 0.999202116 | 0.999208248 | NA | 0 | 0.408 | NA | 0 | NA | 0.204 |
| A5SS | 1257 | NIPA2 | chr15 | + | 22838640 | 22838947 | 22838640 | 22838921 | 22845145 | 22845267 | 0 | 2 | 1 | 0 | 0 | 0 | 0 | 1 | 0 | 1 | 0 | 0 | 75 | 49 | 0.999202116 | 0.999208248 | NA | 1 | 1 | 0 | 1 | NA | 0.5 |
| A5SS | 1813 | NPL | chr1 | + | 182806144 | 182806531 | 182806144 | 182806232 | 182812155 | 182812213 | 0 | 0 | 10 | 0 | 0 | 0 | 0 | 0 | 2 | 0 | 2 | 0 | 98 | 49 | 0.999202116 | 0.999208248 | NA | NA | 1 | NA | 0 | 1 | 0.5 |
| A5SS | 2009 | ABTB1 | chr3 | + | 127674390 | 127674453 | 127674390 | 127674418 | 127674544 | 127674600 | 0 | 2 | 6 | 0 | 0 | 1 | 0 | 6 | 1 | 0 | 0 | 1 | 84 | 49 | 0.999202116 | 0.999208248 | NA | 1 | 0.778 | NA | 1 | 0.368 | 0.205 |
| A5SS | 2108 | DEF8 | chr16 | + | 89949416 | 89949638 | 89949416 | 89949513 | 89954242 | 89954376 | 2 | 0 | 0 | 0 | 0 | 0 | 2 | 1 | 0 | 0 | 1 | 0 | 98 | 49 | 0.999202116 | 0.999208248 | 1 | NA | NA | 1 | 0.333 | NA | 0.334 |
| A5SS | 2486 | BANP | chr16 | + | 87975047 | 87975203 | 87975047 | 87975185 | 87981035 | 87981127 | 0 | 0 | 2 | 0 | 1 | 1 | 0 | 0 | 0 | 0 | 1 | 0 | 67 | 49 | 0.999202116 | 0.999208248 | NA | 0 | 0.594 | NA | 0 | NA | 0.297 |
| A5SS | 2983 | YPEL5 | chr2 | + | 30148244 | 30148541 | 30148244 | 30148305 | 30155849 | 30155910 | 0 | 0 | 8 | 0 | 0 | 0 | 0 | 6 | 1 | 0 | 1 | 0 | 98 | 49 | 0.999202116 | 0.999208248 | NA | NA | 1 | NA | 0.75 | 1 | 0.125 |
| A5SS | 2984 | YPEL5 | chr2 | + | 30148244 | 30148541 | 30148244 | 30148305 | 30156627 | 30156792 | 0 | 0 | 8 | 0 | 0 | 0 | 0 | 6 | 1 | 0 | 1 | 0 | 98 | 49 | 0.999202116 | 0.999208248 | NA | NA | 1 | NA | 0.75 | 1 | 0.125 |
| A5SS | 2989 | NDUFAF7 | chr2 | + | 37241577 | 37241791 | 37241577 | 37241637 | 37242634 | 37242693 | 0 | 0 | 1 | 0 | 0 | 0 | 0 | 2 | 1 | 0 | 1 | 0 | 98 | 49 | 0.999202116 | 0.999208248 | NA | NA | 1 | NA | 0.5 | 1 | 0.25 |
| RI | 91 | CUTA | chr6 | - | 33417480 | 33418288 | 33417480 | 33417695 | 33417958 | 33418288 | 0 | 1 | 1 | 0 | 0 | 1 | 4 | 1 | 0 | 0 | 0 | 0 | 98 | 49 | 0.999202116 | 0.999202116 | NA | 1 | 0.333 | 1 | 1 | NA | -0.334 |
| RI | 250 | TYMP | chr22 | - | 50529495 | 50530085 | 50529495 | 50529719 | 50529903 | 50530085 | 1 | 8 | 20 | 0 | 0 | 13 | 3 | 30 | 5 | 0 | 3 | 2 | 98 | 49 | 0.999202116 | 0.999202116 | 1 | 1 | 0.435 | 1 | 0.833 | 0.556 | 0.015 |
| RI | 270 | POFUT2 | chr21 | - | 45263927 | 45267713 | 45263927 | 45265635 | 45267589 | 45267713 | 0 | 0 | 3 | 0 | 1 | 0 | 3 | 1 | 1 | 0 | 1 | 0 | 98 | 49 | 0.999202116 | 0.999202116 | NA | 0 | 1 | 1 | 0.333 | 1 | -0.278 |
| RI | 293 | SMOX | chr20 | + | 4182088 | 4182848 | 4182088 | 4182323 | 4182482 | 4182848 | 3 | 1 | 0 | 0 | 1 | 0 | 1 | 0 | 0 | 0 | 0 | 0 | 98 | 49 | 0.999202116 | 0.999202116 | 1 | 0.333 | NA | 1 | NA | NA | -0.334 |
| RI | 400 | ULK3 | chr15 | - | 74840233 | 74840641 | 74840233 | 74840316 | 74840497 | 74840641 | 0 | 2 | 1 | 0 | 0 | 3 | 0 | 2 | 0 | 0 | 0 | 0 | 98 | 49 | 0.999202116 | 0.999202116 | NA | 1 | 0.143 | NA | 1 | NA | -0.428 |
| RI | 467 | TP53 | chr17 | - | 7676381 | 7676622 | 7676381 | 7676403 | 7676520 | 7676622 | 3 | 0 | 0 | 0 | 1 | 0 | 1 | 0 | 0 | 0 | 0 | 0 | 98 | 49 | 0.999202116 | 0.999202116 | 1 | 0 | NA | 1 | NA | NA | -0.5 |
| RI | 496 | DNAJC4 | chr11 | + | 64233893 | 64234281 | 64233893 | 64233980 | 64234072 | 64234281 | 0 | 0 | 4 | 0 | 0 | 0 | 2 | 1 | 1 | 0 | 1 | 0 | 98 | 49 | 0.999202116 | 0.999202116 | NA | NA | 1 | 1 | 0.333 | 1 | 0.222 |
| RI | 510 | RAB24 | chr5 | - | 177302127 | 177302496 | 177302127 | 177302178 | 177302396 | 177302496 | 0 | 3 | 14 | 0 | 0 | 12 | 2 | 7 | 0 | 0 | 3 | 2 | 98 | 49 | 0.999202116 | 0.999202116 | NA | 1 | 0.368 | 1 | 0.538 | 0 | 0.171 |
| RI | 511 | RAB24 | chr5 | - | 177303171 | 177303744 | 177303171 | 177303320 | 177303406 | 177303744 | 2 | 6 | 10 | 0 | 2 | 1 | 1 | 3 | 23 | 0 | 1 | 2 | 98 | 49 | 0.999202116 | 0.999202116 | 1 | 0.6 | 0.833 | 1 | 0.6 | 0.852 | -0.006 |
| RI | 582 | CANT1 | chr17 | - | 78996991 | 78997963 | 78996991 | 78997644 | 78997839 | 78997963 | 2 | 0 | 8 | 0 | 0 | 0 | 0 | 3 | 3 | 0 | 1 | 0 | 98 | 49 | 0.999202116 | 0.999202116 | 1 | NA | 1 | NA | 0.6 | 1 | 0.2 |
| RI | 669 | TSTD1 | chr1 | - | 161037631 | 161038075 | 161037631 | 161037826 | 161037912 | 161038075 | 0 | 0 | 0 | 0 | 0 | 1 | 0 | 1 | 0 | 0 | 1 | 1 | 98 | 49 | 0.999202116 | 0.999202116 | NA | NA | 0 | NA | 0.333 | 0 | -0.167 |
| RI | 685 | SAMD8 | chr10 | + | 75176065 | 75182123 | 75176065 | 75176216 | 75176387 | 75182123 | 4 | 0 | 0 | 0 | 0 | 4 | 2 | 6 | 0 | 0 | 2 | 0 | 98 | 49 | 0.999202116 | 0.999202116 | 1 | NA | 0 | 1 | 0.6 | NA | -0.3 |
| RI | 818 | NCF4 | chr22 | + | 36875652 | 36876094 | 36875652 | 36875783 | 36876028 | 36876094 | 1 | 2 | 11 | 0 | 0 | 11 | 0 | 2 | 0 | 0 | 4 | 1 | 98 | 49 | 0.999202116 | 0.999202116 | 1 | 1 | 0.333 | NA | 0.2 | 0 | 0.678 |
| RI | 913 | UBXN1 | chr11 | - | 62676497 | 62677005 | 62676497 | 62676639 | 62676812 | 62677005 | 0 | 0 | 0 | 0 | 0 | 7 | 0 | 3 | 0 | 0 | 2 | 1 | 98 | 49 | 0.999202116 | 0.999202116 | NA | NA | 0 | NA | 0.429 | 0 | -0.214 |
| RI | 919 | ACADVL | chr17 | + | 7219830 | 7220197 | 7219830 | 7220046 | 7220121 | 7220197 | 0 | 2 | 1 | 0 | 2 | 3 | 0 | 0 | 2 | 0 | 2 | 0 | 98 | 49 | 0.999202116 | 0.999202116 | NA | 0.333 | 0.143 | NA | 0 | 1 | -0.262 |
| RI | 1114 | OSCAR | chr19 | - | 54094667 | 54096153 | 54094667 | 54095357 | 54095871 | 54096153 | 3 | 2 | 1 | 0 | 0 | 1 | 4 | 1 | 0 | 0 | 0 | 0 | 98 | 49 | 0.999202116 | 0.999202116 | 1 | 1 | 0.333 | 1 | 1 | NA | -0.222 |
| RI | 1116 | AUP1 | chr2 | - | 74528750 | 74529282 | 74528750 | 74528935 | 74529131 | 74529282 | 0 | 1 | 0 | 0 | 0 | 2 | 2 | 1 | 0 | 0 | 0 | 0 | 98 | 49 | 0.999202116 | 0.999202116 | NA | 1 | 0 | 1 | 1 | NA | -0.5 |
| SE | 172 | PIM2 | chrX | - | 48914394 | 48914571 | 48913181 | 48914294 | 48915019 | 48915392 | 0 | 0 | 2 | 0 | 1 | 2 | 0 | 9 | 1 | 0 | 4 | 0 | 98 | 49 | 0.999202116 | 0.999202116 | NA | 0 | 0.333 | NA | 0.529 | 1 | -0.598 |
| SE | 280 | ASB6 | chr9 | - | 129638572 | 129638659 | 129634603 | 129638457 | 129639201 | 129639310 | 0 | 0 | 1 | 0 | 0 | 0 | 0 | 2 | 1 | 0 | 1 | 0 | 98 | 49 | 0.999202116 | 0.999202116 | NA | NA | 1 | NA | 0.5 | 1 | 0.25 |
| SE | 390 | TMEM2 | chr9 | - | 71734805 | 71734994 | 71732350 | 71732520 | 71740062 | 71740232 | 0 | 0 | 6 | 0 | 0 | 0 | 0 | 4 | 1 | 0 | 1 | 0 | 98 | 49 | 0.999202116 | 0.999202116 | NA | NA | 1 | NA | 0.667 | 1 | 0.166 |
| SE | 626 | LYN | chr8 | + | 55941854 | 55941928 | 55879826 | 55880103 | 55946447 | 55946493 | 0 | 2 | 9 | 0 | 0 | 0 | 0 | 13 | 0 | 0 | 0 | 1 | 98 | 49 | 0.999202116 | 0.999202116 | NA | 1 | 1 | NA | 1 | 0 | 0.5 |
| SE | 627 | LYN | chr8 | + | 55941854 | 55941991 | 55879826 | 55880103 | 55946447 | 55946493 | 0 | 2 | 9 | 0 | 0 | 0 | 0 | 11 | 0 | 0 | 0 | 1 | 98 | 49 | 0.999202116 | 0.999202116 | NA | 1 | 1 | NA | 1 | 0 | 0.5 |
| SE | 694 | DOCK5 | chr8 | + | 25342400 | 25342507 | 25341738 | 25341809 | 25345474 | 25345611 | 0 | 0 | 15 | 0 | 0 | 0 | 0 | 15 | 2 | 0 | 1 | 0 | 98 | 49 | 0.999202116 | 0.999202116 | NA | NA | 1 | NA | 0.882 | 1 | 0.059 |
| SE | 697 | DOCK5 | chr8 | + | 25407982 | 25408154 | 25403557 | 25403724 | 25408801 | 25408940 | 0 | 1 | 23 | 0 | 1 | 1 | 0 | 9 | 3 | 0 | 3 | 0 | 98 | 49 | 0.999202116 | 0.999202116 | NA | 0.333 | 0.92 | NA | 0.6 | 1 | -0.173 |
| SE | 789 | PARP10 | chr8 | - | 143985411 | 143985648 | 143984543 | 143985328 | 143985720 | 143985975 | 2 | 3 | 2 | 0 | 0 | 1 | 0 | 6 | 0 | 1 | 0 | 0 | 98 | 49 | 0.999202116 | 0.999202116 | 1 | 1 | 0.5 | 0 | 1 | NA | 0.333 |
| SE | 925 | SAMD9L | chr7 | - | 93146882 | 93147146 | 93145384 | 93146040 | 93148193 | 93148388 | 0 | 2 | 5 | 0 | 0 | 1 | 0 | 1 | 0 | 0 | 0 | 0 | 98 | 49 | 0.999202116 | 0.999202116 | NA | 1 | 0.714 | NA | 1 | NA | -0.143 |
| SE | 926 | SAMD9L | chr7 | - | 93147087 | 93147146 | 93145384 | 93146040 | 93148193 | 93148388 | 0 | 1 | 7 | 0 | 0 | 1 | 0 | 1 | 0 | 0 | 0 | 0 | 98 | 49 | 0.999202116 | 0.999202116 | NA | 1 | 0.778 | NA | 1 | NA | -0.111 |
| SE | 1015 | IKZF1 | chr7 | + | 50387344 | 50387470 | 50382539 | 50382707 | 50391728 | 50391833 | 0 | 2 | 10 | 0 | 0 | 1 | 0 | 1 | 0 | 0 | 0 | 0 | 98 | 49 | 0.999202116 | 0.999202116 | NA | 1 | 0.833 | NA | 1 | NA | -0.084 |
| SE | 1135 | IGF2R | chr6 | + | 160045744 | 160045882 | 160044513 | 160044657 | 160046497 | 160046645 | 0 | 0 | 35 | 0 | 1 | 1 | 3 | 14 | 1 | 0 | 1 | 0 | 98 | 49 | 0.999202116 | 0.999202116 | NA | 0 | 0.946 | 1 | 0.875 | 1 | -0.485 |
| SE | 1190 | PHACTR2 | chr6 | + | 143765260 | 143765798 | 143753753 | 143753912 | 143772257 | 143772457 | 0 | 1 | 4 | 0 | 0 | 1 | 0 | 1 | 0 | 0 | 0 | 0 | 98 | 49 | 0.999202116 | 0.999202116 | NA | 1 | 0.667 | NA | 1 | NA | -0.166 |
| SE | 1209 | VNN2 | chr6 | - | 132752460 | 132752749 | 132751144 | 132751518 | 132755842 | 132756035 | 0 | 0 | 32 | 0 | 0 | 0 | 0 | 18 | 1 | 0 | 5 | 0 | 98 | 49 | 0.999202116 | 0.999202116 | NA | NA | 1 | NA | 0.643 | 1 | 0.178 |
| SE | 1212 | VNN2 | chr6 | - | 132753417 | 132753491 | 132751144 | 132751518 | 132755842 | 132756035 | 0 | 0 | 3 | 0 | 0 | 0 | 0 | 1 | 1 | 0 | 5 | 0 | 98 | 49 | 0.999202116 | 0.999202116 | NA | NA | 1 | NA | 0.091 | 1 | 0.455 |
| SE | 1508 | HLA-E | chr6 | + | 30492403 | 30492436 | 30491536 | 30491653 | 30492540 | 30492583 | 0 | 26 | 172 | 0 | 0 | 1 | 10 | 116 | 23 | 0 | 1 | 1 | 82 | 49 | 0.999202116 | 0.999202116 | NA | 1 | 0.99 | 1 | 0.986 | 0.932 | 0.022 |
| SE | 1592 | KIF13A | chr6 | - | 17773477 | 17773583 | 17771907 | 17772059 | 17775014 | 17775062 | 0 | 0 | 5 | 0 | 0 | 0 | 2 | 7 | 1 | 0 | 2 | 0 | 98 | 49 | 0.999202116 | 0.999202116 | NA | NA | 1 | 1 | 0.636 | 1 | 0.121 |
| SE | 1593 | KIF13A | chr6 | - | 17789871 | 17789910 | 17787775 | 17787875 | 17794248 | 17794395 | 0 | 0 | 1 | 0 | 2 | 1 | 0 | 0 | 0 | 0 | 2 | 0 | 88 | 49 | 0.999202116 | 0.999202116 | NA | 0 | 0.358 | NA | 0 | NA | 0.179 |
| SE | 1709 | NCF1 | chr7 | + | 74779081 | 74779157 | 74777266 | 74777347 | 74779256 | 74779422 | 0 | 0 | 4 | 0 | 0 | 0 | 2 | 0 | 1 | 0 | 1 | 0 | 98 | 49 | 0.999202116 | 0.999202116 | NA | NA | 1 | 1 | 0 | 1 | 0.333 |
| SE | 1738 | ARHGAP26 | chr5 | + | 143037195 | 143037261 | 143014079 | 143014116 | 143041815 | 143041890 | 0 | 0 | 32 | 0 | 0 | 0 | 0 | 12 | 2 | 0 | 1 | 0 | 98 | 49 | 0.999202116 | 0.999202116 | NA | NA | 1 | NA | 0.857 | 1 | 0.072 |
| SE | 1779 | ASPH | chr8 | - | 61553030 | 61553120 | 61548070 | 61548208 | 61555923 | 61556022 | 0 | 2 | 4 | 0 | 0 | 1 | 0 | 1 | 0 | 0 | 0 | 0 | 98 | 49 | 0.999202116 | 0.999202116 | NA | 1 | 0.667 | NA | 1 | NA | -0.166 |
| SE | 1782 | ASPH | chr8 | - | 61682438 | 61682483 | 61680967 | 61681036 | 61684038 | 61684188 | 0 | 0 | 0 | 0 | 0 | 3 | 0 | 1 | 0 | 0 | 5 | 1 | 94 | 49 | 0.999202116 | 0.999202116 | NA | NA | 0 | NA | 0.094 | 0 | -0.047 |
| SE | 1800 | AP3S1 | chr5 | + | 115902884 | 115902992 | 115895086 | 115895158 | 115913361 | 115914081 | 1 | 0 | 0 | 0 | 0 | 0 | 1 | 1 | 0 | 0 | 1 | 0 | 98 | 49 | 0.999202116 | 0.999202116 | 1 | NA | NA | 1 | 0.333 | NA | 0.334 |
| SE | 1920 | PARP8 | chr5 | + | 50794206 | 50794332 | 50788522 | 50788589 | 50794852 | 50795417 | 0 | 0 | 34 | 0 | 0 | 0 | 7 | 9 | 1 | 0 | 2 | 0 | 98 | 49 | 0.999202116 | 0.999202116 | NA | NA | 1 | 1 | 0.692 | 1 | 0.103 |
| SE | 1987 | DDX60 | chr4 | - | 168283445 | 168283606 | 168280334 | 168280590 | 168284819 | 168284935 | 0 | 0 | 18 | 0 | 0 | 0 | 0 | 9 | 1 | 0 | 2 | 0 | 98 | 49 | 0.999202116 | 0.999202116 | NA | NA | 1 | NA | 0.692 | 1 | 0.154 |
| SE | 2104 | WDFY3 | chr4 | - | 84932269 | 84932363 | 84896910 | 84897010 | 84966208 | 84966391 | 0 | 2 | 13 | 1 | 0 | 0 | 1 | 13 | 9 | 0 | 0 | 5 | 98 | 49 | 0.999202116 | 0.999202116 | 0 | 1 | 1 | 1 | 1 | 0.474 | -0.158 |

| | | | | | | | | | | | | | | | | | | | | | | | | | | | | | | | | | |
|---|---|---|---|---|---|---|---|---|---|---|---|---|---|---|---|---|---|---|---|---|---|---|---|---|---|---|---|---|---|---|---|---|---|
| SE | 2273 | VNN3 | chr6 | - | 132726721 | 132727012 | 132724605 | 132724979 | 132728740 | 132728933 | 0 | 0 | 11 | 0 | 1 | 1 | 0 | 6 | 3 | 0 | 7 | 1 | 98 | 49 | 0.999202116 | 0.999202116 | NA | 0 | 0.846 | NA | 0.3 | 0.6 | -0.027 |
| SE | 2370 | MFSD1 | chr3 | + | 158805361 | 158805474 | 158804318 | 158804371 | 158807039 | 158807082 | 0 | 0 | 10 | 0 | 0 | 0 | 0 | 2 | 2 | 0 | 1 | 0 | 98 | 49 | 0.999202116 | 0.999202116 | NA | NA | 1 | NA | 0.5 | 1 | 0.25 |
| SE | 2467 | HMGA1 | chr6 | + | 34242711 | 34242795 | 34240736 | 34240915 | 34243467 | 34243518 | 0 | 1 | 3 | 0 | 0 | 0 | 0 | 2 | 0 | 0 | 0 | 1 | 98 | 49 | 0.999202116 | 0.999202116 | NA | 1 | 1 | NA | 1 | 0 | 0.5 |
| SE | 2582 | PRKCD | chr3 | + | 53185926 | 53186027 | 53185603 | 53185700 | 53186166 | 53186340 | 0 | 3 | 4 | 0 | 0 | 4 | 0 | 3 | 0 | 0 | 0 | 0 | 98 | 49 | 0.999202116 | 0.999202116 | NA | 1 | 0.333 | NA | 1 | NA | -0.334 |
| SE | 2584 | LCP2 | chr5 | - | 170275319 | 170275351 | 170274300 | 170274338 | 170275794 | 170275860 | 0 | 0 | 23 | 0 | 0 | 0 | 0 | 16 | 3 | 0 | 1 | 0 | 81 | 49 | 0.999202116 | 0.999202116 | NA | NA | 1 | NA | 0.906 | 1 | 0.047 |
| SE | 2714 | TOP2B | chr3 | - | 25624681 | 25624803 | 25624296 | 25624445 | 25626559 | 25626674 | 0 | 0 | 2 | 0 | 0 | 0 | 0 | 1 | 1 | 0 | 1 | 0 | 98 | 49 | 0.999202116 | 0.999202116 | NA | NA | 1 | NA | 0.333 | 1 | 0.334 |
| SE | 2769 | LST1 | chr6 | + | 31587943 | 31587966 | 31587640 | 31587733 | 31588517 | 31588909 | 0 | 0 | 6 | 0 | 0 | 0 | 0 | 0 | 1 | 0 | 1 | 0 | 72 | 49 | 0.999202116 | 0.999202116 | NA | NA | 1 | NA | 0 | 1 | 0.5 |
| SE | 2804 | ARPC4 | chr3 | + | 9801660 | 9801756 | 9800184 | 9800296 | 9803842 | 9804013 | 0 | 3 | 21 | 0 | 0 | 1 | 2 | 29 | 7 | 0 | 0 | 2 | 98 | 49 | 0.999202116 | 0.999202116 | NA | 1 | 0.913 | 1 | 1 | 0.636 | 0.078 |
| SE | 3096 | MX1 | chr21 | + | 41430532 | 41430608 | 41427754 | 41427866 | 41432049 | 41432175 | 0 | 6 | 38 | 0 | 0 | 2 | 0 | 39 | 1 | 0 | 2 | 0 | 98 | 49 | 0.999202116 | 0.999202116 | NA | 1 | 0.905 | NA | 0.907 | 1 | -0.001 |
| SE | 3098 | MX1 | chr21 | + | 41439693 | 41439848 | 41437014 | 41437152 | 41440886 | 41441025 | 0 | 6 | 103 | 0 | 1 | 0 | 4 | 60 | 7 | 0 | 1 | 0 | 98 | 49 | 0.999202116 | 0.999202116 | NA | 0.75 | 1 | 1 | 0.968 | 1 | -0.114 |
| SE | 3100 | MX1 | chr21 | + | 41449136 | 41449295 | 41445999 | 41446141 | 41451166 | 41451243 | 0 | 0 | 30 | 0 | 0 | 0 | 0 | 11 | 1 | 0 | 2 | 0 | 98 | 49 | 0.999202116 | 0.999202116 | NA | NA | 1 | NA | 0.733 | 1 | 0.133 |
| SE | 3104 | MX2 | chr21 | + | 41390564 | 41390703 | 41382409 | 41382564 | 41395586 | 41395785 | 0 | 4 | 23 | 0 | 0 | 13 | 0 | 13 | 0 | 0 | 14 | 1 | 98 | 49 | 0.999202116 | 0.999202116 | NA | 1 | 0.469 | NA | 0.317 | 0 | 0.576 |
| SE | 3172 | SAMSN1 | chr21 | - | 14500528 | 14500735 | 14498441 | 14498592 | 14510309 | 14510461 | 0 | 0 | 43 | 0 | 0 | 0 | 0 | 28 | 2 | 0 | 1 | 0 | 98 | 49 | 0.999202116 | 0.999202116 | NA | NA | 1 | NA | 0.933 | 1 | 0.033 |
| SE | 3173 | SAMSN1 | chr21 | - | 14516891 | 14517041 | 14512443 | 14512573 | 14521149 | 14521221 | 0 | 1 | 12 | 0 | 1 | 3 | 0 | 16 | 2 | 0 | 5 | 0 | 98 | 49 | 0.999202116 | 0.999202116 | NA | 0.333 | 0.667 | NA | 0.615 | 1 | -0.307 |
| SE | 3194 | HELZ2 | chr20 | - | 63562071 | 63562203 | 63561822 | 63561984 | 63562287 | 63562382 | 0 | 0 | 11 | 0 | 0 | 0 | 0 | 6 | 2 | 0 | 2 | 0 | 98 | 49 | 0.999202116 | 0.999202116 | NA | NA | 1 | NA | 0.6 | 1 | 0.2 |
| SE | 3200 | TNIP1 | chr5 | - | 151064959 | 151065127 | 151063612 | 151063747 | 151080879 | 151081088 | 0 | 0 | 4 | 0 | 1 | 2 | 0 | 3 | 1 | 0 | 1 | 0 | 98 | 49 | 0.999202116 | 0.999202116 | NA | 0 | 0.5 | NA | 0.6 | 1 | -0.55 |
| SE | 3201 | TNIP1 | chr5 | - | 151064959 | 151065131 | 151063612 | 151063747 | 151080879 | 151081084 | 0 | 0 | 5 | 0 | 1 | 2 | 0 | 9 | 1 | 0 | 1 | 0 | 98 | 49 | 0.999202116 | 0.999202116 | NA | 0 | 0.556 | NA | 0.818 | 1 | -0.631 |
| SE | 3238 | NUB1 | chr7 | + | 151360140 | 151360247 | 151356127 | 151356222 | 151366938 | 151367125 | 0 | 0 | 3 | 0 | 0 | 0 | 0 | 6 | 1 | 0 | 2 | 0 | 98 | 49 | 0.999202116 | 0.999202116 | NA | NA | 1 | NA | 0.6 | 1 | 0.2 |
| SE | 3247 | SULF2 | chr20 | - | 47702518 | 47702670 | 47690125 | 47690295 | 47736702 | 47736942 | 0 | 1 | 3 | 0 | 0 | 1 | 0 | 1 | 0 | 0 | 0 | 0 | 98 | 49 | 0.999202116 | 0.999202116 | NA | 1 | 0.6 | NA | 1 | NA | -0.2 |
| SE | 3384 | HM13 | chr20 | + | 31549206 | 31549332 | 31549028 | 31549114 | 31550063 | 31550121 | 0 | 3 | 9 | 0 | 0 | 0 | 0 | 3 | 0 | 7 | 0 | 0 | 98 | 49 | 0.999202116 | 0.999202116 | NA | 1 | 1 | 0 | 1 | NA | 0.5 |
| SE | 3452 | GPCPD1 | chr20 | - | 5561464 | 5561530 | 5559939 | 5560076 | 5565016 | 5565078 | 0 | 0 | 22 | 0 | 0 | 0 | 0 | 17 | 2 | 0 | 1 | 0 | 98 | 49 | 0.999202116 | 0.999202116 | NA | NA | 1 | NA | 0.895 | 1 | 0.052 |
| SE | 3613 | NIN | chr14 | - | 50739307 | 50739487 | 50738139 | 50738286 | 50741581 | 50741728 | 0 | 0 | 7 | 0 | 0 | 0 | 4 | 10 | 1 | 0 | 1 | 0 | 98 | 49 | 0.999202116 | 0.999202116 | NA | NA | 1 | 1 | 0.833 | 1 | 0.056 |
| SE | 3635 | USP4 | chr3 | - | 49325718 | 49325845 | 49324893 | 49325039 | 49327685 | 49327816 | 0 | 0 | 17 | 0 | 0 | 0 | 0 | 13 | 3 | 0 | 1 | 0 | 98 | 49 | 0.999202116 | 0.999202116 | NA | NA | 1 | NA | 0.867 | 1 | 0.067 |
| SE | 3660 | ETS2 | chr21 | + | 38810034 | 38810106 | 38805830 | 38806120 | 38813002 | 38813114 | 0 | 0 | 13 | 0 | 0 | 0 | 0 | 10 | 3 | 0 | 2 | 0 | 98 | 49 | 0.999202116 | 0.999202116 | NA | NA | 1 | NA | 0.714 | 1 | 0.143 |
| SE | 3722 | FCHO1 | chr19 | + | 17781230 | 17781343 | 17778608 | 17778884 | 17781451 | 17781539 | 0 | 1 | 3 | 0 | 0 | 3 | 0 | 1 | 0 | 0 | 0 | 0 | 98 | 49 | 0.999202116 | 0.999202116 | NA | 1 | 0.333 | NA | 1 | NA | -0.334 |
| SE | 3749 | STX3 | chr11 | + | 59787036 | 59787136 | 59773210 | 59773294 | 59788872 | 59788947 | 0 | 0 | 14 | 0 | 0 | 0 | 0 | 7 | 2 | 0 | 3 | 0 | 98 | 49 | 0.999202116 | 0.999202116 | NA | NA | 1 | NA | 0.538 | 1 | 0.231 |
| SE | 3890 | MAP7D1 | chr1 | + | 36179261 | 36179315 | 36178920 | 36179025 | 36179511 | 36179557 | 0 | 0 | 9 | 0 | 0 | 0 | 5 | 8 | 1 | 0 | 2 | 0 | 98 | 49 | 0.999202116 | 0.999202116 | NA | NA | 1 | 1 | 0.667 | 1 | 0.111 |
| SE | 3983 | NCOR2 | chr12 | - | 124535564 | 124535611 | 124495146 | 124495368 | 124567307 | 124567464 | 3 | 0 | 3 | 1 | 0 | 0 | 0 | 0 | 1 | 0 | 0 | 0 | 96 | 49 | 0.999202116 | 0.999202116 | 0.605 | NA | 1 | NA | NA | 1 | -0.198 |
| SE | 4021 | LINC01578 | chr15 | + | 92883584 | 92883778 | 92882842 | 92883186 | 92885514 | 92885584 | 0 | 0 | 2 | 0 | 0 | 0 | 0 | 2 | 2 | 0 | 4 | 0 | 98 | 49 | 0.999202116 | 0.999202116 | NA | NA | 1 | NA | 0.2 | 1 | 0.4 |
| SE | 4022 | LINC01578 | chr15 | + | 92885514 | 92885584 | 92882842 | 92883186 | 92886168 | 92886254 | 0 | 1 | 13 | 0 | 0 | 0 | 1 | 14 | 0 | 0 | 0 | 1 | 98 | 49 | 0.999202116 | 0.999202116 | NA | 1 | 1 | 1 | 1 | 0 | 0.333 |
| SE | 4074 | IL1RN | chr2 | + | 113131044 | 113131157 | 113129575 | 113129664 | 113132655 | 113134016 | 0 | 0 | 12 | 0 | 0 | 0 | 0 | 23 | 2 | 0 | 1 | 0 | 98 | 49 | 0.999202116 | 0.999202116 | NA | NA | 1 | NA | 0.92 | 1 | 0.04 |
| SE | 4121 | XPO6 | chr16 | - | 28111816 | 28112006 | 28107521 | 28107677 | 28112903 | 28113050 | 0 | 2 | 40 | 0 | 1 | 1 | 1 | 30 | 3 | 0 | 2 | 0 | 98 | 49 | 0.999202116 | 0.999202116 | NA | 0.5 | 0.952 | 1 | 0.882 | 1 | -0.235 |
| SE | 4123 | XPO6 | chr16 | - | 28121669 | 28121762 | 28117317 | 28117462 | 28125688 | 28125848 | 0 | 6 | 79 | 0 | 0 | 0 | 15 | 61 | 0 | 0 | 0 | 1 | 98 | 49 | 0.999202116 | 0.999202116 | NA | 1 | 1 | 1 | 1 | 0 | 0.333 |
| SE | 4126 | XPO6 | chr16 | - | 28208986 | 28209122 | 28180940 | 28181031 | 28211365 | 28211918 | 0 | 1 | 1 | 0 | 0 | 17 | 0 | 3 | 0 | 3 | 6 | 4 | 98 | 49 | 0.999202116 | 0.999202116 | NA | 1 | 0.029 | 0 | 0.2 | 0 | 0.448 |
| SE | 4141 | TBC1D15 | chr12 | + | 71884810 | 71885021 | 71880468 | 71880607 | 71893221 | 71893324 | 0 | 0 | 3 | 0 | 0 | 0 | 0 | 6 | 1 | 0 | 1 | 0 | 98 | 49 | 0.999202116 | 0.999202116 | NA | NA | 1 | NA | 0.75 | 1 | 0.125 |
| SE | 4170 | RNF10 | chr12 | + | 120571190 | 120571291 | 120566824 | 120566980 | 120575630 | 120575688 | 0 | 3 | 6 | 0 | 0 | 1 | 0 | 3 | 7 | 0 | 6 | 0 | 98 | 49 | 0.999202116 | 0.999202116 | NA | 1 | 0.75 | NA | 0.2 | 1 | 0.275 |
| SE | 4240 | SRCAP | chr16 | + | 30712678 | 30712815 | 30712261 | 30712439 | 30713207 | 30713377 | 0 | 1 | 5 | 0 | 0 | 1 | 0 | 1 | 0 | 0 | 0 | 0 | 98 | 49 | 0.999202116 | 0.999202116 | NA | 1 | 0.714 | NA | 1 | NA | -0.143 |
| SE | 4317 | BIRC6 | chr2 | + | 32392038 | 32392150 | 32388749 | 32388943 | 32395510 | 32395593 | 0 | 0 | 5 | 0 | 0 | 0 | 0 | 2 | 1 | 0 | 1 | 0 | 98 | 49 | 0.999202116 | 0.999202116 | NA | NA | 1 | NA | 0.5 | 1 | 0.25 |
| SE | 4325 | KMT2D | chr12 | - | 49028827 | 49028958 | 49028008 | 49028141 | 49029060 | 49029236 | 0 | 4 | 7 | 0 | 3 | 0 | 0 | 3 | 0 | 0 | 0 | 1 | 98 | 49 | 0.999202116 | 0.999202116 | NA | 0.4 | 1 | NA | 1 | 0 | 0.2 |
| SE | 4422 | PREX1 | chr20 | - | 48644408 | 48644497 | 48642406 | 48642489 | 48645850 | 48646057 | 0 | 5 | 22 | 0 | 0 | 5 | 7 | 18 | 1 | 0 | 3 | 0 | 98 | 49 | 0.999202116 | 0.999202116 | NA | 1 | 0.687 | 1 | 0.75 | 1 | -0.073 |
| SE | 4443 | CORO7 | chr16 | - | 4364596 | 4364689 | 4364275 | 4364413 | 4364774 | 4364920 | 0 | 0 | 8 | 0 | 0 | 0 | 0 | 1 | 3 | 0 | 1 | 0 | 98 | 49 | 0.999202116 | 0.999202116 | NA | NA | 1 | NA | 0.333 | 1 | 0.334 |
| SE | 4504 | PTPN6 | chr12 | + | 6957918 | 6958073 | 6957653 | 6957785 | 6959926 | 6959994 | 0 | 4 | 43 | 0 | 4 | 3 | 2 | 10 | 3 | 0 | 11 | 0 | 98 | 49 | 0.999202116 | 0.999202116 | NA | 0.333 | 0.878 | 1 | 0.312 | 1 | -0.165 |
| SE | 4520 | CHD4 | chr12 | - | 6600925 | 6601053 | 6600533 | 6600669 | 6601288 | 6601530 | 0 | 0 | 7 | 0 | 0 | 0 | 0 | 4 | 1 | 0 | 1 | 0 | 98 | 49 | 0.999202116 | 0.999202116 | NA | NA | 1 | NA | 0.667 | 1 | 0.166 |
| SE | 4534 | TM9SF2 | chr13 | + | 99537738 | 99537863 | 99536607 | 99536737 | 99540713 | 99540793 | 0 | 0 | 4 | 0 | 0 | 0 | 0 | 1 | 1 | 1 | 0 | 0 | 98 | 49 | 0.999202116 | 0.999202116 | NA | NA | 1 | 0 | 1 | 1 | 0.333 |
| SE | 4589 | PLBD1 | chr12 | - | 14553192 | 14553412 | 14542207 | 14542291 | 14567581 | 14567857 | 0 | 2 | 33 | 0 | 0 | 1 | 1 | 1 | 0 | 0 | 0 | 0 | 98 | 49 | 0.999202116 | 0.999202116 | NA | 1 | 0.943 | 1 | 1 | NA | -0.028 |
| SE | 4617 | PTPRJ | chr11 | + | 48144999 | 48145124 | 48144674 | 48144885 | 48146875 | 48146963 | 0 | 11 | 9 | 0 | 0 | 1 | 0 | 1 | 0 | 0 | 0 | 0 | 98 | 49 | 0.999202116 | 0.999202116 | NA | 1 | 0.818 | NA | 1 | NA | -0.091 |
| SE | 4673 | IL10RA | chr11 | + | 117989441 | 117989620 | 117988381 | 117988502 | 117993240 | 117993410 | 0 | 2 | 14 | 0 | 0 | 1 | 0 | 2 | 0 | 0 | 0 | 0 | 98 | 49 | 0.999202116 | 0.999202116 | NA | 1 | 0.875 | NA | 1 | NA | -0.062 |
| SE | 4773 | DDX6 | chr11 | - | 118755401 | 118755503 | 118754704 | 118754887 | 118756259 | 118756323 | 0 | 0 | 12 | 0 | 0 | 0 | 1 | 22 | 1 | 0 | 1 | 0 | 98 | 49 | 0.999202116 | 0.999202116 | NA | NA | 1 | 1 | 0.917 | 1 | 0.028 |
| SE | 4884 | DCAF11 | chr14 | + | 24117265 | 24117393 | 24116916 | 24117044 | 24117667 | 24117732 | 0 | 0 | 4 | 0 | 0 | 0 | 0 | 1 | 2 | 0 | 1 | 0 | 98 | 49 | 0.999202116 | 0.999202116 | NA | NA | 1 | NA | 0.333 | 1 | 0.334 |
| SE | 4939 | ARAP1 | chr11 | - | 72686041 | 72686191 | 72685068 | 72685681 | 72687687 | 72687738 | 0 | 0 | 13 | 0 | 0 | 0 | 0 | 9 | 1 | 0 | 1 | 0 | 98 | 49 | 0.999202116 | 0.999202116 | NA | NA | 1 | NA | 0.818 | 1 | 0.091 |
| SE | 4943 | ARAP1 | chr11 | - | 72697322 | 72697486 | 72696982 | 72697195 | 72697597 | 72697649 | 0 | 2 | 24 | 0 | 1 | 0 | 0 | 16 | 3 | 0 | 1 | 0 | 98 | 49 | 0.999202116 | 0.999202116 | NA | 0.5 | 1 | NA | 0.889 | 1 | -0.195 |
| SE | 4945 | ARAP1 | chr11 | - | 72704151 | 72704334 | 72702904 | 72703079 | 72705804 | 72705890 | 0 | 0 | 25 | 0 | 0 | 0 | 0 | 14 | 10 | 0 | 3 | 0 | 98 | 49 | 0.999202116 | 0.999202116 | NA | NA | 1 | NA | 0.7 | 1 | 0.15 |
| SE | 4947 | ARAP1 | chr11 | - | 72711020 | 72711141 | 72710384 | 72710587 | 72711429 | 72711499 | 0 | 4 | 21 | 0 | 0 | 2 | 0 | 13 | 4 | 0 | 1 | 2 | 98 | 49 | 0.999202116 | 0.999202116 | NA | 1 | 0.84 | NA | 0.867 | 0.5 | 0.236 |
| SE | 4984 | NCOR1 | chr17 | - | 16067893 | 16068121 | 16065484 | 16065694 | 16070164 | 16070525 | 0 | 1 | 2 | 0 | 0 | 1 | 3 | 1 | 0 | 0 | 0 | 0 | 98 | 49 | 0.999202116 | 0.999202116 | NA | 1 | 0.5 | 1 | 1 | NA | -0.25 |
| SE | 5186 | MYO9B | chr19 | + | 17172758 | 17172963 | 17172335 | 17172477 | 17175662 | 17175741 | 0 | 0 | 8 | 0 | 0 | 0 | 0 | 10 | 1 | 0 | 1 | 0 | 98 | 49 | 0.999202116 | 0.999202116 | NA | NA | 1 | NA | 0.833 | 1 | 0.084 |
| SE | 5369 | BAZ1A | chr14 | - | 34862043 | 34862322 | 34826012 | 34826156 | 34874491 | 34874662 | 0 | 0 | 20 | 0 | 0 | 0 | 0 | 4 | 2 | 2 | 1 | 0 | 98 | 49 | 0.999202116 | 0.999202116 | NA | NA | 1 | 0 | 0.667 | 1 | 0.444 |
| SE | 5425 | PRMT3 | chr11 | + | 20397576 | 20397721 | 20395802 | 20395962 | 20402918 | 20402984 | 0 | 3 | 0 | 0 | 0 | 1 | 0 | 1 | 0 | 0 | 0 | 0 | 98 | 49 | 0.999202116 | 0.999202116 | NA | 1 | 0 | NA | 1 | NA | -0.5 |
| SE | 5575 | SAT1 | chrX | + | 23783799 | 23783883 | 23783657 | 23783709 | 23785327 | 23785429 | 0 | 20 | 265 | 0 | 1 | 1 | 2 | 216 | 10 | 0 | 2 | 0 | 98 | 49 | 0.999202116 | 0.999202116 | NA | 0.909 | 0.993 | 1 | 0.982 | 1 | -0.043 |
| SE | 5615 | DOPEY2 | chr21 | + | 36211552 | 36211651 | 36208714 | 36208904 | 36211973 | 36212097 | 0 | 2 | 1 | 0 | 0 | 1 | 0 | 1 | 0 | 0 | 0 | 0 | 98 | 49 | 0.999202116 | 0.999202116 | NA | 1 | 0.333 | NA | 1 | NA | -0.334 |
| SE | 5775 | NCOA2 | chr8 | - | 70213902 | 70214075 | 70174755 | 70174859 | 70216659 | 70216764 | 0 | 0 | 13 | 0 | 0 | 0 | 0 | 18 | 2 | 0 | 2 | 0 | 98 | 49 | 0.999202116 | 0.999202116 | NA | NA | 1 | NA | 0.818 | 1 | 0.091 |
| SE | 5787 | DENND3 | chr8 | + | 141160631 | 141160787 | 141155848 | 141155970 | 141163332 | 141163429 | 0 | 0 | 19 | 0 | 0 | 0 | 0 | 6 | 2 | 0 | 2 | 0 | 98 | 49 | 0.999202116 | 0.999202116 | NA | NA | 1 | NA | 0.6 | 1 | 0.2 |
| SE | 5854 | ITPK1 | chr14 | - | 92993879 | 92993997 | 92962750 | 92962849 | 93016675 | 93016801 | 0 | 2 | 10 | 0 | 0 | 2 | 0 | 2 | 0 | 0 | 0 | 0 | 98 | 49 | 0.999202116 | 0.999202116 | NA | 1 | 0.714 | NA | 1 | NA | -0.143 |
| SE | 5890 | MSN | chrX | + | 65738524 | 65738617 | 65737177 | 65737338 | 65738969 | 65739194 | 0 | 3 | 22 | 0 | 0 | 1 | 4 | 14 | 2 | 0 | 0 | 2 | 98 | 49 | 0.999202116 | 0.999202116 | NA | 1 | 0.917 | 1 | 1 | 0.333 | 0.181 |
| SE | 5968 | CD53 | chr1 | + | 110897808 | 110897892 | 110896652 | 110896733 | 110899123 | 110899936 | 0 | 6 | 27 | 0 | 0 | 0 | 0 | 13 | 0 | 0 | 0 | 1 | 98 | 49 | 0.999202116 | 0.999202116 | NA | 1 | 1 | NA | 1 | 0 | 0.5 |
| SE | 6104 | GLUL | chr1 | - | 182387130 | 182387292 | 182386255 | 182386402 | 182388571 | 182388750 | 0 | 25 | 173 | 0 | 0 | 3 | 4 | 163 | 7 | 0 | 1 | 0 | 98 | 49 | 0.999202116 | 0.999202116 | NA | 1 | 0.966 | 1 | 0.988 | 1 | -0.013 |
| SE | 6179 | ABCA1 | chr9 | - | 104832573 | 104832771 | 104831621 | 104831827 | 104836979 | 104837096 | 0 | 6 | 13 | 0 | 0 | 0 | 0 | 43 | 0 | 0 | 0 | 1 | 98 | 49 | 0.999202116 | 0.999202116 | NA | 1 | 1 | NA | 1 | 0 | 0.5 |
| SE | 6235 | NFKB2 | chr10 | + | 102398384 | 102398523 | 102398211 | 102398297 | 102398738 | 102398864 | 0 | 0 | 7 | 0 | 0 | 0 | 0 | 8 | 1 | 0 | 1 | 0 | 98 | 49 | 0.999202116 | 0.999202116 | NA | NA | 1 | NA | 0.8 | 1 | 0.1 |
| SE | 6236 | RAB8B | chr15 | + | 63249644 | 63249705 | 63244755 | 63244816 | 63255506 | 63255584 | 0 | 0 | 8 | 0 | 0 | 0 | 4 | 8 | 1 | 0 | 1 | 0 | 98 | 49 | 0.999202116 | 0.999202116 | NA | NA | 1 | 1 | 0.8 | 1 | 0.067 |
| SE | 6273 | BAZ2B | chr2 | - | 159448241 | 159448409 | 159446781 | 159446975 | 159478574 | 159478721 | 0 | 0 | 14 | 0 | 0 | 0 | 0 | 3 | 2 | 0 | 2 | 0 | 98 | 49 | 0.999202116 | 0.999202116 | NA | NA | 1 | NA | 0.429 | 1 | 0.285 |
| SE | 6483 | PADI2 | chr1 | - | 17083725 | 17083837 | 17082544 | 17082652 | 17084598 | 17084702 | 0 | 0 | 10 | 0 | 0 | 0 | 0 | 3 | 1 | 0 | 1 | 0 | 98 | 49 | 0.999202116 | 0.999202116 | NA | NA | 1 | NA | 0.6 | 1 | 0.2 |
| SE | 6542 | NRD1 | chr1 | - | 51790532 | 51790649 | 51789567 | 51789657 | 51790899 | 51790990 | 0 | 0 | 17 | 0 | 0 | 0 | 0 | 6 | 1 | 0 | 1 | 0 | 98 | 49 | 0.999202116 | 0.999202116 | NA | NA | 1 | NA | 0.75 | 1 | 0.125 |
| SE | 6573 | CHD8 | chr14 | - | 21393105 | 21393254 | 21392506 | 21392809 | 21393475 | 21394195 | 0 | 2 | 2 | 0 | 0 | 1 | 0 | 2 | 0 | 0 | 0 | 0 | 98 | 49 | 0.999202116 | 0.999202116 | NA | 1 | 0.5 | NA | 1 | NA | -0.25 |
| SE | 6607 | QSOX1 | chr1 | + | 180194212 | 180194392 | 180190432 | 180190580 | 180196261 | 180196599 | 0 | 1 | 3 | 0 | 0 | 1 | 0 | 4 | 0 | 0 | 1 | 1 | 98 | 49 | 0.999202116 | 0.999202116 | NA | 1 | 0.6 | NA | 0.667 | 0 | 0.467 |
| SE | 6616 | AMPD2 | chr1 | + | 109625661 | 109625792 | 109619812 | 109620278 | 109626159 | 109626228 | 0 | 2 | 10 | 0 | 0 | 1 | 0 | 16 | 1 | 0 | 1 | 0 | 98 | 49 | 0.999202116 | 0.999202116 | NA | 1 | 0.833 | NA | 0.889 | 1 | -0.028 |
| SE | 6620 | AMPD2 | chr1 | + | 109630682 | 109630793 | 109630232 | 109630406 | 109630942 | 109632055 | 0 | 0 | 9 | 0 | 0 | 0 | 0 | 30 | 3 | 0 | 5 | 0 | 98 | 49 | 0.999202116 | 0.999202116 | NA | NA | 1 | NA | 0.75 | 1 | 0.125 |
| SE | 6624 | PCNX | chr14 | + | 70995740 | 70995925 | 70976941 | 70978648 | 71009633 | 71009724 | 0 | 0 | 2 | 0 | 0 | 0 | 0 | 1 | 1 | 0 | 1 | 0 | 98 | 49 | 0.999202116 | 0.999202116 | NA | NA | 1 | NA | 0.333 | 1 | 0.334 |
| SE | 6766 | PIK3IP1 | chr22 | - | 31289314 | 31289393 | 31281592 | 31283288 | 31289498 | 31289699 | 0 | 0 | 1 | 0 | 0 | 0 | 0 | 5 | 1 | 0 | 2 | 0 | 98 | 49 | 0.999202116 | 0.999202116 | NA | NA | 1 | NA | 0.556 | 1 | 0.222 |
| SE | 6863 | ATG2A | chr11 | - | 64896461 | 64896616 | 64895289 | 64895442 | 64896747 | 64896869 | 0 | 0 | 5 | 0 | 0 | 0 | 0 | 11 | 2 | 0 | 2 | 0 | 98 | 49 | 0.999202116 | 0.999202116 | NA | NA | 1 | NA | 0.733 | 1 | 0.133 |
| SE | 6939 | CD55 | chr1 | + | 207339396 | 207339417 | 207337328 | 207337409 | 207359545 | 207360966 | 0 | 0 | 7 | 0 | 0 | 0 | 0 | 3 | 1 | 0 | 1 | 0 | 70 | 49 | 0.999202116 | 0.999202116 | NA | NA | 1 | NA | 0.677 | 1 | 0.161 |
| SE | 7066 | EPB41 | chr1 | + | 29058588 | 29058645 | 29053103 | 29053312 | 29064981 | 29065158 | 0 | 0 | 1 | 0 | 2 | 10 | 0 | 0 | 0 | 0 | 2 | 0 | 98 | 49 | 0.999202116 | 0.999202116 | NA | 0 | 0.048 | NA | 0 | NA | 0.024 |
| SE | 7122 | RERE | chr1 | - | 8362682 | 8362844 | 8361762 | 8361876 | 8364055 | 8364255 | 0 | 2 | 6 | 0 | 0 | 1 | 0 | 2 | 0 | 0 | 0 | 0 | 98 | 49 | 0.999202116 | 0.999202116 | NA | 1 | 0.75 | NA | 1 | NA | -0.125 |
| SE | 7139 | KIF21B | chr1 | - | 200999348 | 200999466 | 200998383 | 200998575 | 200999882 | 200999964 | 0 | 2 | 5 | 0 | 0 | 0 | 0 | 3 | 0 | 0 | 0 | 1 | 98 | 49 | 0.999202116 | 0.999202116 | NA | 1 | 1 | NA | 1 | 0 | 0.5 |
| SE | 7142 | CPD | chr17 | + | 30421663 | 30421833 | 30420840 | 30420983 | 30422673 | 30423023 | 0 | 0 | 7 | 0 | 0 | 0 | 0 | 7 | 1 | 0 | 1 | 0 | 98 | 49 | 0.999202116 | 0.999202116 | NA | NA | 1 | NA | 0.778 | 1 | 0.111 |
| SE | 7188 | SNRPB | chr20 | - | 2465707 | 2465819 | 2463746 | 2463899 | 2467606 | 2467758 | 0 | 1 | 10 | 0 | 0 | 1 | 0 | 1 | 0 | 0 | 0 | 0 | 98 | 49 | 0.999202116 | 0.999202116 | NA | 1 | 0.833 | NA | 1 | NA | -0.084 |
| SE | 7341 | EIF4A2 | chr3 | + | 186784961 | 186785101 | 186784563 | 186784696 | 186785882 | 186786051 | 0 | 1 | 6 | 0 | 2 | 3 | 0 | 1 | 0 | 1 | 2 | 0 | 98 | 49 | 0.999202116 | 0.999202116 | NA | 0.2 | 0.5 | 0 | 0.2 | NA | 0.25 |
| SE | 7541 | CD300LF | chr17 | - | 74705203 | 74705377 | 74704477 | 74704816 | 74712823 | 74713000 | 0 | 0 | 4 | 0 | 2 | 0 | 0 | 0 | 0 | 0 | 1 | 0 | 98 | 49 | 0.999202116 | 0.999202116 | NA | 0 | 1 | NA | 0 | NA | 0.5 |
| SE | 7579 | MMP8 | chr11 | - | 102714555 | 102714709 | 102713753 | 102713857 | 102715303 | 102715437 | 0 | 4 | 2 | 0 | 0 | 3 | 0 | 1 | 0 | 0 | 0 | 0 | 98 | 49 | 0.999202116 | 0.999202116 | NA | 1 | 0.25 | NA | 1 | NA | -0.375 |
| SE | 7717 | THEMIS2 | chr1 | + | 27879643 | 27880054 | 27872542 | 27872665 | 27885294 | 27885451 | 0 | 0 | 2 | 0 | 1 | 0 | 0 | 0 | 2 | 3 | 0 | 0 | 98 | 49 | 0.999202116 | 0.999202116 | NA | 0 | 1 | 0 | NA | 1 | 0 |
| SE | 7729 | KIAA1033 | chr12 | + | 105140293 | 105140401 | 105137885 | 105138011 | 105140898 | 105141045 | 0 | 3 | 12 | 0 | 0 | 1 | 0 | 3 | 0 | 0 | 0 | 0 | 98 | 49 | 0.999202116 | 0.999202116 | NA | 1 | 0.857 | NA | 1 | NA | -0.072 |
| SE | 7909 | ATP6V0B | chr1 | + | 43976589 | 43976659 | 43976301 | 43976379 | 43976772 | 43976824 | 0 | 4 | 17 | 0 | 0 | 0 | 0 | 24 | 0 | 0 | 0 | 1 | 98 | 49 | 0.999202116 | 0.999202116 | NA | 1 | 1 | NA | 1 | 0 | 0.5 |
| SE | 7949 | PPP4R1 | chr18 | - | 9549196 | 9549338 | 9546790 | 9547952 | 9550051 | 9550186 | 0 | 1 | 6 | 0 | 0 | 0 | 0 | 2 | 0 | 0 | 0 | 2 | 98 | 49 | 0.999202116 | 0.999202116 | NA | 1 | 1 | NA | 1 | 0 | 0.5 |
| SE | 7967 | OAS3 | chr12 | + | 112965743 | 112966029 | 112964234 | 112964408 | 112967417 | 112967593 | 0 | 0 | 23 | 0 | 0 | 0 | 0 | 14 | 1 | 0 | 1 | 0 | 98 | 49 | 0.999202116 | 0.999202116 | NA | NA | 1 | NA | 0.875 | 1 | 0.062 |
| SE | 8036 | MORF4L1 | chr15 | + | 78884966 | 78885083 | 78880511 | 78880579 | 78886140 | 78886227 | 0 | 0 | 0 | 0 | 0 | 5 | 0 | 1 | 0 | 0 | 3 | 1 | 98 | 49 | 0.999202116 | 0.999202116 | NA | NA | 0 | NA | 0.143 | 0 | -0.071 |
| SE | 8083 | YBX3 | chr12 | - | 10709907 | 10710114 | 10704050 | 10704148 | 10713210 | 10713333 | 0 | 0 | 5 | 0 | 0 | 0 | 0 | 0 | 1 | 0 | 1 | 0 | 98 | 49 | 0.999202116 | 0.999202116 | NA | NA | 1 | NA | 0 | 1 | 0.5 |
| SE | 8096 | JMJD1C | chr10 | - | 63176296 | 63176473 | 63168434 | 63168566 | 63177716 | 63177856 | 0 | 0 | 16 | 0 | 0 | 0 | 1 | 21 | 3 | 0 | 1 | 0 | 98 | 49 | 0.999202116 | 0.999202116 | NA | NA | 1 | 1 | 0.913 | 1 | 0.029 |
| SE | 8125 | ANXA5 | chr4 | - | 121684676 | 121684771 | 121683363 | 121683477 | 121686287 | 121686372 | 0 | 3 | 20 | 0 | 0 | 1 | 0 | 3 | 0 | 0 | 0 | 0 | 98 | 49 | 0.999202116 | 0.999202116 | NA | 1 | 0.909 | NA | 1 | NA | -0.045 |
| SE | 8189 | EPSTI1 | chr13 | - | 42953947 | 42954021 | 42926335 | 42926429 | 42963254 | 42963338 | 0 | 5 | 42 | 0 | 0 | 0 | 0 | 23 | 0 | 0 | 0 | 2 | 98 | 49 | 0.999202116 | 0.999202116 | NA | 1 | 1 | NA | 1 | 0 | 0.5 |
| SE | 8190 | EPSTI1 | chr13 | - | 42970611 | 42970670 | 42969093 | 42969177 | 42991977 | 42992271 | 0 | 0 | 25 | 0 | 1 | 0 | 0 | 7 | 5 | 0 | 1 | 0 | 98 | 49 | 0.999202116 | 0.999202116 | NA | 0 | 1 | NA | 0.778 | 1 | -0.389 |
| SE | 8204 | NDUFS2 | chr1 | + | 161203436 | 161203543 | 161202146 | 161202480 | 161206406 | 161206597 | 0 | 1 | 0 | 0 | 0 | 0 | 0 | 1 | 0 | 3 | 0 | 0 | 98 | 49 | 0.999202116 | 0.999202116 | NA | 1 | NA | 0 | 1 | NA | 0.5 |
| SE | 8326 | COPA | chr1 | - | 160307162 | 160307245 | 160306353 | 160306493 | 160309100 | 160309176 | 0 | 0 | 6 | 0 | 0 | 0 | 0 | 3 | 1 | 0 | 1 | 0 | 98 | 49 | 0.999202116 | 0.999202116 | NA | NA | 1 | NA | 0.6 | 1 | 0.2 |
| SE | 8339 | CHD2 | chr15 | + | 92944414 | 92944515 | 92942842 | 92943068 | 92945820 | 92945865 | 0 | 0 | 5 | 0 | 0 | 0 | 0 | 14 | 2 | 0 | 1 | 0 | 98 | 49 | 0.999202116 | 0.999202116 | NA | NA | 1 | NA | 0.875 | 1 | 0.062 |
| SE | 8385 | CYFIP2 | chr5 | + | 157383264 | 157383359 | 157382589 | 157382662 | 157389188 | 157389427 | 0 | 0 | 4 | 0 | 0 | 0 | 0 | 5 | 1 | 0 | 1 | 0 | 98 | 49 | 0.999202116 | 0.999202116 | NA | NA | 1 | NA | 0.714 | 1 | 0.143 |
| SE | 8481 | MAX | chr14 | - | 65101545 | 65101572 | 65093707 | 65093815 | 65102303 | 65102544 | 0 | 1 | 2 | 0 | 0 | 3 | 0 | 1 | 0 | 0 | 0 | 0 | 76 | 49 | 0.999202116 | 0.999202116 | NA | 1 | 0.301 | NA | 1 | NA | -0.35 |
| SE | 8556 | PIK3AP1 | chr10 | - | 96656797 | 96656934 | 96651508 | 96651651 | 96709566 | 96709983 | 0 | 0 | 16 | 0 | 0 | 0 | 0 | 3 | 3 | 0 | 1 | 0 | 98 | 49 | 0.999202116 | 0.999202116 | NA | NA | 1 | NA | 0.6 | 1 | 0.2 |
| SE | 8657 | DUSP1 | chr5 | - | 172769574 | 172769794 | 172768089 | 172769132 | 172770160 | 172770306 | 0 | 17 | 269 | 0 | 0 | 3 | 7 | 124 | 15 | 0 | 0 | 1 | 98 | 49 | 0.999202116 | 0.999202116 | NA | 1 | 0.978 | 1 | 1 | 0.882 | 0.028 |
| SE | 8659 | DUSP1 | chr5 | - | 172770160 | 172770306 | 172768089 | 172769132 | 172770585 | 172771200 | 4 | 93 | 267 | 0 | 1 | 0 | 23 | 118 | 275 | 0 | 2 | 0 | 98 | 49 | 0.999202116 | 0.999202116 | 1 | 0.979 | 1 | 1 | 0.967 | 1 | 0.004 |
| SE | 8801 | CD46 | chr1 | + | 207790252 | 207790345 | 207785618 | 207785682 | 207793518 | 207795516 | 0 | 0 | 12 | 1 | 0 | 6 | 0 | 12 | 1 | 2 | 2 | 0 | 98 | 49 | 0.999202116 | 0.999202116 | 0 | NA | 0.5 | 0 | 0.75 | 1 | -0.333 |
| SE | 8944 | GUK1 | chr1 | + | 228148370 | 228148456 | 228147614 | 228147699 | 228148664 | 228148954 | 0 | 0 | 8 | 0 | 1 | 3 | 0 | 4 | 1 | 0 | 1 | 0 | 98 | 49 | 0.999202116 | 0.999202116 | NA | 0 | 0.571 | NA | 0.667 | 1 | -0.548 |
| SE | 9046 | FAM65B | chr6 | - | 24832255 | 24832391 | 24830508 | 24830670 | 24839090 | 24839272 | 0 | 1 | 4 | 0 | 0 | 1 | 0 | 1 | 0 | 0 | 0 | 0 | 98 | 49 | 0.999202116 | 0.999202116 | NA | 1 | 0.667 | NA | 1 | NA | -0.166 |
| SE | 9138 | CLK1 | chr2 | - | 200857717 | 200857884 | 200856890 | 200856985 | 200857972 | 200858089 | 0 | 0 | 12 | 0 | 0 | 0 | 0 | 10 | 1 | 0 | 3 | 0 | 98 | 49 | 0.999202116 | 0.999202116 | NA | NA | 1 | NA | 0.625 | 1 | 0.188 |
| SE | 9352 | NABP1 | chr2 | + | 191682491 | 191682595 | 191681945 | 191682017 | 191683728 | 191683804 | 0 | 2 | 4 | 0 | 0 | 5 | 2 | 0 | 1 | 0 | 4 | 0 | 98 | 49 | 0.999202116 | 0.999202116 | NA | 1 | 0.286 | 1 | 0 | 1 | -0.024 |
| SE | 9433 | IMPDH1 | chr7 | - | 128405766 | 128405865 | 128403705 | 128403754 | 128409755 | 128409982 | 0 | 0 | 5 | 0 | 0 | 0 | 0 | 1 | 1 | 0 | 1 | 0 | 98 | 49 | 0.999202116 | 0.999202116 | NA | NA | 1 | NA | 0.333 | 1 | 0.334 |
| SE | 9485 | NAGK | chr2 | + | 71075554 | 71075642 | 71073481 | 71073594 | 71076603 | 71076701 | 0 | 1 | 11 | 0 | 0 | 0 | 0 | 11 | 0 | 0 | 0 | 1 | 98 | 49 | 0.999202116 | 0.999202116 | NA | 1 | 1 | NA | 1 | 0 | 0.5 |
| SE | 9512 | SIPA1 | chr11 | + | 65642249 | 65642377 | 65640800 | 65641600 | 65642462 | 65642639 | 0 | 0 | 14 | 0 | 0 | 0 | 0 | 3 | 1 | 0 | 2 | 0 | 98 | 49 | 0.999202116 | 0.999202116 | NA | NA | 1 | NA | 0.429 | 1 | 0.285 |
| SE | 9563 | OASL | chr12 | - | 121033460 | 121033743 | 121031441 | 121031617 | 121038773 | 121039242 | 0 | 0 | 16 | 0 | 2 | 2 | 2 | 4 | 4 | 0 | 3 | 0 | 98 | 49 | 0.999202116 | 0.999202116 | NA | 0 | 0.8 | 1 | 0.4 | 1 | -0.4 |
| SE | 9755 | LCOR | chr10 | + | 96944112 | 96944245 | 96907667 | 96907747 | 96949007 | 96949295 | 0 | 2 | 0 | 0 | 0 | 1 | 0 | 2 | 0 | 0 | 0 | 0 | 98 | 49 | 0.999202116 | 0.999202116 | NA | 1 | 0 | NA | 1 | NA | -0.5 |
| SE | 9971 | RIOK3 | chr18 | + | 23467398 | 23467526 | 23466132 | 23466276 | 23473428 | 23473626 | 0 | 0 | 6 | 0 | 0 | 0 | 0 | 11 | 2 | 0 | 1 | 0 | 98 | 49 | 0.999202116 | 0.999202116 | NA | NA | 1 | NA | 0.846 | 1 | 0.077 |
| SE | 9990 | TRANK1 | chr3 | - | 36874125 | 36874296 | 36864318 | 36864480 | 36889828 | 36889960 | 0 | 3 | 13 | 0 | 0 | 1 | 0 | 3 | 0 | 0 | 0 | 0 | 98 | 49 | 0.999202116 | 0.999202116 | NA | 1 | 0.867 | NA | 1 | NA | -0.067 |
| SE | 10093 | NUMB | chr14 | - | 73284080 | 73284374 | 73282358 | 73282505 | 73287109 | 73287314 | 0 | 0 | 6 | 0 | 0 | 0 | 0 | 6 | 2 | 0 | 2 | 0 | 98 | 49 | 0.999202116 | 0.999202116 | NA | NA | 1 | NA | 0.6 | 1 | 0.2 |
| SE | 10116 | TRAF3IP3 | chr1 | + | 209762514 | 209762662 | 209759981 | 209760384 | 209762812 | 209762870 | 0 | 0 | 6 | 0 | 0 | 0 | 0 | 0 | 1 | 0 | 1 | 0 | 98 | 49 | 0.999202116 | 0.999202116 | NA | NA | 1 | NA | 0 | 1 | 0.5 |
| SE | 10121 | SPTLC2 | chr14 | - | 77562395 | 77562489 | 77557040 | 77557146 | 77570383 | 77570508 | 0 | 0 | 16 | 0 | 0 | 0 | 0 | 2 | 1 | 0 | 1 | 0 | 98 | 49 | 0.999202116 | 0.999202116 | NA | NA | 1 | NA | 0.5 | 1 | 0.25 |
| SE | 10446 | ELL | chr19 | - | 18458204 | 18458329 | 18451551 | 18451648 | 18461577 | 18461852 | 0 | 0 | 13 | 0 | 0 | 0 | 0 | 1 | 1 | 0 | 3 | 0 | 98 | 49 | 0.999202116 | 0.999202116 | NA | NA | 1 | NA | 0.143 | 1 | 0.428 |
| SE | 10653 | RELA | chr11 | - | 65658717 | 65658822 | 65655854 | 65655935 | 65659665 | 65659788 | 0 | 0 | 2 | 0 | 0 | 0 | 0 | 9 | 1 | 0 | 1 | 0 | 98 | 49 | 0.999202116 | 0.999202116 | NA | NA | 1 | NA | 0.818 | 1 | 0.091 |

| | | | | | | | | | | | | | | | | | | | | | | | | | | | | | | | | | |
|---|---|---|---|---|---|---|---|---|---|---|---|---|---|---|---|---|---|---|---|---|---|---|---|---|---|---|---|---|---|---|---|---|---|
| SE | 10700 | TLE3 | chr15 | - | 70055048 | 70055298 | 70053226 | 70053374 | 70056297 | 70056374 | 0 | 0 | 4 | 0 | 0 | 0 | 0 | 6 | 4 | 0 | 1 | 0 | 98 | 49 | 0.999202116 | 0.999202116 | NA | NA | 1 | NA | 0.75 | 1 | 0.125 |
| SE | 10824 | PSMA4 | chr15 | + | 78542482 | 78542645 | 78542176 | 78542219 | 78544189 | 78544267 | 0 | 1 | 4 | 0 | 0 | 1 | 0 | 1 | 0 | 0 | 0 | 0 | 98 | 49 | 0.999202116 | 0.999202116 | NA | 1 | 0.667 | NA | 1 | NA | -0.166 |
| SE | 10950 | IQGAP1 | chr15 | + | 90472839 | 90473010 | 90467449 | 90467592 | 90473714 | 90473798 | 0 | 0 | 30 | 0 | 0 | 0 | 7 | 16 | 3 | 0 | 1 | 0 | 98 | 49 | 0.999202116 | 0.999202116 | NA | NA | 1 | 1 | 0.889 | 1 | 0.037 |
| SE | 11033 | STAT1 | chr2 | - | 191013524 | 191013678 | 191009875 | 191010004 | 191014017 | 191014250 | 0 | 5 | 34 | 0 | 0 | 2 | 0 | 15 | 4 | 0 | 0 | 2 | 98 | 49 | 0.999202116 | 0.999202116 | NA | 1 | 0.895 | NA | 1 | 0.5 | 0.198 |
| SE | 11193 | PARP14 | chr3 | + | 122699389 | 122701635 | 122695425 | 122695662 | 122703741 | 122703978 | 0 | 8 | 30 | 0 | 0 | 2 | 0 | 39 | 2 | 0 | 1 | 0 | 98 | 49 | 0.999202116 | 0.999202116 | NA | 1 | 0.882 | NA | 0.951 | 1 | -0.034 |
| SE | 11206 | TOP1 | chr20 | + | 41080084 | 41080180 | 41077581 | 41077637 | 41081164 | 41081240 | 0 | 0 | 5 | 0 | 0 | 0 | 0 | 5 | 1 | 0 | 1 | 0 | 98 | 49 | 0.999202116 | 0.999202116 | NA | NA | 1 | NA | 0.714 | 1 | 0.143 |
| SE | 11348 | ADGRE2 | chr19 | - | 14754953 | 14755127 | 14752328 | 14752526 | 14755653 | 14755877 | 0 | 0 | 5 | 0 | 0 | 0 | 0 | 7 | 1 | 0 | 1 | 0 | 98 | 49 | 0.999202116 | 0.999202116 | NA | NA | 1 | NA | 0.778 | 1 | 0.111 |
| SE | 11351 | ADGRE2 | chr19 | - | 14755653 | 14755877 | 14754953 | 14755127 | 14756237 | 14756345 | 0 | 0 | 4 | 0 | 0 | 0 | 0 | 4 | 1 | 0 | 1 | 0 | 98 | 49 | 0.999202116 | 0.999202116 | NA | NA | 1 | NA | 0.667 | 1 | 0.166 |
| SE | 11376 | ARHGEF1 | chr19 | + | 41905174 | 41905261 | 41904948 | 41905036 | 41905759 | 41905827 | 0 | 1 | 10 | 0 | 2 | 0 | 0 | 0 | 3 | 0 | 0 | 0 | 98 | 49 | 0.999202116 | 0.999202116 | NA | 0.2 | 1 | NA | NA | 1 | -0.4 |
| SE | 11391 | CPNE2 | chr16 | + | 57137148 | 57137282 | 57134774 | 57134826 | 57146084 | 57146321 | 0 | 1 | 5 | 0 | 0 | 1 | 0 | 1 | 0 | 0 | 0 | 0 | 98 | 49 | 0.999202116 | 0.999202116 | NA | 1 | 0.714 | NA | 1 | NA | -0.143 |
| SE | 11403 | CSNK2A2 | chr16 | - | 58167206 | 58167308 | 58166583 | 58166684 | 58167684 | 58167795 | 0 | 1 | 0 | 0 | 0 | 1 | 0 | 1 | 0 | 0 | 0 | 0 | 98 | 49 | 0.999202116 | 0.999202116 | NA | 1 | 0 | NA | 1 | NA | -0.5 |
| SE | 11481 | AP1G1 | chr16 | - | 71756018 | 71756159 | 71753832 | 71753887 | 71758807 | 71758921 | 0 | 0 | 7 | 0 | 0 | 0 | 0 | 16 | 3 | 0 | 1 | 0 | 98 | 49 | 0.999202116 | 0.999202116 | NA | NA | 1 | NA | 0.889 | 1 | 0.055 |
| SE | 11563 | NCBP1 | chr9 | + | 97662041 | 97662144 | 97660945 | 97661068 | 97662953 | 97663047 | 0 | 1 | 1 | 0 | 0 | 2 | 0 | 1 | 0 | 0 | 0 | 0 | 98 | 49 | 0.999202116 | 0.999202116 | NA | 1 | 0.2 | NA | 1 | NA | -0.4 |
| SE | 11622 | CIR1 | chr2 | - | 174378922 | 174379040 | 174351618 | 174351725 | 174380201 | 174380250 | 0 | 0 | 8 | 0 | 0 | 0 | 0 | 12 | 1 | 0 | 1 | 0 | 98 | 49 | 0.999202116 | 0.999202116 | NA | NA | 1 | NA | 0.857 | 1 | 0.072 |
| SE | 11641 | ARRB2 | chr17 | + | 4718260 | 4718345 | 4717887 | 4718023 | 4718611 | 4718684 | 0 | 0 | 26 | 0 | 1 | 3 | 4 | 9 | 3 | 0 | 10 | 1 | 98 | 49 | 0.999202116 | 0.999202116 | NA | 0 | 0.812 | 1 | 0.31 | 0.6 | -0.231 |
| SE | 11642 | ARRB2 | chr17 | + | 4720392 | 4720472 | 4720211 | 4720299 | 4720585 | 4720640 | 0 | 2 | 26 | 0 | 1 | 0 | 2 | 12 | 10 | 0 | 4 | 0 | 98 | 49 | 0.999202116 | 0.999202116 | NA | 0.5 | 1 | 1 | 0.6 | 1 | -0.117 |
| SE | 11657 | NLRP1 | chr17 | - | 5520880 | 5521012 | 5517745 | 5517887 | 5521523 | 5521786 | 0 | 1 | 10 | 0 | 0 | 3 | 0 | 5 | 1 | 12 | 1 | 1 | 98 | 49 | 0.999202116 | 0.999202116 | NA | 1 | 0.625 | 0 | 0.714 | 0.333 | 0.464 |
| SE | 11658 | NLRP1 | chr17 | - | 5521523 | 5521786 | 5517745 | 5517887 | 5530480 | 5530704 | 0 | 0 | 9 | 0 | 0 | 0 | 12 | 1 | 1 | 0 | 2 | 0 | 98 | 49 | 0.999202116 | 0.999202116 | NA | NA | 1 | 1 | 0.2 | 1 | 0.267 |
| SE | 11683 | STAT5B | chr17 | - | 42216013 | 42216106 | 42211983 | 42212190 | 42217159 | 42217282 | 0 | 0 | 16 | 0 | 0 | 0 | 0 | 4 | 2 | 0 | 1 | 0 | 98 | 49 | 0.999202116 | 0.999202116 | NA | NA | 1 | NA | 0.667 | 1 | 0.166 |
| SE | 11748 | JTB | chr1 | - | 153976692 | 153976775 | 153975825 | 153975905 | 153976975 | 153977013 | 0 | 0 | 5 | 0 | 0 | 0 | 0 | 8 | 1 | 0 | 1 | 0 | 98 | 49 | 0.999202116 | 0.999202116 | NA | NA | 1 | NA | 0.8 | 1 | 0.1 |
| SE | 11782 | USP22 | chr17 | - | 21015751 | 21015899 | 21012829 | 21012935 | 21017941 | 21018111 | 0 | 4 | 1 | 0 | 0 | 0 | 0 | 2 | 0 | 4 | 0 | 0 | 98 | 49 | 0.999202116 | 0.999202116 | NA | 1 | 1 | 0 | 1 | NA | 0.5 |
| SE | 11805 | KIAA0100 | chr17 | - | 28633548 | 28633751 | 28633282 | 28633401 | 28633894 | 28634086 | 0 | 1 | 6 | 0 | 0 | 0 | 0 | 1 | 0 | 1 | 0 | 0 | 98 | 49 | 0.999202116 | 0.999202116 | NA | 1 | 1 | 0 | 1 | NA | 0.5 |
| SE | 11822 | SSH2 | chr17 | - | 29695458 | 29695523 | 29684562 | 29684684 | 29702958 | 29703062 | 0 | 0 | 20 | 0 | 0 | 0 | 0 | 12 | 1 | 0 | 1 | 0 | 98 | 49 | 0.999202116 | 0.999202116 | NA | NA | 1 | NA | 0.857 | 1 | 0.072 |
| SE | 11876 | MGAM | chr7 | + | 142036822 | 142036977 | 142034669 | 142034841 | 142038530 | 142038615 | 0 | 0 | 14 | 0 | 0 | 0 | 0 | 2 | 6 | 0 | 4 | 0 | 98 | 49 | 0.999202116 | 0.999202116 | NA | NA | 1 | NA | 0.2 | 1 | 0.4 |
| SE | 12165 | USP32 | chr17 | - | 60288522 | 60288682 | 60271349 | 60271481 | 60294682 | 60294801 | 0 | 0 | 12 | 0 | 0 | 0 | 0 | 11 | 3 | 0 | 1 | 0 | 98 | 49 | 0.999202116 | 0.999202116 | NA | NA | 1 | NA | 0.846 | 1 | 0.077 |
| SE | 12220 | BPTF | chr17 | + | 67966571 | 67966656 | 67964211 | 67964404 | 67975771 | 67975958 | 0 | 1 | 3 | 0 | 0 | 1 | 0 | 1 | 0 | 0 | 0 | 0 | 98 | 49 | 0.999202116 | 0.999202116 | NA | 1 | 0.6 | NA | 1 | NA | -0.2 |
| SE | 12394 | ATP5B | chr12 | - | 56642672 | 56642831 | 56642457 | 56642580 | 56643402 | 56643587 | 0 | 0 | 13 | 0 | 0 | 0 | 4 | 3 | 2 | 0 | 2 | 0 | 98 | 49 | 0.999202116 | 0.999202116 | NA | NA | 1 | 1 | 0.429 | 1 | 0.19 |
| SE | 12660 | SURF4 | chr9 | - | 133364839 | 133365026 | 133361448 | 133363759 | 133365984 | 133366028 | 0 | 0 | 2 | 0 | 0 | 0 | 0 | 3 | 1 | 0 | 1 | 0 | 98 | 49 | 0.999202116 | 0.999202116 | NA | NA | 1 | NA | 0.6 | 1 | 0.2 |
| SE | 12704 | ICAM3 | chr19 | - | 10335065 | 10335353 | 10334527 | 10334782 | 10335670 | 10335976 | 0 | 1 | 23 | 0 | 0 | 1 | 3 | 7 | 1 | 0 | 0 | 2 | 98 | 49 | 0.999202116 | 0.999202116 | NA | 1 | 0.92 | 1 | 1 | 0.2 | 0.227 |
| SE | 12712 | SLC44A2 | chr19 | + | 10634973 | 10635073 | 10634755 | 10634887 | 10636322 | 10636585 | 0 | 3 | 8 | 0 | 0 | 1 | 0 | 1 | 0 | 0 | 0 | 0 | 98 | 49 | 0.999202116 | 0.999202116 | NA | 1 | 0.8 | NA | 1 | NA | -0.1 |
| SE | 12820 | MAST3 | chr19 | + | 18124641 | 18124774 | 18124264 | 18124366 | 18128399 | 18128458 | 0 | 1 | 11 | 0 | 0 | 3 | 0 | 2 | 0 | 2 | 0 | 1 | 98 | 49 | 0.999202116 | 0.999202116 | NA | 1 | 0.647 | 0 | 1 | 0 | 0.49 |
| SE | 12825 | TRAPPC10 | chr21 | + | 44074323 | 44074470 | 44063537 | 44063785 | 44075038 | 44075153 | 0 | 0 | 4 | 0 | 0 | 0 | 0 | 3 | 1 | 0 | 1 | 0 | 98 | 49 | 0.999202116 | 0.999202116 | NA | NA | 1 | NA | 0.6 | 1 | 0.2 |
| SE | 12851 | GATAD2A | chr19 | + | 19494293 | 19494383 | 19492580 | 19492712 | 19495753 | 19495885 | 0 | 0 | 4 | 0 | 0 | 0 | 0 | 4 | 2 | 0 | 1 | 0 | 98 | 49 | 0.999202116 | 0.999202116 | NA | NA | 1 | NA | 0.667 | 1 | 0.166 |
| SE | 12950 | GMFG | chr19 | - | 39329543 | 39329626 | 39328999 | 39329073 | 39333076 | 39333126 | 0 | 0 | 53 | 0 | 0 | 0 | 2 | 32 | 1 | 0 | 1 | 0 | 98 | 49 | 0.999202116 | 0.999202116 | NA | NA | 1 | 1 | 0.941 | 1 | 0.02 |
| SE | 12954 | MAN2A2 | chr15 | + | 90910500 | 90910683 | 90909326 | 90909504 | 90910846 | 90910961 | 0 | 0 | 6 | 0 | 0 | 0 | 0 | 1 | 1 | 0 | 1 | 0 | 98 | 49 | 0.999202116 | 0.999202116 | NA | NA | 1 | NA | 0.333 | 1 | 0.334 |
| SE | 12990 | PLAUR | chr19 | - | 43652224 | 43652371 | 43648570 | 43649143 | 43655438 | 43655573 | 0 | 2 | 46 | 0 | 0 | 7 | 0 | 41 | 2 | 0 | 1 | 1 | 98 | 49 | 0.999202116 | 0.999202116 | NA | 1 | 0.767 | NA | 0.953 | 0.5 | 0.157 |
| SE | 13034 | VASP | chr19 | + | 45521321 | 45521406 | 45517928 | 45518094 | 45522167 | 45522217 | 0 | 2 | 37 | 0 | 2 | 3 | 4 | 24 | 4 | 0 | 3 | 0 | 98 | 49 | 0.999202116 | 0.999202116 | NA | 0.333 | 0.86 | 1 | 0.8 | 1 | -0.337 |
| SE | 13052 | NAPA | chr19 | - | 47503422 | 47503502 | 47500632 | 47500749 | 47514842 | 47515258 | 0 | 2 | 18 | 0 | 0 | 0 | 0 | 19 | 1 | 0 | 0 | 4 | 98 | 49 | 0.999202116 | 0.999202116 | NA | 1 | 1 | NA | 1 | 0.111 | 0.445 |
| SE | 13128 | ATXN2 | chr12 | - | 111510384 | 111510582 | 111509890 | 111509998 | 111513356 | 111513539 | 0 | 0 | 2 | 0 | 1 | 0 | 0 | 1 | 1 | 0 | 1 | 0 | 98 | 49 | 0.999202116 | 0.999202116 | NA | 0 | 1 | NA | 0.333 | 1 | -0.166 |
| SE | 13152 | C9orf89 | chr9 | + | 93111878 | 93111938 | 93110567 | 93110721 | 93112217 | 93112289 | 0 | 0 | 6 | 0 | 0 | 0 | 0 | 3 | 2 | 0 | 3 | 0 | 98 | 49 | 0.999202116 | 0.999202116 | NA | NA | 1 | NA | 0.333 | 1 | 0.334 |
| SE | 13179 | NLRP12 | chr19 | - | 53807494 | 53807668 | 53805279 | 53805450 | 53809586 | 53811288 | 0 | 1 | 8 | 0 | 0 | 2 | 0 | 1 | 0 | 0 | 0 | 0 | 98 | 49 | 0.999202116 | 0.999202116 | NA | 1 | 0.667 | NA | 1 | NA | -0.166 |
| SE | 13210 | LILRB2 | chr19 | - | 54280263 | 54280299 | 54279790 | 54280075 | 54280462 | 54280544 | 0 | 0 | 8 | 0 | 0 | 0 | 0 | 1 | 2 | 0 | 1 | 0 | 85 | 49 | 0.999202116 | 0.999202116 | NA | NA | 1 | NA | 0.366 | 1 | 0.317 |
| SE | 13368 | FIP1L1 | chr4 | + | 53414614 | 53414722 | 53399729 | 53399839 | 53425871 | 53425965 | 0 | 1 | 0 | 0 | 0 | 2 | 2 | 1 | 0 | 0 | 0 | 0 | 98 | 49 | 0.999202116 | 0.999202116 | NA | 1 | 0 | 1 | 1 | NA | -0.5 |
| SE | 13635 | CCND3 | chr6 | - | 41940369 | 41940585 | 41937234 | 41937394 | 42048500 | 42048894 | 0 | 1 | 23 | 0 | 0 | 1 | 0 | 0 | 3 | 0 | 1 | 0 | 98 | 49 | 0.999202116 | 0.999202116 | NA | 1 | 0.92 | NA | 0 | 1 | 0.46 |
| SE | 13689 | BAZ2A | chr12 | - | 56609735 | 56609946 | 56606632 | 56606733 | 56610113 | 56610215 | 0 | 0 | 3 | 0 | 0 | 0 | 0 | 1 | 1 | 0 | 1 | 0 | 98 | 49 | 0.999202116 | 0.999202116 | NA | NA | 1 | NA | 0.333 | 1 | 0.334 |
| SE | 13733 | SERPINA1 | chr14 | - | 94388559 | 94388663 | 94382591 | 94383241 | 94390456 | 94390692 | 0 | 0 | 17 | 0 | 2 | 20 | 0 | 6 | 1 | 0 | 5 | 0 | 98 | 49 | 0.999202116 | 0.999202116 | NA | 0 | 0.298 | NA | 0.375 | 1 | -0.538 |
| SE | 13745 | MAP4K4 | chr2 | + | 101887087 | 101887237 | 101885186 | 101885287 | 101887777 | 101887937 | 0 | 0 | 10 | 0 | 0 | 0 | 0 | 2 | 1 | 0 | 1 | 0 | 98 | 49 | 0.999202116 | 0.999202116 | NA | NA | 1 | NA | 0.5 | 1 | 0.25 |
| SE | 13818 | MAP3K2 | chr2 | - | 127325727 | 127325807 | 127324173 | 127324241 | 127326686 | 127326817 | 0 | 0 | 17 | 0 | 0 | 0 | 0 | 15 | 4 | 0 | 1 | 0 | 98 | 49 | 0.999202116 | 0.999202116 | NA | NA | 1 | NA | 0.882 | 1 | 0.059 |
| SE | 13861 | ACSL1 | chr4 | - | 184764852 | 184764925 | 184763166 | 184763255 | 184766621 | 184766756 | 0 | 2 | 44 | 0 | 0 | 2 | 3 | 39 | 5 | 0 | 1 | 1 | 98 | 49 | 0.999202116 | 0.999202116 | NA | 1 | 0.917 | 1 | 0.951 | 0.714 | 0.07 |
| SE | 13862 | ACSL1 | chr4 | - | 184765890 | 184765986 | 184763166 | 184763255 | 184766621 | 184766756 | 0 | 2 | 50 | 0 | 0 | 2 | 0 | 38 | 5 | 0 | 1 | 1 | 98 | 49 | 0.999202116 | 0.999202116 | NA | 1 | 0.926 | NA | 0.95 | 0.714 | 0.131 |
| A5SS | 489 | RCHY1 | chr4 | - | 75491582 | 75491637 | 75491610 | 75491637 | 75490580 | 75490701 | 0 | 0 | 0 | 0 | 1 | 0 | 0 | 0 | 1 | 0 | 1 | 0 | 77 | 49 | 0.999208248 | 0.999208248 | NA | 0 | NA | NA | 0 | 1 | -0.5 |
| | | | | | | | | | | RI | 4 | 197 | 1673 | 1 | 22 | 113 | 78 | 974 | 430 | 28 | 118 | 24 | 10010 | 5047 | 102.917824 | 102.917824 | 1 | 42.212 | 85.167 | 15 | 71.194 | 66.554 | 8.42 |
| | | | | | | | | | | A5SS | 33 | 611 | 5360 | 4 | 71 | 375 | 267 | 3388 | 1071 | 70 | 408 | 84 | 31215 | 15729 | 320.7438914 | 320.7438975 | 12.605 | 132.748 | 259.992 | 64 | 222.913 | 202.991 | 22.352 |
| | | | | | | | | | | A3SS | 80 | 1389 | 11266 | 8 | 190 | 1094 | 583 | 7174 | 2215 | 172 | 1004 | 216 | 71146 | 36162 | 736.0390206 | 737.4113146 | 34.21 | 325.462 | 574.316 | 146.44 | 510.257 | 429.125 | 44.016 |
| A3SS | 13 | LONRF3 | chrX | + | 119012879 | 119013201 | 119013038 | 119013201 | 119011814 | 119011973 | 0 | 0 | 0 | 0 | 0 | 1 | 0 | 1 | 0 | 0 | 0 | 0 | 98 | 49 | NA | NA | NA | NA | 0 | NA | 1 | NA | -1 |
| A3SS | 50 | TFE3 | chrX | - | 49039106 | 49039410 | 49039106 | 49039345 | 49040454 | 49040568 | 0 | 0 | 4 | 0 | 0 | 3 | 0 | 4 | 0 | 0 | 3 | 0 | 98 | 49 | NA | NA | NA | NA | 0.4 | NA | 0.4 | NA | 0 |
| A3SS | 63 | CD99 | chrX | + | 2738199 | 2738256 | 2738202 | 2738256 | 2726259 | 2726373 | 0 | 0 | 10 | 0 | 0 | 1 | 1 | 3 | 0 | 0 | 0 | 0 | 52 | 49 | NA | NA | NA | NA | 0.904 | 1 | 1 | NA | -0.096 |
| A3SS | 71 | EDF1 | chr9 | - | 136862113 | 136862496 | 136862113 | 136862345 | 136862905 | 136862999 | 1 | 0 | 3 | 0 | 0 | 1 | 0 | 1 | 0 | 0 | 2 | 0 | 98 | 49 | NA | NA | 1 | NA | 0.6 | NA | 0.2 | NA | 0.6 |
| A3SS | 95 | CDK5RAP2 | chr9 | - | 120460571 | 120460747 | 120460571 | 120460667 | 120467859 | 120467997 | 3 | 0 | 0 | 0 | 0 | 0 | 0 | 0 | 0 | 0 | 1 | 0 | 98 | 49 | NA | NA | 1 | NA | NA | NA | 0 | NA | 1 |
| A3SS | 117 | TLE4 | chr9 | + | 79652589 | 79652794 | 79652592 | 79652794 | 79627373 | 79627448 | 0 | 0 | 0 | 0 | 0 | 2 | 0 | 1 | 0 | 0 | 1 | 0 | 52 | 49 | NA | NA | NA | NA | 0 | NA | 0.485 | NA | -0.485 |
| A3SS | 213 | CUX1 | chr7 | + | 102178468 | 102178657 | 102178474 | 102178657 | 102170445 | 102170550 | 0 | 0 | 5 | 0 | 0 | 0 | 0 | 0 | 0 | 0 | 2 | 0 | 55 | 49 | NA | NA | NA | NA | 1 | NA | 0 | NA | 1 |
| A3SS | 280 | IPCEF1 | chr6 | - | 154247448 | 154247488 | 154247448 | 154247485 | 154265911 | 154265964 | 0 | 0 | 0 | 0 | 0 | 1 | 0 | 1 | 0 | 0 | 0 | 0 | 52 | 49 | NA | NA | NA | NA | 0 | NA | 1 | NA | -1 |
| A3SS | 287 | PLAGL1 | chr6 | - | 143960468 | 143960580 | 143960468 | 143960542 | 143964786 | 143964818 | 0 | 2 | 0 | 0 | 0 | 1 | 3 | 0 | 0 | 0 | 0 | 0 | 87 | 49 | NA | NA | NA | 1 | 0 | 1 | NA | NA | -0.5 |
| A3SS | 332 | STK38 | chr6 | - | 36540071 | 36540207 | 36540071 | 36540204 | 36547189 | 36547516 | 0 | 0 | 0 | 0 | 0 | 1 | 0 | 1 | 0 | 0 | 0 | 0 | 52 | 49 | NA | NA | NA | NA | 0 | NA | 1 | NA | -1 |
| A3SS | 466 | BTN3A2 | chr6 | + | 26367974 | 26368050 | 26367989 | 26368050 | 26365158 | 26365352 | 0 | 0 | 1 | 0 | 0 | 2 | 0 | 0 | 0 | 0 | 1 | 0 | 64 | 49 | NA | NA | NA | NA | 0.277 | NA | 0 | NA | 0.277 |
| A3SS | 478 | CD83 | chr6 | + | 14135107 | 14136917 | 14135110 | 14136917 | 14133648 | 14133755 | 0 | 0 | 0 | 0 | 0 | 1 | 0 | 14 | 0 | 0 | 0 | 0 | 52 | 49 | NA | NA | NA | NA | 0 | NA | 1 | NA | -1 |
| A3SS | 503 | HARS2 | chr5 | + | 140693490 | 140693665 | 140693590 | 140693665 | 140691425 | 140691756 | 0 | 0 | 2 | 0 | 0 | 0 | 0 | 1 | 0 | 0 | 2 | 0 | 98 | 49 | NA | NA | NA | NA | 1 | NA | 0.2 | NA | 0.8 |
| A3SS | 513 | SRA1 | chr5 | - | 140557146 | 140557332 | 140557146 | 140557272 | 140557427 | 140558093 | 0 | 0 | 1 | 0 | 0 | 1 | 0 | 1 | 0 | 0 | 0 | 0 | 98 | 49 | NA | NA | NA | NA | 0.333 | NA | 1 | NA | -0.667 |
| A3SS | 535 | SSBP2 | chr5 | - | 81466973 | 81467065 | 81466973 | 81467041 | 81473699 | 81473770 | 0 | 0 | 0 | 0 | 0 | 1 | 2 | 0 | 0 | 0 | 0 | 0 | 73 | 49 | NA | NA | NA | NA | 0 | 1 | NA | NA | -1 |
| A3SS | 544 | GAPT | chr5 | + | 58494147 | 58496358 | 58494246 | 58496358 | 58493710 | 58493838 | 0 | 0 | 1 | 0 | 0 | 2 | 5 | 1 | 0 | 0 | 0 | 0 | 98 | 49 | NA | NA | NA | NA | 0.2 | 1 | 1 | NA | -0.8 |
| A3SS | 580 | ALPK1 | chr4 | + | 112377677 | 112377898 | 112377729 | 112377898 | 112315800 | 112315852 | 0 | 0 | 7 | 0 | 0 | 0 | 7 | 2 | 0 | 0 | 4 | 0 | 98 | 49 | NA | NA | NA | NA | 1 | 1 | 0.2 | NA | 0.4 |
| A3SS | 669 | MFSD1 | chr3 | + | 158804318 | 158804371 | 158804322 | 158804371 | 158801925 | 158802315 | 0 | 0 | 5 | 0 | 0 | 0 | 0 | 1 | 0 | 0 | 1 | 0 | 53 | 49 | NA | NA | NA | NA | 1 | NA | 0.48 | NA | 0.52 |
| A3SS | 676 | ARMC8 | chr3 | + | 138188400 | 138188530 | 138188454 | 138188530 | 138187247 | 138187599 | 2 | 0 | 1 | 0 | 0 | 2 | 0 | 1 | 0 | 0 | 0 | 0 | 98 | 49 | NA | NA | 1 | NA | 0.2 | NA | 1 | NA | -0.4 |
| A3SS | 680 | CDV3 | chr3 | + | 133587158 | 133590274 | 133587161 | 133590274 | 133586562 | 133586722 | 0 | 0 | 4 | 0 | 0 | 1 | 0 | 3 | 0 | 0 | 0 | 0 | 52 | 49 | NA | NA | NA | NA | 0.79 | NA | 1 | NA | -0.21 |
| A3SS | 682 | CDV3 | chr3 | + | 133587158 | 133590274 | 133588352 | 133590274 | 133586562 | 133586722 | 0 | 0 | 9 | 0 | 0 | 0 | 0 | 7 | 0 | 0 | 1 | 0 | 98 | 49 | NA | NA | NA | NA | 1 | NA | 0.778 | NA | 0.222 |
| A3SS | 684 | CDV3 | chr3 | + | 133587161 | 133590274 | 133588352 | 133590274 | 133586562 | 133586722 | 0 | 0 | 9 | 0 | 0 | 0 | 0 | 4 | 0 | 0 | 1 | 0 | 98 | 49 | NA | NA | NA | NA | 1 | NA | 0.667 | NA | 0.333 |
| A3SS | 685 | CDV3 | chr3 | + | 133587895 | 133590274 | 133588352 | 133590274 | 133586562 | 133586722 | 0 | 0 | 8 | 0 | 0 | 0 | 0 | 6 | 0 | 0 | 1 | 0 | 98 | 49 | NA | NA | NA | NA | 1 | NA | 0.75 | NA | 0.25 |
| A3SS | 718 | KIAA0922 | chr4 | + | 153584834 | 153584931 | 153584837 | 153584931 | 153583563 | 153583672 | 0 | 0 | 2 | 0 | 0 | 3 | 0 | 0 | 0 | 0 | 2 | 0 | 52 | 49 | NA | NA | NA | NA | 0.386 | NA | 0 | NA | 0.386 |
| A3SS | 726 | USP19 | chr3 | - | 49115723 | 49115944 | 49115723 | 49115938 | 49116046 | 49116162 | 0 | 0 | 1 | 0 | 0 | 0 | 0 | 0 | 0 | 0 | 1 | 0 | 55 | 49 | NA | NA | NA | NA | 1 | NA | 0 | NA | 1 |
| A3SS | 774 | LST1 | chr6 | + | 31588496 | 31588909 | 31588517 | 31588909 | 31587199 | 31587318 | 0 | 0 | 3 | 0 | 0 | 4 | 0 | 0 | 0 | 0 | 4 | 0 | 70 | 49 | NA | NA | NA | NA | 0.344 | NA | 0 | NA | 0.344 |
| A3SS | 813 | POLDIP3 | chr22 | - | 42601969 | 42602107 | 42601969 | 42602056 | 42602769 | 42603160 | 0 | 0 | 0 | 0 | 0 | 1 | 0 | 1 | 0 | 0 | 0 | 0 | 98 | 49 | NA | NA | NA | NA | 0 | NA | 1 | NA | -1 |
| A3SS | 833 | SRP19 | chr5 | + | 112867371 | 112868392 | 112867403 | 112868392 | 112864620 | 112864732 | 0 | 0 | 1 | 0 | 0 | 1 | 0 | 1 | 0 | 0 | 0 | 0 | 81 | 49 | NA | NA | NA | NA | 0.377 | NA | 1 | NA | -0.623 |
| A3SS | 931 | STAU1 | chr20 | - | 49124374 | 49124605 | 49124374 | 49124587 | 49135832 | 49135931 | 0 | 0 | 3 | 0 | 0 | 0 | 0 | 0 | 0 | 0 | 2 | 0 | 67 | 49 | NA | NA | NA | NA | 1 | NA | 0 | NA | 1 |
| A3SS | 938 | ELMO2 | chr20 | - | 46394404 | 46394532 | 46394404 | 46394503 | 46398696 | 46398771 | 0 | 0 | 0 | 0 | 0 | 1 | 2 | 2 | 0 | 0 | 0 | 0 | 78 | 49 | NA | NA | NA | NA | 0 | 1 | 1 | NA | -1 |
| A3SS | 940 | PIGT | chr20 | + | 45419472 | 45419590 | 45419503 | 45419590 | 45419294 | 45419395 | 0 | 0 | 0 | 0 | 0 | 2 | 4 | 1 | 0 | 3 | 1 | 0 | 80 | 49 | NA | NA | NA | NA | 0 | 0.45 | 0.38 | NA | -0.415 |
| A3SS | 952 | TRPC4AP | chr20 | - | 35021189 | 35021356 | 35021189 | 35021332 | 35035122 | 35035308 | 0 | 0 | 3 | 0 | 0 | 1 | 0 | 1 | 0 | 0 | 1 | 0 | 73 | 49 | NA | NA | NA | NA | 0.668 | NA | 0.402 | NA | 0.266 |
| A3SS | 964 | HCK | chr20 | + | 32073715 | 32073818 | 32073718 | 32073818 | 32073318 | 32073361 | 0 | 0 | 6 | 0 | 0 | 0 | 2 | 2 | 0 | 0 | 1 | 0 | 52 | 49 | NA | NA | NA | NA | 1 | 1 | 0.653 | NA | 0.173 |
| A3SS | 1008 | DYRK1A | chr21 | + | 37478180 | 37478300 | 37478207 | 37478300 | 37472683 | 37472880 | 0 | 0 | 0 | 0 | 0 | 3 | 1 | 0 | 0 | 0 | 5 | 0 | 76 | 49 | NA | NA | NA | NA | 0 | 1 | 0 | NA | -0.5 |
| A3SS | 1022 | TRIP12 | chr2 | - | 229805729 | 229805883 | 229805729 | 229805802 | 229807707 | 229807864 | 0 | 0 | 8 | 0 | 0 | 0 | 4 | 9 | 0 | 0 | 1 | 0 | 98 | 49 | NA | NA | NA | NA | 1 | 1 | 0.818 | NA | 0.091 |
| A3SS | 1025 | AGFG1 | chr2 | + | 227553703 | 227553795 | 227553709 | 227553795 | 227551958 | 227552117 | 0 | 0 | 2 | 0 | 0 | 4 | 0 | 1 | 0 | 0 | 0 | 0 | 55 | 49 | NA | NA | NA | NA | 0.308 | NA | 1 | NA | -0.692 |
| A3SS | 1040 | ARHGEF40 | chr14 | + | 21075622 | 21075765 | 21075644 | 21075765 | 21075331 | 21075499 | 0 | 0 | 0 | 0 | 0 | 2 | 2 | 0 | 0 | 0 | 1 | 0 | 71 | 49 | NA | NA | NA | NA | 0 | 1 | 0 | NA | -0.5 |
| A3SS | 1099 | MAP7D1 | chr1 | + | 36179511 | 36179557 | 36179514 | 36179557 | 36179261 | 36179315 | 0 | 0 | 0 | 0 | 0 | 3 | 0 | 1 | 0 | 0 | 3 | 0 | 52 | 49 | NA | NA | NA | NA | 0 | NA | 0.239 | NA | -0.239 |
| A3SS | 1105 | SH3BP5 | chr3 | - | 15304102 | 15304299 | 15304102 | 15304231 | 15330503 | 15330566 | 0 | 0 | 2 | 0 | 0 | 2 | 0 | 1 | 0 | 0 | 2 | 0 | 98 | 49 | NA | NA | NA | NA | 0.333 | NA | 0.2 | NA | 0.133 |
| A3SS | 1109 | ZDHHC20 | chr13 | - | 21381433 | 21381549 | 21381433 | 21381546 | 21382919 | 21383009 | 0 | 0 | 6 | 0 | 0 | 3 | 0 | 1 | 0 | 0 | 0 | 0 | 52 | 49 | NA | NA | NA | NA | 0.653 | NA | 1 | NA | -0.347 |
| A3SS | 1139 | C17orf49 | chr17 | + | 7017256 | 7017524 | 7017260 | 7017524 | 7016909 | 7017014 | 0 | 0 | 0 | 0 | 0 | 2 | 0 | 1 | 0 | 0 | 3 | 0 | 53 | 49 | NA | NA | NA | NA | 0 | NA | 0.236 | NA | -0.236 |
| A3SS | 1193 | SUPT20H | chr13 | - | 37022010 | 37022317 | 37022010 | 37022080 | 37024034 | 37024193 | 0 | 0 | 2 | 0 | 0 | 1 | 0 | 1 | 0 | 0 | 0 | 0 | 98 | 49 | NA | NA | NA | NA | 0.5 | NA | 1 | NA | -0.5 |
| A3SS | 1201 | NFKB1 | chr4 | + | 102533844 | 102533885 | 102533847 | 102533885 | 102529835 | 102529914 | 0 | 0 | 0 | 0 | 0 | 1 | 0 | 1 | 0 | 0 | 1 | 0 | 52 | 49 | NA | NA | NA | NA | 0 | NA | 0.485 | NA | -0.485 |
| A3SS | 1269 | IRF5 | chr7 | + | 128947229 | 128947535 | 128947277 | 128947535 | 128947022 | 128947056 | 0 | 0 | 1 | 0 | 0 | 1 | 2 | 1 | 0 | 0 | 0 | 0 | 97 | 49 | NA | NA | NA | NA | 0.336 | 1 | 1 | NA | -0.664 |
| A3SS | 1279 | ING4 | chr12 | - | 6652661 | 6652770 | 6652661 | 6652767 | 6652935 | 6653050 | 0 | 0 | 2 | 0 | 0 | 2 | 0 | 0 | 0 | 0 | 2 | 0 | 52 | 49 | NA | NA | NA | NA | 0.485 | NA | 0 | NA | 0.485 |
| A3SS | 1301 | GMPR2 | chr14 | + | 24237041 | 24237152 | 24237070 | 24237152 | 24235966 | 24236140 | 0 | 0 | 0 | 0 | 0 | 1 | 0 | 1 | 0 | 0 | 2 | 0 | 78 | 49 | NA | NA | NA | NA | 0 | NA | 0.239 | NA | -0.239 |
| A3SS | 1322 | CASP1 | chr11 | - | 105025509 | 105026356 | 105025509 | 105025600 | 105026841 | 105026951 | 0 | 0 | 29 | 0 | 0 | 0 | 0 | 2 | 0 | 1 | 1 | 0 | 98 | 49 | NA | NA | NA | NA | 1 | 0 | 0.5 | NA | 0.75 |
| A3SS | 1358 | ANAPC15 | chr11 | - | 72109586 | 72109940 | 72109586 | 72109928 | 72110087 | 72110225 | 0 | 0 | 0 | 1 | 0 | 0 | 0 | 1 | 0 | 0 | 1 | 0 | 61 | 49 | NA | NA | 0 | NA | NA | NA | 0.445 | NA | -0.445 |
| A3SS | 1411 | DNM2 | chr19 | + | 10830977 | 10831910 | 10830980 | 10831910 | 10830126 | 10830378 | 0 | 2 | 1 | 0 | 0 | 1 | 1 | 0 | 0 | 0 | 0 | 0 | 52 | 49 | NA | NA | NA | 1 | 0.485 | 1 | NA | NA | -0.258 |
| A3SS | 1533 | TTC23 | chr15 | - | 99139316 | 99139467 | 99139316 | 99139399 | 99156147 | 99156297 | 0 | 0 | 0 | 0 | 0 | 1 | 0 | 1 | 0 | 0 | 0 | 0 | 98 | 49 | NA | NA | NA | NA | 0 | NA | 1 | NA | -1 |
| A3SS | 1537 | CARS | chr11 | - | 3040895 | 3041031 | 3040895 | 3040984 | 3042164 | 3042256 | 2 | 0 | 0 | 0 | 0 | 3 | 0 | 1 | 0 | 0 | 0 | 0 | 96 | 49 | NA | NA | 1 | NA | 0 | NA | 1 | NA | -0.5 |
| A3SS | 1600 | MECR | chr1 | - | 29216587 | 29216812 | 29216587 | 29216685 | 29230730 | 29230958 | 0 | 0 | 0 | 0 | 0 | 1 | 0 | 2 | 0 | 0 | 0 | 0 | 98 | 49 | NA | NA | NA | NA | 0 | NA | 1 | NA | -1 |
| A3SS | 1610 | FAM111A | chr11 | + | 59148796 | 59148953 | 59148835 | 59148953 | 59145826 | 59145856 | 0 | 0 | 0 | 0 | 1 | 0 | 3 | 0 | 0 | 0 | 0 | 1 | 88 | 49 | NA | NA | NA | 0 | NA | 1 | NA | 0 | -0.5 |
| A3SS | 1628 | SRSF10 | chr1 | - | 23964346 | 23967956 | 23964346 | 23967723 | 23971572 | 23971623 | 1 | 0 | 0 | 0 | 0 | 0 | 0 | 0 | 0 | 0 | 2 | 0 | 98 | 49 | NA | NA | 1 | NA | NA | NA | 0 | NA | 1 |
| A3SS | 1629 | SRSF10 | chr1 | - | 23964346 | 23971439 | 23964346 | 23967723 | 23971572 | 23971623 | 1 | 0 | 1 | 0 | 0 | 0 | 0 | 1 | 0 | 0 | 2 | 0 | 98 | 49 | NA | NA | 1 | NA | 1 | NA | 0.2 | NA | 0.8 |
| A3SS | 1635 | METTL9 | chr16 | + | 21655226 | 21657473 | 21655229 | 21657473 | 21624930 | 21625115 | 0 | 0 | 2 | 0 | 0 | 2 | 0 | 1 | 0 | 0 | 1 | 0 | 52 | 49 | NA | NA | NA | NA | 0.485 | NA | 0.485 | NA | 0 |
| A3SS | 1660 | SHC1 | chr1 | - | 154965945 | 154966080 | 154965945 | 154966077 | 154966161 | 154966231 | 0 | 0 | 0 | 0 | 0 | 3 | 0 | 2 | 0 | 0 | 0 | 0 | 52 | 49 | NA | NA | NA | NA | 0 | NA | 1 | NA | -1 |
| A3SS | 1669 | WDR47 | chr1 | - | 109004591 | 109004718 | 109004591 | 109004715 | 109010915 | 109011718 | 0 | 0 | 1 | 0 | 0 | 0 | 0 | 0 | 0 | 0 | 1 | 0 | 52 | 49 | NA | NA | NA | NA | 1 | NA | 0 | NA | 1 |
| A3SS | 1706 | RBMX | chrX | - | 136879016 | 136879158 | 136879016 | 136879123 | 136879318 | 136879453 | 0 | 0 | 0 | 0 | 0 | 4 | 0 | 1 | 0 | 0 | 0 | 0 | 84 | 49 | NA | NA | NA | NA | 0 | NA | 1 | NA | -1 |
| A3SS | 1749 | FAM213B | chr1 | + | 2588335 | 2588453 | 2588389 | 2588453 | 2587740 | 2587792 | 0 | 0 | 0 | 0 | 0 | 1 | 0 | 1 | 0 | 0 | 0 | 0 | 98 | 49 | NA | NA | NA | NA | 0 | NA | 1 | NA | -1 |
| A3SS | 1752 | FAM213B | chr1 | + | 2588921 | 2589040 | 2588953 | 2589040 | 2588549 | 2588625 | 0 | 0 | 1 | 0 | 0 | 0 | 0 | 0 | 0 | 0 | 2 | 0 | 81 | 49 | NA | NA | NA | NA | 1 | NA | 0 | NA | 1 |
| A3SS | 1807 | PCBP2 | chr12 | + | 53468776 | 53468832 | 53468779 | 53468832 | 53467220 | 53467293 | 0 | 0 | 7 | 0 | 0 | 2 | 1 | 3 | 0 | 0 | 0 | 0 | 52 | 49 | NA | NA | NA | NA | 0.767 | 1 | 1 | NA | -0.233 |
| A3SS | 1808 | PCBP2 | chr12 | + | 53468776 | 53468832 | 53468779 | 53468832 | 53467804 | 53467843 | 0 | 0 | 5 | 0 | 0 | 0 | 0 | 1 | 0 | 0 | 2 | 0 | 52 | 49 | NA | NA | NA | NA | 1 | NA | 0.32 | NA | 0.68 |
| A3SS | 1847 | ABI1 | chr10 | - | 26759061 | 26759238 | 26759061 | 26759235 | 26765217 | 26765318 | 0 | 0 | 1 | 0 | 0 | 2 | 0 | 1 | 0 | 0 | 0 | 0 | 52 | 49 | NA | NA | NA | NA | 0.32 | NA | 1 | NA | -0.68 |
| A3SS | 1876 | PSME3 | chr17 | + | 42838691 | 42838799 | 42838730 | 42838799 | 42838092 | 42838205 | 0 | 0 | 1 | 0 | 0 | 4 | 0 | 3 | 0 | 0 | 6 | 0 | 88 | 49 | NA | NA | NA | NA | 0.122 | NA | 0.218 | NA | -0.096 |
| A3SS | 1918 | EPB41 | chr1 | + | 28987430 | 28987905 | 28987447 | 28987905 | 28887090 | 28887210 | 1 | 0 | 0 | 0 | 0 | 0 | 0 | 0 | 0 | 0 | 0 | 1 | 66 | 49 | NA | NA | 1 | NA | NA | NA | NA | 0 | 1 |
| A3SS | 1935 | ATL2 | chr2 | - | 38293956 | 38296590 | 38293956 | 38296113 | 38298143 | 38298575 | 0 | 0 | 1 | 0 | 0 | 0 | 0 | 0 | 0 | 0 | 1 | 0 | 98 | 49 | NA | NA | NA | NA | 1 | NA | 0 | NA | 1 |
| A3SS | 1978 | PCID2 | chr13 | - | 113177538 | 113178008 | 113177538 | 113177976 | 113178173 | 113178287 | 0 | 0 | 3 | 0 | 0 | 0 | 0 | 0 | 0 | 0 | 1 | 0 | 81 | 49 | NA | NA | NA | NA | 1 | NA | 0 | NA | 1 |
| A3SS | 1994 | WNK1 | chr12 | + | 878211 | 878361 | 878214 | 878361 | 862082 | 862270 | 0 | 0 | 1 | 0 | 0 | 2 | 0 | 2 | 0 | 0 | 0 | 0 | 52 | 49 | NA | NA | NA | NA | 0.32 | NA | 1 | NA | -0.68 |

| | | | | | | | | | | | | | | | | | | | | | | | | | | | | | | | | | |
|---|---|---|---|---|---|---|---|---|---|---|---|---|---|---|---|---|---|---|---|---|---|---|---|---|---|---|---|---|---|---|---|---|---|
| A3SS | 1995 | WNK1 | chr12 | + | 878211 | 878361 | 878214 | 878361 | 871264 | 871348 | 0 | 2 | 0 | 0 | 0 | 1 | 3 | 0 | 0 | 0 | 0 | 0 | 52 | 49 | NA | NA | NA | 1 | 0 | 1 | NA | NA | -0.5 |
| A3SS | 2020 | TRPM7 | chr15 | - | 50583088 | 50583159 | 50583088 | 50583156 | 50586391 | 50586488 | 0 | 0 | 1 | 0 | 0 | 0 | 0 | 0 | 0 | 0 | 1 | 0 | 52 | 49 | NA | NA | NA | NA | 1 | NA | 0 | NA | 1 |
| A3SS | 2077 | CD300LF | chr17 | - | 74703034 | 74703143 | 74703034 | 74703098 | 74704477 | 74704816 | 0 | 0 | 2 | 0 | 0 | 4 | 0 | 2 | 0 | 0 | 3 | 0 | 94 | 49 | NA | NA | NA | NA | 0.207 | NA | 0.258 | NA | -0.051 |
| A3SS | 2103 | RALGAPB | chr20 | + | 38565355 | 38565478 | 38565358 | 38565478 | 38562531 | 38562697 | 0 | 0 | 0 | 0 | 0 | 2 | 0 | 1 | 0 | 0 | 0 | 0 | 52 | 49 | NA | NA | NA | NA | 0 | NA | 1 | NA | -1 |
| A3SS | 2128 | ARHGEF2 | chr1 | - | 155964987 | 155965131 | 155964987 | 155965128 | 155965302 | 155965412 | 0 | 0 | 2 | 0 | 0 | 4 | 0 | 1 | 0 | 0 | 1 | 0 | 52 | 49 | NA | NA | NA | NA | 0.32 | NA | 0.485 | NA | -0.165 |
| A3SS | 2171 | TSC2 | chr16 | + | 2079031 | 2079196 | 2079034 | 2079196 | 2076490 | 2076585 | 0 | 0 | 1 | 0 | 0 | 0 | 0 | 1 | 0 | 0 | 2 | 0 | 52 | 49 | NA | NA | NA | NA | 1 | NA | 0.32 | NA | 0.68 |
| A3SS | 2175 | ATP6V0B | chr1 | + | 43977025 | 43978300 | 43977980 | 43978300 | 43976772 | 43976824 | 1 | 0 | 10 | 0 | 0 | 0 | 0 | 4 | 0 | 0 | 1 | 0 | 98 | 49 | NA | NA | 1 | NA | 1 | NA | 0.667 | NA | 0.333 |
| A3SS | 2202 | ATF7IP | chr12 | + | 14434336 | 14434423 | 14434339 | 14434423 | 14423908 | 14425473 | 0 | 0 | 6 | 0 | 0 | 3 | 0 | 1 | 0 | 0 | 0 | 0 | 52 | 49 | NA | NA | NA | NA | 0.653 | NA | 1 | NA | -0.347 |
| A3SS | 2205 | ZNF33A | chr10 | + | 38054374 | 38060067 | 38054377 | 38060067 | 38017290 | 38017386 | 0 | 0 | 1 | 0 | 0 | 1 | 0 | 2 | 0 | 0 | 2 | 0 | 52 | 49 | NA | NA | NA | NA | 0.485 | NA | 0.485 | NA | 0 |
| A3SS | 2214 | ZWILCH | chr15 | + | 66508745 | 66508892 | 66508840 | 66508892 | 66505082 | 66505391 | 0 | 0 | 0 | 0 | 0 | 2 | 6 | 0 | 0 | 0 | 0 | 0 | 98 | 49 | NA | NA | NA | NA | 0 | 1 | NA | NA | -1 |
| A3SS | 2268 | SH2D2A | chr1 | - | 156814204 | 156814324 | 156814204 | 156814294 | 156815036 | 156815221 | 0 | 0 | 2 | 0 | 0 | 1 | 1 | 0 | 0 | 0 | 0 | 0 | 79 | 49 | NA | NA | NA | NA | 0.554 | 1 | NA | NA | -0.446 |
| A3SS | 2286 | COPA | chr1 | - | 160305432 | 160305598 | 160305432 | 160305571 | 160305687 | 160305773 | 0 | 0 | 4 | 0 | 0 | 3 | 0 | 0 | 0 | 0 | 4 | 0 | 76 | 49 | NA | NA | NA | NA | 0.462 | NA | 0 | NA | 0.462 |
| A3SS | 2304 | CAST | chr5 | + | 96765220 | 96765325 | 96765223 | 96765325 | 96762273 | 96762372 | 0 | 0 | 8 | 0 | 0 | 3 | 1 | 0 | 0 | 0 | 2 | 0 | 52 | 49 | NA | NA | NA | NA | 0.715 | 1 | 0 | NA | 0.215 |
| A3SS | 2364 | DGKZ | chr11 | + | 46379453 | 46379568 | 46379468 | 46379568 | 46379202 | 46379236 | 0 | 0 | 0 | 0 | 0 | 2 | 0 | 1 | 0 | 0 | 0 | 0 | 64 | 49 | NA | NA | NA | NA | 0 | NA | 1 | NA | -1 |
| A3SS | 2371 | ZBP1 | chr20 | - | 57613162 | 57613330 | 57613162 | 57613327 | 57614886 | 57615060 | 0 | 0 | 23 | 0 | 0 | 2 | 0 | 3 | 0 | 0 | 0 | 0 | 52 | 49 | NA | NA | NA | NA | 0.916 | NA | 1 | NA | -0.084 |
| A3SS | 2404 | OS9 | chr12 | + | 57716088 | 57716193 | 57716091 | 57716193 | 57715759 | 57715970 | 0 | 0 | 2 | 0 | 0 | 1 | 1 | 2 | 0 | 0 | 0 | 0 | 52 | 49 | NA | NA | NA | NA | 0.653 | 1 | 1 | NA | -0.347 |
| A3SS | 2490 | ESYT1 | chr12 | + | 56134079 | 56134181 | 56134109 | 56134181 | 56133780 | 56133873 | 0 | 0 | 1 | 0 | 0 | 1 | 1 | 0 | 0 | 0 | 1 | 0 | 79 | 49 | NA | NA | NA | NA | 0.383 | 1 | 0 | NA | -0.117 |
| A3SS | 2492 | SON | chr21 | + | 33576364 | 33577514 | 33576417 | 33577514 | 33575777 | 33575893 | 0 | 0 | 5 | 0 | 0 | 0 | 0 | 1 | 0 | 0 | 1 | 0 | 98 | 49 | NA | NA | NA | NA | 1 | NA | 0.333 | NA | 0.667 |
| A3SS | 2500 | FAM65B | chr6 | - | 24839365 | 24840775 | 24839365 | 24839599 | 24842861 | 24843554 | 0 | 0 | 20 | 0 | 0 | 2 | 0 | 13 | 0 | 0 | 1 | 0 | 98 | 49 | NA | NA | NA | NA | 0.833 | NA | 0.867 | NA | -0.034 |
| A3SS | 2521 | ZNF124 | chr1 | - | 247154988 | 247157403 | 247154988 | 247157217 | 247159005 | 247159066 | 0 | 0 | 0 | 0 | 0 | 2 | 1 | 0 | 0 | 0 | 1 | 0 | 98 | 49 | NA | NA | NA | NA | 0 | 1 | 0 | NA | -0.5 |
| A3SS | 2552 | CELF2 | chr10 | + | 11321188 | 11321386 | 11321206 | 11321386 | 11314138 | 11314258 | 0 | 0 | 1 | 0 | 0 | 3 | 0 | 2 | 0 | 0 | 1 | 0 | 67 | 49 | NA | NA | NA | NA | 0.196 | NA | 0.594 | NA | -0.398 |
| A3SS | 2560 | SERPINB1 | chr6 | - | 2835855 | 2836023 | 2835855 | 2836019 | 2836107 | 2836250 | 0 | 0 | 15 | 0 | 0 | 3 | 0 | 12 | 0 | 0 | 1 | 0 | 53 | 49 | NA | NA | NA | NA | 0.822 | NA | 0.917 | NA | -0.095 |
| A3SS | 2588 | APTX | chr9 | - | 32988078 | 32988129 | 32988078 | 32988124 | 32989758 | 32989895 | 0 | 0 | 0 | 0 | 0 | 1 | 0 | 1 | 0 | 0 | 0 | 0 | 54 | 49 | NA | NA | NA | NA | 0 | NA | 1 | NA | -1 |
| A3SS | 2594 | SLC25A45 | chr11 | - | 65380131 | 65380234 | 65380131 | 65380175 | 65380533 | 65380632 | 0 | 0 | 0 | 0 | 0 | 1 | 0 | 1 | 0 | 0 | 0 | 0 | 98 | 49 | NA | NA | NA | NA | 0 | NA | 1 | NA | -1 |
| A3SS | 2616 | RNH1 | chr11 | - | 502061 | 502249 | 502061 | 502181 | 504823 | 504996 | 0 | 0 | 5 | 0 | 0 | 2 | 0 | 2 | 0 | 0 | 0 | 0 | 98 | 49 | NA | NA | NA | NA | 0.556 | NA | 1 | NA | -0.444 |
| A3SS | 2632 | BRD7 | chr16 | - | 50320662 | 50320777 | 50320662 | 50320774 | 50321981 | 50322038 | 0 | 0 | 1 | 0 | 0 | 0 | 0 | 0 | 0 | 0 | 4 | 0 | 52 | 49 | NA | NA | NA | NA | 1 | NA | 0 | NA | 1 |
| A3SS | 2668 | TCF7L2 | chr10 | + | 113150982 | 113151123 | 113150997 | 113151123 | 113146010 | 113146097 | 1 | 0 | 0 | 0 | 0 | 0 | 0 | 0 | 0 | 0 | 1 | 0 | 64 | 49 | NA | NA | 1 | NA | NA | NA | 0 | NA | 1 |
| A3SS | 2709 | ELF2 | chr4 | - | 139071865 | 139072039 | 139071865 | 139072003 | 139073453 | 139073567 | 0 | 0 | 4 | 0 | 0 | 0 | 0 | 4 | 0 | 0 | 1 | 0 | 85 | 49 | NA | NA | NA | NA | 1 | NA | 0.698 | NA | 0.302 |
| A3SS | 2713 | SNRNP70 | chr19 | + | 49107794 | 49108613 | 49107821 | 49108613 | 49107624 | 49107712 | 0 | 0 | 1 | 0 | 0 | 1 | 0 | 3 | 0 | 0 | 0 | 0 | 76 | 49 | NA | NA | NA | NA | 0.392 | NA | 1 | NA | -0.608 |
| A3SS | 2725 | MEF2C | chr5 | - | 88718240 | 88722925 | 88718240 | 88722829 | 88728492 | 88728628 | 0 | 0 | 1 | 0 | 0 | 1 | 3 | 0 | 0 | 0 | 0 | 0 | 98 | 49 | NA | NA | NA | NA | 0.333 | 1 | NA | NA | -0.667 |
| A3SS | 2727 | AFTPH | chr2 | + | 64591884 | 64593004 | 64591887 | 64593004 | 64585421 | 64585545 | 0 | 0 | 1 | 0 | 0 | 0 | 0 | 1 | 0 | 0 | 1 | 0 | 52 | 49 | NA | NA | NA | NA | 1 | NA | 0.485 | NA | 0.515 |
| A3SS | 2728 | SLC43A3 | chr11 | - | 57425540 | 57425709 | 57425540 | 57425670 | 57425988 | 57426389 | 0 | 0 | 0 | 0 | 0 | 2 | 0 | 2 | 0 | 0 | 0 | 0 | 88 | 49 | NA | NA | NA | NA | 0 | NA | 1 | NA | -1 |
| A3SS | 2780 | UPP1 | chr7 | + | 48090187 | 48090364 | 48090249 | 48090364 | 48089211 | 48089418 | 0 | 0 | 1 | 0 | 0 | 0 | 0 | 2 | 0 | 0 | 1 | 0 | 98 | 49 | NA | NA | NA | NA | 1 | NA | 0.5 | NA | 0.5 |
| A3SS | 2861 | TPM3 | chr1 | - | 154170399 | 154170469 | 154170399 | 154170465 | 154170648 | 154170711 | 0 | 0 | 19 | 0 | 0 | 1 | 2 | 14 | 0 | 0 | 0 | 0 | 53 | 49 | NA | NA | NA | NA | 0.946 | 1 | 1 | NA | -0.054 |
| A3SS | 2901 | MEFV | chr16 | - | 3243859 | 3244067 | 3243859 | 3243892 | 3244253 | 3244286 | 0 | 0 | 8 | 0 | 0 | 3 | 0 | 1 | 0 | 0 | 1 | 0 | 98 | 49 | NA | NA | NA | NA | 0.571 | NA | 0.333 | NA | 0.238 |
| A3SS | 2926 | COPS2 | chr15 | - | 49137347 | 49137458 | 49137347 | 49137437 | 49139527 | 49139653 | 0 | 0 | 2 | 0 | 0 | 2 | 0 | 1 | 0 | 0 | 0 | 0 | 70 | 49 | NA | NA | NA | NA | 0.412 | NA | 1 | NA | -0.588 |
| A3SS | 2936 | DMXL2 | chr15 | - | 51476588 | 51476719 | 51476588 | 51476716 | 51478270 | 51478347 | 0 | 0 | 8 | 0 | 0 | 2 | 0 | 2 | 0 | 0 | 0 | 0 | 52 | 49 | NA | NA | NA | NA | 0.79 | NA | 1 | NA | -0.21 |
| A3SS | 2938 | PIK3R5 | chr17 | - | 8888170 | 8888953 | 8888170 | 8888891 | 8889138 | 8889222 | 0 | 0 | 2 | 0 | 0 | 2 | 0 | 1 | 0 | 0 | 1 | 0 | 98 | 49 | NA | NA | NA | NA | 0.333 | NA | 0.333 | NA | 0 |
| A3SS | 2952 | ZNF143 | chr11 | + | 9474549 | 9474633 | 9474552 | 9474633 | 9473940 | 9474024 | 0 | 0 | 1 | 0 | 0 | 3 | 0 | 4 | 0 | 0 | 1 | 0 | 52 | 49 | NA | NA | NA | NA | 0.239 | NA | 0.79 | NA | -0.551 |
| A3SS | 2964 | PTMA | chr2 | + | 231711886 | 231711983 | 231711889 | 231711983 | 231711347 | 231711419 | 0 | 0 | 1 | 0 | 0 | 2 | 0 | 4 | 0 | 0 | 1 | 0 | 52 | 49 | NA | NA | NA | NA | 0.32 | NA | 0.79 | NA | -0.47 |
| A3SS | 2985 | ST20 | chr15 | - | 79907628 | 79907706 | 79907628 | 79907684 | 79914587 | 79915256 | 0 | 0 | 1 | 0 | 0 | 5 | 1 | 0 | 0 | 0 | 0 | 0 | 71 | 49 | NA | NA | NA | NA | 0.121 | 1 | NA | NA | -0.879 |
| A3SS | 2988 | IL16 | chr15 | + | 81306419 | 81306545 | 81306422 | 81306545 | 81305907 | 81306166 | 0 | 0 | 1 | 0 | 0 | 1 | 0 | 0 | 0 | 0 | 4 | 0 | 52 | 49 | NA | NA | NA | NA | 0.485 | NA | 0 | NA | 0.485 |
| A3SS | 3007 | AKAP13 | chr15 | + | 85710578 | 85710645 | 85710581 | 85710645 | 85708018 | 85708086 | 0 | 0 | 13 | 0 | 0 | 0 | 0 | 10 | 0 | 0 | 2 | 0 | 52 | 49 | NA | NA | NA | NA | 1 | NA | 0.825 | NA | 0.175 |
| A3SS | 3057 | ZNF200 | chr16 | - | 3222324 | 3224613 | 3222324 | 3224610 | 3232420 | 3232547 | 0 | 0 | 1 | 0 | 0 | 0 | 0 | 0 | 0 | 0 | 1 | 0 | 52 | 49 | NA | NA | NA | NA | 1 | NA | 0 | NA | 1 |
| A3SS | 3080 | CLEC16A | chr16 | + | 10972931 | 10973061 | 10972937 | 10973061 | 10971124 | 10971230 | 0 | 0 | 2 | 0 | 0 | 0 | 0 | 0 | 0 | 0 | 2 | 0 | 55 | 49 | NA | NA | NA | NA | 1 | NA | 0 | NA | 1 |
| A3SS | 3089 | KIAA0430 | chr16 | - | 15622933 | 15623126 | 15622933 | 15623123 | 15624768 | 15624927 | 0 | 0 | 1 | 0 | 0 | 0 | 0 | 0 | 0 | 0 | 2 | 0 | 52 | 49 | NA | NA | NA | NA | 1 | NA | 0 | NA | 1 |
| A3SS | 3135 | FUS | chr16 | + | 31183857 | 31184002 | 31183860 | 31184002 | 31182512 | 31182664 | 0 | 0 | 5 | 0 | 0 | 3 | 0 | 3 | 0 | 1 | 3 | 0 | 52 | 49 | NA | NA | NA | NA | 0.611 | 0 | 0.485 | NA | 0.368 |
| A3SS | 3251 | LRMP | chr12 | + | 25063719 | 25063816 | 25063722 | 25063816 | 25062819 | 25062900 | 0 | 0 | 3 | 0 | 0 | 2 | 0 | 5 | 0 | 0 | 1 | 0 | 52 | 49 | NA | NA | NA | NA | 0.586 | NA | 0.825 | NA | -0.239 |
| A3SS | 3275 | WDTC1 | chr1 | + | 27297047 | 27297156 | 27297050 | 27297156 | 27296325 | 27296401 | 0 | 0 | 4 | 0 | 0 | 1 | 0 | 1 | 0 | 0 | 0 | 0 | 52 | 49 | NA | NA | NA | NA | 0.79 | NA | 1 | NA | -0.21 |
| A3SS | 3499 | TIAL1 | chr10 | - | 119582458 | 119582608 | 119582458 | 119582557 | 119588151 | 119588248 | 0 | 0 | 0 | 0 | 0 | 1 | 3 | 0 | 0 | 0 | 0 | 0 | 98 | 49 | NA | NA | NA | NA | 0 | 1 | NA | NA | -1 |
| A3SS | 3508 | EXOC7 | chr17 | - | 76090328 | 76090481 | 76090328 | 76090397 | 76091142 | 76091235 | 1 | 0 | 0 | 0 | 0 | 0 | 0 | 0 | 0 | 0 | 1 | 0 | 98 | 49 | NA | NA | 1 | NA | NA | NA | 0 | NA | 1 |
| A3SS | 3511 | RHBDF2 | chr17 | - | 76479732 | 76479941 | 76479732 | 76479854 | 76481374 | 76481545 | 0 | 0 | 1 | 0 | 0 | 5 | 4 | 0 | 0 | 0 | 2 | 0 | 98 | 49 | NA | NA | NA | NA | 0.091 | 1 | 0 | NA | -0.409 |
| A3SS | 3686 | GTPBP3 | chr19 | + | 17341043 | 17341322 | 17341106 | 17341322 | 17339433 | 17339599 | 0 | 0 | 0 | 0 | 2 | 0 | 3 | 0 | 0 | 0 | 0 | 0 | 98 | 49 | NA | NA | NA | 0 | NA | 1 | NA | NA | -1 |
| A3SS | 3695 | TMEM59 | chr1 | - | 54040755 | 54040840 | 54040755 | 54040837 | 54041723 | 54041805 | 0 | 0 | 0 | 0 | 0 | 2 | 0 | 1 | 0 | 0 | 1 | 0 | 52 | 49 | NA | NA | NA | NA | 0 | NA | 0.485 | NA | -0.485 |
| A3SS | 3701 | UXS1 | chr2 | - | 106164735 | 106164799 | 106164735 | 106164784 | 106166055 | 106166083 | 0 | 0 | 1 | 0 | 0 | 0 | 0 | 0 | 0 | 0 | 4 | 0 | 64 | 49 | NA | NA | NA | NA | 1 | NA | 0 | NA | 1 |
| A3SS | 3723 | PITPNM1 | chr11 | - | 67496177 | 67496348 | 67496177 | 67496345 | 67497230 | 67497436 | 0 | 0 | 2 | 0 | 0 | 0 | 0 | 0 | 0 | 0 | 1 | 0 | 52 | 49 | NA | NA | NA | NA | 1 | NA | 0 | NA | 1 |
| A3SS | 3744 | TMEM91 | chr19 | + | 41383714 | 41384082 | 41383862 | 41384082 | 41382771 | 41382921 | 0 | 0 | 2 | 0 | 0 | 3 | 0 | 1 | 0 | 0 | 2 | 0 | 98 | 49 | NA | NA | NA | NA | 0.25 | NA | 0.2 | NA | 0.05 |
| A3SS | 3789 | PTOV1 | chr19 | + | 49860565 | 49860744 | 49860613 | 49860744 | 49860267 | 49860336 | 1 | 0 | 3 | 0 | 0 | 3 | 0 | 2 | 0 | 0 | 0 | 0 | 97 | 49 | NA | NA | 1 | NA | 0.336 | NA | 1 | NA | -0.332 |
| A3SS | 3809 | NFRKB | chr11 | - | 129884744 | 129884882 | 129884744 | 129884846 | 129885434 | 129885609 | 0 | 0 | 1 | 0 | 0 | 0 | 0 | 2 | 0 | 4 | 1 | 0 | 85 | 49 | NA | NA | NA | NA | 1 | 0 | 0.536 | NA | 0.732 |
| A3SS | 3838 | LILRB2 | chr19 | - | 54276806 | 54276979 | 54276806 | 54276929 | 54277549 | 54277597 | 0 | 0 | 1 | 0 | 0 | 3 | 0 | 0 | 0 | 7 | 1 | 0 | 98 | 49 | NA | NA | NA | NA | 0.143 | 0 | 0 | NA | 0.143 |
| A3SS | 3922 | ODC1 | chr2 | - | 10444473 | 10444647 | 10444473 | 10444547 | 10444930 | 10445049 | 0 | 0 | 2 | 0 | 0 | 1 | 3 | 1 | 0 | 0 | 0 | 0 | 98 | 49 | NA | NA | NA | NA | 0.5 | 1 | 1 | NA | -0.5 |
| A3SS | 3939 | SUPT7L | chr2 | - | 27660983 | 27661443 | 27660983 | 27661388 | 27662178 | 27662281 | 0 | 0 | 0 | 0 | 0 | 1 | 0 | 1 | 0 | 0 | 0 | 0 | 98 | 49 | NA | NA | NA | NA | 0 | NA | 1 | NA | -1 |
| A3SS | 3940 | SUPT7L | chr2 | - | 27660983 | 27661601 | 27660983 | 27661388 | 27662178 | 27662281 | 0 | 0 | 0 | 0 | 0 | 1 | 0 | 1 | 0 | 0 | 0 | 0 | 98 | 49 | NA | NA | NA | NA | 0 | NA | 1 | NA | -1 |
| A3SS | 3952 | PPM1B | chr2 | + | 44229992 | 44231473 | 44230412 | 44231473 | 44218479 | 44218537 | 0 | 0 | 3 | 0 | 0 | 2 | 0 | 1 | 0 | 0 | 1 | 0 | 98 | 49 | NA | NA | NA | NA | 0.429 | NA | 0.333 | NA | 0.096 |
| A3SS | 3977 | AUP1 | chr2 | - | 74526940 | 74527121 | 74526940 | 74527059 | 74527247 | 74527363 | 0 | 0 | 0 | 0 | 0 | 3 | 1 | 1 | 0 | 8 | 8 | 0 | 98 | 49 | NA | NA | NA | NA | 0 | 0.059 | 0.059 | NA | -0.059 |
| A3SS | 3997 | SERPINA1 | chr14 | - | 94388559 | 94388663 | 94388559 | 94388660 | 94388800 | 94389010 | 0 | 0 | 3 | 0 | 0 | 2 | 0 | 0 | 0 | 0 | 1 | 0 | 52 | 49 | NA | NA | NA | NA | 0.586 | NA | 0 | NA | 0.586 |
| A3SS | 3998 | SERPINA1 | chr14 | - | 94388559 | 94388663 | 94388559 | 94388660 | 94388818 | 94389010 | 0 | 0 | 3 | 0 | 0 | 1 | 0 | 1 | 0 | 0 | 0 | 0 | 52 | 49 | NA | NA | NA | NA | 0.739 | NA | 1 | NA | -0.261 |
| A5SS | 6 | NAA10 | chrX | - | 153933942 | 153934001 | 153933960 | 153934001 | 153932538 | 153932584 | 0 | 0 | 1 | 0 | 0 | 0 | 0 | 2 | 0 | 0 | 1 | 0 | 67 | 49 | NA | NA | NA | NA | 1 | NA | 0.594 | NA | 0.406 |
| A5SS | 12 | AFF2 | chrX | + | 148651998 | 148652131 | 148651998 | 148652119 | 148661907 | 148662768 | 0 | 0 | 0 | 0 | 0 | 1 | 2 | 0 | 0 | 0 | 1 | 0 | 61 | 49 | NA | NA | NA | NA | 0 | 1 | 0 | NA | -0.5 |
| A5SS | 57 | OTUD5 | chrX | - | 48934695 | 48934867 | 48934710 | 48934867 | 48934463 | 48934612 | 0 | 0 | 3 | 0 | 0 | 2 | 0 | 0 | 0 | 0 | 9 | 0 | 64 | 49 | NA | NA | NA | NA | 0.535 | NA | 0 | NA | 0.535 |
| A5SS | 149 | ENY2 | chr8 | + | 109334322 | 109334474 | 109334322 | 109334402 | 109336127 | 109336204 | 0 | 0 | 1 | 0 | 0 | 2 | 0 | 1 | 0 | 0 | 0 | 0 | 98 | 49 | NA | NA | NA | NA | 0.2 | NA | 1 | NA | -0.8 |
| A5SS | 154 | LYN | chr8 | + | 55941854 | 55941991 | 55941854 | 55941928 | 55946447 | 55946493 | 0 | 0 | 16 | 0 | 0 | 3 | 0 | 4 | 0 | 0 | 6 | 0 | 98 | 49 | NA | NA | NA | NA | 0.727 | NA | 0.25 | NA | 0.477 |
| A5SS | 167 | ADAM28 | chr8 | + | 24343505 | 24343588 | 24343505 | 24343584 | 24349863 | 24349972 | 0 | 0 | 0 | 0 | 0 | 1 | 1 | 0 | 0 | 0 | 0 | 0 | 53 | 49 | NA | NA | NA | NA | 0 | 1 | NA | NA | -1 |
| A5SS | 168 | ENTPD4 | chr8 | - | 23441568 | 23441723 | 23441592 | 23441723 | 23439748 | 23439915 | 0 | 0 | 2 | 0 | 0 | 0 | 0 | 0 | 0 | 0 | 2 | 0 | 73 | 49 | NA | NA | NA | NA | 1 | NA | 0 | NA | 1 |
| A5SS | 223 | IKZF1 | chr7 | + | 50391728 | 50391863 | 50391728 | 50391833 | 50399917 | 50405100 | 0 | 0 | 2 | 0 | 0 | 3 | 3 | 0 | 0 | 0 | 0 | 0 | 79 | 49 | NA | NA | NA | NA | 0.293 | 1 | NA | NA | -0.707 |
| A5SS | 230 | CPVL | chr7 | - | 29146340 | 29146537 | 29146428 | 29146537 | 29120892 | 29121071 | 1 | 0 | 0 | 1 | 0 | 0 | 0 | 0 | 0 | 0 | 2 | 0 | 98 | 49 | NA | NA | 0.333 | NA | NA | NA | 0 | NA | 0.333 |
| A5SS | 232 | SNX10 | chr7 | + | 26291894 | 26292133 | 26291894 | 26292086 | 26346419 | 26346466 | 0 | 0 | 0 | 0 | 0 | 2 | 3 | 0 | 0 | 0 | 0 | 0 | 96 | 49 | NA | NA | NA | NA | 0 | 1 | NA | NA | -1 |
| A5SS | 275 | CUTA | chr6 | - | 33417868 | 33418288 | 33417958 | 33418288 | 33417480 | 33417695 | 0 | 0 | 0 | 0 | 0 | 1 | 4 | 0 | 0 | 0 | 0 | 0 | 98 | 49 | NA | NA | NA | NA | 0 | 1 | NA | NA | -1 |
| A5SS | 383 | USP9X | chrX | + | 41229566 | 41229827 | 41229566 | 41229779 | 41230500 | 41230596 | 0 | 0 | 0 | 0 | 0 | 1 | 0 | 5 | 0 | 0 | 4 | 0 | 97 | 49 | NA | NA | NA | NA | 0 | NA | 0.387 | NA | -0.387 |
| A5SS | 400 | CANX | chr5 | + | 179698928 | 179699102 | 179698928 | 179699049 | 179705678 | 179705852 | 0 | 0 | 0 | 0 | 0 | 1 | 0 | 1 | 0 | 0 | 0 | 0 | 98 | 49 | NA | NA | NA | NA | 0 | NA | 1 | NA | -1 |
| A5SS | 469 | TMEM70 | chr8 | + | 73978755 | 73978875 | 73978755 | 73978861 | 73981154 | 73982783 | 0 | 0 | 0 | 0 | 0 | 1 | 3 | 0 | 0 | 0 | 0 | 0 | 63 | 49 | NA | NA | NA | NA | 0 | 1 | NA | NA | -1 |
| A5SS | 484 | LIN54 | chr4 | - | 82984160 | 82984876 | 82984823 | 82984876 | 82978882 | 82979006 | 1 | 0 | 0 | 0 | 0 | 0 | 0 | 0 | 0 | 0 | 2 | 0 | 98 | 49 | NA | NA | 1 | NA | NA | NA | 0 | NA | 1 |
| A5SS | 496 | KIT | chr4 | + | 54725856 | 54726050 | 54725856 | 54726038 | 54727217 | 54727324 | 0 | 0 | 0 | 0 | 0 | 1 | 1 | 0 | 0 | 0 | 0 | 0 | 61 | 49 | NA | NA | NA | NA | 0 | 1 | NA | NA | -1 |
| A5SS | 507 | CPEB2 | chr4 | + | 15007304 | 15007595 | 15007304 | 15007586 | 15017187 | 15017278 | 0 | 0 | 1 | 0 | 0 | 2 | 0 | 0 | 0 | 0 | 3 | 0 | 58 | 49 | NA | NA | NA | NA | 0.297 | NA | 0 | NA | 0.297 |
| A5SS | 511 | TMEM175 | chr4 | + | 948115 | 948615 | 948115 | 948154 | 950420 | 950518 | 0 | 0 | 0 | 0 | 0 | 1 | 0 | 1 | 0 | 0 | 0 | 0 | 98 | 49 | NA | NA | NA | NA | 0 | NA | 1 | NA | -1 |
| A5SS | 546 | MBD4 | chr3 | - | 129436442 | 129437308 | 129436460 | 129437308 | 129434061 | 129434136 | 0 | 0 | 0 | 0 | 0 | 1 | 0 | 1 | 0 | 0 | 0 | 0 | 67 | 49 | NA | NA | NA | NA | 0 | NA | 1 | NA | -1 |
| A5SS | 549 | CNBP | chr3 | - | 129171633 | 129171771 | 129171654 | 129171771 | 129171445 | 129171538 | 0 | 0 | 4 | 0 | 0 | 5 | 0 | 2 | 0 | 0 | 4 | 0 | 70 | 49 | NA | NA | NA | NA | 0.359 | NA | 0.259 | NA | 0.1 |
| A5SS | 556 | PARP9 | chr3 | - | 122558328 | 122558467 | 122558433 | 122558467 | 122555285 | 122556121 | 0 | 0 | 10 | 0 | 0 | 6 | 1 | 3 | 0 | 0 | 1 | 0 | 98 | 49 | NA | NA | NA | NA | 0.455 | 1 | 0.6 | NA | -0.345 |
| A5SS | 558 | GOLGB1 | chr3 | - | 121730847 | 121730973 | 121730875 | 121730973 | 121729864 | 121730017 | 0 | 0 | 2 | 0 | 0 | 0 | 0 | 1 | 0 | 0 | 1 | 0 | 77 | 49 | NA | NA | NA | NA | 1 | NA | 0.389 | NA | 0.611 |
| A5SS | 572 | CCDC66 | chr3 | + | 56566593 | 56566759 | 56566593 | 56566740 | 56566949 | 56567053 | 0 | 0 | 1 | 0 | 0 | 0 | 0 | 0 | 0 | 6 | 0 | 0 | 68 | 49 | NA | NA | NA | NA | 1 | 0 | NA | NA | 1 |
| A5SS | 603 | FGD5-AS1 | chr3 | - | 14947086 | 14947505 | 14947109 | 14947505 | 14942778 | 14946154 | 0 | 0 | 1 | 0 | 0 | 0 | 0 | 0 | 0 | 0 | 3 | 0 | 72 | 49 | NA | NA | NA | NA | 1 | NA | 0 | NA | 1 |
| A5SS | 607 | BRPF1 | chr3 | + | 9742024 | 9742171 | 9742024 | 9742153 | 9742943 | 9743253 | 0 | 0 | 1 | 0 | 0 | 2 | 0 | 3 | 0 | 0 | 0 | 0 | 67 | 49 | NA | NA | NA | NA | 0.268 | NA | 1 | NA | -0.732 |
| A5SS | 635 | TANGO2 | chr22 | + | 20036759 | 20037094 | 20036759 | 20036854 | 20043354 | 20043443 | 1 | 0 | 2 | 0 | 0 | 7 | 0 | 3 | 0 | 0 | 0 | 0 | 98 | 49 | NA | NA | 1 | NA | 0.125 | NA | 1 | NA | -0.438 |
| A5SS | 684 | NUB1 | chr7 | + | 151374096 | 151374243 | 151374096 | 151374201 | 151375847 | 151375943 | 2 | 0 | 3 | 0 | 0 | 1 | 0 | 1 | 0 | 0 | 2 | 0 | 91 | 49 | NA | NA | 1 | NA | 0.618 | NA | 0.212 | NA | 0.597 |
| A5SS | 702 | C20orf24 | chr20 | + | 36609600 | 36609758 | 36609600 | 36609730 | 36612009 | 36612557 | 0 | 0 | 1 | 0 | 0 | 4 | 0 | 1 | 0 | 0 | 6 | 0 | 77 | 49 | NA | NA | NA | NA | 0.137 | NA | 0.096 | NA | 0.041 |
| A5SS | 716 | NPEPL1 | chr20 | + | 58691723 | 58691830 | 58691723 | 58691780 | 58693736 | 58693922 | 0 | 0 | 5 | 0 | 0 | 1 | 3 | 5 | 0 | 0 | 0 | 0 | 98 | 49 | NA | NA | NA | NA | 0.714 | 1 | 1 | NA | -0.286 |
| A5SS | 738 | DYRK1A | chr21 | + | 37506098 | 37506354 | 37506098 | 37506223 | 37511910 | 37515376 | 0 | 0 | 3 | 0 | 0 | 0 | 0 | 3 | 0 | 0 | 1 | 0 | 98 | 49 | NA | NA | NA | NA | 1 | NA | 0.6 | NA | 0.4 |
| A5SS | 776 | TMEM55B | chr14 | - | 20461162 | 20461612 | 20461183 | 20461612 | 20460654 | 20460845 | 0 | 0 | 2 | 0 | 0 | 0 | 0 | 1 | 0 | 0 | 1 | 0 | 70 | 49 | NA | NA | NA | NA | 1 | NA | 0.412 | NA | 0.588 |
| A5SS | 787 | SUPT4H1 | chr17 | - | 58352064 | 58352238 | 58352066 | 58352238 | 58351401 | 58351508 | 0 | 0 | 5 | 0 | 0 | 0 | 0 | 0 | 0 | 0 | 1 | 0 | 51 | 49 | NA | NA | NA | NA | 1 | NA | 0 | NA | 1 |
| A5SS | 790 | CNOT1 | chr16 | - | 58556846 | 58556993 | 58556861 | 58556993 | 58555783 | 58555908 | 0 | 0 | 1 | 0 | 0 | 1 | 0 | 2 | 0 | 0 | 1 | 0 | 64 | 49 | NA | NA | NA | NA | 0.434 | NA | 0.605 | NA | -0.171 |
| A5SS | 823 | UFM1 | chr13 | + | 38349998 | 38350227 | 38349998 | 38350055 | 38354238 | 38354296 | 0 | 0 | 0 | 0 | 0 | 2 | 1 | 0 | 0 | 0 | 0 | 0 | 98 | 49 | NA | NA | NA | NA | 0 | 1 | NA | NA | -1 |
| A5SS | 852 | HUS1 | chr7 | - | 47979409 | 47979625 | 47979467 | 47979625 | 47978688 | 47978816 | 0 | 0 | 0 | 0 | 0 | 1 | 2 | 1 | 0 | 0 | 1 | 0 | 98 | 49 | NA | NA | NA | NA | 0 | 1 | 0.333 | NA | -0.666 |
| A5SS | 861 | CHST11 | chr12 | + | 104456913 | 104457529 | 104456913 | 104457514 | 104601905 | 104601991 | 0 | 0 | 7 | 0 | 0 | 2 | 0 | 1 | 0 | 0 | 1 | 0 | 64 | 49 | NA | NA | NA | NA | 0.728 | NA | 0.434 | NA | 0.294 |
| A5SS | 990 | SNX19 | chr11 | - | 130914265 | 130916487 | 130916279 | 130916487 | 130911632 | 130911771 | 0 | 0 | 3 | 0 | 0 | 1 | 1 | 0 | 0 | 0 | 0 | 0 | 98 | 49 | NA | NA | NA | NA | 0.6 | 1 | NA | NA | -0.4 |
| A5SS | 1033 | SH3YL1 | chr2 | - | 224863 | 224920 | 224867 | 224920 | 218135 | 219001 | 0 | 0 | 3 | 0 | 0 | 0 | 0 | 0 | 0 | 0 | 1 | 0 | 53 | 49 | NA | NA | NA | NA | 1 | NA | 0 | NA | 1 |
| A5SS | 1118 | IL32 | chr16 | + | 3067376 | 3067613 | 3067376 | 3067415 | 3067983 | 3068010 | 0 | 0 | 1 | 0 | 0 | 0 | 1 | 2 | 0 | 1 | 0 | 0 | 98 | 49 | NA | NA | NA | NA | 1 | 0.333 | 1 | NA | 0.334 |
| A5SS | 1191 | PKM | chr15 | - | 72230899 | 72231386 | 72231115 | 72231386 | 72218943 | 72219110 | 0 | 0 | 0 | 0 | 0 | 1 | 1 | 0 | 0 | 0 | 0 | 0 | 98 | 49 | NA | NA | NA | NA | 0 | 1 | NA | NA | -1 |
| A5SS | 1195 | NUP98 | chr11 | - | 3760487 | 3760626 | 3760538 | 3760626 | 3753315 | 3753408 | 0 | 0 | 4 | 0 | 0 | 2 | 0 | 13 | 0 | 0 | 5 | 0 | 98 | 49 | NA | NA | NA | NA | 0.5 | NA | 0.565 | NA | -0.065 |
| A5SS | 1217 | MAPKAPK5 | chr12 | + | 111888884 | 111889006 | 111888884 | 111889000 | 111890039 | 111890144 | 0 | 0 | 2 | 0 | 0 | 2 | 0 | 0 | 0 | 0 | 1 | 0 | 55 | 49 | NA | NA | NA | NA | 0.471 | NA | 0 | NA | 0.471 |
| A5SS | 1500 | ZNF518A | chr10 | + | 96155325 | 96155398 | 96155325 | 96155376 | 96155923 | 96156047 | 0 | 0 | 0 | 0 | 0 | 3 | 0 | 1 | 0 | 0 | 1 | 0 | 71 | 49 | NA | NA | NA | NA | 0 | NA | 0.408 | NA | -0.408 |
| A5SS | 1514 | AGTRAP | chr1 | + | 11748414 | 11748610 | 11748414 | 11748589 | 11750076 | 11750771 | 0 | 0 | 4 | 0 | 0 | 1 | 1 | 0 | 0 | 0 | 0 | 0 | 70 | 49 | NA | NA | NA | NA | 0.737 | 1 | NA | NA | -0.263 |
| A5SS | 1529 | TMCO1 | chr1 | - | 165768481 | 165768922 | 165768681 | 165768922 | 165768191 | 165768269 | 0 | 0 | 0 | 0 | 1 | 2 | 2 | 0 | 0 | 0 | 0 | 0 | 98 | 49 | NA | NA | NA | 0 | 0 | 1 | NA | NA | -1 |
| A5SS | 1554 | WNK1 | chr12 | + | 890452 | 890513 | 890452 | 890510 | 894561 | 894635 | 0 | 0 | 1 | 0 | 0 | 0 | 0 | 2 | 0 | 0 | 1 | 0 | 52 | 49 | NA | NA | NA | NA | 1 | NA | 0.653 | NA | 0.347 |
| A5SS | 1565 | POLD4 | chr11 | - | 67353293 | 67353596 | 67353302 | 67353596 | 67352987 | 67353077 | 0 | 0 | 0 | 0 | 0 | 1 | 1 | 0 | 0 | 0 | 0 | 0 | 58 | 49 | NA | NA | NA | NA | 0 | 1 | NA | NA | -1 |
| A5SS | 1702 | EIF4B | chr12 | + | 53034611 | 53034724 | 53034611 | 53034709 | 53037408 | 53037622 | 0 | 0 | 0 | 0 | 0 | 3 | 0 | 1 | 0 | 3 | 2 | 0 | 64 | 49 | NA | NA | NA | NA | 0 | 0 | 0.277 | NA | -0.139 |
| A5SS | 1707 | KHDRBS1 | chr1 | + | 32037834 | 32038119 | 32037834 | 32038036 | 32038551 | 32038619 | 0 | 0 | 0 | 0 | 0 | 2 | 0 | 1 | 0 | 0 | 2 | 0 | 98 | 49 | NA | NA | NA | NA | 0 | NA | 0.2 | NA | -0.2 |
| A5SS | 1731 | SSR3 | chr3 | - | 156548865 | 156549003 | 156548904 | 156549003 | 156544307 | 156544439 | 0 | 0 | 0 | 0 | 0 | 1 | 8 | 0 | 0 | 0 | 0 | 0 | 88 | 49 | NA | NA | NA | NA | 0 | 1 | NA | NA | -1 |
| A5SS | 1737 | SUMO3 | chr21 | - | 44813861 | 44814104 | 44813975 | 44814104 | 44809046 | 44809118 | 0 | 0 | 0 | 0 | 0 | 1 | 1 | 0 | 0 | 0 | 1 | 0 | 98 | 49 | NA | NA | NA | NA | 0 | 1 | 0 | NA | -0.5 |
| A5SS | 1792 | MAFF | chr22 | + | 38201931 | 38202238 | 38201931 | 38202212 | 38213822 | 38213889 | 0 | 0 | 0 | 0 | 0 | 2 | 0 | 1 | 0 | 0 | 3 | 0 | 75 | 49 | NA | NA | NA | NA | 0 | NA | 0.179 | NA | -0.179 |
| A5SS | 1889 | CHMP1A | chr16 | - | 89647034 | 89647331 | 89647202 | 89647331 | 89646526 | 89646714 | 0 | 0 | 1 | 0 | 0 | 0 | 0 | 1 | 0 | 0 | 2 | 0 | 98 | 49 | NA | NA | NA | NA | 1 | NA | 0.2 | NA | 0.8 |
| A5SS | 1924 | CELF2 | chr10 | + | 11314138 | 11314270 | 11314138 | 11314258 | 11321206 | 11321386 | 0 | 0 | 1 | 0 | 0 | 3 | 0 | 2 | 0 | 0 | 1 | 0 | 61 | 49 | NA | NA | NA | NA | 0.211 | NA | 0.616 | NA | -0.405 |
| A5SS | 1991 | CAMK2G | chr10 | - | 73825278 | 73825347 | 73825320 | 73825347 | 73824039 | 73824084 | 0 | 0 | 2 | 0 | 0 | 1 | 8 | 2 | 0 | 0 | 0 | 0 | 91 | 49 | NA | NA | NA | NA | 0.519 | 1 | 1 | NA | -0.481 |
| A5SS | 2007 | CPEB3 | chr10 | - | 92192476 | 92192636 | 92192545 | 92192636 | 92180962 | 92181019 | 0 | 0 | 0 | 0 | 0 | 1 | 0 | 1 | 0 | 0 | 0 | 0 | 98 | 49 | NA | NA | NA | NA | 0 | NA | 1 | NA | -1 |

| | | | | | | | | | | | | | | | | | | | | | | | | | | | | | | | | | |
|---|---|---|---|---|---|---|---|---|---|---|---|---|---|---|---|---|---|---|---|---|---|---|---|---|---|---|---|---|---|---|---|---|---|
| A5SS | 2040 | TRAPPC5 | chr19 | + | 7680820 | 7681049 | 7680820 | 7681026 | 7682241 | 7682862 | 1 | 0 | 0 | 0 | 0 | 0 | 0 | 0 | 0 | 0 | 1 | 0 | 72 | 49 | NA | NA | 1 | NA | NA | NA | 0 | NA | 1 |
| A5SS | 2052 | SLC12A9 | chr7 | + | 100852718 | 100852839 | 100852718 | 100852835 | 100854155 | 100854378 | 0 | 0 | 0 | 0 | 0 | 2 | 0 | 2 | 0 | 0 | 0 | 0 | 53 | 49 | NA | NA | NA | NA | 0 | NA | 1 | NA | -1 |
| A5SS | 2078 | CASP8 | chr2 | + | 201276826 | 201277104 | 201276826 | 201276968 | 201284815 | 201285317 | 0 | 0 | 9 | 0 | 0 | 3 | 0 | 9 | 0 | 1 | 1 | 0 | 98 | 49 | NA | NA | NA | NA | 0.6 | 0 | 0.818 | NA | 0.191 |
| A5SS | 2092 | ARID4A | chr14 | + | 58366023 | 58366230 | 58366023 | 58366068 | 58366882 | 58367029 | 1 | 0 | 4 | 0 | 0 | 0 | 0 | 1 | 0 | 0 | 1 | 0 | 98 | 49 | NA | NA | 1 | NA | 1 | NA | 0.333 | NA | 0.667 |
| A5SS | 2122 | ZC3H14 | chr14 | + | 88602827 | 88603060 | 88602827 | 88603045 | 88607242 | 88607363 | 0 | 0 | 2 | 0 | 0 | 0 | 0 | 0 | 0 | 0 | 1 | 0 | 64 | 49 | NA | NA | NA | NA | 1 | NA | 0 | NA | 1 |
| A5SS | 2124 | C14orf159 | chr14 | + | 91114012 | 91114528 | 91114012 | 91114283 | 91145220 | 91145313 | 0 | 0 | 0 | 0 | 0 | 1 | 0 | 1 | 0 | 0 | 0 | 0 | 98 | 49 | NA | NA | NA | NA | 0 | NA | 1 | NA | -1 |
| A5SS | 2125 | C14orf159 | chr14 | + | 91114012 | 91114528 | 91114012 | 91114283 | 91157638 | 91157690 | 0 | 0 | 0 | 0 | 0 | 1 | 0 | 1 | 0 | 0 | 0 | 0 | 98 | 49 | NA | NA | NA | NA | 0 | NA | 1 | NA | -1 |
| A5SS | 2127 | C14orf159 | chr14 | + | 91173288 | 91173439 | 91173288 | 91173403 | 91175933 | 91176056 | 0 | 0 | 12 | 0 | 0 | 2 | 0 | 3 | 0 | 0 | 1 | 0 | 85 | 49 | NA | NA | NA | NA | 0.776 | NA | 0.634 | NA | 0.142 |
| A5SS | 2128 | PPP4R3A | chr14 | - | 91475810 | 91475966 | 91475849 | 91475966 | 91473238 | 91473370 | 0 | 0 | 3 | 0 | 0 | 2 | 0 | 2 | 0 | 0 | 0 | 0 | 88 | 49 | NA | NA | NA | NA | 0.455 | NA | 1 | NA | -0.545 |
| A5SS | 2165 | PACS2 | chr14 | + | 105381913 | 105382058 | 105381913 | 105382046 | 105382476 | 105382581 | 0 | 0 | 0 | 0 | 0 | 2 | 0 | 1 | 0 | 0 | 0 | 0 | 61 | 49 | NA | NA | NA | NA | 0 | NA | 1 | NA | -1 |
| A5SS | 2199 | UQCR10 | chr22 | + | 29767368 | 29767607 | 29767368 | 29767548 | 29769677 | 29770413 | 0 | 0 | 1 | 0 | 0 | 3 | 0 | 1 | 0 | 0 | 1 | 0 | 98 | 49 | NA | NA | NA | NA | 0.143 | NA | 0.333 | NA | -0.19 |
| A5SS | 2205 | PLCB2 | chr15 | - | 40296745 | 40296908 | 40296757 | 40296908 | 40296521 | 40296634 | 0 | 0 | 7 | 0 | 0 | 3 | 0 | 2 | 0 | 0 | 0 | 0 | 61 | 49 | NA | NA | NA | NA | 0.652 | NA | 1 | NA | -0.348 |
| A5SS | 2260 | TLE3 | chr15 | - | 70058149 | 70058291 | 70058185 | 70058291 | 70057458 | 70057658 | 1 | 0 | 2 | 0 | 0 | 1 | 0 | 3 | 0 | 0 | 1 | 0 | 85 | 49 | NA | NA | 1 | NA | 0.536 | NA | 0.634 | NA | 0.134 |
| A5SS | 2273 | PML | chr15 | + | 74033155 | 74033422 | 74033155 | 74033414 | 74034477 | 74034530 | 0 | 0 | 1 | 0 | 0 | 2 | 1 | 0 | 0 | 0 | 0 | 0 | 57 | 49 | NA | NA | NA | NA | 0.301 | 1 | NA | NA | -0.699 |
| A5SS | 2388 | PSEN1 | chr14 | + | 73147966 | 73148106 | 73147966 | 73148094 | 73170796 | 73171047 | 0 | 0 | 2 | 0 | 0 | 2 | 8 | 3 | 0 | 0 | 5 | 0 | 61 | 49 | NA | NA | NA | NA | 0.445 | 1 | 0.325 | NA | -0.217 |
| A5SS | 2404 | CHCHD7 | chr8 | + | 56212761 | 56212909 | 56212761 | 56212889 | 56214597 | 56214667 | 0 | 0 | 0 | 0 | 0 | 1 | 2 | 0 | 0 | 0 | 0 | 0 | 69 | 49 | NA | NA | NA | NA | 0 | 1 | NA | NA | -1 |
| A5SS | 2498 | CRK | chr17 | - | 1436619 | 1437155 | 1436789 | 1437155 | 1421352 | 1423650 | 0 | 0 | 6 | 0 | 0 | 1 | 1 | 0 | 0 | 0 | 1 | 0 | 98 | 49 | NA | NA | NA | NA | 0.75 | 1 | 0 | NA | 0.25 |
| A5SS | 2580 | STAT3 | chr17 | - | 42322281 | 42322494 | 42322284 | 42322494 | 42317181 | 42317224 | 0 | 0 | 19 | 0 | 0 | 4 | 0 | 10 | 0 | 0 | 4 | 0 | 52 | 49 | NA | NA | NA | NA | 0.817 | NA | 0.702 | NA | 0.115 |
| A5SS | 2607 | FASTK | chr7 | - | 151078841 | 151079021 | 151078922 | 151079021 | 151078561 | 151078701 | 0 | 0 | 1 | 0 | 0 | 0 | 0 | 0 | 0 | 0 | 1 | 0 | 98 | 49 | NA | NA | NA | NA | 1 | NA | 0 | NA | 1 |
| A5SS | 2701 | IFI6 | chr1 | - | 27669220 | 27669346 | 27669232 | 27669346 | 27668457 | 27668535 | 0 | 0 | 4 | 0 | 0 | 1 | 0 | 0 | 0 | 0 | 1 | 0 | 61 | 49 | NA | NA | NA | NA | 0.763 | NA | 0 | NA | 0.763 |
| A5SS | 2714 | MLX | chr17 | + | 42567059 | 42567328 | 42567059 | 42567166 | 42567618 | 42567655 | 0 | 0 | 0 | 0 | 0 | 1 | 2 | 0 | 0 | 0 | 0 | 0 | 98 | 49 | NA | NA | NA | NA | 0 | 1 | NA | NA | -1 |
| A5SS | 2723 | ATP5A1 | chr18 | - | 46094979 | 46095131 | 46095052 | 46095131 | 46091681 | 46091851 | 0 | 0 | 0 | 0 | 0 | 2 | 1 | 0 | 0 | 0 | 2 | 0 | 98 | 49 | NA | NA | NA | NA | 0 | 1 | 0 | NA | -0.5 |
| A5SS | 2757 | CAPS | chr19 | + | 5914939 | 5915146 | 5914939 | 5915065 | 5915220 | 5916211 | 0 | 0 | 1 | 0 | 0 | 0 | 0 | 0 | 0 | 0 | 1 | 0 | 98 | 49 | NA | NA | NA | NA | 1 | NA | 0 | NA | 1 |
| A5SS | 2763 | ADGRE1 | chr19 | + | 6928144 | 6928310 | 6928144 | 6928211 | 6934986 | 6935078 | 0 | 0 | 0 | 0 | 0 | 4 | 1 | 0 | 0 | 0 | 0 | 0 | 98 | 49 | NA | NA | NA | NA | 0 | 1 | NA | NA | -1 |
| A5SS | 2772 | LPPR2 | chr19 | + | 11363712 | 11363835 | 11363712 | 11363819 | 11364160 | 11364212 | 0 | 0 | 2 | 0 | 0 | 1 | 0 | 0 | 0 | 0 | 1 | 0 | 65 | 49 | NA | NA | NA | NA | 0.601 | NA | 0 | NA | 0.601 |
| A5SS | 2777 | SGMS2 | chr4 | + | 107893263 | 107893658 | 107893263 | 107893382 | 107895309 | 107896008 | 0 | 0 | 3 | 0 | 0 | 0 | 1 | 0 | 0 | 0 | 1 | 0 | 98 | 49 | NA | NA | NA | NA | 1 | 1 | 0 | NA | 0.5 |
| A5SS | 2801 | EPS15L1 | chr19 | - | 16440861 | 16440909 | 16440884 | 16440909 | 16437769 | 16437865 | 0 | 0 | 2 | 0 | 0 | 0 | 0 | 1 | 0 | 0 | 1 | 0 | 72 | 49 | NA | NA | NA | NA | 1 | NA | 0.405 | NA | 0.595 |
| A5SS | 2950 | HNRNPH3 | chr10 | + | 68338502 | 68338687 | 68338502 | 68338642 | 68339139 | 68339226 | 2 | 0 | 0 | 0 | 0 | 0 | 0 | 0 | 0 | 0 | 2 | 0 | 94 | 49 | NA | NA | 1 | NA | NA | NA | 0 | NA | 1 |
| A5SS | 2996 | VRK2 | chr2 | + | 58046641 | 58046868 | 58046641 | 58046821 | 58048826 | 58048967 | 2 | 0 | 2 | 0 | 0 | 1 | 0 | 0 | 0 | 0 | 1 | 0 | 96 | 49 | NA | NA | 1 | NA | 0.505 | NA | 0 | NA | 0.752 |
| A5SS | 3001 | CCND3 | chr6 | - | 41940951 | 41941848 | 41941451 | 41941848 | 41940369 | 41940585 | 0 | 0 | 0 | 0 | 0 | 5 | 0 | 1 | 0 | 0 | 0 | 0 | 98 | 49 | NA | NA | NA | NA | 0 | NA | 1 | NA | -1 |
| A5SS | 3047 | PLEKHB2 | chr2 | + | 131132901 | 131133015 | 131132901 | 131132991 | 131140169 | 131140275 | 0 | 0 | 2 | 0 | 0 | 0 | 0 | 1 | 0 | 0 | 1 | 0 | 73 | 49 | NA | NA | NA | NA | 1 | NA | 0.402 | NA | 0.598 |
| A5SS | 3074 | HINT1 | chr5 | - | 131165059 | 131165348 | 131165094 | 131165348 | 131162571 | 131162676 | 2 | 0 | 0 | 0 | 0 | 5 | 0 | 0 | 0 | 0 | 1 | 0 | 84 | 49 | NA | NA | 1 | NA | 0 | NA | 0 | NA | 0.5 |
| RI | 33 | VPS28 | chr8 | - | 144423600 | 144424132 | 144423600 | 144423922 | 144424040 | 144424132 | 0 | 0 | 0 | 0 | 0 | 3 | 0 | 2 | 0 | 0 | 2 | 0 | 98 | 49 | NA | NA | NA | NA | 0 | NA | 0.333 | NA | -0.333 |
| RI | 227 | NDUFB4 | chr3 | + | 120601110 | 120602411 | 120601110 | 120601257 | 120602207 | 120602411 | 0 | 0 | 0 | 0 | 0 | 1 | 4 | 0 | 0 | 0 | 1 | 0 | 98 | 49 | NA | NA | NA | NA | 0 | 1 | 0 | NA | -0.5 |
| RI | 288 | NECAB3 | chr20 | - | 33659496 | 33659732 | 33659496 | 33659550 | 33659652 | 33659732 | 0 | 0 | 0 | 0 | 0 | 1 | 2 | 0 | 0 | 0 | 0 | 0 | 98 | 49 | NA | NA | NA | NA | 0 | 1 | NA | NA | -1 |
| RI | 338 | NSD1 | chr5 | + | 177135086 | 177136030 | 177135086 | 177135200 | 177135940 | 177136030 | 0 | 0 | 1 | 0 | 0 | 0 | 0 | 0 | 0 | 0 | 1 | 0 | 98 | 49 | NA | NA | NA | NA | 1 | NA | 0 | NA | 1 |
| RI | 405 | CREBZF | chr11 | - | 85657563 | 85665138 | 85657563 | 85662457 | 85663523 | 85665138 | 0 | 0 | 4 | 0 | 0 | 2 | 2 | 1 | 0 | 0 | 0 | 0 | 98 | 49 | NA | NA | NA | NA | 0.5 | 1 | 1 | NA | -0.5 |
| RI | 429 | IL32 | chr16 | + | 3067376 | 3067613 | 3067376 | 3067415 | 3067553 | 3067613 | 0 | 0 | 1 | 0 | 0 | 3 | 1 | 0 | 0 | 0 | 6 | 0 | 98 | 49 | NA | NA | NA | NA | 0.143 | 1 | 0 | NA | -0.357 |
| RI | 493 | CDK17 | chr12 | - | 96278260 | 96280885 | 96278260 | 96280279 | 96280807 | 96280885 | 0 | 0 | 13 | 0 | 0 | 0 | 3 | 8 | 0 | 0 | 2 | 0 | 98 | 49 | NA | NA | NA | NA | 1 | 1 | 0.667 | NA | 0.166 |
| RI | 514 | DNAJB12 | chr10 | - | 72332829 | 72335931 | 72332829 | 72334617 | 72335779 | 72335931 | 0 | 0 | 1 | 0 | 0 | 3 | 0 | 1 | 0 | 0 | 0 | 0 | 98 | 49 | NA | NA | NA | NA | 0.143 | NA | 1 | NA | -0.857 |
| RI | 530 | OTUB1 | chr11 | + | 63996529 | 63996941 | 63996529 | 63996648 | 63996856 | 63996941 | 0 | 0 | 0 | 0 | 0 | 1 | 2 | 1 | 0 | 0 | 2 | 0 | 98 | 49 | NA | NA | NA | NA | 0 | 1 | 0.2 | NA | -0.6 |
| RI | 585 | PCID2 | chr13 | - | 113177538 | 113178287 | 113177538 | 113177976 | 113178173 | 113178287 | 0 | 0 | 3 | 0 | 0 | 0 | 3 | 0 | 0 | 0 | 1 | 0 | 98 | 49 | NA | NA | NA | NA | 1 | 1 | 0 | NA | 0.5 |
| RI | 619 | IL17RC | chr3 | + | 9923880 | 9924291 | 9923880 | 9924020 | 9924231 | 9924291 | 0 | 1 | 0 | 0 | 0 | 1 | 6 | 0 | 0 | 0 | 0 | 0 | 98 | 49 | NA | NA | NA | 1 | 0 | 1 | NA | NA | -0.5 |
| RI | 729 | CLK1 | chr2 | - | 200859679 | 200861466 | 200859679 | 200859746 | 200861237 | 200861466 | 0 | 0 | 5 | 0 | 0 | 2 | 0 | 18 | 0 | 0 | 3 | 0 | 98 | 49 | NA | NA | NA | NA | 0.556 | NA | 0.75 | NA | -0.194 |
| RI | 834 | CIRBP | chr19 | + | 1271980 | 1273172 | 1271980 | 1272051 | 1272426 | 1273172 | 0 | 0 | 3 | 0 | 0 | 1 | 3 | 0 | 0 | 0 | 4 | 0 | 98 | 49 | NA | NA | NA | NA | 0.6 | 1 | 0 | NA | 0.1 |
| RI | 888 | LAT | chr16 | + | 28986539 | 28986714 | 28986539 | 28986569 | 28986656 | 28986714 | 1 | 0 | 0 | 0 | 0 | 0 | 0 | 1 | 1 | 0 | 1 | 0 | 98 | 49 | NA | NA | 1 | NA | NA | NA | 0.333 | 1 | 0.334 |
| RI | 896 | KAT8 | chr16 | + | 31130745 | 31131393 | 31130745 | 31130900 | 31131194 | 31131393 | 0 | 0 | 0 | 0 | 1 | 0 | 1 | 0 | 0 | 0 | 0 | 0 | 98 | 49 | NA | NA | NA | 0 | NA | 1 | NA | NA | -1 |
| RI | 909 | HNRNPM | chr19 | + | 8465323 | 8465515 | 8465323 | 8465362 | 8465479 | 8465515 | 0 | 0 | 8 | 0 | 0 | 1 | 7 | 0 | 0 | 0 | 0 | 0 | 98 | 49 | NA | NA | NA | NA | 0.8 | 1 | NA | NA | -0.2 |
| RI | 926 | WRAP53 | chr17 | + | 7688348 | 7689079 | 7688348 | 7688561 | 7688647 | 7689079 | 0 | 0 | 0 | 0 | 0 | 2 | 5 | 0 | 0 | 0 | 0 | 0 | 98 | 49 | NA | NA | NA | NA | 0 | 1 | NA | NA | -1 |
| RI | 984 | LUC7L3 | chr17 | + | 50750500 | 50752711 | 50750500 | 50750694 | 50752199 | 50752711 | 0 | 0 | 2 | 0 | 0 | 2 | 0 | 4 | 0 | 0 | 0 | 0 | 98 | 49 | NA | NA | NA | NA | 0.333 | NA | 1 | NA | -0.667 |
| RI | 987 | SRSF1 | chr17 | - | 58000918 | 58005973 | 58000918 | 58005600 | 58005800 | 58005973 | 0 | 0 | 1 | 0 | 0 | 5 | 2 | 0 | 0 | 0 | 0 | 0 | 98 | 49 | NA | NA | NA | NA | 0.091 | 1 | NA | NA | -0.909 |
| RI | 1024 | FAM195B | chr17 | - | 81822360 | 81823311 | 81822360 | 81822907 | 81823208 | 81823311 | 0 | 0 | 3 | 0 | 0 | 0 | 4 | 1 | 0 | 0 | 3 | 0 | 98 | 49 | NA | NA | NA | NA | 1 | 1 | 0.143 | NA | 0.428 |
| RI | 1029 | SNRNP27 | chr2 | + | 69904253 | 69905236 | 69904253 | 69904425 | 69904511 | 69905236 | 1 | 0 | 3 | 0 | 0 | 0 | 0 | 0 | 0 | 0 | 1 | 0 | 98 | 49 | NA | NA | 1 | NA | 1 | NA | 0 | NA | 1 |
| RI | 1037 | CSTF3 | chr11 | - | 33141430 | 33141986 | 33141430 | 33141762 | 33141884 | 33141986 | 0 | 0 | 0 | 0 | 0 | 1 | 8 | 1 | 0 | 0 | 2 | 0 | 98 | 49 | NA | NA | NA | NA | 0 | 1 | 0.2 | NA | -0.6 |
| RI | 1058 | GTPBP3 | chr19 | + | 17338953 | 17339266 | 17338953 | 17339026 | 17339122 | 17339266 | 1 | 0 | 0 | 0 | 0 | 0 | 0 | 0 | 0 | 0 | 1 | 0 | 98 | 49 | NA | NA | 1 | NA | NA | NA | 0 | NA | 1 |
| RI | 1073 | CCPG1 | chr15 | - | 55355222 | 55360944 | 55355222 | 55356409 | 55359538 | 55360944 | 0 | 0 | 1 | 0 | 0 | 1 | 1 | 1 | 0 | 0 | 0 | 0 | 98 | 49 | NA | NA | NA | NA | 0.333 | 1 | 1 | NA | -0.667 |
| RI | 1118 | RPL31 | chr2 | + | 101005958 | 101006423 | 101005958 | 101006071 | 101006349 | 101006423 | 1 | 0 | 4 | 0 | 0 | 5 | 0 | 1 | 0 | 0 | 6 | 0 | 98 | 49 | NA | NA | 1 | NA | 0.286 | NA | 0.077 | NA | 0.566 |
| SE | 37 | FMR1 | chrX | + | 147937465 | 147937600 | 147936503 | 147936613 | 147940575 | 147940662 | 0 | 0 | 1 | 0 | 0 | 0 | 0 | 2 | 0 | 0 | 1 | 0 | 98 | 49 | NA | NA | NA | NA | 1 | NA | 0.5 | NA | 0.5 |
| SE | 38 | FMR1 | chrX | + | 147938098 | 147938161 | 147937465 | 147937600 | 147940575 | 147940662 | 0 | 0 | 0 | 0 | 0 | 1 | 0 | 1 | 0 | 0 | 2 | 0 | 98 | 49 | NA | NA | NA | NA | 0 | NA | 0.2 | NA | -0.2 |
| SE | 57 | SASH3 | chrX | + | 129792626 | 129792836 | 129792327 | 129792476 | 129792988 | 129793139 | 0 | 0 | 20 | 0 | 0 | 0 | 0 | 17 | 0 | 0 | 1 | 0 | 98 | 49 | NA | NA | NA | NA | 1 | NA | 0.895 | NA | 0.105 |
| SE | 59 | STAG2 | chrX | + | 124090853 | 124090964 | 124090574 | 124090764 | 124094017 | 124094144 | 0 | 0 | 0 | 0 | 0 | 3 | 0 | 1 | 0 | 0 | 2 | 0 | 98 | 49 | NA | NA | NA | NA | 0 | NA | 0.2 | NA | -0.2 |
| SE | 129 | XIST | chrX | - | 73833237 | 73833374 | 73831065 | 73831274 | 73837439 | 73837503 | 0 | 0 | 9 | 0 | 0 | 1 | 0 | 2 | 0 | 0 | 0 | 0 | 98 | 49 | NA | NA | NA | NA | 0.818 | NA | 1 | NA | -0.182 |
| SE | 156 | APEX2 | chrX | + | 55002250 | 55002431 | 55001545 | 55001629 | 55002961 | 55003108 | 0 | 0 | 0 | 0 | 0 | 1 | 0 | 1 | 0 | 0 | 0 | 0 | 98 | 49 | NA | NA | NA | NA | 0 | NA | 1 | NA | -1 |
| SE | 173 | PQBP1 | chrX | + | 48898491 | 48898576 | 48897911 | 48898082 | 48901189 | 48901277 | 0 | 0 | 1 | 0 | 0 | 0 | 0 | 1 | 0 | 0 | 1 | 0 | 98 | 49 | NA | NA | NA | NA | 1 | NA | 0.333 | NA | 0.667 |
| SE | 175 | TIMM17B | chrX | - | 48895351 | 48895501 | 48895037 | 48895101 | 48896758 | 48896858 | 0 | 0 | 0 | 0 | 0 | 2 | 0 | 1 | 0 | 0 | 0 | 0 | 98 | 49 | NA | NA | NA | NA | 0 | NA | 1 | NA | -1 |
| SE | 180 | ELK1 | chrX | - | 47637750 | 47638182 | 47637012 | 47637114 | 47638894 | 47639338 | 0 | 0 | 1 | 0 | 0 | 0 | 0 | 0 | 0 | 0 | 1 | 0 | 98 | 49 | NA | NA | NA | NA | 1 | NA | 0 | NA | 1 |
| SE | 182 | UBA1 | chrX | + | 47199208 | 47199377 | 47199047 | 47199106 | 47199479 | 47199614 | 0 | 0 | 5 | 0 | 0 | 0 | 0 | 1 | 0 | 0 | 2 | 0 | 98 | 49 | NA | NA | NA | NA | 1 | NA | 0.2 | NA | 0.8 |
| SE | 184 | RBM10 | chrX | + | 47175018 | 47175092 | 47173127 | 47173197 | 47176499 | 47176586 | 0 | 0 | 1 | 0 | 0 | 0 | 0 | 0 | 0 | 0 | 2 | 0 | 98 | 49 | NA | NA | NA | NA | 1 | NA | 0 | NA | 1 |
| SE | 185 | KDM6A | chrX | + | 45020609 | 45020730 | 45010960 | 45011019 | 45034930 | 45034985 | 0 | 0 | 7 | 0 | 0 | 3 | 0 | 4 | 0 | 0 | 0 | 0 | 98 | 49 | NA | NA | NA | NA | 0.538 | NA | 1 | NA | -0.462 |
| SE | 189 | KDM6A | chrX | + | 45060608 | 45060764 | 45059246 | 45059466 | 45061323 | 45061419 | 0 | 0 | 4 | 0 | 0 | 3 | 0 | 5 | 0 | 0 | 1 | 0 | 98 | 49 | NA | NA | NA | NA | 0.4 | NA | 0.714 | NA | -0.314 |
| SE | 195 | CXorf38 | chrX | - | 40647006 | 40647141 | 40639008 | 40639128 | 40647304 | 40647567 | 0 | 0 | 1 | 0 | 0 | 1 | 0 | 1 | 0 | 0 | 0 | 0 | 98 | 49 | NA | NA | NA | NA | 0.333 | NA | 1 | NA | -0.667 |
| SE | 201 | ACOT9 | chrX | - | 23722669 | 23722753 | 23721880 | 23721984 | 23730526 | 23730564 | 0 | 0 | 7 | 0 | 0 | 0 | 0 | 17 | 0 | 0 | 2 | 0 | 98 | 49 | NA | NA | NA | NA | 1 | NA | 0.81 | NA | 0.19 |
| SE | 202 | ACOT9 | chrX | - | 23734340 | 23734367 | 23733171 | 23733217 | 23735918 | 23736016 | 0 | 0 | 10 | 0 | 0 | 0 | 0 | 4 | 0 | 0 | 1 | 0 | 76 | 49 | NA | NA | NA | NA | 1 | NA | 0.721 | NA | 0.279 |
| SE | 205 | RPS6KA3 | chrX | - | 20187827 | 20187970 | 20186295 | 20186366 | 20188496 | 20188534 | 0 | 0 | 7 | 0 | 0 | 0 | 0 | 2 | 0 | 0 | 1 | 0 | 98 | 49 | NA | NA | NA | NA | 1 | NA | 0.5 | NA | 0.5 |
| SE | 210 | SYAP1 | chrX | + | 16741715 | 16741789 | 16736165 | 16736232 | 16743700 | 16743840 | 0 | 0 | 12 | 0 | 0 | 0 | 0 | 5 | 0 | 0 | 1 | 0 | 98 | 49 | NA | NA | NA | NA | 1 | NA | 0.714 | NA | 0.286 |
| SE | 233 | CSF2RA | chrX | + | 1300490 | 1300626 | 1295426 | 1295456 | 1303922 | 1304019 | 0 | 0 | 19 | 0 | 0 | 4 | 0 | 2 | 0 | 0 | 0 | 0 | 98 | 49 | NA | NA | NA | NA | 0.704 | NA | 1 | NA | -0.296 |
| SE | 234 | CSF2RA | chrX | + | 1303260 | 1303362 | 1300490 | 1300626 | 1303922 | 1304019 | 0 | 0 | 1 | 0 | 0 | 5 | 0 | 0 | 0 | 0 | 1 | 0 | 98 | 49 | NA | NA | NA | NA | 0.091 | NA | 0 | NA | 0.091 |
| SE | 237 | EHMT1 | chr9 | + | 137752330 | 137752408 | 137743901 | 137744090 | 137754170 | 137754291 | 0 | 0 | 4 | 0 | 0 | 0 | 0 | 1 | 0 | 0 | 1 | 0 | 98 | 49 | NA | NA | NA | NA | 1 | NA | 0.333 | NA | 0.667 |
| SE | 252 | SEC16A | chr9 | - | 136445644 | 136445719 | 136443822 | 136443900 | 136446854 | 136446949 | 0 | 0 | 2 | 0 | 0 | 0 | 0 | 0 | 0 | 0 | 1 | 0 | 98 | 49 | NA | NA | NA | NA | 1 | NA | 0 | NA | 1 |
| SE | 253 | SEC16A | chr9 | - | 136457443 | 136457584 | 136456052 | 136456166 | 136459133 | 136459239 | 0 | 0 | 7 | 0 | 0 | 0 | 0 | 2 | 0 | 0 | 1 | 0 | 98 | 49 | NA | NA | NA | NA | 1 | NA | 0.5 | NA | 0.5 |
| SE | 254 | SEC16A | chr9 | - | 136462886 | 136463132 | 136461176 | 136461274 | 136463462 | 136463601 | 0 | 0 | 2 | 0 | 0 | 0 | 0 | 2 | 0 | 0 | 1 | 0 | 98 | 49 | NA | NA | NA | NA | 1 | NA | 0.5 | NA | 0.5 |
| SE | 282 | ASB6 | chr9 | - | 129640540 | 129640722 | 129639401 | 129639508 | 129641886 | 129642169 | 0 | 0 | 1 | 0 | 0 | 1 | 0 | 1 | 0 | 0 | 0 | 0 | 98 | 49 | NA | NA | NA | NA | 0.333 | NA | 1 | NA | -0.667 |
| SE | 283 | PPP2R4 | chr9 | + | 129123051 | 129123138 | 129120512 | 129120610 | 129128984 | 129129110 | 0 | 0 | 1 | 0 | 0 | 0 | 0 | 0 | 0 | 0 | 1 | 0 | 98 | 49 | NA | NA | NA | NA | 1 | NA | 0 | NA | 1 |
| SE | 307 | SH2D3C | chr9 | - | 127761610 | 127761650 | 127751171 | 127751300 | 127773989 | 127774467 | 0 | 0 | 1 | 0 | 0 | 1 | 0 | 3 | 0 | 0 | 0 | 0 | 89 | 49 | NA | NA | NA | NA | 0.355 | NA | 1 | NA | -0.645 |
| SE | 315 | GAPVD1 | chr9 | + | 125360527 | 125360725 | 125359419 | 125359492 | 125362605 | 125365011 | 0 | 0 | 5 | 0 | 0 | 2 | 0 | 3 | 0 | 0 | 0 | 0 | 98 | 49 | NA | NA | NA | NA | 0.556 | NA | 1 | NA | -0.444 |
| SE | 337 | MED12 | chrX | + | 71125350 | 71125495 | 71124975 | 71125146 | 71125662 | 71125713 | 0 | 0 | 3 | 0 | 0 | 0 | 0 | 0 | 0 | 0 | 1 | 0 | 98 | 49 | NA | NA | NA | NA | 1 | NA | 0 | NA | 1 |
| SE | 348 | NR4A3 | chr9 | + | 99826729 | 99826801 | 99825658 | 99825832 | 99828040 | 99828993 | 0 | 0 | 3 | 0 | 0 | 0 | 0 | 1 | 0 | 0 | 5 | 0 | 98 | 49 | NA | NA | NA | NA | 1 | NA | 0.091 | NA | 0.909 |
| SE | 358 | ERCC6L2 | chr9 | + | 95907077 | 95907271 | 95897848 | 95897971 | 95915667 | 95915829 | 0 | 0 | 0 | 0 | 0 | 1 | 0 | 1 | 0 | 0 | 0 | 0 | 98 | 49 | NA | NA | NA | NA | 0 | NA | 1 | NA | -1 |
| SE | 365 | SEMA4D | chr9 | - | 89418362 | 89418437 | 89418055 | 89418118 | 89455887 | 89455953 | 0 | 0 | 2 | 0 | 0 | 0 | 0 | 0 | 0 | 0 | 1 | 0 | 98 | 49 | NA | NA | NA | NA | 1 | NA | 0 | NA | 1 |
| SE | 369 | AGTPBP1 | chr9 | - | 85588297 | 85588478 | 85586830 | 85586960 | 85589527 | 85589681 | 0 | 0 | 12 | 0 | 0 | 1 | 0 | 5 | 0 | 0 | 0 | 0 | 98 | 49 | NA | NA | NA | NA | 0.857 | NA | 1 | NA | -0.143 |
| SE | 377 | TLE4 | chr9 | + | 79612655 | 79612718 | 79576132 | 79576177 | 79627373 | 79627448 | 0 | 0 | 4 | 0 | 0 | 0 | 0 | 5 | 0 | 0 | 1 | 0 | 98 | 49 | NA | NA | NA | NA | 1 | NA | 0.714 | NA | 0.286 |
| SE | 398 | PTAR1 | chr9 | - | 69732138 | 69732352 | 69723325 | 69723630 | 69734149 | 69734254 | 0 | 0 | 1 | 0 | 0 | 0 | 0 | 1 | 0 | 0 | 1 | 0 | 98 | 49 | NA | NA | NA | NA | 1 | NA | 0.333 | NA | 0.667 |
| SE | 430 | UBAP1 | chr9 | + | 34196335 | 34196458 | 34179004 | 34179240 | 34220907 | 34220948 | 0 | 0 | 0 | 0 | 0 | 1 | 0 | 1 | 0 | 0 | 1 | 0 | 98 | 49 | NA | NA | NA | NA | 0 | NA | 0.333 | NA | -0.333 |
| SE | 438 | DNAJA1 | chr9 | + | 33026812 | 33026990 | 33026474 | 33026616 | 33029884 | 33029989 | 0 | 0 | 6 | 0 | 0 | 1 | 0 | 2 | 0 | 0 | 3 | 0 | 98 | 49 | NA | NA | NA | NA | 0.75 | NA | 0.25 | NA | 0.5 |
| SE | 444 | DDX58 | chr9 | - | 32492390 | 32492538 | 32491300 | 32491420 | 32493760 | 32493942 | 0 | 0 | 8 | 0 | 0 | 1 | 0 | 5 | 0 | 0 | 6 | 0 | 98 | 49 | NA | NA | NA | NA | 0.8 | NA | 0.294 | NA | 0.506 |
| SE | 465 | RIC1 | chr9 | + | 5745930 | 5746083 | 5743688 | 5743737 | 5747301 | 5747505 | 0 | 0 | 9 | 0 | 0 | 0 | 0 | 0 | 0 | 0 | 1 | 0 | 98 | 49 | NA | NA | NA | NA | 1 | NA | 0 | NA | 1 |
| SE | 468 | RFX3 | chr9 | - | 3270370 | 3270525 | 3266207 | 3266305 | 3271002 | 3271118 | 0 | 0 | 0 | 0 | 0 | 2 | 0 | 1 | 0 | 0 | 0 | 0 | 98 | 49 | NA | NA | NA | NA | 0 | NA | 1 | NA | -1 |
| SE | 474 | DOCK8 | chr9 | + | 420948 | 421078 | 420400 | 420583 | 422047 | 422135 | 0 | 0 | 11 | 0 | 0 | 1 | 0 | 10 | 0 | 0 | 1 | 0 | 98 | 49 | NA | NA | NA | NA | 0.846 | NA | 0.833 | NA | 0.013 |
| SE | 475 | DOCK8 | chr9 | + | 426884 | 426981 | 422047 | 422135 | 428361 | 428496 | 0 | 0 | 16 | 0 | 0 | 0 | 0 | 5 | 0 | 0 | 1 | 0 | 98 | 49 | NA | NA | NA | NA | 1 | NA | 0.714 | NA | 0.286 |
| SE | 476 | DOCK8 | chr9 | + | 446369 | 446606 | 443426 | 443516 | 449783 | 449927 | 0 | 0 | 19 | 0 | 0 | 0 | 0 | 7 | 0 | 0 | 2 | 0 | 98 | 49 | NA | NA | NA | NA | 1 | NA | 0.636 | NA | 0.364 |
| SE | 487 | GRINA | chr8 | + | 143991877 | 143992078 | 143991691 | 143991804 | 143992244 | 143992373 | 0 | 0 | 5 | 0 | 0 | 0 | 0 | 0 | 0 | 0 | 1 | 0 | 98 | 49 | NA | NA | NA | NA | 1 | NA | 0 | NA | 1 |
| SE | 520 | LRRC6 | chr8 | - | 132632739 | 132632963 | 132625271 | 132625454 | 132637934 | 132638107 | 0 | 0 | 1 | 0 | 0 | 1 | 0 | 0 | 0 | 0 | 3 | 0 | 98 | 49 | NA | NA | NA | NA | 0.333 | NA | 0 | NA | 0.333 |
| SE | 534 | PRRC2B | chr9 | + | 131474453 | 131476535 | 131473507 | 131473724 | 131477743 | 131477949 | 0 | 0 | 3 | 0 | 0 | 0 | 0 | 1 | 0 | 0 | 1 | 0 | 98 | 49 | NA | NA | NA | NA | 1 | NA | 0.333 | NA | 0.667 |
| SE | 553 | UBR5 | chr8 | - | 102278825 | 102278939 | 102276971 | 102277192 | 102279035 | 102279208 | 0 | 0 | 4 | 0 | 0 | 0 | 0 | 2 | 0 | 0 | 1 | 0 | 98 | 49 | NA | NA | NA | NA | 1 | NA | 0.5 | NA | 0.5 |
| SE | 554 | RRM2B | chr8 | - | 102224884 | 102225018 | 102224045 | 102224140 | 102225917 | 102226034 | 0 | 0 | 4 | 0 | 0 | 2 | 0 | 6 | 0 | 0 | 0 | 0 | 98 | 49 | NA | NA | NA | NA | 0.5 | NA | 1 | NA | -0.5 |
| SE | 556 | ZNF706 | chr8 | - | 101202332 | 101202447 | 101201606 | 101201743 | 101205434 | 101205732 | 0 | 0 | 0 | 1 | 0 | 0 | 0 | 2 | 0 | 0 | 0 | 0 | 98 | 49 | NA | NA | 0 | NA | NA | NA | 1 | NA | -1 |
| SE | 560 | RNF19A | chr8 | - | 100264670 | 100264785 | 100259853 | 100259997 | 100268784 | 100268947 | 0 | 0 | 4 | 0 | 0 | 1 | 0 | 6 | 0 | 0 | 0 | 0 | 98 | 49 | NA | NA | NA | NA | 0.667 | NA | 1 | NA | -0.333 |
| SE | 562 | VPS13B | chr8 | + | 99507136 | 99507203 | 99502835 | 99502950 | 99511103 | 99511512 | 0 | 0 | 5 | 0 | 0 | 0 | 0 | 6 | 0 | 0 | 3 | 0 | 98 | 49 | NA | NA | NA | NA | 1 | NA | 0.5 | NA | 0.5 |
| SE | 563 | VPS13B | chr8 | + | 99507769 | 99507911 | 99502835 | 99502950 | 99511103 | 99511512 | 0 | 0 | 2 | 0 | 0 | 0 | 0 | 0 | 0 | 0 | 3 | 0 | 98 | 49 | NA | NA | NA | NA | 1 | NA | 0 | NA | 1 |
| SE | 597 | PAG1 | chr8 | - | 80991478 | 80991530 | 80987369 | 80987466 | 80993102 | 80993307 | 0 | 0 | 3 | 0 | 0 | 0 | 0 | 7 | 0 | 0 | 3 | 0 | 98 | 49 | NA | NA | NA | NA | 1 | NA | 0.538 | NA | 0.462 |
| SE | 598 | PAG1 | chr8 | - | 80993102 | 80993307 | 80991478 | 80991530 | 81029995 | 81030089 | 0 | 0 | 2 | 0 | 0 | 1 | 0 | 4 | 0 | 0 | 0 | 0 | 98 | 49 | NA | NA | NA | NA | 0.5 | NA | 1 | NA | -0.5 |
| SE | 637 | RB1CC1 | chr8 | - | 52676368 | 52676571 | 52673844 | 52674274 | 52683548 | 52683719 | 0 | 0 | 6 | 0 | 0 | 0 | 0 | 9 | 0 | 0 | 1 | 0 | 98 | 49 | NA | NA | NA | NA | 1 | NA | 0.818 | NA | 0.182 |
| SE | 645 | HGSNAT | chr8 | + | 43197839 | 43197952 | 43197671 | 43197742 | 43199387 | 43202827 | 0 | 0 | 2 | 0 | 0 | 0 | 2 | 0 | 0 | 0 | 1 | 0 | 98 | 49 | NA | NA | NA | NA | 1 | 1 | 0 | NA | 0.5 |
| SE | 661 | PLEKHA2 | chr8 | + | 38943788 | 38943837 | 38935993 | 38936050 | 38946123 | 38946221 | 0 | 0 | 2 | 0 | 0 | 0 | 1 | 5 | 0 | 0 | 1 | 0 | 98 | 49 | NA | NA | NA | NA | 1 | 1 | 0.714 | NA | 0.143 |
| SE | 662 | PLEKHA2 | chr8 | + | 38957322 | 38957386 | 38953296 | 38953367 | 38968591 | 38968669 | 0 | 0 | 5 | 0 | 0 | 1 | 2 | 2 | 0 | 0 | 1 | 0 | 98 | 49 | NA | NA | NA | NA | 0.714 | 1 | 0.5 | NA | -0.036 |
| SE | 692 | DOCK5 | chr8 | + | 25340876 | 25340988 | 25336238 | 25336373 | 25341738 | 25341809 | 0 | 0 | 19 | 0 | 0 | 0 | 0 | 10 | 0 | 0 | 1 | 0 | 98 | 49 | NA | NA | NA | NA | 1 | NA | 0.833 | NA | 0.167 |
| SE | 693 | DOCK5 | chr8 | + | 25341738 | 25341809 | 25336238 | 25336373 | 25342400 | 25342507 | 0 | 0 | 6 | 0 | 0 | 0 | 0 | 12 | 0 | 0 | 1 | 0 | 98 | 49 | NA | NA | NA | NA | 1 | NA | 0.857 | NA | 0.143 |
| SE | 696 | DOCK5 | chr8 | + | 25373617 | 25373658 | 25372558 | 25372718 | 25374563 | 25374654 | 0 | 0 | 8 | 0 | 0 | 1 | 0 | 8 | 0 | 0 | 0 | 0 | 90 | 49 | NA | NA | NA | NA | 0.813 | NA | 1 | NA | -0.187 |
| SE | 712 | DOK2 | chr8 | - | 21911900 | 21911988 | 21910672 | 21910857 | 21912228 | 21912510 | 0 | 0 | 7 | 0 | 0 | 2 | 0 | 0 | 0 | 0 | 1 | 0 | 98 | 49 | NA | NA | NA | NA | 0.636 | NA | 0 | NA | 0.636 |
| SE | 735 | CTSB | chr8 | - | 11849045 | 11849164 | 11848066 | 11848152 | 11850865 | 11850980 | 0 | 0 | 32 | 0 | 0 | 2 | 0 | 5 | 0 | 0 | 0 | 0 | 98 | 49 | NA | NA | NA | NA | 0.889 | NA | 1 | NA | -0.111 |
| SE | 793 | SSBP1 | chr7 | + | 141741748 | 141741798 | 141739123 | 141739190 | 141742168 | 141742229 | 0 | 0 | 0 | 0 | 0 | 1 | 0 | 1 | 0 | 0 | 0 | 0 | 98 | 49 | NA | NA | NA | NA | 0 | NA | 1 | NA | -1 |
| SE | 819 | NRF1 | chr7 | + | 129710373 | 129710571 | 129709074 | 129709233 | 129711474 | 129711576 | 0 | 0 | 1 | 0 | 0 | 0 | 0 | 1 | 0 | 0 | 1 | 0 | 98 | 49 | NA | NA | NA | NA | 1 | NA | 0.333 | NA | 0.667 |
| SE | 824 | TNPO3 | chr7 | - | 128993806 | 128993914 | 128991998 | 128992090 | 128997388 | 128997535 | 0 | 0 | 0 | 0 | 0 | 2 | 0 | 2 | 0 | 0 | 0 | 0 | 98 | 49 | NA | NA | NA | NA | 0 | NA | 1 | NA | -1 |

| | | | | | | | | | | | | | | | | | | | | | | | | | | | | | | | | | | | |
|---|---|---|---|---|---|---|---|---|---|---|---|---|---|---|---|---|---|---|---|---|---|---|---|---|---|---|---|---|---|---|---|---|---|---|---|
| SE | 835 | ING3 | chr7 | + | 120973204 | 120973243 | 120970687 | 120970880 | 120974727 | 120975657 | 0 | 0 | 3 | 0 | 0 | 1 | 0 | 3 | 0 | 0 | 0 | 0 | 88 | 49 | NA | NA | NA | NA | | 0.626 | NA | | 1 | NA | -0.374 |
| SE | 864 | NAMPT | chr7 | - | 106269153 | 106269312 | 106263391 | 106263617 | 106272529 | 106272658 | 0 | 0 | 0 | 0 | 0 | 1 | 0 | 2 | 0 | 0 | 0 | 0 | 98 | 49 | NA | NA | NA | NA | | 0 | NA | | 1 | NA | -1 |
| SE | 907 | SMURF1 | chr7 | - | 99040377 | 99040556 | 99038387 | 99038525 | 99042117 | 99042232 | 0 | 0 | 2 | 0 | 0 | 0 | 0 | 2 | 0 | 0 | 1 | 0 | 98 | 49 | NA | NA | NA | NA | | 1 | NA | | 0.5 | NA | 0.5 |
| SE | 910 | TECPR1 | chr7 | - | 98241069 | 98241244 | 98240850 | 98240951 | 98243466 | 98243592 | 0 | 0 | 2 | 0 | 0 | 1 | 0 | 4 | 0 | 0 | 0 | 0 | 98 | 49 | NA | NA | NA | NA | | 0.5 | NA | | 1 | NA | -0.5 |
| SE | 949 | DMTF1 | chr7 | + | 87164933 | 87165050 | 87163494 | 87163617 | 87166482 | 87166605 | 0 | 0 | 2 | 0 | 0 | 0 | 0 | 4 | 0 | 0 | 1 | 0 | 98 | 49 | NA | NA | NA | NA | | 1 | NA | | 0.667 | NA | 0.333 |
| SE | 954 | CD36 | chr7 | + | 80662989 | 80663169 | 80661062 | 80661210 | 80664405 | 80664497 | 0 | 1 | 28 | 0 | 0 | 2 | 15 | 0 | 0 | 0 | 0 | 0 | 98 | 49 | NA | NA | NA | | 1 | 0.875 | | 1 | NA | NA | -0.062 |
| SE | 973 | LAT2 | chr7 | + | 74224638 | 74224760 | 74224017 | 74224197 | 74228943 | 74229834 | 0 | 0 | 7 | 0 | 0 | 1 | 0 | 4 | 0 | 0 | 0 | 0 | 98 | 49 | NA | NA | NA | NA | | 0.778 | NA | | 1 | NA | -0.222 |
| SE | 980 | RABGEF1 | chr7 | + | 66783674 | 66783841 | 66775226 | 66775393 | 66795510 | 66795592 | 0 | 0 | 9 | 0 | 0 | 0 | 0 | 3 | 0 | 0 | 2 | 0 | 98 | 49 | NA | NA | NA | NA | | 1 | NA | | 0.429 | NA | 0.571 |
| SE | 981 | RABGEF1 | chr7 | + | 66795510 | 66795592 | 66775226 | 66775393 | 66797373 | 66797506 | 0 | 0 | 4 | 0 | 0 | 1 | 0 | 7 | 0 | 0 | 0 | 0 | 98 | 49 | NA | NA | NA | NA | | 0.667 | NA | | 1 | NA | -0.333 |
| SE | 993 | SUMF2 | chr7 | + | 56074173 | 56074218 | 56072996 | 56073111 | 56074585 | 56074736 | 0 | 0 | 7 | 0 | 0 | 0 | 0 | 1 | 0 | 0 | 1 | 0 | 94 | 49 | NA | NA | NA | NA | | 1 | NA | | 0.343 | NA | 0.657 |
| SE | 995 | GK | chrX | + | 30694399 | 30694537 | 30691122 | 30691199 | 30696041 | 30696151 | 0 | 0 | 1 | 0 | 0 | 0 | 0 | 2 | 0 | 0 | 2 | 0 | 98 | 49 | NA | NA | NA | NA | | 1 | NA | | 0.333 | NA | 0.667 |
| SE | 996 | GK | chrX | + | 30697731 | 30697749 | 30696616 | 30696683 | 30700413 | 30700449 | 0 | 0 | 1 | 0 | 0 | 2 | 0 | 0 | 0 | 0 | 7 | 0 | 67 | 49 | NA | NA | NA | NA | | 0.268 | NA | | 0 | NA | 0.268 |
| SE | 1008 | IKZF1 | chr7 | + | 50376532 | 50376793 | 50327637 | 50327757 | 50382539 | 50382707 | 0 | 0 | 1 | 0 | 0 | 2 | 2 | 0 | 0 | 0 | 4 | 0 | 98 | 49 | NA | NA | NA | NA | | 0.2 | | 1 | 0 | NA | -0.3 |
| SE | 1034 | ELMO1 | chr7 | - | 36887559 | 36887672 | 36878009 | 36878117 | 36894853 | 36895017 | 0 | 0 | 4 | 0 | 0 | 0 | 0 | 2 | 0 | 0 | 1 | 0 | 98 | 49 | NA | NA | NA | NA | | 1 | NA | | 0.5 | NA | 0.5 |
| SE | 1035 | ELMO1 | chr7 | - | 37216644 | 37216695 | 37213334 | 37213457 | 37222614 | 37222693 | 0 | 0 | 6 | 0 | 0 | 0 | 3 | 3 | 0 | 0 | 1 | 0 | 98 | 49 | NA | NA | NA | NA | | 1 | | 1 | 0.6 | NA | 0.2 |
| SE | 1047 | KBTBD2 | chr7 | - | 32879434 | 32879942 | 32874991 | 32875157 | 32891535 | 32891856 | 0 | 0 | 6 | 0 | 0 | 0 | 0 | 7 | 0 | 0 | 1 | 0 | 98 | 49 | NA | NA | NA | NA | | 1 | NA | | 0.778 | NA | 0.222 |
| SE | 1076 | CCDC126 | chr7 | + | 23597966 | 23598051 | 23597378 | 23597605 | 23611170 | 23611553 | 0 | 0 | 0 | 0 | 0 | 1 | 0 | 1 | 0 | 0 | 0 | 0 | 98 | 49 | NA | NA | NA | NA | | 0 | NA | | 1 | NA | -1 |
| SE | 1090 | HDAC9 | chr7 | + | 18629349 | 18629481 | 18593907 | 18594029 | 18634626 | 18634742 | 0 | 0 | 1 | 0 | 0 | 1 | 0 | 1 | 0 | 0 | 0 | 0 | 98 | 49 | NA | NA | NA | NA | | 0.333 | NA | | 1 | NA | -0.667 |
| SE | 1093 | SNX13 | chr7 | - | 17897333 | 17897446 | 17893331 | 17893434 | 17940283 | 17940508 | 0 | 0 | 2 | 0 | 0 | 0 | 1 | 0 | 0 | 0 | 1 | 0 | 98 | 49 | NA | NA | NA | NA | | 1 | | 1 | 0 | NA | 0.5 |
| SE | 1104 | BRAT1 | chr7 | - | 2544908 | 2545056 | 2543589 | 2543962 | 2547323 | 2547478 | 0 | 0 | 2 | 0 | 0 | 2 | 0 | 2 | 0 | 0 | 0 | 0 | 98 | 49 | NA | NA | NA | NA | | 0.333 | NA | | 1 | NA | -0.667 |
| SE | 1116 | SUN1 | chr7 | + | 838797 | 838986 | 832504 | 832601 | 841945 | 842130 | 0 | 0 | 5 | 0 | 0 | 0 | 0 | 0 | 0 | 0 | 1 | 0 | 98 | 49 | NA | NA | NA | NA | | 1 | NA | | 0 | NA | 1 |
| SE | 1157 | IPCEF1 | chr6 | - | 154212769 | 154212855 | 154199667 | 154200040 | 154214217 | 154214276 | 0 | 0 | 2 | 0 | 0 | 0 | 0 | 4 | 0 | 0 | 1 | 0 | 98 | 49 | NA | NA | NA | NA | | 1 | NA | | 0.667 | NA | 0.333 |
| SE | 1210 | VNN2 | chr6 | - | 132752672 | 132752749 | 132751144 | 132751518 | 132753417 | 132753491 | 0 | 0 | 3 | 0 | 0 | 3 | 0 | 1 | 0 | 0 | 0 | 0 | 98 | 49 | NA | NA | NA | NA | | 0.333 | NA | | 1 | NA | -0.667 |
| SE | 1211 | VNN2 | chr6 | - | 132752672 | 132752749 | 132751144 | 132751518 | 132755842 | 132756035 | 0 | 0 | 10 | 0 | 0 | 0 | 0 | 3 | 0 | 0 | 5 | 0 | 98 | 49 | NA | NA | NA | NA | | 1 | NA | | 0.231 | NA | 0.769 |
| SE | 1302 | SLC17A5 | chr6 | - | 73638411 | 73638499 | 73636620 | 73636707 | 73641690 | 73641924 | 0 | 0 | 0 | 0 | 0 | 1 | 0 | 1 | 0 | 0 | 0 | 0 | 98 | 49 | NA | NA | NA | NA | | 0 | NA | | 1 | NA | -1 |
| SE | 1305 | SMAP1 | chr6 | + | 70791688 | 70791769 | 70773349 | 70773425 | 70798656 | 70798737 | 0 | 0 | 1 | 0 | 0 | 0 | 0 | 0 | 0 | 0 | 1 | 0 | 98 | 49 | NA | NA | NA | NA | | 1 | NA | | 0 | NA | 1 |
| SE | 1314 | RUNX2 | chr6 | + | 45491940 | 45492114 | 45437946 | 45438051 | 45512245 | 45512407 | 0 | 0 | 1 | 0 | 0 | 2 | 0 | 1 | 0 | 0 | 0 | 0 | 98 | 49 | NA | NA | NA | NA | | 0.2 | NA | | 1 | NA | -0.8 |
| SE | 1317 | CDC5L | chr6 | + | 44406322 | 44406467 | 44403808 | 44404027 | 44408443 | 44408632 | 0 | 0 | 1 | 0 | 0 | 0 | 0 | 3 | 0 | 0 | 1 | 0 | 98 | 49 | NA | NA | NA | NA | | 1 | NA | | 0.6 | NA | 0.4 |
| SE | 1323 | YIPF3 | chr6 | - | 43515594 | 43515701 | 43513587 | 43513633 | 43515888 | 43516095 | 0 | 0 | 5 | 0 | 0 | 1 | 0 | 8 | 0 | 0 | 0 | 0 | 98 | 49 | NA | NA | NA | NA | | 0.714 | NA | | 1 | NA | -0.286 |
| SE | 1329 | TRERF1 | chr6 | - | 42263319 | 42263568 | 42259338 | 42259723 | 42264703 | 42264854 | 0 | 0 | 1 | 0 | 1 | 0 | 0 | 0 | 0 | 3 | 0 | 0 | 98 | 49 | NA | NA | NA | | 0 | 1 | | 0 | NA | NA | 0.5 |
| SE | 1338 | CMTR1 | chr6 | + | 37461969 | 37462102 | 37461548 | 37461645 | 37462828 | 37463008 | 0 | 0 | 3 | 0 | 0 | 1 | 0 | 2 | 0 | 0 | 0 | 0 | 98 | 49 | NA | NA | NA | NA | | 0.6 | NA | | 1 | NA | -0.4 |
| SE | 1340 | CMTR1 | chr6 | + | 37476125 | 37476194 | 37475320 | 37475412 | 37477591 | 37477639 | 0 | 0 | 3 | 0 | 0 | 0 | 0 | 3 | 0 | 0 | 1 | 0 | 98 | 49 | NA | NA | NA | NA | | 1 | NA | | 0.6 | NA | 0.4 |
| SE | 1341 | KCTD20 | chr6 | + | 36470051 | 36470257 | 36442766 | 36443111 | 36479590 | 36479711 | 0 | 0 | 1 | 0 | 0 | 0 | 1 | 2 | 0 | 0 | 1 | 0 | 98 | 49 | NA | NA | NA | NA | | 1 | | 1 | 0.5 | NA | 0.25 |
| SE | 1343 | KCTD20 | chr6 | + | 36474788 | 36475062 | 36470051 | 36470257 | 36479120 | 36479223 | 0 | 0 | 2 | 0 | 0 | 4 | 0 | 1 | 0 | 0 | 0 | 0 | 98 | 49 | NA | NA | NA | NA | | 0.2 | NA | | 1 | NA | -0.8 |
| SE | 1344 | KCTD20 | chr6 | + | 36479120 | 36479223 | 36470051 | 36470257 | 36479590 | 36479711 | 0 | 0 | 6 | 0 | 0 | 0 | 0 | 0 | 0 | 0 | 2 | 0 | 98 | 49 | NA | NA | NA | NA | | 1 | NA | | 0 | NA | 1 |
| SE | 1358 | FKBP5 | chr6 | - | 35587033 | 35587117 | 35580035 | 35580221 | 35591129 | 35591220 | 0 | 0 | 12 | 0 | 0 | 1 | 4 | 0 | 0 | 0 | 0 | 0 | 98 | 49 | NA | NA | NA | NA | | 0.857 | | 1 | NA | NA | -0.143 |
| SE | 1372 | HLA-DMB | chr6 | - | 32935535 | 32935652 | 32935341 | 32935377 | 32937171 | 32937456 | 0 | 0 | 4 | 0 | 0 | 1 | 0 | 2 | 0 | 0 | 0 | 0 | 98 | 49 | NA | NA | NA | NA | | 0.667 | NA | | 1 | NA | -0.333 |
| SE | 1445 | BAG6 | chr6 | - | 31639499 | 31639646 | 31639027 | 31639226 | 31640198 | 31640306 | 0 | 0 | 4 | 0 | 0 | 2 | 0 | 2 | 0 | 0 | 2 | 0 | 98 | 49 | NA | NA | NA | NA | | 0.5 | NA | | 0.333 | NA | 0.167 |
| SE | 1450 | BAG6 | chr6 | - | 31647590 | 31647844 | 31646393 | 31646523 | 31648676 | 31648751 | 0 | 0 | 0 | 1 | 0 | 0 | 0 | 1 | 0 | 0 | 0 | 0 | 98 | 49 | NA | NA | 0 | NA | NA | NA | | | 1 | NA | -1 |
| SE | 1456 | ATP6V1G2-DDX39B | chr6 | - | 31538762 | 31538855 | 31536499 | 31536683 | 31540321 | 31540664 | 0 | 0 | 1 | 0 | 0 | 0 | 0 | 2 | 0 | 0 | 3 | 0 | 98 | 49 | NA | NA | NA | NA | | 1 | NA | | 0.25 | NA | 0.75 |
| SE | 1481 | FLOT1 | chr6 | - | 30741791 | 30741867 | 30741189 | 30741333 | 30742146 | 30742203 | 0 | 0 | 2 | 0 | 0 | 2 | 0 | 2 | 0 | 1 | 0 | 0 | 98 | 49 | NA | NA | NA | NA | | 0.333 | | 0 | 1 | NA | -0.167 |
| SE | 1579 | USP9X | chrX | + | 41150920 | 41151057 | 41148368 | 41148575 | 41152947 | 41153081 | 0 | 0 | 10 | 0 | 0 | 0 | 0 | 3 | 0 | 0 | 1 | 0 | 98 | 49 | NA | NA | NA | NA | | 1 | NA | | 0.6 | NA | 0.4 |
| SE | 1580 | USP9X | chrX | + | 41214567 | 41214709 | 41210508 | 41210682 | 41215898 | 41216652 | 0 | 0 | 2 | 0 | 0 | 0 | 0 | 3 | 0 | 0 | 1 | 0 | 98 | 49 | NA | NA | NA | NA | | 1 | NA | | 0.6 | NA | 0.4 |
| SE | 1581 | USP9X | chrX | + | 41219101 | 41219231 | 41218371 | 41218597 | 41223216 | 41223402 | 0 | 0 | 1 | 0 | 0 | 0 | 0 | 6 | 0 | 0 | 1 | 0 | 98 | 49 | NA | NA | NA | NA | | 1 | NA | | 0.75 | NA | 0.25 |
| SE | 1587 | DEK | chr6 | - | 18255730 | 18255851 | 18249650 | 18249839 | 18256360 | 18256455 | 0 | 0 | 8 | 0 | 0 | 0 | 0 | 2 | 0 | 0 | 2 | 0 | 98 | 49 | NA | NA | NA | NA | | 1 | NA | | 0.333 | NA | 0.667 |
| SE | 1594 | KIF13A | chr6 | - | 17817019 | 17817233 | 17808767 | 17808930 | 17825767 | 17825934 | 0 | 0 | 5 | 0 | 0 | 1 | 0 | 3 | 0 | 0 | 0 | 0 | 98 | 49 | NA | NA | NA | NA | | 0.714 | NA | | 1 | NA | -0.286 |
| SE | 1595 | KIF13A | chr6 | - | 17855436 | 17855617 | 17851954 | 17852042 | 17856029 | 17856122 | 0 | 0 | 9 | 0 | 0 | 1 | 0 | 5 | 0 | 0 | 0 | 0 | 98 | 49 | NA | NA | NA | NA | | 0.818 | NA | | 1 | NA | -0.182 |
| SE | 1596 | KIF13A | chr6 | - | 17898167 | 17898180 | 17873376 | 17873437 | 17987053 | 17987144 | 0 | 0 | 6 | 0 | 0 | 1 | 1 | 1 | 0 | 0 | 0 | 0 | 62 | 49 | NA | NA | NA | NA | | 0.826 | | 1 | 1 | NA | -0.174 |
| SE | 1597 | NUP153 | chr6 | - | 17625807 | 17626164 | 17624560 | 17624833 | 17628654 | 17629539 | 0 | 0 | 5 | 0 | 0 | 0 | 1 | 2 | 0 | 0 | 1 | 0 | 98 | 49 | NA | NA | NA | NA | | 1 | | 1 | 0.5 | NA | 0.25 |
| SE | 1600 | NUP153 | chr6 | - | 17661652 | 17661779 | 17649162 | 17649300 | 17662017 | 17662070 | 0 | 0 | 7 | 0 | 0 | 0 | 0 | 9 | 0 | 0 | 1 | 0 | 98 | 49 | NA | NA | NA | NA | | 1 | NA | | 0.818 | NA | 0.182 |
| SE | 1608 | JARID2 | chr6 | + | 15452005 | 15452175 | 15410223 | 15410365 | 15468541 | 15468718 | 0 | 0 | 10 | 0 | 0 | 0 | 0 | 3 | 0 | 0 | 1 | 0 | 98 | 49 | NA | NA | NA | NA | | 1 | NA | | 0.6 | NA | 0.4 |
| SE | 1616 | HIVEP1 | chr6 | + | 12129758 | 12129892 | 12119889 | 12125870 | 12130766 | 12130942 | 0 | 0 | 3 | 0 | 0 | 1 | 0 | 3 | 0 | 0 | 0 | 0 | 98 | 49 | NA | NA | NA | NA | | 0.6 | NA | | 1 | NA | -0.4 |
| SE | 1617 | HIVEP1 | chr6 | + | 12135790 | 12135892 | 12130766 | 12130942 | 12161438 | 12161929 | 0 | 0 | 1 | 0 | 0 | 0 | 0 | 5 | 0 | 0 | 1 | 0 | 98 | 49 | NA | NA | NA | NA | | 1 | NA | | 0.714 | NA | 0.286 |
| SE | 1640 | RIPK1 | chr6 | + | 3085258 | 3085408 | 3083084 | 3083313 | 3089580 | 3089657 | 0 | 0 | 0 | 0 | 0 | 1 | 0 | 1 | 0 | 0 | 0 | 0 | 98 | 49 | NA | NA | NA | NA | | 0 | NA | | 1 | NA | -1 |
| SE | 1652 | HNRNPH1 | chr5 | - | 179615545 | 179615595 | 179614177 | 179614959 | 179616125 | 179616218 | 0 | 0 | 4 | 0 | 0 | 0 | 0 | 8 | 0 | 0 | 1 | 0 | 98 | 49 | NA | NA | NA | NA | | 1 | NA | | 0.8 | NA | 0.2 |
| SE | 1653 | HNRNPH1 | chr5 | - | 179619268 | 179619407 | 179618144 | 179618323 | 179620891 | 179621035 | 0 | 0 | 3 | 0 | 0 | 0 | 0 | 1 | 0 | 0 | 4 | 0 | 98 | 49 | NA | NA | NA | NA | | 1 | NA | | 0.111 | NA | 0.889 |
| SE | 1692 | DOCK2 | chr5 | + | 169983067 | 169983166 | 169840756 | 169840852 | 169985827 | 169985922 | 0 | 0 | 12 | 0 | 0 | 1 | 0 | 5 | 0 | 0 | 0 | 0 | 98 | 49 | NA | NA | NA | NA | | 0.857 | NA | | 1 | NA | -0.143 |
| SE | 1697 | RNF145 | chr5 | - | 159181959 | 159182051 | 159176631 | 159176867 | 159194715 | 159194824 | 0 | 0 | 4 | 0 | 0 | 0 | 0 | 0 | 0 | 0 | 1 | 0 | 98 | 49 | NA | NA | NA | NA | | 1 | NA | | 0 | NA | 1 |
| SE | 1701 | CNOT8 | chr5 | + | 154863206 | 154863395 | 154858518 | 154858768 | 154865191 | 154865385 | 0 | 0 | 2 | 0 | 0 | 0 | 0 | 1 | 0 | 0 | 1 | 0 | 98 | 49 | NA | NA | NA | NA | | 1 | NA | | 0.333 | NA | 0.667 |
| SE | 1703 | CNOT8 | chr5 | + | 154865191 | 154865385 | 154858518 | 154858768 | 154870660 | 154870822 | 0 | 0 | 0 | 0 | 0 | 1 | 0 | 1 | 0 | 0 | 0 | 0 | 98 | 49 | NA | NA | NA | NA | | 0 | NA | | 1 | NA | -1 |
| SE | 1707 | LARP1 | chr5 | + | 154793799 | 154794000 | 154793594 | 154793723 | 154794099 | 154794262 | 0 | 0 | 1 | 0 | 0 | 0 | 0 | 0 | 0 | 0 | 2 | 0 | 98 | 49 | NA | NA | NA | NA | | 1 | NA | | 0 | NA | 1 |
| SE | 1744 | NDFIP1 | chr5 | + | 142137733 | 142137858 | 142135729 | 142135817 | 142140562 | 142140629 | 0 | 0 | 4 | 0 | 0 | 1 | 0 | 3 | 0 | 0 | 0 | 0 | 98 | 49 | NA | NA | NA | NA | | 0.667 | NA | | 1 | NA | -0.333 |
| SE | 1748 | DIAPH1 | chr5 | - | 141529602 | 141529697 | 141529171 | 141529273 | 141534334 | 141534433 | 0 | 0 | 8 | 0 | 0 | 0 | 0 | 1 | 0 | 0 | 1 | 0 | 98 | 49 | NA | NA | NA | NA | | 1 | NA | | 0.333 | NA | 0.667 |
| SE | 1760 | KDM3B | chr5 | + | 138386021 | 138386621 | 138381515 | 138381590 | 138391012 | 138392261 | 0 | 0 | 2 | 0 | 0 | 2 | 0 | 2 | 0 | 0 | 0 | 0 | 98 | 49 | NA | NA | NA | NA | | 0.333 | NA | | 1 | NA | -0.667 |
| SE | 1769 | CAMLG | chr5 | + | 134741062 | 134741523 | 134738479 | 134738792 | 134743986 | 134744052 | 0 | 0 | 0 | 0 | 0 | 1 | 0 | 2 | 0 | 0 | 0 | 0 | 98 | 49 | NA | NA | NA | NA | | 0 | NA | | 1 | NA | -1 |
| SE | 1799 | AP3S1 | chr5 | + | 115870016 | 115870128 | 115866669 | 115866761 | 115895086 | 115895158 | 0 | 0 | 1 | 0 | 0 | 0 | 0 | 4 | 0 | 0 | 1 | 0 | 98 | 49 | NA | NA | NA | NA | | 1 | NA | | 0.667 | NA | 0.333 |
| SE | 1808 | ASAP1 | chr8 | - | 130168991 | 130169067 | 130167535 | 130167622 | 130179263 | 130179349 | 0 | 0 | 15 | 0 | 0 | 0 | 0 | 11 | 0 | 0 | 1 | 0 | 98 | 49 | NA | NA | NA | NA | | 1 | NA | | 0.846 | NA | 0.154 |
| SE | 1815 | TBL1X | chrX | + | 9653544 | 9653689 | 9640272 | 9640360 | 9654214 | 9654322 | 0 | 0 | 1 | 0 | 0 | 3 | 0 | 0 | 0 | 0 | 3 | 0 | 98 | 49 | NA | NA | NA | NA | | 0.143 | NA | | 0 | NA | 0.143 |
| SE | 1816 | CHD1 | chr5 | - | 98858179 | 98858390 | 98855203 | 98856725 | 98858963 | 98859015 | 0 | 0 | 2 | 0 | 0 | 1 | 0 | 19 | 0 | 0 | 5 | 0 | 98 | 49 | NA | NA | NA | NA | | 0.5 | NA | | 0.655 | NA | -0.155 |
| SE | 1820 | ERAP1 | chr5 | - | 96788530 | 96788685 | 96786469 | 96786549 | 96790295 | 96790367 | 0 | 0 | 5 | 0 | 0 | 0 | 0 | 2 | 0 | 0 | 1 | 0 | 98 | 49 | NA | NA | NA | NA | | 1 | NA | | 0.5 | NA | 0.5 |
| SE | 1824 | MCTP1 | chr5 | - | 94799012 | 94799132 | 94779109 | 94779163 | 94868332 | 94868452 | 0 | 0 | 1 | 0 | 0 | 1 | 0 | 3 | 0 | 0 | 0 | 0 | 98 | 49 | NA | NA | NA | NA | | 0.333 | NA | | 1 | NA | -0.667 |
| SE | 1825 | MCTP1 | chr5 | - | 94870871 | 94870973 | 94870416 | 94870491 | 94871314 | 94871417 | 0 | 0 | 2 | 0 | 0 | 0 | 0 | 2 | 0 | 0 | 1 | 0 | 98 | 49 | NA | NA | NA | NA | | 1 | NA | | 0.5 | NA | 0.5 |
| SE | 1828 | MCTP1 | chr5 | - | 94923961 | 94924021 | 94912805 | 94912976 | 94940083 | 94940195 | 0 | 0 | 1 | 0 | 0 | 2 | 0 | 4 | 0 | 0 | 0 | 0 | 98 | 49 | NA | NA | NA | NA | | 0.2 | NA | | 1 | NA | -0.8 |
| SE | 1829 | MCTP1 | chr5 | - | 94931952 | 94931991 | 94912805 | 94912976 | 94940083 | 94940195 | 0 | 0 | 6 | 0 | 0 | 2 | 0 | 4 | 0 | 0 | 0 | 0 | 88 | 49 | NA | NA | NA | NA | | 0.626 | NA | | 1 | NA | -0.374 |
| SE | 1830 | MCTP1 | chr5 | - | 94931952 | 94931991 | 94923961 | 94924021 | 94940083 | 94940195 | 0 | 0 | 9 | 0 | 0 | 1 | 0 | 4 | 0 | 0 | 0 | 0 | 88 | 49 | NA | NA | NA | NA | | 0.834 | NA | | 1 | NA | -0.166 |
| SE | 1831 | FAM172A | chr5 | - | 93958776 | 93958968 | 93881469 | 93881688 | 93964457 | 93964523 | 0 | 0 | 5 | 0 | 0 | 2 | 0 | 4 | 0 | 0 | 0 | 0 | 98 | 49 | NA | NA | NA | NA | | 0.556 | NA | | 1 | NA | -0.444 |
| SE | 1832 | FAM172A | chr5 | - | 94050730 | 94050831 | 93964457 | 93964523 | 94053125 | 94053227 | 0 | 0 | 3 | 0 | 0 | 1 | 0 | 5 | 0 | 0 | 0 | 0 | 98 | 49 | NA | NA | NA | NA | | 0.6 | NA | | 1 | NA | -0.4 |
| SE | 1833 | FAM172A | chr5 | - | 94074645 | 94074769 | 94053125 | 94053227 | 94111510 | 94111699 | 0 | 0 | 2 | 0 | 0 | 1 | 0 | 1 | 0 | 0 | 0 | 0 | 98 | 49 | NA | NA | NA | NA | | 0.5 | NA | | 1 | NA | -0.5 |
| SE | 1841 | VCAN | chr5 | + | 83519348 | 83522309 | 83512102 | 83512396 | 83537006 | 83542268 | 0 | 0 | 1 | 0 | 0 | 19 | 0 | 0 | 0 | 0 | 3 | 0 | 98 | 49 | NA | NA | NA | NA | | 0.026 | NA | | 0 | NA | 0.026 |
| SE | 1847 | SSBP2 | chr5 | - | 81489249 | 81489309 | 81474495 | 81474562 | 81513627 | 81513717 | 0 | 0 | 0 | 0 | 0 | 1 | 1 | 0 | 0 | 0 | 0 | 0 | 98 | 49 | NA | NA | NA | NA | | 0 | | 1 | NA | NA | -1 |
| SE | 1848 | SSBP2 | chr5 | - | 81513627 | 81513717 | 81489249 | 81489309 | 81615472 | 81615557 | 0 | 0 | 4 | 0 | 0 | 3 | 1 | 0 | 0 | 0 | 0 | 0 | 98 | 49 | NA | NA | NA | NA | | 0.4 | | 1 | NA | NA | -0.6 |
| SE | 1850 | PAPD4 | chr5 | + | 79642831 | 79642910 | 79641104 | 79641196 | 79645122 | 79645192 | 0 | 0 | 4 | 0 | 0 | 1 | 0 | 1 | 0 | 0 | 0 | 0 | 98 | 49 | NA | NA | NA | NA | | 0.667 | NA | | 1 | NA | -0.333 |
| SE | 1851 | PAPD4 | chr5 | + | 79642843 | 79642910 | 79641104 | 79641196 | 79645122 | 79645192 | 0 | 0 | 1 | 0 | 0 | 1 | 0 | 1 | 0 | 0 | 0 | 0 | 98 | 49 | NA | NA | NA | NA | | 0.333 | NA | | 1 | NA | -0.667 |
| SE | 1852 | PAPD4 | chr5 | + | 79649061 | 79649190 | 79648616 | 79648693 | 79656957 | 79657001 | 0 | 0 | 0 | 0 | 0 | 1 | 0 | 1 | 0 | 0 | 0 | 0 | 98 | 49 | NA | NA | NA | NA | | 0 | NA | | 1 | NA | -1 |
| SE | 1904 | MAP3K1 | chr5 | + | 56887377 | 56887520 | 56885931 | 56886063 | 56888225 | 56888357 | 0 | 0 | 11 | 0 | 0 | 0 | 0 | 4 | 0 | 0 | 1 | 0 | 98 | 49 | NA | NA | NA | NA | | 1 | NA | | 0.667 | NA | 0.333 |
| SE | 1918 | PARP8 | chr5 | + | 50760291 | 50760362 | 50759642 | 50759732 | 50761820 | 50761898 | 0 | 0 | 17 | 0 | 0 | 0 | 0 | 7 | 0 | 6 | 1 | 0 | 98 | 49 | NA | NA | NA | NA | | 1 | | 0 | 0.778 | NA | 0.611 |
| SE | 1933 | PRKAA1 | chr5 | - | 40769415 | 40769503 | 40767465 | 40767690 | 40771718 | 40771863 | 0 | 0 | 4 | 0 | 0 | 0 | 0 | 6 | 0 | 0 | 1 | 0 | 98 | 49 | NA | NA | NA | NA | | 1 | NA | | 0.75 | NA | 0.25 |
| SE | 1935 | RICTOR | chr5 | - | 38949355 | 38949427 | 38947263 | 38947441 | 38949711 | 38950720 | 0 | 0 | 0 | 0 | 0 | 11 | 0 | 1 | 0 | 0 | 3 | 0 | 98 | 49 | NA | NA | NA | NA | | 0 | NA | | 0.143 | NA | -0.143 |
| SE | 1936 | RICTOR | chr5 | - | 38975536 | 38975604 | 38971876 | 38971959 | 38978582 | 38978648 | 0 | 0 | 12 | 0 | 0 | 1 | 0 | 16 | 0 | 0 | 0 | 0 | 98 | 49 | NA | NA | NA | NA | | 0.857 | NA | | 1 | NA | -0.143 |
| SE | 1946 | ZFR | chr5 | - | 32380074 | 32380172 | 32379114 | 32379210 | 32385507 | 32385649 | 0 | 0 | 4 | 0 | 0 | 2 | 0 | 5 | 0 | 0 | 0 | 0 | 98 | 49 | NA | NA | NA | NA | | 0.5 | NA | | 1 | NA | -0.5 |
| SE | 1952 | 6-Mar | chr5 | + | 10390331 | 10390500 | 10386993 | 10387066 | 10391541 | 10391731 | 0 | 0 | 6 | 0 | 0 | 2 | 0 | 1 | 0 | 0 | 0 | 0 | 98 | 49 | NA | NA | NA | NA | | 0.6 | NA | | 1 | NA | -0.4 |
| SE | 1953 | 6-Mar | chr5 | + | 10426389 | 10426522 | 10423734 | 10423824 | 10429892 | 10430028 | 0 | 0 | 0 | 0 | 0 | 1 | 0 | 3 | 0 | 0 | 2 | 0 | 98 | 49 | NA | NA | NA | NA | | 0 | NA | | 0.429 | NA | -0.429 |
| SE | 1984 | DDX60L | chr4 | - | 168453123 | 168453282 | 168441336 | 168441492 | 168457891 | 168458008 | 0 | 0 | 1 | 0 | 0 | 1 | 0 | 2 | 0 | 0 | 0 | 0 | 98 | 49 | NA | NA | NA | NA | | 0.333 | NA | | 1 | NA | -0.667 |
| SE | 1985 | DDX60L | chr4 | - | 168456038 | 168456152 | 168453123 | 168453282 | 168457891 | 168458008 | 0 | 0 | 35 | 0 | 0 | 0 | 0 | 10 | 0 | 0 | 1 | 0 | 98 | 49 | NA | NA | NA | NA | | 1 | NA | | 0.833 | NA | 0.167 |
| SE | 2000 | UBQLN1 | chr9 | - | 83666349 | 83666433 | 83665029 | 83665145 | 83669184 | 83669327 | 0 | 0 | 4 | 0 | 0 | 1 | 0 | 5 | 0 | 0 | 0 | 0 | 98 | 49 | NA | NA | NA | NA | | 0.667 | NA | | 1 | NA | -0.333 |
| SE | 2008 | ARFIP1 | chr4 | + | 152882722 | 152882880 | 152880962 | 152881184 | 152888132 | 152888307 | 0 | 0 | 1 | 0 | 0 | 0 | 0 | 0 | 0 | 0 | 1 | 0 | 98 | 49 | NA | NA | NA | NA | | 1 | NA | | 0 | NA | 1 |
| SE | 2057 | TET2 | chr4 | + | 105190359 | 105190505 | 105146684 | 105146979 | 105233896 | 105237351 | 0 | 0 | 6 | 0 | 0 | 0 | 0 | 7 | 0 | 0 | 1 | 0 | 98 | 49 | NA | NA | NA | NA | | 1 | NA | | 0.778 | NA | 0.222 |
| SE | 2058 | TET2 | chr4 | + | 105241338 | 105241429 | 105233896 | 105237351 | 105242833 | 105242927 | 0 | 0 | 6 | 0 | 0 | 2 | 0 | 9 | 0 | 0 | 1 | 0 | 98 | 49 | NA | NA | NA | NA | | 0.6 | NA | | 0.818 | NA | -0.218 |
| SE | 2065 | DAPP1 | chr4 | + | 99866556 | 99866618 | 99866033 | 99866121 | 99868116 | 99870189 | 0 | 0 | 3 | 0 | 0 | 6 | 0 | 0 | 0 | 0 | 6 | 0 | 98 | 49 | NA | NA | NA | NA | | 0.2 | NA | | 0 | NA | 0.2 |
| SE | 2079 | HERC5 | chr4 | + | 88486114 | 88486228 | 88479352 | 88479507 | 88489165 | 88489336 | 0 | 0 | 8 | 0 | 0 | 1 | 0 | 9 | 0 | 0 | 2 | 0 | 98 | 49 | NA | NA | NA | NA | | 0.8 | NA | | 0.692 | NA | 0.108 |
| SE | 2083 | HERC6 | chr4 | + | 88417424 | 88417579 | 88413076 | 88413266 | 88423859 | 88423973 | 0 | 0 | 6 | 0 | 0 | 0 | 0 | 2 | 0 | 0 | 2 | 0 | 98 | 49 | NA | NA | NA | NA | | 1 | NA | | 0.333 | NA | 0.667 |
| SE | 2097 | WDFY3 | chr4 | - | 84690505 | 84690664 | 84688085 | 84688265 | 84691630 | 84691785 | 0 | 0 | 5 | 0 | 0 | 0 | 0 | 1 | 0 | 0 | 1 | 0 | 98 | 49 | NA | NA | NA | NA | | 1 | NA | | 0.333 | NA | 0.667 |
| SE | 2098 | WDFY3 | chr4 | - | 84705393 | 84705511 | 84704337 | 84704444 | 84708908 | 84709028 | 0 | 0 | 11 | 0 | 0 | 1 | 2 | 10 | 0 | 0 | 0 | 0 | 98 | 49 | NA | NA | NA | NA | | 0.846 | | 1 | 1 | NA | -0.154 |
| SE | 2100 | WDFY3 | chr4 | - | 84733381 | 84733609 | 84724425 | 84724594 | 84735042 | 84735120 | 0 | 0 | 9 | 0 | 0 | 0 | 0 | 6 | 0 | 0 | 1 | 0 | 98 | 49 | NA | NA | NA | NA | | 1 | NA | | 0.75 | NA | 0.25 |
| SE | 2113 | PPP2CA | chr5 | - | 134200334 | 134200496 | 134199085 | 134199204 | 134200984 | 134201074 | 0 | 0 | 4 | 0 | 0 | 3 | 2 | 1 | 0 | 0 | 1 | 0 | 98 | 49 | NA | NA | NA | NA | | 0.4 | | 1 | 0.333 | NA | -0.266 |
| SE | 2117 | HNRNPD | chr4 | - | 82356536 | 82356683 | 82355303 | 82355401 | 82356795 | 82356895 | 0 | 0 | 1 | 0 | 0 | 0 | 0 | 0 | 0 | 0 | 1 | 0 | 98 | 49 | NA | NA | NA | NA | | 1 | NA | | 0 | NA | 1 |
| SE | 2130 | G3BP2 | chr4 | - | 75653982 | 75654081 | 75648638 | 75648741 | 75655065 | 75655246 | 0 | 0 | 1 | 0 | 0 | 3 | 0 | 0 | 0 | 0 | 6 | 0 | 98 | 49 | NA | NA | NA | NA | | 0.143 | NA | | 0 | NA | 0.143 |
| SE | 2139 | HSPA9 | chr5 | - | 138566625 | 138566718 | 138561579 | 138561789 | 138567000 | 138567163 | 0 | 0 | 2 | 0 | 0 | 0 | 0 | 4 | 0 | 0 | 2 | 0 | 98 | 49 | NA | NA | NA | NA | | 1 | NA | | 0.5 | NA | 0.5 |
| SE | 2142 | ANKRD17 | chr4 | - | 73147240 | 73147432 | 73146763 | 73146873 | 73148812 | 73149050 | 0 | 0 | 10 | 0 | 0 | 0 | 0 | 0 | 0 | 0 | 2 | 0 | 98 | 49 | NA | NA | NA | NA | | 1 | NA | | 0 | NA | 1 |
| SE | 2152 | UBA6 | chr4 | - | 67665188 | 67665292 | 67663884 | 67663947 | 67668550 | 67668674 | 0 | 0 | 7 | 0 | 0 | 0 | 0 | 1 | 0 | 0 | 1 | 0 | 98 | 49 | NA | NA | NA | NA | | 1 | NA | | 0.333 | NA | 0.667 |
| SE | 2165 | EXOC1 | chr4 | + | 55893551 | 55893780 | 55892634 | 55892711 | 55896716 | 55896900 | 0 | 0 | 7 | 0 | 0 | 1 | 0 | 4 | 0 | 0 | 0 | 0 | 98 | 49 | NA | NA | NA | NA | | 0.778 | NA | | 1 | NA | -0.222 |
| SE | 2180 | RBM47 | chr4 | - | 40436440 | 40436647 | 40432650 | 40432862 | 40437770 | 40438924 | 0 | 0 | 6 | 0 | 0 | 0 | 0 | 0 | 0 | 0 | 1 | 0 | 98 | 49 | NA | NA | NA | NA | | 1 | NA | | 0 | NA | 1 |
| SE | 2182 | UBE2K | chr4 | + | 39755656 | 39755739 | 39745751 | 39745810 | 39774833 | 39774933 | 0 | 0 | 2 | 0 | 0 | 1 | 5 | 0 | 0 | 0 | 0 | 0 | 98 | 49 | NA | NA | NA | NA | | 0.5 | | 1 | NA | NA | -0.5 |
| SE | 2199 | CPEB2 | chr4 | + | 15007304 | 15007586 | 15002673 | 15004335 | 15017187 | 15017278 | 0 | 0 | 4 | 0 | 0 | 1 | 0 | 4 | 0 | 0 | 0 | 0 | 98 | 49 | NA | NA | NA | NA | | 0.667 | NA | | 1 | NA | -0.333 |
| SE | 2200 | CPEB2 | chr4 | + | 15007304 | 15007595 | 15002673 | 15004335 | 15017187 | 15017278 | 0 | 0 | 3 | 0 | 0 | 1 | 0 | 1 | 0 | 0 | 0 | 0 | 98 | 49 | NA | NA | NA | NA | | 0.6 | NA | | 1 | NA | -0.4 |
| SE | 2202 | CPEB2 | chr4 | + | 15017187 | 15017278 | 15007304 | 15007586 | 15033160 | 15033211 | 0 | 0 | 5 | 0 | 0 | 0 | 0 | 7 | 0 | 0 | 1 | 0 | 98 | 49 | NA | NA | NA | NA | | 1 | NA | | 0.778 | NA | 0.222 |
| SE | 2208 | IK | chr5 | + | 140655828 | 140655992 | 140654680 | 140654727 | 140657553 | 140657662 | 0 | 0 | 2 | 0 | 0 | 2 | 0 | 1 | 0 | 0 | 0 | 0 | 98 | 49 | NA | NA | NA | NA | | 0.333 | NA | | 1 | NA | -0.667 |
| SE | 2217 | MAN2B2 | chr4 | + | 6598197 | 6598354 | 6597112 | 6597303 | 6600622 | 6600756 | 0 | 0 | 0 | 0 | 0 | 1 | 0 | 1 | 0 | 0 | 0 | 0 | 98 | 49 | NA | NA | NA | NA | | 0 | NA | | 1 | NA | -1 |
| SE | 2256 | KIAA0226 | chr3 | - | 197684156 | 197684217 | 197683306 | 197683439 | 197693714 | 197693816 | 0 | 0 | 1 | 0 | 0 | 0 | 0 | 7 | 0 | 3 | 0 | 0 | 98 | 49 | NA | NA | NA | NA | | 1 | | 0 | 1 | NA | 0.5 |
| SE | 2265 | PCYT1A | chr3 | - | 196283474 | 196283538 | 196270414 | 196270541 | 196287614 | 196287713 | 0 | 0 | 1 | 0 | 0 | 1 | 0 | 1 | 0 | 0 | 0 | 0 | 98 | 49 | NA | NA | NA | NA | | 0.333 | NA | | 1 | NA | -0.667 |
| SE | 2276 | ACAP2 | chr3 | - | 195291705 | 195291815 | 195289120 | 195289231 | 195292264 | 195292452 | 0 | 0 | 12 | 0 | 0 | 2 | 0 | 8 | 0 | 0 | 0 | 0 | 98 | 49 | NA | NA | NA | NA | | 0.75 | NA | | 1 | NA | -0.25 |
| SE | 2279 | OPA1 | chr3 | + | 193617177 | 193617285 | 193615673 | 193615770 | 193618868 | 193618936 | 0 | 0 | 2 | 0 | 0 | 0 | 0 | 0 | 0 | 0 | 1 | 0 | 98 | 49 | NA | NA | NA | NA | | 1 | NA | | 0 | NA | 1 |
| SE | 2288 | IDUA | chr4 | + | 1000881 | 1000989 | 1000611 | 1000697 | 1001467 | 1001563 | 0 | 0 | 0 | 0 | 0 | 2 | 0 | 1 | 0 | 0 | 0 | 0 | 98 | 49 | NA | NA | NA | NA | | 0 | NA | | 1 | NA | -1 |

| | | | | | | | | | | | | | | | | | | | | | | | | | | | | | | | | | |
|---|---|---|---|---|---|---|---|---|---|---|---|---|---|---|---|---|---|---|---|---|---|---|---|---|---|---|---|---|---|---|---|---|---|
| SE | 2304 | PPIP5K2 | chr5 | + | 103133452 | 103133648 | 103129305 | 103129703 | 103136731 | 103136822 | 0 | 0 | 1 | 0 | 0 | 0 | 0 | 1 | 0 | 0 | 1 | 0 | 98 | 49 | NA | NA | NA | NA | 1 | NA | 0.333 | NA | 0.667 |
| SE | 2305 | PPIP5K2 | chr5 | + | 103176865 | 103176910 | 103173857 | 103173972 | 103177666 | 103177774 | 0 | 0 | 2 | 0 | 0 | 1 | 0 | 1 | 0 | 0 | 1 | 0 | 94 | 49 | NA | NA | NA | NA | 0.51 | NA | 0.343 | NA | 0.167 |
| SE | 2314 | TNFAIP3 | chr6 | + | 137880070 | 137880252 | 137878431 | 137879351 | 137881034 | 137883314 | 0 | 0 | 12 | 0 | 0 | 0 | 1 | 33 | 0 | 0 | 1 | 0 | 98 | 49 | NA | NA | NA | NA | 1 | 1 | 0.943 | NA | 0.028 |
| SE | 2342 | ZMAT3 | chr3 | - | 179071326 | 179071447 | 179067482 | 179067809 | 179071594 | 179071868 | 0 | 0 | 0 | 0 | 0 | 1 | 0 | 1 | 0 | 0 | 0 | 0 | 98 | 49 | NA | NA | NA | NA | 0 | NA | 1 | NA | -1 |
| SE | 2347 | FNDC3B | chr3 | + | 172328951 | 172329076 | 172310827 | 172310881 | 172330540 | 172330715 | 0 | 0 | 12 | 0 | 0 | 7 | 0 | 2 | 0 | 0 | 0 | 0 | 98 | 49 | NA | NA | NA | NA | 0.462 | NA | 1 | NA | -0.538 |
| SE | 2348 | FNDC3B | chr3 | + | 172343010 | 172343116 | 172341112 | 172341231 | 172344085 | 172344258 | 0 | 0 | 10 | 0 | 0 | 0 | 0 | 2 | 0 | 0 | 1 | 0 | 98 | 49 | NA | NA | NA | NA | 1 | NA | 0.5 | NA | 0.5 |
| SE | 2382 | MME | chr3 | + | 155168491 | 155168625 | 155166901 | 155167021 | 155168731 | 155168797 | 0 | 0 | 10 | 0 | 0 | 3 | 1 | 10 | 0 | 0 | 2 | 0 | 98 | 49 | NA | NA | NA | NA | 0.625 | 1 | 0.714 | NA | -0.232 |
| SE | 2383 | DHX36 | chr3 | - | 154306215 | 154306295 | 154305093 | 154305168 | 154309652 | 154309823 | 0 | 0 | 1 | 0 | 0 | 1 | 0 | 3 | 0 | 0 | 0 | 0 | 98 | 49 | NA | NA | NA | NA | 0.333 | NA | 1 | NA | -0.667 |
| SE | 2386 | TSC22D2 | chr3 | + | 150423048 | 150423120 | 150408334 | 150411308 | 150457075 | 150457127 | 0 | 0 | 4 | 0 | 0 | 1 | 0 | 0 | 0 | 0 | 1 | 0 | 98 | 49 | NA | NA | NA | NA | 0.667 | NA | 0 | NA | 0.667 |
| SE | 2394 | IMPA1 | chr8 | - | 81670938 | 81671047 | 81660515 | 81660667 | 81673840 | 81673949 | 0 | 0 | 2 | 0 | 0 | 0 | 0 | 0 | 0 | 0 | 1 | 0 | 98 | 49 | NA | NA | NA | NA | 1 | NA | 0 | NA | 1 |
| SE | 2401 | PLSCR1 | chr3 | - | 146528613 | 146528831 | 146525604 | 146525647 | 146533469 | 146533550 | 0 | 0 | 8 | 0 | 0 | 2 | 0 | 2 | 0 | 0 | 3 | 0 | 98 | 49 | NA | NA | NA | NA | 0.667 | NA | 0.25 | NA | 0.417 |
| SE | 2413 | DLD | chr7 | + | 107902324 | 107902393 | 107901737 | 107901817 | 107903477 | 107903547 | 0 | 0 | 7 | 0 | 0 | 0 | 0 | 0 | 0 | 0 | 1 | 0 | 98 | 49 | NA | NA | NA | NA | 1 | NA | 0 | NA | 1 |
| SE | 2418 | PIK3CB | chr3 | - | 138663905 | 138664029 | 138657689 | 138657835 | 138665035 | 138665203 | 0 | 0 | 5 | 0 | 0 | 0 | 0 | 0 | 0 | 0 | 1 | 0 | 98 | 49 | NA | NA | NA | NA | 1 | NA | 0 | NA | 1 |
| SE | 2425 | ARMC8 | chr3 | + | 138223388 | 138223531 | 138221925 | 138221997 | 138223635 | 138223733 | 0 | 0 | 2 | 0 | 0 | 0 | 0 | 4 | 0 | 0 | 2 | 0 | 98 | 49 | NA | NA | NA | NA | 1 | NA | 0.5 | NA | 0.5 |
| SE | 2430 | CEP63 | chr3 | + | 134547334 | 134547472 | 134546148 | 134546288 | 134549061 | 134549176 | 0 | 0 | 4 | 0 | 0 | 6 | 0 | 0 | 0 | 0 | 4 | 0 | 98 | 49 | NA | NA | NA | NA | 0.25 | NA | 0 | NA | 0.25 |
| SE | 2433 | DNAJC13 | chr3 | + | 132495087 | 132495166 | 132494143 | 132494259 | 132496527 | 132496663 | 0 | 0 | 7 | 0 | 0 | 0 | 0 | 3 | 0 | 1 | 0 | 0 | 98 | 49 | NA | NA | NA | NA | 1 | 0 | 1 | NA | 0.5 |
| SE | 2434 | DNAJC13 | chr3 | + | 132511066 | 132511244 | 132507236 | 132507353 | 132513007 | 132513099 | 0 | 0 | 2 | 0 | 0 | 0 | 0 | 1 | 0 | 0 | 1 | 0 | 98 | 49 | NA | NA | NA | NA | 1 | NA | 0.333 | NA | 0.667 |
| SE | 2453 | DOCK11 | chrX | + | 118590209 | 118590302 | 118588412 | 118588478 | 118593213 | 118593337 | 0 | 0 | 15 | 0 | 0 | 1 | 0 | 9 | 0 | 0 | 2 | 0 | 98 | 49 | NA | NA | NA | NA | 0.882 | NA | 0.692 | NA | 0.19 |
| SE | 2457 | MPP1 | chrX | - | 154784027 | 154784108 | 154783426 | 154783507 | 154785050 | 154785157 | 0 | 0 | 10 | 0 | 0 | 0 | 0 | 2 | 0 | 0 | 1 | 0 | 98 | 49 | NA | NA | NA | NA | 1 | NA | 0.5 | NA | 0.5 |
| SE | 2473 | GOLGB1 | chr3 | - | 121699811 | 121699885 | 121693740 | 121698929 | 121702480 | 121702595 | 0 | 0 | 2 | 0 | 0 | 1 | 0 | 7 | 0 | 0 | 0 | 0 | 98 | 49 | NA | NA | NA | NA | 0.5 | NA | 1 | NA | -0.5 |
| SE | 2477 | GSK3B | chr3 | - | 119905754 | 119905852 | 119876412 | 119876508 | 119916043 | 119916174 | 0 | 0 | 2 | 0 | 0 | 0 | 0 | 1 | 0 | 0 | 1 | 0 | 98 | 49 | NA | NA | NA | NA | 1 | NA | 0.333 | NA | 0.667 |
| SE | 2501 | IFT57 | chr3 | - | 108219409 | 108219572 | 108218534 | 108218653 | 108222110 | 108222570 | 0 | 0 | 2 | 0 | 0 | 0 | 0 | 1 | 0 | 0 | 1 | 0 | 98 | 49 | NA | NA | NA | NA | 1 | NA | 0.333 | NA | 0.667 |
| SE | 2503 | CD47 | chr3 | - | 108050577 | 108050602 | 108043093 | 108047292 | 108051938 | 108051970 | 0 | 0 | 1 | 0 | 0 | 1 | 0 | 9 | 0 | 0 | 0 | 0 | 74 | 49 | NA | NA | NA | NA | 0.398 | NA | 1 | NA | -0.602 |
| SE | 2506 | TBC1D23 | chr3 | + | 100295301 | 100295348 | 100295086 | 100295211 | 100296171 | 100296275 | 0 | 0 | 2 | 0 | 0 | 0 | 0 | 0 | 0 | 0 | 1 | 0 | 96 | 49 | NA | NA | NA | NA | 1 | NA | 0 | NA | 1 |
| SE | 2507 | TBC1D23 | chr3 | + | 100310402 | 100310542 | 100306436 | 100306543 | 100316098 | 100316187 | 0 | 0 | 0 | 0 | 0 | 1 | 0 | 3 | 0 | 0 | 0 | 0 | 98 | 49 | NA | NA | NA | NA | 0 | NA | 1 | NA | -1 |
| SE | 2530 | GBE1 | chr3 | - | 81646391 | 81646482 | 81642780 | 81642990 | 81648855 | 81648991 | 0 | 0 | 0 | 0 | 0 | 1 | 0 | 1 | 0 | 0 | 0 | 0 | 98 | 49 | NA | NA | NA | NA | 0 | NA | 1 | NA | -1 |
| SE | 2540 | GRIPAP1 | chrX | - | 48990925 | 48991110 | 48990684 | 48990732 | 48993427 | 48993578 | 0 | 0 | 6 | 0 | 0 | 0 | 0 | 4 | 0 | 0 | 2 | 0 | 98 | 49 | NA | NA | NA | NA | 1 | NA | 0.5 | NA | 0.5 |
| SE | 2552 | ATXN7 | chr3 | + | 63990175 | 63990374 | 63988058 | 63988324 | 63990737 | 63990859 | 0 | 0 | 2 | 0 | 0 | 0 | 0 | 1 | 0 | 0 | 1 | 0 | 98 | 49 | NA | NA | NA | NA | 1 | NA | 0.333 | NA | 0.667 |
| SE | 2553 | ATXN7 | chr3 | + | 63997620 | 63997687 | 63995504 | 63996483 | 63999449 | 64003460 | 0 | 0 | 2 | 0 | 0 | 2 | 0 | 1 | 0 | 0 | 1 | 0 | 98 | 49 | NA | NA | NA | NA | 0.333 | NA | 0.333 | NA | 0 |
| SE | 2569 | FLNB | chr3 | + | 58154790 | 58154928 | 58153374 | 58153641 | 58155959 | 58156075 | 0 | 0 | 1 | 0 | 0 | 0 | 0 | 0 | 0 | 0 | 1 | 0 | 98 | 49 | NA | NA | NA | NA | 1 | NA | 0 | NA | 1 |
| SE | 2570 | SLMAP | chr3 | + | 57860698 | 57860839 | 57858087 | 57858159 | 57861948 | 57862086 | 0 | 0 | 5 | 0 | 0 | 0 | 0 | 6 | 0 | 0 | 1 | 0 | 98 | 49 | NA | NA | NA | NA | 1 | NA | 0.75 | NA | 0.25 |
| SE | 2573 | SLMAP | chr3 | + | 57907883 | 57908006 | 57896872 | 57896932 | 57909075 | 57909150 | 0 | 0 | 1 | 0 | 0 | 0 | 0 | 5 | 0 | 0 | 1 | 0 | 98 | 49 | NA | NA | NA | NA | 1 | NA | 0.714 | NA | 0.286 |
| SE | 2576 | FAM208A | chr3 | - | 56663540 | 56663572 | 56662384 | 56662490 | 56668396 | 56668558 | 0 | 0 | 0 | 0 | 0 | 1 | 0 | 1 | 0 | 0 | 0 | 0 | 81 | 49 | NA | NA | NA | NA | 0 | NA | 1 | NA | -1 |
| SE | 2577 | FAM208A | chr3 | - | 56666259 | 56666384 | 56663540 | 56663572 | 56668396 | 56668558 | 0 | 0 | 1 | 0 | 0 | 0 | 0 | 3 | 0 | 0 | 1 | 0 | 98 | 49 | NA | NA | NA | NA | 1 | NA | 0.6 | NA | 0.4 |
| SE | 2591 | WDR82 | chr3 | - | 52270711 | 52270809 | 52266951 | 52267018 | 52278200 | 52278643 | 0 | 0 | 5 | 0 | 0 | 1 | 4 | 3 | 0 | 0 | 0 | 0 | 98 | 49 | NA | NA | NA | NA | 0.714 | 1 | 1 | NA | -0.286 |
| SE | 2594 | ABHD14B | chr3 | - | 51971459 | 51971698 | 51969942 | 51970184 | 51973964 | 51974630 | 0 | 0 | 0 | 0 | 0 | 1 | 0 | 1 | 0 | 0 | 0 | 0 | 98 | 49 | NA | NA | NA | NA | 0 | NA | 1 | NA | -1 |
| SE | 2595 | ABHD14B | chr3 | - | 51971459 | 51971892 | 51969942 | 51970184 | 51973964 | 51974630 | 0 | 0 | 0 | 0 | 0 | 1 | 0 | 1 | 0 | 0 | 0 | 0 | 98 | 49 | NA | NA | NA | NA | 0 | NA | 1 | NA | -1 |
| SE | 2598 | VPRBP | chr3 | - | 51441384 | 51441897 | 51440969 | 51441071 | 51443765 | 51443903 | 0 | 0 | 1 | 0 | 0 | 0 | 0 | 0 | 0 | 0 | 2 | 0 | 98 | 49 | NA | NA | NA | NA | 1 | NA | 0 | NA | 1 |
| SE | 2606 | RBM5 | chr3 | + | 50100531 | 50100605 | 50099981 | 50100051 | 50103082 | 50103166 | 0 | 0 | 0 | 0 | 0 | 2 | 0 | 2 | 0 | 0 | 4 | 0 | 98 | 49 | NA | NA | NA | NA | 0 | NA | 0.2 | NA | -0.2 |
| SE | 2608 | RBM5 | chr3 | + | 50115905 | 50115980 | 50113382 | 50113544 | 50117073 | 50117171 | 0 | 0 | 5 | 0 | 0 | 0 | 4 | 9 | 0 | 0 | 1 | 0 | 98 | 49 | NA | NA | NA | NA | 1 | 1 | 0.818 | NA | 0.091 |
| SE | 2610 | KIAA0922 | chr4 | + | 153587741 | 153587811 | 153586208 | 153586379 | 153588889 | 153589007 | 0 | 0 | 3 | 0 | 0 | 2 | 0 | 1 | 0 | 0 | 0 | 0 | 98 | 49 | NA | NA | NA | NA | 0.429 | NA | 1 | NA | -0.571 |
| SE | 2615 | APEH | chr3 | + | 49676776 | 49676817 | 49676608 | 49676700 | 49676902 | 49677024 | 0 | 0 | 4 | 0 | 0 | 1 | 0 | 1 | 0 | 0 | 0 | 0 | 90 | 49 | NA | NA | NA | NA | 0.685 | NA | 1 | NA | -0.315 |
| SE | 2661 | SACM1L | chr3 | + | 45735234 | 45735373 | 45732052 | 45732151 | 45737582 | 45737652 | 0 | 0 | 2 | 0 | 0 | 0 | 0 | 2 | 0 | 0 | 1 | 0 | 98 | 49 | NA | NA | NA | NA | 1 | NA | 0.5 | NA | 0.5 |
| SE | 2662 | ABHD5 | chr3 | + | 43717670 | 43717857 | 43714946 | 43715058 | 43718442 | 43722725 | 0 | 0 | 17 | 0 | 0 | 3 | 6 | 3 | 0 | 0 | 1 | 0 | 98 | 49 | NA | NA | NA | NA | 0.739 | 1 | 0.6 | NA | -0.061 |
| SE | 2707 | TGFBR2 | chr3 | + | 30623198 | 30623273 | 30606501 | 30606977 | 30644746 | 30644915 | 0 | 0 | 3 | 0 | 0 | 2 | 0 | 5 | 0 | 0 | 0 | 0 | 98 | 49 | NA | NA | NA | NA | 0.429 | NA | 1 | NA | -0.571 |
| SE | 2713 | TOP2B | chr3 | - | 25604759 | 25604870 | 25601099 | 25601225 | 25606042 | 25606122 | 0 | 0 | 10 | 0 | 0 | 1 | 2 | 0 | 0 | 0 | 0 | 0 | 98 | 49 | NA | NA | NA | NA | 0.833 | 1 | NA | NA | -0.167 |
| SE | 2723 | KAT2B | chr3 | + | 20127422 | 20127549 | 20125904 | 20126113 | 20136941 | 20137052 | 0 | 0 | 12 | 0 | 0 | 0 | 0 | 6 | 0 | 0 | 1 | 0 | 98 | 49 | NA | NA | NA | NA | 1 | NA | 0.75 | NA | 0.25 |
| SE | 2724 | SERPINB9 | chr6 | - | 2893410 | 2893553 | 2891832 | 2891988 | 2895390 | 2895508 | 0 | 0 | 3 | 0 | 0 | 1 | 0 | 5 | 0 | 0 | 1 | 0 | 98 | 49 | NA | NA | NA | NA | 0.6 | NA | 0.714 | NA | -0.114 |
| SE | 2762 | LST1 | chr6 | + | 31587640 | 31587733 | 31587199 | 31587318 | 31587943 | 31587966 | 0 | 0 | 1 | 0 | 0 | 2 | 0 | 1 | 0 | 0 | 0 | 0 | 98 | 49 | NA | NA | NA | NA | 0.2 | NA | 1 | NA | -0.8 |
| SE | 2763 | LST1 | chr6 | + | 31587640 | 31587733 | 31587199 | 31587318 | 31588496 | 31588909 | 0 | 0 | 0 | 0 | 0 | 3 | 0 | 1 | 0 | 0 | 0 | 0 | 98 | 49 | NA | NA | NA | NA | 0 | NA | 1 | NA | -1 |
| SE | 2764 | LST1 | chr6 | + | 31587640 | 31587733 | 31587199 | 31587318 | 31588517 | 31588909 | 0 | 0 | 0 | 0 | 0 | 4 | 0 | 2 | 0 | 0 | 4 | 0 | 98 | 49 | NA | NA | NA | NA | 0 | NA | 0.2 | NA | -0.2 |
| SE | 2773 | XPC | chr3 | - | 14168256 | 14168380 | 14167168 | 14167253 | 14170437 | 14170550 | 0 | 0 | 1 | 0 | 0 | 0 | 0 | 6 | 0 | 0 | 3 | 0 | 98 | 49 | NA | NA | NA | NA | 1 | NA | 0.5 | NA | 0.5 |
| SE | 2776 | POLB | chr8 | + | 42339011 | 42339069 | 42338454 | 42338685 | 42344952 | 42345019 | 0 | 0 | 3 | 0 | 0 | 2 | 0 | 1 | 0 | 0 | 0 | 0 | 98 | 49 | NA | NA | NA | NA | 0.429 | NA | 1 | NA | -0.571 |
| SE | 2777 | POLB | chr8 | + | 42369848 | 42369988 | 42369270 | 42369335 | 42371562 | 42371813 | 0 | 0 | 4 | 0 | 0 | 0 | 0 | 0 | 0 | 0 | 1 | 0 | 98 | 49 | NA | NA | NA | NA | 1 | NA | 0 | NA | 1 |
| SE | 2784 | ATG7 | chr3 | + | 11342134 | 11342279 | 11340644 | 11340735 | 11347876 | 11348035 | 0 | 0 | 16 | 0 | 0 | 1 | 0 | 1 | 0 | 0 | 0 | 0 | 98 | 49 | NA | NA | NA | NA | 0.889 | NA | 1 | NA | -0.111 |
| SE | 2837 | ATRX | chrX | - | 77600433 | 77600564 | 77599731 | 77599820 | 77616612 | 77616730 | 0 | 0 | 2 | 0 | 0 | 2 | 0 | 1 | 0 | 0 | 0 | 0 | 98 | 49 | NA | NA | NA | NA | 0.333 | NA | 1 | NA | -0.667 |
| SE | 2855 | TBC1D22A | chr22 | + | 47037070 | 47037198 | 46997633 | 46997709 | 47111507 | 47111603 | 0 | 0 | 5 | 0 | 0 | 0 | 5 | 4 | 0 | 0 | 2 | 0 | 98 | 49 | NA | NA | NA | NA | 1 | 1 | 0.5 | NA | 0.25 |
| SE | 2888 | TNRC6B | chr22 | + | 40270121 | 40270280 | 40264687 | 40267036 | 40273424 | 40273600 | 0 | 0 | 1 | 0 | 0 | 0 | 0 | 3 | 0 | 0 | 1 | 0 | 98 | 49 | NA | NA | NA | NA | 1 | NA | 0.6 | NA | 0.4 |
| SE | 2901 | TMEM184B | chr22 | - | 38246784 | 38246885 | 38245934 | 38246100 | 38247769 | 38248019 | 0 | 0 | 0 | 0 | 0 | 6 | 3 | 1 | 0 | 0 | 2 | 0 | 98 | 49 | NA | NA | NA | NA | 0 | 1 | 0.2 | NA | -0.6 |
| SE | 2908 | RAF1 | chr3 | - | 12606200 | 12606299 | 12604135 | 12604289 | 12608765 | 12608923 | 0 | 0 | 20 | 0 | 0 | 5 | 0 | 22 | 0 | 0 | 2 | 0 | 98 | 49 | NA | NA | NA | NA | 0.667 | NA | 0.846 | NA | -0.179 |
| SE | 2959 | MTMR3 | chr22 | + | 29970975 | 29971062 | 29957035 | 29957088 | 29978441 | 29978531 | 0 | 0 | 9 | 0 | 0 | 1 | 0 | 3 | 0 | 0 | 0 | 0 | 98 | 49 | NA | NA | NA | NA | 0.818 | NA | 1 | NA | -0.182 |
| SE | 2960 | MTMR3 | chr22 | + | 30012367 | 30012563 | 30009017 | 30009129 | 30013355 | 30013541 | 0 | 0 | 2 | 0 | 0 | 4 | 2 | 8 | 0 | 0 | 0 | 0 | 98 | 49 | NA | NA | NA | NA | 0.2 | 1 | 1 | NA | -0.8 |
| SE | 2962 | MTMR3 | chr22 | + | 30023456 | 30023483 | 30022608 | 30022697 | 30025629 | 30030868 | 0 | 0 | 5 | 0 | 0 | 2 | 0 | 1 | 0 | 3 | 0 | 0 | 76 | 49 | NA | NA | NA | NA | 0.617 | 0 | 1 | NA | 0.117 |
| SE | 2967 | THOC5 | chr22 | - | 29550727 | 29550843 | 29549051 | 29549158 | 29553670 | 29553747 | 0 | 0 | 1 | 0 | 0 | 0 | 0 | 0 | 0 | 0 | 1 | 0 | 98 | 49 | NA | NA | NA | NA | 1 | NA | 0 | NA | 1 |
| SE | 2971 | EWSR1 | chr22 | + | 29282389 | 29282557 | 29278029 | 29278216 | 29286922 | 29287134 | 0 | 0 | 11 | 0 | 0 | 1 | 0 | 6 | 0 | 0 | 0 | 0 | 98 | 49 | NA | NA | NA | NA | 0.846 | NA | 1 | NA | -0.154 |
| SE | 2988 | PPP3CA | chr4 | - | 101063231 | 101063357 | 101061086 | 101061161 | 101080531 | 101080626 | 0 | 0 | 15 | 0 | 0 | 3 | 3 | 0 | 0 | 0 | 0 | 0 | 98 | 49 | NA | NA | NA | NA | 0.714 | 1 | NA | NA | -0.286 |
| SE | 2993 | GUSBP11 | chr22 | - | 23700725 | 23700845 | 23694118 | 23695517 | 23705428 | 23705541 | 0 | 0 | 6 | 0 | 0 | 0 | 0 | 0 | 0 | 0 | 1 | 0 | 98 | 49 | NA | NA | NA | NA | 1 | NA | 0 | NA | 1 |
| SE | 2996 | BCR | chr22 | + | 23260949 | 23261054 | 23253798 | 23253980 | 23261354 | 23261540 | 2 | 0 | 3 | 0 | 0 | 0 | 0 | 3 | 0 | 0 | 1 | 0 | 98 | 49 | NA | NA | 1 | NA | 1 | NA | 0.6 | NA | 0.4 |
| SE | 2997 | BCR | chr22 | + | 23290338 | 23290413 | 23289516 | 23289621 | 23292540 | 23292638 | 0 | 0 | 3 | 0 | 0 | 0 | 0 | 1 | 0 | 0 | 1 | 0 | 98 | 49 | NA | NA | NA | NA | 1 | NA | 0.333 | NA | 0.667 |
| SE | 3026 | TANGO2 | chr22 | + | 20061529 | 20061681 | 20055942 | 20056013 | 20063337 | 20063442 | 0 | 0 | 0 | 0 | 0 | 1 | 0 | 1 | 0 | 0 | 0 | 0 | 98 | 49 | NA | NA | NA | NA | 0 | NA | 1 | NA | -1 |
| SE | 3028 | TANGO2 | chr22 | + | 20061529 | 20061683 | 20055942 | 20056013 | 20063337 | 20063442 | 0 | 0 | 2 | 0 | 0 | 1 | 0 | 1 | 0 | 0 | 0 | 0 | 98 | 49 | NA | NA | NA | NA | 0.5 | NA | 1 | NA | -0.5 |
| SE | 3050 | PRMT2 | chr21 | + | 46658744 | 46658920 | 46649574 | 46649739 | 46664294 | 46665124 | 0 | 0 | 2 | 0 | 0 | 1 | 0 | 2 | 0 | 0 | 0 | 0 | 98 | 49 | NA | NA | NA | NA | 0.5 | NA | 1 | NA | -0.5 |
| SE | 3055 | NQO2 | chr6 | + | 3015529 | 3015643 | 3012543 | 3012674 | 3016883 | 3016985 | 0 | 0 | 4 | 0 | 0 | 1 | 0 | 1 | 0 | 0 | 0 | 0 | 98 | 49 | NA | NA | NA | NA | 0.667 | NA | 1 | NA | -0.333 |
| SE | 3121 | TTC3 | chr21 | + | 37090232 | 37090286 | 37088798 | 37088886 | 37091292 | 37091413 | 0 | 0 | 1 | 0 | 0 | 0 | 0 | 2 | 0 | 0 | 1 | 0 | 98 | 49 | NA | NA | NA | NA | 1 | NA | 0.5 | NA | 0.5 |
| SE | 3135 | IFNGR2 | chr21 | + | 33432176 | 33432336 | 33426883 | 33427032 | 33432713 | 33432871 | 0 | 0 | 16 | 0 | 0 | 1 | 4 | 7 | 0 | 0 | 0 | 0 | 98 | 49 | NA | NA | NA | NA | 0.889 | 1 | 1 | NA | -0.111 |
| SE | 3142 | SYNJ1 | chr21 | - | 32699837 | 32700105 | 32695056 | 32695282 | 32701960 | 32702047 | 0 | 0 | 1 | 0 | 0 | 1 | 0 | 2 | 0 | 0 | 1 | 0 | 98 | 49 | NA | NA | NA | NA | 0.333 | NA | 0.5 | NA | -0.167 |
| SE | 3153 | USP16 | chr21 | + | 29037275 | 29037463 | 29036270 | 29036374 | 29038334 | 29038430 | 0 | 0 | 7 | 0 | 0 | 0 | 0 | 1 | 0 | 0 | 1 | 0 | 98 | 49 | NA | NA | NA | NA | 1 | NA | 0.333 | NA | 0.667 |
| SE | 3154 | USP16 | chr21 | + | 29037278 | 29037463 | 29036270 | 29036374 | 29038334 | 29038430 | 0 | 0 | 4 | 0 | 0 | 0 | 0 | 1 | 0 | 0 | 1 | 0 | 98 | 49 | NA | NA | NA | NA | 1 | NA | 0.333 | NA | 0.667 |
| SE | 3158 | APP | chr21 | - | 25905023 | 25905077 | 25897572 | 25897673 | 25911740 | 25911962 | 0 | 0 | 6 | 0 | 0 | 1 | 0 | 1 | 0 | 0 | 0 | 0 | 98 | 49 | NA | NA | NA | NA | 0.75 | NA | 1 | NA | -0.25 |
| SE | 3159 | APP | chr21 | - | 25997359 | 25997416 | 25982343 | 25982477 | 26000014 | 26000182 | 0 | 0 | 1 | 0 | 0 | 3 | 0 | 0 | 0 | 2 | 1 | 0 | 98 | 49 | NA | NA | NA | NA | 0.143 | 0 | 0 | NA | 0.143 |
| SE | 3160 | APP | chr21 | - | 26000014 | 26000182 | 25982343 | 25982477 | 26021839 | 26022042 | 0 | 0 | 6 | 0 | 0 | 0 | 2 | 2 | 0 | 0 | 1 | 0 | 98 | 49 | NA | NA | NA | NA | 1 | 1 | 0.5 | NA | 0.25 |
| SE | 3171 | NRIP1 | chr21 | - | 15043494 | 15043574 | 15014343 | 15014466 | 15064744 | 15064805 | 2 | 0 | 0 | 0 | 0 | 0 | 0 | 2 | 0 | 0 | 0 | 1 | 98 | 49 | NA | NA | 1 | NA | NA | NA | 1 | 0 | 0.5 |
| SE | 3188 | TPD52L2 | chr20 | + | 63875815 | 63875875 | 63873667 | 63873816 | 63882718 | 63882820 | 0 | 0 | 17 | 0 | 0 | 0 | 0 | 9 | 0 | 0 | 1 | 0 | 98 | 49 | NA | NA | NA | NA | 1 | NA | 0.818 | NA | 0.182 |
| SE | 3191 | TPD52L2 | chr20 | + | 63887563 | 63887605 | 63882718 | 63882820 | 63889189 | 63889238 | 0 | 0 | 1 | 0 | 0 | 8 | 0 | 0 | 0 | 0 | 5 | 0 | 91 | 49 | NA | NA | NA | NA | 0.063 | NA | 0 | NA | 0.063 |
| SE | 3193 | CXorf65 | chrX | - | 71104768 | 71104837 | 71104297 | 71104404 | 71105999 | 71106137 | 0 | 0 | 3 | 0 | 0 | 0 | 0 | 1 | 0 | 0 | 2 | 0 | 98 | 49 | NA | NA | NA | NA | 1 | NA | 0.2 | NA | 0.8 |
| SE | 3197 | TNIP1 | chr5 | - | 151042539 | 151042671 | 151039096 | 151039225 | 151042895 | 151042961 | 0 | 0 | 7 | 0 | 0 | 0 | 0 | 12 | 0 | 0 | 1 | 0 | 98 | 49 | NA | NA | NA | NA | 1 | NA | 0.857 | NA | 0.143 |
| SE | 3198 | TNIP1 | chr5 | - | 151045860 | 151045950 | 151042895 | 151042961 | 151049823 | 151049947 | 0 | 0 | 6 | 0 | 0 | 0 | 0 | 12 | 0 | 0 | 1 | 0 | 98 | 49 | NA | NA | NA | NA | 1 | NA | 0.857 | NA | 0.143 |
| SE | 3209 | STX16-NPEPL1 | chr20 | + | 58673630 | 58673711 | 58671153 | 58671297 | 58691723 | 58691780 | 0 | 0 | 10 | 0 | 0 | 2 | 0 | 5 | 0 | 0 | 0 | 0 | 98 | 49 | NA | NA | NA | NA | 0.714 | NA | 1 | NA | -0.286 |
| SE | 3211 | STX16 | chr20 | + | 58659622 | 58659634 | 58651252 | 58652087 | 58667489 | 58667597 | 0 | 0 | 1 | 0 | 0 | 3 | 0 | 0 | 0 | 0 | 1 | 0 | 61 | 49 | NA | NA | NA | NA | 0.211 | NA | 0 | NA | 0.211 |
| SE | 3216 | PDCD6IP | chr3 | + | 33828837 | 33828969 | 33826479 | 33826580 | 33836043 | 33836266 | 0 | 0 | 1 | 0 | 0 | 1 | 0 | 5 | 0 | 0 | 0 | 0 | 98 | 49 | NA | NA | NA | NA | 0.333 | NA | 1 | NA | -0.667 |
| SE | 3217 | PDCD6IP | chr3 | + | 33828852 | 33828969 | 33826479 | 33826580 | 33836043 | 33836266 | 0 | 0 | 1 | 0 | 0 | 1 | 0 | 5 | 0 | 0 | 0 | 0 | 98 | 49 | NA | NA | NA | NA | 0.333 | NA | 1 | NA | -0.667 |
| SE | 3249 | ZMYND8 | chr20 | - | 47212641 | 47212725 | 47209213 | 47210897 | 47220257 | 47220324 | 0 | 0 | 0 | 0 | 0 | 1 | 0 | 2 | 0 | 0 | 0 | 0 | 98 | 49 | NA | NA | NA | NA | 0 | NA | 1 | NA | -1 |
| SE | 3254 | ZMYND8 | chr20 | - | 47276313 | 47276795 | 47262287 | 47262428 | 47282101 | 47282217 | 0 | 0 | 1 | 0 | 0 | 0 | 0 | 0 | 0 | 0 | 1 | 0 | 98 | 49 | NA | NA | NA | NA | 1 | NA | 0 | NA | 1 |
| SE | 3256 | ZMYND8 | chr20 | - | 47291795 | 47291888 | 47290186 | 47290274 | 47294665 | 47294779 | 0 | 0 | 9 | 0 | 0 | 0 | 0 | 0 | 0 | 2 | 0 | 0 | 98 | 49 | NA | NA | NA | NA | 1 | 0 | NA | NA | 1 |
| SE | 3326 | CTNNBL1 | chr20 | + | 37842338 | 37842419 | 37840101 | 37840199 | 37859898 | 37860036 | 0 | 0 | 2 | 0 | 0 | 1 | 0 | 4 | 0 | 0 | 0 | 0 | 98 | 49 | NA | NA | NA | NA | 0.5 | NA | 1 | NA | -0.5 |
| SE | 3331 | SAMHD1 | chr20 | - | 36898439 | 36898544 | 36897821 | 36897959 | 36904156 | 36904249 | 0 | 0 | 25 | 0 | 0 | 1 | 0 | 5 | 0 | 0 | 0 | 0 | 98 | 49 | NA | NA | NA | NA | 0.926 | NA | 1 | NA | -0.074 |
| SE | 3333 | PHF8 | chrX | - | 53993600 | 53993903 | 53992735 | 53992839 | 53995692 | 53995782 | 0 | 0 | 0 | 0 | 0 | 1 | 0 | 2 | 0 | 0 | 0 | 0 | 98 | 49 | NA | NA | NA | NA | 0 | NA | 1 | NA | -1 |
| SE | 3337 | C20orf24 | chr20 | + | 36607889 | 36608000 | 36607714 | 36607818 | 36609600 | 36609730 | 0 | 0 | 3 | 0 | 0 | 4 | 0 | 3 | 0 | 0 | 2 | 0 | 98 | 49 | NA | NA | NA | NA | 0.273 | NA | 0.429 | NA | -0.156 |
| SE | 3356 | EDEM2 | chr20 | - | 35118597 | 35118719 | 35115356 | 35115933 | 35123889 | 35124034 | 0 | 0 | 6 | 0 | 0 | 1 | 0 | 0 | 0 | 0 | 2 | 0 | 98 | 49 | NA | NA | NA | NA | 0.75 | NA | 0 | NA | 0.75 |
| SE | 3358 | TRPC4AP | chr20 | - | 35021189 | 35021332 | 35016007 | 35016139 | 35035122 | 35035308 | 0 | 0 | 4 | 0 | 0 | 0 | 0 | 3 | 0 | 0 | 1 | 0 | 98 | 49 | NA | NA | NA | NA | 1 | NA | 0.6 | NA | 0.4 |
| SE | 3359 | TRPC4AP | chr20 | - | 35021189 | 35021356 | 35016007 | 35016139 | 35035122 | 35035308 | 0 | 0 | 4 | 0 | 0 | 0 | 0 | 2 | 0 | 0 | 1 | 0 | 98 | 49 | NA | NA | NA | NA | 1 | NA | 0.5 | NA | 0.5 |
| SE | 3360 | TRPC4AP | chr20 | - | 35044504 | 35044712 | 35035122 | 35035308 | 35049865 | 35049994 | 0 | 0 | 19 | 0 | 0 | 0 | 13 | 8 | 0 | 0 | 1 | 0 | 98 | 49 | NA | NA | NA | NA | 1 | 1 | 0.8 | NA | 0.1 |
| SE | 3362 | ACSS2 | chr20 | + | 34926104 | 34926281 | 34925697 | 34925766 | 34926876 | 34926951 | 0 | 0 | 4 | 0 | 0 | 0 | 0 | 0 | 0 | 0 | 1 | 0 | 98 | 49 | NA | NA | NA | NA | 1 | NA | 0 | NA | 1 |
| SE | 3372 | FCHSD1 | chr5 | - | 141646602 | 141646722 | 141646086 | 141646191 | 141647134 | 141647230 | 0 | 0 | 1 | 0 | 0 | 1 | 0 | 2 | 0 | 0 | 0 | 0 | 98 | 49 | NA | NA | NA | NA | 0.333 | NA | 1 | NA | -0.667 |
| SE | 3398 | ELOVL5 | chr6 | - | 53270592 | 53270727 | 53267397 | 53269270 | 53273219 | 53273344 | 0 | 0 | 15 | 0 | 0 | 2 | 0 | 9 | 0 | 0 | 0 | 0 | 98 | 49 | NA | NA | NA | NA | 0.789 | NA | 1 | NA | -0.211 |
| SE | 3411 | XRN2 | chr20 | + | 21356585 | 21356665 | 21356079 | 21356177 | 21357735 | 21357792 | 0 | 0 | 12 | 0 | 0 | 2 | 0 | 2 | 0 | 0 | 0 | 0 | 98 | 49 | NA | NA | NA | NA | 0.75 | NA | 1 | NA | -0.25 |
| SE | 3412 | XRN2 | chr20 | + | 21357735 | 21357792 | 21356585 | 21356665 | 21365420 | 21365489 | 0 | 0 | 9 | 0 | 0 | 1 | 0 | 2 | 0 | 0 | 0 | 0 | 98 | 49 | NA | NA | NA | NA | 0.818 | NA | 1 | NA | -0.182 |
| SE | 3414 | RALGAPA2 | chr20 | - | 20605174 | 20605412 | 20601681 | 20601846 | 20611314 | 20611426 | 0 | 0 | 5 | 0 | 0 | 0 | 0 | 0 | 0 | 0 | 2 | 0 | 98 | 49 | NA | NA | NA | NA | 1 | NA | 0 | NA | 1 |
| SE | 3415 | RALGAPA2 | chr20 | - | 20611314 | 20611426 | 20605174 | 20605412 | 20616042 | 20616191 | 0 | 0 | 6 | 0 | 0 | 1 | 0 | 1 | 0 | 0 | 0 | 0 | 98 | 49 | NA | NA | NA | NA | 0.75 | NA | 1 | NA | -0.25 |
| SE | 3416 | RALGAPA2 | chr20 | - | 20637362 | 20637501 | 20635417 | 20635617 | 20639784 | 20639900 | 0 | 0 | 13 | 0 | 0 | 2 | 0 | 0 | 0 | 0 | 2 | 0 | 98 | 49 | NA | NA | NA | NA | 0.765 | NA | 0 | NA | 0.765 |
| SE | 3419 | SEC23B | chr20 | + | 18525787 | 18525932 | 18524934 | 18525020 | 18526372 | 18526531 | 0 | 0 | 1 | 0 | 0 | 0 | 0 | 1 | 0 | 0 | 1 | 0 | 98 | 49 | NA | NA | NA | NA | 1 | NA | 0.333 | NA | 0.667 |
| SE | 3453 | GPCPD1 | chr20 | - | 5570146 | 5570239 | 5567482 | 5567560 | 5573914 | 5573969 | 0 | 0 | 14 | 0 | 0 | 1 | 0 | 13 | 0 | 0 | 0 | 0 | 98 | 49 | NA | NA | NA | NA | 0.875 | NA | 1 | NA | -0.125 |
| SE | 3454 | GPCPD1 | chr20 | - | 5580007 | 5580131 | 5578379 | 5578611 | 5584280 | 5584322 | 0 | 0 | 17 | 0 | 0 | 1 | 0 | 17 | 0 | 0 | 0 | 0 | 98 | 49 | NA | NA | NA | NA | 0.895 | NA | 1 | NA | -0.105 |
| SE | 3458 | SLC23A2 | chr20 | - | 4885820 | 4885909 | 4884752 | 4884823 | 4899554 | 4899712 | 0 | 0 | 1 | 0 | 0 | 0 | 0 | 4 | 0 | 0 | 1 | 0 | 98 | 49 | NA | NA | NA | NA | 1 | NA | 0.667 | NA | 0.333 |
| SE | 3467 | SIGLEC1 | chr20 | - | 3699201 | 3699459 | 3697797 | 3698133 | 3701341 | 3701641 | 0 | 0 | 8 | 0 | 0 | 1 | 0 | 1 | 0 | 0 | 0 | 0 | 98 | 49 | NA | NA | NA | NA | 0.8 | NA | 1 | NA | -0.2 |
| SE | 3473 | FAM193A | chr4 | + | 2655057 | 2655129 | 2646684 | 2646832 | 2657802 | 2657880 | 0 | 0 | 1 | 0 | 0 | 2 | 0 | 1 | 0 | 0 | 2 | 0 | 98 | 49 | NA | NA | NA | NA | 0.2 | NA | 0.2 | NA | 0 |
| SE | 3479 | STK35 | chr20 | + | 2116665 | 2117415 | 2102767 | 2103365 | 2143783 | 2148552 | 0 | 0 | 1 | 0 | 0 | 0 | 0 | 0 | 0 | 0 | 1 | 0 | 98 | 49 | NA | NA | NA | NA | 1 | NA | 0 | NA | 1 |
| SE | 3509 | HDLBP | chr2 | - | 241238648 | 241238787 | 241236614 | 241236769 | 241239601 | 241239820 | 0 | 0 | 5 | 0 | 0 | 0 | 0 | 4 | 0 | 0 | 1 | 0 | 98 | 49 | NA | NA | NA | NA | 1 | NA | 0.667 | NA | 0.333 |
| SE | 3521 | RNPEPL1 | chr2 | + | 240574112 | 240574348 | 240573774 | 240573891 | 240574514 | 240574628 | 0 | 0 | 6 | 0 | 0 | 1 | 0 | 7 | 0 | 0 | 0 | 0 | 98 | 49 | NA | NA | NA | NA | 0.75 | NA | 1 | NA | -0.25 |
| SE | 3527 | PMPCB | chr7 | + | 103307595 | 103307708 | 103304410 | 103304490 | 103308951 | 103309095 | 0 | 0 | 2 | 0 | 0 | 0 | 0 | 3 | 0 | 0 | 1 | 0 | 98 | 49 | NA | NA | NA | NA | 1 | NA | 0.6 | NA | 0.4 |
| SE | 3531 | LRRFIP1 | chr2 | + | 237708543 | 237708630 | 237692163 | 237692521 | 237748363 | 237748399 | 0 | 0 | 2 | 0 | 0 | 2 | 3 | 2 | 0 | 4 | 1 | 0 | 98 | 49 | NA | NA | NA | NA | 0.333 | 0.273 | 0.5 | NA | -0.053 |
| SE | 3535 | LRRFIP1 | chr2 | + | 237772080 | 237772198 | 237769942 | 237769992 | 237772865 | 237772945 | 0 | 0 | 7 | 0 | 0 | 0 | 0 | 2 | 0 | 5 | 0 | 0 | 98 | 49 | NA | NA | NA | NA | 1 | 0 | 1 | NA | 0.5 |
| SE | 3546 | OSBPL2 | chr20 | + | 62260016 | 62260125 | 62256056 | 62256221 | 62263615 | 62263691 | 0 | 0 | 3 | 0 | 0 | 0 | 0 | 5 | 0 | 0 | 4 | 0 | 98 | 49 | NA | NA | NA | NA | 1 | NA | 0.385 | NA | 0.615 |

| | | | | | | | | | | | | | | | | | | | | | | | | | | | | | | | | | |
|---|---|---|---|---|---|---|---|---|---|---|---|---|---|---|---|---|---|---|---|---|---|---|---|---|---|---|---|---|---|---|---|---|---|
| SE | 3547 | OSBPL2 | chr20 | + | 62264628 | 62264718 | 62263615 | 62263691 | 62272124 | 62272259 | 0 | 0 | 2 | 0 | 0 | 3 | 0 | 5 | 0 | 0 | 5 | 0 | 98 | 49 | NA | NA | NA | NA | 0.25 | NA | 0.333 | NA | -0.083 |
| SE | 3550 | MAPKAPK3 | chr3 | + | 50644408 | 50644532 | 50642252 | 50642332 | 50645709 | 50645785 | 0 | 0 | 11 | 0 | 0 | 5 | 0 | 4 | 0 | 0 | 1 | 0 | 98 | 49 | NA | NA | NA | NA | 0.524 | NA | 0.667 | NA | -0.143 |
| SE | 3555 | TAZ | chrX | + | 154414100 | 154414190 | 154413481 | 154413567 | 154419542 | 154419623 | 0 | 0 | 0 | 0 | 0 | 1 | 0 | 1 | 0 | 0 | 2 | 0 | 98 | 49 | NA | NA | NA | NA | 0 | NA | 0.2 | NA | -0.2 |
| SE | 3558 | DGKD | chr2 | + | 233435817 | 233435924 | 233434768 | 233434901 | 233436315 | 233436441 | 0 | 0 | 12 | 0 | 0 | 1 | 0 | 3 | 0 | 0 | 0 | 0 | 98 | 49 | NA | NA | NA | NA | 0.857 | NA | 1 | NA | -0.143 |
| SE | 3585 | SP110 | chr2 | - | 230170620 | 230170761 | 230168925 | 230169237 | 230172065 | 230172174 | 0 | 0 | 1 | 0 | 0 | 2 | 0 | 1 | 0 | 0 | 1 | 0 | 98 | 49 | NA | NA | NA | NA | 0.2 | NA | 0.333 | NA | -0.133 |
| SE | 3587 | SP110 | chr2 | - | 230172843 | 230172959 | 230172065 | 230172174 | 230177537 | 230177680 | 0 | 0 | 7 | 0 | 0 | 0 | 0 | 2 | 0 | 0 | 1 | 0 | 98 | 49 | NA | NA | NA | NA | 1 | NA | 0.5 | NA | 0.5 |
| SE | 3588 | SP110 | chr2 | - | 230177537 | 230177680 | 230176472 | 230176724 | 230178156 | 230178255 | 0 | 0 | 5 | 0 | 0 | 2 | 0 | 7 | 0 | 0 | 0 | 0 | 98 | 49 | NA | NA | NA | NA | 0.556 | NA | 1 | NA | -0.444 |
| SE | 3592 | TRIP12 | chr2 | - | 229860405 | 229860531 | 229858771 | 229859574 | 229879981 | 229880128 | 0 | 0 | 3 | 0 | 0 | 2 | 0 | 0 | 0 | 0 | 1 | 0 | 98 | 49 | NA | NA | NA | NA | 0.429 | NA | 0 | NA | 0.429 |
| SE | 3608 | DOCK10 | chr2 | - | 224814319 | 224814364 | 224807910 | 224808086 | 224816616 | 224816713 | 0 | 0 | 4 | 0 | 0 | 0 | 0 | 1 | 0 | 0 | 2 | 0 | 94 | 49 | NA | NA | NA | NA | 1 | NA | 0.207 | NA | 0.793 |
| SE | 3614 | NIN | chr14 | - | 50756491 | 50758630 | 50754741 | 50754867 | 50759856 | 50760359 | 0 | 0 | 2 | 0 | 0 | 0 | 0 | 1 | 0 | 0 | 1 | 0 | 98 | 49 | NA | NA | NA | NA | 1 | NA | 0.333 | NA | 0.667 |
| SE | 3615 | NIN | chr14 | - | 50759856 | 50760359 | 50754741 | 50754867 | 50761789 | 50761911 | 0 | 0 | 2 | 0 | 0 | 2 | 0 | 2 | 0 | 0 | 0 | 0 | 98 | 49 | NA | NA | NA | NA | 0.333 | NA | 1 | NA | -0.667 |
| SE | 3616 | NIN | chr14 | - | 50766306 | 50766396 | 50763825 | 50763964 | 50766779 | 50766890 | 0 | 0 | 14 | 0 | 0 | 1 | 0 | 16 | 0 | 0 | 0 | 0 | 98 | 49 | NA | NA | NA | NA | 0.875 | NA | 1 | NA | -0.125 |
| SE | 3623 | ITGAM | chr16 | + | 31321463 | 31321627 | 31321240 | 31321371 | 31324398 | 31324553 | 0 | 0 | 12 | 0 | 0 | 1 | 0 | 0 | 0 | 0 | 1 | 0 | 98 | 49 | NA | NA | NA | NA | 0.857 | NA | 0 | NA | 0.857 |
| SE | 3626 | HECTD1 | chr14 | - | 31113876 | 31114059 | 31113276 | 31113484 | 31114285 | 31114340 | 0 | 0 | 0 | 0 | 0 | 1 | 0 | 1 | 0 | 0 | 0 | 0 | 98 | 49 | NA | NA | NA | NA | 0 | NA | 1 | NA | -1 |
| SE | 3633 | USP4 | chr3 | - | 49310619 | 49310737 | 49305714 | 49305888 | 49324701 | 49324763 | 0 | 0 | 3 | 0 | 0 | 1 | 0 | 8 | 0 | 0 | 0 | 0 | 98 | 49 | NA | NA | NA | NA | 0.6 | NA | 1 | NA | -0.4 |
| SE | 3634 | USP4 | chr3 | - | 49311513 | 49311654 | 49310619 | 49310737 | 49324701 | 49324763 | 0 | 0 | 1 | 0 | 0 | 1 | 0 | 4 | 0 | 0 | 3 | 0 | 98 | 49 | NA | NA | NA | NA | 0.333 | NA | 0.4 | NA | -0.067 |
| SE | 3674 | CALCOCO2 | chr17 | + | 48852505 | 48852628 | 48851558 | 48851628 | 48852925 | 48853012 | 0 | 0 | 3 | 0 | 0 | 0 | 0 | 12 | 0 | 0 | 1 | 0 | 98 | 49 | NA | NA | NA | NA | 1 | NA | 0.857 | NA | 0.143 |
| SE | 3682 | USP15 | chr12 | + | 62396294 | 62396398 | 62393052 | 62393202 | 62401186 | 62401275 | 0 | 0 | 13 | 0 | 0 | 1 | 2 | 11 | 0 | 0 | 0 | 0 | 98 | 49 | NA | NA | NA | NA | 0.867 | 1 | 1 | NA | -0.133 |
| SE | 3686 | NBR1 | chr17 | + | 43196941 | 43197106 | 43196480 | 43196591 | 43200166 | 43200608 | 0 | 0 | 3 | 0 | 0 | 1 | 0 | 3 | 0 | 0 | 0 | 0 | 98 | 49 | NA | NA | NA | NA | 0.6 | NA | 1 | NA | -0.4 |
| SE | 3701 | HBEGF | chr5 | - | 140345910 | 140346084 | 140342634 | 140342812 | 140346282 | 140346603 | 0 | 0 | 5 | 0 | 0 | 1 | 0 | 5 | 0 | 0 | 0 | 0 | 98 | 49 | NA | NA | NA | NA | 0.714 | NA | 1 | NA | -0.286 |
| SE | 3761 | CNOT1 | chr16 | - | 58547565 | 58547682 | 58547185 | 58547296 | 58549718 | 58549898 | 0 | 0 | 13 | 0 | 0 | 0 | 0 | 8 | 0 | 0 | 1 | 0 | 98 | 49 | NA | NA | NA | NA | 1 | NA | 0.8 | NA | 0.2 |
| SE | 3784 | SREK1 | chr5 | + | 66177513 | 66177658 | 66174945 | 66175041 | 66178718 | 66183616 | 0 | 0 | 2 | 0 | 0 | 0 | 0 | 0 | 0 | 0 | 1 | 0 | 98 | 49 | NA | NA | NA | NA | 1 | NA | 0 | NA | 1 |
| SE | 3785 | SREK1 | chr5 | + | 66177516 | 66177658 | 66174945 | 66175041 | 66178718 | 66183616 | 0 | 0 | 1 | 0 | 0 | 0 | 0 | 0 | 0 | 0 | 1 | 0 | 98 | 49 | NA | NA | NA | NA | 1 | NA | 0 | NA | 1 |
| SE | 3793 | STT3B | chr3 | + | 31624913 | 31625085 | 31623673 | 31623861 | 31625953 | 31626127 | 0 | 0 | 3 | 0 | 0 | 0 | 0 | 0 | 0 | 0 | 1 | 0 | 98 | 49 | NA | NA | NA | NA | 1 | NA | 0 | NA | 1 |
| SE | 3802 | ZCCHC2 | chr18 | + | 62556202 | 62556297 | 62550347 | 62550460 | 62558686 | 62558770 | 0 | 0 | 20 | 0 | 0 | 2 | 5 | 12 | 0 | 0 | 0 | 0 | 98 | 49 | NA | NA | NA | NA | 0.833 | 1 | 1 | NA | -0.167 |
| SE | 3804 | ZBTB11 | chr3 | - | 101671129 | 101671361 | 101664963 | 101665808 | 101671977 | 101672213 | 0 | 0 | 3 | 0 | 0 | 0 | 1 | 3 | 0 | 0 | 1 | 0 | 98 | 49 | NA | NA | NA | NA | 1 | 1 | 0.6 | NA | 0.2 |
| SE | 3808 | SNHG6 | chr8 | - | 66922323 | 66922392 | 66921929 | 66922114 | 66922613 | 66922725 | 0 | 0 | 2 | 0 | 0 | 1 | 0 | 4 | 0 | 0 | 0 | 0 | 98 | 49 | NA | NA | NA | NA | 0.5 | NA | 1 | NA | -0.5 |
| SE | 3809 | CPNE1 | chr20 | - | 35655201 | 35655344 | 35632794 | 35632923 | 35658929 | 35658982 | 0 | 0 | 0 | 0 | 0 | 2 | 0 | 1 | 0 | 0 | 0 | 0 | 98 | 49 | NA | NA | NA | NA | 0 | NA | 1 | NA | -1 |
| SE | 3883 | MIS18BP1 | chr14 | - | 45206082 | 45206170 | 45204398 | 45204453 | 45210379 | 45210528 | 0 | 0 | 5 | 0 | 0 | 2 | 5 | 3 | 0 | 0 | 0 | 0 | 98 | 49 | NA | NA | NA | NA | 0.556 | 1 | 1 | NA | -0.444 |
| SE | 3892 | SETX | chr9 | - | 132298079 | 132298312 | 132296886 | 132297054 | 132300629 | 132300803 | 0 | 0 | 16 | 0 | 0 | 1 | 0 | 5 | 0 | 0 | 0 | 0 | 98 | 49 | NA | NA | NA | NA | 0.889 | NA | 1 | NA | -0.111 |
| SE | 3919 | SMG1 | chr16 | - | 18879494 | 18879719 | 18877130 | 18877232 | 18882164 | 18882338 | 0 | 0 | 3 | 0 | 0 | 2 | 0 | 4 | 0 | 0 | 0 | 0 | 98 | 49 | NA | NA | NA | NA | 0.429 | NA | 1 | NA | -0.571 |
| SE | 3920 | SMG1 | chr16 | - | 18889371 | 18889585 | 18885540 | 18885666 | 18890862 | 18890921 | 0 | 0 | 4 | 0 | 0 | 3 | 0 | 2 | 0 | 0 | 0 | 0 | 98 | 49 | NA | NA | NA | NA | 0.4 | NA | 1 | NA | -0.6 |
| SE | 3933 | ATF2 | chr2 | - | 175121443 | 175121540 | 175118250 | 175118369 | 175130137 | 175130207 | 0 | 0 | 4 | 0 | 0 | 1 | 0 | 2 | 0 | 0 | 0 | 0 | 98 | 49 | NA | NA | NA | NA | 0.667 | NA | 1 | NA | -0.333 |
| SE | 3938 | THAP9-AS1 | chr4 | - | 82899712 | 82899737 | 82897988 | 82898062 | 82900076 | 82900223 | 0 | 0 | 3 | 0 | 0 | 0 | 0 | 2 | 0 | 0 | 2 | 0 | 74 | 49 | NA | NA | NA | NA | 1 | NA | 0.398 | NA | 0.602 |
| SE | 3941 | ZDHHC20 | chr13 | - | 21378660 | 21378738 | 21372570 | 21376655 | 21381433 | 21381549 | 0 | 0 | 3 | 0 | 0 | 2 | 0 | 1 | 0 | 0 | 0 | 0 | 98 | 49 | NA | NA | NA | NA | 0.429 | NA | 1 | NA | -0.571 |
| SE | 3942 | ZDHHC20 | chr13 | - | 21381900 | 21381945 | 21381433 | 21381549 | 21382919 | 21383009 | 0 | 0 | 3 | 0 | 0 | 5 | 0 | 4 | 0 | 0 | 1 | 0 | 94 | 49 | NA | NA | NA | NA | 0.238 | NA | 0.676 | NA | -0.438 |
| SE | 3948 | ZMYM2 | chr13 | + | 20019546 | 20019618 | 20006373 | 20006586 | 20027202 | 20027318 | 0 | 0 | 5 | 0 | 0 | 0 | 0 | 4 | 0 | 0 | 1 | 0 | 98 | 49 | NA | NA | NA | NA | 1 | NA | 0.667 | NA | 0.333 |
| SE | 3949 | ZMYM2 | chr13 | + | 20026611 | 20026762 | 20019546 | 20019618 | 20027202 | 20027318 | 0 | 0 | 8 | 0 | 0 | 0 | 0 | 2 | 0 | 0 | 1 | 0 | 98 | 49 | NA | NA | NA | NA | 1 | NA | 0.5 | NA | 0.5 |
| SE | 3971 | HARS | chr5 | - | 140675016 | 140675133 | 140674678 | 140674825 | 140676653 | 140676896 | 0 | 0 | 1 | 0 | 0 | 0 | 0 | 2 | 0 | 1 | 0 | 0 | 98 | 49 | NA | NA | NA | NA | 1 | 0 | 1 | NA | 0.5 |
| SE | 3992 | CLIP1 | chr12 | - | 122336631 | 122336748 | 122334647 | 122334705 | 122340752 | 122341697 | 0 | 0 | 6 | 0 | 0 | 0 | 0 | 3 | 0 | 0 | 1 | 0 | 98 | 49 | NA | NA | NA | NA | 1 | NA | 0.6 | NA | 0.4 |
| SE | 3995 | CLIP1 | chr12 | - | 122355114 | 122355312 | 122354452 | 122354556 | 122360958 | 122361181 | 0 | 0 | 7 | 0 | 0 | 0 | 0 | 0 | 0 | 0 | 1 | 0 | 98 | 49 | NA | NA | NA | NA | 1 | NA | 0 | NA | 1 |
| SE | 4006 | STAB1 | chr3 | + | 52512340 | 52512436 | 52511649 | 52511745 | 52512595 | 52512643 | 0 | 0 | 13 | 0 | 0 | 2 | 3 | 1 | 0 | 0 | 0 | 0 | 98 | 49 | NA | NA | NA | NA | 0.765 | 1 | 1 | NA | -0.235 |
| SE | 4065 | FBXL5 | chr4 | - | 15641523 | 15641627 | 15640787 | 15640883 | 15644492 | 15644708 | 0 | 0 | 1 | 0 | 0 | 31 | 0 | 1 | 0 | 0 | 8 | 0 | 98 | 49 | NA | NA | NA | NA | 0.016 | NA | 0.059 | NA | -0.043 |
| SE | 4071 | TMCO3 | chr13 | + | 113503495 | 113503588 | 113501751 | 113501880 | 113510237 | 113510424 | 0 | 0 | 1 | 0 | 0 | 2 | 0 | 3 | 0 | 0 | 0 | 0 | 98 | 49 | NA | NA | NA | NA | 0.2 | NA | 1 | NA | -0.8 |
| SE | 4087 | TNFRSF1A | chr12 | - | 6330852 | 6330926 | 6330597 | 6330711 | 6333068 | 6333147 | 0 | 0 | 21 | 0 | 0 | 1 | 2 | 3 | 0 | 0 | 1 | 0 | 98 | 49 | NA | NA | NA | NA | 0.913 | 1 | 0.6 | NA | 0.113 |
| SE | 4099 | FAM13B | chr5 | - | 137946227 | 137946311 | 137945901 | 137945997 | 137948954 | 137949184 | 0 | 0 | 5 | 0 | 0 | 0 | 0 | 2 | 0 | 0 | 2 | 0 | 98 | 49 | NA | NA | NA | NA | 1 | NA | 0.333 | NA | 0.667 |
| SE | 4107 | EVI5 | chr1 | - | 92693801 | 92693899 | 92677157 | 92677218 | 92694298 | 92694388 | 0 | 0 | 4 | 0 | 0 | 1 | 1 | 0 | 0 | 0 | 0 | 0 | 98 | 49 | NA | NA | NA | NA | 0.667 | 1 | NA | NA | -0.333 |
| SE | 4112 | ATP2B1 | chr12 | - | 89603711 | 89603925 | 89603042 | 89603254 | 89604154 | 89604346 | 0 | 0 | 6 | 0 | 0 | 0 | 0 | 5 | 0 | 0 | 1 | 0 | 98 | 49 | NA | NA | NA | NA | 1 | NA | 0.714 | NA | 0.286 |
| SE | 4113 | ATP2B1 | chr12 | - | 89616801 | 89617039 | 89611192 | 89611372 | 89619998 | 89620240 | 0 | 0 | 11 | 0 | 0 | 0 | 0 | 17 | 0 | 0 | 3 | 0 | 98 | 49 | NA | NA | NA | NA | 1 | NA | 0.739 | NA | 0.261 |
| SE | 4143 | TBC1D15 | chr12 | + | 71920730 | 71920847 | 71918450 | 71918548 | 71921367 | 71921454 | 0 | 0 | 1 | 0 | 0 | 0 | 0 | 1 | 0 | 0 | 1 | 0 | 98 | 49 | NA | NA | NA | NA | 1 | NA | 0.333 | NA | 0.667 |
| SE | 4151 | MDM2 | chr12 | + | 68816811 | 68816945 | 68813553 | 68813628 | 68820324 | 68820374 | 0 | 0 | 12 | 0 | 0 | 1 | 0 | 1 | 0 | 0 | 0 | 0 | 98 | 49 | NA | NA | NA | NA | 0.857 | NA | 1 | NA | -0.143 |
| SE | 4176 | MARS | chr12 | + | 57489895 | 57489971 | 57489423 | 57489558 | 57490206 | 57490379 | 0 | 0 | 5 | 0 | 0 | 0 | 0 | 2 | 0 | 0 | 1 | 0 | 98 | 49 | NA | NA | NA | NA | 1 | NA | 0.5 | NA | 0.5 |
| SE | 4199 | SUPT20H | chr13 | - | 37048563 | 37048599 | 37047877 | 37047936 | 37051487 | 37051583 | 0 | 0 | 0 | 0 | 0 | 1 | 0 | 1 | 0 | 0 | 0 | 0 | 85 | 49 | NA | NA | NA | NA | 0 | NA | 1 | NA | -1 |
| SE | 4211 | SMARCC2 | chr12 | - | 56172936 | 56173029 | 56172584 | 56172704 | 56173695 | 56173849 | 0 | 0 | 0 | 0 | 0 | 1 | 0 | 1 | 0 | 0 | 0 | 0 | 98 | 49 | NA | NA | NA | NA | 0 | NA | 1 | NA | -1 |
| SE | 4234 | EZH1 | chr17 | - | 42724304 | 42724424 | 42722794 | 42722915 | 42727634 | 42727763 | 0 | 0 | 11 | 0 | 0 | 1 | 0 | 5 | 0 | 0 | 0 | 0 | 98 | 49 | NA | NA | NA | NA | 0.846 | NA | 1 | NA | -0.154 |
| SE | 4243 | ARL8A | chr1 | - | 202135458 | 202135526 | 202135149 | 202135220 | 202135706 | 202135800 | 0 | 0 | 11 | 0 | 0 | 1 | 0 | 9 | 0 | 0 | 0 | 0 | 98 | 49 | NA | NA | NA | NA | 0.846 | NA | 1 | NA | -0.154 |
| SE | 4261 | CLPX | chr15 | - | 65164028 | 65164188 | 65162603 | 65162645 | 65166630 | 65166785 | 0 | 0 | 1 | 0 | 0 | 0 | 0 | 1 | 0 | 2 | 0 | 0 | 98 | 49 | NA | NA | NA | NA | 1 | 0 | 1 | NA | 0.5 |
| SE | 4263 | ITGB7 | chr12 | - | 53192690 | 53192910 | 53192329 | 53192538 | 53193139 | 53193363 | 0 | 0 | 0 | 0 | 0 | 1 | 0 | 1 | 0 | 0 | 0 | 0 | 98 | 49 | NA | NA | NA | NA | 0 | NA | 1 | NA | -1 |
| SE | 4264 | ITGB7 | chr12 | - | 53195625 | 53195721 | 53195373 | 53195463 | 53196040 | 53196199 | 0 | 0 | 2 | 0 | 0 | 0 | 0 | 0 | 0 | 0 | 1 | 0 | 98 | 49 | NA | NA | NA | NA | 1 | NA | 0 | NA | 1 |
| SE | 4275 | TDRD7 | chr9 | + | 97470557 | 97470669 | 97464841 | 97465028 | 97472292 | 97472495 | 0 | 0 | 3 | 0 | 0 | 1 | 0 | 5 | 0 | 0 | 0 | 0 | 98 | 49 | NA | NA | NA | NA | 0.6 | NA | 1 | NA | -0.4 |
| SE | 4278 | SIGLEC10 | chr19 | - | 51415897 | 51416167 | 51415567 | 51415615 | 51416309 | 51416357 | 0 | 0 | 1 | 0 | 0 | 0 | 0 | 0 | 0 | 0 | 1 | 0 | 98 | 49 | NA | NA | NA | NA | 1 | NA | 0 | NA | 1 |
| SE | 4307 | CERS5 | chr12 | - | 50144768 | 50144969 | 50143951 | 50144057 | 50167100 | 50167533 | 0 | 0 | 0 | 0 | 0 | 2 | 0 | 1 | 0 | 0 | 1 | 0 | 98 | 49 | NA | NA | NA | NA | 0 | NA | 0.333 | NA | -0.333 |
| SE | 4311 | SLC39A9 | chr14 | + | 69453240 | 69453309 | 69442068 | 69442266 | 69454811 | 69454897 | 0 | 0 | 0 | 0 | 0 | 1 | 7 | 3 | 0 | 0 | 0 | 0 | 98 | 49 | NA | NA | NA | NA | 0 | 1 | 1 | NA | -1 |
| SE | 4318 | BIRC6 | chr2 | + | 32435495 | 32435585 | 32433643 | 32433804 | 32436052 | 32436184 | 0 | 0 | 1 | 0 | 0 | 0 | 0 | 1 | 0 | 0 | 2 | 0 | 98 | 49 | NA | NA | NA | NA | 1 | NA | 0.2 | NA | 0.8 |
| SE | 4320 | BIRC6 | chr2 | + | 32464508 | 32464823 | 32463193 | 32463381 | 32465064 | 32465164 | 0 | 0 | 1 | 0 | 0 | 1 | 0 | 4 | 0 | 0 | 0 | 0 | 98 | 49 | NA | NA | NA | NA | 0.333 | NA | 1 | NA | -0.667 |
| SE | 4321 | BIRC6 | chr2 | + | 32465064 | 32465164 | 32464508 | 32464823 | 32467524 | 32467739 | 0 | 0 | 2 | 0 | 0 | 0 | 0 | 4 | 0 | 0 | 2 | 0 | 98 | 49 | NA | NA | NA | NA | 1 | NA | 0.5 | NA | 0.5 |
| SE | 4335 | NCF2 | chr1 | - | 183574486 | 183574621 | 183570779 | 183570839 | 183577598 | 183577707 | 0 | 0 | 30 | 0 | 0 | 1 | 0 | 5 | 0 | 0 | 0 | 0 | 98 | 49 | NA | NA | NA | NA | 0.937 | NA | 1 | NA | -0.063 |
| SE | 4336 | NCF2 | chr1 | - | 183574486 | 183574621 | 183573184 | 183573292 | 183577598 | 183577707 | 0 | 0 | 51 | 0 | 0 | 12 | 0 | 15 | 0 | 0 | 7 | 0 | 98 | 49 | NA | NA | NA | NA | 0.68 | NA | 0.517 | NA | 0.163 |
| SE | 4340 | SLC38A2 | chr12 | - | 46363020 | 46363145 | 46362493 | 46362638 | 46363725 | 46363826 | 0 | 0 | 6 | 0 | 0 | 0 | 0 | 4 | 0 | 0 | 1 | 0 | 98 | 49 | NA | NA | NA | NA | 1 | NA | 0.667 | NA | 0.333 |
| SE | 4344 | ARID2 | chr12 | + | 45836740 | 45836991 | 45836588 | 45836655 | 45837320 | 45837417 | 0 | 0 | 2 | 0 | 0 | 1 | 0 | 7 | 0 | 0 | 1 | 0 | 98 | 49 | NA | NA | NA | NA | 0.5 | NA | 0.778 | NA | -0.278 |
| SE | 4353 | APAF1 | chr12 | + | 98665552 | 98665791 | 98662674 | 98662806 | 98666189 | 98666357 | 0 | 0 | 2 | 0 | 0 | 1 | 0 | 2 | 0 | 0 | 0 | 0 | 98 | 49 | NA | NA | NA | NA | 0.5 | NA | 1 | NA | -0.5 |
| SE | 4354 | APAF1 | chr12 | + | 98677424 | 98677551 | 98671534 | 98671719 | 98680276 | 98680402 | 0 | 0 | 7 | 0 | 0 | 1 | 0 | 4 | 0 | 0 | 0 | 0 | 98 | 49 | NA | NA | NA | NA | 0.778 | NA | 1 | NA | -0.222 |
| SE | 4356 | APAF1 | chr12 | + | 98706484 | 98706610 | 98699407 | 98699569 | 98708584 | 98708704 | 0 | 0 | 7 | 0 | 0 | 1 | 0 | 2 | 0 | 0 | 0 | 0 | 98 | 49 | NA | NA | NA | NA | 0.778 | NA | 1 | NA | -0.222 |
| SE | 4360 | LRRK2 | chr12 | + | 40283874 | 40284133 | 40278090 | 40278261 | 40287350 | 40287539 | 0 | 0 | 46 | 0 | 0 | 0 | 0 | 21 | 0 | 0 | 1 | 0 | 98 | 49 | NA | NA | NA | NA | 1 | NA | 0.913 | NA | 0.087 |
| SE | 4375 | FAR2 | chr12 | + | 29297020 | 29297200 | 29293299 | 29293475 | 29307657 | 29307835 | 0 | 0 | 1 | 0 | 0 | 1 | 0 | 2 | 0 | 0 | 0 | 0 | 98 | 49 | NA | NA | NA | NA | 0.333 | NA | 1 | NA | -0.667 |
| SE | 4376 | FAR2 | chr12 | + | 29316840 | 29317012 | 29311882 | 29311950 | 29321794 | 29321924 | 0 | 0 | 1 | 0 | 0 | 1 | 0 | 2 | 0 | 0 | 0 | 0 | 98 | 49 | NA | NA | NA | NA | 0.333 | NA | 1 | NA | -0.667 |
| SE | 4383 | STK38L | chr12 | + | 27315014 | 27315117 | 27314503 | 27314658 | 27315288 | 27315350 | 0 | 0 | 2 | 0 | 0 | 3 | 0 | 1 | 0 | 0 | 0 | 0 | 98 | 49 | NA | NA | NA | NA | 0.25 | NA | 1 | NA | -0.75 |
| SE | 4384 | STK38L | chr12 | + | 27319327 | 27319423 | 27317895 | 27318019 | 27322142 | 27322234 | 0 | 0 | 8 | 0 | 0 | 0 | 0 | 7 | 0 | 0 | 1 | 0 | 98 | 49 | NA | NA | NA | NA | 1 | NA | 0.778 | NA | 0.222 |
| SE | 4394 | TBCEL | chr11 | + | 121055051 | 121055308 | 121053550 | 121053732 | 121058344 | 121058471 | 0 | 0 | 1 | 0 | 0 | 0 | 0 | 0 | 0 | 0 | 1 | 0 | 98 | 49 | NA | NA | NA | NA | 1 | NA | 0 | NA | 1 |
| SE | 4410 | OSM | chr22 | - | 30265001 | 30265144 | 30262829 | 30264464 | 30266765 | 30266840 | 0 | 0 | 6 | 0 | 0 | 0 | 0 | 13 | 0 | 0 | 1 | 0 | 98 | 49 | NA | NA | NA | NA | 1 | NA | 0.867 | NA | 0.133 |
| SE | 4434 | KANSL2 | chr12 | - | 48679035 | 48679150 | 48671798 | 48671962 | 48679654 | 48679833 | 0 | 0 | 6 | 0 | 0 | 2 | 0 | 3 | 0 | 0 | 0 | 0 | 98 | 49 | NA | NA | NA | NA | 0.6 | NA | 1 | NA | -0.4 |
| SE | 4435 | FRS2 | chr12 | + | 69480298 | 69480359 | 69470348 | 69470530 | 69530864 | 69530960 | 0 | 0 | 0 | 0 | 0 | 1 | 0 | 3 | 0 | 0 | 1 | 0 | 98 | 49 | NA | NA | NA | NA | 0 | NA | 0.6 | NA | -0.6 |
| SE | 4460 | CLEC7A | chr12 | - | 10126570 | 10126708 | 10125296 | 10125448 | 10127746 | 10127845 | 0 | 0 | 18 | 0 | 0 | 11 | 0 | 13 | 0 | 2 | 7 | 0 | 98 | 49 | NA | NA | NA | NA | 0.45 | 0 | 0.481 | NA | 0.21 |
| SE | 4469 | MACF1 | chr1 | + | 39378460 | 39378523 | 39372478 | 39372596 | 39379202 | 39379444 | 0 | 0 | 0 | 0 | 0 | 1 | 0 | 1 | 0 | 0 | 3 | 0 | 98 | 49 | NA | NA | NA | NA | 0 | NA | 0.143 | NA | -0.143 |
| SE | 4470 | MACF1 | chr1 | + | 39451051 | 39451211 | 39448593 | 39448763 | 39452155 | 39452350 | 0 | 0 | 2 | 0 | 0 | 0 | 0 | 1 | 0 | 0 | 1 | 0 | 98 | 49 | NA | NA | NA | NA | 1 | NA | 0.333 | NA | 0.667 |
| SE | 4478 | MYO1G | chr7 | - | 44976563 | 44976657 | 44975483 | 44975649 | 44976862 | 44977071 | 0 | 0 | 7 | 0 | 0 | 0 | 0 | 3 | 0 | 0 | 1 | 0 | 98 | 49 | NA | NA | NA | NA | 1 | NA | 0.6 | NA | 0.4 |
| SE | 4505 | PTPN6 | chr12 | + | 6960662 | 6960945 | 6960343 | 6960435 | 6961125 | 6961316 | 0 | 0 | 11 | 0 | 0 | 1 | 4 | 12 | 0 | 0 | 0 | 0 | 98 | 49 | NA | NA | NA | NA | 0.846 | 1 | 1 | NA | -0.154 |
| SE | 4511 | PDE4B | chr1 | + | 66368786 | 66368969 | 66367942 | 66368065 | 66372312 | 66374579 | 0 | 0 | 21 | 0 | 0 | 0 | 0 | 21 | 0 | 0 | 1 | 0 | 98 | 49 | NA | NA | NA | NA | 1 | NA | 0.913 | NA | 0.087 |
| SE | 4518 | CHD4 | chr12 | - | 6582136 | 6582281 | 6581648 | 6581814 | 6582614 | 6582748 | 0 | 0 | 4 | 0 | 0 | 1 | 0 | 5 | 0 | 0 | 1 | 0 | 98 | 49 | NA | NA | NA | NA | 0.667 | NA | 0.714 | NA | -0.047 |
| SE | 4519 | CHD4 | chr12 | - | 6588297 | 6588422 | 6587711 | 6587949 | 6591465 | 6591583 | 0 | 0 | 13 | 0 | 0 | 0 | 0 | 3 | 0 | 0 | 3 | 0 | 98 | 49 | NA | NA | NA | NA | 1 | NA | 0.333 | NA | 0.667 |
| SE | 4521 | GLIPR1 | chr12 | + | 75490405 | 75490518 | 75481833 | 75482079 | 75495576 | 75495662 | 0 | 0 | 15 | 0 | 0 | 1 | 0 | 5 | 0 | 0 | 0 | 0 | 98 | 49 | NA | NA | NA | NA | 0.882 | NA | 1 | NA | -0.118 |
| SE | 4528 | PARP11 | chr12 | - | 3863835 | 3863957 | 3829889 | 3830018 | 3873211 | 3873448 | 0 | 0 | 0 | 0 | 0 | 1 | 0 | 1 | 0 | 0 | 0 | 0 | 98 | 49 | NA | NA | NA | NA | 0 | NA | 1 | NA | -1 |
| SE | 4535 | TM9SF2 | chr13 | + | 99539445 | 99539557 | 99537738 | 99537863 | 99540713 | 99540793 | 0 | 0 | 11 | 0 | 0 | 1 | 0 | 2 | 0 | 0 | 0 | 0 | 98 | 49 | NA | NA | NA | NA | 0.846 | NA | 1 | NA | -0.154 |
| SE | 4553 | MED23 | chr6 | - | 131620627 | 131620729 | 131619826 | 131619896 | 131621880 | 131621979 | 0 | 0 | 4 | 0 | 0 | 0 | 0 | 1 | 0 | 0 | 2 | 0 | 98 | 49 | NA | NA | NA | NA | 1 | NA | 0.2 | NA | 0.8 |
| SE | 4560 | SLC6A6 | chr3 | + | 14468087 | 14468212 | 14467852 | 14467956 | 14472204 | 14472317 | 0 | 0 | 3 | 0 | 0 | 1 | 0 | 1 | 0 | 0 | 0 | 0 | 98 | 49 | NA | NA | NA | NA | 0.6 | NA | 1 | NA | -0.4 |
| SE | 4562 | SNX19 | chr11 | - | 130880621 | 130880806 | 130879623 | 130879711 | 130903254 | 130903384 | 0 | 0 | 2 | 0 | 0 | 0 | 0 | 0 | 0 | 0 | 2 | 0 | 98 | 49 | NA | NA | NA | NA | 1 | NA | 0 | NA | 1 |
| SE | 4584 | SRPR | chr11 | - | 126264375 | 126264539 | 126264190 | 126264289 | 126264958 | 126265172 | 0 | 0 | 3 | 0 | 0 | 0 | 0 | 3 | 0 | 0 | 2 | 0 | 98 | 49 | NA | NA | NA | NA | 1 | NA | 0.429 | NA | 0.571 |
| SE | 4590 | CUL4A | chr13 | + | 113239432 | 113239551 | 113236822 | 113236890 | 113242967 | 113243160 | 0 | 0 | 5 | 0 | 0 | 1 | 0 | 1 | 0 | 0 | 0 | 0 | 98 | 49 | NA | NA | NA | NA | 0.714 | NA | 1 | NA | -0.286 |
| SE | 4594 | KMT2C | chr7 | - | 152146598 | 152146735 | 152145152 | 152145295 | 152148032 | 152149152 | 0 | 0 | 11 | 0 | 0 | 1 | 0 | 13 | 0 | 0 | 1 | 0 | 98 | 49 | NA | NA | NA | NA | 0.846 | NA | 0.867 | NA | -0.021 |
| SE | 4595 | KMT2C | chr7 | - | 152182973 | 152183156 | 152180710 | 152182594 | 152185557 | 152185631 | 0 | 0 | 9 | 0 | 0 | 3 | 1 | 9 | 0 | 0 | 0 | 0 | 98 | 49 | NA | NA | NA | NA | 0.6 | 1 | 1 | NA | -0.4 |
| SE | 4596 | KMT2C | chr7 | - | 152195906 | 152196011 | 152194439 | 152194568 | 152199278 | 152199459 | 0 | 0 | 8 | 0 | 0 | 0 | 0 | 12 | 0 | 0 | 1 | 0 | 98 | 49 | NA | NA | NA | NA | 1 | NA | 0.857 | NA | 0.143 |
| SE | 4597 | KMT2C | chr7 | - | 152224014 | 152224179 | 152222572 | 152222682 | 152224434 | 152224616 | 0 | 0 | 3 | 0 | 0 | 0 | 0 | 3 | 0 | 0 | 1 | 0 | 98 | 49 | NA | NA | NA | NA | 1 | NA | 0.6 | NA | 0.4 |
| SE | 4598 | KMT2C | chr7 | - | 152238706 | 152238826 | 152235816 | 152235933 | 152247901 | 152248620 | 0 | 0 | 9 | 0 | 0 | 0 | 0 | 6 | 0 | 0 | 1 | 0 | 98 | 49 | NA | NA | NA | NA | 1 | NA | 0.75 | NA | 0.25 |
| SE | 4607 | KIAA0319 | chr6 | - | 24556606 | 24556729 | 24554540 | 24554631 | 24559012 | 24559155 | 0 | 0 | 0 | 0 | 0 | 1 | 0 | 1 | 0 | 0 | 0 | 0 | 98 | 49 | NA | NA | NA | NA | 0 | NA | 1 | NA | -1 |
| SE | 4619 | PAN3 | chr13 | + | 28177864 | 28177935 | 28176492 | 28176559 | 28220230 | 28220378 | 0 | 0 | 12 | 0 | 0 | 1 | 0 | 3 | 0 | 0 | 0 | 0 | 98 | 49 | NA | NA | NA | NA | 0.857 | NA | 1 | NA | -0.143 |
| SE | 4620 | PAN3 | chr13 | + | 28197184 | 28197346 | 28177864 | 28177935 | 28220230 | 28220378 | 0 | 0 | 1 | 0 | 0 | 5 | 0 | 0 | 0 | 0 | 1 | 0 | 98 | 49 | NA | NA | NA | NA | 0.091 | NA | 0 | NA | 0.091 |
| SE | 4621 | PAN3 | chr13 | + | 28271980 | 28272071 | 28270700 | 28270866 | 28277236 | 28277376 | 0 | 0 | 0 | 0 | 0 | 1 | 0 | 4 | 0 | 0 | 0 | 0 | 98 | 49 | NA | NA | NA | NA | 0 | NA | 1 | NA | -1 |
| SE | 4630 | SORL1 | chr11 | + | 121608103 | 121608176 | 121607185 | 121607290 | 121611075 | 121611158 | 0 | 0 | 31 | 0 | 0 | 1 | 3 | 18 | 0 | 0 | 0 | 0 | 98 | 49 | NA | NA | NA | NA | 0.939 | 1 | 1 | NA | -0.061 |
| SE | 4672 | IL10RA | chr11 | + | 117988381 | 117988502 | 117986390 | 117986534 | 117993240 | 117993410 | 0 | 0 | 3 | 0 | 0 | 1 | 0 | 2 | 0 | 0 | 0 | 0 | 98 | 49 | NA | NA | NA | NA | 0.6 | NA | 1 | NA | -0.4 |
| SE | 4683 | PITRM1 | chr10 | - | 3156929 | 3157064 | 3155590 | 3155729 | 3157434 | 3157531 | 0 | 0 | 2 | 0 | 0 | 0 | 1 | 1 | 0 | 0 | 1 | 0 | 98 | 49 | NA | NA | NA | NA | 1 | 1 | 0.333 | NA | 0.334 |
| SE | 4689 | IL6ST | chr5 | - | 55959625 | 55959671 | 55957208 | 55957291 | 55960401 | 55960561 | 0 | 0 | 0 | 0 | 0 | 5 | 0 | 2 | 0 | 0 | 2 | 0 | 95 | 49 | NA | NA | NA | NA | 0 | NA | 0.34 | NA | -0.34 |
| SE | 4733 | CDA | chr1 | + | 20613841 | 20613899 | 20604927 | 20605039 | 20618451 | 20618907 | 0 | 0 | 6 | 0 | 0 | 2 | 0 | 3 | 0 | 0 | 1 | 0 | 98 | 49 | NA | NA | NA | NA | 0.6 | NA | 0.6 | NA | 0 |
| SE | 4771 | ATM | chr11 | + | 108248932 | 108249102 | 108246963 | 108247127 | 108250700 | 108251072 | 0 | 0 | 8 | 0 | 0 | 0 | 0 | 3 | 0 | 3 | 0 | 0 | 98 | 49 | NA | NA | NA | NA | 1 | 0 | 1 | NA | 0.5 |
| SE | 4774 | DDX6 | chr11 | - | 118758773 | 118758902 | 118757170 | 118757287 | 118759921 | 118760044 | 0 | 0 | 9 | 0 | 0 | 0 | 0 | 6 | 0 | 0 | 2 | 0 | 98 | 49 | NA | NA | NA | NA | 1 | NA | 0.6 | NA | 0.4 |
| SE | 4775 | DDX6 | chr11 | - | 118763211 | 118763306 | 118759921 | 118760044 | 118765208 | 118765355 | 0 | 0 | 5 | 0 | 0 | 0 | 1 | 15 | 0 | 0 | 1 | 0 | 98 | 49 | NA | NA | NA | NA | 1 | 1 | 0.882 | NA | 0.059 |
| SE | 4791 | OXSR1 | chr3 | + | 38223811 | 38223913 | 38221577 | 38221687 | 38224570 | 38224704 | 0 | 0 | 9 | 0 | 0 | 0 | 0 | 8 | 0 | 0 | 1 | 0 | 98 | 49 | NA | NA | NA | NA | 1 | NA | 0.8 | NA | 0.2 |
| SE | 4795 | CASP1 | chr11 | - | 105033063 | 105033126 | 105031164 | 105031280 | 105034207 | 105034474 | 0 | 0 | 28 | 0 | 0 | 6 | 0 | 7 | 0 | 0 | 4 | 0 | 98 | 49 | NA | NA | NA | NA | 0.7 | NA | 0.467 | NA | 0.233 |
| SE | 4804 | CASP4 | chr11 | - | 104951895 | 104952005 | 104950924 | 104951098 | 104954746 | 104955001 | 0 | 0 | 13 | 0 | 0 | 3 | 0 | 6 | 0 | 0 | 0 | 0 | 98 | 49 | NA | NA | NA | NA | 0.684 | NA | 1 | NA | -0.316 |
| SE | 4810 | CCNH | chr5 | - | 87404843 | 87405007 | 87401701 | 87401772 | 87407975 | 87408186 | 0 | 0 | 11 | 0 | 0 | 0 | 0 | 7 | 0 | 0 | 1 | 0 | 98 | 49 | NA | NA | NA | NA | 1 | NA | 0.778 | NA | 0.222 |
| SE | 4811 | BIRC3 | chr11 | + | 102328051 | 102328130 | 102325465 | 102325565 | 102328896 | 102328945 | 0 | 0 | 3 | 0 | 0 | 0 | 0 | 6 | 0 | 0 | 1 | 0 | 98 | 49 | NA | NA | NA | NA | 1 | NA | 0.75 | NA | 0.25 |
| SE | 4817 | BCAP31 | chrX | - | 153702934 | 153703058 | 153702006 | 153702107 | 153703958 | 153704094 | 1 | 0 | 11 | 0 | 0 | 1 | 0 | 4 | 0 | 0 | 0 | 0 | 98 | 49 | NA | NA | 1 | NA | 0.846 | NA | 1 | NA | -0.077 |
| SE | 4825 | TRIM8 | chr10 | + | 102654652 | 102654748 | 102644494 | 102645187 | 102655079 | 102655313 | 0 | 0 | 8 | 0 | 0 | 1 | 0 | 8 | 0 | 0 | 1 | 0 | 98 | 49 | NA | NA | NA | NA | 0.8 | NA | 0.8 | NA | 0 |
| SE | 4864 | CEP295 | chr11 | + | 93702775 | 93702919 | 93702459 | 93702637 | 93706744 | 93706897 | 0 | 0 | 2 | 0 | 0 | 0 | 0 | 0 | 0 | 0 | 1 | 0 | 98 | 49 | NA | NA | NA | NA | 1 | NA | 0 | NA | 1 |

| | | | | | | | | | | | | | | | | | | | | | | | | | | | | | | | | | | |
|---|---|---|---|---|---|---|---|---|---|---|---|---|---|---|---|---|---|---|---|---|---|---|---|---|---|---|---|---|---|---|---|---|---|---|
| SE | 4876 | TPP1 | chr11 | - | 6616004 | 6616074 | 6615441 | 6615562 | 6616314 | 6616503 | 0 | 0 | 13 | 0 | 0 | 1 | 0 | 5 | 0 | 0 | 0 | 0 | 98 | 49 | NA | NA | NA | NA | 0.867 | NA | | 1 | NA | -0.133 |
| SE | 4885 | DCAF11 | chr14 | + | 24117667 | 24117732 | 24117265 | 24117393 | 24118054 | 24118155 | 0 | 0 | 2 | 0 | 0 | 1 | 0 | 4 | 0 | 0 | 0 | 0 | 98 | 49 | NA | NA | NA | NA | 0.5 | NA | | 1 | NA | -0.5 |
| SE | 4889 | TXNL1 | chr18 | - | 56626161 | 56626266 | 56624287 | 56624461 | 56626360 | 56626457 | 0 | 0 | 3 | 0 | 0 | 1 | 0 | 2 | 0 | 0 | 0 | 0 | 98 | 49 | NA | NA | NA | NA | 0.6 | NA | | 1 | NA | -0.4 |
| SE | 4906 | PAK1 | chr11 | - | 77337323 | 77337423 | 77336085 | 77336282 | 77340645 | 77340763 | 0 | 0 | 12 | 0 | 0 | 0 | 0 | 7 | 0 | 0 | 2 | 0 | 98 | 49 | NA | NA | NA | NA | 1 | NA | | 0.636 | NA | 0.364 |
| SE | 4924 | C2CD3 | chr11 | - | 74161398 | 74161556 | 74139604 | 74139828 | 74168343 | 74168613 | 0 | 0 | 2 | 0 | 0 | 1 | 0 | 2 | 0 | 0 | 0 | 0 | 98 | 49 | NA | NA | NA | NA | 0.5 | NA | | 1 | NA | -0.5 |
| SE | 4949 | RBM6 | chr3 | + | 50059648 | 50059746 | 50058401 | 50058562 | 50060955 | 50060998 | 0 | 0 | 2 | 0 | 0 | 2 | 0 | 2 | 0 | 0 | 1 | 0 | 98 | 49 | NA | NA | NA | NA | 0.333 | NA | | 0.5 | NA | -0.167 |
| SE | 4955 | N4BP1 | chr16 | - | 48546146 | 48546254 | 48538725 | 48543261 | 48548006 | 48548114 | 0 | 0 | 2 | 0 | 0 | 1 | 0 | 1 | 0 | 0 | 0 | 0 | 98 | 49 | NA | NA | NA | NA | 0.5 | NA | | 1 | NA | -0.5 |
| SE | 4965 | DDX3X | chrX | + | 41346858 | 41347012 | 41346504 | 41346622 | 41347314 | 41347451 | 0 | 0 | 33 | 0 | 0 | 0 | 1 | 31 | 0 | 0 | 1 | 0 | 98 | 49 | NA | NA | NA | NA | 1 | | 1 | 0.939 | NA | 0.03 |
| SE | 4981 | RTCB | chr22 | - | 32397940 | 32398100 | 32396073 | 32396249 | 32399602 | 32399759 | 0 | 0 | 4 | 0 | 0 | 1 | 0 | 1 | 0 | 0 | 0 | 0 | 98 | 49 | NA | NA | NA | NA | 0.667 | NA | | 1 | NA | -0.333 |
| SE | 4983 | NCOR1 | chr17 | - | 16048844 | 16048988 | 16046950 | 16047093 | 16057513 | 16057737 | 0 | 0 | 3 | 0 | 0 | 2 | 0 | 3 | 0 | 0 | 0 | 0 | 98 | 49 | NA | NA | NA | NA | 0.429 | NA | | 1 | NA | -0.571 |
| SE | 4988 | HEXB | chr5 | + | 74713505 | 74713635 | 74705218 | 74705320 | 74715509 | 74715690 | 0 | 0 | 12 | 0 | 0 | 2 | 5 | 3 | 0 | 0 | 0 | 0 | 98 | 49 | NA | NA | NA | NA | 0.75 | | 1 | 1 | NA | -0.25 |
| SE | 4991 | TOPBP1 | chr3 | - | 133624051 | 133624175 | 133623310 | 133623457 | 133628361 | 133628471 | 0 | 0 | 2 | 0 | 0 | 2 | 0 | 1 | 0 | 0 | 0 | 0 | 98 | 49 | NA | NA | NA | NA | 0.333 | NA | | 1 | NA | -0.667 |
| SE | 5001 | ABHD2 | chr15 | + | 89104181 | 89104270 | 89103902 | 89104044 | 89113724 | 89113824 | 0 | 0 | 0 | 0 | 0 | 2 | 0 | 1 | 0 | 0 | 2 | 0 | 98 | 49 | NA | NA | NA | NA | 0 | NA | | 0.2 | NA | -0.2 |
| SE | 5007 | RINT1 | chr7 | + | 105550260 | 105550486 | 105550054 | 105550165 | 105551569 | 105551707 | 0 | 0 | 1 | 0 | 0 | 0 | 0 | 0 | 0 | 0 | 2 | 0 | 98 | 49 | NA | NA | NA | NA | 1 | NA | | 0 | NA | 1 |
| SE | 5009 | RPS6KB2 | chr11 | + | 67433125 | 67433216 | 67432951 | 67433042 | 67433339 | 67433447 | 0 | 0 | 0 | 0 | 0 | 1 | 0 | 2 | 0 | 0 | 0 | 0 | 98 | 49 | NA | NA | NA | NA | 0 | NA | | 1 | NA | -1 |
| SE | 5014 | ARHGAP25 | chr2 | + | 68787839 | 68787956 | 68782232 | 68782320 | 68807272 | 68807480 | 0 | 0 | 6 | 0 | 0 | 0 | 0 | 8 | 0 | 0 | 1 | 0 | 98 | 49 | NA | NA | NA | NA | 1 | NA | | 0.8 | NA | 0.2 |
| SE | 5019 | ZFYVE16 | chr5 | + | 80448025 | 80448404 | 80445262 | 80445405 | 80449590 | 80449713 | 0 | 0 | 8 | 0 | 0 | 0 | 0 | 1 | 0 | 0 | 1 | 0 | 98 | 49 | NA | NA | NA | NA | 1 | NA | | 0.333 | NA | 0.667 |
| SE | 5020 | ZFYVE16 | chr5 | + | 80472760 | 80472923 | 80459413 | 80459494 | 80473753 | 80473859 | 0 | 0 | 2 | 0 | 0 | 1 | 0 | 0 | 0 | 0 | 2 | 0 | 98 | 49 | NA | NA | NA | NA | 0.5 | NA | | 0 | NA | 0.5 |
| SE | 5031 | HIAT1 | chr1 | + | 100068472 | 100068586 | 100067976 | 100068195 | 100069613 | 100069685 | 0 | 0 | 2 | 0 | 0 | 0 | 0 | 0 | 0 | 0 | 2 | 0 | 98 | 49 | NA | NA | NA | NA | 1 | NA | | 0 | NA | 1 |
| SE | 5037 | PTS | chr11 | + | 112230207 | 112230230 | 112228593 | 112228673 | 112230625 | 112230682 | 0 | 0 | 0 | 0 | 0 | 4 | 0 | 1 | 0 | 0 | 2 | 0 | 72 | 49 | NA | NA | NA | NA | 0 | NA | | 0.254 | NA | -0.254 |
| SE | 5038 | PTS | chr11 | + | 112230625 | 112230682 | 112228593 | 112228673 | 112233162 | 112233233 | 0 | 0 | 5 | 0 | 0 | 1 | 0 | 2 | 0 | 0 | 0 | 0 | 98 | 49 | NA | NA | NA | NA | 0.714 | NA | | 1 | NA | -0.286 |
| SE | 5040 | RELT | chr11 | + | 73391143 | 73391223 | 73390754 | 73390921 | 73393836 | 73393917 | 0 | 0 | 3 | 0 | 0 | 0 | 0 | 1 | 0 | 2 | 3 | 0 | 98 | 49 | NA | NA | NA | NA | 1 | | 0 | 0.143 | NA | 0.928 |
| SE | 5042 | RELT | chr11 | + | 73392210 | 73392468 | 73391143 | 73391223 | 73393836 | 73393917 | 0 | 0 | 3 | 0 | 0 | 1 | 0 | 10 | 0 | 0 | 0 | 0 | 98 | 49 | NA | NA | NA | NA | 0.6 | NA | | 1 | NA | -0.4 |
| SE | 5071 | CARS2 | chr13 | - | 110651033 | 110651100 | 110647100 | 110647239 | 110663450 | 110663518 | 0 | 0 | 5 | 0 | 0 | 0 | 0 | 0 | 0 | 0 | 1 | 0 | 98 | 49 | NA | NA | NA | NA | 1 | NA | | 0 | NA | 1 |
| SE | 5097 | ARPP19 | chr15 | - | 52564189 | 52564246 | 52557099 | 52557222 | 52568847 | 52568902 | 0 | 0 | 2 | 0 | 0 | 0 | 0 | 1 | 0 | 0 | 2 | 0 | 98 | 49 | NA | NA | NA | NA | 1 | NA | | 0.2 | NA | 0.8 |
| SE | 5110 | WDFY1 | chr2 | - | 223884647 | 223884749 | 223881941 | 223882072 | 223894233 | 223894339 | 0 | 0 | 3 | 0 | 0 | 0 | 0 | 1 | 0 | 0 | 1 | 0 | 98 | 49 | NA | NA | NA | NA | 1 | NA | | 0.333 | NA | 0.667 |
| SE | 5122 | CCM2 | chr7 | + | 45068442 | 45068579 | 45064462 | 45064646 | 45069825 | 45069961 | 0 | 0 | 7 | 0 | 0 | 0 | 0 | 0 | 0 | 0 | 1 | 0 | 98 | 49 | NA | NA | NA | NA | 1 | NA | | 0 | NA | 1 |
| SE | 5125 | DNM2 | chr19 | + | 10808568 | 10808580 | 10805915 | 10805967 | 10812263 | 10812377 | 0 | 0 | 4 | 0 | 0 | 1 | 0 | 3 | 0 | 0 | 1 | 0 | 61 | 49 | NA | NA | NA | NA | 0.763 | NA | | 0.707 | NA | 0.056 |
| SE | 5133 | NXF1 | chr11 | - | 62803418 | 62803572 | 62802176 | 62802260 | 62803791 | 62803978 | 0 | 0 | 9 | 0 | 0 | 0 | 0 | 6 | 0 | 0 | 1 | 0 | 98 | 49 | NA | NA | NA | NA | 1 | NA | | 0.75 | NA | 0.25 |
| SE | 5135 | SKIL | chr3 | + | 170384532 | 170384627 | 170381243 | 170381341 | 170390222 | 170390464 | 0 | 0 | 3 | 0 | 0 | 0 | 0 | 0 | 0 | 0 | 1 | 0 | 98 | 49 | NA | NA | NA | NA | 1 | NA | | 0 | NA | 1 |
| SE | 5136 | SKIL | chr3 | + | 170384532 | 170384765 | 170381243 | 170381341 | 170390222 | 170390464 | 0 | 0 | 4 | 0 | 0 | 0 | 0 | 1 | 0 | 0 | 1 | 0 | 98 | 49 | NA | NA | NA | NA | 1 | NA | | 0.333 | NA | 0.667 |
| SE | 5138 | XRN1 | chr3 | - | 142426743 | 142426841 | 142425428 | 142425538 | 142432660 | 142432893 | 0 | 0 | 8 | 0 | 0 | 1 | 0 | 5 | 0 | 0 | 1 | 0 | 98 | 49 | NA | NA | NA | NA | 0.8 | NA | | 0.714 | NA | 0.086 |
| SE | 5149 | KIAA0226L | chr13 | - | 46377889 | 46378005 | 46371940 | 46372597 | 46387133 | 46387500 | 0 | 0 | 1 | 0 | 0 | 0 | 0 | 1 | 0 | 0 | 2 | 0 | 98 | 49 | NA | NA | NA | NA | 1 | NA | | 0.2 | NA | 0.8 |
| SE | 5159 | IL32 | chr16 | + | 3067553 | 3067613 | 3067376 | 3067415 | 3067983 | 3068010 | 0 | 0 | 4 | 0 | 0 | 0 | 0 | 8 | 0 | 1 | 0 | 0 | 98 | 49 | NA | NA | NA | NA | 1 | | 0 | 1 | NA | 0.5 |
| SE | 5171 | CEP350 | chr1 | + | 180036925 | 180037089 | 180033861 | 180034082 | 180041137 | 180041248 | 0 | 0 | 1 | 0 | 0 | 1 | 0 | 3 | 0 | 0 | 0 | 0 | 98 | 49 | NA | NA | NA | NA | 0.333 | NA | | 1 | NA | -0.667 |
| SE | 5180 | ANKRD13A | chr12 | + | 110018344 | 110018488 | 110016387 | 110016433 | 110019138 | 110019328 | 0 | 0 | 18 | 0 | 0 | 1 | 0 | 4 | 0 | 0 | 0 | 0 | 98 | 49 | NA | NA | NA | NA | 0.9 | NA | | 1 | NA | -0.1 |
| SE | 5191 | PICALM | chr11 | - | 85974707 | 85974812 | 85957170 | 85959060 | 85981128 | 85981228 | 0 | 0 | 13 | 0 | 0 | 0 | 2 | 8 | 0 | 0 | 1 | 0 | 98 | 49 | NA | NA | NA | NA | 1 | | 1 | 0.8 | NA | 0.1 |
| SE | 5197 | PICALM | chr11 | - | 86022366 | 86022469 | 86014869 | 86014963 | 86026291 | 86026367 | 0 | 0 | 36 | 0 | 0 | 1 | 3 | 27 | 0 | 0 | 2 | 0 | 98 | 49 | NA | NA | NA | NA | 0.947 | | 1 | 0.871 | NA | 0.011 |
| SE | 5217 | SUCO | chr1 | + | 172585017 | 172585086 | 172579201 | 172579267 | 172585857 | 172585948 | 0 | 0 | 1 | 0 | 0 | 0 | 0 | 1 | 0 | 0 | 2 | 0 | 98 | 49 | NA | NA | NA | NA | 1 | NA | | 0.2 | NA | 0.8 |
| SE | 5220 | TPM4 | chr19 | + | 16088026 | 16088097 | 16086422 | 16086540 | 16089044 | 16089120 | 0 | 0 | 5 | 0 | 0 | 0 | 0 | 9 | 0 | 0 | 1 | 0 | 98 | 49 | NA | NA | NA | NA | 1 | NA | | 0.818 | NA | 0.182 |
| SE | 5242 | PLXNC1 | chr12 | + | 94259336 | 94259375 | 94255192 | 94255296 | 94259609 | 94259734 | 0 | 0 | 26 | 0 | 0 | 3 | 0 | 15 | 0 | 0 | 0 | 0 | 88 | 49 | NA | NA | NA | NA | 0.828 | NA | | 1 | NA | -0.172 |
| SE | 5244 | PLXNC1 | chr12 | + | 94300909 | 94301057 | 94297315 | 94297423 | 94303755 | 94303896 | 0 | 0 | 11 | 0 | 0 | 0 | 4 | 2 | 0 | 0 | 4 | 0 | 98 | 49 | NA | NA | NA | NA | 1 | | 1 | 0.2 | NA | 0.4 |
| SE | 5256 | PSME4 | chr2 | - | 53899880 | 53900017 | 53898300 | 53898354 | 53901349 | 53901559 | 0 | 0 | 2 | 0 | 0 | 0 | 0 | 1 | 0 | 0 | 1 | 0 | 98 | 49 | NA | NA | NA | NA | 1 | NA | | 0.333 | NA | 0.667 |
| SE | 5290 | PHF21A | chr11 | - | 45938156 | 45938312 | 45936493 | 45936569 | 45945839 | 45946003 | 0 | 0 | 6 | 0 | 0 | 3 | 0 | 5 | 0 | 0 | 1 | 0 | 98 | 49 | NA | NA | NA | NA | 0.5 | NA | | 0.714 | NA | -0.214 |
| SE | 5293 | ENSA | chr1 | - | 150625641 | 150625808 | 150622122 | 150622859 | 150626478 | 150626526 | 0 | 0 | 1 | 0 | 0 | 0 | 0 | 2 | 0 | 0 | 2 | 0 | 98 | 49 | NA | NA | NA | NA | 1 | NA | | 0.333 | NA | 0.667 |
| SE | 5300 | AKAP8L | chr19 | - | 15398629 | 15398766 | 15397713 | 15397855 | 15399301 | 15399410 | 0 | 0 | 0 | 0 | 0 | 4 | 0 | 1 | 0 | 0 | 0 | 0 | 98 | 49 | NA | NA | NA | NA | 0 | NA | | 1 | NA | -1 |
| SE | 5318 | PSMD1 | chr2 | + | 231079535 | 231079614 | 231078658 | 231078747 | 231080140 | 231080314 | 0 | 0 | 5 | 0 | 0 | 0 | 0 | 3 | 0 | 0 | 1 | 0 | 98 | 49 | NA | NA | NA | NA | 1 | NA | | 0.6 | NA | 0.4 |
| SE | 5357 | NLRP3 | chr1 | + | 247434102 | 247434273 | 247423846 | 247425599 | 247443971 | 247444142 | 0 | 0 | 2 | 0 | 0 | 0 | 2 | 1 | 0 | 0 | 1 | 0 | 98 | 49 | NA | NA | NA | NA | 1 | | 1 | 0.333 | NA | 0.334 |
| SE | 5358 | NLRP3 | chr1 | + | 247435969 | 247436140 | 247434102 | 247434273 | 247443971 | 247444142 | 0 | 0 | 1 | 0 | 0 | 2 | 0 | 0 | 0 | 2 | 1 | 0 | 98 | 49 | NA | NA | NA | NA | 0.2 | | 0 | 0 | NA | 0.2 |
| SE | 5367 | BAZ1A | chr14 | - | 34786125 | 34786221 | 34785776 | 34786001 | 34792774 | 34792921 | 0 | 0 | 4 | 0 | 0 | 2 | 0 | 3 | 0 | 0 | 1 | 0 | 98 | 49 | NA | NA | NA | NA | 0.5 | NA | | 0.6 | NA | -0.1 |
| SE | 5388 | CD44 | chr11 | + | 35211245 | 35211449 | 35189834 | 35190065 | 35214851 | 35214914 | 0 | 0 | 1 | 0 | 0 | 3 | 0 | 1 | 0 | 0 | 5 | 0 | 98 | 49 | NA | NA | NA | NA | 0.143 | NA | | 0.091 | NA | 0.052 |
| SE | 5397 | MTMR14 | chr3 | + | 9677983 | 9678058 | 9677316 | 9677387 | 9683177 | 9683244 | 0 | 0 | 2 | 0 | 0 | 1 | 0 | 3 | 0 | 0 | 0 | 0 | 98 | 49 | NA | NA | NA | NA | 0.5 | NA | | 1 | NA | -0.5 |
| SE | 5399 | MTMR14 | chr3 | + | 9697710 | 9697866 | 9689963 | 9690143 | 9701789 | 9702394 | 0 | 0 | 2 | 0 | 0 | 1 | 0 | 0 | 0 | 0 | 1 | 0 | 98 | 49 | NA | NA | NA | NA | 0.5 | NA | | 0 | NA | 0.5 |
| SE | 5406 | RALY | chr20 | + | 34073818 | 34073866 | 34073562 | 34073635 | 34075873 | 34076040 | 0 | 0 | 8 | 0 | 0 | 0 | 0 | 0 | 0 | 0 | 1 | 0 | 97 | 49 | NA | NA | NA | NA | 1 | NA | | 0 | NA | 1 |
| SE | 5417 | VPS39 | chr15 | - | 42173722 | 42173852 | 42169723 | 42169866 | 42178217 | 42178338 | 0 | 0 | 2 | 0 | 0 | 0 | 3 | 5 | 0 | 0 | 1 | 0 | 98 | 49 | NA | NA | NA | NA | 1 | | 1 | 0.714 | NA | 0.143 |
| SE | 5448 | RER1 | chr1 | + | 2399414 | 2399514 | 2397115 | 2397220 | 2400856 | 2400935 | 0 | 0 | 3 | 0 | 0 | 3 | 0 | 1 | 0 | 0 | 1 | 0 | 98 | 49 | NA | NA | NA | NA | 0.333 | NA | | 0.333 | NA | 0 |
| SE | 5471 | ARNTL | chr11 | + | 13355225 | 13355296 | 13354300 | 13354448 | 13356738 | 13356777 | 0 | 0 | 1 | 0 | 0 | 5 | 0 | 0 | 0 | 0 | 2 | 0 | 98 | 49 | NA | NA | NA | NA | 0.091 | NA | | 0 | NA | 0.091 |
| SE | 5472 | ARNTL | chr11 | + | 13355232 | 13355296 | 13354300 | 13354448 | 13356738 | 13356777 | 0 | 0 | 1 | 0 | 0 | 5 | 0 | 0 | 0 | 0 | 2 | 0 | 98 | 49 | NA | NA | NA | NA | 0.091 | NA | | 0 | NA | 0.091 |
| SE | 5487 | CTR9 | chr11 | + | 10755677 | 10755795 | 10754957 | 10755197 | 10756748 | 10756838 | 0 | 0 | 2 | 0 | 0 | 1 | 0 | 4 | 0 | 0 | 0 | 0 | 98 | 49 | NA | NA | NA | NA | 0.5 | NA | | 1 | NA | -0.5 |
| SE | 5500 | MBIP | chr14 | - | 36308091 | 36308189 | 36300784 | 36300823 | 36311572 | 36311725 | 0 | 0 | 0 | 0 | 0 | 1 | 0 | 1 | 0 | 0 | 0 | 0 | 98 | 49 | NA | NA | NA | NA | 0 | NA | | 1 | NA | -1 |
| SE | 5503 | SBF2 | chr11 | - | 9853539 | 9853712 | 9852675 | 9852749 | 9856457 | 9856720 | 0 | 0 | 3 | 0 | 0 | 2 | 0 | 3 | 0 | 0 | 0 | 0 | 98 | 49 | NA | NA | NA | NA | 0.429 | NA | | 1 | NA | -0.571 |
| SE | 5504 | SBF2 | chr11 | - | 9992414 | 9992543 | 9989496 | 9989595 | 9992989 | 9993103 | 0 | 0 | 5 | 0 | 0 | 5 | 0 | 8 | 0 | 0 | 0 | 0 | 98 | 49 | NA | NA | NA | NA | 0.333 | NA | | 1 | NA | -0.667 |
| SE | 5517 | DENND5A | chr11 | - | 9143402 | 9143485 | 9142721 | 9142845 | 9144096 | 9144278 | 0 | 0 | 7 | 0 | 0 | 0 | 0 | 5 | 0 | 0 | 1 | 0 | 98 | 49 | NA | NA | NA | NA | 1 | NA | | 0.714 | NA | 0.286 |
| SE | 5532 | TXLNA | chr1 | + | 32184524 | 32184616 | 32181241 | 32181577 | 32187953 | 32188124 | 0 | 0 | 1 | 0 | 0 | 1 | 0 | 1 | 0 | 0 | 0 | 0 | 98 | 49 | NA | NA | NA | NA | 0.333 | NA | | 1 | NA | -0.667 |
| SE | 5562 | RAPGEF6 | chr5 | - | 131561977 | 131562047 | 131548046 | 131548190 | 131592382 | 131592466 | 0 | 0 | 1 | 0 | 0 | 0 | 0 | 1 | 0 | 0 | 1 | 0 | 98 | 49 | NA | NA | NA | NA | 1 | NA | | 0.333 | NA | 0.667 |
| SE | 5596 | ZNF512 | chr2 | + | 27600690 | 27600815 | 27599969 | 27600053 | 27601355 | 27601442 | 0 | 0 | 0 | 0 | 0 | 1 | 0 | 2 | 0 | 0 | 0 | 0 | 98 | 49 | NA | NA | NA | NA | 0 | NA | | 1 | NA | -1 |
| SE | 5623 | ETF1 | chr5 | - | 138508668 | 138508816 | 138506092 | 138508387 | 138510564 | 138510629 | 0 | 0 | 2 | 0 | 0 | 0 | 0 | 7 | 0 | 0 | 1 | 0 | 98 | 49 | NA | NA | NA | NA | 1 | NA | | 0.778 | NA | 0.222 |
| SE | 5632 | NUP98 | chr11 | - | 3705199 | 3705356 | 3702462 | 3702892 | 3706444 | 3706627 | 0 | 0 | 3 | 0 | 0 | 0 | 0 | 4 | 0 | 0 | 1 | 0 | 98 | 49 | NA | NA | NA | NA | 1 | NA | | 0.667 | NA | 0.333 |
| SE | 5661 | ADD1 | chr4 | + | 2898432 | 2898531 | 2898183 | 2898327 | 2904763 | 2905015 | 0 | 0 | 1 | 0 | 0 | 1 | 0 | 1 | 0 | 0 | 0 | 0 | 98 | 49 | NA | NA | NA | NA | 0.333 | NA | | 1 | NA | -0.667 |
| SE | 5662 | ADD1 | chr4 | + | 2899258 | 2899435 | 2898432 | 2898531 | 2904763 | 2905015 | 0 | 0 | 13 | 0 | 0 | 0 | 1 | 6 | 0 | 0 | 1 | 0 | 98 | 49 | NA | NA | NA | NA | 1 | | 1 | 0.75 | NA | 0.125 |
| SE | 5728 | ZDHHC3 | chr3 | - | 44932906 | 44932990 | 44929305 | 44929436 | 44933117 | 44933199 | 0 | 0 | 2 | 0 | 0 | 0 | 0 | 2 | 0 | 0 | 1 | 0 | 98 | 49 | NA | NA | NA | NA | 1 | NA | | 0.5 | NA | 0.5 |
| SE | 5735 | CCNY | chr10 | + | 35553018 | 35553185 | 35530123 | 35530243 | 35566022 | 35566185 | 0 | 0 | 7 | 0 | 0 | 0 | 8 | 4 | 0 | 0 | 1 | 0 | 98 | 49 | NA | NA | NA | NA | 1 | | 1 | 0.667 | NA | 0.166 |
| SE | 5741 | ENTPD1 | chr10 | + | 95842343 | 95842494 | 95839690 | 95839808 | 95844475 | 95844635 | 0 | 0 | 10 | 0 | 0 | 0 | 0 | 2 | 0 | 0 | 1 | 0 | 98 | 49 | NA | NA | NA | NA | 1 | NA | | 0.5 | NA | 0.5 |
| SE | 5746 | UBE2Q1 | chr1 | - | 154551396 | 154551492 | 154550937 | 154551004 | 154551770 | 154551819 | 0 | 0 | 1 | 0 | 0 | 0 | 0 | 2 | 0 | 0 | 1 | 0 | 98 | 49 | NA | NA | NA | NA | 1 | NA | | 0.5 | NA | 0.5 |
| SE | 5774 | NCOA2 | chr8 | - | 70144641 | 70144848 | 70141183 | 70141399 | 70148272 | 70148483 | 0 | 0 | 5 | 0 | 0 | 1 | 0 | 2 | 0 | 0 | 1 | 0 | 98 | 49 | NA | NA | NA | NA | 0.714 | NA | | 0.5 | NA | 0.214 |
| SE | 5789 | DENND3 | chr8 | + | 141163332 | 141163429 | 141155848 | 141155970 | 141165185 | 141165289 | 0 | 0 | 8 | 0 | 0 | 1 | 3 | 6 | 0 | 0 | 0 | 0 | 98 | 49 | NA | NA | NA | NA | 0.8 | | 1 | 1 | NA | -0.2 |
| SE | 5790 | DENND3 | chr8 | + | 141163332 | 141163429 | 141160631 | 141160787 | 141165185 | 141165289 | 0 | 0 | 16 | 0 | 0 | 0 | 3 | 9 | 0 | 0 | 4 | 0 | 98 | 49 | NA | NA | NA | NA | 1 | | 1 | 0.529 | NA | 0.236 |
| SE | 5792 | AFG3L2 | chr18 | - | 12367275 | 12367382 | 12366964 | 12367117 | 12370848 | 12370926 | 0 | 0 | 0 | 0 | 0 | 1 | 0 | 1 | 0 | 0 | 0 | 0 | 98 | 49 | NA | NA | NA | NA | 0 | NA | | 1 | NA | -1 |
| SE | 5810 | ANXA1 | chr9 | + | 73169031 | 73169154 | 73167496 | 73167555 | 73170050 | 73170394 | 0 | 0 | 72 | 0 | 0 | 0 | 8 | 13 | 0 | 0 | 1 | 0 | 98 | 49 | NA | NA | NA | NA | 1 | | 1 | 0.867 | NA | 0.067 |
| SE | 5820 | HEXA | chr15 | - | 72349078 | 72349259 | 72348047 | 72348134 | 72350517 | 72350650 | 0 | 0 | 3 | 0 | 0 | 2 | 0 | 2 | 0 | 0 | 1 | 0 | 98 | 49 | NA | NA | NA | NA | 0.429 | NA | | 0.5 | NA | -0.071 |
| SE | 5826 | PLEKHO1 | chr1 | + | 150156065 | 150156206 | 150150585 | 150151058 | 150156910 | 150157015 | 0 | 0 | 10 | 0 | 0 | 0 | 0 | 2 | 0 | 0 | 1 | 0 | 98 | 49 | NA | NA | NA | NA | 1 | NA | | 0.5 | NA | 0.5 |
| SE | 5880 | ARID1A | chr1 | + | 26729650 | 26729863 | 26696030 | 26697540 | 26731151 | 26731604 | 0 | 0 | 3 | 0 | 0 | 0 | 0 | 1 | 0 | 1 | 0 | 0 | 98 | 49 | NA | NA | NA | NA | 1 | | 0 | 1 | NA | 0.5 |
| SE | 5896 | FANCM | chr14 | + | 45140631 | 45140709 | 45137068 | 45137241 | 45148836 | 45148995 | 0 | 0 | 0 | 0 | 0 | 1 | 0 | 1 | 0 | 0 | 0 | 0 | 98 | 49 | NA | NA | NA | NA | 0 | NA | | 1 | NA | -1 |
| SE | 5900 | NIPA2 | chr15 | + | 22839654 | 22839790 | 22838640 | 22838921 | 22845145 | 22845267 | 0 | 0 | 1 | 0 | 0 | 0 | 0 | 0 | 0 | 1 | 0 | 0 | 98 | 49 | NA | NA | NA | NA | 1 | | 0 NA | | NA | 1 |
| SE | 5909 | B3GNTL1 | chr17 | - | 82965650 | 82965746 | 82961056 | 82961181 | 83005114 | 83005210 | 0 | 0 | 5 | 0 | 0 | 0 | 0 | 5 | 0 | 0 | 2 | 0 | 98 | 49 | NA | NA | NA | NA | 1 | NA | | 0.556 | NA | 0.444 |
| SE | 5920 | RAP1GAP2 | chr17 | + | 2905283 | 2905368 | 2800514 | 2800550 | 2957758 | 2957794 | 0 | 0 | 4 | 0 | 0 | 0 | 0 | 4 | 0 | 0 | 2 | 0 | 98 | 49 | NA | NA | NA | NA | 1 | NA | | 0.5 | NA | 0.5 |
| SE | 5921 | RAP1GAP2 | chr17 | + | 2962669 | 2962714 | 2957758 | 2957794 | 2963429 | 2963462 | 0 | 0 | 3 | 0 | 0 | 6 | 0 | 2 | 0 | 0 | 1 | 0 | 94 | 49 | NA | NA | NA | NA | 0.207 | NA | | 0.51 | NA | -0.303 |
| SE | 5924 | MKNK1 | chr1 | - | 46574946 | 46575020 | 46572062 | 46572167 | 46576574 | 46576654 | 0 | 0 | 4 | 0 | 0 | 0 | 0 | 4 | 0 | 0 | 1 | 0 | 98 | 49 | NA | NA | NA | NA | 1 | NA | | 0.667 | NA | 0.333 |
| SE | 5928 | ZDHHC5 | chr11 | + | 57695919 | 57696043 | 57693782 | 57693915 | 57696760 | 57696873 | 0 | 0 | 0 | 0 | 0 | 1 | 5 | 5 | 0 | 0 | 0 | 0 | 98 | 49 | NA | NA | NA | NA | 0 | | 1 | 1 | NA | -1 |
| SE | 5937 | SRSF10 | chr1 | - | 23971337 | 23971439 | 23964346 | 23967723 | 23971572 | 23971626 | 0 | 0 | 1 | 0 | 0 | 0 | 0 | 1 | 0 | 0 | 2 | 0 | 98 | 49 | NA | NA | NA | NA | 1 | NA | | 0.2 | NA | 0.8 |
| SE | 5949 | METTL9 | chr16 | + | 21612644 | 21612835 | 21599475 | 21599898 | 21617864 | 21618074 | 0 | 0 | 16 | 0 | 0 | 0 | 0 | 5 | 0 | 2 | 2 | 0 | 98 | 49 | NA | NA | NA | NA | 1 | | 0 | 0.556 | NA | 0.722 |
| SE | 5965 | CD58 | chr1 | - | 116535964 | 116536228 | 116521905 | 116521983 | 116544310 | 116544604 | 0 | 0 | 10 | 0 | 0 | 0 | 0 | 5 | 0 | 0 | 1 | 0 | 98 | 49 | NA | NA | NA | NA | 1 | NA | | 0.714 | NA | 0.286 |
| SE | 5967 | CD53 | chr1 | + | 110894959 | 110895055 | 110891391 | 110891471 | 110896652 | 110896733 | 0 | 0 | 12 | 0 | 0 | 3 | 0 | 3 | 0 | 0 | 0 | 0 | 98 | 49 | NA | NA | NA | NA | 0.667 | NA | | 1 | NA | -0.333 |
| SE | 6044 | GADD45A | chr1 | + | 67686024 | 67686126 | 67685176 | 67685538 | 67687660 | 67688338 | 0 | 0 | 0 | 0 | 0 | 1 | 0 | 1 | 0 | 0 | 0 | 0 | 98 | 49 | NA | NA | NA | NA | 0 | NA | | 1 | NA | -1 |
| SE | 6051 | TROVE2 | chr1 | + | 193069033 | 193069634 | 193059421 | 193059776 | 193075819 | 193076040 | 0 | 0 | 4 | 0 | 0 | 1 | 0 | 0 | 0 | 0 | 1 | 0 | 98 | 49 | NA | NA | NA | NA | 0.667 | NA | | 0 | NA | 0.667 |
| SE | 6063 | PNPLA8 | chr7 | - | 108521475 | 108521521 | 108514435 | 108515574 | 108526028 | 108526318 | 0 | 0 | 3 | 0 | 0 | 0 | 0 | 3 | 0 | 0 | 1 | 0 | 95 | 49 | NA | NA | NA | NA | 1 | NA | | 0.607 | NA | 0.393 |
| SE | 6078 | PTBP2 | chr1 | + | 96813028 | 96813106 | 96812711 | 96812928 | 96813272 | 96815049 | 0 | 0 | 0 | 0 | 0 | 2 | 0 | 1 | 0 | 0 | 0 | 0 | 98 | 49 | NA | NA | NA | NA | 0 | NA | | 1 | NA | -1 |
| SE | 6090 | ZNF207 | chr17 | + | 32366664 | 32366757 | 32365329 | 32365487 | 32367771 | 32368014 | 0 | 0 | 0 | 0 | 0 | 2 | 0 | 3 | 0 | 0 | 0 | 0 | 98 | 49 | NA | NA | NA | NA | 0 | NA | | 1 | NA | -1 |
| SE | 6100 | ZNF217 | chr20 | - | 53571720 | 53571853 | 53567070 | 53569264 | 53575726 | 53577280 | 0 | 0 | 4 | 0 | 0 | 1 | 0 | 2 | 0 | 0 | 0 | 0 | 98 | 49 | NA | NA | NA | NA | 0.667 | NA | | 1 | NA | -0.333 |
| SE | 6101 | CCBL2 | chr1 | - | 88988251 | 88988351 | 88969408 | 88969467 | 88992584 | 88992960 | 0 | 0 | 1 | 0 | 0 | 0 | 0 | 0 | 0 | 0 | 1 | 0 | 98 | 49 | NA | NA | NA | NA | 1 | NA | | 0 | NA | 1 |
| SE | 6126 | ZC3HAV1 | chr7 | - | 139076285 | 139076409 | 139073855 | 139074030 | 139078551 | 139078653 | 0 | 0 | 12 | 0 | 0 | 0 | 4 | 14 | 0 | 0 | 5 | 0 | 98 | 49 | NA | NA | NA | NA | 1 | | 1 | 0.583 | NA | 0.209 |
| SE | 6129 | WDR26 | chr1 | - | 224398888 | 224399034 | 224398514 | 224398593 | 224400949 | 224401069 | 0 | 0 | 5 | 0 | 0 | 1 | 0 | 13 | 0 | 1 | 2 | 0 | 98 | 49 | NA | NA | NA | NA | 0.714 | | 0 | 0.765 | NA | 0.331 |
| SE | 6131 | ANKRD13D | chr11 | + | 67289828 | 67289965 | 67289344 | 67289550 | 67290077 | 67290213 | 0 | 0 | 3 | 0 | 0 | 4 | 0 | 0 | 0 | 0 | 2 | 0 | 98 | 49 | NA | NA | NA | NA | 0.273 | NA | | 0 | NA | 0.273 |
| SE | 6150 | GRASP | chr12 | + | 52011038 | 52011141 | 52010880 | 52010935 | 52013686 | 52013780 | 0 | 0 | 2 | 0 | 0 | 0 | 0 | 0 | 0 | 0 | 1 | 0 | 98 | 49 | NA | NA | NA | NA | 1 | NA | | 0 | NA | 1 |
| SE | 6151 | GRASP | chr12 | + | 52011038 | 52011207 | 52010880 | 52010935 | 52013686 | 52013780 | 0 | 0 | 2 | 0 | 0 | 0 | 0 | 0 | 0 | 0 | 1 | 0 | 98 | 49 | NA | NA | NA | NA | 1 | NA | | 0 | NA | 1 |
| SE | 6165 | EFCAB14 | chr1 | - | 46688352 | 46688544 | 46686783 | 46686870 | 46689586 | 46689691 | 0 | 0 | 0 | 0 | 1 | 0 | 0 | 0 | 1 | 0 | 0 | 0 | 98 | 49 | NA | NA | NA | | 0 NA | NA | | NA | 1 | -1 |
| SE | 6166 | NT5C3A | chr7 | - | 33029628 | 33029714 | 33026816 | 33026915 | 33035933 | 33035988 | 0 | 0 | 0 | 0 | 0 | 1 | 0 | 1 | 0 | 0 | 4 | 0 | 98 | 49 | NA | NA | NA | NA | 0 | NA | | 0.111 | NA | -0.111 |
| SE | 6172 | ABCA1 | chr9 | - | 104786872 | 104786976 | 104786297 | 104786390 | 104787919 | 104788054 | 0 | 0 | 1 | 0 | 0 | 1 | 0 | 3 | 0 | 0 | 1 | 0 | 98 | 49 | NA | NA | NA | NA | 0.333 | NA | | 0.6 | NA | -0.267 |
| SE | 6185 | HERC3 | chr4 | + | 88676217 | 88676241 | 88670138 | 88670252 | 88676333 | 88676423 | 0 | 0 | 6 | 0 | 0 | 3 | 0 | 8 | 0 | 0 | 0 | 0 | 73 | 49 | NA | NA | NA | NA | 0.573 | NA | | 1 | NA | -0.427 |
| SE | 6207 | PPP1R12A | chr12 | - | 79805591 | 79805768 | 79798493 | 79798584 | 79806165 | 79806333 | 0 | 0 | 2 | 0 | 0 | 4 | 0 | 0 | 0 | 0 | 1 | 0 | 98 | 49 | NA | NA | NA | NA | 0.2 | NA | | 0 | NA | 0.2 |
| SE | 6211 | YIPF1 | chr1 | - | 53855000 | 53855051 | 53851718 | 53852270 | 53866199 | 53866382 | 0 | 0 | 2 | 0 | 0 | 0 | 0 | 2 | 0 | 0 | 1 | 0 | 98 | 49 | NA | NA | NA | NA | 1 | NA | | 0.5 | NA | 0.5 |
| SE | 6212 | YIPF1 | chr1 | - | 53860055 | 53860153 | 53851718 | 53852270 | 53866199 | 53866382 | 0 | 0 | 1 | 0 | 0 | 0 | 0 | 0 | 0 | 0 | 1 | 0 | 98 | 49 | NA | NA | NA | NA | 1 | NA | | 0 | NA | 1 |
| SE | 6223 | AHCTF1 | chr1 | - | 246843794 | 246843928 | 246842693 | 246842776 | 246849614 | 246851442 | 0 | 0 | 0 | 0 | 0 | 2 | 0 | 2 | 0 | 0 | 2 | 0 | 98 | 49 | NA | NA | NA | NA | 0 | NA | | 0.333 | NA | -0.333 |
| SE | 6269 | BAZ2B | chr2 | - | 159336941 | 159337077 | 159332539 | 159332686 | 159337566 | 159337772 | 0 | 0 | 4 | 0 | 0 | 0 | 0 | 0 | 0 | 0 | 1 | 0 | 98 | 49 | NA | NA | NA | NA | 1 | NA | | 0 | NA | 1 |
| SE | 6352 | ECD | chr10 | - | 73154255 | 73154448 | 73152292 | 73152421 | 73156274 | 73156453 | 0 | 0 | 2 | 0 | 0 | 1 | 0 | 5 | 0 | 0 | 0 | 0 | 98 | 49 | NA | NA | NA | NA | 0.5 | NA | | 1 | NA | -0.5 |
| SE | 6355 | GATAD2B | chr1 | - | 153818039 | 153818171 | 153817371 | 153817542 | 153818790 | 153818922 | 0 | 0 | 2 | 0 | 0 | 1 | 0 | 4 | 0 | 0 | 0 | 0 | 98 | 49 | NA | NA | NA | NA | 0.5 | NA | | 1 | NA | -0.5 |
| SE | 6363 | FAM91A1 | chr8 | + | 123786494 | 123786610 | 123785628 | 123785741 | 123787260 | 123787373 | 0 | 0 | 4 | 0 | 0 | 1 | 0 | 2 | 0 | 0 | 0 | 0 | 98 | 49 | NA | NA | NA | NA | 0.667 | NA | | 1 | NA | -0.333 |
| SE | 6396 | NDEL1 | chr17 | + | 8445710 | 8445864 | 8444259 | 8444357 | 8446753 | 8446902 | 0 | 0 | 20 | 0 | 0 | 0 | 0 | 8 | 0 | 0 | 2 | 0 | 98 | 49 | NA | NA | NA | NA | 1 | NA | | 0.667 | NA | 0.333 |
| SE | 6435 | R3HDM2 | chr12 | - | 57288856 | 57288910 | 57283823 | 57284056 | 57289008 | 57289040 | 0 | 0 | 0 | 0 | 0 | 3 | 0 | 1 | 0 | 0 | 3 | 0 | 98 | 49 | NA | NA | NA | NA | 0 | NA | | 0.143 | NA | -0.143 |
| SE | 6442 | ATP11A | chr13 | + | 112851036 | 112851218 | 112842275 | 112842379 | 112854278 | 112854530 | 0 | 0 | 8 | 0 | 0 | 0 | 0 | 0 | 0 | 0 | 1 | 0 | 98 | 49 | NA | NA | NA | NA | 1 | NA | | 0 | NA | 1 |
| SE | 6474 | PUM1 | chr1 | - | 30941997 | 30942123 | 30941150 | 30941272 | 30945345 | 30945483 | 0 | 0 | 2 | 0 | 0 | 2 | 3 | 0 | 0 | 0 | 0 | 0 | 98 | 49 | NA | NA | NA | NA | 0.333 | | 1 NA | | NA | -0.667 |
| SE | 6476 | PUM1 | chr1 | - | 30980061 | 30980163 | 30974650 | 30974802 | 30981311 | 30981405 | 0 | 0 | 2 | 0 | 0 | 0 | 0 | 0 | 0 | 0 | 2 | 0 | 98 | 49 | NA | NA | NA | NA | 1 | NA | | 0 | NA | 1 |
| SE | 6495 | PI4KB | chr1 | - | 151326171 | 151326373 | 151315572 | 151316509 | 151327270 | 151327657 | 0 | 0 | 4 | 0 | 0 | 0 | 0 | 2 | 0 | 0 | 1 | 0 | 98 | 49 | NA | NA | NA | NA | 1 | NA | | 0.5 | NA | 0.5 |

| | | | | | | | | | | | | | | | | | | | | | | | | | | | | | | | | | |
|---|---|---|---|---|---|---|---|---|---|---|---|---|---|---|---|---|---|---|---|---|---|---|---|---|---|---|---|---|---|---|---|---|---|
| SE | 6501 | GAS7 | chr17 | - | 9940125 | 9940200 | 9934165 | 9934244 | 9943120 | 9943236 | 0 | 0 | 15 | 0 | 0 | 1 | 0 | 1 | 0 | 0 | 0 | 0 | 98 | 49 | NA | NA | NA | NA | 0.882 | NA | 1 | NA | -0.118 |
| SE | 6505 | RBMS1 | chr2 | - | 160277302 | 160277383 | 160275629 | 160275714 | 160278547 | 160278658 | 0 | 0 | 2 | 0 | 0 | 0 | 0 | 3 | 0 | 0 | 1 | 0 | 98 | 49 | NA | NA | NA | NA | 1 | NA | 0.6 | NA | 0.4 |
| SE | 6514 | HNRNPC | chr14 | - | 21211405 | 21211566 | 21209135 | 21211306 | 21211809 | 21211923 | 0 | 0 | 6 | 0 | 0 | 2 | 0 | 5 | 0 | 0 | 0 | 0 | 98 | 49 | NA | NA | NA | NA | 0.6 | NA | 1 | NA | -0.4 |
| SE | 6516 | PHYKPL | chr5 | - | 178212972 | 178213103 | 178211889 | 178211970 | 178214795 | 178214885 | 0 | 0 | 0 | 0 | 0 | 1 | 0 | 4 | 0 | 0 | 0 | 0 | 98 | 49 | NA | NA | NA | NA | 0 | NA | 1 | NA | -1 |
| SE | 6571 | SENP6 | chr6 | + | 75640683 | 75640704 | 75634706 | 75634811 | 75647730 | 75647801 | 0 | 0 | 1 | 0 | 0 | 1 | 0 | 1 | 0 | 0 | 0 | 0 | 70 | 49 | NA | NA | NA | NA | 0.412 | NA | 1 | NA | -0.588 |
| SE | 6572 | SENP6 | chr6 | + | 75647730 | 75647801 | 75634706 | 75634811 | 75659261 | 75659407 | 0 | 0 | 10 | 0 | 0 | 0 | 0 | 2 | 0 | 1 | 0 | 0 | 98 | 49 | NA | NA | NA | NA | 1 | 0 | 1 | NA | 0.5 |
| SE | 6576 | STX12 | chr1 | + | 27817850 | 27817923 | 27812162 | 27812268 | 27819649 | 27819732 | 0 | 0 | 5 | 0 | 0 | 1 | 0 | 1 | 0 | 0 | 0 | 0 | 98 | 49 | NA | NA | NA | NA | 0.714 | NA | 1 | NA | -0.286 |
| SE | 6579 | RABGGTB | chr1 | + | 75792180 | 75792306 | 75791460 | 75791571 | 75794083 | 75794233 | 0 | 0 | 3 | 0 | 0 | 0 | 0 | 0 | 0 | 0 | 1 | 0 | 98 | 49 | NA | NA | NA | NA | 1 | NA | 0 | NA | 1 |
| SE | 6586 | SP140 | chr2 | + | 230292645 | 230292788 | 230290459 | 230290564 | 230294270 | 230294318 | 0 | 0 | 2 | 0 | 0 | 0 | 0 | 1 | 0 | 0 | 2 | 0 | 98 | 49 | NA | NA | NA | NA | 1 | NA | 0.2 | NA | 0.8 |
| SE | 6596 | ADPGK | chr15 | - | 72755555 | 72755654 | 72751366 | 72752892 | 72771782 | 72771845 | 0 | 0 | 3 | 0 | 0 | 1 | 0 | 1 | 0 | 0 | 0 | 0 | 98 | 49 | NA | NA | NA | NA | 0.6 | NA | 1 | NA | -0.4 |
| SE | 6597 | ADPGK | chr15 | - | 72756250 | 72756447 | 72755555 | 72755654 | 72760406 | 72760527 | 0 | 0 | 5 | 0 | 0 | 0 | 0 | 4 | 0 | 0 | 2 | 0 | 98 | 49 | NA | NA | NA | NA | 1 | NA | 0.5 | NA | 0.5 |
| SE | 6598 | ADPGK | chr15 | - | 72760406 | 72760527 | 72755555 | 72755654 | 72771782 | 72771845 | 0 | 0 | 5 | 0 | 0 | 0 | 0 | 5 | 0 | 0 | 1 | 0 | 98 | 49 | NA | NA | NA | NA | 1 | NA | 0.714 | NA | 0.286 |
| SE | 6599 | ADPGK | chr15 | - | 72774871 | 72775097 | 72771782 | 72771845 | 72783245 | 72783785 | 0 | 0 | 5 | 0 | 0 | 2 | 0 | 3 | 0 | 0 | 0 | 0 | 98 | 49 | NA | NA | NA | NA | 0.556 | NA | 1 | NA | -0.444 |
| SE | 6613 | AMPD2 | chr1 | + | 109620913 | 109621266 | 109619812 | 109620278 | 109625661 | 109625792 | 0 | 0 | 0 | 0 | 0 | 3 | 0 | 5 | 0 | 0 | 7 | 0 | 98 | 49 | NA | NA | NA | NA | 0 | NA | 0.263 | NA | -0.263 |
| SE | 6614 | AMPD2 | chr1 | + | 109625302 | 109625433 | 109619812 | 109620278 | 109625661 | 109625792 | 0 | 0 | 2 | 0 | 0 | 3 | 0 | 3 | 0 | 0 | 7 | 0 | 98 | 49 | NA | NA | NA | NA | 0.25 | NA | 0.176 | NA | 0.074 |
| SE | 6615 | AMPD2 | chr1 | + | 109625302 | 109625433 | 109620580 | 109621266 | 109625661 | 109625792 | 0 | 0 | 2 | 0 | 0 | 0 | 0 | 3 | 0 | 0 | 5 | 0 | 98 | 49 | NA | NA | NA | NA | 1 | NA | 0.231 | NA | 0.769 |
| SE | 6625 | PCNX | chr14 | + | 71011491 | 71011549 | 71009633 | 71009724 | 71012999 | 71013202 | 0 | 0 | 10 | 0 | 0 | 1 | 2 | 1 | 0 | 0 | 0 | 0 | 98 | 49 | NA | NA | NA | NA | 0.833 | 1 | 1 | NA | -0.167 |
| SE | 6629 | PCBP2 | chr12 | + | 53465931 | 53465973 | 53464759 | 53464852 | 53467220 | 53467293 | 0 | 0 | 24 | 0 | 0 | 0 | 0 | 19 | 0 | 0 | 1 | 0 | 91 | 49 | NA | NA | NA | NA | 1 | NA | 0.911 | NA | 0.089 |
| SE | 6665 | THRAP3 | chr1 | + | 36296582 | 36296770 | 36293850 | 36293935 | 36300885 | 36301084 | 0 | 0 | 6 | 0 | 0 | 0 | 0 | 8 | 0 | 0 | 1 | 0 | 98 | 49 | NA | NA | NA | NA | 1 | NA | 0.8 | NA | 0.2 |
| SE | 6666 | THRAP3 | chr1 | + | 36301552 | 36301696 | 36300885 | 36301084 | 36303795 | 36305356 | 0 | 0 | 3 | 0 | 0 | 2 | 0 | 12 | 0 | 0 | 0 | 0 | 98 | 49 | NA | NA | NA | NA | 0.429 | NA | 1 | NA | -0.571 |
| SE | 6714 | ADCK3 | chr1 | + | 226965670 | 226965737 | 226964999 | 226965410 | 226977448 | 226977523 | 0 | 0 | 3 | 0 | 0 | 0 | 0 | 0 | 0 | 0 | 1 | 0 | 98 | 49 | NA | NA | NA | NA | 1 | NA | 0 | NA | 1 |
| SE | 6717 | KIF1B | chr1 | + | 10272240 | 10272306 | 10271501 | 10271579 | 10275427 | 10275503 | 0 | 0 | 6 | 0 | 0 | 0 | 0 | 10 | 0 | 0 | 1 | 0 | 98 | 49 | NA | NA | NA | NA | 1 | NA | 0.833 | NA | 0.167 |
| SE | 6720 | KIF1B | chr1 | + | 10347760 | 10347827 | 10345844 | 10345953 | 10348648 | 10348733 | 0 | 0 | 1 | 0 | 0 | 0 | 0 | 2 | 0 | 0 | 1 | 0 | 98 | 49 | NA | NA | NA | NA | 1 | NA | 0.5 | NA | 0.5 |
| SE | 6735 | DNMT3A | chr2 | - | 25241561 | 25241707 | 25240639 | 25240730 | 25243897 | 25243982 | 0 | 0 | 0 | 0 | 0 | 1 | 0 | 2 | 0 | 0 | 0 | 0 | 98 | 49 | NA | NA | NA | NA | 0 | NA | 1 | NA | -1 |
| SE | 6749 | UBR4 | chr1 | - | 19094905 | 19095025 | 19093948 | 19094139 | 19095544 | 19095652 | 0 | 0 | 3 | 0 | 0 | 1 | 0 | 1 | 0 | 0 | 0 | 0 | 98 | 49 | NA | NA | NA | NA | 0.6 | NA | 1 | NA | -0.4 |
| SE | 6750 | UBR4 | chr1 | - | 19148562 | 19148626 | 19147992 | 19148127 | 19150576 | 19150793 | 0 | 0 | 2 | 0 | 0 | 1 | 0 | 4 | 0 | 0 | 0 | 0 | 98 | 49 | NA | NA | NA | NA | 0.5 | NA | 1 | NA | -0.5 |
| SE | 6751 | UBR4 | chr1 | - | 19153767 | 19153939 | 19153300 | 19153502 | 19154917 | 19155075 | 0 | 0 | 1 | 0 | 0 | 1 | 0 | 2 | 0 | 0 | 0 | 0 | 98 | 49 | NA | NA | NA | NA | 0.333 | NA | 1 | NA | -0.667 |
| SE | 6753 | UBR4 | chr1 | - | 19168026 | 19168184 | 19167021 | 19167231 | 19169434 | 19169532 | 0 | 0 | 11 | 0 | 0 | 0 | 0 | 8 | 0 | 0 | 1 | 0 | 98 | 49 | NA | NA | NA | NA | 1 | NA | 0.8 | NA | 0.2 |
| SE | 6763 | SKIV2L | chr6 | + | 31961195 | 31961386 | 31961040 | 31961094 | 31961546 | 31961675 | 0 | 0 | 1 | 0 | 0 | 1 | 0 | 2 | 0 | 0 | 0 | 0 | 98 | 49 | NA | NA | NA | NA | 0.333 | NA | 1 | NA | -0.667 |
| SE | 6776 | RSAD2 | chr2 | + | 6886934 | 6887164 | 6883370 | 6883532 | 6890175 | 6890325 | 0 | 0 | 36 | 0 | 0 | 0 | 0 | 36 | 0 | 0 | 1 | 0 | 98 | 49 | NA | NA | NA | NA | 1 | NA | 0.947 | NA | 0.053 |
| SE | 6779 | ZFAND6 | chr15 | + | 80098415 | 80098578 | 80059567 | 80059809 | 80120327 | 80120498 | 0 | 0 | 3 | 0 | 0 | 0 | 0 | 1 | 0 | 0 | 2 | 0 | 98 | 49 | NA | NA | NA | NA | 1 | NA | 0.2 | NA | 0.8 |
| SE | 6798 | ABI1 | chr10 | - | 26755654 | 26755741 | 26751597 | 26751783 | 26759061 | 26759235 | 0 | 0 | 1 | 0 | 0 | 10 | 0 | 0 | 0 | 0 | 1 | 0 | 98 | 49 | NA | NA | NA | NA | 0.048 | NA | 0 | NA | 0.048 |
| SE | 6817 | OAS1 | chr12 | + | 112911050 | 112911235 | 112906933 | 112907219 | 112916508 | 112916738 | 0 | 0 | 15 | 0 | 0 | 2 | 0 | 1 | 0 | 0 | 0 | 0 | 98 | 49 | NA | NA | NA | NA | 0.789 | NA | 1 | NA | -0.211 |
| SE | 6825 | PRPF8 | chr17 | - | 1673745 | 1673892 | 1673356 | 1673567 | 1674441 | 1674680 | 0 | 0 | 3 | 0 | 0 | 0 | 0 | 6 | 0 | 0 | 1 | 0 | 98 | 49 | NA | NA | NA | NA | 1 | NA | 0.75 | NA | 0.25 |
| SE | 6826 | PRPF8 | chr17 | - | 1677564 | 1677694 | 1676975 | 1677172 | 1678517 | 1678652 | 0 | 0 | 1 | 0 | 0 | 0 | 0 | 0 | 0 | 0 | 1 | 0 | 98 | 49 | NA | NA | NA | NA | 1 | NA | 0 | NA | 1 |
| SE | 6828 | SH3D21 | chr1 | + | 36307516 | 36307607 | 36307106 | 36307285 | 36307769 | 36307825 | 0 | 0 | 0 | 0 | 0 | 1 | 0 | 1 | 0 | 0 | 0 | 0 | 98 | 49 | NA | NA | NA | NA | 0 | NA | 1 | NA | -1 |
| SE | 6833 | OSBPL9 | chr1 | + | 51669433 | 51669512 | 51651990 | 51652041 | 51714002 | 51714079 | 0 | 0 | 10 | 0 | 0 | 0 | 0 | 8 | 0 | 0 | 1 | 0 | 98 | 49 | NA | NA | NA | NA | 1 | NA | 0.8 | NA | 0.2 |
| SE | 6853 | UBAP2L | chr1 | + | 154234590 | 154234738 | 154228614 | 154228725 | 154235195 | 154235291 | 0 | 0 | 1 | 0 | 0 | 0 | 2 | 2 | 0 | 0 | 1 | 0 | 98 | 49 | NA | NA | NA | NA | 1 | 1 | 0.5 | NA | 0.25 |
| SE | 6854 | UBAP2L | chr1 | + | 154234590 | 154234759 | 154228614 | 154228725 | 154235195 | 154235291 | 0 | 0 | 1 | 0 | 0 | 0 | 2 | 2 | 0 | 0 | 1 | 0 | 98 | 49 | NA | NA | NA | NA | 1 | 1 | 0.5 | NA | 0.25 |
| SE | 6855 | UBAP2L | chr1 | + | 154234590 | 154234792 | 154228614 | 154228725 | 154235195 | 154235291 | 0 | 0 | 2 | 0 | 0 | 0 | 2 | 2 | 0 | 0 | 1 | 0 | 98 | 49 | NA | NA | NA | NA | 1 | 1 | 0.5 | NA | 0.25 |
| SE | 6876 | SOAT1 | chr1 | + | 179344946 | 179345076 | 179343589 | 179343635 | 179347599 | 179347697 | 0 | 0 | 2 | 0 | 0 | 0 | 0 | 0 | 0 | 1 | 0 | 0 | 98 | 49 | NA | NA | NA | NA | 1 | 0 NA | | NA | 1 |
| SE | 6880 | UBN1 | chr16 | + | 4876870 | 4877111 | 4874210 | 4875434 | 4880082 | 4882401 | 0 | 0 | 3 | 0 | 0 | 1 | 0 | 2 | 0 | 0 | 0 | 0 | 98 | 49 | NA | NA | NA | NA | 0.6 | NA | 1 | NA | -0.4 |
| SE | 6881 | UBN1 | chr16 | + | 4877384 | 4877474 | 4876870 | 4877111 | 4880082 | 4882401 | 0 | 0 | 1 | 0 | 0 | 1 | 0 | 0 | 0 | 0 | 2 | 0 | 98 | 49 | NA | NA | NA | NA | 0.333 | NA | 0 | NA | 0.333 |
| SE | 6891 | EP300 | chr22 | + | 41146738 | 41146816 | 41141047 | 41141222 | 41147836 | 41147946 | 0 | 0 | 8 | 0 | 0 | 0 | 0 | 5 | 0 | 0 | 1 | 0 | 98 | 49 | NA | NA | NA | NA | 1 | NA | 0.714 | NA | 0.286 |
| SE | 6907 | ASCC3 | chr6 | - | 100509931 | 100510107 | 100508194 | 100509533 | 100512708 | 100512918 | 0 | 0 | 0 | 0 | 0 | 1 | 0 | 1 | 0 | 0 | 0 | 0 | 98 | 49 | NA | NA | NA | NA | 0 | NA | 1 | NA | -1 |
| SE | 6930 | MADD | chr11 | + | 47293882 | 47293983 | 47290609 | 47290816 | 47295495 | 47295567 | 0 | 0 | 2 | 0 | 0 | 0 | 0 | 0 | 0 | 0 | 1 | 0 | 98 | 49 | NA | NA | NA | NA | 1 | NA | 0 | NA | 1 |
| SE | 6971 | POU2F1 | chr1 | + | 167303473 | 167303574 | 167220885 | 167220958 | 167332469 | 167332535 | 0 | 0 | 1 | 0 | 0 | 0 | 0 | 0 | 0 | 0 | 1 | 0 | 98 | 49 | NA | NA | NA | NA | 1 | NA | 0 | NA | 1 |
| SE | 7032 | IFNAR1 | chr21 | + | 33343267 | 33343422 | 33340998 | 33341174 | 33343534 | 33343676 | 0 | 0 | 6 | 0 | 0 | 1 | 0 | 11 | 0 | 0 | 0 | 0 | 98 | 49 | NA | NA | NA | NA | 0.75 | NA | 1 | NA | -0.25 |
| SE | 7041 | KCNJ15 | chr21 | + | 38298143 | 38298282 | 38296925 | 38297023 | 38299243 | 38301824 | 0 | 0 | 1 | 0 | 0 | 7 | 0 | 1 | 0 | 0 | 2 | 0 | 98 | 49 | NA | NA | NA | NA | 0.067 | NA | 0.2 | NA | -0.133 |
| SE | 7047 | UROD | chr1 | + | 45014939 | 45015006 | 45014735 | 45014836 | 45015336 | 45015669 | 0 | 0 | 4 | 0 | 0 | 0 | 0 | 0 | 0 | 0 | 1 | 0 | 98 | 49 | NA | NA | NA | NA | 1 | NA | 0 | NA | 1 |
| SE | 7060 | AMPD3 | chr11 | + | 10478525 | 10478730 | 10461514 | 10461740 | 10482062 | 10482225 | 0 | 0 | 1 | 0 | 0 | 0 | 0 | 1 | 0 | 0 | 2 | 0 | 98 | 49 | NA | NA | NA | NA | 1 | NA | 0.2 | NA | 0.8 |
| SE | 7073 | NR1D2 | chr3 | + | 23967812 | 23968023 | 23964976 | 23965162 | 23977222 | 23980618 | 0 | 0 | 1 | 0 | 0 | 0 | 0 | 1 | 0 | 0 | 1 | 0 | 98 | 49 | NA | NA | NA | NA | 1 | NA | 0.333 | NA | 0.667 |
| SE | 7074 | ERBB2IP | chr5 | + | 66068876 | 66069020 | 66053405 | 66054951 | 66072168 | 66072291 | 0 | 0 | 0 | 0 | 0 | 2 | 0 | 1 | 0 | 0 | 1 | 0 | 98 | 49 | NA | NA | NA | NA | 0 | NA | 0.333 | NA | -0.333 |
| SE | 7076 | ERBB2IP | chr5 | + | 66072168 | 66072291 | 66053405 | 66054951 | 66075023 | 66075230 | 0 | 0 | 2 | 0 | 0 | 1 | 0 | 1 | 0 | 0 | 4 | 0 | 98 | 49 | NA | NA | NA | NA | 0.5 | NA | 0.111 | NA | 0.389 |
| SE | 7080 | UBR2 | chr6 | + | 42605720 | 42605859 | 42603587 | 42603718 | 42606588 | 42606651 | 0 | 0 | 23 | 0 | 0 | 1 | 1 | 13 | 0 | 0 | 0 | 0 | 98 | 49 | NA | NA | NA | NA | 0.92 | 1 | 1 | NA | -0.08 |
| SE | 7081 | UBR2 | chr6 | + | 42665408 | 42665512 | 42663257 | 42663419 | 42666166 | 42666245 | 0 | 0 | 12 | 0 | 0 | 1 | 0 | 10 | 0 | 0 | 0 | 0 | 98 | 49 | NA | NA | NA | NA | 0.857 | NA | 1 | NA | -0.143 |
| SE | 7083 | RNF38 | chr9 | - | 36342324 | 36342424 | 36336400 | 36339814 | 36344831 | 36344953 | 0 | 0 | 1 | 0 | 0 | 2 | 0 | 2 | 0 | 0 | 0 | 0 | 98 | 49 | NA | NA | NA | NA | 0.2 | NA | 1 | NA | -0.8 |
| SE | 7090 | IL18RAP | chr2 | + | 102443199 | 102443323 | 102441311 | 102441377 | 102445188 | 102445340 | 0 | 0 | 1 | 0 | 0 | 0 | 0 | 1 | 0 | 4 | 0 | 0 | 98 | 49 | NA | NA | NA | NA | 1 | 0 | 1 | NA | 0.5 |
| SE | 7118 | TMEM71 | chr8 | - | 132722039 | 132722115 | 132714153 | 132714215 | 132727797 | 132727986 | 0 | 0 | 14 | 0 | 0 | 0 | 0 | 14 | 0 | 0 | 2 | 0 | 98 | 49 | NA | NA | NA | NA | 1 | NA | 0.778 | NA | 0.222 |
| SE | 7119 | TMEM71 | chr8 | - | 132727797 | 132727986 | 132722039 | 132722115 | 132746941 | 132747057 | 0 | 0 | 9 | 0 | 0 | 0 | 0 | 6 | 0 | 0 | 1 | 0 | 98 | 49 | NA | NA | NA | NA | 1 | NA | 0.75 | NA | 0.25 |
| SE | 7132 | GGA3 | chr17 | - | 75244618 | 75244717 | 75243446 | 75243570 | 75246508 | 75246584 | 0 | 0 | 0 | 0 | 0 | 1 | 0 | 1 | 0 | 0 | 0 | 0 | 98 | 49 | NA | NA | NA | NA | 0 | NA | 1 | NA | -1 |
| SE | 7138 | KIF21B | chr1 | - | 200983055 | 200983094 | 200980959 | 200981096 | 200984858 | 200984972 | 0 | 0 | 1 | 0 | 0 | 0 | 0 | 0 | 0 | 0 | 1 | 0 | 88 | 49 | NA | NA | NA | NA | 1 | NA | 0 | NA | 1 |
| SE | 7144 | CPD | chr17 | + | 30443801 | 30443971 | 30442307 | 30442450 | 30445690 | 30446020 | 0 | 0 | 1 | 0 | 0 | 0 | 0 | 6 | 0 | 0 | 1 | 0 | 98 | 49 | NA | NA | NA | NA | 1 | NA | 0.75 | NA | 0.25 |
| SE | 7187 | LENG8 | chr19 | + | 54453545 | 54453656 | 54452650 | 54452752 | 54454429 | 54454682 | 0 | 0 | 5 | 0 | 0 | 1 | 2 | 1 | 0 | 0 | 0 | 0 | 98 | 49 | NA | NA | NA | NA | 0.714 | 1 | 1 | NA | -0.286 |
| SE | 7203 | CLEC4A | chr12 | + | 8125560 | 8125677 | 8123631 | 8123960 | 8129263 | 8129362 | 0 | 0 | 2 | 0 | 0 | 1 | 0 | 1 | 0 | 0 | 0 | 0 | 98 | 49 | NA | NA | NA | NA | 0.5 | NA | 1 | NA | -0.5 |
| SE | 7251 | GAS5 | chr1 | - | 173865228 | 173865282 | 173864674 | 173864704 | 173865470 | 173865547 | 0 | 0 | 3 | 0 | 0 | 0 | 0 | 2 | 0 | 0 | 1 | 0 | 98 | 49 | NA | NA | NA | NA | 1 | NA | 0.5 | NA | 0.5 |
| SE | 7252 | RYK | chr3 | - | 134175608 | 134175768 | 134159236 | 134159373 | 134175929 | 134176039 | 0 | 0 | 4 | 0 | 0 | 0 | 0 | 0 | 0 | 0 | 1 | 0 | 98 | 49 | NA | NA | NA | NA | 1 | NA | 0 | NA | 1 |
| SE | 7262 | FBXO18 | chr10 | + | 5903019 | 5903175 | 5890202 | 5890346 | 5906036 | 5906632 | 0 | 0 | 3 | 0 | 0 | 0 | 0 | 0 | 0 | 0 | 1 | 0 | 98 | 49 | NA | NA | NA | NA | 1 | NA | 0 | NA | 1 |
| SE | 7269 | CIDEB | chr14 | - | 24305946 | 24306137 | 24305186 | 24305765 | 24306373 | 24306523 | 0 | 0 | 3 | 0 | 0 | 2 | 0 | 2 | 0 | 0 | 0 | 0 | 98 | 49 | NA | NA | NA | NA | 0.429 | NA | 1 | NA | -0.571 |
| SE | 7289 | PGS1 | chr17 | + | 78396307 | 78396385 | 78392475 | 78392665 | 78398251 | 78398351 | 0 | 0 | 5 | 0 | 0 | 1 | 0 | 3 | 0 | 0 | 1 | 0 | 98 | 49 | NA | NA | NA | NA | 0.714 | NA | 0.6 | NA | 0.114 |
| SE | 7314 | WNK1 | chr12 | + | 871264 | 871348 | 862082 | 862270 | 878214 | 878361 | 0 | 0 | 5 | 0 | 0 | 2 | 0 | 1 | 0 | 0 | 0 | 0 | 98 | 49 | NA | NA | NA | NA | 0.556 | NA | 1 | NA | -0.444 |
| SE | 7344 | FAM129A | chr1 | - | 184805945 | 184806056 | 184798078 | 184798190 | 184808073 | 184808235 | 0 | 0 | 4 | 0 | 0 | 0 | 7 | 11 | 0 | 0 | 1 | 0 | 98 | 49 | NA | NA | NA | NA | 1 | 1 | 0.846 | NA | 0.077 |
| SE | 7361 | OXA1L | chr14 | + | 22769790 | 22769934 | 22767957 | 22768171 | 22770192 | 22770278 | 0 | 0 | 5 | 0 | 0 | 0 | 0 | 0 | 0 | 0 | 1 | 0 | 98 | 49 | NA | NA | NA | NA | 1 | NA | 0 | NA | 1 |
| SE | 7362 | NOTCH2 | chr1 | - | 119922235 | 119922446 | 119921712 | 119921809 | 119922635 | 119922778 | 0 | 0 | 10 | 0 | 0 | 0 | 0 | 2 | 0 | 0 | 1 | 0 | 98 | 49 | NA | NA | NA | NA | 1 | NA | 0.5 | NA | 0.5 |
| SE | 7364 | NOTCH2 | chr1 | - | 119959391 | 119959502 | 119955039 | 119955232 | 119963573 | 119963807 | 0 | 0 | 4 | 0 | 0 | 1 | 0 | 5 | 0 | 0 | 0 | 0 | 98 | 49 | NA | NA | NA | NA | 0.667 | NA | 1 | NA | -0.333 |
| SE | 7368 | PSMA3 | chr14 | + | 58270933 | 58270998 | 58270417 | 58270485 | 58271850 | 58272009 | 0 | 0 | 6 | 0 | 0 | 1 | 0 | 3 | 0 | 0 | 0 | 0 | 98 | 49 | NA | NA | NA | NA | 0.75 | NA | 1 | NA | -0.25 |
| SE | 7388 | ZRANB1 | chr10 | + | 124974199 | 124974398 | 124973644 | 124973716 | 124981708 | 124981829 | 0 | 0 | 0 | 0 | 0 | 1 | 0 | 2 | 0 | 0 | 0 | 0 | 98 | 49 | NA | NA | NA | NA | 0 | NA | 1 | NA | -1 |
| SE | 7390 | ZRANB1 | chr10 | + | 124983458 | 124983688 | 124983174 | 124983304 | 124984773 | 124987436 | 0 | 0 | 0 | 0 | 0 | 1 | 0 | 1 | 0 | 0 | 0 | 0 | 98 | 49 | NA | NA | NA | NA | 0 | NA | 1 | NA | -1 |
| SE | 7409 | RBM23 | chr14 | - | 22908332 | 22908380 | 22906194 | 22906368 | 22909482 | 22909595 | 0 | 0 | 2 | 0 | 0 | 0 | 0 | 1 | 0 | 0 | 1 | 0 | 97 | 49 | NA | NA | NA | NA | 1 | NA | 0.336 | NA | 0.664 |
| SE | 7423 | H2AFY | chr5 | - | 135343259 | 135343434 | 135334380 | 135335141 | 135345967 | 135346057 | 0 | 0 | 10 | 0 | 0 | 0 | 2 | 6 | 0 | 0 | 1 | 0 | 98 | 49 | NA | NA | NA | NA | 1 | 1 | 0.75 | NA | 0.125 |
| SE | 7424 | H2AFY | chr5 | - | 135350822 | 135350913 | 135345967 | 135346057 | 135360496 | 135360607 | 0 | 0 | 3 | 0 | 0 | 0 | 0 | 0 | 0 | 0 | 1 | 0 | 98 | 49 | NA | NA | NA | NA | 1 | NA | 0 | NA | 1 |
| SE | 7433 | KCNAB2 | chr1 | + | 6041830 | 6041872 | 6040516 | 6040645 | 6072754 | 6072798 | 0 | 0 | 1 | 0 | 0 | 0 | 0 | 0 | 0 | 2 | 0 | 0 | 91 | 49 | NA | NA | NA | NA | 1 | 0 NA | | NA | 1 |
| SE | 7452 | ARNT | chr1 | - | 150823193 | 150823345 | 150817919 | 150818030 | 150826542 | 150826617 | 0 | 0 | 4 | 0 | 0 | 0 | 0 | 0 | 0 | 0 | 1 | 0 | 98 | 49 | NA | NA | NA | NA | 1 | NA | 0 | NA | 1 |
| SE | 7453 | ARNT | chr1 | - | 150842423 | 150842468 | 150839440 | 150839654 | 150846262 | 150846307 | 0 | 0 | 0 | 0 | 0 | 1 | 0 | 1 | 0 | 0 | 0 | 0 | 94 | 49 | NA | NA | NA | NA | 0 | NA | 1 | NA | -1 |
| SE | 7461 | HNRNPDL | chr4 | - | 82426462 | 82426633 | 82426036 | 82426129 | 82427189 | 82427304 | 0 | 0 | 1 | 0 | 0 | 0 | 0 | 6 | 0 | 0 | 1 | 0 | 98 | 49 | NA | NA | NA | NA | 1 | NA | 0.75 | NA | 0.25 |
| SE | 7467 | SLTM | chr15 | - | 58890280 | 58890461 | 58889429 | 58889554 | 58892896 | 58893060 | 0 | 0 | 2 | 0 | 0 | 1 | 0 | 0 | 0 | 0 | 2 | 0 | 98 | 49 | NA | NA | NA | NA | 0.5 | NA | 0 | NA | 0.5 |
| SE | 7468 | SLTM | chr15 | - | 58894432 | 58894582 | 58894089 | 58894193 | 58897114 | 58897233 | 0 | 0 | 3 | 0 | 0 | 0 | 0 | 1 | 0 | 2 | 0 | 0 | 98 | 49 | NA | NA | NA | NA | 1 | 0 | 1 | NA | 0.5 |
| SE | 7519 | ZNF335 | chr20 | - | 45969450 | 45969691 | 45968284 | 45968362 | 45971209 | 45971460 | 0 | 0 | 0 | 0 | 0 | 1 | 0 | 2 | 0 | 0 | 0 | 0 | 98 | 49 | NA | NA | NA | NA | 0 | NA | 1 | NA | -1 |
| SE | 7542 | ADAR | chr1 | - | 154586180 | 154586363 | 154585752 | 154585865 | 154588124 | 154588258 | 0 | 0 | 38 | 0 | 0 | 2 | 3 | 13 | 0 | 0 | 2 | 0 | 98 | 49 | NA | NA | NA | NA | 0.905 | 1 | 0.765 | NA | 0.022 |
| SE | 7543 | PTPN22 | chr1 | - | 113856547 | 113856619 | 113856381 | 113856441 | 113857737 | 113857776 | 0 | 0 | 1 | 0 | 0 | 0 | 0 | 0 | 0 | 0 | 1 | 0 | 98 | 49 | NA | NA | NA | NA | 1 | NA | 0 | NA | 1 |
| SE | 7567 | ST7L | chr1 | - | 112581991 | 112582106 | 112578344 | 112578417 | 112582374 | 112582472 | 0 | 0 | 2 | 0 | 0 | 0 | 0 | 0 | 0 | 0 | 1 | 0 | 98 | 49 | NA | NA | NA | NA | 1 | NA | 0 | NA | 1 |
| SE | 7569 | ARAP2 | chr4 | - | 36212395 | 36212487 | 36210389 | 36210743 | 36213242 | 36213319 | 0 | 0 | 3 | 0 | 0 | 2 | 1 | 3 | 0 | 0 | 1 | 0 | 98 | 49 | NA | NA | NA | NA | 0.429 | 1 | 0.6 | NA | -0.371 |
| SE | 7585 | STAT2 | chr12 | - | 56350096 | 56350190 | 56349588 | 56349636 | 56350411 | 56350432 | 0 | 0 | 18 | 0 | 0 | 0 | 0 | 12 | 0 | 0 | 2 | 0 | 98 | 49 | NA | NA | NA | NA | 1 | NA | 0.75 | NA | 0.25 |
| SE | 7619 | MAN2B1 | chr19 | - | 12658427 | 12658507 | 12658223 | 12658344 | 12661259 | 12661376 | 0 | 0 | 8 | 0 | 0 | 1 | 0 | 1 | 0 | 0 | 0 | 0 | 98 | 49 | NA | NA | NA | NA | 0.8 | NA | 1 | NA | -0.2 |
| SE | 7620 | MAN2B1 | chr19 | - | 12658427 | 12658510 | 12658223 | 12658344 | 12661259 | 12661376 | 0 | 0 | 10 | 0 | 0 | 1 | 0 | 2 | 0 | 0 | 0 | 0 | 98 | 49 | NA | NA | NA | NA | 0.833 | NA | 1 | NA | -0.167 |
| SE | 7629 | RALGAPB | chr20 | + | 38565355 | 38565478 | 38562531 | 38562697 | 38567095 | 38567232 | 0 | 0 | 0 | 0 | 0 | 4 | 0 | 3 | 0 | 0 | 0 | 0 | 98 | 49 | NA | NA | NA | NA | 0 | NA | 1 | NA | -1 |
| SE | 7630 | RALGAPB | chr20 | + | 38565358 | 38565478 | 38562531 | 38562697 | 38567095 | 38567232 | 0 | 0 | 2 | 0 | 0 | 4 | 0 | 3 | 0 | 0 | 0 | 0 | 98 | 49 | NA | NA | NA | NA | 0.2 | NA | 1 | NA | -0.8 |
| SE | 7632 | BTAF1 | chr10 | + | 92027129 | 92027300 | 92026591 | 92026751 | 92028789 | 92030323 | 0 | 0 | 4 | 0 | 0 | 3 | 0 | 5 | 0 | 0 | 0 | 0 | 98 | 49 | NA | NA | NA | NA | 0.4 | NA | 1 | NA | -0.6 |
| SE | 7685 | EAF1 | chr3 | + | 15432086 | 15432223 | 15429912 | 15430007 | 15434347 | 15434538 | 0 | 0 | 3 | 1 | 0 | 0 | 0 | 2 | 0 | 0 | 0 | 0 | 98 | 49 | NA | NA | 0 | NA | 1 | NA | 1 | NA | -0.5 |
| SE | 7686 | DHX40 | chr17 | + | 59566626 | 59566794 | 59565524 | 59565783 | 59570517 | 59570663 | 0 | 0 | 1 | 0 | 0 | 1 | 0 | 1 | 0 | 0 | 0 | 0 | 98 | 49 | NA | NA | NA | NA | 0.333 | NA | 1 | NA | -0.667 |
| SE | 7707 | CHFR | chr12 | - | 132869618 | 132869834 | 132861466 | 132861634 | 132870723 | 132870783 | 0 | 0 | 0 | 0 | 0 | 2 | 0 | 1 | 0 | 0 | 0 | 0 | 98 | 49 | NA | NA | NA | NA | 0 | NA | 1 | NA | -1 |
| SE | 7722 | ARHGEF2 | chr1 | - | 155965630 | 155965760 | 155965302 | 155965412 | 155966415 | 155966479 | 0 | 0 | 8 | 0 | 0 | 0 | 0 | 0 | 0 | 0 | 1 | 0 | 98 | 49 | NA | NA | NA | NA | 1 | NA | 0 | NA | 1 |
| SE | 7723 | ARHGEF2 | chr1 | - | 155966819 | 155966887 | 155966415 | 155966479 | 155969155 | 155969300 | 0 | 0 | 4 | 0 | 0 | 0 | 0 | 3 | 0 | 0 | 1 | 0 | 98 | 49 | NA | NA | NA | NA | 1 | NA | 0.6 | NA | 0.4 |
| SE | 7728 | KIAA1033 | chr12 | + | 105126234 | 105126362 | 105126003 | 105126127 | 105127128 | 105127289 | 0 | 0 | 12 | 0 | 0 | 1 | 3 | 5 | 0 | 0 | 0 | 0 | 98 | 49 | NA | NA | NA | NA | 0.857 | 1 | 1 | NA | -0.143 |
| SE | 7730 | KIAA1033 | chr12 | + | 105144286 | 105144455 | 105143126 | 105143243 | 105144717 | 105144872 | 0 | 0 | 26 | 0 | 0 | 0 | 0 | 9 | 0 | 0 | 1 | 0 | 98 | 49 | NA | NA | NA | NA | 1 | NA | 0.818 | NA | 0.182 |
| SE | 7732 | FAM32A | chr19 | + | 16190519 | 16190573 | 16185623 | 16185765 | 16190886 | 16192046 | 0 | 0 | 2 | 0 | 0 | 0 | 0 | 2 | 0 | 0 | 1 | 0 | 98 | 49 | NA | NA | NA | NA | 1 | NA | 0.5 | NA | 0.5 |
| SE | 7819 | ADCY10P1 | chr6 | + | 41118923 | 41119107 | 41118128 | 41118184 | 41120522 | 41120657 | 0 | 0 | 1 | 0 | 0 | 0 | 0 | 1 | 0 | 0 | 1 | 0 | 98 | 49 | NA | NA | NA | NA | 1 | NA | 0.333 | NA | 0.667 |
| SE | 7845 | ITPKC | chr19 | + | 40733159 | 40733364 | 40729201 | 40729415 | 40736985 | 40737087 | 0 | 0 | 1 | 0 | 0 | 0 | 0 | 0 | 0 | 0 | 1 | 0 | 98 | 49 | NA | NA | NA | NA | 1 | NA | 0 | NA | 1 |
| SE | 7849 | RPS6KA1 | chr1 | + | 26561504 | 26561663 | 26561044 | 26561134 | 26571448 | 26571610 | 0 | 0 | 12 | 0 | 0 | 1 | 0 | 4 | 0 | 0 | 0 | 0 | 98 | 49 | NA | NA | NA | NA | 0.857 | NA | 1 | NA | -0.143 |
| SE | 7850 | FOXJ3 | chr1 | - | 42199101 | 42199230 | 42194889 | 42195064 | 42205761 | 42205863 | 0 | 0 | 0 | 0 | 0 | 1 | 1 | 7 | 0 | 0 | 0 | 0 | 98 | 49 | NA | NA | NA | NA | 0 | 1 | 1 | NA | -1 |
| SE | 7857 | RBL2 | chr16 | + | 53469915 | 53470185 | 53467057 | 53467169 | 53470382 | 53470663 | 0 | 0 | 3 | 0 | 0 | 0 | 0 | 7 | 0 | 0 | 2 | 0 | 98 | 49 | NA | NA | NA | NA | 1 | NA | 0.636 | NA | 0.364 |
| SE | 7858 | STK24 | chr13 | - | 98457167 | 98457304 | 98450198 | 98453209 | 98460371 | 98460440 | 0 | 0 | 5 | 0 | 0 | 1 | 4 | 3 | 0 | 0 | 0 | 0 | 98 | 49 | NA | NA | NA | NA | 0.714 | 1 | 1 | NA | -0.286 |
| SE | 7870 | CTBS | chr1 | - | 84569930 | 84570139 | 84565840 | 84566012 | 84570581 | 84570720 | 0 | 0 | 4 | 0 | 0 | 0 | 0 | 0 | 0 | 0 | 5 | 0 | 98 | 49 | NA | NA | NA | NA | 1 | NA | 0 | NA | 1 |
| SE | 7872 | MROH6 | chr8 | - | 143567610 | 143567676 | 143566192 | 143567465 | 143567785 | 143567888 | 0 | 0 | 1 | 0 | 0 | 0 | 0 | 0 | 0 | 0 | 2 | 0 | 98 | 49 | NA | NA | NA | NA | 1 | NA | 0 | NA | 1 |
| SE | 7881 | UHRF1BP1L | chr12 | - | 100088914 | 100089095 | 100086274 | 100086358 | 100095666 | 100095837 | 0 | 0 | 11 | 0 | 0 | 3 | 0 | 3 | 0 | 0 | 0 | 0 | 98 | 49 | NA | NA | NA | NA | 0.647 | NA | 1 | NA | -0.353 |
| SE | 7900 | IFIT3 | chr10 | + | 89337424 | 89337542 | 89327818 | 89328078 | 89338660 | 89340968 | 0 | 0 | 0 | 0 | 0 | 1 | 0 | 1 | 0 | 0 | 0 | 0 | 98 | 49 | NA | NA | NA | NA | 0 | NA | 1 | NA | -1 |
| SE | 7906 | IFIT1 | chr10 | + | 89399889 | 89400003 | 89392545 | 89392717 | 89402280 | 89406487 | 0 | 0 | 11 | 0 | 0 | 5 | 0 | 7 | 0 | 4 | 2 | 0 | 98 | 49 | NA | NA | NA | NA | 0.524 | 0 | 0.636 | NA | 0.206 |
| SE | 7913 | DDX21 | chr10 | + | 68977528 | 68977688 | 68974669 | 68974743 | 68978841 | 68978976 | 0 | 0 | 10 | 0 | 0 | 0 | 0 | 8 | 0 | 0 | 1 | 0 | 98 | 49 | NA | NA | NA | NA | 1 | NA | 0.8 | NA | 0.2 |
| SE | 7921 | CSDE1 | chr1 | - | 114730257 | 114730422 | 114726982 | 114727090 | 114730507 | 114730648 | 0 | 0 | 21 | 0 | 0 | 1 | 0 | 16 | 0 | 0 | 0 | 0 | 98 | 49 | NA | NA | NA | NA | 0.913 | NA | 1 | NA | -0.087 |
| SE | 7922 | CSDE1 | chr1 | - | 114734441 | 114734523 | 114733988 | 114734117 | 114736757 | 114736855 | 0 | 0 | 16 | 0 | 0 | 0 | 0 | 13 | 0 | 0 | 1 | 0 | 98 | 49 | NA | NA | NA | NA | 1 | NA | 0.867 | NA | 0.133 |
| SE | 7946 | PPP2R2D | chr10 | + | 131911561 | 131912598 | 131901237 | 131901330 | 131934457 | 131934555 | 0 | 0 | 0 | 0 | 0 | 3 | 0 | 2 | 0 | 0 | 0 | 0 | 98 | 49 | NA | NA | NA | NA | 0 | NA | 1 | NA | -1 |
| SE | 7947 | PPP2R2D | chr10 | + | 131934893 | 131935006 | 131934457 | 131934555 | 131940030 | 131940196 | 0 | 0 | 1 | 0 | 0 | 3 | 0 | 1 | 0 | 0 | 0 | 0 | 98 | 49 | NA | NA | NA | NA | 0.143 | NA | 1 | NA | -0.857 |
| SE | 7960 | CEP104 | chr1 | - | 3826707 | 3826744 | 3826369 | 3826436 | 3829265 | 3829373 | 0 | 0 | 2 | 0 | 0 | 0 | 0 | 0 | 0 | 0 | 1 | 0 | 86 | 49 | NA | NA | NA | NA | 1 | NA | 0 | NA | 1 |

| | | | | | | | | | | | | | | | | | | | | | | | | | | | | | | | | | |
|---|---|---|---|---|---|---|---|---|---|---|---|---|---|---|---|---|---|---|---|---|---|---|---|---|---|---|---|---|---|---|---|---|---|
| SE | 7963 | SP3 | chr2 | - | 173918592 | 173918785 | 173913069 | 173913266 | 173954872 | 173956232 | 0 | 0 | 19 | 0 | 0 | 1 | 0 | 12 | 0 | 0 | 0 | 0 | 98 | 49 | NA | NA | NA | NA | 0.905 | NA | 1 | NA | -0.095 |
| SE | 8007 | ZNF33A | chr10 | + | 38039412 | 38039492 | 38017290 | 38017386 | 38054374 | 38060067 | 0 | 0 | 2 | 0 | 0 | 1 | 0 | 0 | 0 | 0 | 2 | 0 | 98 | 49 | NA | NA | NA | NA | 0.5 | NA | 0 | NA | 0.5 |
| SE | 8010 | DUSP18 | chr22 | - | 30664954 | 30665121 | 30661457 | 30664080 | 30667461 | 30667890 | 0 | 0 | 1 | 0 | 0 | 2 | 0 | 0 | 0 | 0 | 1 | 0 | 98 | 49 | NA | NA | NA | NA | 0.2 | NA | 0 | NA | 0.2 |
| SE | 8016 | HIATL1 | chr9 | + | 94456239 | 94456337 | 94453957 | 94454070 | 94458321 | 94458464 | 0 | 0 | 10 | 0 | 0 | 1 | 0 | 2 | 0 | 0 | 0 | 0 | 98 | 49 | NA | NA | NA | NA | 0.833 | NA | 1 | NA | -0.167 |
| SE | 8018 | MIR22HG | chr17 | - | 1714370 | 1714453 | 1713702 | 1714014 | 1716129 | 1716272 | 0 | 0 | 1 | 0 | 0 | 0 | 0 | 1 | 0 | 0 | 1 | 0 | 98 | 49 | NA | NA | NA | NA | 1 | NA | 0.333 | NA | 0.667 |
| SE | 8037 | MORF4L1 | chr15 | + | 78894057 | 78894230 | 78893538 | 78893627 | 78894819 | 78894904 | 0 | 0 | 7 | 0 | 0 | 3 | 0 | 1 | 0 | 0 | 0 | 0 | 98 | 49 | NA | NA | NA | NA | 0.538 | NA | 1 | NA | -0.462 |
| SE | 8043 | GLYR1 | chr16 | - | 4821546 | 4821597 | 4821379 | 4821453 | 4822874 | 4822931 | 0 | 0 | 9 | 0 | 3 | 0 | 2 | 0 | 0 | 0 | 0 | 0 | 98 | 49 | NA | NA | NA | 0 | 1 | 1 | NA | NA | -0.5 |
| SE | 8044 | GLYR1 | chr16 | - | 4831978 | 4832221 | 4823820 | 4823907 | 4832773 | 4832912 | 0 | 0 | 1 | 0 | 0 | 0 | 0 | 0 | 0 | 0 | 1 | 0 | 98 | 49 | NA | NA | NA | NA | 1 | NA | 0 | NA | 1 |
| SE | 8049 | MARK2 | chr11 | + | 63903985 | 63904147 | 63903060 | 63903158 | 63904785 | 63905043 | 0 | 0 | 1 | 0 | 0 | 0 | 0 | 0 | 0 | 2 | 0 | 0 | 98 | 49 | NA | NA | NA | NA | 1 | 0 | NA | NA | 1 |
| SE | 8066 | P3H1 | chr1 | - | 42755547 | 42755637 | 42755164 | 42755217 | 42757782 | 42757922 | 0 | 0 | 1 | 0 | 0 | 0 | 0 | 0 | 0 | 0 | 1 | 0 | 98 | 49 | NA | NA | NA | NA | 1 | NA | 0 | NA | 1 |
| SE | 8077 | NADK | chr1 | - | 1755373 | 1755476 | 1754543 | 1754698 | 1756257 | 1756343 | 0 | 0 | 12 | 0 | 0 | 3 | 5 | 1 | 0 | 0 | 0 | 0 | 98 | 49 | NA | NA | NA | NA | 0.667 | 1 | 1 | NA | -0.333 |
| SE | 8089 | SCLT1 | chr4 | - | 128942995 | 128943188 | 128936654 | 128936851 | 128946006 | 128946152 | 0 | 0 | 0 | 0 | 0 | 2 | 0 | 1 | 0 | 0 | 0 | 0 | 98 | 49 | NA | NA | NA | NA | 0 | NA | 1 | NA | -1 |
| SE | 8091 | PKN2 | chr1 | + | 88805890 | 88806017 | 88805496 | 88805671 | 88807312 | 88807443 | 0 | 0 | 9 | 0 | 0 | 0 | 0 | 7 | 0 | 0 | 1 | 0 | 98 | 49 | NA | NA | NA | NA | 1 | NA | 0.778 | NA | 0.222 |
| SE | 8093 | TP53INP1 | chr8 | - | 94932069 | 94932107 | 94925971 | 94930728 | 94939859 | 94940220 | 0 | 0 | 4 | 0 | 0 | 1 | 0 | 4 | 0 | 0 | 1 | 0 | 87 | 49 | NA | NA | NA | NA | 0.693 | NA | 0.693 | NA | 0 |
| SE | 8097 | JMJD1C | chr10 | - | 63200475 | 63200677 | 63198512 | 63198727 | 63206594 | 63208801 | 0 | 0 | 13 | 0 | 0 | 1 | 0 | 4 | 0 | 0 | 0 | 0 | 98 | 49 | NA | NA | NA | NA | 0.867 | NA | 1 | NA | -0.133 |
| SE | 8098 | JMJD1C | chr10 | - | 63215552 | 63215696 | 63215262 | 63215455 | 63217206 | 63217331 | 0 | 0 | 21 | 0 | 0 | 1 | 0 | 8 | 0 | 0 | 0 | 0 | 98 | 49 | NA | NA | NA | NA | 0.913 | NA | 1 | NA | -0.087 |
| SE | 8099 | JMJD1C | chr10 | - | 63219877 | 63219983 | 63217206 | 63217331 | 63264650 | 63264764 | 0 | 0 | 14 | 0 | 0 | 2 | 0 | 1 | 0 | 0 | 2 | 0 | 98 | 49 | NA | NA | NA | NA | 0.778 | NA | 0.2 | NA | 0.578 |
| SE | 8111 | COPB2 | chr3 | - | 139373665 | 139373808 | 139373212 | 139373412 | 139374488 | 139374588 | 0 | 0 | 1 | 0 | 0 | 0 | 0 | 0 | 0 | 0 | 2 | 0 | 98 | 49 | NA | NA | NA | NA | 1 | NA | 0 | NA | 1 |
| SE | 8124 | ANXA5 | chr4 | - | 121678414 | 121678494 | 121677893 | 121677950 | 121681670 | 121681761 | 0 | 0 | 22 | 0 | 0 | 1 | 11 | 6 | 0 | 0 | 0 | 0 | 98 | 49 | NA | NA | NA | NA | 0.917 | 1 | 1 | NA | -0.083 |
| SE | 8148 | PPT1 | chr1 | - | 40076841 | 40076913 | 40072709 | 40074183 | 40078559 | 40078658 | 0 | 0 | 9 | 0 | 0 | 5 | 0 | 17 | 0 | 0 | 0 | 0 | 98 | 49 | NA | NA | NA | NA | 0.474 | NA | 1 | NA | -0.526 |
| SE | 8156 | KMT2E | chr7 | + | 105066726 | 105066807 | 105063350 | 105063580 | 105073618 | 105073677 | 0 | 0 | 10 | 0 | 0 | 0 | 0 | 10 | 0 | 0 | 1 | 0 | 98 | 49 | NA | NA | NA | NA | 1 | NA | 0.833 | NA | 0.167 |
| SE | 8178 | MLXIP | chr12 | + | 122130843 | 122130933 | 122129940 | 122130112 | 122132291 | 122132383 | 0 | 0 | 1 | 0 | 0 | 1 | 0 | 1 | 0 | 0 | 0 | 0 | 98 | 49 | NA | NA | NA | NA | 0.333 | NA | 1 | NA | -0.667 |
| SE | 8186 | EPSTI1 | chr13 | - | 42889210 | 42889242 | 42886387 | 42888502 | 42895008 | 42895108 | 0 | 0 | 8 | 0 | 0 | 4 | 0 | 2 | 0 | 4 | 6 | 0 | 81 | 49 | NA | NA | NA | NA | 0.547 | 0 | 0.168 | NA | 0.463 |
| SE | 8192 | ATP5F1 | chr1 | + | 111454210 | 111454356 | 111449836 | 111449873 | 111456085 | 111456249 | 0 | 0 | 8 | 0 | 0 | 1 | 5 | 5 | 0 | 0 | 0 | 0 | 98 | 49 | NA | NA | NA | NA | 0.8 | 1 | 1 | NA | -0.2 |
| SE | 8219 | SLC15A4 | chr12 | - | 128809942 | 128810111 | 128809395 | 128809473 | 128814774 | 128815070 | 0 | 0 | 4 | 0 | 0 | 3 | 5 | 4 | 0 | 0 | 1 | 0 | 98 | 49 | NA | NA | NA | NA | 0.4 | 1 | 0.667 | NA | -0.433 |
| SE | 8237 | LBR | chr1 | - | 225404625 | 225404706 | 225404403 | 225404526 | 225406663 | 225406832 | 0 | 0 | 5 | 0 | 0 | 0 | 0 | 3 | 0 | 0 | 2 | 0 | 98 | 49 | NA | NA | NA | NA | 1 | NA | 0.429 | NA | 0.571 |
| SE | 8249 | PRRC2C | chr1 | + | 171545478 | 171545687 | 171539970 | 171542229 | 171550085 | 171550240 | 0 | 0 | 0 | 0 | 0 | 1 | 0 | 7 | 0 | 0 | 0 | 0 | 98 | 49 | NA | NA | NA | NA | 0 | NA | 1 | NA | -1 |
| SE | 8250 | PRRC2C | chr1 | + | 171571319 | 171571421 | 171568246 | 171568339 | 171574926 | 171575128 | 0 | 0 | 6 | 0 | 0 | 0 | 0 | 2 | 0 | 0 | 1 | 0 | 98 | 49 | NA | NA | NA | NA | 1 | NA | 0.5 | NA | 0.5 |
| SE | 8251 | SMARCA2 | chr9 | + | 2170418 | 2170472 | 2161685 | 2161903 | 2181570 | 2181676 | 0 | 0 | 2 | 0 | 0 | 0 | 0 | 1 | 0 | 0 | 1 | 0 | 98 | 49 | NA | NA | NA | NA | 1 | NA | 0.333 | NA | 0.667 |
| SE | 8260 | ARHGEF11 | chr1 | - | 156947768 | 156947956 | 156947303 | 156947450 | 156948180 | 156948228 | 0 | 0 | 13 | 0 | 0 | 1 | 0 | 3 | 0 | 0 | 0 | 0 | 98 | 49 | NA | NA | NA | NA | 0.867 | NA | 1 | NA | -0.133 |
| SE | 8262 | ARHGEF11 | chr1 | - | 156971696 | 156971816 | 156969997 | 156970043 | 156976982 | 156977054 | 0 | 0 | 1 | 0 | 0 | 0 | 0 | 0 | 0 | 1 | 0 | 0 | 98 | 49 | NA | NA | NA | NA | 1 | 0 | NA | NA | 1 |
| SE | 8268 | DPYD | chr1 | - | 97593217 | 97593387 | 97573759 | 97573970 | 97595058 | 97595166 | 0 | 0 | 31 | 0 | 0 | 0 | 0 | 5 | 0 | 0 | 2 | 0 | 98 | 49 | NA | NA | NA | NA | 1 | NA | 0.556 | NA | 0.444 |
| SE | 8272 | ADAM9 | chr8 | + | 39071297 | 39071403 | 39055576 | 39055772 | 39077227 | 39077411 | 0 | 0 | 4 | 0 | 0 | 0 | 0 | 3 | 0 | 0 | 2 | 0 | 98 | 49 | NA | NA | NA | NA | 1 | NA | 0.429 | NA | 0.571 |
| SE | 8292 | IFI16 | chr1 | + | 159053532 | 159053724 | 159051678 | 159052098 | 159054820 | 159055155 | 0 | 0 | 18 | 0 | 0 | 0 | 1 | 9 | 0 | 0 | 4 | 0 | 98 | 49 | NA | NA | NA | NA | 1 | 1 | 0.529 | NA | 0.236 |
| SE | 8298 | EIF5 | chr14 | + | 103336035 | 103336117 | 103335652 | 103335932 | 103336676 | 103336849 | 0 | 0 | 7 | 0 | 0 | 1 | 0 | 22 | 0 | 0 | 4 | 0 | 98 | 49 | NA | NA | NA | NA | 0.778 | NA | 0.733 | NA | 0.045 |
| SE | 8317 | NUMA1 | chr11 | - | 72004639 | 72004816 | 72004224 | 72004341 | 72005232 | 72005369 | 0 | 0 | 6 | 0 | 0 | 0 | 0 | 1 | 0 | 0 | 1 | 0 | 98 | 49 | NA | NA | NA | NA | 1 | NA | 0.333 | NA | 0.667 |
| SE | 8325 | COPA | chr1 | - | 160294483 | 160294593 | 160293385 | 160293463 | 160294767 | 160294857 | 0 | 0 | 3 | 0 | 0 | 2 | 3 | 3 | 0 | 0 | 0 | 0 | 98 | 49 | NA | NA | NA | NA | 0.429 | 1 | 1 | NA | -0.571 |
| SE | 8327 | COPA | chr1 | - | 160310191 | 160310258 | 160309100 | 160309176 | 160311867 | 160312018 | 0 | 0 | 5 | 0 | 0 | 0 | 0 | 9 | 0 | 0 | 1 | 0 | 98 | 49 | NA | NA | NA | NA | 1 | NA | 0.818 | NA | 0.182 |
| SE | 8328 | COPA | chr1 | - | 160339908 | 160339982 | 160335241 | 160335322 | 160340180 | 160340294 | 0 | 0 | 7 | 0 | 0 | 0 | 1 | 3 | 0 | 0 | 1 | 0 | 98 | 49 | NA | NA | NA | NA | 1 | 1 | 0.6 | NA | 0.2 |
| SE | 8329 | WDR37 | chr10 | + | 1103601 | 1103836 | 1096169 | 1096246 | 1124217 | 1124352 | 0 | 0 | 0 | 0 | 0 | 1 | 0 | 1 | 0 | 0 | 0 | 0 | 98 | 49 | NA | NA | NA | NA | 0 | NA | 1 | NA | -1 |
| SE | 8330 | WDR37 | chr10 | + | 1105125 | 1105267 | 1103601 | 1103836 | 1124217 | 1124352 | 0 | 0 | 4 | 0 | 0 | 0 | 0 | 0 | 0 | 0 | 1 | 0 | 98 | 49 | NA | NA | NA | NA | 1 | NA | 0 | NA | 1 |
| SE | 8335 | NCSTN | chr1 | + | 160356259 | 160356347 | 160354117 | 160354290 | 160356599 | 160356754 | 0 | 0 | 4 | 0 | 0 | 1 | 0 | 5 | 0 | 0 | 0 | 0 | 98 | 49 | NA | NA | NA | NA | 0.667 | NA | 1 | NA | -0.333 |
| SE | 8341 | CHD2 | chr15 | + | 92956458 | 92956649 | 92955422 | 92955512 | 92967324 | 92967513 | 0 | 0 | 13 | 0 | 0 | 1 | 0 | 21 | 0 | 0 | 3 | 0 | 98 | 49 | NA | NA | NA | NA | 0.867 | NA | 0.778 | NA | 0.089 |
| SE | 8342 | CHD2 | chr15 | + | 92978233 | 92978383 | 92974878 | 92974950 | 92979134 | 92979283 | 0 | 0 | 3 | 0 | 0 | 0 | 0 | 4 | 0 | 0 | 1 | 0 | 98 | 49 | NA | NA | NA | NA | 1 | NA | 0.667 | NA | 0.333 |
| SE | 8346 | SF1 | chr11 | - | 64765950 | 64766155 | 64764603 | 64765518 | 64766899 | 64767079 | 0 | 0 | 1 | 0 | 0 | 0 | 0 | 0 | 0 | 0 | 1 | 0 | 98 | 49 | NA | NA | NA | NA | 1 | NA | 0 | NA | 1 |
| SE | 8357 | ARHGAP30 | chr1 | - | 161056387 | 161056532 | 161054622 | 161054705 | 161059613 | 161059716 | 0 | 0 | 26 | 0 | 0 | 3 | 0 | 13 | 0 | 0 | 2 | 0 | 98 | 49 | NA | NA | NA | NA | 0.812 | NA | 0.765 | NA | 0.047 |
| SE | 8377 | UFC1 | chr1 | + | 161158120 | 161158211 | 161157616 | 161157693 | 161158411 | 161158856 | 0 | 0 | 3 | 0 | 0 | 1 | 0 | 1 | 0 | 0 | 0 | 0 | 98 | 49 | NA | NA | NA | NA | 0.6 | NA | 1 | NA | -0.4 |
| SE | 8381 | CYFIP2 | chr5 | + | 157307760 | 157307865 | 157304237 | 157304366 | 157311663 | 157311781 | 0 | 0 | 1 | 0 | 0 | 0 | 4 | 5 | 0 | 0 | 1 | 0 | 98 | 49 | NA | NA | NA | NA | 1 | 1 | 0.714 | NA | 0.143 |
| SE | 8382 | CYFIP2 | chr5 | + | 157309742 | 157309834 | 157307760 | 157307865 | 157311663 | 157311781 | 0 | 0 | 5 | 0 | 0 | 0 | 2 | 1 | 0 | 4 | 0 | 0 | 98 | 49 | NA | NA | NA | NA | 1 | 0.2 | 1 | NA | 0.4 |
| SE | 8384 | CYFIP2 | chr5 | + | 157327972 | 157328049 | 157326170 | 157326267 | 157330741 | 157330850 | 0 | 0 | 3 | 0 | 0 | 0 | 0 | 3 | 0 | 0 | 2 | 0 | 98 | 49 | NA | NA | NA | NA | 1 | NA | 0.429 | NA | 0.571 |
| SE | 8390 | TSNAX-DISC1 | chr1 | + | 231800107 | 231800210 | 231770834 | 231771070 | 231818328 | 231818517 | 0 | 0 | 5 | 0 | 0 | 0 | 0 | 4 | 0 | 0 | 1 | 0 | 98 | 49 | NA | NA | NA | NA | 1 | NA | 0.667 | NA | 0.333 |
| SE | 8395 | ITCH | chr20 | + | 34408650 | 34408792 | 34393790 | 34393881 | 34412514 | 34412639 | 0 | 0 | 5 | 0 | 0 | 1 | 2 | 6 | 0 | 0 | 0 | 0 | 98 | 49 | NA | NA | NA | NA | 0.714 | 1 | 1 | NA | -0.286 |
| SE | 8396 | ITCH | chr20 | + | 34417116 | 34417239 | 34413741 | 34413879 | 34424479 | 34424525 | 0 | 0 | 3 | 0 | 0 | 3 | 0 | 0 | 0 | 0 | 6 | 0 | 98 | 49 | NA | NA | NA | NA | 0.333 | NA | 0 | NA | 0.333 |
| SE | 8397 | ITCH | chr20 | + | 34457389 | 34457474 | 34449410 | 34449480 | 34462092 | 34462221 | 0 | 0 | 3 | 0 | 0 | 1 | 3 | 3 | 0 | 0 | 0 | 0 | 98 | 49 | NA | NA | NA | NA | 0.6 | 1 | 1 | NA | -0.4 |
| SE | 8398 | ITCH | chr20 | + | 34470047 | 34470120 | 34462092 | 34462221 | 34471443 | 34471515 | 0 | 0 | 4 | 0 | 0 | 0 | 1 | 4 | 0 | 0 | 2 | 0 | 98 | 49 | NA | NA | NA | NA | 1 | 1 | 0.5 | NA | 0.25 |
| SE | 8422 | CAST | chr5 | + | 96730779 | 96730860 | 96729611 | 96729725 | 96736171 | 96736240 | 0 | 0 | 8 | 0 | 0 | 1 | 0 | 1 | 0 | 0 | 0 | 0 | 98 | 49 | NA | NA | NA | NA | 0.8 | NA | 1 | NA | -0.2 |
| SE | 8425 | CAST | chr5 | + | 96746341 | 96746425 | 96742654 | 96742756 | 96747344 | 96747392 | 0 | 0 | 11 | 0 | 0 | 0 | 1 | 4 | 0 | 0 | 2 | 0 | 98 | 49 | NA | NA | NA | NA | 1 | 1 | 0.5 | NA | 0.25 |
| SE | 8437 | PIK3C3 | chr18 | + | 41957569 | 41957758 | 41955197 | 41955359 | 41962488 | 41962632 | 0 | 0 | 2 | 0 | 0 | 0 | 0 | 5 | 0 | 0 | 1 | 0 | 98 | 49 | NA | NA | NA | NA | 1 | NA | 0.714 | NA | 0.286 |
| SE | 8451 | MGST3 | chr1 | + | 165651013 | 165651087 | 165649840 | 165649964 | 165651977 | 165652035 | 0 | 0 | 4 | 0 | 0 | 0 | 0 | 0 | 0 | 0 | 1 | 0 | 98 | 49 | NA | NA | NA | NA | 1 | NA | 0 | NA | 1 |
| SE | 8452 | SYF2 | chr1 | - | 25228997 | 25229123 | 25228117 | 25228235 | 25232103 | 25232211 | 0 | 0 | 1 | 0 | 0 | 0 | 0 | 1 | 0 | 0 | 1 | 0 | 98 | 49 | NA | NA | NA | NA | 1 | NA | 0.333 | NA | 0.667 |
| SE | 8470 | SAMD8 | chr10 | + | 75168540 | 75168658 | 75164644 | 75164740 | 75176065 | 75176216 | 0 | 0 | 2 | 0 | 0 | 1 | 0 | 3 | 0 | 0 | 0 | 0 | 98 | 49 | NA | NA | NA | NA | 0.5 | NA | 1 | NA | -0.5 |
| SE | 8471 | EIF3M | chr11 | + | 32587011 | 32587144 | 32583766 | 32583929 | 32588593 | 32588732 | 0 | 0 | 0 | 0 | 0 | 1 | 0 | 1 | 0 | 0 | 0 | 0 | 98 | 49 | NA | NA | NA | NA | 0 | NA | 1 | NA | -1 |
| SE | 8475 | ADCY4 | chr14 | - | 24326298 | 24326342 | 24326078 | 24326165 | 24329060 | 24329234 | 0 | 0 | 1 | 0 | 0 | 0 | 0 | 2 | 0 | 0 | 1 | 0 | 93 | 49 | NA | NA | NA | NA | 1 | NA | 0.513 | NA | 0.487 |
| SE | 8478 | HPSE | chr4 | - | 83310721 | 83310890 | 83310030 | 83310078 | 83319343 | 83319469 | 0 | 0 | 1 | 0 | 0 | 0 | 0 | 1 | 0 | 0 | 1 | 0 | 98 | 49 | NA | NA | NA | NA | 1 | NA | 0.333 | NA | 0.667 |
| SE | 8487 | PELI1 | chr2 | - | 64104700 | 64104830 | 64100397 | 64100499 | 64108239 | 64108379 | 0 | 0 | 13 | 0 | 0 | 0 | 0 | 27 | 0 | 0 | 2 | 0 | 98 | 49 | NA | NA | NA | NA | 1 | NA | 0.871 | NA | 0.129 |
| SE | 8490 | GPBP1 | chr5 | + | 57221375 | 57221396 | 57214073 | 57214193 | 57230845 | 57230969 | 0 | 0 | 3 | 0 | 0 | 5 | 0 | 8 | 0 | 0 | 0 | 0 | 70 | 49 | NA | NA | NA | NA | 0.296 | NA | 1 | NA | -0.704 |
| SE | 8527 | VPS9D1 | chr16 | - | 89708426 | 89708531 | 89707132 | 89707954 | 89708856 | 89708956 | 0 | 0 | 1 | 0 | 0 | 0 | 0 | 0 | 0 | 0 | 1 | 0 | 98 | 49 | NA | NA | NA | NA | 1 | NA | 0 | NA | 1 |
| SE | 8541 | ARAP3 | chr5 | - | 141656503 | 141656637 | 141656193 | 141656276 | 141656717 | 141656846 | 0 | 0 | 2 | 0 | 0 | 1 | 0 | 2 | 0 | 0 | 0 | 0 | 98 | 49 | NA | NA | NA | NA | 0.5 | NA | 1 | NA | -0.5 |
| SE | 8547 | TPP2 | chr13 | + | 102676295 | 102676415 | 102674282 | 102674490 | 102678226 | 102679173 | 0 | 0 | 2 | 0 | 0 | 0 | 0 | 0 | 0 | 0 | 1 | 0 | 98 | 49 | NA | NA | NA | NA | 1 | NA | 0 | NA | 1 |
| SE | 8574 | RC3H1 | chr1 | - | 173943441 | 173943588 | 173941264 | 173941380 | 173946475 | 173946608 | 0 | 0 | 2 | 0 | 0 | 3 | 0 | 0 | 0 | 0 | 3 | 0 | 98 | 49 | NA | NA | NA | NA | 0.25 | NA | 0 | NA | 0.25 |
| SE | 8575 | RC3H1 | chr1 | - | 173943441 | 173943615 | 173941264 | 173941380 | 173946475 | 173946608 | 0 | 0 | 2 | 0 | 0 | 3 | 0 | 0 | 0 | 0 | 3 | 0 | 98 | 49 | NA | NA | NA | NA | 0.25 | NA | 0 | NA | 0.25 |
| SE | 8576 | RC3H1 | chr1 | - | 173951985 | 173952138 | 173947368 | 173947582 | 173961076 | 173961244 | 0 | 0 | 9 | 0 | 0 | 1 | 0 | 5 | 0 | 0 | 0 | 0 | 98 | 49 | NA | NA | NA | NA | 0.818 | NA | 1 | NA | -0.182 |
| SE | 8618 | DGKZ | chr11 | + | 46369940 | 46370009 | 46369493 | 46369550 | 46371312 | 46371384 | 0 | 0 | 5 | 0 | 0 | 0 | 0 | 2 | 0 | 0 | 1 | 0 | 98 | 49 | NA | NA | NA | NA | 1 | NA | 0.5 | NA | 0.5 |
| SE | 8622 | RBM18 | chr9 | - | 122254273 | 122254367 | 122251846 | 122251973 | 122261379 | 122261508 | 0 | 0 | 1 | 0 | 0 | 4 | 0 | 1 | 0 | 0 | 0 | 0 | 98 | 49 | NA | NA | NA | NA | 0.111 | NA | 1 | NA | -0.889 |
| SE | 8632 | SMG7 | chr1 | + | 183538379 | 183538440 | 183537144 | 183537215 | 183540983 | 183541103 | 0 | 0 | 1 | 0 | 0 | 0 | 0 | 2 | 0 | 0 | 1 | 0 | 98 | 49 | NA | NA | NA | NA | 1 | NA | 0.5 | NA | 0.5 |
| SE | 8722 | NOD2 | chr16 | + | 50719924 | 50720008 | 50716890 | 50716974 | 50722621 | 50722705 | 0 | 0 | 3 | 0 | 0 | 1 | 0 | 1 | 0 | 0 | 0 | 0 | 98 | 49 | NA | NA | NA | NA | 0.6 | NA | 1 | NA | -0.4 |
| SE | 8737 | KDM5B | chr1 | - | 202746141 | 202746323 | 202745857 | 202745982 | 202748944 | 202749139 | 0 | 0 | 1 | 0 | 0 | 0 | 0 | 7 | 0 | 0 | 1 | 0 | 98 | 49 | NA | NA | NA | NA | 1 | NA | 0.778 | NA | 0.222 |
| SE | 8744 | BEST1 | chr11 | + | 61958145 | 61958298 | 61957386 | 61957464 | 61959497 | 61959578 | 0 | 0 | 3 | 0 | 0 | 0 | 0 | 1 | 0 | 0 | 1 | 0 | 98 | 49 | NA | NA | NA | NA | 1 | NA | 0.333 | NA | 0.667 |
| SE | 8752 | OS9 | chr12 | + | 57720098 | 57720263 | 57718992 | 57719182 | 57720405 | 57720518 | 0 | 0 | 2 | 0 | 0 | 3 | 2 | 1 | 0 | 1 | 2 | 0 | 98 | 49 | NA | NA | NA | NA | 0.25 | 0.5 | 0.2 | NA | -0.1 |
| SE | 8763 | MITD1 | chr2 | - | 99171342 | 99171429 | 99170536 | 99170652 | 99171509 | 99171646 | 0 | 0 | 2 | 0 | 0 | 0 | 0 | 0 | 0 | 0 | 1 | 0 | 98 | 49 | NA | NA | NA | NA | 1 | NA | 0 | NA | 1 |
| SE | 8770 | YY1 | chr14 | + | 100276489 | 100276648 | 100274697 | 100274758 | 100277417 | 100279034 | 0 | 0 | 2 | 0 | 0 | 1 | 8 | 1 | 0 | 0 | 0 | 0 | 98 | 49 | NA | NA | NA | NA | 0.5 | 1 | 1 | NA | -0.5 |
| SE | 8791 | VPS13D | chr1 | + | 12319496 | 12319630 | 12318071 | 12318337 | 12321808 | 12321964 | 0 | 0 | 2 | 0 | 0 | 0 | 0 | 3 | 0 | 0 | 1 | 0 | 98 | 49 | NA | NA | NA | NA | 1 | NA | 0.6 | NA | 0.4 |
| SE | 8793 | VPS13D | chr1 | + | 12349163 | 12349374 | 12348822 | 12348973 | 12353973 | 12354221 | 0 | 0 | 3 | 0 | 0 | 1 | 0 | 2 | 0 | 0 | 0 | 0 | 98 | 49 | NA | NA | NA | NA | 0.6 | NA | 1 | NA | -0.4 |
| SE | 8794 | VPS13D | chr1 | + | 12358458 | 12358601 | 12356397 | 12356524 | 12362719 | 12362850 | 0 | 0 | 1 | 0 | 0 | 0 | 0 | 1 | 0 | 0 | 2 | 0 | 98 | 49 | NA | NA | NA | NA | 1 | NA | 0.2 | NA | 0.8 |
| SE | 8797 | CD46 | chr1 | + | 207767606 | 207767651 | 207767012 | 207767195 | 207767778 | 207767823 | 0 | 0 | 7 | 0 | 0 | 0 | 0 | 2 | 0 | 0 | 2 | 0 | 94 | 49 | NA | NA | NA | NA | 1 | NA | 0.343 | NA | 0.657 |
| SE | 8814 | KDM2A | chr11 | + | 67121233 | 67121358 | 67119268 | 67120049 | 67180078 | 67180217 | 0 | 0 | 11 | 0 | 0 | 0 | 1 | 5 | 0 | 0 | 2 | 0 | 98 | 49 | NA | NA | NA | NA | 1 | 1 | 0.556 | NA | 0.222 |
| SE | 8815 | KDM2A | chr11 | + | 67181319 | 67181398 | 67180078 | 67180217 | 67181845 | 67181892 | 0 | 0 | 7 | 0 | 0 | 1 | 0 | 1 | 0 | 0 | 0 | 0 | 98 | 49 | NA | NA | NA | NA | 0.778 | NA | 1 | NA | -0.222 |
| SE | 8816 | KDM2A | chr11 | + | 67215855 | 67215949 | 67215339 | 67215446 | 67217730 | 67217884 | 0 | 0 | 5 | 0 | 0 | 1 | 0 | 6 | 0 | 0 | 2 | 0 | 98 | 49 | NA | NA | NA | NA | 0.714 | NA | 0.6 | NA | 0.114 |
| SE | 8826 | HIRA | chr22 | - | 19387710 | 19387816 | 19385520 | 19385736 | 19388483 | 19388554 | 0 | 0 | 8 | 0 | 0 | 2 | 0 | 5 | 0 | 0 | 0 | 0 | 98 | 49 | NA | NA | NA | NA | 0.667 | NA | 1 | NA | -0.333 |
| SE | 8832 | 7-Mar | chr2 | + | 159762879 | 159762993 | 159759225 | 159759335 | 159764625 | 159764674 | 0 | 0 | 8 | 0 | 0 | 0 | 0 | 0 | 0 | 0 | 1 | 0 | 98 | 49 | NA | NA | NA | NA | 1 | NA | 0 | NA | 1 |
| SE | 8844 | GNL2 | chr1 | - | 37574664 | 37574823 | 37574342 | 37574456 | 37575594 | 37575699 | 0 | 0 | 1 | 0 | 0 | 1 | 0 | 3 | 0 | 0 | 0 | 0 | 98 | 49 | NA | NA | NA | NA | 0.333 | NA | 1 | NA | -0.667 |
| SE | 8871 | C1RL | chr12 | - | 7099900 | 7100026 | 7099685 | 7099760 | 7101897 | 7102087 | 1 | 0 | 6 | 0 | 0 | 0 | 0 | 0 | 0 | 0 | 1 | 0 | 98 | 49 | NA | NA | 1 | NA | 1 | NA | 0 | NA | 1 |
| SE | 8883 | FAM160B1 | chr10 | + | 114848646 | 114848737 | 114847089 | 114847233 | 114855196 | 114855340 | 0 | 0 | 3 | 0 | 0 | 0 | 0 | 4 | 0 | 0 | 1 | 0 | 98 | 49 | NA | NA | NA | NA | 1 | NA | 0.667 | NA | 0.333 |
| SE | 8908 | WDR11 | chr10 | + | 120867065 | 120867169 | 120866568 | 120866764 | 120871169 | 120871346 | 0 | 0 | 3 | 0 | 0 | 1 | 0 | 3 | 0 | 0 | 0 | 0 | 98 | 49 | NA | NA | NA | NA | 0.6 | NA | 1 | NA | -0.4 |
| SE | 8922 | SMARCC1 | chr3 | - | 47720665 | 47720735 | 47714414 | 47714490 | 47729024 | 47729094 | 0 | 0 | 2 | 0 | 0 | 0 | 0 | 4 | 0 | 0 | 1 | 0 | 98 | 49 | NA | NA | NA | NA | 1 | NA | 0.667 | NA | 0.333 |
| SE | 8923 | SMARCC1 | chr3 | - | 47729024 | 47729094 | 47720665 | 47720735 | 47736033 | 47736126 | 0 | 0 | 2 | 0 | 0 | 0 | 0 | 5 | 0 | 0 | 1 | 0 | 98 | 49 | NA | NA | NA | NA | 1 | NA | 0.714 | NA | 0.286 |
| SE | 8937 | CD164 | chr6 | - | 109370410 | 109370467 | 109366513 | 109369017 | 109376073 | 109376112 | 0 | 0 | 18 | 0 | 0 | 1 | 0 | 7 | 0 | 0 | 0 | 0 | 98 | 49 | NA | NA | NA | NA | 0.9 | NA | 1 | NA | -0.1 |
| SE | 8955 | REV1 | chr2 | - | 99442316 | 99442469 | 99438600 | 99439310 | 99449335 | 99449504 | 0 | 0 | 5 | 0 | 0 | 0 | 0 | 1 | 0 | 0 | 1 | 0 | 98 | 49 | NA | NA | NA | NA | 1 | NA | 0.333 | NA | 0.667 |
| SE | 8976 | KPNB1 | chr17 | + | 47676408 | 47676491 | 47674637 | 47674782 | 47677019 | 47677127 | 0 | 0 | 13 | 0 | 0 | 0 | 0 | 4 | 0 | 0 | 1 | 0 | 98 | 49 | NA | NA | NA | NA | 1 | NA | 0.667 | NA | 0.333 |
| SE | 8978 | C18orf25 | chr18 | + | 46240004 | 46240187 | 46215810 | 46216595 | 46253696 | 46253823 | 0 | 0 | 2 | 0 | 0 | 0 | 0 | 0 | 0 | 0 | 1 | 0 | 98 | 49 | NA | NA | NA | NA | 1 | NA | 0 | NA | 1 |
| SE | 8992 | BSG | chr19 | + | 579499 | 579656 | 572453 | 572701 | 580378 | 580461 | 0 | 0 | 5 | 0 | 0 | 0 | 0 | 1 | 0 | 1 | 0 | 0 | 98 | 49 | NA | NA | NA | NA | 1 | 0 | 1 | NA | 0.5 |
| SE | 8997 | PANK4 | chr1 | - | 2512887 | 2513039 | 2511627 | 2511683 | 2514001 | 2514089 | 0 | 0 | 4 | 0 | 0 | 0 | 0 | 0 | 0 | 0 | 1 | 0 | 98 | 49 | NA | NA | NA | NA | 1 | NA | 0 | NA | 1 |
| SE | 9002 | AKAP10 | chr17 | - | 19940886 | 19941010 | 19939712 | 19939849 | 19941825 | 19941910 | 0 | 0 | 4 | 0 | 0 | 4 | 0 | 3 | 0 | 0 | 1 | 0 | 98 | 49 | NA | NA | NA | NA | 0.333 | NA | 0.6 | NA | -0.267 |
| SE | 9045 | FNDC3A | chr13 | + | 49191014 | 49191120 | 49188514 | 49188633 | 49191208 | 49191384 | 0 | 0 | 1 | 0 | 0 | 0 | 0 | 0 | 0 | 0 | 1 | 0 | 98 | 49 | NA | NA | NA | NA | 1 | NA | 0 | NA | 1 |
| SE | 9047 | FAM65B | chr6 | - | 24835702 | 24835871 | 24832255 | 24832391 | 24839090 | 24839272 | 0 | 0 | 36 | 0 | 0 | 0 | 0 | 3 | 0 | 0 | 1 | 0 | 98 | 49 | NA | NA | NA | NA | 1 | NA | 0.6 | NA | 0.4 |
| SE | 9048 | FAM65B | chr6 | - | 24839090 | 24839272 | 24835702 | 24835871 | 24842861 | 24843554 | 0 | 0 | 19 | 0 | 0 | 0 | 0 | 4 | 0 | 0 | 1 | 0 | 98 | 49 | NA | NA | NA | NA | 1 | NA | 0.667 | NA | 0.333 |
| SE | 9050 | MPPE1 | chr18 | - | 11884878 | 11885043 | 11883470 | 11884627 | 11885675 | 11885816 | 0 | 0 | 3 | 0 | 0 | 2 | 0 | 1 | 0 | 0 | 0 | 0 | 98 | 49 | NA | NA | NA | NA | 0.429 | NA | 1 | NA | -0.571 |
| SE | 9095 | HPS1 | chr10 | - | 98429790 | 98429889 | 98429145 | 98429642 | 98430570 | 98430670 | 0 | 0 | 1 | 0 | 0 | 1 | 0 | 1 | 0 | 0 | 0 | 0 | 98 | 49 | NA | NA | NA | NA | 0.333 | NA | 1 | NA | -0.667 |
| SE | 9112 | ADGRE3 | chr19 | - | 14658512 | 14658550 | 14654981 | 14655165 | 14661962 | 14662118 | 0 | 0 | 13 | 0 | 0 | 1 | 0 | 6 | 0 | 0 | 0 | 0 | 87 | 49 | NA | NA | NA | NA | 0.88 | NA | 1 | NA | -0.12 |
| SE | 9119 | AGO1 | chr1 | + | 35906934 | 35907119 | 35902203 | 35902337 | 35913841 | 35914001 | 0 | 0 | 2 | 0 | 0 | 1 | 0 | 2 | 0 | 0 | 0 | 0 | 98 | 49 | NA | NA | NA | NA | 0.5 | NA | 1 | NA | -0.5 |
| SE | 9149 | ADD3 | chr10 | + | 110116258 | 110116410 | 110112776 | 110112915 | 110117341 | 110117422 | 0 | 0 | 7 | 0 | 0 | 1 | 0 | 5 | 0 | 0 | 0 | 0 | 98 | 49 | NA | NA | NA | NA | 0.778 | NA | 1 | NA | -0.222 |
| SE | 9150 | ADD3 | chr10 | + | 110132304 | 110132400 | 110130362 | 110130486 | 110133325 | 110135565 | 0 | 0 | 9 | 0 | 0 | 2 | 0 | 7 | 0 | 0 | 0 | 0 | 98 | 49 | NA | NA | NA | NA | 0.692 | NA | 1 | NA | -0.308 |
| SE | 9172 | NDRG1 | chr8 | - | 133258365 | 133258426 | 133256776 | 133256863 | 133259167 | 133259230 | 0 | 0 | 6 | 0 | 0 | 1 | 0 | 6 | 0 | 0 | 0 | 0 | 98 | 49 | NA | NA | NA | NA | 0.75 | NA | 1 | NA | -0.25 |
| SE | 9186 | PPP2R5D | chr6 | + | 43008692 | 43008746 | 43008366 | 43008475 | 43009056 | 43009227 | 0 | 0 | 2 | 0 | 0 | 0 | 0 | 0 | 0 | 0 | 1 | 0 | 98 | 49 | NA | NA | NA | NA | 1 | NA | 0 | NA | 1 |
| SE | 9187 | GIT2 | chr12 | - | 109945259 | 109945349 | 109939164 | 109939247 | 109947255 | 109947504 | 0 | 0 | 0 | 0 | 0 | 1 | 0 | 1 | 0 | 0 | 0 | 0 | 98 | 49 | NA | NA | NA | NA | 0 | NA | 1 | NA | -1 |
| SE | 9200 | PIP5K1A | chr1 | + | 151242122 | 151242269 | 151239954 | 151240039 | 151242437 | 151242567 | 0 | 0 | 0 | 0 | 0 | 1 | 0 | 2 | 0 | 0 | 0 | 0 | 98 | 49 | NA | NA | NA | NA | 0 | NA | 1 | NA | -1 |
| SE | 9202 | ATP5C1 | chr10 | + | 7802757 | 7802854 | 7802269 | 7802425 | 7806973 | 7807010 | 0 | 0 | 6 | 0 | 0 | 1 | 1 | 1 | 0 | 0 | 0 | 0 | 98 | 49 | NA | NA | NA | NA | 0.75 | 1 | 1 | NA | -0.25 |
| SE | 9210 | CELF2 | chr10 | + | 11266597 | 11266677 | 11257737 | 11257872 | 11270665 | 11270824 | 0 | 0 | 15 | 0 | 0 | 5 | 1 | 8 | 0 | 0 | 2 | 0 | 98 | 49 | NA | NA | NA | NA | 0.6 | 1 | 0.667 | NA | -0.234 |
| SE | 9213 | UPF2 | chr10 | - | 12001675 | 12001825 | 11999905 | 12000009 | 12004529 | 12004727 | 0 | 0 | 5 | 0 | 0 | 0 | 0 | 2 | 0 | 0 | 1 | 0 | 98 | 49 | NA | NA | NA | NA | 1 | NA | 0.5 | NA | 0.5 |
| SE | 9215 | PIGG | chr4 | + | 508828 | 508970 | 507404 | 507593 | 515972 | 516185 | 0 | 0 | 2 | 0 | 0 | 0 | 0 | 2 | 0 | 3 | 0 | 0 | 98 | 49 | NA | NA | NA | NA | 1 | 0 | 1 | NA | 0.5 |
| SE | 9227 | SRSF6 | chr20 | + | 43459152 | 43459420 | 43458360 | 43458509 | 43459770 | 43459895 | 0 | 0 | 0 | 0 | 0 | 1 | 0 | 2 | 0 | 0 | 0 | 0 | 98 | 49 | NA | NA | NA | NA | 0 | NA | 1 | NA | -1 |
| SE | 9251 | VIM-AS1 | chr10 | - | 17216166 | 17216274 | 17214238 | 17215364 | 17229344 | 17229985 | 0 | 0 | 1 | 0 | 0 | 0 | 0 | 0 | 0 | 2 | 0 | 0 | 98 | 49 | NA | NA | NA | NA | 1 | 0 | NA | NA | 1 |
| SE | 9257 | SERPINB1 | chr6 | - | 2839334 | 2839466 | 2838548 | 2838686 | 2840418 | 2840594 | 0 | 0 | 2 | 0 | 0 | 7 | 0 | 1 | 0 | 5 | 1 | 0 | 98 | 49 | NA | NA | NA | NA | 0.125 | 0 | 0.333 | NA | -0.042 |

| | | | | | | | | | | | | | | | | | | | | | | | | | | | | | | | | | |
|---|---|---|---|---|---|---|---|---|---|---|---|---|---|---|---|---|---|---|---|---|---|---|---|---|---|---|---|---|---|---|---|---|---|
| SE | 9261 | ATP5L | chr11 | + | 118405656 | 118405709 | 118401388 | 118401717 | 118406936 | 118407097 | 0 | 0 | 2 | 0 | 0 | 0 | 0 | 0 | 0 | 0 | 1 | 0 | 98 | 49 | NA | NA | NA | NA | 1 | NA | 0 | NA | 1 |
| SE | 9264 | PLXDC2 | chr10 | + | 20068169 | 20068239 | 20046868 | 20047015 | 20143294 | 20143417 | 0 | 0 | 21 | 0 | 0 | 1 | 0 | 3 | 0 | 0 | 0 | 0 | 98 | 49 | NA | NA | NA | NA | 0.913 | NA | 1 | NA | -0.087 |
| SE | 9267 | CDC14A | chr1 | + | 100455404 | 100455492 | 100442933 | 100442996 | 100462650 | 100462881 | 0 | 0 | 1 | 0 | 0 | 1 | 5 | 1 | 0 | 0 | 0 | 0 | 98 | 49 | NA | NA | NA | NA | 0.333 | 1 | 1 | NA | -0.667 |
| SE | 9283 | RANBP2 | chr2 | + | 108762100 | 108762195 | 108758412 | 108758548 | 108763236 | 108768388 | 0 | 0 | 1 | 0 | 0 | 0 | 0 | 3 | 0 | 0 | 3 | 0 | 98 | 49 | NA | NA | NA | NA | 1 | NA | 0.333 | NA | 0.667 |
| SE | 9284 | RANBP2 | chr2 | + | 108777129 | 108777231 | 108775731 | 108775936 | 108781268 | 108781429 | 0 | 0 | 1 | 0 | 0 | 0 | 0 | 7 | 0 | 0 | 1 | 0 | 98 | 49 | NA | NA | NA | NA | 1 | NA | 0.778 | NA | 0.222 |
| SE | 9294 | ABCA7 | chr19 | + | 1056906 | 1057084 | 1056329 | 1056499 | 1057313 | 1057429 | 0 | 0 | 4 | 0 | 0 | 0 | 0 | 1 | 0 | 0 | 2 | 0 | 98 | 49 | NA | NA | NA | NA | 1 | NA | 0.2 | NA | 0.8 |
| SE | 9358 | RASGRP1 | chr15 | - | 38503271 | 38503376 | 38502311 | 38502421 | 38505839 | 38505920 | 0 | 0 | 2 | 0 | 0 | 0 | 0 | 1 | 0 | 0 | 1 | 0 | 98 | 49 | NA | NA | NA | NA | 1 | NA | 0.333 | NA | 0.667 |
| SE | 9360 | EIF4G3 | chr1 | - | 20892618 | 20892729 | 20886200 | 20886371 | 20893516 | 20893636 | 0 | 0 | 0 | 0 | 0 | 1 | 0 | 2 | 0 | 0 | 2 | 0 | 98 | 49 | NA | NA | NA | NA | 0 | NA | 0.333 | NA | -0.333 |
| SE | 9382 | PECAM1 | chr17 | - | 64353490 | 64353518 | 64352389 | 64352463 | 64356110 | 64356398 | 0 | 0 | 20 | 0 | 0 | 0 | 0 | 3 | 0 | 0 | 1 | 0 | 77 | 49 | NA | NA | NA | NA | 1 | NA | 0.656 | NA | 0.344 |
| SE | 9388 | NCOA4 | chr10 | - | 46016539 | 46016694 | 46015125 | 46015266 | 46030525 | 46030714 | 0 | 0 | 5 | 0 | 0 | 0 | 5 | 3 | 0 | 4 | 0 | 0 | 98 | 49 | NA | NA | NA | NA | 1 | 0.385 | 1 | NA | 0.307 |
| SE | 9445 | CCNDBP1 | chr15 | + | 43186153 | 43186233 | 43185819 | 43185879 | 43189198 | 43189280 | 0 | 0 | 10 | 0 | 0 | 0 | 0 | 5 | 0 | 0 | 1 | 0 | 98 | 49 | NA | NA | NA | NA | 1 | NA | 0.714 | NA | 0.286 |
| SE | 9446 | CCNDBP1 | chr15 | + | 43190054 | 43190151 | 43189198 | 43189280 | 43190324 | 43190398 | 0 | 0 | 7 | 0 | 0 | 5 | 0 | 2 | 0 | 0 | 3 | 0 | 98 | 49 | NA | NA | NA | NA | 0.412 | NA | 0.25 | NA | 0.162 |
| SE | 9460 | TFAM | chr10 | + | 58390764 | 58390860 | 58388669 | 58388819 | 58394357 | 58394414 | 0 | 0 | 1 | 0 | 0 | 0 | 0 | 1 | 0 | 0 | 1 | 0 | 98 | 49 | NA | NA | NA | NA | 1 | NA | 0.333 | NA | 0.667 |
| SE | 9471 | USP33 | chr1 | - | 77701368 | 77701471 | 77697862 | 77697931 | 77711746 | 77711855 | 0 | 0 | 5 | 0 | 0 | 2 | 0 | 2 | 0 | 0 | 1 | 0 | 98 | 49 | NA | NA | NA | NA | 0.556 | NA | 0.5 | NA | 0.056 |
| SE | 9495 | SAP30BP | chr17 | + | 75690670 | 75690772 | 75671815 | 75671863 | 75693439 | 75693482 | 0 | 0 | 0 | 0 | 0 | 1 | 0 | 1 | 0 | 0 | 6 | 0 | 98 | 49 | NA | NA | NA | NA | 0 | NA | 0.077 | NA | -0.077 |
| SE | 9501 | CCAR1 | chr10 | + | 68771205 | 68771445 | 68765887 | 68766079 | 68772987 | 68773099 | 0 | 0 | 2 | 0 | 0 | 0 | 0 | 0 | 0 | 0 | 1 | 0 | 98 | 49 | NA | NA | NA | NA | 1 | NA | 0 | NA | 1 |
| SE | 9518 | SMG5 | chr1 | - | 156273360 | 156273450 | 156272319 | 156272398 | 156274596 | 156274686 | 0 | 0 | 3 | 0 | 0 | 1 | 0 | 2 | 0 | 0 | 0 | 0 | 98 | 49 | NA | NA | NA | NA | 0.6 | NA | 1 | NA | -0.4 |
| SE | 9531 | PPP3CB | chr10 | - | 73439871 | 73439901 | 73436427 | 73438420 | 73444724 | 73444822 | 0 | 0 | 1 | 0 | 0 | 2 | 0 | 1 | 0 | 0 | 0 | 0 | 79 | 49 | NA | NA | NA | NA | 0.237 | NA | 1 | NA | -0.763 |
| SE | 9534 | CAMK2G | chr10 | - | 73819531 | 73819645 | 73817478 | 73817554 | 73821681 | 73821730 | 0 | 0 | 2 | 0 | 0 | 0 | 0 | 0 | 0 | 0 | 1 | 0 | 98 | 49 | NA | NA | NA | NA | 1 | NA | 0 | NA | 1 |
| SE | 9539 | CAMK2G | chr10 | - | 73848525 | 73848609 | 73847987 | 73848082 | 73849012 | 73849115 | 0 | 0 | 2 | 0 | 0 | 2 | 0 | 4 | 0 | 0 | 4 | 0 | 98 | 49 | NA | NA | NA | NA | 0.333 | NA | 0.333 | NA | 0 |
| SE | 9557 | SENP7 | chr3 | - | 101417592 | 101417790 | 101398860 | 101399055 | 101458954 | 101459052 | 0 | 0 | 0 | 0 | 0 | 1 | 0 | 1 | 0 | 0 | 0 | 0 | 98 | 49 | NA | NA | NA | NA | 0 | NA | 1 | NA | -1 |
| SE | 9572 | FAS | chr10 | + | 89007699 | 89007837 | 89003028 | 89003194 | 89010538 | 89010600 | 0 | 0 | 1 | 0 | 0 | 2 | 0 | 1 | 0 | 0 | 0 | 0 | 98 | 49 | NA | NA | NA | NA | 0.2 | NA | 1 | NA | -0.8 |
| SE | 9652 | BSDC1 | chr1 | - | 32383829 | 32383997 | 32381213 | 32381268 | 32386778 | 32386895 | 0 | 0 | 1 | 0 | 0 | 1 | 0 | 2 | 0 | 0 | 0 | 0 | 98 | 49 | NA | NA | NA | NA | 0.333 | NA | 1 | NA | -0.667 |
| SE | 9659 | TTLL3 | chr3 | + | 9833103 | 9833245 | 9828959 | 9829395 | 9834680 | 9834907 | 0 | 0 | 2 | 0 | 0 | 0 | 0 | 0 | 0 | 5 | 1 | 0 | 98 | 49 | NA | NA | NA | NA | 1 | 0 | 0 | NA | 1 |
| SE | 9667 | PTEN | chr10 | + | 87952117 | 87952259 | 87933012 | 87933251 | 87957852 | 87958019 | 0 | 0 | 4 | 0 | 0 | 1 | 0 | 2 | 0 | 0 | 1 | 0 | 98 | 49 | NA | NA | NA | NA | 0.667 | NA | 0.5 | NA | 0.167 |
| SE | 9688 | IFRD1 | chr7 | + | 112458860 | 112459018 | 112456913 | 112457038 | 112461865 | 112461916 | 0 | 0 | 42 | 0 | 0 | 0 | 0 | 32 | 0 | 0 | 3 | 0 | 98 | 49 | NA | NA | NA | NA | 1 | NA | 0.842 | NA | 0.158 |
| SE | 9697 | ZNF410 | chr14 | + | 73922065 | 73922206 | 73920979 | 73921105 | 73923394 | 73923522 | 0 | 0 | 2 | 0 | 0 | 4 | 0 | 2 | 0 | 0 | 1 | 0 | 98 | 49 | NA | NA | NA | NA | 0.2 | NA | 0.5 | NA | -0.3 |
| SE | 9700 | AAED1 | chr9 | - | 96645892 | 96646024 | 96641250 | 96641886 | 96651389 | 96651495 | 0 | 0 | 1 | 0 | 0 | 1 | 0 | 0 | 0 | 0 | 1 | 0 | 98 | 49 | NA | NA | NA | NA | 0.333 | NA | 0 | NA | 0.333 |
| SE | 9744 | CLEC2D | chr12 | + | 9693033 | 9693115 | 9692827 | 9692931 | 9694759 | 9699555 | 0 | 0 | 1 | 0 | 0 | 2 | 0 | 1 | 0 | 0 | 0 | 0 | 98 | 49 | NA | NA | NA | NA | 0.2 | NA | 1 | NA | -0.8 |
| SE | 9745 | FBXO22 | chr15 | + | 75904490 | 75904629 | 75903858 | 75904103 | 75913202 | 75913290 | 0 | 0 | 3 | 0 | 0 | 0 | 0 | 0 | 0 | 0 | 1 | 0 | 98 | 49 | NA | NA | NA | NA | 1 | NA | 0 | NA | 1 |
| SE | 9766 | SAFB | chr19 | + | 5648988 | 5649499 | 5648015 | 5648043 | 5649925 | 5649975 | 0 | 0 | 1 | 0 | 0 | 3 | 0 | 0 | 0 | 0 | 1 | 0 | 98 | 49 | NA | NA | NA | NA | 0.143 | NA | 0 | NA | 0.143 |
| SE | 9767 | PI4K2A | chr10 | + | 97664884 | 97664984 | 97662906 | 97662968 | 97666437 | 97666571 | 0 | 0 | 1 | 0 | 0 | 0 | 0 | 1 | 0 | 0 | 1 | 0 | 98 | 49 | NA | NA | NA | NA | 1 | NA | 0.333 | NA | 0.667 |
| SE | 9773 | FRYL | chr4 | - | 48549472 | 48549623 | 48548689 | 48548793 | 48550591 | 48550704 | 0 | 0 | 2 | 0 | 0 | 0 | 0 | 0 | 0 | 0 | 1 | 0 | 98 | 49 | NA | NA | NA | NA | 1 | NA | 0 | NA | 1 |
| SE | 9774 | FRYL | chr4 | - | 48609743 | 48609823 | 48608986 | 48609067 | 48619273 | 48619370 | 0 | 0 | 3 | 0 | 0 | 1 | 0 | 1 | 0 | 0 | 0 | 0 | 98 | 49 | NA | NA | NA | NA | 0.6 | NA | 1 | NA | -0.4 |
| SE | 9775 | FRYL | chr4 | - | 48710518 | 48710698 | 48684672 | 48684795 | 48780077 | 48780299 | 0 | 0 | 1 | 0 | 0 | 0 | 0 | 1 | 0 | 0 | 1 | 0 | 98 | 49 | NA | NA | NA | NA | 1 | NA | 0.333 | NA | 0.667 |
| SE | 9776 | FAM188A | chr10 | - | 15838227 | 15838279 | 15837203 | 15837318 | 15841425 | 15841599 | 0 | 0 | 0 | 0 | 0 | 1 | 0 | 1 | 0 | 0 | 1 | 0 | 98 | 49 | NA | NA | NA | NA | 0 | NA | 0.333 | NA | -0.333 |
| SE | 9784 | DCAF6 | chr1 | + | 168038370 | 168038488 | 168004532 | 168004793 | 168043024 | 168043140 | 0 | 0 | 0 | 0 | 0 | 1 | 0 | 1 | 0 | 0 | 0 | 0 | 98 | 49 | NA | NA | NA | NA | 0 | NA | 1 | NA | -1 |
| SE | 9789 | HERC4 | chr10 | - | 68044403 | 68044563 | 68038092 | 68038169 | 68072882 | 68073186 | 0 | 0 | 1 | 0 | 0 | 0 | 0 | 1 | 0 | 0 | 1 | 0 | 98 | 49 | NA | NA | NA | NA | 1 | NA | 0.333 | NA | 0.667 |
| SE | 9795 | CHUK | chr10 | - | 100194424 | 100194521 | 100193983 | 100194131 | 100199970 | 100200020 | 0 | 0 | 2 | 0 | 0 | 2 | 0 | 1 | 0 | 0 | 0 | 0 | 98 | 49 | NA | NA | NA | NA | 0.333 | NA | 1 | NA | -0.667 |
| SE | 9796 | CHUK | chr10 | - | 100219007 | 100219132 | 100218717 | 100218825 | 100219269 | 100219359 | 0 | 0 | 2 | 0 | 0 | 1 | 0 | 0 | 0 | 0 | 2 | 0 | 98 | 49 | NA | NA | NA | NA | 0.5 | NA | 0 | NA | 0.5 |
| SE | 9830 | NFAT5 | chr16 | + | 69568494 | 69568548 | 69565965 | 69566374 | 69626402 | 69626528 | 0 | 0 | 5 | 0 | 0 | 1 | 0 | 1 | 0 | 0 | 0 | 0 | 98 | 49 | NA | NA | NA | NA | 0.714 | NA | 1 | NA | -0.286 |
| SE | 9856 | ZFAND1 | chr8 | - | 81717248 | 81717288 | 81714986 | 81715114 | 81718181 | 81718224 | 0 | 0 | 2 | 0 | 0 | 1 | 0 | 1 | 0 | 0 | 0 | 0 | 89 | 49 | NA | NA | NA | NA | 0.524 | NA | 1 | NA | -0.476 |
| SE | 9899 | C2CD5 | chr12 | - | 22459491 | 22459542 | 22456970 | 22457161 | 22469708 | 22469795 | 0 | 0 | 0 | 0 | 0 | 1 | 0 | 1 | 0 | 0 | 1 | 0 | 98 | 49 | NA | NA | NA | NA | 0 | NA | 0.333 | NA | -0.333 |
| SE | 9910 | IL13RA1 | chrX | + | 118766843 | 118766976 | 118766529 | 118766577 | 118773878 | 118773975 | 0 | 0 | 8 | 0 | 0 | 0 | 0 | 3 | 0 | 0 | 1 | 0 | 98 | 49 | NA | NA | NA | NA | 1 | NA | 0.6 | NA | 0.4 |
| SE | 9912 | CASP8 | chr2 | + | 201271515 | 201271621 | 201266460 | 201266791 | 201272637 | 201272776 | 0 | 0 | 9 | 0 | 0 | 1 | 0 | 17 | 0 | 0 | 3 | 0 | 98 | 49 | NA | NA | NA | NA | 0.818 | NA | 0.739 | NA | 0.079 |
| SE | 9914 | CASP8 | chr2 | + | 201272897 | 201272942 | 201272637 | 201272776 | 201276826 | 201276968 | 0 | 0 | 6 | 0 | 0 | 1 | 0 | 18 | 0 | 0 | 0 | 0 | 94 | 49 | NA | NA | NA | NA | 0.758 | NA | 1 | NA | -0.242 |
| SE | 9915 | CASP8 | chr2 | + | 201274888 | 201274953 | 201272637 | 201272776 | 201276826 | 201276968 | 0 | 0 | 3 | 0 | 0 | 1 | 0 | 3 | 0 | 0 | 0 | 0 | 98 | 49 | NA | NA | NA | NA | 0.6 | NA | 1 | NA | -0.4 |
| SE | 9940 | ZRANB2 | chr1 | - | 71065677 | 71065752 | 71063290 | 71065137 | 71066775 | 71066934 | 0 | 0 | 1 | 0 | 0 | 0 | 5 | 0 | 0 | 0 | 1 | 0 | 98 | 49 | NA | NA | NA | NA | 1 | 1 | 0 | NA | 0.5 |
| SE | 9989 | TRANK1 | chr3 | - | 36855172 | 36858049 | 36852145 | 36852345 | 36858717 | 36858894 | 0 | 0 | 15 | 0 | 0 | 2 | 0 | 3 | 0 | 0 | 0 | 0 | 98 | 49 | NA | NA | NA | NA | 0.789 | NA | 1 | NA | -0.211 |
| SE | 9994 | UPP1 | chr7 | + | 48099669 | 48099787 | 48094762 | 48094827 | 48101823 | 48101982 | 0 | 0 | 1 | 0 | 0 | 0 | 0 | 3 | 0 | 0 | 2 | 0 | 98 | 49 | NA | NA | NA | NA | 1 | NA | 0.429 | NA | 0.571 |
| SE | 10044 | KIAA1468 | chr18 | + | 62280359 | 62280442 | 62275373 | 62275473 | 62280645 | 62280709 | 0 | 0 | 1 | 0 | 0 | 1 | 0 | 1 | 0 | 0 | 0 | 0 | 98 | 49 | NA | NA | NA | NA | 0.333 | NA | 1 | NA | -0.667 |
| SE | 10047 | CHURC1 | chr14 | + | 64926006 | 64926080 | 64923990 | 64924126 | 64932137 | 64935366 | 0 | 0 | 2 | 0 | 0 | 2 | 0 | 1 | 0 | 0 | 1 | 0 | 98 | 49 | NA | NA | NA | NA | 0.333 | NA | 0.333 | NA | 0 |
| SE | 10048 | CHURC1 | chr14 | + | 64926009 | 64926080 | 64923990 | 64924126 | 64932137 | 64935366 | 0 | 0 | 2 | 0 | 0 | 2 | 0 | 2 | 0 | 0 | 1 | 0 | 98 | 49 | NA | NA | NA | NA | 0.333 | NA | 0.5 | NA | -0.167 |
| SE | 10059 | TRRAP | chr7 | + | 98961260 | 98961474 | 98959343 | 98959490 | 98962301 | 98962427 | 0 | 0 | 1 | 0 | 0 | 0 | 0 | 0 | 0 | 0 | 1 | 0 | 98 | 49 | NA | NA | NA | NA | 1 | NA | 0 | NA | 1 |
| SE | 10074 | SRSF5 | chr14 | + | 69768797 | 69768896 | 69768603 | 69768674 | 69769181 | 69769251 | 0 | 0 | 20 | 0 | 0 | 0 | 0 | 26 | 0 | 0 | 2 | 0 | 98 | 49 | NA | NA | NA | NA | 1 | NA | 0.867 | NA | 0.133 |
| SE | 10090 | KIAA0040 | chr1 | - | 175192639 | 175192716 | 175177610 | 175177684 | 175192859 | 175193093 | 0 | 0 | 1 | 0 | 0 | 0 | 0 | 1 | 0 | 0 | 1 | 0 | 98 | 49 | NA | NA | NA | NA | 1 | NA | 0.333 | NA | 0.667 |
| SE | 10118 | TRAF3IP3 | chr1 | + | 209780469 | 209780606 | 209779314 | 209779374 | 209781344 | 209781458 | 0 | 0 | 3 | 0 | 0 | 0 | 0 | 2 | 0 | 0 | 3 | 0 | 98 | 49 | NA | NA | NA | NA | 1 | NA | 0.25 | NA | 0.75 |
| SE | 10134 | CSNK1A1 | chr5 | - | 149511793 | 149511872 | 149509878 | 149509953 | 149513069 | 149513209 | 0 | 0 | 7 | 0 | 0 | 0 | 0 | 7 | 0 | 0 | 1 | 0 | 98 | 49 | NA | NA | NA | NA | 1 | NA | 0.778 | NA | 0.222 |
| SE | 10144 | GALC | chr14 | - | 87945552 | 87945733 | 87941394 | 87941558 | 87947727 | 87947878 | 0 | 0 | 3 | 0 | 0 | 1 | 1 | 3 | 0 | 0 | 0 | 0 | 98 | 49 | NA | NA | NA | NA | 0.6 | 1 | 1 | NA | -0.4 |
| SE | 10145 | GALC | chr14 | - | 87984393 | 87984533 | 87982204 | 87982243 | 87986488 | 87986602 | 0 | 0 | 2 | 0 | 0 | 0 | 0 | 3 | 0 | 0 | 1 | 0 | 98 | 49 | NA | NA | NA | NA | 1 | NA | 0.6 | NA | 0.4 |
| SE | 10160 | C14orf159 | chr14 | + | 91145220 | 91145313 | 91114634 | 91114994 | 91157638 | 91157690 | 0 | 0 | 4 | 0 | 0 | 5 | 0 | 1 | 0 | 0 | 0 | 0 | 98 | 49 | NA | NA | NA | NA | 0.286 | NA | 1 | NA | -0.714 |
| SE | 10161 | C14orf159 | chr14 | + | 91147780 | 91147858 | 91145220 | 91145313 | 91157638 | 91157690 | 0 | 0 | 6 | 0 | 0 | 1 | 0 | 1 | 0 | 0 | 0 | 0 | 98 | 49 | NA | NA | NA | NA | 0.75 | NA | 1 | NA | -0.25 |
| SE | 10181 | ATXN3 | chr14 | - | 92093745 | 92093831 | 92093251 | 92093318 | 92096092 | 92096137 | 0 | 0 | 1 | 0 | 0 | 0 | 0 | 2 | 0 | 0 | 1 | 0 | 98 | 49 | NA | NA | NA | NA | 1 | NA | 0.5 | NA | 0.5 |
| SE | 10195 | PIK3CA | chr3 | + | 179220985 | 179221157 | 179219948 | 179220052 | 179224080 | 179224187 | 0 | 0 | 4 | 0 | 0 | 1 | 0 | 2 | 0 | 0 | 0 | 0 | 98 | 49 | NA | NA | NA | NA | 0.667 | NA | 1 | NA | -0.333 |
| SE | 10234 | PLSCR3 | chr17 | - | 7393143 | 7393364 | 7392790 | 7392952 | 7393466 | 7393509 | 0 | 0 | 1 | 0 | 0 | 1 | 0 | 1 | 0 | 0 | 0 | 0 | 98 | 49 | NA | NA | NA | NA | 0.333 | NA | 1 | NA | -0.667 |
| SE | 10236 | RPS3A | chr4 | + | 151100974 | 151101162 | 151100484 | 151100588 | 151102870 | 151103079 | 1 | 0 | 0 | 0 | 0 | 0 | 0 | 0 | 0 | 0 | 2 | 0 | 98 | 49 | NA | NA | 1 | NA | NA | NA | 0 | NA | 1 |
| SE | 10267 | PBRM1 | chr3 | - | 52615350 | 52615456 | 52609312 | 52609955 | 52617261 | 52617538 | 0 | 0 | 1 | 0 | 0 | 1 | 0 | 1 | 0 | 0 | 0 | 0 | 98 | 49 | NA | NA | NA | NA | 0.333 | NA | 1 | NA | -0.667 |
| SE | 10292 | TECPR2 | chr14 | + | 102376649 | 102376940 | 102362962 | 102363116 | 102407337 | 102407466 | 0 | 0 | 8 | 0 | 0 | 0 | 0 | 5 | 0 | 3 | 0 | 0 | 98 | 49 | NA | NA | NA | NA | 1 | 0 | 1 | NA | 0.5 |
| SE | 10293 | RCOR1 | chr14 | + | 102721319 | 102721377 | 102721006 | 102721084 | 102722186 | 102722416 | 0 | 0 | 7 | 0 | 0 | 2 | 0 | 3 | 0 | 0 | 0 | 0 | 98 | 49 | NA | NA | NA | NA | 0.636 | NA | 1 | NA | -0.364 |
| SE | 10298 | MARK3 | chr14 | + | 103428386 | 103428440 | 103405075 | 103405267 | 103448918 | 103448967 | 0 | 0 | 17 | 0 | 0 | 0 | 0 | 7 | 0 | 0 | 4 | 0 | 98 | 49 | NA | NA | NA | NA | 1 | NA | 0.467 | NA | 0.533 |
| SE | 10300 | MARK3 | chr14 | + | 103465971 | 103466091 | 103465556 | 103465793 | 103466342 | 103466442 | 0 | 0 | 1 | 0 | 0 | 1 | 0 | 3 | 0 | 0 | 0 | 0 | 98 | 49 | NA | NA | NA | NA | 0.333 | NA | 1 | NA | -0.667 |
| SE | 10309 | EIF4E3 | chr3 | - | 71696459 | 71696520 | 71693874 | 71693941 | 71699613 | 71699708 | 0 | 0 | 13 | 0 | 0 | 1 | 0 | 10 | 0 | 0 | 0 | 0 | 98 | 49 | NA | NA | NA | NA | 0.867 | NA | 1 | NA | -0.133 |
| SE | 10320 | GALNS | chr16 | - | 88837621 | 88837765 | 88836200 | 88836267 | 88840991 | 88841094 | 0 | 0 | 1 | 0 | 0 | 2 | 0 | 1 | 0 | 0 | 2 | 0 | 98 | 49 | NA | NA | NA | NA | 0.2 | NA | 0.2 | NA | 0 |
| SE | 10343 | PDCD4 | chr10 | + | 110894411 | 110894522 | 110894090 | 110894198 | 110895947 | 110896087 | 0 | 0 | 6 | 0 | 0 | 0 | 0 | 3 | 0 | 0 | 1 | 0 | 98 | 49 | NA | NA | NA | NA | 1 | NA | 0.6 | NA | 0.4 |
| SE | 10347 | TPM3 | chr1 | - | 154169304 | 154169383 | 154155303 | 154157721 | 154170399 | 154170469 | 0 | 0 | 4 | 0 | 0 | 20 | 0 | 0 | 0 | 0 | 14 | 0 | 98 | 49 | NA | NA | NA | NA | 0.091 | NA | 0 | NA | 0.091 |
| SE | 10362 | SOS2 | chr14 | - | 50138611 | 50138784 | 50134122 | 50134239 | 50139941 | 50140059 | 0 | 0 | 15 | 0 | 0 | 0 | 5 | 8 | 0 | 0 | 1 | 0 | 98 | 49 | NA | NA | NA | NA | 1 | 1 | 0.8 | NA | 0.1 |
| SE | 10373 | LCMT1 | chr16 | + | 25128474 | 25128566 | 25111725 | 25111996 | 25132401 | 25132523 | 0 | 0 | 0 | 0 | 0 | 1 | 0 | 2 | 0 | 0 | 0 | 0 | 98 | 49 | NA | NA | NA | NA | 0 | NA | 1 | NA | -1 |
| SE | 10383 | SIGLEC5 | chr19 | - | 51626031 | 51626113 | 51611502 | 51612422 | 51627148 | 51627248 | 0 | 0 | 13 | 0 | 0 | 1 | 0 | 3 | 0 | 0 | 0 | 0 | 98 | 49 | NA | NA | NA | NA | 0.867 | NA | 1 | NA | -0.133 |
| SE | 10437 | KATNBL1 | chr15 | - | 34147379 | 34147430 | 34147199 | 34147289 | 34148631 | 34148750 | 0 | 0 | 1 | 0 | 0 | 0 | 0 | 0 | 0 | 0 | 1 | 0 | 98 | 49 | NA | NA | NA | NA | 1 | NA | 0 | NA | 1 |
| SE | 10439 | SLC12A6 | chr15 | - | 34242101 | 34242221 | 34241232 | 34241337 | 34243973 | 34244072 | 0 | 0 | 7 | 0 | 0 | 1 | 0 | 4 | 0 | 0 | 2 | 0 | 98 | 49 | NA | NA | NA | NA | 0.778 | NA | 0.5 | NA | 0.278 |
| SE | 10443 | PI4KA | chr22 | - | 20796145 | 20796314 | 20793192 | 20793243 | 20798583 | 20798687 | 0 | 0 | 0 | 0 | 0 | 1 | 0 | 1 | 0 | 0 | 3 | 0 | 98 | 49 | NA | NA | NA | NA | 0 | NA | 0.143 | NA | -0.143 |
| SE | 10455 | TOP3A | chr17 | - | 18280535 | 18280658 | 18277674 | 18278357 | 18282697 | 18282841 | 0 | 0 | 9 | 0 | 0 | 0 | 0 | 1 | 0 | 0 | 1 | 0 | 98 | 49 | NA | NA | NA | NA | 1 | NA | 0.333 | NA | 0.667 |
| SE | 10456 | TOP3A | chr17 | - | 18292644 | 18292852 | 18290841 | 18291027 | 18294702 | 18294785 | 0 | 0 | 1 | 0 | 0 | 1 | 0 | 4 | 0 | 0 | 0 | 0 | 98 | 49 | NA | NA | NA | NA | 0.333 | NA | 1 | NA | -0.667 |
| SE | 10461 | OSBPL11 | chr3 | - | 125552180 | 125552679 | 125547405 | 125547592 | 125560378 | 125560519 | 0 | 0 | 5 | 0 | 0 | 1 | 3 | 3 | 0 | 0 | 0 | 0 | 98 | 49 | NA | NA | NA | NA | 0.714 | 1 | 1 | NA | -0.286 |
| SE | 10468 | PLCB2 | chr15 | - | 40291605 | 40291650 | 40291264 | 40291487 | 40291848 | 40291924 | 0 | 0 | 1 | 0 | 0 | 2 | 0 | 0 | 0 | 0 | 1 | 0 | 94 | 49 | NA | NA | NA | NA | 0.207 | NA | 0 | NA | 0.207 |
| SE | 10491 | DCP1A | chr3 | - | 53312240 | 53312379 | 53292068 | 53292827 | 53319406 | 53319473 | 0 | 0 | 1 | 0 | 0 | 0 | 0 | 2 | 0 | 0 | 1 | 0 | 98 | 49 | NA | NA | NA | NA | 1 | NA | 0.5 | NA | 0.5 |
| SE | 10495 | HSP90AA1 | chr14 | - | 102082110 | 102082444 | 102080737 | 102081821 | 102083033 | 102083302 | 0 | 0 | 5 | 0 | 0 | 1 | 0 | 6 | 0 | 0 | 0 | 0 | 98 | 49 | NA | NA | NA | NA | 0.714 | NA | 1 | NA | -0.286 |
| SE | 10501 | GANC | chr15 | + | 42297610 | 42297656 | 42292734 | 42292917 | 42306545 | 42306612 | 0 | 0 | 2 | 0 | 0 | 1 | 0 | 3 | 0 | 0 | 0 | 0 | 95 | 49 | NA | NA | NA | NA | 0.508 | NA | 1 | NA | -0.492 |
| SE | 10509 | ZNF106 | chr15 | - | 42428017 | 42428134 | 42424833 | 42425025 | 42435383 | 42435518 | 0 | 0 | 7 | 0 | 0 | 1 | 2 | 1 | 0 | 0 | 0 | 0 | 98 | 49 | NA | NA | NA | NA | 0.778 | 1 | 1 | NA | -0.222 |
| SE | 10510 | ZNF106 | chr15 | - | 42449770 | 42451954 | 42448071 | 42448705 | 42466052 | 42466114 | 0 | 0 | 3 | 0 | 0 | 0 | 0 | 0 | 0 | 0 | 2 | 0 | 98 | 49 | NA | NA | NA | NA | 1 | NA | 0 | NA | 1 |
| SE | 10513 | TMEM62 | chr15 | + | 43134256 | 43134368 | 43133523 | 43133982 | 43135511 | 43135649 | 0 | 0 | 2 | 0 | 0 | 0 | 0 | 0 | 0 | 0 | 1 | 0 | 98 | 49 | NA | NA | NA | NA | 1 | NA | 0 | NA | 1 |
| SE | 10514 | TMEM62 | chr15 | + | 43149028 | 43149151 | 43148754 | 43148879 | 43151789 | 43151945 | 0 | 0 | 1 | 0 | 0 | 0 | 0 | 1 | 0 | 0 | 2 | 0 | 98 | 49 | NA | NA | NA | NA | 1 | NA | 0.2 | NA | 0.8 |
| SE | 10517 | TP53BP1 | chr15 | - | 43432193 | 43432677 | 43428015 | 43428168 | 43438323 | 43438416 | 0 | 0 | 1 | 0 | 0 | 0 | 0 | 1 | 0 | 0 | 1 | 0 | 98 | 49 | NA | NA | NA | NA | 1 | NA | 0.333 | NA | 0.667 |
| SE | 10558 | DSC2 | chr18 | - | 31068298 | 31068344 | 31065973 | 31068212 | 31068893 | 31069151 | 0 | 0 | 0 | 0 | 0 | 1 | 0 | 3 | 0 | 0 | 0 | 0 | 95 | 49 | NA | NA | NA | NA | 0 | NA | 1 | NA | -1 |
| SE | 10560 | FAM96A | chr15 | - | 64081101 | 64081151 | 64075491 | 64075537 | 64088686 | 64088851 | 0 | 0 | 5 | 0 | 0 | 0 | 0 | 0 | 0 | 3 | 1 | 0 | 98 | 49 | NA | NA | NA | NA | 1 | 0 | 0 | NA | 1 |
| SE | 10566 | ATP8B4 | chr15 | - | 50002152 | 50002223 | 49987390 | 49987549 | 50010844 | 50010917 | 0 | 0 | 2 | 0 | 0 | 2 | 1 | 2 | 0 | 0 | 0 | 0 | 98 | 49 | NA | NA | NA | NA | 0.333 | 1 | 1 | NA | -0.667 |
| SE | 10570 | LRRC75A-AS1 | chr17 | + | 16439527 | 16439703 | 16438986 | 16439414 | 16440184 | 16440253 | 0 | 0 | 0 | 0 | 0 | 1 | 0 | 3 | 0 | 0 | 1 | 0 | 98 | 49 | NA | NA | NA | NA | 0 | NA | 0.6 | NA | -0.6 |
| SE | 10571 | LRRC75A-AS1 | chr17 | + | 16439580 | 16439703 | 16438986 | 16439414 | 16440184 | 16440253 | 0 | 0 | 0 | 0 | 0 | 1 | 0 | 3 | 0 | 0 | 1 | 0 | 98 | 49 | NA | NA | NA | NA | 0 | NA | 0.6 | NA | -0.6 |
| SE | 10573 | LRRC75A-AS1 | chr17 | + | 16439659 | 16439703 | 16438986 | 16439414 | 16440184 | 16440253 | 0 | 0 | 0 | 0 | 0 | 1 | 0 | 3 | 0 | 0 | 1 | 0 | 93 | 49 | NA | NA | NA | NA | 0 | NA | 0.613 | NA | -0.613 |
| SE | 10576 | DDX47 | chr12 | + | 12823869 | 12824016 | 12823202 | 12823319 | 12824539 | 12824677 | 0 | 0 | 0 | 0 | 0 | 2 | 0 | 1 | 0 | 0 | 0 | 0 | 98 | 49 | NA | NA | NA | NA | 0 | NA | 1 | NA | -1 |
| SE | 10582 | USP8 | chr15 | + | 50484274 | 50484361 | 50481480 | 50482065 | 50489800 | 50489881 | 0 | 0 | 2 | 0 | 0 | 0 | 0 | 1 | 0 | 0 | 2 | 0 | 98 | 49 | NA | NA | NA | NA | 1 | NA | 0.2 | NA | 0.8 |
| SE | 10584 | SPPL2A | chr15 | - | 50732602 | 50732684 | 50730964 | 50731039 | 50736100 | 50736202 | 0 | 0 | 6 | 0 | 0 | 0 | 0 | 4 | 0 | 0 | 1 | 0 | 98 | 49 | NA | NA | NA | NA | 1 | NA | 0.667 | NA | 0.333 |
| SE | 10592 | RNF213 | chr17 | + | 80385038 | 80385171 | 80383676 | 80383928 | 80385537 | 80385621 | 0 | 0 | 23 | 0 | 0 | 2 | 0 | 23 | 0 | 0 | 2 | 0 | 98 | 49 | NA | NA | NA | NA | 0.852 | NA | 0.852 | NA | 0 |
| SE | 10600 | DMXL2 | chr15 | - | 51451644 | 51451697 | 51450128 | 51450346 | 51453549 | 51453641 | 0 | 0 | 8 | 0 | 0 | 0 | 1 | 1 | 0 | 0 | 1 | 0 | 98 | 49 | NA | NA | NA | NA | 1 | 1 | 0.333 | NA | 0.334 |
| SE | 10601 | DMXL2 | chr15 | - | 51464674 | 51464876 | 51463378 | 51463496 | 51465565 | 51465651 | 0 | 0 | 15 | 0 | 0 | 1 | 0 | 2 | 0 | 0 | 0 | 0 | 98 | 49 | NA | NA | NA | NA | 0.882 | NA | 1 | NA | -0.118 |
| SE | 10607 | ANKFY1 | chr17 | - | 4209823 | 4209947 | 4207932 | 4208082 | 4216982 | 4217118 | 0 | 0 | 6 | 0 | 0 | 0 | 0 | 1 | 0 | 0 | 1 | 0 | 98 | 49 | NA | NA | NA | NA | 1 | NA | 0.333 | NA | 0.667 |
| SE | 10610 | MYO5A | chr15 | - | 52348817 | 52348826 | 52346360 | 52346461 | 52351253 | 52351481 | 0 | 0 | 0 | 0 | 0 | 1 | 0 | 1 | 0 | 0 | 0 | 0 | 58 | 49 | NA | NA | NA | NA | 0 | NA | 1 | NA | -1 |
| SE | 10623 | ADAM10 | chr15 | - | 58611807 | 58611991 | 58610998 | 58611107 | 58621470 | 58621621 | 0 | 0 | 11 | 0 | 0 | 1 | 0 | 6 | 0 | 0 | 0 | 0 | 98 | 49 | NA | NA | NA | NA | 0.846 | NA | 1 | NA | -0.154 |
| SE | 10625 | RRP12 | chr10 | - | 97372735 | 97372803 | 97372072 | 97372166 | 97373045 | 97373200 | 0 | 0 | 4 | 0 | 0 | 2 | 0 | 4 | 0 | 0 | 0 | 0 | 98 | 49 | NA | NA | NA | NA | 0.5 | NA | 1 | NA | -0.5 |
| SE | 10629 | BNIP2 | chr15 | - | 59679591 | 59679768 | 59677910 | 59678087 | 59680240 | 59680308 | 0 | 0 | 15 | 0 | 0 | 0 | 0 | 6 | 0 | 0 | 1 | 0 | 98 | 49 | NA | NA | NA | NA | 1 | NA | 0.75 | NA | 0.25 |
| SE | 10640 | USP3 | chr15 | + | 63529013 | 63529166 | 63504510 | 63504830 | 63532646 | 63532707 | 0 | 0 | 0 | 0 | 0 | 2 | 0 | 2 | 0 | 0 | 4 | 0 | 98 | 49 | NA | NA | NA | NA | 0 | NA | 0.2 | NA | -0.2 |
| SE | 10643 | HERC1 | chr15 | - | 63623724 | 63623890 | 63622814 | 63622891 | 63624157 | 63624327 | 0 | 0 | 2 | 0 | 0 | 2 | 0 | 2 | 0 | 0 | 0 | 0 | 98 | 49 | NA | NA | NA | NA | 0.333 | NA | 1 | NA | -0.667 |
| SE | 10644 | HERC1 | chr15 | - | 63632708 | 63632811 | 63630465 | 63630635 | 63633847 | 63633970 | 0 | 0 | 6 | 0 | 0 | 0 | 0 | 1 | 0 | 0 | 1 | 0 | 98 | 49 | NA | NA | NA | NA | 1 | NA | 0.333 | NA | 0.667 |
| SE | 10645 | HERC1 | chr15 | - | 63648068 | 63648199 | 63645482 | 63645682 | 63649724 | 63649925 | 0 | 0 | 2 | 0 | 0 | 0 | 0 | 3 | 0 | 0 | 2 | 0 | 98 | 49 | NA | NA | NA | NA | 1 | NA | 0.429 | NA | 0.571 |
| SE | 10652 | RELA | chr11 | - | 65658286 | 65658499 | 65655854 | 65655935 | 65658717 | 65658822 | 0 | 0 | 4 | 0 | 0 | 0 | 0 | 5 | 0 | 0 | 2 | 0 | 98 | 49 | NA | NA | NA | NA | 1 | NA | 0.556 | NA | 0.444 |
| SE | 10655 | OAZ2 | chr15 | - | 64691433 | 64691588 | 64690373 | 64690549 | 64703028 | 64703281 | 0 | 0 | 2 | 0 | 0 | 0 | 0 | 3 | 0 | 0 | 1 | 0 | 98 | 49 | NA | NA | NA | NA | 1 | NA | 0.6 | NA | 0.4 |
| SE | 10656 | OAZ2 | chr15 | - | 64691433 | 64691591 | 64690373 | 64690549 | 64703028 | 64703281 | 0 | 0 | 3 | 0 | 0 | 0 | 0 | 5 | 0 | 0 | 1 | 0 | 98 | 49 | NA | NA | NA | NA | 1 | NA | 0.714 | NA | 0.286 |
| SE | 10689 | EGLN1 | chr1 | - | 231367568 | 231367636 | 231363750 | 231366475 | 231370561 | 231370698 | 0 | 0 | 7 | 0 | 0 | 1 | 0 | 2 | 0 | 0 | 4 | 0 | 98 | 49 | NA | NA | NA | NA | 0.778 | NA | 0.2 | NA | 0.578 |
| SE | 10703 | TLE3 | chr15 | - | 70095577 | 70095641 | 70094531 | 70094576 | 70096160 | 70096261 | 0 | 0 | 2 | 0 | 0 | 1 | 1 | 8 | 0 | 0 | 0 | 0 | 98 | 49 | NA | NA | NA | NA | 0.5 | 1 | 1 | NA | -0.5 |
| SE | 10709 | ARIH1 | chr15 | + | 72563393 | 72563500 | 72561482 | 72561549 | 72566562 | 72566605 | 0 | 0 | 1 | 0 | 0 | 0 | 0 | 3 | 0 | 0 | 1 | 0 | 98 | 49 | NA | NA | NA | NA | 1 | NA | 0.6 | NA | 0.4 |
| SE | 10726 | PTMA | chr2 | + | 231712442 | 231712516 | 231711347 | 231711419 | 231712803 | 231713540 | 0 | 0 | 1 | 0 | 0 | 0 | 0 | 3 | 0 | 0 | 1 | 0 | 98 | 49 | NA | NA | NA | NA | 1 | NA | 0.6 | NA | 0.4 |
| SE | 10733 | FAM219B | chr15 | - | 74904663 | 74904712 | 74899986 | 74902786 | 74905153 | 74905231 | 0 | 0 | 2 | 0 | 0 | 0 | 0 | 1 | 0 | 0 | 3 | 0 | 98 | 49 | NA | NA | NA | NA | 1 | NA | 0.143 | NA | 0.857 |
| SE | 10745 | MAN2C1 | chr15 | - | 75362641 | 75362748 | 75362342 | 75362453 | 75364487 | 75364665 | 0 | 0 | 4 | 0 | 0 | 1 | 0 | 4 | 0 | 0 | 0 | 0 | 98 | 49 | NA | NA | NA | NA | 0.667 | NA | 1 | NA | -0.333 |

| | | | | | | | | | | | | | | | | | | | | | | | | | | | | | | | | | |
|---|---|---|---|---|---|---|---|---|---|---|---|---|---|---|---|---|---|---|---|---|---|---|---|---|---|---|---|---|---|---|---|---|---|
| SE | 10751 | SIN3A | chr15 | - | 75400729 | 75400940 | 75400039 | 75400156 | 75401851 | 75401970 | 0 | 0 | 4 | 0 | 0 | 0 | 0 | 0 | 0 | 0 | 1 | 0 | 98 | 49 | NA | NA | NA | NA | 1 | NA | 0 | NA | 1 |
| SE | 10827 | AMBRA1 | chr11 | - | 46433473 | 46433628 | 46417912 | 46418052 | 46434848 | 46435037 | 0 | 0 | 2 | 0 | 0 | 0 | 0 | 0 | 0 | 0 | 1 | 0 | 98 | 49 | NA | NA | NA | NA | 1 | NA | 0 | NA | 1 |
| SE | 10846 | IRF1 | chr5 | - | 132484361 | 132484497 | 132481608 | 132484075 | 132486556 | 132486686 | 0 | 0 | 8 | 0 | 0 | 1 | 0 | 5 | 0 | 0 | 0 | 0 | 98 | 49 | NA | NA | NA | NA | 0.8 | NA | 1 | NA | -0.2 |
| SE | 10847 | IRF1 | chr5 | - | 132485666 | 132485716 | 132484361 | 132484497 | 132486556 | 132486686 | 0 | 0 | 11 | 0 | 0 | 0 | 0 | 14 | 0 | 0 | 1 | 0 | 98 | 49 | NA | NA | NA | NA | 1 | NA | 0.875 | NA | 0.125 |
| SE | 10914 | HDAC3 | chr5 | - | 141626012 | 141626071 | 141625684 | 141625764 | 141626193 | 141626283 | 0 | 0 | 6 | 0 | 0 | 2 | 0 | 2 | 0 | 0 | 0 | 0 | 98 | 49 | NA | NA | NA | NA | 0.6 | NA | 1 | NA | -0.4 |
| SE | 10917 | CWF19L1 | chr10 | - | 100256261 | 100256476 | 100253420 | 100253539 | 100260217 | 100260319 | 0 | 0 | 4 | 0 | 0 | 0 | 0 | 1 | 0 | 0 | 1 | 0 | 98 | 49 | NA | NA | NA | NA | 1 | NA | 0.333 | NA | 0.667 |
| SE | 10941 | PLEKHJ1 | chr19 | - | 2234149 | 2234240 | 2233997 | 2234061 | 2235761 | 2235828 | 0 | 0 | 0 | 0 | 0 | 1 | 0 | 1 | 0 | 0 | 0 | 0 | 98 | 49 | NA | NA | NA | NA | 0 | NA | 1 | NA | -1 |
| SE | 10948 | IQGAP1 | chr15 | + | 90466268 | 90466436 | 90466000 | 90466091 | 90467449 | 90467592 | 0 | 0 | 20 | 0 | 0 | 0 | 0 | 18 | 0 | 0 | 3 | 0 | 98 | 49 | NA | NA | NA | NA | 1 | NA | 0.75 | NA | 0.25 |
| SE | 10949 | IQGAP1 | chr15 | + | 90467449 | 90467592 | 90466268 | 90466436 | 90472839 | 90473010 | 0 | 0 | 32 | 0 | 0 | 0 | 1 | 6 | 0 | 0 | 1 | 0 | 98 | 49 | NA | NA | NA | NA | 1 | 1 | 0.75 | NA | 0.125 |
| SE | 10973 | MEF2A | chr15 | + | 99620872 | 99621080 | 99598429 | 99598511 | 99632977 | 99633173 | 0 | 0 | 1 | 0 | 0 | 0 | 0 | 0 | 0 | 0 | 1 | 0 | 98 | 49 | NA | NA | NA | NA | 1 | NA | 0 | NA | 1 |
| SE | 10975 | MEF2A | chr15 | + | 99690240 | 99690428 | 99675398 | 99675458 | 99710633 | 99710760 | 0 | 0 | 7 | 0 | 0 | 1 | 0 | 3 | 0 | 0 | 0 | 0 | 98 | 49 | NA | NA | NA | NA | 0.778 | NA | 1 | NA | -0.222 |
| SE | 10987 | ZCCHC6 | chr9 | - | 86309462 | 86309576 | 86309211 | 86309289 | 86309927 | 86310017 | 0 | 0 | 12 | 0 | 0 | 0 | 0 | 5 | 0 | 0 | 1 | 0 | 98 | 49 | NA | NA | NA | NA | 1 | NA | 0.714 | NA | 0.286 |
| SE | 10988 | ZCCHC6 | chr9 | - | 86318957 | 86319058 | 86317218 | 86317276 | 86319583 | 86319670 | 0 | 0 | 8 | 0 | 0 | 0 | 0 | 4 | 0 | 0 | 1 | 0 | 98 | 49 | NA | NA | NA | NA | 1 | NA | 0.667 | NA | 0.333 |
| SE | 10990 | ZCCHC6 | chr9 | - | 86322324 | 86322475 | 86319583 | 86319670 | 86322872 | 86323960 | 0 | 0 | 6 | 0 | 0 | 0 | 0 | 4 | 0 | 0 | 1 | 0 | 98 | 49 | NA | NA | NA | NA | 1 | NA | 0.667 | NA | 0.333 |
| SE | 11029 | TSR3 | chr16 | - | 1349888 | 1349952 | 1349239 | 1349608 | 1350057 | 1350234 | 0 | 0 | 0 | 0 | 0 | 1 | 0 | 4 | 0 | 0 | 0 | 0 | 98 | 49 | NA | NA | NA | NA | 0 | NA | 1 | NA | -1 |
| SE | 11032 | STAT1 | chr2 | - | 191001073 | 191001163 | 190999625 | 190999704 | 191007562 | 191007661 | 0 | 0 | 49 | 0 | 0 | 1 | 0 | 39 | 0 | 0 | 0 | 0 | 98 | 49 | NA | NA | NA | NA | 0.961 | NA | 1 | NA | -0.039 |
| SE | 11050 | NPEPPS | chr17 | + | 47586147 | 47586244 | 47585499 | 47585700 | 47586364 | 47586398 | 0 | 0 | 6 | 0 | 0 | 1 | 0 | 2 | 0 | 0 | 0 | 0 | 98 | 49 | NA | NA | NA | NA | 0.75 | NA | 1 | NA | -0.25 |
| SE | 11054 | ARFGAP3 | chr22 | - | 42810812 | 42810944 | 42808766 | 42808890 | 42817141 | 42817264 | 0 | 0 | 3 | 0 | 0 | 1 | 0 | 4 | 0 | 0 | 1 | 0 | 98 | 49 | NA | NA | NA | NA | 0.6 | NA | 0.667 | NA | -0.067 |
| SE | 11063 | DENND4C | chr9 | + | 19357964 | 19358160 | 19356971 | 19357154 | 19360243 | 19360489 | 0 | 0 | 3 | 0 | 0 | 2 | 0 | 3 | 0 | 0 | 0 | 0 | 98 | 49 | NA | NA | NA | NA | 0.429 | NA | 1 | NA | -0.571 |
| SE | 11097 | SMU1 | chr9 | - | 33071739 | 33071892 | 33068823 | 33068934 | 33073595 | 33073806 | 0 | 0 | 1 | 0 | 0 | 2 | 0 | 0 | 0 | 0 | 1 | 0 | 98 | 49 | NA | NA | NA | NA | 0.2 | NA | 0 | NA | 0.2 |
| SE | 11098 | SMU1 | chr9 | - | 33073595 | 33073806 | 33068823 | 33068934 | 33076582 | 33076716 | 0 | 0 | 3 | 0 | 0 | 0 | 0 | 2 | 0 | 0 | 1 | 0 | 98 | 49 | NA | NA | NA | NA | 1 | NA | 0.5 | NA | 0.5 |
| SE | 11107 | USP25 | chr21 | + | 15777903 | 15778027 | 15765996 | 15766141 | 15791501 | 15791664 | 0 | 0 | 1 | 0 | 0 | 1 | 0 | 8 | 0 | 0 | 0 | 0 | 98 | 49 | NA | NA | NA | NA | 0.333 | NA | 1 | NA | -0.667 |
| SE | 11108 | USP25 | chr21 | + | 15808808 | 15808885 | 15805120 | 15805258 | 15811136 | 15811210 | 0 | 0 | 6 | 0 | 0 | 6 | 0 | 1 | 0 | 0 | 5 | 0 | 98 | 49 | NA | NA | NA | NA | 0.333 | NA | 0.091 | NA | 0.242 |
| SE | 11111 | USP25 | chr21 | + | 15849776 | 15849872 | 15842397 | 15842540 | 15864267 | 15864446 | 0 | 0 | 0 | 0 | 0 | 3 | 0 | 1 | 0 | 0 | 0 | 0 | 98 | 49 | NA | NA | NA | NA | 0 | NA | 1 | NA | -1 |
| SE | 11141 | TXNDC11 | chr16 | - | 11736016 | 11736233 | 11733981 | 11734079 | 11742476 | 11742878 | 0 | 0 | 1 | 0 | 0 | 0 | 0 | 2 | 0 | 0 | 1 | 0 | 98 | 49 | NA | NA | NA | NA | 1 | NA | 0.5 | NA | 0.5 |
| SE | 11153 | KIAA0430 | chr16 | - | 15625371 | 15625797 | 15625015 | 15625173 | 15630331 | 15630504 | 0 | 0 | 5 | 0 | 0 | 0 | 0 | 4 | 0 | 0 | 2 | 0 | 98 | 49 | NA | NA | NA | NA | 1 | NA | 0.5 | NA | 0.5 |
| SE | 11154 | KIAA0430 | chr16 | - | 15625371 | 15625800 | 15625015 | 15625173 | 15630331 | 15630504 | 0 | 0 | 7 | 0 | 0 | 0 | 0 | 6 | 0 | 0 | 2 | 0 | 98 | 49 | NA | NA | NA | NA | 1 | NA | 0.6 | NA | 0.4 |
| SE | 11207 | TOP1 | chr20 | + | 41116277 | 41116392 | 41115370 | 41115439 | 41118168 | 41118296 | 0 | 0 | 13 | 0 | 0 | 0 | 0 | 2 | 0 | 0 | 1 | 0 | 98 | 49 | NA | NA | NA | NA | 1 | NA | 0.5 | NA | 0.5 |
| SE | 11208 | KIAA1551 | chr12 | + | 31970188 | 31970356 | 31960776 | 31960871 | 31980877 | 31985957 | 0 | 0 | 59 | 0 | 0 | 1 | 0 | 16 | 0 | 0 | 0 | 0 | 98 | 49 | NA | NA | NA | NA | 0.967 | NA | 1 | NA | -0.033 |
| SE | 11218 | RPAIN | chr17 | + | 5425970 | 5426082 | 5422768 | 5422829 | 5432541 | 5433020 | 0 | 0 | 2 | 0 | 0 | 1 | 0 | 1 | 0 | 0 | 0 | 0 | 98 | 49 | NA | NA | NA | NA | 0.5 | NA | 1 | NA | -0.5 |
| SE | 11234 | DCLRE1C | chr10 | - | 14945434 | 14945565 | 14945104 | 14945189 | 14949035 | 14949087 | 0 | 0 | 0 | 0 | 0 | 3 | 0 | 1 | 0 | 0 | 0 | 0 | 98 | 49 | NA | NA | NA | NA | 0 | NA | 1 | NA | -1 |
| SE | 11235 | DCLRE1C | chr10 | - | 14947524 | 14947609 | 14945104 | 14945189 | 14949035 | 14949087 | 0 | 0 | 1 | 0 | 0 | 3 | 0 | 1 | 0 | 0 | 0 | 0 | 98 | 49 | NA | NA | NA | NA | 0.143 | NA | 1 | NA | -0.857 |
| SE | 11253 | IL4R | chr16 | + | 27346466 | 27346618 | 27344868 | 27345020 | 27352539 | 27352696 | 0 | 0 | 8 | 0 | 0 | 1 | 0 | 10 | 0 | 0 | 2 | 0 | 98 | 49 | NA | NA | NA | NA | 0.8 | NA | 0.714 | NA | 0.086 |
| SE | 11254 | IL4R | chr16 | + | 27360765 | 27360815 | 27358915 | 27358994 | 27362251 | 27364778 | 0 | 0 | 4 | 0 | 0 | 2 | 0 | 2 | 0 | 0 | 0 | 0 | 98 | 49 | NA | NA | NA | NA | 0.5 | NA | 1 | NA | -0.5 |
| SE | 11315 | PPP6R3 | chr11 | + | 68551120 | 68551186 | 68548066 | 68548204 | 68554144 | 68554257 | 0 | 0 | 2 | 0 | 0 | 2 | 0 | 3 | 0 | 0 | 0 | 0 | 98 | 49 | NA | NA | NA | NA | 0.333 | NA | 1 | NA | -0.667 |
| SE | 11318 | PPP6R3 | chr11 | + | 68583042 | 68583129 | 68575957 | 68576043 | 68587944 | 68588024 | 0 | 0 | 2 | 0 | 0 | 1 | 0 | 4 | 0 | 0 | 0 | 0 | 98 | 49 | NA | NA | NA | NA | 0.5 | NA | 1 | NA | -0.5 |
| SE | 11323 | ITGAL | chr16 | + | 30479090 | 30479208 | 30475512 | 30475580 | 30481438 | 30481584 | 0 | 0 | 18 | 0 | 0 | 1 | 3 | 3 | 0 | 0 | 0 | 0 | 98 | 49 | NA | NA | NA | NA | 0.9 | 1 | 1 | NA | -0.1 |
| SE | 11352 | ADGRE2 | chr19 | - | 14765657 | 14765804 | 14765523 | 14765570 | 14772341 | 14772497 | 0 | 0 | 0 | 0 | 0 | 1 | 0 | 3 | 0 | 0 | 1 | 0 | 98 | 49 | NA | NA | NA | NA | 0 | NA | 0.6 | NA | -0.6 |
| SE | 11361 | TATDN2 | chr3 | + | 10276360 | 10276488 | 10270130 | 10271015 | 10278278 | 10278462 | 0 | 0 | 2 | 0 | 0 | 0 | 0 | 0 | 0 | 0 | 1 | 0 | 98 | 49 | NA | NA | NA | NA | 1 | NA | 0 | NA | 1 |
| SE | 11385 | NLRC5 | chr16 | + | 57028071 | 57028155 | 57025367 | 57027018 | 57028301 | 57028385 | 0 | 0 | 2 | 0 | 0 | 0 | 0 | 3 | 0 | 0 | 2 | 0 | 98 | 49 | NA | NA | NA | NA | 1 | NA | 0.429 | NA | 0.571 |
| SE | 11388 | NLRC5 | chr16 | + | 57055432 | 57055519 | 57055031 | 57055094 | 57058064 | 57058148 | 0 | 0 | 3 | 0 | 0 | 0 | 0 | 1 | 0 | 0 | 1 | 0 | 98 | 49 | NA | NA | NA | NA | 1 | NA | 0.333 | NA | 0.667 |
| SE | 11389 | NLRC5 | chr16 | + | 57076818 | 57076902 | 57074599 | 57074683 | 57077295 | 57077379 | 0 | 0 | 19 | 0 | 0 | 0 | 0 | 8 | 0 | 0 | 1 | 0 | 98 | 49 | NA | NA | NA | NA | 1 | NA | 0.8 | NA | 0.2 |
| SE | 11390 | NLRC5 | chr16 | + | 57077295 | 57077379 | 57076818 | 57076902 | 57077718 | 57077802 | 0 | 0 | 13 | 0 | 0 | 0 | 0 | 2 | 0 | 0 | 1 | 0 | 98 | 49 | NA | NA | NA | NA | 1 | NA | 0.5 | NA | 0.5 |
| SE | 11392 | RNF24 | chr20 | - | 3974331 | 3974400 | 3963874 | 3964024 | 4015436 | 4015569 | 0 | 0 | 6 | 0 | 0 | 1 | 0 | 1 | 0 | 0 | 0 | 0 | 98 | 49 | NA | NA | NA | NA | 0.75 | NA | 1 | NA | -0.25 |
| SE | 11405 | SETD3 | chr14 | - | 99461191 | 99461340 | 99459112 | 99459185 | 99463485 | 99463578 | 0 | 0 | 4 | 0 | 0 | 1 | 0 | 4 | 0 | 0 | 0 | 0 | 98 | 49 | NA | NA | NA | NA | 0.667 | NA | 1 | NA | -0.333 |
| SE | 11420 | CKLF | chr16 | + | 66563121 | 66563217 | 66558189 | 66558348 | 66565885 | 66566287 | 0 | 0 | 6 | 0 | 0 | 2 | 0 | 0 | 0 | 0 | 2 | 0 | 98 | 49 | NA | NA | NA | NA | 0.6 | NA | 0 | NA | 0.6 |
| SE | 11437 | PLK3 | chr1 | + | 44804649 | 44804779 | 44804338 | 44804501 | 44805265 | 44805379 | 0 | 0 | 2 | 0 | 0 | 3 | 0 | 0 | 0 | 0 | 2 | 0 | 98 | 49 | NA | NA | NA | NA | 0.25 | NA | 0 | NA | 0.25 |
| SE | 11442 | FHOD1 | chr16 | - | 67231187 | 67231349 | 67230600 | 67230791 | 67231429 | 67231549 | 0 | 0 | 3 | 0 | 0 | 0 | 0 | 0 | 0 | 2 | 0 | 0 | 98 | 49 | NA | NA | NA | NA | 1 | 0 | NA | NA | 1 |
| SE | 11449 | GSTO1 | chr10 | + | 104262978 | 104263077 | 104259575 | 104259798 | 104266083 | 104266190 | 0 | 0 | 4 | 0 | 0 | 0 | 1 | 0 | 0 | 0 | 1 | 0 | 98 | 49 | NA | NA | NA | NA | 1 | 1 | 0 | NA | 0.5 |
| SE | 11468 | EMC10 | chr19 | + | 50481860 | 50481947 | 50480883 | 50480977 | 50482148 | 50483526 | 0 | 0 | 1 | 0 | 0 | 2 | 0 | 1 | 0 | 0 | 1 | 0 | 98 | 49 | NA | NA | NA | NA | 0.2 | NA | 0.333 | NA | -0.133 |
| SE | 11472 | ATP8A1 | chr4 | - | 42594288 | 42594362 | 42588259 | 42588329 | 42600477 | 42600518 | 0 | 0 | 1 | 0 | 0 | 0 | 0 | 1 | 0 | 0 | 2 | 0 | 98 | 49 | NA | NA | NA | NA | 1 | NA | 0.2 | NA | 0.8 |
| SE | 11485 | ITM2A | chrX | - | 79363422 | 79363554 | 79362941 | 79363139 | 79367104 | 79367552 | 0 | 0 | 2 | 0 | 0 | 0 | 0 | 0 | 0 | 0 | 1 | 0 | 98 | 49 | NA | NA | NA | NA | 1 | NA | 0 | NA | 1 |
| SE | 11495 | KARS | chr16 | - | 75640183 | 75640349 | 75636453 | 75636547 | 75641563 | 75641723 | 0 | 0 | 2 | 0 | 0 | 0 | 0 | 0 | 0 | 0 | 1 | 0 | 98 | 49 | NA | NA | NA | NA | 1 | NA | 0 | NA | 1 |
| SE | 11520 | NUPL1 | chr13 | + | 25312882 | 25313032 | 25309246 | 25309282 | 25313613 | 25313751 | 0 | 0 | 3 | 0 | 0 | 1 | 0 | 2 | 0 | 0 | 0 | 0 | 98 | 49 | NA | NA | NA | NA | 0.6 | NA | 1 | NA | -0.4 |
| SE | 11521 | PIKFYVE | chr2 | + | 208285725 | 208285933 | 208277536 | 208277708 | 208288728 | 208288818 | 0 | 0 | 6 | 0 | 0 | 0 | 0 | 3 | 0 | 0 | 1 | 0 | 98 | 49 | NA | NA | NA | NA | 1 | NA | 0.6 | NA | 0.4 |
| SE | 11522 | PIKFYVE | chr2 | + | 208304845 | 208305013 | 208304170 | 208304318 | 208312235 | 208312295 | 0 | 0 | 1 | 0 | 0 | 4 | 0 | 1 | 0 | 0 | 0 | 0 | 98 | 49 | NA | NA | NA | NA | 0.111 | NA | 1 | NA | -0.889 |
| SE | 11523 | PIKFYVE | chr2 | + | 208347858 | 208348023 | 208346049 | 208346147 | 208350023 | 208350083 | 0 | 0 | 2 | 0 | 0 | 2 | 0 | 2 | 0 | 0 | 0 | 0 | 98 | 49 | NA | NA | NA | NA | 0.333 | NA | 1 | NA | -0.667 |
| SE | 11524 | PIKFYVE | chr2 | + | 208351351 | 208351455 | 208350770 | 208350947 | 208352653 | 208352782 | 0 | 0 | 3 | 0 | 0 | 1 | 0 | 1 | 0 | 0 | 0 | 0 | 98 | 49 | NA | NA | NA | NA | 0.6 | NA | 1 | NA | -0.4 |
| SE | 11564 | CBLB | chr3 | - | 105734008 | 105734140 | 105720046 | 105720250 | 105737170 | 105737258 | 0 | 0 | 2 | 0 | 0 | 0 | 0 | 2 | 0 | 0 | 1 | 0 | 98 | 49 | NA | NA | NA | NA | 1 | NA | 0.5 | NA | 0.5 |
| SE | 11579 | ANKRD11 | chr16 | - | 89431255 | 89431326 | 89418283 | 89418368 | 89490244 | 89490561 | 0 | 0 | 0 | 0 | 0 | 1 | 0 | 1 | 0 | 0 | 1 | 0 | 98 | 49 | NA | NA | NA | NA | 0 | NA | 0.333 | NA | -0.333 |
| SE | 11621 | IGF1R | chr15 | + | 98934823 | 98935053 | 98930234 | 98930305 | 98935315 | 98935426 | 0 | 0 | 2 | 0 | 0 | 0 | 0 | 0 | 0 | 0 | 1 | 0 | 98 | 49 | NA | NA | NA | NA | 1 | NA | 0 | NA | 1 |
| SE | 11631 | ZZEF1 | chr17 | - | 4056215 | 4056345 | 4054056 | 4054195 | 4057993 | 4058155 | 0 | 0 | 2 | 0 | 0 | 1 | 0 | 2 | 0 | 0 | 0 | 0 | 98 | 49 | NA | NA | NA | NA | 0.5 | NA | 1 | NA | -0.5 |
| SE | 11633 | ZZEF1 | chr17 | - | 4109652 | 4109863 | 4105692 | 4105809 | 4112608 | 4112808 | 0 | 0 | 2 | 0 | 0 | 2 | 0 | 4 | 0 | 0 | 0 | 0 | 98 | 49 | NA | NA | NA | NA | 0.333 | NA | 1 | NA | -0.667 |
| SE | 11667 | NAAA | chr4 | - | 75918760 | 75918789 | 75914867 | 75914985 | 75919908 | 75919975 | 0 | 0 | 0 | 0 | 0 | 2 | 0 | 1 | 0 | 0 | 0 | 0 | 78 | 49 | NA | NA | NA | NA | 0 | NA | 1 | NA | -1 |
| SE | 11676 | ULK1 | chr12 | + | 131919211 | 131919384 | 131918496 | 131918681 | 131919471 | 131919590 | 0 | 0 | 1 | 0 | 0 | 0 | 0 | 2 | 0 | 0 | 2 | 0 | 98 | 49 | NA | NA | NA | NA | 1 | NA | 0.333 | NA | 0.667 |
| SE | 11678 | UBE3B | chr12 | + | 109523977 | 109524115 | 109521440 | 109521551 | 109524437 | 109524503 | 0 | 0 | 2 | 0 | 0 | 0 | 0 | 1 | 0 | 0 | 1 | 0 | 98 | 49 | NA | NA | NA | NA | 1 | NA | 0.333 | NA | 0.667 |
| SE | 11682 | ACADVL | chr17 | + | 7220923 | 7221058 | 7220765 | 7220830 | 7221537 | 7221682 | 0 | 0 | 10 | 0 | 0 | 1 | 0 | 6 | 0 | 0 | 0 | 0 | 98 | 49 | NA | NA | NA | NA | 0.833 | NA | 1 | NA | -0.167 |
| SE | 11686 | HLA-DQB1 | chr6 | - | 32661346 | 32661457 | 32659463 | 32660249 | 32661966 | 32662248 | 0 | 0 | 4 | 0 | 0 | 2 | 0 | 1 | 0 | 0 | 0 | 0 | 98 | 49 | NA | NA | NA | NA | 0.5 | NA | 1 | NA | -0.5 |
| SE | 11705 | TNFSF13 | chr17 | + | 7559845 | 7559893 | 7559623 | 7559702 | 7560048 | 7560167 | 0 | 0 | 11 | 0 | 0 | 1 | 0 | 0 | 0 | 0 | 1 | 0 | 97 | 49 | NA | NA | NA | NA | 0.847 | NA | 0 | NA | 0.847 |
| SE | 11728 | PER1 | chr17 | - | 8145957 | 8146137 | 8144750 | 8144993 | 8146371 | 8146502 | 0 | 0 | 3 | 0 | 0 | 1 | 0 | 1 | 0 | 0 | 6 | 0 | 98 | 49 | NA | NA | NA | NA | 0.6 | NA | 0.077 | NA | 0.523 |
| SE | 11735 | UBE2Q2 | chr15 | + | 75859877 | 75859982 | 75854385 | 75854487 | 75868950 | 75869010 | 0 | 0 | 5 | 0 | 0 | 1 | 0 | 1 | 0 | 0 | 0 | 0 | 98 | 49 | NA | NA | NA | NA | 0.714 | NA | 1 | NA | -0.286 |
| SE | 11745 | MOB1A | chr2 | - | 74159090 | 74159254 | 74156083 | 74156645 | 74165217 | 74165351 | 0 | 0 | 22 | 0 | 0 | 0 | 0 | 15 | 0 | 0 | 1 | 0 | 98 | 49 | NA | NA | NA | NA | 1 | NA | 0.882 | NA | 0.118 |
| SE | 11747 | COPS3 | chr17 | - | 17247479 | 17247560 | 17246623 | 17247151 | 17248925 | 17249039 | 0 | 0 | 1 | 0 | 0 | 2 | 0 | 5 | 0 | 0 | 0 | 0 | 98 | 49 | NA | NA | NA | NA | 0.2 | NA | 1 | NA | -0.8 |
| SE | 11764 | LOC100288778 | chr12 | - | 18372 | 18484 | 18036 | 18183 | 26800 | 26954 | 0 | 0 | 0 | 0 | 0 | 1 | 0 | 2 | 0 | 0 | 0 | 0 | 98 | 49 | NA | NA | NA | NA | 0 | NA | 1 | NA | -1 |
| SE | 11766 | KDM5A | chr12 | - | 306945 | 307089 | 297040 | 297200 | 307453 | 308005 | 0 | 0 | 1 | 0 | 0 | 1 | 0 | 2 | 0 | 0 | 0 | 0 | 98 | 49 | NA | NA | NA | NA | 0.333 | NA | 1 | NA | -0.667 |
| SE | 11780 | PNPLA6 | chr19 | + | 7541983 | 7542067 | 7541521 | 7541684 | 7542560 | 7542670 | 0 | 0 | 5 | 0 | 0 | 0 | 0 | 0 | 0 | 0 | 1 | 0 | 98 | 49 | NA | NA | NA | NA | 1 | NA | 0 | NA | 1 |
| SE | 11802 | KIAA0100 | chr17 | - | 28615663 | 28615798 | 28614439 | 28615246 | 28616095 | 28616197 | 0 | 0 | 5 | 0 | 0 | 0 | 0 | 1 | 0 | 0 | 1 | 0 | 98 | 49 | NA | NA | NA | NA | 1 | NA | 0.333 | NA | 0.667 |
| SE | 11803 | KIAA0100 | chr17 | - | 28616888 | 28616999 | 28616617 | 28616753 | 28617231 | 28617318 | 0 | 0 | 7 | 0 | 0 | 1 | 0 | 2 | 0 | 0 | 0 | 0 | 98 | 49 | NA | NA | NA | NA | 0.778 | NA | 1 | NA | -0.222 |
| SE | 11804 | KIAA0100 | chr17 | - | 28628264 | 28628559 | 28624232 | 28624390 | 28631478 | 28631681 | 0 | 0 | 7 | 0 | 0 | 1 | 0 | 1 | 0 | 0 | 1 | 0 | 98 | 49 | NA | NA | NA | NA | 0.778 | NA | 0.333 | NA | 0.445 |
| SE | 11815 | FLOT2 | chr17 | - | 28884224 | 28884315 | 28883107 | 28883231 | 28888944 | 28889026 | 0 | 0 | 13 | 0 | 0 | 2 | 2 | 3 | 0 | 0 | 0 | 0 | 98 | 49 | NA | NA | NA | NA | 0.765 | 1 | 1 | NA | -0.235 |
| SE | 11816 | DHRS13 | chr17 | - | 28901492 | 28901616 | 28900989 | 28901301 | 28902582 | 28902701 | 0 | 0 | 5 | 0 | 0 | 3 | 0 | 1 | 0 | 0 | 0 | 0 | 98 | 49 | NA | NA | NA | NA | 0.455 | NA | 1 | NA | -0.545 |
| SE | 11817 | PHF12 | chr17 | - | 28908782 | 28908881 | 28907589 | 28907672 | 28910225 | 28910369 | 0 | 0 | 5 | 0 | 0 | 1 | 0 | 5 | 0 | 0 | 2 | 0 | 98 | 49 | NA | NA | NA | NA | 0.714 | NA | 0.556 | NA | 0.158 |
| SE | 11818 | MAP3K3 | chr17 | + | 63634708 | 63634801 | 63632680 | 63632802 | 63646033 | 63646074 | 0 | 0 | 4 | 0 | 0 | 2 | 1 | 3 | 0 | 2 | 0 | 0 | 98 | 49 | NA | NA | NA | NA | 0.5 | 0.2 | 1 | NA | -0.1 |
| SE | 11819 | PROK2 | chr3 | - | 71774444 | 71774507 | 71771654 | 71772828 | 71781466 | 71781592 | 0 | 0 | 2 | 0 | 0 | 4 | 8 | 10 | 0 | 0 | 0 | 0 | 98 | 49 | NA | NA | NA | NA | 0.2 | 1 | 1 | NA | -0.8 |
| SE | 11833 | NCOA1 | chr2 | + | 24693251 | 24693347 | 24691480 | 24691660 | 24697657 | 24697798 | 0 | 0 | 8 | 0 | 0 | 0 | 1 | 3 | 0 | 0 | 1 | 0 | 98 | 49 | NA | NA | NA | NA | 1 | 1 | 0.6 | NA | 0.2 |
| SE | 11843 | RHOT1 | chr17 | + | 32182756 | 32182865 | 32176160 | 32176213 | 32183170 | 32183272 | 0 | 0 | 6 | 0 | 0 | 0 | 7 | 5 | 0 | 0 | 1 | 0 | 98 | 49 | NA | NA | NA | NA | 1 | 1 | 0.714 | NA | 0.143 |
| SE | 11845 | RHOT1 | chr17 | + | 32209349 | 32209445 | 32208106 | 32208309 | 32224615 | 32225771 | 0 | 0 | 1 | 0 | 0 | 2 | 0 | 2 | 0 | 0 | 5 | 0 | 98 | 49 | NA | NA | NA | NA | 0.2 | NA | 0.167 | NA | 0.033 |
| SE | 11846 | RHOT1 | chr17 | + | 32211115 | 32211238 | 32208106 | 32208309 | 32224615 | 32225771 | 0 | 0 | 2 | 0 | 0 | 2 | 0 | 1 | 0 | 0 | 5 | 0 | 98 | 49 | NA | NA | NA | NA | 0.333 | NA | 0.091 | NA | 0.242 |
| SE | 11848 | RHOT1 | chr17 | + | 32221048 | 32221186 | 32211115 | 32211238 | 32224615 | 32225771 | 0 | 0 | 2 | 0 | 0 | 2 | 0 | 0 | 0 | 0 | 1 | 0 | 98 | 49 | NA | NA | NA | NA | 0.333 | NA | 0 | NA | 0.333 |
| SE | 11850 | ARFGAP1 | chr20 | + | 63277204 | 63277305 | 63276479 | 63276651 | 63278116 | 63278203 | 0 | 0 | 1 | 0 | 0 | 0 | 0 | 2 | 0 | 0 | 2 | 0 | 98 | 49 | NA | NA | NA | NA | 1 | NA | 0.333 | NA | 0.667 |
| SE | 11869 | TPGS2 | chr18 | - | 36805373 | 36805397 | 36800197 | 36800311 | 36807846 | 36807934 | 0 | 0 | 1 | 0 | 0 | 0 | 0 | 0 | 0 | 1 | 0 | 0 | 73 | 49 | NA | NA | NA | NA | 1 | 0 | NA | NA | 1 |
| SE | 11870 | TPGS2 | chr18 | - | 36805373 | 36805502 | 36800197 | 36800311 | 36807846 | 36807934 | 0 | 0 | 1 | 0 | 0 | 0 | 0 | 0 | 0 | 1 | 0 | 0 | 98 | 49 | NA | NA | NA | NA | 1 | 0 | NA | NA | 1 |
| SE | 11881 | PHF20 | chr20 | + | 35870954 | 35871134 | 35869437 | 35869551 | 35871649 | 35871829 | 0 | 0 | 1 | 0 | 0 | 0 | 0 | 5 | 0 | 0 | 1 | 0 | 98 | 49 | NA | NA | NA | NA | 1 | NA | 0.714 | NA | 0.286 |
| SE | 11896 | KIAA0319L | chr1 | - | 35448172 | 35448332 | 35444160 | 35444303 | 35449866 | 35450005 | 0 | 0 | 5 | 0 | 0 | 0 | 0 | 1 | 0 | 1 | 0 | 0 | 98 | 49 | NA | NA | NA | NA | 1 | 0 | 1 | NA | 0.5 |
| SE | 11897 | KIAA0319L | chr1 | - | 35451627 | 35451776 | 35450357 | 35450509 | 35453556 | 35453689 | 0 | 0 | 0 | 0 | 0 | 1 | 0 | 1 | 0 | 0 | 0 | 0 | 98 | 49 | NA | NA | NA | NA | 0 | NA | 1 | NA | -1 |
| SE | 11904 | GGNBP2 | chr17 | + | 36557082 | 36557336 | 36554819 | 36554900 | 36560772 | 36560871 | 0 | 0 | 4 | 0 | 0 | 0 | 0 | 4 | 0 | 0 | 1 | 0 | 98 | 49 | NA | NA | NA | NA | 1 | NA | 0.667 | NA | 0.333 |
| SE | 11912 | SYNRG | chr17 | - | 37529782 | 37529851 | 37520537 | 37520648 | 37535978 | 37536127 | 0 | 0 | 0 | 0 | 0 | 2 | 0 | 1 | 0 | 0 | 1 | 0 | 98 | 49 | NA | NA | NA | NA | 0 | NA | 0.333 | NA | -0.333 |
| SE | 11917 | TNRC18 | chr7 | - | 5361593 | 5361722 | 5359397 | 5359569 | 5361896 | 5362033 | 0 | 0 | 4 | 0 | 0 | 2 | 1 | 0 | 0 | 0 | 0 | 0 | 98 | 49 | NA | NA | NA | NA | 0.5 | 1 | NA | NA | -0.5 |
| SE | 11933 | KIAA0355 | chr19 | + | 34333703 | 34333801 | 34330188 | 34330347 | 34339866 | 34339992 | 0 | 0 | 1 | 0 | 0 | 0 | 0 | 2 | 0 | 0 | 1 | 0 | 98 | 49 | NA | NA | NA | NA | 1 | NA | 0.5 | NA | 0.5 |
| SE | 11957 | WSB1 | chr17 | + | 27301787 | 27301956 | 27294079 | 27294435 | 27303366 | 27303635 | 0 | 0 | 12 | 0 | 0 | 1 | 1 | 3 | 0 | 0 | 2 | 0 | 98 | 49 | NA | NA | NA | NA | 0.857 | 1 | 0.429 | NA | 0.142 |
| SE | 11959 | WSB1 | chr17 | + | 27310060 | 27310174 | 27309099 | 27309272 | 27311508 | 27311616 | 0 | 0 | 17 | 0 | 0 | 0 | 0 | 3 | 0 | 0 | 1 | 0 | 98 | 49 | NA | NA | NA | NA | 1 | NA | 0.6 | NA | 0.4 |
| SE | 11978 | MED13 | chr17 | - | 61955679 | 61955838 | 61955381 | 61955567 | 61956338 | 61956481 | 0 | 0 | 2 | 0 | 0 | 1 | 0 | 1 | 0 | 0 | 0 | 0 | 98 | 49 | NA | NA | NA | NA | 0.5 | NA | 1 | NA | -0.5 |
| SE | 11988 | DNAJC7 | chr17 | - | 41989403 | 41989557 | 41988731 | 41988896 | 41990263 | 41990382 | 0 | 0 | 1 | 0 | 0 | 0 | 0 | 3 | 0 | 0 | 1 | 0 | 98 | 49 | NA | NA | NA | NA | 1 | NA | 0.6 | NA | 0.4 |
| SE | 12019 | UBTF | chr17 | - | 44212343 | 44212454 | 44211872 | 44212006 | 44212818 | 44212939 | 0 | 0 | 1 | 0 | 0 | 1 | 1 | 0 | 0 | 0 | 2 | 0 | 98 | 49 | NA | NA | NA | NA | 0.333 | 1 | 0 | NA | -0.167 |
| SE | 12021 | CCDC43 | chr17 | - | 44680584 | 44680643 | 44677436 | 44679043 | 44682002 | 44682138 | 0 | 0 | 1 | 0 | 0 | 0 | 0 | 0 | 0 | 0 | 1 | 0 | 98 | 49 | NA | NA | NA | NA | 1 | NA | 0 | NA | 1 |
| SE | 12025 | KLHL2 | chr4 | + | 165297608 | 165297725 | 165294358 | 165294468 | 165299506 | 165299656 | 0 | 0 | 1 | 0 | 0 | 0 | 0 | 1 | 0 | 0 | 2 | 0 | 98 | 49 | NA | NA | NA | NA | 1 | NA | 0.2 | NA | 0.8 |
| SE | 12114 | ERMARD | chr6 | + | 169766637 | 169766667 | 169762428 | 169762531 | 169768102 | 169768171 | 0 | 0 | 1 | 0 | 0 | 1 | 0 | 1 | 0 | 0 | 0 | 0 | 79 | 49 | NA | NA | NA | NA | 0.383 | NA | 1 | NA | -0.617 |
| SE | 12145 | BZW2 | chr7 | + | 16698047 | 16698186 | 16696914 | 16697061 | 16704546 | 16704669 | 0 | 0 | 1 | 0 | 0 | 0 | 0 | 0 | 0 | 0 | 1 | 0 | 98 | 49 | NA | NA | NA | NA | 1 | NA | 0 | NA | 1 |
| SE | 12154 | CC2D1B | chr1 | - | 52359704 | 52359883 | 52359458 | 52359534 | 52360073 | 52360233 | 0 | 0 | 1 | 0 | 0 | 1 | 0 | 2 | 0 | 0 | 0 | 0 | 98 | 49 | NA | NA | NA | NA | 0.333 | NA | 1 | NA | -0.667 |
| SE | 12158 | VMP1 | chr17 | + | 59738836 | 59738947 | 59735337 | 59735473 | 59764970 | 59765138 | 0 | 0 | 29 | 0 | 0 | 1 | 0 | 7 | 0 | 0 | 0 | 0 | 98 | 49 | NA | NA | NA | NA | 0.935 | NA | 1 | NA | -0.065 |
| SE | 12213 | HELZ | chr17 | - | 67122969 | 67123160 | 67120404 | 67120612 | 67123962 | 67124014 | 0 | 0 | 3 | 0 | 0 | 0 | 0 | 0 | 0 | 0 | 1 | 0 | 98 | 49 | NA | NA | NA | NA | 1 | NA | 0 | NA | 1 |
| SE | 12214 | HELZ | chr17 | - | 67167462 | 67167796 | 67166477 | 67166608 | 67178658 | 67178926 | 0 | 0 | 2 | 0 | 0 | 0 | 1 | 3 | 0 | 2 | 0 | 0 | 98 | 49 | NA | NA | NA | NA | 1 | 0.2 | 1 | NA | 0.4 |
| SE | 12222 | PRKAR1A | chr17 | + | 68514986 | 68515119 | 68512378 | 68512548 | 68515393 | 68515576 | 0 | 0 | 3 | 0 | 0 | 1 | 0 | 0 | 0 | 0 | 1 | 0 | 98 | 49 | NA | NA | NA | NA | 0.6 | NA | 0 | NA | 0.6 |
| SE | 12236 | RNASE4 | chr14 | + | 20693546 | 20693650 | 20684212 | 20684758 | 20699354 | 20700602 | 0 | 0 | 0 | 0 | 0 | 1 | 0 | 1 | 0 | 0 | 0 | 0 | 98 | 49 | NA | NA | NA | NA | 0 | NA | 1 | NA | -1 |
| SE | 12242 | ACOX1 | chr17 | - | 75949717 | 75949897 | 75949494 | 75949600 | 75950773 | 75950964 | 0 | 0 | 3 | 0 | 0 | 0 | 0 | 1 | 0 | 0 | 1 | 0 | 98 | 49 | NA | NA | NA | NA | 1 | NA | 0.333 | NA | 0.667 |
| SE | 12247 | OMA1 | chr1 | - | 58530600 | 58530729 | 58527260 | 58527335 | 58533952 | 58534060 | 0 | 0 | 1 | 0 | 0 | 0 | 0 | 3 | 0 | 0 | 1 | 0 | 98 | 49 | NA | NA | NA | NA | 1 | NA | 0.6 | NA | 0.4 |
| SE | 12269 | SUV420H1 | chr11 | - | 68180131 | 68180200 | 68175017 | 68175183 | 68185780 | 68185928 | 0 | 0 | 8 | 0 | 0 | 0 | 0 | 1 | 0 | 0 | 1 | 0 | 98 | 49 | NA | NA | NA | NA | 1 | NA | 0.333 | NA | 0.667 |
| SE | 12287 | JMJD6 | chr17 | - | 76723771 | 76724058 | 76721797 | 76721933 | 76725466 | 76725855 | 0 | 0 | 4 | 0 | 0 | 1 | 0 | 13 | 0 | 0 | 0 | 0 | 98 | 49 | NA | NA | NA | NA | 0.667 | NA | 1 | NA | -0.333 |
| SE | 12346 | CSNK1D | chr17 | - | 82245975 | 82246039 | 82242660 | 82244831 | 82248874 | 82249014 | 0 | 0 | 2 | 0 | 0 | 1 | 0 | 1 | 0 | 0 | 1 | 0 | 98 | 49 | NA | NA | NA | NA | 0.5 | NA | 0.333 | NA | 0.167 |
| SE | 12384 | VAPA | chr18 | + | 9944923 | 9945058 | 9936985 | 9937066 | 9950394 | 9950568 | 0 | 0 | 1 | 0 | 0 | 3 | 0 | 0 | 0 | 0 | 2 | 0 | 98 | 49 | NA | NA | NA | NA | 0.143 | NA | 0 | NA | 0.143 |
| SE | 12398 | GPS2 | chr17 | - | 7313905 | 7313988 | 7313567 | 7313721 | 7314079 | 7314159 | 0 | 2 | 0 | 0 | 0 | 0 | 0 | 0 | 0 | 0 | 0 | 2 | 98 | 49 | NA | NA | NA | 1 | NA | NA | NA | 0 | 1 |
| SE | 12428 | SS18 | chr18 | - | 26035830 | 26035923 | 26035004 | 26035127 | 26038554 | 26038659 | 0 | 0 | 0 | 0 | 0 | 2 | 0 | 2 | 0 | 0 | 0 | 0 | 98 | 49 | NA | NA | NA | NA | 0 | NA | 1 | NA | -1 |
| SE | 12435 | SPAG9 | chr17 | - | 50985697 | 50985778 | 50982523 | 50982672 | 50987111 | 50987237 | 0 | 0 | 7 | 0 | 0 | 1 | 0 | 8 | 0 | 0 | 0 | 0 | 98 | 49 | NA | NA | NA | NA | 0.778 | NA | 1 | NA | -0.222 |

| | | | | | | | | | | | | | | | | | | | | | | | | | | | | | | | |
|---|---|---|---|---|---|---|---|---|---|---|---|---|---|---|---|---|---|---|---|---|---|---|---|---|---|---|---|---|---|---|---|
| SE | 12438 | SPAG9 | chr17 | - | 51031680 | 51031722 | 51021157 | 51021365 | 51041500 | 51041651 | 0 | 0 | 0 | 0 | 0 | 8 | 0 | 10 | 0 | 0 | 6 | 0 | 91 | 49 | NA | NA | NA | NA | 0 | NA | 0.473 | NA | -0.473 |
| SE | 12446 | TRAPPC8 | chr18 | - | 31866848 | 31866975 | 31864626 | 31864781 | 31867401 | 31867476 | 0 | 0 | 5 | 0 | 0 | 0 | 0 | 0 | 0 | 0 | 2 | 0 | 98 | 49 | NA | NA | NA | NA | 1 | NA | 0 | NA | 1 |
| SE | 12496 | MBD1 | chr18 | - | 50274976 | 50275045 | 50274185 | 50274353 | 50275128 | 50275245 | 0 | 0 | 2 | 0 | 0 | 0 | 0 | 0 | 0 | 0 | 1 | 0 | 98 | 49 | NA | NA | NA | NA | 1 | NA | 0 | NA | 1 |
| SE | 12501 | HELQ | chr4 | - | 83446834 | 83447035 | 83446013 | 83446086 | 83448782 | 83448961 | 0 | 0 | 0 | 0 | 0 | 1 | 0 | 1 | 0 | 0 | 0 | 0 | 98 | 49 | NA | NA | NA | NA | 0 | NA | 1 | NA | -1 |
| SE | 12525 | GUCD1 | chr22 | - | 24547907 | 24548073 | 24546913 | 24547005 | 24554948 | 24555307 | 0 | 0 | 1 | 0 | 0 | 1 | 0 | 5 | 0 | 0 | 0 | 0 | 98 | 49 | NA | NA | NA | NA | 0.333 | NA | 1 | NA | -0.667 |
| SE | 12534 | VPS4B | chr18 | - | 63410289 | 63410446 | 63407431 | 63407499 | 63411466 | 63411578 | 0 | 0 | 7 | 0 | 0 | 0 | 0 | 5 | 0 | 0 | 1 | 0 | 98 | 49 | NA | NA | NA | NA | 1 | NA | 0.714 | NA | 0.286 |
| SE | 12564 | EIF4G1 | chr3 | + | 184315488 | 184315545 | 184314494 | 184314674 | 184315762 | 184315856 | 0 | 0 | 2 | 0 | 0 | 1 | 0 | 1 | 0 | 0 | 0 | 0 | 98 | 49 | NA | NA | NA | NA | 0.5 | NA | 1 | NA | -0.5 |
| SE | 12568 | EIF4G1 | chr3 | + | 184328908 | 184328990 | 184328630 | 184328756 | 184331265 | 184331364 | 0 | 0 | 9 | 0 | 0 | 0 | 2 | 5 | 0 | 0 | 1 | 0 | 98 | 49 | NA | NA | NA | NA | 1 | 1 | 0.714 | NA | 0.143 |
| SE | 12570 | PTBP1 | chr19 | + | 805491 | 805569 | 805012 | 805187 | 806407 | 806556 | 0 | 0 | 1 | 0 | 0 | 0 | 0 | 0 | 0 | 0 | 1 | 0 | 98 | 49 | NA | NA | NA | NA | 1 | NA | 0 | NA | 1 |
| SE | 12571 | PTBP1 | chr19 | + | 805512 | 805569 | 805012 | 805187 | 806407 | 806556 | 0 | 0 | 1 | 0 | 0 | 0 | 0 | 0 | 0 | 0 | 1 | 0 | 98 | 49 | NA | NA | NA | NA | 1 | NA | 0 | NA | 1 |
| SE | 12572 | PTBP1 | chr19 | + | 807868 | 807902 | 806407 | 806556 | 808359 | 808452 | 0 | 0 | 3 | 0 | 0 | 1 | 0 | 1 | 0 | 0 | 0 | 0 | 83 | 49 | NA | NA | NA | NA | 0.639 | NA | 1 | NA | -0.361 |
| SE | 12577 | ARID3A | chr19 | + | 966571 | 966868 | 964247 | 964431 | 968404 | 968503 | 0 | 0 | 0 | 0 | 0 | 1 | 0 | 1 | 0 | 0 | 0 | 0 | 98 | 49 | NA | NA | NA | NA | 0 | NA | 1 | NA | -1 |
| SE | 12596 | GBF1 | chr10 | + | 102369860 | 102369984 | 102369710 | 102369775 | 102370173 | 102370245 | 0 | 0 | 1 | 0 | 0 | 0 | 0 | 1 | 0 | 0 | 2 | 0 | 98 | 49 | NA | NA | NA | NA | 1 | NA | 0.2 | NA | 0.8 |
| SE | 12597 | GBF1 | chr10 | + | 102381126 | 102381255 | 102380505 | 102380686 | 102382055 | 102382899 | 0 | 0 | 1 | 0 | 0 | 0 | 0 | 7 | 0 | 0 | 1 | 0 | 98 | 49 | NA | NA | NA | NA | 1 | NA | 0.778 | NA | 0.222 |
| SE | 12600 | CCT4 | chr2 | - | 61872457 | 61872588 | 61872081 | 61872316 | 61873001 | 61873112 | 0 | 0 | 0 | 0 | 0 | 1 | 0 | 2 | 0 | 0 | 0 | 0 | 98 | 49 | NA | NA | NA | NA | 0 | NA | 1 | NA | -1 |
| SE | 12632 | SAFB2 | chr19 | - | 5611119 | 5611630 | 5610638 | 5610688 | 5612539 | 5612567 | 0 | 0 | 4 | 0 | 0 | 0 | 0 | 1 | 0 | 0 | 1 | 0 | 98 | 49 | NA | NA | NA | NA | 1 | NA | 0.333 | NA | 0.667 |
| SE | 12638 | ZNF638 | chr2 | + | 71426459 | 71427414 | 71424649 | 71424715 | 71428546 | 71428651 | 0 | 0 | 2 | 0 | 0 | 0 | 0 | 2 | 0 | 0 | 1 | 0 | 98 | 49 | NA | NA | NA | NA | 1 | NA | 0.5 | NA | 0.5 |
| SE | 12639 | ZNF638 | chr2 | + | 71428546 | 71428651 | 71426459 | 71427414 | 71431326 | 71431428 | 0 | 0 | 1 | 0 | 0 | 0 | 0 | 0 | 0 | 0 | 1 | 0 | 98 | 49 | NA | NA | NA | NA | 1 | NA | 0 | NA | 1 |
| SE | 12647 | GTF2F1 | chr19 | - | 6383310 | 6383495 | 6381696 | 6381850 | 6387388 | 6387559 | 0 | 0 | 1 | 0 | 0 | 1 | 0 | 1 | 0 | 0 | 0 | 0 | 98 | 49 | NA | NA | NA | NA | 0.333 | NA | 1 | NA | -0.667 |
| SE | 12651 | ITGA4 | chr2 | + | 181530523 | 181530649 | 181529540 | 181529648 | 181531656 | 181531776 | 0 | 0 | 7 | 0 | 0 | 2 | 0 | 1 | 0 | 0 | 0 | 0 | 98 | 49 | NA | NA | NA | NA | 0.636 | NA | 1 | NA | -0.364 |
| SE | 12659 | PPP2R1A | chr19 | + | 52220188 | 52220249 | 52219690 | 52219864 | 52220978 | 52221133 | 0 | 0 | 6 | 0 | 0 | 1 | 0 | 1 | 0 | 0 | 0 | 0 | 98 | 49 | NA | NA | NA | NA | 0.75 | NA | 1 | NA | -0.25 |
| SE | 12663 | PET100 | chr19 | + | 7631231 | 7631321 | 7630822 | 7630846 | 7631472 | 7631624 | 0 | 0 | 0 | 0 | 0 | 8 | 0 | 1 | 0 | 0 | 4 | 0 | 98 | 49 | NA | NA | NA | NA | 0 | NA | 0.111 | NA | -0.111 |
| SE | 12694 | UTRN | chr6 | + | 144437564 | 144437746 | 144435934 | 144436138 | 144438744 | 144438895 | 0 | 0 | 19 | 0 | 0 | 1 | 0 | 10 | 0 | 0 | 0 | 0 | 98 | 49 | NA | NA | NA | NA | 0.905 | NA | 1 | NA | -0.095 |
| SE | 12695 | UTRN | chr6 | + | 144827347 | 144827386 | 144820881 | 144821018 | 144827610 | 144827676 | 0 | 0 | 2 | 0 | 0 | 2 | 0 | 0 | 0 | 0 | 1 | 0 | 88 | 49 | NA | NA | NA | NA | 0.358 | NA | 0 | NA | 0.358 |
| SE | 12702 | ARFGEF1 | chr8 | - | 67277281 | 67277457 | 67271701 | 67271936 | 67287954 | 67288065 | 0 | 0 | 5 | 0 | 0 | 1 | 0 | 2 | 0 | 0 | 3 | 0 | 98 | 49 | NA | NA | NA | NA | 0.714 | NA | 0.25 | NA | 0.464 |
| SE | 12720 | GHITM | chr10 | + | 84148729 | 84148838 | 84144874 | 84145016 | 84150054 | 84150243 | 0 | 0 | 7 | 0 | 0 | 0 | 0 | 9 | 0 | 0 | 1 | 0 | 98 | 49 | NA | NA | NA | NA | 1 | NA | 0.818 | NA | 0.182 |
| SE | 12727 | C19orf43 | chr19 | - | 12731343 | 12731421 | 12730639 | 12731068 | 12734312 | 12734715 | 0 | 0 | 5 | 0 | 0 | 1 | 0 | 9 | 0 | 0 | 0 | 0 | 98 | 49 | NA | NA | NA | NA | 0.714 | NA | 1 | NA | -0.286 |
| SE | 12747 | DYSF | chr2 | + | 71669111 | 71669207 | 71668753 | 71668842 | 71669604 | 71669746 | 0 | 0 | 17 | 0 | 0 | 3 | 0 | 2 | 0 | 0 | 2 | 0 | 98 | 49 | NA | NA | NA | NA | 0.739 | NA | 0.333 | NA | 0.406 |
| SE | 12749 | COL4A3BP | chr5 | - | 75384641 | 75384712 | 75381948 | 75382077 | 75385901 | 75386034 | 0 | 0 | 4 | 0 | 0 | 0 | 0 | 1 | 0 | 0 | 2 | 0 | 98 | 49 | NA | NA | NA | NA | 1 | NA | 0.2 | NA | 0.8 |
| SE | 12751 | COL4A3BP | chr5 | - | 75411010 | 75411103 | 75402971 | 75403058 | 75416875 | 75417033 | 0 | 0 | 1 | 0 | 0 | 0 | 0 | 2 | 0 | 0 | 1 | 0 | 98 | 49 | NA | NA | NA | NA | 1 | NA | 0.5 | NA | 0.5 |
| SE | 12802 | CYP4F3 | chr19 | + | 15643906 | 15644051 | 15641414 | 15641613 | 15647051 | 15647105 | 0 | 0 | 1 | 0 | 0 | 0 | 0 | 0 | 0 | 0 | 3 | 0 | 98 | 49 | NA | NA | NA | NA | 1 | NA | 0 | NA | 1 |
| SE | 12828 | SSBP4 | chr19 | + | 18431352 | 18431418 | 18430840 | 18430930 | 18431646 | 18431706 | 0 | 0 | 0 | 0 | 0 | 1 | 0 | 1 | 0 | 0 | 1 | 0 | 98 | 49 | NA | NA | NA | NA | 0 | NA | 0.333 | NA | -0.333 |
| SE | 12852 | GATAD2A | chr19 | + | 19501968 | 19502043 | 19501117 | 19501416 | 19502330 | 19502526 | 0 | 0 | 3 | 0 | 0 | 0 | 0 | 4 | 0 | 0 | 1 | 0 | 98 | 49 | NA | NA | NA | NA | 1 | NA | 0.667 | NA | 0.333 |
| SE | 12854 | ZNF101 | chr19 | + | 19678725 | 19678786 | 19677863 | 19677990 | 19679180 | 19683506 | 0 | 0 | 1 | 0 | 0 | 1 | 1 | 4 | 0 | 0 | 0 | 0 | 98 | 49 | NA | NA | NA | NA | 0.333 | 1 | 1 | NA | -0.667 |
| SE | 12915 | PITPNM1 | chr11 | - | 67498939 | 67499001 | 67498595 | 67498846 | 67499722 | 67499830 | 0 | 0 | 2 | 0 | 0 | 0 | 0 | 2 | 0 | 0 | 1 | 0 | 98 | 49 | NA | NA | NA | NA | 1 | NA | 0.5 | NA | 0.5 |
| SE | 12934 | EIF3K | chr19 | + | 38620336 | 38620435 | 38619073 | 38619327 | 38624076 | 38624197 | 0 | 0 | 8 | 0 | 0 | 1 | 0 | 10 | 0 | 0 | 0 | 0 | 98 | 49 | NA | NA | NA | NA | 0.8 | NA | 1 | NA | -0.2 |
| SE | 12935 | EIF3K | chr19 | + | 38632600 | 38632678 | 38632429 | 38632496 | 38634992 | 38635118 | 0 | 0 | 5 | 0 | 0 | 1 | 1 | 5 | 0 | 0 | 0 | 0 | 98 | 49 | NA | NA | NA | NA | 0.714 | 1 | 1 | NA | -0.286 |
| SE | 12953 | MAN2A2 | chr15 | + | 90910089 | 90910292 | 90909326 | 90909504 | 90910500 | 90910683 | 0 | 0 | 8 | 0 | 0 | 1 | 0 | 5 | 0 | 0 | 0 | 0 | 98 | 49 | NA | NA | NA | NA | 0.8 | NA | 1 | NA | -0.2 |
| SE | 12982 | CIC | chr19 | + | 42291557 | 42291745 | 42290232 | 42291466 | 42292085 | 42292207 | 0 | 0 | 1 | 0 | 0 | 2 | 0 | 1 | 0 | 0 | 0 | 0 | 98 | 49 | NA | NA | NA | NA | 0.2 | NA | 1 | NA | -0.8 |
| SE | 12983 | ATG13 | chr11 | + | 46618324 | 46618375 | 46617275 | 46617890 | 46644278 | 46644360 | 0 | 0 | 2 | 0 | 0 | 0 | 0 | 0 | 0 | 0 | 1 | 0 | 98 | 49 | NA | NA | NA | NA | 1 | NA | 0 | NA | 1 |
| SE | 12984 | ATG13 | chr11 | + | 46630044 | 46630100 | 46617275 | 46617890 | 46644278 | 46644360 | 0 | 0 | 2 | 0 | 0 | 0 | 0 | 0 | 0 | 0 | 1 | 0 | 98 | 49 | NA | NA | NA | NA | 1 | NA | 0 | NA | 1 |
| SE | 13135 | OCIAD1 | chr4 | + | 48833400 | 48833481 | 48832619 | 48832682 | 48842635 | 48842689 | 0 | 0 | 8 | 0 | 0 | 0 | 0 | 1 | 0 | 1 | 0 | 0 | 98 | 49 | NA | NA | NA | NA | 1 | 0 | 1 | NA | 0.5 |
| SE | 13139 | CNOT10 | chr3 | + | 32713226 | 32713369 | 32708669 | 32708820 | 32716224 | 32716311 | 0 | 0 | 3 | 0 | 0 | 0 | 0 | 0 | 0 | 0 | 2 | 0 | 98 | 49 | NA | NA | NA | NA | 1 | NA | 0 | NA | 1 |
| SE | 13156 | NFRKB | chr11 | - | 129872883 | 129873096 | 129869493 | 129870261 | 129873744 | 129874015 | 0 | 0 | 1 | 0 | 0 | 0 | 0 | 0 | 0 | 0 | 1 | 0 | 98 | 49 | NA | NA | NA | NA | 1 | NA | 0 | NA | 1 |
| SE | 13211 | LILRB2 | chr19 | - | 54280462 | 54280544 | 54279790 | 54280075 | 54280960 | 54281178 | 0 | 0 | 0 | 0 | 0 | 1 | 0 | 1 | 0 | 0 | 1 | 0 | 98 | 49 | NA | NA | NA | NA | 0 | NA | 0.333 | NA | -0.333 |
| SE | 13238 | PAFAH1B2 | chr11 | + | 117161144 | 117161261 | 117159933 | 117160023 | 117163769 | 117163874 | 0 | 0 | 6 | 0 | 0 | 2 | 0 | 10 | 0 | 0 | 0 | 0 | 98 | 49 | NA | NA | NA | NA | 0.6 | NA | 1 | NA | -0.4 |
| SE | 13358 | BROX | chr1 | + | 222715683 | 222715800 | 222713320 | 222713416 | 222718924 | 222719031 | 0 | 0 | 1 | 0 | 0 | 0 | 0 | 0 | 0 | 0 | 1 | 0 | 98 | 49 | NA | NA | NA | NA | 1 | NA | 0 | NA | 1 |
| SE | 13366 | FIP1L1 | chr4 | + | 53379072 | 53379117 | 53377652 | 53377923 | 53379224 | 53379264 | 0 | 0 | 0 | 0 | 0 | 3 | 0 | 1 | 0 | 0 | 0 | 0 | 94 | 49 | NA | NA | NA | NA | 0 | NA | 1 | NA | -1 |
| SE | 13382 | PRKDC | chr8 | - | 47836337 | 47836527 | 47834195 | 47834396 | 47837211 | 47837419 | 0 | 0 | 11 | 0 | 0 | 2 | 0 | 3 | 0 | 0 | 0 | 0 | 98 | 49 | NA | NA | NA | NA | 0.733 | NA | 1 | NA | -0.267 |
| SE | 13385 | HNRNPH3 | chr10 | + | 68337857 | 68337996 | 68337198 | 68337333 | 68338502 | 68338642 | 0 | 0 | 1 | 0 | 0 | 0 | 0 | 1 | 0 | 0 | 1 | 0 | 98 | 49 | NA | NA | NA | NA | 1 | NA | 0.333 | NA | 0.667 |
| SE | 13424 | TBC1D5 | chr3 | - | 17428449 | 17428519 | 17406417 | 17406526 | 17508473 | 17508605 | 0 | 0 | 2 | 0 | 0 | 0 | 5 | 0 | 0 | 0 | 1 | 0 | 98 | 49 | NA | NA | NA | NA | 1 | 1 | 0 | NA | 0.5 |
| SE | 13449 | ADAM17 | chr2 | - | 9535833 | 9535922 | 9527785 | 9527954 | 9536697 | 9536828 | 0 | 0 | 3 | 0 | 0 | 1 | 0 | 1 | 0 | 0 | 0 | 0 | 98 | 49 | NA | NA | NA | NA | 0.6 | NA | 1 | NA | -0.4 |
| SE | 13482 | C10orf128 | chr10 | - | 49165765 | 49165796 | 49161598 | 49161622 | 49166868 | 49167009 | 0 | 0 | 1 | 0 | 0 | 1 | 0 | 1 | 0 | 0 | 0 | 0 | 80 | 49 | NA | NA | NA | NA | 0.38 | NA | 1 | NA | -0.62 |
| SE | 13484 | FAM228B | chr2 | + | 24137908 | 24138100 | 24135118 | 24135187 | 24139369 | 24139450 | 0 | 0 | 1 | 0 | 0 | 0 | 0 | 0 | 0 | 0 | 1 | 0 | 98 | 49 | NA | NA | NA | NA | 1 | NA | 0 | NA | 1 |
| SE | 13487 | ITSN2 | chr2 | - | 24284762 | 24284843 | 24275712 | 24275849 | 24286211 | 24286351 | 0 | 0 | 0 | 0 | 0 | 3 | 2 | 1 | 0 | 0 | 5 | 0 | 98 | 49 | NA | NA | NA | NA | 0 | 1 | 0.091 | NA | -0.545 |
| SE | 13499 | HADHB | chr2 | + | 26260980 | 26261063 | 26254429 | 26254474 | 26263379 | 26263479 | 0 | 0 | 1 | 0 | 0 | 2 | 0 | 0 | 0 | 0 | 1 | 0 | 98 | 49 | NA | NA | NA | NA | 0.2 | NA | 0 | NA | 0.2 |
| SE | 13524 | BRE | chr2 | + | 27892382 | 27892448 | 27890614 | 27890842 | 27894532 | 27894684 | 0 | 0 | 1 | 0 | 0 | 3 | 0 | 2 | 0 | 0 | 0 | 0 | 98 | 49 | NA | NA | NA | NA | 0.143 | NA | 1 | NA | -0.857 |
| SE | 13565 | EML4 | chr2 | + | 42301240 | 42301392 | 42295380 | 42295516 | 42303103 | 42303229 | 0 | 0 | 6 | 0 | 0 | 0 | 0 | 7 | 0 | 0 | 1 | 0 | 98 | 49 | NA | NA | NA | NA | 1 | NA | 0.778 | NA | 0.222 |
| SE | 13566 | RNF144B | chr6 | + | 18457154 | 18457359 | 18439683 | 18439744 | 18459606 | 18459751 | 0 | 0 | 9 | 0 | 0 | 3 | 0 | 1 | 0 | 3 | 0 | 0 | 98 | 49 | NA | NA | NA | NA | 0.6 | 0 | 1 | NA | 0.1 |
| SE | 13567 | THADA | chr2 | - | 43560233 | 43560385 | 43556092 | 43556555 | 43566697 | 43566821 | 0 | 0 | 5 | 0 | 0 | 0 | 0 | 1 | 0 | 4 | 0 | 0 | 98 | 49 | NA | NA | NA | NA | 1 | 0 | 1 | NA | 0.5 |
| SE | 13593 | VRK2 | chr2 | + | 58123100 | 58123233 | 58089630 | 58089723 | 58131807 | 58131928 | 0 | 0 | 2 | 0 | 0 | 0 | 0 | 1 | 0 | 0 | 1 | 0 | 98 | 49 | NA | NA | NA | NA | 1 | NA | 0.333 | NA | 0.667 |
| SE | 13594 | VRK2 | chr2 | + | 58135140 | 58135199 | 58131807 | 58131928 | 58139665 | 58139832 | 0 | 0 | 2 | 0 | 0 | 0 | 0 | 0 | 0 | 0 | 3 | 0 | 98 | 49 | NA | NA | NA | NA | 1 | NA | 0 | NA | 1 |
| SE | 13595 | VRK2 | chr2 | + | 58154753 | 58154802 | 58146315 | 58146474 | 58159348 | 58159920 | 0 | 0 | 2 | 0 | 0 | 1 | 0 | 0 | 0 | 0 | 1 | 0 | 98 | 49 | NA | NA | NA | NA | 0.5 | NA | 0 | NA | 0.5 |
| SE | 13607 | USP34 | chr2 | - | 61192900 | 61192980 | 61190517 | 61190658 | 61203139 | 61203263 | 0 | 0 | 3 | 0 | 0 | 1 | 0 | 1 | 0 | 0 | 0 | 0 | 98 | 49 | NA | NA | NA | NA | 0.6 | NA | 1 | NA | -0.4 |
| SE | 13608 | XPO1 | chr2 | - | 61492914 | 61493053 | 61492566 | 61492748 | 61493893 | 61494091 | 0 | 0 | 4 | 0 | 0 | 2 | 0 | 2 | 0 | 0 | 0 | 0 | 98 | 49 | NA | NA | NA | NA | 0.5 | NA | 1 | NA | -0.5 |
| SE | 13610 | USP47 | chr11 | + | 11891967 | 11892106 | 11884466 | 11884580 | 11897596 | 11897693 | 0 | 0 | 1 | 0 | 0 | 2 | 0 | 7 | 0 | 0 | 0 | 0 | 98 | 49 | NA | NA | NA | NA | 0.2 | NA | 1 | NA | -0.8 |
| SE | 13636 | CCND3 | chr6 | - | 42030008 | 42030190 | 41937234 | 41937394 | 42048500 | 42048894 | 0 | 0 | 1 | 0 | 0 | 1 | 0 | 0 | 0 | 0 | 1 | 0 | 98 | 49 | NA | NA | NA | NA | 0.333 | NA | 0 | NA | 0.333 |
| SE | 13637 | NRBP1 | chr2 | + | 27434470 | 27434560 | 27433988 | 27434090 | 27434721 | 27434762 | 0 | 0 | 5 | 0 | 0 | 1 | 0 | 4 | 0 | 0 | 0 | 0 | 98 | 49 | NA | NA | NA | NA | 0.714 | NA | 1 | NA | -0.286 |
| SE | 13643 | PCBP1-AS1 | chr2 | - | 70083537 | 70083687 | 70059642 | 70059680 | 70085546 | 70085624 | 0 | 0 | 1 | 0 | 0 | 0 | 0 | 2 | 0 | 0 | 1 | 0 | 98 | 49 | NA | NA | NA | NA | 1 | NA | 0.5 | NA | 0.5 |
| SE | 13675 | TP53BP2 | chr1 | - | 223818205 | 223818337 | 223814239 | 223814353 | 223821219 | 223821367 | 0 | 0 | 1 | 0 | 0 | 3 | 0 | 0 | 0 | 0 | 1 | 0 | 98 | 49 | NA | NA | NA | NA | 0.143 | NA | 0 | NA | 0.143 |
| SE | 13704 | LINC00963 | chr9 | + | 129502396 | 129503139 | 129493477 | 129493595 | 129503322 | 129503672 | 0 | 0 | 0 | 0 | 0 | 2 | 0 | 1 | 0 | 0 | 1 | 0 | 98 | 49 | NA | NA | NA | NA | 0 | NA | 0.333 | NA | -0.333 |
| SE | 13714 | MTMR10 | chr15 | - | 30951967 | 30952038 | 30948301 | 30948471 | 30953561 | 30953631 | 0 | 0 | 5 | 0 | 0 | 0 | 0 | 0 | 0 | 0 | 4 | 0 | 98 | 49 | NA | NA | NA | NA | 1 | NA | 0 | NA | 1 |
| SE | 13715 | MTMR10 | chr15 | - | 30976818 | 30976955 | 30974930 | 30975003 | 30990776 | 30990837 | 0 | 0 | 4 | 0 | 0 | 2 | 0 | 2 | 0 | 0 | 0 | 0 | 98 | 49 | NA | NA | NA | NA | 0.5 | NA | 1 | NA | -0.5 |
| SE | 13746 | MAP4K4 | chr2 | + | 101887087 | 101887261 | 101885186 | 101885287 | 101887777 | 101887937 | 0 | 0 | 6 | 0 | 0 | 0 | 0 | 3 | 0 | 0 | 1 | 0 | 98 | 49 | NA | NA | NA | NA | 1 | NA | 0.6 | NA | 0.4 |
| SE | 13747 | MAP4K4 | chr2 | + | 101888795 | 101888935 | 101887777 | 101887937 | 101891165 | 101894690 | 0 | 0 | 8 | 0 | 0 | 0 | 3 | 5 | 0 | 0 | 1 | 0 | 98 | 49 | NA | NA | NA | NA | 1 | 1 | 0.714 | NA | 0.143 |
| SE | 13752 | HMBOX1 | chr8 | + | 28980070 | 28980156 | 28970042 | 28970519 | 29009071 | 29009182 | 0 | 0 | 2 | 0 | 0 | 0 | 0 | 0 | 0 | 0 | 1 | 0 | 98 | 49 | NA | NA | NA | NA | 1 | NA | 0 | NA | 1 |
| SE | 13760 | NCOA6 | chr20 | - | 34727258 | 34727407 | 34714773 | 34715365 | 34732558 | 34732595 | 0 | 0 | 1 | 0 | 0 | 2 | 0 | 1 | 0 | 0 | 0 | 0 | 98 | 49 | NA | NA | NA | NA | 0.2 | NA | 1 | NA | -0.8 |
| SE | 13796 | LGALS8 | chr1 | + | 236542914 | 236543040 | 236542760 | 236542787 | 236543559 | 236543648 | 0 | 0 | 0 | 0 | 0 | 1 | 0 | 1 | 0 | 0 | 0 | 0 | 98 | 49 | NA | NA | NA | NA | 0 | NA | 1 | NA | -1 |
| SE | 13797 | CCDC93 | chr2 | - | 117973907 | 117973994 | 117958364 | 117958481 | 117974849 | 117974900 | 0 | 0 | 2 | 0 | 0 | 0 | 0 | 5 | 0 | 0 | 1 | 0 | 98 | 49 | NA | NA | NA | NA | 1 | NA | 0.714 | NA | 0.286 |
| SE | 13806 | CLASP1 | chr2 | - | 121409023 | 121409047 | 121407470 | 121407715 | 121410865 | 121410969 | 0 | 0 | 0 | 0 | 0 | 2 | 0 | 4 | 0 | 0 | 3 | 0 | 73 | 49 | NA | NA | NA | NA | 0 | NA | 0.472 | NA | -0.472 |
| SE | 13810 | CLASP1 | chr2 | - | 121445448 | 121445496 | 121430072 | 121430177 | 121447336 | 121447507 | 0 | 0 | 0 | 0 | 0 | 1 | 0 | 1 | 0 | 0 | 1 | 0 | 97 | 49 | NA | NA | NA | NA | 0 | NA | 0.336 | NA | -0.336 |
| SE | 13811 | CLASP1 | chr2 | - | 121515664 | 121515762 | 121503166 | 121503234 | 121525844 | 121525920 | 0 | 0 | 1 | 0 | 0 | 2 | 0 | 2 | 0 | 0 | 0 | 0 | 98 | 49 | NA | NA | NA | NA | 0.2 | NA | 1 | NA | -0.8 |
| SE | 13850 | SFT2D1 | chr6 | - | 166331384 | 166331407 | 166330160 | 166330247 | 166342418 | 166342606 | 0 | 0 | 1 | 0 | 0 | 1 | 0 | 0 | 0 | 0 | 2 | 0 | 72 | 49 | NA | NA | NA | NA | 0.405 | NA | 0 | NA | 0.405 |
| SE | 13871 | RAB3GAP1 | chr2 | + | 135135563 | 135135932 | 135135264 | 135135319 | 135150368 | 135150506 | 0 | 0 | 7 | 0 | 0 | 0 | 0 | 3 | 0 | 0 | 1 | 0 | 98 | 49 | NA | NA | NA | NA | 1 | NA | 0.6 | NA | 0.4 |
| SE | 13878 | ZEB2 | chr2 | - | 144398300 | 144400270 | 144396411 | 144396592 | 144401198 | 144401307 | 0 | 0 | 8 | 0 | 0 | 1 | 1 | 3 | 0 | 0 | 0 | 0 | 98 | 49 | NA | NA | NA | NA | 0.8 | 1 | 1 | NA | -0.2 |
| SE | 13880 | ZEB2 | chr2 | - | 144424795 | 144424867 | 144404835 | 144405024 | 144429768 | 144430026 | 0 | 0 | 6 | 0 | 0 | 1 | 0 | 5 | 0 | 0 | 0 | 0 | 98 | 49 | NA | NA | NA | NA | 0.75 | NA | 1 | NA | -0.25 |
| SE | 13881 | ZEB2 | chr2 | - | 144517277 | 144517419 | 144429768 | 144430026 | 144519938 | 144520391 | 0 | 0 | 6 | 0 | 0 | 0 | 0 | 4 | 0 | 0 | 1 | 0 | 98 | 49 | NA | NA | NA | NA | 1 | NA | 0.667 | NA | 0.333 |
| SE | 13890 | CD48 | chr1 | - | 160684886 | 160685189 | 160680421 | 160681468 | 160711681 | 160711851 | 0 | 0 | 22 | 0 | 0 | 1 | 0 | 14 | 0 | 0 | 0 | 0 | 98 | 49 | NA | NA | NA | NA | 0.917 | NA | 1 | NA | -0.083 |
| SE | 13907 | NR4A2 | chr2 | - | 156328403 | 156328533 | 156327850 | 156328014 | 156329322 | 156330188 | 0 | 0 | 1 | 0 | 0 | 1 | 7 | 6 | 0 | 0 | 0 | 0 | 98 | 49 | NA | NA | NA | NA | 0.333 | 1 | 1 | NA | -0.667 |
| SE | 13911 | CSAD | chr12 | - | 53161126 | 53161191 | 53160762 | 53160844 | 53161272 | 53161389 | 0 | 0 | 2 | 0 | 0 | 4 | 0 | 2 | 0 | 0 | 1 | 0 | 98 | 49 | NA | NA | NA | NA | 0.2 | NA | 0.5 | NA | -0.3 |
| SE | 13942 | PDK1 | chr2 | + | 172562740 | 172562800 | 172562219 | 172562291 | 172564502 | 172564687 | 0 | 0 | 1 | 0 | 0 | 0 | 0 | 0 | 0 | 0 | 2 | 0 | 98 | 49 | NA | NA | NA | NA | 1 | NA | 0 | NA | 1 |
| SE | 13944 | PDK1 | chr2 | + | 172564502 | 172564687 | 172562219 | 172562291 | 172564977 | 172565073 | 0 | 0 | 0 | 0 | 0 | 1 | 0 | 2 | 0 | 0 | 1 | 0 | 98 | 49 | NA | NA | NA | NA | 0 | NA | 0.5 | NA | -0.5 |
| SE | 13959 | DIP2B | chr12 | + | 50728547 | 50728678 | 50727702 | 50727812 | 50732365 | 50732536 | 0 | 0 | 1 | 0 | 0 | 0 | 0 | 1 | 0 | 0 | 1 | 0 | 98 | 49 | NA | NA | NA | NA | 1 | NA | 0.333 | NA | 0.667 |
| SE | 13960 | DIP2B | chr12 | + | 50731368 | 50731537 | 50728547 | 50728678 | 50732365 | 50732536 | 0 | 0 | 4 | 0 | 0 | 0 | 0 | 0 | 0 | 0 | 1 | 0 | 98 | 49 | NA | NA | NA | NA | 1 | NA | 0 | NA | 1 |
| SE | 13964 | PARN | chr16 | - | 14627268 | 14627336 | 14627105 | 14627187 | 14628171 | 14628251 | 0 | 0 | 4 | 0 | 0 | 1 | 0 | 2 | 0 | 0 | 0 | 0 | 98 | 49 | NA | NA | NA | NA | 0.667 | NA | 1 | NA | -0.333 |
| SE | 14003 | MS4A7 | chr11 | + | 60392684 | 60392786 | 60389389 | 60389596 | 60393786 | 60395953 | 0 | 0 | 13 | 0 | 0 | 5 | 0 | 0 | 0 | 4 | 0 | 0 | 98 | 49 | NA | NA | NA | NA | 0.565 | 0 | NA | NA | 0.565 |
| SE | 14019 | ANKRD44 | chr2 | - | 196998906 | 196999052 | 196998336 | 196998419 | 197000418 | 197000502 | 0 | 0 | 9 | 0 | 0 | 5 | 0 | 3 | 0 | 0 | 1 | 0 | 98 | 49 | NA | NA | NA | NA | 0.474 | NA | 0.6 | NA | -0.126 |
| SE | 14020 | ANKRD44 | chr2 | - | 197013510 | 197013712 | 197008943 | 197009031 | 197025195 | 197025267 | 0 | 0 | 9 | 0 | 0 | 0 | 0 | 7 | 0 | 0 | 3 | 0 | 98 | 49 | NA | NA | NA | NA | 1 | NA | 0.538 | NA | 0.462 |
| SE | 14029 | FAM126B | chr2 | - | 200992886 | 200993003 | 200992280 | 200992368 | 200997444 | 200997540 | 0 | 0 | 15 | 0 | 0 | 2 | 0 | 3 | 0 | 0 | 0 | 0 | 98 | 49 | NA | NA | NA | NA | 0.789 | NA | 1 | NA | -0.211 |
| SE | 14030 | FAM126B | chr2 | - | 200992886 | 200993003 | 200992280 | 200992368 | 201008972 | 201009088 | 0 | 0 | 10 | 0 | 0 | 0 | 0 | 4 | 0 | 0 | 1 | 0 | 98 | 49 | NA | NA | NA | NA | 1 | NA | 0.667 | NA | 0.333 |
| SE | 14031 | FAM126B | chr2 | - | 200997444 | 200997540 | 200992280 | 200992368 | 201008972 | 201009088 | 0 | 0 | 6 | 0 | 0 | 0 | 0 | 2 | 0 | 0 | 1 | 0 | 98 | 49 | NA | NA | NA | NA | 1 | NA | 0.5 | NA | 0.5 |
| SE | 14032 | FAM126B | chr2 | - | 200997444 | 200997540 | 200992886 | 200993003 | 201008972 | 201009088 | 0 | 0 | 9 | 0 | 0 | 0 | 0 | 2 | 0 | 0 | 1 | 0 | 98 | 49 | NA | NA | NA | NA | 1 | NA | 0.5 | NA | 0.5 |
| SE | 14091 | AREL1 | chr14 | - | 74674033 | 74674111 | 74673076 | 74673218 | 74675698 | 74675946 | 0 | 0 | 2 | 0 | 0 | 0 | 6 | 8 | 0 | 0 | 1 | 0 | 98 | 49 | NA | NA | NA | NA | 1 | 1 | 0.8 | NA | 0.1 |
| SE | 14108 | TMEM9B | chr11 | - | 8962091 | 8962183 | 8956189 | 8956298 | 8964208 | 8964773 | 0 | 0 | 4 | 0 | 0 | 0 | 0 | 0 | 0 | 0 | 1 | 0 | 98 | 49 | NA | NA | NA | NA | 1 | NA | 0 | NA | 1 |

**U12_4_events_filtered**

| | | | | | | | | | | | IJC_SAMPLE_1 | | | SJC_SAMPLE_1 | | | IJC_SAMPLE_2 | | | SJC_SAMPLE_2 | | | | | | | IncLevel_ldg | | | IncLevel2_pmn | | | |
|---|---|---|---|---|---|---|---|---|---|---|---|---|---|---|---|---|---|---|---|---|---|---|---|---|---|---|---|---|---|---|---|---|---|
| **Event** | **ID** | **Gene_ID** | **Chromosome** | ***Strand*** | ***LongExonStart_0base*** | ***LongExonEnd*** | ***ShortES*** | ***ShortEE*** | ***FlankingES*** | ***FlankingEE*** | **ldg_12** | **ldg_14** | **ldg_16** | **ldg_12** | **ldg_14** | **ldg_16** | **pmn_13** | **pmn_15** | **pmn_17** | **pmn_13** | **pmn_15** | **pmn_17** | **IncFormLen** | **SkipFormLen** | **p value** | **FDR** | **ldg_12** | **ldg_14** | **ldg_16** | **pmn_13** | **pmn_15** | **pmn_17** | **IncLevelDifference** |
| A3SS | 1863 | SRPK1 | chr6 | - | 35890894 | 35891041 | 35890894 | 35891013 | 35920467 | 35920528 | 0 | 0 | 0 | 0 | 1 | 2 | 0 | 1 | 1 | 0 | 4 | 1 | 77 | 49 | 0.005762586 | 0.197717869 | NA | 0 | 0 | NA | 0.137 | 0.389 | -0.263 |
| SE | 9353 | GBP5 | chr1 | - | 89262682 | 89262785 | 89262219 | 89262401 | 89263735 | 89263948 | 0 | 3 | 3 | 0 | 0 | 0 | 1 | 6 | 0 | 0 | 2 | 0 | 98 | 49 | 0.019996576 | 0.170760496 | NA | 1 | 1 | 1 | 0.6 | NA | 0.2 |
| SE | 12583 | SBNO2 | chr19 | - | 1109134 | 1109211 | 1108778 | 1108969 | 1109291 | 1109423 | 0 | 4 | 9 | 0 | 0 | 0 | 0 | 25 | 2 | 0 | 3 | 0 | 98 | 49 | 0.024422917 | 0.170760496 | NA | 1 | 1 | NA | 0.806 | 1 | 0.097 |
| SE | 6887 | SRPK1 | chr6 | - | 35870280 | 35870494 | 35869481 | 35869901 | 35870933 | 35870959 | 0 | 0 | 4 | 0 | 1 | 5 | 0 | 2 | 0 | 0 | 3 | 0 | 98 | 49 | 0.027065308 | 0.170760496 | NA | 0 | 0.286 | NA | 0.25 | NA | -0.107 |
| SE | 14102 | SLC12A4 | chr16 | - | 67951822 | 67952037 | 67951139 | 67951304 | 67952183 | 67952425 | 0 | 1 | 0 | 0 | 0 | 0 | 0 | 1 | 0 | 0 | 2 | 0 | 98 | 49 | 0.027259421 | 0.170760496 | NA | 1 | NA | NA | 0.2 | NA | 0.8 |
| SE | 2543 | GAK | chr4 | - | 877614 | 877809 | 877089 | 877207 | 881906 | 882040 | 0 | 2 | 6 | 0 | 0 | 0 | 0 | 3 | 0 | 0 | 1 | 0 | 98 | 49 | 0.032835937 | 0.170760496 | NA | 1 | 1 | NA | 0.6 | NA | 0.4 |
| SE | 691 | DOCK5 | chr8 | + | 25300575 | 25300657 | 25298943 | 25299101 | 25302324 | 25302454 | 0 | 3 | 24 | 0 | 0 | 0 | 0 | 10 | 0 | 0 | 1 | 0 | 98 | 49 | 0.03465568 | 0.170760496 | NA | 1 | 1 | NA | 0.833 | NA | 0.167 |
| SE | 279 | EXOSC2 | chr9 | + | 130698161 | 130698251 | 130697581 | 130697627 | 130699328 | 130699394 | 0 | 1 | 0 | 0 | 0 | 0 | 0 | 1 | 0 | 0 | 1 | 0 | 98 | 49 | 0.037136978 | 0.170760496 | NA | 1 | NA | NA | 0.333 | NA | 0.667 |
| SE | 6888 | SMC3 | chr10 | + | 110590990 | 110591132 | 110590411 | 110590572 | 110593072 | 110593223 | 0 | 1 | 3 | 0 | 0 | 0 | 0 | 1 | 0 | 0 | 1 | 0 | 98 | 49 | 0.037136978 | 0.170760496 | NA | 1 | 1 | NA | 0.333 | NA | 0.667 |
| SE | 5861 | DISC1 | chr1 | + | 231795241 | 231795296 | 231770834 | 231771070 | 231818328 | 231819244 | 0 | 1 | 7 | 0 | 0 | 0 | 0 | 5 | 0 | 0 | 1 | 0 | 98 | 49 | 0.039248064 | 0.170760496 | NA | 1 | 1 | NA | 0.714 | NA | 0.286 |
| SE | 2907 | RAF1 | chr3 | - | 12600151 | 12600279 | 12599690 | 12599808 | 12600387 | 12600415 | 0 | 2 | 19 | 0 | 0 | 0 | 0 | 22 | 3 | 0 | 2 | 0 | 98 | 49 | 0.041304338 | 0.170760496 | NA | 1 | 1 | NA | 0.846 | 1 | 0.077 |
| SE | 1691 | DOCK2 | chr5 | + | 169761518 | 169761625 | 169759704 | 169759775 | 169803057 | 169803206 | 0 | 4 | 19 | 0 | 0 | 0 | 0 | 12 | 1 | 0 | 1 | 0 | 98 | 49 | 0.045620082 | 0.170760496 | NA | 1 | 1 | NA | 0.857 | 1 | 0.072 |
| A3SS | 414 | DCTN3 | chr9 | - | 34614041 | 34614198 | 34614041 | 34614101 | 34614709 | 34614768 | 1 | 1 | 0 | 0 | 0 | 0 | 0 | 0 | 0 | 0 | 3 | 0 | 98 | 49 | 0.059555994 | 0.197717869 | 1 | 1 | NA | NA | 0 | NA | 1 |
| A3SS | 413 | DCTN3 | chr9 | - | 34614041 | 34614132 | 34614041 | 34614101 | 34614709 | 34614768 | 1 | 1 | 0 | 0 | 0 | 0 | 0 | 0 | 0 | 0 | 3 | 0 | 80 | 49 | 0.060211792 | 0.197717869 | 1 | 1 | NA | NA | 0 | NA | 1 |
| SE | 8054 | CUL4B | chrX | - | 120544480 | 120544643 | 120544113 | 120544203 | 120545443 | 120545517 | 0 | 0 | 1 | 0 | 0 | 0 | 0 | 1 | 0 | 0 | 0 | 2 | 98 | 49 | 0.061401334 | 0.170760496 | NA | NA | 1 | NA | 1 | 0 | 0.5 |

| | | | | | | | | | | | | | | | | | | | | | | | | | | | | | | | | |
|---|---|---|---|---|---|---|---|---|---|---|---|---|---|---|---|---|---|---|---|---|---|---|---|---|---|---|---|---|---|---|---|---|
| SE | 5029 | BRMS1 | chr11 | - | 66340120 | 66340213 | 66338720 | 66338785 | 66340773 | 66340870 | 0 | 0 | 2 | 0 | 1 | 0 | 0 | 2 | 0 | 0 | 0 | 0 | 98 | 49 | 0.063770311 | 0.170760496 | NA | 0 | 1 | NA | 1 | NA | -0.5 |
| SE | 8179 | MLXIP | chr12 | + | 122137468 | 122137590 | 122135488 | 122135666 | 122138193 | 122138295 | 0 | 0 | 1 | 0 | 0 | 0 | 0 | 0 | 0 | 0 | 0 | 1 | 98 | 49 | 0.065790241 | 0.170760496 | NA | NA | 1 | NA | NA | 0 | 1 |
| SE | 11719 | CHD3 | chr17 | + | 7907600 | 7907702 | 7907352 | 7907488 | 7907893 | 7908019 | 0 | 0 | 0 | 0 | 0 | 2 | 0 | 0 | 1 | 0 | 0 | 0 | 98 | 49 | 0.0658956 | 0.170760496 | NA | NA | 0 | NA | NA | 1 | -1 |
| SE | 969 | RHBDD2 | chr7 | + | 75881364 | 75881486 | 75878998 | 75879260 | 75881828 | 75882236 | 0 | 0 | 2 | 0 | 1 | 0 | 2 | 1 | 0 | 0 | 0 | 0 | 98 | 49 | 0.068700612 | 0.171142152 | NA | 0 | 1 | 1 | 1 | NA | -0.5 |
| SE | 1355 | MAPK14 | chr6 | + | 36102570 | 36102649 | 36095986 | 36096066 | 36107454 | 36107628 | 0 | 1 | 19 | 0 | 0 | 0 | 0 | 14 | 1 | 0 | 1 | 0 | 98 | 49 | 0.069582416 | 0.17144651 | NA | 1 | 1 | NA | 0.875 | 1 | 0.062 |
| A3SS | 3730 | RASGRP4 | chr19 | - | 38414847 | 38415123 | 38414847 | 38414916 | 38417051 | 38417168 | 0 | 2 | 16 | 0 | 0 | 0 | 1 | 4 | 3 | 0 | 1 | 0 | 98 | 49 | 0.071741814 | 0.198669639 | NA | 1 | 1 | 1 | 0.667 | 1 | 0.111 |
| SE | 4098 | SETD1B | chr12 | + | 121819403 | 121819895 | 121817798 | 121817904 | 121822489 | 121823749 | 0 | 3 | 1 | 0 | 0 | 3 | 1 | 6 | 1 | 0 | 0 | 0 | 98 | 49 | 0.079530423 | 0.185900818 | NA | 1 | 0.143 | 1 | 1 | 1 | -0.428 |
| SE | 12584 | SBNO2 | chr19 | - | 1111018 | 1111093 | 1110744 | 1110888 | 1111505 | 1111614 | 0 | 1 | 6 | 0 | 0 | 1 | 1 | 26 | 2 | 0 | 0 | 0 | 98 | 49 | 0.080803043 | 0.187418168 | NA | 1 | 0.75 | 1 | 1 | 1 | -0.125 |
| SE | 13111 | AP2A1 | chr19 | + | 49800960 | 49801058 | 49799967 | 49800150 | 49801389 | 49801621 | 0 | 1 | 3 | 0 | 1 | 0 | 0 | 4 | 0 | 0 | 0 | 0 | 98 | 49 | 0.115352023 | 0.235883116 | NA | 0.333 | 1 | NA | 1 | NA | -0.334 |
| SE | 6225 | AHCTF1 | chr1 | - | 246916141 | 246916395 | 246913231 | 246913412 | 246918249 | 246918377 | 0 | 0 | 4 | 0 | 0 | 1 | 0 | 0 | 0 | 0 | 0 | 1 | 98 | 49 | 0.133877166 | 0.263718717 | NA | NA | 0.667 | NA | NA | 0 | 0.667 |
| SE | 5934 | RASGRP2 | chr11 | - | 64727295 | 64727360 | 64726910 | 64727131 | 64728862 | 64729042 | 0 | 0 | 7 | 0 | 1 | 0 | 0 | 1 | 1 | 0 | 0 | 0 | 98 | 49 | 0.134979818 | 0.264504128 | NA | 0 | 1 | NA | 1 | 1 | -0.5 |
| A3SS | 1281 | CHD4 | chr12 | - | 6601959 | 6602175 | 6601959 | 6602154 | 6602375 | 6602497 | 0 | 0 | 2 | 0 | 0 | 1 | 0 | 17 | 4 | 4 | 4 | 0 | 70 | 49 | 0.136899463 | 0.324235571 | NA | NA | 0.583 | 0 | 0.748 | 1 | 0 |
| SE | 12586 | SBNO2 | chr19 | - | 1113534 | 1113704 | 1112817 | 1112949 | 1114230 | 1114422 | 0 | 9 | 21 | 0 | 0 | 1 | 0 | 40 | 18 | 0 | 0 | 0 | 98 | 49 | 0.149179961 | 0.284179318 | NA | 1 | 0.913 | NA | 1 | 1 | -0.043 |
| SE | 695 | DOCK5 | chr8 | + | 25368191 | 25368250 | 25366869 | 25366970 | 25368570 | 25368725 | 0 | 0 | 11 | 0 | 0 | 5 | 0 | 5 | 3 | 0 | 1 | 0 | 98 | 49 | 0.166703165 | 0.311886731 | NA | NA | 0.524 | NA | 0.714 | 1 | -0.333 |
| SE | 2487 | ATG3 | chr3 | - | 112553279 | 112553329 | 112550191 | 112550262 | 112558375 | 112558417 | 0 | 0 | 4 | 0 | 0 | 3 | 1 | 4 | 2 | 0 | 0 | 0 | 98 | 49 | 0.167045122 | 0.311886731 | NA | NA | 0.4 | 1 | 1 | 1 | -0.6 |
| SE | 3413 | RALGAPA2 | chr20 | - | 20524405 | 20524543 | 20520916 | 20521100 | 20524829 | 20524898 | 0 | 0 | 2 | 0 | 0 | 1 | 0 | 0 | 2 | 0 | 0 | 0 | 98 | 49 | 0.209881051 | 0.368087771 | NA | NA | 0.5 | NA | NA | 1 | -0.5 |
| SE | 276 | RAPGEF1 | chr9 | - | 131650128 | 131650242 | 131643238 | 131643417 | 131650809 | 131650949 | 0 | 2 | 1 | 0 | 0 | 1 | 3 | 10 | 1 | 0 | 7 | 0 | 98 | 49 | 0.223239446 | 0.388343619 | NA | 1 | 0.333 | 1 | 0.417 | 1 | -0.139 |
| SE | 3236 | SLC9A8 | chr20 | + | 49850761 | 49850844 | 49849578 | 49849680 | 49855437 | 49855581 | 0 | 0 | 4 | 0 | 0 | 4 | 0 | 0 | 0 | 0 | 4 | 1 | 98 | 49 | 0.23228173 | 0.398828525 | NA | NA | 0.333 | NA | 0 | 0 | 0.333 |
| SE | 3237 | SLC9A8 | chr20 | + | 49850809 | 49850844 | 49849578 | 49849680 | 49855437 | 49855581 | 0 | 0 | 4 | 0 | 0 | 4 | 0 | 0 | 0 | 0 | 4 | 1 | 84 | 49 | 0.232676511 | 0.398828525 | NA | NA | 0.368 | NA | 0 | 0 | 0.368 |
| RI | 607 | TAF1C | chr16 | - | 84181327 | 84181663 | 84181327 | 84181463 | 84181591 | 84181663 | 0 | 0 | 3 | 0 | 0 | 1 | 0 | 1 | 2 | 0 | 0 | 0 | 98 | 49 | 0.264638717 | 0.737207854 | NA | NA | 0.6 | NA | 1 | 1 | -0.4 |
| SE | 3872 | WDFY2 | chr13 | + | 51719197 | 51719348 | 51703595 | 51703650 | 51727677 | 51727790 | 0 | 0 | 3 | 0 | 0 | 1 | 3 | 2 | 2 | 0 | 0 | 0 | 98 | 49 | 0.264638717 | 0.435653879 | NA | NA | 0.6 | 1 | 1 | 1 | -0.4 |
| SE | 7725 | ZFC3H1 | chr12 | - | 71636849 | 71637059 | 71636489 | 71636654 | 71638417 | 71638515 | 0 | 0 | 3 | 0 | 0 | 1 | 0 | 9 | 2 | 0 | 0 | 0 | 98 | 49 | 0.264638717 | 0.435653879 | NA | NA | 0.6 | NA | 1 | 1 | -0.4 |
| SE | 5935 | RASGRP2 | chr11 | - | 64735902 | 64735980 | 64735541 | 64735664 | 64736752 | 64737034 | 0 | 1 | 5 | 0 | 0 | 0 | 1 | 3 | 1 | 0 | 0 | 3 | 98 | 49 | 0.311385518 | 0.494727732 | NA | 1 | 1 | 1 | 1 | 0.143 | 0.286 |
| SE | 275 | RAPGEF1 | chr9 | - | 131630235 | 131630324 | 131629101 | 131629254 | 131638634 | 131638791 | 0 | 0 | 6 | 0 | 0 | 2 | 0 | 8 | 1 | 0 | 1 | 0 | 98 | 49 | 0.365054683 | 0.547059405 | NA | NA | 0.6 | NA | 0.8 | 1 | -0.3 |
| SE | 3233 | SLC9A8 | chr20 | + | 49839540 | 49839599 | 49823060 | 49823141 | 49845035 | 49845119 | 0 | 0 | 8 | 0 | 0 | 1 | 0 | 0 | 2 | 0 | 0 | 0 | 98 | 49 | 0.393533929 | 0.579315862 | NA | NA | 0.8 | NA | NA | 1 | -0.2 |
| SE | 2247 | MAEA | chr4 | + | 1327626 | 1327703 | 1315396 | 1315600 | 1332756 | 1332865 | 0 | 3 | 2 | 0 | 1 | 1 | 0 | 1 | 0 | 0 | 1 | 0 | 98 | 49 | 0.429432607 | 0.620451665 | NA | 0.6 | 0.5 | NA | 0.333 | NA | 0.217 |
| RI | 1055 | STX10 | chr19 | - | 13144057 | 13144671 | 13144057 | 13144486 | 13144576 | 13144671 | 0 | 1 | 6 | 0 | 0 | 1 | 0 | 0 | 1 | 0 | 1 | 0 | 98 | 49 | 0.482241355 | 0.94773154 | NA | 1 | 0.75 | NA | 0 | 1 | 0.375 |
| SE | 12585 | SBNO2 | chr19 | - | 1112188 | 1112301 | 1111995 | 1112067 | 1112401 | 1112537 | 0 | 8 | 15 | 0 | 4 | 0 | 0 | 8 | 3 | 0 | 2 | 1 | 98 | 49 | 0.521310981 | 0.720156622 | NA | 0.5 | 1 | NA | 0.667 | 0.6 | 0.117 |
| SE | 3235 | SLC9A8 | chr20 | + | 49849578 | 49849680 | 49845035 | 49845119 | 49855437 | 49855581 | 0 | 0 | 10 | 0 | 0 | 2 | 0 | 6 | 1 | 0 | 2 | 0 | 98 | 49 | 0.539441453 | 0.737743862 | NA | NA | 0.714 | NA | 0.6 | 1 | -0.086 |
| SE | 3417 | RALGAPA2 | chr20 | - | 20676235 | 20676288 | 20653529 | 20653587 | 20680690 | 20680801 | 0 | 0 | 13 | 0 | 0 | 2 | 0 | 5 | 1 | 0 | 0 | 0 | 98 | 49 | 0.588967448 | 0.794630228 | NA | NA | 0.765 | NA | 1 | 1 | -0.235 |
| RI | 230 | ATG3 | chr3 | - | 112532506 | 112534337 | 112532506 | 112532780 | 112534268 | 112534337 | 0 | 0 | 3 | 0 | 3 | 5 | 0 | 0 | 1 | 0 | 4 | 0 | 98 | 49 | 0.654603078 | 0.999202116 | NA | 0 | 0.231 | NA | 0 | 1 | -0.385 |
| SE | 2906 | RAF1 | chr3 | - | 12590797 | 12590974 | 12587590 | 12587637 | 12591707 | 12591792 | 1 | 1 | 24 | 0 | 0 | 2 | 4 | 25 | 0 | 0 | 0 | 0 | 98 | 49 | 0.867186844 | 0.999202116 | 1 | 1 | 0.857 | 1 | 1 | NA | -0.048 |
| SE | 10440 | SLC12A6 | chr15 | - | 34257641 | 34257788 | 34256228 | 34256283 | 34258812 | 34258944 | 0 | 1 | 11 | 0 | 0 | 4 | 0 | 2 | 4 | 0 | 2 | 0 | 98 | 49 | 0.949596688 | 0.999202116 | NA | 1 | 0.579 | NA | 0.333 | 1 | 0.123 |
| SE | 7631 | BTAF1 | chr10 | + | 91953736 | 91953873 | 91951402 | 91951566 | 91956527 | 91956657 | 0 | 1 | 5 | 0 | 0 | 1 | 0 | 4 | 0 | 0 | 0 | 0 | 98 | 49 | 0.952968863 | 0.999202116 | NA | 1 | 0.714 | NA | 1 | NA | -0.143 |
| SE | 11425 | DYNC1LI2 | chr16 | - | 66749196 | 66749313 | 66742437 | 66742668 | 66751272 | 66751346 | 0 | 1 | 6 | 0 | 0 | 1 | 0 | 3 | 0 | 0 | 0 | 0 | 98 | 49 | 0.961336908 | 0.999202116 | NA | 1 | 0.75 | NA | 1 | NA | -0.125 |
| SE | 8808 | WDR1 | chr4 | - | 10103895 | 10103986 | 10098991 | 10099139 | 10116112 | 10116234 | 0 | 5 | 16 | 0 | 0 | 1 | 1 | 9 | 0 | 0 | 0 | 0 | 98 | 49 | 0.976663141 | 0.999202116 | NA | 1 | 0.889 | 1 | 1 | NA | -0.055 |
| SE | 5295 | PEX16 | chr11 | - | 45916226 | 45916303 | 45915702 | 45915836 | 45917457 | 45917493 | 0 | 0 | 3 | 0 | 0 | 0 | 0 | 5 | 2 | 0 | 1 | 0 | 98 | 49 | 0.993205848 | 0.999202116 | NA | NA | 1 | NA | 0.714 | 1 | 0.143 |
| SE | 694 | DOCK5 | chr8 | + | 25342400 | 25342507 | 25341738 | 25341809 | 25345474 | 25345611 | 0 | 0 | 15 | 0 | 0 | 0 | 0 | 15 | 2 | 0 | 1 | 0 | 98 | 49 | 0.999202116 | 0.999202116 | NA | NA | 1 | NA | 0.882 | 1 | 0.059 |
| SE | 697 | DOCK5 | chr8 | + | 25407982 | 25408154 | 25403557 | 25403724 | 25408801 | 25408940 | 0 | 1 | 23 | 0 | 1 | 1 | 0 | 9 | 3 | 0 | 3 | 0 | 98 | 49 | 0.999202116 | 0.999202116 | NA | 0.333 | 0.92 | NA | 0.6 | 1 | -0.173 |
| SE | 3722 | FCHO1 | chr19 | + | 17781230 | 17781343 | 17778608 | 17778884 | 17781451 | 17781539 | 0 | 1 | 3 | 0 | 0 | 3 | 0 | 1 | 0 | 0 | 0 | 0 | 98 | 49 | 0.999202116 | 0.999202116 | NA | 1 | 0.333 | NA | 1 | NA | -0.334 |
| SE | 4520 | CHD4 | chr12 | - | 6600925 | 6601053 | 6600533 | 6600669 | 6601288 | 6601530 | 0 | 0 | 7 | 0 | 0 | 0 | 0 | 4 | 1 | 0 | 1 | 0 | 98 | 49 | 0.999202116 | 0.999202116 | NA | NA | 1 | NA | 0.667 | 1 | 0.166 |
| SE | 11563 | NCBP1 | chr9 | + | 97662041 | 97662144 | 97660945 | 97661068 | 97662953 | 97663047 | 0 | 1 | 1 | 0 | 0 | 2 | 0 | 1 | 0 | 0 | 0 | 0 | 98 | 49 | 0.999202116 | 0.999202116 | NA | 1 | 0.2 | NA | 1 | NA | -0.4 |
| SE | 12825 | TRAPPC10 | chr21 | + | 44074323 | 44074470 | 44063537 | 44063785 | 44075038 | 44075153 | 0 | 0 | 4 | 0 | 0 | 0 | 0 | 3 | 1 | 0 | 1 | 0 | 98 | 49 | 0.999202116 | 0.999202116 | NA | NA | 1 | NA | 0.6 | 1 | 0.2 |
| A3SS | 503 | HARS2 | chr5 | + | 140693490 | 140693665 | 140693590 | 140693665 | 140691425 | 140691756 | 0 | 0 | 2 | 0 | 0 | 0 | 0 | 1 | 0 | 0 | 2 | 0 | 98 | 49 | NA | NA | NA | NA | 1 | NA | 0.2 | NA | 0.8 |
| A3SS | 1533 | TTC23 | chr15 | - | 99139316 | 99139467 | 99139316 | 99139399 | 99156147 | 99156297 | 0 | 0 | 0 | 0 | 0 | 1 | 0 | 1 | 0 | 0 | 0 | 0 | 98 | 49 | NA | NA | NA | NA | 0 | NA | 1 | NA | -1 |
| A3SS | 1628 | SRSF10 | chr1 | - | 23964346 | 23967956 | 23964346 | 23967723 | 23971572 | 23971623 | 1 | 0 | 0 | 0 | 0 | 0 | 0 | 0 | 0 | 0 | 2 | 0 | 98 | 49 | NA | NA | 1 | NA | NA | NA | 0 | NA | 1 |
| A3SS | 1629 | SRSF10 | chr1 | - | 23964346 | 23971439 | 23964346 | 23967723 | 23971572 | 23971623 | 1 | 0 | 1 | 0 | 0 | 0 | 0 | 1 | 0 | 0 | 2 | 0 | 98 | 49 | NA | NA | 1 | NA | 1 | NA | 0.2 | NA | 0.8 |
| A5SS | 1731 | SSR3 | chr3 | - | 156548865 | 156549003 | 156548904 | 156549003 | 156544307 | 156544439 | 0 | 0 | 0 | 0 | 0 | 1 | 8 | 0 | 0 | 0 | 0 | 0 | 88 | 49 | NA | NA | NA | NA | 0 | 1 | NA | NA | -1 |
| A5SS | 2052 | SLC12A9 | chr7 | + | 100852718 | 100852839 | 100852718 | 100852835 | 100854155 | 100854378 | 0 | 0 | 0 | 0 | 0 | 2 | 0 | 2 | 0 | 0 | 0 | 0 | 53 | 49 | NA | NA | NA | NA | 0 | NA | 1 | NA | -1 |
| A5SS | 2205 | PLCB2 | chr15 | - | 40296745 | 40296908 | 40296757 | 40296908 | 40296521 | 40296634 | 0 | 0 | 7 | 0 | 0 | 3 | 0 | 2 | 0 | 0 | 0 | 0 | 61 | 49 | NA | NA | NA | NA | 0.652 | NA | 1 | NA | -0.348 |
| RI | 909 | HNRNPM | chr19 | + | 8465323 | 8465515 | 8465323 | 8465362 | 8465479 | 8465515 | 0 | 0 | 8 | 0 | 0 | 1 | 7 | 0 | 0 | 0 | 0 | 0 | 98 | 49 | NA | NA | NA | NA | 0.8 | 1 | NA | NA | -0.2 |
| SE | 692 | DOCK5 | chr8 | + | 25340876 | 25340988 | 25336238 | 25336373 | 25341738 | 25341809 | 0 | 0 | 19 | 0 | 0 | 0 | 0 | 10 | 0 | 0 | 1 | 0 | 98 | 49 | NA | NA | NA | NA | 1 | NA | 0.833 | NA | 0.167 |
| SE | 693 | DOCK5 | chr8 | + | 25341738 | 25341809 | 25336238 | 25336373 | 25342400 | 25342507 | 0 | 0 | 6 | 0 | 0 | 0 | 0 | 12 | 0 | 0 | 1 | 0 | 98 | 49 | NA | NA | NA | NA | 1 | NA | 0.857 | NA | 0.143 |
| SE | 696 | DOCK5 | chr8 | + | 25373617 | 25373658 | 25372558 | 25372718 | 25374563 | 25374654 | 0 | 0 | 8 | 0 | 0 | 1 | 0 | 8 | 0 | 0 | 1 | 0 | 90 | 49 | NA | NA | NA | NA | 0.813 | NA | 1 | NA | -0.187 |
| SE | 1305 | SMAP1 | chr6 | + | 70791688 | 70791769 | 70773349 | 70773425 | 70798656 | 70798737 | 0 | 0 | 1 | 0 | 0 | 0 | 0 | 0 | 0 | 0 | 1 | 0 | 98 | 49 | NA | NA | NA | NA | 1 | NA | 0 | NA | 1 |
| SE | 1692 | DOCK2 | chr5 | + | 169983067 | 169983166 | 169840756 | 169840852 | 169985827 | 169985922 | 0 | 0 | 12 | 0 | 0 | 1 | 0 | 5 | 0 | 0 | 0 | 0 | 98 | 49 | NA | NA | NA | NA | 0.857 | NA | 1 | NA | -0.143 |
| SE | 1748 | DIAPH1 | chr5 | - | 141529602 | 141529697 | 141529171 | 141529273 | 141534334 | 141534433 | 0 | 0 | 8 | 0 | 0 | 0 | 0 | 1 | 0 | 0 | 1 | 0 | 98 | 49 | NA | NA | NA | NA | 1 | NA | 0.333 | NA | 0.667 |
| SE | 2208 | IK | chr5 | + | 140655828 | 140655992 | 140654680 | 140654727 | 140657553 | 140657662 | 0 | 0 | 2 | 0 | 0 | 2 | 0 | 1 | 0 | 0 | 0 | 0 | 98 | 49 | NA | NA | NA | NA | 0.333 | NA | 1 | NA | -0.667 |
| SE | 2530 | GBE1 | chr3 | - | 81646391 | 81646482 | 81642780 | 81642990 | 81648855 | 81648991 | 0 | 0 | 0 | 0 | 0 | 1 | 0 | 1 | 0 | 0 | 0 | 0 | 98 | 49 | NA | NA | NA | NA | 0 | NA | 1 | NA | -1 |
| SE | 2661 | SACM1L | chr3 | + | 45735234 | 45735373 | 45732052 | 45732151 | 45737582 | 45737652 | 0 | 0 | 2 | 0 | 0 | 0 | 0 | 2 | 0 | 0 | 1 | 0 | 98 | 49 | NA | NA | NA | NA | 1 | NA | 0.5 | NA | 0.5 |
| SE | 2908 | RAF1 | chr3 | - | 12606200 | 12606299 | 12604135 | 12604289 | 12608765 | 12608923 | 0 | 0 | 20 | 0 | 0 | 5 | 0 | 22 | 0 | 0 | 2 | 0 | 98 | 49 | NA | NA | NA | NA | 0.667 | NA | 0.846 | NA | -0.179 |
| SE | 3326 | CTNNBL1 | chr20 | + | 37842338 | 37842419 | 37840101 | 37840199 | 37859898 | 37860036 | 0 | 0 | 2 | 0 | 0 | 1 | 0 | 4 | 0 | 0 | 0 | 0 | 98 | 49 | NA | NA | NA | NA | 0.5 | NA | 1 | NA | -0.5 |
| SE | 3414 | RALGAPA2 | chr20 | - | 20605174 | 20605412 | 20601681 | 20601846 | 20611314 | 20611426 | 0 | 0 | 5 | 0 | 0 | 0 | 0 | 0 | 0 | 0 | 2 | 0 | 98 | 49 | NA | NA | NA | NA | 1 | NA | 0 | NA | 1 |
| SE | 3415 | RALGAPA2 | chr20 | - | 20611314 | 20611426 | 20605174 | 20605412 | 20616042 | 20616191 | 0 | 0 | 6 | 0 | 0 | 1 | 0 | 1 | 0 | 0 | 0 | 0 | 98 | 49 | NA | NA | NA | NA | 0.75 | NA | 1 | NA | -0.25 |
| SE | 3416 | RALGAPA2 | chr20 | - | 20637362 | 20637501 | 20635417 | 20635617 | 20639784 | 20639900 | 0 | 0 | 13 | 0 | 0 | 2 | 0 | 0 | 0 | 0 | 2 | 0 | 98 | 49 | NA | NA | NA | NA | 0.765 | NA | 0 | NA | 0.765 |
| SE | 3971 | HARS | chr5 | - | 140675016 | 140675133 | 140674678 | 140674825 | 140676653 | 140676896 | 0 | 0 | 1 | 0 | 0 | 0 | 0 | 2 | 0 | 1 | 0 | 0 | 98 | 49 | NA | NA | NA | NA | 1 | 0 | 1 | NA | 0.5 |
| SE | 4434 | KANSL2 | chr12 | - | 48679035 | 48679150 | 48671798 | 48671962 | 48679654 | 48679833 | 0 | 0 | 6 | 0 | 0 | 2 | 0 | 3 | 0 | 0 | 0 | 0 | 98 | 49 | NA | NA | NA | NA | 0.6 | NA | 1 | NA | -0.4 |
| SE | 4518 | CHD4 | chr12 | - | 6582136 | 6582281 | 6581648 | 6581814 | 6582614 | 6582748 | 0 | 0 | 4 | 0 | 0 | 1 | 0 | 5 | 0 | 0 | 1 | 0 | 98 | 49 | NA | NA | NA | NA | 0.667 | NA | 0.714 | NA | -0.047 |
| SE | 4519 | CHD4 | chr12 | - | 6588297 | 6588422 | 6587711 | 6587949 | 6591465 | 6591583 | 0 | 0 | 13 | 0 | 0 | 0 | 0 | 3 | 0 | 0 | 3 | 0 | 98 | 49 | NA | NA | NA | NA | 1 | NA | 0.333 | NA | 0.667 |
| SE | 4553 | MED23 | chr6 | - | 131620627 | 131620729 | 131619826 | 131619896 | 131621880 | 131621979 | 0 | 0 | 4 | 0 | 0 | 0 | 0 | 1 | 0 | 0 | 2 | 0 | 98 | 49 | NA | NA | NA | NA | 1 | NA | 0.2 | NA | 0.8 |
| SE | 4590 | CUL4A | chr13 | + | 113239432 | 113239551 | 113236822 | 113236890 | 113242967 | 113243160 | 0 | 0 | 5 | 0 | 0 | 1 | 0 | 1 | 0 | 0 | 0 | 0 | 98 | 49 | NA | NA | NA | NA | 0.714 | NA | 1 | NA | -0.286 |
| SE | 4924 | C2CD3 | chr11 | - | 74161398 | 74161556 | 74139604 | 74139828 | 74168343 | 74168613 | 0 | 0 | 2 | 0 | 0 | 1 | 0 | 2 | 0 | 0 | 0 | 0 | 98 | 49 | NA | NA | NA | NA | 0.5 | NA | 1 | NA | -0.5 |
| SE | 5007 | RINT1 | chr7 | + | 105550260 | 105550486 | 105550054 | 105550165 | 105551569 | 105551707 | 0 | 0 | 1 | 0 | 0 | 0 | 0 | 0 | 0 | 0 | 2 | 0 | 98 | 49 | NA | NA | NA | NA | 1 | NA | 0 | NA | 1 |
| SE | 5110 | WDFY1 | chr2 | - | 223884647 | 223884749 | 223881941 | 223882072 | 223894233 | 223894339 | 0 | 0 | 3 | 0 | 0 | 0 | 0 | 1 | 0 | 0 | 1 | 0 | 98 | 49 | NA | NA | NA | NA | 1 | NA | 0.333 | NA | 0.667 |
| SE | 5928 | ZDHHC5 | chr11 | + | 57695919 | 57696043 | 57693782 | 57693915 | 57696760 | 57696873 | 0 | 0 | 0 | 0 | 0 | 1 | 5 | 5 | 0 | 0 | 0 | 0 | 98 | 49 | NA | NA | NA | NA | 0 | 1 | 1 | NA | -1 |
| SE | 5937 | SRSF10 | chr1 | - | 23971337 | 23971439 | 23964346 | 23967723 | 23971572 | 23971626 | 0 | 0 | 1 | 0 | 0 | 0 | 0 | 1 | 0 | 0 | 2 | 0 | 98 | 49 | NA | NA | NA | NA | 1 | NA | 0.2 | NA | 0.8 |
| SE | 6090 | ZNF207 | chr17 | + | 32366664 | 32366757 | 32365329 | 32365487 | 32367771 | 32368014 | 0 | 0 | 0 | 0 | 0 | 2 | 0 | 3 | 0 | 0 | 0 | 0 | 98 | 49 | NA | NA | NA | NA | 0 | NA | 1 | NA | -1 |
| SE | 6223 | AHCTF1 | chr1 | - | 246843794 | 246843928 | 246842693 | 246842776 | 246849614 | 246851442 | 0 | 0 | 0 | 0 | 0 | 2 | 0 | 2 | 0 | 0 | 2 | 0 | 98 | 49 | NA | NA | NA | NA | 0 | NA | 0.333 | NA | -0.333 |
| SE | 7632 | BTAF1 | chr10 | + | 92027129 | 92027300 | 92026591 | 92026751 | 92028789 | 92030323 | 0 | 0 | 4 | 0 | 0 | 3 | 0 | 5 | 0 | 0 | 0 | 0 | 98 | 49 | NA | NA | NA | NA | 0.4 | NA | 1 | NA | -0.6 |
| SE | 7946 | PPP2R2D | chr10 | + | 131911561 | 131912598 | 131901237 | 131901330 | 131934457 | 131934555 | 0 | 0 | 0 | 0 | 0 | 3 | 0 | 2 | 0 | 0 | 0 | 0 | 98 | 49 | NA | NA | NA | NA | 0 | NA | 1 | NA | -1 |
| SE | 7947 | PPP2R2D | chr10 | + | 131934893 | 131935006 | 131934457 | 131934555 | 131940030 | 131940196 | 0 | 0 | 1 | 0 | 0 | 3 | 0 | 1 | 0 | 0 | 0 | 0 | 98 | 49 | NA | NA | NA | NA | 0.143 | NA | 1 | NA | -0.857 |
| SE | 8178 | MLXIP | chr12 | + | 122130843 | 122130933 | 122129940 | 122130112 | 122132291 | 122132383 | 0 | 0 | 1 | 0 | 0 | 1 | 0 | 1 | 0 | 0 | 0 | 0 | 98 | 49 | NA | NA | NA | NA | 0.333 | NA | 1 | NA | -0.667 |
| SE | 8908 | WDR11 | chr10 | + | 120867065 | 120867169 | 120866568 | 120866764 | 120871169 | 120871346 | 0 | 0 | 3 | 0 | 0 | 1 | 0 | 3 | 0 | 0 | 0 | 0 | 98 | 49 | NA | NA | NA | NA | 0.6 | NA | 1 | NA | -0.4 |
| SE | 9095 | HPS1 | chr10 | - | 98429790 | 98429889 | 98429145 | 98429642 | 98430570 | 98430670 | 0 | 0 | 1 | 0 | 0 | 1 | 0 | 1 | 0 | 0 | 0 | 0 | 98 | 49 | NA | NA | NA | NA | 0.333 | NA | 1 | NA | -0.667 |
| SE | 9358 | RASGRP1 | chr15 | - | 38503271 | 38503376 | 38502311 | 38502421 | 38505839 | 38505920 | 0 | 0 | 2 | 0 | 0 | 0 | 0 | 1 | 0 | 0 | 1 | 0 | 98 | 49 | NA | NA | NA | NA | 1 | NA | 0.333 | NA | 0.667 |
| SE | 9667 | PTEN | chr10 | + | 87952117 | 87952259 | 87933012 | 87933251 | 87957852 | 87958019 | 0 | 0 | 4 | 0 | 0 | 1 | 0 | 2 | 0 | 0 | 1 | 0 | 98 | 49 | NA | NA | NA | NA | 0.667 | NA | 0.5 | NA | 0.167 |
| SE | 10439 | SLC12A6 | chr15 | - | 34242101 | 34242221 | 34241232 | 34241337 | 34243973 | 34244072 | 0 | 0 | 7 | 0 | 0 | 1 | 0 | 4 | 0 | 0 | 2 | 0 | 98 | 49 | NA | NA | NA | NA | 0.778 | NA | 0.5 | NA | 0.278 |
| SE | 10468 | PLCB2 | chr15 | - | 40291605 | 40291650 | 40291264 | 40291487 | 40291848 | 40291924 | 0 | 0 | 1 | 0 | 0 | 2 | 0 | 0 | 0 | 0 | 1 | 0 | 94 | 49 | NA | NA | NA | NA | 0.207 | NA | 0 | NA | 0.207 |
| SE | 12021 | CCDC43 | chr17 | - | 44680584 | 44680643 | 44677436 | 44679043 | 44682002 | 44682138 | 0 | 0 | 1 | 0 | 0 | 0 | 0 | 0 | 0 | 0 | 1 | 0 | 98 | 49 | NA | NA | NA | NA | 1 | NA | 0 | NA | 1 |
| SE | 12446 | TRAPPC8 | chr18 | - | 31866848 | 31866975 | 31864626 | 31864781 | 31867401 | 31867476 | 0 | 0 | 5 | 0 | 0 | 0 | 0 | 0 | 0 | 0 | 2 | 0 | 98 | 49 | NA | NA | NA | NA | 1 | NA | 0 | NA | 1 |
| SE | 12934 | EIF3K | chr19 | + | 38620336 | 38620435 | 38619073 | 38619327 | 38624076 | 38624197 | 0 | 0 | 8 | 0 | 0 | 1 | 0 | 10 | 0 | 0 | 0 | 0 | 98 | 49 | NA | NA | NA | NA | 0.8 | NA | 1 | NA | -0.2 |
| SE | 12935 | EIF3K | chr19 | + | 38632600 | 38632678 | 38632429 | 38632496 | 38634992 | 38635118 | 0 | 0 | 5 | 0 | 0 | 1 | 1 | 5 | 0 | 0 | 0 | 0 | 98 | 49 | NA | NA | NA | NA | 0.714 | 1 | 1 | NA | -0.286 |
| SE | 13565 | EML4 | chr2 | + | 42301240 | 42301392 | 42295380 | 42295516 | 42303103 | 42303229 | 0 | 0 | 6 | 0 | 0 | 0 | 0 | 7 | 0 | 0 | 1 | 0 | 98 | 49 | NA | NA | NA | NA | 1 | NA | 0.778 | NA | 0.222 |

**MXE_all**

| | | | | | | | | | | | | Inclusion_counts | | | Skipping_counts | | | Inclusion_counts | | | Skipping_counts | | | | | | | Inclusion level(normalized) | | | | | | |
|---|---|---|---|---|---|---|---|---|---|---|---|---|---|---|---|---|---|---|---|---|---|---|---|---|---|---|---|---|---|---|---|---|---|---|
| Splicing event | ID | Gene_ID | Chromosome | Strand | 1stExonStart_0base | 1stExonEnd | 2ndExonSta | 2ndExonEn | upstreamES | upstreamEE | downstr | downstr | Idg_12 | Idg_14 | Idg_16 | Idg_12 | Idg_14 | Idg_16 | pmn_13 | pmn_15 | pmn_17 | pmn_13 | pmn_15 | pmn_17 | IncFormLen | SkipFormLen | p value | FDR | Idg_12 | Idg_14 | Idg_16 | pmn_13 | pmn_15 | pmn_17 | IncLevelDifference |
| MXE | 631 | IRF7 | chr11 | - | 613950 | 614037 | 614173 | 614399 | 613784 | 613865 | 614475 | 614534 | 0 | 0 | 0 | 0 | 8 | 14 | 0 | 4 | 1 | 0 | 7 | 3 | 98 | 98 | 0.0009 | 0.11226 | NA | 0 | 0 | NA | 0.364 | 0.25 | -0.307 |
| MXE | 401 | DDX3X | chrX | + | 41346858 | 41347012 | 41347314 | 41347451 | 41346504 | 41346622 | 4E+07 | 4E+07 | 0 | 0 | 27 | 0 | 1 | 11 | 1 | 32 | 0 | 0 | 19 | 0 | 98 | 98 | 0.0056 | 0.19855 | NA | 0 | 0.711 | 1 | 0.627 | NA | -0.458 |
| MXE | 716 | CTSS | chr1 | - | 150750005 | 150750171 | 150751780 | 150752008 | 150747776 | 150747879 | 2E+08 | 2E+08 | 0 | 2 | 110 | 0 | 0 | 58 | 2 | 11 | 0 | 0 | 11 | 0 | 98 | 98 | 0.0085 | 0.19855 | NA | 1 | 0.655 | 1 | 0.5 | NA | 0.078 |
| MXE | 170 | MAEA | chr4 | + | 1315396 | 1315600 | 1322380 | 1322503 | 1311978 | 1312161 | 1E+06 | 1E+06 | 0 | 2 | 1 | 0 | 0 | 0 | 0 | 2 | 7 | 0 | 3 | 1 | 98 | 98 | 0.0098 | 0.19855 | NA | 1 | 1 | NA | 0.4 | 0.875 | 0.363 |
| MXE | 344 | NCF2 | chr1 | - | 183573184 | 183573292 | 183574486 | 183574621 | 183570779 | 183570839 | 2E+08 | 2E+08 | 0 | 0 | 30 | 2 | 5 | 53 | 0 | 5 | 0 | 0 | 44 | 0 | 98 | 98 | 0.0116 | 0.19855 | 0 | 0 | 0.361 | NA | 0.102 | NA | 0.018 |
| MXE | 294 | USP15 | chr12 | + | 62325871 | 62325933 | 62349220 | 62349307 | 62321463 | 62321609 | 6E+07 | 6E+07 | 0 | 2 | 22 | 0 | 0 | 6 | 0 | 7 | 0 | 2 | 6 | 0 | 98 | 98 | 0.0117 | 0.19855 | NA | 1 | 0.786 | 0 | 0.538 | NA | 0.624 |
| MXE | 518 | FCAR | chr19 | + | 54885234 | 54885525 | 54888006 | 54888294 | 54874092 | 54874323 | 5E+07 | 5E+07 | 0 | 0 | 1 | 0 | 4 | 17 | 0 | 3 | 0 | 0 | 9 | 0 | 98 | 98 | 0.0151 | 0.19855 | NA | 0 | 0.056 | NA | 0.25 | NA | -0.222 |
| MXE | 948 | PTPN12 | chr7 | + | 77583554 | 77583650 | 77585542 | 77585581 | 77581426 | 77581503 | 8E+07 | 8E+07 | 0 | 0 | 9 | 0 | 1 | 1 | 0 | 5 | 0 | 0 | 4 | 0 | 98 | 88 | 0.0153 | 0.19855 | NA | 0 | 0.89 | NA | 0.529 | NA | -0.084 |
| MXE | 541 | PCBP2 | chr12 | + | 53464759 | 53464852 | 53465931 | 53465973 | 53462480 | 53462567 | 5E+07 | 5E+07 | 0 | 0 | 13 | 0 | 2 | 30 | 0 | 3 | 0 | 0 | 11 | 0 | 98 | 91 | 0.0171 | 0.19855 | NA | 0 | 0.287 | NA | 0.202 | NA | -0.059 |
| MXE | 227 | TNRC6B | chr22 | + | 40264687 | 40267036 | 40270121 | 40270280 | 40261831 | 40262173 | 4E+07 | 4E+07 | 0 | 1 | 4 | 0 | 0 | 1 | 0 | 4 | 0 | 0 | 3 | 0 | 98 | 98 | 0.019 | 0.19855 | NA | 1 | 0.8 | NA | 0.571 | NA | 0.329 |
| MXE | 563 | OSBPL9 | chr1 | + | 51651990 | 51652041 | 51669433 | 51669512 | 51616873 | 51617221 | 5E+07 | 5E+07 | 0 | 1 | 1 | 0 | 0 | 8 | 0 | 1 | 0 | 0 | 6 | 0 | 98 | 98 | 0.0201 | 0.19855 | NA | 1 | 0.111 | NA | 0.143 | NA | 0.412 |
| MXE | 771 | PLXDC2 | chr10 | + | 20046868 | 20047015 | 20068169 | 20068239 | 20001774 | 20001986 | 2E+07 | 2E+07 | 0 | 1 | 6 | 0 | 0 | 11 | 2 | 2 | 0 | 0 | 3 | 0 | 98 | 98 | 0.0206 | 0.19855 | NA | 1 | 0.353 | 1 | 0.4 | NA | -0.023 |
| MXE | 601 | NOTCH2 | chr1 | - | 119922235 | 119922446 | 119922635 | 119922778 | 119921712 | 119921809 | 1E+08 | 1E+08 | 0 | 2 | 2 | 0 | 0 | 8 | 0 | 6 | 0 | 0 | 2 | 0 | 98 | 98 | 0.0213 | 0.19855 | NA | 1 | 0.2 | NA | 0.75 | NA | -0.15 |
| MXE | 919 | ARFGAP3 | chr22 | - | 42834241 | 42834325 | 42835361 | 42835493 | 42831548 | 42831636 | 4E+07 | 4E+07 | 0 | 1 | 0 | 0 | 0 | 1 | 0 | 1 | 0 | 0 | 3 | 0 | 98 | 98 | 0.0238 | 0.19855 | NA | 1 | 0 | NA | 0.25 | NA | 0.25 |
| MXE | 559 | PLEKHM2 | chr1 | + | 15719733 | 15719920 | 15721328 | 15721388 | 15718537 | 15718625 | 2E+07 | 2E+07 | 0 | 1 | 2 | 0 | 0 | 0 | 0 | 3 | 0 | 0 | 2 | 0 | 98 | 98 | 0.0239 | 0.19855 | NA | 1 | 1 | NA | 0.6 | NA | 0.4 |
| MXE | 1080 | RASGRP4 | chr19 | - | 38417051 | 38417168 | 38419859 | 38420013 | 38413393 | 38413474 | 4E+07 | 4E+07 | 0 | 4 | 7 | 0 | 0 | 0 | 1 | 1 | 0 | 0 | 1 | 0 | 98 | 98 | 0.0304 | 0.19855 | NA | 1 | 1 | 1 | 0.5 | NA | 0.25 |
| MXE | 566 | PSME3 | chr17 | + | 42838092 | 42838205 | 42838730 | 42838799 | 42837648 | 42837697 | 4E+07 | 4E+07 | 0 | 0 | 1 | 0 | 2 | 1 | 0 | 1 | 0 | 0 | 2 | 0 | 98 | 98 | 0.0326 | 0.19855 | NA | 0 | 0.5 | NA | 0.333 | NA | -0.083 |
| MXE | 606 | RBM23 | chr14 | - | 22908332 | 22908380 | 22909482 | 22909595 | 22906194 | 22906368 | 2E+07 | 2E+07 | 0 | 2 | 3 | 0 | 0 | 2 | 0 | 7 | 0 | 0 | 1 | 0 | 98 | 97 | 0.0356 | 0.19855 | NA | 1 | 0.598 | NA | 0.874 | NA | -0.075 |
| MXE | 853 | SNX17 | chr2 | + | 27372622 | 27372740 | 27373039 | 27373113 | 27371268 | 27371343 | 3E+07 | 3E+07 | 0 | 1 | 4 | 0 | 0 | 0 | 0 | 2 | 0 | 0 | 1 | 0 | 98 | 98 | 0.0365 | 0.19855 | NA | 1 | 1 | NA | 0.667 | NA | 0.333 |
| MXE | 603 | ZRANB1 | chr10 | + | 124974199 | 124974398 | 124981708 | 124981829 | 124973644 | 124973716 | 1E+08 | 1E+08 | 0 | 0 | 1 | 0 | 1 | 2 | 0 | 1 | 0 | 0 | 2 | 0 | 98 | 98 | 0.0367 | 0.19855 | NA | 0 | 0.333 | NA | 0.333 | NA | -0.167 |
| MXE | 692 | ADAM9 | chr8 | + | 39082967 | 39083073 | 39090046 | 39090188 | 39082640 | 39082721 | 4E+07 | 4E+07 | 0 | 0 | 1 | 0 | 1 | 3 | 0 | 1 | 0 | 0 | 3 | 0 | 98 | 98 | 0.0374 | 0.19855 | NA | 0 | 0.25 | NA | 0.25 | NA | -0.125 |
| MXE | 767 | ATP5C1 | chr10 | + | 7802757 | 7802854 | 7806973 | 7807010 | 7802269 | 7802425 | 8E+06 | 8E+06 | 0 | 0 | 0 | 0 | 1 | 8 | 1 | 1 | 0 | 0 | 2 | 0 | 98 | 86 | 0.0375 | 0.19855 | NA | 0 | 0 | 1 | 0.305 | NA | -0.652 |
| MXE | 443 | ADAM8 | chr10 | - | 133269444 | 133269529 | 133269896 | 133269974 | 133267928 | 133268118 | 1E+08 | 1E+08 | 0 | 5 | 18 | 0 | 0 | 0 | 0 | 34 | 6 | 0 | 2 | 0 | 98 | 98 | 0.0376 | 0.19855 | NA | 1 | 1 | NA | 0.944 | 1 | 0.028 |
| MXE | 722 | BEST1 | chr11 | + | 61959497 | 61959578 | 61959891 | 61960043 | 61958145 | 61958298 | 6E+07 | 6E+07 | 0 | 0 | 2 | 0 | 1 | 3 | 0 | 1 | 0 | 0 | 4 | 0 | 98 | 98 | 0.0384 | 0.19855 | NA | 0 | 0.4 | NA | 0.2 | NA | 0 |
| MXE | 1143 | ACSL1 | chr4 | - | 184770254 | 184770332 | 184770398 | 184770476 | 184768315 | 184768450 | 2E+08 | 2E+08 | 0 | 1 | 55 | 0 | 2 | 3 | 4 | 29 | 0 | 0 | 3 | 0 | 98 | 98 | 0.0391 | 0.19855 | NA | 0.333 | 0.948 | 1 | 0.906 | NA | -0.313 |
| MXE | 740 | CD164 | chr6 | - | 109370410 | 109370467 | 109376073 | 109376112 | 109366513 | 109369017 | 1E+08 | 1E+08 | 0 | 1 | 6 | 0 | 0 | 5 | 1 | 6 | 0 | 0 | 1 | 0 | 88 | 98 | 0.0413 | 0.20153 | NA | 1 | 0.572 | 1 | 0.87 | NA | -0.149 |

| | | | | | | | | | | | | | | | | | | | | | | | | | | | | | | | | | | | |
|---|---|---|---|---|---|---|---|---|---|---|---|---|---|---|---|---|---|---|---|---|---|---|---|---|---|---|---|---|---|---|---|---|---|---|---|
| MXE | 320 | ZDHHC20 | chr13 | - | 21421060 | 21421164 | 21425651 | 21425678 | 21413651 | 21413772 | 2E+07 | 2E+07 | 0 | 0 | 0 | 0 | 0 | 3 | 0 | 1 | 2 | 0 | 0 | 0 | 76 | 98 | 0.0581 | 0.24305 | NA | NA | 0 | NA | 1 | 1 | -1 |
| MXE | 425 | PHF21A | chr11 | - | 45945839 | 45946003 | 45946075 | 45946098 | 45938156 | 45938312 | 5E+07 | 5E+07 | 0 | 1 | 0 | 0 | 0 | 4 | 0 | 0 | 0 | 0 | 4 | 0 | 72 | 98 | 0.0595 | 0.24305 | NA | 1 | 0 | NA | 0 | NA | 0.5 |
| MXE | 872 | PIK3R5 | chr17 | - | 8889972 | 8890126 | 8890737 | 8890912 | 8889138 | 8889222 | 9E+06 | 9E+06 | 0 | 0 | 6 | 0 | 1 | 1 | 0 | 3 | 0 | 0 | 0 | 0 | 98 | 98 | 0.0596 | 0.24305 | NA | 0 | 0.857 | NA | 1 | NA | -0.572 |
| MXE | 906 | DEDD2 | chr19 | - | 42202050 | 42202072 | 42204461 | 42204559 | 42198592 | 42199829 | 4E+07 | 4E+07 | 0 | 1 | 0 | 0 | 0 | 3 | 0 | 0 | 0 | 0 | 2 | 0 | 98 | 71 | 0.0614 | 0.24305 | NA | 1 | 0 | NA | 0 | NA | 0.5 |
| MXE | 1148 | PDK1 | chr2 | + | 172562740 | 172562800 | 172564502 | 172564687 | 172562219 | 172562291 | 2E+08 | 2E+08 | 0 | 1 | 0 | 0 | 0 | 0 | 0 | 0 | 0 | 0 | 2 | 0 | 98 | 98 | 0.0614 | 0.24305 | NA | 1 | NA | NA | 0 | NA | 1 |
| MXE | 1052 | ARID3A | chr19 | + | 964832 | 965080 | 966571 | 966868 | 964247 | 964431 | 968404 | 968503 | 0 | 1 | 1 | 0 | 0 | 0 | 0 | 0 | 0 | 0 | 1 | 0 | 98 | 98 | 0.0659 | 0.24305 | NA | 1 | 1 | NA | 0 | NA | 1 |
| MXE | 3 | ALG13 | chrX | + | 111682131 | 111682235 | 111682235 | 111682294 | 111681117 | 111681299 | 1E+08 | 1E+08 | 0 | 0 | 0 | 0 | 1 | 0 | 0 | 1 | 0 | 0 | 0 | 0 | 98 | 98 | 0.0659 | 0.24305 | NA | 0 | NA | NA | 1 | NA | -1 |
| MXE | 611 | PTGES3 | chr12 | - | 56670274 | 56670364 | 56671748 | 56671847 | 56666203 | 56666266 | 6E+07 | 6E+07 | 0 | 0 | 2 | 0 | 1 | 2 | 0 | 1 | 0 | 0 | 0 | 0 | 98 | 98 | 0.0659 | 0.24305 | NA | 0 | 0.5 | NA | 1 | NA | -0.75 |
| MXE | 216 | LST1 | chr6 | + | 31587640 | 31587733 | 31587943 | 31587966 | 31587199 | 31587318 | 3E+07 | 3E+07 | 0 | 0 | 0 | 0 | 1 | 1 | 0 | 1 | 0 | 0 | 0 | 0 | 98 | 72 | 0.067 | 0.24305 | NA | 0 | 0 | NA | 1 | NA | -1 |
| MXE | 290 | NIN | chr14 | - | 50821873 | 50822077 | 50830463 | 50830517 | 50806736 | 50806818 | 5E+07 | 5E+07 | 0 | 0 | 1 | 0 | 0 | 6 | 2 | 0 | 13 | 0 | 7 | 0 | 98 | 98 | 0.0791 | 0.2761 | NA | NA | 0.143 | 1 | 0 | 1 | -0.524 |
| MXE | 327 | FOS | chr14 | + | 75279876 | 75280128 | 75280559 | 75280667 | 75278777 | 75279123 | 8E+07 | 8E+07 | 0 | 6 | 55 | 0 | 12 | 92 | 11 | 47 | 3 | 3 | 119 | 9 | 98 | 98 | 0.0804 | 0.2761 | NA | 0.333 | 0.374 | 0.786 | 0.283 | 0.25 | -0.086 |
| MXE | 111 | HLA-E | chr6 | + | 30491536 | 30491653 | 30492540 | 30492583 | 30491136 | 30491412 | 3E+07 | 3E+07 | 0 | 13 | 184 | 0 | 3 | 54 | 11 | 69 | 9 | 0 | 33 | 3 | 98 | 92 | 0.0886 | 0.29606 | NA | 0.803 | 0.762 | 1 | 0.662 | 0.738 | -0.017 |
| MXE | 1102 | NLRP12 | chr19 | - | 53807494 | 53807665 | 53809586 | 53811288 | 53805279 | 53805450 | 5E+07 | 5E+07 | 0 | 1 | 8 | 0 | 1 | 8 | 0 | 3 | 2 | 0 | 1 | 0 | 98 | 98 | 0.1035 | 0.337 | NA | 0.5 | 0.5 | NA | 0.75 | 1 | -0.375 |
| MXE | 918 | HLA-B | chr6 | - | 31354632 | 31354665 | 31355106 | 31355223 | 31354478 | 31354526 | 3E+07 | 3E+07 | 0 | 40 | 238 | 1 | 12 | 113 | 10 | 193 | 14 | 8 | 112 | 7 | 98 | 82 | 0.1085 | 0.33916 | 0 | 0.736 | 0.638 | 0.511 | 0.59 | 0.626 | -0.118 |
| MXE | 176 | BCL6 | chr3 | - | 187726730 | 187726898 | 187728359 | 187728544 | 187725498 | 187725629 | 2E+08 | 2E+08 | 0 | 2 | 13 | 0 | 1 | 13 | 0 | 5 | 1 | 0 | 1 | 0 | 98 | 98 | 0.1105 | 0.33916 | NA | 0.667 | 0.5 | NA | 0.833 | 1 | -0.333 |
| MXE | 964 | CKLF | chr16 | + | 66558189 | 66558348 | 66563121 | 66563217 | 66552562 | 66552793 | 7E+07 | 7E+07 | 0 | 0 | 9 | 0 | 0 | 7 | 0 | 5 | 5 | 0 | 0 | 0 | 98 | 98 | 0.1148 | 0.33916 | NA | NA | 0.562 | NA | 1 | 1 | -0.438 |
| MXE | 1081 | RASGRP4 | chr19 | - | 38417051 | 38417168 | 38419859 | 38420013 | 38414847 | 38415123 | 4E+07 | 4E+07 | 0 | 4 | 7 | 0 | 2 | 13 | 1 | 1 | 0 | 0 | 4 | 0 | 98 | 98 | 0.1166 | 0.33916 | NA | 0.667 | 0.35 | 1 | 0.2 | NA | -0.092 |
| MXE | 417 | SUCO | chr1 | + | 172551511 | 172551626 | 172553259 | 172553370 | 172533117 | 172533497 | 2E+08 | 2E+08 | 0 | 1 | 0 | 0 | 0 | 1 | 0 | 0 | 0 | 0 | 2 | 1 | 98 | 98 | 0.1192 | 0.33916 | NA | 1 | 0 | NA | 0 | 0 | 0.5 |
| MXE | 909 | IQGAP1 | chr15 | + | 90467449 | 90467592 | 90472839 | 90473010 | 90466268 | 90466436 | 9E+07 | 9E+07 | 0 | 0 | 16 | 0 | 0 | 14 | 0 | 4 | 0 | 6 | 14 | 3 | 98 | 98 | 0.1202 | 0.33916 | NA | NA | 0.533 | 0 | 0.222 | 0 | 0.459 |
| MXE | 332 | STAT6 | chr12 | - | 57107604 | 57107743 | 57108162 | 57108299 | 57107230 | 57107314 | 6E+07 | 6E+07 | 0 | 1 | 5 | 0 | 1 | 6 | 0 | 3 | 0 | 0 | 0 | 0 | 98 | 98 | 0.1246 | 0.34414 | NA | 0.5 | 0.455 | NA | 1 | NA | -0.522 |
| MXE | 97 | VNN2 | chr6 | - | 132752672 | 132752749 | 132753417 | 132753491 | 132751144 | 132751518 | 1E+08 | 1E+08 | 0 | 0 | 3 | 0 | 0 | 10 | 0 | 1 | 1 | 0 | 3 | 0 | 98 | 98 | 0.1412 | 0.38142 | NA | NA | 0.231 | NA | 0.25 | 1 | -0.394 |
| MXE | 405 | RELT | chr11 | + | 73391143 | 73391223 | 73392210 | 73392468 | 73390754 | 73390921 | 7E+07 | 7E+07 | 0 | 0 | 3 | 0 | 0 | 3 | 0 | 1 | 0 | 0 | 6 | 2 | 98 | 98 | 0.1464 | 0.38743 | NA | NA | 0.5 | NA | 0.143 | 0 | 0.428 |
| MXE | 366 | PTPRJ | chr11 | + | 48112746 | 48112983 | 48121002 | 48121266 | 48110057 | 48110076 | 5E+07 | 5E+07 | 2 | 11 | 21 | 0 | 4 | 13 | 0 | 6 | 5 | 0 | 2 | 0 | 98 | 98 | 0.16 | 0.3991 | 1 | 0.733 | 0.618 | NA | 0.75 | 1 | -0.091 |
| MXE | 986 | TNFSF13 | chr17 | + | 7559623 | 7559702 | 7559850 | 7559893 | 7558291 | 7559297 | 8E+06 | 8E+06 | 0 | 0 | 4 | 0 | 0 | 9 | 0 | 1 | 1 | 0 | 0 | 0 | 98 | 92 | 0.1617 | 0.3991 | NA | NA | 0.294 | NA | 1 | 1 | -0.706 |
| MXE | 951 | APLP2 | chr11 | + | 130109428 | 130109505 | 130120705 | 130120818 | 130069820 | 130070082 | 1E+08 | 1E+08 | 0 | 0 | 9 | 0 | 0 | 20 | 0 | 1 | 1 | 0 | 8 | 0 | 98 | 98 | 0.1628 | 0.3991 | NA | NA | 0.31 | NA | 0.111 | 1 | -0.245 |
| MXE | 96 | VNN2 | chr6 | - | 132752460 | 132752749 | 132753417 | 132753491 | 132751144 | 132751518 | 1E+08 | 1E+08 | 0 | 0 | 3 | 0 | 0 | 32 | 0 | 1 | 1 | 0 | 18 | 1 | 98 | 98 | 0.1663 | 0.3991 | NA | NA | 0.086 | NA | 0.053 | 0.5 | -0.191 |
| MXE | 985 | TNFSF13 | chr17 | + | 7559623 | 7559702 | 7559847 | 7559893 | 7558291 | 7559297 | 8E+06 | 8E+06 | 0 | 0 | 4 | 0 | 0 | 8 | 0 | 1 | 1 | 0 | 0 | 0 | 98 | 95 | 0.169 | 0.3991 | NA | NA | 0.326 | NA | 1 | 1 | -0.674 |
| MXE | 710 | CKLF-CMTM1 | chr16 | + | 66558189 | 66558348 | 66563121 | 66563217 | 66552562 | 66552793 | 7E+07 | 7E+07 | 0 | 0 | 7 | 0 | 0 | 2 | 0 | 3 | 5 | 0 | 0 | 0 | 98 | 98 | 0.1724 | 0.3991 | NA | NA | 0.778 | NA | 1 | 1 | -0.222 |
| MXE | 478 | DENND3 | chr8 | + | 141160631 | 141160787 | 141163332 | 141163429 | 141155848 | 141155970 | 1E+08 | 1E+08 | 0 | 0 | 11 | 0 | 0 | 8 | 0 | 5 | 2 | 3 | 6 | 0 | 98 | 98 | 0.1751 | 0.3991 | NA | NA | 0.579 | 0 | 0.455 | 1 | 0.094 |
| MXE | 299 | ZCCHC2 | chr18 | + | 62556202 | 62556297 | 62558686 | 62558770 | 62550347 | 62550460 | 6E+07 | 6E+07 | 0 | 0 | 16 | 0 | 0 | 9 | 0 | 9 | 0 | 0 | 8 | 1 | 98 | 98 | 0.176 | 0.3991 | NA | NA | 0.64 | NA | 0.529 | 0 | 0.376 |
| MXE | 1140 | PLEKHB2 | chr2 | + | 131126683 | 131126786 | 131132901 | 131132991 | 131125752 | 131125905 | 1E+08 | 1E+08 | 0 | 1 | 0 | 0 | 1 | 3 | 0 | 1 | 0 | 0 | 4 | 0 | 98 | 98 | 0.1953 | 0.43511 | NA | 0.5 | 0 | NA | 0.2 | NA | 0.05 |
| MXE | 877 | LAPTM5 | chr1 | - | 30741639 | 30741716 | 30742455 | 30742549 | 30739808 | 30739937 | 3E+07 | 3E+07 | 0 | 0 | 4 | 0 | 1 | 20 | 0 | 2 | 0 | 5 | 28 | 6 | 98 | 98 | 0.219 | 0.47953 | NA | 0 | 0.167 | 0 | 0.067 | 0 | 0.061 |
| MXE | 905 | DEDD2 | chr19 | - | 42201988 | 42202072 | 42204461 | 42204559 | 42198592 | 42199829 | 4E+07 | 4E+07 | 0 | 1 | 0 | 0 | 1 | 2 | 0 | 0 | 0 | 0 | 1 | 0 | 98 | 98 | 0.2322 | 0.49986 | NA | 0.5 | 0 | NA | 0 | NA | 0.25 |
| MXE | 942 | PARP14 | chr3 | + | 122699389 | 122701635 | 122703741 | 122703978 | 122695425 | 122695662 | 1E+08 | 1E+08 | 0 | 7 | 16 | 0 | 4 | 9 | 0 | 27 | 2 | 0 | 11 | 0 | 98 | 98 | 0.2868 | 0.607 | NA | 0.636 | 0.64 | NA | 0.711 | 1 | -0.217 |
| MXE | 392 | DCAF11 | chr14 | + | 24117265 | 24117393 | 24117667 | 24117732 | 24116916 | 24117044 | 2E+07 | 2E+07 | 0 | 0 | 4 | 0 | 0 | 1 | 0 | 0 | 2 | 0 | 4 | 0 | 98 | 98 | 0.3035 | 0.62254 | NA | NA | 0.8 | NA | 0 | 1 | 0.3 |
| MXE | 748 | MYD88 | chr3 | + | 38139863 | 38139998 | 38140387 | 38140568 | 38138477 | 38139028 | 4E+07 | 4E+07 | 0 | 6 | 12 | 0 | 3 | 7 | 0 | 11 | 3 | 0 | 2 | 2 | 98 | 98 | 0.3039 | 0.62254 | NA | 0.667 | 0.632 | NA | 0.846 | 0.6 | -0.074 |
| MXE | 367 | SORL1 | chr11 | + | 121543547 | 121543726 | 121545242 | 121545429 | 121532463 | 121532552 | 1E+08 | 1E+08 | 0 | 0 | 18 | 0 | 2 | 13 | 0 | 11 | 0 | 0 | 15 | 2 | 98 | 98 | 0.3121 | 0.62925 | NA | 0 | 0.581 | NA | 0.423 | 0 | 0.079 |
| MXE | 717 | CTSS | chr1 | - | 150751780 | 150752008 | 150755000 | 150755150 | 150750005 | 150750171 | 2E+08 | 2E+08 | 0 | 2 | 126 | 0 | 1 | 73 | 4 | 28 | 7 | 5 | 13 | 1 | 98 | 98 | 0.3255 | 0.64594 | NA | 0.667 | 0.633 | 0.444 | 0.683 | 0.875 | -0.017 |
| MXE | 110 | HLA-E | chr6 | + | 30491536 | 30491653 | 30492403 | 30492436 | 30491136 | 30491412 | 3E+07 | 3E+07 | 0 | 12 | 185 | 0 | 7 | 64 | 11 | 70 | 10 | 0 | 48 | 3 | 98 | 82 | 0.3366 | 0.65006 | NA | 0.589 | 0.707 | 1 | 0.55 | 0.736 | -0.114 |
| MXE | 727 | 7-Mar | chr2 | + | 159762879 | 159762993 | 159764625 | 159764674 | 159759225 | 159759335 | 2E+08 | 2E+08 | 0 | 0 | 2 | 0 | 0 | 5 | 0 | 0 | 0 | 0 | 5 | 1 | 98 | 98 | 0.3401 | 0.65006 | NA | NA | 0.286 | NA | 0 | 0 | 0.286 |
| MXE | 683 | ITGAX | chr16 | + | 31355892 | 31355998 | 31356624 | 31356728 | 31355133 | 31355291 | 3E+07 | 3E+07 | 0 | 0 | 4 | 0 | 9 | 38 | 0 | 5 | 0 | 0 | 34 | 6 | 98 | 98 | 0.3429 | 0.65006 | NA | 0 | 0.095 | NA | 0.128 | 0 | -0.017 |
| MXE | 800 | TAPBP | chr6 | - | 33313216 | 33313477 | 33313693 | 33313864 | 33304988 | 33305387 | 3E+07 | 3E+07 | 0 | 0 | 1 | 0 | 5 | 4 | 0 | 0 | 1 | 0 | 1 | 1 | 98 | 98 | 0.3696 | 0.69031 | NA | 0 | 0.2 | NA | 0 | 0.5 | -0.15 |
| MXE | 959 | CREBBP | chr16 | - | 3782683 | 3782926 | 3791980 | 3792094 | 3781203 | 3781306 | 4E+06 | 4E+06 | 0 | 0 | 5 | 0 | 0 | 2 | 0 | 3 | 1 | 0 | 4 | 0 | 98 | 98 | 0.4066 | 0.74842 | NA | NA | 0.714 | NA | 0.429 | 1 | -0.001 |
| MXE | 416 | PICALM | chr11 | - | 86022366 | 86022469 | 86026291 | 86026367 | 86014869 | 86014963 | 9E+07 | 9E+07 | 0 | 1 | 14 | 0 | 0 | 16 | 0 | 20 | 1 | 0 | 24 | 0 | 98 | 98 | 0.4149 | 0.75274 | NA | 1 | 0.467 | NA | 0.455 | 1 | 0.006 |
| MXE | 1118 | CCND3 | chr6 | - | 41940369 | 41940585 | 42030008 | 42030190 | 41937234 | 41937394 | 4E+07 | 4E+07 | 0 | 0 | 1 | 0 | 1 | 23 | 0 | 0 | 0 | 0 | 0 | 3 | 98 | 98 | 0.4635 | 0.82909 | NA | 0 | 0.042 | NA | NA | 0 | 0.021 |
| MXE | 645 | THEMIS2 | chr1 | + | 27879643 | 27880054 | 27881970 | 27883043 | 27876587 | 27876728 | 3E+07 | 3E+07 | 0 | 6 | 25 | 0 | 0 | 2 | 1 | 12 | 8 | 0 | 2 | 0 | 98 | 98 | 0.4758 | 0.83779 | NA | 1 | 0.926 | 1 | 0.857 | 1 | 0.011 |
| MXE | 348 | PREX1 | chr20 | - | 48726289 | 48726391 | 48734545 | 48734650 | 48708259 | 48708421 | 5E+07 | 5E+07 | 0 | 7 | 26 | 1 | 18 | 53 | 3 | 42 | 5 | 8 | 31 | 14 | 98 | 98 | 0.4816 | 0.83779 | 0 | 0.28 | 0.329 | 0.273 | 0.575 | 0.263 | -0.167 |
| MXE | 1061 | MYO1F | chr19 | - | 8539946 | 8540028 | 8541905 | 8541991 | 8536948 | 8537055 | 9E+06 | 9E+06 | 0 | 6 | 17 | 0 | 1 | 42 | 0 | 11 | 2 | 1 | 20 | 3 | 98 | 98 | 0.4957 | 0.85071 | NA | 0.857 | 0.288 | 0 | 0.355 | 0.4 | 0.321 |
| MXE | 253 | SLC9A8 | chr20 | + | 49845035 | 49845119 | 49849578 | 49849680 | 49839540 | 49839599 | 5E+07 | 5E+07 | 0 | 0 | 4 | 0 | 0 | 4 | 0 | 2 | 2 | 0 | 4 | 1 | 98 | 98 | 0.5531 | 0.91543 | NA | NA | 0.5 | NA | 0.333 | 0.667 | 0 |
| MXE | 495 | HCLS1 | chr3 | - | 121632100 | 121632184 | 121632331 | 121632563 | 121631397 | 121631982 | 1E+08 | 1E+08 | 0 | 8 | 38 | 0 | 1 | 28 | 0 | 21 | 12 | 1 | 8 | 4 | 98 | 98 | 0.5547 | 0.91543 | NA | 0.889 | 0.576 | 0 | 0.724 | 0.75 | 0.241 |
| MXE | 20 | ASB6 | chr9 | - | 129638572 | 129638659 | 129639201 | 129639310 | 129634603 | 129638457 | 1E+08 | 1E+08 | 0 | 0 | 1 | 0 | 0 | 1 | 0 | 1 | 2 | 0 | 1 | 1 | 98 | 98 | 0.555 | 0.91543 | NA | NA | 0.5 | NA | 0.5 | 0.667 | -0.084 |
| MXE | 144 | DBNL | chr7 | + | 44050224 | 44050280 | 44052866 | 44052941 | 44044639 | 44044820 | 4E+07 | 4E+07 | 0 | 1 | 1 | 0 | 2 | 5 | 0 | 1 | 0 | 0 | 1 | 6 | 98 | 98 | 0.5928 | 0.95849 | NA | 0.333 | 0.167 | NA | 0.5 | 0 | 0 |
| MXE | 321 | GLT1D1 | chr12 | + | 128875913 | 128876062 | 128888638 | 128888744 | 128853535 | 128853649 | 1E+08 | 1E+08 | 0 | 2 | 9 | 0 | 5 | 8 | 0 | 4 | 2 | 0 | 3 | 2 | 98 | 98 | 0.5962 | 0.95849 | NA | 0.286 | 0.529 | NA | 0.571 | 0.5 | -0.128 |
| MXE | 809 | IFRD1 | chr7 | + | 112456913 | 112457038 | 112458860 | 112459018 | 112456001 | 112456086 | 1E+08 | 1E+08 | 0 | 6 | 38 | 0 | 0 | 28 | 0 | 72 | 4 | 0 | 20 | 0 | 98 | 98 | 0.6106 | 0.96816 | NA | 1 | 0.576 | NA | 0.783 | 1 | -0.103 |
| MXE | 157 | HERC5 | chr4 | + | 88470613 | 88470673 | 88472408 | 88472502 | 88469156 | 88469260 | 9E+07 | 9E+07 | 0 | 0 | 9 | 0 | 0 | 16 | 0 | 20 | 1 | 0 | 15 | 3 | 98 | 98 | 0.6214 | 0.96816 | NA | NA | 0.36 | NA | 0.571 | 0.25 | -0.05 |
| MXE | 354 | CLEC7A | chr12 | - | 10123244 | 10123363 | 10125086 | 10125173 | 10116780 | 10118590 | 1E+07 | 1E+07 | 0 | 3 | 18 | 0 | 0 | 38 | 0 | 4 | 3 | 0 | 16 | 2 | 98 | 98 | 0.6251 | 0.96816 | NA | 1 | 0.321 | NA | 0.2 | 0.6 | 0.26 |
| MXE | 1142 | ACSL1 | chr4 | - | 184764852 | 184764925 | 184765890 | 184765986 | 184763166 | 184763255 | 2E+08 | 2E+08 | 0 | 2 | 50 | 0 | 2 | 44 | 0 | 38 | 5 | 3 | 39 | 5 | 98 | 98 | 0.6342 | 0.97045 | NA | 0.5 | 0.532 | 0 | 0.494 | 0.5 | 0.185 |
| MXE | 1083 | GMFG | chr19 | - | 39335260 | 39335310 | 39335434 | 39335531 | 39333076 | 39333126 | 4E+07 | 4E+07 | 0 | 0 | 1 | 0 | 1 | 12 | 0 | 1 | 0 | 13 | 12 | 1 | 98 | 98 | 0.659 | 0.99468 | NA | 0 | 0.077 | 0 | 0.077 | 0 | 0.013 |
| MXE | 139 | ARHGAP26 | chr5 | + | 143014079 | 143014116 | 143037195 | 143037261 | 142932046 | 142932125 | 1E+08 | 1E+08 | 0 | 1 | 28 | 0 | 0 | 13 | 0 | 18 | 2 | 0 | 9 | 0 | 86 | 98 | 0.6657 | 0.99468 | NA | 1 | 0.711 | NA | 0.695 | 1 | 0.008 |
| MXE | 69 | TBXAS1 | chr7 | + | 139955458 | 139955607 | 139957633 | 139957764 | 139953367 | 139953456 | 1E+08 | 1E+08 | 0 | 1 | 5 | 0 | 2 | 12 | 0 | 1 | 0 | 2 | 3 | 0 | 98 | 98 | 0.703 | 0.9992 | NA | 0.333 | 0.294 | 0 | 0.25 | NA | 0.189 |
| MXE | 963 | ADGRG3 | chr16 | + | 57680504 | 57680617 | 57684389 | 57684483 | 57680264 | 57680365 | 6E+07 | 6E+07 | 0 | 5 | 10 | 0 | 0 | 6 | 0 | 9 | 14 | 0 | 6 | 1 | 98 | 98 | 0.7158 | 0.9992 | NA | 1 | 0.625 | NA | 0.6 | 0.933 | 0.046 |
| MXE | 1087 | PLAUR | chr19 | - | 43655438 | 43655573 | 43656478 | 43656640 | 43652224 | 43652371 | 4E+07 | 4E+07 | 0 | 6 | 38 | 0 | 0 | 29 | 0 | 62 | 6 | 0 | 26 | 2 | 98 | 98 | 0.7567 | 0.9992 | NA | 1 | 0.567 | NA | 0.705 | 0.75 | 0.056 |
| MXE | 614 | IFI44 | chr1 | + | 78660553 | 78660654 | 78662703 | 78662878 | 78659311 | 78659483 | 8E+07 | 8E+07 | 0 | 0 | 31 | 0 | 6 | 51 | 0 | 20 | 1 | 0 | 87 | 6 | 98 | 98 | 0.7641 | 0.9992 | NA | 0 | 0.378 | NA | 0.187 | 0.143 | 0.024 |
| MXE | 644 | THEMIS2 | chr1 | + | 27876587 | 27876728 | 27879643 | 27880054 | 27872542 | 27872665 | 3E+07 | 3E+07 | 0 | 10 | 8 | 0 | 0 | 2 | 6 | 6 | 15 | 0 | 0 | 2 | 98 | 98 | 0.7725 | 0.9992 | NA | 1 | 0.8 | 1 | 1 | 0.882 | -0.061 |
| MXE | 400 | DDX3X | chrX | + | 41343215 | 41343351 | 41343736 | 41343822 | 41342736 | 41342836 | 4E+07 | 4E+07 | 0 | 2 | 15 | 0 | 2 | 21 | 0 | 19 | 2 | 0 | 28 | 1 | 98 | 98 | 0.7867 | 0.9992 | NA | 0.5 | 0.417 | NA | 0.404 | 0.667 | -0.077 |
| MXE | 714 | DUSP1 | chr5 | - | 172769574 | 172769794 | 172770160 | 172770306 | 172768089 | 172769132 | 2E+08 | 2E+08 | 4 | 93 | 267 | 0 | 6 | 110 | 23 | 118 | 275 | 3 | 47 | 10 | 98 | 98 | 0.7892 | 0.9992 | 1 | 0.939 | 0.708 | 0.885 | 0.715 | 0.965 | 0.027 |
| MXE | 269 | ELOVL5 | chr6 | - | 53270592 | 53270727 | 53273219 | 53273344 | 53267397 | 53269270 | 5E+07 | 5E+07 | 0 | 1 | 16 | 0 | 0 | 8 | 3 | 6 | 4 | 0 | 7 | 0 | 98 | 98 | 0.868 | 0.9992 | NA | 1 | 0.667 | 1 | 0.462 | 1 | 0.013 |
| MXE | 540 | AMPD2 | chr1 | + | 109629326 | 109629490 | 109629795 | 109629916 | 109629108 | 109629235 | 1E+08 | 1E+08 | 0 | 2 | 5 | 0 | 1 | 4 | 0 | 8 | 1 | 0 | 6 | 1 | 98 | 98 | 0.887 | 0.9992 | NA | 0.667 | 0.556 | NA | 0.571 | 0.5 | 0.076 |
| MXE | 1034 | CSNK1D | chr17 | - | 82253015 | 82253244 | 82255428 | 82255577 | 82252433 | 82252604 | 8E+07 | 8E+07 | 0 | 1 | 2 | 0 | 2 | 4 | 0 | 3 | 1 | 0 | 4 | 3 | 98 | 98 | 0.8881 | 0.9992 | NA | 0.333 | 0.333 | NA | 0.429 | 0.25 | -0.007 |
| MXE | 1077 | PTTG1IP | chr21 | - | 44861162 | 44861271 | 44865394 | 44865447 | 44856192 | 44856364 | 4E+07 | 4E+07 | 0 | 1 | 3 | 0 | 0 | 7 | 0 | 1 | 1 | 0 | 3 | 0 | 98 | 98 | 0.922 | 0.9992 | NA | 1 | 0.3 | NA | 0.25 | 1 | 0.025 |
| MXE | 436 | TAGAP | chr6 | - | 159042077 | 159042244 | 159043588 | 159043655 | 159041353 | 159041515 | 2E+08 | 2E+08 | 0 | 0 | 4 | 0 | 3 | 8 | 0 | 5 | 0 | 0 | 15 | 2 | 98 | 98 | 0.9267 | 0.9992 | NA | 0 | 0.333 | NA | 0.25 | 0 | 0.042 |
| MXE | 643 | THEMIS2 | chr1 | + | 27876587 | 27876728 | 27879643 | 27880054 | 27872542 | 27872665 | 3E+07 | 3E+07 | 0 | 7 | 8 | 0 | 1 | 19 | 6 | 5 | 15 | 0 | 13 | 5 | 98 | 98 | 0.9303 | 0.9992 | NA | 0.875 | 0.296 | 1 | 0.278 | 0.75 | -0.091 |
| MXE | 142 | VCAN | chr5 | + | 83519348 | 83522309 | 83537006 | 83542268 | 83512102 | 83512396 | 8E+07 | 8E+07 | 0 | 0 | 1 | 0 | 2 | 51 | 0 | 0 | 0 | 0 | 10 | 0 | 98 | 98 | 0.9435 | 0.9992 | NA | 0 | 0.019 | NA | 0 | NA | 0.009 |
| MXE | 1010 | KLHL2 | chr4 | + | 165228806 | 165228913 | 165238777 | 165238899 | 165219933 | 165220059 | 2E+08 | 2E+08 | 0 | 1 | 6 | 0 | 0 | 4 | 0 | 5 | 0 | 0 | 0 | 0 | 98 | 98 | 0.9442 | 0.9992 | NA | 1 | 0.6 | NA | 1 | NA | -0.2 |
| MXE | 1076 | CYP4F3 | chr19 | + | 15643906 | 15644051 | 15645718 | 15645863 | 15641414 | 15641613 | 2E+07 | 2E+07 | 0 | 0 | 1 | 0 | 5 | 21 | 0 | 0 | 0 | 7 | 13 | 0 | 98 | 98 | 0.9631 | 0.9992 | NA | 0 | 0.045 | 0 | 0 | NA | 0.022 |
| MXE | 1023 | APH1A | chr1 | - | 150267715 | 150267789 | 150267956 | 150268127 | 150267355 | 150267478 | 2E+08 | 2E+08 | 0 | 0 | 3 | 0 | 1 | 1 | 0 | 0 | 0 | 0 | 2 | 0 | 98 | 98 | 0.9714 | 0.9992 | NA | 0 | 0.75 | NA | 0 | NA | 0.375 |
| MXE | 358 | GZMH | chr14 | - | 24607148 | 24607406 | 24607611 | 24607747 | 24606479 | 24606746 | 2E+07 | 2E+07 | 0 | 1 | 0 | 0 | 0 | 3 | 0 | 2 | 0 | 0 | 0 | 0 | 98 | 98 | 0.9717 | 0.9992 | NA | 1 | 0 | NA | 1 | NA | -0.5 |
| MXE | 359 | GZMH | chr14 | - | 24607340 | 24607406 | 24607611 | 24607747 | 24606479 | 24606746 | 2E+07 | 2E+07 | 0 | 1 | 0 | 0 | 0 | 3 | 0 | 2 | 0 | 0 | 0 | 0 | 98 | 98 | 0.9717 | 0.9992 | NA | 1 | 0 | NA | 1 | NA | -0.5 |
| MXE | 911 | MEF2A | chr15 | + | 99671322 | 99671454 | 99671519 | 99671657 | 99645560 | 99645764 | 1E+08 | 1E+08 | 0 | 1 | 5 | 0 | 0 | 3 | 0 | 2 | 0 | 0 | 0 | 0 | 98 | 98 | 0.9717 | 0.9992 | NA | 1 | 0.625 | NA | 1 | NA | -0.188 |
| MXE | 18 | CSF2RA | chrX | + | 1300490 | 1300626 | 1303922 | 1304019 | 1295426 | 1295456 | 1E+06 | 1E+06 | 0 | 0 | 15 | 0 | 1 | 10 | 0 | 6 | 1 | 0 | 1 | 0 | 98 | 98 | 0.9992 | 0.9992 | NA | 0 | 0.6 | NA | 0.857 | 1 | -0.629 |
| MXE | 21 | PPP2R4 | chr9 | + | 129123051 | 129123138 | 129128984 | 129129110 | 129120512 | 129120610 | 1E+08 | 1E+08 | 0 | 0 | 1 | 0 | 2 | 1 | 0 | 0 | 0 | 0 | 1 | 0 | 98 | 98 | 0.9992 | 0.9992 | NA | 0 | 0.5 | NA | 0 | NA | 0.25 |
| MXE | 125 | PDLIM7 | chr5 | - | 177491403 | 177491420 | 177491806 | 177491925 | 177491009 | 177491146 | 2E+08 | 2E+08 | 0 | 0 | 10 | 0 | 0 | 0 | 0 | 6 | 4 | 0 | 1 | 0 | 98 | 66 | 0.9992 | 0.9992 | NA | NA | 1 | NA | 0.802 | 1 | 0.099 |
| MXE | 179 | NDUFB5 | chr3 | + | 179616982 | 179617044 | 179618414 | 179618521 | 179615982 | 179616049 | 2E+08 | 2E+08 | 0 | 0 | 5 | 0 | 1 | 5 | 0 | 0 | 0 | 0 | 1 | 0 | 98 | 98 | 0.9992 | 0.9992 | NA | 0 | 0.5 | NA | 0 | NA | 0.25 |
| MXE | 217 | LST1 | chr6 | + | 31587640 | 31587733 | 31587943 | 31587966 | 31587199 | 31587318 | 3E+07 | 3E+07 | 0 | 0 | 0 | 0 | 0 | 7 | 0 | 2 | 0 | 0 | 0 | 1 | 98 | 72 | 0.9992 | 0.9992 | NA | NA | 0 | NA | 1 | 0 | -0.5 |
| MXE | 245 | MX1 | chr21 | + | 41437014 | 41437152 | 41439693 | 41439848 | 41435836 | 41436029 | 4E+07 | 4E+07 | 0 | 5 | 29 | 0 | 6 | 76 | 4 | 18 | 2 | 1 | 47 | 6 | 98 | 98 | 0.9992 | 0.9992 | NA | 0.455 | 0.276 | 0.8 | 0.277 | 0.25 | -0.077 |
| MXE | 331 | XPO6 | chr16 | - | 28117317 | 28117462 | 28121669 | 28121762 | 28112903 | 28113050 | 3E+07 | 3E+07 | 0 | 5 | 50 | 0 | 0 | 25 | 7 | 34 | 0 | 0 | 19 | 1 | 98 | 98 | 0.9992 | 0.9992 | NA | 1 | 0.667 | 1 | 0.642 | 0 | 0.286 |
| MXE | 390 | DCAF11 | chr14 | + | 24116916 | 24117044 | 24117265 | 24117393 | 24115381 | 24115749 | 2E+07 | 2E+07 | 0 | 0 | 0 | 0 | 0 | 2 | 0 | 3 | 0 | 0 | 3 | 2 | 98 | 98 | 0.9992 | 0.9992 | NA | NA | 0 | NA | 0.5 | 0 | -0.25 |
| MXE | 391 | DCAF11 | chr14 | + | 24116971 | 24117044 | 24117265 | 24117393 | 24115282 | 24115749 | 2E+07 | 2E+07 | 0 | 0 | 0 | 0 | 0 | 2 | 0 | 1 | 0 | 0 | 3 | 2 | 98 | 98 | 0.9992 | 0.9992 | NA | NA | 0 | NA | 0.25 | 0 | -0.125 |
| MXE | 422 | BID | chr22 | - | 17743802 | 17744013 | 17750104 | 17750174 | 17739348 | 17739488 | 2E+07 | 2E+07 | 0 | 0 | 0 | 0 | 0 | 9 | 0 | 2 | 0 | 0 | 5 | 2 | 98 | 98 | 0.9992 | 0.9992 | NA | NA | 0 | NA | 0.286 | 0 | -0.143 |
| MXE | 484 | NFKBID | chr19 | - | 35896912 | 35897058 | 35897650 | 35897856 | 35896725 | 35896831 | 4E+07 | 4E+07 | 0 | 6 | 4 | 0 | 7 | 1 | 0 | 4 | 1 | 0 | 4 | 1 | 98 | 98 | 0.9992 | 0.9992 | NA | 0.462 | 0.8 | NA | 0.5 | 0.5 | 0.131 |
| MXE | 607 | H2AFY | chr5 | - | 135350822 | 135350913 | 135352945 | 135353045 | 135345967 | 135346057 | 1E+08 | 1E+08 | 0 | 2 | 10 | 0 | 0 | 3 | 0 | 1 | 0 | 0 | 0 | 0 | 98 | 98 | 0.9992 | 0.9992 | NA | 1 | 0.769 | NA | 1 | NA | -0.115 |
| MXE | 656 | HIF1A | chr14 | + | 61738086 | 61738373 | 61740504 | 61740627 | 61736888 | 61737109 | 6E+07 | 6E+07 | 0 | 2 | 6 | 0 | 0 | 11 | 0 | 22 | 1 | 0 | 6 | 1 | 98 | 98 | 0.9992 | 0.9992 | NA | 1 | 0.353 | NA | 0.786 | 0.5 | 0.033 |
| MXE | 700 | ARHGAP30 | chr1 | - | 161056387 | 161056532 | 161059613 | 161059716 | 161054622 | 161054705 | 2E+08 | 2E+08 | 0 | 1 | 18 | 0 | 0 | 18 | 0 | 7 | 0 | 0 | 11 | 1 | 98 | 98 | 0.9992 | 0.9992 | NA | 1 | 0.5 | NA | 0.389 | 0 | 0.555 |
| MXE | 766 | GIT2 | chr12 | - | 109947255 | 109947504 | 109951211 | 109951316 | 109939164 | 109939247 | 1E+08 | 1E+08 | 0 | 1 | 2 | 0 | 0 | 2 | 0 | 1 | 0 | 0 | 0 | 0 | 98 | 98 | 0.9992 | 0.9992 | NA | 1 | 0.5 | NA | 1 | NA | -0.25 |
| MXE | 828 | PTPRE | chr10 | + | 128056122 | 128056213 | 128060938 | 128061015 | 128049529 | 128049666 | 1E+08 | 1E+08 | 1 | 1 | 25 | 0 | 3 | 5 | 0 | 34 | 1 | 0 | 19 | 0 | 98 | 98 | 0.9992 | 0.9992 | 1 | 0.25 | 0.833 | NA | 0.642 | 1 | -0.127 |
| MXE | 998 | MGAM | chr7 | + | 142036168 | 142036285 | 142036822 | 142036977 | 142034669 | 142034841 | 1E+08 | 1E+08 | 0 | 2 | 18 | 0 | 0 | 14 | 2 | 5 | 1 | 0 | 2 | 6 | 98 | 98 | 0.9992 | 0.9992 | NA | 1 | 0.562 | 1 | 0.714 | 0.143 | 0.162 |
| MXE | 1021 | TRIM25 | chr17 | - | 56895520 | 56895604 | 56895925 | 56895952 | 56895342 | 56895441 | 6E+07 | 6E+07 | 0 | 1 | 4 | 0 | 0 | 14 | 0 | 5 | 0 | 0 | 11 | 4 | 76 | 98 | 0.9992 | 0.9992 | NA | 1 | 0.269 | NA | 0.37 | 0 | 0.45 |
| MXE | 1082 | SMCHD1 | chr18 | + | 2722518 | 2722663 | 2724898 | 2724995 | 2718314 | 2718434 | 3E+06 | 3E+06 | 0 | 0 | 26 | 0 | 1 | 40 | 1 | 25 | 1 | 0 | 48 | 1 | 98 | 98 | 0.9992 | 0.9992 | NA | 0 | 0.394 | 1 | 0.342 | 0.5 | -0.417 |
| MXE | 1089 | C17orf62 | chr17 | - | 82446622 | 82446696 | 82447579 | 82447621 | 82445863 | 82445960 | 8E+07 | 8E+07 | 0 | 2 | 4 | 0 | 2 | 5 | 0 | 0 | 3 | 0 | 6 | 0 | 91 | 98 | 0.9992 | 0.9992 | NA | 0.519 | 0.463 | NA | 0 | 1 | -0.009 |
| MXE | 1114 | ALOX5 | chr10 | + | 45382482 | 45382681 | 45395854 | 45395936 | 45374175 | 45374429 | 5E+07 | 5E+07 | 1 | 1 | 30 | 0 | 0 | 31 | 0 | 3 | 0 | 0 | 4 | 1 | 98 | 98 | 0.9992 | 0.9992 | 1 | 1 | 0.492 | NA | 0.429 | 0 | 0.616 |
| MXE | 1158 | NLRP6 | chr11 | + | 279833 | 279872 | 280083 | 281839 | 279326 | 279607 | 282704 | 282797 | 0 | 1 | 0 | 0 | 1 | 1 | 0 | 1 | 2 | 0 | 1 | 4 | 88 | 98 | 0.9992 | 0.9992 | NA | 0.527 | 0 | NA | 0.527 | 0.358 | -0.179 |
| MXE | 22 | TLE4 | chr9 | + | 79612655 | 79612718 | 79627373 | 79627448 | 79576132 | 79576177 | 8E+07 | 8E+07 | 0 | 0 | 0 | 0 | 0 | 2 | 0 | 4 | 0 | 0 | 2 | 0 | 98 | 98 | NA | NA | NA | NA | 0 | NA | 0.667 | NA | -0.667 |
| MXE | 31 | UBAP1 | chr9 | + | 34220907 | 34220948 | 34234215 | 34234340 | 34179004 | 34179240 | 3E+07 | 3E+07 | 0 | 0 | 1 | 0 | 0 | 0 | 0 | 1 | 0 | 0 | 2 | 0 | 90 | 98 | NA | NA | NA | NA | 1 | NA | 0.353 | NA | 0.647 |
| MXE | 40 | VPS13B | chr8 | + | 99507136 | 99507203 | 99507769 | 99507911 | 99502835 | 99502950 | 1E+08 | 1E+08 | 0 | 0 | 5 | 0 | 0 | 2 | 0 | 6 | 0 | 0 | 0 | 0 | 98 | 98 | NA | NA | NA | NA | 0.714 | NA | 1 | NA | -0.286 |
| MXE | 51 | PAG1 | chr8 | - | 80991478 | 80991530 | 80993102 | 80993307 | 80987369 | 80987466 | 8E+07 | 8E+07 | 0 | 0 | 0 | 0 | 0 | 2 | 0 | 3 | 0 | 0 | 3 | 0 | 98 | 98 | NA | NA | NA | NA | 0 | NA | 0.5 | NA | -0.5 |
| MXE | 54 | YTHDF3 | chr8 | + | 63175330 | 63175416 | 63186146 | 63187745 | 63169386 | 63169411 | 6E+07 | 6E+07 | 0 | 0 | 3 | 0 | 0 | 5 | 0 | 0 | 0 | 0 | 6 | 0 | 98 | 98 | NA | NA | NA | NA | 0.375 | NA | 0 | NA | 0.375 |
| MXE | 57 | GSR | chr8 | - | 30689160 | 30689319 | 30692968 | 30693055 | 30684087 | 30684199 | 3E+07 | 3E+07 | 0 | 0 | 2 | 0 | 0 | 3 | 0 | 2 | 0 | 0 | 0 | 0 | 98 | 98 | NA | NA | NA | NA | 0.4 | NA | 1 | NA | -0.6 |
| MXE | 70 | LINC-PINT | chr7 | - | 130959868 | 130959970 | 130967388 | 130967517 | 130944159 | 130945835 | 1E+08 | 1E+08 | 0 | 0 | 5 | 0 | 0 | 1 | 0 | 1 | 0 | 0 | 1 | 0 | 98 | 98 | NA | NA | NA | NA | 0.833 | NA | 0.5 | NA | 0.333 |
| MXE | 80 | IKZF1 | chr7 | + | 50382539 | 50382707 | 50387344 | 50387470 | 50327637 | 50327757 | 5E+07 | 5E+07 | 0 | 0 | 3 | 0 | 0 | 3 | 0 | 4 | 0 | 0 | 0 | 0 | 98 | 98 | NA | NA | NA | NA | 0.5 | NA | 1 | NA | -0.5 |
| MXE | 87 | SUN1 | chr7 | + | 832504 | 832601 | 838797 | 838986 | 816614 | 816673 | 841945 | 842130 | 0 | 0 | 0 | 0 | 0 | 5 | 0 | 1 | 0 | 0 | 0 | 0 | 98 | 98 | NA | NA | NA | NA | 0 | NA | 1 | NA | -1 |
| MXE | 101 | KCTD20 | chr6 | + | 36470051 | 36470257 | 36479590 | 36479711 | 36442766 | 36443111 | 4E+07 | 4E+07 | 0 | 0 | 1 | 0 | 0 | 1 | 1 | 0 | 0 | 0 | 2 | 0 | 98 | 98 | NA | NA | NA | NA | 0.5 | 1 | 0 | NA | 0 |
| MXE | 102 | KCTD20 | chr6 | + | 36479120 | 36479223 | 36479590 | 36479711 | 36470051 | 36470257 | 4E+07 | 4E+07 | 0 | 0 | 4 | 0 | 0 | 1 | 0 | 0 | 0 | 0 | 3 | 0 | 98 | 98 | NA | NA | NA | NA | 0.8 | NA | 0 | NA | 0.8 |
| MXE | 104 | MAPK14 | chr6 | + | 36095986 | 36096066 | 36100187 | 36100267 | 36076536 | 36076608 | 4E+07 | 4E+07 | 0 | 0 | 8 | 0 | 0 | 1 | 0 | 6 | 0 | 0 | 0 | 0 | 98 | 98 | NA | NA | NA | NA | 0.889 | NA | 1 | NA | -0.111 |
| MXE | 134 | CNOT8 | chr5 | + | 154865191 | 154865385 | 154870660 | 154870822 | 154863206 | 154863395 | 2E+08 | 2E+08 | 0 | 0 | 0 | 0 | 0 | 3 | 0 | 1 | 0 | 0 | 1 | 0 | 98 | 98 | NA | NA | NA | NA | 0 | NA | 0.5 | NA | -0.5 |
| MXE | 160 | MRPS18C | chr4 | + | 83458345 | 83458429 | 83459739 | 83459797 | 83456908 | 83456958 | 8E+07 | 8E+07 | 0 | 0 | 3 | 0 | 0 | 5 | 0 | 0 | 0 | 0 | 1 | 0 | 98 | 98 | NA | NA | NA | NA | 0.375 | NA | 0 | NA | 0.375 |
| MXE | 168 | CPEB2 | chr4 | + | 15007304 | 15007586 | 15017187 | 15017278 | 15002673 | 15004335 | 2E+07 | 2E+07 | 0 | 0 | 2 | 0 | 0 | 4 | 0 | 2 | 0 | 0 | 4 | 0 | 98 | 98 | NA | NA | NA | NA | 0.333 | NA | 0.333 | NA | 0 |
| MXE | 172 | DLG1 | chr3 | - | 197075836 | 197075870 | 197076585 | 197076685 | 197069218 | 197069260 | 2E+08 | 2E+08 | 0 | 0 | 4 | 0 | 0 | 0 | 0 | 0 | 0 | 0 | 1 | 0 | 98 | 83 | NA | NA | NA | NA | 1 | NA | 0 | NA | 1 |
| MXE | 175 | OPA1 | chr3 | + | 193617177 | 193617285 | 193617783 | 193617837 | 193615673 | 193615770 | 2E+08 | 2E+08 | 0 | 0 | 2 | 0 | 0 | 0 | 0 | 0 | 0 | 0 | 1 | 0 | 98 | 98 | NA | NA | NA | NA | 1 | NA | 0 | NA | 1 |
| MXE | 201 | MBNL1 | chr3 | + | 152455541 | 152455577 | 152456266 | 152456361 | 152447619 | 152447773 | 2E+08 | 2E+08 | 0 | 0 | 10 | 0 | 0 | 5 | 0 | 0 | 0 | 2 | 7 | 0 | 85 | 98 | NA | NA | NA | NA | 0.698 | 0 | 0 | NA | 0.698 |
| MXE | 231 | TANGO2 | chr22 | + | 20053436 | 20053551 | 20055942 | 20056013 | 20052464 | 20052584 | 2E+07 | 2E+07 | 0 | 0 | 0 | 0 | 0 | 1 | 5 | 0 | 0 | 0 | 0 | 0 | 98 | 98 | NA | NA | NA | NA | 0 | 1 | NA | NA | -1 |
| MXE | 232 | TANGO2 | chr22 | + | 20053436 | 20053551 | 20061529 | 20061683 | 20052464 | 20052584 | 2E+07 | 2E+07 | 0 | 0 | 0 | 0 | 0 | 2 | 5 | 0 | 0 | 0 | 0 | 0 | 98 | 98 | NA | NA | NA | NA | 0 | 1 | NA | NA | -1 |
| MXE | 234 | TANGO2 | chr22 | + | 20055942 | 20056013 | 20063337 | 20063442 | 20053436 | 20053551 | 2E+07 | 2E+07 | 0 | 0 | 4 | 0 | 0 | 3 | 0 | 0 | 0 | 0 | 1 | 0 | 98 | 98 | NA | NA | NA | NA | 0.571 | NA | 0 | NA | 0.571 |
| MXE | 235 | PRMT2 | chr21 | + | 46658744 | 46658920 | 46663382 | 46663554 | 46649574 | 46649739 | 5E+07 | 5E+07 | 0 | 0 | 2 | 0 | 0 | 1 | 0 | 2 | 0 | 0 | 0 | 0 | 98 | 98 | NA | NA | NA | NA | 0.667 | NA | 1 | NA | -0.333 |

| | | | | | | | | | | | | | | | | | | | | | | | | | | | | | | | | | | | |
|---|---|---|---|---|---|---|---|---|---|---|---|---|---|---|---|---|---|---|---|---|---|---|---|---|---|---|---|---|---|---|---|---|---|---|---|
| MXE | 252 | SLC9A8 | chr20 | + | 49839540 | 49839599 | 49845035 | 49845119 | 49823060 | 49823141 | 5E+07 | 5E+07 | 0 | 0 | 6 | 0 | 0 | 7 | 0 | 0 | 0 | 0 | 2 | 0 | 98 | 98 | NA | NA | NA | NA | 0.462 | NA | 0 | NA | 0.462 |
| MXE | 254 | ZMYND8 | chr20 | - | 47262287 | 47262428 | 47276313 | 47276795 | 47249286 | 47249439 | 5E+07 | 5E+07 | 0 | 0 | 1 | 0 | 0 | 1 | 0 | 0 | 0 | 0 | 1 | 0 | 98 | 98 | NA | NA | NA | NA | 0.5 | NA | 0 | NA | 0.5 |
| MXE | 272 | XRN2 | chr20 | + | 21356585 | 21356665 | 21357735 | 21357792 | 21356079 | 21356177 | 2E+07 | 2E+07 | 0 | 0 | 8 | 0 | 0 | 6 | 0 | 2 | 0 | 0 | 2 | 0 | 98 | 98 | NA | NA | NA | NA | 0.571 | NA | 0.5 | NA | 0.071 |
| MXE | 273 | RALGAPA2 | chr20 | - | 20605174 | 20605412 | 20611314 | 20611426 | 20601681 | 20601846 | 2E+07 | 2E+07 | 0 | 0 | 5 | 0 | 0 | 5 | 0 | 3 | 0 | 0 | 0 | 0 | 98 | 98 | NA | NA | NA | NA | 0.5 | NA | 1 | NA | -0.5 |
| MXE | 291 | CASP10 | chr2 | + | 201203729 | 201203766 | 201208074 | 201208183 | 201195841 | 201195948 | 2E+08 | 2E+08 | 0 | 0 | 1 | 0 | 0 | 1 | 0 | 0 | 0 | 0 | 1 | 0 | 86 | 98 | NA | NA | NA | NA | 0.533 | NA | 0 | NA | 0.533 |
| MXE | 292 | CALCOCO2 | chr17 | + | 48841697 | 48841887 | 48848063 | 48848166 | 48830987 | 48831078 | 5E+07 | 5E+07 | 0 | 0 | 0 | 0 | 0 | 3 | 0 | 1 | 0 | 0 | 4 | 0 | 98 | 98 | NA | NA | NA | NA | 0 | NA | 0.2 | NA | -0.2 |
| MXE | 328 | SLC25A3 | chr12 | + | 98595432 | 98595557 | 98595726 | 98595848 | 98593974 | 98594135 | 1E+08 | 1E+08 | 0 | 0 | 0 | 0 | 0 | 15 | 0 | 2 | 0 | 0 | 5 | 0 | 98 | 98 | NA | NA | NA | NA | 0 | NA | 0.286 | NA | -0.286 |
| MXE | 340 | HNRNPR | chr1 | - | 23323555 | 23323732 | 23333517 | 23333631 | 23321527 | 23321663 | 2E+07 | 2E+07 | 0 | 0 | 0 | 0 | 0 | 1 | 5 | 0 | 0 | 0 | 1 | 0 | 98 | 98 | NA | NA | NA | NA | 0 | 1 | 0 | NA | -0.5 |
| MXE | 343 | BIRC6 | chr2 | + | 32464508 | 32464823 | 32465064 | 32465164 | 32463193 | 32463381 | 3E+07 | 3E+07 | 0 | 0 | 0 | 0 | 0 | 2 | 0 | 5 | 0 | 0 | 3 | 0 | 98 | 98 | NA | NA | NA | NA | 0 | NA | 0.625 | NA | -0.625 |
| MXE | 357 | PTPN6 | chr12 | + | 6954804 | 6954994 | 6955150 | 6955267 | 6951982 | 6952177 | 7E+06 | 7E+06 | 0 | 0 | 15 | 0 | 0 | 11 | 0 | 14 | 0 | 0 | 17 | 0 | 98 | 98 | NA | NA | NA | NA | 0.577 | NA | 0.452 | NA | 0.125 |
| MXE | 362 | BLOC1S6 | chr15 | + | 45603099 | 45603187 | 45605427 | 45605514 | 45592134 | 45592276 | 5E+07 | 5E+07 | 0 | 0 | 0 | 0 | 0 | 4 | 0 | 1 | 0 | 0 | 1 | 0 | 98 | 98 | NA | NA | NA | NA | 0 | NA | 0.5 | NA | -0.5 |
| MXE | 376 | TXNRD1 | chr12 | + | 104288930 | 104289040 | 104311289 | 104311412 | 104286681 | 104287109 | 1E+08 | 1E+08 | 0 | 0 | 0 | 0 | 0 | 2 | 0 | 1 | 0 | 0 | 14 | 0 | 98 | 98 | NA | NA | NA | NA | 0 | NA | 0.067 | NA | -0.067 |
| MXE | 408 | DNM2 | chr19 | + | 10796060 | 10796199 | 10797379 | 10797518 | 10795371 | 10795439 | 1E+07 | 1E+07 | 0 | 0 | 2 | 0 | 0 | 4 | 0 | 4 | 0 | 1 | 3 | 0 | 98 | 98 | NA | NA | NA | NA | 0.333 | 0 | 0.571 | NA | 0.048 |
| MXE | 413 | IL32 | chr16 | + | 3067553 | 3067613 | 3067983 | 3068010 | 3067376 | 3067415 | 3E+06 | 3E+06 | 0 | 0 | 3 | 0 | 0 | 1 | 0 | 6 | 0 | 1 | 3 | 0 | 98 | 76 | NA | NA | NA | NA | 0.699 | 0 | 0.608 | NA | 0.395 |
| MXE | 421 | PRR5L | chr11 | + | 36446299 | 36446440 | 36451208 | 36451335 | 36437384 | 36437476 | 4E+07 | 4E+07 | 0 | 0 | 1 | 0 | 0 | 0 | 0 | 0 | 0 | 0 | 1 | 0 | 98 | 98 | NA | NA | NA | NA | 1 | NA | 0 | NA | 1 |
| MXE | 426 | ENSA | chr1 | - | 150625641 | 150625808 | 150626478 | 150626526 | 150622122 | 150622859 | 2E+08 | 2E+08 | 0 | 0 | 0 | 0 | 0 | 1 | 0 | 2 | 0 | 0 | 2 | 0 | 97 | 98 | NA | NA | NA | NA | 0 | NA | 0.503 | NA | -0.503 |
| MXE | 455 | ZNF512 | chr2 | + | 27599969 | 27600053 | 27600690 | 27600815 | 27599582 | 27599678 | 3E+07 | 3E+07 | 0 | 0 | 1 | 0 | 0 | 0 | 0 | 0 | 0 | 0 | 2 | 0 | 98 | 98 | NA | NA | NA | NA | 1 | NA | 0 | NA | 1 |
| MXE | 486 | NIPA2 | chr15 | + | 22839654 | 22839790 | 22845145 | 22845267 | 22838640 | 22838921 | 2E+07 | 2E+07 | 0 | 0 | 1 | 0 | 0 | 0 | 0 | 0 | 0 | 1 | 1 | 0 | 98 | 98 | NA | NA | NA | NA | 1 | 0 | 0 | NA | 1 |
| MXE | 496 | CLN3 | chr16 | - | 28487661 | 28487741 | 28488590 | 28488662 | 28487455 | 28487541 | 3E+07 | 3E+07 | 0 | 0 | 4 | 0 | 0 | 1 | 0 | 1 | 0 | 0 | 0 | 0 | 98 | 98 | NA | NA | NA | NA | 0.8 | NA | 1 | NA | -0.2 |
| MXE | 504 | YIPF1 | chr1 | - | 53855000 | 53855051 | 53860055 | 53860153 | 53851718 | 53852270 | 5E+07 | 5E+07 | 0 | 0 | 1 | 0 | 0 | 2 | 0 | 0 | 0 | 0 | 2 | 0 | 98 | 98 | NA | NA | NA | NA | 0.333 | NA | 0 | NA | 0.333 |
| MXE | 520 | ECD | chr10 | - | 73152292 | 73152421 | 73154255 | 73154448 | 73148275 | 73148404 | 7E+07 | 7E+07 | 0 | 0 | 1 | 0 | 0 | 1 | 0 | 4 | 0 | 0 | 0 | 0 | 98 | 98 | NA | NA | NA | NA | 0.5 | NA | 1 | NA | -0.5 |
| MXE | 527 | AGO4 | chr1 | + | 35832436 | 35832570 | 35833989 | 35834174 | 35832056 | 35832185 | 4E+07 | 4E+07 | 0 | 0 | 15 | 0 | 0 | 2 | 0 | 3 | 0 | 0 | 2 | 0 | 98 | 98 | NA | NA | NA | NA | 0.882 | NA | 0.6 | NA | 0.282 |
| MXE | 534 | FRMD8 | chr11 | + | 65389360 | 65389528 | 65393572 | 65393674 | 65387036 | 65387121 | 7E+07 | 7E+07 | 0 | 0 | 0 | 0 | 0 | 1 | 0 | 3 | 0 | 0 | 0 | 0 | 98 | 98 | NA | NA | NA | NA | 0 | NA | 1 | NA | -1 |
| MXE | 560 | ABI1 | chr10 | - | 26755654 | 26755741 | 26759061 | 26759235 | 26751597 | 26751783 | 3E+07 | 3E+07 | 0 | 0 | 12 | 0 | 0 | 1 | 0 | 1 | 0 | 0 | 0 | 0 | 98 | 98 | NA | NA | NA | NA | 0.923 | NA | 1 | NA | -0.077 |
| MXE | 561 | ABI1 | chr10 | - | 26755654 | 26755741 | 26759061 | 26759238 | 26751597 | 26751783 | 3E+07 | 3E+07 | 0 | 0 | 11 | 0 | 0 | 1 | 0 | 2 | 0 | 0 | 0 | 0 | 98 | 98 | NA | NA | NA | NA | 0.917 | NA | 1 | NA | -0.083 |
| MXE | 562 | CPEB4 | chr5 | + | 173932449 | 173932500 | 173943025 | 173943049 | 173910522 | 173910604 | 2E+08 | 2E+08 | 0 | 0 | 5 | 0 | 0 | 0 | 0 | 2 | 0 | 0 | 1 | 0 | 98 | 73 | NA | NA | NA | NA | 1 | NA | 0.598 | NA | 0.402 |
| MXE | 564 | SOAT1 | chr1 | + | 179302676 | 179302802 | 179323436 | 179323495 | 179293713 | 179293936 | 2E+08 | 2E+08 | 0 | 0 | 3 | 0 | 0 | 0 | 0 | 0 | 0 | 0 | 2 | 0 | 98 | 98 | NA | NA | NA | NA | 1 | NA | 0 | NA | 1 |
| MXE | 583 | TK2 | chr16 | - | 66536963 | 66537017 | 66541878 | 66541953 | 66531379 | 66531469 | 7E+07 | 7E+07 | 0 | 0 | 2 | 0 | 0 | 2 | 0 | 1 | 0 | 0 | 2 | 0 | 98 | 98 | NA | NA | NA | NA | 0.5 | NA | 0.333 | NA | 0.167 |
| MXE | 594 | H2AFV | chr7 | - | 44840898 | 44841012 | 44843276 | 44843354 | 44835528 | 44835658 | 4E+07 | 4E+07 | 0 | 0 | 0 | 0 | 0 | 3 | 0 | 1 | 0 | 0 | 0 | 0 | 98 | 98 | NA | NA | NA | NA | 0 | NA | 1 | NA | -1 |
| MXE | 595 | PGS1 | chr17 | + | 78392475 | 78392665 | 78396307 | 78396385 | 78378617 | 78378808 | 8E+07 | 8E+07 | 0 | 0 | 4 | 0 | 0 | 4 | 0 | 3 | 0 | 0 | 0 | 0 | 98 | 98 | NA | NA | NA | NA | 0.5 | NA | 1 | NA | -0.5 |
| MXE | 596 | PGS1 | chr17 | + | 78392513 | 78392665 | 78396307 | 78396385 | 78378617 | 78378808 | 8E+07 | 8E+07 | 0 | 0 | 1 | 0 | 0 | 4 | 0 | 1 | 0 | 0 | 0 | 0 | 98 | 98 | NA | NA | NA | NA | 0.2 | NA | 1 | NA | -0.8 |
| MXE | 610 | PTGES3 | chr12 | - | 56666203 | 56666266 | 56670274 | 56670364 | 56664775 | 56664800 | 6E+07 | 6E+07 | 0 | 0 | 0 | 0 | 0 | 5 | 0 | 2 | 0 | 0 | 1 | 0 | 98 | 98 | NA | NA | NA | NA | 0 | NA | 0.667 | NA | -0.667 |
| MXE | 615 | CD300LF | chr17 | - | 74696194 | 74696217 | 74698368 | 74698481 | 74695724 | 74695859 | 7E+07 | 7E+07 | 0 | 0 | 0 | 0 | 0 | 1 | 0 | 3 | 0 | 0 | 0 | 0 | 98 | 72 | NA | NA | NA | NA | 0 | NA | 1 | NA | -1 |
| MXE | 636 | DHX40 | chr17 | + | 59566626 | 59566709 | 59570517 | 59570663 | 59565524 | 59565783 | 6E+07 | 6E+07 | 0 | 0 | 1 | 0 | 0 | 1 | 0 | 0 | 0 | 0 | 1 | 0 | 98 | 98 | NA | NA | NA | NA | 0.5 | NA | 0 | NA | 0.5 |
| MXE | 653 | FOXJ3 | chr1 | - | 42199101 | 42199230 | 42205761 | 42205863 | 42194889 | 42195064 | 4E+07 | 4E+07 | 0 | 0 | 1 | 0 | 0 | 0 | 0 | 1 | 0 | 1 | 4 | 0 | 98 | 98 | NA | NA | NA | NA | 1 | 0 | 0.2 | NA | 0.9 |
| MXE | 721 | BEST1 | chr11 | + | 61958145 | 61958298 | 61959497 | 61959578 | 61957386 | 61957464 | 6E+07 | 6E+07 | 0 | 0 | 2 | 0 | 0 | 0 | 0 | 0 | 0 | 0 | 2 | 0 | 98 | 98 | NA | NA | NA | NA | 1 | NA | 0 | NA | 1 |
| MXE | 736 | FBXO28 | chr1 | + | 224134073 | 224134212 | 224153141 | 224153337 | 224130471 | 224130581 | 2E+08 | 2E+08 | 0 | 0 | 3 | 0 | 0 | 0 | 0 | 3 | 0 | 0 | 4 | 0 | 98 | 98 | NA | NA | NA | NA | 1 | NA | 0.429 | NA | 0.571 |
| MXE | 738 | SMARCC1 | chr3 | - | 47720665 | 47720735 | 47729024 | 47729094 | 47714414 | 47714490 | 5E+07 | 5E+07 | 0 | 0 | 2 | 0 | 0 | 2 | 0 | 2 | 0 | 0 | 1 | 0 | 98 | 98 | NA | NA | NA | NA | 0.5 | NA | 0.667 | NA | -0.167 |
| MXE | 751 | FAM65B | chr6 | - | 24835702 | 24835871 | 24839090 | 24839272 | 24832255 | 24832391 | 2E+07 | 2E+07 | 0 | 0 | 4 | 0 | 0 | 21 | 0 | 5 | 0 | 0 | 4 | 0 | 98 | 98 | NA | NA | NA | NA | 0.16 | NA | 0.556 | NA | -0.396 |
| MXE | 760 | ADGRE3 | chr19 | - | 14658512 | 14658550 | 14661962 | 14662118 | 14654981 | 14655165 | 1E+07 | 1E+07 | 0 | 0 | 5 | 0 | 0 | 5 | 5 | 1 | 0 | 0 | 3 | 0 | 98 | 87 | NA | NA | NA | NA | 0.47 | 1 | 0.228 | NA | -0.144 |
| MXE | 791 | SAP30BP | chr17 | + | 75668515 | 75668625 | 75671815 | 75671863 | 75667115 | 75667478 | 8E+07 | 8E+07 | 0 | 0 | 1 | 0 | 0 | 1 | 0 | 5 | 0 | 0 | 7 | 0 | 98 | 97 | NA | NA | NA | NA | 0.497 | NA | 0.414 | NA | 0.083 |
| MXE | 821 | CLTA | chr9 | + | 36197550 | 36197588 | 36198978 | 36199096 | 36190855 | 36191273 | 4E+07 | 4E+07 | 0 | 0 | 0 | 0 | 0 | 4 | 0 | 2 | 0 | 0 | 2 | 0 | 87 | 98 | NA | NA | NA | NA | 0 | NA | 0.53 | NA | -0.53 |
| MXE | 823 | ATE1 | chr10 | - | 121898840 | 121898969 | 121899865 | 121899994 | 121870005 | 121870038 | 1E+08 | 1E+08 | 0 | 0 | 1 | 0 | 0 | 0 | 0 | 0 | 0 | 0 | 1 | 0 | 98 | 98 | NA | NA | NA | NA | 1 | NA | 0 | NA | 1 |
| MXE | 826 | CASP8 | chr2 | + | 201272897 | 201272942 | 201274888 | 201274953 | 201272637 | 201272776 | 2E+08 | 2E+08 | 0 | 0 | 6 | 0 | 0 | 3 | 0 | 18 | 0 | 0 | 3 | 0 | 94 | 98 | NA | NA | NA | NA | 0.676 | NA | 0.862 | NA | -0.186 |
| MXE | 844 | ATXN3 | chr14 | - | 92093251 | 92093318 | 92096092 | 92096137 | 92088729 | 92088817 | 9E+07 | 9E+07 | 0 | 0 | 1 | 0 | 0 | 1 | 0 | 0 | 0 | 3 | 0 | 0 | 94 | 98 | NA | NA | NA | NA | 0.51 | 0 | NA | NA | 0.51 |
| MXE | 845 | ATXN3 | chr14 | - | 92093745 | 92093831 | 92096092 | 92096137 | 92093251 | 92093318 | 9E+07 | 9E+07 | 0 | 0 | 1 | 0 | 0 | 1 | 0 | 1 | 0 | 0 | 0 | 0 | 94 | 98 | NA | NA | NA | NA | 0.51 | NA | 1 | NA | -0.49 |
| MXE | 846 | ATXN3 | chr14 | - | 92093745 | 92093831 | 92096092 | 92096137 | 92093251 | 92093318 | 9E+07 | 9E+07 | 0 | 0 | 0 | 0 | 0 | 1 | 0 | 1 | 0 | 0 | 0 | 0 | 94 | 98 | NA | NA | NA | NA | 0 | NA | 1 | NA | -1 |
| MXE | 857 | MARK3 | chr14 | + | 103465556 | 103465793 | 103465971 | 103466091 | 103462404 | 103462461 | 1E+08 | 1E+08 | 0 | 0 | 5 | 0 | 0 | 1 | 0 | 1 | 0 | 0 | 2 | 0 | 98 | 98 | NA | NA | NA | NA | 0.833 | NA | 0.333 | NA | 0.5 |
| MXE | 908 | IQGAP1 | chr15 | + | 90466268 | 90466436 | 90467449 | 90467592 | 90466000 | 90466091 | 9E+07 | 9E+07 | 0 | 0 | 4 | 0 | 0 | 16 | 0 | 16 | 0 | 1 | 6 | 0 | 98 | 98 | NA | NA | NA | NA | 0.2 | 0 | 0.727 | NA | -0.163 |
| MXE | 910 | VPS33B | chr15 | - | 91016962 | 91017024 | 91017804 | 91017885 | 91014383 | 91014433 | 9E+07 | 9E+07 | 0 | 0 | 2 | 0 | 0 | 1 | 0 | 0 | 0 | 0 | 1 | 0 | 98 | 98 | NA | NA | NA | NA | 0.667 | NA | 0 | NA | 0.667 |
| MXE | 949 | DCLRE1C | chr10 | - | 14945104 | 14945189 | 14949035 | 14949087 | 14939809 | 14939869 | 1E+07 | 1E+07 | 0 | 0 | 2 | 0 | 0 | 0 | 0 | 0 | 0 | 0 | 1 | 0 | 98 | 98 | NA | NA | NA | NA | 1 | NA | 0 | NA | 1 |
| MXE | 950 | DCLRE1C | chr10 | - | 14945434 | 14945565 | 14947524 | 14947609 | 14945104 | 14945189 | 1E+07 | 1E+07 | 0 | 0 | 1 | 0 | 0 | 0 | 0 | 1 | 0 | 0 | 1 | 0 | 98 | 98 | NA | NA | NA | NA | 1 | NA | 0.5 | NA | 0.5 |
| MXE | 960 | ADGRE2 | chr19 | - | 14754953 | 14755127 | 14755653 | 14755877 | 14752328 | 14752526 | 1E+07 | 1E+07 | 0 | 0 | 2 | 0 | 0 | 3 | 0 | 2 | 0 | 0 | 5 | 0 | 98 | 98 | NA | NA | NA | NA | 0.4 | NA | 0.286 | NA | 0.114 |
| MXE | 962 | NLRC5 | chr16 | + | 57076818 | 57076902 | 57077295 | 57077379 | 57074599 | 57074683 | 6E+07 | 6E+07 | 0 | 0 | 7 | 0 | 0 | 1 | 0 | 7 | 0 | 0 | 1 | 0 | 98 | 98 | NA | NA | NA | NA | 0.875 | NA | 0.875 | NA | 0 |
| MXE | 965 | DYNC1LI2 | chr16 | - | 66742437 | 66742668 | 66749196 | 66749313 | 66736074 | 66736244 | 7E+07 | 7E+07 | 0 | 0 | 2 | 0 | 0 | 1 | 0 | 2 | 0 | 0 | 0 | 0 | 98 | 98 | NA | NA | NA | NA | 0.667 | NA | 1 | NA | -0.333 |
| MXE | 976 | NUPL1 | chr13 | + | 25309246 | 25309282 | 25312882 | 25313032 | 25307805 | 25307948 | 3E+07 | 3E+07 | 0 | 0 | 5 | 0 | 0 | 2 | 0 | 2 | 0 | 0 | 2 | 0 | 85 | 98 | NA | NA | NA | NA | 0.742 | NA | 0.536 | NA | 0.206 |
| MXE | 995 | RHOT1 | chr17 | + | 32209349 | 32209445 | 32211115 | 32211238 | 32208106 | 32208309 | 3E+07 | 3E+07 | 0 | 0 | 1 | 0 | 0 | 2 | 0 | 2 | 0 | 0 | 1 | 0 | 98 | 98 | NA | NA | NA | NA | 0.333 | NA | 0.667 | NA | -0.334 |
| MXE | 1049 | MBD1 | chr18 | - | 50274185 | 50274353 | 50274976 | 50275045 | 50273563 | 50273863 | 5E+07 | 5E+07 | 0 | 0 | 1 | 0 | 0 | 0 | 0 | 0 | 0 | 0 | 2 | 0 | 98 | 98 | NA | NA | NA | NA | 1 | NA | 0 | NA | 1 |
| MXE | 1060 | ZNF638 | chr2 | + | 71426459 | 71427414 | 71428546 | 71428651 | 71424649 | 71424715 | 7E+07 | 7E+07 | 0 | 0 | 1 | 0 | 0 | 0 | 0 | 3 | 0 | 0 | 1 | 0 | 98 | 98 | NA | NA | NA | NA | 1 | NA | 0.75 | NA | 0.25 |
| MXE | 1124 | BCL2L11 | chr2 | + | 111123732 | 111123869 | 111150043 | 111150147 | 111120913 | 111121188 | 1E+08 | 1E+08 | 0 | 0 | 1 | 0 | 0 | 1 | 0 | 0 | 0 | 0 | 1 | 0 | 98 | 98 | NA | NA | NA | NA | 0.5 | NA | 0 | NA | 0.5 |
| MXE | 1126 | BCL2L11 | chr2 | + | 111123732 | 111124139 | 111150043 | 111150147 | 111120913 | 111121188 | 1E+08 | 1E+08 | 0 | 0 | 1 | 0 | 0 | 1 | 0 | 0 | 0 | 0 | 1 | 0 | 98 | 98 | NA | NA | NA | NA | 0.5 | NA | 0 | NA | 0.5 |
| MXE | 1139 | SMPD4 | chr2 | - | 130173278 | 130173354 | 130173513 | 130173656 | 130172786 | 130172895 | 1E+08 | 1E+08 | 0 | 0 | 2 | 0 | 0 | 0 | 0 | 0 | 0 | 0 | 1 | 0 | 98 | 98 | NA | NA | NA | NA | 1 | NA | 0 | NA | 1 |
| MXE | 1155 | FAM126B | chr2 | - | 200992886 | 200993003 | 200997444 | 200997540 | 200992280 | 200992368 | 2E+08 | 2E+08 | 0 | 0 | 6 | 0 | 0 | 10 | 0 | 2 | 0 | 0 | 4 | 0 | 98 | 98 | NA | NA | NA | NA | 0.375 | NA | 0.333 | NA | 0.042 |

**U12_MXE_filtered**

| | | | | | | | | | | | | | Inclusion_counts | | | Skipping_counts | | | Inclusion_counts | | | Skipping_counts | | | | | | | Inclusion level(normalized) | | | | | | |
|---|---|---|---|---|---|---|---|---|---|---|---|---|---|---|---|---|---|---|---|---|---|---|---|---|---|---|---|---|---|---|---|---|---|---|---|
| **Splicing event** | **ID** | **Gene_ID** | **Chromosome** | **Strand** | **1stExonStart_0base** | **1stExonEnd** | **2ndExonSta** | **2ndExonEn** | **upstreamES** | **upstreamEE** | **downstr** | **downstr** | **ldg_12** | **ldg_14** | **ldg_16** | **ldg_12** | **ldg_14** | **ldg_16** | **pmn_13** | **pmn_15** | **pmn_17** | **pmn_13** | **pmn_15** | **pmn_17** | **IncFormLen** | **SkipFormLen** | **p value** | **FDR** | **ldg_12** | **ldg_14** | **ldg_16** | **pmn_13** | **pmn_15** | **pmn_17** | **IncLevelDifference** |
| MXE | 170 | MAEA | chr4 | + | 1315396 | 1315600 | 1322380 | 1322503 | 1311978 | 1312161 | 1E+06 | 1E+06 | 0 | 2 | 1 | 0 | 0 | 0 | 0 | 2 | 7 | 0 | 3 | 1 | 98 | 98 | 0.0098 | 0.19855 | NA | 1 | 1 | NA | 0.4 | 0.875 | 0.363 |
| MXE | 1080 | RASGRP4 | chr19 | - | 38417051 | 38417168 | 38419859 | 38420013 | 38413393 | 38413474 | 4E+07 | 4E+07 | 0 | 4 | 7 | 0 | 0 | 0 | 1 | 1 | 0 | 0 | 1 | 0 | 98 | 98 | 0.0304 | 0.19855 | NA | 1 | 1 | 1 | 0.5 | NA | 0.25 |
| MXE | 1081 | RASGRP4 | chr19 | - | 38417051 | 38417168 | 38419859 | 38420013 | 38414847 | 38415123 | 4E+07 | 4E+07 | 0 | 4 | 7 | 0 | 2 | 13 | 1 | 1 | 0 | 0 | 4 | 0 | 98 | 98 | 0.1166 | 0.33916 | NA | 0.667 | 0.35 | 1 | 0.2 | NA | -0.092 |
| MXE | 253 | SLC9A8 | chr20 | + | 49845035 | 49845119 | 49849578 | 49849680 | 49839540 | 49839599 | 5E+07 | 5E+07 | 0 | 0 | 4 | 0 | 0 | 4 | 0 | 2 | 2 | 0 | 4 | 1 | 98 | 98 | 0.5531 | 0.91543 | NA | NA | 0.5 | NA | 0.333 | 0.667 | 0 |
| MXE | 104 | MAPK14 | chr6 | + | 36095986 | 36096066 | 36100187 | 36100267 | 36076536 | 36076608 | 4E+07 | 4E+07 | 0 | 0 | 8 | 0 | 0 | 1 | 0 | 6 | 0 | 0 | 0 | 0 | 98 | 98 | NA | NA | NA | NA | 0.889 | NA | 1 | NA | -0.111 |
| MXE | 252 | SLC9A8 | chr20 | + | 49839540 | 49839599 | 49845035 | 49845119 | 49823060 | 49823141 | 5E+07 | 5E+07 | 0 | 0 | 6 | 0 | 0 | 7 | 0 | 0 | 0 | 0 | 2 | 0 | 98 | 98 | NA | NA | NA | NA | 0.462 | NA | 0 | NA | 0.462 |
| MXE | 273 | RALGAPA2 | chr20 | - | 20605174 | 20605412 | 20611314 | 20611426 | 20601681 | 20601846 | 2E+07 | 2E+07 | 0 | 0 | 5 | 0 | 0 | 5 | 0 | 3 | 0 | 0 | 0 | 0 | 98 | 98 | NA | NA | NA | NA | 0.5 | NA | 1 | NA | -0.5 |
| MXE | 965 | DYNC1LI2 | chr16 | - | 66742437 | 66742668 | 66749196 | 66749313 | 66736074 | 66736244 | 7E+07 | 7E+07 | 0 | 0 | 2 | 0 | 0 | 1 | 0 | 2 | 0 | 0 | 0 | 0 | 98 | 98 | NA | NA | NA | NA | 0.667 | NA | 1 | NA | -0.333 |
| MXE | 1139 | SMPD4 | chr2 | - | 130173278 | 130173354 | 130173513 | 130173656 | 130172786 | 130172895 | 1E+08 | 1E+08 | 0 | 0 | 2 | 0 | 0 | 0 | 0 | 0 | 0 | 0 | 1 | 0 | 98 | 98 | NA | NA | NA | NA | 1 | NA | 0 | NA | 1 |

**Supplementary_Data_3S**
**4_events_all**
***HD NDG versus SLE NDG***

| | | | | | | | | | | | | Inclusion_count | | | Skipping_count | | | Inclusion_count | | | Skipping_count | | | | | | | Inclusion_level (normalized) | | | | | | |
|---|---|---|---|---|---|---|---|---|---|---|---|---|---|---|---|---|---|---|---|---|---|---|---|---|---|---|---|---|---|---|---|---|---|---|
| *Splice_type* | *ID* | *Gene_ID* | *Gene_Symbol* | *Chromosome* | *Strand* | *LongExonStart_0base* | *LongExonEnd* | *ShortES* | *ShortEE* | *FlankingES* | *FlankingEE* | *control_3* | *control_4* | *control_10* | *control_3* | *control_4* | *control_10* | *pmn_13* | *pmn_15* | *pmn_17* | *pmn_13* | *pmn_15* | *pmn_17* | *IncFormLen* | *SkipFormLen* | *p value* | *FDR* | *control_3* | *control_4* | *control_10* | *pmn_13* | *pmn_15* | *pmn_17* | *IncLevelDifference* |
| SE | 13845 | C10orf54 | C10orf54 | chr10 | - | 71761597 | 71762026 | 71760867 | 71760924 | 71773357 | 71773580 | 0 | 4 | 72 | 0 | 0 | 0 | 1 | 16 | 24 | 0 | 1 | 3 | 98 | 49 | 0.001516497 | 0.170760496 | NA | 1 | 1 | 1 | 0.889 | 0.8 | 0.104 |
| SE | 1661 | FAM193B | FAM193B | chr5 | - | 177531281 | 177531521 | 177524184 | 177525205 | 177532442 | 177532641 | 0 | 3 | 7 | 0 | 0 | 0 | 0 | 1 | 2 | 0 | 1 | 2 | 98 | 49 | 0.001603576 | 0.170760496 | NA | 1 | 1 | NA | 0.333 | 0.333 | 0.667 |
| SE | 7639 | IRF7 | IRF7 | chr11 | - | 614173 | 614399 | 613784 | 613865 | 614475 | 614534 | 0 | 0 | 0 | 0 | 8 | 3 | 0 | 4 | 1 | 0 | 3 | 2 | 98 | 49 | 0.001756837 | 0.170760496 | NA | 0 | 0 | NA | 0.4 | 0.2 | -0.3 |
| SE | 6866 | ATG2A | ATG2A | chr11 | - | 64906664 | 64906815 | 64906333 | 64906533 | 64907254 | 64907439 | 0 | 2 | 5 | 0 | 0 | 0 | 0 | 6 | 5 | 0 | 2 | 1 | 98 | 49 | 0.001882581 | 0.170760496 | NA | 1 | 1 | NA | 0.6 | 0.714 | 0.343 |
| SE | 9706 | REC8 | REC8 | chr14 | + | 24175542 | 24175624 | 24173290 | 24173411 | 24176821 | 24176901 | 0 | 3 | 10 | 0 | 0 | 0 | 1 | 3 | 1 | 0 | 2 | 4 | 98 | 49 | 0.002515847 | 0.170760496 | NA | 1 | 1 | 1 | 0.429 | 0.111 | 0.487 |
| SE | 13860 | ACSL1 | ACSL1 | chr4 | - | 184764852 | 184764925 | 184763166 | 184763255 | 184765890 | 184765986 | 0 | 2 | 50 | 0 | 0 | 0 | 7 | 51 | 14 | 0 | 2 | 2 | 98 | 49 | 0.003078802 | 0.170760496 | NA | 1 | 1 | 1 | 0.927 | 0.778 | 0.098 |
| SE | 8555 | PIK3AP1 | PIK3AP1 | chr10 | - | 96652697 | 96652842 | 96651508 | 96651651 | 96656797 | 96656934 | 0 | 1 | 15 | 0 | 0 | 0 | 0 | 4 | 7 | 0 | 2 | 1 | 98 | 49 | 0.003471069 | 0.170760496 | NA | 1 | 1 | NA | 0.5 | 0.778 | 0.361 |
| SE | 11710 | SENP3-EIF4A1 | SENP3-EIF4A1 | chr17 | + | 7575118 | 7575258 | 7574545 | 7574678 | 7576523 | 7576692 | 0 | 2 | 33 | 0 | 0 | 0 | 3 | 11 | 2 | 0 | 1 | 1 | 98 | 49 | 0.004779846 | 0.170760496 | NA | 1 | 1 | 1 | 0.846 | 0.5 | 0.218 |
| SE | 12939 | SMCHD1 | SMCHD1 | chr18 | + | 2666156 | 2666232 | 2655886 | 2656261 | 2666869 | 2667031 | 0 | 20 | 98 | 0 | 0 | 0 | 4 | 61 | 8 | 0 | 1 | 1 | 98 | 49 | 0.00491362 | 0.170760496 | NA | 1 | 1 | 1 | 0.968 | 0.8 | 0.077 |
| A3SS | 1863 | SRPK1 | SRPK1 | chr6 | - | 35890894 | 35891041 | 35890894 | 35891013 | 35920467 | 35920528 | 0 | 0 | 0 | 0 | 1 | 2 | 0 | 1 | 1 | 0 | 4 | 1 | 77 | 49 | 0.005762586 | 0.197717869 | NA | 0 | 0 | NA | 0.137 | 0.389 | -0.263 |
| SE | 12824 | SEC14L1 | SEC14L1 | chr17 | + | 77196201 | 77196311 | 77194676 | 77194911 | 77200483 | 77200673 | 0 | 11 | 34 | 0 | 0 | 0 | 0 | 20 | 0 | 0 | 11 | 2 | 98 | 49 | 0.005971369 | 0.170760496 | NA | 1 | 1 | NA | 0.476 | 0 | 0.762 |
| A5SS | 72 | CSF2RA | CSF2RA | chrX | + | 1274754 | 1274842 | 1274754 | 1274818 | 1282677 | 1282779 | 0 | 1 | 3 | 0 | 0 | 0 | 0 | 2 | 0 | 0 | 2 | 1 | 73 | 49 | 0.006411688 | 0.170414363 | NA | 1 | 1 | NA | 0.402 | 0 | 0.799 |
| SE | 4425 | PREX1 | PREX1 | chr20 | - | 48659918 | 48660061 | 48658135 | 48658228 | 48666282 | 48666355 | 0 | 1 | 31 | 0 | 0 | 0 | 0 | 16 | 2 | 0 | 1 | 1 | 98 | 49 | 0.008019007 | 0.170760496 | NA | 1 | 1 | NA | 0.889 | 0.5 | 0.305 |
| SE | 11393 | ADGRG3 | ADGRG3 | chr16 | + | 57676199 | 57676338 | 57673320 | 57673468 | 57678169 | 57678316 | 0 | 2 | 14 | 0 | 0 | 0 | 0 | 25 | 2 | 0 | 3 | 3 | 98 | 49 | 0.008245186 | 0.170760496 | NA | 1 | 1 | NA | 0.806 | 0.25 | 0.472 |
| SE | 9930 | PTPRE | PTPRE | chr10 | + | 128060938 | 128061015 | 128056122 | 128056213 | 128061678 | 128061715 | 0 | 3 | 11 | 0 | 0 | 0 | 0 | 22 | 0 | 0 | 7 | 0 | 98 | 49 | 0.009650972 | 0.170760496 | NA | 1 | 1 | NA | 0.611 | NA | 0.389 |
| SE | 5497 | ADAM8 | ADAM8 | chr10 | - | 133270880 | 133271070 | 133270735 | 133270805 | 133271199 | 133271289 | 1 | 20 | 40 | 0 | 0 | 0 | 4 | 55 | 21 | 0 | 1 | 2 | 98 | 49 | 0.010036062 | 0.170760496 | 1 | 1 | 1 | 1 | 0.965 | 0.84 | 0.065 |
| SE | 11923 | RBM39 | RBM39 | chr20 | - | 35740524 | 35740597 | 35738967 | 35739017 | 35740823 | 35740887 | 0 | 2 | 8 | 0 | 0 | 1 | 0 | 8 | 0 | 0 | 8 | 0 | 98 | 49 | 0.010118399 | 0.170760496 | NA | 1 | 0.8 | NA | 0.333 | NA | 0.567 |
| SE | 14093 | NLRP6 | NLRP6 | chr11 | + | 280083 | 281839 | 279833 | 279872 | 282704 | 282797 | 0 | 1 | 2 | 0 | 0 | 0 | 0 | 1 | 0 | 0 | 1 | 2 | 98 | 49 | 0.010649553 | 0.170760496 | NA | 1 | 1 | NA | 0.333 | 0 | 0.834 |
| SE | 10848 | IRF1 | IRF1 | chr5 | - | 132486250 | 132486373 | 132485666 | 132485716 | 132486556 | 132486686 | 0 | 2 | 36 | 0 | 0 | 1 | 7 | 13 | 0 | 0 | 6 | 0 | 98 | 49 | 0.011205299 | 0.170760496 | NA | 1 | 0.947 | 1 | 0.52 | NA | 0.214 |
| SE | 2075 | HERC5 | HERC5 | chr4 | + | 88462134 | 88462356 | 88460094 | 88460171 | 88463531 | 88463623 | 0 | 4 | 25 | 0 | 0 | 2 | 0 | 11 | 0 | 0 | 5 | 0 | 98 | 49 | 0.011241877 | 0.170760496 | NA | 1 | 0.862 | NA | 0.524 | NA | 0.407 |
| SE | 8188 | EPSTI1 | EPSTI1 | chr13 | - | 42919299 | 42919332 | 42917540 | 42917624 | 42926335 | 42926429 | 0 | 0 | 4 | 0 | 4 | 14 | 0 | 6 | 0 | 0 | 10 | 0 | 82 | 49 | 0.011354222 | 0.170760496 | NA | 0 | 0.146 | NA | 0.264 | NA | -0.191 |
| SE | 4426 | PREX1 | PREX1 | chr20 | - | 48726289 | 48726391 | 48708259 | 48708421 | 48734545 | 48734650 | 1 | 23 | 82 | 0 | 0 | 0 | 8 | 65 | 15 | 0 | 2 | 3 | 98 | 49 | 0.01202775 | 0.170760496 | 1 | 1 | 1 | 1 | 0.942 | 0.714 | 0.115 |
| SE | 3738 | BRD2 | BRD2 | chr6 | + | 32975020 | 32975112 | 32974461 | 32974765 | 32975383 | 32975521 | 0 | 1 | 3 | 0 | 0 | 3 | 0 | 4 | 0 | 0 | 8 | 0 | 98 | 49 | 0.012603077 | 0.170760496 | NA | 1 | 0.333 | NA | 0.2 | NA | 0.466 |
| SE | 3103 | MX2 | MX2 | chr21 | + | 41380016 | 41380151 | 41377788 | 41377981 | 41382409 | 41382564 | 0 | 4 | 25 | 0 | 0 | 2 | 0 | 16 | 5 | 0 | 11 | 2 | 98 | 49 | 0.013190421 | 0.170760496 | NA | 1 | 0.862 | NA | 0.421 | 0.556 | 0.443 |
| SE | 10602 | DMXL2 | DMXL2 | chr15 | - | 51495023 | 51495134 | 51491577 | 51491747 | 51498551 | 51500231 | 0 | 1 | 12 | 0 | 0 | 0 | 0 | 7 | 0 | 0 | 5 | 0 | 98 | 49 | 0.014099979 | 0.170760496 | NA | 1 | 1 | NA | 0.412 | NA | 0.588 |
| SE | 3545 | OSBPL2 | OSBPL2 | chr20 | + | 62259980 | 62260125 | 62256056 | 62256221 | 62263615 | 62263691 | 0 | 2 | 3 | 0 | 0 | 0 | 0 | 5 | 0 | 0 | 4 | 0 | 98 | 49 | 0.014587636 | 0.170760496 | NA | 1 | 1 | NA | 0.385 | NA | 0.615 |
| SE | 12803 | CYP4F3 | CYP4F3 | chr19 | + | 15645718 | 15645863 | 15641414 | 15641613 | 15647051 | 15647105 | 0 | 5 | 21 | 0 | 0 | 0 | 7 | 13 | 0 | 0 | 3 | 0 | 98 | 49 | 0.014732939 | 0.170760496 | NA | 1 | 1 | 1 | 0.684 | NA | 0.158 |
| SE | 1826 | MCTP1 | MCTP1 | chr5 | - | 94917895 | 94917973 | 94912805 | 94912976 | 94923961 | 94924021 | 0 | 2 | 3 | 0 | 0 | 0 | 0 | 4 | 0 | 0 | 4 | 0 | 98 | 49 | 0.014963682 | 0.170760496 | NA | 1 | 1 | NA | 0.333 | NA | 0.667 |
| SE | 6336 | FCAR | FCAR | chr19 | + | 54888006 | 54888294 | 54885234 | 54885525 | 54889648 | 54891420 | 0 | 5 | 28 | 0 | 0 | 1 | 0 | 19 | 0 | 0 | 3 | 0 | 98 | 49 | 0.01511042 | 0.170760496 | NA | 1 | 0.933 | NA | 0.76 | NA | 0.207 |
| SE | 2203 | CPEB2 | CPEB2 | chr4 | + | 15040463 | 15040487 | 15033160 | 15033211 | 15052413 | 15052584 | 0 | 0 | 1 | 0 | 5 | 1 | 0 | 5 | 0 | 0 | 2 | 0 | 73 | 49 | 0.015421435 | 0.170760496 | NA | 0 | 0.402 | NA | 0.627 | NA | -0.426 |
| SE | 12815 | BST2 | BST2 | chr19 | - | 17404370 | 17404437 | 17404128 | 17404189 | 17405290 | 17405648 | 0 | 4 | 23 | 0 | 0 | 0 | 0 | 4 | 0 | 0 | 3 | 0 | 98 | 49 | 0.015554946 | 0.170760496 | NA | 1 | 1 | NA | 0.4 | NA | 0.6 |
| SE | 11842 | RHOT1 | RHOT1 | chr17 | + | 32172936 | 32173060 | 32171042 | 32171101 | 32173830 | 32173912 | 0 | 5 | 3 | 0 | 0 | 0 | 0 | 3 | 0 | 0 | 3 | 0 | 98 | 49 | 0.015643271 | 0.170760496 | NA | 1 | 1 | NA | 0.333 | NA | 0.667 |
| SE | 6335 | FCAR | FCAR | chr19 | + | 54888006 | 54888228 | 54885234 | 54885525 | 54889648 | 54891420 | 0 | 3 | 17 | 0 | 0 | 1 | 0 | 12 | 0 | 0 | 3 | 0 | 98 | 49 | 0.015872783 | 0.170760496 | NA | 1 | 0.895 | NA | 0.667 | NA | 0.28 |
| SE | 5621 | ANKRD12 | ANKRD12 | chr18 | + | 9204475 | 9204544 | 9195550 | 9195698 | 9208656 | 9208803 | 0 | 1 | 3 | 0 | 0 | 2 | 0 | 5 | 0 | 0 | 4 | 0 | 98 | 49 | 0.016160337 | 0.170760496 | NA | 1 | 0.429 | NA | 0.385 | NA | 0.33 |
| SE | 10536 | SGK1 | SGK1 | chr6 | - | 134170825 | 134170915 | 134169245 | 134170435 | 134171022 | 134171178 | 0 | 1 | 11 | 0 | 0 | 0 | 1 | 16 | 0 | 0 | 4 | 0 | 98 | 49 | 0.016182238 | 0.170760496 | NA | 1 | 1 | 1 | 0.667 | NA | 0.166 |
| SE | 6177 | ABCA1 | ABCA1 | chr9 | - | 104825682 | 104825887 | 104824464 | 104824578 | 104826947 | 104827169 | 0 | 4 | 6 | 0 | 0 | 0 | 0 | 21 | 1 | 0 | 5 | 0 | 98 | 49 | 0.016495032 | 0.170760496 | NA | 1 | 1 | NA | 0.677 | 1 | 0.161 |
| SE | 799 | TBXAS1 | TBXAS1 | chr7 | + | 139911224 | 139911321 | 139875584 | 139875637 | 139936190 | 139936307 | 0 | 2 | 22 | 0 | 0 | 0 | 0 | 7 | 0 | 0 | 3 | 0 | 98 | 49 | 0.016691194 | 0.170760496 | NA | 1 | 1 | NA | 0.538 | NA | 0.462 |
| A5SS | 1875 | N4BP2L2 | N4BP2L2 | chr13 | - | 32527407 | 32527532 | 32527457 | 32527532 | 32522181 | 32522270 | 0 | 2 | 17 | 0 | 0 | 2 | 0 | 9 | 0 | 0 | 3 | 0 | 98 | 49 | 0.016742666 | 0.170414363 | NA | 1 | 0.81 | NA | 0.6 | NA | 0.305 |
| SE | 7593 | ATG16L2 | ATG16L2 | chr11 | + | 72822847 | 72822961 | 72822477 | 72822543 | 72824059 | 72824122 | 0 | 9 | 41 | 0 | 0 | 1 | 4 | 19 | 16 | 0 | 4 | 2 | 98 | 49 | 0.017045614 | 0.170760496 | NA | 1 | 0.953 | 1 | 0.704 | 0.8 | 0.142 |
| A3SS | 2805 | TPT1 | TPT1 | chr13 | - | 45333470 | 45337553 | 45333470 | 45337388 | 45338659 | 45338776 | 0 | 0 | 2 | 1 | 3 | 72 | 0 | 4 | 0 | 0 | 47 | 0 | 98 | 49 | 0.017197054 | 0.197717869 | 0 | 0 | 0.014 | NA | 0.041 | NA | -0.036 |
| SE | 883 | SH2B2 | SH2B2 | chr7 | + | 102300521 | 102301279 | 102285061 | 102285258 | 102306720 | 102306822 | 0 | 4 | 0 | 0 | 0 | 0 | 0 | 2 | 1 | 0 | 3 | 0 | 98 | 49 | 0.017202771 | 0.170760496 | NA | 1 | NA | NA | 0.25 | 1 | 0.375 |
| SE | 1452 | PRRC2A | PRRC2A | chr6 | + | 31625791 | 31625871 | 31625459 | 31625611 | 31626019 | 31626162 | 0 | 4 | 4 | 0 | 0 | 1 | 0 | 1 | 0 | 0 | 5 | 0 | 98 | 49 | 0.017880599 | 0.170760496 | NA | 1 | 0.667 | NA | 0.091 | NA | 0.743 |
| SE | 6626 | PCBP2 | PCBP2 | chr12 | + | 53464759 | 53464852 | 53462480 | 53462567 | 53465931 | 53465973 | 0 | 0 | 23 | 0 | 2 | 16 | 0 | 11 | 0 | 0 | 1 | 0 | 98 | 49 | 0.017987559 | 0.170760496 | NA | 0 | 0.418 | NA | 0.846 | NA | -0.637 |
| SE | 6174 | ABCA1 | ABCA1 | chr9 | - | 104810799 | 104810924 | 104809465 | 104809564 | 104812573 | 104812722 | 0 | 0 | 3 | 0 | 1 | 1 | 0 | 4 | 0 | 2 | 4 | 0 | 98 | 49 | 0.018518561 | 0.170760496 | NA | 0 | 0.6 | 0 | 0.333 | NA | 0.133 |
| SE | 11731 | TAOK3 | TAOK3 | chr12 | - | 118212913 | 118212995 | 118201295 | 118201463 | 118214016 | 118214110 | 1 | 2 | 4 | 0 | 0 | 1 | 0 | 2 | 0 | 0 | 3 | 0 | 98 | 49 | 0.018766874 | 0.170760496 | 1 | 1 | 0.667 | NA | 0.25 | NA | 0.639 |
| SE | 3208 | STX16-NPEPL1 | STX16-NPEPL1 | chr20 | + | 58671153 | 58671297 | 58670511 | 58670603 | 58673630 | 58673711 | 0 | 1 | 16 | 0 | 0 | 1 | 0 | 5 | 0 | 0 | 3 | 0 | 98 | 49 | 0.018766876 | 0.170760496 | NA | 1 | 0.889 | NA | 0.455 | NA | 0.489 |
| SE | 1982 | DDX60L | DDX60L | chr4 | - | 168419301 | 168419397 | 168416681 | 168416797 | 168420260 | 168420380 | 0 | 1 | 32 | 0 | 0 | 1 | 0 | 8 | 0 | 0 | 3 | 0 | 98 | 49 | 0.018767562 | 0.170760496 | NA | 1 | 0.941 | NA | 0.571 | NA | 0.399 |
| SE | 6334 | FCAR | FCAR | chr19 | + | 54888006 | 54888220 | 54885234 | 54885525 | 54889648 | 54891420 | 0 | 1 | 11 | 0 | 0 | 1 | 0 | 10 | 0 | 0 | 3 | 0 | 98 | 49 | 0.018937654 | 0.170760496 | NA | 1 | 0.846 | NA | 0.625 | NA | 0.298 |
| SE | 4057 | APBB1IP | APBB1IP | chr10 | + | 26536073 | 26536217 | 26533438 | 26533525 | 26541581 | 26541692 | 0 | 1 | 12 | 0 | 0 | 0 | 3 | 4 | 0 | 0 | 3 | 0 | 98 | 49 | 0.018984944 | 0.170760496 | NA | 1 | 1 | 1 | 0.4 | NA | 0.3 |
| SE | 3889 | MAP7D1 | MAP7D1 | chr1 | + | 36174897 | 36175008 | 36173363 | 36173478 | 36176198 | 36176321 | 0 | 0 | 4 | 0 | 2 | 2 | 0 | 8 | 0 | 0 | 1 | 0 | 98 | 49 | 0.019005986 | 0.170760496 | NA | 0 | 0.5 | NA | 0.8 | NA | -0.55 |
| A3SS | 1378 | OSBPL8 | OSBPL8 | chr12 | - | 76450850 | 76450988 | 76450850 | 76450979 | 76487509 | 76487618 | 0 | 0 | 3 | 0 | 1 | 1 | 0 | 5 | 0 | 0 | 2 | 0 | 58 | 49 | 0.019031697 | 0.197717869 | NA | 0 | 0.717 | NA | 0.679 | NA | -0.321 |
| SE | 5821 | MGEA5 | MGEA5 | chr10 | - | 101793912 | 101793998 | 101792838 | 101792943 | 101797979 | 101798154 | 0 | 0 | 3 | 0 | 2 | 1 | 0 | 3 | 0 | 0 | 4 | 0 | 98 | 49 | 0.019064109 | 0.170760496 | NA | 0 | 0.6 | NA | 0.273 | NA | 0.027 |
| SE | 5241 | PLXNC1 | PLXNC1 | chr12 | + | 94209589 | 94209704 | 94186372 | 94186473 | 94220015 | 94220163 | 0 | 4 | 27 | 0 | 0 | 1 | 0 | 5 | 0 | 0 | 2 | 0 | 98 | 49 | 0.019110193 | 0.170760496 | NA | 1 | 0.931 | NA | 0.556 | NA | 0.409 |
| A3SS | 3389 | RBM39 | RBM39 | chr20 | - | 35709223 | 35709274 | 35709223 | 35709256 | 35713018 | 35713096 | 0 | 0 | 7 | 0 | 1 | 1 | 0 | 4 | 0 | 0 | 3 | 0 | 67 | 49 | 0.019156561 | 0.197717869 | NA | 0 | 0.837 | NA | 0.494 | NA | -0.076 |
| SE | 10591 | RNF213 | RNF213 | chr17 | + | 80383676 | 80383928 | 80382978 | 80383070 | 80385537 | 80385621 | 0 | 1 | 30 | 0 | 0 | 0 | 1 | 12 | 0 | 0 | 3 | 0 | 98 | 49 | 0.019182317 | 0.170760496 | NA | 1 | 1 | 1 | 0.667 | NA | 0.166 |
| SE | 11022 | TAP1 | TAP1 | chr6 | - | 32847918 | 32848092 | 32847512 | 32847675 | 32848651 | 32848840 | 0 | 4 | 21 | 0 | 0 | 0 | 0 | 7 | 0 | 0 | 2 | 0 | 98 | 49 | 0.019207741 | 0.170760496 | NA | 1 | 1 | NA | 0.636 | NA | 0.364 |
| A5SS | 2325 | DEDD2 | DEDD2 | chr19 | - | 42201988 | 42202072 | 42202050 | 42202072 | 42198592 | 42199829 | 0 | 4 | 0 | 0 | 0 | 1 | 2 | 3 | 0 | 0 | 2 | 0 | 98 | 49 | 0.019484918 | 0.170414363 | NA | 1 | 0 | 1 | 0.429 | NA | -0.215 |
| SE | 5270 | BID | BID | chr22 | - | 17750104 | 17750174 | 17743802 | 17744013 | 17774380 | 17774495 | 0 | 4 | 1 | 0 | 0 | 0 | 0 | 3 | 0 | 0 | 2 | 0 | 98 | 49 | 0.019484918 | 0.170760496 | NA | 1 | 1 | NA | 0.429 | NA | 0.571 |
| SE | 7455 | FPR1 | FPR1 | chr19 | - | 51750202 | 51750253 | 51745769 | 51747005 | 51751813 | 51751897 | 0 | 1 | 2 | 0 | 0 | 1 | 0 | 3 | 0 | 0 | 3 | 0 | 98 | 49 | 0.019485123 | 0.170760496 | NA | 1 | 0.5 | NA | 0.333 | NA | 0.417 |
| SE | 7415 | VAV1 | VAV1 | chr19 | + | 6821790 | 6821859 | 6821621 | 6821680 | 6822220 | 6822329 | 0 | 4 | 4 | 0 | 0 | 0 | 3 | 11 | 0 | 0 | 2 | 0 | 98 | 49 | 0.01972669 | 0.170760496 | NA | 1 | 1 | 1 | 0.733 | NA | 0.133 |
| SE | 9526 | EIF4G2 | EIF4G2 | chr11 | - | 10801660 | 10801774 | 10800961 | 10801087 | 10802048 | 10802209 | 0 | 0 | 6 | 0 | 1 | 1 | 2 | 10 | 0 | 0 | 1 | 0 | 98 | 49 | 0.01979002 | 0.170760496 | NA | 0 | 0.75 | 1 | 0.833 | NA | -0.541 |
| SE | 9353 | GBP5 | GBP5 | chr1 | - | 89262682 | 89262785 | 89262219 | 89262401 | 89263735 | 89263948 | 0 | 3 | 3 | 0 | 0 | 0 | 1 | 6 | 0 | 0 | 2 | 0 | 98 | 49 | 0.019996576 | 0.170760496 | NA | 1 | 1 | 1 | 0.6 | NA | 0.2 |
| SE | 12946 | SMCHD1 | SMCHD1 | chr18 | + | 2750042 | 2750122 | 2747521 | 2747647 | 2750349 | 2750507 | 0 | 2 | 54 | 0 | 0 | 8 | 0 | 46 | 0 | 0 | 3 | 0 | 98 | 49 | 0.020053524 | 0.170760496 | NA | 1 | 0.771 | NA | 0.885 | NA | 0 |
| SE | 496 | EEF1D | EEF1D | chr8 | - | 143586218 | 143586290 | 143581228 | 143581328 | 143586728 | 143586849 | 0 | 3 | 12 | 0 | 0 | 0 | 0 | 8 | 0 | 0 | 2 | 0 | 98 | 49 | 0.020218801 | 0.170760496 | NA | 1 | 1 | NA | 0.667 | NA | 0.333 |
| SE | 4780 | TXNRD1 | TXNRD1 | chr12 | + | 104321090 | 104321316 | 104319469 | 104319585 | 104325336 | 104325429 | 0 | 3 | 2 | 0 | 0 | 0 | 0 | 8 | 0 | 0 | 2 | 0 | 98 | 49 | 0.020218801 | 0.170760496 | NA | 1 | 1 | NA | 0.667 | NA | 0.333 |
| SE | 4967 | DDX3X | DDX3X | chrX | + | 41347314 | 41347451 | 41346858 | 41347012 | 41347639 | 41350287 | 0 | 1 | 17 | 0 | 0 | 0 | 0 | 20 | 0 | 0 | 3 | 0 | 98 | 49 | 0.020293915 | 0.170760496 | NA | 1 | 1 | NA | 0.769 | NA | 0.231 |
| SE | 1577 | USP9X | USP9X | chrX | + | 41123470 | 41123724 | 41085634 | 41086109 | 41128999 | 41129145 | 0 | 0 | 4 | 0 | 0 | 0 | 0 | 1 | 2 | 0 | 0 | 2 | 98 | 49 | 0.020294338 | 0.170760496 | NA | NA | 1 | NA | 1 | 0.333 | 0.334 |
| A5SS | 976 | CTAGE5 | CTAGE5 | chr14 | + | 39276933 | 39277065 | 39276933 | 39276955 | 39279336 | 39279358 | 0 | 1 | 6 | 0 | 0 | 0 | 0 | 2 | 0 | 0 | 3 | 0 | 98 | 49 | 0.020627966 | 0.170414363 | NA | 1 | 1 | NA | 0.25 | NA | 0.75 |
| SE | 9058 | FAM107B | FAM107B | chr10 | - | 14553321 | 14553387 | 14530331 | 14530515 | 14604239 | 14604418 | 0 | 1 | 0 | 0 | 0 | 0 | 0 | 2 | 0 | 0 | 3 | 0 | 98 | 49 | 0.020627966 | 0.170760496 | NA | 1 | NA | NA | 0.25 | NA | 0.75 |
| SE | 11109 | USP25 | USP25 | chr21 | + | 15824038 | 15824166 | 15818697 | 15818846 | 15824965 | 15825061 | 0 | 1 | 3 | 0 | 0 | 2 | 0 | 2 | 0 | 0 | 3 | 0 | 98 | 49 | 0.020627966 | 0.170760496 | NA | 1 | 0.429 | NA | 0.25 | NA | 0.465 |
| SE | 7818 | DPEP2 | DPEP2 | chr16 | - | 67990046 | 67990131 | 67989322 | 67989398 | 67990820 | 67990997 | 0 | 1 | 15 | 0 | 0 | 0 | 0 | 5 | 0 | 0 | 1 | 1 | 98 | 49 | 0.020811238 | 0.170760496 | NA | 1 | 1 | NA | 0.714 | 0 | 0.643 |
| SE | 3457 | CDS2 | CDS2 | chr20 | + | 5186686 | 5186839 | 5185757 | 5185826 | 5189066 | 5189186 | 0 | 3 | 8 | 0 | 0 | 0 | 0 | 1 | 0 | 0 | 3 | 0 | 98 | 49 | 0.020882905 | 0.170760496 | NA | 1 | 1 | NA | 0.143 | NA | 0.857 |
| A5SS | 2345 | HN1 | HN1 | chr17 | - | 75147519 | 75147653 | 75147555 | 75147653 | 75146665 | 75146684 | 2 | 0 | 1 | 0 | 1 | 2 | 0 | 0 | 0 | 4 | 5 | 0 | 85 | 49 | 0.020913098 | 0.170414363 | 1 | 0 | 0.224 | 0 | 0 | NA | 0.408 |
| SE | 6630 | PCBP2 | PCBP2 | chr12 | + | 53467804 | 53467843 | 53467220 | 53467293 | 53468776 | 53468832 | 0 | 1 | 9 | 0 | 0 | 7 | 8 | 2 | 0 | 1 | 3 | 0 | 98 | 49 | 0.020946855 | 0.170760496 | NA | 1 | 0.417 | 0.817 | 0.271 | NA | 0.164 |
| SE | 1457 | ATP6V1G2-DDX39B | ATP6V1G2-DDX39B | chr6 | - | 31539146 | 31539274 | 31538762 | 31538855 | 31540321 | 31540664 | 0 | 2 | 20 | 0 | 0 | 0 | 0 | 4 | 0 | 0 | 2 | 0 | 98 | 49 | 0.021248483 | 0.170760496 | NA | 1 | 1 | NA | 0.5 | NA | 0.5 |
| SE | 6867 | ATG2A | ATG2A | chr11 | - | 64908990 | 64909150 | 64907747 | 64907890 | 64909270 | 64909367 | 0 | 0 | 2 | 0 | 0 | 0 | 0 | 10 | 4 | 0 | 1 | 2 | 98 | 49 | 0.021248483 | 0.170760496 | NA | NA | 1 | NA | 0.833 | 0.5 | 0.334 |
| A5SS | 2264 | MTCH1 | MTCH1 | chr6 | - | 36972651 | 36972796 | 36972702 | 36972796 | 36970646 | 36970694 | 0 | 2 | 9 | 0 | 0 | 8 | 0 | 6 | 0 | 0 | 2 | 0 | 98 | 49 | 0.021315983 | 0.170414363 | NA | 1 | 0.36 | NA | 0.6 | NA | 0.08 |
| SE | 12526 | GUCD1 | GUCD1 | chr22 | - | 24548916 | 24549001 | 24547907 | 24548073 | 24554948 | 24555307 | 0 | 2 | 0 | 0 | 0 | 0 | 0 | 6 | 2 | 0 | 2 | 0 | 98 | 49 | 0.021315983 | 0.170760496 | NA | 1 | NA | NA | 0.6 | 1 | 0.2 |
| SE | 5012 | ADRBK1 | ADRBK1 | chr11 | + | 67281092 | 67281184 | 67280731 | 67280783 | 67281458 | 67281558 | 0 | 2 | 32 | 0 | 0 | 0 | 0 | 7 | 0 | 0 | 2 | 0 | 98 | 49 | 0.021459804 | 0.170760496 | NA | 1 | 1 | NA | 0.636 | NA | 0.364 |
| SE | 2371 | MFSD1 | MFSD1 | chr3 | + | 158821983 | 158822140 | 158821596 | 158821653 | 158823427 | 158823525 | 0 | 2 | 16 | 0 | 0 | 0 | 0 | 3 | 0 | 0 | 2 | 0 | 98 | 49 | 0.021489596 | 0.170760496 | NA | 1 | 1 | NA | 0.429 | NA | 0.571 |
| SE | 10746 | MAN2C1 | MAN2C1 | chr15 | - | 75363998 | 75364188 | 75362641 | 75362748 | 75364487 | 75364665 | 0 | 2 | 0 | 0 | 0 | 2 | 0 | 3 | 0 | 0 | 2 | 0 | 98 | 49 | 0.021489596 | 0.170760496 | NA | 1 | 0 | NA | 0.429 | NA | 0.071 |
| SE | 13957 | DIP2B | DIP2B | chr12 | + | 50671185 | 50671398 | 50660193 | 50660319 | 50674473 | 50674629 | 0 | 2 | 5 | 0 | 0 | 0 | 0 | 3 | 0 | 0 | 2 | 0 | 98 | 49 | 0.021489596 | 0.170760496 | NA | 1 | 1 | NA | 0.429 | NA | 0.571 |
| SE | 12161 | HSH2D | HSH2D | chr19 | + | 16151446 | 16151578 | 16148723 | 16148875 | 16152551 | 16153208 | 0 | 0 | 1 | 0 | 4 | 15 | 0 | 2 | 0 | 0 | 9 | 0 | 98 | 49 | 0.021888757 | 0.170760496 | NA | 0 | 0.032 | NA | 0.1 | NA | -0.084 |
| SE | 6970 | MFN2 | MFN2 | chr1 | + | 12007052 | 12007249 | 12006537 | 12006693 | 12009591 | 12009726 | 0 | 0 | 5 | 0 | 1 | 0 | 0 | 6 | 0 | 0 | 1 | 0 | 98 | 49 | 0.021892715 | 0.170760496 | NA | 0 | 1 | NA | 0.75 | NA | -0.25 |
| A3SS | 689 | CNBP | CNBP | chr3 | - | 129171078 | 129171280 | 129171078 | 129171277 | 129171445 | 129171538 | 0 | 0 | 2 | 0 | 1 | 4 | 0 | 3 | 0 | 0 | 11 | 0 | 52 | 49 | 0.021942316 | 0.197717869 | NA | 0 | 0.32 | NA | 0.204 | NA | -0.044 |
| A3SS | 490 | GRK6 | GRK6 | chr5 | + | 177441734 | 177442849 | 177441736 | 177442849 | 177440918 | 177441053 | 0 | 1 | 6 | 0 | 0 | 2 | 0 | 0 | 0 | 0 | 2 | 2 | 51 | 49 | 0.021987328 | 0.197717869 | NA | 1 | 0.742 | NA | 0 | 0 | 0.871 |
| SE | 4999 | WIPI2 | WIPI2 | chr7 | + | 5193117 | 5193171 | 5190203 | 5190493 | 5199575 | 5199658 | 0 | 1 | 3 | 0 | 0 | 1 | 0 | 0 | 0 | 0 | 2 | 2 | 98 | 49 | 0.021999349 | 0.170760496 | NA | 1 | 0.6 | NA | 0 | 0 | 0.8 |
| SE | 2204 | CPEB2 | CPEB2 | chr4 | + | 15059186 | 15059301 | 15058420 | 15058539 | 15062078 | 15062260 | 0 | 1 | 0 | 0 | 1 | 2 | 0 | 3 | 3 | 0 | 0 | 0 | 98 | 49 | 0.022049079 | 0.170760496 | NA | 0.333 | 0 | NA | 1 | 1 | -0.834 |
| A5SS | 2821 | LILRA6 | LILRA6 | chr19 | - | 54241575 | 54241878 | 54241661 | 54241878 | 54240830 | 54241127 | 0 | 0 | 0 | 0 | 2 | 1 | 0 | 4 | 0 | 0 | 1 | 0 | 98 | 49 | 0.022093415 | 0.170414363 | NA | 0 | 0 | NA | 0.667 | NA | -0.667 |
| SE | 9285 | VIM | VIM | chr10 | + | 17235845 | 17235889 | 17235168 | 17235389 | 17236293 | 17236379 | 0 | 12 | 171 | 0 | 0 | 0 | 3 | 36 | 3 | 0 | 3 | 0 | 93 | 49 | 0.022225644 | 0.170760496 | NA | 1 | 1 | 1 | 0.863 | 1 | 0.046 |
| SE | 13688 | BAZ2A | BAZ2A | chr12 | - | 56605829 | 56606063 | 56605072 | 56605327 | 56606246 | 56606312 | 0 | 2 | 2 | 0 | 0 | 0 | 0 | 2 | 0 | 0 | 2 | 0 | 98 | 49 | 0.022262485 | 0.170760496 | NA | 1 | 1 | NA | 0.333 | NA | 0.667 |
| SE | 14090 | AREL1 | AREL1 | chr14 | - | 74669946 | 74670126 | 74669648 | 74669774 | 74670761 | 74670871 | 0 | 2 | 2 | 0 | 0 | 3 | 0 | 2 | 0 | 0 | 2 | 0 | 98 | 49 | 0.022262485 | 0.170760496 | NA | 1 | 0.25 | NA | 0.333 | NA | 0.292 |
| SE | 6878 | UBN1 | UBN1 | chr16 | + | 4868832 | 4868903 | 4860663 | 4861102 | 4870211 | 4870341 | 0 | 0 | 4 | 0 | 1 | 0 | 0 | 5 | 0 | 0 | 1 | 0 | 98 | 49 | 0.022832961 | 0.170760496 | NA | 0 | 1 | NA | 0.714 | NA | -0.214 |
| SE | 11349 | ADGRE2 | ADGRE2 | chr19 | - | 14754953 | 14755127 | 14752328 | 14752526 | 14756237 | 14756345 | 0 | 0 | 3 | 0 | 1 | 1 | 0 | 5 | 0 | 0 | 1 | 0 | 98 | 49 | 0.022832961 | 0.170760496 | NA | 0 | 0.6 | NA | 0.714 | NA | -0.414 |
| SE | 3680 | USP15 | USP15 | chr12 | + | 62325871 | 62325933 | 62321463 | 62321609 | 62349220 | 62349307 | 0 | 2 | 32 | 0 | 0 | 1 | 0 | 15 | 0 | 0 | 2 | 0 | 98 | 49 | 0.023010064 | 0.170760496 | NA | 1 | 0.941 | NA | 0.789 | NA | 0.181 |
| SE | 10210 | RFWD2 | RFWD2 | chr1 | - | 176175909 | 176176007 | 176163814 | 176163891 | 176184632 | 176184692 | 0 | 2 | 13 | 0 | 0 | 0 | 0 | 17 | 0 | 0 | 2 | 0 | 98 | 49 | 0.023408389 | 0.170760496 | NA | 1 | 1 | NA | 0.81 | NA | 0.19 |
| SE | 558 | PABPC1 | PABPC1 | chr8 | - | 100713086 | 100713181 | 100712651 | 100712789 | 100715461 | 100715601 | 0 | 6 | 63 | 0 | 0 | 0 | 5 | 32 | 2 | 0 | 3 | 0 | 98 | 49 | 0.0234244 | 0.170760496 | NA | 1 | 1 | 1 | 0.842 | 1 | 0.053 |
| SE | 17 | FLNA | FLNA | chrX | - | 154353553 | 154353727 | 154353295 | 154353457 | 154353914 | 154354043 | 0 | 2 | 35 | 0 | 0 | 1 | 0 | 18 | 0 | 2 | 2 | 0 | 98 | 49 | 0.023592394 | 0.170760496 | NA | 1 | 0.946 | 0 | 0.818 | NA | 0.564 |
| A5SS | 93 | GAPVD1 | GAPVD1 | chr9 | + | 125321432 | 125321562 | 125321432 | 125321499 | 125323797 | 125323923 | 0 | 3 | 3 | 0 | 0 | 0 | 0 | 1 | 0 | 0 | 2 | 0 | 98 | 49 | 0.023673881 | 0.170414363 | NA | 1 | 1 | NA | 0.2 | NA | 0.8 |
| SE | 2652 | SETD2 | SETD2 | chr3 | - | 47101457 | 47101555 | 47097954 | 47098081 | 47103345 | 47103423 | 0 | 1 | 10 | 0 | 0 | 0 | 0 | 4 | 0 | 0 | 2 | 0 | 98 | 49 | 0.023775553 | 0.170760496 | NA | 1 | 1 | NA | 0.5 | NA | 0.5 |
| SE | 4064 | FBXL5 | FBXL5 | chr4 | - | 15638507 | 15638694 | 15636493 | 15636676 | 15640787 | 15640883 | 0 | 3 | 71 | 0 | 0 | 0 | 6 | 32 | 0 | 0 | 2 | 0 | 98 | 49 | 0.0238404 | 0.170760496 | NA | 1 | 1 | 1 | 0.889 | NA | 0.055 |
| A3SS | 681 | CDV3 | CDV3 | chr3 | + | 133587158 | 133590274 | 133587895 | 133590274 | 133586562 | 133586722 | 0 | 1 | 2 | 0 | 0 | 0 | 0 | 5 | 0 | 0 | 2 | 0 | 98 | 49 | 0.023845162 | 0.197717869 | NA | 1 | 1 | NA | 0.556 | NA | 0.444 |
| SE | 3215 | PDCD6IP | PDCD6IP | chr3 | + | 33812071 | 33812126 | 33798570 | 33798937 | 33813558 | 33813628 | 0 | 1 | 6 | 0 | 0 | 0 | 0 | 1 | 0 | 0 | 3 | 0 | 98 | 49 | 0.023847213 | 0.170760496 | NA | 1 | 1 | NA | 0.143 | NA | 0.857 |
| SE | 4847 | TINF2 | TINF2 | chr14 | - | 24241889 | 24241994 | 24241674 | 24241776 | 24242140 | 24242674 | 0 | 1 | 0 | 0 | 0 | 0 | 0 | 1 | 0 | 0 | 3 | 0 | 98 | 49 | 0.023847213 | 0.170760496 | NA | 1 | NA | NA | 0.143 | NA | 0.857 |
| SE | 2082 | HERC6 | HERC6 | chr4 | + | 88393487 | 88393582 | 88390651 | 88390879 | 88396014 | 88396142 | 0 | 1 | 12 | 0 | 0 | 0 | 0 | 3 | 0 | 0 | 2 | 0 | 98 | 49 | 0.023922591 | 0.170760496 | NA | 1 | 1 | NA | 0.429 | NA | 0.571 |
| SE | 7201 | MDM4 | MDM4 | chr1 | + | 204537429 | 204537497 | 204532190 | 204532246 | 204538208 | 204538308 | 0 | 1 | 6 | 0 | 0 | 0 | 0 | 3 | 0 | 0 | 2 | 0 | 98 | 49 | 0.023922591 | 0.170760496 | NA | 1 | 1 | NA | 0.429 | NA | 0.571 |
| SE | 10725 | PTMA | PTMA | chr2 | + | 231711889 | 231711983 | 231711347 | 231711419 | 231712442 | 231712516 | 0 | 1 | 3 | 0 | 0 | 0 | 0 | 3 | 0 | 0 | 2 | 0 | 98 | 49 | 0.023922591 | 0.170760496 | NA | 1 | 1 | NA | 0.429 | NA | 0.571 |
| SE | 6590 | UQCRH | UQCRH | chr1 | + | 46309896 | 46310044 | 46309100 | 46309127 | 46310154 | 46310316 | 0 | 0 | 1 | 0 | 2 | 0 | 0 | 2 | 0 | 3 | 3 | 0 | 98 | 49 | 0.024004538 | 0.170760496 | NA | 0 | 1 | 0 | 0.25 | NA | 0.375 |
| SE | 4111 | ATP2B1 | ATP2B1 | chr12 | - | 89601325 | 89601433 | 89599116 | 89599299 | 89603042 | 89603254 | 0 | 1 | 2 | 0 | 0 | 0 | 2 | 6 | 0 | 0 | 2 | 0 | 98 | 49 | 0.024023827 | 0.170760496 | NA | 1 | 1 | 1 | 0.6 | NA | 0.2 |
| SE | 5016 | ARHGAP25 | ARHGAP25 | chr2 | + | 68813286 | 68813419 | 68807272 | 68807480 | 68816288 | 68816362 | 0 | 1 | 15 | 0 | 0 | 0 | 0 | 6 | 0 | 0 | 2 | 0 | 98 | 49 | 0.024023827 | 0.170760496 | NA | 1 | 1 | NA | 0.6 | NA | 0.4 |
| SE | 7011 | HIPK3 | HIPK3 | chr11 | + | 33348521 | 33348818 | 33347851 | 33348013 | 33349146 | 33349287 | 0 | 1 | 4 | 0 | 0 | 0 | 0 | 6 | 0 | 0 | 2 | 0 | 98 | 49 | 0.024023827 | 0.170760496 | NA | 1 | 1 | NA | 0.6 | NA | 0.4 |
| SE | 10363 | SOS2 | SOS2 | chr14 | - | 50145169 | 50145332 | 50139941 | 50140059 | 50145476 | 50145596 | 0 | 1 | 9 | 0 | 0 | 1 | 0 | 6 | 0 | 0 | 2 | 0 | 98 | 49 | 0.024023827 | 0.170760496 | NA | 1 | 0.818 | NA | 0.6 | NA | 0.309 |
| SE | 10724 | PTMA | PTMA | chr2 | + | 231711886 | 231711983 | 231711347 | 231711419 | 231712442 | 231712516 | 0 | 1 | 2 | 0 | 0 | 0 | 0 | 6 | 0 | 0 | 2 | 0 | 98 | 49 | 0.024023827 | 0.170760496 | NA | 1 | 1 | NA | 0.6 | NA | 0.4 |
| SE | 11386 | NLRC5 | NLRC5 | chr16 | + | 57046551 | 57046641 | 57045447 | 57045492 | 57047544 | 57047628 | 0 | 1 | 10 | 0 | 0 | 0 | 0 | 6 | 0 | 0 | 2 | 0 | 98 | 49 | 0.024023827 | 0.170760496 | NA | 1 | 1 | NA | 0.6 | NA | 0.4 |
| SE | 690 | CDK16 | CDK16 | chrX | + | 47225771 | 47225866 | 47224987 | 47225102 | 47225964 | 47226027 | 0 | 1 | 4 | 0 | 0 | 0 | 0 | 7 | 0 | 0 | 2 | 0 | 98 | 49 | 0.02426018 | 0.170760496 | NA | 1 | 1 | NA | 0.636 | NA | 0.364 |
| SE | 7907 | IFIT1 | IFIT1 | chr10 | + | 89399894 | 89400003 | 89392545 | 89392717 | 89402280 | 89406487 | 0 | 1 | 12 | 0 | 0 | 5 | 0 | 7 | 0 | 4 | 2 | 0 | 98 | 49 | 0.02426018 | 0.170760496 | NA | 1 | 0.545 | 0 | 0.636 | NA | 0.454 |
| SE | 8359 | ARHGAP30 | ARHGAP30 | chr1 | - | 161059613 | 161059716 | 161054622 | 161054705 | 161069527 | 161069970 | 0 | 1 | 18 | 0 | 0 | 2 | 0 | 7 | 0 | 0 | 2 | 0 | 98 | 49 | 0.02426018 | 0.170760496 | NA | 1 | 0.818 | NA | 0.636 | NA | 0.273 |
| SE | 9209 | CELF2 | CELF2 | chr10 | + | 11217424 | 11217507 | 11165485 | 11165682 | 11249152 | 11249201 | 0 | 1 | 31 | 0 | 0 | 2 | 5 | 7 | 0 | 0 | 2 | 0 | 98 | 49 | 0.02426018 | 0.170760496 | NA | 1 | 0.886 | 1 | 0.636 | NA | 0.125 |
| SE | 8140 | ITGAX | ITGAX | chr16 | + | 31373248 | 31373390 | 31372596 | 31372670 | 31376798 | 31376915 | 0 | 12 | 40 | 0 | 0 | 5 | 6 | 46 | 13 | 0 | 0 | 0 | 98 | 49 | 0.024390083 | 0.170760496 | NA | 1 | 0.8 | 1 | 1 | 1 | -0.1 |
| SE | 12583 | SBNO2 | SBNO2 | chr19 | - | 1109134 | 1109211 | 1108778 | 1108969 | 1109291 | 1109423 | 0 | 4 | 9 | 0 | 0 | 0 | 0 | 25 | 2 | 0 | 3 | 0 | 98 | 49 | 0.024422917 | 0.170760496 | NA | 1 | 1 | NA | 0.806 | 1 | 0.097 |
| SE | 13765 | ABCG1 | ABCG1 | chr21 | + | 42276894 | 42276945 | 42273302 | 42273435 | 42282273 | 42282419 | 0 | 1 | 11 | 0 | 0 | 0 | 0 | 8 | 0 | 0 | 2 | 0 | 98 | 49 | 0.024517355 | 0.170760496 | NA | 1 | 1 | NA | 0.667 | NA | 0.333 |
| A3SS | 683 | CDV3 | CDV3 | chr3 | + | 133587161 | 133590274 | 133587895 | 133590274 | 133586562 | 133586722 | 0 | 1 | 2 | 0 | 0 | 0 | 0 | 2 | 0 | 0 | 2 | 0 | 98 | 49 | 0.024627853 | 0.197717869 | NA | 1 | 1 | NA | 0.333 | NA | 0.667 |
| SE | 1480 | FLOT1 | FLOT1 | chr6 | - | 30741613 | 30741704 | 30741189 | 30741333 | 30741791 | 30741867 | 0 | 1 | 6 | 0 | 0 | 1 | 0 | 2 | 0 | 0 | 2 | 0 | 98 | 49 | 0.024627853 | 0.170760496 | NA | 1 | 0.75 | NA | 0.333 | NA | 0.542 |
| SE | 3632 | USP4 | USP4 | chr3 | - | 49300466 | 49300691 | 49298551 | 49298635 | 49302383 | 49302542 | 0 | 1 | 0 | 0 | 0 | 0 | 0 | 2 | 0 | 0 | 2 | 0 | 98 | 49 | 0.024627853 | 0.170760496 | NA | 1 | NA | NA | 0.333 | NA | 0.667 |
| SE | 4078 | PHF1 | PHF1 | chr6 | + | 33414954 | 33415144 | 33414724 | 33414829 | 33415234 | 33415329 | 0 | 1 | 1 | 0 | 0 | 0 | 0 | 2 | 0 | 0 | 2 | 0 | 98 | 49 | 0.024627853 | 0.170760496 | NA | 1 | 1 | NA | 0.333 | NA | 0.667 |
| SE | 4079 | PHF1 | PHF1 | chr6 | + | 33414965 | 33415144 | 33414724 | 33414829 | 33415234 | 33415329 | 0 | 1 | 1 | 0 | 0 | 0 | 0 | 2 | 0 | 0 | 2 | 0 | 98 | 49 | 0.024627853 | 0.170760496 | NA | 1 | 1 | NA | 0.333 | NA | 0.667 |
| SE | 6133 | STXBP3 | STXBP3 | chr1 | + | 108779285 | 108779410 | 108776332 | 108776423 | 108782421 | 108782517 | 0 | 2 | 7 | 0 | 0 | 0 | 0 | 1 | 0 | 0 | 2 | 0 | 98 | 49 | 0.02492004 | 0.170760496 | NA | 1 | 1 | NA | 0.2 | NA | 0.8 |
| SE | 7313 | WNK1 | WNK1 | chr12 | + | 871264 | 871348 | 862082 | 862270 | 878211 | 878361 | 0 | 2 | 4 | 0 | 0 | 1 | 3 | 1 | 0 | 0 | 2 | 0 | 98 | 49 | 0.02492004 | 0.170760496 | NA | 1 | 0.667 | 1 | 0.2 | NA | 0.234 |
| SE | 8790 | VPS13D | VPS13D | chr1 | + | 12311453 | 12311625 | 12308430 | 12308641 | 12311812 | 12311925 | 0 | 2 | 6 | 0 | 0 | 0 | 0 | 1 | 0 | 0 | 2 | 0 | 98 | 49 | 0.02492004 | 0.170760496 | NA | 1 | 1 | NA | 0.2 | NA | 0.8 |
| A5SS | 1139 | TBC1D10C | TBC1D10C | chr11 | + | 67406626 | 67406738 | 67406626 | 67406692 | 67406826 | 67407016 | 0 | 1 | 2 | 0 | 0 | 4 | 0 | 3 | 3 | 0 | 0 | 0 | 95 | 49 | 0.024974929 | 0.170414363 | NA | 1 | 0.205 | NA | 1 | 1 | -0.397 |
| SE | 5184 | MYO9B | MYO9B | chr19 | + | 17145396 | 17145491 | 17101659 | 17102557 | 17152643 | 17152706 | 0 | 3 | 23 | 0 | 0 | 0 | 0 | 19 | 2 | 0 | 4 | 0 | 98 | 49 | 0.024989397 | 0.170760496 | NA | 1 | 1 | NA | 0.704 | 1 | 0.148 |
| SE | 13164 | PPP1CB | PPP1CB | chr2 | + | 28788657 | 28788809 | 28783906 | 28783978 | 28793862 | 28793997 | 0 | 1 | 6 | 0 | 0 | 0 | 0 | 10 | 0 | 0 | 2 | 0 | 98 | 49 | 0.025074234 | 0.170760496 | NA | 1 | 1 | NA | 0.714 | NA | 0.286 |
| SE | 2167 | FYB | FYB | chr5 | - | 39201825 | 39202987 | 39153447 | 39153604 | 39219442 | 39219578 | 0 | 8 | 22 | 0 | 0 | 0 | 1 | 10 | 1 | 0 | 2 | 0 | 98 | 49 | 0.025260336 | 0.170760496 | NA | 1 | 1 | 1 | 0.714 | 1 | 0.095 |
| SE | 2076 | HERC5 | HERC5 | chr4 | + | 88463854 | 88463985 | 88463531 | 88463623 | 88467058 | 88467204 | 0 | 2 | 27 | 0 | 0 | 1 | 0 | 28 | 0 | 0 | 2 | 0 | 98 | 49 | 0.02534026 | 0.170760496 | NA | 1 | 0.931 | NA | 0.875 | NA | 0.091 |
| SE | 4015 | RPLP0 | RPLP0 | chr12 | - | 120198553 | 120198739 | 120197321 | 120197462 | 120198853 | 120199000 | 0 | 1 | 3 | 0 | 0 | 0 | 0 | 12 | 0 | 0 | 2 | 0 | 98 | 49 | 0.025646206 | 0.170760496 | NA | 1 | 1 | NA | 0.75 | NA | 0.25 |
| SE | 5192 | PICALM | PICALM | chr11 | - | 85976622 | 85976682 | 85974707 | 85974812 | 85981128 | 85981228 | 0 | 2 | 30 | 0 | 0 | 0 | 3 | 30 | 0 | 0 | 2 | 0 | 98 | 49 | 0.025657506 | 0.170760496 | NA | 1 | 1 | 1 | 0.882 | NA | 0.059 |
| A3SS | 932 | CXCL1 | CXCL1 | chr4 | + | 73869645 | 73869792 | 73869668 | 73869792 | 73869391 | 73869570 | 0 | 0 | 0 | 0 | 0 | 3 | 3 | 1 | 2 | 0 | 1 | 1 | 72 | 49 | 0.025891313 | 0.197717869 | NA | NA | 0 | 1 | 0.405 | 0.576 | -0.66 |
| SE | 7720 | THEMIS2 | THEMIS2 | chr1 | + | 27881970 | 27883043 | 27876587 | 27876728 | 27885294 | 27885451 | 0 | 0 | 2 | 0 | 3 | 1 | 0 | 2 | 0 | 0 | 1 | 0 | 98 | 49 | 0.025897097 | 0.170760496 | NA | 0 | 0.5 | NA | 0.5 | NA | -0.25 |

| | | | | | | | | | | | | | | | | | | | | | | | | | | | | | | | | | | |
|---|---|---|---|---|---|---|---|---|---|---|---|---|---|---|---|---|---|---|---|---|---|---|---|---|---|---|---|---|---|---|---|---|---|---|
| SE | 13194 | MBOAT7 | MBOAT7 | chr19 | - | 54188432 | 54188511 | 54188216 | 54188346 | 54189337 | 54189882 | 0 | 9 | 20 | 0 | 0 | 1 | 1 | 12 | 31 | 0 | 2 | 6 | 98 | 49 | 0.025921935 | 0.170760496 | NA | 1 | 0.909 | 1 | 0.75 | 0.721 | 0.131 |
| SE | 6180 | ABCA1 | ABCA1 | chr9 | - | 104884426 | 104884568 | 104883038 | 104883157 | 104889101 | 104889195 | 0 | 3 | 20 | 0 | 1 | 1 | 0 | 16 | 2 | 0 | 7 | 3 | 98 | 49 | 0.026099399 | 0.170760496 | NA | 0.6 | 0.909 | NA | 0.533 | 0.25 | 0.363 |
| SE | 2583 | LCP2 | LCP2 | chr5 | - | 170270718 | 170270917 | 170268384 | 170268482 | 170274300 | 170274338 | 1 | 1 | 19 | 0 | 0 | 0 | 0 | 14 | 0 | 0 | 2 | 0 | 98 | 49 | 0.026208434 | 0.170760496 | 1 | 1 | 1 | NA | 0.778 | NA | 0.222 |
| A5SS | 2038 | TRAPPC5 | TRAPPC5 | chr19 | + | 7680820 | 7681026 | 7680820 | 7680878 | 7682241 | 7682862 | 0 | 0 | 0 | 0 | 0 | 1 | 0 | 4 | 3 | 0 | 0 | 1 | 98 | 49 | 0.026257417 | 0.170414363 | NA | NA | 0 | NA | 1 | 0.6 | -0.8 |
| A5SS | 2039 | TRAPPC5 | TRAPPC5 | chr19 | + | 7680820 | 7681049 | 7680820 | 7680878 | 7682241 | 7682862 | 0 | 0 | 0 | 0 | 0 | 1 | 0 | 3 | 3 | 0 | 0 | 1 | 98 | 49 | 0.026257417 | 0.170414363 | NA | NA | 0 | NA | 1 | 0.6 | -0.8 |
| SE | 6627 | PCBP2 | PCBP2 | chr12 | + | 53464759 | 53464852 | 53462480 | 53462567 | 53467220 | 53467293 | 0 | 0 | 13 | 0 | 1 | 3 | 0 | 3 | 0 | 0 | 1 | 0 | 98 | 49 | 0.026257417 | 0.170760496 | NA | 0 | 0.684 | NA | 0.6 | NA | -0.258 |
| SE | 7010 | HIPK3 | HIPK3 | chr11 | + | 33348165 | 33348228 | 33347851 | 33348013 | 33348521 | 33348818 | 0 | 0 | 7 | 0 | 1 | 1 | 0 | 3 | 0 | 0 | 1 | 0 | 98 | 49 | 0.026257417 | 0.170760496 | NA | 0 | 0.778 | NA | 0.6 | NA | -0.211 |
| SE | 12300 | CYTH1 | CYTH1 | chr17 | - | 78702537 | 78702604 | 78702121 | 78702240 | 78708196 | 78708261 | 0 | 0 | 6 | 0 | 1 | 3 | 0 | 3 | 0 | 0 | 1 | 0 | 98 | 49 | 0.026257417 | 0.170760496 | NA | 0 | 0.5 | NA | 0.6 | NA | -0.35 |
| SE | 6175 | ABCA1 | ABCA1 | chr9 | - | 104812573 | 104812722 | 104809465 | 104809564 | 104814117 | 104814231 | 0 | 1 | 3 | 0 | 0 | 0 | 2 | 15 | 0 | 0 | 2 | 0 | 98 | 49 | 0.026481025 | 0.170760496 | NA | 1 | 1 | 1 | 0.789 | NA | 0.105 |
| SE | 5194 | PICALM | PICALM | chr11 | - | 85990249 | 85990378 | 85983865 | 85983973 | 85996825 | 85996929 | 0 | 1 | 18 | 0 | 0 | 21 | 0 | 16 | 0 | 0 | 2 | 0 | 98 | 49 | 0.026734839 | 0.170760496 | NA | 1 | 0.3 | NA | 0.8 | NA | -0.15 |
| SE | 5975 | TAP2 | TAP2 | chr6 | - | 32835153 | 32835359 | 32832626 | 32832824 | 32835642 | 32835773 | 0 | 0 | 11 | 0 | 0 | 0 | 0 | 11 | 1 | 0 | 0 | 1 | 98 | 49 | 0.026878049 | 0.170760496 | NA | NA | 1 | NA | 1 | 0.333 | 0.334 |
| SE | 9687 | IFRD1 | IFRD1 | chr7 | + | 112456913 | 112457038 | 112456001 | 112456086 | 112458860 | 112459018 | 0 | 6 | 52 | 0 | 0 | 0 | 0 | 84 | 4 | 0 | 3 | 0 | 98 | 49 | 0.026897321 | 0.170760496 | NA | 1 | 1 | NA | 0.933 | 1 | 0.033 |
| SE | 6887 | SRPK1 | SRPK1 | chr6 | - | 35870280 | 35870494 | 35869481 | 35869901 | 35870933 | 35870959 | 0 | 0 | 4 | 0 | 1 | 5 | 0 | 2 | 0 | 0 | 3 | 0 | 98 | 49 | 0.027065308 | 0.170760496 | NA | 0 | 0.286 | NA | 0.25 | NA | -0.107 |
| A3SS | 3741 | TMEM91 | TMEM91 | chr19 | + | 41383560 | 41384082 | 41383862 | 41384082 | 41382771 | 41382921 | 0 | 1 | 3 | 0 | 0 | 3 | 0 | 1 | 0 | 0 | 2 | 0 | 98 | 49 | 0.027259421 | 0.197717869 | NA | 1 | 0.333 | NA | 0.2 | NA | 0.466 |
| SE | 2726 | SATB1 | SATB1 | chr3 | - | 18351991 | 18352195 | 18347640 | 18349682 | 18378169 | 18378325 | 0 | 1 | 3 | 0 | 0 | 0 | 0 | 1 | 0 | 0 | 2 | 0 | 98 | 49 | 0.027259421 | 0.170760496 | NA | 1 | 1 | NA | 0.2 | NA | 0.8 |
| SE | 9126 | HGS | HGS | chr17 | + | 81690181 | 81690234 | 81688703 | 81688827 | 81690673 | 81690742 | 0 | 1 | 1 | 0 | 0 | 0 | 0 | 1 | 0 | 0 | 2 | 0 | 98 | 49 | 0.027259421 | 0.170760496 | NA | 1 | 1 | NA | 0.2 | NA | 0.8 |
| SE | 14102 | SLC12A4 | SLC12A4 | chr16 | - | 67951822 | 67952037 | 67951139 | 67951304 | 67952183 | 67952425 | 0 | 1 | 0 | 0 | 0 | 0 | 0 | 1 | 0 | 0 | 2 | 0 | 98 | 49 | 0.027259421 | 0.170760496 | NA | 1 | NA | NA | 0.2 | NA | 0.8 |
| A5SS | 1672 | PTBP3 | PTBP3 | chr9 | - | 112297831 | 112297916 | 112297840 | 112297916 | 112275843 | 112276013 | 0 | 0 | 3 | 0 | 2 | 2 | 0 | 2 | 0 | 0 | 1 | 0 | 58 | 49 | 0.027334905 | 0.170414363 | NA | 0 | 0.559 | NA | 0.628 | NA | -0.348 |
| SE | 8906 | FBXW7 | FBXW7 | chr4 | - | 152412479 | 152412529 | 152411159 | 152411872 | 152534940 | 152535033 | 0 | 0 | 0 | 0 | 2 | 0 | 0 | 2 | 0 | 0 | 1 | 1 | 98 | 49 | 0.02734801 | 0.170760496 | NA | 0 | NA | NA | 0.5 | 0 | -0.25 |
| SE | 3622 | ITGAM | ITGAM | chr16 | + | 31266029 | 31266147 | 31265810 | 31265881 | 31270953 | 31271084 | 0 | 6 | 27 | 0 | 0 | 0 | 0 | 3 | 0 | 0 | 1 | 0 | 98 | 49 | 0.027600753 | 0.170760496 | NA | 1 | 1 | NA | 0.6 | NA | 0.4 |
| A3SS | 405 | BAG6 | BAG6 | chr6 | - | 31647590 | 31647844 | 31647590 | 31647826 | 31648676 | 31648751 | 0 | 0 | 0 | 0 | 1 | 2 | 0 | 2 | 0 | 0 | 2 | 0 | 67 | 49 | 0.027623127 | 0.197717869 | NA | 0 | 0 | NA | 0.422 | NA | -0.422 |
| SE | 3281 | RAB11FIP1 | RAB11FIP1 | chr8 | - | 37871277 | 37873179 | 37870419 | 37870528 | 37874514 | 37875322 | 0 | 0 | 4 | 0 | 1 | 4 | 0 | 2 | 0 | 0 | 2 | 0 | 98 | 49 | 0.027623276 | 0.170760496 | NA | 0 | 0.333 | NA | 0.333 | NA | -0.167 |
| SE | 5922 | MKNK1 | MKNK1 | chr1 | - | 46560233 | 46560277 | 46557406 | 46558800 | 46561477 | 46561642 | 0 | 0 | 2 | 0 | 1 | 2 | 0 | 2 | 0 | 0 | 2 | 0 | 93 | 49 | 0.027624169 | 0.170760496 | NA | 0 | 0.345 | NA | 0.345 | NA | -0.172 |
| SE | 472 | DOCK8 | DOCK8 | chr9 | + | 372184 | 372286 | 371427 | 371566 | 376209 | 376305 | 0 | 10 | 33 | 0 | 0 | 0 | 0 | 9 | 0 | 0 | 1 | 0 | 98 | 49 | 0.027770234 | 0.170760496 | NA | 1 | 1 | NA | 0.818 | NA | 0.182 |
| SE | 20 | FLNA | FLNA | chrX | - | 154362417 | 154362578 | 154362241 | 154362332 | 154362660 | 154362784 | 0 | 5 | 37 | 0 | 0 | 0 | 2 | 18 | 1 | 0 | 2 | 0 | 98 | 49 | 0.027864407 | 0.170760496 | NA | 1 | 1 | 1 | 0.818 | 1 | 0.061 |
| SE | 5181 | ANKRD13A | ANKRD13A | chr12 | + | 110024045 | 110024112 | 110019138 | 110019328 | 110025741 | 110025823 | 0 | 6 | 10 | 0 | 0 | 0 | 0 | 4 | 0 | 0 | 1 | 0 | 98 | 49 | 0.027908407 | 0.170760496 | NA | 1 | 1 | NA | 0.667 | NA | 0.333 |
| SE | 6868 | ATG2A | ATG2A | chr11 | - | 64909270 | 64909367 | 64907747 | 64907890 | 64909680 | 64909924 | 0 | 0 | 6 | 0 | 0 | 0 | 0 | 7 | 4 | 0 | 0 | 1 | 98 | 49 | 0.027908407 | 0.170760496 | NA | NA | 1 | NA | 1 | 0.667 | 0.166 |
| SE | 4661 | AMICA1 | AMICA1 | chr11 | - | 118205881 | 118205991 | 118203427 | 118203665 | 118210486 | 118210712 | 0 | 1 | 9 | 0 | 0 | 0 | 0 | 21 | 0 | 0 | 2 | 0 | 98 | 49 | 0.028047628 | 0.170760496 | NA | 1 | 1 | NA | 0.84 | NA | 0.16 |
| SE | 5195 | PICALM | PICALM | chr11 | - | 85990249 | 85990399 | 85983865 | 85983973 | 85996825 | 85996929 | 0 | 1 | 25 | 0 | 0 | 21 | 0 | 21 | 0 | 0 | 2 | 0 | 98 | 49 | 0.028047628 | 0.170760496 | NA | 1 | 0.373 | NA | 0.84 | NA | -0.153 |
| SE | 13787 | PSD4 | PSD4 | chr2 | + | 113198739 | 113198884 | 113197746 | 113197913 | 113199082 | 113199226 | 0 | 5 | 2 | 0 | 0 | 0 | 0 | 7 | 1 | 0 | 3 | 0 | 98 | 49 | 0.028063223 | 0.170760496 | NA | 1 | 1 | NA | 0.538 | 1 | 0.231 |
| SE | 16 | FLNA | FLNA | chrX | - | 154352552 | 154352675 | 154352180 | 154352447 | 154352771 | 154352924 | 0 | 9 | 44 | 0 | 0 | 1 | 0 | 9 | 0 | 0 | 1 | 0 | 98 | 49 | 0.028070044 | 0.170760496 | NA | 1 | 0.957 | NA | 0.818 | NA | 0.16 |
| SE | 688 | PTK2B | PTK2B | chr8 | + | 27397547 | 27397788 | 27325478 | 27325681 | 27419894 | 27420073 | 0 | 7 | 27 | 0 | 0 | 0 | 0 | 6 | 0 | 0 | 1 | 0 | 98 | 49 | 0.028077309 | 0.170760496 | NA | 1 | 1 | NA | 0.75 | NA | 0.25 |
| SE | 8356 | ARHGAP30 | ARHGAP30 | chr1 | - | 161052625 | 161052797 | 161052439 | 161052543 | 161053257 | 161053385 | 0 | 7 | 26 | 0 | 0 | 0 | 0 | 6 | 0 | 0 | 1 | 0 | 98 | 49 | 0.028077309 | 0.170760496 | NA | 1 | 1 | NA | 0.75 | NA | 0.25 |
| SE | 12701 | ARFGEF1 | ARFGEF1 | chr8 | - | 67275975 | 67276109 | 67271701 | 67271936 | 67277281 | 67277457 | 0 | 5 | 2 | 0 | 0 | 0 | 7 | 2 | 0 | 0 | 1 | 0 | 98 | 49 | 0.028306385 | 0.170760496 | NA | 1 | 1 | 1 | 0.5 | NA | 0.25 |
| SE | 4456 | CLEC7A | CLEC7A | chr12 | - | 10123244 | 10123363 | 10116780 | 10118590 | 10125086 | 10125173 | 0 | 0 | 38 | 0 | 2 | 1 | 0 | 15 | 3 | 0 | 0 | 0 | 98 | 49 | 0.028599911 | 0.170760496 | NA | 0 | 0.95 | NA | 1 | 1 | -0.525 |
| A5SS | 672 | STX16 | STX16 | chr20 | + | 58651252 | 58652138 | 58651252 | 58652087 | 58667489 | 58667597 | 0 | 4 | 7 | 0 | 0 | 3 | 1 | 2 | 0 | 0 | 1 | 0 | 98 | 49 | 0.029221508 | 0.170414363 | NA | 1 | 0.538 | 1 | 0.5 | NA | 0.019 |
| SE | 5228 | NFKBIA | NFKBIA | chr14 | - | 35402770 | 35402859 | 35402393 | 35402663 | 35403149 | 35403360 | 0 | 10 | 63 | 0 | 0 | 0 | 7 | 167 | 19 | 0 | 4 | 0 | 98 | 49 | 0.029442582 | 0.170760496 | NA | 1 | 1 | 1 | 0.954 | 1 | 0.015 |
| SE | 5000 | WIPI2 | WIPI2 | chr7 | + | 5193117 | 5193345 | 5190203 | 5190493 | 5199575 | 5199658 | 0 | 1 | 1 | 0 | 0 | 1 | 0 | 0 | 0 | 0 | 2 | 2 | 98 | 49 | 0.029568334 | 0.170760496 | NA | 1 | 0.333 | NA | 0 | 0 | 0.666 |
| SE | 4834 | ULK3 | ULK3 | chr15 | - | 74838265 | 74838344 | 74838151 | 74838192 | 74838439 | 74838504 | 0 | 5 | 0 | 0 | 0 | 0 | 0 | 1 | 0 | 0 | 1 | 0 | 98 | 49 | 0.029576123 | 0.170760496 | NA | 1 | NA | NA | 0.333 | NA | 0.667 |
| SE | 4835 | ULK3 | ULK3 | chr15 | - | 74838269 | 74838344 | 74838151 | 74838192 | 74838439 | 74838504 | 0 | 5 | 0 | 0 | 0 | 0 | 0 | 1 | 0 | 0 | 1 | 0 | 98 | 49 | 0.029576123 | 0.170760496 | NA | 1 | NA | NA | 0.333 | NA | 0.667 |
| SE | 7841 | B4GALT5 | B4GALT5 | chr20 | - | 49639677 | 49639800 | 49637340 | 49637442 | 49640477 | 49640665 | 0 | 5 | 13 | 0 | 0 | 1 | 0 | 6 | 0 | 0 | 1 | 0 | 98 | 49 | 0.029628726 | 0.170760496 | NA | 1 | 0.867 | NA | 0.75 | NA | 0.183 |
| SE | 11675 | ULK1 | ULK1 | chr12 | + | 131915334 | 131915421 | 131915082 | 131915231 | 131915890 | 131916159 | 0 | 0 | 5 | 0 | 0 | 0 | 0 | 3 | 6 | 0 | 0 | 1 | 98 | 49 | 0.029628726 | 0.170760496 | NA | NA | 1 | NA | 1 | 0.75 | 0.125 |
| SE | 6193 | FCER1G | FCER1G | chr1 | + | 161217985 | 161218077 | 161215296 | 161215370 | 161218240 | 161218276 | 0 | 7 | 28 | 0 | 0 | 0 | 4 | 10 | 0 | 0 | 1 | 0 | 98 | 49 | 0.029643012 | 0.170760496 | NA | 1 | 1 | 1 | 0.833 | NA | 0.084 |
| A5SS | 1841 | OS9 | OS9 | chr12 | + | 57718145 | 57718421 | 57718145 | 57718376 | 57718992 | 57719182 | 0 | 6 | 6 | 0 | 0 | 0 | 3 | 8 | 0 | 0 | 1 | 0 | 94 | 49 | 0.029835799 | 0.170414363 | NA | 1 | 1 | 1 | 0.807 | NA | 0.097 |
| SE | 11769 | FLII | FLII | chr17 | - | 18255182 | 18255263 | 18254768 | 18254854 | 18256525 | 18256597 | 0 | 6 | 12 | 0 | 0 | 0 | 0 | 9 | 0 | 0 | 1 | 0 | 98 | 49 | 0.03004548 | 0.170760496 | NA | 1 | 1 | NA | 0.818 | NA | 0.182 |
| A3SS | 3631 | SURF4 | SURF4 | chr9 | - | 133367258 | 133367543 | 133367258 | 133367445 | 133375921 | 133376166 | 0 | 0 | 0 | 0 | 1 | 0 | 1 | 2 | 0 | 0 | 1 | 0 | 98 | 49 | 0.030221083 | 0.197717869 | NA | 0 | NA | 1 | 0.5 | NA | -0.75 |
| SE | 388 | ZFAND5 | ZFAND5 | chr9 | - | 72363469 | 72363606 | 72360627 | 72360787 | 72364695 | 72365247 | 0 | 0 | 4 | 0 | 1 | 0 | 0 | 2 | 0 | 0 | 1 | 0 | 98 | 49 | 0.030221083 | 0.170760496 | NA | 0 | 1 | NA | 0.5 | NA | 0 |
| SE | 11350 | ADGRE2 | ADGRE2 | chr19 | - | 14755653 | 14755877 | 14752328 | 14752526 | 14756237 | 14756345 | 0 | 0 | 2 | 0 | 1 | 1 | 0 | 2 | 0 | 0 | 1 | 0 | 98 | 49 | 0.030221083 | 0.170760496 | NA | 0 | 0.5 | NA | 0.5 | NA | -0.25 |
| SE | 5198 | PICALM | PICALM | chr11 | - | 86026291 | 86026367 | 86022366 | 86022469 | 86031468 | 86031611 | 0 | 1 | 33 | 0 | 0 | 0 | 3 | 24 | 1 | 0 | 3 | 0 | 98 | 49 | 0.03042033 | 0.170760496 | NA | 1 | 1 | 1 | 0.8 | 1 | 0.067 |
| SE | 1451 | BAG6 | BAG6 | chr6 | - | 31648910 | 31648964 | 31648676 | 31648751 | 31649198 | 31649395 | 0 | 3 | 2 | 0 | 0 | 0 | 0 | 2 | 0 | 0 | 1 | 0 | 98 | 49 | 0.030496725 | 0.170760496 | NA | 1 | 1 | NA | 0.5 | NA | 0.5 |
| SE | 8360 | ARHGAP30 | ARHGAP30 | chr1 | - | 161059613 | 161059716 | 161056387 | 161056532 | 161069527 | 161069970 | 0 | 1 | 23 | 0 | 0 | 0 | 0 | 8 | 0 | 0 | 1 | 1 | 98 | 49 | 0.030607676 | 0.170760496 | NA | 1 | 1 | NA | 0.8 | 0 | 0.6 |
| SE | 143 | NONO | NONO | chrX | + | 71297835 | 71297938 | 71297376 | 71297461 | 71298468 | 71298508 | 0 | 3 | 2 | 0 | 0 | 0 | 0 | 3 | 0 | 0 | 1 | 0 | 98 | 49 | 0.030693475 | 0.170760496 | NA | 1 | 1 | NA | 0.6 | NA | 0.4 |
| SE | 9279 | PIK3CD | PIK3CD | chr1 | + | 9723968 | 9724092 | 9723124 | 9723292 | 9724275 | 9724421 | 0 | 3 | 12 | 0 | 0 | 0 | 0 | 3 | 0 | 0 | 1 | 0 | 98 | 49 | 0.030693475 | 0.170760496 | NA | 1 | 1 | NA | 0.6 | NA | 0.4 |
| SE | 9304 | SNRNP200 | SNRNP200 | chr2 | - | 96298283 | 96298420 | 96297636 | 96297720 | 96298602 | 96298702 | 0 | 3 | 3 | 0 | 0 | 0 | 0 | 3 | 0 | 0 | 1 | 0 | 98 | 49 | 0.030693475 | 0.170760496 | NA | 1 | 1 | NA | 0.6 | NA | 0.4 |
| SE | 6670 | FGR | FGR | chr1 | - | 27616856 | 27617006 | 27615688 | 27615844 | 27621558 | 27621657 | 0 | 2 | 21 | 0 | 0 | 0 | 0 | 9 | 0 | 0 | 1 | 1 | 98 | 49 | 0.030846049 | 0.170760496 | NA | 1 | 1 | NA | 0.818 | 0 | 0.591 |
| A3SS | 2290 | SF1 | SF1 | chr11 | - | 64764603 | 64766155 | 64764603 | 64765518 | 64766899 | 64767079 | 0 | 3 | 7 | 0 | 0 | 0 | 0 | 4 | 0 | 0 | 1 | 0 | 98 | 49 | 0.03121249 | 0.197717869 | NA | 1 | 1 | NA | 0.667 | NA | 0.333 |
| SE | 6445 | RALGDS | RALGDS | chr9 | - | 133102475 | 133102571 | 133101937 | 133102139 | 133102778 | 133102900 | 0 | 3 | 0 | 0 | 0 | 0 | 0 | 4 | 0 | 0 | 1 | 0 | 98 | 49 | 0.03121249 | 0.170760496 | NA | 1 | NA | NA | 0.667 | NA | 0.333 |
| SE | 54 | ELF4 | ELF4 | chrX | - | 130071039 | 130071232 | 130069299 | 130069677 | 130071335 | 130071419 | 0 | 3 | 7 | 0 | 0 | 0 | 0 | 1 | 0 | 0 | 1 | 0 | 98 | 49 | 0.031636388 | 0.170760496 | NA | 1 | 1 | NA | 0.333 | NA | 0.667 |
| SE | 8271 | ADAM9 | ADAM9 | chr8 | + | 39025802 | 39025884 | 39023155 | 39023325 | 39026676 | 39026810 | 0 | 3 | 4 | 0 | 0 | 0 | 0 | 5 | 0 | 0 | 1 | 0 | 98 | 49 | 0.031777706 | 0.170760496 | NA | 1 | 1 | NA | 0.714 | NA | 0.286 |
| SE | 11767 | KDM5A | KDM5A | chr12 | - | 354075 | 354234 | 352204 | 352324 | 355157 | 355249 | 0 | 3 | 7 | 0 | 0 | 1 | 0 | 5 | 0 | 0 | 1 | 0 | 98 | 49 | 0.031777706 | 0.170760496 | NA | 1 | 1 | 1 | 0.714 | NA | 0.143 |
| SE | 12649 | TLN1 | TLN1 | chr9 | - | 35722860 | 35722921 | 35722118 | 35722223 | 35723951 | 35724079 | 0 | 3 | 26 | 0 | 0 | 2 | 0 | 5 | 0 | 0 | 1 | 0 | 98 | 49 | 0.031777706 | 0.170760496 | NA | 1 | 1 | 1 | 0.714 | NA | 0.143 |
| SE | 13848 | GNLY | GNLY | chr2 | + | 85697505 | 85697677 | 85695957 | 85696056 | 85698563 | 85698851 | 0 | 3 | 2 | 0 | 0 | 0 | 0 | 5 | 0 | 0 | 1 | 0 | 98 | 49 | 0.031777706 | 0.170760496 | NA | 1 | 1 | NA | 0.714 | NA | 0.286 |
| SE | 8338 | CHD2 | CHD2 | chr15 | + | 92901166 | 92901299 | 92900320 | 92900824 | 92924320 | 92924552 | 0 | 4 | 13 | 0 | 0 | 2 | 0 | 8 | 0 | 0 | 1 | 0 | 98 | 49 | 0.031803979 | 0.170760496 | NA | 1 | 0.765 | NA | 0.8 | NA | 0.083 |
| SE | 11875 | MGAM | MGAM | chr7 | + | 142036168 | 142036285 | 142034669 | 142034841 | 142038530 | 142038615 | 0 | 2 | 18 | 0 | 0 | 0 | 2 | 5 | 1 | 0 | 4 | 0 | 98 | 49 | 0.032046501 | 0.170760496 | NA | 1 | 1 | 1 | 0.385 | 1 | 0.205 |
| A5SS | 2853 | CD37 | CD37 | chr19 | + | 49335419 | 49335609 | 49335419 | 49335575 | 49335713 | 49335786 | 0 | 3 | 10 | 0 | 0 | 0 | 0 | 4 | 0 | 0 | 1 | 0 | 83 | 49 | 0.032094074 | 0.170414363 | NA | 1 | 1 | NA | 0.703 | NA | 0.297 |
| SE | 2478 | GSK3B | GSK3B | chr3 | - | 119912703 | 119912810 | 119905754 | 119905852 | 119916043 | 119916174 | 0 | 2 | 8 | 0 | 0 | 2 | 0 | 2 | 0 | 0 | 1 | 0 | 98 | 49 | 0.032484896 | 0.170760496 | NA | 1 | 0.667 | NA | 0.5 | NA | 0.334 |
| SE | 6900 | PSME3 | PSME3 | chr17 | + | 42838691 | 42838799 | 42838092 | 42838205 | 42838944 | 42839012 | 0 | 2 | 1 | 0 | 0 | 0 | 0 | 2 | 0 | 0 | 1 | 0 | 98 | 49 | 0.032484896 | 0.170760496 | NA | 1 | 1 | NA | 0.5 | NA | 0.5 |
| SE | 9190 | GIT2 | GIT2 | chr12 | - | 109980951 | 109981046 | 109967457 | 109967503 | 109983372 | 109983503 | 0 | 2 | 20 | 0 | 0 | 0 | 0 | 2 | 0 | 0 | 1 | 0 | 98 | 49 | 0.032484896 | 0.170760496 | NA | 1 | 1 | NA | 0.5 | NA | 0.5 |
| SE | 13501 | HADHB | HADHB | chr2 | + | 26285406 | 26285571 | 26284882 | 26284957 | 26289917 | 26290465 | 0 | 2 | 11 | 0 | 0 | 0 | 0 | 2 | 0 | 0 | 1 | 0 | 98 | 49 | 0.032484896 | 0.170760496 | NA | 1 | 1 | NA | 0.5 | NA | 0.5 |
| SE | 13782 | SLC20A1 | SLC20A1 | chr2 | + | 112647323 | 112647464 | 112646562 | 112647162 | 112647652 | 112647738 | 0 | 2 | 0 | 0 | 0 | 0 | 0 | 2 | 0 | 0 | 1 | 0 | 98 | 49 | 0.032484896 | 0.170760496 | NA | 1 | NA | NA | 0.5 | NA | 0.5 |
| SE | 4966 | DDX3X | DDX3X | chrX | + | 41347311 | 41347451 | 41346858 | 41347012 | 41347639 | 41347915 | 0 | 1 | 19 | 0 | 0 | 0 | 0 | 34 | 1 | 0 | 3 | 0 | 98 | 49 | 0.032593127 | 0.170760496 | NA | 1 | 1 | NA | 0.85 | 1 | 0.075 |
| SE | 5243 | PLXNC1 | PLXNC1 | chr12 | + | 94298631 | 94298795 | 94297315 | 94297423 | 94300909 | 94301057 | 0 | 4 | 20 | 0 | 0 | 0 | 1 | 10 | 0 | 0 | 1 | 0 | 98 | 49 | 0.03278556 | 0.170760496 | NA | 1 | 1 | 1 | 0.833 | NA | 0.084 |
| A5SS | 1502 | MADD | MADD | chr11 | + | 47284940 | 47285194 | 47284940 | 47285065 | 47285450 | 47285590 | 0 | 2 | 5 | 0 | 0 | 0 | 3 | 3 | 0 | 0 | 1 | 0 | 98 | 49 | 0.032835937 | 0.170414363 | NA | 1 | 1 | 1 | 0.6 | NA | 0.2 |
| SE | 1141 | IGF2R | IGF2R | chr6 | + | 160064803 | 160064901 | 160064400 | 160064531 | 160068248 | 160068385 | 0 | 2 | 17 | 0 | 0 | 3 | 0 | 3 | 0 | 0 | 1 | 0 | 98 | 49 | 0.032835937 | 0.170760496 | NA | 1 | 0.739 | NA | 0.6 | NA | 0.269 |
| SE | 2318 | FAM105A | FAM105A | chr5 | + | 14601350 | 14601442 | 14601212 | 14601244 | 14602182 | 14602332 | 0 | 2 | 3 | 0 | 0 | 2 | 0 | 3 | 0 | 0 | 1 | 0 | 98 | 49 | 0.032835937 | 0.170760496 | NA | 1 | 0.429 | NA | 0.6 | NA | 0.115 |
| SE | 2543 | GAK | GAK | chr4 | - | 877614 | 877809 | 877089 | 877207 | 881906 | 882040 | 0 | 2 | 6 | 0 | 0 | 0 | 0 | 3 | 0 | 0 | 1 | 0 | 98 | 49 | 0.032835937 | 0.170760496 | NA | 1 | 1 | NA | 0.6 | NA | 0.4 |
| SE | 11659 | NLRP1 | NLRP1 | chr17 | - | 5536850 | 5536940 | 5533896 | 5533988 | 5539414 | 5539585 | 0 | 2 | 30 | 0 | 0 | 0 | 0 | 3 | 0 | 0 | 1 | 0 | 98 | 49 | 0.032835937 | 0.170760496 | NA | 1 | 1 | NA | 0.6 | NA | 0.4 |
| SE | 652 | KAT6A | KAT6A | chr8 | - | 41942792 | 41943000 | 41940841 | 41941444 | 41943747 | 41943979 | 0 | 3 | 9 | 0 | 0 | 1 | 0 | 7 | 0 | 0 | 1 | 0 | 98 | 49 | 0.032969828 | 0.170760496 | NA | 1 | 0.818 | NA | 0.778 | NA | 0.131 |
| SE | 1651 | CANX | CANX | chr5 | + | 179708977 | 179709059 | 179708238 | 179708380 | 179709872 | 179710065 | 1 | 3 | 6 | 0 | 0 | 0 | 0 | 7 | 0 | 0 | 1 | 0 | 98 | 49 | 0.032969828 | 0.170760496 | 1 | 1 | 1 | NA | 0.778 | NA | 0.222 |
| A3SS | 3558 | TMEM127 | TMEM127 | chr2 | - | 96265137 | 96265512 | 96265137 | 96265489 | 96265868 | 96266013 | 0 | 4 | 2 | 0 | 3 | 4 | 0 | 6 | 1 | 0 | 0 | 0 | 72 | 49 | 0.033183051 | 0.197717869 | NA | 0.476 | 0.254 | NA | 1 | 1 | -0.635 |
| SE | 13035 | VASP | VASP | chr19 | + | 45522339 | 45522581 | 45522167 | 45522217 | 45522717 | 45522818 | 0 | 4 | 13 | 0 | 0 | 0 | 0 | 11 | 0 | 0 | 1 | 0 | 98 | 49 | 0.033237301 | 0.170760496 | NA | 1 | 1 | NA | 0.846 | NA | 0.154 |
| SE | 10971 | ARHGAP15 | ARHGAP15 | chr2 | + | 143436912 | 143437042 | 143435600 | 143435699 | 143487372 | 143487495 | 0 | 0 | 19 | 0 | 2 | 0 | 0 | 21 | 2 | 0 | 0 | 0 | 98 | 49 | 0.033416613 | 0.170760496 | NA | 0 | 1 | NA | 1 | 1 | -0.5 |
| SE | 2607 | RBM5 | RBM5 | chr3 | + | 50104247 | 50104308 | 50103082 | 50103166 | 50105076 | 50105142 | 0 | 2 | 6 | 0 | 0 | 1 | 0 | 1 | 0 | 3 | 1 | 0 | 98 | 49 | 0.033473796 | 0.170760496 | NA | 1 | 0.75 | 0 | 0.333 | NA | 0.709 |
| SE | 7140 | KIF21B | KIF21B | chr1 | - | 201004765 | 201004933 | 201004339 | 201004455 | 201005307 | 201005442 | 0 | 2 | 3 | 0 | 0 | 0 | 0 | 1 | 0 | 0 | 1 | 0 | 98 | 49 | 0.033473796 | 0.170760496 | NA | 1 | 1 | NA | 0.333 | NA | 0.667 |
| SE | 7425 | H2AFY | H2AFY | chr5 | - | 135352945 | 135353045 | 135345967 | 135346057 | 135360496 | 135360604 | 0 | 2 | 10 | 0 | 0 | 0 | 0 | 1 | 0 | 0 | 1 | 0 | 98 | 49 | 0.033473796 | 0.170760496 | NA | 1 | 1 | NA | 0.333 | NA | 0.667 |
| SE | 7587 | DHX9 | DHX9 | chr1 | + | 182855576 | 182855713 | 182854029 | 182854178 | 182856531 | 182856578 | 0 | 2 | 1 | 0 | 0 | 5 | 0 | 1 | 0 | 0 | 1 | 0 | 98 | 49 | 0.033473796 | 0.170760496 | NA | 1 | 0.091 | NA | 0.333 | NA | 0.212 |
| SE | 9970 | SCYL3 | SCYL3 | chr1 | - | 169870254 | 169870357 | 169868927 | 169869039 | 169873695 | 169873752 | 0 | 2 | 1 | 0 | 0 | 0 | 0 | 1 | 0 | 0 | 1 | 0 | 98 | 49 | 0.033473796 | 0.170760496 | NA | 1 | 1 | NA | 0.333 | NA | 0.667 |
| SE | 12042 | TMC8 | TMC8 | chr17 | + | 78137234 | 78137358 | 78134869 | 78135009 | 78137716 | 78137814 | 0 | 2 | 4 | 0 | 0 | 0 | 0 | 1 | 0 | 0 | 1 | 0 | 98 | 49 | 0.033473796 | 0.170760496 | NA | 1 | 1 | NA | 0.333 | NA | 0.667 |
| SE | 12085 | DCAF8 | DCAF8 | chr1 | - | 160225061 | 160225119 | 160224441 | 160224549 | 160225590 | 160225663 | 0 | 2 | 9 | 0 | 0 | 0 | 2 | 1 | 0 | 0 | 1 | 0 | 98 | 49 | 0.033473796 | 0.170760496 | NA | 1 | 1 | 1 | 0.333 | NA | 0.334 |
| SE | 12285 | ST6GALNAC2 | ST6GALNAC2 | chr17 | - | 76578755 | 76578816 | 76574364 | 76574539 | 76585683 | 76586063 | 0 | 2 | 2 | 0 | 0 | 4 | 0 | 1 | 0 | 0 | 1 | 0 | 98 | 49 | 0.033473796 | 0.170760496 | NA | 1 | 0.2 | NA | 0.333 | NA | 0.267 |
| SE | 12756 | ADGRE5 | ADGRE5 | chr19 | + | 14397076 | 14397223 | 14396341 | 14396473 | 14397657 | 14397804 | 0 | 2 | 8 | 0 | 0 | 0 | 1 | 1 | 0 | 5 | 1 | 0 | 98 | 49 | 0.033473796 | 0.170760496 | NA | 1 | 1 | 0.091 | 0.333 | NA | 0.788 |
| SE | 13958 | DIP2B | DIP2B | chr12 | + | 50692948 | 50693013 | 50691048 | 50691151 | 50695266 | 50695360 | 0 | 2 | 5 | 0 | 0 | 0 | 0 | 1 | 0 | 0 | 1 | 0 | 98 | 49 | 0.033473796 | 0.170760496 | NA | 1 | 1 | NA | 0.333 | NA | 0.667 |
| SE | 10454 | THBS1 | THBS1 | chr15 | + | 39588525 | 39588699 | 39588041 | 39588218 | 39589958 | 39590086 | 0 | 2 | 8 | 0 | 0 | 4 | 0 | 4 | 0 | 0 | 1 | 0 | 98 | 49 | 0.033475981 | 0.170760496 | NA | 1 | 0.5 | NA | 0.667 | NA | 0.083 |
| SE | 13548 | STRN | STRN | chr2 | - | 36855211 | 36855352 | 36850999 | 36851107 | 36857855 | 36858023 | 0 | 2 | 0 | 0 | 0 | 0 | 0 | 4 | 0 | 0 | 1 | 0 | 98 | 49 | 0.033475981 | 0.170760496 | NA | 1 | NA | NA | 0.667 | NA | 0.333 |
| RI | 452 | BIN2 | BIN2 | chr12 | - | 51302019 | 51302780 | 51302019 | 51302115 | 51302685 | 51302780 | 0 | 0 | 1 | 0 | 4 | 5 | 1 | 1 | 0 | 4 | 1 | 0 | 98 | 49 | 0.033604201 | 0.297702651 | NA | 0 | 0.091 | 1 | 0.111 | NA | -0.51 |
| SE | 8340 | CHD2 | CHD2 | chr15 | + | 92945820 | 92945865 | 92942842 | 92943068 | 92946037 | 92946216 | 0 | 2 | 8 | 0 | 0 | 0 | 0 | 4 | 0 | 0 | 1 | 0 | 94 | 49 | 0.033727313 | 0.170760496 | NA | 1 | 1 | NA | 0.676 | NA | 0.324 |
| SE | 4559 | SLC6A6 | SLC6A6 | chr3 | + | 14457949 | 14458082 | 14447581 | 14447816 | 14466515 | 14466650 | 0 | 3 | 36 | 0 | 0 | 0 | 0 | 9 | 2 | 0 | 3 | 0 | 98 | 49 | 0.033971646 | 0.170760496 | NA | 1 | 1 | NA | 0.6 | 1 | 0.2 |
| SE | 2911 | PPP6C | PPP6C | chr9 | - | 125153532 | 125153742 | 125146572 | 125149921 | 125153905 | 125153985 | 0 | 3 | 7 | 0 | 0 | 0 | 0 | 9 | 0 | 0 | 1 | 0 | 98 | 49 | 0.034107533 | 0.170760496 | NA | 1 | 1 | NA | 0.818 | NA | 0.182 |
| SE | 8654 | SVIL | SVIL | chr10 | - | 29555010 | 29555108 | 29554782 | 29554934 | 29563200 | 29563292 | 0 | 3 | 13 | 0 | 0 | 0 | 0 | 9 | 0 | 0 | 1 | 0 | 98 | 49 | 0.034107533 | 0.170760496 | NA | 1 | 1 | NA | 0.818 | NA | 0.182 |
| SE | 5187 | MYO9B | MYO9B | chr19 | + | 17192745 | 17193062 | 17191096 | 17191219 | 17194555 | 17195473 | 0 | 2 | 5 | 0 | 0 | 0 | 0 | 5 | 0 | 0 | 1 | 0 | 98 | 49 | 0.03419934 | 0.170760496 | NA | 1 | 1 | NA | 0.714 | NA | 0.286 |
| SE | 1137 | IGF2R | IGF2R | chr6 | + | 160058022 | 160058124 | 160056423 | 160056525 | 160058905 | 160059098 | 0 | 2 | 35 | 0 | 0 | 0 | 4 | 21 | 1 | 0 | 2 | 0 | 98 | 49 | 0.034257364 | 0.170760496 | NA | 1 | 1 | 1 | 0.84 | 1 | 0.053 |
| SE | 6669 | FGR | FGR | chr1 | - | 27614429 | 27614583 | 27613218 | 27613350 | 27614849 | 27614926 | 0 | 25 | 47 | 0 | 0 | 0 | 2 | 22 | 2 | 0 | 1 | 0 | 98 | 49 | 0.034275909 | 0.170760496 | NA | 1 | 1 | 1 | 0.917 | 1 | 0.028 |
| SE | 8631 | SMG7 | SMG7 | chr1 | + | 183512836 | 183512868 | 183472370 | 183472649 | 183515873 | 183515991 | 0 | 0 | 2 | 0 | 0 | 0 | 0 | 1 | 1 | 0 | 0 | 1 | 81 | 49 | 0.034410483 | 0.170760496 | NA | NA | 1 | NA | 1 | 0.377 | 0.311 |
| SE | 4362 | LRRK2 | LRRK2 | chr12 | + | 40305784 | 40305966 | 40303947 | 40304134 | 40308466 | 40308696 | 0 | 2 | 32 | 0 | 0 | 0 | 6 | 23 | 1 | 0 | 2 | 0 | 98 | 49 | 0.034647673 | 0.170760496 | NA | 1 | 1 | 1 | 0.852 | 1 | 0.049 |
| SE | 691 | DOCK5 | DOCK5 | chr8 | + | 25300575 | 25300657 | 25298943 | 25299101 | 25302324 | 25302454 | 0 | 3 | 24 | 0 | 0 | 0 | 0 | 10 | 0 | 0 | 1 | 0 | 98 | 49 | 0.03465568 | 0.170760496 | NA | 1 | 1 | NA | 0.833 | NA | 0.167 |
| A3SS | 806 | TYMP | TYMP | chr22 | - | 50529495 | 50529754 | 50529495 | 50529731 | 50529903 | 50530085 | 0 | 2 | 4 | 0 | 0 | 1 | 3 | 3 | 0 | 0 | 1 | 0 | 72 | 49 | 0.034683351 | 0.197717869 | NA | 1 | 0.731 | 1 | 0.671 | NA | 0.03 |
| SE | 4962 | DDX3X | DDX3X | chrX | + | 41339035 | 41339083 | 41337407 | 41337465 | 41341483 | 41341616 | 0 | 2 | 15 | 0 | 0 | 0 | 0 | 19 | 3 | 0 | 3 | 0 | 97 | 49 | 0.034732632 | 0.170760496 | NA | 1 | 1 | NA | 0.762 | 1 | 0.119 |
| SE | 9028 | MYD88 | MYD88 | chr3 | + | 38139863 | 38139998 | 38138477 | 38139028 | 38140756 | 38140848 | 0 | 6 | 12 | 0 | 0 | 0 | 0 | 11 | 3 | 0 | 2 | 0 | 98 | 49 | 0.03475926 | 0.170760496 | NA | 1 | 1 | NA | 0.733 | 1 | 0.133 |
| A3SS | 2212 | GNAS | GNAS | chr20 | + | 58903527 | 58903585 | 58903530 | 58903585 | 58895611 | 58895684 | 0 | 3 | 4 | 0 | 0 | 2 | 0 | 4 | 0 | 0 | 1 | 0 | 52 | 49 | 0.034903266 | 0.197717869 | NA | 1 | 0.653 | NA | 0.79 | NA | 0.036 |
| SE | 3591 | TRIP12 | TRIP12 | chr2 | - | 229778875 | 229778990 | 229778432 | 229778587 | 229785756 | 229785855 | 0 | 2 | 11 | 0 | 0 | 0 | 0 | 6 | 0 | 0 | 1 | 0 | 98 | 49 | 0.034934183 | 0.170760496 | NA | 1 | 1 | NA | 0.75 | NA | 0.25 |
| SE | 863 | NAMPT | NAMPT | chr7 | - | 106268463 | 106268600 | 106263391 | 106263617 | 106269153 | 106269312 | 0 | 9 | 124 | 0 | 0 | 0 | 0 | 88 | 7 | 0 | 2 | 0 | 98 | 49 | 0.035105524 | 0.170760496 | NA | 1 | 1 | NA | 0.957 | 1 | 0.022 |
| SE | 7287 | HDAC5 | HDAC5 | chr17 | - | 44078794 | 44078879 | 44078499 | 44078665 | 44079143 | 44079277 | 0 | 1 | 2 | 0 | 0 | 0 | 0 | 5 | 1 | 0 | 3 | 0 | 98 | 49 | 0.035115165 | 0.170760496 | NA | 1 | 1 | NA | 0.455 | 1 | 0.272 |
| SE | 9359 | FERMT3 | FERMT3 | chr11 | + | 64219523 | 64219658 | 64219250 | 64219358 | 64219739 | 64219789 | 0 | 3 | 13 | 0 | 0 | 0 | 0 | 11 | 0 | 0 | 1 | 0 | 98 | 49 | 0.03520334 | 0.170760496 | NA | 1 | 1 | NA | 0.846 | NA | 0.154 |
| SE | 7828 | SRGN | SRGN | chr10 | + | 69097083 | 69097231 | 69088071 | 69088236 | 69103870 | 69104811 | 0 | 28 | 397 | 0 | 0 | 0 | 31 | 320 | 22 | 0 | 2 | 0 | 98 | 49 | 0.035427964 | 0.170760496 | NA | 1 | 1 | 1 | 0.988 | 1 | 0.004 |
| SE | 8959 | N4BP2L2 | N4BP2L2 | chr13 | - | 32527407 | 32527532 | 32522181 | 32522270 | 32535768 | 32537027 | 0 | 2 | 9 | 0 | 0 | 1 | 0 | 7 | 0 | 0 | 1 | 0 | 98 | 49 | 0.0356372 | 0.170760496 | NA | 1 | 0.818 | NA | 0.778 | NA | 0.131 |
| A3SS | 1851 | OAS1 | OAS1 | chr12 | + | 112919388 | 112919907 | 112919486 | 112919907 | 112917546 | 112917700 | 0 | 3 | 5 | 0 | 0 | 4 | 0 | 15 | 2 | 0 | 0 | 0 | 98 | 49 | 0.035922391 | 0.197717869 | NA | 1 | 0.385 | NA | 1 | 1 | -0.307 |
| A5SS | 478 | UBE2D3 | UBE2D3 | chr4 | - | 102827426 | 102827948 | 102827591 | 102827948 | 102826484 | 102826636 | 0 | 2 | 2 | 0 | 0 | 2 | 1 | 9 | 3 | 0 | 0 | 0 | 98 | 49 | 0.036267739 | 0.170414363 | NA | 1 | 0.333 | 1 | 1 | 1 | -0.334 |
| SE | 14036 | CFLAR | CFLAR | chr2 | + | 201135971 | 201136107 | 201132964 | 201133134 | 201140356 | 201140439 | 1 | 6 | 52 | 0 | 0 | 0 | 0 | 29 | 0 | 0 | 1 | 0 | 98 | 49 | 0.0362875 | 0.170760496 | 1 | 1 | 1 | NA | 0.935 | NA | 0.065 |
| SE | 1368 | RGL2 | RGL2 | chr6 | - | 33294687 | 33294762 | 33294033 | 33294066 | 33294976 | 33295045 | 0 | 2 | 17 | 0 | 0 | 0 | 2 | 8 | 0 | 0 | 1 | 0 | 98 | 49 | 0.036328243 | 0.170760496 | NA | 1 | 1 | 1 | 0.8 | NA | 0.1 |
| SE | 4793 | CASP1 | CASP1 | chr11 | - | 105026225 | 105026356 | 105025509 | 105025600 | 105026841 | 105026951 | 0 | 2 | 29 | 0 | 0 | 0 | 0 | 8 | 0 | 1 | 1 | 0 | 98 | 49 | 0.036328243 | 0.170760496 | NA | 1 | 1 | 0 | 0.8 | NA | 0.6 |
| SE | 6901 | PSME3 | PSME3 | chr17 | + | 42838730 | 42838799 | 42838092 | 42838205 | 42838944 | 42839012 | 0 | 2 | 5 | 0 | 0 | 0 | 0 | 8 | 0 | 0 | 1 | 0 | 98 | 49 | 0.036328243 | 0.170760496 | NA | 1 | 1 | NA | 0.8 | NA | 0.2 |
| SE | 338 | DNAJC25 | DNAJC25 | chr9 | + | 111642856 | 111643094 | 111631351 | 111631743 | 111647106 | 111647259 | 0 | 1 | 1 | 0 | 0 | 0 | 0 | 2 | 0 | 0 | 1 | 0 | 98 | 49 | 0.036515846 | 0.170760496 | NA | 1 | 1 | NA | 0.5 | NA | 0.5 |
| SE | 2653 | CCDC12 | CCDC12 | chr3 | - | 46923606 | 46923668 | 46923328 | 46923363 | 46925455 | 46925535 | 0 | 1 | 1 | 0 | 0 | 0 | 0 | 2 | 0 | 0 | 1 | 0 | 98 | 49 | 0.036515846 | 0.170760496 | NA | 1 | 1 | NA | 0.5 | NA | 0.5 |
| SE | 3010 | MED15 | MED15 | chr22 | + | 20584854 | 20585015 | 20584358 | 20584425 | 20585100 | 20585267 | 0 | 1 | 1 | 0 | 0 | 0 | 0 | 2 | 0 | 0 | 1 | 0 | 98 | 49 | 0.036515846 | 0.170760496 | NA | 1 | 1 | NA | 0.5 | NA | 0.5 |
| SE | 6389 | SUPT6H | SUPT6H | chr17 | + | 28690920 | 28691063 | 28690081 | 28690229 | 28693695 | 28693836 | 0 | 1 | 2 | 0 | 0 | 0 | 0 | 2 | 0 | 0 | 1 | 0 | 98 | 49 | 0.036515846 | 0.170760496 | NA | 1 | 1 | NA | 0.5 | NA | 0.5 |
| SE | 6793 | PLEKHM2 | PLEKHM2 | chr1 | + | 15719733 | 15719920 | 15718537 | 15718625 | 15721328 | 15721388 | 0 | 1 | 2 | 0 | 0 | 0 | 0 | 2 | 0 | 0 | 1 | 0 | 98 | 49 | 0.036515846 | 0.170760496 | NA | 1 | 1 | NA | 0.5 | NA | 0.5 |
| SE | 7794 | G6PD | G6PD | chrX | - | 154532941 | 154533128 | 154532566 | 154532802 | 154533575 | 154533669 | 0 | 1 | 4 | 0 | 0 | 0 | 0 | 2 | 0 | 0 | 1 | 0 | 98 | 49 | 0.036515846 | 0.170760496 | NA | 1 | 1 | NA | 0.5 | NA | 0.5 |
| SE | 8149 | TTC39C | TTC39C | chr18 | + | 24069156 | 24069271 | 24066011 | 24066140 | 24080584 | 24080939 | 0 | 1 | 2 | 0 | 0 | 0 | 0 | 2 | 0 | 0 | 1 | 0 | 98 | 49 | 0.036515846 | 0.170760496 | NA | 1 | 1 | NA | 0.5 | NA | 0.5 |
| SE | 8182 | LSM14A | LSM14A | chr19 | + | 34215123 | 34215300 | 34208928 | 34209051 | 34215595 | 34215661 | 0 | 1 | 7 | 0 | 0 | 0 | 0 | 2 | 0 | 0 | 1 | 0 | 98 | 49 | 0.036515846 | 0.170760496 | NA | 1 | 1 | NA | 0.5 | NA | 0.5 |
| SE | 9120 | CR1 | CR1 | chr1 | + | 207609289 | 207609688 | 207607250 | 207607336 | 207611676 | 207611853 | 0 | 1 | 20 | 0 | 0 | 0 | 0 | 2 | 0 | 0 | 1 | 0 | 98 | 49 | 0.036515846 | 0.170760496 | NA | 1 | 1 | NA | 0.5 | NA | 0.5 |
| SE | 9391 | ZNF346 | ZNF346 | chr5 | + | 177023159 | 177023234 | 177022679 | 177022913 | 177041125 | 177041229 | 0 | 1 | 0 | 0 | 0 | 0 | 0 | 2 | 0 | 0 | 1 | 0 | 98 | 49 | 0.036515846 | 0.170760496 | NA | 1 | NA | NA | 0.5 | NA | 0.5 |
| SE | 10262 | SNX17 | SNX17 | chr2 | + | 27372622 | 27372740 | 27371268 | 27371343 | 27373039 | 27373113 | 0 | 1 | 3 | 0 | 0 | 0 | 0 | 2 | 0 | 0 | 1 | 0 | 98 | 49 | 0.036515846 | 0.170760496 | NA | 1 | 1 | NA | 0.5 | NA | 0.5 |
| SE | 10380 | NFKBIE | NFKBIE | chr6 | - | 44262559 | 44262662 | 44261625 | 44261848 | 44264981 | 44265788 | 0 | 1 | 2 | 0 | 0 | 0 | 0 | 2 | 0 | 0 | 1 | 0 | 98 | 49 | 0.036515846 | 0.170760496 | NA | 1 | 1 | NA | 0.5 | NA | 0.5 |
| SE | 12595 | GBF1 | GBF1 | chr10 | + | 102363255 | 102363396 | 102362474 | 102362664 | 102363709 | 102363798 | 0 | 1 | 0 | 0 | 0 | 0 | 0 | 2 | 0 | 0 | 1 | 0 | 98 | 49 | 0.036515846 | 0.170760496 | NA | 1 | NA | NA | 0.5 | NA | 0.5 |
| SE | 12839 | DENND1B | DENND1B | chr1 | - | 197647054 | 197647114 | 197645689 | 197645743 | 197652234 | 197652315 | 0 | 1 | 0 | 0 | 0 | 0 | 0 | 2 | 0 | 0 | 1 | 0 | 98 | 49 | 0.036515846 | 0.170760496 | NA | 1 | NA | NA | 0.5 | NA | 0.5 |
| SE | 13738 | SERPINA1 | SERPINA1 | chr14 | - | 94388818 | 94389010 | 94388559 | 94388663 | 94390456 | 94390692 | 0 | 1 | 17 | 0 | 0 | 4 | 0 | 2 | 0 | 0 | 1 | 0 | 98 | 49 | 0.036515846 | 0.170760496 | NA | 1 | 0.68 | NA | 0.5 | NA | 0.34 |
| SE | 347 | OGT | OGT | chrX | + | 71561774 | 71561900 | 71559587 | 71559677 | 71562846 | 71563017 | 0 | 3 | 24 | 0 | 0 | 0 | 5 | 14 | 0 | 0 | 1 | 0 | 98 | 49 | 0.036639257 | 0.170760496 | NA | 1 | 1 | 1 | 0.875 | NA | 0.062 |
| SE | 7594 | ATG16L2 | ATG16L2 | chr11 | + | 72826517 | 72826589 | 72826172 | 72826243 | 72826702 | 72826823 | 0 | 12 | 63 | 0 | 0 | 0 | 11 | 29 | 8 | 0 | 2 | 0 | 98 | 49 | 0.036924565 | 0.170760496 | NA | 1 | 1 | 1 | 0.879 | 1 | 0.04 |
| SE | 3590 | TRIP12 | TRIP12 | chr2 | - | 229771518 | 229771632 | 229769230 | 229769325 | 229774096 | 229774261 | 0 | 2 | 2 | 0 | 0 | 0 | 0 | 9 | 0 | 0 | 1 | 0 | 98 | 49 | 0.037002078 | 0.170760496 | NA | 1 | 1 | NA | 0.818 | NA | 0.182 |
| SE | 8756 | ZC3H11A | ZC3H11A | chr1 | + | 203848330 | 203848407 | 203847183 | 203847687 | 203849710 | 203850026 | 0 | 2 | 10 | 0 | 0 | 0 | 0 | 9 | 0 | 0 | 1 | 0 | 98 | 49 | 0.037002078 | 0.170760496 | NA | 1 | 1 | NA | 0.818 | NA | 0.182 |
| SE | 279 | EXOSC2 | EXOSC2 | chr9 | + | 130698161 | 130698251 | 130697581 | 130697627 | 130699328 | 130699394 | 0 | 1 | 0 | 0 | 0 | 0 | 0 | 1 | 0 | 0 | 1 | 0 | 98 | 49 | 0.037136978 | 0.170760496 | NA | 1 | NA | NA | 0.333 | NA | 0.667 |
| SE | 3510 | HDLBP | HDLBP | chr2 | - | 241255373 | 241255580 | 241255050 | 241255158 | 241256183 | 241256399 | 0 | 1 | 1 | 0 | 0 | 0 | 0 | 1 | 0 | 0 | 1 | 0 | 98 | 49 | 0.037136978 | 0.170760496 | NA | 1 | 1 | NA | 0.333 | NA | 0.667 |

| | | | | | | | | | | | | | | | | | | | | | | | | | | | | | | | | | | |
|---|---|---|---|---|---|---|---|---|---|---|---|---|---|---|---|---|---|---|---|---|---|---|---|---|---|---|---|---|---|---|---|---|---|---|
| SE | 4017 | AGAP2 | AGAP2 | chr12 | - | 57731808 | 57731967 | 57731555 | 57731642 | 57732402 | 57732512 | 0 | 1 | 1 | 0 | 0 | 0 | 4 | 1 | 0 | 0 | 1 | 0 | 98 | 49 | 0.037136978 | 0.170760496 | NA | 1 | 1 | 1 | 0.333 | NA | 0.334 |
| SE | 4033 | TRAFD1 | TRAFD1 | chr12 | + | 112151800 | 112152140 | 112149750 | 112149871 | 112152426 | 112152499 | 0 | 1 | 5 | 0 | 0 | 0 | 0 | 1 | 0 | 0 | 1 | 0 | 98 | 49 | 0.037136978 | 0.170760496 | NA | 1 | 1 | NA | 0.333 | NA | 0.667 |
| SE | 5121 | CCM2 | CCM2 | chr7 | + | 45038252 | 45038426 | 45027633 | 45027797 | 45063917 | 45064001 | 0 | 1 | 0 | 0 | 0 | 0 | 0 | 1 | 0 | 0 | 1 | 0 | 98 | 49 | 0.037136978 | 0.170760496 | NA | 1 | NA | NA | 0.333 | NA | 0.667 |
| SE | 6443 | ATP11A | ATP11A | chr13 | + | 112858144 | 112858290 | 112857817 | 112857920 | 112859392 | 112859452 | 0 | 1 | 1 | 0 | 0 | 1 | 0 | 1 | 0 | 0 | 1 | 0 | 98 | 49 | 0.037136978 | 0.170760496 | NA | 1 | 0.333 | NA | 0.333 | NA | 0.333 |
| SE | 6888 | SMC3 | SMC3 | chr10 | + | 110590990 | 110591132 | 110590411 | 110590572 | 110593072 | 110593223 | 0 | 1 | 3 | 0 | 0 | 0 | 0 | 1 | 0 | 0 | 1 | 0 | 98 | 49 | 0.037136978 | 0.170760496 | NA | 1 | 1 | NA | 0.333 | NA | 0.667 |
| SE | 10311 | KLC1 | KLC1 | chr14 | + | 103657545 | 103657776 | 103654563 | 103654825 | 103662115 | 103662194 | 0 | 1 | 8 | 0 | 0 | 0 | 0 | 1 | 0 | 0 | 1 | 0 | 98 | 49 | 0.037136978 | 0.170760496 | NA | 1 | 1 | NA | 0.333 | NA | 0.667 |
| SE | 11210 | GYPC | GYPC | chr2 | + | 126690254 | 126690311 | 126655934 | 126656312 | 126693863 | 126693947 | 0 | 1 | 0 | 0 | 0 | 0 | 0 | 1 | 0 | 0 | 1 | 0 | 98 | 49 | 0.037136978 | 0.170760496 | NA | 1 | NA | NA | 0.333 | NA | 0.667 |
| SE | 13735 | SERPINA1 | SERPINA1 | chr14 | - | 94388800 | 94389010 | 94388559 | 94388663 | 94390456 | 94390692 | 0 | 1 | 17 | 0 | 0 | 4 | 0 | 1 | 0 | 0 | 1 | 0 | 98 | 49 | 0.037136978 | 0.170760496 | NA | 1 | 0.68 | NA | 0.333 | NA | 0.507 |
| SE | 13855 | FBXW11 | FBXW11 | chr5 | - | 171914342 | 171914405 | 171910571 | 171910797 | 172006457 | 172006873 | 0 | 0 | 1 | 0 | 0 | 0 | 0 | 1 | 1 | 0 | 0 | 1 | 98 | 49 | 0.037136978 | 0.170760496 | NA | NA | 1 | NA | 1 | 0.333 | 0.334 |
| SE | 1807 | ASAP1 | ASAP1 | chr8 | - | 130091972 | 130092143 | 130079901 | 130079971 | 130112093 | 130112322 | 0 | 1 | 0 | 0 | 0 | 1 | 0 | 3 | 0 | 0 | 1 | 0 | 98 | 49 | 0.03722674 | 0.170760496 | NA | 1 | 0 | NA | 0.6 | NA | -0.1 |
| SE | 3240 | ZFAS1 | ZFAS1 | chr20 | + | 49280902 | 49280964 | 49280484 | 49280570 | 49289044 | 49289258 | 0 | 1 | 12 | 0 | 0 | 1 | 0 | 3 | 0 | 0 | 1 | 0 | 98 | 49 | 0.03722674 | 0.170760496 | NA | 1 | 0.857 | NA | 0.6 | NA | 0.329 |
| SE | 5144 | KIAA0226L | KIAA0226L | chr13 | - | 46371940 | 46372210 | 46368732 | 46368815 | 46387133 | 46387500 | 0 | 1 | 1 | 0 | 0 | 0 | 0 | 3 | 0 | 0 | 1 | 0 | 98 | 49 | 0.03722674 | 0.170760496 | NA | 1 | 1 | NA | 0.6 | NA | 0.4 |
| SE | 5505 | SBF2 | SBF2 | chr11 | - | 10031047 | 10031170 | 10029764 | 10029875 | 10042843 | 10042981 | 0 | 1 | 12 | 0 | 0 | 0 | 0 | 3 | 0 | 0 | 1 | 0 | 98 | 49 | 0.03722674 | 0.170760496 | NA | 1 | 1 | NA | 0.6 | NA | 0.4 |
| SE | 6391 | KMT2B | KMT2B | chr19 | + | 35736689 | 35736827 | 35733762 | 35733872 | 35736911 | 35736986 | 0 | 1 | 1 | 0 | 0 | 0 | 0 | 3 | 0 | 0 | 1 | 0 | 98 | 49 | 0.03722674 | 0.170760496 | NA | 1 | 1 | NA | 0.6 | NA | 0.4 |
| SE | 8166 | ARL5B | ARL5B | chr10 | + | 18672621 | 18672705 | 18668529 | 18668677 | 18673983 | 18674135 | 0 | 1 | 5 | 0 | 0 | 0 | 0 | 3 | 0 | 0 | 1 | 0 | 98 | 49 | 0.03722674 | 0.170760496 | NA | 1 | 1 | NA | 0.6 | NA | 0.4 |
| SE | 9178 | SH3BP1 | SH3BP1 | chr22 | + | 37644869 | 37644960 | 37644636 | 37644705 | 37645364 | 37645510 | 0 | 1 | 4 | 0 | 0 | 0 | 2 | 3 | 0 | 0 | 1 | 0 | 98 | 49 | 0.03722674 | 0.170760496 | NA | 1 | 1 | 1 | 0.6 | NA | 0.2 |
| SE | 10631 | VPS13C | VPS13C | chr15 | - | 61951823 | 61951980 | 61950944 | 61951024 | 61954420 | 61954554 | 0 | 1 | 3 | 0 | 0 | 0 | 0 | 3 | 0 | 0 | 1 | 0 | 98 | 49 | 0.03722674 | 0.170760496 | NA | 1 | 1 | NA | 0.6 | NA | 0.4 |
| SE | 10953 | FES | FES | chr15 | + | 90889516 | 90889636 | 90889305 | 90889443 | 90889839 | 90889962 | 0 | 1 | 7 | 0 | 0 | 0 | 0 | 3 | 0 | 0 | 1 | 0 | 98 | 49 | 0.03722674 | 0.170760496 | NA | 1 | 1 | NA | 0.6 | NA | 0.4 |
| SE | 11935 | STK40 | STK40 | chr1 | - | 36367847 | 36368084 | 36361220 | 36361340 | 36385722 | 36385927 | 0 | 1 | 1 | 0 | 0 | 0 | 0 | 3 | 0 | 0 | 1 | 0 | 98 | 49 | 0.03722674 | 0.170760496 | NA | 1 | 1 | NA | 0.6 | NA | 0.4 |
| SE | 13687 | SIRPA | SIRPA | chr20 | + | 1924763 | 1924877 | 1915098 | 1915455 | 1927874 | 1927899 | 0 | 1 | 10 | 0 | 0 | 0 | 0 | 3 | 0 | 0 | 1 | 0 | 98 | 49 | 0.03722674 | 0.170760496 | NA | 1 | 1 | NA | 0.6 | NA | 0.4 |
| SE | 331 | AKNA | AKNA | chr9 | - | 114343707 | 114343803 | 114342008 | 114342125 | 114345862 | 114346009 | 0 | 0 | 10 | 0 | 2 | 2 | 0 | 1 | 0 | 0 | 1 | 0 | 98 | 49 | 0.037261148 | 0.170760496 | NA | 0 | 0.714 | NA | 0.333 | NA | 0.024 |
| SE | 6388 | SUPT6H | SUPT6H | chr17 | + | 28682941 | 28683092 | 28682726 | 28682856 | 28683267 | 28683422 | 0 | 0 | 1 | 0 | 2 | 0 | 0 | 1 | 0 | 0 | 1 | 0 | 98 | 49 | 0.037261148 | 0.170760496 | NA | 0 | 1 | NA | 0.333 | NA | 0.167 |
| SE | 7325 | SREBF1 | SREBF1 | chr17 | - | 17814614 | 17814747 | 17814244 | 17814410 | 17814834 | 17814944 | 0 | 0 | 0 | 0 | 2 | 0 | 0 | 1 | 0 | 0 | 1 | 0 | 98 | 49 | 0.037261148 | 0.170760496 | NA | 0 | NA | NA | 0.333 | NA | -0.333 |
| SE | 559 | PABPC1 | PABPC1 | chr8 | - | 100718086 | 100718280 | 100717772 | 100717888 | 100721390 | 100722087 | 0 | 0 | 29 | 0 | 1 | 0 | 1 | 19 | 6 | 0 | 0 | 0 | 98 | 49 | 0.037496585 | 0.170760496 | NA | 0 | 1 | 1 | 1 | 1 | -0.5 |
| SE | 371 | AGTPBP1 | AGTPBP1 | chr9 | - | 85678334 | 85678398 | 85677435 | 85677582 | 85681267 | 85681335 | 0 | 2 | 31 | 0 | 0 | 0 | 0 | 10 | 0 | 0 | 1 | 0 | 98 | 49 | 0.037649757 | 0.170760496 | NA | 1 | 1 | NA | 0.833 | NA | 0.167 |
| SE | 5702 | PPP4C | PPP4C | chr16 | + | 30083654 | 30083781 | 30083393 | 30083567 | 30084665 | 30084855 | 0 | 2 | 14 | 0 | 0 | 0 | 0 | 10 | 0 | 0 | 1 | 0 | 98 | 49 | 0.037649757 | 0.170760496 | NA | 1 | 1 | NA | 0.833 | NA | 0.167 |
| SE | 11023 | TAP1 | TAP1 | chr6 | - | 32848989 | 32849118 | 32848651 | 32848840 | 32850319 | 32850517 | 1 | 2 | 34 | 0 | 0 | 0 | 0 | 10 | 0 | 0 | 1 | 0 | 98 | 49 | 0.037649757 | 0.170760496 | 1 | 1 | 1 | NA | 0.833 | NA | 0.167 |
| A3SS | 1947 | SNRPB | SNRPB | chr20 | - | 2461634 | 2461939 | 2461634 | 2461793 | 2462635 | 2462761 | 0 | 0 | 3 | 0 | 1 | 2 | 7 | 5 | 1 | 0 | 0 | 0 | 98 | 49 | 0.037878306 | 0.197717869 | NA | 0 | 0.429 | 1 | 1 | 1 | -0.785 |
| SE | 370 | AGTPBP1 | AGTPBP1 | chr9 | - | 85660935 | 85660973 | 85657434 | 85657643 | 85669484 | 85669578 | 0 | 1 | 9 | 0 | 0 | 0 | 0 | 3 | 0 | 0 | 1 | 0 | 87 | 49 | 0.038097905 | 0.170760496 | NA | 1 | 1 | NA | 0.628 | NA | 0.372 |
| SE | 5887 | SIPA1L1 | SIPA1L1 | chr14 | + | 71650334 | 71650509 | 71624047 | 71624236 | 71658332 | 71658436 | 0 | 1 | 14 | 0 | 0 | 0 | 0 | 4 | 0 | 0 | 1 | 0 | 98 | 49 | 0.038206571 | 0.170760496 | NA | 1 | 1 | NA | 0.667 | NA | 0.333 |
| SE | 8614 | TM9SF3 | TM9SF3 | chr10 | - | 96533050 | 96533190 | 96530539 | 96530608 | 96544075 | 96544206 | 0 | 1 | 11 | 0 | 0 | 0 | 0 | 4 | 0 | 0 | 1 | 0 | 98 | 49 | 0.038206571 | 0.170760496 | NA | 1 | 1 | NA | 0.667 | NA | 0.333 |
| SE | 11155 | NDE1 | NDE1 | chr16 | + | 15694164 | 15694256 | 15691143 | 15691323 | 15696708 | 15696860 | 0 | 1 | 16 | 0 | 0 | 0 | 0 | 4 | 0 | 0 | 1 | 0 | 98 | 49 | 0.038206571 | 0.170760496 | NA | 1 | 1 | NA | 0.667 | NA | 0.333 |
| SE | 11295 | CREBBP | CREBBP | chr16 | - | 3782683 | 3782926 | 3781203 | 3781306 | 3791980 | 3792094 | 0 | 1 | 6 | 0 | 0 | 0 | 0 | 4 | 0 | 0 | 1 | 0 | 98 | 49 | 0.038206571 | 0.170760496 | NA | 1 | 1 | NA | 0.667 | NA | 0.333 |
| SE | 12432 | SPAG9 | SPAG9 | chr17 | - | 50974770 | 50974947 | 50970706 | 50970856 | 50977107 | 50977221 | 0 | 1 | 3 | 0 | 0 | 0 | 0 | 4 | 0 | 0 | 1 | 0 | 98 | 49 | 0.038206571 | 0.170760496 | NA | 1 | 1 | NA | 0.667 | NA | 0.333 |
| SE | 3199 | TNIP1 | TNIP1 | chr5 | - | 151060317 | 151060395 | 151056765 | 151056957 | 151062126 | 151062212 | 0 | 2 | 8 | 0 | 0 | 0 | 0 | 11 | 0 | 0 | 1 | 0 | 98 | 49 | 0.038274566 | 0.170760496 | NA | 1 | 1 | NA | 0.846 | NA | 0.154 |
| SE | 349 | NR4A3 | NR4A3 | chr9 | + | 99832688 | 99832818 | 99828040 | 99828993 | 99833281 | 99833454 | 0 | 1 | 3 | 0 | 0 | 0 | 0 | 7 | 1 | 0 | 2 | 0 | 98 | 49 | 0.038321227 | 0.170760496 | NA | 1 | 1 | NA | 0.636 | 1 | 0.182 |
| SE | 11639 | ARRB2 | ARRB2 | chr17 | + | 4716146 | 4716191 | 4715972 | 4716033 | 4716411 | 4716608 | 0 | 6 | 32 | 0 | 0 | 1 | 0 | 13 | 1 | 0 | 5 | 2 | 94 | 49 | 0.038650672 | 0.170760496 | NA | 1 | 0.943 | NA | 0.575 | 0.207 | 0.581 |
| SE | 9931 | PTPRE | PTPRE | chr10 | + | 128066074 | 128066194 | 128063082 | 128063180 | 128068122 | 128068286 | 0 | 4 | 51 | 0 | 0 | 0 | 0 | 47 | 9 | 0 | 3 | 0 | 98 | 49 | 0.038824113 | 0.170760496 | NA | 1 | 1 | NA | 0.887 | 1 | 0.056 |
| SE | 11065 | CORO1A | CORO1A | chr16 | + | 30187941 | 30188087 | 30187724 | 30187829 | 30188191 | 30188249 | 0 | 18 | 59 | 0 | 0 | 0 | 5 | 24 | 3 | 0 | 1 | 0 | 98 | 49 | 0.038917622 | 0.170760496 | NA | 1 | 1 | 1 | 0.923 | 1 | 0.026 |
| A5SS | 3018 | SERPINA1 | SERPINA1 | chr14 | - | 94388800 | 94389010 | 94388818 | 94389010 | 94388559 | 94388663 | 0 | 1 | 6 | 0 | 0 | 2 | 0 | 2 | 0 | 0 | 1 | 0 | 67 | 49 | 0.039150683 | 0.170414363 | NA | 1 | 0.687 | NA | 0.594 | NA | 0.25 |
| A3SS | 3771 | STRN4 | STRN4 | chr19 | - | 46727451 | 46727567 | 46727451 | 46727546 | 46727893 | 46728007 | 0 | 1 | 1 | 0 | 0 | 4 | 0 | 1 | 0 | 0 | 1 | 0 | 70 | 49 | 0.039208323 | 0.197717869 | NA | 1 | 0.149 | NA | 0.412 | NA | 0.163 |
| SE | 2205 | BOD1L1 | BOD1L1 | chr4 | - | 13595859 | 13595944 | 13591922 | 13591966 | 13597103 | 13597168 | 0 | 1 | 12 | 0 | 0 | 0 | 0 | 5 | 0 | 0 | 1 | 0 | 98 | 49 | 0.039248064 | 0.170760496 | NA | 1 | 1 | NA | 0.714 | NA | 0.286 |
| SE | 2551 | ATXN7 | ATXN7 | chr3 | + | 63913156 | 63913225 | 63912587 | 63912923 | 63952378 | 63952483 | 0 | 1 | 7 | 0 | 0 | 0 | 0 | 5 | 0 | 0 | 1 | 0 | 98 | 49 | 0.039248064 | 0.170760496 | NA | 1 | 1 | NA | 0.714 | NA | 0.286 |
| SE | 4158 | SH3BP2 | SH3BP2 | chr4 | + | 2831922 | 2831978 | 2831570 | 2831679 | 2832330 | 2832412 | 0 | 1 | 12 | 0 | 0 | 1 | 1 | 5 | 0 | 0 | 1 | 0 | 98 | 49 | 0.039248064 | 0.170760496 | NA | 1 | 0.857 | 1 | 0.714 | NA | 0.072 |
| SE | 4319 | BIRC6 | BIRC6 | chr2 | + | 32441328 | 32441462 | 32439507 | 32439686 | 32442064 | 32442226 | 0 | 1 | 5 | 0 | 0 | 0 | 0 | 5 | 0 | 0 | 1 | 0 | 98 | 49 | 0.039248064 | 0.170760496 | NA | 1 | 1 | NA | 0.714 | NA | 0.286 |
| SE | 5146 | KIAA0226L | KIAA0226L | chr13 | - | 46371940 | 46372597 | 46368732 | 46368815 | 46387133 | 46387500 | 0 | 1 | 1 | 0 | 0 | 0 | 0 | 5 | 0 | 0 | 1 | 0 | 98 | 49 | 0.039248064 | 0.170760496 | NA | 1 | 1 | NA | 0.714 | NA | 0.286 |
| SE | 5222 | CLEC4E | CLEC4E | chr12 | - | 8537114 | 8537266 | 8536089 | 8536205 | 8539216 | 8539306 | 0 | 1 | 35 | 0 | 0 | 0 | 0 | 5 | 0 | 0 | 1 | 0 | 98 | 49 | 0.039248064 | 0.170760496 | NA | 1 | 1 | NA | 0.714 | NA | 0.286 |
| SE | 5861 | DISC1 | DISC1 | chr1 | + | 231795241 | 231795296 | 231770834 | 231771070 | 231818328 | 231819244 | 0 | 1 | 7 | 0 | 0 | 0 | 0 | 5 | 0 | 0 | 1 | 0 | 98 | 49 | 0.039248064 | 0.170760496 | NA | 1 | 1 | NA | 0.714 | NA | 0.286 |
| SE | 6191 | FOXN2 | FOXN2 | chr2 | + | 48328560 | 48328702 | 48314655 | 48314814 | 48346200 | 48346751 | 0 | 1 | 2 | 0 | 0 | 1 | 0 | 5 | 0 | 0 | 1 | 0 | 98 | 49 | 0.039248064 | 0.170760496 | NA | 1 | 0.5 | NA | 0.714 | NA | 0.036 |
| SE | 8721 | NOD2 | NOD2 | chr16 | + | 50707854 | 50707960 | 50699349 | 50699954 | 50710557 | 50712373 | 0 | 1 | 3 | 0 | 0 | 0 | 0 | 5 | 0 | 0 | 1 | 0 | 98 | 49 | 0.039248064 | 0.170760496 | NA | 1 | 1 | NA | 0.714 | NA | 0.286 |
| SE | 8817 | KDM2A | KDM2A | chr11 | + | 67243008 | 67243092 | 67231565 | 67231960 | 67245188 | 67245458 | 0 | 1 | 11 | 0 | 0 | 0 | 2 | 5 | 0 | 0 | 1 | 0 | 98 | 49 | 0.039248064 | 0.170760496 | NA | 1 | 1 | 1 | 0.714 | NA | 0.143 |
| SE | 9383 | PECAM1 | PECAM1 | chr17 | - | 64354932 | 64355040 | 64353490 | 64353518 | 64356110 | 64356398 | 0 | 1 | 38 | 0 | 0 | 8 | 0 | 5 | 0 | 0 | 1 | 0 | 98 | 49 | 0.039248064 | 0.170760496 | NA | 1 | 0.704 | NA | 0.714 | NA | 0.138 |
| SE | 11702 | POLR2A | POLR2A | chr17 | + | 7496209 | 7496346 | 7495940 | 7496073 | 7496439 | 7496613 | 0 | 1 | 9 | 0 | 0 | 0 | 0 | 5 | 0 | 0 | 1 | 0 | 98 | 49 | 0.039248064 | 0.170760496 | NA | 1 | 1 | NA | 0.714 | NA | 0.286 |
| SE | 11979 | TSEN34 | TSEN34 | chr19 | + | 54191720 | 54191964 | 54191213 | 54191607 | 54192115 | 54192373 | 0 | 5 | 17 | 0 | 0 | 1 | 0 | 8 | 4 | 0 | 3 | 1 | 98 | 49 | 0.039311924 | 0.170760496 | NA | 1 | 0.895 | NA | 0.571 | 0.667 | 0.329 |
| SE | 7521 | AQP9 | AQP9 | chr15 | + | 58173067 | 58173205 | 58166672 | 58166799 | 58179127 | 58179345 | 0 | 1 | 40 | 0 | 0 | 0 | 0 | 21 | 1 | 0 | 2 | 0 | 98 | 49 | 0.039675176 | 0.170760496 | NA | 1 | 1 | NA | 0.84 | 1 | 0.08 |
| A3SS | 462 | BTN3A1 | BTN3A1 | chr6 | + | 26412704 | 26415216 | 26413168 | 26415216 | 26411554 | 26411581 | 0 | 1 | 5 | 0 | 0 | 2 | 0 | 6 | 0 | 0 | 1 | 0 | 98 | 49 | 0.040279671 | 0.197717869 | NA | 1 | 0.556 | NA | 0.75 | NA | 0.028 |
| SE | 1761 | FAM53C | FAM53C | chr5 | + | 138341808 | 138341866 | 138341183 | 138341413 | 138344824 | 138345609 | 0 | 1 | 7 | 0 | 0 | 0 | 0 | 6 | 0 | 0 | 1 | 0 | 98 | 49 | 0.040279671 | 0.170760496 | NA | 1 | 1 | NA | 0.75 | NA | 0.25 |
| SE | 3245 | SULF2 | SULF2 | chr20 | - | 47665198 | 47665293 | 47664129 | 47664189 | 47665856 | 47665953 | 0 | 1 | 14 | 0 | 0 | 0 | 0 | 6 | 0 | 0 | 1 | 0 | 98 | 49 | 0.040279671 | 0.170760496 | NA | 1 | 1 | NA | 0.75 | NA | 0.25 |
| SE | 8677 | MANBA | MANBA | chr4 | - | 102714437 | 102714561 | 102690595 | 102690771 | 102722870 | 102723041 | 0 | 1 | 4 | 0 | 0 | 0 | 0 | 6 | 0 | 0 | 1 | 0 | 98 | 49 | 0.040279671 | 0.170760496 | NA | 1 | 1 | NA | 0.75 | NA | 0.25 |
| SE | 12664 | STXBP2 | STXBP2 | chr19 | + | 7643164 | 7643245 | 7642982 | 7643048 | 7644613 | 7644752 | 0 | 1 | 4 | 0 | 0 | 0 | 1 | 6 | 0 | 0 | 1 | 0 | 98 | 49 | 0.040279671 | 0.170760496 | NA | 1 | 1 | 1 | 0.75 | NA | 0.125 |
| SE | 19 | FLNA | FLNA | chrX | - | 154357250 | 154357274 | 154354824 | 154355072 | 154357433 | 154357623 | 0 | 1 | 4 | 0 | 7 | 5 | 2 | 10 | 0 | 0 | 4 | 0 | 73 | 49 | 0.040614468 | 0.170760496 | NA | 0.087 | 0.349 | 1 | 0.627 | NA | -0.596 |
| SE | 12838 | PTTG1IP | PTTG1IP | chr21 | - | 44865394 | 44865447 | 44861162 | 44861271 | 44873501 | 44873903 | 0 | 2 | 4 | 0 | 0 | 0 | 0 | 3 | 1 | 0 | 2 | 0 | 98 | 49 | 0.040702408 | 0.170760496 | NA | 1 | 1 | NA | 0.429 | 1 | 0.285 |
| SE | 4624 | SORL1 | SORL1 | chr11 | + | 121543547 | 121543726 | 121532463 | 121532552 | 121545242 | 121545429 | 0 | 4 | 58 | 0 | 0 | 0 | 0 | 34 | 0 | 0 | 1 | 0 | 98 | 49 | 0.04109126 | 0.170760496 | NA | 1 | 1 | NA | 0.944 | NA | 0.056 |
| SE | 1140 | IGF2R | IGF2R | chr6 | + | 160063414 | 160063630 | 160062531 | 160062619 | 160064400 | 160064531 | 0 | 2 | 31 | 0 | 0 | 0 | 0 | 16 | 0 | 0 | 1 | 0 | 98 | 49 | 0.041121367 | 0.170760496 | NA | 1 | 1 | NA | 0.889 | NA | 0.111 |
| SE | 5495 | ADAM8 | ADAM8 | chr10 | - | 133269444 | 133269529 | 133267928 | 133268118 | 133269896 | 133269974 | 0 | 2 | 12 | 0 | 0 | 0 | 0 | 16 | 0 | 0 | 1 | 0 | 98 | 49 | 0.041121367 | 0.170760496 | NA | 1 | 1 | NA | 0.889 | NA | 0.111 |
| SE | 2907 | RAF1 | RAF1 | chr3 | - | 12600151 | 12600279 | 12599690 | 12599808 | 12600387 | 12600415 | 0 | 2 | 19 | 0 | 0 | 0 | 0 | 22 | 3 | 0 | 2 | 0 | 98 | 49 | 0.041304338 | 0.170760496 | NA | 1 | 1 | NA | 0.846 | 1 | 0.077 |
| SE | 5124 | DNM2 | DNM2 | chr19 | + | 10759737 | 10759811 | 10718052 | 10718403 | 10772478 | 10772628 | 0 | 11 | 23 | 0 | 0 | 0 | 0 | 12 | 3 | 0 | 1 | 0 | 98 | 49 | 0.041331042 | 0.170760496 | NA | 1 | 1 | NA | 0.857 | 1 | 0.072 |
| A3SS | 3207 | ANKRD11 | ANKRD11 | chr16 | - | 89316932 | 89317078 | 89316932 | 89317075 | 89418283 | 89418368 | 0 | 1 | 2 | 0 | 0 | 0 | 0 | 1 | 0 | 0 | 1 | 0 | 52 | 49 | 0.041336327 | 0.197717869 | NA | 1 | 1 | NA | 0.485 | NA | 0.515 |
| SE | 8601 | PYGL | PYGL | chr14 | - | 50912155 | 50912303 | 50911977 | 50912036 | 50913028 | 50913130 | 0 | 1 | 14 | 0 | 0 | 1 | 0 | 7 | 0 | 0 | 1 | 0 | 98 | 49 | 0.041336891 | 0.170760496 | NA | 1 | 0.875 | NA | 0.778 | NA | 0.159 |
| SE | 9493 | SAP30BP | SAP30BP | chr17 | + | 75668515 | 75668625 | 75667115 | 75667478 | 75671815 | 75671863 | 0 | 1 | 1 | 0 | 0 | 0 | 0 | 7 | 0 | 0 | 1 | 0 | 98 | 49 | 0.041336891 | 0.170760496 | NA | 1 | 1 | NA | 0.778 | NA | 0.222 |
| SE | 10364 | SOS2 | SOS2 | chr14 | - | 50158564 | 50158646 | 50156998 | 50157121 | 50159430 | 50160086 | 0 | 1 | 14 | 0 | 0 | 2 | 0 | 7 | 0 | 0 | 1 | 0 | 98 | 49 | 0.041336891 | 0.170760496 | NA | 1 | 0.778 | NA | 0.778 | NA | 0.111 |
| SE | 11078 | TCEB2 | TCEB2 | chr16 | - | 2775450 | 2775556 | 2771413 | 2772102 | 2776992 | 2777127 | 0 | 1 | 12 | 0 | 0 | 0 | 1 | 7 | 0 | 0 | 1 | 0 | 98 | 49 | 0.041336891 | 0.170760496 | NA | 1 | 1 | 1 | 0.778 | NA | 0.111 |
| SE | 10594 | RNF213 | RNF213 | chr17 | + | 80390011 | 80390196 | 80389827 | 80389917 | 80393344 | 80398781 | 0 | 0 | 79 | 0 | 1 | 1 | 0 | 30 | 3 | 0 | 0 | 0 | 98 | 49 | 0.041591024 | 0.170760496 | NA | 0 | 0.975 | NA | 1 | 1 | -0.512 |
| SE | 1662 | FAM193B | FAM193B | chr5 | - | 177531952 | 177532200 | 177531281 | 177531521 | 177532442 | 177532641 | 0 | 2 | 0 | 0 | 0 | 1 | 0 | 0 | 1 | 0 | 0 | 2 | 98 | 49 | 0.041785228 | 0.170760496 | NA | 1 | 0 | NA | NA | 0.2 | 0.3 |
| SE | 9538 | CAMK2G | CAMK2G | chr10 | - | 73839538 | 73839601 | 73837467 | 73837511 | 73842168 | 73842211 | 0 | 0 | 0 | 0 | 1 | 4 | 0 | 1 | 0 | 3 | 2 | 0 | 98 | 49 | 0.041785228 | 0.170760496 | NA | 0 | 0 | 0 | 0.2 | NA | -0.1 |
| A5SS | 2638 | BPTF | BPTF | chr17 | + | 67945408 | 67946325 | 67945408 | 67945896 | 67947725 | 67947808 | 0 | 0 | 0 | 0 | 1 | 1 | 0 | 1 | 0 | 0 | 1 | 0 | 98 | 49 | 0.041803869 | 0.170414363 | NA | 0 | 0 | NA | 0.333 | NA | -0.333 |
| RI | 287 | EDEM2 | EDEM2 | chr20 | - | 35146824 | 35147358 | 35146824 | 35146935 | 35147151 | 35147358 | 0 | 0 | 0 | 0 | 0 | 1 | 0 | 0 | 1 | 0 | 0 | 1 | 98 | 49 | 0.041803869 | 0.297702651 | NA | NA | 0 | NA | NA | 0.333 | -0.333 |
| SE | 888 | GIGYF1 | GIGYF1 | chr7 | - | 100683549 | 100683632 | 100683303 | 100683444 | 100683817 | 100683918 | 0 | 0 | 0 | 0 | 1 | 0 | 0 | 1 | 0 | 0 | 1 | 0 | 98 | 49 | 0.041803869 | 0.170760496 | NA | 0 | NA | NA | 0.333 | NA | -0.333 |
| SE | 6999 | AGTRAP | AGTRAP | chr1 | + | 11746126 | 11746223 | 11745802 | 11745837 | 11747439 | 11747545 | 0 | 0 | 0 | 0 | 1 | 2 | 0 | 1 | 0 | 0 | 1 | 0 | 98 | 49 | 0.041803869 | 0.170760496 | NA | 0 | 0 | NA | 0.333 | NA | -0.333 |
| SE | 8430 | GS1-124K5.11 | GS1-124K5.11 | chr7 | - | 66560578 | 66560675 | 66553467 | 66554837 | 66573384 | 66573550 | 0 | 0 | 1 | 0 | 1 | 0 | 0 | 1 | 0 | 0 | 1 | 0 | 98 | 49 | 0.041803869 | 0.170760496 | NA | 0 | 1 | NA | 0.333 | NA | 0.167 |
| SE | 9926 | PPP4R3B | PPP4R3B | chr2 | - | 55578246 | 55578342 | 55577314 | 55577356 | 55579678 | 55579781 | 0 | 0 | 0 | 0 | 1 | 1 | 0 | 1 | 0 | 0 | 1 | 0 | 98 | 49 | 0.041803869 | 0.170760496 | NA | 0 | 0 | NA | 0.333 | NA | -0.333 |
| SE | 7544 | TKT | TKT | chr3 | - | 53237796 | 53237820 | 53234982 | 53235174 | 53240250 | 53240348 | 0 | 2 | 1 | 0 | 1 | 33 | 0 | 0 | 0 | 0 | 10 | 6 | 73 | 49 | 0.041838974 | 0.170760496 | NA | 0.573 | 0.02 | NA | 0 | 0 | 0.296 |
| SE | 2150 | YTHDC1 | YTHDC1 | chr4 | - | 68322748 | 68322915 | 68320122 | 68320205 | 68324138 | 68324223 | 0 | 1 | 3 | 0 | 0 | 0 | 0 | 8 | 0 | 0 | 1 | 0 | 98 | 49 | 0.04224465 | 0.170760496 | NA | 1 | 1 | NA | 0.8 | NA | 0.2 |
| SE | 2915 | LPCAT1 | LPCAT1 | chr5 | - | 1483427 | 1483486 | 1480941 | 1480976 | 1488390 | 1488451 | 0 | 1 | 2 | 0 | 0 | 0 | 0 | 8 | 0 | 0 | 1 | 0 | 98 | 49 | 0.04224465 | 0.170760496 | NA | 1 | 1 | NA | 0.8 | NA | 0.2 |
| SE | 3672 | CALCOCO2 | CALCOCO2 | chr17 | + | 48848063 | 48848166 | 48841697 | 48841887 | 48848321 | 48848455 | 0 | 1 | 8 | 0 | 0 | 0 | 0 | 8 | 0 | 0 | 1 | 0 | 98 | 49 | 0.04224465 | 0.170760496 | NA | 1 | 1 | NA | 0.8 | NA | 0.2 |
| SE | 1138 | IGF2R | IGF2R | chr6 | + | 160061502 | 160061646 | 160060546 | 160060717 | 160062531 | 160062619 | 0 | 5 | 19 | 0 | 0 | 0 | 2 | 9 | 3 | 0 | 4 | 0 | 98 | 49 | 0.042663026 | 0.170760496 | NA | 1 | 1 | 1 | 0.529 | 1 | 0.157 |
| SE | 5043 | CFL1 | CFL1 | chr11 | - | 65855653 | 65855730 | 65854813 | 65855448 | 65855934 | 65856242 | 0 | 9 | 66 | 0 | 0 | 0 | 10 | 26 | 1 | 0 | 1 | 0 | 98 | 49 | 0.042685592 | 0.170760496 | NA | 1 | 1 | 1 | 0.929 | 1 | 0.024 |
| SE | 7672 | SF3B1 | SF3B1 | chr2 | - | 197392971 | 197393188 | 197391973 | 197392461 | 197396055 | 197396328 | 0 | 5 | 39 | 0 | 0 | 0 | 0 | 9 | 1 | 0 | 1 | 0 | 98 | 49 | 0.042920223 | 0.170760496 | NA | 1 | 1 | NA | 0.818 | 1 | 0.091 |
| SE | 9384 | PECAM1 | PECAM1 | chr17 | - | 64375050 | 64375356 | 64369749 | 64370025 | 64377823 | 64378117 | 0 | 2 | 51 | 0 | 0 | 0 | 8 | 20 | 0 | 0 | 1 | 0 | 98 | 49 | 0.043140749 | 0.170760496 | NA | 1 | 1 | 1 | 0.909 | NA | 0.045 |
| SE | 8137 | ITGAX | ITGAX | chr16 | + | 31356624 | 31356728 | 31355892 | 31355998 | 31357030 | 31357101 | 0 | 26 | 76 | 0 | 0 | 0 | 8 | 82 | 30 | 0 | 2 | 0 | 98 | 49 | 0.043231591 | 0.170760496 | NA | 1 | 1 | 1 | 0.953 | 1 | 0.016 |
| SE | 22 | ARHGAP4 | ARHGAP4 | chrX | - | 153919566 | 153919686 | 153919154 | 153919283 | 153920625 | 153920808 | 0 | 2 | 0 | 2 | 2 | 2 | 0 | 0 | 0 | 0 | 6 | 3 | 98 | 49 | 0.043694816 | 0.170760496 | 0 | 0.333 | 0 | NA | 0 | 0 | 0.111 |
| A3SS | 4015 | PLEKHB2 | PLEKHB2 | chr2 | + | 131140166 | 131140275 | 131140169 | 131140275 | 131132901 | 131132991 | 0 | 1 | 3 | 0 | 0 | 0 | 0 | 4 | 0 | 0 | 1 | 0 | 52 | 49 | 0.043807749 | 0.197717869 | NA | 1 | 1 | NA | 0.79 | NA | 0.21 |
| A3SS | 461 | BTN3A1 | BTN3A1 | chr6 | + | 26412411 | 26415216 | 26413168 | 26415216 | 26411554 | 26411581 | 0 | 1 | 5 | 0 | 0 | 2 | 0 | 10 | 0 | 0 | 1 | 0 | 98 | 49 | 0.044092288 | 0.197717869 | NA | 1 | 0.556 | NA | 0.833 | NA | -0.055 |
| SE | 7268 | IVNS1ABP | IVNS1ABP | chr1 | - | 185309002 | 185309172 | 185308799 | 185308875 | 185309382 | 185309511 | 0 | 1 | 10 | 0 | 0 | 0 | 0 | 10 | 0 | 0 | 1 | 0 | 98 | 49 | 0.044092288 | 0.170760496 | NA | 1 | 1 | NA | 0.833 | NA | 0.167 |
| SE | 5494 | ADAM8 | ADAM8 | chr10 | - | 133268747 | 133268862 | 133267928 | 133268118 | 133269444 | 133269529 | 0 | 2 | 19 | 0 | 0 | 0 | 0 | 25 | 4 | 0 | 2 | 0 | 98 | 49 | 0.044237698 | 0.170760496 | NA | 1 | 1 | NA | 0.862 | 1 | 0.069 |
| SE | 1818 | CHD1 | CHD1 | chr5 | - | 98881974 | 98882123 | 98881278 | 98881375 | 98883087 | 98883237 | 0 | 2 | 14 | 0 | 0 | 0 | 0 | 23 | 0 | 0 | 1 | 0 | 98 | 49 | 0.044532178 | 0.170760496 | NA | 1 | 1 | NA | 0.92 | NA | 0.08 |
| SE | 814 | LINC-PINT | LINC-PINT | chr7 | - | 130959868 | 130959970 | 130944159 | 130945835 | 130984052 | 130984109 | 0 | 0 | 1 | 0 | 1 | 2 | 0 | 1 | 0 | 0 | 4 | 0 | 98 | 49 | 0.044611886 | 0.170760496 | NA | 0 | 0.2 | NA | 0.111 | NA | -0.011 |
| SE | 815 | LINC-PINT | LINC-PINT | chr7 | - | 130967388 | 130967517 | 130944159 | 130945835 | 130984052 | 130984109 | 0 | 0 | 5 | 0 | 1 | 2 | 0 | 1 | 0 | 0 | 4 | 0 | 98 | 49 | 0.044611886 | 0.170760496 | NA | 0 | 0.556 | NA | 0.111 | NA | 0.167 |
| A5SS | 3046 | PLEKHB2 | PLEKHB2 | chr2 | + | 131132901 | 131133015 | 131132901 | 131132991 | 131140166 | 131140275 | 0 | 0 | 4 | 0 | 1 | 3 | 0 | 1 | 0 | 0 | 4 | 0 | 73 | 49 | 0.044612593 | 0.170414363 | NA | 0 | 0.472 | NA | 0.144 | NA | 0.092 |
| SE | 11394 | ADGRG3 | ADGRG3 | chr16 | + | 57679176 | 57679311 | 57678169 | 57678316 | 57679815 | 57679855 | 0 | 0 | 15 | 0 | 1 | 0 | 0 | 11 | 3 | 0 | 0 | 0 | 98 | 49 | 0.044729667 | 0.170760496 | NA | 0 | 1 | NA | 1 | 1 | -0.5 |
| SE | 2103 | WDFY3 | WDFY3 | chr4 | - | 84836928 | 84837090 | 84831412 | 84831605 | 84841153 | 84841263 | 0 | 1 | 20 | 0 | 0 | 0 | 0 | 11 | 0 | 0 | 1 | 0 | 98 | 49 | 0.044972537 | 0.170760496 | NA | 1 | 1 | NA | 0.846 | NA | 0.154 |
| SE | 7942 | IFI44L | IFI44L | chr1 | + | 78629719 | 78629915 | 78628950 | 78628999 | 78635336 | 78635489 | 0 | 3 | 61 | 0 | 0 | 2 | 4 | 38 | 0 | 0 | 1 | 0 | 98 | 49 | 0.045281167 | 0.170760496 | NA | 1 | 0.938 | 1 | 0.95 | NA | -0.006 |
| SE | 7117 | MED13L | MED13L | chr12 | - | 116007410 | 116007636 | 116006305 | 116006411 | 116008400 | 116009132 | 0 | 2 | 11 | 0 | 0 | 0 | 0 | 25 | 0 | 0 | 1 | 0 | 98 | 49 | 0.045416917 | 0.170760496 | NA | 1 | 1 | NA | 0.926 | NA | 0.074 |
| SE | 4382 | STK38L | STK38L | chr12 | + | 27308338 | 27308461 | 27302136 | 27302188 | 27309113 | 27309197 | 0 | 3 | 8 | 0 | 0 | 0 | 0 | 5 | 1 | 0 | 1 | 0 | 98 | 49 | 0.045453113 | 0.170760496 | NA | 1 | 1 | NA | 0.714 | 1 | 0.143 |
| SE | 1691 | DOCK2 | DOCK2 | chr5 | + | 169761518 | 169761625 | 169759704 | 169759775 | 169803057 | 169803206 | 0 | 4 | 19 | 0 | 0 | 0 | 0 | 12 | 1 | 0 | 1 | 0 | 98 | 49 | 0.045620082 | 0.170760496 | NA | 1 | 1 | NA | 0.857 | 1 | 0.072 |
| SE | 6818 | OAS1 | OAS1 | chr12 | + | 112917546 | 112917700 | 112916508 | 112916738 | 112919388 | 112919907 | 0 | 3 | 8 | 0 | 0 | 0 | 0 | 8 | 1 | 0 | 1 | 0 | 98 | 49 | 0.045786327 | 0.170760496 | NA | 1 | 1 | NA | 0.8 | 1 | 0.1 |
| SE | 7899 | IFIT3 | IFIT3 | chr10 | + | 89337420 | 89337542 | 89332478 | 89332627 | 89338660 | 89340968 | 0 | 0 | 0 | 0 | 1 | 1 | 0 | 1 | 0 | 0 | 5 | 0 | 98 | 49 | 0.046158728 | 0.170760496 | NA | 0 | 0 | NA | 0.091 | NA | -0.091 |
| SE | 9005 | LYST | LYST | chr1 | - | 235709090 | 235709308 | 235702746 | 235702977 | 235712056 | 235712197 | 0 | 3 | 10 | 0 | 0 | 0 | 0 | 4 | 1 | 0 | 1 | 0 | 98 | 49 | 0.046507952 | 0.170760496 | NA | 1 | 1 | NA | 0.667 | 1 | 0.166 |
| A5SS | 2803 | TYROBP | TYROBP | chr19 | - | 35908167 | 35908309 | 35908174 | 35908309 | 35907445 | 35907580 | 0 | 0 | 5 | 0 | 3 | 1 | 0 | 1 | 2 | 0 | 0 | 0 | 56 | 49 | 0.046582587 | 0.170414363 | NA | 0 | 0.814 | NA | 1 | 1 | -0.593 |
| SE | 5273 | NIPBL | NIPBL | chr5 | + | 37036378 | 37036487 | 37027358 | 37027412 | 37038601 | 37038738 | 0 | 1 | 11 | 0 | 0 | 0 | 2 | 13 | 0 | 0 | 1 | 0 | 98 | 49 | 0.046638092 | 0.170760496 | NA | 1 | 1 | 1 | 0.867 | NA | 0.067 |
| SE | 9385 | CLTC | CLTC | chr17 | + | 59682270 | 59682428 | 59681646 | 59681839 | 59682628 | 59682793 | 0 | 1 | 12 | 0 | 0 | 0 | 0 | 13 | 0 | 0 | 1 | 0 | 98 | 49 | 0.046638092 | 0.170760496 | NA | 1 | 1 | NA | 0.867 | NA | 0.133 |
| SE | 4948 | ARAP1 | ARAP1 | chr11 | - | 72712195 | 72712339 | 72711429 | 72711499 | 72712437 | 72712568 | 0 | 7 | 9 | 0 | 0 | 0 | 0 | 6 | 3 | 0 | 1 | 0 | 98 | 49 | 0.047039397 | 0.170760496 | NA | 1 | 1 | NA | 0.75 | 1 | 0.125 |
| SE | 9916 | CASP8 | CASP8 | chr2 | + | 201274888 | 201274953 | 201272897 | 201272942 | 201276826 | 201276968 | 0 | 1 | 6 | 0 | 0 | 2 | 0 | 14 | 0 | 0 | 1 | 0 | 98 | 49 | 0.047441935 | 0.170760496 | NA | 1 | 0.6 | NA | 0.875 | NA | -0.075 |
| SE | 11995 | STAT3 | STAT3 | chr17 | - | 42348388 | 42348517 | 42346568 | 42346713 | 42388278 | 42388495 | 0 | 1 | 28 | 0 | 0 | 0 | 0 | 14 | 0 | 0 | 1 | 0 | 98 | 49 | 0.047441935 | 0.170760496 | NA | 1 | 1 | NA | 0.875 | NA | 0.125 |
| SE | 9286 | VIM | VIM | chr10 | + | 17236293 | 17236379 | 17235168 | 17235389 | 17237229 | 17237593 | 0 | 7 | 135 | 0 | 0 | 0 | 15 | 37 | 1 | 0 | 1 | 0 | 98 | 49 | 0.0475366 | 0.170760496 | NA | 1 | 1 | 1 | 0.949 | 1 | 0.017 |
| SE | 510 | ST3GAL1 | ST3GAL1 | chr8 | - | 133475718 | 133476600 | 133465893 | 133466090 | 133499134 | 133499189 | 0 | 1 | 1 | 0 | 0 | 3 | 0 | 3 | 1 | 0 | 0 | 0 | 98 | 49 | 0.047617 | 0.170760496 | NA | 1 | 0.143 | NA | 1 | 1 | -0.428 |
| SE | 12755 | ADGRE5 | ADGRE5 | chr19 | + | 14396341 | 14396473 | 14390923 | 14391079 | 14397657 | 14397804 | 0 | 0 | 4 | 0 | 1 | 11 | 5 | 1 | 0 | 0 | 6 | 0 | 98 | 49 | 0.047680582 | 0.170760496 | NA | 0 | 0.154 | 1 | 0.077 | NA | -0.461 |
| SE | 5193 | PICALM | PICALM | chr11 | - | 85978069 | 85978093 | 85976622 | 85976682 | 85981128 | 85981228 | 0 | 0 | 12 | 0 | 2 | 14 | 0 | 1 | 0 | 0 | 14 | 0 | 73 | 49 | 0.047713059 | 0.170760496 | NA | 0 | 0.365 | NA | 0.046 | NA | 0.137 |
| SE | 12573 | R3HDM4 | R3HDM4 | chr19 | - | 899600 | 899686 | 899439 | 899495 | 900060 | 900146 | 0 | 6 | 13 | 0 | 0 | 0 | 0 | 32 | 2 | 0 | 1 | 0 | 98 | 49 | 0.047912713 | 0.170760496 | NA | 1 | 1 | NA | 0.941 | 1 | 0.03 |
| SE | 6173 | ABCA1 | ABCA1 | chr9 | - | 104804625 | 104804720 | 104803283 | 104803316 | 104806240 | 104806430 | 0 | 4 | 7 | 0 | 0 | 0 | 0 | 25 | 1 | 0 | 1 | 0 | 98 | 49 | 0.048012831 | 0.170760496 | NA | 1 | 1 | NA | 0.926 | 1 | 0.037 |
| SE | 11306 | SP100 | SP100 | chr2 | + | 230461261 | 230461414 | 230450171 | 230450255 | 230462434 | 230462518 | 0 | 3 | 35 | 0 | 0 | 0 | 8 | 15 | 5 | 0 | 2 | 0 | 98 | 49 | 0.048106877 | 0.170760496 | NA | 1 | 1 | 1 | 0.789 | 1 | 0.07 |
| A3SS | 1145 | FBXL5 | FBXL5 | chr4 | - | 15644492 | 15644708 | 15644492 | 15644657 | 15655203 | 15655412 | 0 | 1 | 40 | 0 | 0 | 0 | 0 | 15 | 0 | 0 | 1 | 0 | 98 | 49 | 0.048230843 | 0.197717869 | NA | 1 | 1 | NA | 0.882 | NA | 0.118 |
| SE | 8157 | KMT2E | KMT2E | chr7 | + | 105078845 | 105078963 | 105077302 | 105077433 | 105081687 | 105081797 | 0 | 1 | 12 | 0 | 0 | 0 | 0 | 15 | 0 | 0 | 1 | 0 | 98 | 49 | 0.048230843 | 0.170760496 | NA | 1 | 1 | NA | 0.882 | NA | 0.118 |
| SE | 10588 | RNF213 | RNF213 | chr17 | + | 80363608 | 80363790 | 80363101 | 80363314 | 80364432 | 80364553 | 0 | 15 | 83 | 0 | 0 | 0 | 0 | 31 | 8 | 0 | 1 | 0 | 98 | 49 | 0.048276428 | 0.170760496 | NA | 1 | 1 | NA | 0.939 | 1 | 0.03 |
| SE | 12401 | CSF2RB | CSF2RB | chr22 | + | 36936548 | 36936652 | 36935629 | 36935687 | 36937376 | 36940437 | 0 | 7 | 34 | 0 | 0 | 0 | 4 | 32 | 2 | 0 | 1 | 0 | 98 | 49 | 0.048373702 | 0.170760496 | NA | 1 | 1 | 1 | 0.941 | 1 | 0.02 |
| SE | 7883 | HIF1A | HIF1A | chr14 | + | 61740504 | 61740627 | 61738086 | 61738373 | 61740754 | 61741188 | 0 | 0 | 14 | 0 | 1 | 0 | 0 | 18 | 1 | 0 | 0 | 0 | 98 | 49 | 0.048559241 | 0.170760496 | NA | 0 | 1 | NA | 1 | 1 | -0.5 |
| SE | 4361 | LRRK2 | LRRK2 | chr12 | + | 40293544 | 40293663 | 40287350 | 40287539 | 40294844 | 40294914 | 0 | 2 | 74 | 0 | 0 | 0 | 0 | 33 | 0 | 0 | 1 | 0 | 98 | 49 | 0.048704034 | 0.170760496 | NA | 1 | 1 | NA | 0.943 | NA | 0.057 |
| SE | 11375 | ARHGEF1 | ARHGEF1 | chr19 | + | 41894625 | 41894661 | 41894450 | 41894547 | 41895348 | 41895486 | 0 | 5 | 22 | 0 | 0 | 0 | 0 | 16 | 2 | 0 | 1 | 0 | 85 | 49 | 0.048992317 | 0.170760496 | NA | 1 | 1 | NA | 0.902 | 1 | 0.049 |
| A3SS | 555 | RICTOR | RICTOR | chr5 | - | 38945490 | 38945724 | 38945490 | 38945610 | 38946467 | 38946552 | 0 | 1 | 15 | 0 | 0 | 1 | 0 | 16 | 0 | 0 | 1 | 0 | 98 | 49 | 0.048997106 | 0.197717869 | NA | 1 | 0.882 | NA | 0.889 | NA | 0.052 |
| SE | 9006 | LYST | LYST | chr1 | - | 235728075 | 235728131 | 235724027 | 235724180 | 235729595 | 235729657 | 0 | 1 | 35 | 0 | 0 | 0 | 2 | 16 | 0 | 0 | 1 | 0 | 98 | 49 | 0.048997106 | 0.170760496 | NA | 1 | 1 | 1 | 0.889 | NA | 0.055 |
| SE | 6618 | AMPD2 | AMPD2 | chr1 | + | 109629326 | 109629490 | 109629108 | 109629235 | 109629795 | 109629916 | 0 | 2 | 5 | 0 | 0 | 0 | 0 | 6 | 1 | 0 | 1 | 0 | 98 | 49 | 0.049093709 | 0.170760496 | NA | 1 | 1 | NA | 0.75 | 1 | 0.125 |
| SE | 6896 | ITGB2 | ITGB2 | chr21 | - | 44889275 | 44889495 | 44888692 | 44888895 | 44889977 | 44890222 | 0 | 4 | 78 | 0 | 0 | 0 | 3 | 11 | 2 | 0 | 1 | 0 | 98 | 49 | 0.049219655 | 0.170760496 | NA | 1 | 1 | 1 | 0.846 | 1 | 0.051 |
| SE | 11046 | HLA-B | HLA-B | chr6 | - | 31355106 | 31355223 | 31354478 | 31354526 | 31355316 | 31355592 | 0 | 40 | 238 | 0 | 0 | 0 | 10 | 193 | 14 | 0 | 1 | 0 | 98 | 49 | 0.049293349 | 0.170760496 | NA | 1 | 1 | 1 | 0.99 | 1 | 0.003 |
| SE | 9374 | FMNL1 | FMNL1 | chr17 | + | 45233647 | 45233731 | 45233223 | 45233297 | 45234071 | 45234200 | 0 | 4 | 23 | 0 | 0 | 0 | 0 | 21 | 1 | 0 | 1 | 0 | 98 | 49 | 0.049821364 | 0.170760496 | NA | 1 | 1 | NA | 0.913 | 1 | 0.043 |
| SE | 7884 | HIF1A | HIF1A | chr14 | + | 61740754 | 61741188 | 61738086 | 61738373 | 61744704 | 61744813 | 0 | 1 | 13 | 0 | 0 | 3 | 0 | 34 | 1 | 0 | 0 | 0 | 98 | 49 | 0.049963886 | 0.170760496 | NA | 1 | 0.684 | NA | 1 | 1 | -0.158 |
| SE | 9513 | SIPA1 | SIPA1 | chr11 | + | 65644954 | 65645129 | 65642462 | 65642639 | 65646220 | 65646378 | 0 | 1 | 11 | 0 | 2 | 0 | 0 | 7 | 7 | 0 | 0 | 0 | 98 | 49 | 0.050052195 | 0.170760496 | NA | 0.2 | 1 | NA | 1 | 1 | -0.4 |
| SE | 6178 | ABCA1 | ABCA1 | chr9 | - | 104828915 | 104829138 | 104826947 | 104827169 | 104830924 | 104831101 | 0 | 4 | 14 | 0 | 0 | 0 | 0 | 25 | 1 | 0 | 1 | 0 | 98 | 49 | 0.050181472 | 0.170760496 | NA | 1 | 1 | NA | 0.926 | 1 | 0.037 |
| SE | 1146 | IGF2R | IGF2R | chr6 | + | 160102518 | 160102671 | 160096438 | 160096625 | 160103745 | 160103815 | 0 | 6 | 13 | 0 | 0 | 0 | 0 | 14 | 7 | 0 | 2 | 0 | 98 | 49 | 0.050333626 | 0.170760496 | NA | 1 | 1 | NA | 0.778 | 1 | 0.111 |
| SE | 10067 | ACTN1 | ACTN1 | chr14 | - | 68884774 | 68884883 | 68884167 | 68884308 | 68885424 | 68885575 | 0 | 1 | 15 | 0 | 0 | 0 | 0 | 18 | 0 | 0 | 1 | 0 | 98 | 49 | 0.050460196 | 0.170760496 | NA | 1 | 1 | NA | 0.9 | NA | 0.1 |
| SE | 4427 | PREX1 | PREX1 | chr20 | - | 48734545 | 48734650 | 48726289 | 48726391 | 48745024 | 48745147 | 0 | 12 | 55 | 0 | 0 | 0 | 3 | 75 | 3 | 0 | 1 | 0 | 98 | 49 | 0.050469046 | 0.170760496 | NA | 1 | 1 | 1 | 0.974 | 1 | 0.009 |
| SE | 12942 | SMCHD1 | SMCHD1 | chr18 | + | 2703691 | 2703886 | 2700734 | 2700918 | 2705693 | 2705807 | 1 | 4 | 139 | 0 | 0 | 0 | 0 | 77 | 0 | 0 | 1 | 0 | 98 | 49 | 0.05052881 | 0.170760496 | 1 | 1 | 1 | NA | 0.975 | NA | 0.025 |
| SE | 7030 | ARPC2 | ARPC2 | chr2 | + | 218217462 | 218217544 | 218217093 | 218217254 | 218225919 | 218225954 | 0 | 4 | 9 | 0 | 0 | 0 | 0 | 24 | 2 | 0 | 1 | 0 | 98 | 49 | 0.050622824 | 0.170760496 | NA | 1 | 1 | NA | 0.923 | 1 | 0.038 |

| | | | | | | | | | | | | | | | | | | | | | | | | | | | | | | | | | | |
|---|---|---|---|---|---|---|---|---|---|---|---|---|---|---|---|---|---|---|---|---|---|---|---|---|---|---|---|---|---|---|---|---|---|---|
| SE | 12325 | MAPKAPK2 | MAPKAPK2 | chr1 | + | 206731838 | 206731919 | 206731639 | 206731725 | 206732574 | 206734285 | 0 | 1 | 14 | 0 | 2 | 0 | 5 | 16 | 3 | 0 | 0 | 0 | 98 | 49 | 0.050687516 | 0.170760496 | NA | 0.2 | 1 | 1 | 1 | 1 | -0.4 |
| SE | 11256 | SRSF7 | SRSF7 | chr2 | - | 38746143 | 38746179 | 38743598 | 38745187 | 38746693 | 38746747 | 0 | 2 | 5 | 0 | 0 | 0 | 0 | 7 | 1 | 0 | 1 | 0 | 85 | 49 | 0.05078876 | 0.170760496 | NA | 1 | 1 | NA | 0.801 | 1 | 0.099 |
| SE | 5864 | NFKBID | NFKBID | chr19 | - | 35896912 | 35897058 | 35896725 | 35896831 | 35897650 | 35897856 | 0 | 6 | 7 | 0 | 0 | 0 | 0 | 4 | 7 | 0 | 0 | 1 | 98 | 49 | 0.050792748 | 0.170760496 | NA | 1 | 1 | NA | 1 | 0.778 | 0.111 |
| SE | 9455 | PFKFB3 | PFKFB3 | chr10 | + | 6216124 | 6216191 | 6215220 | 6215317 | 6216705 | 6216780 | 0 | 19 | 68 | 0 | 1 | 1 | 1 | 72 | 24 | 0 | 0 | 0 | 98 | 49 | 0.050850888 | 0.170760496 | NA | 0.905 | 0.971 | 1 | 1 | 1 | -0.062 |
| SE | 8552 | EHBP1L1 | EHBP1L1 | chr11 | + | 65581538 | 65583765 | 65581210 | 65581373 | 65584240 | 65584398 | 0 | 2 | 12 | 0 | 0 | 0 | 0 | 5 | 1 | 0 | 1 | 0 | 98 | 49 | 0.051092621 | 0.170760496 | NA | 1 | 1 | NA | 0.714 | 1 | 0.143 |
| SE | 5410 | TAGAP | TAGAP | chr6 | - | 159043588 | 159043655 | 159042077 | 159042244 | 159043977 | 159044031 | 0 | 1 | 14 | 0 | 0 | 0 | 0 | 19 | 0 | 0 | 1 | 0 | 98 | 49 | 0.051175801 | 0.170760496 | NA | 1 | 1 | NA | 0.905 | NA | 0.095 |
| RI | 1060 | SEC14L1 | SEC14L1 | chr17 | + | 77213917 | 77217099 | 77213917 | 77214019 | 77216548 | 77217099 | 0 | 13 | 22 | 0 | 0 | 0 | 6 | 16 | 9 | 0 | 1 | 0 | 98 | 49 | 0.051358786 | 0.297702651 | NA | 1 | 1 | 1 | 0.889 | 1 | 0.037 |
| SE | 4670 | TNFRSF1B | TNFRSF1B | chr1 | + | 12201966 | 12202171 | 12194583 | 12194618 | 12206739 | 12209220 | 0 | 5 | 27 | 0 | 0 | 0 | 0 | 27 | 2 | 0 | 1 | 0 | 98 | 49 | 0.051369383 | 0.170760496 | NA | 1 | 1 | NA | 0.931 | 1 | 0.034 |
| SE | 4501 | PTPN6 | PTPN6 | chr12 | + | 6951608 | 6951731 | 6951270 | 6951520 | 6951982 | 6952177 | 0 | 5 | 25 | 0 | 0 | 0 | 0 | 8 | 3 | 0 | 1 | 0 | 98 | 49 | 0.051433279 | 0.170760496 | NA | 1 | 1 | NA | 0.8 | 1 | 0.1 |
| SE | 7417 | VAV1 | VAV1 | chr19 | + | 6836984 | 6837050 | 6836431 | 6836568 | 6843134 | 6843166 | 0 | 1 | 12 | 0 | 0 | 0 | 3 | 20 | 0 | 0 | 1 | 0 | 98 | 49 | 0.051859277 | 0.170760496 | NA | 1 | 1 | 1 | 0.909 | NA | 0.045 |
| SE | 12684 | HNRNPK | HNRNPK | chr9 | - | 83973901 | 83973973 | 83973285 | 83973399 | 83974516 | 83974589 | 0 | 1 | 53 | 0 | 0 | 3 | 1 | 20 | 0 | 0 | 1 | 0 | 98 | 49 | 0.051859277 | 0.170760496 | NA | 1 | 0.898 | 1 | 0.909 | NA | -0.005 |
| SE | 3134 | IFNGR2 | IFNGR2 | chr21 | + | 33414887 | 33415020 | 33402895 | 33403616 | 33421479 | 33421685 | 9 | 14 | 31 | 0 | 0 | 0 | 0 | 15 | 47 | 0 | 0 | 1 | 98 | 49 | 0.052010728 | 0.170760496 | 1 | 1 | 1 | NA | 1 | 0.959 | 0.02 |
| SE | 7031 | ARPC2 | ARPC2 | chr2 | + | 218239390 | 218239484 | 218238663 | 218238850 | 218245419 | 218245546 | 0 | 0 | 56 | 0 | 0 | 1 | 1 | 33 | 2 | 0 | 1 | 2 | 98 | 49 | 0.052035219 | 0.170760496 | NA | NA | 0.966 | 1 | 0.943 | 0.333 | 0.207 |
| SE | 1737 | ARHGAP26 | ARHGAP26 | chr5 | + | 143014079 | 143014116 | 142932046 | 142932125 | 143037195 | 143037261 | 0 | 1 | 47 | 0 | 0 | 0 | 0 | 23 | 4 | 0 | 2 | 0 | 86 | 49 | 0.052066191 | 0.170760496 | NA | 1 | 1 | NA | 0.868 | 1 | 0.066 |
| SE | 13867 | ACSL1 | ACSL1 | chr4 | - | 184776483 | 184776662 | 184773842 | 184773875 | 184776883 | 184776983 | 0 | 3 | 33 | 0 | 0 | 1 | 1 | 36 | 1 | 0 | 6 | 0 | 98 | 49 | 0.052282739 | 0.170760496 | NA | 1 | 0.943 | 1 | 0.75 | 1 | 0.055 |
| SE | 4669 | TNFRSF1B | TNFRSF1B | chr1 | + | 12188795 | 12188895 | 12167002 | 12167169 | 12190956 | 12191085 | 0 | 7 | 37 | 0 | 2 | 3 | 5 | 25 | 32 | 0 | 0 | 1 | 98 | 49 | 0.052384975 | 0.170760496 | NA | 0.636 | 0.86 | 1 | 1 | 0.941 | -0.232 |
| SE | 9376 | FMNL1 | FMNL1 | chr17 | + | 45245252 | 45245416 | 45244978 | 45245108 | 45245877 | 45245973 | 0 | 3 | 24 | 0 | 0 | 0 | 0 | 14 | 0 | 0 | 1 | 2 | 98 | 49 | 0.052499618 | 0.170760496 | NA | 1 | 1 | NA | 0.875 | 0 | 0.562 |
| SE | 10927 | DEDD2 | DEDD2 | chr19 | - | 42201988 | 42202072 | 42198592 | 42199829 | 42209699 | 42209840 | 0 | 1 | 2 | 0 | 3 | 5 | 0 | 1 | 0 | 0 | 16 | 3 | 98 | 49 | 0.052585138 | 0.170760496 | NA | 0.143 | 0.167 | NA | 0.03 | 0 | 0.14 |
| SE | 1854 | DBNL | DBNL | chr7 | + | 44058431 | 44058480 | 44058128 | 44058280 | 44058901 | 44058983 | 0 | 4 | 5 | 0 | 0 | 0 | 0 | 3 | 2 | 0 | 1 | 0 | 98 | 49 | 0.052858149 | 0.170760496 | NA | 1 | 1 | NA | 0.6 | 1 | 0.2 |
| RI | 576 | SRSF2 | SRSF2 | chr17 | - | 76734114 | 76736464 | 76734114 | 76735158 | 76736153 | 76736464 | 0 | 7 | 12 | 0 | 2 | 3 | 0 | 16 | 1 | 0 | 12 | 1 | 98 | 49 | 0.052909456 | 0.297702651 | NA | 0.636 | 0.667 | NA | 0.4 | 0.333 | 0.285 |
| A3SS | 459 | BTN3A1 | BTN3A1 | chr6 | + | 26407670 | 26407952 | 26407826 | 26407952 | 26405908 | 26406256 | 0 | 2 | 3 | 0 | 0 | 0 | 2 | 2 | 1 | 0 | 1 | 0 | 98 | 49 | 0.05291102 | 0.197717869 | NA | 1 | 1 | 1 | 0.5 | 1 | 0.167 |
| SE | 8417 | GRAMD1A | GRAMD1A | chr19 | + | 35021950 | 35022038 | 35021690 | 35021864 | 35023235 | 35023343 | 0 | 4 | 17 | 0 | 0 | 0 | 0 | 4 | 12 | 0 | 0 | 1 | 98 | 49 | 0.052978196 | 0.170760496 | NA | 1 | 1 | NA | 1 | 0.857 | 0.072 |
| SE | 7910 | ATP6V0B | ATP6V0B | chr1 | + | 43977025 | 43977216 | 43976772 | 43976824 | 43977980 | 43978300 | 0 | 4 | 22 | 0 | 0 | 0 | 1 | 11 | 3 | 0 | 1 | 0 | 98 | 49 | 0.053042143 | 0.170760496 | NA | 1 | 1 | 1 | 0.846 | 1 | 0.051 |
| SE | 10586 | RNF213 | RNF213 | chr17 | + | 80360060 | 80360206 | 80358287 | 80358479 | 80361733 | 80361888 | 0 | 1 | 46 | 0 | 0 | 0 | 2 | 41 | 4 | 0 | 2 | 0 | 98 | 49 | 0.053139356 | 0.170760496 | NA | 1 | 1 | 1 | 0.911 | 1 | 0.03 |
| SE | 6938 | CD55 | CD55 | chr1 | + | 207331107 | 207331296 | 207326751 | 207326837 | 207336692 | 207336818 | 0 | 2 | 28 | 0 | 0 | 0 | 0 | 46 | 0 | 0 | 1 | 0 | 98 | 49 | 0.053445903 | 0.170760496 | NA | 1 | 1 | NA | 0.958 | NA | 0.042 |
| SE | 7392 | IRAK3 | IRAK3 | chr12 | + | 66211445 | 66211597 | 66210146 | 66210201 | 66217170 | 66217235 | 0 | 2 | 30 | 0 | 0 | 0 | 2 | 10 | 1 | 0 | 1 | 0 | 98 | 49 | 0.053461329 | 0.170760496 | NA | 1 | 1 | 1 | 0.833 | 1 | 0.056 |
| SE | 9740 | PXN | PXN | chr12 | - | 120222548 | 120222750 | 120221622 | 120221758 | 120222862 | 120222999 | 0 | 3 | 23 | 0 | 0 | 0 | 0 | 6 | 2 | 0 | 1 | 0 | 98 | 49 | 0.053502796 | 0.170760496 | NA | 1 | 1 | NA | 0.75 | 1 | 0.125 |
| SE | 4088 | TNFRSF1A | TNFRSF1A | chr12 | - | 6334090 | 6334244 | 6333736 | 6333865 | 6341775 | 6342117 | 0 | 2 | 17 | 0 | 3 | 1 | 0 | 5 | 3 | 0 | 0 | 0 | 98 | 49 | 0.053505644 | 0.170760496 | NA | 0.25 | 0.895 | NA | 1 | 1 | -0.427 |
| SE | 13863 | ACSL1 | ACSL1 | chr4 | - | 184765890 | 184765986 | 184764852 | 184764925 | 184766621 | 184766756 | 0 | 2 | 59 | 0 | 0 | 3 | 4 | 48 | 12 | 0 | 0 | 0 | 98 | 49 | 0.053534738 | 0.170760496 | NA | 1 | 0.908 | 1 | 1 | 1 | -0.046 |
| SE | 1144 | IGF2R | IGF2R | chr6 | + | 160084994 | 160085131 | 160083949 | 160084184 | 160088032 | 160088147 | 0 | 4 | 17 | 0 | 0 | 0 | 1 | 20 | 3 | 0 | 1 | 0 | 98 | 49 | 0.05376828 | 0.170760496 | NA | 1 | 1 | 1 | 0.909 | 1 | 0.03 |
| SE | 8590 | LTB | LTB | chr6 | - | 31581813 | 31581859 | 31581558 | 31581630 | 31582255 | 31582425 | 0 | 0 | 5 | 0 | 3 | 0 | 0 | 11 | 0 | 0 | 0 | 0 | 95 | 49 | 0.054362012 | 0.170760496 | NA | 0 | 1 | NA | 1 | NA | -0.5 |
| SE | 4241 | ITGA5 | ITGA5 | chr12 | - | 54403624 | 54403779 | 54403186 | 54403324 | 54403910 | 54403966 | 0 | 0 | 12 | 0 | 0 | 0 | 0 | 7 | 0 | 0 | 0 | 2 | 98 | 49 | 0.054536123 | 0.170760496 | NA | NA | 1 | NA | 1 | 0 | 0.5 |
| A3SS | 690 | CNBP | CNBP | chr3 | - | 129171078 | 129171283 | 129171078 | 129171277 | 129171445 | 129171538 | 0 | 0 | 0 | 0 | 1 | 4 | 0 | 1 | 0 | 0 | 11 | 0 | 55 | 49 | 0.054675117 | 0.197717869 | NA | 0 | 0 | NA | 0.075 | NA | -0.075 |
| SE | 5196 | PICALM | PICALM | chr11 | - | 86001034 | 86001158 | 86000642 | 86000779 | 86003365 | 86003451 | 0 | 6 | 50 | 0 | 0 | 0 | 2 | 55 | 1 | 0 | 1 | 0 | 98 | 49 | 0.054703796 | 0.170760496 | NA | 1 | 1 | 1 | 0.965 | 1 | 0.012 |
| SE | 11996 | STAT3 | STAT3 | chr17 | - | 42348388 | 42348539 | 42346568 | 42346713 | 42388278 | 42388387 | 0 | 2 | 29 | 0 | 0 | 0 | 0 | 12 | 1 | 0 | 1 | 0 | 98 | 49 | 0.054806328 | 0.170760496 | NA | 1 | 1 | NA | 0.857 | 1 | 0.072 |
| SE | 7497 | IFI44 | IFI44 | chr1 | + | 78660553 | 78660654 | 78659311 | 78659483 | 78663764 | 78664078 | 0 | 0 | 31 | 0 | 1 | 1 | 0 | 20 | 1 | 0 | 0 | 0 | 98 | 49 | 0.054840236 | 0.170760496 | NA | 0 | 0.939 | NA | 1 | 1 | -0.53 |
| SE | 7726 | FGD3 | FGD3 | chr9 | + | 93002924 | 93003014 | 92976207 | 92976709 | 93004000 | 93004137 | 0 | 10 | 22 | 0 | 0 | 0 | 0 | 16 | 9 | 0 | 1 | 0 | 98 | 49 | 0.054857277 | 0.170760496 | NA | 1 | 1 | NA | 0.889 | 1 | 0.055 |
| SE | 4940 | ARAP1 | ARAP1 | chr11 | - | 72687438 | 72687502 | 72686041 | 72686191 | 72687687 | 72687738 | 0 | 3 | 10 | 0 | 0 | 0 | 0 | 4 | 2 | 0 | 1 | 0 | 98 | 49 | 0.055117945 | 0.170760496 | NA | 1 | 1 | NA | 0.667 | 1 | 0.166 |
| SE | 12992 | PLAUR | PLAUR | chr19 | - | 43656478 | 43656640 | 43655438 | 43655573 | 43665315 | 43665459 | 0 | 14 | 45 | 0 | 0 | 0 | 0 | 84 | 14 | 0 | 1 | 0 | 98 | 49 | 0.055245993 | 0.170760496 | NA | 1 | 1 | NA | 0.977 | 1 | 0.012 |
| SE | 11304 | SP100 | SP100 | chr2 | + | 230449560 | 230449710 | 230449087 | 230449150 | 230450171 | 230450255 | 0 | 7 | 33 | 0 | 0 | 0 | 0 | 28 | 7 | 0 | 1 | 0 | 98 | 49 | 0.055357429 | 0.170760496 | NA | 1 | 1 | NA | 0.933 | 1 | 0.033 |
| SE | 3739 | BRD2 | BRD2 | chr6 | + | 32976561 | 32976936 | 32976249 | 32976464 | 32977441 | 32977570 | 0 | 0 | 8 | 0 | 0 | 0 | 0 | 8 | 0 | 0 | 0 | 2 | 98 | 49 | 0.055397548 | 0.170760496 | NA | NA | 1 | NA | 1 | 0 | 0.5 |
| A3SS | 1302 | GMPR2 | GMPR2 | chr14 | + | 24238245 | 24239241 | 24238588 | 24239241 | 24237519 | 24237562 | 1 | 0 | 3 | 0 | 2 | 0 | 0 | 10 | 0 | 0 | 0 | 0 | 98 | 49 | 0.055620739 | 0.197717869 | 1 | 0 | 1 | NA | 1 | NA | -0.333 |
| SE | 2096 | WDFY3 | WDFY3 | chr4 | - | 84678918 | 84679242 | 84678167 | 84678279 | 84682373 | 84682470 | 0 | 2 | 8 | 0 | 0 | 0 | 0 | 2 | 1 | 0 | 1 | 0 | 98 | 49 | 0.055836208 | 0.170760496 | NA | 1 | 1 | NA | 0.5 | 1 | 0.25 |
| SE | 6103 | GLUL | GLUL | chr1 | - | 182385759 | 182385887 | 182385356 | 182385556 | 182386255 | 182386402 | 0 | 16 | 127 | 0 | 0 | 0 | 6 | 106 | 11 | 0 | 1 | 0 | 98 | 49 | 0.055843452 | 0.170760496 | NA | 1 | 1 | 1 | 0.981 | 1 | 0.006 |
| SE | 10624 | RRP12 | RRP12 | chr10 | - | 97366107 | 97366233 | 97363853 | 97363903 | 97366445 | 97366621 | 0 | 0 | 2 | 0 | 0 | 0 | 0 | 1 | 0 | 0 | 0 | 5 | 98 | 49 | 0.055847068 | 0.170760496 | NA | NA | 1 | NA | 1 | 0 | 0.5 |
| SE | 2782 | ATG7 | ATG7 | chr3 | + | 11299361 | 11299416 | 11298685 | 11298855 | 11306942 | 11307060 | 0 | 0 | 7 | 0 | 1 | 0 | 0 | 9 | 1 | 0 | 0 | 0 | 98 | 49 | 0.055867249 | 0.170760496 | NA | 0 | 1 | NA | 1 | 1 | -0.5 |
| SE | 11397 | ADGRG3 | ADGRG3 | chr16 | + | 57684389 | 57684483 | 57680504 | 57680617 | 57685642 | 57685926 | 0 | 0 | 7 | 0 | 2 | 0 | 0 | 9 | 0 | 0 | 0 | 0 | 98 | 49 | 0.055954714 | 0.170760496 | NA | 0 | 1 | NA | 1 | NA | -0.5 |
| SE | 2283 | IL1RAP | IL1RAP | chr3 | + | 190550341 | 190550505 | 190514050 | 190514219 | 190556129 | 190556216 | 0 | 0 | 3 | 0 | 3 | 3 | 0 | 3 | 0 | 0 | 4 | 2 | 98 | 49 | 0.05597701 | 0.170760496 | NA | 0 | 0 | NA | 0.273 | 0 | -0.137 |
| SE | 1506 | HLA-E | HLA-E | chr6 | + | 30491536 | 30491653 | 30491136 | 30491412 | 30492540 | 30492583 | 0 | 12 | 185 | 0 | 0 | 0 | 11 | 70 | 10 | 0 | 1 | 0 | 98 | 49 | 0.056110759 | 0.170760496 | NA | 1 | 1 | 1 | 0.972 | 1 | 0.009 |
| SE | 3382 | HCK | HCK | chr20 | + | 32084390 | 32084543 | 32083893 | 32084043 | 32086627 | 32086807 | 0 | 3 | 55 | 0 | 1 | 5 | 5 | 17 | 4 | 0 | 0 | 0 | 98 | 49 | 0.056127965 | 0.170760496 | NA | 0.6 | 0.846 | 1 | 1 | 1 | -0.277 |
| SE | 3057 | COL18A1 | COL18A1 | chr21 | + | 45505133 | 45505278 | 45504521 | 45504556 | 45505357 | 45505431 | 0 | 0 | 6 | 0 | 0 | 0 | 0 | 2 | 0 | 0 | 0 | 2 | 98 | 49 | 0.056129395 | 0.170760496 | NA | NA | 1 | NA | 1 | 0 | 0.5 |
| SE | 10626 | RRP12 | RRP12 | chr10 | - | 97379292 | 97379414 | 97373829 | 97373894 | 97379627 | 97379770 | 0 | 2 | 7 | 0 | 0 | 0 | 0 | 9 | 2 | 0 | 1 | 0 | 98 | 49 | 0.056251461 | 0.170760496 | NA | 1 | 1 | NA | 0.818 | 1 | 0.091 |
| SE | 1507 | HLA-E | HLA-E | chr6 | + | 30492403 | 30492436 | 30491136 | 30491412 | 30492540 | 30492583 | 0 | 7 | 64 | 0 | 0 | 0 | 0 | 48 | 3 | 0 | 1 | 0 | 82 | 49 | 0.05627258 | 0.170760496 | NA | 1 | 1 | NA | 0.966 | 1 | 0.017 |
| SE | 9587 | ZNF467 | ZNF467 | chr7 | - | 149770439 | 149770556 | 149769089 | 149769200 | 149770998 | 149771074 | 0 | 0 | 1 | 0 | 1 | 0 | 0 | 2 | 3 | 0 | 0 | 0 | 98 | 49 | 0.056299876 | 0.170760496 | NA | 0 | 1 | NA | 1 | 1 | -0.5 |
| SE | 3981 | NCOR2 | NCOR2 | chr12 | - | 124374412 | 124374463 | 124372021 | 124372610 | 124378236 | 124378384 | 0 | 0 | 2 | 0 | 0 | 0 | 0 | 0 | 0 | 0 | 0 | 4 | 98 | 49 | 0.056348041 | 0.170760496 | NA | NA | 1 | NA | NA | 0 | 1 |
| SE | 6199 | RPRD2 | RPRD2 | chr1 | + | 150441880 | 150441958 | 150440922 | 150441023 | 150443230 | 150443283 | 0 | 2 | 0 | 0 | 0 | 0 | 0 | 0 | 0 | 0 | 4 | 0 | 98 | 49 | 0.056348041 | 0.170760496 | NA | 1 | NA | NA | 0 | NA | 1 |
| SE | 636 | RB1CC1 | RB1CC1 | chr8 | - | 52660595 | 52660657 | 52658872 | 52658976 | 52660925 | 52661007 | 0 | 0 | 3 | 0 | 0 | 0 | 0 | 4 | 0 | 0 | 0 | 3 | 98 | 49 | 0.056460428 | 0.170760496 | NA | NA | 1 | NA | 1 | 0 | 0.5 |
| SE | 12437 | SPAG9 | SPAG9 | chr17 | - | 51014231 | 51014353 | 51007268 | 51007326 | 51020158 | 51020258 | 0 | 0 | 8 | 0 | 1 | 0 | 0 | 14 | 0 | 0 | 0 | 0 | 98 | 49 | 0.056479947 | 0.170760496 | NA | 0 | 1 | NA | 1 | NA | -0.5 |
| SE | 6672 | FGR | FGR | chr1 | - | 27621558 | 27621657 | 27615688 | 27615844 | 27623041 | 27623144 | 0 | 3 | 15 | 0 | 0 | 0 | 0 | 10 | 3 | 0 | 1 | 0 | 98 | 49 | 0.0565412 | 0.170760496 | NA | 1 | 1 | NA | 0.833 | 1 | 0.084 |
| SE | 3944 | ZDHHC20 | ZDHHC20 | chr13 | - | 21421060 | 21421164 | 21413651 | 21413772 | 21425651 | 21425678 | 0 | 0 | 5 | 0 | 0 | 0 | 0 | 0 | 0 | 0 | 1 | 2 | 98 | 49 | 0.056609827 | 0.170760496 | NA | NA | 1 | NA | 0 | 0 | 1 |
| SE | 10040 | SYNE2 | SYNE2 | chr14 | + | 64029894 | 64030059 | 64027483 | 64027793 | 64031015 | 64031357 | 0 | 0 | 5 | 0 | 2 | 0 | 0 | 7 | 0 | 0 | 0 | 0 | 98 | 49 | 0.056662448 | 0.170760496 | NA | 0 | 1 | NA | 1 | NA | -0.5 |
| SE | 3801 | ZCCHC2 | ZCCHC2 | chr18 | + | 62544801 | 62544873 | 62542497 | 62542574 | 62550347 | 62550460 | 0 | 0 | 29 | 0 | 1 | 2 | 3 | 13 | 0 | 0 | 0 | 0 | 98 | 49 | 0.056686045 | 0.170760496 | NA | 0 | 0.879 | 1 | 1 | NA | -0.56 |
| SE | 1817 | CHD1 | CHD1 | chr5 | - | 98876397 | 98876558 | 98875071 | 98875113 | 98879551 | 98879728 | 0 | 1 | 2 | 0 | 0 | 0 | 0 | 8 | 2 | 0 | 9 | 0 | 98 | 49 | 0.056740038 | 0.170760496 | NA | 1 | 1 | NA | 0.308 | 1 | 0.346 |
| SE | 2275 | VNN3 | VNN3 | chr6 | - | 132728740 | 132728979 | 132724605 | 132724979 | 132734252 | 132734383 | 0 | 1 | 2 | 0 | 0 | 0 | 0 | 7 | 1 | 0 | 1 | 0 | 98 | 49 | 0.056900121 | 0.170760496 | NA | 1 | 1 | NA | 0.778 | 1 | 0.111 |
| A5SS | 2964 | USF1 | USF1 | chr1 | - | 161042554 | 161042670 | 161042585 | 161042670 | 161042115 | 161042217 | 0 | 0 | 9 | 0 | 2 | 2 | 0 | 6 | 0 | 0 | 0 | 0 | 80 | 49 | 0.057135855 | 0.170414363 | NA | 0 | 0.734 | NA | 1 | NA | -0.633 |
| SE | 3101 | MX1 | MX1 | chr21 | + | 41451166 | 41451243 | 41445999 | 41446141 | 41452620 | 41452869 | 0 | 2 | 24 | 0 | 0 | 0 | 1 | 15 | 1 | 0 | 1 | 0 | 98 | 49 | 0.057325035 | 0.170760496 | NA | 1 | 1 | 1 | 0.882 | 1 | 0.039 |
| SE | 12164 | USP32 | USP32 | chr17 | - | 60180544 | 60180637 | 60177329 | 60179428 | 60181323 | 60181748 | 0 | 0 | 12 | 0 | 1 | 0 | 0 | 10 | 0 | 0 | 0 | 0 | 98 | 49 | 0.057462192 | 0.170760496 | NA | 0 | 1 | NA | 1 | NA | -0.5 |
| SE | 6865 | ATG2A | ATG2A | chr11 | - | 64901961 | 64902176 | 64900493 | 64900629 | 64902259 | 64902386 | 0 | 0 | 2 | 0 | 0 | 0 | 0 | 2 | 0 | 0 | 1 | 3 | 98 | 49 | 0.057547789 | 0.170760496 | NA | NA | 1 | NA | 0.5 | 0 | 0.75 |
| SE | 8248 | PRRC2C | PRRC2C | chr1 | + | 171513000 | 171513172 | 171485526 | 171485735 | 171514535 | 171514645 | 0 | 1 | 1 | 0 | 0 | 0 | 2 | 2 | 1 | 0 | 1 | 0 | 98 | 49 | 0.057579548 | 0.170760496 | NA | 1 | 1 | 1 | 0.5 | 1 | 0.167 |
| SE | 8660 | DUSP1 | DUSP1 | chr5 | - | 172770160 | 172770306 | 172769574 | 172769794 | 172770585 | 172771200 | 4 | 105 | 424 | 0 | 1 | 1 | 27 | 197 | 285 | 0 | 2 | 6 | 98 | 49 | 0.057728535 | 0.170760496 | 1 | 0.981 | 0.995 | 1 | 0.98 | 0.96 | 0.012 |
| SE | 5304 | CD82 | CD82 | chr11 | + | 44605354 | 44605429 | 44605057 | 44605182 | 44615271 | 44615373 | 0 | 0 | 1 | 0 | 2 | 0 | 0 | 5 | 0 | 0 | 0 | 0 | 98 | 49 | 0.057729 | 0.170760496 | NA | 0 | 1 | NA | 1 | NA | -0.5 |
| SE | 4803 | CASP4 | CASP4 | chr11 | - | 104950924 | 104951098 | 104949542 | 104949777 | 104954746 | 104955001 | 0 | 0 | 17 | 0 | 1 | 2 | 4 | 9 | 0 | 0 | 0 | 0 | 98 | 49 | 0.057774487 | 0.170760496 | NA | 0 | 0.81 | 1 | 1 | NA | -0.595 |
| SE | 5518 | DENND5A | DENND5A | chr11 | - | 9145669 | 9145815 | 9144994 | 9145113 | 9147029 | 9147151 | 0 | 0 | 13 | 0 | 1 | 0 | 2 | 9 | 0 | 0 | 0 | 0 | 98 | 49 | 0.057774487 | 0.170760496 | NA | 0 | 1 | 1 | 1 | NA | -0.5 |
| SE | 5966 | 15-Sep | 15-Sep | chr1 | - | 86868052 | 86868102 | 86862444 | 86863605 | 86880661 | 86880725 | 0 | 0 | 12 | 0 | 1 | 0 | 0 | 9 | 0 | 0 | 0 | 0 | 98 | 49 | 0.057774487 | 0.170760496 | NA | 0 | 1 | NA | 1 | NA | -0.5 |
| SE | 6091 | SRRM2 | SRRM2 | chr16 | + | 2756333 | 2756606 | 2752328 | 2752846 | 2757471 | 2757579 | 0 | 0 | 7 | 0 | 1 | 0 | 0 | 9 | 0 | 0 | 0 | 0 | 98 | 49 | 0.057774487 | 0.170760496 | NA | 0 | 1 | NA | 1 | NA | -0.5 |
| SE | 11055 | ARFGAP3 | ARFGAP3 | chr22 | - | 42834241 | 42834325 | 42831548 | 42831636 | 42835361 | 42835493 | 0 | 0 | 8 | 0 | 1 | 0 | 0 | 9 | 0 | 0 | 0 | 0 | 98 | 49 | 0.057774487 | 0.170760496 | NA | 0 | 1 | NA | 1 | NA | -0.5 |
| SE | 15 | FLNA | FLNA | chrX | - | 154352180 | 154352447 | 154351883 | 154352021 | 154352771 | 154352924 | 0 | 1 | 15 | 0 | 0 | 0 | 5 | 8 | 3 | 0 | 2 | 0 | 98 | 49 | 0.058033181 | 0.170760496 | NA | 1 | 1 | 1 | 0.667 | 1 | 0.111 |
| SE | 9987 | TRANK1 | TRANK1 | chr3 | - | 36838371 | 36838504 | 36834761 | 36834907 | 36838612 | 36838716 | 0 | 0 | 3 | 0 | 0 | 0 | 0 | 3 | 0 | 0 | 0 | 2 | 98 | 49 | 0.058049942 | 0.170760496 | NA | NA | 1 | NA | 1 | 0 | 0.5 |
| SE | 8136 | ITGAX | ITGAX | chr16 | + | 31355892 | 31355998 | 31355133 | 31355291 | 31356624 | 31356728 | 0 | 17 | 42 | 0 | 0 | 0 | 8 | 52 | 24 | 0 | 1 | 0 | 98 | 49 | 0.058165611 | 0.170760496 | NA | 1 | 1 | 1 | 0.963 | 1 | 0.012 |
| SE | 4886 | DCAF11 | DCAF11 | chr14 | + | 24120837 | 24120991 | 24119710 | 24119896 | 24121364 | 24121517 | 0 | 0 | 4 | 0 | 1 | 0 | 0 | 8 | 0 | 0 | 0 | 0 | 98 | 49 | 0.058181256 | 0.170760496 | NA | 0 | 1 | NA | 1 | NA | -0.5 |
| RI | 523 | DDAH2 | DDAH2 | chr6 | - | 31728864 | 31730265 | 31728864 | 31729225 | 31730050 | 31730265 | 0 | 0 | 0 | 0 | 0 | 3 | 0 | 0 | 3 | 0 | 0 | 0 | 98 | 49 | 0.058431081 | 0.297702651 | NA | 1 | 0 | NA | NA | 1 | -0.5 |
| SE | 4634 | GMPR2 | GMPR2 | chr14 | + | 24238245 | 24238405 | 24237519 | 24237562 | 24238588 | 24239241 | 0 | 0 | 1 | 0 | 2 | 0 | 0 | 4 | 0 | 0 | 0 | 0 | 98 | 49 | 0.058521801 | 0.170760496 | NA | 0 | 1 | NA | 1 | NA | -0.5 |
| SE | 11517 | CMIP | CMIP | chr16 | + | 81664268 | 81664349 | 81660883 | 81660946 | 81670141 | 81670245 | 0 | 0 | 5 | 0 | 2 | 0 | 0 | 4 | 0 | 0 | 0 | 0 | 98 | 49 | 0.058521801 | 0.170760496 | NA | 0 | 1 | NA | 1 | NA | -0.5 |
| SE | 3630 | USP4 | USP4 | chr3 | - | 49286097 | 49286325 | 49284848 | 49284919 | 49292509 | 49292598 | 0 | 2 | 22 | 0 | 0 | 0 | 6 | 10 | 2 | 0 | 1 | 0 | 98 | 49 | 0.058749844 | 0.170760496 | NA | 1 | 1 | 1 | 0.833 | 1 | 0.056 |
| SE | 14092 | NLRP6 | NLRP6 | chr11 | + | 279833 | 279872 | 279326 | 279607 | 280083 | 281839 | 0 | 1 | 1 | 0 | 0 | 0 | 0 | 0 | 0 | 0 | 0 | 4 | 88 | 49 | 0.058768543 | 0.170760496 | NA | 1 | 1 | NA | NA | 0 | 1 |
| SE | 13371 | CCT7 | CCT7 | chr2 | + | 73243996 | 73244049 | 73243003 | 73243129 | 73244543 | 73244715 | 1 | 0 | 0 | 0 | 1 | 0 | 6 | 1 | 0 | 0 | 0 | 0 | 98 | 49 | 0.058830341 | 0.170760496 | 1 | 0 | NA | 1 | 1 | NA | -0.5 |
| SE | 1853 | DBNL | DBNL | chr7 | + | 44058428 | 44058480 | 44058128 | 44058280 | 44058901 | 44058983 | 0 | 1 | 3 | 0 | 0 | 0 | 0 | 3 | 1 | 0 | 1 | 0 | 98 | 49 | 0.058903155 | 0.170760496 | NA | 1 | 1 | NA | 0.6 | 1 | 0.2 |
| SE | 3097 | MX1 | MX1 | chr21 | + | 41437014 | 41437152 | 41435836 | 41436029 | 41439693 | 41439848 | 0 | 4 | 56 | 0 | 0 | 0 | 7 | 31 | 3 | 0 | 1 | 0 | 98 | 49 | 0.05891182 | 0.170760496 | NA | 1 | 1 | 1 | 0.939 | 1 | 0.02 |
| SE | 2611 | KIAA0922 | KIAA0922 | chr4 | + | 153592474 | 153592584 | 153591052 | 153591194 | 153593798 | 153593871 | 0 | 0 | 7 | 0 | 0 | 0 | 0 | 0 | 0 | 0 | 0 | 1 | 98 | 49 | 0.05894622 | 0.170760496 | NA | NA | 1 | NA | NA | 0 | 1 |
| SE | 6096 | CLN3 | CLN3 | chr16 | - | 28491481 | 28491560 | 28489289 | 28489386 | 28491713 | 28491835 | 0 | 0 | 7 | 0 | 0 | 0 | 0 | 0 | 0 | 0 | 0 | 1 | 98 | 49 | 0.05894622 | 0.170760496 | NA | NA | 1 | NA | NA | 0 | 1 |
| SE | 9258 | MOV10 | MOV10 | chr1 | + | 112694748 | 112694896 | 112694452 | 112694629 | 112695415 | 112695574 | 0 | 5 | 7 | 0 | 0 | 0 | 0 | 0 | 0 | 0 | 0 | 1 | 98 | 49 | 0.05894622 | 0.170760496 | NA | 1 | 1 | NA | NA | 0 | 1 |
| SE | 12991 | PLAUR | PLAUR | chr19 | - | 43655438 | 43655573 | 43652224 | 43652371 | 43656478 | 43656640 | 0 | 8 | 42 | 0 | 0 | 6 | 0 | 48 | 10 | 0 | 1 | 0 | 98 | 49 | 0.059088669 | 0.170760496 | NA | 1 | 0.778 | NA | 0.96 | 1 | -0.091 |
| SE | 13915 | CD300A | CD300A | chr17 | + | 74473535 | 74473874 | 74466382 | 74466743 | 74474531 | 74474685 | 0 | 0 | 1 | 0 | 1 | 1 | 0 | 6 | 0 | 0 | 0 | 0 | 98 | 49 | 0.05910855 | 0.170760496 | NA | 0 | 0.333 | NA | 1 | NA | -0.834 |
| SE | 4341 | SLC38A2 | SLC38A2 | chr12 | - | 46370511 | 46370627 | 46367266 | 46367340 | 46370775 | 46370857 | 0 | 2 | 6 | 0 | 0 | 1 | 0 | 0 | 0 | 0 | 2 | 0 | 98 | 49 | 0.05919998 | 0.170760496 | NA | 1 | 0.75 | NA | 0 | NA | 0.875 |
| SE | 5039 | RELT | RELT | chr11 | + | 73391143 | 73391223 | 73390754 | 73390921 | 73392210 | 73392468 | 0 | 0 | 2 | 0 | 0 | 0 | 0 | 6 | 0 | 0 | 1 | 2 | 98 | 49 | 0.05919998 | 0.170760496 | NA | NA | 1 | NA | 0.75 | 0 | 0.625 |
| SE | 7302 | PPP6R2 | PPP6R2 | chr22 | + | 50436987 | 50437068 | 50436366 | 50436452 | 50437505 | 50437603 | 0 | 0 | 2 | 0 | 0 | 0 | 0 | 0 | 0 | 0 | 0 | 2 | 98 | 49 | 0.05919998 | 0.170760496 | NA | NA | 1 | NA | NA | 0 | 1 |
| SE | 12511 | WDR7 | WDR7 | chr18 | + | 56717963 | 56718159 | 56696241 | 56696462 | 56731382 | 56731597 | 0 | 0 | 2 | 0 | 0 | 0 | 0 | 3 | 0 | 0 | 0 | 2 | 98 | 49 | 0.05919998 | 0.170760496 | NA | NA | 1 | NA | 1 | 0 | 0.5 |
| SE | 8184 | EVI2A | EVI2A | chr17 | - | 31320372 | 31320513 | 31316409 | 31319023 | 31321494 | 31321749 | 0 | 0 | 0 | 0 | 0 | 4 | 0 | 0 | 2 | 0 | 0 | 0 | 98 | 49 | 0.059276699 | 0.170760496 | NA | NA | 0 | NA | NA | 1 | -1 |
| SE | 9171 | KLF6 | KLF6 | chr10 | - | 3780105 | 3780229 | 3775995 | 3779590 | 3781640 | 3782214 | 0 | 3 | 14 | 1 | 0 | 0 | 0 | 6 | 3 | 0 | 1 | 0 | 98 | 49 | 0.059302042 | 0.170760496 | 0 | 1 | 1 | NA | 0.75 | 1 | -0.208 |
| SE | 7343 | FAM129A | FAM129A | chr1 | - | 184803584 | 184803692 | 184798078 | 184798190 | 184805945 | 184806056 | 0 | 1 | 38 | 0 | 0 | 0 | 0 | 7 | 1 | 0 | 1 | 0 | 98 | 49 | 0.059353638 | 0.170760496 | NA | 1 | 1 | NA | 0.778 | 1 | 0.111 |
| A5SS | 553 | HMGA1 | HMGA1 | chr6 | + | 34240736 | 34240915 | 34240736 | 34240882 | 34242711 | 34242795 | 0 | 1 | 4 | 0 | 0 | 3 | 0 | 5 | 1 | 0 | 0 | 0 | 82 | 49 | 0.059398863 | 0.170414363 | NA | 1 | 0.443 | NA | 1 | 1 | -0.278 |
| A3SS | 414 | DCTN3 | DCTN3 | chr9 | - | 34614041 | 34614198 | 34614041 | 34614101 | 34614709 | 34614768 | 1 | 1 | 0 | 0 | 0 | 0 | 0 | 0 | 0 | 0 | 3 | 0 | 98 | 49 | 0.059555994 | 0.197717869 | 1 | 1 | NA | NA | 0 | NA | 1 |
| A5SS | 2673 | RNF31 | RNF31 | chr14 | + | 24148257 | 24148477 | 24148257 | 24148413 | 24148641 | 24148701 | 0 | 1 | 1 | 0 | 0 | 3 | 0 | 0 | 0 | 0 | 3 | 0 | 98 | 49 | 0.059555994 | 0.170414363 | NA | 1 | 0.143 | NA | 0 | NA | 0.572 |
| SE | 6361 | ZSWIM8 | ZSWIM8 | chr10 | + | 73789366 | 73789539 | 73789095 | 73789190 | 73789716 | 73789824 | 0 | 1 | 4 | 0 | 0 | 0 | 0 | 0 | 0 | 0 | 3 | 0 | 98 | 49 | 0.059555994 | 0.170760496 | NA | 1 | 1 | NA | 0 | NA | 1 |
| SE | 7469 | SLTM | SLTM | chr15 | - | 58899468 | 58899883 | 58898802 | 58898852 | 58901259 | 58901287 | 0 | 1 | 0 | 0 | 0 | 0 | 0 | 0 | 0 | 0 | 3 | 0 | 98 | 49 | 0.059555994 | 0.170760496 | NA | 1 | NA | NA | 0 | NA | 1 |
| SE | 7470 | SLTM | SLTM | chr15 | - | 58899468 | 58899937 | 58898802 | 58898852 | 58901259 | 58901287 | 0 | 1 | 0 | 0 | 0 | 0 | 0 | 0 | 0 | 0 | 3 | 0 | 98 | 49 | 0.059555994 | 0.170760496 | NA | 1 | NA | NA | 0 | NA | 1 |
| SE | 9472 | USP33 | USP33 | chr1 | - | 77745420 | 77745599 | 77741616 | 77741748 | 77759642 | 77759879 | 0 | 0 | 1 | 0 | 0 | 0 | 0 | 0 | 0 | 0 | 0 | 3 | 98 | 49 | 0.059555994 | 0.170760496 | NA | NA | 1 | NA | NA | 0 | 1 |
| SE | 13888 | RIF1 | RIF1 | chr2 | + | 151410413 | 151410527 | 151409882 | 151410033 | 151411259 | 151411338 | 0 | 0 | 1 | 0 | 0 | 0 | 0 | 2 | 0 | 0 | 0 | 3 | 98 | 49 | 0.059555994 | 0.170760496 | NA | NA | 1 | NA | 1 | 0 | 0.5 |
| SE | 4181 | DDIT3 | DDIT3 | chr12 | - | 57517268 | 57517595 | 57516587 | 57517180 | 57517705 | 57517753 | 0 | 1 | 5 | 0 | 0 | 0 | 0 | 4 | 1 | 0 | 1 | 0 | 98 | 49 | 0.059568296 | 0.170760496 | NA | 1 | 1 | NA | 0.667 | 1 | 0.166 |
| SE | 12744 | DYSF | DYSF | chr2 | + | 71561751 | 71561944 | 71555964 | 71556071 | 71564057 | 71564213 | 0 | 5 | 41 | 0 | 0 | 0 | 2 | 10 | 26 | 0 | 0 | 1 | 98 | 49 | 0.059610573 | 0.170760496 | NA | 1 | 1 | 1 | 1 | 0.929 | 0.024 |
| SE | 2865 | POLDIP3 | POLDIP3 | chr22 | - | 42591954 | 42592084 | 42587505 | 42587572 | 42595536 | 42595603 | 0 | 0 | 0 | 0 | 0 | 2 | 0 | 3 | 3 | 0 | 0 | 0 | 98 | 49 | 0.059642855 | 0.170760496 | NA | NA | 0 | NA | 1 | 1 | -1 |
| SE | 7379 | RNF19B | RNF19B | chr1 | - | 32949568 | 32949774 | 32948221 | 32948363 | 32964050 | 32964813 | 0 | 0 | 7 | 0 | 2 | 0 | 0 | 3 | 0 | 0 | 0 | 0 | 98 | 49 | 0.059642855 | 0.170760496 | NA | 0 | 1 | NA | 1 | NA | -0.5 |
| SE | 7408 | RBM23 | RBM23 | chr14 | - | 22905605 | 22905659 | 22905335 | 22905453 | 22906194 | 22906368 | 0 | 0 | 2 | 0 | 2 | 1 | 0 | 3 | 0 | 0 | 0 | 0 | 98 | 49 | 0.059642855 | 0.170760496 | NA | 0 | 0.5 | NA | 1 | NA | -0.75 |
| SE | 9772 | FRYL | FRYL | chr4 | - | 48512480 | 48512688 | 48510834 | 48510984 | 48515027 | 48515275 | 0 | 0 | 2 | 0 | 2 | 0 | 7 | 3 | 0 | 0 | 0 | 0 | 98 | 49 | 0.059642855 | 0.170760496 | NA | 0 | 1 | 1 | 1 | NA | -0.5 |
| SE | 11703 | POLR2A | POLR2A | chr17 | + | 7511810 | 7512014 | 7511405 | 7511588 | 7512157 | 7512344 | 0 | 2 | 19 | 0 | 0 | 0 | 0 | 5 | 2 | 0 | 1 | 0 | 98 | 49 | 0.059650539 | 0.170760496 | NA | 1 | 1 | NA | 0.714 | 1 | 0.143 |
| SE | 6864 | ATG2A | ATG2A | chr11 | - | 64900883 | 64901092 | 64900493 | 64900629 | 64901961 | 64902176 | 0 | 0 | 0 | 0 | 0 | 1 | 0 | 5 | 5 | 0 | 0 | 0 | 98 | 49 | 0.059786145 | 0.170760496 | NA | NA | 0 | NA | 1 | 1 | -1 |
| A5SS | 2457 | SCARF1 | SCARF1 | chr17 | - | 1645559 | 1645789 | 1645596 | 1645789 | 1645177 | 1645239 | 0 | 0 | 0 | 0 | 0 | 1 | 0 | 3 | 5 | 0 | 3 | 0 | 86 | 49 | 0.059797532 | 0.170414363 | NA | NA | 0 | NA | 0.363 | 1 | -0.681 |
| A3SS | 1375 | ARHGAP25 | ARHGAP25 | chr2 | + | 68807272 | 68807480 | 68807275 | 68807480 | 68787839 | 68787956 | 0 | 0 | 5 | 0 | 1 | 0 | 0 | 5 | 0 | 0 | 0 | 0 | 52 | 49 | 0.059800864 | 0.197717869 | NA | 0 | 1 | NA | 1 | NA | -0.5 |
| SE | 12948 | SMCHD1 | SMCHD1 | chr18 | + | 2778168 | 2778239 | 2777805 | 2777915 | 2784449 | 2784621 | 0 | 2 | 47 | 0 | 0 | 1 | 2 | 28 | 1 | 0 | 4 | 0 | 98 | 49 | 0.059897198 | 0.170760496 | NA | 1 | 0.959 | 1 | 0.778 | 1 | 0.053 |
| SE | 5027 | OSBPL8 | OSBPL8 | chr12 | - | 76459858 | 76459895 | 76450850 | 76450988 | 76487509 | 76487618 | 0 | 1 | 0 | 0 | 0 | 2 | 0 | 0 | 0 | 0 | 3 | 0 | 86 | 49 | 0.059993631 | 0.170760496 | NA | 1 | 0 | NA | 0 | NA | 0.5 |
| SE | 9705 | ABTB1 | ABTB1 | chr3 | + | 127676966 | 127677083 | 127676535 | 127676581 | 127677167 | 127677286 | 0 | 1 | 20 | 0 | 0 | 0 | 5 | 4 | 1 | 0 | 1 | 0 | 98 | 49 | 0.060016261 | 0.170760496 | NA | 1 | 1 | 1 | 0.667 | 1 | 0.111 |
| SE | 4178 | DDIT3 | DDIT3 | chr12 | - | 57517268 | 57517438 | 57516587 | 57517180 | 57517705 | 57517753 | 0 | 1 | 6 | 0 | 0 | 0 | 0 | 5 | 1 | 0 | 1 | 0 | 98 | 49 | 0.060034631 | 0.170760496 | NA | 1 | 1 | NA | 0.714 | 1 | 0.143 |
| SE | 364 | SEMA4D | SEMA4D | chr9 | - | 89418055 | 89418118 | 89405350 | 89405699 | 89455887 | 89455953 | 0 | 0 | 5 | 0 | 0 | 0 | 0 | 7 | 0 | 1 | 0 | 1 | 98 | 49 | 0.060038159 | 0.170760496 | NA | NA | 1 | 0 | 1 | 0 | 0.667 |
| A5SS | 1404 | RBMS1 | RBMS1 | chr2 | - | 160286968 | 160287084 | 160286977 | 160287084 | 160284994 | 160285044 | 1 | 0 | 2 | 0 | 0 | 2 | 0 | 1 | 0 | 4 | 0 | 0 | 58 | 49 | 0.060055861 | 0.170414363 | 1 | NA | 0.458 | 0 | 1 | NA | 0.229 |
| A3SS | 413 | DCTN3 | DCTN3 | chr9 | - | 34614041 | 34614132 | 34614041 | 34614101 | 34614709 | 34614768 | 1 | 1 | 0 | 0 | 0 | 0 | 0 | 0 | 0 | 0 | 3 | 0 | 80 | 49 | 0.060211792 | 0.197717869 | 1 | 1 | NA | NA | 0 | NA | 1 |
| SE | 12010 | GPR108 | GPR108 | chr19 | - | 6731890 | 6731934 | 6731472 | 6731522 | 6732024 | 6732155 | 0 | 0 | 5 | 0 | 0 | 0 | 0 | 5 | 0 | 0 | 0 | 1 | 93 | 49 | 0.060224876 | 0.170760496 | NA | NA | 1 | NA | 1 | 0 | 0.5 |
| SE | 8079 | NADK | NADK | chr1 | - | 1757180 | 1757310 | 1756257 | 1756343 | 1761951 | 1762035 | 0 | 1 | 24 | 0 | 0 | 0 | 0 | 6 | 1 | 0 | 1 | 0 | 98 | 49 | 0.060230863 | 0.170760496 | NA | 1 | 1 | NA | 0.75 | 1 | 0.125 |
| SE | 2349 | FNDC3B | FNDC3B | chr3 | + | 172346326 | 172346440 | 172344085 | 172344258 | 172347211 | 172347361 | 0 | 1 | 6 | 0 | 0 | 0 | 0 | 7 | 3 | 0 | 2 | 0 | 98 | 49 | 0.060484453 | 0.170760496 | NA | 1 | 1 | NA | 0.636 | 1 | 0.182 |
| SE | 4177 | DDIT3 | DDIT3 | chr12 | - | 57517268 | 57517424 | 57516587 | 57517180 | 57517705 | 57517753 | 0 | 1 | 5 | 0 | 0 | 0 | 0 | 3 | 1 | 0 | 1 | 0 | 98 | 49 | 0.060546566 | 0.170760496 | NA | 1 | 1 | NA | 0.6 | 1 | 0.2 |
| SE | 4179 | DDIT3 | DDIT3 | chr12 | - | 57517268 | 57517519 | 57516587 | 57517180 | 57517705 | 57517753 | 0 | 1 | 5 | 0 | 0 | 0 | 0 | 3 | 1 | 0 | 1 | 0 | 98 | 49 | 0.060546566 | 0.170760496 | NA | 1 | 1 | NA | 0.6 | 1 | 0.2 |
| SE | 4180 | DDIT3 | DDIT3 | chr12 | - | 57517268 | 57517542 | 57516587 | 57517180 | 57517705 | 57517753 | 0 | 1 | 5 | 0 | 0 | 0 | 0 | 3 | 1 | 0 | 1 | 0 | 98 | 49 | 0.060546566 | 0.170760496 | NA | 1 | 1 | NA | 0.6 | 1 | 0.2 |
| SE | 13577 | STK17B | STK17B | chr2 | - | 196163261 | 196163427 | 196156438 | 196156651 | 196171332 | 196171612 | 0 | 21 | 136 | 0 | 0 | 0 | 6 | 68 | 29 | 0 | 1 | 0 | 98 | 49 | 0.060570323 | 0.170760496 | NA | 1 | 1 | 1 | 0.971 | 1 | 0.01 |
| SE | 1540 | HLA-F | HLA-F | chr6 | + | 29725030 | 29725306 | 29724172 | 29724448 | 29725446 | 29725563 | 0 | 4 | 52 | 0 | 0 | 0 | 0 | 23 | 5 | 0 | 1 | 0 | 98 | 49 | 0.06060798 | 0.170760496 | NA | 1 | 1 | NA | 0.92 | 1 | 0.04 |
| SE | 11556 | KIAA0513 | KIAA0513 | chr16 | + | 85078414 | 85078455 | 85077424 | 85077632 | 85078924 | 85079003 | 0 | 0 | 9 | 0 | 1 | 0 | 0 | 4 | 0 | 0 | 0 | 0 | 90 | 49 | 0.060658984 | 0.170760496 | NA | 0 | 1 | NA | 1 | NA | -0.5 |
| A3SS | 3134 | STX4 | STX4 | chr16 | + | 31034203 | 31034325 | 31034225 | 31034325 | 31034012 | 31034114 | 0 | 0 | 3 | 0 | 1 | 2 | 1 | 4 | 0 | 0 | 0 | 0 | 71 | 49 | 0.06066392 | 0.197717869 | NA | 0 | 0.509 | 1 | 1 | NA | -0.746 |
| SE | 2842 | HDAC10 | HDAC10 | chr22 | - | 50247691 | 50247776 | 50246874 | 50246966 | 50247889 | 50248145 | 0 | 0 | 0 | 0 | 0 | 1 | 0 | 0 | 4 | 0 | 2 | 0 | 98 | 49 | 0.060672385 | 0.170760496 | NA | NA | 0 | NA | 0 | 1 | -0.5 |
| SE | 6993 | MATR3 | MATR3 | chr5 | + | 139307238 | 139308327 | 139293643 | 139293805 | 139314674 | 139314736 | 0 | 0 | 0 | 0 | 1 | 0 | 0 | 4 | 0 | 0 | 0 | 0 | 98 | 49 | 0.060672385 | 0.170760496 | NA | 0 | NA | | 1 | NA | -1 |
| SE | 7078 | ERBB2IP | ERBB2IP | chr5 | + | 66075023 | 66075230 | 66053405 | 66054951 | 66076315 | 66076408 | 0 | 0 | 2 | 0 | 1 | 0 | 0 | 4 | 0 | 0 | 0 | 0 | 98 | 49 | 0.060672385 | 0.170760496 | NA | 0 | 1 | NA | 1 | NA | -0.5 |
| SE | 1858 | AP3B1 | AP3B1 | chr5 | - | 78228143 | 78228239 | 78227371 | 78227532 | 78240861 | 78240936 | 0 | 1 | 13 | 0 | 0 | 0 | 0 | 5 | 1 | 0 | 1 | 0 | 98 | 49 | 0.060716183 | 0.170760496 | NA | 1 | 1 | NA | 0.714 | 1 | 0.143 |
| A3SS | 2183 | PPP4R1 | PPP4R1 | chr18 | - | 9595017 | 9595153 | 9595017 | 9595102 | 9614225 | 9614270 | 0 | 0 | 4 | 0 | 0 | 0 | 0 | 2 | 0 | 0 | 2 | 1 | 98 | 49 | 0.060734355 | 0.197717869 | NA | NA | 1 | NA | 0.333 | 0 | 0.834 |
| SE | 788 | PARP10 | PARP10 | chr8 | - | 143984007 | 143984104 | 143983166 | 143983811 | 143984209 | 143984431 | 0 | 0 | 4 | 0 | 0 | 0 | 0 | 4 | 0 | 0 | 0 | 1 | 98 | 49 | 0.060734355 | 0.170760496 | NA | NA | 1 | NA | 1 | 0 | 0.5 |
| SE | 5889 | SIPA1L1 | SIPA1L1 | chr14 | + | 71723646 | 71723886 | 71709221 | 71709664 | 71724669 | 71724835 | 0 | 0 | 4 | 0 | 0 | 0 | 0 | 0 | 0 | 0 | 0 | 1 | 98 | 49 | 0.060734355 | 0.170760496 | NA | NA | 1 | NA | NA | 0 | 1 |

| | | | | | | | | | | | | | | | | | | | | | | | | | | | | | | | | | | |
|---|---|---|---|---|---|---|---|---|---|---|---|---|---|---|---|---|---|---|---|---|---|---|---|---|---|---|---|---|---|---|---|---|---|---|
| A5SS | 3026 | ABCG1 | ABCG1 | chr21 | + | 42287888 | 42288073 | 42287888 | 42288037 | 42288210 | 42288312 | 0 | 2 | 3 | 0 | 0 | 2 | 0 | 8 | 2 | 0 | 0 | 0 | 85 | 49 | 0.060789913 | 0.170414363 | NA | 1 | 0.464 | NA | 1 | 1 | -0.268 |
| SE | 4502 | PTPN6 | PTPN6 | chr12 | + | 6954804 | 6954994 | 6951982 | 6952177 | 6955150 | 6955267 | 0 | 2 | 31 | 0 | 0 | 0 | 0 | 21 | 1 | 0 | 1 | 0 | 98 | 49 | 0.061131927 | 0.170760496 | NA | 1 | 1 | NA | 0.913 | 1 | 0.043 |
| SE | 10395 | HES4 | HES4 | chr1 | - | 999691 | 999787 | 999525 | 999613 | 999865 | 1000172 | 0 | 0 | 0 | 0 | 1 | 1 | 0 | 1 | 0 | 0 | 4 | 1 | 98 | 49 | 0.061306069 | 0.170760496 | NA | 0 | 0 | NA | 0.111 | 0 | -0.056 |
| RI | 672 | PDE4D | PDE4D | chr5 | - | 59038858 | 59039512 | 59038858 | 59038971 | 59039057 | 59039512 | 0 | 0 | 1 | 0 | 0 | 0 | 3 | 3 | 0 | 0 | 0 | 2 | 98 | 49 | 0.061401334 | 0.297702651 | NA | NA | 1 | 1 | 1 | 0 | 0.333 |
| SE | 7868 | VRK3 | VRK3 | chr19 | - | 49989638 | 49989771 | 49988371 | 49988492 | 49992859 | 49992952 | 0 | 1 | 2 | 0 | 0 | 0 | 0 | 0 | 0 | 0 | 2 | 0 | 98 | 49 | 0.061401334 | 0.170760496 | NA | 1 | 1 | NA | 0 | NA | 1 |
| SE | 8054 | CUL4B | CUL4B | chrX | - | 120544480 | 120544643 | 120544113 | 120544203 | 120545443 | 120545517 | 0 | 0 | 1 | 0 | 0 | 0 | 0 | 1 | 0 | 0 | 0 | 2 | 98 | 49 | 0.061401334 | 0.170760496 | NA | NA | 1 | NA | 1 | 0 | 0.5 |
| SE | 11362 | RSRC2 | RSRC2 | chr12 | - | 122520503 | 122520581 | 122518838 | 122519029 | 122521384 | 122521428 | 0 | 1 | 3 | 0 | 0 | 1 | 0 | 0 | 0 | 0 | 2 | 0 | 98 | 49 | 0.061401334 | 0.170760496 | NA | 1 | 0.6 | NA | 0 | NA | 0.8 |
| SE | 12007 | DHX8 | DHX8 | chr17 | + | 43508338 | 43508520 | 43507808 | 43508019 | 43513361 | 43513502 | 0 | 1 | 1 | 0 | 0 | 0 | 0 | 0 | 0 | 0 | 2 | 0 | 98 | 49 | 0.061401334 | 0.170760496 | NA | 1 | 1 | NA | 0 | NA | 1 |
| SE | 9457 | PFKFB3 | PFKFB3 | chr10 | + | 6221640 | 6221745 | 6221380 | 6221527 | 6222854 | 6222984 | 0 | 16 | 35 | 0 | 0 | 0 | 0 | 44 | 26 | 0 | 1 | 0 | 98 | 49 | 0.0614161 | 0.170760496 | NA | 1 | 1 | NA | 0.957 | 1 | 0.022 |
| A3SS | 3638 | MAP2K7 | MAP2K7 | chr19 | + | 7911414 | 7911578 | 7911435 | 7911578 | 7911249 | 7911330 | 0 | 0 | 0 | 0 | 1 | 2 | 0 | 0 | 2 | 0 | 0 | 0 | 70 | 49 | 0.061460457 | 0.197717869 | NA | 0 | 0 | NA | NA | 1 | -1 |
| A5SS | 725 | FAM193A | FAM193A | chr4 | + | 2694945 | 2695142 | 2694945 | 2695129 | 2696362 | 2696593 | 0 | 0 | 0 | 0 | 2 | 0 | 0 | 2 | 0 | 0 | 0 | 0 | 62 | 49 | 0.061467502 | 0.170414363 | NA | 0 | NA | NA | 1 | NA | -1 |
| A3SS | 16 | ALG13 | ALG13 | chrX | + | 111682131 | 111682294 | 111682235 | 111682294 | 111681169 | 111681424 | 0 | 0 | 0 | 0 | 2 | 0 | 0 | 2 | 0 | 0 | 0 | 0 | 98 | 49 | 0.061489123 | 0.197717869 | NA | 0 | NA | NA | 1 | NA | -1 |
| A5SS | 1081 | WIPI2 | WIPI2 | chr7 | + | 5193117 | 5193345 | 5193117 | 5193171 | 5199575 | 5199658 | 0 | 0 | 0 | 0 | 0 | 2 | 0 | 0 | 2 | 0 | 0 | 0 | 98 | 49 | 0.061489123 | 0.170414363 | NA | NA | 0 | NA | NA | 1 | -1 |
| RI | 843 | FAM96A | FAM96A | chr15 | - | 64072558 | 64075537 | 64072558 | 64073028 | 64075491 | 64075537 | 0 | 0 | 0 | 0 | 0 | 2 | 1 | 0 | 2 | 0 | 1 | 0 | 98 | 49 | 0.061489123 | 0.297702651 | NA | NA | 0 | 1 | 0 | 1 | -0.667 |
| SE | 4053 | CASC3 | CASC3 | chr17 | + | 40167849 | 40167948 | 40167497 | 40167612 | 40168202 | 40168417 | 0 | 0 | 5 | 0 | 2 | 0 | 0 | 2 | 0 | 0 | 0 | 0 | 98 | 49 | 0.061489123 | 0.170760496 | NA | 0 | 1 | NA | 1 | NA | -0.5 |
| SE | 10886 | AKAP13 | AKAP13 | chr15 | + | 85662387 | 85662453 | 85658536 | 85658590 | 85664562 | 85664755 | 0 | 0 | 2 | 0 | 2 | 11 | 0 | 2 | 0 | 0 | 0 | 0 | 98 | 49 | 0.061489123 | 0.170760496 | NA | 0 | 0.083 | NA | 1 | NA | -0.959 |
| SE | 11013 | RHOT2 | RHOT2 | chr16 | + | 669552 | 669606 | 668655 | 668699 | 670122 | 670175 | 0 | 0 | 1 | 0 | 2 | 0 | 0 | 2 | 0 | 0 | 0 | 0 | 98 | 49 | 0.061489123 | 0.170760496 | NA | 0 | 1 | NA | 1 | NA | -0.5 |
| SE | 11296 | CREBBP | CREBBP | chr16 | - | 3791980 | 3792094 | 3782683 | 3782926 | 3793385 | 3793626 | 0 | 1 | 9 | 0 | 0 | 0 | 0 | 3 | 1 | 0 | 1 | 0 | 98 | 49 | 0.061515403 | 0.170760496 | NA | 1 | 1 | NA | 0.6 | 1 | 0.2 |
| A3SS | 3506 | DYRK1B | DYRK1B | chr19 | - | 39826671 | 39826987 | 39826671 | 39826867 | 39827284 | 39827425 | 0 | 0 | 3 | 0 | 0 | 0 | 0 | 0 | 0 | 0 | 2 | 1 | 98 | 49 | 0.06170155 | 0.197717869 | NA | NA | 1 | NA | 0 | 0 | 1 |
| A5SS | 1533 | BTF3 | BTF3 | chr5 | + | 73498424 | 73498799 | 73498424 | 73498558 | 73499133 | 73499202 | 3 | 0 | 2 | 0 | 0 | 2 | 0 | 2 | 0 | 1 | 3 | 0 | 98 | 49 | 0.06170155 | 0.170414363 | 1 | NA | 0.333 | 0 | 0.25 | NA | 0.541 |
| SE | 7580 | MMP8 | MMP8 | chr11 | - | 102722990 | 102723078 | 102722428 | 102722673 | 102724753 | 102724954 | 0 | 3 | 3 | 0 | 0 | 1 | 0 | 0 | 0 | 0 | 1 | 0 | 98 | 49 | 0.06170155 | 0.170760496 | NA | 1 | 0.6 | NA | 0 | NA | 0.8 |
| SE | 7581 | MMP8 | MMP8 | chr11 | - | 102722990 | 102723081 | 102722428 | 102722673 | 102724753 | 102724954 | 0 | 3 | 3 | 0 | 0 | 1 | 0 | 0 | 0 | 0 | 1 | 0 | 98 | 49 | 0.06170155 | 0.170760496 | NA | 1 | 0.6 | NA | 0 | NA | 0.8 |
| SE | 8154 | CD302 | CD302 | chr2 | - | 159783358 | 159783469 | 159780881 | 159780998 | 159798131 | 159798255 | 0 | 0 | 3 | 0 | 0 | 0 | 0 | 3 | 0 | 0 | 0 | 1 | 98 | 49 | 0.06170155 | 0.170760496 | NA | NA | 1 | NA | 1 | 0 | 0.5 |
| SE | 9121 | KDM4B | KDM4B | chr19 | + | 5071009 | 5071059 | 5047475 | 5047669 | 5077366 | 5077470 | 0 | 0 | 3 | 0 | 0 | 0 | 0 | 4 | 0 | 0 | 0 | 1 | 98 | 49 | 0.06170155 | 0.170760496 | NA | NA | 1 | NA | 1 | 0 | 0.5 |
| SE | 10516 | ZNF316 | ZNF316 | chr7 | + | 6642381 | 6642764 | 6641824 | 6641962 | 6642963 | 6643073 | 0 | 3 | 0 | 0 | 0 | 0 | 0 | 0 | 0 | 0 | 1 | 0 | 98 | 49 | 0.06170155 | 0.170760496 | NA | 1 | NA | NA | 0 | NA | 1 |
| SE | 11106 | MGRN1 | MGRN1 | chr16 | + | 4680031 | 4680097 | 4677462 | 4677572 | 4681549 | 4681776 | 0 | 0 | 3 | 0 | 0 | 0 | 0 | 1 | 0 | 0 | 0 | 1 | 98 | 49 | 0.06170155 | 0.170760496 | NA | NA | 1 | NA | 1 | 0 | 0.5 |
| A3SS | 910 | LIME1 | LIME1 | chr20 | + | 63737972 | 63738060 | 63738007 | 63738060 | 63737820 | 63737902 | 0 | 0 | 1 | 0 | 1 | 0 | 0 | 3 | 0 | 0 | 0 | 0 | 84 | 49 | 0.061899751 | 0.197717869 | NA | 0 | 1 | NA | 1 | NA | -0.5 |
| A3SS | 2475 | ZBTB17 | ZBTB17 | chr1 | - | 15946153 | 15946316 | 15946153 | 15946294 | 15946934 | 15947123 | 0 | 0 | 1 | 0 | 1 | 2 | 0 | 3 | 0 | 0 | 0 | 0 | 71 | 49 | 0.061903 | 0.197717869 | NA | 0 | 0.257 | NA | 1 | NA | -0.871 |
| A3SS | 122 | FAM219A | FAM219A | chr9 | - | 34398183 | 34401122 | 34398183 | 34401119 | 34401665 | 34401720 | 0 | 0 | 0 | 0 | 1 | 0 | 0 | 3 | 0 | 0 | 0 | 0 | 52 | 49 | 0.061903774 | 0.197717869 | NA | 0 | NA | NA | 1 | NA | -1 |
| A3SS | 3899 | PPP1R12C | PPP1R12C | chr19 | - | 55092782 | 55092868 | 55092782 | 55092865 | 55093015 | 55093076 | 0 | 0 | 0 | 0 | 0 | 1 | 0 | 0 | 3 | 0 | 1 | 0 | 52 | 49 | 0.061903774 | 0.197717869 | NA | NA | 0 | NA | 0 | 1 | -0.5 |
| RI | 528 | NFKB2 | NFKB2 | chr10 | + | 102395595 | 102395980 | 102395595 | 102395796 | 102395887 | 102395980 | 0 | 0 | 0 | 0 | 0 | 1 | 0 | 5 | 3 | 0 | 0 | 0 | 98 | 49 | 0.061911422 | 0.297702651 | NA | NA | 0 | NA | 1 | 1 | -1 |
| SE | 4260 | SLC22A4 | SLC22A4 | chr5 | + | 132312160 | 132312264 | 132294451 | 132295009 | 132313613 | 132313768 | 0 | 0 | 4 | 0 | 1 | 0 | 0 | 3 | 0 | 0 | 0 | 0 | 98 | 49 | 0.061911422 | 0.170760496 | NA | 0 | 1 | NA | 1 | NA | -0.5 |
| SE | 5368 | BAZ1A | BAZ1A | chr14 | - | 34826012 | 34826156 | 34810934 | 34811036 | 34874491 | 34874662 | 0 | 0 | 13 | 0 | 1 | 0 | 2 | 3 | 0 | 0 | 0 | 0 | 98 | 49 | 0.061911422 | 0.170760496 | NA | 0 | 1 | 1 | 1 | NA | -0.5 |
| SE | 6045 | GADD45A | GADD45A | chr1 | + | 67686349 | 67686587 | 67685176 | 67685538 | 67687660 | 67688338 | 0 | 0 | 0 | 0 | 0 | 1 | 0 | 1 | 3 | 0 | 0 | 0 | 98 | 49 | 0.061911422 | 0.170760496 | NA | NA | 0 | NA | 1 | 1 | -1 |
| SE | 12169 | HDAC7 | HDAC7 | chr12 | - | 47786584 | 47786703 | 47785751 | 47785885 | 47787711 | 47787809 | 0 | 0 | 4 | 0 | 1 | 1 | 0 | 3 | 0 | 0 | 0 | 0 | 98 | 49 | 0.061911422 | 0.170760496 | NA | 0 | 0.667 | NA | 1 | NA | -0.666 |
| SE | 12235 | FAM104A | FAM104A | chr17 | - | 73227164 | 73227264 | 73207352 | 73209768 | 73232085 | 73232394 | 0 | 0 | 1 | 0 | 1 | 0 | 0 | 3 | 0 | 0 | 0 | 0 | 98 | 49 | 0.061911422 | 0.170760496 | NA | 0 | 1 | NA | 1 | NA | -0.5 |
| A5SS | 2118 | CSNK1A1 | CSNK1A1 | chr5 | - | 149505446 | 149505595 | 149505482 | 149505595 | 149495893 | 149496860 | 3 | 0 | 11 | 0 | 0 | 1 | 0 | 5 | 0 | 1 | 1 | 0 | 85 | 49 | 0.062188913 | 0.170414363 | 1 | NA | 0.864 | 0 | 0.742 | NA | 0.561 |
| SE | 12245 | DYRK1B | DYRK1B | chr19 | - | 39826951 | 39826987 | 39826671 | 39826867 | 39827284 | 39827425 | 0 | 0 | 3 | 0 | 0 | 0 | 0 | 0 | 0 | 0 | 2 | 1 | 85 | 49 | 0.062188913 | 0.170760496 | NA | NA | 1 | NA | 0 | 0 | 1 |
| A3SS | 2080 | AP2S1 | AP2S1 | chr19 | - | 46839464 | 46839620 | 46839464 | 46839578 | 46845992 | 46846142 | 0 | 0 | 0 | 0 | 0 | 5 | 0 | 0 | 1 | 5 | 2 | 0 | 91 | 49 | 0.062394347 | 0.197717869 | NA | NA | 0 | 0 | 0 | 1 | -0.333 |
| SE | 4424 | PREX1 | PREX1 | chr20 | - | 48658135 | 48658228 | 48657039 | 48657188 | 48666282 | 48666355 | 0 | 2 | 10 | 0 | 0 | 0 | 0 | 20 | 2 | 0 | 1 | 0 | 98 | 49 | 0.062524268 | 0.170760496 | NA | 1 | 1 | NA | 0.909 | 1 | 0.045 |
| SE | 2274 | VNN3 | VNN3 | chr6 | - | 132728740 | 132728933 | 132724605 | 132724979 | 132734252 | 132734383 | 0 | 1 | 5 | 0 | 0 | 0 | 0 | 9 | 1 | 0 | 1 | 0 | 98 | 49 | 0.062727245 | 0.170760496 | NA | 1 | 1 | NA | 0.818 | 1 | 0.091 |
| SE | 13163 | PPP1CB | PPP1CB | chr2 | + | 28752037 | 28752176 | 28751747 | 28751820 | 28776850 | 28776982 | 0 | 2 | 2 | 0 | 0 | 0 | 1 | 4 | 3 | 0 | 1 | 0 | 98 | 49 | 0.06302451 | 0.170760496 | NA | 1 | 1 | 1 | 0.667 | 1 | 0.111 |
| A3SS | 2776 | MYADM | MYADM | chr19 | + | 53869689 | 53869838 | 53869753 | 53869838 | 53867864 | 53867943 | 0 | 0 | 2 | 0 | 0 | 0 | 3 | 0 | 0 | 0 | 0 | 1 | 98 | 49 | 0.063117334 | 0.197717869 | NA | NA | 1 | 1 | NA | 0 | 0.5 |
| A5SS | 1568 | POLD4 | POLD4 | chr11 | - | 67353302 | 67353596 | 67353363 | 67353596 | 67352987 | 67353077 | 0 | 1 | 2 | 0 | 0 | 0 | 0 | 0 | 0 | 0 | 0 | 1 | 98 | 49 | 0.063117334 | 0.170414363 | NA | 1 | 1 | NA | NA | 0 | 1 |
| SE | 498 | NAPRT | NAPRT | chr8 | - | 143577061 | 143577177 | 143576645 | 143576842 | 143577268 | 143577399 | 0 | 2 | 4 | 0 | 0 | 0 | 0 | 0 | 0 | 0 | 1 | 0 | 98 | 49 | 0.063117334 | 0.170760496 | NA | 1 | 1 | NA | 0 | NA | 1 |
| SE | 3230 | TMEM189 | TMEM189 | chr20 | - | 50130855 | 50130947 | 50129545 | 50129690 | 50143501 | 50143621 | 0 | 2 | 1 | 0 | 0 | 0 | 0 | 0 | 0 | 0 | 1 | 0 | 98 | 49 | 0.063117334 | 0.170760496 | NA | 1 | 1 | NA | 0 | NA | 1 |
| SE | 3231 | TMEM189 | TMEM189 | chr20 | - | 50130864 | 50130947 | 50129545 | 50129690 | 50143501 | 50143621 | 0 | 2 | 0 | 0 | 0 | 0 | 0 | 0 | 0 | 0 | 1 | 0 | 98 | 49 | 0.063117334 | 0.170760496 | NA | 1 | NA | NA | 0 | NA | 1 |
| SE | 3810 | CPNE1 | CPNE1 | chr20 | - | 35658929 | 35658982 | 35632794 | 35632923 | 35664759 | 35664937 | 0 | 0 | 2 | 0 | 0 | 0 | 0 | 0 | 0 | 0 | 0 | 1 | 98 | 49 | 0.063117334 | 0.170760496 | NA | NA | 1 | NA | NA | 0 | 1 |
| SE | 3811 | CPNE1 | CPNE1 | chr20 | - | 35658929 | 35659014 | 35632794 | 35632923 | 35664759 | 35664956 | 0 | 0 | 2 | 0 | 0 | 0 | 0 | 0 | 0 | 0 | 0 | 1 | 98 | 49 | 0.063117334 | 0.170760496 | NA | NA | 1 | NA | NA | 0 | 1 |
| SE | 6736 | MTOR | MTOR | chr1 | - | 11256932 | 11257165 | 11255991 | 11256192 | 11258484 | 11258593 | 0 | 2 | 4 | 0 | 0 | 0 | 0 | 0 | 0 | 0 | 1 | 0 | 98 | 49 | 0.063117334 | 0.170760496 | NA | 1 | 1 | NA | 0 | NA | 1 |
| SE | 6925 | DMTN | DMTN | chr8 | + | 22067084 | 22067159 | 22066704 | 22066893 | 22067526 | 22067682 | 0 | 2 | 1 | 0 | 0 | 0 | 0 | 0 | 0 | 0 | 1 | 0 | 98 | 49 | 0.063117334 | 0.170760496 | NA | 1 | 1 | NA | 0 | NA | 1 |
| SE | 9827 | SHISA5 | SHISA5 | chr3 | - | 48503094 | 48503193 | 48501136 | 48501293 | 48504422 | 48504814 | 0 | 0 | 2 | 0 | 0 | 0 | 0 | 0 | 0 | 0 | 0 | 1 | 98 | 49 | 0.063117334 | 0.170760496 | NA | NA | 1 | NA | NA | 0 | 1 |
| SE | 10041 | SYNE2 | SYNE2 | chr14 | + | 64031015 | 64031357 | 64027483 | 64027793 | 64047999 | 64048155 | 0 | 2 | 0 | 0 | 0 | 0 | 0 | 0 | 0 | 0 | 1 | 0 | 98 | 49 | 0.063117334 | 0.170760496 | NA | 1 | NA | NA | 0 | NA | 1 |
| SE | 10233 | SYNE3 | SYNE3 | chr14 | - | 95467794 | 95467967 | 95465930 | 95466240 | 95475677 | 95475836 | 0 | 2 | 0 | 0 | 0 | 0 | 0 | 0 | 0 | 0 | 1 | 0 | 98 | 49 | 0.063117334 | 0.170760496 | NA | 1 | NA | NA | 0 | NA | 1 |
| SE | 12575 | ARID3A | ARID3A | chr19 | + | 964832 | 965080 | 964247 | 964431 | 966571 | 966868 | 0 | 2 | 4 | 0 | 0 | 0 | 0 | 0 | 0 | 0 | 1 | 0 | 98 | 49 | 0.063117334 | 0.170760496 | NA | 1 | 1 | NA | 0 | NA | 1 |
| SE | 13045 | CNOT7 | CNOT7 | chr8 | - | 17232426 | 17232537 | 17229230 | 17230848 | 17234715 | 17234860 | 0 | 2 | 2 | 0 | 0 | 0 | 0 | 0 | 0 | 0 | 1 | 0 | 98 | 49 | 0.063117334 | 0.170760496 | NA | 1 | 1 | NA | 0 | NA | 1 |
| A3SS | 1438 | FGD4 | FGD4 | chr12 | + | 32607937 | 32608095 | 32607956 | 32608095 | 32602160 | 32602317 | 1 | 0 | 0 | 0 | 0 | 4 | 0 | 2 | 1 | 0 | 0 | 0 | 68 | 49 | 0.063176479 | 0.197717869 | 1 | NA | 0 | NA | 1 | 1 | -0.5 |
| A5SS | 1600 | TRAPPC3 | TRAPPC3 | chr1 | - | 36149091 | 36149514 | 36149336 | 36149514 | 36140068 | 36140166 | 0 | 0 | 0 | 0 | 4 | 0 | 8 | 1 | 0 | 0 | 0 | 0 | 98 | 49 | 0.063182848 | 0.170414363 | NA | 0 | NA | 1 | 1 | NA | -1 |
| SE | 5519 | DENND5A | DENND5A | chr11 | - | 9203659 | 9204317 | 9193493 | 9193681 | 9206672 | 9206782 | 0 | 2 | 20 | 0 | 0 | 0 | 0 | 18 | 2 | 0 | 1 | 0 | 98 | 49 | 0.063221519 | 0.170760496 | NA | 1 | 1 | NA | 0.9 | 1 | 0.05 |
| SE | 7498 | IFI44 | IFI44 | chr1 | + | 78662703 | 78662878 | 78659311 | 78659483 | 78663764 | 78664078 | 0 | 6 | 51 | 0 | 1 | 1 | 0 | 87 | 6 | 0 | 0 | 0 | 98 | 49 | 0.063259916 | 0.170760496 | NA | 0.75 | 0.962 | NA | 1 | 1 | -0.144 |
| SE | 12385 | JAK3 | JAK3 | chr19 | - | 17831227 | 17831400 | 17830502 | 17830620 | 17831673 | 17831798 | 0 | 2 | 10 | 0 | 0 | 0 | 0 | 7 | 3 | 0 | 1 | 0 | 98 | 49 | 0.063527186 | 0.170760496 | NA | 1 | 1 | NA | 0.778 | 1 | 0.111 |
| A3SS | 3618 | CYB561D1 | CYB561D1 | chr1 | + | 109495678 | 109500441 | 109495755 | 109500441 | 109495142 | 109495180 | 0 | 0 | 0 | 0 | 0 | 1 | 0 | 0 | 2 | 0 | 0 | 0 | 98 | 49 | 0.063770311 | 0.197717869 | NA | NA | 0 | NA | NA | 1 | -1 |
| A5SS | 1161 | GAPDH | GAPDH | chr12 | + | 6534404 | 6534661 | 6534404 | 6534569 | 6534809 | 6534861 | 0 | 0 | 0 | 0 | 0 | 1 | 0 | 0 | 2 | 0 | 1 | 0 | 98 | 49 | 0.063770311 | 0.170414363 | NA | NA | 0 | NA | 0 | 1 | -0.5 |
| A5SS | 2679 | OXLD1 | OXLD1 | chr17 | - | 81666145 | 81666637 | 81666517 | 81666637 | 81665035 | 81665584 | 0 | 0 | 0 | 0 | 0 | 1 | 0 | 0 | 2 | 0 | 0 | 0 | 98 | 49 | 0.063770311 | 0.170414363 | NA | NA | 0 | NA | NA | 1 | -1 |
| A5SS | 2683 | OXLD1 | OXLD1 | chr17 | - | 81666149 | 81666637 | 81666517 | 81666637 | 81665035 | 81665584 | 0 | 0 | 0 | 0 | 0 | 1 | 0 | 0 | 2 | 0 | 0 | 0 | 98 | 49 | 0.063770311 | 0.170414363 | NA | NA | 0 | NA | NA | 1 | -1 |
| A5SS | 2688 | OXLD1 | OXLD1 | chr17 | - | 81666163 | 81666637 | 81666517 | 81666637 | 81665035 | 81665584 | 0 | 0 | 0 | 0 | 0 | 1 | 0 | 0 | 2 | 0 | 0 | 0 | 98 | 49 | 0.063770311 | 0.170414363 | NA | NA | 0 | NA | NA | 1 | -1 |
| A5SS | 2689 | OXLD1 | OXLD1 | chr17 | - | 81666276 | 81666637 | 81666517 | 81666637 | 81665035 | 81665584 | 0 | 0 | 0 | 0 | 0 | 1 | 0 | 0 | 2 | 0 | 0 | 0 | 98 | 49 | 0.063770311 | 0.170414363 | NA | NA | 0 | NA | NA | 1 | -1 |
| A5SS | 2944 | UNKL | UNKL | chr16 | - | 1369933 | 1370357 | 1370129 | 1370357 | 1367655 | 1367858 | 0 | 1 | 0 | 0 | 0 | 1 | 0 | 0 | 2 | 0 | 0 | 0 | 98 | 49 | 0.063770311 | 0.170414363 | NA | 1 | 0 | NA | NA | 1 | -0.5 |
| RI | 746 | PTPMT1 | PTPMT1 | chr11 | + | 47565429 | 47565986 | 47565429 | 47565796 | 47565905 | 47565986 | 1 | 0 | 2 | 0 | 1 | 0 | 0 | 2 | 0 | 0 | 0 | 0 | 98 | 49 | 0.063770311 | 0.297702651 | 1 | 0 | 1 | NA | 1 | NA | -0.333 |
| RI | 790 | CCDC125 | CCDC125 | chr5 | - | 69320236 | 69320580 | 69320236 | 69320345 | 69320464 | 69320580 | 0 | 0 | 0 | 0 | 0 | 1 | 1 | 1 | 2 | 0 | 0 | 0 | 98 | 49 | 0.063770311 | 0.297702651 | NA | NA | 0 | 1 | 1 | 1 | -1 |
| SE | 255 | SEC16A | SEC16A | chr9 | - | 136468414 | 136468512 | 136466956 | 136467083 | 136471974 | 136472111 | 0 | 0 | 1 | 0 | 1 | 0 | 0 | 2 | 0 | 0 | 0 | 0 | 98 | 49 | 0.063770311 | 0.170760496 | NA | 0 | 1 | NA | 1 | NA | -0.5 |
| SE | 506 | SLC45A4 | SLC45A4 | chr8 | - | 141221576 | 141221765 | 141219649 | 141219829 | 141253988 | 141254629 | 0 | 0 | 5 | 0 | 1 | 0 | 2 | 2 | 0 | 0 | 0 | 0 | 98 | 49 | 0.063770311 | 0.170760496 | NA | 0 | 1 | 1 | 1 | NA | -0.5 |
| SE | 3157 | XBP1 | XBP1 | chr22 | - | 28796046 | 28796192 | 28794559 | 28795732 | 28797076 | 28797205 | 0 | 0 | 6 | 0 | 1 | 0 | 0 | 2 | 0 | 0 | 0 | 0 | 98 | 49 | 0.063770311 | 0.170760496 | NA | 0 | 1 | NA | 1 | NA | -0.5 |
| SE | 3781 | MYCBP2 | MYCBP2 | chr13 | - | 77126317 | 77126542 | 77125335 | 77125468 | 77139195 | 77139336 | 0 | 0 | 2 | 0 | 1 | 0 | 0 | 2 | 0 | 0 | 0 | 0 | 98 | 49 | 0.063770311 | 0.170760496 | NA | 0 | 1 | NA | 1 | NA | -0.5 |
| SE | 5029 | BRMS1 | BRMS1 | chr11 | - | 66340120 | 66340213 | 66338720 | 66338785 | 66340773 | 66340870 | 0 | 0 | 2 | 0 | 1 | 0 | 0 | 2 | 0 | 0 | 0 | 0 | 98 | 49 | 0.063770311 | 0.170760496 | NA | 0 | 1 | NA | 1 | NA | -0.5 |
| SE | 10606 | ANKFY1 | ANKFY1 | chr17 | - | 4189381 | 4189479 | 4184817 | 4185046 | 4194977 | 4195177 | 0 | 0 | 0 | 0 | 0 | 1 | 0 | 0 | 2 | 0 | 0 | 0 | 98 | 49 | 0.063770311 | 0.170760496 | NA | NA | 0 | NA | NA | 1 | -1 |
| SE | 13383 | ZNF805 | ZNF805 | chr19 | + | 57243922 | 57244049 | 57240684 | 57240921 | 57248604 | 57248700 | 0 | 0 | 0 | 0 | 1 | 0 | 0 | 2 | 0 | 0 | 0 | 0 | 98 | 49 | 0.063770311 | 0.170760496 | NA | 0 | NA | NA | 1 | NA | -1 |
| SE | 12368 | NBEAL2 | NBEAL2 | chr3 | + | 47007065 | 47007155 | 47006332 | 47006449 | 47007240 | 47007350 | 0 | 2 | 16 | 0 | 0 | 0 | 0 | 8 | 3 | 0 | 1 | 0 | 98 | 49 | 0.063873063 | 0.170760496 | NA | 1 | 1 | NA | 0.8 | 1 | 0.1 |
| SE | 8540 | ARAP3 | ARAP3 | chr5 | - | 141655361 | 141655400 | 141653400 | 141654435 | 141655620 | 141655758 | 0 | 0 | 0 | 0 | 1 | 2 | 0 | 2 | 0 | 0 | 0 | 0 | 88 | 49 | 0.063945674 | 0.170760496 | NA | 0 | 0 | NA | 1 | NA | -1 |
| A3SS | 530 | GLRX | GLRX | chr5 | - | 95813848 | 95814807 | 95813848 | 95814389 | 95816506 | 95816626 | 0 | 0 | 0 | 0 | 0 | 3 | 0 | 0 | 1 | 0 | 1 | 0 | 98 | 49 | 0.064268453 | 0.197717869 | NA | NA | 0 | NA | 0 | 1 | -0.5 |
| A3SS | 531 | GLRX | GLRX | chr5 | - | 95813848 | 95814957 | 95813848 | 95814389 | 95816506 | 95816626 | 0 | 0 | 0 | 0 | 0 | 3 | 0 | 0 | 1 | 0 | 1 | 0 | 98 | 49 | 0.064268453 | 0.197717869 | NA | NA | 0 | NA | 0 | 1 | -0.5 |
| SE | 5261 | PCSK7 | PCSK7 | chr11 | - | 117208896 | 117209053 | 117207969 | 117208064 | 117218465 | 117218568 | 0 | 0 | 0 | 0 | 0 | 3 | 0 | 0 | 1 | 0 | 0 | 0 | 98 | 49 | 0.064268453 | 0.170760496 | NA | NA | 0 | NA | NA | 1 | -1 |
| SE | 6387 | PSRC1 | PSRC1 | chr1 | - | 109280395 | 109280489 | 109279553 | 109280156 | 109280776 | 109280916 | 0 | 0 | 1 | 0 | 3 | 0 | 0 | 1 | 0 | 0 | 0 | 0 | 98 | 49 | 0.064268453 | 0.170760496 | NA | 0 | 1 | NA | 1 | NA | -0.5 |
| A5SS | 1969 | SLC35A2 | SLC35A2 | chrX | - | 48911506 | 48911958 | 48911545 | 48911958 | 48909813 | 48909996 | 0 | 0 | 0 | 0 | 0 | 3 | 0 | 0 | 1 | 0 | 0 | 0 | 88 | 49 | 0.064272896 | 0.170414363 | NA | NA | 0 | NA | NA | 1 | -1 |
| A5SS | 1774 | SF1 | SF1 | chr11 | - | 64767566 | 64767844 | 64767570 | 64767844 | 64767191 | 64767251 | 0 | 0 | 0 | 0 | 0 | 3 | 6 | 0 | 1 | 0 | 2 | 0 | 53 | 49 | 0.064284969 | 0.170414363 | NA | NA | 0 | 1 | 0 | 1 | -0.667 |
| SE | 10332 | CYTIP | CYTIP | chr2 | - | 157434369 | 157434424 | 157430859 | 157430962 | 157434697 | 157434747 | 0 | 9 | 93 | 0 | 0 | 0 | 18 | 103 | 8 | 0 | 1 | 0 | 98 | 49 | 0.064528457 | 0.170760496 | NA | 1 | 1 | 1 | 0.981 | 1 | 0.006 |
| SE | 11921 | RBM39 | RBM39 | chr20 | - | 35729461 | 35729527 | 35729311 | 35729365 | 35731940 | 35732135 | 0 | 3 | 27 | 0 | 0 | 0 | 2 | 37 | 2 | 0 | 1 | 0 | 98 | 49 | 0.064578009 | 0.170760496 | NA | 1 | 1 | 1 | 0.949 | 1 | 0.017 |
| SE | 8709 | CD74 | CD74 | chr5 | - | 150403120 | 150403312 | 150402562 | 150402625 | 150404679 | 150404767 | 0 | 2 | 7 | 0 | 2 | 32 | 1 | 1 | 0 | 0 | 20 | 0 | 98 | 49 | 0.064910289 | 0.170760496 | NA | 0.333 | 0.099 | 1 | 0.024 | NA | -0.296 |
| SE | 250 | NOTCH1 | NOTCH1 | chr9 | - | 136515980 | 136516094 | 136515482 | 136515716 | 136517271 | 136517385 | 1 | 1 | 10 | 0 | 0 | 0 | 0 | 2 | 1 | 0 | 1 | 0 | 98 | 49 | 0.065113604 | 0.170760496 | 1 | 1 | 1 | NA | 0.5 | 1 | 0.25 |
| SE | 6697 | FCGR2A | FCGR2A | chr1 | + | 161506333 | 161506591 | 161505986 | 161506007 | 161509819 | 161510074 | 0 | 2 | 42 | 0 | 0 | 0 | 1 | 17 | 3 | 0 | 1 | 0 | 98 | 49 | 0.065339874 | 0.170760496 | NA | 1 | 1 | 1 | 0.895 | 1 | 0.035 |
| SE | 12941 | SMCHD1 | SMCHD1 | chr18 | + | 2694526 | 2694693 | 2688627 | 2688747 | 2697031 | 2697122 | 0 | 3 | 97 | 0 | 0 | 0 | 0 | 55 | 1 | 0 | 1 | 0 | 98 | 49 | 0.065369701 | 0.170760496 | NA | 1 | 1 | NA | 0.965 | 1 | 0.018 |
| A3SS | 2057 | FBXO7 | FBXO7 | chr22 | + | 32478980 | 32479275 | 32479132 | 32479275 | 32474719 | 32475124 | 0 | 1 | 1 | 0 | 0 | 0 | 0 | 0 | 0 | 0 | 1 | 0 | 98 | 49 | 0.065790241 | 0.197717869 | NA | 1 | 1 | NA | 0 | NA | 1 |
| A3SS | 2058 | FBXO7 | FBXO7 | chr22 | + | 32478980 | 32479275 | 32479132 | 32479275 | 32475236 | 32475398 | 0 | 1 | 1 | 0 | 0 | 0 | 0 | 0 | 0 | 0 | 1 | 0 | 98 | 49 | 0.065790241 | 0.197717869 | NA | 1 | 1 | NA | 0 | NA | 1 |
| A3SS | 2827 | APOL3 | APOL3 | chr22 | - | 36149045 | 36149371 | 36149045 | 36149142 | 36160668 | 36160929 | 1 | 1 | 0 | 0 | 0 | 0 | 0 | 0 | 0 | 0 | 1 | 0 | 98 | 49 | 0.065790241 | 0.197717869 | 1 | 1 | NA | NA | 0 | NA | 1 |
| A3SS | 3029 | POLM | POLM | chr7 | - | 44072061 | 44073430 | 44072061 | 44073377 | 44073624 | 44073708 | 1 | 0 | 1 | 0 | 0 | 0 | 0 | 3 | 0 | 0 | 0 | 1 | 98 | 49 | 0.065790241 | 0.197717869 | 1 | NA | 1 | NA | 1 | 0 | 0.5 |
| A5SS | 1566 | POLD4 | POLD4 | chr11 | - | 67353293 | 67353596 | 67353363 | 67353596 | 67352987 | 67353077 | 0 | 1 | 1 | 0 | 0 | 0 | 0 | 0 | 0 | 0 | 0 | 1 | 98 | 49 | 0.065790241 | 0.170414363 | NA | 1 | 1 | NA | NA | 0 | 1 |
| A5SS | 2228 | USP8 | USP8 | chr15 | + | 50424376 | 50424654 | 50424376 | 50424514 | 50439008 | 50439177 | 0 | 0 | 1 | 0 | 0 | 0 | 0 | 0 | 0 | 0 | 3 | 1 | 98 | 49 | 0.065790241 | 0.170414363 | NA | NA | 1 | NA | 0 | 0 | 1 |
| A5SS | 2719 | NBPF11 | NBPF11 | chr1 | - | 148110377 | 148110541 | 148110457 | 148110541 | 148109283 | 148109335 | 0 | 1 | 0 | 0 | 0 | 0 | 0 | 0 | 0 | 0 | 1 | 1 | 98 | 49 | 0.065790241 | 0.170414363 | NA | 1 | NA | NA | 0 | 0 | 1 |
| SE | 1021 | PPIA | PPIA | chr7 | + | 44798747 | 44798904 | 44796635 | 44796793 | 44799246 | 44799277 | 0 | 0 | 1 | 0 | 0 | 0 | 0 | 0 | 0 | 0 | 1 | 1 | 98 | 49 | 0.065790241 | 0.170760496 | NA | NA | 1 | NA | 0 | 0 | 1 |
| SE | 2879 | ACO2 | ACO2 | chr22 | + | 41522829 | 41522987 | 41520170 | 41520276 | 41523204 | 41523278 | 0 | 1 | 2 | 0 | 0 | 0 | 0 | 0 | 0 | 0 | 1 | 0 | 98 | 49 | 0.065790241 | 0.170760496 | NA | 1 | 1 | NA | 0 | NA | 1 |
| SE | 4101 | FAM13B | FAM13B | chr5 | - | 138018954 | 138019146 | 138018301 | 138018514 | 138021030 | 138021197 | 0 | 0 | 1 | 0 | 0 | 0 | 0 | 1 | 0 | 3 | 0 | 1 | 98 | 49 | 0.065790241 | 0.170760496 | NA | NA | 1 | 0 | 1 | 0 | 0.667 |
| SE | 4610 | HSPA8 | HSPA8 | chr11 | - | 123060592 | 123060798 | 123060115 | 123060268 | 123061119 | 123061329 | 0 | 0 | 1 | 0 | 0 | 0 | 0 | 0 | 0 | 0 | 2 | 1 | 98 | 49 | 0.065790241 | 0.170760496 | NA | NA | 1 | NA | 0 | 0 | 1 |
| SE | 6531 | FRMD8 | FRMD8 | chr11 | + | 65393572 | 65393674 | 65389360 | 65389528 | 65394040 | 65394099 | 0 | 1 | 3 | 0 | 0 | 0 | 0 | 0 | 0 | 0 | 1 | 0 | 98 | 49 | 0.065790241 | 0.170760496 | NA | 1 | 1 | NA | 0 | NA | 1 |
| SE | 7131 | GGA3 | GGA3 | chr17 | - | 75243062 | 75243166 | 75242830 | 75242911 | 75243446 | 75243570 | 0 | 1 | 5 | 0 | 0 | 0 | 0 | 0 | 0 | 0 | 1 | 0 | 98 | 49 | 0.065790241 | 0.170760496 | NA | 1 | 1 | NA | 0 | NA | 1 |
| SE | 7595 | ATG16L2 | ATG16L2 | chr11 | + | 72826702 | 72826823 | 72826172 | 72826243 | 72828358 | 72828508 | 0 | 0 | 1 | 0 | 0 | 0 | 0 | 2 | 0 | 0 | 1 | 1 | 98 | 49 | 0.065790241 | 0.170760496 | NA | NA | 1 | NA | 0.5 | 0 | 0.75 |
| SE | 8118 | POLE | POLE | chr12 | - | 132648928 | 132649072 | 132643836 | 132643977 | 132649305 | 132649515 | 0 | 1 | 1 | 0 | 0 | 0 | 0 | 0 | 0 | 0 | 1 | 0 | 98 | 49 | 0.065790241 | 0.170760496 | NA | 1 | 1 | NA | 0 | NA | 1 |
| SE | 8179 | MLXIP | MLXIP | chr12 | + | 122137468 | 122137590 | 122135488 | 122135666 | 122138193 | 122138295 | 0 | 0 | 1 | 0 | 0 | 0 | 0 | 0 | 0 | 0 | 0 | 1 | 98 | 49 | 0.065790241 | 0.170760496 | NA | NA | 1 | NA | NA | 0 | 1 |
| SE | 8364 | STRAP | STRAP | chr12 | + | 15883540 | 15883676 | 15882353 | 15882819 | 15889927 | 15890009 | 0 | 0 | 1 | 0 | 0 | 0 | 0 | 2 | 0 | 0 | 1 | 1 | 98 | 49 | 0.065790241 | 0.170760496 | NA | NA | 1 | NA | 0.5 | 0 | 0.75 |
| SE | 10100 | GGA1 | GGA1 | chr22 | + | 37625796 | 37625949 | 37624968 | 37625076 | 37629461 | 37629526 | 0 | 1 | 1 | 0 | 0 | 0 | 0 | 0 | 0 | 0 | 1 | 0 | 98 | 49 | 0.065790241 | 0.170760496 | NA | 1 | 1 | NA | 0 | NA | 1 |
| SE | 10460 | OSBPL11 | OSBPL11 | chr3 | - | 125547405 | 125547592 | 125538450 | 125538633 | 125560378 | 125560519 | 0 | 1 | 2 | 0 | 0 | 0 | 0 | 0 | 0 | 0 | 1 | 0 | 98 | 49 | 0.065790241 | 0.170760496 | NA | 1 | 1 | NA | 0 | NA | 1 |
| SE | 10539 | P4HA1 | P4HA1 | chr10 | - | 73051052 | 73051249 | 73046924 | 73047101 | 73053350 | 73053590 | 0 | 0 | 1 | 0 | 0 | 0 | 0 | 0 | 0 | 0 | 0 | 1 | 98 | 49 | 0.065790241 | 0.170760496 | NA | NA | 1 | NA | NA | 0 | 1 |
| SE | 11584 | FANCA | FANCA | chr16 | - | 89742799 | 89742938 | 89740803 | 89740866 | 89744958 | 89745071 | 0 | 1 | 1 | 0 | 0 | 0 | 0 | 0 | 0 | 0 | 1 | 0 | 98 | 49 | 0.065790241 | 0.170760496 | NA | 1 | 1 | NA | 0 | NA | 1 |
| SE | 13943 | PDK1 | PDK1 | chr2 | + | 172562740 | 172562800 | 172562219 | 172562291 | 172564977 | 172565073 | 0 | 1 | 0 | 0 | 0 | 1 | 0 | 0 | 0 | 0 | 1 | 0 | 98 | 49 | 0.065790241 | 0.170760496 | NA | 1 | 0 | NA | 0 | NA | 0.5 |
| A3SS | 2853 | TGOLN2 | TGOLN2 | chr2 | - | 85318017 | 85322799 | 85318017 | 85322741 | 85324914 | 85324998 | 0 | 0 | 0 | 0 | 0 | 2 | 0 | 2 | 1 | 2 | 4 | 0 | 98 | 49 | 0.0658956 | 0.197717869 | NA | NA | 0 | 0 | 0.2 | 1 | -0.4 |
| A5SS | 1409 | PHYKPL | PHYKPL | chr5 | - | 178231668 | 178232822 | 178232491 | 178232822 | 178231404 | 178231523 | 0 | 0 | 3 | 0 | 2 | 0 | 0 | 1 | 0 | 0 | 0 | 0 | 98 | 49 | 0.0658956 | 0.170414363 | NA | 0 | 1 | NA | 1 | NA | -0.5 |
| SE | 145 | SNX12 | SNX12 | chrX | - | 71061842 | 71061967 | 71059246 | 71061118 | 71062853 | 71062949 | 0 | 0 | 3 | 0 | 2 | 0 | 0 | 1 | 0 | 0 | 0 | 0 | 98 | 49 | 0.0658956 | 0.170760496 | NA | 0 | 1 | NA | 1 | NA | -0.5 |
| SE | 239 | NDOR1 | NDOR1 | chr9 | + | 137215406 | 137215521 | 137215094 | 137215202 | 137215658 | 137215805 | 0 | 0 | 0 | 0 | 2 | 0 | 0 | 1 | 0 | 0 | 0 | 0 | 98 | 49 | 0.0658956 | 0.170760496 | NA | 0 | NA | NA | 1 | NA | -1 |
| SE | 436 | UBE2R2 | UBE2R2 | chr9 | + | 33911963 | 33912098 | 33900173 | 33900271 | 33917017 | 33920403 | 0 | 0 | 6 | 0 | 2 | 1 | 0 | 1 | 0 | 0 | 0 | 0 | 98 | 49 | 0.0658956 | 0.170760496 | NA | 0 | 0.75 | NA | 1 | NA | -0.625 |
| SE | 4582 | ST3GAL4 | ST3GAL4 | chr11 | + | 126409267 | 126409411 | 126408306 | 126408496 | 126413504 | 126413648 | 0 | 0 | 3 | 0 | 2 | 0 | 0 | 1 | 0 | 0 | 0 | 0 | 98 | 49 | 0.0658956 | 0.170760496 | NA | 0 | 1 | NA | 1 | NA | -0.5 |
| SE | 7326 | SREBF1 | SREBF1 | chr17 | - | 17819012 | 17819234 | 17818259 | 17818374 | 17819319 | 17819454 | 0 | 0 | 0 | 0 | 2 | 0 | 0 | 1 | 0 | 0 | 0 | 0 | 98 | 49 | 0.0658956 | 0.170760496 | NA | 0 | NA | NA | 1 | NA | -1 |
| SE | 8223 | MAPK3 | MAPK3 | chr16 | - | 30118046 | 30118163 | 30117669 | 30117784 | 30118348 | 30118538 | 0 | 0 | 0 | 0 | 0 | 2 | 0 | 0 | 1 | 0 | 0 | 0 | 98 | 49 | 0.0658956 | 0.170760496 | NA | NA | 0 | NA | NA | 1 | -1 |
| SE | 11719 | CHD3 | CHD3 | chr17 | + | 7907600 | 7907702 | 7907352 | 7907488 | 7907893 | 7908019 | 0 | 0 | 0 | 0 | 0 | 2 | 0 | 0 | 1 | 0 | 0 | 0 | 98 | 49 | 0.0658956 | 0.170760496 | NA | NA | 0 | NA | NA | 1 | -1 |
| A3SS | 81 | GBGT1 | GBGT1 | chr9 | - | 133156014 | 133156102 | 133156014 | 133156065 | 133161466 | 133161532 | 0 | 0 | 0 | 0 | 0 | 2 | 0 | 0 | 1 | 0 | 1 | 0 | 86 | 49 | 0.065901144 | 0.197717869 | NA | NA | 0 | NA | 0 | 1 | -0.5 |
| SE | 4442 | CORO7 | CORO7 | chr16 | - | 4357967 | 4358103 | 4357167 | 4357259 | 4358366 | 4358483 | 0 | 1 | 4 | 0 | 0 | 0 | 5 | 8 | 2 | 0 | 1 | 0 | 98 | 49 | 0.067252723 | 0.171142152 | NA | 1 | 1 | 1 | 0.8 | 1 | 0.067 |
| SE | 8028 | GNAS | GNAS | chr20 | + | 58898940 | 58898985 | 58895611 | 58895684 | 58903530 | 58903585 | 0 | 1 | 3 | 0 | 0 | 2 | 0 | 0 | 0 | 0 | 1 | 0 | 94 | 49 | 0.067458351 | 0.171142152 | NA | 1 | 0.439 | NA | 0 | NA | 0.72 |
| SE | 911 | LMTK2 | LMTK2 | chr7 | + | 98171540 | 98171674 | 98159337 | 98159425 | 98185050 | 98185135 | 0 | 1 | 2 | 0 | 0 | 0 | 0 | 3 | 2 | 0 | 2 | 0 | 98 | 49 | 0.068013122 | 0.171142152 | NA | 1 | 1 | NA | 0.429 | 1 | 0.285 |
| SE | 2940 | LIMK2 | LIMK2 | chr22 | + | 31273451 | 31273507 | 31272529 | 31272704 | 31275150 | 31275308 | 0 | 13 | 40 | 0 | 0 | 0 | 3 | 93 | 36 | 0 | 1 | 0 | 98 | 49 | 0.068060835 | 0.171142152 | NA | 1 | 1 | 1 | 0.979 | 1 | 0.007 |
| SE | 2968 | AP1B1 | AP1B1 | chr22 | - | 29329711 | 29329720 | 29327679 | 29328895 | 29330377 | 29330532 | 0 | 0 | 1 | 0 | 0 | 0 | 0 | 0 | 0 | 0 | 0 | 1 | 58 | 49 | 0.068063814 | 0.171142152 | NA | NA | 1 | NA | NA | 0 | 1 |
| SE | 12329 | ARHGDIA | ARHGDIA | chr17 | - | 81869329 | 81869406 | 81869172 | 81869236 | 81869541 | 81869625 | 0 | 0 | 21 | 0 | 0 | 1 | 0 | 11 | 1 | 0 | 0 | 2 | 98 | 49 | 0.068249658 | 0.171142152 | NA | NA | 0.913 | NA | 1 | 0.2 | 0.313 |
| SE | 5783 | DENND3 | DENND3 | chr8 | + | 141138021 | 141138137 | 141136557 | 141136791 | 141151618 | 141151837 | 0 | 3 | 15 | 0 | 0 | 0 | 0 | 19 | 7 | 0 | 1 | 0 | 98 | 49 | 0.068459154 | 0.171142152 | NA | 1 | 1 | NA | 0.905 | 1 | 0.047 |
| A3SS | 1319 | P2RX7 | P2RX7 | chr12 | + | 121162270 | 121162520 | 121162423 | 121162520 | 121160901 | 121160974 | 0 | 0 | 0 | 0 | 0 | 1 | 0 | 0 | 1 | 0 | 0 | 0 | 98 | 49 | 0.068700612 | 0.197717869 | NA | NA | 0 | NA | NA | 1 | -1 |
| A3SS | 3364 | CCL4L2 | CCL4L2 | chr17 | + | 36212292 | 36212878 | 36212496 | 36212878 | 36211775 | 36211890 | 0 | 0 | 0 | 0 | 0 | 1 | 2 | 4 | 1 | 0 | 0 | 0 | 98 | 49 | 0.068700612 | 0.197717869 | NA | NA | 0 | 1 | 1 | 1 | -1 |
| A3SS | 3368 | CCL4L2 | CCL4L2 | chr17 | + | 36212302 | 36212878 | 36212496 | 36212878 | 36211775 | 36211890 | 0 | 0 | 0 | 0 | 0 | 1 | 2 | 4 | 1 | 0 | 0 | 0 | 98 | 49 | 0.068700612 | 0.197717869 | NA | NA | 0 | 1 | 1 | 1 | -1 |
| A3SS | 3371 | CCL4L2 | CCL4L2 | chr17 | + | 36212312 | 36212878 | 36212496 | 36212878 | 36211775 | 36211890 | 0 | 0 | 0 | 0 | 0 | 1 | 2 | 4 | 1 | 0 | 0 | 0 | 98 | 49 | 0.068700612 | 0.197717869 | NA | NA | 0 | 1 | 1 | 1 | -1 |
| A3SS | 3373 | CCL4L2 | CCL4L2 | chr17 | + | 36212333 | 36212878 | 36212496 | 36212878 | 36211775 | 36211890 | 0 | 0 | 0 | 0 | 0 | 1 | 2 | 4 | 1 | 0 | 0 | 0 | 98 | 49 | 0.068700612 | 0.197717869 | NA | NA | 0 | 1 | 1 | 1 | -1 |
| A3SS | 3374 | CCL4L2 | CCL4L2 | chr17 | + | 36212363 | 36212878 | 36212496 | 36212878 | 36211775 | 36211890 | 0 | 0 | 0 | 0 | 0 | 1 | 2 | 4 | 1 | 0 | 0 | 0 | 98 | 49 | 0.068700612 | 0.197717869 | NA | NA | 0 | 1 | 1 | 1 | -1 |
| A3SS | 3842 | CARKD | CARKD | chr13 | + | 110638241 | 110639995 | 110638377 | 110639995 | 110637128 | 110637249 | 0 | 0 | 0 | 0 | 0 | 1 | 0 | 0 | 1 | 0 | 0 | 0 | 98 | 49 | 0.068700612 | 0.197717869 | NA | NA | 0 | NA | NA | 1 | -1 |
| A3SS | 3925 | C10orf128 | C10orf128 | chr10 | - | 49154723 | 49156259 | 49154723 | 49155909 | 49161598 | 49161622 | 0 | 0 | 0 | 0 | 0 | 1 | 7 | 0 | 1 | 0 | 0 | 0 | 98 | 49 | 0.068700612 | 0.197717869 | NA | NA | 0 | 1 | NA | 1 | -1 |
| A5SS | 407 | ARHGAP26 | ARHGAP26 | chr5 | + | 143207197 | 143207473 | 143207197 | 143207308 | 143213996 | 143214088 | 0 | 0 | 0 | 0 | 0 | 1 | 0 | 0 | 1 | 0 | 1 | 0 | 98 | 49 | 0.068700612 | 0.170414363 | NA | NA | 0 | NA | 0 | 1 | -0.5 |
| A5SS | 1221 | C6orf203 | C6orf203 | chr6 | + | 107028171 | 107028797 | 107028171 | 107028271 | 107039748 | 107040230 | 0 | 0 | 0 | 0 | 0 | 1 | 0 | 0 | 1 | 0 | 0 | 0 | 98 | 49 | 0.068700612 | 0.170414363 | NA | NA | 0 | NA | NA | 1 | -1 |
| A5SS | 1816 | POLR2F | POLR2F | chr22 | + | 37956772 | 37956901 | 37956772 | 37956842 | 37959345 | 37959476 | 0 | 0 | 0 | 0 | 0 | 1 | 0 | 0 | 1 | 2 | 0 | 0 | 98 | 49 | 0.068700612 | 0.170414363 | NA | NA | 0 | 0 | NA | 1 | -0.5 |
| A5SS | 2218 | TRIM69 | TRIM69 | chr15 | + | 44758612 | 44758854 | 44758612 | 44758803 | 44759639 | 44759662 | 0 | 0 | 0 | 0 | 0 | 1 | 0 | 2 | 1 | 0 | 0 | 0 | 98 | 49 | 0.068700612 | 0.170414363 | NA | NA | 0 | NA | 1 | 1 | -1 |

| | | | | | | | | | | | | | | | | | | | | | | | | | | | | | | | | | | |
|---|---|---|---|---|---|---|---|---|---|---|---|---|---|---|---|---|---|---|---|---|---|---|---|---|---|---|---|---|---|---|---|---|---|---|
| RI | 334 | ARL6IP4 | ARL6IP4 | chr12 | + | 122982468 | 122982913 | 122982468 | 122982538 | 122982619 | 122982913 | 0 | 0 | 0 | 0 | 0 | 1 | 0 | 0 | 1 | 0 | 1 | 0 | 98 | 49 | 0.068700612 | 0.297702651 | NA | NA | 0 | NA | 0 | 1 | -0.5 |
| RI | 369 | RECQL | RECQL | chr12 | - | 21501169 | 21501669 | 21501169 | 21501215 | 21501412 | 21501669 | 0 | 0 | 0 | 0 | 0 | 1 | 2 | 1 | 1 | 0 | 1 | 0 | 98 | 49 | 0.068700612 | 0.297702651 | NA | NA | 0 | 1 | 0.333 | 1 | -0.778 |
| RI | 581 | TK2 | TK2 | chr16 | - | 66548977 | 66550412 | 66548977 | 66549009 | 66549937 | 66550412 | 0 | 0 | 0 | 0 | 0 | 1 | 0 | 3 | 1 | 0 | 0 | 0 | 98 | 49 | 0.068700612 | 0.297702651 | NA | NA | 0 | NA | 1 | 1 | -1 |
| RI | 682 | NAP1L1 | NAP1L1 | chr12 | - | 76045382 | 76048464 | 76045382 | 76047718 | 76048392 | 76048464 | 0 | 0 | 0 | 0 | 0 | 1 | 0 | 2 | 1 | 0 | 2 | 0 | 98 | 49 | 0.068700612 | 0.297702651 | NA | NA | 0 | NA | 0.333 | 1 | -0.666 |
| RI | 968 | CCL4L2 | CCL4L2 | chr17 | + | 36211775 | 36212878 | 36211775 | 36211890 | 36212496 | 36212878 | 0 | 0 | 0 | 0 | 0 | 1 | 2 | 4 | 1 | 0 | 0 | 0 | 98 | 49 | 0.068700612 | 0.297702651 | NA | NA | 0 | 1 | 1 | 1 | -1 |
| RI | 1065 | CPT1B | CPT1B | chr22 | - | 50568860 | 50569421 | 50568860 | 50569081 | 50569335 | 50569421 | 0 | 0 | 0 | 0 | 2 | 1 | 0 | 0 | 1 | 2 | 0 | 0 | 98 | 49 | 0.068700612 | 0.297702651 | NA | 0 | 0 | 0 | NA | 1 | -0.5 |
| RI | 1071 | NFKBIB | NFKBIB | chr19 | + | 38907398 | 38908894 | 38907398 | 38907659 | 38908730 | 38908894 | 0 | 0 | 0 | 0 | 0 | 1 | 0 | 2 | 1 | 0 | 0 | 0 | 98 | 49 | 0.068700612 | 0.297702651 | NA | NA | 0 | NA | 1 | 1 | -1 |
| RI | 1115 | DCTN1 | DCTN1 | chr2 | - | 74366243 | 74366620 | 74366243 | 74366375 | 74366458 | 74366620 | 0 | 0 | 1 | 0 | 1 | 1 | 1 | 1 | 0 | 0 | 0 | 0 | 98 | 49 | 0.068700612 | 0.297702651 | NA | 0 | 0.333 | 1 | 1 | NA | -0.834 |
| SE | 689 | PTK2B | PTK2B | chr8 | + | 27445793 | 27445919 | 27444205 | 27444271 | 27450748 | 27450895 | 0 | 0 | 0 | 0 | 1 | 2 | 0 | 1 | 0 | 0 | 0 | 0 | 98 | 49 | 0.068700612 | 0.171142152 | NA | 0 | 0 | NA | 1 | NA | -1 |
| SE | 969 | RHBDD2 | RHBDD2 | chr7 | + | 75881364 | 75881486 | 75878998 | 75879260 | 75881828 | 75882236 | 0 | 0 | 2 | 0 | 1 | 0 | 2 | 1 | 0 | 0 | 0 | 0 | 98 | 49 | 0.068700612 | 0.171142152 | NA | 0 | 1 | 1 | 1 | NA | -0.5 |
| SE | 3923 | HAGH | HAGH | chr16 | - | 1826506 | 1826602 | 1822864 | 1823037 | 1826711 | 1827194 | 0 | 0 | 0 | 0 | 1 | 0 | 0 | 1 | 0 | 0 | 0 | 0 | 98 | 49 | 0.068700612 | 0.171142152 | NA | 0 | NA | NA | 1 | NA | -1 |
| SE | 3999 | ASL | ASL | chr7 | + | 66086743 | 66086821 | 66086584 | 66086662 | 66087333 | 66087386 | 0 | 0 | 2 | 0 | 1 | 0 | 0 | 1 | 0 | 0 | 0 | 0 | 98 | 49 | 0.068700612 | 0.171142152 | NA | 0 | 1 | NA | 1 | NA | -0.5 |
| SE | 4928 | FCRL6 | FCRL6 | chr1 | + | 159808960 | 159809245 | 159808177 | 159808444 | 159809401 | 159809683 | 0 | 0 | 0 | 0 | 0 | 1 | 0 | 0 | 1 | 0 | 0 | 0 | 98 | 49 | 0.068700612 | 0.171142152 | NA | NA | 0 | NA | NA | 1 | -1 |
| SE | 5379 | SYNE1 | SYNE1 | chr6 | - | 152455885 | 152456044 | 152455425 | 152455590 | 152458756 | 152458930 | 0 | 0 | 1 | 0 | 1 | 0 | 0 | 1 | 0 | 0 | 0 | 0 | 98 | 49 | 0.068700612 | 0.171142152 | NA | 0 | 1 | NA | 1 | NA | -0.5 |
| SE | 6602 | CCDC88B | CCDC88B | chr11 | + | 64351159 | 64351255 | 64349550 | 64349668 | 64351475 | 64351616 | 0 | 0 | 0 | 0 | 0 | 1 | 0 | 5 | 1 | 0 | 0 | 0 | 98 | 49 | 0.068700612 | 0.171142152 | NA | NA | 0 | NA | 1 | 1 | -1 |
| SE | 6680 | TTN-AS1 | TTN-AS1 | chr2 | + | 178571299 | 178571449 | 178542221 | 178542361 | 178597595 | 178597681 | 0 | 0 | 2 | 0 | 1 | 0 | 0 | 1 | 0 | 0 | 0 | 0 | 98 | 49 | 0.068700612 | 0.171142152 | NA | 0 | 1 | NA | 1 | NA | -0.5 |
| SE | 7590 | UBE4B | UBE4B | chr1 | + | 10158355 | 10158482 | 10151325 | 10151561 | 10161141 | 10161286 | 0 | 0 | 0 | 0 | 0 | 1 | 0 | 0 | 1 | 0 | 1 | 0 | 98 | 49 | 0.068700612 | 0.171142152 | NA | NA | 0 | NA | 0 | 1 | -0.5 |
| SE | 10301 | MARK3 | MARK3 | chr14 | + | 103491776 | 103492034 | 103480386 | 103480490 | 103502881 | 103503829 | 0 | 0 | 0 | 0 | 0 | 1 | 0 | 1 | 1 | 0 | 0 | 0 | 98 | 49 | 0.068700612 | 0.171142152 | NA | NA | 0 | NA | 1 | 1 | -1 |
| SE | 11137 | CLEC16A | CLEC16A | chr16 | + | 10977224 | 10977399 | 10972931 | 10973061 | 10979328 | 10979382 | 0 | 0 | 0 | 0 | 1 | 0 | 0 | 1 | 0 | 0 | 0 | 0 | 98 | 49 | 0.068700612 | 0.171142152 | NA | 0 | NA | NA | 1 | NA | -1 |
| SE | 11367 | LONP2 | LONP2 | chr16 | + | 48256609 | 48256741 | 48252130 | 48252365 | 48258617 | 48258740 | 0 | 0 | 0 | 0 | 0 | 1 | 0 | 0 | 1 | 0 | 1 | 0 | 98 | 49 | 0.068700612 | 0.171142152 | NA | NA | 0 | NA | 0 | 1 | -0.5 |
| SE | 12842 | SMARCA4 | SMARCA4 | chr19 | + | 11033766 | 11033865 | 11033289 | 11033517 | 11034122 | 11034200 | 0 | 0 | 0 | 0 | 0 | 1 | 0 | 0 | 1 | 0 | 0 | 0 | 98 | 49 | 0.068700612 | 0.171142152 | NA | NA | 0 | NA | NA | 1 | -1 |
| SE | 13356 | TMEM86B | TMEM86B | chr19 | - | 55228190 | 55228437 | 55226633 | 55227563 | 55228690 | 55229264 | 0 | 0 | 1 | 0 | 1 | 0 | 0 | 1 | 0 | 0 | 0 | 0 | 98 | 49 | 0.068700612 | 0.171142152 | NA | 0 | 1 | NA | 1 | NA | -0.5 |
| SE | 13996 | QRICH1 | QRICH1 | chr3 | - | 49092396 | 49092456 | 49076708 | 49077038 | 49093911 | 49094071 | 0 | 0 | 0 | 0 | 1 | 0 | 0 | 1 | 0 | 0 | 0 | 0 | 98 | 49 | 0.068700612 | 0.171142152 | NA | 0 | NA | NA | 1 | NA | -1 |
| SE | 14058 | ATG2B | ATG2B | chr14 | - | 96322111 | 96322254 | 96317697 | 96317855 | 96322539 | 96322735 | 0 | 0 | 0 | 0 | 1 | 0 | 0 | 1 | 0 | 0 | 0 | 0 | 98 | 49 | 0.068700612 | 0.171142152 | NA | 0 | NA | NA | 1 | NA | -1 |
| A5SS | 1424 | C7orf49 | C7orf49 | chr7 | - | 135168748 | 135169060 | 135168785 | 135169060 | 135167731 | 135167807 | 0 | 0 | 0 | 0 | 0 | 1 | 0 | 6 | 1 | 0 | 0 | 0 | 86 | 49 | 0.068703243 | 0.170414363 | NA | NA | 0 | NA | 1 | 1 | -1 |
| A5SS | 3076 | NCEH1 | NCEH1 | chr3 | - | 172647861 | 172648114 | 172647885 | 172648114 | 172645622 | 172645692 | 0 | 0 | 0 | 0 | 0 | 1 | 0 | 0 | 1 | 0 | 0 | 0 | 73 | 49 | 0.06870403 | 0.170414363 | NA | NA | 0 | NA | NA | 1 | -1 |
| A5SS | 903 | ATF7 | ATF7 | chr12 | - | 53543296 | 53543448 | 53543329 | 53543448 | 53537414 | 53537552 | 1 | 0 | 0 | 0 | 0 | 1 | 0 | 0 | 1 | 0 | 0 | 0 | 82 | 49 | 0.068705526 | 0.170414363 | 1 | NA | 0 | NA | NA | 1 | -0.5 |
| A3SS | 2808 | SRRT | SRRT | chr7 | + | 100885212 | 100885370 | 100885215 | 100885370 | 100884922 | 100885040 | 0 | 0 | 0 | 0 | 0 | 1 | 0 | 4 | 1 | 0 | 2 | 0 | 52 | 49 | 0.068726673 | 0.197717869 | NA | NA | 0 | NA | 0.653 | 1 | -0.827 |
| A3SS | 3999 | SERPINA1 | SERPINA1 | chr14 | - | 94388559 | 94388663 | 94388559 | 94388660 | 94390456 | 94390692 | 0 | 0 | 5 | 0 | 1 | 3 | 0 | 1 | 0 | 4 | 0 | 0 | 52 | 49 | 0.068726673 | 0.197717869 | NA | 0 | 0.611 | 0 | 1 | NA | -0.195 |
| SE | 9686 | IFRD1 | IFRD1 | chr7 | + | 112450507 | 112450782 | 112423143 | 112423432 | 112455762 | 112455867 | 0 | 3 | 58 | 0 | 0 | 0 | 0 | 33 | 5 | 0 | 1 | 0 | 98 | 49 | 0.068775688 | 0.171142152 | NA | 1 | 1 | NA | 0.943 | 1 | 0.028 |
| A5SS | 2089 | CLASRP | CLASRP | chr19 | + | 45068014 | 45068058 | 45068014 | 45068054 | 45068419 | 45068480 | 0 | 0 | 0 | 0 | 0 | 1 | 0 | 1 | 1 | 0 | 0 | 0 | 53 | 49 | 0.068946269 | 0.170414363 | NA | NA | 0 | NA | 1 | 1 | -1 |
| SE | 11969 | RARA | RARA | chr17 | + | 40348315 | 40348464 | 40342018 | 40342867 | 40349783 | 40349925 | 0 | 2 | 18 | 0 | 0 | 0 | 0 | 9 | 4 | 0 | 1 | 0 | 98 | 49 | 0.069039705 | 0.17126162 | NA | 1 | 1 | NA | 0.818 | 1 | 0.091 |
| SE | 3617 | NIN | NIN | chr14 | - | 50772300 | 50772468 | 50771331 | 50771468 | 50772948 | 50773095 | 0 | 1 | 12 | 0 | 0 | 0 | 0 | 13 | 1 | 0 | 1 | 0 | 98 | 49 | 0.069097631 | 0.17126162 | NA | 1 | 1 | NA | 0.867 | 1 | 0.067 |
| SE | 784 | ZYX | ZYX | chr7 | + | 143388218 | 143388339 | 143382800 | 143383322 | 143388488 | 143388658 | 0 | 5 | 34 | 0 | 0 | 0 | 0 | 9 | 7 | 0 | 1 | 0 | 98 | 49 | 0.069165538 | 0.17126162 | NA | 1 | 1 | NA | 0.818 | 1 | 0.091 |
| A3SS | 2597 | FERMT3 | FERMT3 | chr11 | + | 64219878 | 64220015 | 64219890 | 64220015 | 64219739 | 64219789 | 0 | 2 | 1 | 2 | 8 | 9 | 0 | 0 | 0 | 3 | 6 | 6 | 61 | 49 | 0.069201254 | 0.197717869 | 0 | 0.167 | 0.082 | 0 | 0 | 0 | 0.083 |
| SE | 2329 | ATP11B | ATP11B | chr3 | + | 182842074 | 182842122 | 182837070 | 182837174 | 182845457 | 182845522 | 0 | 1 | 22 | 0 | 0 | 0 | 0 | 6 | 2 | 0 | 1 | 0 | 97 | 49 | 0.069320212 | 0.171362302 | NA | 1 | 1 | NA | 0.752 | 1 | 0.124 |
| SE | 5624 | ETF1 | ETF1 | chr5 | - | 138517560 | 138517700 | 138513567 | 138513706 | 138518691 | 138518867 | 0 | 0 | 12 | 0 | 2 | 0 | 0 | 10 | 1 | 0 | 1 | 0 | 98 | 49 | 0.069479149 | 0.17144651 | NA | 0 | 1 | NA | 0.833 | 1 | -0.416 |
| SE | 1355 | MAPK14 | MAPK14 | chr6 | + | 36102570 | 36102649 | 36095986 | 36096066 | 36107454 | 36107628 | 0 | 1 | 19 | 0 | 0 | 0 | 0 | 14 | 1 | 0 | 1 | 0 | 98 | 49 | 0.069582416 | 0.17144651 | NA | 1 | 1 | NA | 0.875 | 1 | 0.062 |
| SE | 4942 | ARAP1 | ARAP1 | chr11 | - | 72695717 | 72695865 | 72695541 | 72695628 | 72696548 | 72696654 | 0 | 1 | 24 | 0 | 0 | 0 | 0 | 15 | 1 | 0 | 1 | 0 | 98 | 49 | 0.06991611 | 0.171507349 | NA | 1 | 1 | NA | 0.882 | 1 | 0.059 |
| SE | 10851 | IRF1 | IRF1 | chr5 | - | 132489391 | 132489483 | 132486953 | 132487130 | 132490544 | 132490773 | 0 | 0 | 6 | 0 | 0 | 2 | 0 | 8 | 2 | 0 | 2 | 7 | 98 | 49 | 0.069995497 | 0.171507349 | NA | NA | 0.6 | NA | 0.667 | 0.125 | 0.204 |
| SE | 4877 | TPP1 | TPP1 | chr11 | - | 6618775 | 6618915 | 6617625 | 6617776 | 6619195 | 6619267 | 1 | 2 | 13 | 0 | 0 | 0 | 0 | 1 | 1 | 0 | 1 | 0 | 98 | 49 | 0.07001504 | 0.171507349 | 1 | 1 | 1 | NA | 0.333 | 1 | 0.334 |
| SE | 6551 | SESN2 | SESN2 | chr1 | + | 28271673 | 28271871 | 28269182 | 28269248 | 28272283 | 28272466 | 0 | 1 | 1 | 0 | 0 | 0 | 0 | 7 | 4 | 0 | 1 | 0 | 98 | 49 | 0.070207347 | 0.171507349 | NA | 1 | 1 | NA | 0.778 | 1 | 0.111 |
| SE | 8748 | ADIPOR1 | ADIPOR1 | chr1 | - | 202954240 | 202954307 | 202950929 | 202951164 | 202958184 | 202958396 | 0 | 3 | 1 | 0 | 1 | 1 | 0 | 2 | 2 | 0 | 0 | 0 | 98 | 49 | 0.070257706 | 0.171507349 | NA | 0.6 | 0.333 | NA | 1 | 1 | -0.533 |
| SE | 13036 | VASP | VASP | chr19 | + | 45523840 | 45523877 | 45523643 | 45523695 | 45524096 | 45524142 | 0 | 5 | 37 | 0 | 0 | 3 | 0 | 33 | 3 | 0 | 0 | 0 | 86 | 49 | 0.070291768 | 0.171507349 | NA | 1 | 0.875 | NA | 1 | 1 | -0.062 |
| SE | 7499 | IFI44 | IFI44 | chr1 | + | 78662703 | 78662878 | 78660553 | 78660654 | 78663764 | 78664078 | 0 | 6 | 52 | 0 | 0 | 2 | 0 | 85 | 6 | 0 | 0 | 0 | 98 | 49 | 0.070496338 | 0.171727708 | NA | 1 | 0.929 | NA | 1 | 1 | -0.035 |
| SE | 12903 | TYROBP | TYROBP | chr19 | - | 35907729 | 35907762 | 35907445 | 35907580 | 35908174 | 35908309 | 0 | 3 | 6 | 0 | 3 | 1 | 0 | 5 | 1 | 0 | 0 | 0 | 82 | 49 | 0.070693174 | 0.171928544 | NA | 0.374 | 0.782 | NA | 1 | 1 | -0.422 |
| A3SS | 581 | LRCH4 | LRCH4 | chr7 | - | 100574010 | 100575304 | 100574010 | 100574356 | 100575704 | 100575782 | 0 | 1 | 2 | 0 | 0 | 4 | 0 | 3 | 9 | 0 | 1 | 1 | 98 | 49 | 0.070727262 | 0.198669639 | NA | 1 | 0.2 | NA | 0.6 | 0.818 | -0.109 |
| SE | 12640 | CYTH4 | CYTH4 | chr22 | + | 37310987 | 37311064 | 37309211 | 37309323 | 37311455 | 37311527 | 0 | 2 | 30 | 0 | 0 | 0 | 0 | 38 | 1 | 0 | 1 | 0 | 98 | 49 | 0.07087784 | 0.172099182 | NA | 1 | 1 | NA | 0.95 | 1 | 0.025 |
| SE | 12746 | DYSF | DYSF | chr2 | + | 71656161 | 71656290 | 71643964 | 71644063 | 71658877 | 71659033 | 0 | 2 | 17 | 0 | 2 | 0 | 0 | 7 | 6 | 0 | 0 | 0 | 98 | 49 | 0.071270681 | 0.172773926 | NA | 0.333 | 1 | NA | 1 | 1 | -0.334 |
| SE | 333 | AKNA | AKNA | chr9 | - | 114356862 | 114356969 | 114355924 | 114356136 | 114357920 | 114358167 | 0 | 1 | 3 | 0 | 0 | 0 | 6 | 8 | 3 | 0 | 1 | 0 | 98 | 49 | 0.071722008 | 0.173368597 | NA | 1 | 1 | 1 | 0.8 | 1 | 0.067 |
| A3SS | 3730 | RASGRP4 | RASGRP4 | chr19 | - | 38414847 | 38415123 | 38414847 | 38414916 | 38417051 | 38417168 | 0 | 2 | 16 | 0 | 0 | 0 | 1 | 4 | 3 | 0 | 1 | 0 | 98 | 49 | 0.071741814 | 0.198669639 | NA | 1 | 1 | 1 | 0.667 | 1 | 0.111 |
| SE | 4363 | LRRK2 | LRRK2 | chr12 | + | 40313971 | 40314173 | 40310430 | 40310649 | 40315211 | 40315300 | 0 | 3 | 44 | 0 | 0 | 1 | 1 | 19 | 1 | 0 | 3 | 0 | 98 | 49 | 0.071746685 | 0.173368597 | NA | 1 | 0.957 | 1 | 0.76 | 1 | 0.058 |
| A3SS | 590 | UBE2D3 | UBE2D3 | chr4 | - | 102826484 | 102826844 | 102826484 | 102826636 | 102827591 | 102827948 | 0 | 5 | 24 | 0 | 0 | 2 | 2 | 32 | 7 | 0 | 0 | 0 | 98 | 49 | 0.073395949 | 0.200170771 | NA | 1 | 0.857 | 1 | 1 | 1 | -0.072 |
| SE | 4359 | LRRK2 | LRRK2 | chr12 | + | 40274582 | 40274727 | 40263788 | 40263901 | 40274853 | 40274993 | 0 | 1 | 45 | 0 | 0 | 0 | 0 | 7 | 3 | 0 | 1 | 0 | 98 | 49 | 0.073501365 | 0.177323519 | NA | 1 | 1 | NA | 0.778 | 1 | 0.111 |
| SE | 6869 | ATG2A | ATG2A | chr11 | - | 64910615 | 64910708 | 64910039 | 64910195 | 64910806 | 64910954 | 0 | 8 | 7 | 0 | 0 | 0 | 8 | 26 | 23 | 0 | 1 | 0 | 98 | 49 | 0.07395786 | 0.178138883 | NA | 1 | 1 | 1 | 0.929 | 1 | 0.024 |
| SE | 1736 | ARHGAP26 | ARHGAP26 | chr5 | + | 142901934 | 142902039 | 142894237 | 142894348 | 142903539 | 142903669 | 0 | 2 | 101 | 0 | 0 | 0 | 0 | 61 | 1 | 0 | 1 | 0 | 98 | 49 | 0.074241382 | 0.178535676 | NA | 1 | 1 | NA | 0.968 | 1 | 0.016 |
| SE | 8074 | BCL2A1 | BCL2A1 | chr15 | - | 79967636 | 79967692 | 79960889 | 79961174 | 79970699 | 79971301 | 0 | 0 | 2 | 0 | 1 | 19 | 0 | 0 | 0 | 1 | 40 | 1 | 98 | 49 | 0.074407684 | 0.178649759 | NA | 0 | 0.05 | 0 | 0 | 0 | 0.025 |
| SE | 13869 | ACSL1 | ACSL1 | chr4 | - | 184788378 | 184788513 | 184783926 | 184783991 | 184788616 | 184788731 | 0 | 0 | 0 | 0 | 1 | 55 | 0 | 2 | 0 | 1 | 37 | 9 | 98 | 49 | 0.074750662 | 0.178951805 | NA | 0 | 0 | 0 | 0.026 | 0 | -0.009 |
| SE | 11135 | ECE1 | ECE1 | chr1 | - | 21279190 | 21279332 | 21272698 | 21272911 | 21290069 | 21290156 | 0 | 2 | 12 | 0 | 0 | 0 | 0 | 8 | 5 | 0 | 1 | 0 | 98 | 49 | 0.074771613 | 0.178951805 | NA | 1 | 1 | NA | 0.8 | 1 | 0.1 |
| SE | 2078 | HERC5 | HERC5 | chr4 | + | 88472408 | 88472502 | 88470613 | 88470673 | 88475840 | 88476030 | 0 | 2 | 29 | 0 | 0 | 0 | 0 | 34 | 3 | 0 | 1 | 0 | 98 | 49 | 0.075329785 | 0.180001061 | NA | 1 | 1 | NA | 0.944 | 1 | 0.028 |
| A5SS | 2702 | IFI6 | IFI6 | chr1 | - | 27669220 | 27669346 | 27669244 | 27669346 | 27668457 | 27668535 | 0 | 0 | 5 | 0 | 1 | 20 | 0 | 0 | 0 | 0 | 9 | 1 | 73 | 49 | 0.075687381 | 0.183611979 | NA | 0 | 0.144 | NA | 0 | 0 | 0.072 |
| SE | 801 | TBXAS1 | TBXAS1 | chr7 | + | 139955458 | 139955607 | 139953367 | 139953456 | 139957633 | 139957764 | 0 | 1 | 15 | 0 | 0 | 0 | 0 | 3 | 2 | 0 | 1 | 0 | 98 | 49 | 0.07582917 | 0.180642255 | NA | 1 | 1 | NA | 0.6 | 1 | 0.2 |
| SE | 12943 | SMCHD1 | SMCHD1 | chr18 | + | 2722518 | 2722663 | 2718314 | 2718434 | 2724898 | 2724995 | 0 | 1 | 43 | 0 | 0 | 5 | 1 | 37 | 1 | 0 | 12 | 0 | 98 | 49 | 0.075958015 | 0.180642255 | NA | 1 | 0.811 | 1 | 0.607 | 1 | 0.036 |
| SE | 6058 | HCLS1 | HCLS1 | chr3 | - | 121637145 | 121637256 | 121636433 | 121636489 | 121642926 | 121642981 | 0 | 0 | 9 | 0 | 1 | 0 | 1 | 4 | 1 | 0 | 0 | 0 | 98 | 49 | 0.075958686 | 0.180642255 | NA | 0 | 1 | 1 | 1 | 1 | -0.5 |
| RI | 461 | WTAP | WTAP | chr6 | + | 159748190 | 159749421 | 159748190 | 159748369 | 159748589 | 159749421 | 0 | 1 | 13 | 0 | 0 | 0 | 3 | 9 | 5 | 0 | 2 | 0 | 98 | 49 | 0.076502744 | 0.314063896 | NA | 1 | 1 | 1 | 0.692 | 1 | 0.103 |
| SE | 11482 | AP1G1 | AP1G1 | chr16 | - | 71789278 | 71789482 | 71774467 | 71774592 | 71808762 | 71809073 | 0 | 1 | 7 | 0 | 0 | 0 | 0 | 7 | 3 | 0 | 1 | 0 | 98 | 49 | 0.076633988 | 0.181960321 | NA | 1 | 1 | NA | 0.778 | 1 | 0.111 |
| SE | 4413 | MEGF6 | MEGF6 | chr1 | - | 3496987 | 3497119 | 3496654 | 3496783 | 3497232 | 3497361 | 0 | 3 | 0 | 0 | 0 | 2 | 0 | 1 | 1 | 0 | 0 | 0 | 98 | 49 | 0.076772626 | 0.182001983 | NA | 1 | 0 | NA | 1 | 1 | -0.5 |
| SE | 5396 | MTMR14 | MTMR14 | chr3 | + | 9662266 | 9662375 | 9653620 | 9653769 | 9668718 | 9668794 | 0 | 0 | 1 | 0 | 1 | 0 | 0 | 1 | 3 | 0 | 0 | 0 | 98 | 49 | 0.077074472 | 0.182348728 | NA | 0 | 1 | NA | 1 | 1 | -0.5 |
| SE | 5680 | GMIP | GMIP | chr19 | - | 19641809 | 19641867 | 19640445 | 19640571 | 19641975 | 19642051 | 0 | 1 | 27 | 0 | 0 | 3 | 0 | 15 | 6 | 0 | 0 | 0 | 98 | 49 | 0.077161537 | 0.182348728 | NA | 1 | 0.818 | NA | 1 | 1 | -0.091 |
| SE | 6871 | ATG2A | ATG2A | chr11 | - | 64914080 | 64914233 | 64913820 | 64913923 | 64914337 | 64914500 | 0 | 2 | 8 | 0 | 0 | 0 | 4 | 20 | 8 | 0 | 1 | 0 | 98 | 49 | 0.077718556 | 0.183327985 | NA | 1 | 1 | 1 | 0.909 | 1 | 0.03 |
| SE | 10716 | GADD45B | GADD45B | chr19 | + | 2477028 | 2477251 | 2476528 | 2476630 | 2477487 | 2478259 | 0 | 1 | 6 | 0 | 0 | 1 | 0 | 22 | 4 | 0 | 0 | 0 | 98 | 49 | 0.077819863 | 0.183327985 | NA | 1 | 0.75 | NA | 1 | 1 | -0.125 |
| SE | 10342 | PDCD4 | PDCD4 | chr10 | + | 110887664 | 110887886 | 110885252 | 110885366 | 110889532 | 110889630 | 0 | 1 | 4 | 0 | 0 | 0 | 0 | 21 | 2 | 0 | 1 | 0 | 98 | 49 | 0.078019098 | 0.183509709 | NA | 1 | 1 | NA | 0.913 | 1 | 0.043 |
| A3SS | 1541 | GMIP | GMIP | chr19 | - | 19636706 | 19636796 | 19636706 | 19636787 | 19636916 | 19637029 | 0 | 1 | 9 | 0 | 0 | 0 | 0 | 5 | 2 | 0 | 1 | 0 | 58 | 49 | 0.078029018 | 0.208872272 | NA | 1 | 1 | NA | 0.809 | 1 | 0.095 |
| SE | 11436 | PLK3 | PLK3 | chr1 | + | 44803267 | 44803391 | 44802954 | 44803153 | 44803599 | 44803691 | 0 | 2 | 8 | 0 | 0 | 0 | 0 | 6 | 5 | 0 | 1 | 0 | 98 | 49 | 0.078722929 | 0.184875878 | NA | 1 | 1 | NA | 0.75 | 1 | 0.125 |
| A3SS | 1062 | ATXN2L | ATXN2L | chr16 | + | 28832186 | 28832399 | 28832204 | 28832399 | 28830961 | 28831072 | 0 | 0 | 1 | 0 | 1 | 0 | 0 | 0 | 0 | 0 | 1 | 2 | 67 | 49 | 0.078907303 | 0.208872272 | NA | 0 | 1 | NA | 0 | 0 | 0.5 |
| SE | 9029 | MYD88 | MYD88 | chr3 | + | 38140387 | 38140568 | 38138477 | 38139028 | 38140756 | 38140848 | 0 | 3 | 7 | 0 | 0 | 0 | 0 | 2 | 2 | 0 | 2 | 0 | 98 | 49 | 0.079080192 | 0.185334725 | NA | 1 | 1 | NA | 0.333 | 1 | 0.334 |
| SE | 9375 | FMNL1 | FMNL1 | chr17 | + | 45236135 | 45236244 | 45234071 | 45234200 | 45237280 | 45237357 | 0 | 8 | 42 | 0 | 2 | 1 | 3 | 17 | 7 | 0 | 0 | 0 | 98 | 49 | 0.079164932 | 0.185334725 | NA | 0.667 | 0.955 | 1 | 1 | 1 | -0.189 |
| SE | 4098 | SETD1B | SETD1B | chr12 | + | 121819403 | 121819895 | 121817798 | 121817904 | 121822489 | 121823749 | 0 | 3 | 1 | 0 | 0 | 3 | 1 | 6 | 1 | 0 | 0 | 0 | 98 | 49 | 0.079530423 | 0.185900818 | NA | 1 | 0.143 | 1 | 1 | 1 | -0.428 |
| SE | 5185 | MYO9B | MYO9B | chr19 | + | 17159394 | 17159484 | 17156908 | 17157038 | 17162349 | 17162466 | 0 | 4 | 24 | 0 | 0 | 0 | 1 | 16 | 12 | 0 | 1 | 0 | 98 | 49 | 0.079811629 | 0.186229709 | NA | 1 | 1 | 1 | 0.889 | 1 | 0.037 |
| SE | 6698 | FCGR2A | FCGR2A | chr1 | + | 161506336 | 161506591 | 161505986 | 161506007 | 161509819 | 161510074 | 0 | 1 | 45 | 0 | 0 | 0 | 1 | 23 | 2 | 0 | 1 | 0 | 98 | 49 | 0.079918937 | 0.186229709 | NA | 1 | 1 | 1 | 0.92 | 1 | 0.027 |
| SE | 7345 | FAM129A | FAM129A | chr1 | - | 184823637 | 184823742 | 184823166 | 184823329 | 184831846 | 184831962 | 0 | 2 | 32 | 0 | 0 | 4 | 1 | 19 | 1 | 0 | 0 | 0 | 98 | 49 | 0.080074419 | 0.186303176 | NA | 1 | 0.8 | 1 | 1 | 1 | -0.1 |
| SE | 7416 | VAV1 | VAV1 | chr19 | + | 6822418 | 6822514 | 6822220 | 6822329 | 6825052 | 6825121 | 0 | 0 | 15 | 0 | 2 | 0 | 0 | 11 | 4 | 0 | 2 | 0 | 98 | 49 | 0.080433731 | 0.18684992 | NA | 0 | 1 | NA | 0.733 | 1 | -0.367 |
| SE | 12584 | SBNO2 | SBNO2 | chr19 | - | 1111018 | 1111093 | 1110744 | 1110888 | 1111505 | 1111614 | 0 | 1 | 6 | 0 | 0 | 1 | 1 | 26 | 2 | 0 | 0 | 0 | 98 | 49 | 0.080803043 | 0.187418168 | NA | 1 | 0.75 | 1 | 1 | 1 | -0.125 |
| SE | 10297 | TNFAIP2 | TNFAIP2 | chr14 | + | 103127004 | 103127629 | 103126326 | 103126692 | 103129739 | 103129854 | 0 | 2 | 36 | 0 | 0 | 0 | 0 | 35 | 5 | 0 | 1 | 0 | 98 | 49 | 0.081018716 | 0.187601066 | NA | 1 | 1 | NA | 0.946 | 1 | 0.027 |
| SE | 4657 | PRKD2 | PRKD2 | chr19 | - | 46681649 | 46681748 | 46678395 | 46678663 | 46689536 | 46689698 | 0 | 1 | 4 | 0 | 0 | 0 | 1 | 17 | 4 | 0 | 1 | 0 | 98 | 49 | 0.081231413 | 0.187601066 | NA | 1 | 1 | 1 | 0.895 | 1 | 0.035 |
| SE | 11395 | ADGRG3 | ADGRG3 | chr16 | + | 57680504 | 57680617 | 57680264 | 57680365 | 57684389 | 57684483 | 0 | 3 | 11 | 0 | 0 | 0 | 0 | 12 | 14 | 0 | 0 | 1 | 98 | 49 | 0.081296035 | 0.187601066 | NA | 1 | 1 | NA | 1 | 0.875 | 0.062 |
| SE | 5698 | RPLP2 | RPLP2 | chr11 | + | 812534 | 812633 | 811596 | 811645 | 812759 | 812876 | 0 | 2 | 43 | 0 | 0 | 0 | 0 | 41 | 4 | 0 | 1 | 0 | 98 | 49 | 0.081381168 | 0.187601066 | NA | 1 | 1 | NA | 0.953 | 1 | 0.024 |
| SE | 10951 | IQGAP1 | IQGAP1 | chr15 | + | 90482196 | 90482281 | 90481959 | 90482100 | 90483360 | 90483593 | 0 | 1 | 25 | 0 | 0 | 0 | 0 | 12 | 4 | 0 | 1 | 0 | 98 | 49 | 0.083074196 | 0.191210592 | NA | 1 | 1 | NA | 0.857 | 1 | 0.072 |
| SE | 3399 | ELOVL5 | ELOVL5 | chr6 | - | 53273219 | 53273344 | 53270592 | 53270727 | 53275089 | 53275261 | 0 | 1 | 21 | 0 | 0 | 0 | 3 | 9 | 4 | 0 | 1 | 0 | 98 | 49 | 0.083349562 | 0.191551057 | NA | 1 | 1 | 1 | 0.818 | 1 | 0.061 |
| SE | 5156 | PSAP | PSAP | chr10 | - | 71819030 | 71819111 | 71818616 | 71818724 | 71819464 | 71819622 | 1 | 15 | 181 | 0 | 0 | 0 | 10 | 44 | 9 | 0 | 1 | 0 | 98 | 49 | 0.084300652 | 0.193441038 | 1 | 1 | 1 | 1 | 0.957 | 1 | 0.014 |
| SE | 9929 | PTPRE | PTPRE | chr10 | + | 128056122 | 128056213 | 128049529 | 128049666 | 128060938 | 128061015 | 1 | 1 | 31 | 0 | 0 | 0 | 0 | 31 | 1 | 0 | 1 | 0 | 98 | 49 | 0.084566773 | 0.193502494 | 1 | 1 | 1 | NA | 0.939 | 1 | 0.03 |
| SE | 6870 | ATG2A | ATG2A | chr11 | - | 64912326 | 64912423 | 64912084 | 64912249 | 64913037 | 64913136 | 0 | 2 | 5 | 0 | 0 | 1 | 4 | 16 | 7 | 0 | 0 | 0 | 98 | 49 | 0.084584923 | 0.193502494 | NA | 1 | 0.714 | 1 | 1 | 1 | -0.143 |
| A5SS | 886 | DDIT3 | DDIT3 | chr12 | - | 57517268 | 57517753 | 57517705 | 57517753 | 57516587 | 57517180 | 1 | 3 | 7 | 0 | 0 | 0 | 2 | 20 | 3 | 0 | 1 | 0 | 98 | 49 | 0.084700702 | 0.201741673 | 1 | 1 | 1 | 1 | 0.909 | 1 | 0.03 |
| SE | 10384 | SIGLEC5 | SIGLEC5 | chr19 | - | 51627461 | 51627746 | 51627148 | 51627248 | 51627833 | 51628091 | 0 | 2 | 8 | 0 | 2 | 0 | 0 | 4 | 5 | 0 | 0 | 0 | 98 | 49 | 0.084994123 | 0.19414311 | NA | 0.333 | 1 | NA | 1 | 1 | -0.334 |
| SE | 12730 | PILRA | PILRA | chr7 | + | 100397878 | 100397912 | 100374043 | 100374433 | 100399290 | 100399340 | 0 | 1 | 14 | 0 | 0 | 0 | 0 | 3 | 2 | 0 | 3 | 0 | 83 | 49 | 0.085434257 | 0.194790215 | NA | 1 | 1 | NA | 0.371 | 1 | 0.315 |
| SE | 4324 | KMT2D | KMT2D | chr12 | - | 49026181 | 49027322 | 49024809 | 49024946 | 49027802 | 49027930 | 0 | 1 | 7 | 0 | 0 | 0 | 0 | 2 | 2 | 0 | 1 | 0 | 98 | 49 | 0.085536621 | 0.194790215 | NA | 1 | 1 | NA | 0.5 | 1 | 0.25 |
| SE | 11837 | RAB11FIP4 | RAB11FIP4 | chr17 | + | 31517650 | 31517877 | 31488000 | 31488272 | 31521165 | 31521360 | 0 | 1 | 2 | 0 | 0 | 0 | 0 | 6 | 5 | 0 | 1 | 0 | 98 | 49 | 0.086321731 | 0.196280728 | NA | 1 | 1 | NA | 0.75 | 1 | 0.125 |
| RI | 359 | DDIT3 | DDIT3 | chr12 | - | 57517268 | 57517753 | 57517268 | 57517595 | 57517705 | 57517753 | 4 | 2 | 10 | 0 | 0 | 0 | 8 | 27 | 2 | 0 | 1 | 0 | 98 | 49 | 0.086408407 | 0.336992786 | 1 | 1 | 1 | 1 | 0.931 | 1 | 0.023 |
| SE | 10863 | ARHGAP9 | ARHGAP9 | chr12 | - | 57473602 | 57473708 | 57472254 | 57472688 | 57474041 | 57474176 | 0 | 0 | 32 | 0 | 0 | 3 | 0 | 6 | 1 | 0 | 2 | 2 | 98 | 49 | 0.086690204 | 0.196820811 | NA | NA | 0.842 | NA | 0.6 | 0.2 | 0.442 |
| SE | 9932 | PTPRE | PTPRE | chr10 | + | 128072137 | 128072214 | 128070807 | 128070901 | 128073336 | 128073471 | 0 | 1 | 29 | 0 | 0 | 0 | 0 | 27 | 2 | 0 | 1 | 0 | 98 | 49 | 0.087167679 | 0.197606366 | NA | 1 | 1 | NA | 0.931 | 1 | 0.034 |
| SE | 10447 | ELL | ELL | chr19 | - | 18465796 | 18465918 | 18465411 | 18465575 | 18472834 | 18472882 | 0 | 2 | 13 | 0 | 4 | 0 | 0 | 10 | 0 | 0 | 0 | 0 | 98 | 49 | 0.088641831 | 0.200598284 | NA | 0.2 | 1 | NA | 1 | NA | -0.4 |
| SE | 10822 | C16orf72 | C16orf72 | chr16 | + | 9092920 | 9093001 | 9091679 | 9092270 | 9102983 | 9103259 | 0 | 1 | 22 | 0 | 1 | 1 | 0 | 20 | 0 | 0 | 0 | 0 | 98 | 49 | 0.088754397 | 0.200598284 | NA | 0.333 | 0.917 | NA | 1 | NA | -0.375 |
| SE | 13628 | SUN2 | SUN2 | chr22 | - | 38754641 | 38754706 | 38752506 | 38752665 | 38755762 | 38756019 | 0 | 0 | 7 | 0 | 1 | 2 | 0 | 0 | 0 | 0 | 0 | 6 | 98 | 49 | 0.088928004 | 0.200688873 | NA | 0 | 0.636 | NA | NA | 0 | 0.318 |
| SE | 1903 | MAP3K1 | MAP3K1 | chr5 | + | 56875031 | 56875310 | 56872824 | 56873005 | 56878979 | 56879101 | 0 | 0 | 5 | 0 | 1 | 1 | 1 | 3 | 2 | 0 | 0 | 0 | 98 | 49 | 0.089518138 | 0.201717783 | NA | 0 | 0.714 | 1 | 1 | 1 | -0.643 |
| SE | 7640 | IRF7 | IRF7 | chr11 | - | 614173 | 614399 | 613950 | 614037 | 614475 | 614534 | 0 | 1 | 2 | 0 | 7 | 10 | 0 | 5 | 1 | 0 | 3 | 3 | 98 | 49 | 0.089929548 | 0.202173556 | NA | 0.067 | 0.091 | NA | 0.455 | 0.143 | -0.22 |
| SE | 5219 | TPM4 | TPM4 | chr19 | + | 16081912 | 16082046 | 16076324 | 16076697 | 16086422 | 16086540 | 0 | 4 | 6 | 0 | 0 | 0 | 0 | 10 | 7 | 0 | 0 | 1 | 98 | 49 | 0.09016239 | 0.202173556 | NA | 1 | 1 | NA | 1 | 0.778 | 0.111 |
| SE | 6816 | OAS1 | OAS1 | chr12 | + | 112908535 | 112908824 | 112906933 | 112907219 | 112911050 | 112911235 | 0 | 1 | 25 | 0 | 0 | 0 | 5 | 3 | 3 | 0 | 1 | 0 | 98 | 49 | 0.090232466 | 0.202173556 | NA | 1 | 1 | 1 | 0.6 | 1 | 0.133 |
| SE | 2081 | HERC5 | HERC5 | chr4 | + | 88499925 | 88499992 | 88494164 | 88494331 | 88500914 | 88500985 | 0 | 3 | 48 | 0 | 1 | 1 | 0 | 33 | 0 | 0 | 2 | 0 | 98 | 49 | 0.090258454 | 0.202173556 | NA | 0.6 | 0.96 | NA | 0.892 | NA | -0.112 |
| A3SS | 872 | DGCR2 | DGCR2 | chr22 | - | 19064847 | 19065075 | 19064847 | 19065067 | 19068099 | 19068225 | 0 | 0 | 3 | 0 | 1 | 4 | 0 | 0 | 0 | 0 | 2 | 2 | 57 | 49 | 0.092162455 | 0.240423795 | NA | 0 | 0.392 | NA | 0 | 0 | 0.196 |
| RI | 832 | HES4 | HES4 | chr1 | - | 999691 | 1000172 | 999691 | 999787 | 999865 | 1000172 | 1 | 2 | 1 | 0 | 0 | 0 | 3 | 13 | 4 | 0 | 1 | 0 | 98 | 49 | 0.092178476 | 0.342377196 | 1 | 1 | 1 | 1 | 0.867 | 1 | 0.044 |
| SE | 1266 | MICAL1 | MICAL1 | chr6 | - | 109447356 | 109447440 | 109447072 | 109447229 | 109447680 | 109447722 | 0 | 3 | 5 | 0 | 0 | 0 | 0 | 2 | 1 | 0 | 0 | 1 | 98 | 49 | 0.094107379 | 0.210481236 | NA | 1 | 1 | NA | 1 | 0.333 | 0.334 |
| SE | 4124 | XPO6 | XPO6 | chr16 | - | 28135215 | 28135324 | 28133840 | 28133933 | 28146093 | 28146203 | 0 | 5 | 50 | 0 | 0 | 2 | 3 | 60 | 4 | 0 | 0 | 0 | 98 | 49 | 0.094351496 | 0.210563613 | NA | 1 | 0.926 | 1 | 1 | 1 | -0.037 |
| SE | 3589 | SP110 | SP110 | chr2 | - | 230202578 | 230202728 | 230200884 | 230200965 | 230207990 | 230208059 | 0 | 1 | 8 | 0 | 0 | 0 | 0 | 10 | 6 | 0 | 1 | 0 | 98 | 49 | 0.094424401 | 0.210563613 | NA | 1 | 1 | NA | 0.833 | 1 | 0.084 |
| SE | 10651 | SLC43A2 | SLC43A2 | chr17 | - | 1616561 | 1616769 | 1614978 | 1615034 | 1627714 | 1627920 | 0 | 0 | 9 | 0 | 0 | 1 | 0 | 9 | 0 | 0 | 0 | 2 | 98 | 49 | 0.094824344 | 0.211028906 | NA | NA | 0.818 | NA | 1 | 0 | 0.318 |
| SE | 7418 | VAV1 | VAV1 | chr19 | + | 6847997 | 6848114 | 6843134 | 6843166 | 6850669 | 6850757 | 0 | 3 | 13 | 0 | 4 | 0 | 3 | 10 | 0 | 0 | 0 | 0 | 98 | 49 | 0.095128464 | 0.211028906 | NA | 0.273 | 1 | 1 | 1 | NA | -0.363 |
| SE | 10849 | IRF1 | IRF1 | chr5 | - | 132486794 | 132486844 | 132486556 | 132486686 | 132486953 | 132487130 | 0 | 2 | 21 | 0 | 0 | 1 | 0 | 36 | 5 | 0 | 8 | 0 | 98 | 49 | 0.095196923 | 0.211028906 | NA | 1 | 0.913 | NA | 0.692 | 1 | 0.111 |
| SE | 4963 | DDX3X | DDX3X | chrX | + | 41343215 | 41343351 | 41342736 | 41342836 | 41343736 | 41343822 | 0 | 2 | 31 | 0 | 0 | 1 | 0 | 24 | 1 | 0 | 3 | 0 | 98 | 49 | 0.095320071 | 0.211028906 | NA | 1 | 0.939 | NA | 0.8 | 1 | 0.07 |
| SE | 1143 | IGF2R | IGF2R | chr6 | + | 160079579 | 160079787 | 160078200 | 160078362 | 160080128 | 160080275 | 0 | 4 | 40 | 0 | 0 | 1 | 0 | 6 | 2 | 0 | 3 | 0 | 98 | 49 | 0.095335081 | 0.211028906 | NA | 1 | 0.952 | NA | 0.5 | 1 | 0.226 |
| SE | 10701 | TLE3 | TLE3 | chr15 | - | 70059409 | 70059460 | 70058662 | 70058815 | 70060529 | 70060649 | 0 | 4 | 31 | 0 | 2 | 0 | 0 | 21 | 0 | 0 | 0 | 0 | 98 | 49 | 0.095684167 | 0.211266553 | NA | 0.5 | 1 | NA | 1 | NA | -0.25 |
| SE | 6176 | ABCA1 | ABCA1 | chr9 | - | 104819585 | 104819723 | 104818662 | 104818883 | 104819926 | 104820069 | 0 | 1 | 17 | 0 | 0 | 0 | 0 | 29 | 3 | 0 | 1 | 0 | 98 | 49 | 0.095723568 | 0.211266553 | NA | 1 | 1 | NA | 0.935 | 1 | 0.032 |
| SE | 8187 | EPSTI1 | EPSTI1 | chr13 | - | 42895008 | 42895108 | 42886387 | 42888502 | 42900309 | 42900383 | 0 | 1 | 9 | 0 | 0 | 2 | 4 | 11 | 1 | 0 | 0 | 0 | 98 | 49 | 0.096411782 | 0.212473472 | NA | 1 | 0.692 | 1 | 1 | 1 | -0.154 |
| SE | 12365 | NBEAL2 | NBEAL2 | chr3 | + | 46995931 | 46996051 | 46995713 | 46995846 | 46996270 | 46996592 | 0 | 8 | 19 | 0 | 0 | 0 | 0 | 5 | 4 | 0 | 0 | 2 | 98 | 49 | 0.09699006 | 0.213434934 | NA | 1 | 1 | NA | 1 | 0.5 | 0.25 |
| SE | 8710 | CD74 | CD74 | chr5 | - | 150406880 | 150406960 | 150406258 | 150406321 | 150407151 | 150407324 | 0 | 7 | 68 | 0 | 0 | 1 | 2 | 12 | 4 | 0 | 3 | 0 | 98 | 49 | 0.097385214 | 0.213991193 | NA | 1 | 0.971 | 1 | 0.667 | 1 | 0.097 |
| SE | 11126 | PSEN1 | PSEN1 | chr14 | + | 73198030 | 73198129 | 73192643 | 73192864 | 73206385 | 73206472 | 0 | 1 | 8 | 0 | 2 | 1 | 0 | 5 | 0 | 0 | 0 | 0 | 98 | 49 | 0.098181291 | 0.215425518 | NA | 0.2 | 0.8 | NA | 1 | NA | -0.5 |
| A3SS | 1986 | PGS1 | PGS1 | chr17 | + | 78414878 | 78415027 | 78414951 | 78415027 | 78403567 | 78404089 | 0 | 1 | 5 | 0 | 0 | 0 | 3 | 5 | 5 | 0 | 1 | 0 | 98 | 49 | 0.098812002 | 0.253724157 | NA | 1 | 1 | 1 | 0.714 | 1 | 0.095 |
| SE | 13029 | FOSB | FOSB | chr19 | + | 45471193 | 45471301 | 45470628 | 45470949 | 45472550 | 45475179 | 0 | 3 | 9 | 0 | 2 | 0 | 0 | 13 | 0 | 0 | 0 | 0 | 98 | 49 | 0.099118539 | 0.217164962 | NA | 0.429 | 1 | NA | 1 | NA | -0.285 |
| SE | 13040 | MAP3K8 | MAP3K8 | chr10 | + | 30437175 | 30437406 | 30434020 | 30434378 | 30438915 | 30439274 | 0 | 0 | 6 | 0 | 0 | 1 | 0 | 2 | 0 | 0 | 2 | 2 | 98 | 49 | 0.099502086 | 0.217687969 | NA | NA | 0.75 | NA | 0.333 | 0 | 0.584 |
| SE | 12128 | LRRC59 | LRRC59 | chr17 | - | 50392738 | 50392897 | 50392397 | 50392502 | 50394928 | 50394988 | 1 | 1 | 3 | 0 | 0 | 0 | 0 | 1 | 1 | 0 | 3 | 0 | 98 | 49 | 0.099753465 | 0.217812352 | 1 | 1 | 1 | NA | 0.143 | 1 | 0.428 |
| SE | 55 | ELF4 | ELF4 | chrX | - | 130081255 | 130081539 | 130074580 | 130074752 | 130110324 | 130110500 | 0 | 2 | 5 | 0 | 1 | 0 | 0 | 0 | 1 | 0 | 1 | 4 | 98 | 49 | 0.100072297 | 0.217812352 | NA | 0.5 | 1 | NA | 0 | 0.111 | 0.695 |
| A3SS | 2860 | TPM3 | TPM3 | chr1 | - | 154155303 | 154159048 | 154155303 | 154157721 | 154170399 | 154170469 | 1 | 0 | 1 | 0 | 0 | 20 | 0 | 2 | 1 | 0 | 14 | 0 | 98 | 49 | 0.100080084 | 0.253724157 | 1 | NA | 0.024 | NA | 0.067 | 1 | -0.021 |
| SE | 9378 | FMNL1 | FMNL1 | chr17 | + | 45246504 | 45246604 | 45246209 | 45246330 | 45246866 | 45247316 | 0 | 1 | 0 | 0 | 1 | 15 | 5 | 0 | 0 | 0 | 13 | 2 | 98 | 49 | 0.100134734 | 0.217812352 | NA | 0.333 | 0 | 1 | 0 | 0 | -0.167 |
| SE | 6719 | KIF1B | KIF1B | chr1 | + | 10345844 | 10345953 | 10343231 | 10343287 | 10348648 | 10348733 | 0 | 4 | 2 | 0 | 0 | 0 | 0 | 1 | 2 | 0 | 1 | 0 | 98 | 49 | 0.100138613 | 0.217812352 | NA | 1 | 1 | NA | 0.333 | 1 | 0.334 |
| SE | 13866 | ACSL1 | ACSL1 | chr4 | - | 184773080 | 184773154 | 184768315 | 184768450 | 184773662 | 184773714 | 0 | 2 | 38 | 0 | 0 | 1 | 3 | 28 | 2 | 0 | 3 | 0 | 98 | 49 | 0.100289399 | 0.217825096 | NA | 1 | 0.95 | 1 | 0.824 | 1 | 0.034 |
| SE | 13870 | ACSL1 | ACSL1 | chr4 | - | 184803319 | 184803546 | 184788616 | 184788731 | 184825915 | 184826114 | 0 | 4 | 91 | 0 | 0 | 0 | 8 | 83 | 44 | 0 | 3 | 0 | 98 | 49 | 0.100452402 | 0.217864301 | NA | 1 | 1 | 1 | 0.933 | 1 | 0.022 |
| SE | 13051 | NAPA | NAPA | chr19 | - | 47500632 | 47500749 | 47495549 | 47495596 | 47514842 | 47515258 | 0 | 1 | 13 | 0 | 0 | 0 | 0 | 11 | 7 | 0 | 1 | 0 | 98 | 49 | 0.100911104 | 0.218543788 | NA | 1 | 1 | NA | 0.846 | 1 | 0.077 |
| SE | 7577 | SPATS2L | SPATS2L | chr2 | + | 200439121 | 200439328 | 200419249 | 200419496 | 200440648 | 200440784 | 0 | 0 | 3 | 0 | 0 | 1 | 0 | 1 | 0 | 0 | 0 | 3 | 98 | 49 | 0.101162648 | 0.218773323 | NA | NA | 0.6 | NA | 1 | 0 | 0.1 |
| SE | 12742 | DYSF | DYSF | chr2 | + | 71520177 | 71520208 | 71516988 | 71517039 | 71520788 | 71520904 | 0 | 1 | 12 | 0 | 1 | 0 | 2 | 7 | 0 | 0 | 0 | 0 | 80 | 49 | 0.10199177 | 0.220129306 | NA | 0.38 | 1 | 1 | 1 | NA | -0.31 |
| SE | 10929 | DEDD2 | DEDD2 | chr19 | - | 42204461 | 42204559 | 42198592 | 42199829 | 42209699 | 42209840 | 0 | 1 | 0 | 0 | 3 | 5 | 0 | 0 | 0 | 0 | 16 | 3 | 98 | 49 | 0.102192668 | 0.220129306 | NA | 0.143 | 0 | NA | 0 | 0 | 0.071 |
| SE | 3914 | ALOX5AP | ALOX5AP | chr13 | + | 30755943 | 30756025 | 30752051 | 30752122 | 30763943 | 30764428 | 0 | 2 | 74 | 0 | 0 | 0 | 0 | 36 | 17 | 0 | 1 | 0 | 98 | 49 | 0.102229046 | 0.220129306 | NA | 1 | 1 | NA | 0.947 | 1 | 0.026 |
| SE | 3886 | LCP1 | LCP1 | chr13 | - | 46146907 | 46147103 | 46144441 | 46144520 | 46148351 | 46148447 | 0 | 9 | 167 | 0 | 0 | 1 | 8 | 67 | 3 | 0 | 2 | 0 | 98 | 49 | 0.102410835 | 0.220205272 | NA | 1 | 0.988 | 1 | 0.944 | 1 | 0.013 |
| SE | 3619 | NIN | NIN | chr14 | - | 50830463 | 50830517 | 50821873 | 50822077 | 50831005 | 50831121 | 0 | 3 | 3 | 0 | 0 | 1 | 2 | 0 | 18 | 0 | 0 | 0 | 98 | 49 | 0.10387195 | 0.223027916 | NA | 1 | 0.6 | 1 | NA | 1 | -0.2 |
| SE | 803 | TBXAS1 | TBXAS1 | chr7 | + | 140017670 | 140017833 | 140015722 | 140015860 | 140020024 | 140020325 | 0 | 0 | 14 | 0 | 0 | 1 | 0 | 2 | 0 | 0 | 1 | 1 | 98 | 49 | 0.104215706 | 0.223185153 | NA | NA | 0.875 | NA | 0.5 | 0 | 0.625 |

| | | | | | | | | | | | | | | | | | | | | | | | | | | | | | | | | | | |
|---|---|---|---|---|---|---|---|---|---|---|---|---|---|---|---|---|---|---|---|---|---|---|---|---|---|---|---|---|---|---|---|---|---|---|
| SE | 10226 | ACAP1 | ACAP1 | chr17 | + | 7337311 | 7337369 | 7336528 | 7336787 | 7341947 | 7342067 | 2 | 1 | 3 | 0 | 0 | 0 | 0 | 2 | 3 | 0 | 1 | 0 | 98 | 49 | 0.104242167 | 0.223185153 | 1 | 1 | 1 | NA | 0.5 | 1 | 0.25 |
| SE | 6858 | SELL | SELL | chr1 | - | 169701559 | 169701688 | 169696534 | 169696553 | 169703254 | 169703440 | 0 | 2 | 95 | 0 | 1 | 0 | 12 | 29 | 3 | 0 | 0 | 0 | 98 | 49 | 0.105494926 | 0.225138011 | NA | 0.5 | 1 | 1 | 1 | 1 | -0.25 |
| SE | 796 | PARP12 | PARP12 | chr7 | - | 140057898 | 140058034 | 140056855 | 140057153 | 140062521 | 140063721 | 0 | 1 | 8 | 0 | 1 | 0 | 0 | 6 | 0 | 0 | 0 | 0 | 98 | 49 | 0.105498898 | 0.225138011 | NA | 0.333 | 1 | NA | 1 | NA | -0.334 |
| SE | 4946 | ARAP1 | ARAP1 | chr11 | - | 72705804 | 72705890 | 72702904 | 72703079 | 72707174 | 72707374 | 0 | 2 | 13 | 0 | 0 | 0 | 0 | 5 | 7 | 0 | 1 | 0 | 98 | 49 | 0.105603658 | 0.225138011 | NA | 1 | 1 | NA | 0.714 | 1 | 0.143 |
| SE | 7968 | OAS3 | OAS3 | chr12 | + | 112967417 | 112967593 | 112964234 | 112964408 | 112967935 | 112968174 | 0 | 0 | 27 | 0 | 0 | 1 | 0 | 7 | 1 | 0 | 2 | 1 | 98 | 49 | 0.106158162 | 0.225817599 | NA | NA | 0.931 | NA | 0.636 | 0.333 | 0.447 |
| SE | 4056 | FOS | FOS | chr14 | + | 75280559 | 75280667 | 75279876 | 75280128 | 75280782 | 75282234 | 0 | 25 | 189 | 0 | 1 | 0 | 3 | 248 | 16 | 0 | 0 | 0 | 98 | 49 | 0.106325146 | 0.225817599 | NA | 0.926 | 1 | 1 | 1 | 1 | -0.037 |
| SE | 1509 | HLA-E | HLA-E | chr6 | + | 30492540 | 30492583 | 30491536 | 30491653 | 30492748 | 30494205 | 0 | 3 | 55 | 0 | 1 | 0 | 0 | 33 | 4 | 0 | 0 | 0 | 92 | 49 | 0.106659671 | 0.225817599 | NA | 0.615 | 1 | NA | 1 | 1 | -0.193 |
| SE | 11045 | HLA-B | HLA-B | chr6 | - | 31354632 | 31354665 | 31354478 | 31354526 | 31355316 | 31355592 | 1 | 12 | 113 | 0 | 0 | 0 | 8 | 112 | 7 | 0 | 1 | 0 | 82 | 49 | 0.106709745 | 0.225817599 | 1 | 1 | 1 | 1 | 0.985 | 1 | 0.005 |
| SE | 7902 | PACSIN2 | PACSIN2 | chr22 | - | 42890946 | 42891182 | 42888642 | 42888798 | 42893456 | 42893613 | 0 | 1 | 8 | 0 | 0 | 0 | 0 | 10 | 8 | 0 | 1 | 0 | 98 | 49 | 0.106828817 | 0.225817599 | NA | 1 | 1 | NA | 0.833 | 1 | 0.084 |
| SE | 7882 | HIF1A | HIF1A | chr14 | + | 61738086 | 61738373 | 61736888 | 61737109 | 61740504 | 61740627 | 0 | 1 | 10 | 0 | 0 | 1 | 0 | 34 | 1 | 0 | 0 | 0 | 98 | 49 | 0.106940257 | 0.225817599 | NA | 1 | 0.833 | NA | 1 | 1 | -0.084 |
| SE | 7143 | CPD | CPD | chr17 | + | 30438974 | 30439077 | 30431771 | 30431881 | 30442307 | 30442450 | 0 | 1 | 10 | 0 | 0 | 2 | 0 | 11 | 1 | 0 | 0 | 0 | 98 | 49 | 0.107022882 | 0.225817599 | NA | 1 | 0.714 | NA | 1 | 1 | -0.143 |
| SE | 5742 | ENTPD1 | ENTPD1 | chr10 | + | 95860468 | 95860582 | 95847445 | 95847706 | 95864723 | 95864861 | 0 | 1 | 5 | 0 | 0 | 0 | 0 | 2 | 3 | 0 | 1 | 0 | 98 | 49 | 0.107124383 | 0.225817599 | NA | 1 | 1 | NA | 0.5 | 1 | 0.25 |
| A3SS | 166 | SDCBP | SDCBP | chr8 | + | 58575899 | 58576061 | 58575902 | 58576061 | 58572204 | 58572314 | 0 | 3 | 20 | 0 | 0 | 2 | 0 | 9 | 1 | 0 | 4 | 0 | 52 | 49 | 0.10775843 | 0.269396074 | NA | 1 | 0.904 | NA | 0.68 | 1 | 0.112 |
| SE | 4628 | SORL1 | SORL1 | chr11 | + | 121577280 | 121577400 | 121574240 | 121574363 | 121583457 | 121583583 | 0 | 8 | 61 | 0 | 0 | 3 | 0 | 33 | 7 | 0 | 0 | 0 | 98 | 49 | 0.107936364 | 0.227210581 | NA | 1 | 0.91 | NA | 1 | 1 | -0.045 |
| SE | 9584 | ANXA11 | ANXA11 | chr10 | - | 80176106 | 80176155 | 80172806 | 80172869 | 80205342 | 80205444 | 0 | 0 | 4 | 0 | 0 | 2 | 0 | 4 | 0 | 0 | 0 | 3 | 98 | 49 | 0.108478756 | 0.228032965 | NA | NA | 0.5 | NA | 1 | 0 | 0 |
| SE | 10587 | RNF213 | RNF213 | chr17 | + | 80361733 | 80361888 | 80358287 | 80358479 | 80363101 | 80363314 | 0 | 1 | 25 | 0 | 0 | 0 | 0 | 32 | 6 | 0 | 1 | 0 | 98 | 49 | 0.108649764 | 0.228073457 | NA | 1 | 1 | NA | 0.941 | 1 | 0.03 |
| SE | 9787 | HERC4 | HERC4 | chr10 | - | 67990210 | 67990400 | 67988662 | 67988835 | 67990903 | 67991015 | 0 | 0 | 1 | 0 | 0 | 2 | 0 | 2 | 3 | 0 | 0 | 0 | 98 | 49 | 0.108971078 | 0.228428912 | NA | NA | 0.2 | NA | 1 | 1 | -0.8 |
| A5SS | 1408 | HNRNPC | HNRNPC | chr14 | - | 21230957 | 21231072 | 21230996 | 21231072 | 21230318 | 21230366 | 0 | 6 | 6 | 0 | 1 | 5 | 4 | 15 | 0 | 1 | 7 | 0 | 88 | 49 | 0.108976627 | 0.254927467 | NA | 0.77 | 0.401 | 0.69 | 0.544 | NA | -0.031 |
| SE | 11640 | ARRB2 | ARRB2 | chr17 | + | 4716411 | 4716608 | 4715972 | 4716033 | 4717216 | 4717276 | 0 | 0 | 2 | 0 | 0 | 3 | 0 | 5 | 4 | 0 | 0 | 0 | 98 | 49 | 0.1093759 | 0.228660806 | NA | NA | 0.25 | NA | 1 | 1 | -0.75 |
| SE | 7570 | PGD | PGD | chr1 | + | 10403070 | 10403136 | 10400392 | 10400572 | 10404160 | 10404279 | 0 | 7 | 13 | 0 | 4 | 0 | 0 | 8 | 0 | 0 | 0 | 0 | 98 | 49 | 0.109641955 | 0.228660806 | NA | 0.467 | 1 | NA | 1 | NA | -0.266 |
| SE | 5385 | CD44 | CD44 | chr11 | + | 35180273 | 35180407 | 35176574 | 35176740 | 35186831 | 35186900 | 1 | 4 | 16 | 0 | 2 | 1 | 0 | 9 | 0 | 0 | 0 | 0 | 98 | 49 | 0.109674831 | 0.228660806 | 1 | 0.5 | 0.889 | NA | 1 | NA | -0.204 |
| SE | 1986 | DDX60 | DDX60 | chr4 | - | 168272042 | 168272138 | 168268853 | 168268969 | 168273278 | 168273398 | 0 | 1 | 7 | 0 | 1 | 0 | 0 | 5 | 0 | 0 | 0 | 0 | 98 | 49 | 0.109690247 | 0.228660806 | NA | 0.333 | 1 | NA | 1 | NA | -0.334 |
| SE | 1038 | AOAH | AOAH | chr7 | - | 36659165 | 36659265 | 36637850 | 36637910 | 36673942 | 36674009 | 0 | 0 | 10 | 0 | 0 | 1 | 1 | 8 | 0 | 0 | 1 | 1 | 98 | 49 | 0.110142389 | 0.229285332 | NA | NA | 0.833 | 1 | 0.8 | 0 | 0.233 |
| SE | 5785 | DENND3 | DENND3 | chr8 | + | 141144147 | 141144259 | 141138021 | 141138137 | 141150833 | 141150953 | 0 | 0 | 6 | 0 | 0 | 2 | 1 | 0 | 0 | 0 | 4 | 2 | 98 | 49 | 0.111042654 | 0.230839709 | NA | NA | 0.6 | 1 | 0 | 0 | 0.267 |
| SE | 5585 | WTAP | WTAP | chr6 | + | 159736257 | 159736295 | 159726997 | 159727333 | 159738989 | 159739045 | 0 | 3 | 10 | 0 | 0 | 0 | 0 | 6 | 3 | 0 | 0 | 1 | 87 | 49 | 0.111205631 | 0.230859204 | NA | 1 | 1 | NA | 1 | 0.628 | 0.186 |
| SE | 13176 | ZNF274 | ZNF274 | chr19 | + | 58209960 | 58210073 | 58206719 | 58207202 | 58211559 | 58211686 | 0 | 1 | 0 | 0 | 3 | 0 | 0 | 2 | 0 | 0 | 0 | 0 | 98 | 49 | 0.111677163 | 0.231518312 | NA | 0.143 | NA | NA | 1 | NA | -0.857 |
| SE | 422 | FAM214B | FAM214B | chr9 | - | 35106537 | 35106643 | 35106251 | 35106410 | 35106832 | 35106891 | 0 | 1 | 10 | 0 | 0 | 0 | 0 | 4 | 4 | 0 | 2 | 0 | 98 | 49 | 0.112112283 | 0.232100222 | NA | 1 | 1 | NA | 0.5 | 1 | 0.25 |
| SE | 13719 | HMHA1 | HMHA1 | chr19 | + | 1073673 | 1073746 | 1073505 | 1073590 | 1073947 | 1074014 | 0 | 0 | 23 | 0 | 0 | 1 | 0 | 6 | 1 | 0 | 3 | 1 | 98 | 49 | 0.112614145 | 0.232818515 | NA | NA | 0.92 | NA | 0.5 | 0.333 | 0.504 |
| SE | 8977 | RPS9 | RPS9 | chr19 | + | 54206275 | 54206462 | 54201486 | 54201609 | 54207397 | 54207647 | 0 | 3 | 56 | 0 | 0 | 1 | 0 | 22 | 2 | 0 | 2 | 0 | 98 | 49 | 0.113459253 | 0.234243486 | NA | 1 | 0.966 | NA | 0.846 | 1 | 0.06 |
| SE | 12370 | NBEAL2 | NBEAL2 | chr3 | + | 47007524 | 47007697 | 47006332 | 47006449 | 47007815 | 47007910 | 0 | 14 | 33 | 0 | 0 | 0 | 2 | 7 | 17 | 0 | 1 | 0 | 98 | 49 | 0.114232158 | 0.235336193 | NA | 1 | 1 | 1 | 0.778 | 1 | 0.074 |
| SE | 4364 | LRRK2 | LRRK2 | chr12 | + | 40319987 | 40320175 | 40315211 | 40315300 | 40321033 | 40321188 | 0 | 2 | 64 | 0 | 4 | 2 | 0 | 16 | 1 | 0 | 3 | 0 | 98 | 49 | 0.114457785 | 0.235336193 | NA | 0.2 | 0.941 | NA | 0.727 | 1 | -0.293 |
| SE | 11194 | PARP14 | PARP14 | chr3 | + | 122703741 | 122703978 | 122699389 | 122701635 | 122704526 | 122704748 | 0 | 6 | 21 | 0 | 1 | 0 | 0 | 22 | 0 | 0 | 0 | 0 | 98 | 49 | 0.114458255 | 0.235336193 | NA | 0.75 | 1 | NA | 1 | NA | -0.125 |
| A5SS | 1123 | MYO9B | MYO9B | chr19 | + | 17211646 | 17211786 | 17211646 | 17211774 | 17211894 | 17213295 | 0 | 0 | 2 | 0 | 0 | 2 | 0 | 0 | 0 | 0 | 0 | 4 | 61 | 49 | 0.11468766 | 0.263580411 | NA | NA | 0.445 | NA | NA | 0 | 0.445 |
| SE | 10702 | TLE3 | TLE3 | chr15 | - | 70064453 | 70064470 | 70060529 | 70060649 | 70066013 | 70066218 | 0 | 3 | 13 | 0 | 0 | 1 | 0 | 11 | 1 | 0 | 3 | 0 | 66 | 49 | 0.114960195 | 0.235829003 | NA | 1 | 0.906 | NA | 0.731 | 1 | 0.088 |
| SE | 2899 | GTPBP1 | GTPBP1 | chr22 | + | 38708844 | 38708956 | 38705801 | 38706147 | 38715906 | 38716087 | 0 | 0 | 8 | 0 | 0 | 1 | 0 | 9 | 0 | 0 | 6 | 1 | 98 | 49 | 0.115011749 | 0.235829003 | NA | NA | 0.8 | NA | 0.429 | 0 | 0.586 |
| SE | 7731 | KIAA1033 | KIAA1033 | chr12 | + | 105162748 | 105162845 | 105160000 | 105160148 | 105164110 | 105164307 | 0 | 1 | 7 | 0 | 1 | 0 | 0 | 4 | 0 | 0 | 0 | 0 | 98 | 49 | 0.115352023 | 0.235883116 | NA | 0.333 | 1 | NA | 1 | NA | -0.334 |
| SE | 13111 | AP2A1 | AP2A1 | chr19 | + | 49800960 | 49801058 | 49799967 | 49800150 | 49801389 | 49801621 | 0 | 1 | 3 | 0 | 1 | 0 | 0 | 4 | 0 | 0 | 0 | 0 | 98 | 49 | 0.115352023 | 0.235883116 | NA | 0.333 | 1 | NA | 1 | NA | -0.334 |
| SE | 3524 | DYRK1A | DYRK1A | chr21 | + | 37420298 | 37420384 | 37367556 | 37367628 | 37472683 | 37472880 | 0 | 0 | 2 | 0 | 0 | 2 | 0 | 2 | 4 | 0 | 0 | 0 | 98 | 49 | 0.115941495 | 0.236766395 | NA | NA | 0.333 | NA | 1 | 1 | -0.667 |
| SE | 7378 | RNF19B | RNF19B | chr1 | - | 32945513 | 32945628 | 32944018 | 32944159 | 32946401 | 32946564 | 0 | 1 | 24 | 0 | 0 | 0 | 1 | 30 | 9 | 0 | 1 | 0 | 98 | 49 | 0.116142758 | 0.236855585 | NA | 1 | 1 | 1 | 0.937 | 1 | 0.021 |
| SE | 6360 | ZSWIM8 | ZSWIM8 | chr10 | + | 73788669 | 73788823 | 73785623 | 73786086 | 73789095 | 73789190 | 0 | 2 | 3 | 0 | 5 | 0 | 0 | 2 | 0 | 0 | 0 | 0 | 98 | 49 | 0.116399553 | 0.237028277 | NA | 0.167 | 1 | NA | 1 | NA | -0.416 |
| SE | 12686 | MYO1F | MYO1F | chr19 | - | 8541905 | 8541991 | 8539946 | 8540028 | 8544296 | 8544464 | 0 | 11 | 31 | 0 | 0 | 3 | 0 | 21 | 4 | 0 | 0 | 0 | 98 | 49 | 0.116542846 | 0.237028277 | NA | 1 | 0.838 | NA | 1 | 1 | -0.081 |
| SE | 6861 | CARD8 | CARD8 | chr19 | - | 48238382 | 48238532 | 48234402 | 48234543 | 48240961 | 48241063 | 0 | 0 | 7 | 0 | 0 | 1 | 0 | 0 | 0 | 0 | 0 | 1 | 98 | 49 | 0.11681427 | 0.237259253 | NA | NA | 0.778 | NA | NA | 0 | 0.778 |
| A3SS | 412 | PRRC2A | PRRC2A | chr6 | + | 31622689 | 31622901 | 31622726 | 31622901 | 31620672 | 31620858 | 0 | 0 | 2 | 0 | 0 | 2 | 0 | 0 | 0 | 0 | 3 | 4 | 86 | 49 | 0.117982531 | 0.29091583 | NA | NA | 0.363 | NA | 0 | 0 | 0.363 |
| SE | 11993 | STAT5A | STAT5A | chr17 | + | 42291971 | 42292061 | 42289865 | 42290022 | 42295618 | 42295793 | 0 | 1 | 2 | 0 | 2 | 0 | 0 | 4 | 0 | 0 | 1 | 0 | 98 | 49 | 0.118118135 | 0.239583747 | NA | 0.2 | 1 | NA | 0.667 | NA | -0.067 |
| SE | 12348 | CSNK1D | CSNK1D | chr17 | - | 82255428 | 82255577 | 82253015 | 82253244 | 82265685 | 82265796 | 0 | 0 | 3 | 0 | 0 | 2 | 0 | 2 | 5 | 0 | 0 | 0 | 98 | 49 | 0.118962034 | 0.240970265 | NA | NA | 0.429 | NA | 1 | 1 | -0.571 |
| SE | 4455 | GABARAPL1 | GABARAPL1 | chr12 | + | 10218062 | 10218141 | 10212889 | 10213219 | 10220439 | 10220558 | 0 | 0 | 8 | 0 | 2 | 0 | 0 | 6 | 3 | 0 | 2 | 0 | 98 | 49 | 0.119908107 | 0.242559736 | NA | 0 | 1 | NA | 0.6 | 1 | -0.3 |
| SE | 11089 | IQSEC1 | IQSEC1 | chr3 | - | 12902772 | 12902822 | 12897042 | 12901522 | 12908348 | 12908525 | 0 | 0 | 2 | 0 | 0 | 3 | 0 | 0 | 0 | 0 | 5 | 5 | 98 | 49 | 0.120556389 | 0.24354335 | NA | NA | 0.25 | NA | 0 | 0 | 0.25 |
| SE | 7230 | NFKBIZ | NFKBIZ | chr3 | + | 101855394 | 101855458 | 101855061 | 101855208 | 101855732 | 101855902 | 0 | 1 | 28 | 0 | 0 | 0 | 5 | 65 | 2 | 0 | 1 | 0 | 98 | 49 | 0.120755712 | 0.243618571 | NA | 1 | 1 | 1 | 0.97 | 1 | 0.01 |
| SE | 6272 | BAZ2B | BAZ2B | chr2 | - | 159430862 | 159431156 | 159429199 | 159429260 | 159432756 | 159433363 | 0 | 0 | 18 | 0 | 0 | 3 | 0 | 5 | 0 | 0 | 0 | 1 | 98 | 49 | 0.12105318 | 0.243891326 | NA | NA | 0.75 | NA | 1 | 0 | 0.25 |
| SE | 4944 | ARAP1 | ARAP1 | chr11 | - | 72699004 | 72699107 | 72697910 | 72698106 | 72699416 | 72699552 | 2 | 8 | 34 | 0 | 0 | 2 | 0 | 30 | 7 | 0 | 0 | 0 | 98 | 49 | 0.121497957 | 0.24431631 | 1 | 1 | 0.895 | NA | 1 | 1 | -0.035 |
| SE | 11629 | ATP2A3 | ATP2A3 | chr17 | - | 3953346 | 3953429 | 3951580 | 3951685 | 3953692 | 3953710 | 0 | 2 | 4 | 0 | 0 | 0 | 0 | 8 | 4 | 0 | 0 | 1 | 98 | 49 | 0.121589221 | 0.24431631 | NA | 1 | 1 | NA | 1 | 0.667 | 0.166 |
| SE | 3974 | GLT1D1 | GLT1D1 | chr12 | + | 128875913 | 128876062 | 128853535 | 128853649 | 128888638 | 128888744 | 0 | 3 | 18 | 0 | 0 | 0 | 0 | 7 | 3 | 0 | 0 | 1 | 98 | 49 | 0.122907287 | 0.246635051 | NA | 1 | 1 | NA | 1 | 0.6 | 0.2 |
| SE | 3455 | GPCPD1 | GPCPD1 | chr20 | - | 5586193 | 5586269 | 5584280 | 5584322 | 5593326 | 5593411 | 0 | 0 | 11 | 0 | 0 | 2 | 0 | 7 | 0 | 0 | 6 | 1 | 98 | 49 | 0.123901232 | 0.24829807 | NA | NA | 0.733 | NA | 0.368 | 0 | 0.549 |
| SE | 812 | MKLN1 | MKLN1 | chr7 | + | 131445773 | 131445903 | 131443480 | 131443702 | 131463216 | 131463364 | 0 | 1 | 9 | 0 | 1 | 0 | 2 | 3 | 0 | 0 | 0 | 0 | 98 | 49 | 0.124958526 | 0.24977771 | NA | 0.333 | 1 | 1 | 1 | NA | -0.334 |
| SE | 8076 | NADK | NADK | chr1 | - | 1754050 | 1754208 | 1753566 | 1753649 | 1754283 | 1754383 | 0 | 4 | 21 | 0 | 0 | 2 | 4 | 17 | 5 | 0 | 0 | 0 | 98 | 49 | 0.125009625 | 0.24977771 | NA | 1 | 0.84 | 1 | 1 | 1 | -0.08 |
| SE | 6057 | HCLS1 | HCLS1 | chr3 | - | 121632331 | 121632563 | 121632100 | 121632184 | 121633066 | 121633171 | 0 | 10 | 51 | 0 | 0 | 2 | 0 | 41 | 11 | 0 | 0 | 0 | 98 | 49 | 0.125175468 | 0.24977771 | NA | 1 | 0.927 | NA | 1 | 1 | -0.036 |
| SE | 11932 | FBRS | FBRS | chr16 | + | 30661303 | 30661333 | 30661179 | 30661215 | 30662419 | 30662468 | 0 | 0 | 1 | 0 | 0 | 1 | 0 | 0 | 3 | 0 | 0 | 0 | 79 | 49 | 0.12530432 | 0.24977771 | NA | NA | 0.383 | NA | NA | 1 | -0.617 |
| A3SS | 1733 | ZSWIM8 | ZSWIM8 | chr10 | + | 73794540 | 73794639 | 73794555 | 73794639 | 73794146 | 73794330 | 0 | 1 | 7 | 0 | 1 | 0 | 0 | 3 | 0 | 5 | 0 | 0 | 64 | 49 | 0.125310145 | 0.30480846 | NA | 0.434 | 1 | 0 | 1 | NA | 0.217 |
| SE | 7879 | ALPL | ALPL | chr1 | + | 21573664 | 21573799 | 21570304 | 21570374 | 21575732 | 21575924 | 0 | 0 | 18 | 0 | 0 | 1 | 0 | 14 | 1 | 0 | 1 | 1 | 98 | 49 | 0.125632388 | 0.250099972 | NA | NA | 0.9 | NA | 0.875 | 0.333 | 0.296 |
| SE | 8247 | PRRC2C | PRRC2C | chr1 | + | 171512031 | 171512200 | 171485526 | 171485735 | 171513000 | 171513172 | 0 | 1 | 4 | 0 | 1 | 0 | 0 | 8 | 2 | 0 | 0 | 0 | 98 | 49 | 0.127922639 | 0.254322389 | NA | 0.333 | 1 | NA | 1 | 1 | -0.334 |
| SE | 10699 | TLE3 | TLE3 | chr15 | - | 70054437 | 70054685 | 70053226 | 70053374 | 70055048 | 70055298 | 0 | 4 | 15 | 0 | 0 | 4 | 0 | 21 | 6 | 0 | 1 | 0 | 98 | 49 | 0.128234493 | 0.254605606 | NA | 1 | 0.652 | NA | 0.913 | 1 | -0.13 |
| SE | 3888 | LCP1 | LCP1 | chr13 | - | 46158825 | 46158989 | 46158521 | 46158651 | 46159598 | 46159686 | 0 | 1 | 110 | 0 | 0 | 5 | 0 | 59 | 2 | 0 | 6 | 0 | 98 | 49 | 0.129117546 | 0.256020675 | NA | 1 | 0.917 | NA | 0.831 | 1 | 0.043 |
| SE | 6628 | PCBP2 | PCBP2 | chr12 | + | 53465931 | 53465973 | 53462480 | 53462567 | 53467220 | 53467293 | 0 | 2 | 30 | 0 | 1 | 3 | 0 | 11 | 0 | 0 | 1 | 0 | 91 | 49 | 0.129961836 | 0.257034707 | NA | 0.519 | 0.843 | NA | 0.856 | NA | -0.175 |
| SE | 7727 | FGD3 | FGD3 | chr9 | + | 93019830 | 93019861 | 93018135 | 93018215 | 93020316 | 93020424 | 0 | 3 | 10 | 0 | 0 | 1 | 0 | 10 | 1 | 0 | 3 | 0 | 80 | 49 | 0.129970976 | 0.257034707 | NA | 1 | 0.86 | NA | 0.671 | 1 | 0.094 |
| SE | 9562 | OASL | OASL | chr12 | - | 121027575 | 121027817 | 121023989 | 121024137 | 121031441 | 121031617 | 0 | 0 | 26 | 0 | 1 | 2 | 1 | 10 | 1 | 0 | 0 | 0 | 98 | 49 | 0.130217062 | 0.257182975 | NA | 0 | 0.867 | 1 | 1 | 1 | -0.567 |
| SE | 8139 | ITGAX | ITGAX | chr16 | + | 31372377 | 31372509 | 31371629 | 31371784 | 31372596 | 31372670 | 0 | 16 | 38 | 0 | 0 | 2 | 0 | 24 | 19 | 0 | 0 | 0 | 98 | 49 | 0.132864223 | 0.262066834 | NA | 1 | 0.905 | NA | 1 | 1 | -0.047 |
| SE | 6225 | AHCTF1 | AHCTF1 | chr1 | - | 246916141 | 246916395 | 246913231 | 246913412 | 246918249 | 246918377 | 0 | 0 | 4 | 0 | 0 | 1 | 0 | 0 | 0 | 0 | 0 | 1 | 98 | 49 | 0.133877166 | 0.263718717 | NA | NA | 0.667 | NA | NA | 0 | 0.667 |
| SE | 5499 | ADAM8 | ADAM8 | chr10 | - | 133275483 | 133275587 | 133274158 | 133274235 | 133276771 | 133276903 | 0 | 7 | 13 | 0 | 0 | 0 | 1 | 20 | 14 | 0 | 0 | 2 | 98 | 49 | 0.134548885 | 0.264504128 | NA | 1 | 1 | 1 | 1 | 0.778 | 0.074 |
| SE | 12648 | TLN1 | TLN1 | chr9 | - | 35704668 | 35704815 | 35704331 | 35704498 | 35705550 | 35705670 | 0 | 3 | 28 | 0 | 0 | 0 | 1 | 6 | 8 | 0 | 2 | 0 | 98 | 49 | 0.134879669 | 0.264504128 | NA | 1 | 1 | 1 | 0.6 | 1 | 0.133 |
| SE | 5934 | RASGRP2 | RASGRP2 | chr11 | - | 64727295 | 64727360 | 64726910 | 64727131 | 64728862 | 64729042 | 0 | 0 | 7 | 0 | 1 | 0 | 0 | 1 | 1 | 0 | 0 | 0 | 98 | 49 | 0.134979818 | 0.264504128 | NA | 0 | 1 | NA | 1 | 1 | -0.5 |
| SE | 9259 | MOV10 | MOV10 | chr1 | + | 112695415 | 112695574 | 112694452 | 112694629 | 112696147 | 112696251 | 0 | 0 | 7 | 0 | 1 | 0 | 0 | 1 | 1 | 0 | 0 | 0 | 98 | 49 | 0.134979818 | 0.264504128 | NA | 0 | 1 | NA | 1 | 1 | -0.5 |
| SE | 7950 | PPP4R1 | PPP4R1 | chr18 | - | 9562921 | 9563046 | 9561979 | 9562075 | 9563377 | 9563550 | 0 | 0 | 4 | 0 | 0 | 5 | 0 | 2 | 2 | 0 | 1 | 0 | 98 | 49 | 0.135910558 | 0.265981209 | NA | NA | 0.286 | NA | 0.5 | 1 | -0.464 |
| A3SS | 2930 | LRRC75A-AS1 | LRRC75A-AS1 | chr17 | + | 16440110 | 16440253 | 16440184 | 16440253 | 16439326 | 16439414 | 0 | 0 | 2 | 0 | 0 | 1 | 0 | 1 | 4 | 0 | 1 | 0 | 98 | 49 | 0.136565787 | 0.324235571 | NA | NA | 0.5 | NA | 0.333 | 1 | -0.166 |
| SE | 865 | NAMPT | NAMPT | chr7 | - | 106274945 | 106275049 | 106272529 | 106272658 | 106277022 | 106277179 | 0 | 15 | 313 | 0 | 0 | 2 | 3 | 467 | 4 | 0 | 6 | 0 | 98 | 49 | 0.136632841 | 0.267047022 | NA | 1 | 0.987 | 1 | 0.975 | 1 | 0.002 |
| A3SS | 1281 | CHD4 | CHD4 | chr12 | - | 6601959 | 6602175 | 6601959 | 6602154 | 6602375 | 6602497 | 0 | 0 | 2 | 0 | 0 | 1 | 0 | 17 | 4 | 4 | 4 | 0 | 70 | 49 | 0.136899463 | 0.324235571 | NA | NA | 0.583 | 0 | 0.748 | 1 | 0 |
| A5SS | 1981 | C12orf57 | C12orf57 | chr12 | + | 6944379 | 6944652 | 6944379 | 6944565 | 6945770 | 6946003 | 0 | 2 | 1 | 0 | 0 | 1 | 0 | 4 | 3 | 0 | 0 | 0 | 98 | 49 | 0.137146559 | 0.309762055 | NA | 1 | 0.333 | NA | 1 | 1 | -0.334 |
| SE | 8155 | KMT2E | KMT2E | chr7 | + | 105038125 | 105038199 | 105014189 | 105014535 | 105040838 | 105041023 | 0 | 0 | 7 | 0 | 0 | 2 | 0 | 12 | 0 | 0 | 2 | 1 | 98 | 49 | 0.137637656 | 0.268661555 | NA | NA | 0.636 | NA | 0.75 | 0 | 0.261 |
| SE | 11184 | MEF2D | MEF2D | chr1 | - | 156480833 | 156480971 | 156479585 | 156479796 | 156482436 | 156482640 | 1 | 4 | 3 | 0 | 0 | 0 | 0 | 3 | 5 | 0 | 2 | 0 | 98 | 49 | 0.139889993 | 0.272539469 | 1 | 1 | 1 | NA | 0.429 | 1 | 0.285 |
| SE | 1447 | BAG6 | BAG6 | chr6 | - | 31643889 | 31643974 | 31643100 | 31643115 | 31644081 | 31644194 | 0 | 1 | 2 | 0 | 0 | 1 | 0 | 6 | 1 | 0 | 0 | 0 | 98 | 49 | 0.139987006 | 0.272539469 | NA | 1 | 0.5 | NA | 1 | 1 | -0.25 |
| SE | 1213 | VNN2 | VNN2 | chr6 | - | 132753417 | 132753491 | 132752672 | 132752749 | 132755842 | 132756035 | 0 | 0 | 3 | 0 | 0 | 10 | 0 | 2 | 1 | 0 | 3 | 0 | 98 | 49 | 0.140487467 | 0.273115403 | NA | NA | 0.13 | NA | 0.25 | 1 | -0.495 |
| SE | 4623 | SORL1 | SORL1 | chr11 | + | 121470006 | 121470123 | 121452202 | 121452616 | 121478117 | 121478243 | 0 | 9 | 67 | 0 | 0 | 1 | 1 | 40 | 3 | 0 | 2 | 0 | 98 | 49 | 0.140646255 | 0.273115403 | NA | 1 | 0.971 | 1 | 0.909 | 1 | 0.016 |
| SE | 2848 | PIM3 | PIM3 | chr22 | + | 49961124 | 49961234 | 49960494 | 49961032 | 49961317 | 49961368 | 0 | 8 | 15 | 0 | 1 | 0 | 0 | 32 | 7 | 0 | 0 | 0 | 98 | 49 | 0.14293986 | 0.277211109 | NA | 0.8 | 1 | NA | 1 | 1 | -0.1 |
| RI | 341 | CDKN1C | CDKN1C | chr11 | - | 2884669 | 2885765 | 2884669 | 2885466 | 2885633 | 2885765 | 1 | 1 | 2 | 0 | 0 | 2 | 2 | 0 | 1 | 0 | 0 | 0 | 98 | 49 | 0.143188949 | 0.496482453 | 1 | 1 | 0.333 | 1 | NA | 1 | -0.222 |
| SE | 2268 | TNK2 | TNK2 | chr3 | - | 195882050 | 195882328 | 195879048 | 195879175 | 195883156 | 195883309 | 0 | 0 | 3 | 0 | 0 | 1 | 2 | 2 | 0 | 0 | 2 | 1 | 98 | 49 | 0.143961591 | 0.278473967 | NA | NA | 0.6 | 1 | 0.333 | 0 | 0.156 |
| SE | 8090 | SCLT1 | SCLT1 | chr4 | - | 128965226 | 128965318 | 128959599 | 128959777 | 128970377 | 128970468 | 0 | 0 | 3 | 0 | 0 | 1 | 0 | 6 | 0 | 0 | 0 | 1 | 98 | 49 | 0.143961591 | 0.278473967 | NA | NA | 0.6 | NA | 1 | 0 | 0.1 |
| SE | 9704 | ABTB1 | ABTB1 | chr3 | + | 127676271 | 127676431 | 127675969 | 127676114 | 127676535 | 127676581 | 0 | 3 | 20 | 0 | 0 | 0 | 1 | 11 | 4 | 0 | 0 | 1 | 98 | 49 | 0.145196396 | 0.280100163 | NA | 1 | 1 | 1 | 1 | 0.667 | 0.111 |
| A5SS | 1928 | PIGG | PIGG | chr4 | + | 499199 | 499725 | 499199 | 499489 | 500395 | 500601 | 0 | 0 | 1 | 0 | 0 | 3 | 0 | 0 | 1 | 0 | 0 | 0 | 98 | 49 | 0.145772881 | 0.312928566 | NA | NA | 0.143 | NA | NA | 1 | -0.857 |
| A5SS | 1929 | PIGG | PIGG | chr4 | + | 499199 | 500047 | 499199 | 499489 | 500395 | 500601 | 0 | 0 | 1 | 0 | 0 | 3 | 0 | 0 | 1 | 0 | 0 | 0 | 98 | 49 | 0.145772881 | 0.312928566 | NA | NA | 0.143 | NA | NA | 1 | -0.857 |
| SE | 7876 | TLR4 | TLR4 | chr9 | + | 117708562 | 117708729 | 117704174 | 117704565 | 117712388 | 117717491 | 0 | 0 | 10 | 0 | 0 | 10 | 0 | 2 | 0 | 0 | 6 | 2 | 98 | 49 | 0.146224332 | 0.280100163 | NA | NA | 0.333 | NA | 0.143 | 0 | 0.262 |
| SE | 324 | STOM | STOM | chr9 | - | 121366115 | 121366245 | 121356052 | 121356156 | 121370126 | 121370304 | 0 | 2 | 0 | 0 | 6 | 0 | 0 | 1 | 0 | 0 | 0 | 0 | 98 | 49 | 0.146368947 | 0.280100163 | NA | 0.143 | NA | NA | 1 | NA | -0.857 |
| A5SS | 468 | ARFIP1 | ARFIP1 | chr4 | + | 152779936 | 152780226 | 152779936 | 152780091 | 152829624 | 152829726 | 0 | 0 | 1 | 0 | 0 | 1 | 0 | 2 | 2 | 0 | 0 | 0 | 98 | 49 | 0.146398672 | 0.312928566 | NA | NA | 0.333 | NA | 1 | 1 | -0.667 |
| RI | 1023 | OXLD1 | OXLD1 | chr17 | - | 81665035 | 81666637 | 81665035 | 81665584 | 81666517 | 81666637 | 0 | 0 | 1 | 0 | 0 | 1 | 0 | 0 | 2 | 0 | 0 | 0 | 98 | 49 | 0.146398672 | 0.496482453 | NA | NA | 0.333 | NA | NA | 1 | -0.667 |
| SE | 3770 | KIAA1109 | KIAA1109 | chr4 | + | 122226706 | 122226835 | 122224497 | 122224677 | 122229120 | 122229240 | 0 | 0 | 1 | 0 | 0 | 1 | 0 | 9 | 2 | 0 | 1 | 0 | 98 | 49 | 0.146398672 | 0.280100163 | NA | NA | 0.333 | NA | 0.818 | 1 | -0.576 |
| SE | 6502 | GAS7 | GAS7 | chr17 | - | 9969676 | 9969762 | 9959201 | 9959255 | 9981803 | 9981884 | 0 | 1 | 9 | 0 | 1 | 0 | 0 | 2 | 0 | 0 | 0 | 0 | 98 | 49 | 0.146398672 | 0.280100163 | NA | 0.333 | 1 | NA | 1 | NA | -0.334 |
| SE | 11396 | ADGRG3 | ADGRG3 | chr16 | + | 57683931 | 57684212 | 57680504 | 57680617 | 57684389 | 57684483 | 0 | 0 | 1 | 0 | 0 | 1 | 0 | 3 | 2 | 0 | 3 | 0 | 98 | 49 | 0.146398672 | 0.280100163 | NA | NA | 0.333 | NA | 0.333 | 1 | -0.333 |
| SE | 13361 | SMARCA5 | SMARCA5 | chr4 | + | 143540362 | 143540495 | 143538785 | 143538938 | 143543508 | 143543657 | 0 | 0 | 1 | 0 | 0 | 1 | 0 | 3 | 2 | 0 | 0 | 0 | 98 | 49 | 0.146398672 | 0.280100163 | NA | NA | 0.333 | NA | 1 | 1 | -0.667 |
| SE | 13734 | SERPINA1 | SERPINA1 | chr14 | - | 94388800 | 94389010 | 94388559 | 94388660 | 94390456 | 94390692 | 0 | 1 | 17 | 0 | 1 | 3 | 0 | 2 | 0 | 4 | 0 | 0 | 98 | 49 | 0.146398672 | 0.280100163 | NA | 0.333 | 0.739 | 0 | 1 | NA | 0.036 |
| SE | 4332 | NCF2 | NCF2 | chr1 | - | 183567203 | 183567345 | 183566919 | 183566988 | 183569141 | 183569185 | 0 | 1 | 39 | 0 | 0 | 2 | 2 | 30 | 2 | 0 | 0 | 0 | 98 | 49 | 0.146479526 | 0.280100163 | NA | 1 | 0.907 | 1 | 1 | 1 | -0.046 |
| SE | 9574 | FAS | FAS | chr10 | + | 89010752 | 89010815 | 89010538 | 89010600 | 89011998 | 89012081 | 0 | 0 | 2 | 0 | 0 | 2 | 0 | 6 | 2 | 0 | 0 | 0 | 98 | 49 | 0.147583451 | 0.281852512 | NA | NA | 0.333 | NA | 1 | 1 | -0.667 |
| A5SS | 2262 | TLE3 | TLE3 | chr15 | - | 70074502 | 70074607 | 70074532 | 70074607 | 70066013 | 70066218 | 0 | 0 | 7 | 0 | 0 | 4 | 0 | 2 | 3 | 0 | 2 | 0 | 79 | 49 | 0.148103596 | 0.312928566 | NA | NA | 0.52 | NA | 0.383 | 1 | -0.171 |
| SE | 2581 | PRKCD | PRKCD | chr3 | + | 53165103 | 53165215 | 53161206 | 53161428 | 53178403 | 53178537 | 0 | 4 | 3 | 0 | 2 | 1 | 0 | 1 | 2 | 0 | 1 | 3 | 98 | 49 | 0.148640738 | 0.283511458 | NA | 0.5 | 0.6 | NA | 0.333 | 0.25 | 0.259 |
| SE | 12586 | SBNO2 | SBNO2 | chr19 | - | 1113534 | 1113704 | 1112817 | 1112949 | 1114230 | 1114422 | 0 | 9 | 21 | 0 | 0 | 1 | 0 | 40 | 18 | 0 | 0 | 0 | 98 | 49 | 0.149179961 | 0.284179318 | NA | 1 | 0.913 | NA | 1 | 1 | -0.043 |
| SE | 8708 | CTSS | CTSS | chr1 | - | 150764637 | 150764764 | 150757857 | 150757980 | 150765697 | 150765957 | 0 | 1 | 129 | 0 | 0 | 0 | 20 | 18 | 6 | 0 | 0 | 1 | 98 | 49 | 0.150561932 | 0.28644884 | NA | 1 | 1 | 1 | 1 | 0.75 | 0.083 |
| SE | 759 | RBM33 | RBM33 | chr7 | + | 155665174 | 155665253 | 155644508 | 155644919 | 155672866 | 155672915 | 0 | 4 | 8 | 0 | 0 | 0 | 0 | 7 | 2 | 0 | 0 | 1 | 98 | 49 | 0.15316134 | 0.291025909 | NA | 1 | 1 | NA | 1 | 0.5 | 0.25 |
| SE | 6532 | CARD16 | CARD16 | chr11 | - | 105043482 | 105043545 | 105041325 | 105041719 | 105044391 | 105044658 | 0 | 1 | 14 | 0 | 1 | 5 | 4 | 1 | 0 | 2 | 5 | 0 | 98 | 49 | 0.15394085 | 0.292137749 | NA | 0.333 | 0.583 | 0.5 | 0.091 | NA | 0.162 |
| SE | 7042 | SRSF2 | SRSF2 | chr17 | - | 76735771 | 76735875 | 76734114 | 76735158 | 76736153 | 76736464 | 0 | 1 | 2 | 0 | 2 | 3 | 0 | 3 | 0 | 0 | 12 | 1 | 98 | 49 | 0.154541762 | 0.292908283 | NA | 0.2 | 0.25 | NA | 0.111 | 0 | 0.17 |
| SE | 9278 | PIK3CD | PIK3CD | chr1 | + | 9721443 | 9721587 | 9721126 | 9721248 | 9721760 | 9721860 | 0 | 1 | 19 | 0 | 0 | 3 | 0 | 8 | 1 | 0 | 0 | 0 | 98 | 49 | 0.156377277 | 0.296013914 | NA | 1 | 0.76 | NA | 1 | 1 | -0.12 |
| SE | 6507 | TRIM33 | TRIM33 | chr1 | - | 114397939 | 114397990 | 114392776 | 114397860 | 114399456 | 114399609 | 0 | 0 | 7 | 0 | 0 | 2 | 0 | 4 | 5 | 0 | 0 | 0 | 98 | 49 | 0.156576702 | 0.296018595 | NA | NA | 0.636 | NA | 1 | 1 | -0.364 |
| A3SS | 3032 | NME4 | NME4 | chr16 | + | 400108 | 400754 | 400218 | 400754 | 399626 | 399739 | 2 | 0 | 0 | 1 | 0 | 0 | 3 | 0 | 0 | 0 | 0 | 0 | 98 | 49 | 0.156977715 | 0.366960892 | 0.5 | NA | NA | 1 | NA | NA | -0.5 |
| SE | 5476 | MICAL2 | MICAL2 | chr11 | + | 12138389 | 12138460 | 12110575 | 12110726 | 12162078 | 12162419 | 0 | 0 | 2 | 0 | 0 | 1 | 1 | 2 | 3 | 0 | 2 | 0 | 98 | 49 | 0.156977715 | 0.296403901 | NA | NA | 0.5 | 1 | 0.333 | 1 | -0.278 |
| SE | 4367 | LRRK2 | LRRK2 | chr12 | + | 40367005 | 40367077 | 40364841 | 40365050 | 40367643 | 40369284 | 0 | 0 | 13 | 0 | 0 | 3 | 0 | 13 | 6 | 0 | 3 | 0 | 98 | 49 | 0.157632777 | 0.297267333 | NA | NA | 0.684 | NA | 0.684 | 1 | -0.158 |
| SE | 9609 | TAPBP | TAPBP | chr6 | - | 33303954 | 33303989 | 33299694 | 33301771 | 33304127 | 33304217 | 0 | 2 | 23 | 0 | 1 | 0 | 4 | 10 | 1 | 0 | 0 | 0 | 84 | 49 | 0.158787547 | 0.299069779 | NA | 0.538 | 1 | 1 | 1 | 1 | -0.231 |
| A5SS | 3020 | SERPINA1 | SERPINA1 | chr14 | - | 94390419 | 94390692 | 94390456 | 94390692 | 94388818 | 94389010 | 0 | 4 | 5 | 0 | 1 | 15 | 3 | 3 | 2 | 0 | 1 | 0 | 86 | 49 | 0.159752672 | 0.332184127 | NA | 0.695 | 0.16 | 1 | 0.631 | 1 | -0.45 |
| SE | 8027 | GNAS | GNAS | chr20 | + | 58898940 | 58898985 | 58895611 | 58895684 | 58903527 | 58903585 | 0 | 1 | 2 | 0 | 3 | 2 | 0 | 0 | 0 | 0 | 4 | 0 | 94 | 49 | 0.16039283 | 0.301490206 | NA | 0.148 | 0.343 | NA | 0 | NA | 0.245 |
| SE | 1448 | BAG6 | BAG6 | chr6 | - | 31643889 | 31643977 | 31642828 | 31643115 | 31644081 | 31644194 | 0 | 1 | 3 | 0 | 0 | 1 | 0 | 7 | 1 | 0 | 0 | 0 | 98 | 49 | 0.160473829 | 0.301490206 | NA | 1 | 0.6 | NA | 1 | 1 | -0.2 |
| SE | 4626 | SORL1 | SORL1 | chr11 | + | 121557313 | 121557405 | 121555186 | 121555318 | 121558590 | 121558837 | 0 | 10 | 73 | 0 | 0 | 2 | 0 | 39 | 5 | 0 | 3 | 0 | 98 | 49 | 0.162347432 | 0.304629451 | NA | 1 | 0.948 | NA | 0.867 | 1 | 0.04 |
| A3SS | 1989 | PPP6R2 | PPP6R2 | chr22 | + | 50419345 | 50419462 | 50419348 | 50419462 | 50418866 | 50418979 | 0 | 1 | 0 | 0 | 0 | 1 | 0 | 2 | 1 | 0 | 0 | 0 | 52 | 49 | 0.163314358 | 0.376879289 | NA | 1 | 0 | NA | 1 | 1 | -0.5 |
| SE | 8529 | CKLF | CKLF | chr16 | + | 66558189 | 66558348 | 66552562 | 66552793 | 66563121 | 66563217 | 0 | 0 | 8 | 0 | 0 | 2 | 0 | 3 | 5 | 0 | 0 | 0 | 98 | 49 | 0.164123685 | 0.307578427 | NA | NA | 0.667 | NA | 1 | 1 | -0.333 |
| SE | 695 | DOCK5 | DOCK5 | chr8 | + | 25368191 | 25368250 | 25366869 | 25366970 | 25368570 | 25368725 | 0 | 0 | 11 | 0 | 0 | 5 | 0 | 5 | 3 | 0 | 1 | 0 | 98 | 49 | 0.166703165 | 0.311886731 | NA | NA | 0.524 | NA | 0.714 | 1 | -0.333 |
| SE | 7636 | IRF7 | IRF7 | chr11 | - | 612998 | 613117 | 612554 | 612800 | 613205 | 613595 | 0 | 9 | 20 | 0 | 0 | 0 | 0 | 20 | 5 | 0 | 0 | 1 | 98 | 49 | 0.16703421 | 0.311886731 | NA | 1 | 1 | NA | 1 | 0.714 | 0.143 |
| SE | 2487 | ATG3 | ATG3 | chr3 | - | 112553279 | 112553329 | 112550191 | 112550262 | 112558375 | 112558417 | 0 | 0 | 4 | 0 | 0 | 3 | 1 | 4 | 2 | 0 | 0 | 0 | 98 | 49 | 0.167045122 | 0.311886731 | NA | NA | 0.4 | 1 | 1 | 1 | -0.6 |
| SE | 471 | DOCK8 | DOCK8 | chr9 | + | 317042 | 317128 | 311953 | 312166 | 325670 | 325737 | 0 | 10 | 33 | 0 | 0 | 0 | 2 | 15 | 7 | 0 | 0 | 2 | 98 | 49 | 0.167661893 | 0.311930048 | NA | 1 | 1 | 1 | 1 | 0.636 | 0.121 |
| SE | 4637 | ARID5A | ARID5A | chr2 | + | 96549320 | 96549459 | 96547401 | 96547517 | 96549752 | 96549805 | 0 | 2 | 2 | 0 | 0 | 3 | 0 | 1 | 0 | 0 | 6 | 4 | 98 | 49 | 0.167831186 | 0.311930048 | NA | 1 | 0.25 | NA | 0.077 | 0 | 0.587 |
| RI | 566 | SPRTN | SPRTN | chr1 | + | 231351303 | 231355027 | 231351303 | 231351571 | 231352609 | 231355027 | 0 | 0 | 1 | 0 | 0 | 2 | 2 | 2 | 1 | 0 | 0 | 0 | 98 | 49 | 0.167898476 | 0.545670046 | NA | NA | 0.2 | 1 | 1 | 1 | -0.8 |
| SE | 5473 | ARNTL | ARNTL | chr11 | + | 13365499 | 13365592 | 13360344 | 13360424 | 13366664 | 13366782 | 0 | 0 | 1 | 0 | 0 | 2 | 2 | 4 | 1 | 0 | 3 | 0 | 98 | 49 | 0.167898476 | 0.311930048 | NA | NA | 0.2 | 1 | 0.4 | 1 | -0.6 |
| SE | 11688 | CDK11B | CDK11B | chr1 | - | 1637080 | 1637202 | 1636896 | 1637004 | 1637407 | 1637513 | 0 | 1 | 0 | 0 | 2 | 0 | 0 | 1 | 1 | 0 | 0 | 0 | 98 | 49 | 0.167898476 | 0.311930048 | NA | 0.2 | NA | NA | 1 | 1 | -0.8 |
| A3SS | 3932 | OTOF | OTOF | chr2 | - | 26472518 | 26472649 | 26472518 | 26472589 | 26473131 | 26473294 | 0 | 0 | 1 | 0 | 1 | 1 | 0 | 0 | 0 | 0 | 2 | 2 | 98 | 49 | 0.167980116 | 0.379975466 | NA | 0 | 0.333 | NA | 0 | 0 | 0.167 |
| A3SS | 2018 | RNF19B | RNF19B | chr1 | - | 32948221 | 32948366 | 32948221 | 32948363 | 32949568 | 32949774 | 0 | 0 | 1 | 0 | 0 | 2 | 0 | 1 | 1 | 0 | 1 | 0 | 52 | 49 | 0.168877985 | 0.379975466 | NA | NA | 0.32 | NA | 0.485 | 1 | -0.422 |
| SE | 7215 | LSP1 | LSP1 | chr11 | + | 1880086 | 1880224 | 1852969 | 1853197 | 1881431 | 1881596 | 0 | 7 | 39 | 0 | 0 | 0 | 2 | 37 | 11 | 0 | 0 | 1 | 98 | 49 | 0.17099036 | 0.31689431 | NA | 1 | 1 | 1 | 1 | 0.846 | 0.051 |
| SE | 1069 | HNRNPA2B1 | HNRNPA2B1 | chr7 | - | 26197830 | 26197866 | 26197621 | 26197732 | 26200571 | 26200793 | 0 | 0 | 1 | 0 | 3 | 7 | 0 | 2 | 0 | 0 | 2 | 1 | 85 | 49 | 0.170992206 | 0.31689431 | NA | 0 | 0.076 | NA | 0.366 | 0 | -0.145 |
| SE | 4615 | PTPRJ | PTPRJ | chr11 | + | 48112746 | 48112983 | 48110057 | 48110076 | 48121002 | 48121266 | 2 | 11 | 32 | 0 | 2 | 0 | 0 | 9 | 5 | 0 | 0 | 0 | 98 | 49 | 0.171767102 | 0.317938368 | 1 | 0.733 | 1 | NA | 1 | 1 | -0.089 |
| A3SS | 502 | ARG1 | ARG1 | chr6 | + | 131579086 | 131579285 | 131579110 | 131579285 | 131576662 | 131576735 | 0 | 0 | 2 | 0 | 0 | 4 | 0 | 0 | 1 | 0 | 0 | 0 | 73 | 49 | 0.173440158 | 0.385422574 | NA | NA | 0.251 | NA | NA | 1 | -0.749 |
| SE | 11874 | MGAM | MGAM | chr7 | + | 142036168 | 142036285 | 142034669 | 142034841 | 142036822 | 142036977 | 0 | 2 | 23 | 0 | 0 | 0 | 6 | 6 | 3 | 0 | 0 | 2 | 98 | 49 | 0.173570524 | 0.3208813 | NA | 1 | 1 | 1 | 1 | 0.429 | 0.19 |
| A5SS | 1004 | GMPR2 | GMPR2 | chr14 | + | 24232421 | 24232847 | 24232421 | 24232788 | 24233218 | 24233340 | 3 | 0 | 2 | 0 | 0 | 4 | 0 | 4 | 1 | 0 | 1 | 0 | 98 | 49 | 0.174787188 | 0.357767525 | 1 | NA | 0.2 | NA | 0.667 | 1 | -0.234 |
| SE | 7320 | IL18R1 | IL18R1 | chr2 | + | 102373936 | 102373993 | 102371952 | 102372118 | 102375906 | 102376063 | 0 | 0 | 2 | 0 | 0 | 4 | 0 | 0 | 1 | 0 | 0 | 0 | 98 | 49 | 0.174787188 | 0.322733591 | NA | NA | 0.2 | NA | NA | 1 | -0.8 |
| SE | 4120 | XPO6 | XPO6 | chr16 | - | 28104545 | 28104707 | 28101846 | 28101945 | 28106042 | 28106214 | 0 | 7 | 58 | 0 | 2 | 0 | 1 | 60 | 4 | 0 | 2 | 0 | 98 | 49 | 0.176059763 | 0.324652462 | NA | 0.636 | 1 | 1 | 0.937 | 1 | -0.161 |
| SE | 235 | CSF2RA | CSF2RA | chrX | + | 1303922 | 1304019 | 1300490 | 1300626 | 1305445 | 1305527 | 0 | 1 | 11 | 0 | 0 | 1 | 0 | 2 | 0 | 0 | 5 | 1 | 98 | 49 | 0.176258423 | 0.324652462 | NA | 1 | 0.846 | NA | 0.167 | 0 | 0.84 |
| SE | 12687 | MYO1F | MYO1F | chr19 | - | 8548035 | 8548122 | 8545649 | 8545736 | 8548236 | 8548317 | 0 | 9 | 61 | 0 | 0 | 2 | 0 | 38 | 5 | 0 | 0 | 0 | 98 | 49 | 0.177681081 | 0.326872294 | NA | 1 | 0.938 | NA | 1 | 1 | -0.031 |
| SE | 10039 | SYNE2 | SYNE2 | chr14 | + | 64009965 | 64010116 | 64007042 | 64007222 | 64016472 | 64016631 | 0 | 0 | 3 | 0 | 0 | 2 | 0 | 7 | 2 | 0 | 0 | 0 | 98 | 49 | 0.179363678 | 0.329564313 | NA | NA | 0.429 | NA | 1 | 1 | -0.571 |
| SE | 5557 | SPATA13 | SPATA13 | chr13 | + | 24278613 | 24278821 | 24270686 | 24270912 | 24284134 | 24284271 | 0 | 0 | 1 | 0 | 1 | 3 | 0 | 0 | 0 | 0 | 3 | 4 | 98 | 49 | 0.18048442 | 0.331070043 | NA | 0 | 0.143 | NA | 0 | 0 | 0.071 |
| SE | 10589 | RNF213 | RNF213 | chr17 | + | 80369767 | 80369867 | 80369501 | 80369671 | 80371873 | 80371985 | 0 | 10 | 63 | 0 | 0 | 2 | 3 | 33 | 13 | 0 | 0 | 0 | 98 | 49 | 0.180942085 | 0.331070043 | NA | 1 | 0.94 | 1 | 1 | 1 | -0.03 |
| SE | 10850 | IRF1 | IRF1 | chr5 | - | 132487925 | 132488025 | 132486953 | 132487130 | 132489391 | 132489483 | 0 | 7 | 14 | 0 | 0 | 1 | 0 | 12 | 1 | 0 | 2 | 0 | 98 | 49 | 0.181005184 | 0.331070043 | NA | 1 | 0.875 | NA | 0.75 | 1 | 0.062 |

| | | | | | | | | | | | | | | | | | | | | | | | | | | | | | | | | | | |
|---|---|---|---|---|---|---|---|---|---|---|---|---|---|---|---|---|---|---|---|---|---|---|---|---|---|---|---|---|---|---|---|---|---|---|
| SE | 4941 | ARAP1 | ARAP1 | chr11 | - | 72692752 | 72692785 | 72688454 | 72688537 | 72693324 | 72693470 | 0 | 0 | 2 | 0 | 1 | 16 | 0 | 2 | 0 | 0 | 2 | 1 | 82 | 49 | 0.181064255 | 0.331070043 | NA | 0 | 0.07 | NA | 0.374 | 0 | -0.152 |
| SE | 6699 | FCGR2A | FCGR2A | chr1 | + | 161510833 | 161510956 | 161509819 | 161510074 | 161513894 | 161513932 | 0 | 1 | 66 | 0 | 0 | 2 | 0 | 40 | 2 | 0 | 0 | 0 | 98 | 49 | 0.181721355 | 0.331867797 | NA | 1 | 0.943 | NA | 1 | 1 | -0.028 |
| SE | 7908 | ATP6V0B | ATP6V0B | chr1 | + | 43975799 | 43975848 | 43974647 | 43975107 | 43976089 | 43976173 | 0 | 5 | 19 | 0 | 1 | 4 | 1 | 10 | 5 | 0 | 3 | 3 | 98 | 49 | 0.183348939 | 0.334433805 | NA | 0.714 | 0.704 | 1 | 0.625 | 0.455 | 0.016 |
| A3SS | 457 | BTN2A1 | BTN2A1 | chr6 | + | 26467718 | 26469653 | 26467947 | 26469653 | 26466061 | 26466088 | 0 | 2 | 4 | 0 | 0 | 2 | 0 | 3 | 1 | 0 | 0 | 0 | 98 | 49 | 0.184424674 | 0.403832852 | NA | 1 | 0.5 | NA | 1 | 1 | -0.25 |
| SE | 2077 | HERC5 | HERC5 | chr4 | + | 88470613 | 88470673 | 88469156 | 88469260 | 88472408 | 88472502 | 0 | 2 | 23 | 0 | 0 | 1 | 0 | 41 | 1 | 0 | 3 | 0 | 98 | 49 | 0.18526689 | 0.337522588 | NA | 1 | 0.92 | NA | 0.872 | 1 | 0.024 |
| SE | 8553 | EHBP1L1 | EHBP1L1 | chr11 | + | 65589935 | 65589991 | 65589750 | 65589820 | 65590086 | 65590210 | 0 | 1 | 10 | 0 | 0 | 0 | 3 | 7 | 2 | 0 | 0 | 1 | 98 | 49 | 0.185747044 | 0.337684951 | NA | 1 | 1 | 1 | 1 | 0.5 | 0.167 |
| SE | 5679 | GMIP | GMIP | chr19 | - | 19640295 | 19640360 | 19640084 | 19640192 | 19640445 | 19640571 | 0 | 9 | 25 | 0 | 0 | 2 | 2 | 12 | 3 | 0 | 3 | 0 | 98 | 49 | 0.185805359 | 0.337684951 | NA | 1 | 0.862 | 1 | 0.667 | 1 | 0.042 |
| SE | 9611 | TAPBP | TAPBP | chr6 | - | 33313693 | 33313864 | 33313216 | 33313477 | 33314004 | 33314387 | 0 | 4 | 1 | 0 | 0 | 1 | 0 | 9 | 3 | 0 | 0 | 0 | 98 | 49 | 0.186099817 | 0.337811625 | NA | 1 | 0.333 | NA | 1 | 1 | -0.334 |
| SE | 6577 | RILPL2 | RILPL2 | chr12 | - | 123430507 | 123430659 | 123423043 | 123423157 | 123436081 | 123436733 | 0 | 1 | 12 | 0 | 4 | 4 | 0 | 13 | 6 | 0 | 3 | 3 | 98 | 49 | 0.189327441 | 0.343255903 | NA | 0.111 | 0.6 | NA | 0.684 | 0.5 | -0.237 |
| SE | 6105 | GLUL | GLUL | chr1 | - | 182390496 | 182390852 | 182388571 | 182388750 | 182391678 | 182392206 | 0 | 2 | 1 | 0 | 5 | 18 | 0 | 1 | 1 | 3 | 24 | 22 | 98 | 49 | 0.189877016 | 0.343454188 | NA | 0.167 | 0.027 | 0 | 0.02 | 0.022 | 0.083 |
| SE | 2138 | CXCL8 | CXCL8 | chr4 | + | 73741541 | 73741677 | 73740505 | 73740722 | 73741948 | 73742032 | 1 | 26 | 133 | 0 | 1 | 0 | 6 | 985 | 18 | 0 | 1 | 0 | 98 | 49 | 0.189893833 | 0.343454188 | 1 | 0.929 | 1 | 1 | 0.998 | 1 | -0.023 |
| RI | 624 | MCL1 | MCL1 | chr1 | - | 150578842 | 150579738 | 150578842 | 150578963 | 150579422 | 150579738 | 0 | 41 | 264 | 0 | 0 | 2 | 13 | 148 | 29 | 0 | 0 | 0 | 98 | 49 | 0.190371362 | 0.586344893 | NA | 1 | 0.985 | 1 | 1 | 1 | -0.008 |
| A3SS | 504 | NDUFA2 | NDUFA2 | chr5 | - | 140645362 | 140645725 | 140645362 | 140645678 | 140647255 | 140647362 | 0 | 0 | 3 | 0 | 1 | 2 | 0 | 0 | 0 | 0 | 0 | 1 | 96 | 49 | 0.190583999 | 0.403832852 | NA | 0 | 0.434 | NA | NA | 0 | 0.217 |
| A5SS | 1894 | MYD88 | MYD88 | chr3 | + | 38140387 | 38140592 | 38140387 | 38140568 | 38140756 | 38140848 | 0 | 0 | 4 | 0 | 1 | 7 | 0 | 0 | 0 | 0 | 0 | 2 | 73 | 49 | 0.190693809 | 0.381630589 | NA | 0 | 0.277 | NA | NA | 0 | 0.139 |
| SE | 12366 | NBEAL2 | NBEAL2 | chr3 | + | 46996750 | 46996833 | 46996270 | 46996592 | 46996953 | 46997046 | 0 | 7 | 12 | 0 | 0 | 0 | 0 | 4 | 6 | 0 | 4 | 0 | 98 | 49 | 0.19095347 | 0.344955607 | NA | 1 | 1 | NA | 0.333 | 1 | 0.334 |
| A3SS | 2130 | FGD3 | FGD3 | chr9 | + | 93034540 | 93034681 | 93034543 | 93034681 | 93032768 | 93032873 | 0 | 6 | 5 | 0 | 0 | 1 | 0 | 11 | 5 | 0 | 0 | 0 | 52 | 49 | 0.19141247 | 0.403832852 | NA | 1 | 0.825 | NA | 1 | 1 | -0.088 |
| SE | 5865 | NFKBID | NFKBID | chr19 | - | 35897650 | 35897856 | 35896912 | 35897058 | 35898471 | 35898532 | 0 | 7 | 10 | 0 | 2 | 0 | 0 | 4 | 6 | 0 | 0 | 0 | 98 | 49 | 0.192464506 | 0.34726789 | NA | 0.636 | 1 | NA | 1 | 1 | -0.182 |
| A3SS | 4000 | MAP4K4 | MAP4K4 | chr2 | + | 101859642 | 101859864 | 101859735 | 101859864 | 101858995 | 101859082 | 0 | 0 | 4 | 0 | 0 | 1 | 0 | 1 | 4 | 0 | 0 | 0 | 98 | 49 | 0.193038976 | 0.403832852 | NA | NA | 0.667 | NA | 1 | 1 | -0.333 |
| SE | 3456 | GPCPD1 | GPCPD1 | chr20 | - | 5593326 | 5593411 | 5584280 | 5584322 | 5598724 | 5598821 | 0 | 1 | 27 | 0 | 0 | 1 | 0 | 38 | 4 | 0 | 0 | 0 | 98 | 49 | 0.193523387 | 0.348759773 | NA | 1 | 0.931 | NA | 1 | 1 | -0.034 |
| A3SS | 1806 | PCBP2 | PCBP2 | chr12 | + | 53462480 | 53462567 | 53462492 | 53462567 | 53461014 | 53461143 | 1 | 1 | 8 | 0 | 1 | 19 | 0 | 9 | 0 | 4 | 7 | 0 | 61 | 49 | 0.194077531 | 0.403832852 | 1 | 0.445 | 0.253 | 0 | 0.508 | NA | 0.312 |
| SE | 8686 | PTPRC | PTPRC | chr1 | + | 198702386 | 198702530 | 198699563 | 198699704 | 198703297 | 198703372 | 1 | 0 | 4 | 0 | 3 | 6 | 4 | 3 | 2 | 1 | 11 | 0 | 98 | 49 | 0.195041125 | 0.351074025 | 1 | 0 | 0.25 | 0.667 | 0.12 | 1 | -0.179 |
| A5SS | 1245 | TUBA1A | TUBA1A | chr12 | - | 49188416 | 49189324 | 49188976 | 49189324 | 49186610 | 49186833 | 0 | 1 | 1 | 0 | 1 | 3 | 0 | 1 | 0 | 0 | 4 | 0 | 98 | 49 | 0.195185111 | 0.381630589 | NA | 0.333 | 0.143 | NA | 0.111 | NA | 0.127 |
| A5SS | 1246 | TUBA1A | TUBA1A | chr12 | - | 49188888 | 49189324 | 49188976 | 49189324 | 49186610 | 49186833 | 0 | 1 | 1 | 0 | 1 | 3 | 0 | 1 | 0 | 0 | 4 | 0 | 98 | 49 | 0.195185111 | 0.381630589 | NA | 0.333 | 0.143 | NA | 0.111 | NA | 0.127 |
| A3SS | 570 | DDX60L | DDX60L | chr4 | - | 168395958 | 168396127 | 168395958 | 168396124 | 168400825 | 168400978 | 0 | 0 | 9 | 0 | 2 | 12 | 0 | 16 | 0 | 0 | 5 | 1 | 52 | 49 | 0.195185878 | 0.403832852 | NA | 0 | 0.414 | NA | 0.751 | 0 | -0.169 |
| RI | 52 | GSTK1 | GSTK1 | chr7 | + | 143264991 | 143265296 | 143264991 | 143265092 | 143265260 | 143265296 | 0 | 2 | 3 | 0 | 1 | 6 | 2 | 4 | 1 | 9 | 2 | 0 | 98 | 49 | 0.195448298 | 0.586344893 | NA | 0.5 | 0.2 | 0.1 | 0.5 | 1 | -0.183 |
| SE | 5839 | TUBA1A | TUBA1A | chr12 | - | 49186309 | 49186458 | 49184794 | 49185990 | 49186610 | 49186833 | 0 | 5 | 29 | 0 | 1 | 2 | 6 | 17 | 0 | 0 | 1 | 0 | 98 | 49 | 0.195483437 | 0.35144929 | NA | 0.714 | 0.879 | 1 | 0.895 | NA | -0.151 |
| SE | 12944 | SMCHD1 | SMCHD1 | chr18 | + | 2724898 | 2724995 | 2722518 | 2722663 | 2726451 | 2726524 | 0 | 2 | 53 | 0 | 0 | 1 | 0 | 50 | 1 | 0 | 2 | 0 | 98 | 49 | 0.196703531 | 0.353220319 | NA | 1 | 0.964 | NA | 0.926 | 1 | 0.019 |
| SE | 2289 | BCL6 | BCL6 | chr3 | - | 187726730 | 187726898 | 187725498 | 187725629 | 187728359 | 187728544 | 0 | 1 | 25 | 0 | 0 | 2 | 3 | 7 | 3 | 0 | 3 | 0 | 98 | 49 | 0.197315011 | 0.353895539 | NA | 1 | 0.862 | 1 | 0.538 | 1 | 0.085 |
| SE | 12685 | MYO1F | MYO1F | chr19 | - | 8539946 | 8540028 | 8536948 | 8537055 | 8541905 | 8541991 | 0 | 6 | 55 | 0 | 0 | 2 | 1 | 30 | 5 | 0 | 0 | 0 | 98 | 49 | 0.197961525 | 0.354631909 | NA | 1 | 0.932 | 1 | 1 | 1 | -0.034 |
| SE | 5477 | MICAL2 | MICAL2 | chr11 | + | 12138389 | 12138464 | 12110575 | 12110726 | 12162078 | 12162419 | 0 | 0 | 3 | 0 | 0 | 1 | 1 | 3 | 3 | 0 | 2 | 0 | 98 | 49 | 0.199033101 | 0.354859729 | NA | NA | 0.6 | 1 | 0.429 | 1 | -0.21 |
| SE | 5479 | MICAL2 | MICAL2 | chr11 | + | 12220200 | 12220458 | 12216218 | 12216319 | 12221643 | 12221759 | 0 | 0 | 3 | 0 | 0 | 1 | 0 | 1 | 3 | 0 | 0 | 0 | 98 | 49 | 0.199033101 | 0.354859729 | NA | NA | 0.6 | NA | 1 | 1 | -0.4 |
| SE | 7788 | CLK3 | CLK3 | chr15 | + | 74627969 | 74628052 | 74627538 | 74627668 | 74628603 | 74628683 | 0 | 0 | 3 | 0 | 0 | 1 | 0 | 0 | 3 | 0 | 0 | 0 | 98 | 49 | 0.199033101 | 0.354859729 | NA | NA | 0.6 | NA | NA | 1 | -0.4 |
| SE | 8840 | C1orf43 | C1orf43 | chr1 | - | 154213892 | 154213946 | 154212457 | 154212624 | 154214456 | 154214574 | 0 | 0 | 3 | 0 | 0 | 1 | 0 | 5 | 3 | 0 | 0 | 0 | 98 | 49 | 0.199033101 | 0.354859729 | NA | NA | 0.6 | NA | 1 | 1 | -0.4 |
| SE | 9582 | ANXA11 | ANXA11 | chr10 | - | 80168968 | 80169358 | 80167225 | 80167313 | 80170799 | 80170915 | 0 | 0 | 6 | 0 | 0 | 2 | 0 | 2 | 3 | 0 | 0 | 0 | 98 | 49 | 0.200611555 | 0.357250198 | NA | NA | 0.6 | NA | 1 | 1 | -0.4 |
| SE | 10668 | LAPTM5 | LAPTM5 | chr1 | - | 30741639 | 30741716 | 30739808 | 30739937 | 30742455 | 30742549 | 0 | 4 | 37 | 0 | 0 | 1 | 6 | 52 | 6 | 0 | 0 | 0 | 98 | 49 | 0.201003059 | 0.357523784 | NA | 1 | 0.949 | 1 | 1 | 1 | -0.026 |
| SE | 1136 | IGF2R | IGF2R | chr6 | + | 160048374 | 160048543 | 160047791 | 160047907 | 160050472 | 160050652 | 0 | 3 | 42 | 0 | 0 | 2 | 0 | 21 | 4 | 0 | 0 | 0 | 98 | 49 | 0.201533872 | 0.357884259 | NA | 1 | 0.913 | NA | 1 | 1 | -0.043 |
| SE | 5119 | SLC3A2 | SLC3A2 | chr11 | + | 62883988 | 62884112 | 62882907 | 62882999 | 62884456 | 62884525 | 0 | 0 | 1 | 0 | 0 | 4 | 0 | 0 | 0 | 0 | 5 | 4 | 98 | 49 | 0.201681948 | 0.357884259 | NA | NA | 0.111 | NA | 0 | 0 | 0.111 |
| A3SS | 3948 | ALOX5 | ALOX5 | chr10 | + | 45443037 | 45443216 | 45443133 | 45443216 | 45441343 | 45441430 | 2 | 7 | 33 | 0 | 0 | 0 | 1 | 6 | 7 | 0 | 3 | 0 | 98 | 49 | 0.202413488 | 0.410636839 | 1 | 1 | 1 | 1 | 0.5 | 1 | 0.167 |
| SE | 13916 | CD300A | CD300A | chr17 | + | 74481288 | 74481326 | 74477435 | 74477530 | 74484000 | 74484798 | 0 | 2 | 17 | 0 | 0 | 2 | 0 | 5 | 6 | 0 | 0 | 0 | 87 | 49 | 0.203130618 | 0.359943165 | NA | 1 | 0.827 | NA | 1 | 1 | -0.087 |
| SE | 11717 | KDM6B | KDM6B | chr17 | + | 7844900 | 7845020 | 7839916 | 7840024 | 7845313 | 7845456 | 0 | 3 | 14 | 0 | 0 | 1 | 0 | 15 | 9 | 0 | 0 | 0 | 98 | 49 | 0.203321189 | 0.359943165 | NA | 1 | 0.875 | NA | 1 | 1 | -0.062 |
| A5SS | 519 | TNFAIP3 | TNFAIP3 | chr6 | + | 137867187 | 137867518 | 137867187 | 137867494 | 137871212 | 137871522 | 0 | 3 | 1 | 0 | 2 | 0 | 0 | 8 | 4 | 0 | 1 | 0 | 73 | 49 | 0.204508982 | 0.388363884 | NA | 0.502 | 1 | NA | 0.843 | 1 | -0.17 |
| A3SS | 1274 | PTPN6 | PTPN6 | chr12 | + | 6960662 | 6960945 | 6960805 | 6960945 | 6960343 | 6960435 | 0 | 1 | 8 | 0 | 1 | 17 | 0 | 2 | 0 | 3 | 6 | 3 | 98 | 49 | 0.204565043 | 0.410636839 | NA | 0.333 | 0.19 | 0 | 0.143 | 0 | 0.214 |
| SE | 13219 | LILRA5 | LILRA5 | chr19 | - | 54312334 | 54312370 | 54311863 | 54312148 | 54312536 | 54312621 | 0 | 0 | 15 | 0 | 0 | 2 | 1 | 2 | 1 | 0 | 3 | 1 | 85 | 49 | 0.205560222 | 0.36347884 | NA | NA | 0.812 | 1 | 0.278 | 0.366 | 0.264 |
| SE | 5498 | ADAM8 | ADAM8 | chr10 | - | 133274158 | 133274235 | 133273950 | 133274029 | 133276771 | 133276903 | 0 | 8 | 19 | 0 | 0 | 0 | 1 | 26 | 10 | 0 | 0 | 2 | 98 | 49 | 0.206600072 | 0.364888259 | NA | 1 | 1 | 1 | 1 | 0.714 | 0.095 |
| A3SS | 3540 | MAPKAPK2 | MAPKAPK2 | chr1 | + | 206732037 | 206734285 | 206732574 | 206734285 | 206731838 | 206731919 | 0 | 0 | 4 | 0 | 1 | 9 | 0 | 8 | 9 | 5 | 10 | 2 | 98 | 49 | 0.208792326 | 0.410636839 | NA | 0 | 0.182 | 0 | 0.286 | 0.692 | -0.235 |
| A3SS | 1990 | PPP6R2 | PPP6R2 | chr22 | + | 50439697 | 50439857 | 50439700 | 50439857 | 50438598 | 50438762 | 0 | 0 | 1 | 0 | 0 | 2 | 0 | 0 | 0 | 0 | 3 | 2 | 52 | 49 | 0.209323809 | 0.410636839 | NA | NA | 0.32 | NA | 0 | 0 | 0.32 |
| A5SS | 1572 | ACAD9 | ACAD9 | chr3 | + | 128879489 | 128880017 | 128879489 | 128879841 | 128884652 | 128884746 | 0 | 0 | 4 | 0 | 0 | 2 | 0 | 0 | 2 | 0 | 0 | 0 | 98 | 49 | 0.209418448 | 0.388363884 | NA | NA | 0.5 | NA | NA | 1 | -0.5 |
| SE | 3047 | FAM157A | FAM157A | chr3 | + | 198169784 | 198169872 | 198167675 | 198167871 | 198180754 | 198180857 | 0 | 4 | 48 | 0 | 0 | 3 | 0 | 7 | 4 | 0 | 1 | 4 | 98 | 49 | 0.209675253 | 0.368087771 | NA | 1 | 0.889 | NA | 0.778 | 0.333 | 0.389 |
| A3SS | 2544 | TRIM26 | TRIM26 | chr6 | - | 30198665 | 30199264 | 30198665 | 30199153 | 30201034 | 30201138 | 0 | 0 | 2 | 0 | 0 | 1 | 1 | 2 | 2 | 0 | 2 | 0 | 98 | 49 | 0.209881051 | 0.410636839 | NA | NA | 0.5 | 1 | 0.333 | 1 | -0.278 |
| A5SS | 2686 | OXLD1 | OXLD1 | chr17 | - | 81666154 | 81666637 | 81666517 | 81666637 | 81665035 | 81665584 | 0 | 0 | 2 | 0 | 0 | 1 | 0 | 0 | 2 | 0 | 0 | 0 | 98 | 49 | 0.209881051 | 0.388363884 | NA | NA | 0.5 | NA | NA | 1 | -0.5 |
| SE | 533 | PRRC2B | PRRC2B | chr9 | + | 131446400 | 131446642 | 131444184 | 131444328 | 131447084 | 131447206 | 0 | 0 | 2 | 0 | 0 | 1 | 0 | 1 | 2 | 0 | 0 | 0 | 98 | 49 | 0.209881051 | 0.368087771 | NA | NA | 0.5 | NA | 1 | 1 | -0.5 |
| SE | 3379 | TM9SF4 | TM9SF4 | chr20 | + | 32159991 | 32160111 | 32158450 | 32158514 | 32161275 | 32161365 | 0 | 2 | 4 | 0 | 1 | 0 | 0 | 2 | 0 | 0 | 0 | 0 | 98 | 49 | 0.209881051 | 0.368087771 | NA | 0.5 | 1 | NA | 1 | NA | -0.25 |
| SE | 3413 | RALGAPA2 | RALGAPA2 | chr20 | - | 20524405 | 20524543 | 20520916 | 20521100 | 20524829 | 20524898 | 0 | 0 | 2 | 0 | 0 | 1 | 0 | 0 | 2 | 0 | 0 | 0 | 98 | 49 | 0.209881051 | 0.368087771 | NA | NA | 0.5 | NA | NA | 1 | -0.5 |
| SE | 10410 | RBM42 | RBM42 | chr19 | + | 35633686 | 35634019 | 35633076 | 35633252 | 35634255 | 35634373 | 0 | 2 | 0 | 0 | 1 | 0 | 0 | 2 | 2 | 0 | 0 | 0 | 98 | 49 | 0.209881051 | 0.368087771 | NA | 0.5 | NA | NA | 1 | 1 | -0.5 |
| SE | 12955 | MAN2A2 | MAN2A2 | chr15 | + | 90912541 | 90912664 | 90912042 | 90912279 | 90912876 | 90912991 | 0 | 0 | 2 | 0 | 0 | 1 | 0 | 2 | 2 | 0 | 0 | 0 | 98 | 49 | 0.209881051 | 0.368087771 | NA | NA | 0.5 | NA | 1 | 1 | -0.5 |
| A5SS | 2442 | RNASEK | RNASEK | chr17 | + | 7012416 | 7012761 | 7012416 | 7012757 | 7013665 | 7013742 | 0 | 0 | 4 | 0 | 0 | 2 | 1 | 2 | 2 | 0 | 1 | 0 | 53 | 49 | 0.210487296 | 0.388363884 | NA | NA | 0.649 | 1 | 0.649 | 1 | -0.234 |
| SE | 5409 | TAGAP | TAGAP | chr6 | - | 159042077 | 159042244 | 159041353 | 159041515 | 159043588 | 159043655 | 0 | 4 | 19 | 0 | 0 | 1 | 0 | 28 | 2 | 0 | 0 | 0 | 98 | 49 | 0.21465984 | 0.37603 | NA | 1 | 0.905 | NA | 1 | 1 | -0.047 |
| SE | 4197 | CPPED1 | CPPED1 | chr16 | - | 12704623 | 12705049 | 12659798 | 12665115 | 12781184 | 12781403 | 0 | 1 | 12 | 0 | 0 | 3 | 0 | 2 | 1 | 0 | 3 | 0 | 98 | 49 | 0.215298398 | 0.376709536 | NA | 1 | 0.667 | NA | 0.25 | 1 | 0.209 |
| SE | 3187 | RGS19 | RGS19 | chr20 | - | 64076856 | 64076954 | 64076524 | 64076646 | 64079293 | 64079492 | 0 | 0 | 7 | 0 | 0 | 2 | 0 | 1 | 3 | 0 | 0 | 0 | 98 | 49 | 0.220448488 | 0.384909961 | NA | NA | 0.636 | NA | 1 | 1 | -0.364 |
| SE | 8747 | ADIPOR1 | ADIPOR1 | chr1 | - | 202943757 | 202943945 | 202942024 | 202942218 | 202944982 | 202945169 | 0 | 1 | 9 | 0 | 0 | 0 | 0 | 5 | 1 | 0 | 0 | 1 | 98 | 49 | 0.220497323 | 0.384909961 | NA | 1 | 1 | NA | 1 | 0.333 | 0.334 |
| SE | 6274 | BAZ2B | BAZ2B | chr2 | - | 159453612 | 159453801 | 159448241 | 159448409 | 159478574 | 159478721 | 0 | 0 | 11 | 0 | 0 | 5 | 0 | 3 | 1 | 0 | 0 | 2 | 98 | 49 | 0.220936482 | 0.385229157 | NA | NA | 0.524 | NA | 1 | 0.2 | -0.076 |
| SE | 838 | CAPZA2 | CAPZA2 | chr7 | + | 116906262 | 116906342 | 116904176 | 116904383 | 116910232 | 116910311 | 0 | 0 | 23 | 0 | 0 | 3 | 0 | 10 | 6 | 0 | 1 | 0 | 98 | 49 | 0.221735441 | 0.386174239 | NA | NA | 0.793 | NA | 0.833 | 1 | -0.123 |
| SE | 276 | RAPGEF1 | RAPGEF1 | chr9 | - | 131650128 | 131650242 | 131643238 | 131643417 | 131650809 | 131650949 | 0 | 2 | 1 | 0 | 0 | 1 | 3 | 10 | 1 | 0 | 7 | 0 | 98 | 49 | 0.223239446 | 0.388343619 | NA | 1 | 0.333 | 1 | 0.417 | 1 | -0.139 |
| SE | 7716 | THEMIS2 | THEMIS2 | chr1 | + | 27876587 | 27876728 | 27872542 | 27872665 | 27885294 | 27885451 | 0 | 10 | 8 | 0 | 1 | 0 | 6 | 6 | 15 | 3 | 0 | 0 | 98 | 49 | 0.223535788 | 0.388409582 | NA | 0.833 | 1 | 0.5 | 1 | 1 | 0.083 |
| SE | 7875 | TLR4 | TLR4 | chr9 | + | 117708198 | 117708318 | 117704174 | 117704565 | 117708562 | 117708729 | 0 | 0 | 18 | 0 | 0 | 5 | 0 | 9 | 3 | 0 | 0 | 0 | 98 | 49 | 0.224282921 | 0.389257772 | NA | NA | 0.643 | NA | 1 | 1 | -0.357 |
| SE | 4169 | RNF10 | RNF10 | chr12 | + | 120552498 | 120552698 | 120546404 | 120546601 | 120554717 | 120554808 | 0 | 1 | 5 | 0 | 0 | 1 | 0 | 6 | 2 | 0 | 0 | 0 | 98 | 49 | 0.225232695 | 0.390455294 | NA | 1 | 0.714 | NA | 1 | 1 | -0.143 |
| RI | 991 | HSH2D | HSH2D | chr19 | + | 16152551 | 16153208 | 16152551 | 16152641 | 16153042 | 16153208 | 0 | 5 | 21 | 0 | 1 | 1 | 4 | 8 | 1 | 0 | 0 | 0 | 98 | 49 | 0.226524618 | 0.654404451 | NA | 0.714 | 0.913 | 1 | 1 | 1 | -0.186 |
| A5SS | 1015 | AMICA1 | AMICA1 | chr11 | - | 118197967 | 118198091 | 118197997 | 118198091 | 118196734 | 118196821 | 0 | 0 | 14 | 0 | 0 | 6 | 0 | 9 | 2 | 0 | 1 | 0 | 79 | 49 | 0.228281084 | 0.409690921 | NA | NA | 0.591 | NA | 0.848 | 1 | -0.333 |
| SE | 12396 | TMEM127 | TMEM127 | chr2 | - | 96265137 | 96265512 | 96254832 | 96254997 | 96265868 | 96266013 | 0 | 3 | 11 | 0 | 0 | 1 | 0 | 13 | 4 | 0 | 0 | 0 | 98 | 49 | 0.229343062 | 0.397122837 | NA | 1 | 0.846 | NA | 1 | 1 | -0.077 |
| SE | 10331 | CYTIP | CYTIP | chr2 | - | 157418522 | 157418589 | 157414618 | 157416143 | 157427350 | 157427420 | 0 | 5 | 36 | 0 | 0 | 1 | 4 | 50 | 1 | 0 | 0 | 0 | 98 | 49 | 0.231211633 | 0.398828525 | NA | 1 | 0.947 | 1 | 1 | 1 | -0.026 |
| SE | 6671 | FGR | FGR | chr1 | - | 27617192 | 27617296 | 27616856 | 27617006 | 27621558 | 27621657 | 0 | 3 | 47 | 0 | 0 | 1 | 5 | 12 | 2 | 0 | 1 | 0 | 98 | 49 | 0.231633345 | 0.398828525 | NA | 1 | 0.959 | 1 | 0.857 | 1 | 0.027 |
| A3SS | 2782 | GCH1 | GCH1 | chr14 | - | 54842005 | 54844143 | 54842005 | 54843054 | 54845767 | 54845852 | 0 | 0 | 2 | 0 | 0 | 2 | 0 | 2 | 0 | 1 | 0 | 1 | 98 | 49 | 0.232250281 | 0.43959786 | NA | NA | 0.333 | 0 | 1 | 0 | 0 |
| SE | 3236 | SLC9A8 | SLC9A8 | chr20 | + | 49850761 | 49850844 | 49849578 | 49849680 | 49855437 | 49855581 | 0 | 0 | 4 | 0 | 0 | 4 | 0 | 0 | 0 | 0 | 4 | 1 | 98 | 49 | 0.23228173 | 0.398828525 | NA | NA | 0.333 | NA | 0 | 0 | 0.333 |
| SE | 10641 | USP3 | USP3 | chr15 | + | 63533802 | 63533918 | 63532646 | 63532707 | 63537024 | 63537156 | 0 | 0 | 6 | 0 | 0 | 6 | 0 | 3 | 0 | 0 | 2 | 1 | 98 | 49 | 0.232454602 | 0.398828525 | NA | NA | 0.333 | NA | 0.429 | 0 | 0.119 |
| SE | 4196 | FAM49B | FAM49B | chr8 | - | 129904498 | 129904585 | 129903311 | 129903350 | 129970942 | 129970995 | 0 | 0 | 3 | 0 | 0 | 3 | 0 | 1 | 0 | 0 | 3 | 1 | 98 | 49 | 0.232595516 | 0.398828525 | NA | NA | 0.333 | NA | 0.143 | 0 | 0.262 |
| SE | 3237 | SLC9A8 | SLC9A8 | chr20 | + | 49850809 | 49850844 | 49849578 | 49849680 | 49855437 | 49855581 | 0 | 0 | 4 | 0 | 0 | 4 | 0 | 0 | 0 | 0 | 4 | 1 | 84 | 49 | 0.232676511 | 0.398828525 | NA | NA | 0.368 | NA | 0 | 0 | 0.368 |
| SE | 4286 | RASSF3 | RASSF3 | chr12 | + | 64688215 | 64688453 | 64684786 | 64684894 | 64691469 | 64691579 | 0 | 1 | 21 | 0 | 0 | 2 | 1 | 10 | 1 | 0 | 0 | 0 | 98 | 49 | 0.233640332 | 0.398828525 | NA | 1 | 0.84 | 1 | 1 | 1 | -0.08 |
| SE | 10884 | AKAP13 | AKAP13 | chr15 | + | 85658536 | 85658590 | 85655416 | 85655787 | 85664562 | 85664755 | 0 | 2 | 13 | 0 | 0 | 1 | 0 | 4 | 3 | 0 | 2 | 0 | 98 | 49 | 0.233666262 | 0.398828525 | NA | 1 | 0.867 | NA | 0.5 | 1 | 0.183 |
| A3SS | 650 | TMEM175 | TMEM175 | chr4 | + | 951645 | 951717 | 951681 | 951717 | 951206 | 951258 | 0 | 0 | 1 | 0 | 0 | 1 | 0 | 0 | 1 | 0 | 0 | 0 | 85 | 49 | 0.234374295 | 0.43959786 | NA | NA | 0.366 | NA | NA | 1 | -0.634 |
| A3SS | 1341 | FAM89B | FAM89B | chr11 | + | 65573352 | 65574198 | 65573362 | 65574198 | 65572348 | 65572960 | 0 | 0 | 1 | 0 | 0 | 1 | 0 | 0 | 1 | 0 | 1 | 0 | 59 | 49 | 0.234434278 | 0.43959786 | NA | NA | 0.454 | NA | 0 | 1 | -0.046 |
| A3SS | 193 | ZYX | ZYX | chr7 | + | 143381556 | 143381779 | 143381559 | 143381779 | 143381266 | 143381409 | 2 | 1 | 1 | 0 | 1 | 0 | 0 | 1 | 0 | 0 | 0 | 0 | 52 | 49 | 0.234452192 | 0.43959786 | 1 | 0.485 | 1 | NA | 1 | NA | -0.172 |
| A5SS | 696 | PDCD6 | PDCD6 | chr5 | + | 306601 | 306760 | 306601 | 306754 | 311292 | 311402 | 0 | 0 | 1 | 0 | 0 | 1 | 0 | 0 | 1 | 0 | 0 | 0 | 55 | 49 | 0.234460292 | 0.409690921 | NA | NA | 0.471 | NA | NA | 1 | -0.529 |
| A5SS | 2321 | ISG20 | ISG20 | chr15 | + | 88638742 | 88639178 | 88638742 | 88639076 | 88639342 | 88639594 | 0 | 0 | 1 | 0 | 0 | 1 | 0 | 1 | 1 | 0 | 2 | 0 | 98 | 49 | 0.234545893 | 0.409690921 | NA | NA | 0.333 | NA | 0.2 | 1 | -0.267 |
| SE | 739 | CTSB | CTSB | chr8 | - | 11864375 | 11864463 | 11853328 | 11853479 | 11868000 | 11868137 | 0 | 0 | 1 | 0 | 2 | 1 | 0 | 0 | 1 | 0 | 0 | 0 | 98 | 49 | 0.234545893 | 0.398828525 | NA | 0 | 0.333 | NA | NA | 1 | -0.834 |
| SE | 5272 | NIPBL | NIPBL | chr5 | + | 37000372 | 37000570 | 36995621 | 36995804 | 37000816 | 37000888 | 0 | 0 | 1 | 0 | 0 | 1 | 0 | 2 | 1 | 0 | 0 | 0 | 98 | 49 | 0.234545893 | 0.398828525 | NA | NA | 0.333 | NA | 1 | 1 | -0.667 |
| SE | 7330 | NEDD9 | NEDD9 | chr6 | - | 11305065 | 11305159 | 11213280 | 11213727 | 11305991 | 11306155 | 0 | 0 | 1 | 0 | 0 | 1 | 0 | 2 | 1 | 0 | 0 | 0 | 98 | 49 | 0.234545893 | 0.398828525 | NA | NA | 0.333 | NA | 1 | 1 | -0.667 |
| SE | 9136 | TRIM5 | TRIM5 | chr11 | - | 5665980 | 5666081 | 5663194 | 5665682 | 5667688 | 5667711 | 0 | 0 | 1 | 0 | 0 | 1 | 0 | 0 | 1 | 0 | 2 | 0 | 98 | 49 | 0.234545893 | 0.398828525 | NA | NA | 0.333 | NA | 0 | 1 | -0.167 |
| SE | 10646 | HERC1 | HERC1 | chr15 | - | 63662983 | 63663204 | 63661752 | 63662021 | 63664469 | 63664594 | 0 | 0 | 1 | 0 | 0 | 1 | 0 | 2 | 1 | 0 | 1 | 0 | 98 | 49 | 0.234545893 | 0.398828525 | NA | NA | 0.333 | NA | 0.5 | 1 | -0.417 |
| SE | 11342 | BCKDK | BCKDK | chr16 | + | 31110687 | 31110761 | 31110400 | 31110499 | 31111090 | 31111219 | 0 | 0 | 1 | 0 | 0 | 1 | 0 | 0 | 1 | 0 | 0 | 0 | 98 | 49 | 0.234545893 | 0.398828525 | NA | NA | 0.333 | NA | NA | 1 | -0.667 |
| SE | 13736 | SERPINA1 | SERPINA1 | chr14 | - | 94388818 | 94389010 | 94388559 | 94388660 | 94390456 | 94390692 | 0 | 1 | 16 | 0 | 1 | 3 | 0 | 1 | 0 | 4 | 0 | 0 | 98 | 49 | 0.234545893 | 0.398828525 | NA | 0.333 | 0.727 | 0 | 1 | NA | 0.03 |
| A5SS | 2440 | SMARCB1 | SMARCB1 | chr22 | + | 23791755 | 23791894 | 23791755 | 23791867 | 23793558 | 23793688 | 2 | 0 | 1 | 0 | 0 | 1 | 0 | 2 | 1 | 0 | 0 | 0 | 76 | 49 | 0.234555871 | 0.409690921 | 1 | NA | 0.392 | NA | 1 | 1 | -0.304 |
| SE | 11000 | TBC1D14 | TBC1D14 | chr4 | + | 7004843 | 7004924 | 7001144 | 7001251 | 7010652 | 7010781 | 0 | 0 | 3 | 0 | 0 | 3 | 0 | 1 | 1 | 0 | 0 | 0 | 98 | 49 | 0.234573797 | 0.398828525 | NA | NA | 0.333 | NA | 1 | 1 | -0.667 |
| SE | 2249 | PCGF3 | PCGF3 | chr4 | + | 764983 | 765064 | 761278 | 761416 | 766031 | 770639 | 0 | 0 | 2 | 0 | 1 | 0 | 0 | 1 | 1 | 0 | 0 | 0 | 98 | 49 | 0.236755697 | 0.402083405 | NA | 0 | 1 | NA | 1 | 1 | -0.5 |
| SE | 12650 | TLN1 | TLN1 | chr9 | - | 35725564 | 35725727 | 35725223 | 35725321 | 35732074 | 35732395 | 0 | 1 | 14 | 0 | 0 | 1 | 0 | 15 | 3 | 0 | 0 | 0 | 98 | 49 | 0.239010624 | 0.405454817 | NA | 1 | 0.875 | NA | 1 | 1 | -0.062 |
| SE | 9492 | SLC15A3 | SLC15A3 | chr11 | - | 60941121 | 60941290 | 60939479 | 60939638 | 60942034 | 60942145 | 0 | 2 | 5 | 0 | 1 | 1 | 0 | 2 | 3 | 0 | 2 | 2 | 98 | 49 | 0.239519623 | 0.405735216 | NA | 0.5 | 0.714 | NA | 0.333 | 0.429 | 0.226 |
| SE | 13178 | NLRP12 | NLRP12 | chr19 | - | 53807494 | 53807665 | 53805279 | 53805450 | 53809586 | 53811288 | 0 | 1 | 13 | 0 | 0 | 2 | 0 | 3 | 4 | 0 | 0 | 0 | 98 | 49 | 0.23984512 | 0.405735216 | NA | 1 | 0.765 | NA | 1 | 1 | -0.117 |
| SE | 6632 | MRVI1 | MRVI1 | chr11 | - | 10600917 | 10601059 | 10594145 | 10594195 | 10603119 | 10603251 | 0 | 0 | 4 | 0 | 0 | 1 | 0 | 6 | 3 | 0 | 0 | 0 | 98 | 49 | 0.239985767 | 0.405735216 | NA | NA | 0.667 | NA | 1 | 1 | -0.333 |
| SE | 13627 | SUN2 | SUN2 | chr22 | - | 38750897 | 38751035 | 38750224 | 38750320 | 38751209 | 38751373 | 0 | 1 | 6 | 0 | 0 | 0 | 0 | 1 | 3 | 0 | 2 | 0 | 98 | 49 | 0.241094036 | 0.407150939 | NA | 1 | 1 | NA | 0.2 | 1 | 0.4 |
| SE | 3099 | MX1 | MX1 | chr21 | + | 41443787 | 41443866 | 41441715 | 41441914 | 41445447 | 41445570 | 0 | 4 | 62 | 0 | 0 | 2 | 18 | 50 | 6 | 0 | 3 | 0 | 98 | 49 | 0.244079828 | 0.411554843 | NA | 1 | 0.939 | 1 | 0.893 | 1 | 0.005 |
| SE | 11814 | FLOT2 | FLOT2 | chr17 | - | 28881191 | 28881375 | 28880712 | 28880862 | 28881813 | 28882028 | 0 | 3 | 27 | 0 | 0 | 2 | 0 | 12 | 1 | 0 | 2 | 0 | 98 | 49 | 0.244249447 | 0.411554843 | NA | 1 | 0.871 | NA | 0.75 | 1 | 0.06 |
| A5SS | 1373 | FCAR | FCAR | chr19 | + | 54888006 | 54888294 | 54888006 | 54888228 | 54889648 | 54891420 | 1 | 5 | 40 | 0 | 2 | 6 | 9 | 17 | 4 | 0 | 2 | 0 | 98 | 49 | 0.244859012 | 0.422059613 | 1 | 0.556 | 0.769 | 1 | 0.81 | 1 | -0.162 |
| SE | 1039 | AOAH | AOAH | chr7 | - | 36686698 | 36686794 | 36673942 | 36674009 | 36724021 | 36724549 | 0 | 5 | 37 | 0 | 0 | 1 | 0 | 10 | 2 | 0 | 1 | 0 | 98 | 49 | 0.245974583 | 0.413770577 | NA | 1 | 0.949 | NA | 0.833 | 1 | 0.058 |
| SE | 12364 | NBEAL2 | NBEAL2 | chr3 | + | 46992474 | 46992555 | 46991839 | 46991946 | 46993936 | 46994020 | 0 | 0 | 10 | 0 | 0 | 9 | 0 | 2 | 1 | 0 | 1 | 0 | 98 | 49 | 0.246115034 | 0.413770577 | NA | NA | 0.357 | NA | 0.5 | 1 | -0.393 |
| SE | 12574 | R3HDM4 | R3HDM4 | chr19 | - | 901421 | 901546 | 900828 | 900952 | 901975 | 902130 | 0 | 6 | 19 | 0 | 0 | 1 | 5 | 27 | 1 | 0 | 0 | 0 | 98 | 49 | 0.247240838 | 0.41489777 | NA | 1 | 0.905 | 1 | 1 | 1 | -0.047 |
| SE | 516 | SLA | SLA | chr8 | - | 133044983 | 133045115 | 133039997 | 133040130 | 133047829 | 133047933 | 0 | 0 | 35 | 0 | 0 | 6 | 0 | 8 | 4 | 0 | 1 | 0 | 98 | 49 | 0.247337593 | 0.41489777 | NA | NA | 0.745 | NA | 0.8 | 1 | -0.155 |
| SE | 4618 | PTPRJ | PTPRJ | chr11 | + | 48164379 | 48164515 | 48163457 | 48163618 | 48167203 | 48170842 | 0 | 1 | 10 | 0 | 0 | 1 | 0 | 10 | 3 | 0 | 0 | 0 | 98 | 49 | 0.248230356 | 0.415931131 | NA | 1 | 0.833 | NA | 1 | 1 | -0.084 |
| SE | 4055 | FOS | FOS | chr14 | + | 75279876 | 75280128 | 75278777 | 75279123 | 75280559 | 75280667 | 0 | 18 | 153 | 0 | 0 | 1 | 11 | 176 | 10 | 0 | 0 | 0 | 98 | 49 | 0.249706448 | 0.417938521 | NA | 1 | 0.987 | 1 | 1 | 1 | -0.006 |
| A3SS | 3814 | NLRP12 | NLRP12 | chr19 | - | 53807494 | 53807668 | 53807494 | 53807665 | 53809586 | 53811288 | 0 | 0 | 1 | 0 | 0 | 6 | 0 | 0 | 0 | 0 | 2 | 4 | 52 | 49 | 0.250107194 | 0.464116443 | NA | NA | 0.136 | NA | 0 | 0 | 0.136 |
| SE | 2700 | MBNL1 | MBNL1 | chr3 | + | 152432716 | 152432920 | 152414940 | 152415111 | 152445281 | 152445539 | 0 | 1 | 24 | 0 | 0 | 1 | 7 | 7 | 1 | 0 | 1 | 0 | 98 | 49 | 0.252076189 | 0.421435498 | NA | 1 | 0.923 | 1 | 0.778 | 1 | 0.035 |
| SE | 12940 | SMCHD1 | SMCHD1 | chr18 | + | 2674014 | 2674145 | 2673280 | 2673363 | 2688393 | 2688508 | 0 | 7 | 143 | 0 | 0 | 2 | 5 | 100 | 1 | 0 | 3 | 0 | 98 | 49 | 0.254061338 | 0.424282435 | NA | 1 | 0.973 | 1 | 0.943 | 1 | 0.005 |
| A3SS | 3504 | FCGR3A | FCGR3A | chr1 | - | 161548420 | 161548678 | 161548420 | 161548675 | 161549010 | 161549031 | 0 | 1 | 5 | 0 | 0 | 2 | 0 | 2 | 1 | 0 | 0 | 0 | 52 | 49 | 0.25427504 | 0.467035788 | NA | 1 | 0.702 | NA | 1 | 1 | -0.149 |
| SE | 4164 | NT5C2 | NT5C2 | chr10 | - | 103090610 | 103090787 | 103088016 | 103089908 | 103090935 | 103090996 | 0 | 0 | 14 | 0 | 0 | 5 | 0 | 2 | 2 | 0 | 0 | 0 | 98 | 49 | 0.254640733 | 0.424778049 | NA | NA | 0.583 | NA | 1 | 1 | -0.417 |
| A5SS | 2258 | TLE3 | TLE3 | chr15 | - | 70057458 | 70057658 | 70057473 | 70057658 | 70056297 | 70056374 | 0 | 1 | 9 | 0 | 0 | 3 | 0 | 3 | 1 | 0 | 3 | 0 | 64 | 49 | 0.254834903 | 0.430953855 | NA | 1 | 0.697 | NA | 0.434 | 1 | 0.132 |
| SE | 13858 | MMP25 | MMP25 | chr16 | + | 3050253 | 3050546 | 3050008 | 3050144 | 3057032 | 3057209 | 0 | 7 | 19 | 0 | 0 | 1 | 0 | 17 | 10 | 0 | 0 | 0 | 98 | 49 | 0.255901705 | 0.426408273 | NA | 1 | 0.905 | NA | 1 | 1 | -0.047 |
| A5SS | 403 | CNOT8 | CNOT8 | chr5 | + | 154858518 | 154858816 | 154858518 | 154858768 | 154863206 | 154863395 | 1 | 0 | 3 | 0 | 0 | 2 | 1 | 0 | 1 | 0 | 0 | 0 | 97 | 49 | 0.256598479 | 0.430953855 | 1 | NA | 0.431 | 1 | NA | 1 | -0.284 |
| SE | 12369 | NBEAL2 | NBEAL2 | chr3 | + | 47007240 | 47007350 | 47006332 | 47006449 | 47007524 | 47007697 | 0 | 1 | 7 | 0 | 1 | 0 | 0 | 5 | 0 | 0 | 2 | 0 | 98 | 49 | 0.256741576 | 0.427007803 | NA | 0.333 | 1 | NA | 0.556 | NA | 0.11 |
| SE | 12947 | SMCHD1 | SMCHD1 | chr18 | + | 2751277 | 2751393 | 2750349 | 2750507 | 2752487 | 2752552 | 1 | 8 | 40 | 0 | 0 | 1 | 0 | 46 | 4 | 0 | 0 | 0 | 98 | 49 | 0.25682971 | 0.427007803 | 1 | 1 | 0.952 | NA | 1 | 1 | -0.016 |
| SE | 2166 | FYB | FYB | chr5 | - | 39125997 | 39126135 | 39124252 | 39124278 | 39127740 | 39127807 | 0 | 0 | 17 | 0 | 0 | 9 | 6 | 25 | 1 | 0 | 6 | 2 | 98 | 49 | 0.257579272 | 0.427780824 | NA | NA | 0.486 | 1 | 0.676 | 0.2 | -0.139 |
| SE | 7217 | LSP1 | LSP1 | chr11 | + | 1887236 | 1887314 | 1886731 | 1886866 | 1887473 | 1887576 | 0 | 6 | 18 | 0 | 0 | 1 | 0 | 22 | 2 | 0 | 0 | 0 | 98 | 49 | 0.260907854 | 0.432830579 | NA | 1 | 0.9 | NA | 1 | 1 | -0.05 |
| SE | 7231 | NFKBIZ | NFKBIZ | chr3 | + | 101857072 | 101857183 | 101855732 | 101855902 | 101857291 | 101857459 | 0 | 0 | 19 | 0 | 0 | 3 | 5 | 34 | 4 | 0 | 7 | 0 | 98 | 49 | 0.261478977 | 0.432929688 | NA | NA | 0.76 | 1 | 0.708 | 1 | -0.143 |
| SE | 5891 | CSF1 | CSF1 | chr1 | + | 109914258 | 109914381 | 109910610 | 109911062 | 109915633 | 109915696 | 0 | 1 | 1 | 0 | 3 | 0 | 0 | 0 | 0 | 0 | 2 | 0 | 98 | 49 | 0.261543684 | 0.432929688 | NA | 0.143 | 1 | NA | 0 | NA | 0.572 |
| SE | 6056 | HCLS1 | HCLS1 | chr3 | - | 121632100 | 121632184 | 121631397 | 121631982 | 121632331 | 121632563 | 0 | 3 | 39 | 0 | 0 | 0 | 1 | 28 | 4 | 0 | 0 | 1 | 98 | 49 | 0.261910932 | 0.433060651 | NA | 1 | 1 | 1 | 1 | 0.667 | 0.111 |
| SE | 1667 | GRK6 | GRK6 | chr5 | + | 177431994 | 177432107 | 177430871 | 177430967 | 177432232 | 177432310 | 0 | 7 | 9 | 0 | 0 | 1 | 0 | 4 | 3 | 0 | 3 | 0 | 98 | 49 | 0.26379167 | 0.435653879 | NA | 1 | 0.818 | NA | 0.4 | 1 | 0.209 |
| SE | 10928 | DEDD2 | DEDD2 | chr19 | - | 42202050 | 42202072 | 42198592 | 42199829 | 42209699 | 42209840 | 0 | 0 | 3 | 0 | 3 | 5 | 0 | 2 | 0 | 0 | 16 | 3 | 71 | 49 | 0.264511909 | 0.435653879 | NA | 0 | 0.293 | NA | 0.079 | 0 | 0.107 |
| RI | 607 | TAF1C | TAF1C | chr16 | - | 84181327 | 84181663 | 84181327 | 84181463 | 84181591 | 84181663 | 0 | 0 | 3 | 0 | 0 | 1 | 0 | 1 | 2 | 0 | 0 | 0 | 98 | 49 | 0.264638717 | 0.737207854 | NA | NA | 0.6 | NA | 1 | 1 | -0.4 |
| SE | 3872 | WDFY2 | WDFY2 | chr13 | + | 51719197 | 51719348 | 51703595 | 51703650 | 51727677 | 51727790 | 0 | 0 | 3 | 0 | 0 | 1 | 3 | 2 | 2 | 0 | 0 | 0 | 98 | 49 | 0.264638717 | 0.435653879 | NA | NA | 0.6 | 1 | 1 | 1 | -0.4 |
| SE | 7725 | ZFC3H1 | ZFC3H1 | chr12 | - | 71636849 | 71637059 | 71636489 | 71636654 | 71638417 | 71638515 | 0 | 0 | 3 | 0 | 0 | 1 | 0 | 9 | 2 | 0 | 0 | 0 | 98 | 49 | 0.264638717 | 0.435653879 | NA | NA | 0.6 | NA | 1 | 1 | -0.4 |
| A5SS | 1432 | ADPGK | ADPGK | chr15 | - | 72783245 | 72783785 | 72783458 | 72783785 | 72774871 | 72775097 | 0 | 3 | 7 | 0 | 0 | 1 | 2 | 2 | 8 | 0 | 0 | 0 | 98 | 49 | 0.265624158 | 0.440465375 | NA | 1 | 0.778 | 1 | 1 | 1 | -0.111 |
| SE | 5576 | SAT1 | SAT1 | chrX | + | 23784293 | 23784403 | 23783799 | 23783883 | 23785327 | 23785429 | 0 | 5 | 44 | 0 | 10 | 78 | 0 | 46 | 0 | 0 | 89 | 9 | 98 | 49 | 0.267669304 | 0.440160792 | NA | 0.2 | 0.22 | NA | 0.205 | 0 | 0.108 |
| A3SS | 1360 | DDX3X | DDX3X | chrX | + | 41347311 | 41347451 | 41347314 | 41347451 | 41346858 | 41347012 | 0 | 0 | 8 | 0 | 0 | 6 | 0 | 16 | 1 | 0 | 2 | 0 | 52 | 49 | 0.267844114 | 0.486989299 | NA | NA | 0.557 | NA | 0.883 | 1 | -0.384 |
| SE | 7628 | RALGAPB | RALGAPB | chr20 | + | 38548688 | 38548795 | 38546242 | 38546430 | 38551070 | 38551223 | 0 | 0 | 4 | 0 | 0 | 3 | 0 | 0 | 1 | 0 | 1 | 0 | 98 | 49 | 0.268677998 | 0.441336646 | NA | NA | 0.4 | NA | 0 | 1 | -0.1 |
| SE | 6197 | TPCN2 | TPCN2 | chr11 | + | 69072626 | 69072708 | 69071922 | 69072023 | 69072914 | 69073001 | 0 | 2 | 0 | 0 | 0 | 1 | 0 | 3 | 1 | 0 | 0 | 0 | 98 | 49 | 0.27131636 | 0.444793103 | NA | 1 | 0 | NA | 1 | 1 | -0.5 |
| A5SS | 435 | PAPD4 | PAPD4 | chr5 | + | 79612419 | 79612769 | 79612419 | 79612735 | 79613010 | 79613075 | 0 | 2 | 1 | 0 | 0 | 1 | 0 | 2 | 1 | 0 | 0 | 0 | 83 | 49 | 0.271326485 | 0.444297119 | NA | 1 | 0.371 | NA | 1 | 1 | -0.315 |
| SE | 1339 | CMTR1 | CMTR1 | chr6 | + | 37462828 | 37463008 | 37461548 | 37461645 | 37471020 | 37471077 | 0 | 0 | 3 | 0 | 1 | 0 | 0 | 2 | 5 | 0 | 0 | 0 | 98 | 49 | 0.271849638 | 0.444793103 | NA | 0 | 1 | NA | 1 | 1 | -0.5 |
| SE | 9456 | PFKFB3 | PFKFB3 | chr10 | + | 6217134 | 6217191 | 6216705 | 6216780 | 6219568 | 6219693 | 0 | 2 | 31 | 0 | 0 | 1 | 0 | 32 | 1 | 0 | 0 | 0 | 98 | 49 | 0.271902353 | 0.444793103 | NA | 1 | 0.939 | NA | 1 | 1 | -0.03 |
| SE | 9007 | LYST | LYST | chr1 | - | 235804503 | 235804665 | 235802907 | 235803064 | 235805742 | 235806772 | 0 | 4 | 25 | 0 | 0 | 2 | 0 | 11 | 1 | 0 | 0 | 0 | 98 | 49 | 0.271988796 | 0.444793103 | NA | 1 | 0.862 | NA | 1 | 1 | -0.069 |
| SE | 12740 | DYSF | DYSF | chr2 | + | 71511806 | 71511921 | 71503213 | 71503319 | 71513715 | 71513921 | 0 | 0 | 5 | 0 | 0 | 1 | 0 | 4 | 3 | 0 | 0 | 0 | 98 | 49 | 0.272557849 | 0.444793103 | NA | NA | 0.714 | NA | 1 | 1 | -0.286 |
| SE | 13209 | LILRB2 | LILRB2 | chr19 | - | 54276380 | 54276456 | 54276263 | 54276301 | 54276806 | 54276929 | 0 | 0 | 5 | 0 | 0 | 1 | 0 | 2 | 3 | 0 | 0 | 0 | 98 | 49 | 0.272557849 | 0.444793103 | NA | NA | 0.714 | NA | 1 | 1 | -0.286 |
| A3SS | 831 | TOM1 | TOM1 | chr22 | + | 35322998 | 35323177 | 35323027 | 35323177 | 35317876 | 35317961 | 0 | 1 | 5 | 0 | 0 | 1 | 0 | 0 | 3 | 0 | 0 | 0 | 78 | 49 | 0.272864991 | 0.491156984 | NA | 1 | 0.759 | NA | NA | 1 | -0.121 |
| SE | 12823 | SEC14L1 | SEC14L1 | chr17 | + | 77193420 | 77193549 | 77191180 | 77191312 | 77194676 | 77194911 | 0 | 5 | 49 | 0 | 0 | 1 | 2 | 50 | 3 | 0 | 0 | 0 | 98 | 49 | 0.274578151 | 0.44760408 | NA | 1 | 0.961 | 1 | 1 | 1 | -0.02 |
| SE | 4882 | DCAF11 | DCAF11 | chr14 | + | 24116916 | 24117044 | 24115381 | 24115749 | 24117265 | 24117393 | 0 | 0 | 3 | 0 | 0 | 1 | 0 | 2 | 2 | 0 | 2 | 2 | 98 | 49 | 0.277220408 | 0.450933196 | NA | NA | 0.6 | NA | 0.333 | 0.333 | 0.267 |
| SE | 4883 | DCAF11 | DCAF11 | chr14 | + | 24116971 | 24117044 | 24115282 | 24115749 | 24117265 | 24117393 | 0 | 0 | 3 | 0 | 0 | 1 | 0 | 0 | 2 | 0 | 2 | 2 | 98 | 49 | 0.277220408 | 0.450933196 | NA | NA | 0.6 | NA | 0 | 0.333 | 0.433 |

| | | | | | | | | | | | | | | | | | | | | | | | | | | | | | | | | | | |
|---|---|---|---|---|---|---|---|---|---|---|---|---|---|---|---|---|---|---|---|---|---|---|---|---|---|---|---|---|---|---|---|---|---|---|
| A3SS | 3597 | EIF4G1 | EIF4G1 | chr3 | + | 184323404 | 184323593 | 184323407 | 184323593 | 184323082 | 184323241 | 0 | 0 | 2 | 0 | 0 | 3 | 0 | 0 | 0 | 0 | 1 | 1 | 52 | 49 | 0.278939638 | 0.491656631 | NA | NA | 0.386 | NA | 0 | 0 | 0.386 |
| SE | 8081 | CSF3R | CSF3R | chr1 | - | 36472065 | 36472139 | 36471432 | 36471646 | 36472237 | 36472391 | 0 | 94 | 256 | 0 | 0 | 1 | 16 | 183 | 131 | 0 | 0 | 0 | 98 | 49 | 0.278963874 | 0.453278598 | NA | 1 | 0.992 | 1 | 1 | 1 | -0.004 |
| A3SS | 1548 | PPP4C | PPP4C | chr16 | + | 30081199 | 30081310 | 30081258 | 30081310 | 30076314 | 30076475 | 0 | 0 | 3 | 0 | 0 | 2 | 0 | 2 | 1 | 0 | 5 | 0 | 98 | 49 | 0.280935793 | 0.491656631 | NA | NA | 0.429 | NA | 0.167 | 1 | -0.155 |
| SE | 3522 | HDAC4 | HDAC4 | chr2 | - | 239144582 | 239144714 | 239139683 | 239139796 | 239156651 | 239156773 | 0 | 0 | 3 | 0 | 0 | 2 | 0 | 0 | 1 | 0 | 0 | 0 | 98 | 49 | 0.280935793 | 0.454548838 | NA | NA | 0.429 | NA | NA | 1 | -0.571 |
| SE | 4254 | MAPKAP1 | MAPKAP1 | chr9 | - | 125672315 | 125672643 | 125669817 | 125669907 | 125706970 | 125707234 | 0 | 0 | 3 | 0 | 0 | 2 | 8 | 0 | 1 | 0 | 0 | 0 | 98 | 49 | 0.280935793 | 0.454548838 | NA | NA | 0.429 | 1 | NA | 1 | -0.571 |
| SE | 4531 | DUSP6 | DUSP6 | chr12 | - | 89350587 | 89351025 | 89347824 | 89349561 | 89351639 | 89352859 | 0 | 3 | 21 | 0 | 2 | 1 | 0 | 1 | 0 | 0 | 0 | 0 | 98 | 49 | 0.280935793 | 0.454548838 | NA | 0.429 | 0.913 | NA | 1 | NA | -0.329 |
| SE | 10627 | RRP12 | RRP12 | chr10 | - | 97390422 | 97390539 | 97388488 | 97388624 | 97390738 | 97390844 | 0 | 7 | 7 | 0 | 0 | 1 | 5 | 9 | 3 | 0 | 0 | 0 | 98 | 49 | 0.281057395 | 0.454548838 | NA | 1 | 0.778 | 1 | 1 | 1 | -0.111 |
| SE | 13626 | SUN2 | SUN2 | chr22 | - | 38738872 | 38738988 | 38738586 | 38738754 | 38739341 | 38739426 | 0 | 0 | 6 | 0 | 0 | 4 | 0 | 6 | 1 | 0 | 0 | 0 | 98 | 49 | 0.281257764 | 0.454548838 | NA | NA | 0.429 | NA | 1 | 1 | -0.571 |
| A3SS | 1827 | FCGR2A | FCGR2A | chr1 | + | 161506333 | 161506591 | 161506336 | 161506591 | 161505986 | 161506007 | 0 | 1 | 20 | 0 | 0 | 17 | 0 | 6 | 2 | 0 | 10 | 1 | 52 | 49 | 0.28133685 | 0.491656631 | NA | 1 | 0.526 | NA | 0.361 | 0.653 | 0.256 |
| SE | 12951 | GMFG | GMFG | chr19 | - | 39335260 | 39335310 | 39333076 | 39333126 | 39335434 | 39335531 | 0 | 1 | 18 | 0 | 0 | 1 | 13 | 16 | 2 | 0 | 0 | 0 | 98 | 49 | 0.283872459 | 0.458281747 | NA | 1 | 0.9 | 1 | 1 | 1 | -0.05 |
| SE | 3214 | RTFDC1 | RTFDC1 | chr20 | + | 56516934 | 56516989 | 56513314 | 56513428 | 56517105 | 56517201 | 0 | 0 | 7 | 0 | 0 | 2 | 0 | 13 | 2 | 0 | 0 | 0 | 98 | 49 | 0.286074243 | 0.461340759 | NA | NA | 0.636 | NA | 1 | 1 | -0.364 |
| SE | 9377 | FMNL1 | FMNL1 | chr17 | + | 45245631 | 45245733 | 45245252 | 45245416 | 45245877 | 45245973 | 0 | 5 | 21 | 0 | 3 | 6 | 0 | 24 | 7 | 0 | 7 | 0 | 98 | 49 | 0.287117418 | 0.462526773 | NA | 0.455 | 0.636 | NA | 0.632 | 1 | -0.271 |
| SE | 9458 | PFKFB3 | PFKFB3 | chr10 | + | 6222854 | 6222984 | 6221380 | 6221527 | 6223957 | 6224020 | 0 | 3 | 22 | 0 | 0 | 1 | 0 | 21 | 7 | 0 | 2 | 0 | 98 | 49 | 0.28811154 | 0.463631312 | NA | 1 | 0.917 | NA | 0.84 | 1 | 0.039 |
| SE | 10611 | PIK3R5 | PIK3R5 | chr17 | - | 8889972 | 8890126 | 8889138 | 8889222 | 8890737 | 8890912 | 0 | 5 | 10 | 0 | 0 | 1 | 0 | 3 | 10 | 0 | 0 | 0 | 98 | 49 | 0.288581572 | 0.463891019 | NA | 1 | 0.833 | NA | 1 | 1 | -0.084 |
| SE | 816 | LINC-PINT | LINC-PINT | chr7 | - | 131002196 | 131002320 | 130984052 | 130984109 | 131052424 | 131052555 | 0 | 0 | 0 | 0 | 2 | 9 | 0 | 0 | 1 | 0 | 8 | 1 | 98 | 49 | 0.290996266 | 0.46727285 | NA | 0 | 0 | NA | 0 | 0.333 | -0.167 |
| A3SS | 2039 | H2AFY | H2AFY | chr5 | - | 135360496 | 135360607 | 135360496 | 135360604 | 135369405 | 135369603 | 0 | 0 | 3 | 0 | 0 | 5 | 6 | 1 | 0 | 0 | 0 | 1 | 52 | 49 | 0.292512865 | 0.506272265 | NA | NA | 0.361 | 1 | 1 | 0 | -0.306 |
| SE | 7721 | THEMIS2 | THEMIS2 | chr1 | + | 27881970 | 27883043 | 27879643 | 27880054 | 27885294 | 27885451 | 0 | 1 | 19 | 0 | 0 | 1 | 0 | 15 | 3 | 0 | 0 | 0 | 98 | 49 | 0.292531456 | 0.469236689 | NA | 1 | 0.905 | NA | 1 | 1 | -0.047 |
| SE | 11044 | HLA-B | HLA-B | chr6 | - | 31354632 | 31354665 | 31354478 | 31354526 | 31355106 | 31355223 | 1 | 16 | 132 | 0 | 0 | 1 | 8 | 135 | 6 | 0 | 0 | 0 | 82 | 49 | 0.294858598 | 0.472465323 | 1 | 1 | 0.987 | 1 | 1 | 1 | -0.004 |
| SE | 2178 | WAS | WAS | chrX | + | 48685733 | 48685836 | 48685546 | 48685633 | 48685945 | 48685987 | 0 | 5 | 20 | 0 | 0 | 0 | 0 | 20 | 2 | 0 | 0 | 1 | 98 | 49 | 0.297924203 | 0.476869091 | NA | 1 | 1 | NA | 1 | 0.5 | 0.25 |
| RI | 186 | PAPD4 | PAPD4 | chr5 | + | 79612419 | 79613075 | 79612419 | 79612735 | 79613010 | 79613075 | 2 | 3 | 7 | 0 | 0 | 1 | 2 | 7 | 2 | 0 | 0 | 0 | 98 | 49 | 0.299346388 | 0.80513856 | 1 | 1 | 0.778 | 1 | 1 | 1 | -0.074 |
| A3SS | 2683 | PXN | PXN | chr12 | - | 120215102 | 120215417 | 120215102 | 120215273 | 120221622 | 120221758 | 0 | 1 | 7 | 0 | 1 | 3 | 0 | 5 | 3 | 0 | 2 | 0 | 98 | 49 | 0.29950383 | 0.513435137 | NA | 0.333 | 0.538 | NA | 0.556 | 1 | -0.343 |
| SE | 8361 | TNFRSF10C | TNFRSF10C | chr8 | + | 23111719 | 23111825 | 23102813 | 23103181 | 23114656 | 23114770 | 0 | 0 | 51 | 0 | 0 | 4 | 0 | 12 | 6 | 0 | 1 | 0 | 98 | 49 | 0.302014034 | 0.482901162 | NA | NA | 0.864 | NA | 0.857 | 1 | -0.065 |
| SE | 9507 | FOXP1 | FOXP1 | chr3 | - | 71359149 | 71359244 | 71299819 | 71299880 | 71493425 | 71493555 | 0 | 0 | 4 | 0 | 0 | 1 | 0 | 5 | 2 | 0 | 0 | 0 | 98 | 49 | 0.303158096 | 0.484215322 | NA | NA | 0.667 | NA | 1 | 1 | -0.333 |
| A5SS | 2161 | SIRPB2 | SIRPB2 | chr20 | - | 1479699 | 1480065 | 1479993 | 1480065 | 1478265 | 1478607 | 0 | 0 | 8 | 0 | 0 | 2 | 0 | 1 | 2 | 0 | 0 | 0 | 98 | 49 | 0.304701638 | 0.492789069 | NA | NA | 0.667 | NA | 1 | 1 | -0.333 |
| SE | 7847 | RPS6KA1 | RPS6KA1 | chr1 | + | 26555536 | 26555625 | 26555150 | 26555221 | 26556653 | 26556718 | 0 | 0 | 27 | 0 | 1 | 4 | 4 | 8 | 0 | 0 | 3 | 1 | 98 | 49 | 0.305107208 | 0.48651819 | NA | 0 | 0.771 | 1 | 0.571 | 0 | -0.138 |
| SE | 4904 | PAK1 | PAK1 | chr11 | - | 77325327 | 77325376 | 77322014 | 77323360 | 77332729 | 77332867 | 0 | 0 | 3 | 0 | 1 | 12 | 0 | 0 | 0 | 0 | 2 | 1 | 98 | 49 | 0.305247274 | 0.48651819 | NA | 0 | 0.111 | NA | 0 | 0 | 0.056 |
| A3SS | 3600 | CNN2 | CNN2 | chr19 | + | 1036066 | 1036246 | 1036129 | 1036246 | 1032558 | 1032696 | 0 | 2 | 9 | 0 | 1 | 10 | 11 | 5 | 10 | 0 | 5 | 1 | 98 | 49 | 0.309557191 | 0.525663154 | NA | 0.5 | 0.31 | 1 | 0.333 | 0.833 | -0.317 |
| SE | 8217 | FURIN | FURIN | chr15 | + | 90877526 | 90877615 | 90877134 | 90877211 | 90878131 | 90878304 | 1 | 1 | 6 | 0 | 0 | 1 | 0 | 2 | 4 | 0 | 0 | 0 | 98 | 49 | 0.310862931 | 0.494727732 | 1 | 1 | 0.75 | NA | 1 | 1 | -0.083 |
| SE | 12945 | SMCHD1 | SMCHD1 | chr18 | + | 2732264 | 2732492 | 2729274 | 2729409 | 2738396 | 2738545 | 0 | 1 | 52 | 0 | 0 | 1 | 1 | 43 | 2 | 0 | 0 | 0 | 98 | 49 | 0.311165715 | 0.494727732 | NA | 1 | 0.963 | 1 | 1 | 1 | -0.018 |
| SE | 5935 | RASGRP2 | RASGRP2 | chr11 | - | 64735902 | 64735980 | 64735541 | 64735664 | 64736752 | 64737034 | 0 | 1 | 5 | 0 | 0 | 0 | 1 | 3 | 1 | 0 | 0 | 3 | 98 | 49 | 0.311385518 | 0.494727732 | NA | 1 | 1 | 1 | 1 | 0.143 | 0.286 |
| SE | 13180 | NLRP12 | NLRP12 | chr19 | - | 53809586 | 53811288 | 53807494 | 53807665 | 53814907 | 53814988 | 0 | 1 | 12 | 0 | 0 | 1 | 0 | 5 | 6 | 0 | 0 | 0 | 98 | 49 | 0.311762966 | 0.49480437 | NA | 1 | 0.857 | NA | 1 | 1 | -0.072 |
| SE | 3534 | LRRFIP1 | LRRFIP1 | chr2 | + | 237751199 | 237751271 | 237749198 | 237749324 | 237753308 | 237753479 | 0 | 0 | 2 | 0 | 1 | 13 | 0 | 7 | 0 | 0 | 4 | 2 | 98 | 49 | 0.314453037 | 0.498547379 | NA | 0 | 0.071 | NA | 0.467 | 0 | -0.198 |
| SE | 6694 | RBCK1 | RBCK1 | chr20 | + | 420870 | 421031 | 419557 | 419731 | 422126 | 422238 | 0 | 2 | 3 | 0 | 0 | 1 | 0 | 1 | 3 | 0 | 0 | 0 | 98 | 49 | 0.315518625 | 0.499709688 | NA | 1 | 0.6 | NA | 1 | 1 | -0.2 |
| SE | 1664 | PDLIM7 | PDLIM7 | chr5 | - | 177489392 | 177489627 | 177488067 | 177488248 | 177489770 | 177489832 | 0 | 3 | 11 | 0 | 0 | 1 | 0 | 6 | 5 | 0 | 0 | 0 | 98 | 49 | 0.31710227 | 0.50058381 | NA | 1 | 0.846 | NA | 1 | 1 | -0.077 |
| SE | 3246 | SULF2 | SULF2 | chr20 | - | 47666259 | 47666488 | 47665856 | 47665953 | 47672197 | 47672393 | 0 | 1 | 5 | 0 | 0 | 1 | 0 | 3 | 2 | 0 | 0 | 0 | 98 | 49 | 0.317618048 | 0.50058381 | NA | 1 | 0.714 | NA | 1 | 1 | -0.143 |
| A5SS | 1014 | PRKD2 | PRKD2 | chr19 | - | 46716814 | 46717127 | 46716950 | 46717127 | 46716130 | 46716535 | 0 | 0 | 2 | 0 | 0 | 1 | 0 | 0 | 1 | 0 | 0 | 0 | 98 | 49 | 0.318735 | 0.509198598 | NA | NA | 0.5 | NA | NA | 1 | -0.5 |
| RI | 727 | TRIM5 | TRIM5 | chr11 | - | 5663194 | 5665682 | 5663194 | 5665395 | 5665655 | 5665682 | 0 | 0 | 2 | 0 | 0 | 1 | 4 | 0 | 1 | 0 | 0 | 0 | 98 | 49 | 0.318735 | 0.805848647 | NA | NA | 0.5 | 1 | NA | 1 | -0.5 |
| SE | 702 | ENTPD4 | ENTPD4 | chr8 | - | 23442006 | 23442066 | 23441568 | 23441723 | 23443849 | 23443953 | 0 | 0 | 2 | 0 | 0 | 1 | 0 | 2 | 1 | 0 | 1 | 0 | 98 | 49 | 0.318735 | 0.50058381 | NA | NA | 0.5 | NA | 0.5 | 1 | -0.25 |
| SE | 2612 | UBA7 | UBA7 | chr3 | - | 49809389 | 49809464 | 49808975 | 49809159 | 49809541 | 49809725 | 0 | 0 | 2 | 0 | 0 | 1 | 0 | 1 | 1 | 0 | 0 | 0 | 98 | 49 | 0.318735 | 0.50058381 | NA | NA | 0.5 | NA | 1 | 1 | -0.5 |
| SE | 2722 | KAT2B | KAT2B | chr3 | + | 20125904 | 20126113 | 20122667 | 20122804 | 20136941 | 20137052 | 0 | 0 | 2 | 0 | 0 | 1 | 2 | 1 | 1 | 0 | 0 | 0 | 98 | 49 | 0.318735 | 0.50058381 | NA | NA | 0.5 | 1 | 1 | 1 | -0.5 |
| SE | 4806 | GNB1 | GNB1 | chr1 | - | 1839189 | 1839238 | 1825396 | 1825499 | 1890819 | 1891117 | 0 | 2 | 5 | 0 | 1 | 0 | 0 | 1 | 0 | 0 | 0 | 0 | 98 | 49 | 0.318735 | 0.50058381 | NA | 0.5 | 1 | NA | 1 | NA | -0.25 |
| SE | 6222 | DNAJB12 | DNAJB12 | chr10 | - | 72338201 | 72338311 | 72336523 | 72336696 | 72340788 | 72340868 | 0 | 0 | 2 | 0 | 0 | 1 | 0 | 0 | 1 | 0 | 0 | 0 | 98 | 49 | 0.318735 | 0.50058381 | NA | NA | 0.5 | NA | NA | 1 | -0.5 |
| SE | 12813 | SRP54 | SRP54 | chr14 | + | 35013345 | 35013494 | 35011508 | 35011659 | 35013801 | 35013902 | 0 | 0 | 2 | 0 | 0 | 1 | 0 | 1 | 1 | 0 | 0 | 0 | 98 | 49 | 0.318735 | 0.50058381 | NA | NA | 0.5 | NA | 1 | 1 | -0.5 |
| SE | 10593 | RNF213 | RNF213 | chr17 | + | 80389172 | 80389367 | 80388611 | 80388689 | 80389827 | 80389917 | 0 | 7 | 56 | 0 | 0 | 1 | 2 | 23 | 1 | 0 | 1 | 0 | 98 | 49 | 0.320121262 | 0.502236176 | NA | 1 | 0.966 | 1 | 0.92 | 1 | 0.01 |
| RI | 389 | GMPR2 | GMPR2 | chr14 | + | 24238245 | 24239241 | 24238245 | 24238405 | 24238588 | 24239241 | 1 | 0 | 7 | 0 | 0 | 1 | 0 | 8 | 3 | 0 | 0 | 0 | 98 | 49 | 0.32027318 | 0.805848647 | 1 | NA | 0.778 | NA | 1 | 1 | -0.111 |
| A3SS | 3968 | SUN2 | SUN2 | chr22 | - | 38750224 | 38750383 | 38750224 | 38750320 | 38750897 | 38751035 | 0 | 0 | 2 | 0 | 1 | 1 | 0 | 2 | 1 | 0 | 0 | 1 | 98 | 49 | 0.320307546 | 0.538835124 | NA | 0 | 0.5 | NA | 1 | 0.333 | -0.416 |
| SE | 7718 | THEMIS2 | THEMIS2 | chr1 | + | 27879643 | 27880054 | 27876587 | 27876728 | 27881970 | 27882125 | 0 | 7 | 42 | 0 | 0 | 1 | 1 | 25 | 11 | 0 | 0 | 0 | 98 | 49 | 0.322009305 | 0.504671517 | NA | 1 | 0.955 | 1 | 1 | 1 | -0.022 |
| SE | 372 | AGTPBP1 | AGTPBP1 | chr9 | - | 85692688 | 85692813 | 85681267 | 85681335 | 85741774 | 85742029 | 0 | 3 | 9 | 0 | 0 | 1 | 3 | 8 | 1 | 0 | 0 | 0 | 98 | 49 | 0.326777752 | 0.511611417 | NA | 1 | 0.818 | 1 | 1 | 1 | -0.091 |
| SE | 13193 | MBOAT7 | MBOAT7 | chr19 | - | 54187160 | 54187287 | 54183520 | 54183680 | 54188216 | 54188346 | 0 | 10 | 24 | 0 | 0 | 1 | 2 | 13 | 11 | 0 | 0 | 0 | 98 | 49 | 0.327855035 | 0.51276391 | NA | 1 | 0.923 | 1 | 1 | 1 | -0.038 |
| SE | 2938 | LIMK2 | LIMK2 | chr22 | + | 31225719 | 31225819 | 31212263 | 31212424 | 31258290 | 31258426 | 0 | 14 | 29 | 0 | 0 | 0 | 3 | 19 | 8 | 0 | 0 | 5 | 98 | 49 | 0.328408932 | 0.512945188 | NA | 1 | 1 | 1 | 1 | 0.444 | 0.185 |
| SE | 3975 | GLT1D1 | GLT1D1 | chr12 | + | 128888638 | 128888744 | 128875913 | 128876062 | 128899235 | 128899287 | 0 | 6 | 18 | 0 | 0 | 1 | 0 | 7 | 2 | 0 | 1 | 0 | 98 | 49 | 0.328653504 | 0.512945188 | NA | 1 | 0.9 | NA | 0.778 | 1 | 0.061 |
| SE | 2767 | LST1 | LST1 | chr6 | + | 31587943 | 31587966 | 31587199 | 31587318 | 31588517 | 31588909 | 0 | 0 | 7 | 0 | 0 | 4 | 0 | 0 | 1 | 0 | 4 | 0 | 72 | 49 | 0.329153667 | 0.513192906 | NA | NA | 0.544 | NA | 0 | 1 | 0.044 |
| SE | 4457 | CLEC7A | CLEC7A | chr12 | - | 10123244 | 10123363 | 10116780 | 10118590 | 10125296 | 10125448 | 0 | 0 | 38 | 0 | 0 | 8 | 0 | 16 | 2 | 0 | 1 | 0 | 98 | 49 | 0.329686827 | 0.513491504 | NA | NA | 0.704 | NA | 0.889 | 1 | -0.241 |
| SE | 10068 | ACTN1 | ACTN1 | chr14 | - | 68909954 | 68910042 | 68909317 | 68909396 | 68912155 | 68912242 | 0 | 9 | 25 | 0 | 2 | 0 | 0 | 18 | 1 | 0 | 2 | 1 | 98 | 49 | 0.331082125 | 0.515130884 | NA | 0.692 | 1 | NA | 0.818 | 0.333 | 0.27 |
| A3SS | 1213 | CIB1 | CIB1 | chr15 | - | 90232218 | 90232447 | 90232218 | 90232327 | 90233668 | 90233703 | 0 | 0 | 5 | 0 | 0 | 1 | 2 | 0 | 2 | 0 | 0 | 0 | 98 | 49 | 0.332444824 | 0.545043142 | NA | NA | 0.714 | 1 | NA | 1 | -0.286 |
| A3SS | 1850 | OAS1 | OAS1 | chr12 | + | 112919388 | 112919907 | 112919486 | 112919907 | 112916508 | 112916738 | 0 | 0 | 5 | 0 | 0 | 1 | 0 | 12 | 2 | 0 | 1 | 0 | 98 | 49 | 0.332444824 | 0.545043142 | NA | NA | 0.714 | NA | 0.857 | 1 | -0.215 |
| RI | 641 | ATPIF1 | ATPIF1 | chr1 | + | 28236360 | 28238105 | 28236360 | 28236452 | 28237836 | 28238105 | 0 | 0 | 5 | 0 | 0 | 1 | 0 | 0 | 2 | 0 | 0 | 0 | 98 | 49 | 0.332444824 | 0.810334259 | NA | NA | 0.714 | NA | NA | 1 | -0.286 |
| SE | 7846 | RPS6KA1 | RPS6KA1 | chr1 | + | 26546866 | 26546983 | 26545851 | 26546035 | 26547188 | 26547270 | 0 | 3 | 19 | 0 | 0 | 1 | 0 | 10 | 6 | 0 | 0 | 0 | 98 | 49 | 0.332491859 | 0.516037235 | NA | 1 | 0.905 | NA | 1 | 1 | -0.047 |
| SE | 9991 | TRANK1 | TRANK1 | chr3 | - | 36892900 | 36892984 | 36892201 | 36892340 | 36899108 | 36899259 | 0 | 0 | 15 | 0 | 0 | 2 | 1 | 3 | 3 | 0 | 0 | 0 | 98 | 49 | 0.332635761 | 0.516037235 | NA | NA | 0.789 | 1 | 1 | 1 | -0.211 |
| SE | 10092 | NUMB | NUMB | chr14 | - | 73279280 | 73279424 | 73275209 | 73277293 | 73282358 | 73282505 | 0 | 0 | 2 | 0 | 0 | 10 | 0 | 0 | 0 | 0 | 0 | 2 | 98 | 49 | 0.332694665 | 0.516037235 | NA | NA | 0.091 | NA | NA | 0 | 0.091 |
| A3SS | 653 | VNN3 | VNN3 | chr6 | - | 132728740 | 132728979 | 132728740 | 132728933 | 132734252 | 132734383 | 0 | 0 | 9 | 0 | 0 | 4 | 0 | 23 | 1 | 0 | 2 | 0 | 95 | 49 | 0.33308192 | 0.545043142 | NA | NA | 0.537 | NA | 0.856 | 1 | -0.391 |
| SE | 13633 | CCND3 | CCND3 | chr6 | - | 41936558 | 41936695 | 41934932 | 41936107 | 41937234 | 41937394 | 0 | 3 | 19 | 0 | 2 | 0 | 0 | 2 | 2 | 0 | 0 | 2 | 98 | 49 | 0.333586462 | 0.516887065 | NA | 0.429 | 1 | NA | 1 | 0.333 | 0.048 |
| SE | 9234 | RPS24 | RPS24 | chr10 | + | 78040203 | 78040225 | 78037193 | 78037304 | 78040614 | 78040715 | 0 | 0 | 11 | 0 | 0 | 26 | 5 | 22 | 0 | 0 | 6 | 1 | 71 | 49 | 0.336551286 | 0.520943957 | NA | NA | 0.226 | 1 | 0.717 | 0 | -0.346 |
| SE | 2900 | DDX17 | DDX17 | chr22 | - | 38495795 | 38495937 | 38494885 | 38495046 | 38498084 | 38498150 | 0 | 1 | 31 | 0 | 0 | 1 | 3 | 22 | 2 | 0 | 0 | 0 | 98 | 49 | 0.337825803 | 0.522378788 | NA | 1 | 0.939 | 1 | 1 | 1 | -0.03 |
| SE | 12582 | CNN2 | CNN2 | chr19 | + | 1036129 | 1036246 | 1032558 | 1032696 | 1036415 | 1036562 | 0 | 1 | 25 | 0 | 0 | 1 | 0 | 17 | 2 | 0 | 0 | 0 | 98 | 49 | 0.338322411 | 0.522609027 | NA | 1 | 0.926 | NA | 1 | 1 | -0.037 |
| SE | 7966 | OAS3 | OAS3 | chr12 | + | 112941569 | 112941852 | 112938432 | 112938707 | 112944475 | 112944651 | 0 | 7 | 34 | 0 | 0 | 1 | 0 | 17 | 11 | 0 | 0 | 0 | 98 | 49 | 0.338676564 | 0.52261897 | NA | 1 | 0.944 | NA | 1 | 1 | -0.028 |
| SE | 12901 | TYROBP | TYROBP | chr19 | - | 35907217 | 35907264 | 35904400 | 35904634 | 35907445 | 35907580 | 0 | 18 | 77 | 0 | 0 | 1 | 0 | 30 | 10 | 0 | 1 | 0 | 96 | 49 | 0.342984435 | 0.527529386 | NA | 1 | 0.975 | NA | 0.939 | 1 | 0.018 |
| SE | 4627 | SORL1 | SORL1 | chr11 | + | 121570156 | 121570270 | 121566939 | 121567113 | 121574240 | 121574363 | 0 | 0 | 56 | 0 | 1 | 12 | 0 | 27 | 2 | 4 | 1 | 1 | 98 | 49 | 0.343393791 | 0.527529386 | NA | 0 | 0.7 | 0 | 0.931 | 0.5 | -0.127 |
| SE | 5558 | SPATA13 | SPATA13 | chr13 | + | 24286764 | 24286950 | 24286213 | 24286393 | 24288998 | 24289178 | 0 | 0 | 5 | 0 | 0 | 2 | 0 | 1 | 1 | 0 | 1 | 0 | 98 | 49 | 0.34342743 | 0.527529386 | NA | NA | 0.556 | NA | 0.333 | 1 | -0.11 |
| SE | 11729 | PER1 | PER1 | chr17 | - | 8146896 | 8147002 | 8146593 | 8146765 | 8147249 | 8147381 | 0 | 0 | 5 | 0 | 0 | 2 | 0 | 1 | 1 | 0 | 2 | 0 | 98 | 49 | 0.34342743 | 0.527529386 | NA | NA | 0.556 | NA | 0.2 | 1 | -0.044 |
| SE | 12395 | TMEM127 | TMEM127 | chr2 | - | 96265137 | 96265489 | 96254832 | 96254997 | 96265868 | 96266013 | 0 | 3 | 13 | 0 | 0 | 1 | 0 | 7 | 4 | 0 | 0 | 0 | 98 | 49 | 0.343613619 | 0.527529386 | NA | 1 | 0.867 | NA | 1 | 1 | -0.067 |
| SE | 13844 | C10orf54 | C10orf54 | chr10 | - | 71755358 | 71755466 | 71752974 | 71753002 | 71760867 | 71760924 | 0 | 3 | 48 | 0 | 0 | 1 | 2 | 32 | 4 | 0 | 0 | 0 | 98 | 49 | 0.344174583 | 0.527680329 | NA | 1 | 0.96 | 1 | 1 | 1 | -0.02 |
| SE | 13536 | ALOX5 | ALOX5 | chr10 | + | 45382482 | 45382681 | 45374175 | 45374429 | 45395854 | 45395936 | 1 | 6 | 67 | 0 | 0 | 2 | 1 | 16 | 6 | 0 | 0 | 0 | 98 | 49 | 0.344414107 | 0.527680329 | 1 | 1 | 0.944 | 1 | 1 | 1 | -0.019 |
| SE | 12400 | CSF2RB | CSF2RB | chr22 | + | 36925986 | 36926177 | 36923243 | 36923367 | 36929401 | 36929559 | 0 | 1 | 39 | 0 | 0 | 1 | 1 | 24 | 4 | 0 | 0 | 0 | 98 | 49 | 0.345199125 | 0.528344486 | NA | 1 | 0.951 | 1 | 1 | 1 | -0.024 |
| SE | 4506 | PTPN6 | PTPN6 | chr12 | + | 6960805 | 6960945 | 6960343 | 6960435 | 6961125 | 6961316 | 0 | 1 | 28 | 0 | 0 | 1 | 7 | 18 | 3 | 0 | 0 | 0 | 98 | 49 | 0.346027303 | 0.528422573 | NA | 1 | 0.933 | 1 | 1 | 1 | -0.033 |
| SE | 9933 | PTPRE | PTPRE | chr10 | + | 128076602 | 128076728 | 128073336 | 128073471 | 128077616 | 128077783 | 0 | 3 | 39 | 0 | 0 | 0 | 0 | 24 | 2 | 0 | 0 | 1 | 98 | 49 | 0.346119905 | 0.528422573 | NA | 1 | 1 | NA | 1 | 0.5 | 0.25 |
| SE | 7719 | THEMIS2 | THEMIS2 | chr1 | + | 27879643 | 27880054 | 27876587 | 27876728 | 27885294 | 27885451 | 0 | 6 | 25 | 0 | 3 | 1 | 1 | 12 | 8 | 0 | 1 | 0 | 98 | 49 | 0.346439311 | 0.528422573 | NA | 0.5 | 0.926 | 1 | 0.857 | 1 | -0.239 |
| SE | 12745 | DYSF | DYSF | chr2 | + | 71620546 | 71620609 | 71613333 | 71613410 | 71643964 | 71644063 | 0 | 0 | 13 | 0 | 0 | 3 | 0 | 3 | 2 | 0 | 2 | 0 | 98 | 49 | 0.346656459 | 0.528422573 | NA | NA | 0.684 | NA | 0.429 | 1 | -0.03 |
| SE | 13437 | HBP1 | HBP1 | chr7 | + | 107189291 | 107189448 | 107186568 | 107186681 | 107190172 | 107190317 | 0 | 1 | 1 | 0 | 0 | 1 | 0 | 1 | 1 | 0 | 3 | 0 | 98 | 49 | 0.348964996 | 0.531353191 | NA | 1 | 0.333 | NA | 0.143 | 1 | 0.095 |
| SE | 9997 | GCH1 | GCH1 | chr14 | - | 54844000 | 54844143 | 54842005 | 54843054 | 54845767 | 54845852 | 0 | 0 | 1 | 0 | 0 | 2 | 0 | 0 | 0 | 1 | 0 | 1 | 98 | 49 | 0.349286063 | 0.531353191 | NA | NA | 0.2 | 0 | NA | 0 | 0.2 |
| SE | 3661 | ETS2 | ETS2 | chr21 | + | 38821585 | 38821704 | 38819502 | 38819766 | 38822673 | 38824954 | 0 | 0 | 8 | 0 | 0 | 3 | 3 | 13 | 1 | 0 | 0 | 0 | 98 | 49 | 0.35298525 | 0.535895789 | NA | NA | 0.571 | 1 | 1 | 1 | -0.429 |
| SE | 9992 | TRANK1 | TRANK1 | chr3 | - | 36895639 | 36895758 | 36892900 | 36892984 | 36899108 | 36899259 | 0 | 0 | 8 | 0 | 0 | 3 | 0 | 4 | 1 | 0 | 1 | 0 | 98 | 49 | 0.35298525 | 0.535895789 | NA | NA | 0.571 | NA | 0.667 | 1 | -0.263 |
| SE | 11324 | ITGAL | ITGAL | chr16 | + | 30479330 | 30479461 | 30479090 | 30479208 | 30481438 | 30481584 | 0 | 0 | 11 | 0 | 0 | 4 | 0 | 6 | 1 | 0 | 0 | 0 | 98 | 49 | 0.355559836 | 0.539081642 | NA | NA | 0.579 | NA | 1 | 1 | -0.421 |
| SE | 2458 | MPP1 | MPP1 | chrX | - | 154799744 | 154799893 | 154792141 | 154792285 | 154805271 | 154805527 | 0 | 0 | 1 | 0 | 0 | 6 | 0 | 0 | 0 | 0 | 0 | 2 | 98 | 49 | 0.355801057 | 0.539081642 | NA | NA | 0.077 | NA | NA | 0 | 0.077 |
| A3SS | 564 | PHF20L1 | PHF20L1 | chr8 | + | 132777791 | 132777911 | 132777795 | 132777911 | 132775357 | 132775645 | 0 | 0 | 1 | 0 | 3 | 3 | 0 | 3 | 0 | 2 | 5 | 3 | 53 | 49 | 0.358798866 | 0.579927566 | NA | 0 | 0.236 | 0 | 0.357 | 0 | -0.001 |
| SE | 2884 | MKL1 | MKL1 | chr22 | - | 40594673 | 40594735 | 40552105 | 40552367 | 40636477 | 40636719 | 0 | 0 | 4 | 0 | 0 | 2 | 0 | 0 | 1 | 0 | 0 | 0 | 98 | 49 | 0.359545146 | 0.544205795 | NA | NA | 0.5 | NA | NA | 1 | -0.5 |
| SE | 3751 | ARHGAP27 | ARHGAP27 | chr17 | - | 45396486 | 45396585 | 45396206 | 45396284 | 45396667 | 45396790 | 0 | 4 | 14 | 0 | 0 | 1 | 0 | 8 | 3 | 0 | 0 | 0 | 98 | 49 | 0.360380165 | 0.544920914 | NA | 1 | 0.875 | NA | 1 | 1 | -0.062 |
| A3SS | 140 | LY6E | LY6E | chr8 | + | 143020882 | 143020991 | 143020887 | 143020991 | 143018484 | 143018586 | 0 | 0 | 2 | 0 | 0 | 1 | 2 | 1 | 1 | 0 | 1 | 0 | 54 | 49 | 0.360843819 | 0.579927566 | NA | NA | 0.645 | 1 | 0.476 | 1 | -0.18 |
| SE | 18 | FLNA | FLNA | chrX | - | 154354615 | 154354711 | 154354380 | 154354483 | 154354824 | 154355072 | 0 | 2 | 25 | 0 | 0 | 1 | 0 | 15 | 2 | 0 | 0 | 0 | 98 | 49 | 0.361294614 | 0.545754578 | NA | 1 | 0.926 | NA | 1 | 1 | -0.037 |
| SE | 12741 | DYSF | DYSF | chr2 | + | 71513239 | 71513332 | 71511806 | 71511921 | 71513715 | 71513921 | 0 | 1 | 10 | 0 | 0 | 1 | 0 | 3 | 2 | 0 | 1 | 0 | 98 | 49 | 0.362088102 | 0.546301307 | NA | 1 | 0.833 | NA | 0.6 | 1 | 0.116 |
| SE | 12142 | TRIM25 | TRIM25 | chr17 | - | 56895520 | 56895604 | 56895342 | 56895441 | 56895925 | 56895952 | 0 | 0 | 13 | 0 | 0 | 1 | 0 | 15 | 4 | 0 | 0 | 0 | 98 | 49 | 0.362383502 | 0.546301307 | NA | NA | 0.867 | NA | 1 | 1 | -0.133 |
| SE | 275 | RAPGEF1 | RAPGEF1 | chr9 | - | 131630235 | 131630324 | 131629101 | 131629254 | 131638634 | 131638791 | 0 | 0 | 6 | 0 | 0 | 2 | 0 | 8 | 1 | 0 | 1 | 0 | 98 | 49 | 0.365054683 | 0.547059405 | NA | NA | 0.6 | NA | 0.8 | 1 | -0.3 |
| SE | 5253 | SEC31A | SEC31A | chr4 | - | 82848803 | 82848977 | 82844385 | 82844509 | 82851430 | 82851604 | 0 | 0 | 3 | 0 | 0 | 1 | 0 | 4 | 1 | 0 | 0 | 0 | 98 | 49 | 0.365118555 | 0.547059405 | NA | NA | 0.6 | NA | 1 | 1 | -0.4 |
| SE | 6132 | ANKRD13D | ANKRD13D | chr11 | + | 67299057 | 67299124 | 67291990 | 67292180 | 67299529 | 67299611 | 0 | 0 | 3 | 0 | 0 | 1 | 0 | 8 | 1 | 0 | 0 | 0 | 98 | 49 | 0.365118555 | 0.547059405 | NA | NA | 0.6 | NA | 1 | 1 | -0.4 |
| SE | 7202 | MDM4 | MDM4 | chr1 | + | 204544534 | 204544684 | 204542783 | 204542944 | 204546796 | 204546877 | 0 | 0 | 3 | 0 | 0 | 1 | 0 | 7 | 1 | 0 | 0 | 0 | 98 | 49 | 0.365118555 | 0.547059405 | NA | NA | 0.6 | NA | 1 | 1 | -0.4 |
| SE | 13919 | LUCAT1 | LUCAT1 | chr5 | - | 91313077 | 91313229 | 91303872 | 91303971 | 91313637 | 91313777 | 0 | 2 | 3 | 0 | 0 | 1 | 0 | 0 | 1 | 0 | 0 | 0 | 98 | 49 | 0.365118555 | 0.547059405 | NA | 1 | 0.6 | NA | NA | 1 | -0.2 |
| SE | 1533 | HLA-A | HLA-A | chr6 | + | 29944499 | 29944616 | 29944121 | 29944397 | 29945058 | 29945091 | 1 | 41 | 187 | 0 | 1 | 2 | 5 | 230 | 19 | 0 | 6 | 0 | 98 | 49 | 0.3653902 | 0.547059405 | 1 | 0.953 | 0.979 | 1 | 0.95 | 1 | -0.006 |
| SE | 3397 | APMAP | APMAP | chr20 | - | 24978766 | 24978882 | 24973644 | 24973737 | 24983902 | 24984019 | 0 | 1 | 5 | 0 | 0 | 1 | 0 | 2 | 2 | 0 | 0 | 0 | 98 | 49 | 0.365434227 | 0.547059405 | NA | 1 | 0.714 | NA | 1 | 1 | -0.143 |
| SE | 11877 | MGAM | MGAM | chr7 | + | 142036822 | 142036977 | 142036168 | 142036285 | 142038530 | 142038615 | 0 | 0 | 20 | 0 | 0 | 1 | 4 | 3 | 6 | 0 | 0 | 0 | 98 | 49 | 0.366623914 | 0.548294271 | NA | NA | 0.909 | 1 | 1 | 1 | -0.091 |
| SE | 2284 | IL1RAP | IL1RAP | chr3 | + | 190556129 | 190556216 | 190514050 | 190514219 | 190564288 | 190564353 | 0 | 3 | 7 | 0 | 0 | 1 | 0 | 4 | 2 | 0 | 0 | 0 | 98 | 49 | 0.367396868 | 0.548904067 | NA | 1 | 0.778 | NA | 1 | 1 | -0.111 |
| SE | 12976 | RPS19 | RPS19 | chr19 | + | 41869030 | 41869214 | 41861111 | 41861212 | 41869698 | 41869753 | 1 | 6 | 58 | 0 | 0 | 1 | 3 | 38 | 4 | 0 | 0 | 0 | 98 | 49 | 0.368911011 | 0.550618918 | 1 | 1 | 0.967 | 1 | 1 | 1 | -0.011 |
| SE | 8375 | OAS2 | OAS2 | chr12 | + | 112987037 | 112987308 | 112978468 | 112978785 | 112995295 | 112995474 | 0 | 6 | 42 | 0 | 0 | 3 | 3 | 7 | 1 | 0 | 0 | 0 | 98 | 49 | 0.370767489 | 0.552840809 | NA | 1 | 0.875 | 1 | 1 | 1 | -0.062 |
| SE | 6279 | IDS | IDS | chrX | - | 149491537 | 149491667 | 149490313 | 149490440 | 149496345 | 149496516 | 0 | 0 | 1 | 0 | 0 | 7 | 0 | 0 | 0 | 0 | 2 | 2 | 98 | 49 | 0.37479931 | 0.558298674 | NA | NA | 0.067 | NA | 0 | 0 | 0.067 |
| SE | 6619 | AMPD2 | AMPD2 | chr1 | + | 109629795 | 109629916 | 109629326 | 109629490 | 109630232 | 109630406 | 0 | 1 | 7 | 0 | 0 | 3 | 0 | 7 | 1 | 0 | 4 | 0 | 98 | 49 | 0.376779701 | 0.560692961 | NA | 1 | 0.538 | NA | 0.467 | 1 | 0.035 |
| SE | 12898 | TXNIP | TXNIP | chr1 | - | 145994543 | 145994800 | 145994280 | 145994437 | 145994928 | 145995031 | 0 | 17 | 121 | 0 | 0 | 1 | 4 | 61 | 19 | 0 | 0 | 0 | 98 | 49 | 0.378921937 | 0.563323118 | NA | 1 | 0.984 | 1 | 1 | 1 | -0.008 |
| A3SS | 3632 | STXBP2 | STXBP2 | chr19 | + | 7640697 | 7640809 | 7640730 | 7640809 | 7639730 | 7639807 | 0 | 0 | 7 | 0 | 1 | 1 | 2 | 0 | 2 | 0 | 0 | 0 | 82 | 49 | 0.379879433 | 0.605117681 | NA | 0 | 0.807 | 1 | NA | 1 | -0.597 |
| RI | 545 | CDK5RAP3 | CDK5RAP3 | chr17 | + | 47974399 | 47975337 | 47974399 | 47974448 | 47975158 | 47975337 | 0 | 7 | 10 | 0 | 0 | 1 | 0 | 4 | 5 | 0 | 0 | 0 | 98 | 49 | 0.38169658 | 0.878475733 | NA | 1 | 0.833 | NA | 1 | 1 | -0.084 |
| SE | 8578 | RC3H1 | RC3H1 | chr1 | - | 173983417 | 173983657 | 173982726 | 173982902 | 173984498 | 173984619 | 0 | 1 | 7 | 0 | 0 | 1 | 0 | 3 | 1 | 0 | 1 | 0 | 98 | 49 | 0.382084955 | 0.567464117 | NA | 1 | 0.778 | NA | 0.6 | 1 | 0.089 |
| SE | 11360 | TATDN2 | TATDN2 | chr3 | + | 10249194 | 10249614 | 10248492 | 10249067 | 10260136 | 10260670 | 0 | 3 | 6 | 0 | 0 | 1 | 2 | 4 | 1 | 0 | 0 | 0 | 98 | 49 | 0.382895649 | 0.568106772 | NA | 1 | 0.75 | 1 | 1 | 1 | -0.125 |
| RI | 660 | CSF3R | CSF3R | chr1 | - | 36466042 | 36466827 | 36466042 | 36466200 | 36466620 | 36466827 | 1 | 19 | 165 | 0 | 0 | 1 | 7 | 83 | 24 | 0 | 0 | 0 | 98 | 49 | 0.382925319 | 0.878475733 | 1 | 1 | 0.988 | 1 | 1 | 1 | -0.004 |
| SE | 7522 | AQP9 | AQP9 | chr15 | + | 58174917 | 58175036 | 58173067 | 58173205 | 58179127 | 58179345 | 0 | 1 | 43 | 0 | 0 | 1 | 3 | 20 | 6 | 0 | 0 | 0 | 98 | 49 | 0.383614897 | 0.568612613 | NA | 1 | 0.956 | 1 | 1 | 1 | -0.022 |
| SE | 802 | TBXAS1 | TBXAS1 | chr7 | + | 139957633 | 139957764 | 139955458 | 139955607 | 139961918 | 139962233 | 0 | 3 | 22 | 0 | 1 | 0 | 2 | 4 | 2 | 0 | 0 | 0 | 98 | 49 | 0.385851284 | 0.571364019 | NA | 0.6 | 1 | 1 | 1 | 1 | -0.2 |
| SE | 8267 | DPYD | DPYD | chr1 | - | 97549559 | 97549744 | 97515725 | 97515941 | 97573759 | 97573970 | 0 | 4 | 25 | 0 | 0 | 1 | 0 | 11 | 4 | 0 | 1 | 0 | 98 | 49 | 0.389051763 | 0.57553622 | NA | 1 | 0.926 | NA | 0.846 | 1 | 0.04 |
| A5SS | 1166 | IRF2BP2 | IRF2BP2 | chr1 | - | 234608446 | 234609525 | 234608494 | 234609525 | 234604268 | 234607852 | 0 | 0 | 5 | 0 | 1 | 5 | 0 | 4 | 0 | 0 | 10 | 1 | 97 | 49 | 0.389300363 | 0.614437923 | NA | 0 | 0.336 | NA | 0.168 | 0 | 0.084 |
| SE | 306 | SH2D3C | SH2D3C | chr9 | - | 127741787 | 127741959 | 127740257 | 127740369 | 127742848 | 127742964 | 0 | 2 | 7 | 0 | 1 | 0 | 0 | 2 | 2 | 0 | 0 | 0 | 98 | 49 | 0.390828967 | 0.577596791 | NA | 0.5 | 1 | NA | 1 | 1 | -0.25 |
| SE | 4458 | CLEC7A | CLEC7A | chr12 | - | 10125086 | 10125173 | 10116780 | 10118590 | 10125296 | 10125448 | 0 | 3 | 18 | 0 | 0 | 8 | 0 | 4 | 3 | 0 | 1 | 0 | 98 | 49 | 0.391525682 | 0.578058056 | NA | 1 | 0.529 | NA | 0.667 | 1 | -0.069 |
| SE | 6890 | EP300 | EP300 | chr22 | + | 41135812 | 41135906 | 41131387 | 41131633 | 41137652 | 41137790 | 0 | 2 | 14 | 0 | 0 | 1 | 0 | 8 | 1 | 0 | 0 | 0 | 98 | 49 | 0.392944473 | 0.579315862 | NA | 1 | 0.875 | NA | 1 | 1 | -0.062 |
| SE | 3233 | SLC9A8 | SLC9A8 | chr20 | + | 49839540 | 49839599 | 49823060 | 49823141 | 49845035 | 49845119 | 0 | 0 | 8 | 0 | 0 | 1 | 0 | 0 | 2 | 0 | 0 | 0 | 98 | 49 | 0.393533929 | 0.579315862 | NA | NA | 0.8 | NA | NA | 1 | -0.2 |
| SE | 11134 | ECE1 | ECE1 | chr1 | - | 21225249 | 21225440 | 21221746 | 21221842 | 21227158 | 21227226 | 1 | 0 | 8 | 0 | 0 | 1 | 0 | 12 | 2 | 0 | 0 | 0 | 98 | 49 | 0.393533929 | 0.579315862 | 1 | NA | 0.8 | NA | 1 | 1 | -0.1 |
| SE | 11656 | NLRP1 | NLRP1 | chr17 | - | 5517745 | 5517887 | 5515472 | 5515517 | 5530480 | 5530704 | 0 | 0 | 7 | 0 | 1 | 0 | 0 | 2 | 3 | 0 | 0 | 0 | 98 | 49 | 0.39950276 | 0.587426728 | NA | 0 | 1 | NA | 1 | 1 | -0.5 |
| SE | 1505 | HLA-E | HLA-E | chr6 | + | 30491536 | 30491653 | 30491136 | 30491412 | 30492403 | 30492436 | 0 | 33 | 293 | 0 | 0 | 1 | 21 | 137 | 29 | 0 | 0 | 0 | 98 | 49 | 0.399825378 | 0.587426728 | NA | 1 | 0.993 | 1 | 1 | 1 | -0.004 |
| SE | 5153 | FTH1 | FTH1 | chr11 | - | 61964986 | 61965112 | 61964284 | 61964891 | 61965368 | 61965515 | 0 | 8 | 114 | 0 | 0 | 1 | 7 | 66 | 2 | 0 | 0 | 0 | 98 | 49 | 0.400407723 | 0.587707819 | NA | 1 | 0.983 | 1 | 1 | 1 | -0.008 |
| SE | 11835 | NCOA1 | NCOA1 | chr2 | + | 24768078 | 24768135 | 24762686 | 24762776 | 24768220 | 24770701 | 0 | 0 | 8 | 0 | 0 | 2 | 0 | 2 | 1 | 0 | 0 | 0 | 98 | 49 | 0.401159139 | 0.58823628 | NA | NA | 0.667 | NA | 1 | 1 | -0.333 |
| A3SS | 909 | TPD52L2 | TPD52L2 | chr20 | + | 63882718 | 63882820 | 63882749 | 63882820 | 63875815 | 63875875 | 0 | 2 | 11 | 0 | 0 | 1 | 3 | 5 | 2 | 0 | 0 | 0 | 80 | 49 | 0.401186798 | 0.63162252 | NA | 1 | 0.871 | 1 | 1 | 1 | -0.065 |
| SE | 9689 | IFRD1 | IFRD1 | chr7 | + | 112462269 | 112462378 | 112462000 | 112462179 | 112467980 | 112468115 | 0 | 0 | 10 | 0 | 3 | 37 | 4 | 3 | 0 | 2 | 22 | 2 | 98 | 49 | 0.40162165 | 0.588340487 | NA | 0 | 0.119 | 0.5 | 0.064 | 0 | -0.129 |
| A3SS | 4073 | CFLAR | CFLAR | chr2 | + | 201162995 | 201172688 | 201163834 | 201172688 | 201160431 | 201160942 | 0 | 0 | 4 | 0 | 3 | 38 | 2 | 3 | 2 | 7 | 14 | 3 | 98 | 49 | 0.40353661 | 0.63162252 | NA | 0 | 0.05 | 0.125 | 0.097 | 0.25 | -0.132 |
| SE | 13946 | AP3D1 | AP3D1 | chr19 | - | 2108686 | 2108766 | 2100987 | 2102268 | 2109085 | 2109207 | 0 | 2 | 2 | 0 | 0 | 1 | 0 | 1 | 1 | 0 | 0 | 0 | 98 | 49 | 0.409859466 | 0.599823542 | NA | 1 | 0.5 | NA | 1 | 1 | -0.25 |
| SE | 13218 | LILRA5 | LILRA5 | chr19 | - | 54311863 | 54312148 | 54311164 | 54311716 | 54312536 | 54312621 | 0 | 1 | 18 | 0 | 0 | 1 | 0 | 9 | 1 | 0 | 0 | 0 | 98 | 49 | 0.410993926 | 0.600898707 | NA | 1 | 0.9 | NA | 1 | 1 | -0.05 |
| SE | 6789 | RAB4B-EGLN2 | RAB4B-EGLN2 | chr19 | + | 40806554 | 40806674 | 40800338 | 40801415 | 40807137 | 40807274 | 0 | 2 | 3 | 0 | 0 | 0 | 0 | 1 | 0 | 0 | 0 | 1 | 98 | 49 | 0.412150794 | 0.601420042 | NA | 1 | 1 | NA | 1 | 0 | 0.5 |
| SE | 7891 | FHL3 | FHL3 | chr1 | - | 37999256 | 37999432 | 37998973 | 37999148 | 38005356 | 38005515 | 0 | 2 | 3 | 0 | 0 | 0 | 0 | 1 | 0 | 0 | 0 | 1 | 98 | 49 | 0.412150794 | 0.601420042 | NA | 1 | 1 | NA | 1 | 0 | 0.5 |
| SE | 12902 | TYROBP | TYROBP | chr19 | - | 35907729 | 35907762 | 35907445 | 35907580 | 35908167 | 35908309 | 0 | 4 | 15 | 0 | 0 | 5 | 0 | 7 | 3 | 0 | 1 | 0 | 82 | 49 | 0.416321308 | 0.606916515 | NA | 1 | 0.642 | NA | 0.807 | 1 | -0.083 |
| RI | 981 | ACAA1 | ACAA1 | chr3 | - | 38126161 | 38126700 | 38126161 | 38126341 | 38126509 | 38126700 | 0 | 0 | 9 | 0 | 0 | 1 | 0 | 1 | 3 | 0 | 1 | 0 | 98 | 49 | 0.421251047 | 0.934008656 | NA | NA | 0.818 | NA | 0.333 | 1 | 0.151 |
| SE | 5079 | WDR13 | WDR13 | chrX | + | 48599586 | 48599717 | 48599352 | 48599462 | 48600318 | 48600626 | 0 | 1 | 4 | 0 | 0 | 0 | 0 | 1 | 0 | 0 | 0 | 1 | 98 | 49 | 0.421888846 | 0.613929698 | NA | 1 | 1 | NA | 1 | 0 | 0.5 |
| SE | 11377 | ARHGEF1 | ARHGEF1 | chr19 | + | 41906456 | 41906620 | 41905938 | 41906025 | 41906702 | 41906803 | 0 | 0 | 18 | 0 | 0 | 2 | 0 | 5 | 3 | 0 | 0 | 0 | 98 | 49 | 0.422105527 | 0.613929698 | NA | NA | 0.818 | NA | 1 | 1 | -0.182 |
| SE | 3692 | DOK3 | DOK3 | chr5 | - | 177508236 | 177508542 | 177505010 | 177505110 | 177509474 | 177509575 | 0 | 7 | 32 | 0 | 0 | 1 | 0 | 13 | 5 | 0 | 0 | 0 | 98 | 49 | 0.42235749 | 0.613929698 | NA | 1 | 0.941 | NA | 1 | 1 | -0.03 |
| SE | 13638 | RNF114 | RNF114 | chr20 | + | 49941560 | 49941711 | 49936376 | 49936552 | 49945381 | 49945488 | 0 | 1 | 2 | 0 | 0 | 0 | 0 | 0 | 1 | 0 | 1 | 0 | 98 | 49 | 0.423875417 | 0.615287988 | NA | 1 | 1 | NA | 0 | 1 | 0.5 |
| SE | 5214 | SUCO | SUCO | chr1 | + | 172551511 | 172551626 | 172533117 | 172533497 | 172553259 | 172553370 | 0 | 1 | 1 | 0 | 0 | 0 | 1 | 1 | 0 | 0 | 0 | 1 | 98 | 49 | 0.424110682 | 0.615287988 | NA | 1 | 1 | 1 | 1 | 0 | 0.333 |
| SE | 5013 | ADRBK1 | ADRBK1 | chr11 | + | 67283706 | 67283773 | 67283127 | 67283228 | 67283853 | 67283949 | 0 | 13 | 34 | 0 | 5 | 0 | 4 | 7 | 17 | 0 | 1 | 0 | 98 | 49 | 0.425222253 | 0.6162202 | NA | 0.565 | 1 | 1 | 0.778 | 1 | -0.144 |
| SE | 2290 | BCL6 | BCL6 | chr3 | - | 187728359 | 187728544 | 187726730 | 187726898 | 187729049 | 187730021 | 0 | 2 | 24 | 0 | 0 | 1 | 3 | 8 | 4 | 0 | 0 | 0 | 98 | 49 | 0.425573232 | 0.6162202 | NA | 1 | 0.923 | 1 | 1 | 1 | -0.038 |
| A5SS | 1299 | WDR26 | WDR26 | chr1 | - | 224431476 | 224431581 | 224431524 | 224431581 | 224424517 | 224424654 | 0 | 0 | 11 | 0 | 0 | 4 | 4 | 5 | 1 | 0 | 2 | 0 | 97 | 49 | 0.426432576 | 0.665031756 | NA | NA | 0.581 | 1 | 0.558 | 1 | -0.272 |
| SE | 6186 | HERC3 | HERC3 | chr4 | + | 88681158 | 88681325 | 88680092 | 88680236 | 88686735 | 88686802 | 0 | 0 | 12 | 0 | 0 | 2 | 0 | 8 | 2 | 0 | 0 | 0 | 98 | 49 | 0.429038403 | 0.620451665 | NA | NA | 0.75 | NA | 1 | 1 | -0.25 |
| SE | 2247 | MAEA | MAEA | chr4 | + | 1327626 | 1327703 | 1315396 | 1315600 | 1332756 | 1332865 | 0 | 3 | 2 | 0 | 1 | 1 | 0 | 1 | 0 | 0 | 1 | 0 | 98 | 49 | 0.429432607 | 0.620451665 | NA | 0.6 | 0.5 | NA | 0.333 | NA | 0.217 |

| | | | | | | | | | | | | | | | | | | | | | | | | | | | | | | | | | | |
|---|---|---|---|---|---|---|---|---|---|---|---|---|---|---|---|---|---|---|---|---|---|---|---|---|---|---|---|---|---|---|---|---|---|---|
| SE | 12900 | TYROBP | TYROBP | chr19 | - | 35907217 | 35907261 | 35904400 | 35904634 | 35907445 | 35907580 | 0 | 7 | 39 | 0 | 0 | 1 | 0 | 22 | 1 | 0 | 1 | 0 | 93 | 49 | 0.429733987 | 0.620451665 | NA | 1 | 0.954 | NA | 0.921 | 1 | 0.016 |
| SE | 679 | SARAF | SARAF | chr8 | - | 30073876 | 30074055 | 30069641 | 30070059 | 30082846 | 30083208 | 0 | 3 | 13 | 0 | 1 | 5 | 0 | 17 | 4 | 0 | 5 | 6 | 98 | 49 | 0.430454253 | 0.620561858 | NA | 0.6 | 0.565 | NA | 0.63 | 0.25 | 0.143 |
| SE | 5660 | KCNQ1 | KCNQ1 | chr11 | + | 2585211 | 2585307 | 2583434 | 2583545 | 2587569 | 2587692 | 0 | 0 | 4 | 0 | 1 | 0 | 0 | 1 | 1 | 0 | 0 | 0 | 98 | 49 | 0.430636073 | 0.620561858 | NA | 0 | 1 | NA | 1 | 1 | -0.5 |
| RI | 694 | UNC119 | UNC119 | chr17 | - | 28546706 | 28547849 | 28546706 | 28547409 | 28547676 | 28547849 | 0 | 1 | 10 | 0 | 0 | 1 | 2 | 3 | 2 | 1 | 0 | 0 | 98 | 49 | 0.431080918 | 0.934008656 | NA | 1 | 0.833 | 0.5 | 1 | 1 | 0.083 |
| A3SS | 3011 | ISG20 | ISG20 | chr15 | + | 88650234 | 88652310 | 88652109 | 88652310 | 88639342 | 88639594 | 0 | 0 | 2 | 0 | 0 | 11 | 0 | 3 | 1 | 0 | 1 | 2 | 98 | 49 | 0.432135137 | 0.670554523 | NA | NA | 0.083 | NA | 0.6 | 0.2 | -0.317 |
| SE | 13731 | SERPINA1 | SERPINA1 | chr14 | - | 94388559 | 94388660 | 94382591 | 94383241 | 94390456 | 94390692 | 0 | 1 | 16 | 0 | 2 | 20 | 4 | 5 | 1 | 0 | 5 | 0 | 98 | 49 | 0.432495196 | 0.621784006 | NA | 0.2 | 0.286 | 1 | 0.333 | 1 | -0.535 |
| SE | 1921 | PARP8 | PARP8 | chr5 | + | 50821212 | 50821338 | 50815431 | 50815524 | 50822334 | 50822400 | 0 | 2 | 9 | 0 | 0 | 1 | 0 | 4 | 1 | 0 | 0 | 0 | 98 | 49 | 0.432725263 | 0.621784006 | NA | 1 | 0.818 | NA | 1 | 1 | -0.091 |
| SE | 2181 | RBM47 | RBM47 | chr4 | - | 40437770 | 40438924 | 40432650 | 40432862 | 40466576 | 40466699 | 0 | 2 | 9 | 0 | 0 | 1 | 0 | 4 | 1 | 0 | 0 | 0 | 98 | 49 | 0.432725263 | 0.621784006 | NA | 1 | 0.818 | NA | 1 | 1 | -0.091 |
| A5SS | 1440 | MLKL | MLKL | chr16 | - | 74675323 | 74675404 | 74675354 | 74675404 | 74674959 | 74675100 | 1 | 1 | 1 | 0 | 5 | 3 | 0 | 3 | 0 | 1 | 1 | 0 | 80 | 49 | 0.432994858 | 0.667321487 | 1 | 0.109 | 0.17 | 0 | 0.648 | NA | 0.102 |
| SE | 4557 | SLC6A6 | SLC6A6 | chr3 | + | 14443623 | 14443863 | 14416411 | 14416453 | 14445716 | 14445851 | 0 | 5 | 30 | 0 | 0 | 1 | 0 | 16 | 5 | 0 | 1 | 0 | 98 | 49 | 0.434955535 | 0.624391756 | NA | 1 | 0.937 | NA | 0.889 | 1 | 0.024 |
| SE | 5786 | DENND3 | DENND3 | chr8 | + | 141150833 | 141150953 | 141138021 | 141138137 | 141151618 | 141151837 | 0 | 0 | 22 | 0 | 0 | 1 | 0 | 10 | 4 | 0 | 0 | 0 | 98 | 49 | 0.435694354 | 0.624855547 | NA | NA | 0.917 | NA | 1 | 1 | -0.083 |
| SE | 3704 | LYZ | LYZ | chr12 | + | 69352219 | 69352298 | 69350107 | 69350272 | 69353152 | 69354233 | 0 | 40 | 456 | 0 | 0 | 1 | 43 | 267 | 1 | 0 | 0 | 0 | 98 | 49 | 0.441484717 | 0.632034884 | NA | 1 | 0.996 | 1 | 1 | 1 | -0.002 |
| SE | 2699 | MBNL1 | MBNL1 | chr3 | + | 152299404 | 152300367 | 152268039 | 152269092 | 152414940 | 152415111 | 0 | 0 | 6 | 0 | 0 | 1 | 5 | 3 | 1 | 0 | 1 | 0 | 98 | 49 | 0.441541336 | 0.632034884 | NA | NA | 0.75 | 1 | 0.6 | 1 | -0.117 |
| SE | 13538 | ALOX5 | ALOX5 | chr10 | + | 45428617 | 45428764 | 45424959 | 45425132 | 45440429 | 45440633 | 0 | 9 | 46 | 0 | 0 | 1 | 0 | 19 | 4 | 0 | 0 | 0 | 98 | 49 | 0.442803979 | 0.633239183 | NA | 1 | 0.958 | NA | 1 | 1 | -0.021 |
| SE | 3207 | STX16 | STX16 | chr20 | + | 58669290 | 58669453 | 58667986 | 58668127 | 58670511 | 58670603 | 0 | 0 | 9 | 0 | 0 | 3 | 0 | 9 | 1 | 0 | 4 | 0 | 98 | 49 | 0.44356605 | 0.633726019 | NA | NA | 0.6 | NA | 0.529 | 1 | -0.164 |
| SE | 10590 | RNF213 | RNF213 | chr17 | + | 80379619 | 80379714 | 80377761 | 80377796 | 80380830 | 80380987 | 0 | 3 | 53 | 0 | 0 | 1 | 0 | 22 | 3 | 0 | 0 | 0 | 98 | 49 | 0.44486502 | 0.634978277 | NA | 1 | 0.964 | NA | 1 | 1 | -0.018 |
| A5SS | 2190 | HES4 | HES4 | chr1 | - | 999691 | 1000172 | 999865 | 1000172 | 999525 | 999613 | 1 | 1 | 1 | 0 | 1 | 1 | 3 | 11 | 4 | 0 | 4 | 1 | 98 | 49 | 0.445770867 | 0.679023064 | 1 | 0.333 | 0.333 | 1 | 0.579 | 0.667 | -0.193 |
| SE | 1983 | DDX60L | DDX60L | chr4 | - | 168448637 | 168448779 | 168441336 | 168441492 | 168453123 | 168453282 | 0 | 2 | 53 | 0 | 0 | 1 | 0 | 35 | 2 | 0 | 1 | 0 | 98 | 49 | 0.445818676 | 0.63573574 | NA | 1 | 0.964 | NA | 0.946 | 1 | 0.009 |
| SE | 4191 | FAM49B | FAM49B | chr8 | - | 129879388 | 129879471 | 129871374 | 129871496 | 129903311 | 129903350 | 0 | 0 | 14 | 0 | 0 | 1 | 4 | 13 | 4 | 0 | 4 | 0 | 98 | 49 | 0.449034561 | 0.639399819 | NA | NA | 0.875 | 1 | 0.619 | 1 | 0.002 |
| SE | 13835 | SPG21 | SPG21 | chr15 | - | 64983506 | 64983593 | 64980863 | 64981025 | 64989664 | 64989946 | 0 | 1 | 6 | 0 | 0 | 1 | 0 | 2 | 1 | 0 | 0 | 0 | 98 | 49 | 0.449609977 | 0.639399819 | NA | 1 | 0.75 | NA | 1 | 1 | -0.125 |
| SE | 11826 | ARHGDIB | ARHGDIB | chr12 | - | 14949801 | 14949885 | 14947872 | 14947949 | 14950531 | 14950724 | 0 | 13 | 90 | 0 | 0 | 1 | 12 | 34 | 7 | 0 | 0 | 0 | 98 | 49 | 0.44966441 | 0.639399819 | NA | 1 | 0.978 | 1 | 1 | 1 | -0.011 |
| RI | 780 | SHISA5 | SHISA5 | chr3 | - | 48467797 | 48469573 | 48467797 | 48469186 | 48469360 | 48469573 | 0 | 3 | 1 | 0 | 5 | 13 | 5 | 0 | 1 | 0 | 10 | 2 | 98 | 49 | 0.455153121 | 0.94773154 | NA | 0.231 | 0.037 | 1 | 0 | 0.2 | -0.266 |
| SE | 2102 | WDFY3 | WDFY3 | chr4 | - | 84817391 | 84817585 | 84809886 | 84810344 | 84820084 | 84820186 | 0 | 1 | 19 | 0 | 1 | 0 | 1 | 6 | 3 | 0 | 0 | 0 | 98 | 49 | 0.458220849 | 0.650950791 | NA | 0.333 | 1 | 1 | 1 | 1 | -0.334 |
| SE | 13718 | HMHA1 | HMHA1 | chr19 | + | 1068413 | 1068744 | 1067165 | 1067495 | 1073148 | 1073292 | 0 | 8 | 14 | 0 | 0 | 1 | 0 | 2 | 6 | 0 | 0 | 0 | 98 | 49 | 0.460039403 | 0.652917113 | NA | 1 | 0.875 | NA | 1 | 1 | -0.062 |
| SE | 8333 | NCSTN | NCSTN | chr1 | + | 160355654 | 160355757 | 160354117 | 160354290 | 160355862 | 160355958 | 0 | 1 | 4 | 0 | 0 | 1 | 0 | 1 | 1 | 0 | 0 | 0 | 98 | 49 | 0.46501066 | 0.659350021 | NA | 1 | 0.667 | NA | 1 | 1 | -0.166 |
| A3SS | 2675 | ABTB1 | ABTB1 | chr3 | + | 127674344 | 127674418 | 127674390 | 127674418 | 127672937 | 127673081 | 0 | 0 | 8 | 0 | 3 | 0 | 0 | 2 | 0 | 0 | 3 | 2 | 95 | 49 | 0.468058768 | 0.720090412 | NA | 0 | 1 | NA | 0.256 | 0 | 0.372 |
| SE | 1195 | MAP3K5 | MAP3K5 | chr6 | - | 136720449 | 136720589 | 136705109 | 136705133 | 136791709 | 136792518 | 0 | 1 | 12 | 0 | 0 | 1 | 0 | 4 | 1 | 0 | 0 | 0 | 98 | 49 | 0.46879248 | 0.66408586 | NA | 1 | 0.857 | NA | 1 | 1 | -0.072 |
| A5SS | 3016 | SERPINA1 | SERPINA1 | chr14 | - | 94390419 | 94390692 | 94390456 | 94390692 | 94382591 | 94383241 | 0 | 4 | 5 | 0 | 2 | 20 | 3 | 3 | 2 | 0 | 5 | 0 | 86 | 49 | 0.47090828 | 0.709068789 | NA | 0.533 | 0.125 | 1 | 0.255 | 1 | -0.423 |
| SE | 9545 | PRKCB | PRKCB | chr16 | + | 24172269 | 24172361 | 24154683 | 24154857 | 24174517 | 24174580 | 0 | 9 | 27 | 0 | 0 | 1 | 0 | 11 | 2 | 0 | 0 | 0 | 98 | 49 | 0.472851524 | 0.669205122 | NA | 1 | 0.931 | NA | 1 | 1 | -0.034 |
| A3SS | 1443 | TBC1D10C | TBC1D10C | chr11 | + | 67405902 | 67406017 | 67405906 | 67406017 | 67405594 | 67405701 | 0 | 0 | 7 | 0 | 0 | 2 | 0 | 0 | 1 | 0 | 0 | 0 | 53 | 49 | 0.472854846 | 0.721304002 | NA | NA | 0.764 | NA | NA | 1 | -0.236 |
| RI | 556 | CDC16 | CDC16 | chr13 | + | 114272183 | 114272675 | 114272183 | 114272481 | 114272563 | 114272675 | 0 | 0 | 7 | 0 | 0 | 2 | 0 | 1 | 1 | 0 | 0 | 0 | 98 | 49 | 0.47352014 | 0.94773154 | NA | NA | 0.636 | NA | 1 | 1 | -0.364 |
| SE | 11966 | MSL1 | MSL1 | chr17 | + | 40131536 | 40131584 | 40129244 | 40129627 | 40132033 | 40132098 | 0 | 0 | 7 | 0 | 0 | 2 | 0 | 4 | 1 | 0 | 1 | 0 | 97 | 49 | 0.474814393 | 0.671350925 | NA | NA | 0.639 | NA | 0.669 | 1 | -0.196 |
| SE | 2939 | LIMK2 | LIMK2 | chr22 | + | 31266983 | 31267070 | 31265945 | 31266132 | 31267775 | 31267907 | 0 | 0 | 20 | 0 | 0 | 1 | 0 | 34 | 5 | 0 | 0 | 0 | 98 | 49 | 0.475908434 | 0.671934183 | NA | NA | 0.909 | NA | 1 | 1 | -0.091 |
| SE | 1139 | IGF2R | IGF2R | chr6 | + | 160061752 | 160061928 | 160061502 | 160061646 | 160062531 | 160062619 | 0 | 5 | 27 | 0 | 0 | 1 | 0 | 13 | 8 | 0 | 1 | 0 | 98 | 49 | 0.476121028 | 0.671934183 | NA | 1 | 0.931 | NA | 0.867 | 1 | 0.032 |
| SE | 3270 | CTSA | CTSA | chr20 | + | 45893219 | 45893311 | 45892724 | 45892880 | 45893987 | 45894072 | 0 | 1 | 14 | 0 | 1 | 0 | 0 | 4 | 2 | 2 | 0 | 0 | 98 | 49 | 0.476656296 | 0.672058549 | NA | 0.333 | 1 | 0 | 1 | 1 | 0 |
| SE | 5812 | HAUS4 | HAUS4 | chr14 | - | 22951554 | 22951689 | 22950313 | 22950410 | 22952327 | 22952459 | 0 | 0 | 16 | 0 | 0 | 1 | 0 | 3 | 2 | 0 | 0 | 0 | 98 | 49 | 0.480086309 | 0.676260283 | NA | NA | 0.889 | NA | 1 | 1 | -0.111 |
| RI | 1055 | STX10 | STX10 | chr19 | - | 13144057 | 13144671 | 13144057 | 13144486 | 13144576 | 13144671 | 0 | 1 | 6 | 0 | 0 | 1 | 0 | 0 | 1 | 0 | 1 | 0 | 98 | 49 | 0.482241355 | 0.94773154 | NA | 1 | 0.75 | NA | 0 | 1 | 0.375 |
| A5SS | 2804 | TYROBP | TYROBP | chr19 | - | 35908167 | 35908309 | 35908174 | 35908309 | 35907729 | 35907762 | 0 | 1 | 10 | 0 | 0 | 1 | 0 | 3 | 4 | 0 | 1 | 0 | 56 | 49 | 0.482508271 | 0.718279358 | NA | 1 | 0.897 | NA | 0.724 | 1 | 0.087 |
| SE | 8138 | ITGAX | ITGAX | chr16 | + | 31371333 | 31371497 | 31371083 | 31371214 | 31371629 | 31371784 | 0 | 1 | 13 | 0 | 1 | 0 | 0 | 10 | 6 | 0 | 3 | 1 | 98 | 49 | 0.482836455 | 0.67949737 | NA | 0.333 | 1 | NA | 0.625 | 0.75 | -0.021 |
| SE | 12743 | DYSF | DYSF | chr2 | + | 71549349 | 71549391 | 71539156 | 71539239 | 71551040 | 71551156 | 0 | 2 | 2 | 0 | 1 | 11 | 0 | 2 | 1 | 0 | 4 | 0 | 91 | 49 | 0.489990933 | 0.688920834 | NA | 0.519 | 0.089 | NA | 0.212 | 1 | -0.302 |
| A3SS | 1013 | INPP5D | INPP5D | chr2 | + | 233125744 | 233125919 | 233125747 | 233125919 | 233122106 | 233122257 | 1 | 0 | 7 | 0 | 1 | 5 | 0 | 5 | 0 | 1 | 2 | 0 | 52 | 49 | 0.490380444 | 0.74107425 | 1 | 0 | 0.569 | 0 | 0.702 | NA | 0.172 |
| SE | 10208 | RFWD2 | RFWD2 | chr1 | - | 176135009 | 176135086 | 176116623 | 176116681 | 176149005 | 176149074 | 0 | 1 | 21 | 0 | 0 | 1 | 0 | 7 | 1 | 0 | 0 | 0 | 98 | 49 | 0.493056216 | 0.692582704 | NA | 1 | 0.913 | NA | 1 | 1 | -0.043 |
| RI | 322 | MAP7D1 | MAP7D1 | chr1 | + | 36176198 | 36176581 | 36176198 | 36176321 | 36176417 | 36176581 | 0 | 1 | 8 | 0 | 1 | 1 | 3 | 5 | 6 | 0 | 1 | 0 | 98 | 49 | 0.496928876 | 0.94773154 | NA | 0.333 | 0.8 | 1 | 0.714 | 1 | -0.338 |
| SE | 4467 | CLEC12A | CLEC12A | chr12 | + | 9978965 | 9979064 | 9971408 | 9971687 | 9979335 | 9979524 | 0 | 0 | 25 | 0 | 0 | 3 | 0 | 0 | 2 | 0 | 0 | 0 | 98 | 49 | 0.497638055 | 0.696506389 | NA | NA | 0.806 | NA | NA | 1 | -0.194 |
| SE | 4625 | SORL1 | SORL1 | chr11 | + | 121545242 | 121545429 | 121543547 | 121543726 | 121549959 | 121550088 | 0 | 6 | 54 | 0 | 0 | 1 | 0 | 38 | 2 | 0 | 1 | 0 | 98 | 49 | 0.497995567 | 0.696506389 | NA | 1 | 0.964 | NA | 0.95 | 1 | 0.007 |
| A3SS | 1731 | SECISBP2 | SECISBP2 | chr9 | + | 89325426 | 89325676 | 89325547 | 89325676 | 89319651 | 89319797 | 0 | 0 | 4 | 0 | 0 | 1 | 0 | 4 | 1 | 0 | 0 | 0 | 98 | 49 | 0.498166579 | 0.74107425 | NA | NA | 0.667 | NA | 1 | 1 | -0.333 |
| A3SS | 3445 | PHOSPHO1 | PHOSPHO1 | chr17 | - | 49223369 | 49225131 | 49223369 | 49225004 | 49226646 | 49226758 | 0 | 0 | 4 | 0 | 0 | 1 | 0 | 2 | 1 | 0 | 0 | 0 | 98 | 49 | 0.498166579 | 0.74107425 | NA | NA | 0.667 | NA | 1 | 1 | -0.333 |
| RI | 392 | PRKD2 | PRKD2 | chr19 | - | 46716130 | 46717127 | 46716130 | 46716535 | 46716950 | 46717127 | 0 | 0 | 4 | 0 | 0 | 1 | 0 | 0 | 1 | 0 | 0 | 0 | 98 | 49 | 0.498166579 | 0.94773154 | NA | NA | 0.667 | NA | NA | 1 | -0.333 |
| SE | 3691 | ARHGEF40 | ARHGEF40 | chr14 | + | 21087319 | 21087463 | 21087000 | 21087105 | 21087967 | 21088098 | 0 | 0 | 4 | 0 | 0 | 1 | 0 | 0 | 1 | 0 | 0 | 0 | 98 | 49 | 0.498166579 | 0.696506389 | NA | NA | 0.667 | NA | NA | 1 | -0.333 |
| SE | 6752 | UBR4 | UBR4 | chr1 | - | 19165655 | 19165757 | 19165248 | 19165349 | 19167021 | 19167231 | 0 | 0 | 4 | 0 | 0 | 1 | 0 | 4 | 1 | 0 | 1 | 0 | 98 | 49 | 0.498166579 | 0.696506389 | NA | NA | 0.667 | NA | 0.667 | 1 | -0.166 |
| SE | 7087 | PRDX5 | PRDX5 | chr11 | + | 64320660 | 64320792 | 64319733 | 64319868 | 64320864 | 64320903 | 0 | 0 | 4 | 0 | 0 | 1 | 0 | 5 | 1 | 0 | 0 | 0 | 98 | 49 | 0.498166579 | 0.696506389 | NA | NA | 0.667 | NA | 1 | 1 | -0.333 |
| SE | 4125 | XPO6 | XPO6 | chr16 | - | 28177219 | 28177332 | 28175897 | 28176095 | 28180940 | 28181031 | 0 | 3 | 53 | 0 | 0 | 1 | 0 | 35 | 8 | 0 | 1 | 0 | 98 | 49 | 0.499162452 | 0.69697095 | NA | 1 | 0.964 | NA | 0.946 | 1 | 0.009 |
| SE | 13537 | ALOX5 | ALOX5 | chr10 | + | 45395854 | 45395936 | 45382482 | 45382681 | 45412190 | 45412313 | 0 | 5 | 67 | 0 | 0 | 1 | 1 | 17 | 7 | 0 | 0 | 0 | 98 | 49 | 0.49942629 | 0.69697095 | NA | 1 | 0.971 | 1 | 1 | 1 | -0.014 |
| A5SS | 2491 | ZNF276 | ZNF276 | chr16 | + | 89727278 | 89727357 | 89727278 | 89727325 | 89729234 | 89729318 | 0 | 0 | 8 | 0 | 0 | 2 | 0 | 3 | 1 | 0 | 4 | 0 | 81 | 49 | 0.50024699 | 0.736318604 | NA | NA | 0.708 | NA | 0.312 | 1 | 0.052 |
| SE | 10650 | SLC43A2 | SLC43A2 | chr17 | - | 1591268 | 1591471 | 1590801 | 1590948 | 1591565 | 1591699 | 0 | 5 | 19 | 0 | 0 | 1 | 0 | 5 | 8 | 0 | 1 | 0 | 98 | 49 | 0.500994407 | 0.698510756 | NA | 1 | 0.905 | NA | 0.714 | 1 | 0.096 |
| A3SS | 1434 | PICALM | PICALM | chr11 | - | 85990249 | 85990399 | 85990249 | 85990378 | 85996825 | 85996929 | 0 | 0 | 18 | 0 | 0 | 1 | 0 | 21 | 4 | 0 | 1 | 0 | 70 | 49 | 0.504274672 | 0.744011811 | NA | NA | 0.926 | NA | 0.936 | 1 | -0.042 |
| SE | 12757 | PKN1 | PKN1 | chr19 | + | 14469751 | 14469814 | 14469340 | 14469517 | 14469911 | 14469988 | 0 | 1 | 5 | 0 | 0 | 1 | 2 | 1 | 1 | 0 | 0 | 0 | 98 | 49 | 0.507272183 | 0.706608055 | NA | 1 | 0.714 | 1 | 1 | 1 | -0.143 |
| SE | 6543 | NRD1 | NRD1 | chr1 | - | 51800555 | 51800683 | 51798248 | 51798411 | 51803813 | 51803964 | 0 | 0 | 18 | 0 | 0 | 1 | 0 | 10 | 4 | 0 | 0 | 0 | 98 | 49 | 0.507742839 | 0.706608784 | NA | NA | 0.9 | NA | 1 | 1 | -0.1 |
| SE | 1150 | SOD2 | SOD2 | chr6 | - | 159688125 | 159688242 | 159684853 | 159685033 | 159692660 | 159692863 | 3 | 7 | 82 | 0 | 2 | 0 | 18 | 158 | 13 | 0 | 13 | 1 | 98 | 49 | 0.509005729 | 0.707237006 | 1 | 0.636 | 1 | 1 | 0.859 | 0.867 | -0.03 |
| SE | 13859 | MMP25 | MMP25 | chr16 | + | 3057309 | 3057394 | 3057032 | 3057209 | 3057530 | 3057613 | 0 | 1 | 33 | 0 | 0 | 1 | 3 | 8 | 3 | 0 | 0 | 0 | 98 | 49 | 0.509135356 | 0.707237006 | NA | 1 | 0.943 | 1 | 1 | 1 | -0.028 |
| SE | 8685 | PTPRC | PTPRC | chr1 | + | 198699563 | 198699704 | 198692346 | 198692373 | 198703297 | 198703372 | 0 | 3 | 18 | 0 | 0 | 25 | 2 | 15 | 1 | 0 | 11 | 6 | 98 | 49 | 0.509721028 | 0.707396773 | NA | 1 | 0.265 | 1 | 0.405 | 0.077 | 0.139 |
| SE | 8376 | OAS2 | OAS2 | chr12 | + | 113004933 | 113005222 | 113002931 | 113003102 | 113006412 | 113006600 | 0 | 2 | 43 | 0 | 0 | 1 | 0 | 11 | 3 | 0 | 0 | 0 | 98 | 49 | 0.512035658 | 0.709568442 | NA | 1 | 0.956 | NA | 1 | 1 | -0.022 |
| SE | 4905 | PAK1 | PAK1 | chr11 | - | 77332729 | 77332867 | 77322014 | 77323360 | 77336085 | 77336282 | 0 | 2 | 22 | 0 | 1 | 0 | 0 | 7 | 1 | 0 | 0 | 0 | 98 | 49 | 0.512585144 | 0.709568442 | NA | 0.5 | 1 | NA | 1 | 1 | -0.25 |
| SE | 5056 | SRSF3 | SRSF3 | chr6 | + | 36599820 | 36600276 | 36598848 | 36598983 | 36601151 | 36601190 | 0 | 1 | 11 | 0 | 0 | 4 | 0 | 7 | 1 | 0 | 1 | 0 | 98 | 49 | 0.512702147 | 0.709568442 | NA | 1 | 0.579 | NA | 0.778 | 1 | -0.1 |
| A5SS | 3021 | MAP4K4 | MAP4K4 | chr2 | + | 101887087 | 101887261 | 101887087 | 101887237 | 101887777 | 101887937 | 0 | 0 | 2 | 0 | 0 | 5 | 0 | 5 | 1 | 0 | 2 | 1 | 73 | 49 | 0.51637545 | 0.751613155 | NA | NA | 0.212 | NA | 0.627 | 0.402 | -0.302 |
| SE | 2564 | HK3 | HK3 | chr5 | - | 176887200 | 176887337 | 176887001 | 176887121 | 176887450 | 176887746 | 0 | 10 | 17 | 0 | 1 | 0 | 2 | 5 | 2 | 0 | 0 | 0 | 98 | 49 | 0.517806019 | 0.715972812 | NA | 0.833 | 1 | 1 | 1 | 1 | -0.084 |
| SE | 12585 | SBNO2 | SBNO2 | chr19 | - | 1112188 | 1112301 | 1111995 | 1112067 | 1112401 | 1112537 | 0 | 8 | 15 | 0 | 4 | 0 | 0 | 8 | 3 | 0 | 2 | 1 | 98 | 49 | 0.521310981 | 0.720156622 | NA | 0.5 | 1 | NA | 0.667 | 0.6 | 0.117 |
| SE | 4558 | SLC6A6 | SLC6A6 | chr3 | + | 14443627 | 14443863 | 14416411 | 14416453 | 14445716 | 14445851 | 0 | 5 | 14 | 0 | 0 | 1 | 0 | 9 | 4 | 0 | 1 | 0 | 98 | 49 | 0.523615544 | 0.722675999 | NA | 1 | 0.875 | NA | 0.818 | 1 | 0.028 |
| SE | 5960 | LILRA2 | LILRA2 | chr19 | + | 54574075 | 54574111 | 54573697 | 54573912 | 54574300 | 54574582 | 0 | 0 | 5 | 0 | 0 | 1 | 1 | 1 | 1 | 0 | 0 | 0 | 85 | 49 | 0.5248053 | 0.723653547 | NA | NA | 0.742 | 1 | 1 | 1 | -0.258 |
| SE | 7023 | HPCAL1 | HPCAL1 | chr2 | + | 10408561 | 10408734 | 10396834 | 10396920 | 10419733 | 10420135 | 0 | 0 | 2 | 0 | 2 | 7 | 0 | 2 | 0 | 0 | 1 | 2 | 98 | 49 | 0.527686141 | 0.726959 | NA | 0 | 0.125 | NA | 0.5 | 0 | -0.188 |
| SE | 9067 | ERGIC1 | ERGIC1 | chr5 | + | 172909666 | 172909761 | 172897001 | 172897074 | 172914713 | 172914838 | 0 | 1 | 13 | 0 | 0 | 1 | 0 | 1 | 3 | 0 | 0 | 0 | 98 | 49 | 0.528398477 | 0.727273728 | NA | 1 | 0.867 | NA | 1 | 1 | -0.067 |
| SE | 11047 | HLA-B | HLA-B | chr6 | - | 31355106 | 31355223 | 31354632 | 31354665 | 31355316 | 31355592 | 0 | 46 | 261 | 0 | 2 | 4 | 10 | 220 | 15 | 0 | 4 | 2 | 98 | 49 | 0.529614267 | 0.728280186 | NA | 0.92 | 0.97 | 1 | 0.965 | 0.789 | 0.027 |
| SE | 7715 | THEMIS2 | THEMIS2 | chr1 | + | 27876587 | 27876728 | 27872542 | 27872665 | 27879643 | 27879963 | 0 | 13 | 31 | 0 | 0 | 1 | 7 | 17 | 23 | 0 | 0 | 2 | 98 | 49 | 0.530434112 | 0.728740832 | NA | 1 | 0.939 | 1 | 1 | 0.852 | 0.019 |
| A5SS | 2259 | TLE3 | TLE3 | chr15 | - | 70058149 | 70058291 | 70058158 | 70058291 | 70057458 | 70057658 | 0 | 0 | 2 | 0 | 1 | 4 | 0 | 2 | 0 | 0 | 3 | 1 | 58 | 49 | 0.531185703 | 0.764673924 | NA | 0 | 0.297 | NA | 0.36 | 0 | -0.032 |
| A3SS | 1208 | SDF4 | SDF4 | chr1 | - | 1216907 | 1217804 | 1216907 | 1217688 | 1218457 | 1218633 | 1 | 0 | 0 | 0 | 0 | 4 | 1 | 0 | 1 | 2 | 0 | 0 | 98 | 49 | 0.53213915 | 0.762573929 | 1 | NA | 0 | 0.2 | NA | 1 | -0.1 |
| SE | 4662 | AMICA1 | AMICA1 | chr11 | - | 118214823 | 118214886 | 118212406 | 118212561 | 118224940 | 118225094 | 0 | 7 | 17 | 0 | 1 | 5 | 0 | 13 | 2 | 0 | 2 | 0 | 98 | 49 | 0.537451897 | 0.737707946 | NA | 0.778 | 0.63 | NA | 0.765 | 1 | -0.179 |
| A3SS | 1184 | DDIT3 | DDIT3 | chr12 | - | 57517268 | 57517519 | 57517268 | 57517438 | 57517705 | 57517753 | 0 | 0 | 5 | 0 | 0 | 1 | 0 | 11 | 1 | 0 | 2 | 0 | 98 | 49 | 0.538906714 | 0.762573929 | NA | NA | 0.714 | NA | 0.733 | 1 | -0.153 |
| A3SS | 1185 | DDIT3 | DDIT3 | chr12 | - | 57517268 | 57517542 | 57517268 | 57517438 | 57517705 | 57517753 | 0 | 0 | 5 | 0 | 0 | 1 | 0 | 11 | 1 | 0 | 2 | 0 | 98 | 49 | 0.538906714 | 0.762573929 | NA | NA | 0.714 | NA | 0.733 | 1 | -0.153 |
| A3SS | 1186 | DDIT3 | DDIT3 | chr12 | - | 57517268 | 57517595 | 57517268 | 57517438 | 57517705 | 57517753 | 0 | 0 | 5 | 0 | 0 | 1 | 0 | 12 | 1 | 0 | 2 | 0 | 98 | 49 | 0.538906714 | 0.762573929 | NA | NA | 0.714 | NA | 0.75 | 1 | -0.161 |
| SE | 8577 | RC3H1 | RC3H1 | chr1 | - | 173964838 | 173965120 | 173963972 | 173964187 | 173970504 | 173970617 | 0 | 0 | 5 | 0 | 0 | 1 | 0 | 5 | 1 | 0 | 0 | 0 | 98 | 49 | 0.538906714 | 0.737743862 | NA | NA | 0.714 | NA | 1 | 1 | -0.286 |
| SE | 1576 | TRIM38 | TRIM38 | chr6 | + | 25973053 | 25973076 | 25971868 | 25972099 | 25973172 | 25973285 | 0 | 0 | 5 | 0 | 0 | 1 | 0 | 9 | 1 | 0 | 1 | 0 | 72 | 49 | 0.539026433 | 0.737743862 | NA | NA | 0.773 | NA | 0.86 | 1 | -0.157 |
| SE | 3235 | SLC9A8 | SLC9A8 | chr20 | + | 49849578 | 49849680 | 49845035 | 49845119 | 49855437 | 49855581 | 0 | 0 | 10 | 0 | 0 | 2 | 0 | 6 | 1 | 0 | 2 | 0 | 98 | 49 | 0.539441453 | 0.737743862 | NA | NA | 0.714 | NA | 0.6 | 1 | -0.086 |
| SE | 5886 | RASSF5 | RASSF5 | chr1 | + | 206585179 | 206585295 | 206584386 | 206584684 | 206586825 | 206589284 | 0 | 0 | 10 | 0 | 0 | 2 | 0 | 10 | 1 | 0 | 6 | 0 | 98 | 49 | 0.539441453 | 0.737743862 | NA | NA | 0.714 | NA | 0.455 | 1 | -0.014 |
| A3SS | 3256 | FLII | FLII | chr17 | - | 18251679 | 18251816 | 18251679 | 18251813 | 18251998 | 18252146 | 0 | 2 | 6 | 0 | 2 | 3 | 0 | 1 | 0 | 0 | 2 | 0 | 52 | 49 | 0.542191316 | 0.762573929 | NA | 0.485 | 0.653 | NA | 0.32 | NA | 0.249 |
| A3SS | 116 | AGTPBP1 | AGTPBP1 | chr9 | - | 85655142 | 85655320 | 85655142 | 85655200 | 85657434 | 85657643 | 0 | 0 | 20 | 0 | 0 | 4 | 0 | 8 | 1 | 0 | 2 | 0 | 98 | 49 | 0.542274794 | 0.762573929 | NA | NA | 0.714 | NA | 0.667 | 1 | -0.12 |
| SE | 4629 | SORL1 | SORL1 | chr11 | + | 121605401 | 121605571 | 121605112 | 121605239 | 121606844 | 121606957 | 0 | 11 | 57 | 0 | 0 | 1 | 0 | 45 | 4 | 0 | 1 | 0 | 98 | 49 | 0.544440604 | 0.743903843 | NA | 1 | 0.966 | NA | 0.957 | 1 | 0.005 |
| SE | 2099 | WDFY3 | WDFY3 | chr4 | - | 84726860 | 84726911 | 84724425 | 84724594 | 84733381 | 84733609 | 0 | 5 | 2 | 0 | 0 | 3 | 0 | 5 | 3 | 0 | 1 | 0 | 98 | 49 | 0.545895829 | 0.745214742 | NA | 1 | 0.25 | NA | 0.714 | 1 | -0.232 |
| SE | 6349 | FLI1 | FLI1 | chr11 | + | 128807179 | 128807239 | 128805365 | 128805431 | 128809156 | 128809204 | 0 | 1 | 11 | 0 | 0 | 1 | 1 | 2 | 1 | 0 | 0 | 0 | 98 | 49 | 0.547491166 | 0.746714358 | NA | 1 | 0.846 | 1 | 1 | 1 | -0.077 |
| A5SS | 1194 | ETF1 | ETF1 | chr5 | - | 138542633 | 138542936 | 138542832 | 138542936 | 138518691 | 138518867 | 0 | 0 | 2 | 0 | 2 | 4 | 0 | 4 | 3 | 0 | 3 | 0 | 98 | 49 | 0.547780994 | 0.779992502 | NA | 0 | 0.2 | NA | 0.4 | 1 | -0.6 |
| SE | 1134 | IGF2R | IGF2R | chr6 | + | 160029549 | 160029655 | 160027184 | 160027314 | 160032550 | 160032713 | 0 | 14 | 45 | 0 | 0 | 4 | 0 | 27 | 6 | 0 | 1 | 0 | 98 | 49 | 0.549943569 | 0.749379133 | NA | 1 | 0.849 | NA | 0.931 | 1 | -0.041 |
| SE | 11237 | APLP2 | APLP2 | chr11 | + | 130123611 | 130123779 | 130122304 | 130122513 | 130126699 | 130126830 | 0 | 0 | 33 | 0 | 0 | 3 | 0 | 5 | 2 | 0 | 0 | 0 | 98 | 49 | 0.550569813 | 0.749552925 | NA | NA | 0.846 | NA | 1 | 1 | -0.154 |
| SE | 8917 | IRF9 | IRF9 | chr14 | + | 24163877 | 24163959 | 24163377 | 24163508 | 24164062 | 24164134 | 0 | 3 | 25 | 0 | 1 | 5 | 0 | 7 | 3 | 0 | 8 | 0 | 98 | 49 | 0.551132735 | 0.749640272 | NA | 0.6 | 0.714 | NA | 0.304 | 1 | 0.005 |
| SE | 5788 | DENND3 | DENND3 | chr8 | + | 141160631 | 141160787 | 141155848 | 141155970 | 141165185 | 141165289 | 0 | 0 | 11 | 0 | 0 | 1 | 0 | 5 | 2 | 0 | 0 | 0 | 98 | 49 | 0.551645974 | 0.749659945 | NA | NA | 0.846 | NA | 1 | 1 | -0.154 |
| RI | 675 | PSME1 | PSME1 | chr14 | + | 24138473 | 24138967 | 24138473 | 24138560 | 24138735 | 24138967 | 0 | 0 | 2 | 0 | 0 | 11 | 3 | 0 | 0 | 0 | 6 | 1 | 98 | 49 | 0.557006469 | 0.999202116 | NA | NA | 0.083 | 1 | 0 | 0 | -0.25 |
| A3SS | 2017 | RNF19B | RNF19B | chr1 | - | 32936445 | 32937309 | 32936445 | 32937259 | 32938396 | 32938528 | 0 | 1 | 9 | 0 | 0 | 5 | 0 | 3 | 2 | 0 | 5 | 0 | 98 | 49 | 0.558523697 | 0.779335391 | NA | 1 | 0.474 | NA | 0.231 | 1 | 0.121 |
| SE | 13539 | ALOX5 | ALOX5 | chr10 | + | 45444115 | 45444286 | 45443727 | 45443828 | 45445507 | 45446119 | 0 | 2 | 41 | 0 | 0 | 2 | 1 | 4 | 7 | 0 | 0 | 1 | 98 | 49 | 0.559152751 | 0.759174873 | NA | 1 | 0.911 | 1 | 1 | 0.778 | 0.029 |
| SE | 4964 | DDX3X | DDX3X | chrX | + | 41343736 | 41343822 | 41343215 | 41343351 | 41344029 | 41344128 | 0 | 4 | 36 | 0 | 2 | 0 | 0 | 34 | 1 | 0 | 4 | 1 | 98 | 49 | 0.565342278 | 0.76688578 | NA | 0.5 | 1 | NA | 0.81 | 0.333 | 0.178 |
| A5SS | 520 | TNFAIP3 | TNFAIP3 | chr6 | + | 137867187 | 137867542 | 137867187 | 137867494 | 137871212 | 137871522 | 0 | 3 | 6 | 0 | 2 | 0 | 0 | 9 | 4 | 0 | 1 | 0 | 97 | 49 | 0.570399439 | 0.803465876 | NA | 0.431 | 1 | NA | 0.82 | 1 | -0.194 |
| A3SS | 1102 | FKBP8 | FKBP8 | chr19 | - | 18538215 | 18538436 | 18538215 | 18538433 | 18539370 | 18539459 | 0 | 2 | 2 | 0 | 0 | 5 | 0 | 2 | 1 | 2 | 5 | 0 | 52 | 49 | 0.570761829 | 0.79028561 | NA | 1 | 0.274 | 0 | 0.274 | 1 | 0.212 |
| SE | 7496 | IFI44 | IFI44 | chr1 | + | 78660553 | 78660654 | 78659311 | 78659483 | 78662703 | 78662878 | 0 | 0 | 36 | 0 | 0 | 6 | 0 | 23 | 1 | 0 | 5 | 0 | 98 | 49 | 0.575224839 | 0.778766917 | NA | NA | 0.75 | NA | 0.697 | 1 | -0.099 |
| SE | 8238 | LBR | LBR | chr1 | - | 225410290 | 225410416 | 225406663 | 225406832 | 225411336 | 225411440 | 0 | 0 | 18 | 0 | 0 | 3 | 1 | 10 | 1 | 0 | 1 | 0 | 98 | 49 | 0.575643233 | 0.778766917 | NA | NA | 0.75 | 1 | 0.833 | 1 | -0.194 |
| SE | 11307 | SP100 | SP100 | chr2 | + | 230464066 | 230464150 | 230462434 | 230462518 | 230466300 | 230466354 | 0 | 0 | 25 | 0 | 0 | 2 | 0 | 12 | 2 | 0 | 2 | 0 | 98 | 49 | 0.575655386 | 0.778766917 | NA | NA | 0.862 | NA | 0.75 | 1 | -0.013 |
| A5SS | 3019 | SERPINA1 | SERPINA1 | chr14 | - | 94390419 | 94390692 | 94390456 | 94390692 | 94388559 | 94388663 | 0 | 4 | 5 | 0 | 0 | 4 | 3 | 3 | 2 | 0 | 1 | 0 | 86 | 49 | 0.578375981 | 0.804333654 | NA | 1 | 0.416 | 1 | 0.631 | 1 | -0.169 |
| A5SS | 1552 | PGS1 | PGS1 | chr17 | + | 78419545 | 78419725 | 78419545 | 78419675 | 78424060 | 78424659 | 0 | 10 | 32 | 0 | 0 | 1 | 0 | 27 | 6 | 0 | 1 | 0 | 98 | 49 | 0.583295398 | 0.804333654 | NA | 1 | 0.941 | NA | 0.931 | 1 | 0.005 |
| SE | 6362 | ZSWIM8 | ZSWIM8 | chr10 | + | 73791323 | 73791499 | 73790974 | 73791176 | 73791858 | 73792852 | 0 | 0 | 6 | 0 | 1 | 0 | 0 | 1 | 0 | 0 | 0 | 1 | 98 | 49 | 0.584480703 | 0.789995051 | NA | 0 | 1 | NA | 1 | 0 | 0 |
| SE | 3417 | RALGAPA2 | RALGAPA2 | chr20 | - | 20676235 | 20676288 | 20653529 | 20653587 | 20680690 | 20680801 | 0 | 0 | 13 | 0 | 0 | 2 | 0 | 5 | 1 | 0 | 0 | 0 | 98 | 49 | 0.588967448 | 0.794630228 | NA | NA | 0.765 | NA | 1 | 1 | -0.235 |
| SE | 3803 | ZCCHC2 | ZCCHC2 | chr18 | + | 62558686 | 62558770 | 62556202 | 62556297 | 62560586 | 62560644 | 0 | 0 | 13 | 0 | 0 | 2 | 5 | 11 | 1 | 0 | 0 | 0 | 98 | 49 | 0.588967448 | 0.794630228 | NA | NA | 0.765 | 1 | 1 | 1 | -0.235 |
| SE | 1922 | PARP8 | PARP8 | chr5 | + | 50824907 | 50824975 | 50822334 | 50822400 | 50826754 | 50826803 | 0 | 0 | 14 | 0 | 0 | 1 | 0 | 5 | 2 | 0 | 0 | 0 | 98 | 49 | 0.592731575 | 0.797560929 | NA | NA | 0.875 | NA | 1 | 1 | -0.125 |
| SE | 3771 | KIAA1109 | KIAA1109 | chr4 | + | 122356620 | 122356755 | 122355795 | 122355990 | 122359556 | 122359722 | 0 | 0 | 14 | 0 | 0 | 1 | 0 | 7 | 2 | 0 | 0 | 0 | 98 | 49 | 0.592731575 | 0.797560929 | NA | NA | 0.875 | NA | 1 | 1 | -0.125 |
| SE | 4732 | CDA | CDA | chr1 | + | 20604927 | 20605039 | 20588950 | 20589283 | 20618451 | 20618907 | 0 | 0 | 14 | 0 | 0 | 1 | 0 | 5 | 2 | 0 | 0 | 0 | 98 | 49 | 0.592731575 | 0.797560929 | NA | NA | 0.875 | NA | 1 | 1 | -0.125 |
| SE | 7937 | TMC6 | TMC6 | chr17 | - | 78124888 | 78124985 | 78124523 | 78124781 | 78125157 | 78125263 | 1 | 4 | 2 | 0 | 1 | 0 | 0 | 3 | 3 | 0 | 2 | 0 | 98 | 49 | 0.596766039 | 0.802271339 | 1 | 0.667 | 1 | NA | 0.429 | 1 | 0.174 |
| A5SS | 2516 | NLRP1 | NLRP1 | chr17 | - | 5533291 | 5533384 | 5533303 | 5533384 | 5532821 | 5532984 | 0 | 0 | 1 | 0 | 0 | 7 | 0 | 5 | 0 | 0 | 4 | 1 | 61 | 49 | 0.597621998 | 0.815505018 | NA | NA | 0.103 | NA | 0.501 | 0 | -0.148 |
| SE | 11312 | ALDOA | ALDOA | chr16 | + | 30066884 | 30067038 | 30065672 | 30065927 | 30067233 | 30067366 | 0 | 0 | 1 | 0 | 0 | 7 | 0 | 0 | 0 | 0 | 5 | 1 | 98 | 49 | 0.59822268 | 0.803510892 | NA | NA | 0.067 | NA | 0 | 0 | 0.067 |
| A3SS | 1568 | ENTPD1 | ENTPD1 | chr10 | + | 95847445 | 95847706 | 95847568 | 95847706 | 95845356 | 95845596 | 0 | 4 | 26 | 0 | 0 | 1 | 2 | 4 | 1 | 0 | 0 | 0 | 98 | 49 | 0.600161541 | 0.824649445 | NA | 1 | 0.929 | 1 | 1 | 1 | -0.035 |
| SE | 4503 | PTPN6 | PTPN6 | chr12 | + | 6955150 | 6955267 | 6954804 | 6954994 | 6955371 | 6955485 | 0 | 2 | 28 | 0 | 0 | 1 | 0 | 24 | 1 | 0 | 1 | 0 | 98 | 49 | 0.600780878 | 0.805924887 | NA | 1 | 0.933 | NA | 0.923 | 1 | 0.005 |
| SE | 2258 | KIAA0226 | KIAA0226 | chr3 | - | 197717976 | 197718130 | 197705091 | 197705175 | 197736654 | 197736926 | 0 | 0 | 2 | 0 | 1 | 0 | 0 | 3 | 1 | 0 | 0 | 1 | 98 | 49 | 0.601092347 | 0.805924887 | NA | 0 | 1 | NA | 1 | 0.333 | -0.166 |
| SE | 2636 | PFKFB4 | PFKFB4 | chr3 | - | 48539253 | 48539310 | 48538497 | 48538619 | 48539696 | 48539771 | 0 | 0 | 7 | 0 | 0 | 1 | 0 | 1 | 1 | 0 | 0 | 0 | 98 | 49 | 0.602141516 | 0.806612031 | NA | NA | 0.778 | NA | 1 | 1 | -0.222 |
| SE | 9030 | MYD88 | MYD88 | chr3 | + | 38140387 | 38140568 | 38139863 | 38139998 | 38140756 | 38140848 | 0 | 1 | 15 | 0 | 1 | 1 | 0 | 0 | 2 | 0 | 0 | 0 | 98 | 49 | 0.608707381 | 0.814681384 | NA | 0.333 | 0.882 | NA | NA | 1 | -0.392 |
| SE | 4423 | PREX1 | PREX1 | chr20 | - | 48650893 | 48651055 | 48649995 | 48650206 | 48651395 | 48651583 | 0 | 3 | 32 | 0 | 0 | 2 | 0 | 37 | 5 | 0 | 1 | 0 | 98 | 49 | 0.611249391 | 0.817355724 | NA | 1 | 0.889 | NA | 0.949 | 1 | -0.03 |
| SE | 8705 | CTSS | CTSS | chr1 | - | 150750005 | 150750171 | 150747776 | 150747879 | 150751780 | 150752008 | 0 | 1 | 129 | 0 | 0 | 1 | 5 | 24 | 1 | 0 | 0 | 0 | 98 | 49 | 0.612568689 | 0.818391768 | NA | 1 | 0.985 | 1 | 1 | 1 | -0.008 |
| SE | 8707 | CTSS | CTSS | chr1 | - | 150755000 | 150755150 | 150751780 | 150752008 | 150757857 | 150757980 | 0 | 4 | 234 | 0 | 0 | 1 | 6 | 39 | 7 | 0 | 0 | 0 | 98 | 49 | 0.614855148 | 0.820716951 | NA | 1 | 0.992 | 1 | 1 | 1 | -0.004 |
| SE | 56 | SASH3 | SASH3 | chrX | + | 129788430 | 129788577 | 129787974 | 129788070 | 129790939 | 129791081 | 0 | 0 | 8 | 0 | 0 | 1 | 0 | 12 | 1 | 0 | 1 | 0 | 98 | 49 | 0.622026252 | 0.829552314 | NA | NA | 0.8 | NA | 0.857 | 1 | -0.128 |
| SE | 11031 | STAT1 | STAT1 | chr2 | - | 190998216 | 190998308 | 190997855 | 190998007 | 190999625 | 190999704 | 0 | 0 | 33 | 0 | 0 | 2 | 0 | 14 | 2 | 0 | 4 | 0 | 98 | 49 | 0.624060418 | 0.831527312 | NA | NA | 0.892 | NA | 0.636 | 1 | 0.074 |
| RI | 30 | PLIN2 | PLIN2 | chr9 | - | 19126113 | 19126448 | 19126113 | 19126309 | 19126396 | 19126448 | 2 | 0 | 1 | 0 | 1 | 0 | 4 | 1 | 0 | 0 | 1 | 0 | 98 | 49 | 0.627465042 | 0.999202116 | 1 | 0 | 1 | 1 | 0.333 | NA | 0 |
| SE | 12347 | CSNK1D | CSNK1D | chr17 | - | 82253015 | 82253244 | 82252433 | 82252604 | 82255428 | 82255577 | 0 | 2 | 3 | 0 | 1 | 0 | 0 | 4 | 7 | 0 | 1 | 0 | 98 | 49 | 0.630476403 | 0.839332182 | NA | 0.5 | 1 | NA | 0.667 | 1 | -0.084 |
| SE | 11030 | STAT1 | STAT1 | chr2 | - | 190979771 | 190979866 | 190978855 | 190979001 | 190980619 | 190980669 | 0 | 1 | 20 | 0 | 1 | 2 | 0 | 2 | 0 | 0 | 3 | 0 | 98 | 49 | 0.633284155 | 0.842323969 | NA | 0.333 | 0.833 | NA | 0.25 | NA | 0.333 |
| SE | 381 | GNAQ | GNAQ | chr9 | - | 77794462 | 77794592 | 77728513 | 77728667 | 77797519 | 77797648 | 0 | 0 | 9 | 0 | 0 | 1 | 0 | 2 | 1 | 0 | 0 | 0 | 98 | 49 | 0.642008309 | 0.852419159 | NA | NA | 0.818 | NA | 1 | 1 | -0.182 |
| SE | 8334 | NCSTN | NCSTN | chr1 | + | 160355862 | 160355958 | 160354117 | 160354290 | 160356259 | 160356347 | 0 | 0 | 9 | 0 | 0 | 1 | 0 | 2 | 1 | 0 | 2 | 0 | 98 | 49 | 0.642008309 | 0.852419159 | NA | NA | 0.818 | NA | 0.333 | 1 | 0.151 |
| SE | 5678 | GMIP | GMIP | chr19 | - | 19635643 | 19635721 | 19635414 | 19635569 | 19636706 | 19636796 | 0 | 9 | 8 | 1 | 1 | 0 | 0 | 5 | 1 | 0 | 0 | 1 | 98 | 49 | 0.642808803 | 0.852728712 | 0 | 0.818 | 1 | NA | 1 | 0.333 | -0.06 |
| SE | 8358 | ARHGAP30 | ARHGAP30 | chr1 | - | 161056387 | 161056532 | 161054622 | 161054705 | 161069527 | 161069970 | 0 | 0 | 18 | 0 | 0 | 2 | 0 | 11 | 1 | 0 | 2 | 0 | 98 | 49 | 0.643507131 | 0.852902309 | NA | NA | 0.818 | NA | 0.733 | 1 | -0.049 |
| SE | 4961 | DDX3X | DDX3X | chrX | + | 41337407 | 41337465 | 41333307 | 41334297 | 41341483 | 41341616 | 0 | 2 | 10 | 0 | 0 | 2 | 0 | 23 | 2 | 0 | 2 | 0 | 98 | 49 | 0.648092299 | 0.858222665 | NA | 1 | 0.714 | NA | 0.852 | 1 | -0.069 |
| SE | 11305 | SP100 | SP100 | chr2 | + | 230450171 | 230450255 | 230449087 | 230449150 | 230461261 | 230461414 | 0 | 2 | 23 | 0 | 1 | 0 | 8 | 15 | 5 | 0 | 1 | 0 | 98 | 49 | 0.652122993 | 0.862079722 | NA | 0.5 | 1 | 1 | 0.882 | 1 | -0.211 |
| SE | 4428 | PREX1 | PREX1 | chr20 | - | 48747808 | 48747880 | 48745024 | 48745147 | 48827641 | 48827883 | 0 | 20 | 45 | 0 | 0 | 1 | 2 | 45 | 9 | 0 | 1 | 0 | 98 | 49 | 0.652532646 | 0.862079722 | NA | 1 | 0.957 | 1 | 0.957 | 1 | -0.007 |
| SE | 9514 | SIPA1 | SIPA1 | chr11 | + | 65645853 | 65645957 | 65644954 | 65645129 | 65646220 | 65646378 | 0 | 0 | 19 | 0 | 0 | 2 | 0 | 5 | 1 | 0 | 0 | 0 | 98 | 49 | 0.652725698 | 0.862079722 | NA | NA | 0.826 | NA | 1 | 1 | -0.174 |
| RI | 230 | ATG3 | ATG3 | chr3 | - | 112532506 | 112534337 | 112532506 | 112532780 | 112534268 | 112534337 | 0 | 0 | 3 | 0 | 3 | 5 | 0 | 1 | 0 | 0 | 4 | 0 | 98 | 49 | 0.654603078 | 0.999202116 | NA | 0 | 0.231 | NA | 0 | 1 | -0.385 |
| SE | 6617 | AMPD2 | AMPD2 | chr1 | + | 109628642 | 109628806 | 109628363 | 109628495 | 109629108 | 109629235 | 0 | 1 | 2 | 0 | 1 | 0 | 0 | 7 | 2 | 0 | 5 | 1 | 98 | 49 | 0.656411303 | 0.866186294 | NA | 0.333 | 1 | NA | 0.412 | 0.5 | 0.211 |
| RI | 644 | ATP6V0B | ATP6V0B | chr1 | + | 43977025 | 43978300 | 43977025 | 43977216 | 43977980 | 43978300 | 3 | 0 | 3 | 0 | 4 | 15 | 0 | 0 | 4 | 1 | 7 | 3 | 98 | 49 | 0.657605162 | 0.999202116 | 1 | 0 | 0.091 | 0 | 0 | 0.4 | 0.23 |
| SE | 2839 | TYMP | TYMP | chr22 | - | 50526000 | 50526141 | 50525752 | 50525918 | 50526245 | 50526476 | 1 | 8 | 30 | 0 | 1 | 0 | 2 | 8 | 14 | 0 | 0 | 1 | 98 | 49 | 0.661103457 | 0.871612716 | 1 | 0.8 | 1 | 1 | 1 | 0.875 | -0.025 |
| SE | 3332 | SAMHD1 | SAMHD1 | chr20 | - | 36935028 | 36935189 | 36930759 | 36930875 | 36941038 | 36941111 | 0 | 0 | 21 | 0 | 0 | 1 | 0 | 3 | 2 | 0 | 1 | 0 | 98 | 49 | 0.661813092 | 0.871783591 | NA | NA | 0.913 | NA | 0.6 | 1 | 0.113 |
| SE | 8706 | CTSS | CTSS | chr1 | - | 150751780 | 150752008 | 150750005 | 150750171 | 150755000 | 150755150 | 0 | 3 | 181 | 0 | 0 | 1 | 7 | 24 | 1 | 0 | 0 | 0 | 98 | 49 | 0.66301558 | 0.872602817 | NA | 1 | 0.989 | 1 | 1 | 1 | -0.006 |
| SE | 11325 | ITGAL | ITGAL | chr16 | + | 30510880 | 30510960 | 30510360 | 30510471 | 30511049 | 30511136 | 0 | 2 | 13 | 0 | 0 | 1 | 0 | 1 | 1 | 0 | 0 | 0 | 98 | 49 | 0.665452763 | 0.875044185 | NA | 1 | 0.867 | NA | 1 | 1 | -0.067 |
| SE | 7596 | ATG16L2 | ATG16L2 | chr11 | + | 72827187 | 72827293 | 72826702 | 72826823 | 72828358 | 72828508 | 0 | 1 | 41 | 0 | 0 | 1 | 1 | 4 | 1 | 0 | 0 | 0 | 98 | 49 | 0.668079787 | 0.8777307 | NA | 1 | 0.953 | 1 | 1 | 1 | -0.024 |

| | | | | | | | | | | | | | | | | | | | | | | | | | | | | | | | | | | |
|---|---|---|---|---|---|---|---|---|---|---|---|---|---|---|---|---|---|---|---|---|---|---|---|---|---|---|---|---|---|---|---|---|---|---|
| SE | 9411 | WDFY4 | WDFY4 | chr10 | + | 48760346 | 48760440 | 48742967 | 48743548 | 48774457 | 48774672 | 0 | 0 | 11 | 0 | 0 | 1 | 0 | 1 | 1 | 0 | 0 | 1 | 98 | 49 | 0.668749011 | 0.877842588 | NA | NA | 0.846 | NA | 1 | 1 | -0.154 |
| A5SS | 1756 | PSME1 | PSME1 | chr14 | + | 24138473 | 24138568 | 24138473 | 24138560 | 24138735 | 24138967 | 0 | 0 | 1 | 0 | 0 | 11 | 0 | 0 | 0 | 0 | 6 | 1 | 57 | 49 | 0.670811132 | 0.905940807 | NA | NA | 0.072 | NA | 0 | 0 | 0.072 |
| SE | 4459 | CLEC7A | CLEC7A | chr12 | - | 10125086 | 10125173 | 10123244 | 10123363 | 10125296 | 10125448 | 0 | 1 | 25 | 0 | 0 | 8 | 0 | 7 | 4 | 0 | 4 | 0 | 98 | 49 | 0.674199264 | 0.884224689 | NA | 1 | 0.61 | NA | 0.467 | 1 | 0.071 |
| SE | 7535 | MCL1 | MCL1 | chr1 | - | 150578243 | 150578491 | 150574550 | 150577491 | 150578842 | 150578963 | 0 | 5 | 123 | 1 | 4 | 15 | 1 | 70 | 5 | 0 | 30 | 0 | 98 | 49 | 0.678459838 | 0.889036737 | 0 | 0.385 | 0.804 | 1 | 0.538 | 1 | -0.45 |
| SE | 8082 | CSF3R | CSF3R | chr1 | - | 36473434 | 36473622 | 36472516 | 36472686 | 36473763 | 36473887 | 0 | 19 | 197 | 0 | 1 | 3 | 20 | 148 | 32 | 0 | 2 | 1 | 98 | 49 | 0.679743226 | 0.889942569 | NA | 0.905 | 0.97 | 1 | 0.974 | 0.941 | -0.034 |
| A5SS | 2261 | TLE3 | TLE3 | chr15 | - | 70058158 | 70058291 | 70058185 | 70058291 | 70057458 | 70057658 | 1 | 1 | 4 | 0 | 0 | 1 | 0 | 4 | 1 | 0 | 1 | 0 | 76 | 49 | 0.682465604 | 0.91227545 | 1 | 1 | 0.721 | NA | 0.721 | 1 | 0.046 |
| SE | 6482 | PADI2 | PADI2 | chr1 | - | 17079263 | 17079415 | 17075678 | 17075823 | 17082544 | 17082652 | 0 | 0 | 12 | 0 | 0 | 1 | 0 | 0 | 1 | 0 | 0 | 0 | 98 | 49 | 0.685696462 | 0.895643369 | NA | NA | 0.857 | NA | NA | 1 | -0.143 |
| SE | 6819 | OAS1 | OAS1 | chr12 | + | 112917546 | 112917700 | 112916508 | 112916738 | 112919486 | 112919907 | 0 | 0 | 12 | 0 | 0 | 1 | 0 | 4 | 1 | 0 | 1 | 0 | 98 | 49 | 0.685696462 | 0.895643369 | NA | NA | 0.857 | NA | 0.667 | 1 | 0.023 |
| SE | 8684 | PTPRC | PTPRC | chr1 | + | 198696711 | 198696909 | 198692346 | 198692373 | 198699563 | 198699704 | 0 | 0 | 1 | 0 | 0 | 12 | 0 | 3 | 0 | 1 | 4 | 1 | 98 | 49 | 0.685885241 | 0.895643369 | NA | NA | 0.04 | 0 | 0.273 | 0 | -0.051 |
| SE | 3887 | LCP1 | LCP1 | chr13 | - | 46148351 | 46148447 | 46144441 | 46144520 | 46150935 | 46151078 | 0 | 0 | 37 | 0 | 0 | 3 | 0 | 8 | 1 | 0 | 0 | 0 | 98 | 49 | 0.687307489 | 0.89672149 | NA | NA | 0.86 | NA | 1 | 1 | -0.14 |
| A3SS | 803 | TYMP | TYMP | chr22 | - | 50526000 | 50526156 | 50526000 | 50526141 | 50526245 | 50526476 | 0 | 2 | 0 | 0 | 5 | 13 | 1 | 0 | 6 | 2 | 5 | 5 | 64 | 49 | 0.687426061 | 0.937399174 | NA | 0.234 | 0 | 0.277 | 0 | 0.479 | -0.135 |
| SE | 13917 | CD300A | CD300A | chr17 | + | 74481725 | 74481833 | 74481288 | 74481326 | 74484000 | 74484798 | 0 | 4 | 15 | 0 | 0 | 1 | 0 | 6 | 0 | 0 | 0 | 1 | 98 | 49 | 0.688861707 | 0.89796977 | NA | 1 | 0.882 | NA | 1 | 0 | 0.441 |
| SE | 2765 | LST1 | LST1 | chr6 | + | 31587640 | 31587733 | 31587199 | 31587318 | 31588562 | 31588909 | 0 | 0 | 2 | 0 | 1 | 6 | 0 | 1 | 0 | 0 | 2 | 1 | 98 | 49 | 0.690552858 | 0.899394233 | NA | 0 | 0.143 | NA | 0.2 | 0 | -0.029 |
| SE | 1145 | IGF2R | IGF2R | chr6 | + | 160088032 | 160088147 | 160083949 | 160084184 | 160089106 | 160089253 | 0 | 0 | 13 | 0 | 0 | 1 | 0 | 2 | 1 | 0 | 0 | 0 | 98 | 49 | 0.699215366 | 0.909888048 | NA | NA | 0.867 | NA | 1 | 1 | -0.133 |
| SE | 13865 | ACSL1 | ACSL1 | chr4 | - | 184770398 | 184770476 | 184768315 | 184768450 | 184773080 | 184773154 | 0 | 1 | 55 | 0 | 0 | 1 | 4 | 29 | 0 | 0 | 0 | 0 | 98 | 49 | 0.700770989 | 0.911123526 | NA | 1 | 0.965 | 1 | 1 | NA | -0.018 |
| SE | 12581 | CNN2 | CNN2 | chr19 | + | 1036066 | 1036246 | 1032558 | 1032696 | 1036415 | 1036562 | 0 | 0 | 15 | 0 | 0 | 1 | 0 | 12 | 1 | 0 | 0 | 0 | 98 | 49 | 0.713986072 | 0.927503083 | NA | NA | 0.882 | NA | 1 | 1 | -0.118 |
| SE | 8658 | DUSP1 | DUSP1 | chr5 | - | 172769574 | 172769794 | 172768089 | 172769132 | 172770585 | 172771200 | 0 | 6 | 110 | 0 | 1 | 0 | 3 | 47 | 10 | 0 | 2 | 0 | 98 | 49 | 0.7163398 | 0.929757097 | NA | 0.75 | 1 | 1 | 0.922 | 1 | -0.099 |
| A3SS | 775 | LST1 | LST1 | chr6 | + | 31588496 | 31588909 | 31588562 | 31588909 | 31587199 | 31587318 | 0 | 3 | 13 | 0 | 1 | 6 | 4 | 11 | 0 | 0 | 2 | 1 | 98 | 49 | 0.718836501 | 0.97286143 | NA | 0.6 | 0.52 | 1 | 0.733 | 0 | -0.018 |
| SE | 11238 | APLP2 | APLP2 | chr11 | + | 130137255 | 130137291 | 130135562 | 130135715 | 130140397 | 130140483 | 0 | 1 | 13 | 0 | 1 | 11 | 0 | 3 | 3 | 0 | 10 | 0 | 85 | 49 | 0.719023331 | 0.932434915 | NA | 0.366 | 0.405 | NA | 0.147 | 1 | -0.188 |
| SE | 1142 | IGF2R | IGF2R | chr6 | + | 160072764 | 160072884 | 160071909 | 160072036 | 160073212 | 160073469 | 0 | 1 | 16 | 0 | 0 | 1 | 0 | 18 | 2 | 0 | 1 | 0 | 98 | 49 | 0.724649909 | 0.938921391 | NA | 1 | 0.889 | NA | 0.9 | 1 | -0.005 |
| SE | 1037 | AOAH | AOAH | chr7 | - | 36548611 | 36548686 | 36540318 | 36540491 | 36549438 | 36549475 | 0 | 0 | 16 | 0 | 0 | 1 | 0 | 3 | 1 | 0 | 0 | 0 | 98 | 49 | 0.727128809 | 0.941321792 | NA | NA | 0.889 | NA | 1 | 1 | -0.111 |
| RI | 251 | TYMP | TYMP | chr22 | - | 50529495 | 50530085 | 50529495 | 50529731 | 50529903 | 50530085 | 1 | 7 | 17 | 0 | 0 | 1 | 7 | 16 | 4 | 0 | 1 | 0 | 98 | 49 | 0.727141068 | 0.999202116 | 1 | 1 | 0.895 | 1 | 0.889 | 1 | 0.002 |
| SE | 5784 | DENND3 | DENND3 | chr8 | + | 141141202 | 141141324 | 141138021 | 141138137 | 141144147 | 141144259 | 0 | 0 | 17 | 0 | 0 | 1 | 2 | 10 | 1 | 0 | 0 | 0 | 98 | 49 | 0.73240269 | 0.947333256 | NA | NA | 0.895 | 1 | 1 | 1 | -0.105 |
| SE | 5221 | CLEC4E | CLEC4E | chr12 | - | 8536089 | 8536205 | 8533304 | 8534809 | 8539216 | 8539306 | 0 | 0 | 35 | 0 | 0 | 1 | 0 | 8 | 2 | 0 | 0 | 0 | 98 | 49 | 0.736969319 | 0.952420366 | NA | NA | 0.946 | NA | 1 | 1 | -0.054 |
| A5SS | 2690 | ARHGDIA | ARHGDIA | chr17 | - | 81869329 | 81869625 | 81869541 | 81869625 | 81869172 | 81869236 | 1 | 3 | 16 | 0 | 0 | 1 | 0 | 7 | 3 | 0 | 0 | 2 | 98 | 49 | 0.739861649 | 0.968908208 | 1 | 1 | 0.889 | NA | 1 | 0.429 | 0.249 |
| A5SS | 2132 | RFWD2 | RFWD2 | chr1 | - | 176163814 | 176163891 | 176163826 | 176163891 | 176162868 | 176162988 | 0 | 1 | 17 | 0 | 1 | 5 | 0 | 5 | 0 | 0 | 6 | 0 | 61 | 49 | 0.745584037 | 0.968908208 | NA | 0.445 | 0.732 | NA | 0.401 | NA | 0.188 |
| A3SS | 1291 | SLC6A6 | SLC6A6 | chr3 | + | 14443623 | 14443863 | 14443627 | 14443863 | 14416411 | 14416453 | 0 | 0 | 19 | 0 | 0 | 1 | 0 | 9 | 1 | 0 | 0 | 0 | 53 | 49 | 0.746035627 | 0.999202116 | NA | NA | 0.946 | NA | 1 | 1 | -0.054 |
| A5SS | 2703 | IFI6 | IFI6 | chr1 | - | 27669232 | 27669346 | 27669244 | 27669346 | 27668457 | 27668535 | 0 | 0 | 2 | 0 | 1 | 20 | 0 | 1 | 0 | 0 | 9 | 1 | 61 | 49 | 0.747020832 | 0.968908208 | NA | 0 | 0.074 | NA | 0.082 | 0 | -0.004 |
| SE | 862 | NAMPT | NAMPT | chr7 | - | 106254363 | 106254504 | 106253016 | 106253151 | 106261587 | 106261707 | 0 | 24 | 228 | 0 | 0 | 4 | 14 | 336 | 15 | 0 | 3 | 0 | 98 | 49 | 0.748257683 | 0.966178091 | NA | 1 | 0.966 | 1 | 0.982 | 1 | -0.011 |
| SE | 7082 | UBR2 | UBR2 | chr6 | + | 42676055 | 42676191 | 42674125 | 42674193 | 42676782 | 42676873 | 0 | 0 | 20 | 0 | 0 | 1 | 0 | 6 | 1 | 0 | 0 | 0 | 98 | 49 | 0.751735186 | 0.96983518 | NA | NA | 0.909 | NA | 1 | 1 | -0.091 |
| SE | 8416 | GRAMD1A | GRAMD1A | chr19 | + | 35014188 | 35014387 | 35013540 | 35013691 | 35015823 | 35015967 | 0 | 3 | 13 | 0 | 0 | 0 | 0 | 1 | 0 | 0 | 0 | 2 | 98 | 49 | 0.752841225 | 0.970277462 | NA | 1 | 1 | NA | 1 | 0 | 0.5 |
| SE | 4184 | STAT6 | STAT6 | chr12 | - | 57107604 | 57107743 | 57107230 | 57107314 | 57108162 | 57108299 | 0 | 1 | 13 | 0 | 1 | 5 | 0 | 3 | 1 | 0 | 3 | 0 | 98 | 49 | 0.753369127 | 0.970277462 | NA | 0.333 | 0.565 | NA | 0.333 | 1 | -0.218 |
| SE | 3752 | ARHGAP27 | ARHGAP27 | chr17 | - | 45404268 | 45404334 | 45404028 | 45404096 | 45404444 | 45404528 | 0 | 2 | 4 | 0 | 0 | 3 | 0 | 0 | 1 | 0 | 5 | 0 | 98 | 49 | 0.759645396 | 0.976832296 | NA | 1 | 0.4 | NA | 0 | 1 | 0.2 |
| SE | 9027 | MYD88 | MYD88 | chr3 | + | 38139863 | 38139998 | 38138477 | 38139028 | 38140387 | 38140568 | 0 | 5 | 19 | 0 | 2 | 0 | 0 | 11 | 3 | 0 | 2 | 0 | 98 | 49 | 0.759758452 | 0.976832296 | NA | 0.556 | 1 | NA | 0.733 | 1 | -0.089 |
| SE | 2840 | TYMP | TYMP | chr22 | - | 50526000 | 50526156 | 50525752 | 50525918 | 50526245 | 50526476 | 1 | 3 | 17 | 0 | 1 | 0 | 0 | 3 | 9 | 0 | 0 | 1 | 98 | 49 | 0.762114536 | 0.979024058 | 1 | 0.6 | 1 | NA | 1 | 0.818 | -0.042 |
| A3SS | 3416 | STAT3 | STAT3 | chr17 | - | 42348388 | 42348539 | 42348388 | 42348517 | 42388278 | 42388495 | 0 | 2 | 14 | 0 | 1 | 6 | 0 | 2 | 1 | 0 | 2 | 0 | 71 | 49 | 0.764799772 | 0.999202116 | NA | 0.58 | 0.617 | NA | 0.408 | 1 | -0.105 |
| RI | 356 | DDIT3 | DDIT3 | chr12 | - | 57517268 | 57517753 | 57517268 | 57517438 | 57517705 | 57517753 | 1 | 2 | 7 | 0 | 0 | 1 | 2 | 28 | 3 | 0 | 2 | 0 | 98 | 49 | 0.777903504 | 0.999202116 | 1 | 1 | 0.778 | 1 | 0.875 | 1 | -0.032 |
| SE | 9233 | RPS24 | RPS24 | chr10 | + | 78040200 | 78040225 | 78037193 | 78037304 | 78040614 | 78040715 | 0 | 0 | 1 | 0 | 0 | 26 | 0 | 8 | 0 | 0 | 6 | 1 | 74 | 49 | 0.781756708 | 0.999202116 | NA | NA | 0.025 | NA | 0.469 | 0 | -0.209 |
| RI | 733 | CELF2 | CELF2 | chr10 | + | 11328925 | 11336673 | 11328925 | 11329097 | 11330455 | 11336673 | 0 | 0 | 29 | 0 | 0 | 1 | 1 | 10 | 1 | 0 | 1 | 0 | 98 | 49 | 0.794160768 | 0.999202116 | NA | NA | 0.935 | 1 | 0.833 | 1 | -0.009 |
| SE | 8415 | GRAMD1A | GRAMD1A | chr19 | + | 35013255 | 35013368 | 35011473 | 35011554 | 35013540 | 35013691 | 0 | 1 | 5 | 0 | 0 | 4 | 0 | 0 | 1 | 0 | 1 | 0 | 98 | 49 | 0.795544674 | 0.999202116 | NA | 1 | 0.385 | NA | 0 | 1 | 0.193 |
| RI | 1027 | ARHGDIA | ARHGDIA | chr17 | - | 81869329 | 81869625 | 81869329 | 81869406 | 81869541 | 81869625 | 1 | 5 | 15 | 0 | 0 | 13 | 5 | 7 | 6 | 0 | 8 | 1 | 98 | 49 | 0.797337051 | 0.999202116 | 1 | 1 | 0.366 | 1 | 0.304 | 0.75 | 0.104 |
| SE | 11387 | NLRC5 | NLRC5 | chr16 | + | 57054750 | 57054840 | 57051537 | 57051621 | 57055031 | 57055094 | 0 | 3 | 12 | 0 | 0 | 1 | 0 | 9 | 11 | 0 | 1 | 0 | 98 | 49 | 0.799295763 | 0.999202116 | NA | 1 | 0.857 | NA | 0.818 | 1 | 0.019 |
| A5SS | 2652 | ACOX1 | ACOX1 | chr17 | - | 75978790 | 75979434 | 75978964 | 75979434 | 75978533 | 75978693 | 0 | 0 | 0 | 0 | 2 | 2 | 1 | 2 | 0 | 0 | 5 | 1 | 98 | 49 | 0.808907745 | 0.999208248 | NA | 0 | 0 | 1 | 1 | 0 | -0.667 |
| A3SS | 3230 | ARRB2 | ARRB2 | chr17 | + | 4720549 | 4720640 | 4720585 | 4720640 | 4720392 | 4720472 | 0 | 0 | 2 | 0 | 2 | 13 | 0 | 1 | 0 | 2 | 5 | 8 | 85 | 49 | 0.814988864 | 0.999202116 | NA | 0 | 0.081 | 0 | 0.103 | 0 | 0.006 |
| SE | 1331 | TREM1 | TREM1 | chr6 | - | 41280960 | 41281153 | 41275260 | 41276230 | 41282394 | 41282751 | 0 | 0 | 37 | 0 | 0 | 1 | 0 | 38 | 1 | 0 | 3 | 0 | 98 | 49 | 0.817902743 | 0.999202116 | NA | NA | 0.949 | NA | 0.864 | 1 | 0.017 |
| SE | 8687 | PTPRC | PTPRC | chr1 | + | 198749415 | 198749549 | 198748108 | 198748199 | 198750491 | 198750626 | 0 | 1 | 72 | 0 | 0 | 1 | 9 | 51 | 0 | 0 | 0 | 0 | 98 | 49 | 0.818606032 | 0.999202116 | NA | 1 | 0.973 | 1 | 1 | NA | -0.014 |
| SE | 6603 | CCDC88B | CCDC88B | chr11 | + | 64354003 | 64354170 | 64353714 | 64353813 | 64355193 | 64355400 | 0 | 4 | 4 | 0 | 1 | 2 | 0 | 2 | 2 | 0 | 0 | 1 | 98 | 49 | 0.823294092 | 0.999202116 | NA | 0.667 | 0.5 | NA | 1 | 0.5 | -0.166 |
| A5SS | 1623 | ADAR | ADAR | chr1 | - | 154590183 | 154590409 | 154590261 | 154590409 | 154589756 | 154589928 | 0 | 4 | 24 | 0 | 0 | 3 | 0 | 17 | 2 | 0 | 1 | 0 | 98 | 49 | 0.823441962 | 0.999208248 | NA | 1 | 0.8 | NA | 0.895 | 1 | -0.047 |
| SE | 5377 | SYNE1 | SYNE1 | chr6 | - | 152300640 | 152300781 | 152293959 | 152294127 | 152301868 | 152302063 | 0 | 0 | 1 | 0 | 1 | 1 | 0 | 5 | 0 | 0 | 0 | 1 | 98 | 49 | 0.827502283 | 0.999202116 | NA | 0 | 0.333 | NA | 1 | 0 | -0.334 |
| SE | 4122 | XPO6 | XPO6 | chr16 | - | 28117317 | 28117462 | 28112903 | 28113050 | 28121669 | 28121762 | 0 | 1 | 55 | 0 | 0 | 1 | 8 | 46 | 0 | 0 | 0 | 0 | 98 | 49 | 0.827647446 | 0.999202116 | NA | 1 | 0.965 | 1 | 1 | NA | -0.018 |
| SE | 12729 | PILRA | PILRA | chr7 | + | 100389887 | 100390106 | 100374043 | 100374433 | 100397878 | 100397912 | 0 | 4 | 21 | 0 | 0 | 5 | 7 | 11 | 3 | 0 | 2 | 0 | 98 | 49 | 0.837221542 | 0.999202116 | NA | 1 | 0.677 | 1 | 0.733 | 1 | -0.073 |
| SE | 11878 | MGAM | MGAM | chr7 | + | 142064383 | 142064522 | 142063498 | 142063586 | 142065334 | 142065468 | 0 | 4 | 8 | 0 | 0 | 2 | 3 | 8 | 2 | 0 | 1 | 0 | 98 | 49 | 0.840992744 | 0.999202116 | NA | 1 | 0.667 | 1 | 0.8 | 1 | -0.1 |
| A3SS | 1500 | ADAM8 | ADAM8 | chr10 | - | 133273253 | 133273443 | 133273253 | 133273430 | 133273761 | 133273838 | 0 | 6 | 11 | 0 | 0 | 2 | 2 | 5 | 3 | 0 | 1 | 0 | 62 | 49 | 0.846278337 | 0.999202116 | NA | 1 | 0.813 | 1 | 0.798 | 1 | -0.026 |
| A3SS | 537 | DBNL | DBNL | chr7 | + | 44058428 | 44058480 | 44058431 | 44058480 | 44058128 | 44058280 | 0 | 0 | 6 | 0 | 3 | 2 | 0 | 1 | 0 | 0 | 1 | 1 | 52 | 49 | 0.847104365 | 0.999202116 | NA | 0 | 0.739 | NA | 0.485 | 0 | 0.127 |
| SE | 9515 | SIPA1 | SIPA1 | chr11 | + | 65650400 | 65650479 | 65650137 | 65650192 | 65650568 | 65650920 | 0 | 0 | 9 | 0 | 1 | 0 | 0 | 4 | 2 | 0 | 0 | 2 | 98 | 49 | 0.852675535 | 0.999202116 | NA | 0 | 1 | NA | 1 | 0.333 | -0.166 |
| SE | 6906 | FCGR3B | FCGR3B | chr1 | - | 161626144 | 161626402 | 161623197 | 161624639 | 161629777 | 161630032 | 0 | 0 | 59 | 0 | 0 | 1 | 2 | 19 | 1 | 0 | 2 | 0 | 98 | 49 | 0.85537265 | 0.999202116 | NA | NA | 0.967 | 1 | 0.826 | 1 | 0.025 |
| A3SS | 3415 | STAT3 | STAT3 | chr17 | - | 42316788 | 42316901 | 42316788 | 42316851 | 42317181 | 42317224 | 3 | 2 | 12 | 0 | 0 | 4 | 0 | 8 | 2 | 0 | 2 | 0 | 98 | 49 | 0.866784393 | 0.999202116 | 1 | 1 | 0.6 | NA | 0.667 | 1 | 0.033 |
| SE | 2906 | RAF1 | RAF1 | chr3 | - | 12590797 | 12590974 | 12587590 | 12587637 | 12591707 | 12591792 | 1 | 1 | 24 | 0 | 0 | 2 | 4 | 25 | 0 | 0 | 0 | 0 | 98 | 49 | 0.867186844 | 0.999202116 | 1 | 1 | 0.857 | 1 | 1 | NA | -0.048 |
| SE | 144 | IL2RG | IL2RG | chrX | - | 71108598 | 71108695 | 71108276 | 71108346 | 71109227 | 71109390 | 0 | 1 | 4 | 0 | 2 | 0 | 0 | 12 | 1 | 0 | 10 | 0 | 98 | 49 | 0.876241607 | 0.999202116 | NA | 0.2 | 1 | NA | 0.375 | 1 | -0.088 |
| SE | 4365 | LRRK2 | LRRK2 | chr12 | + | 40322034 | 40322181 | 40321033 | 40321188 | 40322318 | 40322510 | 0 | 1 | 45 | 0 | 0 | 2 | 0 | 21 | 0 | 0 | 0 | 0 | 98 | 49 | 0.881723484 | 0.999202116 | NA | 1 | 0.918 | NA | 1 | NA | -0.041 |
| RI | 809 | ZFP36L1 | ZFP36L1 | chr14 | - | 68787654 | 68790492 | 68787654 | 68788809 | 68789526 | 68790492 | 0 | 4 | 17 | 0 | 0 | 2 | 8 | 27 | 5 | 0 | 2 | 0 | 98 | 49 | 0.883440896 | 0.999202116 | NA | 1 | 0.81 | 1 | 0.871 | 1 | -0.052 |
| SE | 9293 | ABCA7 | ABCA7 | chr19 | + | 1054569 | 1054694 | 1054192 | 1054341 | 1054779 | 1054878 | 0 | 1 | 4 | 0 | 0 | 3 | 0 | 4 | 1 | 0 | 2 | 0 | 98 | 49 | 0.891129146 | 0.999202116 | NA | 1 | 0.4 | NA | 0.5 | 1 | -0.05 |
| SE | 6059 | HCLS1 | HCLS1 | chr3 | - | 121647318 | 121647448 | 121644817 | 121644928 | 121657278 | 121657352 | 0 | 1 | 33 | 0 | 0 | 1 | 3 | 18 | 0 | 0 | 0 | 0 | 98 | 49 | 0.89175776 | 0.999202116 | NA | 1 | 0.943 | 1 | 1 | NA | -0.028 |
| SE | 9137 | CLK1 | CLK1 | chr2 | - | 200854615 | 200854695 | 200853902 | 200853993 | 200855003 | 200855086 | 0 | 1 | 8 | 0 | 0 | 2 | 0 | 18 | 0 | 0 | 0 | 0 | 98 | 49 | 0.89175776 | 0.999202116 | NA | 1 | 0.667 | NA | 1 | NA | -0.166 |
| A3SS | 1625 | RPS3 | RPS3 | chr11 | + | 75402303 | 75402446 | 75402351 | 75402446 | 75401639 | 75401733 | 0 | 0 | 2 | 0 | 2 | 7 | 3 | 0 | 0 | 0 | 15 | 0 | 97 | 49 | 0.906255181 | 0.999202116 | NA | 0 | 0.126 | 1 | 0 | NA | -0.437 |
| SE | 1126 | RNASET2 | RNASET2 | chr6 | - | 166948569 | 166948625 | 166946681 | 166946739 | 166952487 | 166952548 | 0 | 8 | 51 | 0 | 1 | 0 | 0 | 18 | 2 | 0 | 1 | 0 | 98 | 49 | 0.909429153 | 0.999202116 | NA | 0.8 | 1 | NA | 0.9 | 1 | -0.05 |
| SE | 11303 | SP100 | SP100 | chr2 | + | 230440475 | 230440557 | 230417590 | 230417665 | 230442936 | 230443099 | 0 | 0 | 1 | 0 | 1 | 3 | 0 | 0 | 0 | 0 | 7 | 0 | 98 | 49 | 0.909814497 | 0.999202116 | NA | 0 | 0.143 | NA | 0 | NA | 0.071 |
| SE | 1834 | ARRDC3 | ARRDC3 | chr5 | - | 91376620 | 91376768 | 91375510 | 91375613 | 91378693 | 91378775 | 0 | 1 | 13 | 0 | 0 | 6 | 0 | 13 | 0 | 0 | 0 | 0 | 98 | 49 | 0.910129019 | 0.999202116 | NA | 1 | 0.52 | NA | 1 | NA | -0.24 |
| SE | 11821 | SSH2 | SSH2 | chr17 | - | 29650653 | 29650800 | 29648143 | 29648344 | 29655560 | 29655607 | 0 | 1 | 32 | 0 | 0 | 3 | 0 | 13 | 0 | 0 | 0 | 0 | 98 | 49 | 0.910129019 | 0.999202116 | NA | 1 | 0.842 | NA | 1 | NA | -0.079 |
| A3SS | 3715 | TYROBP | TYROBP | chr19 | - | 35907217 | 35907264 | 35907217 | 35907261 | 35907445 | 35907580 | 0 | 11 | 42 | 0 | 0 | 4 | 0 | 9 | 9 | 0 | 1 | 0 | 52 | 49 | 0.912303287 | 0.999202116 | NA | 1 | 0.908 | NA | 0.895 | 1 | 0.006 |
| SE | 3136 | IFNAR2 | IFNAR2 | chr21 | + | 33248708 | 33248854 | 33246717 | 33246890 | 33252661 | 33252830 | 0 | 1 | 7 | 0 | 0 | 1 | 0 | 12 | 0 | 0 | 0 | 0 | 98 | 49 | 0.91423345 | 0.999202116 | NA | 1 | 0.778 | NA | 1 | NA | -0.111 |
| SE | 6130 | WDR26 | WDR26 | chr1 | - | 224431681 | 224431781 | 224431476 | 224431581 | 224433683 | 224434299 | 0 | 1 | 6 | 0 | 0 | 1 | 0 | 12 | 0 | 0 | 0 | 0 | 98 | 49 | 0.91423345 | 0.999202116 | NA | 1 | 0.75 | NA | 1 | NA | -0.125 |
| SE | 12143 | TRIM25 | TRIM25 | chr17 | - | 56895925 | 56895952 | 56895520 | 56895604 | 56899114 | 56899180 | 0 | 1 | 3 | 0 | 0 | 1 | 0 | 9 | 0 | 0 | 0 | 0 | 76 | 49 | 0.917397557 | 0.999202116 | NA | 1 | 0.659 | NA | 1 | NA | -0.17 |
| A3SS | 1626 | RPS3 | RPS3 | chr11 | + | 75405582 | 75406913 | 75405613 | 75406913 | 75404671 | 75404868 | 1 | 0 | 4 | 0 | 1 | 4 | 0 | 0 | 0 | 0 | 6 | 0 | 80 | 49 | 0.917875723 | 0.999202116 | 1 | 0 | 0.38 | NA | 0 | NA | 0.46 |
| SE | 3618 | NIN | NIN | chr14 | - | 50821873 | 50822077 | 50806736 | 50806818 | 50830463 | 50830517 | 0 | 3 | 8 | 0 | 0 | 1 | 0 | 7 | 6 | 0 | 0 | 1 | 98 | 49 | 0.918267371 | 0.999202116 | NA | 1 | 0.8 | NA | 1 | 0.75 | 0.025 |
| SE | 8078 | NADK | NADK | chr1 | - | 1756502 | 1756608 | 1756257 | 1756343 | 1757180 | 1757310 | 0 | 1 | 24 | 0 | 0 | 3 | 0 | 11 | 0 | 0 | 0 | 0 | 98 | 49 | 0.918460091 | 0.999202116 | NA | 1 | 0.8 | NA | 1 | NA | -0.1 |
| A3SS | 2809 | SRRT | SRRT | chr7 | + | 100888041 | 100888143 | 100888053 | 100888143 | 100887702 | 100887859 | 0 | 1 | 1 | 0 | 0 | 3 | 0 | 7 | 0 | 0 | 0 | 0 | 61 | 49 | 0.919108663 | 0.999202116 | NA | 1 | 0.211 | NA | 1 | NA | -0.394 |
| SE | 10061 | TPT1 | TPT1 | chr13 | - | 45339496 | 45339602 | 45338659 | 45338776 | 45339993 | 45340184 | 0 | 6 | 79 | 0 | 0 | 3 | 10 | 61 | 0 | 0 | 0 | 0 | 98 | 49 | 0.922183161 | 0.999202116 | NA | 1 | 0.929 | 1 | 1 | NA | -0.035 |
| RI | 894 | STX4 | STX4 | chr16 | + | 31034225 | 31034536 | 31034225 | 31034325 | 31034461 | 31034536 | 0 | 1 | 5 | 0 | 0 | 3 | 6 | 10 | 0 | 0 | 0 | 0 | 98 | 49 | 0.922793683 | 0.999202116 | NA | 1 | 0.455 | 1 | 1 | NA | -0.272 |
| SE | 5493 | ADAM8 | ADAM8 | chr10 | - | 133263687 | 133263765 | 133262415 | 133263233 | 133267351 | 133267417 | 0 | 4 | 10 | 0 | 2 | 2 | 3 | 19 | 13 | 0 | 5 | 4 | 98 | 49 | 0.923886696 | 0.999202116 | NA | 0.5 | 0.714 | 1 | 0.655 | 0.619 | -0.151 |
| SE | 1583 | SYTL3 | SYTL3 | chr6 | + | 158663204 | 158663378 | 158662749 | 158662812 | 158665394 | 158665613 | 0 | 1 | 1 | 0 | 0 | 1 | 0 | 2 | 0 | 0 | 0 | 1 | 98 | 49 | 0.925071736 | 0.999202116 | NA | 1 | 0.333 | NA | 1 | 0 | 0.166 |
| SE | 4333 | NCF2 | NCF2 | chr1 | - | 183573184 | 183573292 | 183570779 | 183570839 | 183574486 | 183574621 | 2 | 5 | 63 | 0 | 0 | 1 | 0 | 47 | 0 | 0 | 0 | 0 | 98 | 49 | 0.925576519 | 0.999202116 | 1 | 1 | 0.969 | NA | 1 | NA | -0.01 |
| SE | 8261 | ARHGEF11 | ARHGEF11 | chr1 | - | 156957791 | 156957815 | 156956419 | 156956564 | 156958741 | 156958864 | 0 | 2 | 7 | 0 | 0 | 0 | 0 | 2 | 0 | 0 | 0 | 1 | 73 | 49 | 0.92640979 | 0.999202116 | NA | 1 | 1 | NA | 1 | 0 | 0.5 |
| A5SS | 2405 | SP100 | SP100 | chr2 | + | 230467119 | 230467215 | 230467119 | 230467206 | 230469042 | 230469096 | 0 | 3 | 13 | 0 | 0 | 1 | 0 | 15 | 0 | 0 | 0 | 0 | 58 | 49 | 0.926498131 | 0.999208248 | NA | 1 | 0.917 | NA | 1 | NA | -0.041 |
| RI | 1099 | RPL28 | RPL28 | chr19 | + | 55387929 | 55392083 | 55387929 | 55388048 | 55388242 | 55392083 | 1 | 0 | 1 | 0 | 0 | 13 | 0 | 0 | 1 | 4 | 0 | 0 | 98 | 49 | 0.927341297 | 0.999202116 | 1 | NA | 0.037 | 0 | NA | 1 | 0.018 |
| SE | 11228 | PTPN12 | PTPN12 | chr7 | + | 77585542 | 77585581 | 77583554 | 77583650 | 77592184 | 77592256 | 0 | 1 | 9 | 0 | 0 | 1 | 0 | 8 | 0 | 0 | 0 | 0 | 88 | 49 | 0.9284209 | 0.999202116 | NA | 1 | 0.834 | NA | 1 | NA | -0.083 |
| SE | 10669 | LAPTM5 | LAPTM5 | chr1 | - | 30742455 | 30742549 | 30741639 | 30741716 | 30757658 | 30757836 | 0 | 3 | 22 | 0 | 0 | 2 | 1 | 26 | 0 | 0 | 0 | 0 | 98 | 49 | 0.928473035 | 0.999202116 | NA | 1 | 0.846 | 1 | 1 | NA | -0.077 |
| SE | 4334 | NCF2 | NCF2 | chr1 | - | 183573184 | 183573292 | 183570779 | 183570839 | 183577598 | 183577707 | 2 | 5 | 53 | 0 | 0 | 1 | 0 | 44 | 0 | 0 | 0 | 0 | 98 | 49 | 0.928484533 | 0.999202116 | 1 | 1 | 0.964 | NA | 1 | NA | -0.012 |
| SE | 11373 | CYLD | CYLD | chr16 | + | 50744729 | 50744866 | 50742761 | 50742841 | 50749575 | 50750202 | 0 | 0 | 0 | 0 | 2 | 1 | 2 | 0 | 0 | 0 | 7 | 2 | 98 | 49 | 0.928492447 | 0.999202116 | NA | 0 | 0 | 1 | 0 | 0 | -0.333 |
| A5SS | 2858 | RPL13A | RPL13A | chr19 | + | 49489849 | 49489922 | 49489849 | 49489912 | 49490231 | 49490297 | 0 | 3 | 11 | 0 | 0 | 1 | 4 | 17 | 0 | 0 | 0 | 0 | 59 | 49 | 0.928715497 | 0.999208248 | NA | 1 | 0.901 | 1 | 1 | NA | -0.049 |
| SE | 1923 | EMB | EMB | chr5 | - | 50410876 | 50410965 | 50405724 | 50405852 | 50411196 | 50411383 | 1 | 1 | 22 | 0 | 0 | 1 | 1 | 9 | 0 | 0 | 0 | 0 | 98 | 49 | 0.928729465 | 0.999202116 | 1 | 1 | 0.917 | 1 | 1 | NA | -0.028 |
| SE | 3671 | CALCOCO2 | CALCOCO2 | chr17 | + | 48846442 | 48846514 | 48841697 | 48841887 | 48848063 | 48848166 | 0 | 0 | 1 | 0 | 1 | 5 | 0 | 0 | 0 | 0 | 4 | 0 | 98 | 49 | 0.930162674 | 0.999202116 | NA | 0 | 0.091 | NA | 0 | NA | 0.045 |
| SE | 10146 | GALC | GALC | chr14 | - | 87988454 | 87988523 | 87988143 | 87988207 | 87992969 | 87993271 | 0 | 1 | 5 | 0 | 1 | 0 | 0 | 5 | 12 | 0 | 0 | 2 | 98 | 49 | 0.93135784 | 0.999202116 | NA | 0.333 | 1 | NA | 1 | 0.75 | -0.209 |
| A3SS | 538 | DBNL | DBNL | chr7 | + | 44058877 | 44058983 | 44058901 | 44058983 | 44058428 | 44058480 | 0 | 0 | 2 | 0 | 1 | 3 | 3 | 1 | 0 | 0 | 2 | 1 | 73 | 49 | 0.931399982 | 0.999202116 | NA | 0 | 0.309 | 1 | 0.251 | 0 | -0.262 |
| SE | 10209 | RFWD2 | RFWD2 | chr1 | - | 176136487 | 176136547 | 176135009 | 176135086 | 176149005 | 176149074 | 0 | 1 | 16 | 0 | 0 | 4 | 3 | 15 | 3 | 0 | 3 | 0 | 98 | 49 | 0.931793753 | 0.999202116 | NA | 1 | 0.667 | 1 | 0.714 | 1 | -0.071 |
| SE | 5478 | MICAL2 | MICAL2 | chr11 | + | 12204249 | 12204457 | 12162078 | 12162419 | 12208022 | 12208139 | 0 | 1 | 17 | 0 | 0 | 1 | 0 | 8 | 0 | 0 | 0 | 0 | 98 | 49 | 0.932280727 | 0.999202116 | NA | 1 | 0.895 | NA | 1 | NA | -0.052 |
| SE | 11480 | AP1G1 | AP1G1 | chr16 | - | 71748250 | 71748378 | 71746587 | 71746692 | 71749893 | 71749983 | 0 | 1 | 4 | 0 | 0 | 1 | 0 | 8 | 0 | 0 | 0 | 0 | 98 | 49 | 0.932280727 | 0.999202116 | NA | 1 | 0.667 | NA | 1 | NA | -0.166 |
| SE | 13003 | HNRNPU | HNRNPU | chr1 | - | 244858010 | 244858274 | 244857597 | 244857717 | 244858728 | 244858841 | 0 | 2 | 18 | 0 | 0 | 2 | 2 | 13 | 0 | 0 | 0 | 0 | 98 | 49 | 0.932958395 | 0.999202116 | NA | 1 | 0.818 | 1 | 1 | NA | -0.091 |
| SE | 7216 | LSP1 | LSP1 | chr11 | + | 1884279 | 1884323 | 1883931 | 1884024 | 1884499 | 1884581 | 0 | 5 | 71 | 0 | 0 | 1 | 0 | 36 | 0 | 0 | 0 | 0 | 93 | 49 | 0.933472812 | 0.999202116 | NA | 1 | 0.974 | NA | 1 | NA | -0.013 |
| SE | 4366 | LRRK2 | LRRK2 | chr12 | + | 40354298 | 40354492 | 40351538 | 40351733 | 40356114 | 40356187 | 0 | 2 | 26 | 0 | 0 | 2 | 0 | 15 | 0 | 0 | 0 | 0 | 98 | 49 | 0.934689043 | 0.999202116 | NA | 1 | 0.867 | NA | 1 | NA | -0.067 |
| SE | 13049 | DHX34 | DHX34 | chr19 | + | 47357865 | 47358120 | 47355038 | 47355350 | 47359967 | 47360070 | 0 | 2 | 24 | 0 | 0 | 1 | 0 | 15 | 0 | 0 | 0 | 0 | 98 | 49 | 0.934689043 | 0.999202116 | NA | 1 | 0.923 | NA | 1 | NA | -0.038 |
| SE | 3908 | MYL6 | MYL6 | chr12 | + | 56160625 | 56160670 | 56160242 | 56160320 | 56161386 | 56161582 | 0 | 0 | 1 | 0 | 3 | 15 | 0 | 0 | 0 | 0 | 11 | 0 | 94 | 49 | 0.935467898 | 0.999202116 | NA | 0 | 0.034 | NA | 0 | NA | 0.017 |
| A3SS | 2959 | MTCH1 | MTCH1 | chr6 | - | 36977633 | 36977784 | 36977633 | 36977691 | 36978077 | 36978155 | 0 | 1 | 1 | 0 | 0 | 3 | 2 | 6 | 0 | 0 | 0 | 0 | 98 | 49 | 0.936414148 | 0.999202116 | NA | 1 | 0.143 | 1 | 1 | NA | -0.428 |
| RI | 864 | SUMO1 | SUMO1 | chr2 | - | 202206179 | 202207321 | 202206179 | 202206803 | 202207252 | 202207321 | 0 | 1 | 10 | 0 | 0 | 1 | 0 | 6 | 0 | 0 | 0 | 0 | 98 | 49 | 0.936414148 | 0.999202116 | NA | 1 | 0.833 | NA | 1 | NA | -0.084 |
| SE | 11834 | NCOA1 | NCOA1 | chr2 | + | 24705085 | 24705233 | 24697657 | 24697798 | 24706567 | 24707888 | 0 | 1 | 11 | 0 | 0 | 1 | 0 | 6 | 0 | 0 | 0 | 0 | 98 | 49 | 0.936414148 | 0.999202116 | NA | 1 | 0.846 | NA | 1 | NA | -0.077 |
| SE | 12434 | SPAG9 | SPAG9 | chr17 | - | 50984922 | 50984990 | 50982523 | 50982672 | 50985697 | 50985778 | 0 | 1 | 8 | 0 | 0 | 1 | 0 | 6 | 0 | 0 | 0 | 0 | 98 | 49 | 0.936414148 | 0.999202116 | NA | 1 | 0.8 | NA | 1 | NA | -0.1 |
| SE | 13879 | ZEB2 | ZEB2 | chr2 | - | 144404835 | 144405024 | 144403915 | 144404130 | 144429768 | 144430026 | 0 | 1 | 3 | 0 | 0 | 1 | 0 | 6 | 0 | 0 | 0 | 0 | 98 | 49 | 0.936414148 | 0.999202116 | NA | 1 | 0.6 | NA | 1 | NA | -0.2 |
| A3SS | 3220 | LILRB3 | LILRB3 | chr19 | - | 54217318 | 54217477 | 54217318 | 54217474 | 54218360 | 54218413 | 0 | 2 | 8 | 0 | 3 | 11 | 0 | 1 | 3 | 4 | 6 | 1 | 52 | 49 | 0.937019423 | 0.999202116 | NA | 0.386 | 0.407 | 0 | 0.136 | 0.739 | 0.105 |
| RI | 712 | CD164 | CD164 | chr6 | - | 109366513 | 109369017 | 109366513 | 109368325 | 109368921 | 109369017 | 0 | 1 | 31 | 0 | 0 | 1 | 4 | 7 | 0 | 0 | 0 | 0 | 98 | 49 | 0.937374675 | 0.999202116 | NA | 1 | 0.939 | 1 | 1 | NA | -0.03 |
| SE | 2080 | HERC5 | HERC5 | chr4 | + | 88487068 | 88487179 | 88486114 | 88486228 | 88489165 | 88489336 | 0 | 1 | 19 | 0 | 0 | 1 | 0 | 7 | 0 | 0 | 0 | 0 | 98 | 49 | 0.937374675 | 0.999202116 | NA | 1 | 0.905 | NA | 1 | NA | -0.047 |
| SE | 2091 | HSD17B11 | HSD17B11 | chr4 | - | 87374698 | 87374830 | 87372708 | 87372815 | 87382254 | 87382362 | 0 | 3 | 26 | 0 | 0 | 1 | 2 | 20 | 0 | 0 | 0 | 0 | 98 | 49 | 0.939243248 | 0.999202116 | NA | 1 | 0.929 | 1 | 1 | NA | -0.035 |
| RI | 861 | JUP | JUP | chr17 | - | 41754606 | 41755895 | 41754606 | 41755432 | 41755729 | 41755895 | 1 | 0 | 0 | 0 | 0 | 0 | 5 | 1 | 0 | 0 | 1 | 0 | 98 | 49 | 0.94318207 | 0.999202116 | 1 | NA | NA | 1 | 0.333 | NA | 0.334 |
| SE | 8746 | BEST1 | BEST1 | chr11 | + | 61959891 | 61960043 | 61959497 | 61959578 | 61962254 | 61962893 | 0 | 1 | 3 | 0 | 0 | 1 | 0 | 5 | 0 | 0 | 0 | 0 | 98 | 49 | 0.94318207 | 0.999202116 | NA | 1 | 0.6 | NA | 1 | NA | -0.2 |
| SE | 9188 | GIT2 | GIT2 | chr12 | - | 109951166 | 109951316 | 109947255 | 109947504 | 109953091 | 109953234 | 0 | 1 | 5 | 0 | 0 | 1 | 0 | 5 | 0 | 0 | 0 | 0 | 98 | 49 | 0.94318207 | 0.999202116 | NA | 1 | 0.714 | NA | 1 | NA | -0.143 |
| SE | 1843 | VCAN | VCAN | chr5 | + | 83537006 | 83542268 | 83512102 | 83512396 | 83545536 | 83545650 | 0 | 2 | 51 | 0 | 0 | 1 | 0 | 10 | 0 | 0 | 0 | 0 | 98 | 49 | 0.944027198 | 0.999202116 | NA | 1 | 0.962 | NA | 1 | NA | -0.019 |
| SE | 3459 | RASSF2 | RASSF2 | chr20 | - | 4795814 | 4795966 | 4792538 | 4792627 | 4798009 | 4798085 | 0 | 4 | 29 | 0 | 0 | 1 | 0 | 20 | 0 | 0 | 0 | 0 | 98 | 49 | 0.944343057 | 0.999202116 | NA | 1 | 0.935 | NA | 1 | NA | -0.032 |
| SE | 3950 | ARID4B | ARID4B | chr1 | - | 235213768 | 235214026 | 235196030 | 235196115 | 235219792 | 235219968 | 0 | 0 | 1 | 0 | 1 | 2 | 0 | 0 | 0 | 0 | 3 | 0 | 98 | 49 | 0.94451324 | 0.999202116 | NA | 0 | 0.2 | NA | 0 | NA | 0.1 |
| SE | 8841 | C1orf43 | C1orf43 | chr1 | - | 154219835 | 154219937 | 154214456 | 154214574 | 154220341 | 154220797 | 0 | 0 | 1 | 0 | 1 | 1 | 0 | 0 | 0 | 0 | 3 | 0 | 98 | 49 | 0.94451324 | 0.999202116 | NA | 0 | 0.333 | NA | 0 | NA | 0.167 |
| SE | 12024 | KLHL2 | KLHL2 | chr4 | + | 165238777 | 165238899 | 165228806 | 165228913 | 165263196 | 165263359 | 0 | 0 | 4 | 0 | 1 | 2 | 0 | 0 | 0 | 0 | 3 | 0 | 98 | 49 | 0.94451324 | 0.999202116 | NA | 0 | 0.5 | NA | 0 | NA | 0.25 |
| A5SS | 192 | TBXAS1 | TBXAS1 | chr7 | + | 139875584 | 139875666 | 139875584 | 139875637 | 139911224 | 139911321 | 0 | 0 | 1 | 0 | 1 | 15 | 5 | 0 | 0 | 0 | 3 | 0 | 78 | 49 | 0.945027512 | 0.999208248 | NA | 0 | 0.04 | 1 | 0 | NA | -0.48 |
| RI | 396 | CASP1 | CASP1 | chr11 | - | 105025509 | 105026356 | 105025509 | 105025600 | 105026225 | 105026356 | 0 | 0 | 15 | 0 | 2 | 6 | 0 | 0 | 0 | 0 | 6 | 0 | 98 | 49 | 0.945187435 | 0.999202116 | NA | 0 | 0.556 | NA | 0 | NA | 0.278 |
| SE | 11313 | ALDOA | ALDOA | chr16 | + | 30068645 | 30068700 | 30067449 | 30067661 | 30068817 | 30068978 | 0 | 3 | 41 | 0 | 0 | 1 | 4 | 14 | 0 | 0 | 0 | 0 | 98 | 49 | 0.946228808 | 0.999202116 | NA | 1 | 0.953 | 1 | 1 | NA | -0.024 |
| SE | 10440 | SLC12A6 | SLC12A6 | chr15 | - | 34257641 | 34257788 | 34256228 | 34256283 | 34258812 | 34258944 | 0 | 1 | 11 | 0 | 0 | 4 | 0 | 2 | 4 | 0 | 2 | 0 | 98 | 49 | 0.949596688 | 0.999202116 | NA | 1 | 0.579 | NA | 0.333 | 1 | 0.123 |
| SE | 10864 | ARHGAP9 | ARHGAP9 | chr12 | - | 57476870 | 57476963 | 57476589 | 57476651 | 57477155 | 57477269 | 0 | 2 | 25 | 0 | 0 | 2 | 0 | 9 | 0 | 0 | 0 | 0 | 98 | 49 | 0.950138287 | 0.999202116 | NA | 1 | 0.862 | NA | 1 | NA | -0.069 |
| SE | 7901 | PACSIN2 | PACSIN2 | chr22 | - | 42876887 | 42877010 | 42876136 | 42876333 | 42879047 | 42879169 | 0 | 0 | 5 | 0 | 3 | 0 | 0 | 0 | 0 | 0 | 8 | 0 | 98 | 49 | 0.950176987 | 0.999202116 | NA | 0 | 1 | NA | 0 | NA | 0.5 |
| SE | 7638 | IRF7 | IRF7 | chr11 | - | 613950 | 614037 | 613784 | 613865 | 614475 | 614534 | 0 | 8 | 14 | 0 | 8 | 3 | 0 | 7 | 3 | 0 | 3 | 2 | 98 | 49 | 0.950918257 | 0.999202116 | NA | 0.333 | 0.7 | NA | 0.538 | 0.429 | 0.033 |
| A3SS | 779 | LST1 | LST1 | chr6 | + | 31588517 | 31588909 | 31588562 | 31588909 | 31587640 | 31587733 | 0 | 3 | 10 | 0 | 0 | 2 | 4 | 12 | 0 | 0 | 0 | 0 | 94 | 49 | 0.951788859 | 0.999202116 | NA | 1 | 0.723 | 1 | 1 | NA | -0.139 |
| A5SS | 1230 | STRADA | STRADA | chr17 | - | 63704340 | 63704582 | 63704431 | 63704582 | 63704004 | 63704047 | 1 | 0 | 0 | 0 | 0 | 0 | 4 | 1 | 0 | 0 | 1 | 0 | 98 | 49 | 0.952968863 | 0.999208248 | 1 | NA | NA | 1 | 0.333 | NA | 0.334 |
| RI | 174 | NDUFA2 | NDUFA2 | chr5 | - | 140647255 | 140647785 | 140647255 | 140647362 | 140647482 | 140647785 | 1 | 0 | 0 | 0 | 0 | 2 | 4 | 3 | 0 | 0 | 1 | 0 | 98 | 49 | 0.952968863 | 0.999202116 | 1 | NA | 0 | 1 | 0.6 | NA | -0.3 |
| SE | 1981 | DDX60L | DDX60L | chr4 | - | 168379725 | 168379830 | 168379362 | 168379504 | 168384611 | 168384812 | 0 | 1 | 18 | 0 | 0 | 1 | 0 | 4 | 0 | 0 | 0 | 0 | 98 | 49 | 0.952968863 | 0.999202116 | NA | 1 | 0.9 | NA | 1 | NA | -0.05 |
| SE | 2114 | KDM5C | KDM5C | chrX | - | 53218275 | 53218398 | 53217795 | 53217966 | 53220838 | 53220916 | 0 | 1 | 12 | 0 | 0 | 2 | 0 | 4 | 0 | 0 | 0 | 0 | 98 | 49 | 0.952968863 | 0.999202116 | NA | 1 | 0.75 | NA | 1 | NA | -0.125 |
| SE | 7631 | BTAF1 | BTAF1 | chr10 | + | 91953736 | 91953873 | 91951402 | 91951566 | 91956527 | 91956657 | 0 | 1 | 5 | 0 | 0 | 1 | 0 | 4 | 0 | 0 | 0 | 0 | 98 | 49 | 0.952968863 | 0.999202116 | NA | 1 | 0.714 | NA | 1 | NA | -0.143 |
| SE | 10952 | FES | FES | chr15 | + | 90885411 | 90885585 | 90885036 | 90885258 | 90886960 | 90887057 | 0 | 1 | 3 | 0 | 0 | 1 | 0 | 4 | 0 | 0 | 0 | 0 | 98 | 49 | 0.952968863 | 0.999202116 | NA | 1 | 0.6 | NA | 1 | NA | -0.2 |
| SE | 11294 | CREBBP | CREBBP | chr16 | - | 3780731 | 3780878 | 3778699 | 3778817 | 3781203 | 3781306 | 0 | 1 | 9 | 0 | 0 | 1 | 2 | 4 | 0 | 0 | 0 | 0 | 98 | 49 | 0.952968863 | 0.999202116 | NA | 1 | 0.818 | 1 | 1 | NA | -0.091 |
| SE | 11440 | PSMB3 | PSMB3 | chr17 | + | 38760430 | 38760608 | 38755882 | 38755990 | 38762410 | 38762505 | 0 | 1 | 12 | 0 | 0 | 1 | 0 | 4 | 0 | 0 | 0 | 0 | 98 | 49 | 0.952968863 | 0.999202116 | NA | 1 | 0.857 | NA | 1 | NA | -0.072 |
| A5SS | 3017 | SERPINA1 | SERPINA1 | chr14 | - | 94388800 | 94389010 | 94388818 | 94389010 | 94388559 | 94388660 | 0 | 1 | 6 | 0 | 0 | 1 | 0 | 3 | 0 | 0 | 0 | 0 | 67 | 49 | 0.954109788 | 0.999208248 | NA | 1 | 0.814 | NA | 1 | NA | -0.093 |
| A5SS | 399 | BTNL8 | BTNL8 | chr5 | + | 180947511 | 180947755 | 180947511 | 180947625 | 180948354 | 180948375 | 0 | 2 | 2 | 0 | 0 | 1 | 0 | 7 | 0 | 0 | 0 | 0 | 98 | 49 | 0.957490202 | 0.999208248 | NA | 1 | 0.5 | NA | 1 | NA | -0.25 |
| SE | 5182 | ANKRD13A | ANKRD13A | chr12 | + | 110033796 | 110033957 | 110030644 | 110030758 | 110036260 | 110036328 | 0 | 2 | 4 | 0 | 0 | 1 | 0 | 7 | 0 | 0 | 0 | 0 | 98 | 49 | 0.957490202 | 0.999202116 | NA | 1 | 0.667 | NA | 1 | NA | -0.166 |
| SE | 6877 | UBN1 | UBN1 | chr16 | + | 4857989 | 4858076 | 4852878 | 4853166 | 4858567 | 4858663 | 0 | 2 | 11 | 0 | 0 | 1 | 1 | 7 | 0 | 0 | 0 | 0 | 98 | 49 | 0.957490202 | 0.999202116 | NA | 1 | 0.846 | 1 | 1 | NA | -0.077 |
| SE | 9478 | JAK1 | JAK1 | chr1 | - | 64839602 | 64839795 | 64838464 | 64838589 | 64841244 | 64841339 | 0 | 2 | 17 | 0 | 0 | 1 | 0 | 7 | 0 | 0 | 0 | 0 | 98 | 49 | 0.957490202 | 0.999202116 | NA | 1 | 0.895 | NA | 1 | NA | -0.052 |
| SE | 11195 | PARP14 | PARP14 | chr3 | + | 122704526 | 122704748 | 122699389 | 122701635 | 122708189 | 122708268 | 0 | 2 | 3 | 0 | 0 | 1 | 0 | 7 | 0 | 0 | 0 | 0 | 98 | 49 | 0.957490202 | 0.999202116 | NA | 1 | 0.6 | NA | 1 | NA | -0.2 |
| A3SS | 778 | LST1 | LST1 | chr6 | + | 31588496 | 31588909 | 31588562 | 31588909 | 31587640 | 31587733 | 0 | 3 | 10 | 0 | 0 | 2 | 4 | 11 | 0 | 0 | 0 | 0 | 98 | 49 | 0.957822245 | 0.999202116 | NA | 1 | 0.714 | 1 | 1 | NA | -0.143 |

| | | | | | | | | | | | | | | | | | | | | | | | | | | | | | | | | | | | |
|---|---|---|---|---|---|---|---|---|---|---|---|---|---|---|---|---|---|---|---|---|---|---|---|---|---|---|---|---|---|---|---|---|---|---|---|
| SE | 11576 | PPIF | PPIF | chr10 | + | 79349075 | 79349106 | 79347463 | 79347743 | 79349664 | 79349753 | 0 | 0 | 2 | 0 | 0 | 0 | 0 | 13 | 6 | 0 | 2 | 0 | 80 | 49 | 0.958077329 | 0.999202116 | NA | | NA | 1 | NA | 0.799 | 1 | 0.101 |
| SE | 6631 | PCBP2 | PCBP2 | chr12 | + | 53467804 | 53467843 | 53467220 | 53467293 | 53468779 | 53468832 | 0 | 1 | 4 | 0 | 0 | 2 | 8 | 3 | 0 | 0 | 0 | 0 | 88 | 49 | 0.960459131 | 0.999202116 | NA | | 1 | 0.527 | 1 | 1 | NA | -0.236 |
| RI | 557 | QSOX1 | QSOX1 | chr1 | + | 180196261 | 180198034 | 180196261 | 180196599 | 180197332 | 180198034 | 1 | 0 | 11 | 0 | 0 | 3 | 3 | 8 | 0 | 0 | 1 | 0 | 98 | 49 | 0.961336908 | 0.999202116 | | 1 | NA | 0.647 | 1 | 0.8 | NA | -0.077 |
| RI | 1064 | FXYD5 | FXYD5 | chr19 | + | 35154768 | 35155611 | 35154768 | 35154869 | 35155550 | 35155611 | 1 | 0 | 0 | 0 | 0 | 0 | 3 | 1 | 0 | 0 | 1 | 0 | 98 | 49 | 0.961336908 | 0.999202116 | | 1 | NA | NA | 1 | 0.333 | NA | 0.334 |
| SE | 1449 | BAG6 | BAG6 | chr6 | - | 31647590 | 31647826 | 31646393 | 31646523 | 31648676 | 31648751 | 0 | 1 | 2 | 1 | 0 | 0 | 0 | 3 | 0 | 0 | 0 | 0 | 98 | 49 | 0.961336908 | 0.999202116 | | 0 | 1 | 1 | NA | 1 | NA | -0.333 |
| SE | 3433 | RRBP1 | RRBP1 | chr20 | - | 17616731 | 17616839 | 17615925 | 17616009 | 17618595 | 17618679 | 0 | 1 | 0 | 0 | 0 | 1 | 0 | 3 | 0 | 0 | 0 | 0 | 98 | 49 | 0.961336908 | 0.999202116 | NA | | 1 | 0 | NA | 1 | NA | -0.5 |
| SE | 4548 | GZMH | GZMH | chr14 | - | 24607611 | 24607747 | 24607148 | 24607406 | 24608264 | 24608412 | 0 | 1 | 0 | 0 | 0 | 3 | 0 | 3 | 0 | 0 | 0 | 0 | 98 | 49 | 0.961336908 | 0.999202116 | NA | | 1 | 0 | NA | 1 | NA | -0.5 |
| SE | 5312 | AFF1 | AFF1 | chr4 | + | 87089983 | 87090070 | 87084119 | 87084164 | 87091792 | 87091829 | 0 | 1 | 12 | 0 | 0 | 1 | 0 | 3 | 0 | 0 | 0 | 0 | 98 | 49 | 0.961336908 | 0.999202116 | NA | | 1 | 0.857 | NA | 1 | NA | -0.072 |
| SE | 6569 | HADHA | HADHA | chr2 | - | 26193576 | 26193772 | 26192309 | 26192424 | 26194569 | 26194638 | 0 | 1 | 11 | 0 | 0 | 1 | 0 | 3 | 0 | 0 | 0 | 0 | 98 | 49 | 0.961336908 | 0.999202116 | NA | | 1 | 0.846 | NA | 1 | NA | -0.077 |
| SE | 7389 | ZRANB1 | ZRANB1 | chr10 | + | 124981708 | 124981829 | 124974199 | 124974398 | 124983174 | 124983304 | 0 | 1 | 1 | 0 | 0 | 1 | 0 | 3 | 0 | 0 | 0 | 0 | 98 | 49 | 0.961336908 | 0.999202116 | NA | | 1 | 0.333 | NA | 1 | NA | -0.334 |
| SE | 7848 | RPS6KA1 | RPS6KA1 | chr1 | + | 26560725 | 26560851 | 26558806 | 26558937 | 26561044 | 26561134 | 0 | 1 | 11 | 0 | 0 | 2 | 0 | 3 | 0 | 0 | 0 | 0 | 98 | 49 | 0.961336908 | 0.999202116 | NA | | 1 | 0.733 | NA | 1 | NA | -0.133 |
| SE | 7862 | C1orf162 | C1orf162 | chr1 | + | 111477725 | 111477775 | 111477333 | 111477428 | 111477982 | 111478512 | 0 | 1 | 9 | 0 | 0 | 1 | 0 | 3 | 0 | 0 | 0 | 0 | 98 | 49 | 0.961336908 | 0.999202116 | NA | | 1 | 0.818 | NA | 1 | NA | -0.091 |
| SE | 11425 | DYNC1LI2 | DYNC1LI2 | chr16 | - | 66749196 | 66749313 | 66742437 | 66742668 | 66751272 | 66751346 | 0 | 1 | 6 | 0 | 0 | 1 | 0 | 3 | 0 | 0 | 0 | 0 | 98 | 49 | 0.961336908 | 0.999202116 | NA | | 1 | 0.75 | NA | 1 | NA | -0.125 |
| SE | 12705 | GANAB | GANAB | chr11 | - | 62632564 | 62632745 | 62631029 | 62631183 | 62633004 | 62633101 | 0 | 1 | 3 | 0 | 0 | 1 | 0 | 3 | 0 | 0 | 0 | 0 | 98 | 49 | 0.961336908 | 0.999202116 | NA | | 1 | 0.6 | NA | 1 | NA | -0.2 |
| SE | 10094 | NUMB | NUMB | chr14 | - | 73316389 | 73316422 | 73297210 | 73297285 | 73323129 | 73323204 | 0 | 0 | 2 | 0 | 3 | 8 | 0 | 0 | 0 | 0 | 6 | 0 | 82 | 49 | 0.961535291 | 0.999202116 | NA | | 0 | 0.13 | NA | 0 | NA | 0.065 |
| A3SS | 3132 | ITGAL | ITGAL | chr16 | + | 30505243 | 30505300 | 30505246 | 30505300 | 30504174 | 30504264 | 1 | 0 | 8 | 0 | 0 | 2 | 2 | 3 | 0 | 0 | 0 | 0 | 52 | 49 | 0.961575457 | 0.999202116 | | 1 | NA | 0.79 | 1 | 1 | NA | -0.105 |
| A5SS | 2154 | SNX17 | SNX17 | chr2 | + | 27371268 | 27371468 | 27371268 | 27371343 | 27372622 | 27372740 | 1 | 0 | 0 | 0 | 1 | 3 | 0 | 0 | 0 | 0 | 2 | 0 | 98 | 49 | 0.962033383 | 0.999208248 | | 1 | 0 | 0 | NA | 0 | NA | 0.333 |
| SE | 9739 | PXN | PXN | chr12 | - | 120215559 | 120215661 | 120215102 | 120215273 | 120221622 | 120221758 | 0 | 0 | 1 | 0 | 1 | 3 | 0 | 0 | 0 | 0 | 2 | 0 | 98 | 49 | 0.962033383 | 0.999202116 | NA | | 0 | 0.143 | NA | 0 | NA | 0.071 |
| SE | 11604 | INPP5K | INPP5K | chr17 | - | 1515426 | 1515585 | 1513871 | 1513979 | 1516455 | 1516888 | 0 | 1 | 0 | 0 | 0 | 1 | 0 | 0 | 0 | 0 | 0 | 2 | 98 | 49 | 0.962033383 | 0.999202116 | NA | | 1 | 0 | NA | NA | 0 | 0.5 |
| SE | 13744 | MAP4K4 | MAP4K4 | chr2 | + | 101871493 | 101871685 | 101870294 | 101870415 | 101873646 | 101873764 | 0 | 0 | 7 | 0 | 1 | 4 | 0 | 0 | 0 | 0 | 2 | 0 | 98 | 49 | 0.962033383 | 0.999202116 | NA | | 0 | 0.467 | NA | 0 | NA | 0.234 |
| SE | 6797 | BHLHE40 | BHLHE40 | chr3 | + | 4981391 | 4981515 | 4980300 | 4980408 | 4982835 | 4985180 | 0 | 3 | 3 | 0 | 0 | 1 | 0 | 9 | 0 | 0 | 0 | 0 | 98 | 49 | 0.962139286 | 0.999202116 | NA | | 1 | 0.6 | NA | 1 | NA | -0.2 |
| SE | 2917 | TOM1 | TOM1 | chr22 | + | 35321958 | 35322037 | 35317876 | 35317961 | 35323027 | 35323177 | 0 | 4 | 11 | 0 | 0 | 1 | 0 | 12 | 0 | 0 | 0 | 0 | 98 | 49 | 0.962709327 | 0.999202116 | NA | | 1 | 0.846 | NA | 1 | NA | -0.077 |
| SE | 1267 | MICAL1 | MICAL1 | chr6 | - | 109450299 | 109450557 | 109449969 | 109450085 | 109451599 | 109451700 | 0 | 2 | 4 | 0 | 0 | 1 | 0 | 6 | 0 | 0 | 0 | 0 | 98 | 49 | 0.963061436 | 0.999202116 | NA | | 1 | 0.667 | NA | 1 | NA | -0.166 |
| SE | 4007 | FRY | FRY | chr13 | + | 32175545 | 32175632 | 32173366 | 32173549 | 32178176 | 32178436 | 0 | 2 | 7 | 0 | 0 | 2 | 0 | 6 | 0 | 0 | 0 | 0 | 98 | 49 | 0.963061436 | 0.999202116 | NA | | 1 | 0.636 | NA | 1 | NA | -0.182 |
| SE | 6402 | AGO4 | AGO4 | chr1 | + | 35832436 | 35832570 | 35832056 | 35832185 | 35833989 | 35834174 | 0 | 2 | 14 | 0 | 0 | 1 | 0 | 6 | 0 | 0 | 0 | 0 | 98 | 49 | 0.963061436 | 0.999202116 | NA | | 1 | 0.875 | NA | 1 | NA | -0.062 |
| SE | 7412 | RBM23 | RBM23 | chr14 | - | 22909482 | 22909595 | 22908332 | 22908380 | 22911327 | 22911403 | 0 | 2 | 4 | 0 | 0 | 1 | 0 | 6 | 0 | 0 | 0 | 0 | 98 | 49 | 0.963061436 | 0.999202116 | NA | | 1 | 0.667 | NA | 1 | NA | -0.166 |
| SE | 1919 | PARP8 | PARP8 | chr5 | + | 50763147 | 50763242 | 50761820 | 50761898 | 50778068 | 50778129 | 0 | 3 | 15 | 0 | 0 | 1 | 0 | 8 | 0 | 0 | 0 | 0 | 98 | 49 | 0.964425677 | 0.999202116 | NA | | 1 | 0.882 | NA | 1 | NA | -0.059 |
| SE | 12367 | NBEAL2 | NBEAL2 | chr3 | + | 47003173 | 47003309 | 47002956 | 47003081 | 47003815 | 47003976 | 0 | 14 | 31 | 0 | 0 | 2 | 2 | 12 | 14 | 0 | 1 | 0 | 98 | 49 | 0.965210998 | 0.999202116 | NA | | 1 | 0.886 | 1 | 0.857 | 1 | -0.009 |
| SE | 12157 | VMP1 | VMP1 | chr17 | + | 59737452 | 59737543 | 59735337 | 59735473 | 59738836 | 59738947 | 0 | 4 | 43 | 0 | 0 | 4 | 0 | 10 | 0 | 0 | 0 | 0 | 98 | 49 | 0.967766702 | 0.999202116 | NA | | 1 | 0.843 | NA | 1 | NA | -0.079 |
| SE | 1590 | KDM1B | KDM1B | chr6 | + | 18162834 | 18162924 | 18161326 | 18161454 | 18166266 | 18166378 | 0 | 2 | 3 | 0 | 0 | 1 | 0 | 5 | 0 | 0 | 0 | 0 | 98 | 49 | 0.967767969 | 0.999202116 | NA | | 1 | 0.6 | NA | 1 | NA | -0.2 |
| SE | 4616 | PTPRJ | PTPRJ | chr11 | + | 48121002 | 48121266 | 48112746 | 48112983 | 48123612 | 48123870 | 0 | 2 | 25 | 0 | 0 | 1 | 0 | 5 | 0 | 0 | 0 | 0 | 98 | 49 | 0.967767969 | 0.999202116 | NA | | 1 | 0.926 | NA | 1 | NA | -0.037 |
| SE | 6403 | AGO4 | AGO4 | chr1 | + | 35833989 | 35834174 | 35832436 | 35832570 | 35835833 | 35835993 | 0 | 2 | 1 | 0 | 0 | 1 | 0 | 5 | 0 | 0 | 0 | 0 | 98 | 49 | 0.967767969 | 0.999202116 | NA | | 1 | 0.333 | NA | 1 | NA | -0.334 |
| SE | 6446 | RALGDS | RALGDS | chr9 | - | 133107084 | 133107300 | 133106644 | 133106748 | 133107987 | 133108406 | 0 | 2 | 1 | 0 | 0 | 1 | 0 | 5 | 0 | 0 | 0 | 0 | 98 | 49 | 0.967767969 | 0.999202116 | NA | | 1 | 0.333 | NA | 1 | NA | -0.334 |
| SE | 7363 | NOTCH2 | NOTCH2 | chr1 | - | 119922635 | 119922778 | 119922235 | 119922446 | 119923636 | 119923984 | 0 | 2 | 5 | 0 | 0 | 1 | 0 | 5 | 0 | 0 | 0 | 0 | 98 | 49 | 0.967767969 | 0.999202116 | NA | | 1 | 0.714 | NA | 1 | NA | -0.143 |
| SE | 13360 | BROX | BROX | chr1 | + | 222731356 | 222731516 | 222730026 | 222730177 | 222732627 | 222735196 | 0 | 2 | 6 | 0 | 0 | 4 | 0 | 5 | 0 | 0 | 0 | 0 | 98 | 49 | 0.967767969 | 0.999202116 | NA | | 1 | 0.429 | NA | 1 | NA | -0.285 |
| A5SS | 2323 | POLG | POLG | chr15 | - | 89334672 | 89334795 | 89334684 | 89334795 | 89333095 | 89333913 | 0 | 0 | 0 | 0 | 0 | 3 | 0 | 1 | 0 | 0 | 0 | 4 | 61 | 49 | 0.969523928 | 0.999208248 | NA | | NA | 0 | NA | 1 | 0 | -0.5 |
| SE | 3907 | MYL6 | MYL6 | chr12 | + | 56159586 | 56159730 | 56158683 | 56158711 | 56159974 | 56160148 | 0 | 7 | 24 | 0 | 0 | 1 | 0 | 11 | 0 | 0 | 0 | 0 | 98 | 49 | 0.969652412 | 0.999202116 | NA | | 1 | 0.923 | NA | 1 | NA | -0.038 |
| SE | 9995 | UPP1 | UPP1 | chr7 | + | 48101823 | 48101870 | 48094762 | 48094827 | 48106872 | 48107082 | 0 | 0 | 2 | 0 | 0 | 0 | 0 | 3 | 3 | 0 | 3 | 0 | 96 | 49 | 0.970230495 | 0.999202116 | NA | | NA | 1 | NA | 0.338 | 1 | 0.331 |
| RI | 401 | CTNNB1 | CTNNB1 | chr3 | + | 41239133 | 41240448 | 41239133 | 41239354 | 41239659 | 41240448 | 4 | 0 | 6 | 0 | 0 | 3 | 6 | 8 | 0 | 0 | 2 | 0 | 98 | 49 | 0.970780408 | 0.999202116 | | 1 | NA | 0.5 | 1 | 0.667 | NA | -0.084 |
| SE | 813 | LINC-PINT | LINC-PINT | chr7 | - | 130912109 | 130912152 | 130884316 | 130884396 | 130945719 | 130945835 | 0 | 0 | 1 | 0 | 0 | 0 | 0 | 4 | 2 | 0 | 1 | 0 | 92 | 49 | 0.970840435 | 0.999202116 | NA | | NA | 1 | NA | 0.681 | 1 | 0.159 |
| SE | 3380 | HCK | HCK | chr20 | + | 32054215 | 32054281 | 32052187 | 32052486 | 32071661 | 32071782 | 0 | 0 | 5 | 0 | 9 | 33 | 0 | 0 | 0 | 2 | 14 | 0 | 98 | 49 | 0.971752688 | 0.999202116 | NA | | 0 | 0.07 | 0 | 0 | NA | 0.035 |
| A3SS | 80 | GBGT1 | GBGT1 | chr9 | - | 133155177 | 133155349 | 133155177 | 133155312 | 133155900 | 133155936 | 1 | 0 | 1 | 0 | 0 | 1 | 2 | 0 | 0 | 0 | 0 | 0 | 86 | 49 | 0.972021626 | 0.999202116 | | 1 | NA | 0.363 | 1 | NA | NA | -0.319 |
| A3SS | 2931 | LRRC75A-AS1 | LRRC75A-AS1 | chr17 | + | 16440110 | 16440253 | 16440184 | 16440253 | 16439659 | 16439703 | 0 | 0 | 2 | 0 | 0 | 0 | 0 | 1 | 4 | 0 | 3 | 0 | 98 | 49 | 0.97202508 | 0.999202116 | NA | | NA | 1 | NA | 0.143 | 1 | 0.428 |
| RI | 848 | ARSA | ARSA | chr22 | - | 50627555 | 50628173 | 50627555 | 50627792 | 50627897 | 50628173 | 0 | 0 | 2 | 0 | 0 | 0 | 0 | 0 | 4 | 0 | 2 | 0 | 98 | 49 | 0.97202508 | 0.999202116 | NA | | NA | 1 | NA | 0 | 1 | 0.5 |
| SE | 3390 | ANKHD1-EIF4EBP3 | ANKHD1-EIF4EBP3 | chr5 | + | 140496519 | 140497278 | 140486957 | 140487060 | 140504820 | 140504966 | 0 | 2 | 6 | 0 | 0 | 3 | 0 | 4 | 0 | 0 | 0 | 0 | 98 | 49 | 0.97202508 | 0.999202116 | NA | | 1 | 0.5 | NA | 1 | NA | -0.25 |
| SE | 4583 | BECN1 | BECN1 | chr17 | - | 42815907 | 42816054 | 42814523 | 42814673 | 42818220 | 42818415 | 0 | 2 | 7 | 0 | 0 | 3 | 0 | 4 | 0 | 0 | 0 | 0 | 98 | 49 | 0.97202508 | 0.999202116 | NA | | 1 | 0.538 | NA | 1 | NA | -0.231 |
| A5SS | 2532 | TTC19 | TTC19 | chr17 | + | 16000117 | 16000382 | 16000117 | 16000245 | 16001914 | 16002025 | 1 | 0 | 0 | 0 | 0 | 1 | 2 | 0 | 0 | 0 | 0 | 0 | 98 | 49 | 0.9723413 | 0.999208248 | | 1 | NA | 0 | 1 | NA | NA | -0.5 |
| SE | 1578 | USP9X | USP9X | chrX | + | 41140655 | 41140771 | 41136803 | 41137022 | 41140965 | 41141217 | 0 | 1 | 4 | 0 | 0 | 1 | 0 | 2 | 0 | 0 | 0 | 0 | 98 | 49 | 0.9723413 | 0.999202116 | NA | | 1 | 0.667 | NA | 1 | NA | -0.166 |
| SE | 3631 | USP4 | USP4 | chr3 | - | 49294406 | 49294598 | 49292509 | 49292598 | 49297869 | 49297964 | 0 | 1 | 7 | 0 | 0 | 2 | 4 | 2 | 0 | 0 | 0 | 0 | 98 | 49 | 0.9723413 | 0.999202116 | NA | | 1 | 0.636 | 1 | 1 | NA | -0.182 |
| SE | 3782 | MYCBP2 | MYCBP2 | chr13 | - | 77278757 | 77278911 | 77273471 | 77273668 | 77288160 | 77288376 | 0 | 1 | 2 | 0 | 0 | 0 | 0 | 2 | 0 | 5 | 0 | 0 | 98 | 49 | 0.9723413 | 0.999202116 | NA | | 1 | 1 | 0 | 1 | NA | 0.5 |
| SE | 6270 | BAZ2B | BAZ2B | chr2 | - | 159337566 | 159337772 | 159332539 | 159332686 | 159347485 | 159347646 | 0 | 1 | 9 | 0 | 0 | 1 | 0 | 2 | 0 | 0 | 0 | 0 | 98 | 49 | 0.9723413 | 0.999202116 | NA | | 1 | 0.818 | NA | 1 | NA | -0.091 |
| SE | 9517 | SMG5 | SMG5 | chr1 | - | 156266540 | 156266678 | 156265780 | 156266380 | 156267469 | 156267678 | 0 | 1 | 3 | 0 | 0 | 1 | 0 | 2 | 0 | 0 | 0 | 0 | 98 | 49 | 0.9723413 | 0.999202116 | NA | | 1 | 0.6 | NA | 1 | NA | -0.2 |
| SE | 9988 | TRANK1 | TRANK1 | chr3 | - | 36838612 | 36838716 | 36834761 | 36834907 | 36842621 | 36842710 | 0 | 0 | 1 | 0 | 0 | 0 | 0 | 1 | 2 | 0 | 1 | 0 | 98 | 49 | 0.9723413 | 0.999202116 | NA | | NA | 1 | NA | 0.333 | 1 | 0.334 |
| SE | 10095 | NUMB | NUMB | chr14 | - | 73366896 | 73366981 | 73355625 | 73355766 | 73409936 | 73410068 | 0 | 1 | 10 | 0 | 0 | 1 | 2 | 2 | 0 | 0 | 0 | 0 | 98 | 49 | 0.9723413 | 0.999202116 | NA | | 1 | 0.833 | 1 | 1 | NA | -0.084 |
| SE | 11441 | PSMB3 | PSMB3 | chr17 | + | 38760507 | 38760608 | 38755882 | 38755990 | 38762410 | 38762505 | 0 | 1 | 7 | 0 | 0 | 1 | 0 | 2 | 0 | 0 | 0 | 0 | 98 | 49 | 0.9723413 | 0.999202116 | NA | | 1 | 0.778 | NA | 1 | NA | -0.111 |
| SE | 13354 | TFIP11 | TFIP11 | chr22 | - | 26494138 | 26494304 | 26491928 | 26492368 | 26494796 | 26494939 | 0 | 1 | 0 | 0 | 0 | 1 | 0 | 2 | 0 | 0 | 0 | 0 | 98 | 49 | 0.9723413 | 0.999202116 | NA | | 1 | 0 | NA | 1 | NA | -0.5 |
| SE | 1601 | NUP153 | NUP153 | chr6 | - | 17668974 | 17669028 | 17665238 | 17665385 | 17669292 | 17669337 | 0 | 3 | 7 | 0 | 0 | 1 | 0 | 6 | 0 | 0 | 0 | 0 | 98 | 49 | 0.973770068 | 0.999202116 | NA | | 1 | 0.778 | NA | 1 | NA | -0.111 |
| SE | 332 | AKNA | AKNA | chr9 | - | 114355924 | 114356136 | 114350858 | 114351021 | 114357920 | 114358167 | 0 | 2 | 12 | 0 | 0 | 1 | 0 | 3 | 0 | 0 | 0 | 0 | 98 | 49 | 0.974386813 | 0.999202116 | NA | | 1 | 0.857 | NA | 1 | NA | -0.072 |
| SE | 3924 | MTMR6 | MTMR6 | chr13 | - | 25251648 | 25251775 | 25246200 | 25249492 | 25251852 | 25251984 | 0 | 2 | 5 | 0 | 0 | 2 | 0 | 3 | 0 | 0 | 0 | 0 | 98 | 49 | 0.974386813 | 0.999202116 | NA | | 1 | 0.556 | NA | 1 | NA | -0.222 |
| SE | 11516 | PDXDC1 | PDXDC1 | chr16 | + | 15026642 | 15026706 | 15022703 | 15022754 | 15028877 | 15028966 | 0 | 2 | 0 | 0 | 0 | 2 | 0 | 3 | 0 | 0 | 0 | 0 | 98 | 49 | 0.974386813 | 0.999202116 | NA | | 1 | 0 | NA | 1 | NA | -0.5 |
| SE | 13864 | ACSL1 | ACSL1 | chr4 | - | 184770254 | 184770332 | 184768315 | 184768450 | 184773080 | 184773154 | 0 | 2 | 3 | 0 | 0 | 1 | 0 | 3 | 0 | 0 | 0 | 0 | 98 | 49 | 0.974386813 | 0.999202116 | NA | | 1 | 0.6 | NA | 1 | NA | -0.2 |
| A3SS | 1029 | ITGAM | ITGAM | chr16 | + | 31297741 | 31297954 | 31297744 | 31297954 | 31297513 | 31297654 | 0 | 2 | 2 | 0 | 1 | 7 | 0 | 3 | 0 | 6 | 1 | 2 | 52 | 49 | 0.975810876 | 0.999202116 | NA | | 0.653 | 0.212 | 0 | 0.739 | 0 | 0.186 |
| SE | 7116 | MED13L | MED13L | chr12 | - | 115968939 | 115969097 | 115966081 | 115966243 | 115970593 | 115970770 | 0 | 6 | 8 | 0 | 0 | 1 | 0 | 10 | 0 | 0 | 0 | 0 | 98 | 49 | 0.976427731 | 0.999202116 | NA | | 1 | 0.8 | NA | 1 | NA | -0.1 |
| SE | 12588 | MIDN | MIDN | chr19 | + | 1251561 | 1251649 | 1249889 | 1250529 | 1251838 | 1251901 | 0 | 6 | 6 | 0 | 0 | 1 | 0 | 10 | 0 | 0 | 0 | 0 | 98 | 49 | 0.976427731 | 0.999202116 | NA | | 1 | 0.75 | NA | 1 | NA | -0.125 |
| SE | 9816 | LDB1 | LDB1 | chr10 | - | 102110528 | 102110701 | 102109920 | 102110043 | 102110868 | 102110971 | 0 | 3 | 5 | 0 | 0 | 2 | 3 | 5 | 0 | 0 | 0 | 0 | 98 | 49 | 0.976610097 | 0.999202116 | NA | | 1 | 0.556 | 1 | 1 | NA | -0.222 |
| SE | 8808 | WDR1 | WDR1 | chr4 | - | 10103895 | 10103986 | 10098991 | 10099139 | 10116112 | 10116234 | 0 | 5 | 16 | 0 | 0 | 1 | 1 | 9 | 0 | 0 | 0 | 0 | 98 | 49 | 0.976663141 | 0.999202116 | NA | | 1 | 0.889 | 1 | 1 | NA | -0.055 |
| SE | 3259 | ELMO2 | ELMO2 | chr20 | - | 46387337 | 46387437 | 46386123 | 46386275 | 46389038 | 46389220 | 0 | 0 | 3 | 0 | 0 | 0 | 0 | 0 | 4 | 0 | 2 | 0 | 98 | 49 | 0.977411915 | 0.999202116 | NA | | NA | 1 | NA | 0 | 1 | 0.5 |
| SE | 186 | KDM6A | KDM6A | chrX | + | 45059005 | 45059104 | 45053828 | 45053955 | 45059246 | 45059466 | 0 | 4 | 4 | 0 | 0 | 1 | 0 | 3 | 0 | 0 | 0 | 0 | 98 | 49 | 0.99285348 | 0.999202116 | NA | | 1 | 0.667 | NA | 1 | NA | -0.166 |
| SE | 1798 | FBXO38 | FBXO38 | chr5 | + | 148427212 | 148427722 | 148425521 | 148425701 | 148433423 | 148433524 | 0 | 3 | 2 | 0 | 0 | 2 | 0 | 2 | 0 | 0 | 0 | 0 | 98 | 49 | 0.993205848 | 0.999202116 | NA | | 1 | 0.333 | NA | 1 | NA | -0.334 |
| SE | 5041 | RELT | RELT | chr11 | + | 73392210 | 73392468 | 73390754 | 73390921 | 73393836 | 73393917 | 0 | 0 | 3 | 0 | 0 | 0 | 0 | 6 | 2 | 2 | 3 | 0 | 98 | 49 | 0.993205848 | 0.999202116 | NA | | NA | 1 | 0 | 0.5 | 1 | 0.5 |
| SE | 5295 | PEX16 | PEX16 | chr11 | - | 45916226 | 45916303 | 45915702 | 45915836 | 45917457 | 45917493 | 0 | 0 | 3 | 0 | 0 | 0 | 0 | 5 | 2 | 0 | 1 | 0 | 98 | 49 | 0.993205848 | 0.999202116 | NA | | NA | 1 | NA | 0.714 | 1 | 0.143 |
| A3SS | 107 | ZNF189 | ZNF189 | chr9 | + | 101399883 | 101400010 | 101399925 | 101400010 | 101398853 | 101399189 | 0 | 0 | 1 | 0 | 0 | 1 | 3 | 0 | 1 | 0 | 0 | 1 | 91 | 49 | 0.997361002 | 0.999202116 | NA | | NA | 0.35 | 1 | NA | 0.35 | -0.325 |
| A5SS | 2698 | ANAPC11 | ANAPC11 | chr17 | + | 81894466 | 81894624 | 81894466 | 81894586 | 81899919 | 81900533 | 1 | 0 | 0 | 0 | 1 | 0 | 0 | 0 | 0 | 0 | 1 | 0 | 87 | 49 | 0.997625179 | 0.999208248 | | 1 | 0 | NA | NA | 0 | NA | 0.5 |
| A3SS | 70 | MAN1B1 | MAN1B1 | chr9 | + | 137107249 | 137107447 | 137107274 | 137107447 | 137106688 | 137106809 | 0 | 0 | 3 | 0 | 0 | 0 | 0 | 0 | 2 | 0 | 1 | 0 | 74 | 49 | 0.999202116 | 0.999202116 | NA | | NA | 1 | NA | 0 | 1 | 0.5 |
| A3SS | 338 | CUTA | CUTA | chr6 | - | 33417480 | 33417816 | 33417480 | 33417695 | 33417958 | 33418288 | 0 | 1 | 1 | 0 | 0 | 1 | 0 | 1 | 0 | 0 | 0 | 0 | 98 | 49 | 0.999202116 | 0.999202116 | NA | | 1 | 0.333 | NA | 1 | NA | -0.334 |
| A3SS | 776 | LST1 | LST1 | chr6 | + | 31588517 | 31588909 | 31588562 | 31588909 | 31587199 | 31587318 | 0 | 3 | 14 | 0 | 1 | 6 | 4 | 15 | 0 | 0 | 2 | 1 | 94 | 49 | 0.999202116 | 0.999202116 | NA | | 0.61 | 0.549 | 1 | 0.796 | 0 | -0.019 |
| A3SS | 780 | LST1 | LST1 | chr6 | + | 31588496 | 31588909 | 31588517 | 31588909 | 31587943 | 31587966 | 0 | 1 | 0 | 0 | 0 | 6 | 0 | 0 | 0 | 0 | 0 | 1 | 70 | 49 | 0.999202116 | 0.999202116 | NA | | 1 | 0 | NA | NA | 0 | 0.5 |
| A3SS | 804 | TYMP | TYMP | chr22 | - | 50529495 | 50529731 | 50529495 | 50529719 | 50529903 | 50530085 | 0 | 1 | 6 | 0 | 0 | 13 | 0 | 20 | 1 | 0 | 3 | 2 | 61 | 49 | 0.999202116 | 0.999202116 | NA | | 1 | 0.27 | NA | 0.843 | 0.287 | 0.07 |
| A3SS | 805 | TYMP | TYMP | chr22 | - | 50529495 | 50529754 | 50529495 | 50529719 | 50529903 | 50530085 | 0 | 3 | 9 | 0 | 0 | 13 | 3 | 22 | 1 | 0 | 3 | 2 | 84 | 49 | 0.999202116 | 0.999202116 | NA | | 1 | 0.288 | 1 | 0.811 | 0.226 | -0.035 |
| A3SS | 832 | TOM1 | TOM1 | chr22 | + | 35322998 | 35323177 | 35323027 | 35323177 | 35321958 | 35322037 | 0 | 1 | 5 | 0 | 4 | 6 | 0 | 0 | 3 | 0 | 4 | 0 | 78 | 49 | 0.999202116 | 0.999202116 | NA | | 0.136 | 0.344 | NA | 0 | 1 | -0.26 |
| A3SS | 837 | TUG1 | TUG1 | chr22 | + | 30972854 | 30973601 | 30972923 | 30973601 | 30969210 | 30971779 | 0 | 0 | 1 | 0 | 0 | 0 | 3 | 2 | 1 | 0 | 2 | 0 | 98 | 49 | 0.999202116 | 0.999202116 | NA | | NA | 1 | 1 | 0.333 | 1 | 0.222 |
| A3SS | 845 | GAS2L1 | GAS2L1 | chr22 | + | 29308058 | 29308738 | 29308086 | 29308738 | 29306994 | 29307153 | 2 | 0 | 0 | 0 | 1 | 0 | 2 | 0 | 0 | 0 | 0 | 0 | 77 | 49 | 0.999202116 | 0.999202116 | | 1 | 0 | NA | 1 | NA | NA | -0.5 |
| A3SS | 1313 | BRD9 | BRD9 | chr5 | - | 889586 | 889766 | 889586 | 889647 | 891154 | 891287 | 0 | 2 | 2 | 0 | 0 | 2 | 0 | 2 | 0 | 0 | 0 | 0 | 98 | 49 | 0.999202116 | 0.999202116 | NA | | 1 | 0.333 | NA | 1 | NA | -0.334 |
| A3SS | 1412 | GLIPR2 | GLIPR2 | chr9 | + | 36162231 | 36163913 | 36162361 | 36163913 | 36150871 | 36150949 | 0 | 1 | 6 | 0 | 0 | 5 | 0 | 2 | 0 | 0 | 5 | 2 | 98 | 49 | 0.999202116 | 0.999202116 | NA | | 1 | 0.375 | NA | 0.167 | 0 | 0.604 |
| A3SS | 1450 | SPI1 | SPI1 | chr11 | - | 47375632 | 47375732 | 47375632 | 47375729 | 47378308 | 47378576 | 0 | 8 | 16 | 0 | 4 | 19 | 0 | 9 | 1 | 0 | 9 | 2 | 52 | 49 | 0.999202116 | 0.999202116 | NA | | 0.653 | 0.442 | NA | 0.485 | 0.32 | 0.145 |
| A3SS | 1515 | KRCC1 | KRCC1 | chr2 | - | 88034133 | 88034292 | 88034133 | 88034231 | 88036942 | 88037051 | 0 | 0 | 1 | 0 | 0 | 0 | 0 | 0 | 1 | 0 | 1 | 0 | 98 | 49 | 0.999202116 | 0.999202116 | NA | | NA | 1 | NA | 0 | 1 | 0.5 |
| A3SS | 1711 | NFKB2 | NFKB2 | chr10 | + | 102402251 | 102402529 | 102402254 | 102402529 | 102402047 | 102402159 | 0 | 1 | 3 | 0 | 0 | 3 | 0 | 6 | 0 | 0 | 0 | 1 | 52 | 49 | 0.999202116 | 0.999202116 | NA | | 1 | 0.485 | NA | 1 | 0 | 0.242 |
| A3SS | 1716 | YY1AP1 | YY1AP1 | chr1 | - | 155668626 | 155668777 | 155668626 | 155668717 | 155670319 | 155670464 | 0 | 2 | 1 | 0 | 0 | 3 | 0 | 2 | 0 | 0 | 0 | 0 | 98 | 49 | 0.999202116 | 0.999202116 | NA | | 1 | 0.143 | NA | 1 | NA | -0.428 |
| A3SS | 1761 | ATP11A | ATP11A | chr13 | + | 112881815 | 112887168 | 112881875 | 112887168 | 112875775 | 112875941 | 1 | 3 | 1 | 0 | 0 | 1 | 0 | 1 | 0 | 0 | 0 | 0 | 98 | 49 | 0.999202116 | 0.999202116 | | 1 | 1 | 0.333 | NA | 1 | NA | -0.222 |
| A3SS | 1775 | PHYKPL | PHYKPL | chr5 | - | 178222354 | 178222580 | 178222354 | 178222492 | 178222851 | 178222934 | 0 | 0 | 5 | 0 | 0 | 0 | 0 | 5 | 1 | 0 | 1 | 0 | 98 | 49 | 0.999202116 | 0.999202116 | NA | | NA | 1 | NA | 0.714 | 1 | 0.143 |
| A3SS | 1875 | ITGB2 | ITGB2 | chr21 | - | 44910283 | 44910415 | 44910283 | 44910372 | 44910724 | 44910785 | 0 | 1 | 3 | 0 | 1 | 16 | 0 | 1 | 1 | 2 | 6 | 1 | 92 | 49 | 0.999202116 | 0.999202116 | NA | | 0.348 | 0.091 | 0 | 0.082 | 0.348 | 0.076 |
| A3SS | 1896 | ADAT1 | ADAT1 | chr16 | - | 75618585 | 75618698 | 75618585 | 75618640 | 75620265 | 75620334 | 0 | 0 | 0 | 0 | 1 | 0 | 0 | 0 | 1 | 0 | 1 | 0 | 98 | 49 | 0.999202116 | 0.999202116 | NA | | 0 | NA | NA | 0 | 1 | -0.5 |
| A3SS | 2090 | STAT2 | STAT2 | chr12 | - | 56355707 | 56355803 | 56355707 | 56355791 | 56356131 | 56356285 | 0 | 4 | 17 | 0 | 0 | 3 | 0 | 6 | 2 | 0 | 1 | 0 | 61 | 49 | 0.999202116 | 0.999202116 | NA | | 1 | 0.82 | NA | 0.828 | 1 | -0.004 |
| A3SS | 2169 | HIF1A | HIF1A | chr14 | + | 61720378 | 61720572 | 61720381 | 61720572 | 61695400 | 61695839 | 0 | 1 | 4 | 0 | 1 | 14 | 0 | 11 | 0 | 1 | 25 | 3 | 52 | 49 | 0.999202116 | 0.999202116 | NA | | 0.485 | 0.212 | 0 | 0.293 | 0 | 0.251 |
| A3SS | 2223 | CSF3R | CSF3R | chr1 | - | 36466042 | 36466908 | 36466042 | 36466827 | 36467229 | 36467311 | 1 | 3 | 22 | 0 | 17 | 96 | 3 | 23 | 5 | 11 | 73 | 20 | 98 | 49 | 0.999202116 | 0.999202116 | | 1 | 0.081 | 0.103 | 0.12 | 0.136 | 0.111 | 0.272 |
| A3SS | 2815 | DEF8 | DEF8 | chr16 | + | 89964169 | 89964310 | 89964220 | 89964310 | 89963362 | 89963443 | 0 | 3 | 3 | 0 | 0 | 1 | 0 | 1 | 0 | 0 | 0 | 0 | 98 | 49 | 0.999202116 | 0.999202116 | NA | | 1 | 0.6 | NA | 1 | NA | -0.2 |
| A3SS | 3095 | MEF2D | MEF2D | chr1 | - | 156483238 | 156483430 | 156483238 | 156483418 | 156500485 | 156500842 | 0 | 5 | 4 | 0 | 0 | 0 | 0 | 10 | 0 | 0 | 0 | 1 | 61 | 49 | 0.999202116 | 0.999202116 | NA | | 1 | 1 | NA | 1 | 0 | 0.5 |
| A3SS | 3209 | ZNF276 | ZNF276 | chr16 | + | 89737720 | 89737905 | 89737805 | 89737905 | 89733920 | 89734038 | 0 | 1 | 1 | 0 | 0 | 5 | 0 | 1 | 0 | 0 | 0 | 0 | 98 | 49 | 0.999202116 | 0.999202116 | NA | | 1 | 0.091 | NA | 1 | NA | -0.455 |
| A3SS | 3689 | DDX49 | DDX49 | chr19 | + | 18920579 | 18920703 | 18920650 | 18920703 | 18919674 | 18919856 | 0 | 0 | 1 | 0 | 0 | 0 | 0 | 1 | 1 | 0 | 1 | 0 | 98 | 49 | 0.999202116 | 0.999202116 | NA | | NA | 1 | NA | 0.333 | 1 | 0.334 |
| A3SS | 3839 | LILRB2 | LILRB2 | chr19 | - | 54277549 | 54277600 | 54277549 | 54277597 | 54277888 | 54277939 | 0 | 1 | 2 | 0 | 0 | 6 | 1 | 1 | 0 | 2 | 1 | 1 | 52 | 49 | 0.999202116 | 0.999202116 | NA | | 1 | 0.239 | 0.32 | 0.485 | 0 | 0.351 |
| A5SS | 59 | RBM10 | RBM10 | chrX | + | 47145217 | 47145516 | 47145217 | 47145485 | 47147356 | 47147498 | 0 | 0 | 2 | 0 | 2 | 3 | 0 | 0 | 0 | 0 | 2 | 0 | 80 | 49 | 0.999202116 | 0.999208248 | NA | | 0 | 0.29 | NA | 0 | NA | 0.145 |
| A5SS | 140 | GRINA | GRINA | chr8 | + | 143990057 | 143990311 | 143990057 | 143990199 | 143991199 | 143991602 | 0 | 1 | 0 | 0 | 0 | 0 | 0 | 1 | 8 | 0 | 0 | 1 | 98 | 49 | 0.999202116 | 0.999208248 | NA | | 1 | NA | NA | 1 | 0.8 | 0.1 |
| A5SS | 211 | SAMD9L | SAMD9L | chr7 | - | 93146882 | 93147146 | 93147087 | 93147146 | 93145384 | 93146040 | 0 | 4 | 10 | 0 | 0 | 6 | 2 | 1 | 0 | 0 | 0 | 0 | 98 | 49 | 0.999202116 | 0.999208248 | NA | | 1 | 0.455 | 1 | 1 | NA | -0.272 |
| A5SS | 388 | SYTL3 | SYTL3 | chr6 | + | 158662749 | 158663378 | 158662749 | 158662812 | 158665394 | 158665613 | 0 | 1 | 2 | 0 | 0 | 1 | 2 | 2 | 1 | 0 | 0 | 1 | 98 | 49 | 0.999202116 | 0.999208248 | NA | | 1 | 0.5 | 1 | 1 | 0.333 | -0.028 |
| A5SS | 615 | TOM1 | TOM1 | chr22 | + | 35346929 | 35346972 | 35346929 | 35346969 | 35347054 | 35347994 | 0 | 0 | 0 | 0 | 0 | 5 | 0 | 2 | 0 | 0 | 4 | 1 | 52 | 49 | 0.999202116 | 0.999208248 | NA | | NA | 0 | NA | 0.32 | 0 | -0.16 |
| A5SS | 671 | STX16 | STX16 | chr20 | + | 58651252 | 58652138 | 58651252 | 58652087 | 58659622 | 58659634 | 0 | 4 | 7 | 0 | 0 | 1 | 1 | 1 | 0 | 0 | 0 | 0 | 98 | 49 | 0.999202116 | 0.999208248 | NA | | 1 | 0.778 | 1 | 1 | NA | -0.111 |
| A5SS | 745 | TAZ | TAZ | chrX | + | 154413206 | 154413358 | 154413206 | 154413252 | 154413481 | 154413567 | 0 | 0 | 4 | 0 | 2 | 1 | 0 | 0 | 0 | 0 | 1 | 0 | 98 | 49 | 0.999202116 | 0.999208248 | NA | | 0 | 0.667 | NA | 0 | NA | 0.334 |
| A5SS | 791 | SLAIN1 | SLAIN1 | chr13 | + | 77744282 | 77744516 | 77744282 | 77744432 | 77746513 | 77746855 | 1 | 0 | 0 | 0 | 0 | 1 | 1 | 0 | 0 | 0 | 0 | 0 | 98 | 49 | 0.999202116 | 0.999208248 | | 1 | NA | 0 | 1 | NA | NA | -0.5 |
| A5SS | 1115 | GLIPR2 | GLIPR2 | chr9 | + | 36147785 | 36147916 | 36147785 | 36147894 | 36148546 | 36148650 | 0 | 0 | 3 | 0 | 2 | 3 | 0 | 0 | 0 | 0 | 1 | 0 | 71 | 49 | 0.999202116 | 0.999208248 | NA | | 0 | 0.408 | NA | 0 | NA | 0.204 |
| A5SS | 1257 | NIPA2 | NIPA2 | chr15 | + | 22838640 | 22838947 | 22838640 | 22838921 | 22845145 | 22845267 | 0 | 2 | 1 | 0 | 0 | 0 | 0 | 1 | 0 | 1 | 0 | 0 | 75 | 49 | 0.999202116 | 0.999208248 | NA | | 1 | 1 | 0 | 1 | NA | 0.5 |
| A5SS | 1813 | NPL | NPL | chr1 | + | 182806144 | 182806531 | 182806144 | 182806232 | 182812155 | 182812213 | 0 | 0 | 10 | 0 | 0 | 0 | 0 | 0 | 2 | 0 | 2 | 0 | 98 | 49 | 0.999202116 | 0.999208248 | NA | | NA | 1 | NA | 0 | 1 | 0.5 |
| A5SS | 2009 | ABTB1 | ABTB1 | chr3 | + | 127674390 | 127674453 | 127674390 | 127674418 | 127674544 | 127674600 | 0 | 2 | 6 | 0 | 0 | 1 | 0 | 6 | 1 | 0 | 0 | 1 | 84 | 49 | 0.999202116 | 0.999208248 | NA | | 1 | 0.778 | NA | 1 | 0.368 | 0.205 |
| A5SS | 2108 | DEF8 | DEF8 | chr16 | + | 89949416 | 89949638 | 89949416 | 89949513 | 89954242 | 89954376 | 2 | 0 | 0 | 0 | 0 | 0 | 2 | 1 | 0 | 0 | 1 | 0 | 98 | 49 | 0.999202116 | 0.999208248 | | 1 | NA | NA | 1 | 0.333 | NA | 0.334 |
| A5SS | 2486 | BANP | BANP | chr16 | + | 87975047 | 87975203 | 87975047 | 87975185 | 87981035 | 87981127 | 0 | 0 | 2 | 0 | 1 | 1 | 0 | 0 | 0 | 0 | 1 | 0 | 67 | 49 | 0.999202116 | 0.999208248 | NA | | 0 | 0.594 | NA | 0 | NA | 0.297 |
| A5SS | 2983 | YPEL5 | YPEL5 | chr2 | + | 30148244 | 30148541 | 30148244 | 30148305 | 30155849 | 30155910 | 0 | 0 | 8 | 0 | 0 | 0 | 0 | 6 | 1 | 0 | 1 | 0 | 98 | 49 | 0.999202116 | 0.999208248 | NA | | NA | 1 | NA | 0.75 | 1 | 0.125 |
| A5SS | 2984 | YPEL5 | YPEL5 | chr2 | + | 30148244 | 30148541 | 30148244 | 30148305 | 30156627 | 30156792 | 0 | 0 | 8 | 0 | 0 | 0 | 0 | 6 | 1 | 0 | 1 | 0 | 98 | 49 | 0.999202116 | 0.999208248 | NA | | NA | 1 | NA | 0.75 | 1 | 0.125 |
| A5SS | 2989 | NDUFAF7 | NDUFAF7 | chr2 | + | 37241577 | 37241791 | 37241577 | 37241637 | 37242634 | 37242693 | 0 | 0 | 1 | 0 | 0 | 0 | 0 | 2 | 1 | 0 | 1 | 0 | 98 | 49 | 0.999202116 | 0.999208248 | NA | | NA | 1 | NA | 0.5 | 1 | 0.25 |
| RI | 91 | CUTA | CUTA | chr6 | - | 33417480 | 33418288 | 33417480 | 33417695 | 33417958 | 33418288 | 0 | 1 | 1 | 0 | 0 | 1 | 4 | 1 | 0 | 0 | 0 | 0 | 98 | 49 | 0.999202116 | 0.999202116 | NA | | 1 | 0.333 | 1 | 1 | NA | -0.334 |
| RI | 250 | TYMP | TYMP | chr22 | - | 50529495 | 50530085 | 50529495 | 50529719 | 50529903 | 50530085 | 1 | 8 | 20 | 0 | 0 | 13 | 3 | 30 | 5 | 0 | 3 | 2 | 98 | 49 | 0.999202116 | 0.999202116 | | 1 | 1 | 0.435 | 1 | 0.833 | 0.556 | 0.015 |
| RI | 270 | POFUT2 | POFUT2 | chr21 | - | 45263927 | 45267713 | 45263927 | 45265635 | 45267589 | 45267713 | 0 | 0 | 3 | 0 | 1 | 0 | 3 | 1 | 1 | 0 | 1 | 0 | 98 | 49 | 0.999202116 | 0.999202116 | NA | | 0 | 1 | 1 | 0.333 | 1 | -0.278 |
| RI | 293 | SMOX | SMOX | chr20 | + | 4182088 | 4182848 | 4182088 | 4182323 | 4182482 | 4182848 | 3 | 1 | 0 | 0 | 1 | 0 | 1 | 0 | 0 | 0 | 0 | 0 | 98 | 49 | 0.999202116 | 0.999202116 | | 1 | 0.333 | NA | 1 | NA | NA | -0.334 |
| RI | 400 | ULK3 | ULK3 | chr15 | - | 74840233 | 74840641 | 74840233 | 74840316 | 74840497 | 74840641 | 0 | 2 | 1 | 0 | 0 | 3 | 0 | 2 | 0 | 0 | 0 | 0 | 98 | 49 | 0.999202116 | 0.999202116 | NA | | 1 | 0.143 | NA | 1 | NA | -0.428 |
| RI | 467 | TP53 | TP53 | chr17 | - | 7676381 | 7676622 | 7676381 | 7676403 | 7676520 | 7676622 | 3 | 0 | 0 | 0 | 1 | 0 | 1 | 0 | 0 | 0 | 0 | 0 | 98 | 49 | 0.999202116 | 0.999202116 | | 1 | 0 | NA | 1 | NA | NA | -0.5 |
| RI | 496 | DNAJC4 | DNAJC4 | chr11 | + | 64233893 | 64234281 | 64233893 | 64233980 | 64234072 | 64234281 | 0 | 0 | 4 | 0 | 0 | 0 | 2 | 1 | 1 | 0 | 1 | 0 | 98 | 49 | 0.999202116 | 0.999202116 | NA | | NA | 1 | 1 | 0.333 | 1 | 0.222 |
| RI | 510 | RAB24 | RAB24 | chr5 | - | 177302127 | 177302496 | 177302127 | 177302178 | 177302396 | 177302496 | 0 | 3 | 14 | 0 | 0 | 12 | 2 | 7 | 0 | 0 | 3 | 2 | 98 | 49 | 0.999202116 | 0.999202116 | NA | | 1 | 0.368 | 1 | 0.538 | 0 | 0.171 |
| RI | 511 | RAB24 | RAB24 | chr5 | - | 177303171 | 177303744 | 177303171 | 177303320 | 177303406 | 177303744 | 2 | 6 | 10 | 0 | 2 | 1 | 1 | 3 | 23 | 0 | 1 | 2 | 98 | 49 | 0.999202116 | 0.999202116 | | 1 | 0.6 | 0.833 | 1 | 0.6 | 0.852 | -0.006 |
| RI | 582 | CANT1 | CANT1 | chr17 | - | 78996991 | 78997963 | 78996991 | 78997644 | 78997839 | 78997963 | 2 | 0 | 8 | 0 | 0 | 0 | 0 | 3 | 3 | 0 | 1 | 0 | 98 | 49 | 0.999202116 | 0.999202116 | | 1 | NA | 1 | NA | 0.6 | 1 | 0.2 |
| RI | 669 | TSTD1 | TSTD1 | chr1 | - | 161037631 | 161038075 | 161037631 | 161037826 | 161037912 | 161038075 | 0 | 0 | 0 | 0 | 0 | 1 | 0 | 1 | 0 | 0 | 1 | 1 | 98 | 49 | 0.999202116 | 0.999202116 | NA | | NA | 0 | NA | 0.333 | 0 | -0.167 |
| RI | 685 | SAMD8 | SAMD8 | chr10 | + | 75176065 | 75182123 | 75176065 | 75176216 | 75176387 | 75182123 | 4 | 0 | 0 | 0 | 0 | 4 | 2 | 6 | 0 | 0 | 2 | 0 | 98 | 49 | 0.999202116 | 0.999202116 | | 1 | NA | 0 | 1 | 0.6 | NA | -0.3 |
| RI | 818 | NCF4 | NCF4 | chr22 | + | 36875652 | 36876094 | 36875652 | 36875783 | 36876028 | 36876094 | 1 | 2 | 11 | 0 | 0 | 11 | 0 | 2 | 0 | 0 | 4 | 1 | 98 | 49 | 0.999202116 | 0.999202116 | | 1 | 1 | 0.333 | NA | 0.2 | 0 | 0.678 |
| RI | 913 | UBXN1 | UBXN1 | chr11 | - | 62676497 | 62677005 | 62676497 | 62676639 | 62676812 | 62677005 | 0 | 0 | 0 | 0 | 0 | 7 | 0 | 3 | 0 | 0 | 2 | 1 | 98 | 49 | 0.999202116 | 0.999202116 | NA | | NA | 0 | NA | 0.429 | 0 | -0.214 |
| RI | 919 | ACADVL | ACADVL | chr17 | + | 7219830 | 7220197 | 7219830 | 7220046 | 7220121 | 7220197 | 0 | 2 | 1 | 0 | 2 | 3 | 0 | 0 | 2 | 0 | 2 | 0 | 98 | 49 | 0.999202116 | 0.999202116 | NA | | 0.333 | 0.143 | NA | 0 | 1 | -0.262 |
| RI | 1114 | OSCAR | OSCAR | chr19 | - | 54094667 | 54096153 | 54094667 | 54095357 | 54095871 | 54096153 | 3 | 2 | 1 | 0 | 0 | 1 | 4 | 1 | 0 | 0 | 0 | 0 | 98 | 49 | 0.999202116 | 0.999202116 | | 1 | 1 | 0.333 | 1 | 1 | NA | -0.222 |
| RI | 1116 | AUP1 | AUP1 | chr2 | - | 74528750 | 74529282 | 74528750 | 74528935 | 74529131 | 74529282 | 0 | 1 | 0 | 0 | 0 | 2 | 2 | 1 | 0 | 0 | 0 | 0 | 98 | 49 | 0.999202116 | 0.999202116 | NA | | 1 | 0 | 1 | 1 | NA | -0.5 |
| SE | 172 | PIM2 | PIM2 | chrX | - | 48914394 | 48914571 | 48913181 | 48914294 | 48915019 | 48915392 | 0 | 0 | 2 | 0 | 1 | 2 | 0 | 9 | 1 | 0 | 4 | 0 | 98 | 49 | 0.999202116 | 0.999202116 | NA | | 0 | 0.333 | NA | 0.529 | 1 | -0.598 |
| SE | 280 | ASB6 | ASB6 | chr9 | - | 129638572 | 129638659 | 129634603 | 129638457 | 129639201 | 129639310 | 0 | 0 | 1 | 0 | 0 | 0 | 0 | 2 | 1 | 0 | 1 | 0 | 98 | 49 | 0.999202116 | 0.999202116 | NA | | NA | 1 | NA | 0.5 | 1 | 0.25 |
| SE | 390 | TMEM2 | TMEM2 | chr9 | - | 71734805 | 71734994 | 71732350 | 71732520 | 71740062 | 71740232 | 0 | 0 | 6 | 0 | 0 | 0 | 0 | 4 | 1 | 0 | 1 | 0 | 98 | 49 | 0.999202116 | 0.999202116 | NA | | NA | 1 | NA | 0.667 | 1 | 0.166 |
| SE | 626 | LYN | LYN | chr8 | + | 55941854 | 55941928 | 55879826 | 55880103 | 55946447 | 55946493 | 0 | 2 | 9 | 0 | 0 | 0 | 0 | 13 | 0 | 0 | 0 | 1 | 98 | 49 | 0.999202116 | 0.999202116 | NA | | 1 | 1 | NA | 1 | 0 | 0.5 |
| SE | 627 | LYN | LYN | chr8 | + | 55941854 | 55941991 | 55879826 | 55880103 | 55946447 | 55946493 | 0 | 2 | 9 | 0 | 0 | 0 | 0 | 11 | 0 | 0 | 0 | 1 | 98 | 49 | 0.999202116 | 0.999202116 | NA | | 1 | 1 | NA | 1 | 0 | 0.5 |
| SE | 694 | DOCK5 | DOCK5 | chr8 | + | 25342400 | 25342507 | 25341738 | 25341809 | 25345474 | 25345611 | 0 | 0 | 15 | 0 | 0 | 0 | 0 | 15 | 2 | 0 | 1 | 0 | 98 | 49 | 0.999202116 | 0.999202116 | NA | | NA | 1 | NA | 0.882 | 1 | 0.059 |
| SE | 697 | DOCK5 | DOCK5 | chr8 | + | 25407982 | 25408154 | 25403557 | 25403724 | 25408801 | 25408940 | 0 | 1 | 23 | 0 | 1 | 1 | 0 | 9 | 3 | 0 | 3 | 0 | 98 | 49 | 0.999202116 | 0.999202116 | NA | | 0.333 | 0.92 | NA | 0.6 | 1 | -0.173 |
| SE | 789 | PARP10 | PARP10 | chr8 | - | 143985411 | 143985648 | 143984543 | 143985328 | 143985720 | 143985975 | 2 | 3 | 2 | 0 | 0 | 1 | 0 | 6 | 0 | 1 | 0 | 0 | 98 | 49 | 0.999202116 | 0.999202116 | | 1 | 1 | 0.5 | 0 | 1 | NA | 0.333 |

| | | | | | | | | | | | | | | | | | | | | | | | | | | | | | | | | | | |
|---|---|---|---|---|---|---|---|---|---|---|---|---|---|---|---|---|---|---|---|---|---|---|---|---|---|---|---|---|---|---|---|---|---|---|
| SE | 925 | SAMD9L | SAMD9L | chr7 | - | 93146882 | 93147146 | 93145384 | 93146040 | 93148193 | 93148388 | 0 | 2 | 5 | 0 | 0 | 1 | 0 | 1 | 0 | 0 | 0 | 0 | 98 | 49 | 0.999202116 | 0.999202116 | NA | 1 | 0.714 | NA | 1 | NA | -0.143 |
| SE | 926 | SAMD9L | SAMD9L | chr7 | - | 93147087 | 93147146 | 93145384 | 93146040 | 93148193 | 93148388 | 0 | 1 | 7 | 0 | 0 | 1 | 0 | 1 | 0 | 0 | 0 | 0 | 98 | 49 | 0.999202116 | 0.999202116 | NA | 1 | 0.778 | NA | 1 | NA | -0.111 |
| SE | 1015 | IKZF1 | IKZF1 | chr7 | + | 50387344 | 50387470 | 50382539 | 50382707 | 50391728 | 50391833 | 0 | 2 | 10 | 0 | 0 | 1 | 0 | 1 | 0 | 0 | 0 | 0 | 98 | 49 | 0.999202116 | 0.999202116 | NA | 1 | 0.833 | NA | 1 | NA | -0.084 |
| SE | 1135 | IGF2R | IGF2R | chr6 | + | 160045744 | 160045882 | 160044513 | 160044657 | 160046497 | 160046645 | 0 | 0 | 35 | 0 | 1 | 1 | 3 | 14 | 1 | 0 | 1 | 0 | 98 | 49 | 0.999202116 | 0.999202116 | NA | 0 | 0.946 | 1 | 0.875 | 1 | -0.485 |
| SE | 1190 | PHACTR2 | PHACTR2 | chr6 | + | 143765260 | 143765798 | 143753753 | 143753912 | 143772257 | 143772457 | 0 | 1 | 4 | 0 | 0 | 1 | 0 | 1 | 0 | 0 | 0 | 0 | 98 | 49 | 0.999202116 | 0.999202116 | NA | 1 | 0.667 | NA | 1 | NA | -0.166 |
| SE | 1209 | VNN2 | VNN2 | chr6 | - | 132752460 | 132752749 | 132751144 | 132751518 | 132755842 | 132756035 | 0 | 0 | 32 | 0 | 0 | 0 | 0 | 18 | 1 | 0 | 5 | 0 | 98 | 49 | 0.999202116 | 0.999202116 | NA | NA | 1 | NA | 0.643 | 1 | 0.178 |
| SE | 1212 | VNN2 | VNN2 | chr6 | - | 132753417 | 132753491 | 132751144 | 132751518 | 132755842 | 132756035 | 0 | 0 | 3 | 0 | 0 | 0 | 0 | 1 | 1 | 0 | 5 | 0 | 98 | 49 | 0.999202116 | 0.999202116 | NA | NA | 1 | NA | 0.091 | 1 | 0.455 |
| SE | 1508 | HLA-E | HLA-E | chr6 | + | 30492403 | 30492436 | 30491536 | 30491653 | 30492540 | 30492583 | 0 | 26 | 172 | 0 | 0 | 1 | 10 | 116 | 23 | 0 | 1 | 1 | 82 | 49 | 0.999202116 | 0.999202116 | NA | 1 | 0.99 | 1 | 0.986 | 0.932 | 0.022 |
| SE | 1592 | KIF13A | KIF13A | chr6 | - | 17773477 | 17773583 | 17771907 | 17772059 | 17775014 | 17775062 | 0 | 0 | 5 | 0 | 0 | 1 | 2 | 7 | 1 | 0 | 2 | 0 | 98 | 49 | 0.999202116 | 0.999202116 | NA | NA | 1 | 1 | 0.636 | 1 | 0.121 |
| SE | 1593 | KIF13A | KIF13A | chr6 | - | 17789871 | 17789910 | 17787775 | 17787875 | 17794248 | 17794395 | 0 | 0 | 1 | 0 | 2 | 1 | 0 | 0 | 0 | 0 | 2 | 0 | 88 | 49 | 0.999202116 | 0.999202116 | NA | 0 | 0.358 | NA | 0 | NA | 0.179 |
| SE | 1709 | NCF1 | NCF1 | chr7 | + | 74779081 | 74779157 | 74777266 | 74777347 | 74779256 | 74779422 | 0 | 0 | 4 | 0 | 0 | 0 | 2 | 0 | 1 | 0 | 1 | 0 | 98 | 49 | 0.999202116 | 0.999202116 | NA | NA | 1 | 1 | 0 | 1 | 0.333 |
| SE | 1738 | ARHGAP26 | ARHGAP26 | chr5 | + | 143037195 | 143037261 | 143014079 | 143014116 | 143041815 | 143041890 | 0 | 0 | 32 | 0 | 0 | 0 | 0 | 12 | 2 | 0 | 1 | 0 | 98 | 49 | 0.999202116 | 0.999202116 | NA | | 1 | NA | 0.857 | 1 | 0.072 |
| SE | 1779 | ASPH | ASPH | chr8 | - | 61553030 | 61553120 | 61548070 | 61548208 | 61555923 | 61556022 | 0 | 2 | 4 | 0 | 0 | 1 | 0 | 1 | 0 | 0 | 0 | 0 | 98 | 49 | 0.999202116 | 0.999202116 | NA | 1 | 0.667 | NA | 1 | NA | -0.166 |
| SE | 1782 | ASPH | ASPH | chr8 | - | 61682438 | 61682483 | 61680967 | 61681036 | 61684038 | 61684188 | 0 | 0 | 0 | 0 | 0 | 3 | 0 | 1 | 0 | 0 | 5 | 1 | 94 | 49 | 0.999202116 | 0.999202116 | NA | NA | 0 | NA | 0.094 | 0 | -0.047 |
| SE | 1800 | AP3S1 | AP3S1 | chr5 | + | 115902884 | 115902992 | 115895086 | 115895158 | 115913361 | 115914081 | 1 | 0 | 0 | 0 | 0 | 0 | 1 | 1 | 0 | 0 | 1 | 0 | 98 | 49 | 0.999202116 | 0.999202116 | 1 | NA | NA | 1 | 0.333 | NA | 0.334 |
| SE | 1920 | PARP8 | PARP8 | chr5 | + | 50794206 | 50794332 | 50788522 | 50788589 | 50794852 | 50795417 | 0 | 0 | 34 | 0 | 0 | 0 | 7 | 9 | 1 | 0 | 2 | 0 | 98 | 49 | 0.999202116 | 0.999202116 | NA | NA | 1 | 1 | 0.692 | 1 | 0.103 |
| SE | 1987 | DDX60 | DDX60 | chr4 | - | 168283445 | 168283606 | 168280334 | 168280590 | 168284819 | 168284935 | 0 | 0 | 18 | 0 | 0 | 0 | 0 | 9 | 1 | 0 | 2 | 0 | 98 | 49 | 0.999202116 | 0.999202116 | NA | NA | 1 | NA | 0.692 | 1 | 0.154 |
| SE | 2104 | WDFY3 | WDFY3 | chr4 | - | 84932269 | 84932363 | 84896910 | 84897010 | 84966208 | 84966391 | 0 | 2 | 13 | 1 | 0 | 0 | 1 | 13 | 9 | 0 | 0 | 5 | 98 | 49 | 0.999202116 | 0.999202116 | 0 | 1 | 1 | 1 | 1 | 0.474 | -0.158 |
| SE | 2273 | VNN3 | VNN3 | chr6 | - | 132726721 | 132727012 | 132724605 | 132724979 | 132728740 | 132728933 | 0 | 0 | 11 | 0 | 1 | 1 | 0 | 6 | 3 | 0 | 7 | 1 | 98 | 49 | 0.999202116 | 0.999202116 | NA | 0 | 0.846 | NA | 0.3 | 0.6 | -0.027 |
| SE | 2370 | MFSD1 | MFSD1 | chr3 | + | 158805361 | 158805474 | 158804318 | 158804371 | 158807039 | 158807082 | 0 | 0 | 10 | 0 | 0 | 0 | 0 | 2 | 2 | 0 | 1 | 0 | 98 | 49 | 0.999202116 | 0.999202116 | NA | NA | 1 | NA | 0.5 | 1 | 0.25 |
| SE | 2467 | HMGA1 | HMGA1 | chr6 | + | 34242711 | 34242795 | 34240736 | 34240915 | 34243467 | 34243518 | 0 | 1 | 3 | 0 | 0 | 0 | 0 | 2 | 0 | 0 | 0 | 1 | 98 | 49 | 0.999202116 | 0.999202116 | NA | 1 | 1 | NA | 1 | 0 | 0.5 |
| SE | 2582 | PRKCD | PRKCD | chr3 | + | 53185926 | 53186027 | 53185603 | 53185700 | 53186166 | 53186340 | 0 | 3 | 4 | 0 | 0 | 4 | 0 | 3 | 0 | 0 | 0 | 0 | 98 | 49 | 0.999202116 | 0.999202116 | NA | 1 | 0.333 | NA | 1 | NA | -0.334 |
| SE | 2584 | LCP2 | LCP2 | chr5 | - | 170275319 | 170275351 | 170274300 | 170274338 | 170275794 | 170275860 | 0 | 0 | 23 | 0 | 0 | 0 | 0 | 16 | 3 | 0 | 1 | 0 | 81 | 49 | 0.999202116 | 0.999202116 | NA | NA | 1 | NA | 0.906 | 1 | 0.047 |
| SE | 2714 | TOP2B | TOP2B | chr3 | - | 25624681 | 25624803 | 25624296 | 25624445 | 25626559 | 25626674 | 0 | 0 | 2 | 0 | 0 | 0 | 0 | 1 | 1 | 0 | 1 | 0 | 98 | 49 | 0.999202116 | 0.999202116 | NA | NA | 1 | NA | 0.333 | 1 | 0.334 |
| SE | 2769 | LST1 | LST1 | chr6 | + | 31587943 | 31587966 | 31587640 | 31587733 | 31588517 | 31588909 | 0 | 0 | 6 | 0 | 0 | 0 | 0 | 1 | 0 | 0 | 1 | 0 | 72 | 49 | 0.999202116 | 0.999202116 | NA | NA | 1 | NA | 0 | 1 | 0.5 |
| SE | 2804 | ARPC4 | ARPC4 | chr3 | + | 9801660 | 9801756 | 9800184 | 9800296 | 9803842 | 9804013 | 0 | 3 | 21 | 0 | 0 | 1 | 2 | 29 | 7 | 0 | 0 | 2 | 98 | 49 | 0.999202116 | 0.999202116 | NA | 1 | 0.913 | 1 | 1 | 0.636 | 0.078 |
| SE | 3096 | MX1 | MX1 | chr21 | + | 41430532 | 41430608 | 41427754 | 41427866 | 41432049 | 41432175 | 0 | 6 | 38 | 0 | 0 | 2 | 0 | 39 | 1 | 0 | 2 | 0 | 98 | 49 | 0.999202116 | 0.999202116 | NA | 1 | 0.905 | NA | 0.907 | 1 | -0.001 |
| SE | 3098 | MX1 | MX1 | chr21 | + | 41439693 | 41439848 | 41437014 | 41437152 | 41440886 | 41441025 | 0 | 6 | 103 | 0 | 1 | 0 | 4 | 60 | 7 | 0 | 1 | 0 | 98 | 49 | 0.999202116 | 0.999202116 | NA | 0.75 | 1 | 1 | 0.968 | 1 | -0.114 |
| SE | 3100 | MX1 | MX1 | chr21 | + | 41449136 | 41449295 | 41445999 | 41446141 | 41451166 | 41451243 | 0 | 0 | 30 | 0 | 0 | 0 | 0 | 11 | 1 | 0 | 2 | 0 | 98 | 49 | 0.999202116 | 0.999202116 | NA | NA | 1 | NA | 0.733 | 1 | 0.133 |
| SE | 3104 | MX2 | MX2 | chr21 | + | 41390564 | 41390703 | 41382409 | 41382564 | 41395586 | 41395785 | 0 | 4 | 23 | 0 | 0 | 13 | 0 | 13 | 0 | 0 | 14 | 1 | 98 | 49 | 0.999202116 | 0.999202116 | NA | 1 | 0.469 | NA | 0.317 | 0 | 0.576 |
| SE | 3172 | SAMSN1 | SAMSN1 | chr21 | - | 14500528 | 14500735 | 14498441 | 14498592 | 14510309 | 14510461 | 0 | 0 | 43 | 0 | 0 | 0 | 0 | 28 | 2 | 0 | 1 | 0 | 98 | 49 | 0.999202116 | 0.999202116 | NA | NA | 1 | NA | 0.933 | 1 | 0.033 |
| SE | 3173 | SAMSN1 | SAMSN1 | chr21 | - | 14516891 | 14517041 | 14512443 | 14512573 | 14521149 | 14521221 | 0 | 1 | 12 | 0 | 1 | 3 | 0 | 16 | 2 | 0 | 5 | 0 | 98 | 49 | 0.999202116 | 0.999202116 | NA | 0.333 | 0.667 | NA | 0.615 | 1 | -0.307 |
| SE | 3194 | HELZ2 | HELZ2 | chr20 | - | 63562071 | 63562203 | 63561822 | 63561984 | 63562287 | 63562382 | 0 | 0 | 11 | 0 | 0 | 0 | 0 | 6 | 2 | 0 | 2 | 0 | 98 | 49 | 0.999202116 | 0.999202116 | NA | NA | 1 | NA | 0.6 | 1 | 0.2 |
| SE | 3200 | TNIP1 | TNIP1 | chr5 | - | 151064959 | 151065127 | 151063612 | 151063747 | 151080879 | 151081088 | 0 | 0 | 4 | 0 | 1 | 2 | 0 | 3 | 1 | 0 | 1 | 0 | 98 | 49 | 0.999202116 | 0.999202116 | NA | 0 | 0.5 | NA | 0.6 | 1 | -0.55 |
| SE | 3201 | TNIP1 | TNIP1 | chr5 | - | 151064959 | 151065131 | 151063612 | 151063747 | 151080879 | 151081084 | 0 | 0 | 5 | 0 | 1 | 2 | 0 | 9 | 1 | 0 | 1 | 0 | 98 | 49 | 0.999202116 | 0.999202116 | NA | 0 | 0.556 | NA | 0.818 | 1 | -0.631 |
| SE | 3238 | NUB1 | NUB1 | chr7 | + | 151360140 | 151360247 | 151356127 | 151356222 | 151366938 | 151367125 | 0 | 0 | 3 | 0 | 0 | 0 | 0 | 6 | 1 | 0 | 2 | 0 | 98 | 49 | 0.999202116 | 0.999202116 | NA | NA | 1 | NA | 0.6 | 1 | 0.2 |
| SE | 3247 | SULF2 | SULF2 | chr20 | - | 47702518 | 47702670 | 47690125 | 47690295 | 47736702 | 47736942 | 0 | 1 | 3 | 0 | 0 | 1 | 0 | 1 | 0 | 0 | 0 | 0 | 98 | 49 | 0.999202116 | 0.999202116 | NA | 1 | 0.6 | NA | 1 | NA | -0.2 |
| SE | 3384 | HM13 | HM13 | chr20 | + | 31549206 | 31549332 | 31549028 | 31549114 | 31550063 | 31550121 | 0 | 3 | 9 | 0 | 0 | 0 | 0 | 3 | 0 | 7 | 0 | 0 | 98 | 49 | 0.999202116 | 0.999202116 | NA | 1 | 1 | 0 | 1 | NA | 0.5 |
| SE | 3452 | GPCPD1 | GPCPD1 | chr20 | - | 5561464 | 5561530 | 5559939 | 5560076 | 5565016 | 5565078 | 0 | 0 | 22 | 0 | 0 | 0 | 0 | 17 | 2 | 0 | 1 | 0 | 98 | 49 | 0.999202116 | 0.999202116 | NA | NA | 1 | NA | 0.895 | 1 | 0.052 |
| SE | 3613 | NIN | NIN | chr14 | - | 50739307 | 50739487 | 50738139 | 50738286 | 50741581 | 50741728 | 0 | 0 | 7 | 0 | 0 | 0 | 4 | 10 | 1 | 0 | 1 | 0 | 98 | 49 | 0.999202116 | 0.999202116 | NA | NA | 1 | 1 | 0.833 | 1 | 0.056 |
| SE | 3635 | USP4 | USP4 | chr3 | - | 49325718 | 49325845 | 49324893 | 49325039 | 49327685 | 49327816 | 0 | 0 | 17 | 0 | 0 | 0 | 0 | 13 | 3 | 0 | 1 | 0 | 98 | 49 | 0.999202116 | 0.999202116 | NA | NA | 1 | NA | 0.867 | 1 | 0.067 |
| SE | 3660 | ETS2 | ETS2 | chr21 | + | 38810034 | 38810106 | 38805830 | 38806120 | 38813002 | 38813114 | 0 | 0 | 13 | 0 | 0 | 0 | 0 | 10 | 3 | 0 | 2 | 0 | 98 | 49 | 0.999202116 | 0.999202116 | NA | NA | 1 | NA | 0.714 | 1 | 0.143 |
| SE | 3722 | FCHO1 | FCHO1 | chr19 | + | 17781230 | 17781343 | 17778608 | 17778884 | 17781451 | 17781539 | 0 | 1 | 3 | 0 | 0 | 3 | 0 | 1 | 0 | 0 | 0 | 0 | 98 | 49 | 0.999202116 | 0.999202116 | NA | 1 | 0.333 | NA | 1 | NA | -0.334 |
| SE | 3749 | STX3 | STX3 | chr11 | + | 59787036 | 59787136 | 59773210 | 59773294 | 59788872 | 59788947 | 0 | 0 | 14 | 0 | 0 | 0 | 0 | 7 | 2 | 0 | 3 | 0 | 98 | 49 | 0.999202116 | 0.999202116 | NA | NA | 1 | NA | 0.538 | 1 | 0.231 |
| SE | 3890 | MAP7D1 | MAP7D1 | chr1 | + | 36179261 | 36179315 | 36178920 | 36179025 | 36179511 | 36179557 | 0 | 0 | 9 | 0 | 0 | 0 | 5 | 8 | 1 | 0 | 2 | 0 | 98 | 49 | 0.999202116 | 0.999202116 | NA | NA | 1 | 1 | 0.667 | 1 | 0.111 |
| SE | 3983 | NCOR2 | NCOR2 | chr12 | - | 124535564 | 124535611 | 124495146 | 124495368 | 124567307 | 124567464 | 3 | 0 | 3 | 1 | 0 | 0 | 0 | 0 | 1 | 0 | 0 | 0 | 96 | 49 | 0.999202116 | 0.999202116 | 0.605 | NA | 1 | NA | NA | 1 | -0.198 |
| SE | 4021 | LINC01578 | LINC01578 | chr15 | + | 92883584 | 92883778 | 92882842 | 92883186 | 92885514 | 92885584 | 0 | 0 | 2 | 0 | 0 | 0 | 0 | 2 | 2 | 0 | 4 | 0 | 98 | 49 | 0.999202116 | 0.999202116 | NA | NA | 1 | NA | 0.2 | 1 | 0.4 |
| SE | 4022 | LINC01578 | LINC01578 | chr15 | + | 92885514 | 92885584 | 92882842 | 92883186 | 92886168 | 92886254 | 0 | 1 | 13 | 0 | 0 | 0 | 1 | 14 | 0 | 0 | 0 | 1 | 98 | 49 | 0.999202116 | 0.999202116 | NA | 1 | 1 | 1 | 1 | 0 | 0.333 |
| SE | 4074 | IL1RN | IL1RN | chr2 | + | 113131044 | 113131157 | 113129575 | 113129664 | 113132655 | 113134016 | 0 | 0 | 12 | 0 | 0 | 0 | 0 | 23 | 2 | 0 | 1 | 0 | 98 | 49 | 0.999202116 | 0.999202116 | NA | NA | 1 | NA | 0.92 | 1 | 0.04 |
| SE | 4121 | XPO6 | XPO6 | chr16 | - | 28111816 | 28112006 | 28107521 | 28107677 | 28112903 | 28113050 | 0 | 2 | 40 | 0 | 1 | 1 | 1 | 30 | 3 | 0 | 2 | 0 | 98 | 49 | 0.999202116 | 0.999202116 | NA | 0.5 | 0.952 | 1 | 0.882 | 1 | -0.235 |
| SE | 4123 | XPO6 | XPO6 | chr16 | - | 28121669 | 28121762 | 28117317 | 28117462 | 28125688 | 28125848 | 0 | 6 | 79 | 0 | 0 | 0 | 15 | 61 | 0 | 0 | 0 | 1 | 98 | 49 | 0.999202116 | 0.999202116 | NA | 1 | 1 | 1 | 1 | 0 | 0.333 |
| SE | 4126 | XPO6 | XPO6 | chr16 | - | 28208986 | 28209122 | 28180940 | 28181031 | 28211365 | 28211918 | 0 | 1 | 1 | 0 | 0 | 17 | 0 | 3 | 0 | 3 | 6 | 4 | 98 | 49 | 0.999202116 | 0.999202116 | NA | 1 | 0.029 | 0 | 0.2 | 0 | 0.448 |
| SE | 4141 | TBC1D15 | TBC1D15 | chr12 | + | 71884810 | 71885021 | 71880468 | 71880607 | 71893221 | 71893324 | 0 | 0 | 3 | 0 | 0 | 0 | 0 | 6 | 1 | 0 | 1 | 0 | 98 | 49 | 0.999202116 | 0.999202116 | NA | NA | 1 | NA | 0.75 | 1 | 0.125 |
| SE | 4170 | RNF10 | RNF10 | chr12 | + | 120571190 | 120571291 | 120566824 | 120566980 | 120575630 | 120575688 | 0 | 3 | 6 | 0 | 0 | 1 | 0 | 3 | 7 | 0 | 6 | 0 | 98 | 49 | 0.999202116 | 0.999202116 | NA | 1 | 0.75 | NA | 0.2 | 1 | 0.275 |
| SE | 4240 | SRCAP | SRCAP | chr16 | + | 30712678 | 30712815 | 30712261 | 30712439 | 30713207 | 30713377 | 0 | 1 | 5 | 0 | 0 | 1 | 0 | 1 | 0 | 0 | 0 | 0 | 98 | 49 | 0.999202116 | 0.999202116 | NA | 1 | 0.714 | NA | 1 | NA | -0.143 |
| SE | 4317 | BIRC6 | BIRC6 | chr2 | + | 32392038 | 32392150 | 32388749 | 32388943 | 32395510 | 32395593 | 0 | 0 | 5 | 0 | 0 | 0 | 0 | 2 | 1 | 0 | 1 | 0 | 98 | 49 | 0.999202116 | 0.999202116 | NA | NA | 1 | NA | 0.5 | 1 | 0.25 |
| SE | 4325 | KMT2D | KMT2D | chr12 | - | 49028827 | 49028958 | 49028008 | 49028141 | 49029060 | 49029236 | 0 | 4 | 7 | 0 | 3 | 0 | 0 | 3 | 0 | 0 | 0 | 1 | 98 | 49 | 0.999202116 | 0.999202116 | NA | 0.4 | 1 | NA | 1 | 0 | 0.2 |
| SE | 4422 | PREX1 | PREX1 | chr20 | - | 48644408 | 48644497 | 48642406 | 48642489 | 48645850 | 48646057 | 0 | 5 | 22 | 0 | 0 | 5 | 7 | 18 | 1 | 0 | 3 | 0 | 98 | 49 | 0.999202116 | 0.999202116 | NA | 1 | 0.687 | 1 | 0.75 | 1 | -0.073 |
| SE | 4443 | CORO7 | CORO7 | chr16 | - | 4364596 | 4364689 | 4364275 | 4364413 | 4364774 | 4364920 | 0 | 0 | 8 | 0 | 0 | 0 | 0 | 1 | 3 | 0 | 1 | 0 | 98 | 49 | 0.999202116 | 0.999202116 | NA | NA | 1 | NA | 0.333 | 1 | 0.334 |
| SE | 4504 | PTPN6 | PTPN6 | chr12 | + | 6957918 | 6958073 | 6957653 | 6957785 | 6959926 | 6959994 | 0 | 4 | 43 | 0 | 4 | 3 | 2 | 10 | 3 | 0 | 11 | 0 | 98 | 49 | 0.999202116 | 0.999202116 | NA | 0.333 | 0.878 | 1 | 0.312 | 1 | -0.165 |
| SE | 4520 | CHD4 | CHD4 | chr12 | - | 6600925 | 6601053 | 6600533 | 6600669 | 6601288 | 6601530 | 0 | 0 | 7 | 0 | 0 | 0 | 0 | 4 | 1 | 0 | 1 | 0 | 98 | 49 | 0.999202116 | 0.999202116 | NA | NA | 1 | NA | 0.667 | 1 | 0.166 |
| SE | 4534 | TM9SF2 | TM9SF2 | chr13 | + | 99537738 | 99537863 | 99536607 | 99536737 | 99540713 | 99540793 | 0 | 0 | 4 | 0 | 0 | 0 | 0 | 1 | 1 | 1 | 0 | 0 | 98 | 49 | 0.999202116 | 0.999202116 | NA | NA | 1 | 0 | 1 | 1 | 0.333 |
| SE | 4589 | PLBD1 | PLBD1 | chr12 | - | 14553192 | 14553412 | 14542207 | 14542291 | 14567581 | 14567857 | 0 | 2 | 33 | 0 | 0 | 1 | 1 | 1 | 0 | 0 | 0 | 0 | 98 | 49 | 0.999202116 | 0.999202116 | NA | 1 | 0.943 | 1 | 1 | NA | -0.028 |
| SE | 4617 | PTPRJ | PTPRJ | chr11 | + | 48144999 | 48145124 | 48144674 | 48144885 | 48146875 | 48146963 | 0 | 11 | 9 | 0 | 0 | 1 | 0 | 1 | 0 | 0 | 0 | 0 | 98 | 49 | 0.999202116 | 0.999202116 | NA | 1 | 0.818 | NA | 1 | NA | -0.091 |
| SE | 4673 | IL10RA | IL10RA | chr11 | + | 117989441 | 117989620 | 117988381 | 117988502 | 117993240 | 117993410 | 0 | 2 | 14 | 0 | 0 | 1 | 0 | 2 | 0 | 0 | 0 | 0 | 98 | 49 | 0.999202116 | 0.999202116 | NA | 1 | 0.875 | NA | 1 | NA | -0.062 |
| SE | 4773 | DDX6 | DDX6 | chr11 | - | 118755401 | 118755503 | 118754704 | 118754887 | 118756259 | 118756323 | 0 | 0 | 12 | 0 | 0 | 0 | 1 | 22 | 1 | 0 | 1 | 0 | 98 | 49 | 0.999202116 | 0.999202116 | NA | NA | 1 | 1 | 0.917 | 1 | 0.028 |
| SE | 4884 | DCAF11 | DCAF11 | chr14 | + | 24117265 | 24117393 | 24116916 | 24117044 | 24117667 | 24117732 | 0 | 0 | 4 | 0 | 0 | 0 | 0 | 1 | 2 | 0 | 1 | 0 | 98 | 49 | 0.999202116 | 0.999202116 | NA | NA | 1 | NA | 0.333 | 1 | 0.334 |
| SE | 4939 | ARAP1 | ARAP1 | chr11 | - | 72686041 | 72686191 | 72685068 | 72685681 | 72687687 | 72687738 | 0 | 0 | 13 | 0 | 0 | 0 | 0 | 9 | 1 | 0 | 1 | 0 | 98 | 49 | 0.999202116 | 0.999202116 | NA | NA | 1 | NA | 0.818 | 1 | 0.091 |
| SE | 4943 | ARAP1 | ARAP1 | chr11 | - | 72697322 | 72697486 | 72696982 | 72697195 | 72697597 | 72697649 | 0 | 2 | 24 | 0 | 1 | 0 | 0 | 16 | 3 | 0 | 1 | 0 | 98 | 49 | 0.999202116 | 0.999202116 | NA | 0.5 | 1 | NA | 0.889 | 1 | -0.195 |
| SE | 4945 | ARAP1 | ARAP1 | chr11 | - | 72704151 | 72704334 | 72702904 | 72703079 | 72705804 | 72705890 | 0 | 0 | 25 | 0 | 0 | 0 | 0 | 14 | 10 | 0 | 3 | 0 | 98 | 49 | 0.999202116 | 0.999202116 | NA | NA | 1 | NA | 0.7 | 1 | 0.15 |
| SE | 4947 | ARAP1 | ARAP1 | chr11 | - | 72711020 | 72711141 | 72710384 | 72710587 | 72711429 | 72711499 | 0 | 4 | 21 | 0 | 0 | 2 | 0 | 13 | 4 | 0 | 1 | 2 | 98 | 49 | 0.999202116 | 0.999202116 | NA | 1 | 0.84 | NA | 0.867 | 0.5 | 0.236 |
| SE | 4984 | NCOR1 | NCOR1 | chr17 | - | 16067893 | 16068121 | 16065484 | 16065694 | 16070164 | 16070525 | 0 | 1 | 2 | 0 | 0 | 1 | 3 | 1 | 0 | 0 | 0 | 0 | 98 | 49 | 0.999202116 | 0.999202116 | NA | 1 | 0.5 | 1 | 1 | NA | -0.25 |
| SE | 5186 | MYO9B | MYO9B | chr19 | + | 17172758 | 17172963 | 17172335 | 17172477 | 17175662 | 17175741 | 0 | 0 | 8 | 0 | 0 | 0 | 0 | 10 | 1 | 0 | 1 | 0 | 98 | 49 | 0.999202116 | 0.999202116 | NA | NA | 1 | NA | 0.833 | 1 | 0.084 |
| SE | 5369 | BAZ1A | BAZ1A | chr14 | - | 34862043 | 34862322 | 34826012 | 34826156 | 34874491 | 34874662 | 0 | 0 | 20 | 0 | 0 | 0 | 0 | 4 | 2 | 2 | 1 | 0 | 98 | 49 | 0.999202116 | 0.999202116 | NA | NA | 1 | 0 | 0.667 | 1 | 0.444 |
| SE | 5425 | PRMT3 | PRMT3 | chr11 | + | 20397576 | 20397721 | 20395802 | 20395962 | 20402918 | 20402984 | 0 | 3 | 0 | 0 | 0 | 1 | 0 | 1 | 0 | 0 | 0 | 0 | 98 | 49 | 0.999202116 | 0.999202116 | NA | 1 | 0 | NA | 1 | NA | -0.5 |
| SE | 5575 | SAT1 | SAT1 | chrX | + | 23783799 | 23783883 | 23783657 | 23783709 | 23785327 | 23785429 | 0 | 20 | 265 | 0 | 1 | 1 | 2 | 216 | 10 | 0 | 2 | 0 | 98 | 49 | 0.999202116 | 0.999202116 | NA | 0.909 | 0.993 | 1 | 0.982 | 1 | -0.043 |
| SE | 5615 | DOPEY2 | DOPEY2 | chr21 | + | 36211552 | 36211651 | 36208714 | 36208904 | 36211973 | 36212097 | 0 | 2 | 1 | 0 | 0 | 1 | 0 | 1 | 0 | 0 | 0 | 0 | 98 | 49 | 0.999202116 | 0.999202116 | NA | 1 | 0.333 | NA | 1 | NA | -0.334 |
| SE | 5775 | NCOA2 | NCOA2 | chr8 | - | 70213902 | 70214075 | 70174755 | 70174859 | 70216659 | 70216764 | 0 | 0 | 13 | 0 | 0 | 0 | 0 | 18 | 2 | 0 | 2 | 0 | 98 | 49 | 0.999202116 | 0.999202116 | NA | NA | 1 | NA | 0.818 | 1 | 0.091 |
| SE | 5787 | DENND3 | DENND3 | chr8 | + | 141160631 | 141160787 | 141155848 | 141155970 | 141163332 | 141163429 | 0 | 0 | 19 | 0 | 0 | 0 | 0 | 6 | 2 | 0 | 2 | 0 | 98 | 49 | 0.999202116 | 0.999202116 | NA | NA | 1 | NA | 0.6 | 1 | 0.2 |
| SE | 5854 | ITPK1 | ITPK1 | chr14 | - | 92993879 | 92993997 | 92962750 | 92962849 | 93016675 | 93016801 | 0 | 2 | 10 | 0 | 0 | 2 | 0 | 2 | 0 | 0 | 0 | 0 | 98 | 49 | 0.999202116 | 0.999202116 | NA | 1 | 0.714 | NA | 1 | NA | -0.143 |
| SE | 5890 | MSN | MSN | chrX | + | 65738524 | 65738617 | 65737177 | 65737338 | 65738969 | 65739194 | 0 | 3 | 22 | 0 | 0 | 1 | 4 | 14 | 2 | 0 | 0 | 2 | 98 | 49 | 0.999202116 | 0.999202116 | NA | 1 | 0.917 | 1 | 1 | 0.333 | 0.181 |
| SE | 5968 | CD53 | CD53 | chr1 | + | 110897808 | 110897892 | 110896652 | 110896733 | 110899123 | 110899936 | 0 | 6 | 27 | 0 | 0 | 0 | 0 | 13 | 0 | 0 | 0 | 1 | 98 | 49 | 0.999202116 | 0.999202116 | NA | 1 | 1 | NA | 1 | 0 | 0.5 |
| SE | 6104 | GLUL | GLUL | chr1 | - | 182387130 | 182387292 | 182386255 | 182386402 | 182388571 | 182388750 | 0 | 25 | 173 | 0 | 0 | 3 | 4 | 163 | 7 | 0 | 1 | 0 | 98 | 49 | 0.999202116 | 0.999202116 | NA | 1 | 0.966 | 1 | 0.988 | 1 | -0.013 |
| SE | 6179 | ABCA1 | ABCA1 | chr9 | - | 104832573 | 104832771 | 104831621 | 104831827 | 104836979 | 104837096 | 0 | 6 | 13 | 0 | 0 | 0 | 0 | 43 | 0 | 0 | 0 | 1 | 98 | 49 | 0.999202116 | 0.999202116 | NA | 1 | 1 | NA | 1 | 0 | 0.5 |
| SE | 6235 | NFKB2 | NFKB2 | chr10 | + | 102398384 | 102398523 | 102398211 | 102398297 | 102398738 | 102398864 | 0 | 0 | 7 | 0 | 0 | 0 | 0 | 8 | 1 | 0 | 1 | 0 | 98 | 49 | 0.999202116 | 0.999202116 | NA | NA | 1 | NA | 0.8 | 1 | 0.1 |
| SE | 6236 | RAB8B | RAB8B | chr15 | + | 63249644 | 63249705 | 63244755 | 63244816 | 63255506 | 63255584 | 0 | 0 | 8 | 0 | 0 | 0 | 4 | 8 | 1 | 0 | 1 | 0 | 98 | 49 | 0.999202116 | 0.999202116 | NA | NA | 1 | 1 | 0.8 | 1 | 0.067 |
| SE | 6273 | BAZ2B | BAZ2B | chr2 | - | 159448241 | 159448409 | 159446781 | 159446975 | 159478574 | 159478721 | 0 | 0 | 14 | 0 | 0 | 0 | 0 | 3 | 2 | 0 | 2 | 0 | 98 | 49 | 0.999202116 | 0.999202116 | NA | NA | 1 | NA | 0.429 | 1 | 0.285 |
| SE | 6483 | PADI2 | PADI2 | chr1 | - | 17083725 | 17083837 | 17082544 | 17082652 | 17084598 | 17084702 | 0 | 0 | 10 | 0 | 0 | 0 | 0 | 3 | 1 | 0 | 1 | 0 | 98 | 49 | 0.999202116 | 0.999202116 | NA | NA | 1 | NA | 0.6 | 1 | 0.2 |
| SE | 6542 | NRD1 | NRD1 | chr1 | - | 51790532 | 51790649 | 51789567 | 51789657 | 51790899 | 51790990 | 0 | 0 | 17 | 0 | 0 | 0 | 0 | 6 | 1 | 0 | 1 | 0 | 98 | 49 | 0.999202116 | 0.999202116 | NA | NA | 1 | NA | 0.75 | 1 | 0.125 |
| SE | 6573 | CHD8 | CHD8 | chr14 | - | 21393105 | 21393254 | 21392506 | 21392809 | 21393475 | 21394195 | 0 | 2 | 2 | 0 | 0 | 1 | 0 | 2 | 0 | 0 | 0 | 0 | 98 | 49 | 0.999202116 | 0.999202116 | NA | 1 | 0.5 | NA | 1 | NA | -0.25 |
| SE | 6607 | QSOX1 | QSOX1 | chr1 | + | 180194212 | 180194392 | 180190432 | 180190580 | 180196261 | 180196599 | 0 | 1 | 3 | 0 | 0 | 1 | 0 | 4 | 0 | 0 | 1 | 1 | 98 | 49 | 0.999202116 | 0.999202116 | NA | 1 | 0.6 | NA | 0.667 | 0 | 0.467 |
| SE | 6616 | AMPD2 | AMPD2 | chr1 | + | 109625661 | 109625792 | 109619812 | 109620278 | 109626159 | 109626228 | 0 | 2 | 10 | 0 | 0 | 1 | 0 | 16 | 1 | 0 | 1 | 0 | 98 | 49 | 0.999202116 | 0.999202116 | NA | 1 | 0.833 | NA | 0.889 | 1 | -0.028 |
| SE | 6620 | AMPD2 | AMPD2 | chr1 | + | 109630682 | 109630793 | 109630232 | 109630406 | 109630942 | 109632055 | 0 | 0 | 9 | 0 | 0 | 0 | 0 | 30 | 3 | 0 | 5 | 0 | 98 | 49 | 0.999202116 | 0.999202116 | NA | NA | 1 | NA | 0.75 | 1 | 0.125 |
| SE | 6624 | PCNX | PCNX | chr14 | + | 70995740 | 70995925 | 70976941 | 70978648 | 71009633 | 71009724 | 0 | 0 | 2 | 0 | 0 | 0 | 0 | 1 | 1 | 0 | 1 | 0 | 98 | 49 | 0.999202116 | 0.999202116 | NA | NA | 1 | NA | 0.333 | 1 | 0.334 |
| SE | 6766 | PIK3IP1 | PIK3IP1 | chr22 | - | 31289314 | 31289393 | 31281592 | 31283288 | 31289498 | 31289699 | 0 | 0 | 1 | 0 | 0 | 0 | 0 | 5 | 1 | 0 | 2 | 0 | 98 | 49 | 0.999202116 | 0.999202116 | NA | NA | 1 | NA | 0.556 | 1 | 0.222 |
| SE | 6863 | ATG2A | ATG2A | chr11 | - | 64896461 | 64896616 | 64895289 | 64895442 | 64896747 | 64896869 | 0 | 0 | 5 | 0 | 0 | 0 | 0 | 11 | 2 | 0 | 2 | 0 | 98 | 49 | 0.999202116 | 0.999202116 | NA | NA | 1 | NA | 0.733 | 1 | 0.133 |
| SE | 6939 | CD55 | CD55 | chr1 | + | 207339396 | 207339417 | 207337328 | 207337409 | 207359545 | 207360966 | 0 | 0 | 7 | 0 | 0 | 0 | 0 | 3 | 1 | 0 | 1 | 0 | 70 | 49 | 0.999202116 | 0.999202116 | NA | NA | 1 | NA | 0.677 | 1 | 0.161 |
| SE | 7066 | EPB41 | EPB41 | chr1 | + | 29058588 | 29058645 | 29053103 | 29053312 | 29064981 | 29065158 | 0 | 0 | 1 | 0 | 2 | 10 | 0 | 0 | 0 | 0 | 2 | 0 | 98 | 49 | 0.999202116 | 0.999202116 | NA | 0 | 0.048 | NA | 0 | NA | 0.024 |
| SE | 7122 | RERE | RERE | chr1 | - | 8362682 | 8362844 | 8361762 | 8361876 | 8364055 | 8364255 | 0 | 2 | 6 | 0 | 0 | 1 | 0 | 2 | 0 | 0 | 0 | 0 | 98 | 49 | 0.999202116 | 0.999202116 | NA | 1 | 0.75 | NA | 1 | NA | -0.125 |
| SE | 7139 | KIF21B | KIF21B | chr1 | - | 200999348 | 200999466 | 200998383 | 200998575 | 200999882 | 200999964 | 0 | 2 | 5 | 0 | 0 | 0 | 0 | 3 | 0 | 0 | 0 | 1 | 98 | 49 | 0.999202116 | 0.999202116 | NA | 1 | 1 | NA | 1 | 0 | 0.5 |
| SE | 7142 | CPD | CPD | chr17 | + | 30421663 | 30421833 | 30420840 | 30420983 | 30422673 | 30423023 | 0 | 0 | 7 | 0 | 0 | 0 | 0 | 7 | 1 | 0 | 1 | 0 | 98 | 49 | 0.999202116 | 0.999202116 | NA | NA | 1 | NA | 0.778 | 1 | 0.111 |
| SE | 7188 | SNRPB | SNRPB | chr20 | - | 2465707 | 2465819 | 2463746 | 2463899 | 2467606 | 2467758 | 0 | 1 | 10 | 0 | 0 | 1 | 0 | 1 | 0 | 0 | 0 | 0 | 98 | 49 | 0.999202116 | 0.999202116 | NA | 1 | 0.833 | NA | 1 | NA | -0.084 |
| SE | 7341 | EIF4A2 | EIF4A2 | chr3 | + | 186784961 | 186785101 | 186784563 | 186784696 | 186785882 | 186786051 | 0 | 1 | 6 | 0 | 2 | 3 | 0 | 1 | 0 | 1 | 2 | 0 | 98 | 49 | 0.999202116 | 0.999202116 | NA | 0.2 | 0.5 | 0 | 0.2 | NA | 0.25 |
| SE | 7541 | CD300LF | CD300LF | chr17 | - | 74705203 | 74705377 | 74704477 | 74704816 | 74712823 | 74713000 | 0 | 0 | 4 | 0 | 2 | 0 | 0 | 0 | 0 | 0 | 1 | 0 | 98 | 49 | 0.999202116 | 0.999202116 | NA | 0 | 1 | NA | 0 | NA | 0.5 |
| SE | 7579 | MMP8 | MMP8 | chr11 | - | 102714555 | 102714709 | 102713753 | 102713857 | 102715303 | 102715437 | 0 | 4 | 2 | 0 | 0 | 3 | 0 | 1 | 0 | 0 | 0 | 0 | 98 | 49 | 0.999202116 | 0.999202116 | NA | 1 | 0.25 | NA | 1 | NA | -0.375 |
| SE | 7717 | THEMIS2 | THEMIS2 | chr1 | + | 27879643 | 27880054 | 27872542 | 27872665 | 27885294 | 27885451 | 0 | 0 | 2 | 0 | 1 | 0 | 0 | 0 | 2 | 3 | 0 | 0 | 98 | 49 | 0.999202116 | 0.999202116 | NA | 0 | 1 | 0 | NA | 1 | 0 |
| SE | 7729 | KIAA1033 | KIAA1033 | chr12 | + | 105140293 | 105140401 | 105137885 | 105138011 | 105140898 | 105141045 | 0 | 3 | 12 | 0 | 0 | 1 | 0 | 3 | 0 | 0 | 0 | 0 | 98 | 49 | 0.999202116 | 0.999202116 | NA | 1 | 0.857 | NA | 1 | NA | -0.072 |
| SE | 7909 | ATP6V0B | ATP6V0B | chr1 | + | 43976569 | 43976659 | 43976301 | 43976379 | 43976772 | 43976824 | 0 | 4 | 17 | 0 | 0 | 0 | 0 | 24 | 0 | 0 | 0 | 1 | 98 | 49 | 0.999202116 | 0.999202116 | NA | 1 | 1 | NA | 1 | 0 | 0.5 |
| SE | 7949 | PPP4R1 | PPP4R1 | chr18 | - | 9549196 | 9549338 | 9546790 | 9547952 | 9550051 | 9550186 | 0 | 1 | 6 | 0 | 0 | 0 | 0 | 2 | 0 | 0 | 0 | 2 | 98 | 49 | 0.999202116 | 0.999202116 | NA | 1 | 1 | NA | 1 | 0 | 0.5 |
| SE | 7967 | OAS3 | OAS3 | chr12 | + | 112965743 | 112966029 | 112964234 | 112964408 | 112967417 | 112967593 | 0 | 0 | 23 | 0 | 0 | 0 | 0 | 14 | 1 | 0 | 1 | 0 | 98 | 49 | 0.999202116 | 0.999202116 | NA | NA | 1 | NA | 0.875 | 1 | 0.062 |
| SE | 8036 | MORF4L1 | MORF4L1 | chr15 | + | 78884966 | 78885083 | 78880511 | 78880579 | 78886140 | 78886227 | 0 | 0 | 0 | 0 | 0 | 5 | 0 | 1 | 0 | 0 | 3 | 1 | 98 | 49 | 0.999202116 | 0.999202116 | NA | NA | 0 | NA | 0.143 | 0 | -0.071 |
| SE | 8083 | YBX3 | YBX3 | chr12 | - | 10709907 | 10710114 | 10704050 | 10704148 | 10713210 | 10713333 | 0 | 0 | 5 | 0 | 0 | 0 | 0 | 0 | 1 | 0 | 1 | 0 | 98 | 49 | 0.999202116 | 0.999202116 | NA | NA | 1 | NA | 0 | 1 | 0.5 |
| SE | 8096 | JMJD1C | JMJD1C | chr10 | - | 63176296 | 63176473 | 63168434 | 63168566 | 63177716 | 63177856 | 0 | 0 | 16 | 0 | 0 | 0 | 1 | 21 | 3 | 0 | 1 | 0 | 98 | 49 | 0.999202116 | 0.999202116 | NA | NA | 1 | 1 | 0.913 | 1 | 0.029 |
| SE | 8125 | ANXA5 | ANXA5 | chr4 | - | 121684676 | 121684771 | 121683363 | 121683477 | 121686287 | 121686372 | 0 | 3 | 20 | 0 | 0 | 1 | 0 | 3 | 0 | 0 | 0 | 0 | 98 | 49 | 0.999202116 | 0.999202116 | NA | 1 | 0.909 | NA | 1 | NA | -0.045 |
| SE | 8189 | EPSTI1 | EPSTI1 | chr13 | - | 42953947 | 42954021 | 42926335 | 42926429 | 42963254 | 42963338 | 0 | 5 | 42 | 0 | 0 | 0 | 0 | 23 | 0 | 0 | 0 | 2 | 98 | 49 | 0.999202116 | 0.999202116 | NA | 1 | 1 | NA | 1 | 0 | 0.5 |
| SE | 8190 | EPSTI1 | EPSTI1 | chr13 | - | 42970611 | 42970670 | 42969093 | 42969177 | 42991977 | 42992271 | 0 | 0 | 25 | 0 | 1 | 0 | 0 | 7 | 5 | 0 | 1 | 0 | 98 | 49 | 0.999202116 | 0.999202116 | NA | 0 | 1 | NA | 0.778 | 1 | -0.389 |
| SE | 8204 | NDUFS2 | NDUFS2 | chr1 | + | 161203436 | 161203543 | 161202146 | 161202480 | 161206406 | 161206597 | 0 | 1 | 0 | 0 | 0 | 0 | 0 | 1 | 0 | 3 | 0 | 0 | 98 | 49 | 0.999202116 | 0.999202116 | NA | 1 | NA | 0 | 1 | NA | 0.5 |
| SE | 8326 | COPA | COPA | chr1 | - | 160307162 | 160307245 | 160306353 | 160306493 | 160309100 | 160309176 | 0 | 0 | 6 | 0 | 0 | 0 | 0 | 3 | 1 | 0 | 1 | 0 | 98 | 49 | 0.999202116 | 0.999202116 | NA | NA | 1 | NA | 0.6 | 1 | 0.2 |
| SE | 8339 | CHD2 | CHD2 | chr15 | + | 92944414 | 92944515 | 92942842 | 92943068 | 92945820 | 92945865 | 0 | 0 | 5 | 0 | 0 | 0 | 0 | 14 | 2 | 0 | 1 | 0 | 98 | 49 | 0.999202116 | 0.999202116 | NA | NA | 1 | NA | 0.875 | 1 | 0.062 |
| SE | 8385 | CYFIP2 | CYFIP2 | chr5 | + | 157383264 | 157383359 | 157382589 | 157382662 | 157389188 | 157389427 | 0 | 0 | 4 | 0 | 0 | 0 | 0 | 5 | 1 | 0 | 1 | 0 | 98 | 49 | 0.999202116 | 0.999202116 | NA | NA | 1 | NA | 0.714 | 1 | 0.143 |
| SE | 8481 | MAX | MAX | chr14 | - | 65101545 | 65101572 | 65093707 | 65093815 | 65102303 | 65102544 | 0 | 1 | 2 | 0 | 0 | 3 | 0 | 1 | 0 | 0 | 0 | 0 | 76 | 49 | 0.999202116 | 0.999202116 | NA | 1 | 0.301 | NA | 1 | NA | -0.35 |
| SE | 8556 | PIK3AP1 | PIK3AP1 | chr10 | - | 96656797 | 96656934 | 96651508 | 96651651 | 96709566 | 96709983 | 0 | 0 | 16 | 0 | 0 | 0 | 0 | 3 | 3 | 0 | 1 | 0 | 98 | 49 | 0.999202116 | 0.999202116 | NA | NA | 1 | NA | 0.6 | 1 | 0.2 |
| SE | 8657 | DUSP1 | DUSP1 | chr5 | - | 172769574 | 172769794 | 172768089 | 172769132 | 172770160 | 172770306 | 0 | 17 | 269 | 0 | 0 | 3 | 7 | 124 | 15 | 0 | 0 | 1 | 98 | 49 | 0.999202116 | 0.999202116 | NA | 1 | 0.978 | 1 | 1 | 0.882 | 0.028 |
| SE | 8659 | DUSP1 | DUSP1 | chr5 | - | 172770160 | 172770306 | 172768089 | 172769132 | 172770585 | 172771200 | 4 | 93 | 267 | 0 | 1 | 0 | 23 | 118 | 275 | 0 | 2 | 0 | 98 | 49 | 0.999202116 | 0.999202116 | 1 | 0.979 | 1 | 1 | 0.967 | 1 | 0.004 |
| SE | 8801 | CD46 | CD46 | chr1 | + | 207790252 | 207790345 | 207785618 | 207785682 | 207793518 | 207795516 | 0 | 0 | 12 | 1 | 0 | 6 | 0 | 12 | 1 | 2 | 2 | 0 | 98 | 49 | 0.999202116 | 0.999202116 | 0 | NA | 0.5 | 0 | 0.75 | 1 | -0.333 |
| SE | 8944 | GUK1 | GUK1 | chr1 | + | 228148370 | 228148456 | 228147614 | 228147699 | 228148664 | 228148954 | 0 | 0 | 8 | 0 | 1 | 3 | 0 | 4 | 1 | 0 | 1 | 0 | 98 | 49 | 0.999202116 | 0.999202116 | NA | 0 | 0.571 | NA | 0.667 | 1 | -0.548 |
| SE | 9046 | FAM65B | FAM65B | chr6 | - | 24832255 | 24832391 | 24830508 | 24830670 | 24839090 | 24839272 | 0 | 1 | 4 | 0 | 0 | 1 | 0 | 1 | 0 | 0 | 0 | 0 | 98 | 49 | 0.999202116 | 0.999202116 | NA | 1 | 0.667 | NA | 1 | NA | -0.166 |
| SE | 9138 | CLK1 | CLK1 | chr2 | - | 200857717 | 200857884 | 200856890 | 200856985 | 200857972 | 200858089 | 0 | 0 | 12 | 0 | 0 | 0 | 0 | 10 | 1 | 0 | 3 | 0 | 98 | 49 | 0.999202116 | 0.999202116 | NA | NA | 1 | NA | 0.625 | 1 | 0.188 |
| SE | 9352 | NABP1 | NABP1 | chr2 | + | 191682491 | 191682595 | 191681945 | 191682017 | 191683728 | 191683804 | 0 | 2 | 4 | 0 | 0 | 5 | 2 | 0 | 1 | 0 | 4 | 0 | 98 | 49 | 0.999202116 | 0.999202116 | NA | 1 | 0.286 | 1 | 0 | 1 | -0.024 |
| SE | 9433 | IMPDH1 | IMPDH1 | chr7 | - | 128405766 | 128405865 | 128403705 | 128403754 | 128409755 | 128409982 | 0 | 0 | 5 | 0 | 0 | 0 | 0 | 1 | 1 | 0 | 1 | 0 | 98 | 49 | 0.999202116 | 0.999202116 | NA | NA | 1 | NA | 0.333 | 1 | 0.334 |
| SE | 9485 | NAGK | NAGK | chr2 | + | 71075554 | 71075642 | 71073481 | 71073594 | 71076603 | 71076701 | 0 | 1 | 11 | 0 | 0 | 0 | 0 | 11 | 0 | 0 | 0 | 1 | 98 | 49 | 0.999202116 | 0.999202116 | NA | 1 | 1 | NA | 1 | 0 | 0.5 |
| SE | 9512 | SIPA1 | SIPA1 | chr11 | + | 65642249 | 65642377 | 65640800 | 65641600 | 65642462 | 65642639 | 0 | 0 | 14 | 0 | 0 | 0 | 0 | 3 | 1 | 0 | 2 | 0 | 98 | 49 | 0.999202116 | 0.999202116 | NA | NA | 1 | NA | 0.429 | 1 | 0.285 |
| SE | 9563 | OASL | OASL | chr12 | - | 121033460 | 121033743 | 121031441 | 121031617 | 121038773 | 121039242 | 0 | 0 | 16 | 0 | 2 | 2 | 2 | 4 | 4 | 0 | 3 | 0 | 98 | 49 | 0.999202116 | 0.999202116 | NA | 0 | 0.8 | 1 | 0.4 | 1 | -0.4 |
| SE | 9755 | LCOR | LCOR | chr10 | + | 96944112 | 96944245 | 96907667 | 96907747 | 96949007 | 96949295 | 0 | 2 | 0 | 0 | 0 | 1 | 0 | 2 | 0 | 0 | 0 | 0 | 98 | 49 | 0.999202116 | 0.999202116 | NA | 1 | 0 | NA | 1 | NA | -0.5 |
| SE | 9971 | RIOK3 | RIOK3 | chr18 | + | 23467398 | 23467526 | 23466132 | 23466276 | 23473428 | 23473626 | 0 | 0 | 6 | 0 | 0 | 0 | 0 | 11 | 2 | 0 | 1 | 0 | 98 | 49 | 0.999202116 | 0.999202116 | NA | NA | 1 | NA | 0.846 | 1 | 0.077 |
| SE | 9990 | TRANK1 | TRANK1 | chr3 | - | 36874125 | 36874296 | 36864318 | 36864480 | 36889828 | 36889960 | 0 | 3 | 13 | 0 | 0 | 1 | 0 | 3 | 0 | 0 | 0 | 0 | 98 | 49 | 0.999202116 | 0.999202116 | NA | 1 | 0.867 | NA | 1 | NA | -0.067 |
| SE | 10093 | NUMB | NUMB | chr14 | - | 73284080 | 73284374 | 73282358 | 73282505 | 73287109 | 73287314 | 0 | 0 | 6 | 0 | 0 | 0 | 0 | 6 | 2 | 0 | 2 | 0 | 98 | 49 | 0.999202116 | 0.999202116 | NA | NA | 1 | NA | 0.6 | 1 | 0.2 |
| SE | 10116 | TRAF3IP3 | TRAF3IP3 | chr1 | + | 209762514 | 209762662 | 209759981 | 209760384 | 209762812 | 209762870 | 0 | 0 | 6 | 0 | 0 | 0 | 0 | 0 | 1 | 0 | 1 | 0 | 98 | 49 | 0.999202116 | 0.999202116 | NA | NA | 1 | NA | 0 | 1 | 0.5 |
| SE | 10121 | SPTLC2 | SPTLC2 | chr14 | - | 77562395 | 77562489 | 77557040 | 77557146 | 77570383 | 77570508 | 0 | 0 | 16 | 0 | 0 | 0 | 0 | 2 | 1 | 0 | 1 | 0 | 98 | 49 | 0.999202116 | 0.999202116 | NA | NA | 1 | NA | 0.5 | 1 | 0.25 |
| SE | 10446 | ELL | ELL | chr19 | - | 18458204 | 18458329 | 18451551 | 18451648 | 18461577 | 18461852 | 0 | 0 | 13 | 0 | 0 | 0 | 0 | 1 | 1 | 0 | 3 | 0 | 98 | 49 | 0.999202116 | 0.999202116 | NA | NA | 1 | NA | 0.143 | 1 | 0.428 |
| SE | 10653 | RELA | RELA | chr11 | - | 65658717 | 65658822 | 65655854 | 65655935 | 65659665 | 65659788 | 0 | 0 | 2 | 0 | 0 | 0 | 0 | 9 | 1 | 0 | 1 | 0 | 98 | 49 | 0.999202116 | 0.999202116 | NA | NA | 1 | NA | 0.818 | 1 | 0.091 |
| SE | 10700 | TLE3 | TLE3 | chr15 | - | 70055048 | 70055298 | 70053226 | 70053374 | 70056297 | 70056374 | 0 | 0 | 4 | 0 | 0 | 0 | 0 | 6 | 4 | 0 | 1 | 0 | 98 | 49 | 0.999202116 | 0.999202116 | NA | NA | 1 | NA | 0.75 | 1 | 0.125 |

| | | | | | | | | | | | | | | | | | | | | | | | | | | | | | | | | | | | | |
|---|---|---|---|---|---|---|---|---|---|---|---|---|---|---|---|---|---|---|---|---|---|---|---|---|---|---|---|---|---|---|---|---|---|---|---|---|
| SE | 10824 | PSMA4 | PSMA4 | chr15 | + | 78542482 | 78542645 | 78542176 | 78542219 | 78544189 | 78544267 | 0 | 1 | 4 | 0 | 0 | 1 | 0 | 1 | 0 | 1 | 0 | 0 | 0 | 0 | 98 | 49 | 0.999202116 | 0.999202116 | NA | 1 | 0.667 | NA | 1 | NA | -0.166 |
| SE | 10950 | IQGAP1 | IQGAP1 | chr15 | + | 90472839 | 90473010 | 90467449 | 90467592 | 90473714 | 90473798 | 0 | 0 | 30 | 0 | 0 | 0 | 0 | 0 | 7 | 16 | 3 | 0 | 1 | 0 | 98 | 49 | 0.999202116 | 0.999202116 | NA | NA | 1 | 1 | 0.889 | 1 | 0.037 |
| SE | 11033 | STAT1 | STAT1 | chr2 | - | 191013524 | 191013678 | 191009875 | 191010004 | 191014017 | 191014250 | 0 | 5 | 34 | 0 | 0 | 2 | 0 | 2 | 0 | 15 | 4 | 0 | 0 | 2 | 98 | 49 | 0.999202116 | 0.999202116 | NA | 1 | 0.895 | NA | 1 | 0.5 | 0.198 |
| SE | 11193 | PARP14 | PARP14 | chr3 | + | 122699389 | 122701635 | 122695425 | 122695662 | 122703741 | 122703978 | 0 | 8 | 30 | 0 | 0 | 2 | 0 | 2 | 0 | 39 | 2 | 0 | 1 | 0 | 98 | 49 | 0.999202116 | 0.999202116 | NA | 1 | 0.882 | NA | 0.951 | 1 | -0.034 |
| SE | 11206 | TOP1 | TOP1 | chr20 | + | 41080084 | 41080180 | 41077581 | 41077637 | 41081164 | 41081240 | 0 | 0 | 5 | 0 | 0 | 0 | 0 | 0 | 0 | 5 | 1 | 0 | 1 | 0 | 98 | 49 | 0.999202116 | 0.999202116 | NA | NA | 1 | NA | 0.714 | 1 | 0.143 |
| SE | 11348 | ADGRE2 | ADGRE2 | chr19 | - | 14754953 | 14755127 | 14752328 | 14752526 | 14755653 | 14755877 | 0 | 0 | 5 | 0 | 0 | 0 | 0 | 0 | 0 | 7 | 1 | 0 | 1 | 0 | 98 | 49 | 0.999202116 | 0.999202116 | NA | NA | 1 | NA | 0.778 | 1 | 0.111 |
| SE | 11351 | ADGRE2 | ADGRE2 | chr19 | - | 14755653 | 14755877 | 14754953 | 14755127 | 14756237 | 14756345 | 0 | 0 | 4 | 0 | 0 | 0 | 0 | 0 | 0 | 4 | 1 | 0 | 1 | 0 | 98 | 49 | 0.999202116 | 0.999202116 | NA | NA | 1 | NA | 0.667 | 1 | 0.166 |
| SE | 11376 | ARHGEF1 | ARHGEF1 | chr19 | + | 41905174 | 41905261 | 41904948 | 41905036 | 41905759 | 41905827 | 0 | 1 | 10 | 0 | 2 | 0 | 2 | 0 | 0 | 0 | 3 | 0 | 0 | 0 | 98 | 49 | 0.999202116 | 0.999202116 | NA | 0.2 | 1 | NA | NA | 1 | -0.4 |
| SE | 11391 | CPNE2 | CPNE2 | chr16 | + | 57137148 | 57137282 | 57134774 | 57134826 | 57146084 | 57146321 | 0 | 1 | 5 | 0 | 0 | 1 | 0 | 1 | 0 | 1 | 0 | 0 | 0 | 0 | 98 | 49 | 0.999202116 | 0.999202116 | NA | 1 | 0.714 | NA | 1 | NA | -0.143 |
| SE | 11403 | CSNK2A2 | CSNK2A2 | chr16 | - | 58167206 | 58167308 | 58166583 | 58166684 | 58167684 | 58167795 | 0 | 1 | 0 | 0 | 0 | 1 | 0 | 1 | 0 | 1 | 0 | 0 | 0 | 0 | 98 | 49 | 0.999202116 | 0.999202116 | NA | 1 | 0 | NA | 1 | NA | -0.5 |
| SE | 11481 | AP1G1 | AP1G1 | chr16 | - | 71756018 | 71756159 | 71753832 | 71753887 | 71758807 | 71758921 | 0 | 0 | 7 | 0 | 0 | 0 | 0 | 0 | 0 | 16 | 3 | 0 | 1 | 0 | 98 | 49 | 0.999202116 | 0.999202116 | NA | NA | 1 | NA | 0.889 | 1 | 0.055 |
| SE | 11563 | NCBP1 | NCBP1 | chr9 | + | 97662041 | 97662144 | 97660945 | 97661068 | 97662953 | 97663047 | 0 | 1 | 1 | 0 | 0 | 2 | 0 | 2 | 0 | 1 | 0 | 0 | 0 | 0 | 98 | 49 | 0.999202116 | 0.999202116 | NA | 1 | 0.2 | NA | 1 | NA | -0.4 |
| SE | 11622 | CIR1 | CIR1 | chr2 | - | 174378922 | 174379040 | 174351618 | 174351725 | 174380201 | 174380250 | 0 | 0 | 8 | 0 | 0 | 0 | 0 | 0 | 0 | 12 | 1 | 0 | 1 | 0 | 98 | 49 | 0.999202116 | 0.999202116 | NA | NA | 1 | NA | 0.857 | 1 | 0.072 |
| SE | 11641 | ARRB2 | ARRB2 | chr17 | + | 4718260 | 4718345 | 4717887 | 4718023 | 4718611 | 4718684 | 0 | 0 | 26 | 0 | 1 | 3 | 1 | 3 | 4 | 9 | 3 | 0 | 10 | 1 | 98 | 49 | 0.999202116 | 0.999202116 | NA | 0 | 0.812 | 1 | 0.31 | 0.6 | -0.231 |
| SE | 11642 | ARRB2 | ARRB2 | chr17 | + | 4720392 | 4720472 | 4720211 | 4720299 | 4720585 | 4720640 | 0 | 2 | 26 | 0 | 1 | 0 | 1 | 0 | 2 | 12 | 10 | 0 | 4 | 0 | 98 | 49 | 0.999202116 | 0.999202116 | NA | 0.5 | 1 | 1 | 0.6 | 1 | -0.117 |
| SE | 11657 | NLRP1 | NLRP1 | chr17 | - | 5520880 | 5521012 | 5517745 | 5517887 | 5521523 | 5521786 | 0 | 1 | 10 | 0 | 0 | 3 | 0 | 3 | 0 | 5 | 1 | 12 | 1 | 1 | 98 | 49 | 0.999202116 | 0.999202116 | NA | 1 | 0.625 | 0 | 0.714 | 0.333 | 0.464 |
| SE | 11658 | NLRP1 | NLRP1 | chr17 | - | 5521523 | 5521786 | 5517745 | 5517887 | 5530480 | 5530704 | 0 | 0 | 9 | 0 | 0 | 0 | 0 | 0 | 12 | 1 | 1 | 0 | 2 | 0 | 98 | 49 | 0.999202116 | 0.999202116 | NA | NA | 1 | 1 | 0.2 | 1 | 0.267 |
| SE | 11683 | STAT5B | STAT5B | chr17 | - | 42216013 | 42216106 | 42211983 | 42212190 | 42217159 | 42217282 | 0 | 0 | 16 | 0 | 0 | 0 | 0 | 0 | 0 | 4 | 2 | 0 | 1 | 0 | 98 | 49 | 0.999202116 | 0.999202116 | NA | NA | 1 | NA | 0.667 | 1 | 0.166 |
| SE | 11748 | JTB | JTB | chr1 | - | 153976692 | 153976775 | 153975825 | 153975905 | 153976975 | 153977013 | 0 | 0 | 5 | 0 | 0 | 0 | 0 | 0 | 0 | 8 | 1 | 0 | 1 | 0 | 98 | 49 | 0.999202116 | 0.999202116 | NA | NA | 1 | NA | 0.8 | 1 | 0.1 |
| SE | 11782 | USP22 | USP22 | chr17 | - | 21015751 | 21015899 | 21012829 | 21012935 | 21017941 | 21018111 | 0 | 4 | 1 | 0 | 0 | 0 | 0 | 0 | 0 | 2 | 0 | 4 | 0 | 0 | 98 | 49 | 0.999202116 | 0.999202116 | NA | 1 | 1 | 0 | 1 | NA | 0.5 |
| SE | 11805 | KIAA0100 | KIAA0100 | chr17 | - | 28633548 | 28633751 | 28633282 | 28633401 | 28633894 | 28634086 | 0 | 1 | 6 | 0 | 0 | 0 | 0 | 0 | 0 | 1 | 0 | 1 | 0 | 0 | 98 | 49 | 0.999202116 | 0.999202116 | NA | 1 | 1 | 0 | 1 | NA | 0.5 |
| SE | 11822 | SSH2 | SSH2 | chr17 | - | 29695458 | 29695523 | 29684562 | 29684684 | 29702958 | 29703062 | 0 | 0 | 20 | 0 | 0 | 0 | 0 | 0 | 0 | 12 | 1 | 0 | 1 | 0 | 98 | 49 | 0.999202116 | 0.999202116 | NA | NA | 1 | NA | 0.857 | 1 | 0.072 |
| SE | 11876 | MGAM | MGAM | chr7 | + | 142036822 | 142036977 | 142034669 | 142034841 | 142038530 | 142038615 | 0 | 0 | 14 | 0 | 0 | 0 | 0 | 0 | 0 | 2 | 6 | 0 | 4 | 0 | 98 | 49 | 0.999202116 | 0.999202116 | NA | NA | 1 | NA | 0.2 | 1 | 0.4 |
| SE | 12165 | USP32 | USP32 | chr17 | - | 60288522 | 60288682 | 60271349 | 60271481 | 60294682 | 60294801 | 0 | 0 | 12 | 0 | 0 | 0 | 0 | 0 | 0 | 11 | 3 | 0 | 1 | 0 | 98 | 49 | 0.999202116 | 0.999202116 | NA | NA | 1 | NA | 0.846 | 1 | 0.077 |
| SE | 12220 | BPTF | BPTF | chr17 | + | 67966571 | 67966656 | 67964211 | 67964404 | 67975771 | 67975958 | 0 | 1 | 3 | 0 | 0 | 1 | 0 | 1 | 0 | 1 | 0 | 0 | 0 | 0 | 98 | 49 | 0.999202116 | 0.999202116 | NA | 1 | 0.6 | NA | 1 | NA | -0.2 |
| SE | 12394 | ATP5B | ATP5B | chr12 | - | 56642672 | 56642831 | 56642457 | 56642580 | 56643402 | 56643587 | 0 | 0 | 13 | 0 | 0 | 0 | 0 | 0 | 4 | 3 | 2 | 0 | 2 | 0 | 98 | 49 | 0.999202116 | 0.999202116 | NA | NA | 1 | 1 | 0.429 | 1 | 0.19 |
| SE | 12660 | SURF4 | SURF4 | chr9 | - | 133364839 | 133365026 | 133361448 | 133363759 | 133365984 | 133366028 | 0 | 0 | 2 | 0 | 0 | 0 | 0 | 0 | 0 | 3 | 1 | 0 | 1 | 0 | 98 | 49 | 0.999202116 | 0.999202116 | NA | NA | 1 | NA | 0.6 | 1 | 0.2 |
| SE | 12704 | ICAM3 | ICAM3 | chr19 | - | 10335065 | 10335353 | 10334527 | 10334782 | 10335670 | 10335976 | 0 | 1 | 23 | 0 | 0 | 1 | 0 | 1 | 3 | 7 | 1 | 0 | 0 | 2 | 98 | 49 | 0.999202116 | 0.999202116 | NA | 1 | 0.92 | 1 | 1 | 0.2 | 0.227 |
| SE | 12712 | SLC44A2 | SLC44A2 | chr19 | + | 10634973 | 10635073 | 10634755 | 10634887 | 10636322 | 10636585 | 0 | 3 | 8 | 0 | 0 | 1 | 0 | 1 | 0 | 1 | 0 | 0 | 0 | 0 | 98 | 49 | 0.999202116 | 0.999202116 | NA | 1 | 0.8 | NA | 1 | NA | -0.1 |
| SE | 12820 | MAST3 | MAST3 | chr19 | + | 18124641 | 18124774 | 18124264 | 18124366 | 18128399 | 18128458 | 0 | 1 | 11 | 0 | 0 | 3 | 0 | 3 | 0 | 2 | 0 | 2 | 0 | 1 | 98 | 49 | 0.999202116 | 0.999202116 | NA | 1 | 0.647 | 0 | 1 | 0 | 0.49 |
| SE | 12825 | TRAPPC10 | TRAPPC10 | chr21 | + | 44074323 | 44074470 | 44063537 | 44063785 | 44075038 | 44075153 | 0 | 0 | 4 | 0 | 0 | 0 | 0 | 0 | 0 | 3 | 1 | 0 | 1 | 0 | 98 | 49 | 0.999202116 | 0.999202116 | NA | NA | 1 | NA | 0.6 | 1 | 0.2 |
| SE | 12851 | GATAD2A | GATAD2A | chr19 | + | 19494293 | 19494383 | 19492580 | 19492712 | 19495753 | 19495885 | 0 | 0 | 4 | 0 | 0 | 0 | 0 | 0 | 0 | 4 | 2 | 0 | 1 | 0 | 98 | 49 | 0.999202116 | 0.999202116 | NA | NA | 1 | NA | 0.667 | 1 | 0.166 |
| SE | 12950 | GMFG | GMFG | chr19 | - | 39329543 | 39329626 | 39328999 | 39329073 | 39333076 | 39333126 | 0 | 0 | 53 | 0 | 0 | 0 | 0 | 0 | 2 | 32 | 1 | 0 | 1 | 0 | 98 | 49 | 0.999202116 | 0.999202116 | NA | NA | 1 | 1 | 0.941 | 1 | 0.02 |
| SE | 12954 | MAN2A2 | MAN2A2 | chr15 | + | 90910500 | 90910683 | 90909326 | 90909504 | 90910846 | 90910961 | 0 | 0 | 6 | 0 | 0 | 0 | 0 | 0 | 0 | 1 | 1 | 0 | 1 | 0 | 98 | 49 | 0.999202116 | 0.999202116 | NA | NA | 1 | NA | 0.333 | 1 | 0.334 |
| SE | 12990 | PLAUR | PLAUR | chr19 | - | 43652224 | 43652371 | 43648570 | 43649143 | 43655438 | 43655573 | 0 | 2 | 46 | 0 | 0 | 7 | 0 | 7 | 0 | 41 | 2 | 0 | 1 | 1 | 98 | 49 | 0.999202116 | 0.999202116 | NA | 1 | 0.767 | NA | 0.953 | 0.5 | 0.157 |
| SE | 13034 | VASP | VASP | chr19 | + | 45521321 | 45521406 | 45517928 | 45518094 | 45522167 | 45522217 | 0 | 2 | 37 | 0 | 2 | 3 | 2 | 3 | 4 | 24 | 4 | 0 | 3 | 0 | 98 | 49 | 0.999202116 | 0.999202116 | NA | 0.333 | 0.86 | 1 | 0.8 | 1 | -0.337 |
| SE | 13052 | NAPA | NAPA | chr19 | - | 47503422 | 47503502 | 47500632 | 47500749 | 47514842 | 47515258 | 0 | 2 | 18 | 0 | 0 | 0 | 0 | 0 | 0 | 19 | 1 | 0 | 0 | 4 | 98 | 49 | 0.999202116 | 0.999202116 | NA | 1 | 1 | NA | 1 | 0.111 | 0.445 |
| SE | 13128 | ATXN2 | ATXN2 | chr12 | - | 111510384 | 111510582 | 111509890 | 111509998 | 111513356 | 111513539 | 0 | 0 | 2 | 0 | 1 | 0 | 1 | 0 | 0 | 1 | 1 | 0 | 1 | 0 | 98 | 49 | 0.999202116 | 0.999202116 | NA | 0 | 1 | NA | 0.333 | 1 | -0.166 |
| SE | 13152 | C9orf89 | C9orf89 | chr9 | + | 93111878 | 93111938 | 93110567 | 93110721 | 93112217 | 93112289 | 0 | 0 | 6 | 0 | 0 | 0 | 0 | 0 | 0 | 3 | 2 | 0 | 3 | 0 | 98 | 49 | 0.999202116 | 0.999202116 | NA | NA | 1 | NA | 0.333 | 1 | 0.334 |
| SE | 13179 | NLRP12 | NLRP12 | chr19 | - | 53807494 | 53807668 | 53805279 | 53805450 | 53809586 | 53811288 | 0 | 1 | 8 | 0 | 0 | 2 | 0 | 2 | 0 | 1 | 0 | 0 | 0 | 0 | 98 | 49 | 0.999202116 | 0.999202116 | NA | 1 | 0.667 | NA | 1 | NA | -0.166 |
| SE | 13210 | LILRB2 | LILRB2 | chr19 | - | 54280263 | 54280299 | 54279790 | 54280075 | 54280462 | 54280544 | 0 | 0 | 8 | 0 | 0 | 0 | 0 | 0 | 0 | 1 | 2 | 0 | 1 | 0 | 85 | 49 | 0.999202116 | 0.999202116 | NA | NA | 1 | NA | 0.366 | 1 | 0.317 |
| SE | 13368 | FIP1L1 | FIP1L1 | chr4 | + | 53414614 | 53414722 | 53399729 | 53399839 | 53425871 | 53425965 | 0 | 1 | 0 | 0 | 0 | 2 | 0 | 2 | 2 | 1 | 0 | 0 | 0 | 0 | 98 | 49 | 0.999202116 | 0.999202116 | NA | 1 | 0 | 1 | 1 | NA | -0.5 |
| SE | 13635 | CCND3 | CCND3 | chr6 | - | 41940369 | 41940585 | 41937234 | 41937394 | 42048500 | 42048894 | 0 | 1 | 23 | 0 | 0 | 1 | 0 | 1 | 0 | 0 | 3 | 0 | 1 | 0 | 98 | 49 | 0.999202116 | 0.999202116 | NA | 1 | 0.92 | NA | 0 | 1 | 0.46 |
| SE | 13689 | BAZ2A | BAZ2A | chr12 | - | 56609735 | 56609946 | 56606632 | 56606733 | 56610113 | 56610215 | 0 | 0 | 3 | 0 | 0 | 0 | 0 | 0 | 0 | 1 | 1 | 0 | 1 | 0 | 98 | 49 | 0.999202116 | 0.999202116 | NA | NA | 1 | NA | 0.333 | 1 | 0.334 |
| SE | 13733 | SERPINA1 | SERPINA1 | chr14 | - | 94388559 | 94388663 | 94382591 | 94383241 | 94390456 | 94390692 | 0 | 0 | 17 | 0 | 2 | 20 | 2 | 20 | 0 | 6 | 1 | 0 | 5 | 0 | 98 | 49 | 0.999202116 | 0.999202116 | NA | 0 | 0.298 | NA | 0.375 | 1 | -0.538 |
| SE | 13745 | MAP4K4 | MAP4K4 | chr2 | + | 101887087 | 101887237 | 101885186 | 101885287 | 101887777 | 101887937 | 0 | 0 | 10 | 0 | 0 | 0 | 0 | 0 | 0 | 2 | 1 | 0 | 1 | 0 | 98 | 49 | 0.999202116 | 0.999202116 | NA | NA | 1 | NA | 0.5 | 1 | 0.25 |
| SE | 13818 | MAP3K2 | MAP3K2 | chr2 | - | 127325727 | 127325807 | 127324173 | 127324241 | 127326686 | 127326817 | 0 | 0 | 17 | 0 | 0 | 0 | 0 | 0 | 0 | 15 | 4 | 0 | 1 | 0 | 98 | 49 | 0.999202116 | 0.999202116 | NA | NA | 1 | NA | 0.882 | 1 | 0.059 |
| SE | 13861 | ACSL1 | ACSL1 | chr4 | - | 184764852 | 184764925 | 184763166 | 184763255 | 184766621 | 184766756 | 0 | 2 | 44 | 0 | 0 | 2 | 0 | 2 | 3 | 39 | 5 | 0 | 1 | 1 | 98 | 49 | 0.999202116 | 0.999202116 | NA | 1 | 0.917 | 1 | 0.951 | 0.714 | 0.07 |
| SE | 13862 | ACSL1 | ACSL1 | chr4 | - | 184765890 | 184765986 | 184763166 | 184763255 | 184766621 | 184766756 | 0 | 2 | 50 | 0 | 0 | 2 | 0 | 2 | 0 | 38 | 5 | 0 | 1 | 1 | 98 | 49 | 0.999202116 | 0.999202116 | NA | 1 | 0.926 | NA | 0.95 | 0.714 | 0.131 |
| A5SS | 489 | RCHY1 | RCHY1 | chr4 | - | 75491582 | 75491637 | 75491610 | 75491637 | 75490580 | 75490701 | 0 | 0 | 0 | 0 | 1 | 0 | 1 | 0 | 0 | 0 | 1 | 0 | 1 | 0 | 77 | 49 | 0.999208248 | 0.999208248 | NA | 0 | NA | NA | 0 | 1 | -0.5 |
| A3SS | 13 | LONRF3 | LONRF3 | chrX | + | 119012879 | 119013201 | 119013038 | 119013201 | 119011814 | 119011973 | 0 | 0 | 0 | 0 | 0 | 1 | 0 | 1 | 0 | 1 | 0 | 0 | 0 | 0 | 98 | 49 | NA | NA | NA | NA | 0 | NA | 1 | NA | -1 |
| A3SS | 50 | TFE3 | TFE3 | chrX | - | 49039106 | 49039410 | 49039106 | 49039345 | 49040454 | 49040568 | 0 | 0 | 4 | 0 | 0 | 3 | 0 | 3 | 0 | 4 | 0 | 0 | 3 | 0 | 98 | 49 | NA | NA | NA | NA | 0.4 | NA | 0.4 | NA | 0 |
| A3SS | 63 | CD99 | CD99 | chrX | + | 2738199 | 2738256 | 2738202 | 2738256 | 2726259 | 2726373 | 0 | 0 | 10 | 0 | 0 | 1 | 0 | 1 | 1 | 3 | 0 | 0 | 0 | 0 | 52 | 49 | NA | NA | NA | NA | 0.904 | 1 | 1 | NA | -0.096 |
| A3SS | 71 | EDF1 | EDF1 | chr9 | - | 136862113 | 136862496 | 136862113 | 136862345 | 136862905 | 136862999 | 1 | 0 | 3 | 0 | 0 | 1 | 0 | 1 | 0 | 1 | 0 | 0 | 2 | 0 | 98 | 49 | NA | NA | 1 | NA | 0.6 | NA | 0.2 | NA | 0.6 |
| A3SS | 95 | CDK5RAP2 | CDK5RAP2 | chr9 | - | 120460571 | 120460747 | 120460571 | 120460667 | 120467859 | 120467997 | 3 | 0 | 0 | 0 | 0 | 0 | 0 | 0 | 0 | 0 | 0 | 0 | 1 | 0 | 98 | 49 | NA | NA | 1 | NA | NA | NA | 0 | NA | 1 |
| A3SS | 117 | TLE4 | TLE4 | chr9 | + | 79652589 | 79652794 | 79652592 | 79652794 | 79627373 | 79627448 | 0 | 0 | 0 | 0 | 0 | 2 | 0 | 2 | 0 | 1 | 0 | 0 | 1 | 0 | 52 | 49 | NA | NA | NA | NA | 0 | NA | 0.485 | NA | -0.485 |
| A3SS | 213 | CUX1 | CUX1 | chr7 | + | 102178468 | 102178657 | 102178474 | 102178657 | 102170445 | 102170550 | 0 | 0 | 5 | 0 | 0 | 0 | 0 | 0 | 0 | 0 | 0 | 0 | 2 | 0 | 55 | 49 | NA | NA | NA | NA | 1 | NA | 0 | NA | 1 |
| A3SS | 280 | IPCEF1 | IPCEF1 | chr6 | - | 154247448 | 154247488 | 154247448 | 154247485 | 154265911 | 154265964 | 0 | 0 | 0 | 0 | 0 | 1 | 0 | 1 | 0 | 1 | 0 | 0 | 0 | 0 | 52 | 49 | NA | NA | NA | NA | 0 | NA | 1 | NA | -1 |
| A3SS | 287 | PLAGL1 | PLAGL1 | chr6 | - | 143960468 | 143960580 | 143960468 | 143960542 | 143964786 | 143964818 | 0 | 2 | 0 | 0 | 0 | 1 | 0 | 1 | 3 | 0 | 0 | 0 | 0 | 0 | 87 | 49 | NA | NA | NA | 1 | 0 | 1 | NA | NA | -0.5 |
| A3SS | 332 | STK38 | STK38 | chr6 | - | 36540071 | 36540207 | 36540071 | 36540204 | 36547189 | 36547516 | 0 | 0 | 0 | 0 | 0 | 1 | 0 | 1 | 0 | 1 | 0 | 0 | 0 | 0 | 52 | 49 | NA | NA | NA | NA | 0 | NA | 1 | NA | -1 |
| A3SS | 466 | BTN3A2 | BTN3A2 | chr6 | + | 26367974 | 26368050 | 26367989 | 26368050 | 26365158 | 26365352 | 0 | 0 | 1 | 0 | 0 | 2 | 0 | 2 | 0 | 0 | 0 | 0 | 1 | 0 | 64 | 49 | NA | NA | NA | NA | 0.277 | NA | 0 | NA | 0.277 |
| A3SS | 478 | CD83 | CD83 | chr6 | + | 14135107 | 14136917 | 14135110 | 14136917 | 14133648 | 14133755 | 0 | 0 | 0 | 0 | 0 | 1 | 0 | 1 | 0 | 14 | 0 | 0 | 0 | 0 | 52 | 49 | NA | NA | NA | NA | 0 | NA | 1 | NA | -1 |
| A3SS | 503 | HARS2 | HARS2 | chr5 | + | 140693490 | 140693665 | 140693590 | 140693665 | 140691425 | 140691756 | 0 | 0 | 2 | 0 | 0 | 0 | 0 | 0 | 0 | 1 | 0 | 0 | 2 | 0 | 98 | 49 | NA | NA | NA | NA | 1 | NA | 0.2 | NA | 0.8 |
| A3SS | 513 | SRA1 | SRA1 | chr5 | - | 140557146 | 140557332 | 140557146 | 140557272 | 140557427 | 140558093 | 0 | 0 | 1 | 0 | 0 | 1 | 0 | 1 | 0 | 1 | 0 | 0 | 0 | 0 | 98 | 49 | NA | NA | NA | NA | 0.333 | NA | 1 | NA | -0.667 |
| A3SS | 535 | SSBP2 | SSBP2 | chr5 | - | 81466973 | 81467065 | 81466973 | 81467041 | 81473699 | 81473770 | 0 | 0 | 0 | 0 | 0 | 1 | 0 | 1 | 2 | 0 | 0 | 0 | 0 | 0 | 73 | 49 | NA | NA | NA | NA | 0 | 1 | NA | NA | -1 |
| A3SS | 544 | GAPT | GAPT | chr5 | + | 58494147 | 58496358 | 58494246 | 58496358 | 58493710 | 58493838 | 0 | 0 | 1 | 0 | 0 | 2 | 0 | 2 | 5 | 1 | 0 | 0 | 0 | 0 | 98 | 49 | NA | NA | NA | NA | 0.2 | 1 | 1 | NA | -0.8 |
| A3SS | 580 | ALPK1 | ALPK1 | chr4 | + | 112377677 | 112377898 | 112377729 | 112377898 | 112315800 | 112315852 | 0 | 0 | 7 | 0 | 0 | 0 | 0 | 0 | 7 | 2 | 0 | 0 | 4 | 0 | 98 | 49 | NA | NA | NA | NA | 1 | 1 | 0.2 | NA | 0.4 |
| A3SS | 669 | MFSD1 | MFSD1 | chr3 | + | 158804318 | 158804371 | 158804322 | 158804371 | 158801925 | 158802315 | 0 | 0 | 5 | 0 | 0 | 0 | 0 | 0 | 0 | 1 | 0 | 0 | 1 | 0 | 53 | 49 | NA | NA | NA | NA | 1 | NA | 0.48 | NA | 0.52 |
| A3SS | 676 | ARMC8 | ARMC8 | chr3 | + | 138188400 | 138188530 | 138188454 | 138188530 | 138187247 | 138187599 | 2 | 0 | 1 | 0 | 0 | 2 | 0 | 2 | 0 | 1 | 0 | 0 | 0 | 0 | 98 | 49 | NA | NA | 1 | NA | 0.2 | NA | 1 | NA | -0.4 |
| A3SS | 680 | CDV3 | CDV3 | chr3 | + | 133587158 | 133590274 | 133587161 | 133590274 | 133586562 | 133586722 | 0 | 0 | 4 | 0 | 0 | 1 | 0 | 1 | 0 | 3 | 0 | 0 | 0 | 0 | 52 | 49 | NA | NA | NA | NA | 0.79 | NA | 1 | NA | -0.21 |
| A3SS | 682 | CDV3 | CDV3 | chr3 | + | 133587158 | 133590274 | 133588352 | 133590274 | 133586562 | 133586722 | 0 | 0 | 9 | 0 | 0 | 0 | 0 | 0 | 0 | 7 | 0 | 0 | 1 | 0 | 98 | 49 | NA | NA | NA | NA | 1 | NA | 0.778 | NA | 0.222 |
| A3SS | 684 | CDV3 | CDV3 | chr3 | + | 133587161 | 133590274 | 133588352 | 133590274 | 133586562 | 133586722 | 0 | 0 | 9 | 0 | 0 | 0 | 0 | 0 | 0 | 4 | 0 | 0 | 1 | 0 | 98 | 49 | NA | NA | NA | NA | 1 | NA | 0.667 | NA | 0.333 |
| A3SS | 685 | CDV3 | CDV3 | chr3 | + | 133587895 | 133590274 | 133588352 | 133590274 | 133586562 | 133586722 | 0 | 0 | 8 | 0 | 0 | 0 | 0 | 0 | 0 | 6 | 0 | 0 | 1 | 0 | 98 | 49 | NA | NA | NA | NA | 1 | NA | 0.75 | NA | 0.25 |
| A3SS | 718 | KIAA0922 | KIAA0922 | chr4 | + | 153584834 | 153584931 | 153584837 | 153584931 | 153583563 | 153583672 | 0 | 0 | 2 | 0 | 0 | 3 | 0 | 3 | 0 | 0 | 0 | 0 | 2 | 0 | 52 | 49 | NA | NA | NA | NA | 0.386 | NA | 0 | NA | 0.386 |
| A3SS | 726 | USP19 | USP19 | chr3 | - | 49115723 | 49115944 | 49115723 | 49115938 | 49116046 | 49116162 | 0 | 0 | 1 | 0 | 0 | 0 | 0 | 0 | 0 | 0 | 0 | 0 | 1 | 0 | 55 | 49 | NA | NA | NA | NA | 1 | NA | 0 | NA | 1 |
| A3SS | 774 | LST1 | LST1 | chr6 | + | 31588496 | 31588909 | 31588517 | 31588909 | 31587199 | 31587318 | 0 | 0 | 3 | 0 | 0 | 4 | 0 | 4 | 0 | 0 | 0 | 0 | 4 | 0 | 70 | 49 | NA | NA | NA | NA | 0.344 | NA | 0 | NA | 0.344 |
| A3SS | 813 | POLDIP3 | POLDIP3 | chr22 | - | 42601969 | 42602107 | 42601969 | 42602056 | 42602769 | 42603160 | 0 | 0 | 0 | 0 | 0 | 1 | 0 | 1 | 0 | 1 | 0 | 0 | 0 | 0 | 98 | 49 | NA | NA | NA | NA | 0 | NA | 1 | NA | -1 |
| A3SS | 833 | SRP19 | SRP19 | chr5 | + | 112867371 | 112868392 | 112867403 | 112868392 | 112864620 | 112864732 | 0 | 0 | 1 | 0 | 0 | 1 | 0 | 1 | 0 | 1 | 0 | 0 | 0 | 0 | 81 | 49 | NA | NA | NA | NA | 0.377 | NA | 1 | NA | -0.623 |
| A3SS | 931 | STAU1 | STAU1 | chr20 | - | 49124374 | 49124605 | 49124374 | 49124587 | 49135832 | 49135931 | 0 | 0 | 3 | 0 | 0 | 0 | 0 | 0 | 0 | 0 | 0 | 0 | 2 | 0 | 67 | 49 | NA | NA | NA | NA | 1 | NA | 0 | NA | 1 |
| A3SS | 938 | ELMO2 | ELMO2 | chr20 | - | 46394404 | 46394532 | 46394404 | 46394503 | 46398696 | 46398771 | 0 | 0 | 0 | 0 | 0 | 1 | 0 | 1 | 2 | 2 | 0 | 0 | 0 | 0 | 78 | 49 | NA | NA | NA | NA | 0 | 1 | 1 | NA | -1 |
| A3SS | 940 | PIGT | PIGT | chr20 | + | 45419472 | 45419590 | 45419503 | 45419590 | 45419294 | 45419395 | 0 | 0 | 0 | 0 | 0 | 2 | 0 | 2 | 4 | 1 | 0 | 3 | 1 | 0 | 80 | 49 | NA | NA | NA | NA | 0 | 0.45 | 0.38 | NA | -0.415 |
| A3SS | 952 | TRPC4AP | TRPC4AP | chr20 | - | 35021189 | 35021356 | 35021189 | 35021332 | 35035122 | 35035308 | 0 | 0 | 3 | 0 | 0 | 1 | 0 | 1 | 0 | 1 | 0 | 0 | 1 | 0 | 73 | 49 | NA | NA | NA | NA | 0.668 | NA | 0.402 | NA | 0.266 |
| A3SS | 964 | HCK | HCK | chr20 | + | 32073715 | 32073818 | 32073718 | 32073818 | 32073318 | 32073361 | 0 | 0 | 6 | 0 | 0 | 0 | 0 | 0 | 2 | 2 | 0 | 0 | 1 | 0 | 52 | 49 | NA | NA | NA | NA | 1 | 1 | 0.653 | NA | 0.173 |
| A3SS | 1008 | DYRK1A | DYRK1A | chr21 | + | 37478180 | 37478300 | 37478207 | 37478300 | 37472683 | 37472880 | 0 | 0 | 0 | 0 | 0 | 3 | 0 | 3 | 1 | 0 | 0 | 0 | 5 | 0 | 76 | 49 | NA | NA | NA | NA | 0 | 1 | 0 | NA | -0.5 |
| A3SS | 1022 | TRIP12 | TRIP12 | chr2 | - | 229805729 | 229805883 | 229805729 | 229805802 | 229807707 | 229807864 | 0 | 0 | 8 | 0 | 0 | 0 | 0 | 0 | 4 | 9 | 0 | 0 | 1 | 0 | 98 | 49 | NA | NA | NA | NA | 1 | 1 | 0.818 | NA | 0.091 |
| A3SS | 1025 | AGFG1 | AGFG1 | chr2 | + | 227553703 | 227553795 | 227553709 | 227553795 | 227551958 | 227552117 | 0 | 0 | 2 | 0 | 0 | 4 | 0 | 4 | 0 | 1 | 0 | 0 | 0 | 0 | 55 | 49 | NA | NA | NA | NA | 0.308 | NA | 1 | NA | -0.692 |
| A3SS | 1040 | ARHGEF40 | ARHGEF40 | chr14 | + | 21075622 | 21075765 | 21075644 | 21075765 | 21075331 | 21075499 | 0 | 0 | 0 | 0 | 0 | 2 | 0 | 2 | 2 | 0 | 0 | 0 | 1 | 0 | 71 | 49 | NA | NA | NA | NA | 0 | 1 | 0 | NA | -0.5 |
| A3SS | 1099 | MAP7D1 | MAP7D1 | chr1 | + | 36179511 | 36179557 | 36179514 | 36179557 | 36179261 | 36179315 | 0 | 0 | 0 | 0 | 0 | 3 | 0 | 3 | 0 | 1 | 0 | 0 | 3 | 0 | 52 | 49 | NA | NA | NA | NA | 0 | NA | 0.239 | NA | -0.239 |
| A3SS | 1105 | SH3BP5 | SH3BP5 | chr3 | - | 15304102 | 15304299 | 15304102 | 15304231 | 15330503 | 15330566 | 0 | 0 | 2 | 0 | 0 | 2 | 0 | 2 | 0 | 1 | 0 | 0 | 2 | 0 | 98 | 49 | NA | NA | NA | NA | 0.333 | NA | 0.2 | NA | 0.133 |
| A3SS | 1109 | ZDHHC20 | ZDHHC20 | chr13 | - | 21381433 | 21381549 | 21381433 | 21381546 | 21382919 | 21383009 | 0 | 0 | 6 | 0 | 0 | 3 | 0 | 3 | 0 | 1 | 0 | 0 | 0 | 0 | 52 | 49 | NA | NA | NA | NA | 0.653 | NA | 1 | NA | -0.347 |
| A3SS | 1139 | C17orf49 | C17orf49 | chr17 | + | 7017256 | 7017524 | 7017260 | 7017524 | 7016909 | 7017014 | 0 | 0 | 0 | 0 | 0 | 2 | 0 | 2 | 0 | 1 | 0 | 0 | 3 | 0 | 53 | 49 | NA | NA | NA | NA | 0 | NA | 0.236 | NA | -0.236 |
| A3SS | 1193 | SUPT20H | SUPT20H | chr13 | - | 37022010 | 37022317 | 37022010 | 37022080 | 37024034 | 37024193 | 0 | 0 | 2 | 0 | 0 | 1 | 0 | 1 | 0 | 1 | 0 | 0 | 0 | 0 | 98 | 49 | NA | NA | NA | NA | 0.5 | NA | 1 | NA | -0.5 |
| A3SS | 1201 | NFKB1 | NFKB1 | chr4 | + | 102533844 | 102533885 | 102533847 | 102533885 | 102529835 | 102529914 | 0 | 0 | 0 | 0 | 0 | 1 | 0 | 1 | 0 | 1 | 0 | 0 | 1 | 0 | 52 | 49 | NA | NA | NA | NA | 0 | NA | 0.485 | NA | -0.485 |
| A3SS | 1269 | IRF5 | IRF5 | chr7 | + | 128947229 | 128947535 | 128947277 | 128947535 | 128947022 | 128947056 | 0 | 0 | 1 | 0 | 0 | 1 | 0 | 1 | 2 | 1 | 0 | 0 | 0 | 0 | 97 | 49 | NA | NA | NA | NA | 0.336 | 1 | 1 | NA | -0.664 |
| A3SS | 1279 | ING4 | ING4 | chr12 | - | 6652661 | 6652770 | 6652661 | 6652767 | 6652935 | 6653050 | 0 | 0 | 2 | 0 | 0 | 2 | 0 | 2 | 0 | 0 | 0 | 0 | 2 | 0 | 52 | 49 | NA | NA | NA | NA | 0.485 | NA | 0 | NA | 0.485 |
| A3SS | 1301 | GMPR2 | GMPR2 | chr14 | + | 24237041 | 24237152 | 24237070 | 24237152 | 24235966 | 24236140 | 0 | 0 | 0 | 0 | 0 | 1 | 0 | 1 | 0 | 1 | 0 | 0 | 2 | 0 | 78 | 49 | NA | NA | NA | NA | 0 | NA | 0.239 | NA | -0.239 |
| A3SS | 1322 | CASP1 | CASP1 | chr11 | - | 105025509 | 105026356 | 105025509 | 105025600 | 105026841 | 105026951 | 0 | 0 | 29 | 0 | 0 | 0 | 0 | 0 | 0 | 2 | 0 | 1 | 1 | 0 | 98 | 49 | NA | NA | NA | NA | 1 | 0 | 0.5 | NA | 0.75 |
| A3SS | 1358 | ANAPC15 | ANAPC15 | chr11 | - | 72109586 | 72109940 | 72109586 | 72109928 | 72110087 | 72110225 | 0 | 0 | 0 | 1 | 0 | 0 | 0 | 0 | 0 | 1 | 0 | 0 | 1 | 0 | 61 | 49 | NA | NA | 0 | NA | NA | NA | 0.445 | NA | -0.445 |
| A3SS | 1411 | DNM2 | DNM2 | chr19 | + | 10830977 | 10831910 | 10830980 | 10831910 | 10830126 | 10830378 | 0 | 2 | 1 | 0 | 0 | 1 | 0 | 1 | 1 | 0 | 0 | 0 | 0 | 0 | 52 | 49 | NA | NA | NA | 1 | 0.485 | 1 | NA | NA | -0.258 |
| A3SS | 1533 | TTC23 | TTC23 | chr15 | - | 99139316 | 99139467 | 99139316 | 99139399 | 99156147 | 99156297 | 0 | 0 | 0 | 0 | 0 | 1 | 0 | 1 | 0 | 1 | 0 | 0 | 0 | 0 | 98 | 49 | NA | NA | NA | NA | 0 | NA | 1 | NA | -1 |
| A3SS | 1537 | CARS | CARS | chr11 | - | 3040895 | 3041031 | 3040895 | 3040984 | 3042164 | 3042256 | 2 | 0 | 0 | 0 | 0 | 3 | 0 | 3 | 0 | 1 | 0 | 0 | 0 | 0 | 96 | 49 | NA | NA | 1 | NA | 0 | NA | 1 | NA | -0.5 |
| A3SS | 1600 | MECR | MECR | chr1 | - | 29216587 | 29216812 | 29216587 | 29216685 | 29230730 | 29230958 | 0 | 0 | 0 | 0 | 0 | 1 | 0 | 1 | 0 | 2 | 0 | 0 | 0 | 0 | 98 | 49 | NA | NA | NA | NA | 0 | NA | 1 | NA | -1 |
| A3SS | 1610 | FAM111A | FAM111A | chr11 | + | 59148796 | 59148953 | 59148835 | 59148953 | 59145826 | 59145856 | 0 | 0 | 0 | 0 | 1 | 0 | 1 | 0 | 3 | 0 | 0 | 0 | 0 | 1 | 88 | 49 | NA | NA | NA | 0 | NA | 1 | NA | 0 | -0.5 |
| A3SS | 1628 | SRSF10 | SRSF10 | chr1 | - | 23964346 | 23967956 | 23964346 | 23967723 | 23971572 | 23971623 | 1 | 0 | 0 | 0 | 0 | 0 | 0 | 0 | 0 | 0 | 0 | 0 | 2 | 0 | 98 | 49 | NA | NA | 1 | NA | NA | NA | 0 | NA | 1 |
| A3SS | 1629 | SRSF10 | SRSF10 | chr1 | - | 23964346 | 23971439 | 23964346 | 23967723 | 23971572 | 23971623 | 1 | 0 | 1 | 0 | 0 | 0 | 0 | 0 | 0 | 1 | 0 | 0 | 2 | 0 | 98 | 49 | NA | NA | 1 | NA | 1 | NA | 0.2 | NA | 0.8 |
| A3SS | 1635 | METTL9 | METTL9 | chr16 | + | 21655226 | 21657473 | 21655229 | 21657473 | 21624930 | 21625115 | 0 | 0 | 2 | 0 | 0 | 2 | 0 | 2 | 0 | 1 | 0 | 0 | 1 | 0 | 52 | 49 | NA | NA | NA | NA | 0.485 | NA | 0.485 | NA | 0 |
| A3SS | 1660 | SHC1 | SHC1 | chr1 | - | 154965945 | 154966080 | 154965945 | 154966077 | 154966161 | 154966231 | 0 | 0 | 0 | 0 | 0 | 3 | 0 | 3 | 0 | 2 | 0 | 0 | 0 | 0 | 52 | 49 | NA | NA | NA | NA | 0 | NA | 1 | NA | -1 |
| A3SS | 1669 | WDR47 | WDR47 | chr1 | - | 109004591 | 109004718 | 109004591 | 109004715 | 109010915 | 109011718 | 0 | 0 | 1 | 0 | 0 | 0 | 0 | 0 | 0 | 0 | 0 | 0 | 1 | 0 | 52 | 49 | NA | NA | NA | NA | 1 | NA | 0 | NA | 1 |
| A3SS | 1706 | RBMX | RBMX | chrX | - | 136879016 | 136879158 | 136879016 | 136879123 | 136879318 | 136879453 | 0 | 0 | 0 | 0 | 0 | 4 | 0 | 4 | 0 | 1 | 0 | 0 | 0 | 0 | 84 | 49 | NA | NA | NA | NA | 0 | NA | 1 | NA | -1 |
| A3SS | 1749 | FAM213B | FAM213B | chr1 | + | 2588335 | 2588453 | 2588389 | 2588453 | 2587740 | 2587792 | 0 | 0 | 0 | 0 | 0 | 1 | 0 | 1 | 0 | 1 | 0 | 0 | 0 | 0 | 98 | 49 | NA | NA | NA | NA | 0 | NA | 1 | NA | -1 |
| A3SS | 1752 | FAM213B | FAM213B | chr1 | + | 2588921 | 2589040 | 2588953 | 2589040 | 2588549 | 2588625 | 0 | 0 | 1 | 0 | 0 | 0 | 0 | 0 | 0 | 0 | 0 | 0 | 2 | 0 | 81 | 49 | NA | NA | NA | NA | 1 | NA | 0 | NA | 1 |
| A3SS | 1807 | PCBP2 | PCBP2 | chr12 | + | 53468776 | 53468832 | 53468779 | 53468832 | 53467220 | 53467293 | 0 | 0 | 7 | 0 | 0 | 2 | 0 | 2 | 1 | 3 | 0 | 0 | 0 | 0 | 52 | 49 | NA | NA | NA | NA | 0.767 | 1 | 1 | NA | -0.233 |
| A3SS | 1808 | PCBP2 | PCBP2 | chr12 | + | 53468776 | 53468832 | 53468779 | 53468832 | 53467804 | 53467843 | 0 | 0 | 5 | 0 | 0 | 0 | 0 | 0 | 0 | 1 | 0 | 0 | 2 | 0 | 52 | 49 | NA | NA | NA | NA | 1 | NA | 0.32 | NA | 0.68 |
| A3SS | 1847 | ABI1 | ABI1 | chr10 | - | 26759061 | 26759238 | 26759061 | 26759235 | 26765217 | 26765318 | 0 | 0 | 1 | 0 | 0 | 2 | 0 | 2 | 0 | 1 | 0 | 0 | 0 | 0 | 52 | 49 | NA | NA | NA | NA | 0.32 | NA | 1 | NA | -0.68 |
| A3SS | 1876 | PSME3 | PSME3 | chr17 | + | 42838691 | 42838799 | 42838730 | 42838799 | 42838092 | 42838205 | 0 | 0 | 1 | 0 | 0 | 4 | 0 | 4 | 0 | 3 | 0 | 0 | 6 | 0 | 88 | 49 | NA | NA | NA | NA | 0.122 | NA | 0.218 | NA | -0.096 |
| A3SS | 1918 | EPB41 | EPB41 | chr1 | + | 28987430 | 28987905 | 28987447 | 28987905 | 28887090 | 28887210 | 1 | 0 | 0 | 0 | 0 | 0 | 0 | 0 | 0 | 0 | 0 | 0 | 0 | 1 | 66 | 49 | NA | NA | 1 | NA | NA | NA | NA | 0 | 1 |
| A3SS | 1935 | ATL2 | ATL2 | chr2 | - | 38293956 | 38296590 | 38293956 | 38296113 | 38298143 | 38298575 | 0 | 0 | 1 | 0 | 0 | 0 | 0 | 0 | 0 | 0 | 0 | 0 | 1 | 0 | 98 | 49 | NA | NA | NA | NA | 1 | NA | 0 | NA | 1 |
| A3SS | 1978 | PCID2 | PCID2 | chr13 | - | 113177538 | 113178008 | 113177538 | 113177976 | 113178173 | 113178287 | 0 | 0 | 3 | 0 | 0 | 0 | 0 | 0 | 0 | 0 | 0 | 0 | 1 | 0 | 81 | 49 | NA | NA | NA | NA | 1 | NA | 0 | NA | 1 |
| A3SS | 1994 | WNK1 | WNK1 | chr12 | + | 878211 | 878361 | 878214 | 878361 | 862082 | 862270 | 0 | 0 | 1 | 0 | 0 | 2 | 0 | 2 | 0 | 2 | 0 | 0 | 0 | 0 | 52 | 49 | NA | NA | NA | NA | 0.32 | NA | 1 | NA | -0.68 |
| A3SS | 1995 | WNK1 | WNK1 | chr12 | + | 878211 | 878361 | 878214 | 878361 | 871264 | 871348 | 0 | 2 | 0 | 0 | 0 | 1 | 0 | 1 | 3 | 0 | 0 | 0 | 0 | 0 | 52 | 49 | NA | NA | NA | 1 | 0 | 1 | NA | NA | -0.5 |
| A3SS | 2020 | TRPM7 | TRPM7 | chr15 | - | 50583088 | 50583159 | 50583088 | 50583156 | 50586391 | 50586488 | 0 | 0 | 1 | 0 | 0 | 0 | 0 | 0 | 0 | 0 | 0 | 0 | 1 | 0 | 52 | 49 | NA | NA | NA | NA | 1 | NA | 0 | NA | 1 |
| A3SS | 2077 | CD300LF | CD300LF | chr17 | - | 74703034 | 74703143 | 74703034 | 74703098 | 74704477 | 74704816 | 0 | 0 | 2 | 0 | 0 | 4 | 0 | 4 | 0 | 2 | 0 | 0 | 3 | 0 | 94 | 49 | NA | NA | NA | NA | 0.207 | NA | 0.258 | NA | -0.051 |
| A3SS | 2103 | RALGAPB | RALGAPB | chr20 | + | 38565355 | 38565478 | 38565358 | 38565478 | 38562531 | 38562697 | 0 | 0 | 0 | 0 | 0 | 2 | 0 | 2 | 0 | 1 | 0 | 0 | 0 | 0 | 52 | 49 | NA | NA | NA | NA | 0 | NA | 1 | NA | -1 |
| A3SS | 2128 | ARHGEF2 | ARHGEF2 | chr1 | - | 155964987 | 155965131 | 155964987 | 155965128 | 155965302 | 155965412 | 0 | 0 | 2 | 0 | 0 | 4 | 0 | 4 | 0 | 1 | 0 | 0 | 1 | 0 | 52 | 49 | NA | NA | NA | NA | 0.32 | NA | 0.485 | NA | -0.165 |
| A3SS | 2171 | TSC2 | TSC2 | chr16 | + | 2079031 | 2079196 | 2079034 | 2079196 | 2076490 | 2076585 | 0 | 0 | 1 | 0 | 0 | 0 | 0 | 0 | 0 | 1 | 0 | 0 | 2 | 0 | 52 | 49 | NA | NA | NA | NA | 1 | NA | 0.32 | NA | 0.68 |
| A3SS | 2175 | ATP6V0B | ATP6V0B | chr1 | + | 43977025 | 43978300 | 43977980 | 43978300 | 43976772 | 43976824 | 1 | 0 | 10 | 0 | 0 | 0 | 0 | 0 | 0 | 4 | 0 | 0 | 1 | 0 | 98 | 49 | NA | NA | 1 | NA | 1 | NA | 0.667 | NA | 0.333 |
| A3SS | 2202 | ATF7IP | ATF7IP | chr12 | + | 14434336 | 14434423 | 14434339 | 14434423 | 14423908 | 14425473 | 0 | 0 | 6 | 0 | 0 | 3 | 0 | 3 | 0 | 1 | 0 | 0 | 0 | 0 | 52 | 49 | NA | NA | NA | NA | 0.653 | NA | 1 | NA | -0.347 |
| A3SS | 2205 | ZNF33A | ZNF33A | chr10 | + | 38054374 | 38060067 | 38054377 | 38060067 | 38017290 | 38017386 | 0 | 0 | 1 | 0 | 0 | 1 | 0 | 1 | 0 | 2 | 0 | 0 | 2 | 0 | 52 | 49 | NA | NA | NA | NA | 0.485 | NA | 0.485 | NA | 0 |
| A3SS | 2214 | ZWILCH | ZWILCH | chr15 | + | 66508745 | 66508892 | 66508840 | 66508892 | 66505082 | 66505391 | 0 | 0 | 0 | 0 | 0 | 2 | 0 | 2 | 6 | 0 | 0 | 0 | 0 | 0 | 98 | 49 | NA | NA | NA | NA | 0 | 1 | NA | NA | -1 |
| A3SS | 2268 | SH2D2A | SH2D2A | chr1 | - | 156814204 | 156814324 | 156814204 | 156814294 | 156815036 | 156815221 | 0 | 0 | 2 | 0 | 0 | 1 | 0 | 1 | 1 | 0 | 0 | 0 | 0 | 0 | 79 | 49 | NA | NA | NA | NA | 0.554 | 1 | NA | NA | -0.446 |
| A3SS | 2286 | COPA | COPA | chr1 | - | 160305432 | 160305598 | 160305432 | 160305571 | 160305687 | 160305773 | 0 | 0 | 4 | 0 | 0 | 3 | 0 | 3 | 0 | 0 | 0 | 0 | 4 | 0 | 76 | 49 | NA | NA | NA | NA | 0.462 | NA | 0 | NA | 0.462 |
| A3SS | 2304 | CAST | CAST | chr5 | + | 96765220 | 96765325 | 96765223 | 96765325 | 96762273 | 96762372 | 0 | 0 | 8 | 0 | 0 | 3 | 0 | 3 | 1 | 0 | 0 | 0 | 2 | 0 | 52 | 49 | NA | NA | NA | NA | 0.715 | 1 | 0 | NA | 0.215 |
| A3SS | 2364 | DGKZ | DGKZ | chr11 | + | 46379453 | 46379568 | 46379468 | 46379568 | 46379202 | 46379236 | 0 | 0 | 0 | 0 | 0 | 2 | 0 | 2 | 0 | 1 | 0 | 0 | 0 | 0 | 64 | 49 | NA | NA | NA | NA | 0 | NA | 1 | NA | -1 |
| A3SS | 2371 | ZBP1 | ZBP1 | chr20 | - | 57613162 | 57613330 | 57613162 | 57613327 | 57614886 | 57615060 | 0 | 0 | 23 | 0 | 0 | 2 | 0 | 2 | 0 | 3 | 0 | 0 | 0 | 0 | 52 | 49 | NA | NA | NA | NA | 0.916 | NA | 1 | NA | -0.084 |
| A3SS | 2404 | OS9 | OS9 | chr12 | + | 57716088 | 57716193 | 57716091 | 57716193 | 57715759 | 57715970 | 0 | 0 | 2 | 0 | 0 | 1 | 0 | 1 | 1 | 2 | 0 | 0 | 0 | 0 | 52 | 49 | NA | NA | NA | NA | 0.653 | 1 | 1 | NA | -0.347 |
| A3SS | 2490 | ESYT1 | ESYT1 | chr12 | + | 56134079 | 56134181 | 56134109 | 56134181 | 56133780 | 56133873 | 0 | 0 | 1 | 0 | 0 | 1 | 0 | 1 | 1 | 0 | 0 | 0 | 1 | 0 | 79 | 49 | NA | NA | NA | NA | 0.383 | 1 | 0 | NA | -0.117 |
| A3SS | 2492 | SON | SON | chr21 | + | 33576364 | 33577514 | 33576417 | 33577514 | 33575777 | 33575893 | 0 | 0 | 5 | 0 | 0 | 0 | 0 | 0 | 0 | 1 | 0 | 0 | 1 | 0 | 98 | 49 | NA | NA | NA | NA | 1 | NA | 0.333 | NA | 0.667 |
| A3SS | 2500 | FAM65B | FAM65B | chr6 | - | 24839365 | 24840775 | 24839365 | 24839599 | 24842861 | 24843554 | 0 | 0 | 20 | 0 | 0 | 2 | 0 | 2 | 0 | 13 | 0 | 0 | 1 | 0 | 98 | 49 | NA | NA | NA | NA | 0.833 | NA | 0.867 | NA | -0.034 |
| A3SS | 2521 | ZNF124 | ZNF124 | chr1 | - | 247154988 | 247157403 | 247154988 | 247157217 | 247159005 | 247159066 | 0 | 0 | 0 | 0 | 0 | 2 | 0 | 2 | 1 | 0 | 0 | 0 | 1 | 0 | 98 | 49 | NA | NA | NA | NA | 0 | 1 | 0 | NA | -0.5 |
| A3SS | 2552 | CELF2 | CELF2 | chr10 | + | 11321188 | 11321386 | 11321206 | 11321386 | 11314138 | 11314258 | 0 | 0 | 1 | 0 | 0 | 3 | 0 | 3 | 0 | 2 | 0 | 0 | 1 | 0 | 67 | 49 | NA | NA | NA | NA | 0.196 | NA | 0.594 | NA | -0.398 |
| A3SS | 2560 | SERPINB1 | SERPINB1 | chr6 | - | 2835855 | 2836023 | 2835855 | 2836019 | 2836107 | 2836250 | 0 | 0 | 15 | 0 | 0 | 3 | 0 | 3 | 0 | 12 | 0 | 0 | 1 | 0 | 53 | 49 | NA | NA | NA | NA | 0.822 | NA | 0.917 | NA | -0.095 |
| A3SS | 2588 | APTX | APTX | chr9 | - | 32988078 | 32988129 | 32988078 | 32988124 | 32989758 | 32989895 | 0 | 0 | 0 | 0 | 0 | 1 | 0 | 1 | 0 | 1 | 0 | 0 | 0 | 0 | 54 | 49 | NA | NA | NA | NA | 0 | NA | 1 | NA | -1 |

| | | | | | | | | | | | | | | | | | | | | | | | | | | | | | | | | | | |
|---|---|---|---|---|---|---|---|---|---|---|---|---|---|---|---|---|---|---|---|---|---|---|---|---|---|---|---|---|---|---|---|---|---|---|
| A3SS | 2594 | SLC25A45 | SLC25A45 | chr11 | - | 65380131 | 65380234 | 65380131 | 65380175 | 65380533 | 65380632 | 0 | 0 | 0 | 0 | 0 | 1 | 0 | 1 | 0 | 0 | 0 | 0 | 98 | 49 | NA | NA | NA | NA | 0 | NA | 1 | NA | -1 |
| A3SS | 2616 | RNH1 | RNH1 | chr11 | - | 502061 | 502249 | 502061 | 502181 | 504823 | 504996 | 0 | 0 | 5 | 0 | 0 | 2 | 0 | 2 | 0 | 0 | 0 | 0 | 98 | 49 | NA | NA | NA | NA | 0.556 | NA | 1 | NA | -0.444 |
| A3SS | 2632 | BRD7 | BRD7 | chr16 | - | 50320662 | 50320777 | 50320662 | 50320774 | 50321981 | 50322038 | 0 | 0 | 1 | 0 | 0 | 0 | 0 | 0 | 0 | 0 | 4 | 0 | 52 | 49 | NA | NA | NA | NA | 1 | NA | 0 | NA | 1 |
| A3SS | 2668 | TCF7L2 | TCF7L2 | chr10 | + | 113150982 | 113151123 | 113150997 | 113151123 | 113146010 | 113146097 | 1 | 0 | 0 | 0 | 0 | 0 | 0 | 0 | 0 | 0 | 1 | 0 | 64 | 49 | NA | NA | 1 | NA | NA | NA | 0 | NA | 1 |
| A3SS | 2709 | ELF2 | ELF2 | chr4 | - | 139071865 | 139072039 | 139071865 | 139072003 | 139073453 | 139073567 | 0 | 0 | 4 | 0 | 0 | 0 | 0 | 4 | 0 | 0 | 1 | 0 | 85 | 49 | NA | NA | NA | NA | 1 | NA | 0.698 | NA | 0.302 |
| A3SS | 2713 | SNRNP70 | SNRNP70 | chr19 | + | 49107794 | 49108613 | 49107821 | 49108613 | 49107624 | 49107712 | 0 | 0 | 1 | 0 | 0 | 1 | 0 | 3 | 0 | 0 | 0 | 0 | 76 | 49 | NA | NA | NA | NA | 0.392 | NA | 1 | NA | -0.608 |
| A3SS | 2725 | MEF2C | MEF2C | chr5 | - | 88718240 | 88722925 | 88718240 | 88722829 | 88728492 | 88728628 | 0 | 0 | 1 | 0 | 0 | 1 | 3 | 0 | 0 | 0 | 0 | 0 | 98 | 49 | NA | NA | NA | NA | 0.333 | 1 | NA | NA | -0.667 |
| A3SS | 2727 | AFTPH | AFTPH | chr2 | + | 64591884 | 64593004 | 64591887 | 64593004 | 64585421 | 64585545 | 0 | 0 | 1 | 0 | 0 | 0 | 0 | 1 | 0 | 0 | 1 | 0 | 52 | 49 | NA | NA | NA | NA | 1 | NA | 0.485 | NA | 0.515 |
| A3SS | 2728 | SLC43A3 | SLC43A3 | chr11 | - | 57425540 | 57425709 | 57425540 | 57425670 | 57425988 | 57426389 | 0 | 0 | 0 | 0 | 0 | 2 | 0 | 2 | 0 | 0 | 0 | 0 | 88 | 49 | NA | NA | NA | NA | 0 | NA | 1 | NA | -1 |
| A3SS | 2780 | UPP1 | UPP1 | chr7 | + | 48090187 | 48090364 | 48090249 | 48090364 | 48089211 | 48089418 | 0 | 0 | 1 | 0 | 0 | 0 | 0 | 2 | 0 | 0 | 1 | 0 | 98 | 49 | NA | NA | NA | NA | 1 | NA | 0.5 | NA | 0.5 |
| A3SS | 2861 | TPM3 | TPM3 | chr1 | - | 154170399 | 154170469 | 154170399 | 154170465 | 154170648 | 154170711 | 0 | 0 | 19 | 0 | 0 | 1 | 2 | 14 | 0 | 0 | 0 | 0 | 53 | 49 | NA | NA | NA | NA | 0.946 | 1 | 1 | NA | -0.054 |
| A3SS | 2901 | MEFV | MEFV | chr16 | - | 3243859 | 3244067 | 3243859 | 3243892 | 3244253 | 3244286 | 0 | 0 | 8 | 0 | 0 | 3 | 0 | 1 | 0 | 0 | 1 | 0 | 98 | 49 | NA | NA | NA | NA | 0.571 | NA | 0.333 | NA | 0.238 |
| A3SS | 2926 | COPS2 | COPS2 | chr15 | - | 49137347 | 49137458 | 49137347 | 49137437 | 49139527 | 49139653 | 0 | 0 | 2 | 0 | 0 | 2 | 0 | 1 | 0 | 0 | 0 | 0 | 70 | 49 | NA | NA | NA | NA | 0.412 | NA | 1 | NA | -0.588 |
| A3SS | 2936 | DMXL2 | DMXL2 | chr15 | - | 51476588 | 51476719 | 51476588 | 51476716 | 51478270 | 51478347 | 0 | 0 | 8 | 0 | 0 | 2 | 0 | 2 | 0 | 0 | 0 | 0 | 52 | 49 | NA | NA | NA | NA | 0.79 | NA | 1 | NA | -0.21 |
| A3SS | 2938 | PIK3R5 | PIK3R5 | chr17 | - | 8888170 | 8888953 | 8888170 | 8888891 | 8889138 | 8889222 | 0 | 0 | 2 | 0 | 0 | 2 | 0 | 1 | 0 | 0 | 1 | 0 | 98 | 49 | NA | NA | NA | NA | 0.333 | NA | 0.333 | NA | 0 |
| A3SS | 2952 | ZNF143 | ZNF143 | chr11 | + | 9474549 | 9474633 | 9474552 | 9474633 | 9473940 | 9474024 | 0 | 0 | 1 | 0 | 0 | 3 | 0 | 4 | 0 | 0 | 1 | 0 | 52 | 49 | NA | NA | NA | NA | 0.239 | NA | 0.79 | NA | -0.551 |
| A3SS | 2964 | PTMA | PTMA | chr2 | + | 231711886 | 231711983 | 231711889 | 231711983 | 231711347 | 231711419 | 0 | 0 | 1 | 0 | 0 | 2 | 0 | 4 | 0 | 0 | 1 | 0 | 52 | 49 | NA | NA | NA | NA | 0.32 | NA | 0.79 | NA | -0.47 |
| A3SS | 2985 | ST20 | ST20 | chr15 | - | 79907628 | 79907706 | 79907628 | 79907684 | 79914587 | 79915256 | 0 | 0 | 1 | 0 | 0 | 5 | 1 | 0 | 0 | 0 | 0 | 0 | 71 | 49 | NA | NA | NA | NA | 0.121 | 1 | NA | NA | -0.879 |
| A3SS | 2988 | IL16 | IL16 | chr15 | + | 81306419 | 81306545 | 81306422 | 81306545 | 81305907 | 81306166 | 0 | 0 | 1 | 0 | 0 | 1 | 0 | 0 | 0 | 0 | 4 | 0 | 52 | 49 | NA | NA | NA | NA | 0.485 | NA | 0 | NA | 0.485 |
| A3SS | 3007 | AKAP13 | AKAP13 | chr15 | + | 85710578 | 85710645 | 85710581 | 85710645 | 85708018 | 85708086 | 0 | 0 | 13 | 0 | 0 | 0 | 0 | 10 | 0 | 0 | 2 | 0 | 52 | 49 | NA | NA | NA | NA | 1 | NA | 0.825 | NA | 0.175 |
| A3SS | 3057 | ZNF200 | ZNF200 | chr16 | - | 3222324 | 3224613 | 3222324 | 3224610 | 3232420 | 3232547 | 0 | 0 | 1 | 0 | 0 | 0 | 0 | 0 | 0 | 0 | 1 | 0 | 52 | 49 | NA | NA | NA | NA | 1 | NA | 0 | NA | 1 |
| A3SS | 3080 | CLEC16A | CLEC16A | chr16 | + | 10972931 | 10973061 | 10972937 | 10973061 | 10971124 | 10971230 | 0 | 0 | 2 | 0 | 0 | 0 | 0 | 0 | 0 | 0 | 2 | 0 | 55 | 49 | NA | NA | NA | NA | 1 | NA | 0 | NA | 1 |
| A3SS | 3089 | KIAA0430 | KIAA0430 | chr16 | - | 15622933 | 15623126 | 15622933 | 15623123 | 15624768 | 15624927 | 0 | 0 | 1 | 0 | 0 | 0 | 0 | 0 | 0 | 0 | 2 | 0 | 52 | 49 | NA | NA | NA | NA | 1 | NA | 0 | NA | 1 |
| A3SS | 3135 | FUS | FUS | chr16 | + | 31183857 | 31184002 | 31183860 | 31184002 | 31182512 | 31182664 | 0 | 0 | 5 | 0 | 0 | 3 | 0 | 3 | 0 | 1 | 3 | 0 | 52 | 49 | NA | NA | NA | NA | 0.611 | 0 | 0.485 | NA | 0.368 |
| A3SS | 3251 | LRMP | LRMP | chr12 | + | 25063719 | 25063816 | 25063722 | 25063816 | 25062819 | 25062900 | 0 | 0 | 3 | 0 | 0 | 2 | 0 | 5 | 0 | 0 | 1 | 0 | 52 | 49 | NA | NA | NA | NA | 0.586 | NA | 0.825 | NA | -0.239 |
| A3SS | 3275 | WDTC1 | WDTC1 | chr1 | + | 27297047 | 27297156 | 27297050 | 27297156 | 27296325 | 27296401 | 0 | 0 | 4 | 0 | 0 | 1 | 0 | 1 | 0 | 0 | 0 | 0 | 52 | 49 | NA | NA | NA | NA | 0.79 | NA | 1 | NA | -0.21 |
| A3SS | 3499 | TIAL1 | TIAL1 | chr10 | - | 119582458 | 119582608 | 119582458 | 119582557 | 119588151 | 119588248 | 0 | 0 | 0 | 0 | 0 | 1 | 3 | 0 | 0 | 0 | 0 | 0 | 98 | 49 | NA | NA | NA | NA | 0 | 1 | NA | NA | -1 |
| A3SS | 3508 | EXOC7 | EXOC7 | chr17 | - | 76090328 | 76090481 | 76090328 | 76090397 | 76091142 | 76091235 | 1 | 0 | 0 | 0 | 0 | 0 | 0 | 0 | 0 | 0 | 1 | 0 | 98 | 49 | NA | NA | 1 | NA | NA | NA | 0 | NA | 1 |
| A3SS | 3511 | RHBDF2 | RHBDF2 | chr17 | - | 76479732 | 76479941 | 76479732 | 76479854 | 76481374 | 76481545 | 0 | 0 | 1 | 0 | 0 | 5 | 4 | 0 | 0 | 0 | 2 | 0 | 98 | 49 | NA | NA | NA | NA | 0.091 | 1 | 0 | NA | -0.409 |
| A3SS | 3686 | GTPBP3 | GTPBP3 | chr19 | + | 17341043 | 17341322 | 17341106 | 17341322 | 17339433 | 17339599 | 0 | 0 | 0 | 0 | 2 | 0 | 3 | 0 | 0 | 0 | 0 | 0 | 98 | 49 | NA | NA | NA | 0 | NA | 1 | NA | NA | -1 |
| A3SS | 3695 | TMEM59 | TMEM59 | chr1 | - | 54040755 | 54040840 | 54040755 | 54040837 | 54041723 | 54041805 | 0 | 0 | 0 | 0 | 0 | 2 | 0 | 1 | 0 | 0 | 1 | 0 | 52 | 49 | NA | NA | NA | NA | 0 | NA | 0.485 | NA | -0.485 |
| A3SS | 3701 | UXS1 | UXS1 | chr2 | - | 106164735 | 106164799 | 106164735 | 106164784 | 106166055 | 106166083 | 0 | 0 | 1 | 0 | 0 | 0 | 0 | 0 | 0 | 0 | 4 | 0 | 64 | 49 | NA | NA | NA | NA | 1 | NA | 0 | NA | 1 |
| A3SS | 3723 | PITPNM1 | PITPNM1 | chr11 | - | 67496177 | 67496348 | 67496177 | 67496345 | 67497230 | 67497436 | 0 | 0 | 2 | 0 | 0 | 0 | 0 | 0 | 0 | 0 | 1 | 0 | 52 | 49 | NA | NA | NA | NA | 1 | NA | 0 | NA | 1 |
| A3SS | 3744 | TMEM91 | TMEM91 | chr19 | + | 41383714 | 41384082 | 41383862 | 41384082 | 41382771 | 41382921 | 0 | 0 | 2 | 0 | 0 | 3 | 0 | 1 | 0 | 0 | 2 | 0 | 98 | 49 | NA | NA | NA | NA | 0.25 | NA | 0.2 | NA | 0.05 |
| A3SS | 3789 | PTOV1 | PTOV1 | chr19 | + | 49860565 | 49860744 | 49860613 | 49860744 | 49860267 | 49860336 | 1 | 0 | 3 | 0 | 0 | 3 | 0 | 2 | 0 | 0 | 0 | 0 | 97 | 49 | NA | NA | 1 | NA | 0.336 | NA | 1 | NA | -0.332 |
| A3SS | 3809 | NFRKB | NFRKB | chr11 | - | 129884744 | 129884882 | 129884744 | 129884846 | 129885434 | 129885609 | 0 | 0 | 1 | 0 | 0 | 0 | 0 | 2 | 0 | 4 | 1 | 0 | 85 | 49 | NA | NA | NA | NA | 1 | 0 | 0.536 | NA | 0.732 |
| A3SS | 3838 | LILRB2 | LILRB2 | chr19 | - | 54276806 | 54276979 | 54276806 | 54276929 | 54277549 | 54277597 | 0 | 0 | 1 | 0 | 0 | 3 | 0 | 0 | 0 | 7 | 1 | 0 | 98 | 49 | NA | NA | NA | NA | 0.143 | 0 | 0 | NA | 0.143 |
| A3SS | 3922 | ODC1 | ODC1 | chr2 | - | 10444473 | 10444647 | 10444473 | 10444547 | 10444930 | 10445049 | 0 | 0 | 2 | 0 | 0 | 1 | 3 | 1 | 0 | 0 | 0 | 0 | 98 | 49 | NA | NA | NA | NA | 0.5 | 1 | 1 | NA | -0.5 |
| A3SS | 3939 | SUPT7L | SUPT7L | chr2 | - | 27660983 | 27661443 | 27660983 | 27661388 | 27662178 | 27662281 | 0 | 0 | 0 | 0 | 0 | 1 | 0 | 1 | 0 | 0 | 0 | 0 | 98 | 49 | NA | NA | NA | NA | 0 | NA | 1 | NA | -1 |
| A3SS | 3940 | SUPT7L | SUPT7L | chr2 | - | 27660983 | 27661601 | 27660983 | 27661388 | 27662178 | 27662281 | 0 | 0 | 0 | 0 | 0 | 1 | 0 | 1 | 0 | 0 | 0 | 0 | 98 | 49 | NA | NA | NA | NA | 0 | NA | 1 | NA | -1 |
| A3SS | 3952 | PPM1B | PPM1B | chr2 | + | 44229992 | 44231473 | 44230412 | 44231473 | 44218479 | 44218537 | 0 | 0 | 3 | 0 | 0 | 2 | 0 | 1 | 0 | 0 | 1 | 0 | 98 | 49 | NA | NA | NA | NA | 0.429 | NA | 0.333 | NA | 0.096 |
| A3SS | 3977 | AUP1 | AUP1 | chr2 | - | 74526940 | 74527121 | 74526940 | 74527059 | 74527247 | 74527363 | 0 | 0 | 0 | 0 | 0 | 3 | 1 | 1 | 0 | 8 | 8 | 0 | 98 | 49 | NA | NA | NA | NA | 0 | 0.059 | 0.059 | NA | -0.059 |
| A3SS | 3997 | SERPINA1 | SERPINA1 | chr14 | - | 94388559 | 94388663 | 94388559 | 94388660 | 94388800 | 94389010 | 0 | 0 | 3 | 0 | 0 | 2 | 0 | 0 | 0 | 0 | 1 | 0 | 52 | 49 | NA | NA | NA | NA | 0.586 | NA | 0 | NA | 0.586 |
| A3SS | 3998 | SERPINA1 | SERPINA1 | chr14 | - | 94388559 | 94388663 | 94388559 | 94388660 | 94388818 | 94389010 | 0 | 0 | 3 | 0 | 0 | 1 | 0 | 1 | 0 | 0 | 0 | 0 | 52 | 49 | NA | NA | NA | NA | 0.739 | NA | 1 | NA | -0.261 |
| A5SS | 6 | NAA10 | NAA10 | chrX | - | 153933942 | 153934001 | 153933960 | 153934001 | 153932538 | 153932584 | 0 | 0 | 1 | 0 | 0 | 0 | 0 | 2 | 0 | 0 | 1 | 0 | 67 | 49 | NA | NA | NA | NA | 1 | NA | 0.594 | NA | 0.406 |
| A5SS | 12 | AFF2 | AFF2 | chrX | + | 148651998 | 148652131 | 148651998 | 148652119 | 148661907 | 148662768 | 0 | 0 | 0 | 0 | 0 | 1 | 2 | 0 | 0 | 0 | 1 | 0 | 61 | 49 | NA | NA | NA | NA | 0 | 1 | 0 | NA | -0.5 |
| A5SS | 57 | OTUD5 | OTUD5 | chrX | - | 48934695 | 48934867 | 48934710 | 48934867 | 48934463 | 48934612 | 0 | 0 | 3 | 0 | 0 | 2 | 0 | 0 | 0 | 0 | 9 | 0 | 64 | 49 | NA | NA | NA | NA | 0.535 | NA | 0 | NA | 0.535 |
| A5SS | 149 | ENY2 | ENY2 | chr8 | + | 109334322 | 109334474 | 109334322 | 109334402 | 109336127 | 109336204 | 0 | 0 | 1 | 0 | 0 | 2 | 0 | 1 | 0 | 0 | 0 | 0 | 98 | 49 | NA | NA | NA | NA | 0.2 | NA | 1 | NA | -0.8 |
| A5SS | 154 | LYN | LYN | chr8 | + | 55941854 | 55941991 | 55941854 | 55941928 | 55946447 | 55946493 | 0 | 0 | 16 | 0 | 0 | 3 | 0 | 4 | 0 | 0 | 6 | 0 | 98 | 49 | NA | NA | NA | NA | 0.727 | NA | 0.25 | NA | 0.477 |
| A5SS | 167 | ADAM28 | ADAM28 | chr8 | + | 24343505 | 24343588 | 24343505 | 24343584 | 24349863 | 24349972 | 0 | 0 | 0 | 0 | 0 | 1 | 1 | 0 | 0 | 0 | 0 | 0 | 53 | 49 | NA | NA | NA | NA | 0 | 1 | NA | NA | -1 |
| A5SS | 168 | ENTPD4 | ENTPD4 | chr8 | - | 23441568 | 23441723 | 23441592 | 23441723 | 23439748 | 23439915 | 0 | 0 | 2 | 0 | 0 | 0 | 0 | 0 | 0 | 0 | 2 | 0 | 73 | 49 | NA | NA | NA | NA | 1 | NA | 0 | NA | 1 |
| A5SS | 223 | IKZF1 | IKZF1 | chr7 | + | 50391728 | 50391863 | 50391728 | 50391833 | 50399917 | 50405100 | 0 | 0 | 2 | 0 | 0 | 3 | 3 | 0 | 0 | 0 | 0 | 0 | 79 | 49 | NA | NA | NA | NA | 0.293 | 1 | NA | NA | -0.707 |
| A5SS | 230 | CPVL | CPVL | chr7 | - | 29146340 | 29146537 | 29146428 | 29146537 | 29120892 | 29121071 | 1 | 0 | 0 | 1 | 0 | 0 | 0 | 0 | 0 | 0 | 2 | 0 | 98 | 49 | NA | NA | 0.333 | NA | NA | NA | 0 | NA | 0.333 |
| A5SS | 232 | SNX10 | SNX10 | chr7 | + | 26291894 | 26292133 | 26291894 | 26292086 | 26346419 | 26346466 | 0 | 0 | 0 | 0 | 0 | 2 | 3 | 0 | 0 | 0 | 0 | 0 | 96 | 49 | NA | NA | NA | NA | 0 | 1 | NA | NA | -1 |
| A5SS | 275 | CUTA | CUTA | chr6 | - | 33417868 | 33418288 | 33417958 | 33418288 | 33417480 | 33417695 | 0 | 0 | 0 | 0 | 0 | 1 | 4 | 0 | 0 | 0 | 0 | 0 | 98 | 49 | NA | NA | NA | NA | 0 | 1 | NA | NA | -1 |
| A5SS | 383 | USP9X | USP9X | chrX | + | 41229566 | 41229827 | 41229566 | 41229779 | 41230500 | 41230596 | 0 | 0 | 0 | 0 | 0 | 1 | 0 | 5 | 0 | 0 | 4 | 0 | 97 | 49 | NA | NA | NA | NA | 0 | NA | 0.387 | NA | -0.387 |
| A5SS | 400 | CANX | CANX | chr5 | + | 179698928 | 179699102 | 179698928 | 179699049 | 179705678 | 179705852 | 0 | 0 | 0 | 0 | 0 | 1 | 0 | 1 | 0 | 0 | 0 | 0 | 98 | 49 | NA | NA | NA | NA | 0 | NA | 1 | NA | -1 |
| A5SS | 469 | TMEM70 | TMEM70 | chr8 | + | 73978755 | 73978875 | 73978755 | 73978861 | 73981154 | 73982783 | 0 | 0 | 0 | 0 | 0 | 1 | 3 | 0 | 0 | 0 | 0 | 0 | 63 | 49 | NA | NA | NA | NA | 0 | 1 | NA | NA | -1 |
| A5SS | 484 | LIN54 | LIN54 | chr4 | - | 82984160 | 82984876 | 82984823 | 82984876 | 82978882 | 82979006 | 1 | 0 | 0 | 0 | 0 | 0 | 0 | 0 | 0 | 0 | 2 | 0 | 98 | 49 | NA | NA | 1 | NA | NA | NA | 0 | NA | 1 |
| A5SS | 496 | KIT | KIT | chr4 | + | 54725856 | 54726050 | 54725856 | 54726038 | 54727217 | 54727324 | 0 | 0 | 0 | 0 | 0 | 1 | 1 | 0 | 0 | 0 | 0 | 0 | 61 | 49 | NA | NA | NA | NA | 0 | 1 | NA | NA | -1 |
| A5SS | 507 | CPEB2 | CPEB2 | chr4 | + | 15007304 | 15007595 | 15007304 | 15007586 | 15017187 | 15017278 | 0 | 0 | 1 | 0 | 0 | 2 | 0 | 0 | 0 | 0 | 3 | 0 | 58 | 49 | NA | NA | NA | NA | 0.297 | NA | 0 | NA | 0.297 |
| A5SS | 511 | TMEM175 | TMEM175 | chr4 | + | 948115 | 948615 | 948115 | 948154 | 950420 | 950518 | 0 | 0 | 0 | 0 | 0 | 1 | 0 | 1 | 0 | 0 | 0 | 0 | 98 | 49 | NA | NA | NA | NA | 0 | NA | 1 | NA | -1 |
| A5SS | 546 | MBD4 | MBD4 | chr3 | - | 129436442 | 129437308 | 129436460 | 129437308 | 129434061 | 129434136 | 0 | 0 | 0 | 0 | 0 | 1 | 0 | 1 | 0 | 0 | 0 | 0 | 67 | 49 | NA | NA | NA | NA | 0 | NA | 1 | NA | -1 |
| A5SS | 549 | CNBP | CNBP | chr3 | - | 129171633 | 129171771 | 129171654 | 129171771 | 129171445 | 129171538 | 0 | 0 | 4 | 0 | 0 | 5 | 0 | 2 | 0 | 0 | 4 | 0 | 70 | 49 | NA | NA | NA | NA | 0.359 | NA | 0.259 | NA | 0.1 |
| A5SS | 556 | PARP9 | PARP9 | chr3 | - | 122558328 | 122558467 | 122558433 | 122558467 | 122555285 | 122556121 | 0 | 0 | 10 | 0 | 0 | 6 | 1 | 3 | 0 | 0 | 1 | 0 | 98 | 49 | NA | NA | NA | NA | 0.455 | 1 | 0.6 | NA | -0.345 |
| A5SS | 558 | GOLGB1 | GOLGB1 | chr3 | - | 121730847 | 121730973 | 121730875 | 121730973 | 121729864 | 121730017 | 0 | 0 | 2 | 0 | 0 | 0 | 0 | 1 | 0 | 0 | 1 | 0 | 77 | 49 | NA | NA | NA | NA | 1 | NA | 0.389 | NA | 0.611 |
| A5SS | 572 | CCDC66 | CCDC66 | chr3 | + | 56566593 | 56566759 | 56566593 | 56566740 | 56566949 | 56567053 | 0 | 0 | 1 | 0 | 0 | 0 | 0 | 0 | 0 | 6 | 0 | 0 | 68 | 49 | NA | NA | NA | NA | 1 | 0 | NA | NA | 1 |
| A5SS | 603 | FGD5-AS1 | FGD5-AS1 | chr3 | - | 14947086 | 14947505 | 14947109 | 14947505 | 14942778 | 14946154 | 0 | 0 | 1 | 0 | 0 | 0 | 0 | 0 | 0 | 0 | 3 | 0 | 72 | 49 | NA | NA | NA | NA | 1 | NA | 0 | NA | 1 |
| A5SS | 607 | BRPF1 | BRPF1 | chr3 | + | 9742024 | 9742171 | 9742024 | 9742153 | 9742943 | 9743253 | 0 | 0 | 1 | 0 | 0 | 2 | 0 | 3 | 0 | 0 | 0 | 0 | 67 | 49 | NA | NA | NA | NA | 0.268 | NA | 1 | NA | -0.732 |
| A5SS | 635 | TANGO2 | TANGO2 | chr22 | + | 20036759 | 20037094 | 20036759 | 20036854 | 20043354 | 20043443 | 1 | 0 | 2 | 0 | 0 | 7 | 0 | 3 | 0 | 0 | 0 | 0 | 98 | 49 | NA | NA | 1 | NA | 0.125 | NA | 1 | NA | -0.438 |
| A5SS | 684 | NUB1 | NUB1 | chr7 | + | 151374096 | 151374243 | 151374096 | 151374201 | 151375847 | 151375943 | 2 | 0 | 3 | 0 | 0 | 1 | 0 | 1 | 0 | 0 | 2 | 0 | 91 | 49 | NA | NA | 1 | NA | 0.618 | NA | 0.212 | NA | 0.597 |
| A5SS | 702 | C20orf24 | C20orf24 | chr20 | + | 36609600 | 36609758 | 36609600 | 36609730 | 36612009 | 36612557 | 0 | 0 | 1 | 0 | 0 | 4 | 0 | 1 | 0 | 0 | 6 | 0 | 77 | 49 | NA | NA | NA | NA | 0.137 | NA | 0.096 | NA | 0.041 |
| A5SS | 716 | NPEPL1 | NPEPL1 | chr20 | + | 58691723 | 58691830 | 58691723 | 58691780 | 58693736 | 58693922 | 0 | 0 | 5 | 0 | 0 | 1 | 3 | 5 | 0 | 0 | 0 | 0 | 98 | 49 | NA | NA | NA | NA | 0.714 | 1 | 1 | NA | -0.286 |
| A5SS | 738 | DYRK1A | DYRK1A | chr21 | + | 37506098 | 37506354 | 37506098 | 37506223 | 37511910 | 37515376 | 0 | 0 | 3 | 0 | 0 | 0 | 0 | 3 | 0 | 0 | 1 | 0 | 98 | 49 | NA | NA | NA | NA | 1 | NA | 0.6 | NA | 0.4 |
| A5SS | 776 | TMEM55B | TMEM55B | chr14 | - | 20461162 | 20461612 | 20461183 | 20461612 | 20460654 | 20460845 | 0 | 0 | 2 | 0 | 0 | 0 | 0 | 1 | 0 | 0 | 1 | 0 | 70 | 49 | NA | NA | NA | NA | 1 | NA | 0.412 | NA | 0.588 |
| A5SS | 787 | SUPT4H1 | SUPT4H1 | chr17 | - | 58352064 | 58352238 | 58352066 | 58352238 | 58351401 | 58351508 | 0 | 0 | 5 | 0 | 0 | 0 | 0 | 0 | 0 | 0 | 1 | 0 | 51 | 49 | NA | NA | NA | NA | 1 | NA | 0 | NA | 1 |
| A5SS | 790 | CNOT1 | CNOT1 | chr16 | - | 58556846 | 58556993 | 58556861 | 58556993 | 58555783 | 58555908 | 0 | 0 | 1 | 0 | 0 | 1 | 0 | 2 | 0 | 0 | 1 | 0 | 64 | 49 | NA | NA | NA | NA | 0.434 | NA | 0.605 | NA | -0.171 |
| A5SS | 823 | UFM1 | UFM1 | chr13 | + | 38349998 | 38350227 | 38349998 | 38350055 | 38354238 | 38354296 | 0 | 0 | 0 | 0 | 0 | 2 | 1 | 0 | 0 | 0 | 0 | 0 | 98 | 49 | NA | NA | NA | NA | 0 | 1 | NA | NA | -1 |
| A5SS | 852 | HUS1 | HUS1 | chr7 | - | 47979409 | 47979625 | 47979467 | 47979625 | 47978688 | 47978816 | 0 | 0 | 0 | 0 | 0 | 1 | 2 | 1 | 0 | 0 | 1 | 0 | 98 | 49 | NA | NA | NA | NA | 0 | 1 | 0.333 | NA | -0.666 |
| A5SS | 861 | CHST11 | CHST11 | chr12 | + | 104456913 | 104457529 | 104456913 | 104457514 | 104601905 | 104601991 | 0 | 0 | 7 | 0 | 0 | 2 | 0 | 1 | 0 | 0 | 1 | 0 | 64 | 49 | NA | NA | NA | NA | 0.728 | NA | 0.434 | NA | 0.294 |
| A5SS | 990 | SNX19 | SNX19 | chr11 | - | 130914265 | 130916487 | 130916279 | 130916487 | 130911632 | 130911771 | 0 | 0 | 3 | 0 | 0 | 1 | 1 | 0 | 0 | 0 | 0 | 0 | 98 | 49 | NA | NA | NA | NA | 0.6 | 1 | NA | NA | -0.4 |
| A5SS | 1033 | SH3YL1 | SH3YL1 | chr2 | - | 224863 | 224920 | 224867 | 224920 | 218135 | 219001 | 0 | 0 | 3 | 0 | 0 | 0 | 0 | 0 | 0 | 0 | 1 | 0 | 53 | 49 | NA | NA | NA | NA | 1 | NA | 0 | NA | 1 |
| A5SS | 1118 | IL32 | IL32 | chr16 | + | 3067376 | 3067613 | 3067376 | 3067415 | 3067983 | 3068010 | 0 | 0 | 1 | 0 | 0 | 0 | 1 | 2 | 0 | 1 | 0 | 0 | 98 | 49 | NA | NA | NA | NA | 1 | 0.333 | 1 | NA | 0.334 |
| A5SS | 1191 | PKM | PKM | chr15 | - | 72230899 | 72231386 | 72231115 | 72231386 | 72218943 | 72219110 | 0 | 0 | 0 | 0 | 0 | 1 | 1 | 0 | 0 | 0 | 0 | 0 | 98 | 49 | NA | NA | NA | NA | 0 | 1 | NA | NA | -1 |
| A5SS | 1195 | NUP98 | NUP98 | chr11 | - | 3760487 | 3760626 | 3760538 | 3760626 | 3753315 | 3753408 | 0 | 0 | 4 | 0 | 0 | 2 | 0 | 13 | 0 | 0 | 5 | 0 | 98 | 49 | NA | NA | NA | NA | 0.5 | NA | 0.565 | NA | -0.065 |
| A5SS | 1217 | MAPKAPK5 | MAPKAPK5 | chr12 | + | 111888884 | 111889006 | 111888884 | 111889000 | 111890039 | 111890144 | 0 | 0 | 2 | 0 | 0 | 2 | 0 | 0 | 0 | 0 | 1 | 0 | 55 | 49 | NA | NA | NA | NA | 0.471 | NA | 0 | NA | 0.471 |
| A5SS | 1500 | ZNF518A | ZNF518A | chr10 | + | 96155325 | 96155398 | 96155325 | 96155376 | 96155923 | 96156047 | 0 | 0 | 0 | 0 | 0 | 3 | 0 | 1 | 0 | 0 | 1 | 0 | 71 | 49 | NA | NA | NA | NA | 0 | NA | 0.408 | NA | -0.408 |
| A5SS | 1514 | AGTRAP | AGTRAP | chr1 | + | 11748414 | 11748610 | 11748414 | 11748589 | 11750076 | 11750771 | 0 | 0 | 4 | 0 | 0 | 1 | 1 | 0 | 0 | 0 | 0 | 0 | 70 | 49 | NA | NA | NA | NA | 0.737 | 1 | NA | NA | -0.263 |
| A5SS | 1529 | TMCO1 | TMCO1 | chr1 | - | 165768481 | 165768922 | 165768681 | 165768922 | 165768191 | 165768269 | 0 | 0 | 0 | 0 | 1 | 2 | 2 | 0 | 0 | 0 | 0 | 0 | 98 | 49 | NA | NA | NA | 0 | 0 | 1 | NA | NA | -1 |
| A5SS | 1554 | WNK1 | WNK1 | chr12 | + | 890452 | 890513 | 890452 | 890510 | 894561 | 894635 | 0 | 0 | 1 | 0 | 0 | 0 | 0 | 2 | 0 | 0 | 1 | 0 | 52 | 49 | NA | NA | NA | NA | 1 | NA | 0.653 | NA | 0.347 |
| A5SS | 1565 | POLD4 | POLD4 | chr11 | - | 67353293 | 67353596 | 67353302 | 67353596 | 67352987 | 67353077 | 0 | 0 | 0 | 0 | 0 | 1 | 1 | 0 | 0 | 0 | 0 | 0 | 58 | 49 | NA | NA | NA | NA | 0 | 1 | NA | NA | -1 |
| A5SS | 1702 | EIF4B | EIF4B | chr12 | + | 53034611 | 53034724 | 53034611 | 53034709 | 53037408 | 53037622 | 0 | 0 | 0 | 0 | 0 | 3 | 0 | 1 | 0 | 3 | 2 | 0 | 64 | 49 | NA | NA | NA | NA | 0 | 0 | 0.277 | NA | -0.139 |
| A5SS | 1707 | KHDRBS1 | KHDRBS1 | chr1 | + | 32037834 | 32038119 | 32037834 | 32038036 | 32038551 | 32038619 | 0 | 0 | 0 | 0 | 0 | 2 | 0 | 1 | 0 | 0 | 2 | 0 | 98 | 49 | NA | NA | NA | NA | 0 | NA | 0.2 | NA | -0.2 |
| A5SS | 1731 | SSR3 | SSR3 | chr3 | - | 156548865 | 156549003 | 156548904 | 156549003 | 156544307 | 156544439 | 0 | 0 | 0 | 0 | 0 | 1 | 8 | 0 | 0 | 0 | 0 | 0 | 88 | 49 | NA | NA | NA | NA | 0 | 1 | NA | NA | -1 |
| A5SS | 1737 | SUMO3 | SUMO3 | chr21 | - | 44813861 | 44814104 | 44813975 | 44814104 | 44809046 | 44809118 | 0 | 0 | 0 | 0 | 0 | 1 | 1 | 0 | 0 | 0 | 1 | 0 | 98 | 49 | NA | NA | NA | NA | 0 | 1 | 0 | NA | -0.5 |
| A5SS | 1792 | MAFF | MAFF | chr22 | + | 38201931 | 38202238 | 38201931 | 38202212 | 38213822 | 38213889 | 0 | 0 | 0 | 0 | 0 | 2 | 0 | 1 | 0 | 0 | 3 | 0 | 75 | 49 | NA | NA | NA | NA | 0 | NA | 0.179 | NA | -0.179 |
| A5SS | 1889 | CHMP1A | CHMP1A | chr16 | - | 89647034 | 89647331 | 89647202 | 89647331 | 89646526 | 89646714 | 0 | 0 | 1 | 0 | 0 | 0 | 0 | 1 | 0 | 0 | 2 | 0 | 98 | 49 | NA | NA | NA | NA | 1 | NA | 0.2 | NA | 0.8 |
| A5SS | 1924 | CELF2 | CELF2 | chr10 | + | 11314138 | 11314270 | 11314138 | 11314258 | 11321206 | 11321386 | 0 | 0 | 1 | 0 | 0 | 3 | 0 | 2 | 0 | 0 | 1 | 0 | 61 | 49 | NA | NA | NA | NA | 0.211 | NA | 0.616 | NA | -0.405 |
| A5SS | 1991 | CAMK2G | CAMK2G | chr10 | - | 73825278 | 73825347 | 73825320 | 73825347 | 73824039 | 73824084 | 0 | 0 | 2 | 0 | 0 | 1 | 8 | 2 | 0 | 0 | 0 | 0 | 91 | 49 | NA | NA | NA | NA | 0.519 | 1 | 1 | NA | -0.481 |
| A5SS | 2007 | CPEB3 | CPEB3 | chr10 | - | 92192476 | 92192636 | 92192545 | 92192636 | 92180962 | 92181019 | 0 | 0 | 0 | 0 | 0 | 1 | 0 | 1 | 0 | 0 | 0 | 0 | 98 | 49 | NA | NA | NA | NA | 0 | NA | 1 | NA | -1 |
| A5SS | 2040 | TRAPPC5 | TRAPPC5 | chr19 | + | 7680820 | 7681049 | 7680820 | 7681026 | 7682241 | 7682862 | 1 | 0 | 0 | 0 | 0 | 0 | 0 | 0 | 0 | 0 | 1 | 0 | 72 | 49 | NA | NA | 1 | NA | NA | NA | 0 | NA | 1 |
| A5SS | 2052 | SLC12A9 | SLC12A9 | chr7 | + | 100852718 | 100852839 | 100852718 | 100852835 | 100854155 | 100854378 | 0 | 0 | 0 | 0 | 0 | 2 | 0 | 2 | 0 | 0 | 0 | 0 | 53 | 49 | NA | NA | NA | NA | 0 | NA | 1 | NA | -1 |
| A5SS | 2078 | CASP8 | CASP8 | chr2 | + | 201276826 | 201277104 | 201276826 | 201276968 | 201284815 | 201285317 | 0 | 0 | 9 | 0 | 0 | 3 | 0 | 9 | 0 | 1 | 1 | 0 | 98 | 49 | NA | NA | NA | NA | 0.6 | 0 | 0.818 | NA | 0.191 |
| A5SS | 2092 | ARID4A | ARID4A | chr14 | + | 58366023 | 58366230 | 58366023 | 58366068 | 58366882 | 58367029 | 1 | 0 | 4 | 0 | 0 | 0 | 0 | 1 | 0 | 0 | 1 | 0 | 98 | 49 | NA | NA | 1 | NA | 1 | NA | 0.333 | NA | 0.667 |
| A5SS | 2122 | ZC3H14 | ZC3H14 | chr14 | + | 88602827 | 88603060 | 88602827 | 88603045 | 88607242 | 88607363 | 0 | 0 | 2 | 0 | 0 | 0 | 0 | 0 | 0 | 0 | 1 | 0 | 64 | 49 | NA | NA | NA | NA | 1 | NA | 0 | NA | 1 |
| A5SS | 2124 | C14orf159 | C14orf159 | chr14 | + | 91114012 | 91114528 | 91114012 | 91114283 | 91145220 | 91145313 | 0 | 0 | 0 | 0 | 0 | 1 | 0 | 1 | 0 | 0 | 0 | 0 | 98 | 49 | NA | NA | NA | NA | 0 | NA | 1 | NA | -1 |
| A5SS | 2125 | C14orf159 | C14orf159 | chr14 | + | 91114012 | 91114528 | 91114012 | 91114283 | 91157638 | 91157690 | 0 | 0 | 0 | 0 | 0 | 1 | 0 | 1 | 0 | 0 | 0 | 0 | 98 | 49 | NA | NA | NA | NA | 0 | NA | 1 | NA | -1 |
| A5SS | 2127 | C14orf159 | C14orf159 | chr14 | + | 91173288 | 91173439 | 91173288 | 91173403 | 91175933 | 91176056 | 0 | 0 | 12 | 0 | 0 | 2 | 0 | 3 | 0 | 0 | 1 | 0 | 85 | 49 | NA | NA | NA | NA | 0.776 | NA | 0.634 | NA | 0.142 |
| A5SS | 2128 | PPP4R3A | PPP4R3A | chr14 | - | 91475810 | 91475966 | 91475849 | 91475966 | 91473238 | 91473370 | 0 | 0 | 3 | 0 | 0 | 2 | 0 | 2 | 0 | 0 | 0 | 0 | 88 | 49 | NA | NA | NA | NA | 0.455 | NA | 1 | NA | -0.545 |
| A5SS | 2165 | PACS2 | PACS2 | chr14 | + | 105381913 | 105382058 | 105381913 | 105382046 | 105382476 | 105382581 | 0 | 0 | 0 | 0 | 0 | 2 | 0 | 1 | 0 | 0 | 0 | 0 | 61 | 49 | NA | NA | NA | NA | 0 | NA | 1 | NA | -1 |
| A5SS | 2199 | UQCR10 | UQCR10 | chr22 | + | 29767368 | 29767607 | 29767368 | 29767548 | 29769677 | 29770413 | 0 | 0 | 1 | 0 | 0 | 3 | 0 | 1 | 0 | 0 | 1 | 0 | 98 | 49 | NA | NA | NA | NA | 0.143 | NA | 0.333 | NA | -0.19 |
| A5SS | 2205 | PLCB2 | PLCB2 | chr15 | - | 40296745 | 40296908 | 40296757 | 40296908 | 40296521 | 40296634 | 0 | 0 | 7 | 0 | 0 | 3 | 0 | 2 | 0 | 0 | 0 | 0 | 61 | 49 | NA | NA | NA | NA | 0.652 | NA | 1 | NA | -0.348 |
| A5SS | 2260 | TLE3 | TLE3 | chr15 | - | 70058149 | 70058291 | 70058185 | 70058291 | 70057458 | 70057658 | 1 | 0 | 2 | 0 | 0 | 1 | 0 | 3 | 0 | 0 | 1 | 0 | 85 | 49 | NA | NA | 1 | NA | 0.536 | NA | 0.634 | NA | 0.134 |
| A5SS | 2273 | PML | PML | chr15 | + | 74033155 | 74033422 | 74033155 | 74033414 | 74034477 | 74034530 | 0 | 0 | 1 | 0 | 0 | 2 | 1 | 0 | 0 | 0 | 0 | 0 | 57 | 49 | NA | NA | NA | NA | 0.301 | 1 | NA | NA | -0.699 |
| A5SS | 2388 | PSEN1 | PSEN1 | chr14 | + | 73147966 | 73148106 | 73147966 | 73148094 | 73170796 | 73171047 | 0 | 0 | 2 | 0 | 0 | 2 | 8 | 3 | 0 | 0 | 5 | 0 | 61 | 49 | NA | NA | NA | NA | 0.445 | 1 | 0.325 | NA | -0.217 |
| A5SS | 2404 | CHCHD7 | CHCHD7 | chr8 | + | 56212761 | 56212909 | 56212761 | 56212889 | 56214597 | 56214667 | 0 | 0 | 0 | 0 | 0 | 1 | 2 | 0 | 0 | 0 | 0 | 0 | 69 | 49 | NA | NA | NA | NA | 0 | 1 | NA | NA | -1 |
| A5SS | 2498 | CRK | CRK | chr17 | - | 1436619 | 1437155 | 1436789 | 1437155 | 1421352 | 1423650 | 0 | 0 | 6 | 0 | 0 | 1 | 1 | 0 | 0 | 0 | 1 | 0 | 98 | 49 | NA | NA | NA | NA | 0.75 | 1 | 0 | NA | 0.25 |
| A5SS | 2580 | STAT3 | STAT3 | chr17 | - | 42322281 | 42322494 | 42322284 | 42322494 | 42317181 | 42317224 | 0 | 0 | 19 | 0 | 0 | 4 | 0 | 10 | 0 | 0 | 4 | 0 | 52 | 49 | NA | NA | NA | NA | 0.817 | NA | 0.702 | NA | 0.115 |
| A5SS | 2607 | FASTK | FASTK | chr7 | - | 151078841 | 151079021 | 151078922 | 151079021 | 151078561 | 151078701 | 0 | 0 | 1 | 0 | 0 | 0 | 0 | 0 | 0 | 0 | 1 | 0 | 98 | 49 | NA | NA | NA | NA | 1 | NA | 0 | NA | 1 |
| A5SS | 2701 | IFI6 | IFI6 | chr1 | - | 27669220 | 27669346 | 27669232 | 27669346 | 27668457 | 27668535 | 0 | 0 | 4 | 0 | 0 | 1 | 0 | 0 | 0 | 0 | 1 | 0 | 61 | 49 | NA | NA | NA | NA | 0.763 | NA | 0 | NA | 0.763 |
| A5SS | 2714 | MLX | MLX | chr17 | + | 42567059 | 42567328 | 42567059 | 42567166 | 42567618 | 42567655 | 0 | 0 | 0 | 0 | 0 | 1 | 2 | 0 | 0 | 0 | 0 | 0 | 98 | 49 | NA | NA | NA | NA | 0 | 1 | NA | NA | -1 |
| A5SS | 2723 | ATP5A1 | ATP5A1 | chr18 | - | 46094979 | 46095131 | 46095052 | 46095131 | 46091681 | 46091851 | 0 | 0 | 0 | 0 | 0 | 2 | 1 | 0 | 0 | 0 | 2 | 0 | 98 | 49 | NA | NA | NA | NA | 0 | 1 | 0 | NA | -0.5 |
| A5SS | 2757 | CAPS | CAPS | chr19 | + | 5914939 | 5915146 | 5914939 | 5915065 | 5915220 | 5916211 | 0 | 0 | 1 | 0 | 0 | 0 | 0 | 0 | 0 | 0 | 1 | 0 | 98 | 49 | NA | NA | NA | NA | 1 | NA | 0 | NA | 1 |
| A5SS | 2763 | ADGRE1 | ADGRE1 | chr19 | + | 6928144 | 6928310 | 6928144 | 6928211 | 6934986 | 6935078 | 0 | 0 | 0 | 0 | 0 | 4 | 1 | 0 | 0 | 0 | 0 | 0 | 98 | 49 | NA | NA | NA | NA | 0 | 1 | NA | NA | -1 |
| A5SS | 2772 | LPPR2 | LPPR2 | chr19 | + | 11363712 | 11363835 | 11363712 | 11363819 | 11364160 | 11364212 | 0 | 0 | 2 | 0 | 0 | 1 | 0 | 0 | 0 | 0 | 1 | 0 | 65 | 49 | NA | NA | NA | NA | 0.601 | NA | 0 | NA | 0.601 |
| A5SS | 2777 | SGMS2 | SGMS2 | chr4 | + | 107893263 | 107893658 | 107893263 | 107893382 | 107895309 | 107896008 | 0 | 0 | 3 | 0 | 0 | 0 | 1 | 0 | 0 | 0 | 1 | 0 | 98 | 49 | NA | NA | NA | NA | 1 | 1 | 0 | NA | 0.5 |
| A5SS | 2801 | EPS15L1 | EPS15L1 | chr19 | - | 16440861 | 16440909 | 16440884 | 16440909 | 16437769 | 16437865 | 0 | 0 | 2 | 0 | 0 | 0 | 0 | 1 | 0 | 0 | 1 | 0 | 72 | 49 | NA | NA | NA | NA | 1 | NA | 0.405 | NA | 0.595 |
| A5SS | 2950 | HNRNPH3 | HNRNPH3 | chr10 | + | 68338502 | 68338687 | 68338502 | 68338642 | 68339139 | 68339226 | 2 | 0 | 0 | 0 | 0 | 0 | 0 | 0 | 0 | 0 | 2 | 0 | 94 | 49 | NA | NA | 1 | NA | NA | NA | 0 | NA | 1 |
| A5SS | 2996 | VRK2 | VRK2 | chr2 | + | 58046641 | 58046868 | 58046641 | 58046821 | 58048826 | 58048967 | 2 | 0 | 2 | 0 | 0 | 1 | 0 | 0 | 0 | 0 | 1 | 0 | 96 | 49 | NA | NA | 1 | NA | 0.505 | NA | 0 | NA | 0.752 |
| A5SS | 3001 | CCND3 | CCND3 | chr6 | - | 41940951 | 41941848 | 41941451 | 41941848 | 41940369 | 41940585 | 0 | 0 | 0 | 0 | 0 | 5 | 0 | 1 | 0 | 0 | 0 | 0 | 98 | 49 | NA | NA | NA | NA | 0 | NA | 1 | NA | -1 |
| A5SS | 3047 | PLEKHB2 | PLEKHB2 | chr2 | + | 131132901 | 131133015 | 131132901 | 131132991 | 131140169 | 131140275 | 0 | 0 | 2 | 0 | 0 | 0 | 0 | 1 | 0 | 0 | 1 | 0 | 73 | 49 | NA | NA | NA | NA | 1 | NA | 0.402 | NA | 0.598 |
| A5SS | 3074 | HINT1 | HINT1 | chr5 | - | 131165059 | 131165348 | 131165094 | 131165348 | 131162571 | 131162676 | 2 | 0 | 0 | 0 | 0 | 5 | 0 | 0 | 0 | 0 | 1 | 0 | 84 | 49 | NA | NA | 1 | NA | 0 | NA | 0 | NA | 0.5 |
| RI | 33 | VPS28 | VPS28 | chr8 | - | 144423600 | 144424132 | 144423600 | 144423922 | 144424040 | 144424132 | 0 | 0 | 0 | 0 | 0 | 3 | 0 | 2 | 0 | 0 | 2 | 0 | 98 | 49 | NA | NA | NA | NA | 0 | NA | 0.333 | NA | -0.333 |
| RI | 227 | NDUFB4 | NDUFB4 | chr3 | + | 120601110 | 120602411 | 120601110 | 120601257 | 120602207 | 120602411 | 0 | 0 | 0 | 0 | 0 | 1 | 4 | 0 | 0 | 0 | 1 | 0 | 98 | 49 | NA | NA | NA | NA | 0 | 1 | 0 | NA | -0.5 |
| RI | 288 | NECAB3 | NECAB3 | chr20 | - | 33659496 | 33659732 | 33659496 | 33659550 | 33659652 | 33659732 | 0 | 0 | 0 | 0 | 0 | 1 | 2 | 0 | 0 | 0 | 0 | 0 | 98 | 49 | NA | NA | NA | NA | 0 | 1 | NA | NA | -1 |
| RI | 338 | NSD1 | NSD1 | chr5 | + | 177135086 | 177136030 | 177135086 | 177135200 | 177135940 | 177136030 | 0 | 0 | 1 | 0 | 0 | 0 | 0 | 0 | 0 | 0 | 1 | 0 | 98 | 49 | NA | NA | NA | NA | 1 | NA | 0 | NA | 1 |
| RI | 405 | CREBZF | CREBZF | chr11 | - | 85657563 | 85665138 | 85657563 | 85662457 | 85663523 | 85665138 | 0 | 0 | 4 | 0 | 0 | 2 | 2 | 1 | 0 | 0 | 0 | 0 | 98 | 49 | NA | NA | NA | NA | 0.5 | 1 | 1 | NA | -0.5 |
| RI | 429 | IL32 | IL32 | chr16 | + | 3067376 | 3067613 | 3067376 | 3067415 | 3067553 | 3067613 | 0 | 0 | 1 | 0 | 0 | 3 | 1 | 0 | 0 | 0 | 6 | 0 | 98 | 49 | NA | NA | NA | NA | 0.143 | 1 | 0 | NA | -0.357 |
| RI | 493 | CDK17 | CDK17 | chr12 | - | 96278260 | 96280885 | 96278260 | 96280279 | 96280807 | 96280885 | 0 | 0 | 13 | 0 | 0 | 0 | 3 | 8 | 0 | 0 | 2 | 0 | 98 | 49 | NA | NA | NA | NA | 1 | 1 | 0.667 | NA | 0.166 |
| RI | 514 | DNAJB12 | DNAJB12 | chr10 | - | 72332829 | 72335931 | 72332829 | 72334617 | 72335779 | 72335931 | 0 | 0 | 1 | 0 | 0 | 3 | 0 | 1 | 0 | 0 | 0 | 0 | 98 | 49 | NA | NA | NA | NA | 0.143 | NA | 1 | NA | -0.857 |
| RI | 530 | OTUB1 | OTUB1 | chr11 | + | 63996529 | 63996941 | 63996529 | 63996648 | 63996856 | 63996941 | 0 | 0 | 0 | 0 | 0 | 1 | 2 | 1 | 0 | 0 | 2 | 0 | 98 | 49 | NA | NA | NA | NA | 0 | 1 | 0.2 | NA | -0.6 |
| RI | 585 | PCID2 | PCID2 | chr13 | - | 113177538 | 113178287 | 113177538 | 113177976 | 113178173 | 113178287 | 0 | 0 | 3 | 0 | 0 | 0 | 3 | 0 | 0 | 0 | 1 | 0 | 98 | 49 | NA | NA | NA | NA | 1 | 1 | 0 | NA | 0.5 |

| | | | | | | | | | | | | | | | | | | | | | | | | | | | | | | | | | | |
|---|---|---|---|---|---|---|---|---|---|---|---|---|---|---|---|---|---|---|---|---|---|---|---|---|---|---|---|---|---|---|---|---|---|---|
| RI | 619 | IL17RC | IL17RC | chr3 | + | 9923880 | 9924291 | 9923880 | 9924020 | 9924231 | 9924291 | 0 | 1 | 0 | 0 | 0 | 1 | 6 | 0 | 0 | 0 | 0 | 0 | 98 | 49 | NA | NA | NA | 1 | 0 | 1 | NA | NA | -0.5 |
| RI | 729 | CLK1 | CLK1 | chr2 | - | 200859679 | 200861466 | 200859679 | 200859746 | 200861237 | 200861466 | 0 | 0 | 5 | 0 | 0 | 2 | 0 | 18 | 0 | 0 | 3 | 0 | 98 | 49 | NA | NA | NA | NA | 0.556 | NA | 0.75 | NA | -0.194 |
| RI | 834 | CIRBP | CIRBP | chr19 | + | 1271980 | 1273172 | 1271980 | 1272051 | 1272426 | 1273172 | 0 | 0 | 3 | 0 | 0 | 1 | 3 | 0 | 0 | 0 | 4 | 0 | 98 | 49 | NA | NA | NA | NA | 0.6 | 1 | 0 | NA | 0.1 |
| RI | 888 | LAT | LAT | chr16 | + | 28986539 | 28986714 | 28986539 | 28986569 | 28986656 | 28986714 | 1 | 0 | 0 | 0 | 0 | 0 | 0 | 1 | 1 | 0 | 1 | 0 | 98 | 49 | NA | NA | 1 | NA | NA | NA | 0.333 | 1 | 0.334 |
| RI | 896 | KAT8 | KAT8 | chr16 | + | 31130745 | 31131393 | 31130745 | 31130900 | 31131194 | 31131393 | 0 | 0 | 0 | 0 | 1 | 0 | 1 | 0 | 0 | 0 | 0 | 0 | 98 | 49 | NA | NA | NA | 0 | NA | 1 | NA | NA | -1 |
| RI | 909 | HNRNPM | HNRNPM | chr19 | + | 8465323 | 8465515 | 8465323 | 8465362 | 8465479 | 8465515 | 0 | 0 | 8 | 0 | 0 | 1 | 7 | 0 | 0 | 0 | 0 | 0 | 98 | 49 | NA | NA | NA | NA | 0.8 | 1 | NA | NA | -0.2 |
| RI | 926 | WRAP53 | WRAP53 | chr17 | + | 7688348 | 7689079 | 7688348 | 7688561 | 7688647 | 7689079 | 0 | 0 | 0 | 0 | 0 | 2 | 5 | 0 | 0 | 0 | 0 | 0 | 98 | 49 | NA | NA | NA | NA | 0 | 1 | NA | NA | -1 |
| RI | 984 | LUC7L3 | LUC7L3 | chr17 | + | 50750500 | 50752711 | 50750500 | 50750694 | 50752199 | 50752711 | 0 | 0 | 2 | 0 | 0 | 2 | 0 | 4 | 0 | 0 | 0 | 0 | 98 | 49 | NA | NA | NA | NA | 0.333 | NA | 1 | NA | -0.667 |
| RI | 987 | SRSF1 | SRSF1 | chr17 | - | 58000918 | 58005973 | 58000918 | 58005600 | 58005800 | 58005973 | 0 | 0 | 1 | 0 | 0 | 5 | 2 | 0 | 0 | 0 | 0 | 0 | 98 | 49 | NA | NA | NA | NA | 0.091 | 1 | NA | NA | -0.909 |
| RI | 1024 | FAM195B | FAM195B | chr17 | - | 81822360 | 81823311 | 81822360 | 81822907 | 81823208 | 81823311 | 0 | 0 | 3 | 0 | 0 | 0 | 4 | 1 | 0 | 0 | 3 | 0 | 98 | 49 | NA | NA | NA | NA | 1 | 1 | 0.143 | NA | 0.428 |
| RI | 1029 | SNRNP27 | SNRNP27 | chr2 | + | 69904253 | 69905236 | 69904253 | 69904425 | 69904511 | 69905236 | 1 | 0 | 3 | 0 | 0 | 0 | 0 | 0 | 0 | 0 | 1 | 0 | 98 | 49 | NA | NA | 1 | NA | 1 | NA | 0 | NA | 1 |
| RI | 1037 | CSTF3 | CSTF3 | chr11 | - | 33141430 | 33141986 | 33141430 | 33141762 | 33141884 | 33141986 | 0 | 0 | 0 | 0 | 0 | 1 | 8 | 1 | 0 | 0 | 2 | 0 | 98 | 49 | NA | NA | NA | NA | 0 | 1 | 0.2 | NA | -0.6 |
| RI | 1058 | GTPBP3 | GTPBP3 | chr19 | + | 17338953 | 17339266 | 17338953 | 17339026 | 17339122 | 17339266 | 1 | 0 | 0 | 0 | 0 | 0 | 0 | 0 | 0 | 0 | 1 | 0 | 98 | 49 | NA | NA | 1 | NA | NA | NA | 0 | NA | 1 |
| RI | 1073 | CCPG1 | CCPG1 | chr15 | - | 55355222 | 55360944 | 55355222 | 55356409 | 55359538 | 55360944 | 0 | 0 | 1 | 0 | 0 | 1 | 1 | 1 | 0 | 0 | 0 | 0 | 98 | 49 | NA | NA | NA | NA | 0.333 | 1 | 1 | NA | -0.667 |
| RI | 1118 | RPL31 | RPL31 | chr2 | + | 101005958 | 101006423 | 101005958 | 101006071 | 101006349 | 101006423 | 1 | 0 | 4 | 0 | 0 | 5 | 0 | 1 | 0 | 0 | 6 | 0 | 98 | 49 | NA | NA | 1 | NA | 0.286 | NA | 0.077 | NA | 0.566 |
| SE | 37 | FMR1 | FMR1 | chrX | + | 147937465 | 147937600 | 147936503 | 147936613 | 147940575 | 147940662 | 0 | 0 | 1 | 0 | 0 | 0 | 0 | 2 | 0 | 0 | 1 | 0 | 98 | 49 | NA | NA | NA | NA | 1 | NA | 0.5 | NA | 0.5 |
| SE | 38 | FMR1 | FMR1 | chrX | + | 147938098 | 147938161 | 147937465 | 147937600 | 147940575 | 147940662 | 0 | 0 | 0 | 0 | 0 | 1 | 0 | 1 | 0 | 0 | 2 | 0 | 98 | 49 | NA | NA | NA | NA | 0 | NA | 0.2 | NA | -0.2 |
| SE | 57 | SASH3 | SASH3 | chrX | + | 129792626 | 129792836 | 129792327 | 129792476 | 129792988 | 129793139 | 0 | 0 | 20 | 0 | 0 | 0 | 0 | 17 | 0 | 0 | 1 | 0 | 98 | 49 | NA | NA | NA | NA | 1 | NA | 0.895 | NA | 0.105 |
| SE | 59 | STAG2 | STAG2 | chrX | + | 124090853 | 124090964 | 124090574 | 124090764 | 124094017 | 124094144 | 0 | 0 | 0 | 0 | 0 | 3 | 0 | 1 | 0 | 0 | 2 | 0 | 98 | 49 | NA | NA | NA | NA | 0 | NA | 0.2 | NA | -0.2 |
| SE | 129 | XIST | XIST | chrX | - | 73833237 | 73833374 | 73831065 | 73831274 | 73837439 | 73837503 | 0 | 0 | 9 | 0 | 0 | 1 | 0 | 2 | 0 | 0 | 0 | 0 | 98 | 49 | NA | NA | NA | NA | 0.818 | NA | 1 | NA | -0.182 |
| SE | 156 | APEX2 | APEX2 | chrX | + | 55002250 | 55002431 | 55001545 | 55001629 | 55002961 | 55003108 | 0 | 0 | 0 | 0 | 0 | 1 | 0 | 1 | 0 | 0 | 0 | 0 | 98 | 49 | NA | NA | NA | NA | 0 | NA | 1 | NA | -1 |
| SE | 173 | PQBP1 | PQBP1 | chrX | + | 48898491 | 48898576 | 48897911 | 48898082 | 48901189 | 48901277 | 0 | 0 | 1 | 0 | 0 | 0 | 0 | 1 | 0 | 0 | 1 | 0 | 98 | 49 | NA | NA | NA | NA | 1 | NA | 0.333 | NA | 0.667 |
| SE | 175 | TIMM17B | TIMM17B | chrX | - | 48895351 | 48895501 | 48895037 | 48895101 | 48896758 | 48896858 | 0 | 0 | 0 | 0 | 0 | 2 | 0 | 1 | 0 | 0 | 0 | 0 | 98 | 49 | NA | NA | NA | NA | 0 | NA | 1 | NA | -1 |
| SE | 180 | ELK1 | ELK1 | chrX | - | 47637750 | 47638182 | 47637012 | 47637114 | 47638894 | 47639338 | 0 | 0 | 1 | 0 | 0 | 0 | 0 | 0 | 0 | 0 | 1 | 0 | 98 | 49 | NA | NA | NA | NA | 1 | NA | 0 | NA | 1 |
| SE | 182 | UBA1 | UBA1 | chrX | + | 47199208 | 47199377 | 47199047 | 47199106 | 47199479 | 47199614 | 0 | 0 | 5 | 0 | 0 | 0 | 0 | 1 | 0 | 0 | 2 | 0 | 98 | 49 | NA | NA | NA | NA | 1 | NA | 0.2 | NA | 0.8 |
| SE | 184 | RBM10 | RBM10 | chrX | + | 47175018 | 47175092 | 47173127 | 47173197 | 47176499 | 47176586 | 0 | 0 | 1 | 0 | 0 | 0 | 0 | 0 | 0 | 0 | 2 | 0 | 98 | 49 | NA | NA | NA | NA | 1 | NA | 0 | NA | 1 |
| SE | 185 | KDM6A | KDM6A | chrX | + | 45020609 | 45020730 | 45010960 | 45011019 | 45034930 | 45034985 | 0 | 0 | 7 | 0 | 0 | 3 | 0 | 4 | 0 | 0 | 0 | 0 | 98 | 49 | NA | NA | NA | NA | 0.538 | NA | 1 | NA | -0.462 |
| SE | 189 | KDM6A | KDM6A | chrX | + | 45060608 | 45060764 | 45059246 | 45059466 | 45061323 | 45061419 | 0 | 0 | 4 | 0 | 0 | 3 | 0 | 5 | 0 | 0 | 1 | 0 | 98 | 49 | NA | NA | NA | NA | 0.4 | NA | 0.714 | NA | -0.314 |
| SE | 195 | CXorf38 | CXorf38 | chrX | - | 40647006 | 40647141 | 40639008 | 40639128 | 40647304 | 40647567 | 0 | 0 | 1 | 0 | 0 | 1 | 0 | 1 | 0 | 0 | 0 | 0 | 98 | 49 | NA | NA | NA | NA | 0.333 | NA | 1 | NA | -0.667 |
| SE | 201 | ACOT9 | ACOT9 | chrX | - | 23722669 | 23722753 | 23721880 | 23721984 | 23730526 | 23730564 | 0 | 0 | 7 | 0 | 0 | 0 | 0 | 17 | 0 | 0 | 2 | 0 | 98 | 49 | NA | NA | NA | NA | 1 | NA | 0.81 | NA | 0.19 |
| SE | 202 | ACOT9 | ACOT9 | chrX | - | 23734340 | 23734367 | 23733171 | 23733217 | 23735918 | 23736016 | 0 | 0 | 10 | 0 | 0 | 0 | 0 | 4 | 0 | 0 | 1 | 0 | 76 | 49 | NA | NA | NA | NA | 1 | NA | 0.721 | NA | 0.279 |
| SE | 205 | RPS6KA3 | RPS6KA3 | chrX | - | 20187827 | 20187970 | 20186295 | 20186366 | 20188496 | 20188534 | 0 | 0 | 7 | 0 | 0 | 0 | 0 | 2 | 0 | 0 | 1 | 0 | 98 | 49 | NA | NA | NA | NA | 1 | NA | 0.5 | NA | 0.5 |
| SE | 210 | SYAP1 | SYAP1 | chrX | + | 16741715 | 16741789 | 16736165 | 16736232 | 16743700 | 16743840 | 0 | 0 | 12 | 0 | 0 | 0 | 0 | 5 | 0 | 0 | 1 | 0 | 98 | 49 | NA | NA | NA | NA | 1 | NA | 0.714 | NA | 0.286 |
| SE | 233 | CSF2RA | CSF2RA | chrX | + | 1300490 | 1300626 | 1295426 | 1295456 | 1303922 | 1304019 | 0 | 0 | 19 | 0 | 0 | 4 | 0 | 2 | 0 | 0 | 0 | 0 | 98 | 49 | NA | NA | NA | NA | 0.704 | NA | 1 | NA | -0.296 |
| SE | 234 | CSF2RA | CSF2RA | chrX | + | 1303260 | 1303362 | 1300490 | 1300626 | 1303922 | 1304019 | 0 | 0 | 1 | 0 | 0 | 5 | 0 | 0 | 0 | 0 | 1 | 0 | 98 | 49 | NA | NA | NA | NA | 0.091 | NA | 0 | NA | 0.091 |
| SE | 237 | EHMT1 | EHMT1 | chr9 | + | 137752330 | 137752408 | 137743901 | 137744090 | 137754170 | 137754291 | 0 | 0 | 4 | 0 | 0 | 0 | 0 | 1 | 0 | 0 | 1 | 0 | 98 | 49 | NA | NA | NA | NA | 1 | NA | 0.333 | NA | 0.667 |
| SE | 252 | SEC16A | SEC16A | chr9 | - | 136445644 | 136445719 | 136443822 | 136443900 | 136446854 | 136446949 | 0 | 0 | 2 | 0 | 0 | 0 | 0 | 0 | 0 | 0 | 1 | 0 | 98 | 49 | NA | NA | NA | NA | 1 | NA | 0 | NA | 1 |
| SE | 253 | SEC16A | SEC16A | chr9 | - | 136457443 | 136457584 | 136456052 | 136456166 | 136459133 | 136459239 | 0 | 0 | 7 | 0 | 0 | 0 | 0 | 2 | 0 | 0 | 1 | 0 | 98 | 49 | NA | NA | NA | NA | 1 | NA | 0.5 | NA | 0.5 |
| SE | 254 | SEC16A | SEC16A | chr9 | - | 136462886 | 136463132 | 136461176 | 136461274 | 136463462 | 136463601 | 0 | 0 | 2 | 0 | 0 | 0 | 0 | 2 | 0 | 0 | 1 | 0 | 98 | 49 | NA | NA | NA | NA | 1 | NA | 0.5 | NA | 0.5 |
| SE | 282 | ASB6 | ASB6 | chr9 | - | 129640540 | 129640722 | 129639401 | 129639508 | 129641886 | 129642169 | 0 | 0 | 1 | 0 | 0 | 1 | 0 | 1 | 0 | 0 | 0 | 0 | 98 | 49 | NA | NA | NA | NA | 0.333 | NA | 1 | NA | -0.667 |
| SE | 283 | PPP2R4 | PPP2R4 | chr9 | + | 129123051 | 129123138 | 129120512 | 129120610 | 129128984 | 129129110 | 0 | 0 | 1 | 0 | 0 | 0 | 0 | 0 | 0 | 0 | 1 | 0 | 98 | 49 | NA | NA | NA | NA | 1 | NA | 0 | NA | 1 |
| SE | 307 | SH2D3C | SH2D3C | chr9 | - | 127761610 | 127761650 | 127751171 | 127751300 | 127773989 | 127774467 | 0 | 0 | 1 | 0 | 0 | 1 | 0 | 3 | 0 | 0 | 0 | 0 | 89 | 49 | NA | NA | NA | NA | 0.355 | NA | 1 | NA | -0.645 |
| SE | 315 | GAPVD1 | GAPVD1 | chr9 | + | 125360527 | 125360725 | 125359419 | 125359492 | 125362605 | 125365011 | 0 | 0 | 5 | 0 | 0 | 2 | 0 | 3 | 0 | 0 | 0 | 0 | 98 | 49 | NA | NA | NA | NA | 0.556 | NA | 1 | NA | -0.444 |
| SE | 337 | MED12 | MED12 | chrX | + | 71125350 | 71125495 | 71124975 | 71125146 | 71125662 | 71125713 | 0 | 0 | 3 | 0 | 0 | 0 | 0 | 0 | 0 | 0 | 1 | 0 | 98 | 49 | NA | NA | NA | NA | 1 | NA | 0 | NA | 1 |
| SE | 348 | NR4A3 | NR4A3 | chr9 | + | 99826729 | 99826801 | 99825658 | 99825832 | 99828040 | 99828993 | 0 | 0 | 3 | 0 | 0 | 0 | 0 | 1 | 0 | 0 | 5 | 0 | 98 | 49 | NA | NA | NA | NA | 1 | NA | 0.091 | NA | 0.909 |
| SE | 358 | ERCC6L2 | ERCC6L2 | chr9 | + | 95907077 | 95907271 | 95897848 | 95897971 | 95915667 | 95915829 | 0 | 0 | 0 | 0 | 0 | 1 | 0 | 1 | 0 | 0 | 0 | 0 | 98 | 49 | NA | NA | NA | NA | 0 | NA | 1 | NA | -1 |
| SE | 365 | SEMA4D | SEMA4D | chr9 | - | 89418362 | 89418437 | 89418055 | 89418118 | 89455887 | 89455953 | 0 | 0 | 2 | 0 | 0 | 0 | 0 | 0 | 0 | 0 | 1 | 0 | 98 | 49 | NA | NA | NA | NA | 1 | NA | 0 | NA | 1 |
| SE | 369 | AGTPBP1 | AGTPBP1 | chr9 | - | 85588297 | 85588478 | 85586830 | 85586960 | 85589527 | 85589681 | 0 | 0 | 12 | 0 | 0 | 1 | 0 | 5 | 0 | 0 | 0 | 0 | 98 | 49 | NA | NA | NA | NA | 0.857 | NA | 1 | NA | -0.143 |
| SE | 377 | TLE4 | TLE4 | chr9 | + | 79612655 | 79612718 | 79576132 | 79576177 | 79627373 | 79627448 | 0 | 0 | 4 | 0 | 0 | 0 | 0 | 5 | 0 | 0 | 1 | 0 | 98 | 49 | NA | NA | NA | NA | 1 | NA | 0.714 | NA | 0.286 |
| SE | 398 | PTAR1 | PTAR1 | chr9 | - | 69732138 | 69732352 | 69723325 | 69723630 | 69734149 | 69734254 | 0 | 0 | 1 | 0 | 0 | 0 | 0 | 1 | 0 | 0 | 1 | 0 | 98 | 49 | NA | NA | NA | NA | 1 | NA | 0.333 | NA | 0.667 |
| SE | 430 | UBAP1 | UBAP1 | chr9 | + | 34196335 | 34196458 | 34179004 | 34179240 | 34220907 | 34220948 | 0 | 0 | 0 | 0 | 0 | 1 | 0 | 1 | 0 | 0 | 1 | 0 | 98 | 49 | NA | NA | NA | NA | 0 | NA | 0.333 | NA | -0.333 |
| SE | 438 | DNAJA1 | DNAJA1 | chr9 | + | 33026812 | 33026990 | 33026474 | 33026616 | 33029884 | 33029989 | 0 | 0 | 6 | 0 | 0 | 1 | 0 | 2 | 0 | 0 | 3 | 0 | 98 | 49 | NA | NA | NA | NA | 0.75 | NA | 0.25 | NA | 0.5 |
| SE | 444 | DDX58 | DDX58 | chr9 | - | 32492390 | 32492538 | 32491300 | 32491420 | 32493760 | 32493942 | 0 | 0 | 8 | 0 | 0 | 1 | 0 | 5 | 0 | 0 | 6 | 0 | 98 | 49 | NA | NA | NA | NA | 0.8 | NA | 0.294 | NA | 0.506 |
| SE | 465 | RIC1 | RIC1 | chr9 | + | 5745930 | 5746083 | 5743688 | 5743737 | 5747301 | 5747505 | 0 | 0 | 9 | 0 | 0 | 0 | 0 | 0 | 0 | 0 | 1 | 0 | 98 | 49 | NA | NA | NA | NA | 1 | NA | 0 | NA | 1 |
| SE | 468 | RFX3 | RFX3 | chr9 | - | 3270370 | 3270525 | 3266207 | 3266305 | 3271002 | 3271118 | 0 | 0 | 0 | 0 | 0 | 2 | 0 | 1 | 0 | 0 | 0 | 0 | 98 | 49 | NA | NA | NA | NA | 0 | NA | 1 | NA | -1 |
| SE | 474 | DOCK8 | DOCK8 | chr9 | + | 420948 | 421078 | 420400 | 420583 | 422047 | 422135 | 0 | 0 | 11 | 0 | 0 | 1 | 0 | 10 | 0 | 0 | 1 | 0 | 98 | 49 | NA | NA | NA | NA | 0.846 | NA | 0.833 | NA | 0.013 |
| SE | 475 | DOCK8 | DOCK8 | chr9 | + | 426884 | 426981 | 422047 | 422135 | 428361 | 428496 | 0 | 0 | 16 | 0 | 0 | 0 | 0 | 5 | 0 | 0 | 1 | 0 | 98 | 49 | NA | NA | NA | NA | 1 | NA | 0.714 | NA | 0.286 |
| SE | 476 | DOCK8 | DOCK8 | chr9 | + | 446369 | 446606 | 443426 | 443516 | 449783 | 449927 | 0 | 0 | 19 | 0 | 0 | 0 | 0 | 7 | 0 | 0 | 2 | 0 | 98 | 49 | NA | NA | NA | NA | 1 | NA | 0.636 | NA | 0.364 |
| SE | 487 | GRINA | GRINA | chr8 | + | 143991877 | 143992078 | 143991691 | 143991804 | 143992244 | 143992373 | 0 | 0 | 5 | 0 | 0 | 0 | 0 | 0 | 0 | 0 | 1 | 0 | 98 | 49 | NA | NA | NA | NA | 1 | NA | 0 | NA | 1 |
| SE | 520 | LRRC6 | LRRC6 | chr8 | - | 132632739 | 132632963 | 132625271 | 132625454 | 132637934 | 132638107 | 0 | 0 | 1 | 0 | 0 | 1 | 0 | 0 | 0 | 0 | 3 | 0 | 98 | 49 | NA | NA | NA | NA | 0.333 | NA | 0 | NA | 0.333 |
| SE | 534 | PRRC2B | PRRC2B | chr9 | + | 131474453 | 131476535 | 131473507 | 131473724 | 131477743 | 131477949 | 0 | 0 | 3 | 0 | 0 | 0 | 0 | 1 | 0 | 0 | 1 | 0 | 98 | 49 | NA | NA | NA | NA | 1 | NA | 0.333 | NA | 0.667 |
| SE | 553 | UBR5 | UBR5 | chr8 | - | 102278825 | 102278939 | 102276971 | 102277192 | 102279035 | 102279208 | 0 | 0 | 4 | 0 | 0 | 0 | 0 | 2 | 0 | 0 | 1 | 0 | 98 | 49 | NA | NA | NA | NA | 1 | NA | 0.5 | NA | 0.5 |
| SE | 554 | RRM2B | RRM2B | chr8 | - | 102224884 | 102225018 | 102224045 | 102224140 | 102225917 | 102226034 | 0 | 0 | 4 | 0 | 0 | 2 | 0 | 6 | 0 | 0 | 0 | 0 | 98 | 49 | NA | NA | NA | NA | 0.5 | NA | 1 | NA | -0.5 |
| SE | 556 | ZNF706 | ZNF706 | chr8 | - | 101202332 | 101202447 | 101201606 | 101201743 | 101205434 | 101205732 | 0 | 1 | 0 | 0 | 0 | 0 | 0 | 2 | 0 | 0 | 0 | 0 | 98 | 49 | NA | NA | 0 | NA | NA | NA | 1 | NA | -1 |
| SE | 560 | RNF19A | RNF19A | chr8 | - | 100264670 | 100264785 | 100259853 | 100259997 | 100268784 | 100268947 | 0 | 0 | 4 | 0 | 0 | 1 | 0 | 6 | 0 | 0 | 0 | 0 | 98 | 49 | NA | NA | NA | NA | 0.667 | NA | 1 | NA | -0.333 |
| SE | 562 | VPS13B | VPS13B | chr8 | + | 99507136 | 99507203 | 99502835 | 99502950 | 99511103 | 99511512 | 0 | 0 | 5 | 0 | 0 | 0 | 0 | 6 | 0 | 0 | 3 | 0 | 98 | 49 | NA | NA | NA | NA | 1 | NA | 0.5 | NA | 0.5 |
| SE | 563 | VPS13B | VPS13B | chr8 | + | 99507769 | 99507911 | 99502835 | 99502950 | 99511103 | 99511512 | 0 | 0 | 2 | 0 | 0 | 0 | 0 | 0 | 0 | 0 | 3 | 0 | 98 | 49 | NA | NA | NA | NA | 1 | NA | 0 | NA | 1 |
| SE | 597 | PAG1 | PAG1 | chr8 | - | 80991478 | 80991530 | 80987369 | 80987466 | 80993102 | 80993307 | 0 | 0 | 3 | 0 | 0 | 0 | 0 | 7 | 0 | 0 | 3 | 0 | 98 | 49 | NA | NA | NA | NA | 1 | NA | 0.538 | NA | 0.462 |
| SE | 598 | PAG1 | PAG1 | chr8 | - | 80993102 | 80993307 | 80991478 | 80991530 | 81029995 | 81030089 | 0 | 0 | 2 | 0 | 0 | 1 | 0 | 4 | 0 | 0 | 0 | 0 | 98 | 49 | NA | NA | NA | NA | 0.5 | NA | 1 | NA | -0.5 |
| SE | 637 | RB1CC1 | RB1CC1 | chr8 | - | 52676368 | 52676571 | 52673844 | 52674274 | 52683548 | 52683719 | 0 | 0 | 6 | 0 | 0 | 0 | 0 | 9 | 0 | 0 | 1 | 0 | 98 | 49 | NA | NA | NA | NA | 1 | NA | 0.818 | NA | 0.182 |
| SE | 645 | HGSNAT | HGSNAT | chr8 | + | 43197839 | 43197952 | 43197671 | 43197742 | 43199387 | 43202827 | 0 | 0 | 2 | 0 | 0 | 0 | 2 | 0 | 0 | 0 | 1 | 0 | 98 | 49 | NA | NA | NA | NA | 1 | 1 | 0 | NA | 0.5 |
| SE | 661 | PLEKHA2 | PLEKHA2 | chr8 | + | 38943788 | 38943837 | 38935993 | 38936050 | 38946123 | 38946221 | 0 | 0 | 2 | 0 | 0 | 0 | 1 | 5 | 0 | 0 | 1 | 0 | 98 | 49 | NA | NA | NA | NA | 1 | 1 | 0.714 | NA | 0.143 |
| SE | 662 | PLEKHA2 | PLEKHA2 | chr8 | + | 38957322 | 38957386 | 38953296 | 38953367 | 38968591 | 38968669 | 0 | 0 | 5 | 0 | 0 | 1 | 2 | 2 | 0 | 0 | 1 | 0 | 98 | 49 | NA | NA | NA | NA | 0.714 | 1 | 0.5 | NA | -0.036 |
| SE | 692 | DOCK5 | DOCK5 | chr8 | + | 25340876 | 25340988 | 25336238 | 25336373 | 25341738 | 25341809 | 0 | 0 | 19 | 0 | 0 | 0 | 0 | 10 | 0 | 0 | 1 | 0 | 98 | 49 | NA | NA | NA | NA | 1 | NA | 0.833 | NA | 0.167 |
| SE | 693 | DOCK5 | DOCK5 | chr8 | + | 25341738 | 25341809 | 25336238 | 25336373 | 25342400 | 25342507 | 0 | 0 | 6 | 0 | 0 | 0 | 0 | 12 | 0 | 0 | 1 | 0 | 98 | 49 | NA | NA | NA | NA | 1 | NA | 0.857 | NA | 0.143 |
| SE | 696 | DOCK5 | DOCK5 | chr8 | + | 25373617 | 25373658 | 25372558 | 25372718 | 25374563 | 25374654 | 0 | 0 | 8 | 0 | 0 | 1 | 0 | 8 | 0 | 0 | 0 | 0 | 90 | 49 | NA | NA | NA | NA | 0.813 | NA | 1 | NA | -0.187 |
| SE | 712 | DOK2 | DOK2 | chr8 | - | 21911900 | 21911988 | 21910672 | 21910857 | 21912228 | 21912510 | 0 | 0 | 7 | 0 | 0 | 2 | 0 | 0 | 0 | 0 | 1 | 0 | 98 | 49 | NA | NA | NA | NA | 0.636 | NA | 0 | NA | 0.636 |
| SE | 735 | CTSB | CTSB | chr8 | - | 11849045 | 11849164 | 11848066 | 11848152 | 11850865 | 11850980 | 0 | 0 | 32 | 0 | 0 | 2 | 0 | 5 | 0 | 0 | 0 | 0 | 98 | 49 | NA | NA | NA | NA | 0.889 | NA | 1 | NA | -0.111 |
| SE | 793 | SSBP1 | SSBP1 | chr7 | + | 141741748 | 141741798 | 141739123 | 141739190 | 141742168 | 141742229 | 0 | 0 | 0 | 0 | 0 | 1 | 0 | 1 | 0 | 0 | 0 | 0 | 98 | 49 | NA | NA | NA | NA | 0 | NA | 1 | NA | -1 |
| SE | 819 | NRF1 | NRF1 | chr7 | + | 129710373 | 129710571 | 129709074 | 129709233 | 129711474 | 129711576 | 0 | 0 | 1 | 0 | 0 | 0 | 0 | 1 | 0 | 0 | 1 | 0 | 98 | 49 | NA | NA | NA | NA | 1 | NA | 0.333 | NA | 0.667 |
| SE | 824 | TNPO3 | TNPO3 | chr7 | - | 128993806 | 128993914 | 128991998 | 128992090 | 128997388 | 128997535 | 0 | 0 | 0 | 0 | 0 | 2 | 0 | 2 | 0 | 0 | 0 | 0 | 98 | 49 | NA | NA | NA | NA | 0 | NA | 1 | NA | -1 |
| SE | 835 | ING3 | ING3 | chr7 | + | 120973204 | 120973243 | 120970687 | 120970880 | 120974727 | 120975657 | 0 | 0 | 3 | 0 | 0 | 1 | 0 | 3 | 0 | 0 | 0 | 0 | 88 | 49 | NA | NA | NA | NA | 0.626 | NA | 1 | NA | -0.374 |
| SE | 864 | NAMPT | NAMPT | chr7 | - | 106269153 | 106269312 | 106263391 | 106263617 | 106272529 | 106272658 | 0 | 0 | 0 | 0 | 0 | 1 | 0 | 2 | 0 | 0 | 0 | 0 | 98 | 49 | NA | NA | NA | NA | 0 | NA | 1 | NA | -1 |
| SE | 907 | SMURF1 | SMURF1 | chr7 | - | 99040377 | 99040556 | 99038387 | 99038525 | 99042117 | 99042232 | 0 | 0 | 2 | 0 | 0 | 0 | 0 | 2 | 0 | 0 | 1 | 0 | 98 | 49 | NA | NA | NA | NA | 1 | NA | 0.5 | NA | 0.5 |
| SE | 910 | TECPR1 | TECPR1 | chr7 | - | 98241069 | 98241244 | 98240850 | 98240951 | 98243466 | 98243592 | 0 | 0 | 2 | 0 | 0 | 1 | 0 | 4 | 0 | 0 | 0 | 0 | 98 | 49 | NA | NA | NA | NA | 0.5 | NA | 1 | NA | -0.5 |
| SE | 949 | DMTF1 | DMTF1 | chr7 | + | 87164933 | 87165050 | 87163494 | 87163617 | 87166482 | 87166605 | 0 | 0 | 2 | 0 | 0 | 0 | 0 | 4 | 0 | 0 | 1 | 0 | 98 | 49 | NA | NA | NA | NA | 1 | NA | 0.667 | NA | 0.333 |
| SE | 954 | CD36 | CD36 | chr7 | + | 80662989 | 80663169 | 80661062 | 80661210 | 80664405 | 80664497 | 0 | 1 | 28 | 0 | 0 | 2 | 15 | 0 | 0 | 0 | 0 | 0 | 98 | 49 | NA | NA | NA | 1 | 0.875 | 1 | NA | NA | -0.062 |
| SE | 973 | LAT2 | LAT2 | chr7 | + | 74224638 | 74224760 | 74224017 | 74224197 | 74228943 | 74229834 | 0 | 0 | 7 | 0 | 0 | 1 | 0 | 4 | 0 | 0 | 0 | 0 | 98 | 49 | NA | NA | NA | NA | 0.778 | NA | 1 | NA | -0.222 |
| SE | 980 | RABGEF1 | RABGEF1 | chr7 | + | 66783674 | 66783841 | 66775226 | 66775393 | 66795510 | 66795592 | 0 | 0 | 9 | 0 | 0 | 0 | 0 | 3 | 0 | 0 | 2 | 0 | 98 | 49 | NA | NA | NA | NA | 1 | NA | 0.429 | NA | 0.571 |
| SE | 981 | RABGEF1 | RABGEF1 | chr7 | + | 66795510 | 66795592 | 66775226 | 66775393 | 66797373 | 66797506 | 0 | 0 | 4 | 0 | 0 | 1 | 0 | 7 | 0 | 0 | 0 | 0 | 98 | 49 | NA | NA | NA | NA | 0.667 | NA | 1 | NA | -0.333 |
| SE | 993 | SUMF2 | SUMF2 | chr7 | + | 56074173 | 56074218 | 56072996 | 56073111 | 56074585 | 56074736 | 0 | 0 | 7 | 0 | 0 | 0 | 0 | 1 | 0 | 0 | 1 | 0 | 94 | 49 | NA | NA | NA | NA | 1 | NA | 0.343 | NA | 0.657 |
| SE | 995 | GK | GK | chrX | + | 30694399 | 30694537 | 30691122 | 30691199 | 30696041 | 30696151 | 0 | 0 | 1 | 0 | 0 | 0 | 0 | 2 | 0 | 0 | 2 | 0 | 98 | 49 | NA | NA | NA | NA | 1 | NA | 0.333 | NA | 0.667 |
| SE | 996 | GK | GK | chrX | + | 30697731 | 30697749 | 30696616 | 30696683 | 30700413 | 30700449 | 0 | 0 | 1 | 0 | 0 | 2 | 0 | 0 | 0 | 0 | 7 | 0 | 67 | 49 | NA | NA | NA | NA | 0.268 | NA | 0 | NA | 0.268 |
| SE | 1008 | IKZF1 | IKZF1 | chr7 | + | 50376532 | 50376793 | 50327637 | 50327757 | 50382539 | 50382707 | 0 | 0 | 1 | 0 | 0 | 2 | 2 | 0 | 0 | 0 | 4 | 0 | 98 | 49 | NA | NA | NA | NA | 0.2 | 1 | 0 | NA | -0.3 |
| SE | 1034 | ELMO1 | ELMO1 | chr7 | - | 36887559 | 36887672 | 36878009 | 36878117 | 36894853 | 36895017 | 0 | 0 | 4 | 0 | 0 | 0 | 0 | 2 | 0 | 0 | 1 | 0 | 98 | 49 | NA | NA | NA | NA | 1 | NA | 0.5 | NA | 0.5 |
| SE | 1035 | ELMO1 | ELMO1 | chr7 | - | 37216644 | 37216695 | 37213334 | 37213457 | 37222614 | 37222693 | 0 | 0 | 6 | 0 | 0 | 0 | 3 | 3 | 0 | 0 | 1 | 0 | 98 | 49 | NA | NA | NA | NA | 1 | 1 | 0.6 | NA | 0.2 |
| SE | 1047 | KBTBD2 | KBTBD2 | chr7 | - | 32879434 | 32879942 | 32874991 | 32875157 | 32891535 | 32891856 | 0 | 0 | 6 | 0 | 0 | 0 | 0 | 7 | 0 | 0 | 1 | 0 | 98 | 49 | NA | NA | NA | NA | 1 | NA | 0.778 | NA | 0.222 |
| SE | 1076 | CCDC126 | CCDC126 | chr7 | + | 23597966 | 23598051 | 23597378 | 23597605 | 23611170 | 23611553 | 0 | 0 | 0 | 0 | 0 | 1 | 0 | 1 | 0 | 0 | 0 | 0 | 98 | 49 | NA | NA | NA | NA | 0 | NA | 1 | NA | -1 |
| SE | 1090 | HDAC9 | HDAC9 | chr7 | + | 18629349 | 18629481 | 18593907 | 18594029 | 18634626 | 18634742 | 0 | 0 | 1 | 0 | 0 | 1 | 0 | 1 | 0 | 0 | 1 | 0 | 98 | 49 | NA | NA | NA | NA | 0.333 | NA | 1 | NA | -0.667 |
| SE | 1093 | SNX13 | SNX13 | chr7 | - | 17897333 | 17897446 | 17893331 | 17893434 | 17940283 | 17940508 | 0 | 0 | 2 | 0 | 0 | 0 | 1 | 0 | 0 | 0 | 1 | 0 | 98 | 49 | NA | NA | NA | NA | 1 | 1 | 0 | NA | 0.5 |
| SE | 1104 | BRAT1 | BRAT1 | chr7 | - | 2544908 | 2545056 | 2543589 | 2543962 | 2547323 | 2547478 | 0 | 0 | 2 | 0 | 0 | 2 | 0 | 2 | 0 | 0 | 0 | 0 | 98 | 49 | NA | NA | NA | NA | 0.333 | NA | 1 | NA | -0.667 |
| SE | 1116 | SUN1 | SUN1 | chr7 | + | 838797 | 838986 | 832504 | 832601 | 841945 | 842130 | 0 | 0 | 5 | 0 | 0 | 0 | 0 | 0 | 0 | 0 | 1 | 0 | 98 | 49 | NA | NA | NA | NA | 1 | NA | 0 | NA | 1 |
| SE | 1157 | IPCEF1 | IPCEF1 | chr6 | - | 154212769 | 154212855 | 154199667 | 154200040 | 154214217 | 154214276 | 0 | 0 | 2 | 0 | 0 | 0 | 0 | 4 | 0 | 0 | 1 | 0 | 98 | 49 | NA | NA | NA | NA | 1 | NA | 0.667 | NA | 0.333 |
| SE | 1210 | VNN2 | VNN2 | chr6 | - | 132752672 | 132752749 | 132751144 | 132751518 | 132753417 | 132753491 | 0 | 0 | 3 | 0 | 0 | 3 | 0 | 1 | 0 | 0 | 0 | 0 | 98 | 49 | NA | NA | NA | NA | 0.333 | NA | 1 | NA | -0.667 |
| SE | 1211 | VNN2 | VNN2 | chr6 | - | 132752672 | 132752749 | 132751144 | 132751518 | 132755842 | 132756035 | 0 | 0 | 10 | 0 | 0 | 0 | 0 | 3 | 0 | 0 | 5 | 0 | 98 | 49 | NA | NA | NA | NA | 1 | NA | 0.231 | NA | 0.769 |
| SE | 1302 | SLC17A5 | SLC17A5 | chr6 | - | 73638411 | 73638499 | 73636620 | 73636707 | 73641690 | 73641924 | 0 | 0 | 0 | 0 | 0 | 1 | 0 | 1 | 0 | 0 | 0 | 0 | 98 | 49 | NA | NA | NA | NA | 0 | NA | 1 | NA | -1 |
| SE | 1305 | SMAP1 | SMAP1 | chr6 | + | 70791688 | 70791769 | 70773349 | 70773425 | 70798656 | 70798737 | 0 | 0 | 1 | 0 | 0 | 0 | 0 | 0 | 0 | 0 | 1 | 0 | 98 | 49 | NA | NA | NA | NA | 1 | NA | 0 | NA | 1 |
| SE | 1314 | RUNX2 | RUNX2 | chr6 | + | 45491940 | 45492114 | 45437946 | 45438051 | 45512245 | 45512407 | 0 | 0 | 1 | 0 | 0 | 2 | 0 | 1 | 0 | 0 | 0 | 0 | 98 | 49 | NA | NA | NA | NA | 0.2 | NA | 1 | NA | -0.8 |
| SE | 1317 | CDC5L | CDC5L | chr6 | + | 44406322 | 44406467 | 44403808 | 44404027 | 44408443 | 44408632 | 0 | 0 | 1 | 0 | 0 | 0 | 0 | 3 | 0 | 0 | 1 | 0 | 98 | 49 | NA | NA | NA | NA | 1 | NA | 0.6 | NA | 0.4 |
| SE | 1323 | YIPF3 | YIPF3 | chr6 | - | 43515594 | 43515701 | 43513587 | 43513633 | 43515888 | 43516095 | 0 | 0 | 5 | 0 | 0 | 1 | 0 | 8 | 0 | 0 | 0 | 0 | 98 | 49 | NA | NA | NA | NA | 0.714 | NA | 1 | NA | -0.286 |
| SE | 1329 | TRERF1 | TRERF1 | chr6 | - | 42263319 | 42263568 | 42259338 | 42259723 | 42264703 | 42264854 | 0 | 0 | 1 | 0 | 1 | 0 | 0 | 0 | 0 | 3 | 0 | 0 | 98 | 49 | NA | NA | NA | 0 | 1 | 0 | NA | NA | 0.5 |
| SE | 1338 | CMTR1 | CMTR1 | chr6 | + | 37461969 | 37462102 | 37461548 | 37461645 | 37462828 | 37463008 | 0 | 0 | 3 | 0 | 0 | 1 | 0 | 2 | 0 | 0 | 0 | 0 | 98 | 49 | NA | NA | NA | NA | 0.6 | NA | 1 | NA | -0.4 |
| SE | 1340 | CMTR1 | CMTR1 | chr6 | + | 37476125 | 37476194 | 37475320 | 37475412 | 37477591 | 37477639 | 0 | 0 | 3 | 0 | 0 | 0 | 0 | 3 | 0 | 0 | 1 | 0 | 98 | 49 | NA | NA | NA | NA | 1 | NA | 0.6 | NA | 0.4 |
| SE | 1341 | KCTD20 | KCTD20 | chr6 | + | 36470051 | 36470257 | 36442766 | 36443111 | 36479590 | 36479711 | 0 | 0 | 1 | 0 | 0 | 0 | 1 | 2 | 0 | 0 | 1 | 0 | 98 | 49 | NA | NA | NA | NA | 1 | 1 | 0.5 | NA | 0.25 |
| SE | 1343 | KCTD20 | KCTD20 | chr6 | + | 36474788 | 36475062 | 36470051 | 36470257 | 36479120 | 36479223 | 0 | 0 | 2 | 0 | 0 | 4 | 0 | 1 | 0 | 0 | 0 | 0 | 98 | 49 | NA | NA | NA | NA | 0.2 | NA | 1 | NA | -0.8 |
| SE | 1344 | KCTD20 | KCTD20 | chr6 | + | 36479120 | 36479223 | 36470051 | 36470257 | 36479590 | 36479711 | 0 | 0 | 6 | 0 | 0 | 0 | 0 | 0 | 0 | 0 | 2 | 0 | 98 | 49 | NA | NA | NA | NA | 1 | NA | 0 | NA | 1 |
| SE | 1358 | FKBP5 | FKBP5 | chr6 | - | 35587033 | 35587117 | 35580035 | 35580221 | 35591129 | 35591220 | 0 | 0 | 12 | 0 | 0 | 1 | 4 | 0 | 0 | 0 | 0 | 0 | 98 | 49 | NA | NA | NA | NA | 0.857 | 1 | NA | NA | -0.143 |
| SE | 1372 | HLA-DMB | HLA-DMB | chr6 | - | 32935535 | 32935652 | 32935341 | 32935377 | 32937171 | 32937456 | 0 | 0 | 4 | 0 | 0 | 1 | 0 | 2 | 0 | 0 | 0 | 0 | 98 | 49 | NA | NA | NA | NA | 0.667 | NA | 1 | NA | -0.333 |
| SE | 1445 | BAG6 | BAG6 | chr6 | - | 31639499 | 31639646 | 31639027 | 31639226 | 31640198 | 31640306 | 0 | 0 | 4 | 0 | 0 | 2 | 0 | 2 | 0 | 0 | 2 | 0 | 98 | 49 | NA | NA | NA | NA | 0.5 | NA | 0.333 | NA | 0.167 |
| SE | 1450 | BAG6 | BAG6 | chr6 | - | 31647590 | 31647844 | 31646393 | 31646523 | 31648676 | 31648751 | 0 | 1 | 0 | 0 | 0 | 0 | 0 | 1 | 0 | 0 | 0 | 0 | 98 | 49 | NA | NA | 0 | NA | NA | NA | 1 | NA | -1 |
| SE | 1456 | ATP6V1G2-DDX39B | ATP6V1G2-DDX39B | chr6 | - | 31538762 | 31538855 | 31536499 | 31536683 | 31540321 | 31540664 | 0 | 0 | 1 | 0 | 0 | 0 | 0 | 2 | 0 | 0 | 3 | 0 | 98 | 49 | NA | NA | NA | NA | 1 | NA | 0.25 | NA | 0.75 |
| SE | 1481 | FLOT1 | FLOT1 | chr6 | - | 30741791 | 30741867 | 30741189 | 30741333 | 30742146 | 30742203 | 0 | 0 | 2 | 0 | 0 | 2 | 0 | 2 | 0 | 1 | 0 | 0 | 98 | 49 | NA | NA | NA | NA | 0.333 | 0 | 1 | NA | -0.167 |
| SE | 1579 | USP9X | USP9X | chrX | + | 41150920 | 41151057 | 41148368 | 41148575 | 41152947 | 41153081 | 0 | 0 | 10 | 0 | 0 | 0 | 0 | 3 | 0 | 0 | 1 | 0 | 98 | 49 | NA | NA | NA | NA | 1 | NA | 0.6 | NA | 0.4 |
| SE | 1580 | USP9X | USP9X | chrX | + | 41214567 | 41214709 | 41210508 | 41210682 | 41215898 | 41216652 | 0 | 0 | 2 | 0 | 0 | 0 | 0 | 3 | 0 | 0 | 1 | 0 | 98 | 49 | NA | NA | NA | NA | 1 | NA | 0.6 | NA | 0.4 |
| SE | 1581 | USP9X | USP9X | chrX | + | 41219101 | 41219231 | 41218371 | 41218597 | 41223216 | 41223402 | 0 | 0 | 1 | 0 | 0 | 0 | 0 | 6 | 0 | 0 | 1 | 0 | 98 | 49 | NA | NA | NA | NA | 1 | NA | 0.75 | NA | 0.25 |
| SE | 1587 | DEK | DEK | chr6 | - | 18255730 | 18255851 | 18249650 | 18249839 | 18256360 | 18256455 | 0 | 0 | 8 | 0 | 0 | 0 | 0 | 2 | 0 | 0 | 2 | 0 | 98 | 49 | NA | NA | NA | NA | 1 | NA | 0.333 | NA | 0.667 |
| SE | 1594 | KIF13A | KIF13A | chr6 | - | 17817019 | 17817233 | 17808767 | 17808930 | 17825767 | 17825934 | 0 | 0 | 5 | 0 | 0 | 1 | 0 | 3 | 0 | 0 | 0 | 0 | 98 | 49 | NA | NA | NA | NA | 0.714 | NA | 1 | NA | -0.286 |
| SE | 1595 | KIF13A | KIF13A | chr6 | - | 17855436 | 17855617 | 17851954 | 17852042 | 17856029 | 17856122 | 0 | 0 | 9 | 0 | 0 | 1 | 0 | 5 | 0 | 0 | 0 | 0 | 98 | 49 | NA | NA | NA | NA | 0.818 | NA | 1 | NA | -0.182 |
| SE | 1596 | KIF13A | KIF13A | chr6 | - | 17898167 | 17898180 | 17873376 | 17873437 | 17987053 | 17987144 | 0 | 0 | 6 | 0 | 0 | 1 | 1 | 1 | 0 | 0 | 0 | 0 | 62 | 49 | NA | NA | NA | NA | 0.826 | 1 | 1 | NA | -0.174 |
| SE | 1597 | NUP153 | NUP153 | chr6 | - | 17625807 | 17626164 | 17624560 | 17624833 | 17628654 | 17629539 | 0 | 0 | 5 | 0 | 0 | 0 | 1 | 2 | 0 | 0 | 1 | 0 | 98 | 49 | NA | NA | NA | NA | 1 | 1 | 0.5 | NA | 0.25 |
| SE | 1600 | NUP153 | NUP153 | chr6 | - | 17661652 | 17661779 | 17649162 | 17649300 | 17662017 | 17662070 | 0 | 0 | 7 | 0 | 0 | 0 | 0 | 9 | 0 | 0 | 1 | 0 | 98 | 49 | NA | NA | NA | NA | 1 | NA | 0.818 | NA | 0.182 |
| SE | 1608 | JARID2 | JARID2 | chr6 | + | 15452005 | 15452175 | 15410223 | 15410365 | 15468541 | 15468718 | 0 | 0 | 10 | 0 | 0 | 0 | 0 | 3 | 0 | 0 | 1 | 0 | 98 | 49 | NA | NA | NA | NA | 1 | NA | 0.6 | NA | 0.4 |
| SE | 1616 | HIVEP1 | HIVEP1 | chr6 | + | 12129758 | 12129892 | 12119889 | 12125870 | 12130766 | 12130942 | 0 | 0 | 3 | 0 | 0 | 1 | 0 | 3 | 0 | 0 | 0 | 0 | 98 | 49 | NA | NA | NA | NA | 0.6 | NA | 1 | NA | -0.4 |
| SE | 1617 | HIVEP1 | HIVEP1 | chr6 | + | 12135790 | 12135892 | 12130766 | 12130942 | 12161438 | 12161929 | 0 | 0 | 1 | 0 | 0 | 0 | 0 | 5 | 0 | 0 | 1 | 0 | 98 | 49 | NA | NA | NA | NA | 1 | NA | 0.714 | NA | 0.286 |
| SE | 1640 | RIPK1 | RIPK1 | chr6 | + | 3085258 | 3085408 | 3083084 | 3083313 | 3089580 | 3089657 | 0 | 0 | 0 | 0 | 0 | 1 | 0 | 1 | 0 | 0 | 0 | 0 | 98 | 49 | NA | NA | NA | NA | 0 | NA | 1 | NA | -1 |
| SE | 1652 | HNRNPH1 | HNRNPH1 | chr5 | - | 179615545 | 179615595 | 179614177 | 179614959 | 179616125 | 179616218 | 0 | 0 | 4 | 0 | 0 | 0 | 0 | 8 | 0 | 0 | 1 | 0 | 98 | 49 | NA | NA | NA | NA | 1 | NA | 0.8 | NA | 0.2 |
| SE | 1653 | HNRNPH1 | HNRNPH1 | chr5 | - | 179619268 | 179619407 | 179618144 | 179618323 | 179620891 | 179621035 | 0 | 0 | 3 | 0 | 0 | 0 | 0 | 1 | 0 | 0 | 4 | 0 | 98 | 49 | NA | NA | NA | NA | 1 | NA | 0.111 | NA | 0.889 |
| SE | 1692 | DOCK2 | DOCK2 | chr5 | + | 169983067 | 169983166 | 169840756 | 169840852 | 169985827 | 169985922 | 0 | 0 | 12 | 0 | 0 | 1 | 0 | 5 | 0 | 0 | 0 | 0 | 98 | 49 | NA | NA | NA | NA | 0.857 | NA | 1 | NA | -0.143 |
| SE | 1697 | RNF145 | RNF145 | chr5 | - | 159181959 | 159182051 | 159176631 | 159176867 | 159194715 | 159194824 | 0 | 0 | 4 | 0 | 0 | 0 | 0 | 0 | 0 | 0 | 1 | 0 | 98 | 49 | NA | NA | NA | NA | 1 | NA | 0 | NA | 1 |
| SE | 1701 | CNOT8 | CNOT8 | chr5 | + | 154863206 | 154863395 | 154858518 | 154858768 | 154865191 | 154865385 | 0 | 0 | 2 | 0 | 0 | 0 | 0 | 1 | 0 | 0 | 1 | 0 | 98 | 49 | NA | NA | NA | NA | 1 | NA | 0.333 | NA | 0.667 |
| SE | 1703 | CNOT8 | CNOT8 | chr5 | + | 154865191 | 154865385 | 154858518 | 154858768 | 154870660 | 154870822 | 0 | 0 | 0 | 0 | 0 | 1 | 0 | 1 | 0 | 0 | 0 | 0 | 98 | 49 | NA | NA | NA | NA | 0 | NA | 1 | NA | -1 |
| SE | 1707 | LARP1 | LARP1 | chr5 | + | 154793799 | 154794000 | 154793594 | 154793723 | 154794099 | 154794262 | 0 | 0 | 1 | 0 | 0 | 0 | 0 | 0 | 0 | 0 | 2 | 0 | 98 | 49 | NA | NA | NA | NA | 1 | NA | 0 | NA | 1 |

| | | | | | | | | | | | | | | | | | | | | | | | | | | | | | | | | | | | |
|---|---|---|---|---|---|---|---|---|---|---|---|---|---|---|---|---|---|---|---|---|---|---|---|---|---|---|---|---|---|---|---|---|---|---|---|
| SE | 1744 | NDFIP1 | NDFIP1 | chr5 | + | 142137733 | 142137858 | 142135729 | 142135817 | 142140562 | 142140629 | 0 | 0 | 4 | 0 | 0 | 1 | 0 | 3 | 0 | 0 | 0 | 0 | 98 | 49 | NA | NA | NA | NA | 0.667 | NA | | 1 | NA | -0.333 |
| SE | 1748 | DIAPH1 | DIAPH1 | chr5 | - | 141529602 | 141529697 | 141529171 | 141529273 | 141534334 | 141534433 | 0 | 0 | 8 | 0 | 0 | 0 | 0 | 1 | 0 | 0 | 1 | 0 | 98 | 49 | NA | NA | NA | NA | 1 | NA | | 0.333 | NA | 0.667 |
| SE | 1760 | KDM3B | KDM3B | chr5 | + | 138386021 | 138386621 | 138381515 | 138381590 | 138391012 | 138392261 | 0 | 0 | 2 | 0 | 0 | 2 | 0 | 2 | 0 | 0 | 0 | 0 | 98 | 49 | NA | NA | NA | NA | 0.333 | NA | | 1 | NA | -0.667 |
| SE | 1769 | CAMLG | CAMLG | chr5 | + | 134741062 | 134741523 | 134738479 | 134738792 | 134743986 | 134744052 | 0 | 0 | 0 | 0 | 0 | 1 | 0 | 2 | 0 | 0 | 0 | 0 | 98 | 49 | NA | NA | NA | NA | 0 | NA | | 1 | NA | -1 |
| SE | 1799 | AP3S1 | AP3S1 | chr5 | + | 115870016 | 115870128 | 115866669 | 115866761 | 115895086 | 115895158 | 0 | 0 | 1 | 0 | 0 | 0 | 0 | 4 | 0 | 0 | 1 | 0 | 98 | 49 | NA | NA | NA | NA | 1 | NA | | 0.667 | NA | 0.333 |
| SE | 1808 | ASAP1 | ASAP1 | chr8 | - | 130168991 | 130169067 | 130167535 | 130167622 | 130179263 | 130179349 | 0 | 0 | 15 | 0 | 0 | 0 | 0 | 11 | 0 | 0 | 1 | 0 | 98 | 49 | NA | NA | NA | NA | 1 | NA | | 0.846 | NA | 0.154 |
| SE | 1815 | TBL1X | TBL1X | chrX | + | 9653544 | 9653689 | 9640272 | 9640360 | 9654214 | 9654322 | 0 | 0 | 1 | 0 | 0 | 3 | 0 | 0 | 0 | 0 | 3 | 0 | 98 | 49 | NA | NA | NA | NA | 0.143 | NA | | 0 | NA | 0.143 |
| SE | 1816 | CHD1 | CHD1 | chr5 | - | 98858179 | 98858390 | 98855203 | 98856725 | 98858963 | 98859015 | 0 | 0 | 2 | 0 | 0 | 1 | 0 | 19 | 0 | 0 | 5 | 0 | 98 | 49 | NA | NA | NA | NA | 0.5 | NA | | 0.655 | NA | -0.155 |
| SE | 1820 | ERAP1 | ERAP1 | chr5 | - | 96788530 | 96788685 | 96786469 | 96786549 | 96790295 | 96790367 | 0 | 0 | 5 | 0 | 0 | 0 | 0 | 2 | 0 | 0 | 1 | 0 | 98 | 49 | NA | NA | NA | NA | 1 | NA | | 0.5 | NA | 0.5 |
| SE | 1824 | MCTP1 | MCTP1 | chr5 | - | 94799012 | 94799132 | 94779109 | 94779163 | 94868332 | 94868452 | 0 | 0 | 1 | 0 | 0 | 1 | 0 | 3 | 0 | 0 | 0 | 0 | 98 | 49 | NA | NA | NA | NA | 0.333 | NA | | 1 | NA | -0.667 |
| SE | 1825 | MCTP1 | MCTP1 | chr5 | - | 94870871 | 94870973 | 94870416 | 94870491 | 94871314 | 94871417 | 0 | 0 | 2 | 0 | 0 | 0 | 0 | 2 | 0 | 0 | 1 | 0 | 98 | 49 | NA | NA | NA | NA | 1 | NA | | 0.5 | NA | 0.5 |
| SE | 1828 | MCTP1 | MCTP1 | chr5 | - | 94923961 | 94924021 | 94912805 | 94912976 | 94940083 | 94940195 | 0 | 0 | 1 | 0 | 0 | 2 | 0 | 4 | 0 | 0 | 0 | 0 | 98 | 49 | NA | NA | NA | NA | 0.2 | NA | | 1 | NA | -0.8 |
| SE | 1829 | MCTP1 | MCTP1 | chr5 | - | 94931952 | 94931991 | 94912805 | 94912976 | 94940083 | 94940195 | 0 | 0 | 6 | 0 | 0 | 2 | 0 | 4 | 0 | 0 | 0 | 0 | 88 | 49 | NA | NA | NA | NA | 0.626 | NA | | 1 | NA | -0.374 |
| SE | 1830 | MCTP1 | MCTP1 | chr5 | - | 94931952 | 94931991 | 94923961 | 94924021 | 94940083 | 94940195 | 0 | 0 | 9 | 0 | 0 | 1 | 0 | 4 | 0 | 0 | 0 | 0 | 88 | 49 | NA | NA | NA | NA | 0.834 | NA | | 1 | NA | -0.166 |
| SE | 1831 | FAM172A | FAM172A | chr5 | - | 93958776 | 93958968 | 93881469 | 93881688 | 93964457 | 93964523 | 0 | 0 | 5 | 0 | 0 | 2 | 0 | 4 | 0 | 0 | 0 | 0 | 98 | 49 | NA | NA | NA | NA | 0.556 | NA | | 1 | NA | -0.444 |
| SE | 1832 | FAM172A | FAM172A | chr5 | - | 94050730 | 94050831 | 93964457 | 93964523 | 94053125 | 94053227 | 0 | 0 | 3 | 0 | 0 | 1 | 0 | 5 | 0 | 0 | 0 | 0 | 98 | 49 | NA | NA | NA | NA | 0.6 | NA | | 1 | NA | -0.4 |
| SE | 1833 | FAM172A | FAM172A | chr5 | - | 94074645 | 94074769 | 94053125 | 94053227 | 94111510 | 94111699 | 0 | 0 | 2 | 0 | 0 | 1 | 0 | 1 | 0 | 0 | 0 | 0 | 98 | 49 | NA | NA | NA | NA | 0.5 | NA | | 1 | NA | -0.5 |
| SE | 1841 | VCAN | VCAN | chr5 | + | 83519348 | 83522309 | 83512102 | 83512396 | 83537006 | 83542268 | 0 | 0 | 1 | 0 | 0 | 19 | 0 | 0 | 0 | 0 | 3 | 0 | 98 | 49 | NA | NA | NA | NA | 0.026 | NA | | 0 | NA | 0.026 |
| SE | 1847 | SSBP2 | SSBP2 | chr5 | - | 81489249 | 81489309 | 81474495 | 81474562 | 81513627 | 81513717 | 0 | 0 | 0 | 0 | 0 | 1 | 1 | 0 | 0 | 0 | 0 | 0 | 98 | 49 | NA | NA | NA | NA | 0 | | 1 | NA | NA | -1 |
| SE | 1848 | SSBP2 | SSBP2 | chr5 | - | 81513627 | 81513717 | 81489249 | 81489309 | 81615472 | 81615557 | 0 | 0 | 4 | 0 | 0 | 3 | 1 | 0 | 0 | 0 | 0 | 0 | 98 | 49 | NA | NA | NA | NA | 0.4 | | 1 | NA | NA | -0.6 |
| SE | 1850 | PAPD4 | PAPD4 | chr5 | + | 79642831 | 79642910 | 79641104 | 79641196 | 79645122 | 79645192 | 0 | 0 | 4 | 0 | 0 | 1 | 0 | 1 | 0 | 0 | 0 | 0 | 98 | 49 | NA | NA | NA | NA | 0.667 | NA | | 1 | NA | -0.333 |
| SE | 1851 | PAPD4 | PAPD4 | chr5 | + | 79642843 | 79642910 | 79641104 | 79641196 | 79645122 | 79645192 | 0 | 0 | 1 | 0 | 0 | 1 | 0 | 1 | 0 | 0 | 0 | 0 | 98 | 49 | NA | NA | NA | NA | 0.333 | NA | | 1 | NA | -0.667 |
| SE | 1852 | PAPD4 | PAPD4 | chr5 | + | 79649061 | 79649190 | 79648616 | 79648693 | 79656957 | 79657001 | 0 | 0 | 0 | 0 | 0 | 1 | 0 | 1 | 0 | 0 | 0 | 0 | 98 | 49 | NA | NA | NA | NA | 0 | NA | | 1 | NA | -1 |
| SE | 1904 | MAP3K1 | MAP3K1 | chr5 | + | 56887377 | 56887520 | 56885931 | 56886063 | 56888225 | 56888357 | 0 | 0 | 11 | 0 | 0 | 0 | 0 | 4 | 0 | 0 | 1 | 0 | 98 | 49 | NA | NA | NA | NA | 1 | NA | | 0.667 | NA | 0.333 |
| SE | 1918 | PARP8 | PARP8 | chr5 | + | 50760291 | 50760362 | 50759642 | 50759732 | 50761820 | 50761898 | 0 | 0 | 17 | 0 | 0 | 0 | 0 | 7 | 0 | 6 | 1 | 0 | 98 | 49 | NA | NA | NA | NA | 1 | | 0 | 0.778 | NA | 0.611 |
| SE | 1933 | PRKAA1 | PRKAA1 | chr5 | - | 40769415 | 40769503 | 40767465 | 40767690 | 40771718 | 40771863 | 0 | 0 | 4 | 0 | 0 | 0 | 0 | 6 | 0 | 0 | 1 | 0 | 98 | 49 | NA | NA | NA | NA | 1 | NA | | 0.75 | NA | 0.25 |
| SE | 1935 | RICTOR | RICTOR | chr5 | - | 38949355 | 38949427 | 38947263 | 38947441 | 38949711 | 38950720 | 0 | 0 | 0 | 0 | 0 | 11 | 0 | 1 | 0 | 0 | 3 | 0 | 98 | 49 | NA | NA | NA | NA | 0 | NA | | 0.143 | NA | -0.143 |
| SE | 1936 | RICTOR | RICTOR | chr5 | - | 38975536 | 38975604 | 38971876 | 38971959 | 38978582 | 38978648 | 0 | 0 | 12 | 0 | 0 | 1 | 0 | 16 | 0 | 0 | 0 | 0 | 98 | 49 | NA | NA | NA | NA | 0.857 | NA | | 1 | NA | -0.143 |
| SE | 1946 | ZFR | ZFR | chr5 | - | 32380074 | 32380172 | 32379114 | 32379210 | 32385507 | 32385649 | 0 | 0 | 4 | 0 | 0 | 2 | 0 | 5 | 0 | 0 | 0 | 0 | 98 | 49 | NA | NA | NA | NA | 0.5 | NA | | 1 | NA | -0.5 |
| SE | 1952 | 6-Mar | 6-Mar | chr5 | + | 10390331 | 10390500 | 10386993 | 10387066 | 10391541 | 10391731 | 0 | 0 | 6 | 0 | 0 | 2 | 0 | 1 | 0 | 0 | 0 | 0 | 98 | 49 | NA | NA | NA | NA | 0.6 | NA | | 1 | NA | -0.4 |
| SE | 1953 | 6-Mar | 6-Mar | chr5 | + | 10426389 | 10426522 | 10423734 | 10423824 | 10429892 | 10430028 | 0 | 0 | 0 | 0 | 0 | 1 | 0 | 3 | 0 | 0 | 2 | 0 | 98 | 49 | NA | NA | NA | NA | 0 | NA | | 0.429 | NA | -0.429 |
| SE | 1984 | DDX60L | DDX60L | chr4 | - | 168453123 | 168453282 | 168441336 | 168441492 | 168457891 | 168458008 | 0 | 0 | 1 | 0 | 0 | 1 | 0 | 2 | 0 | 0 | 0 | 0 | 98 | 49 | NA | NA | NA | NA | 0.333 | NA | | 1 | NA | -0.667 |
| SE | 1985 | DDX60L | DDX60L | chr4 | - | 168456038 | 168456152 | 168453123 | 168453282 | 168457891 | 168458008 | 0 | 0 | 35 | 0 | 0 | 0 | 0 | 10 | 0 | 0 | 1 | 0 | 98 | 49 | NA | NA | NA | NA | 1 | NA | | 0.833 | NA | 0.167 |
| SE | 2000 | UBQLN1 | UBQLN1 | chr9 | - | 83666349 | 83666433 | 83665029 | 83665145 | 83669184 | 83669327 | 0 | 0 | 4 | 0 | 0 | 1 | 0 | 5 | 0 | 0 | 0 | 0 | 98 | 49 | NA | NA | NA | NA | 0.667 | NA | | 1 | NA | -0.333 |
| SE | 2008 | ARFIP1 | ARFIP1 | chr4 | + | 152882722 | 152882880 | 152880962 | 152881184 | 152888132 | 152888307 | 0 | 0 | 1 | 0 | 0 | 0 | 0 | 0 | 0 | 0 | 1 | 0 | 98 | 49 | NA | NA | NA | NA | 1 | NA | | 0 | NA | 1 |
| SE | 2057 | TET2 | TET2 | chr4 | + | 105190359 | 105190505 | 105146684 | 105146979 | 105233896 | 105237351 | 0 | 0 | 6 | 0 | 0 | 0 | 0 | 7 | 0 | 0 | 1 | 0 | 98 | 49 | NA | NA | NA | NA | 1 | NA | | 0.778 | NA | 0.222 |
| SE | 2058 | TET2 | TET2 | chr4 | + | 105241338 | 105241429 | 105233896 | 105237351 | 105242833 | 105242927 | 0 | 0 | 6 | 0 | 0 | 2 | 0 | 9 | 0 | 0 | 1 | 0 | 98 | 49 | NA | NA | NA | NA | 0.6 | NA | | 0.818 | NA | -0.218 |
| SE | 2065 | DAPP1 | DAPP1 | chr4 | + | 99866556 | 99866618 | 99866033 | 99866121 | 99868116 | 99870189 | 0 | 0 | 3 | 0 | 0 | 6 | 0 | 0 | 0 | 0 | 6 | 0 | 98 | 49 | NA | NA | NA | NA | 0.2 | NA | | 0 | NA | 0.2 |
| SE | 2079 | HERC5 | HERC5 | chr4 | + | 88486114 | 88486228 | 88479352 | 88479507 | 88489165 | 88489336 | 0 | 0 | 8 | 0 | 0 | 1 | 0 | 9 | 0 | 0 | 2 | 0 | 98 | 49 | NA | NA | NA | NA | 0.8 | NA | | 0.692 | NA | 0.108 |
| SE | 2083 | HERC6 | HERC6 | chr4 | + | 88417424 | 88417579 | 88413076 | 88413266 | 88423859 | 88423973 | 0 | 0 | 6 | 0 | 0 | 0 | 0 | 2 | 0 | 0 | 2 | 0 | 98 | 49 | NA | NA | NA | NA | 1 | NA | | 0.333 | NA | 0.667 |
| SE | 2097 | WDFY3 | WDFY3 | chr4 | - | 84690505 | 84690664 | 84688085 | 84688265 | 84691630 | 84691785 | 0 | 0 | 5 | 0 | 0 | 0 | 0 | 1 | 0 | 0 | 1 | 0 | 98 | 49 | NA | NA | NA | NA | 1 | NA | | 0.333 | NA | 0.667 |
| SE | 2098 | WDFY3 | WDFY3 | chr4 | - | 84705393 | 84705511 | 84704337 | 84704444 | 84708908 | 84709028 | 0 | 0 | 11 | 0 | 0 | 1 | 2 | 10 | 0 | 0 | 0 | 0 | 98 | 49 | NA | NA | NA | NA | 0.846 | | 1 | 1 | NA | -0.154 |
| SE | 2100 | WDFY3 | WDFY3 | chr4 | - | 84733381 | 84733609 | 84724425 | 84724594 | 84735042 | 84735120 | 0 | 0 | 9 | 0 | 0 | 0 | 0 | 6 | 0 | 0 | 1 | 0 | 98 | 49 | NA | NA | NA | NA | 1 | NA | | 0.75 | NA | 0.25 |
| SE | 2113 | PPP2CA | PPP2CA | chr5 | - | 134200334 | 134200496 | 134199085 | 134199204 | 134200984 | 134201074 | 0 | 0 | 4 | 0 | 0 | 3 | 2 | 1 | 0 | 0 | 1 | 0 | 98 | 49 | NA | NA | NA | NA | 0.4 | | 1 | 0.333 | NA | -0.266 |
| SE | 2117 | HNRNPD | HNRNPD | chr4 | - | 82356536 | 82356683 | 82355303 | 82355401 | 82356795 | 82356895 | 0 | 0 | 1 | 0 | 0 | 0 | 0 | 0 | 0 | 0 | 1 | 0 | 98 | 49 | NA | NA | NA | NA | 1 | NA | | 0 | NA | 1 |
| SE | 2130 | G3BP2 | G3BP2 | chr4 | - | 75653982 | 75654081 | 75648638 | 75648741 | 75655065 | 75655246 | 0 | 0 | 1 | 0 | 0 | 3 | 0 | 0 | 0 | 0 | 6 | 0 | 98 | 49 | NA | NA | NA | NA | 0.143 | NA | | 0 | NA | 0.143 |
| SE | 2139 | HSPA9 | HSPA9 | chr5 | - | 138566625 | 138566718 | 138561579 | 138561789 | 138567000 | 138567163 | 0 | 0 | 2 | 0 | 0 | 0 | 0 | 4 | 0 | 0 | 2 | 0 | 98 | 49 | NA | NA | NA | NA | 1 | NA | | 0.5 | NA | 0.5 |
| SE | 2142 | ANKRD17 | ANKRD17 | chr4 | - | 73147240 | 73147432 | 73146763 | 73146873 | 73148812 | 73149050 | 0 | 0 | 10 | 0 | 0 | 0 | 0 | 0 | 0 | 0 | 2 | 0 | 98 | 49 | NA | NA | NA | NA | 1 | NA | | 0 | NA | 1 |
| SE | 2152 | UBA6 | UBA6 | chr4 | - | 67665188 | 67665292 | 67663884 | 67663947 | 67668550 | 67668674 | 0 | 0 | 7 | 0 | 0 | 0 | 0 | 1 | 0 | 0 | 1 | 0 | 98 | 49 | NA | NA | NA | NA | 1 | NA | | 0.333 | NA | 0.667 |
| SE | 2165 | EXOC1 | EXOC1 | chr4 | + | 55893551 | 55893780 | 55892634 | 55892711 | 55896716 | 55896900 | 0 | 0 | 7 | 0 | 0 | 1 | 0 | 4 | 0 | 0 | 0 | 0 | 98 | 49 | NA | NA | NA | NA | 0.778 | NA | | 1 | NA | -0.222 |
| SE | 2180 | RBM47 | RBM47 | chr4 | - | 40436440 | 40436647 | 40432650 | 40432862 | 40437770 | 40438924 | 0 | 0 | 6 | 0 | 0 | 0 | 0 | 0 | 0 | 0 | 1 | 0 | 98 | 49 | NA | NA | NA | NA | 1 | NA | | 0 | NA | 1 |
| SE | 2182 | UBE2K | UBE2K | chr4 | + | 39755656 | 39755739 | 39745751 | 39745810 | 39774833 | 39774933 | 0 | 0 | 2 | 0 | 0 | 1 | 5 | 0 | 0 | 0 | 0 | 0 | 98 | 49 | NA | NA | NA | NA | 0.5 | | 1 | NA | NA | -0.5 |
| SE | 2199 | CPEB2 | CPEB2 | chr4 | + | 15007304 | 15007586 | 15002673 | 15004335 | 15017187 | 15017278 | 0 | 0 | 4 | 0 | 0 | 1 | 0 | 4 | 0 | 0 | 0 | 0 | 98 | 49 | NA | NA | NA | NA | 0.667 | NA | | 1 | NA | -0.333 |
| SE | 2200 | CPEB2 | CPEB2 | chr4 | + | 15007304 | 15007595 | 15002673 | 15004335 | 15017187 | 15017278 | 0 | 0 | 3 | 0 | 0 | 1 | 0 | 1 | 0 | 0 | 0 | 0 | 98 | 49 | NA | NA | NA | NA | 0.6 | NA | | 1 | NA | -0.4 |
| SE | 2202 | CPEB2 | CPEB2 | chr4 | + | 15017187 | 15017278 | 15007304 | 15007586 | 15033160 | 15033211 | 0 | 0 | 5 | 0 | 0 | 0 | 0 | 7 | 0 | 0 | 1 | 0 | 98 | 49 | NA | NA | NA | NA | 1 | NA | | 0.778 | NA | 0.222 |
| SE | 2208 | IK | IK | chr5 | + | 140655828 | 140655992 | 140654680 | 140654727 | 140657553 | 140657662 | 0 | 0 | 2 | 0 | 0 | 2 | 0 | 1 | 0 | 0 | 0 | 0 | 98 | 49 | NA | NA | NA | NA | 0.333 | NA | | 1 | NA | -0.667 |
| SE | 2217 | MAN2B2 | MAN2B2 | chr4 | + | 6598197 | 6598354 | 6597112 | 6597303 | 6600622 | 6600756 | 0 | 0 | 0 | 0 | 0 | 1 | 0 | 1 | 0 | 0 | 0 | 0 | 98 | 49 | NA | NA | NA | NA | 0 | NA | | 1 | NA | -1 |
| SE | 2256 | KIAA0226 | KIAA0226 | chr3 | - | 197684156 | 197684217 | 197683306 | 197683439 | 197693714 | 197693816 | 0 | 0 | 1 | 0 | 0 | 0 | 0 | 7 | 0 | 3 | 0 | 0 | 98 | 49 | NA | NA | NA | NA | 1 | | 0 | 1 | NA | 0.5 |
| SE | 2265 | PCYT1A | PCYT1A | chr3 | - | 196283474 | 196283538 | 196270414 | 196270541 | 196287614 | 196287713 | 0 | 0 | 1 | 0 | 0 | 1 | 0 | 1 | 0 | 0 | 0 | 0 | 98 | 49 | NA | NA | NA | NA | 0.333 | NA | | 1 | NA | -0.667 |
| SE | 2276 | ACAP2 | ACAP2 | chr3 | - | 195291705 | 195291815 | 195289120 | 195289231 | 195292264 | 195292452 | 0 | 0 | 12 | 0 | 0 | 2 | 0 | 8 | 0 | 0 | 0 | 0 | 98 | 49 | NA | NA | NA | NA | 0.75 | NA | | 1 | NA | -0.25 |
| SE | 2279 | OPA1 | OPA1 | chr3 | + | 193617177 | 193617285 | 193615673 | 193615770 | 193618868 | 193618936 | 0 | 0 | 2 | 0 | 0 | 0 | 0 | 0 | 0 | 0 | 1 | 0 | 98 | 49 | NA | NA | NA | NA | 1 | NA | | 0 | NA | 1 |
| SE | 2288 | IDUA | IDUA | chr4 | + | 1000881 | 1000989 | 1000611 | 1000697 | 1001467 | 1001563 | 0 | 0 | 0 | 0 | 0 | 2 | 0 | 1 | 0 | 0 | 0 | 0 | 98 | 49 | NA | NA | NA | NA | 0 | NA | | 1 | NA | -1 |
| SE | 2304 | PPIP5K2 | PPIP5K2 | chr5 | + | 103133452 | 103133648 | 103129305 | 103129703 | 103136731 | 103136822 | 0 | 0 | 1 | 0 | 0 | 0 | 0 | 1 | 0 | 0 | 1 | 0 | 98 | 49 | NA | NA | NA | NA | 1 | NA | | 0.333 | NA | 0.667 |
| SE | 2305 | PPIP5K2 | PPIP5K2 | chr5 | + | 103176865 | 103176910 | 103173857 | 103173972 | 103177666 | 103177774 | 0 | 0 | 2 | 0 | 0 | 1 | 0 | 1 | 0 | 0 | 1 | 0 | 94 | 49 | NA | NA | NA | NA | 0.51 | NA | | 0.343 | NA | 0.167 |
| SE | 2314 | TNFAIP3 | TNFAIP3 | chr6 | + | 137880070 | 137880252 | 137878431 | 137879351 | 137881034 | 137883314 | 0 | 0 | 12 | 0 | 0 | 0 | 1 | 33 | 0 | 0 | 1 | 0 | 98 | 49 | NA | NA | NA | NA | 1 | | 1 | 0.943 | NA | 0.028 |
| SE | 2342 | ZMAT3 | ZMAT3 | chr3 | - | 179071326 | 179071447 | 179067482 | 179067809 | 179071594 | 179071868 | 0 | 0 | 0 | 0 | 0 | 1 | 0 | 1 | 0 | 0 | 0 | 0 | 98 | 49 | NA | NA | NA | NA | 0 | NA | | 1 | NA | -1 |
| SE | 2347 | FNDC3B | FNDC3B | chr3 | + | 172328951 | 172329076 | 172310827 | 172310881 | 172330540 | 172330715 | 0 | 0 | 12 | 0 | 0 | 7 | 0 | 2 | 0 | 0 | 0 | 0 | 98 | 49 | NA | NA | NA | NA | 0.462 | NA | | 1 | NA | -0.538 |
| SE | 2348 | FNDC3B | FNDC3B | chr3 | + | 172343010 | 172343116 | 172341112 | 172341231 | 172344085 | 172344258 | 0 | 0 | 10 | 0 | 0 | 0 | 0 | 2 | 0 | 0 | 1 | 0 | 98 | 49 | NA | NA | NA | NA | 1 | NA | | 0.5 | NA | 0.5 |
| SE | 2382 | MME | MME | chr3 | + | 155168491 | 155168625 | 155166901 | 155167021 | 155168731 | 155168797 | 0 | 0 | 10 | 0 | 0 | 3 | 1 | 10 | 0 | 0 | 2 | 0 | 98 | 49 | NA | NA | NA | NA | 0.625 | | 1 | 0.714 | NA | -0.232 |
| SE | 2383 | DHX36 | DHX36 | chr3 | - | 154306215 | 154306295 | 154305093 | 154305168 | 154309652 | 154309823 | 0 | 0 | 1 | 0 | 0 | 1 | 0 | 3 | 0 | 0 | 0 | 0 | 98 | 49 | NA | NA | NA | NA | 0.333 | NA | | 1 | NA | -0.667 |
| SE | 2386 | TSC22D2 | TSC22D2 | chr3 | + | 150423048 | 150423120 | 150408334 | 150411308 | 150457075 | 150457127 | 0 | 0 | 4 | 0 | 0 | 1 | 0 | 0 | 0 | 0 | 1 | 0 | 98 | 49 | NA | NA | NA | NA | 0.667 | NA | | 0 | NA | 0.667 |
| SE | 2394 | IMPA1 | IMPA1 | chr8 | - | 81670938 | 81671047 | 81660515 | 81660667 | 81673840 | 81673949 | 0 | 0 | 2 | 0 | 0 | 0 | 0 | 0 | 0 | 0 | 1 | 0 | 98 | 49 | NA | NA | NA | NA | 1 | NA | | 0 | NA | 1 |
| SE | 2401 | PLSCR1 | PLSCR1 | chr3 | - | 146528613 | 146528831 | 146525604 | 146525647 | 146533469 | 146533550 | 0 | 0 | 8 | 0 | 0 | 2 | 0 | 2 | 0 | 0 | 3 | 0 | 98 | 49 | NA | NA | NA | NA | 0.667 | NA | | 0.25 | NA | 0.417 |
| SE | 2413 | DLD | DLD | chr7 | + | 107902324 | 107902393 | 107901737 | 107901817 | 107903477 | 107903547 | 0 | 0 | 7 | 0 | 0 | 0 | 0 | 0 | 0 | 0 | 1 | 0 | 98 | 49 | NA | NA | NA | NA | 1 | NA | | 0 | NA | 1 |
| SE | 2418 | PIK3CB | PIK3CB | chr3 | - | 138663905 | 138664029 | 138657689 | 138657835 | 138665035 | 138665203 | 0 | 0 | 5 | 0 | 0 | 0 | 0 | 0 | 0 | 0 | 1 | 0 | 98 | 49 | NA | NA | NA | NA | 1 | NA | | 0 | NA | 1 |
| SE | 2425 | ARMC8 | ARMC8 | chr3 | + | 138223388 | 138223531 | 138221925 | 138221997 | 138223635 | 138223733 | 0 | 0 | 2 | 0 | 0 | 0 | 0 | 4 | 0 | 0 | 2 | 0 | 98 | 49 | NA | NA | NA | NA | 1 | NA | | 0.5 | NA | 0.5 |
| SE | 2430 | CEP63 | CEP63 | chr3 | + | 134547334 | 134547472 | 134546148 | 134546288 | 134549061 | 134549176 | 0 | 0 | 4 | 0 | 0 | 6 | 0 | 0 | 0 | 0 | 4 | 0 | 98 | 49 | NA | NA | NA | NA | 0.25 | NA | | 0 | NA | 0.25 |
| SE | 2433 | DNAJC13 | DNAJC13 | chr3 | + | 132495087 | 132495166 | 132494143 | 132494259 | 132496527 | 132496663 | 0 | 0 | 7 | 0 | 0 | 0 | 0 | 3 | 0 | 1 | 0 | 0 | 98 | 49 | NA | NA | NA | NA | 1 | | 0 | 1 | NA | 0.5 |
| SE | 2434 | DNAJC13 | DNAJC13 | chr3 | + | 132511066 | 132511244 | 132507236 | 132507353 | 132513007 | 132513099 | 0 | 0 | 2 | 0 | 0 | 0 | 0 | 1 | 0 | 0 | 1 | 0 | 98 | 49 | NA | NA | NA | NA | 1 | NA | | 0.333 | NA | 0.667 |
| SE | 2453 | DOCK11 | DOCK11 | chrX | + | 118590209 | 118590302 | 118588412 | 118588478 | 118593213 | 118593337 | 0 | 0 | 15 | 0 | 0 | 1 | 0 | 9 | 0 | 0 | 2 | 0 | 98 | 49 | NA | NA | NA | NA | 0.882 | NA | | 0.692 | NA | 0.19 |
| SE | 2457 | MPP1 | MPP1 | chrX | - | 154784027 | 154784108 | 154783426 | 154783507 | 154785050 | 154785157 | 0 | 0 | 10 | 0 | 0 | 0 | 0 | 2 | 0 | 0 | 1 | 0 | 98 | 49 | NA | NA | NA | NA | 1 | NA | | 0.5 | NA | 0.5 |
| SE | 2473 | GOLGB1 | GOLGB1 | chr3 | - | 121699811 | 121699885 | 121693740 | 121698929 | 121702480 | 121702595 | 0 | 0 | 2 | 0 | 0 | 1 | 0 | 7 | 0 | 0 | 0 | 0 | 98 | 49 | NA | NA | NA | NA | 0.5 | NA | | 1 | NA | -0.5 |
| SE | 2477 | GSK3B | GSK3B | chr3 | - | 119905754 | 119905852 | 119876412 | 119876508 | 119916043 | 119916174 | 0 | 0 | 2 | 0 | 0 | 0 | 0 | 1 | 0 | 0 | 1 | 0 | 98 | 49 | NA | NA | NA | NA | 1 | NA | | 0.333 | NA | 0.667 |
| SE | 2501 | IFT57 | IFT57 | chr3 | - | 108219409 | 108219572 | 108218534 | 108218653 | 108222110 | 108222570 | 0 | 0 | 2 | 0 | 0 | 0 | 0 | 1 | 0 | 0 | 1 | 0 | 98 | 49 | NA | NA | NA | NA | 1 | NA | | 0.333 | NA | 0.667 |
| SE | 2503 | CD47 | CD47 | chr3 | - | 108050577 | 108050602 | 108043093 | 108047292 | 108051938 | 108051970 | 0 | 0 | 1 | 0 | 0 | 1 | 0 | 9 | 0 | 0 | 0 | 0 | 74 | 49 | NA | NA | NA | NA | 0.398 | NA | | 1 | NA | -0.602 |
| SE | 2506 | TBC1D23 | TBC1D23 | chr3 | + | 100295301 | 100295348 | 100295086 | 100295211 | 100296171 | 100296275 | 0 | 0 | 2 | 0 | 0 | 0 | 0 | 0 | 0 | 0 | 1 | 0 | 96 | 49 | NA | NA | NA | NA | 1 | NA | | 0 | NA | 1 |
| SE | 2507 | TBC1D23 | TBC1D23 | chr3 | + | 100310402 | 100310542 | 100306436 | 100306543 | 100316098 | 100316187 | 0 | 0 | 0 | 0 | 0 | 1 | 0 | 3 | 0 | 0 | 0 | 0 | 98 | 49 | NA | NA | NA | NA | 0 | NA | | 1 | NA | -1 |
| SE | 2530 | GBE1 | GBE1 | chr3 | - | 81646391 | 81646482 | 81642780 | 81642990 | 81648855 | 81648991 | 0 | 0 | 0 | 0 | 0 | 1 | 0 | 1 | 0 | 0 | 0 | 0 | 98 | 49 | NA | NA | NA | NA | 0 | NA | | 1 | NA | -1 |
| SE | 2540 | GRIPAP1 | GRIPAP1 | chrX | - | 48990925 | 48991110 | 48990684 | 48990732 | 48993427 | 48993578 | 0 | 0 | 6 | 0 | 0 | 0 | 0 | 4 | 0 | 0 | 2 | 0 | 98 | 49 | NA | NA | NA | NA | 1 | NA | | 0.5 | NA | 0.5 |
| SE | 2552 | ATXN7 | ATXN7 | chr3 | + | 63990175 | 63990374 | 63988058 | 63988324 | 63990737 | 63990859 | 0 | 0 | 2 | 0 | 0 | 0 | 0 | 1 | 0 | 0 | 1 | 0 | 98 | 49 | NA | NA | NA | NA | 1 | NA | | 0.333 | NA | 0.667 |
| SE | 2553 | ATXN7 | ATXN7 | chr3 | + | 63997620 | 63997687 | 63995504 | 63996483 | 63999449 | 64003460 | 0 | 0 | 2 | 0 | 0 | 2 | 0 | 1 | 0 | 0 | 1 | 0 | 98 | 49 | NA | NA | NA | NA | 0.333 | NA | | 0.333 | NA | 0 |
| SE | 2569 | FLNB | FLNB | chr3 | + | 58154790 | 58154928 | 58153374 | 58153641 | 58155959 | 58156075 | 0 | 0 | 1 | 0 | 0 | 0 | 0 | 0 | 0 | 0 | 1 | 0 | 98 | 49 | NA | NA | NA | NA | 1 | NA | | 0 | NA | 1 |
| SE | 2570 | SLMAP | SLMAP | chr3 | + | 57860698 | 57860839 | 57858087 | 57858159 | 57861948 | 57862086 | 0 | 0 | 5 | 0 | 0 | 0 | 0 | 6 | 0 | 0 | 1 | 0 | 98 | 49 | NA | NA | NA | NA | 1 | NA | | 0.75 | NA | 0.25 |
| SE | 2573 | SLMAP | SLMAP | chr3 | + | 57907883 | 57908006 | 57896872 | 57896932 | 57909075 | 57909150 | 0 | 0 | 1 | 0 | 0 | 0 | 0 | 5 | 0 | 0 | 1 | 0 | 98 | 49 | NA | NA | NA | NA | 1 | NA | | 0.714 | NA | 0.286 |
| SE | 2576 | FAM208A | FAM208A | chr3 | - | 56663540 | 56663572 | 56662384 | 56662490 | 56668396 | 56668558 | 0 | 0 | 0 | 0 | 0 | 1 | 0 | 1 | 0 | 0 | 0 | 0 | 81 | 49 | NA | NA | NA | NA | 0 | NA | | 1 | NA | -1 |
| SE | 2577 | FAM208A | FAM208A | chr3 | - | 56666259 | 56666384 | 56663540 | 56663572 | 56668396 | 56668558 | 0 | 0 | 1 | 0 | 0 | 0 | 0 | 3 | 0 | 0 | 1 | 0 | 98 | 49 | NA | NA | NA | NA | 1 | NA | | 0.6 | NA | 0.4 |
| SE | 2591 | WDR82 | WDR82 | chr3 | - | 52270711 | 52270809 | 52266951 | 52267018 | 52278200 | 52278643 | 0 | 0 | 5 | 0 | 0 | 1 | 4 | 3 | 0 | 0 | 0 | 0 | 98 | 49 | NA | NA | NA | NA | 0.714 | | 1 | 1 | NA | -0.286 |
| SE | 2594 | ABHD14B | ABHD14B | chr3 | - | 51971459 | 51971698 | 51969942 | 51970184 | 51973964 | 51974630 | 0 | 0 | 0 | 0 | 0 | 1 | 0 | 1 | 0 | 0 | 0 | 0 | 98 | 49 | NA | NA | NA | NA | 0 | NA | | 1 | NA | -1 |
| SE | 2595 | ABHD14B | ABHD14B | chr3 | - | 51971459 | 51971892 | 51969942 | 51970184 | 51973964 | 51974630 | 0 | 0 | 0 | 0 | 0 | 1 | 0 | 1 | 0 | 0 | 0 | 0 | 98 | 49 | NA | NA | NA | NA | 0 | NA | | 1 | NA | -1 |
| SE | 2598 | VPRBP | VPRBP | chr3 | - | 51441384 | 51441897 | 51440969 | 51441071 | 51443765 | 51443903 | 0 | 0 | 1 | 0 | 0 | 0 | 0 | 0 | 0 | 0 | 2 | 0 | 98 | 49 | NA | NA | NA | NA | 1 | NA | | 0 | NA | 1 |
| SE | 2606 | RBM5 | RBM5 | chr3 | + | 50100531 | 50100605 | 50099981 | 50100051 | 50103082 | 50103166 | 0 | 0 | 0 | 0 | 0 | 2 | 0 | 2 | 0 | 0 | 4 | 0 | 98 | 49 | NA | NA | NA | NA | 0 | NA | | 0.2 | NA | -0.2 |
| SE | 2608 | RBM5 | RBM5 | chr3 | + | 50115905 | 50115980 | 50113382 | 50113544 | 50117073 | 50117171 | 0 | 0 | 5 | 0 | 0 | 0 | 4 | 9 | 0 | 0 | 1 | 0 | 98 | 49 | NA | NA | NA | NA | 1 | | 1 | 0.818 | NA | 0.091 |
| SE | 2610 | KIAA0922 | KIAA0922 | chr4 | + | 153587741 | 153587811 | 153586208 | 153586379 | 153588889 | 153589007 | 0 | 0 | 3 | 0 | 0 | 2 | 0 | 1 | 0 | 0 | 0 | 0 | 98 | 49 | NA | NA | NA | NA | 0.429 | NA | | 1 | NA | -0.571 |
| SE | 2615 | APEH | APEH | chr3 | + | 49676776 | 49676817 | 49676608 | 49676700 | 49676902 | 49677024 | 0 | 0 | 4 | 0 | 0 | 1 | 0 | 1 | 0 | 0 | 0 | 0 | 90 | 49 | NA | NA | NA | NA | 0.685 | NA | | 1 | NA | -0.315 |
| SE | 2661 | SACM1L | SACM1L | chr3 | + | 45735234 | 45735373 | 45732052 | 45732151 | 45737582 | 45737652 | 0 | 0 | 2 | 0 | 0 | 0 | 0 | 2 | 0 | 0 | 1 | 0 | 98 | 49 | NA | NA | NA | NA | 1 | NA | | 0.5 | NA | 0.5 |
| SE | 2662 | ABHD5 | ABHD5 | chr3 | + | 43717670 | 43717857 | 43714946 | 43715058 | 43718442 | 43722725 | 0 | 0 | 17 | 0 | 0 | 3 | 6 | 3 | 0 | 0 | 1 | 0 | 98 | 49 | NA | NA | NA | NA | 0.739 | | 1 | 0.6 | NA | -0.061 |
| SE | 2707 | TGFBR2 | TGFBR2 | chr3 | + | 30623198 | 30623273 | 30606501 | 30606977 | 30644746 | 30644915 | 0 | 0 | 3 | 0 | 0 | 2 | 0 | 5 | 0 | 0 | 0 | 0 | 98 | 49 | NA | NA | NA | NA | 0.429 | NA | | 1 | NA | -0.571 |
| SE | 2713 | TOP2B | TOP2B | chr3 | - | 25604759 | 25604870 | 25601099 | 25601225 | 25606042 | 25606122 | 0 | 0 | 10 | 0 | 0 | 1 | 2 | 0 | 0 | 0 | 0 | 0 | 98 | 49 | NA | NA | NA | NA | 0.833 | | 1 | NA | NA | -0.167 |
| SE | 2723 | KAT2B | KAT2B | chr3 | + | 20127422 | 20127549 | 20125904 | 20126113 | 20136941 | 20137052 | 0 | 0 | 12 | 0 | 0 | 0 | 0 | 6 | 0 | 0 | 1 | 0 | 98 | 49 | NA | NA | NA | NA | 1 | NA | | 0.75 | NA | 0.25 |
| SE | 2724 | SERPINB9 | SERPINB9 | chr6 | - | 2893410 | 2893553 | 2891832 | 2891988 | 2895390 | 2895508 | 0 | 0 | 3 | 0 | 0 | 1 | 0 | 5 | 0 | 0 | 1 | 0 | 98 | 49 | NA | NA | NA | NA | 0.6 | NA | | 0.714 | NA | -0.114 |
| SE | 2762 | LST1 | LST1 | chr6 | + | 31587640 | 31587733 | 31587199 | 31587318 | 31587943 | 31587966 | 0 | 0 | 1 | 0 | 0 | 2 | 0 | 1 | 0 | 0 | 0 | 0 | 98 | 49 | NA | NA | NA | NA | 0.2 | NA | | 1 | NA | -0.8 |
| SE | 2763 | LST1 | LST1 | chr6 | + | 31587640 | 31587733 | 31587199 | 31587318 | 31588496 | 31588909 | 0 | 0 | 0 | 0 | 0 | 3 | 0 | 1 | 0 | 0 | 0 | 0 | 98 | 49 | NA | NA | NA | NA | 0 | NA | | 1 | NA | -1 |
| SE | 2764 | LST1 | LST1 | chr6 | + | 31587640 | 31587733 | 31587199 | 31587318 | 31588517 | 31588909 | 0 | 0 | 0 | 0 | 0 | 4 | 0 | 2 | 0 | 0 | 4 | 0 | 98 | 49 | NA | NA | NA | NA | 0 | NA | | 0.2 | NA | -0.2 |
| SE | 2773 | XPC | XPC | chr3 | - | 14168256 | 14168380 | 14167168 | 14167253 | 14170437 | 14170550 | 0 | 0 | 1 | 0 | 0 | 0 | 0 | 6 | 0 | 0 | 3 | 0 | 98 | 49 | NA | NA | NA | NA | 1 | NA | | 0.5 | NA | 0.5 |
| SE | 2776 | POLB | POLB | chr8 | + | 42339011 | 42339069 | 42338454 | 42338685 | 42344952 | 42345019 | 0 | 0 | 3 | 0 | 0 | 2 | 0 | 1 | 0 | 0 | 0 | 0 | 98 | 49 | NA | NA | NA | NA | 0.429 | NA | | 1 | NA | -0.571 |
| SE | 2777 | POLB | POLB | chr8 | + | 42369848 | 42369988 | 42369270 | 42369335 | 42371562 | 42371813 | 0 | 0 | 4 | 0 | 0 | 0 | 0 | 0 | 0 | 0 | 1 | 0 | 98 | 49 | NA | NA | NA | NA | 1 | NA | | 0 | NA | 1 |
| SE | 2784 | ATG7 | ATG7 | chr3 | + | 11342134 | 11342279 | 11340644 | 11340735 | 11347876 | 11348035 | 0 | 0 | 16 | 0 | 0 | 1 | 0 | 1 | 0 | 0 | 0 | 0 | 98 | 49 | NA | NA | NA | NA | 0.889 | NA | | 1 | NA | -0.111 |
| SE | 2837 | ATRX | ATRX | chrX | - | 77600433 | 77600564 | 77599731 | 77599820 | 77616612 | 77616730 | 0 | 0 | 2 | 0 | 0 | 2 | 0 | 1 | 0 | 0 | 0 | 0 | 98 | 49 | NA | NA | NA | NA | 0.333 | NA | | 1 | NA | -0.667 |
| SE | 2855 | TBC1D22A | TBC1D22A | chr22 | + | 47037070 | 47037198 | 46997633 | 46997709 | 47111507 | 47111603 | 0 | 0 | 5 | 0 | 0 | 0 | 5 | 4 | 0 | 0 | 2 | 0 | 98 | 49 | NA | NA | NA | NA | 1 | | 1 | 0.5 | NA | 0.25 |
| SE | 2888 | TNRC6B | TNRC6B | chr22 | + | 40270121 | 40270280 | 40264687 | 40267036 | 40273424 | 40273600 | 0 | 0 | 1 | 0 | 0 | 0 | 0 | 3 | 0 | 0 | 1 | 0 | 98 | 49 | NA | NA | NA | NA | 1 | NA | | 0.6 | NA | 0.4 |
| SE | 2901 | TMEM184B | TMEM184B | chr22 | - | 38246784 | 38246885 | 38245934 | 38246100 | 38247769 | 38248019 | 0 | 0 | 0 | 0 | 0 | 6 | 3 | 1 | 0 | 0 | 2 | 0 | 98 | 49 | NA | NA | NA | NA | 0 | | 1 | 0.2 | NA | -0.6 |
| SE | 2908 | RAF1 | RAF1 | chr3 | - | 12606200 | 12606299 | 12604135 | 12604289 | 12608765 | 12608923 | 0 | 0 | 20 | 0 | 0 | 5 | 0 | 22 | 0 | 0 | 2 | 0 | 98 | 49 | NA | NA | NA | NA | 0.667 | NA | | 0.846 | NA | -0.179 |
| SE | 2959 | MTMR3 | MTMR3 | chr22 | + | 29970975 | 29971062 | 29957035 | 29957088 | 29978441 | 29978531 | 0 | 0 | 9 | 0 | 0 | 1 | 0 | 3 | 0 | 0 | 0 | 0 | 98 | 49 | NA | NA | NA | NA | 0.818 | NA | | 1 | NA | -0.182 |
| SE | 2960 | MTMR3 | MTMR3 | chr22 | + | 30012367 | 30012563 | 30009017 | 30009129 | 30013355 | 30013541 | 0 | 0 | 2 | 0 | 0 | 4 | 2 | 8 | 0 | 0 | 0 | 0 | 98 | 49 | NA | NA | NA | NA | 0.2 | | 1 | 1 | NA | -0.8 |
| SE | 2962 | MTMR3 | MTMR3 | chr22 | + | 30023456 | 30023483 | 30022608 | 30022697 | 30025629 | 30030868 | 0 | 0 | 5 | 0 | 0 | 2 | 0 | 1 | 0 | 3 | 0 | 0 | 76 | 49 | NA | NA | NA | NA | 0.617 | | 0 | 1 | NA | 0.117 |
| SE | 2967 | THOC5 | THOC5 | chr22 | - | 29550727 | 29550843 | 29549051 | 29549158 | 29553670 | 29553747 | 0 | 0 | 1 | 0 | 0 | 0 | 0 | 0 | 0 | 0 | 1 | 0 | 98 | 49 | NA | NA | NA | NA | 1 | NA | | 0 | NA | 1 |
| SE | 2971 | EWSR1 | EWSR1 | chr22 | + | 29282389 | 29282557 | 29278029 | 29278216 | 29286922 | 29287134 | 0 | 0 | 11 | 0 | 0 | 1 | 0 | 6 | 0 | 0 | 0 | 0 | 98 | 49 | NA | NA | NA | NA | 0.846 | NA | | 1 | NA | -0.154 |
| SE | 2988 | PPP3CA | PPP3CA | chr4 | - | 101063231 | 101063357 | 101061086 | 101061161 | 101080531 | 101080626 | 0 | 0 | 15 | 0 | 0 | 3 | 3 | 0 | 0 | 0 | 0 | 0 | 98 | 49 | NA | NA | NA | NA | 0.714 | | 1 | NA | NA | -0.286 |
| SE | 2993 | GUSBP11 | GUSBP11 | chr22 | - | 23700725 | 23700845 | 23694118 | 23695517 | 23705428 | 23705541 | 0 | 0 | 6 | 0 | 0 | 0 | 0 | 0 | 0 | 0 | 1 | 0 | 98 | 49 | NA | NA | NA | NA | 1 | NA | | 0 | NA | 1 |
| SE | 2996 | BCR | BCR | chr22 | + | 23260949 | 23261054 | 23253798 | 23253980 | 23261354 | 23261540 | 2 | 0 | 3 | 0 | 0 | 0 | 0 | 3 | 0 | 0 | 1 | 0 | 98 | 49 | NA | NA | 1 | NA | 1 | NA | | 0.6 | NA | 0.4 |
| SE | 2997 | BCR | BCR | chr22 | + | 23290338 | 23290413 | 23289516 | 23289621 | 23292540 | 23292638 | 0 | 0 | 3 | 0 | 0 | 0 | 0 | 1 | 0 | 0 | 1 | 0 | 98 | 49 | NA | NA | NA | NA | 1 | NA | | 0.333 | NA | 0.667 |
| SE | 3026 | TANGO2 | TANGO2 | chr22 | + | 20061529 | 20061681 | 20055942 | 20056013 | 20063337 | 20063442 | 0 | 0 | 0 | 0 | 0 | 1 | 0 | 1 | 0 | 0 | 0 | 0 | 98 | 49 | NA | NA | NA | NA | 0 | NA | | 1 | NA | -1 |
| SE | 3028 | TANGO2 | TANGO2 | chr22 | + | 20061529 | 20061683 | 20055942 | 20056013 | 20063337 | 20063442 | 0 | 0 | 2 | 0 | 0 | 1 | 0 | 1 | 0 | 0 | 0 | 0 | 98 | 49 | NA | NA | NA | NA | 0.5 | NA | | 1 | NA | -0.5 |
| SE | 3050 | PRMT2 | PRMT2 | chr21 | + | 46658744 | 46658920 | 46649574 | 46649739 | 46664294 | 46665124 | 0 | 0 | 2 | 0 | 0 | 1 | 0 | 2 | 0 | 0 | 0 | 0 | 98 | 49 | NA | NA | NA | NA | 0.5 | NA | | 1 | NA | -0.5 |
| SE | 3055 | NQO2 | NQO2 | chr6 | + | 3015529 | 3015643 | 3012543 | 3012674 | 3016883 | 3016985 | 0 | 0 | 4 | 0 | 0 | 1 | 0 | 1 | 0 | 0 | 0 | 0 | 98 | 49 | NA | NA | NA | NA | 0.667 | NA | | 1 | NA | -0.333 |
| SE | 3121 | TTC3 | TTC3 | chr21 | + | 37090232 | 37090286 | 37088798 | 37088886 | 37091292 | 37091413 | 0 | 0 | 1 | 0 | 0 | 0 | 0 | 2 | 0 | 0 | 1 | 0 | 98 | 49 | NA | NA | NA | NA | 1 | NA | | 0.5 | NA | 0.5 |
| SE | 3135 | IFNGR2 | IFNGR2 | chr21 | + | 33432176 | 33432336 | 33426883 | 33427032 | 33432713 | 33432871 | 0 | 0 | 16 | 0 | 0 | 1 | 4 | 7 | 0 | 0 | 0 | 0 | 98 | 49 | NA | NA | NA | NA | 0.889 | | 1 | 1 | NA | -0.111 |
| SE | 3142 | SYNJ1 | SYNJ1 | chr21 | - | 32699837 | 32700105 | 32695056 | 32695282 | 32701960 | 32702047 | 0 | 0 | 1 | 0 | 0 | 1 | 0 | 2 | 0 | 0 | 1 | 0 | 98 | 49 | NA | NA | NA | NA | 0.333 | NA | | 0.5 | NA | -0.167 |
| SE | 3153 | USP16 | USP16 | chr21 | + | 29037275 | 29037463 | 29036270 | 29036374 | 29038334 | 29038430 | 0 | 0 | 7 | 0 | 0 | 0 | 0 | 1 | 0 | 0 | 1 | 0 | 98 | 49 | NA | NA | NA | NA | 1 | NA | | 0.333 | NA | 0.667 |
| SE | 3154 | USP16 | USP16 | chr21 | + | 29037278 | 29037463 | 29036270 | 29036374 | 29038334 | 29038430 | 0 | 0 | 4 | 0 | 0 | 0 | 0 | 1 | 0 | 0 | 1 | 0 | 98 | 49 | NA | NA | NA | NA | 1 | NA | | 0.333 | NA | 0.667 |
| SE | 3158 | APP | APP | chr21 | - | 25905023 | 25905077 | 25897572 | 25897673 | 25911740 | 25911962 | 0 | 0 | 6 | 0 | 0 | 1 | 0 | 1 | 0 | 0 | 0 | 0 | 98 | 49 | NA | NA | NA | NA | 0.75 | NA | | 1 | NA | -0.25 |
| SE | 3159 | APP | APP | chr21 | - | 25997359 | 25997416 | 25982343 | 25982477 | 26000014 | 26000182 | 0 | 0 | 1 | 0 | 0 | 3 | 0 | 0 | 0 | 2 | 1 | 0 | 98 | 49 | NA | NA | NA | NA | 0.143 | | 0 | 0 | NA | 0.143 |

| | | | | | | | | | | | | | | | | | | | | | | | | | | | | | | | | | | | | | | | |
|---|---|---|---|---|---|---|---|---|---|---|---|---|---|---|---|---|---|---|---|---|---|---|---|---|---|---|---|---|---|---|---|---|---|---|---|---|---|---|---|
| SE | 3160 | APP | APP | chr21 | - | 26000014 | 26000182 | 25982343 | 25982477 | 26021839 | 26022042 | 0 | 0 | 6 | 0 | 0 | 0 | 2 | 2 | 0 | 0 | 1 | 0 | 98 | 49 | NA | NA | NA | | NA | | 1 | | 1 | | 0.5 | NA | | 0.25 |
| SE | 3171 | NRIP1 | NRIP1 | chr21 | - | 15043494 | 15043574 | 15014343 | 15014466 | 15064744 | 15064805 | 2 | 0 | 0 | 0 | 0 | 0 | 0 | 2 | 0 | 0 | 0 | 1 | 98 | 49 | NA | NA | | 1 | NA | NA | | NA | | | 1 | | 0 | 0.5 |
| SE | 3188 | TPD52L2 | TPD52L2 | chr20 | + | 63875815 | 63875875 | 63873667 | 63873816 | 63882718 | 63882820 | 0 | 0 | 17 | 0 | 0 | 0 | 0 | 9 | 0 | 0 | 1 | 0 | 98 | 49 | NA | NA | NA | | NA | | 1 | NA | | | 0.818 | NA | | 0.182 |
| SE | 3191 | TPD52L2 | TPD52L2 | chr20 | + | 63887563 | 63887605 | 63882718 | 63882820 | 63889189 | 63889238 | 0 | 0 | 1 | 0 | 0 | 8 | 0 | 0 | 0 | 0 | 5 | 0 | 91 | 49 | NA | NA | NA | | NA | | 0.063 | NA | | | 0 | NA | | 0.063 |
| SE | 3193 | CXorf65 | CXorf65 | chrX | - | 71104768 | 71104837 | 71104297 | 71104404 | 71105999 | 71106137 | 0 | 0 | 3 | 0 | 0 | 0 | 0 | 1 | 0 | 0 | 2 | 0 | 98 | 49 | NA | NA | NA | | NA | | 1 | NA | | | 0.2 | NA | | 0.8 |
| SE | 3197 | TNIP1 | TNIP1 | chr5 | - | 151042539 | 151042671 | 151039096 | 151039225 | 151042895 | 151042961 | 0 | 0 | 7 | 0 | 0 | 0 | 0 | 12 | 0 | 0 | 1 | 0 | 98 | 49 | NA | NA | NA | | NA | | 1 | NA | | | 0.857 | NA | | 0.143 |
| SE | 3198 | TNIP1 | TNIP1 | chr5 | - | 151045860 | 151045950 | 151042895 | 151042961 | 151049823 | 151049947 | 0 | 0 | 6 | 0 | 0 | 0 | 0 | 12 | 0 | 0 | 1 | 0 | 98 | 49 | NA | NA | NA | | NA | | 1 | NA | | | 0.857 | NA | | 0.143 |
| SE | 3209 | STX16-NPEPL1 | STX16-NPEPL1 | chr20 | + | 58673630 | 58673711 | 58671153 | 58671297 | 58691723 | 58691780 | 0 | 0 | 10 | 0 | 0 | 2 | 0 | 5 | 0 | 0 | 0 | 0 | 98 | 49 | NA | NA | NA | | NA | | 0.714 | NA | | | 1 | NA | | -0.286 |
| SE | 3211 | STX16 | STX16 | chr20 | + | 58659622 | 58659634 | 58651252 | 58652087 | 58667489 | 58667597 | 0 | 0 | 1 | 0 | 0 | 3 | 0 | 0 | 0 | 0 | 1 | 0 | 61 | 49 | NA | NA | NA | | NA | | 0.211 | NA | | | 0 | NA | | 0.211 |
| SE | 3216 | PDCD6IP | PDCD6IP | chr3 | + | 33828837 | 33828969 | 33826479 | 33826580 | 33836043 | 33836266 | 0 | 0 | 1 | 0 | 0 | 1 | 0 | 5 | 0 | 0 | 0 | 0 | 98 | 49 | NA | NA | NA | | NA | | 0.333 | NA | | | 1 | NA | | -0.667 |
| SE | 3217 | PDCD6IP | PDCD6IP | chr3 | + | 33828852 | 33828969 | 33826479 | 33826580 | 33836043 | 33836266 | 0 | 0 | 1 | 0 | 0 | 1 | 0 | 5 | 0 | 0 | 0 | 0 | 98 | 49 | NA | NA | NA | | NA | | 0.333 | NA | | | 1 | NA | | -0.667 |
| SE | 3249 | ZMYND8 | ZMYND8 | chr20 | - | 47212641 | 47212725 | 47209213 | 47210897 | 47220257 | 47220324 | 0 | 0 | 0 | 0 | 0 | 1 | 0 | 2 | 0 | 0 | 0 | 0 | 98 | 49 | NA | NA | NA | | NA | | 0 | NA | | | 1 | NA | | -1 |
| SE | 3254 | ZMYND8 | ZMYND8 | chr20 | - | 47276313 | 47276795 | 47262287 | 47262428 | 47282101 | 47282217 | 0 | 0 | 1 | 0 | 0 | 0 | 0 | 0 | 0 | 0 | 1 | 0 | 98 | 49 | NA | NA | NA | | NA | | 1 | NA | | | 0 | NA | | 1 |
| SE | 3256 | ZMYND8 | ZMYND8 | chr20 | - | 47291795 | 47291888 | 47290186 | 47290274 | 47294665 | 47294779 | 0 | 0 | 9 | 0 | 0 | 0 | 0 | 0 | 0 | 2 | 0 | 0 | 98 | 49 | NA | NA | NA | | NA | | 1 | | 0 | NA | | NA | | 1 |
| SE | 3326 | CTNNBL1 | CTNNBL1 | chr20 | + | 37842338 | 37842419 | 37840101 | 37840199 | 37859898 | 37860036 | 0 | 0 | 2 | 0 | 0 | 1 | 0 | 4 | 0 | 0 | 0 | 0 | 98 | 49 | NA | NA | NA | | NA | | 0.5 | NA | | | 1 | NA | | -0.5 |
| SE | 3331 | SAMHD1 | SAMHD1 | chr20 | - | 36898439 | 36898544 | 36897821 | 36897959 | 36904156 | 36904249 | 0 | 0 | 25 | 0 | 0 | 1 | 0 | 5 | 0 | 0 | 0 | 0 | 98 | 49 | NA | NA | NA | | NA | | 0.926 | NA | | | 1 | NA | | -0.074 |
| SE | 3333 | PHF8 | PHF8 | chrX | - | 53993600 | 53993903 | 53992735 | 53992839 | 53995692 | 53995782 | 0 | 0 | 0 | 0 | 0 | 1 | 0 | 2 | 0 | 0 | 0 | 0 | 98 | 49 | NA | NA | NA | | NA | | 0 | NA | | | 1 | NA | | -1 |
| SE | 3337 | C20orf24 | C20orf24 | chr20 | + | 36607889 | 36608000 | 36607714 | 36607818 | 36609600 | 36609730 | 0 | 0 | 3 | 0 | 0 | 4 | 0 | 3 | 0 | 0 | 2 | 0 | 98 | 49 | NA | NA | NA | | NA | | 0.273 | NA | | | 0.429 | NA | | -0.156 |
| SE | 3356 | EDEM2 | EDEM2 | chr20 | - | 35118597 | 35118719 | 35115356 | 35115933 | 35123889 | 35124034 | 0 | 0 | 6 | 0 | 0 | 1 | 0 | 0 | 0 | 0 | 2 | 0 | 98 | 49 | NA | NA | NA | | NA | | 0.75 | NA | | | 0 | NA | | 0.75 |
| SE | 3358 | TRPC4AP | TRPC4AP | chr20 | - | 35021189 | 35021332 | 35016007 | 35016139 | 35035122 | 35035308 | 0 | 0 | 4 | 0 | 0 | 0 | 0 | 3 | 0 | 0 | 1 | 0 | 98 | 49 | NA | NA | NA | | NA | | 1 | NA | | | 0.6 | NA | | 0.4 |
| SE | 3359 | TRPC4AP | TRPC4AP | chr20 | - | 35021189 | 35021356 | 35016007 | 35016139 | 35035122 | 35035308 | 0 | 0 | 4 | 0 | 0 | 0 | 0 | 2 | 0 | 0 | 1 | 0 | 98 | 49 | NA | NA | NA | | NA | | 1 | NA | | | 0.5 | NA | | 0.5 |
| SE | 3360 | TRPC4AP | TRPC4AP | chr20 | - | 35044504 | 35044712 | 35035122 | 35035308 | 35049865 | 35049994 | 0 | 0 | 19 | 0 | 0 | 0 | 13 | 8 | 0 | 0 | 1 | 0 | 98 | 49 | NA | NA | NA | | NA | | 1 | | 1 | | 0.8 | NA | | 0.1 |
| SE | 3362 | ACSS2 | ACSS2 | chr20 | + | 34926104 | 34926281 | 34925697 | 34925766 | 34926876 | 34926951 | 0 | 0 | 4 | 0 | 0 | 0 | 0 | 0 | 0 | 0 | 1 | 0 | 98 | 49 | NA | NA | NA | | NA | | 1 | NA | | | 0 | NA | | 1 |
| SE | 3372 | FCHSD1 | FCHSD1 | chr5 | - | 141646602 | 141646722 | 141646086 | 141646191 | 141647134 | 141647230 | 0 | 0 | 1 | 0 | 0 | 1 | 0 | 2 | 0 | 0 | 0 | 0 | 98 | 49 | NA | NA | NA | | NA | | 0.333 | NA | | | 1 | NA | | -0.667 |
| SE | 3398 | ELOVL5 | ELOVL5 | chr6 | - | 53270592 | 53270727 | 53267397 | 53269270 | 53273219 | 53273344 | 0 | 0 | 15 | 0 | 0 | 2 | 0 | 9 | 0 | 0 | 0 | 0 | 98 | 49 | NA | NA | NA | | NA | | 0.789 | NA | | | 1 | NA | | -0.211 |
| SE | 3411 | XRN2 | XRN2 | chr20 | + | 21356585 | 21356665 | 21356079 | 21356177 | 21357735 | 21357792 | 0 | 0 | 12 | 0 | 0 | 2 | 0 | 2 | 0 | 0 | 0 | 0 | 98 | 49 | NA | NA | NA | | NA | | 0.75 | NA | | | 1 | NA | | -0.25 |
| SE | 3412 | XRN2 | XRN2 | chr20 | + | 21357735 | 21357792 | 21356585 | 21356665 | 21365420 | 21365489 | 0 | 0 | 9 | 0 | 0 | 1 | 0 | 2 | 0 | 0 | 0 | 0 | 98 | 49 | NA | NA | NA | | NA | | 0.818 | NA | | | 1 | NA | | -0.182 |
| SE | 3414 | RALGAPA2 | RALGAPA2 | chr20 | - | 20605174 | 20605412 | 20601681 | 20601846 | 20611314 | 20611426 | 0 | 0 | 5 | 0 | 0 | 0 | 0 | 0 | 0 | 0 | 2 | 0 | 98 | 49 | NA | NA | NA | | NA | | 1 | NA | | | 0 | NA | | 1 |
| SE | 3415 | RALGAPA2 | RALGAPA2 | chr20 | - | 20611314 | 20611426 | 20605174 | 20605412 | 20616042 | 20616191 | 0 | 0 | 6 | 0 | 0 | 1 | 0 | 1 | 0 | 0 | 0 | 0 | 98 | 49 | NA | NA | NA | | NA | | 0.75 | NA | | | 1 | NA | | -0.25 |
| SE | 3416 | RALGAPA2 | RALGAPA2 | chr20 | - | 20637362 | 20637501 | 20635417 | 20635617 | 20639784 | 20639900 | 0 | 0 | 13 | 0 | 0 | 2 | 0 | 0 | 0 | 0 | 2 | 0 | 98 | 49 | NA | NA | NA | | NA | | 0.765 | NA | | | 0 | NA | | 0.765 |
| SE | 3419 | SEC23B | SEC23B | chr20 | + | 18525787 | 18525932 | 18524934 | 18525020 | 18526372 | 18526531 | 0 | 0 | 1 | 0 | 0 | 0 | 0 | 1 | 0 | 0 | 1 | 0 | 98 | 49 | NA | NA | NA | | NA | | 1 | NA | | | 0.333 | NA | | 0.667 |
| SE | 3453 | GPCPD1 | GPCPD1 | chr20 | - | 5570146 | 5570239 | 5567482 | 5567560 | 5573914 | 5573969 | 0 | 0 | 14 | 0 | 0 | 1 | 0 | 13 | 0 | 0 | 0 | 0 | 98 | 49 | NA | NA | NA | | NA | | 0.875 | NA | | | 1 | NA | | -0.125 |
| SE | 3454 | GPCPD1 | GPCPD1 | chr20 | - | 5580007 | 5580131 | 5578379 | 5578611 | 5584280 | 5584322 | 0 | 0 | 17 | 0 | 0 | 1 | 0 | 17 | 0 | 0 | 0 | 0 | 98 | 49 | NA | NA | NA | | NA | | 0.895 | NA | | | 1 | NA | | -0.105 |
| SE | 3458 | SLC23A2 | SLC23A2 | chr20 | - | 4885820 | 4885909 | 4884752 | 4884823 | 4899554 | 4899712 | 0 | 0 | 1 | 0 | 0 | 0 | 0 | 4 | 0 | 0 | 1 | 0 | 98 | 49 | NA | NA | NA | | NA | | 1 | NA | | | 0.667 | NA | | 0.333 |
| SE | 3467 | SIGLEC1 | SIGLEC1 | chr20 | - | 3699201 | 3699459 | 3697797 | 3698133 | 3701341 | 3701641 | 0 | 0 | 8 | 0 | 0 | 1 | 0 | 1 | 0 | 0 | 0 | 0 | 98 | 49 | NA | NA | NA | | NA | | 0.8 | NA | | | 1 | NA | | -0.2 |
| SE | 3473 | FAM193A | FAM193A | chr4 | + | 2655057 | 2655129 | 2646684 | 2646832 | 2657802 | 2657880 | 0 | 0 | 1 | 0 | 0 | 2 | 0 | 1 | 0 | 0 | 2 | 0 | 98 | 49 | NA | NA | NA | | NA | | 0.2 | NA | | | 0.2 | NA | | 0 |
| SE | 3479 | STK35 | STK35 | chr20 | + | 2116665 | 2117415 | 2102767 | 2103365 | 2143783 | 2148552 | 0 | 0 | 1 | 0 | 0 | 0 | 0 | 0 | 0 | 0 | 1 | 0 | 98 | 49 | NA | NA | NA | | NA | | 1 | NA | | | 0 | NA | | 1 |
| SE | 3509 | HDLBP | HDLBP | chr2 | - | 241238648 | 241238787 | 241236614 | 241236769 | 241239601 | 241239820 | 0 | 0 | 5 | 0 | 0 | 0 | 0 | 4 | 0 | 0 | 1 | 0 | 98 | 49 | NA | NA | NA | | NA | | 1 | NA | | | 0.667 | NA | | 0.333 |
| SE | 3521 | RNPEPL1 | RNPEPL1 | chr2 | + | 240574112 | 240574348 | 240573774 | 240573891 | 240574514 | 240574628 | 0 | 0 | 6 | 0 | 0 | 1 | 0 | 7 | 0 | 0 | 0 | 0 | 98 | 49 | NA | NA | NA | | NA | | 0.75 | NA | | | 1 | NA | | -0.25 |
| SE | 3527 | PMPCB | PMPCB | chr7 | + | 103307595 | 103307708 | 103304410 | 103304490 | 103308951 | 103309095 | 0 | 0 | 2 | 0 | 0 | 0 | 0 | 3 | 0 | 0 | 1 | 0 | 98 | 49 | NA | NA | NA | | NA | | 1 | NA | | | 0.6 | NA | | 0.4 |
| SE | 3531 | LRRFIP1 | LRRFIP1 | chr2 | + | 237708543 | 237708630 | 237692163 | 237692521 | 237748363 | 237748399 | 0 | 0 | 2 | 0 | 0 | 2 | 3 | 2 | 0 | 4 | 1 | 0 | 98 | 49 | NA | NA | NA | | NA | | 0.333 | | 0.273 | | 0.5 | NA | | -0.053 |
| SE | 3535 | LRRFIP1 | LRRFIP1 | chr2 | + | 237772080 | 237772198 | 237769942 | 237769992 | 237772865 | 237772945 | 0 | 0 | 7 | 0 | 0 | 0 | 0 | 2 | 0 | 5 | 0 | 0 | 98 | 49 | NA | NA | NA | | NA | | 1 | | 0 | | 1 | NA | | 0.5 |
| SE | 3546 | OSBPL2 | OSBPL2 | chr20 | + | 62260016 | 62260125 | 62256056 | 62256221 | 62263615 | 62263691 | 0 | 0 | 3 | 0 | 0 | 0 | 0 | 5 | 0 | 0 | 4 | 0 | 98 | 49 | NA | NA | NA | | NA | | 1 | NA | | | 0.385 | NA | | 0.615 |
| SE | 3547 | OSBPL2 | OSBPL2 | chr20 | + | 62264628 | 62264718 | 62263615 | 62263691 | 62272124 | 62272259 | 0 | 0 | 2 | 0 | 0 | 3 | 0 | 5 | 0 | 0 | 5 | 0 | 98 | 49 | NA | NA | NA | | NA | | 0.25 | NA | | | 0.333 | NA | | -0.083 |
| SE | 3550 | MAPKAPK3 | MAPKAPK3 | chr3 | + | 50644408 | 50644532 | 50642252 | 50642332 | 50645709 | 50645785 | 0 | 0 | 11 | 0 | 0 | 5 | 0 | 4 | 0 | 0 | 1 | 0 | 98 | 49 | NA | NA | NA | | NA | | 0.524 | NA | | | 0.667 | NA | | -0.143 |
| SE | 3555 | TAZ | TAZ | chrX | + | 154414100 | 154414190 | 154413481 | 154413567 | 154419542 | 154419623 | 0 | 0 | 0 | 0 | 0 | 1 | 0 | 1 | 0 | 0 | 2 | 0 | 98 | 49 | NA | NA | NA | | NA | | 0 | NA | | | 0.2 | NA | | -0.2 |
| SE | 3558 | DGKD | DGKD | chr2 | + | 233435817 | 233435924 | 233434768 | 233434901 | 233436315 | 233436441 | 0 | 0 | 12 | 0 | 0 | 1 | 0 | 3 | 0 | 0 | 0 | 0 | 98 | 49 | NA | NA | NA | | NA | | 0.857 | NA | | | 1 | NA | | -0.143 |
| SE | 3585 | SP110 | SP110 | chr2 | - | 230170620 | 230170761 | 230168925 | 230169237 | 230172065 | 230172174 | 0 | 0 | 1 | 0 | 0 | 2 | 0 | 1 | 0 | 0 | 1 | 0 | 98 | 49 | NA | NA | NA | | NA | | 0.2 | NA | | | 0.333 | NA | | -0.133 |
| SE | 3587 | SP110 | SP110 | chr2 | - | 230172843 | 230172959 | 230172065 | 230172174 | 230177537 | 230177680 | 0 | 0 | 7 | 0 | 0 | 0 | 0 | 2 | 0 | 0 | 1 | 0 | 98 | 49 | NA | NA | NA | | NA | | 1 | NA | | | 0.5 | NA | | 0.5 |
| SE | 3588 | SP110 | SP110 | chr2 | - | 230177537 | 230177680 | 230176472 | 230176724 | 230178156 | 230178255 | 0 | 0 | 5 | 0 | 0 | 2 | 0 | 7 | 0 | 0 | 0 | 0 | 98 | 49 | NA | NA | NA | | NA | | 0.556 | NA | | | 1 | NA | | -0.444 |
| SE | 3592 | TRIP12 | TRIP12 | chr2 | - | 229860405 | 229860531 | 229858771 | 229859574 | 229879981 | 229880128 | 0 | 0 | 3 | 0 | 0 | 2 | 0 | 0 | 0 | 0 | 1 | 0 | 98 | 49 | NA | NA | NA | | NA | | 0.429 | NA | | | 0 | NA | | 0.429 |
| SE | 3608 | DOCK10 | DOCK10 | chr2 | - | 224814319 | 224814364 | 224807910 | 224808086 | 224816616 | 224816713 | 0 | 0 | 4 | 0 | 0 | 0 | 0 | 1 | 0 | 0 | 2 | 0 | 94 | 49 | NA | NA | NA | | NA | | 1 | NA | | | 0.207 | NA | | 0.793 |
| SE | 3614 | NIN | NIN | chr14 | - | 50756491 | 50758630 | 50754741 | 50754867 | 50759856 | 50760359 | 0 | 0 | 2 | 0 | 0 | 0 | 0 | 1 | 0 | 0 | 1 | 0 | 98 | 49 | NA | NA | NA | | NA | | 1 | NA | | | 0.333 | NA | | 0.667 |
| SE | 3615 | NIN | NIN | chr14 | - | 50759856 | 50760359 | 50754741 | 50754867 | 50761789 | 50761911 | 0 | 0 | 2 | 0 | 0 | 2 | 0 | 2 | 0 | 0 | 0 | 0 | 98 | 49 | NA | NA | NA | | NA | | 0.333 | NA | | | 1 | NA | | -0.667 |
| SE | 3616 | NIN | NIN | chr14 | - | 50766306 | 50766396 | 50763825 | 50763964 | 50766779 | 50766890 | 0 | 0 | 14 | 0 | 0 | 1 | 0 | 16 | 0 | 0 | 0 | 0 | 98 | 49 | NA | NA | NA | | NA | | 0.875 | NA | | | 1 | NA | | -0.125 |
| SE | 3623 | ITGAM | ITGAM | chr16 | + | 31321463 | 31321627 | 31321240 | 31321371 | 31324398 | 31324553 | 0 | 0 | 12 | 0 | 0 | 1 | 0 | 0 | 0 | 0 | 1 | 0 | 98 | 49 | NA | NA | NA | | NA | | 0.857 | NA | | | 0 | NA | | 0.857 |
| SE | 3626 | HECTD1 | HECTD1 | chr14 | - | 31113876 | 31114059 | 31113276 | 31113484 | 31114285 | 31114340 | 0 | 0 | 0 | 0 | 0 | 1 | 0 | 1 | 0 | 0 | 0 | 0 | 98 | 49 | NA | NA | NA | | NA | | 0 | NA | | | 1 | NA | | -1 |
| SE | 3633 | USP4 | USP4 | chr3 | - | 49310619 | 49310737 | 49305714 | 49305888 | 49324701 | 49324763 | 0 | 0 | 3 | 0 | 0 | 1 | 0 | 8 | 0 | 0 | 0 | 0 | 98 | 49 | NA | NA | NA | | NA | | 0.6 | NA | | | 1 | NA | | -0.4 |
| SE | 3634 | USP4 | USP4 | chr3 | - | 49311513 | 49311654 | 49310619 | 49310737 | 49324701 | 49324763 | 0 | 0 | 1 | 0 | 0 | 1 | 0 | 4 | 0 | 0 | 3 | 0 | 98 | 49 | NA | NA | NA | | NA | | 0.333 | NA | | | 0.4 | NA | | -0.067 |
| SE | 3674 | CALCOCO2 | CALCOCO2 | chr17 | + | 48852505 | 48852628 | 48851558 | 48851628 | 48852925 | 48853012 | 0 | 0 | 3 | 0 | 0 | 0 | 0 | 12 | 0 | 0 | 1 | 0 | 98 | 49 | NA | NA | NA | | NA | | 1 | NA | | | 0.857 | NA | | 0.143 |
| SE | 3682 | USP15 | USP15 | chr12 | + | 62396294 | 62396398 | 62393052 | 62393202 | 62401186 | 62401275 | 0 | 0 | 13 | 0 | 0 | 1 | 2 | 11 | 0 | 0 | 0 | 0 | 98 | 49 | NA | NA | NA | | NA | | 0.867 | | 1 | | 1 | NA | | -0.133 |
| SE | 3686 | NBR1 | NBR1 | chr17 | + | 43196941 | 43197106 | 43196480 | 43196591 | 43200166 | 43200608 | 0 | 0 | 3 | 0 | 0 | 1 | 0 | 3 | 0 | 0 | 0 | 0 | 98 | 49 | NA | NA | NA | | NA | | 0.6 | NA | | | 1 | NA | | -0.4 |
| SE | 3701 | HBEGF | HBEGF | chr5 | - | 140345910 | 140346084 | 140342634 | 140342812 | 140346282 | 140346603 | 0 | 0 | 5 | 0 | 0 | 1 | 0 | 5 | 0 | 0 | 0 | 0 | 98 | 49 | NA | NA | NA | | NA | | 0.714 | NA | | | 1 | NA | | -0.286 |
| SE | 3761 | CNOT1 | CNOT1 | chr16 | - | 58547565 | 58547682 | 58547185 | 58547296 | 58549718 | 58549898 | 0 | 0 | 13 | 0 | 0 | 0 | 0 | 8 | 0 | 0 | 1 | 0 | 98 | 49 | NA | NA | NA | | NA | | 1 | NA | | | 0.8 | NA | | 0.2 |
| SE | 3784 | SREK1 | SREK1 | chr5 | + | 66177513 | 66177658 | 66174945 | 66175041 | 66178718 | 66183616 | 0 | 0 | 2 | 0 | 0 | 0 | 0 | 0 | 0 | 0 | 1 | 0 | 98 | 49 | NA | NA | NA | | NA | | 1 | NA | | | 0 | NA | | 1 |
| SE | 3785 | SREK1 | SREK1 | chr5 | + | 66177516 | 66177658 | 66174945 | 66175041 | 66178718 | 66183616 | 0 | 0 | 1 | 0 | 0 | 0 | 0 | 0 | 0 | 0 | 1 | 0 | 98 | 49 | NA | NA | NA | | NA | | 1 | NA | | | 0 | NA | | 1 |
| SE | 3793 | STT3B | STT3B | chr3 | + | 31624913 | 31625085 | 31623673 | 31623861 | 31625953 | 31626127 | 0 | 0 | 3 | 0 | 0 | 0 | 0 | 0 | 0 | 0 | 1 | 0 | 98 | 49 | NA | NA | NA | | NA | | 1 | NA | | | 0 | NA | | 1 |
| SE | 3802 | ZCCHC2 | ZCCHC2 | chr18 | + | 62556202 | 62556297 | 62550347 | 62550460 | 62558686 | 62558770 | 0 | 0 | 20 | 0 | 0 | 2 | 5 | 12 | 0 | 0 | 0 | 0 | 98 | 49 | NA | NA | NA | | NA | | 0.833 | | 1 | | 1 | NA | | -0.167 |
| SE | 3804 | ZBTB11 | ZBTB11 | chr3 | - | 101671129 | 101671361 | 101664963 | 101665808 | 101671977 | 101672213 | 0 | 0 | 3 | 0 | 0 | 0 | 1 | 3 | 0 | 0 | 1 | 0 | 98 | 49 | NA | NA | NA | | NA | | 1 | | 1 | | 0.6 | NA | | 0.2 |
| SE | 3808 | SNHG6 | SNHG6 | chr8 | - | 66922323 | 66922392 | 66921929 | 66922114 | 66922613 | 66922725 | 0 | 0 | 2 | 0 | 0 | 1 | 0 | 4 | 0 | 0 | 0 | 0 | 98 | 49 | NA | NA | NA | | NA | | 0.5 | NA | | | 1 | NA | | -0.5 |
| SE | 3809 | CPNE1 | CPNE1 | chr20 | - | 35655201 | 35655344 | 35632794 | 35632923 | 35658929 | 35658982 | 0 | 0 | 0 | 0 | 0 | 2 | 0 | 1 | 0 | 0 | 0 | 0 | 98 | 49 | NA | NA | NA | | NA | | 0 | NA | | | 1 | NA | | -1 |
| SE | 3883 | MIS18BP1 | MIS18BP1 | chr14 | - | 45206082 | 45206170 | 45204398 | 45204453 | 45210379 | 45210528 | 0 | 0 | 5 | 0 | 0 | 2 | 5 | 3 | 0 | 0 | 0 | 0 | 98 | 49 | NA | NA | NA | | NA | | 0.556 | | 1 | | 1 | NA | | -0.444 |
| SE | 3892 | SETX | SETX | chr9 | - | 132298079 | 132298312 | 132296886 | 132297054 | 132300629 | 132300803 | 0 | 0 | 16 | 0 | 0 | 1 | 0 | 5 | 0 | 0 | 0 | 0 | 98 | 49 | NA | NA | NA | | NA | | 0.889 | NA | | | 1 | NA | | -0.111 |
| SE | 3919 | SMG1 | SMG1 | chr16 | - | 18879494 | 18879719 | 18877130 | 18877232 | 18882164 | 18882338 | 0 | 0 | 3 | 0 | 0 | 2 | 0 | 4 | 0 | 0 | 0 | 0 | 98 | 49 | NA | NA | NA | | NA | | 0.429 | NA | | | 1 | NA | | -0.571 |
| SE | 3920 | SMG1 | SMG1 | chr16 | - | 18889371 | 18889585 | 18885540 | 18885666 | 18890862 | 18890921 | 0 | 0 | 4 | 0 | 0 | 3 | 0 | 2 | 0 | 0 | 0 | 0 | 98 | 49 | NA | NA | NA | | NA | | 0.4 | NA | | | 1 | NA | | -0.6 |
| SE | 3933 | ATF2 | ATF2 | chr2 | - | 175121443 | 175121540 | 175118250 | 175118369 | 175130137 | 175130207 | 0 | 0 | 4 | 0 | 0 | 1 | 0 | 2 | 0 | 0 | 0 | 0 | 98 | 49 | NA | NA | NA | | NA | | 0.667 | NA | | | 1 | NA | | -0.333 |
| SE | 3938 | THAP9-AS1 | THAP9-AS1 | chr4 | - | 82899712 | 82899737 | 82897988 | 82898062 | 82900076 | 82900223 | 0 | 0 | 3 | 0 | 0 | 0 | 0 | 2 | 0 | 0 | 2 | 0 | 74 | 49 | NA | NA | NA | | NA | | 1 | NA | | | 0.398 | NA | | 0.602 |
| SE | 3941 | ZDHHC20 | ZDHHC20 | chr13 | - | 21378660 | 21378738 | 21372570 | 21376655 | 21381433 | 21381549 | 0 | 0 | 3 | 0 | 0 | 2 | 0 | 1 | 0 | 0 | 0 | 0 | 98 | 49 | NA | NA | NA | | NA | | 0.429 | NA | | | 1 | NA | | -0.571 |
| SE | 3942 | ZDHHC20 | ZDHHC20 | chr13 | - | 21381900 | 21381945 | 21381433 | 21381549 | 21382919 | 21383009 | 0 | 0 | 3 | 0 | 0 | 5 | 0 | 4 | 0 | 0 | 1 | 0 | 94 | 49 | NA | NA | NA | | NA | | 0.238 | NA | | | 0.676 | NA | | -0.438 |
| SE | 3948 | ZMYM2 | ZMYM2 | chr13 | + | 20019546 | 20019618 | 20006373 | 20006586 | 20027202 | 20027318 | 0 | 0 | 5 | 0 | 0 | 0 | 0 | 4 | 0 | 0 | 1 | 0 | 98 | 49 | NA | NA | NA | | NA | | 1 | NA | | | 0.667 | NA | | 0.333 |
| SE | 3949 | ZMYM2 | ZMYM2 | chr13 | + | 20026611 | 20026762 | 20019546 | 20019618 | 20027202 | 20027318 | 0 | 0 | 8 | 0 | 0 | 0 | 0 | 2 | 0 | 0 | 1 | 0 | 98 | 49 | NA | NA | NA | | NA | | 1 | NA | | | 0.5 | NA | | 0.5 |
| SE | 3971 | HARS | HARS | chr5 | - | 140675016 | 140675133 | 140674678 | 140674825 | 140676653 | 140676896 | 0 | 0 | 1 | 0 | 0 | 0 | 0 | 2 | 0 | 1 | 0 | 0 | 98 | 49 | NA | NA | NA | | NA | | 1 | | 0 | | 1 | NA | | 0.5 |
| SE | 3992 | CLIP1 | CLIP1 | chr12 | - | 122336631 | 122336748 | 122334647 | 122334705 | 122340752 | 122341697 | 0 | 0 | 6 | 0 | 0 | 0 | 0 | 3 | 0 | 0 | 1 | 0 | 98 | 49 | NA | NA | NA | | NA | | 1 | NA | | | 0.6 | NA | | 0.4 |
| SE | 3995 | CLIP1 | CLIP1 | chr12 | - | 122355114 | 122355312 | 122354452 | 122354556 | 122360958 | 122361181 | 0 | 0 | 7 | 0 | 0 | 0 | 0 | 0 | 0 | 0 | 1 | 0 | 98 | 49 | NA | NA | NA | | NA | | 1 | NA | | | 0 | NA | | 1 |
| SE | 4006 | STAB1 | STAB1 | chr3 | + | 52512340 | 52512436 | 52511649 | 52511745 | 52512595 | 52512643 | 0 | 0 | 13 | 0 | 0 | 2 | 3 | 1 | 0 | 0 | 0 | 0 | 98 | 49 | NA | NA | NA | | NA | | 0.765 | | 1 | | 1 | NA | | -0.235 |
| SE | 4065 | FBXL5 | FBXL5 | chr4 | - | 15641523 | 15641627 | 15640787 | 15640883 | 15644492 | 15644708 | 0 | 0 | 1 | 0 | 0 | 31 | 0 | 1 | 0 | 0 | 8 | 0 | 98 | 49 | NA | NA | NA | | NA | | 0.016 | NA | | | 0.059 | NA | | -0.043 |
| SE | 4071 | TMCO3 | TMCO3 | chr13 | + | 113503495 | 113503588 | 113501751 | 113501880 | 113510237 | 113510424 | 0 | 0 | 1 | 0 | 0 | 2 | 0 | 3 | 0 | 0 | 0 | 0 | 98 | 49 | NA | NA | NA | | NA | | 0.2 | NA | | | 1 | NA | | -0.8 |
| SE | 4087 | TNFRSF1A | TNFRSF1A | chr12 | - | 6330852 | 6330926 | 6330597 | 6330711 | 6333068 | 6333147 | 0 | 0 | 21 | 0 | 0 | 1 | 2 | 3 | 0 | 0 | 1 | 0 | 98 | 49 | NA | NA | NA | | NA | | 0.913 | | 1 | | 0.6 | NA | | 0.113 |
| SE | 4099 | FAM13B | FAM13B | chr5 | - | 137946227 | 137946311 | 137945901 | 137945997 | 137948954 | 137949184 | 0 | 0 | 5 | 0 | 0 | 0 | 0 | 2 | 0 | 0 | 2 | 0 | 98 | 49 | NA | NA | NA | | NA | | 1 | NA | | | 0.333 | NA | | 0.667 |
| SE | 4107 | EVI5 | EVI5 | chr1 | - | 92693801 | 92693899 | 92677157 | 92677218 | 92694298 | 92694388 | 0 | 0 | 4 | 0 | 0 | 1 | 1 | 0 | 0 | 0 | 0 | 0 | 98 | 49 | NA | NA | NA | | NA | | 0.667 | | 1 | NA | | NA | | -0.333 |
| SE | 4112 | ATP2B1 | ATP2B1 | chr12 | - | 89603711 | 89603925 | 89603042 | 89603254 | 89604154 | 89604346 | 0 | 0 | 6 | 0 | 0 | 0 | 0 | 5 | 0 | 0 | 1 | 0 | 98 | 49 | NA | NA | NA | | NA | | 1 | NA | | | 0.714 | NA | | 0.286 |
| SE | 4113 | ATP2B1 | ATP2B1 | chr12 | - | 89616801 | 89617039 | 89611192 | 89611372 | 89619998 | 89620240 | 0 | 0 | 11 | 0 | 0 | 0 | 0 | 17 | 0 | 0 | 3 | 0 | 98 | 49 | NA | NA | NA | | NA | | 1 | NA | | | 0.739 | NA | | 0.261 |
| SE | 4143 | TBC1D15 | TBC1D15 | chr12 | + | 71920730 | 71920847 | 71918450 | 71918548 | 71921367 | 71921454 | 0 | 0 | 1 | 0 | 0 | 0 | 0 | 1 | 0 | 0 | 1 | 0 | 98 | 49 | NA | NA | NA | | NA | | 1 | NA | | | 0.333 | NA | | 0.667 |
| SE | 4151 | MDM2 | MDM2 | chr12 | + | 68816811 | 68816945 | 68813553 | 68813628 | 68820324 | 68820374 | 0 | 0 | 12 | 0 | 0 | 1 | 0 | 1 | 0 | 0 | 0 | 0 | 98 | 49 | NA | NA | NA | | NA | | 0.857 | NA | | | 1 | NA | | -0.143 |
| SE | 4176 | MARS | MARS | chr12 | + | 57489895 | 57489971 | 57489423 | 57489558 | 57490206 | 57490379 | 0 | 0 | 5 | 0 | 0 | 0 | 0 | 2 | 0 | 0 | 1 | 0 | 98 | 49 | NA | NA | NA | | NA | | 1 | NA | | | 0.5 | NA | | 0.5 |
| SE | 4199 | SUPT20H | SUPT20H | chr13 | - | 37048563 | 37048599 | 37047877 | 37047936 | 37051487 | 37051583 | 0 | 0 | 0 | 0 | 0 | 1 | 0 | 1 | 0 | 0 | 0 | 0 | 85 | 49 | NA | NA | NA | | NA | | 0 | NA | | | 1 | NA | | -1 |
| SE | 4211 | SMARCC2 | SMARCC2 | chr12 | - | 56172936 | 56173029 | 56172584 | 56172704 | 56173695 | 56173849 | 0 | 0 | 0 | 0 | 0 | 1 | 0 | 1 | 0 | 0 | 0 | 0 | 98 | 49 | NA | NA | NA | | NA | | 0 | NA | | | 1 | NA | | -1 |
| SE | 4234 | EZH1 | EZH1 | chr17 | - | 42724304 | 42724424 | 42722794 | 42722915 | 42727634 | 42727763 | 0 | 0 | 11 | 0 | 0 | 1 | 0 | 5 | 0 | 0 | 0 | 0 | 98 | 49 | NA | NA | NA | | NA | | 0.846 | NA | | | 1 | NA | | -0.154 |
| SE | 4243 | ARL8A | ARL8A | chr1 | - | 202135458 | 202135526 | 202135149 | 202135220 | 202135706 | 202135800 | 0 | 0 | 11 | 0 | 0 | 1 | 0 | 9 | 0 | 0 | 0 | 0 | 98 | 49 | NA | NA | NA | | NA | | 0.846 | NA | | | 1 | NA | | -0.154 |
| SE | 4261 | CLPX | CLPX | chr15 | - | 65164028 | 65164188 | 65162603 | 65162645 | 65166630 | 65166785 | 0 | 0 | 1 | 0 | 0 | 0 | 0 | 1 | 0 | 2 | 0 | 0 | 98 | 49 | NA | NA | NA | | NA | | 1 | | 0 | | 1 | NA | | 0.5 |
| SE | 4263 | ITGB7 | ITGB7 | chr12 | - | 53192690 | 53192910 | 53192329 | 53192538 | 53193139 | 53193363 | 0 | 0 | 0 | 0 | 0 | 1 | 0 | 1 | 0 | 0 | 0 | 0 | 98 | 49 | NA | NA | NA | | NA | | 0 | NA | | | 1 | NA | | -1 |
| SE | 4264 | ITGB7 | ITGB7 | chr12 | - | 53195625 | 53195721 | 53195373 | 53195463 | 53196040 | 53196199 | 0 | 0 | 2 | 0 | 0 | 0 | 0 | 0 | 0 | 0 | 1 | 0 | 98 | 49 | NA | NA | NA | | NA | | 1 | NA | | | 0 | NA | | 1 |
| SE | 4275 | TDRD7 | TDRD7 | chr9 | + | 97470557 | 97470669 | 97464841 | 97465028 | 97472292 | 97472495 | 0 | 0 | 3 | 0 | 0 | 1 | 0 | 5 | 0 | 0 | 0 | 0 | 98 | 49 | NA | NA | NA | | NA | | 0.6 | NA | | | 1 | NA | | -0.4 |
| SE | 4278 | SIGLEC10 | SIGLEC10 | chr19 | - | 51415897 | 51416167 | 51415567 | 51415615 | 51416309 | 51416357 | 0 | 0 | 1 | 0 | 0 | 0 | 0 | 0 | 0 | 0 | 1 | 0 | 98 | 49 | NA | NA | NA | | NA | | 1 | NA | | | 0 | NA | | 1 |
| SE | 4307 | CERS5 | CERS5 | chr12 | - | 50144768 | 50144969 | 50143951 | 50144057 | 50167100 | 50167533 | 0 | 0 | 0 | 0 | 0 | 2 | 0 | 1 | 0 | 0 | 1 | 0 | 98 | 49 | NA | NA | NA | | NA | | 0 | NA | | | 0.333 | NA | | -0.333 |
| SE | 4311 | SLC39A9 | SLC39A9 | chr14 | + | 69453240 | 69453309 | 69442068 | 69442266 | 69454811 | 69454897 | 0 | 0 | 0 | 0 | 0 | 1 | 7 | 3 | 0 | 0 | 0 | 0 | 98 | 49 | NA | NA | NA | | NA | | 0 | | 1 | | 1 | NA | | -1 |
| SE | 4318 | BIRC6 | BIRC6 | chr2 | + | 32435495 | 32435585 | 32433643 | 32433804 | 32436052 | 32436184 | 0 | 0 | 1 | 0 | 0 | 0 | 0 | 1 | 0 | 0 | 2 | 0 | 98 | 49 | NA | NA | NA | | NA | | 1 | NA | | | 0.2 | NA | | 0.8 |
| SE | 4320 | BIRC6 | BIRC6 | chr2 | + | 32464508 | 32464823 | 32463193 | 32463381 | 32465064 | 32465164 | 0 | 0 | 1 | 0 | 0 | 1 | 0 | 4 | 0 | 0 | 0 | 0 | 98 | 49 | NA | NA | NA | | NA | | 0.333 | NA | | | 1 | NA | | -0.667 |
| SE | 4321 | BIRC6 | BIRC6 | chr2 | + | 32465064 | 32465164 | 32464508 | 32464823 | 32467524 | 32467739 | 0 | 0 | 2 | 0 | 0 | 0 | 0 | 4 | 0 | 0 | 2 | 0 | 98 | 49 | NA | NA | NA | | NA | | 1 | NA | | | 0.5 | NA | | 0.5 |
| SE | 4335 | NCF2 | NCF2 | chr1 | - | 183574486 | 183574621 | 183570779 | 183570839 | 183577598 | 183577707 | 0 | 0 | 30 | 0 | 0 | 1 | 0 | 5 | 0 | 0 | 0 | 0 | 98 | 49 | NA | NA | NA | | NA | | 0.937 | NA | | | 1 | NA | | -0.063 |
| SE | 4336 | NCF2 | NCF2 | chr1 | - | 183574486 | 183574621 | 183573184 | 183573292 | 183577598 | 183577707 | 0 | 0 | 51 | 0 | 0 | 12 | 0 | 15 | 0 | 0 | 7 | 0 | 98 | 49 | NA | NA | NA | | NA | | 0.68 | NA | | | 0.517 | NA | | 0.163 |
| SE | 4340 | SLC38A2 | SLC38A2 | chr12 | - | 46363020 | 46363145 | 46362493 | 46362638 | 46363725 | 46363826 | 0 | 0 | 6 | 0 | 0 | 0 | 0 | 4 | 0 | 0 | 1 | 0 | 98 | 49 | NA | NA | NA | | NA | | 1 | NA | | | 0.667 | NA | | 0.333 |
| SE | 4344 | ARID2 | ARID2 | chr12 | + | 45836740 | 45836991 | 45836588 | 45836655 | 45837320 | 45837417 | 0 | 0 | 2 | 0 | 0 | 1 | 0 | 7 | 0 | 0 | 1 | 0 | 98 | 49 | NA | NA | NA | | NA | | 0.5 | NA | | | 0.778 | NA | | -0.278 |
| SE | 4353 | APAF1 | APAF1 | chr12 | + | 98665552 | 98665791 | 98662674 | 98662806 | 98666189 | 98666357 | 0 | 0 | 2 | 0 | 0 | 1 | 0 | 2 | 0 | 0 | 0 | 0 | 98 | 49 | NA | NA | NA | | NA | | 0.5 | NA | | | 1 | NA | | -0.5 |
| SE | 4354 | APAF1 | APAF1 | chr12 | + | 98677424 | 98677551 | 98671534 | 98671719 | 98680276 | 98680402 | 0 | 0 | 7 | 0 | 0 | 1 | 0 | 4 | 0 | 0 | 0 | 0 | 98 | 49 | NA | NA | NA | | NA | | 0.778 | NA | | | 1 | NA | | -0.222 |
| SE | 4356 | APAF1 | APAF1 | chr12 | + | 98706484 | 98706610 | 98699407 | 98699569 | 98708584 | 98708704 | 0 | 0 | 7 | 0 | 0 | 1 | 0 | 2 | 0 | 0 | 0 | 0 | 98 | 49 | NA | NA | NA | | NA | | 0.778 | NA | | | 1 | NA | | -0.222 |
| SE | 4360 | LRRK2 | LRRK2 | chr12 | + | 40283874 | 40284133 | 40278090 | 40278261 | 40287350 | 40287539 | 0 | 0 | 46 | 0 | 0 | 0 | 0 | 21 | 0 | 0 | 1 | 0 | 98 | 49 | NA | NA | NA | | NA | | 1 | NA | | | 0.913 | NA | | 0.087 |
| SE | 4375 | FAR2 | FAR2 | chr12 | + | 29297020 | 29297200 | 29293299 | 29293475 | 29307657 | 29307835 | 0 | 0 | 1 | 0 | 0 | 1 | 0 | 2 | 0 | 0 | 0 | 0 | 98 | 49 | NA | NA | NA | | NA | | 0.333 | NA | | | 1 | NA | | -0.667 |
| SE | 4376 | FAR2 | FAR2 | chr12 | + | 29316840 | 29317012 | 29311882 | 29311950 | 29321794 | 29321924 | 0 | 0 | 1 | 0 | 0 | 1 | 0 | 2 | 0 | 0 | 0 | 0 | 98 | 49 | NA | NA | NA | | NA | | 0.333 | NA | | | 1 | NA | | -0.667 |
| SE | 4383 | STK38L | STK38L | chr12 | + | 27315014 | 27315117 | 27314503 | 27314658 | 27315288 | 27315350 | 0 | 0 | 2 | 0 | 0 | 3 | 0 | 1 | 0 | 0 | 0 | 0 | 98 | 49 | NA | NA | NA | | NA | | 0.25 | NA | | | 1 | NA | | -0.75 |
| SE | 4384 | STK38L | STK38L | chr12 | + | 27319327 | 27319423 | 27317895 | 27318019 | 27322142 | 27322234 | 0 | 0 | 8 | 0 | 0 | 0 | 0 | 7 | 0 | 0 | 1 | 0 | 98 | 49 | NA | NA | NA | | NA | | 1 | NA | | | 0.778 | NA | | 0.222 |
| SE | 4394 | TBCEL | TBCEL | chr11 | + | 121055051 | 121055308 | 121053550 | 121053732 | 121058344 | 121058471 | 0 | 0 | 1 | 0 | 0 | 0 | 0 | 0 | 0 | 0 | 1 | 0 | 98 | 49 | NA | NA | NA | | NA | | 1 | NA | | | 0 | NA | | 1 |
| SE | 4410 | OSM | OSM | chr22 | - | 30265001 | 30265144 | 30262829 | 30264464 | 30266765 | 30266840 | 0 | 0 | 6 | 0 | 0 | 0 | 0 | 13 | 0 | 0 | 1 | 0 | 98 | 49 | NA | NA | NA | | NA | | 1 | NA | | | 0.867 | NA | | 0.133 |
| SE | 4434 | KANSL2 | KANSL2 | chr12 | - | 48679035 | 48679150 | 48671798 | 48671962 | 48679654 | 48679833 | 0 | 0 | 6 | 0 | 0 | 2 | 0 | 3 | 0 | 0 | 0 | 0 | 98 | 49 | NA | NA | NA | | NA | | 0.6 | NA | | | 1 | NA | | -0.4 |
| SE | 4435 | FRS2 | FRS2 | chr12 | + | 69480298 | 69480359 | 69470348 | 69470530 | 69530864 | 69530960 | 0 | 0 | 0 | 0 | 0 | 1 | 0 | 3 | 0 | 0 | 1 | 0 | 98 | 49 | NA | NA | NA | | NA | | 0 | NA | | | 0.6 | NA | | -0.6 |
| SE | 4460 | CLEC7A | CLEC7A | chr12 | - | 10126570 | 10126708 | 10125296 | 10125448 | 10127746 | 10127845 | 0 | 0 | 18 | 0 | 0 | 11 | 0 | 13 | 0 | 2 | 7 | 0 | 98 | 49 | NA | NA | NA | | NA | | 0.45 | | 0 | | 0.481 | NA | | 0.21 |
| SE | 4469 | MACF1 | MACF1 | chr1 | + | 39378460 | 39378523 | 39372478 | 39372596 | 39379202 | 39379444 | 0 | 0 | 0 | 0 | 0 | 1 | 0 | 1 | 0 | 0 | 3 | 0 | 98 | 49 | NA | NA | NA | | NA | | 0 | NA | | | 0.143 | NA | | -0.143 |
| SE | 4470 | MACF1 | MACF1 | chr1 | + | 39451051 | 39451211 | 39448593 | 39448763 | 39452155 | 39452350 | 0 | 0 | 2 | 0 | 0 | 0 | 0 | 1 | 0 | 0 | 1 | 0 | 98 | 49 | NA | NA | NA | | NA | | 1 | NA | | | 0.333 | NA | | 0.667 |
| SE | 4478 | MYO1G | MYO1G | chr7 | - | 44976563 | 44976657 | 44975483 | 44975649 | 44976862 | 44977071 | 0 | 0 | 7 | 0 | 0 | 0 | 0 | 3 | 0 | 0 | 1 | 0 | 98 | 49 | NA | NA | NA | | NA | | 1 | NA | | | 0.6 | NA | | 0.4 |
| SE | 4505 | PTPN6 | PTPN6 | chr12 | + | 6960662 | 6960945 | 6960343 | 6960435 | 6961125 | 6961316 | 0 | 0 | 11 | 0 | 0 | 1 | 4 | 12 | 0 | 0 | 0 | 0 | 98 | 49 | NA | NA | NA | | NA | | 0.846 | | 1 | | 1 | NA | | -0.154 |
| SE | 4511 | PDE4B | PDE4B | chr1 | + | 66368786 | 66368969 | 66367942 | 66368065 | 66372312 | 66374579 | 0 | 0 | 21 | 0 | 0 | 0 | 0 | 21 | 0 | 0 | 1 | 0 | 98 | 49 | NA | NA | NA | | NA | | 1 | NA | | | 0.913 | NA | | 0.087 |
| SE | 4518 | CHD4 | CHD4 | chr12 | - | 6582136 | 6582281 | 6581648 | 6581814 | 6582614 | 6582748 | 0 | 0 | 4 | 0 | 0 | 1 | 0 | 5 | 0 | 0 | 1 | 0 | 98 | 49 | NA | NA | NA | | NA | | 0.667 | NA | | | 0.714 | NA | | -0.047 |
| SE | 4519 | CHD4 | CHD4 | chr12 | - | 6588297 | 6588422 | 6587711 | 6587949 | 6591465 | 6591583 | 0 | 0 | 13 | 0 | 0 | 0 | 0 | 3 | 0 | 0 | 3 | 0 | 98 | 49 | NA | NA | NA | | NA | | 1 | NA | | | 0.333 | NA | | 0.667 |
| SE | 4521 | GLIPR1 | GLIPR1 | chr12 | + | 75490405 | 75490518 | 75481833 | 75482079 | 75495576 | 75495662 | 0 | 0 | 15 | 0 | 0 | 1 | 0 | 5 | 0 | 0 | 0 | 0 | 98 | 49 | NA | NA | NA | | NA | | 0.882 | NA | | | 1 | NA | | -0.118 |
| SE | 4528 | PARP11 | PARP11 | chr12 | - | 3863835 | 3863957 | 3829889 | 3830018 | 3873211 | 3873448 | 0 | 0 | 0 | 0 | 0 | 1 | 0 | 1 | 0 | 0 | 0 | 0 | 98 | 49 | NA | NA | NA | | NA | | 0 | NA | | | 1 | NA | | -1 |
| SE | 4535 | TM9SF2 | TM9SF2 | chr13 | + | 99539445 | 99539557 | 99537738 | 99537863 | 99540713 | 99540793 | 0 | 0 | 11 | 0 | 0 | 1 | 0 | 2 | 0 | 0 | 0 | 0 | 98 | 49 | NA | NA | NA | | NA | | 0.846 | NA | | | 1 | NA | | -0.154 |
| SE | 4553 | MED23 | MED23 | chr6 | - | 131620627 | 131620729 | 131619826 | 131619896 | 131621880 | 131621979 | 0 | 0 | 4 | 0 | 0 | 0 | 0 | 1 | 0 | 0 | 2 | 0 | 98 | 49 | NA | NA | NA | | NA | | 1 | NA | | | 0.2 | NA | | 0.8 |
| SE | 4560 | SLC6A6 | SLC6A6 | chr3 | + | 14468087 | 14468212 | 14467852 | 14467956 | 14472204 | 14472317 | 0 | 0 | 3 | 0 | 0 | 1 | 0 | 1 | 0 | 0 | 0 | 0 | 98 | 49 | NA | NA | NA | | NA | | 0.6 | NA | | | 1 | NA | | -0.4 |
| SE | 4562 | SNX19 | SNX19 | chr11 | - | 130880621 | 130880806 | 130879623 | 130879711 | 130903254 | 130903384 | 0 | 0 | 2 | 0 | 0 | 0 | 0 | 0 | 0 | 0 | 2 | 0 | 98 | 49 | NA | NA | NA | | NA | | 1 | NA | | | 0 | NA | | 1 |
| SE | 4584 | SRPR | SRPR | chr11 | - | 126264375 | 126264539 | 126264190 | 126264289 | 126264958 | 126265172 | 0 | 0 | 3 | 0 | 0 | 0 | 0 | 3 | 0 | 0 | 2 | 0 | 98 | 49 | NA | NA | NA | | NA | | 1 | NA | | | 0.429 | NA | | 0.571 |
| SE | 4590 | CUL4A | CUL4A | chr13 | + | 113239432 | 113239551 | 113236822 | 113236890 | 113242967 | 113243160 | 0 | 0 | 5 | 0 | 0 | 1 | 0 | 1 | 0 | 0 | 0 | 0 | 98 | 49 | NA | NA | NA | | NA | | 0.714 | NA | | | 1 | NA | | -0.286 |
| SE | 4594 | KMT2C | KMT2C | chr7 | - | 152146598 | 152146735 | 152145152 | 152145295 | 152148032 | 152149152 | 0 | 0 | 11 | 0 | 0 | 1 | 0 | 13 | 0 | 0 | 1 | 0 | 98 | 49 | NA | NA | NA | | NA | | 0.846 | NA | | | 0.867 | NA | | -0.021 |

| | | | | | | | | | | | | | | | | | | | | | | | | | | | | | | | | | | | | |
|---|---|---|---|---|---|---|---|---|---|---|---|---|---|---|---|---|---|---|---|---|---|---|---|---|---|---|---|---|---|---|---|---|---|---|---|---|
| SE | 4595 | KMT2C | KMT2C | chr7 | - | 152182973 | 152183156 | 152180710 | 152182594 | 152185557 | 152185631 | 0 | 0 | 9 | 0 | 0 | 3 | 1 | 9 | 0 | 0 | 0 | 0 | 98 | 49 | NA | NA | NA | | NA | 0.6 | | 1 | 1 | NA | -0.4 |
| SE | 4596 | KMT2C | KMT2C | chr7 | - | 152195906 | 152196011 | 152194439 | 152194568 | 152199278 | 152199459 | 0 | 0 | 8 | 0 | 0 | 0 | 0 | 12 | 0 | 0 | 1 | 0 | 98 | 49 | NA | NA | NA | | NA | 1 | NA | | 0.857 | NA | 0.143 |
| SE | 4597 | KMT2C | KMT2C | chr7 | - | 152224014 | 152224179 | 152222572 | 152222682 | 152224434 | 152224616 | 0 | 0 | 3 | 0 | 0 | 0 | 0 | 3 | 0 | 0 | 1 | 0 | 98 | 49 | NA | NA | NA | | NA | 1 | NA | | 0.6 | NA | 0.4 |
| SE | 4598 | KMT2C | KMT2C | chr7 | - | 152238706 | 152238826 | 152235816 | 152235933 | 152247901 | 152248620 | 0 | 0 | 9 | 0 | 0 | 0 | 0 | 6 | 0 | 0 | 1 | 0 | 98 | 49 | NA | NA | NA | | NA | 1 | NA | | 0.75 | NA | 0.25 |
| SE | 4607 | KIAA0319 | KIAA0319 | chr6 | - | 24556606 | 24556729 | 24554540 | 24554631 | 24559012 | 24559155 | 0 | 0 | 0 | 0 | 0 | 1 | 0 | 1 | 0 | 0 | 0 | 0 | 98 | 49 | NA | NA | NA | | NA | 0 | NA | | 1 | NA | -1 |
| SE | 4619 | PAN3 | PAN3 | chr13 | + | 28177864 | 28177935 | 28176492 | 28176559 | 28220230 | 28220378 | 0 | 0 | 12 | 0 | 0 | 1 | 0 | 3 | 0 | 0 | 0 | 0 | 98 | 49 | NA | NA | NA | | NA | 0.857 | NA | | 1 | NA | -0.143 |
| SE | 4620 | PAN3 | PAN3 | chr13 | + | 28197184 | 28197346 | 28177864 | 28177935 | 28220230 | 28220378 | 0 | 0 | 1 | 0 | 0 | 5 | 0 | 0 | 0 | 0 | 1 | 0 | 98 | 49 | NA | NA | NA | | NA | 0.091 | NA | | 0 | NA | 0.091 |
| SE | 4621 | PAN3 | PAN3 | chr13 | + | 28271980 | 28272071 | 28270700 | 28270866 | 28277236 | 28277376 | 0 | 0 | 0 | 0 | 0 | 1 | 0 | 4 | 0 | 0 | 0 | 0 | 98 | 49 | NA | NA | NA | | NA | 0 | NA | | 1 | NA | -1 |
| SE | 4630 | SORL1 | SORL1 | chr11 | + | 121608103 | 121608176 | 121607185 | 121607290 | 121611075 | 121611158 | 0 | 0 | 31 | 0 | 0 | 1 | 3 | 18 | 0 | 0 | 0 | 0 | 98 | 49 | NA | NA | NA | | NA | 0.939 | | 1 | 1 | NA | -0.061 |
| SE | 4672 | IL10RA | IL10RA | chr11 | + | 117988381 | 117988502 | 117986390 | 117986534 | 117993240 | 117993410 | 0 | 0 | 3 | 0 | 0 | 1 | 0 | 2 | 0 | 0 | 0 | 0 | 98 | 49 | NA | NA | NA | | NA | 0.6 | NA | | 1 | NA | -0.4 |
| SE | 4683 | PITRM1 | PITRM1 | chr10 | - | 3156929 | 3157064 | 3155590 | 3155729 | 3157434 | 3157531 | 0 | 0 | 2 | 0 | 0 | 0 | 1 | 1 | 0 | 0 | 1 | 0 | 98 | 49 | NA | NA | NA | | NA | 1 | | 1 | 0.333 | NA | 0.334 |
| SE | 4689 | IL6ST | IL6ST | chr5 | - | 55959625 | 55959671 | 55957208 | 55957291 | 55960401 | 55960561 | 0 | 0 | 0 | 0 | 0 | 5 | 0 | 2 | 0 | 0 | 2 | 0 | 95 | 49 | NA | NA | NA | | NA | 0 | NA | | 0.34 | NA | -0.34 |
| SE | 4733 | CDA | CDA | chr1 | + | 20613841 | 20613899 | 20604927 | 20605039 | 20618451 | 20618907 | 0 | 0 | 6 | 0 | 0 | 2 | 0 | 3 | 0 | 0 | 1 | 0 | 98 | 49 | NA | NA | NA | | NA | 0.6 | NA | | 0.6 | NA | 0 |
| SE | 4771 | ATM | ATM | chr11 | + | 108248932 | 108249102 | 108246963 | 108247127 | 108250700 | 108251072 | 0 | 0 | 8 | 0 | 0 | 0 | 0 | 3 | 0 | 3 | 0 | 0 | 98 | 49 | NA | NA | NA | | NA | 1 | | 0 | 1 | NA | 0.5 |
| SE | 4774 | DDX6 | DDX6 | chr11 | - | 118758773 | 118758902 | 118757170 | 118757287 | 118759921 | 118760044 | 0 | 0 | 9 | 0 | 0 | 0 | 0 | 6 | 0 | 0 | 2 | 0 | 98 | 49 | NA | NA | NA | | NA | 1 | NA | | 0.6 | NA | 0.4 |
| SE | 4775 | DDX6 | DDX6 | chr11 | - | 118763211 | 118763306 | 118759921 | 118760044 | 118765208 | 118765355 | 0 | 0 | 5 | 0 | 0 | 0 | 1 | 15 | 0 | 0 | 1 | 0 | 98 | 49 | NA | NA | NA | | NA | 1 | | 1 | 0.882 | NA | 0.059 |
| SE | 4791 | OXSR1 | OXSR1 | chr3 | + | 38223811 | 38223913 | 38221577 | 38221687 | 38224570 | 38224704 | 0 | 0 | 9 | 0 | 0 | 0 | 0 | 8 | 0 | 0 | 1 | 0 | 98 | 49 | NA | NA | NA | | NA | 1 | NA | | 0.8 | NA | 0.2 |
| SE | 4795 | CASP1 | CASP1 | chr11 | - | 105033063 | 105033126 | 105031164 | 105031280 | 105034207 | 105034474 | 0 | 0 | 28 | 0 | 0 | 6 | 0 | 7 | 0 | 0 | 4 | 0 | 98 | 49 | NA | NA | NA | | NA | 0.7 | NA | | 0.467 | NA | 0.233 |
| SE | 4804 | CASP4 | CASP4 | chr11 | - | 104951895 | 104952005 | 104950924 | 104951098 | 104954746 | 104955001 | 0 | 0 | 13 | 0 | 0 | 3 | 0 | 6 | 0 | 0 | 0 | 0 | 98 | 49 | NA | NA | NA | | NA | 0.684 | NA | | 1 | NA | -0.316 |
| SE | 4810 | CCNH | CCNH | chr5 | - | 87404843 | 87405007 | 87401701 | 87401772 | 87407975 | 87408186 | 0 | 0 | 11 | 0 | 0 | 0 | 0 | 7 | 0 | 0 | 1 | 0 | 98 | 49 | NA | NA | NA | | NA | 1 | NA | | 0.778 | NA | 0.222 |
| SE | 4811 | BIRC3 | BIRC3 | chr11 | + | 102328051 | 102328130 | 102325465 | 102325565 | 102328896 | 102328945 | 0 | 0 | 3 | 0 | 0 | 0 | 0 | 6 | 0 | 0 | 1 | 0 | 98 | 49 | NA | NA | NA | | NA | 1 | NA | | 0.75 | NA | 0.25 |
| SE | 4817 | BCAP31 | BCAP31 | chrX | - | 153702934 | 153703058 | 153702006 | 153702107 | 153703958 | 153704094 | 1 | 0 | 11 | 0 | 0 | 1 | 0 | 4 | 0 | 0 | 0 | 0 | 98 | 49 | NA | NA | | 1 | NA | 0.846 | NA | | 1 | NA | -0.077 |
| SE | 4825 | TRIM8 | TRIM8 | chr10 | + | 102654652 | 102654748 | 102644494 | 102645187 | 102655079 | 102655313 | 0 | 0 | 8 | 0 | 0 | 1 | 0 | 8 | 0 | 0 | 1 | 0 | 98 | 49 | NA | NA | NA | | NA | 0.8 | NA | | 0.8 | NA | 0 |
| SE | 4864 | CEP295 | CEP295 | chr11 | + | 93702775 | 93702919 | 93702459 | 93702637 | 93706744 | 93706897 | 0 | 0 | 2 | 0 | 0 | 0 | 0 | 0 | 0 | 0 | 1 | 0 | 98 | 49 | NA | NA | NA | | NA | 1 | NA | | 0 | NA | 1 |
| SE | 4876 | TPP1 | TPP1 | chr11 | - | 6616004 | 6616074 | 6615441 | 6615562 | 6616314 | 6616503 | 0 | 0 | 13 | 0 | 0 | 1 | 0 | 5 | 0 | 0 | 0 | 0 | 98 | 49 | NA | NA | NA | | NA | 0.867 | NA | | 1 | NA | -0.133 |
| SE | 4885 | DCAF11 | DCAF11 | chr14 | + | 24117667 | 24117732 | 24117265 | 24117393 | 24118054 | 24118155 | 0 | 0 | 2 | 0 | 0 | 1 | 0 | 4 | 0 | 0 | 0 | 0 | 98 | 49 | NA | NA | NA | | NA | 0.5 | NA | | 1 | NA | -0.5 |
| SE | 4889 | TXNL1 | TXNL1 | chr18 | - | 56626161 | 56626266 | 56624287 | 56624461 | 56626360 | 56626457 | 0 | 0 | 3 | 0 | 0 | 1 | 0 | 2 | 0 | 0 | 0 | 0 | 98 | 49 | NA | NA | NA | | NA | 0.6 | NA | | 1 | NA | -0.4 |
| SE | 4906 | PAK1 | PAK1 | chr11 | - | 77337323 | 77337423 | 77336085 | 77336282 | 77340645 | 77340763 | 0 | 0 | 12 | 0 | 0 | 0 | 0 | 7 | 0 | 0 | 2 | 0 | 98 | 49 | NA | NA | NA | | NA | 1 | NA | | 0.636 | NA | 0.364 |
| SE | 4924 | C2CD3 | C2CD3 | chr11 | - | 74161398 | 74161556 | 74139604 | 74139828 | 74168343 | 74168613 | 0 | 0 | 2 | 0 | 0 | 1 | 0 | 2 | 0 | 0 | 0 | 0 | 98 | 49 | NA | NA | NA | | NA | 0.5 | NA | | 1 | NA | -0.5 |
| SE | 4949 | RBM6 | RBM6 | chr3 | + | 50059648 | 50059746 | 50058401 | 50058562 | 50060955 | 50060998 | 0 | 0 | 2 | 0 | 0 | 2 | 0 | 2 | 0 | 0 | 0 | 0 | 98 | 49 | NA | NA | NA | | NA | 0.333 | NA | | 0.5 | NA | -0.167 |
| SE | 4955 | N4BP1 | N4BP1 | chr16 | - | 48546146 | 48546254 | 48538725 | 48543261 | 48548006 | 48548114 | 0 | 0 | 2 | 0 | 0 | 1 | 0 | 1 | 0 | 0 | 0 | 0 | 98 | 49 | NA | NA | NA | | NA | 0.5 | NA | | 1 | NA | -0.5 |
| SE | 4965 | DDX3X | DDX3X | chrX | + | 41346858 | 41347012 | 41346504 | 41346622 | 41347314 | 41347451 | 0 | 0 | 33 | 0 | 0 | 0 | 1 | 31 | 0 | 0 | 1 | 0 | 98 | 49 | NA | NA | NA | | NA | 1 | | 1 | 0.939 | NA | 0.03 |
| SE | 4981 | RTCB | RTCB | chr22 | - | 32397940 | 32398100 | 32396073 | 32396249 | 32399602 | 32399759 | 0 | 0 | 4 | 0 | 0 | 1 | 0 | 1 | 0 | 0 | 0 | 0 | 98 | 49 | NA | NA | NA | | NA | 0.667 | NA | | 1 | NA | -0.333 |
| SE | 4983 | NCOR1 | NCOR1 | chr17 | - | 16048844 | 16048988 | 16046950 | 16047093 | 16057513 | 16057737 | 0 | 0 | 3 | 0 | 0 | 2 | 0 | 3 | 0 | 0 | 0 | 0 | 98 | 49 | NA | NA | NA | | NA | 0.429 | NA | | 1 | NA | -0.571 |
| SE | 4988 | HEXB | HEXB | chr5 | + | 74713505 | 74713635 | 74705218 | 74705320 | 74715509 | 74715690 | 0 | 0 | 12 | 0 | 0 | 2 | 5 | 3 | 0 | 0 | 0 | 0 | 98 | 49 | NA | NA | NA | | NA | 0.75 | | 1 | 1 | NA | -0.25 |
| SE | 4991 | TOPBP1 | TOPBP1 | chr3 | - | 133624051 | 133624175 | 133623310 | 133623457 | 133628361 | 133628471 | 0 | 0 | 2 | 0 | 0 | 2 | 0 | 1 | 0 | 0 | 0 | 0 | 98 | 49 | NA | NA | NA | | NA | 0.333 | NA | | 1 | NA | -0.667 |
| SE | 5001 | ABHD2 | ABHD2 | chr15 | + | 89104181 | 89104270 | 89103902 | 89104044 | 89113724 | 89113824 | 0 | 0 | 0 | 0 | 0 | 2 | 0 | 1 | 0 | 0 | 2 | 0 | 98 | 49 | NA | NA | NA | | NA | 0 | NA | | 0.2 | NA | -0.2 |
| SE | 5007 | RINT1 | RINT1 | chr7 | + | 105550260 | 105550486 | 105550054 | 105550165 | 105551569 | 105551707 | 0 | 0 | 1 | 0 | 0 | 0 | 0 | 0 | 0 | 0 | 2 | 0 | 98 | 49 | NA | NA | NA | | NA | 1 | NA | | 0 | NA | 1 |
| SE | 5009 | RPS6KB2 | RPS6KB2 | chr11 | + | 67433125 | 67433216 | 67432951 | 67433042 | 67433339 | 67433447 | 0 | 0 | 0 | 0 | 0 | 1 | 0 | 2 | 0 | 0 | 0 | 0 | 98 | 49 | NA | NA | NA | | NA | 0 | NA | | 1 | NA | -1 |
| SE | 5014 | ARHGAP25 | ARHGAP25 | chr2 | + | 68787839 | 68787956 | 68782232 | 68782320 | 68807272 | 68807480 | 0 | 0 | 6 | 0 | 0 | 0 | 0 | 8 | 0 | 0 | 1 | 0 | 98 | 49 | NA | NA | NA | | NA | 1 | NA | | 0.8 | NA | 0.2 |
| SE | 5019 | ZFYVE16 | ZFYVE16 | chr5 | + | 80448025 | 80448404 | 80445262 | 80445405 | 80449590 | 80449713 | 0 | 0 | 8 | 0 | 0 | 0 | 0 | 1 | 0 | 0 | 1 | 0 | 98 | 49 | NA | NA | NA | | NA | 1 | NA | | 0.333 | NA | 0.667 |
| SE | 5020 | ZFYVE16 | ZFYVE16 | chr5 | + | 80472760 | 80472923 | 80459413 | 80459494 | 80473753 | 80473859 | 0 | 0 | 2 | 0 | 0 | 1 | 0 | 0 | 0 | 0 | 2 | 0 | 98 | 49 | NA | NA | NA | | NA | 0.5 | NA | | 0 | NA | 0.5 |
| SE | 5031 | HIAT1 | HIAT1 | chr1 | + | 100068472 | 100068586 | 100067976 | 100068195 | 100069613 | 100069685 | 0 | 0 | 2 | 0 | 0 | 0 | 0 | 0 | 0 | 0 | 2 | 0 | 98 | 49 | NA | NA | NA | | NA | 1 | NA | | 0 | NA | 1 |
| SE | 5037 | PTS | PTS | chr11 | + | 112230207 | 112230230 | 112228593 | 112228673 | 112230625 | 112230682 | 0 | 0 | 0 | 0 | 0 | 4 | 0 | 1 | 0 | 0 | 2 | 0 | 72 | 49 | NA | NA | NA | | NA | 0 | NA | | 0.254 | NA | -0.254 |
| SE | 5038 | PTS | PTS | chr11 | + | 112230625 | 112230682 | 112228593 | 112228673 | 112233162 | 112233233 | 0 | 0 | 5 | 0 | 0 | 1 | 0 | 2 | 0 | 0 | 0 | 0 | 98 | 49 | NA | NA | NA | | NA | 0.714 | NA | | 1 | NA | -0.286 |
| SE | 5040 | RELT | RELT | chr11 | + | 73391143 | 73391223 | 73390754 | 73390921 | 73393836 | 73393917 | 0 | 0 | 3 | 0 | 0 | 0 | 0 | 1 | 0 | 2 | 3 | 0 | 98 | 49 | NA | NA | NA | | NA | 1 | | 0 | 0.143 | NA | 0.928 |
| SE | 5042 | RELT | RELT | chr11 | + | 73392210 | 73392468 | 73391143 | 73391223 | 73393836 | 73393917 | 0 | 0 | 3 | 0 | 0 | 1 | 0 | 10 | 0 | 0 | 0 | 0 | 98 | 49 | NA | NA | NA | | NA | 0.6 | NA | | 1 | NA | -0.4 |
| SE | 5071 | CARS2 | CARS2 | chr13 | - | 110651033 | 110651100 | 110647100 | 110647239 | 110663450 | 110663518 | 0 | 0 | 5 | 0 | 0 | 0 | 0 | 0 | 0 | 0 | 1 | 0 | 98 | 49 | NA | NA | NA | | NA | 1 | NA | | 0 | NA | 1 |
| SE | 5097 | ARPP19 | ARPP19 | chr15 | - | 52564189 | 52564246 | 52557099 | 52557222 | 52568847 | 52568902 | 0 | 0 | 2 | 0 | 0 | 0 | 0 | 1 | 0 | 0 | 2 | 0 | 98 | 49 | NA | NA | NA | | NA | 1 | NA | | 0.2 | NA | 0.8 |
| SE | 5110 | WDFY1 | WDFY1 | chr2 | - | 223884647 | 223884749 | 223881941 | 223882072 | 223894233 | 223894339 | 0 | 0 | 3 | 0 | 0 | 0 | 0 | 1 | 0 | 0 | 1 | 0 | 98 | 49 | NA | NA | NA | | NA | 1 | NA | | 0.333 | NA | 0.667 |
| SE | 5122 | CCM2 | CCM2 | chr7 | + | 45068442 | 45068579 | 45064462 | 45064646 | 45069825 | 45069961 | 0 | 0 | 7 | 0 | 0 | 0 | 0 | 0 | 0 | 0 | 1 | 0 | 98 | 49 | NA | NA | NA | | NA | 1 | NA | | 0 | NA | 1 |
| SE | 5125 | DNM2 | DNM2 | chr19 | + | 10808568 | 10808580 | 10805915 | 10805967 | 10812263 | 10812377 | 0 | 0 | 4 | 0 | 0 | 1 | 0 | 3 | 0 | 0 | 1 | 0 | 61 | 49 | NA | NA | NA | | NA | 0.763 | NA | | 0.707 | NA | 0.056 |
| SE | 5133 | NXF1 | NXF1 | chr11 | - | 62803418 | 62803572 | 62802176 | 62802260 | 62803791 | 62803978 | 0 | 0 | 9 | 0 | 0 | 0 | 0 | 6 | 0 | 0 | 1 | 0 | 98 | 49 | NA | NA | NA | | NA | 1 | NA | | 0.75 | NA | 0.25 |
| SE | 5135 | SKIL | SKIL | chr3 | + | 170384532 | 170384627 | 170381243 | 170381341 | 170390222 | 170390464 | 0 | 0 | 3 | 0 | 0 | 0 | 0 | 0 | 0 | 0 | 1 | 0 | 98 | 49 | NA | NA | NA | | NA | 1 | NA | | 0 | NA | 1 |
| SE | 5136 | SKIL | SKIL | chr3 | + | 170384532 | 170384765 | 170381243 | 170381341 | 170390222 | 170390464 | 0 | 0 | 4 | 0 | 0 | 0 | 0 | 1 | 0 | 0 | 1 | 0 | 98 | 49 | NA | NA | NA | | NA | 1 | NA | | 0.333 | NA | 0.667 |
| SE | 5138 | XRN1 | XRN1 | chr3 | - | 142426743 | 142426841 | 142425428 | 142425538 | 142432660 | 142432893 | 0 | 0 | 8 | 0 | 0 | 1 | 0 | 5 | 0 | 0 | 1 | 0 | 98 | 49 | NA | NA | NA | | NA | 0.8 | NA | | 0.714 | NA | 0.086 |
| SE | 5149 | KIAA0226L | KIAA0226L | chr13 | - | 46377889 | 46378005 | 46371940 | 46372597 | 46387133 | 46387500 | 0 | 0 | 1 | 0 | 0 | 0 | 0 | 1 | 0 | 0 | 2 | 0 | 98 | 49 | NA | NA | NA | | NA | 1 | NA | | 0.2 | NA | 0.8 |
| SE | 5159 | IL32 | IL32 | chr16 | + | 3067553 | 3067613 | 3067376 | 3067415 | 3067983 | 3068010 | 0 | 0 | 4 | 0 | 0 | 0 | 0 | 8 | 0 | 1 | 0 | 0 | 98 | 49 | NA | NA | NA | | NA | 1 | | 0 | 1 | NA | 0.5 |
| SE | 5171 | CEP350 | CEP350 | chr1 | + | 180036925 | 180037089 | 180033861 | 180034082 | 180041137 | 180041248 | 0 | 0 | 1 | 0 | 0 | 0 | 0 | 3 | 0 | 0 | 0 | 0 | 98 | 49 | NA | NA | NA | | NA | 0.333 | NA | | 1 | NA | -0.667 |
| SE | 5180 | ANKRD13A | ANKRD13A | chr12 | + | 110018344 | 110018488 | 110016387 | 110016433 | 110019138 | 110019328 | 0 | 0 | 18 | 0 | 0 | 1 | 0 | 4 | 0 | 0 | 0 | 0 | 98 | 49 | NA | NA | NA | | NA | 0.9 | NA | | 1 | NA | -0.1 |
| SE | 5191 | PICALM | PICALM | chr11 | - | 85974707 | 85974812 | 85957170 | 85959060 | 85981128 | 85981228 | 0 | 0 | 13 | 0 | 0 | 0 | 2 | 8 | 0 | 0 | 1 | 0 | 98 | 49 | NA | NA | NA | | NA | 1 | | 1 | 0.8 | NA | 0.1 |
| SE | 5197 | PICALM | PICALM | chr11 | - | 86022366 | 86022469 | 86014869 | 86014963 | 86026291 | 86026367 | 0 | 0 | 36 | 0 | 0 | 1 | 3 | 27 | 0 | 0 | 2 | 0 | 98 | 49 | NA | NA | NA | | NA | 0.947 | | 1 | 0.871 | NA | 0.011 |
| SE | 5217 | SUCO | SUCO | chr1 | + | 172585017 | 172585086 | 172579201 | 172579267 | 172585857 | 172585948 | 0 | 0 | 1 | 0 | 0 | 0 | 0 | 0 | 0 | 0 | 2 | 0 | 98 | 49 | NA | NA | NA | | NA | 1 | NA | | 0.2 | NA | 0.8 |
| SE | 5220 | TPM4 | TPM4 | chr19 | + | 16088026 | 16088097 | 16086422 | 16086540 | 16089044 | 16089120 | 0 | 0 | 5 | 0 | 0 | 0 | 0 | 9 | 0 | 0 | 1 | 0 | 98 | 49 | NA | NA | NA | | NA | 1 | NA | | 0.818 | NA | 0.182 |
| SE | 5242 | PLXNC1 | PLXNC1 | chr12 | + | 94259336 | 94259375 | 94255192 | 94255296 | 94259609 | 94259734 | 0 | 0 | 26 | 0 | 0 | 3 | 0 | 15 | 0 | 0 | 0 | 0 | 88 | 49 | NA | NA | NA | | NA | 0.828 | NA | | 1 | NA | -0.172 |
| SE | 5244 | PLXNC1 | PLXNC1 | chr12 | + | 94300909 | 94301057 | 94297315 | 94297423 | 94303755 | 94303896 | 0 | 0 | 11 | 0 | 0 | 0 | 4 | 2 | 0 | 0 | 4 | 0 | 98 | 49 | NA | NA | NA | | NA | 1 | | 1 | 0.2 | NA | 0.4 |
| SE | 5256 | PSME4 | PSME4 | chr2 | - | 53899880 | 53900017 | 53898300 | 53898354 | 53901349 | 53901559 | 0 | 0 | 2 | 0 | 0 | 0 | 0 | 1 | 0 | 0 | 1 | 0 | 98 | 49 | NA | NA | NA | | NA | 1 | NA | | 0.333 | NA | 0.667 |
| SE | 5290 | PHF21A | PHF21A | chr11 | - | 45938156 | 45938312 | 45936493 | 45936569 | 45945839 | 45946003 | 0 | 0 | 6 | 0 | 0 | 3 | 0 | 5 | 0 | 0 | 1 | 0 | 98 | 49 | NA | NA | NA | | NA | 0.5 | NA | | 0.714 | NA | -0.214 |
| SE | 5293 | ENSA | ENSA | chr1 | - | 150625641 | 150625808 | 150622122 | 150622859 | 150626478 | 150626526 | 0 | 0 | 1 | 0 | 0 | 0 | 0 | 2 | 0 | 0 | 2 | 0 | 98 | 49 | NA | NA | NA | | NA | 1 | NA | | 0.333 | NA | 0.667 |
| SE | 5300 | AKAP8L | AKAP8L | chr19 | - | 15398629 | 15398766 | 15397713 | 15397855 | 15399301 | 15399410 | 0 | 0 | 0 | 0 | 0 | 4 | 0 | 1 | 0 | 0 | 0 | 0 | 98 | 49 | NA | NA | NA | | NA | 0 | NA | | 1 | NA | -1 |
| SE | 5318 | PSMD1 | PSMD1 | chr2 | + | 231079535 | 231079614 | 231078658 | 231078747 | 231080140 | 231080314 | 0 | 0 | 5 | 0 | 0 | 0 | 0 | 3 | 0 | 0 | 1 | 0 | 98 | 49 | NA | NA | NA | | NA | 1 | NA | | 0.6 | NA | 0.4 |
| SE | 5357 | NLRP3 | NLRP3 | chr1 | + | 247434102 | 247434273 | 247423846 | 247425599 | 247443971 | 247444142 | 0 | 0 | 2 | 0 | 0 | 0 | 2 | 1 | 0 | 0 | 1 | 0 | 98 | 49 | NA | NA | NA | | NA | 1 | | 1 | 0.333 | NA | 0.334 |
| SE | 5358 | NLRP3 | NLRP3 | chr1 | + | 247435969 | 247436140 | 247434102 | 247434273 | 247443971 | 247444142 | 0 | 0 | 1 | 0 | 0 | 2 | 0 | 0 | 0 | 2 | 1 | 0 | 98 | 49 | NA | NA | NA | | NA | 0.2 | | 0 | 0 | NA | 0.2 |
| SE | 5367 | BAZ1A | BAZ1A | chr14 | - | 34786125 | 34786221 | 34785776 | 34786001 | 34792774 | 34792921 | 0 | 0 | 4 | 0 | 0 | 2 | 0 | 3 | 0 | 0 | 1 | 0 | 98 | 49 | NA | NA | NA | | NA | 0.5 | NA | | 0.6 | NA | -0.1 |
| SE | 5388 | CD44 | CD44 | chr11 | + | 35211245 | 35211449 | 35189834 | 35190065 | 35214851 | 35214914 | 0 | 0 | 1 | 0 | 0 | 3 | 0 | 1 | 0 | 0 | 5 | 0 | 98 | 49 | NA | NA | NA | | NA | 0.143 | NA | | 0.091 | NA | 0.052 |
| SE | 5397 | MTMR14 | MTMR14 | chr3 | + | 9677983 | 9678058 | 9677316 | 9677387 | 9683177 | 9683244 | 0 | 0 | 2 | 0 | 0 | 1 | 0 | 3 | 0 | 0 | 0 | 0 | 98 | 49 | NA | NA | NA | | NA | 0.5 | NA | | 1 | NA | -0.5 |
| SE | 5399 | MTMR14 | MTMR14 | chr3 | + | 9697710 | 9697866 | 9689963 | 9690143 | 9701789 | 9702394 | 0 | 0 | 2 | 0 | 0 | 1 | 0 | 0 | 0 | 0 | 0 | 0 | 98 | 49 | NA | NA | NA | | NA | 0.5 | NA | | 0 | NA | 0.5 |
| SE | 5406 | RALY | RALY | chr20 | + | 34073818 | 34073866 | 34073562 | 34073635 | 34075873 | 34076040 | 0 | 0 | 8 | 0 | 0 | 0 | 0 | 0 | 0 | 0 | 1 | 0 | 97 | 49 | NA | NA | NA | | NA | 1 | NA | | 0 | NA | 1 |
| SE | 5417 | VPS39 | VPS39 | chr15 | - | 42173722 | 42173852 | 42169723 | 42169866 | 42178217 | 42178338 | 0 | 0 | 2 | 0 | 0 | 0 | 3 | 5 | 0 | 0 | 1 | 0 | 98 | 49 | NA | NA | NA | | NA | 1 | | 1 | 0.714 | NA | 0.143 |
| SE | 5448 | RER1 | RER1 | chr1 | + | 2399414 | 2399514 | 2397115 | 2397220 | 2400856 | 2400935 | 0 | 0 | 3 | 0 | 0 | 3 | 0 | 1 | 0 | 0 | 1 | 0 | 98 | 49 | NA | NA | NA | | NA | 0.333 | NA | | 0.333 | NA | 0 |
| SE | 5471 | ARNTL | ARNTL | chr11 | + | 13355225 | 13355296 | 13354300 | 13354448 | 13356738 | 13356777 | 0 | 0 | 1 | 0 | 0 | 5 | 0 | 0 | 0 | 0 | 2 | 0 | 98 | 49 | NA | NA | NA | | NA | 0.091 | NA | | 0 | NA | 0.091 |
| SE | 5472 | ARNTL | ARNTL | chr11 | + | 13355232 | 13355296 | 13354300 | 13354448 | 13356738 | 13356777 | 0 | 0 | 1 | 0 | 0 | 5 | 0 | 0 | 0 | 0 | 2 | 0 | 98 | 49 | NA | NA | NA | | NA | 0.091 | NA | | 0 | NA | 0.091 |
| SE | 5487 | CTR9 | CTR9 | chr11 | + | 10755677 | 10755795 | 10754957 | 10755197 | 10756748 | 10756838 | 0 | 0 | 2 | 0 | 0 | 1 | 0 | 4 | 0 | 0 | 0 | 0 | 98 | 49 | NA | NA | NA | | NA | 0.5 | NA | | 1 | NA | -0.5 |
| SE | 5500 | MBIP | MBIP | chr14 | - | 36308091 | 36308189 | 36300784 | 36300823 | 36311572 | 36311725 | 0 | 0 | 0 | 0 | 0 | 1 | 0 | 1 | 0 | 0 | 0 | 0 | 98 | 49 | NA | NA | NA | | NA | 0 | NA | | 1 | NA | -1 |
| SE | 5503 | SBF2 | SBF2 | chr11 | - | 9853539 | 9853712 | 9852675 | 9852749 | 9856457 | 9856720 | 0 | 0 | 3 | 0 | 0 | 2 | 0 | 3 | 0 | 0 | 0 | 0 | 98 | 49 | NA | NA | NA | | NA | 0.429 | NA | | 1 | NA | -0.571 |
| SE | 5504 | SBF2 | SBF2 | chr11 | - | 9992414 | 9992543 | 9989496 | 9989595 | 9992989 | 9993103 | 0 | 0 | 5 | 0 | 0 | 5 | 0 | 8 | 0 | 0 | 0 | 0 | 98 | 49 | NA | NA | NA | | NA | 0.333 | NA | | 1 | NA | -0.667 |
| SE | 5517 | DENND5A | DENND5A | chr11 | - | 9143402 | 9143485 | 9142721 | 9142845 | 9144096 | 9144278 | 0 | 0 | 7 | 0 | 0 | 0 | 0 | 5 | 0 | 0 | 1 | 0 | 98 | 49 | NA | NA | NA | | NA | 1 | NA | | 0.714 | NA | 0.286 |
| SE | 5532 | TXLNA | TXLNA | chr1 | + | 32184524 | 32184616 | 32181241 | 32181577 | 32187953 | 32188124 | 0 | 0 | 1 | 0 | 0 | 1 | 0 | 1 | 0 | 0 | 0 | 0 | 98 | 49 | NA | NA | NA | | NA | 0.333 | NA | | 1 | NA | -0.667 |
| SE | 5562 | RAPGEF6 | RAPGEF6 | chr5 | - | 131561977 | 131562047 | 131548046 | 131548190 | 131592382 | 131592466 | 0 | 0 | 1 | 0 | 0 | 0 | 0 | 1 | 0 | 0 | 1 | 0 | 98 | 49 | NA | NA | NA | | NA | 1 | NA | | 0.333 | NA | 0.667 |
| SE | 5596 | ZNF512 | ZNF512 | chr2 | + | 27600690 | 27600815 | 27599969 | 27600053 | 27601355 | 27601442 | 0 | 0 | 0 | 0 | 0 | 1 | 0 | 2 | 0 | 0 | 0 | 0 | 98 | 49 | NA | NA | NA | | NA | 0 | NA | | 1 | NA | -1 |
| SE | 5623 | ETF1 | ETF1 | chr5 | - | 138508668 | 138508816 | 138506092 | 138508387 | 138510564 | 138510629 | 0 | 0 | 2 | 0 | 0 | 0 | 0 | 7 | 0 | 0 | 1 | 0 | 98 | 49 | NA | NA | NA | | NA | 1 | NA | | 0.778 | NA | 0.222 |
| SE | 5632 | NUP98 | NUP98 | chr11 | - | 3705199 | 3705356 | 3702462 | 3702892 | 3706444 | 3706627 | 0 | 0 | 3 | 0 | 0 | 0 | 0 | 4 | 0 | 0 | 1 | 0 | 98 | 49 | NA | NA | NA | | NA | 1 | NA | | 0.667 | NA | 0.333 |
| SE | 5661 | ADD1 | ADD1 | chr4 | + | 2898432 | 2898531 | 2898183 | 2898327 | 2904763 | 2905015 | 0 | 0 | 1 | 0 | 0 | 1 | 0 | 1 | 0 | 0 | 0 | 0 | 98 | 49 | NA | NA | NA | | NA | 0.333 | NA | | 1 | NA | -0.667 |
| SE | 5662 | ADD1 | ADD1 | chr4 | + | 2899258 | 2899435 | 2898432 | 2898531 | 2904763 | 2905015 | 0 | 0 | 13 | 0 | 0 | 0 | 1 | 6 | 0 | 0 | 1 | 0 | 98 | 49 | NA | NA | NA | | NA | 1 | | 1 | 0.75 | NA | 0.125 |
| SE | 5728 | ZDHHC3 | ZDHHC3 | chr3 | - | 44932906 | 44932990 | 44929305 | 44929436 | 44933117 | 44933199 | 0 | 0 | 2 | 0 | 0 | 0 | 0 | 2 | 0 | 0 | 1 | 0 | 98 | 49 | NA | NA | NA | | NA | 1 | NA | | 0.5 | NA | 0.5 |
| SE | 5735 | CCNY | CCNY | chr10 | + | 35553018 | 35553185 | 35530123 | 35530243 | 35566022 | 35566185 | 0 | 0 | 7 | 0 | 0 | 0 | 8 | 4 | 0 | 0 | 1 | 0 | 98 | 49 | NA | NA | NA | | NA | 1 | | 1 | 0.667 | NA | 0.166 |
| SE | 5741 | ENTPD1 | ENTPD1 | chr10 | + | 95842343 | 95842494 | 95839690 | 95839808 | 95844475 | 95844635 | 0 | 0 | 10 | 0 | 0 | 0 | 0 | 2 | 0 | 0 | 1 | 0 | 98 | 49 | NA | NA | NA | | NA | 1 | NA | | 0.5 | NA | 0.5 |
| SE | 5746 | UBE2Q1 | UBE2Q1 | chr1 | - | 154551396 | 154551492 | 154550937 | 154551004 | 154551770 | 154551819 | 0 | 0 | 1 | 0 | 0 | 0 | 0 | 2 | 0 | 0 | 1 | 0 | 98 | 49 | NA | NA | NA | | NA | 1 | NA | | 0.5 | NA | 0.5 |
| SE | 5774 | NCOA2 | NCOA2 | chr8 | - | 70144641 | 70144848 | 70141183 | 70141399 | 70148272 | 70148483 | 0 | 0 | 5 | 0 | 0 | 1 | 0 | 2 | 0 | 0 | 1 | 0 | 98 | 49 | NA | NA | NA | | NA | 0.714 | NA | | 0.5 | NA | 0.214 |
| SE | 5789 | DENND3 | DENND3 | chr8 | + | 141163332 | 141163429 | 141155848 | 141155970 | 141165185 | 141165289 | 0 | 0 | 8 | 0 | 0 | 1 | 3 | 6 | 0 | 0 | 0 | 0 | 98 | 49 | NA | NA | NA | | NA | 0.8 | | 1 | 1 | NA | -0.2 |
| SE | 5790 | DENND3 | DENND3 | chr8 | + | 141163332 | 141163429 | 141160631 | 141160787 | 141165185 | 141165289 | 0 | 0 | 16 | 0 | 0 | 0 | 3 | 9 | 0 | 0 | 4 | 0 | 98 | 49 | NA | NA | NA | | NA | 1 | | 1 | 0.529 | NA | 0.236 |
| SE | 5792 | AFG3L2 | AFG3L2 | chr18 | - | 12367275 | 12367382 | 12366964 | 12367117 | 12370848 | 12370926 | 0 | 0 | 0 | 0 | 0 | 1 | 0 | 1 | 0 | 0 | 0 | 0 | 98 | 49 | NA | NA | NA | | NA | 0 | NA | | 1 | NA | -1 |
| SE | 5810 | ANXA1 | ANXA1 | chr9 | + | 73169031 | 73169154 | 73167496 | 73167555 | 73170050 | 73170394 | 0 | 0 | 72 | 0 | 0 | 0 | 8 | 13 | 0 | 0 | 1 | 0 | 98 | 49 | NA | NA | NA | | NA | 1 | | 1 | 0.867 | NA | 0.067 |
| SE | 5820 | HEXA | HEXA | chr15 | - | 72349078 | 72349259 | 72348047 | 72348134 | 72350517 | 72350650 | 0 | 0 | 3 | 0 | 0 | 2 | 0 | 2 | 0 | 0 | 1 | 0 | 98 | 49 | NA | NA | NA | | NA | 0.429 | NA | | 0.5 | NA | -0.071 |
| SE | 5826 | PLEKHO1 | PLEKHO1 | chr1 | + | 150156065 | 150156206 | 150150585 | 150151058 | 150156910 | 150157015 | 0 | 0 | 10 | 0 | 0 | 0 | 0 | 2 | 0 | 0 | 1 | 0 | 98 | 49 | NA | NA | NA | | NA | 1 | NA | | 0.5 | NA | 0.5 |
| SE | 5880 | ARID1A | ARID1A | chr1 | + | 26729650 | 26729863 | 26696030 | 26697540 | 26731151 | 26731604 | 0 | 0 | 3 | 0 | 0 | 0 | 0 | 1 | 0 | 1 | 0 | 0 | 98 | 49 | NA | NA | NA | | NA | 1 | | 0 | 1 | NA | 0.5 |
| SE | 5896 | FANCM | FANCM | chr14 | + | 45140631 | 45140709 | 45137068 | 45137241 | 45148836 | 45148995 | 0 | 0 | 0 | 0 | 0 | 1 | 0 | 1 | 0 | 0 | 0 | 0 | 98 | 49 | NA | NA | NA | | NA | 0 | NA | | 1 | NA | -1 |
| SE | 5900 | NIPA2 | NIPA2 | chr15 | - | 22839654 | 22839790 | 22838640 | 22838921 | 22845145 | 22845267 | 0 | 0 | 1 | 0 | 0 | 0 | 0 | 0 | 0 | 1 | 0 | 0 | 98 | 49 | NA | NA | NA | | NA | 1 | | 0 | NA | NA | 1 |
| SE | 5909 | B3GNTL1 | B3GNTL1 | chr17 | - | 82965650 | 82965746 | 82961056 | 82961181 | 83005114 | 83005210 | 0 | 0 | 5 | 0 | 0 | 0 | 0 | 5 | 0 | 0 | 2 | 0 | 98 | 49 | NA | NA | NA | | NA | 1 | NA | | 0.556 | NA | 0.444 |
| SE | 5920 | RAP1GAP2 | RAP1GAP2 | chr17 | + | 2905283 | 2905368 | 2800514 | 2800550 | 2957758 | 2957794 | 0 | 0 | 4 | 0 | 0 | 0 | 0 | 4 | 0 | 0 | 2 | 0 | 98 | 49 | NA | NA | NA | | NA | 1 | NA | | 0.5 | NA | 0.5 |
| SE | 5921 | RAP1GAP2 | RAP1GAP2 | chr17 | + | 2962669 | 2962714 | 2957758 | 2957794 | 2963429 | 2963462 | 0 | 0 | 3 | 0 | 0 | 6 | 0 | 2 | 0 | 0 | 1 | 0 | 94 | 49 | NA | NA | NA | | NA | 0.207 | NA | | 0.51 | NA | -0.303 |
| SE | 5924 | MKNK1 | MKNK1 | chr1 | - | 46574946 | 46575020 | 46572062 | 46572167 | 46576574 | 46576654 | 0 | 0 | 3 | 0 | 0 | 0 | 0 | 4 | 0 | 0 | 1 | 0 | 98 | 49 | NA | NA | NA | | NA | 1 | NA | | 0.667 | NA | 0.333 |
| SE | 5928 | ZDHHC5 | ZDHHC5 | chr11 | + | 57695919 | 57696043 | 57693782 | 57693915 | 57696760 | 57696873 | 0 | 0 | 0 | 0 | 0 | 1 | 5 | 5 | 0 | 0 | 0 | 0 | 98 | 49 | NA | NA | NA | | NA | 0 | | 1 | 1 | NA | -1 |
| SE | 5937 | SRSF10 | SRSF10 | chr1 | - | 23971337 | 23971439 | 23964346 | 23967723 | 23971572 | 23971626 | 0 | 0 | 1 | 0 | 0 | 0 | 0 | 1 | 0 | 0 | 2 | 0 | 98 | 49 | NA | NA | NA | | NA | 1 | NA | | 0.2 | NA | 0.8 |
| SE | 5949 | METTL9 | METTL9 | chr16 | + | 21612644 | 21612835 | 21599475 | 21599898 | 21617864 | 21618074 | 0 | 0 | 16 | 0 | 0 | 0 | 0 | 5 | 0 | 2 | 2 | 0 | 98 | 49 | NA | NA | NA | | NA | 1 | | 0 | 0.556 | NA | 0.722 |
| SE | 5965 | CD58 | CD58 | chr1 | - | 116535964 | 116536228 | 116521905 | 116521983 | 116544310 | 116544604 | 0 | 0 | 10 | 0 | 0 | 0 | 0 | 5 | 0 | 0 | 1 | 0 | 98 | 49 | NA | NA | NA | | NA | 1 | NA | | 0.714 | NA | 0.286 |
| SE | 5967 | CD53 | CD53 | chr1 | + | 110894959 | 110895055 | 110891391 | 110891471 | 110896652 | 110896733 | 0 | 0 | 12 | 0 | 0 | 3 | 0 | 3 | 0 | 0 | 0 | 0 | 98 | 49 | NA | NA | NA | | NA | 0.667 | NA | | 1 | NA | -0.333 |
| SE | 6044 | GADD45A | GADD45A | chr1 | + | 67686024 | 67686126 | 67685176 | 67685538 | 67687660 | 67688338 | 0 | 0 | 0 | 0 | 0 | 1 | 0 | 1 | 0 | 0 | 0 | 0 | 98 | 49 | NA | NA | NA | | NA | 0 | NA | | 1 | NA | -1 |
| SE | 6051 | TROVE2 | TROVE2 | chr1 | + | 193069033 | 193069634 | 193059421 | 193059776 | 193075819 | 193076040 | 0 | 0 | 4 | 0 | 0 | 1 | 0 | 0 | 0 | 0 | 1 | 0 | 98 | 49 | NA | NA | NA | | NA | 0.667 | NA | | 0 | NA | 0.667 |
| SE | 6063 | PNPLA8 | PNPLA8 | chr7 | - | 108521475 | 108521521 | 108514435 | 108515574 | 108526028 | 108526318 | 0 | 0 | 3 | 0 | 0 | 0 | 0 | 3 | 0 | 0 | 1 | 0 | 95 | 49 | NA | NA | NA | | NA | 1 | NA | | 0.607 | NA | 0.393 |
| SE | 6078 | PTBP2 | PTBP2 | chr1 | + | 96813028 | 96813106 | 96812711 | 96812928 | 96813272 | 96815049 | 0 | 0 | 0 | 0 | 0 | 2 | 0 | 1 | 0 | 0 | 0 | 0 | 98 | 49 | NA | NA | NA | | NA | 0 | NA | | 1 | NA | -1 |
| SE | 6090 | ZNF207 | ZNF207 | chr17 | + | 32366664 | 32366757 | 32365329 | 32365487 | 32367771 | 32368014 | 0 | 0 | 0 | 0 | 0 | 2 | 0 | 3 | 0 | 0 | 0 | 0 | 98 | 49 | NA | NA | NA | | NA | 0 | NA | | 1 | NA | -1 |
| SE | 6100 | ZNF217 | ZNF217 | chr20 | - | 53571720 | 53571853 | 53567070 | 53569264 | 53575726 | 53577280 | 0 | 0 | 4 | 0 | 0 | 1 | 0 | 2 | 0 | 0 | 0 | 0 | 98 | 49 | NA | NA | NA | | NA | 0.667 | NA | | 1 | NA | -0.333 |
| SE | 6101 | CCBL2 | CCBL2 | chr1 | - | 88988251 | 88988351 | 88969408 | 88969467 | 88992584 | 88992960 | 0 | 0 | 1 | 0 | 0 | 0 | 0 | 0 | 0 | 0 | 1 | 0 | 98 | 49 | NA | NA | NA | | NA | 1 | NA | | 0 | NA | 1 |
| SE | 6126 | ZC3HAV1 | ZC3HAV1 | chr7 | - | 139076285 | 139076409 | 139073855 | 139074030 | 139078551 | 139078653 | 0 | 0 | 12 | 0 | 0 | 0 | 4 | 14 | 0 | 0 | 5 | 0 | 98 | 49 | NA | NA | NA | | NA | 1 | | 1 | 0.583 | NA | 0.209 |
| SE | 6129 | WDR26 | WDR26 | chr1 | - | 224398888 | 224399034 | 224398514 | 224398593 | 224400949 | 224401069 | 0 | 0 | 5 | 0 | 0 | 1 | 0 | 13 | 0 | 1 | 2 | 0 | 98 | 49 | NA | NA | NA | | NA | 0.714 | | 0 | 0.765 | NA | 0.331 |
| SE | 6131 | ANKRD13D | ANKRD13D | chr11 | + | 67289828 | 67289965 | 67289344 | 67289550 | 67290077 | 67290213 | 0 | 0 | 3 | 0 | 0 | 4 | 0 | 0 | 0 | 0 | 2 | 0 | 98 | 49 | NA | NA | NA | | NA | 0.273 | NA | | 0 | NA | 0.273 |
| SE | 6150 | GRASP | GRASP | chr12 | + | 52011038 | 52011141 | 52010880 | 52010935 | 52013686 | 52013780 | 0 | 0 | 2 | 0 | 0 | 0 | 0 | 0 | 0 | 0 | 1 | 0 | 98 | 49 | NA | NA | NA | | NA | 1 | NA | | 0 | NA | 1 |
| SE | 6151 | GRASP | GRASP | chr12 | + | 52011038 | 52011207 | 52010880 | 52010935 | 52013686 | 52013780 | 0 | 0 | 2 | 0 | 0 | 0 | 0 | 0 | 0 | 0 | 1 | 0 | 98 | 49 | NA | NA | NA | | NA | 1 | NA | | 0 | NA | 1 |
| SE | 6165 | EFCAB14 | EFCAB14 | chr1 | - | 46688352 | 46688544 | 46686783 | 46686870 | 46689586 | 46689691 | 0 | 0 | 0 | 0 | 1 | 0 | 0 | 0 | 1 | 0 | 0 | 0 | 98 | 49 | NA | NA | NA | | | 0 NA | NA | | NA | 1 | -1 |
| SE | 6166 | NT5C3A | NT5C3A | chr7 | - | 33029628 | 33029714 | 33026816 | 33026915 | 33035933 | 33035988 | 0 | 0 | 0 | 0 | 0 | 1 | 0 | 1 | 0 | 0 | 4 | 0 | 98 | 49 | NA | NA | NA | | NA | 0 | NA | | 0.111 | NA | -0.111 |
| SE | 6172 | ABCA1 | ABCA1 | chr9 | - | 104786872 | 104786976 | 104786297 | 104786390 | 104787919 | 104788054 | 0 | 0 | 1 | 0 | 0 | 1 | 0 | 3 | 0 | 0 | 1 | 0 | 98 | 49 | NA | NA | NA | | NA | 0.333 | NA | | 0.6 | NA | -0.267 |
| SE | 6185 | HERC3 | HERC3 | chr4 | + | 88676217 | 88676241 | 88670138 | 88670252 | 88676333 | 88676423 | 0 | 0 | 6 | 0 | 0 | 3 | 0 | 8 | 0 | 0 | 0 | 0 | 73 | 49 | NA | NA | NA | | NA | 0.573 | NA | | 1 | NA | -0.427 |
| SE | 6207 | PPP1R12A | PPP1R12A | chr12 | - | 79805591 | 79805768 | 79798493 | 79798584 | 79806165 | 79806333 | 0 | 0 | 2 | 0 | 0 | 4 | 0 | 0 | 0 | 0 | 1 | 0 | 98 | 49 | NA | NA | NA | | NA | 0.2 | NA | | 0 | NA | 0.2 |
| SE | 6211 | YIPF1 | YIPF1 | chr1 | - | 53855000 | 53855051 | 53851718 | 53852270 | 53866199 | 53866382 | 0 | 0 | 2 | 0 | 0 | 0 | 0 | 2 | 0 | 0 | 1 | 0 | 98 | 49 | NA | NA | NA | | NA | 1 | NA | | 0.5 | NA | 0.5 |
| SE | 6212 | YIPF1 | YIPF1 | chr1 | - | 53860055 | 53860153 | 53851718 | 53852270 | 53866199 | 53866382 | 0 | 0 | 1 | 0 | 0 | 0 | 0 | 0 | 0 | 0 | 1 | 0 | 98 | 49 | NA | NA | NA | | NA | 1 | NA | | 0 | NA | 1 |
| SE | 6223 | AHCTF1 | AHCTF1 | chr1 | - | 246843794 | 246843928 | 246842693 | 246842776 | 246849614 | 246851442 | 0 | 0 | 0 | 0 | 0 | 2 | 0 | 2 | 0 | 0 | 2 | 0 | 98 | 49 | NA | NA | NA | | NA | 0 | NA | | 0.333 | NA | -0.333 |
| SE | 6269 | BAZ2B | BAZ2B | chr2 | - | 159336941 | 159337077 | 159332539 | 159332686 | 159337566 | 159337772 | 0 | 0 | 4 | 0 | 0 | 0 | 0 | 0 | 0 | 0 | 1 | 0 | 98 | 49 | NA | NA | NA | | NA | 1 | NA | | 0 | NA | 1 |
| SE | 6352 | ECD | ECD | chr10 | - | 73154255 | 73154448 | 73152292 | 73152421 | 73156274 | 73156453 | 0 | 0 | 2 | 0 | 0 | 1 | 0 | 5 | 0 | 0 | 0 | 0 | 98 | 49 | NA | NA | NA | | NA | 0.5 | NA | | 1 | NA | -0.5 |
| SE | 6355 | GATAD2B | GATAD2B | chr1 | - | 153818039 | 153818171 | 153817371 | 153817542 | 153818790 | 153818922 | 0 | 0 | 2 | 0 | 0 | 1 | 0 | 4 | 0 | 0 | 0 | 0 | 98 | 49 | NA | NA | NA | | NA | 0.5 | NA | | 1 | NA | -0.5 |
| SE | 6363 | FAM91A1 | FAM91A1 | chr8 | + | 123786494 | 123786610 | 123785628 | 123785741 | 123787260 | 123787373 | 0 | 0 | 4 | 0 | 0 | 1 | 0 | 2 | 0 | 0 | 0 | 0 | 98 | 49 | NA | NA | NA | | NA | 0.667 | NA | | 1 | NA | -0.333 |
| SE | 6396 | NDEL1 | NDEL1 | chr17 | + | 8445710 | 8445864 | 8444259 | 8444357 | 8446753 | 8446902 | 0 | 0 | 20 | 0 | 0 | 0 | 0 | 8 | 0 | 0 | 2 | 0 | 98 | 49 | NA | NA | NA | | NA | 1 | NA | | 0.667 | NA | 0.333 |

| | | | | | | | | | | | | | | | | | | | | | | | | | | | | | | | | | | | | | |
|---|---|---|---|---|---|---|---|---|---|---|---|---|---|---|---|---|---|---|---|---|---|---|---|---|---|---|---|---|---|---|---|---|---|---|---|---|---|
| SE | 6435 | R3HDM2 | R3HDM2 | chr12 | - | 57288856 | 57288910 | 57283823 | 57284056 | 57289008 | 57289040 | 0 | 0 | 0 | 0 | 0 | 3 | 0 | 1 | 0 | 0 | 3 | 0 | 98 | 49 | NA | NA | NA | NA | | 0 | NA | | | 0.143 | NA | -0.143 |
| SE | 6442 | ATP11A | ATP11A | chr13 | + | 112851036 | 112851218 | 112842275 | 112842379 | 112854278 | 112854530 | 0 | 0 | 8 | 0 | 0 | 0 | 0 | 0 | 0 | 0 | 1 | 0 | 98 | 49 | NA | NA | NA | NA | | 1 | NA | | | 0 | NA | 1 |
| SE | 6474 | PUM1 | PUM1 | chr1 | - | 30941997 | 30942123 | 30941150 | 30941272 | 30945345 | 30945483 | 0 | 0 | 2 | 0 | 0 | 2 | 3 | 0 | 0 | 0 | 0 | 0 | 98 | 49 | NA | NA | NA | NA | | 0.333 | | 1 | NA | | NA | -0.667 |
| SE | 6476 | PUM1 | PUM1 | chr1 | - | 30980061 | 30980163 | 30974650 | 30974802 | 30981311 | 30981405 | 0 | 0 | 2 | 0 | 0 | 0 | 0 | 0 | 0 | 0 | 2 | 0 | 98 | 49 | NA | NA | NA | NA | | 1 | NA | | | 0 | NA | 1 |
| SE | 6495 | PI4KB | PI4KB | chr1 | - | 151326171 | 151326373 | 151315572 | 151316509 | 151327270 | 151327657 | 0 | 0 | 4 | 0 | 0 | 0 | 0 | 2 | 0 | 0 | 1 | 0 | 98 | 49 | NA | NA | NA | NA | | 1 | NA | | | 0.5 | NA | 0.5 |
| SE | 6501 | GAS7 | GAS7 | chr17 | - | 9940125 | 9940200 | 9934165 | 9934244 | 9943120 | 9943236 | 0 | 0 | 15 | 0 | 0 | 1 | 0 | 1 | 0 | 0 | 0 | 0 | 98 | 49 | NA | NA | NA | NA | | 0.882 | NA | | | 1 | NA | -0.118 |
| SE | 6505 | RBMS1 | RBMS1 | chr2 | - | 160277302 | 160277383 | 160275629 | 160275714 | 160278547 | 160278658 | 0 | 0 | 2 | 0 | 0 | 0 | 0 | 3 | 0 | 0 | 1 | 0 | 98 | 49 | NA | NA | NA | NA | | 1 | NA | | | 0.6 | NA | 0.4 |
| SE | 6514 | HNRNPC | HNRNPC | chr14 | - | 21211405 | 21211566 | 21209135 | 21211306 | 21211809 | 21211923 | 0 | 0 | 6 | 0 | 0 | 2 | 0 | 5 | 0 | 0 | 0 | 0 | 98 | 49 | NA | NA | NA | NA | | 0.6 | NA | | | 1 | NA | -0.4 |
| SE | 6516 | PHYKPL | PHYKPL | chr5 | - | 178212972 | 178213103 | 178211889 | 178211970 | 178214795 | 178214885 | 0 | 0 | 0 | 0 | 0 | 1 | 0 | 4 | 0 | 0 | 0 | 0 | 98 | 49 | NA | NA | NA | NA | | 0 | NA | | | 1 | NA | -1 |
| SE | 6571 | SENP6 | SENP6 | chr6 | + | 75640683 | 75640704 | 75634706 | 75634811 | 75647730 | 75647801 | 0 | 0 | 1 | 0 | 0 | 1 | 0 | 1 | 0 | 0 | 0 | 0 | 70 | 49 | NA | NA | NA | NA | | 0.412 | NA | | | 1 | NA | -0.588 |
| SE | 6572 | SENP6 | SENP6 | chr6 | + | 75647730 | 75647801 | 75634706 | 75634811 | 75659261 | 75659407 | 0 | 0 | 10 | 0 | 0 | 0 | 0 | 2 | 0 | 1 | 0 | 0 | 98 | 49 | NA | NA | NA | NA | | 1 | | 0 | | 1 | NA | 0.5 |
| SE | 6576 | STX12 | STX12 | chr1 | + | 27817850 | 27817923 | 27812162 | 27812268 | 27819649 | 27819732 | 0 | 0 | 5 | 0 | 0 | 1 | 0 | 1 | 0 | 0 | 0 | 0 | 98 | 49 | NA | NA | NA | NA | | 0.714 | NA | | | 1 | NA | -0.286 |
| SE | 6579 | RABGGTB | RABGGTB | chr1 | + | 75792180 | 75792306 | 75791460 | 75791571 | 75794083 | 75794233 | 0 | 0 | 3 | 0 | 0 | 0 | 0 | 0 | 0 | 0 | 1 | 0 | 98 | 49 | NA | NA | NA | NA | | 1 | NA | | | 0 | NA | 1 |
| SE | 6586 | SP140 | SP140 | chr2 | + | 230292645 | 230292788 | 230290459 | 230290564 | 230294270 | 230294318 | 0 | 0 | 2 | 0 | 0 | 0 | 0 | 1 | 0 | 0 | 2 | 0 | 98 | 49 | NA | NA | NA | NA | | 1 | NA | | | 0.2 | NA | 0.8 |
| SE | 6596 | ADPGK | ADPGK | chr15 | - | 72755555 | 72755654 | 72751366 | 72752892 | 72771782 | 72771845 | 0 | 0 | 3 | 0 | 0 | 1 | 0 | 1 | 0 | 0 | 0 | 0 | 98 | 49 | NA | NA | NA | NA | | 0.6 | NA | | | 1 | NA | -0.4 |
| SE | 6597 | ADPGK | ADPGK | chr15 | - | 72756250 | 72756447 | 72755555 | 72755654 | 72760406 | 72760527 | 0 | 0 | 5 | 0 | 0 | 0 | 0 | 4 | 0 | 0 | 2 | 0 | 98 | 49 | NA | NA | NA | NA | | 1 | NA | | | 0.5 | NA | 0.5 |
| SE | 6598 | ADPGK | ADPGK | chr15 | - | 72760406 | 72760527 | 72755555 | 72755654 | 72771782 | 72771845 | 0 | 0 | 5 | 0 | 0 | 0 | 0 | 5 | 0 | 0 | 1 | 0 | 98 | 49 | NA | NA | NA | NA | | 1 | NA | | | 0.714 | NA | 0.286 |
| SE | 6599 | ADPGK | ADPGK | chr15 | - | 72774871 | 72775097 | 72771782 | 72771845 | 72783245 | 72783785 | 0 | 0 | 5 | 0 | 0 | 2 | 0 | 3 | 0 | 0 | 0 | 0 | 98 | 49 | NA | NA | NA | NA | | 0.556 | NA | | | 1 | NA | -0.444 |
| SE | 6613 | AMPD2 | AMPD2 | chr1 | + | 109620913 | 109621266 | 109619812 | 109620278 | 109625661 | 109625792 | 0 | 0 | 0 | 0 | 0 | 3 | 0 | 5 | 0 | 0 | 7 | 0 | 98 | 49 | NA | NA | NA | NA | | 0 | NA | | | 0.263 | NA | -0.263 |
| SE | 6614 | AMPD2 | AMPD2 | chr1 | + | 109625302 | 109625433 | 109619812 | 109620278 | 109625661 | 109625792 | 0 | 0 | 2 | 0 | 0 | 3 | 0 | 3 | 0 | 0 | 7 | 0 | 98 | 49 | NA | NA | NA | NA | | 0.25 | NA | | | 0.176 | NA | 0.074 |
| SE | 6615 | AMPD2 | AMPD2 | chr1 | + | 109625302 | 109625433 | 109620580 | 109621266 | 109625661 | 109625792 | 0 | 0 | 2 | 0 | 0 | 0 | 0 | 3 | 0 | 0 | 5 | 0 | 98 | 49 | NA | NA | NA | NA | | 1 | NA | | | 0.231 | NA | 0.769 |
| SE | 6625 | PCNX | PCNX | chr14 | + | 71011491 | 71011549 | 71009633 | 71009724 | 71012999 | 71013202 | 0 | 0 | 10 | 0 | 0 | 1 | 2 | 1 | 0 | 0 | 0 | 0 | 98 | 49 | NA | NA | NA | NA | | 0.833 | | 1 | | 1 | NA | -0.167 |
| SE | 6629 | PCBP2 | PCBP2 | chr12 | + | 53465931 | 53465973 | 53464759 | 53464852 | 53467220 | 53467293 | 0 | 0 | 24 | 0 | 0 | 0 | 0 | 19 | 0 | 0 | 1 | 0 | 91 | 49 | NA | NA | NA | NA | | 1 | NA | | | 0.911 | NA | 0.089 |
| SE | 6665 | THRAP3 | THRAP3 | chr1 | + | 36296582 | 36296770 | 36293850 | 36293935 | 36300885 | 36301084 | 0 | 0 | 6 | 0 | 0 | 0 | 0 | 8 | 0 | 0 | 1 | 0 | 98 | 49 | NA | NA | NA | NA | | 1 | NA | | | 0.8 | NA | 0.2 |
| SE | 6666 | THRAP3 | THRAP3 | chr1 | + | 36301552 | 36301696 | 36300885 | 36301084 | 36303795 | 36305356 | 0 | 0 | 3 | 0 | 0 | 2 | 0 | 12 | 0 | 0 | 0 | 0 | 98 | 49 | NA | NA | NA | NA | | 0.429 | NA | | | 1 | NA | -0.571 |
| SE | 6714 | ADCK3 | ADCK3 | chr1 | + | 226965670 | 226965737 | 226964999 | 226965410 | 226977448 | 226977523 | 0 | 0 | 3 | 0 | 0 | 0 | 0 | 0 | 0 | 0 | 1 | 0 | 98 | 49 | NA | NA | NA | NA | | 1 | NA | | | 0 | NA | 1 |
| SE | 6717 | KIF1B | KIF1B | chr1 | + | 10272240 | 10272306 | 10271501 | 10271579 | 10275427 | 10275503 | 0 | 0 | 6 | 0 | 0 | 0 | 0 | 10 | 0 | 0 | 1 | 0 | 98 | 49 | NA | NA | NA | NA | | 1 | NA | | | 0.833 | NA | 0.167 |
| SE | 6720 | KIF1B | KIF1B | chr1 | + | 10347760 | 10347827 | 10345844 | 10345953 | 10348648 | 10348733 | 0 | 0 | 1 | 0 | 0 | 0 | 0 | 2 | 0 | 0 | 1 | 0 | 98 | 49 | NA | NA | NA | NA | | 1 | NA | | | 0.5 | NA | 0.5 |
| SE | 6735 | DNMT3A | DNMT3A | chr2 | - | 25241561 | 25241707 | 25240639 | 25240730 | 25243897 | 25243982 | 0 | 0 | 0 | 0 | 0 | 1 | 0 | 2 | 0 | 0 | 0 | 0 | 98 | 49 | NA | NA | NA | NA | | 0 | NA | | | 1 | NA | -1 |
| SE | 6749 | UBR4 | UBR4 | chr1 | - | 19094905 | 19095025 | 19093948 | 19094139 | 19095544 | 19095652 | 0 | 0 | 3 | 0 | 0 | 1 | 0 | 1 | 0 | 0 | 0 | 0 | 98 | 49 | NA | NA | NA | NA | | 0.6 | NA | | | 1 | NA | -0.4 |
| SE | 6750 | UBR4 | UBR4 | chr1 | - | 19148562 | 19148626 | 19147992 | 19148127 | 19150576 | 19150793 | 0 | 0 | 2 | 0 | 0 | 1 | 0 | 4 | 0 | 0 | 0 | 0 | 98 | 49 | NA | NA | NA | NA | | 0.5 | NA | | | 1 | NA | -0.5 |
| SE | 6751 | UBR4 | UBR4 | chr1 | - | 19153767 | 19153939 | 19153300 | 19153502 | 19154917 | 19155075 | 0 | 0 | 1 | 0 | 0 | 1 | 0 | 2 | 0 | 0 | 0 | 0 | 98 | 49 | NA | NA | NA | NA | | 0.333 | NA | | | 1 | NA | -0.667 |
| SE | 6753 | UBR4 | UBR4 | chr1 | - | 19168026 | 19168184 | 19167021 | 19167231 | 19169434 | 19169532 | 0 | 0 | 11 | 0 | 0 | 0 | 0 | 8 | 0 | 0 | 1 | 0 | 98 | 49 | NA | NA | NA | NA | | 1 | NA | | | 0.8 | NA | 0.2 |
| SE | 6763 | SKIV2L | SKIV2L | chr6 | + | 31961195 | 31961386 | 31961040 | 31961094 | 31961546 | 31961675 | 0 | 0 | 1 | 0 | 0 | 1 | 0 | 2 | 0 | 0 | 0 | 0 | 98 | 49 | NA | NA | NA | NA | | 0.333 | NA | | | 1 | NA | -0.667 |
| SE | 6776 | RSAD2 | RSAD2 | chr2 | + | 6886934 | 6887164 | 6883370 | 6883532 | 6890175 | 6890325 | 0 | 0 | 36 | 0 | 0 | 0 | 0 | 36 | 0 | 0 | 1 | 0 | 98 | 49 | NA | NA | NA | NA | | 1 | NA | | | 0.947 | NA | 0.053 |
| SE | 6779 | ZFAND6 | ZFAND6 | chr15 | + | 80098415 | 80098578 | 80059567 | 80059809 | 80120327 | 80120498 | 0 | 0 | 3 | 0 | 0 | 0 | 0 | 1 | 0 | 0 | 2 | 0 | 98 | 49 | NA | NA | NA | NA | | 1 | NA | | | 0.2 | NA | 0.8 |
| SE | 6798 | ABI1 | ABI1 | chr10 | - | 26755654 | 26755741 | 26751597 | 26751783 | 26759061 | 26759235 | 0 | 0 | 1 | 0 | 0 | 10 | 0 | 0 | 0 | 0 | 1 | 0 | 98 | 49 | NA | NA | NA | NA | | 0.048 | NA | | | 0 | NA | 0.048 |
| SE | 6817 | OAS1 | OAS1 | chr12 | + | 112911050 | 112911235 | 112906933 | 112907219 | 112916508 | 112916738 | 0 | 0 | 15 | 0 | 0 | 2 | 0 | 1 | 0 | 0 | 0 | 0 | 98 | 49 | NA | NA | NA | NA | | 0.789 | NA | | | 1 | NA | -0.211 |
| SE | 6825 | PRPF8 | PRPF8 | chr17 | - | 1673745 | 1673892 | 1673356 | 1673567 | 1674441 | 1674680 | 0 | 0 | 3 | 0 | 0 | 0 | 0 | 6 | 0 | 0 | 1 | 0 | 98 | 49 | NA | NA | NA | NA | | 1 | NA | | | 0.75 | NA | 0.25 |
| SE | 6826 | PRPF8 | PRPF8 | chr17 | - | 1677564 | 1677694 | 1676975 | 1677172 | 1678517 | 1678652 | 0 | 0 | 1 | 0 | 0 | 0 | 0 | 0 | 0 | 0 | 1 | 0 | 98 | 49 | NA | NA | NA | NA | | 1 | NA | | | 0 | NA | 1 |
| SE | 6828 | SH3D21 | SH3D21 | chr1 | + | 36307516 | 36307607 | 36307106 | 36307285 | 36307769 | 36307825 | 0 | 0 | 0 | 0 | 0 | 1 | 0 | 1 | 0 | 0 | 0 | 0 | 98 | 49 | NA | NA | NA | NA | | 0 | NA | | | 1 | NA | -1 |
| SE | 6833 | OSBPL9 | OSBPL9 | chr1 | + | 51669433 | 51669512 | 51651990 | 51652041 | 51714002 | 51714079 | 0 | 0 | 10 | 0 | 0 | 0 | 0 | 8 | 0 | 0 | 1 | 0 | 98 | 49 | NA | NA | NA | NA | | 1 | NA | | | 0.8 | NA | 0.2 |
| SE | 6853 | UBAP2L | UBAP2L | chr1 | + | 154234590 | 154234738 | 154228614 | 154228725 | 154235195 | 154235291 | 0 | 0 | 1 | 0 | 0 | 0 | 2 | 2 | 0 | 0 | 1 | 0 | 98 | 49 | NA | NA | NA | NA | | 1 | | 1 | | 0.5 | NA | 0.25 |
| SE | 6854 | UBAP2L | UBAP2L | chr1 | + | 154234590 | 154234759 | 154228614 | 154228725 | 154235195 | 154235291 | 0 | 0 | 1 | 0 | 0 | 0 | 2 | 2 | 0 | 0 | 1 | 0 | 98 | 49 | NA | NA | NA | NA | | 1 | | 1 | | 0.5 | NA | 0.25 |
| SE | 6855 | UBAP2L | UBAP2L | chr1 | + | 154234590 | 154234792 | 154228614 | 154228725 | 154235195 | 154235291 | 0 | 0 | 2 | 0 | 0 | 0 | 2 | 2 | 0 | 0 | 1 | 0 | 98 | 49 | NA | NA | NA | NA | | 1 | | 1 | | 0.5 | NA | 0.25 |
| SE | 6876 | SOAT1 | SOAT1 | chr1 | + | 179344946 | 179345076 | 179343589 | 179343635 | 179347599 | 179347697 | 0 | 0 | 2 | 0 | 0 | 0 | 0 | 0 | 0 | 1 | 0 | 0 | 98 | 49 | NA | NA | NA | NA | | 1 | | 0 | NA | | NA | 1 |
| SE | 6880 | UBN1 | UBN1 | chr16 | + | 4876870 | 4877111 | 4874210 | 4875434 | 4880082 | 4882401 | 0 | 0 | 3 | 0 | 0 | 1 | 0 | 2 | 0 | 0 | 0 | 0 | 98 | 49 | NA | NA | NA | NA | | 0.6 | NA | | | 1 | NA | -0.4 |
| SE | 6881 | UBN1 | UBN1 | chr16 | + | 4877384 | 4877474 | 4876870 | 4877111 | 4880082 | 4882401 | 0 | 0 | 1 | 0 | 0 | 1 | 0 | 0 | 0 | 0 | 2 | 0 | 98 | 49 | NA | NA | NA | NA | | 0.333 | NA | | | 0 | NA | 0.333 |
| SE | 6891 | EP300 | EP300 | chr22 | + | 41146738 | 41146816 | 41141047 | 41141222 | 41147836 | 41147946 | 0 | 0 | 8 | 0 | 0 | 0 | 0 | 5 | 0 | 0 | 1 | 0 | 98 | 49 | NA | NA | NA | NA | | 1 | NA | | | 0.714 | NA | 0.286 |
| SE | 6907 | ASCC3 | ASCC3 | chr6 | - | 100509931 | 100510107 | 100508194 | 100509533 | 100512708 | 100512918 | 0 | 0 | 0 | 0 | 0 | 1 | 0 | 1 | 0 | 0 | 0 | 0 | 98 | 49 | NA | NA | NA | NA | | 0 | NA | | | 1 | NA | -1 |
| SE | 6930 | MADD | MADD | chr11 | + | 47293882 | 47293983 | 47290609 | 47290816 | 47295495 | 47295567 | 0 | 0 | 2 | 0 | 0 | 0 | 0 | 0 | 0 | 0 | 1 | 0 | 98 | 49 | NA | NA | NA | NA | | 1 | NA | | | 0 | NA | 1 |
| SE | 6971 | POU2F1 | POU2F1 | chr1 | + | 167303473 | 167303574 | 167220885 | 167220958 | 167332469 | 167332535 | 0 | 0 | 1 | 0 | 0 | 0 | 0 | 0 | 0 | 0 | 1 | 0 | 98 | 49 | NA | NA | NA | NA | | 1 | NA | | | 0 | NA | 1 |
| SE | 7032 | IFNAR1 | IFNAR1 | chr21 | + | 33343267 | 33343422 | 33340998 | 33341174 | 33343534 | 33343676 | 0 | 0 | 6 | 0 | 0 | 1 | 0 | 11 | 0 | 0 | 0 | 0 | 98 | 49 | NA | NA | NA | NA | | 0.75 | NA | | | 1 | NA | -0.25 |
| SE | 7041 | KCNJ15 | KCNJ15 | chr21 | + | 38298143 | 38298282 | 38296925 | 38297023 | 38299243 | 38301824 | 0 | 0 | 1 | 0 | 0 | 7 | 0 | 1 | 0 | 0 | 2 | 0 | 98 | 49 | NA | NA | NA | NA | | 0.067 | NA | | | 0.2 | NA | -0.133 |
| SE | 7047 | UROD | UROD | chr1 | + | 45014939 | 45015006 | 45014735 | 45014836 | 45015336 | 45015669 | 0 | 0 | 4 | 0 | 0 | 0 | 0 | 0 | 0 | 0 | 1 | 0 | 98 | 49 | NA | NA | NA | NA | | 1 | NA | | | 0 | NA | 1 |
| SE | 7060 | AMPD3 | AMPD3 | chr11 | + | 10478525 | 10478730 | 10461514 | 10461740 | 10482062 | 10482225 | 0 | 0 | 1 | 0 | 0 | 0 | 0 | 1 | 0 | 0 | 2 | 0 | 98 | 49 | NA | NA | NA | NA | | 1 | NA | | | 0.2 | NA | 0.8 |
| SE | 7073 | NR1D2 | NR1D2 | chr3 | + | 23967812 | 23968023 | 23964976 | 23965162 | 23977222 | 23980618 | 0 | 0 | 1 | 0 | 0 | 0 | 0 | 1 | 0 | 0 | 1 | 0 | 98 | 49 | NA | NA | NA | NA | | 1 | NA | | | 0.333 | NA | 0.667 |
| SE | 7074 | ERBB2IP | ERBB2IP | chr5 | + | 66068876 | 66069020 | 66053405 | 66054951 | 66072168 | 66072291 | 0 | 0 | 0 | 0 | 0 | 2 | 0 | 1 | 0 | 0 | 1 | 0 | 98 | 49 | NA | NA | NA | NA | | 0 | NA | | | 0.333 | NA | -0.333 |
| SE | 7076 | ERBB2IP | ERBB2IP | chr5 | + | 66072168 | 66072291 | 66053405 | 66054951 | 66075023 | 66075230 | 0 | 0 | 2 | 0 | 0 | 1 | 0 | 1 | 0 | 0 | 4 | 0 | 98 | 49 | NA | NA | NA | NA | | 0.5 | NA | | | 0.111 | NA | 0.389 |
| SE | 7080 | UBR2 | UBR2 | chr6 | + | 42605720 | 42605859 | 42603587 | 42603718 | 42606588 | 42606651 | 0 | 0 | 23 | 0 | 0 | 1 | 1 | 13 | 0 | 0 | 0 | 0 | 98 | 49 | NA | NA | NA | NA | | 0.92 | | 1 | | 1 | NA | -0.08 |
| SE | 7081 | UBR2 | UBR2 | chr6 | + | 42665408 | 42665512 | 42663257 | 42663419 | 42666166 | 42666245 | 0 | 0 | 12 | 0 | 0 | 1 | 0 | 10 | 0 | 0 | 0 | 0 | 98 | 49 | NA | NA | NA | NA | | 0.857 | NA | | | 1 | NA | -0.143 |
| SE | 7083 | RNF38 | RNF38 | chr9 | - | 36342324 | 36342424 | 36336400 | 36339814 | 36344831 | 36344953 | 0 | 0 | 1 | 0 | 0 | 2 | 0 | 2 | 0 | 0 | 0 | 0 | 98 | 49 | NA | NA | NA | NA | | 0.2 | NA | | | 1 | NA | -0.8 |
| SE | 7090 | IL18RAP | IL18RAP | chr2 | + | 102443199 | 102443323 | 102441311 | 102441377 | 102445188 | 102445340 | 0 | 0 | 1 | 0 | 0 | 0 | 0 | 1 | 0 | 4 | 0 | 0 | 98 | 49 | NA | NA | NA | NA | | 1 | | 0 | | 1 | NA | 0.5 |
| SE | 7118 | TMEM71 | TMEM71 | chr8 | - | 132722039 | 132722115 | 132714153 | 132714215 | 132727797 | 132727986 | 0 | 0 | 14 | 0 | 0 | 0 | 0 | 14 | 0 | 0 | 2 | 0 | 98 | 49 | NA | NA | NA | NA | | 1 | NA | | | 0.778 | NA | 0.222 |
| SE | 7119 | TMEM71 | TMEM71 | chr8 | - | 132727797 | 132727986 | 132722039 | 132722115 | 132746941 | 132747057 | 0 | 0 | 9 | 0 | 0 | 0 | 0 | 6 | 0 | 0 | 1 | 0 | 98 | 49 | NA | NA | NA | NA | | 1 | NA | | | 0.75 | NA | 0.25 |
| SE | 7132 | GGA3 | GGA3 | chr17 | - | 75244618 | 75244717 | 75243446 | 75243570 | 75246508 | 75246584 | 0 | 0 | 0 | 0 | 0 | 1 | 0 | 1 | 0 | 0 | 0 | 0 | 98 | 49 | NA | NA | NA | NA | | 0 | NA | | | 1 | NA | -1 |
| SE | 7138 | KIF21B | KIF21B | chr1 | - | 200983055 | 200983094 | 200980959 | 200981096 | 200984858 | 200984972 | 0 | 0 | 1 | 0 | 0 | 0 | 0 | 0 | 0 | 0 | 1 | 0 | 88 | 49 | NA | NA | NA | NA | | 1 | NA | | | 0 | NA | 1 |
| SE | 7144 | CPD | CPD | chr17 | + | 30443801 | 30443971 | 30442307 | 30442450 | 30445690 | 30446020 | 0 | 0 | 1 | 0 | 0 | 0 | 0 | 6 | 0 | 0 | 1 | 0 | 98 | 49 | NA | NA | NA | NA | | 1 | NA | | | 0.75 | NA | 0.25 |
| SE | 7187 | LENG8 | LENG8 | chr19 | + | 54453545 | 54453656 | 54452650 | 54452752 | 54454429 | 54454682 | 0 | 0 | 5 | 0 | 0 | 1 | 2 | 1 | 0 | 0 | 0 | 0 | 98 | 49 | NA | NA | NA | NA | | 0.714 | | 1 | | 1 | NA | -0.286 |
| SE | 7203 | CLEC4A | CLEC4A | chr12 | + | 8125560 | 8125677 | 8123631 | 8123960 | 8129263 | 8129362 | 0 | 0 | 2 | 0 | 0 | 1 | 0 | 1 | 0 | 0 | 0 | 0 | 98 | 49 | NA | NA | NA | NA | | 0.5 | NA | | | 1 | NA | -0.5 |
| SE | 7251 | GAS5 | GAS5 | chr1 | - | 173865228 | 173865282 | 173864674 | 173864704 | 173865470 | 173865547 | 0 | 0 | 3 | 0 | 0 | 0 | 0 | 2 | 0 | 0 | 1 | 0 | 98 | 49 | NA | NA | NA | NA | | 1 | NA | | | 0.5 | NA | 0.5 |
| SE | 7252 | RYK | RYK | chr3 | - | 134175608 | 134175768 | 134159236 | 134159373 | 134175929 | 134176039 | 0 | 0 | 4 | 0 | 0 | 0 | 0 | 0 | 0 | 0 | 1 | 0 | 98 | 49 | NA | NA | NA | NA | | 1 | NA | | | 0 | NA | 1 |
| SE | 7262 | FBXO18 | FBXO18 | chr10 | + | 5903019 | 5903175 | 5890202 | 5890346 | 5906036 | 5906632 | 0 | 0 | 3 | 0 | 0 | 0 | 0 | 0 | 0 | 0 | 1 | 0 | 98 | 49 | NA | NA | NA | NA | | 1 | NA | | | 0 | NA | 1 |
| SE | 7269 | CIDEB | CIDEB | chr14 | - | 24305946 | 24306137 | 24305186 | 24305765 | 24306373 | 24306523 | 0 | 0 | 3 | 0 | 0 | 2 | 0 | 2 | 0 | 0 | 0 | 0 | 98 | 49 | NA | NA | NA | NA | | 0.429 | NA | | | 1 | NA | -0.571 |
| SE | 7289 | PGS1 | PGS1 | chr17 | + | 78396307 | 78396385 | 78392475 | 78392665 | 78398251 | 78398351 | 0 | 0 | 5 | 0 | 0 | 1 | 0 | 3 | 0 | 0 | 1 | 0 | 98 | 49 | NA | NA | NA | NA | | 0.714 | NA | | | 0.6 | NA | 0.114 |
| SE | 7314 | WNK1 | WNK1 | chr12 | + | 871264 | 871348 | 862082 | 862270 | 878214 | 878361 | 0 | 0 | 5 | 0 | 0 | 2 | 0 | 1 | 0 | 0 | 0 | 0 | 98 | 49 | NA | NA | NA | NA | | 0.556 | NA | | | 1 | NA | -0.444 |
| SE | 7344 | FAM129A | FAM129A | chr1 | - | 184805945 | 184806056 | 184798078 | 184798190 | 184808073 | 184808235 | 0 | 0 | 4 | 0 | 0 | 0 | 7 | 11 | 0 | 0 | 1 | 0 | 98 | 49 | NA | NA | NA | NA | | 1 | | 1 | | 0.846 | NA | 0.077 |
| SE | 7361 | OXA1L | OXA1L | chr14 | + | 22769790 | 22769934 | 22767957 | 22768171 | 22770192 | 22770278 | 0 | 0 | 5 | 0 | 0 | 0 | 0 | 0 | 0 | 0 | 1 | 0 | 98 | 49 | NA | NA | NA | NA | | 1 | NA | | | 0 | NA | 1 |
| SE | 7362 | NOTCH2 | NOTCH2 | chr1 | - | 119922235 | 119922446 | 119921712 | 119921809 | 119922635 | 119922778 | 0 | 0 | 10 | 0 | 0 | 0 | 0 | 2 | 0 | 0 | 1 | 0 | 98 | 49 | NA | NA | NA | NA | | 1 | NA | | | 0.5 | NA | 0.5 |
| SE | 7364 | NOTCH2 | NOTCH2 | chr1 | - | 119959391 | 119959502 | 119955039 | 119955232 | 119963573 | 119963807 | 0 | 0 | 4 | 0 | 0 | 1 | 0 | 5 | 0 | 0 | 0 | 0 | 98 | 49 | NA | NA | NA | NA | | 0.667 | NA | | | 1 | NA | -0.333 |
| SE | 7368 | PSMA3 | PSMA3 | chr14 | + | 58270933 | 58270998 | 58270417 | 58270485 | 58271850 | 58272009 | 0 | 0 | 6 | 0 | 0 | 1 | 0 | 3 | 0 | 0 | 0 | 0 | 98 | 49 | NA | NA | NA | NA | | 0.75 | NA | | | 1 | NA | -0.25 |
| SE | 7388 | ZRANB1 | ZRANB1 | chr10 | + | 124974199 | 124974398 | 124973644 | 124973716 | 124981708 | 124981829 | 0 | 0 | 0 | 0 | 0 | 1 | 0 | 2 | 0 | 0 | 0 | 0 | 98 | 49 | NA | NA | NA | NA | | 0 | NA | | | 1 | NA | -1 |
| SE | 7390 | ZRANB1 | ZRANB1 | chr10 | + | 124983458 | 124983688 | 124983174 | 124983304 | 124984773 | 124987436 | 0 | 0 | 0 | 0 | 0 | 1 | 0 | 1 | 0 | 0 | 0 | 0 | 98 | 49 | NA | NA | NA | NA | | 0 | NA | | | 1 | NA | -1 |
| SE | 7409 | RBM23 | RBM23 | chr14 | - | 22908332 | 22908380 | 22906194 | 22906368 | 22909482 | 22909595 | 0 | 0 | 2 | 0 | 0 | 0 | 0 | 1 | 0 | 0 | 1 | 0 | 97 | 49 | NA | NA | NA | NA | | 1 | NA | | | 0.336 | NA | 0.664 |
| SE | 7423 | H2AFY | H2AFY | chr5 | - | 135343259 | 135343434 | 135334380 | 135335141 | 135345967 | 135346057 | 0 | 0 | 10 | 0 | 0 | 0 | 2 | 6 | 0 | 0 | 1 | 0 | 98 | 49 | NA | NA | NA | NA | | 1 | | 1 | | 0.75 | NA | 0.125 |
| SE | 7424 | H2AFY | H2AFY | chr5 | - | 135350822 | 135350913 | 135345967 | 135346057 | 135360496 | 135360607 | 0 | 0 | 3 | 0 | 0 | 0 | 0 | 0 | 0 | 0 | 1 | 0 | 98 | 49 | NA | NA | NA | NA | | 1 | NA | | | 0 | NA | 1 |
| SE | 7433 | KCNAB2 | KCNAB2 | chr1 | + | 6041830 | 6041872 | 6040516 | 6040645 | 6072754 | 6072798 | 0 | 0 | 1 | 0 | 0 | 0 | 0 | 0 | 0 | 2 | 0 | 0 | 91 | 49 | NA | NA | NA | NA | | 1 | | 0 | NA | | NA | 1 |
| SE | 7452 | ARNT | ARNT | chr1 | - | 150823193 | 150823345 | 150817919 | 150818030 | 150826542 | 150826617 | 0 | 0 | 4 | 0 | 0 | 0 | 0 | 0 | 0 | 0 | 1 | 0 | 98 | 49 | NA | NA | NA | NA | | 1 | NA | | | 0 | NA | 1 |
| SE | 7453 | ARNT | ARNT | chr1 | - | 150842423 | 150842468 | 150839440 | 150839654 | 150846262 | 150846307 | 0 | 0 | 0 | 0 | 0 | 1 | 0 | 1 | 0 | 0 | 0 | 0 | 94 | 49 | NA | NA | NA | NA | | 0 | NA | | | 1 | NA | -1 |
| SE | 7461 | HNRNPDL | HNRNPDL | chr4 | - | 82426462 | 82426633 | 82426036 | 82426129 | 82427189 | 82427304 | 0 | 0 | 1 | 0 | 0 | 0 | 0 | 6 | 0 | 0 | 1 | 0 | 98 | 49 | NA | NA | NA | NA | | 1 | NA | | | 0.75 | NA | 0.25 |
| SE | 7467 | SLTM | SLTM | chr15 | - | 58890280 | 58890461 | 58889429 | 58889554 | 58892896 | 58893060 | 0 | 0 | 2 | 0 | 0 | 1 | 0 | 0 | 0 | 0 | 2 | 0 | 98 | 49 | NA | NA | NA | NA | | 0.5 | NA | | | 0 | NA | 0.5 |
| SE | 7468 | SLTM | SLTM | chr15 | - | 58894432 | 58894582 | 58894089 | 58894193 | 58897114 | 58897233 | 0 | 0 | 3 | 0 | 0 | 0 | 0 | 1 | 0 | 2 | 0 | 0 | 98 | 49 | NA | NA | NA | NA | | 1 | | 0 | | 1 | NA | 0.5 |
| SE | 7519 | ZNF335 | ZNF335 | chr20 | - | 45969450 | 45969691 | 45968284 | 45968362 | 45971209 | 45971460 | 0 | 0 | 0 | 0 | 0 | 1 | 0 | 2 | 0 | 0 | 0 | 0 | 98 | 49 | NA | NA | NA | NA | | 0 | NA | | | 1 | NA | -1 |
| SE | 7542 | ADAR | ADAR | chr1 | - | 154586180 | 154586363 | 154585752 | 154585865 | 154588124 | 154588258 | 0 | 0 | 38 | 0 | 0 | 2 | 3 | 13 | 0 | 0 | 2 | 0 | 98 | 49 | NA | NA | NA | NA | | 0.905 | | 1 | | 0.765 | NA | 0.022 |
| SE | 7543 | PTPN22 | PTPN22 | chr1 | - | 113856547 | 113856619 | 113856381 | 113856441 | 113857737 | 113857776 | 0 | 0 | 1 | 0 | 0 | 0 | 0 | 0 | 0 | 0 | 1 | 0 | 98 | 49 | NA | NA | NA | NA | | 1 | NA | | | 0 | NA | 1 |
| SE | 7567 | ST7L | ST7L | chr1 | - | 112581991 | 112582106 | 112578344 | 112578417 | 112582374 | 112582472 | 0 | 0 | 2 | 0 | 0 | 0 | 0 | 0 | 0 | 0 | 1 | 0 | 98 | 49 | NA | NA | NA | NA | | 1 | NA | | | 0 | NA | 1 |
| SE | 7569 | ARAP2 | ARAP2 | chr4 | - | 36212395 | 36212487 | 36210389 | 36210743 | 36213242 | 36213319 | 0 | 0 | 3 | 0 | 0 | 2 | 1 | 3 | 0 | 0 | 1 | 0 | 98 | 49 | NA | NA | NA | NA | | 0.429 | | 1 | | 0.6 | NA | -0.371 |
| SE | 7585 | STAT2 | STAT2 | chr12 | - | 56350096 | 56350190 | 56349588 | 56349636 | 56350411 | 56350432 | 0 | 0 | 18 | 0 | 0 | 0 | 0 | 12 | 0 | 0 | 2 | 0 | 98 | 49 | NA | NA | NA | NA | | 1 | NA | | | 0.75 | NA | 0.25 |
| SE | 7619 | MAN2B1 | MAN2B1 | chr19 | - | 12658427 | 12658507 | 12658223 | 12658344 | 12661259 | 12661376 | 0 | 0 | 8 | 0 | 0 | 1 | 0 | 1 | 0 | 0 | 0 | 0 | 98 | 49 | NA | NA | NA | NA | | 0.8 | NA | | | 1 | NA | -0.2 |
| SE | 7620 | MAN2B1 | MAN2B1 | chr19 | - | 12658427 | 12658510 | 12658223 | 12658344 | 12661259 | 12661376 | 0 | 0 | 10 | 0 | 0 | 1 | 0 | 2 | 0 | 0 | 0 | 0 | 98 | 49 | NA | NA | NA | NA | | 0.833 | NA | | | 1 | NA | -0.167 |
| SE | 7629 | RALGAPB | RALGAPB | chr20 | + | 38565355 | 38565478 | 38562531 | 38562697 | 38567095 | 38567232 | 0 | 0 | 0 | 0 | 0 | 4 | 0 | 3 | 0 | 0 | 0 | 0 | 98 | 49 | NA | NA | NA | NA | | 0 | NA | | | 1 | NA | -1 |
| SE | 7630 | RALGAPB | RALGAPB | chr20 | + | 38565358 | 38565478 | 38562531 | 38562697 | 38567095 | 38567232 | 0 | 0 | 2 | 0 | 0 | 4 | 0 | 3 | 0 | 0 | 0 | 0 | 98 | 49 | NA | NA | NA | NA | | 0.2 | NA | | | 1 | NA | -0.8 |
| SE | 7632 | BTAF1 | BTAF1 | chr10 | + | 92027129 | 92027300 | 92026591 | 92026751 | 92028789 | 92030323 | 0 | 0 | 4 | 0 | 0 | 3 | 0 | 5 | 0 | 0 | 0 | 0 | 98 | 49 | NA | NA | NA | NA | | 0.4 | NA | | | 1 | NA | -0.6 |
| SE | 7685 | EAF1 | EAF1 | chr3 | + | 15432086 | 15432223 | 15429912 | 15430007 | 15434347 | 15434538 | 0 | 0 | 3 | 1 | 0 | 0 | 0 | 2 | 0 | 0 | 0 | 0 | 98 | 49 | NA | NA | 0 | NA | | 1 | NA | | | 1 | NA | -0.5 |
| SE | 7686 | DHX40 | DHX40 | chr17 | + | 59566626 | 59566794 | 59565524 | 59565783 | 59570517 | 59570663 | 0 | 0 | 1 | 0 | 0 | 1 | 0 | 1 | 0 | 0 | 0 | 0 | 98 | 49 | NA | NA | NA | NA | | 0.333 | NA | | | 1 | NA | -0.667 |
| SE | 7707 | CHFR | CHFR | chr12 | - | 132869618 | 132869834 | 132861466 | 132861634 | 132870723 | 132870783 | 0 | 0 | 0 | 0 | 0 | 2 | 0 | 1 | 0 | 0 | 0 | 0 | 98 | 49 | NA | NA | NA | NA | | 0 | NA | | | 1 | NA | -1 |
| SE | 7722 | ARHGEF2 | ARHGEF2 | chr1 | - | 155965630 | 155965760 | 155965302 | 155965412 | 155966415 | 155966479 | 0 | 0 | 8 | 0 | 0 | 0 | 0 | 0 | 0 | 0 | 1 | 0 | 98 | 49 | NA | NA | NA | NA | | 1 | NA | | | 0 | NA | 1 |
| SE | 7723 | ARHGEF2 | ARHGEF2 | chr1 | - | 155966819 | 155966887 | 155966415 | 155966479 | 155969155 | 155969300 | 0 | 0 | 4 | 0 | 0 | 0 | 0 | 3 | 0 | 0 | 1 | 0 | 98 | 49 | NA | NA | NA | NA | | 1 | NA | | | 0.6 | NA | 0.4 |
| SE | 7728 | KIAA1033 | KIAA1033 | chr12 | + | 105126234 | 105126362 | 105126003 | 105126127 | 105127128 | 105127289 | 0 | 0 | 12 | 0 | 0 | 1 | 3 | 5 | 0 | 0 | 0 | 0 | 98 | 49 | NA | NA | NA | NA | | 0.857 | | 1 | | 1 | NA | -0.143 |
| SE | 7730 | KIAA1033 | KIAA1033 | chr12 | + | 105144286 | 105144455 | 105143126 | 105143243 | 105144717 | 105144872 | 0 | 0 | 26 | 0 | 0 | 0 | 0 | 9 | 0 | 0 | 1 | 0 | 98 | 49 | NA | NA | NA | NA | | 1 | NA | | | 0.818 | NA | 0.182 |
| SE | 7732 | FAM32A | FAM32A | chr19 | + | 16190519 | 16190573 | 16185623 | 16185765 | 16190886 | 16192046 | 0 | 0 | 2 | 0 | 0 | 0 | 0 | 2 | 0 | 0 | 1 | 0 | 98 | 49 | NA | NA | NA | NA | | 1 | NA | | | 0.5 | NA | 0.5 |
| SE | 7819 | ADCY10P1 | ADCY10P1 | chr6 | + | 41118923 | 41119107 | 41118128 | 41118184 | 41120522 | 41120657 | 0 | 0 | 1 | 0 | 0 | 0 | 0 | 1 | 0 | 0 | 1 | 0 | 98 | 49 | NA | NA | NA | NA | | 1 | NA | | | 0.333 | NA | 0.667 |
| SE | 7845 | ITPKC | ITPKC | chr19 | + | 40733159 | 40733364 | 40729201 | 40729415 | 40736985 | 40737087 | 0 | 0 | 1 | 0 | 0 | 0 | 0 | 0 | 0 | 0 | 1 | 0 | 98 | 49 | NA | NA | NA | NA | | 1 | NA | | | 0 | NA | 1 |
| SE | 7849 | RPS6KA1 | RPS6KA1 | chr1 | + | 26561504 | 26561663 | 26561044 | 26561134 | 26571448 | 26571610 | 0 | 0 | 12 | 0 | 0 | 1 | 0 | 4 | 0 | 0 | 0 | 0 | 98 | 49 | NA | NA | NA | NA | | 0.857 | NA | | | 1 | NA | -0.143 |
| SE | 7850 | FOXJ3 | FOXJ3 | chr1 | - | 42199101 | 42199230 | 42194889 | 42195064 | 42205761 | 42205863 | 0 | 0 | 0 | 0 | 0 | 1 | 1 | 7 | 0 | 0 | 0 | 0 | 98 | 49 | NA | NA | NA | NA | | 0 | | 1 | | 1 | NA | -1 |
| SE | 7857 | RBL2 | RBL2 | chr16 | + | 53469915 | 53470185 | 53467057 | 53467169 | 53470382 | 53470663 | 0 | 0 | 3 | 0 | 0 | 0 | 0 | 7 | 0 | 0 | 2 | 0 | 98 | 49 | NA | NA | NA | NA | | 1 | NA | | | 0.636 | NA | 0.364 |
| SE | 7858 | STK24 | STK24 | chr13 | - | 98457167 | 98457304 | 98450198 | 98453209 | 98460371 | 98460440 | 0 | 0 | 5 | 0 | 0 | 1 | 4 | 3 | 0 | 0 | 0 | 0 | 98 | 49 | NA | NA | NA | NA | | 0.714 | | 1 | | 1 | NA | -0.286 |
| SE | 7870 | CTBS | CTBS | chr1 | - | 84569930 | 84570139 | 84565840 | 84566012 | 84570581 | 84570720 | 0 | 0 | 4 | 0 | 0 | 0 | 0 | 0 | 0 | 0 | 5 | 0 | 98 | 49 | NA | NA | NA | NA | | 1 | NA | | | 0 | NA | 1 |
| SE | 7872 | MROH6 | MROH6 | chr8 | - | 143567610 | 143567676 | 143566192 | 143567465 | 143567785 | 143567888 | 0 | 0 | 1 | 0 | 0 | 0 | 0 | 0 | 0 | 0 | 2 | 0 | 98 | 49 | NA | NA | NA | NA | | 1 | NA | | | 0 | NA | 1 |
| SE | 7881 | UHRF1BP1L | UHRF1BP1L | chr12 | - | 100088914 | 100089095 | 100086274 | 100086358 | 100095666 | 100095837 | 0 | 0 | 11 | 0 | 0 | 3 | 0 | 3 | 0 | 0 | 0 | 0 | 98 | 49 | NA | NA | NA | NA | | 0.647 | NA | | | 1 | NA | -0.353 |
| SE | 7900 | IFIT3 | IFIT3 | chr10 | + | 89337424 | 89337542 | 89327818 | 89328078 | 89338660 | 89340968 | 0 | 0 | 0 | 0 | 0 | 1 | 0 | 1 | 0 | 0 | 0 | 0 | 98 | 49 | NA | NA | NA | NA | | 0 | NA | | | 1 | NA | -1 |
| SE | 7906 | IFIT1 | IFIT1 | chr10 | + | 89399889 | 89400003 | 89392545 | 89392717 | 89402280 | 89406487 | 0 | 0 | 11 | 0 | 0 | 5 | 0 | 7 | 0 | 4 | 2 | 0 | 98 | 49 | NA | NA | NA | NA | | 0.524 | | 0 | | 0.636 | NA | 0.206 |
| SE | 7913 | DDX21 | DDX21 | chr10 | + | 68977528 | 68977688 | 68974669 | 68974743 | 68978841 | 68978976 | 0 | 0 | 10 | 0 | 0 | 0 | 0 | 8 | 0 | 0 | 1 | 0 | 98 | 49 | NA | NA | NA | NA | | 1 | NA | | | 0.8 | NA | 0.2 |
| SE | 7921 | CSDE1 | CSDE1 | chr1 | - | 114730257 | 114730422 | 114726982 | 114727090 | 114730507 | 114730648 | 0 | 0 | 21 | 0 | 0 | 1 | 0 | 16 | 0 | 0 | 0 | 0 | 98 | 49 | NA | NA | NA | NA | | 0.913 | NA | | | 1 | NA | -0.087 |
| SE | 7922 | CSDE1 | CSDE1 | chr1 | - | 114734441 | 114734523 | 114733988 | 114734117 | 114736757 | 114736855 | 0 | 0 | 16 | 0 | 0 | 0 | 0 | 13 | 0 | 0 | 1 | 0 | 98 | 49 | NA | NA | NA | NA | | 1 | NA | | | 0.867 | NA | 0.133 |
| SE | 7946 | PPP2R2D | PPP2R2D | chr10 | + | 131911561 | 131912598 | 131901237 | 131901330 | 131934457 | 131934555 | 0 | 0 | 0 | 0 | 0 | 3 | 0 | 2 | 0 | 0 | 0 | 0 | 98 | 49 | NA | NA | NA | NA | | 0 | NA | | | 1 | NA | -1 |
| SE | 7947 | PPP2R2D | PPP2R2D | chr10 | + | 131934893 | 131935006 | 131934457 | 131934555 | 131940030 | 131940196 | 0 | 0 | 1 | 0 | 0 | 3 | 0 | 1 | 0 | 0 | 0 | 0 | 98 | 49 | NA | NA | NA | NA | | 0.143 | NA | | | 1 | NA | -0.857 |
| SE | 7960 | CEP104 | CEP104 | chr1 | - | 3826707 | 3826744 | 3826369 | 3826436 | 3829265 | 3829373 | 0 | 0 | 2 | 0 | 0 | 0 | 0 | 0 | 0 | 0 | 1 | 0 | 86 | 49 | NA | NA | NA | NA | | 1 | NA | | | 0 | NA | 1 |
| SE | 7963 | SP3 | SP3 | chr2 | - | 173918592 | 173918785 | 173913069 | 173913266 | 173954872 | 173956232 | 0 | 0 | 19 | 0 | 0 | 1 | 0 | 12 | 0 | 0 | 0 | 0 | 98 | 49 | NA | NA | NA | NA | | 0.905 | NA | | | 1 | NA | -0.095 |
| SE | 8007 | ZNF33A | ZNF33A | chr10 | + | 38039412 | 38039492 | 38017290 | 38017386 | 38054374 | 38060067 | 0 | 0 | 2 | 0 | 0 | 1 | 0 | 0 | 0 | 0 | 2 | 0 | 98 | 49 | NA | NA | NA | NA | | 0.5 | NA | | | 0 | NA | 0.5 |
| SE | 8010 | DUSP18 | DUSP18 | chr22 | - | 30664954 | 30665121 | 30661457 | 30664080 | 30667461 | 30667890 | 0 | 0 | 1 | 0 | 0 | 2 | 0 | 0 | 0 | 0 | 1 | 0 | 98 | 49 | NA | NA | NA | NA | | 0.2 | NA | | | 0 | NA | 0.2 |
| SE | 8016 | HIATL1 | HIATL1 | chr9 | + | 94456239 | 94456337 | 94453957 | 94454070 | 94458321 | 94458464 | 0 | 0 | 10 | 0 | 0 | 1 | 0 | 2 | 0 | 0 | 0 | 0 | 98 | 49 | NA | NA | NA | NA | | 0.833 | NA | | | 1 | NA | -0.167 |
| SE | 8018 | MIR22HG | MIR22HG | chr17 | - | 1714370 | 1714453 | 1713702 | 1714014 | 1716129 | 1716272 | 0 | 0 | 1 | 0 | 0 | 0 | 0 | 1 | 0 | 0 | 1 | 0 | 98 | 49 | NA | NA | NA | NA | | 1 | NA | | | 0.333 | NA | 0.667 |
| SE | 8037 | MORF4L1 | MORF4L1 | chr15 | + | 78894057 | 78894230 | 78893538 | 78893627 | 78894819 | 78894904 | 0 | 0 | 7 | 0 | 0 | 3 | 0 | 1 | 0 | 0 | 0 | 0 | 98 | 49 | NA | NA | NA | NA | | 0.538 | NA | | | 1 | NA | -0.462 |
| SE | 8043 | GLYR1 | GLYR1 | chr16 | - | 4821546 | 4821597 | 4821379 | 4821453 | 4822874 | 4822931 | 0 | 0 | 9 | 0 | 3 | 0 | 2 | 0 | 0 | 0 | 0 | 0 | 98 | 49 | NA | NA | NA | NA | 0 | 1 | | 1 | NA | | NA | -0.5 |
| SE | 8044 | GLYR1 | GLYR1 | chr16 | - | 4831978 | 4832221 | 4823820 | 4823907 | 4832773 | 4832912 | 0 | 0 | 1 | 0 | 0 | 0 | 0 | 0 | 0 | 0 | 1 | 0 | 98 | 49 | NA | NA | NA | NA | | 1 | NA | | | 0 | NA | 1 |
| SE | 8049 | MARK2 | MARK2 | chr11 | + | 63903985 | 63904147 | 63903060 | 63903158 | 63904785 | 63905043 | 0 | 0 | 1 | 0 | 0 | 0 | 0 | 0 | 0 | 2 | 0 | 0 | 98 | 49 | NA | NA | NA | NA | | 1 | | 0 | NA | | NA | 1 |
| SE | 8066 | P3H1 | P3H1 | chr1 | - | 42755547 | 42755637 | 42755164 | 42755217 | 42757782 | 42757922 | 0 | 0 | 1 | 0 | 0 | 0 | 0 | 0 | 0 | 0 | 1 | 0 | 98 | 49 | NA | NA | NA | NA | | 1 | NA | | | 0 | NA | 1 |
| SE | 8077 | NADK | NADK | chr1 | - | 1755373 | 1755476 | 1754543 | 1754698 | 1756257 | 1756343 | 0 | 0 | 12 | 0 | 0 | 3 | 5 | 1 | 0 | 0 | 0 | 0 | 98 | 49 | NA | NA | NA | NA | | 0.667 | | 1 | | 1 | NA | -0.333 |
| SE | 8089 | SCLT1 | SCLT1 | chr4 | - | 128942995 | 128943188 | 128936654 | 128936851 | 128946006 | 128946152 | 0 | 0 | 0 | 0 | 0 | 2 | 0 | 1 | 0 | 0 | 0 | 0 | 98 | 49 | NA | NA | NA | NA | | 0 | NA | | | 1 | NA | -1 |
| SE | 8091 | PKN2 | PKN2 | chr1 | + | 88805890 | 88806017 | 88805496 | 88805671 | 88807312 | 88807443 | 0 | 0 | 9 | 0 | 0 | 0 | 0 | 7 | 0 | 0 | 1 | 0 | 98 | 49 | NA | NA | NA | NA | | 1 | NA | | | 0.778 | NA | 0.222 |
| SE | 8093 | TP53INP1 | TP53INP1 | chr8 | - | 94932069 | 94932107 | 94925971 | 94930728 | 94939859 | 94940220 | 0 | 0 | 4 | 0 | 0 | 1 | 0 | 4 | 0 | 0 | 1 | 0 | 87 | 49 | NA | NA | NA | NA | | 0.693 | NA | | | 0.693 | NA | 0 |

| | | | | | | | | | | | | | | | | | | | | | | | | | | | | | | | | | | |
|---|---|---|---|---|---|---|---|---|---|---|---|---|---|---|---|---|---|---|---|---|---|---|---|---|---|---|---|---|---|---|---|---|---|---|
| SE | 8097 | JMJD1C | JMJD1C | chr10 | - | 63200475 | 63200677 | 63198512 | 63198727 | 63206594 | 63208801 | 0 | 0 | 13 | 0 | 0 | 1 | 0 | 4 | 0 | 0 | 0 | 0 | 98 | 49 | NA | NA | NA | NA | 0.867 | NA | 1 | NA | -0.133 |
| SE | 8098 | JMJD1C | JMJD1C | chr10 | - | 63215552 | 63215696 | 63215262 | 63215455 | 63217206 | 63217331 | 0 | 0 | 21 | 0 | 0 | 1 | 0 | 8 | 0 | 0 | 0 | 0 | 98 | 49 | NA | NA | NA | NA | 0.913 | NA | 1 | NA | -0.087 |
| SE | 8099 | JMJD1C | JMJD1C | chr10 | - | 63219877 | 63219983 | 63217206 | 63217331 | 63264650 | 63264764 | 0 | 0 | 14 | 0 | 0 | 2 | 0 | 1 | 0 | 0 | 2 | 0 | 98 | 49 | NA | NA | NA | NA | 0.778 | NA | 0.2 | NA | 0.578 |
| SE | 8111 | COPB2 | COPB2 | chr3 | - | 139373665 | 139373808 | 139373212 | 139373412 | 139374488 | 139374588 | 0 | 0 | 1 | 0 | 0 | 0 | 0 | 0 | 0 | 0 | 2 | 0 | 98 | 49 | NA | NA | NA | NA | 1 | NA | 0 | NA | 1 |
| SE | 8124 | ANXA5 | ANXA5 | chr4 | - | 121678414 | 121678494 | 121677893 | 121677950 | 121681670 | 121681761 | 0 | 0 | 22 | 0 | 0 | 1 | 11 | 6 | 0 | 0 | 0 | 0 | 98 | 49 | NA | NA | NA | NA | 0.917 | 1 | 1 | NA | -0.083 |
| SE | 8148 | PPT1 | PPT1 | chr1 | - | 40076841 | 40076913 | 40072709 | 40074183 | 40078559 | 40078658 | 0 | 0 | 9 | 0 | 0 | 5 | 0 | 17 | 0 | 0 | 0 | 0 | 98 | 49 | NA | NA | NA | NA | 0.474 | NA | 1 | NA | -0.526 |
| SE | 8156 | KMT2E | KMT2E | chr7 | + | 105066726 | 105066807 | 105063350 | 105063580 | 105073618 | 105073677 | 0 | 0 | 10 | 0 | 0 | 0 | 0 | 10 | 0 | 0 | 1 | 0 | 98 | 49 | NA | NA | NA | NA | 1 | NA | 0.833 | NA | 0.167 |
| SE | 8178 | MLXIP | MLXIP | chr12 | + | 122130843 | 122130933 | 122129940 | 122130112 | 122132291 | 122132383 | 0 | 0 | 1 | 0 | 0 | 1 | 0 | 1 | 0 | 0 | 0 | 0 | 98 | 49 | NA | NA | NA | NA | 0.333 | NA | 1 | NA | -0.667 |
| SE | 8186 | EPSTI1 | EPSTI1 | chr13 | - | 42889210 | 42889242 | 42886387 | 42888502 | 42895008 | 42895108 | 0 | 0 | 8 | 0 | 0 | 4 | 0 | 2 | 0 | 4 | 6 | 0 | 81 | 49 | NA | NA | NA | NA | 0.547 | 0 | 0.168 | NA | 0.463 |
| SE | 8192 | ATP5F1 | ATP5F1 | chr1 | + | 111454210 | 111454356 | 111449836 | 111449873 | 111456085 | 111456249 | 0 | 0 | 8 | 0 | 0 | 1 | 5 | 5 | 0 | 0 | 0 | 0 | 98 | 49 | NA | NA | NA | NA | 0.8 | 1 | 1 | NA | -0.2 |
| SE | 8219 | SLC15A4 | SLC15A4 | chr12 | - | 128809942 | 128810111 | 128809395 | 128809473 | 128814774 | 128815070 | 0 | 0 | 4 | 0 | 0 | 3 | 5 | 4 | 0 | 0 | 1 | 0 | 98 | 49 | NA | NA | NA | NA | 0.4 | 1 | 0.667 | NA | -0.433 |
| SE | 8237 | LBR | LBR | chr1 | - | 225404625 | 225404706 | 225404403 | 225404526 | 225406663 | 225406832 | 0 | 0 | 5 | 0 | 0 | 0 | 0 | 3 | 0 | 0 | 2 | 0 | 98 | 49 | NA | NA | NA | NA | 1 | NA | 0.429 | NA | 0.571 |
| SE | 8249 | PRRC2C | PRRC2C | chr1 | + | 171545478 | 171545687 | 171539970 | 171542229 | 171550085 | 171550240 | 0 | 0 | 0 | 0 | 0 | 1 | 0 | 7 | 0 | 0 | 0 | 0 | 98 | 49 | NA | NA | NA | NA | 0 | NA | 1 | NA | -1 |
| SE | 8250 | PRRC2C | PRRC2C | chr1 | + | 171571319 | 171571421 | 171568246 | 171568339 | 171574926 | 171575128 | 0 | 0 | 6 | 0 | 0 | 0 | 0 | 2 | 0 | 0 | 1 | 0 | 98 | 49 | NA | NA | NA | NA | 1 | NA | 0.5 | NA | 0.5 |
| SE | 8251 | SMARCA2 | SMARCA2 | chr9 | + | 2170418 | 2170472 | 2161685 | 2161903 | 2181570 | 2181676 | 0 | 0 | 2 | 0 | 0 | 0 | 0 | 1 | 0 | 0 | 1 | 0 | 98 | 49 | NA | NA | NA | NA | 1 | NA | 0.333 | NA | 0.667 |
| SE | 8260 | ARHGEF11 | ARHGEF11 | chr1 | - | 156947768 | 156947956 | 156947303 | 156947450 | 156948180 | 156948228 | 0 | 0 | 13 | 0 | 0 | 1 | 0 | 3 | 0 | 0 | 0 | 0 | 98 | 49 | NA | NA | NA | NA | 0.867 | NA | 1 | NA | -0.133 |
| SE | 8262 | ARHGEF11 | ARHGEF11 | chr1 | - | 156971696 | 156971816 | 156969997 | 156970043 | 156976982 | 156977054 | 0 | 0 | 1 | 0 | 0 | 0 | 0 | 0 | 0 | 1 | 0 | 0 | 98 | 49 | NA | NA | NA | NA | 1 | 0 | NA | NA | 1 |
| SE | 8268 | DPYD | DPYD | chr1 | - | 97593217 | 97593387 | 97573759 | 97573970 | 97595058 | 97595166 | 0 | 0 | 31 | 0 | 0 | 0 | 0 | 5 | 0 | 0 | 2 | 0 | 98 | 49 | NA | NA | NA | NA | 1 | NA | 0.556 | NA | 0.444 |
| SE | 8272 | ADAM9 | ADAM9 | chr8 | + | 39071297 | 39071403 | 39055576 | 39055772 | 39077227 | 39077411 | 0 | 0 | 4 | 0 | 0 | 0 | 0 | 3 | 0 | 0 | 2 | 0 | 98 | 49 | NA | NA | NA | NA | 1 | NA | 0.429 | NA | 0.571 |
| SE | 8292 | IFI16 | IFI16 | chr1 | + | 159053532 | 159053724 | 159051678 | 159052098 | 159054820 | 159055155 | 0 | 0 | 18 | 0 | 0 | 0 | 1 | 9 | 0 | 0 | 4 | 0 | 98 | 49 | NA | NA | NA | NA | 1 | 1 | 0.529 | NA | 0.236 |
| SE | 8298 | EIF5 | EIF5 | chr14 | + | 103336035 | 103336117 | 103335652 | 103335932 | 103336676 | 103336849 | 0 | 0 | 7 | 0 | 0 | 1 | 0 | 22 | 0 | 0 | 4 | 0 | 98 | 49 | NA | NA | NA | NA | 0.778 | NA | 0.733 | NA | 0.045 |
| SE | 8317 | NUMA1 | NUMA1 | chr11 | - | 72004639 | 72004816 | 72004224 | 72004341 | 72005232 | 72005369 | 0 | 0 | 6 | 0 | 0 | 0 | 0 | 1 | 0 | 0 | 1 | 0 | 98 | 49 | NA | NA | NA | NA | 1 | NA | 0.333 | NA | 0.667 |
| SE | 8325 | COPA | COPA | chr1 | - | 160294483 | 160294593 | 160293385 | 160293463 | 160294767 | 160294857 | 0 | 0 | 3 | 0 | 0 | 2 | 3 | 3 | 0 | 0 | 0 | 0 | 98 | 49 | NA | NA | NA | NA | 0.429 | 1 | 1 | NA | -0.571 |
| SE | 8327 | COPA | COPA | chr1 | - | 160310191 | 160310258 | 160309100 | 160309176 | 160311867 | 160312018 | 0 | 0 | 5 | 0 | 0 | 0 | 0 | 9 | 0 | 0 | 1 | 0 | 98 | 49 | NA | NA | NA | NA | 1 | NA | 0.818 | NA | 0.182 |
| SE | 8328 | COPA | COPA | chr1 | - | 160339908 | 160339982 | 160335241 | 160335322 | 160340180 | 160340294 | 0 | 0 | 7 | 0 | 0 | 0 | 1 | 3 | 0 | 0 | 1 | 0 | 98 | 49 | NA | NA | NA | NA | 1 | 1 | 0.6 | NA | 0.2 |
| SE | 8329 | WDR37 | WDR37 | chr10 | + | 1103601 | 1103836 | 1096169 | 1096246 | 1124217 | 1124352 | 0 | 0 | 0 | 0 | 0 | 1 | 0 | 1 | 0 | 0 | 0 | 0 | 98 | 49 | NA | NA | NA | NA | 0 | NA | 1 | NA | -1 |
| SE | 8330 | WDR37 | WDR37 | chr10 | + | 1105125 | 1105267 | 1103601 | 1103836 | 1124217 | 1124352 | 0 | 0 | 4 | 0 | 0 | 0 | 0 | 0 | 0 | 0 | 1 | 0 | 98 | 49 | NA | NA | NA | NA | 1 | NA | 0 | NA | 1 |
| SE | 8335 | NCSTN | NCSTN | chr1 | + | 160356259 | 160356347 | 160354117 | 160354290 | 160356599 | 160356754 | 0 | 0 | 4 | 0 | 0 | 1 | 0 | 5 | 0 | 0 | 0 | 0 | 98 | 49 | NA | NA | NA | NA | 0.667 | NA | 1 | NA | -0.333 |
| SE | 8341 | CHD2 | CHD2 | chr15 | + | 92956458 | 92956649 | 92955422 | 92955512 | 92967324 | 92967513 | 0 | 0 | 13 | 0 | 0 | 1 | 0 | 21 | 0 | 0 | 3 | 0 | 98 | 49 | NA | NA | NA | NA | 0.867 | NA | 0.778 | NA | 0.089 |
| SE | 8342 | CHD2 | CHD2 | chr15 | + | 92978233 | 92978383 | 92974878 | 92974950 | 92979134 | 92979283 | 0 | 0 | 3 | 0 | 0 | 0 | 0 | 4 | 0 | 0 | 1 | 0 | 98 | 49 | NA | NA | NA | NA | 1 | NA | 0.667 | NA | 0.333 |
| SE | 8346 | SF1 | SF1 | chr11 | - | 64765950 | 64766155 | 64764603 | 64765518 | 64766899 | 64767079 | 0 | 0 | 1 | 0 | 0 | 0 | 0 | 0 | 0 | 0 | 1 | 0 | 98 | 49 | NA | NA | NA | NA | 1 | NA | 0 | NA | 1 |
| SE | 8357 | ARHGAP30 | ARHGAP30 | chr1 | - | 161056387 | 161056532 | 161054622 | 161054705 | 161059613 | 161059716 | 0 | 0 | 26 | 0 | 0 | 3 | 0 | 13 | 0 | 0 | 2 | 0 | 98 | 49 | NA | NA | NA | NA | 0.812 | NA | 0.765 | NA | 0.047 |
| SE | 8377 | UFC1 | UFC1 | chr1 | + | 161158120 | 161158211 | 161157616 | 161157693 | 161158411 | 161158856 | 0 | 0 | 3 | 0 | 0 | 1 | 0 | 1 | 0 | 0 | 0 | 0 | 98 | 49 | NA | NA | NA | NA | 0.6 | NA | 1 | NA | -0.4 |
| SE | 8381 | CYFIP2 | CYFIP2 | chr5 | + | 157307760 | 157307865 | 157304237 | 157304366 | 157311663 | 157311781 | 0 | 0 | 1 | 0 | 0 | 0 | 4 | 5 | 0 | 0 | 1 | 0 | 98 | 49 | NA | NA | NA | NA | 1 | 1 | 0.714 | NA | 0.143 |
| SE | 8382 | CYFIP2 | CYFIP2 | chr5 | + | 157309742 | 157309834 | 157307760 | 157307865 | 157311663 | 157311781 | 0 | 0 | 5 | 0 | 0 | 0 | 2 | 1 | 0 | 4 | 0 | 0 | 98 | 49 | NA | NA | NA | NA | 1 | 0.2 | 1 | NA | 0.4 |
| SE | 8384 | CYFIP2 | CYFIP2 | chr5 | + | 157327972 | 157328049 | 157326170 | 157326267 | 157330741 | 157330850 | 0 | 0 | 3 | 0 | 0 | 0 | 0 | 3 | 0 | 0 | 2 | 0 | 98 | 49 | NA | NA | NA | NA | 1 | NA | 0.429 | NA | 0.571 |
| SE | 8390 | TSNAX-DISC1 | TSNAX-DISC1 | chr1 | + | 231800107 | 231800210 | 231770834 | 231771070 | 231818328 | 231818517 | 0 | 0 | 5 | 0 | 0 | 0 | 0 | 4 | 0 | 0 | 1 | 0 | 98 | 49 | NA | NA | NA | NA | 1 | NA | 0.667 | NA | 0.333 |
| SE | 8395 | ITCH | ITCH | chr20 | + | 34408650 | 34408792 | 34393790 | 34393881 | 34412514 | 34412639 | 0 | 0 | 5 | 0 | 0 | 1 | 2 | 6 | 0 | 0 | 0 | 0 | 98 | 49 | NA | NA | NA | NA | 0.714 | 1 | 1 | NA | -0.286 |
| SE | 8396 | ITCH | ITCH | chr20 | + | 34417116 | 34417239 | 34413741 | 34413879 | 34424479 | 34424525 | 0 | 0 | 3 | 0 | 0 | 3 | 0 | 0 | 0 | 0 | 6 | 0 | 98 | 49 | NA | NA | NA | NA | 0.333 | NA | 0 | NA | 0.333 |
| SE | 8397 | ITCH | ITCH | chr20 | + | 34457389 | 34457474 | 34449410 | 34449480 | 34462092 | 34462221 | 0 | 0 | 3 | 0 | 0 | 1 | 3 | 3 | 0 | 0 | 0 | 0 | 98 | 49 | NA | NA | NA | NA | 0.6 | 1 | 1 | NA | -0.4 |
| SE | 8398 | ITCH | ITCH | chr20 | + | 34470047 | 34470120 | 34462092 | 34462221 | 34471443 | 34471515 | 0 | 0 | 4 | 0 | 0 | 0 | 1 | 4 | 0 | 0 | 2 | 0 | 98 | 49 | NA | NA | NA | NA | 1 | 1 | 0.5 | NA | 0.25 |
| SE | 8422 | CAST | CAST | chr5 | + | 96730779 | 96730860 | 96729611 | 96729725 | 96736171 | 96736240 | 0 | 0 | 8 | 0 | 0 | 1 | 0 | 1 | 0 | 0 | 0 | 0 | 98 | 49 | NA | NA | NA | NA | 0.8 | NA | 1 | NA | -0.2 |
| SE | 8425 | CAST | CAST | chr5 | + | 96746341 | 96746425 | 96742654 | 96742756 | 96747344 | 96747392 | 0 | 0 | 11 | 0 | 0 | 0 | 1 | 4 | 0 | 0 | 2 | 0 | 98 | 49 | NA | NA | NA | NA | 1 | 1 | 0.5 | NA | 0.25 |
| SE | 8437 | PIK3C3 | PIK3C3 | chr18 | + | 41957569 | 41957758 | 41955197 | 41955359 | 41962488 | 41962632 | 0 | 0 | 2 | 0 | 0 | 0 | 0 | 5 | 0 | 0 | 1 | 0 | 98 | 49 | NA | NA | NA | NA | 1 | NA | 0.714 | NA | 0.286 |
| SE | 8451 | MGST3 | MGST3 | chr1 | + | 165651013 | 165651087 | 165649840 | 165649964 | 165651977 | 165652035 | 0 | 0 | 4 | 0 | 0 | 0 | 0 | 0 | 0 | 0 | 1 | 0 | 98 | 49 | NA | NA | NA | NA | 1 | NA | 0 | NA | 1 |
| SE | 8452 | SYF2 | SYF2 | chr1 | - | 25228997 | 25229123 | 25228117 | 25228235 | 25232103 | 25232211 | 0 | 0 | 1 | 0 | 0 | 0 | 0 | 1 | 0 | 0 | 1 | 0 | 98 | 49 | NA | NA | NA | NA | 1 | NA | 0.333 | NA | 0.667 |
| SE | 8470 | SAMD8 | SAMD8 | chr10 | + | 75168540 | 75168658 | 75164644 | 75164740 | 75176065 | 75176216 | 0 | 0 | 2 | 0 | 0 | 1 | 0 | 3 | 0 | 0 | 0 | 0 | 98 | 49 | NA | NA | NA | NA | 0.5 | NA | 1 | NA | -0.5 |
| SE | 8471 | EIF3M | EIF3M | chr11 | + | 32587011 | 32587144 | 32583766 | 32583929 | 32588593 | 32588732 | 0 | 0 | 0 | 0 | 0 | 1 | 0 | 1 | 0 | 0 | 0 | 0 | 98 | 49 | NA | NA | NA | NA | 0 | NA | 1 | NA | -1 |
| SE | 8475 | ADCY4 | ADCY4 | chr14 | - | 24326298 | 24326342 | 24326078 | 24326165 | 24329060 | 24329234 | 0 | 0 | 1 | 0 | 0 | 0 | 0 | 2 | 0 | 0 | 1 | 0 | 93 | 49 | NA | NA | NA | NA | 1 | NA | 0.513 | NA | 0.487 |
| SE | 8478 | HPSE | HPSE | chr4 | - | 83310721 | 83310890 | 83310030 | 83310078 | 83319343 | 83319469 | 0 | 0 | 1 | 0 | 0 | 0 | 0 | 1 | 0 | 0 | 1 | 0 | 98 | 49 | NA | NA | NA | NA | 1 | NA | 0.333 | NA | 0.667 |
| SE | 8487 | PELI1 | PELI1 | chr2 | - | 64104700 | 64104830 | 64100397 | 64100499 | 64108239 | 64108379 | 0 | 0 | 13 | 0 | 0 | 0 | 0 | 27 | 0 | 0 | 2 | 0 | 98 | 49 | NA | NA | NA | NA | 1 | NA | 0.871 | NA | 0.129 |
| SE | 8490 | GPBP1 | GPBP1 | chr5 | + | 57221375 | 57221396 | 57214073 | 57214193 | 57230845 | 57230969 | 0 | 0 | 3 | 0 | 0 | 5 | 0 | 8 | 0 | 0 | 0 | 0 | 70 | 49 | NA | NA | NA | NA | 0.296 | NA | 1 | NA | -0.704 |
| SE | 8527 | VPS9D1 | VPS9D1 | chr16 | - | 89708426 | 89708531 | 89707132 | 89707954 | 89708856 | 89708956 | 0 | 0 | 1 | 0 | 0 | 0 | 0 | 0 | 0 | 0 | 1 | 0 | 98 | 49 | NA | NA | NA | NA | 1 | NA | 0 | NA | 1 |
| SE | 8541 | ARAP3 | ARAP3 | chr5 | - | 141656503 | 141656637 | 141656193 | 141656276 | 141656717 | 141656846 | 0 | 0 | 2 | 0 | 0 | 1 | 0 | 2 | 0 | 0 | 0 | 0 | 98 | 49 | NA | NA | NA | NA | 0.5 | NA | 1 | NA | -0.5 |
| SE | 8547 | TPP2 | TPP2 | chr13 | + | 102676295 | 102676415 | 102674282 | 102674490 | 102678226 | 102679173 | 0 | 0 | 2 | 0 | 0 | 0 | 0 | 0 | 0 | 0 | 1 | 0 | 98 | 49 | NA | NA | NA | NA | 1 | NA | 0 | NA | 1 |
| SE | 8574 | RC3H1 | RC3H1 | chr1 | - | 173943441 | 173943588 | 173941264 | 173941380 | 173946475 | 173946608 | 0 | 0 | 2 | 0 | 0 | 3 | 0 | 0 | 0 | 0 | 3 | 0 | 98 | 49 | NA | NA | NA | NA | 0.25 | NA | 0 | NA | 0.25 |
| SE | 8575 | RC3H1 | RC3H1 | chr1 | - | 173943441 | 173943615 | 173941264 | 173941380 | 173946475 | 173946608 | 0 | 0 | 2 | 0 | 0 | 3 | 0 | 0 | 0 | 0 | 3 | 0 | 98 | 49 | NA | NA | NA | NA | 0.25 | NA | 0 | NA | 0.25 |
| SE | 8576 | RC3H1 | RC3H1 | chr1 | - | 173951985 | 173952138 | 173947368 | 173947582 | 173961076 | 173961244 | 0 | 0 | 9 | 0 | 0 | 1 | 0 | 5 | 0 | 0 | 0 | 0 | 98 | 49 | NA | NA | NA | NA | 0.818 | NA | 1 | NA | -0.182 |
| SE | 8618 | DGKZ | DGKZ | chr11 | + | 46369940 | 46370009 | 46369493 | 46369550 | 46371312 | 46371384 | 0 | 0 | 5 | 0 | 0 | 0 | 0 | 2 | 0 | 0 | 1 | 0 | 98 | 49 | NA | NA | NA | NA | 1 | NA | 0.5 | NA | 0.5 |
| SE | 8622 | RBM18 | RBM18 | chr9 | - | 122254273 | 122254367 | 122251846 | 122251973 | 122261379 | 122261508 | 0 | 0 | 1 | 0 | 0 | 4 | 0 | 1 | 0 | 0 | 0 | 0 | 98 | 49 | NA | NA | NA | NA | 0.111 | NA | 1 | NA | -0.889 |
| SE | 8632 | SMG7 | SMG7 | chr1 | + | 183538379 | 183538440 | 183537144 | 183537215 | 183540983 | 183541103 | 0 | 0 | 1 | 0 | 0 | 0 | 0 | 2 | 0 | 0 | 1 | 0 | 98 | 49 | NA | NA | NA | NA | 1 | NA | 0.5 | NA | 0.5 |
| SE | 8722 | NOD2 | NOD2 | chr16 | + | 50719924 | 50720008 | 50716890 | 50716974 | 50722621 | 50722705 | 0 | 0 | 3 | 0 | 0 | 1 | 0 | 1 | 0 | 0 | 0 | 0 | 98 | 49 | NA | NA | NA | NA | 0.6 | NA | 1 | NA | -0.4 |
| SE | 8737 | KDM5B | KDM5B | chr1 | - | 202746141 | 202746323 | 202745857 | 202745982 | 202748944 | 202749139 | 0 | 0 | 1 | 0 | 0 | 0 | 0 | 7 | 0 | 0 | 1 | 0 | 98 | 49 | NA | NA | NA | NA | 1 | NA | 0.778 | NA | 0.222 |
| SE | 8744 | BEST1 | BEST1 | chr11 | + | 61958145 | 61958298 | 61957386 | 61957464 | 61959497 | 61959578 | 0 | 0 | 3 | 0 | 0 | 0 | 0 | 1 | 0 | 0 | 1 | 0 | 98 | 49 | NA | NA | NA | NA | 1 | NA | 0.333 | NA | 0.667 |
| SE | 8752 | OS9 | OS9 | chr12 | + | 57720098 | 57720263 | 57718992 | 57719182 | 57720405 | 57720518 | 0 | 0 | 2 | 0 | 0 | 3 | 2 | 1 | 0 | 1 | 2 | 0 | 98 | 49 | NA | NA | NA | NA | 0.25 | 0.5 | 0.2 | NA | -0.1 |
| SE | 8763 | MITD1 | MITD1 | chr2 | - | 99171342 | 99171429 | 99170536 | 99170652 | 99171509 | 99171646 | 0 | 0 | 2 | 0 | 0 | 0 | 0 | 0 | 0 | 0 | 1 | 0 | 98 | 49 | NA | NA | NA | NA | 1 | NA | 0 | NA | 1 |
| SE | 8770 | YY1 | YY1 | chr14 | + | 100276489 | 100276648 | 100274697 | 100274758 | 100277417 | 100279034 | 0 | 0 | 2 | 0 | 0 | 1 | 8 | 1 | 0 | 0 | 0 | 0 | 98 | 49 | NA | NA | NA | NA | 0.5 | 1 | 1 | NA | -0.5 |
| SE | 8791 | VPS13D | VPS13D | chr1 | + | 12319496 | 12319630 | 12318071 | 12318337 | 12321808 | 12321964 | 0 | 0 | 2 | 0 | 0 | 0 | 0 | 3 | 0 | 0 | 1 | 0 | 98 | 49 | NA | NA | NA | NA | 1 | NA | 0.6 | NA | 0.4 |
| SE | 8793 | VPS13D | VPS13D | chr1 | + | 12349163 | 12349374 | 12348822 | 12348973 | 12353973 | 12354221 | 0 | 0 | 3 | 0 | 0 | 1 | 0 | 2 | 0 | 0 | 0 | 0 | 98 | 49 | NA | NA | NA | NA | 0.6 | NA | 1 | NA | -0.4 |
| SE | 8794 | VPS13D | VPS13D | chr1 | + | 12358458 | 12358601 | 12356397 | 12356524 | 12362719 | 12362850 | 0 | 0 | 1 | 0 | 0 | 0 | 0 | 1 | 0 | 0 | 2 | 0 | 98 | 49 | NA | NA | NA | NA | 1 | NA | 0.2 | NA | 0.8 |
| SE | 8797 | CD46 | CD46 | chr1 | + | 207767606 | 207767651 | 207767012 | 207767195 | 207767778 | 207767823 | 0 | 0 | 7 | 0 | 0 | 0 | 0 | 2 | 0 | 0 | 2 | 0 | 94 | 49 | NA | NA | NA | NA | 1 | NA | 0.343 | NA | 0.657 |
| SE | 8814 | KDM2A | KDM2A | chr11 | + | 67121233 | 67121358 | 67119268 | 67120049 | 67180078 | 67180217 | 0 | 0 | 11 | 0 | 0 | 0 | 1 | 5 | 0 | 0 | 2 | 0 | 98 | 49 | NA | NA | NA | NA | 1 | 1 | 0.556 | NA | 0.222 |
| SE | 8815 | KDM2A | KDM2A | chr11 | + | 67181319 | 67181398 | 67180078 | 67180217 | 67181845 | 67181892 | 0 | 0 | 7 | 0 | 0 | 1 | 0 | 1 | 0 | 0 | 0 | 0 | 98 | 49 | NA | NA | NA | NA | 0.778 | NA | 1 | NA | -0.222 |
| SE | 8816 | KDM2A | KDM2A | chr11 | + | 67215855 | 67215949 | 67215339 | 67215446 | 67217730 | 67217884 | 0 | 0 | 5 | 0 | 0 | 1 | 0 | 6 | 0 | 0 | 2 | 0 | 98 | 49 | NA | NA | NA | NA | 0.714 | NA | 0.6 | NA | 0.114 |
| SE | 8826 | HIRA | HIRA | chr22 | - | 19387710 | 19387816 | 19385520 | 19385736 | 19388483 | 19388554 | 0 | 0 | 8 | 0 | 0 | 2 | 0 | 5 | 0 | 0 | 0 | 0 | 98 | 49 | NA | NA | NA | NA | 0.667 | NA | 1 | NA | -0.333 |
| SE | 8832 | 7-Mar | 7-Mar | chr2 | + | 159762879 | 159762993 | 159759225 | 159759335 | 159764625 | 159764674 | 0 | 0 | 8 | 0 | 0 | 0 | 0 | 0 | 0 | 0 | 1 | 0 | 98 | 49 | NA | NA | NA | NA | 1 | NA | 0 | NA | 1 |
| SE | 8844 | GNL2 | GNL2 | chr1 | - | 37574664 | 37574823 | 37574342 | 37574456 | 37575594 | 37575699 | 0 | 0 | 1 | 0 | 0 | 1 | 0 | 3 | 0 | 0 | 0 | 0 | 98 | 49 | NA | NA | NA | NA | 0.333 | NA | 1 | NA | -0.667 |
| SE | 8871 | C1RL | C1RL | chr12 | - | 7099900 | 7100026 | 7099685 | 7099760 | 7101897 | 7102087 | 1 | 0 | 6 | 0 | 0 | 0 | 0 | 0 | 0 | 0 | 1 | 0 | 98 | 49 | NA | NA | 1 | NA | 1 | NA | 0 | NA | 1 |
| SE | 8883 | FAM160B1 | FAM160B1 | chr10 | + | 114848646 | 114848737 | 114847089 | 114847233 | 114855196 | 114855340 | 0 | 0 | 3 | 0 | 0 | 0 | 0 | 4 | 0 | 0 | 1 | 0 | 98 | 49 | NA | NA | NA | NA | 1 | NA | 0.667 | NA | 0.333 |
| SE | 8908 | WDR11 | WDR11 | chr10 | + | 120867065 | 120867169 | 120866568 | 120866764 | 120871169 | 120871346 | 0 | 0 | 3 | 0 | 0 | 1 | 0 | 3 | 0 | 0 | 0 | 0 | 98 | 49 | NA | NA | NA | NA | 0.6 | NA | 1 | NA | -0.4 |
| SE | 8922 | SMARCC1 | SMARCC1 | chr3 | - | 47720665 | 47720735 | 47714414 | 47714490 | 47729024 | 47729094 | 0 | 0 | 2 | 0 | 0 | 0 | 0 | 4 | 0 | 0 | 1 | 0 | 98 | 49 | NA | NA | NA | NA | 1 | NA | 0.667 | NA | 0.333 |
| SE | 8923 | SMARCC1 | SMARCC1 | chr3 | - | 47729024 | 47729094 | 47720665 | 47720735 | 47736033 | 47736126 | 0 | 0 | 2 | 0 | 0 | 0 | 0 | 5 | 0 | 0 | 1 | 0 | 98 | 49 | NA | NA | NA | NA | 1 | NA | 0.714 | NA | 0.286 |
| SE | 8937 | CD164 | CD164 | chr6 | - | 109370410 | 109370467 | 109366513 | 109369017 | 109376073 | 109376112 | 0 | 0 | 18 | 0 | 0 | 1 | 0 | 7 | 0 | 0 | 0 | 0 | 98 | 49 | NA | NA | NA | NA | 0.9 | NA | 1 | NA | -0.1 |
| SE | 8955 | REV1 | REV1 | chr2 | - | 99442316 | 99442469 | 99438600 | 99439310 | 99449335 | 99449504 | 0 | 0 | 5 | 0 | 0 | 0 | 0 | 1 | 0 | 0 | 1 | 0 | 98 | 49 | NA | NA | NA | NA | 1 | NA | 0.333 | NA | 0.667 |
| SE | 8976 | KPNB1 | KPNB1 | chr17 | + | 47676408 | 47676491 | 47674637 | 47674782 | 47677019 | 47677127 | 0 | 0 | 13 | 0 | 0 | 0 | 0 | 4 | 0 | 0 | 1 | 0 | 98 | 49 | NA | NA | NA | NA | 1 | NA | 0.667 | NA | 0.333 |
| SE | 8978 | C18orf25 | C18orf25 | chr18 | + | 46240004 | 46240187 | 46215810 | 46216595 | 46253696 | 46253823 | 0 | 0 | 2 | 0 | 0 | 0 | 0 | 0 | 0 | 0 | 1 | 0 | 98 | 49 | NA | NA | NA | NA | 1 | NA | 0 | NA | 1 |
| SE | 8992 | BSG | BSG | chr19 | + | 579499 | 579656 | 572453 | 572701 | 580378 | 580461 | 0 | 0 | 5 | 0 | 0 | 0 | 0 | 1 | 0 | 1 | 0 | 0 | 98 | 49 | NA | NA | NA | NA | 1 | 0 | 1 | NA | 0.5 |
| SE | 8997 | PANK4 | PANK4 | chr1 | - | 2512887 | 2513039 | 2511627 | 2511683 | 2514001 | 2514089 | 0 | 0 | 4 | 0 | 0 | 0 | 0 | 0 | 0 | 0 | 1 | 0 | 98 | 49 | NA | NA | NA | NA | 1 | NA | 0 | NA | 1 |
| SE | 9002 | AKAP10 | AKAP10 | chr17 | - | 19940886 | 19941010 | 19939712 | 19939849 | 19941825 | 19941910 | 0 | 0 | 4 | 0 | 0 | 4 | 0 | 3 | 0 | 0 | 1 | 0 | 98 | 49 | NA | NA | NA | NA | 0.333 | NA | 0.6 | NA | -0.267 |
| SE | 9045 | FNDC3A | FNDC3A | chr13 | + | 49191014 | 49191120 | 49188514 | 49188633 | 49191208 | 49191384 | 0 | 0 | 1 | 0 | 0 | 0 | 0 | 0 | 0 | 0 | 1 | 0 | 98 | 49 | NA | NA | NA | NA | 1 | NA | 0 | NA | 1 |
| SE | 9047 | FAM65B | FAM65B | chr6 | - | 24835702 | 24835871 | 24832255 | 24832391 | 24839090 | 24839272 | 0 | 0 | 36 | 0 | 0 | 0 | 0 | 3 | 0 | 0 | 1 | 0 | 98 | 49 | NA | NA | NA | NA | 1 | NA | 0.6 | NA | 0.4 |
| SE | 9048 | FAM65B | FAM65B | chr6 | - | 24839090 | 24839272 | 24835702 | 24835871 | 24842861 | 24843554 | 0 | 0 | 19 | 0 | 0 | 0 | 0 | 4 | 0 | 0 | 1 | 0 | 98 | 49 | NA | NA | NA | NA | 1 | NA | 0.667 | NA | 0.333 |
| SE | 9050 | MPPE1 | MPPE1 | chr18 | - | 11884878 | 11885043 | 11883470 | 11884627 | 11885675 | 11885816 | 0 | 0 | 3 | 0 | 0 | 2 | 0 | 1 | 0 | 0 | 0 | 0 | 98 | 49 | NA | NA | NA | NA | 0.429 | NA | 1 | NA | -0.571 |
| SE | 9095 | HPS1 | HPS1 | chr10 | - | 98429790 | 98429889 | 98429145 | 98429642 | 98430570 | 98430670 | 0 | 0 | 1 | 0 | 0 | 1 | 0 | 1 | 0 | 0 | 0 | 0 | 98 | 49 | NA | NA | NA | NA | 0.333 | NA | 1 | NA | -0.667 |
| SE | 9112 | ADGRE3 | ADGRE3 | chr19 | - | 14658512 | 14658550 | 14654981 | 14655165 | 14661962 | 14662118 | 0 | 0 | 13 | 0 | 0 | 1 | 0 | 6 | 0 | 0 | 0 | 0 | 87 | 49 | NA | NA | NA | NA | 0.88 | NA | 1 | NA | -0.12 |
| SE | 9119 | AGO1 | AGO1 | chr1 | + | 35906934 | 35907119 | 35902203 | 35902337 | 35913841 | 35914001 | 0 | 0 | 2 | 0 | 0 | 1 | 0 | 2 | 0 | 0 | 0 | 0 | 98 | 49 | NA | NA | NA | NA | 0.5 | NA | 1 | NA | -0.5 |
| SE | 9149 | ADD3 | ADD3 | chr10 | + | 110116258 | 110116410 | 110112776 | 110112915 | 110117341 | 110117422 | 0 | 0 | 7 | 0 | 0 | 1 | 0 | 5 | 0 | 0 | 0 | 0 | 98 | 49 | NA | NA | NA | NA | 0.778 | NA | 1 | NA | -0.222 |
| SE | 9150 | ADD3 | ADD3 | chr10 | + | 110132304 | 110132400 | 110130362 | 110130486 | 110133325 | 110135565 | 0 | 0 | 9 | 0 | 0 | 2 | 0 | 7 | 0 | 0 | 0 | 0 | 98 | 49 | NA | NA | NA | NA | 0.692 | NA | 1 | NA | -0.308 |
| SE | 9172 | NDRG1 | NDRG1 | chr8 | - | 133258365 | 133258426 | 133256776 | 133256863 | 133259167 | 133259230 | 0 | 0 | 6 | 0 | 0 | 1 | 0 | 6 | 0 | 0 | 0 | 0 | 98 | 49 | NA | NA | NA | NA | 0.75 | NA | 1 | NA | -0.25 |
| SE | 9186 | PPP2R5D | PPP2R5D | chr6 | + | 43008692 | 43008746 | 43008366 | 43008475 | 43009056 | 43009227 | 0 | 0 | 2 | 0 | 0 | 0 | 0 | 0 | 0 | 0 | 1 | 0 | 98 | 49 | NA | NA | NA | NA | 1 | NA | 0 | NA | 1 |
| SE | 9187 | GIT2 | GIT2 | chr12 | - | 109945259 | 109945349 | 109939164 | 109939247 | 109947255 | 109947504 | 0 | 0 | 0 | 0 | 0 | 1 | 0 | 1 | 0 | 0 | 0 | 0 | 98 | 49 | NA | NA | NA | NA | 0 | NA | 1 | NA | -1 |
| SE | 9200 | PIP5K1A | PIP5K1A | chr1 | + | 151242122 | 151242269 | 151239954 | 151240039 | 151242437 | 151242567 | 0 | 0 | 0 | 0 | 0 | 1 | 0 | 2 | 0 | 0 | 0 | 0 | 98 | 49 | NA | NA | NA | NA | 0 | NA | 1 | NA | -1 |
| SE | 9202 | ATP5C1 | ATP5C1 | chr10 | + | 7802757 | 7802854 | 7802269 | 7802425 | 7806973 | 7807010 | 0 | 0 | 6 | 0 | 0 | 1 | 1 | 1 | 0 | 0 | 0 | 0 | 98 | 49 | NA | NA | NA | NA | 0.75 | 1 | 1 | NA | -0.25 |
| SE | 9210 | CELF2 | CELF2 | chr10 | + | 11266597 | 11266677 | 11257737 | 11257872 | 11270665 | 11270824 | 0 | 0 | 15 | 0 | 0 | 5 | 1 | 8 | 0 | 0 | 2 | 0 | 98 | 49 | NA | NA | NA | NA | 0.6 | 1 | 0.667 | NA | -0.234 |
| SE | 9213 | UPF2 | UPF2 | chr10 | - | 12001675 | 12001825 | 11999905 | 12000009 | 12004529 | 12004727 | 0 | 0 | 5 | 0 | 0 | 0 | 0 | 2 | 0 | 0 | 1 | 0 | 98 | 49 | NA | NA | NA | NA | 1 | NA | 0.5 | NA | 0.5 |
| SE | 9215 | PIGG | PIGG | chr4 | + | 508828 | 508970 | 507404 | 507593 | 515972 | 516185 | 0 | 0 | 2 | 0 | 0 | 0 | 0 | 2 | 0 | 3 | 0 | 0 | 98 | 49 | NA | NA | NA | NA | 1 | 0 | 1 | NA | 0.5 |
| SE | 9227 | SRSF6 | SRSF6 | chr20 | + | 43459152 | 43459420 | 43458360 | 43458509 | 43459770 | 43459895 | 0 | 0 | 0 | 0 | 0 | 1 | 0 | 2 | 0 | 0 | 0 | 0 | 98 | 49 | NA | NA | NA | NA | 0 | NA | 1 | NA | -1 |
| SE | 9251 | VIM-AS1 | VIM-AS1 | chr10 | - | 17216166 | 17216274 | 17214238 | 17215364 | 17229344 | 17229985 | 0 | 0 | 1 | 0 | 0 | 0 | 0 | 0 | 0 | 2 | 0 | 0 | 98 | 49 | NA | NA | NA | NA | 1 | 0 | NA | NA | 1 |
| SE | 9257 | SERPINB1 | SERPINB1 | chr6 | - | 2839334 | 2839466 | 2838548 | 2838686 | 2840418 | 2840594 | 0 | 0 | 2 | 0 | 0 | 7 | 0 | 1 | 0 | 5 | 1 | 0 | 98 | 49 | NA | NA | NA | NA | 0.125 | 0 | 0.333 | NA | -0.042 |
| SE | 9261 | ATP5L | ATP5L | chr11 | + | 118405656 | 118405709 | 118401388 | 118401717 | 118406936 | 118407097 | 0 | 0 | 2 | 0 | 0 | 0 | 0 | 0 | 0 | 0 | 1 | 0 | 98 | 49 | NA | NA | NA | NA | 1 | NA | 0 | NA | 1 |
| SE | 9264 | PLXDC2 | PLXDC2 | chr10 | + | 20068169 | 20068239 | 20046868 | 20047015 | 20143294 | 20143417 | 0 | 0 | 21 | 0 | 0 | 1 | 0 | 3 | 0 | 0 | 0 | 0 | 98 | 49 | NA | NA | NA | NA | 0.913 | NA | 1 | NA | -0.087 |
| SE | 9267 | CDC14A | CDC14A | chr1 | + | 100455404 | 100455492 | 100442933 | 100442996 | 100462650 | 100462881 | 0 | 0 | 1 | 0 | 0 | 1 | 5 | 1 | 0 | 0 | 0 | 0 | 98 | 49 | NA | NA | NA | NA | 0.333 | 1 | 1 | NA | -0.667 |
| SE | 9283 | RANBP2 | RANBP2 | chr2 | + | 108762100 | 108762195 | 108758412 | 108758548 | 108763236 | 108768388 | 0 | 0 | 1 | 0 | 0 | 0 | 0 | 3 | 0 | 0 | 3 | 0 | 98 | 49 | NA | NA | NA | NA | 1 | NA | 0.333 | NA | 0.667 |
| SE | 9284 | RANBP2 | RANBP2 | chr2 | + | 108777129 | 108777231 | 108775731 | 108775936 | 108781268 | 108781429 | 0 | 0 | 1 | 0 | 0 | 0 | 0 | 7 | 0 | 0 | 1 | 0 | 98 | 49 | NA | NA | NA | NA | 1 | NA | 0.778 | NA | 0.222 |
| SE | 9294 | ABCA7 | ABCA7 | chr19 | + | 1056906 | 1057084 | 1056329 | 1056499 | 1057313 | 1057429 | 0 | 0 | 4 | 0 | 0 | 0 | 0 | 1 | 0 | 0 | 2 | 0 | 98 | 49 | NA | NA | NA | NA | 1 | NA | 0.2 | NA | 0.8 |
| SE | 9358 | RASGRP1 | RASGRP1 | chr15 | - | 38503271 | 38503376 | 38502311 | 38502421 | 38505839 | 38505920 | 0 | 0 | 2 | 0 | 0 | 0 | 0 | 1 | 0 | 0 | 1 | 0 | 98 | 49 | NA | NA | NA | NA | 1 | NA | 0.333 | NA | 0.667 |
| SE | 9360 | EIF4G3 | EIF4G3 | chr1 | - | 20892618 | 20892729 | 20886200 | 20886371 | 20893516 | 20893636 | 0 | 0 | 0 | 0 | 0 | 1 | 0 | 2 | 0 | 0 | 2 | 0 | 98 | 49 | NA | NA | NA | NA | 0 | NA | 0.333 | NA | -0.333 |
| SE | 9382 | PECAM1 | PECAM1 | chr17 | - | 64353490 | 64353518 | 64352389 | 64352463 | 64356110 | 64356398 | 0 | 0 | 20 | 0 | 0 | 0 | 0 | 3 | 0 | 0 | 1 | 0 | 77 | 49 | NA | NA | NA | NA | 1 | NA | 0.656 | NA | 0.344 |
| SE | 9388 | NCOA4 | NCOA4 | chr10 | - | 46016539 | 46016694 | 46015125 | 46015266 | 46030525 | 46030714 | 0 | 0 | 5 | 0 | 0 | 0 | 5 | 3 | 0 | 4 | 0 | 0 | 98 | 49 | NA | NA | NA | NA | 1 | 0.385 | 1 | NA | 0.307 |
| SE | 9445 | CCNDBP1 | CCNDBP1 | chr15 | + | 43186153 | 43186233 | 43185819 | 43185879 | 43189198 | 43189280 | 0 | 0 | 10 | 0 | 0 | 0 | 0 | 5 | 0 | 0 | 1 | 0 | 98 | 49 | NA | NA | NA | NA | 1 | NA | 0.714 | NA | 0.286 |
| SE | 9446 | CCNDBP1 | CCNDBP1 | chr15 | + | 43190054 | 43190151 | 43189198 | 43189280 | 43190324 | 43190398 | 0 | 0 | 7 | 0 | 0 | 5 | 0 | 2 | 0 | 0 | 3 | 0 | 98 | 49 | NA | NA | NA | NA | 0.412 | NA | 0.25 | NA | 0.162 |
| SE | 9460 | TFAM | TFAM | chr10 | + | 58390764 | 58390860 | 58388669 | 58388819 | 58394357 | 58394414 | 0 | 0 | 1 | 0 | 0 | 0 | 0 | 1 | 0 | 0 | 1 | 0 | 98 | 49 | NA | NA | NA | NA | 1 | NA | 0.333 | NA | 0.667 |
| SE | 9471 | USP33 | USP33 | chr1 | - | 77701368 | 77701471 | 77697862 | 77697931 | 77711746 | 77711855 | 0 | 0 | 5 | 0 | 0 | 2 | 0 | 2 | 0 | 0 | 1 | 0 | 98 | 49 | NA | NA | NA | NA | 0.556 | NA | 0.5 | NA | 0.056 |
| SE | 9495 | SAP30BP | SAP30BP | chr17 | + | 75690670 | 75690772 | 75671815 | 75671863 | 75693439 | 75693482 | 0 | 0 | 0 | 0 | 0 | 1 | 0 | 1 | 0 | 0 | 6 | 0 | 98 | 49 | NA | NA | NA | NA | 0 | NA | 0.077 | NA | -0.077 |
| SE | 9501 | CCAR1 | CCAR1 | chr10 | + | 68771205 | 68771445 | 68765887 | 68766079 | 68772987 | 68773099 | 0 | 0 | 2 | 0 | 0 | 0 | 0 | 0 | 0 | 0 | 1 | 0 | 98 | 49 | NA | NA | NA | NA | 1 | NA | 0 | NA | 1 |
| SE | 9518 | SMG5 | SMG5 | chr1 | - | 156273360 | 156273450 | 156272319 | 156272398 | 156274596 | 156274686 | 0 | 0 | 3 | 0 | 0 | 1 | 0 | 2 | 0 | 0 | 0 | 0 | 98 | 49 | NA | NA | NA | NA | 0.6 | NA | 1 | NA | -0.4 |
| SE | 9531 | PPP3CB | PPP3CB | chr10 | - | 73439871 | 73439901 | 73436427 | 73438420 | 73444724 | 73444822 | 0 | 0 | 1 | 0 | 0 | 2 | 0 | 1 | 0 | 0 | 0 | 0 | 79 | 49 | NA | NA | NA | NA | 0.237 | NA | 1 | NA | -0.763 |
| SE | 9534 | CAMK2G | CAMK2G | chr10 | - | 73819531 | 73819645 | 73817478 | 73817554 | 73821681 | 73821730 | 0 | 0 | 2 | 0 | 0 | 0 | 0 | 0 | 0 | 0 | 1 | 0 | 98 | 49 | NA | NA | NA | NA | 1 | NA | 0 | NA | 1 |
| SE | 9539 | CAMK2G | CAMK2G | chr10 | - | 73848525 | 73848609 | 73847987 | 73848082 | 73849012 | 73849115 | 0 | 0 | 2 | 0 | 0 | 2 | 0 | 4 | 0 | 0 | 4 | 0 | 98 | 49 | NA | NA | NA | NA | 0.333 | NA | 0.333 | NA | 0 |
| SE | 9557 | SENP7 | SENP7 | chr3 | - | 101417592 | 101417790 | 101398860 | 101399055 | 101458954 | 101459052 | 0 | 0 | 0 | 0 | 0 | 1 | 0 | 1 | 0 | 0 | 0 | 0 | 98 | 49 | NA | NA | NA | NA | 0 | NA | 1 | NA | -1 |
| SE | 9572 | FAS | FAS | chr10 | + | 89007699 | 89007837 | 89003028 | 89003194 | 89010538 | 89010600 | 0 | 0 | 1 | 0 | 0 | 2 | 0 | 1 | 0 | 0 | 0 | 0 | 98 | 49 | NA | NA | NA | NA | 0.2 | NA | 1 | NA | -0.8 |
| SE | 9652 | BSDC1 | BSDC1 | chr1 | - | 32383829 | 32383997 | 32381213 | 32381268 | 32386778 | 32386895 | 0 | 0 | 1 | 0 | 0 | 1 | 0 | 2 | 0 | 0 | 0 | 0 | 98 | 49 | NA | NA | NA | NA | 0.333 | NA | 1 | NA | -0.667 |
| SE | 9659 | TTLL3 | TTLL3 | chr3 | + | 9833103 | 9833245 | 9828959 | 9829395 | 9834680 | 9834907 | 0 | 0 | 2 | 0 | 0 | 0 | 0 | 0 | 0 | 5 | 1 | 0 | 98 | 49 | NA | NA | NA | NA | 1 | 0 | 0 | NA | 1 |
| SE | 9667 | PTEN | PTEN | chr10 | + | 87952117 | 87952259 | 87933012 | 87933251 | 87957852 | 87958019 | 0 | 0 | 4 | 0 | 0 | 1 | 0 | 2 | 0 | 0 | 1 | 0 | 98 | 49 | NA | NA | NA | NA | 0.667 | NA | 0.5 | NA | 0.167 |
| SE | 9688 | IFRD1 | IFRD1 | chr7 | + | 112458860 | 112459018 | 112456913 | 112457038 | 112461865 | 112461916 | 0 | 0 | 42 | 0 | 0 | 0 | 0 | 32 | 0 | 0 | 3 | 0 | 98 | 49 | NA | NA | NA | NA | 1 | NA | 0.842 | NA | 0.158 |
| SE | 9697 | ZNF410 | ZNF410 | chr14 | + | 73922065 | 73922206 | 73920979 | 73921105 | 73923394 | 73923522 | 0 | 0 | 2 | 0 | 0 | 4 | 0 | 2 | 0 | 0 | 1 | 0 | 98 | 49 | NA | NA | NA | NA | 0.2 | NA | 0.5 | NA | -0.3 |
| SE | 9700 | AAED1 | AAED1 | chr9 | - | 96645892 | 96646024 | 96641250 | 96641886 | 96651389 | 96651495 | 0 | 0 | 1 | 0 | 0 | 1 | 0 | 0 | 0 | 0 | 1 | 0 | 98 | 49 | NA | NA | NA | NA | 0.333 | NA | 0 | NA | 0.333 |
| SE | 9744 | CLEC2D | CLEC2D | chr12 | + | 9693033 | 9693115 | 9692827 | 9692931 | 9694759 | 9699555 | 0 | 0 | 1 | 0 | 0 | 2 | 0 | 1 | 0 | 0 | 0 | 0 | 98 | 49 | NA | NA | NA | NA | 0.2 | NA | 1 | NA | -0.8 |
| SE | 9745 | FBXO22 | FBXO22 | chr15 | + | 75904490 | 75904629 | 75903858 | 75904103 | 75913202 | 75913290 | 0 | 0 | 3 | 0 | 0 | 0 | 0 | 0 | 0 | 0 | 1 | 0 | 98 | 49 | NA | NA | NA | NA | 1 | NA | 0 | NA | 1 |
| SE | 9766 | SAFB | SAFB | chr19 | + | 5648988 | 5649499 | 5648015 | 5648043 | 5649925 | 5649975 | 0 | 0 | 1 | 0 | 0 | 3 | 0 | 0 | 0 | 0 | 1 | 0 | 98 | 49 | NA | NA | NA | NA | 0.143 | NA | 0 | NA | 0.143 |
| SE | 9767 | PI4K2A | PI4K2A | chr10 | + | 97664884 | 97664984 | 97662906 | 97662968 | 97666437 | 97666571 | 0 | 0 | 1 | 0 | 0 | 0 | 0 | 1 | 0 | 0 | 1 | 0 | 98 | 49 | NA | NA | NA | NA | 1 | NA | 0.333 | NA | 0.667 |
| SE | 9773 | FRYL | FRYL | chr4 | - | 48549472 | 48549623 | 48548689 | 48548793 | 48550591 | 48550704 | 0 | 0 | 2 | 0 | 0 | 0 | 0 | 0 | 0 | 0 | 1 | 0 | 98 | 49 | NA | NA | NA | NA | 1 | NA | 0 | NA | 1 |

| | | | | | | | | | | | | | | | | | | | | | | | | | | | | | | | | | | | | |
|---|---|---|---|---|---|---|---|---|---|---|---|---|---|---|---|---|---|---|---|---|---|---|---|---|---|---|---|---|---|---|---|---|---|---|---|---|
| SE | 9774 | FRYL | FRYL | chr4 | - | 48609743 | 48609823 | 48608986 | 48609067 | 48619273 | 48619370 | 0 | 0 | 3 | 0 | 0 | 1 | 0 | 1 | 0 | 0 | 0 | 0 | 98 | 49 | NA | NA | NA | NA | | 0.6 | NA | | 1 | NA | -0.4 |
| SE | 9775 | FRYL | FRYL | chr4 | - | 48710518 | 48710698 | 48684672 | 48684795 | 48780077 | 48780299 | 0 | 0 | 1 | 0 | 0 | 0 | 0 | 1 | 0 | 0 | 1 | 0 | 98 | 49 | NA | NA | NA | NA | | 1 | NA | | 0.333 | NA | 0.667 |
| SE | 9776 | FAM188A | FAM188A | chr10 | - | 15838227 | 15838279 | 15837203 | 15837318 | 15841425 | 15841599 | 0 | 0 | 0 | 0 | 0 | 1 | 0 | 1 | 0 | 0 | 1 | 0 | 98 | 49 | NA | NA | NA | NA | | 0 | NA | | 0.333 | NA | -0.333 |
| SE | 9784 | DCAF6 | DCAF6 | chr1 | + | 168038370 | 168038488 | 168004532 | 168004793 | 168043024 | 168043140 | 0 | 0 | 0 | 0 | 0 | 1 | 0 | 1 | 0 | 0 | 0 | 0 | 98 | 49 | NA | NA | NA | NA | | 0 | NA | | 1 | NA | -1 |
| SE | 9789 | HERC4 | HERC4 | chr10 | - | 68044403 | 68044563 | 68038092 | 68038169 | 68072882 | 68073186 | 0 | 0 | 1 | 0 | 0 | 0 | 0 | 1 | 0 | 0 | 1 | 0 | 98 | 49 | NA | NA | NA | NA | | 1 | NA | | 0.333 | NA | 0.667 |
| SE | 9795 | CHUK | CHUK | chr10 | - | 100194424 | 100194521 | 100193983 | 100194131 | 100199970 | 100200020 | 0 | 0 | 2 | 0 | 0 | 2 | 0 | 1 | 0 | 0 | 0 | 0 | 98 | 49 | NA | NA | NA | NA | | 0.333 | NA | | 1 | NA | -0.667 |
| SE | 9796 | CHUK | CHUK | chr10 | - | 100219007 | 100219132 | 100218717 | 100218825 | 100219269 | 100219359 | 0 | 0 | 2 | 0 | 0 | 1 | 0 | 0 | 0 | 0 | 2 | 0 | 98 | 49 | NA | NA | NA | NA | | 0.5 | NA | | 0 | NA | 0.5 |
| SE | 9830 | NFAT5 | NFAT5 | chr16 | + | 69568494 | 69568548 | 69565965 | 69566374 | 69626402 | 69626528 | 0 | 0 | 5 | 0 | 0 | 1 | 0 | 1 | 0 | 0 | 0 | 0 | 98 | 49 | NA | NA | NA | NA | | 0.714 | NA | | 1 | NA | -0.286 |
| SE | 9856 | ZFAND1 | ZFAND1 | chr8 | - | 81717248 | 81717288 | 81714986 | 81715114 | 81718181 | 81718224 | 0 | 0 | 2 | 0 | 0 | 1 | 0 | 1 | 0 | 0 | 0 | 0 | 89 | 49 | NA | NA | NA | NA | | 0.524 | NA | | 1 | NA | -0.476 |
| SE | 9899 | C2CD5 | C2CD5 | chr12 | - | 22459491 | 22459542 | 22456970 | 22457161 | 22469708 | 22469795 | 0 | 0 | 0 | 0 | 0 | 1 | 0 | 1 | 0 | 0 | 1 | 0 | 98 | 49 | NA | NA | NA | NA | | 0 | NA | | 0.333 | NA | -0.333 |
| SE | 9910 | IL13RA1 | IL13RA1 | chrX | + | 118766843 | 118766976 | 118766529 | 118766577 | 118773878 | 118773975 | 0 | 0 | 8 | 0 | 0 | 0 | 0 | 3 | 0 | 0 | 1 | 0 | 98 | 49 | NA | NA | NA | NA | | 1 | NA | | 0.6 | NA | 0.4 |
| SE | 9912 | CASP8 | CASP8 | chr2 | + | 201271515 | 201271621 | 201266460 | 201266791 | 201272637 | 201272776 | 0 | 0 | 9 | 0 | 0 | 1 | 0 | 17 | 0 | 0 | 3 | 0 | 98 | 49 | NA | NA | NA | NA | | 0.818 | NA | | 0.739 | NA | 0.079 |
| SE | 9914 | CASP8 | CASP8 | chr2 | + | 201272897 | 201272942 | 201272637 | 201272776 | 201276826 | 201276968 | 0 | 0 | 6 | 0 | 0 | 1 | 0 | 18 | 0 | 0 | 0 | 0 | 94 | 49 | NA | NA | NA | NA | | 0.758 | NA | | 1 | NA | -0.242 |
| SE | 9915 | CASP8 | CASP8 | chr2 | + | 201274888 | 201274953 | 201272637 | 201272776 | 201276826 | 201276968 | 0 | 0 | 3 | 0 | 0 | 1 | 0 | 3 | 0 | 0 | 0 | 0 | 98 | 49 | NA | NA | NA | NA | | 0.6 | NA | | 1 | NA | -0.4 |
| SE | 9940 | ZRANB2 | ZRANB2 | chr1 | - | 71065677 | 71065752 | 71063290 | 71065137 | 71066775 | 71066934 | 0 | 0 | 1 | 0 | 0 | 0 | 5 | 0 | 0 | 0 | 1 | 0 | 98 | 49 | NA | NA | NA | NA | | 1 | | 1 | 0 | NA | 0.5 |
| SE | 9989 | TRANK1 | TRANK1 | chr3 | - | 36855172 | 36858049 | 36852145 | 36852345 | 36858717 | 36858894 | 0 | 0 | 15 | 0 | 0 | 2 | 0 | 3 | 0 | 0 | 0 | 0 | 98 | 49 | NA | NA | NA | NA | | 0.789 | NA | | 1 | NA | -0.211 |
| SE | 9994 | UPP1 | UPP1 | chr7 | + | 48099669 | 48099787 | 48094762 | 48094827 | 48101823 | 48101982 | 0 | 0 | 1 | 0 | 0 | 0 | 0 | 3 | 0 | 0 | 2 | 0 | 98 | 49 | NA | NA | NA | NA | | 1 | NA | | 0.429 | NA | 0.571 |
| SE | 10044 | KIAA1468 | KIAA1468 | chr18 | + | 62280359 | 62280442 | 62275373 | 62275473 | 62280645 | 62280709 | 0 | 0 | 1 | 0 | 0 | 1 | 0 | 1 | 0 | 0 | 0 | 0 | 98 | 49 | NA | NA | NA | NA | | 0.333 | NA | | 1 | NA | -0.667 |
| SE | 10047 | CHURC1 | CHURC1 | chr14 | + | 64926006 | 64926080 | 64923990 | 64924126 | 64932137 | 64935366 | 0 | 0 | 2 | 0 | 0 | 2 | 0 | 1 | 0 | 0 | 1 | 0 | 98 | 49 | NA | NA | NA | NA | | 0.333 | NA | | 0.333 | NA | 0 |
| SE | 10048 | CHURC1 | CHURC1 | chr14 | + | 64926009 | 64926080 | 64923990 | 64924126 | 64932137 | 64935366 | 0 | 0 | 2 | 0 | 0 | 2 | 0 | 2 | 0 | 0 | 1 | 0 | 98 | 49 | NA | NA | NA | NA | | 0.333 | NA | | 0.5 | NA | -0.167 |
| SE | 10059 | TRRAP | TRRAP | chr7 | + | 98961260 | 98961474 | 98959343 | 98959490 | 98962301 | 98962427 | 0 | 0 | 1 | 0 | 0 | 0 | 0 | 0 | 0 | 0 | 1 | 0 | 98 | 49 | NA | NA | NA | NA | | 1 | NA | | 0 | NA | 1 |
| SE | 10074 | SRSF5 | SRSF5 | chr14 | + | 69768797 | 69768896 | 69768603 | 69768674 | 69769181 | 69769251 | 0 | 0 | 20 | 0 | 0 | 0 | 0 | 26 | 0 | 0 | 2 | 0 | 98 | 49 | NA | NA | NA | NA | | 1 | NA | | 0.867 | NA | 0.133 |
| SE | 10090 | KIAA0040 | KIAA0040 | chr1 | - | 175192639 | 175192716 | 175177610 | 175177684 | 175192859 | 175193093 | 0 | 0 | 1 | 0 | 0 | 0 | 0 | 1 | 0 | 0 | 1 | 0 | 98 | 49 | NA | NA | NA | NA | | 1 | NA | | 0.333 | NA | 0.667 |
| SE | 10118 | TRAF3IP3 | TRAF3IP3 | chr1 | + | 209780469 | 209780606 | 209779314 | 209779374 | 209781344 | 209781458 | 0 | 0 | 3 | 0 | 0 | 0 | 0 | 2 | 0 | 0 | 3 | 0 | 98 | 49 | NA | NA | NA | NA | | 1 | NA | | 0.25 | NA | 0.75 |
| SE | 10134 | CSNK1A1 | CSNK1A1 | chr5 | - | 149511793 | 149511872 | 149509878 | 149509953 | 149513069 | 149513209 | 0 | 0 | 7 | 0 | 0 | 0 | 0 | 7 | 0 | 0 | 1 | 0 | 98 | 49 | NA | NA | NA | NA | | 1 | NA | | 0.778 | NA | 0.222 |
| SE | 10144 | GALC | GALC | chr14 | - | 87945552 | 87945733 | 87941394 | 87941558 | 87947727 | 87947878 | 0 | 0 | 3 | 0 | 0 | 1 | 1 | 3 | 0 | 0 | 0 | 0 | 98 | 49 | NA | NA | NA | NA | | 0.6 | | 1 | 1 | NA | -0.4 |
| SE | 10145 | GALC | GALC | chr14 | - | 87984393 | 87984533 | 87982204 | 87982243 | 87986488 | 87986602 | 0 | 0 | 2 | 0 | 0 | 0 | 0 | 3 | 0 | 0 | 1 | 0 | 98 | 49 | NA | NA | NA | NA | | 1 | NA | | 0.6 | NA | 0.4 |
| SE | 10160 | C14orf159 | C14orf159 | chr14 | + | 91145220 | 91145313 | 91114634 | 91114994 | 91157638 | 91157690 | 0 | 0 | 4 | 0 | 0 | 5 | 0 | 1 | 0 | 0 | 0 | 0 | 98 | 49 | NA | NA | NA | NA | | 0.286 | NA | | 1 | NA | -0.714 |
| SE | 10161 | C14orf159 | C14orf159 | chr14 | + | 91147780 | 91147858 | 91145220 | 91145313 | 91157638 | 91157690 | 0 | 0 | 6 | 0 | 0 | 1 | 0 | 1 | 0 | 0 | 0 | 0 | 98 | 49 | NA | NA | NA | NA | | 0.75 | NA | | 1 | NA | -0.25 |
| SE | 10181 | ATXN3 | ATXN3 | chr14 | - | 92093745 | 92093831 | 92093251 | 92093318 | 92096092 | 92096137 | 0 | 0 | 1 | 0 | 0 | 0 | 0 | 2 | 0 | 0 | 1 | 0 | 98 | 49 | NA | NA | NA | NA | | 1 | NA | | 0.5 | NA | 0.5 |
| SE | 10195 | PIK3CA | PIK3CA | chr3 | + | 179220985 | 179221157 | 179219948 | 179220052 | 179224080 | 179224187 | 0 | 0 | 4 | 0 | 0 | 1 | 0 | 2 | 0 | 0 | 0 | 0 | 98 | 49 | NA | NA | NA | NA | | 0.667 | NA | | 1 | NA | -0.333 |
| SE | 10234 | PLSCR3 | PLSCR3 | chr17 | - | 7393143 | 7393364 | 7392790 | 7392952 | 7393466 | 7393509 | 0 | 0 | 1 | 0 | 0 | 1 | 0 | 1 | 0 | 0 | 0 | 0 | 98 | 49 | NA | NA | NA | NA | | 0.333 | NA | | 1 | NA | -0.667 |
| SE | 10236 | RPS3A | RPS3A | chr4 | + | 151100974 | 151101162 | 151100484 | 151100588 | 151102870 | 151103079 | 1 | 0 | 0 | 0 | 0 | 0 | 0 | 0 | 0 | 0 | 2 | 0 | 98 | 49 | NA | NA | 1 | NA | NA | | NA | | 0 | NA | 1 |
| SE | 10267 | PBRM1 | PBRM1 | chr3 | - | 52615350 | 52615456 | 52609312 | 52609955 | 52617261 | 52617538 | 0 | 0 | 1 | 0 | 0 | 1 | 0 | 1 | 0 | 0 | 0 | 0 | 98 | 49 | NA | NA | NA | NA | | 0.333 | NA | | 1 | NA | -0.667 |
| SE | 10292 | TECPR2 | TECPR2 | chr14 | + | 102376649 | 102376940 | 102362962 | 102363116 | 102407337 | 102407466 | 0 | 0 | 8 | 0 | 0 | 0 | 0 | 5 | 0 | 3 | 0 | 0 | 98 | 49 | NA | NA | NA | NA | | 1 | | 0 | 1 | NA | 0.5 |
| SE | 10293 | RCOR1 | RCOR1 | chr14 | + | 102721319 | 102721377 | 102721006 | 102721084 | 102722186 | 102722416 | 0 | 0 | 7 | 0 | 0 | 2 | 0 | 3 | 0 | 0 | 0 | 0 | 98 | 49 | NA | NA | NA | NA | | 0.636 | NA | | 1 | NA | -0.364 |
| SE | 10298 | MARK3 | MARK3 | chr14 | + | 103428386 | 103428440 | 103405075 | 103405267 | 103448918 | 103448967 | 0 | 0 | 17 | 0 | 0 | 0 | 0 | 7 | 0 | 0 | 4 | 0 | 98 | 49 | NA | NA | NA | NA | | 1 | NA | | 0.467 | NA | 0.533 |
| SE | 10300 | MARK3 | MARK3 | chr14 | + | 103465971 | 103466091 | 103465556 | 103465793 | 103466342 | 103466442 | 0 | 0 | 1 | 0 | 0 | 1 | 0 | 3 | 0 | 0 | 0 | 0 | 98 | 49 | NA | NA | NA | NA | | 0.333 | NA | | 1 | NA | -0.667 |
| SE | 10309 | EIF4E3 | EIF4E3 | chr3 | - | 71696459 | 71696520 | 71693874 | 71693941 | 71699613 | 71699708 | 0 | 0 | 13 | 0 | 0 | 1 | 0 | 10 | 0 | 0 | 0 | 0 | 98 | 49 | NA | NA | NA | NA | | 0.867 | NA | | 1 | NA | -0.133 |
| SE | 10320 | GALNS | GALNS | chr16 | - | 88837621 | 88837765 | 88836200 | 88836267 | 88840991 | 88841094 | 0 | 0 | 1 | 0 | 0 | 2 | 0 | 1 | 0 | 0 | 2 | 0 | 98 | 49 | NA | NA | NA | NA | | 0.2 | NA | | 0.2 | NA | 0 |
| SE | 10343 | PDCD4 | PDCD4 | chr10 | + | 110894411 | 110894522 | 110894090 | 110894198 | 110895947 | 110896087 | 0 | 0 | 6 | 0 | 0 | 0 | 0 | 3 | 0 | 0 | 1 | 0 | 98 | 49 | NA | NA | NA | NA | | 1 | NA | | 0.6 | NA | 0.4 |
| SE | 10347 | TPM3 | TPM3 | chr1 | - | 154169304 | 154169383 | 154155303 | 154157721 | 154170399 | 154170469 | 0 | 0 | 4 | 0 | 0 | 20 | 0 | 0 | 0 | 0 | 14 | 0 | 98 | 49 | NA | NA | NA | NA | | 0.091 | NA | | 0 | NA | 0.091 |
| SE | 10362 | SOS2 | SOS2 | chr14 | - | 50138611 | 50138784 | 50134122 | 50134239 | 50139941 | 50140059 | 0 | 0 | 15 | 0 | 0 | 0 | 5 | 8 | 0 | 0 | 1 | 0 | 98 | 49 | NA | NA | NA | NA | | 1 | | 1 | 0.8 | NA | 0.1 |
| SE | 10373 | LCMT1 | LCMT1 | chr16 | + | 25128474 | 25128566 | 25111725 | 25111996 | 25132401 | 25132523 | 0 | 0 | 0 | 0 | 0 | 1 | 0 | 2 | 0 | 0 | 0 | 0 | 98 | 49 | NA | NA | NA | NA | | 0 | NA | | 1 | NA | -1 |
| SE | 10383 | SIGLEC5 | SIGLEC5 | chr19 | - | 51626031 | 51626113 | 51611502 | 51612422 | 51627148 | 51627248 | 0 | 0 | 13 | 0 | 0 | 1 | 0 | 3 | 0 | 0 | 0 | 0 | 98 | 49 | NA | NA | NA | NA | | 0.867 | NA | | 1 | NA | -0.133 |
| SE | 10437 | KATNBL1 | KATNBL1 | chr15 | - | 34147379 | 34147430 | 34147199 | 34147289 | 34148631 | 34148750 | 0 | 0 | 1 | 0 | 0 | 0 | 0 | 0 | 0 | 0 | 1 | 0 | 98 | 49 | NA | NA | NA | NA | | 1 | NA | | 0 | NA | 1 |
| SE | 10439 | SLC12A6 | SLC12A6 | chr15 | - | 34242101 | 34242221 | 34241232 | 34241337 | 34243973 | 34244072 | 0 | 0 | 7 | 0 | 0 | 1 | 0 | 4 | 0 | 0 | 2 | 0 | 98 | 49 | NA | NA | NA | NA | | 0.778 | NA | | 0.5 | NA | 0.278 |
| SE | 10443 | PI4KA | PI4KA | chr22 | - | 20796145 | 20796314 | 20793192 | 20793243 | 20798583 | 20798687 | 0 | 0 | 0 | 0 | 0 | 1 | 0 | 1 | 0 | 0 | 3 | 0 | 98 | 49 | NA | NA | NA | NA | | 0 | NA | | 0.143 | NA | -0.143 |
| SE | 10455 | TOP3A | TOP3A | chr17 | - | 18280535 | 18280658 | 18277674 | 18278357 | 18282697 | 18282841 | 0 | 0 | 9 | 0 | 0 | 0 | 0 | 1 | 0 | 0 | 1 | 0 | 98 | 49 | NA | NA | NA | NA | | 1 | NA | | 0.333 | NA | 0.667 |
| SE | 10456 | TOP3A | TOP3A | chr17 | - | 18292644 | 18292852 | 18290841 | 18291027 | 18294702 | 18294785 | 0 | 0 | 1 | 0 | 0 | 1 | 0 | 4 | 0 | 0 | 0 | 0 | 98 | 49 | NA | NA | NA | NA | | 0.333 | NA | | 1 | NA | -0.667 |
| SE | 10461 | OSBPL11 | OSBPL11 | chr3 | - | 125552180 | 125552679 | 125547405 | 125547592 | 125560378 | 125560519 | 0 | 0 | 5 | 0 | 0 | 1 | 3 | 3 | 0 | 0 | 0 | 0 | 98 | 49 | NA | NA | NA | NA | | 0.714 | | 1 | 1 | NA | -0.286 |
| SE | 10468 | PLCB2 | PLCB2 | chr15 | - | 40291605 | 40291650 | 40291264 | 40291487 | 40291848 | 40291924 | 0 | 0 | 1 | 0 | 0 | 2 | 0 | 0 | 0 | 0 | 1 | 0 | 94 | 49 | NA | NA | NA | NA | | 0.207 | NA | | 0 | NA | 0.207 |
| SE | 10491 | DCP1A | DCP1A | chr3 | - | 53312240 | 53312379 | 53292068 | 53292827 | 53319406 | 53319473 | 0 | 0 | 1 | 0 | 0 | 0 | 0 | 2 | 0 | 0 | 1 | 0 | 98 | 49 | NA | NA | NA | NA | | 1 | NA | | 0.5 | NA | 0.5 |
| SE | 10495 | HSP90AA1 | HSP90AA1 | chr14 | - | 102082110 | 102082444 | 102080737 | 102081821 | 102083033 | 102083302 | 0 | 0 | 5 | 0 | 0 | 1 | 0 | 6 | 0 | 0 | 0 | 0 | 98 | 49 | NA | NA | NA | NA | | 0.714 | NA | | 1 | NA | -0.286 |
| SE | 10501 | GANC | GANC | chr15 | + | 42297610 | 42297656 | 42292734 | 42292917 | 42306545 | 42306612 | 0 | 0 | 2 | 0 | 0 | 1 | 0 | 3 | 0 | 0 | 0 | 0 | 95 | 49 | NA | NA | NA | NA | | 0.508 | NA | | 1 | NA | -0.492 |
| SE | 10509 | ZNF106 | ZNF106 | chr15 | - | 42428017 | 42428134 | 42424833 | 42425025 | 42435383 | 42435518 | 0 | 0 | 7 | 0 | 0 | 1 | 2 | 1 | 0 | 0 | 0 | 0 | 98 | 49 | NA | NA | NA | NA | | 0.778 | | 1 | 1 | NA | -0.222 |
| SE | 10510 | ZNF106 | ZNF106 | chr15 | - | 42449770 | 42451954 | 42448071 | 42448705 | 42466052 | 42466114 | 0 | 0 | 3 | 0 | 0 | 0 | 0 | 0 | 0 | 0 | 2 | 0 | 98 | 49 | NA | NA | NA | NA | | 1 | NA | | 0 | NA | 1 |
| SE | 10513 | TMEM62 | TMEM62 | chr15 | + | 43134256 | 43134368 | 43133523 | 43133982 | 43135511 | 43135649 | 0 | 0 | 2 | 0 | 0 | 0 | 0 | 0 | 0 | 0 | 1 | 0 | 98 | 49 | NA | NA | NA | NA | | 1 | NA | | 0 | NA | 1 |
| SE | 10514 | TMEM62 | TMEM62 | chr15 | + | 43149028 | 43149151 | 43148754 | 43148879 | 43151789 | 43151945 | 0 | 0 | 1 | 0 | 0 | 0 | 0 | 1 | 0 | 0 | 2 | 0 | 98 | 49 | NA | NA | NA | NA | | 1 | NA | | 0.2 | NA | 0.8 |
| SE | 10517 | TP53BP1 | TP53BP1 | chr15 | - | 43432193 | 43432677 | 43428015 | 43428168 | 43438323 | 43438416 | 0 | 0 | 1 | 0 | 0 | 0 | 0 | 1 | 0 | 0 | 1 | 0 | 98 | 49 | NA | NA | NA | NA | | 1 | NA | | 0.333 | NA | 0.667 |
| SE | 10558 | DSC2 | DSC2 | chr18 | - | 31068298 | 31068344 | 31065973 | 31068212 | 31068893 | 31069151 | 0 | 0 | 0 | 0 | 0 | 1 | 0 | 3 | 0 | 0 | 0 | 0 | 95 | 49 | NA | NA | NA | NA | | 0 | NA | | 1 | NA | -1 |
| SE | 10560 | FAM96A | FAM96A | chr15 | - | 64081101 | 64081151 | 64075491 | 64075537 | 64088686 | 64088851 | 0 | 0 | 5 | 0 | 0 | 0 | 0 | 0 | 0 | 3 | 1 | 0 | 98 | 49 | NA | NA | NA | NA | | 1 | | 0 | 0 | NA | 1 |
| SE | 10566 | ATP8B4 | ATP8B4 | chr15 | - | 50002152 | 50002223 | 49987390 | 49987549 | 50010844 | 50010917 | 0 | 0 | 2 | 0 | 0 | 2 | 1 | 2 | 0 | 0 | 0 | 0 | 98 | 49 | NA | NA | NA | NA | | 0.333 | | 1 | 1 | NA | -0.667 |
| SE | 10570 | LRRC75A-AS1 | LRRC75A-AS1 | chr17 | + | 16439527 | 16439703 | 16438986 | 16439414 | 16440184 | 16440253 | 0 | 0 | 0 | 0 | 0 | 1 | 0 | 3 | 0 | 0 | 1 | 0 | 98 | 49 | NA | NA | NA | NA | | 0 | NA | | 0.6 | NA | -0.6 |
| SE | 10571 | LRRC75A-AS1 | LRRC75A-AS1 | chr17 | + | 16439580 | 16439703 | 16438986 | 16439414 | 16440184 | 16440253 | 0 | 0 | 0 | 0 | 0 | 1 | 0 | 3 | 0 | 0 | 1 | 0 | 98 | 49 | NA | NA | NA | NA | | 0 | NA | | 0.6 | NA | -0.6 |
| SE | 10573 | LRRC75A-AS1 | LRRC75A-AS1 | chr17 | + | 16439659 | 16439703 | 16438986 | 16439414 | 16440184 | 16440253 | 0 | 0 | 0 | 0 | 0 | 1 | 0 | 3 | 0 | 0 | 1 | 0 | 93 | 49 | NA | NA | NA | NA | | 0 | NA | | 0.613 | NA | -0.613 |
| SE | 10576 | DDX47 | DDX47 | chr12 | + | 12823869 | 12824016 | 12823202 | 12823319 | 12824539 | 12824677 | 0 | 0 | 0 | 0 | 0 | 2 | 0 | 1 | 0 | 0 | 0 | 0 | 98 | 49 | NA | NA | NA | NA | | 0 | NA | | 1 | NA | -1 |
| SE | 10582 | USP8 | USP8 | chr15 | + | 50484274 | 50484361 | 50481480 | 50482065 | 50489800 | 50489881 | 0 | 0 | 2 | 0 | 0 | 0 | 0 | 1 | 0 | 0 | 2 | 0 | 98 | 49 | NA | NA | NA | NA | | 1 | NA | | 0.2 | NA | 0.8 |
| SE | 10584 | SPPL2A | SPPL2A | chr15 | - | 50732602 | 50732684 | 50730964 | 50731039 | 50736100 | 50736202 | 0 | 0 | 6 | 0 | 0 | 0 | 0 | 4 | 0 | 0 | 1 | 0 | 98 | 49 | NA | NA | NA | NA | | 1 | NA | | 0.667 | NA | 0.333 |
| SE | 10592 | RNF213 | RNF213 | chr17 | + | 80385038 | 80385171 | 80383676 | 80383928 | 80385537 | 80385621 | 0 | 0 | 23 | 0 | 0 | 2 | 0 | 23 | 0 | 0 | 2 | 0 | 98 | 49 | NA | NA | NA | NA | | 0.852 | NA | | 0.852 | NA | 0 |
| SE | 10600 | DMXL2 | DMXL2 | chr15 | - | 51451644 | 51451697 | 51450128 | 51450346 | 51453549 | 51453641 | 0 | 0 | 8 | 0 | 0 | 0 | 1 | 1 | 0 | 0 | 1 | 0 | 98 | 49 | NA | NA | NA | NA | | 1 | | 1 | 0.333 | NA | 0.334 |
| SE | 10601 | DMXL2 | DMXL2 | chr15 | - | 51464674 | 51464876 | 51463378 | 51463496 | 51465565 | 51465651 | 0 | 0 | 15 | 0 | 0 | 1 | 0 | 2 | 0 | 0 | 0 | 0 | 98 | 49 | NA | NA | NA | NA | | 0.882 | NA | | 1 | NA | -0.118 |
| SE | 10607 | ANKFY1 | ANKFY1 | chr17 | - | 4209823 | 4209947 | 4207932 | 4208082 | 4216982 | 4217118 | 0 | 0 | 6 | 0 | 0 | 0 | 0 | 1 | 0 | 0 | 1 | 0 | 98 | 49 | NA | NA | NA | NA | | 1 | NA | | 0.333 | NA | 0.667 |
| SE | 10610 | MYO5A | MYO5A | chr15 | - | 52348817 | 52348826 | 52346360 | 52346461 | 52351253 | 52351481 | 0 | 0 | 0 | 0 | 0 | 1 | 0 | 1 | 0 | 0 | 0 | 0 | 58 | 49 | NA | NA | NA | NA | | 0 | NA | | 1 | NA | -1 |
| SE | 10623 | ADAM10 | ADAM10 | chr15 | - | 58611807 | 58611991 | 58610998 | 58611107 | 58621470 | 58621621 | 0 | 0 | 11 | 0 | 0 | 1 | 0 | 6 | 0 | 0 | 0 | 0 | 98 | 49 | NA | NA | NA | NA | | 0.846 | NA | | 1 | NA | -0.154 |
| SE | 10625 | RRP12 | RRP12 | chr10 | - | 97372735 | 97372803 | 97372072 | 97372166 | 97373045 | 97373200 | 0 | 0 | 4 | 0 | 0 | 2 | 0 | 4 | 0 | 0 | 0 | 0 | 98 | 49 | NA | NA | NA | NA | | 0.5 | NA | | 1 | NA | -0.5 |
| SE | 10629 | BNIP2 | BNIP2 | chr15 | - | 59679591 | 59679768 | 59677910 | 59678087 | 59680240 | 59680308 | 0 | 0 | 15 | 0 | 0 | 0 | 0 | 6 | 0 | 0 | 1 | 0 | 98 | 49 | NA | NA | NA | NA | | 1 | NA | | 0.75 | NA | 0.25 |
| SE | 10640 | USP3 | USP3 | chr15 | + | 63529013 | 63529166 | 63504510 | 63504830 | 63532646 | 63532707 | 0 | 0 | 0 | 0 | 0 | 2 | 0 | 2 | 0 | 0 | 4 | 0 | 98 | 49 | NA | NA | NA | NA | | 0 | NA | | 0.2 | NA | -0.2 |
| SE | 10643 | HERC1 | HERC1 | chr15 | - | 63623724 | 63623890 | 63622814 | 63622891 | 63624157 | 63624327 | 0 | 0 | 2 | 0 | 0 | 2 | 0 | 2 | 0 | 0 | 0 | 0 | 98 | 49 | NA | NA | NA | NA | | 0.333 | NA | | 1 | NA | -0.667 |
| SE | 10644 | HERC1 | HERC1 | chr15 | - | 63632708 | 63632811 | 63630465 | 63630635 | 63633847 | 63633970 | 0 | 0 | 6 | 0 | 0 | 0 | 0 | 1 | 0 | 0 | 1 | 0 | 98 | 49 | NA | NA | NA | NA | | 1 | NA | | 0.333 | NA | 0.667 |
| SE | 10645 | HERC1 | HERC1 | chr15 | - | 63648068 | 63648199 | 63645482 | 63645682 | 63649724 | 63649925 | 0 | 0 | 2 | 0 | 0 | 0 | 0 | 3 | 0 | 0 | 2 | 0 | 98 | 49 | NA | NA | NA | NA | | 1 | NA | | 0.429 | NA | 0.571 |
| SE | 10652 | RELA | RELA | chr11 | - | 65658286 | 65658499 | 65655854 | 65655935 | 65658717 | 65658822 | 0 | 0 | 4 | 0 | 0 | 0 | 0 | 5 | 0 | 0 | 2 | 0 | 98 | 49 | NA | NA | NA | NA | | 1 | NA | | 0.556 | NA | 0.444 |
| SE | 10655 | OAZ2 | OAZ2 | chr15 | - | 64691433 | 64691588 | 64690373 | 64690549 | 64703028 | 64703281 | 0 | 0 | 2 | 0 | 0 | 0 | 0 | 3 | 0 | 0 | 1 | 0 | 98 | 49 | NA | NA | NA | NA | | 1 | NA | | 0.6 | NA | 0.4 |
| SE | 10656 | OAZ2 | OAZ2 | chr15 | - | 64691433 | 64691591 | 64690373 | 64690549 | 64703028 | 64703281 | 0 | 0 | 3 | 0 | 0 | 0 | 0 | 5 | 0 | 0 | 1 | 0 | 98 | 49 | NA | NA | NA | NA | | 1 | NA | | 0.714 | NA | 0.286 |
| SE | 10689 | EGLN1 | EGLN1 | chr1 | - | 231367568 | 231367636 | 231363750 | 231366475 | 231370561 | 231370698 | 0 | 0 | 7 | 0 | 0 | 1 | 0 | 2 | 0 | 0 | 4 | 0 | 98 | 49 | NA | NA | NA | NA | | 0.778 | NA | | 0.2 | NA | 0.578 |
| SE | 10703 | TLE3 | TLE3 | chr15 | - | 70095577 | 70095641 | 70094531 | 70094576 | 70096160 | 70096261 | 0 | 0 | 2 | 0 | 0 | 1 | 1 | 8 | 0 | 0 | 0 | 0 | 98 | 49 | NA | NA | NA | NA | | 0.5 | | 1 | 1 | NA | -0.5 |
| SE | 10709 | ARIH1 | ARIH1 | chr15 | + | 72563393 | 72563500 | 72561482 | 72561549 | 72566562 | 72566605 | 0 | 0 | 1 | 0 | 0 | 0 | 0 | 3 | 0 | 0 | 1 | 0 | 98 | 49 | NA | NA | NA | NA | | 1 | NA | | 0.6 | NA | 0.4 |
| SE | 10726 | PTMA | PTMA | chr2 | + | 231712442 | 231712516 | 231711347 | 231711419 | 231712803 | 231713540 | 0 | 0 | 1 | 0 | 0 | 0 | 0 | 3 | 0 | 0 | 1 | 0 | 98 | 49 | NA | NA | NA | NA | | 1 | NA | | 0.6 | NA | 0.4 |
| SE | 10733 | FAM219B | FAM219B | chr15 | - | 74904663 | 74904712 | 74899986 | 74902786 | 74905153 | 74905231 | 0 | 0 | 2 | 0 | 0 | 0 | 0 | 1 | 0 | 0 | 3 | 0 | 98 | 49 | NA | NA | NA | NA | | 1 | NA | | 0.143 | NA | 0.857 |
| SE | 10745 | MAN2C1 | MAN2C1 | chr15 | - | 75362641 | 75362748 | 75362342 | 75362453 | 75364487 | 75364665 | 0 | 0 | 4 | 0 | 0 | 1 | 0 | 4 | 0 | 0 | 0 | 0 | 98 | 49 | NA | NA | NA | NA | | 0.667 | NA | | 1 | NA | -0.333 |
| SE | 10751 | SIN3A | SIN3A | chr15 | - | 75400729 | 75400940 | 75400039 | 75400156 | 75401851 | 75401970 | 0 | 0 | 4 | 0 | 0 | 0 | 0 | 0 | 0 | 0 | 1 | 0 | 98 | 49 | NA | NA | NA | NA | | 1 | NA | | 0 | NA | 1 |
| SE | 10827 | AMBRA1 | AMBRA1 | chr11 | - | 46433473 | 46433628 | 46417912 | 46418052 | 46434848 | 46435037 | 0 | 0 | 2 | 0 | 0 | 0 | 0 | 0 | 0 | 0 | 1 | 0 | 98 | 49 | NA | NA | NA | NA | | 1 | NA | | 0 | NA | 1 |
| SE | 10846 | IRF1 | IRF1 | chr5 | - | 132484361 | 132484497 | 132481608 | 132484075 | 132486556 | 132486686 | 0 | 0 | 8 | 0 | 0 | 1 | 0 | 5 | 0 | 0 | 0 | 0 | 98 | 49 | NA | NA | NA | NA | | 0.8 | NA | | 1 | NA | -0.2 |
| SE | 10847 | IRF1 | IRF1 | chr5 | - | 132485666 | 132485716 | 132484361 | 132484497 | 132486556 | 132486686 | 0 | 0 | 11 | 0 | 0 | 0 | 0 | 14 | 0 | 0 | 1 | 0 | 98 | 49 | NA | NA | NA | NA | | 1 | NA | | 0.875 | NA | 0.125 |
| SE | 10914 | HDAC3 | HDAC3 | chr5 | - | 141626012 | 141626071 | 141625684 | 141625764 | 141626193 | 141626283 | 0 | 0 | 6 | 0 | 0 | 2 | 0 | 2 | 0 | 0 | 0 | 0 | 98 | 49 | NA | NA | NA | NA | | 0.6 | NA | | 1 | NA | -0.4 |
| SE | 10917 | CWF19L1 | CWF19L1 | chr10 | - | 100256261 | 100256476 | 100253420 | 100253539 | 100260217 | 100260319 | 0 | 0 | 4 | 0 | 0 | 0 | 0 | 1 | 0 | 0 | 1 | 0 | 98 | 49 | NA | NA | NA | NA | | 1 | NA | | 0.333 | NA | 0.667 |
| SE | 10941 | PLEKHJ1 | PLEKHJ1 | chr19 | - | 2234149 | 2234240 | 2233997 | 2234061 | 2235761 | 2235828 | 0 | 0 | 0 | 0 | 0 | 1 | 0 | 1 | 0 | 0 | 0 | 0 | 98 | 49 | NA | NA | NA | NA | | 0 | NA | | 1 | NA | -1 |
| SE | 10948 | IQGAP1 | IQGAP1 | chr15 | + | 90466268 | 90466436 | 90466000 | 90466091 | 90467449 | 90467592 | 0 | 0 | 20 | 0 | 0 | 0 | 0 | 18 | 0 | 0 | 3 | 0 | 98 | 49 | NA | NA | NA | NA | | 1 | NA | | 0.75 | NA | 0.25 |
| SE | 10949 | IQGAP1 | IQGAP1 | chr15 | + | 90467449 | 90467592 | 90466268 | 90466436 | 90472839 | 90473010 | 0 | 0 | 32 | 0 | 0 | 0 | 1 | 6 | 0 | 0 | 1 | 0 | 98 | 49 | NA | NA | NA | NA | | 1 | | 1 | 0.75 | NA | 0.125 |
| SE | 10973 | MEF2A | MEF2A | chr15 | + | 99620872 | 99621080 | 99598429 | 99598511 | 99632977 | 99633173 | 0 | 0 | 1 | 0 | 0 | 0 | 0 | 0 | 0 | 0 | 1 | 0 | 98 | 49 | NA | NA | NA | NA | | 1 | NA | | 0 | NA | 1 |
| SE | 10975 | MEF2A | MEF2A | chr15 | + | 99690240 | 99690428 | 99675398 | 99675458 | 99710633 | 99710760 | 0 | 0 | 7 | 0 | 0 | 1 | 0 | 3 | 0 | 0 | 0 | 0 | 98 | 49 | NA | NA | NA | NA | | 0.778 | NA | | 1 | NA | -0.222 |
| SE | 10987 | ZCCHC6 | ZCCHC6 | chr9 | - | 86309462 | 86309576 | 86309211 | 86309289 | 86309927 | 86310017 | 0 | 0 | 12 | 0 | 0 | 0 | 0 | 5 | 0 | 0 | 1 | 0 | 98 | 49 | NA | NA | NA | NA | | 1 | NA | | 0.714 | NA | 0.286 |
| SE | 10988 | ZCCHC6 | ZCCHC6 | chr9 | - | 86318957 | 86319058 | 86317218 | 86317276 | 86319583 | 86319670 | 0 | 0 | 8 | 0 | 0 | 0 | 0 | 4 | 0 | 0 | 1 | 0 | 98 | 49 | NA | NA | NA | NA | | 1 | NA | | 0.667 | NA | 0.333 |
| SE | 10990 | ZCCHC6 | ZCCHC6 | chr9 | - | 86322324 | 86322475 | 86319583 | 86319670 | 86322872 | 86323960 | 0 | 0 | 6 | 0 | 0 | 0 | 0 | 4 | 0 | 0 | 1 | 0 | 98 | 49 | NA | NA | NA | NA | | 1 | NA | | 0.667 | NA | 0.333 |
| SE | 11029 | TSR3 | TSR3 | chr16 | - | 1349888 | 1349952 | 1349239 | 1349608 | 1350057 | 1350234 | 0 | 0 | 0 | 0 | 0 | 1 | 0 | 4 | 0 | 0 | 0 | 0 | 98 | 49 | NA | NA | NA | NA | | 0 | NA | | 1 | NA | -1 |
| SE | 11032 | STAT1 | STAT1 | chr2 | - | 191001073 | 191001163 | 190999625 | 190999704 | 191007562 | 191007661 | 0 | 0 | 49 | 0 | 0 | 1 | 0 | 39 | 0 | 0 | 0 | 0 | 98 | 49 | NA | NA | NA | NA | | 0.961 | NA | | 1 | NA | -0.039 |
| SE | 11050 | NPEPPS | NPEPPS | chr17 | + | 47586147 | 47586244 | 47585499 | 47585700 | 47586364 | 47586398 | 0 | 0 | 6 | 0 | 0 | 1 | 0 | 2 | 0 | 0 | 0 | 0 | 98 | 49 | NA | NA | NA | NA | | 0.75 | NA | | 1 | NA | -0.25 |
| SE | 11054 | ARFGAP3 | ARFGAP3 | chr22 | - | 42810812 | 42810944 | 42808766 | 42808890 | 42817141 | 42817264 | 0 | 0 | 3 | 0 | 0 | 1 | 0 | 4 | 0 | 0 | 1 | 0 | 98 | 49 | NA | NA | NA | NA | | 0.6 | NA | | 0.667 | NA | -0.067 |
| SE | 11063 | DENND4C | DENND4C | chr9 | + | 19357964 | 19358160 | 19356971 | 19357154 | 19360243 | 19360489 | 0 | 0 | 3 | 0 | 0 | 2 | 0 | 3 | 0 | 0 | 0 | 0 | 98 | 49 | NA | NA | NA | NA | | 0.429 | NA | | 1 | NA | -0.571 |
| SE | 11097 | SMU1 | SMU1 | chr9 | - | 33071739 | 33071892 | 33068823 | 33068934 | 33073595 | 33073806 | 0 | 0 | 1 | 0 | 0 | 2 | 0 | 0 | 0 | 0 | 1 | 0 | 98 | 49 | NA | NA | NA | NA | | 0.2 | NA | | 0 | NA | 0.2 |
| SE | 11098 | SMU1 | SMU1 | chr9 | - | 33073595 | 33073806 | 33068823 | 33068934 | 33076582 | 33076716 | 0 | 0 | 3 | 0 | 0 | 0 | 0 | 2 | 0 | 0 | 1 | 0 | 98 | 49 | NA | NA | NA | NA | | 1 | NA | | 0.5 | NA | 0.5 |
| SE | 11107 | USP25 | USP25 | chr21 | + | 15777903 | 15778027 | 15765996 | 15766141 | 15791501 | 15791664 | 0 | 0 | 1 | 0 | 0 | 1 | 0 | 8 | 0 | 0 | 0 | 0 | 98 | 49 | NA | NA | NA | NA | | 0.333 | NA | | 1 | NA | -0.667 |
| SE | 11108 | USP25 | USP25 | chr21 | + | 15808808 | 15808885 | 15805120 | 15805258 | 15811136 | 15811210 | 0 | 0 | 6 | 0 | 0 | 6 | 0 | 1 | 0 | 0 | 5 | 0 | 98 | 49 | NA | NA | NA | NA | | 0.333 | NA | | 0.091 | NA | 0.242 |
| SE | 11111 | USP25 | USP25 | chr21 | + | 15849776 | 15849872 | 15842397 | 15842540 | 15864267 | 15864446 | 0 | 0 | 0 | 0 | 0 | 3 | 0 | 1 | 0 | 0 | 0 | 0 | 98 | 49 | NA | NA | NA | NA | | 0 | NA | | 1 | NA | -1 |
| SE | 11141 | TXNDC11 | TXNDC11 | chr16 | - | 11736016 | 11736233 | 11733981 | 11734079 | 11742476 | 11742878 | 0 | 0 | 1 | 0 | 0 | 0 | 0 | 2 | 0 | 0 | 1 | 0 | 98 | 49 | NA | NA | NA | NA | | 1 | NA | | 0.5 | NA | 0.5 |
| SE | 11153 | KIAA0430 | KIAA0430 | chr16 | - | 15625371 | 15625797 | 15625015 | 15625173 | 15630331 | 15630504 | 0 | 0 | 5 | 0 | 0 | 0 | 0 | 4 | 0 | 0 | 2 | 0 | 98 | 49 | NA | NA | NA | NA | | 1 | NA | | 0.5 | NA | 0.5 |
| SE | 11154 | KIAA0430 | KIAA0430 | chr16 | - | 15625371 | 15625800 | 15625015 | 15625173 | 15630331 | 15630504 | 0 | 0 | 7 | 0 | 0 | 0 | 0 | 6 | 0 | 0 | 2 | 0 | 98 | 49 | NA | NA | NA | NA | | 1 | NA | | 0.6 | NA | 0.4 |
| SE | 11207 | TOP1 | TOP1 | chr20 | + | 41116277 | 41116392 | 41115370 | 41115439 | 41118168 | 41118296 | 0 | 0 | 13 | 0 | 0 | 0 | 0 | 2 | 0 | 0 | 1 | 0 | 98 | 49 | NA | NA | NA | NA | | 1 | NA | | 0.5 | NA | 0.5 |
| SE | 11208 | KIAA1551 | KIAA1551 | chr12 | + | 31970188 | 31970356 | 31960776 | 31960871 | 31980877 | 31985957 | 0 | 0 | 59 | 0 | 0 | 1 | 0 | 16 | 0 | 0 | 0 | 0 | 98 | 49 | NA | NA | NA | NA | | 0.967 | NA | | 1 | NA | -0.033 |
| SE | 11218 | RPAIN | RPAIN | chr17 | + | 5425970 | 5426082 | 5422768 | 5422829 | 5432541 | 5433020 | 0 | 0 | 2 | 0 | 0 | 1 | 0 | 1 | 0 | 0 | 0 | 0 | 98 | 49 | NA | NA | NA | NA | | 0.5 | NA | | 1 | NA | -0.5 |
| SE | 11234 | DCLRE1C | DCLRE1C | chr10 | - | 14945434 | 14945565 | 14945104 | 14945189 | 14949035 | 14949087 | 0 | 0 | 0 | 0 | 0 | 3 | 0 | 1 | 0 | 0 | 0 | 0 | 98 | 49 | NA | NA | NA | NA | | 0 | NA | | 1 | NA | -1 |
| SE | 11235 | DCLRE1C | DCLRE1C | chr10 | - | 14947524 | 14947609 | 14945104 | 14945189 | 14949035 | 14949087 | 0 | 0 | 1 | 0 | 0 | 3 | 0 | 1 | 0 | 0 | 0 | 0 | 98 | 49 | NA | NA | NA | NA | | 0.143 | NA | | 1 | NA | -0.857 |
| SE | 11253 | IL4R | IL4R | chr16 | + | 27346466 | 27346618 | 27344868 | 27345020 | 27352539 | 27352696 | 0 | 0 | 8 | 0 | 0 | 1 | 0 | 10 | 0 | 0 | 2 | 0 | 98 | 49 | NA | NA | NA | NA | | 0.8 | NA | | 0.714 | NA | 0.086 |
| SE | 11254 | IL4R | IL4R | chr16 | + | 27360765 | 27360815 | 27358915 | 27358994 | 27362251 | 27364778 | 0 | 0 | 4 | 0 | 0 | 2 | 0 | 2 | 0 | 0 | 0 | 0 | 98 | 49 | NA | NA | NA | NA | | 0.5 | NA | | 1 | NA | -0.5 |
| SE | 11315 | PPP6R3 | PPP6R3 | chr11 | + | 68551120 | 68551186 | 68548066 | 68548204 | 68554144 | 68554257 | 0 | 0 | 2 | 0 | 0 | 2 | 0 | 3 | 0 | 0 | 0 | 0 | 98 | 49 | NA | NA | NA | NA | | 0.333 | NA | | 1 | NA | -0.667 |
| SE | 11318 | PPP6R3 | PPP6R3 | chr11 | + | 68583042 | 68583129 | 68575957 | 68576043 | 68587944 | 68588024 | 0 | 0 | 2 | 0 | 0 | 1 | 0 | 4 | 0 | 0 | 0 | 0 | 98 | 49 | NA | NA | NA | NA | | 0.5 | NA | | 1 | NA | -0.5 |
| SE | 11323 | ITGAL | ITGAL | chr16 | + | 30479090 | 30479208 | 30475512 | 30475580 | 30481438 | 30481584 | 0 | 0 | 18 | 0 | 0 | 1 | 3 | 3 | 0 | 0 | 0 | 0 | 98 | 49 | NA | NA | NA | NA | | 0.9 | | 1 | 1 | NA | -0.1 |
| SE | 11352 | ADGRE2 | ADGRE2 | chr19 | - | 14765657 | 14765804 | 14765523 | 14765570 | 14772341 | 14772497 | 0 | 0 | 0 | 0 | 0 | 1 | 0 | 3 | 0 | 0 | 1 | 0 | 98 | 49 | NA | NA | NA | NA | | 0 | NA | | 0.6 | NA | -0.6 |
| SE | 11361 | TATDN2 | TATDN2 | chr3 | + | 10276360 | 10276488 | 10270130 | 10271015 | 10278278 | 10278462 | 0 | 0 | 2 | 0 | 0 | 0 | 0 | 0 | 0 | 0 | 1 | 0 | 98 | 49 | NA | NA | NA | NA | | 1 | NA | | 0 | NA | 1 |
| SE | 11385 | NLRC5 | NLRC5 | chr16 | + | 57028071 | 57028155 | 57025367 | 57027018 | 57028301 | 57028385 | 0 | 0 | 2 | 0 | 0 | 0 | 0 | 3 | 0 | 0 | 2 | 0 | 98 | 49 | NA | NA | NA | NA | | 1 | NA | | 0.429 | NA | 0.571 |
| SE | 11388 | NLRC5 | NLRC5 | chr16 | + | 57055432 | 57055519 | 57055031 | 57055094 | 57058064 | 57058148 | 0 | 0 | 3 | 0 | 0 | 0 | 0 | 1 | 0 | 0 | 1 | 0 | 98 | 49 | NA | NA | NA | NA | | 1 | NA | | 0.333 | NA | 0.667 |
| SE | 11389 | NLRC5 | NLRC5 | chr16 | + | 57076818 | 57076902 | 57074599 | 57074683 | 57077295 | 57077379 | 0 | 0 | 19 | 0 | 0 | 0 | 0 | 8 | 0 | 0 | 1 | 0 | 98 | 49 | NA | NA | NA | NA | | 1 | NA | | 0.8 | NA | 0.2 |
| SE | 11390 | NLRC5 | NLRC5 | chr16 | + | 57077295 | 57077379 | 57076818 | 57076902 | 57077718 | 57077802 | 0 | 0 | 13 | 0 | 0 | 0 | 0 | 2 | 0 | 0 | 1 | 0 | 98 | 49 | NA | NA | NA | NA | | 1 | NA | | 0.5 | NA | 0.5 |
| SE | 11392 | RNF24 | RNF24 | chr20 | - | 3974331 | 3974400 | 3963874 | 3964024 | 4015436 | 4015569 | 0 | 0 | 6 | 0 | 0 | 1 | 0 | 1 | 0 | 0 | 0 | 0 | 98 | 49 | NA | NA | NA | NA | | 0.75 | NA | | 1 | NA | -0.25 |
| SE | 11405 | SETD3 | SETD3 | chr14 | - | 99461191 | 99461340 | 99459112 | 99459185 | 99463485 | 99463578 | 0 | 0 | 4 | 0 | 0 | 1 | 0 | 4 | 0 | 0 | 0 | 0 | 98 | 49 | NA | NA | NA | NA | | 0.667 | NA | | 1 | NA | -0.333 |
| SE | 11420 | CKLF | CKLF | chr16 | + | 66563121 | 66563217 | 66558189 | 66558348 | 66565885 | 66566287 | 0 | 0 | 6 | 0 | 0 | 2 | 0 | 0 | 0 | 0 | 2 | 0 | 98 | 49 | NA | NA | NA | NA | | 0.6 | NA | | 0 | NA | 0.6 |
| SE | 11437 | PLK3 | PLK3 | chr1 | + | 44804649 | 44804779 | 44804338 | 44804501 | 44805265 | 44805379 | 0 | 0 | 2 | 0 | 0 | 3 | 0 | 0 | 0 | 0 | 2 | 0 | 98 | 49 | NA | NA | NA | NA | | 0.25 | NA | | 0 | NA | 0.25 |
| SE | 11442 | FHOD1 | FHOD1 | chr16 | - | 67231187 | 67231349 | 67230600 | 67230791 | 67231429 | 67231549 | 0 | 0 | 3 | 0 | 0 | 0 | 0 | 0 | 0 | 2 | 0 | 0 | 98 | 49 | NA | NA | NA | NA | | 1 | | 0 | NA | NA | 1 |
| SE | 11449 | GSTO1 | GSTO1 | chr10 | + | 104262978 | 104263077 | 104259575 | 104259798 | 104266083 | 104266190 | 0 | 0 | 4 | 0 | 0 | 0 | 1 | 0 | 0 | 0 | 1 | 0 | 98 | 49 | NA | NA | NA | NA | | 1 | | 1 | 0 | NA | 0.5 |
| SE | 11468 | EMC10 | EMC10 | chr19 | + | 50481860 | 50481947 | 50480883 | 50480977 | 50482148 | 50483526 | 0 | 0 | 1 | 0 | 0 | 2 | 0 | 1 | 0 | 0 | 1 | 0 | 98 | 49 | NA | NA | NA | NA | | 0.2 | NA | | 0.333 | NA | -0.133 |
| SE | 11472 | ATP8A1 | ATP8A1 | chr4 | - | 42594288 | 42594362 | 42588259 | 42588329 | 42600477 | 42600518 | 0 | 0 | 1 | 0 | 0 | 0 | 0 | 1 | 0 | 0 | 2 | 0 | 98 | 49 | NA | NA | NA | NA | | 1 | NA | | 0.2 | NA | 0.8 |
| SE | 11485 | ITM2A | ITM2A | chrX | - | 79363422 | 79363554 | 79362941 | 79363139 | 79367104 | 79367552 | 0 | 0 | 2 | 0 | 0 | 0 | 0 | 0 | 0 | 0 | 1 | 0 | 98 | 49 | NA | NA | NA | NA | | 1 | NA | | 0 | NA | 1 |

| | | | | | | | | | | | | | | | | | | | | | | | | | | | | | | | | | | | | |
|---|---|---|---|---|---|---|---|---|---|---|---|---|---|---|---|---|---|---|---|---|---|---|---|---|---|---|---|---|---|---|---|---|---|---|---|---|
| SE | 11495 | KARS | KARS | chr16 | - | 75640183 | 75640349 | 75636453 | 75636547 | 75641563 | 75641723 | 0 | 0 | 2 | 0 | 0 | 0 | 0 | 0 | 0 | 0 | 1 | 0 | 98 | 49 | NA | NA | NA | NA | 1 | NA | | | 0 | NA | 1 |
| SE | 11520 | NUPL1 | NUPL1 | chr13 | + | 25312882 | 25313032 | 25309246 | 25309282 | 25313613 | 25313751 | 0 | 0 | 3 | 0 | 0 | 1 | 0 | 2 | 0 | 0 | 0 | 0 | 98 | 49 | NA | NA | NA | NA | 0.6 | NA | | | 1 | NA | -0.4 |
| SE | 11521 | PIKFYVE | PIKFYVE | chr2 | + | 208285725 | 208285933 | 208277536 | 208277708 | 208288728 | 208288818 | 0 | 0 | 6 | 0 | 0 | 0 | 0 | 3 | 0 | 0 | 1 | 0 | 98 | 49 | NA | NA | NA | NA | 1 | NA | | | 0.6 | NA | 0.4 |
| SE | 11522 | PIKFYVE | PIKFYVE | chr2 | + | 208304845 | 208305013 | 208304170 | 208304318 | 208312235 | 208312295 | 0 | 0 | 1 | 0 | 0 | 4 | 0 | 1 | 0 | 0 | 0 | 0 | 98 | 49 | NA | NA | NA | NA | 0.111 | NA | | | 1 | NA | -0.889 |
| SE | 11523 | PIKFYVE | PIKFYVE | chr2 | + | 208347858 | 208348023 | 208346049 | 208346147 | 208350023 | 208350083 | 0 | 0 | 2 | 0 | 0 | 2 | 0 | 2 | 0 | 0 | 0 | 0 | 98 | 49 | NA | NA | NA | NA | 0.333 | NA | | | 1 | NA | -0.667 |
| SE | 11524 | PIKFYVE | PIKFYVE | chr2 | + | 208351351 | 208351455 | 208350770 | 208350947 | 208352653 | 208352782 | 0 | 0 | 3 | 0 | 0 | 1 | 0 | 1 | 0 | 0 | 0 | 0 | 98 | 49 | NA | NA | NA | NA | 0.6 | NA | | | 1 | NA | -0.4 |
| SE | 11564 | CBLB | CBLB | chr3 | - | 105734008 | 105734140 | 105720046 | 105720250 | 105737170 | 105737258 | 0 | 0 | 2 | 0 | 0 | 0 | 0 | 2 | 0 | 0 | 1 | 0 | 98 | 49 | NA | NA | NA | NA | 1 | NA | | | 0.5 | NA | 0.5 |
| SE | 11579 | ANKRD11 | ANKRD11 | chr16 | - | 89431255 | 89431326 | 89418283 | 89418368 | 89490244 | 89490561 | 0 | 0 | 0 | 0 | 0 | 1 | 0 | 1 | 0 | 0 | 1 | 0 | 98 | 49 | NA | NA | NA | NA | 0 | NA | | | 0.333 | NA | -0.333 |
| SE | 11621 | IGF1R | IGF1R | chr15 | + | 98934823 | 98935053 | 98930234 | 98930305 | 98935315 | 98935426 | 0 | 0 | 2 | 0 | 0 | 0 | 0 | 0 | 0 | 0 | 1 | 0 | 98 | 49 | NA | NA | NA | NA | 1 | NA | | | 0 | NA | 1 |
| SE | 11631 | ZZEF1 | ZZEF1 | chr17 | - | 4056215 | 4056345 | 4054056 | 4054195 | 4057993 | 4058155 | 0 | 0 | 2 | 0 | 0 | 1 | 0 | 2 | 0 | 0 | 0 | 0 | 98 | 49 | NA | NA | NA | NA | 0.5 | NA | | | 1 | NA | -0.5 |
| SE | 11633 | ZZEF1 | ZZEF1 | chr17 | - | 4109652 | 4109863 | 4105692 | 4105809 | 4112608 | 4112808 | 0 | 0 | 2 | 0 | 0 | 2 | 0 | 4 | 0 | 0 | 0 | 0 | 98 | 49 | NA | NA | NA | NA | 0.333 | NA | | | 1 | NA | -0.667 |
| SE | 11667 | NAAA | NAAA | chr4 | - | 75918760 | 75918789 | 75914867 | 75914985 | 75919908 | 75919975 | 0 | 0 | 0 | 0 | 0 | 2 | 0 | 1 | 0 | 0 | 0 | 0 | 78 | 49 | NA | NA | NA | NA | 0 | NA | | | 1 | NA | -1 |
| SE | 11676 | ULK1 | ULK1 | chr12 | + | 131919211 | 131919384 | 131918496 | 131918681 | 131919471 | 131919590 | 0 | 0 | 1 | 0 | 0 | 0 | 0 | 2 | 0 | 0 | 2 | 0 | 98 | 49 | NA | NA | NA | NA | 1 | NA | | | 0.333 | NA | 0.667 |
| SE | 11678 | UBE3B | UBE3B | chr12 | + | 109523977 | 109524115 | 109521440 | 109521551 | 109524437 | 109524503 | 0 | 0 | 2 | 0 | 0 | 0 | 0 | 1 | 0 | 0 | 1 | 0 | 98 | 49 | NA | NA | NA | NA | 1 | NA | | | 0.333 | NA | 0.667 |
| SE | 11682 | ACADVL | ACADVL | chr17 | + | 7220923 | 7221058 | 7220765 | 7220830 | 7221537 | 7221682 | 0 | 0 | 10 | 0 | 0 | 1 | 0 | 6 | 0 | 0 | 0 | 0 | 98 | 49 | NA | NA | NA | NA | 0.833 | NA | | | 1 | NA | -0.167 |
| SE | 11686 | HLA-DQB1 | HLA-DQB1 | chr6 | - | 32661346 | 32661457 | 32659463 | 32660249 | 32661966 | 32662248 | 0 | 0 | 4 | 0 | 0 | 2 | 0 | 1 | 0 | 0 | 0 | 0 | 98 | 49 | NA | NA | NA | NA | 0.5 | NA | | | 1 | NA | -0.5 |
| SE | 11705 | TNFSF13 | TNFSF13 | chr17 | + | 7559845 | 7559893 | 7559623 | 7559702 | 7560048 | 7560167 | 0 | 0 | 11 | 0 | 0 | 1 | 0 | 0 | 0 | 0 | 1 | 0 | 97 | 49 | NA | NA | NA | NA | 0.847 | NA | | | 0 | NA | 0.847 |
| SE | 11728 | PER1 | PER1 | chr17 | - | 8145957 | 8146137 | 8144750 | 8144993 | 8146371 | 8146502 | 0 | 0 | 3 | 0 | 0 | 1 | 0 | 1 | 0 | 0 | 6 | 0 | 98 | 49 | NA | NA | NA | NA | 0.6 | NA | | | 0.077 | NA | 0.523 |
| SE | 11735 | UBE2Q2 | UBE2Q2 | chr15 | + | 75859877 | 75859982 | 75854385 | 75854487 | 75868950 | 75869010 | 0 | 0 | 5 | 0 | 0 | 1 | 0 | 1 | 0 | 0 | 0 | 0 | 98 | 49 | NA | NA | NA | NA | 0.714 | NA | | | 1 | NA | -0.286 |
| SE | 11745 | MOB1A | MOB1A | chr2 | - | 74159090 | 74159254 | 74156083 | 74156645 | 74165217 | 74165351 | 0 | 0 | 22 | 0 | 0 | 0 | 0 | 15 | 0 | 0 | 1 | 0 | 98 | 49 | NA | NA | NA | NA | 1 | NA | | | 0.882 | NA | 0.118 |
| SE | 11747 | COPS3 | COPS3 | chr17 | - | 17247479 | 17247560 | 17246623 | 17247151 | 17248925 | 17249039 | 0 | 0 | 1 | 0 | 0 | 2 | 0 | 5 | 0 | 0 | 0 | 0 | 98 | 49 | NA | NA | NA | NA | 0.2 | NA | | | 1 | NA | -0.8 |
| SE | 11764 | LOC100288778 | LOC100288778 | chr12 | - | 18372 | 18484 | 18036 | 18183 | 26800 | 26954 | 0 | 0 | 0 | 0 | 0 | 1 | 0 | 2 | 0 | 0 | 0 | 0 | 98 | 49 | NA | NA | NA | NA | 0 | NA | | | 1 | NA | -1 |
| SE | 11766 | KDM5A | KDM5A | chr12 | - | 306945 | 307089 | 297040 | 297200 | 307453 | 308005 | 0 | 0 | 1 | 0 | 0 | 1 | 0 | 2 | 0 | 0 | 0 | 0 | 98 | 49 | NA | NA | NA | NA | 0.333 | NA | | | 1 | NA | -0.667 |
| SE | 11780 | PNPLA6 | PNPLA6 | chr19 | + | 7541983 | 7542067 | 7541521 | 7541684 | 7542560 | 7542670 | 0 | 0 | 5 | 0 | 0 | 0 | 0 | 0 | 0 | 0 | 1 | 0 | 98 | 49 | NA | NA | NA | NA | 1 | NA | | | 0 | NA | 1 |
| SE | 11802 | KIAA0100 | KIAA0100 | chr17 | - | 28615663 | 28615798 | 28614439 | 28615246 | 28616095 | 28616197 | 0 | 0 | 5 | 0 | 0 | 0 | 0 | 1 | 0 | 0 | 1 | 0 | 98 | 49 | NA | NA | NA | NA | 1 | NA | | | 0.333 | NA | 0.667 |
| SE | 11803 | KIAA0100 | KIAA0100 | chr17 | - | 28616888 | 28616999 | 28616617 | 28616753 | 28617231 | 28617318 | 0 | 0 | 7 | 0 | 0 | 1 | 0 | 2 | 0 | 0 | 0 | 0 | 98 | 49 | NA | NA | NA | NA | 0.778 | NA | | | 1 | NA | -0.222 |
| SE | 11804 | KIAA0100 | KIAA0100 | chr17 | - | 28628264 | 28628559 | 28624232 | 28624390 | 28631478 | 28631681 | 0 | 0 | 7 | 0 | 0 | 1 | 0 | 1 | 0 | 0 | 1 | 0 | 98 | 49 | NA | NA | NA | NA | 0.778 | NA | | | 0.333 | NA | 0.445 |
| SE | 11815 | FLOT2 | FLOT2 | chr17 | - | 28884224 | 28884315 | 28883107 | 28883231 | 28888944 | 28889026 | 0 | 0 | 13 | 0 | 0 | 2 | 2 | 3 | 0 | 0 | 0 | 0 | 98 | 49 | NA | NA | NA | NA | 0.765 | | 1 | | 1 | NA | -0.235 |
| SE | 11816 | DHRS13 | DHRS13 | chr17 | - | 28901492 | 28901616 | 28900989 | 28901301 | 28902582 | 28902701 | 0 | 0 | 5 | 0 | 0 | 3 | 0 | 1 | 0 | 0 | 0 | 0 | 98 | 49 | NA | NA | NA | NA | 0.455 | NA | | | 1 | NA | -0.545 |
| SE | 11817 | PHF12 | PHF12 | chr17 | - | 28908782 | 28908881 | 28907589 | 28907672 | 28910225 | 28910369 | 0 | 0 | 5 | 0 | 0 | 1 | 0 | 5 | 0 | 0 | 2 | 0 | 98 | 49 | NA | NA | NA | NA | 0.714 | NA | | | 0.556 | NA | 0.158 |
| SE | 11818 | MAP3K3 | MAP3K3 | chr17 | + | 63634708 | 63634801 | 63632680 | 63632802 | 63646033 | 63646074 | 0 | 0 | 4 | 0 | 0 | 2 | 1 | 3 | 0 | 2 | 0 | 0 | 98 | 49 | NA | NA | NA | NA | 0.5 | | 0.2 | | 1 | NA | -0.1 |
| SE | 11819 | PROK2 | PROK2 | chr3 | - | 71774444 | 71774507 | 71771654 | 71772828 | 71781466 | 71781592 | 0 | 0 | 2 | 0 | 0 | 4 | 8 | 10 | 0 | 0 | 0 | 0 | 98 | 49 | NA | NA | NA | NA | 0.2 | | 1 | | 1 | NA | -0.8 |
| SE | 11833 | NCOA1 | NCOA1 | chr2 | + | 24693251 | 24693347 | 24691480 | 24691660 | 24697657 | 24697798 | 0 | 0 | 8 | 0 | 0 | 0 | 1 | 3 | 0 | 0 | 1 | 0 | 98 | 49 | NA | NA | NA | NA | 1 | | 1 | | 0.6 | NA | 0.2 |
| SE | 11843 | RHOT1 | RHOT1 | chr17 | + | 32182756 | 32182865 | 32176160 | 32176213 | 32183170 | 32183272 | 0 | 0 | 6 | 0 | 0 | 0 | 7 | 5 | 0 | 0 | 1 | 0 | 98 | 49 | NA | NA | NA | NA | 1 | | 1 | | 0.714 | NA | 0.143 |
| SE | 11845 | RHOT1 | RHOT1 | chr17 | + | 32209349 | 32209445 | 32208106 | 32208309 | 32224615 | 32225771 | 0 | 0 | 1 | 0 | 0 | 2 | 0 | 2 | 0 | 0 | 5 | 0 | 98 | 49 | NA | NA | NA | NA | 0.2 | NA | | | 0.167 | NA | 0.033 |
| SE | 11846 | RHOT1 | RHOT1 | chr17 | + | 32211115 | 32211238 | 32208106 | 32208309 | 32224615 | 32225771 | 0 | 0 | 2 | 0 | 0 | 2 | 0 | 1 | 0 | 0 | 5 | 0 | 98 | 49 | NA | NA | NA | NA | 0.333 | NA | | | 0.091 | NA | 0.242 |
| SE | 11848 | RHOT1 | RHOT1 | chr17 | + | 32221048 | 32221186 | 32211115 | 32211238 | 32224615 | 32225771 | 0 | 0 | 2 | 0 | 0 | 2 | 0 | 0 | 0 | 0 | 1 | 0 | 98 | 49 | NA | NA | NA | NA | 0.333 | NA | | | 0 | NA | 0.333 |
| SE | 11850 | ARFGAP1 | ARFGAP1 | chr20 | + | 63277204 | 63277305 | 63276479 | 63276651 | 63278116 | 63278203 | 0 | 0 | 1 | 0 | 0 | 0 | 0 | 2 | 0 | 0 | 2 | 0 | 98 | 49 | NA | NA | NA | NA | 1 | NA | | | 0.333 | NA | 0.667 |
| SE | 11869 | TPGS2 | TPGS2 | chr18 | - | 36805373 | 36805397 | 36800197 | 36800311 | 36807846 | 36807934 | 0 | 0 | 1 | 0 | 0 | 0 | 0 | 0 | 0 | 1 | 0 | 0 | 73 | 49 | NA | NA | NA | NA | 1 | | 0 | NA | | NA | 1 |
| SE | 11870 | TPGS2 | TPGS2 | chr18 | - | 36805373 | 36805502 | 36800197 | 36800311 | 36807846 | 36807934 | 0 | 0 | 1 | 0 | 0 | 0 | 0 | 0 | 0 | 1 | 0 | 0 | 98 | 49 | NA | NA | NA | NA | 1 | | 0 | NA | | NA | 1 |
| SE | 11881 | PHF20 | PHF20 | chr20 | + | 35870954 | 35871134 | 35869437 | 35869551 | 35871649 | 35871829 | 0 | 0 | 1 | 0 | 0 | 0 | 0 | 5 | 0 | 0 | 1 | 0 | 98 | 49 | NA | NA | NA | NA | 1 | NA | | | 0.714 | NA | 0.286 |
| SE | 11896 | KIAA0319L | KIAA0319L | chr1 | - | 35448172 | 35448332 | 35444160 | 35444303 | 35449866 | 35450005 | 0 | 0 | 5 | 0 | 0 | 0 | 0 | 1 | 0 | 1 | 0 | 0 | 98 | 49 | NA | NA | NA | NA | 1 | | 0 | | 1 | NA | 0.5 |
| SE | 11897 | KIAA0319L | KIAA0319L | chr1 | - | 35451627 | 35451776 | 35450357 | 35450509 | 35453556 | 35453689 | 0 | 0 | 0 | 0 | 0 | 1 | 0 | 1 | 0 | 0 | 0 | 0 | 98 | 49 | NA | NA | NA | NA | 0 | NA | | | 1 | NA | -1 |
| SE | 11904 | GGNBP2 | GGNBP2 | chr17 | + | 36557082 | 36557336 | 36554819 | 36554900 | 36560772 | 36560871 | 0 | 0 | 4 | 0 | 0 | 0 | 0 | 4 | 0 | 0 | 1 | 0 | 98 | 49 | NA | NA | NA | NA | 1 | NA | | | 0.667 | NA | 0.333 |
| SE | 11912 | SYNRG | SYNRG | chr17 | - | 37529782 | 37529851 | 37520537 | 37520648 | 37535978 | 37536127 | 0 | 0 | 0 | 0 | 0 | 2 | 0 | 1 | 0 | 0 | 1 | 0 | 98 | 49 | NA | NA | NA | NA | 0 | NA | | | 0.333 | NA | -0.333 |
| SE | 11917 | TNRC18 | TNRC18 | chr7 | - | 5361593 | 5361722 | 5359397 | 5359569 | 5361896 | 5362033 | 0 | 0 | 4 | 0 | 0 | 2 | 1 | 0 | 0 | 0 | 0 | 0 | 98 | 49 | NA | NA | NA | NA | 0.5 | | 1 | NA | | NA | -0.5 |
| SE | 11933 | KIAA0355 | KIAA0355 | chr19 | + | 34333703 | 34333801 | 34330188 | 34330347 | 34339866 | 34339992 | 0 | 0 | 1 | 0 | 0 | 0 | 0 | 2 | 0 | 0 | 1 | 0 | 98 | 49 | NA | NA | NA | NA | 1 | NA | | | 0.5 | NA | 0.5 |
| SE | 11957 | WSB1 | WSB1 | chr17 | + | 27301787 | 27301956 | 27294079 | 27294435 | 27303366 | 27303635 | 0 | 0 | 12 | 0 | 0 | 1 | 1 | 3 | 0 | 0 | 2 | 0 | 98 | 49 | NA | NA | NA | NA | 0.857 | | 1 | | 0.429 | NA | 0.142 |
| SE | 11959 | WSB1 | WSB1 | chr17 | + | 27310060 | 27310174 | 27309099 | 27309272 | 27311508 | 27311616 | 0 | 0 | 17 | 0 | 0 | 0 | 0 | 3 | 0 | 0 | 1 | 0 | 98 | 49 | NA | NA | NA | NA | 1 | NA | | | 0.6 | NA | 0.4 |
| SE | 11978 | MED13 | MED13 | chr17 | - | 61955679 | 61955838 | 61955381 | 61955567 | 61956338 | 61956481 | 0 | 0 | 2 | 0 | 0 | 1 | 0 | 1 | 0 | 0 | 0 | 0 | 98 | 49 | NA | NA | NA | NA | 0.5 | NA | | | 1 | NA | -0.5 |
| SE | 11988 | DNAJC7 | DNAJC7 | chr17 | - | 41989403 | 41989557 | 41988731 | 41988896 | 41990263 | 41990382 | 0 | 0 | 1 | 0 | 0 | 0 | 0 | 3 | 0 | 0 | 1 | 0 | 98 | 49 | NA | NA | NA | NA | 1 | NA | | | 0.6 | NA | 0.4 |
| SE | 12019 | UBTF | UBTF | chr17 | - | 44212343 | 44212454 | 44211872 | 44212006 | 44212818 | 44212939 | 0 | 0 | 1 | 0 | 0 | 1 | 1 | 0 | 0 | 0 | 2 | 0 | 98 | 49 | NA | NA | NA | NA | 0.333 | | 1 | | 0 | NA | -0.167 |
| SE | 12021 | CCDC43 | CCDC43 | chr17 | - | 44680584 | 44680643 | 44677436 | 44679043 | 44682002 | 44682138 | 0 | 0 | 1 | 0 | 0 | 0 | 0 | 0 | 0 | 0 | 1 | 0 | 98 | 49 | NA | NA | NA | NA | 1 | NA | | | 0 | NA | 1 |
| SE | 12025 | KLHL2 | KLHL2 | chr4 | + | 165297608 | 165297725 | 165294358 | 165294468 | 165299506 | 165299656 | 0 | 0 | 1 | 0 | 0 | 0 | 0 | 1 | 0 | 0 | 2 | 0 | 98 | 49 | NA | NA | NA | NA | 1 | NA | | | 0.2 | NA | 0.8 |
| SE | 12114 | ERMARD | ERMARD | chr6 | + | 169766637 | 169766667 | 169762428 | 169762531 | 169768102 | 169768171 | 0 | 0 | 1 | 0 | 0 | 1 | 0 | 1 | 0 | 0 | 0 | 0 | 79 | 49 | NA | NA | NA | NA | 0.383 | NA | | | 1 | NA | -0.617 |
| SE | 12145 | BZW2 | BZW2 | chr7 | + | 16698047 | 16698186 | 16696914 | 16697061 | 16704546 | 16704669 | 0 | 0 | 1 | 0 | 0 | 0 | 0 | 0 | 0 | 0 | 1 | 0 | 98 | 49 | NA | NA | NA | NA | 1 | NA | | | 0 | NA | 1 |
| SE | 12154 | CC2D1B | CC2D1B | chr1 | - | 52359704 | 52359883 | 52359458 | 52359534 | 52360073 | 52360233 | 0 | 0 | 1 | 0 | 0 | 1 | 0 | 2 | 0 | 0 | 0 | 0 | 98 | 49 | NA | NA | NA | NA | 0.333 | NA | | | 1 | NA | -0.667 |
| SE | 12158 | VMP1 | VMP1 | chr17 | + | 59738836 | 59738947 | 59735337 | 59735473 | 59764970 | 59765138 | 0 | 0 | 29 | 0 | 0 | 1 | 0 | 7 | 0 | 0 | 0 | 0 | 98 | 49 | NA | NA | NA | NA | 0.935 | NA | | | 1 | NA | -0.065 |
| SE | 12213 | HELZ | HELZ | chr17 | - | 67122969 | 67123160 | 67120404 | 67120612 | 67123962 | 67124014 | 0 | 0 | 3 | 0 | 0 | 0 | 0 | 0 | 0 | 0 | 1 | 0 | 98 | 49 | NA | NA | NA | NA | 1 | NA | | | 0 | NA | 1 |
| SE | 12214 | HELZ | HELZ | chr17 | - | 67167462 | 67167796 | 67166477 | 67166608 | 67178658 | 67178926 | 0 | 0 | 2 | 0 | 0 | 0 | 1 | 3 | 0 | 2 | 0 | 0 | 98 | 49 | NA | NA | NA | NA | 1 | | 0.2 | | 1 | NA | 0.4 |
| SE | 12222 | PRKAR1A | PRKAR1A | chr17 | + | 68514986 | 68515119 | 68512378 | 68512548 | 68515393 | 68515576 | 0 | 0 | 3 | 0 | 0 | 1 | 0 | 0 | 0 | 0 | 1 | 0 | 98 | 49 | NA | NA | NA | NA | 0.6 | NA | | | 0 | NA | 0.6 |
| SE | 12236 | RNASE4 | RNASE4 | chr14 | + | 20693546 | 20693650 | 20684212 | 20684758 | 20699354 | 20700602 | 0 | 0 | 0 | 0 | 0 | 1 | 0 | 1 | 0 | 0 | 0 | 0 | 98 | 49 | NA | NA | NA | NA | 0 | NA | | | 1 | NA | -1 |
| SE | 12242 | ACOX1 | ACOX1 | chr17 | - | 75949717 | 75949897 | 75949494 | 75949600 | 75950773 | 75950964 | 0 | 0 | 3 | 0 | 0 | 0 | 0 | 1 | 0 | 0 | 1 | 0 | 98 | 49 | NA | NA | NA | NA | 1 | NA | | | 0.333 | NA | 0.667 |
| SE | 12247 | OMA1 | OMA1 | chr1 | - | 58530600 | 58530729 | 58527260 | 58527335 | 58533952 | 58534060 | 0 | 0 | 1 | 0 | 0 | 0 | 0 | 3 | 0 | 0 | 1 | 0 | 98 | 49 | NA | NA | NA | NA | 1 | NA | | | 0.6 | NA | 0.4 |
| SE | 12269 | SUV420H1 | SUV420H1 | chr11 | - | 68180131 | 68180200 | 68175017 | 68175183 | 68185780 | 68185928 | 0 | 0 | 8 | 0 | 0 | 0 | 0 | 1 | 0 | 0 | 1 | 0 | 98 | 49 | NA | NA | NA | NA | 1 | NA | | | 0.333 | NA | 0.667 |
| SE | 12287 | JMJD6 | JMJD6 | chr17 | - | 76723771 | 76724058 | 76721797 | 76721933 | 76725466 | 76725855 | 0 | 0 | 4 | 0 | 0 | 1 | 0 | 13 | 0 | 0 | 0 | 0 | 98 | 49 | NA | NA | NA | NA | 0.667 | NA | | | 1 | NA | -0.333 |
| SE | 12346 | CSNK1D | CSNK1D | chr17 | - | 82245975 | 82246039 | 82242660 | 82244831 | 82248874 | 82249014 | 0 | 0 | 2 | 0 | 0 | 1 | 0 | 1 | 0 | 0 | 1 | 0 | 98 | 49 | NA | NA | NA | NA | 0.5 | NA | | | 0.333 | NA | 0.167 |
| SE | 12384 | VAPA | VAPA | chr18 | + | 9944923 | 9945058 | 9936985 | 9937066 | 9950394 | 9950568 | 0 | 0 | 1 | 0 | 0 | 3 | 0 | 0 | 0 | 0 | 2 | 0 | 98 | 49 | NA | NA | NA | NA | 0.143 | NA | | | 0 | NA | 0.143 |
| SE | 12398 | GPS2 | GPS2 | chr17 | - | 7313905 | 7313988 | 7313567 | 7313721 | 7314079 | 7314159 | 0 | 2 | 0 | 0 | 0 | 0 | 0 | 0 | 0 | 0 | 0 | 2 | 98 | 49 | NA | NA | NA | | 1 | NA | NA | NA | 0 | | 1 |
| SE | 12428 | SS18 | SS18 | chr18 | - | 26035830 | 26035923 | 26035004 | 26035127 | 26038554 | 26038659 | 0 | 0 | 0 | 0 | 0 | 2 | 0 | 2 | 0 | 0 | 0 | 0 | 98 | 49 | NA | NA | NA | NA | 0 | NA | | | 1 | NA | -1 |
| SE | 12435 | SPAG9 | SPAG9 | chr17 | - | 50985697 | 50985778 | 50982523 | 50982672 | 50987111 | 50987237 | 0 | 0 | 7 | 0 | 0 | 1 | 0 | 8 | 0 | 0 | 0 | 0 | 98 | 49 | NA | NA | NA | NA | 0.778 | NA | | | 1 | NA | -0.222 |
| SE | 12438 | SPAG9 | SPAG9 | chr17 | - | 51031680 | 51031722 | 51021157 | 51021365 | 51041500 | 51041651 | 0 | 0 | 0 | 0 | 0 | 8 | 0 | 10 | 0 | 0 | 6 | 0 | 91 | 49 | NA | NA | NA | NA | 0 | NA | | | 0.473 | NA | -0.473 |
| SE | 12446 | TRAPPC8 | TRAPPC8 | chr18 | - | 31866848 | 31866975 | 31864626 | 31864781 | 31867401 | 31867476 | 0 | 0 | 5 | 0 | 0 | 0 | 0 | 0 | 0 | 0 | 2 | 0 | 98 | 49 | NA | NA | NA | NA | 1 | NA | | | 0 | NA | 1 |
| SE | 12496 | MBD1 | MBD1 | chr18 | - | 50274976 | 50275045 | 50274185 | 50274353 | 50275128 | 50275245 | 0 | 0 | 2 | 0 | 0 | 0 | 0 | 0 | 0 | 0 | 1 | 0 | 98 | 49 | NA | NA | NA | NA | 1 | NA | | | 0 | NA | 1 |
| SE | 12501 | HELQ | HELQ | chr4 | - | 83446834 | 83447035 | 83446013 | 83446086 | 83448782 | 83448961 | 0 | 0 | 0 | 0 | 0 | 1 | 0 | 1 | 0 | 0 | 0 | 0 | 98 | 49 | NA | NA | NA | NA | 0 | NA | | | 1 | NA | -1 |
| SE | 12525 | GUCD1 | GUCD1 | chr22 | - | 24547907 | 24548073 | 24546913 | 24547005 | 24554948 | 24555307 | 0 | 0 | 1 | 0 | 0 | 1 | 0 | 5 | 0 | 0 | 0 | 0 | 98 | 49 | NA | NA | NA | NA | 0.333 | NA | | | 1 | NA | -0.667 |
| SE | 12534 | VPS4B | VPS4B | chr18 | - | 63410289 | 63410446 | 63407431 | 63407499 | 63411466 | 63411578 | 0 | 0 | 7 | 0 | 0 | 0 | 0 | 5 | 0 | 0 | 1 | 0 | 98 | 49 | NA | NA | NA | NA | 1 | NA | | | 0.714 | NA | 0.286 |
| SE | 12564 | EIF4G1 | EIF4G1 | chr3 | + | 184315488 | 184315545 | 184314494 | 184314674 | 184315762 | 184315856 | 0 | 0 | 2 | 0 | 0 | 1 | 0 | 1 | 0 | 0 | 0 | 0 | 98 | 49 | NA | NA | NA | NA | 0.5 | NA | | | 1 | NA | -0.5 |
| SE | 12568 | EIF4G1 | EIF4G1 | chr3 | + | 184328908 | 184328990 | 184328630 | 184328756 | 184331265 | 184331364 | 0 | 0 | 9 | 0 | 0 | 0 | 2 | 5 | 0 | 0 | 1 | 0 | 98 | 49 | NA | NA | NA | NA | 1 | | 1 | | 0.714 | NA | 0.143 |
| SE | 12570 | PTBP1 | PTBP1 | chr19 | + | 805491 | 805569 | 805012 | 805187 | 806407 | 806556 | 0 | 0 | 1 | 0 | 0 | 0 | 0 | 0 | 0 | 0 | 1 | 0 | 98 | 49 | NA | NA | NA | NA | 1 | NA | | | 0 | NA | 1 |
| SE | 12571 | PTBP1 | PTBP1 | chr19 | + | 805512 | 805569 | 805012 | 805187 | 806407 | 806556 | 0 | 0 | 1 | 0 | 0 | 0 | 0 | 0 | 0 | 0 | 1 | 0 | 98 | 49 | NA | NA | NA | NA | 1 | NA | | | 0 | NA | 1 |
| SE | 12572 | PTBP1 | PTBP1 | chr19 | + | 807868 | 807902 | 806407 | 806556 | 808359 | 808452 | 0 | 0 | 3 | 0 | 0 | 1 | 0 | 1 | 0 | 0 | 0 | 0 | 83 | 49 | NA | NA | NA | NA | 0.639 | NA | | | 1 | NA | -0.361 |
| SE | 12577 | ARID3A | ARID3A | chr19 | + | 966571 | 966868 | 964247 | 964431 | 968404 | 968503 | 0 | 0 | 0 | 0 | 0 | 1 | 0 | 1 | 0 | 0 | 0 | 0 | 98 | 49 | NA | NA | NA | NA | 0 | NA | | | 1 | NA | -1 |
| SE | 12596 | GBF1 | GBF1 | chr10 | + | 102369860 | 102369984 | 102369710 | 102369775 | 102370173 | 102370245 | 0 | 0 | 1 | 0 | 0 | 0 | 0 | 1 | 0 | 0 | 2 | 0 | 98 | 49 | NA | NA | NA | NA | 1 | NA | | | 0.2 | NA | 0.8 |
| SE | 12597 | GBF1 | GBF1 | chr10 | + | 102381126 | 102381255 | 102380505 | 102380686 | 102382055 | 102382899 | 0 | 0 | 1 | 0 | 0 | 0 | 0 | 7 | 0 | 0 | 1 | 0 | 98 | 49 | NA | NA | NA | NA | 1 | NA | | | 0.778 | NA | 0.222 |
| SE | 12600 | CCT4 | CCT4 | chr2 | - | 61872457 | 61872588 | 61872081 | 61872316 | 61873001 | 61873112 | 0 | 0 | 0 | 0 | 0 | 1 | 0 | 2 | 0 | 0 | 0 | 0 | 98 | 49 | NA | NA | NA | NA | 0 | NA | | | 1 | NA | -1 |
| SE | 12632 | SAFB2 | SAFB2 | chr19 | - | 5611119 | 5611630 | 5610638 | 5610688 | 5612539 | 5612567 | 0 | 0 | 4 | 0 | 0 | 0 | 0 | 1 | 0 | 0 | 1 | 0 | 98 | 49 | NA | NA | NA | NA | 1 | NA | | | 0.333 | NA | 0.667 |
| SE | 12638 | ZNF638 | ZNF638 | chr2 | + | 71426459 | 71427414 | 71424649 | 71424715 | 71428546 | 71428651 | 0 | 0 | 2 | 0 | 0 | 0 | 0 | 2 | 0 | 0 | 1 | 0 | 98 | 49 | NA | NA | NA | NA | 1 | NA | | | 0.5 | NA | 0.5 |
| SE | 12639 | ZNF638 | ZNF638 | chr2 | + | 71428546 | 71428651 | 71426459 | 71427414 | 71431326 | 71431428 | 0 | 0 | 1 | 0 | 0 | 0 | 0 | 0 | 0 | 0 | 1 | 0 | 98 | 49 | NA | NA | NA | NA | 1 | NA | | | 0 | NA | 1 |
| SE | 12647 | GTF2F1 | GTF2F1 | chr19 | - | 6383310 | 6383495 | 6381696 | 6381850 | 6387388 | 6387559 | 0 | 0 | 1 | 0 | 0 | 1 | 0 | 1 | 0 | 0 | 0 | 0 | 98 | 49 | NA | NA | NA | NA | 0.333 | NA | | | 1 | NA | -0.667 |
| SE | 12651 | ITGA4 | ITGA4 | chr2 | + | 181530523 | 181530649 | 181529540 | 181529648 | 181531656 | 181531776 | 0 | 0 | 7 | 0 | 0 | 2 | 0 | 1 | 0 | 0 | 0 | 0 | 98 | 49 | NA | NA | NA | NA | 0.636 | NA | | | 1 | NA | -0.364 |
| SE | 12659 | PPP2R1A | PPP2R1A | chr19 | + | 52220188 | 52220249 | 52219690 | 52219864 | 52220978 | 52221133 | 0 | 0 | 6 | 0 | 0 | 1 | 0 | 1 | 0 | 0 | 0 | 0 | 98 | 49 | NA | NA | NA | NA | 0.75 | NA | | | 1 | NA | -0.25 |
| SE | 12663 | PET100 | PET100 | chr19 | + | 7631231 | 7631321 | 7630822 | 7630846 | 7631472 | 7631624 | 0 | 0 | 0 | 0 | 0 | 8 | 0 | 1 | 0 | 0 | 4 | 0 | 98 | 49 | NA | NA | NA | NA | 0 | NA | | | 0.111 | NA | -0.111 |
| SE | 12694 | UTRN | UTRN | chr6 | + | 144437564 | 144437746 | 144435934 | 144436138 | 144438744 | 144438895 | 0 | 0 | 19 | 0 | 0 | 1 | 0 | 10 | 0 | 0 | 0 | 0 | 98 | 49 | NA | NA | NA | NA | 0.905 | NA | | | 1 | NA | -0.095 |
| SE | 12695 | UTRN | UTRN | chr6 | + | 144827347 | 144827386 | 144820881 | 144821018 | 144827610 | 144827676 | 0 | 0 | 2 | 0 | 0 | 2 | 0 | 0 | 0 | 0 | 1 | 0 | 88 | 49 | NA | NA | NA | NA | 0.358 | NA | | | 0 | NA | 0.358 |
| SE | 12702 | ARFGEF1 | ARFGEF1 | chr8 | - | 67277281 | 67277457 | 67271701 | 67271936 | 67287954 | 67288065 | 0 | 0 | 5 | 0 | 0 | 1 | 0 | 2 | 0 | 0 | 3 | 0 | 98 | 49 | NA | NA | NA | NA | 0.714 | NA | | | 0.25 | NA | 0.464 |
| SE | 12720 | GHITM | GHITM | chr10 | + | 84148729 | 84148838 | 84144874 | 84145016 | 84150054 | 84150243 | 0 | 0 | 7 | 0 | 0 | 0 | 0 | 9 | 0 | 0 | 1 | 0 | 98 | 49 | NA | NA | NA | NA | 1 | NA | | | 0.818 | NA | 0.182 |
| SE | 12727 | C19orf43 | C19orf43 | chr19 | - | 12731343 | 12731421 | 12730639 | 12731068 | 12734312 | 12734715 | 0 | 0 | 5 | 0 | 0 | 1 | 0 | 9 | 0 | 0 | 0 | 0 | 98 | 49 | NA | NA | NA | NA | 0.714 | NA | | | 1 | NA | -0.286 |
| SE | 12747 | DYSF | DYSF | chr2 | + | 71669111 | 71669207 | 71668753 | 71668842 | 71669604 | 71669746 | 0 | 0 | 17 | 0 | 0 | 3 | 0 | 2 | 0 | 0 | 2 | 0 | 98 | 49 | NA | NA | NA | NA | 0.739 | NA | | | 0.333 | NA | 0.406 |
| SE | 12749 | COL4A3BP | COL4A3BP | chr5 | - | 75384641 | 75384712 | 75381948 | 75382077 | 75385901 | 75386034 | 0 | 0 | 4 | 0 | 0 | 0 | 0 | 1 | 0 | 0 | 2 | 0 | 98 | 49 | NA | NA | NA | NA | 1 | NA | | | 0.2 | NA | 0.8 |
| SE | 12751 | COL4A3BP | COL4A3BP | chr5 | - | 75411010 | 75411103 | 75402971 | 75403058 | 75416875 | 75417033 | 0 | 0 | 1 | 0 | 0 | 0 | 0 | 2 | 0 | 0 | 1 | 0 | 98 | 49 | NA | NA | NA | NA | 1 | NA | | | 0.5 | NA | 0.5 |
| SE | 12802 | CYP4F3 | CYP4F3 | chr19 | + | 15643906 | 15644051 | 15641414 | 15641613 | 15647051 | 15647105 | 0 | 0 | 1 | 0 | 0 | 0 | 0 | 0 | 0 | 0 | 3 | 0 | 98 | 49 | NA | NA | NA | NA | 1 | NA | | | 0 | NA | 1 |
| SE | 12828 | SSBP4 | SSBP4 | chr19 | + | 18431352 | 18431418 | 18430840 | 18430930 | 18431646 | 18431706 | 0 | 0 | 0 | 0 | 0 | 1 | 0 | 1 | 0 | 0 | 1 | 0 | 98 | 49 | NA | NA | NA | NA | 0 | NA | | | 0.333 | NA | -0.333 |
| SE | 12852 | GATAD2A | GATAD2A | chr19 | + | 19501968 | 19502043 | 19501117 | 19501416 | 19502330 | 19502526 | 0 | 0 | 3 | 0 | 0 | 0 | 0 | 4 | 0 | 0 | 1 | 0 | 98 | 49 | NA | NA | NA | NA | 1 | NA | | | 0.667 | NA | 0.333 |
| SE | 12854 | ZNF101 | ZNF101 | chr19 | + | 19678725 | 19678786 | 19677863 | 19677990 | 19679180 | 19683506 | 0 | 0 | 1 | 0 | 0 | 1 | 1 | 4 | 0 | 0 | 0 | 0 | 98 | 49 | NA | NA | NA | NA | 0.333 | | 1 | | 1 | NA | -0.667 |
| SE | 12915 | PITPNM1 | PITPNM1 | chr11 | - | 67498939 | 67499001 | 67498595 | 67498846 | 67499722 | 67499830 | 0 | 0 | 2 | 0 | 0 | 0 | 0 | 2 | 0 | 0 | 1 | 0 | 98 | 49 | NA | NA | NA | NA | 1 | NA | | | 0.5 | NA | 0.5 |
| SE | 12934 | EIF3K | EIF3K | chr19 | + | 38620336 | 38620435 | 38619073 | 38619327 | 38624076 | 38624197 | 0 | 0 | 8 | 0 | 0 | 1 | 0 | 10 | 0 | 0 | 0 | 0 | 98 | 49 | NA | NA | NA | NA | 0.8 | NA | | | 1 | NA | -0.2 |
| SE | 12935 | EIF3K | EIF3K | chr19 | + | 38632600 | 38632678 | 38632429 | 38632496 | 38634992 | 38635118 | 0 | 0 | 5 | 0 | 0 | 1 | 1 | 5 | 0 | 0 | 0 | 0 | 98 | 49 | NA | NA | NA | NA | 0.714 | | 1 | | 1 | NA | -0.286 |
| SE | 12953 | MAN2A2 | MAN2A2 | chr15 | + | 90910089 | 90910292 | 90909326 | 90909504 | 90910500 | 90910683 | 0 | 0 | 8 | 0 | 0 | 1 | 0 | 5 | 0 | 0 | 0 | 0 | 98 | 49 | NA | NA | NA | NA | 0.8 | NA | | | 1 | NA | -0.2 |
| SE | 12982 | CIC | CIC | chr19 | + | 42291557 | 42291745 | 42290232 | 42291466 | 42292085 | 42292207 | 0 | 0 | 1 | 0 | 0 | 2 | 0 | 1 | 0 | 0 | 0 | 0 | 98 | 49 | NA | NA | NA | NA | 0.2 | NA | | | 1 | NA | -0.8 |
| SE | 12983 | ATG13 | ATG13 | chr11 | + | 46618324 | 46618375 | 46617275 | 46617890 | 46644278 | 46644360 | 0 | 0 | 2 | 0 | 0 | 0 | 0 | 0 | 0 | 0 | 1 | 0 | 98 | 49 | NA | NA | NA | NA | 1 | NA | | | 0 | NA | 1 |
| SE | 12984 | ATG13 | ATG13 | chr11 | + | 46630044 | 46630100 | 46617275 | 46617890 | 46644278 | 46644360 | 0 | 0 | 2 | 0 | 0 | 0 | 0 | 0 | 0 | 0 | 1 | 0 | 98 | 49 | NA | NA | NA | NA | 1 | NA | | | 0 | NA | 1 |
| SE | 13135 | OCIAD1 | OCIAD1 | chr4 | + | 48833400 | 48833481 | 48832619 | 48832682 | 48842635 | 48842689 | 0 | 0 | 8 | 0 | 0 | 0 | 0 | 1 | 0 | 1 | 0 | 0 | 98 | 49 | NA | NA | NA | NA | 1 | | 0 | | 1 | NA | 0.5 |
| SE | 13139 | CNOT10 | CNOT10 | chr3 | + | 32713226 | 32713369 | 32708669 | 32708820 | 32716224 | 32716311 | 0 | 0 | 3 | 0 | 0 | 0 | 0 | 0 | 0 | 0 | 2 | 0 | 98 | 49 | NA | NA | NA | NA | 1 | NA | | | 0 | NA | 1 |
| SE | 13156 | NFRKB | NFRKB | chr11 | - | 129872883 | 129873096 | 129869493 | 129870261 | 129873744 | 129874015 | 0 | 0 | 1 | 0 | 0 | 0 | 0 | 0 | 0 | 0 | 1 | 0 | 98 | 49 | NA | NA | NA | NA | 1 | NA | | | 0 | NA | 1 |
| SE | 13211 | LILRB2 | LILRB2 | chr19 | - | 54280462 | 54280544 | 54279790 | 54280075 | 54280960 | 54281178 | 0 | 0 | 0 | 0 | 0 | 1 | 0 | 1 | 0 | 0 | 1 | 0 | 98 | 49 | NA | NA | NA | NA | 0 | NA | | | 0.333 | NA | -0.333 |
| SE | 13238 | PAFAH1B2 | PAFAH1B2 | chr11 | + | 117161144 | 117161261 | 117159933 | 117160023 | 117163769 | 117163874 | 0 | 0 | 6 | 0 | 0 | 2 | 0 | 10 | 0 | 0 | 0 | 0 | 98 | 49 | NA | NA | NA | NA | 0.6 | NA | | | 1 | NA | -0.4 |
| SE | 13358 | BROX | BROX | chr1 | + | 222715683 | 222715800 | 222713320 | 222713416 | 222718924 | 222719031 | 0 | 0 | 1 | 0 | 0 | 0 | 0 | 0 | 0 | 0 | 1 | 0 | 98 | 49 | NA | NA | NA | NA | 1 | NA | | | 0 | NA | 1 |
| SE | 13366 | FIP1L1 | FIP1L1 | chr4 | + | 53379072 | 53379117 | 53377652 | 53377923 | 53379224 | 53379264 | 0 | 0 | 0 | 0 | 0 | 3 | 0 | 1 | 0 | 0 | 0 | 0 | 94 | 49 | NA | NA | NA | NA | 0 | NA | | | 1 | NA | -1 |
| SE | 13382 | PRKDC | PRKDC | chr8 | - | 47836337 | 47836527 | 47834195 | 47834396 | 47837211 | 47837419 | 0 | 0 | 11 | 0 | 0 | 2 | 0 | 3 | 0 | 0 | 0 | 0 | 98 | 49 | NA | NA | NA | NA | 0.733 | NA | | | 1 | NA | -0.267 |
| SE | 13385 | HNRNPH3 | HNRNPH3 | chr10 | + | 68337857 | 68337996 | 68337198 | 68337333 | 68338502 | 68338642 | 0 | 0 | 1 | 0 | 0 | 0 | 0 | 1 | 0 | 0 | 1 | 0 | 98 | 49 | NA | NA | NA | NA | 1 | NA | | | 0.333 | NA | 0.667 |
| SE | 13424 | TBC1D5 | TBC1D5 | chr3 | - | 17428449 | 17428519 | 17406417 | 17406526 | 17508473 | 17508605 | 0 | 0 | 2 | 0 | 0 | 0 | 5 | 0 | 0 | 0 | 1 | 0 | 98 | 49 | NA | NA | NA | NA | 1 | | 1 | | 0 | NA | 0.5 |
| SE | 13449 | ADAM17 | ADAM17 | chr2 | - | 9535833 | 9535922 | 9527785 | 9527954 | 9536697 | 9536828 | 0 | 0 | 3 | 0 | 0 | 1 | 0 | 1 | 0 | 0 | 0 | 0 | 98 | 49 | NA | NA | NA | NA | 0.6 | NA | | | 1 | NA | -0.4 |
| SE | 13482 | C10orf128 | C10orf128 | chr10 | - | 49165765 | 49165796 | 49161598 | 49161622 | 49166868 | 49167009 | 0 | 0 | 1 | 0 | 0 | 1 | 0 | 1 | 0 | 0 | 0 | 0 | 80 | 49 | NA | NA | NA | NA | 0.38 | NA | | | 1 | NA | -0.62 |
| SE | 13484 | FAM228B | FAM228B | chr2 | + | 24137908 | 24138100 | 24135118 | 24135187 | 24139369 | 24139450 | 0 | 0 | 1 | 0 | 0 | 0 | 0 | 0 | 0 | 0 | 1 | 0 | 98 | 49 | NA | NA | NA | NA | 1 | NA | | | 0 | NA | 1 |
| SE | 13487 | ITSN2 | ITSN2 | chr2 | - | 24284762 | 24284843 | 24275712 | 24275849 | 24286211 | 24286351 | 0 | 0 | 0 | 0 | 0 | 3 | 2 | 1 | 0 | 0 | 5 | 0 | 98 | 49 | NA | NA | NA | NA | 0 | | 1 | | 0.091 | NA | -0.545 |
| SE | 13499 | HADHB | HADHB | chr2 | + | 26260980 | 26261063 | 26254429 | 26254474 | 26263379 | 26263479 | 0 | 0 | 1 | 0 | 0 | 2 | 0 | 0 | 0 | 0 | 1 | 0 | 98 | 49 | NA | NA | NA | NA | 0.2 | NA | | | 0 | NA | 0.2 |
| SE | 13524 | BRE | BRE | chr2 | + | 27892382 | 27892448 | 27890614 | 27890842 | 27894532 | 27894684 | 0 | 0 | 1 | 0 | 0 | 3 | 0 | 2 | 0 | 0 | 0 | 0 | 98 | 49 | NA | NA | NA | NA | 0.143 | NA | | | 1 | NA | -0.857 |
| SE | 13565 | EML4 | EML4 | chr2 | + | 42301240 | 42301392 | 42295380 | 42295516 | 42303103 | 42303229 | 0 | 0 | 6 | 0 | 0 | 0 | 0 | 7 | 0 | 0 | 1 | 0 | 98 | 49 | NA | NA | NA | NA | 1 | NA | | | 0.778 | NA | 0.222 |
| SE | 13566 | RNF144B | RNF144B | chr6 | + | 18457154 | 18457359 | 18439683 | 18439744 | 18459606 | 18459751 | 0 | 0 | 9 | 0 | 0 | 3 | 0 | 1 | 0 | 3 | 0 | 0 | 98 | 49 | NA | NA | NA | NA | 0.6 | | 0 | | 1 | NA | 0.1 |
| SE | 13567 | THADA | THADA | chr2 | - | 43560233 | 43560385 | 43556092 | 43556555 | 43566697 | 43566821 | 0 | 0 | 5 | 0 | 0 | 0 | 0 | 1 | 0 | 4 | 0 | 0 | 98 | 49 | NA | NA | NA | NA | 1 | | 0 | | 1 | NA | 0.5 |
| SE | 13593 | VRK2 | VRK2 | chr2 | + | 58123100 | 58123233 | 58089630 | 58089723 | 58131807 | 58131928 | 0 | 0 | 2 | 0 | 0 | 0 | 0 | 1 | 0 | 0 | 1 | 0 | 98 | 49 | NA | NA | NA | NA | 1 | NA | | | 0.333 | NA | 0.667 |
| SE | 13594 | VRK2 | VRK2 | chr2 | + | 58135140 | 58135199 | 58131807 | 58131928 | 58139665 | 58139832 | 0 | 0 | 2 | 0 | 0 | 0 | 0 | 0 | 0 | 0 | 3 | 0 | 98 | 49 | NA | NA | NA | NA | 1 | NA | | | 0 | NA | 1 |
| SE | 13595 | VRK2 | VRK2 | chr2 | + | 58154753 | 58154802 | 58146315 | 58146474 | 58159348 | 58159920 | 0 | 0 | 2 | 0 | 0 | 1 | 0 | 0 | 0 | 0 | 1 | 0 | 98 | 49 | NA | NA | NA | NA | 0.5 | NA | | | 0 | NA | 0.5 |
| SE | 13607 | USP34 | USP34 | chr2 | - | 61192900 | 61192980 | 61190517 | 61190658 | 61203139 | 61203263 | 0 | 0 | 3 | 0 | 0 | 1 | 0 | 1 | 0 | 0 | 0 | 0 | 98 | 49 | NA | NA | NA | NA | 0.6 | NA | | | 1 | NA | -0.4 |
| SE | 13608 | XPO1 | XPO1 | chr2 | - | 61492914 | 61493053 | 61492566 | 61492748 | 61493893 | 61494091 | 0 | 0 | 4 | 0 | 0 | 2 | 0 | 2 | 0 | 0 | 0 | 0 | 98 | 49 | NA | NA | NA | NA | 0.5 | NA | | | 1 | NA | -0.5 |
| SE | 13610 | USP47 | USP47 | chr11 | + | 11891967 | 11892106 | 11884466 | 11884580 | 11897596 | 11897693 | 0 | 0 | 1 | 0 | 0 | 2 | 0 | 7 | 0 | 0 | 0 | 0 | 98 | 49 | NA | NA | NA | NA | 0.2 | NA | | | 1 | NA | -0.8 |
| SE | 13636 | CCND3 | CCND3 | chr6 | - | 42030008 | 42030190 | 41937234 | 41937394 | 42048500 | 42048894 | 0 | 0 | 1 | 0 | 0 | 1 | 0 | 0 | 0 | 0 | 1 | 0 | 98 | 49 | NA | NA | NA | NA | 0.333 | NA | | | 0 | NA | 0.333 |
| SE | 13637 | NRBP1 | NRBP1 | chr2 | + | 27434470 | 27434560 | 27433988 | 27434090 | 27434721 | 27434762 | 0 | 0 | 5 | 0 | 0 | 1 | 0 | 4 | 0 | 0 | 0 | 0 | 98 | 49 | NA | NA | NA | NA | 0.714 | NA | | | 1 | NA | -0.286 |
| SE | 13643 | PCBP1-AS1 | PCBP1-AS1 | chr2 | - | 70083537 | 70083687 | 70059642 | 70059680 | 70085546 | 70085624 | 0 | 0 | 1 | 0 | 0 | 0 | 0 | 2 | 0 | 0 | 1 | 0 | 98 | 49 | NA | NA | NA | NA | 1 | NA | | | 0.5 | NA | 0.5 |
| SE | 13675 | TP53BP2 | TP53BP2 | chr1 | - | 223818205 | 223818337 | 223814239 | 223814353 | 223821219 | 223821367 | 0 | 0 | 1 | 0 | 0 | 3 | 0 | 0 | 0 | 0 | 1 | 0 | 98 | 49 | NA | NA | NA | NA | 0.143 | NA | | | 0 | NA | 0.143 |
| SE | 13704 | LINC00963 | LINC00963 | chr9 | + | 129502396 | 129503139 | 129493477 | 129493595 | 129503322 | 129503672 | 0 | 0 | 0 | 0 | 0 | 2 | 0 | 1 | 0 | 0 | 1 | 0 | 98 | 49 | NA | NA | NA | NA | 0 | NA | | | 0.333 | NA | -0.333 |

| | | | | | | | | | | | | | | | | | | | | | | | | | | | | | | | | | | |
|---|---|---|---|---|---|---|---|---|---|---|---|---|---|---|---|---|---|---|---|---|---|---|---|---|---|---|---|---|---|---|---|---|---|---|
| SE | 13714 | MTMR10 | MTMR10 | chr15 | - | 30951967 | 30952038 | 30948301 | 30948471 | 30953561 | 30953631 | 0 | 0 | 5 | 0 | 0 | 0 | 0 | 0 | 0 | 0 | 4 | 0 | 98 | 49 | NA | NA | NA | NA | 1 | NA | 0 | NA | 1 |
| SE | 13715 | MTMR10 | MTMR10 | chr15 | - | 30976818 | 30976955 | 30974930 | 30975003 | 30990776 | 30990837 | 0 | 0 | 4 | 0 | 0 | 2 | 0 | 2 | 0 | 0 | 0 | 0 | 98 | 49 | NA | NA | NA | NA | 0.5 | NA | 1 | NA | -0.5 |
| SE | 13746 | MAP4K4 | MAP4K4 | chr2 | + | 101887087 | 101887261 | 101885186 | 101885287 | 101887777 | 101887937 | 0 | 0 | 6 | 0 | 0 | 0 | 0 | 3 | 0 | 0 | 1 | 0 | 98 | 49 | NA | NA | NA | NA | 1 | NA | 0.6 | NA | 0.4 |
| SE | 13747 | MAP4K4 | MAP4K4 | chr2 | + | 101888795 | 101888935 | 101887777 | 101887937 | 101891165 | 101894690 | 0 | 0 | 8 | 0 | 0 | 0 | 3 | 5 | 0 | 0 | 1 | 0 | 98 | 49 | NA | NA | NA | NA | 1 | 1 | 0.714 | NA | 0.143 |
| SE | 13752 | HMBOX1 | HMBOX1 | chr8 | + | 28980070 | 28980156 | 28970042 | 28970519 | 29009071 | 29009182 | 0 | 0 | 2 | 0 | 0 | 0 | 0 | 0 | 0 | 0 | 1 | 0 | 98 | 49 | NA | NA | NA | NA | 1 | NA | 0 | NA | 1 |
| SE | 13760 | NCOA6 | NCOA6 | chr20 | - | 34727258 | 34727407 | 34714773 | 34715365 | 34732558 | 34732595 | 0 | 0 | 1 | 0 | 0 | 2 | 0 | 1 | 0 | 0 | 0 | 0 | 98 | 49 | NA | NA | NA | NA | 0.2 | NA | 1 | NA | -0.8 |
| SE | 13796 | LGALS8 | LGALS8 | chr1 | + | 236542914 | 236543040 | 236542760 | 236542787 | 236543559 | 236543648 | 0 | 0 | 0 | 0 | 0 | 1 | 0 | 1 | 0 | 0 | 0 | 0 | 98 | 49 | NA | NA | NA | NA | 0 | NA | 1 | NA | -1 |
| SE | 13797 | CCDC93 | CCDC93 | chr2 | - | 117973907 | 117973994 | 117958364 | 117958481 | 117974849 | 117974900 | 0 | 0 | 2 | 0 | 0 | 0 | 0 | 5 | 0 | 0 | 1 | 0 | 98 | 49 | NA | NA | NA | NA | 1 | NA | 0.714 | NA | 0.286 |
| SE | 13806 | CLASP1 | CLASP1 | chr2 | - | 121409023 | 121409047 | 121407470 | 121407715 | 121410865 | 121410969 | 0 | 0 | 0 | 0 | 0 | 2 | 0 | 4 | 0 | 0 | 3 | 0 | 73 | 49 | NA | NA | NA | NA | 0 | NA | 0.472 | NA | -0.472 |
| SE | 13810 | CLASP1 | CLASP1 | chr2 | - | 121445448 | 121445496 | 121430072 | 121430177 | 121447336 | 121447507 | 0 | 0 | 0 | 0 | 0 | 1 | 0 | 1 | 0 | 0 | 1 | 0 | 97 | 49 | NA | NA | NA | NA | 0 | NA | 0.336 | NA | -0.336 |
| SE | 13811 | CLASP1 | CLASP1 | chr2 | - | 121515664 | 121515762 | 121503166 | 121503234 | 121525844 | 121525920 | 0 | 0 | 1 | 0 | 0 | 2 | 0 | 2 | 0 | 0 | 0 | 0 | 98 | 49 | NA | NA | NA | NA | 0.2 | NA | 1 | NA | -0.8 |
| SE | 13850 | SFT2D1 | SFT2D1 | chr6 | - | 166331384 | 166331407 | 166330160 | 166330247 | 166342418 | 166342606 | 0 | 0 | 1 | 0 | 0 | 1 | 0 | 0 | 0 | 0 | 2 | 0 | 72 | 49 | NA | NA | NA | NA | 0.405 | NA | 0 | NA | 0.405 |
| SE | 13871 | RAB3GAP1 | RAB3GAP1 | chr2 | + | 135135563 | 135135932 | 135135264 | 135135319 | 135150368 | 135150506 | 0 | 0 | 7 | 0 | 0 | 0 | 0 | 3 | 0 | 0 | 1 | 0 | 98 | 49 | NA | NA | NA | NA | 1 | NA | 0.6 | NA | 0.4 |
| SE | 13878 | ZEB2 | ZEB2 | chr2 | - | 144398300 | 144400270 | 144396411 | 144396592 | 144401198 | 144401307 | 0 | 0 | 8 | 0 | 0 | 1 | 1 | 3 | 0 | 0 | 0 | 0 | 98 | 49 | NA | NA | NA | NA | 0.8 | 1 | 1 | NA | -0.2 |
| SE | 13880 | ZEB2 | ZEB2 | chr2 | - | 144424795 | 144424867 | 144404835 | 144405024 | 144429768 | 144430026 | 0 | 0 | 6 | 0 | 0 | 1 | 0 | 5 | 0 | 0 | 0 | 0 | 98 | 49 | NA | NA | NA | NA | 0.75 | NA | 1 | NA | -0.25 |
| SE | 13881 | ZEB2 | ZEB2 | chr2 | - | 144517277 | 144517419 | 144429768 | 144430026 | 144519938 | 144520391 | 0 | 0 | 6 | 0 | 0 | 0 | 0 | 4 | 0 | 0 | 1 | 0 | 98 | 49 | NA | NA | NA | NA | 1 | NA | 0.667 | NA | 0.333 |
| SE | 13890 | CD48 | CD48 | chr1 | - | 160684886 | 160685189 | 160680421 | 160681468 | 160711681 | 160711851 | 0 | 0 | 22 | 0 | 0 | 1 | 0 | 14 | 0 | 0 | 0 | 0 | 98 | 49 | NA | NA | NA | NA | 0.917 | NA | 1 | NA | -0.083 |
| SE | 13907 | NR4A2 | NR4A2 | chr2 | - | 156328403 | 156328533 | 156327850 | 156328014 | 156329322 | 156330188 | 0 | 0 | 1 | 0 | 0 | 1 | 7 | 6 | 0 | 0 | 0 | 0 | 98 | 49 | NA | NA | NA | NA | 0.333 | 1 | 1 | NA | -0.667 |
| SE | 13911 | CSAD | CSAD | chr12 | - | 53161126 | 53161191 | 53160762 | 53160844 | 53161272 | 53161389 | 0 | 0 | 2 | 0 | 0 | 4 | 0 | 2 | 0 | 0 | 1 | 0 | 98 | 49 | NA | NA | NA | NA | 0.2 | NA | 0.5 | NA | -0.3 |
| SE | 13942 | PDK1 | PDK1 | chr2 | + | 172562740 | 172562800 | 172562219 | 172562291 | 172564502 | 172564687 | 0 | 0 | 1 | 0 | 0 | 0 | 0 | 0 | 0 | 0 | 2 | 0 | 98 | 49 | NA | NA | NA | NA | 1 | NA | 0 | NA | 1 |
| SE | 13944 | PDK1 | PDK1 | chr2 | + | 172564502 | 172564687 | 172562219 | 172562291 | 172564977 | 172565073 | 0 | 0 | 0 | 0 | 0 | 1 | 0 | 2 | 0 | 0 | 1 | 0 | 98 | 49 | NA | NA | NA | NA | 0 | NA | 0.5 | NA | -0.5 |
| SE | 13959 | DIP2B | DIP2B | chr12 | + | 50728547 | 50728678 | 50727702 | 50727812 | 50732365 | 50732536 | 0 | 0 | 1 | 0 | 0 | 0 | 0 | 1 | 0 | 0 | 1 | 0 | 98 | 49 | NA | NA | NA | NA | 1 | NA | 0.333 | NA | 0.667 |
| SE | 13960 | DIP2B | DIP2B | chr12 | + | 50731368 | 50731537 | 50728547 | 50728678 | 50732365 | 50732536 | 0 | 0 | 4 | 0 | 0 | 0 | 0 | 0 | 0 | 0 | 1 | 0 | 98 | 49 | NA | NA | NA | NA | 1 | NA | 0 | NA | 1 |
| SE | 13964 | PARN | PARN | chr16 | - | 14627268 | 14627336 | 14627105 | 14627187 | 14628171 | 14628251 | 0 | 0 | 4 | 0 | 0 | 1 | 0 | 2 | 0 | 0 | 0 | 0 | 98 | 49 | NA | NA | NA | NA | 0.667 | NA | 1 | NA | -0.333 |
| SE | 14003 | MS4A7 | MS4A7 | chr11 | + | 60392684 | 60392786 | 60389389 | 60389596 | 60393786 | 60395953 | 0 | 0 | 13 | 0 | 0 | 5 | 0 | 0 | 0 | 4 | 0 | 0 | 98 | 49 | NA | NA | NA | NA | 0.565 | 0 | NA | NA | 0.565 |
| SE | 14019 | ANKRD44 | ANKRD44 | chr2 | - | 196998906 | 196999052 | 196998336 | 196998419 | 197000418 | 197000502 | 0 | 0 | 9 | 0 | 0 | 5 | 0 | 3 | 0 | 0 | 1 | 0 | 98 | 49 | NA | NA | NA | NA | 0.474 | NA | 0.6 | NA | -0.126 |
| SE | 14020 | ANKRD44 | ANKRD44 | chr2 | - | 197013510 | 197013712 | 197008943 | 197009031 | 197025195 | 197025267 | 0 | 0 | 9 | 0 | 0 | 0 | 0 | 7 | 0 | 0 | 3 | 0 | 98 | 49 | NA | NA | NA | NA | 1 | NA | 0.538 | NA | 0.462 |
| SE | 14029 | FAM126B | FAM126B | chr2 | - | 200992886 | 200993003 | 200992280 | 200992368 | 200997444 | 200997540 | 0 | 0 | 15 | 0 | 0 | 2 | 0 | 3 | 0 | 0 | 0 | 0 | 98 | 49 | NA | NA | NA | NA | 0.789 | NA | 1 | NA | -0.211 |
| SE | 14030 | FAM126B | FAM126B | chr2 | - | 200992886 | 200993003 | 200992280 | 200992368 | 201008972 | 201009088 | 0 | 0 | 10 | 0 | 0 | 0 | 0 | 4 | 0 | 0 | 1 | 0 | 98 | 49 | NA | NA | NA | NA | 1 | NA | 0.667 | NA | 0.333 |
| SE | 14031 | FAM126B | FAM126B | chr2 | - | 200997444 | 200997540 | 200992280 | 200992368 | 201008972 | 201009088 | 0 | 0 | 6 | 0 | 0 | 0 | 0 | 2 | 0 | 0 | 1 | 0 | 98 | 49 | NA | NA | NA | NA | 1 | NA | 0.5 | NA | 0.5 |
| SE | 14032 | FAM126B | FAM126B | chr2 | - | 200997444 | 200997540 | 200992886 | 200993003 | 201008972 | 201009088 | 0 | 0 | 9 | 0 | 0 | 0 | 0 | 2 | 0 | 0 | 1 | 0 | 98 | 49 | NA | NA | NA | NA | 1 | NA | 0.5 | NA | 0.5 |
| SE | 14091 | AREL1 | AREL1 | chr14 | - | 74674033 | 74674111 | 74673076 | 74673218 | 74675698 | 74675946 | 0 | 0 | 2 | 0 | 0 | 0 | 6 | 8 | 0 | 0 | 1 | 0 | 98 | 49 | NA | NA | NA | NA | 1 | 1 | 0.8 | NA | 0.1 |
| SE | 14108 | TMEM9B | TMEM9B | chr11 | - | 8962091 | 8962183 | 8956189 | 8956298 | 8964208 | 8964773 | 0 | 0 | 4 | 0 | 0 | 0 | 0 | 0 | 0 | 0 | 1 | 0 | 98 | 49 | NA | NA | NA | NA | 1 | NA | 0 | NA | 1 |

**U12_4_events_filtered**

| | | | | | | | | | | | | Inclusion_count | | | Skipping_count | | | Inclusion_count | | | Skipping_count | | | | | | | Inclusion_level (normalized) | | | | | | |
|---|---|---|---|---|---|---|---|---|---|---|---|---|---|---|---|---|---|---|---|---|---|---|---|---|---|---|---|---|---|---|---|---|---|---|
| **Splice_type** | **ID** | **Gene_ID** | **Gene_Symbol** | **Chromosome** | **Strand** | **LongExonStart_0base** | **LongExonEnd** | **ShortES** | **ShortEE** | **FlankingES** | **FlankingEE** | **control_3** | **control_4** | **control_10** | **control_3** | **control_4** | **control_10** | **pmn_13** | **pmn_15** | **pmn_17** | **pmn_13** | **pmn_15** | **pmn_17** | **IncFormLen** | **SkipFormLen** | **p value** | **FDR** | **control_3** | **control_4** | **control_10** | **pmn_13** | **pmn_15** | **pmn_17** | **IncLevelDifference** |
| A3SS | 1863 | SRPK1 | SRPK1 | chr6 | - | 35890894 | 35891041 | 35890894 | 35891013 | 35920467 | 35920528 | 0 | 0 | 0 | 0 | 1 | 2 | 0 | 1 | 1 | 0 | 4 | 1 | 77 | 49 | 0.005762586 | 0.197717869 | NA | 0 | 0 | NA | 0.137 | 0.389 | -0.263 |
| SE | 9353 | GBP5 | GBP5 | chr1 | - | 89262682 | 89262785 | 89262219 | 89262401 | 89263735 | 89263948 | 0 | 3 | 3 | 0 | 0 | 0 | 1 | 6 | 0 | 0 | 2 | 0 | 98 | 49 | 0.019996576 | 0.170760496 | NA | 1 | 1 | 1 | 0.6 | NA | 0.2 |
| SE | 12583 | SBNO2 | SBNO2 | chr19 | - | 1109134 | 1109211 | 1108778 | 1108969 | 1109291 | 1109423 | 0 | 4 | 9 | 0 | 0 | 0 | 0 | 25 | 2 | 0 | 3 | 0 | 98 | 49 | 0.024422917 | 0.170760496 | NA | 1 | 1 | NA | 0.806 | 1 | 0.097 |
| SE | 6887 | SRPK1 | SRPK1 | chr6 | - | 35870280 | 35870494 | 35869481 | 35869901 | 35870933 | 35870959 | 0 | 0 | 4 | 0 | 1 | 5 | 0 | 2 | 0 | 0 | 3 | 0 | 98 | 49 | 0.027065308 | 0.170760496 | NA | 0 | 0.286 | NA | 0.25 | NA | -0.107 |
| SE | 14102 | SLC12A4 | SLC12A4 | chr16 | - | 67951822 | 67952037 | 67951139 | 67951304 | 67952183 | 67952425 | 0 | 1 | 0 | 0 | 0 | 0 | 0 | 1 | 0 | 0 | 2 | 0 | 98 | 49 | 0.027259421 | 0.170760496 | NA | 1 | NA | NA | 0.2 | NA | 0.8 |
| SE | 2543 | GAK | GAK | chr4 | - | 877614 | 877809 | 877089 | 877207 | 881906 | 882040 | 0 | 2 | 6 | 0 | 0 | 0 | 0 | 3 | 0 | 0 | 1 | 0 | 98 | 49 | 0.032835937 | 0.170760496 | NA | 1 | 1 | NA | 0.6 | NA | 0.4 |
| SE | 691 | DOCK5 | DOCK5 | chr8 | + | 25300575 | 25300657 | 25298943 | 25299101 | 25302324 | 25302454 | 0 | 3 | 24 | 0 | 0 | 0 | 0 | 10 | 0 | 0 | 1 | 0 | 98 | 49 | 0.03465568 | 0.170760496 | NA | 1 | 1 | NA | 0.833 | NA | 0.167 |
| SE | 279 | EXOSC2 | EXOSC2 | chr9 | + | 130698161 | 130698251 | 130697581 | 130697627 | 130699328 | 130699394 | 0 | 1 | 0 | 0 | 0 | 0 | 0 | 1 | 0 | 0 | 1 | 0 | 98 | 49 | 0.037136978 | 0.170760496 | NA | 1 | NA | NA | 0.333 | NA | 0.667 |
| SE | 6888 | SMC3 | SMC3 | chr10 | + | 110590990 | 110591132 | 110590411 | 110590572 | 110593072 | 110593223 | 0 | 1 | 3 | 0 | 0 | 0 | 0 | 1 | 0 | 0 | 1 | 0 | 98 | 49 | 0.037136978 | 0.170760496 | NA | 1 | 1 | NA | 0.333 | NA | 0.667 |
| SE | 5861 | DISC1 | DISC1 | chr1 | + | 231795241 | 231795296 | 231770834 | 231771070 | 231818328 | 231819244 | 0 | 1 | 7 | 0 | 0 | 0 | 0 | 5 | 0 | 0 | 1 | 0 | 98 | 49 | 0.039248064 | 0.170760496 | NA | 1 | 1 | NA | 0.714 | NA | 0.286 |
| SE | 2907 | RAF1 | RAF1 | chr3 | - | 12600151 | 12600279 | 12599690 | 12599808 | 12600387 | 12600415 | 0 | 2 | 19 | 0 | 0 | 0 | 0 | 22 | 3 | 0 | 2 | 0 | 98 | 49 | 0.041304338 | 0.170760496 | NA | 1 | 1 | NA | 0.846 | 1 | 0.077 |
| SE | 1691 | DOCK2 | DOCK2 | chr5 | + | 169761518 | 169761625 | 169759704 | 169759775 | 169803057 | 169803206 | 0 | 4 | 19 | 0 | 0 | 0 | 0 | 12 | 1 | 0 | 1 | 0 | 98 | 49 | 0.045620082 | 0.170760496 | NA | 1 | 1 | NA | 0.857 | 1 | 0.072 |
| A3SS | 414 | DCTN3 | DCTN3 | chr9 | - | 34614041 | 34614198 | 34614041 | 34614101 | 34614709 | 34614768 | 1 | 1 | 0 | 0 | 0 | 0 | 0 | 0 | 0 | 0 | 3 | 0 | 98 | 49 | 0.059555994 | 0.197717869 | 1 | 1 | NA | NA | 0 | NA | 1 |
| A3SS | 413 | DCTN3 | DCTN3 | chr9 | - | 34614041 | 34614132 | 34614041 | 34614101 | 34614709 | 34614768 | 1 | 1 | 0 | 0 | 0 | 0 | 0 | 0 | 0 | 0 | 3 | 0 | 80 | 49 | 0.060211792 | 0.197717869 | 1 | 1 | NA | NA | 0 | NA | 1 |
| SE | 8054 | CUL4B | CUL4B | chrX | - | 120544480 | 120544643 | 120544113 | 120544203 | 120545443 | 120545517 | 0 | 0 | 1 | 0 | 0 | 0 | 0 | 1 | 0 | 0 | 0 | 2 | 98 | 49 | 0.061401334 | 0.170760496 | NA | NA | 1 | NA | 1 | 0 | 0.5 |
| SE | 5029 | BRMS1 | BRMS1 | chr11 | - | 66340120 | 66340213 | 66338720 | 66338785 | 66340773 | 66340870 | 0 | 0 | 2 | 0 | 1 | 0 | 0 | 2 | 0 | 0 | 0 | 0 | 98 | 49 | 0.063770311 | 0.170760496 | NA | 0 | 1 | NA | 1 | NA | -0.5 |
| SE | 8179 | MLXIP | MLXIP | chr12 | + | 122137468 | 122137590 | 122135488 | 122135666 | 122138193 | 122138295 | 0 | 0 | 1 | 0 | 0 | 0 | 0 | 0 | 0 | 0 | 0 | 1 | 98 | 49 | 0.065790241 | 0.170760496 | NA | NA | 1 | NA | NA | 0 | 1 |
| SE | 11719 | CHD3 | CHD3 | chr17 | + | 7907600 | 7907702 | 7907352 | 7907488 | 7907893 | 7908019 | 0 | 0 | 0 | 0 | 0 | 2 | 0 | 0 | 1 | 0 | 0 | 0 | 98 | 49 | 0.0658956 | 0.170760496 | NA | NA | 0 | NA | NA | 1 | -1 |
| SE | 969 | RHBDD2 | RHBDD2 | chr7 | + | 75881364 | 75881486 | 75878998 | 75879260 | 75881828 | 75882236 | 0 | 0 | 2 | 0 | 1 | 0 | 2 | 1 | 0 | 0 | 0 | 0 | 98 | 49 | 0.068700612 | 0.171142152 | NA | 0 | 1 | 1 | 1 | NA | -0.5 |
| SE | 1355 | MAPK14 | MAPK14 | chr6 | + | 36102570 | 36102649 | 36095986 | 36096066 | 36107454 | 36107628 | 0 | 1 | 19 | 0 | 0 | 0 | 0 | 14 | 1 | 0 | 1 | 0 | 98 | 49 | 0.069582416 | 0.17144651 | NA | 1 | 1 | NA | 0.875 | 1 | 0.062 |
| A3SS | 3730 | RASGRP4 | RASGRP4 | chr19 | - | 38414847 | 38415123 | 38414847 | 38414916 | 38417051 | 38417168 | 0 | 2 | 16 | 0 | 0 | 0 | 1 | 4 | 3 | 0 | 1 | 0 | 98 | 49 | 0.071741814 | 0.198669639 | NA | 1 | 1 | 1 | 0.667 | 1 | 0.111 |
| SE | 4098 | SETD1B | SETD1B | chr12 | + | 121819403 | 121819895 | 121817798 | 121817904 | 121822489 | 121823749 | 0 | 3 | 1 | 0 | 0 | 3 | 1 | 6 | 1 | 0 | 0 | 0 | 98 | 49 | 0.079530423 | 0.185900818 | NA | 1 | 0.143 | 1 | 1 | 1 | -0.428 |
| SE | 12584 | SBNO2 | SBNO2 | chr19 | - | 1111018 | 1111093 | 1110744 | 1110888 | 1111505 | 1111614 | 0 | 1 | 6 | 0 | 0 | 1 | 1 | 26 | 2 | 0 | 0 | 0 | 98 | 49 | 0.080803043 | 0.187418168 | NA | 1 | 0.75 | 1 | 1 | 1 | -0.125 |
| SE | 13111 | AP2A1 | AP2A1 | chr19 | + | 49800960 | 49801058 | 49799967 | 49800150 | 49801389 | 49801621 | 0 | 1 | 3 | 0 | 1 | 0 | 0 | 4 | 0 | 0 | 0 | 0 | 98 | 49 | 0.115352023 | 0.235883116 | NA | 0.333 | 1 | NA | 1 | NA | -0.334 |
| SE | 6225 | AHCTF1 | AHCTF1 | chr1 | - | 246916141 | 246916395 | 246913231 | 246913412 | 246918249 | 246918377 | 0 | 0 | 4 | 0 | 0 | 1 | 0 | 0 | 0 | 0 | 0 | 1 | 98 | 49 | 0.133877166 | 0.263718717 | NA | NA | 0.667 | NA | NA | 0 | 0.667 |
| SE | 5934 | RASGRP2 | RASGRP2 | chr11 | - | 64727295 | 64727360 | 64726910 | 64727131 | 64728862 | 64729042 | 0 | 0 | 7 | 0 | 1 | 0 | 0 | 1 | 1 | 0 | 0 | 0 | 98 | 49 | 0.134979818 | 0.264504128 | NA | 0 | 1 | NA | 1 | 1 | -0.5 |
| A3SS | 1281 | CHD4 | CHD4 | chr12 | - | 6601959 | 6602175 | 6601959 | 6602154 | 6602375 | 6602497 | 0 | 0 | 2 | 0 | 0 | 1 | 0 | 17 | 4 | 4 | 4 | 0 | 70 | 49 | 0.136899463 | 0.324235571 | NA | NA | 0.583 | 0 | 0.748 | 1 | 0 |
| SE | 12586 | SBNO2 | SBNO2 | chr19 | - | 1113534 | 1113704 | 1112817 | 1112949 | 1114230 | 1114422 | 0 | 9 | 21 | 0 | 0 | 1 | 0 | 40 | 18 | 0 | 0 | 0 | 98 | 49 | 0.149179961 | 0.284179318 | NA | 1 | 0.913 | NA | 1 | 1 | -0.043 |
| SE | 695 | DOCK5 | DOCK5 | chr8 | + | 25368191 | 25368250 | 25366869 | 25366970 | 25368570 | 25368725 | 0 | 0 | 11 | 0 | 0 | 5 | 0 | 5 | 3 | 0 | 1 | 0 | 98 | 49 | 0.166703165 | 0.311886731 | NA | NA | 0.524 | NA | 0.714 | 1 | -0.333 |
| SE | 2487 | ATG3 | ATG3 | chr3 | - | 112553279 | 112553329 | 112550191 | 112550262 | 112558375 | 112558417 | 0 | 0 | 4 | 0 | 0 | 3 | 1 | 4 | 2 | 0 | 0 | 0 | 98 | 49 | 0.167045122 | 0.311886731 | NA | NA | 0.4 | 1 | 1 | 1 | -0.6 |
| SE | 3413 | RALGAPA2 | RALGAPA2 | chr20 | - | 20524405 | 20524543 | 20520916 | 20521100 | 20524829 | 20524898 | 0 | 0 | 2 | 0 | 0 | 1 | 0 | 0 | 2 | 0 | 0 | 0 | 98 | 49 | 0.209881051 | 0.368087771 | NA | NA | 0.5 | NA | NA | 1 | -0.5 |
| SE | 276 | RAPGEF1 | RAPGEF1 | chr9 | - | 131650128 | 131650242 | 131643238 | 131643417 | 131650809 | 131650949 | 0 | 2 | 1 | 0 | 0 | 1 | 3 | 10 | 1 | 0 | 7 | 0 | 98 | 49 | 0.223239446 | 0.388343619 | NA | 1 | 0.333 | 1 | 0.417 | 1 | -0.139 |
| SE | 3236 | SLC9A8 | SLC9A8 | chr20 | + | 49850761 | 49850844 | 49849578 | 49849680 | 49855437 | 49855581 | 0 | 0 | 4 | 0 | 0 | 4 | 0 | 0 | 0 | 0 | 4 | 1 | 98 | 49 | 0.23228173 | 0.398828525 | NA | NA | 0.333 | NA | 0 | 0 | 0.333 |
| SE | 3237 | SLC9A8 | SLC9A8 | chr20 | + | 49850809 | 49850844 | 49849578 | 49849680 | 49855437 | 49855581 | 0 | 0 | 4 | 0 | 0 | 4 | 0 | 0 | 0 | 0 | 4 | 1 | 84 | 49 | 0.232676511 | 0.398828525 | NA | NA | 0.368 | NA | 0 | 0 | 0.368 |
| RI | 607 | TAF1C | TAF1C | chr16 | - | 84181327 | 84181663 | 84181327 | 84181463 | 84181591 | 84181663 | 0 | 0 | 3 | 0 | 0 | 1 | 0 | 1 | 2 | 0 | 0 | 0 | 98 | 49 | 0.264638717 | 0.737207854 | NA | NA | 0.6 | NA | 1 | 1 | -0.4 |
| SE | 3872 | WDFY2 | WDFY2 | chr13 | + | 51719197 | 51719348 | 51703595 | 51703650 | 51727677 | 51727790 | 0 | 0 | 3 | 0 | 0 | 1 | 3 | 2 | 2 | 0 | 0 | 0 | 98 | 49 | 0.264638717 | 0.435653879 | NA | NA | 0.6 | 1 | 1 | 1 | -0.4 |
| SE | 7725 | ZFC3H1 | ZFC3H1 | chr12 | - | 71636849 | 71637059 | 71636489 | 71636654 | 71638417 | 71638515 | 0 | 0 | 3 | 0 | 0 | 1 | 0 | 9 | 2 | 0 | 0 | 0 | 98 | 49 | 0.264638717 | 0.435653879 | NA | NA | 0.6 | NA | 1 | 1 | -0.4 |
| SE | 5935 | RASGRP2 | RASGRP2 | chr11 | - | 64735902 | 64735980 | 64735541 | 64735664 | 64736752 | 64737034 | 0 | 1 | 5 | 0 | 0 | 0 | 1 | 3 | 1 | 0 | 0 | 3 | 98 | 49 | 0.311385518 | 0.494727732 | NA | 1 | 1 | 1 | 1 | 0.143 | 0.286 |
| SE | 275 | RAPGEF1 | RAPGEF1 | chr9 | - | 131630235 | 131630324 | 131629101 | 131629254 | 131638634 | 131638791 | 0 | 0 | 6 | 0 | 0 | 2 | 0 | 8 | 1 | 0 | 1 | 0 | 98 | 49 | 0.365054683 | 0.547059405 | NA | NA | 0.6 | NA | 0.8 | 1 | -0.3 |
| SE | 3233 | SLC9A8 | SLC9A8 | chr20 | + | 49839540 | 49839599 | 49823060 | 49823141 | 49845035 | 49845119 | 0 | 0 | 8 | 0 | 0 | 1 | 0 | 0 | 2 | 0 | 0 | 0 | 98 | 49 | 0.393533929 | 0.579315862 | NA | NA | 0.8 | NA | NA | 1 | -0.2 |
| SE | 2247 | MAEA | MAEA | chr4 | + | 1327626 | 1327703 | 1315396 | 1315600 | 1332756 | 1332865 | 0 | 3 | 2 | 0 | 1 | 1 | 0 | 1 | 0 | 0 | 1 | 0 | 98 | 49 | 0.429432607 | 0.620451665 | NA | 0.6 | 0.5 | NA | 0.333 | NA | 0.217 |
| RI | 1055 | STX10 | STX10 | chr19 | - | 13144057 | 13144671 | 13144057 | 13144486 | 13144576 | 13144671 | 0 | 1 | 6 | 0 | 0 | 1 | 0 | 0 | 1 | 0 | 1 | 0 | 98 | 49 | 0.482241355 | 0.94773154 | NA | 1 | 0.75 | NA | 0 | 1 | 0.375 |
| SE | 12585 | SBNO2 | SBNO2 | chr19 | - | 1112188 | 1112301 | 1111995 | 1112067 | 1112401 | 1112537 | 0 | 8 | 15 | 0 | 4 | 0 | 0 | 8 | 3 | 0 | 2 | 1 | 98 | 49 | 0.521310981 | 0.720156622 | NA | 0.5 | 1 | NA | 0.667 | 0.6 | 0.117 |
| SE | 3235 | SLC9A8 | SLC9A8 | chr20 | + | 49849578 | 49849680 | 49845035 | 49845119 | 49855437 | 49855581 | 0 | 0 | 10 | 0 | 0 | 2 | 0 | 6 | 1 | 0 | 2 | 0 | 98 | 49 | 0.539441453 | 0.737743862 | NA | NA | 0.714 | NA | 0.6 | 1 | -0.086 |
| SE | 3417 | RALGAPA2 | RALGAPA2 | chr20 | - | 20676235 | 20676288 | 20653529 | 20653587 | 20680690 | 20680801 | 0 | 0 | 13 | 0 | 0 | 2 | 0 | 5 | 1 | 0 | 0 | 0 | 98 | 49 | 0.588967448 | 0.794630228 | NA | NA | 0.765 | NA | 1 | 1 | -0.235 |
| RI | 230 | ATG3 | ATG3 | chr3 | - | 112532506 | 112534337 | 112532506 | 112532780 | 112534268 | 112534337 | 0 | 0 | 3 | 0 | 3 | 5 | 0 | 0 | 1 | 0 | 4 | 0 | 98 | 49 | 0.654603078 | 0.999202116 | NA | 0 | 0.231 | NA | 0 | 1 | -0.385 |
| SE | 2906 | RAF1 | RAF1 | chr3 | - | 12590797 | 12590974 | 12587590 | 12587637 | 12591707 | 12591792 | 1 | 1 | 24 | 0 | 0 | 2 | 4 | 25 | 0 | 0 | 0 | 0 | 98 | 49 | 0.867186844 | 0.999202116 | 1 | 1 | 0.857 | 1 | 1 | NA | -0.048 |
| SE | 10440 | SLC12A6 | SLC12A6 | chr15 | - | 34257641 | 34257788 | 34256228 | 34256283 | 34258812 | 34258944 | 0 | 1 | 11 | 0 | 0 | 4 | 0 | 2 | 4 | 0 | 2 | 0 | 98 | 49 | 0.949596688 | 0.999202116 | NA | 1 | 0.579 | NA | 0.333 | 1 | 0.123 |
| SE | 7631 | BTAF1 | BTAF1 | chr10 | + | 91953736 | 91953873 | 91951402 | 91951566 | 91956527 | 91956657 | 0 | 1 | 5 | 0 | 0 | 1 | 0 | 4 | 0 | 0 | 0 | 0 | 98 | 49 | 0.952968863 | 0.999202116 | NA | 1 | 0.714 | NA | 1 | NA | -0.143 |
| SE | 11425 | DYNC1LI2 | DYNC1LI2 | chr16 | - | 66749196 | 66749313 | 66742437 | 66742668 | 66751272 | 66751346 | 0 | 1 | 6 | 0 | 0 | 1 | 0 | 3 | 0 | 0 | 0 | 0 | 98 | 49 | 0.961336908 | 0.999202116 | NA | 1 | 0.75 | NA | 1 | NA | -0.125 |
| SE | 8808 | WDR1 | WDR1 | chr4 | - | 10103895 | 10103986 | 10098991 | 10099139 | 10116112 | 10116234 | 0 | 5 | 16 | 0 | 0 | 1 | 1 | 9 | 0 | 0 | 0 | 0 | 98 | 49 | 0.976663141 | 0.999202116 | NA | 1 | 0.889 | 1 | 1 | NA | -0.055 |
| SE | 5295 | PEX16 | PEX16 | chr11 | - | 45916226 | 45916303 | 45915702 | 45915836 | 45917457 | 45917493 | 0 | 0 | 3 | 0 | 0 | 0 | 0 | 5 | 2 | 0 | 1 | 0 | 98 | 49 | 0.993205848 | 0.999202116 | NA | NA | 1 | NA | 0.714 | 1 | 0.143 |
| SE | 694 | DOCK5 | DOCK5 | chr8 | + | 25342400 | 25342507 | 25341738 | 25341809 | 25345474 | 25345611 | 0 | 0 | 15 | 0 | 0 | 0 | 0 | 15 | 2 | 0 | 1 | 0 | 98 | 49 | 0.999202116 | 0.999202116 | NA | NA | 1 | NA | 0.882 | 1 | 0.059 |
| SE | 697 | DOCK5 | DOCK5 | chr8 | + | 25407982 | 25408154 | 25403557 | 25403724 | 25408801 | 25408940 | 0 | 1 | 23 | 0 | 1 | 1 | 0 | 9 | 3 | 0 | 3 | 0 | 98 | 49 | 0.999202116 | 0.999202116 | NA | 0.333 | 0.92 | NA | 0.6 | 1 | -0.173 |
| SE | 3722 | FCHO1 | FCHO1 | chr19 | + | 17781230 | 17781343 | 17778608 | 17778884 | 17781451 | 17781539 | 0 | 1 | 3 | 0 | 0 | 3 | 0 | 1 | 0 | 0 | 0 | 0 | 98 | 49 | 0.999202116 | 0.999202116 | NA | 1 | 0.333 | NA | 1 | NA | -0.334 |
| SE | 4520 | CHD4 | CHD4 | chr12 | - | 6600925 | 6601053 | 6600533 | 6600669 | 6601288 | 6601530 | 0 | 0 | 7 | 0 | 0 | 0 | 0 | 4 | 1 | 0 | 1 | 0 | 98 | 49 | 0.999202116 | 0.999202116 | NA | NA | 1 | NA | 0.667 | 1 | 0.166 |
| SE | 11563 | NCBP1 | NCBP1 | chr9 | + | 97662041 | 97662144 | 97660945 | 97661068 | 97662953 | 97663047 | 0 | 1 | 1 | 0 | 0 | 2 | 0 | 1 | 0 | 0 | 0 | 0 | 98 | 49 | 0.999202116 | 0.999202116 | NA | 1 | 0.2 | NA | 1 | NA | -0.4 |
| SE | 12825 | TRAPPC10 | TRAPPC10 | chr21 | + | 44074323 | 44074470 | 44063537 | 44063785 | 44075038 | 44075153 | 0 | 0 | 4 | 0 | 0 | 0 | 0 | 3 | 1 | 0 | 1 | 0 | 98 | 49 | 0.999202116 | 0.999202116 | NA | NA | 1 | NA | 0.6 | 1 | 0.2 |
| A3SS | 503 | HARS2 | HARS2 | chr5 | + | 140693490 | 140693665 | 140693590 | 140693665 | 140691425 | 140691756 | 0 | 0 | 2 | 0 | 0 | 0 | 0 | 1 | 0 | 0 | 2 | 0 | 98 | 49 | NA | NA | NA | NA | 1 | NA | 0.2 | NA | 0.8 |
| A3SS | 1533 | TTC23 | TTC23 | chr15 | - | 99139316 | 99139467 | 99139316 | 99139399 | 99156147 | 99156297 | 0 | 0 | 0 | 0 | 0 | 1 | 0 | 1 | 0 | 0 | 0 | 0 | 98 | 49 | NA | NA | NA | NA | 0 | NA | 1 | NA | -1 |
| A3SS | 1628 | SRSF10 | SRSF10 | chr1 | - | 23964346 | 23967956 | 23964346 | 23967723 | 23971572 | 23971623 | 1 | 0 | 0 | 0 | 0 | 0 | 0 | 0 | 0 | 0 | 2 | 0 | 98 | 49 | NA | NA | 1 | NA | NA | NA | 0 | NA | 1 |
| A3SS | 1629 | SRSF10 | SRSF10 | chr1 | - | 23964346 | 23971439 | 23964346 | 23967723 | 23971572 | 23971623 | 1 | 0 | 1 | 0 | 0 | 0 | 0 | 1 | 0 | 0 | 2 | 0 | 98 | 49 | NA | NA | 1 | NA | 1 | NA | 0.2 | NA | 0.8 |
| A5SS | 1731 | SSR3 | SSR3 | chr3 | - | 156548865 | 156549003 | 156548904 | 156549003 | 156544307 | 156544439 | 0 | 0 | 0 | 0 | 0 | 1 | 8 | 0 | 0 | 0 | 0 | 0 | 88 | 49 | NA | NA | NA | NA | 0 | 1 | NA | NA | -1 |
| A5SS | 2052 | SLC12A9 | SLC12A9 | chr7 | + | 100852718 | 100852839 | 100852718 | 100852835 | 100854155 | 100854378 | 0 | 0 | 0 | 0 | 0 | 2 | 0 | 2 | 0 | 0 | 0 | 0 | 53 | 49 | NA | NA | NA | NA | 0 | NA | 1 | NA | -1 |
| A5SS | 2205 | PLCB2 | PLCB2 | chr15 | - | 40296745 | 40296908 | 40296757 | 40296908 | 40296521 | 40296634 | 0 | 0 | 7 | 0 | 0 | 3 | 0 | 2 | 0 | 0 | 0 | 0 | 61 | 49 | NA | NA | NA | NA | 0.652 | NA | 1 | NA | -0.348 |
| RI | 909 | HNRNPM | HNRNPM | chr19 | + | 8465323 | 8465515 | 8465323 | 8465362 | 8465479 | 8465515 | 0 | 0 | 8 | 0 | 0 | 1 | 7 | 0 | 0 | 0 | 0 | 0 | 98 | 49 | NA | NA | NA | NA | 0.8 | 1 | NA | NA | -0.2 |
| SE | 692 | DOCK5 | DOCK5 | chr8 | + | 25340876 | 25340988 | 25336238 | 25336373 | 25341738 | 25341809 | 0 | 0 | 19 | 0 | 0 | 0 | 0 | 10 | 0 | 0 | 1 | 0 | 98 | 49 | NA | NA | NA | NA | 1 | NA | 0.833 | NA | 0.167 |
| SE | 693 | DOCK5 | DOCK5 | chr8 | + | 25341738 | 25341809 | 25336238 | 25336373 | 25342400 | 25342507 | 0 | 0 | 6 | 0 | 0 | 0 | 0 | 12 | 0 | 0 | 1 | 0 | 98 | 49 | NA | NA | NA | NA | 1 | NA | 0.857 | NA | 0.143 |
| SE | 696 | DOCK5 | DOCK5 | chr8 | + | 25373617 | 25373658 | 25372558 | 25372718 | 25374563 | 25374654 | 0 | 0 | 8 | 0 | 0 | 1 | 0 | 8 | 0 | 0 | 0 | 0 | 90 | 49 | NA | NA | NA | NA | 0.813 | NA | 1 | NA | -0.187 |
| SE | 1305 | SMAP1 | SMAP1 | chr6 | + | 70791688 | 70791769 | 70773349 | 70773425 | 70798656 | 70798737 | 0 | 0 | 1 | 0 | 0 | 0 | 0 | 0 | 0 | 0 | 1 | 0 | 98 | 49 | NA | NA | NA | NA | 1 | NA | 0 | NA | 1 |
| SE | 1692 | DOCK2 | DOCK2 | chr5 | + | 169983067 | 169983166 | 169840756 | 169840852 | 169985827 | 169985922 | 0 | 0 | 12 | 0 | 0 | 1 | 0 | 5 | 0 | 0 | 0 | 0 | 98 | 49 | NA | NA | NA | NA | 0.857 | NA | 1 | NA | -0.143 |
| SE | 1748 | DIAPH1 | DIAPH1 | chr5 | - | 141529602 | 141529697 | 141529171 | 141529273 | 141534334 | 141534433 | 0 | 0 | 8 | 0 | 0 | 0 | 0 | 1 | 0 | 0 | 1 | 0 | 98 | 49 | NA | NA | NA | NA | 1 | NA | 0.333 | NA | 0.667 |
| SE | 2208 | IK | IK | chr5 | + | 140655828 | 140655992 | 140654680 | 140654727 | 140657553 | 140657662 | 0 | 0 | 2 | 0 | 0 | 2 | 0 | 1 | 0 | 0 | 0 | 0 | 98 | 49 | NA | NA | NA | NA | 0.333 | NA | 1 | NA | -0.667 |
| SE | 2530 | GBE1 | GBE1 | chr3 | - | 81646391 | 81646482 | 81642780 | 81642990 | 81648855 | 81648991 | 0 | 0 | 0 | 0 | 0 | 1 | 0 | 1 | 0 | 0 | 0 | 0 | 98 | 49 | NA | NA | NA | NA | 0 | NA | 1 | NA | -1 |
| SE | 2661 | SACM1L | SACM1L | chr3 | + | 45735234 | 45735373 | 45732052 | 45732151 | 45737582 | 45737652 | 0 | 0 | 2 | 0 | 0 | 0 | 0 | 2 | 0 | 0 | 1 | 0 | 98 | 49 | NA | NA | NA | NA | 1 | NA | 0.5 | NA | 0.5 |
| SE | 2908 | RAF1 | RAF1 | chr3 | - | 12606200 | 12606299 | 12604135 | 12604289 | 12608765 | 12608923 | 0 | 0 | 20 | 0 | 0 | 5 | 0 | 22 | 0 | 0 | 2 | 0 | 98 | 49 | NA | NA | NA | NA | 0.667 | NA | 0.846 | NA | -0.179 |
| SE | 3326 | CTNNBL1 | CTNNBL1 | chr20 | + | 37842338 | 37842419 | 37840101 | 37840199 | 37859898 | 37860036 | 0 | 0 | 2 | 0 | 0 | 1 | 0 | 4 | 0 | 0 | 0 | 0 | 98 | 49 | NA | NA | NA | NA | 0.5 | NA | 1 | NA | -0.5 |
| SE | 3414 | RALGAPA2 | RALGAPA2 | chr20 | - | 20605174 | 20605412 | 20601681 | 20601846 | 20611314 | 20611426 | 0 | 0 | 5 | 0 | 0 | 0 | 0 | 0 | 0 | 0 | 2 | 0 | 98 | 49 | NA | NA | NA | NA | 1 | NA | 0 | NA | 1 |
| SE | 3415 | RALGAPA2 | RALGAPA2 | chr20 | - | 20611314 | 20611426 | 20605174 | 20605412 | 20616042 | 20616191 | 0 | 0 | 6 | 0 | 0 | 1 | 0 | 1 | 0 | 0 | 0 | 0 | 98 | 49 | NA | NA | NA | NA | 0.75 | NA | 1 | NA | -0.25 |
| SE | 3416 | RALGAPA2 | RALGAPA2 | chr20 | - | 20637362 | 20637501 | 20635417 | 20635617 | 20639784 | 20639900 | 0 | 0 | 13 | 0 | 0 | 2 | 0 | 0 | 0 | 0 | 2 | 0 | 98 | 49 | NA | NA | NA | NA | 0.765 | NA | 0 | NA | 0.765 |
| SE | 3971 | HARS | HARS | chr5 | - | 140675016 | 140675133 | 140674678 | 140674825 | 140676653 | 140676896 | 0 | 0 | 1 | 0 | 0 | 0 | 0 | 2 | 0 | 1 | 0 | 0 | 98 | 49 | NA | NA | NA | NA | 1 | 0 | 1 | NA | 0.5 |
| SE | 4434 | KANSL2 | KANSL2 | chr12 | - | 48679035 | 48679150 | 48671798 | 48671962 | 48679654 | 48679833 | 0 | 0 | 6 | 0 | 0 | 2 | 0 | 3 | 0 | 0 | 0 | 0 | 98 | 49 | NA | NA | NA | NA | 0.6 | NA | 1 | NA | -0.4 |
| SE | 4518 | CHD4 | CHD4 | chr12 | - | 6582136 | 6582281 | 6581648 | 6581814 | 6582614 | 6582748 | 0 | 0 | 4 | 0 | 0 | 1 | 0 | 5 | 0 | 0 | 1 | 0 | 98 | 49 | NA | NA | NA | NA | 0.667 | NA | 0.714 | NA | -0.047 |
| SE | 4519 | CHD4 | CHD4 | chr12 | - | 6588297 | 6588422 | 6587711 | 6587949 | 6591465 | 6591583 | 0 | 0 | 13 | 0 | 0 | 0 | 0 | 3 | 0 | 0 | 3 | 0 | 98 | 49 | NA | NA | NA | NA | 1 | NA | 0.333 | NA | 0.667 |
| SE | 4553 | MED23 | MED23 | chr6 | - | 131620627 | 131620729 | 131619826 | 131619896 | 131621880 | 131621979 | 0 | 0 | 4 | 0 | 0 | 0 | 0 | 1 | 0 | 0 | 2 | 0 | 98 | 49 | NA | NA | NA | NA | 1 | NA | 0.2 | NA | 0.8 |
| SE | 4590 | CUL4A | CUL4A | chr13 | + | 113239432 | 113239551 | 113236822 | 113236890 | 113242967 | 113243160 | 0 | 0 | 5 | 0 | 0 | 1 | 0 | 1 | 0 | 0 | 0 | 0 | 98 | 49 | NA | NA | NA | NA | 0.714 | NA | 1 | NA | -0.286 |
| SE | 4924 | C2CD3 | C2CD3 | chr11 | - | 74161398 | 74161556 | 74139604 | 74139828 | 74168343 | 74168613 | 0 | 0 | 2 | 0 | 0 | 1 | 0 | 2 | 0 | 0 | 0 | 0 | 98 | 49 | NA | NA | NA | NA | 0.5 | NA | 1 | NA | -0.5 |
| SE | 5007 | RINT1 | RINT1 | chr7 | + | 105550260 | 105550486 | 105550054 | 105550165 | 105551569 | 105551707 | 0 | 0 | 1 | 0 | 0 | 0 | 0 | 0 | 0 | 0 | 2 | 0 | 98 | 49 | NA | NA | NA | NA | 1 | NA | 0 | NA | 1 |
| SE | 5110 | WDFY1 | WDFY1 | chr2 | - | 223884647 | 223884749 | 223881941 | 223882072 | 223894233 | 223894339 | 0 | 0 | 3 | 0 | 0 | 0 | 0 | 1 | 0 | 0 | 1 | 0 | 98 | 49 | NA | NA | NA | NA | 1 | NA | 0.333 | NA | 0.667 |
| SE | 5928 | ZDHHC5 | ZDHHC5 | chr11 | + | 57695919 | 57696043 | 57693782 | 57693915 | 57696760 | 57696873 | 0 | 0 | 0 | 0 | 0 | 1 | 5 | 5 | 0 | 0 | 0 | 0 | 98 | 49 | NA | NA | NA | NA | 0 | 1 | 1 | NA | -1 |
| SE | 5937 | SRSF10 | SRSF10 | chr1 | - | 23971337 | 23971439 | 23964346 | 23967723 | 23971572 | 23971626 | 0 | 0 | 1 | 0 | 0 | 0 | 0 | 1 | 0 | 0 | 2 | 0 | 98 | 49 | NA | NA | NA | NA | 1 | NA | 0.2 | NA | 0.8 |
| SE | 6090 | ZNF207 | ZNF207 | chr17 | + | 32366664 | 32366757 | 32365329 | 32365487 | 32367771 | 32368014 | 0 | 0 | 0 | 0 | 0 | 2 | 0 | 3 | 0 | 0 | 0 | 0 | 98 | 49 | NA | NA | NA | NA | 0 | NA | 1 | NA | -1 |
| SE | 6223 | AHCTF1 | AHCTF1 | chr1 | - | 246843794 | 246843928 | 246842693 | 246842776 | 246849614 | 246851442 | 0 | 0 | 0 | 0 | 0 | 2 | 0 | 2 | 0 | 0 | 2 | 0 | 98 | 49 | NA | NA | NA | NA | 0 | NA | 0.333 | NA | -0.333 |
| SE | 7632 | BTAF1 | BTAF1 | chr10 | + | 92027129 | 92027300 | 92026591 | 92026751 | 92028789 | 92030323 | 0 | 0 | 4 | 0 | 0 | 3 | 0 | 5 | 0 | 0 | 0 | 0 | 98 | 49 | NA | NA | NA | NA | 0.4 | NA | 1 | NA | -0.6 |
| SE | 7946 | PPP2R2D | PPP2R2D | chr10 | + | 131911561 | 131912598 | 131901237 | 131901330 | 131934457 | 131934555 | 0 | 0 | 0 | 0 | 0 | 3 | 0 | 2 | 0 | 0 | 0 | 0 | 98 | 49 | NA | NA | NA | NA | 0 | NA | 1 | NA | -1 |
| SE | 7947 | PPP2R2D | PPP2R2D | chr10 | + | 131934893 | 131935006 | 131934457 | 131934555 | 131940030 | 131940196 | 0 | 0 | 1 | 0 | 0 | 3 | 0 | 1 | 0 | 0 | 0 | 0 | 98 | 49 | NA | NA | NA | NA | 0.143 | NA | 1 | NA | -0.857 |
| SE | 8178 | MLXIP | MLXIP | chr12 | + | 122130843 | 122130933 | 122129940 | 122130112 | 122132291 | 122132383 | 0 | 0 | 1 | 0 | 0 | 1 | 0 | 1 | 0 | 0 | 0 | 0 | 98 | 49 | NA | NA | NA | NA | 0.333 | NA | 1 | NA | -0.667 |
| SE | 8908 | WDR11 | WDR11 | chr10 | + | 120867065 | 120867169 | 120866568 | 120866764 | 120871169 | 120871346 | 0 | 0 | 3 | 0 | 0 | 1 | 0 | 3 | 0 | 0 | 0 | 0 | 98 | 49 | NA | NA | NA | NA | 0.6 | NA | 1 | NA | -0.4 |
| SE | 9095 | HPS1 | HPS1 | chr10 | - | 98429790 | 98429889 | 98429145 | 98429642 | 98430570 | 98430670 | 0 | 0 | 1 | 0 | 0 | 1 | 0 | 1 | 0 | 0 | 0 | 0 | 98 | 49 | NA | NA | NA | NA | 0.333 | NA | 1 | NA | -0.667 |
| SE | 9358 | RASGRP1 | RASGRP1 | chr15 | - | 38503271 | 38503376 | 38502311 | 38502421 | 38505839 | 38505920 | 0 | 0 | 2 | 0 | 0 | 0 | 0 | 1 | 0 | 0 | 1 | 0 | 98 | 49 | NA | NA | NA | NA | 1 | NA | 0.333 | NA | 0.667 |
| SE | 9667 | PTEN | PTEN | chr10 | + | 87952117 | 87952259 | 87933012 | 87933251 | 87957852 | 87958019 | 0 | 0 | 4 | 0 | 0 | 1 | 0 | 2 | 0 | 0 | 1 | 0 | 98 | 49 | NA | NA | NA | NA | 0.667 | NA | 0.5 | NA | 0.167 |
| SE | 10439 | SLC12A6 | SLC12A6 | chr15 | - | 34242101 | 34242221 | 34241232 | 34241337 | 34243973 | 34244072 | 0 | 0 | 7 | 0 | 0 | 1 | 0 | 4 | 0 | 0 | 2 | 0 | 98 | 49 | NA | NA | NA | NA | 0.778 | NA | 0.5 | NA | 0.278 |
| SE | 10468 | PLCB2 | PLCB2 | chr15 | - | 40291605 | 40291650 | 40291264 | 40291487 | 40291848 | 40291924 | 0 | 0 | 1 | 0 | 0 | 2 | 0 | 0 | 0 | 0 | 1 | 0 | 94 | 49 | NA | NA | NA | NA | 0.207 | NA | 0 | NA | 0.207 |
| SE | 12021 | CCDC43 | CCDC43 | chr17 | - | 44680584 | 44680643 | 44677436 | 44679043 | 44682002 | 44682138 | 0 | 0 | 1 | 0 | 0 | 0 | 0 | 0 | 0 | 0 | 1 | 0 | 98 | 49 | NA | NA | NA | NA | 1 | NA | 0 | NA | 1 |
| SE | 12446 | TRAPPC8 | TRAPPC8 | chr18 | - | 31866848 | 31866975 | 31864626 | 31864781 | 31867401 | 31867476 | 0 | 0 | 5 | 0 | 0 | 0 | 0 | 0 | 0 | 0 | 2 | 0 | 98 | 49 | NA | NA | NA | NA | 1 | NA | 0 | NA | 1 |

| | | | | | | | | | | | | | | | | | | | | | | | | | | | | | | | | | | |
|---|---|---|---|---|---|---|---|---|---|---|---|---|---|---|---|---|---|---|---|---|---|---|---|---|---|---|---|---|---|---|---|---|---|---|
| SE | 12934 | EIF3K | EIF3K | chr19 | + | 38620336 | 38620435 | 38619073 | 38619327 | 38624076 | 38624197 | 0 | 0 | 8 | 0 | 0 | 1 | 0 | 10 | 0 | 0 | 0 | 0 | 98 | 49 | NA | NA | NA | NA | 0.8 | NA | 1 | NA | -0.2 |
| SE | 12935 | EIF3K | EIF3K | chr19 | + | 38632600 | 38632678 | 38632429 | 38632496 | 38634992 | 38635118 | 0 | 0 | 5 | 0 | 0 | 1 | 1 | 5 | 0 | 0 | 0 | 0 | 98 | 49 | NA | NA | NA | NA | 0.714 | 1 | 1 | NA | -0.286 |
| SE | 13565 | EML4 | EML4 | chr2 | + | 42301240 | 42301392 | 42295380 | 42295516 | 42303103 | 42303229 | 0 | 0 | 6 | 0 | 0 | 0 | 0 | 7 | 0 | 0 | 1 | 0 | 98 | 49 | NA | NA | NA | NA | 1 | NA | 0.778 | NA | 0.222 |

**MXE_all**

| | | | | | | | | | | | | | Inclusion_count | | | Skipping_count | | | Inclusion_count | | | Skipping_count | | | | | | | Inclusion_level (normalized) | | | | | | |
|---|---|---|---|---|---|---|---|---|---|---|---|---|---|---|---|---|---|---|---|---|---|---|---|---|---|---|---|---|---|---|---|---|---|---|---|
| Splice_type | ID | Gene_name | Chromosome | Strand | 1stExon | 1stExonEnd | 2ndExonStart_0t | 2ndExonEn | UpstreamES | UpstreamEE | DownstreamE | Downstrear | control_3 | control_4 | control_10 | control_3 | control_4 | control_10 | pmn_13 | pmn_15 | pmn_17 | pmn_13 | pmn_15 | pmn_17 | IncFormLen | SkipFormLen | p value | FDR | control_3 | control_4 | control_10 | pmn_13 | pmn_15 | pmn_17 | IncLevelDifference |
| MXE | 955 | DUSP1 | chr5 | - | 2E+08 | 172769794 | 172770160 | 172770306 | 172768089 | 172769132 | 172770585 | 172771200 | 561 | 303 | 269 | 438 | 200 | 178 | 23 | 118 | 275 | 3 | 47 | 10 | 98 | 98 | 2.30E-10 | 1.43E-07 | 0.562 | 0.602 | 0.602 | 0.885 | 0.715 | 0.965 | -0.266 |
| MXE | 1418 | SLC44A2 | chr19 | + | 1E+07 | 10635073 | 10636322 | 10636585 | 10634755 | 10634887 | 10636661 | 10636756 | 29 | 14 | 12 | 53 | 28 | 9 | 0 | 0 | 0 | 1 | 7 | 4 | 98 | 98 | 7.63E-08 | 2.24E-05 | 0.354 | 0.333 | 0.571 | 0 | 0 | 0 | 0.419 |
| MXE | 1132 | RFWD2 | chr1 | - | 2E+08 | 176136547 | 176175909 | 176176007 | 176135009 | 176135086 | 176184632 | 176184692 | 32 | 16 | 3 | 89 | 32 | 10 | 0 | 0 | 0 | 3 | 13 | 3 | 98 | 98 | 1.08E-07 | 2.24E-05 | 0.264 | 0.333 | 0.231 | 0 | 0 | 0 | 0.276 |
| MXE | 761 | CPEB4 | chr5 | + | 2E+08 | 173932500 | 173944966 | 173945140 | 173910522 | 173910604 | 173949507 | 173949597 | 8 | 3 | 7 | 28 | 12 | 10 | 0 | 0 | 0 | 5 | 5 | 3 | 98 | 98 | 1.22E-06 | 0.0001899 | 0.222 | 0.2 | 0.412 | 0 | 0 | 0 | 0.278 |
| MXE | 1306 | NLRP1 | chr17 | - | 6E+06 | 5532984 | 5533896 | 5533988 | 5530480 | 5530704 | 5536850 | 5536940 | 55 | 19 | 11 | 47 | 39 | 5 | 0 | 2 | 0 | 1 | 16 | 1 | 98 | 98 | 2.03E-06 | 0.0002533 | 0.539 | 0.328 | 0.687 | 0 | 0.111 | 0 | 0.481 |
| MXE | 1223 | IQGAP1 | chr15 | + | 9E+07 | 90467592 | 90472839 | 90473010 | 90466268 | 90466436 | 90473714 | 90473798 | 109 | 32 | 24 | 82 | 41 | 22 | 0 | 4 | 0 | 6 | 14 | 3 | 98 | 98 | 1.59E-05 | 0.0014171 | 0.571 | 0.438 | 0.522 | 0 | 0.222 | 0 | 0.436 |
| MXE | 964 | CTSS | chr1 | - | 2E+08 | 150757980 | 150764637 | 150764764 | 150755000 | 150755150 | 150765697 | 150765957 | 62 | 24 | 13 | 412 | 132 | 122 | 0 | 1 | 0 | 4 | 28 | 8 | 98 | 98 | 1.59E-05 | 0.0014171 | 0.131 | 0.154 | 0.096 | 0 | 0.034 | 0 | 0.116 |
| MXE | 1022 | CR1 | chr1 | + | 2E+08 | 207623068 | 207630516 | 207630621 | 207621972 | 207621996 | 207639396 | 207641765 | 16 | 5 | 8 | 41 | 25 | 5 | 4 | 3 | 1 | 0 | 2 | 0 | 98 | 98 | 4.26E-05 | 0.00332 | 0.281 | 0.167 | 0.615 | 1 | 0.6 | 1 | -0.512 |
| MXE | 873 | THEMIS2 | chr1 | + | 3E+07 | 27880054 | 27881970 | 27883043 | 27876587 | 27876728 | 27885294 | 27885451 | 111 | 23 | 15 | 25 | 14 | 1 | 1 | 12 | 8 | 0 | 2 | 0 | 98 | 98 | 0.00053316 | 0.0369065 | 0.816 | 0.622 | 0.937 | 1 | 0.857 | 1 | -0.161 |
| MXE | 113 | IGF2R | chr6 | + | 2E+08 | 160085131 | 160088032 | 160088147 | 160083949 | 160084184 | 160089106 | 160089253 | 119 | 27 | 21 | 77 | 19 | 6 | 1 | 14 | 3 | 0 | 2 | 1 | 98 | 98 | 0.000955554 | 0.059531 | 0.607 | 0.587 | 0.778 | 1 | 0.875 | 0.75 | -0.218 |
| MXE | 435 | XPO6 | chr16 | - | 3E+07 | 28177332 | 28180940 | 28181031 | 28175897 | 28176095 | 28211365 | 28211918 | 50 | 24 | 9 | 18 | 34 | 6 | 3 | 7 | 4 | 0 | 3 | 0 | 98 | 98 | 0.001633586 | 0.0925204 | 0.735 | 0.414 | 0.6 | 1 | 0.7 | 1 | -0.317 |
| MXE | 963 | CTSS | chr1 | - | 2E+08 | 150755150 | 150757857 | 150757980 | 150751780 | 150752008 | 150764637 | 150764764 | 410 | 160 | 108 | 262 | 144 | 131 | 20 | 17 | 6 | 2 | 11 | 0 | 98 | 98 | 0.002339859 | 0.1214777 | 0.61 | 0.526 | 0.452 | 0.909 | 0.607 | 1 | -0.309 |
| MXE | 672 | CD53 | chr1 | + | 1E+08 | 110892533 | 110897808 | 110897892 | 110891391 | 110891471 | 110899123 | 110899936 | 141 | 54 | 53 | 98 | 50 | 56 | 4 | 22 | 3 | 0 | 9 | 0 | 98 | 98 | 0.0040486 | 0.1940214 | 0.59 | 0.519 | 0.486 | 1 | 0.71 | 1 | -0.372 |
| MXE | 1034 | VIM | chr10 | + | 2E+07 | 17235889 | 17236293 | 17236379 | 17235168 | 17235389 | 17237229 | 17237593 | 146 | 40 | 21 | 218 | 59 | 45 | 0 | 15 | 0 | 15 | 37 | 1 | 93 | 98 | 0.005383209 | 0.2367954 | 0.414 | 0.417 | 0.33 | 0 | 0.299 | 0 | 0.287 |
| MXE | 956 | PTPRC | chr1 | + | 2E+08 | 198742026 | 198742231 | 198742367 | 198735126 | 198735252 | 198744053 | 198744203 | 196 | 110 | 52 | 207 | 86 | 77 | 0 | 33 | 1 | 8 | 49 | 9 | 98 | 98 | 0.005701333 | 0.2367954 | 0.486 | 0.561 | 0.403 | 0 | 0.402 | 0.1 | 0.316 |
| MXE | 973 | WDR1 | chr4 | - | 1E+07 | 10097891 | 10103895 | 10103986 | 10088663 | 10088741 | 10116112 | 10116234 | 36 | 16 | 13 | 37 | 7 | 7 | 1 | 7 | 0 | 0 | 1 | 1 | 98 | 98 | 0.00681759 | 0.2654599 | 0.493 | 0.696 | 0.65 | 1 | 0.875 | 0 | -0.012 |
| MXE | 428 | TNFRSF1A | chr12 | - | 6E+06 | 6333865 | 6334090 | 6334244 | 6333366 | 6333516 | 6341775 | 6342117 | 62 | 26 | 20 | 64 | 36 | 11 | 0 | 2 | 3 | 1 | 1 | 0 | 98 | 98 | 0.00983149 | 0.3602952 | 0.492 | 0.419 | 0.645 | 0 | 0.667 | 1 | -0.037 |
| MXE | 1444 | GMFG | chr19 | - | 4E+07 | 39329626 | 39333076 | 39333126 | 39328999 | 39329073 | 39335260 | 39335310 | 99 | 64 | 46 | 185 | 118 | 124 | 13 | 13 | 1 | 2 | 18 | 0 | 98 | 98 | 0.012757035 | 0.4415352 | 0.349 | 0.352 | 0.271 | 0.867 | 0.419 | 1 | -0.438 |
| MXE | 1131 | RFWD2 | chr1 | - | 2E+08 | 176136547 | 176149005 | 176149074 | 176135009 | 176135086 | 176184632 | 176184692 | 29 | 7 | 0 | 89 | 32 | 10 | 0 | 3 | 0 | 3 | 13 | 3 | 98 | 98 | 0.020043077 | 0.6497286 | 0.246 | 0.179 | 0 | 0 | 0.187 | 0 | 0.079 |
| MXE | 568 | MYO9B | chr19 | + | 2E+07 | 17157038 | 17159394 | 17159484 | 17154314 | 17154415 | 17162349 | 17162466 | 57 | 25 | 4 | 57 | 54 | 5 | 0 | 6 | 0 | 1 | 9 | 9 | 98 | 98 | 0.021827882 | 0.6497286 | 0.5 | 0.316 | 0.444 | 0 | 0.4 | 0 | 0.287 |
| MXE | 505 | SORL1 | chr11 | + | 1E+08 | 121545429 | 121549959 | 121550088 | 121543547 | 121543726 | 121550584 | 121550670 | 254 | 80 | 55 | 196 | 49 | 36 | 0 | 24 | 0 | 8 | 11 | 2 | 98 | 98 | 0.021900965 | 0.6497286 | 0.564 | 0.62 | 0.604 | 0 | 0.686 | 0 | 0.367 |
| MXE | 254 | ATXN7 | chr3 | + | 6E+07 | 63913225 | 63952378 | 63952483 | 63912587 | 63912923 | 63979914 | 63980167 | 13 | 7 | 7 | 29 | 5 | 3 | 0 | 3 | 0 | 1 | 1 | 2 | 98 | 98 | 0.025627133 | 0.7257138 | 0.31 | 0.583 | 0.7 | 0 | 0.75 | 0 | 0.281 |
| MXE | 1257 | KIAA1551 | chr12 | + | 3E+07 | 31960871 | 31970188 | 31970356 | 31959418 | 31959491 | 31980877 | 31985957 | 45 | 20 | 11 | 204 | 96 | 57 | 1 | 2 | 2 | 0 | 13 | 0 | 98 | 98 | 0.035676022 | 0.9651035 | 0.181 | 0.172 | 0.162 | 1 | 0.133 | 1 | -0.539 |
| MXE | 172 | DOCK2 | chr5 | + | 2E+08 | 169684350 | 169689251 | 169689333 | 169681743 | 169681879 | 169695802 | 169695938 | 109 | 35 | 17 | 83 | 48 | 5 | 0 | 13 | 2 | 4 | 19 | 4 | 98 | 98 | 0.038579299 | 0.9651035 | 0.568 | 0.422 | 0.773 | 0 | 0.406 | 0.333 | 0.341 |
| MXE | 436 | XPO6 | chr16 | - | 3E+07 | 28181031 | 28208986 | 28209122 | 28177219 | 28177332 | 28211365 | 28211918 | 8 | 6 | 1 | 338 | 139 | 60 | 0 | 0 | 0 | 3 | 38 | 12 | 98 | 98 | 0.038728069 | 0.9651035 | 0.023 | 0.041 | 0.016 | 0 | 0 | 0 | 0.027 |
| MXE | 1210 | ARHGAP9 | chr12 | - | 6E+07 | 57473708 | 57474041 | 57474176 | 57472254 | 57472688 | 57474422 | 57474476 | 214 | 71 | 70 | 116 | 35 | 41 | 2 | 11 | 2 | 0 | 4 | 0 | 98 | 98 | 0.043656592 | 1 | 0.648 | 0.67 | 0.631 | 1 | 0.733 | 1 | -0.261 |
| MXE | 110 | RNASET2 | chr6 | - | 2E+08 | 166934136 | 166938894 | 166939008 | 166931043 | 166931118 | 166943018 | 166943089 | 162 | 68 | 52 | 119 | 30 | 59 | 3 | 15 | 2 | 0 | 8 | 0 | 98 | 95 | 0.04769713 | 1 | 0.569 | 0.687 | 0.461 | 1 | 0.645 | 1 | -0.309 |
| MXE | 572 | NFKBIA | chr14 | - | 4E+07 | 35402859 | 35403149 | 35403360 | 35402393 | 35402663 | 35403689 | 35403798 | 77 | 218 | 23 | 78 | 178 | 23 | 0 | 83 | 4 | 7 | 108 | 6 | 98 | 98 | 0.051787064 | 1 | 0.497 | 0.551 | 0.5 | 0 | 0.435 | 0.4 | 0.238 |
| MXE | 390 | STX3 | chr11 | + | 6E+07 | 59787136 | 59788872 | 59788947 | 59773210 | 59773294 | 59790518 | 59790586 | 38 | 20 | 10 | 57 | 26 | 14 | 0 | 2 | 1 | 3 | 5 | 0 | 98 | 98 | 0.054812008 | 1 | 0.4 | 0.435 | 0.417 | 0 | 0.286 | 1 | -0.011 |
| MXE | 680 | GLUL | chr1 | - | 2E+08 | 182385556 | 182385759 | 182385887 | 182378092 | 182384723 | 182386255 | 182386402 | 662 | 192 | 81 | 1299 | 443 | 148 | 2 | 59 | 8 | 4 | 163 | 26 | 98 | 98 | 0.067852504 | 1 | 0.338 | 0.302 | 0.354 | 0.333 | 0.266 | 0.235 | 0.053 |
| MXE | 1145 | CYTIP | chr2 | - | 2E+08 | 157430962 | 157434369 | 157434424 | 157430558 | 157430652 | 157434697 | 157434747 | 184 | 88 | 26 | 187 | 85 | 27 | 6 | 55 | 6 | 5 | 32 | 1 | 98 | 98 | 0.074560925 | 1 | 0.496 | 0.509 | 0.491 | 0.545 | 0.632 | 0.857 | -0.179 |
| MXE | 114 | SOD2 | chr6 | - | 2E+08 | 159682638 | 159684853 | 159685033 | 159679115 | 159679298 | 159688125 | 159688242 | 413 | 247 | 217 | 148 | 110 | 100 | 8 | 78 | 4 | 0 | 31 | 0 | 98 | 98 | 0.095051109 | 1 | 0.736 | 0.692 | 0.685 | 1 | 0.716 | 1 | -0.201 |
| MXE | 125 | VNN2 | chr6 | - | 1E+08 | 132751518 | 132755842 | 132756035 | 132749694 | 132749865 | 132757415 | 132757546 | 299 | 172 | 111 | 126 | 82 | 68 | 1 | 34 | 5 | 2 | 6 | 0 | 98 | 98 | 0.09769198 | 1 | 0.704 | 0.677 | 0.62 | 0.333 | 0.85 | 1 | -0.061 |
| MXE | 221 | FYB | chr5 | - | 4E+07 | 39126135 | 39127740 | 39127807 | 39124252 | 39124278 | 39130589 | 39130612 | 91 | 29 | 17 | 59 | 19 | 8 | 0 | 12 | 3 | 6 | 16 | 1 | 98 | 98 | 0.109894645 | 1 | 0.607 | 0.604 | 0.68 | 0 | 0.429 | 0.75 | 0.237 |
| MXE | 431 | XPO6 | chr16 | - | 3E+07 | 28121762 | 28125688 | 28125848 | 28117317 | 28117462 | 28132333 | 28132403 | 129 | 71 | 20 | 153 | 88 | 40 | 0 | 12 | 1 | 8 | 27 | 0 | 98 | 98 | 0.112227384 | 1 | 0.457 | 0.447 | 0.333 | 0 | 0.308 | 1 | -0.024 |
| MXE | 1528 | ACSL1 | chr4 | - | 2E+08 | 184770476 | 184773080 | 184773154 | 184768315 | 184768450 | 184773662 | 184773714 | 163 | 95 | 53 | 85 | 32 | 78 | 3 | 28 | 2 | 0 | 15 | 0 | 98 | 98 | 0.127001838 | 1 | 0.657 | 0.748 | 0.405 | 1 | 0.651 | 1 | -0.28 |
| MXE | 822 | FAM129A | chr1 | - | 2E+08 | 184823742 | 184831846 | 184831962 | 184823166 | 184823329 | 184884632 | 184884800 | 136 | 33 | 28 | 122 | 66 | 18 | 0 | 6 | 1 | 1 | 13 | 1 | 98 | 98 | 0.129302592 | 1 | 0.527 | 0.333 | 0.609 | 0 | 0.316 | 0.5 | 0.218 |
| MXE | 910 | NADK | chr1 | - | 2E+06 | 1753649 | 1754050 | 1754208 | 1751231 | 1753060 | 1754283 | 1754383 | 60 | 35 | 36 | 54 | 19 | 38 | 4 | 14 | 3 | 0 | 4 | 5 | 98 | 98 | 0.157678336 | 1 | 0.526 | 0.648 | 0.486 | 1 | 0.778 | 0.375 | -0.164 |
| MXE | 115 | SOD2 | chr6 | - | 2E+08 | 159685033 | 159688125 | 159688242 | 159682473 | 159682638 | 159692660 | 159692863 | 370 | 266 | 195 | 468 | 299 | 215 | 10 | 80 | 9 | 5 | 105 | 2 | 98 | 98 | 0.163187452 | 1 | 0.442 | 0.471 | 0.476 | 0.667 | 0.432 | 0.818 | -0.176 |
| MXE | 456 | NCF2 | chr1 | - | 2E+08 | 183573292 | 183577598 | 183577707 | 183570779 | 183570839 | 183586894 | 183586977 | 161 | 90 | 98 | 280 | 132 | 80 | 2 | 18 | 1 | 0 | 37 | 0 | 98 | 98 | 0.174332737 | 1 | 0.365 | 0.405 | 0.551 | 1 | 0.327 | 1 | -0.335 |
| MXE | 1529 | ACSL1 | chr4 | - | 2E+08 | 184773875 | 184776483 | 184776662 | 184773662 | 184773714 | 184776883 | 184776983 | 82 | 24 | 41 | 106 | 47 | 33 | 0 | 20 | 1 | 2 | 39 | 0 | 98 | 82 | 0.182090854 | 1 | 0.393 | 0.299 | 0.51 | 0 | 0.3 | 1 | -0.033 |
| MXE | 145 | HLA-E | chr6 | + | 3E+07 | 30491653 | 30492403 | 30492436 | 30491136 | 30491412 | 30492540 | 30492583 | 302 | 247 | 202 | 143 | 135 | 125 | 11 | 70 | 10 | 0 | 48 | 3 | 98 | 82 | 0.184229858 | 1 | 0.639 | 0.605 | 0.575 | 1 | 0.55 | 0.736 | -0.156 |
| MXE | 962 | CTSS | chr1 | - | 2E+08 | 150752008 | 150755000 | 150755150 | 150750005 | 150750171 | 150757857 | 150757980 | 412 | 132 | 123 | 268 | 149 | 97 | 4 | 28 | 7 | 5 | 13 | 1 | 98 | 98 | 0.199307564 | 1 | 0.606 | 0.47 | 0.559 | 0.444 | 0.683 | 0.875 | -0.122 |
| MXE | 327 | MX1 | chr21 | + | 4E+07 | 41437152 | 41439693 | 41439848 | 41435836 | 41436029 | 41440886 | 41441025 | 18 | 10 | 0 | 55 | 40 | 2 | 4 | 18 | 2 | 1 | 47 | 6 | 98 | 98 | 0.224680315 | 1 | 0.247 | 0.2 | 0 | 0.8 | 0.277 | 0.25 | -0.293 |
| MXE | 1268 | IL4R | chr16 | + | 3E+07 | 27330198 | 27340185 | 27340273 | 27313908 | 27314020 | 27342120 | 27342259 | 0 | 1 | 4 | 54 | 21 | 12 | 0 | 0 | 0 | 1 | 12 | 18 | 98 | 98 | 0.228796443 | 1 | 0 | 0.045 | 0.25 | 0 | 0 | 0 | 0.098 |
| MXE | 430 | XPO6 | chr16 | - | 3E+07 | 28117462 | 28121669 | 28121762 | 28112903 | 28113050 | 28125688 | 28125848 | 186 | 130 | 42 | 115 | 75 | 26 | 7 | 34 | 0 | 0 | 19 | 1 | 98 | 98 | 0.228892606 | 1 | 0.618 | 0.634 | 0.618 | 1 | 0.642 | 0 | 0.076 |
| MXE | 786 | ARPC2 | chr2 | + | 2E+08 | 218239484 | 218245419 | 218245546 | 218238663 | 218238850 | 218249363 | 218249464 | 144 | 58 | 65 | 163 | 76 | 46 | 1 | 14 | 2 | 3 | 22 | 2 | 98 | 98 | 0.241047021 | 1 | 0.469 | 0.433 | 0.586 | 0.25 | 0.389 | 0.5 | 0.116 |
| MXE | 1526 | ACSL1 | chr4 | - | 2E+08 | 184764925 | 184765890 | 184765986 | 184763166 | 184763255 | 184766621 | 184766756 | 186 | 79 | 41 | 147 | 62 | 37 | 0 | 38 | 5 | 3 | 39 | 5 | 98 | 98 | 0.246337284 | 1 | 0.559 | 0.56 | 0.526 | 0 | 0.494 | 0.5 | 0.217 |
| MXE | 467 | LRRK2 | chr12 | + | 4E+07 | 40305966 | 40308466 | 40308696 | 40303947 | 40304134 | 40309105 | 40309233 | 213 | 78 | 30 | 288 | 118 | 21 | 6 | 11 | 1 | 0 | 13 | 2 | 98 | 98 | 0.253469383 | 1 | 0.425 | 0.398 | 0.588 | 1 | 0.458 | 0.333 | -0.127 |
| MXE | 827 | RNF19B | chr1 | - | 3E+07 | 32945628 | 32946401 | 32946564 | 32944018 | 32944159 | 32948221 | 32948363 | 67 | 25 | 18 | 53 | 35 | 4 | 0 | 9 | 1 | 1 | 15 | 0 | 98 | 98 | 0.284668519 | 1 | 0.558 | 0.417 | 0.818 | 0 | 0.375 | 1 | 0.139 |
| MXE | 432 | XPO6 | chr16 | - | 3E+07 | 28133933 | 28135215 | 28135324 | 28132333 | 28132403 | 28146093 | 28146203 | 125 | 73 | 21 | 134 | 92 | 25 | 0 | 19 | 0 | 1 | 21 | 1 | 98 | 98 | 0.294003081 | 1 | 0.483 | 0.442 | 0.457 | 0 | 0.475 | 0 | 0.302 |
| MXE | 1416 | MYO1F | chr19 | - | 9E+06 | 8530573 | 8536251 | 8536396 | 8530195 | 8530365 | 8536498 | 8536597 | 221 | 148 | 54 | 56 | 48 | 22 | 5 | 18 | 4 | 0 | 9 | 0 | 98 | 98 | 0.294584837 | 1 | 0.798 | 0.755 | 0.711 | 1 | 0.667 | 1 | -0.134 |
| MXE | 185 | CDC42SE2 | chr5 | + | 1E+08 | 131316144 | 131359208 | 131359547 | 131264008 | 131264166 | 131385542 | 131385644 | 9 | 2 | 0 | 50 | 14 | 10 | 1 | 1 | 1 | 0 | 9 | 3 | 98 | 98 | 0.294720446 | 1 | 0.153 | 0.125 | 0 | 1 | 0.1 | 0.25 | -0.357 |
| MXE | 441 | FAM49B | chr8 | - | 1E+08 | 129879471 | 129903311 | 129903350 | 129871374 | 129871496 | 129970942 | 129970995 | 33 | 27 | 14 | 92 | 55 | 26 | 0 | 7 | 1 | 4 | 13 | 4 | 88 | 98 | 0.29592073 | 1 | 0.285 | 0.353 | 0.375 | 0 | 0.375 | 0.218 | 0.14 |
| MXE | 1168 | RNF213 | chr17 | + | 8E+07 | 80361888 | 80363101 | 80363314 | 80360060 | 80360206 | 80363608 | 80363790 | 43 | 13 | 9 | 66 | 30 | 7 | 2 | 22 | 3 | 0 | 20 | 5 | 98 | 98 | 0.31350776 | 1 | 0.394 | 0.302 | 0.562 | 1 | 0.524 | 0.375 | -0.214 |
| MXE | 181 | ARHGAP26 | chr5 | + | 1E+08 | 142885399 | 142901934 | 142902039 | 142879373 | 142879445 | 142903539 | 142903669 | 267 | 142 | 17 | 160 | 74 | 22 | 1 | 64 | 1 | 0 | 30 | 1 | 98 | 98 | 0.314970171 | 1 | 0.625 | 0.657 | 0.436 | 1 | 0.681 | 0.5 | -0.154 |
| MXE | 454 | NCF2 | chr1 | - | 2E+08 | 183565779 | 183566919 | 183566988 | 183564004 | 183564030 | 183567203 | 183567345 | 55 | 44 | 53 | 135 | 63 | 66 | 0 | 10 | 1 | 2 | 15 | 0 | 98 | 98 | 0.432725555 | 1 | 0.289 | 0.411 | 0.445 | 0 | 0.4 | 1 | -0.085 |
| MXE | 797 | MED13L | chr12 | - | 1E+08 | 116096752 | 116111427 | 116111512 | 116022455 | 116022601 | 116237467 | 116237705 | 84 | 36 | 16 | 56 | 27 | 5 | 0 | 9 | 1 | 3 | 8 | 0 | 98 | 98 | 0.434851809 | 1 | 0.6 | 0.571 | 0.762 | 0 | 0.529 | 1 | 0.135 |
| MXE | 1458 | VASP | chr19 | + | 5E+07 | 45518094 | 45521321 | 45521406 | 45517662 | 45517834 | 45522167 | 45522217 | 121 | 49 | 30 | 98 | 38 | 44 | 0 | 18 | 2 | 3 | 11 | 3 | 98 | 98 | 0.440960111 | 1 | 0.553 | 0.563 | 0.405 | 0 | 0.621 | 0.4 | 0.167 |
| MXE | 872 | THEMIS2 | chr1 | + | 3E+07 | 27880054 | 27881970 | 27882125 | 27876587 | 27876728 | 27882512 | 27883043 | 110 | 23 | 14 | 0 | 5 | 0 | 1 | 12 | 8 | 0 | 0 | 0 | 98 | 98 | 0.469758481 | 1 | 1 | 0.821 | 1 | 1 | 1 | 1 | -0.06 |
| MXE | 1453 | PLAUR | chr19 | - | 4E+07 | 43656640 | 43665315 | 43665459 | 43655438 | 43655573 | 43667580 | 43667691 | 44 | 41 | 9 | 27 | 17 | 3 | 3 | 62 | 14 | 0 | 23 | 8 | 98 | 98 | 0.478260994 | 1 | 0.62 | 0.707 | 0.75 | 1 | 0.729 | 0.636 | -0.096 |
| MXE | 823 | MNDA | chr1 | + | 2E+08 | 158844122 | 158845586 | 158846003 | 158843278 | 158843415 | 158847727 | 158847916 | 358 | 171 | 135 | 428 | 329 | 232 | 0 | 32 | 6 | 1 | 36 | 1 | 98 | 98 | 0.481004194 | 1 | 0.455 | 0.342 | 0.368 | 0 | 0.471 | 0.857 | -0.054 |
| MXE | 474 | PREX1 | chr20 | - | 5E+07 | 48650206 | 48650893 | 48651055 | 48649299 | 48649576 | 48651395 | 48651583 | 31 | 22 | 13 | 84 | 37 | 35 | 0 | 9 | 3 | 6 | 22 | 3 | 98 | 98 | 0.486082517 | 1 | 0.27 | 0.373 | 0.271 | 0 | 0.29 | 0.5 | 0.041 |
| MXE | 502 | SORL1 | chr11 | + | 1E+08 | 121470123 | 121478117 | 121478243 | 121452202 | 121452616 | 121488031 | 121488193 | 174 | 85 | 67 | 166 | 80 | 53 | 1 | 24 | 2 | 0 | 22 | 1 | 98 | 98 | 0.488900024 | 1 | 0.512 | 0.515 | 0.558 | 1 | 0.522 | 0.667 | -0.201 |
| MXE | 491 | SLC6A6 | chr3 | + | 1E+07 | 14416453 | 14443623 | 14443863 | 14402575 | 14402847 | 14445716 | 14445851 | 95 | 41 | 9 | 75 | 39 | 28 | 1 | 10 | 3 | 0 | 9 | 4 | 91 | 98 | 0.548356538 | 1 | 0.577 | 0.531 | 0.257 | 1 | 0.545 | 0.447 | -0.209 |
| MXE | 1380 | NBEAL2 | chr3 | + | 5E+07 | 47003309 | 47003815 | 47003976 | 47002956 | 47003081 | 47004076 | 47004393 | 47 | 20 | 6 | 13 | 4 | 4 | 2 | 5 | 2 | 0 | 5 | 0 | 98 | 98 | 0.574951688 | 1 | 0.783 | 0.833 | 0.6 | 1 | 0.5 | 1 | -0.095 |
| MXE | 871 | THEMIS2 | chr1 | + | 3E+07 | 27876728 | 27879643 | 27880054 | 27872542 | 27872665 | 27885294 | 27885451 | 75 | 50 | 16 | 3 | 0 | 1 | 6 | 6 | 15 | 0 | 0 | 2 | 98 | 98 | 0.579301345 | 1 | 0.962 | 1 | 0.941 | 1 | 1 | 0.882 | 0.007 |
| MXE | 433 | XPO6 | chr16 | - | 3E+07 | 28135324 | 28146093 | 28146203 | 28133840 | 28133933 | 28152658 | 28152785 | 127 | 43 | 20 | 198 | 116 | 46 | 3 | 19 | 5 | 3 | 41 | 4 | 98 | 98 | 0.593249562 | 1 | 0.391 | 0.27 | 0.303 | 0.5 | 0.317 | 0.556 | -0.136 |
| MXE | 690 | RAB8B | chr15 | + | 6E+07 | 63249705 | 63255506 | 63255584 | 63244755 | 63244816 | 63256504 | 63256594 | 19 | 14 | 5 | 12 | 2 | 8 | 4 | 1 | 1 | 0 | 3 | 1 | 98 | 98 | 0.608491204 | 1 | 0.613 | 0.875 | 0.385 | 1 | 0.25 | 0.5 | 0.041 |
| MXE | 607 | ADAM8 | chr10 | - | 1E+08 | 133274235 | 133275483 | 133275587 | 133273950 | 133274029 | 133276771 | 133276903 | 18 | 15 | 1 | 84 | 46 | 27 | 1 | 6 | 4 | 1 | 26 | 10 | 98 | 98 | 0.649266983 | 1 | 0.176 | 0.246 | 0.036 | 0.5 | 0.187 | 0.286 | -0.172 |
| MXE | 202 | PARP8 | chr5 | + | 5E+07 | 50794332 | 50794852 | 50795417 | 50788522 | 50788589 | 50796981 | 50797032 | 104 | 16 | 13 | 81 | 16 | 1 | 4 | 3 | 0 | 0 | 7 | 1 | 98 | 98 | 0.67128589 | 1 | 0.562 | 0.5 | 0.929 | 1 | 0.3 | 0 | 0.23 |
| MXE | 870 | THEMIS2 | chr1 | + | 3E+07 | 27876728 | 27879643 | 27880054 | 27872542 | 27872665 | 27881970 | 27882125 | 72 | 53 | 10 | 47 | 9 | 18 | 6 | 5 | 15 | 0 | 13 | 5 | 98 | 98 | 0.679724192 | 1 | 0.605 | 0.855 | 0.357 | 1 | 0.278 | 0.75 | -0.07 |
| MXE | 1334 | MGAM | chr7 | + | 1E+08 | 142036285 | 142036822 | 142036977 | 142034669 | 142034841 | 142038530 | 142038615 | 110 | 29 | 10 | 97 | 19 | 10 | 2 | 5 | 1 | 0 | 2 | 6 | 98 | 98 | 0.688352365 | 1 | 0.531 | 0.604 | 0.5 | 1 | 0.714 | 0.143 | -0.074 |
| MXE | 1414 | CYTH4 | chr22 | + | 4E+07 | 37311064 | 37311455 | 37311527 | 37309211 | 37309323 | 37312019 | 37312174 | 82 | 23 | 19 | 55 | 41 | 44 | 0 | 11 | 1 | 3 | 10 | 0 | 98 | 98 | 0.744744769 | 1 | 0.599 | 0.359 | 0.302 | 0 | 0.524 | 1 | -0.088 |
| MXE | 677 | HCLS1 | chr3 | - | 1E+08 | 121632184 | 121632331 | 121632563 | 121631397 | 121631982 | 121633066 | 121633171 | 102 | 63 | 71 | 82 | 51 | 19 | 0 | 21 | 12 | 1 | 8 | 4 | 98 | 98 | 0.764021251 | 1 | 0.554 | 0.553 | 0.789 | 0 | 0.724 | 0.75 | 0.141 |
| MXE | 1447 | GMFG | chr19 | - | 4E+07 | 39335310 | 39335434 | 39335531 | 39333076 | 39333126 | 39335973 | 39336086 | 5 | 0 | 0 | 97 | 63 | 46 | 0 | 1 | 0 | 13 | 12 | 1 | 98 | 98 | 0.773168367 | 1 | 0.049 | 0 | 0 | 0 | 0.077 | 0 | -0.009 |
| MXE | 174 | DOCK2 | chr5 | + | 2E+08 | 169761625 | 169803057 | 169803206 | 169759704 | 169759775 | 169840756 | 169840852 | 45 | 19 | 11 | 79 | 34 | 16 | 0 | 8 | 1 | 1 | 9 | 2 | 98 | 98 | 0.779672199 | 1 | 0.363 | 0.358 | 0.407 | 0 | 0.471 | 0.333 | 0.108 |
| MXE | 461 | LRRK2 | chr12 | + | 4E+07 | 40240617 | 40243549 | 40243681 | 40237968 | 40238103 | 40249825 | 40249945 | 491 | 161 | 45 | 171 | 64 | 23 | 2 | 27 | 1 | 0 | 6 | 4 | 98 | 98 | 0.791444635 | 1 | 0.742 | 0.716 | 0.662 | 1 | 0.818 | 0.2 | 0.034 |
| MXE | 1152 | MEFV | chr16 | - | 3E+06 | 3247246 | 3248908 | 3249004 | 3246524 | 3246547 | 3249430 | 3249780 | 23 | 10 | 12 | 84 | 58 | 39 | 0 | 7 | 0 | 2 | 12 | 2 | 98 | 98 | 0.804538892 | 1 | 0.215 | 0.147 | 0.235 | 0 | 0.368 | 0 | 0.076 |
| MXE | 654 | DENND3 | chr8 | + | 1E+08 | 141160787 | 141163332 | 141163429 | 141155848 | 141155970 | 141165185 | 141165289 | 52 | 20 | 10 | 60 | 17 | 13 | 0 | 5 | 2 | 3 | 6 | 0 | 98 | 98 | 0.80764641 | 1 | 0.464 | 0.541 | 0.435 | 0 | 0.455 | 1 | -0.005 |
| MXE | 1442 | SMCHD1 | chr18 | + | 3E+06 | 2722663 | 2724898 | 2724995 | 2718314 | 2718434 | 2726451 | 2726524 | 103 | 62 | 32 | 217 | 108 | 24 | 1 | 25 | 1 | 0 | 48 | 1 | 98 | 98 | 0.865049573 | 1 | 0.322 | 0.365 | 0.571 | 1 | 0.342 | 0.5 | -0.195 |
| MXE | 483 | CLEC7A | chr12 | - | 1E+07 | 10123363 | 10125296 | 10125448 | 10116780 | 10118590 | 10127746 | 10127845 | 48 | 19 | 15 | 92 | 36 | 25 | 2 | 8 | 0 | 0 | 12 | 2 | 98 | 98 | 0.866957778 | 1 | 0.343 | 0.345 | 0.375 | 1 | 0.4 | 0 | -0.112 |
| MXE | 476 | PREX1 | chr20 | - | 5E+07 | 48726391 | 48734545 | 48734650 | 48708259 | 48708421 | 48745024 | 48745147 | 112 | 69 | 46 | 182 | 100 | 47 | 3 | 42 | 5 | 8 | 31 | 14 | 98 | 98 | 0.884198696 | 1 | 0.381 | 0.408 | 0.495 | 0.273 | 0.575 | 0.263 | 0.058 |
| MXE | 1153 | MEFV | chr16 | - | 3E+06 | 3249004 | 3249430 | 3249780 | 3247015 | 3247246 | 3254157 | 3254790 | 46 | 22 | 18 | 48 | 18 | 35 | 0 | 11 | 2 | 3 | 13 | 0 | 98 | 98 | 0.884277074 | 1 | 0.489 | 0.55 | 0.34 | 0 | 0.458 | 1 | -0.026 |
| MXE | 746 | CDC42SE1 | chr1 | - | 2E+08 | 151055126 | 151055676 | 151055993 | 151054230 | 151054321 | 151059478 | 151059649 | 49 | 16 | 7 | 128 | 59 | 33 | 2 | 4 | 1 | 7 | 13 | 0 | 98 | 98 | 0.915529711 | 1 | 0.277 | 0.213 | 0.175 | 0.222 | 0.235 | 1 | -0.264 |
| MXE | 503 | SORL1 | chr11 | + | 1E+08 | 121478243 | 121488031 | 121488193 | 121470006 | 121470123 | 121490042 | 121490110 | 184 | 118 | 36 | 296 | 153 | 134 | 0 | 16 | 1 | 1 | 21 | 1 | 98 | 98 | 0.916016225 | 1 | 0.383 | 0.435 | 0.212 | 0 | 0.432 | 0.5 | 0.033 |
| MXE | 272 | ABHD5 | chr3 | + | 4E+07 | 43715058 | 43717670 | 43717857 | 43711708 | 43711863 | 43718442 | 43722725 | 61 | 17 | 23 | 40 | 7 | 7 | 0 | 4 | 1 | 6 | 1 | 0 | 98 | 98 | 0.926381298 | 1 | 0.604 | 0.708 | 0.767 | 0 | 0.8 | 1 | 0.093 |
| MXE | 455 | NCF2 | chr1 | - | 2E+08 | 183566988 | 183567203 | 183567345 | 183565703 | 183565779 | 183569141 | 183569185 | 111 | 80 | 66 | 180 | 77 | 88 | 2 | 20 | 1 | 6 | 7 | 4 | 98 | 98 | 0.955623146 | 1 | 0.381 | 0.51 | 0.429 | 0.25 | 0.741 | 0.2 | 0.043 |
| MXE | 804 | LSP1 | chr11 | + | 2E+06 | 1880224 | 1881431 | 1881596 | 1852969 | 1853197 | 1883418 | 1883560 | 149 | 52 | 33 | 111 | 83 | 45 | 0 | 21 | 11 | 3 | 15 | 5 | 98 | 98 | 0.97674093 | 1 | 0.573 | 0.385 | 0.423 | 0 | 0.583 | 0.687 | 0.037 |
| MXE | 4 | SASH3 | chrX | + | 1E+08 | 129792676 | 129792626 | 129792836 | 129790939 | 129791081 | 129792988 | 129793139 | 33 | 18 | 3 | 24 | 12 | 2 | 0 | 5 | 1 | 0 | 1 | 0 | 98 | 98 | 1 | 1 | 0.579 | 0.6 | 0.6 | NA | 0.833 | 1 | -0.324 |
| MXE | 6 | ALG13 | chrX | + | 1E+08 | 111682235 | 111682235 | 111682294 | 111681117 | 111681299 | 111684964 | 111685103 | 0 | 0 | 0 | 14 | 0 | 2 | 0 | 1 | 0 | 0 | 0 | 0 | 98 | 98 | 1 | 1 | 0 | NA | 0 | NA | 1 | NA | -1 |
| MXE | 9 | MORF4L2 | chrX | - | 1E+08 | 103678600 | 103684680 | 103684729 | 103675497 | 103677051 | 103685170 | 103685260 | 1 | 2 | 0 | 10 | 2 | 3 | 0 | 0 | 0 | 0 | 1 | 0 | 98 | 98 | 1 | 1 | 0.091 | 0.5 | 0 | NA | 0 | NA | 0.197 |
| MXE | 10 | MORF4L2 | chrX | - | 1E+08 | 103678620 | 103684680 | 103684729 | 103675497 | 103677051 | 103685170 | 103685260 | 1 | 2 | 0 | 10 | 2 | 3 | 0 | 0 | 0 | 0 | 1 | 0 | 98 | 98 | 1 | 1 | 0.091 | 0.5 | 0 | NA | 0 | NA | 0.197 |
| MXE | 11 | MORF4L2 | chrX | - | 1E+08 | 103678651 | 103684680 | 103684729 | 103675497 | 103677051 | 103685170 | 103685260 | 1 | 2 | 0 | 12 | 2 | 3 | 0 | 0 | 0 | 0 | 1 | 0 | 98 | 98 | 1 | 1 | 0.077 | 0.5 | 0 | NA | 0 | NA | 0.192 |
| MXE | 19 | KDM6A | chrX | + | 5E+07 | 45060156 | 45060608 | 45060764 | 45059246 | 45059466 | 45061323 | 45061419 | 7 | 0 | 0 | 11 | 8 | 2 | 0 | 0 | 0 | 0 | 5 | 0 | 98 | 98 | 1 | 1 | 0.389 | 0 | 0 | NA | 0 | NA | 0.13 |
| MXE | 21 | ASMTL | chrX | - | 1E+06 | 1439144 | 1442185 | 1442317 | 1435693 | 1435758 | 1452747 | 1452977 | 2 | 0 | 0 | 3 | 2 | 0 | 0 | 0 | 0 | 0 | 3 | 0 | 98 | 97 | 1 | 1 | 0.398 | 0 | NA | NA | 0 | NA | 0.199 |
| MXE | 22 | CSF2RA | chrX | + | 1E+06 | 1300626 | 1303922 | 1304019 | 1295426 | 1295456 | 1305445 | 1305527 | 15 | 11 | 8 | 8 | 3 | 10 | 0 | 6 | 1 | 0 | 1 | 0 | 98 | 98 | 1 | 1 | 0.652 | 0.786 | 0.444 | NA | 0.857 | 1 | -0.301 |
| MXE | 24 | ASB6 | chr9 | - | 1E+08 | 129638659 | 129639201 | 129639310 | 129634603 | 129638457 | 129639401 | 129639508 | 2 | 0 | 4 | 4 | 2 | 0 | 0 | 1 | 2 | 0 | 1 | 1 | 98 | 98 | 1 | 1 | 0.333 | 0 | 1 | NA | 0.5 | 0.667 | -0.139 |
| MXE | 25 | PPP2R4 | chr9 | + | 1E+08 | 129123138 | 129128984 | 129129110 | 129120512 | 129120610 | 129131521 | 129131639 | 1 | 0 | 0 | 2 | 0 | 0 | 0 | 0 | 0 | 0 | 1 | 0 | 98 | 98 | 1 | 1 | 0.333 | NA | NA | | 0 | NA | 0.333 |
| MXE | 26 | AKNA | chr9 | - | 1E+08 | 114377532 | 114381059 | 114381446 | 114374092 | 114374167 | 114394356 | 114394405 | 13 | 9 | 1 | 56 | 16 | 8 | 0 | 1 | 0 | 0 | 7 | 0 | 98 | 98 | 1 | 1 | 0.188 | 0.36 | 0.111 | NA | 0.125 | NA | 0.095 |
| MXE | 27 | TLE4 | chr9 | + | 8E+07 | 79612718 | 79627373 | 79627448 | 79576132 | 79576177 | 79652589 | 79652794 | 23 | 4 | 3 | 4 | 0 | 0 | 0 | 4 | 0 | 0 | 2 | 0 | 98 | 98 | 1 | 1 | 0.852 | 1 | 1 | NA | 0.667 | NA | 0.284 |
| MXE | 28 | TLE4 | chr9 | + | 8E+07 | 79612718 | 79627373 | 79627448 | 79576132 | 79576177 | 79652592 | 79652794 | 22 | 4 | 3 | 7 | 1 | 0 | 0 | 4 | 0 | 0 | 2 | 0 | 98 | 98 | 1 | 1 | 0.759 | 0.8 | 1 | NA | 0.667 | NA | 0.186 |
| MXE | 29 | ZFAND5 | chr9 | - | 7E+07 | 72359521 | 72360109 | 72360221 | 72356930 | 72357056 | 72360627 | 72360784 | 2 | 0 | 0 | 42 | 6 | 4 | 0 | 0 | 0 | 0 | 2 | 0 | 98 | 98 | 1 | 1 | 0.045 | 0 | 0 | NA | 0 | NA | 0.015 |
| MXE | 38 | UBAP1 | chr9 | + | 3E+07 | 34220948 | 34234215 | 34234340 | 34179004 | 34179240 | 34241184 | 34242108 | 7 | 3 | 4 | 4 | 0 | 4 | 0 | 1 | 0 | 0 | 2 | 0 | 90 | 98 | 1 | 1 | 0.656 | 1 | 0.521 | NA | 0.353 | NA | 0.373 |
| MXE | 41 | UHRF2 | chr9 | + | 6E+06 | 6421142 | 6433913 | 6434173 | 6413150 | 6413643 | 6460572 | 6460791 | 4 | 3 | 0 | 3 | 6 | 1 | 0 | 2 | 0 | 0 | 1 | 0 | 98 | 98 | 1 | 1 | 0.571 | 0.333 | 0 | NA | 0.667 | NA | -0.366 |
| MXE | 49 | VPS13B | chr8 | + | 1E+08 | 99507203 | 99507769 | 99507911 | 99502835 | 99502950 | 99511103 | 99511512 | 38 | 11 | 0 | 4 | 1 | 0 | 0 | 6 | 0 | 0 | 0 | 0 | 98 | 98 | 1 | 1 | 0.905 | 0.917 | NA | NA | 1 | NA | -0.089 |
| MXE | 50 | CPQ | chr8 | + | 1E+08 | 96835180 | 96879797 | 96880005 | 96784863 | 96785330 | 96965934 | 96966046 | 33 | 16 | 7 | 24 | 15 | 5 | 0 | 4 | 0 | 0 | 3 | 1 | 98 | 98 | 1 | 1 | 0.579 | 0.516 | 0.583 | NA | 0.571 | 0 | 0.274 |
| MXE | 63 | YTHDF3 | chr8 | + | 6E+07 | 63175416 | 63186146 | 63187745 | 63169386 | 63169411 | 63209682 | 63212788 | 3 | 6 | 0 | 19 | 3 | 0 | 0 | 0 | 0 | 0 | 6 | 0 | 98 | 98 | 1 | 1 | 0.136 | 0.667 | NA | NA | 0 | NA | 0.402 |
| MXE | 66 | GSR | chr8 | - | 3E+07 | 30689319 | 30692968 | 30693055 | 30684087 | 30684199 | 30696379 | 30696479 | 11 | 1 | 1 | 6 | 5 | 7 | 0 | 2 | 0 | 0 | 0 | 0 | 98 | 98 | 1 | 1 | 0.647 | 0.167 | 0.125 | NA | 1 | NA | -0.687 |
| MXE | 79 | PARP10 | chr8 | - | 1E+08 | 143983811 | 143984007 | 143984104 | 143982931 | 143983065 | 143984209 | 143984431 | 8 | 3 | 1 | 10 | 1 | 0 | 0 | 3 | 0 | 0 | 3 | 1 | 98 | 98 | 1 | 1 | 0.444 | 0.75 | 1 | NA | 0.5 | 0 | 0.481 |
| MXE | 80 | LINC-PINT | chr7 | - | 1E+08 | 130959970 | 130967388 | 130967517 | 130944159 | 130945835 | 130984052 | 130984109 | 36 | 7 | 0 | 21 | 1 | 0 | 0 | 1 | 0 | 0 | 1 | 0 | 98 | 98 | 1 | 1 | 0.632 | 0.875 | NA | NA | 0.5 | NA | 0.254 |
| MXE | 82 | NAMPT | chr7 | - | 1E+08 | 106254504 | 106263391 | 106263617 | 106253016 | 106253151 | 106268463 | 106268600 | 0 | 1 | 0 | 1558 | 790 | 321 | 0 | 0 | 0 | 14 | 339 | 15 | 98 | 98 | 1 | 1 | 0 | 0.001 | 0 | 0 | | 0 | 0 |
| MXE | 83 | NAMPT | chr7 | - | 1E+08 | 106263617 | 106268463 | 106268600 | 106261587 | 106261707 | 106269153 | 106269312 | 680 | 331 | 111 | 1 | 1 | 1 | 0 | 88 | 7 | 0 | 2 | 0 | 98 | 98 | 1 | 1 | 0.999 | 0.997 | 0.991 | NA | 0.978 | 1 | 0.007 |
| MXE | 88 | LMTK2 | chr7 | + | 1E+08 | 98171674 | 98185050 | 98185135 | 98159337 | 98159425 | 98186876 | 98186998 | 15 | 15 | 4 | 4 | 1 | 2 | 0 | 2 | 2 | 0 | 7 | 0 | 98 | 98 | 1 | 1 | 0.789 | 0.937 | 0.667 | NA | 0.222 | 1 | 0.187 |
| MXE | 90 | DMTF1 | chr7 | + | 9E+07 | 87165050 | 87166482 | 87166605 | 87163494 | 87163617 | 87170994 | 87171089 | 9 | 7 | 4 | 15 | 11 | 0 | 0 | 0 | 0 | 0 | 5 | 0 | 98 | 98 | 1 | 1 | 0.375 | 0.389 | 1 | NA | 0 | NA | 0.588 |
| MXE | 91 | GUSB | chr7 | - | 7E+07 | 65976202 | 65979833 | 65979911 | 65974918 | 65975071 | 65980223 | 65980409 | 1 | 0 | 0 | 0 | 0 | 1 | 0 | 2 | 0 | 0 | 0 | 0 | 98 | 98 | 1 | 1 | 1 | NA | 0 | NA | 1 | NA | -0.5 |
| MXE | 92 | GK | chrX | + | 3E+07 | 30691199 | 30694399 | 30694537 | 30677374 | 30677452 | 30696041 | 30696151 | 9 | 2 | 0 | 13 | 4 | 10 | 0 | 2 | 0 | 0 | 2 | 0 | 98 | 98 | 1 | 1 | 0.409 | 0.333 | 0 | NA | 0.5 | NA | -0.253 |
| MXE | 96 | IKZF1 | chr7 | + | 5E+07 | 50382707 | 50387344 | 50387470 | 50327637 | 50327757 | 50391728 | 50391833 | 15 | 1 | 3 | 5 | 2 | 5 | 0 | 4 | 0 | 0 | 0 | 0 | 98 | 98 | 1 | 1 | 0.75 | 0.333 | 0.375 | NA | 1 | NA | -0.514 |
| MXE | 99 | IKZF1 | chr7 | + | 5E+07 | 50387470 | 50391728 | 50391833 | 50382539 | 50382707 | 50399917 | 50405100 | 25 | 3 | 4 | 5 | 0 | 2 | 0 | 1 | 0 | 0 | 0 | 0 | 98 | 98 | 1 | 1 | 0.833 | 1 | 0.667 | NA | 1 | NA | -0.167 |
| MXE | 100 | IKZF1 | chr7 | + | 5E+07 | 50387470 | 50391728 | 50391863 | 50382539 | 50382707 | 50399917 | 50405100 | 25 | 3 | 4 | 6 | 2 | 0 | 0 | 1 | 0 | 0 | 0 | 0 | 98 | 98 | 1 | 1 | 0.806 | 0.6 | 1 | NA | 1 | NA | -0.198 |
| MXE | 101 | CALU | chr7 | + | 1E+08 | 128754455 | 128754528 | 128754722 | 128748572 | 128748804 | 128758870 | 128759037 | 0 | 0 | 3 | 0 | 0 | 0 | 0 | 0 | 0 | 0 | 1 | 0 | 98 | 98 | 1 | 1 | NA | NA | 1 | NA | 0 | NA | 1 |
| MXE | 103 | AOAH | chr7 | - | 4E+07 | 36637910 | 36659165 | 36659265 | 36632035 | 36632106 | 36673942 | 36674009 | 18 | 4 | 0 | 32 | 13 | 4 | 0 | 4 | 0 | 0 | 6 | 1 | 98 | 98 | 1 | 1 | 0.36 | 0.235 | 0 | NA | 0.4 | 0 | -0.002 |
| MXE | 109 | SUN1 | chr7 | + | 832504 | 832601 | 838797 | 838986 | 816614 | 816673 | 841945 | 842130 | 0 | 0 | 0 | 2 | 0 | 0 | 0 | 1 | 0 | 0 | 0 | 0 | 98 | 98 | 1 | 1 | 0 | NA | NA | NA | 1 | NA | -1 |
| MXE | 112 | IGF2R | chr6 | + | 2E+08 | 160027314 | 160029549 | 160029655 | 160024571 | 160024704 | 160032550 | 160032713 | 252 | 84 | 46 | 147 | 63 | 26 | 0 | 13 | 0 | 0 | 9 | 1 | 98 | 98 | 1 | 1 | 0.632 | 0.571 | 0.639 | NA | 0.591 | 0 | 0.319 |
| MXE | 116 | SCAF8 | chr6 | + | 2E+08 | 154822409 | 154824233 | 154824378 | 154820176 | 154820333 | 154827171 | 154827240 | 16 | 4 | 8 | 13 | 0 | 0 | 0 | 1 | 0 | 2 | 0 | 0 | 98 | 98 | 1 | 1 | 0.552 | 1 | 1 | 0 | 1 | NA | 0.351 |
| MXE | 126 | VNN2 | chr6 | - | 1E+08 | 132752749 | 132753417 | 132753491 | 132751144 | 132751518 | 132755842 | 132756035 | 34 | 5 | 4 | 330 | 178 | 186 | 0 | 1 | 1 | 0 | 18 | 1 | 98 | 98 | 1 | 1 | 0.093 | 0.027 | 0.021 | NA | 0.053 | 0.5 | -0.23 |
| MXE | 127 | VNN2 | chr6 | - | 1E+08 | 132752749 | 132753417 | 132753491 | 132751144 | 132751518 | 132755842 | 132756035 | 34 | 5 | 4 | 115 | 82 | 84 | 0 | 1 | 1 | 0 | 3 | 0 | 98 | 98 | 1 | 1 | 0.228 | 0.057 | 0.045 | NA | 0.25 | 1 | -0.515 |
| MXE | 130 | SNX14 | chr6 | - | 9E+07 | 85558060 | 85565331 | 85565419 | 85549722 | 85549879 | 85567533 | 85567577 | 9 | 1 | 3 | 4 | 2 | 0 | 0 | 0 | 1 | 0 | 1 | 0 | 98 | 98 | 1 | 1 | 0.692 | 0.333 | 1 | NA | 0 | 1 | 0.175 |
| MXE | 132 | KCTD20 | chr6 | + | 4E+07 | 36470257 | 36479120 | 36479223 | 36442766 | 36443111 | 36479590 | 36479711 | 5 | 3 | 0 | 4 | 1 | 0 | 1 | 2 | 0 | 0 | 0 | 0 | 98 | 98 | 1 | 1 | 0.556 | 0.75 | NA | 1 | 1 | NA | -0.347 |
| MXE | 133 | KCTD20 | chr6 | + | 4E+07 | 36470257 | 36479120 | 36479223 | 36442766 | 36443111 | 36481561 | 36481759 | 5 | 0 | 0 | 0 | 1 | 0 | 1 | 0 | 0 | 0 | 0 | 0 | 98 | 98 | 1 | 1 | 1 | 0 | NA | 1 | NA | NA | -0.5 |
| MXE | 134 | KCTD20 | chr6 | + | 4E+07 | 36470257 | 36479590 | 36479711 | 36442766 | 36443111 | 36481561 | 36481759 | 5 | 0 | 0 | 8 | 2 | 2 | 1 | 0 | 0 | 0 | 2 | 0 | 98 | 98 | 1 | 1 | 0.385 | 0 | 0 | 1 | 0 | NA | -0.372 |
| MXE | 135 | KCTD20 | chr6 | + | 4E+07 | 36479223 | 36479590 | 36479711 | 36442766 | 36443111 | 36481561 | 36481759 | 0 | 1 | 0 | 8 | 2 | 2 | 0 | 0 | 0 | 0 | 2 | 0 | 98 | 98 | 1 | 1 | 0 | 0.333 | 0 | NA | 0 | NA | 0.111 |
| MXE | 136 | KCTD20 | chr6 | + | 4E+07 | 36479223 | 36479590 | 36479711 | 36470051 | 36470257 | 36481561 | 36481759 | 1 | 0 | 0 | 8 | 5 | 2 | 0 | 0 | 0 | 0 | 3 | 0 | 98 | 98 | 1 | 1 | 0.111 | 0 | 0 | NA | 0 | NA | 0.037 |
| MXE | 138 | MAPK14 | chr6 | + | 4E+07 | 36096066 | 36100187 | 36100267 | 36076536 | 36076608 | 36102570 | 36102649 | 31 | 21 | 14 | 18 | 11 | 4 | 0 | 6 | 0 | 0 | 0 | 0 | 98 | 98 | 1 | 1 | 0.633 | 0.656 | 0.778 | NA | 1 | NA | -0.311 |
| MXE | 144 | FLOT1 | chr6 | - | 3E+07 | 30741333 | 30741613 | 30741704 | 30740678 | 30740798 | 30741791 | 30741867 | 24 | 14 | 13 | 39 | 14 | 17 | 0 | 1 | 0 | 0 | 5 | 1 | 98 | 98 | 1 | 1 | 0.381 | 0.5 | 0.433 | NA | 0.167 | 0 | 0.354 |

| | | | | | | | | | | | | | | | | | | | | | | | | | | | | | | | | | | | | |
|---|---|---|---|---|---|---|---|---|---|---|---|---|---|---|---|---|---|---|---|---|---|---|---|---|---|---|---|---|---|---|---|---|---|---|---|---|
| MXE | 146 | HLA-F | chr6 | + | 3E+07 | 29725306 | 29725446 | 29725563 | 29724172 | 29724448 | 29726402 | 29726526 | 120 | 51 | 32 | 2 | 0 | 3 | 0 | 13 | 0 | 0 | 1 | 0 | 98 | 98 | 1 | 1 | 0.984 | 1 | 0.914 | NA | 0.929 | NA | | 0.037 |
| MXE | 151 | BTN3A2 | chr6 | + | 3E+07 | 26370603 | 26372896 | 26373097 | 26368564 | 26368912 | 26373275 | 26373296 | 8 | 2 | 6 | 0 | 0 | 1 | 0 | 0 | 0 | 2 | 0 | 0 | 98 | 98 | 1 | 1 | 1 | 1 | 1 | 0 | NA | NA | | 1 |
| MXE | 154 | USP9X | chrX | + | 4E+07 | 41137022 | 41140655 | 41140771 | 41134724 | 41134837 | 41140965 | 41141217 | 19 | 2 | 4 | 6 | 9 | 1 | 0 | 3 | 0 | 0 | 0 | 0 | 98 | 98 | 1 | 1 | 0.76 | 0.182 | 0.8 | NA | 1 | NA | | -0.419 |
| MXE | 162 | PDLIM7 | chr5 | - | 2E+08 | 177491420 | 177491806 | 177491925 | 177491009 | 177491146 | 177492404 | 177492435 | 30 | 14 | 17 | 0 | 0 | 0 | 0 | 6 | 4 | 0 | 1 | 0 | 98 | 66 | 1 | 1 | 1 | 1 | 1 | NA | 0.802 | | 1 | 0.099 |
| MXE | 171 | DOCK2 | chr5 | + | 2E+08 | 169670597 | 169671077 | 169671174 | 169669287 | 169669328 | 169674296 | 169674445 | 32 | 11 | 5 | 142 | 34 | 5 | 0 | 6 | 0 | 0 | 15 | 1 | 98 | 98 | 1 | 1 | 0.184 | 0.244 | 0.5 | NA | 0.286 | | 0 | 0.166 |
| MXE | 173 | DOCK2 | chr5 | + | 2E+08 | 169759775 | 169761518 | 169761625 | 169747395 | 169747504 | 169803057 | 169803206 | 58 | 31 | 6 | 67 | 41 | 4 | 0 | 19 | 0 | 0 | 4 | 0 | 98 | 98 | 1 | 1 | 0.464 | 0.431 | 0.6 | NA | 0.826 | NA | | -0.328 |
| MXE | 175 | CNOT8 | chr5 | + | 2E+08 | 154865385 | 154870660 | 154870822 | 154863206 | 154863395 | 154871729 | 154871874 | 6 | 3 | 4 | 9 | 2 | 2 | 0 | 1 | 0 | 0 | 1 | 0 | 98 | 98 | 1 | 1 | 0.4 | 0.6 | 0.667 | NA | 0.5 | NA | | 0.056 |
| MXE | 176 | CNOT8 | chr5 | + | 2E+08 | 154867792 | 154870660 | 154870822 | 154865191 | 154865385 | 154871729 | 154871874 | 0 | 0 | 1 | 13 | 4 | 7 | 0 | 0 | 0 | 0 | 1 | 0 | 98 | 98 | 1 | 1 | 0 | 0 | 0.125 | NA | 0 | NA | | 0.042 |
| MXE | 182 | ARHGAP26 | chr5 | + | 1E+08 | 143014116 | 143037195 | 143037261 | 142932046 | 142932125 | 143041815 | 143041890 | 134 | 37 | 12 | 50 | 33 | 4 | 0 | 18 | 2 | 0 | 9 | 0 | 86 | 98 | 1 | 1 | 0.753 | 0.561 | 0.774 | NA | 0.695 | | 1 | -0.151 |
| MXE | 184 | FNIP1 | chr5 | - | 1E+08 | 131677872 | 131679028 | 131679175 | 131671504 | 131672924 | 131698916 | 131699002 | 16 | 2 | 2 | 34 | 15 | 2 | 0 | 2 | 0 | 0 | 4 | 0 | 98 | 98 | 1 | 1 | 0.32 | 0.118 | 0.5 | NA | 0.333 | NA | | -0.02 |
| MXE | 187 | ASAP1 | chr8 | - | 1E+08 | 130092143 | 130112093 | 130112322 | 130079901 | 130079971 | 130115627 | 130115735 | 37 | 7 | 0 | 5 | 5 | 2 | 0 | 4 | 1 | 0 | 2 | 0 | 98 | 98 | 1 | 1 | 0.881 | 0.583 | 0 | NA | 0.667 | | 1 | -0.346 |
| MXE | 188 | DCP2 | chr5 | + | 1E+08 | 113004077 | 113007937 | 113008042 | 113001566 | 113001674 | 113010755 | 113010807 | 12 | 8 | 7 | 26 | 7 | 2 | 0 | 1 | 0 | 0 | 0 | 0 | 98 | 98 | 1 | 1 | 0.316 | 0.533 | 0.778 | NA | 1 | NA | | -0.458 |
| MXE | 190 | MCTP1 | chr5 | - | 9E+07 | 94917973 | 94931952 | 94931991 | 94912805 | 94912976 | 94940083 | 94940195 | 25 | 9 | 6 | 8 | 15 | 2 | 0 | 4 | 0 | 0 | 2 | 0 | 88 | 98 | 1 | 1 | 0.777 | 0.401 | 0.77 | NA | 0.69 | NA | | -0.041 |
| MXE | 191 | VCAN | chr5 | + | 8E+07 | 83522309 | 83537006 | 83542268 | 83512102 | 83512396 | 83545536 | 83545650 | 1 | 0 | 0 | 56 | 0 | 0 | 0 | 0 | 0 | 0 | 10 | 0 | 98 | 98 | 1 | 1 | 0.018 | NA | NA | NA | 0 | NA | | 0.018 |
| MXE | 193 | PAPD4 | chr5 | + | 8E+07 | 79649190 | 79656957 | 79657001 | 79648616 | 79648693 | 79668891 | 79669028 | 18 | 4 | 0 | 13 | 3 | 0 | 0 | 1 | 0 | 0 | 2 | 0 | 98 | 93 | 1 | 1 | 0.568 | 0.559 | NA | NA | 0.322 | NA | | 0.241 |
| MXE | 194 | DBNL | chr7 | + | 4E+07 | 44050280 | 44052866 | 44052941 | 44044639 | 44044820 | 44056756 | 44056903 | 11 | 3 | 5 | 24 | 6 | 9 | 0 | 1 | 0 | 0 | 1 | 6 | 98 | 98 | 1 | 1 | 0.314 | 0.333 | 0.357 | NA | 0.5 | | 0 | 0.085 |
| MXE | 195 | FCHO2 | chr5 | + | 7E+07 | 73006549 | 73015625 | 73015724 | 72990711 | 72990864 | 73017211 | 73017308 | 15 | 3 | 0 | 22 | 9 | 1 | 0 | 1 | 0 | 0 | 0 | 0 | 98 | 98 | 1 | 1 | 0.405 | 0.25 | 0 | NA | 1 | NA | | -0.782 |
| MXE | 204 | DDX60L | chr4 | - | 2E+08 | 168441492 | 168448637 | 168448779 | 168433009 | 168433115 | 168453123 | 168453282 | 30 | 22 | 3 | 44 | 27 | 0 | 0 | 17 | 0 | 0 | 15 | 0 | 98 | 98 | 1 | 1 | 0.405 | 0.449 | 1 | NA | 0.531 | NA | | 0.087 |
| MXE | 206 | MGST2 | chr4 | + | 1E+08 | 139678642 | 139695196 | 139695267 | 139665767 | 139666077 | 139703454 | 139703536 | 0 | 0 | 0 | 4 | 0 | 0 | 0 | 1 | 0 | 0 | 0 | 0 | 98 | 98 | 1 | 1 | 0 | NA | NA | NA | 1 | NA | | -1 |
| MXE | 210 | HERC5 | chr4 | + | 9E+07 | 88470673 | 88472408 | 88472502 | 88469156 | 88469260 | 88475840 | 88476030 | 7 | 18 | 1 | 4 | 11 | 1 | 0 | 20 | 1 | 0 | 15 | 3 | 98 | 98 | 1 | 1 | 0.636 | 0.621 | 0.5 | NA | 0.571 | | 0.25 | 0.175 |
| MXE | 213 | MRPS18C | chr4 | + | 8E+07 | 83458429 | 83459739 | 83459797 | 83456908 | 83456958 | 83460972 | 83461032 | 3 | 0 | 1 | 2 | 2 | 5 | 0 | 0 | 0 | 0 | 1 | 0 | 98 | 98 | 1 | 1 | 0.6 | 0 | 0.167 | NA | 0 | NA | | 0.256 |
| MXE | 222 | FYB | chr5 | - | 4E+07 | 39153604 | 39201825 | 39202987 | 39141094 | 39141141 | 39219442 | 39219578 | 52 | 17 | 0 | 90 | 43 | 10 | 0 | 6 | 0 | 14 | 9 | 0 | 98 | 98 | 1 | 1 | 0.366 | 0.283 | 0 | 0 | 0.4 | NA | | 0.016 |
| MXE | 225 | MAEA | chr4 | + | 1E+06 | 1315600 | 1322380 | 1322503 | 1311978 | 1312161 | 1327626 | 1327703 | 15 | 13 | 2 | 1 | 4 | 1 | 0 | 2 | 7 | 0 | 3 | 1 | 98 | 98 | 1 | 1 | 0.937 | 0.765 | 0.667 | NA | 0.4 | | 0.875 | 0.152 |
| MXE | 226 | MAEA | chr4 | + | 1E+06 | 1315600 | 1322380 | 1322503 | 1311978 | 1312161 | 1332756 | 1332865 | 15 | 16 | 2 | 0 | 3 | 1 | 0 | 2 | 7 | 0 | 0 | 0 | 98 | 98 | 1 | 1 | 1 | 0.842 | 0.667 | NA | 1 | | 1 | -0.164 |
| MXE | 227 | MAEA | chr4 | + | 1E+06 | 1322503 | 1327626 | 1327703 | 1315396 | 1315600 | 1332756 | 1332865 | 11 | 5 | 2 | 12 | 5 | 7 | 0 | 1 | 0 | 0 | 1 | 0 | 98 | 98 | 1 | 1 | 0.478 | 0.5 | 0.222 | NA | 0.5 | NA | | -0.1 |
| MXE | 228 | DLG1 | chr3 | - | 2E+08 | 197075870 | 197076585 | 197076682 | 197069218 | 197069260 | 197081050 | 197081117 | 1 | 0 | 0 | 2 | 0 | 0 | 0 | 0 | 0 | 0 | 1 | 0 | 98 | 83 | 1 | 1 | 0.297 | NA | NA | NA | 0 | NA | | 0.297 |
| MXE | 229 | DLG1 | chr3 | - | 2E+08 | 197075870 | 197076585 | 197076685 | 197069218 | 197069260 | 197081050 | 197081117 | 1 | 0 | 0 | 2 | 0 | 0 | 0 | 0 | 0 | 0 | 1 | 0 | 98 | 83 | 1 | 1 | 0.297 | NA | NA | NA | 0 | NA | | 0.297 |
| MXE | 232 | VNN3 | chr6 | - | 1E+08 | 132727012 | 132728740 | 132728933 | 132724605 | 132724979 | 132734252 | 132734383 | 51 | 21 | 4 | 39 | 26 | 1 | 0 | 9 | 1 | 0 | 2 | 0 | 98 | 98 | 1 | 1 | 0.567 | 0.447 | 0.8 | NA | 0.818 | | 1 | -0.304 |
| MXE | 233 | VNN3 | chr6 | - | 1E+08 | 132727012 | 132728740 | 132728979 | 132724605 | 132724979 | 132734252 | 132734383 | 44 | 14 | 6 | 39 | 26 | 1 | 0 | 7 | 1 | 0 | 2 | 0 | 98 | 98 | 1 | 1 | 0.53 | 0.35 | 0.857 | NA | 0.778 | | 1 | -0.31 |
| MXE | 234 | ACAP2 | chr3 | - | 2E+08 | 195289231 | 195291705 | 195291815 | 195285795 | 195285857 | 195292264 | 195292452 | 20 | 13 | 2 | 31 | 13 | 9 | 0 | 3 | 0 | 3 | 10 | 0 | 98 | 98 | 1 | 1 | 0.392 | 0.5 | 0.182 | 0 | 0.231 | NA | | 0.243 |
| MXE | 235 | OPA1 | chr3 | + | 2E+08 | 193617285 | 193617783 | 193617837 | 193615673 | 193615770 | 193618868 | 193618936 | 11 | 0 | 1 | 3 | 0 | 0 | 0 | 0 | 0 | 0 | 1 | 0 | 98 | 98 | 1 | 1 | 0.786 | NA | 1 | NA | 0 | NA | | 0.893 |
| MXE | 236 | IL1RAP | chr3 | + | 2E+08 | 190550505 | 190556129 | 190556216 | 190514050 | 190514219 | 190564288 | 190564353 | 0 | 1 | 0 | 22 | 20 | 1 | 0 | 0 | 0 | 0 | 4 | 2 | 98 | 98 | 1 | 1 | 0 | 0.048 | 0 | NA | 0 | | 0 | 0.016 |
| MXE | 240 | FNDC3B | chr3 | + | 2E+08 | 172247776 | 172251259 | 172251541 | 172226870 | 172226947 | 172285925 | 172285984 | 20 | 6 | 8 | 18 | 4 | 7 | 0 | 1 | 0 | 0 | 2 | 0 | 98 | 98 | 1 | 1 | 0.526 | 0.6 | 0.533 | NA | 0.333 | NA | | 0.22 |
| MXE | 242 | FYN | chr6 | - | 1E+08 | 111699670 | 111700103 | 111700268 | 111696276 | 111696456 | 111702884 | 111703034 | 0 | 0 | 1 | 5 | 3 | 0 | 0 | 0 | 0 | 2 | 1 | 0 | 98 | 98 | 1 | 1 | 0 | 0 | 1 | 0 | 0 | NA | | 0.333 |
| MXE | 243 | MME | chr3 | + | 2E+08 | 155142127 | 155142236 | 155142330 | 155140190 | 155140292 | 155143442 | 155143571 | 67 | 18 | 23 | 95 | 36 | 23 | 0 | 2 | 0 | 0 | 6 | 0 | 98 | 98 | 1 | 1 | 0.414 | 0.333 | 0.5 | NA | 0.25 | NA | | 0.166 |
| MXE | 244 | EIF2A | chr3 | + | 2E+08 | 150572529 | 150575648 | 150575762 | 150568175 | 150568292 | 150581617 | 150581746 | 6 | 0 | 2 | 6 | 1 | 0 | 0 | 2 | 0 | 0 | 0 | 0 | 98 | 98 | 1 | 1 | 0.5 | 0 | 1 | NA | 1 | NA | | -0.5 |
| MXE | 256 | SLMAP | chr3 | + | 6E+07 | 57864857 | 57871635 | 57871698 | 57864547 | 57864716 | 57890040 | 57890100 | 6 | 3 | 3 | 4 | 10 | 0 | 0 | 1 | 0 | 0 | 1 | 0 | 98 | 98 | 1 | 1 | 0.6 | 0.231 | 1 | NA | 0.5 | NA | | 0.11 |
| MXE | 258 | VPRBP | chr3 | - | 5E+07 | 51441897 | 51443765 | 51443903 | 51440969 | 51441071 | 51463113 | 51463227 | 5 | 1 | 0 | 4 | 0 | 3 | 0 | 3 | 1 | 0 | 0 | 0 | 98 | 98 | 1 | 1 | 0.556 | 1 | 0 | NA | 1 | | 1 | -0.481 |
| MXE | 259 | KIAA0922 | chr4 | + | 2E+08 | 153591194 | 153592474 | 153592584 | 153588889 | 153589007 | 153593798 | 153593871 | 19 | 14 | 1 | 9 | 13 | 0 | 0 | 2 | 1 | 0 | 0 | 0 | 98 | 98 | 1 | 1 | 0.679 | 0.519 | 1 | NA | 1 | | 1 | -0.267 |
| MXE | 274 | MBNL1 | chr3 | + | 2E+08 | 152455577 | 152456266 | 152456361 | 152447619 | 152447773 | 152459270 | 152459345 | 17 | 1 | 8 | 35 | 7 | 9 | 0 | 0 | 0 | 2 | 7 | 0 | 85 | 98 | 1 | 1 | 0.359 | 0.141 | 0.506 | 0 | 0 | NA | | 0.335 |
| MXE | 289 | LST1 | chr6 | + | 3E+07 | 31587733 | 31587943 | 31587966 | 31587199 | 31587318 | 31588496 | 31588909 | 39 | 24 | 22 | 9 | 9 | 2 | 0 | 1 | 0 | 0 | 0 | 0 | 98 | 72 | 1 | 1 | 0.761 | 0.662 | 0.89 | NA | 1 | NA | | -0.229 |
| MXE | 290 | LST1 | chr6 | + | 3E+07 | 31587733 | 31587943 | 31587966 | 31587199 | 31587318 | 31588517 | 31588909 | 42 | 28 | 26 | 17 | 17 | 14 | 0 | 2 | 0 | 0 | 0 | 1 | 98 | 72 | 1 | 1 | 0.645 | 0.548 | 0.577 | NA | 1 | | 0 | 0.09 |
| MXE | 292 | FBXL13 | chr7 | - | 1E+08 | 102822440 | 102832839 | 102832974 | 102822039 | 102822203 | 102854776 | 102854860 | 8 | 1 | 0 | 1 | 5 | 1 | 0 | 3 | 0 | 0 | 1 | 0 | 98 | 98 | 1 | 1 | 0.889 | 0.167 | 0 | NA | 0.75 | NA | | -0.398 |
| MXE | 293 | FBXL13 | chr7 | - | 1E+08 | 102832974 | 102854776 | 102854860 | 102822039 | 102822203 | 102877466 | 102877593 | 18 | 7 | 0 | 6 | 0 | 0 | 0 | 1 | 0 | 0 | 0 | 0 | 98 | 98 | 1 | 1 | 0.75 | 1 | NA | NA | 1 | NA | | -0.125 |
| MXE | 294 | FBXL13 | chr7 | - | 1E+08 | 102931933 | 102963532 | 102963665 | 102926273 | 102926374 | 102968021 | 102968117 | 26 | 1 | 0 | 30 | 18 | 2 | 0 | 2 | 0 | 6 | 1 | 0 | 98 | 98 | 1 | 1 | 0.464 | 0.053 | 0 | 0 | 0.667 | NA | | -0.161 |
| MXE | 300 | TBC1D22A | chr22 | + | 5E+07 | 46797620 | 46878652 | 46878723 | 46793500 | 46793841 | 46891265 | 46891394 | 24 | 2 | 2 | 15 | 6 | 0 | 0 | 0 | 0 | 0 | 0 | 0 | 98 | 98 | 1 | 1 | 0.615 | 0.25 | 1 | NA | 1 | NA | | -0.378 |
| MXE | 302 | TBC1D22A | chr22 | + | 5E+07 | 46974399 | 46997633 | 46997709 | 46912073 | 46912188 | 47037070 | 47037198 | 11 | 2 | 0 | 13 | 8 | 4 | 0 | 0 | 0 | 5 | 2 | 0 | 98 | 98 | 1 | 1 | 0.458 | 0.2 | 0 | 0 | 0 | NA | | 0.219 |
| MXE | 303 | TBC1D22A | chr22 | + | 5E+07 | 46997709 | 47037070 | 47037198 | 46974289 | 46974399 | 47111507 | 47111603 | 6 | 3 | 10 | 12 | 10 | 1 | 0 | 2 | 0 | 0 | 2 | 0 | 98 | 98 | 1 | 1 | 0.333 | 0.231 | 0.909 | NA | 0.5 | NA | | -0.009 |
| MXE | 304 | MKL1 | chr22 | - | 4E+07 | 40423685 | 40424205 | 40424381 | 40420846 | 40421100 | 40429605 | 40429767 | 10 | 3 | 1 | 5 | 6 | 0 | 0 | 1 | 0 | 0 | 1 | 0 | 98 | 98 | 1 | 1 | 0.667 | 0.333 | 1 | NA | 0.5 | NA | | 0.167 |
| MXE | 305 | TNRC6B | chr22 | + | 4E+07 | 40267036 | 40270121 | 40270280 | 40261831 | 40262173 | 40273424 | 40273600 | 27 | 14 | 0 | 6 | 2 | 1 | 0 | 4 | 0 | 0 | 3 | 0 | 98 | 98 | 1 | 1 | 0.818 | 0.875 | 0 | NA | 0.571 | NA | | -0.007 |
| MXE | 306 | GTPBP1 | chr22 | + | 4E+07 | 38727348 | 38727982 | 38728161 | 38726257 | 38726440 | 38729461 | 38729662 | 16 | 7 | 5 | 27 | 12 | 0 | 0 | 7 | 1 | 0 | 5 | 0 | 98 | 98 | 1 | 1 | 0.372 | 0.368 | 1 | NA | 0.583 | | 1 | -0.212 |
| MXE | 309 | PGM2 | chr4 | + | 4E+07 | 37848651 | 37850183 | 37850373 | 37847201 | 37847295 | 37855607 | 37855741 | 7 | 18 | 3 | 23 | 10 | 2 | 0 | 0 | 0 | 0 | 1 | 0 | 98 | 98 | 1 | 1 | 0.233 | 0.643 | 0.6 | NA | 0 | NA | | 0.492 |
| MXE | 312 | TANGO2 | chr22 | + | 2E+07 | 20053551 | 20055942 | 20056013 | 20052464 | 20052584 | 20063337 | 20063442 | 1 | 6 | 7 | 2 | 3 | 1 | 5 | 0 | 0 | 0 | 0 | 0 | 98 | 98 | 1 | 1 | 0.333 | 0.667 | 0.875 | 1 | NA | NA | | -0.375 |
| MXE | 313 | TANGO2 | chr22 | + | 2E+07 | 20053551 | 20061529 | 20061683 | 20052464 | 20052584 | 20063337 | 20063442 | 1 | 6 | 7 | 0 | 3 | 2 | 5 | 0 | 0 | 0 | 0 | 0 | 98 | 98 | 1 | 1 | 1 | 0.667 | 0.778 | 1 | NA | NA | | -0.185 |
| MXE | 315 | TANGO2 | chr22 | + | 2E+07 | 20056013 | 20063337 | 20063442 | 20053436 | 20053551 | 20064541 | 20065926 | 4 | 7 | 1 | 5 | 9 | 5 | 0 | 0 | 0 | 0 | 1 | 0 | 98 | 98 | 1 | 1 | 0.444 | 0.437 | 0.167 | NA | 0 | NA | | 0.349 |
| MXE | 316 | PRMT2 | chr21 | + | 5E+07 | 46658920 | 46663382 | 46663554 | 46649574 | 46649739 | 46664294 | 46665124 | 9 | 2 | 5 | 4 | 9 | 2 | 0 | 2 | 0 | 0 | 0 | 0 | 98 | 98 | 1 | 1 | 0.692 | 0.182 | 0.714 | NA | 1 | NA | | -0.471 |
| MXE | 332 | NRIP1 | chr21 | - | 2E+07 | 15014466 | 15043494 | 15043574 | 14961234 | 14968526 | 15064744 | 15064805 | 3 | 4 | 0 | 0 | 1 | 0 | 0 | 2 | 0 | 0 | 1 | 1 | 98 | 98 | 1 | 1 | 1 | 0.8 | NA | NA | 0.667 | | 0 | 0.567 |
| MXE | 333 | SAMSN1 | chr21 | - | 1E+07 | 14500735 | 14510309 | 14510461 | 14498441 | 14498592 | 14512443 | 14512573 | 10 | 5 | 0 | 22 | 5 | 12 | 0 | 6 | 0 | 0 | 11 | 2 | 98 | 98 | 1 | 1 | 0.312 | 0.5 | 0 | NA | 0.353 | | 0 | 0.094 |
| MXE | 336 | SLC9A8 | chr20 | + | 5E+07 | 49845119 | 49849578 | 49849680 | 49839540 | 49839599 | 49855437 | 49855581 | 13 | 2 | 0 | 14 | 9 | 0 | 0 | 2 | 2 | 0 | 4 | 1 | 98 | 98 | 1 | 1 | 0.481 | 0.182 | NA | NA | 0.333 | | 0.667 | -0.168 |
| MXE | 337 | NUB1 | chr7 | + | 2E+08 | 151356222 | 151360140 | 151360247 | 151355767 | 151355950 | 151366938 | 151367125 | 17 | 6 | 0 | 9 | 7 | 7 | 0 | 4 | 0 | 0 | 3 | 1 | 98 | 98 | 1 | 1 | 0.654 | 0.462 | 0 | NA | 0.571 | | 0 | 0.087 |
| MXE | 338 | SULF2 | chr20 | - | 5E+07 | 47702670 | 47736702 | 47736942 | 47690125 | 47690295 | 47757188 | 47757463 | 32 | 7 | 4 | 7 | 5 | 3 | 0 | 10 | 4 | 0 | 1 | 0 | 98 | 98 | 1 | 1 | 0.821 | 0.583 | 0.571 | NA | 0.909 | | 1 | -0.296 |
| MXE | 339 | ZMYND8 | chr20 | - | 5E+07 | 47262428 | 47276313 | 47276795 | 47249286 | 47249439 | 47282101 | 47282217 | 5 | 7 | 1 | 1 | 3 | 0 | 0 | 0 | 0 | 0 | 1 | 0 | 98 | 98 | 1 | 1 | 0.833 | 0.7 | 1 | NA | 0 | NA | | 0.844 |
| MXE | 351 | C20orf24 | chr20 | + | 4E+07 | 36608000 | 36609600 | 36609730 | 36607714 | 36607818 | 36612009 | 36612557 | 3 | 3 | 1 | 29 | 41 | 54 | 0 | 2 | 0 | 0 | 8 | 0 | 98 | 98 | 1 | 1 | 0.094 | 0.068 | 0.018 | NA | 0.2 | NA | | -0.14 |
| MXE | 352 | C20orf24 | chr20 | + | 4E+07 | 36608000 | 36609600 | 36609758 | 36607714 | 36607818 | 36612009 | 36612557 | 3 | 3 | 1 | 9 | 11 | 26 | 0 | 2 | 0 | 0 | 6 | 0 | 98 | 98 | 1 | 1 | 0.25 | 0.214 | 0.037 | NA | 0.5 | NA | | -0.333 |
| MXE | 354 | EDEM2 | chr20 | - | 4E+07 | 35145018 | 35146824 | 35146935 | 35142372 | 35142478 | 35147151 | 35147358 | 5 | 6 | 0 | 0 | 5 | 5 | 0 | 0 | 1 | 0 | 2 | 0 | 98 | 89 | 1 | 1 | 1 | 0.521 | 0 | NA | 0 | | 1 | 0.007 |
| MXE | 366 | GPCPD1 | chr20 | - | 6E+06 | 5560076 | 5561464 | 5561530 | 5558683 | 5558819 | 5565016 | 5565078 | 72 | 14 | 11 | 86 | 39 | 10 | 0 | 15 | 0 | 0 | 11 | 0 | 98 | 98 | 1 | 1 | 0.456 | 0.264 | 0.524 | NA | 0.577 | NA | | -0.162 |
| MXE | 376 | MAPKAPK3 | chr3 | + | 5E+07 | 50644532 | 50645709 | 50645785 | 50642252 | 50642332 | 50646139 | 50646264 | 29 | 17 | 9 | 15 | 14 | 4 | 0 | 4 | 0 | 0 | 3 | 0 | 98 | 98 | 1 | 1 | 0.659 | 0.548 | 0.692 | NA | 0.571 | NA | | 0.062 |
| MXE | 378 | MFF | chr2 | + | 2E+08 | 227342819 | 227352513 | 227352573 | 227340291 | 227340380 | 227355676 | 227355761 | 0 | 0 | 2 | 2 | 2 | 0 | 0 | 2 | 0 | 0 | 1 | 0 | 98 | 98 | 1 | 1 | 0 | NA | 1 | NA | 0.667 | NA | | -0.167 |
| MXE | 380 | CASP10 | chr2 | + | 2E+08 | 201203766 | 201208074 | 201208183 | 201195841 | 201195948 | 201209069 | 201209562 | 2 | 0 | 0 | 2 | 0 | 0 | 0 | 0 | 0 | 0 | 0 | 0 | 86 | 98 | 1 | 1 | 0.533 | NA | NA | NA | 0 | NA | | 0.533 |
| MXE | 381 | CALCOCO2 | chr17 | + | 5E+07 | 48841887 | 48848063 | 48848166 | 48830987 | 48831078 | 48848321 | 48848455 | 0 | 0 | 0 | 7 | 4 | 4 | 0 | 1 | 0 | 0 | 4 | 0 | 98 | 98 | 1 | 1 | 0 | 0 | 0 | NA | 0.2 | NA | | -0.2 |
| MXE | 383 | CALCOCO2 | chr17 | + | 5E+07 | 48849377 | 48851088 | 48851177 | 48848321 | 48848455 | 48851558 | 48851628 | 15 | 15 | 10 | 11 | 5 | 3 | 0 | 2 | 0 | 0 | 4 | 0 | 98 | 98 | 1 | 1 | 0.577 | 0.75 | 0.769 | NA | 0.333 | NA | | 0.366 |
| MXE | 384 | USP15 | chr12 | + | 6E+07 | 62325933 | 62349220 | 62349307 | 62321463 | 62321609 | 62355330 | 62355475 | 193 | 56 | 26 | 80 | 33 | 23 | 0 | 7 | 0 | 2 | 6 | 0 | 98 | 98 | 1 | 1 | 0.707 | 0.629 | 0.531 | 0 | 0.538 | NA | | 0.353 |
| MXE | 387 | ARHGEF7 | chr13 | + | 1E+08 | 111205373 | 111209871 | 111210002 | 111153713 | 111153991 | 111217678 | 111217880 | 4 | 2 | 0 | 6 | 1 | 1 | 4 | 1 | 0 | 0 | 1 | 0 | 98 | 98 | 1 | 1 | 0.4 | 0.667 | 0 | 1 | 0.5 | NA | | -0.394 |
| MXE | 388 | ELP2 | chr18 | + | 4E+07 | 36138872 | 36139418 | 36139613 | 36138269 | 36138426 | 36141136 | 36141201 | 7 | 2 | 0 | 0 | 5 | 0 | 0 | 1 | 0 | 0 | 0 | 0 | 98 | 98 | 1 | 1 | 1 | 0.286 | NA | NA | 1 | NA | | -0.357 |
| MXE | 389 | ELP2 | chr18 | + | 4E+07 | 36146048 | 36146249 | 36146381 | 36144938 | 36145034 | 36154849 | 36154999 | 3 | 0 | 0 | 3 | 0 | 0 | 0 | 0 | 0 | 0 | 2 | 0 | 98 | 98 | 1 | 1 | 0.5 | NA | NA | NA | 0 | NA | | 0.5 |
| MXE | 407 | NRM | chr6 | - | 3E+07 | 30689452 | 30690024 | 30690243 | 30688046 | 30688942 | 30690841 | 30691129 | 1 | 0 | 0 | 1 | 0 | 0 | 0 | 0 | 1 | 0 | 0 | 0 | 98 | 98 | 1 | 1 | 0.5 | NA | NA | NA | NA | | 1 | -0.5 |
| MXE | 408 | NRM | chr6 | - | 3E+07 | 30689452 | 30690046 | 30690243 | 30688046 | 30688942 | 30690841 | 30691129 | 1 | 0 | 0 | 1 | 0 | 0 | 0 | 0 | 1 | 0 | 0 | 0 | 98 | 98 | 1 | 1 | 0.5 | NA | NA | NA | NA | | 1 | -0.5 |
| MXE | 409 | LDLR | chr19 | + | 1E+07 | 11102786 | 11105219 | 11105600 | 11100222 | 11100345 | 11106564 | 11106687 | 2 | 0 | 0 | 1 | 0 | 0 | 0 | 1 | 0 | 0 | 0 | 0 | 98 | 98 | 1 | 1 | 0.667 | NA | NA | NA | 1 | NA | | -0.333 |
| MXE | 410 | LCP1 | chr13 | - | 5E+07 | 46158651 | 46158825 | 46158989 | 46156437 | 46156570 | 46159598 | 46159686 | 50 | 39 | 15 | 250 | 100 | 86 | 0 | 6 | 0 | 0 | 43 | 0 | 98 | 98 | 1 | 1 | 0.167 | 0.281 | 0.149 | NA | 0.122 | NA | | 0.077 |
| MXE | 411 | SETX | chr9 | - | 1E+08 | 132334727 | 132336295 | 132336515 | 132331276 | 132331448 | 132342689 | 132342799 | 116 | 29 | 8 | 80 | 20 | 2 | 0 | 6 | 0 | 0 | 2 | 0 | 98 | 98 | 1 | 1 | 0.592 | 0.592 | 0.8 | NA | 0.75 | NA | | -0.089 |
| MXE | 412 | SETX | chr9 | - | 1E+08 | 132336515 | 132342689 | 132342799 | 132334607 | 132334727 | 132346260 | 132346471 | 72 | 17 | 2 | 91 | 20 | 4 | 0 | 5 | 1 | 0 | 8 | 0 | 98 | 98 | 1 | 1 | 0.442 | 0.459 | 0.333 | NA | 0.385 | | 1 | -0.281 |
| MXE | 416 | THAP9-AS1 | chr4 | - | 8E+07 | 82895774 | 82897988 | 82898062 | 82893451 | 82894847 | 82900076 | 82900223 | 3 | 0 | 0 | 1 | 0 | 0 | 0 | 3 | 0 | 0 | 1 | 0 | 98 | 98 | 1 | 1 | 0.75 | NA | NA | NA | 0.75 | NA | | 0 |
| MXE | 417 | ZDHHC20 | chr13 | - | 2E+07 | 21421164 | 21425651 | 21425678 | 21413651 | 21413772 | 21459053 | 21459370 | 0 | 0 | 0 | 7 | 4 | 2 | 0 | 1 | 2 | 0 | 0 | 0 | 76 | 98 | 1 | 1 | 0 | 0 | 0 | NA | 1 | | 1 | -1 |
| MXE | 418 | GLT1D1 | chr12 | + | 1E+08 | 128876062 | 128888638 | 128888744 | 128853535 | 128853649 | 128899235 | 128899287 | 31 | 12 | 3 | 51 | 29 | 7 | 0 | 4 | 2 | 0 | 3 | 2 | 98 | 98 | 1 | 1 | 0.378 | 0.293 | 0.3 | NA | 0.571 | | 0.5 | -0.212 |
| MXE | 419 | GLT1D1 | chr12 | + | 1E+08 | 128888744 | 128899235 | 128899287 | 128875913 | 128876062 | 128945325 | 128945369 | 29 | 15 | 11 | 3 | 0 | 0 | 0 | 4 | 1 | 0 | 2 | 0 | 98 | 98 | 1 | 1 | 0.906 | 1 | 1 | NA | 0.667 | | 1 | 0.135 |
| MXE | 425 | SLC25A3 | chr12 | + | 1E+08 | 98595557 | 98595726 | 98595848 | 98593974 | 98594135 | 98597855 | 98598035 | 0 | 0 | 0 | 36 | 19 | 9 | 0 | 2 | 0 | 0 | 5 | 0 | 98 | 98 | 1 | 1 | 0 | 0 | 0 | NA | 0.286 | NA | | -0.286 |
| MXE | 427 | PHF1 | chr6 | + | 3E+07 | 33415144 | 33415234 | 33415329 | 33414724 | 33414829 | 33415589 | 33415670 | 1 | 2 | 0 | 4 | 3 | 3 | 0 | 0 | 0 | 1 | 4 | 0 | 98 | 98 | 1 | 1 | 0.2 | 0.4 | 0 | 0 | 0 | NA | | 0.2 |
| MXE | 434 | XPO6 | chr16 | - | 3E+07 | 28177332 | 28180940 | 28181031 | 28175897 | 28176095 | 28208986 | 28209122 | 4 | 6 | 3 | 18 | 25 | 6 | 0 | 4 | 0 | 0 | 3 | 0 | 98 | 98 | 1 | 1 | 0.182 | 0.194 | 0.333 | NA | 0.571 | NA | | -0.335 |
| MXE | 437 | TBC1D15 | chr12 | + | 7E+07 | 71880607 | 71884810 | 71885021 | 71872928 | 71873003 | 71893221 | 71893324 | 7 | 4 | 0 | 4 | 9 | 0 | 0 | 2 | 0 | 0 | 1 | 0 | 98 | 98 | 1 | 1 | 0.636 | 0.308 | NA | NA | 1 | NA | | -0.528 |
| MXE | 438 | NT5C2 | chr10 | - | 1E+08 | 103106706 | 103139405 | 103139479 | 103105705 | 103105801 | 103174857 | 103174982 | 36 | 30 | 8 | 43 | 15 | 11 | 0 | 0 | 0 | 0 | 4 | 0 | 98 | 98 | 1 | 1 | 0.456 | 0.667 | 0.421 | NA | 0 | NA | | 0.515 |
| MXE | 439 | RNF10 | chr12 | + | 1E+08 | 120566980 | 120571190 | 120571291 | 120565427 | 120565529 | 120575630 | 120575688 | 19 | 19 | 11 | 14 | 16 | 5 | 0 | 10 | 0 | 0 | 1 | 4 | 98 | 98 | 1 | 1 | 0.576 | 0.543 | 0.687 | NA | 0.909 | | 0 | 0.147 |
| MXE | 440 | STAT6 | chr12 | - | 6E+07 | 57107743 | 57108162 | 57108299 | 57107230 | 57107314 | 57111128 | 57111413 | 28 | 11 | 3 | 36 | 20 | 11 | 0 | 3 | 0 | 0 | 0 | 0 | 98 | 98 | 1 | 1 | 0.437 | 0.355 | 0.214 | NA | 1 | NA | | -0.665 |
| MXE | 443 | DAPK1 | chr9 | + | 9E+07 | 87686739 | 87697006 | 87697204 | 87681403 | 87681626 | 87698655 | 87698794 | 4 | 4 | 5 | 5 | 5 | 0 | 0 | 1 | 0 | 0 | 0 | 0 | 98 | 98 | 1 | 1 | 0.444 | 0.444 | 1 | NA | 1 | NA | | -0.371 |
| MXE | 447 | YIPF5 | chr5 | - | 1E+08 | 144165604 | 144169845 | 144169965 | 144164110 | 144164256 | 144170283 | 144170714 | 0 | 0 | 0 | 2 | 0 | 4 | 0 | 1 | 0 | 0 | 0 | 0 | 98 | 98 | 1 | 1 | 0 | NA | 0 | NA | 1 | NA | | -1 |
| MXE | 448 | ITGA5 | chr12 | - | 5E+07 | 54409597 | 54411833 | 54411964 | 54409231 | 54409352 | 54418980 | 54419266 | 6 | 5 | 2 | 20 | 15 | 8 | 0 | 2 | 0 | 0 | 1 | 0 | 98 | 98 | 1 | 1 | 0.231 | 0.25 | 0.2 | NA | 0.667 | NA | | -0.44 |
| MXE | 450 | MAPKAP1 | chr9 | - | 1E+08 | 125484583 | 125506309 | 125506417 | 125467971 | 125468109 | 125543058 | 125543168 | 3 | 0 | 1 | 9 | 3 | 0 | 0 | 0 | 0 | 0 | 2 | 0 | 98 | 98 | 1 | 1 | 0.25 | 0 | 1 | NA | 0 | NA | | 0.417 |
| MXE | 451 | HNRNPR | chr1 | - | 2E+07 | 23323732 | 23333517 | 23333631 | 23321527 | 23321663 | 23337753 | 23337861 | 7 | 0 | 0 | 6 | 3 | 0 | 5 | 0 | 0 | 0 | 1 | 0 | 98 | 98 | 1 | 1 | 0.538 | 0 | NA | 1 | 0 | NA | | -0.231 |
| MXE | 457 | NCF2 | chr1 | - | 2E+08 | 183577707 | 183586894 | 183586977 | 183573184 | 183573292 | 183590155 | 183590604 | 310 | 137 | 123 | 25 | 16 | 0 | 0 | 47 | 6 | 0 | 7 | 0 | 98 | 98 | 1 | 1 | 0.925 | 0.895 | 1 | NA | 0.87 | | 1 | 0.005 |
| MXE | 458 | NDUFA5 | chr7 | - | 1E+08 | 123550586 | 123557403 | 123557448 | 123545610 | 123545676 | 123557774 | 123557904 | 1 | 0 | 0 | 0 | 0 | 0 | 0 | 1 | 0 | 0 | 0 | 0 | 94 | 98 | 1 | 1 | 1 | NA | NA | NA | 0.51 | NA | | 0.49 |
| MXE | 460 | APAF1 | chr12 | + | 1E+08 | 98725540 | 98727172 | 98727316 | 98723638 | 98723764 | 98732419 | 98735433 | 15 | 4 | 5 | 32 | 10 | 4 | 0 | 2 | 0 | 0 | 2 | 0 | 98 | 98 | 1 | 1 | 0.319 | 0.286 | 0.556 | NA | 0.5 | NA | | -0.113 |
| MXE | 462 | LRRK2 | chr12 | + | 4E+07 | 40257377 | 40259479 | 40259604 | 40252909 | 40253016 | 40263788 | 40263901 | 175 | 69 | 34 | 372 | 93 | 25 | 8 | 15 | 0 | 9 | 29 | 0 | 98 | 98 | 1 | 1 | 0.32 | 0.426 | 0.576 | 0.471 | 0.341 | NA | | 0.035 |
| MXE | 463 | LRRK2 | chr12 | + | 4E+07 | 40278261 | 40283874 | 40284133 | 40277887 | 40278016 | 40287350 | 40287539 | 102 | 91 | 48 | 177 | 94 | 32 | 5 | 7 | 0 | 0 | 8 | 0 | 98 | 98 | 1 | 1 | 0.366 | 0.492 | 0.6 | 1 | 0.467 | NA | | -0.248 |
| MXE | 464 | LRRK2 | chr12 | + | 4E+07 | 40295644 | 40298242 | 40298493 | 40294844 | 40294914 | 40299108 | 40299257 | 319 | 82 | 30 | 215 | 48 | 19 | 0 | 15 | 0 | 0 | 7 | 0 | 98 | 98 | 1 | 1 | 0.597 | 0.631 | 0.612 | NA | 0.682 | NA | | -0.069 |
| MXE | 465 | LRRK2 | chr12 | + | 4E+07 | 40298493 | 40299108 | 40299257 | 40295426 | 40295644 | 40302788 | 40302882 | 192 | 59 | 33 | 390 | 124 | 39 | 0 | 14 | 0 | 0 | 27 | 2 | 98 | 98 | 1 | 1 | 0.33 | 0.322 | 0.458 | NA | 0.341 | | 0 | 0.2 |
| MXE | 466 | LRRK2 | chr12 | + | 4E+07 | 40304134 | 40305784 | 40305966 | 40302788 | 40302882 | 40308466 | 40308696 | 208 | 58 | 33 | 91 | 86 | 28 | 0 | 8 | 0 | 0 | 12 | 0 | 98 | 98 | 1 | 1 | 0.696 | 0.403 | 0.541 | NA | 0.4 | NA | | 0.147 |
| MXE | 468 | LRRK2 | chr12 | + | 4E+07 | 40348509 | 40351538 | 40351733 | 40346752 | 40346923 | 40354298 | 40354492 | 144 | 58 | 52 | 83 | 38 | 10 | 0 | 21 | 0 | 0 | 7 | 0 | 98 | 98 | 1 | 1 | 0.634 | 0.604 | 0.839 | NA | 0.75 | NA | | -0.058 |
| MXE | 469 | LRRK2 | chr12 | + | 4E+07 | 40351733 | 40354298 | 40354492 | 40348408 | 40348509 | 40356114 | 40356187 | 164 | 62 | 21 | 126 | 40 | 21 | 2 | 5 | 0 | 0 | 8 | 0 | 98 | 98 | 1 | 1 | 0.566 | 0.608 | 0.5 | 1 | 0.385 | NA | | -0.135 |
| MXE | 470 | FAR2 | chr12 | + | 3E+07 | 29270638 | 29293299 | 29293475 | 29223622 | 29223894 | 29297020 | 29297200 | 1 | 0 | 0 | 18 | 3 | 9 | 0 | 0 | 0 | 0 | 1 | 0 | 98 | 98 | 1 | 1 | 0.053 | 0 | 0 | NA | 0 | NA | | 0.018 |
| MXE | 471 | STK38L | chr12 | + | 3E+07 | 27315117 | 27315288 | 27315350 | 27314503 | 27314658 | 27317335 | 27317453 | 2 | 0 | 1 | 16 | 5 | 0 | 0 | 1 | 0 | 0 | 1 | 0 | 98 | 98 | 1 | 1 | 0.111 | 0 | 1 | NA | 0.5 | NA | | -0.13 |
| MXE | 472 | STK38L | chr12 | + | 3E+07 | 27319423 | 27322142 | 27322234 | 27317895 | 27318019 | 27322327 | 27325957 | 31 | 10 | 8 | 17 | 8 | 0 | 0 | 3 | 0 | 0 | 2 | 0 | 98 | 98 | 1 | 1 | 0.646 | 0.556 | 1 | NA | 0.6 | NA | | 0.134 |
| MXE | 475 | PREX1 | chr20 | - | 5E+07 | 48676268 | 48679359 | 48679409 | 48666282 | 48666355 | 48679650 | 48679754 | 40 | 25 | 28 | 109 | 83 | 47 | 0 | 19 | 1 | 0 | 43 | 4 | 98 | 98 | 1 | 1 | 0.268 | 0.231 | 0.373 | NA | 0.306 | | 0.2 | 0.038 |
| MXE | 478 | FRS2 | chr12 | + | 7E+07 | 69480359 | 69486201 | 69486297 | 69470348 | 69470530 | 69530864 | 69530960 | 2 | 1 | 0 | 1 | 0 | 2 | 0 | 3 | 0 | 0 | 0 | 0 | 98 | 98 | 1 | 1 | 0.667 | 1 | 0 | NA | 1 | NA | | -0.444 |
| MXE | 482 | CLEC7A | chr12 | - | 1E+07 | 10123363 | 10125086 | 10125173 | 10116780 | 10118590 | 10125296 | 10125448 | 26 | 2 | 10 | 141 | 62 | 38 | 0 | 4 | 3 | 0 | 16 | 2 | 98 | 98 | 1 | 1 | 0.156 | 0.031 | 0.208 | NA | 0.2 | | 0.6 | -0.268 |
| MXE | 484 | CLEC7A | chr12 | - | 1E+07 | 10125448 | 10127746 | 10127845 | 10123244 | 10123363 | 10129979 | 10130269 | 199 | 74 | 36 | 52 | 26 | 13 | 9 | 20 | 0 | 0 | 4 | 0 | 98 | 98 | 1 | 1 | 0.793 | 0.74 | 0.735 | 1 | 0.833 | NA | | -0.161 |
| MXE | 487 | PTPN6 | chr12 | + | 7E+06 | 6954994 | 6955150 | 6955267 | 6951982 | 6952177 | 6955371 | 6955485 | 49 | 28 | 32 | 59 | 30 | 14 | 0 | 14 | 0 | 0 | 17 | 0 | 98 | 98 | 1 | 1 | 0.454 | 0.483 | 0.696 | NA | 0.452 | NA | | 0.092 |
| MXE | 488 | PTPN6 | chr12 | + | 7E+06 | 6960435 | 6960805 | 6960945 | 6960087 | 6960239 | 6961125 | 6961316 | 84 | 34 | 19 | 105 | 37 | 13 | 2 | 8 | 0 | 4 | 12 | 0 | 98 | 98 | 1 | 1 | 0.444 | 0.479 | 0.594 | 0.333 | 0.4 | NA | | 0.139 |
| MXE | 490 | ZNF266 | chr19 | - | 9E+06 | 9434235 | 9434797 | 9434859 | 9420064 | 9420218 | 9435107 | 9435193 | 6 | 3 | 0 | 1 | 0 | 0 | 0 | 1 | 0 | 0 | 0 | 0 | 98 | 98 | 1 | 1 | 0.857 | 1 | NA | NA | 1 | NA | | -0.072 |
| MXE | 492 | SLC6A6 | chr3 | + | 1E+07 | 14478568 | 14479084 | 14479185 | 14477204 | 14477342 | 14481670 | 14481841 | 59 | 20 | 15 | 97 | 39 | 18 | 0 | 6 | 4 | 0 | 2 | 0 | 98 | 98 | 1 | 1 | 0.378 | 0.339 | 0.455 | NA | 0.75 | | 1 | -0.484 |
| MXE | 493 | BLOC1S6 | chr15 | + | 5E+07 | 45592276 | 45603099 | 45603187 | 45587122 | 45587525 | 45605427 | 45605514 | 6 | 0 | 0 | 9 | 3 | 0 | 0 | 1 | 0 | 0 | 2 | 0 | 98 | 98 | 1 | 1 | 0.4 | 0 | NA | NA | 0.333 | NA | | -0.133 |
| MXE | 494 | BLOC1S6 | chr15 | + | 5E+07 | 45592276 | 45603103 | 45603187 | 45587122 | 45587525 | 45605427 | 45605514 | 6 | 0 | 0 | 7 | 3 | 0 | 0 | 1 | 0 | 0 | 2 | 0 | 98 | 98 | 1 | 1 | 0.462 | 0 | NA | NA | 0.333 | NA | | -0.102 |
| MXE | 495 | BLOC1S6 | chr15 | + | 5E+07 | 45603187 | 45605427 | 45605514 | 45592134 | 45592276 | 45606394 | 45609716 | 1 | 0 | 1 | 2 | 3 | 0 | 0 | 1 | 0 | 0 | 1 | 0 | 98 | 98 | 1 | 1 | 0.333 | 0 | 1 | NA | 0.5 | NA | | -0.056 |
| MXE | 496 | ST3GAL4 | chr11 | + | 1E+08 | 126407634 | 126408306 | 126408496 | 126407251 | 126407349 | 126409267 | 126409411 | 16 | 3 | 5 | 7 | 2 | 1 | 0 | 2 | 0 | 0 | 2 | 0 | 98 | 98 | 1 | 1 | 0.696 | 0.6 | 0.833 | NA | 0.5 | NA | | 0.21 |
| MXE | 499 | KIAA0319 | chr6 | - | 2E+07 | 24551525 | 24556606 | 24556729 | 24544103 | 24547343 | 24559012 | 24559155 | 0 | 0 | 0 | 3 | 0 | 0 | 0 | 1 | 0 | 0 | 0 | 0 | 98 | 98 | 1 | 1 | 0 | NA | NA | NA | 1 | NA | | -1 |
| MXE | 501 | PTPRJ | chr11 | + | 5E+07 | 48112983 | 48121002 | 48121266 | 48110057 | 48110076 | 48123612 | 48123870 | 63 | 43 | 22 | 43 | 18 | 17 | 0 | 6 | 5 | 0 | 2 | 0 | 98 | 98 | 1 | 1 | 0.594 | 0.705 | 0.564 | NA | 0.75 | | 1 | -0.254 |
| MXE | 504 | SORL1 | chr11 | + | 1E+08 | 121543726 | 121545242 | 121545429 | 121532463 | 121532552 | 121549959 | 121550088 | 120 | 67 | 33 | 137 | 71 | 36 | 0 | 11 | 0 | 0 | 15 | 2 | 98 | 98 | 1 | 1 | 0.467 | 0.486 | 0.478 | NA | 0.423 | | 0 | 0.266 |
| MXE | 506 | SORL1 | chr11 | + | 1E+08 | 121608176 | 121611075 | 121611158 | 121607185 | 121607290 | 121612735 | 121612832 | 73 | 47 | 30 | 239 | 98 | 59 | 0 | 7 | 0 | 0 | 21 | 3 | 98 | 98 | 1 | 1 | 0.234 | 0.324 | 0.337 | NA | 0.25 | | 0 | 0.173 |
| MXE | 512 | CDA | chr1 | + | 2E+07 | 20605039 | 20613841 | 20613899 | 20588950 | 20589283 | 20618451 | 20618907 | 15 | 12 | 15 | 26 | 7 | 10 | 0 | 5 | 2 | 0 | 3 | 0 | 98 | 98 | 1 | 1 | 0.366 | 0.632 | 0.6 | NA | 0.625 | | 1 | -0.28 |
| MXE | 515 | GPSM3 | chr6 | - | 3E+07 | 32191908 | 32192147 | 32192250 | 32190765 | 32191503 | 32192471 | 32192530 | 20 | 13 | 8 | 50 | 23 | 25 | 0 | 2 | 0 | 0 | 7 | 0 | 98 | 98 | 1 | 1 | 0.286 | 0.361 | 0.242 | NA | 0.222 | NA | | 0.074 |
| MXE | 517 | DDX6 | chr11 | - | 1E+08 | 118763306 | 118765208 | 118765355 | 118759921 | 118760044 | 118768222 | 118768352 | 15 | 12 | 7 | 29 | 11 | 6 | 0 | 8 | 0 | 1 | 9 | 0 | 98 | 98 | 1 | 1 | 0.341 | 0.522 | 0.538 | 0 | 0.471 | NA | | 0.232 |
| MXE | 518 | TXNRD1 | chr12 | + | 1E+08 | 104289040 | 104311289 | 104311412 | 104286681 | 104287109 | 104313244 | 104313317 | 0 | 1 | 0 | 27 | 20 | 2 | 0 | 1 | 0 | 0 | 14 | 0 | 98 | 98 | 1 | 1 | 0 | 0.048 | 0 | NA | 0.067 | NA | | -0.051 |
| MXE | 520 | CASP5 | chr11 | - | 1E+08 | 105007334 | 105008806 | 105008980 | 105003273 | 105003383 | 105023129 | 105023168 | 1 | 0 | 0 | 10 | 0 | 0 | 0 | 0 | 0 | 0 | 4 | 0 | 98 | 98 | 1 | 1 | 0.091 | NA | NA | NA | 0 | NA | | 0.091 |
| MXE | 521 | CASP5 | chr11 | - | 1E+08 | 105007334 | 105008806 | 105009019 | 105003273 | 105003383 | 105023129 | 105023168 | 1 | 0 | 0 | 10 | 0 | 0 | 0 | 0 | 0 | 0 | 4 | 0 | 98 | 98 | 1 | 1 | 0.091 | NA | NA | NA | 0 | NA | | 0.091 |
| MXE | 534 | DCAF11 | chr14 | + | 2E+07 | 24117044 | 24117265 | 24117393 | 24115381 | 24115749 | 24117667 | 24117732 | 3 | 6 | 0 | 9 | 8 | 3 | 0 | 3 | 0 | 0 | 3 | 2 | 98 | 98 | 1 | 1 | 0.25 | 0.429 | 0 | NA | 0.5 | | 0 | -0.024 |
| MXE | 535 | DCAF11 | chr14 | + | 2E+07 | 24117044 | 24117265 | 24117393 | 24115282 | 24115749 | 24117667 | 24117732 | 0 | 0 | 0 | 9 | 8 | 3 | 0 | 1 | 0 | 0 | 3 | 2 | 98 | 98 | 1 | 1 | 0 | 0 | 0 | NA | 0.25 | | 0 | -0.125 |
| MXE | 536 | DCAF11 | chr14 | + | 2E+07 | 24117393 | 24117667 | 24117732 | 24116916 | 24117044 | 24118054 | 24118155 | 7 | 7 | 2 | 7 | 3 | 2 | 0 | 0 | 2 | 0 | 4 | 0 | 98 | 98 | 1 | 1 | 0.5 | 0.7 | 0.5 | NA | 0 | | 1 | 0.067 |
| MXE | 540 | MILR1 | chr17 | + | 6E+07 | 64457684 | 64460821 | 64460932 | 64452596 | 64452866 | 64465451 | 64465541 | 6 | 2 | 0 | 6 | 0 | 0 | 0 | 0 | 0 | 0 | 2 | 0 | 98 | 98 | 1 | 1 | 0.5 | 1 | NA | NA | 0 | NA | | 0.75 |
| MXE | 544 | ARAP1 | chr11 | - | 7E+07 | 72686191 | 72687438 | 72687502 | 72685068 | 72685681 | 72687687 | 72687738 | 29 | 18 | 25 | 65 | 28 | 16 | 0 | 0 | 0 | 0 | 9 | 1 | 98 | 98 | 1 | 1 | 0.309 | 0.391 | 0.61 | NA | 0 | | 0 | 0.437 |
| MXE | 545 | ARAP1 | chr11 | - | 7E+07 | 72698106 | 72699004 | 72699107 | 72697597 | 72697649 | 72699416 | 72699552 | 97 | 76 | 23 | 26 | 29 | 15 | 0 | 18 | 5 | 0 | 6 | 2 | 98 | 98 | 1 | 1 | 0.789 | 0.724 | 0.605 | NA | 0.75 | | 0.714 | -0.026 |
| MXE | 547 | DDX3X | chrX | + | 4E+07 | 41343351 | 41343736 | 41343822 | 41342736 | 41342836 | 41344029 | 41344128 | 147 | 42 | 21 | 166 | 36 | 32 | 0 | 19 | 2 | 0 | 28 | 1 | 98 | 98 | 1 | 1 | 0.47 | 0.538 | 0.396 | NA | 0.404 | | 0.667 | -0.068 |
| MXE | 548 | DDX3X | chrX | + | 4E+07 | 41347012 | 41347311 | 41347451 | 41346504 | 41346622 | 41347639 | 41347915 | 108 | 47 | 22 | 108 | 30 | 29 | 1 | 32 | 0 | 0 | 18 | 0 | 98 | 98 | 1 | 1 | 0.5 | 0.61 | 0.431 | 1 | 0.64 | NA | | -0.306 |
| MXE | 549 | DDX3X | chrX | + | 4E+07 | 41347012 | 41347314 | 41347451 | 41346504 | 41346622 | 41347639 | 41350287 | 108 | 47 | 22 | 102 | 29 | 29 | 1 | 32 | 0 | 0 | 19 | 0 | 98 | 98 | 1 | 1 | 0.514 | 0.618 | 0.431 | 1 | 0.627 | NA | | -0.292 |
| MXE | 551 | PTCD3 | chr2 | + | 9E+07 | 86125515 | 86125794 | 86125880 | 86124994 | 86125082 | 86127160 | 86127305 | 11 | 3 | 0 | 3 | 0 | 0 | 0 | 1 | 0 | 0 | 3 | 0 | 98 | 98 | 1 | 1 | 0.786 | 1 | NA | NA | 0.25 | NA | | 0.643 |

| | | | | | | | | | | | | | | | | | | | | | | | | | | | | | | | | | | | | |
|---|---|---|---|---|---|---|---|---|---|---|---|---|---|---|---|---|---|---|---|---|---|---|---|---|---|---|---|---|---|---|---|---|---|---|---|---|
| MXE | 556 | CCM2 | chr7 | + | 5E+07 | 45068579 | 45069825 | 45069961 | 45064462 | 45064646 | 45072725 | 45072783 | 9 | 8 | 6 | 11 | 11 | 10 | 0 | 0 | 0 | 2 | 3 | 0 | 98 | 98 | 1 | 1 | 0.45 | 0.421 | 0.375 | 0 | 0 | NA | | 0.415 |
| MXE | 557 | DNM2 | chr19 | + | 1E+07 | 10796199 | 10797379 | 10797518 | 10795371 | 10795439 | 10798485 | 10798572 | 12 | 5 | 1 | 10 | 6 | 8 | 0 | 4 | 0 | 1 | 3 | 0 | 98 | 98 | 1 | 1 | 0.545 | 0.455 | 0.111 | 0 | 0.571 | NA | | 0.085 |
| MXE | 559 | GLIPR2 | chr9 | + | 4E+07 | 36148650 | 36150871 | 36150949 | 36147785 | 36147894 | 36162361 | 36163913 | 20 | 9 | 9 | 52 | 13 | 20 | 0 | 1 | 0 | 0 | 5 | 2 | 98 | 98 | 1 | 1 | 0.278 | 0.409 | 0.31 | NA | 0.167 | | 0 | 0.249 |
| MXE | 560 | XRN1 | chr3 | - | 1E+08 | 142412663 | 142414134 | 142414291 | 142404906 | 142405076 | 142417139 | 142417229 | 21 | 11 | 2 | 16 | 14 | 1 | 0 | 2 | 0 | 7 | 8 | 0 | 98 | 98 | 1 | 1 | 0.568 | 0.44 | 0.667 | 0 | 0.2 | NA | | 0.458 |
| MXE | 561 | KIAA0226L | chr13 | - | 5E+07 | 46361573 | 46362537 | 46362598 | 46359485 | 46359631 | 46363114 | 46363213 | 42 | 11 | 10 | 64 | 9 | 9 | 0 | 4 | 0 | 0 | 13 | 5 | 98 | 98 | 1 | 1 | 0.396 | 0.55 | 0.526 | NA | 0.235 | | 0 | 0.373 |
| MXE | 565 | IL32 | chr16 | + | 3E+06 | 3067613 | 3067983 | 3068010 | 3067376 | 3067415 | 3068179 | 3068239 | 14 | 13 | 2 | 11 | 2 | 0 | 0 | 6 | 0 | 1 | 3 | 0 | 98 | 76 | 1 | 1 | 0.497 | 0.834 | 1 | 0 | 0.608 | NA | | 0.473 |
| MXE | 567 | CD6 | chr11 | + | 6E+07 | 61014014 | 61015712 | 61015835 | 61013422 | 61013563 | 61017478 | 61017550 | 7 | 2 | 0 | 13 | 3 | 0 | 0 | 0 | 0 | 0 | 2 | 2 | 98 | 98 | 1 | 1 | 0.35 | 0.4 | NA | NA | 0 | | 0 | 0.375 |
| MXE | 569 | PICALM | chr11 | - | 9E+07 | 86022469 | 86026291 | 86026367 | 86014869 | 86014963 | 86031468 | 86031611 | 144 | 66 | 56 | 188 | 89 | 37 | 0 | 20 | 1 | 0 | 24 | 0 | 98 | 98 | 1 | 1 | 0.434 | 0.426 | 0.602 | NA | 0.455 | | 1 | -0.24 |
| MXE | 570 | SUCO | chr1 | + | 2E+08 | 172551626 | 172553259 | 172553370 | 172533117 | 172533497 | 172555868 | 172556023 | 5 | 1 | 0 | 7 | 22 | 3 | 0 | 0 | 0 | 0 | 2 | 1 | 98 | 98 | 1 | 1 | 0.417 | 0.043 | 0 | NA | 0 | | 0 | 0.153 |
| MXE | 574 | PLXNC1 | chr12 | + | 9E+07 | 94298795 | 94300909 | 94301057 | 94297315 | 94297423 | 94303755 | 94303896 | 31 | 26 | 12 | 68 | 30 | 19 | 1 | 8 | 0 | 4 | 2 | 0 | 98 | 98 | 1 | 1 | 0.313 | 0.464 | 0.387 | 0.2 | 0.8 | NA | | -0.112 |
| MXE | 576 | PRR5L | chr11 | + | 4E+07 | 36446440 | 36451208 | 36451335 | 36437384 | 36437476 | 36462341 | 36465204 | 4 | 5 | 1 | 1 | 0 | 0 | 0 | 0 | 0 | 0 | 1 | 0 | 98 | 98 | 1 | 1 | 0.8 | 1 | 1 | NA | 0 | NA | | 0.933 |
| MXE | 577 | TRIP11 | chr14 | - | 9E+07 | 91975286 | 91976107 | 91976189 | 91974626 | 91974743 | 91988283 | 91988383 | 1 | 7 | 0 | 2 | 0 | 0 | 0 | 0 | 0 | 0 | 1 | 0 | 98 | 98 | 1 | 1 | 0.333 | 1 | NA | NA | 0 | NA | | 0.666 |
| MXE | 578 | SHKBP1 | chr19 | + | 4E+07 | 40582466 | 40583397 | 40583485 | 40580745 | 40580936 | 40583600 | 40583717 | 65 | 23 | 14 | 107 | 42 | 23 | 0 | 8 | 0 | 2 | 10 | 0 | 98 | 98 | 1 | 1 | 0.378 | 0.354 | 0.378 | 0 | 0.444 | NA | | 0.148 |
| MXE | 579 | BID | chr22 | - | 2E+07 | 17744013 | 17750104 | 17750174 | 17739348 | 17739488 | 17774380 | 17774495 | 7 | 0 | 0 | 11 | 15 | 12 | 0 | 2 | 0 | 0 | 5 | 2 | 98 | 98 | 1 | 1 | 0.389 | 0 | 0 | NA | 0.286 | | 0 | -0.013 |
| MXE | 580 | NIPBL | chr5 | + | 4E+07 | 37057332 | 37060843 | 37061018 | 37052365 | 37052566 | 37063789 | 37063978 | 16 | 6 | 0 | 4 | 2 | 1 | 0 | 0 | 1 | 0 | 1 | 0 | 98 | 98 | 1 | 1 | 0.8 | 0.75 | 0 | NA | 0 | | 1 | 0.017 |
| MXE | 582 | NF2 | chr22 | + | 3E+07 | 29636876 | 29639089 | 29639212 | 29603555 | 29604112 | 29642201 | 29642285 | 2 | 0 | 0 | 1 | 0 | 0 | 0 | 0 | 1 | 0 | 0 | 0 | 98 | 98 | 1 | 1 | 0.667 | NA | NA | NA | NA | | 1 | -0.333 |
| MXE | 583 | PHF21A | chr11 | - | 5E+07 | 45938312 | 45945839 | 45946003 | 45936493 | 45936569 | 45948885 | 45948946 | 8 | 3 | 0 | 21 | 2 | 0 | 0 | 2 | 0 | 0 | 2 | 0 | 98 | 98 | 1 | 1 | 0.276 | 0.6 | NA | NA | 0.5 | NA | | -0.062 |
| MXE | 584 | PHF21A | chr11 | - | 5E+07 | 45946003 | 45946075 | 45946098 | 45938156 | 45938312 | 45948885 | 45948946 | 7 | 6 | 0 | 16 | 2 | 1 | 0 | 0 | 0 | 0 | 4 | 0 | 72 | 98 | 1 | 1 | 0.373 | 0.803 | 0 | NA | 0 | NA | | 0.392 |
| MXE | 585 | ENSA | chr1 | - | 2E+08 | 150625808 | 150626478 | 150626526 | 150622122 | 150622859 | 150627466 | 150627592 | 3 | 0 | 0 | 10 | 1 | 1 | 0 | 2 | 0 | 0 | 2 | 0 | 97 | 98 | 1 | 1 | 0.233 | 0 | 0 | NA | 0.503 | NA | | -0.425 |
| MXE | 592 | NLRP3 | chr1 | + | 2E+08 | 247429755 | 247434102 | 247434273 | 247423846 | 247425599 | 247443971 | 247444142 | 8 | 0 | 0 | 2 | 0 | 0 | 0 | 0 | 0 | 2 | 1 | 0 | 98 | 98 | 1 | 1 | 0.8 | NA | NA | 0 | 0 | NA | | 0.8 |
| MXE | 594 | BAZ1A | chr14 | - | 3E+07 | 34786001 | 34786125 | 34786221 | 34783761 | 34783927 | 34792774 | 34792921 | 13 | 6 | 4 | 38 | 16 | 6 | 0 | 1 | 0 | 0 | 2 | 0 | 98 | 98 | 1 | 1 | 0.255 | 0.273 | 0.4 | NA | 0.333 | NA | | -0.024 |
| MXE | 596 | CD44 | chr11 | + | 4E+07 | 35210054 | 35211245 | 35211449 | 35208104 | 35208206 | 35214851 | 35214914 | 5 | 0 | 0 | 6 | 0 | 0 | 0 | 0 | 0 | 0 | 1 | 0 | 98 | 98 | 1 | 1 | 0.455 | NA | NA | NA | 0 | NA | | 0.455 |
| MXE | 597 | CRELD2 | chr22 | + | 5E+07 | 49923317 | 49924359 | 49924455 | 49922611 | 49922707 | 49925416 | 49925557 | 1 | 0 | 2 | 2 | 0 | 0 | 0 | 0 | 0 | 0 | 1 | 0 | 98 | 98 | 1 | 1 | 0.333 | NA | 1 | NA | 0 | NA | | 0.666 |
| MXE | 598 | TAGAP | chr6 | - | 2E+08 | 159042244 | 159043588 | 159043655 | 159041353 | 159041515 | 159043977 | 159044031 | 36 | 17 | 15 | 82 | 41 | 21 | 0 | 5 | 0 | 0 | 15 | 2 | 98 | 98 | 1 | 1 | 0.305 | 0.293 | 0.417 | NA | 0.25 | | 0 | 0.213 |
| MXE | 599 | TAGAP | chr6 | - | 2E+08 | 159043655 | 159043977 | 159044031 | 159042077 | 159042244 | 159044119 | 159044204 | 149 | 47 | 12 | 62 | 25 | 19 | 0 | 18 | 0 | 0 | 14 | 0 | 98 | 98 | 1 | 1 | 0.706 | 0.653 | 0.387 | NA | 0.562 | NA | | 0.02 |
| MXE | 603 | ARNTL | chr11 | + | 1E+07 | 13350070 | 13354300 | 13354448 | 13309986 | 13310059 | 13356738 | 13356777 | 8 | 5 | 5 | 10 | 2 | 1 | 0 | 1 | 0 | 0 | 2 | 0 | 98 | 98 | 1 | 1 | 0.444 | 0.714 | 0.833 | NA | 0.333 | NA | | 0.331 |
| MXE | 604 | MICAL2 | chr11 | + | 1E+07 | 12244112 | 12249183 | 12249246 | 12239435 | 12239585 | 12255642 | 12255750 | 0 | 0 | 0 | 4 | 2 | 1 | 0 | 1 | 0 | 0 | 0 | 0 | 98 | 98 | 1 | 1 | 0 | 0 | 0 | NA | 1 | NA | | -1 |
| MXE | 606 | ADAM8 | chr10 | - | 1E+08 | 133269529 | 133269896 | 133269974 | 133267928 | 133268118 | 133270359 | 133270510 | 159 | 65 | 43 | 3 | 0 | 0 | 0 | 34 | 6 | 0 | 2 | 0 | 98 | 98 | 1 | 1 | 0.981 | 1 | 1 | NA | 0.944 | | 1 | 0.022 |
| MXE | 608 | DENND5A | chr11 | - | 9E+06 | 9142845 | 9143402 | 9143485 | 9141939 | 9142108 | 9144096 | 9144278 | 36 | 15 | 7 | 65 | 20 | 10 | 0 | 5 | 0 | 0 | 5 | 1 | 98 | 98 | 1 | 1 | 0.356 | 0.429 | 0.412 | NA | 0.5 | | 0 | 0.149 |
| MXE | 609 | DENND5A | chr11 | - | 9E+06 | 9145815 | 9147029 | 9147151 | 9144994 | 9145113 | 9150080 | 9150209 | 26 | 15 | 10 | 43 | 12 | 8 | 0 | 2 | 0 | 0 | 3 | 0 | 98 | 98 | 1 | 1 | 0.377 | 0.556 | 0.556 | NA | 0.4 | NA | | 0.096 |
| MXE | 610 | RAB6A | chr11 | - | 7E+07 | 73718718 | 73718784 | 73718890 | 73716250 | 73716362 | 73720845 | 73720899 | 0 | 0 | 1 | 11 | 3 | 4 | 0 | 0 | 0 | 2 | 1 | 0 | 98 | 98 | 1 | 1 | 0 | 0 | 0.2 | 0 | 0 | NA | | 0.067 |
| MXE | 619 | FAM53B | chr10 | - | 1E+08 | 124696212 | 124706635 | 124706887 | 124681606 | 124682379 | 124744012 | 124744361 | 12 | 3 | 0 | 4 | 6 | 3 | 0 | 3 | 2 | 0 | 1 | 1 | 98 | 98 | 1 | 1 | 0.75 | 0.333 | 0 | NA | 0.75 | 0.667 | | -0.348 |
| MXE | 620 | RAPGEF6 | chr5 | - | 1E+08 | 131562047 | 131592382 | 131592466 | 131548046 | 131548190 | 131603270 | 131603327 | 4 | 3 | 0 | 8 | 6 | 1 | 0 | 1 | 0 | 0 | 0 | 0 | 98 | 98 | 1 | 1 | 0.333 | 0.333 | 0 | NA | 0.5 | NA | | -0.278 |
| MXE | 621 | QKI | chr6 | + | 2E+08 | 163563719 | 163566720 | 163566795 | 163561981 | 163562069 | 163570693 | 163578596 | 28 | 11 | 10 | 10 | 4 | 5 | 0 | 4 | 0 | 0 | 3 | 0 | 98 | 98 | 1 | 1 | 0.737 | 0.733 | 0.667 | NA | 0.571 | NA | | 0.141 |
| MXE | 622 | QKI | chr6 | + | 2E+08 | 163563719 | 163566720 | 163566795 | 163561981 | 163562069 | 163570693 | 163578596 | 2 | 2 | 0 | 10 | 4 | 5 | 0 | 0 | 0 | 0 | 3 | 0 | 98 | 98 | 1 | 1 | 0.167 | 0.333 | 0 | NA | 0 | NA | | 0.167 |
| MXE | 625 | PKM | chr15 | - | 7E+07 | 72202620 | 72203021 | 72203188 | 72200473 | 72200655 | 72206727 | 72206880 | 3 | 0 | 0 | 36 | 6 | 7 | 0 | 0 | 0 | 0 | 5 | 0 | 98 | 98 | 1 | 1 | 0.077 | 0 | 0 | NA | 0 | NA | | 0.026 |
| MXE | 636 | ZNF195 | chr11 | - | 3E+06 | 3373656 | 3377616 | 3377802 | 3371576 | 3371703 | 3379037 | 3379222 | 0 | 0 | 0 | 1 | 0 | 0 | 0 | 1 | 0 | 0 | 0 | 0 | 98 | 98 | 1 | 1 | 0 | NA | NA | NA | 1 | NA | | -1 |
| MXE | 637 | ZNF195 | chr11 | - | 3E+06 | 3373675 | 3377616 | 3377802 | 3371576 | 3371703 | 3379037 | 3379222 | 0 | 0 | 0 | 1 | 0 | 0 | 0 | 1 | 0 | 0 | 0 | 0 | 98 | 98 | 1 | 1 | 0 | NA | NA | NA | 1 | NA | | -1 |
| MXE | 638 | CARS | chr11 | - | 3E+06 | 3026797 | 3028995 | 3029084 | 3020219 | 3020332 | 3029302 | 3029443 | 27 | 10 | 3 | 12 | 8 | 3 | 0 | 4 | 3 | 0 | 3 | 0 | 98 | 98 | 1 | 1 | 0.692 | 0.556 | 0.5 | NA | 0.571 | | 1 | -0.203 |
| MXE | 649 | ZDHHC3 | chr3 | - | 4E+07 | 44929436 | 44932906 | 44932990 | 44915260 | 44926847 | 44933117 | 44933199 | 1 | 1 | 1 | 15 | 3 | 2 | 0 | 0 | 0 | 0 | 1 | 0 | 98 | 98 | 1 | 1 | 0.062 | 0.25 | 0.333 | NA | 0 | NA | | 0.215 |
| MXE | 650 | NCOA2 | chr8 | - | 7E+07 | 70174859 | 70213902 | 70214075 | 70170201 | 70170379 | 70216659 | 70216764 | 61 | 27 | 8 | 14 | 16 | 4 | 0 | 10 | 2 | 0 | 3 | 0 | 98 | 98 | 1 | 1 | 0.813 | 0.628 | 0.667 | NA | 0.769 | | 1 | -0.182 |
| MXE | 651 | DENND3 | chr8 | + | 1E+08 | 141136791 | 141150833 | 141150953 | 141128620 | 141128809 | 141151618 | 141151837 | 4 | 2 | 0 | 83 | 42 | 10 | 0 | 2 | 0 | 0 | 6 | 2 | 98 | 98 | 1 | 1 | 0.046 | 0.045 | 0 | NA | 0.25 | | 0 | -0.095 |
| MXE | 652 | DENND3 | chr8 | + | 1E+08 | 141138137 | 141150833 | 141150953 | 141136557 | 141136791 | 141151618 | 141151837 | 53 | 54 | 11 | 84 | 39 | 10 | 0 | 19 | 7 | 0 | 6 | 2 | 98 | 98 | 1 | 1 | 0.387 | 0.581 | 0.524 | NA | 0.76 | 0.778 | | -0.272 |
| MXE | 653 | DENND3 | chr8 | + | 1E+08 | 141138137 | 141151618 | 141151837 | 141136557 | 141136791 | 141155848 | 141155970 | 54 | 53 | 11 | 74 | 50 | 17 | 0 | 19 | 7 | 0 | 14 | 0 | 98 | 98 | 1 | 1 | 0.422 | 0.515 | 0.393 | NA | 0.576 | | 1 | -0.345 |
| MXE | 655 | DENND3 | chr8 | + | 1E+08 | 141190417 | 141192330 | 141192449 | 141188985 | 141189146 | 141192525 | 141192663 | 38 | 11 | 3 | 29 | 8 | 6 | 0 | 10 | 0 | 0 | 11 | 2 | 98 | 98 | 1 | 1 | 0.567 | 0.579 | 0.333 | NA | 0.476 | | 0 | 0.255 |
| MXE | 657 | EPG5 | chr18 | - | 5E+07 | 45879214 | 45880074 | 45880223 | 45878375 | 45878448 | 45882273 | 45882487 | 1 | 5 | 0 | 4 | 1 | 6 | 0 | 2 | 0 | 0 | 1 | 0 | 98 | 98 | 1 | 1 | 0.2 | 0.833 | 0 | NA | 0.667 | NA | | -0.323 |
| MXE | 660 | ANP32E | chr1 | - | 2E+08 | 150229237 | 150230570 | 150230693 | 150226607 | 150226795 | 150231776 | 150231926 | 3 | 3 | 2 | 2 | 0 | 1 | 0 | 2 | 0 | 0 | 0 | 0 | 98 | 98 | 1 | 1 | 0.6 | 1 | 0.667 | NA | 1 | NA | | -0.244 |
| MXE | 661 | ZNF627 | chr19 | + | 1E+07 | 11614653 | 11614826 | 11614887 | 11597419 | 11597630 | 11616694 | 11619159 | 2 | 2 | 0 | 1 | 6 | 0 | 0 | 4 | 0 | 0 | 2 | 0 | 98 | 98 | 1 | 1 | 0.667 | 0.25 | NA | 0 NA | 0.667 | NA | | -0.209 |
| MXE | 664 | NFKBID | chr19 | - | 4E+07 | 35897058 | 35897650 | 35897856 | 35896725 | 35896831 | 35898471 | 35898532 | 5 | 0 | 1 | 7 | 1 | 4 | 0 | 4 | 1 | 0 | 4 | 1 | 98 | 98 | 1 | 1 | 0.417 | 0 | 0.2 | NA | 0.5 | 0.5 | | -0.294 |
| MXE | 665 | TUBA1C | chr12 | + | 5E+07 | 49256511 | 49257883 | 49258001 | 49227925 | 49228166 | 49269464 | 49269687 | 0 | 1 | 0 | 0 | 0 | 0 | 0 | 0 | 0 | 0 | 4 | 0 | 98 | 98 | 1 | 1 | NA | 1 | NA | NA | 0 | NA | | 1 |
| MXE | 673 | CD53 | chr1 | + | 1E+08 | 110896733 | 110897808 | 110897892 | 110894959 | 110895055 | 110899123 | 110899936 | 33 | 29 | 35 | 99 | 49 | 56 | 0 | 2 | 1 | 0 | 9 | 0 | 98 | 98 | 1 | 1 | 0.25 | 0.372 | 0.385 | NA | 0.182 | | 1 | -0.255 |
| MXE | 675 | GADD45A | chr1 | + | 7E+07 | 67686126 | 67686349 | 67686587 | 67685176 | 67685538 | 67687660 | 67688338 | 4 | 0 | 2 | 3 | 4 | 1 | 0 | 1 | 0 | 0 | 1 | 3 | 98 | 98 | 1 | 1 | 0.571 | 0 | 0.667 | NA | 0.5 | | 0 | 0.163 |
| MXE | 678 | CLN3 | chr16 | - | 3E+07 | 28487741 | 28488590 | 28488662 | 28487455 | 28487541 | 28489289 | 28489386 | 4 | 2 | 0 | 8 | 6 | 2 | 0 | 1 | 0 | 0 | 0 | 0 | 98 | 98 | 1 | 1 | 0.333 | 0.25 | 0 | NA | 1 | NA | | -0.806 |
| MXE | 687 | PPP1R12A | chr12 | - | 8E+07 | 79805768 | 79806165 | 79806333 | 79798493 | 79798583 | 79807225 | 79807330 | 22 | 12 | 3 | 8 | 2 | 0 | 0 | 1 | 0 | 0 | 0 | 0 | 98 | 98 | 1 | 1 | 0.733 | 0.857 | 1 | NA | 1 | NA | | -0.137 |
| MXE | 688 | YIPF1 | chr1 | - | 5E+07 | 53860153 | 53866199 | 53866382 | 53851718 | 53852270 | 53866757 | 53866924 | 10 | 2 | 5 | 1 | 0 | 0 | 0 | 1 | 0 | 0 | 0 | 0 | 98 | 98 | 1 | 1 | 0.909 | 1 | 1 | NA | 1 | NA | | -0.03 |
| MXE | 691 | BAZ2B | chr2 | - | 2E+08 | 159448403 | 159453612 | 159453801 | 159446781 | 159446975 | 159478574 | 159478721 | 36 | 15 | 2 | 37 | 22 | 5 | 0 | 1 | 1 | 0 | 3 | 0 | 98 | 98 | 1 | 1 | 0.493 | 0.405 | 0.286 | NA | 0.25 | | 1 | -0.23 |
| MXE | 692 | BAZ2B | chr2 | - | 2E+08 | 159448409 | 159453612 | 159453801 | 159446781 | 159446975 | 159478574 | 159478721 | 36 | 15 | 2 | 43 | 28 | 5 | 0 | 1 | 1 | 0 | 3 | 2 | 98 | 98 | 1 | 1 | 0.456 | 0.349 | 0.286 | NA | 0.25 | 0.333 | | 0.072 |
| MXE | 695 | BST1 | chr4 | + | 2E+07 | 15715361 | 15715706 | 15715799 | 15711806 | 15711889 | 15718906 | 15718993 | 30 | 11 | 11 | 37 | 19 | 16 | 0 | 1 | 3 | 0 | 6 | 0 | 98 | 98 | 1 | 1 | 0.448 | 0.367 | 0.407 | NA | 0.143 | | 1 | -0.164 |
| MXE | 706 | FCAR | chr19 | + | 5E+07 | 54885525 | 54888006 | 54888294 | 54874092 | 54874323 | 54889648 | 54891420 | 4 | 10 | 0 | 28 | 6 | 2 | 0 | 3 | 0 | 0 | 9 | 0 | 98 | 98 | 1 | 1 | 0.125 | 0.625 | 0 | NA | 0.25 | NA | | 0 |
| MXE | 708 | ECD | chr10 | - | 7E+07 | 73152421 | 73154255 | 73154448 | 73148275 | 73148404 | 73156274 | 73156453 | 3 | 0 | 0 | 4 | 0 | 0 | 0 | 4 | 0 | 0 | 0 | 0 | 98 | 98 | 1 | 1 | 0.429 | NA | NA | NA | 1 | NA | | -0.571 |
| MXE | 715 | NDEL1 | chr17 | + | 8E+06 | 8445864 | 8446753 | 8446902 | 8444259 | 8444357 | 8448549 | 8448686 | 28 | 12 | 1 | 69 | 38 | 14 | 0 | 5 | 0 | 0 | 10 | 0 | 98 | 98 | 1 | 1 | 0.289 | 0.24 | 0.067 | NA | 0.333 | NA | | -0.134 |
| MXE | 717 | RAB5C | chr17 | - | 4E+07 | 42131822 | 42137761 | 42137878 | 42130336 | 42130590 | 42154892 | 42155044 | 3 | 1 | 3 | 1 | 1 | 1 | 0 | 0 | 0 | 0 | 1 | 0 | 98 | 98 | 1 | 1 | 0.75 | 0.5 | 0.75 | NA | 0 | NA | | 0.667 |
| MXE | 722 | PI4KB | chr1 | - | 2E+08 | 151324917 | 151326171 | 151326373 | 151315572 | 151316509 | 151327270 | 151327657 | 3 | 1 | 0 | 1 | 2 | 0 | 0 | 2 | 0 | 0 | 0 | 0 | 98 | 98 | 1 | 1 | 0.75 | 0.333 | NA | NA | 1 | NA | | -0.459 |
| MXE | 724 | FRMD8 | chr11 | + | 7E+07 | 65389528 | 65393572 | 65393674 | 65387036 | 65387121 | 65394040 | 65394099 | 4 | 3 | 0 | 4 | 3 | 6 | 0 | 3 | 0 | 0 | 0 | 0 | 98 | 98 | 1 | 1 | 0.5 | 0.5 | 0 | NA | 1 | NA | | -0.667 |
| MXE | 725 | IST1 | chr16 | + | 7E+07 | 71923380 | 71924768 | 71924817 | 71922473 | 71922680 | 71927613 | 71930637 | 20 | 8 | 6 | 4 | 5 | 3 | 0 | 1 | 0 | 0 | 0 | 1 | 98 | 98 | 1 | 1 | 0.833 | 0.615 | 0.667 | NA | 1 | | 0 | 0.205 |
| MXE | 726 | STAG3L4 | chr7 | + | 7E+07 | 67308652 | 67308934 | 67309139 | 67305986 | 67306083 | 67309563 | 67309715 | 1 | 0 | 0 | 0 | 2 | 1 | 0 | 1 | 0 | 0 | 0 | 0 | 98 | 98 | 1 | 1 | 1 | 0 | 0 | NA | 1 | NA | | -0.667 |
| MXE | 729 | EIF2AK2 | chr2 | - | 4E+07 | 37122664 | 37126288 | 37126411 | 37119958 | 37120139 | 37135483 | 37135546 | 17 | 7 | 0 | 12 | 13 | 2 | 0 | 2 | 0 | 0 | 4 | 1 | 98 | 98 | 1 | 1 | 0.586 | 0.35 | 0 | NA | 0.333 | | 0 | 0.145 |
| MXE | 730 | SP140 | chr2 | + | 2E+08 | 230253417 | 230255451 | 230255532 | 230250980 | 230251061 | 230269531 | 230269618 | 1 | 0 | 0 | 4 | 0 | 0 | 0 | 0 | 0 | 0 | 3 | 0 | 98 | 98 | 1 | 1 | 0.2 | NA | NA | NA | 0.25 | NA | | -0.05 |
| MXE | 732 | AMPD2 | chr1 | + | 1E+08 | 109629490 | 109629795 | 109629916 | 109629108 | 109629235 | 109630232 | 109630406 | 19 | 25 | 0 | 31 | 8 | 12 | 0 | 8 | 1 | 0 | 6 | 1 | 98 | 98 | 1 | 1 | 0.38 | 0.758 | 0 | NA | 0.571 | | 0.5 | -0.156 |
| MXE | 733 | PCBP2 | chr12 | + | 5E+07 | 53464852 | 53465931 | 53465973 | 53462480 | 53462567 | 53467220 | 53467293 | 54 | 17 | 19 | 72 | 31 | 6 | 0 | 3 | 0 | 0 | 11 | 0 | 98 | 91 | 1 | 1 | 0.411 | 0.337 | 0.746 | NA | 0.202 | NA | | 0.296 |
| MXE | 734 | MRVI1 | chr11 | - | 1E+07 | 10594195 | 10600917 | 10601059 | 10593491 | 10593599 | 10603119 | 10603251 | 27 | 9 | 9 | 10 | 2 | 4 | 0 | 5 | 3 | 0 | 0 | 0 | 98 | 98 | 1 | 1 | 0.73 | 0.818 | 0.692 | NA | 1 | | 1 | -0.253 |
| MXE | 739 | MLKL | chr16 | - | 7E+07 | 74678980 | 74682650 | 74682786 | 74675612 | 74675764 | 74685485 | 74685583 | 17 | 2 | 0 | 6 | 1 | 8 | 0 | 1 | 1 | 0 | 1 | 0 | 98 | 98 | 1 | 1 | 0.739 | 0.667 | 0 | NA | 0.5 | | 1 | -0.281 |
| MXE | 740 | FGR | chr1 | - | 3E+07 | 27614583 | 27614849 | 27614926 | 27613218 | 27613350 | 27615433 | 27615613 | 67 | 28 | 22 | 87 | 22 | 38 | 0 | 8 | 1 | 0 | 14 | 1 | 98 | 98 | 1 | 1 | 0.435 | 0.56 | 0.367 | NA | 0.364 | | 0.5 | 0.022 |
| MXE | 751 | UBR4 | chr1 | - | 2E+07 | 19148626 | 19150576 | 19150793 | 19147992 | 19148127 | 19151642 | 19151859 | 5 | 8 | 0 | 8 | 4 | 3 | 0 | 3 | 0 | 0 | 4 | 0 | 98 | 98 | 1 | 1 | 0.385 | 0.667 | 0 | NA | 0.429 | NA | | -0.078 |
| MXE | 756 | PLEKHM2 | chr1 | + | 2E+07 | 15719920 | 15721328 | 15721388 | 15718537 | 15718625 | 15725316 | 15725545 | 10 | 6 | 5 | 2 | 2 | 1 | 0 | 3 | 0 | 0 | 2 | 0 | 98 | 98 | 1 | 1 | 0.833 | 0.75 | 0.833 | NA | 0.6 | NA | | 0.205 |
| MXE | 760 | CPEB4 | chr5 | + | 2E+08 | 173932500 | 173943025 | 173943049 | 173910522 | 173910604 | 173944966 | 173945140 | 11 | 2 | 9 | 1 | 0 | 0 | 0 | 2 | 0 | 0 | 1 | 0 | 98 | 73 | 1 | 1 | 0.891 | 1 | 1 | NA | 0.598 | NA | | 0.366 |
| MXE | 762 | OSBPL9 | chr1 | + | 5E+07 | 51652041 | 51669433 | 51669512 | 51616873 | 51617221 | 51714002 | 51714079 | 4 | 7 | 0 | 11 | 8 | 3 | 0 | 1 | 0 | 0 | 6 | 0 | 98 | 98 | 1 | 1 | 0.267 | 0.467 | 0 | NA | 0.143 | NA | | 0.102 |
| MXE | 763 | RCBTB2 | chr13 | - | 5E+07 | 48521962 | 48522307 | 48522403 | 48515585 | 48515741 | 48524658 | 48524757 | 2 | 1 | 0 | 6 | 3 | 0 | 0 | 0 | 0 | 0 | 1 | 0 | 98 | 98 | 1 | 1 | 0.25 | 0.25 | NA | NA | 0 | NA | | 0.25 |
| MXE | 764 | SELL | chr1 | - | 2E+08 | 169696553 | 169701559 | 169701688 | 169690664 | 169691802 | 169703254 | 169703440 | 124 | 55 | 47 | 38 | 26 | 7 | 2 | 8 | 0 | 4 | 3 | 0 | 98 | 68 | 1 | 1 | 0.694 | 0.595 | 0.823 | 0.258 | 0.649 | NA | | 0.251 |
| MXE | 765 | SOAT1 | chr1 | + | 2E+08 | 179302802 | 179323436 | 179323495 | 179293713 | 179293936 | 179335505 | 179335657 | 1 | 1 | 0 | 8 | 6 | 0 | 0 | 0 | 0 | 0 | 2 | 0 | 98 | 98 | 1 | 1 | 0.111 | 0.143 | NA | NA | 0 | NA | | 0.127 |
| MXE | 767 | PSME3 | chr17 | + | 4E+07 | 42838205 | 42838730 | 42838799 | 42837648 | 42837697 | 42838944 | 42839012 | 17 | 9 | 7 | 24 | 14 | 10 | 0 | 1 | 0 | 0 | 2 | 0 | 98 | 98 | 1 | 1 | 0.415 | 0.391 | 0.412 | NA | 0.333 | NA | | 0.073 |
| MXE | 769 | ZNF518A | chr10 | + | 1E+08 | 96151566 | 96155325 | 96155398 | 96133555 | 96133648 | 96155923 | 96156047 | 2 | 0 | 0 | 2 | 1 | 0 | 0 | 0 | 0 | 0 | 1 | 0 | 98 | 98 | 1 | 1 | 0.5 | 0 | NA | NA | 0 | NA | | 0.25 |
| MXE | 781 | MFN2 | chr1 | + | 1E+07 | 11982114 | 11989164 | 11989343 | 11980180 | 11980484 | 11992554 | 11992690 | 7 | 3 | 0 | 29 | 10 | 0 | 0 | 0 | 0 | 0 | 3 | 0 | 98 | 98 | 1 | 1 | 0.194 | 0.231 | NA | NA | 0 | NA | | 0.213 |
| MXE | 782 | HIPK3 | chr11 | + | 3E+07 | 33348228 | 33348521 | 33348818 | 33347851 | 33348013 | 33349146 | 33349287 | 14 | 4 | 2 | 54 | 19 | 5 | 0 | 2 | 0 | 0 | 6 | 0 | 98 | 98 | 1 | 1 | 0.206 | 0.174 | 0.286 | NA | 0.25 | NA | | -0.028 |
| MXE | 783 | HIPK3 | chr11 | + | 3E+07 | 33348818 | 33349146 | 33349287 | 33347851 | 33348013 | 33351607 | 33351843 | 16 | 2 | 2 | 31 | 16 | 4 | 0 | 1 | 0 | 0 | 2 | 0 | 98 | 98 | 1 | 1 | 0.34 | 0.111 | 0.333 | NA | 0.333 | NA | | -0.072 |
| MXE | 784 | HIPK3 | chr11 | + | 3E+07 | 33348818 | 33349146 | 33349287 | 33348165 | 33348228 | 33351607 | 33351843 | 14 | 6 | 2 | 32 | 17 | 4 | 0 | 1 | 0 | 0 | 0 | 0 | 98 | 98 | 1 | 1 | 0.304 | 0.261 | 0.333 | NA | 1 | NA | | -0.701 |
| MXE | 785 | RAB2A | chr8 | + | 6E+07 | 60558923 | 60572045 | 60572113 | 60516909 | 60517253 | 60584207 | 60584290 | 14 | 7 | 0 | 9 | 6 | 4 | 1 | 2 | 0 | 0 | 0 | 0 | 98 | 98 | 1 | 1 | 0.609 | 0.538 | 0 | 1 | 1 | NA | | -0.618 |
| MXE | 787 | AMPD3 | chr11 | + | 1E+07 | 10478730 | 10482062 | 10482225 | 10461514 | 10461740 | 10484819 | 10485039 | 3 | 0 | 0 | 18 | 5 | 0 | 0 | 0 | 0 | 0 | 0 | 0 | 98 | 98 | 1 | 1 | 0.143 | 0 | NA | NA | 0 | NA | | 0.071 |
| MXE | 788 | EPB41 | chr1 | + | 3E+07 | 28993542 | 28997214 | 28997319 | 28987430 | 28987905 | 29011864 | 29011907 | 46 | 19 | 5 | 27 | 7 | 7 | 0 | 3 | 0 | 0 | 5 | 0 | 98 | 98 | 1 | 1 | 0.63 | 0.731 | 0.417 | NA | 0.375 | NA | | 0.218 |
| MXE | 790 | EPB41 | chr1 | + | 3E+07 | 29097935 | 29109335 | 29109437 | 29064981 | 29065158 | 29112367 | 29112448 | 31 | 14 | 5 | 13 | 12 | 5 | 0 | 0 | 0 | 0 | 3 | 0 | 98 | 98 | 1 | 1 | 0.705 | 0.538 | 0.5 | NA | 0 | NA | | 0.581 |
| MXE | 791 | ERBB2IP | chr5 | + | 7E+07 | 66069020 | 66072168 | 66072291 | 66053405 | 66054951 | 66075023 | 66075230 | 2 | 0 | 0 | 6 | 9 | 0 | 0 | 0 | 0 | 0 | 1 | 0 | 98 | 98 | 1 | 1 | 0.25 | 0 | NA | NA | 0 | NA | | 0.125 |
| MXE | 793 | RFFL | chr17 | - | 4E+07 | 35017606 | 35021370 | 35021781 | 35016369 | 35016580 | 35026373 | 35026561 | 9 | 1 | 0 | 15 | 4 | 5 | 0 | 0 | 0 | 0 | 1 | 0 | 98 | 98 | 1 | 1 | 0.375 | 0.2 | 0 | NA | 0 | NA | | 0.192 |
| MXE | 794 | PPP2R3C | chr14 | - | 4E+07 | 35091207 | 35095047 | 35095184 | 35087950 | 35088010 | 35096557 | 35096633 | 3 | 1 | 1 | 9 | 5 | 4 | 0 | 0 | 0 | 0 | 1 | 0 | 98 | 98 | 1 | 1 | 0.25 | 0.167 | 0.2 | NA | 0 | NA | | 0.206 |
| MXE | 795 | CYB5R4 | chr6 | + | 8E+07 | 83914448 | 83918004 | 83918065 | 83909008 | 83909090 | 83919396 | 83919454 | 65 | 27 | 25 | 68 | 29 | 20 | 0 | 3 | 0 | 0 | 5 | 0 | 82 | 98 | 1 | 1 | 0.533 | 0.527 | 0.599 | NA | 0.418 | NA | | 0.135 |
| MXE | 796 | TK2 | chr16 | - | 7E+07 | 66537017 | 66541878 | 66541953 | 66531379 | 66531469 | 66548977 | 66549009 | 5 | 0 | 0 | 23 | 4 | 0 | 0 | 1 | 0 | 0 | 2 | 0 | 98 | 98 | 1 | 1 | 0.179 | 0 | NA | NA | 0.333 | NA | | -0.244 |
| MXE | 798 | MED13L | chr12 | - | 1E+08 | 116111512 | 116237467 | 116237705 | 116096668 | 116096752 | 116277059 | 116277186 | 56 | 19 | 3 | 47 | 30 | 16 | 0 | 11 | 3 | 0 | 9 | 0 | 98 | 98 | 1 | 1 | 0.544 | 0.388 | 0.158 | NA | 0.55 | | 1 | -0.412 |
| MXE | 801 | CLEC4A | chr12 | + | 8E+06 | 8125677 | 8129263 | 8129362 | 8123631 | 8123960 | 8135584 | 8135736 | 3 | 4 | 1 | 8 | 1 | 3 | 0 | 0 | 0 | 0 | 1 | 0 | 98 | 98 | 1 | 1 | 0.273 | 0.8 | 1 | NA | 0 | NA | | 0.691 |
| MXE | 802 | CLEC4A | chr12 | + | 8E+06 | 8129362 | 8135584 | 8135736 | 8123631 | 8123960 | 8136787 | 8136903 | 0 | 0 | 9 | 4 | 1 | 3 | 0 | 0 | 0 | 0 | 1 | 0 | 98 | 98 | 1 | 1 | 0 | 0 | 0.75 | NA | 0 | NA | | 0.25 |
| MXE | 803 | CLEC4A | chr12 | + | 8E+06 | 8129362 | 8135584 | 8135736 | 8125560 | 8125677 | 8136787 | 8136903 | 5 | 2 | 22 | 4 | 1 | 3 | 0 | 1 | 0 | 0 | 1 | 0 | 98 | 98 | 1 | 1 | 0.556 | 0.667 | 0.88 | NA | 0.5 | NA | | 0.201 |
| MXE | 810 | FBXO18 | chr10 | + | 6E+06 | 5918478 | 5921257 | 5921357 | 5917589 | 5917676 | 5921447 | 5921569 | 4 | 4 | 1 | 6 | 5 | 0 | 0 | 3 | 0 | 0 | 2 | 0 | 98 | 98 | 1 | 1 | 0.4 | 0.444 | 1 | NA | 0.6 | NA | | 0.015 |
| MXE | 812 | FDFT1 | chr8 | + | 1E+07 | 11802931 | 11808793 | 11808891 | 11795572 | 11796011 | 11809666 | 11809850 | 0 | 0 | 0 | 7 | 1 | 1 | 0 | 1 | 0 | 0 | 0 | 0 | 98 | 98 | 1 | 1 | 0 | 0 | 0 | NA | 1 | NA | | -1 |
| MXE | 814 | H2AFV | chr7 | - | 4E+07 | 44841012 | 44843276 | 44843354 | 44835528 | 44835658 | 44847968 | 44848126 | 4 | 8 | 1 | 7 | 1 | 10 | 0 | 1 | 0 | 0 | 0 | 0 | 98 | 98 | 1 | 1 | 0.364 | 0.889 | 0.091 | NA | 1 | NA | | -0.552 |
| MXE | 815 | PGS1 | chr17 | + | 8E+07 | 78392665 | 78396307 | 78396385 | 78378617 | 78378808 | 78398251 | 78398351 | 12 | 14 | 1 | 11 | 19 | 1 | 0 | 3 | 0 | 0 | 0 | 0 | 98 | 98 | 1 | 1 | 0.522 | 0.424 | 0.5 | NA | 1 | NA | | -0.518 |
| MXE | 816 | PGS1 | chr17 | + | 8E+07 | 78392665 | 78396307 | 78396385 | 78378617 | 78378808 | 78398251 | 78398351 | 4 | 5 | 1 | 11 | 19 | 1 | 0 | 1 | 0 | 0 | 0 | 0 | 98 | 98 | 1 | 1 | 0.267 | 0.208 | 0.5 | NA | 1 | NA | | -0.675 |
| MXE | 821 | FAM129A | chr1 | - | 2E+08 | 184803692 | 184805945 | 184806056 | 184798078 | 184798190 | 184808073 | 184808235 | 59 | 34 | 14 | 104 | 42 | 32 | 7 | 11 | 0 | 0 | 5 | 0 | 98 | 98 | 1 | 1 | 0.362 | 0.447 | 0.304 | 1 | 0.687 | NA | | -0.473 |
| MXE | 824 | PSMA3 | chr14 | + | 6E+07 | 58247832 | 58252118 | 58252242 | 58244804 | 58244941 | 58257744 | 58257846 | 0 | 1 | 1 | 11 | 4 | 0 | 1 | 0 | 0 | 0 | 1 | 0 | 98 | 98 | 1 | 1 | 0 | 0.2 | 1 | 1 | 0 | NA | | -0.1 |
| MXE | 826 | RNF19B | chr1 | - | 3E+07 | 32938528 | 32942251 | 32942459 | 32936445 | 32937259 | 32944018 | 32944159 | 55 | 25 | 38 | 43 | 11 | 11 | 0 | 10 | 2 | 0 | 5 | 0 | 98 | 98 | 1 | 1 | 0.561 | 0.694 | 0.776 | NA | 0.667 | | 1 | -0.157 |
| MXE | 830 | RBM23 | chr14 | - | 2E+07 | 22908380 | 22909482 | 22909595 | 22906194 | 22906368 | 22911327 | 22911403 | 67 | 35 | 36 | 7 | 10 | 0 | 0 | 7 | 0 | 0 | 1 | 0 | 98 | 97 | 1 | 1 | 0.905 | 0.776 | 1 | NA | 0.874 | NA | | 0.02 |
| MXE | 831 | RBM23 | chr14 | - | 2E+07 | 22908380 | 22911327 | 22911403 | 22906194 | 22906368 | 22918998 | 22919187 | 7 | 13 | 3 | 5 | 8 | 0 | 0 | 0 | 1 | 0 | 1 | 0 | 98 | 97 | 1 | 1 | 0.581 | 0.617 | 1 | NA | 0 | | 1 | 0.233 |
| MXE | 832 | NUP50 | chr22 | + | 5E+07 | 45176080 | 45178237 | 45178900 | 45171599 | 45171683 | 45181285 | 45181367 | 19 | 3 | 3 | 12 | 12 | 3 | 0 | 5 | 0 | 0 | 0 | 0 | 98 | 98 | 1 | 1 | 0.613 | 0.2 | 0.5 | NA | 0.5 | NA | | -0.062 |
| MXE | 833 | H2AFY | chr5 | - | 1E+08 | 135350913 | 135352945 | 135353045 | 135345967 | 135346057 | 135360496 | 135360607 | 16 | 12 | 14 | 5 | 3 | 3 | 0 | 1 | 0 | 0 | 0 | 0 | 98 | 98 | 1 | 1 | 0.762 | 0.8 | 0.824 | NA | 1 | NA | | -0.205 |
| MXE | 837 | PTGES3 | chr12 | - | 6E+07 | 56666266 | 56670274 | 56670364 | 56664775 | 56664800 | 56671748 | 56671847 | 6 | 3 | 0 | 6 | 13 | 1 | 0 | 2 | 0 | 0 | 0 | 0 | 98 | 98 | 1 | 1 | 0.5 | 0.187 | 0 | NA | 0.667 | NA | | -0.438 |
| MXE | 838 | PTGES3 | chr12 | - | 6E+07 | 56670364 | 56671748 | 56671847 | 56666203 | 56666266 | 56672739 | 56672809 | 13 | 2 | 0 | 17 | 6 | 14 | 0 | 1 | 0 | 0 | 0 | 0 | 98 | 98 | 1 | 1 | 0.433 | 0.25 | 0 | NA | 1 | NA | | -0.772 |
| MXE | 841 | CD300LF | chr17 | - | 7E+07 | 74696217 | 74698368 | 74698481 | 74695724 | 74695859 | 74703034 | 74703098 | 15 | 7 | 10 | 1 | 1 | 2 | 0 | 3 | 0 | 0 | 0 | 0 | 98 | 72 | 1 | 1 | 0.917 | 0.837 | 0.786 | NA | 1 | NA | | -0.153 |
| MXE | 842 | DHDDS | chr1 | + | 3E+07 | 26446432 | 26447558 | 26447660 | 26442730 | 26442873 | 26457790 | 26457905 | 2 | 0 | 0 | 3 | 0 | 0 | 0 | 0 | 0 | 0 | 0 | 0 | 98 | 98 | 1 | 1 | 0.4 | NA | NA | NA | 1 | NA | | -0.6 |
| MXE | 846 | DHX9 | chr1 | + | 2E+08 | 182855713 | 182856531 | 182856578 | 182854029 | 182854178 | 182858103 | 182858240 | 9 | 4 | 0 | 28 | 10 | 3 | 0 | 0 | 0 | 0 | 3 | 0 | 98 | 96 | 1 | 1 | 0.239 | 0.282 | 0 | NA | 0 | NA | | 0.174 |
| MXE | 858 | IRF7 | chr11 | - | 613950 | 614037 | 614173 | 614399 | 613784 | 613865 | 614475 | 614534 | 2 | 3 | 7 | 13 | 11 | 0 | 0 | 4 | 1 | 0 | 7 | 3 | 98 | 98 | 1 | 1 | 0.133 | 0.214 | 1 | NA | 0.364 | 0.25 | | 0.142 |
| MXE | 859 | RREB1 | chr6 | + | 7E+06 | 7240602 | 7246423 | 7247221 | 7228996 | 7231907 | 7248510 | 7251980 | 2 | 0 | 0 | 1 | 0 | 0 | 0 | 0 | 0 | 0 | 0 | 0 | 98 | 98 | 1 | 1 | 0.667 | NA | NA | NA | 0 | NA | | 0.667 |
| MXE | 864 | ZNF480 | chr19 | + | 5E+07 | 52314279 | 52315833 | 52315962 | 52300393 | 52300484 | 52321578 | 52325927 | 0 | 0 | 0 | 4 | 0 | 0 | 0 | 0 | 0 | 0 | 1 | 0 | 98 | 98 | 1 | 1 | 0 | NA | NA | NA | 1 | NA | | -1 |
| MXE | 874 | ARHGEF2 | chr1 | - | 2E+08 | 155954969 | 155957712 | 155957882 | 155952627 | 155952828 | 155958319 | 155958396 | 9 | 15 | 12 | 25 | 14 | 2 | 0 | 4 | 0 | 0 | 0 | 1 | 98 | 98 | 1 | 1 | 0.265 | 0.517 | 0.857 | NA | 1 | | 0 | 0.046 |
| MXE | 879 | C11orf54 | chr11 | + | 9E+07 | 93755386 | 93757315 | 93757465 | 93753935 | 93754037 | 93759741 | 93759858 | 7 | 1 | 0 | 11 | 3 | 0 | 0 | 2 | 0 | 0 | 0 | 0 | 98 | 98 | 1 | 1 | 0.389 | 0.25 | NA | NA | 1 | NA | | -0.68 |
| MXE | 883 | TLR4 | chr9 | + | 1E+08 | 117708318 | 117708562 | 117708729 | 117704174 | 117704565 | 117712388 | 117717491 | 80 | 13 | 10 | 72 | 13 | 36 | 0 | 7 | 3 | 0 | 2 | 0 | 98 | 98 | 1 | 1 | 0.526 | 0.5 | 0.217 | NA | 0.778 | | 1 | -0.475 |
| MXE | 884 | ALPL | chr1 | + | 2E+07 | 21554142 | 21560625 | 21560745 | 21509364 | 21509517 | 21561096 | 21561212 | 25 | 7 | 6 | 57 | 25 | 15 | 0 | 1 | 3 | 0 | 2 | 0 | 98 | 98 | 1 | 1 | 0.305 | 0.219 | 0.286 | NA | 0.333 | | 1 | -0.396 |
| MXE | 885 | ALPL | chr1 | + | 2E+07 | 21554142 | 21560625 | 21560745 | 21509364 | 21509517 | 21563109 | 21563284 | 25 | 7 | 6 | 5 | 4 | 0 | 0 | 1 | 3 | 0 | 0 | 0 | 98 | 98 | 1 | 1 | 0.833 | 0.636 | 1 | NA | 1 | | 1 | -0.177 |
| MXE | 892 | CSDE1 | chr1 | - | 1E+08 | 114739890 | 114749820 | 114750207 | 114737962 | 114738072 | 114757924 | 114758050 | 3 | 3 | 0 | 41 | 19 | 5 | 0 | 0 | 0 | 0 | 11 | 0 | 98 | 98 | 1 | 1 | 0.068 | 0.136 | 0 | NA | 0 | NA | | 0.068 |
| MXE | 912 | JMJD1C | chr10 | - | 6E+07 | 63186383 | 63189167 | 63189446 | 63185562 | 63185653 | 63190893 | 63191108 | 140 | 33 | 16 | 63 | 23 | 6 | 1 | 18 | 0 | 0 | 11 | 0 | 98 | 98 | 1 | 1 | 0.69 | 0.589 | 0.727 | 1 | 0.621 | NA | | -0.142 |
| MXE | 913 | JMJD1C | chr10 | - | 6E+07 | 63219983 | 63264650 | 63264764 | 63217206 | 63217331 | 63268662 | 63269223 | 152 | 34 | 22 | 28 | 10 | 1 | 0 | 18 | 5 | 0 | 0 | 0 | 98 | 98 | 1 | 1 | 0.844 | 0.773 | 0.957 | NA | 1 | | 1 | -0.142 |
| MXE | 914 | JMJD1C | chr10 | - | 6E+07 | 63219983 | 63264650 | 63264764 | 63217206 | 63217331 | 63380317 | 63380482 | 21 | 6 | 5 | 29 | 10 | 1 | 0 | 3 | 0 | 0 | 0 | 0 | 98 | 98 | 1 | 1 | 0.42 | 0.375 | 0.833 | NA | 1 | NA | | -0.457 |
| MXE | 917 | ITGAX | chr16 | + | 3E+07 | 31355998 | 31356624 | 31356728 | 31355133 | 31355291 | 31357030 | 31357101 | 12 | 18 | 3 | 112 | 81 | 16 | 0 | 5 | 0 | 0 | 34 | 6 | 98 | 98 | 1 | 1 | 0.097 | 0.182 | 0.158 | NA | 0.128 | | 0 | 0.082 |
| MXE | 919 | MED8 | chr1 | - | 4E+07 | 43386226 | 43386588 | 43386670 | 43383907 | 43385106 | 43386857 | 43386998 | 5 | 2 | 0 | 3 | 1 | 2 | 0 | 0 | 0 | 0 | 1 | 0 | 98 | 98 | 1 | 1 | 0.625 | 0.667 | 0 | NA | 0 | NA | | 0.431 |
| MXE | 928 | ADAM9 | chr8 | + | 4E+07 | 39083073 | 39090046 | 39090188 | 39082640 | 39082721 | 39091258 | 39091346 | 2 | 3 | 0 | 11 | 0 | 0 | 0 | 1 | 0 | 0 | 3 | 0 | 98 | 98 | 1 | 1 | 0.154 | 1 | NA | NA | 0.25 | NA | | 0.327 |
| MXE | 930 | SCP2 | chr1 | + | 5E+07 | 52941853 | 52948008 | 52948080 | 52927228 | 52927465 | 52950754 | 52950886 | 0 | 0 | 0 | 1 | 1 | 0 | 0 | 1 | 0 | 0 | 0 | 0 | 98 | 98 | 1 | 1 | 0 | 0 | NA | NA | 1 | NA | | -1 |
| MXE | 935 | WDR37 | chr10 | + | 1E+06 | 1072293 | 1077906 | 1078003 | 1056835 | 1056968 | 1080010 | 1080106 | 2 | 4 | 2 | 9 | 2 | 6 | 0 | 0 | 0 | 0 | 0 | 0 | 98 | 98 | 1 | 1 | 0.182 | 0.667 | 0.25 | NA | 0 | NA | | 0.366 |
| MXE | 936 | CHD2 | chr15 | + | 9E+07 | 92901299 | 92924320 | 92924552 | 92900320 | 92900824 | 92927243 | 92927330 | 30 | 20 | 8 | 45 | 15 | 9 | 0 | 7 | 0 | 0 | 13 | 0 | 98 | 98 | 1 | 1 | 0.4 | 0.571 | 0.471 | NA | 0.35 | NA | | 0.131 |
| MXE | 937 | CHD2 | chr15 | + | 9E+07 | 92944515 | 92945820 | 92945865 | 92942842 | 92943068 | 92946037 | 92946216 | 23 | 3 | 0 | 44 | 13 | 8 | 0 | 7 | 2 | 0 | 4 | 0 | 98 | 94 | 1 | 1 | 0.334 | 0.181 | 0 | NA | 0.627 | | 1 | -0.642 |
| MXE | 939 | ARHGAP30 | chr1 | - | 2E+08 | 161056532 | 161059613 | 161059716 | 161054622 | 161054705 | 161069527 | 161069970 | 40 | 18 | 10 | 54 | 33 | 7 | 0 | 7 | 0 | 0 | 11 | 1 | 98 | 98 | 1 | 1 | 0.426 | 0.353 | 0.588 | NA | 0.389 | | 0 | 0.261 |
| MXE | 942 | ITCH | chr20 | + | 3E+07 | 34393881 | 34408650 | 34408792 | 34369393 | 34369470 | 34412514 | 34412639 | 9 | 6 | 0 | 28 | 28 | 16 | 0 | 1 | 0 | 2 | 2 | 0 | 98 | 98 | 1 | 1 | 0.243 | 0.176 | 0 | 0 | 0.333 | NA | | -0.027 |
| MXE | 945 | GS1-124K5.11 | chr7 | - | 7E+07 | 66576932 | 66584344 | 66584466 | 66573384 | 66573550 | 66588511 | 66588580 | 0 | 1 | 0 | 4 | 2 | 0 | 0 | 0 | 0 | 0 | 3 | 0 | 98 | 98 | 1 | 1 | 0 | 0.333 | NA | NA | 0 | NA | | 0.167 |
| MXE | 950 | CKLF-CMTM1 | chr16 | + | 7E+07 | 66558348 | 66563121 | 66563217 | 66552562 | 66552793 | 66577103 | 66577202 | 69 | 49 | 18 | 17 | 11 | 4 | 0 | 3 | 5 | 0 | 0 | 0 | 98 | 98 | 1 | 1 | 0.802 | 0.817 | 0.818 | NA | 1 | | 1 | -0.188 |
| MXE | 953 | PYGL | chr14 | - | 5E+07 | 50914815 | 50915335 | 50915499 | 50913028 | 50913130 | 50915824 | 50915971 | 46 | 46 | 19 | 68 | 25 | 23 | 0 | 1 | 0 | 0 | 8 | 0 | 98 | 98 | 1 | 1 | 0.404 | 0.648 | 0.452 | NA | 0.111 | NA | | 0.39 |
| MXE | 954 | MED24 | chr17 | - | 4E+07 | 40035795 | 40036115 | 40036154 | 40035116 | 40035349 | 40053297 | 40053380 | 1 | 0 | 0 | 1 | 0 | 0 | 0 | 0 | 0 | 0 | 3 | 0 | 88 | 98 | 1 | 1 | 0.527 | NA | NA | NA | 0 | NA | | 0.527 |
| MXE | 958 | PAM | chr5 | + | 1E+08 | 103025334 | 103028184 | 103028238 | 103019789 | 103019843 | 103028886 | 103031104 | 2 | 3 | 0 | 2 | 2 | 0 | 0 | 0 | 0 | 0 | 2 | 0 | 98 | 98 | 1 | 1 | 0.5 | 0.6 | NA | NA | 0 | NA | | 0.55 |

| | | | | | | | | | | | | | | | | | | | | | | | | | | | | | | | | | | | | |
|---|---|---|---|---|---|---|---|---|---|---|---|---|---|---|---|---|---|---|---|---|---|---|---|---|---|---|---|---|---|---|---|---|---|---|---|---|
| MXE | 959 | PAM | chr5 | + | 1E+08 | 103025334 | 103028184 | 103028238 | 103019789 | 103019843 | 103028886 | 103031104 | 1 | 0 | 0 | 2 | 2 | 0 | 0 | 0 | 0 | 0 | 2 | 0 | 98 | 98 | 1 | 1 | 0.333 | 0 | NA | NA | | 0 | NA | 0.167 |
| MXE | 960 | CTSS | chr1 | - | 2E+08 | 150750171 | 150751780 | 150752008 | 150747776 | 150747879 | 150755000 | 150755150 | 261 | 144 | 134 | 104 | 71 | 67 | 2 | 11 | 0 | 0 | 11 | 0 | 98 | 98 | 1 | 1 | 0.715 | 0.67 | 0.667 | | 1 | 0.5 | NA | -0.066 |
| MXE | 961 | CTSS | chr1 | - | 2E+08 | 150752008 | 150755000 | 150755150 | 150747776 | 150747879 | 150757857 | 150757980 | 414 | 132 | 122 | 2 | 0 | 5 | 4 | 28 | 7 | 0 | 0 | 0 | 98 | 98 | 1 | 1 | 0.995 | 1 | 0.961 | | 1 | 1 | 1 | -0.015 |
| MXE | 965 | SNX27 | chr1 | + | 2E+08 | 151662270 | 151665932 | 151666011 | 151660797 | 151660862 | 151668471 | 151668635 | 17 | 0 | 6 | 21 | 11 | 3 | 0 | 1 | 0 | 0 | 4 | 0 | 98 | 98 | 1 | 1 | 0.447 | 0 | 0.667 | NA | | 0.2 | NA | 0.171 |
| MXE | 969 | BEST1 | chr11 | + | 6E+07 | 61958298 | 61959497 | 61959578 | 61957386 | 61957464 | 61959891 | 61960043 | 14 | 7 | 2 | 18 | 6 | 3 | 0 | 0 | 0 | 0 | 2 | 0 | 98 | 98 | 1 | 1 | 0.437 | 0.538 | 0.4 | NA | | 0 | NA | 0.458 |
| MXE | 970 | BEST1 | chr11 | + | 6E+07 | 61959578 | 61959891 | 61960043 | 61958145 | 61958298 | 61962254 | 61962893 | 7 | 1 | 5 | 26 | 7 | 17 | 0 | 1 | 0 | 0 | 4 | 0 | 98 | 98 | 1 | 1 | 0.212 | 0.125 | 0.227 | NA | | 0.2 | NA | -0.012 |
| MXE | 972 | PAQR6 | chr1 | - | 2E+08 | 156245238 | 156245781 | 156245987 | 156244760 | 156244911 | 156246122 | 156246250 | 0 | 0 | 6 | 0 | 0 | 1 | 0 | 0 | 0 | 0 | 1 | 0 | 98 | 98 | 1 | 1 | NA | NA | 0.857 | NA | | 0 | NA | 0.857 |
| MXE | 976 | 7-Mar | chr2 | + | 2E+08 | 159762993 | 159764625 | 159764674 | 159759225 | 159759335 | 159767283 | 159768583 | 16 | 10 | 0 | 26 | 16 | 15 | 0 | 0 | 0 | 0 | 5 | 1 | 98 | 98 | 1 | 1 | 0.381 | 0.385 | 0 | NA | | 0 | 0 | 0.255 |
| MXE | 977 | C1orf43 | chr1 | - | 2E+08 | 154213946 | 154214456 | 154214574 | 154212457 | 154212624 | 154220341 | 154220797 | 4 | 2 | 0 | 7 | 6 | 7 | 0 | 3 | 0 | 0 | 5 | 3 | 98 | 98 | 1 | 1 | 0.364 | 0.25 | 0 | NA | | 0.375 | 0 | 0.017 |
| MXE | 983 | RCAN3 | chr1 | + | 2E+07 | 24531391 | 24533082 | 24533254 | 24514313 | 24514567 | 24535092 | 24537020 | 5 | 0 | 0 | 4 | 0 | 0 | 0 | 1 | 0 | 0 | 0 | 0 | 98 | 98 | 1 | 1 | 0.556 | NA | NA | NA | | 1 | NA | -0.444 |
| MXE | 986 | FBXO28 | chr1 | + | 2E+08 | 224134212 | 224153141 | 224153337 | 224130471 | 224130581 | 224157351 | 224162047 | 13 | 3 | 5 | 1 | 1 | 0 | 0 | 3 | 0 | 0 | 4 | 0 | 98 | 98 | 1 | 1 | 0.929 | 0.75 | 1 | NA | | 0.429 | NA | 0.464 |
| MXE | 988 | SMARCC1 | chr3 | - | 5E+07 | 47675588 | 47676628 | 47676782 | 47670657 | 47670717 | 47678197 | 47678311 | 3 | 1 | 0 | 2 | 0 | 0 | 0 | 0 | 0 | 0 | 1 | 0 | 98 | 98 | 1 | 1 | 0.6 | 1 | NA | NA | | 0 | NA | 0.8 |
| MXE | 989 | SMARCC1 | chr3 | - | 5E+07 | 47720735 | 47729024 | 47729094 | 47714414 | 47714490 | 47736033 | 47736126 | 5 | 5 | 4 | 10 | 4 | 1 | 0 | 2 | 0 | 0 | 1 | 0 | 98 | 98 | 1 | 1 | 0.333 | 0.556 | 0.8 | NA | | 0.667 | NA | -0.104 |
| MXE | 991 | CD164 | chr6 | - | 1E+08 | 109370467 | 109376073 | 109376112 | 109366513 | 109369017 | 109377899 | 109377971 | 34 | 17 | 3 | 20 | 10 | 0 | 1 | 6 | 0 | 0 | 1 | 0 | 88 | 98 | 1 | 1 | 0.654 | 0.654 | 1 | | 1 | 0.87 | NA | -0.166 |
| MXE | 994 | UBE2D1 | chr10 | + | 6E+07 | 58361526 | 58363608 | 58363686 | 58361337 | 58361401 | 58364770 | 58364876 | 12 | 0 | 5 | 33 | 4 | 13 | 0 | 2 | 0 | 0 | 1 | 0 | 81 | 98 | 1 | 1 | 0.306 | 0 | 0.318 | NA | | 0.708 | NA | -0.5 |
| MXE | 999 | MYD88 | chr3 | + | 4E+07 | 38139998 | 38140387 | 38140568 | 38138477 | 38139028 | 38140756 | 38140848 | 50 | 12 | 5 | 12 | 1 | 3 | 0 | 11 | 3 | 0 | 2 | 2 | 98 | 98 | 1 | 1 | 0.806 | 0.923 | 0.625 | NA | | 0.846 | 0.6 | 0.062 |
| MXE | 1002 | FAM65B | chr6 | - | 2E+07 | 24835871 | 24839090 | 24839272 | 24832255 | 24832391 | 24842861 | 24843554 | 81 | 33 | 16 | 61 | 28 | 18 | 0 | 5 | 0 | 0 | 4 | 0 | 98 | 98 | 1 | 1 | 0.57 | 0.541 | 0.471 | NA | | 0.556 | NA | -0.029 |
| MXE | 1006 | RBM27 | chr5 | + | 1E+08 | 146251875 | 146254942 | 146255092 | 146237297 | 146237432 | 146258448 | 146258593 | 6 | 1 | 0 | 7 | 7 | 0 | 0 | 0 | 0 | 0 | 4 | 0 | 98 | 98 | 1 | 1 | 0.462 | 0.125 | NA | NA | | 0 | NA | 0.293 |
| MXE | 1009 | SLAMF7 | chr1 | + | 2E+08 | 160750423 | 160751344 | 160751448 | 160749820 | 160750093 | 160752185 | 160752248 | 6 | 1 | 0 | 1 | 0 | 0 | 2 | 0 | 0 | 0 | 0 | 0 | 98 | 98 | 1 | 1 | 0.857 | 1 | NA | NA | 1 | NA | NA | -0.072 |
| MXE | 1015 | ADGRE3 | chr19 | - | 1E+07 | 14655165 | 14658512 | 14658550 | 14651084 | 14651204 | 14661962 | 14662118 | 98 | 26 | 12 | 77 | 34 | 21 | 0 | 3 | 0 | 0 | 3 | 0 | 87 | 98 | 1 | 1 | 0.589 | 0.463 | 0.392 | NA | | 0.53 | NA | -0.049 |
| MXE | 1016 | ADGRE3 | chr19 | - | 1E+07 | 14658550 | 14661962 | 14662118 | 14654981 | 14655165 | 14663417 | 14663540 | 72 | 31 | 17 | 66 | 13 | 15 | 5 | 1 | 0 | 0 | 3 | 0 | 98 | 87 | 1 | 1 | 0.492 | 0.679 | 0.502 | | 1 | 0.228 | NA | -0.056 |
| MXE | 1017 | ADGRE3 | chr19 | - | 1E+07 | 14662118 | 14663417 | 14663540 | 14658512 | 14658550 | 14668801 | 14668852 | 96 | 25 | 4 | 102 | 33 | 12 | 0 | 4 | 1 | 0 | 3 | 0 | 98 | 98 | 1 | 1 | 0.485 | 0.431 | 0.25 | NA | | 0.571 | 1 | -0.397 |
| MXE | 1018 | ADGRE3 | chr19 | - | 1E+07 | 14663540 | 14668801 | 14668852 | 14661962 | 14662118 | 14674745 | 14674918 | 11 | 5 | 0 | 72 | 32 | 17 | 0 | 1 | 0 | 5 | 1 | 0 | 98 | 98 | 1 | 1 | 0.133 | 0.135 | 0 | | 0 | 0.5 | NA | -0.161 |
| MXE | 1021 | HACL1 | chr3 | - | 2E+07 | 15589612 | 15591599 | 15591680 | 15586524 | 15586602 | 15596383 | 15596424 | 1 | 3 | 0 | 0 | 0 | 1 | 0 | 0 | 0 | 0 | 1 | 0 | 98 | 98 | 1 | 1 | 1 | 1 | 0 | NA | | 0 | NA | 0.667 |
| MXE | 1026 | GIT2 | chr12 | - | 1E+08 | 109947504 | 109951211 | 109951316 | 109939164 | 109939247 | 109953091 | 109953234 | 17 | 5 | 5 | 1 | 3 | 1 | 0 | 1 | 0 | 0 | 0 | 0 | 98 | 98 | 1 | 1 | 0.944 | 0.625 | 0.833 | NA | | 1 | NA | -0.199 |
| MXE | 1027 | ATP5C1 | chr10 | + | 8E+06 | 7802854 | 7806973 | 7807010 | 7802269 | 7802425 | 7807658 | 7807799 | 20 | 7 | 2 | 3 | 1 | 3 | 1 | 1 | 0 | 0 | 2 | 0 | 98 | 86 | 1 | 1 | 0.854 | 0.86 | 0.369 | | 1 | 0.305 | NA | 0.042 |
| MXE | 1029 | CELF2 | chr10 | + | 1E+07 | 11266677 | 11270665 | 11270824 | 11257737 | 11257872 | 11275056 | 11275120 | 20 | 6 | 1 | 104 | 45 | 14 | 0 | 4 | 0 | 0 | 11 | 3 | 98 | 98 | 1 | 1 | 0.161 | 0.118 | 0.067 | NA | | 0.267 | 0 | -0.018 |
| MXE | 1032 | PIK3CD | chr1 | + | 1E+07 | 9724092 | 9724275 | 9724421 | 9723124 | 9723292 | 9724803 | 9724936 | 34 | 17 | 14 | 43 | 22 | 17 | 0 | 0 | 0 | 0 | 6 | 0 | 98 | 98 | 1 | 1 | 0.442 | 0.436 | 0.452 | NA | | 0 | NA | 0.443 |
| MXE | 1033 | RANBP2 | chr2 | + | 1E+08 | 108775936 | 108777129 | 108777231 | 108772867 | 108773046 | 108781268 | 108781429 | 6 | 0 | 0 | 5 | 4 | 1 | 5 | 7 | 0 | 0 | 4 | 0 | 98 | 98 | 1 | 1 | 0.545 | 0 | 0 | | 1 | 0.636 | NA | -0.636 |
| MXE | 1039 | WAC | chr10 | + | 3E+07 | 28596041 | 28608185 | 28608431 | 28590719 | 28590832 | 28610698 | 28610821 | 26 | 13 | 2 | 26 | 9 | 9 | 0 | 3 | 0 | 0 | 4 | 1 | 98 | 98 | 1 | 1 | 0.5 | 0.591 | 0.182 | NA | | 0.429 | 0 | 0.21 |
| MXE | 1042 | GBP5 | chr1 | - | 9E+07 | 89267526 | 89268728 | 89268856 | 89266956 | 89267153 | 89269365 | 89269574 | 27 | 6 | 4 | 91 | 27 | 18 | 0 | 11 | 0 | 0 | 15 | 0 | 98 | 98 | 1 | 1 | 0.229 | 0.182 | 0.182 | NA | | 0.423 | NA | -0.225 |
| MXE | 1043 | GBP5 | chr1 | - | 9E+07 | 89268856 | 89269365 | 89269574 | 89267416 | 89267526 | 89272451 | 89272861 | 5 | 4 | 1 | 48 | 11 | 5 | 0 | 0 | 0 | 0 | 4 | 0 | 98 | 98 | 1 | 1 | 0.094 | 0.267 | 0.167 | NA | | 0 | NA | 0.176 |
| MXE | 1046 | PECAM1 | chr17 | - | 6E+07 | 64329722 | 64341633 | 64341690 | 64319415 | 64323845 | 64348259 | 64348322 | 21 | 26 | 28 | 56 | 21 | 12 | 0 | 1 | 0 | 0 | 3 | 0 | 98 | 72 | 1 | 1 | 0.216 | 0.476 | 0.632 | NA | | 0.197 | NA | 0.244 |
| MXE | 1047 | PECAM1 | chr17 | - | 6E+07 | 64341690 | 64348259 | 64348322 | 64329699 | 64329722 | 64350379 | 64350433 | 42 | 38 | 40 | 104 | 68 | 31 | 0 | 2 | 0 | 2 | 6 | 0 | 98 | 98 | 1 | 1 | 0.288 | 0.358 | 0.563 | | 0 | 0.25 | NA | 0.278 |
| MXE | 1048 | PECAM1 | chr17 | - | 6E+07 | 64353518 | 64356110 | 64356398 | 64352389 | 64352463 | 64360139 | 64360415 | 118 | 59 | 42 | 103 | 51 | 21 | 0 | 3 | 0 | 0 | 2 | 0 | 98 | 77 | 1 | 1 | 0.474 | 0.476 | 0.611 | NA | | 0.541 | NA | -0.021 |
| MXE | 1057 | PFKFB3 | chr10 | + | 6E+06 | 6221745 | 6222854 | 6222984 | 6221380 | 6221527 | 6223957 | 6224020 | 56 | 41 | 5 | 69 | 41 | 7 | 0 | 28 | 22 | 0 | 21 | 7 | 98 | 98 | 1 | 1 | 0.448 | 0.5 | 0.417 | NA | | 0.571 | 0.759 | -0.21 |
| MXE | 1061 | SAP30BP | chr17 | + | 8E+07 | 75668625 | 75671815 | 75671863 | 75667115 | 75667478 | 75693439 | 75693482 | 10 | 8 | 1 | 18 | 8 | 1 | 0 | 5 | 0 | 0 | 7 | 0 | 98 | 97 | 1 | 1 | 0.355 | 0.497 | 0.497 | NA | | 0.414 | NA | 0.036 |
| MXE | 1062 | SAP30BP | chr17 | + | 8E+07 | 75671863 | 75693439 | 75693482 | 75668515 | 75668625 | 75699782 | 75699871 | 25 | 6 | 5 | 15 | 5 | 5 | 0 | 2 | 0 | 0 | 5 | 0 | 97 | 92 | 1 | 1 | 0.613 | 0.532 | 0.487 | NA | | 0.275 | NA | 0.269 |
| MXE | 1067 | CAMK2G | chr10 | - | 7E+07 | 73847347 | 73847987 | 73848082 | 73842457 | 73842541 | 73848525 | 73848609 | 13 | 8 | 5 | 15 | 1 | 2 | 0 | 0 | 0 | 0 | 1 | 0 | 98 | 98 | 1 | 1 | 0.464 | 0.889 | 0.714 | NA | | 0 | NA | 0.689 |
| MXE | 1068 | CAMK2G | chr10 | - | 7E+07 | 73848082 | 73848525 | 73848609 | 73847224 | 73847347 | 73849012 | 73849115 | 23 | 16 | 17 | 13 | 15 | 1 | 0 | 4 | 0 | 0 | 4 | 1 | 98 | 98 | 1 | 1 | 0.639 | 0.516 | 0.944 | NA | | 0.5 | 0 | 0.45 |
| MXE | 1070 | SENP7 | chr3 | - | 1E+08 | 101399055 | 101417592 | 101417790 | 101372007 | 101372126 | 101458954 | 101459052 | 11 | 8 | 2 | 13 | 4 | 0 | 0 | 0 | 0 | 0 | 3 | 0 | 98 | 98 | 1 | 1 | 0.458 | 0.667 | 1 | NA | | 0 | NA | 0.708 |
| MXE | 1077 | SNW1 | chr14 | - | 8E+07 | 77720828 | 77723180 | 77723277 | 77718366 | 77718530 | 77730987 | 77731129 | 9 | 4 | 2 | 6 | 11 | 0 | 0 | 2 | 0 | 0 | 1 | 0 | 98 | 98 | 1 | 1 | 0.6 | 0.267 | 1 | NA | | 0.667 | NA | -0.045 |
| MXE | 1082 | IFRD1 | chr7 | + | 1E+08 | 112457038 | 112458860 | 112459018 | 112456001 | 112456086 | 112461865 | 112461916 | 216 | 63 | 15 | 61 | 44 | 24 | 0 | 72 | 4 | 0 | 20 | 0 | 98 | 98 | 1 | 1 | 0.78 | 0.589 | 0.385 | NA | | 0.783 | 1 | -0.307 |
| MXE | 1089 | LCOR | chr10 | + | 1E+08 | 96944245 | 96949007 | 96949295 | 96907667 | 96907747 | 96952102 | 96952196 | 7 | 2 | 3 | 11 | 4 | 1 | 0 | 2 | 0 | 0 | 2 | 0 | 98 | 98 | 1 | 1 | 0.389 | 0.333 | 0.75 | NA | | 0.5 | NA | -0.009 |
| MXE | 1091 | ZFYVE27 | chr10 | + | 1E+08 | 97738674 | 97744728 | 97744915 | 97737120 | 97737321 | 97748268 | 97748364 | 3 | 5 | 0 | 1 | 1 | 0 | 0 | 1 | 0 | 0 | 0 | 0 | 98 | 98 | 1 | 1 | 0.75 | 0.833 | NA | NA | | 1 | NA | -0.209 |
| MXE | 1093 | ELF2 | chr4 | - | 1E+08 | 139062057 | 139067683 | 139067770 | 139060323 | 139060674 | 139071865 | 139072003 | 33 | 3 | 2 | 49 | 12 | 2 | 0 | 4 | 0 | 0 | 7 | 0 | 98 | 98 | 1 | 1 | 0.402 | 0.2 | 0.5 | NA | | 0.364 | NA | 0.003 |
| MXE | 1098 | CLTA | chr9 | + | 4E+07 | 36197588 | 36198978 | 36199096 | 36190855 | 36191273 | 36204067 | 36204179 | 11 | 0 | 2 | 3 | 1 | 4 | 0 | 2 | 0 | 0 | 2 | 0 | 87 | 98 | 1 | 1 | 0.805 | 0 | 0.36 | NA | | 0.53 | NA | -0.142 |
| MXE | 1100 | ATE1 | chr10 | - | 1E+08 | 121898969 | 121899865 | 121899994 | 121870005 | 121870038 | 121902390 | 121902620 | 4 | 0 | 0 | 3 | 0 | 0 | 0 | 0 | 0 | 0 | 1 | 0 | 98 | 98 | 1 | 1 | 0.571 | NA | NA | NA | | 0 | NA | 0.571 |
| MXE | 1103 | CASP8 | chr2 | + | 2E+08 | 201272942 | 201274888 | 201274953 | 201272637 | 201272776 | 201276826 | 201276968 | 106 | 49 | 12 | 89 | 36 | 17 | 0 | 18 | 0 | 0 | 3 | 0 | 94 | 98 | 1 | 1 | 0.554 | 0.587 | 0.424 | NA | | 0.862 | NA | -0.34 |
| MXE | 1105 | PTPRE | chr10 | + | 1E+08 | 128056213 | 128060938 | 128061015 | 128049529 | 128049666 | 128061678 | 128061715 | 78 | 52 | 7 | 71 | 33 | 15 | 0 | 34 | 1 | 0 | 19 | 0 | 98 | 98 | 1 | 1 | 0.523 | 0.612 | 0.318 | NA | | 0.642 | 1 | -0.337 |
| MXE | 1115 | ACTN1 | chr14 | - | 7E+07 | 68878523 | 68878988 | 68879069 | 68877081 | 68877240 | 68879961 | 68880108 | 85 | 49 | 27 | 0 | 1 | 0 | 0 | 14 | 2 | 0 | 0 | 0 | 98 | 98 | 1 | 1 | 1 | 0.98 | 1 | NA | | 1 | 1 | -0.007 |
| MXE | 1119 | TRAF3IP3 | chr1 | + | 2E+08 | 209779374 | 209780469 | 209780606 | 209778110 | 209778173 | 209781344 | 209781458 | 23 | 6 | 0 | 8 | 2 | 0 | 0 | 9 | 0 | 0 | 2 | 0 | 98 | 98 | 1 | 1 | 0.742 | 0.75 | NA | NA | | 0.818 | NA | -0.072 |
| MXE | 1123 | ATXN3 | chr14 | - | 9E+07 | 92093318 | 92096092 | 92096137 | 92088729 | 92088817 | 92096673 | 92096838 | 7 | 0 | 2 | 9 | 1 | 0 | 0 | 0 | 0 | 3 | 0 | 0 | 94 | 98 | 1 | 1 | 0.448 | 0 | 1 | | 0 | NA | NA | 0.483 |
| MXE | 1124 | ATXN3 | chr14 | - | 9E+07 | 92093831 | 92096092 | 92096137 | 92093251 | 92093318 | 92096673 | 92096838 | 10 | 1 | 2 | 4 | 2 | 0 | 0 | 1 | 0 | 0 | 0 | 0 | 94 | 98 | 1 | 1 | 0.723 | 0.343 | 1 | NA | | 1 | NA | -0.311 |
| MXE | 1125 | ATXN3 | chr14 | - | 9E+07 | 92093831 | 92096092 | 92096137 | 92093251 | 92093318 | 92106528 | 92106621 | 3 | 1 | 0 | 4 | 2 | 0 | 0 | 1 | 0 | 0 | 0 | 0 | 94 | 98 | 1 | 1 | 0.439 | 0.343 | NA | NA | | 1 | NA | -0.609 |
| MXE | 1130 | RFWD2 | chr1 | - | 2E+08 | 176135086 | 176149005 | 176149074 | 176116623 | 176116681 | 176175909 | 176176007 | 3 | 1 | 0 | 118 | 57 | 9 | 0 | 0 | 0 | 0 | 4 | 1 | 98 | 98 | 1 | 1 | 0.025 | 0.017 | 0 | NA | | 0 | 0 | 0.014 |
| MXE | 1133 | RFWD2 | chr1 | - | 2E+08 | 176149074 | 176175909 | 176176007 | 176135009 | 176135086 | 176184632 | 176184692 | 32 | 16 | 3 | 29 | 7 | 0 | 0 | 0 | 0 | 0 | 3 | 0 | 98 | 98 | 1 | 1 | 0.525 | 0.696 | 1 | NA | | 0 | NA | 0.74 |
| MXE | 1136 | SNX17 | chr2 | + | 3E+07 | 27372740 | 27373039 | 27373113 | 27371268 | 27371343 | 27373246 | 27373311 | 6 | 1 | 1 | 1 | 0 | 0 | 0 | 2 | 0 | 0 | 1 | 0 | 98 | 98 | 1 | 1 | 0.857 | 1 | 1 | NA | | 0.667 | NA | 0.285 |
| MXE | 1140 | TECPR2 | chr14 | + | 1E+08 | 102425291 | 102431795 | 102432128 | 102414635 | 102414793 | 102434234 | 102435211 | 36 | 8 | 2 | 19 | 5 | 3 | 0 | 3 | 0 | 0 | 2 | 0 | 98 | 98 | 1 | 1 | 0.655 | 0.615 | 0.4 | NA | | 0.6 | NA | -0.043 |
| MXE | 1141 | MARK3 | chr14 | + | 1E+08 | 103428440 | 103448918 | 103448967 | 103405075 | 103405267 | 103451917 | 103451983 | 7 | 4 | 1 | 19 | 9 | 2 | 0 | 1 | 0 | 0 | 6 | 0 | 98 | 98 | 1 | 1 | 0.269 | 0.308 | 0.333 | NA | | 0.143 | NA | 0.16 |
| MXE | 1143 | EIF4E3 | chr3 | - | 7E+07 | 71696520 | 71699613 | 71699708 | 71693874 | 71693941 | 71710411 | 71710484 | 11 | 12 | 8 | 29 | 15 | 15 | 0 | 1 | 0 | 0 | 6 | 0 | 98 | 98 | 1 | 1 | 0.275 | 0.444 | 0.348 | NA | | 0.143 | NA | 0.213 |
| MXE | 1144 | KIZ | chr20 | + | 2E+07 | 21132159 | 21145564 | 21145654 | 21125982 | 21126204 | 21161870 | 21162507 | 7 | 4 | 2 | 2 | 4 | 0 | 0 | 0 | 0 | 0 | 1 | 0 | 98 | 98 | 1 | 1 | 0.778 | 0.5 | 1 | NA | | 0 | NA | 0.759 |
| MXE | 1146 | TPM3 | chr1 | - | 2E+08 | 154171488 | 154172028 | 154172104 | 154170648 | 154170711 | 154172907 | 154172978 | 159 | 96 | 73 | 3 | 0 | 0 | 0 | 21 | 0 | 0 | 0 | 0 | 98 | 98 | 1 | 1 | 0.981 | 1 | 1 | NA | | 1 | NA | -0.006 |
| MXE | 1147 | ZNF92 | chr7 | + | 7E+07 | 65388028 | 65388805 | 65388901 | 65373798 | 65374000 | 65398340 | 65401135 | 0 | 0 | 4 | 3 | 1 | 0 | 0 | 2 | 0 | 1 | 0 | 0 | 98 | 98 | 1 | 1 | 0 | 0 | 1 | | 0 | 1 | NA | -0.167 |
| MXE | 1148 | SOS2 | chr14 | - | 5E+07 | 50174552 | 50180571 | 50180682 | 50161481 | 50161609 | 50182462 | 50182606 | 70 | 30 | 7 | 37 | 13 | 5 | 9 | 10 | 0 | 0 | 3 | 0 | 98 | 98 | 1 | 1 | 0.654 | 0.698 | 0.583 | | 1 | 0.769 | NA | -0.24 |
| MXE | 1154 | MEFV | chr16 | - | 3E+06 | 3249780 | 3254157 | 3254790 | 3247015 | 3247246 | 3256310 | 3256627 | 81 | 26 | 30 | 0 | 2 | 1 | 9 | 22 | 0 | 0 | 0 | 0 | 98 | 98 | 1 | 1 | 1 | 0.929 | 0.968 | | 1 | 1 | NA | -0.034 |
| MXE | 1155 | MEFV | chr16 | - | 3E+06 | 3249780 | 3254157 | 3254790 | 3248908 | 3249004 | 3256310 | 3256627 | 81 | 26 | 30 | 23 | 10 | 11 | 9 | 22 | 0 | 0 | 7 | 0 | 98 | 98 | 1 | 1 | 0.779 | 0.722 | 0.732 | | 1 | 0.759 | NA | -0.135 |
| MXE | 1157 | RDH11 | chr14 | - | 7E+07 | 67685204 | 67690211 | 67690421 | 67676800 | 67678423 | 67691139 | 67691244 | 9 | 0 | 0 | 1 | 0 | 0 | 0 | 0 | 0 | 0 | 1 | 0 | 98 | 98 | 1 | 1 | 0.9 | NA | NA | NA | | 0 | NA | 0.9 |
| MXE | 1158 | SGK1 | chr6 | - | 1E+08 | 134171732 | 134172192 | 134172316 | 134171022 | 134171178 | 134172661 | 134172774 | 82 | 52 | 12 | 90 | 38 | 6 | 0 | 13 | 0 | 0 | 7 | 0 | 98 | 98 | 1 | 1 | 0.477 | 0.578 | 0.667 | NA | | 0.65 | NA | -0.076 |
| MXE | 1159 | P4HA1 | chr10 | - | 7E+07 | 73043957 | 73044980 | 73045051 | 73030266 | 73030370 | 73046924 | 73047101 | 7 | 2 | 0 | 5 | 0 | 3 | 0 | 1 | 0 | 0 | 1 | 0 | 98 | 98 | 1 | 1 | 0.583 | 1 | 0 | NA | | 0.5 | NA | 0.028 |
| MXE | 1165 | USP8 | chr15 | + | 5E+07 | 50482065 | 50484274 | 50484361 | 50477275 | 50477499 | 50489800 | 50489881 | 13 | 6 | 1 | 4 | 2 | 2 | 0 | 3 | 0 | 0 | 1 | 0 | 98 | 98 | 1 | 1 | 0.765 | 0.75 | 0.333 | NA | | 0.75 | NA | -0.134 |
| MXE | 1167 | RNF213 | chr17 | + | 8E+07 | 80360206 | 80361733 | 80361888 | 80358287 | 80358479 | 80363101 | 80363314 | 31 | 19 | 2 | 32 | 41 | 4 | 0 | 19 | 1 | 0 | 32 | 6 | 98 | 98 | 1 | 1 | 0.492 | 0.317 | 0.333 | NA | | 0.373 | 0.143 | 0.123 |
| MXE | 1169 | RNF213 | chr17 | + | 8E+07 | 80363314 | 80363608 | 80363790 | 80361733 | 80361888 | 80364432 | 80364553 | 32 | 40 | 4 | 24 | 19 | 5 | 0 | 31 | 6 | 0 | 11 | 3 | 98 | 98 | 1 | 1 | 0.571 | 0.678 | 0.444 | NA | | 0.738 | 0.667 | -0.138 |
| MXE | 1170 | DMXL2 | chr15 | - | 5E+07 | 51464876 | 51465565 | 51465651 | 51463378 | 51463496 | 51466183 | 51466311 | 33 | 5 | 3 | 25 | 6 | 3 | 0 | 0 | 0 | 0 | 1 | 0 | 98 | 98 | 1 | 1 | 0.569 | 0.455 | 0.5 | NA | | 0 | NA | 0.508 |
| MXE | 1171 | PIK3R5 | chr17 | - | 9E+06 | 8890126 | 8890737 | 8890912 | 8889138 | 8889222 | 8893585 | 8893655 | 16 | 11 | 14 | 4 | 4 | 1 | 0 | 3 | 0 | 0 | 0 | 0 | 98 | 98 | 1 | 1 | 0.8 | 0.733 | 0.933 | NA | | 1 | NA | -0.178 |
| MXE | 1173 | RRP12 | chr10 | - | 1E+08 | 97379414 | 97379627 | 97379770 | 97373829 | 97373894 | 97380798 | 97380913 | 22 | 3 | 0 | 18 | 0 | 0 | 0 | 1 | 1 | 0 | 5 | 0 | 98 | 98 | 1 | 1 | 0.55 | 1 | NA | NA | | 0.167 | 1 | 0.192 |
| MXE | 1174 | BNIP2 | chr15 | - | 6E+07 | 59668990 | 59669275 | 59669362 | 59662862 | 59664120 | 59671182 | 59671314 | 27 | 15 | 3 | 44 | 31 | 10 | 0 | 8 | 0 | 0 | 3 | 0 | 98 | 98 | 1 | 1 | 0.38 | 0.326 | 0.231 | NA | | 0.727 | NA | -0.415 |
| MXE | 1179 | DPP8 | chr15 | - | 7E+07 | 65452102 | 65456224 | 65456371 | 65450998 | 65451110 | 65463760 | 65463906 | 5 | 3 | 3 | 3 | 0 | 0 | 0 | 1 | 0 | 0 | 0 | 0 | 98 | 98 | 1 | 1 | 0.625 | 1 | 1 | NA | | 1 | NA | -0.125 |
| MXE | 1185 | PPCDC | chr15 | + | 8E+07 | 75043536 | 75044385 | 75044514 | 75028246 | 75028453 | 75048552 | 75048721 | 0 | 2 | 0 | 16 | 0 | 4 | 0 | 0 | 0 | 0 | 4 | 2 | 98 | 98 | 1 | 1 | 0 | 1 | 0 | NA | | 0 | 0 | 0.333 |
| MXE | 1204 | IRF1 | chr5 | - | 1E+08 | 132484497 | 132485666 | 132485716 | 132481608 | 132484075 | 132486250 | 132486373 | 53 | 34 | 26 | 43 | 18 | 10 | 0 | 5 | 0 | 0 | 4 | 0 | 98 | 98 | 1 | 1 | 0.552 | 0.654 | 0.722 | NA | | 0.556 | NA | 0.087 |
| MXE | 1205 | IRF1 | chr5 | - | 1E+08 | 132484497 | 132486250 | 132486373 | 132481608 | 132484075 | 132486556 | 132486686 | 65 | 21 | 29 | 44 | 24 | 13 | 7 | 8 | 0 | 0 | 5 | 0 | 98 | 98 | 1 | 1 | 0.596 | 0.467 | 0.69 | | 1 | 0.615 | NA | -0.223 |
| MXE | 1206 | IRF1 | chr5 | - | 1E+08 | 132485716 | 132486250 | 132486373 | 132484361 | 132484497 | 132486556 | 132486686 | 65 | 18 | 29 | 72 | 36 | 16 | 7 | 8 | 0 | 0 | 14 | 0 | 98 | 98 | 1 | 1 | 0.474 | 0.333 | 0.644 | | 1 | 0.364 | NA | -0.198 |
| MXE | 1207 | IRF1 | chr5 | - | 1E+08 | 132488025 | 132489391 | 132489483 | 132486953 | 132487130 | 132490544 | 132490773 | 43 | 25 | 13 | 75 | 60 | 22 | 0 | 8 | 2 | 0 | 11 | 1 | 98 | 98 | 1 | 1 | 0.364 | 0.294 | 0.371 | NA | | 0.421 | 0.667 | -0.201 |
| MXE | 1208 | BTBD1 | chr15 | - | 8E+07 | 83018853 | 83020674 | 83020762 | 83016428 | 83018225 | 83030135 | 83030328 | 3 | 4 | 0 | 5 | 2 | 0 | 0 | 0 | 0 | 0 | 2 | 0 | 98 | 98 | 1 | 1 | 0.375 | 0.667 | NA | NA | | 0 | NA | 0.521 |
| MXE | 1213 | SEC11A | chr15 | - | 8E+07 | 84680832 | 84687624 | 84687774 | 84670724 | 84670782 | 84691534 | 84691644 | 15 | 4 | 20 | 3 | 2 | 4 | 0 | 3 | 0 | 0 | 0 | 0 | 98 | 98 | 1 | 1 | 0.833 | 0.667 | 0.833 | NA | | 1 | NA | -0.222 |
| MXE | 1214 | SEC11A | chr15 | - | 8E+07 | 84687774 | 84691534 | 84691644 | 84680712 | 84680832 | 84716024 | 84716460 | 5 | 2 | 0 | 0 | 2 | 0 | 0 | 2 | 0 | 0 | 0 | 0 | 98 | 98 | 1 | 1 | 1 | 0.5 | NA | NA | | 1 | NA | -0.25 |
| MXE | 1215 | AKAP13 | chr15 | + | 9E+07 | 85658590 | 85662387 | 85662453 | 85655416 | 85655787 | 85664562 | 85664755 | 80 | 39 | 4 | 3 | 1 | 0 | 0 | 4 | 3 | 0 | 0 | 0 | 98 | 98 | 1 | 1 | 0.964 | 0.975 | 1 | NA | | 1 | 1 | -0.02 |
| MXE | 1216 | POLG | chr15 | - | 9E+07 | 89322902 | 89323403 | 89323511 | 89321961 | 89322015 | 89323814 | 89323901 | 7 | 3 | 0 | 2 | 0 | 0 | 0 | 1 | 0 | 0 | 0 | 0 | 98 | 98 | 1 | 1 | 0.778 | 1 | NA | NA | | 1 | NA | -0.111 |
| MXE | 1219 | WWC3 | chrX | + | 1E+07 | 10125110 | 10126113 | 10126306 | 10124264 | 10124339 | 10127987 | 10128161 | 16 | 3 | 0 | 9 | 1 | 6 | 0 | 1 | 0 | 0 | 1 | 0 | 98 | 98 | 1 | 1 | 0.64 | 0.75 | 0 | NA | | 0.5 | NA | -0.037 |
| MXE | 1220 | WWC3 | chrX | + | 1E+07 | 10126306 | 10127987 | 10128161 | 10125023 | 10125110 | 10128556 | 10128708 | 11 | 5 | 2 | 15 | 8 | 10 | 0 | 0 | 0 | 0 | 1 | 0 | 98 | 98 | 1 | 1 | 0.423 | 0.385 | 0.167 | NA | | 0 | NA | 0.325 |
| MXE | 1222 | IQGAP1 | chr15 | + | 9E+07 | 90466436 | 90467449 | 90467592 | 90466000 | 90466091 | 90472839 | 90473010 | 63 | 42 | 35 | 71 | 42 | 17 | 0 | 16 | 0 | 1 | 6 | 0 | 98 | 98 | 1 | 1 | 0.47 | 0.5 | 0.673 | | 0 | 0.727 | NA | 0.184 |
| MXE | 1224 | VPS33B | chr15 | - | 9E+07 | 91017024 | 91017804 | 91017885 | 91014383 | 91014433 | 91022153 | 91022621 | 2 | 0 | 0 | 1 | 0 | 0 | 0 | 0 | 0 | 0 | 1 | 0 | 98 | 98 | 1 | 1 | 0.667 | NA | NA | NA | | 0 | NA | 0.667 |
| MXE | 1225 | ARHGAP15 | chr2 | + | 1E+08 | 143487495 | 143519265 | 143519364 | 143436912 | 143437042 | 143556407 | 143556485 | 128 | 34 | 12 | 98 | 12 | 10 | 0 | 15 | 2 | 0 | 13 | 0 | 98 | 98 | 1 | 1 | 0.566 | 0.739 | 0.545 | NA | | 0.536 | 1 | -0.151 |
| MXE | 1226 | MEF2A | chr15 | + | 1E+08 | 99671454 | 99671519 | 99671657 | 99645560 | 99645764 | 99674392 | 99674612 | 20 | 3 | 2 | 6 | 0 | 0 | 0 | 2 | 0 | 0 | 0 | 0 | 98 | 98 | 1 | 1 | 0.769 | 1 | 1 | NA | | 1 | NA | -0.077 |
| MXE | 1245 | USP25 | chr21 | + | 2E+07 | 15849872 | 15864267 | 15864446 | 15842397 | 15842540 | 15866265 | 15866344 | 1 | 0 | 0 | 8 | 8 | 0 | 0 | 0 | 0 | 0 | 1 | 0 | 98 | 98 | 1 | 1 | 0.111 | 0 | NA | NA | | 0 | NA | 0.056 |
| MXE | 1251 | NDE1 | chr16 | + | 2E+07 | 15694256 | 15696708 | 15696860 | 15691143 | 15691323 | 15724190 | 15726351 | 16 | 25 | 7 | 11 | 11 | 3 | 0 | 3 | 0 | 0 | 1 | 0 | 98 | 98 | 1 | 1 | 0.593 | 0.694 | 0.7 | NA | | 0.75 | NA | -0.088 |
| MXE | 1258 | KIAA1551 | chr12 | + | 3E+07 | 31970356 | 31980877 | 31985957 | 31960776 | 31960871 | 31987238 | 31987322 | 65 | 52 | 41 | 95 | 43 | 25 | 0 | 3 | 0 | 0 | 9 | 0 | 98 | 98 | 1 | 1 | 0.406 | 0.547 | 0.621 | NA | | 0.25 | NA | 0.275 |
| MXE | 1259 | KIAA1551 | chr12 | + | 3E+07 | 31985957 | 31987238 | 31987322 | 31970188 | 31970356 | 31992377 | 31993107 | 203 | 97 | 57 | 69 | 25 | 16 | 0 | 13 | 0 | 0 | 0 | 1 | 98 | 98 | 1 | 1 | 0.746 | 0.795 | 0.781 | NA | | 1 | 0 | 0.274 |
| MXE | 1262 | RPAIN | chr17 | + | 5E+06 | 5426082 | 5428070 | 5428211 | 5422768 | 5422829 | 5432541 | 5433020 | 0 | 2 | 0 | 2 | 0 | 0 | 0 | 1 | 0 | 0 | 0 | 0 | 98 | 98 | 1 | 1 | 0 | 1 | NA | NA | | 1 | NA | -0.5 |
| MXE | 1265 | DCLRE1C | chr10 | - | 1E+07 | 14945189 | 14949035 | 14949087 | 14939809 | 14939869 | 14953901 | 14954432 | 4 | 1 | 0 | 6 | 1 | 0 | 0 | 0 | 0 | 0 | 1 | 0 | 98 | 98 | 1 | 1 | 0.4 | 0.5 | NA | NA | | 0 | NA | 0.45 |
| MXE | 1266 | DCLRE1C | chr10 | - | 1E+07 | 14945565 | 14947524 | 14947609 | 14945104 | 14945189 | 14949035 | 14949087 | 3 | 0 | 0 | 0 | 0 | 0 | 0 | 1 | 0 | 0 | 1 | 0 | 98 | 98 | 1 | 1 | 1 | NA | NA | NA | | 0.5 | NA | 0.5 |
| MXE | 1267 | APLP2 | chr11 | + | 1E+08 | 130109505 | 130120705 | 130120818 | 130069820 | 130070082 | 130121613 | 130121810 | 21 | 12 | 5 | 53 | 34 | 10 | 0 | 1 | 1 | 0 | 8 | 0 | 98 | 98 | 1 | 1 | 0.284 | 0.261 | 0.333 | NA | | 0.111 | 1 | -0.263 |
| MXE | 1272 | BCL2L13 | chr22 | + | 2E+07 | 17689142 | 17702242 | 17702386 | 17683213 | 17683321 | 17726676 | 17730855 | 3 | 0 | 0 | 1 | 0 | 0 | 0 | 1 | 0 | 0 | 0 | 0 | 98 | 98 | 1 | 1 | 0.75 | NA | NA | NA | | 1 | NA | -0.25 |
| MXE | 1273 | BCL2L13 | chr22 | + | 2E+07 | 17696210 | 17702242 | 17702386 | 17688985 | 17689142 | 17726676 | 17730855 | 7 | 2 | 0 | 1 | 0 | 0 | 0 | 1 | 0 | 0 | 0 | 0 | 98 | 98 | 1 | 1 | 0.875 | 1 | NA | NA | | 1 | NA | -0.062 |
| MXE | 1276 | CREBBP | chr16 | - | 4E+06 | 3782926 | 3791980 | 3792094 | 3781203 | 3781306 | 3793385 | 3793626 | 32 | 8 | 4 | 18 | 3 | 8 | 0 | 3 | 1 | 0 | 4 | 0 | 98 | 98 | 1 | 1 | 0.64 | 0.727 | 0.333 | NA | | 0.429 | 1 | -0.148 |
| MXE | 1277 | MVP | chr16 | + | 3E+07 | 29842112 | 29844492 | 29844879 | 29841595 | 29841840 | 29845862 | 29845979 | 31 | 8 | 7 | 30 | 16 | 18 | 0 | 3 | 2 | 0 | 7 | 0 | 98 | 98 | 1 | 1 | 0.508 | 0.333 | 0.28 | NA | | 0.3 | 1 | -0.276 |
| MXE | 1278 | DEDD | chr1 | - | 2E+08 | 161124526 | 161130814 | 161130847 | 161123838 | 161123946 | 161132550 | 161132688 | 0 | 1 | 0 | 2 | 3 | 1 | 0 | 0 | 0 | 0 | 3 | 0 | 82 | 98 | 1 | 1 | 0 | 0.285 | 0 | NA | | 0 | NA | 0.095 |
| MXE | 1279 | ADGRE2 | chr19 | - | 1E+07 | 14755127 | 14755653 | 14755877 | 14752328 | 14752526 | 14756237 | 14756345 | 6 | 6 | 12 | 20 | 5 | 0 | 0 | 2 | 0 | 0 | 5 | 0 | 98 | 98 | 1 | 1 | 0.231 | 0.545 | 1 | NA | | 0.286 | NA | 0.306 |
| MXE | 1281 | ARHGEF1 | chr19 | + | 4E+07 | 41905036 | 41905174 | 41905261 | 41904215 | 41904383 | 41905759 | 41905827 | 19 | 9 | 0 | 35 | 10 | 14 | 0 | 4 | 0 | 0 | 0 | 2 | 98 | 98 | 1 | 1 | 0.352 | 0.474 | 0 | NA | | 1 | 0 | -0.225 |
| MXE | 1282 | NLRC5 | chr16 | + | 6E+07 | 57076902 | 57077295 | 57077379 | 57074599 | 57074683 | 57077718 | 57077802 | 4 | 6 | 1 | 6 | 1 | 1 | 0 | 7 | 0 | 0 | 1 | 0 | 98 | 98 | 1 | 1 | 0.4 | 0.857 | 0.5 | NA | | 0.875 | NA | -0.289 |
| MXE | 1283 | ADGRG3 | chr16 | + | 6E+07 | 57680617 | 57684389 | 57684483 | 57680264 | 57680365 | 57685642 | 57685926 | 60 | 24 | 15 | 36 | 10 | 17 | 0 | 9 | 14 | 0 | 6 | 1 | 98 | 98 | 1 | 1 | 0.625 | 0.706 | 0.469 | NA | | 0.6 | 0.933 | -0.166 |
| MXE | 1284 | USB1 | chr16 | + | 6E+07 | 58014326 | 58017333 | 58017439 | 58009928 | 58010112 | 58018971 | 58019055 | 14 | 4 | 1 | 6 | 1 | 0 | 0 | 0 | 0 | 0 | 2 | 0 | 98 | 98 | 1 | 1 | 0.7 | 0.8 | 1 | NA | | 0 | NA | 0.833 |
| MXE | 1285 | CKLF | chr16 | + | 7E+07 | 66558348 | 66563121 | 66563217 | 66552562 | 66552793 | 66565885 | 66566287 | 71 | 53 | 18 | 47 | 24 | 20 | 0 | 5 | 5 | 0 | 0 | 0 | 98 | 98 | 1 | 1 | 0.602 | 0.688 | 0.474 | NA | | 1 | 1 | -0.412 |
| MXE | 1286 | ZMYM5 | chr13 | - | 2E+07 | 19838985 | 19851354 | 19851448 | 19837452 | 19837821 | 19851688 | 19852190 | 6 | 1 | 0 | 2 | 2 | 0 | 0 | 0 | 0 | 0 | 1 | 0 | 98 | 98 | 1 | 1 | 0.75 | 0.333 | NA | NA | | 0 | NA | 0.541 |
| MXE | 1296 | SCMH1 | chr1 | - | 4E+07 | 41151684 | 41159724 | 41159933 | 41142877 | 41143112 | 41160874 | 41160898 | 0 | 0 | 0 | 0 | 2 | 0 | 0 | 0 | 1 | 0 | 0 | 0 | 98 | 98 | 1 | 1 | NA | 0 | NA | NA | | NA | 1 | -1 |
| MXE | 1297 | PIKFYVE | chr2 | + | 2E+08 | 208276830 | 208285725 | 208285933 | 208273583 | 208273733 | 208288728 | 208288818 | 10 | 1 | 0 | 12 | 9 | 0 | 0 | 3 | 0 | 0 | 2 | 0 | 98 | 98 | 1 | 1 | 0.455 | 0.1 | NA | NA | | 0.6 | NA | -0.322 |
| MXE | 1298 | PIKFYVE | chr2 | + | 2E+08 | 208277708 | 208285725 | 208285933 | 208276711 | 208276830 | 208288728 | 208288818 | 8 | 2 | 3 | 13 | 9 | 0 | 0 | 3 | 0 | 0 | 2 | 0 | 98 | 98 | 1 | 1 | 0.381 | 0.182 | 1 | NA | | 0.6 | NA | -0.079 |
| MXE | 1304 | ATP2A3 | chr17 | - | 4E+06 | 3951685 | 3953346 | 3953429 | 3951250 | 3951389 | 3953692 | 3953710 | 18 | 10 | 8 | 27 | 24 | 13 | 0 | 5 | 3 | 0 | 6 | 4 | 98 | 98 | 1 | 1 | 0.4 | 0.294 | 0.381 | NA | | 0.455 | 0.429 | -0.084 |
| MXE | 1305 | ATP2A3 | chr17 | - | 4E+06 | 3953429 | 3953692 | 3953710 | 3951580 | 3951685 | 3964173 | 3964464 | 6 | 4 | 5 | 19 | 13 | 6 | 0 | 0 | 1 | 0 | 3 | 1 | 67 | 98 | 1 | 1 | 0.316 | 0.31 | 0.549 | NA | | 0 | 0.594 | 0.095 |
| MXE | 1307 | NLRP1 | chr17 | - | 6E+06 | 5533988 | 5536850 | 5536940 | 5533303 | 5533384 | 5539414 | 5539585 | 36 | 25 | 5 | 31 | 12 | 6 | 0 | 1 | 0 | 0 | 3 | 0 | 98 | 98 | 1 | 1 | 0.537 | 0.676 | 0.455 | NA | | 0.25 | NA | 0.306 |
| MXE | 1308 | XAF1 | chr17 | + | 7E+06 | 6759718 | 6760405 | 6760601 | 6758088 | 6758224 | 6762154 | 6762240 | 11 | 14 | 1 | 3 | 6 | 1 | 0 | 8 | 0 | 0 | 6 | 0 | 98 | 98 | 1 | 1 | 0.786 | 0.7 | 0.5 | NA | | 0.571 | NA | 0.091 |
| MXE | 1309 | XAF1 | chr17 | + | 7E+06 | 6760601 | 6762154 | 6762240 | 6758088 | 6758224 | 6770642 | 6770984 | 0 | 2 | 0 | 5 | 5 | 0 | 0 | 0 | 0 | 0 | 6 | 0 | 98 | 98 | 1 | 1 | 0 | 0.286 | NA | NA | | 0 | NA | 0.143 |
| MXE | 1310 | XAF1 | chr17 | + | 7E+06 | 6760601 | 6762154 | 6762240 | 6759661 | 6759718 | 6770642 | 6770984 | 3 | 7 | 0 | 5 | 6 | 0 | 0 | 10 | 0 | 0 | 6 | 0 | 98 | 98 | 1 | 1 | 0.375 | 0.538 | NA | NA | | 0.625 | NA | -0.168 |
| MXE | 1313 | TNFSF13 | chr17 | + | 8E+06 | 7559702 | 7559845 | 7559893 | 7558291 | 7559297 | 7560048 | 7560167 | 6 | 3 | 0 | 12 | 3 | 1 | 0 | 1 | 1 | 0 | 0 | 0 | 98 | 97 | 1 | 1 | 0.331 | 0.497 | 0 | NA | | 1 | 1 | -0.724 |
| MXE | 1314 | TNFSF13 | chr17 | + | 8E+06 | 7559702 | 7559847 | 7559893 | 7558291 | 7559297 | 7560048 | 7560167 | 6 | 3 | 0 | 11 | 3 | 1 | 0 | 1 | 1 | 0 | 0 | 0 | 98 | 95 | 1 | 1 | 0.346 | 0.492 | 0 | NA | | 1 | 1 | -0.721 |
| MXE | 1315 | TNFSF13 | chr17 | + | 8E+06 | 7559702 | 7559850 | 7559893 | 7558291 | 7559297 | 7560048 | 7560167 | 6 | 3 | 0 | 13 | 2 | 1 | 0 | 1 | 1 | 0 | 0 | 0 | 98 | 92 | 1 | 1 | 0.302 | 0.585 | 0 | NA | | 1 | 1 | -0.704 |
| MXE | 1321 | FLII | chr17 | - | 2E+07 | 18255263 | 18256525 | 18256597 | 18254768 | 18254854 | 18256908 | 18257019 | 18 | 3 | 5 | 18 | 3 | 2 | 0 | 5 | 2 | 0 | 5 | 0 | 98 | 98 | 1 | 1 | 0.5 | 0.5 | 0.714 | NA | | 0.5 | 1 | -0.179 |
| MXE | 1322 | ASAH1 | chr8 | - | 2E+07 | 18062423 | 18063184 | 18063230 | 18061685 | 18061740 | 18064456 | 18064531 | 40 | 11 | 20 | 39 | 29 | 15 | 0 | 3 | 0 | 0 | 6 | 0 | 95 | 98 | 1 | 1 | 0.514 | 0.281 | 0.579 | NA | | 0.34 | NA | 0.118 |
| MXE | 1323 | KIAA0100 | chr17 | - | 3E+07 | 28633401 | 28633548 | 28633751 | 28632941 | 28633162 | 28633894 | 28634086 | 6 | 2 | 0 | 5 | 1 | 0 | 0 | 1 | 0 | 1 | 1 | 0 | 98 | 98 | 1 | 1 | 0.545 | 0.667 | NA | | 0 | 0.5 | NA | 0.356 |
| MXE | 1324 | KIAA0100 | chr17 | - | 3E+07 | 28642078 | 28642274 | 28642335 | 28640559 | 28640683 | 28642868 | 28642987 | 9 | 7 | 0 | 6 | 3 | 0 | 0 | 2 | 0 | 0 | 1 | 0 | 98 | 98 | 1 | 1 | 0.6 | 0.7 | NA | NA | | 0.667 | NA | -0.017 |
| MXE | 1328 | FLOT2 | chr17 | - | 3E+07 | 28884315 | 28888944 | 28889026 | 28883107 | 28883231 | 28897525 | 28897697 | 26 | 7 | 14 | 33 | 28 | 26 | 0 | 1 | 2 | 0 | 2 | 0 | 98 | 98 | 1 | 1 | 0.441 | 0.2 | 0.35 | NA | | 0.333 | 1 | -0.336 |
| MXE | 1329 | SSH2 | chr17 | - | 3E+07 | 29695523 | 29702958 | 29703062 | 29684562 | 29684684 | 29761149 | 29761432 | 41 | 20 | 13 | 48 | 33 | 12 | 0 | 1 | 0 | 0 | 5 | 0 | 98 | 98 | 1 | 1 | 0.461 | 0.377 | 0.52 | NA | | 0.167 | NA | 0.286 |
| MXE | 1330 | NCOA1 | chr2 | + | 2E+07 | 24762776 | 24768078 | 24768135 | 24757972 | 24758156 | 24768220 | 24770701 | 37 | 9 | 5 | 18 | 5 | 0 | 2 | 2 | 0 | 0 | 0 | 0 | 98 | 98 | 1 | 1 | 0.673 | 0.643 | 1 | | 1 | 1 | NA | -0.228 |
| MXE | 1331 | RHOT1 | chr17 | + | 3E+07 | 32209445 | 32211115 | 32211238 | 32208106 | 32208309 | 32224615 | 32225771 | 3 | 0 | 0 | 4 | 0 | 5 | 0 | 2 | 0 | 0 | 1 | 0 | 98 | 98 | 1 | 1 | 0.429 | NA | 0 | NA | | 0.667 | NA | -0.453 |
| MXE | 1335 | MGAM | chr7 | + | 1E+08 | 142047873 | 142050234 | 142050284 | 142040721 | 142040846 | 142050696 | 142050864 | 85 | 20 | 17 | 124 | 20 | 11 | 1 | 3 | 0 | 0 | 10 | 0 | 98 | 98 | 1 | 1 | 0.407 | 0.5 | 0.607 | | 1 | 0.231 | NA | -0.111 |
| MXE | 1337 | STK40 | chr1 | - | 4E+07 | 36358822 | 36361220 | 36361340 | 36358238 | 36358382 | 36385722 | 36385927 | 19 | 8 | 3 | 30 | 18 | 9 | 0 | 1 | 0 | 0 | 3 | 0 | 98 | 98 | 1 | 1 | 0.388 | 0.308 | 0.25 | NA | | 0.25 | NA | 0.065 |
| MXE | 1346 | TSEN34 | chr19 | + | 5E+07 | 54191607 | 54191720 | 54191964 | 54190267 | 54190450 | 54192115 | 54192373 | 3 | 7 | 3 | 44 | 21 | 17 | 0 | 3 | 1 | 0 | 3 | 1 | 98 | 98 | 1 | 1 | 0.064 | 0.25 | 0.15 | NA | | 0.5 | 0.5 | -0.345 |
| MXE | 1347 | STAT3 | chr17 | - | 4E+07 | 42324846 | 42324962 | 42325061 | 42323572 | 42323625 | 42326115 | 42326199 | 27 | 18 | 26 | 49 | 25 | 19 | 0 | 3 | 0 | 0 | 7 | 0 | 98 | 98 | 1 | 1 | 0.355 | 0.419 | 0.578 | NA | | 0.3 | NA | 0.151 |
| MXE | 1349 | DHX8 | chr17 | + | 4E+07 | 43493040 | 43493444 | 43493589 | 43492182 | 43492292 | 43493682 | 43493886 | 21 | 9 | 0 | 15 | 11 | 4 | 0 | 2 | 0 | 0 | 0 | 0 | 98 | 98 | 1 | 1 | 0.583 | 0.45 | 0 | NA | | 1 | NA | -0.656 |
| MXE | 1351 | KLHL2 | chr4 | + | 2E+08 | 165228913 | 165238777 | 165238899 | 165219933 | 165220059 | 165263196 | 165263359 | 12 | 5 | 9 | 17 | 4 | 9 | 0 | 5 | 0 | 0 | 0 | 0 | 98 | 98 | 1 | 1 | 0.414 | 0.556 | 0.5 | NA | | 1 | NA | -0.51 |
| MXE | 1352 | SNX3 | chr6 | - | 1E+08 | 108214622 | 108222949 | 108223045 | 108211216 | 108212254 | 108260759 | 108261260 | 6 | 5 | 3 | 13 | 14 | 5 | 0 | 0 | 0 | 0 | 2 | 0 | 98 | 98 | 1 | 1 | 0.316 | 0.263 | 0.375 | NA | | 0 | NA | 0.318 |
| MXE | 1357 | DCAF8 | chr1 | - | 2E+08 | 160244034 | 160261284 | 160261358 | 160239696 | 160240370 | 160262116 | 160262528 | 1 | 0 | 0 | 3 | 2 | 0 | 0 | 0 | 0 | 0 | 1 | 0 | 98 | 98 | 1 | 1 | 0.25 | 0 | NA | NA | | 0 | NA | 0.125 |

| | | | | | | | | | | | | | | | | | | | | | | | | | | | | | | | | | | | |
|---|---|---|---|---|---|---|---|---|---|---|---|---|---|---|---|---|---|---|---|---|---|---|---|---|---|---|---|---|---|---|---|---|---|---|---|
| MXE | 1358 | DCAF8 | chr1 | - | 2E+08 | 160244034 | 160261284 | 160261358 | 160239696 | 160240370 | 160262448 | 160262560 | 1 | 2 | 0 | 4 | 2 | 0 | 0 | 0 | 0 | 0 | 1 | 0 | 98 | 98 | 1 | 1 | 0.2 | 0.5 | NA | NA | 0 | NA | 0.35 |
| MXE | 1360 | ACAA1 | chr3 | - | 4E+07 | 38129388 | 38131595 | 38131638 | 38127785 | 38127866 | 38131925 | 38132005 | 3 | 0 | 3 | 9 | 8 | 2 | 0 | 0 | 0 | 0 | 1 | 0 | 92 | 98 | 1 | 1 | 0.262 | 0 | 0.615 | NA | 0 | NA | 0.292 |
| MXE | 1363 | KAT7 | chr17 | + | 5E+07 | 49805445 | 49809118 | 49809208 | 49798318 | 49798558 | 49811475 | 49811574 | 5 | 2 | 0 | 15 | 8 | 0 | 0 | 2 | 0 | 0 | 1 | 0 | 98 | 98 | 1 | 1 | 0.25 | 0.2 | NA | NA | 0.667 | NA | -0.442 |
| MXE | 1365 | C18orf8 | chr18 | + | 2E+07 | 23504447 | 23509192 | 23509279 | 23503469 | 23503720 | 23515855 | 23515996 | 1 | 7 | 0 | 15 | 5 | 2 | 0 | 0 | 0 | 0 | 5 | 0 | 98 | 98 | 1 | 1 | 0.062 | 0.583 | 0 | NA | 0 | NA | 0.215 |
| MXE | 1366 | APH1A | chr1 | - | 2E+08 | 150267789 | 150267956 | 150267991 | 150267355 | 150267478 | 150268697 | 150269209 | 2 | 3 | 1 | 11 | 2 | 6 | 0 | 0 | 0 | 0 | 2 | 0 | 84 | 98 | 1 | 1 | 0.175 | 0.636 | 0.163 | NA | 0 | NA | 0.325 |
| MXE | 1367 | APH1A | chr1 | - | 2E+08 | 150267789 | 150267956 | 150268127 | 150267355 | 150267478 | 150268697 | 150269209 | 9 | 0 | 0 | 11 | 2 | 6 | 0 | 0 | 0 | 0 | 2 | 0 | 98 | 98 | 1 | 1 | 0.45 | 0 | 0 | NA | 0 | NA | 0.15 |
| MXE | 1368 | VMP1 | chr17 | + | 6E+07 | 59735473 | 59737452 | 59737543 | 59731420 | 59731522 | 59738836 | 59738947 | 65 | 35 | 48 | 124 | 101 | 62 | 0 | 3 | 0 | 0 | 7 | 0 | 98 | 98 | 1 | 1 | 0.344 | 0.257 | 0.436 | NA | 0.3 | NA | 0.046 |
| MXE | 1384 | CSF2RB | chr22 | + | 4E+07 | 36926177 | 36929401 | 36929559 | 36923243 | 36923367 | 36929638 | 36929807 | 46 | 34 | 37 | 94 | 42 | 44 | 0 | 6 | 1 | 0 | 13 | 0 | 98 | 98 | 1 | 1 | 0.329 | 0.447 | 0.457 | NA | 0.316 | 1 | -0.247 |
| MXE | 1392 | SPAG9 | chr17 | - | 5E+07 | 50984990 | 50985697 | 50985778 | 50982523 | 50982672 | 50987111 | 50987237 | 21 | 10 | 4 | 27 | 19 | 5 | 0 | 8 | 0 | 0 | 5 | 0 | 98 | 98 | 1 | 1 | 0.438 | 0.345 | 0.444 | NA | 0.615 | NA | -0.206 |
| MXE | 1397 | MBD1 | chr18 | - | 5E+07 | 50274353 | 50274976 | 50275045 | 50273563 | 50273863 | 50275128 | 50275245 | 7 | 0 | 1 | 0 | 4 | 0 | 0 | 0 | 0 | 0 | 2 | 0 | 98 | 98 | 1 | 1 | 1 | 0 | 1 | NA | 0 | NA | 0.667 |
| MXE | 1399 | GUCD1 | chr22 | - | 2E+07 | 24548073 | 24548916 | 24549001 | 24546913 | 24547005 | 24554948 | 24555307 | 5 | 4 | 0 | 13 | 7 | 0 | 0 | 6 | 0 | 0 | 5 | 0 | 98 | 98 | 1 | 1 | 0.278 | 0.364 | NA | NA | 0.545 | NA | -0.224 |
| MXE | 1400 | GUCD1 | chr22 | - | 2E+07 | 24548073 | 24548916 | 24549001 | 24546913 | 24547005 | 24555614 | 24555935 | 2 | 1 | 0 | 11 | 7 | 0 | 0 | 0 | 0 | 0 | 3 | 0 | 98 | 98 | 1 | 1 | 0.154 | 0.125 | NA | NA | 0 | NA | 0.14 |
| MXE | 1403 | CTDP1 | chr18 | + | 8E+07 | 79729069 | 79736354 | 79736521 | 79717809 | 79718016 | 79753651 | 79754510 | 5 | 0 | 1 | 3 | 3 | 0 | 0 | 1 | 0 | 0 | 1 | 3 | 98 | 98 | 1 | 1 | 0.625 | 0 | 1 | NA | 0.5 | 0 | 0.292 |
| MXE | 1404 | TXNL4A | chr18 | - | 8E+07 | 79977701 | 79986601 | 79986750 | 79972866 | 79973856 | 79988239 | 79988592 | 2 | 0 | 0 | 8 | 0 | 10 | 0 | 0 | 1 | 0 | 0 | 0 | 98 | 98 | 1 | 1 | 0.2 | NA | 0 | NA | NA | 1 | -0.9 |
| MXE | 1413 | ZNF638 | chr2 | + | 7E+07 | 71427414 | 71428546 | 71428651 | 71424649 | 71424715 | 71431326 | 71431428 | 3 | 0 | 0 | 8 | 1 | 2 | 0 | 3 | 0 | 0 | 1 | 0 | 98 | 98 | 1 | 1 | 0.273 | 0 | 0 | NA | 0.75 | NA | -0.659 |
| MXE | 1415 | TLN1 | chr9 | - | 4E+07 | 35699461 | 35699973 | 35700081 | 35699031 | 35699156 | 35700190 | 35700376 | 85 | 21 | 17 | 74 | 21 | 10 | 0 | 11 | 1 | 0 | 4 | 0 | 98 | 98 | 1 | 1 | 0.535 | 0.5 | 0.63 | NA | 0.733 | 1 | -0.311 |
| MXE | 1417 | MAP3K11 | chr11 | - | 7E+07 | 65608070 | 65608267 | 65608448 | 65607640 | 65607816 | 65613017 | 65614249 | 25 | 18 | 11 | 17 | 10 | 11 | 0 | 2 | 0 | 0 | 0 | 0 | 98 | 98 | 1 | 1 | 0.595 | 0.643 | 0.5 | NA | 1 | NA | -0.421 |
| MXE | 1420 | GPATCH2L | chr14 | + | 8E+07 | 76177931 | 76177987 | 76178042 | 76176622 | 76176690 | 76180763 | 76180849 | 2 | 0 | 0 | 9 | 13 | 0 | 0 | 0 | 0 | 0 | 0 | 3 | 89 | 98 | 1 | 1 | 0.197 | 0 | NA | NA | NA | 0 | 0.099 |
| MXE | 1422 | ADGRE5 | chr19 | + | 1E+07 | 14407229 | 14407907 | 14408009 | 14406868 | 14406960 | 14408091 | 14408725 | 45 | 51 | 44 | 59 | 32 | 34 | 0 | 9 | 0 | 0 | 4 | 0 | 98 | 98 | 1 | 1 | 0.433 | 0.614 | 0.564 | NA | 0.692 | NA | -0.155 |
| MXE | 1434 | CYP4F3 | chr19 | + | 2E+07 | 15644051 | 15645718 | 15645863 | 15641414 | 15641613 | 15647051 | 15647105 | 2 | 3 | 5 | 161 | 45 | 36 | 0 | 0 | 0 | 7 | 13 | 0 | 98 | 98 | 1 | 1 | 0.012 | 0.062 | 0.122 | 0 | 0 | NA | 0.065 |
| MXE | 1437 | PTTG1IP | chr21 | - | 4E+07 | 44861271 | 44865394 | 44865447 | 44856192 | 44856364 | 44873501 | 44873903 | 10 | 3 | 6 | 27 | 15 | 4 | 0 | 1 | 1 | 0 | 3 | 0 | 98 | 98 | 1 | 1 | 0.27 | 0.167 | 0.6 | NA | 0.25 | 1 | -0.279 |
| MXE | 1440 | RASGRP4 | chr19 | - | 4E+07 | 38417168 | 38419859 | 38420013 | 38413393 | 38413474 | 38420130 | 38420262 | 20 | 18 | 4 | 1 | 0 | 1 | 1 | 1 | 0 | 0 | 1 | 0 | 98 | 98 | 1 | 1 | 0.952 | 1 | 0.8 | 1 | 0.5 | NA | 0.167 |
| MXE | 1441 | RASGRP4 | chr19 | - | 4E+07 | 38417168 | 38419859 | 38420013 | 38414847 | 38415123 | 38420130 | 38420262 | 20 | 18 | 4 | 39 | 11 | 9 | 1 | 1 | 0 | 0 | 4 | 0 | 98 | 98 | 1 | 1 | 0.339 | 0.621 | 0.308 | 1 | 0.2 | NA | -0.177 |
| MXE | 1443 | SMCHD1 | chr18 | + | 3E+06 | 2760739 | 2762104 | 2762236 | 2752487 | 2752552 | 2763636 | 2763789 | 110 | 56 | 32 | 120 | 94 | 33 | 0 | 18 | 0 | 5 | 28 | 0 | 98 | 98 | 1 | 1 | 0.478 | 0.373 | 0.492 | 0 | 0.391 | NA | 0.252 |
| MXE | 1445 | GMFG | chr19 | - | 4E+07 | 39333126 | 39335260 | 39335310 | 39329543 | 39329626 | 39335434 | 39335531 | 60 | 42 | 35 | 107 | 88 | 99 | 0 | 4 | 1 | 0 | 14 | 1 | 98 | 98 | 1 | 1 | 0.359 | 0.323 | 0.261 | NA | 0.222 | 0.5 | -0.047 |
| MXE | 1446 | GMFG | chr19 | - | 4E+07 | 39333126 | 39335260 | 39335310 | 39329543 | 39329626 | 39335973 | 39336086 | 0 | 1 | 0 | 104 | 88 | 99 | 0 | 0 | 0 | 0 | 14 | 1 | 98 | 98 | 1 | 1 | 0 | 0.011 | 0 | NA | 0 | 0 | 0.004 |
| MXE | 1451 | PLAUR | chr19 | - | 4E+07 | 43655573 | 43656478 | 43656640 | 43648566 | 43649143 | 43665315 | 43665459 | 76 | 31 | 10 | 10 | 2 | 1 | 0 | 61 | 6 | 0 | 2 | 1 | 98 | 98 | 1 | 1 | 0.884 | 0.939 | 0.909 | NA | 0.968 | 0.857 | -0.002 |
| MXE | 1452 | PLAUR | chr19 | - | 4E+07 | 43655573 | 43656478 | 43656640 | 43652224 | 43652371 | 43665315 | 43665459 | 78 | 27 | 10 | 40 | 12 | 15 | 0 | 62 | 6 | 0 | 26 | 2 | 98 | 98 | 1 | 1 | 0.661 | 0.692 | 0.4 | NA | 0.705 | 0.75 | -0.143 |
| MXE | 1455 | C17orf62 | chr17 | - | 8E+07 | 82446696 | 82447579 | 82447621 | 82445863 | 82445960 | 82449169 | 82449292 | 28 | 15 | 1 | 47 | 20 | 15 | 0 | 0 | 3 | 0 | 6 | 0 | 91 | 98 | 1 | 1 | 0.391 | 0.447 | 0.067 | NA | 0 | 1 | -0.198 |
| MXE | 1459 | CNOT7 | chr8 | - | 2E+07 | 17232537 | 17234715 | 17234860 | 17229230 | 17230848 | 17237211 | 17237373 | 6 | 11 | 1 | 6 | 1 | 0 | 0 | 2 | 0 | 0 | 0 | 0 | 98 | 98 | 1 | 1 | 0.5 | 0.917 | 1 | NA | 1 | NA | -0.194 |
| MXE | 1464 | POC1B | chr12 | - | 9E+07 | 89425379 | 89459637 | 89459718 | 89419720 | 89421257 | 89466769 | 89466922 | 0 | 3 | 3 | 0 | 0 | 2 | 0 | 1 | 0 | 0 | 1 | 0 | 98 | 98 | 1 | 1 | NA | 1 | 0.6 | NA | 0.5 | NA | 0.3 |
| MXE | 1465 | KLHDC3 | chr6 | + | 4E+07 | 43017695 | 43017852 | 43017968 | 43017133 | 43017346 | 43018144 | 43018216 | 7 | 1 | 4 | 2 | 4 | 6 | 0 | 0 | 0 | 4 | 0 | 0 | 98 | 98 | 1 | 1 | 0.778 | 0.2 | 0.4 | 0 | NA | NA | 0.459 |
| MXE | 1466 | KLHDC3 | chr6 | + | 4E+07 | 43017968 | 43018144 | 43018216 | 43017518 | 43017695 | 43018342 | 43018556 | 7 | 7 | 0 | 3 | 2 | 0 | 0 | 0 | 0 | 0 | 1 | 0 | 98 | 98 | 1 | 1 | 0.7 | 0.778 | NA | NA | 0 | NA | 0.739 |
| MXE | 1475 | MBOAT7 | chr19 | - | 5E+07 | 54187287 | 54188216 | 54188346 | 54183520 | 54183680 | 54188432 | 54188511 | 15 | 16 | 8 | 62 | 24 | 33 | 0 | 4 | 3 | 0 | 8 | 5 | 98 | 98 | 1 | 1 | 0.195 | 0.4 | 0.195 | NA | 0.333 | 0.375 | -0.091 |
| MXE | 1476 | LILRA5 | chr19 | - | 5E+07 | 54312370 | 54312536 | 54312621 | 54311863 | 54312148 | 54313016 | 54313139 | 21 | 14 | 12 | 25 | 14 | 23 | 0 | 5 | 3 | 0 | 2 | 0 | 98 | 85 | 1 | 1 | 0.421 | 0.464 | 0.312 | NA | 0.684 | 1 | -0.443 |
| MXE | 1477 | CD28 | chr2 | + | 2E+08 | 203726698 | 203729647 | 203729772 | 203706474 | 203706748 | 203734783 | 203738912 | 3 | 0 | 1 | 3 | 0 | 0 | 0 | 0 | 0 | 0 | 1 | 0 | 98 | 98 | 1 | 1 | 0.5 | NA | 1 | NA | 0 | NA | 0.75 |
| MXE | 1482 | R3HCC1L | chr10 | + | 1E+08 | 98209899 | 98231511 | 98231687 | 98163292 | 98163397 | 98234445 | 98234516 | 4 | 2 | 0 | 2 | 3 | 0 | 0 | 0 | 0 | 0 | 1 | 0 | 98 | 98 | 1 | 1 | 0.667 | 0.4 | NA | NA | 0 | NA | 0.534 |
| MXE | 1484 | SART3 | chr12 | - | 1E+08 | 108543152 | 108544426 | 108544478 | 108538933 | 108539089 | 108545138 | 108545323 | 5 | 6 | 1 | 6 | 0 | 8 | 0 | 1 | 0 | 0 | 1 | 0 | 98 | 98 | 1 | 1 | 0.455 | 1 | 0.111 | NA | 0.5 | NA | 0.022 |
| MXE | 1490 | THADA | chr2 | - | 4E+07 | 43556555 | 43560233 | 43560385 | 43552067 | 43552339 | 43566697 | 43566821 | 8 | 2 | 3 | 1 | 6 | 0 | 0 | 0 | 0 | 4 | 0 | 0 | 98 | 98 | 1 | 1 | 0.889 | 0.25 | 1 | 0 | NA | NA | 0.713 |
| MXE | 1491 | PPM1B | chr2 | + | 4E+07 | 44202045 | 44209209 | 44209327 | 44168802 | 44169274 | 44217966 | 44218078 | 4 | 7 | 3 | 19 | 7 | 7 | 0 | 2 | 0 | 0 | 0 | 0 | 98 | 98 | 1 | 1 | 0.174 | 0.5 | 0.3 | NA | 1 | NA | -0.675 |
| MXE | 1496 | PCBP1-AS1 | chr2 | - | 7E+07 | 70059680 | 70083537 | 70083687 | 70051202 | 70051305 | 70085546 | 70085624 | 8 | 0 | 0 | 5 | 3 | 0 | 0 | 0 | 0 | 0 | 1 | 0 | 98 | 87 | 1 | 1 | 0.587 | 0 | NA | NA | 0 | NA | 0.293 |
| MXE | 1501 | SERPINA1 | chr14 | - | 9E+07 | 94379611 | 94382591 | 94383241 | 94376746 | 94378640 | 94388559 | 94388660 | 70 | 15 | 37 | 142 | 66 | 104 | 0 | 5 | 1 | 0 | 26 | 1 | 98 | 98 | 1 | 1 | 0.33 | 0.185 | 0.262 | NA | 0.161 | 0.5 | -0.072 |
| MXE | 1502 | SERPINA1 | chr14 | - | 9E+07 | 94379611 | 94382591 | 94383241 | 94376746 | 94378640 | 94390456 | 94390692 | 108 | 35 | 38 | 142 | 65 | 104 | 0 | 5 | 0 | 0 | 26 | 1 | 98 | 98 | 1 | 1 | 0.432 | 0.35 | 0.268 | NA | 0.161 | 0 | 0.27 |
| MXE | 1503 | SERPINA1 | chr14 | - | 9E+07 | 94379611 | 94388800 | 94389010 | 94376746 | 94378640 | 94390456 | 94390692 | 47 | 13 | 5 | 142 | 65 | 104 | 0 | 1 | 0 | 0 | 26 | 1 | 98 | 98 | 1 | 1 | 0.249 | 0.167 | 0.046 | NA | 0.037 | 0 | 0.136 |
| MXE | 1504 | SERPINA1 | chr14 | - | 9E+07 | 94383241 | 94388800 | 94389010 | 94376746 | 94378640 | 94390456 | 94390692 | 47 | 13 | 5 | 108 | 35 | 38 | 0 | 1 | 0 | 0 | 5 | 0 | 98 | 98 | 1 | 1 | 0.303 | 0.271 | 0.116 | NA | 0.167 | NA | 0.063 |
| MXE | 1506 | ORAI2 | chr7 | + | 1E+08 | 102436333 | 102438943 | 102439181 | 102433529 | 102433661 | 102446512 | 102456821 | 1 | 4 | 0 | 11 | 6 | 5 | 0 | 0 | 0 | 0 | 1 | 3 | 98 | 98 | 1 | 1 | 0.083 | 0.4 | 0 | NA | 0 | 0 | 0.161 |
| MXE | 1507 | BCL2L11 | chr2 | + | 1E+08 | 111123869 | 111150043 | 111150147 | 111120913 | 111121188 | 111164132 | 111168445 | 3 | 0 | 0 | 3 | 2 | 0 | 0 | 0 | 0 | 0 | 1 | 0 | 98 | 98 | 1 | 1 | 0.5 | 0 | NA | NA | 0 | NA | 0.25 |
| MXE | 1509 | BCL2L11 | chr2 | + | 1E+08 | 111124139 | 111150043 | 111150147 | 111120913 | 111121188 | 111164132 | 111168445 | 3 | 0 | 0 | 3 | 2 | 0 | 0 | 0 | 0 | 0 | 1 | 0 | 98 | 98 | 1 | 1 | 0.5 | 0 | NA | NA | 0 | NA | 0.25 |
| MXE | 1512 | BCL2L11 | chr2 | + | 1E+08 | 111124139 | 111150043 | 111150147 | 111123732 | 111123869 | 111164132 | 111168445 | 6 | 0 | 3 | 1 | 3 | 0 | 0 | 0 | 0 | 0 | 1 | 0 | 98 | 98 | 1 | 1 | 0.857 | 0 | 1 | NA | 0 | NA | 0.619 |
| MXE | 1513 | BCL2L11 | chr2 | + | 1E+08 | 111150147 | 111153779 | 111153904 | 111123732 | 111124139 | 111164132 | 111168445 | 7 | 2 | 1 | 0 | 1 | 0 | 0 | 1 | 0 | 0 | 0 | 0 | 98 | 98 | 1 | 1 | 1 | 0.667 | 1 | NA | 1 | NA | -0.111 |
| MXE | 1514 | BCL2L11 | chr2 | + | 1E+08 | 111153904 | 111161418 | 111161550 | 111150043 | 111150147 | 111164132 | 111168445 | 3 | 1 | 0 | 0 | 0 | 0 | 0 | 0 | 0 | 0 | 1 | 0 | 98 | 98 | 1 | 1 | 1 | 1 | NA | NA | 0 | NA | 1 |
| MXE | 1523 | SMIM7 | chr19 | - | 2E+07 | 16654125 | 16659394 | 16659447 | 16646147 | 16647261 | 16659958 | 16660000 | 1 | 3 | 0 | 5 | 0 | 0 | 0 | 0 | 0 | 0 | 1 | 0 | 98 | 98 | 1 | 1 | 0.167 | 1 | NA | NA | 0 | NA | 0.584 |
| MXE | 1524 | PLEKHB2 | chr2 | + | 1E+08 | 131126786 | 131132901 | 131132991 | 131125752 | 131125905 | 131140166 | 131140275 | 24 | 9 | 2 | 18 | 5 | 2 | 0 | 1 | 0 | 0 | 4 | 0 | 98 | 98 | 1 | 1 | 0.571 | 0.643 | 0.5 | NA | 0.2 | NA | 0.371 |
| MXE | 1527 | ACSL1 | chr4 | - | 2E+08 | 184770332 | 184770398 | 184770476 | 184768315 | 184768450 | 184773080 | 184773154 | 230 | 105 | 146 | 18 | 3 | 6 | 4 | 29 | 0 | 0 | 3 | 0 | 98 | 98 | 1 | 1 | 0.927 | 0.972 | 0.961 | 1 | 0.906 | NA | 0 |
| MXE | 1534 | PDK1 | chr2 | + | 2E+08 | 172562800 | 172564502 | 172564687 | 172562219 | 172562291 | 172564977 | 172565073 | 3 | 3 | 0 | 8 | 0 | 2 | 0 | 0 | 0 | 0 | 2 | 0 | 98 | 98 | 1 | 1 | 0.273 | 1 | 0 | NA | 0 | NA | 0.424 |
| MXE | 1540 | CCDC117 | chr22 | + | 3E+07 | 28773778 | 28780947 | 28781172 | 28772673 | 28773034 | 28783507 | 28783645 | 1 | 0 | 0 | 2 | 1 | 0 | 0 | 1 | 0 | 0 | 0 | 0 | 98 | 98 | 1 | 1 | 0.333 | 0 | NA | NA | 1 | NA | -0.834 |
| MXE | 1541 | ANKRD44 | chr2 | - | 2E+08 | 196999052 | 197000418 | 197000502 | 196998336 | 196998419 | 197001752 | 197001840 | 47 | 11 | 6 | 25 | 7 | 13 | 0 | 5 | 0 | 0 | 1 | 0 | 98 | 98 | 1 | 1 | 0.653 | 0.611 | 0.316 | NA | 0.833 | NA | -0.306 |
| MXE | 1542 | FAM126B | chr2 | - | 2E+08 | 200993003 | 200997444 | 200997540 | 200992280 | 200992368 | 201008972 | 201009088 | 39 | 9 | 6 | 39 | 11 | 3 | 0 | 2 | 0 | 0 | 4 | 0 | 98 | 98 | 1 | 1 | 0.5 | 0.45 | 0.667 | NA | 0.333 | NA | 0.206 |
| MXE | 1543 | NDUFS1 | chr2 | - | 2E+08 | 206149925 | 206152418 | 206152510 | 206149019 | 206149096 | 206153617 | 206153682 | 3 | 4 | 2 | 10 | 6 | 0 | 0 | 2 | 0 | 0 | 0 | 0 | 98 | 98 | 1 | 1 | 0.231 | 0.4 | 1 | NA | 1 | NA | -0.456 |
| MXE | 1545 | NLRP6 | chr11 | + | 279833 | 279872 | 280083 | 281839 | 279326 | 279607 | 282701 | 282797 | 5 | 7 | 1 | 1 | 0 | 3 | 0 | 0 | 0 | 0 | 0 | 4 | 88 | 98 | 1 | 1 | 0.848 | 1 | 0.271 | NA | NA | 0 | 0.706 |
| MXE | 1546 | NLRP6 | chr11 | + | 279833 | 279872 | 280083 | 281839 | 279326 | 279607 | 282704 | 282797 | 5 | 7 | 4 | 7 | 8 | 2 | 0 | 1 | 2 | 0 | 1 | 4 | 88 | 98 | 1 | 1 | 0.443 | 0.494 | 0.69 | NA | 0.527 | 0.358 | 0.1 |
| MXE | 1547 | SLC11A1 | chr2 | + | 2E+08 | 218383102 | 218384242 | 218384365 | 218382028 | 218382375 | 218385146 | 218385266 | 27 | 17 | 0 | 106 | 58 | 23 | 0 | 0 | 0 | 0 | 14 | 5 | 98 | 98 | 1 | 1 | 0.203 | 0.227 | 0 | NA | 0 | 0 | 0.143 |

**U12_MXE_filtered**

| | | | | | | | | | | | | | Inclusion_count | | | Skipping_count | | | Inclusion_count | | | Skipping_count | | | | | | | Inclusion_level (normalized) | | | | | | |
|---|---|---|---|---|---|---|---|---|---|---|---|---|---|---|---|---|---|---|---|---|---|---|---|---|---|---|---|---|---|---|---|---|---|---|---|
| Splice_type | ID | Gene_name | Chromosome | Strand | 1stExon | 1stExonEnd | 2ndExonStart_0t | 2ndExonEn | UpstreamES | UpstreamEE | DownstreamE | Downstrear | control_3 | control_4 | control_10 | control_3 | control_4 | control_10 | pmn_13 | pmn_15 | pmn_17 | pmn_13 | pmn_15 | pmn_17 | IncFormLen | SkipFormLen | p value | FDR | control_3 | control_4 | control_10 | pmn_13 | pmn_15 | pmn_17 | IncLevelDifference |
| MXE | 973 | WDR1 | chr4 | - | 1E+07 | 10097891 | 10103895 | 10103986 | 10088663 | 10088741 | 10116112 | 10116234 | 36 | 16 | 13 | 37 | 7 | 7 | 1 | 7 | 0 | 0 | 1 | 1 | 98 | 98 | 0.00681759 | 0.2654599 | 0.493 | 0.696 | 0.65 | 1 | 0.875 | 0 | -0.012 |
| MXE | 172 | DOCK2 | chr5 | + | 2E+08 | 169684350 | 169689251 | 169689333 | 169681743 | 169681879 | 169695802 | 169695938 | 109 | 35 | 17 | 83 | 48 | 5 | 0 | 13 | 2 | 4 | 19 | 4 | 98 | 98 | 0.0385793 | 0.9651035 | 0.568 | 0.422 | 0.773 | 0 | 0.406 | 0.333 | 0.341 |
| MXE | 174 | DOCK2 | chr5 | + | 2E+08 | 169761625 | 169803057 | 169803206 | 169759704 | 169759775 | 169840756 | 169840852 | 45 | 19 | 11 | 79 | 34 | 16 | 0 | 8 | 1 | 1 | 9 | 2 | 98 | 98 | 0.7796722 | 1 | 0.363 | 0.358 | 0.407 | 0 | 0.471 | 0.333 | 0.108 |
| MXE | 138 | MAPK14 | chr6 | + | 4E+07 | 36096066 | 36100187 | 36100267 | 36076536 | 36076608 | 36102570 | 36102649 | 31 | 21 | 14 | 18 | 11 | 4 | 0 | 6 | 0 | 0 | 0 | 0 | 98 | 98 | 1 | 1 | 0.633 | 0.656 | 0.778 | NA | 1 | NA | -0.311 |
| MXE | 171 | DOCK2 | chr5 | + | 2E+08 | 169670597 | 169671077 | 169671174 | 169669287 | 169669328 | 169674296 | 169674445 | 32 | 11 | 5 | 142 | 34 | 5 | 0 | 6 | 0 | 0 | 15 | 1 | 98 | 98 | 1 | 1 | 0.184 | 0.244 | 0.5 | NA | 0.286 | 0 | 0.166 |
| MXE | 173 | DOCK2 | chr5 | + | 2E+08 | 169759775 | 169761518 | 169761625 | 169747395 | 169747504 | 169803057 | 169803206 | 58 | 31 | 6 | 67 | 41 | 4 | 0 | 19 | 0 | 0 | 4 | 0 | 98 | 98 | 1 | 1 | 0.464 | 0.431 | 0.6 | NA | 0.826 | NA | -0.328 |
| MXE | 188 | DCP2 | chr5 | + | 1E+08 | 113004077 | 113007937 | 113008042 | 113001566 | 113001674 | 113010755 | 113010807 | 12 | 8 | 7 | 26 | 7 | 2 | 0 | 1 | 0 | 0 | 0 | 0 | 98 | 98 | 1 | 1 | 0.316 | 0.533 | 0.778 | NA | 1 | NA | -0.458 |
| MXE | 195 | FCHO2 | chr5 | + | 7E+07 | 73006549 | 73015625 | 73015724 | 72990711 | 72990864 | 73017211 | 73017308 | 15 | 3 | 0 | 22 | 9 | 1 | 0 | 1 | 0 | 0 | 0 | 0 | 98 | 98 | 1 | 1 | 0.405 | 0.25 | 0 | NA | 1 | NA | -0.782 |
| MXE | 225 | MAEA | chr4 | + | 1E+06 | 1315600 | 1322380 | 1322503 | 1311978 | 1312161 | 1327626 | 1327703 | 15 | 13 | 2 | 1 | 4 | 1 | 0 | 2 | 7 | 0 | 3 | 1 | 98 | 98 | 1 | 1 | 0.937 | 0.765 | 0.667 | NA | 0.4 | 0.875 | 0.152 |
| MXE | 226 | MAEA | chr4 | + | 1E+06 | 1315600 | 1322380 | 1322503 | 1311978 | 1312161 | 1332756 | 1332865 | 15 | 16 | 2 | 0 | 3 | 1 | 0 | 2 | 7 | 0 | 0 | 0 | 98 | 98 | 1 | 1 | 1 | 0.842 | 0.667 | NA | 1 | 1 | -0.164 |
| MXE | 227 | MAEA | chr4 | + | 1E+06 | 1322503 | 1327626 | 1327703 | 1315396 | 1315600 | 1332756 | 1332865 | 11 | 5 | 2 | 12 | 5 | 7 | 0 | 1 | 0 | 0 | 1 | 0 | 98 | 98 | 1 | 1 | 0.478 | 0.5 | 0.222 | NA | 0.5 | NA | -0.1 |
| MXE | 336 | SLC9A8 | chr20 | + | 5E+07 | 49845119 | 49849578 | 49849680 | 49839540 | 49839599 | 49855437 | 49855581 | 13 | 2 | 0 | 14 | 9 | 0 | 0 | 2 | 2 | 0 | 4 | 1 | 98 | 98 | 1 | 1 | 0.481 | 0.182 | NA | NA | 0.333 | 0.667 | -0.168 |
| MXE | 610 | RAB6A | chr11 | - | 7E+07 | 73718718 | 73718784 | 73718890 | 73716250 | 73716362 | 73720845 | 73720899 | 0 | 0 | 1 | 11 | 3 | 4 | 0 | 0 | 0 | 2 | 1 | 0 | 98 | 98 | 1 | 1 | 0 | 0 | 0.2 | 0 | 0 | NA | 0.067 |
| MXE | 1042 | GBP5 | chr1 | - | 9E+07 | 89267526 | 89268728 | 89268856 | 89266956 | 89267153 | 89269365 | 89269574 | 27 | 6 | 4 | 91 | 27 | 18 | 0 | 11 | 0 | 0 | 15 | 0 | 98 | 98 | 1 | 1 | 0.229 | 0.182 | 0.182 | NA | 0.423 | NA | -0.225 |
| MXE | 1043 | GBP5 | chr1 | - | 9E+07 | 89268856 | 89269365 | 89269574 | 89267416 | 89267526 | 89272451 | 89272861 | 5 | 4 | 1 | 48 | 11 | 5 | 0 | 0 | 0 | 0 | 4 | 0 | 98 | 98 | 1 | 1 | 0.094 | 0.267 | 0.167 | NA | 0 | NA | 0.176 |
| MXE | 1440 | RASGRP4 | chr19 | - | 4E+07 | 38417168 | 38419859 | 38420013 | 38413393 | 38413474 | 38420130 | 38420262 | 20 | 18 | 4 | 1 | 0 | 1 | 1 | 1 | 0 | 0 | 1 | 0 | 98 | 98 | 1 | 1 | 0.952 | 1 | 0.8 | 1 | 0.5 | NA | 0.167 |
| MXE | 1441 | RASGRP4 | chr19 | - | 4E+07 | 38417168 | 38419859 | 38420013 | 38414847 | 38415123 | 38420130 | 38420262 | 20 | 18 | 4 | 39 | 11 | 9 | 1 | 1 | 0 | 0 | 4 | 0 | 98 | 98 | 1 | 1 | 0.339 | 0.621 | 0.308 | 1 | 0.2 | NA | -0.177 |

**Supplementary_Data_4S**

**A3SS**

***HC PBMC versus SLE PBMC***

| | | | | | | | | | | | | *Inclusion_counts_HC* | | | | | *Skipping_counts_HC* | | | | | *Inclusion counts_SLE* | | | | | *Skipping_counts_SLE* | | | | | | | | | | | | | *Inclusion_level(normalized)* | | | | | | |
|---|---|---|---|---|---|---|---|---|---|---|---|---|---|---|---|---|---|---|---|---|---|---|---|---|---|---|---|---|---|---|---|---|---|---|---|---|---|---|---|---|---|---|---|---|---|---|
| *Splice_typ* | *ID* | *GeneID* | *Gene_Symbol* | *Chromosome* | *Strand* | *LongExonStart_0base* | *LongExonEnd* | *ShortES* | *ShortEE* | *FlankingES* | *FlankingEE* | *SRR13214146* | *SRR13214147* | *SRR13214148* | *SRR13214149* | *SRR13214150* | *SRR13214146* | *SRR13214147* | *SRR13214148* | *SRR13214149* | *SRR13214150* | *SRR13214141* | *SRR13214242* | *SRR13214143* | *SRR13214144* | *SRR13214145* | *SRR13214141* | *SRR13214242* | *SRR13214143* | *SRR13214144* | *SRR13214145* | *IncFormLen* | *SkipFormLen* | *p value* | *FDR* | *SRR13214146* | *SRR13214147* | *SRR13214148* | *SRR13214149* | *SRR13214150* | *SRR13214141* | *SRR13214242* | *SRR13214143* | *SRR13214144* | *SRR13214145* | *Inclevel_diff* |
| A3SS | 289 | ENSG00000170889 | RPS9 | chr19 | + | 54206552 | 54207647 | 54207397 | 54207647 | 54206275 | 54206462 | 256 | 150 | 267 | 206 | 275 | 2265 | 1226 | 2895 | 2512 | 2463 | 718 | 474 | 350 | 445 | 758 | 2132 | 2377 | 1630 | 1587 | 3063 | 298 | 149 | 0.000000 | 0 | 0.053 | 0.058 | 0.044 | 0.039 | 0.053 | 0.144 | 0.091 | 0.097 | 0.123 | 0.11 | -0.064 |
| A3SS | 7099 | ENSG00000019582 | CD74 | chr5 | - | 150405084 | 150405671 | 150405084 | 150405180 | 150406258 | 150406321 | 29 | 36 | 26 | 31 | 47 | 3501 | 1525 | 2441 | 3131 | 1791 | 308 | 85 | 43 | 166 | 428 | 5579 | 3547 | 1674 | 3067 | 8990 | 298 | 149 | 0.000000 | 0 | 0.004 | 0.012 | 0.005 | 0.005 | 0.013 | 0.027 | 0.012 | 0.013 | 0.026 | 0.023 | -0.012 |
| A3SS | 13094 | ENSG00000063046 | EIF4B | chr12 | + | 53021715 | 53021860 | 53021805 | 53021860 | 53019909 | 53020026 | 68 | 32 | 72 | 75 | 66 | 3021 | 1875 | 3073 | 2767 | 3862 | 145 | 101 | 186 | 196 | 43 | 1960 | 1494 | 1784 | 1149 | 817 | 239 | 149 | 0.000000 | 0 | 0.014 | 0.011 | 0.014 | 0.017 | 0.011 | 0.044 | 0.04 | 0.061 | 0.096 | 0.032 | -0.041 |
| A3SS | 2907 | ENSG00000175061 | SNHG29 | chr17 | + | 16439580 | 16440253 | 16440184 | 16440253 | 16439326 | 16439414 | 49 | 52 | 73 | 63 | 50 | 617 | 324 | 818 | 518 | 513 | 84 | 50 | 75 | 117 | 121 | 272 | 214 | 229 | 323 | 462 | 298 | 149 | 0.000000 | 0 | 0.038 | 0.074 | 0.043 | 0.057 | 0.046 | 0.134 | 0.105 | 0.141 | 0.153 | 0.116 | -0.078 |
| A3SS | 2908 | ENSG00000175061 | SNHG29 | chr17 | + | 16440110 | 16440253 | 16440184 | 16440253 | 16439326 | 16439414 | 14 | 13 | 13 | 21 | 36 | 617 | 324 | 818 | 518 | 513 | 32 | 18 | 24 | 74 | 60 | 272 | 214 | 229 | 323 | 462 | 223 | 149 | 0.000000 | 0 | 0.015 | 0.026 | 0.011 | 0.026 | 0.045 | 0.073 | 0.053 | 0.065 | 0.133 | 0.08 | -0.056 |
| A3SS | 4361 | ENSG00000156976 | EIF4A2 | chr3 | + | 186787478 | 186787584 | 186787494 | 186787584 | 186787126 | 186787264 | 42 | 30 | 40 | 34 | 25 | 1528 | 908 | 1431 | 1475 | 4405 | 23 | 57 | 41 | 40 | 0 | 276 | 819 | 949 | 552 | 73 | 165 | 149 | 0.000000 | 0 | 0.024 | 0.029 | 0.025 | 0.02 | 0.005 | 0.07 | 0.059 | 0.038 | 0.061 | 0 | -0.025 |
| A3SS | 3924 | ENSG00000100650 | SRSF5 | chr14 | + | 69768797 | 69769251 | 69769181 | 69769251 | 69768603 | 69768674 | 2591 | 2084 | 2699 | 3220 | 3492 | 26 | 26 | 21 | 55 | 32 | 3166 | 3026 | 2799 | 3179 | 1823 | 277 | 216 | 76 | 561 | 95 | 298 | 149 | 0.000000 | 0 | 0.98 | 0.976 | 0.985 | 0.967 | 0.982 | 0.851 | 0.875 | 0.948 | 0.739 | 0.906 | 0.114 |
| A3SS | 6809 | ENSG00000169554 | ZEB2 | chr2 | - | 144424795 | 144424953 | 144424795 | 144424867 | 144429768 | 144430026 | 59 | 85 | 36 | 57 | 50 | 365 | 285 | 193 | 280 | 258 | 6 | 17 | 40 | 5 | 6 | 217 | 376 | 440 | 146 | 76 | 235 | 149 | 0.000000 | 0 | 0.093 | 0.159 | 0.106 | 0.114 | 0.109 | 0.017 | 0.028 | 0.054 | 0.021 | 0.048 | 0.083 |
| A3SS | 5020 | ENSG00000080822 | CLDND1 | chr3 | - | 98515482 | 98516879 | 98515482 | 98515832 | 98517051 | 98517189 | 334 | 229 | 424 | 423 | 668 | 16 | 4 | 29 | 24 | 67 | 57 | 130 | 185 | 74 | 26 | 44 | 36 | 118 | 53 | 18 | 298 | 149 | 0.000000 | 0 | 0.913 | 0.966 | 0.88 | 0.898 | 0.833 | 0.393 | 0.644 | 0.439 | 0.411 | 0.419 | 0.437 |
| A3SS | 10061 | ENSG00000122566 | HNRNPA2B1 | chr7 | - | 26193250 | 26193694 | 26193250 | 26193373 | 26195846 | 26195909 | 1499 | 1890 | 1233 | 1814 | 1494 | 8 | 10 | 11 | 37 | 35 | 645 | 1588 | 2198 | 1379 | 210 | 35 | 38 | 100 | 73 | 8 | 298 | 149 | 0.000000 | 0 | 0.989 | 0.99 | 0.982 | 0.961 | 0.955 | 0.902 | 0.954 | 0.917 | 0.904 | 0.929 | 0.054 |
| A3SS | 1525 | ENSG00000168259 | DNAJC7 | chr17 | - | 41982254 | 41982601 | 41982254 | 41982401 | 41983562 | 41983636 | 6 | 6 | 6 | 14 | 11 | 171 | 147 | 139 | 176 | 311 | 81 | 28 | 37 | 53 | 44 | 451 | 215 | 282 | 338 | 132 | 298 | 149 | 0.000000 | 8.15E-14 | 0.017 | 0.02 | 0.021 | 0.038 | 0.017 | 0.082 | 0.061 | 0.062 | 0.073 | 0.143 | -0.062 |
| A3SS | 13270 | ENSG00000197111 | PCBP2 | chr12 | + | 53459799 | 53461143 | 53461014 | 53461143 | 53459271 | 53459403 | 191 | 145 | 174 | 214 | 228 | 1224 | 657 | 1117 | 1066 | 1879 | 653 | 399 | 382 | 433 | 308 | 1119 | 1084 | 884 | 863 | 2088 | 298 | 149 | 0.000000 | 1.49E-13 | 0.072 | 0.099 | 0.072 | 0.091 | 0.057 | 0.226 | 0.155 | 0.178 | 0.201 | 0.069 | -0.088 |
| A3SS | 4761 | ENSG00000004534 | RBM6 | chr3 | + | 50059631 | 50059746 | 50059648 | 50059746 | 50058401 | 50058562 | 6 | 2 | 5 | 3 | 5 | 245 | 197 | 210 | 214 | 247 | 15 | 24 | 9 | 23 | 11 | 200 | 163 | 113 | 154 | 83 | 166 | 149 | 0.000000 | 2.76E-13 | 0.022 | 0.009 | 0.021 | 0.012 | 0.018 | 0.063 | 0.117 | 0.067 | 0.118 | 0.106 | -0.078 |
| A3SS | 5790 | ENSG00000108654 | DDX5 | chr17 | - | 64500548 | 64500882 | 64500548 | 64500773 | 64501056 | 64501143 | 1627 | 1028 | 1574 | 1804 | 1306 | 126 | 80 | 88 | 99 | 101 | 226 | 1391 | 1482 | 875 | 85 | 9 | 40 | 12 | 9 | 10 | 258 | 149 | 0.000000 | 1.73E-12 | 0.882 | 0.881 | 0.912 | 0.913 | 0.882 | 0.935 | 0.953 | 0.986 | 0.983 | 0.831 | -0.044 |
| A3SS | 10713 | ENSG00000215301 | DDX3X | chrX | + | 41337690 | 41339083 | 41339035 | 41339083 | 41337407 | 41337465 | 69 | 80 | 47 | 94 | 74 | 1127 | 591 | 871 | 1034 | 437 | 68 | 80 | 112 | 170 | 14 | 177 | 530 | 614 | 413 | 178 | 298 | 149 | 0.000000 | 3.02E-11 | 0.03 | 0.063 | 0.026 | 0.043 | 0.078 | 0.161 | 0.07 | 0.084 | 0.171 | 0.038 | -0.057 |
| A3SS | 4251 | ENSG00000150995 | ITPR1 | chr3 | + | 4727122 | 4727173 | 4727125 | 4727173 | 4711756 | 4711868 | 5 | 2 | 1 | 1 | 4 | 1 | 0 | 1 | 0 | 2 | 2 | 0 | 1 | 0 | 0 | 6 | 12 | 9 | 4 | 5 | 152 | 149 | 0.000000 | 7.36E-11 | 0.831 | 1 | 0.495 | 1 | 0.662 | 0.246 | 0 | 0.098 | 0 | 0 | 0.729 |
| A3SS | 3940 | ENSG00000203875 | SNHG5 | chr6 | - | 85677423 | 85677875 | 85677423 | 85677492 | 85677953 | 85678032 | 305 | 636 | 392 | 320 | 618 | 32 | 22 | 29 | 20 | 98 | 249 | 438 | 537 | 635 | 1003 | 0 | 16 | 16 | 0 | 40 | 298 | 149 | 0.000000 | 8.57E-11 | 0.827 | 0.935 | 0.871 | 0.889 | 0.759 | 1 | 0.932 | 0.944 | 1 | 0.926 | -0.104 |
| A3SS | 1173 | ENSG00000126561 | STAT5A | chr17 | + | 42309376 | 42311903 | 42310506 | 42311903 | 42308177 | 42308333 | 6 | 6 | 12 | 9 | 3 | 5 | 3 | 9 | 20 | 13 | 44 | 32 | 6 | 41 | 55 | 0 | 6 | 2 | 3 | 1 | 298 | 149 | 0.000000 | 8.86E-10 | 0.375 | 0.5 | 0.4 | 0.184 | 0.103 | 1 | 0.727 | 0.6 | 0.872 | 0.965 | -0.52 |
| A3SS | 4736 | ENSG00000151726 | ACSL1 | chr4 | - | 184762406 | 184762523 | 184762406 | 184762493 | 184763166 | 184763255 | 148 | 81 | 99 | 121 | 181 | 0 | 0 | 1 | 0 | 1 | 295 | 365 | 861 | 430 | 127 | 0 | 17 | 61 | 30 | 5 | 179 | 149 | 0.000000 | 1.04E-09 | 1 | 1 | 0.988 | 1 | 0.993 | 1 | 0.947 | 0.922 | 0.923 | 0.955 | 0.047 |
| A3SS | 9778 | ENSG00000142657 | PGD | chr1 | + | 10400375 | 10400572 | 10400392 | 10400572 | 10399628 | 10399704 | 16 | 8 | 8 | 10 | 13 | 369 | 215 | 204 | 263 | 295 | 127 | 58 | 75 | 159 | 133 | 1011 | 787 | 480 | 1053 | 1993 | 166 | 149 | 0.000000 | 1.22E-09 | 0.037 | 0.032 | 0.034 | 0.033 | 0.038 | 0.101 | 0.062 | 0.123 | 0.119 | 0.057 | -0.058 |
| A3SS | 6424 | ENSG00000165458 | INPPL1 | chr11 | + | 72233440 | 72233744 | 72233654 | 72233744 | 72233074 | 72233163 | 175 | 76 | 128 | 127 | 121 | 0 | 0 | 3 | 0 | 0 | 260 | 272 | 124 | 146 | 485 | 13 | 24 | 7 | 9 | 15 | 298 | 149 | 0.000000 | 1.39E-09 | 1 | 1 | 0.955 | 1 | 1 | 0.909 | 0.85 | 0.899 | 0.89 | 0.942 | 0.093 |
| A3SS | 9952 | ENSG00000141480 | ARRB2 | chr17 | + | 4720549 | 4720640 | 4720585 | 4720640 | 4720215 | 4720299 | 24 | 16 | 16 | 37 | 11 | 0 | 1 | 0 | 0 | 0 | 66 | 47 | 46 | 54 | 117 | 23 | 10 | 6 | 15 | 13 | 185 | 149 | 0.000000 | 1.50E-09 | 1 | 0.928 | 1 | 1 | 1 | 0.698 | 0.791 | 0.861 | 0.744 | 0.879 | 0.191 |
| A3SS | 6967 | ENSG00000137275 | RIPK1 | chr6 | + | 3082546 | 3083313 | 3083084 | 3083313 | 3080978 | 3081116 | 28 | 46 | 26 | 16 | 23 | 144 | 78 | 82 | 99 | 103 | 96 | 91 | 53 | 114 | 18 | 47 | 90 | 67 | 102 | 31 | 298 | 149 | 0.000000 | 2.44E-09 | 0.089 | 0.228 | 0.137 | 0.075 | 0.1 | 0.505 | 0.336 | 0.283 | 0.358 | 0.225 | -0.216 |
| A3SS | 6822 | ENSG00000142089 | IFITM3 | chr11 | - | 319675 | 320030 | 319675 | 319990 | 320564 | 320860 | 24 | 23 | 14 | 41 | 68 | 534 | 397 | 268 | 1089 | 405 | 34 | 81 | 55 | 133 | 23 | 2696 | 2714 | 1527 | 3441 | 1123 | 189 | 149 | 0.000000 | 2.75E-09 | 0.034 | 0.044 | 0.04 | 0.029 | 0.117 | 0.01 | 0.023 | 0.028 | 0.03 | 0.016 | 0.031 |
| A3SS | 1867 | ENSG00000197912 | SPG7 | chr16 | + | 89528631 | 89529579 | 89529476 | 89529579 | 89526328 | 89526468 | 3 | 3 | 2 | 1 | 3 | 51 | 42 | 65 | 48 | 39 | 21 | 13 | 4 | 14 | 36 | 67 | 56 | 25 | 59 | 114 | 298 | 149 | 0.000000 | 3.22E-09 | 0.029 | 0.034 | 0.015 | 0.01 | 0.037 | 0.135 | 0.104 | 0.074 | 0.106 | 0.136 | -0.086 |
| A3SS | 8248 | ENSG00000124151 | NCOA3 | chr20 | + | 47639576 | 47639822 | 47639621 | 47639822 | 47637647 | 47637783 | 31 | 24 | 43 | 55 | 76 | 0 | 0 | 0 | 0 | 0 | 26 | 34 | 32 | 23 | 4 | 0 | 12 | 6 | 11 | 0 | 194 | 149 | 0.000000 | 3.22E-09 | 1 | 1 | 1 | 1 | 1 | 1 | 0.685 | 0.804 | 0.616 | 1 | 0.179 |
| A3SS | 4104 | ENSG00000087266 | SH3BP2 | chr4 | + | 2831922 | 2832412 | 2832330 | 2832412 | 2831570 | 2831679 | 231 | 114 | 135 | 153 | 93 | 7 | 6 | 0 | 11 | 2 | 219 | 183 | 109 | 214 | 386 | 36 | 20 | 14 | 47 | 39 | 298 | 149 | 0.000000 | 4.70E-09 | 0.943 | 0.905 | 1 | 0.874 | 0.959 | 0.753 | 0.821 | 0.796 | 0.695 | 0.832 | 0.157 |
| A3SS | 10591 | ENSG00000102103 | PQBP1 | chrX | + | 48902232 | 48902795 | 48902731 | 48902795 | 48901929 | 48902042 | 192 | 118 | 136 | 175 | 153 | 3 | 2 | 1 | 2 | 0 | 277 | 143 | 161 | 144 | 246 | 37 | 5 | 14 | 3 | 12 | 298 | 149 | 0.000000 | 4.70E-09 | 0.97 | 0.967 | 0.986 | 0.978 | 1 | 0.789 | 0.935 | 0.852 | 0.96 | 0.911 | 0.091 |
| A3SS | 3085 | ENSG00000130811 | EIF3G | chr19 | - | 10119087 | 10119255 | 10119087 | 10119171 | 10119653 | 10119700 | 5 | 2 | 10 | 6 | 7 | 336 | 151 | 350 | 306 | 242 | 58 | 38 | 3 | 7 | 46 | 512 | 408 | 248 | 348 | 479 | 233 | 149 | 0.000000 | 5.50E-09 | 0.009 | 0.008 | 0.018 | 0.012 | 0.018 | 0.068 | 0.056 | 0.008 | 0.013 | 0.058 | -0.028 |
| A3SS | 11790 | ENSG00000149182 | ARFGAP2 | chr11 | - | 47175181 | 47175413 | 47175181 | 47175313 | 47175850 | 47175923 | 12 | 14 | 12 | 15 | 7 | 259 | 139 | 194 | 191 | 343 | 39 | 23 | 17 | 24 | 51 | 220 | 272 | 94 | 114 | 276 | 249 | 149 | 0.000000 | 6.02E-09 | 0.027 | 0.057 | 0.036 | 0.045 | 0.012 | 0.096 | 0.048 | 0.098 | 0.112 | 0.1 | -0.055 |
| A3SS | 3357 | ENSG00000145715 | RASA1 | chr5 | + | 87389302 | 87389527 | 87389392 | 87389527 | 87386825 | 87386903 | 18 | 32 | 9 | 17 | 29 | 101 | 74 | 100 | 140 | 101 | 0 | 0 | 1 | 4 | 0 | 35 | 59 | 37 | 22 | 24 | 239 | 149 | 0.000000 | 8.74E-09 | 0.1 | 0.212 | 0.053 | 0.07 | 0.152 | 0 | 0 | 0.017 | 0.102 | 0 | 0.094 |
| A3SS | 7190 | ENSG00000001497 | LAS1L | chrX | - | 65517986 | 65522285 | 65517986 | 65518465 | 65523559 | 65523707 | 8 | 1 | 8 | 7 | 2 | 41 | 35 | 60 | 48 | 68 | 36 | 23 | 19 | 10 | 21 | 44 | 40 | 55 | 46 | 42 | 298 | 149 | 0.000000 | 8.81E-09 | 0.089 | 0.014 | 0.062 | 0.068 | 0.014 | 0.29 | 0.223 | 0.147 | 0.098 | 0.2 | -0.142 |
| A3SS | 12515 | ENSG00000120438 | TCP1 | chr6 | - | 159787742 | 159787902 | 159787742 | 159787871 | 159788057 | 159788143 | 12 | 8 | 13 | 32 | 18 | 171 | 91 | 142 | 148 | 196 | 20 | 33 | 45 | 24 | 7 | 79 | 56 | 63 | 99 | 13 | 180 | 149 | 0.000000 | 1.63E-08 | 0.055 | 0.068 | 0.07 | 0.152 | 0.071 | 0.173 | 0.328 | 0.372 | 0.167 | 0.308 | -0.186 |
| A3SS | 4356 | ENSG00000135870 | RC3H1 | chr1 | - | 173943441 | 173943615 | 173943441 | 173943588 | 173946475 | 173946608 | 62 | 56 | 49 | 55 | 57 | 32 | 14 | 15 | 39 | 45 | 96 | 50 | 77 | 52 | 12 | 0 | 4 | 20 | 2 | 0 | 176 | 149 | 0.000000 | 1.70E-08 | 0.621 | 0.772 | 0.734 | 0.544 | 0.517 | 1 | 0.914 | 0.765 | 0.957 | 1 | -0.29 |
| A3SS | 9352 | ENSG00000170919 | TPT1-AS1 | chr13 | + | 45377111 | 45377215 | 45377139 | 45377215 | 45369958 | 45370182 | 4 | 2 | 0 | 5 | 0 | 2 | 4 | 1 | 5 | 1 | 2 | 4 | 1 | 8 | 5 | 0 | 1 | 0 | 0 | 0 | 177 | 149 | 0.000000 | 2.24E-08 | 0.627 | 0.296 | 0 | 0.457 | 0 | 1 | 0.771 | 1 | 1 | 1 | -0.678 |
| A3SS | 9126 | ENSG00000105393 | BABAM1 | chr19 | + | 17275776 | 17275825 | 17275788 | 17275825 | 17274106 | 17274185 | 3 | 1 | 1 | 1 | 5 | 0 | 0 | 0 | 0 | 0 | 8 | 0 | 2 | 3 | 10 | 11 | 2 | 2 | 0 | 8 | 161 | 149 | 0.000000 | 2.90E-08 | 1 | 1 | 1 | 1 | 1 | 0.402 | 0 | 0.481 | 1 | 0.536 | 0.516 |
| A3SS | 4380 | ENSG00000009790 | TRAF3IP3 | chr1 | + | 209779314 | 209779655 | 209779430 | 209779655 | 209778110 | 209778173 | 846 | 616 | 935 | 867 | 792 | 24 | 14 | 23 | 26 | 24 | 469 | 716 | 421 | 441 | 542 | 24 | 56 | 31 | 24 | 37 | 265 | 149 | 0.000000 | 3.51E-08 | 0.952 | 0.961 | 0.958 | 0.949 | 0.949 | 0.917 | 0.878 | 0.884 | 0.912 | 0.892 | 0.057 |
| A3SS | 10918 | ENSG00000171988 | JMJD1C | chr10 | - | 63215552 | 63215696 | 63215552 | 63215617 | 63217206 | 63217331 | 384 | 364 | 325 | 445 | 862 | 18 | 19 | 17 | 19 | 11 | 105 | 251 | 189 | 35 | 50 | 0 | 0 | 0 | 0 | 0 | 228 | 149 | 0.000000 | 3.88E-08 | 0.933 | 0.926 | 0.926 | 0.939 | 0.981 | 1 | 1 | 1 | 1 | 1 | -0.059 |
| A3SS | 4395 | ENSG00000155975 | VPS37A | chr8 | + | 17279721 | 17280155 | 17280027 | 17280155 | 17276396 | 17276467 | 5 | 26 | 10 | 14 | 14 | 165 | 141 | 106 | 182 | 220 | 0 | 1 | 7 | 1 | 0 | 128 | 173 | 125 | 97 | 160 | 298 | 149 | 0.000000 | 6.83E-08 | 0.015 | 0.084 | 0.045 | 0.037 | 0.031 | 0 | 0.003 | 0.027 | 0.005 | 0 | 0.035 |
| A3SS | 6522 | ENSG00000113812 | ACTR8 | chr3 | - | 53873031 | 53873146 | 53873031 | 53873127 | 53874210 | 53874364 | 1 | 5 | 9 | 13 | 19 | 87 | 43 | 77 | 65 | 152 | 0 | 0 | 2 | 0 | 0 | 41 | 67 | 37 | 43 | 51 | 168 | 149 | 0.000000 | 1.06E-07 | 0.01 | 0.093 | 0.094 | 0.151 | 0.1 | 0 | 0 | 0.046 | 0 | 0 | 0.08 |
| [illegible] | [illegible] | [illegible] | [illegible] | [illegible] | [illegible] | [illegible] | [illegible] | [illegible] | [illegible] | [illegible] | [illegible] | [illegible] | [illegible] | [illegible] | [illegible] | [illegible] | [illegible] | [illegible] | [illegible] | [illegible] | [illegible] | [illegible] | [illegible] | [illegible] | [illegible] | [illegible] | [illegible] | [illegible] | [illegible] | [illegible] | [illegible] | [illegible] | [illegible] | [illegible] | [illegible] | [illegible] | [illegible] | [illegible] | [illegible] | [illegible] | [illegible] | [illegible] | [illegible] | [illegible] | [illegible] | [illegible] |

| | | | | | | | | | | | | | | | |
|---|---|---|---|---|---|---|---|---|---|---|---|---|---|---|---|
| A3SS | 10814 | ENSG00000142046 | TMEM91 | chr19 | + | 41383862 | 41384083 | 41383885 | 41384083 | 41382771 | 41382921 | 9 | 4 | 2 | [illegible] |
| A3SS | 6375 | ENSG00000115568 | ZNF142 | chr2 | - | 218658700 | 218658876 | 218658700 | 218658801 | 218658979 | 218659130 | 28 | 25 | 45 | [illegible] |
| A3SS | 7195 | ENSG00000001497 | LAS1L | chrX | - | 65523559 | 65523876 | 65523559 | 65523707 | 65524055 | 65524614 | 2 | 2 | 4 | [illegible] |
| A3SS | 12998 | ENSG00000092531 | SNAP23 | chr15 | + | 42513173 | 42513447 | 42513398 | 42513447 | 42512954 | 42512996 | 74 | 65 | 84 | [illegible] |
| A3SS | 13134 | ENSG00000157538 | VPS26C | chr21 | - | 37232376 | 37232645 | 37232376 | 37232451 | 37233361 | 37233442 | 25 | 19 | 14 | [illegible] |
| A3SS | 12829 | ENSG00000123200 | ZC3H13 | chr13 | - | 46045390 | 46045516 | 46045390 | 46045513 | 46052697 | 46052759 | 21 | 18 | 31 | [illegible] |
| A3SS | 6407 | ENSG00000132781 | MUTYH | chr1 | - | 45332762 | 45333170 | 45332762 | 45332834 | 45333284 | 45333324 | 22 | 6 | 28 | [illegible] |
| A3SS | 7140 | ENSG00000124571 | XPO5 | chr6 | - | 43525838 | 43525954 | 43525838 | 43525921 | 43526684 | 43526747 | 0 | 1 | 2 | [illegible] |
| A3SS | 8642 | ENSG00000160767 | FAM189B | chr1 | - | 155251732 | 155251908 | 155251732 | 155251905 | 155253624 | 155253732 | 0 | 0 | 0 | [illegible] |
| A3SS | 7251 | ENSG00000065413 | ANKRD44 | chr2 | - | 197088710 | 197088828 | 197088710 | 197088774 | 197089949 | 197090032 | 27 | 15 | 27 | [illegible] |
| A3SS | 3034 | ENSG00000125459 | MSTO1 | chr1 | + | 155613424 | 155613566 | 155613461 | 155613566 | 155613048 | 155613233 | 0 | 0 | 2 | [illegible] |
| A3SS | 6910 | ENSG00000168610 | STAT3 | chr17 | - | 42329746 | 42329791 | 42329746 | 42329776 | 42331471 | 42331531 | 8 | 4 | 3 | [illegible] |
| A3SS | 10232 | ENSG00000196998 | WDR45 | chrX | - | 49075363 | 49075453 | 49075363 | 49075423 | 49075544 | 49075753 | 12 | 9 | 10 | [illegible] |
| A3SS | 1673 | ENSG00000140829 | DHX38 | chr16 | + | 72098926 | 72099280 | 72099203 | 72099280 | 72098644 | 72098792 | 100 | 46 | 92 | [illegible] |
| A3SS | 895 | ENSG00000080189 | SLC35C2 | chr20 | - | 46355072 | 46355241 | 46355072 | 46355178 | 46355801 | 46355874 | 51 | 21 | 53 | [illegible] |
| A3SS | 8154 | ENSG00000131051 | RBM39 | chr20 | - | 35734185 | 35735081 | 35734185 | 35734257 | 35738967 | 35739017 | 130 | 161 | 115 | [illegible] |
| A3SS | 10722 | ENSG00000215301 | DDX3X | chrX | + | 41346858 | 41347451 | 41347311 | 41347451 | 41346504 | 41346622 | 1105 | 731 | 950 | [illegible] |
| A3SS | 5742 | ENSG00000204475 | NCR3 | chr6 | - | 31588894 | 31589231 | 31588894 | 31589176 | 31589525 | 31589633 | 63 | 28 | 87 | [illegible] |
| A3SS | 3306 | ENSG00000143774 | GUK1 | chr1 | + | 228145470 | 228145624 | 228145510 | 228145624 | 228140227 | 228140363 | 2 | 1 | 2 | [illegible] |
| A3SS | 1208 | ENSG00000185624 | P4HB | chr17 | - | 81845143 | 81845303 | 81845143 | 81845230 | 81845560 | 81845742 | 10 | 7 | 13 | [illegible] |
| A3SS | 9241 | ENSG00000100316 | RPL3 | chr22 | - | 39316705 | 39317021 | 39316705 | 39316841 | 39317460 | 39317629 | 59 | 14 | 27 | [illegible] |
| A3SS | 3167 | ENSG00000146094 | DOK3 | chr5 | - | 177509474 | 177509657 | 177509474 | 177509584 | 177509757 | 177509826 | 20 | 7 | 12 | [illegible] |
| A3SS | 11864 | ENSG00000198879 | SFMBT2 | chr10 | - | 7283903 | 7284150 | 7283903 | 7283959 | 7285865 | 7285954 | 42 | 34 | 56 | [illegible] |
| A3SS | 5539 | ENSG00000138002 | IFT172 | chr2 | - | 27481045 | 27481260 | 27481045 | 27481126 | 27483288 | 27483376 | 14 | 7 | 21 | [illegible] |
| A3SS | 1435 | ENSG00000141030 | COPS3 | chr17 | - | 17267884 | 17268045 | 17267884 | 17267977 | 17270757 | 17270807 | 30 | 22 | 14 | [illegible] |

[illegible]

| | | | | | | | | | | | | | | | | | | | | | | | | | | | | | | | | | | | | | | | | | | | | | | |
|---|---|---|---|---|---|---|---|---|---|---|---|---|---|---|---|---|---|---|---|---|---|---|---|---|---|---|---|---|---|---|---|---|---|---|---|---|---|---|---|---|---|---|---|---|---|---|
| A3SS | 360 | ENSG00000167604 | NFKBID | chr19 | - | 35896912 | 35897103 | 35896912 | 35897058 | 35897650 | 35897856 | 1 | 0 | 0 | 2 | 2 | 7 | 4 | 8 | 8 | 5 | 4 | 3 | 2 | 12 | 10 | 21 | 4 | 2 | 16 | 23 | 194 | 149 | 0.000125 | 0.002421907 | 0.099 | 0 | 0 | 0.161 | 0.235 | 0.128 | 0.365 | 0.434 | 0.365 | 0.25 | -0.209 |
| A3SS | 106 | ENSG00000100417 | PMM1 | chr22 | - | 41583958 | 41584372 | 41583958 | 41584058 | 41584526 | 41584603 | 31 | 12 | 38 | 36 | 37 | 0 | 0 | 0 | 0 | 4 | 53 | 44 | 35 | 22 | 80 | 0 | 13 | 4 | 3 | 0 | 298 | 149 | 0.000126 | 0.002453624 | 1 | 1 | 1 | 1 | 0.822 | 1 | 0.629 | 0.814 | 0.786 | 1 | 0.119 |
| A3SS | 1515 | ENSG00000198231 | DDX42 | chr17 | + | 63805043 | 63805175 | 63805070 | 63805175 | 63800467 | 63800617 | 3 | 6 | 9 | 6 | 6 | 101 | 70 | 111 | 94 | 142 | 0 | 4 | 2 | 0 | 0 | 68 | 122 | 76 | 70 | 8 | 176 | 149 | 0.000129 | 0.002503103 | 0.025 | 0.068 | 0.064 | 0.051 | 0.035 | 0 | 0.027 | 0.022 | 0 | 0 | 0.039 |
| A3SS | 2701 | ENSG00000099797 | TECR | chr19 | + | 14563140 | 14563257 | 14563205 | 14563257 | 14562524 | 14562575 | 1 | 0 | 4 | 0 | 0 | 73 | 33 | 109 | 71 | 51 | 0 | 4 | 0 | 6 | 15 | 20 | 77 | 87 | 66 | 123 | 214 | 149 | 0.000131 | 0.00252093 | 0.009 | 0 | 0.025 | 0 | 0 | 0 | 0.035 | 0 | 0.06 | 0.078 | -0.028 |
| A3SS | 9677 | ENSG00000026950 | BTN3A1 | chr6 | + | 26409893 | 26410032 | 26410005 | 26410032 | 26409532 | 26409733 | 277 | 180 | 284 | 270 | 334 | 12 | 9 | 11 | 7 | 7 | 211 | 327 | 198 | 281 | 198 | 11 | 43 | 9 | 16 | 10 | 261 | 149 | 0.000131 | 0.00252315 | 0.929 | 0.919 | 0.936 | 0.957 | 0.965 | 0.916 | 0.813 | 0.926 | 0.909 | 0.919 | 0.045 |
| A3SS | 11833 | ENSG00000139626 | ITGB7 | chr12 | - | 53191322 | 53191731 | 53191322 | 53191636 | 53191858 | 53192019 | 43 | 19 | 40 | 36 | 41 | 434 | 190 | 425 | 339 | 357 | 19 | 47 | 10 | 10 | 12 | 301 | 428 | 209 | 241 | 432 | 244 | 149 | 0.000132 | 0.002540947 | 0.057 | 0.058 | 0.054 | 0.061 | 0.066 | 0.037 | 0.063 | 0.028 | 0.025 | 0.017 | 0.025 |
| A3SS | 9835 | ENSG00000006534 | ALDH3B1 | chr11 | + | 68019065 | 68019255 | 68019169 | 68019255 | 68018772 | 68018893 | 2 | 0 | 2 | 0 | 2 | 53 | 20 | 37 | 31 | 17 | 44 | 6 | 1 | 15 | 0 | 12 | 62 | 31 | 58 | 56 | 253 | 149 | 0.000133 | 0.002540947 | 0.022 | 0 | 0.031 | 0 | 0.065 | 0.683 | 0.054 | 0.019 | 0.132 | 0 | -0.154 |
| A3SS | 1812 | ENSG00000103275 | UBE2I | chr16 | + | 1319701 | 1320283 | 1320173 | 1320283 | 1315653 | 1315726 | 69 | 40 | 68 | 75 | 90 | 659 | 334 | 504 | 582 | 1004 | 167 | 102 | 109 | 172 | 214 | 1234 | 821 | 654 | 813 | 1237 | 298 | 149 | 0.000134 | 0.002560137 | 0.05 | 0.056 | 0.063 | 0.061 | 0.043 | 0.063 | 0.058 | 0.077 | 0.096 | 0.08 | -0.02 |
| A3SS | 11866 | ENSG00000149823 | VPS51 | chr11 | + | 65107186 | 65107727 | 65107580 | 65107727 | 65096997 | 65097127 | 25 | 20 | 23 | 22 | 11 | 266 | 119 | 257 | 185 | 232 | 37 | 12 | 7 | 25 | 52 | 175 | 205 | 101 | 83 | 172 | 298 | 149 | 0.000136 | 0.002601416 | 0.045 | 0.078 | 0.043 | 0.056 | 0.023 | 0.096 | 0.028 | 0.033 | 0.131 | 0.131 | -0.035 |
| A3SS | 6800 | ENSG00000189266 | PNRC2 | chr1 | + | 23960928 | 23961134 | 23960932 | 23961134 | 23959827 | 23959998 | 877 | 512 | 706 | 932 | 961 | 25 | 7 | 20 | 16 | 15 | 157 | 637 | 584 | 319 | 61 | 4 | 21 | 10 | 22 | 16 | 153 | 149 | 0.000137 | 0.002607037 | 0.972 | 0.986 | 0.972 | 0.983 | 0.984 | 0.975 | 0.967 | 0.983 | 0.934 | 0.788 | 0.05 |
| A3SS | 6657 | ENSG00000083544 | TDRD3 | chr13 | + | 60444682 | 60444748 | 60444686 | 60444748 | 60439687 | 60439772 | 90 | 69 | 65 | 102 | 149 | 13 | 7 | 6 | 16 | 12 | 74 | 68 | 36 | 51 | 91 | 0 | 2 | 5 | 0 | 7 | 153 | 149 | 0.000138 | 0.002615168 | 0.871 | 0.906 | 0.913 | 0.861 | 0.924 | 1 | 0.971 | 0.875 | 1 | 0.927 | -0.06 |
| A3SS | 2948 | ENSG00000135506 | OS9 | chr12 | + | 57694705 | 57694926 | 57694749 | 57694926 | 57694131 | 57694323 | 39 | 14 | 16 | 43 | 36 | 381 | 200 | 227 | 332 | 248 | 50 | 29 | 26 | 34 | 41 | 741 | 494 | 270 | 599 | 790 | 193 | 149 | 0.000138 | 0.002619647 | 0.073 | 0.051 | 0.052 | 0.091 | 0.101 | 0.05 | 0.043 | 0.069 | 0.042 | 0.039 | 0.025 |
| A3SS | 13146 | ENSG00000123349 | PFDN5 | chr12 | + | 53296218 | 53296275 | 53296243 | 53296275 | 53295838 | 53295941 | 26 | 20 | 35 | 20 | 32 | 1022 | 540 | 881 | 838 | 1024 | 32 | 37 | 18 | 30 | 93 | 1143 | 630 | 468 | 705 | 1141 | 174 | 149 | 0.000144 | 0.002715578 | 0.021 | 0.031 | 0.033 | 0.02 | 0.026 | 0.023 | 0.048 | 0.032 | 0.035 | 0.065 | -0.014 |
| A3SS | 2430 | ENSG00000140474 | ULK3 | chr15 | - | 74842074 | 74842195 | 74842074 | 74842118 | 74842279 | 74842420 | 59 | 49 | 75 | 91 | 127 | 3 | 1 | 2 | 8 | 9 | 55 | 104 | 31 | 37 | 123 | 0 | 0 | 1 | 3 | 2 | 226 | 149 | 0.000146 | 0.002758164 | 0.928 | 0.97 | 0.961 | 0.882 | 0.903 | 1 | 1 | 0.953 | 0.89 | 0.976 | -0.035 |
| A3SS | 123 | ENSG00000100330 | MTMR3 | chr22 | + | 29978424 | 29978531 | 29978441 | 29978531 | 29970975 | 29971062 | 4 | 2 | 3 | 0 | 3 | 190 | 85 | 120 | 140 | 151 | 12 | 6 | 13 | 21 | 8 | 307 | 281 | 245 | 313 | 247 | 166 | 149 | 0.000147 | 0.002758164 | 0.019 | 0.021 | 0.022 | 0 | 0.018 | 0.034 | 0.019 | 0.045 | 0.057 | 0.028 | -0.021 |
| A3SS | 1068 | ENSG00000154889 | MPPE1 | chr18 | - | 11885675 | 11885912 | 11885675 | 11885816 | 11886916 | 11887025 | 37 | 21 | 28 | 25 | 34 | 2 | 0 | 2 | 0 | 2 | 60 | 34 | 31 | 73 | 48 | 14 | 12 | 0 | 8 | 2 | 245 | 149 | 0.000148 | 0.002769874 | 0.918 | 1 | 0.895 | 1 | 0.912 | 0.723 | 0.633 | 1 | 0.847 | 0.936 | 0.117 |
| A3SS | 984 | ENSG00000124155 | PIGT | chr20 | + | 45419277 | 45419395 | 45419294 | 45419395 | 45416516 | 45416694 | 0 | 0 | 3 | 0 | 0 | 7 | 3 | 10 | 14 | 4 | 0 | 4 | 0 | 6 | 4 | 21 | 6 | 20 | 18 | 13 | 166 | 149 | 0.000148 | 0.002769874 | 0 | 0 | 0.212 | 0 | 0 | 0 | 0.374 | 0 | 0.23 | 0.216 | -0.122 |
| A3SS | 8733 | ENSG00000077380 | DYNC1I2 | chr2 | + | 171747775 | 171748411 | 171747778 | 171748411 | 171745801 | 171745927 | 202 | 133 | 156 | 175 | 222 | 7 | 2 | 9 | 6 | 8 | 89 | 171 | 91 | 117 | 12 | 0 | 3 | 1 | 0 | 0 | 152 | 149 | 0.000149 | 0.002783629 | 0.966 | 0.985 | 0.944 | 0.966 | 0.965 | 1 | 0.982 | 0.989 | 1 | 1 | -0.029 |
| A3SS | 2232 | ENSG00000166783 | MARF1 | chr16 | - | 15622933 | 15623126 | 15622933 | 15623057 | 15624768 | 15624927 | 148 | 84 | 143 | 131 | 191 | 0 | 0 | 0 | 2 | 0 | 77 | 121 | 97 | 135 | 16 | 0 | 2 | 16 | 0 | 0 | 218 | 149 | 0.000149 | 0.002783629 | 1 | 1 | 1 | 0.978 | 1 | 1 | 0.976 | 0.806 | 1 | 1 | 0.039 |
| A3SS | 11018 | ENSG00000172269 | DPAGT1 | chr11 | - | 119097141 | 119097551 | 119097141 | 119097297 | 119097854 | 119098043 | 30 | 16 | 19 | 39 | 19 | 0 | 1 | 0 | 3 | 0 | 15 | 12 | 10 | 19 | 32 | 3 | 6 | 0 | 2 | 0 | 298 | 149 | 0.000150 | 0.002790455 | 1 | 0.889 | 1 | 0.867 | 1 | 0.714 | 0.5 | 1 | 0.826 | 1 | 0.143 |
| A3SS | 13027 | ENSG00000166012 | TAF1D | chr11 | - | 93733349 | 93733505 | 93733349 | 93733397 | 93734624 | 93734660 | 80 | 51 | 60 | 67 | 55 | 443 | 423 | 418 | 397 | 868 | 16 | 20 | 61 | 11 | 17 | 19 | 195 | 183 | 55 | 11 | 257 | 149 | 0.000156 | 0.002895429 | 0.095 | 0.065 | 0.077 | 0.089 | 0.035 | 0.328 | 0.056 | 0.162 | 0.104 | 0.473 | -0.152 |
| A3SS | 6529 | ENSG00000143771 | CNIH4 | chr1 | + | 224375794 | 224379452 | 224379068 | 224379452 | 224371282 | 224371423 | 263 | 145 | 209 | 202 | 307 | 10 | 2 | 5 | 1 | 5 | 194 | 339 | 237 | 279 | 254 | 10 | 7 | 3 | 12 | 35 | 298 | 149 | 0.000156 | 0.002895429 | 0.929 | 0.973 | 0.954 | 0.99 | 0.968 | 0.907 | 0.96 | 0.975 | 0.921 | 0.784 | 0.053 |
| A3SS | 2498 | ENSG00000275835 | TUBGCP5 | chr15 | - | 23027191 | 23027309 | 23027191 | 23027306 | 23030884 | 23031020 | 3 | 1 | 2 | 5 | 2 | 16 | 18 | 26 | 21 | 34 | 0 | 1 | 0 | 0 | 0 | 22 | 4 | 23 | 7 | 4 | 152 | 149 | 0.000157 | 0.002898547 | 0.155 | 0.052 | 0.07 | 0.189 | 0.055 | 0 | 0.197 | 0 | 0 | 0 | 0.065 |
| A3SS | 5821 | ENSG00000160058 | BSDC1 | chr1 | - | 32383829 | 32384048 | 32383829 | 32383997 | 32386778 | 32386895 | 12 | 10 | 24 | 27 | 10 | 89 | 56 | 46 | 62 | 113 | 43 | 21 | 26 | 31 | 40 | 35 | 39 | 37 | 62 | 141 | 200 | 149 | 0.000159 | 0.00293661 | 0.091 | 0.117 | 0.28 | 0.245 | 0.062 | 0.478 | 0.286 | 0.344 | 0.271 | 0.174 | -0.152 |
| A3SS | 7818 | ENSG00000112855 | HARS2 | chr5 | + | 140696028 | 140696201 | 140696102 | 140696201 | 140695737 | 140695845 | 18 | 12 | 12 | 12 | 19 | 44 | 31 | 56 | 42 | 52 | 54 | 21 | 31 | 18 | 9 | 60 | 45 | 28 | 11 | 20 | 223 | 149 | 0.000160 | 0.002946001 | 0.215 | 0.205 | 0.125 | 0.16 | 0.196 | 0.376 | 0.238 | 0.425 | 0.522 | 0.231 | -0.178 |
| A3SS | 253 | ENSG00000131351 | HAUS8 | chr19 | - | 17059996 | 17060092 | 17059996 | 17060089 | 17062697 | 17062779 | 31 | 12 | 14 | 17 | 20 | 9 | 0 | 3 | 1 | 5 | 24 | 3 | 21 | 4 | 18 | 0 | 3 | 0 | 0 | 0 | 152 | 149 | 0.000162 | 0.002966754 | 0.772 | 1 | 0.821 | 0.943 | 0.797 | 1 | 0.495 | 1 | 1 | 1 | -0.032 |
| A3SS | 5892 | ENSG00000058272 | PPP1R12A | chr12 | - | 79806165 | 79806333 | 79806165 | 79806297 | 79807225 | 79807330 | 348 | 227 | 229 | 255 | 497 | 23 | 28 | 6 | 25 | 11 | 64 | 160 | 177 | 56 | 27 | 0 | 7 | 4 | 2 | 0 | 185 | 149 | 0.000163 | 0.00296793 | 0.924 | 0.867 | 0.968 | 0.891 | 0.973 | 1 | 0.948 | 0.973 | 0.958 | 1 | -0.051 |
| A3SS | 9135 | ENSG00000168000 | BSCL2 | chr11 | - | 62690786 | 62690873 | 62690786 | 62690867 | 62691074 | 62691141 | 0 | 2 | 2 | 8 | 3 | 51 | 19 | 34 | 32 | 72 | 0 | 0 | 1 | 0 | 1 | 97 | 42 | 27 | 33 | 202 | 155 | 149 | 0.000163 | 0.00296793 | 0 | 0.092 | 0.054 | 0.194 | 0.039 | 0 | 0 | 0.034 | 0 | 0.005 | 0.068 |
| A3SS | 1833 | ENSG00000103051 | COG4 | chr16 | - | 70496265 | 70496431 | 70496265 | 70496379 | 70497220 | 70497387 | 60 | 53 | 53 | 53 | 35 | 5 | 1 | 4 | 4 | 4 | 26 | 57 | 14 | 36 | 74 | 30 | 11 | 2 | 3 | 19 | 201 | 149 | 0.000163 | 0.00296793 | 0.899 | 0.975 | 0.908 | 0.908 | 0.866 | 0.391 | 0.793 | 0.838 | 0.899 | 0.743 | 0.178 |
| A3SS | 9985 | ENSG00000106348 | IMPDH1 | chr7 | - | 128400094 | 128400204 | 128400094 | 128400182 | 128400332 | 128400536 | 17 | 12 | 20 | 22 | 7 | 125 | 74 | 105 | 94 | 97 | 77 | 31 | 20 | 28 | 121 | 270 | 128 | 81 | 144 | 180 | 171 | 149 | 0.000164 | 0.00296793 | 0.106 | 0.124 | 0.142 | 0.169 | 0.059 | 0.199 | 0.174 | 0.177 | 0.145 | 0.369 | -0.093 |
| A3SS | 2381 | ENSG00000140553 | UNC45A | chr15 | + | 90936136 | 90936460 | 90936284 | 90936460 | 90935945 | 90935982 | 8 | 3 | 8 | 11 | 2 | 50 | 19 | 43 | 47 | 51 | 19 | 11 | 7 | 15 | 21 | 17 | 27 | 20 | 38 | 103 | 297 | 149 | 0.000164 | 0.00296793 | 0.074 | 0.073 | 0.085 | 0.105 | 0.019 | 0.359 | 0.17 | 0.149 | 0.165 | 0.093 | -0.116 |
| A3SS | 12240 | ENSG00000172183 | ISG20 | chr15 | + | 88650234 | 88652310 | 88652109 | 88652310 | 88639342 | 88639594 | 50 | 25 | 46 | 41 | 38 | 194 | 98 | 225 | 168 | 71 | 37 | 98 | 82 | 88 | 132 | 77 | 345 | 234 | 138 | 109 | 298 | 149 | 0.000164 | 0.002970062 | 0.114 | 0.113 | 0.093 | 0.109 | 0.211 | 0.194 | 0.124 | 0.149 | 0.242 | 0.377 | -0.089 |
| A3SS | 7279 | ENSG00000125633 | CCDC93 | chr2 | - | 117958364 | 117958481 | 117958364 | 117958478 | 117973907 | 117973994 | 60 | 37 | 47 | 62 | 70 | 2 | 1 | 4 | 4 | 6 | 18 | 40 | 41 | 38 | 21 | 0 | 0 | 1 | 0 | 0 | 152 | 149 | 0.000165 | 0.002977994 | 0.967 | 0.973 | 0.92 | 0.938 | 0.92 | 1 | 1 | 0.976 | 1 | 1 | -0.052 |
| A3SS | 13393 | ENSG00000148362 | PAXX | chr9 | + | 136992924 | 136993243 | 136993052 | 136993243 | 136992639 | 136992700 | 39 | 13 | 30 | 38 | 21 | 0 | 0 | 0 | 0 | 0 | 4 | 17 | 11 | 4 | 30 | 9 | 5 | 0 | 0 | 0 | 277 | 149 | 0.000166 | 0.002986732 | 1 | 1 | 1 | 1 | 1 | 0.193 | 0.647 | 1 | 1 | 1 | 0.232 |
| A3SS | 6709 | ENSG00000145979 | TBC1D7 | chr6 | - | 13326786 | 13326963 | 13326786 | 13326906 | 13328455 | 13328531 | 8 | 2 | 3 | 8 | 8 | 4 | 2 | 3 | 7 | 6 | 9 | 7 | 0 | 3 | 0 | 9 | 17 | 5 | 11 | 23 | 206 | 149 | 0.000167 | 0.003004476 | 0.591 | 0.42 | 0.42 | 0.453 | 0.491 | 0.42 | 0.229 | 0 | 0.165 | 0 | 0.312 |
| A3SS | 8912 | ENSG00000136699 | SMPD4 | chr2 | - | 130156034 | 130156376 | 130156034 | 130156135 | 130156584 | 130156675 | 4 | 4 | 12 | 8 | 2 | 39 | 9 | 27 | 30 | 29 | 9 | 19 | 7 | 27 | 19 | 28 | 36 | 18 | 21 | 25 | 298 | 149 | 0.000172 | 0.003077514 | 0.049 | 0.182 | 0.182 | 0.118 | 0.033 | 0.138 | 0.209 | 0.163 | 0.391 | 0.275 | -0.122 |
| A3SS | 10407 | ENSG00000139436 | GIT2 | chr12 | - | 109965525 | 109965577 | 109965525 | 109965574 | 109967457 | 109967503 | 315 | 262 | 262 | 257 | 282 | 20 | 10 | 26 | 19 | 36 | 548 | 349 | 286 | 420 | 264 | 60 | 52 | 36 | 39 | 54 | 152 | 149 | 0.000178 | 0.003189467 | 0.939 | 0.963 | 0.908 | 0.93 | 0.885 | 0.9 | 0.868 | 0.886 | 0.913 | 0.827 | 0.046 |
| A3SS | 11425 | ENSG00000167118 | URM1 | chr9 | + | 128377994 | 128378106 | 128378035 | 128378106 | 128371360 | 128371415 | 0 | 0 | 1 | 2 | 0 | 22 | 14 | 39 | 42 | 5 | 0 | 0 | 2 | 4 | 12 | 30 | 29 | 33 | 37 | 55 | 190 | 149 | 0.000181 | 0.003227448 | 0 | 0 | 0.02 | 0.036 | 0 | 0 | 0 | 0.045 | 0.078 | 0.146 | -0.043 |
| A3SS | 10888 | ENSG00000205581 | HMGN1 | chr21 | - | 39345829 | 39347292 | 39345829 | 39345958 | 39347378 | 39347483 | 221 | 216 | 227 | 229 | 308 | 2 | 3 | 12 | 8 | 18 | 10 | 116 | 148 | 67 | 30 | 0 | 8 | 8 | 22 | 13 | 298 | 149 | 0.000182 | 0.003246832 | 0.982 | 0.973 | 0.904 | 0.935 | 0.895 | 1 | 0.879 | 0.902 | 0.604 | 0.536 | 0.154 |
| A3SS | 8247 | ENSG00000124151 | NCOA3 | chr20 | + | 47634017 | 47634195 | 47634047 | 47634195 | 47633495 | 47633636 | 17 | 10 | 30 | 29 | 24 | 187 | 101 | 190 | 147 | 415 | 11 | 46 | 27 | 41 | 24 | 68 | 192 | 154 | 106 | 13 | 179 | 149 | 0.000183 | 0.003250846 | 0.07 | 0.076 | 0.116 | 0.141 | 0.046 | 0.119 | 0.166 | 0.127 | 0.244 | 0.606 | -0.163 |
| A3SS | 7569 | ENSG00000185619 | PCGF3 | chr4 | + | 730951 | 731110 | 730969 | 731110 | 730629 | 730668 | 6 | 1 | 0 | 0 | 5 | 18 | 14 | 18 | 23 | 10 | 0 | 0 | 0 | 0 | 0 | 35 | 31 | 11 | 55 | 26 | 167 | 149 | 0.000183 | 0.003250846 | 0.229 | 0.06 | 0 | 0 | 0.308 | 0 | 0 | 0 | 0 | 0 | 0.119 |
| A3SS | 2552 | ENSG00000197102 | DYNC1H1 | chr14 | + | 101979648 | 101979974 | 101979718 | 101979974 | 101979318 | 101979492 | 21 | 16 | 13 | 20 | 29 | 193 | 115 | 169 | 167 | 168 | 84 | 26 | 52 | 52 | 30 | 322 | 212 | 136 | 214 | 266 | 219 | 149 | 0.000190 | 0.003364936 | 0.069 | 0.086 | 0.05 | 0.075 | 0.105 | 0.151 | 0.077 | 0.206 | 0.142 | 0.071 | -0.052 |
| A3SS | 11914 | ENSG00000205758 | CRYZL1 | chr21 | - | 33599149 | 33599285 | 33599149 | 33599248 | 33602233 | 33602345 | 19 | 23 | 17 | 25 | 16 | 46 | 27 | 53 | 36 | 56 | 0 | 6 | 3 | 10 | 2 | 8 | 52 | 20 | 32 | 8 | 186 | 149 | 0.000191 | 0.003364936 | 0.249 | 0.406 | 0.204 | 0.357 | 0.186 | 0 | 0.085 | 0.107 | 0.2 | 0.167 | 0.169 |
| A3SS | 3621 | ENSG00000125089 | SH3TC1 | chr4 | + | 8236277 | 8237670 | 8237473 | 8237670 | 8235432 | 8235555 | 50 | 27 | 27 | 32 | 19 | 5 | 1 | 0 | 6 | 1 | 39 | 37 | 14 | 29 | 44 | 5 | 22 | 18 | 13 | 4 | 298 | 149 | 0.000193 | 0.003399146 | 0.833 | 0.931 | 1 | 0.727 | 0.905 | 0.796 | 0.457 | 0.28 | 0.527 | 0.846 | 0.298 |
| A3SS | 10582 | ENSG00000123562 | MORF4L2 | chrX | - | 103678498 | 103678620 | 103678498 | 103678600 | 103685170 | 103685260 | 23 | 16 | 32 | 30 | 27 | 177 | 93 | 131 | 133 | 227 | 32 | 8 | 11 | 3 | 0 | 234 | 93 | 86 | 105 | 69 | 169 | 149 | 0.000195 | 0.003428464 | 0.103 | 0.132 | 0.177 | 0.166 | 0.095 | 0.108 | 0.07 | 0.101 | 0.025 | 0 | 0.074 |
| A3SS | 7796 | ENSG00000122507 | BBS9 | chr7 | + | 33351026 | 33351323 | 33351218 | 33351323 | 33349067 | 33349170 | 11 | 8 | 3 | 15 | 6 | 68 | 47 | 59 | 78 | 145 | 0 | 4 | 1 | 0 | 0 | 17 | 30 | 24 | 32 | 10 | 298 | 149 | 0.000195 | 0.003428464 | 0.075 | 0.078 | 0.025 | 0.088 | 0.02 | 0 | 0.062 | 0.02 | 0 | 0 | 0.041 |
| A3SS | 10518 | ENSG00000102178 | UBL4A | chrX | - | 154485770 | 154486083 | 154485770 | 154485979 | 154486227 | 154486333 | 2 | 0 | 0 | 1 | 1 | 14 | 9 | 10 | 23 | 1 | 0 | 8 | 6 | 3 | 3 | 1 | 20 | 14 | 4 | 44 | 253 | 149 | 0.000197 | 0.003442691 | 0.078 | 0 | 0 | 0.025 | 0.371 | 0 | 0.191 | 0.202 | 0.306 | 0.039 | -0.053 |
| A3SS | 10479 | ENSG00000101199 | ARFGAP1 | chr20 | + | 63286365 | 63286442 | 63286371 | 63286442 | 63285653 | 63285713 | 61 | 32 | 48 | 59 | 27 | 1 | 2 | 0 | 0 | 0 | 102 | 54 | 31 | 57 | 87 | 0 | 0 | 7 | 0 | 13 | 155 | 149 | 0.000201 | 0.003504904 | 0.983 | 0.939 | 1 | 1 | 1 | 1 | 1 | 0.81 | 1 | 0.865 | 0.049 |
| A3SS | 12054 | ENSG00000135365 | PHF21A | chr11 | - | 46076753 | 46076840 | 46076753 | 46076819 | 46079133 | 46079166 | 30 | 21 | 21 | 10 | 26 | 239 | 218 | 175 | 249 | 584 | 12 | 24 | 18 | 14 | 7 | 461 | 299 | 293 | 447 | 343 | 170 | 149 | 0.000203 | 0.003532823 | 0.099 | 0.078 | 0.095 | 0.034 | 0.038 | 0.022 | 0.066 | 0.051 | 0.027 | 0.018 | 0.032 |
| A3SS | 9678 | ENSG00000169660 | HEXD | chr17 | + | 82424393 | 82424503 | 82424418 | 82424503 | 82419748 | 82419883 | 78 | 53 | 86 | 83 | 143 | 4 | 3 | 9 | 5 | 7 | 89 | 47 | 28 | 38 | 73 | 0 | 2 | 0 | 4 | 0 | 174 | 149 | 0.000204 | 0.003544167 | 0.943 | 0.938 | 0.891 | 0.934 | 0.946 | 1 | 0.953 | 1 | 0.891 | 1 | -0.038 |
| A3SS | 8903 | ENSG00000143499 | SMYD2 | chr1 | + | 214330949 | 214334308 | 214334199 | 214334308 | 214330167 | 214330278 | 82 | 48 | 96 | 84 | 115 | 3 | 1 | 1 | 2 | 0 | 56 | 30 | 40 | 53 | 71 | 26 | 4 | 2 | 0 | 0 | 298 | 149 | 0.000211 | 0.00366794 | 0.932 | 0.96 | 0.98 | 0.955 | 1 | 0.519 | 0.789 | 0.909 | 1 | 1 | 0.122 |
| A3SS | 4455 | ENSG00000114867 | EIF4G1 | chr3 | + | 184320842 | 184320993 | 184320926 | 184320993 | 184320629 | 184320722 | 12 | 3 | 13 | 15 | 2 | 168 | 106 | 127 | 190 | 181 | 11 | 27 | 7 | 34 | 43 | 270 | 215 | 120 | 188 | 302 | 233 | 149 | 0.000213 | 0.00368125 | 0.044 | 0.018 | 0.061 | 0.048 | 0.007 | 0.025 | 0.074 | 0.036 | 0.104 | 0.083 | -0.029 |
| A3SS | 10575 | ENSG00000035681 | NSMAF | chr8 | - | 58589451 | 58589575 | 58589451 | 58589558 | 58590036 | 58590074 | 216 | 145 | 207 | 180 | 250 | 2 | 2 | 3 | 3 | 3 | 277 | 150 | 83 | 105 | 95 | 7 | 4 | 10 | 8 | 0 | 166 | 149 | 0.000215 | 0.00370923 | 0.99 | 0.985 | 0.984 | 0.982 | 0.987 | 0.973 | 0.971 | 0.882 | 0.922 | 1 | 0.036 |
| A3SS | 11611 | ENSG00000172716 | SLFN11 | chr17 | - | 35366946 | 35367063 | 35366946 | 35367043 | 35373581 | 35373620 | 23 | 21 | 7 | 10 | 15 | 0 | 0 | 0 | 0 | 0 | 25 | 12 | 17 | 15 | 0 | 14 | 0 | 1 | 0 | 7 | 169 | 149 | 0.000229 | 0.003945179 | 1 | 1 | 1 | 1 | 1 | 0.612 | 1 | 0.937 | 1 | 0 | 0.29 |
| A3SS | 8355 | ENSG00000135838 | NPL | chr1 | + | 182828723 | 182830384 | 182829602 | 182830384 | 182825780 | 182825820 | 149 | 72 | 105 | 132 | 202 | 0 | 3 | 0 | 2 | 2 | 171 | 127 | 106 | 211 | 59 | 0 | 3 | 13 | 8 | 0 | 298 | 149 | 0.000232 | 0.003993674 | 1 | 0.923 | 1 | 0.971 | 0.981 | 1 | 0.955 | 0.803 | 0.93 | 1 | 0.037 |
| A3SS | 3887 | ENSG00000101109 | STK4 | chr20 | + | 44973206 | 44973433 | 44973300 | 44973433 | 44972077 | 44972158 | 62 | 44 | 48 | 59 | 104 | 4 | 1 | 4 | 2 | 17 | 27 | 29 | 67 | 19 | 3 | 0 | 0 | 5 | 0 | 0 | 243 | 149 | 0.000236 | 0.004060985 | 0.905 | 0.964 | 0.88 | 0.948 | 0.79 | 1 | 1 | 0.891 | 1 | 1 | -0.081 |
| A3SS | 4823 | ENSG00000197386 | HTT | chr4 | + | 3213916 | 3214135 | 3213957 | 3214135 | 3212563 | 3212709 | 2 | 0 | 8 | 3 | 4 | 152 | 72 | 101 | 116 | 118 | 12 | 14 | 7 | 11 | 18 | 176 | 198 | 90 | 127 | 291 | 190 | 149 | 0.000238 | 0.004084763 | 0.01 | 0 | 0.058 | 0.02 | 0.026 | 0.051 | 0.053 | 0.057 | 0.064 | 0.046 | -0.031 |
| A3SS | 13083 | ENSG00000134285 | FKBP11 | chr12 | - | 48923781 | 48923974 | 48923781 | 48923852 | 48924222 | 48924256 | 28 | 18 | 31 | 30 | 26 | 126 | 37 | 128 | 87 | 121 | 32 | 41 | 37 | 27 | 98 | 26 | 115 | 110 | 59 | 115 | 271 | 149 | 0.000242 | 0.004144937 | 0.109 | 0.211 | 0.118 | 0.159 | 0.106 | 0.404 | 0.164 | 0.156 | 0.201 | 0.319 | -0.108 |
| A3SS | 9549 | ENSG00000214765 | SEPTIN7P2 | chr7 | - | 45724352 | 45725715 | 45724352 | 45724480 | 45726183 | 45726236 | 11 | 8 | 11 | 16 | 10 | 4 | 7 | 3 | 9 | 10 | 0 | 7 | 12 | 22 | 4 | 9 | 0 | 0 | 0 | 0 | 298 | 149 | 0.000243 | 0.004152576 | 0.579 | 0.364 | 0.647 | 0.471 | 0.333 | 0 | 1 | 1 | 1 | 1 | -0.321 |
| A3SS | 7426 | ENSG00000103168 | TAF1C | chr16 | - | 84184850 | 84185060 | 84184850 | 84184899 | 84186900 | 84187057 | 43 | 9 | 23 | 25 | 11 | 5 | 2 | 8 | 7 | 0 | 5 | 44 | 19 | 19 | 12 | 9 | 8 | 23 | 15 | 22 | 298 | 149 | 0.000245 | 0.004178353 | 0.811 | 0.692 | 0.59 | 0.641 | 1 | 0.217 | 0.733 | 0.292 | 0.388 | 0.214 | 0.378 |
| A3SS | 4841 | ENSG00000064961 | HMG20B | chr19 | + | 3575501 | 3575660 | 3575539 | 3575660 | 3574369 | 3574586 | 1 | 0 | 0 | 1 | 0 | 43 | 31 | 35 | 58 | 12 | 5 | 4 | 2 | 1 | 12 | 84 | 53 | 36 | 64 | 178 | 187 | 149 | 0.000256 | 0.004353133 | 0.018 | 0 | 0 | 0.014 | 0 | 0.045 | 0.057 | 0.042 | 0.012 | 0.051 | -0.035 |
| A3SS | 12516 | ENSG00000175470 | PPP2R2D | chr10 | + | 131943967 | 131944145 | 131944094 | 131944145 | 131940030 | 131940196 | 165 | 146 | 119 | 98 | 248 | 0 | 0 | 0 | 0 | 0 | 131 | 90 | 91 | 110 | 105 | 16 | 0 | 1 | 0 | 0 | 276 | 149 | 0.000258 | 0.004379656 | 1 | 1 | 1 | 1 | 1 | 0.816 | 1 | 0.98 | 1 | 1 | 0.041 |
| A3SS | 8874 | ENSG00000116793 | PHTF1 | chr1 | - | 113726417 | 113726574 | 113726417 | 113726458 | 113738109 | 113738268 | 52 | 63 | 37 | 51 | 63 | 2 | 2 | 4 | 2 | 0 | 64 | 127 | 93 | 89 | 107 | 0 | 17 | 22 | 23 | 2 | 265 | 149 | 0.000260 | 0.004392755 | 0.936 | 0.947 | 0.839 | 0.935 | 1 | 1 | 0.808 | 0.704 | 0.685 | 0.968 | 0.098 |
| A3SS | 8686 | ENSG00000064012 | CASP8 | chr2 | + | 201266460 | 201266791 | 201266486 | 201266791 | 201234015 | 201234112 | 194 | 83 | 156 | 153 | 273 | 8 | 8 | 16 | 9 | 2 | 118 | 197 | 227 | 174 | 92 | 0 | 3 | 7 | 2 | 4 | 175 | 149 | 0.000261 | 0.004403286 | 0.954 | 0.898 | 0.892 | 0.935 | 0.991 | 1 | 0.982 | 0.965 | 0.987 | 0.951 | -0.043 |
| A3SS | 4637 | ENSG00000073111 | MCM2 | chr3 | + | 127604876 | 127605156 | 127604895 | 127605156 | 127604607 | 127604783 | 1 | 0 | 0 | 0 | 0 | 23 | 7 | 25 | 34 | 27 | 0 | 2 | 1 | 10 | 2 | 46 | 35 | 34 | 37 | 27 | 168 | 149 | 0.000263 | 0.004439447 | 0.037 | 0 | 0 | 0 | 0 | 0 | 0.048 | 0.025 | 0.193 | 0.062 | -0.058 |
| A3SS | 7529 | ENSG00000237298 | TTN-AS1 | chr2 | + | 178600475 | 178600667 | 178600557 | 178600667 | 178599593 | 178599750 | 7 | 10 | 4 | 12 | 30 | 13 | 4 | 8 | 12 | 10 | 4 | 6 | 1 | 13 | 10 | 129 | 6 | 24 | 13 | 21 | 231 | 149 | 0.000265 | 0.004449854 | 0.258 | 0.617 | 0.244 | 0.392 | 0.659 | 0.02 | 0.392 | 0.026 | 0.392 | 0.235 | 0.221 |
| A3SS | 7667 | ENSG00000106992 | AK1 | chr9 | - | 127873025 | 127873180 | 127873025 | 127873061 | 127874610 | 127874649 | 1 | 0 | 0 | 0 | 1 | 44 | 21 | 28 | 31 | 22 | 28 | 0 | 0 | 2 | 0 | 26 | 37 | 40 | 34 | 33 | 268 | 149 | 0.000269 | 0.004510794 | 0.012 | 0 | 0 | 0 | 0.025 | 0.375 | 0 | 0 | 0.032 | 0 | -0.074 |
| A3SS | 11312 | ENSG00000132334 | PTPRE | chr10 | + | 128047763 | 128047837 | 128047769 | 128047837 | 128047630 | 128047683 | 169 | 93 | 82 | 122 | 49 | 1 | 2 | 1 | 4 | 5 | 161 | 88 | 150 | 245 | 73 | 28 | 8 | 10 | 2 | 0 | 155 | 149 | 0.000271 | 0.004531475 | 0.994 | 0.978 | 0.987 | 0.967 | 0.904 | 0.847 | 0.914 | 0.935 | 0.992 | 1 | 0.028 |
| A3SS | 1922 | ENSG00000157353 | FCSK | chr16 | + | 70478796 | 70479403 | 70479179 | 70479403 | 70478550 | 70478650 | 1 | 0 | 0 | 2 | 3 | 9 | 4 | 6 | 5 | 2 | 4 | 7 | 0 | 10 | 6 | 0 | 2 | 2 | 11 | 3 | 298 | 149 | 0.000272 | 0.0045375 | 0.053 | 0 | 0 | 0.167 | 0.429 | 1 | 0.636 | 0 | 0.312 | 0.5 | -0.36 |
| A3SS | 11590 | ENSG00000213930 | GALT | chr9 | + | 34647474 | 34647567 | 34647540 | 34647567 | 34647088 | 34647258 | 41 | 29 | 43 | 34 | 41 | 3 | 0 | 5 | 4 | 1 | 16 | 37 | 25 | 41 | 40 | 0 | 0 | 2 | 0 | 0 | 215 | 149 | 0.000275 | 0.004578257 | 0.905 | 1 | 0.856 | 0.855 | 0.966 | 1 | 1 | 0.897 | 1 | 1 | -0.063 |
| A3SS | 6843 | ENSG00000151694 | ADAM17 | chr2 | - | 9543152 | 9543285 | 9543152 | 9543226 | 9543695 | 9543760 | 74 | 79 | 41 | 61 | 19 | 1 | 0 | 5 | 1 | 0 | 160 | 148 | 119 | 172 | 29 | 0 | 15 | 5 | 15 | 7 | 208 | 149 | 0.000278 | 0.004627557 | 0.981 | 1 | 0.855 | 0.978 | 1 | 1 | 0.876 | 0.945 | 0.891 | 0.748 | 0.071 |
| A3SS | 1124 | ENSG00000170004 | CHD3 | chr17 | + | 7907890 | 7908019 | 7907893 | 7908019 | 7907352 | 7907488 | 7 | 4 | 15 | 13 | 3 | 63 | 54 | 100 | 72 | 75 | 0 | 0 | 6 | 6 | 5 | 59 | 61 | 87 | 91 | 88 | 152 | 149 | 0.000280 | 0.004657605 | 0.098 | 0.068 | 0.128 | 0.15 | 0.038 | 0 | 0 | 0.063 | 0.061 | 0.053 | 0.061 |
| A3SS | 6324 | ENSG00000137575 | SDCBP | chr8 | + | 58575899 | 58576061 | 58575902 | 58576061 | 58572204 | 58572314 | 571 | 414 | 392 | 423 | 529 | 202 | 118 | 127 | 171 | 255 | 49 | 254 | 510 | 160 | 96 | 11 | 62 | 152 | 36 | 18 | 152 | 149 | 0.000287 | 0.004751759 | 0.735 | 0.775 | 0.752 | 0.708 | 0.67 | 0.814 | 0.801 | 0.767 | 0.813 | 0.839 | -0.079 |
| A3SS | 7991 | ENSG00000152689 | RASGRP3 | chr2 | + | 33539093 | 33539210 | 33539096 | 33539210 | 33534322 | 33534400 | 17 | 15 | 22 | 31 | 47 | 0 | 0 | 4 | 3 | 7 | 25 | 49 | 17 | 20 | 11 | 14 | 25 | 1 | 12 | 1 | 152 | 149 | 0.000287 | 0.00475504 | 1 | 1 | 0.844 | 0.91 | 0.868 | 0.636 | 0.658 | 0.943 | 0.62 | 0.915 | 0.17 |
| A3SS | 3638 | ENSG00000026297 | RNASET2 | chr6 | - | 166943018 | 166943809 | 166943018 | 166943089 | 166946681 | 166946739 | 116 | 131 | 101 | 186 | 121 | 375 | 207 | 275 | 455 | 625 | 341 | 181 | 131 | 173 | 258 | 630 | 272 | 245 | 418 | 573 | 298 | 149 | 0.000288 | 0.0047602 | 0.134 | 0.24 | 0.155 | 0.17 | 0.088 | 0.213 | 0.25 | 0.211 | 0.171 | 0.184 | -0.048 |
| A3SS | 3520 | ENSG00000140350 | ANP32A | chr15 | - | 68787412 | 68787535 | 68787412 | 68787495 | 68787769 | 68787919 | 1427 | 1101 | 1239 | 1387 | 2164 | 14 | 6 | 21 | 10 | 8 | 2766 | 1433 | 1280 | 2113 | 1117 | 34 | 33 | 5 | 25 | 31 | 189 | 149 | 0.000293 | 0.004827055 | 0.988 | 0.993 | 0.979 | 0.991 | 0.995 | 0.985 | 0.972 | 0.995 | 0.985 | 0.966 | 0.009 |
| A3SS | 8560 | ENSG00000115282 | TTC31 | chr2 | + | 74490655 | 74491184 | 74491127 | 74491184 | 74490243 | 74490473 | 90 | 64 | 79 | 99 | 108 | 1 | 0 | 3 | 5 | 0 | 65 | 73 | 93 | 106 | 148 | 0 | 4 | 10 | 6 | 10 | 298 | 149 | 0.000294 | 0.004839567 | 0.978 | 1 | 0.929 | 0.908 | 1 | 1 | 0.901 | 0.823 | 0.898 | 0.881 | 0.062 |
| A3SS | 5655 | ENSG00000203667 | COX20 | chr1 | + | 244842194 | 244842258 | 244842198 | 244842258 | 244841943 | 244842058 | 160 | 118 | 137 | 93 | 205 | 13 | 4 | 3 | 6 | 2 | 70 | 97 | 131 | 137 | 89 | 0 | 0 | 0 | 0 | 6 | 153 | 149 | 0.000297 | 0.004866843 | 0.923 | 0.966 | 0.978 | 0.938 | 0.99 | 1 | 1 | 1 | 1 | 0.935 | -0.028 |
| A3SS | 11076 | ENSG00000079459 | FDFT1 | chr8 | + | 11826023 | 11826215 | 11826078 | 11826215 | 11821749 | 11821878 | 505 | 235 | 408 | 389 | 530 | 3 | 10 | 12 | 11 | 10 | 344 | 337 | 264 | 472 | 455 | 14 | 9 | 13 | 25 | 22 | 204 | 149 | 0.000298 | 0.004881526 | 0.992 | 0.945 | 0.961 | 0.963 | 0.975 | 0.947 | 0.965 | 0.937 | 0.932 | 0.938 | 0.023 |
| A3SS | 2386 | ENSG00000140400 | MAN2C1 | chr15 | - | 75367510 | 75367645 | 75367510 | 75367634 | 75368072 | 75368198 | 2 | 2 | 5 | 0 | 2 | 132 | 35 | 123 | 92 | 28 | 7 | 6 | 16 | 0 | 16 | 248 | 112 | 112 | 81 | 104 | 160 | 149 | 0.000300 | 0.004894128 | 0.014 | 0.051 | 0.036 | 0 | 0.062 | 0.026 | 0.048 | 0.117 | 0 | 0.125 | -0.031 |
| A3SS | 10689 | ENSG00000102312 | PORCN | chrX | + | 48512834 | 48514365 | 48514239 | 48514365 | 48512588 | 48512719 | 2 | 1 | 1 | 1 | 0 | 6 | 2 | 3 | 8 | 7 | 8 | 4 | 2 | 0 | 5 | 0 | 7 | 1 | 3 | 4 | 298 | 149 | 0.000301 | 0.004905954 | 0.143 | 0.2 | 0.143 | 0.059 | 0 | 1 | 0.222 | 0.5 | 0 | 0.385 | -0.312 |
| A3SS | 3661 | ENSG00000077232 | DNAJC10 | chr2 | + | 182739544 | 182740388 | 182740298 | 182740388 | 182736248 | 182736386 | 59 | 60 | 80 | 48 | 133 | 89 | 74 | 71 | 89 | 184 | 64 | 37 | 27 | 26 | 12 | 4 | 39 | 30 | 4 | 0 | 298 | 149 | 0.000305 | 0.004964156 | 0.249 | 0.288 | 0.36 | 0.212 | 0.265 | 0.889 | 0.322 | 0.31 | 0.765 | 1 | -0.382 |
| A3SS | 2909 | ENSG00000175061 | SNHG29 | chr17 | + | 16440110 | 16440253 | 16440184 | 16440253 | 16439527 | 16439703 | 14 | 14 | 14 | 22 | 36 | 132 | 65 | 180 | 131 | 95 | 32 | 18 | 26 | 75 | 60 | 72 | 114 | 177 | 90 | 111 | 223 | 149 | 0.000309 | 0.005011645 | 0.066 | 0.126 | 0.049 | 0.101 | 0.202 | 0.229 | 0.095 | 0.089 | 0.358 | 0.265 | -0.098 |
| A3SS | 9790 | ENSG00000122515 | ZMIZ2 | chr7 | + | 44756974 | 44757149 | 44757042 | 44757149 | 44756424 | 44756539 | 15 | 3 | 10 | 10 | 5 | 5 | 2 | 3 | 6 | 0 | 10 | 20 | 0 | 0 | 16 | 26 | 12 | 4 | 7 | 20 | 217 | 149 | 0.000310 | 0.005011645 | 0.673 | 0.507 | 0.696 | 0.534 | 1 | 0.209 | 0.534 | 0 | 0 | 0.355 | 0.462 |
| A3SS | 4060 | ENSG00000205726 | ITSN1 | chr21 | + | 33750142 | 33750322 | 33750253 | 33750322 | 33735043 | 33735204 | 14 | 14 | 8 | 12 | 14 | 0 | 0 | 0 | 0 | 3 | 16 | 24 | 9 | 12 | 0 | 0 | 0 | 2 | 2 | 12 | 260 | 149 | 0.000310 | 0.005011645 | 1 | 1 | 1 | 1 | 0.728 | 1 | 1 | 0.721 | 0.775 | 0 | 0.246 |
| A3SS | 5243 | ENSG00000183258 | DDX41 | chr5 | - | 177516287 | 177516516 | 177516287 | 177516447 | 177516724 | 177516835 | 10 | 3 | 2 | 5 | 0 | 142 | 75 | 90 | 101 | 65 | 20 | 11 | 3 | 5 | 18 | 97 | 126 | 65 | 105 | 189 | 218 | 149 | 0.000312 | 0.005026611 | 0.046 | 0.027 | 0.015 | 0.033 | 0 | 0.124 | 0.056 | 0.031 | 0.032 | 0.061 | -0.037 |
| A3SS | 11929 | ENSG00000126705 | AHDC1 | chr1 | - | 27553073 | 27553304 | 27553073 | 27553185 | 27558705 | 27558883 | 11 | 3 | 5 | 6 | 4 | 0 | 0 | 1 | 1 | 0 | 7 | 10 | 14 | 4 | 16 | 0 | 0 | 2 | 3 | 5 | 268 | 149 | 0.000312 | 0.005026611 | 1 | 1 | 0.735 | 0.769 | 1 | 1 | 1 | 0.796 | 0.426 | 0.64 | 0.128 |
| A3SS | 11542 | ENSG00000157693 | TMEM268 | chr9 | + | 114626895 | 114627006 | 114626898 | 114627006 | 114624349 | 114624459 | 1 | 3 | 5 | 1 | 2 | 33 | 18 | 22 | 26 | 15 | 0 | 1 | 0 | 0 | 0 | 57 | 6 | 16 | 26 | 32 | 152 | 149 | 0.000313 | 0.005033993 | 0.029 | 0.14 | 0.182 | 0.036 | 0.116 | 0 | 0.14 | 0 | 0 | 0 | 0.073 |
| A3SS | 11447 | ENSG00000132740 | IGHMBP2 | chr11 | + | 68929070 | 68929357 | 68929182 | 68929357 | 68917735 | 68917883 | 1 | 2 | 0 | 1 | 4 | 49 | 37 | 15 | 29 | 45 | 5 | 5 | 0 | 8 | 6 | 6 | 17 | 32 | 34 | 59 | 261 | 149 | 0.000327 | 0.005244046 | 0.012 | 0.03 | 0 | 0.019 | 0.048 | 0.322 | 0.144 | 0 | 0.118 | 0.055 | -0.106 |
| A3SS | 13145 | ENSG00000123349 | PFDN5 | chr12 | + | 53295838 | 53296275 | 53296218 | 53296275 | 53295524 | 53295639 | 995 | 597 | 863 | 853 | 686 | 6 | 8 | 10 | 12 | 2 | 1158 | 749 | 634 | 705 | 1203 | 36 | 8 | 9 | 6 | 26 | 298 | 149 | 0.000330 | 0.005285138 | 0.988 | 0.974 | 0.977 | 0.973 | 0.994 | 0.941 | 0.979 | 0.972 | 0.983 | 0.959 | 0.014 |
| A3SS | 681 | ENSG00000105397 | TYK2 | chr19 | - | 10368294 | 10368418 | 10368294 | 10368371 | 10378213 | 10378508 | 161 | 77 | 148 | 139 | 111 | 0 | 1 | 0 | 1 | 0 | 307 | 195 | 109 | 201 | 443 | 13 | 11 | 1 | 0 | 5 | 196 | 149 | 0.000337 | 0.005383256 | 1 | 0.983 | 1 | 0.991 | 1 | 0.947 | 0.931 | 0.988 | 1 | 0.985 | 0.025 |
| A3SS | 4957 | ENSG00000158864 | NDUFS2 | chr1 | + | 161209842 | 161209931 | 161209856 | 161209931 | 161209482 | 161209595 | 24 | 15 | 10 | 11 | 17 | 192 | 106 | 210 | 180 | 240 | 74 | 15 | 22 | 16 | 57 | 178 | 242 | 149 | 186 | 308 | 163 | 149 | 0.000340 | 0.005417856 | 0.103 | 0.115 | 0.042 | 0.053 | 0.061 | 0.275 | 0.054 | 0.119 | 0.073 | 0.145 | -0.058 |
| A3SS | 8330 | ENSG00000112130 | RNF8 | chr6 | + | 37368377 | 37369218 | 37368483 | 37369218 | 37360445 | 37360574 | 9 | 13 | 6 | 14 | 10 | 69 | 43 | 43 | 56 | 52 | 23 | 41 | 5 | 46 | 31 | 47 | 53 | 63 | 71 | 86 | 255 | 149 | 0.000341 | 0.005432555 | 0.071 | 0.15 | 0.075 | 0.127 | 0.101 | 0.222 | 0.311 | 0.044 | 0.275 | 0.174 | -0.1 |
| A3SS | 9780 | ENSG00000198198 | SZT2 | chr1 | + | 43426367 | 43426538 | 43426437 | 43426538 | 43426037 | 43426151 | 25 | 5 | 19 | 22 | 10 | 5 | 5 | 7 | 6 | 14 | 8 | 30 | 10 | 21 | 23 | 28 | 27 | 24 | 34 | 39 | 219 | 149 | 0.000343 | 0.005442426 | 0.773 | 0.405 | 0.649 | 0.714 | 0.327 | 0.163 | 0.431 | 0.221 | 0.296 | 0.286 | 0.294 |
| A3SS | 11901 | ENSG00000149547 | EI24 | chr11 | + | 125575250 | 125575408 | 125575304 | 125575408 | 125572457 | 125572569 | 38 | 15 | 25 | 51 | 14 | 0 | 2 | 8 | 3 | 0 | 12 | 30 | 18 | 37 | 44 | 0 | 0 | 0 | 4 | 0 | 203 | 149 | 0.000344 | 0.005462931 | 1 | 0.846 | 0.696 | 0.926 | 1 | 1 | 1 | 1 | 0.872 | 1 | -0.081 |
| A3SS | 13033 | ENSG00000139178 | C1RL | chr12 | - | 7099685 | 7099807 | 7099685 | 7099760 | 7099900 | 7100026 | 5 | 2 | 2 | 10 | 6 | 52 | 25 | 28 | 27 | 49 | 53 | 7 | 24 | 27 | 11 | 84 | 40 | 52 | 108 | 63 | 196 | 149 | 0.000351 | 0.00554931 | 0.068 | 0.057 | 0.052 | 0.22 | 0.085 | 0.324 | 0.117 | 0.26 | 0.16 | 0.117 | -0.099 |
| A3SS | 7930 | ENSG00000167220 | HDHD2 | chr18 | - | 47134495 | 47134704 | 47134495 | 47134677 | 47136338 | 47136449 | 64 | 51 | 42 | 53 | 92 | 3 | 3 | 7 | 4 | 4 | 19 | 45 | 37 | 28 | 16 | 0 | 4 | 0 | 0 | 0 | 176 | 149 | 0.000353 | 0.005579519 | 0.948 | 0.935 | 0.836 | 0.918 | 0.951 | 1 | 0.905 | 1 | 1 | 1 | -0.063 |
| A3SS | 9669 | ENSG00000128534 | LSM8 | chr7 | + | 118185646 | 118185694 | 118185653 | 118185694 | 118184163 | 118184254 | 26 | 8 | 12 | 22 | 18 | 118 | 62 | 103 | 111 | 43 | 0 | 21 | 20 | 1 | 0 | 83 | 84 | 74 | 131 | 85 | 156 | 149 | 0.000360 | 0.00565874 | 0.174 | 0.11 | 0.1 | 0.159 | 0.286 | 0 | 0.193 | 0.205 | 0.007 | 0 | 0.085 |
| A3SS | 5299 | ENSG00000143110 | C1orf162 | chr1 | + | 111477650 | 111477775 | 111477725 | 111477775 | 111476797 | 111476867 | 42 | 30 | 30 | 44 | 39 | 2 | 2 | 1 | 13 | 5 | 120 | 49 | 46 | 74 | 126 | 7 | 1 | 0 | 5 | 0 | 224 | 149 | 0.000361 | 0.00565874 | 0.933 | 0.909 | 0.952 | 0.692 | 0.838 | 0.919 | 0.97 | 1 | 0.908 | 1 | -0.095 |
| A3SS | 4202 | ENSG00000114745 | GORASP1 | chr3 | - | 39100304 | 39100503 | 39100304 | 39100454 | 39101015 | 39101102 | 7 | 8 | 5 | 10 | 6 | 0 | 0 | 0 | 1 | 0 | 27 | 15 | 10 | 10 | 40 | 0 | 9 | 2 | 1 | 0 | 198 | 149 | 0.000361 | 0.00565874 | 1 | 1 | 1 | 0.883 | 1 | 1 | 0.556 | 0.79 | 0.883 | 1 | 0.131 |
| A3SS | 10200 | ENSG00000091073 | DTX2 | chr7 | + | 76476942 | 76477178 | 76477113 | 76477178 | 76463544 | 76463709 | 0 | 2 | 1 | 2 | 4 | 1 | 1 | 0 | 1 | 0 | 2 | 5 | 0 | 0 | 10 | 12 | 0 | 5 | 1 | 0 | 298 | 149 | 0.000362 | 0.00565874 | 0 | 0.5 | 1 | 0.5 | 1 | 0.077 | 1 | 0 | 0 | 1 | 0.185 |
| A3SS | 1413 | ENSG00000178971 | CTC1 | chr17 | - | 8229890 | 8229968 | 8229890 | 8229922 | 8230293 | 8230468 | 71 | 56 | 74 | 91 | 101 | 1 | 5 | 3 | 3 | 1 | 44 | 40 | 28 | 68 | 118 | 11 | 5 | 2 | 3 | 9 | 195 | 149 | 0.000362 | 0.00565874 | 0.982 | 0.895 | 0.95 | 0.959 | 0.987 | 0.753 | 0.859 | 0.915 | 0.945 | 0.909 | 0.078 |
| A3SS | 9349 | ENSG00000170919 | TPT1-AS1 | chr13 | + | 45369958 | 45370182 | 45370079 | 45370182 | 45344328 | 45344446 | 32 | 12 | 27 | 31 | 12 | 3 | 1 | 2 | 1 | 3 | 43 | 45 | 35 | 35 | 28 | 0 | 0 | 1 | 0 | 0 | 270 | 149 | 0.000362 | 0.00565874 | 0.855 | 0.869 | 0.882 | 0.945 | 0.688 | 1 | 1 | 0.951 | 1 | 1 | -0.142 |
| A3SS | 6699 | ENSG00000143951 | WDPCP | chr2 | - | 63439756 | 63439871 | 63439756 | 63439855 | 63484603 | 63484663 | 26 | 21 | 38 | 26 | 30 | 3 | 0 | 0 | 1 | 1 | 5 | 4 | 29 | 14 | 1 | 0 | 11 | 2 | 5 | 0 | 165 | 149 | 0.000363 | 0.005661976 | 0.887 | 1 | 1 | 0.959 | 0.964 | 1 | 0.247 | 0.929 | 0.717 | 1 | 0.183 |
| A3SS | 12135 | ENSG00000132199 | ENOSF1 | chr18 | - | 690548 | 690745 | 690548 | 690631 | 691067 | 691106 | 13 | 1 | 20 | 15 | 7 | 8 | 12 | 33 | 16 | 39 | 4 | 0 | 8 | 1 | 0 | 25 | 21 | 13 | 19 | 12 | 263 | 149 | 0.000366 | 0.005696855 | 0.479 | 0.045 | 0.256 | 0.347 | 0.092 | 0.083 | 0 | 0.259 | 0.029 | 0 | 0.17 |
| A3SS | 11928 | ENSG00000126705 | AHDC1 | chr1 | - | 27553073 | 27553290 | 27553073 | 27553185 | 27558705 | 27558883 | 11 | 3 | 5 | 6 | 4 | 0 | 0 | 1 | 1 | 0 | 7 | 10 | 18 | 6 | 16 | 0 | 0 | 2 | 3 | 5 | 254 | 149 | 0.000369 | 0.005730741 | 1 | 1 | 0.746 | 0.779 | 1 | 1 | 1 | 0.841 | 0.54 | 0.652 | 0.098 |
| A3SS | 2867 | ENSG00000138081 | FBXO11 | chr2 | - | 47838858 | 47839003 | 47838858 | 47839000 | 47839418 | 47839500 | 168 | 165 | 166 | 188 | 234 | 5 | 5 | 7 | 6 | 28 | 142 | 138 | 168 | 112 | 59 | 0 | 1 | 1 | 5 | 7 | 152 | 149 | 0.000370 | 0.005730741 | 0.971 | 0.97 | 0.959 | 0.968 | 0.891 | 1 | 0.993 | 0.994 | 0.956 | 0.892 | -0.015 |
| A3SS | 9191 | ENSG00000204386 | NEU1 | chr6 | - | 31860041 | 31860264 | 31860041 | 31860178 | 31860438 | 31860621 | 106 | 46 | 85 | 127 | 143 | 0 | 0 | 2 | 2 | 0 | 209 | 125 | 79 | 145 | 166 | 5 | 5 | 1 | 1 | 10 | 235 | 149 | 0.000371 | 0.005742833 | 1 | 1 | 0.964 | 0.976 | 1 | 0.964 | 0.941 | 0.98 | 0.989 | 0.913 | 0.031 |
| A3SS | 4317 | ENSG00000172053 | QARS1 | chr3 | - | 49103862 | 49104098 | 49103862 | 49103972 | 49104323 | 49104471 | 11 | 3 | 8 | 22 | 0 | 156 | 81 | 149 | 194 | 86 | 0 | 11 | 1 | 5 | 9 | 199 | 182 | 139 | 105 | 356 | 275 | 149 | 0.000374 | 0.00577324 | 0.037 | 0.02 | 0.028 | 0.058 | 0 | 0 | 0.032 | 0.004 | 0.025 | 0.014 | 0.014 |
| A3SS | 6162 | ENSG00000142864 | SERBP1 | chr1 | - | 67424199 | 67424322 | 67424199 | 67424277 | 67424887 | 67424959 | 326 | 309 | 278 | 366 | 491 | 804 | 712 | 787 | 901 | 1742 | 520 | 204 | 266 | 334 | 123 | 1238 | 1118 | 1339 | 1107 | 403 | 194 | 149 | 0.000376 | 0.005788722 | 0.237 | 0.25 | 0.213 | 0.238 | 0.178 | 0.244 | 0.123 | 0.132 | 0.188 | 0.19 | 0.048 |
| A3SS | 10440 | ENSG00000140332 | TLE3 | chr15 | - | 70066013 | 70066218 | 70066013 | 70066215 | 70074532 | 70074607 | 108 | 55 | 117 | 127 | 194 | 13 | 7 | 8 | 1 | 6 | 183 | 107 | 160 | 165 | 241 | 36 | 20 | 19 | 12 | 18 | 152 | 149 | 0.000377 | 0.005788722 | 0.891 | 0.885 | 0.935 | 0.992 | 0.96 | 0.833 | 0.84 | 0.892 | 0.931 | 0.929 | 0.048 |
| A3SS | 5102 | ENSG00000085433 | WDR47 | chr1 | - | 108982608 | 108982785 | 108982608 | 108982779 | 108983281 | 108983451 | 10 | 3 | 2 | 3 | 1 | 80 | 50 | 38 | 78 | 69 | 0 | 3 | 0 | 0 | 0 | 24 | 70 | 44 | 53 | 10 | 155 | 149 | 0.000378 | 0.005788722 | 0.107 | 0.055 | 0.048 | 0.036 | 0.014 | 0 | 0.04 | 0 | 0 | 0 | 0.044 |
| A3SS | 2609 | ENSG00000092199 | HNRNPC | chr14 | - | 21263310 | 21263582 | 21263310 | 21263336 | 21263666 | 21263829 | 230 | 239 | 208 | 283 | 349 | 11 | 7 | 11 | 15 | 14 | 334 | 381 | 496 | 437 | 63 | 0 | 11 | 6 | 15 | 3 | 298 | 149 | 0.000378 | 0.005788722 | 0.913 | 0.945 | 0.904 | 0.904 | 0.926 | 1 | 0.945 | 0.976 | 0.936 | 0.913 | -0.036 |
| A3SS | 4056 | ENSG00000119922 | IFIT2 | chr10 | + | 89305936 | 89309271 | 89305961 | 89309271 | 89302045 | 89302128 | 0 | 0 | 0 | 2 | 8 | 63 | 68 | 39 | 121 | 44 | 23 | 0 | 1 | 12 | 0 | 28 | 53 | 53 | 94 | 6 | 174 | 149 | 0.000388 | 0.005933546 | 0 | 0 | 0 | 0.014 | 0.135 | 0.413 | 0 | 0.016 | 0.099 | 0 | -0.076 |
| A3SS | 11958 | ENSG00000168071 | CCDC88B | chr11 | + | 64342501 | 64342680 | 64342521 | 64342680 | 64342293 | 64342375 | 5 | 3 | 1 | 4 | 0 | 37 | 27 | 35 | 32 | 13 | 0 | 0 | 0 | 0 | 0 | 20 | 18 | 14 | 29 | 57 | 169 | 149 | 0.000391 | 0.005964154 | 0.106 | 0.089 | 0.025 | 0.099 | 0 | 0 | 0 | 0 | 0 | 0 | 0.064 |
| A3SS | 9830 | ENSG00000142303 | ADAMTS10 | chr19 | - | 8597233 | 8597322 | 8597233 | 8597317 | 8600927 | 8601145 | 32 | 7 | 23 | 39 | 42 | 14 | 3 | 9 | 9 | 15 | 0 | 7 | 4 | 4 | 3 | 12 | 2 | 4 | 9 | 6 | 154 | 149 | 0.000392 | 0.005969291 | 0.689 | 0.693 | 0.712 | 0.807 | 0.73 | 0 | 0.772 | 0.492 | 0.301 | 0.326 | 0.348 |
| A3SS | 3292 | ENSG00000161960 | EIF4A1 | chr17 | + | 7578119 | 7578244 | 7578164 | 7578244 | 7577826 | 7577916 | 47 | 35 | 50 | 59 | 54 | 475 | 269 | 429 | 485 | 715 | 108 | 97 | 36 | 79 | 115 | 714 | 569 | 371 | 514 | 669 | 194 | 149 | 0.000393 | 0.005969291 | 0.071 | 0.091 | 0.082 | 0.085 | 0.055 | 0.104 | 0.116 | 0.069 | 0.106 | 0.117 | -0.026 |
| A3SS | 12027 | ENSG00000175166 | PSMD2 | chr3 | + | 184301793 | 184302071 | 184301846 | 184302071 | 184301536 | 184301658 | 8 | 9 | 3 | 11 | 2 | 291 | 214 | 275 | 308 | 308 | 24 | 22 | 11 | 24 | 20 | 544 | 454 | 260 | 308 | 672 | 202 | 149 | 0.000397 | 0.006019367 | 0.02 | 0.03 | 0.008 | 0.026 | 0.005 | 0.032 | 0.035 | 0.03 | 0.054 | 0.021 | -0.017 |
| A3SS | 4839 | ENSG00000064961 | HMG20B | chr19 | + | 3574340 | 3574586 | 3574382 | 3574586 | 3573691 | 3573800 | 2 | 8 | 3 | 3 | 0 | 22 | 20 | 20 | 44 | 8 | 7 | 8 | 5 | 9 | 22 | 10 | 29 | 6 | 32 | 46 | 191 | 149 | 0.000401 | 0.006075345 | 0.066 | 0.238 | 0.105 | 0.051 | 0 | 0.353 | 0.177 | 0.394 | 0.18 | 0.272 | -0.183 |
| A3SS | 10989 | ENSG00000078487 | ZCWPW1 | chr7 | - | 100419629 | 100419883 | 100419629 | 100419813 | 100420621 | 100420678 | 18 | 15 | 40 | 12 | 12 | 0 | 1 | 0 | 1 | 1 | 36 | 8 | 13 | 35 | 20 | 11 | 0 | 0 | 3 | 0 | 219 | 149 | 0.000404 | 0.006107228 | 1 | 0.911 | 1 | 0.891 | 0.891 | 0.69 | 1 | 1 | 0.888 | 1 | 0.023 |
| A3SS | 13377 | ENSG00000102710 | SUPT20H | chr13 | - | 37022010 | 37022317 | 37022010 | 37022080 | 37024034 | 37024193 | 64 | 54 | 73 | 74 | 71 | 38 | 31 | 62 | 51 | 54 | 32 | 41 | 64 | 25 | 19 | 51 | 104 | 50 | 56 | 13 | 298 | 149 | 0.000419 | 0.006328684 | 0.457 | 0.466 | 0.371 | 0.42 | 0.397 | 0.239 | 0.165 | 0.39 | 0.182 | 0.422 | 0.143 |
| A3SS | 4943 | ENSG00000197948 | FCHSD1 | chr5 | - | 141644858 | 141644942 | 141644858 | 141644938 | 141645019 | 141645148 | 77 | 30 | 55 | 61 | 33 | 5 | 1 | 1 | 0 | 3 | 81 | 83 | 32 | 75 | 36 | 18 | 9 | 2 | 6 | 0 | 153 | 149 | 0.000423 | 0.006370041 | 0.937 | 0.967 | 0.982 | 1 | 0.915 | 0.814 | 0.9 | 0.94 | 0.924 | 1 | 0.045 |
| A3SS | 7833 | ENSG00000131748 | STARD3 | chr17 | + | 39660159 | 39660273 | 39660210 | 39660273 | 39659460 | 39659553 | 14 | 4 | 5 | 6 | 1 | 69 | 32 | 38 | 65 | 45 | 0 | 2 | 5 | 9 | 0 | 83 | 78 | 29 | 60 | 165 | 200 | 149 | 0.000426 | 0.006402185 | 0.131 | 0.085 | 0.089 | 0.064 | 0.016 | 0 | 0.019 | 0.114 | 0.101 | 0 | 0.03 |
| A3SS | 9056 | ENSG00000116539 | ASH1L | chr1 | - | 155338088 | 155338405 | 155338088 | 155338390 | 155339327 | 155339368 | 3 | 4 | 4 | 8 | 2 | 103 | 86 | 140 | 104 | 142 | 11 | 7 | 3 | 10 | 3 | 37 | 116 | 109 | 42 | 38 | 164 | 149 | 0.000433 | 0.006479462 | 0.026 | 0.041 | 0.025 | 0.065 | 0.013 | 0.213 | 0.052 | 0.024 | 0.178 | 0.067 | -0.073 |
| A3SS | 2491 | ENSG00000206190 | ATP10A | chr15 | - | 25687702 | 25687996 | 25687702 | 25687828 | 25691714 | 25691791 | 7 | 11 | 4 | 11 | 6 | 30 | 24 | 48 | 37 | 23 | 45 | 15 | 3 | 13 | 13 | 32 | 32 | 20 | 21 | 17 | 298 | 149 | 0.000434 | 0.006479462 | 0.104 | 0.186 | 0.04 | 0.129 | 0.115 | 0.413 | 0.19 | 0.07 | 0.236 | 0.277 | -0.122 |
| A3SS | 342 | ENSG00000129351 | ILF3 | chr19 | + | 10683124 | 10683262 | 10683136 | 10683262 | 10681992 | 10682192 | 19 | 6 | 23 | 25 | 29 | 0 | 0 | 0 | 0 | 0 | 35 | 23 | 24 | 54 | 33 | 0 | 3 | 2 | 8 | 0 | 161 | 149 | 0.000434 | 0.006479462 | 1 | 1 | 1 | 1 | 1 | 1 | 0.876 | 0.917 | 0.862 | 1 | 0.069 |
| A3SS | 1364 | ENSG00000161526 | SAP30BP | chr17 | + | 75705610 | 75706092 | 75706007 | 75706092 | 75704755 | 75704814 | 55 | 45 | 38 | 57 | 69 | 198 | 110 | 143 | 104 | 379 | 77 | 61 | 66 | 112 | 128 | 181 | 237 | 135 | 150 | 209 | 298 | 149 | 0.000434 | 0.006479462 | 0.122 | 0.17 | 0.117 | 0.215 | 0.083 | 0.175 | 0.114 | 0.196 | 0.272 | 0.234 | -0.057 |
| A3SS | 12173 | ENSG00000149187 | CELF1 | chr11 | - | 47478876 | 47478952 | 47478876 | 47478940 | 47482694 | 47482856 | 114 | 71 | 75 | 99 | 228 | 14 | 8 | 11 | 7 | 4 | 86 | 50 | 47 | 67 | 45 | 0 | 20 | 21 | 11 | 10 | 161 | 149 | 0.000436 | 0.006482828 | 0.883 | 0.891 | 0.863 | 0.929 | 0.981 | 1 | 0.698 | 0.674 | 0.849 | 0.806 | 0.104 |
| A3SS | 7412 | ENSG00000174748 | RPL15 | chr3 | + | 23917946 | 23918031 | 23917972 | 23918031 | 23917147 | 23917173 | 389 | 124 | 388 | 291 | 296 | 5 | 8 | 10 | 7 | 8 | 185 | 183 | 113 | 107 | 308 | 21 | 1 | 4 | 14 | 12 | 175 | 149 | 0.000436 | 0.006482828 | 0.985 | 0.93 | 0.971 | 0.973 | 0.969 | 0.882 | 0.994 | 0.96 | 0.867 | 0.956 | 0.034 |
| A3SS | 13247 | ENSG00000075089 | ACTR6 | chr12 | + | 100220007 | 100220146 | 100220067 | 100220146 | 100218414 | 100218586 | 114 | 89 | 110 | 118 | 180 | 2 | 0 | 1 | 6 | 11 | 29 | 67 | 56 | 48 | 5 | 0 | 0 | 0 | 0 | 0 | 209 | 149 | 0.000439 | 0.006509021 | 0.976 | 1 | 0.987 | 0.933 | 0.921 | 1 | 1 | 1 | 1 | 1 | -0.037 |
| A3SS | 6072 | ENSG00000157193 | LRP8 | chr1 | - | 53255116 | 53257464 | 53255116 | 53255185 | 53258318 | 53258471 | 8 | 9 | 7 | 0 | 0 | 0 | 3 | 0 | 2 | 2 | 8 | 19 | 19 | 12 | 40 | 0 | 3 | 0 | 0 | 0 | 298 | 149 | 0.000439 | 0.006509021 | 1 | 0.6 | 1 | 0 | 0 | 1 | 0.76 | 1 | 1 | 1 | -0.432 |
| A3SS | 9585 | ENSG00000106244 | PDAP1 | chr7 | - | 99403397 | 99403525 | 99403397 | 99403505 | 99404861 | 99404953 | 8 | 9 | 3 | 8 | 4 | 67 | 40 | 67 | 64 | 72 | 49 | 10 | 17 | 30 | 44 | 210 | 78 | 58 | 141 | 262 | 169 | 149 | 0.000443 | 0.006544714 | 0.095 | 0.166 | 0.038 | 0.099 | 0.047 | 0.171 | 0.102 | 0.205 | 0.158 | 0.129 | -0.064 |
| A3SS | 3192 | ENSG00000113240 | CLK4 | chr5 | - | 178619837 | 178620673 | 178619837 | 178619896 | 178623255 | 178623416 | 51 | 23 | 43 | 55 | 18 | 28 | 13 | 10 | 17 | 8 | 15 | 34 | 35 | 12 | 8 | 0 | 5 | 11 | 0 | 0 | 298 | 149 | 0.000449 | 0.00662217 | 0.477 | 0.469 | 0.683 | 0.618 | 0.529 | 1 | 0.773 | 0.614 | 1 | 1 | -0.322 |
| A3SS | 9536 | ENSG00000196923 | PDLIM7 | chr5 | - | 177489770 | 177490031 | 177489770 | 177489832 | 177490869 | 177490906 | 8 | 0 | 1 | 3 | 1 | 11 | 6 | 10 | 8 | 4 | 18 | 15 | 27 | 36 | 45 | 25 | 18 | 6 | 36 | 70 | 298 | 149 | 0.000459 | 0.006755076 | 0.267 | 0 | 0.048 | 0.158 | 0.111 | 0.265 | 0.294 | 0.692 | 0.333 | 0.243 | -0.249 |
| A3SS | 3989 | ENSG00000178104 | PDE4DIP | chr1 | + | 149003610 | 149003730 | 149003613 | 149003730 | 149002849 | 149003042 | 3 | 6 | 16 | 17 | 17 | 3 | 4 | 9 | 3 | 4 | 0 | 3 | 1 | 6 | 16 | 7 | 8 | 5 | 11 | 7 | 152 | 149 | 0.000464 | 0.006817955 | 0.495 | 0.595 | 0.635 | 0.847 | 0.806 | 0 | 0.269 | 0.164 | 0.348 | 0.691 | 0.381 |
| A3SS | 6862 | ENSG00000133056 | PIK3C2B | chr1 | - | 204433315 | 204433949 | 204433315 | 204433425 | 204434438 | 204434608 | 21 | 9 | 35 | 27 | 45 | 1 | 1 | 0 | 2 | 0 | 17 | 34 | 18 | 12 | 41 | 0 | 3 | 5 | 5 | 0 | 298 | 149 | 0.000470 | 0.006891803 | 0.913 | 0.818 | 1 | 0.871 | 1 | 1 | 0.85 | 0.643 | 0.545 | 1 | 0.113 |
| A3SS | 3499 | ENSG00000247774 | PCED1B-AS1 | chr12 | - | 47209280 | 47209482 | 47209280 | 47209458 | 47216164 | 47216264 | 17 | 9 | 26 | 13 | 42 | 51 | 31 | 83 | 78 | 240 | 0 | 1 | 9 | 6 | 0 | 19 | 44 | 51 | 21 | 22 | 173 | 149 | 0.000471 | 0.0069005 | 0.223 | 0.2 | 0.212 | 0.126 | 0.131 | 0 | 0.019 | 0.132 | 0.197 | 0 | 0.109 |
| A3SS | 7613 | ENSG00000138757 | G3BP2 | chr4 | - | 75646337 | 75647157 | 75646337 | 75646456 | 75648638 | 75648741 | 897 | 744 | 766 | 1059 | 955 | 3 | 1 | 4 | 5 | 4 | 761 | 944 | 1508 | 1121 | 250 | 10 | 24 | 4 | 12 | 0 | 298 | 149 | 0.000472 | 0.006904338 | 0.993 | 0.997 | 0.99 | 0.991 | 0.992 | 0.974 | 0.952 | 0.995 | 0.979 | 1 | 0.013 |
| A3SS | 1653 | ENSG00000168488 | ATXN2L | chr16 | + | 28835545 | 28835758 | 28835548 | 28835758 | 28835277 | 28835399 | 5 | 0 | 6 | 7 | 0 | 11 | 18 | 5 | 14 | 4 | 5 | 2 | 0 | 0 | 0 | 61 | 4 | 14 | 25 | 4 | 152 | 149 | 0.000475 | 0.006926216 | 0.308 | 0 | 0.541 | 0.329 | 0 | 0.074 | 0.329 | 0 | 0 | 0 | 0.155 |
| A3SS | 3155 | ENSG00000105698 | USF2 | chr19 | + | 35278945 | 35279074 | 35278969 | 35279074 | 35278697 | 35278792 | 729 | 578 | 549 | 768 | 927 | 7 | 15 | 3 | 15 | 4 | 976 | 800 | 463 | 625 | 1658 | 34 | 14 | 5 | 34 | 15 | 173 | 149 | 0.000479 | 0.006983321 | 0.989 | 0.971 | 0.994 | 0.978 | 0.995 | 0.961 | 0.98 | 0.988 | 0.941 | 0.99 | 0.013 |
| A3SS | 5049 | ENSG00000125347 | IRF1 | chr5 | - | 132487925 | 132488052 | 132487925 | 132488025 | 132489391 | 132489483 | 66 | 28 | 68 | 56 | 31 | 294 | 231 | 341 | 381 | 258 | 263 | 109 | 40 | 150 | 178 | 1037 | 204 | 349 | 597 | 622 | 176 | 149 | 0.000495 | 0.007184957 | 0.16 | 0.093 | 0.144 | 0.111 | 0.092 | 0.177 | 0.311 | 0.088 | 0.175 | 0.195 | -0.069 |
| A3SS | 3907 | ENSG00000127419 | TMEM175 | chr4 | + | 950420 | 950518 | 950423 | 950518 | 948410 | 948615 | 14 | 10 | 21 | 29 | 16 | 0 | 0 | 3 | 0 | 0 | 25 | 20 | 10 | 34 | 13 | 0 | 2 | 1 | 0 | 16 | 152 | 149 | 0.000496 | 0.007184957 | 1 | 1 | 0.873 | 1 | 1 | 1 | 0.907 | 0.907 | 1 | 0.443 | 0.123 |
| A3SS | 12989 | ENSG00000198563 | DDX39B | chr6 | - | 31538762 | 31538939 | 31538762 | 31538855 | 31539146 | 31539274 | 17 | 12 | 21 | 17 | 15 | 371 | 278 | 404 | 375 | 233 | 37 | 27 | 15 | 34 | 47 | 414 | 334 | 263 | 406 | 514 | 233 | 149 | 0.000496 | 0.007184957 | 0.028 | 0.027 | 0.032 | 0.028 | 0.04 | 0.054 | 0.049 | 0.035 | 0.051 | 0.055 | -0.018 |
| A3SS | 332 | ENSG00000105723 | GSK3A | chr19 | - | 42232056 | 42232170 | 42232056 | 42232149 | 42232495 | 42232682 | 0 | 0 | 3 | 0 | 1 | 146 | 77 | 94 | 145 | 190 | 0 | 7 | 0 | 8 | 18 | 250 | 234 | 119 | 84 | 341 | 170 | 149 | 0.000499 | 0.007225742 | 0 | 0 | 0.027 | 0 | 0.005 | 0 | 0.026 | 0 | 0.077 | 0.044 | -0.023 |
| A3SS | 3151 | ENSG00000140853 | NLRC5 | chr16 | + | 57069771 | 57069919 | 57069835 | 57069919 | 57067735 | 57067828 | 10 | 0 | 14 | 7 | 0 | 165 | 64 | 136 | 141 | 77 | 26 | 4 | 5 | 12 | 105 | 179 | 205 | 134 | 280 | 314 | 213 | 149 | 0.000502 | 0.007247458 | 0.041 | 0 | 0.067 | 0.034 | 0 | 0.092 | 0.013 | 0.025 | 0.029 | 0.19 | -0.041 |
| A3SS | 12698 | ENSG00000184988 | TMEM106A | chr17 | + | 43213020 | 43213252 | 43213114 | 43213252 | 43212226 | 43212394 | 40 | 23 | 44 | 40 | 47 | 0 | 0 | 0 | 1 | 0 | 51 | 91 | 22 | 57 | 137 | 0 | 12 | 2 | 1 | 0 | 243 | 149 | 0.000510 | 0.007344636 | 1 | 1 | 1 | 0.961 | 1 | 1 | 0.823 | 0.871 | 0.972 | 1 | 0.059 |
| A3SS | 11589 | ENSG00000213930 | GALT | chr9 | + | 34647474 | 34647567 | 34647491 | 34647567 | 34647088 | 34647258 | 13 | 4 | 6 | 6 | 13 | 59 | 33 | 97 | 65 | 33 | 0 | 2 | 1 | 11 | 0 | 60 | 56 | 41 | 37 | 65 | 166 | 149 | 0.000519 | 0.007471175 | 0.165 | 0.098 | 0.053 | 0.077 | 0.261 | 0 | 0.031 | 0.021 | 0.211 | 0 | 0.078 |
| A3SS | 2780 | ENSG00000100461 | RBM23 | chr14 | - | 22908332 | 22908399 | 22908332 | 22908380 | 22909482 | 22909595 | 2 | 0 | 2 | 3 | 0 | 60 | 37 | 33 | 63 | 68 | 23 | 0 | 2 | 6 | 1 | 95 | 82 | 51 | 88 | 114 | 168 | 149 | 0.000530 | 0.007609325 | 0.029 | 0 | 0.051 | 0.041 | 0 | 0.177 | 0 | 0.034 | 0.057 | 0.008 | -0.031 |
| A3SS | 384 | ENSG00000076944 | STXBP2 | chr19 | + | 7646171 | 7646344 | 7646248 | 7646344 | 7645196 | 7645306 | 27 | 8 | 5 | 9 | 4 | 108 | 61 | 59 | 129 | 55 | 56 | 35 | 31 | 43 | 58 | 217 | 78 | 63 | 224 | 560 | 226 | 149 | 0.000543 | 0.007789964 | 0.142 | 0.08 | 0.053 | 0.044 | 0.046 | 0.145 | 0.228 | 0.245 | 0.112 | 0.064 | -0.086 |
| A3SS | 4217 | ENSG00000188010 | MORN2 | chr2 | + | 38880581 | 38880706 | 38880599 | 38880706 | 38880178 | 38880229 | 0 | 1 | 0 | 0 | 0 | 20 | 18 | 9 | 14 | 7 | 13 | 6 | 0 | 0 | 0 | 108 | 12 | 14 | 53 | 49 | 167 | 149 | 0.000554 | 0.007926519 | 0 | 0.047 | 0 | 0 | 0 | 0.097 | 0.308 | 0 | 0 | 0 | -0.072 |
| A3SS | 9702 | ENSG00000231721 | LINC-PINT | chr7 | - | 130945719 | 130945855 | 130945719 | 130945835 | 130984052 | 130984109 | 22 | 22 | 27 | 21 | 34 | 52 | 64 | 56 | 49 | 60 | 39 | 21 | 52 | 35 | 15 | 22 | 27 | 79 | 35 | 3 | 169 | 149 | 0.000555 | 0.007926519 | 0.272 | 0.233 | 0.298 | 0.274 | 0.333 | 0.61 | 0.407 | 0.367 | 0.469 | 0.815 | -0.252 |
| A3SS | 6880 | ENSG00000104894 | CD37 | chr19 | + | 49339329 | 49340606 | 49340250 | 49340606 | 49338699 | 49338936 | 760 | 261 | 488 | 546 | 461 | 1 | 0 | 3 | 4 | 0 | 620 | 703 | 400 | 570 | 1438 | 17 | 15 | 3 | 8 | 0 | 298 | 149 | 0.000556 | 0.007926519 | 0.997 | 1 | 0.988 | 0.986 | 1 | 0.948 | 0.959 | 0.985 | 0.973 | 1 | 0.021 |
| A3SS | 7110 | ENSG00000117616 | RSRP1 | chr1 | - | 25243549 | 25244224 | 25243549 | 25243633 | 25244453 | 25244498 | 484 | 486 | 438 | 479 | 755 | 10 | 11 | 9 | 10 | 31 | 301 | 211 | 294 | 282 | 114 | 12 | 19 | 13 | 22 | 0 | 298 | 149 | 0.000560 | 0.007966601 | 0.96 | 0.957 | 0.961 | 0.96 | 0.924 | 0.926 | 0.847 | 0.919 | 0.865 | 1 | 0.041 |
| A3SS | 8315 | ENSG00000135912 | TTLL4 | chr2 | + | 218754133 | 218754693 | 218754230 | 218754693 | 218753114 | 218753185 | 20 | 9 | 13 | 22 | 26 | 0 | 0 | 0 | 0 | 0 | 27 | 11 | 8 | 10 | 19 | 0 | 0 | 0 | 6 | 4 | 246 | 149 | 0.000565 | 0.008031824 | 1 | 1 | 1 | 1 | 1 | 1 | 1 | 1 | 0.502 | 0.742 | 0.151 |
| A3SS | 628 | ENSG00000187994 | RINL | chr19 | - | 38876330 | 38876490 | 38876330 | 38876474 | 38876692 | 38876781 | 22 | 11 | 6 | 26 | 5 | 5 | 0 | 6 | 1 | 0 | 22 | 36 | 2 | 18 | 43 | 0 | 4 | 0 | 0 | 0 | 165 | 149 | 0.000567 | 0.008045587 | 0.799 | 1 | 0.475 | 0.959 | 1 | 1 | 0.89 | 1 | 1 | 1 | -0.131 |
| A3SS | 8301 | ENSG00000068024 | HDAC4 | chr2 | - | 239126455 | 239126694 | 239126455 | 239126679 | 239134244 | 239134443 | 8 | 5 | 10 | 12 | 2 | 1 | 1 | 3 | 2 | 0 | 0 | 8 | 12 | 2 | 5 | 3 | 12 | 2 | 1 | 9 | 164 | 149 | 0.000575 | 0.008118318 | 0.879 | 0.82 | 0.752 | 0.845 | 1 | 0 | 0.377 | 0.845 | 0.645 | 0.335 | 0.419 |
| A3SS | 11731 | ENSG00000166405 | RIC3 | chr11 | - | 8126658 | 8126891 | 8126658 | 8126804 | 8128218 | 8128354 | 3 | 7 | 5 | 2 | 9 | 4 | 0 | 6 | 6 | 0 | 12 | 3 | 0 | 1 | 0 | 0 | 0 | 1 | 0 | 1 | 236 | 149 | 0.000575 | 0.008118318 | 0.321 | 1 | 0.345 | 0.174 | 1 | 1 | 1 | 0 | 1 | 0 | -0.032 |
| A3SS | 6133 | ENSG00000102805 | CLN5 | chr13 | + | 76995842 | 76996127 | 76995901 | 76996127 | 76995062 | 76995228 | 1 | 8 | 5 | 2 | 7 | 34 | 40 | 45 | 37 | 59 | 0 | 4 | 0 | 1 | 0 | 47 | 19 | 10 | 29 | 19 | 208 | 149 | 0.000576 | 0.008118318 | 0.021 | 0.125 | 0.074 | 0.037 | 0.078 | 0 | 0.131 | 0 | 0.024 | 0 | 0.036 |
| A3SS | 3495 | ENSG00000247774 | PCED1B-AS1 | chr12 | - | 47205897 | 47206145 | 47205897 | 47206069 | 47210639 | 47210761 | 258 | 179 | 258 | 251 | 636 | 21 | 7 | 17 | 13 | 42 | 110 | 122 | 152 | 108 | 95 | 0 | 1 | 4 | 13 | 0 | 225 | 149 | 0.000576 | 0.008118318 | 0.891 | 0.944 | 0.91 | 0.927 | 0.909 | 1 | 0.988 | 0.962 | 0.846 | 1 | -0.043 |
| A3SS | 11830 | ENSG00000073921 | PICALM | chr11 | - | 85990249 | 85990399 | 85990249 | 85990378 | 85996825 | 85996929 | 323 | 184 | 210 | 204 | 192 | 23 | 11 | 4 | 21 | 7 | 97 | 260 | 376 | 260 | 56 | 0 | 33 | 48 | 29 | 11 | 170 | 149 | 0.000581 | 0.008163568 | 0.925 | 0.936 | 0.979 | 0.895 | 0.96 | 1 | 0.874 | 0.873 | 0.887 | 0.817 | 0.049 |
| A3SS | 8077 | ENSG00000123240 | OPTN | chr10 | + | 13109096 | 13109288 | 13109111 | 13109288 | 13100172 | 13100302 | 5 | 0 | 2 | 2 | 5 | 35 | 10 | 14 | 36 | 5 | 0 | 0 | 0 | 5 | 0 | 29 | 23 | 26 | 33 | 23 | 164 | 149 | 0.000589 | 0.008272859 | 0.115 | 0 | 0.115 | 0.048 | 0.476 | 0 | 0 | 0 | 0.121 | 0 | 0.127 |
| A3SS | 4811 | ENSG00000184014 | DENND5A | chr11 | - | 9141939 | 9143485 | 9141939 | 9142108 | 9144096 | 9144278 | 96 | 79 | 89 | 96 | 63 | 1 | 4 | 2 | 1 | 0 | 239 | 112 | 95 | 142 | 171 | 7 | 3 | 6 | 16 | 4 | 298 | 149 | 0.000593 | 0.008313714 | 0.98 | 0.908 | 0.957 | 0.98 | 1 | 0.945 | 0.949 | 0.888 | 0.816 | 0.955 | 0.054 |
| A3SS | 11857 | ENSG00000142186 | SCYL1 | chr11 | + | 65538441 | 65538704 | 65538446 | 65538704 | 65538269 | 65538324 | 104 | 40 | 84 | 74 | 46 | 2 | 0 | 4 | 1 | 0 | 132 | 114 | 101 | 88 | 257 | 0 | 5 | 4 | 11 | 14 | 154 | 149 | 0.000597 | 0.008348092 | 0.981 | 1 | 0.953 | 0.986 | 1 | 1 | 0.957 | 0.961 | 0.886 | 0.947 | 0.034 |
| A3SS | 8440 | ENSG00000135926 | TMBIM1 | chr2 | - | 218277934 | 218278165 | 218277934 | 218277974 | 218278514 | 218278565 | 13 | 19 | 9 | 26 | 24 | 288 | 125 | 100 | 201 | 256 | 69 | 29 | 36 | 33 | 66 | 268 | 209 | 97 | 179 | 1044 | 298 | 149 | 0.000601 | 0.008399939 | 0.022 | 0.071 | 0.043 | 0.061 | 0.045 | 0.114 | 0.065 | 0.157 | 0.084 | 0.031 | -0.042 |
| A3SS | 12215 | ENSG00000185187 | SIGIRR | chr11 | - | 407424 | 407568 | 407424 | 407534 | 407816 | 407957 | 86 | 40 | 69 | 63 | 52 | 6 | 1 | 1 | 2 | 0 | 26 | 54 | 20 | 27 | 66 | 0 | 0 | 6 | 2 | 19 | 183 | 149 | 0.000605 | 0.008432849 | 0.921 | 0.97 | 0.983 | 0.962 | 1 | 1 | 1 | 0.731 | 0.917 | 0.739 | 0.09 |
| A3SS | 6328 | ENSG00000139636 | LMBR1L | chr12 | - | 49101249 | 49101323 | 49101249 | 49101263 | 49101471 | 49101549 | 20 | 10 | 21 | 18 | 2 | 3 | 5 | 7 | 10 | 0 | 13 | 24 | 14 | 20 | 33 | 35 | 18 | 17 | 23 | 27 | 209 | 149 | 0.000606 | 0.00843985 | 0.826 | 0.588 | 0.681 | 0.562 | 1 | 0.209 | 0.487 | 0.37 | 0.383 | 0.466 | 0.348 |
| A3SS | 12820 | ENSG00000109685 | NSD2 | chr4 | + | 1951998 | 1952231 | 1952107 | 1952231 | 1951071 | 1951203 | 1 | 4 | 7 | 0 | 0 | 34 | 30 | 25 | 39 | 40 | 12 | 8 | 3 | 1 | 14 | 36 | 25 | 34 | 69 | 35 | 258 | 149 | 0.000611 | 0.008482863 | 0.017 | 0.071 | 0.139 | 0 | 0 | 0.161 | 0.156 | 0.048 | 0.008 | 0.188 | -0.067 |
| A3SS | 1792 | ENSG00000140931 | CMTM3 | chr16 | + | 66609758 | 66610003 | 66609882 | 66610003 | 66609434 | 66609530 | 3 | 8 | 7 | 6 | 15 | 74 | 25 | 72 | 71 | 28 | 30 | 17 | 13 | 19 | 25 | 86 | 58 | 58 | 32 | 89 | 273 | 149 | 0.000615 | 0.00853058 | 0.022 | 0.149 | 0.05 | 0.044 | 0.226 | 0.16 | 0.138 | 0.109 | 0.245 | 0.133 | -0.059 |
| A3SS | 12044 | ENSG00000149091 | DGKZ | chr11 | + | 46367651 | 46367747 | 46367654 | 46367747 | 46367290 | 46367399 | 233 | 96 | 126 | 161 | 85 | 10 | 5 | 12 | 4 | 0 | 228 | 141 | 85 | 136 | 337 | 0 | 0 | 2 | 1 | 12 | 152 | 149 | 0.000621 | 0.008596473 | 0.958 | 0.95 | 0.911 | 0.975 | 1 | 1 | 1 | 0.977 | 0.993 | 0.965 | -0.028 |
| A3SS | 8830 | ENSG00000072415 | MPP5 | chr14 | + | 67279017 | 67279537 | 67279249 | 67279537 | 67269700 | 67269783 | 69 | 104 | 76 | 99 | 174 | 1 | 2 | 3 | 4 | 6 | 45 | 101 | 88 | 36 | 26 | 0 | 0 | 0 | 2 | 0 | 298 | 149 | 0.000623 | 0.008607646 | 0.972 | 0.963 | 0.927 | 0.925 | 0.935 | 1 | 1 | 1 | 0.9 | 1 | -0.036 |
| A3SS | 11357 | ENSG00000152763 | DNAI4 | chr1 | - | 66893228 | 66893413 | 66893228 | 66893361 | 66905200 | 66905375 | 40 | 23 | 27 | 40 | 108 | 2 | 0 | 0 | 14 | 0 | 11 | 32 | 16 | 14 | 16 | 0 | 2 | 0 | 0 | 0 | 201 | 149 | 0.000627 | 0.008652983 | 0.937 | 1 | 1 | 0.679 | 1 | 1 | 0.922 | 1 | 1 | 1 | -0.061 |
| A3SS | 13192 | ENSG00000177575 | CD163 | chr12 | - | 7479859 | 7479996 | 7479859 | 7479913 | 7482642 | 7482762 | 30 | 19 | 9 | 31 | 33 | 0 | 0 | 0 | 2 | 0 | 102 | 129 | 60 | 84 | 7 | 0 | 8 | 4 | 3 | 2 | 232 | 149 | 0.000646 | 0.008885341 | 1 | 1 | 1 | 0.909 | 1 | 1 | 0.912 | 0.906 | 0.947 | 0.692 | 0.09 |
| A3SS | 9969 | ENSG00000011275 | RNF216 | chr7 | - | 5739275 | 5739621 | 5739275 | 5739352 | 5740972 | 5741644 | 71 | 59 | 86 | 110 | 103 | 109 | 96 | 89 | 98 | 170 | 111 | 104 | 136 | 145 | 99 | 99 | 53 | 60 | 55 | 146 | 298 | 149 | 0.000646 | 0.008885341 | 0.246 | 0.235 | 0.326 | 0.359 | 0.233 | 0.359 | 0.495 | 0.531 | 0.569 | 0.253 | -0.162 |
| A3SS | 2418 | ENSG00000140365 | COMMD4 | chr15 | + | 75338984 | 75339104 | 75339036 | 75339104 | 75338645 | 75338685 | 73 | 30 | 50 | 57 | 55 | 0 | 0 | 0 | 0 | 0 | 45 | 50 | 49 | 69 | 142 | 0 | 16 | 1 | 0 | 0 | 201 | 149 | 0.000653 | 0.008960909 | 1 | 1 | 1 | 1 | 1 | 1 | 0.698 | 0.973 | 1 | 1 | 0.066 |
| A3SS | 8212 | ENSG00000172493 | AFF1 | chr4 | + | 87134470 | 87134694 | 87134473 | 87134694 | 87131792 | 87131864 | 10 | 18 | 9 | 10 | 17 | 1 | 0 | 1 | 0 | 0 | 4 | 11 | 38 | 42 | 40 | 6 | 0 | 8 | 2 | 0 | 152 | 149 | 0.000660 | 0.009044203 | 0.907 | 1 | 0.898 | 1 | 1 | 0.395 | 1 | 0.823 | 0.954 | 1 | 0.127 |
| A3SS | 10208 | ENSG00000156831 | NSMCE2 | chr8 | + | 125102013 | 125102146 | 125102050 | 125102146 | 125091843 | 125091958 | 16 | 7 | 8 | 5 | 1 | 7 | 7 | 6 | 8 | 4 | 8 | 6 | 13 | 22 | 4 | 0 | 4 | 7 | 1 | 0 | 186 | 149 | 0.000664 | 0.009084154 | 0.647 | 0.445 | 0.516 | 0.334 | 0.167 | 1 | 0.546 | 0.598 | 0.946 | 1 | -0.396 |
| A3SS | 10155 | ENSG00000184863 | RBM33 | chr7 | + | 155711172 | 155711455 | 155711202 | 155711455 | 155706859 | 155707068 | 37 | 15 | 73 | 53 | 60 | 46 | 26 | 38 | 53 | 24 | 32 | 25 | 22 | 21 | 4 | 36 | 78 | 45 | 24 | 58 | 179 | 149 | 0.000671 | 0.009161646 | 0.401 | 0.324 | 0.615 | 0.454 | 0.675 | 0.425 | 0.211 | 0.289 | 0.421 | 0.054 | 0.214 |
| A3SS | 8562 | ENSG00000115282 | TTC31 | chr2 | + | 74491943 | 74492055 | 74492003 | 74492055 | 74491583 | 74491672 | 3 | 9 | 2 | 2 | 5 | 28 | 44 | 24 | 29 | 49 | 18 | 4 | 7 | 7 | 17 | 31 | 40 | 39 | 19 | 43 | 209 | 149 | 0.000673 | 0.009175651 | 0.071 | 0.127 | 0.056 | 0.047 | 0.068 | 0.293 | 0.067 | 0.113 | 0.208 | 0.22 | -0.106 |
| A3SS | 3930 | ENSG00000100650 | SRSF5 | chr14 | + | 69771193 | 69772005 | 69771196 | 69772005 | 69770994 | 69771105 | 1143 | 802 | 1304 | 1291 | 1820 | 80 | 43 | 69 | 80 | 123 | 984 | 939 | 750 | 964 | 387 | 17 | 47 | 25 | 47 | 33 | 152 | 149 | 0.000674 | 0.009175651 | 0.933 | 0.948 | 0.949 | 0.941 | 0.936 | 0.983 | 0.951 | 0.967 | 0.953 | 0.92 | -0.013 |
| A3SS | 715 | ENSG00000196365 | LONP1 | chr19 | - | 5711770 | 5712070 | 5711770 | 5712002 | 5713133 | 5713253 | 6 | 3 | 2 | 1 | 0 | 128 | 49 | 116 | 111 | 98 | 13 | 4 | 3 | 2 | 16 | 157 | 103 | 72 | 148 | 153 | 217 | 149 | 0.000698 | 0.009490804 | 0.031 | 0.04 | 0.012 | 0.006 | 0 | 0.054 | 0.026 | 0.028 | 0.009 | 0.067 | -0.019 |
| A3SS | 8575 | ENSG00000116750 | UCHL5 | chr1 | - | 193022925 | 193023039 | 193022925 | 193023036 | 193023843 | 193023946 | 99 | 70 | 68 | 78 | 168 | 145 | 109 | 109 | 124 | 199 | 19 | 37 | 75 | 11 | 21 | 2 | 53 | 54 | 9 | 0 | 152 | 149 | 0.000706 | 0.009580937 | 0.401 | 0.386 | 0.379 | 0.381 | 0.453 | 0.903 | 0.406 | 0.577 | 0.545 | 1 | -0.286 |
| A3SS | 11394 | ENSG00000116580 | GON4L | chr1 | - | 155751959 | 155752590 | 155751959 | 155752587 | 155753203 | 155753414 | 1 | 0 | 0 | 1 | 0 | 44 | 22 | 41 | 50 | 26 | 8 | 5 | 0 | 0 | 0 | 14 | 51 | 14 | 34 | 10 | 152 | 149 | 0.000708 | 0.009586147 | 0.022 | 0 | 0 | 0.019 | 0 | 0.359 | 0.088 | 0 | 0 | 0 | -0.081 |
| A3SS | 9099 | ENSG00000248592 | STIMATE-MUSTN1 | chr3 | - | 52834939 | 52835302 | 52834939 | 52835003 | 52835788 | 52840262 | 44 | 29 | 36 | 51 | 34 | 5 | 3 | 2 | 4 | 0 | 162 | 112 | 49 | 127 | 106 | 9 | 17 | 17 | 26 | 27 | 298 | 149 | 0.000709 | 0.009592104 | 0.815 | 0.829 | 0.9 | 0.864 | 1 | 0.9 | 0.767 | 0.59 | 0.709 | 0.662 | 0.156 |
| A3SS | 7242 | ENSG00000089127 | OAS1 | chr12 | + | 112911050 | 112911235 | 112911097 | 112911235 | 112906972 | 112907219 | 83 | 36 | 34 | 71 | 29 | 1 | 1 | 3 | 6 | 6 | 149 | 135 | 42 | 204 | 61 | 18 | 24 | 6 | 25 | 15 | 196 | 149 | 0.000713 | 0.009619398 | 0.984 | 0.965 | 0.896 | 0.9 | 0.786 | 0.863 | 0.81 | 0.842 | 0.861 | 0.756 | 0.08 |
| A3SS | 9128 | ENSG00000105393 | BABAM1 | chr19 | + | 17275788 | 17275825 | 17275800 | 17275825 | 17274106 | 17274185 | 2 | 0 | 3 | 0 | 0 | 126 | 58 | 135 | 111 | 192 | 11 | 2 | 2 | 1 | 16 | 344 | 149 | 114 | 176 | 218 | 161 | 149 | 0.000714 | 0.009628208 | 0.014 | 0 | 0.02 | 0 | 0 | 0.029 | 0.012 | 0.016 | 0.005 | 0.064 | -0.018 |
| A3SS | 2654 | ENSG00000119638 | NEK9 | chr14 | - | 75107342 | 75107755 | 75107342 | 75107487 | 75109684 | 75109877 | 10 | 4 | 12 | 9 | 2 | 77 | 66 | 69 | 86 | 58 | 26 | 27 | 11 | 2 | 0 | 59 | 40 | 25 | 85 | 29 | 298 | 149 | 0.000717 | 0.009643233 | 0.061 | 0.029 | 0.08 | 0.05 | 0.017 | 0.181 | 0.252 | 0.18 | 0.012 | 0 | -0.078 |
| A3SS | 10031 | ENSG00000104133 | SPG11 | chr15 | - | 44583813 | 44584558 | 44583813 | 44584414 | 44585635 | 44585850 | 309 | 160 | 190 | 185 | 272 | 6 | 3 | 9 | 2 | 2 | 172 | 237 | 203 | 175 | 93 | 0 | 3 | 0 | 1 | 0 | 293 | 149 | 0.000719 | 0.009654656 | 0.963 | 0.964 | 0.915 | 0.979 | 0.986 | 1 | 0.976 | 1 | 0.989 | 1 | -0.032 |
| A3SS | 5618 | ENSG00000163346 | PBXIP1 | chr1 | - | 154951470 | 154951921 | 154951470 | 154951535 | 154953670 | 154953757 | 268 | 158 | 231 | 304 | 142 | 1 | 0 | 9 | 5 | 0 | 257 | 222 | 240 | 156 | 360 | 16 | 12 | 3 | 6 | 3 | 298 | 149 | 0.000725 | 0.009727475 | 0.993 | 1 | 0.928 | 0.968 | 1 | 0.889 | 0.902 | 0.976 | 0.929 | 0.984 | 0.042 |
| A3SS | 923 | ENSG00000105576 | TNPO2 | chr19 | - | 12715632 | 12715739 | 12715632 | 12715710 | 12719028 | 12719178 | 55 | 24 | 58 | 51 | 107 | 0 | 4 | 0 | 1 | 0 | 53 | 80 | 54 | 98 | 206 | 2 | 5 | 2 | 0 | 15 | 178 | 149 | 0.000729 | 0.009760334 | 1 | 0.834 | 1 | 0.977 | 1 | 0.957 | 0.931 | 0.958 | 1 | 0.92 | 0.009 |
| A3SS | 5226 | ENSG00000084072 | PPIE | chr1 | + | 39745374 | 39745498 | 39745413 | 39745498 | 39743823 | 39743924 | 120 | 68 | 124 | 138 | 171 | 8 | 6 | 18 | 8 | 32 | 123 | 49 | 45 | 84 | 190 | 42 | 10 | 13 | 14 | 27 | 188 | 149 | 0.000731 | 0.009766795 | 0.922 | 0.9 | 0.845 | 0.932 | 0.809 | 0.699 | 0.795 | 0.733 | 0.826 | 0.848 | 0.101 |
| A3SS | 1855 | ENSG00000205336 | ADGRG1 | chr16 | + | 57651242 | 57651622 | 57651355 | 57651622 | 57650252 | 57650351 | 84 | 8 | 12 | 25 | 13 | 9 | 0 | 1 | 3 | 0 | 33 | 71 | 42 | 16 | 25 | 0 | 5 | 0 | 0 | 0 | 262 | 149 | 0.000736 | 0.009814679 | 0.841 | 1 | 0.872 | 0.826 | 1 | 1 | 0.89 | 1 | 1 | 1 | -0.07 |
| A3SS | 3024 | ENSG00000133706 | LARS1 | chr5 | - | 146140203 | 146143084 | 146140203 | 146140261 | 146143411 | 146143550 | 61 | 55 | 73 | 76 | 74 | 1 | 1 | 0 | 1 | 0 | 8 | 63 | 23 | 41 | 6 | 0 | 2 | 8 | 1 | 0 | 298 | 149 | 0.000745 | 0.009925311 | 0.968 | 0.965 | 1 | 0.974 | 1 | 1 | 0.94 | 0.59 | 0.953 | 1 | 0.085 |
| A3SS | 5490 | ENSG00000099622 | CIRBP | chr19 | + | 1270866 | 1271036 | 1270927 | 1271036 | 1269330 | 1269410 | 63 | 21 | 69 | 59 | 48 | 34 | 23 | 18 | 12 | 8 | 115 | 113 | 92 | 80 | 185 | 9 | 17 | 14 | 10 | 32 | 210 | 149 | 0.000747 | 0.009936573 | 0.568 | 0.393 | 0.731 | 0.777 | 0.81 | 0.901 | 0.825 | 0.823 | 0.85 | 0.804 | -0.185 |
| A3SS | 12792 | ENSG00000069493 | CLEC2D | chr12 | + | 9693415 | 9693948 | 9693824 | 9693948 | 9692827 | 9692931 | 111 | 108 | 116 | 123 | 117 | 3 | 0 | 1 | 5 | 20 | 17 | 7 | 30 | 9 | 16 | 0 | 1 | 2 | 0 | 0 | 298 | 149 | 0.000749 | 0.009938611 | 0.949 | 1 | 0.983 | 0.925 | 0.745 | 1 | 0.778 | 0.882 | 1 | 1 | -0.012 |
| A3SS | 5394 | ENSG00000126088 | UROD | chr1 | + | 45013574 | 45013791 | 45013593 | 45013791 | 45013135 | 45013354 | 9 | 4 | 7 | 11 | 18 | 50 | 33 | 40 | 61 | 46 | 18 | 18 | 12 | 32 | 31 | 60 | 56 | 22 | 46 | 72 | 168 | 149 | 0.000755 | 0.010005531 | 0.138 | 0.097 | 0.134 | 0.138 | 0.258 | 0.21 | 0.222 | 0.326 | 0.382 | 0.276 | -0.13 |
| A3SS | 10995 | ENSG00000156787 | TBC1D31 | chr8 | + | 123129065 | 123129218 | 123129069 | 123129218 | 123128280 | 123128513 | 160 | 150 | 143 | 177 | 230 | 5 | 8 | 3 | 5 | 14 | 33 | 93 | 49 | 11 | 39 | 0 | 2 | 2 | 0 | 0 | 153 | 149 | 0.000775 | 0.01025776 | 0.969 | 0.948 | 0.979 | 0.972 | 0.941 | 1 | 0.978 | 0.96 | 1 | 1 | -0.026 |
| A3SS | 12761 | ENSG00000155256 | ZFYVE27 | chr10 | + | 97751375 | 97751462 | 97751390 | 97751462 | 97750330 | 97750470 | 9 | 7 | 11 | 26 | 11 | 52 | 23 | 46 | 38 | 41 | 43 | 28 | 19 | 30 | 13 | 48 | 39 | 32 | 47 | 31 | 164 | 149 | 0.000788 | 0.010408018 | 0.136 | 0.217 | 0.178 | 0.383 | 0.196 | 0.449 | 0.395 | 0.35 | 0.367 | 0.276 | -0.145 |
| A3SS | 2904 | ENSG00000175061 | SNHG29 | chr17 | + | 16439531 | 16439703 | 16439659 | 16439703 | 16439326 | 16439414 | 129 | 72 | 169 | 146 | 97 | 17 | 10 | 43 | 20 | 7 | 173 | 137 | 156 | 186 | 225 | 0 | 10 | 31 | 26 | 1 | 277 | 149 | 0.000793 | 0.01045604 | 0.803 | 0.795 | 0.679 | 0.797 | 0.882 | 1 | 0.881 | 0.73 | 0.794 | 0.992 | -0.088 |
| A3SS | 8413 | ENSG00000239779 | WBP1 | chr2 | + | 74459437 | 74459745 | 74459642 | 74459745 | 74458996 | 74459098 | 11 | 7 | 25 | 24 | 17 | 0 | 0 | 3 | 3 | 0 | 19 | 11 | 22 | 3 | 9 | 3 | 7 | 6 | 0 | 2 | 298 | 149 | 0.000794 | 0.01045604 | 1 | 1 | 0.806 | 0.8 | 1 | 0.76 | 0.44 | 0.647 | 1 | 0.692 | 0.213 |
| A3SS | 2676 | ENSG00000254505 | CHMP4A | chr14 | - | 24211410 | 24211592 | 24211410 | 24211512 | 24211679 | 24211829 | 378 | 243 | 262 | 354 | 201 | 0 | 4 | 3 | 3 | 17 | 552 | 262 | 259 | 321 | 538 | 0 | 2 | 3 | 0 | 0 | 229 | 149 | 0.000798 | 0.010488411 | 1 | 0.975 | 0.983 | 0.987 | 0.885 | 1 | 0.988 | 0.983 | 1 | 1 | -0.028 |
| A3SS | 12465 | ENSG00000166987 | MBD6 | chr12 | + | 57525347 | 57526388 | 57526310 | 57526388 | 57524952 | 57525115 | 49 | 19 | 36 | 42 | 18 | 0 | 0 | 0 | 2 | 0 | 96 | 59 | 67 | 97 | 166 | 10 | 4 | 0 | 2 | 0 | 298 | 149 | 0.000808 | 0.010604763 | 1 | 1 | 1 | 0.913 | 1 | 0.828 | 0.881 | 1 | 0.96 | 1 | 0.049 |
| A3SS | 5206 | ENSG00000137776 | SLTM | chr15 | - | 58899468 | 58899937 | 58899468 | 58899883 | 58901259 | 58901287 | 143 | 137 | 106 | 141 | 162 | 49 | 35 | 46 | 48 | 71 | 58 | 77 | 72 | 34 | 27 | 4 | 22 | 30 | 1 | 0 | 203 | 149 | 0.000821 | 0.01075527 | 0.682 | 0.742 | 0.628 | 0.683 | 0.626 | 0.914 | 0.72 | 0.638 | 0.961 | 1 | -0.174 |
| A3SS | 6485 | ENSG00000181523 | SGSH | chr17 | - | 80214614 | 80214783 | 80214614 | 80214765 | 80215032 | 80215138 | 0 | 0 | 0 | 0 | 0 | 16 | 6 | 21 | 11 | 14 | 0 | 0 | 4 | 5 | 2 | 22 | 22 | 12 | 16 | 16 | 167 | 149 | 0.000830 | 0.010856632 | 0 | 0 | 0 | 0 | 0 | 0 | 0 | 0.229 | 0.218 | 0.1 | -0.109 |
| A3SS | 904 | ENSG00000062598 | ELMO2 | chr20 | - | 46387337 | 46387437 | 46387337 | 46387431 | 46389038 | 46389220 | 70 | 39 | 47 | 61 | 49 | 1 | 0 | 2 | 0 | 1 | 41 | 59 | 33 | 31 | 98 | 5 | 7 | 0 | 0 | 4 | 155 | 149 | 0.000833 | 0.010881329 | 0.985 | 1 | 0.958 | 1 | 0.979 | 0.887 | 0.89 | 1 | 1 | 0.959 | 0.037 |
| A3SS | 329 | ENSG00000028277 | POU2F2 | chr19 | - | 42092068 | 42092270 | 42092068 | 42092134 | 42093828 | 42093895 | 102 | 50 | 61 | 83 | 44 | 2 | 2 | 2 | 1 | 3 | 147 | 66 | 38 | 50 | 138 | 0 | 12 | 7 | 8 | 6 | 285 | 149 | 0.000836 | 0.010896782 | 0.964 | 0.929 | 0.941 | 0.977 | 0.885 | 1 | 0.742 | 0.739 | 0.766 | 0.923 | 0.105 |
| A3SS | 10429 | ENSG00000285171 | NA | chrX | - | 71106531 | 71106770 | 71106531 | 71106590 | 71108276 | 71108346 | 21 | 10 | 23 | 17 | 11 | 0 | 3 | 0 | 1 | 0 | 21 | 24 | 32 | 27 | 54 | 0 | 0 | 11 | 1 | 9 | 298 | 149 | 0.000838 | 0.010906815 | 1 | 0.625 | 1 | 0.895 | 1 | 1 | 1 | 0.593 | 0.931 | 0.75 | 0.049 |
| A3SS | 131 | ENSG00000100395 | L3MBTL2 | chr22 | + | 41213850 | 41214026 | 41213892 | 41214026 | 41209695 | 41209933 | 6 | 6 | 11 | 9 | 12 | 82 | 42 | 75 | 86 | 54 | 6 | 21 | 11 | 16 | 5 | 14 | 26 | 28 | 38 | 95 | 191 | 149 | 0.000848 | 0.011016394 | 0.054 | 0.1 | 0.103 | 0.075 | 0.148 | 0.251 | 0.387 | 0.235 | 0.247 | 0.039 | -0.136 |
| A3SS | 7306 | ENSG00000164828 | SUN1 | chr7 | + | 852809 | 852952 | 852812 | 852952 | 852608 | 852667 | 22 | 24 | 24 | 22 | 12 | 10 | 6 | 13 | 4 | 13 | 30 | 30 | 19 | 34 | 43 | 0 | 3 | 9 | 3 | 7 | 152 | 149 | 0.000858 | 0.011134009 | 0.683 | 0.797 | 0.644 | 0.844 | 0.475 | 1 | 0.907 | 0.674 | 0.917 | 0.858 | -0.183 |
| A3SS | 5475 | ENSG00000111325 | OGFOD2 | chr12 | + | 122978359 | 122978569 | 122978441 | 122978569 | 122976870 | 122976970 | 14 | 9 | 20 | 21 | 7 | 1 | 1 | 4 | 1 | 3 | 24 | 15 | 17 | 2 | 16 | 0 | 0 | 3 | 0 | 0 | 231 | 149 | 0.000861 | 0.011139341 | 0.9 | 0.853 | 0.763 | 0.931 | 0.601 | 1 | 1 | 0.785 | 1 | 1 | -0.147 |

| | | | | | | | | | | | | |
|---|---|---|---|---|---|---|---|---|---|---|---|---|
| A3SS | 1424 | ENSG00000170037 | CNTROB | chr17 | + | 7949081 | 7949157 | 7949084 | 7949157 | 7948486 | 7948619 | 11 |
| A3SS | 7384 | ENSG00000048162 | NOP16 | chr5 | - | 176386839 | 176386943 | 176386839 | 176386909 | 176388234 | 176388343 | 2 |
| A3SS | 11565 | ENSG00000135047 | CTSL | chr9 | + | 87727957 | 87728149 | 87728026 | 87728149 | 87727593 | 87727729 | 0 |
| A3SS | 2069 | ENSG00000069345 | DNAJA2 | chr16 | - | 46971348 | 46971659 | 46971348 | 46971572 | 46971895 | 46971955 | 39 |
| A3SS | 4452 | ENSG00000114867 | EIF4G1 | chr3 | + | 184317716 | 184317816 | 184317746 | 184317816 | 184317320 | 184317497 | 206 |
| A3SS | 2553 | ENSG00000197102 | DYNC1H1 | chr14 | + | 102018450 | 102018616 | 102018511 | 102018616 | 102017382 | 102017504 | 280 |
| A3SS | 5351 | ENSG00000116001 | TIA1 | chr2 | - | 70227734 | 70227822 | 70227734 | 70227791 | 70229058 | 70229091 | 149 |
| A3SS | 12419 | ENSG00000011478 | QPCTL | chr19 | + | 45695400 | 45695718 | 45695436 | 45695718 | 45693412 | 45693556 | 0 |
| A3SS | 2237 | ENSG00000166783 | MARF1 | chr16 | - | 15633616 | 15633843 | 15633616 | 15633834 | 15634756 | 15634931 | 190 |
| A3SS | 10724 | ENSG00000215301 | DDX3X | chrX | + | 41347311 | 41347451 | 41347314 | 41347451 | 41346504 | 41346622 | 16 |
| A3SS | 11853 | ENSG00000142186 | SCYL1 | chr11 | + | 65536985 | 65537128 | 65537036 | 65537128 | 65536585 | 65536750 | 45 |
| A3SS | 8688 | ENSG00000165280 | VCP | chr9 | - | 35056063 | 35057222 | 35056063 | 35057211 | 35057375 | 35057530 | 334 |
| A3SS | 1619 | ENSG00000083454 | P2RX5 | chr17 | - | 3688625 | 3688759 | 3688625 | 3688676 | 3690069 | 3690150 | 73 |
| A3SS | 6808 | ENSG00000169554 | ZEB2 | chr2 | - | 144424795 | 144424953 | 144424795 | 144424864 | 144429768 | 144430026 | 63 |
| A3SS | 8323 | ENSG00000018280 | SLC11A1 | chr2 | + | 218384242 | 218384365 | 218384318 | 218384365 | 218382959 | 218383102 | 147 |
| A3SS | 11223 | ENSG00000196150 | ZNF250 | chr8 | - | 144889932 | 144890074 | 144889932 | 144890059 | 144890307 | 144890403 | 3 |
| A3SS | 2101 | ENSG00000140983 | RHOT2 | chr16 | + | 670888 | 671000 | 670891 | 671000 | 670674 | 670773 | 8 |
| A3SS | 12163 | ENSG00000188690 | UROS | chr10 | - | 125789282 | 125789528 | 125789282 | 125789363 | 125794879 | 125794978 | 14 |
| A3SS | 12871 | ENSG00000241370 | RPP21 | chr6 | + | 30345466 | 30345573 | 30345490 | 30345573 | 30345297 | 30345398 | 0 |
| A3SS | 10041 | ENSG00000087077 | TRIP6 | chr7 | + | 100867860 | 100867988 | 100867910 | 100867988 | 100867386 | 100867606 | 8 |
| A3SS | 2732 | ENSG00000100722 | ZC3H14 | chr14 | + | 88609711 | 88609803 | 88609714 | 88609803 | 88609266 | 88609403 | 72 |
| A3SS | 4351 | ENSG00000187742 | SECISBP2 | chr9 | + | 89334521 | 89334730 | 89334635 | 89334730 | 89334024 | 89334155 | 147 |
| A3SS | 1212 | ENSG00000185624 | P4HB | chr17 | - | 81846428 | 81847072 | 81846428 | 81847068 | 81847242 | 81847347 | 702 |
| A3SS | 13165 | ENSG00000134283 | PPHLN1 | chr12 | + | 42445972 | 42446290 | 42445990 | 42446290 | 42442640 | 42442741 | 7 |
| A3SS | 5144 | ENSG00000064687 | ABCA7 | chr19 | + | 1056906 | 1058747 | 1058617 | 1058747 | 1056329 | 1056499 | 33 |
| A3SS | 2046 | ENSG00000167261 | DPEP2 | chr16 | - | 67987851 | 67988057 | 67987851 | 67987987 | 67989322 | 67989398 | 13 |
| A3SS | 2724 | ENSG00000105647 | PIK3R2 | chr19 | + | 18166022 | 18166302 | 18166159 | 18166302 | 18163262 | 18163388 | 1 |
| A3SS | 3679 | ENSG00000126522 | ASL | chr7 | + | 66086743 | 66087791 | 66087728 | 66087791 | 66086584 | 66086662 | 45 |
| A3SS | 3537 | ENSG00000131503 | ANKHD1 | chr5 | + | 140537337 | 140537589 | 140537388 | 140537589 | 140528183 | 140529796 | 36 |
| A3SS | 11827 | ENSG00000119392 | GLE1 | chr9 | + | 128525191 | 128525423 | 128525196 | 128525423 | 128523591 | 128523846 | 25 |
| A3SS | 10446 | ENSG00000183337 | BCOR | chrX | - | 40063607 | 40063952 | 40063607 | 40063850 | 40064335 | 40064599 | 51 |
| A3SS | 9057 | ENSG00000116539 | ASH1L | chr1 | - | 155347655 | 155347904 | 155347655 | 155347901 | 155349326 | 155349459 | 135 |
| A3SS | 10427 | ENSG00000285171 | NA | chrX | - | 71106531 | 71106662 | 71106531 | 71106590 | 71108276 | 71108346 | 10 |
| A3SS | 11925 | ENSG00000066926 | FECH | chr18 | - | 57571391 | 57571603 | 57571391 | 57571540 | 57573245 | 57573365 | 13 |
| A3SS | 1861 | ENSG00000103313 | MEFV | chr16 | - | 3243859 | 3244067 | 3243859 | 3243892 | 3244253 | 3244286 | 5 |
| A3SS | 7731 | ENSG00000115525 | ST3GAL5 | chr2 | - | 85847860 | 85850834 | 85847860 | 85848204 | 85851263 | 85851703 | 71 |
| A3SS | 6766 | ENSG00000158796 | DEDD | chr1 | - | 161123074 | 161123221 | 161123074 | 161123174 | 161123838 | 161123946 | 82 |
| A3SS | 8328 | ENSG00000039650 | PNKP | chr19 | - | 49865126 | 49865426 | 49865126 | 49865242 | 49866398 | 49866445 | 138 |
| A3SS | 1327 | ENSG00000109113 | RAB34 | chr17 | - | 28717212 | 28717620 | 28717212 | 28717425 | 28718121 | 28718167 | 2 |
| A3SS | 1328 | ENSG00000109113 | RAB34 | chr17 | - | 28717212 | 28717647 | 28717212 | 28717425 | 28718121 | 28718167 | 2 |
| A3SS | 2886 | ENSG00000162711 | NLRP3 | chr1 | + | 247418052 | 247419077 | 247418756 | 247419077 | 247416155 | 247416193 | 83 |
| A3SS | 5947 | ENSG00000095787 | WAC | chr10 | + | 28595694 | 28596041 | 28595732 | 28596041 | 28590719 | 28590832 | 24 |
| A3SS | 4618 | ENSG00000143554 | SLC27A3 | chr1 | + | 153779825 | 153780157 | 153779930 | 153780157 | 153779342 | 153779473 | 117 |
| A3SS | 6090 | ENSG00000079313 | REXO1 | chr19 | - | 1819933 | 1820180 | 1819933 | 1820057 | 1820263 | 1820395 | 8 |
| A3SS | 10230 | ENSG00000029363 | BCLAF1 | chr6 | - | 136277864 | 136278776 | 136277864 | 136278770 | 136279762 | 136279876 | 676 |
| A3SS | 10796 | ENSG00000168066 | SF1 | chr11 | - | 64767191 | 64767275 | 64767191 | 64767251 | 64767570 | 64767844 | 7 |
| A3SS | 7730 | ENSG00000115525 | ST3GAL5 | chr2 | - | 85847860 | 85848381 | 85847860 | 85848204 | 85851263 | 85851703 | 72 |
| A3SS | 2436 | ENSG00000140382 | HMG20A | chr15 | + | 77467094 | 77467307 | 77467182 | 77467307 | 77464239 | 77464387 | 135 |
| A3SS | 9408 | ENSG00000167483 | NIBAN3 | chr19 | + | 17545222 | 17546797 | 17546685 | 17546797 | 17543523 | 17543631 | 1 |
| A3SS | 10428 | ENSG00000285171 | NA | chrX | - | 71106531 | 71106703 | 71106531 | 71106590 | 71108276 | 71108346 | 13 |
| A3SS | 6685 | ENSG00000183918 | SH2D1A | chrX | + | 124370175 | 124370320 | 124370230 | 124370320 | 124365760 | 124365824 | 142 |
| A3SS | 7179 | ENSG00000101361 | NOP56 | chr20 | + | 2655237 | 2655512 | 2655324 | 2655512 | 2654748 | 2654947 | 6 |
| A3SS | 1702 | ENSG00000033011 | ALG1 | chr16 | + | 5074878 | 5075536 | 5075387 | 5075536 | 5073152 | 5073256 | 2 |
| A3SS | 10602 | ENSG00000215021 | PHB2 | chr12 | - | 6969955 | 6970068 | 6969955 | 6970063 | 6970195 | 6970280 | 2 |
| A3SS | 9100 | ENSG00000163939 | PBRM1 | chr3 | - | 52587352 | 52587510 | 52587352 | 52587435 | 52589069 | 52589255 | 141 |
| A3SS | 6396 | ENSG00000143353 | LYPLAL1 | chr1 | + | 219210514 | 219210647 | 219210531 | 219210647 | 219193129 | 219193251 | 3 |
| A3SS | 90 | ENSG00000138867 | GUCD1 | chr22 | - | 24540437 | 24543100 | 24540437 | 24543097 | 24543841 | 24544083 | 13 |
| A3SS | 4055 | ENSG00000003056 | M6PR | chr12 | - | 8946228 | 8946405 | 8946228 | 8946387 | 8946715 | 8946824 | 145 |
| A3SS | 12087 | ENSG00000087053 | MTMR2 | chr11 | - | 95887611 | 95887759 | 95887611 | 95887684 | 95888155 | 95888261 | 1 |
| A3SS | 127 | ENSG00000100031 | GGT1 | chr22 | + | 24611074 | 24611245 | 24611077 | 24611245 | 24610249 | 24610531 | 12 |
| A3SS | 7455 | ENSG00000122223 | CD244 | chr1 | - | 160841209 | 160841500 | 160841209 | 160841485 | 160841583 | 160841901 | 13 |
| A3SS | 5830 | ENSG00000170322 | NFRKB | chr11 | - | 129884744 | 129884882 | 129884744 | 129884846 | 129885434 | 129885609 | 38 |
| A3SS | 6393 | ENSG00000143353 | LYPLAL1 | chr1 | + | 219210514 | 219210647 | 219210531 | 219210647 | 219179146 | 219179246 | 4 |
| A3SS | 2913 | ENSG00000175061 | SNHG29 | chr17 | + | 16441356 | 16442018 | 16441367 | 16442018 | 16441073 | 16441130 | 41 |
| A3SS | 4451 | ENSG00000114867 | EIF4G1 | chr3 | + | 184316071 | 184316218 | 184316131 | 184316218 | 184315762 | 184315856 | 9 |
| A3SS | 11849 | ENSG00000173120 | KDM2A | chr11 | + | 67217730 | 67217884 | 67217755 | 67217884 | 67215855 | 67215949 | 249 |
| A3SS | 6773 | ENSG00000092847 | AGO1 | chr1 | + | 35893591 | 35893810 | 35893673 | 35893810 | 35893096 | 35893278 | 5 |
| A3SS | 6147 | ENSG00000178498 | DTX3 | chr12 | + | 57605988 | 57606252 | 57606189 | 57606252 | 57605570 | 57605731 | 2 |
| A3SS | 4169 | ENSG00000100003 | SEC14L2 | chr22 | + | 30407414 | 30407603 | 30407523 | 30407603 | 30407094 | 30407154 | 12 |
| A3SS | 13326 | ENSG00000081189 | MEF2C | chr5 | - | 88823734 | 88823930 | 88823734 | 88823927 | 88887501 | 88887598 | 7 |
| A3SS | 8521 | ENSG00000173559 | NABP1 | chr2 | + | 191683289 | 191683804 | 191683728 | 191683804 | 191682491 | 191682595 | 141 |
| A3SS | 6598 | ENSG00000133805 | AMPD3 | chr11 | + | 10504548 | 10504659 | 10504579 | 10504659 | 10502720 | 10502894 | 96 |
| A3SS | 8494 | ENSG00000106799 | TGFBR1 | chr9 | + | 99132496 | 99132739 | 99132508 | 99132739 | 99128854 | 99129100 | 28 |
| A3SS | 3220 | ENSG00000153113 | CAST | chr5 | + | 96726753 | 96726859 | 96726793 | 96726859 | 96722638 | 96722698 | 19 |
| A3SS | 1978 | ENSG00000126602 | TRAP1 | chr16 | - | 3674338 | 3674597 | 3674338 | 3674494 | 3675323 | 3675397 | 3 |
| A3SS | 861 | ENSG00000160410 | SHKBP1 | chr19 | + | 40578447 | 40578542 | 40578461 | 40578542 | 40578153 | 40578212 | 2 |
| A3SS | 8297 | ENSG00000162923 | WDR26 | chr1 | - | 224418259 | 224418416 | 224418259 | 224418398 | 224419517 | 224419615 | 444 |
| A3SS | 8200 | ENSG00000131653 | TRAF7 | chr16 | + | 2165856 | 2165936 | 2165878 | 2165936 | 2163882 | 2164001 | 0 |
| A3SS | 8826 | ENSG00000118495 | PLAGL1 | chr6 | - | 143966157 | 143966285 | 143966157 | 143966198 | 143968906 | 143968978 | 11 |
| A3SS | 3941 | ENSG00000203875 | SNHG5 | chr6 | - | 85677794 | 85677875 | 85677794 | 85677868 | 85677953 | 85678032 | 250 |
| A3SS | 615 | ENSG00000105204 | DYRK1B | chr19 | - | 39826671 | 39826987 | 39826671 | 39826867 | 39827284 | 39827425 | 17 |
| A3SS | 5134 | ENSG00000123473 | STIL | chr1 | - | 47262902 | 47263116 | 47262902 | 47263113 | 47269634 | 47269866 | 2 |
| A3SS | 11164 | ENSG00000176623 | RMDN1 | chr8 | - | 86472348 | 86474358 | 86472348 | 86472479 | 86474819 | 86474953 | 92 |
| A3SS | 9143 | ENSG00000130023 | ERMARD | chr6 | + | 169755282 | 169755422 | 169755287 | 169755422 | 169753863 | 169754032 | 30 |
| A3SS | 437 | ENSG00000161265 | U2AF1L4 | chr19 | - | 35744322 | 35744738 | 35744322 | 35744421 | 35745124 | 35745187 | 64 |
| A3SS | 3578 | ENSG00000158290 | CUL4B | chrX | - | 120543726 | 120543964 | 120543726 | 120543809 | 120544113 | 120544203 | 6 |
| A3SS | 11599 | ENSG00000213930 | GALT | chr9 | + | 34648091 | 34648171 | 34648114 | 34648171 | 34647831 | 34647961 | 11 |
| A3SS | 10795 | ENSG00000168066 | SF1 | chr11 | - | 64767191 | 64767275 | 64767191 | 64767246 | 64767570 | 64767844 | 26 |
| A3SS | 8902 | ENSG00000115307 | AUP1 | chr2 | - | 74526940 | 74527121 | 74526940 | 74527059 | 74527247 | 74527363 | 93 |
| A3SS | 5504 | ENSG00000105323 | HNRNPUL1 | chr19 | + | 41305675 | 41305867 | 41305831 | 41305867 | 41303971 | 41304261 | 90 |
| A3SS | 6686 | ENSG00000135953 | MFSD9 | chr2 | - | 102723699 | 102723909 | 102723699 | 102723878 | 102726844 | 102726974 | 11 |
| A3SS | 5252 | ENSG00000247828 | TMEM161B-AS1 | chr5 | + | 88285740 | 88285851 | 88285743 | 88285851 | 88283009 | 88283086 | 11 |
| A3SS | 7367 | ENSG00000119004 | CYP20A1 | chr2 | + | 203272645 | 203272748 | 203272669 | 203272748 | 203266513 | 203266681 | 10 |
| A3SS | 4668 | ENSG00000179262 | RAD23A | chr19 | + | 12948496 | 12948552 | 12948513 | 12948552 | 12948176 | 12948358 | 152 |
| A3SS | 4609 | ENSG00000138246 | DNAJC13 | chr3 | + | 132465950 | 132466070 | 132465994 | 132466070 | 132463695 | 132463817 | 19 |
| A3SS | 2119 | ENSG00000167965 | MLST8 | chr16 | + | 2207192 | 2207345 | 2207196 | 2207345 | 2207034 | 2207110 | 24 |
| A3SS | 11117 | ENSG00000120533 | ENY2 | chr8 | + | 109340443 | 109340563 | 109340488 | 109340563 | 109339319 | 109339390 | 11 |
| A3SS | 8954 | ENSG00000114770 | ABCC5 | chr3 | - | 183985303 | 183985455 | 183985303 | 183985378 | 183987769 | 183987917 | 15 |
| A3SS | 1791 | ENSG00000188603 | CLN3 | chr16 | - | 28487455 | 28488662 | 28487455 | 28487541 | 28489289 | 28489386 | 72 |
| A3SS | 3304 | ENSG00000178338 | ZNF354B | chr5 | + | 178887608 | 178888122 | 178887620 | 178888122 | 178885613 | 178885733 | 9 |
| A3SS | 438 | ENSG00000161265 | U2AF1L4 | chr19 | - | 35744322 | 35744981 | 35744322 | 35744421 | 35745124 | 35745212 | 56 |
| A3SS | 3014 | ENSG00000129515 | SNX6 | chr14 | - | 34604134 | 34604268 | 34604134 | 34604174 | 34605595 | 34605717 | 22 |
| A3SS | 11825 | ENSG00000106290 | TAF6 | chr7 | - | 100113348 | 100113769 | 100113348 | 100113405 | 100113867 | 100113954 | 62 |
| A3SS | 5859 | ENSG00000028310 | BRD9 | chr5 | - | 889586 | 889766 | 889586 | 889647 | 891154 | 891287 | 31 |
| A3SS | 1894 | ENSG00000132600 | PRMT7 | chr16 | + | 68353394 | 68353566 | 68353491 | 68353566 | 68352247 | 68352409 | 10 |
| A3SS | 5702 | ENSG00000104419 | NDRG1 | chr8 | - | 133250439 | 133250676 | 133250439 | 133250543 | 133254538 | 133254595 | 32 |
| A3SS | 1953 | ENSG00000149923 | PPP4C | chr16 | + | 30082745 | 30083567 | 30083393 | 30083567 | 30082483 | 30082534 | 158 |
| A3SS | 5350 | ENSG00000116001 | TIA1 | chr2 | - | 70227734 | 70227822 | 70227734 | 70227791 | 70228343 | 70228462 | 88 |
| A3SS | 3334 | ENSG00000115758 | ODC1 | chr2 | - | 10444473 | 10444647 | 10444473 | 10444547 | 10444930 | 10445049 | 227 |
| A3SS | 286 | ENSG00000104880 | ARHGEF18 | chr19 | + | 7440188 | 7440482 | 7440343 | 7440482 | 7383061 | 7383203 | 28 |
| A3SS | 2048 | ENSG00000167261 | DPEP2 | chr16 | - | 67990820 | 67990997 | 67990820 | 67990939 | 67991837 | 67991979 | 35 |
| A3SS | 1140 | ENSG00000278053 | DDX52 | chr17 | - | 37619767 | 37621277 | 37619767 | 37619839 | 37621397 | 37621520 | 76 |
| A3SS | 2647 | ENSG00000075413 | MARK3 | chr14 | + | 103468032 | 103468186 | 103468085 | 103468186 | 103467078 | 103467191 | 136 |
| A3SS | 4227 | ENSG00000131370 | SH3BP5 | chr3 | - | 15304102 | 15304299 | 15304102 | 15304231 | 15330503 | 15330566 | 41 |
| A3SS | 12025 | ENSG00000175166 | PSMD2 | chr3 | + | 184301770 | 184302071 | 184301846 | 184302071 | 184301536 | 184301658 | 11 |
| A3SS | 1680 | ENSG00000125124 | BBS2 | chr16 | - | 56496966 | 56497124 | 56496966 | 56497079 | 56497742 | 56497880 | 7 |
| A3SS | 8128 | ENSG00000091527 | CDV3 | chr3 | + | 133584001 | 133584150 | 133584004 | 133584150 | 133575038 | 133575115 | 288 |
| A3SS | 8141 | ENSG00000102081 | FMR1 | chrX | + | 147944904 | 147945051 | 147944943 | 147945051 | 147943130 | 147943326 | 26 |
| A3SS | 2989 | ENSG00000154309 | DISP1 | chr1 | + | 222836922 | 222837384 | 222836995 | 222837384 | 222835118 | 222835215 | 4 |
| A3SS | 11541 | ENSG00000157693 | TMEM268 | chr9 | + | 114624135 | 114624459 | 114624349 | 114624459 | 114617117 | 114617301 | 10 |
| A3SS | 5282 | ENSG00000127054 | INTS11 | chr1 | - | 1313731 | 1314105 | 1313731 | 1313921 | 1314300 | 1314365 | 12 |
| A3SS | 10256 | ENSG00000168807 | SNTB2 | chr16 | + | 69260049 | 69260260 | 69260170 | 69260260 | 69245601 | 69245815 | 133 |
| A3SS | 4874 | ENSG00000186074 | CD300LF | chr17 | - | 74703034 | 74703143 | 74703034 | 74703098 | 74704477 | 74704816 | 56 |
| A3SS | 4203 | ENSG00000114745 | GORASP1 | chr3 | - | 39100304 | 39100877 | 39100304 | 39100454 | 39101015 | 39101102 | 24 |
| A3SS | 9030 | ENSG00000137478 | FCHSD2 | chr11 | - | 72849754 | 72849889 | 72849754 | 72849879 | 72867864 | 72868026 | 97 |
| A3SS | 2674 | ENSG00000100888 | CHD8 | chr14 | - | 21400150 | 21400331 | 21400150 | 21400307 | 21400412 | 21400612 | 8 |
| A3SS | 6532 | ENSG00000123636 | BAZ2B | chr2 | - | 159448241 | 159448409 | 159448241 | 159448403 | 159453612 | 159453801 | 58 |
| A3SS | 7304 | ENSG00000077254 | USP33 | chr1 | - | 77721830 | 77721925 | 77721830 | 77721901 | 77722023 | 77722196 | 117 |
| A3SS | 5980 | ENSG00000156873 | PHKG2 | chr16 | + | 30756354 | 30756520 | 30756366 | 30756520 | 30756181 | 30756272 | 5 |
| A3SS | 8600 | ENSG00000188092 | GPR89B | chr1 | + | 147968874 | 147968963 | 147968894 | 147968963 | 147968331 | 147968428 | 22 |
| A3SS | 6934 | ENSG00000135241 | PNPLA8 | chr7 | - | 108514435 | 108515574 | 108514435 | 108515447 | 108521475 | 108521521 | 236 |
| A3SS | 8233 | ENSG00000162910 | MRPL55 | chr1 | - | 228108234 | 228108321 | 228108234 | 228108318 | 228108436 | 228108474 | 1 |
| A3SS | 10486 | ENSG00000177646 | ACAD9 | chr3 | + | 128907907 | 128908264 | 128908184 | 128908264 | 128906120 | 128906249 | 3 |
| A3SS | 8134 | ENSG00000161813 | LARP4 | chr12 | + | 50430494 | 50430570 | 50430497 | 50430570 | 50428934 | 50429090 | 95 |
| A3SS | 6392 | ENSG00000143353 | LYPLAL1 | chr1 | + | 219193081 | 219193251 | 219193129 | 219193251 | 219179146 | 219179246 | 183 |
| A3SS | 9385 | ENSG00000168036 | CTNNB1 | chr3 | + | 41239640 | 41239900 | 41239659 | 41239900 | 41239133 | 41239354 | 5 |
| A3SS | 5909 | ENSG00000055609 | KMT2C | chr7 | - | 152156204 | 152156346 | 152156204 | 152156274 | 152157760 | 152157922 | 173 |
| A3SS | 10719 | ENSG00000215301 | DDX3X | chrX | + | 41343736 | 41344128 | 41344029 | 41344128 | 41343215 | 41343351 | 494 |
| A3SS | 5145 | ENSG00000064687 | ABCA7 | chr19 | + | 1057914 | 1058059 | 1057968 | 1058059 | 1057313 | 1057429 | 77 |
| A3SS | 1323 | ENSG00000109113 | RAB34 | chr17 | - | 28715992 | 28716061 | 28715992 | 28716058 | 28716902 | 28716994 | 4 |
| A3SS | 7136 | ENSG00000127124 | HIVEP3 | chr1 | - | 41506364 | 41511266 | 41506364 | 41511263 | 41512815 | 41513750 | 19 |
| A3SS | 6010 | ENSG00000145687 | SSBP2 | chr5 | - | 81466973 | 81467065 | 81466973 | 81467041 | 81473699 | 81473770 | 4 |
| A3SS | 4325 | ENSG00000163930 | BAP1 | chr3 | - | 52402778 | 52402940 | 52402778 | 52402871 | 52403137 | 52403298 | 20 |
| A3SS | 4478 | ENSG00000136560 | TANK | chr2 | + | 161203483 | 161203595 | 161203486 | 161203595 | 161179613 | 161179761 | 4 |
| A3SS | 11788 | ENSG00000149182 | ARFGAP2 | chr11 | - | 47173758 | 47173915 | 47173758 | 47173840 | 47175014 | 47175098 | 31 |
| A3SS | 7087 | ENSG00000136108 | CKAP2 | chr13 | + | 52460895 | 52460974 | 52460898 | 52460974 | 52456522 | 52456607 | 16 |
| A3SS | 7584 | ENSG00000124772 | CPNE5 | chr6 | - | 36744267 | 36744526 | 36744267 | 36744325 | 36745047 | 36745150 | 3 |
| A3SS | 4029 | ENSG00000196954 | CASP4 | chr11 | - | 104951895 | 104952063 | 104951895 | 104952005 | 104954746 | 104955001 | 53 |
| A3SS | 8007 | ENSG00000149499 | EML3 | chr11 | - | 62602758 | 62602889 | 62602758 | 62602779 | 62603148 | 62603247 | 102 |
| A3SS | 7379 | ENSG00000145246 | ATP10D | chr4 | + | 47536685 | 47536938 | 47536730 | 47536938 | 47536436 | 47536564 | 86 |
| A3SS | 9472 | ENSG00000089820 | ARHGAP4 | chrX | - | 153921096 | 153921223 | 153921096 | 153921159 | 153921364 | 153921527 | 0 |
| A3SS | 5985 | ENSG00000066322 | ELOVL1 | chr1 | - | 43364927 | 43365097 | 43364927 | 43365008 | 43365185 | 43365376 | 7 |
| A3SS | 7005 | ENSG00000144218 | AFF3 | chr2 | - | 99554310 | 99554534 | 99554310 | 99554486 | 99554682 | 99554732 | 7 |
| A3SS | 7717 | ENSG00000049167 | ERCC8 | chr5 | - | 60924290 | 60924508 | 60924290 | 60924367 | 60928863 | 60928959 | 14 |
| A3SS | 1739 | ENSG00000179583 | CIITA | chr16 | + | 10901797 | 10902184 | 10902037 | 10902184 | 10901513 | 10901558 | 42 |
| A3SS | 10262 | ENSG00000139218 | SCAF11 | chr12 | - | 45919130 | 45922194 | 45919130 | 45919309 | 45922462 | 45922582 | 519 |
| A3SS | 10961 | ENSG00000095485 | CWF19L1 | chr10 | - | 100260965 | 100261044 | 100260965 | 100261028 | 100261978 | 100262063 | 71 |
| A3SS | 8081 | ENSG00000123240 | OPTN | chr10 | + | 13118887 | 13119040 | 13118905 | 13119040 | 13116266 | 13116340 | 163 |
| A3SS | 13417 | ENSG00000071564 | TCF3 | chr19 | - | 1612206 | 1612433 | 1612206 | 1612430 | 1615685 | 1615821 | 27 |
| A3SS | 11775 | ENSG00000214413 | BBIP1 | chr10 | - | 110901025 | 110901279 | 110901025 | 110901122 | 110901537 | 110901612 | 41 |
| A3SS | 8341 | ENSG00000172890 | NADSYN1 | chr11 | + | 71476032 | 71477438 | 71477344 | 71477438 | 71474394 | 71474526 | 16 |
| A3SS | 11606 | ENSG00000122565 | CBX3 | chr7 | + | 26208392 | 26208555 | 26208465 | 26208555 | 26206367 | 26206510 | 968 |
| A3SS | 6549 | ENSG00000127463 | EMC1 | chr1 | - | 19219568 | 19219698 | 19219568 | 19219669 | 19220763 | 19220848 | 39 |
| A3SS | 10040 | ENSG00000146828 | SLC12A9 | chr7 | + | 100861011 | 100861262 | 100861137 | 100861262 | 100860149 | 100860232 | 3 |
| A3SS | 12898 | ENSG00000135720 | DYNC1LI2 | chr16 | - | 66751272 | 66751386 | 66751272 | 66751346 | 66751484 | 66751601 | 0 |
| A3SS | 2921 | ENSG00000130779 | CLIP1 | chr12 | - | 122274037 | 122274162 | 122274037 | 122274143 | 122278153 | 122278203 | 148 |
| A3SS | 2368 | ENSG00000185787 | MORF4L1 | chr15 | + | 78884966 | 78885083 | 78884969 | 78885083 | 78880511 | 78880579 | 15 |
| A3SS | 11677 | ENSG00000137073 | UBAP2 | chr9 | - | 33971650 | 33971754 | 33971650 | 33971737 | 33973182 | 33973237 | 71 |
| A3SS | 9260 | ENSG00000065029 | ZNF76 | chr6 | + | 35290252 | 35290382 | 35290265 | 35290382 | 35287645 | 35287845 | 3 |
| A3SS | 2288 | ENSG00000102901 | CENPT | chr16 | - | 67832011 | 67832136 | 67832011 | 67832108 | 67832227 | 67832315 | 2 |
| A3SS | 3613 | ENSG00000125089 | SH3TC1 | chr4 | + | 8209664 | 8209822 | 8209747 | 8209822 | 8205166 | 8205366 | 29 |
| A3SS | 9363 | ENSG00000141642 | ELAC1 | chr18 | + | 50984095 | 50984563 | 50984486 | 50984563 | 50974396 | 50974561 | 50 |
| A3SS | 5637 | ENSG00000119402 | FBXW2 | chr9 | - | 120787768 | 120788278 | 120787768 | 120788186 | 120793148 | 120793257 | 151 |
| A3SS | 8258 | ENSG00000082269 | FAM135A | chr6 | + | 70477158 | 70477332 | 70477292 | 70477332 | 70475662 | 70475733 | 99 |
| A3SS | 9817 | ENSG00000168137 | SETD5 | chr3 | + | 9434276 | 9434485 | 9434333 | 9434485 | 9433844 | 9433950 | 64 |
| A3SS | 178 | ENSG00000198089 | SFI1 | chr22 | + | 31604868 | 31606430 | 31606327 | 31606430 | 31604308 | 31604404 | 46 |
| A3SS | 5359 | ENSG00000048740 | CELF2 | chr10 | + | 11321188 | 11321386 | 11321206 | 11321386 | 11314138 | 11314270 | 11 |
| A3SS | 3317 | ENSG00000104731 | KLHDC4 | chr16 | - | 87707811 | 87708075 | 87707811 | 87708072 | 87708349 | 87708466 | 19 |
| A3SS | 11349 | ENSG00000165525 | NEMF | chr14 | - | 49814753 | 49814857 | 49814753 | 49814807 | 49825866 | 49825955 | 145 |
| A3SS | 11552 | ENSG00000266714 | MYO15B | chr17 | + | 75592427 | 75592541 | 75592463 | 75592541 | 75592237 | 75592301 | 4 |
| A3SS | 1969 | ENSG00000197818 | SLC9A8 | chr20 | + | 49850761 | 49850844 | 49850809 | 49850844 | 49849578 | 49849680 | 20 |
| A3SS | 2897 | ENSG00000143379 | SETDB1 | chr1 | + | 150929966 | 150930118 | 150929997 | 150930118 | 150927703 | 150927974 | 152 |
| A3SS | 8711 | ENSG00000123983 | ACSL3 | chr2 | + | 222918045 | 222918155 | 222918069 | 222918155 | 222916318 | 222916496 | 143 |
| A3SS | 5774 | ENSG00000049541 | RFC2 | chr7 | - | 74249738 | 74249829 | 74249738 | 74249780 | 74252428 | 74252498 | 7 |
| A3SS | 7907 | ENSG00000166801 | FAM111A | chr11 | + | 59144537 | 59144896 | 59144808 | 59144896 | 59143496 | 59143695 | 43 |
| A3SS | 9816 | ENSG00000077454 | LRCH4 | chr7 | - | 100577085 | 100577389 | 100577085 | 100577154 | 100577663 | 100577740 | 4 |
| A3SS | 3909 | ENSG00000153234 | NR4A2 | chr2 | - | 156326149 | 156326328 | 156326149 | 156326207 | 156326717 | 156326920 | 44 |
| A3SS | 13266 | ENSG00000137100 | DCTN3 | chr9 | - | 34614041 | 34614198 | 34614041 | 34614101 | 34614709 | 34614768 | 15 |
| A3SS | 7170 | ENSG00000135913 | USP37 | chr2 | - | 218558497 | 218558717 | 218558497 | 218558677 | 218560819 | 218560883 | 10 |
| A3SS | 1848 | ENSG00000205336 | ADGRG1 | chr16 | + | 57651184 | 57651474 | 57651199 | 57651474 | 57650252 | 57650351 | 4 |
| A3SS | 4001 | ENSG00000108591 | DRG2 | chr17 | + | 18104559 | 18104681 | 18104622 | 18104681 | 18103800 | 18103889 | 2 |
| A3SS | 10490 | ENSG00000177646 | ACAD9 | chr3 | + | 128910390 | 128910813 | 128910740 | 128910813 | 128910020 | 128910149 | 4 |
| A3SS | 13180 | ENSG00000256683 | ZNF350 | chr19 | - | 51969004 | 51969145 | 51969004 | 51969131 | 51974345 | 51974531 | 6 |
| A3SS | 6634 | ENSG00000159023 | EPB41 | chr1 | + | 28987430 | 28987905 | 28987447 | 28987905 | 28887123 | 28887210 | 47 |
| A3SS | 9593 | ENSG00000088038 | CNOT3 | chr19 | + | 54145541 | 54145817 | 54145597 | 54145817 | 54144236 | 54144332 | 17 |
| A3SS | 8416 | ENSG00000085978 | ATG16L1 | chr2 | + | 233289853 | 233290353 | 233290247 | 233290353 | 233282681 | 233282753 | 92 |
| A3SS | 4499 | ENSG00000163755 | HPS3 | chr3 | + | 149167033 | 149167240 | 149167064 | 149167240 | 149163841 | 149163949 | 180 |
| A3SS | 2917 | ENSG00000196367 | TRRAP | chr7 | + | 98910509 | 98910607 | 98910512 | 98910607 | 98910055 | 98910419 | 199 |
| A3SS | 12141 | ENSG00000255717 | SNHG1 | chr11 | - | 62853800 | 62853849 | 62853800 | 62853833 | 62854017 | 62854064 | 2 |
| A3SS | 5720 | ENSG00000116030 | SUMO1 | chr2 | - | 202214356 | 202214434 | 202214356 | 202214429 | 202220031 | 202220106 | 347 |
| A3SS | 3945 | ENSG00000203875 | SNHG5 | chr6 | - | 85677423 | 85678032 | 85677423 | 85677492 | 85678135 | 85678188 | 259 |
| A3SS | 9730 | ENSG00000272333 | KMT2B | chr19 | + | 35737588 | 35737743 | 35737635 | 35737743 | 35737085 | 35737263 | 0 |
| A3SS | 5532 | ENSG00000148019 | CEP78 | chr9 | + | 78254835 | 78254964 | 78254842 | 78254964 | 78253231 | 78253277 | 67 |
| A3SS | 3662 | ENSG00000077232 | DNAJC10 | chr2 | + | 182743597 | 182743712 | 182743657 | 182743712 | 182741242 | 182741356 | 156 |
| A3SS | 2297 | ENSG00000140199 | SLC12A6 | chr15 | - | 34238231 | 34238401 | 34238231 | 34238331 | 34238964 | 34239160 | 182 |
| A3SS | 387 | ENSG00000167685 | ZNF444 | chr19 | + | 56159623 | 56160893 | 56159626 | 56160893 | 56158493 | 56158602 | 9 |
| A3SS | 10418 | ENSG00000103037 | SETD6 | chr16 | + | 58518019 | 58518231 | 58518050 | 58518231 | 58516807 | 58516928 | 11 |
| A3SS | 4853 | ENSG00000137074 | APTX | chr9 | - | 32987543 | 32988129 | 32987543 | 32987846 | 32989758 | 32989895 | 28 |
| A3SS | 11800 | ENSG00000089159 | PXN | chr12 | - | 120215102 | 120215417 | 120215102 | 120215273 | 120215559 | 120215661 | 37 |

| | | | | | | | | | | | | | | | | | | | | | | | | | | | | | | | | | | | | | | | | | | | | | | |
|---|---|---|---|---|---|---|---|---|---|---|---|---|---|---|---|---|---|---|---|---|---|---|---|---|---|---|---|---|---|---|---|---|---|---|---|---|---|---|---|---|---|---|---|---|---|---|
| A3SS | 1070 | ENSG00000154889 | MPPE1 | chr18 | - | 11886712 | 11886781 | 11886712 | 11886778 | 11886916 | 11887025 | 11 | 2 | 5 | 9 | 6 | 22 | 11 | 12 | 11 | 12 | 0 | 1 | 7 | 10 | 0 | 32 | 21 | 23 | 12 | 17 | 152 | 149 | 0.003272 | 0.031711255 | 0.329 | 0.151 | 0.29 | 0.445 | 0.329 | 0 | 0.045 | 0.23 | 0.45 | 0 | 0.164 |
| A3SS | 4636 | ENSG00000187862 | TTC24 | chr1 | + | 156583796 | 156583895 | 156583799 | 156583895 | 156583337 | 156583450 | 16 | 17 | 15 | 26 | 3 | 2 | 0 | 0 | 2 | 1 | 7 | 14 | 6 | 4 | 16 | 0 | 13 | 5 | 0 | 0 | 152 | 149 | 0.003278 | 0.031724341 | 0.887 | 1 | 1 | 0.927 | 0.746 | 1 | 0.514 | 0.541 | 1 | 1 | 0.101 |
| A3SS | 5677 | ENSG00000117682 | DHDDS | chr1 | + | 26442707 | 26442873 | 26442730 | 26442873 | 26438167 | 26438284 | 5 | 5 | 8 | 11 | 12 | 68 | 39 | 100 | 68 | 85 | 0 | 9 | 3 | 12 | 5 | 178 | 77 | 55 | 101 | 138 | 172 | 149 | 0.003292 | 0.031806729 | 0.06 | 0.1 | 0.065 | 0.123 | 0.109 | 0 | 0.092 | 0.045 | 0.093 | 0.03 | 0.039 |
| A3SS | 9791 | ENSG00000012124 | CD22 | chr19 | + | 35331926 | 35332074 | 35332018 | 35332074 | 35329175 | 35329230 | 13 | 12 | 15 | 39 | 15 | 42 | 14 | 29 | 41 | 40 | 36 | 70 | 21 | 39 | 66 | 62 | 69 | 6 | 17 | 58 | 241 | 149 | 0.003294 | 0.031806729 | 0.161 | 0.346 | 0.242 | 0.37 | 0.188 | 0.264 | 0.385 | 0.684 | 0.586 | 0.413 | -0.205 |
| A3SS | 5004 | ENSG00000117385 | P3H1 | chr1 | - | 42757782 | 42757945 | 42757782 | 42757922 | 42758851 | 42758983 | 2 | 0 | 6 | 1 | 0 | 93 | 62 | 73 | 82 | 53 | 15 | 0 | 4 | 0 | 9 | 144 | 89 | 87 | 46 | 128 | 172 | 149 | 0.003304 | 0.031826135 | 0.018 | 0 | 0.066 | 0.01 | 0 | 0.083 | 0 | 0.038 | 0 | 0.057 | -0.017 |
| A3SS | 10154 | ENSG00000184863 | RBM33 | chr7 | + | 155711172 | 155711455 | 155711202 | 155711455 | 155700772 | 155700944 | 12 | 9 | 10 | 13 | 10 | 3 | 0 | 0 | 1 | 5 | 4 | 5 | 14 | 2 | 0 | 5 | 0 | 4 | 3 | 9 | 179 | 149 | 0.003304 | 0.031826135 | 0.769 | 1 | 1 | 0.915 | 0.625 | 0.4 | 1 | 0.744 | 0.357 | 0 | 0.362 |
| A3SS | 7126 | ENSG00000117360 | PRPF3 | chr1 | + | 150344375 | 150344547 | 150344433 | 150344547 | 150344161 | 150344261 | 21 | 23 | 50 | 29 | 23 | 76 | 70 | 95 | 95 | 93 | 19 | 56 | 88 | 97 | 30 | 97 | 85 | 96 | 99 | 56 | 207 | 149 | 0.003347 | 0.032202676 | 0.166 | 0.191 | 0.275 | 0.18 | 0.151 | 0.124 | 0.322 | 0.398 | 0.414 | 0.278 | -0.115 |
| A3SS | 12107 | ENSG00000137760 | ALKBH8 | chr11 | - | 107553846 | 107553978 | 107553846 | 107553973 | 107556765 | 107557003 | 38 | 53 | 39 | 40 | 68 | 13 | 9 | 14 | 10 | 7 | 16 | 45 | 43 | 22 | 24 | 0 | 8 | 5 | 0 | 3 | 154 | 149 | 0.003358 | 0.032266316 | 0.739 | 0.851 | 0.729 | 0.795 | 0.904 | 1 | 0.845 | 0.893 | 1 | 0.886 | -0.121 |
| A3SS | 10922 | ENSG00000075856 | SART3 | chr12 | - | 108544426 | 108544532 | 108544426 | 108544478 | 108545138 | 108545323 | 16 | 22 | 27 | 19 | 18 | 113 | 46 | 106 | 126 | 135 | 17 | 19 | 5 | 3 | 2 | 99 | 69 | 101 | 58 | 89 | 203 | 149 | 0.003380 | 0.032441103 | 0.094 | 0.26 | 0.158 | 0.1 | 0.089 | 0.112 | 0.168 | 0.035 | 0.037 | 0.016 | 0.067 |
| A3SS | 12194 | ENSG00000186104 | CYP2R1 | chr11 | - | 14879113 | 14879443 | 14879113 | 14879207 | 14880135 | 14880768 | 100 | 73 | 72 | 65 | 115 | 9 | 4 | 10 | 11 | 12 | 29 | 51 | 40 | 23 | 0 | 0 | 2 | 0 | 0 | 2 | 298 | 149 | 0.003385 | 0.032447823 | 0.847 | 0.901 | 0.783 | 0.747 | 0.827 | 1 | 0.927 | 1 | 1 | 0 | 0.036 |
| A3SS | 7269 | ENSG00000107223 | EDF1 | chr9 | - | 136862118 | 136862496 | 136862118 | 136862345 | 136862905 | 136862999 | 50 | 22 | 54 | 60 | 58 | 116 | 79 | 148 | 142 | 138 | 103 | 45 | 38 | 67 | 61 | 115 | 87 | 62 | 81 | 193 | 298 | 149 | 0.003395 | 0.032507384 | 0.177 | 0.122 | 0.154 | 0.174 | 0.174 | 0.309 | 0.205 | 0.235 | 0.293 | 0.136 | -0.075 |
| A3SS | 7738 | ENSG00000115525 | ST3GAL5 | chr2 | - | 85847860 | 85848381 | 85847860 | 85848084 | 85861180 | 85861292 | 133 | 137 | 114 | 186 | 168 | 3 | 5 | 8 | 0 | 2 | 314 | 220 | 107 | 202 | 136 | 23 | 8 | 2 | 0 | 21 | 298 | 149 | 0.003420 | 0.032709454 | 0.957 | 0.932 | 0.877 | 1 | 0.977 | 0.872 | 0.932 | 0.964 | 1 | 0.764 | 0.042 |
| A3SS | 4749 | ENSG00000153551 | CMTM7 | chr3 | + | 32452065 | 32452473 | 32452391 | 32452473 | 32449453 | 32449552 | 19 | 15 | 20 | 28 | 6 | 46 | 28 | 71 | 76 | 37 | 13 | 18 | 22 | 3 | 20 | 83 | 77 | 54 | 74 | 114 | 298 | 149 | 0.003430 | 0.032757358 | 0.171 | 0.211 | 0.123 | 0.156 | 0.075 | 0.073 | 0.105 | 0.169 | 0.02 | 0.081 | 0.058 |
| A3SS | 1207 | ENSG00000033627 | ATP6V0A1 | chr17 | + | 42477638 | 42477742 | 42477659 | 42477742 | 42470089 | 42470218 | 9 | 11 | 10 | 15 | 41 | 51 | 28 | 27 | 30 | 95 | 69 | 46 | 14 | 26 | 94 | 75 | 36 | 49 | 54 | 83 | 170 | 149 | 0.003434 | 0.032757358 | 0.134 | 0.256 | 0.245 | 0.305 | 0.274 | 0.446 | 0.528 | 0.2 | 0.297 | 0.498 | -0.151 |
| A3SS | 1850 | ENSG00000205336 | ADGRG1 | chr16 | + | 57651184 | 57651622 | 57651355 | 57651622 | 57650252 | 57650351 | 139 | 22 | 19 | 56 | 14 | 9 | 0 | 1 | 3 | 0 | 39 | 95 | 53 | 33 | 42 | 0 | 5 | 0 | 0 | 0 | 298 | 149 | 0.003441 | 0.032757358 | 0.885 | 1 | 0.905 | 0.903 | 1 | 1 | 0.905 | 1 | 1 | 1 | -0.042 |
| A3SS | 2902 | ENSG00000175061 | SNHG29 | chr17 | + | 16439527 | 16439703 | 16439659 | 16439703 | 16439326 | 16439414 | 209 | 98 | 228 | 205 | 152 | 17 | 10 | 43 | 20 | 7 | 225 | 183 | 223 | 217 | 268 | 0 | 10 | 31 | 26 | 1 | 281 | 149 | 0.003443 | 0.032757358 | 0.867 | 0.839 | 0.738 | 0.845 | 0.92 | 1 | 0.907 | 0.792 | 0.816 | 0.993 | -0.06 |
| A3SS | 11563 | ENSG00000096996 | IL12RB1 | chr19 | - | 18082149 | 18082276 | 18082149 | 18082264 | 18083431 | 18083491 | 5 | 0 | 6 | 4 | 0 | 35 | 11 | 28 | 20 | 7 | 7 | 19 | 0 | 3 | 30 | 19 | 31 | 10 | 10 | 70 | 161 | 149 | 0.003445 | 0.032757358 | 0.117 | 0 | 0.165 | 0.156 | 0 | 0.254 | 0.362 | 0 | 0.217 | 0.284 | -0.136 |
| A3SS | 7034 | ENSG00000160679 | CHTOP | chr1 | + | 153642242 | 153642429 | 153642245 | 153642429 | 153638294 | 153638448 | 143 | 84 | 117 | 144 | 98 | 139 | 89 | 110 | 133 | 76 | 165 | 99 | 76 | 65 | 121 | 187 | 121 | 93 | 171 | 149 | 152 | 149 | 0.003474 | 0.03299233 | 0.502 | 0.481 | 0.51 | 0.515 | 0.558 | 0.464 | 0.445 | 0.445 | 0.271 | 0.443 | 0.1 |
| A3SS | 4391 | ENSG00000172046 | USP19 | chr3 | - | 49115723 | 49115944 | 49115723 | 49115938 | 49116046 | 49116162 | 100 | 46 | 92 | 81 | 108 | 35 | 19 | 23 | 22 | 29 | 138 | 118 | 54 | 41 | 135 | 13 | 34 | 3 | 14 | 20 | 155 | 149 | 0.003487 | 0.033075037 | 0.733 | 0.699 | 0.794 | 0.78 | 0.782 | 0.911 | 0.769 | 0.945 | 0.738 | 0.866 | -0.088 |
| A3SS | 2854 | ENSG00000214900 | LINC01588 | chr14 | - | 50010933 | 50011158 | 50010933 | 50011026 | 50039014 | 50039257 | 34 | 22 | 27 | 7 | 18 | 4 | 0 | 6 | 10 | 6 | 53 | 31 | 16 | 35 | 31 | 0 | 7 | 3 | 0 | 0 | 281 | 149 | 0.003514 | 0.033283221 | 0.818 | 1 | 0.705 | 0.271 | 0.614 | 1 | 0.701 | 0.739 | 1 | 1 | -0.206 |
| A3SS | 2758 | ENSG00000119684 | MLH3 | chr14 | - | 75030542 | 75030715 | 75030542 | 75030702 | 75032067 | 75032179 | 2 | 1 | 1 | 0 | 8 | 37 | 16 | 26 | 21 | 31 | 0 | 2 | 0 | 0 | 0 | 8 | 7 | 16 | 2 | 12 | 162 | 149 | 0.003517 | 0.033283221 | 0.047 | 0.054 | 0.034 | 0 | 0.192 | 0 | 0.208 | 0 | 0 | 0 | 0.024 |
| A3SS | 3080 | ENSG00000141279 | NPEPPS | chr17 | + | 47591903 | 47592060 | 47591955 | 47592060 | 47590716 | 47590881 | 9 | 6 | 2 | 7 | 0 | 122 | 106 | 84 | 120 | 140 | 8 | 1 | 14 | 23 | 0 | 78 | 124 | 80 | 77 | 169 | 201 | 149 | 0.003564 | 0.033683971 | 0.052 | 0.04 | 0.017 | 0.041 | 0 | 0.071 | 0.006 | 0.115 | 0.181 | 0 | -0.045 |
| A3SS | 7575 | ENSG00000131408 | NR1H2 | chr19 | + | 50379091 | 50379181 | 50379133 | 50379181 | 50378521 | 50378796 | 10 | 9 | 13 | 20 | 20 | 1 | 2 | 0 | 6 | 6 | 2 | 23 | 16 | 16 | 34 | 0 | 11 | 11 | 12 | 37 | 191 | 149 | 0.003619 | 0.034166703 | 0.886 | 0.778 | 1 | 0.722 | 0.722 | 1 | 0.62 | 0.532 | 0.51 | 0.418 | 0.206 |
| A3SS | 10442 | ENSG00000099204 | ABLIM1 | chr10 | - | 114447879 | 114448026 | 114447879 | 114448020 | 114451623 | 114451671 | 19 | 10 | 4 | 11 | 20 | 60 | 36 | 89 | 68 | 105 | 0 | 14 | 22 | 20 | 87 | 24 | 35 | 36 | 8 | 16 | 155 | 149 | 0.003624 | 0.034176498 | 0.233 | 0.211 | 0.041 | 0.135 | 0.155 | 0 | 0.278 | 0.37 | 0.706 | 0.839 | -0.284 |
| A3SS | 13104 | ENSG00000090686 | USP48 | chr1 | - | 21706743 | 21706907 | 21706743 | 21706868 | 21715388 | 21715457 | 18 | 7 | 4 | 10 | 10 | 151 | 172 | 108 | 147 | 272 | 6 | 30 | 16 | 15 | 0 | 65 | 226 | 201 | 99 | 36 | 188 | 149 | 0.003657 | 0.034421997 | 0.086 | 0.031 | 0.029 | 0.051 | 0.028 | 0.068 | 0.095 | 0.059 | 0.107 | 0 | -0.021 |
| A3SS | 6057 | ENSG00000117543 | DPH5 | chr1 | - | 101001466 | 101001587 | 101001466 | 101001544 | 101021640 | 101021765 | 55 | 37 | 37 | 36 | 98 | 7 | 2 | 9 | 8 | 15 | 17 | 7 | 21 | 31 | 16 | 1 | 12 | 21 | 11 | 6 | 192 | 149 | 0.003659 | 0.034421997 | 0.859 | 0.935 | 0.761 | 0.777 | 0.835 | 0.93 | 0.312 | 0.437 | 0.686 | 0.674 | 0.226 |
| A3SS | 2641 | ENSG00000100453 | GZMB | chr14 | - | 24631857 | 24632119 | 24631857 | 24632118 | 24632914 | 24633062 | 8 | 1 | 3 | 5 | 3 | 56 | 12 | 12 | 47 | 11 | 0 | 0 | 5 | 0 | 5 | 15 | 38 | 125 | 23 | 12 | 150 | 149 | 0.003683 | 0.034609864 | 0.124 | 0.076 | 0.199 | 0.096 | 0.213 | 0 | 0 | 0.038 | 0 | 0.293 | 0.075 |
| A3SS | 6726 | ENSG00000143851 | PTPN7 | chr1 | - | 202159280 | 202159454 | 202159280 | 202159450 | 202160544 | 202161581 | 55 | 14 | 40 | 42 | 19 | 1 | 0 | 0 | 1 | 0 | 10 | 53 | 36 | 23 | 41 | 0 | 0 | 13 | 0 | 0 | 153 | 149 | 0.003691 | 0.034642285 | 0.982 | 1 | 1 | 0.976 | 1 | 1 | 1 | 0.729 | 1 | 1 | 0.046 |
| A3SS | 7827 | ENSG00000166986 | MARS1 | chr12 | + | 57512235 | 57512353 | 57512302 | 57512353 | 57512007 | 57512103 | 135 | 79 | 133 | 169 | 167 | 12 | 4 | 8 | 22 | 2 | 169 | 198 | 112 | 186 | 131 | 20 | 14 | 13 | 15 | 43 | 216 | 149 | 0.003704 | 0.034724177 | 0.886 | 0.932 | 0.92 | 0.841 | 0.983 | 0.854 | 0.907 | 0.856 | 0.895 | 0.678 | 0.074 |
| A3SS | 3898 | ENSG00000121210 | TMEM131L | chr4 | + | 153584834 | 153584931 | 153584837 | 153584931 | 153583189 | 153583248 | 7 | 7 | 13 | 16 | 42 | 0 | 4 | 0 | 1 | 0 | 3 | 25 | 8 | 14 | 5 | 0 | 0 | 4 | 7 | 0 | 152 | 149 | 0.003755 | 0.035161826 | 1 | 0.632 | 1 | 0.94 | 1 | 1 | 1 | 0.662 | 0.662 | 1 | 0.05 |
| A3SS | 3956 | ENSG00000154743 | TSEN2 | chr3 | + | 12531538 | 12531659 | 12531569 | 12531659 | 12529761 | 12529873 | 2 | 2 | 2 | 1 | 4 | 23 | 16 | 32 | 32 | 38 | 0 | 0 | 0 | 0 | 0 | 8 | 13 | 27 | 4 | 37 | 180 | 149 | 0.003773 | 0.035287782 | 0.067 | 0.094 | 0.049 | 0.025 | 0.08 | 0 | 0 | 0 | 0 | 0 | 0.063 |
| A3SS | 4293 | ENSG00000114127 | XRN1 | chr3 | - | 142332374 | 142332531 | 142332374 | 142332424 | 142332966 | 142333089 | 237 | 229 | 257 | 190 | 477 | 4 | 7 | 2 | 3 | 13 | 12 | 105 | 106 | 18 | 14 | 0 | 0 | 3 | 0 | 0 | 256 | 149 | 0.003780 | 0.035319594 | 0.972 | 0.95 | 0.987 | 0.974 | 0.955 | 1 | 1 | 0.954 | 1 | 1 | -0.023 |
| A3SS | 7780 | ENSG00000106105 | GARS1 | chr7 | + | 30631447 | 30632437 | 30632246 | 30632437 | 30628559 | 30628669 | 169 | 149 | 160 | 162 | 367 | 1 | 3 | 9 | 0 | 0 | 168 | 223 | 150 | 185 | 161 | 4 | 14 | 10 | 0 | 0 | 298 | 149 | 0.003787 | 0.035343853 | 0.988 | 0.961 | 0.899 | 1 | 1 | 0.955 | 0.888 | 0.882 | 1 | 1 | 0.025 |
| A3SS | 2475 | ENSG00000172575 | RASGRP1 | chr15 | - | 38498793 | 38499184 | 38498793 | 38498946 | 38500102 | 38500139 | 40 | 36 | 44 | 44 | 41 | 272 | 177 | 304 | 324 | 299 | 18 | 23 | 59 | 11 | 2 | 30 | 174 | 102 | 59 | 32 | 298 | 149 | 0.003796 | 0.035382635 | 0.068 | 0.092 | 0.067 | 0.064 | 0.064 | 0.231 | 0.062 | 0.224 | 0.085 | 0.03 | -0.055 |
| A3SS | 711 | ENSG00000141968 | VAV1 | chr19 | + | 6829785 | 6829918 | 6829872 | 6829918 | 6828814 | 6828900 | 299 | 175 | 301 | 338 | 206 | 13 | 4 | 2 | 6 | 0 | 510 | 372 | 252 | 382 | 535 | 0 | 6 | 8 | 0 | 0 | 236 | 149 | 0.003804 | 0.035382635 | 0.936 | 0.965 | 0.99 | 0.973 | 1 | 1 | 0.975 | 0.952 | 1 | 1 | -0.013 |
| A3SS | 9814 | ENSG00000077454 | LRCH4 | chr7 | - | 100574010 | 100575304 | 100574010 | 100574356 | 100575704 | 100575782 | 26 | 16 | 17 | 12 | 12 | 159 | 62 | 155 | 196 | 197 | 33 | 13 | 34 | 12 | 3 | 99 | 115 | 72 | 53 | 161 | 298 | 149 | 0.003805 | 0.035382635 | 0.076 | 0.114 | 0.052 | 0.03 | 0.03 | 0.143 | 0.053 | 0.191 | 0.102 | 0.009 | -0.039 |
| A3SS | 3431 | ENSG00000126456 | IRF3 | chr19 | - | 49660712 | 49660844 | 49660712 | 49660828 | 49661947 | 49662328 | 18 | 4 | 23 | 15 | 7 | 65 | 17 | 55 | 84 | 88 | 42 | 15 | 9 | 13 | 32 | 64 | 54 | 30 | 26 | 93 | 165 | 149 | 0.003835 | 0.035617271 | 0.2 | 0.175 | 0.274 | 0.139 | 0.067 | 0.372 | 0.201 | 0.213 | 0.311 | 0.237 | -0.096 |
| A3SS | 2652 | ENSG00000075413 | MARK3 | chr14 | + | 103490951 | 103492034 | 103491776 | 103492034 | 103480386 | 103480490 | 24 | 59 | 25 | 16 | 34 | 192 | 182 | 159 | 220 | 403 | 75 | 42 | 76 | 38 | 11 | 327 | 219 | 202 | 262 | 143 | 298 | 149 | 0.003839 | 0.035617271 | 0.059 | 0.139 | 0.073 | 0.035 | 0.04 | 0.103 | 0.087 | 0.158 | 0.068 | 0.037 | -0.021 |
| A3SS | 11027 | ENSG00000172269 | DPAGT1 | chr11 | - | 119100629 | 119100877 | 119100629 | 119100843 | 119101017 | 119101138 | 2 | 2 | 0 | 4 | 2 | 9 | 8 | 1 | 16 | 6 | 0 | 4 | 0 | 0 | 0 | 16 | 7 | 4 | 5 | 15 | 183 | 149 | 0.003850 | 0.035619393 | 0.153 | 0.169 | 0 | 0.169 | 0.213 | 0 | 0.318 | 0 | 0 | 0 | 0.077 |
| A3SS | 10552 | ENSG00000068438 | FTSJ1 | chrX | + | 48482402 | 48482506 | 48482408 | 48482506 | 48481631 | 48481715 | 22 | 5 | 7 | 9 | 11 | 8 | 6 | 5 | 14 | 3 | 35 | 30 | 16 | 18 | 12 | 0 | 2 | 5 | 2 | 15 | 155 | 149 | 0.003851 | 0.035619393 | 0.726 | 0.445 | 0.574 | 0.382 | 0.779 | 1 | 0.935 | 0.755 | 0.896 | 0.435 | -0.223 |
| A3SS | 3314 | ENSG00000103496 | STX4 | chr16 | + | 31034203 | 31034325 | 31034225 | 31034325 | 31034012 | 31034114 | 2 | 5 | 1 | 1 | 0 | 69 | 32 | 58 | 54 | 9 | 0 | 3 | 7 | 12 | 10 | 98 | 40 | 54 | 74 | 104 | 171 | 149 | 0.003852 | 0.035619393 | 0.025 | 0.12 | 0.015 | 0.016 | 0 | 0 | 0.061 | 0.101 | 0.124 | 0.077 | -0.037 |
| A3SS | 8411 | ENSG00000140368 | PSTPIP1 | chr15 | + | 77035801 | 77035935 | 77035810 | 77035935 | 77035507 | 77035563 | 122 | 54 | 92 | 108 | 63 | 2 | 1 | 2 | 8 | 0 | 165 | 67 | 69 | 196 | 94 | 14 | 0 | 5 | 5 | 13 | 158 | 149 | 0.003864 | 0.035684218 | 0.983 | 0.981 | 0.977 | 0.927 | 1 | 0.917 | 1 | 0.929 | 0.974 | 0.872 | 0.035 |
| A3SS | 12286 | ENSG00000110011 | DNAJC4 | chr11 | + | 64232393 | 64232614 | 64232429 | 64232614 | 64231877 | 64231964 | 6 | 1 | 7 | 1 | 4 | 76 | 37 | 62 | 74 | 68 | 7 | 5 | 4 | 5 | 33 | 80 | 92 | 65 | 73 | 114 | 185 | 149 | 0.003935 | 0.036261364 | 0.06 | 0.021 | 0.083 | 0.011 | 0.045 | 0.066 | 0.042 | 0.047 | 0.052 | 0.189 | -0.035 |
| A3SS | 713 | ENSG00000160633 | SAFB | chr19 | + | 5661517 | 5661808 | 5661520 | 5661808 | 5657240 | 5657347 | 171 | 72 | 98 | 104 | 103 | 138 | 81 | 104 | 121 | 61 | 158 | 65 | 70 | 69 | 57 | 163 | 91 | 98 | 105 | 110 | 152 | 149 | 0.003935 | 0.036261364 | 0.548 | 0.466 | 0.48 | 0.457 | 0.623 | 0.487 | 0.412 | 0.412 | 0.392 | 0.337 | 0.107 |
| A3SS | 756 | ENSG00000101347 | SAMHD1 | chr20 | - | 36916948 | 36917096 | 36916948 | 36917049 | 36919363 | 36919519 | 64 | 59 | 72 | 45 | 72 | 1172 | 866 | 780 | 1028 | 1652 | 102 | 65 | 71 | 54 | 60 | 2155 | 714 | 384 | 959 | 696 | 196 | 149 | 0.004068 | 0.037444425 | 0.04 | 0.049 | 0.066 | 0.032 | 0.032 | 0.035 | 0.065 | 0.123 | 0.041 | 0.062 | -0.021 |
| A3SS | 10597 | ENSG00000215021 | PHB2 | chr12 | - | 6967170 | 6967779 | 6967170 | 6967335 | 6967891 | 6968021 | 334 | 203 | 359 | 297 | 359 | 9 | 1 | 3 | 2 | 2 | 397 | 240 | 247 | 301 | 500 | 5 | 4 | 4 | 18 | 7 | 298 | 149 | 0.004111 | 0.0377975 | 0.949 | 0.99 | 0.984 | 0.987 | 0.989 | 0.975 | 0.968 | 0.969 | 0.893 | 0.973 | 0.024 |
| A3SS | 6765 | ENSG00000180353 | HCLS1 | chr3 | - | 121647318 | 121647482 | 121647318 | 121647448 | 121657278 | 121657352 | 23 | 25 | 8 | 32 | 14 | 1222 | 793 | 913 | 1011 | 1171 | 6 | 15 | 30 | 29 | 0 | 2374 | 1587 | 1397 | 1768 | 2092 | 183 | 149 | 0.004133 | 0.03795782 | 0.015 | 0.025 | 0.007 | 0.025 | 0.01 | 0.002 | 0.008 | 0.017 | 0.013 | 0 | 0.008 |
| A3SS | 9723 | ENSG00000139641 | ESYT1 | chr12 | + | 56134079 | 56134181 | 56134109 | 56134181 | 56133780 | 56133873 | 91 | 62 | 75 | 83 | 182 | 281 | 138 | 233 | 241 | 365 | 48 | 55 | 26 | 29 | 70 | 183 | 224 | 105 | 141 | 164 | 179 | 149 | 0.004138 | 0.037960931 | 0.212 | 0.272 | 0.211 | 0.223 | 0.293 | 0.179 | 0.17 | 0.171 | 0.146 | 0.262 | 0.057 |
| A3SS | 5924 | ENSG00000055609 | KMT2C | chr7 | - | 152330171 | 152330742 | 152330171 | 152330739 | 152358586 | 152358675 | 13 | 12 | 6 | 5 | 23 | 164 | 131 | 160 | 164 | 196 | 0 | 7 | 10 | 2 | 0 | 80 | 117 | 106 | 104 | 86 | 152 | 149 | 0.004167 | 0.038179184 | 0.072 | 0.082 | 0.035 | 0.029 | 0.103 | 0 | 0.055 | 0.085 | 0.019 | 0 | 0.032 |
| A3SS | 10488 | ENSG00000177646 | ACAD9 | chr3 | + | 128908922 | 128909099 | 128908972 | 128909099 | 128908184 | 128908264 | 6 | 0 | 1 | 1 | 0 | 20 | 12 | 19 | 19 | 23 | 12 | 0 | 6 | 8 | 0 | 20 | 38 | 8 | 16 | 34 | 199 | 149 | 0.004199 | 0.038426129 | 0.183 | 0 | 0.038 | 0.038 | 0 | 0.31 | 0 | 0.36 | 0.272 | 0 | -0.137 |
| A3SS | 1864 | ENSG00000185324 | CDK10 | chr16 | + | 89695594 | 89696354 | 89695728 | 89696354 | 89695292 | 89695345 | 92 | 45 | 83 | 90 | 80 | 0 | 0 | 0 | 4 | 11 | 38 | 22 | 34 | 48 | 98 | 0 | 2 | 0 | 0 | 2 | 283 | 149 | 0.004209 | 0.038458035 | 1 | 1 | 1 | 0.922 | 0.793 | 1 | 0.853 | 1 | 1 | 0.963 | -0.02 |
| A3SS | 12864 | ENSG00000082898 | XPO1 | chr2 | - | 61482956 | 61483122 | 61482956 | 61483091 | 61483936 | 61484105 | 1 | 2 | 4 | 7 | 10 | 246 | 177 | 239 | 231 | 520 | 11 | 0 | 2 | 6 | 0 | 37 | 126 | 76 | 54 | 59 | 180 | 149 | 0.004212 | 0.038458035 | 0.003 | 0.009 | 0.014 | 0.024 | 0.016 | 0.197 | 0 | 0.021 | 0.084 | 0 | -0.047 |
| A3SS | 7173 | ENSG00000160917 | CPSF4 | chr7 | + | 99453965 | 99454136 | 99454040 | 99454136 | 99452367 | 99452440 | 20 | 2 | 17 | 28 | 7 | 15 | 5 | 10 | 14 | 6 | 9 | 46 | 28 | 22 | 27 | 0 | 18 | 8 | 0 | 15 | 224 | 149 | 0.004219 | 0.038458035 | 0.47 | 0.21 | 0.531 | 0.571 | 0.437 | 1 | 0.63 | 0.7 | 1 | 0.545 | -0.331 |
| A3SS | 7521 | ENSG00000141524 | TMC6 | chr17 | - | 78120176 | 78121164 | 78120176 | 78120832 | 78122604 | 78122749 | 4 | 5 | 17 | 21 | 2 | 0 | 0 | 0 | 0 | 0 | 10 | 25 | 16 | 3 | 31 | 0 | 11 | 0 | 0 | 0 | 298 | 149 | 0.004222 | 0.038458035 | 1 | 1 | 1 | 1 | 1 | 1 | 0.532 | 1 | 1 | 1 | 0.094 |
| A3SS | 10723 | ENSG00000215301 | DDX3X | chrX | + | 41346858 | 41347451 | 41347314 | 41347451 | 41346504 | 41346622 | 1115 | 733 | 956 | 1135 | 1503 | 1 | 0 | 3 | 2 | 0 | 108 | 426 | 797 | 414 | 49 | 3 | 4 | 0 | 7 | 8 | 298 | 149 | 0.004226 | 0.038458035 | 0.998 | 1 | 0.994 | 0.996 | 1 | 0.947 | 0.982 | 1 | 0.967 | 0.754 | 0.068 |
| A3SS | 5503 | ENSG00000105323 | HNRNPUL1 | chr19 | + | 41305645 | 41305867 | 41305831 | 41305867 | 41303971 | 41304261 | 50 | 25 | 33 | 34 | 15 | 0 | 0 | 4 | 1 | 0 | 115 | 74 | 42 | 77 | 102 | 20 | 5 | 0 | 0 | 0 | 298 | 149 | 0.004294 | 0.039027699 | 1 | 1 | 0.805 | 0.944 | 1 | 0.742 | 0.881 | 1 | 1 | 1 | 0.025 |
| A3SS | 1065 | ENSG00000152234 | ATP5F1A | chr18 | - | 46095052 | 46095147 | 46095052 | 46095131 | 46098171 | 46098296 | 11 | 11 | 20 | 14 | 6 | 246 | 126 | 226 | 225 | 94 | 21 | 27 | 12 | 27 | 39 | 288 | 196 | 182 | 150 | 362 | 165 | 149 | 0.004298 | 0.039027699 | 0.039 | 0.073 | 0.074 | 0.053 | 0.054 | 0.062 | 0.111 | 0.056 | 0.14 | 0.089 | -0.033 |
| A3SS | 4585 | ENSG00000111642 | CHD4 | chr12 | - | 6578028 | 6578135 | 6578028 | 6578117 | 6578408 | 6578546 | 21 | 14 | 34 | 22 | 22 | 1 | 1 | 4 | 2 | 16 | 69 | 23 | 13 | 15 | 6 | 16 | 9 | 27 | 5 | 5 | 167 | 149 | 0.004341 | 0.039367651 | 0.949 | 0.926 | 0.884 | 0.908 | 0.551 | 0.794 | 0.695 | 0.3 | 0.728 | 0.517 | 0.237 |
| A3SS | 493 | ENSG00000063241 | ISOC2 | chr19 | - | 55455259 | 55455378 | 55455259 | 55455330 | 55455635 | 55455845 | 1 | 0 | 1 | 1 | 2 | 5 | 1 | 11 | 1 | 4 | 0 | 4 | 1 | 7 | 6 | 4 | 5 | 10 | 9 | 7 | 197 | 149 | 0.004396 | 0.039828844 | 0.131 | 0 | 0.064 | 0.431 | 0.274 | 0 | 0.377 | 0.07 | 0.37 | 0.393 | -0.062 |
| A3SS | 6255 | ENSG00000171793 | CTPS1 | chr1 | + | 40995916 | 40997526 | 40997393 | 40997526 | 40991764 | 40991845 | 36 | 41 | 60 | 55 | 59 | 0 | 0 | 0 | 0 | 0 | 163 | 59 | 102 | 139 | 96 | 11 | 0 | 3 | 0 | 0 | 298 | 149 | 0.004430 | 0.040084661 | 1 | 1 | 1 | 1 | 1 | 0.881 | 1 | 0.944 | 1 | 1 | 0.035 |
| A3SS | 4266 | ENSG00000168291 | PDHB | chr3 | - | 58429707 | 58429941 | 58429707 | 58429799 | 58430127 | 58430238 | 21 | 13 | 10 | 13 | 13 | 120 | 76 | 118 | 98 | 202 | 16 | 19 | 34 | 31 | 5 | 98 | 86 | 85 | 141 | 135 | 291 | 149 | 0.004484 | 0.040532361 | 0.082 | 0.081 | 0.042 | 0.064 | 0.032 | 0.077 | 0.102 | 0.17 | 0.101 | 0.019 | -0.034 |
| A3SS | 7945 | ENSG00000163006 | CCDC138 | chr2 | + | 108804888 | 108805008 | 108804891 | 108805008 | 108798427 | 108798586 | 13 | 12 | 23 | 15 | 40 | 0 | 5 | 1 | 0 | 7 | 22 | 12 | 19 | 13 | 6 | 0 | 3 | 2 | 0 | 0 | 152 | 149 | 0.004490 | 0.040541773 | 1 | 0.702 | 0.958 | 1 | 0.849 | 1 | 0.797 | 0.903 | 1 | 1 | -0.038 |
| A3SS | 12696 | ENSG00000119977 | TCTN3 | chr10 | - | 95683100 | 95683195 | 95683100 | 95683180 | 95683521 | 95683629 | 46 | 22 | 44 | 45 | 82 | 1 | 2 | 4 | 1 | 4 | 87 | 38 | 21 | 19 | 24 | 0 | 0 | 0 | 0 | 0 | 164 | 149 | 0.004510 | 0.040649939 | 0.977 | 0.909 | 0.909 | 0.976 | 0.949 | 1 | 1 | 1 | 1 | 1 | -0.056 |
| A3SS | 12590 | ENSG00000028116 | VRK2 | chr2 | + | 58048826 | 58048967 | 58048880 | 58048967 | 58046874 | 58047128 | 57 | 55 | 44 | 52 | 45 | 0 | 0 | 0 | 0 | 0 | 24 | 23 | 22 | 8 | 14 | 0 | 11 | 0 | 0 | 0 | 203 | 149 | 0.004512 | 0.040649939 | 1 | 1 | 1 | 1 | 1 | 1 | 0.605 | 1 | 1 | 1 | 0.079 |
| A3SS | 2836 | ENSG00000119599 | DCAF4 | chr14 | + | 72940219 | 72940377 | 72940251 | 72940377 | 72939801 | 72939902 | 63 | 27 | 36 | 44 | 79 | 4 | 0 | 2 | 1 | 5 | 118 | 48 | 54 | 56 | 66 | 0 | 0 | 0 | 0 | 0 | 181 | 149 | 0.004520 | 0.040674543 | 0.928 | 1 | 0.937 | 0.973 | 0.929 | 1 | 1 | 1 | 1 | 1 | -0.047 |
| A3SS | 10596 | ENSG00000215021 | PHB2 | chr12 | - | 6967170 | 6967779 | 6967170 | 6967238 | 6967891 | 6968021 | 359 | 233 | 394 | 347 | 428 | 14 | 6 | 2 | 5 | 13 | 504 | 275 | 279 | 370 | 552 | 31 | 14 | 22 | 11 | 0 | 298 | 149 | 0.004584 | 0.041203187 | 0.928 | 0.951 | 0.99 | 0.972 | 0.943 | 0.89 | 0.908 | 0.864 | 0.944 | 1 | 0.036 |
| A3SS | 1818 | ENSG00000170100 | ZNF778 | chr16 | + | 89220996 | 89221152 | 89221003 | 89221152 | 89217709 | 89217910 | 6 | 5 | 4 | 10 | 3 | 0 | 0 | 0 | 2 | 0 | 21 | 2 | 6 | 5 | 22 | 0 | 4 | 0 | 0 | 9 | 156 | 149 | 0.004592 | 0.04123152 | 1 | 1 | 1 | 0.827 | 1 | 1 | 0.323 | 1 | 1 | 0.7 | 0.161 |
| A3SS | 7853 | ENSG00000186281 | GPAT2 | chr2 | - | 96022957 | 96023440 | 96022957 | 96023023 | 96023922 | 96024000 | 3 | 3 | 13 | 3 | 3 | 0 | 0 | 0 | 1 | 0 | 15 | 0 | 6 | 14 | 14 | 0 | 7 | 0 | 0 | 3 | 298 | 149 | 0.004616 | 0.041397681 | 1 | 1 | 1 | 0.6 | 1 | 1 | 0 | 1 | 1 | 0.7 | 0.18 |
| A3SS | 4352 | ENSG00000187742 | SECISBP2 | chr9 | + | 89357284 | 89357565 | 89357410 | 89357565 | 89350631 | 89350852 | 29 | 41 | 42 | 50 | 15 | 108 | 91 | 124 | 98 | 163 | 22 | 75 | 54 | 41 | 21 | 84 | 112 | 59 | 86 | 60 | 275 | 149 | 0.004641 | 0.041519055 | 0.127 | 0.196 | 0.155 | 0.217 | 0.047 | 0.124 | 0.266 | 0.332 | 0.205 | 0.159 | -0.069 |
| A3SS | 7083 | ENSG00000116560 | SFPQ | chr1 | - | 35176377 | 35176474 | 35176377 | 35176471 | 35177318 | 35177399 | 78 | 95 | 129 | 104 | 90 | 52 | 66 | 72 | 58 | 87 | 97 | 122 | 193 | 80 | 88 | 24 | 75 | 127 | 21 | 29 | 152 | 149 | 0.004644 | 0.041519055 | 0.595 | 0.585 | 0.637 | 0.637 | 0.503 | 0.798 | 0.615 | 0.598 | 0.789 | 0.748 | -0.118 |
| A3SS | 12893 | ENSG00000179889 | PDXDC1 | chr16 | + | 14998339 | 14998405 | 14998342 | 14998405 | 14997752 | 14997826 | 23 | 23 | 44 | 23 | 39 | 2 | 2 | 0 | 2 | 4 | 40 | 35 | 21 | 36 | 25 | 0 | 4 | 5 | 5 | 13 | 152 | 149 | 0.004645 | 0.041519055 | 0.919 | 0.919 | 1 | 0.919 | 0.905 | 1 | 0.896 | 0.805 | 0.876 | 0.653 | 0.086 |
| A3SS | 9793 | ENSG00000012124 | CD22 | chr19 | + | 35331905 | 35332074 | 35332018 | 35332074 | 35329186 | 35329230 | 14 | 15 | 16 | 42 | 13 | 42 | 14 | 29 | 41 | 40 | 37 | 71 | 21 | 47 | 69 | 62 | 69 | 6 | 17 | 58 | 262 | 149 | 0.004654 | 0.041555503 | 0.159 | 0.379 | 0.239 | 0.368 | 0.156 | 0.253 | 0.369 | 0.666 | 0.611 | 0.404 | -0.2 |
| A3SS | 10493 | ENSG00000147955 | SIGMAR1 | chr9 | - | 34636996 | 34637112 | 34636996 | 34637089 | 34637219 | 34637420 | 0 | 0 | 5 | 5 | 0 | 14 | 10 | 25 | 30 | 3 | 0 | 1 | 1 | 0 | 0 | 17 | 9 | 15 | 4 | 1 | 172 | 149 | 0.004660 | 0.04156771 | 0 | 0 | 0.148 | 0.126 | 0 | 0 | 0.088 | 0.055 | 0 | 0 | 0.026 |
| A3SS | 4702 | ENSG00000175792 | RUVBL1 | chr3 | - | 128087705 | 128087844 | 128087705 | 128087808 | 128097299 | 128097498 | 1 | 3 | 4 | 2 | 7 | 77 | 46 | 93 | 58 | 89 | 0 | 0 | 1 | 2 | 0 | 103 | 86 | 46 | 59 | 95 | 185 | 149 | 0.004675 | 0.041618155 | 0.01 | 0.05 | 0.033 | 0.027 | 0.06 | 0 | 0 | 0.017 | 0.027 | 0 | 0.027 |
| A3SS | 2833 | ENSG00000119688 | ABCD4 | chr14 | - | 74293153 | 74293310 | 74293153 | 74293248 | 74295147 | 74295198 | 3 | 6 | 5 | 9 | 4 | 11 | 5 | 14 | 31 | 23 | 2 | 8 | 5 | 13 | 33 | 7 | 6 | 4 | 21 | 18 | 211 | 149 | 0.004676 | 0.041618155 | 0.161 | 0.459 | 0.201 | 0.17 | 0.109 | 0.168 | 0.485 | 0.469 | 0.304 | 0.564 | -0.178 |
| A3SS | 8311 | ENSG00000181381 | DDX60L | chr4 | - | 168395958 | 168396127 | 168395958 | 168396124 | 168400825 | 168400978 | 322 | 0 | 155 | 130 | 128 | 0 | 263 | 2 | 145 | 184 | 0 | 99 | 117 | 0 | 20 | 103 | 109 | 126 | 134 | 2 | 152 | 149 | 0.004717 | 0.041900991 | 1 | 0 | 0.987 | 0.468 | 0.405 | 0 | 0.471 | 0.477 | 0 | 0.907 | 0.201 |
| A3SS | 7185 | ENSG00000001497 | LAS1L | chrX | - | 65514822 | 65514973 | 65514822 | 65514970 | 65517986 | 65518465 | 35 | 24 | 62 | 26 | 48 | 4 | 10 | 1 | 8 | 1 | 41 | 77 | 40 | 44 | 59 | 0 | 3 | 2 | 4 | 3 | 152 | 149 | 0.004719 | 0.041900991 | 0.896 | 0.702 | 0.984 | 0.761 | 0.979 | 1 | 0.962 | 0.951 | 0.915 | 0.951 | -0.091 |
| A3SS | 3461 | ENSG00000011243 | AKAP8L | chr19 | - | 15397713 | 15399984 | 15397713 | 15397855 | 15400294 | 15400358 | 28 | 13 | 17 | 20 | 5 | 0 | 0 | 0 | 1 | 0 | 18 | 37 | 48 | 74 | 64 | 25 | 1 | 0 | 1 | 0 | 298 | 149 | 0.004745 | 0.042085984 | 1 | 1 | 1 | 0.909 | 1 | 0.265 | 0.949 | 1 | 0.974 | 1 | 0.144 |
| A3SS | 2397 | ENSG00000196547 | MAN2A2 | chr15 | + | 90911366 | 90911550 | 90911384 | 90911550 | 90911170 | 90911238 | 2 | 0 | 1 | 2 | 0 | 33 | 13 | 20 | 31 | 14 | 0 | 5 | 3 | 7 | 8 | 51 | 69 | 19 | 37 | 127 | 167 | 149 | 0.004762 | 0.042195913 | 0.051 | 0 | 0.043 | 0.054 | 0 | 0 | 0.061 | 0.123 | 0.144 | 0.053 | -0.047 |
| A3SS | 5087 | ENSG00000184432 | COPB2 | chr3 | - | 139373665 | 139373833 | 139373665 | 139373808 | 139374488 | 139374588 | 7 | 5 | 9 | 4 | 3 | 460 | 399 | 405 | 369 | 521 | 7 | 15 | 17 | 0 | 0 | 435 | 314 | 186 | 477 | 35 | 174 | 149 | 0.004769 | 0.042205642 | 0.013 | 0.011 | 0.019 | 0.009 | 0.005 | 0.014 | 0.039 | 0.073 | 0 | 0 | -0.014 |
| A3SS | 6750 | ENSG00000173575 | CHD2 | chr15 | + | 92978147 | 92978383 | 92978233 | 92978383 | 92974878 | 92974950 | 37 | 37 | 39 | 50 | 43 | 308 | 263 | 335 | 314 | 353 | 25 | 71 | 74 | 51 | 34 | 122 | 432 | 573 | 268 | 60 | 235 | 149 | 0.004791 | 0.042356299 | 0.071 | 0.082 | 0.069 | 0.092 | 0.072 | 0.115 | 0.094 | 0.076 | 0.108 | 0.264 | -0.054 |
| A3SS | 565 | ENSG00000167613 | LAIR1 | chr19 | - | 54364294 | 54364486 | 54364294 | 54364330 | 54365029 | 54365099 | 46 | 16 | 28 | 30 | 19 | 2 | 2 | 3 | 0 | 4 | 14 | 30 | 11 | 51 | 36 | 16 | 7 | 2 | 23 | 0 | 298 | 149 | 0.004815 | 0.042527927 | 0.92 | 0.8 | 0.824 | 1 | 0.704 | 0.304 | 0.682 | 0.733 | 0.526 | 1 | 0.201 |
| A3SS | 6920 | ENSG00000135905 | DOCK10 | chr2 | - | 224886458 | 224886531 | 224886458 | 224886526 | 224896294 | 224896377 | 139 | 116 | 151 | 147 | 216 | 4 | 7 | 4 | 3 | 5 | 61 | 61 | 102 | 35 | 38 | 2 | 9 | 6 | 0 | 7 | 154 | 149 | 0.004823 | 0.042549387 | 0.971 | 0.941 | 0.973 | 0.979 | 0.977 | 0.967 | 0.868 | 0.943 | 1 | 0.84 | 0.045 |
| A3SS | 11927 | ENSG00000126705 | AHDC1 | chr1 | - | 27553073 | 27553290 | 27553073 | 27553185 | 27558304 | 27558530 | 11 | 4 | 7 | 7 | 4 | 16 | 0 | 10 | 11 | 7 | 7 | 16 | 18 | 6 | 16 | 0 | 17 | 2 | 0 | 4 | 254 | 149 | 0.004879 | 0.042990927 | 0.287 | 1 | 0.291 | 0.272 | 0.251 | 1 | 0.356 | 0.841 | 1 | 0.701 | -0.359 |
| A3SS | 8324 | ENSG00000018280 | SLC11A1 | chr2 | + | 218392114 | 218393130 | 218392980 | 218393130 | 218391375 | 218391495 | 49 | 43 | 46 | 19 | 20 | 37 | 10 | 8 | 26 | 16 | 93 | 145 | 224 | 56 | 113 | 10 | 26 | 12 | 18 | 58 | 298 | 149 | 0.004888 | 0.043026182 | 0.398 | 0.683 | 0.742 | 0.268 | 0.385 | 0.823 | 0.736 | 0.903 | 0.609 | 0.493 | -0.218 |
| A3SS | 4528 | ENSG00000156709 | AIFM1 | chrX | - | 130147751 | 130147935 | 130147751 | 130147876 | 130149468 | 130149568 | 2 | 0 | 2 | 0 | 0 | 74 | 48 | 45 | 51 | 88 | 4 | 0 | 4 | 11 | 0 | 96 | 56 | 51 | 84 | 103 | 208 | 149 | 0.004940 | 0.043399206 | 0.019 | 0 | 0.031 | 0 | 0 | 0.029 | 0 | 0.053 | 0.086 | 0 | -0.024 |
| A3SS | 10004 | ENSG00000113013 | HSPA9 | chr5 | - | 138574067 | 138574169 | 138574067 | 138574126 | 138575237 | 138575401 | 17 | 18 | 21 | 13 | 18 | 79 | 37 | 57 | 60 | 20 | 45 | 19 | 27 | 51 | 103 | 35 | 71 | 64 | 47 | 4 | 192 | 149 | 0.004941 | 0.043399206 | 0.143 | 0.274 | 0.222 | 0.144 | 0.411 | 0.499 | 0.172 | 0.247 | 0.457 | 0.952 | -0.227 |
| A3SS | 10128 | ENSG00000181513 | ACBD4 | chr17 | + | 45137721 | 45137830 | 45137759 | 45137830 | 45137367 | 45137454 | 12 | 8 | 5 | 16 | 9 | 12 | 2 | 5 | 23 | 10 | 2 | 4 | 4 | 4 | 2 | 20 | 5 | 10 | 2 | 28 | 187 | 149 | 0.004957 | 0.043495516 | 0.443 | 0.761 | 0.443 | 0.357 | 0.418 | 0.074 | 0.389 | 0.242 | 0.614 | 0.054 | 0.21 |
| A3SS | 9688 | ENSG00000145191 | EIF2B5 | chr3 | + | 184143430 | 184143565 | 184143441 | 184143565 | 184143051 | 184143142 | 2 | 8 | 4 | 1 | 4 | 105 | 58 | 92 | 81 | 119 | 18 | 4 | 11 | 6 | 0 | 114 | 105 | 33 | 50 | 130 | 160 | 149 | 0.004971 | 0.043564836 | 0.017 | 0.114 | 0.039 | 0.011 | 0.03 | 0.128 | 0.034 | 0.237 | 0.101 | 0 | -0.058 |
| A3SS | 8195 | ENSG00000129292 | PHF20L1 | chr8 | + | 132804580 | 132804740 | 132804614 | 132804740 | 132803818 | 132804032 | 8 | 10 | 2 | 8 | 16 | 157 | 172 | 87 | 158 | 238 | 0 | 3 | 8 | 0 | 0 | 78 | 112 | 112 | 59 | 27 | 183 | 149 | 0.004988 | 0.043660372 | 0.04 | 0.045 | 0.018 | 0.04 | 0.052 | 0 | 0.021 | 0.055 | 0 | 0 | 0.024 |
| A3SS | 11201 | ENSG00000104529 | EEF1D | chr8 | - | 143586728 | 143586852 | 143586728 | 143586849 | 143596878 | 143597388 | 12 | 7 | 14 | 18 | 15 | 0 | 0 | 0 | 4 | 0 | 2 | 8 | 17 | 1 | 26 | 0 | 0 | 0 | 0 | 10 | 152 | 149 | 0.004996 | 0.043660372 | 1 | 1 | 1 | 0.815 | 1 | 1 | 1 | 1 | 1 | 0.718 | 0.019 |
| A3SS | 9246 | ENSG00000213654 | GPSM3 | chr6 | - | 32191708 | 32191908 | 32191708 | 32191870 | 32192147 | 32192250 | 303 | 105 | 234 | 274 | 404 | 28 | 10 | 15 | 23 | 24 | 531 | 269 | 289 | 333 | 877 | 85 | 25 | 29 | 46 | 64 | 187 | 149 | 0.005002 | 0.043660372 | 0.896 | 0.893 | 0.926 | 0.905 | 0.931 | 0.833 | 0.896 | 0.888 | 0.852 | 0.916 | 0.033 |
| A3SS | 678 | ENSG00000127507 | ADGRE2 | chr19 | - | 14765657 | 14766381 | 14765657 | 14765804 | 14766977 | 14767109 | 48 | 32 | 18 | 30 | 42 | 24 | 19 | 5 | 20 | 41 | 53 | 50 | 48 | 97 | 76 | 0 | 31 | 13 | 13 | 36 | 298 | 149 | 0.005008 | 0.043660372 | 0.5 | 0.457 | 0.643 | 0.429 | 0.339 | 1 | 0.446 | 0.649 | 0.789 | 0.514 | -0.206 |
| A3SS | 9804 | ENSG00000012124 | CD22 | chr19 | + | 35332546 | 35332924 | 35332593 | 35332924 | 35332018 | 35332074 | 45 | 22 | 44 | 67 | 121 | 2 | 0 | 0 | 0 | 8 | 92 | 93 | 2 | 56 | 99 | 8 | 1 | 0 | 0 | 20 | 196 | 149 | 0.005009 | 0.043660372 | 0.945 | 1 | 1 | 1 | 0.92 | 0.897 | 0.986 | 1 | 1 | 0.79 | 0.038 |
| A3SS | 9181 | ENSG00000004139 | SARM1 | chr17 | + | 28388173 | 28388539 | 28388349 | 28388539 | 28385430 | 28385466 | 4 | 3 | 6 | 13 | 8 | 1 | 1 | 4 | 2 | 0 | 9 | 15 | 10 | 5 | 16 | 3 | 1 | 0 | 0 | 0 | 298 | 149 | 0.005042 | 0.043902881 | 0.667 | 0.6 | 0.429 | 0.765 | 1 | 0.6 | 0.882 | 1 | 1 | 1 | -0.204 |
| A3SS | 8702 | ENSG00000165280 | VCP | chr9 | - | 35067890 | 35068166 | 35067890 | 35068063 | 35068250 | 35068362 | 0 | 3 | 4 | 5 | 0 | 131 | 63 | 116 | 109 | 145 | 7 | 0 | 4 | 10 | 13 | 170 | 124 | 91 | 139 | 202 | 252 | 149 | 0.005097 | 0.044335097 | 0 | 0.027 | 0.02 | 0.026 | 0 | 0.024 | 0 | 0.025 | 0.041 | 0.037 | -0.011 |
| A3SS | 4458 | ENSG00000114867 | EIF4G1 | chr3 | + | 184323404 | 184323593 | 184323407 | 184323593 | 184323082 | 184323241 | 75 | 58 | 74 | 105 | 121 | 213 | 123 | 134 | 202 | 254 | 207 | 131 | 96 | 146 | 233 | 357 | 190 | 202 | 238 | 312 | 152 | 149 | 0.005115 | 0.044440769 | 0.257 | 0.316 | 0.351 | 0.338 | 0.318 | 0.362 | 0.403 | 0.318 | 0.376 | 0.423 | -0.06 |
| A3SS | 11175 | ENSG00000133794 | ARNTL | chr11 | + | 13376544 | 13376728 | 13376639 | 13376728 | 13375611 | 13375762 | 14 | 6 | 6 | 9 | 10 | 92 | 49 | 74 | 66 | 104 | 5 | 2 | 48 | 19 | 4 | 47 | 64 | 122 | 67 | 11 | 244 | 149 | 0.005212 | 0.045233726 | 0.085 | 0.07 | 0.047 | 0.077 | 0.055 | 0.061 | 0.019 | 0.194 | 0.148 | 0.182 | -0.054 |
| A3SS | 12183 | ENSG00000110075 | PPP6R3 | chr11 | + | 68587926 | 68588024 | 68587944 | 68588024 | 68583042 | 68583129 | 333 | 276 | 270 | 298 | 472 | 50 | 66 | 46 | 90 | 83 | 267 | 361 | 309 | 253 | 44 | 81 | 127 | 128 | 128 | 3 | 167 | 149 | 0.005226 | 0.045312227 | 0.856 | 0.789 | 0.84 | 0.747 | 0.835 | 0.746 | 0.717 | 0.683 | 0.638 | 0.929 | 0.071 |
| A3SS | 10497 | ENSG00000072778 | ACADVL | chr17 | + | 7222666 | 7223237 | 7223132 | 7223237 | 7222176 | 7222302 | 220 | 153 | 155 | 222 | 170 | 0 | 0 | 1 | 2 | 0 | 296 | 352 | 192 | 284 | 504 | 14 | 3 | 3 | 0 | 3 | 298 | 149 | 0.005279 | 0.045720657 | 1 | 1 | 0.987 | 0.982 | 1 | 0.914 | 0.983 | 0.97 | 1 | 0.988 | 0.023 |
| A3SS | 8376 | ENSG00000122085 | MTERF4 | chr2 | - | 241094276 | 241094432 | 241094276 | 241094429 | 241097242 | 241097427 | 8 | 8 | 18 | 4 | 19 | 2 | 1 | 3 | 10 | 6 | 14 | 8 | 1 | 6 | 15 | 0 | 2 | 0 | 4 | 0 | 152 | 149 | 0.005328 | 0.046049568 | 0.797 | 0.887 | 0.855 | 0.282 | 0.756 | 1 | 0.797 | 1 | 0.595 | 1 | -0.163 |
| A3SS | 7755 | ENSG00000114120 | SLC25A36 | chr3 | + | 140966933 | 140967036 | 140966999 | 140967036 | 140963126 | 140963227 | 19 | 17 | 28 | 27 | 30 | 0 | 1 | 0 | 3 | 10 | 9 | 6 | 7 | 8 | 5 | 0 | 0 | 2 | 2 | 0 | 215 | 149 | 0.005328 | 0.046049568 | 1 | 0.922 | 1 | 0.862 | 0.675 | 1 | 1 | 0.708 | 0.735 | 1 | 0.003 |
| A3SS | 2531 | ENSG00000079134 | THOC1 | chr18 | - | 224083 | 224200 | 224083 | 224179 | 224923 | 224994 | 5 | 3 | 4 | 2 | 6 | 81 | 81 | 89 | 73 | 132 | 0 | 0 | 2 | 0 | 0 | 21 | 24 | 34 | 3 | 10 | 170 | 149 | 0.005345 | 0.046147077 | 0.051 | 0.031 | 0.038 | 0.023 | 0.038 | 0 | 0 | 0.049 | 0 | 0 | 0.026 |
| A3SS | 12640 | ENSG00000107862 | GBF1 | chr10 | + | 102365382 | 102365599 | 102365396 | 102365599 | 102363709 | 102363798 | 2 | 1 | 1 | 0 | 0 | 90 | 71 | 66 | 127 | 113 | 12 | 2 | 0 | 2 | 9 | 193 | 128 | 86 | 189 | 226 | 163 | 149 | 0.005425 | 0.046781927 | 0.02 | 0.013 | 0.014 | 0 | 0 | 0.054 | 0.014 | 0 | 0.01 | 0.035 | -0.013 |
| A3SS | 358 | ENSG00000011600 | TYROBP | chr19 | - | 35907217 | 35907357 | 35907217 | 35907264 | 35907445 | 35907580 | 38 | 25 | 6 | 28 | 16 | 984 | 322 | 413 | 590 | 354 | 15 | 19 | 42 | 41 | 54 | 1690 | 943 | 500 | 1298 | 1856 | 242 | 149 | 0.005437 | 0.046839306 | 0.023 | 0.046 | 0.009 | 0.028 | 0.027 | 0.005 | 0.012 | 0.049 | 0.019 | 0.018 | 0.006 |
| A3SS | 8214 | ENSG00000162645 | GBP2 | chr1 | - | 89106131 | 89108291 | 89106131 | 89107052 | 89109676 | 89109870 | 338 | 243 | 288 | 325 | 503 | 7 | 6 | 6 | 6 | 25 | 332 | 165 | 181 | 254 | 161 | 7 | 0 | 6 | 4 | 0 | 298 | 149 | 0.005486 | 0.047206647 | 0.96 | 0.953 | 0.96 | 0.964 | 0.91 | 0.96 | 1 | 0.938 | 0.969 | 1 | -0.024 |
| A3SS | 7706 | ENSG00000113761 | ZNF346 | chr5 | + | 177050702 | 177050936 | 177050750 | 177050936 | 177044388 | 177044533 | 12 | 6 | 9 | 6 | 3 | 37 | 27 | 42 | 51 | 47 | 23 | 12 | 28 | 25 | 26 | 64 | 42 | 56 | 66 | 151 | 197 | 149 | 0.005495 | 0.047239857 | 0.197 | 0.144 | 0.139 | 0.082 | 0.046 | 0.214 | 0.178 | 0.274 | 0.223 | 0.115 | -0.079 |
| A3SS | 7586 | ENSG00000142082 | SIRT3 | chr11 | - | 230451 | 230641 | 230451 | 230552 | 232982 | 233215 | 0 | 3 | 0 | 0 | 0 | 9 | 11 | 16 | 11 | 5 | 16 | 0 | 0 | 0 | 7 | 16 | 17 | 8 | 10 | 0 | 238 | 149 | 0.005545 | 0.047590532 | 0 | 0.146 | 0 | 0 | 0 | 0.385 | 0 | 0 | 0 | 1 | -0.248 |
| A3SS | 986 | ENSG00000124155 | PIGT | chr20 | + | 45419472 | 45419590 | 45419503 | 45419590 | 45419294 | 45419395 | 21 | 3 | 15 | 10 | 10 | 106 | 39 | 88 | 69 | 74 | 12 | 22 | 4 | 4 | 13 | 78 | 180 | 111 | 103 | 166 | 180 | 149 | 0.005548 | 0.047590532 | 0.141 | 0.06 | 0.124 | 0.107 | 0.101 | 0.113 | 0.092 | 0.029 | 0.031 | 0.061 | 0.041 |
| A3SS | 10610 | ENSG00000185485 | SDHAP1 | chr3 | - | 195982161 | 195982352 | 195982161 | 195982310 | 195984111 | 195984276 | 2 | 2 | 3 | 3 | 0 | 14 | 19 | 48 | 42 | 26 | 1 | 4 | 0 | 8 | 14 | 18 | 46 | 33 | 29 | 27 | 191 | 149 | 0.005560 | 0.047644821 | 0.1 | 0.076 | 0.046 | 0.053 | 0 | 0.042 | 0.064 | 0 | 0.177 | 0.288 | -0.059 |
| A3SS | 2544 | ENSG00000100813 | ACIN1 | chr14 | - | 23064104 | 23064257 | 23064104 | 23064221 | 23064354 | 23064485 | 326 | 210 | 305 | 389 | 238 | 110 | 92 | 103 | 99 | 67 | 526 | 376 | 264 | 495 | 296 | 111 | 83 | 53 | 124 | 101 | 185 | 149 | 0.005596 | 0.047900637 | 0.705 | 0.648 | 0.705 | 0.76 | 0.741 | 0.792 | 0.785 | 0.8 | 0.763 | 0.702 | -0.057 |
| A3SS | 7006 | ENSG00000188186 | LAMTOR4 | chr7 | + | 100153849 | 100154203 | 100153866 | 100154203 | 100153399 | 100153517 | 10 | 2 | 4 | 3 | 2 | 114 | 75 | 100 | 106 | 58 | 0 | 0 | 5 | 4 | 3 | 146 | 100 | 85 | 140 | 214 | 166 | 149 | 0.005620 | 0.048054308 | 0.073 | 0.023 | 0.035 | 0.025 | 0.03 | 0 | 0 | 0.05 | 0.025 | 0.012 | 0.02 |
| A3SS | 5255 | ENSG00000116497 | S100PBP | chr1 | + | 32827992 | 32828081 | 32827995 | 32828081 | 32826097 | 32826930 | 49 | 36 | 62 | 58 | 106 | 25 | 24 | 27 | 15 | 72 | 59 | 116 | 60 | 53 | 49 | 0 | 51 | 15 | 41 | 3 | 152 | 149 | 0.005645 | 0.048221751 | 0.658 | 0.595 | 0.692 | 0.791 | 0.591 | 1 | 0.69 | 0.797 | 0.559 | 0.941 | -0.132 |
| A3SS | 8748 | ENSG00000074054 | CLASP1 | chr2 | - | 121407470 | 121407715 | 121407470 | 121407712 | 121410865 | 121410969 | 69 | 31 | 44 | 63 | 64 | 3 | 0 | 1 | 0 | 3 | 41 | 50 | 67 | 63 | 16 | 0 | 2 | 5 | 0 | 16 | 152 | 149 | 0.005657 | 0.048255841 | 0.958 | 1 | 0.977 | 1 | 0.954 | 1 | 0.961 | 0.929 | 1 | 0.495 | 0.101 |
| A3SS | 4012 | ENSG00000055955 | ITIH4 | chr3 | - | 52819753 | 52819990 | 52819753 | 52819807 | 52820290 | 52820317 | 51 | 18 | 32 | 30 | 31 | 0 | 0 | 0 | 0 | 0 | 43 | 103 | 46 | 75 | 143 | 16 | 0 | 0 | 0 | 0 | 298 | 149 | 0.005661 | 0.048255841 | 1 | 1 | 1 | 1 | 1 | 0.573 | 1 | 1 | 1 | 1 | 0.085 |
| A3SS | 12124 | ENSG00000070047 | PHRF1 | chr11 | + | 581961 | 582081 | 581964 | 582081 | 581491 | 581606 | 66 | 20 | 41 | 35 | 20 | 14 | 10 | 10 | 19 | 5 | 8 | 37 | 19 | 28 | 37 | 31 | 11 | 21 | 21 | 18 | 152 | 149 | 0.005698 | 0.048519696 | 0.822 | 0.662 | 0.801 | 0.644 | 0.797 | 0.202 | 0.767 | 0.47 | 0.567 | 0.668 | 0.21 |
| A3SS | 8362 | ENSG00000020922 | MRE11 | chr11 | - | 94445809 | 94445893 | 94445809 | 94445890 | 94447218 | 94447438 | 51 | 53 | 55 | 44 | 42 | 1 | 3 | 2 | 3 | 5 | 10 | 22 | 36 | 16 | 15 | 0 | 2 | 1 | 0 | 0 | 152 | 149 | 0.005708 | 0.04855141 | 0.98 | 0.945 | 0.964 | 0.935 | 0.892 | 1 | 0.915 | 0.972 | 1 | 1 | -0.034 |
| A3SS | 3818 | ENSG00000104972 | LILRB1 | chr19 | + | 54636493 | 54636652 | 54636496 | 54636652 | 54635556 | 54635609 | 35 | 19 | 9 | 16 | 10 | 43 | 20 | 8 | 25 | 24 | 57 | 21 | 27 | 38 | 72 | 39 | 20 | 10 | 29 | 58 | 152 | 149 | 0.005716 | 0.048569207 | 0.444 | 0.482 | 0.524 | 0.386 | 0.29 | 0.589 | 0.507 | 0.726 | 0.562 | 0.549 | -0.161 |
| A3SS | 12458 | ENSG00000122033 | MTIF3 | chr13 | - | 27445087 | 27445156 | 27445087 | 27445137 | 27449851 | 27449918 | 24 | 9 | 17 | 6 | 20 | 0 | 0 | 0 | 0 | 3 | 21 | 28 | 5 | 6 | 5 | 12 | 0 | 0 | 0 | 0 | 168 | 149 | 0.005740 | 0.048721866 | 1 | 1 | 1 | 1 | 0.855 | 0.608 | 1 | 1 | 1 | 1 | 0.049 |
| A3SS | 6007 | ENSG00000162869 | PPP1R21 | chr2 | + | 48458001 | 48458227 | 48458125 | 48458227 | 48454594 | 48454741 | 1 | 6 | 11 | 2 | 1 | 181 | 149 | 125 | 179 | 306 | 0 | 11 | 16 | 2 | 7 | 239 | 171 | 88 | 153 | 116 | 273 | 149 | 0.005922 | 0.050210567 | 0.003 | 0.022 | 0.046 | 0.006 | 0.002 | 0 | 0.034 | 0.09 | 0.007 | 0.032 | -0.017 |
| A3SS | 1110 | ENSG00000040633 | PHF23 | chr17 | - | 7235929 | 7236767 | 7235929 | 7236566 | 7237384 | 7237477 | 190 | 123 | 166 | 154 | 226 | 16 | 3 | 17 | 2 | 13 | 104 | 139 | 148 | 85 | 100 | 0 | 3 | 0 | 1 | 13 | 298 | 149 | 0.006011 | 0.050910259 | 0.856 | 0.953 | 0.83 | 0.975 | 0.897 | 1 | 0.959 | 1 | 0.977 | 0.794 | -0.044 |
| A3SS | 3438 | ENSG00000134644 | PUM1 | chr1 | - | 30980061 | 30980166 | 30980061 | 30980163 | 30981311 | 30981405 | 35 | 30 | 34 | 29 | 22 | 44 | 23 | 54 | 62 | 61 | 15 | 22 | 14 | 15 | 5 | 96 | 36 | 33 | 30 | 46 | 152 | 149 | 0.006099 | 0.051605876 | 0.438 | 0.561 | 0.382 | 0.314 | 0.261 | 0.133 | 0.375 | 0.294 | 0.329 | 0.096 | 0.146 |
| A3SS | 8754 | ENSG00000160710 | ADAR | chr1 | - | 154588124 | 154588673 | 154588124 | 154588258 | 154589368 | 154589462 | 430 | 319 | 393 | 401 | 592 | 0 | 2 | 0 | 0 | 0 | 791 | 454 | 316 | 642 | 321 | 26 | 0 | 0 | 4 | 0 | 298 | 149 | 0.006140 | 0.051896748 | 1 | 0.988 | 1 | 1 | 1 | 0.938 | 1 | 1 | 0.988 | 1 | 0.012 |
| A3SS | 1160 | ENSG00000141580 | WDR45B | chr17 | - | 82619042 | 82619128 | 82619042 | 82619094 | 82621608 | 82621799 | 157 | 111 | 187 | 175 | 148 | 9 | 5 | 10 | 11 | 16 | 381 | 228 | 249 | 280 | 416 | 0 | 14 | 11 | 15 | 15 | 183 | 149 | 0.006152 | 0.05194398 | 0.934 | 0.948 | 0.938 | 0.928 | 0.883 | 1 | 0.93 | 0.949 | 0.938 | 0.958 | -0.029 |
| A3SS | 3630 | ENSG00000101182 | PSMA7 | chr20 | - | 62137363 | 62137465 | 62137363 | 62137426 | 62138170 | 62138290 | 1 | 5 | 9 | 3 | 5 | 297 | 98 | 216 | 276 | 478 | 14 | 12 | 7 | 8 | 0 | 241 | 275 | 169 | 248 | 278 | 188 | 149 | 0.006197 | 0.052273219 | 0.003 | 0.039 | 0.032 | 0.009 | 0.008 | 0.044 | 0.033 | 0.032 | 0.025 | 0 | -0.009 |
| A3SS | 2400 | ENSG00000196547 | MAN2A2 | chr15 | + | 90912516 | 90912664 | 90912541 | 90912664 | 90912042 | 90912279 | 2 | 5 | 2 | 3 | 0 | 20 | 12 | 13 | 23 | 3 | 4 | 7 | 11 | 22 | 5 | 0 | 19 | 16 | 15 | 63 | 174 | 149 | 0.006335 | 0.053374016 | 0.079 | 0.263 | 0.116 | 0.1 | 0 | 1 | 0.24 | 0.371 | 0.557 | 0.064 | -0.335 |
| A3SS | 2702 | ENSG00000099797 | TECR | chr19 | + | 14563173 | 14563257 | 14563205 | 14563257 | 14562524 | 14562575 | 2 | 4 | 10 | 1 | 0 | 73 | 33 | 109 | 71 | 51 | 0 | 8 | 2 | 12 | 17 | 20 | 77 | 87 | 66 | 123 | 181 | 149 | 0.006361 | 0.053493793 | 0.022 | 0.091 | 0.07 | 0.011 | 0 | 0 | 0.079 | 0.019 | 0.13 | 0.102 | -0.027 |
| A3SS | 12555 | ENSG00000040341 | STAU2 | chr8 | - | 73615674 | 73615807 | 73615674 | 73615782 | 73617291 | 73617451 | 0 | 0 | 0 | 0 | 0 | 122 | 88 | 109 | 127 | 192 | 0 | 0 | 0 | 0 | 16 | 44 | 57 | 100 | 60 | 60 | 174 | 149 | 0.006362 | 0.053493793 | 0 | 0 | 0 | 0 | 0 | 0 | 0 | 0 | 0 | 0.186 | -0.037 |
| A3SS | 2231 | ENSG00000166783 | MARF1 | chr16 | - | 15622933 | 15623123 | 15622933 | 15623057 | 15624768 | 15624927 | 211 | 154 | 209 | 221 | 304 | 0 | 0 | 0 | 2 | 0 | 115 | 217 | 188 | 178 | 38 | 0 | 2 | 16 | 0 | 0 | 215 | 149 | 0.006385 | 0.053631264 | 1 | 1 | 1 | 0.987 | 1 | 1 | 0.987 | 0.891 | 1 | 1 | 0.022 |
| A3SS | 4022 | ENSG00000131042 | LILRB2 | chr19 | - | 54279790 | 54280075 | 54279790 | 54279960 | 54280462 | 54280544 | 173 | 167 | 91 | 248 | 257 | 5 | 3 | 0 | 2 | 0 | 607 | 347 | 161 | 442 | 500 | 0 | 6 | 0 | 10 | 25 | 264 | 149 | 0.006447 | 0.054049492 | 0.951 | 0.969 | 1 | 0.986 | 1 | 1 | 0.97 | 1 | 0.961 | 0.919 | 0.011 |
| A3SS | 3879 | ENSG00000061918 | GUCY1B1 | chr4 | + | 155802341 | 155802579 | 155802440 | 155802579 | 155799876 | 155800074 | 89 | 76 | 19 | 163 | 194 | 1 | 2 | 0 | 2 | 2 | 99 | 89 | 90 | 58 | 228 | 0 | 0 | 2 | 13 | 4 | 248 | 149 | 0.006448 | 0.054049492 | 0.982 | 0.958 | 1 | 0.98 | 0.983 | 1 | 1 | 0.964 | 0.728 | 0.972 | 0.048 |
| A3SS | 5514 | ENSG00000196628 | TCF4 | chr18 | - | 55257314 | 55257391 | 55257314 | 55257388 | 55259948 | 55260027 | 29 | 27 | 40 | 32 | 25 | 2 | 2 | 0 | 4 | 3 | 31 | 29 | 21 | 4 | 2 | 0 | 0 | 0 | 0 | 0 | 152 | 149 | 0.006475 | 0.054214543 | 0.934 | 0.93 | 1 | 0.887 | 0.891 | 1 | 1 | 1 | 1 | 1 | -0.072 |
| A3SS | 1002 | ENSG00000100342 | APOL1 | chr22 | + | 36254897 | 36254999 | 36254936 | 36254999 | 36253837 | 36253991 | 6 | 2 | 2 | 0 | 2 | 6 | 1 | 2 | 1 | 0 | 0 | 0 | 0 | 2 | 10 | 10 | 5 | 5 | 0 | 14 | 188 | 149 | 0.006499 | 0.054359783 | 0.442 | 0.613 | 0.442 | 0 | 1 | 0 | 0 | 0 | 1 | 0.361 | 0.227 |
| A3SS | 11920 | ENSG00000139697 | SBNO1 | chr12 | - | 123345257 | 123345570 | 123345257 | 123345567 | 123348028 | 123348133 | 88 | 64 | 90 | 67 | 180 | 86 | 45 | 77 | 93 | 127 | 54 | 73 | 55 | 38 | 14 | 72 | 68 | 78 | 75 | 30 | 152 | 149 | 0.006543 | 0.054676337 | 0.501 | 0.582 | 0.534 | 0.414 | 0.581 | 0.424 | 0.513 | 0.409 | 0.332 | 0.314 | 0.124 |
| A3SS | 9905 | ENSG00000157764 | BRAF | chr7 | - | 140800361 | 140800481 | 140800361 | 140800462 | 140801411 | 140801560 | 201 | 170 | 192 | 197 | 544 | 10 | 13 | 6 | 6 | 53 | 177 | 179 | 152 | 94 | 172 | 27 | 18 | 26 | 7 | 6 | 168 | 149 | 0.006649 | 0.055502567 | 0.947 | 0.921 | 0.966 | 0.967 | 0.901 | 0.853 | 0.898 | 0.838 | 0.923 | 0.962 | 0.046 |
| A3SS | 10221 | ENSG00000166579 | NDEL1 | chr17 | + | 8443926 | 8444357 | 8444259 | 8444357 | 8413194 | 8413269 | 38 | 37 | 38 | 41 | 40 | 15 | 8 | 21 | 36 | 32 | 12 | 76 | 72 | 78 | 10 | 36 | 41 | 114 | 55 | 19 | 298 | 149 | 0.006666 | 0.055586983 | 0.559 | 0.698 | 0.475 | 0.363 | 0.385 | 0.143 | 0.481 | 0.24 | 0.415 | 0.208 | 0.199 |
| A3SS | 1108 | ENSG00000184792 | OSBP2 | chr22 | + | 30889176 | 30889234 | 30889179 | 30889234 | 30888222 | 30888340 | 37 | 6 | 4 | 13 | 32 | 4 | 0 | 0 | 1 | 0 | 14 | 96 | 18 | 11 | 1626 | 0 | 8 | 0 | 0 | 99 | 152 | 149 | 0.006675 | 0.055606549 | 0.901 | 1 | 1 | 0.927 | 1 | 1 | 0.922 | 1 | 1 | 0.942 | -0.007 |
| A3SS | 5657 | ENSG00000198265 | HELZ | chr17 | - | 67160260 | 67160365 | 67160260 | 67160362 | 67160896 | 67161076 | 50 | 27 | 39 | 56 | 32 | 139 | 104 | 98 | 110 | 100 | 0 | 26 | 39 | 9 | 3 | 25 | 131 | 112 | 46 | 12 | 152 | 149 | 0.006687 | 0.055645923 | 0.261 | 0.203 | 0.281 | 0.333 | 0.239 | 0 | 0.163 | 0.254 | 0.161 | 0.197 | 0.108 |
| A3SS | 4127 | ENSG00000163913 | IFT122 | chr3 | + | 129463544 | 129463626 | 129463559 | 129463626 | 129451913 | 129451998 | 0 | 1 | 2 | 1 | 0 | 10 | 5 | 6 | 1 | 11 | 12 | 0 | 1 | 0 | 0 | 22 | 9 | 10 | 16 | 11 | 164 | 149 | 0.006727 | 0.055869643 | 0 | 0.154 | 0.232 | 0.476 | 0 | 0.331 | 0 | 0.083 | 0 | 0 | 0.09 |
| A3SS | 3350 | ENSG00000055163 | CYFIP2 | chr5 | + | 157304237 | 157304366 | 157304249 | 157304366 | 157302793 | 157302890 | 398 | 154 | 354 | 326 | 241 | 4 | 1 | 0 | 0 | 5 | 227 | 343 | 252 | 320 | 332 | 7 | 3 | 3 | 6 | 12 | 161 | 149 | 0.006728 | 0.055869643 | 0.989 | 0.993 | 1 | 1 | 0.978 | 0.968 | 0.991 | 0.987 | 0.98 | 0.962 | 0.014 |
| A3SS | 1799 | ENSG00000005844 | ITGAL | chr16 | + | 30475288 | 30475400 | 30475305 | 30475400 | 30474195 | 30474287 | 30 | 18 | 16 | 28 | 17 | 4 | 0 | 3 | 3 | 0 | 36 | 31 | 20 | 34 | 60 | 0 | 4 | 0 | 0 | 0 | 166 | 149 | 0.006755 | 0.056031387 | 0.871 | 1 | 0.827 | 0.893 | 1 | 1 | 0.874 | 1 | 1 | 1 | -0.057 |
| A3SS | 2905 | ENSG00000175061 | SNHG29 | chr17 | + | 16439580 | 16439703 | 16439659 | 16439703 | 16439326 | 16439414 | 93 | 56 | 150 | 103 | 76 | 17 | 10 | 43 | 20 | 7 | 87 | 105 | 122 | 101 | 149 | 0 | 10 | 31 | 26 | 1 | 228 | 149 | 0.006761 | 0.056031387 | 0.781 | 0.785 | 0.695 | 0.771 | 0.876 | 1 | 0.873 | 0.72 | 0.717 | 0.99 | -0.078 |
| A3SS | 8564 | ENSG00000115282 | TTC31 | chr2 | + | 74492921 | 74494559 | 74492924 | 74494559 | 74492645 | 74492747 | 35 | 11 | 17 | 14 | 17 | 4 | 0 | 1 | 2 | 4 | 17 | 19 | 8 | 24 | 60 | 0 | 2 | 0 | 3 | 0 | 152 | 149 | 0.006804 | 0.056332086 | 0.896 | 1 | 0.943 | 0.873 | 0.806 | 1 | 0.903 | 1 | 0.887 | 1 | -0.054 |
| A3SS | 4038 | ENSG00000100412 | ACO2 | chr22 | + | 41499545 | 41499862 | 41499725 | 41499862 | 41469116 | 41469182 | 24 | 6 | 19 | 26 | 2 | 77 | 20 | 54 | 55 | 39 | 59 | 32 | 33 | 25 | 35 | 59 | 77 | 31 | 81 | 100 | 298 | 149 | 0.006829 | 0.05647806 | 0.135 | 0.13 | 0.15 | 0.191 | 0.025 | 0.333 | 0.172 | 0.347 | 0.134 | 0.149 | -0.101 |
| A3SS | 1952 | ENSG00000149923 | PPP4C | chr16 | + | 30081199 | 30081310 | 30081258 | 30081310 | 30076314 | 30076475 | 10 | 10 | 7 | 8 | 4 | 145 | 55 | 98 | 78 | 106 | 30 | 14 | 10 | 29 | 20 | 154 | 159 | 108 | 154 | 202 | 208 | 149 | 0.006870 | 0.056759349 | 0.047 | 0.115 | 0.049 | 0.068 | 0.026 | 0.122 | 0.059 | 0.062 | 0.119 | 0.066 | -0.025 |
| A3SS | 1724 | ENSG00000167522 | ANKRD11 | chr16 | - | 89316932 | 89317078 | 89316932 | 89317075 | 89418283 | 89418368 | 49 | 32 | 34 | 19 | 25 | 35 | 22 | 27 | 32 | 28 | 104 | 34 | 54 | 71 | 128 | 74 | 19 | 22 | 37 | 64 | 152 | 149 | 0.006919 | 0.057107024 | 0.578 | 0.588 | 0.552 | 0.368 | 0.467 | 0.579 | 0.637 | 0.706 | 0.653 | 0.662 | -0.137 |
| A3SS | 2175 | ENSG00000140263 | SORD | chr15 | + | 45060981 | 45061226 | 45061066 | 45061226 | 45043256 | 45043421 | 4 | 0 | 0 | 1 | 8 | 37 | 11 | 23 | 23 | 58 | 0 | 0 | 0 | 0 | 0 | 18 | 13 | 18 | 38 | 41 | 234 | 149 | 0.006961 | 0.057390517 | 0.064 | 0 | 0 | 0.027 | 0.081 | 0 | 0 | 0 | 0 | 0 | 0.034 |
| A3SS | 7029 | ENSG00000166311 | SMPD1 | chr11 | + | 6394177 | 6394778 | 6394197 | 6394778 | 6393895 | 6394041 | 7 | 9 | 21 | 12 | 13 | 95 | 61 | 96 | 101 | 111 | 23 | 23 | 10 | 18 | 97 | 109 | 95 | 64 | 68 | 510 | 169 | 149 | 0.007001 | 0.057619361 | 0.061 | 0.115 | 0.162 | 0.095 | 0.094 | 0.157 | 0.176 | 0.121 | 0.189 | 0.144 | -0.052 |
| A3SS | 4469 | ENSG00000169714 | CNBP | chr3 | - | 129171078 | 129171283 | 129171078 | 129171280 | 129171445 | 129171538 | 15 | 11 | 19 | 11 | 28 | 371 | 220 | 368 | 333 | 354 | 48 | 19 | 16 | 39 | 17 | 439 | 274 | 410 | 338 | 350 | 152 | 149 | 0.007006 | 0.057619361 | 0.038 | 0.047 | 0.048 | 0.031 | 0.072 | 0.097 | 0.064 | 0.037 | 0.102 | 0.045 | -0.022 |
| A3SS | 1083 | ENSG00000073605 | GSDMB | chr17 | - | 39906110 | 39906987 | 39906110 | 39906271 | 39908957 | 39909042 | 34 | 27 | 43 | 39 | 12 | 0 | 3 | 0 | 1 | 0 | 13 | 22 | 32 | 16 | 17 | 7 | 0 | 0 | 3 | 0 | 298 | 149 | 0.007010 | 0.057619361 | 1 | 0.818 | 1 | 0.951 | 1 | 0.481 | 1 | 1 | 0.727 | 1 | 0.112 |
| A3SS | 12872 | ENSG00000105221 | AKT2 | chr19 | - | 40236023 | 40236142 | 40236023 | 40236104 | 40236256 | 40236385 | 5 | 3 | 2 | 8 | 0 | 79 | 50 | 94 | 83 | 98 | 0 | 1 | 0 | 6 | 0 | 104 | 121 | 86 | 82 | 129 | 187 | 149 | 0.007038 | 0.057789907 | 0.048 | 0.046 | 0.017 | 0.071 | 0 | 0 | 0.007 | 0 | 0.055 | 0 | 0.024 |
| A3SS | 10658 | ENSG00000131269 | ABCB7 | chrX | - | 75071508 | 75071704 | 75071508 | 75071683 | 75073688 | 75073776 | 0 | 3 | 0 | 4 | 8 | 44 | 33 | 44 | 33 | 100 | 0 | 0 | 3 | 0 | 0 | 33 | 10 | 35 | 20 | 7 | 170 | 149 | 0.007052 | 0.057846922 | 0 | 0.074 | 0 | 0.096 | 0.066 | 0 | 0 | 0.07 | 0 | 0 | 0.033 |
| A3SS | 12726 | ENSG00000130640 | TUBGCP2 | chr10 | - | 133283077 | 133283221 | 133283077 | 133283184 | 133283881 | 133284002 | 100 | 62 | 79 | 72 | 48 | 2 | 1 | 6 | 10 | 1 | 104 | 112 | 57 | 97 | 194 | 0 | 0 | 2 | 0 | 10 | 186 | 149 | 0.007092 | 0.058077834 | 0.976 | 0.98 | 0.913 | 0.852 | 0.975 | 1 | 1 | 0.958 | 1 | 0.94 | -0.04 |
| A3SS | 2903 | ENSG00000175061 | SNHG29 | chr17 | + | 16439531 | 16439703 | 16439580 | 16439703 | 16439326 | 16439414 | 54 | 29 | 55 | 61 | 32 | 18 | 13 | 36 | 18 | 11 | 97 | 50 | 53 | 110 | 92 | 11 | 18 | 19 | 25 | 16 | 198 | 149 | 0.007094 | 0.058077834 | 0.693 | 0.627 | 0.535 | 0.718 | 0.686 | 0.869 | 0.676 | 0.677 | 0.768 | 0.812 | -0.109 |
| A3SS | 7631 | ENSG00000231500 | RPS18 | chr6 | + | 33272586 | 33272726 | 33272627 | 33272726 | 33272074 | 33272122 | 47 | 13 | 26 | 32 | 15 | 1195 | 526 | 1341 | 1104 | 868 | 37 | 12 | 28 | 29 | 53 | 723 | 845 | 742 | 661 | 1200 | 190 | 149 | 0.007123 | 0.058251543 | 0.03 | 0.019 | 0.015 | 0.022 | 0.013 | 0.039 | 0.011 | 0.029 | 0.033 | 0.033 | -0.009 |
| A3SS | 12823 | ENSG00000142166 | IFNAR1 | chr21 | + | 33340998 | 33341174 | 33341077 | 33341174 | 33335523 | 33335647 | 254 | 205 | 263 | 289 | 429 | 1 | 4 | 1 | 1 | 1 | 146 | 184 | 246 | 221 | 22 | 9 | 0 | 7 | 2 | 0 | 228 | 149 | 0.007148 | 0.058397207 | 0.994 | 0.971 | 0.994 | 0.995 | 0.996 | 0.914 | 1 | 0.958 | 0.986 | 1 | 0.018 |
| A3SS | 11749 | ENSG00000149428 | HYOU1 | chr11 | - | 119049017 | 119049370 | 119049017 | 119049203 | 119049555 | 119049635 | 9 | 4 | 9 | 15 | 5 | 424 | 213 | 361 | 399 | 345 | 24 | 12 | 16 | 28 | 18 | 687 | 461 | 426 | 400 | 567 | 298 | 149 | 0.007172 | 0.058537277 | 0.011 | 0.009 | 0.012 | 0.018 | 0.007 | 0.017 | 0.013 | 0.018 | 0.034 | 0.016 | -0.008 |
| A3SS | 11271 | ENSG00000132386 | SERPINF1 | chr17 | + | 1775057 | 1775200 | 1775062 | 1775200 | 1771718 | 1772075 | 33 | 32 | 39 | 61 | 30 | 0 | 1 | 13 | 8 | 0 | 47 | 19 | 8 | 36 | 29 | 0 | 0 | 0 | 0 | 7 | 154 | 149 | 0.007191 | 0.058628267 | 1 | 0.969 | 0.744 | 0.881 | 1 | 1 | 1 | 1 | 1 | 0.8 | -0.041 |
| A3SS | 5862 | ENSG00000178028 | DMAP1 | chr1 | + | 44213662 | 44213858 | 44213708 | 44213858 | 44213472 | 44213527 | 6 | 1 | 7 | 3 | 2 | 13 | 8 | 12 | 7 | 3 | 16 | 6 | 3 | 4 | 22 | 3 | 17 | 1 | 7 | 7 | 195 | 149 | 0.007239 | 0.058926203 | 0.261 | 0.087 | 0.308 | 0.247 | 0.337 | 0.803 | 0.212 | 0.696 | 0.304 | 0.706 | -0.296 |
| A3SS | 7207 | ENSG00000093010 | COMT | chr22 | + | 19961696 | 19962815 | 19962526 | 19962815 | 19961198 | 19961289 | 10 | 3 | 3 | 4 | 0 | 89 | 49 | 49 | 70 | 64 | 2 | 5 | 1 | 0 | 0 | 95 | 135 | 45 | 54 | 187 | 298 | 149 | 0.007243 | 0.058926203 | 0.053 | 0.03 | 0.03 | 0.028 | 0 | 0.01 | 0.018 | 0.011 | 0 | 0 | 0.02 |
| A3SS | 4595 | ENSG00000161217 | PCYT1A | chr3 | - | 196247366 | 196247643 | 196247366 | 196247518 | 196248206 | 196248323 | 25 | 15 | 19 | 24 | 15 | 126 | 77 | 100 | 110 | 123 | 22 | 33 | 26 | 45 | 27 | 149 | 113 | 66 | 107 | 120 | 274 | 149 | 0.007249 | 0.058926203 | 0.097 | 0.096 | 0.094 | 0.106 | 0.062 | 0.074 | 0.137 | 0.176 | 0.186 | 0.109 | -0.045 |
| A3SS | 1625 | ENSG00000108559 | NUP88 | chr17 | - | 5387778 | 5387952 | 5387778 | 5387904 | 5388801 | 5388960 | 7 | 0 | 3 | 1 | 14 | 108 | 64 | 103 | 107 | 236 | 0 | 0 | 2 | 4 | 0 | 72 | 62 | 88 | 91 | 42 | 197 | 149 | 0.007284 | 0.059108456 | 0.047 | 0 | 0.022 | 0.007 | 0.043 | 0 | 0 | 0.017 | 0.032 | 0 | 0.014 |
| A3SS | 2568 | ENSG00000092036 | HAUS4 | chr14 | - | 22947867 | 22948033 | 22947867 | 22948013 | 22950313 | 22950410 | 1 | 2 | 9 | 2 | 3 | 99 | 54 | 62 | 79 | 108 | 0 | 0 | 2 | 3 | 0 | 61 | 83 | 49 | 75 | 133 | 169 | 149 | 0.007286 | 0.059108456 | 0.009 | 0.032 | 0.113 | 0.022 | 0.024 | 0 | 0 | 0.035 | 0.034 | 0 | 0.026 |
| A3SS | 2403 | ENSG00000176700 | SCAND2P | chr15 | + | 84638353 | 84638477 | 84638395 | 84638477 | 84637346 | 84637420 | 2 | 7 | 8 | 2 | 14 | 19 | 12 | 19 | 18 | 33 | 37 | 11 | 12 | 31 | 5 | 5 | 28 | 18 | 15 | 20 | 191 | 149 | 0.007293 | 0.059108548 | 0.076 | 0.313 | 0.247 | 0.08 | 0.249 | 0.852 | 0.235 | 0.342 | 0.617 | 0.163 | -0.249 |
| A3SS | 6665 | ENSG00000138688 | KIAA1109 | chr4 | + | 122224497 | 122224677 | 122224500 | 122224677 | 122220332 | 122220470 | 136 | 95 | 153 | 166 | 118 | 22 | 12 | 25 | 31 | 18 | 27 | 41 | 53 | 20 | 21 | 23 | 2 | 12 | 9 | 16 | 152 | 149 | 0.007304 | 0.05913886 | 0.858 | 0.886 | 0.857 | 0.84 | 0.865 | 0.535 | 0.953 | 0.812 | 0.685 | 0.563 | 0.152 |
| A3SS | 8601 | ENSG00000164548 | TRA2A | chr7 | - | 23505745 | 23505813 | 23505745 | 23505810 | 23506137 | 23506266 | 494 | 387 | 433 | 554 | 1228 | 11 | 15 | 14 | 5 | 16 | 106 | 250 | 446 | 186 | 59 | 22 | 3 | 1 | 14 | 0 | 152 | 149 | 0.007366 | 0.05957847 | 0.978 | 0.962 | 0.968 | 0.991 | 0.987 | 0.825 | 0.988 | 0.998 | 0.929 | 1 | 0.029 |
| A3SS | 9167 | ENSG00000175575 | PAAF1 | chr11 | + | 73886928 | 73887099 | 73887039 | 73887099 | 73878778 | 73878819 | 1 | 4 | 9 | 1 | 5 | 0 | 1 | 1 | 0 | 0 | 11 | 15 | 15 | 17 | 5 | 0 | 0 | 2 | 0 | 16 | 260 | 149 | 0.007439 | 0.060108119 | 1 | 0.696 | 0.838 | 1 | 1 | 1 | 1 | 0.811 | 1 | 0.152 | 0.114 |
| A3SS | 7848 | ENSG00000109332 | UBE2D3 | chr4 | - | 102826484 | 102826844 | 102826484 | 102826636 | 102827591 | 102827948 | 212 | 134 | 201 | 261 | 109 | 16 | 3 | 7 | 12 | 14 | 89 | 282 | 326 | 270 | 206 | 12 | 4 | 10 | 7 | 0 | 298 | 149 | 0.007577 | 0.061102401 | 0.869 | 0.957 | 0.935 | 0.916 | 0.796 | 0.788 | 0.972 | 0.942 | 0.951 | 1 | -0.036 |
| A3SS | 11235 | ENSG00000099810 | MTAP | chr9 | + | 21859285 | 21859425 | 21859302 | 21859425 | 21854630 | 21854870 | 3 | 1 | 4 | 1 | 4 | 48 | 43 | 44 | 55 | 18 | 0 | 3 | 0 | 0 | 0 | 9 | 39 | 44 | 18 | 10 | 166 | 149 | 0.007577 | 0.061102401 | 0.053 | 0.02 | 0.075 | 0.016 | 0.166 | 0 | 0.065 | 0 | 0 | 0 | 0.053 |
| A3SS | 7784 | ENSG00000013810 | TACC3 | chr4 | + | 1738347 | 1739778 | 1739701 | 1739778 | 1737597 | 1737702 | 11 | 5 | 10 | 11 | 0 | 94 | 66 | 90 | 92 | 107 | 27 | 22 | 9 | 12 | 42 | 156 | 134 | 115 | 183 | 255 | 298 | 149 | 0.007664 | 0.061743116 | 0.055 | 0.036 | 0.053 | 0.056 | 0 | 0.08 | 0.076 | 0.038 | 0.032 | 0.076 | -0.02 |
| A3SS | 5455 | ENSG00000186470 | BTN3A2 | chr6 | + | 26367974 | 26368050 | 26367989 | 26368050 | 26365230 | 26365352 | 8 | 6 | 4 | 7 | 13 | 152 | 114 | 94 | 75 | 213 | 7 | 3 | 1 | 0 | 0 | 91 | 52 | 36 | 37 | 146 | 164 | 149 | 0.007739 | 0.06227928 | 0.046 | 0.046 | 0.037 | 0.078 | 0.053 | 0.065 | 0.05 | 0.025 | 0 | 0 | 0.024 |
| A3SS | 1256 | ENSG00000141503 | MINK1 | chr17 | + | 4891455 | 4891716 | 4891515 | 4891716 | 4890950 | 4891124 | 32 | 19 | 27 | 34 | 62 | 12 | 7 | 10 | 26 | 18 | 100 | 61 | 57 | 42 | 309 | 18 | 34 | 11 | 6 | 38 | 209 | 149 | 0.007759 | 0.062384143 | 0.655 | 0.659 | 0.658 | 0.482 | 0.711 | 0.798 | 0.561 | 0.787 | 0.833 | 0.853 | -0.133 |
| A3SS | 5508 | ENSG00000058673 | ZC3H11A | chr1 | + | 203850494 | 203850681 | 203850514 | 203850681 | 203849710 | 203850026 | 88 | 74 | 91 | 92 | 157 | 269 | 231 | 279 | 299 | 459 | 7 | 79 | 74 | 32 | 2 | 68 | 294 | 265 | 128 | 31 | 169 | 149 | 0.007812 | 0.062747317 | 0.224 | 0.22 | 0.223 | 0.213 | 0.232 | 0.083 | 0.192 | 0.198 | 0.181 | 0.054 | 0.081 |
| A3SS | 5921 | ENSG00000055609 | KMT2C | chr7 | - | 152207299 | 152207428 | 152207299 | 152207425 | 152220522 | 152220735 | 297 | 345 | 281 | 392 | 386 | 24 | 21 | 12 | 15 | 21 | 36 | 323 | 297 | 235 | 21 | 11 | 21 | 36 | 18 | 0 | 152 | 149 | 0.007823 | 0.062770644 | 0.924 | 0.942 | 0.958 | 0.962 | 0.947 | 0.762 | 0.938 | 0.89 | 0.928 | 1 | 0.043 |
| A3SS | 12875 | ENSG00000105221 | AKT2 | chr19 | - | 40255157 | 40255299 | 40255157 | 40255269 | 40256925 | 40257054 | 2 | 1 | 1 | 0 | 0 | 77 | 31 | 64 | 40 | 37 | 0 | 10 | 1 | 0 | 7 | 62 | 79 | 33 | 90 | 107 | 179 | 149 | 0.007840 | 0.062846297 | 0.021 | 0.026 | 0.013 | 0.02 | 0 | 0 | 0.095 | 0.025 | 0 | 0.052 | -0.018 |
| A3SS | 8230 | ENSG00000204681 | GABBR1 | chr6 | - | 29629086 | 29629247 | 29629086 | 29629107 | 29630457 | 29630643 | 37 | 36 | 29 | 29 | 42 | 1 | 5 | 7 | 4 | 3 | 48 | 9 | 20 | 35 | 38 | 0 | 3 | 0 | 4 | 0 | 289 | 149 | 0.007895 | 0.063218706 | 0.95 | 0.788 | 0.681 | 0.789 | 0.878 | 1 | 0.607 | 1 | 0.819 | 1 | -0.068 |
| A3SS | 4509 | ENSG00000166501 | PRKCB | chr16 | + | 24214657 | 24220611 | 24219960 | 24220611 | 24191089 | 24191230 | 515 | 313 | 420 | 427 | 681 | 5 | 5 | 9 | 14 | 11 | 367 | 429 | 467 | 421 | 621 | 11 | 4 | 13 | 3 | 44 | 298 | 149 | 0.007960 | 0.063681584 | 0.981 | 0.969 | 0.959 | 0.938 | 0.969 | 0.943 | 0.982 | 0.947 | 0.986 | 0.876 | 0.016 |
| A3SS | 11185 | ENSG00000104365 | IKBKB | chr8 | + | 42317656 | 42317771 | 42317709 | 42317771 | 42316709 | 42316904 | 233 | 118 | 138 | 182 | 189 | 9 | 7 | 10 | 14 | 2 | 212 | 155 | 121 | 204 | 232 | 7 | 25 | 22 | 12 | 11 | 202 | 149 | 0.007973 | 0.063722638 | 0.95 | 0.926 | 0.911 | 0.906 | 0.986 | 0.957 | 0.821 | 0.802 | 0.926 | 0.94 | 0.047 |
| A3SS | 10034 | ENSG00000184216 | IRAK1 | chrX | - | 154013042 | 154013433 | 154013042 | 154013343 | 154014041 | 154014278 | 92 | 42 | 57 | 79 | 73 | 9 | 4 | 2 | 2 | 10 | 80 | 100 | 48 | 62 | 97 | 21 | 18 | 0 | 7 | 21 | 239 | 149 | 0.007994 | 0.063803175 | 0.864 | 0.867 | 0.947 | 0.961 | 0.82 | 0.704 | 0.776 | 1 | 0.847 | 0.742 | 0.078 |
| A3SS | 5829 | ENSG00000174175 | SELP | chr1 | - | 169588848 | 169589565 | 169588848 | 169589461 | 169590146 | 169590202 | 98 | 150 | 33 | 145 | 194 | 1 | 1 | 0 | 12 | 4 | 443 | 151 | 119 | 118 | 2921 | 25 | 14 | 12 | 10 | 91 | 253 | 149 | 0.007999 | 0.063803175 | 0.983 | 0.989 | 1 | 0.877 | 0.966 | 0.913 | 0.864 | 0.854 | 0.874 | 0.95 | 0.072 |
| A3SS | 1874 | ENSG00000197912 | SPG7 | chr16 | + | 89548979 | 89549336 | 89549080 | 89549336 | 89548002 | 89548113 | 11 | 4 | 14 | 9 | 9 | 0 | 0 | 0 | 1 | 0 | 29 | 23 | 28 | 33 | 41 | 12 | 0 | 0 | 0 | 0 | 250 | 149 | 0.008066 | 0.064271801 | 1 | 1 | 1 | 0.843 | 1 | 0.59 | 1 | 1 | 1 | 1 | 0.051 |
| A3SS | 6950 | ENSG00000185885 | IFITM1 | chr11 | + | 314881 | 315272 | 314921 | 315272 | 314039 | 314356 | 55 | 27 | 32 | 42 | 92 | 1289 | 463 | 877 | 871 | 636 | 51 | 34 | 39 | 54 | 40 | 852 | 1173 | 897 | 1229 | 755 | 189 | 149 | 0.008153 | 0.064844973 | 0.033 | 0.044 | 0.028 | 0.037 | 0.102 | 0.045 | 0.022 | 0.033 | 0.033 | 0.04 | 0.014 |
| A3SS | 2994 | ENSG00000164180 | TMEM161B | chr5 | - | 88207028 | 88207180 | 88207028 | 88207142 | 88220562 | 88220719 | 55 | 29 | 38 | 51 | 102 | 1 | 5 | 3 | 3 | 0 | 2 | 18 | 12 | 1 | 5 | 0 | 0 | 0 | 0 | 0 | 187 | 149 | 0.008154 | 0.064844973 | 0.978 | 0.822 | 0.91 | 0.931 | 1 | 1 | 1 | 1 | 1 | 1 | -0.072 |
| A3SS | 11651 | ENSG00000130714 | POMT1 | chr9 | + | 131507367 | 131509022 | 131508910 | 131509022 | 131506402 | 131506453 | 28 | 9 | 33 | 15 | 20 | 0 | 0 | 0 | 2 | 0 | 15 | 36 | 11 | 11 | 26 | 0 | 0 | 2 | 0 | 7 | 298 | 149 | 0.008171 | 0.064913702 | 1 | 1 | 1 | 0.789 | 1 | 1 | 1 | 0.733 | 1 | 0.65 | 0.081 |
| A3SS | 7896 | ENSG00000163219 | ARHGAP25 | chr2 | + | 68807272 | 68807480 | 68807275 | 68807480 | 68787839 | 68787956 | 317 | 95 | 166 | 182 | 218 | 56 | 33 | 55 | 37 | 48 | 88 | 158 | 110 | 82 | 160 | 23 | 27 | 23 | 15 | 5 | 152 | 149 | 0.008222 | 0.065238181 | 0.847 | 0.738 | 0.747 | 0.828 | 0.817 | 0.789 | 0.852 | 0.824 | 0.843 | 0.969 | -0.06 |
| A3SS | 4077 | ENSG00000085982 | USP40 | chr2 | - | 233540661 | 233540769 | 233540661 | 233540766 | 233542267 | 233542363 | 47 | 21 | 18 | 40 | 101 | 1 | 1 | 0 | 0 | 11 | 40 | 35 | 23 | 9 | 1 | 0 | 0 | 0 | 2 | 0 | 152 | 149 | 0.008234 | 0.065238181 | 0.979 | 0.954 | 1 | 1 | 0.9 | 1 | 1 | 1 | 0.815 | 1 | 0.004 |
| A3SS | 6842 | ENSG00000151694 | ADAM17 | chr2 | - | 9526110 | 9526264 | 9526110 | 9526244 | 9527785 | 9527954 | 8 | 10 | 21 | 26 | 17 | 193 | 182 | 127 | 160 | 218 | 4 | 4 | 11 | 17 | 24 | 285 | 182 | 285 | 371 | 75 | 169 | 149 | 0.008236 | 0.065238181 | 0.035 | 0.046 | 0.127 | 0.125 | 0.064 | 0.012 | 0.019 | 0.033 | 0.039 | 0.22 | 0.015 |
| A3SS | 9779 | ENSG00000198198 | SZT2 | chr1 | + | 43403600 | 43403774 | 43403606 | 43403774 | 43403176 | 43403302 | 54 | 16 | 45 | 57 | 28 | 0 | 0 | 0 | 0 | 0 | 52 | 62 | 43 | 37 | 70 | 0 | 1 | 0 | 8 | 4 | 155 | 149 | 0.008253 | 0.065314906 | 1 | 1 | 1 | 1 | 1 | 1 | 0.983 | 1 | 0.816 | 0.944 | 0.051 |
| A3SS | 13204 | ENSG00000050426 | LETMD1 | chr12 | + | 51053777 | 51053860 | 51053811 | 51053860 | 51052091 | 51052207 | 83 | 45 | 75 | 50 | 151 | 5 | 0 | 0 | 2 | 1 | 39 | 40 | 48 | 36 | 50 | 0 | 0 | 2 | 15 | 0 | 183 | 149 | 0.008344 | 0.065969505 | 0.931 | 1 | 1 | 0.953 | 0.992 | 1 | 1 | 0.951 | 0.661 | 1 | 0.053 |
| A3SS | 4300 | ENSG00000285774 | NA | chr14 | - | 54684752 | 54685001 | 54684752 | 54684985 | 54685736 | 54685875 | 7 | 7 | 1 | 10 | 16 | 0 | 1 | 0 | 1 | 0 | 24 | 13 | 2 | 2 | 26 | 10 | 0 | 0 | 0 | 0 | 165 | 149 | 0.008384 | 0.066221839 | 1 | 0.863 | 1 | 0.9 | 1 | 0.684 | 1 | 1 | 1 | 1 | 0.016 |
| A3SS | 9458 | ENSG00000242802 | AP5Z1 | chr7 | + | 4790388 | 4790591 | 4790458 | 4790591 | 4789831 | 4789929 | 7 | 1 | 2 | 3 | 0 | 1 | 0 | 10 | 3 | 1 | 0 | 4 | 1 | 0 | 0 | 6 | 4 | 5 | 14 | 2 | 219 | 149 | 0.008485 | 0.066953995 | 0.826 | 1 | 0.12 | 0.405 | 0 | 0 | 0.405 | 0.12 | 0 | 0 | 0.365 |
| A3SS | 9259 | ENSG00000136044 | APPL2 | chr12 | - | 105207970 | 105208047 | 105207970 | 105208029 | 105208157 | 105208199 | 5 | 1 | 3 | 5 | 11 | 49 | 24 | 45 | 57 | 40 | 3 | 0 | 6 | 1 | 0 | 37 | 33 | 35 | 48 | 33 | 167 | 149 | 0.008534 | 0.067269339 | 0.083 | 0.036 | 0.056 | 0.073 | 0.197 | 0.067 | 0 | 0.133 | 0.018 | 0 | 0.045 |
| A3SS | 10239 | ENSG00000196998 | WDR45 | chrX | - | 49076644 | 49076825 | 49076644 | 49076750 | 49077642 | 49077747 | 13 | 1 | 11 | 12 | 0 | 50 | 15 | 40 | 35 | 49 | 45 | 0 | 23 | 33 | 26 | 83 | 76 | 21 | 36 | 73 | 224 | 149 | 0.008545 | 0.067290722 | 0.147 | 0.042 | 0.155 | 0.186 | 0 | 0.265 | 0 | 0.421 | 0.379 | 0.192 | -0.145 |
| A3SS | 6778 | ENSG00000175376 | EIF1AD | chr11 | - | 66000302 | 66000476 | 66000302 | 66000406 | 66002039 | 66002166 | 46 | 16 | 52 | 51 | 25 | 15 | 6 | 10 | 11 | 5 | 73 | 72 | 54 | 101 | 117 | 48 | 29 | 18 | 40 | 41 | 219 | 149 | 0.008560 | 0.067348666 | 0.676 | 0.645 | 0.78 | 0.759 | 0.773 | 0.509 | 0.628 | 0.671 | 0.632 | 0.66 | 0.107 |
| A3SS | 13296 | ENSG00000136104 | RNASEH2B | chr13 | + | 50929460 | 50929582 | 50929515 | 50929582 | 50927406 | 50927478 | 148 | 102 | 111 | 129 | 268 | 0 | 1 | 3 | 10 | 3 | 58 | 96 | 87 | 67 | 116 | 5 | 1 | 3 | 10 | 4 | 204 | 149 | 0.008643 | 0.067936048 | 1 | 0.987 | 0.964 | 0.904 | 0.985 | 0.894 | 0.986 | 0.955 | 0.83 | 0.955 | 0.044 |
| A3SS | 8618 | ENSG00000095059 | DHPS | chr19 | - | 12679800 | 12680030 | 12679800 | 12679922 | 12680160 | 12680325 | 26 | 13 | 14 | 42 | 15 | 97 | 44 | 85 | 80 | 58 | 80 | 37 | 26 | 36 | 66 | 108 | 77 | 79 | 84 | 193 | 257 | 149 | 0.008802 | 0.069117978 | 0.134 | 0.146 | 0.087 | 0.233 | 0.13 | 0.3 | 0.218 | 0.16 | 0.199 | 0.165 | -0.062 |
| A3SS | 7400 | ENSG00000111801 | BTN3A3 | chr6 | + | 26443569 | 26443659 | 26443640 | 26443659 | 26443381 | 26443442 | 106 | 61 | 104 | 97 | 87 | 4 | 5 | 2 | 8 | 4 | 59 | 45 | 37 | 31 | 52 | 11 | 2 | 3 | 7 | 4 | 220 | 149 | 0.008824 | 0.069223063 | 0.947 | 0.892 | 0.972 | 0.891 | 0.936 | 0.784 | 0.938 | 0.893 | 0.75 | 0.898 | 0.075 |
| A3SS | 2040 | ENSG00000159593 | NAE1 | chr16 | - | 66805776 | 66805862 | 66805776 | 66805826 | 66805911 | 66806026 | 9 | 7 | 7 | 1 | 9 | 99 | 69 | 92 | 112 | 195 | 3 | 2 | 3 | 1 | 10 | 0 | 53 | 36 | 3 | 1 | 185 | 149 | 0.008911 | 0.06972305 | 0.068 | 0.076 | 0.058 | 0.007 | 0.036 | 1 | 0.029 | 0.063 | 0.212 | 0.89 | -0.39 |
| A3SS | 10642 | ENSG00000124275 | MTRR | chr5 | + | 7877943 | 7878322 | 7877948 | 7878322 | 7875257 | 7875375 | 137 | 108 | 170 | 168 | 156 | 8 | 7 | 6 | 15 | 8 | 72 | 171 | 164 | 75 | 38 | 7 | 9 | 20 | 15 | 4 | 154 | 149 | 0.008913 | 0.06972305 | 0.943 | 0.937 | 0.965 | 0.916 | 0.95 | 0.909 | 0.948 | 0.888 | 0.829 | 0.902 | 0.047 |
| A3SS | 11265 | ENSG00000165233 | CARD19 | chr9 | + | 93110567 | 93112289 | 93112217 | 93112289 | 93107673 | 93107816 | 29 | 19 | 12 | 23 | 4 | 0 | 0 | 1 | 0 | 0 | 41 | 35 | 25 | 36 | 35 | 0 | 0 | 0 | 0 | 11 | 298 | 149 | 0.008917 | 0.06972305 | 1 | 1 | 0.857 | 1 | 1 | 1 | 1 | 1 | 1 | 0.614 | 0.049 |
| A3SS | 9262 | ENSG00000135486 | HNRNPA1 | chr12 | + | 54281352 | 54281502 | 54281385 | 54281502 | 54280725 | 54280822 | 29 | 15 | 14 | 32 | 16 | 1299 | 857 | 1234 | 1175 | 1024 | 37 | 24 | 12 | 51 | 17 | 887 | 1034 | 1505 | 1337 | 406 | 182 | 149 | 0.008931 | 0.06972305 | 0.018 | 0.014 | 0.009 | 0.022 | 0.013 | 0.033 | 0.019 | 0.006 | 0.03 | 0.033 | -0.009 |
| A3SS | 5796 | ENSG00000108654 | DDX5 | chr17 | - | 64500548 | 64501891 | 64500548 | 64501143 | 64502009 | 64502069 | 1170 | 697 | 1166 | 1205 | 1180 | 10 | 12 | 4 | 14 | 21 | 185 | 812 | 760 | 425 | 116 | 4 | 7 | 4 | 0 | 0 | 298 | 149 | 0.008931 | 0.06972305 | 0.983 | 0.967 | 0.993 | 0.977 | 0.966 | 0.959 | 0.983 | 0.99 | 1 | 1 | -0.009 |
| A3SS | 6981 | ENSG00000163463 | KRTCAP2 | chr1 | - | 155169790 | 155172489 | 155169790 | 155169857 | 155172564 | 155172892 | 12 | 9 | 13 | 1 | 2 | 162 | 54 | 121 | 141 | 86 | 11 | 17 | 8 | 3 | 33 | 97 | 162 | 87 | 122 | 173 | 298 | 149 | 0.008953 | 0.069826812 | 0.036 | 0.077 | 0.051 | 0.004 | 0.011 | 0.054 | 0.05 | 0.044 | 0.012 | 0.087 | -0.014 |
| A3SS | 12474 | ENSG00000196531 | NACA | chr12 | - | 56724451 | 56724549 | 56724451 | 56724523 | 56725262 | 56725299 | 27 | 14 | 36 | 24 | 25 | 1083 | 558 | 1050 | 932 | 510 | 24 | 5 | 7 | 17 | 8 | 95 | 479 | 442 | 232 | 61 | 175 | 149 | 0.009004 | 0.070152136 | 0.021 | 0.021 | 0.028 | 0.021 | 0.04 | 0.177 | 0.009 | 0.013 | 0.059 | 0.1 | -0.045 |
| A3SS | 13126 | ENSG00000168010 | ATG16L2 | chr11 | + | 72821198 | 72821741 | 72821547 | 72821741 | 72817755 | 72817855 | 63 | 37 | 59 | 64 | 20 | 9 | 6 | 9 | 2 | 6 | 93 | 115 | 65 | 104 | 179 | 7 | 6 | 3 | 6 | 11 | 298 | 149 | 0.009017 | 0.070189711 | 0.778 | 0.755 | 0.766 | 0.941 | 0.625 | 0.869 | 0.906 | 0.915 | 0.897 | 0.891 | -0.123 |
| A3SS | 9496 | ENSG00000214021 | TTLL3 | chr3 | + | 9818554 | 9818920 | 9818821 | 9818920 | 9817644 | 9817759 | 63 | 38 | 62 | 55 | 39 | 2 | 2 | 12 | 15 | 6 | 310 | 68 | 90 | 145 | 287 | 63 | 3 | 0 | 1 | 11 | 298 | 149 | 0.009069 | 0.070458307 | 0.94 | 0.905 | 0.721 | 0.647 | 0.765 | 0.711 | 0.919 | 1 | 0.986 | 0.929 | -0.113 |
| A3SS | 6818 | ENSG00000112303 | VNN2 | chr6 | - | 132757670 | 132757902 | 132757670 | 132757759 | 132760631 | 132760803 | 508 | 184 | 395 | 317 | 654 | 12 | 12 | 10 | 18 | 17 | 197 | 784 | 767 | 464 | 131 | 0 | 19 | 7 | 22 | 0 | 292 | 149 | 0.009069 | 0.070458307 | 0.956 | 0.887 | 0.953 | 0.9 | 0.952 | 1 | 0.955 | 0.982 | 0.915 | 1 | -0.041 |
| A3SS | 434 | ENSG00000161265 | U2AF1L4 | chr19 | - | 35742530 | 35742803 | 35742530 | 35742619 | 35743808 | 35743904 | 44 | 21 | 81 | 53 | 74 | 3 | 3 | 2 | 1 | 2 | 86 | 44 | 38 | 43 | 42 | 0 | 2 | 2 | 0 | 0 | 298 | 149 | 0.009088 | 0.07053359 | 0.88 | 0.778 | 0.953 | 0.964 | 0.949 | 1 | 0.917 | 0.905 | 1 | 1 | -0.06 |
| A3SS | 7325 | ENSG00000159348 | CYB5R1 | chr1 | - | 202965886 | 202966109 | 202965886 | 202965993 | 202966527 | 202966600 | 8 | 0 | 3 | 7 | 7 | 75 | 14 | 31 | 39 | 47 | 10 | 34 | 11 | 4 | 18 | 51 | 60 | 35 | 30 | 202 | 265 | 149 | 0.009116 | 0.070682583 | 0.057 | 0 | 0.052 | 0.092 | 0.077 | 0.099 | 0.242 | 0.15 | 0.07 | 0.048 | -0.066 |

| | | | | | | | | | | | | | | | | | | | | | | | | | | | | | | | | | | | | | | | | | | | | | | |
|---|---|---|---|---|---|---|---|---|---|---|---|---|---|---|---|---|---|---|---|---|---|---|---|---|---|---|---|---|---|---|---|---|---|---|---|---|---|---|---|---|---|---|---|---|---|---|
| A3SS | 7646 | ENSG00000153029 | MR1 | chr1 | + | 181052234 | 181052510 | 181052321 | 181052510 | 181050010 | 181050286 | 41 | 23 | 39 | 27 | 46 | 3 | 1 | 3 | 3 | 4 | 43 | 27 | 19 | 40 | 39 | 0 | 0 | 0 | 0 | 6 | 236 | 149 | 0.009251 | 0.071663275 | 0.896 | 0.936 | 0.891 | 0.85 | 0.879 | 1 | 1 | 1 | 1 | 0.804 | -0.07 |
| A3SS | 6018 | ENSG00000014138 | POLA2 | chr11 | + | 65295863 | 65295990 | 65295933 | 65295990 | 65295539 | 65295599 | 43 | 27 | 48 | 37 | 25 | 6 | 1 | 8 | 0 | 0 | 18 | 41 | 7 | 50 | 25 | 3 | 5 | 8 | 8 | 4 | 219 | 149 | 0.009284 | 0.071853418 | 0.83 | 0.948 | 0.803 | 1 | 1 | 0.803 | 0.848 | 0.373 | 0.81 | 0.81 | 0.187 |
| A3SS | 8139 | ENSG00000102081 | FMR1 | chrX | + | 147944868 | 147945051 | 147944904 | 147945051 | 147943130 | 147943326 | 96 | 102 | 91 | 125 | 170 | 15 | 19 | 20 | 22 | 40 | 46 | 67 | 90 | 101 | 5 | 11 | 0 | 13 | 12 | 0 | 185 | 149 | 0.009342 | 0.072229112 | 0.838 | 0.812 | 0.786 | 0.821 | 0.774 | 0.771 | 1 | 0.848 | 0.871 | 1 | -0.092 |
| A3SS | 6011 | ENSG00000100029 | PES1 | chr22 | - | 30578836 | 30578998 | 30578836 | 30578995 | 30579136 | 30579303 | 68 | 38 | 90 | 72 | 118 | 7 | 4 | 4 | 2 | 0 | 174 | 77 | 63 | 52 | 136 | 10 | 10 | 3 | 13 | 6 | 152 | 149 | 0.009389 | 0.072523245 | 0.905 | 0.903 | 0.957 | 0.972 | 1 | 0.945 | 0.883 | 0.954 | 0.797 | 0.957 | 0.04 |
| A3SS | 6230 | ENSG00000263956 | NBPF11 | chr1 | - | 148109283 | 148109411 | 148109283 | 148109335 | 148110377 | 148110541 | 0 | 3 | 3 | 7 | 7 | 26 | 17 | 18 | 24 | 14 | 0 | 7 | 0 | 2 | 0 | 60 | 25 | 6 | 3 | 25 | 225 | 149 | 0.009458 | 0.072986681 | 0 | 0.105 | 0.099 | 0.162 | 0.249 | 0 | 0.156 | 0 | 0.306 | 0 | 0.031 |
| A3SS | 9957 | ENSG00000135164 | DMTF1 | chr7 | + | 87182055 | 87182337 | 87182227 | 87182337 | 87181308 | 87181341 | 232 | 222 | 235 | 270 | 241 | 127 | 125 | 120 | 115 | 119 | 36 | 75 | 207 | 50 | 24 | 27 | 78 | 109 | 31 | 22 | 298 | 149 | 0.009485 | 0.073129318 | 0.477 | 0.47 | 0.495 | 0.54 | 0.503 | 0.4 | 0.325 | 0.487 | 0.446 | 0.353 | 0.095 |
| A3SS | 5746 | ENSG00000082146 | STRADB | chr2 | + | 201480007 | 201480846 | 201480031 | 201480846 | 201479488 | 201479531 | 3 | 2 | 6 | 0 | 2 | 47 | 22 | 35 | 56 | 67 | 0 | 0 | 0 | 0 | 0 | 60 | 55 | 28 | 47 | 44 | 173 | 149 | 0.009547 | 0.073534813 | 0.052 | 0.073 | 0.129 | 0 | 0.025 | 0 | 0 | 0 | 0 | 0 | 0.056 |
| A3SS | 12604 | ENSG00000119953 | SMNDC1 | chr10 | - | 110297566 | 110297745 | 110297566 | 110297728 | 110298647 | 110298790 | 14 | 20 | 15 | 19 | 12 | 227 | 166 | 170 | 228 | 268 | 0 | 27 | 8 | 19 | 9 | 324 | 317 | 280 | 242 | 170 | 166 | 149 | 0.009582 | 0.073732241 | 0.052 | 0.098 | 0.073 | 0.07 | 0.039 | 0 | 0.071 | 0.025 | 0.066 | 0.045 | 0.025 |
| A3SS | 12226 | ENSG00000166333 | ILK | chr11 | + | 6608693 | 6608967 | 6608883 | 6608967 | 6608393 | 6608489 | 428 | 258 | 272 | 472 | 343 | 2 | 0 | 1 | 0 | 2 | 899 | 460 | 260 | 418 | 3064 | 10 | 5 | 2 | 26 | 0 | 298 | 149 | 0.009606 | 0.073809721 | 0.991 | 1 | 0.993 | 1 | 0.988 | 0.978 | 0.979 | 0.985 | 0.889 | 1 | 0.028 |
| A3SS | 4844 | ENSG00000138750 | NUP54 | chr4 | - | 76117663 | 76117774 | 76117663 | 76117742 | 76118074 | 76118194 | 147 | 72 | 104 | 88 | 148 | 0 | 2 | 3 | 4 | 12 | 15 | 57 | 60 | 6 | 7 | 0 | 0 | 4 | 0 | 0 | 181 | 149 | 0.009610 | 0.073809721 | 1 | 0.967 | 0.966 | 0.948 | 0.91 | 1 | 1 | 0.925 | 1 | 1 | -0.027 |
| A3SS | 4611 | ENSG00000161204 | ABCF3 | chr3 | + | 184187396 | 184187761 | 184187663 | 184187761 | 184186795 | 184186875 | 153 | 71 | 97 | 116 | 98 | 3 | 0 | 1 | 1 | 0 | 156 | 125 | 114 | 153 | 180 | 6 | 6 | 0 | 0 | 4 | 298 | 149 | 0.009627 | 0.073829383 | 0.962 | 1 | 0.98 | 0.983 | 1 | 0.929 | 0.912 | 1 | 1 | 0.957 | 0.025 |
| A3SS | 6515 | ENSG00000042429 | MED17 | chr11 | + | 93809716 | 93812378 | 93811852 | 93812378 | 93807517 | 93807635 | 97 | 75 | 70 | 55 | 91 | 0 | 0 | 4 | 7 | 9 | 73 | 44 | 68 | 63 | 10 | 0 | 0 | 7 | 0 | 0 | 298 | 149 | 0.009631 | 0.073829383 | 1 | 1 | 0.897 | 0.797 | 0.835 | 1 | 1 | 0.829 | 1 | 1 | -0.06 |
| A3SS | 11389 | ENSG00000168003 | SLC3A2 | chr11 | + | 62888112 | 62888218 | 62888134 | 62888218 | 62885464 | 62885608 | 1 | 2 | 1 | 1 | 0 | 41 | 15 | 39 | 30 | 7 | 0 | 5 | 1 | 0 | 9 | 24 | 52 | 19 | 48 | 25 | 171 | 149 | 0.009732 | 0.074530312 | 0.021 | 0.104 | 0.022 | 0.028 | 0 | 0 | 0.077 | 0.044 | 0 | 0.239 | -0.037 |
| A3SS | 13297 | ENSG00000136104 | RNASEH2B | chr13 | + | 50929474 | 50929582 | 50929515 | 50929582 | 50927406 | 50927478 | 292 | 200 | 228 | 282 | 512 | 0 | 1 | 3 | 10 | 3 | 143 | 220 | 154 | 140 | 199 | 5 | 1 | 3 | 10 | 4 | 190 | 149 | 0.009781 | 0.074765559 | 1 | 0.994 | 0.983 | 0.957 | 0.993 | 0.957 | 0.994 | 0.976 | 0.917 | 0.975 | 0.022 |
| A3SS | 6387 | ENSG00000112182 | BACH2 | chr6 | - | 90206568 | 90206681 | 90206568 | 90206676 | 90271848 | 90271941 | 12 | 17 | 35 | 30 | 30 | 0 | 0 | 0 | 1 | 0 | 8 | 14 | 2 | 9 | 6 | 0 | 1 | 0 | 0 | 16 | 154 | 149 | 0.009783 | 0.074765559 | 1 | 1 | 1 | 0.967 | 1 | 1 | 0.931 | 1 | 1 | 0.266 | 0.154 |
| A3SS | 1430 | ENSG00000126351 | THRA | chr17 | + | 40093019 | 40093867 | 40093136 | 40093867 | 40089205 | 40089333 | 40 | 21 | 35 | 41 | 27 | 6 | 0 | 0 | 0 | 0 | 2 | 34 | 14 | 23 | 49 | 0 | 0 | 6 | 0 | 6 | 266 | 149 | 0.009790 | 0.074765559 | 0.789 | 1 | 1 | 1 | 1 | 1 | 1 | 0.567 | 1 | 0.821 | 0.08 |
| A3SS | 7228 | ENSG00000082701 | GSK3B | chr3 | - | 119912703 | 119912800 | 119912703 | 119912750 | 119916043 | 119916174 | 102 | 82 | 103 | 134 | 98 | 0 | 0 | 0 | 0 | 1 | 115 | 68 | 118 | 66 | 55 | 0 | 0 | 5 | 9 | 0 | 199 | 149 | 0.009863 | 0.075184428 | 1 | 1 | 1 | 1 | 0.987 | 1 | 1 | 0.946 | 0.846 | 1 | 0.039 |
| A3SS | 9075 | ENSG00000128284 | APOL3 | chr22 | - | 36149045 | 36149185 | 36149045 | 36149142 | 36160668 | 36160929 | 30 | 25 | 21 | 24 | 22 | 114 | 71 | 133 | 100 | 59 | 35 | 31 | 18 | 31 | 40 | 81 | 88 | 32 | 100 | 99 | 192 | 149 | 0.009864 | 0.075184428 | 0.17 | 0.215 | 0.109 | 0.157 | 0.224 | 0.251 | 0.215 | 0.304 | 0.194 | 0.239 | -0.066 |
| A3SS | 2100 | ENSG00000140983 | RHOT2 | chr16 | + | 670248 | 670357 | 670265 | 670357 | 670122 | 670175 | 217 | 120 | 147 | 147 | 162 | 32 | 5 | 25 | 13 | 9 | 241 | 165 | 171 | 134 | 245 | 49 | 41 | 23 | 17 | 22 | 166 | 149 | 0.009894 | 0.075347211 | 0.859 | 0.956 | 0.841 | 0.91 | 0.942 | 0.815 | 0.783 | 0.87 | 0.876 | 0.909 | 0.051 |
| A3SS | 7059 | ENSG00000131165 | CHMP1A | chr16 | - | 89647202 | 89648478 | 89647202 | 89647331 | 89649350 | 89649497 | 11 | 0 | 2 | 2 | 0 | 76 | 47 | 57 | 42 | 85 | 12 | 8 | 0 | 0 | 19 | 89 | 66 | 45 | 64 | 152 | 298 | 149 | 0.009937 | 0.075586097 | 0.067 | 0 | 0.017 | 0.023 | 0 | 0.063 | 0.057 | 0 | 0 | 0.059 | -0.014 |
| A3SS | 12212 | ENSG00000142102 | PGGHG | chr11 | + | 294266 | 294478 | 294320 | 294478 | 294098 | 294196 | 156 | 64 | 153 | 98 | 33 | 2 | 0 | 4 | 2 | 0 | 32 | 66 | 82 | 58 | 80 | 0 | 0 | 3 | 3 | 9 | 203 | 149 | 0.009945 | 0.075586097 | 0.983 | 1 | 0.966 | 0.973 | 1 | 1 | 1 | 0.953 | 0.934 | 0.867 | 0.034 |
| A3SS | 10093 | ENSG00000061938 | TNK2 | chr3 | - | 195867168 | 195867264 | 195867168 | 195867258 | 195867360 | 195868709 | 31 | 6 | 17 | 21 | 2 | 0 | 1 | 3 | 2 | 4 | 12 | 12 | 2 | 35 | 13 | 0 | 2 | 2 | 0 | 0 | 155 | 149 | 0.009965 | 0.075667409 | 1 | 0.852 | 0.845 | 0.91 | 0.325 | 1 | 0.852 | 0.49 | 1 | 1 | -0.082 |
| A3SS | 5074 | ENSG00000170581 | STAT2 | chr12 | - | 56348751 | 56348851 | 56348751 | 56348804 | 56349344 | 56349509 | 12 | 11 | 11 | 11 | 16 | 1 | 0 | 6 | 0 | 7 | 89 | 28 | 21 | 65 | 43 | 41 | 10 | 10 | 19 | 29 | 196 | 149 | 0.010018 | 0.076003734 | 0.901 | 1 | 0.582 | 1 | 0.635 | 0.623 | 0.68 | 0.615 | 0.722 | 0.53 | 0.19 |
| A3SS | 5639 | ENSG00000183020 | AP2A2 | chr11 | + | 985431 | 985582 | 985434 | 985582 | 984644 | 984753 | 3 | 1 | 6 | 9 | 10 | 27 | 18 | 31 | 32 | 19 | 0 | 4 | 6 | 3 | 0 | 54 | 17 | 11 | 26 | 56 | 152 | 149 | 0.010127 | 0.076736052 | 0.098 | 0.052 | 0.159 | 0.216 | 0.34 | 0 | 0.187 | 0.348 | 0.102 | 0 | 0.046 |
| A3SS | 9596 | ENSG00000088038 | CNOT3 | chr19 | + | 54153496 | 54153840 | 54153714 | 54153840 | 54152866 | 54152999 | 3 | 9 | 8 | 5 | 5 | 73 | 22 | 60 | 53 | 80 | 2 | 0 | 6 | 7 | 5 | 77 | 61 | 56 | 122 | 94 | 298 | 149 | 0.010141 | 0.076736052 | 0.02 | 0.17 | 0.062 | 0.045 | 0.03 | 0.013 | 0 | 0.051 | 0.028 | 0.026 | 0.042 |
| A3SS | 7850 | ENSG00000115464 | USP34 | chr2 | - | 61214059 | 61214278 | 61214059 | 61214218 | 61220309 | 61220457 | 206 | 186 | 147 | 227 | 285 | 4 | 0 | 9 | 6 | 2 | 39 | 122 | 124 | 36 | 15 | 0 | 0 | 0 | 1 | 0 | 209 | 149 | 0.010153 | 0.076736052 | 0.973 | 1 | 0.921 | 0.964 | 0.99 | 1 | 1 | 1 | 0.962 | 1 | -0.023 |
| A3SS | 576 | ENSG00000090372 | STRN4 | chr19 | - | 46727451 | 46727691 | 46727451 | 46727546 | 46727893 | 46728007 | 11 | 11 | 9 | 3 | 6 | 70 | 34 | 57 | 59 | 77 | 34 | 15 | 5 | 10 | 22 | 78 | 79 | 31 | 74 | 113 | 294 | 149 | 0.010161 | 0.076736052 | 0.074 | 0.141 | 0.074 | 0.025 | 0.038 | 0.181 | 0.088 | 0.076 | 0.064 | 0.09 | -0.029 |
| A3SS | 4907 | ENSG00000148399 | DPH7 | chr9 | - | 137574751 | 137574846 | 137574751 | 137574843 | 137576079 | 137576167 | 3 | 0 | 0 | 3 | 2 | 7 | 13 | 14 | 15 | 15 | 8 | 6 | 4 | 1 | 0 | 0 | 10 | 4 | 10 | 11 | 152 | 149 | 0.010171 | 0.076736052 | 0.296 | 0 | 0 | 0.164 | 0.116 | 1 | 0.37 | 0.495 | 0.089 | 0 | -0.276 |
| A3SS | 2056 | ENSG00000129636 | ITFG1 | chr16 | - | 47162539 | 47162664 | 47162539 | 47162601 | 47218867 | 47218946 | 220 | 148 | 115 | 221 | 367 | 2 | 0 | 0 | 2 | 6 | 242 | 218 | 204 | 186 | 217 | 0 | 11 | 5 | 0 | 7 | 212 | 149 | 0.010172 | 0.076736052 | 0.987 | 1 | 1 | 0.987 | 0.977 | 1 | 0.933 | 0.966 | 1 | 0.956 | 0.019 |
| A3SS | 3112 | ENSG00000159753 | CARMIL2 | chr16 | + | 67656800 | 67656881 | 67656833 | 67656881 | 67656423 | 67656645 | 48 | 20 | 45 | 27 | 37 | 6 | 2 | 9 | 5 | 3 | 13 | 43 | 18 | 17 | 48 | 0 | 3 | 2 | 4 | 0 | 182 | 149 | 0.010201 | 0.076883302 | 0.868 | 0.891 | 0.804 | 0.816 | 0.91 | 1 | 0.921 | 0.88 | 0.777 | 1 | -0.058 |
| A3SS | 1379 | ENSG00000215041 | NEURL4 | chr17 | - | 7324270 | 7324107 | 7324266 | 7324394 | 7324480 | [illegible] | 19 | 26 | 15 | 16 | 3 | 2 | 0 | 2 | 0 | 0 | 9 | 25 | 4 | 10 | 28 | 0 | 2 | 0 | 0 | 8 | 153 | 149 | 0.010220 | 0.076957325 | 0.902 | 1 | 0.88 | 1 | 1 | 1 | 0.924 | 1 | 1 | 0.773 | 0.017 |
| A3SS | 12657 | ENSG00000107862 | GBF1 | chr10 | + | 102379852 | 102379954 | 102379876 | 102379954 | 102379520 | 102379651 | 45 | 10 | 22 | 21 | 26 | 3 | 0 | 0 | 0 | 0 | 32 | 39 | 12 | 46 | 93 | 0 | 5 | 0 | 7 | 0 | 173 | 149 | 0.010300 | 0.077485371 | 0.928 | 1 | 1 | 1 | 1 | 1 | 0.87 | 1 | 0.85 | 1 | 0.042 |
| A3SS | 1392 | ENSG00000129657 | SEC14L1 | chr17 | + | 77142645 | 77142750 | 77142649 | 77142750 | 77141069 | 77141107 | 163 | 105 | 122 | 185 | 68 | 0 | 1 | 3 | 5 | 1 | 407 | 249 | 326 | 250 | 940 | 11 | 1 | 5 | 5 | 35 | 153 | 149 | 0.010330 | 0.077641931 | 1 | 0.99 | 0.975 | 0.973 | 0.985 | 0.973 | 0.996 | 0.984 | 0.98 | 0.963 | 0.005 |
| A3SS | 4546 | ENSG00000076108 | BAZ2A | chr12 | - | 56610408 | 56610539 | 56610408 | 56610517 | 56611572 | 56611633 | 19 | 7 | 12 | 17 | 14 | 152 | 124 | 228 | 206 | 158 | 19 | 8 | 12 | 25 | 23 | 396 | 329 | 284 | 374 | 395 | 171 | 149 | 0.010482 | 0.078705869 | 0.098 | 0.047 | 0.044 | 0.067 | 0.072 | 0.04 | 0.021 | 0.036 | 0.055 | 0.048 | 0.026 |
| A3SS | 1241 | ENSG00000108788 | MLX | chr17 | + | 42568469 | 42568943 | 42568836 | 42568943 | 42567618 | 42567655 | 47 | 24 | 52 | 55 | 37 | 6 | 3 | 10 | 12 | 2 | 100 | 39 | 34 | 41 | 137 | 36 | 11 | 16 | 17 | 16 | 298 | 149 | 0.010525 | 0.078956768 | 0.797 | 0.8 | 0.722 | 0.696 | 0.902 | 0.581 | 0.639 | 0.515 | 0.547 | 0.811 | 0.165 |
| A3SS | 8488 | ENSG00000167674 | HDGFL2 | chr19 | + | 4496918 | 4498031 | 4497957 | 4498031 | 4496301 | 4496405 | 0 | 0 | 3 | 2 | 7 | 29 | 20 | 18 | 19 | 4 | 0 | 0 | 3 | 0 | 0 | 42 | 17 | 16 | 11 | 4 | 298 | 149 | 0.010589 | 0.079365016 | 0 | 0 | 0.077 | 0.05 | 0.467 | 0 | 0 | 0.086 | 0 | 0 | 0.102 |
| A3SS | 13376 | ENSG00000102710 | SUPT20H | chr13 | - | 37017244 | 37017364 | 37017244 | 37017328 | 37019341 | 37019397 | 74 | 51 | 72 | 89 | 107 | 11 | 16 | 11 | 8 | 22 | 25 | 38 | 48 | 36 | 17 | 0 | 10 | 5 | 1 | 0 | 185 | 149 | 0.010696 | 0.080088972 | 0.844 | 0.72 | 0.841 | 0.9 | 0.797 | 1 | 0.754 | 0.885 | 0.967 | 1 | -0.101 |
| A3SS | 12991 | ENSG00000076513 | ANKRD13A | chr12 | + | 110027704 | 110027766 | 110027707 | 110027766 | 110025741 | 110025823 | 568 | 439 | 498 | 533 | 861 | 96 | 69 | 90 | 103 | 209 | 1422 | 605 | 627 | 754 | 1126 | 193 | 86 | 103 | 105 | 201 | 152 | 149 | 0.010759 | 0.08048727 | 0.853 | 0.862 | 0.844 | 0.835 | 0.802 | 0.878 | 0.873 | 0.856 | 0.876 | 0.846 | -0.027 |
| A3SS | 9409 | ENSG00000131018 | SYNE1 | chr6 | - | 152283977 | 152284184 | 152283977 | 152284172 | 152293587 | 152293749 | 27 | 16 | 25 | 15 | 13 | 65 | 37 | 49 | 76 | 46 | 0 | 12 | 13 | 7 | 1 | 28 | 59 | 32 | 31 | 14 | 161 | 149 | 0.010907 | 0.081520205 | 0.278 | 0.286 | 0.321 | 0.154 | 0.207 | 0 | 0.158 | 0.273 | 0.173 | 0.062 | 0.116 |
| A3SS | 5938 | ENSG00000175309 | PHYKPL | chr5 | - | 178222354 | 178222545 | 178222354 | 178222492 | 178222851 | 178222934 | 36 | 23 | 22 | 34 | 13 | 3 | 7 | 2 | 3 | 5 | 87 | 38 | 10 | 42 | 74 | 0 | 2 | 0 | 2 | 11 | 202 | 149 | 0.010993 | 0.082087308 | 0.898 | 0.708 | 0.89 | 0.893 | 0.657 | 1 | 0.933 | 1 | 0.939 | 0.832 | -0.132 |
| A3SS | 12393 | ENSG00000107485 | GATA3 | chr10 | + | 8063992 | 8064138 | 8063995 | 8064138 | 8058304 | 8058841 | 40 | 11 | 32 | 39 | 26 | 20 | 2 | 14 | 8 | 0 | 0 | 17 | 31 | 5 | 1 | 6 | 7 | 7 | 5 | 4 | 152 | 149 | 0.011111 | 0.082894349 | 0.662 | 0.844 | 0.691 | 0.827 | 1 | 0 | 0.704 | 0.813 | 0.495 | 0.197 | 0.363 |
| A3SS | 12244 | ENSG00000166471 | TMEM41B | chr11 | - | 9288441 | 9288542 | 9288441 | 9288535 | 9295258 | 9295387 | 7 | 6 | 11 | 17 | 4 | 59 | 50 | 72 | 69 | 121 | 0 | 12 | 10 | 16 | 4 | 11 | 32 | 32 | 15 | 28 | 156 | 149 | 0.011170 | 0.083249438 | 0.102 | 0.103 | 0.127 | 0.19 | 0.031 | 0 | 0.264 | 0.23 | 0.505 | 0.12 | -0.113 |
| A3SS | 3604 | ENSG00000124279 | FASTKD3 | chr5 | - | 7861582 | 7861773 | 7861582 | 7861652 | 7862822 | 7862997 | 16 | 10 | 28 | 13 | 19 | 36 | 31 | 35 | 55 | 44 | 0 | 0 | 26 | 6 | 0 | 21 | 37 | 34 | 9 | 9 | 270 | 149 | 0.011180 | 0.083249438 | 0.197 | 0.151 | 0.306 | 0.115 | 0.192 | 0 | 0 | 0.297 | 0.269 | 0 | 0.079 |
| A3SS | 2760 | ENSG00000119718 | EIF2B2 | chr14 | + | 75005840 | 75005961 | 75005896 | 75005961 | 75004736 | 75004900 | 19 | 14 | 11 | 23 | 52 | 1 | 3 | 3 | 1 | 1 | 3 | 8 | 19 | 12 | 30 | 0 | 0 | 1 | 1 | 0 | 205 | 149 | 0.011225 | 0.083511861 | 0.932 | 0.772 | 0.727 | 0.944 | 0.974 | 1 | 1 | 0.932 | 0.897 | 1 | -0.096 |
| A3SS | 2898 | ENSG00000143379 | SETDB1 | chr1 | + | 150949366 | 150949525 | 150949369 | 150949525 | 150949121 | 150949278 | 44 | 24 | 24 | 28 | 31 | 9 | 5 | 10 | 7 | 18 | 7 | 62 | 16 | 14 | 89 | 7 | 2 | 0 | 10 | 0 | 152 | 149 | 0.011436 | 0.084999231 | 0.827 | 0.825 | 0.702 | 0.797 | 0.628 | 0.495 | 0.968 | 1 | 0.578 | 1 | -0.052 |
| A3SS | 981 | ENSG00000124155 | PIGT | chr20 | + | 45418771 | 45418979 | 45418851 | 45418979 | 45416516 | 45416694 | 15 | 9 | 20 | 24 | 24 | 109 | 37 | 61 | 70 | 86 | 7 | 33 | 10 | 27 | 35 | 124 | 131 | 88 | 103 | 249 | 229 | 149 | 0.011455 | 0.085066612 | 0.082 | 0.137 | 0.176 | 0.182 | 0.154 | 0.035 | 0.141 | 0.069 | 0.146 | 0.084 | 0.051 |
| A3SS | 1925 | ENSG00000168872 | DDX19A | chr16 | + | 70365932 | 70366262 | 70366084 | 70366262 | 70365016 | 70365131 | 23 | 18 | 12 | 23 | 12 | 44 | 27 | 45 | 39 | 69 | 37 | 30 | 21 | 17 | 50 | 60 | 31 | 35 | 48 | 57 | 298 | 149 | 0.011493 | 0.085269095 | 0.207 | 0.25 | 0.118 | 0.228 | 0.08 | 0.236 | 0.326 | 0.231 | 0.15 | 0.305 | -0.073 |
| A3SS | 4301 | ENSG00000183323 | CCDC125 | chr5 | - | 69313984 | 69314046 | 69313984 | 69314043 | 69320236 | 69320580 | 11 | 8 | 10 | 7 | 14 | 59 | 50 | 37 | 37 | 46 | 12 | 3 | 1 | 23 | 0 | 67 | 41 | 70 | 117 | 17 | 152 | 149 | 0.011555 | 0.085652368 | 0.155 | 0.136 | 0.209 | 0.156 | 0.23 | 0.149 | 0.067 | 0.014 | 0.162 | 0 | 0.099 |
| A3SS | 6462 | ENSG00000143621 | ILF2 | chr1 | - | 153668452 | 153668576 | 153668452 | 153668557 | 153669835 | 153669878 | 8 | 3 | 5 | 9 | 10 | 355 | 227 | 322 | 346 | 261 | 13 | 3 | 2 | 3 | 0 | 560 | 372 | 343 | 508 | 545 | 168 | 149 | 0.011610 | 0.085939234 | 0.02 | 0.012 | 0.014 | 0.023 | 0.033 | 0.02 | 0.007 | 0.005 | 0.005 | 0 | 0.013 |
| A3SS | 264 | ENSG00000079432 | CIC | chr19 | + | 42292565 | 42292859 | 42292568 | 42292859 | 42292299 | 42292466 | 46 | 33 | 23 | 39 | 12 | 6 | 4 | 3 | 10 | 1 | 51 | 27 | 28 | 32 | 113 | 10 | 2 | 4 | 0 | 0 | 152 | 149 | 0.011615 | 0.085939234 | 0.883 | 0.89 | 0.883 | 0.793 | 0.922 | 0.833 | 0.93 | 0.873 | 1 | 1 | -0.053 |
| A3SS | 5238 | ENSG00000183258 | DDX41 | chr5 | - | 177515185 | 177515361 | 177515185 | 177515258 | 177515684 | 177515821 | 2 | 4 | 16 | 13 | 2 | 52 | 19 | 46 | 60 | 60 | 14 | 10 | 24 | 10 | 20 | 71 | 38 | 22 | 41 | 111 | 252 | 149 | 0.011652 | 0.086065027 | 0.022 | 0.111 | 0.171 | 0.114 | 0.019 | 0.104 | 0.135 | 0.392 | 0.126 | 0.096 | -0.083 |
| A3SS | 12821 | ENSG00000109685 | NSD2 | chr4 | + | 1952751 | 1953524 | 1953323 | 1953524 | 1952104 | 1952231 | 6 | 2 | 5 | 5 | 10 | 24 | 10 | 18 | 16 | 9 | 30 | 8 | 10 | 12 | 13 | 10 | 8 | 26 | 24 | 16 | 298 | 149 | 0.011654 | 0.086065027 | 0.111 | 0.091 | 0.122 | 0.135 | 0.357 | 0.6 | 0.333 | 0.161 | 0.2 | 0.289 | -0.153 |
| A3SS | 11174 | ENSG00000133794 | ARNTL | chr11 | + | 13372162 | 13372409 | 13372165 | 13372409 | 13369617 | 13369767 | 70 | 34 | 79 | 59 | 68 | 44 | 22 | 24 | 55 | 26 | 40 | 63 | 83 | 36 | 13 | 0 | 27 | 47 | 14 | 0 | 152 | 149 | 0.011709 | 0.086390767 | 0.609 | 0.602 | 0.763 | 0.513 | 0.719 | 1 | 0.696 | 0.634 | 0.716 | 1 | -0.168 |
| A3SS | 6272 | ENSG00000154222 | CC2D1B | chr1 | - | 52359704 | 52359883 | 52359704 | 52359836 | 52360073 | 52360233 | 81 | 63 | 104 | 58 | 92 | 3 | 2 | 9 | 10 | 3 | 72 | 63 | 47 | 39 | 45 | 0 | 8 | 4 | 10 | 25 | 196 | 149 | 0.011786 | 0.086883715 | 0.954 | 0.96 | 0.898 | 0.815 | 0.959 | 1 | 0.857 | 0.899 | 0.748 | 0.578 | 0.101 |
| A3SS | 573 | ENSG00000105486 | LIG1 | chr19 | - | 48151231 | 48151339 | 48151231 | 48151336 | 48153871 | 48153967 | 23 | 13 | 21 | 28 | 24 | 16 | 6 | 11 | 10 | 11 | 63 | 45 | 31 | 39 | 83 | 5 | 14 | 11 | 9 | 31 | 152 | 149 | 0.011835 | 0.087165748 | 0.585 | 0.68 | 0.652 | 0.733 | 0.681 | 0.925 | 0.759 | 0.734 | 0.809 | 0.724 | -0.124 |
| A3SS | 8079 | ENSG00000123240 | OPTN | chr10 | + | 13110179 | 13110476 | 13110273 | 13110476 | 13109111 | 13109288 | 36 | 13 | 32 | 32 | 28 | 83 | 54 | 81 | 65 | 48 | 70 | 63 | 48 | 50 | 49 | 91 | 97 | 105 | 98 | 26 | 243 | 149 | 0.011883 | 0.087437742 | 0.21 | 0.129 | 0.195 | 0.232 | 0.263 | 0.32 | 0.285 | 0.219 | 0.238 | 0.536 | -0.114 |
| A3SS | 12622 | ENSG00000107959 | PITRM1 | chr10 | - | 3143388 | 3143775 | 3143388 | 3143501 | 3144291 | 3144366 | 18 | 9 | 32 | 23 | 16 | 60 | 34 | 60 | 52 | 119 | 40 | 21 | 46 | 39 | 24 | 59 | 60 | 60 | 74 | 97 | 298 | 149 | 0.011939 | 0.087772835 | 0.13 | 0.117 | 0.211 | 0.181 | 0.063 | 0.253 | 0.149 | 0.277 | 0.209 | 0.11 | -0.059 |
| A3SS | 11828 | ENSG00000073921 | PICALM | chr11 | - | 85981871 | 85982003 | 85981871 | 85981988 | 85983865 | 85983973 | 543 | 340 | 280 | 418 | 740 | 414 | 185 | 244 | 323 | 534 | 181 | 403 | 412 | 340 | 102 | 159 | 369 | 355 | 287 | 119 | 164 | 149 | 0.011962 | 0.087859683 | 0.544 | 0.625 | 0.51 | 0.54 | 0.557 | 0.508 | 0.498 | 0.513 | 0.518 | 0.438 | 0.06 |
| A3SS | 1030 | ENSG00000175354 | PTPN2 | chr18 | - | 12817155 | 12817365 | 12817155 | 12817265 | 12821335 | 12821425 | 244 | 170 | 173 | 206 | 539 | 2 | 1 | 0 | 0 | 1 | 54 | 153 | 175 | 133 | 21 | 0 | 0 | 7 | 7 | 0 | 249 | 149 | 0.011978 | 0.087888492 | 0.986 | 0.99 | 1 | 1 | 0.997 | 1 | 1 | 0.937 | 0.919 | 1 | 0.023 |
| A3SS | 3711 | ENSG00000177058 | SLC38A9 | chr5 | - | 55672562 | 55672822 | 55672562 | 55672695 | 55697845 | 55697994 | 11 | 9 | 5 | 14 | 3 | 102 | 71 | 73 | 90 | 128 | 0 | 8 | 9 | 16 | 1 | 31 | 72 | 33 | 30 | 4 | 276 | 149 | 0.011992 | 0.087888492 | 0.055 | 0.064 | 0.036 | 0.077 | 0.012 | 0 | 0.057 | 0.128 | 0.224 | 0.119 | -0.057 |
| A3SS | 3429 | ENSG00000160783 | PMF1 | chr1 | + | 156236287 | 156236483 | 156236348 | 156236483 | 156233627 | 156233728 | 204 | 189 | 184 | 201 | 241 | 13 | 0 | 0 | 0 | 0 | 762 | 398 | 284 | 510 | 985 | 0 | 15 | 18 | 0 | 28 | 210 | 149 | 0.011999 | 0.087888492 | 0.918 | 1 | 1 | 1 | 1 | 1 | 0.95 | 0.918 | 1 | 0.961 | 0.018 |
| A3SS | 8764 | ENSG00000079785 | DDX1 | chr2 | + | 15595457 | 15595553 | 15595489 | 15595553 | 15595144 | 15595196 | 6 | 5 | 10 | 4 | 11 | 108 | 66 | 130 | 103 | 86 | 0 | 1 | 4 | 5 | 0 | 41 | 51 | 71 | 64 | 11 | 181 | 149 | 0.012051 | 0.088191034 | 0.044 | 0.059 | 0.06 | 0.031 | 0.095 | 0 | 0.016 | 0.044 | 0.06 | 0 | 0.034 |
| A3SS | 5758 | ENSG00000144283 | PKP4 | chr2 | + | 158624877 | 158625427 | 158624954 | 158625427 | 158621230 | 158621421 | 51 | 60 | 65 | 69 | 177 | 1 | 0 | 0 | 3 | 0 | 19 | 67 | 105 | 178 | 171 | 0 | 13 | 0 | 0 | 2 | 226 | 149 | 0.012109 | 0.088534295 | 0.971 | 1 | 1 | 0.938 | 1 | 1 | 0.773 | 1 | 1 | 0.983 | 0.031 |
| A3SS | 1491 | ENSG00000082641 | NFE2L1 | chr17 | + | 48056385 | 48056598 | 48056418 | 48056598 | 48050606 | 48051628 | 133 | 96 | 112 | 168 | 107 | 18 | 12 | 11 | 14 | 32 | 323 | 214 | 119 | 160 | 274 | 12 | 11 | 13 | 18 | 40 | 182 | 149 | 0.012160 | 0.088830248 | 0.858 | 0.868 | 0.893 | 0.908 | 0.732 | 0.957 | 0.941 | 0.882 | 0.879 | 0.849 | -0.05 |
| A3SS | 11109 | ENSG00000104164 | BLOC1S6 | chr15 | + | 45603099 | 45603187 | 45603103 | 45603187 | 45592134 | 45592276 | 141 | 128 | 96 | 113 | 160 | 14 | 15 | 8 | 24 | 12 | 62 | 126 | 142 | 70 | 68 | 3 | 20 | 34 | 19 | 14 | 153 | 149 | 0.012344 | 0.0900918 | 0.907 | 0.893 | 0.921 | 0.821 | 0.928 | 0.953 | 0.86 | 0.803 | 0.782 | 0.825 | 0.049 |
| A3SS | 5311 | ENSG00000172113 | NME6 | chr3 | - | 48296118 | 48296236 | 48296118 | 48296158 | 48296726 | 48296829 | 12 | 1 | 8 | 10 | 1 | 17 | 11 | 18 | 26 | 28 | 5 | 2 | 3 | 0 | 3 | 8 | 43 | 15 | 19 | 18 | 227 | 149 | 0.012402 | 0.090436055 | 0.317 | 0.056 | 0.226 | 0.202 | 0.023 | 0.291 | 0.03 | 0.116 | 0 | 0.099 | 0.058 |
| A3SS | 6423 | ENSG00000165458 | INPPL1 | chr11 | + | 72228347 | 72228498 | 72228453 | 72228498 | 72227880 | 72228253 | 37 | 22 | 48 | 62 | 69 | 0 | 0 | 0 | 0 | 0 | 116 | 65 | 55 | 79 | 124 | 0 | 0 | 0 | 8 | 13 | 255 | 149 | 0.012416 | 0.090454775 | 1 | 1 | 1 | 1 | 1 | 1 | 1 | 1 | 1 | 0.848 | 0.03 |
| A3SS | 6425 | ENSG00000197785 | ATAD3A | chr1 | + | 1517166 | 1517412 | 1517310 | 1517412 | 1516011 | 1516088 | 1 | 0 | 3 | 0 | 3 | 17 | 17 | 3 | 10 | 2 | 12 | 0 | 0 | 5 | 21 | 11 | 24 | 25 | 25 | 25 | 293 | 149 | 0.012490 | 0.090912139 | 0.029 | 0 | 0.337 | 0 | 0.433 | 0.357 | 0 | 0 | 0.241 | 0.299 | -0.02 |
| [illegible] | [illegible] | [illegible] | [illegible] | [illegible] | [illegible] | [illegible] | [illegible] | [illegible] | [illegible] | [illegible] | [illegible] | [illegible] | [illegible] | [illegible] | [illegible] | [illegible] | [illegible] | [illegible] | [illegible] | [illegible] | [illegible] | [illegible] | [illegible] | [illegible] | [illegible] | [illegible] | [illegible] | [illegible] | [illegible] | [illegible] | [illegible] | [illegible] | [illegible] | [illegible] | [illegible] | [illegible] | [illegible] | [illegible] | [illegible] | [illegible] | [illegible] | [illegible] | [illegible] | [illegible] | [illegible] | [illegible] |

| | | | | | | |
|---|---|---|---|---|---|---|
| A3SS | 7739 | ENSG00000115525 | ST3GAL5 | chr2 | - | [illegible] |
| A3SS | 12202 | ENSG00000115368 | WDR75 | chr2 | + | [illegible] |
| A3SS | 9867 | ENSG00000121716 | PILRB | chr7 | + | [illegible] |
| A3SS | 9454 | ENSG00000242802 | AP5Z1 | chr7 | + | [illegible] |
| A3SS | 809 | ENSG00000185513 | L3MBTL1 | chr20 | + | [illegible] |
| A3SS | 9887 | ENSG00000126883 | NUP214 | chr9 | + | [illegible] |
| A3SS | 10238 | ENSG00000196998 | WDR45 | chrX | - | [illegible] |
| A3SS | 10976 | ENSG00000167925 | GHDC | chr17 | - | [illegible] |
| A3SS | 11610 | ENSG00000172716 | SLFN11 | chr17 | - | [illegible] |
| A3SS | 6137 | ENSG00000198821 | CD247 | chr1 | - | [illegible] |
| A3SS | 4294 | ENSG00000114127 | XRN1 | chr3 | - | [illegible] |
| A3SS | 4945 | ENSG00000197948 | FCHSD1 | chr5 | - | [illegible] |
| A3SS | 13221 | ENSG00000111335 | OAS2 | chr12 | + | [illegible] |
| A3SS | 13128 | ENSG00000168010 | ATG16L2 | chr11 | + | [illegible] |
| A3SS | 9059 | ENSG00000116539 | ASH1L | chr1 | - | [illegible] |
| A3SS | 5505 | ENSG00000110172 | CHORDC1 | chr11 | - | [illegible] |
| A3SS | 9673 | ENSG00000160219 | GAB3 | chrX | - | [illegible] |
| A3SS | 9394 | ENSG00000135317 | SNX14 | chr6 | - | [illegible] |
| A3SS | 9732 | ENSG00000134905 | CARS2 | chr13 | - | [illegible] |
| A3SS | 399 | ENSG00000142546 | NOSIP | chr19 | - | [illegible] |
| A3SS | 13026 | ENSG00000166012 | TAF1D | chr11 | - | [illegible] |
| A3SS | 2923 | ENSG00000071994 | PDCD2 | chr6 | - | [illegible] |
| A3SS | 4117 | ENSG00000161013 | MGAT4B | chr5 | - | [illegible] |
| A3SS | 8409 | ENSG00000140368 | PSTPIP1 | chr15 | + | [illegible] |
| A3SS | 18 | ENSG00000185917 | SETD4 | chr21 | - | [illegible] |
| A3SS | 9292 | ENSG00000204351 | SKIV2L | chr6 | + | [illegible] |
| A3SS | 1198 | ENSG00000141252 | VPS53 | chr17 | - | [illegible] |
| A3SS | 3242 | ENSG00000254996 | ANKHD1-EIF4EBP3 | chr5 | + | [illegible] |
| A3SS | 5132 | ENSG00000123473 | STIL | chr1 | - | [illegible] |
| A3SS | 13351 | ENSG00000137822 | TUBGCP4 | chr15 | + | [illegible] |
| A3SS | 5997 | ENSG00000110619 | CARS1 | chr11 | - | [illegible] |
| A3SS | 12048 | ENSG00000149091 | DGKZ | chr11 | + | [illegible] |
| A3SS | 12178 | ENSG00000110075 | PPP6R3 | chr11 | + | [illegible] |
| A3SS | 3614 | ENSG00000125089 | SH3TC1 | chr4 | + | [illegible] |
| A3SS | 738 | ENSG00000196562 | SULF2 | chr20 | - | [illegible] |
| A3SS | 435 | ENSG00000161265 | U2AF1L4 | chr19 | - | [illegible] |
| A3SS | 6037 | ENSG00000170476 | MZB1 | chr5 | - | [illegible] |
| A3SS | 3293 | ENSG00000161960 | EIF4A1 | chr17 | + | [illegible] |
| A3SS | 2200 | ENSG00000182511 | FES | chr15 | + | [illegible] |
| A3SS | 10605 | ENSG00000071889 | FAM3A | chrX | - | [illegible] |
| A3SS | 9807 | ENSG00000012124 | CD22 | chr19 | + | [illegible] |
| A3SS | 2975 | ENSG00000119900 | OGFRL1 | chr6 | + | [illegible] |
| A3SS | 10563 | ENSG00000182858 | ALG12 | chr22 | - | [illegible] |
| A3SS | 3796 | ENSG00000070831 | CDC42 | chr1 | + | [illegible] |
| A3SS | 7572 | ENSG00000131408 | NR1H2 | chr19 | + | [illegible] |
| A3SS | 3906 | ENSG00000127419 | TMEM175 | chr4 | + | [illegible] |
| A3SS | 6554 | ENSG00000155363 | MOV10 | chr1 | + | [illegible] |
| A3SS | 3589 | ENSG00000214655 | ZSWIM8 | chr10 | + | [illegible] |
| A3SS | 13242 | ENSG00000155957 | TMBIM4 | chr12 | - | [illegible] |
| A3SS | 7439 | ENSG00000188976 | NOC2L | chr1 | - | [illegible] |
| A3SS | 3430 | ENSG00000138640 | FAM13A | chr4 | - | [illegible] |
| A3SS | 963 | ENSG00000101751 | POLI | chr18 | + | [illegible] |
| A3SS | 7537 | ENSG00000138398 | PPIG | chr2 | + | [illegible] |
| A3SS | 8032 | ENSG00000147526 | TACC1 | chr8 | + | [illegible] |
| A3SS | 4045 | ENSG00000187621 | TCL6 | chr14 | + | [illegible] |
| A3SS | 563 | ENSG00000167613 | LAIR1 | chr19 | - | [illegible] |
| A3SS | 13011 | ENSG00000135297 | MTO1 | chr6 | + | [illegible] |
| A3SS | 11799 | ENSG00000100731 | PCNX1 | chr14 | + | [illegible] |
| A3SS | 7550 | ENSG00000078808 | SDF4 | chr1 | - | [illegible] |
| A3SS | 3311 | ENSG00000124357 | NAGK | chr2 | + | [illegible] |
| A3SS | 8091 | ENSG00000116809 | ZBTB17 | chr1 | - | [illegible] |
| A3SS | 12826 | ENSG00000123200 | ZC3H13 | chr13 | - | [illegible] |
| A3SS | 3439 | ENSG00000134644 | PUM1 | chr1 | - | [illegible] |
| A3SS | 12051 | ENSG00000135365 | PHF21A | chr11 | - | [illegible] |
| A3SS | 8836 | ENSG00000034152 | MAP2K3 | chr17 | + | [illegible] |
| A3SS | 10269 | ENSG00000111581 | NUP107 | chr12 | + | [illegible] |
| A3SS | 10529 | ENSG00000005302 | MSL3 | chrX | + | [illegible] |
| A3SS | 11860 | ENSG00000118363 | SPCS2 | chr11 | + | [illegible] |
| A3SS | 4810 | ENSG00000184014 | DENND5A | chr11 | - | [illegible] |
| A3SS | 5207 | ENSG00000137776 | SLTM | chr15 | - | [illegible] |
| A3SS | 3415 | ENSG00000113272 | THG1L | chr5 | + | [illegible] |
| A3SS | 11180 | ENSG00000156738 | MS4A1 | chr11 | + | [illegible] |
| A3SS | 4419 | ENSG00000109065 | NAT9 | chr17 | - | [illegible] |
| A3SS | 10496 | ENSG00000072778 | ACADVL | chr17 | + | [illegible] |
| A3SS | 3003 | ENSG00000093134 | VNN3 | chr6 | - | [illegible] |
| A3SS | 11162 | ENSG00000134539 | KLRD1 | chr12 | + | [illegible] |
| A3SS | 2529 | ENSG00000258289 | CHURC1 | chr14 | + | [illegible] |
| A3SS | 3571 | ENSG00000105875 | WDR91 | chr7 | - | [illegible] |
| A3SS | 2387 | ENSG00000140400 | MAN2C1 | chr15 | - | [illegible] |
| A3SS | 2951 | ENSG00000196182 | STK40 | chr1 | - | [illegible] |
| A3SS | 3119 | ENSG00000131844 | MCCC2 | chr5 | + | [illegible] |
| A3SS | 381 | ENSG00000076944 | STXBP2 | chr19 | + | [illegible] |
| A3SS | 7063 | ENSG00000131165 | CHMP1A | chr16 | - | [illegible] |
| A3SS | 10023 | ENSG00000051825 | MPHOSPH9 | chr12 | - | [illegible] |
| A3SS | 13007 | ENSG00000135297 | MTO1 | chr6 | + | [illegible] |
| A3SS | 1798 | ENSG00000005844 | ITGAL | chr16 | + | [illegible] |
| A3SS | 9188 | ENSG00000197081 | IGF2R | chr6 | + | [illegible] |
| A3SS | 10978 | ENSG00000147576 | ADHFE1 | chr8 | + | [illegible] |
| A3SS | 13051 | ENSG00000005379 | TSPOAP1 | chr17 | - | [illegible] |
| A3SS | 11034 | ENSG00000110057 | UNC93B1 | chr11 | - | [illegible] |
| A3SS | 13130 | ENSG00000168010 | ATG16L2 | chr11 | + | [illegible] |
| A3SS | 5930 | ENSG00000141027 | NCOR1 | chr17 | - | [illegible] |
| A3SS | 6720 | ENSG00000143851 | PTPN7 | chr1 | - | [illegible] |
| A3SS | 8148 | ENSG00000131051 | RBM39 | chr20 | - | [illegible] |
| A3SS | 227 | ENSG00000099331 | MYO9B | chr19 | + | [illegible] |
| A3SS | 10699 | ENSG00000046651 | OFD1 | chrX | + | [illegible] |
| A3SS | 12477 | ENSG00000100109 | TFIP11 | chr22 | - | [illegible] |
| A3SS | 1914 | ENSG00000090861 | AARS1 | chr16 | - | [illegible] |
| A3SS | 1631 | ENSG00000185722 | ANKFY1 | chr17 | - | [illegible] |
| A3SS | 13210 | ENSG00000166598 | HSP90B1 | chr12 | + | [illegible] |
| A3SS | 7232 | ENSG00000117523 | PRRC2C | chr1 | + | [illegible] |
| A3SS | 9298 | ENSG00000124508 | BTN2A2 | chr6 | + | [illegible] |
| A3SS | 3654 | ENSG00000073331 | ALPK1 | chr4 | + | [illegible] |
| A3SS | 5084 | ENSG00000008952 | SEC62 | chr3 | + | [illegible] |
| A3SS | 2624 | ENSG00000171469 | ZNF561 | chr19 | - | [illegible] |
| A3SS | 9450 | ENSG00000168944 | CEP120 | chr5 | - | [illegible] |
| A3SS | 1935 | ENSG00000089280 | FUS | chr16 | + | [illegible] |
| A3SS | 11591 | ENSG00000213930 | GALT | chr9 | + | [illegible] |
| A3SS | 9413 | ENSG00000173511 | VEGFB | chr11 | + | [illegible] |
| A3SS | 7791 | ENSG00000177084 | POLE | chr12 | - | [illegible] |
| A3SS | 10712 | ENSG00000172732 | MUS81 | chr11 | + | [illegible] |
| A3SS | 7823 | ENSG00000162882 | HAAO | chr2 | - | [illegible] |
| A3SS | 5766 | ENSG00000163608 | NEPRO | chr3 | - | [illegible] |
| A3SS | 1829 | ENSG00000135723 | FHOD1 | chr16 | - | [illegible] |
| A3SS | 1456 | ENSG00000087157 | PGS1 | chr17 | + | [illegible] |
| A3SS | 4420 | ENSG00000109065 | NAT9 | chr17 | - | [illegible] |
| A3SS | 10549 | ENSG00000197713 | RPE | chr2 | + | [illegible] |
| A3SS | 11161 | ENSG00000134539 | KLRD1 | chr12 | + | [illegible] |
| A3SS | 13211 | ENSG00000166598 | HSP90B1 | chr12 | + | [illegible] |
| A3SS | 3150 | ENSG00000140853 | NLRC5 | chr16 | + | [illegible] |
| A3SS | 9076 | ENSG00000128284 | APOL3 | chr22 | - | [illegible] |
| A3SS | 408 | ENSG00000104872 | PIH1D1 | chr19 | - | [illegible] |
| A3SS | 8467 | ENSG00000115694 | STK25 | chr2 | - | [illegible] |
| A3SS | 2090 | ENSG00000102858 | MGRN1 | chr16 | + | [illegible] |
| A3SS | 12580 | ENSG00000023191 | RNH1 | chr11 | - | [illegible] |
| A3SS | 4205 | ENSG00000114745 | GORASP1 | chr3 | - | [illegible] |
| A3SS | 2419 | ENSG00000140395 | WDR61 | chr15 | - | [illegible] |
| A3SS | 10258 | ENSG00000130985 | UBA1 | chrX | + | [illegible] |
| A3SS | 6836 | ENSG00000151694 | ADAM17 | chr2 | - | [illegible] |
| A3SS | 2353 | ENSG00000167491 | GATAD2A | chr19 | + | [illegible] |
| A3SS | 1505 | ENSG00000108433 | GOSR2 | chr17 | + | [illegible] |
| A3SS | 5227 | ENSG00000118308 | IRAG2 | chr12 | + | [illegible] |
| A3SS | 9601 | ENSG00000168014 | C2CD3 | chr11 | - | [illegible] |
| A3SS | 4926 | ENSG00000160298 | C21orf58 | chr21 | - | [illegible] |
| A3SS | 3261 | ENSG00000280987 | MATR3 | chr5 | + | [illegible] |
| A3SS | 12605 | ENSG00000119953 | SMNDC1 | chr10 | - | [illegible] |
| A3SS | 951 | ENSG00000125827 | TMX4 | chr20 | - | [illegible] |
| A3SS | 7168 | ENSG00000154845 | PPP4R1 | chr18 | - | [illegible] |
| A3SS | 9464 | ENSG00000167380 | ZNF226 | chr19 | + | [illegible] |
| A3SS | 6891 | ENSG00000072110 | ACTN1 | chr14 | - | [illegible] |
| A3SS | 4421 | ENSG00000109065 | NAT9 | chr17 | - | [illegible] |
| A3SS | 6191 | ENSG00000107551 | RASSF4 | chr10 | + | [illegible] |
| A3SS | 10009 | ENSG00000089006 | SNX5 | chr20 | - | [illegible] |
| A3SS | 7174 | ENSG00000160917 | CPSF4 | chr7 | + | [illegible] |
| A3SS | 10085 | ENSG00000105835 | NAMPT | chr7 | - | [illegible] |
| A3SS | 2611 | ENSG00000092199 | HNRNPC | chr14 | - | [illegible] |
| A3SS | 1909 | ENSG00000087263 | OGFOD1 | chr16 | + | [illegible] |
| A3SS | 10545 | ENSG00000169359 | SLC33A1 | chr3 | - | [illegible] |
| A3SS | 10489 | ENSG00000177646 | ACAD9 | chr3 | + | [illegible] |
| A3SS | 7342 | ENSG00000115204 | MPV17 | chr2 | - | [illegible] |
| A3SS | 1119 | ENSG00000161939 | RNASEK-C17orf49 | chr17 | + | [illegible] |
| A3SS | 10307 | ENSG00000172613 | RAD9A | chr11 | + | [illegible] |
| A3SS | 6620 | ENSG00000087191 | PSMC5 | chr17 | + | [illegible] |
| A3SS | 4888 | ENSG00000130024 | PHF10 | chr6 | - | [illegible] |
| A3SS | 2562 | ENSG00000197102 | DYNC1H1 | chr14 | + | [illegible] |
| A3SS | 12079 | ENSG00000172932 | ANKRD13D | chr11 | + | [illegible] |
| A3SS | 12143 | ENSG00000255717 | SNHG1 | chr11 | - | [illegible] |
| A3SS | 6893 | ENSG00000119725 | ZNF410 | chr14 | + | [illegible] |
| A3SS | 4264 | ENSG00000163349 | HIPK1 | chr1 | + | [illegible] |
| A3SS | 12239 | ENSG00000172183 | ISG20 | chr15 | + | [illegible] |
| A3SS | 12647 | ENSG00000107862 | GBF1 | chr10 | + | [illegible] |
| A3SS | 1961 | ENSG00000261740 | BOLA2-SMG1P6 | chr16 | - | [illegible] |
| A3SS | 2199 | ENSG00000182511 | FES | chr15 | + | [illegible] |
| A3SS | 9968 | ENSG00000129103 | SUMF2 | chr7 | + | [illegible] |
| A3SS | 8938 | ENSG00000138380 | CARF | chr2 | + | [illegible] |
| A3SS | 5379 | ENSG00000176444 | CLK2 | chr1 | - | [illegible] |
| A3SS | 12060 | ENSG00000175550 | DRAP1 | chr11 | + | [illegible] |
| A3SS | 346 | ENSG00000129353 | SLC44A2 | chr19 | + | [illegible] |
| A3SS | 11912 | ENSG00000205758 | CRYZL1 | chr21 | - | [illegible] |
| A3SS | 3708 | ENSG00000083896 | YTHDC1 | chr4 | - | [illegible] |
| A3SS | 2941 | ENSG00000135596 | MICAL1 | chr6 | - | [illegible] |
| A3SS | 4355 | ENSG00000096063 | SRPK1 | chr6 | - | [illegible] |
| A3SS | 2639 | ENSG00000139990 | DCAF5 | chr14 | - | [illegible] |
| A3SS | 1394 | ENSG00000129657 | SEC14L1 | chr17 | + | [illegible] |
| A3SS | 12950 | ENSG00000110801 | PSMD9 | chr12 | + | [illegible] |
| A3SS | 892 | ENSG00000124228 | DDX27 | chr20 | + | [illegible] |
| A3SS | 4114 | ENSG00000178950 | GAK | chr4 | - | [illegible] |
| A3SS | 12269 | ENSG00000186166 | CENATAC | chr11 | + | [illegible] |
| A3SS | 7496 | ENSG00000115211 | EIF2B4 | chr2 | - | [illegible] |
| A3SS | 3878 | ENSG00000061918 | GUCY1B1 | chr4 | + | [illegible] |
| A3SS | 13272 | ENSG00000197111 | PCBP2 | chr12 | + | [illegible] |
| A3SS | 1449 | ENSG00000266967 | AARSD1 | chr17 | - | [illegible] |
| A3SS | 11683 | ENSG00000185920 | PTCH1 | chr9 | - | [illegible] |
| A3SS | 1306 | ENSG00000256525 | POLG2 | chr17 | - | [illegible] |
| A3SS | 13416 | ENSG00000071564 | TCF3 | chr19 | - | [illegible] |
| A3SS | 5568 | ENSG00000114626 | ABTB1 | chr3 | + | [illegible] |
| A3SS | 1974 | ENSG00000196118 | CCDC189 | chr16 | - | [illegible] |
| A3SS | 13100 | ENSG00000127328 | RAB3IP | chr12 | + | [illegible] |
| A3SS | 339 | ENSG00000105723 | GSK3A | chr19 | - | [illegible] |
| A3SS | 13158 | ENSG00000205352 | PRR13 | chr12 | + | [illegible] |
| A3SS | 2607 | ENSG00000155465 | SLC7A7 | chr14 | - | [illegible] |
| A3SS | 12506 | ENSG00000146826 | TRAPPC14 | chr7 | - | [illegible] |
| A3SS | 12615 | ENSG00000221817 | PPP3CB-AS1 | chr10 | + | [illegible] |
| A3SS | 6248 | ENSG00000015153 | YAF2 | chr12 | - | [illegible] |
| A3SS | 4975 | ENSG00000142794 | NBPF3 | chr1 | + | [illegible] |
| A3SS | 5493 | ENSG00000099622 | CIRBP | chr19 | + | [illegible] |
| A3SS | 4326 | ENSG00000163930 | BAP1 | chr3 | - | [illegible] |
| A3SS | 2004 | ENSG00000174177 | CTU2 | chr16 | + | [illegible] |
| A3SS | 1092 | ENSG00000169718 | DUS1L | chr17 | - | [illegible] |
| A3SS | 13098 | ENSG00000135108 | FBXO21 | chr12 | - | [illegible] |
| A3SS | 6064 | ENSG00000138821 | SLC39A8 | chr4 | - | [illegible] |
| A3SS | 5383 | ENSG00000169231 | THBS3 | chr1 | - | [illegible] |
| A3SS | 12538 | ENSG00000175197 | DDIT3 | chr12 | - | [illegible] |
| A3SS | 13052 | ENSG00000205885 | C1RL-AS1 | chr12 | + | [illegible] |
| A3SS | 6060 | ENSG00000196204 | RNF216P1 | chr7 | + | [illegible] |
| A3SS | 774 | ENSG00000088833 | NSFL1C | chr20 | - | [illegible] |
| A3SS | 5984 | ENSG00000143363 | PRUNE1 | chr1 | + | [illegible] |
| A3SS | 4385 | ENSG00000010295 | IFFO1 | chr12 | - | [illegible] |
| A3SS | 12452 | ENSG00000123992 | DNPEP | chr2 | - | [illegible] |

| | | | | | | | | | | | | | | |
|---|---|---|---|---|---|---|---|---|---|---|---|---|---|---|
| A3SS | 6311 | ENSG00000114933 | INO80D | chr2 | - | 206062798 | 206063045 | 206062798 | 206062954 | 206063150 | 206063244 | 65 | 71 | 103 |
| A3SS | 2508 | ENSG00000197555 | SIPA1L1 | chr14 | + | 71650289 | 71650509 | 71650334 | 71650509 | 71624047 | 71624236 | 2 | 0 | 1 |
| A3SS | 13411 | ENSG00000134899 | ERCC5 | chr13 | + | 102854142 | 102854374 | 102854287 | 102854374 | 102853756 | 102853872 | 69 | 71 | 81 |
| A3SS | 5741 | ENSG00000169994 | MYO7B | chr2 | + | 127634175 | 127634683 | 127634595 | 127634683 | 127633257 | 127633363 | 2 | 5 | 3 |
| A3SS | 6340 | ENSG00000176463 | SLCO3A1 | chr15 | + | 92150949 | 92151014 | 92150954 | 92151014 | 92146983 | 92147159 | 57 | 30 | 52 |
| A3SS | 156 | ENSG00000100150 | DEPDC5 | chr22 | + | 31758526 | 31758633 | 31758545 | 31758633 | 31754861 | 31754979 | 0 | 0 | 5 |
| A3SS | 11856 | ENSG00000142186 | SCYL1 | chr11 | + | 65538269 | 65538704 | 65538441 | 65538704 | 65537966 | 65538182 | 122 | 45 | 107 |
| A3SS | 9871 | ENSG00000085514 | PILRA | chr7 | + | 100399784 | 100400096 | 100399822 | 100400096 | 100399290 | 100399340 | 26 | 7 | 12 |
| A3SS | 13327 | ENSG00000121310 | ECHDC2 | chr1 | - | 52905033 | 52906611 | 52905033 | 52905090 | 52907867 | 52907954 | 27 | 20 | 28 |
| A3SS | 5852 | ENSG00000198301 | SDAD1 | chr4 | - | 75981371 | 75981545 | 75981371 | 75981470 | 75981932 | 75982037 | 32 | 22 | 35 |
| A3SS | 3456 | ENSG00000011243 | AKAP8L | chr19 | - | 15397713 | 15398766 | 15397713 | 15397855 | 15399301 | 15399410 | 38 | 23 | 24 |
| A3SS | 12451 | ENSG00000123992 | DNPEP | chr2 | - | 219385630 | 219385751 | 219385630 | 219385706 | 219385967 | 219386098 | 6 | 8 | 1 |
| A3SS | 3994 | ENSG00000100298 | APOBEC3H | chr22 | + | 39103688 | 39104067 | 39103691 | 39104067 | 39101917 | 39102042 | 8 | 4 | 5 |
| A3SS | 13233 | ENSG00000135473 | PAN2 | chr12 | - | 56324036 | 56324185 | 56324036 | 56324182 | 56324293 | 56324493 | 53 | 60 | 43 |
| A3SS | 4353 | ENSG00000213523 | SRA1 | chr5 | - | 140551060 | 140552184 | 140551060 | 140551169 | 140557146 | 140557446 | 48 | 20 | 51 |
| A3SS | 10579 | ENSG00000123562 | MORF4L2 | chrX | - | 103678498 | 103678620 | 103678498 | 103678600 | 103684680 | 103684729 | 20 | 8 | 25 |
| A3SS | 7630 | ENSG00000187098 | MITF | chr3 | + | 69951811 | 69951886 | 69951829 | 69951886 | 69949050 | 69949168 | 8 | 3 | 2 |
| A3SS | 8561 | ENSG00000115282 | TTC31 | chr2 | + | 74491943 | 74492055 | 74492003 | 74492055 | 74491480 | 74491666 | 4 | 10 | 0 |
| A3SS | 10090 | ENSG00000067182 | TNFRSF1A | chr12 | - | 6333736 | 6333890 | 6333736 | 6333865 | 6334090 | 6334244 | 16 | 3 | 3 |
| A3SS | 1351 | ENSG00000180891 | CUEDC1 | chr17 | - | 57862975 | 57863357 | 57862975 | 57863285 | 57866473 | 57866544 | 6 | 5 | 1 |
| [illegible] | | | | | | | | | | | | | | |

| | | | | | | | | | | | | | | | | | | | | | | | | | | | | | | | | | | | | | | | | | | | | | | |
|---|---|---|---|---|---|---|---|---|---|---|---|---|---|---|---|---|---|---|---|---|---|---|---|---|---|---|---|---|---|---|---|---|---|---|---|---|---|---|---|---|---|---|---|---|---|---|
| A3SS | 8196 | ENSG00000129292 | PHF20L1 | chr8 | + | 132804611 | 132804740 | 132804614 | 132804740 | 132803818 | 132804032 | 40 | 29 | 17 | 33 | 47 | 157 | 172 | 87 | 158 | 238 | 0 | 36 | 19 | 3 | 7 | 78 | 112 | 112 | 59 | 27 | 152 | 149 | 0.074024 | 0.358012203 | 0.2 | 0.142 | 0.161 | 0.17 | 0.162 | 0 | 0.24 | 0.143 | 0.047 | 0.203 | 0.04 |
| A3SS | 12089 | ENSG00000149577 | SIDT2 | chr11 | + | 117188707 | 117188826 | 117188728 | 117188826 | 117187627 | 117187699 | 146 | 109 | 69 | 259 | 77 | 6 | 1 | 2 | 15 | 0 | 160 | 177 | 33 | 97 | 101 | 8 | 13 | 3 | 3 | 6 | 170 | 149 | 0.074910 | 0.362079011 | 0.955 | 0.99 | 0.968 | 0.938 | 1 | 0.946 | 0.923 | 0.906 | 0.966 | 0.937 | 0.035 |
| A3SS | 5073 | ENSG00000170581 | STAT2 | chr12 | - | 56348923 | 56349083 | 56348923 | 56349059 | 56349162 | 56349261 | 2 | 2 | 4 | 1 | 2 | 48 | 25 | 38 | 56 | 33 | 6 | 9 | 2 | 5 | 13 | 147 | 31 | 18 | 95 | 149 | 173 | 149 | 0.075294 | 0.363720422 | 0.035 | 0.064 | 0.083 | 0.015 | 0.05 | 0.034 | 0.2 | 0.087 | 0.043 | 0.07 | -0.037 |
| A3SS | 4674 | ENSG00000162302 | RPS6KA4 | chr11 | + | 64368467 | 64368601 | 64368485 | 64368601 | 64368131 | 64368260 | 54 | 25 | 23 | 29 | 12 | 6 | 3 | 3 | 2 | 0 | 12 | 19 | 13 | 37 | 101 | 0 | 1 | 3 | 0 | 5 | 167 | 149 | 0.075482 | 0.364409043 | 0.889 | 0.881 | 0.872 | 0.928 | 1 | 1 | 0.944 | 0.795 | 1 | 0.947 | -0.023 |
| A3SS | 2713 | ENSG00000203485 | INF2 | chr14 | + | 104710881 | 104711007 | 104710936 | 104711007 | 104710087 | 104710188 | 2 | 7 | 1 | 0 | 0 | 34 | 24 | 32 | 40 | 13 | 0 | 0 | 0 | 0 | 0 | 49 | 36 | 20 | 32 | 115 | 204 | 149 | 0.075551 | 0.364522674 | 0.041 | 0.176 | 0.022 | 0 | 0 | 0 | | 0 | 0 | 0 | 0.048 |
| A3SS | 2395 | ENSG00000196547 | MAN2A2 | chr15 | + | 90905844 | 90906016 | 90905921 | 90906016 | 90905578 | 90905723 | 55 | 32 | 46 | 100 | 27 | 0 | 0 | 0 | 1 | 0 | 88 | 138 | 125 | 103 | 412 | 0 | 0 | 0 | 1 | 15 | 226 | 149 | 0.075748 | 0.36525385 | 1 | 1 | 1 | 0.985 | 1 | 1 | 1 | 1 | 0.985 | 0.948 | 0.01 |
| A3SS | 11654 | ENSG00000130714 | POMT1 | chr9 | + | 131510259 | 131510415 | 131510356 | 131510415 | 131508910 | 131509022 | 6 | 10 | 11 | 13 | 4 | 1 | 0 | 1 | 0 | 0 | 13 | 31 | 3 | 7 | 22 | 0 | 0 | 0 | 0 | 8 | 246 | 149 | 0.076160 | 0.366989682 | 0.784 | 1 | 0.869 | 1 | 1 | 1 | 1 | 1 | 1 | 0.625 | 0.006 |
| A3SS | 439 | ENSG00000161265 | U2AF1L4 | chr19 | - | 35745124 | 35745212 | 35745124 | 35745187 | 35745347 | 35745418 | 14 | 1 | 12 | 7 | 3 | 9 | 3 | 9 | 7 | 42 | 10 | 0 | 3 | 12 | 11 | 14 | 41 | 22 | 30 | 23 | 174 | 149 | 0.076199 | 0.366989682 | 0.571 | 0.222 | 0.533 | 0.461 | 0.058 | 0.38 | 0 | 0.105 | 0.255 | 0.291 | 0.163 |
| A3SS | 10512 | ENSG00000041988 | THAP3 | chr1 | + | 6632369 | 6632495 | 6632390 | 6632495 | 6630287 | 6630353 | 1 | 0 | 0 | 0 | 0 | 33 | 2 | 8 | 14 | 14 | 0 | 3 | 0 | 2 | 3 | 6 | 7 | 11 | 3 | 31 | 170 | 149 | 0.076345 | 0.367474858 | 0.026 | 0 | 0 | 0 | 0 | 0 | 0.273 | 0 | 0.369 | 0.078 | -0.139 |
| A3SS | 8153 | ENSG00000131051 | RBM39 | chr20 | - | 35729461 | 35729551 | 35729461 | 35729527 | 35731940 | 35732135 | 3 | 9 | 11 | 19 | 15 | 672 | 515 | 639 | 750 | 825 | 0 | 8 | 14 | 20 | 10 | 209 | 512 | 793 | 442 | 153 | 173 | 149 | 0.076506 | 0.368031229 | 0.004 | 0.015 | 0.015 | 0.021 | 0.015 | 0 | 0.013 | 0.015 | 0.038 | 0.053 | -0.01 |
| A3SS | 2301 | ENSG00000140199 | SLC12A6 | chr15 | - | 34336409 | 34336752 | 34336409 | 34336686 | 34337960 | 34338057 | 53 | 61 | 63 | 62 | 132 | 4 | 1 | 2 | 4 | 4 | 185 | 85 | 159 | 146 | 117 | 15 | 5 | 12 | 9 | 0 | 215 | 149 | 0.076899 | 0.369703527 | 0.902 | 0.977 | 0.956 | 0.915 | 0.958 | 0.895 | 0.922 | 0.902 | 0.918 | 1 | 0.014 |
| A3SS | 11967 | ENSG00000068831 | RASGRP2 | chr11 | - | 64727295 | 64727363 | 64727295 | 64727360 | 64728862 | 64729042 | 6 | 2 | 8 | 8 | 3 | 122 | 67 | 141 | 165 | 43 | 5 | 11 | 4 | 4 | 12 | 18 | 123 | 104 | 111 | 90 | 251 | 152 | 149 | 0.076949 | 0.369722731 | 0.046 | 0.028 | 0.053 | 0.045 | 0.064 | 0.038 | 0.094 | 0.034 | 0.116 | 0.066 | -0.022 |
| A3SS | 5891 | ENSG00000058272 | PPP1R12A | chr12 | - | 79778549 | 79779351 | 79778549 | 79778600 | 79781814 | 79781862 | 53 | 49 | 29 | 37 | 45 | 392 | 279 | 267 | 390 | 666 | 18 | 8 | 4 | 2 | 0 | 21 | 189 | 233 | 45 | 12 | 298 | 149 | 0.077020 | 0.369766613 | 0.063 | 0.081 | 0.052 | 0.045 | 0.033 | 0.3 | 0.021 | 0.009 | 0.022 | 0 | -0.016 |
| A3SS | 544 | ENSG00000105483 | CARD8 | chr19 | - | 48238382 | 48238532 | 48238382 | 48238526 | 48240961 | 48241063 | 396 | 232 | 300 | 365 | 366 | 22 | 9 | 15 | 7 | 18 | 319 | 241 | 238 | 139 | 207 | 8 | 6 | 10 | 11 | 0 | 155 | 149 | 0.077050 | 0.369766613 | 0.945 | 0.961 | 0.951 | 0.98 | 0.951 | 0.975 | 0.975 | 0.958 | 0.924 | 1 | -0.009 |
| A3SS | 9810 | ENSG00000012124 | CD22 | chr19 | + | 35332577 | 35332924 | 35332652 | 35332924 | 35332018 | 35332074 | 50 | 36 | 46 | 50 | 59 | 9 | 2 | 0 | 2 | 9 | 151 | 166 | 22 | 91 | 113 | 16 | 1 | 0 | 8 | 0 | 224 | 149 | 0.077231 | 0.370416127 | 0.787 | 0.923 | 1 | 0.943 | 0.813 | 0.863 | 0.991 | 1 | 0.883 | 1 | -0.054 |
| A3SS | 5908 | ENSG00000055609 | KMT2C | chr7 | - | 152156204 | 152156343 | 152156204 | 152156274 | 152157760 | 152157922 | 93 | 96 | 85 | 133 | 164 | 20 | 9 | 8 | 12 | 31 | 34 | 75 | 113 | 48 | 10 | 0 | 11 | 31 | 31 | 11 | 218 | 149 | 0.077788 | 0.372772624 | 0.761 | 0.879 | 0.879 | 0.883 | 0.783 | 1 | 0.823 | 0.714 | 0.514 | 0.383 | 0.15 |
| A3SS | 4075 | ENSG00000145247 | OCIAD2 | chr4 | - | 48892771 | 48894053 | 48892771 | 48892889 | 48897803 | 48897857 | 39 | 22 | 56 | 59 | 79 | 0 | 0 | 0 | 0 | 0 | 11 | 35 | 40 | 18 | 48 | 0 | 0 | 3 | 5 | 0 | 298 | 149 | 0.077815 | 0.372772624 | 1 | 1 | 1 | 1 | 1 | 1 | 1 | 0.87 | 0.643 | 1 | 0.097 |
| A3SS | 12185 | ENSG00000110075 | PPP6R3 | chr11 | + | 68587944 | 68588024 | 68587977 | 68588024 | 68583042 | 68583129 | 123 | 114 | 85 | 160 | 130 | 69 | 54 | 44 | 74 | 65 | 128 | 162 | 196 | 163 | 15 | 16 | 55 | 50 | 44 | 20 | 182 | 149 | 0.077891 | 0.372913732 | 0.593 | 0.633 | 0.613 | 0.639 | 0.621 | 0.868 | 0.707 | 0.762 | 0.752 | 0.38 | -0.074 |
| A3SS | 11780 | ENSG00000198561 | CTNND1 | chr11 | + | 57791384 | 57791673 | 57791491 | 57791673 | 57789036 | 57789155 | 45 | 42 | 33 | 43 | 70 | 13 | 4 | 4 | 5 | 16 | 62 | 30 | 14 | 62 | 61 | 33 | 7 | 0 | 18 | 38 | 256 | 149 | 0.078160 | 0.373801766 | 0.668 | 0.859 | 0.828 | 0.833 | 0.718 | 0.522 | 0.714 | 1 | 0.667 | 0.483 | 0.104 |
| A3SS | 12196 | ENSG00000214787 | MS4A4E | chr11 | - | 60214570 | 60214627 | 60214570 | 60214614 | 60224964 | 60225093 | 5 | 1 | 0 | 1 | 0 | 3 | 11 | 8 | 14 | 2 | 0 | 0 | 0 | 0 | 0 | 17 | 2 | 17 | 14 | 5 | 162 | 149 | 0.078169 | 0.373801766 | 0.605 | 0.077 | 0 | 0.062 | 0 | 0 | 0 | 0 | 0 | 0 | 0.149 |
| A3SS | 1860 | ENSG00000059122 | FLYWCH1 | chr16 | + | 2930409 | 2930880 | 2930412 | 2930880 | 2929612 | 2930010 | 20 | 11 | 20 | 31 | 27 | 9 | 2 | 8 | 15 | 0 | 39 | 16 | 32 | 23 | 37 | 0 | 2 | 3 | 3 | 17 | 152 | 149 | 0.078516 | 0.375238199 | 0.685 | 0.844 | 0.71 | 0.67 | 1 | 1 | 0.887 | 0.913 | 0.883 | 0.681 | -0.091 |
| A3SS | 12509 | ENSG00000148824 | MTG1 | chr10 | + | 133401528 | 133402245 | 133402148 | 133402245 | 133399528 | 133399619 | 35 | 15 | 54 | 52 | 48 | 8 | 1 | 5 | 6 | 5 | 31 | 38 | 17 | 26 | 67 | 0 | 10 | 7 | 5 | 26 | 298 | 149 | 0.078764 | 0.37620303 | 0.686 | 0.882 | 0.844 | 0.812 | 0.828 | 1 | 0.655 | 0.548 | 0.722 | 0.563 | 0.113 |
| A3SS | 11747 | ENSG00000166181 | API5 | chr11 | + | 43342427 | 43344529 | 43342432 | 43344529 | 43335857 | 43335994 | 175 | 136 | 132 | 146 | 184 | 132 | 116 | 111 | 147 | 136 | 47 | 81 | 73 | 116 | 26 | 5 | 93 | 67 | 102 | 3 | 154 | 149 | 0.079149 | 0.377777476 | 0.562 | 0.531 | 0.535 | 0.49 | 0.567 | 0.901 | 0.457 | 0.513 | 0.524 | 0.893 | -0.121 |
| A3SS | 7224 | ENSG00000175634 | RPS6KB2 | chr11 | + | 67433125 | 67433447 | 67433339 | 67433447 | 67432951 | 67433042 | 62 | 30 | 72 | 60 | 49 | 5 | 2 | 3 | 6 | 2 | 68 | 114 | 70 | 54 | 126 | 5 | 9 | 14 | 4 | 11 | 298 | 149 | 0.079187 | 0.377777476 | 0.861 | 0.882 | 0.923 | 0.833 | 0.925 | 0.872 | 0.864 | 0.714 | 0.871 | 0.851 | 0.05 |
| A3SS | 1849 | ENSG00000205336 | ADGRG1 | chr16 | + | 57651184 | 57651622 | 57651242 | 57651622 | 57650252 | 57650351 | 65 | 16 | 8 | 37 | 5 | 2 | 0 | 1 | 0 | 4 | 6 | 28 | 19 | 18 | 21 | 0 | 4 | 7 | 1 | 0 | 207 | 149 | 0.079607 | 0.379553353 | 0.959 | 1 | 0.852 | 1 | 0.474 | 1 | 0.834 | 0.661 | 0.928 | 1 | -0.028 |
| A3SS | 7908 | ENSG00000166801 | FAM111A | chr11 | + | 59148703 | 59148953 | 59148796 | 59148953 | 59145826 | 59145856 | 46 | 37 | 36 | 33 | 39 | 71 | 95 | 54 | 134 | 137 | 0 | 27 | 53 | 9 | 0 | 34 | 61 | 51 | 64 | 12 | 242 | 149 | 0.079702 | 0.379693445 | 0.285 | 0.193 | 0.291 | 0.132 | 0.149 | 0 | 0.214 | 0.39 | 0.08 | 0 | 0.073 |
| A3SS | 5015 | ENSG00000105171 | POP4 | chr19 | + | 29610319 | 29610632 | 29610408 | 29610632 | 29608656 | 29608709 | 0 | 1 | 5 | 4 | 7 | 75 | 35 | 62 | 82 | 41 | 0 | 0 | 0 | 3 | 6 | 76 | 44 | 19 | 46 | 84 | 238 | 149 | 0.079763 | 0.379693445 | 0 | 0.018 | 0.048 | 0.03 | 0.097 | 0 | 0 | 0 | 0.039 | 0.043 | 0.022 |
| A3SS | 9291 | ENSG00000204351 | SKIV2L | chr6 | + | 31961195 | 31961386 | 31961340 | 31961386 | 31961040 | 31961094 | 187 | 103 | 176 | 180 | 276 | 7 | 0 | 0 | 2 | 5 | 160 | 167 | 107 | 137 | 223 | 0 | 2 | 2 | 0 | 0 | 294 | 149 | 0.079777 | 0.379693445 | 0.931 | 1 | 1 | 0.979 | 0.965 | 1 | 0.977 | 0.964 | 1 | 1 | -0.013 |
| A3SS | 12725 | ENSG00000130640 | TUBGCP2 | chr10 | - | 133281272 | 133282106 | 133281272 | 133281436 | 133282222 | 133282342 | 13 | 5 | 17 | 2 | 12 | 75 | 30 | 48 | 56 | 25 | 25 | 18 | 12 | 14 | 32 | 54 | 89 | 42 | 50 | 109 | 298 | 149 | 0.080149 | 0.381240596 | 0.08 | 0.077 | 0.15 | 0.018 | 0.194 | 0.188 | 0.092 | 0.125 | 0.123 | 0.128 | -0.027 |
| A3SS | 9478 | ENSG00000008405 | CRY1 | chr12 | - | 106997293 | 106997406 | 106997293 | 106997386 | 106997487 | 106997690 | 0 | 0 | 0 | 0 | 2 | 77 | 69 | 76 | 73 | 69 | 10 | 0 | 0 | 0 | 0 | 15 | 44 | 73 | 27 | 12 | 169 | 149 | 0.080247 | 0.381482157 | 0 | 0 | 0 | 0 | 0.025 | 0.37 | 0 | 0 | 0 | 0 | -0.069 |
| A3SS | 9537 | ENSG00000196923 | PDLIM7 | chr5 | - | 177491806 | 177491976 | 177491806 | 177491925 | 177492404 | 177492435 | 2 | 0 | 0 | 2 | 5 | 25 | 3 | 7 | 14 | 18 | 0 | 4 | 0 | 3 | 2 | 40 | 34 | 11 | 52 | 150 | 200 | 149 | 0.080791 | 0.3838411 | 0.056 | 0 | 0 | 0.096 | 0.171 | 0 | 0.081 | 0 | 0.041 | 0.01 | 0.038 |
| A3SS | 111 | ENSG00000015475 | BID | chr22 | - | 17739348 | 17740176 | 17739348 | 17739488 | 17743802 | 17744013 | 4 | 9 | 2 | 7 | 2 | 76 | 45 | 26 | 64 | 49 | 8 | 8 | 5 | 10 | 29 | 92 | 73 | 42 | 83 | 165 | 298 | 149 | 0.080928 | 0.384263899 | 0.026 | 0.091 | 0.037 | 0.052 | 0.02 | 0.042 | 0.052 | 0.056 | 0.057 | 0.081 | -0.012 |
| A3SS | 3341 | ENSG00000125846 | ZNF133 | chr20 | + | 18315068 | 18316994 | 18315071 | 18316994 | 18306297 | 18306393 | 10 | 0 | 5 | 2 | 18 | 3 | 2 | 2 | 3 | 0 | 18 | 3 | 1 | 2 | 3 | 0 | 1 | 0 | 7 | 0 | 152 | 149 | 0.080996 | 0.384361445 | 0.766 | 0 | 0.71 | 0.395 | 1 | 1 | 0.746 | 1 | 0.219 | 1 | -0.219 |
| A3SS | 11353 | ENSG00000187239 | FNBP1 | chr9 | - | 129923843 | 129923996 | 129923843 | 129923939 | 129924959 | 129925157 | 13 | 13 | 7 | 13 | 16 | 5 | 3 | 1 | 6 | 7 | 36 | 10 | 7 | 6 | 12 | 0 | 0 | 6 | 8 | 0 | 206 | 149 | 0.081414 | 0.386120549 | 0.653 | 0.758 | 0.835 | 0.61 | 0.623 | 1 | 1 | 0.458 | 0.352 | 1 | -0.066 |
| A3SS | 8395 | ENSG00000087087 | SRRT | chr7 | + | 100885212 | 100885370 | 100885215 | 100885370 | 100884922 | 100885040 | 43 | 23 | 56 | 54 | 55 | 31 | 31 | 29 | 61 | 38 | 96 | 80 | 53 | 38 | 67 | 76 | 68 | 59 | 94 | 97 | 152 | 149 | 0.081744 | 0.387458247 | 0.576 | 0.421 | 0.654 | 0.465 | 0.587 | 0.553 | 0.536 | 0.468 | 0.284 | 0.404 | 0.092 |
| A3SS | 13408 | ENSG00000005810 | MYCBP2 | chr13 | - | 77118339 | 77118519 | 77118339 | 77118495 | 77121372 | 77121495 | 15 | 25 | 7 | 18 | 21 | 21 | 21 | 14 | 29 | 29 | 0 | 7 | 3 | 1 | 7 | 12 | 18 | 18 | 9 | 0 | 173 | 149 | 0.082101 | 0.388919013 | 0.381 | 0.506 | 0.301 | 0.348 | 0.384 | 0 | 0.251 | 0.126 | 0.087 | 1 | 0.091 |
| A3SS | 3359 | ENSG00000146067 | FAM193B | chr5 | - | 177531952 | 177532200 | 177531952 | 177532131 | 177532442 | 177532641 | 45 | 50 | 51 | 56 | 47 | 5 | 6 | 2 | 11 | 2 | 133 | 65 | 47 | 87 | 127 | 19 | 11 | 7 | 13 | 18 | 218 | 149 | 0.082540 | 0.390770747 | 0.86 | 0.851 | 0.946 | 0.777 | 0.941 | 0.827 | 0.802 | 0.821 | 0.821 | 0.828 | 0.055 |
| A3SS | 5083 | ENSG00000008952 | SEC62 | chr3 | + | 169982636 | 169982911 | 169982706 | 169982911 | 169976945 | 169977051 | 10 | 14 | 9 | 19 | 13 | 279 | 242 | 318 | 326 | 445 | 0 | 16 | 16 | 2 | 5 | 14 | 232 | 249 | 54 | 44 | 219 | 149 | 0.082599 | 0.390770747 | 0.024 | 0.038 | 0.019 | 0.038 | 0.019 | 0 | 0.045 | 0.042 | 0.025 | 0.072 | -0.009 |
| A3SS | 11424 | ENSG00000175854 | SWI5 | chr9 | + | 128276586 | 128276755 | 128276706 | 128276755 | 128276221 | 128276402 | 9 | 4 | 7 | 5 | 13 | 59 | 41 | 36 | 83 | 11 | 2 | 22 | 22 | 7 | 44 | 77 | 43 | 44 | 29 | 123 | 269 | 149 | 0.082637 | 0.390770747 | 0.078 | 0.051 | 0.097 | 0.032 | 0.396 | 0.014 | 0.221 | 0.217 | 0.118 | 0.165 | -0.016 |
| A3SS | 133 | ENSG00000100241 | SBF1 | chr22 | - | 50467531 | 50467693 | 50467531 | 50467690 | 50467785 | 50467923 | 11 | 3 | 15 | 7 | 0 | 33 | 18 | 39 | 45 | 26 | 0 | 10 | 5 | 7 | 28 | 5 | 20 | 13 | 31 | 40 | 152 | 149 | 0.082926 | 0.391909926 | 0.246 | 0.14 | 0.274 | 0.132 | 0 | 0 | 0.329 | 0.274 | 0.181 | 0.407 | -0.08 |
| A3SS | 3905 | ENSG00000127419 | TMEM175 | chr4 | + | 948410 | 948615 | 948510 | 948615 | 948115 | 948154 | 38 | 13 | 33 | 30 | 14 | 0 | 0 | 0 | 0 | 0 | 9 | 20 | 8 | 27 | 27 | 0 | 0 | 0 | 0 | 8 | 249 | 149 | 0.083093 | 0.392467041 | 1 | 1 | 1 | 1 | 1 | 1 | 1 | 1 | 1 | 0.669 | 0.066 |
| A3SS | 1029 | ENSG00000175354 | PTPN2 | chr18 | - | 12817155 | 12817365 | 12817155 | 12817265 | 12819213 | 12819282 | 288 | 213 | 212 | 237 | 649 | 8 | 1 | 1 | 6 | 6 | 61 | 189 | 260 | 205 | 26 | 2 | 6 | 6 | 6 | 0 | 249 | 149 | 0.083170 | 0.39255054 | 0.956 | 0.992 | 0.992 | 0.959 | 0.985 | 0.948 | 0.95 | 0.963 | 0.953 | 1 | 0.014 |
| A3SS | 6711 | ENSG00000188529 | SRSF10 | chr1 | - | 23971572 | 23971626 | 23971572 | 23971623 | 23971849 | 23972012 | 281 | 250 | 237 | 338 | 421 | 66 | 53 | 95 | 90 | 127 | 71 | 191 | 284 | 96 | 30 | 33 | 58 | 79 | 55 | 8 | 152 | 149 | 0.083249 | 0.39255054 | 0.807 | 0.822 | 0.71 | 0.786 | 0.765 | 0.678 | 0.763 | 0.779 | 0.631 | 0.786 | 0.051 |
| A3SS | 1013 | ENSG00000133313 | CNDP2 | chr18 | + | 74499881 | 74500033 | 74499936 | 74500033 | 74496306 | 74496431 | 139 | 101 | 101 | 124 | 75 | 28 | 22 | 15 | 26 | 7 | 345 | 193 | 80 | 118 | 311 | 31 | 23 | 6 | 24 | 54 | 204 | 149 | 0.083257 | 0.39255054 | 0.784 | 0.77 | 0.831 | 0.777 | 0.887 | 0.89 | 0.86 | 0.907 | 0.782 | 0.808 | -0.04 |
| A3SS | 12539 | ENSG00000175197 | DDIT3 | chr12 | - | 57517268 | 57517595 | 57517268 | 57517438 | 57517705 | 57517753 | 32 | 23 | 42 | 43 | 40 | 18 | 10 | 10 | 12 | 21 | 103 | 115 | 113 | 71 | 168 | 26 | 11 | 63 | 29 | 18 | 298 | 149 | 0.083331 | 0.392671765 | 0.471 | 0.535 | 0.677 | 0.642 | 0.488 | 0.665 | 0.839 | 0.473 | 0.55 | 0.824 | -0.108 |
| A3SS | 9527 | ENSG00000173200 | PARP15 | chr3 | + | 122617014 | 122617164 | 122617085 | 122617164 | 122613040 | 122613268 | 100 | 88 | 59 | 123 | 143 | 2 | 1 | 0 | 2 | 1 | 23 | 76 | 70 | 31 | 19 | 0 | 2 | 1 | 6 | 0 | 220 | 149 | 0.083407 | 0.392729771 | 0.971 | 0.983 | 1 | 0.977 | 0.99 | 1 | 0.963 | 0.979 | 0.778 | 1 | 0.04 |
| A3SS | 4459 | ENSG00000114867 | EIF4G1 | chr3 | + | 184331696 | 184331809 | 184331727 | 184331809 | 184331471 | 184331606 | 1 | 4 | 3 | 3 | 2 | 242 | 132 | 192 | 216 | 145 | 9 | 15 | 4 | 3 | 3 | 523 | 269 | 144 | 247 | 358 | 180 | 149 | 0.083440 | 0.392729771 | 0.003 | 0.024 | 0.013 | 0.011 | 0.011 | 0.014 | 0.044 | 0.022 | 0.01 | 0.007 | -0.007 |
| A3SS | 11275 | ENSG00000150776 | NKAPD1 | chr11 | + | 112082464 | 112085150 | 112082467 | 112085150 | 112081981 | 112082035 | 183 | 114 | 139 | 157 | 487 | 10 | 17 | 12 | 17 | 54 | 146 | 125 | 111 | 90 | 22 | 14 | 10 | 18 | 10 | 22 | 152 | 149 | 0.083830 | 0.394334219 | 0.947 | 0.868 | 0.919 | 0.901 | 0.898 | 0.911 | 0.925 | 0.858 | 0.898 | 0.495 | 0.089 |
| A3SS | 3455 | ENSG00000011243 | AKAP8L | chr19 | - | 15397149 | 15397320 | 15397149 | 15397280 | 15397519 | 15397625 | 3 | 0 | 1 | 8 | 3 | 183 | 107 | 188 | 210 | 169 | 20 | 4 | 5 | 2 | 7 | 350 | 317 | 275 | 238 | 309 | 189 | 149 | 0.084222 | 0.395946207 | 0.013 | 0 | 0.004 | 0.029 | 0.014 | 0.043 | 0.01 | 0.014 | 0.007 | 0.018 | -0.006 |
| A3SS | 12482 | ENSG00000100109 | TFIP11 | chr22 | - | 26511118 | 26511195 | 26511118 | 26511153 | 26512098 | 26512175 | 13 | 8 | 20 | 17 | 6 | 1 | 3 | 0 | 2 | 1 | 27 | 17 | 21 | 16 | 20 | 0 | 8 | 0 | 2 | 8 | 191 | 149 | 0.084322 | 0.396184375 | 0.91 | 0.675 | 1 | 0.869 | 0.824 | 1 | 0.624 | 1 | 0.862 | 0.661 | 0.026 |
| A3SS | 5208 | ENSG00000137776 | SLTM | chr15 | - | 58916934 | 58917020 | 58916934 | 58916999 | 58932355 | 58932443 | 2 | 3 | 4 | 3 | 10 | 45 | 28 | 36 | 52 | 41 | 13 | 5 | 7 | 4 | 5 | 10 | 39 | 44 | 39 | 49 | 170 | 149 | 0.084386 | 0.396254451 | 0.037 | 0.086 | 0.089 | 0.048 | 0.176 | 0.533 | 0.101 | 0.122 | 0.082 | 0.082 | -0.097 |
| A3SS | 7855 | ENSG00000110066 | KMT5B | chr11 | - | 68171231 | 68171296 | 68171231 | 68171251 | 68171542 | 68171709 | 9 | 10 | 2 | 8 | 4 | 120 | 87 | 82 | 128 | 230 | 0 | 7 | 6 | 9 | 0 | 27 | 69 | 55 | 53 | 12 | 194 | 149 | 0.084476 | 0.396449012 | 0.054 | 0.081 | 0.018 | 0.046 | 0.013 | 0 | 0.072 | 0.077 | 0.115 | 0 | -0.01 |
| A3SS | 2668 | ENSG00000100938 | GMPR2 | chr14 | + | 24237041 | 24237152 | 24237070 | 24237152 | 24235966 | 24236140 | 5 | 5 | 10 | 16 | 1 | 84 | 42 | 87 | 92 | 59 | 9 | 0 | 10 | 14 | 0 | 124 | 120 | 72 | 69 | 155 | 178 | 149 | 0.084561 | 0.396451436 | 0.047 | 0.091 | 0.088 | 0.127 | 0.014 | 0.057 | 0 | 0.104 | 0.145 | 0 | 0.012 |
| A3SS | 3401 | ENSG00000185627 | PSMD13 | chr11 | + | 244040 | 244216 | 244160 | 244216 | 238997 | 239076 | 82 | 77 | 88 | 166 | 46 | 2 | 1 | 0 | 0 | 0 | 307 | 135 | 109 | 155 | 283 | 0 | 5 | 0 | 2 | 8 | 269 | 149 | 0.084575 | 0.396451436 | 0.958 | 0.977 | 1 | 1 | 1 | 1 | 0.937 | 1 | 0.977 | 0.951 | 0.014 |
| A3SS | 7249 | ENSG00000089127 | OAS1 | chr12 | + | 112917546 | 112919903 | 112919486 | 112919903 | 112916508 | 112916738 | 231 | 282 | 187 | 279 | 427 | 1 | 3 | 3 | 14 | 4 | 1155 | 470 | 153 | 664 | 352 | 12 | 9 | 15 | 18 | 10 | 298 | 149 | 0.084914 | 0.397583228 | 0.991 | 0.979 | 0.969 | 0.909 | 0.982 | 0.98 | 0.963 | 0.836 | 0.949 | 0.946 | 0.031 |
| A3SS | 12454 | ENSG00000138326 | RPS24 | chr10 | + | 78037438 | 78040225 | 78040203 | 78040225 | 78037193 | 78037304 | 27 | 10 | 21 | 9 | 66 | 390 | 271 | 423 | 320 | 713 | 8 | 10 | 4 | 13 | 11 | 178 | 198 | 163 | 186 | 253 | 298 | 149 | 0.084949 | 0.397583228 | 0.033 | 0.018 | 0.024 | 0.014 | 0.044 | 0.022 | 0.025 | 0.012 | 0.034 | 0.021 | 0.004 |
| A3SS | 8127 | ENSG00000185480 | PARPBP | chr12 | + | 102148229 | 102148263 | 102148234 | 102148263 | 102123885 | 102124041 | 7 | 11 | 7 | 13 | 14 | 1 | 0 | 5 | 1 | 0 | 12 | 8 | 1 | 19 | 17 | 0 | 0 | 0 | 0 | 0 | 154 | 149 | 0.084964 | 0.397583228 | 0.871 | 1 | 0.575 | 0.926 | 1 | 1 | 1 | 1 | 1 | 1 | -0.126 |
| A3SS | 219 | ENSG00000198131 | ZNF544 | chr19 | + | 58246710 | 58246794 | 58246745 | 58246794 | 58246300 | 58246427 | 25 | 7 | 40 | 17 | 6 | 18 | 13 | 18 | 9 | 11 | 2 | 34 | 12 | 18 | 46 | 3 | 7 | 18 | 4 | 7 | 184 | 149 | 0.085266 | 0.398681529 | 0.529 | 0.304 | 0.643 | 0.605 | 0.306 | 0.351 | 0.797 | 0.351 | 0.785 | 0.842 | -0.148 |
| A3SS | 12704 | ENSG00000151651 | ADAM8 | chr10 | - | 133273253 | 133273443 | 133273253 | 133273430 | 133273761 | 133273838 | 190 | 76 | 76 | 100 | 34 | 15 | 2 | 17 | 11 | 10 | 208 | 113 | 57 | 52 | 133 | 9 | 10 | 7 | 5 | 13 | 162 | 149 | 0.085298 | 0.398681529 | 0.921 | 0.972 | 0.804 | 0.893 | 0.758 | 0.955 | 0.912 | 0.882 | 0.905 | 0.904 | -0.042 |
| A3SS | 8453 | ENSG00000180902 | D2HGDH | chr2 | + | 241743621 | 241743815 | 241743638 | 241743815 | 241742434 | 241742574 | 18 | 10 | 20 | 16 | 2 | 2 | 7 | 3 | 7 | 2 | 0 | 2 | 26 | 35 | 61 | 11 | 0 | 7 | 0 | 0 | 166 | 149 | 0.085359 | 0.398738775 | 0.89 | 0.562 | 0.857 | 0.672 | 0.473 | 0 | 1 | 0.769 | 1 | 1 | -0.063 |
| A3SS | 12707 | ENSG00000151923 | TIAL1 | chr10 | - | 119580470 | 119580846 | 119580470 | 119580538 | 119581921 | 119582009 | 125 | 151 | 129 | 130 | 166 | 21 | 7 | 13 | 21 | 41 | 61 | 175 | 263 | 95 | 45 | 12 | 9 | 12 | 9 | 10 | 298 | 149 | 0.085703 | 0.399934605 | 0.749 | 0.915 | 0.832 | 0.756 | 0.669 | 0.718 | 0.907 | 0.916 | 0.841 | 0.692 | -0.031 |
| A3SS | 12874 | ENSG00000105221 | AKT2 | chr19 | - | 40240044 | 40240121 | 40240044 | 40240110 | 40241937 | 40242069 | 5 | 0 | 6 | 15 | 22 | 294 | 210 | 324 | 334 | 284 | 0 | 17 | 8 | 11 | 5 | 380 | 442 | 200 | 332 | 501 | 160 | 149 | 0.085714 | 0.399934605 | 0.016 | 0 | 0.017 | 0.04 | 0.067 | 0 | 0.035 | 0.036 | 0.03 | 0.009 | 0.006 |
| A3SS | 7822 | ENSG00000112242 | E2F3 | chr6 | + | 20481205 | 20481425 | 20481223 | 20481425 | 20479845 | 20479957 | 58 | 39 | 51 | 81 | 81 | 1 | 10 | 19 | 12 | 14 | 69 | 109 | 74 | 116 | 76 | 16 | 7 | 12 | 9 | 6 | 167 | 149 | 0.085822 | 0.400203733 | 0.981 | 0.777 | 0.705 | 0.858 | 0.838 | 0.794 | 0.933 | 0.846 | 0.92 | 0.919 | -0.051 |
| A3SS | 4010 | ENSG00000055955 | ITIH4 | chr3 | - | 52819753 | 52819807 | 52819753 | 52819792 | 52820290 | 52820317 | 8 | 2 | 8 | 2 | 12 | 0 | 0 | 0 | 0 | 0 | 22 | 4 | 5 | 8 | 5 | 0 | 0 | 0 | 8 | 0 | 164 | 149 | 0.086046 | 0.401016801 | 1 | 1 | 1 | 1 | 1 | 1 | 1 | 1 | 0.476 | 1 | 0.105 |
| A3SS | 9372 | ENSG00000112511 | PHF1 | chr6 | + | 33414954 | 33415144 | 33414965 | 33415144 | 33414724 | 33414829 | 77 | 53 | 73 | 58 | 108 | 48 | 21 | 62 | 39 | 70 | 60 | 87 | 31 | 34 | 40 | 43 | 40 | 44 | 35 | 58 | 160 | 149 | 0.086286 | 0.401902234 | 0.599 | 0.702 | 0.523 | 0.581 | 0.59 | 0.565 | 0.669 | 0.396 | 0.475 | 0.391 | 0.1 |
| A3SS | 3143 | ENSG00000043143 | JADE2 | chr5 | + | 134578493 | 134579317 | 134578496 | 134579317 | 134576767 | 134576896 | 80 | 47 | 74 | 84 | 164 | 64 | 16 | 41 | 62 | 81 | 47 | 108 | 37 | 44 | 87 | 6 | 24 | 26 | 21 | 58 | 152 | 149 | 0.086507 | 0.402701766 | 0.551 | 0.742 | 0.639 | 0.57 | 0.665 | 0.885 | 0.815 | 0.582 | 0.673 | 0.595 | -0.077 |
| A3SS | 2352 | ENSG00000167491 | GATAD2A | chr19 | + | 19465339 | 19465614 | 19465344 | 19465614 | 19427695 | 19427992 | 3 | 10 | 13 | 12 | 5 | 0 | 3 | 0 | 1 | 0 | 40 | 23 | 14 | 14 | 20 | 7 | 2 | 0 | 0 | 0 | 154 | 149 | 0.086651 | 0.403137567 | 1 | 0.763 | 1 | 0.921 | 1 | 0.847 | 0.918 | 1 | 1 | 1 | -0.016 |
| A3SS | 836 | ENSG00000125991 | ERGIC3 | chr20 | + | 35556876 | 35557109 | 35556972 | 35557109 | 35556206 | 35556271 | 10 | 6 | 6 | 11 | 10 | 265 | 138 | 248 | 233 | 366 | 6 | 4 | 7 | 0 | 7 | 262 | 208 | 136 | 154 | 269 | 245 | 149 | 0.086824 | 0.403713498 | 0.022 | 0.026 | 0.015 | 0.028 | 0.016 | 0.014 | 0.012 | 0.03 | 0 | 0.016 | 0.007 |
| A3SS | 3375 | ENSG00000105223 | PLD3 | chr19 | + | 40366383 | 40366510 | 40366418 | 40366510 | 40365717 | 40365930 | 3 | 2 | 4 | 2 | 0 | 24 | 20 | 15 | 25 | 15 | 0 | 4 | 2 | 3 | 0 | 65 | 58 | 16 | 30 | 46 | 184 | 149 | 0.086888 | 0.403775149 | 0.092 | 0.075 | 0.178 | 0.061 | 0 | 0 | 0.053 | 0.092 | 0.075 | 0 | 0.037 |
| A3SS | 11325 | ENSG00000127084 | FGD3 | chr9 | + | 93034540 | 93034681 | 93034543 | 93034681 | 93032768 | 93032873 | 104 | 47 | 81 | 72 | 46 | 17 | 13 | 25 | 20 | 12 | 71 | 88 | 57 | 65 | 73 | 16 | 33 | 2 | 4 | 13 | 152 | 149 | 0.087315 | 0.405525457 | 0.857 | 0.78 | 0.761 | 0.779 | 0.79 | 0.813 | 0.723 | 0.965 | 0.941 | 0.846 | -0.064 |
| A3SS | 4084 | ENSG00000108509 | CAMTA2 | chr17 | - | 4981677 | 4981831 | 4981677 | 4981742 | 4982756 | 4982892 | 35 | 23 | 33 | 32 | 14 | 8 | 5 | 0 | 0 | 9 | 12 | 52 | 23 | 24 | 61 | 0 | 0 | 4 | 0 | 9 | 238 | 149 | 0.087379 | 0.405590705 | 0.733 | 0.742 | 1 | 1 | 0.493 | 1 | 1 | 0.783 | 1 | 0.809 | -0.125 |
| A3SS | 9087 | ENSG00000114388 | NPRL2 | chr3 | - | 50349664 | 50349833 | 50349664 | 50349828 | 50349930 | 50350022 | 72 | 28 | 50 | 77 | 43 | 24 | 9 | 18 | 13 | 21 | 55 | 59 | 28 | 24 | 52 | 12 | 20 | 15 | 15 | 46 | 154 | 149 | 0.087597 | 0.406369476 | 0.744 | 0.751 | 0.729 | 0.851 | 0.665 | 0.816 | 0.741 | 0.644 | 0.608 | 0.522 | 0.082 |
| A3SS | 2893 | ENSG00000006744 | ELAC2 | chr17 | - | 13004988 | 13005110 | 13004988 | 13005101 | 13005752 | 13005825 | 4 | 0 | 0 | 1 | 0 | 59 | 20 | 54 | 41 | 85 | 0 | 0 | 0 | 7 | 2 | 38 | 19 | 20 | 29 | 32 | 158 | 149 | 0.088080 | 0.408043024 | 0.06 | 0 | 0 | 0.022 | 0 | 0 | 0 | 0 | 0.185 | 0.056 | -0.032 |
| A3SS | 10792 | ENSG00000168066 | SF1 | chr11 | - | 64764605 | 64766155 | 64764605 | 64765518 | 64767191 | 64767251 | 133 | 75 | 159 | 105 | 101 | 2 | 1 | 5 | 7 | 0 | 225 | 228 | 135 | 166 | 108 | 0 | 3 | 6 | 2 | 0 | 298 | 149 | 0.088081 | 0.408043024 | 0.971 | 0.974 | 0.941 | 0.882 | 1 | 1 | 0.974 | 0.918 | 0.976 | 1 | -0.02 |
| A3SS | 5142 | ENSG00000064687 | ABCA7 | chr19 | + | 1055082 | 1055351 | 1055096 | 1055351 | 1054779 | 1054878 | 2 | 0 | 0 | 0 | 0 | 14 | 4 | 7 | 11 | 5 | 0 | 5 | 0 | 0 | 3 | 4 | 18 | 5 | 7 | 45 | 163 | 149 | 0.088109 | 0.408043024 | 0.116 | 0 | 0 | 0 | 0 | 0 | 0.203 | 0 | 0 | 0.057 | -0.029 |
| A3SS | 5233 | ENSG00000117640 | MTFR1L | chr1 | + | 25829508 | 25829830 | 25829544 | 25829830 | 25826614 | 25826826 | 60 | 41 | 58 | 68 | 63 | 6 | 3 | 1 | 1 | 0 | 93 | 90 | 19 | 35 | 152 | 0 | 0 | 0 | 3 | 4 | 185 | 149 | 0.088260 | 0.408506667 | 0.89 | 0.917 | 0.979 | 0.982 | 1 | 1 | 1 | 1 | 0.904 | 0.968 | -0.021 |
| A3SS | 10498 | ENSG00000072778 | ACADVL | chr17 | + | 7224479 | 7224552 | 7224483 | 7224552 | 7224320 | 7224393 | 184 | 83 | 114 | 152 | 63 | 23 | 8 | 10 | 13 | 2 | 130 | 276 | 135 | 192 | 300 | 21 | 7 | 19 | 6 | 10 | 153 | 149 | 0.088382 | 0.40883705 | 0.886 | 0.91 | 0.917 | 0.919 | 0.968 | 0.858 | 0.975 | 0.874 | 0.969 | 0.967 | -0.009 |
| A3SS | 9243 | ENSG00000213676 | ATF6B | chr6 | - | 32117036 | 32117130 | 32117036 | 32117107 | 32117322 | 32117404 | 16 | 8 | 19 | 11 | 22 | 160 | 75 | 126 | 131 | 97 | 14 | 22 | 18 | 7 | 32 | 207 | 177 | 106 | 155 | 350 | 172 | 149 | 0.088554 | 0.409397593 | 0.08 | 0.085 | 0.116 | 0.068 | 0.164 | 0.055 | 0.097 | 0.128 | 0.038 | 0.073 | 0.024 |
| A3SS | 9065 | ENSG00000090975 | PITPNM2 | chr12 | - | 123004393 | 123004603 | 123004393 | 123004489 | 123005239 | 123005548 | 1 | 2 | 0 | 0 | 0 | 27 | 16 | 28 | 32 | 14 | 0 | 1 | 4 | 0 | 12 | 46 | 42 | 25 | 25 | 243 | 263 | 149 | 0.088614 | 0.409442477 | 0.021 | 0.066 | 0 | 0 | 0 | 0 | 0.013 | 0.083 | 0 | 0.027 | -0.007 |
| A3SS | 5466 | ENSG00000171853 | TRAPPC12 | chr2 | + | 3477691 | 3477795 | 3477694 | 3477795 | 3465596 | 3465695 | 4 | 2 | 3 | 3 | 3 | 79 | 84 | 86 | 126 | 104 | 17 | 5 | 0 | 12 | 0 | 224 | 88 | 44 | 121 | 245 | 152 | 149 | 0.088749 | 0.409831638 | 0.047 | 0.023 | 0.033 | 0.023 | 0.027 | 0.069 | 0.053 | 0 | 0.089 | 0 | -0.012 |
| A3SS | 8042 | ENSG00000138795 | LEF1 | chr4 | - | 108063622 | 108063663 | 108063622 | 108063657 | 108064335 | 108064384 | 319 | 353 | 861 | 560 | 1549 | 10 | 7 | 15 | 17 | 25 | 66 | 173 | 212 | 148 | 44 | 0 | 0 | 4 | 0 | 7 | 155 | 149 | 0.089057 | 0.411015723 | 0.968 | 0.98 | 0.982 | 0.969 | 0.983 | 1 | 1 | 0.981 | 1 | 0.858 | 0.009 |
| A3SS | 9285 | ENSG00000132424 | PNISR | chr6 | - | 99414571 | 99414690 | 99414571 | 99414686 | 99416348 | 99416428 | 390 | 416 | 416 | 478 | 837 | 69 | 75 | 106 | 82 | 148 | 201 | 197 | 238 | 154 | 125 | 43 | 41 | 35 | 53 | 42 | 153 | 149 | 0.089895 | 0.414645713 | 0.846 | 0.844 | 0.793 | 0.85 | 0.846 | 0.82 | 0.824 | 0.869 | 0.739 | 0.743 | 0.037 |
| A3SS | 4815 | ENSG00000148843 | PDCD11 | chr10 | + | 103442209 | 103442460 | 103442212 | 103442460 | 103441825 | 103441975 | 5 | 9 | 16 | 13 | 5 | 9 | 3 | 10 | 19 | 12 | 0 | 1 | 13 | 4 | 9 | 42 | 3 | 6 | 5 | 15 | 152 | 149 | 0.090004 | 0.414911413 | 0.353 | 0.746 | 0.611 | 0.401 | 0.29 | 0 | 0.246 | 0.68 | 0.44 | 0.37 | 0.133 |
| A3SS | 8189 | ENSG00000084623 | EIF3I | chr1 | + | 32231114 | 32231604 | 32231120 | 32231604 | 32230926 | 32231019 | 104 | 67 | 136 | 151 | 110 | 0 | 0 | 0 | 0 | 0 | 184 | 111 | 90 | 122 | 279 | 13 | 0 | 5 | 0 | 0 | 155 | 149 | 0.090181 | 0.415493173 | 1 | 1 | 1 | 1 | 1 | 0.932 | 1 | 0.945 | 1 | 1 | 0.025 |
| A3SS | 2832 | ENSG00000119688 | ABCD4 | chr14 | - | 74293153 | 74293279 | 74293153 | 74293248 | 74295147 | 74295198 | 2 | 0 | 2 | 2 | 4 | 11 | 5 | 14 | 31 | 23 | 0 | 1 | 0 | 2 | 2 | 7 | 6 | 4 | 21 | 18 | 180 | 149 | 0.090380 | 0.416172332 | 0.131 | 0 | 0.106 | 0.051 | 0.126 | 0 | 0.121 | 0 | 0.073 | 0.084 | 0.027 |
| A3SS | 5193 | ENSG00000269893 | SNHG8 | chr4 | + | 118278933 | 118279137 | 118279106 | 118279137 | 118278754 | 118278792 | 156 | 57 | 230 | 156 | 161 | 11 | 2 | 6 | 10 | 0 | 64 | 126 | 116 | 72 | 123 | 0 | 0 | 5 | 2 | 6 | 298 | 149 | 0.090607 | 0.416979238 | 0.876 | 0.934 | 0.95 | 0.886 | 1 | 1 | 1 | 0.921 | 0.947 | 0.911 | -0.027 |
| A3SS | 5634 | ENSG00000101940 | WDR13 | chrX | + | 48598716 | 48598957 | 48598750 | 48598957 | 48597957 | 48598037 | 38 | 8 | 21 | 40 | 18 | 4 | 0 | 2 | 6 | 12 | 34 | 28 | 24 | 27 | 74 | 0 | 4 | 1 | 0 | 13 | 183 | 149 | 0.090683 | 0.417091948 | 0.886 | 1 | 0.895 | 0.844 | 0.55 | 1 | 0.851 | 0.951 | 1 | 0.823 | -0.09 |
| A3SS | 168 | ENSG00000100307 | CBX7 | chr22 | - | 39134400 | 39134752 | 39134400 | 39134473 | 39138635 | 39138702 | 119 | 44 | 101 | 77 | 61 | 11 | 2 | 6 | 4 | 8 | 8 | 75 | 47 | 22 | 118 | 0 | 8 | 0 | 3 | 2 | 298 | 149 | 0.090902 | 0.417861548 | 0.844 | 0.917 | 0.894 | 0.906 | 0.792 | 1 | 0.824 | 1 | 0.786 | 0.967 | -0.045 |
| A3SS | 4543 | ENSG00000076108 | BAZ2A | chr12 | - | 56600942 | 56601098 | 56600942 | 56601086 | 56601179 | 56601402 | 179 | 79 | 185 | 209 | 148 | 8 | 4 | 2 | 2 | 2 | 385 | 362 | 236 | 349 | 449 | 2 | 0 | 5 | 3 | 8 | 161 | 149 | 0.091289 | 0.419401718 | 0.954 | 0.948 | 0.988 | 0.99 | 0.986 | 0.994 | 1 | 0.978 | 0.991 | 0.981 | -0.016 |
| A3SS | 9189 | ENSG00000197081 | IGF2R | chr6 | + | 160088032 | 160089253 | 160089106 | 160089253 | 160084994 | 160085131 | 356 | 159 | 228 | 262 | 226 | 0 | 0 | 2 | 2 | 1 | 303 | 277 | 335 | 253 | 354 | 0 | 8 | 4 | 6 | 0 | 298 | 149 | 0.091630 | 0.420662983 | 1 | 1 | 0.983 | 0.985 | 0.991 | 1 | 0.945 | 0.977 | 0.955 | 1 | 0.016 |
| A3SS | 7456 | ENSG00000162373 | BEND5 | chr1 | - | 48758899 | 48759366 | 48758899 | 48759284 | 48761336 | 48761470 | 8 | 5 | 8 | 4 | 0 | 2 | 4 | 10 | 6 | 4 | 8 | 5 | 2 | 8 | 0 | 0 | 13 | 0 | 1 | 2 | 231 | 149 | 0.091678 | 0.420662983 | 0.721 | 0.446 | 0.34 | 0.301 | 0 | 1 | 0.199 | 1 | 0.838 | 0 | -0.246 |
| A3SS | 3325 | ENSG00000155660 | PDIA4 | chr7 | - | 149005896 | 149006056 | 149005896 | 149006053 | 149008158 | 149008310 | 17 | 12 | 27 | 12 | 8 | 302 | 144 | 390 | 346 | 415 | 0 | 9 | 13 | 6 | 20 | 348 | 288 | 301 | 173 | 259 | 152 | 149 | 0.091747 | 0.420662983 | 0.052 | 0.076 | 0.064 | 0.033 | 0.019 | 0 | 0.03 | 0.041 | 0.033 | 0.07 | 0.014 |
| A3SS | 2315 | ENSG00000184508 | HDDC3 | chr15 | - | 90930924 | 90931556 | 90930924 | 90931405 | 90931703 | 90931944 | 14 | 16 | 23 | 31 | 18 | 54 | 20 | 19 | 24 | 14 | 5 | 17 | 15 | 12 | 26 | 7 | 33 | 14 | 26 | 67 | 298 | 149 | 0.091772 | 0.420662983 | 0.115 | 0.286 | 0.377 | 0.392 | 0.391 | 0.263 | 0.205 | 0.349 | 0.188 | 0.163 | 0.079 |
| A3SS | 6717 | ENSG00000183479 | TREX2 | chrX | - | 153455541 | 153455766 | 153455541 | 153455648 | 153456264 | 153456364 | 33 | 8 | 16 | 20 | 14 | 2 | 2 | 1 | 1 | 1 | 20 | 49 | 5 | 22 | 56 | 0 | 0 | 0 | 0 | 0 | 267 | 149 | 0.092186 | 0.422320454 | 0.902 | 0.691 | 0.899 | 0.918 | 0.887 | 1 | 1 | 1 | 1 | 1 | -0.141 |
| A3SS | 3472 | ENSG00000189067 | LITAF | chr16 | - | 11553532 | 11553748 | 11553532 | 11553689 | 11556510 | 11556735 | 10 | 1 | 3 | 5 | 4 | 225 | 61 | 200 | 190 | 129 | 0 | 6 | 4 | 2 | 0 | 99 | 182 | 246 | 184 | 155 | 208 | 149 | 0.092922 | 0.425447454 | 0.031 | 0.012 | 0.011 | 0.019 | 0.022 | 0 | 0.023 | 0.012 | 0.008 | 0 | 0.01 |
| A3SS | 11383 | ENSG00000124444 | ZNF576 | chr19 | + | 43597093 | 43597193 | 43597096 | 43597193 | 43596594 | 43596743 | 27 | 21 | 18 | 26 | 53 | 4 | 0 | 4 | 0 | 17 | 0 | 5 | 6 | 8 | 17 | 5 | 1 | 0 | 0 | 0 | 152 | 149 | 0.093066 | 0.425865349 | 0.869 | 1 | 0.815 | 1 | 0.753 | 0 | 0.831 | 1 | 1 | 1 | 0.121 |
| A3SS | 5641 | ENSG00000026508 | CD44 | chr11 | + | [illegible] | 35206243 | 35206114 | 35206243 | 35204511 | 35204640 | 3 | 8 | 4 | 5 | 8 | 1 | 0 | 2 | 0 | 3 | 26 | 4 | 2 | 18 | 5 | 7 | 0 | 0 | 0 | 0 | 152 | 149 | 0.093394 | 0.427123963 | 0.746 | 1 | 0.662 | 1 | 0.723 | 0.785 | 1 | 1 | 1 | 1 | -0.131 |
| A3SS | 6059 | ENSG00000196204 | RNF216P1 | chr7 | + | 4984909 | 4985075 | 4984961 | 4985075 | 4983670 | 4983773 | 108 | 93 | 105 | 103 | 217 | 9 | 6 | 14 | 17 | 9 | 114 | 39 | 74 | 50 | 79 | 19 | 60 | 10 | 7 | 0 | 201 | 149 | 0.094103 | 0.429886424 | 0.899 | 0.92 | 0.848 | 0.818 | 0.947 | 0.816 | 0.325 | 0.846 | 0.841 | 1 | 0.121 |
| A3SS | 10931 | ENSG00000094914 | AAAS | chr12 | - | 53315334 | 53315486 | 53315334 | 53315426 | 53315726 | 53315782 | 3 | 1 | 2 | 2 | 5 | 81 | 46 | 89 | 72 | 124 | 14 | 2 | 2 | 1 | 0 | 55 | 60 | 46 | 81 | 77 | 209 | 149 | 0.094113 | 0.429886424 | 0.026 | 0.015 | 0.016 | 0.019 | 0.028 | 0.154 | 0.023 | 0.03 | 0.009 | 0 | -0.022 |
| A3SS | 8780 | ENSG00000115053 | NCL | chr2 | - | 231458265 | 231458389 | 231458265 | 231458389 | 231459000 | 231459125 | 61 | 66 | 72 | 103 | 52 | 1841 | 1584 | 1802 | 1962 | 2837 | 28 | 79 | 54 | 48 | 0 | 1511 | 1478 | 1675 | 1874 | 325 | 298 | 149 | 0.094157 | 0.429886424 | 0.016 | 0.02 | 0.02 | 0.026 | 0.009 | 0.009 | 0.026 | 0.016 | 0.013 | 0 | 0.005 |
| A3SS | 541 | ENSG00000006712 | PAF1 | chr19 | - | 39390068 | 39390289 | 39390068 | 39390161 | 39390817 | 39391154 | 42 | 20 | 47 | 50 | 35 | 0 | 0 | 0 | 0 | 0 | 50 | 58 | 43 | 74 | 0 | 1 | 1 | 0 | 0 | 7 | 277 | 149 | 0.094371 | 0.430617933 | 1 | 1 | 1 | 1 | 1 | 1 | 0.969 | 0.959 | 1 | 0.85 | 0.044 |
| A3SS | 8700 | ENSG00000165280 | VCP | chr9 | - | 35067890 | 35068084 | 35067890 | 35068063 | 35068250 | 35068362 | 0 | 2 | 1 | 4 | 0 | 131 | 63 | 116 | 109 | 145 | 7 | 0 | 3 | 4 | 5 | 170 | 124 | 91 | 139 | 202 | 170 | 149 | 0.094534 | 0.431117191 | 0 | 0.027 | 0.007 | 0.031 | 0 | 0.035 | 0 | 0.028 | 0.025 | 0.021 | -0.009 |
| A3SS | 11682 | ENSG00000165140 | FBP1 | chr9 | - | 94606814 | 94607112 | 94606814 | 94606952 | 94609920 | 94610061 | 2 | 1 | 5 | 3 | 2 | 162 | 98 | 83 | 76 | 181 | 18 | 0 | 15 | 2 | 17 | 366 | 391 | 207 | 145 | 523 | 298 | 149 | 0.095253 | 0.434152252 | 0.006 | 0.005 | 0.029 | 0.019 | 0.005 | 0.024 | 0 | 0.035 | 0.007 | 0.016 | -0.004 |
| A3SS | 9581 | ENSG00000177156 | TALDO1 | chr11 | + | 758949 | 759057 | 759008 | 759057 | 755878 | 756002 | 642 | 344 | 438 | 560 | 614 | 2 | 6 | 2 | 3 | 2 | 1550 | 800 | 572 | 1341 | 2267 | 10 | 10 | 11 | 19 | 16 | 208 | 149 | 0.095389 | 0.434524231 | 0.996 | 0.976 | 0.994 | 0.993 | 0.995 | 0.991 | 0.983 | 0.974 | 0.981 | 0.99 | 0.007 |
| A3SS | 1341 | ENSG00000011143 | MKS1 | chr17 | - | 58214740 | 58214898 | 58214740 | 58214838 | 58216087 | 58216243 | 1 | 1 | 1 | 0 | 9 | 20 | 4 | 17 | 15 | 20 | 9 | 0 | 2 | 0 | 12 | 0 | 13 | 3 | 4 | 11 | 209 | 149 | 0.095710 | 0.435741206 | 0.034 | 0.151 | 0.04 | 0 | 0.243 | 1 | 0 | 0.322 | 0 | 0.437 | -0.258 |
| A3SS | 7832 | ENSG00000131748 | STARD3 | chr17 | + | 39658708 | 39658820 | 39658721 | 39658820 | 39658404 | 39658522 | 0 | 2 | 2 | 0 | 4 | 70 | 23 | 81 | 60 | 69 | 5 | 0 | 5 | 0 | 6 | 66 | 83 | 66 | 68 | 89 | 162 | 149 | 0.096027 | 0.436938502 | 0 | 0.074 | 0.022 | 0 | 0.051 | 0.065 | 0 | 0.065 | 0 | 0.058 | -0.008 |
| A3SS | 8389 | ENSG00000198589 | LRBA | chr4 | - | 150321190 | 150321383 | 150321190 | 150321368 | 150325808 | 150325898 | 13 | 17 | 22 | 34 | 32 | 144 | 119 | 105 | 168 | 255 | 0 | 28 | 19 | 1 | 0 | 15 | 132 | 133 | 70 | 32 | 164 | 149 | 0.096097 | 0.437010371 | 0.076 | 0.115 | 0.16 | 0.155 | 0.102 | 0 | 0.162 | 0.115 | 0.013 | 0 | 0.064 |
| A3SS | 12116 | ENSG00000086848 | ALG9 | chr11 | - | 111838248 | 111838399 | 111838248 | 111838378 | 111844600 | 111844723 | 17 | 7 | 11 | 11 | 17 | 4 | 1 | 1 | 7 | 8 | 55 | 10 | 24 | 10 | 15 | 5 | 8 | 22 | 10 | 13 | 170 | 149 | 0.096259 | 0.43750242 | 0.788 | 0.86 | 0.906 | 0.579 | 0.651 | 0.906 | 0.523 | 0.489 | 0.467 | 0.503 | 0.179 |
| A3SS | 2818 | ENSG00000161558 | TMEM143 | chr19 | - | 48360071 | 48360183 | 48360071 | 48360176 | 48363290 | 48363674 | 3 | 1 | 2 | 4 | 0 | 8 | 3 | 11 | 4 | 2 | 9 | 0 | 0 | 1 | 0 | 5 | 20 | 10 | 7 | 11 | 156 | 149 | 0.096378 | 0.437795365 | 0.264 | 0.241 | 0.148 | 0.489 | 0 | 0.632 | 0 | 0 | 0.12 | 0 | 0.078 |
| A3SS | 12624 | ENSG00000107959 | PITRM1 | chr10 | - | 3159847 | 3160001 | 3159847 | 3159936 | 3160203 | 3160330 | 9 | 8 | 3 | 2 | 1 | 75 | 60 | 78 | 71 | 93 | 2 | 5 | 15 | 11 | 0 | 51 | 66 | 49 | 72 | 110 | 214 | 149 | 0.096551 | 0.438336746 | 0.077 | 0.085 | 0.026 | 0.019 | 0.007 | 0.027 | 0.05 | 0.176 | 0.096 | 0 | -0.027 |
| A3SS | 10921 | ENSG00000064115 | TM7SF3 | chr12 | - | 26999525 | 26999766 | 26999525 | 26999676 | 27003235 | 27003390 | 7 | 2 | 1 | 6 | 1 | 112 | 64 | 85 | 76 | 198 | 0 | 10 | 3 | 0 | 9 | 104 | 87 | 38 | 59 | 103 | 239 | 149 | 0.096650 | 0.438538941 | 0.038 | 0.019 | 0.007 | 0.047 | 0.003 | 0 | 0.067 | 0.047 | 0 | 0.052 | -0.01 |
| A3SS | 11676 | ENSG00000137073 | UBAP2 | chr9 | - | 33933489 | 33933646 | 33933489 | 33933628 | 33935838 | 33935878 | 1 | 0 | 1 | 0 | 1 | 40 | 24 | 40 | 30 | 66 | 0 | 0 | 6 | 6 | 0 | 55 | 62 | 50 | 83 | 58 | 167 | 149 | 0.096834 | 0.439127684 | 0.022 | 0 | 0.022 | 0 | 0.013 | 0 | 0 | 0.097 | 0.061 | 0 | -0.02 |
| A3SS | 11658 | ENSG00000130714 | POMT1 | chr9 | + | 131519083 | 131519486 | 131519388 | 131519486 | 131518836 | 131518957 | 0 | 0 | 3 | 10 | 3 | 33 | 20 | 33 | 37 | 32 | 0 | 0 | 6 | 3 | 16 | 21 | 24 | 25 | 26 | 43 | 298 | 149 | 0.097289 | 0.440502193 | 0 | 0 | 0.043 | 0.119 | 0.045 | 0 | 0 | 0.107 | 0.055 | 0.157 | -0.022 |
| A3SS | 12349 | ENSG00000149311 | ATM | chr11 | + | 108227775 | 108227888 | 108227779 | 108227888 | 108227594 | 108227696 | 785 | 547 | 732 | 695 | 1091 | 18 | 14 | 21 | 16 | 62 | 143 | 200 | 126 | 87 | 61 | 11 | 7 | 14 | 2 | 0 | 153 | 149 | 0.097305 | 0.440502193 | 0.977 | 0.974 | 0.971 | 0.977 | 0.945 | 0.927 | 0.965 | 0.898 | 0.977 | 1 | 0.015 |
| A3SS | 10758 | ENSG00000008838 | MED24 | chr17 | - | 40053468 | 40053657 | 40053468 | 40053635 | 40054360 | 40054405 | 21 | 13 | 21 | 10 | 14 | 3 | 13 | 11 | 2 | 14 | 28 | 25 | 3 | 18 | 33 | 16 | 6 | 3 | 1 | 3 | 171 | 149 | 0.097309 | 0.440502193 | 0.859 | 0.466 | 0.625 | 0.813 | 0.466 | 0.604 | 0.784 | 0.466 | 0.94 | 0.906 | -0.094 |
| A3SS | 3004 | ENSG00000093134 | VNN3 | chr6 | - | 132728740 | 132728979 | 132728740 | 132728933 | 132734252 | 132734383 | 5 | 7 | 11 | 13 | 14 | 4 | 3 | 6 | 9 | 11 | 0 | 17 | 75 | 13 | 3 | 27 | 8 | 44 | 7 | 8 | 195 | 149 | 0.097356 | 0.440502193 | 0.489 | 0.641 | 0.583 | 0.525 | 0.493 | 0 | 0.619 | 0.566 | 0.587 | 0.223 | 0.147 |
| A3SS | 10680 | ENSG00000071553 | ATP6AP1 | chrX | + | 154433590 | 154433720 | 154433634 | 154433720 | 154432930 | 154432971 | 1 | 2 | 0 | 2 | 0 | 102 | 33 | 69 | 98 | 57 | 0 | 14 | 4 | 1 | 0 | 146 | 121 | 70 | 106 | 212 | 193 | 149 | 0.097517 | 0.440985303 | 0.008 | 0.045 | 0 | 0.016 | 0 | 0 | 0.082 | 0.042 | 0.007 | 0 | -0.012 |
| A3SS | 10612 | ENSG00000085224 | ATRX | chrX | - | 77688817 | 77688927 | 77688817 | 77688924 | 77696576 | 77696704 | 27 | 27 | 25 | 25 | 53 | 35 | 24 | 26 | 41 | 49 | 32 | 13 | 3 | 0 | 0 | 0 | 2 | 0 | 5 | 2 | 152 | 149 | 0.098832 | 0.446681778 | 0.431 | 0.524 | 0.485 | 0.374 | 0.515 | 1 | 0.864 | 1 | 0 | 0 | -0.107 |
| A3SS | 11841 | ENSG00000113163 | CERT1 | chr5 | - | 75385901 | 75386057 | 75385901 | 75386034 | 75389591 | 75389687 | 1 | 3 | 2 | 4 | 8 | 124 | 111 | 135 | 133 | 156 | 0 | 0 | 1 | 0 | 0 | 49 | 89 | 113 | 81 | 59 | 172 | 149 | 0.098889 | 0.446687454 | 0.007 | 0.023 | 0.013 | 0.025 | 0.043 | 0 | 0 | 0.008 | 0 | 0 | 0.021 |
| A3SS | 12045 | ENSG00000149091 | DGKZ | chr11 | + | 46368001 | 46368079 | 46368004 | 46368079 | 46367651 | 46367747 | 140 | 51 | 73 | 99 | 70 | 53 | 40 | 41 | 65 | 51 | 109 | 123 | 47 | 44 | 178 | 24 | 63 | 28 | 26 | 80 | 152 | 149 | 0.098958 | 0.446749885 | 0.721 | 0.556 | 0.636 | 0.599 | 0.574 | 0.817 | 0.657 | 0.622 | 0.624 | 0.686 | -0.064 |
| A3SS | 3176 | ENSG00000111641 | NOP2 | chr12 | - | 6566100 | 6566336 | 6566100 | 6566324 | 6566528 | 6566617 | 101 | 72 | 87 | 84 | 83 | 49 | 38 | 44 | 44 | 19 | 115 | 56 | 75 | 81 | 150 | 33 | 29 | 18 | 37 | 40 | 161 | 149 | 0.099264 | 0.447879758 | 0.656 | 0.637 | 0.647 | 0.639 | 0.802 | 0.763 | 0.641 | 0.794 | 0.67 | 0.776 | -0.053 |
| A3SS | 3444 | ENSG00000112992 | NNT | chr5 | + | 43647923 | 43649308 | 43649146 | 43649308 | 43645356 | 43645510 | 20 | 19 | 14 | 6 | 14 | 100 | 43 | 60 | 79 | 102 | 1 | 18 | 18 | 6 | 10 | 114 | 86 | 63 | 64 | 61 | 298 | 149 | 0.099771 | 0.449902821 | 0.091 | 0.181 | 0.104 | 0.037 | 0.064 | 0.004 | 0.095 | 0.125 | 0.045 | 0.076 | 0.026 |
| A3SS | 7664 | ENSG00000135766 | EGLN1 | chr1 | - | 231367568 | 231368187 | 231367568 | 231367636 | 231370561 | 231370698 | 32 | 43 | 31 | 25 | 21 | 131 | 98 | 128 | 138 | 177 | 0 | 11 | 31 | 31 | 14 | 201 | 102 | 105 | 85 | 35 | 298 | 149 | 0.099824 | 0.449902821 | 0.109 | 0.18 | 0.108 | 0.083 | 0.056 | 0 | 0.051 | 0.129 | 0.154 | 0.167 | 0.007 |
| A3SS | 2177 | ENSG00000260916 | CCPG1 | chr15 | - | 55371792 | 55372044 | 55371792 | 55372030 | 55376948 | 55377150 | 264 | 153 | 105 | 157 | 268 | 4 | 0 | 1 | 1 | 0 | 39 | 88 | 42 | 57 | 73 | 0 | 0 | 0 | 11 | 0 | 163 | 149 | 0.099918 | 0.450076613 | 0.984 | 1 | 0.99 | 0.993 | 1 | 1 | 1 | 1 | 0.826 | 1 | 0.028 |
| A3SS | 6190 | ENSG00000107551 | RASSF4 | chr10 | + | 44982520 | 44982663 | 44982524 | 44982663 | 44971772 | 44971848 | 134 | 57 | 74 | 98 | 71 | 0 | 2 | 0 | 1 | 2 | 222 | 118 | 54 | 91 | 158 | 0 | 4 | 3 | 0 | 9 | 153 | 149 | 0.100020 | 0.450284324 | 1 | 0.965 | 1 | 0.99 | 0.972 | 1 | 0.966 | 0.946 | 1 | 0.945 | 0.014 |
| A3SS | 3042 | ENSG00000068796 | KIF2A | chr5 | + | 62353256 | 62353375 | 62353274 | 62353375 | 62352587 | 62352710 | 4 | 6 | 2 | 0 | 14 | 189 | 213 | 115 | 315 | 369 | 0 | 0 | 0 | 0 | 13 | 318 | 243 | 215 | 178 | 446 | 167 | 149 | 0.100231 | 0.450811101 | 0.019 | 0.025 | 0.015 | 0 | 0.033 | 0 | 0 | 0 | 0 | 0.025 | 0.013 |
| A3SS | 5194 | ENSG00000269893 | SNHG8 | chr4 | + | 118278943 | 118279137 | 118279106 | 118279137 | 118278754 | 118278792 | 163 | 60 | 238 | 181 | 164 | 11 | 2 | 6 | 10 | 0 | 65 | 130 | 120 | 79 | 127 | 0 | 0 | 5 | 2 | 6 | 298 | 149 | 0.100249 | 0.450811101 | 0.881 | 0.938 | 0.952 | 0.9 | 1 | 1 | 1 | 0.923 | 0.952 | 0.914 | -0.024 |
| A3SS | 8304 | ENSG00000175215 | CTDSP2 | chr12 | - | 57826995 | 57827123 | 57826995 | 57827097 | 57827551 | 57827590 | 48 | 40 | 43 | 52 | 44 | 304 | 136 | 231 | 246 | 113 | 71 | 64 | 40 | 53 | 129 | 241 | 220 | 144 | 147 | 562 | 175 | 149 | 0.100409 | 0.451243169 | 0.119 | 0.2 | 0.137 | 0.153 | 0.249 | 0.201 | 0.199 | 0.191 | 0.235 | 0.163 | -0.026 |
| A3SS | 4620 | ENSG00000185507 | IRF7 | chr11 | - | 614475 | 614534 | 614475 | 614531 | 614796 | 615007 | 30 | 10 | 31 | 59 | 3 | 5 | 2 | 9 | 7 | 0 | 27 | 57 | 14 | 63 | 58 | 0 | 4 | 3 | 12 | 2 | 152 | 149 | 0.100485 | 0.451243169 | 0.855 | 0.831 | 0.772 | 0.892 | 1 | 1 | 0.933 | 0.821 | 0.837 | 0.966 | -0.041 |
| A3SS | 7865 | ENSG00000032444 | PNPLA6 | chr19 | + | 7535788 | 7536020 | 7535932 | 7536020 | 7535508 | 7535577 | 27 | 22 | 37 | 24 | 26 | 4 | 2 | 5 | 8 | 3 | 34 | 73 | 30 | 33 | 158 | 22 | 2 | 0 | 1 | 8 | 293 | 149 | 0.100531 | 0.451243169 | 0.774 | 0.848 | 0.79 | 0.604 | 0.815 | 0.44 | 0.949 | 1 | 0.944 | 0.909 | -0.082 |
| A3SS | 3075 | ENSG00000124784 | RIOK1 | chr6 | + | 7395020 | 7395143 | 7395052 | 7395143 | 7393098 | 7393303 | 16 | 17 | 18 | 20 | 11 | 64 | 45 | 56 | 55 | 65 | 25 | 25 | 3 | 9 | 6 | 12 | 35 | 36 | 34 | 6 | 181 | 149 | 0.100568 | 0.451243169 | 0.171 | 0.237 | 0.209 | 0.23 | 0.122 | 0.632 | 0.37 | 0.064 | 0.179 | 0.452 | -0.146 |
| A3SS | 6249 | ENSG00000015153 | YAF2 | chr12 | - | 42210547 | 42210680 | 42210547 | 42210619 | 42237598 | 42237724 | 8 | 9 | 3 | 6 | 6 | 1 | 0 | 0 | 4 | 2 | 8 | 3 | 10 | 12 | 2 | 0 | 0 | 0 | 4 | 0 | 210 | 149 | 0.100746 | 0.451788213 | 0.85 | 1 | 1 | 0.516 | 0.68 | 1 | 1 | 1 | 0.68 | 1 | -0.127 |
| A3SS | 8408 | ENSG00000140368 | PSTPIP1 | chr15 | + | 77027745 | 77027914 | 77027799 | 77027914 | 77025497 | 77025604 | 8 | 6 | 3 | 5 | 0 | 16 | 19 | 29 | 21 | 7 | 16 | 7 | 8 | 4 | 9 | 18 | 33 | 15 | 5 | 50 | 203 | 149 | 0.101270 | 0.453889001 | 0.268 | 0.188 | 0.071 | 0.149 | 0 | 0.395 | 0.135 | 0.281 | 0.37 | 0.117 | -0.124 |
| A3SS | 11700 | ENSG00000188566 | NDOR1 | chr9 | + | 137216066 | 137216188 | 137216093 | 137216188 | 137215898 | 137216017 | 3 | 0 | 0 | 0 | 4 | 15 | 2 | 16 | 16 | 3 | 29 | 0 | 0 | 0 | 9 | 23 | 9 | 13 | 12 | 29 | 176 | 149 | 0.101534 | 0.454817378 | 0.145 | 0 | 0 | 0 | 0.53 | 0.516 | 0 | 0 | 0 | 0.208 | -0.01 |
| A3SS | 3354 | ENSG00000114107 | CEP70 | chr3 | - | 138572858 | 138572984 | 138572858 | 138572914 | 138591853 | 138591956 | 26 | 22 | 14 | 23 | 50 | 1 | 2 | 0 | 6 | 0 | 14 | 9 | 3 | 5 | 1 | 0 | 0 | 0 | 3 | 0 | 219 | 149 | 0.102417 | 0.458520698 | 0.946 | 0.882 | 1 | 0.723 | 1 | 1 | 1 | 1 | 0.531 | 1 | 0.004 |
| A3SS | 4142 | ENSG00000112701 | SENP6 | chr6 | + | 75640683 | 75640704 | 75640686 | 75640704 | 75634706 | 75634811 | 65 | 48 | 50 | 57 | 78 | 63 | 60 | 44 | 47 | 107 | 0 | 18 | 14 | 10 | 11 | 10 | 25 | 23 | 25 | 6 | 152 | 149 | 0.102731 | 0.459664534 | 0.503 | 0.44 | 0.527 | 0.543 | 0.417 | 0 | 0.414 | 0.374 | 0.282 | 0.642 | 0.144 |
| A3SS | 7851 | ENSG00000115464 | USP34 | chr2 | - | 61214059 | 61214694 | 61214059 | 61214218 | 61220309 | 61220457 | 419 | 310 | 326 | 427 | 516 | 4 | 0 | 9 | 6 | 2 | 66 | 226 | 267 | 52 | 15 | 0 | 0 | 0 | 1 | 0 | 298 | 149 | 0.102828 | 0.459664534 | 0.981 | 1 | 0.948 | 0.973 | 0.992 | 1 | 1 | 1 | 0.963 | 1 | -0.014 |
| A3SS | 4028 | ENSG00000132466 | ANKRD17 | chr4 | - | 73135116 | 73135265 | 73135116 | 73135262 | 73141740 | 73141843 | 40 | 32 | 32 | 45 | 24 | 6 | 0 | 3 | 7 | 2 | 41 | 69 | 47 | 49 | 3 | 8 | 1 | 1 | 2 | 0 | 152 | 149 | 0.102844 | 0.459664534 | 0.867 | 1 | 0.913 | 0.863 | 0.922 | 0.834 | 0.985 | 0.979 | 0.96 | 1 | -0.039 |
| A3SS | 4068 | ENSG00000017260 | ATP2C1 | chr3 | + | 130996679 | 130996796 | 130996752 | 130996796 | 130996042 | 130996111 | 191 | 190 | 213 | 199 | 301 | 1 | 0 | 1 | 4 | 3 | 98 | 133 | 145 | 84 | 64 | 0 | 6 | 6 | 0 | 0 | 222 | 149 | 0.102996 | 0.460089609 | 0.992 | 1 | 0.993 | 0.971 | 0.985 | 1 | 0.937 | 0.942 | 1 | 1 | 0.012 |
| A3SS | 13235 | ENSG00000004700 | RECQL | chr12 | - | 21501169 | 21501372 | 21501169 | 21501215 | 21501462 | 21501669 | 106 | 72 | 93 | 94 | 50 | 9 | 4 | 5 | 24 | 4 | 18 | 39 | 33 | 40 | 18 | 0 | 14 | 1 | 0 | 0 | 298 | 149 | 0.103176 | 0.460641366 | 0.855 | 0.9 | 0.903 | 0.662 | 0.862 | 1 | 0.582 | 0.943 | 1 | 1 | -0.069 |
| A3SS | 7707 | ENSG00000120265 | PCMT1 | chr6 | + | 149772013 | 149772247 | 149772047 | 149772247 | 149771161 | 149771266 | 8 | 4 | 5 | 3 | 3 | 1 | 3 | 3 | 2 | 2 | 12 | 8 | 4 | 10 | 10 | 0 | 0 | 8 | 4 | 0 | 183 | 149 | 0.103607 | 0.46230584 | 0.867 | 0.521 | 0.576 | 0.55 | 0.55 | 1 | 1 | 0.289 | 0.671 | 1 | -0.179 |
| A3SS | 9870 | ENSG00000121716 | PILRB | chr7 | + | 100358689 | 100359008 | 100358932 | 100359008 | 100358219 | 100358366 | 272 | 123 | 217 | 227 | 259 | 3 | 2 | 6 | 4 | 0 | 186 | 328 | 164 | 298 | 235 | 0 | 2 | 3 | 3 | 0 | 298 | 149 | 0.103664 | 0.46230584 | 0.978 | 0.969 | 0.948 | 0.966 | 1 | 1 | 0.988 | 0.965 | 0.98 | 1 | -0.014 |
| A3SS | 5851 | ENSG00000198301 | SDAD1 | chr4 | - | 75981371 | 75981514 | 75981371 | 75981470 | 75981932 | 75982037 | 2 | 2 | 3 | 13 | 22 | 55 | 40 | 74 | 60 | 91 | 0 | 2 | 7 | 0 | 1 | 32 | 26 | 20 | 13 | 0 | 193 | 149 | 0.103753 | 0.462450707 | 0.027 | 0.037 | 0.03 | 0.143 | 0.157 | 0 | 0.056 | 0.213 | 0 | 1 | -0.175 |
| A3SS | 100 | ENSG00000184702 | SEPTIN5 | chr22 | + | 19721821 | 19721957 | 19721832 | 19721957 | 19721639 | 19721736 | 57 | 61 | 22 | 178 | 49 | 2 | 7 | 0 | 13 | 14 | 246 | 110 | 75 | 120 | 1419 | 0 | 22 | 1 | 11 | 75 | 160 | 149 | 0.104035 | 0.463450796 | 0.964 | 0.89 | 1 | 0.927 | 0.765 | 1 | 0.823 | 0.986 | 0.91 | 0.946 | -0.024 |
| A3SS | 3975 | ENSG00000070423 | RNF126 | chr19 | - | 651610 | 651915 | 651610 | 651855 | 652232 | 652296 | 10 | 6 | 6 | 6 | 0 | 87 | 33 | 110 | 48 | 60 | 0 | 2 | 7 | 6 | 0 | 22 | 72 | 52 | 37 | 96 | 209 | 149 | 0.104204 | 0.463947602 | 0.076 | 0.115 | 0.037 | 0.082 | 0 | 0 | 0.019 | 0.088 | 0.104 | 0 | 0.02 |
| A3SS | 613 | ENSG00000105738 | SIPA1L3 | chr19 | + | 38106288 | 38106640 | 38106536 | 38106640 | 38101051 | 38101226 | 0 | 2 | 4 | 3 | 1 | 28 | 9 | 32 | 38 | 33 | 11 | 16 | 1 | 1 | 3 | 52 | 105 | 24 | 76 | 76 | 298 | 149 | 0.104421 | 0.464655168 | 0 | 0.1 | 0.059 | 0.038 | 0.015 | 0.096 | 0.071 | 0.02 | 0.007 | 0.019 | 0 |
| A3SS | 458 | ENSG00000104960 | PTOV1 | chr19 | + | 49859985 | 49860183 | 49860055 | 49860183 | 49858548 | 49858653 | 252 | 165 | 214 | 254 | 198 | 3 | 1 | 3 | 0 | 0 | 363 | 195 | 123 | 127 | 349 | 0 | 6 | 10 | 0 | 2 | 219 | 149 | 0.104653 | 0.465430283 | 0.983 | 0.991 | 0.98 | 1 | 1 | 1 | 0.957 | 0.893 | 1 | 0.992 | 0.022 |
| A3SS | 4267 | ENSG00000168297 | PXK | chr3 | + | 58412900 | 58412963 | 58412903 | 58412963 | 58410089 | 58410159 | 93 | 54 | 63 | 89 | 237 | 39 | 21 | 29 | 26 | 57 | 160 | 99 | 104 | 86 | 40 | 82 | 56 | 37 | 54 | 7 | 152 | 149 | 0.105003 | 0.466731683 | 0.7 | 0.716 | 0.68 | 0.77 | 0.803 | 0.657 | 0.634 | 0.734 | 0.61 | 0.849 | 0.037 |
| A3SS | 1743 | ENSG00000179583 | CIITA | chr16 | + | 10908982 | 10909187 | 10909028 | 10909187 | 10906498 | 10908149 | 14 | 10 | 13 | 17 | 16 | 36 | 16 | 18 | 32 | 17 | 9 | 25 | 5 | 9 | 17 | 100 | 39 | 9 | 17 | 26 | 195 | 149 | 0.105151 | 0.467132804 | 0.229 | 0.323 | 0.356 | 0.289 | 0.418 | 0.064 | 0.329 | 0.298 | 0.288 | 0.333 | 0.061 |
| A3SS | 10715 | ENSG00000215301 | DDX3X | chrX | + | 41339035 | 41339083 | 41339038 | 41339083 | 41337407 | 41337465 | 1134 | 602 | 877 | 1047 | 440 | 21 | 19 | 27 | 21 | 30 | 184 | 530 | 621 | 413 | 178 | 13 | 31 | 24 | 7 | 2 | 152 | 149 | 0.105322 | 0.467636057 | 0.981 | 0.969 | 0.97 | 0.98 | 0.935 | 0.933 | 0.944 | 0.962 | 0.983 | 0.989 | 0.005 |
| A3SS | 3015 | ENSG00000129515 | SNX6 | chr14 | - | 34629906 | 34629954 | 34629906 | 34629950 | 34630110 | 34630158 | 99 | 65 | 97 | 96 | 67 | 2 | 0 | 3 | 0 | 8 | 49 | 66 | 43 | 46 | 33 | 0 | 2 | 0 | 0 | 0 | 153 | 149 | 0.105437 | 0.467889174 | 0.98 | 1 | 0.969 | 1 | 0.891 | 1 | 0.97 | 1 | 1 | 1 | -0.026 |
| A3SS | 7711 | ENSG00000049167 | ERCC8 | chr5 | - | 60887439 | 60891086 | 60887439 | 60887520 | 60898275 | 60898400 | 48 | 28 | 30 | 25 | 41 | 0 | 0 | 0 | 0 | 0 | 41 | 45 | 17 | 17 | 4 | 8 | 0 | 0 | 0 | 0 | 298 | 149 | 0.105544 | 0.468106223 | 1 | 1 | 1 | 1 | 1 | 0.719 | 1 | 1 | 1 | 1 | 0.056 |
| A3SS | 6197 | ENSG00000115020 | PIKFYVE | chr2 | + | 208274017 | 208274053 | 208274020 | 208274053 | 208273583 | 208273733 | 5 | 6 | 8 | 12 | 8 | 3 | 5 | 3 | 9 | 0 | 32 | 0 | 4 | 4 | 4 | 1 | 5 | 7 | 5 | 11 | 152 | 149 | 0.106231 | 0.470892981 | 0.62 | 0.541 | 0.723 | 0.567 | 1 | 0.969 | 0 | 0.359 | 0.44 | 0.263 | 0.284 |
| A3SS | 8307 | ENSG00000126768 | TIMM17B | chrX | - | 48895351 | 48895972 | 48895351 | 48895501 | 48896758 | 48896858 | 13 | 6 | 12 | 15 | 10 | 0 | 0 | 4 | 2 | 1 | 23 | 22 | 10 | 9 | 17 | 0 | 0 | 1 | 0 | 0 | 298 | 149 | 0.106359 | 0.47120213 | 1 | 1 | 0.6 | 0.789 | 0.833 | 1 | 1 | 0.833 | 1 | 1 | -0.122 |
| A3SS | 12512 | ENSG00000171307 | ZDHHC16 | chr10 | + | 97455638 | 97455783 | 97455659 | 97455783 | 97454713 | 97454799 | 5 | 2 | 5 | 3 | 2 | 41 | 17 | 30 | 28 | 36 | 8 | 8 | 4 | 6 | 0 | 63 | 25 | 11 | 19 | 18 | 170 | 149 | 0.106463 | 0.471404734 | 0.097 | 0.093 | 0.127 | 0.086 | 0.046 | 0.1 | 0.219 | 0.242 | 0.217 | 0 | -0.066 |
| A3SS | 5092 | ENSG00000008128 | CDK11A | chr1 | - | 1712263 | 1712427 | 1712263 | 1712400 | 1716345 | 1716478 | 1 | 1 | 3 | 2 | 4 | 23 | 44 | 16 | 27 | 10 | 9 | 1 | 1 | 0 | 17 | 58 | 4 | 20 | 55 | 41 | 176 | 149 | 0.106585 | 0.471684507 | 0.036 | 0.019 | 0.137 | 0.059 | 0.253 | 0.116 | 0.175 | 0.041 | 0 | 0.26 | -0.018 |
| A3SS | 11600 | ENSG00000213930 | GALT | chr9 | + | 34648333 | 34648894 | 34648761 | 34648894 | 34648114 | 34648171 | 61 | 50 | 65 | 44 | 49 | 3 | 0 | 4 | 2 | 0 | 48 | 55 | 38 | 42 | 52 | 0 | 0 | 1 | 4 | 0 | 298 | 149 | 0.106741 | 0.472115017 | 0.91 | 1 | 0.89 | 0.917 | 1 | 1 | 1 | 0.95 | 0.84 | 1 | -0.015 |
| A3SS | 11807 | ENSG00000153339 | TRAPPC8 | chr18 | - | 31857539 | 31857982 | 31857539 | 31857720 | 31864626 | 31864781 | 127 | 73 | 87 | 89 | 162 | 0 | 1 | 2 | 1 | 0 | 0 | 70 | 38 | 3 | 14 | 4 | 0 | 0 | 0 | 0 | 298 | 149 | 0.107455 | 0.474900999 | 1 | 0.973 | 0.956 | 0.978 | 1 | 0 | 1 | 1 | 1 | 1 | 0.181 |
| A3SS | 11077 | ENSG00000173064 | HECTD4 | chr12 | - | 112171116 | 112171347 | 112171116 | 112171263 | 112172670 | 112172861 | 2 | 0 | 3 | 11 | 4 | 21 | 21 | 19 | 27 | 23 | 20 | 17 | 7 | 1 | 5 | 17 | 40 | 32 | 29 | 43 | 233 | 149 | 0.107488 | 0.474900999 | 0.057 | 0 | 0.092 | 0.207 | 0.1 | 0.429 | 0.214 | 0.123 | 0.022 | 0.069 | -0.08 |
| A3SS | 1965 | ENSG00000177455 | CD19 | chr16 | + | 28938672 | 28938768 | 28938675 | 28938768 | 28937768 | 28937825 | 8 | 4 | 3 | 8 | 4 | 46 | 21 | 42 | 40 | 54 | 5 | 2 | 2 | 8 | 10 | 79 | 113 | 12 | 31 | 122 | 152 | 149 | 0.108050 | 0.477106629 | 0.146 | 0.157 | 0.065 | 0.164 | 0.068 | 0.058 | 0.017 | 0.14 | 0.202 | 0.074 | 0.022 |
| A3SS | 2116 | ENSG00000167965 | MLST8 | chr16 | + | 2206483 | 2206659 | 2206496 | 2206659 | 2206357 | 2206409 | 2 | 0 | 0 | 0 | 0 | 6 | 4 | 7 | 8 | 5 | 0 | 0 | 0 | 0 | 6 | 12 | 3 | 8 | 2 | 0 | 162 | 149 | 0.108106 | 0.477106629 | 0.235 | 0 | 0 | 0 | 0 | 0 | 0 | 0 | 0 | 1 | -0.153 |
| A3SS | 7538 | ENSG00000138398 | PPIG | chr2 | + | 169631765 | 169631933 | 169631774 | 169631933 | 169630773 | 169630987 | 72 | 50 | 68 | 62 | 179 | 0 | 0 | 5 | 2 | 6 | 19 | 45 | 69 | 23 | 11 | 0 | 0 | 0 | 0 | 0 | 158 | 149 | 0.108773 | 0.479789786 | 1 | 1 | 0.928 | 0.967 | 0.966 | 1 | 1 | 1 | 1 | 1 | -0.028 |
| A3SS | 9695 | ENSG00000133606 | MKRN1 | chr7 | - | 140456866 | 140456919 | 140456651 | 140456866 | 140459006 | 140459233 | 3 | 3 | 7 | 3 | 1 | 283 | 284 | 255 | 373 | 346 | 33 | 0 | 9 | 27 | 5 | 974 | 469 | 467 | 701 | 1294 | 202 | 149 | 0.109485 | 0.482664771 | 0.008 | 0.008 | 0.02 | 0.006 | 0.006 | 0.024 | 0 | 0.014 | 0.028 | 0.003 | -0.004 |
| A3SS | 5679 | ENSG00000117682 | DHDDS | chr1 | + | 26468891 | 26471294 | 26468894 | 26471294 | 26460036 | 26460144 | 7 | 1 | 8 | 5 | 1 | 12 | 9 | 19 | 6 | 13 | 5 | 7 | 1 | 3 | 8 | 35 | 17 | 24 | 17 | 44 | 152 | 149 | 0.109545 | 0.482669359 | 0.364 | 0.098 | 0.292 | 0.45 | 0.07 | 0.123 | 0.288 | 0.039 | 0.147 | 0.151 | 0.105 |
| A3SS | 11558 | ENSG00000266714 | MYO15B | chr17 | + | 75625765 | 75625977 | 75625843 | 75625977 | 75625526 | 75625660 | 5 | 1 | 1 | 2 | 0 | 34 | 11 | 15 | 36 | 31 | 0 | 1 | 0 | 0 | 0 | 12 | 22 | 9 | 48 | 47 | 227 | 149 | 0.109837 | 0.483691937 | 0.088 | 0.056 | 0.042 | 0.035 | 0 | 0 | 0.029 | 0 | 0 | 0 | 0.038 |
| A3SS | 8563 | ENSG00000115282 | TTC31 | chr2 | + | 74492645 | 74492747 | 74492675 | 74492747 | 74492303 | 74492445 | 43 | 16 | 30 | 31 | 47 | 1 | 0 | 1 | 0 | 3 | 25 | 32 | 14 | 32 | 69 | 5 | 0 | 2 | 3 | 0 | 179 | 149 | 0.110198 | 0.485014006 | 0.973 | 1 | 0.961 | 1 | 0.929 | 0.806 | 1 | 0.854 | 0.899 | 1 | 0.061 |
| A3SS | 5736 | ENSG00000100991 | TRPC4AP | chr20 | - | 35021189 | 35021356 | 35021189 | 35021332 | 35035122 | 35035308 | 99 | 48 | 64 | 85 | 76 | 25 | 6 | 29 | 30 | 49 | 112 | 98 | 67 | 54 | 117 | 52 | 36 | 19 | 38 | 92 | 173 | 149 | 0.110482 | 0.485979305 | 0.773 | 0.873 | 0.655 | 0.709 | 0.572 | 0.65 | 0.701 | 0.752 | 0.55 | 0.523 | 0.081 |
| A3SS | 9192 | ENSG00000204386 | NEU1 | chr6 | - | 31860438 | 31861450 | 31860438 | 31860621 | 31861998 | 31862191 | 34 | 17 | 32 | 47 | 9 | 3 | 1 | 0 | 1 | 5 | 54 | 35 | 21 | 37 | 55 | 20 | 0 | 0 | 5 | 12 | 298 | 149 | 0.110537 | 0.485979305 | 0.85 | 0.895 | 1 | 0.959 | 0.474 | 0.574 | 1 | 1 | 0.787 | 0.696 | 0.024 |
| A3SS | 2265 | ENSG00000114062 | UBE3A | chr15 | - | 25407066 | 25407262 | 25407066 | 25407137 | 25408619 | 25408684 | 49 | 21 | 28 | 28 | 59 | 44 | 19 | 24 | 25 | 45 | 40 | 58 | 9 | 38 | 33 | 29 | 10 | 21 | 4 | 21 | 274 | 149 | 0.111238 | 0.488794123 | 0.377 | 0.375 | 0.388 | 0.379 | 0.416 | 0.429 | 0.759 | 0.189 | 0.838 | 0.461 | -0.148 |
| A3SS | 11286 | ENSG00000147162 | OGT | chrX | + | 71536177 | 71536358 | 71536207 | 71536358 | 71533103 | 71533336 | 1041 | 741 | 988 | 1078 | 309 | 67 | 59 | 72 | 50 | 20 | 644 | 803 | 788 | 831 | 431 | 21 | 33 | 46 | 51 | 37 | 179 | 149 | 0.111615 | 0.490182102 | 0.928 | 0.913 | 0.92 | 0.947 | 0.928 | 0.962 | 0.953 | 0.934 | 0.931 | 0.907 | -0.01 |
| A3SS | 2233 | ENSG00000166783 | MARF1 | chr16 | - | 15622933 | 15623126 | 15622933 | 15623123 | 15624768 | 15624927 | 32 | 15 | 24 | 22 | 39 | 95 | 85 | 90 | 112 | 152 | 17 | 16 | 13 | 16 | 4 | 55 | 112 | 104 | 59 | 26 | 152 | 149 | 0.112094 | 0.492017988 | 0.248 | 0.147 | 0.207 | 0.161 | 0.201 | 0.233 | 0.123 | 0.109 | 0.21 | 0.131 | 0.032 |
| A3SS | 3241 | ENSG00000204574 | ABCF1 | chr6 | + | 30578072 | 30578202 | 30578075 | 30578202 | 30577817 | 30577913 | 38 | 30 | 33 | 35 | 28 | 28 | 12 | 19 | 20 | 14 | 130 | 23 | 17 | 56 | 45 | 59 | 16 | 46 | 39 | 38 | 152 | 149 | 0.112449 | 0.493307134 | 0.571 | 0.71 | 0.63 | 0.632 | 0.662 | 0.684 | 0.585 | 0.266 | 0.585 | 0.537 | 0.11 |
| A3SS | 2023 | ENSG00000039523 | RIPOR1 | chr16 | + | 67540551 | 67540704 | 67540578 | 67540704 | 67540457 | 67540501 | 2 | 4 | 1 | 3 | 2 | 87 | 47 | 63 | 75 | 21 | 0 | 3 | 8 | 10 | 3 | 99 | 75 | 14 | 56 | 157 | 176 | 149 | 0.113012 | 0.495508185 | 0.019 | 0.067 | 0.013 | 0.033 | 0.075 | 0 | 0.033 | 0.326 | 0.131 | 0.016 | -0.06 |
| A3SS | 241 | ENSG00000127616 | SMARCA4 | chr19 | + | 11058254 | 11058363 | 11058257 | 11058363 | 11041297 | 11041560 | 56 | 43 | 54 | 57 | 85 | 29 | 18 | 31 | 29 | 22 | 153 | 56 | 37 | 69 | 111 | 37 | 18 | 18 | 21 | 45 | 152 | 149 | 0.113310 | 0.496545754 | 0.654 | 0.701 | 0.631 | 0.658 | 0.791 | 0.802 | 0.753 | 0.668 | 0.763 | 0.707 | -0.052 |
| A3SS | 8589 | ENSG00000117984 | CTSD | chr11 | - | 1761308 | 1761611 | 1761308 | 1761468 | 1763791 | 1763927 | 10 | 7 | 6 | 17 | 6 | 58 | 17 | 28 | 39 | 2 | 0 | 9 | 8 | 9 | 26 | 36 | 76 | 18 | 55 | 63 | 292 | 149 | 0.114895 | 0.503021506 | 0.081 | 0.174 | 0.099 | 0.182 | 0.605 | 0 | 0.057 | 0.185 | 0.077 | 0.174 | 0.13 |
| A3SS | 6039 | ENSG00000266028 | SRGAP2 | chr1 | + | 206401420 | 206401645 | 206401423 | 206401645 | 206393544 | 206393673 | 14 | 9 | 12 | 11 | 25 | 4 | 5 | 2 | 5 | 18 | 23 | 39 | 15 | 39 | 35 | 11 | 4 | 10 | 13 | 0 | 152 | 149 | 0.114912 | 0.503021506 | 0.774 | 0.638 | 0.855 | 0.683 | 0.577 | 0.672 | 0.905 | 0.595 | 0.746 | 1 | -0.078 |
| A3SS | 8209 | ENSG00000172493 | AFF1 | chr4 | + | 87084119 | 87084164 | 87084122 | 87084164 | 87046694 | 87047594 | 84 | 70 | 84 | 65 | 171 | 20 | 25 | 19 | 13 | 46 | 62 | 93 | 52 | 103 | 33 | 38 | 31 | 27 | 22 | 6 | 152 | 149 | 0.115441 | 0.50506128 | 0.805 | 0.733 | 0.813 | 0.831 | 0.785 | 0.615 | 0.746 | 0.654 | 0.821 | 0.844 | 0.057 |
| A3SS | 4694 | ENSG00000158062 | UBXN11 | chr1 | - | 26284141 | 26284281 | 26284141 | 26284245 | 26284361 | 26285541 | 4 | 1 | 3 | 4 | 6 | 153 | 120 | 125 | 146 | 77 | 0 | 0 | 4 | 5 | 0 | 338 | 185 | 150 | 185 | 288 | 185 | 149 | 0.116060 | 0.507497219 | 0.021 | 0.007 | 0.019 | 0.022 | 0.059 | 0 | 0 | 0.021 | 0.021 | 0 | 0.017 |
| A3SS | 6264 | ENSG00000162298 | SYVN1 | chr11 | - | 65132247 | 65132780 | 65132247 | 65132351 | 65132921 | 65133074 | 106 | 34 | 86 | 92 | 72 | 0 | 0 | 0 | 0 | 3 | 44 | 134 | 59 | 37 | 186 | 0 | 5 | 1 | 4 | 0 | 298 | 149 | 0.116672 | 0.509672424 | 1 | 1 | 1 | 1 | 0.923 | 1 | 0.931 | 0.967 | 0.822 | 1 | 0.041 |
| A3SS | 6919 | ENSG00000135905 | DOCK10 | chr2 | - | 224805407 | 224805529 | 224805407 | 224805502 | 224806125 | 224806237 | 203 | 237 | 254 | 250 | 381 | 28 | 30 | 60 | 69 | 72 | 11 | 152 | 131 | 27 | 11 | 9 | 48 | 29 | 15 | 0 | 176 | 149 | 0.116684 | 0.509672424 | 0.86 | 0.87 | 0.782 | 0.754 | 0.818 | 0.509 | 0.728 | 0.793 | 0.604 | 1 | 0.09 |
| A3SS | 9313 | ENSG00000136861 | CDK5RAP2 | chr9 | - | 120470110 | 120470220 | 120470110 | 120470217 | 120471747 | 120471878 | 43 | 76 | 49 | 46 | 55 | 11 | 8 | 14 | 4 | 6 | 18 | 60 | 96 | 64 | 15 | 0 | 9 | 13 | 6 | 0 | 152 | 149 | 0.117044 | 0.510966195 | 0.793 | 0.903 | 0.774 | 0.919 | 0.9 | 1 | 0.867 | 0.879 | 0.913 | 1 | -0.074 |
| A3SS | 8503 | ENSG00000135390 | ATP5MC2 | chr12 | - | 53669870 | 53670114 | 53669870 | 53669948 | 53672575 | 53672645 | 33 | 22 | 44 | 40 | 42 | 207 | 68 | 138 | 172 | 99 | 88 | 56 | 46 | 63 | 64 | 172 | 228 | 148 | 175 | 256 | 298 | 149 | 0.117424 | 0.51234837 | 0.074 | 0.139 | 0.138 | 0.104 | 0.175 | 0.204 | 0.109 | 0.135 | 0.153 | 0.111 | -0.016 |
| A3SS | 4261 | ENSG00000058056 | USP13 | chr3 | + | 179763936 | 179764168 | 179764001 | 179764168 | 179761111 | 179761255 | 0 | 0 | 0 | 0 | 0 | 14 | 5 | 15 | 9 | 9 | 0 | 4 | 0 | 0 | 2 | 12 | 8 | 4 | 8 | 0 | 214 | 149 | 0.117587 | 0.512749861 | 0 | 0 | 0 | 0 | 0 | 0 | 0.258 | 0 | 0 | 1 | -0.252 |
| A3SS | 10584 | ENSG00000123562 | MORF4L2 | chrX | - | 103678498 | 103678651 | 103678498 | 103678620 | 103685170 | 103685260 | 53 | 43 | 36 | 52 | 139 | 6 | 8 | 11 | 19 | 12 | 62 | 33 | 63 | 73 | 17 | 25 | 7 | 2 | 2 | 0 | 180 | 149 | 0.117643 | 0.512749861 | 0.88 | 0.816 | 0.73 | 0.694 | 0.906 | 0.672 | 0.796 | 0.963 | 0.968 | 1 | -0.075 |
| A3SS | 3308 | ENSG00000143774 | GUK1 | chr1 | + | 228146841 | 228146938 | 228146860 | 228146938 | 228146012 | 228146067 | 187 | 61 | 129 | 140 | 126 | 5 | 2 | 4 | 1 | 0 | 159 | 219 | 130 | 138 | 320 | 9 | 4 | 2 | 6 | 5 | 168 | 149 | 0.118147 | 0.514670316 | 0.971 | 0.964 | 0.966 | 0.992 | 1 | 0.94 | 0.98 | 0.983 | 0.953 | 0.983 | 0.011 |
| A3SS | 5333 | ENSG00000197343 | ZNF655 | chr7 | + | 99562343 | 99562494 | 99562348 | 99562494 | 99561887 | 99561973 | 4 | 2 | 0 | 6 | 1 | 0 | 0 | 2 | 0 | 2 | 9 | 16 | 14 | 14 | 5 | 0 | 2 | 5 | 0 | 6 | 154 | 149 | 0.118300 | 0.515059569 | 1 | 1 | 0 | 1 | 0.326 | 1 | 0.886 | 0.73 | 1 | 0.446 | -0.147 |
| A3SS | 8222 | ENSG00000170906 | NDUFA3 | chr19 | + | 54106763 | 54106961 | 54106810 | 54106961 | 54106352 | 54106446 | 2 | 2 | 3 | 5 | 6 | 8 | 7 | 5 | 5 | 8 | 0 | 6 | 8 | 0 | 0 | 2 | 10 | 3 | 9 | 16 | 196 | 149 | 0.118534 | 0.515660519 | 0.16 | 0.178 | 0.313 | 0.432 | 0.363 | 0 | 0.313 | 0.67 | 0 | 0 | 0.093 |
| A3SS | 1125 | ENSG00000170004 | CHD3 | chr17 | + | 7907890 | 7908019 | 7907893 | 7908019 | 7907600 | 7907702 | 23 | 13 | 45 | 44 | 45 | 64 | 64 | 75 | 76 | 81 | 7 | 16 | 17 | 29 | 33 | 47 | 74 | 48 | 53 | 81 | 152 | 149 | 0.118566 | 0.515660519 | 0.261 | 0.166 | 0.37 | 0.362 | 0.353 | 0.127 | 0.175 | 0.258 | 0.349 | 0.285 | 0.064 |
| A3SS | 859 | ENSG00000160410 | SHKBP1 | chr19 | + | 40578096 | 40578212 | 40578153 | 40578212 | 40577556 | 40577630 | 5 | 4 | 3 | 2 | 3 | 101 | 43 | 79 | 67 | 51 | 14 | 0 | 11 | 12 | 11 | 135 | 129 | 75 | 111 | 275 | 206 | 149 | 0.118757 | 0.516213593 | 0.035 | 0.063 | 0.027 | 0.021 | 0.041 | 0.07 | 0 | 0.096 | 0.073 | 0.028 | -0.016 |
| A3SS | 9238 | ENSG00000160185 | UBASH3A | chr21 | + | 42443153 | 42443418 | 42443311 | 42443418 | 42442451 | 42442596 | 7 | 9 | 17 | 11 | 12 | 13 | 15 | 24 | 36 | 72 | 8 | 15 | 5 | 1 | 14 | 7 | 31 | 4 | 10 | 14 | 298 | 149 | 0.118872 | 0.516434786 | 0.212 | 0.231 | 0.262 | 0.133 | 0.077 | 0.364 | 0.195 | 0.385 | 0.048 | 0.333 | -0.082 |
| A3SS | 12074 | ENSG00000174669 | SLC29A2 | chr11 | - | 66366125 | 66366231 | 66366125 | 66366191 | 66366430 | 66366564 | 5 | 3 | 14 | 3 | 6 | 3 | 1 | 4 | 1 | 1 | 0 | 2 | 8 | 3 | 0 | 12 | 0 | 2 | 0 | 10 | 189 | 149 | 0.118940 | 0.516453725 | 0.568 | 0.703 | 0.734 | 0.703 | 0.825 | 0 | 1 | 0.759 | 1 | 0 | 0.155 |
| A3SS | 5470 | ENSG00000181666 | ZNF875 | chr19 | + | 37347189 | 37347316 | 37347231 | 37347316 | 37346422 | 37346492 | 3 | 10 | 11 | 11 | 13 | 1 | 3 | 0 | 3 | 0 | 13 | 25 | 14 | 3 | 7 | 9 | 0 | 2 | 7 | 0 | 191 | 149 | 0.119259 | 0.517558566 | 0.701 | 0.722 | 1 | 0.741 | 1 | 0.53 | 1 | 0.845 | 0.251 | 1 | 0.108 |
| A3SS | 255 | ENSG00000131351 | HAUS8 | chr19 | - | 17069030 | 17069099 | 17069030 | 17069086 | 17073273 | 17073335 | 0 | 0 | 0 | 0 | 0 | 25 | 4 | 16 | 9 | 18 | 6 | 0 | 0 | 0 | 0 | 2 | 18 | 10 | 10 | 12 | 162 | 149 | 0.119719 | 0.519250168 | 0 | 0 | 0 | 0 | 0 | 0.734 | 0 | 0 | 0 | 0 | -0.147 |
| A3SS | 2216 | ENSG00000137804 | NUSAP1 | chr15 | + | 41350987 | 41351129 | 41350990 | 41351129 | 41349097 | 41349241 | 8 | 7 | 20 | 21 | 23 | 1 | 3 | 0 | 2 | 2 | 5 | 29 | 15 | 37 | 8 | 0 | 0 | 0 | 5 | 0 | 152 | 149 | 0.119777 | 0.519250168 | 0.887 | 0.696 | 1 | 0.911 | 0.919 | 1 | 1 | 1 | 0.879 | 1 | -0.093 |
| A3SS | 1470 | ENSG00000169738 | DCXR | chr17 | - | 82036190 | 82036322 | 82036190 | 82036308 | 82036383 | 82036448 | 3 | 2 | 2 | 4 | 0 | 65 | 36 | 63 | 53 | 41 | 0 | 0 | 0 | 0 | 5 | 34 | 27 | 30 | 34 | 48 | 163 | 149 | 0.120009 | 0.519972764 | 0.04 | 0.048 | 0.028 | 0.065 | 0 | 0 | 0 | 0 | 0 | 0.087 | 0.019 |
| A3SS | 10185 | ENSG00000150637 | CD226 | chr18 | - | 69947360 | 69947532 | 69947360 | 69947433 | 69961591 | 69961803 | 16 | 51 | 7 | 34 | 29 | 27 | 57 | 14 | 82 | 127 | 41 | 12 | 69 | 42 | 121 | 82 | 51 | 37 | 72 | 123 | 248 | 149 | 0.120719 | 0.522771596 | 0.263 | 0.35 | 0.231 | 0.199 | 0.121 | 0.231 | 0.124 | 0.528 | 0.26 | 0.371 | -0.07 |
| A3SS | 6322 | ENSG00000115685 | PPP1R7 | chr2 | + | 241158465 | 241158549 | 241158483 | 241158549 | 241157806 | 241157862 | 6 | 1 | 7 | 2 | 6 | 241 | 171 | 204 | 252 | 323 | 15 | 4 | 1 | 4 | 9 | 369 | 235 | 139 | 221 | 216 | 167 | 149 | 0.121090 | 0.524095954 | 0.022 | 0.005 | 0.03 | 0.007 | 0.016 | 0.035 | 0.015 | 0.006 | 0.016 | 0.036 | -0.006 |
| A3SS | 1868 | ENSG00000197912 | SPG7 | chr16 | + | 89529476 | 89530808 | 89530682 | 89530808 | 89526328 | 89526468 | 61 | 46 | 71 | 51 | 41 | 1 | 4 | 2 | 1 | 0 | 67 | 62 | 43 | 79 | 146 | 8 | 4 | 5 | 0 | 0 | 298 | 149 | 0.121224 | 0.524308003 | 0.968 | 0.852 | 0.947 | 0.962 | 1 | 0.807 | 0.886 | 0.811 | 1 | 1 | 0.045 |
| A3SS | 3534 | ENSG00000131503 | ANKHD1 | chr5 | + | 140438460 | 140438617 | 140438502 | 140438617 | 140436103 | 140436257 | 435 | 367 | 360 | 399 | 508 | 41 | 25 | 27 | 49 | 38 | 580 | 464 | 621 | 468 | 319 | 30 | 45 | 31 | 49 | 19 | 191 | 149 | 0.121269 | 0.524308003 | 0.892 | 0.92 | 0.912 | 0.864 | 0.913 | 0.938 | 0.889 | 0.94 | 0.882 | 0.929 | -0.015 |
| A3SS | 4204 | ENSG00000114745 | GORASP1 | chr3 | - | 39100304 | 39100877 | 39100304 | 39100503 | 39101015 | 39101102 | 21 | 20 | 34 | 14 | 28 | 1 | 0 | 0 | 1 | 0 | 32 | 49 | 11 | 34 | 100 | 5 | 3 | 0 | 0 | 0 | 298 | 149 | 0.121436 | 0.524747813 | 0.913 | 1 | 1 | 0.875 | 1 | 0.762 | 0.891 | 1 | 1 | 1 | 0.027 |
| A3SS | 548 | ENSG00000186818 | LILRB4 | chr19 | + | 54666693 | 54666749 | 54666696 | 54666749 | 54666398 | 54666436 | 13 | 4 | 4 | 14 | 10 | 38 | 47 | 10 | 25 | 18 | 12 | 29 | 9 | 8 | 22 | 119 | 93 | 36 | 44 | 88 | 152 | 149 | 0.121976 | 0.52680081 | 0.251 | 0.077 | 0.282 | 0.354 | 0.353 | 0.09 | 0.234 | 0.197 | 0.151 | 0.197 | 0.09 |
| A3SS | 5447 | ENSG00000239900 | ADSL | chr22 | + | 40361087 | 40361342 | 40361272 | 40361342 | 40360401 | 40360492 | 10 | 3 | 16 | 0 | 17 | 121 | 78 | 118 | 129 | 174 | 25 | 18 | 5 | 2 | 10 | 110 | 122 | 100 | 100 | 156 | 298 | 149 | 0.122137 | 0.52721406 | 0.04 | 0.019 | 0.063 | 0 | 0.047 | 0.102 | 0.069 | 0.024 | 0.01 | 0.031 | -0.013 |
| A3SS | [illegible] | [illegible] | [illegible] | [illegible] | [illegible] | [illegible] | [illegible] | [illegible] | [illegible] | [illegible] | [illegible] | [illegible] | [illegible] | [illegible] | [illegible] | [illegible] | [illegible] | [illegible] | [illegible] | [illegible] | [illegible] | [illegible] | [illegible] | [illegible] | [illegible] | [illegible] | [illegible] | [illegible] | [illegible] | [illegible] | [illegible] | [illegible] | [illegible] | [illegible] | [illegible] | [illegible] | [illegible] | [illegible] | [illegible] | [illegible] | [illegible] | [illegible] | [illegible] | [illegible] | [illegible] | [illegible] |

| | | | | | | | | | | | | | | |
|---|---|---|---|---|---|---|---|---|---|---|---|---|---|---|
| A3SS | 11748 | ENSG00000149428 | HYOU1 | chr11 | - | 119045780 | 119045834 | 119045780 | 119045831 | 119046416 | 119046467 | 17 | 7 | 24 | [illegible] |
| A3SS | 10473 | ENSG00000008294 | SPAG9 | chr17 | - | 50989676 | 50989872 | 50989676 | 50989827 | 50990449 | 50990668 | 63 | 54 | 56 | [illegible] |
| A3SS | 4295 | ENSG00000114127 | XRN1 | chr3 | - | 142332374 | 142332534 | 142332374 | 142332531 | 142332966 | 142333089 | 183 | 145 | 144 | [illegible] |
| A3SS | 2471 | ENSG00000171606 | ZNF274 | chr19 | + | 58185696 | 58185838 | 58185711 | 58185838 | 58183920 | 58183998 | 0 | 0 | 0 | [illegible] |
| A3SS | 2695 | ENSG00000100897 | DCAF11 | chr14 | + | 24116971 | 24117044 | 24116987 | 24117044 | 24115381 | 24115749 | 30 | 14 | 24 | [illegible] |
| A3SS | 7345 | ENSG00000115204 | MPV17 | chr2 | - | 27312993 | 27313109 | 27312993 | 27313074 | 27322447 | 27322522 | 50 | 22 | 39 | [illegible] |
| A3SS | 3033 | ENSG00000125459 | MSTO1 | chr1 | + | 155611920 | 155612100 | 155611982 | 155612100 | 155611688 | 155611827 | 21 | 5 | 15 | [illegible] |
| A3SS | 564 | ENSG00000167613 | LAIR1 | chr19 | - | 54364294 | 54364486 | 54364294 | 54364330 | 54364770 | 54364931 | 51 | 22 | 29 | [illegible] |
| A3SS | 10494 | ENSG00000175463 | TBC1D10C | chr11 | + | 67405902 | 67406017 | 67405906 | 67406017 | 67405594 | 67405701 | 200 | 52 | 148 | [illegible] |
| A3SS | 12181 | ENSG00000110075 | PPP6R3 | chr11 | + | 68587797 | 68588024 | 68587944 | 68588024 | 68583042 | 68583129 | 93 | 67 | 87 | [illegible] |
| A3SS | 12609 | ENSG00000196233 | LCOR | chr10 | + | 96949007 | 96952196 | 96952102 | 96952196 | 96944112 | 96944245 | 218 | 193 | 198 | [illegible] |
| A3SS | 1469 | ENSG00000062716 | VMP1 | chr17 | + | 59838294 | 59838397 | 59838311 | 59838397 | 59837854 | 59838212 | 38 | 20 | 6 | [illegible] |
| A3SS | 3278 | ENSG00000133835 | HSD17B4 | chr5 | + | 119529893 | 119529980 | 119529900 | 119529980 | 119527132 | 119527219 | 243 | 129 | 100 | [illegible] |
| A3SS | 1218 | ENSG00000185624 | P4HB | chr17 | - | 81855452 | 81855586 | 81855452 | 81855579 | 81859180 | 81859387 | 364 | 180 | 324 | [illegible] |
| A3SS | 10547 | ENSG00000120860 | WASHC3 | chr12 | - | 102039867 | 102039978 | 102039867 | 102039975 | 102044104 | 102044212 | 140 | 115 | 129 | [illegible] |
| A3SS | 8487 | ENSG00000196576 | PLXNB2 | chr22 | - | 50284131 | 50284244 | 50284131 | 50284213 | 50284572 | 50284665 | 2 | 0 | 1 | [illegible] |
| A3SS | 11854 | ENSG00000142186 | SCYL1 | chr11 | + | 65537808 | 65537880 | 65537811 | 65537880 | 65536985 | 65537128 | 58 | 21 | 53 | [illegible] |
| A3SS | 8231 | ENSG00000204681 | GABBR1 | chr6 | - | 29629086 | 29629247 | 29629086 | 29629107 | 29631395 | 29631599 | 37 | 36 | 29 | [illegible] |
| A3SS | 7403 | ENSG00000152683 | SLC30A6 | chr2 | + | 32197317 | 32197392 | 32197343 | 32197392 | 32193888 | 32193983 | 3 | 5 | 0 | [illegible] |
| A3SS | 13273 | ENSG00000197111 | PCBP2 | chr12 | + | 53468776 | 53468832 | 53468779 | 53468832 | 53467804 | 53467843 | 395 | 176 | 391 | [illegible] |
| A3SS | 10668 | ENSG00000258890 | CEP95 | chr17 | + | 64514228 | 64514358 | 64514247 | 64514358 | 64510172 | 64510280 | 1 | 7 | 6 | [illegible] |
| A3SS | 11687 | ENSG00000125944 | HNRNPR | chr1 | - | 23318482 | 23318697 | 23318482 | 23318688 | 23321527 | 23321663 | 122 | 99 | 104 | [illegible] |
| A3SS | 163 | ENSG00000100150 | DEPDC5 | chr22 | + | 31876086 | 31876265 | 31876156 | 31876265 | 31874272 | 31874405 | 3 | 0 | 3 | [illegible] |
| A3SS | 130 | ENSG00000100359 | SGSM3 | chr22 | + | 40409249 | 40409372 | 40409339 | 40409372 | 40408932 | 40409018 | 32 | 21 | 30 | [illegible] |
| A3SS | 2759 | ENSG00000119718 | EIF2B2 | chr14 | + | 75005840 | 75005961 | 75005865 | 75005961 | 75004736 | 75004900 | 2 | 0 | 2 | [illegible] |
| A3SS | 3748 | ENSG00000138698 | RAP1GDS1 | chr4 | + | 98293404 | 98293515 | 98293407 | 98293515 | 98261485 | 98261569 | 19 | 8 | 9 | [illegible] |
| A3SS | 13121 | ENSG00000110906 | KCTD10 | chr12 | - | 109457991 | 109458184 | 109457991 | 109458081 | 109460635 | 109460805 | 10 | 8 | 7 | [illegible] |
| A3SS | 10705 | ENSG00000046651 | OFD1 | chrX | + | 13767126 | 13768224 | 13768053 | 13768224 | 13763744 | 13763855 | 77 | 85 | 94 | [illegible] |
| A3SS | 10400 | ENSG00000164074 | ABHD18 | chr4 | + | 128017357 | 128017501 | 128017362 | 128017501 | 128011674 | 128011700 | 7 | 2 | 2 | [illegible] |
| A3SS | 12125 | ENSG00000070047 | PHRF1 | chr11 | + | 605604 | 605724 | 605607 | 605724 | 605118 | 605300 | 129 | 67 | 122 | [illegible] |
| A3SS | 8479 | ENSG00000168958 | MFF | chr2 | + | 227330625 | 227330846 | 227330780 | 227330846 | 227329605 | 227329783 | 233 | 110 | 238 | [illegible] |
| A3SS | 8810 | ENSG00000169045 | HNRNPH1 | chr5 | - | 179623036 | 179623187 | 179623036 | 179623164 | 179623635 | 179623670 | 33 | 24 | 9 | [illegible] |
| A3SS | 12754 | ENSG00000139620 | KANSL2 | chr12 | - | 48679035 | 48679238 | 48679035 | 48679150 | 48679654 | 48679833 | 37 | 33 | 25 | [illegible] |
| A3SS | 2792 | ENSG00000100784 | RPS6KA5 | chr14 | - | 90923112 | 90923233 | 90923112 | 90923196 | 90943077 | 90943185 | 3 | 17 | 1 | [illegible] |
| A3SS | 12064 | ENSG00000175467 | SART1 | chr11 | + | 65964005 | 65964131 | 65964073 | 65964131 | 65961728 | 65962093 | 3 | 0 | 4 | [illegible] |
| A3SS | 4851 | ENSG00000137074 | APTX | chr9 | - | 32987543 | 32987846 | 32987543 | 32987731 | 32989758 | 32989895 | 24 | 24 | 19 | [illegible] |
| A3SS | 8602 | ENSG00000164548 | TRA2A | chr7 | - | 23521706 | 23522432 | 23521706 | 23521840 | 23531788 | 23531981 | 299 | 277 | 317 | [illegible] |
| A3SS | 12295 | ENSG00000179532 | DNHD1 | chr11 | + | 6544791 | 6547666 | 6544876 | 6547666 | 6544573 | 6544671 | 15 | 8 | 9 | [illegible] |
| A3SS | 13135 | ENSG00000166037 | CEP57 | chr11 | + | 95812903 | 95813111 | 95812931 | 95813111 | 95799231 | 95799388 | 5 | 5 | 5 | [illegible] |
| A3SS | 11612 | ENSG00000148400 | NOTCH1 | chr9 | - | 136502271 | 136502775 | 136502271 | 136502488 | 136503181 | 136503330 | 9 | 6 | 3 | [illegible] |
| A3SS | 13255 | ENSG00000121073 | SLC35B1 | chr17 | - | 49706203 | 49706334 | 49706203 | 49706309 | 49706964 | 49707068 | 28 | 19 | 53 | [illegible] |
| A3SS | 8314 | ENSG00000135912 | TTLL4 | chr2 | + | 218753114 | 218753185 | 218753120 | 218753185 | 218752762 | 218752973 | 20 | 13 | 16 | [illegible] |
| A3SS | 10652 | ENSG00000116754 | SRSF11 | chr1 | + | 70249930 | 70250047 | 70249951 | 70250047 | 70246817 | 70246907 | 10 | 12 | 28 | [illegible] |
| A3SS | 10223 | ENSG00000166579 | NDEL1 | chr17 | + | 8443926 | 8444357 | 8444259 | 8444357 | 8436650 | 8436694 | 40 | 37 | 37 | [illegible] |
| A3SS | 1687 | ENSG00000125124 | BBS2 | chr16 | - | 56501352 | 56502076 | 56501352 | 56501497 | 56502316 | 56502456 | 17 | 7 | 23 | [illegible] |
| A3SS | 1852 | ENSG00000205336 | ADGRG1 | chr16 | + | 57651199 | 57651457 | 57651242 | 57651457 | 57650252 | 57650351 | 215 | 41 | 29 | [illegible] |
| A3SS | 5744 | ENSG00000139370 | SLC15A4 | chr12 | - | 128809942 | 128810111 | 128809942 | 128810053 | 128814774 | 128815070 | 189 | 83 | 122 | [illegible] |
| A3SS | 6277 | ENSG00000025708 | TYMP | chr22 | - | 50529495 | 50529747 | 50529495 | 50529719 | 50529903 | 50530012 | 17 | 9 | 14 | [illegible] |
| A3SS | 1748 | ENSG00000103042 | SLC38A7 | chr16 | - | 58683954 | 58684178 | 58683954 | 58684083 | 58684716 | 58684770 | 17 | 11 | 6 | [illegible] |
| A3SS | 5205 | ENSG00000284879 | NA | chr2 | + | 87719806 | 87720021 | 87719831 | 87720021 | 87716611 | 87716790 | 12 | 5 | 13 | [illegible] |
| A3SS | 5229 | ENSG00000117640 | MTFR1L | chr1 | + | 25822995 | 25823128 | 25823018 | 25823128 | 25819928 | 25820029 | 0 | 0 | 0 | [illegible] |
| A3SS | 4248 | ENSG00000177963 | RIC8A | chr11 | + | 212596 | 212759 | 212614 | 212759 | 212415 | 212511 | 3 | 3 | 8 | [illegible] |
| A3SS | 35 | ENSG00000141959 | PFKL | chr21 | + | 44325731 | 44326060 | 44325960 | 44326060 | 44325152 | 44325264 | 16 | 15 | 15 | [illegible] |
| A3SS | 10052 | ENSG00000128604 | IRF5 | chr7 | + | 128947229 | 128947535 | 128947277 | 128947535 | 128947022 | 128947056 | 52 | 10 | 21 | [illegible] |
| A3SS | 20 | ENSG00000160216 | AGPAT3 | chr21 | + | 43967868 | 43968115 | 43967945 | 43968115 | 43960631 | 43960779 | 7 | 4 | 3 | [illegible] |
| A3SS | 901 | ENSG00000170471 | RALGAPB | chr20 | + | 38535073 | 38535207 | 38535085 | 38535207 | 38532729 | 38532859 | 20 | 12 | 17 | [illegible] |
| A3SS | 6828 | ENSG00000100092 | SH3BP1 | chr22 | + | 37650085 | 37650249 | 37650151 | 37650249 | 37648318 | 37648435 | 7 | 3 | 3 | [illegible] |
| A3SS | 12741 | ENSG00000023041 | ZDHHC6 | chr10 | - | 112443514 | 112443606 | 112443514 | 112443594 | 112445169 | 112445650 | 76 | 62 | 94 | [illegible] |
| A3SS | 2045 | ENSG00000102908 | NFAT5 | chr16 | + | 69684886 | 69684970 | 69684889 | 69684970 | 69677202 | 69677335 | 45 | 35 | 34 | [illegible] |
| A3SS | 9606 | ENSG00000075975 | MKRN2 | chr3 | + | 12576610 | 12576741 | 12576630 | 12576741 | 12574791 | 12575006 | 0 | 0 | 0 | [illegible] |
| A3SS | 8984 | ENSG00000123600 | METTL8 | chr2 | - | 171330558 | 171330698 | 171330558 | 171330616 | 171331803 | 171331867 | 66 | 62 | 65 | [illegible] |
| A3SS | 13358 | ENSG00000013364 | MVP | chr16 | + | 29830473 | 29830674 | 29830514 | 29830674 | 29820472 | 29820510 | 33 | 16 | 16 | [illegible] |
| A3SS | 6131 | ENSG00000198408 | OGA | chr10 | - | 101792838 | 101792943 | 101792838 | 101792901 | 101793912 | 101793998 | 216 | 180 | 234 | [illegible] |
| A3SS | 3073 | ENSG00000205133 | TRIQK | chr8 | - | 92916928 | 92917010 | 92916928 | 92917007 | 92954405 | 92954564 | 41 | 34 | 27 | [illegible] |
| A3SS | 8237 | ENSG00000123136 | DDX39A | chr19 | - | 14409741 | 14409981 | 14409741 | 14409873 | 14410215 | 14410334 | 14 | 7 | 7 | [illegible] |
| A3SS | 12253 | ENSG00000187049 | TMEM216 | chr11 | + | 61397727 | 61397975 | 61397773 | 61397975 | 61393883 | 61393976 | 6 | 2 | 3 | [illegible] |
| A3SS | 9621 | ENSG00000173598 | NUDT4 | chr12 | + | 93398767 | 93398855 | 93398770 | 93398855 | 93395488 | 93395533 | 39 | 28 | 21 | [illegible] |
| A3SS | 2579 | ENSG00000119682 | AREL1 | chr14 | - | 74669946 | 74670142 | 74669946 | 74670126 | 74670761 | 74670871 | 6 | 6 | 8 | [illegible] |
| A3SS | 150 | ENSG00000100239 | PPP6R2 | chr22 | + | 50419345 | 50419462 | 50419348 | 50419462 | 50416091 | 50416157 | 0 | 0 | 4 | [illegible] |
| A3SS | 11443 | ENSG00000048828 | FAM120A | chr9 | + | 93561108 | 93561250 | 93561111 | 93561250 | 93558580 | 93558718 | 55 | 17 | 29 | [illegible] |
| A3SS | 1330 | ENSG00000109113 | RAB34 | chr17 | - | 28717212 | 28717425 | 28717212 | 28717291 | 28718121 | 28718268 | 5 | 6 | 5 | [illegible] |
| A3SS | 865 | ENSG00000160410 | SHKBP1 | chr19 | + | 40590729 | 40590853 | 40590732 | 40590853 | 40590243 | 40590422 | 29 | 11 | 14 | [illegible] |
| A3SS | 3238 | ENSG00000100284 | TOM1 | chr22 | + | 35333174 | 35333497 | 35333403 | 35333497 | 35332980 | 35333014 | 17 | 2 | 10 | [illegible] |
| A3SS | 5308 | ENSG00000265241 | RBM8A | chr1 | - | 145926808 | 145926886 | 145926808 | 145926883 | 145927017 | 145927077 | 372 | 224 | 321 | [illegible] |
| A3SS | 3568 | ENSG00000105875 | WDR91 | chr7 | - | 135186971 | 135188545 | 135186971 | 135187169 | 135189343 | 135189452 | 22 | 20 | 32 | [illegible] |
| A3SS | 172 | ENSG00000100242 | SUN2 | chr22 | - | 38754641 | 38754706 | 38754641 | 38754690 | 38755762 | 38756000 | 97 | 19 | 50 | [illegible] |
| A3SS | 10778 | ENSG00000156504 | PABIR2 | chrX | - | 134785885 | 134785950 | 134785885 | 134785947 | 134787471 | 134787533 | 87 | 77 | 98 | [illegible] |
| A3SS | 2831 | ENSG00000119688 | ABCD4 | chr14 | - | 74292747 | 74293248 | 74292747 | 74292869 | 74295147 | 74295198 | 31 | 15 | 27 | [illegible] |
| A3SS | 3861 | ENSG00000145495 | MARCHF6 | chr5 | + | 10415472 | 10415669 | 10415487 | 10415669 | 10414432 | 10414502 | 10 | 14 | 7 | [illegible] |
| A3SS | 1376 | ENSG00000215041 | NEURL4 | chr17 | - | 7317460 | 7317684 | 7317460 | 7317573 | 7317787 | 7317932 | 5 | 6 | 7 | [illegible] |
| A3SS | 12980 | ENSG00000116198 | CEP104 | chr1 | - | 3847474 | 3847631 | 3847474 | 3847613 | 3848607 | 3848781 | 0 | 3 | 4 | [illegible] |
| A3SS | 3268 | ENSG00000197603 | CPLANE1 | chr5 | - | 37179360 | 37179443 | 37179360 | 37179440 | 37180016 | 37180183 | 25 | 21 | 30 | [illegible] |
| A3SS | 2402 | ENSG00000140598 | EFL1 | chr15 | - | 82227449 | 82227579 | 82227449 | 82227572 | 82228190 | 82228327 | 6 | 3 | 1 | [illegible] |
| A3SS | 12267 | ENSG00000204356 | NELFE | chr6 | - | 31954554 | 31954892 | 31954554 | 31954877 | 31955058 | 31955096 | 93 | 43 | 75 | [illegible] |
| A3SS | 7404 | ENSG00000143633 | C1orf131 | chr1 | - | 231229149 | 231229209 | 231229149 | 231229206 | 231238856 | 231239188 | 63 | 27 | 47 | [illegible] |
| A3SS | 4879 | ENSG00000132549 | VPS13B | chr8 | + | 99391556 | 99391704 | 99391559 | 99391704 | 99384207 | 99384317 | 135 | 123 | 118 | [illegible] |
| A3SS | 660 | ENSG00000105705 | SUGP1 | chr19 | - | 19303347 | 19303448 | 19303347 | 19303406 | 19305848 | 19306076 | 22 | 18 | 30 | [illegible] |
| A3SS | 7716 | ENSG00000049167 | ERCC8 | chr5 | - | 60924290 | 60924429 | 60924290 | 60924367 | 60928863 | 60928959 | 3 | 13 | 3 | [illegible] |
| A3SS | 2648 | ENSG00000075413 | MARK3 | chr14 | + | 103468080 | 103468186 | 103468085 | 103468186 | 103467078 | 103467191 | 24 | 18 | 26 | [illegible] |
| A3SS | 9457 | ENSG00000242802 | AP5Z1 | chr7 | + | 4790213 | 4790591 | 4790458 | 4790591 | 4789831 | 4789929 | 18 | 7 | 15 | [illegible] |
| A3SS | 11653 | ENSG00000130714 | POMT1 | chr9 | + | 131509742 | 131510415 | 131510356 | 131510415 | 131508910 | 131509022 | 25 | 21 | 34 | [illegible] |
| A3SS | 11910 | ENSG00000197798 | FAM118B | chr11 | + | 126261424 | 126261484 | 126261427 | 126261484 | 126256566 | 126256852 | 30 | 20 | 21 | [illegible] |
| A3SS | 2025 | ENSG00000103248 | MTHFSD | chr16 | - | 86554644 | 86554751 | 86554644 | 86554748 | 86555168 | 86555203 | 7 | 5 | 12 | [illegible] |
| A3SS | 3585 | ENSG00000214655 | ZSWIM8 | chr10 | + | 73794540 | 73794639 | 73794555 | 73794639 | 73794146 | 73794330 | 70 | 38 | 50 | [illegible] |
| A3SS | 4482 | ENSG00000108219 | TSPAN14 | chr10 | + | 80507198 | 80507374 | 80507227 | 80507374 | 80504727 | 80504778 | 3 | 1 | 0 | [illegible] |
| A3SS | 7469 | ENSG00000163534 | FCRL1 | chr1 | - | 157801914 | 157802193 | 157801914 | 157802162 | 157802376 | 157802664 | 66 | 15 | 31 | [illegible] |
| A3SS | 10157 | ENSG00000184863 | RBM33 | chr7 | + | 155763808 | 155764018 | 155763811 | 155764018 | 155744960 | 155745602 | 30 | 18 | 17 | [illegible] |
| A3SS | 176 | ENSG00000198089 | SFI1 | chr22 | + | 31604868 | 31606430 | 31606327 | 31606430 | 31589446 | 31589577 | 19 | 23 | 43 | [illegible] |
| A3SS | 3727 | ENSG00000128609 | NDUFA5 | chr7 | - | 123557403 | 123557507 | 123557403 | 123557448 | 123557774 | 123557804 | 6 | 4 | 10 | [illegible] |
| A3SS | 4468 | ENSG00000169714 | CNBP | chr3 | - | 129171078 | 129171283 | 129171078 | 129171277 | 129171445 | 129171538 | 27 | 13 | 28 | [illegible] |
| A3SS | 6904 | ENSG00000168610 | STAT3 | chr17 | - | 42316788 | 42316889 | 42316788 | 42316851 | 42317181 | 42317224 | 84 | 61 | 96 | [illegible] |
| A3SS | 12018 | ENSG00000149179 | C11orf49 | chr11 | + | 47161445 | 47162247 | 47161463 | 47162247 | 47161126 | 47161190 | 6 | 3 | 1 | [illegible] |
| A3SS | 12610 | ENSG00000150760 | DOCK1 | chr10 | + | 127012231 | 127012374 | 127012294 | 127012374 | 127008731 | 127008804 | 4 | 2 | 0 | [illegible] |
| A3SS | 5940 | ENSG00000175309 | PHYKPL | chr5 | - | 178222354 | 178222580 | 178222354 | 178222545 | 178222851 | 178222934 | 43 | 14 | 40 | [illegible] |
| A3SS | 6278 | ENSG00000025708 | TYMP | chr22 | - | 50529495 | 50529754 | 50529495 | 50529719 | 50529903 | 50530012 | 12 | 11 | 15 | [illegible] |
| A3SS | 6776 | ENSG00000164300 | SERINC5 | chr5 | - | 80158835 | 80158962 | 80158835 | 80158956 | 80166382 | 80166478 | 132 | 87 | 187 | [illegible] |
| A3SS | 3728 | ENSG00000163950 | SLBP | chr4 | - | 1700010 | 1700091 | 1700010 | 1700070 | 1703595 | 1703700 | 3 | 3 | 0 | [illegible] |
| A3SS | 8849 | ENSG00000172765 | TMCC1 | chr3 | - | 129827802 | 129828508 | 129827802 | 129828077 | 129832773 | 129832826 | 82 | 70 | 83 | [illegible] |
| A3SS | 12067 | ENSG00000087365 | SF3B2 | chr11 | + | 66057265 | 66057375 | 66057268 | 66057375 | 66056837 | 66056955 | 296 | 193 | 294 | [illegible] |
| A3SS | 2530 | ENSG00000133961 | NUMB | chr14 | - | 73323129 | 73323229 | 73323129 | 73323204 | 73355625 | 73355766 | 5 | 3 | 2 | [illegible] |
| A3SS | 649 | ENSG00000130479 | MAP1S | chr19 | + | 17725029 | 17725189 | 17725048 | 17725189 | 17724125 | 17724208 | 10 | 0 | 2 | [illegible] |
| A3SS | 12059 | ENSG00000175550 | DRAP1 | chr11 | + | 65920342 | 65920462 | 65920415 | 65920462 | 65919947 | 65919997 | 105 | 59 | 68 | [illegible] |
| A3SS | 876 | ENSG00000188785 | ZNF548 | chr19 | + | 57394042 | 57394223 | 57394187 | 57394223 | 57389874 | 57390114 | 4 | 4 | 3 | [illegible] |
| A3SS | 12796 | ENSG00000079974 | RABL2B | chr22 | - | 50769454 | 50769552 | 50769454 | 50769549 | 50769904 | 50770016 | 32 | 20 | 20 | [illegible] |
| A3SS | 10235 | ENSG00000196998 | WDR45 | chrX | - | 49075544 | 49075753 | 49075544 | 49075663 | 49075865 | 49075945 | 220 | 84 | 127 | [illegible] |
| A3SS | 6445 | ENSG00000144136 | SLC20A1 | chr2 | + | 112658824 | 112659094 | 112658973 | 112659094 | 112657121 | 112657241 | 341 | 204 | 306 | [illegible] |
| A3SS | 3583 | ENSG00000083312 | TNPO1 | chr5 | + | 72851237 | 72851319 | 72851243 | 72851319 | 72848384 | 72848498 | 19 | 0 | 7 | [illegible] |
| A3SS | 2312 | ENSG00000184056 | VPS33B | chr15 | - | 91016962 | 91017064 | 91016962 | 91017024 | 91017804 | 91017885 | 2 | 2 | 3 | [illegible] |
| A3SS | 11385 | ENSG00000168003 | SLC3A2 | chr11 | + | 62881880 | 62882066 | 62881892 | 62882066 | 62880895 | 62881447 | 2 | 0 | 0 | [illegible] |
| A3SS | 5827 | ENSG00000170248 | PDCD6IP | chr3 | + | 33828837 | 33828969 | 33828852 | 33828969 | 33826479 | 33826580 | 42 | 33 | 37 | [illegible] |
| A3SS | 4269 | ENSG00000157601 | MX1 | chr21 | + | 41432030 | 41432175 | 41432049 | 41432175 | 41427754 | 41427866 | 5 | 8 | 0 | [illegible] |
| A3SS | 7298 | ENSG00000177628 | GBA | chr1 | - | 155239885 | 155240077 | 155239885 | 155239962 | 155240629 | 155240717 | 196 | 75 | 118 | [illegible] |
| A3SS | 8701 | ENSG00000165280 | VCP | chr9 | - | 35067890 | 35068099 | 35067890 | 35068063 | 35068250 | 35068362 | 0 | 2 | 2 | [illegible] |
| A3SS | 11101 | ENSG00000160948 | VPS28 | chr8 | - | 144424945 | 144425051 | 144424945 | 144425000 | 144425682 | 144425772 | 196 | 95 | 144 | [illegible] |
| A3SS | 3224 | ENSG00000215068 | NA | chr5 | + | 43044392 | 43045268 | 43044698 | 43045268 | 43042132 | 43043567 | 25 | 14 | 14 | [illegible] |
| A3SS | 451 | ENSG00000115268 | RPS15 | chr19 | + | 1439967 | 1440253 | 1440025 | 1440253 | 1438806 | 1438892 | 67 | 21 | 61 | [illegible] |
| A3SS | 10908 | ENSG00000158669 | GPAT4 | chr8 | + | 41612785 | 41612960 | 41612844 | 41612960 | 41612179 | 41612273 | 2 | 1 | 4 | [illegible] |
| A3SS | 10455 | ENSG00000147394 | ZNF185 | chrX | + | 152920706 | 152920748 | 152920709 | 152920748 | 152920327 | 152920411 | 52 | 55 | 13 | [illegible] |
| A3SS | 763 | ENSG00000064601 | CTSA | chr20 | + | 45896455 | 45897040 | 45896964 | 45897040 | 45894993 | 45895133 | 9 | 2 | 8 | [illegible] |
| A3SS | 5103 | ENSG00000085433 | WDR47 | chr1 | - | 109004591 | 109004718 | 109004591 | 109004715 | 109010915 | 109011718 | 13 | 10 | 8 | [illegible] |
| A3SS | 420 | ENSG00000105520 | PLPPR2 | chr19 | + | 11359531 | 11359720 | 11359606 | 11359720 | 11357659 | 11357739 | 13 | 4 | 5 | [illegible] |
| A3SS | 7247 | ENSG00000089127 | OAS1 | chr12 | + | 112911050 | 112911235 | 112911097 | 112911235 | 112908535 | 112908824 | 173 | 125 | 94 | [illegible] |
| A3SS | 897 | ENSG00000080189 | SLC35C2 | chr20 | - | 46355072 | 46355279 | 46355072 | 46355241 | 46355801 | 46355874 | 2 | 0 | 4 | [illegible] |
| A3SS | 449 | ENSG00000115268 | RPS15 | chr19 | + | 1439960 | 1440253 | 1440025 | 1440253 | 1438806 | 1438892 | 70 | 21 | 59 | [illegible] |
| A3SS | 8329 | ENSG00000039650 | PNKP | chr19 | - | 49866398 | 49866528 | 49866398 | 49866445 | 49867053 | 49867217 | 8 | 7 | 11 | [illegible] |
| A3SS | 343 | ENSG00000129351 | ILF3 | chr19 | + | 10683124 | 10683262 | 10683136 | 10683262 | 10682536 | 10682704 | 97 | 49 | 133 | [illegible] |
| A3SS | 13000 | ENSG00000089041 | P2RX7 | chr12 | + | 121162270 | 121162520 | 121162423 | 121162520 | 121156078 | 121156147 | 3 | 5 | 1 | [illegible] |
| A3SS | 11069 | ENSG00000100023 | PPIL2 | chr22 | + | 21695866 | 21698248 | 21696746 | 21698248 | 21695393 | 21695496 | 23 | 11 | 12 | [illegible] |
| A3SS | 292 | ENSG00000105618 | PRPF31 | chr19 | + | 54126499 | 54126617 | 54126527 | 54126617 | 54124498 | 54124656 | 1 | 1 | 5 | [illegible] |
| A3SS | 4139 | ENSG00000204463 | BAG6 | chr6 | - | 31644081 | 31644227 | 31644081 | 31644194 | 31644306 | 31644414 | 12 | 0 | 1 | [illegible] |
| A3SS | 5449 | ENSG00000177728 | TMEM94 | chr17 | + | 75494626 | 75494808 | 75494709 | 75494808 | 75493698 | 75493916 | 21 | 11 | 19 | [illegible] |
| A3SS | 8419 | ENSG00000223960 | CHROMR | chr2 | + | 178430677 | 178430889 | 178430769 | 178430889 | 178426365 | 178426505 | 15 | 17 | 14 | [illegible] |
| A3SS | 2219 | ENSG00000137804 | NUSAP1 | chr15 | + | 41365401 | 41365589 | 41365404 | 41365589 | 41358148 | 41358258 | 9 | 11 | 20 | [illegible] |
| A3SS | 13155 | ENSG00000205352 | PRR13 | chr12 | + | 53443390 | 53443773 | 53443540 | 53443773 | 53442694 | 53442733 | 260 | 141 | 178 | [illegible] |
| A3SS | 10065 | ENSG00000181444 | ZNF467 | chr7 | - | 149764181 | 149766239 | 149764181 | 149764678 | 149769089 | 149769200 | 118 | 97 | 75 | [illegible] |
| A3SS | 12437 | ENSG00000090621 | PABPC4 | chr1 | - | 39563613 | 39563922 | 39563613 | 39563741 | 39564422 | 39564542 | 51 | 31 | 50 | [illegible] |
| A3SS | 2837 | ENSG00000119599 | DCAF4 | chr14 | + | 72945883 | 72946027 | 72945887 | 72946027 | 72942993 | 72943096 | 28 | 17 | 28 | [illegible] |
| A3SS | 492 | ENSG00000063245 | EPN1 | chr19 | + | 55692950 | 55693037 | 55692953 | 55693037 | 55692685 | 55692796 | 15 | 10 | 8 | [illegible] |
| A3SS | 9992 | ENSG00000161036 | LRWD1 | chr7 | + | 102472881 | 102473168 | 102472908 | 102473168 | 102472691 | 102472804 | 3 | 3 | 2 | [illegible] |
| A3SS | 5286 | ENSG00000127054 | INTS11 | chr1 | - | 1319295 | 1319524 | 1319295 | 1319433 | 1320455 | 1320529 | 67 | 30 | 63 | [illegible] |
| A3SS | 6314 | ENSG00000196369 | SRGAP2B | chr1 | - | 144897115 | 144897340 | 144897115 | 144897337 | 144905090 | 144905219 | 13 | 4 | 3 | [illegible] |
| A3SS | 2039 | ENSG00000226232 | NPIPB14P | chr16 | - | 69982349 | 69982545 | 69982349 | 69982494 | 69982657 | 69982757 | 11 | 1 | 1 | [illegible] |
| A3SS | 6632 | ENSG00000237877 | LINC01473 | chr2 | - | 186083153 | 186083224 | 186083153 | 186083221 | 186086300 | 186086370 | 3 | 2 | 7 | [illegible] |
| A3SS | 8322 | ENSG00000018280 | SLC11A1 | chr2 | + | 218384242 | 218384365 | 218384318 | 218384365 | 218382959 | 218383070 | 68 | 32 | 16 | [illegible] |
| A3SS | 12811 | ENSG00000130119 | GNL3L | chrX | + | 54543145 | 54543342 | 54543206 | 54543342 | 54542954 | 54543038 | 4 | 0 | 1 | [illegible] |
| A3SS | 1638 | ENSG00000178188 | SH2B1 | chr16 | + | 28867330 | 28867432 | 28867373 | 28867432 | 28866011 | 28867033 | 92 | 58 | 116 | [illegible] |
| A3SS | 11295 | ENSG00000105879 | CBLL1 | chr7 | + | 107753410 | 107753511 | 107753413 | 107753511 | 107752544 | 107752617 | 18 | 15 | 24 | [illegible] |
| A3SS | 3434 | ENSG00000119685 | TTLL5 | chr14 | + | 75792912 | 75793100 | 75792915 | 75793100 | 75783146 | 75783530 | 14 | 9 | 5 | [illegible] |
| A3SS | 2696 | ENSG00000100897 | DCAF11 | chr14 | + | 24116916 | 24117044 | 24116971 | 24117044 | 24115514 | 24115726 | 55 | 24 | 37 | [illegible] |
| A3SS | 5411 | ENSG00000133226 | SRRM1 | chr1 | + | 24661309 | 24661396 | 24661312 | 24661396 | 24660718 | 24660799 | 189 | 170 | 153 | [illegible] |
| A3SS | 3177 | ENSG00000065357 | DGKA | chr12 | + | 55940081 | 55940433 | 55940313 | 55940433 | 55939414 | 55939529 | 122 | 87 | 224 | [illegible] |
| A3SS | 12960 | ENSG00000204482 | LST1 | chr6 | + | 31588496 | 31588909 | 31588517 | 31588909 | 31587943 | 31587966 | 26 | 19 | 7 | [illegible] |
| A3SS | 1032 | ENSG00000100983 | GSS | chr20 | - | 34928472 | 34929237 | 34928472 | 34928951 | 34929400 | 34929590 | 16 | 8 | 9 | [illegible] |
| A3SS | 3797 | ENSG00000145907 | G3BP1 | chr5 | + | 151786571 | 151786715 | 151786602 | 151786715 | 151771951 | 151772036 | 88 | 42 | 85 | [illegible] |
| A3SS | 8653 | ENSG00000178719 | GRINA | chr8 | + | 143991877 | 143992373 | 143992244 | 143992373 | 143991691 | 143991804 | 214 | 94 | 120 | [illegible] |
| A3SS | 3286 | ENSG00000161960 | EIF4A1 | chr17 | + | 7574521 | 7574678 | 7574545 | 7574678 | 7574259 | 7574308 | 8 | 2 | 11 | [illegible] |
| A3SS | 9686 | ENSG00000145191 | EIF2B5 | chr3 | + | 184141920 | 184142070 | 184141924 | 184142070 | 184140417 | 184140730 | 4 | 4 | 3 | [illegible] |
| A3SS | 12653 | ENSG00000107862 | GBF1 | chr10 | + | 102375319 | 102375584 | 102375358 | 102375584 | 102370706 | 102370860 | 2 | 0 | 0 | [illegible] |
| A3SS | 9904 | ENSG00000157764 | BRAF | chr7 | - | 140794307 | 140794467 | 140794307 | 140794415 | 140800361 | 140800481 | 160 | 114 | 155 | [illegible] |
| A3SS | 4315 | ENSG00000172053 | QARS1 | chr3 | - | 49103344 | 49103466 | 49103344 | 49103409 | 49103630 | 49103706 | 16 | 12 | 20 | [illegible] |
| A3SS | 13109 | ENSG00000130396 | AFDN | chr6 | + | 167870382 | 167870498 | 167870385 | 167870498 | 167864550 | 167864746 | 3 | 1 | 1 | [illegible] |
| A3SS | 12627 | ENSG00000107959 | PITRM1 | chr10 | - | 3165412 | 3165527 | 3165412 | 3165483 | 3166228 | 3166380 | 140 | 73 | 93 | [illegible] |
| A3SS | 2692 | ENSG00000100897 | DCAF11 | chr14 | + | 24115514 | 24115726 | 24115542 | 24115726 | 24114776 | 24114924 | 13 | 9 | 9 | [illegible] |
| A3SS | 3148 | ENSG00000140853 | NLRC5 | chr16 | + | 57055432 | 57055519 | 57055435 | 57055519 | 57055031 | 57055094 | 46 | 13 | 47 | [illegible] |
| A3SS | 9676 | ENSG00000026950 | BTN3A1 | chr6 | + | 26407670 | 26407952 | 26407826 | 26407952 | 26405908 | 26406256 | 455 | 218 | 460 | [illegible] |
| A3SS | 2513 | ENSG00000176438 | SYNE3 | chr14 | - | 95436819 | 95436981 | 95436819 | 95436966 | 95439032 | 95439162 | 32 | 16 | 18 | [illegible] |
| A3SS | 12780 | ENSG00000171792 | RHNO1 | chr12 | + | 2885161 | 2885534 | 2885282 | 2885534 | 2877222 | 2877282 | 5 | 3 | 22 | [illegible] |
| A3SS | 13123 | ENSG00000139428 | MMAB | chr12 | - | 109561039 | 109561329 | 109561039 | 109561104 | 109561419 | 109561517 | 3 | 0 | 4 | [illegible] |
| A3SS | 3936 | ENSG00000146350 | TBC1D32 | chr6 | - | 121317494 | 121317690 | 121317494 | 121317672 | 121321632 | 121321794 | 4 | 3 | 3 | [illegible] |
| A3SS | 7621 | ENSG00000061273 | HDAC7 | chr12 | - | 47789812 | 47791308 | 47789812 | 47789920 | 47791585 | 47791706 | 69 | 32 | 60 | [illegible] |
| A3SS | 11903 | ENSG00000149547 | EI24 | chr11 | + | 125575262 | 125575408 | 125575304 | 125575408 | 125572457 | 125572569 | 161 | 64 | 110 | [illegible] |
| A3SS | 10329 | ENSG00000166432 | ZMAT1 | chrX | - | 101895813 | 101895882 | 101895813 | 101895879 | 101897867 | 101898042 | 22 | 33 | 21 | [illegible] |
| A3SS | 8274 | ENSG00000106636 | YKT6 | chr7 | + | 44208406 | 44211124 | 44211022 | 44211124 | 44208132 | 44208198 | 15 | 12 | 9 | [illegible] |
| A3SS | 2289 | ENSG00000102901 | CENPT | chr16 | - | 67832227 | 67832350 | 67832227 | 67832315 | 67832454 | 67832545 | 3 | 6 | 5 | [illegible] |
| A3SS | 11631 | ENSG00000148356 | LRSAM1 | chr9 | + | 127487536 | 127487763 | 127487675 | 127487763 | 127485735 | 127485835 | 1 | 5 | 5 | [illegible] |
| A3SS | 9580 | ENSG00000255036 | SUGT1P4-STRA6LP-CCDC180 | chr9 | + | 97364017 | 97364128 | 97364050 | 97364128 | 97362362 | 97362441 | 0 | 5 | 4 | [illegible] |
| A3SS | 3512 | ENSG00000013306 | SLC25A39 | chr17 | - | 44322418 | 44322552 | 44322418 | 44322528 | 44322807 | 44322852 | 12 | 4 | 12 | [illegible] |
| A3SS | 4182 | ENSG00000242086 | MUC20-OT1 | chr3 | + | 195708428 | 195708841 | 195708746 | 195708841 | 195708177 | 195708268 | 7 | 0 | 7 | [illegible] |
| A3SS | 8152 | ENSG00000131051 | RBM39 | chr20 | - | 35729311 | 35729365 | 35729311 | 35729362 | 35729461 | 35729527 | 728 | 623 | 727 | [illegible] |
| A3SS | 12708 | ENSG00000151923 | TIAL1 | chr10 | - | 119582458 | 119582608 | 119582458 | 119582557 | 119588151 | 119588248 | 301 | 195 | 241 | [illegible] |
| A3SS | 4542 | ENSG00000159433 | STARD9 | chr15 | + | 42663280 | 42663917 | 42663819 | 42663917 | 42662793 | 42662891 | 10 | 19 | 13 | [illegible] |
| A3SS | 5675 | ENSG00000101096 | NFATC2 | chr20 | - | 51523080 | 51524110 | 51523080 | 51523596 | 51542369 | 51542719 | 79 | 18 | 40 | [illegible] |
| A3SS | 10824 | ENSG00000111364 | DDX55 | chr12 | + | 123607599 | 123607662 | 123607603 | 123607662 | 123607431 | 123607523 | 42 | 35 | 43 | [illegible] |
| A3SS | 9249 | ENSG00000137312 | FLOT1 | chr6 | - | 30741189 | 30741704 | 30741189 | 30741333 | 30741791 | 30741867 | 92 | 44 | 73 | [illegible] |
| A3SS | 1308 | ENSG00000141295 | SCRN2 | chr17 | - | 47839443 | 47839858 | 47839443 | 47839643 | 47840190 | 47840372 | 6 | 5 | 3 | [illegible] |
| A3SS | 1975 | ENSG00000217930 | PAM16 | chr16 | - | 4341367 | 4341583 | 4341367 | 4341504 | 4343206 | 4343291 | 3 | 0 | 3 | [illegible] |
| A3SS | 7062 | ENSG00000131165 | CHMP1A | chr16 | - | 89649350 | 89649520 | 89649350 | 89649497 | 89651568 | 89651646 | 9 | 7 | 8 | [illegible] |
| A3SS | 10028 | ENSG00000136451 | VEZF1 | chr17 | - | 57980602 | 57980804 | 57980602 | 57980786 | 57981872 | 57981936 | 18 | 13 | 18 | [illegible] |
| A3SS | 11704 | ENSG00000148343 | MIGA2 | chr9 | + | 129068018 | 129068332 | 129068197 | 129068332 | 129067772 | 129067871 | 0 | 0 | 0 | [illegible] |
| A3SS | 12374 | ENSG00000162241 | SLC25A45 | chr11 | - | 65380131 | 65380234 | 65380131 | 65380175 | 65381914 | 65382373 | 0 | 0 | 7 | [illegible] |
| A3SS | 10068 | ENSG00000204257 | HLA-DMA | chr6 | - | 32949610 | 32949889 | 32949610 | 32949787 | 32950518 | 32950803 | 504 | 282 | 403 | [illegible] |
| A3SS | 13250 | ENSG00000186908 | ZDHHC17 | chr12 | + | 76815856 | 76816019 | 76815897 | 76816019 | 76815145 | 76815210 | 120 | 88 | 85 | [illegible] |
| A3SS | 655 | ENSG00000130475 | FCHO1 | chr19 | + | 17788269 | 17788557 | 17788283 | 17788557 | 17787681 | 17787846 | 1 | 0 | 0 | [illegible] |
| A3SS | 4074 | ENSG00000163694 | RBM47 | chr4 | - | 40436440 | 40436885 | 40436440 | 40436647 | 40437770 | 40438924 | 9 | 15 | 12 | [illegible] |
| A3SS | 10469 | ENSG00000100802 | C14orf93 | chr14 | - | 22999053 | 22999402 | 22999053 | 22999156 | 23000958 | 23001037 | 24 | 8 | 23 | [illegible] |
| A3SS | 1536 | ENSG00000131467 | PSME3 | chr17 | + | 42838691 | 42838799 | 42838730 | 42838799 | 42838092 | 42838205 | 1 | 3 | 1 | [illegible] |
| A3SS | 6413 | ENSG00000132781 | MUTYH | chr1 | - | 45333412 | 45333594 | 45333412 | 45333561 | 45334390 | 45334511 | 9 | 8 | 11 | [illegible] |
| A3SS | 11354 | ENSG00000187239 | FNBP1 | chr9 | - | 129923843 | 129924026 | 129923843 | 129923939 | 129924959 | 129925157 | 28 | 29 | 17 | [illegible] |
| A3SS | 11630 | ENSG00000148356 | LRSAM1 | chr9 | + | 127487494 | 127487763 | 127487675 | 127487763 | 127485735 | 127485835 | 1 | 5 | 4 | [illegible] |

| | | | | | | | | | | | | | | |
|---|---|---|---|---|---|---|---|---|---|---|---|---|---|---|
| A3SS | 5983 | ENSG00000183023 | SLC8A1 | chr2 | - | 40428472 | 40430304 | 40428472 | 40430301 | 40451903 | 40452069 | 34 | 27 | 20 |
| A3SS | 8835 | ENSG00000135426 | TESPA1 | chr12 | - | 54967842 | 54968013 | 54967842 | 54967892 | 54973476 | 54973519 | 8 | 14 | 38 |
| A3SS | 4519 | ENSG00000156709 | AIFM1 | chrX | - | 130129969 | 130130193 | 130129969 | 130130166 | 130131674 | 130131799 | 3 | 0 | 2 |
| A3SS | 4973 | ENSG00000213024 | NUP62 | chr19 | - | 49927693 | 49927848 | 49927693 | 49927815 | 49929416 | 49929504 | 3 | 6 | 8 |
| A3SS | 8300 | ENSG00000120159 | CAAP1 | chr9 | - | 26884809 | 26884902 | 26884809 | 26884885 | 26886103 | 26886188 | 2 | 0 | 1 |
| A3SS | 8454 | ENSG00000180902 | D2HGDH | chr2 | + | 241755044 | 241756014 | 241755848 | 241756014 | 241751245 | 241751388 | 13 | 4 | 14 |
| A3SS | 10746 | ENSG00000147050 | KDM6A | chrX | + | 45106541 | 45107536 | 45107409 | 45107536 | 45090722 | 45090864 | 27 | 33 | 20 |
| A3SS | 12948 | ENSG00000111144 | LTA4H | chr12 | - | 96003837 | 96003981 | 96003837 | 96003920 | 96006313 | 96006409 | 17 | 7 | 7 |
| A3SS | 6552 | ENSG00000127463 | EMC1 | chr1 | - | 19238794 | 19238857 | 19238794 | 19238854 | 19239230 | 19239302 | 37 | 13 | 35 |
| A3SS | 1888 | ENSG00000130175 | PRKCSH | chr19 | + | 11447692 | 11447789 | 11447722 | 11447789 | 11447438 | 11447618 | 39 | 17 | 63 |
| A3SS | 9268 | ENSG00000135486 | HNRNPA1 | chr12 | + | 54283791 | 54283967 | 54283811 | 54283967 | 54282799 | 54282874 | 37 | 46 | 57 |
| A3SS | 12870 | ENSG00000161638 | ITGA5 | chr12 | - | 54403186 | 54403779 | 54403186 | 54403324 | 54403910 | 54403966 | 129 | 60 | 115 |
| A3SS | 11273 | ENSG00000180776 | ZDHHC20 | chr13 | - | 21381433 | 21381549 | 21381433 | 21381546 | 21382919 | 21383009 | 212 | 161 | 163 |
| A3SS | 577 | ENSG00000090372 | STRN4 | chr19 | - | 46727451 | 46727691 | 46727451 | 46727567 | 46727893 | 46728007 | 8 | 11 | 8 |
| A3SS | 2885 | ENSG00000111913 | RIPOR2 | chr6 | - | 24839358 | 24840775 | 24839358 | 24839599 | 24842861 | 24843554 | 773 | 449 | 784 |
| A3SS | 9367 | ENSG00000237441 | RGL2 | chr6 | - | 33293412 | 33293699 | 33293412 | 33293524 | 33293794 | 33293916 | 74 | 38 | 31 |
| A3SS | 12140 | ENSG00000255717 | SNHG1 | chr11 | - | 62852810 | 62852993 | 62852810 | 62852839 | 62853074 | 62853115 | 211 | 160 | 270 |
| A3SS | 2226 | ENSG00000128915 | ICE2 | chr15 | - | 60468060 | 60468322 | 60468060 | 60468162 | 60476062 | 60476167 | 140 | 94 | 93 |
| A3SS | 10112 | ENSG00000145375 | SPATA5 | chr4 | + | 122927633 | 122927758 | 122927636 | 122927758 | 122923077 | 122923305 | 12 | 10 | 15 |
| A3SS | 1875 | ENSG00000197912 | SPG7 | chr16 | + | 89552851 | 89553135 | 89552978 | 89553135 | 89550493 | 89550609 | 2 | 0 | 2 |
| A3SS | 13348 | ENSG00000085872 | CHERP | chr19 | - | 16530768 | 16530913 | 16530768 | 16530880 | 16532597 | 16532749 | 8 | 3 | 5 |
| A3SS | 1803 | ENSG00000005844 | ITGAL | chr16 | + | 30505243 | 30505300 | 30505246 | 30505300 | 30499337 | 30499489 | 6 | 4 | 7 |
| A3SS | 3366 | ENSG00000172785 | CBWD1 | chr9 | - | 121088 | 122090 | 121088 | 121573 | 123216 | 123282 | 53 | 126 | 48 |
| A3SS | 13306 | ENSG00000168883 | USP39 | chr2 | + | 85623625 | 85623782 | 85623645 | 85623782 | 85621484 | 85621579 | 15 | 1 | 6 |
| A3SS | 13379 | ENSG00000102710 | SUPT20H | chr13 | - | 37022010 | 37023044 | 37022010 | 37022317 | 37024034 | 37024193 | 126 | 84 | 119 |
| A3SS | 2248 | ENSG00000128739 | SNRPN | chr15 | + | 24974310 | 24974456 | 24974386 | 24974456 | 24967931 | 24968082 | 418 | 226 | 409 |
| A3SS | 12167 | ENSG00000174483 | BBS1 | chr11 | + | 66515539 | 66515731 | 66515692 | 66515731 | 66514405 | 66514678 | 10 | 17 | 23 |
| A3SS | 10806 | ENSG00000142046 | TMEM91 | chr19 | + | 41383560 | 41384083 | 41383714 | 41384083 | 41382771 | 41382921 | 15 | 19 | 19 |
| A3SS | 4489 | ENSG00000134802 | SLC43A3 | chr11 | - | 57425540 | 57425709 | 57425540 | 57425670 | 57425988 | 57426316 | 18 | 6 | 11 |
| A3SS | 2327 | ENSG00000092439 | TRPM7 | chr15 | - | 50583088 | 50583159 | 50583088 | 50583156 | 50586391 | 50586488 | 153 | 111 | 113 |
| A3SS | 6344 | ENSG00000142784 | WDTC1 | chr1 | + | 27305000 | 27305193 | 27305058 | 27305193 | 27303620 | 27303795 | 158 | 67 | 134 |
| A3SS | 8346 | ENSG00000197858 | GPAA1 | chr8 | + | 144084110 | 144084330 | 144084133 | 144084330 | 144083938 | 144084040 | 3 | 2 | 4 |
| A3SS | 9187 | ENSG00000135049 | AGTPBP1 | chr9 | - | 85655142 | 85655320 | 85655142 | 85655200 | 85657434 | 85657643 | 417 | 250 | 300 |
| A3SS | 2130 | ENSG00000008710 | PKD1 | chr16 | - | 2105321 | 2105707 | 2105321 | 2105474 | 2105864 | 2106024 | 12 | 0 | 3 |
| A3SS | 12250 | ENSG00000167986 | DDB1 | chr11 | - | 61314046 | 61314210 | 61314046 | 61314202 | 61314307 | 61314486 | 306 | 137 | 259 |
| [illegible] | | | | | | | | | | | | | | |

| | | | | | | | | | | | | | |
|---|---|---|---|---|---|---|---|---|---|---|---|---|---|
| A3SS | 9218 | ENSG00000204231 | RXRB | chr6 | - | 33195362 | 33195466 | 33195362 | 33195454 | 33195569 | 33195702 | 37 | 11 | 26 | [illegible] |
| A3SS | 2385 | ENSG00000140400 | MAN2C1 | chr15 | - | 75361854 | 75362025 | 75361854 | 75361947 | 75362342 | 75362453 | 5 | 1 | 4 | [illegible] |
| A3SS | 1069 | ENSG00000154889 | MPPE1 | chr18 | - | 11886498 | 11886778 | 11886498 | 11886621 | 11886916 | 11887025 | 41 | 19 | 25 | [illegible] |
| A3SS | 3102 | ENSG00000151612 | ZNF827 | chr4 | - | 145892242 | 145892415 | 145892242 | 145892409 | 145902165 | 145903215 | 23 | 12 | 18 | [illegible] |
| A3SS | 8306 | ENSG00000126768 | TIMM17B | chrX | - | 48895351 | 48895972 | 48895351 | 48895454 | 48896758 | 48896858 | 14 | 8 | 13 | [illegible] |
| A3SS | 8582 | ENSG00000144306 | SCRN3 | chr2 | + | 174398274 | 174398442 | 174398365 | 174398442 | 174397335 | 174397404 | 45 | 62 | 54 | [illegible] |
| A3SS | 12483 | ENSG00000100109 | TFIP11 | chr22 | - | 26511118 | 26511317 | 26511118 | 26511153 | 26512098 | 26512175 | 15 | 14 | 21 | [illegible] |
| A3SS | 9737 | ENSG00000168255 | POLR2J3 | chr7 | - | 102543524 | 102543702 | 102543524 | 102543697 | 102544705 | 102544776 | 6 | 2 | 7 | [illegible] |
| A3SS | 2172 | ENSG00000159322 | ADPGK | chr15 | - | 72751368 | 72752895 | 72751368 | 72752892 | 72755555 | 72755654 | 58 | 46 | 49 | [illegible] |
| A3SS | 11594 | ENSG00000213930 | GALT | chr9 | + | 34647540 | 34648171 | 34648114 | 34648171 | 34647088 | 34647258 | 27 | 12 | 22 | [illegible] |
| A3SS | 9009 | ENSG00000111011 | RSRC2 | chr12 | - | 122518838 | 122519051 | 122518838 | 122519029 | 122520503 | 122520581 | 60 | 57 | 62 | [illegible] |
| A3SS | 10560 | ENSG00000172534 | HCFC1 | chrX | - | 153951840 | 153952161 | 153951840 | 153952158 | 153952513 | 153952958 | 2 | 3 | 3 | [illegible] |
| A3SS | 8520 | ENSG00000177674 | AGTRAP | chr1 | + | 11745987 | 11746223 | 11746126 | 11746223 | 11745802 | 11745837 | 12 | 6 | 29 | [illegible] |
| A3SS | 4759 | ENSG00000099330 | OCEL1 | chr19 | + | 17228802 | 17229219 | 17229008 | 17229219 | 17228255 | 17228309 | 51 | 48 | 61 | [illegible] |
| A3SS | 2669 | ENSG00000257621 | PSMA3-AS1 | chr14 | - | 58273978 | 58274135 | 58273978 | 58274010 | 58291634 | 58291707 | 17 | 18 | 25 | [illegible] |
| A3SS | 5224 | ENSG00000084072 | PPIE | chr1 | + | 39740088 | 39740263 | 39740164 | 39740263 | 39738900 | 39738931 | 2 | 0 | 0 | [illegible] |
| A3SS | 12695 | ENSG00000152778 | IFIT5 | chr10 | + | 89417187 | 89420997 | 89417204 | 89420997 | 89414567 | 89414803 | 5 | 7 | 5 | [illegible] |
| A3SS | 5811 | ENSG00000162434 | JAK1 | chr1 | - | 64866865 | 64867079 | 64866865 | 64867076 | 64869310 | 64869474 | 32 | 16 | 44 | [illegible] |
| A3SS | 717 | ENSG00000031823 | RANBP3 | chr19 | - | 5933413 | 5933656 | 5933413 | 5933479 | 5941620 | 5941707 | 14 | 10 | 16 | [illegible] |
| A3SS | 3413 | ENSG00000150593 | PDCD4 | chr10 | + | 110881232 | 110881535 | 110881274 | 110881535 | 110875965 | 110876070 | 689 | 664 | 709 | [illegible] |
| A3SS | 316 | ENSG00000141867 | BRD4 | chr19 | - | 15268153 | 15268281 | 15268153 | 15268240 | 15268904 | 15269042 | 21 | 32 | 31 | [illegible] |
| A3SS | 1322 | ENSG00000109113 | RAB34 | chr17 | - | 28715194 | 28715276 | 28715194 | 28715214 | 28715455 | 28715507 | 51 | 34 | 46 | [illegible] |
| A3SS | 13328 | ENSG00000121310 | ECHDC2 | chr1 | - | 52911722 | 52912021 | 52911722 | 52911790 | 52913894 | 52914095 | 14 | 23 | 25 | [illegible] |
| A3SS | 4695 | ENSG00000158062 | UBXN11 | chr1 | - | 26300925 | 26301053 | 26300925 | 26301024 | 26301693 | 26301722 | 20 | 8 | 4 | [illegible] |
| A3SS | 12951 | ENSG00000127314 | RAP1B | chr12 | + | 68624507 | 68624873 | 68624670 | 68624873 | 68610907 | 68611043 | 32 | 38 | 25 | [illegible] |
| A3SS | 5152 | ENSG00000092841 | MYL6 | chr12 | + | 56159586 | 56160148 | 56159974 | 56160148 | 56158683 | 56158711 | 409 | 312 | 284 | [illegible] |
| A3SS | 6871 | ENSG00000117859 | OSBPL9 | chr1 | + | 51760665 | 51760780 | 51760689 | 51760780 | 51756319 | 51756358 | 1 | 3 | 7 | [illegible] |
| A3SS | 410 | ENSG00000104872 | PIH1D1 | chr19 | - | 49446550 | 49446794 | 49446550 | 49446694 | 49447023 | 49447099 | 3 | 2 | 3 | [illegible] |
| A3SS | 13087 | ENSG00000187109 | NAP1L1 | chr12 | - | 76056032 | 76056161 | 76056032 | 76056158 | 76059797 | 76059878 | 1685 | 1913 | 1666 | [illegible] |
| A3SS | 9350 | ENSG00000170919 | TPT1-AS1 | chr13 | + | 45377060 | 45377215 | 45377111 | 45377215 | 45369958 | 45370182 | 7 | 7 | 11 | [illegible] |
| A3SS | 1154 | ENSG00000100425 | BRD1 | chr22 | - | 49787389 | 49787522 | 49787389 | 49787494 | 49794033 | 49794294 | 18 | 6 | 9 | [illegible] |
| A3SS | 1940 | ENSG00000213658 | LAT | chr16 | + | 28986354 | 28986446 | 28986381 | 28986446 | 28986134 | 28986216 | 7 | 1 | 7 | [illegible] |
| A3SS | 6863 | ENSG00000189362 | NEMP2 | chr2 | - | 190518951 | 190519183 | 190518951 | 190519022 | 190525262 | 190525378 | 54 | 36 | 61 | [illegible] |
| A3SS | 5478 | ENSG00000135597 | REPS1 | chr6 | - | 138917554 | 138917627 | 138917554 | 138917624 | 138920214 | 138920316 | 123 | 113 | 120 | [illegible] |
| A3SS | 9151 | ENSG00000072121 | ZFYVE26 | chr14 | - | 67775015 | 67775132 | 67775015 | 67775114 | 67775859 | 67776106 | 4 | 4 | 0 | [illegible] |
| A3SS | 2509 | ENSG00000197555 | SIPA1L1 | chr14 | + | 71704720 | 71705340 | 71705221 | 71705340 | 71702380 | 71702505 | 6 | 10 | 13 | [illegible] |
| A3SS | 12563 | ENSG00000078668 | VDAC3 | chr8 | + | 42396655 | 42396680 | 42396677 | 42396680 | 42395083 | 42395133 | 0 | 6 | 0 | [illegible] |
| A3SS | 3345 | ENSG00000156603 | MED19 | chr11 | - | 57704301 | 57704635 | 57704301 | 57704396 | 57704718 | 57704815 | 16 | 10 | 3 | [illegible] |
| A3SS | 10672 | ENSG00000248323 | LUCAT1 | chr5 | - | 91310626 | 91312495 | 91310626 | 91312413 | 91313077 | 91313229 | 11 | 16 | 9 | [illegible] |
| A3SS | 12881 | ENSG00000172819 | RARG | chr12 | - | 53215292 | 53215462 | 53215292 | 53215434 | 53215645 | 53215794 | 2 | 0 | 3 | [illegible] |
| A3SS | 9440 | ENSG00000064419 | TNPO3 | chr7 | - | 128989960 | 128990202 | 128989960 | 128990100 | 128991998 | 128992090 | 5 | 6 | 6 | [illegible] |
| A3SS | 2134 | ENSG00000100726 | TELO2 | chr16 | + | 1506224 | 1506329 | 1506237 | 1506329 | 1505409 | 1505601 | 0 | 0 | 4 | [illegible] |
| A3SS | 4592 | ENSG00000111642 | CHD4 | chr12 | - | 6600216 | 6600395 | 6600216 | 6600356 | 6600533 | 6600669 | 156 | 98 | 162 | [illegible] |
| A3SS | 5520 | ENSG00000116584 | ARHGEF2 | chr1 | - | 155964987 | 155965131 | 155964987 | 155965128 | 155965302 | 155965412 | 97 | 77 | 96 | [illegible] |
| A3SS | 9884 | ENSG00000126883 | NUP214 | chr9 | + | 131144279 | 131144754 | 131144567 | 131144754 | 131140548 | 131140710 | 75 | 46 | 68 | [illegible] |
| A3SS | 8427 | ENSG00000196295 | GARS1-DT | chr7 | - | 30563730 | 30563998 | 30563730 | 30563893 | 30568832 | 30568989 | 127 | 119 | 59 | [illegible] |
| A3SS | 1921 | ENSG00000157353 | FCSK | chr16 | + | 70474527 | 70474694 | 70474625 | 70474694 | 70474128 | 70474339 | 6 | 4 | 3 | [illegible] |
| A3SS | 11697 | ENSG00000148396 | SEC16A | chr9 | - | 136446854 | 136447015 | 136446854 | 136446949 | 136447226 | 136447364 | 10 | 1 | 3 | [illegible] |
| A3SS | 10199 | ENSG00000087274 | ADD1 | chr4 | + | 2926638 | 2926675 | 2926641 | 2926675 | 2926013 | 2926112 | 0 | 0 | 0 | [illegible] |
| A3SS | 8 | ENSG00000160255 | ITGB2 | chr21 | - | 44910283 | 44910415 | 44910283 | 44910372 | 44910599 | 44910785 | 7 | 4 | 8 | [illegible] |
| A3SS | 4948 | ENSG00000113810 | SMC4 | chr3 | + | 160404327 | 160404504 | 160404377 | 160404504 | 160402675 | 160402867 | 281 | 263 | 249 | [illegible] |
| A3SS | 1633 | ENSG00000172530 | BANP | chr16 | + | 88035322 | 88035394 | 88035331 | 88035394 | 88033108 | 88033245 | 23 | 5 | 10 | [illegible] |
| A3SS | 8445 | ENSG00000147454 | SLC25A37 | chr8 | + | 23568258 | 23568378 | 23568321 | 23568378 | 23566107 | 23566336 | 24 | 23 | 19 | [illegible] |
| A3SS | 8352 | ENSG00000162999 | DUSP19 | chr2 | + | 183087039 | 183087192 | 183087157 | 183087192 | 183083507 | 183083554 | 10 | 13 | 9 | [illegible] |
| A3SS | 11671 | ENSG00000142396 | ERVK3-1 | chr19 | + | 58312027 | 58312462 | 58312196 | 58312462 | 58306087 | 58306216 | 76 | 69 | 114 | [illegible] |
| A3SS | 42 | ENSG00000159131 | GART | chr21 | - | 33539170 | 33539356 | 33539170 | 33539323 | 33542547 | 33542820 | 13 | 6 | 9 | [illegible] |
| A3SS | 12961 | ENSG00000130177 | CDC16 | chr13 | + | 114236798 | 114236896 | 114236801 | 114236896 | 114236644 | 114236699 | 69 | 57 | 87 | [illegible] |
| A3SS | 3420 | ENSG00000171206 | TRIM8 | chr10 | + | 102656244 | 102656385 | 102656269 | 102656385 | 102656105 | 102656137 | 3 | 4 | 4 | [illegible] |
| A3SS | 5745 | ENSG00000116455 | WDR77 | chr1 | - | 111442024 | 111442129 | 111442024 | 111442093 | 111442652 | 111442761 | 0 | 0 | 0 | [illegible] |
| A3SS | 8659 | ENSG00000163516 | ANKZF1 | chr2 | + | 219234132 | 219235312 | 219234825 | 219235312 | 219233285 | 219233433 | 205 | 132 | 171 | [illegible] |
| A3SS | 10484 | ENSG00000177646 | ACAD9 | chr3 | + | 128893518 | 128893656 | 128893554 | 128893656 | 128884652 | 128884746 | 0 | 1 | 1 | [illegible] |
| A3SS | 5159 | ENSG00000122483 | CCDC18 | chr1 | + | 93214742 | 93214966 | 93214745 | 93214966 | 93212100 | 93212261 | 26 | 22 | 17 | [illegible] |
| A3SS | 9006 | ENSG00000111011 | RSRC2 | chr12 | - | 122508217 | 122508450 | 122508217 | 122508447 | 122511108 | 122511188 | 194 | 181 | 169 | [illegible] |
| A3SS | 10038 | ENSG00000065000 | AP3D1 | chr19 | - | 2112859 | 2112967 | 2112859 | 2112931 | 2113335 | 2113413 | 35 | 13 | 23 | [illegible] |
| A3SS | 10807 | ENSG00000142046 | TMEM91 | chr19 | + | 41383560 | 41384083 | 41383862 | 41384083 | 41382771 | 41382921 | 30 | 20 | 23 | [illegible] |
| A3SS | 9267 | ENSG00000135486 | HNRNPA1 | chr12 | + | 54283778 | 54283967 | 54283811 | 54283967 | 54282799 | 54282874 | 42 | 58 | 81 | [illegible] |
| A3SS | 9841 | ENSG00000099901 | RANBP1 | chr22 | + | 20126951 | 20127355 | 20126954 | 20127355 | 20126302 | 20126368 | 128 | 78 | 131 | [illegible] |
| A3SS | 9159 | ENSG00000204410 | MSH5 | chr6 | + | 31761817 | 31761955 | 31761865 | 31761955 | 31761471 | 31761615 | 14 | 8 | 11 | [illegible] |
| A3SS | 13148 | ENSG00000123349 | PFDN5 | chr12 | + | 53296429 | 53297924 | 53297849 | 53297924 | 53295838 | 53295941 | 47 | 41 | 68 | [illegible] |
| A3SS | 9722 | ENSG00000119616 | FCF1 | chr14 | + | 74715950 | 74716099 | 74715993 | 74716099 | 74714871 | 74714943 | 254 | 189 | 231 | [illegible] |
| A3SS | 10102 | ENSG00000106263 | EIF3B | chr7 | + | 2380203 | 2380740 | 2380278 | 2380740 | 2379393 | 2379511 | 97 | 26 | 68 | [illegible] |
| A3SS | 10173 | ENSG00000160408 | ST6GALNAC6 | chr9 | - | 127894511 | 127894691 | 127894511 | 127894629 | 127896241 | 127896332 | 122 | 32 | 93 | [illegible] |
| A3SS | 4332 | ENSG00000247596 | TWF2 | chr3 | - | 52231943 | 52232122 | 52231943 | 52232117 | 52235028 | 52235106 | 164 | 58 | 132 | [illegible] |
| A3SS | 10890 | ENSG00000205581 | HMGN1 | chr21 | - | 39347378 | 39347483 | 39347378 | 39347478 | 39348291 | 39348339 | 18 | 14 | 16 | [illegible] |
| A3SS | 10240 | ENSG00000196998 | WDR45 | chrX | - | 49076644 | 49076804 | 49076644 | 49076753 | 49077642 | 49077747 | 4 | 0 | 8 | [illegible] |
| A3SS | 8519 | ENSG00000177674 | AGTRAP | chr1 | + | 11745929 | 11746223 | 11746126 | 11746223 | 11745802 | 11745837 | 12 | 5 | 29 | [illegible] |
| A3SS | 1956 | ENSG00000169592 | INO80E | chr16 | + | 30001413 | 30001530 | 30001464 | 30001530 | 30000928 | 30001040 | 112 | 59 | 104 | [illegible] |
| A3SS | 5578 | ENSG00000182923 | CEP63 | chr3 | + | 134545572 | 134545819 | 134545585 | 134545819 | 134537154 | 134537268 | 1 | 1 | 2 | [illegible] |
| A3SS | 2844 | ENSG00000103811 | CTSH | chr15 | - | 78931450 | 78932041 | 78931450 | 78932038 | 78932371 | 78932458 | 5 | 3 | 4 | [illegible] |
| A3SS | 11842 | ENSG00000113163 | CERT1 | chr5 | - | 75400204 | 75400297 | 75400204 | 75400291 | 75402971 | 75403058 | 176 | 193 | 164 | [illegible] |
| A3SS | 8254 | ENSG00000135632 | SMYD5 | chr2 | + | 73219951 | 73220190 | 73220050 | 73220190 | 73218860 | 73218969 | 9 | 7 | 12 | [illegible] |
| A3SS | 10546 | ENSG00000102054 | RBBP7 | chrX | - | 16858675 | 16858849 | 16858675 | 16858822 | 16862954 | 16863100 | 408 | 275 | 419 | [illegible] |
| A3SS | 8399 | ENSG00000131236 | CAP1 | chr1 | + | 40059333 | 40059458 | 40059336 | 40059458 | 40040728 | 40040801 | 151 | 70 | 87 | [illegible] |
| A3SS | 1202 | ENSG00000120071 | KANSL1 | chr17 | - | 46034160 | 46034285 | 46034160 | 46034282 | 46038537 | 46038686 | 72 | 24 | 55 | [illegible] |
| A3SS | 1942 | ENSG00000213658 | LAT | chr16 | + | 28986381 | 28986446 | 28986384 | 28986446 | 28986134 | 28986216 | 63 | 55 | 72 | [illegible] |
| A3SS | 4887 | ENSG00000116871 | MAP7D1 | chr1 | + | 36179511 | 36179557 | 36179514 | 36179557 | 36179261 | 36179315 | 33 | 16 | 52 | [illegible] |
| A3SS | 685 | ENSG00000167468 | GPX4 | chr19 | + | 1106377 | 1106459 | 1106399 | 1106459 | 1106241 | 1106266 | 15 | 7 | 14 | [illegible] |
| A3SS | 3964 | ENSG00000171298 | GAA | chr17 | + | 80112844 | 80113027 | 80112875 | 80113027 | 80112577 | 80112711 | 5 | 2 | 6 | [illegible] |
| A3SS | 921 | ENSG00000105576 | TNPO2 | chr19 | - | 12702071 | 12702177 | 12702071 | 12702147 | 12702822 | 12702918 | 4 | 4 | 6 | [illegible] |
| A3SS | 5420 | ENSG00000125637 | PSD4 | chr2 | + | 113185661 | 113186255 | 113185876 | 113186255 | 113185364 | 113185440 | 64 | 13 | 27 | [illegible] |
| A3SS | 2761 | ENSG00000119718 | EIF2B2 | chr14 | + | 75005865 | 75005961 | 75005896 | 75005961 | 75004736 | 75004900 | 68 | 39 | 41 | [illegible] |
| A3SS | 10633 | ENSG00000059728 | MXD1 | chr2 | + | 69937018 | 69937394 | 69937234 | 69937394 | 69935350 | 69935465 | 14 | 11 | 6 | [illegible] |
| A3SS | 6578 | ENSG00000048707 | VPS13D | chr1 | + | 12341779 | 12341885 | 12341782 | 12341885 | 12335704 | 12335827 | 28 | 16 | 21 | [illegible] |
| A3SS | 2679 | ENSG00000196497 | IPO4 | chr14 | - | 24182277 | 24182423 | 24182277 | 24182403 | 24182791 | 24182842 | 0 | 0 | 0 | [illegible] |
| A3SS | 11962 | ENSG00000160226 | CFAP410 | chr21 | - | 44333032 | 44333262 | 44333032 | 44333253 | 44335757 | 44335804 | 9 | 10 | 6 | [illegible] |
| A3SS | 7308 | ENSG00000196154 | S100A4 | chr1 | - | 153543612 | 153543939 | 153543612 | 153543923 | 153544653 | 153544809 | 76 | 25 | 16 | [illegible] |
| A3SS | 12007 | ENSG00000136527 | TRA2B | chr3 | - | 185923795 | 185924008 | 185923795 | 185923984 | 185925463 | 185925626 | 25 | 24 | 26 | [illegible] |
| A3SS | 116 | ENSG00000189306 | RRP7A | chr22 | - | 42516010 | 42516296 | 42516010 | 42516136 | 42518004 | 42518147 | 27 | 20 | 31 | [illegible] |
| A3SS | 489 | ENSG00000071626 | DAZAP1 | chr19 | + | 1421147 | 1421258 | 1421150 | 1421258 | 1418665 | 1418731 | 236 | 149 | 252 | [illegible] |
| A3SS | 8512 | ENSG00000004897 | CDC27 | chr17 | - | 47141852 | 47142025 | 47141852 | 47142022 | 47142228 | 47142436 | 35 | 27 | 31 | [illegible] |
| A3SS | 7989 | ENSG00000152689 | RASGRP3 | chr2 | + | 33511709 | 33511842 | 33511811 | 33511842 | 33476648 | 33476707 | 18 | 21 | 28 | [illegible] |
| A3SS | 4270 | ENSG00000157601 | MX1 | chr21 | + | 41432030 | 41432175 | 41432049 | 41432175 | 41430532 | 41430608 | 9 | 14 | 5 | [illegible] |
| A3SS | 4067 | ENSG00000017260 | ATP2C1 | chr3 | + | 130992930 | 130993001 | 130992950 | 130993001 | 130980581 | 130980679 | 8 | 11 | 6 | [illegible] |
| A3SS | 9514 | ENSG00000152457 | DCLRE1C | chr10 | - | 14942113 | 14942435 | 14942113 | 14942240 | 14945104 | 14945189 | 20 | 13 | 12 | [illegible] |
| A3SS | 10309 | ENSG00000164062 | APEH | chr3 | + | 49676055 | 49676515 | 49676377 | 49676515 | 49675890 | 49675966 | 320 | 179 | 229 | [illegible] |
| A3SS | 11898 | ENSG00000129255 | MPDU1 | chr17 | + | 7586655 | 7586777 | 7586691 | 7586777 | 7585945 | 7586078 | 15 | 11 | 16 | [illegible] |
| A3SS | 496 | ENSG00000180061 | TMEM150B | chr19 | - | 55320968 | 55321093 | 55320968 | 55321044 | 55322647 | 55322743 | 2 | 12 | 0 | [illegible] |
| A3SS | 2389 | ENSG00000140400 | MAN2C1 | chr15 | - | 75368072 | 75368198 | 75368072 | 75368159 | 75368482 | 75368612 | 74 | 30 | 97 | [illegible] |
| A3SS | 1858 | ENSG00000205336 | ADGRG1 | chr16 | + | 57659394 | 57659681 | 57659412 | 57659681 | 57657372 | 57657491 | 68 | 8 | 14 | [illegible] |
| A3SS | 12184 | ENSG00000110075 | PPP6R3 | chr11 | + | 68587926 | 68588024 | 68587977 | 68588024 | 68583042 | 68583129 | 406 | 324 | 309 | [illegible] |
| A3SS | 1843 | ENSG00000141002 | TCF25 | chr16 | + | 89895037 | 89895137 | 89895082 | 89895137 | 89893727 | 89893858 | 225 | 99 | 139 | [illegible] |
| A3SS | 7549 | ENSG00000078808 | SDF4 | chr1 | - | 1218768 | 1218943 | 1218768 | 1218927 | 1223243 | 1223357 | 4 | 7 | 2 | [illegible] |
| A3SS | 10708 | ENSG00000147133 | TAF1 | chrX | + | 71397252 | 71397466 | 71397255 | 71397466 | 71394066 | 71394245 | 175 | 141 | 159 | [illegible] |
| A3SS | 13357 | ENSG00000144567 | RETREG2 | chr2 | + | 219180109 | 219180245 | 219180197 | 219180245 | 219179732 | 219179763 | 62 | 26 | 94 | [illegible] |
| A3SS | 6973 | ENSG00000149474 | KAT14 | chr20 | + | 18161819 | 18162239 | 18161822 | 18162239 | 18159083 | 18159265 | 54 | 44 | 69 | [illegible] |
| A3SS | 6530 | ENSG00000071054 | MAP4K4 | chr2 | + | 101859642 | 101859864 | 101859735 | 101859864 | 101858995 | 101859082 | 90 | 79 | 85 | [illegible] |
| A3SS | 7676 | ENSG00000092529 | CAPN3 | chr15 | + | 42410794 | 42411000 | 42410883 | 42411000 | 42410587 | 42410666 | 5 | 1 | 5 | [illegible] |
| A3SS | 7041 | ENSG00000120948 | TARDBP | chr1 | + | 11024063 | 11024335 | 11024301 | 11024335 | 11023192 | 11023329 | 123 | 113 | 142 | [illegible] |
| A3SS | 12287 | ENSG00000110011 | DNAJC4 | chr11 | + | 64232700 | 64232865 | 64232703 | 64232865 | 64232429 | 64232614 | 3 | 1 | 14 | [illegible] |
| A3SS | 3712 | ENSG00000109814 | UGDH | chr4 | - | 39514082 | 39514184 | 39514082 | 39514180 | 39521350 | 39521519 | 71 | 40 | 53 | [illegible] |
| A3SS | 13355 | ENSG00000164649 | CDCA7L | chr7 | - | 21911616 | 21911754 | 21911616 | 21911751 | 21916753 | 21916894 | 30 | 16 | 48 | [illegible] |
| A3SS | 11805 | ENSG00000062822 | POLD1 | chr19 | + | 50408706 | 50408901 | 50408784 | 50408901 | 50407326 | 50407415 | 1 | 0 | 2 | [illegible] |
| A3SS | 3877 | ENSG00000061918 | GUCY1B1 | chr4 | + | 155802212 | 155802579 | 155802341 | 155802579 | 155799876 | 155800074 | 1 | 2 | 2 | [illegible] |
| A3SS | 4950 | ENSG00000141293 | SKAP1 | chr17 | - | 48162468 | 48162569 | 48162468 | 48162566 | 48170608 | 48170659 | 188 | 116 | 201 | [illegible] |
| A3SS | 9751 | ENSG00000106617 | PRKAG2 | chr7 | - | 151565719 | 151566699 | 151565719 | 151565885 | 151568715 | 151568842 | 23 | 16 | 15 | [illegible] |
| A3SS | 4731 | ENSG00000182481 | KPNA2 | chr17 | + | 68037329 | 68037495 | 68037357 | 68037495 | 68037109 | 68037207 | 1 | 0 | 3 | [illegible] |
| A3SS | 9026 | ENSG00000116698 | SMG7 | chr1 | + | 183517625 | 183517820 | 183517687 | 183517820 | 183515873 | 183515991 | 29 | 17 | 15 | [illegible] |
| A3SS | 5071 | ENSG00000170581 | STAT2 | chr12 | - | 56348923 | 56349059 | 56348923 | 56349029 | 56349162 | 56349261 | 73 | 42 | 57 | [illegible] |
| A3SS | 4666 | ENSG00000173889 | PHC3 | chr3 | - | 170172556 | 170172712 | 170172556 | 170172700 | 170178772 | 170178938 | 131 | 67 | 88 | [illegible] |
| A3SS | 4470 | ENSG00000169714 | CNBP | chr3 | - | 129171445 | 129171538 | 129171445 | 129171508 | 129171633 | 129171771 | 608 | 354 | 630 | [illegible] |
| A3SS | 5216 | ENSG00000138674 | SEC31A | chr4 | - | 82881857 | 82881940 | 82881857 | 82881937 | 82891087 | 82891178 | 50 | 29 | 26 | [illegible] |
| A3SS | 1401 | ENSG00000154957 | ZNF18 | chr17 | - | 11983296 | 11983407 | 11983296 | 11983404 | 11984112 | 11984197 | 21 | 17 | 28 | [illegible] |
| A3SS | 9127 | ENSG00000105393 | BABAM1 | chr19 | + | 17275776 | 17275825 | 17275800 | 17275825 | 17274106 | 17274185 | 5 | 1 | 4 | [illegible] |
| A3SS | 7732 | ENSG00000115525 | ST3GAL5 | chr2 | - | 85847860 | 85850834 | 85847860 | 85848381 | 85851263 | 85851703 | 20 | 19 | 23 | [illegible] |
| A3SS | 12891 | ENSG00000145901 | TNIP1 | chr5 | - | 151032286 | 151032400 | 151032286 | 151032383 | 151033607 | 151033799 | 10 | 6 | 13 | [illegible] |
| A3SS | 3683 | ENSG00000104517 | UBR5 | chr8 | - | 102342472 | 102342684 | 102342472 | 102342666 | 102345395 | 102345545 | 66 | 46 | 73 | [illegible] |
| A3SS | 11877 | ENSG00000155158 | TTC39B | chr9 | - | 15185279 | 15185528 | 15185279 | 15185406 | 15186943 | 15187035 | 13 | 26 | 27 | [illegible] |
| A3SS | 10339 | ENSG00000185753 | CXorf38 | chrX | - | 40639008 | 40640320 | 40639008 | 40639128 | 40647006 | 40647141 | 5 | 9 | 4 | [illegible] |
| A3SS | 10064 | ENSG00000122566 | HNRNPA2B1 | chr7 | - | 26196806 | 26197461 | 26196806 | 26197017 | 26197621 | 26197732 | 1894 | 1657 | 1729 | [illegible] |
| A3SS | 9703 | ENSG00000054523 | KIF1B | chr1 | + | 10282321 | 10282533 | 10282399 | 10282533 | 10277985 | 10278128 | 0 | 1 | 6 | [illegible] |
| A3SS | 6455 | ENSG00000066427 | ATXN3 | chr14 | - | 92083125 | 92083258 | 92083125 | 92083235 | 92088729 | 92088817 | 159 | 115 | 113 | [illegible] |
| A3SS | 7561 | ENSG00000028528 | SNX1 | chr15 | + | 64118116 | 64118244 | 64118202 | 64118244 | 64112572 | 64112684 | 775 | 442 | 570 | [illegible] |
| A3SS | 281 | ENSG00000142002 | DPP9 | chr19 | - | 4700215 | 4700277 | 4700215 | 4700273 | 4702026 | 4702155 | 79 | 35 | 36 | [illegible] |
| A3SS | 11885 | ENSG00000116786 | PLEKHM2 | chr1 | + | 15731888 | 15732048 | 15731949 | 15732048 | 15731191 | 15731257 | 60 | 25 | 52 | [illegible] |
| A3SS | 7888 | ENSG00000138363 | ATIC | chr2 | + | 215325179 | 215325329 | 215325240 | 215325329 | 215319664 | 215319731 | 10 | 3 | 3 | [illegible] |
| A3SS | 5192 | ENSG00000269893 | SNHG8 | chr4 | + | 118278933 | 118279137 | 118278943 | 118279137 | 118278754 | 118278792 | 1 | 2 | 0 | [illegible] |
| A3SS | 1436 | ENSG00000141030 | COPS3 | chr17 | - | 17276034 | 17276183 | 17276034 | 17276164 | 17281131 | 17281246 | 4 | 3 | 8 | [illegible] |
| A3SS | 5999 | ENSG00000288701 | PRRC2B | chr9 | + | 131482370 | 131482562 | 131482373 | 131482562 | 131481725 | 131481808 | 94 | 57 | 109 | [illegible] |
| A3SS | 22 | ENSG00000160209 | PDXK | chr21 | + | 43755476 | 43755764 | 43755697 | 43755764 | 43753582 | 43753719 | 0 | 2 | 1 | [illegible] |
| A3SS | 13156 | ENSG00000205352 | PRR13 | chr12 | + | 53443390 | 53443773 | 53443566 | 53443773 | 53442694 | 53442733 | 311 | 169 | 215 | [illegible] |
| A3SS | 13178 | ENSG00000111596 | CNOT2 | chr12 | + | 70332766 | 70332846 | 70332790 | 70332846 | 70330286 | 70330469 | 208 | 136 | 163 | [illegible] |
| A3SS | 12225 | ENSG00000171202 | TMEM126A | chr11 | + | 85654040 | 85654234 | 85654062 | 85654234 | 85650248 | 85650341 | 11 | 2 | 9 | [illegible] |
| A3SS | 4392 | ENSG00000172046 | USP19 | chr3 | - | 49117656 | 49117830 | 49117656 | 49117785 | 49117946 | 49118120 | 29 | 21 | 28 | [illegible] |
| A3SS | 8075 | ENSG00000143537 | ADAM15 | chr1 | + | 155055786 | 155055852 | 155055789 | 155055852 | 155054313 | 155054506 | 4 | 9 | 1 | [illegible] |
| A3SS | 9815 | ENSG00000077454 | LRCH4 | chr7 | - | 100577085 | 100577389 | 100577085 | 100577154 | 100577496 | 100577558 | 185 | 145 | 187 | [illegible] |
| A3SS | 3908 | ENSG00000127419 | TMEM175 | chr4 | + | 951645 | 951717 | 951681 | 951717 | 951206 | 951258 | 4 | 9 | 6 | [illegible] |
| A3SS | 2634 | ENSG00000092108 | SCFD1 | chr14 | + | 30673234 | 30673347 | 30673256 | 30673347 | 30670255 | 30670395 | 12 | 17 | 15 | [illegible] |
| A3SS | 7841 | ENSG00000181163 | NPM1 | chr5 | + | 171406396 | 171407161 | 171407128 | 171407161 | 171405301 | 171405403 | 194 | 181 | 227 | [illegible] |
| A3SS | 4171 | ENSG00000100003 | SEC14L2 | chr22 | + | 30415758 | 30416087 | 30415947 | 30416087 | 30410595 | 30410679 | 9 | 6 | 19 | [illegible] |
| A3SS | 8381 | ENSG00000101104 | PABPC1L | chr20 | + | 44930459 | 44930726 | 44930517 | 44930726 | 44924160 | 44924256 | 0 | 2 | 1 | [illegible] |
| A3SS | 13129 | ENSG00000168010 | ATG16L2 | chr11 | + | 72823543 | 72823797 | 72823765 | 72823797 | 72822847 | 72822961 | 40 | 15 | 15 | [illegible] |
| A3SS | 1620 | ENSG00000083454 | P2RX5 | chr17 | - | 3689491 | 3689630 | 3689491 | 3689627 | 3690069 | 3690150 | 92 | 32 | 81 | [illegible] |
| A3SS | 7868 | ENSG00000143387 | CTSK | chr1 | - | 150799167 | 150799285 | 150799167 | 150799273 | 150799543 | 150799709 | 0 | 0 | 3 | [illegible] |
| A3SS | 6427 | ENSG00000227372 | TP73-AS1 | chr1 | - | 3742803 | 3742957 | 3742803 | 3742911 | 3743060 | 3743190 | 3 | 1 | 3 | [illegible] |
| A3SS | 9312 | ENSG00000136861 | CDK5RAP2 | chr9 | - | 120460571 | 120460747 | 120460571 | 120460667 | 120467859 | 120467997 | 6 | 1 | 6 | [illegible] |
| A3SS | 378 | ENSG00000104980 | TIMM44 | chr19 | - | 7934088 | 7934241 | 7934088 | 7934238 | 7935064 | 7935145 | 8 | 10 | 5 | [illegible] |
| A3SS | 2478 | ENSG00000141956 | PRDM15 | chr21 | - | 41836467 | 41836649 | 41836467 | 41836589 | 41837933 | 41838063 | 34 | 20 | 34 | [illegible] |
| A3SS | 4999 | ENSG00000132581 | SDF2 | chr17 | - | 28651485 | 28651577 | 28651485 | 28651542 | 28655286 | 28655483 | 12 | 13 | 9 | [illegible] |
| A3SS | 10895 | ENSG00000140632 | GLYR1 | chr16 | - | 4821379 | 4821664 | 4821379 | 4821453 | 4822874 | 4822931 | 6 | 19 | 14 | [illegible] |
| A3SS | 4936 | ENSG00000109320 | NFKB1 | chr4 | + | 102533844 | 102533885 | 102533847 | 102533885 | 102529835 | 102529914 | 82 | 52 | 56 | [illegible] |
| A3SS | 1872 | ENSG00000197912 | SPG7 | chr16 | + | 89545871 | 89546760 | 89546657 | 89546760 | 89544647 | 89544772 | 4 | 1 | 1 | [illegible] |
| A3SS | 686 | ENSG00000167468 | GPX4 | chr19 | + | 1106539 | 1106779 | 1106572 | 1106779 | 1106399 | 1106459 | 769 | 467 | 584 | [illegible] |
| A3SS | 11795 | ENSG00000109911 | ELP4 | chr11 | + | 31603764 | 31603907 | 31603767 | 31603907 | 31594769 | 31594901 | 14 | 11 | 11 | [illegible] |
| A3SS | 2646 | ENSG00000075413 | MARK3 | chr14 | + | 103468032 | 103468186 | 103468080 | 103468186 | 103467078 | 103467191 | 135 | 76 | 140 | [illegible] |
| A3SS | 3539 | ENSG00000049192 | ADAMTS6 | chr5 | - | 65452132 | 65452221 | 65452132 | 65452216 | 65452706 | 65452918 | 21 | 23 | 22 | [illegible] |
| A3SS | 12886 | ENSG00000050438 | SLC4A8 | chr12 | + | 51450641 | 51451022 | 51450875 | 51451022 | 51440707 | 51440789 | 15 | 9 | 10 | [illegible] |
| A3SS | 9955 | ENSG00000135164 | DMTF1 | chr7 | + | 87182016 | 87182337 | 87182055 | 87182337 | 87181308 | 87181341 | 40 | 42 | 33 | [illegible] |
| A3SS | 561 | ENSG00000167613 | LAIR1 | chr19 | - | 54360915 | 54361209 | 54360915 | 54361206 | 54364294 | 54364330 | 113 | 47 | 79 | [illegible] |
| A3SS | 4867 | ENSG00000138172 | CALHM2 | chr10 | - | 103451082 | 103451285 | 103451082 | 103451182 | 103451878 | 103452036 | 44 | 28 | 21 | [illegible] |
| A3SS | 6737 | ENSG00000143575 | HAX1 | chr1 | + | 154275077 | 154275260 | 154275153 | 154275260 | 154274949 | 154275001 | 13 | 6 | 7 | [illegible] |
| A3SS | 2610 | ENSG00000092199 | HNRNPC | chr14 | - | 21263310 | 21263582 | 21263310 | 21263336 | 21269297 | 21269425 | 230 | 239 | 208 | [illegible] |
| A3SS | 3938 | ENSG00000203875 | SNHG5 | chr6 | - | 85677081 | 85677215 | 85677081 | 85677191 | 85677423 | 85677492 | 4 | 6 | 7 | [illegible] |
| A3SS | 7307 | ENSG00000182552 | RWDD4 | chr4 | - | 183651217 | 183651327 | 183651217 | 183651323 | 183655880 | 183655961 | 50 | 37 | 49 | [illegible] |
| A3SS | 10039 | ENSG00000065000 | AP3D1 | chr19 | - | 2116604 | 2116796 | 2116604 | 2116746 | 2117221 | 2117367 | 1 | 1 | 5 | [illegible] |
| A3SS | 12758 | ENSG00000132305 | IMMT | chr2 | - | 86171207 | 86171381 | 86171207 | 86171345 | 86173649 | 86173761 | 7 | 1 | 2 | [illegible] |
| A3SS | 10822 | ENSG00000111364 | DDX55 | chr12 | + | 123605864 | 123605981 | 123605930 | 123605981 | 123602127 | 123602256 | 0 | 1 | 6 | [illegible] |
| A3SS | 8383 | ENSG00000101104 | PABPC1L | chr20 | + | 44933056 | 44933185 | 44933071 | 44933185 | 44931832 | 44932026 | 5 | 1 | 3 | [illegible] |
| A3SS | 400 | ENSG00000142546 | NOSIP | chr19 | - | 49557089 | 49558199 | 49557089 | 49557249 | 49558896 | 49558978 | 36 | 15 | 51 | [illegible] |
| A3SS | 3635 | ENSG00000161912 | ADCY10P1 | chr6 | + | 41108948 | 41109904 | 41109712 | 41109904 | 41108304 | 41108509 | 15 | 26 | 19 | [illegible] |
| A3SS | 9407 | ENSG00000167483 | NIBAN3 | chr19 | + | 17533524 | 17533701 | 17533586 | 17533701 | 17532262 | 17532388 | 6 | 4 | 6 | [illegible] |
| A3SS | 9805 | ENSG00000012124 | CD22 | chr19 | + | 35332546 | 35332924 | 35332652 | 35332924 | 35332018 | 35332074 | 83 | 52 | 84 | [illegible] |
| A3SS | 12679 | ENSG00000078403 | MLLT10 | chr10 | + | 21740029 | 21740236 | 21740153 | 21740236 | 21735138 | 21735235 | 50 | 40 | 69 | [illegible] |
| A3SS | 4107 | ENSG00000124535 | WRNIP1 | chr6 | + | 2770119 | 2770361 | 2770194 | 2770361 | 2768690 | 2768882 | 228 | 122 | 218 | [illegible] |
| A3SS | 9060 | ENSG00000123219 | CENPK | chr5 | - | 65528918 | 65529049 | 65528918 | 65529017 | 65529116 | 65529199 | 20 | 1 | 12 | [illegible] |
| A3SS | 1611 | ENSG00000109046 | WSB1 | chr17 | + | 27307743 | 27309272 | 27309099 | 27309272 | 27306781 | 27306882 | 214 | 199 | 231 | [illegible] |
| A3SS | 5769 | ENSG00000109323 | MANBA | chr4 | - | 102672008 | 102672146 | 102672008 | 102672054 | 102673918 | 102674070 | 1 | 3 | 4 | [illegible] |
| A3SS | 825 | ENSG00000101473 | ACOT8 | chr20 | - | 45843526 | 45843772 | 45843526 | 45843721 | 45844262 | 45844420 | 4 | 9 | 14 | [illegible] |
| A3SS | 5899 | ENSG00000055070 | SZRD1 | chr1 | + | 16393227 | 16393482 | 16393230 | 16393482 | 16391374 | 16391424 | 59 | 23 | 46 | [illegible] |
| A3SS | 1439 | ENSG00000105053 | VRK3 | chr19 | - | 50009235 | 50009385 | 50009235 | 50009289 | 50016023 | 50016163 | 143 | 77 | 110 | [illegible] |
| A3SS | 7915 | ENSG00000213185 | FAM24B | chr10 | - | 122850423 | 122850550 | 122850423 | 122850520 | 122855644 | 122855786 | 11 | 7 | 17 | [illegible] |
| A3SS | 9319 | ENSG00000047579 | DTNBP1 | chr6 | - | 15651312 | 15651408 | 15651312 | 15651363 | 15652086 | 15652140 | 2 | 7 | 3 | [illegible] |
| A3SS | 6373 | ENSG00000048052 | HDAC9 | chr7 | + | 18590335 | 18590486 | 18590344 | 18590486 | 18585280 | 18585522 | 116 | 124 | 47 | [illegible] |
| A3SS | 1800 | ENSG00000005844 | ITGAL | chr16 | + | 30475288 | 30475400 | 30475305 | 30475400 | 30474195 | 30474298 | 30 | 26 | 19 | [illegible] |
| A3SS | 6412 | ENSG00000132781 | MUTYH | chr1 | - | 45333412 | 45333564 | 45333412 | 45333561 | 45334390 | 45334511 | 5 | 5 | 4 | [illegible] |
| A3SS | 2146 | ENSG00000103197 | TSC2 | chr16 | + | 2062496 | 2062600 | 2062528 | 2062600 | 2061870 | 2062008 | 75 | 28 | 53 | [illegible] |
| A3SS | 6129 | ENSG00000011295 | TTC19 | chr17 | + | 16002656 | 16002831 | 16002792 | 16002831 | 16001914 | 16002025 | 22 | 17 | 23 | [illegible] |
| A3SS | 2670 | ENSG00000100567 | PSMA3 | chr14 | + | 58252118 | 58252242 | 58252123 | 58252242 | 58247749 | 58247832 | 201 | 124 | 185 | [illegible] |
| A3SS | 2626 | ENSG00000171469 | ZNF561 | chr19 | - | 9618090 | 9618179 | 9618090 | 9618166 | 9619431 | 9619582 | 73 | 46 | 73 | [illegible] |
| A3SS | 1901 | ENSG00000214078 | CPNE1 | chr20 | - | 35626566 | 35626826 | 35626566 | 35626803 | 35627279 | 35627413 | 0 | 2 | 3 | [illegible] |

| | | | | | | | | | | | | | | | | | | | | | | | | | | | | | | | | | | | | | | | | | | | | | | |
|---|---|---|---|---|---|---|---|---|---|---|---|---|---|---|---|---|---|---|---|---|---|---|---|---|---|---|---|---|---|---|---|---|---|---|---|---|---|---|---|---|---|---|---|---|---|---|
| A3SS | 11592 | ENSG00000213930 | GALT | chr9 | + | 34647491 | 34647961 | 34647831 | 34647961 | 34647088 | 34647258 | 77 | 44 | 113 | 84 | 47 | 3 | 0 | 4 | 0 | 0 | 81 | 88 | 45 | 49 | 99 | 10 | 1 | 0 | 0 | 0 | 298 | 149 | 0.329140 | 1 | 0.928 | 1 | 0.934 | 1 | 1 | 0.802 | 0.978 | 1 | 1 | 1 | 0.016 |
| A3SS | 12956 | ENSG00000102225 | CDK16 | chrX | + | 47227159 | 47227223 | 47227180 | 47227223 | 47226993 | 47227099 | 4 | 4 | 2 | 7 | 5 | 53 | 18 | 52 | 59 | 25 | 23 | 6 | 4 | 2 | 12 | 59 | 36 | 62 | 71 | 121 | 170 | 149 | 0.329157 | 1 | 0.062 | 0.163 | 0.033 | 0.094 | 0.149 | 0.255 | 0.127 | 0.054 | 0.024 | 0.08 | -0.008 |
| A3SS | 8059 | ENSG00000224078 | SNHG14 | chr15 | + | 25101822 | 25103970 | 25103858 | 25103970 | 25099144 | 25099278 | 5 | 7 | 0 | 2 | 6 | 16 | 23 | 43 | 12 | 29 | 7 | 1 | 6 | 5 | 0 | 16 | 12 | 26 | 5 | 10 | 298 | 149 | 0.329334 | 1 | 0.135 | 0.132 | 0 | 0.077 | 0.094 | 0.179 | 0.04 | 0.103 | 0.333 | 0 | -0.043 |
| A3SS | 9495 | ENSG00000214021 | TTLL3 | chr3 | + | 9817644 | 9820741 | 9820545 | 9820741 | 9816073 | 9816202 | 64 | 45 | 56 | 64 | 47 | 0 | 1 | 0 | 0 | 0 | 86 | 69 | 88 | 125 | 160 | 3 | 0 | 2 | 0 | 4 | 298 | 149 | 0.329489 | 1 | 1 | 0.957 | 1 | 1 | 1 | 0.935 | 1 | 0.957 | 1 | 0.952 | 0.023 |
| A3SS | 11517 | ENSG00000244187 | TMEM141 | chr9 | + | 136791946 | 136792358 | 136792250 | 136792358 | 136791710 | 136791777 | 75 | 25 | 33 | 38 | 21 | 5 | 2 | 1 | 4 | 4 | 25 | 47 | 17 | 35 | 89 | 0 | 11 | 4 | 13 | 0 | 298 | 149 | 0.330423 | 1 | 0.882 | 0.862 | 0.943 | 0.826 | 0.724 | 1 | 0.681 | 0.68 | 0.574 | 1 | 0.06 |
| A3SS | 1863 | ENSG00000185324 | CDK10 | chr16 | + | 89694568 | 89694788 | 89694664 | 89694788 | 89694172 | 89694232 | 8 | 10 | 16 | 8 | 5 | 18 | 6 | 8 | 10 | 9 | 44 | 14 | 4 | 8 | 55 | 22 | 8 | 12 | 16 | 9 | 245 | 149 | 0.330443 | 1 | 0.213 | 0.503 | 0.549 | 0.327 | 0.253 | 0.549 | 0.516 | 0.169 | 0.233 | 0.788 | -0.082 |
| A3SS | 1897 | ENSG00000103066 | PLA2G15 | chr16 | + | 68255765 | 68255990 | 68255890 | 68255990 | 68255281 | 68255380 | 24 | 10 | 21 | 11 | 30 | 15 | 3 | 7 | 0 | 0 | 24 | 32 | 14 | 45 | 35 | 18 | 2 | 4 | 10 | 10 | 274 | 149 | 0.330597 | 1 | 0.465 | 0.644 | 0.62 | 1 | 1 | 0.42 | 0.897 | 0.656 | 0.71 | 0.656 | 0.078 |
| A3SS | 7895 | ENSG00000163219 | ARHGAP25 | chr2 | + | 68807272 | 68807480 | 68807275 | 68807480 | 68782232 | 68782320 | 18 | 6 | 17 | 7 | 40 | 2 | 1 | 3 | 0 | 0 | 15 | 31 | 54 | 38 | 28 | 0 | 6 | 3 | 2 | 0 | 152 | 149 | 0.331576 | 1 | 0.898 | 0.855 | 0.847 | 1 | 1 | 1 | 0.835 | 0.946 | 0.949 | 1 | -0.026 |
| A3SS | 13325 | ENSG00000081189 | MEF2C | chr5 | - | 88823734 | 88823930 | 88823734 | 88823927 | 88882954 | 88883184 | 48 | 36 | 32 | 41 | 92 | 85 | 79 | 66 | 99 | 106 | 178 | 43 | 34 | 72 | 99 | 250 | 131 | 54 | 129 | 52 | 152 | 149 | 0.332084 | 1 | 0.356 | 0.309 | 0.322 | 0.289 | 0.46 | 0.411 | 0.243 | 0.382 | 0.354 | 0.651 | -0.061 |
| A3SS | 7887 | ENSG00000138363 | ATIC | chr2 | + | 215318130 | 215318233 | 215318156 | 215318233 | 215312497 | 215312624 | 0 | 0 | 0 | 0 | 0 | 79 | 24 | 72 | 69 | 30 | 3 | 0 | 4 | 5 | 0 | 104 | 71 | 49 | 67 | 90 | 175 | 149 | 0.332291 | 1 | 0 | 0 | 0 | 0 | 0 | 0.024 | 0 | 0.065 | 0.06 | 0 | -0.03 |
| A3SS | 5321 | ENSG00000163956 | LRPAP1 | chr4 | - | 3520071 | 3520229 | 3520071 | 3520193 | 3524906 | 3525051 | 6 | 5 | 6 | 9 | 6 | 204 | 119 | 148 | 196 | 184 | 37 | 2 | 2 | 16 | 19 | 440 | 339 | 198 | 259 | 509 | 185 | 149 | 0.333049 | 1 | 0.023 | 0.033 | 0.032 | 0.036 | 0.026 | 0.063 | 0.005 | 0.008 | 0.047 | 0.029 | 0 |
| A3SS | 297 | ENSG00000239382 | ALKBH6 | chr19 | - | 36010566 | 36010683 | 36010566 | 36010645 | 36010893 | 36011045 | 11 | 4 | 14 | 4 | 9 | 1 | 2 | 3 | 3 | 10 | 0 | 20 | 4 | 2 | 24 | 9 | 5 | 1 | 0 | 0 | 187 | 149 | 0.333202 | 1 | 0.898 | 0.614 | 0.788 | 0.515 | 0.418 | 0 | 0.761 | 0.761 | 1 | 1 | -0.058 |
| A3SS | 10507 | ENSG00000223959 | AFG3L1P | chr16 | + | 89979301 | 89980336 | 89980241 | 89980336 | 89978809 | 89978878 | 22 | 5 | 13 | 15 | 27 | 15 | 20 | 18 | 24 | 9 | 99 | 47 | 22 | 30 | 59 | 49 | 26 | 33 | 58 | 28 | 298 | 149 | 0.333214 | 1 | 0.423 | 0.111 | 0.265 | 0.238 | 0.6 | 0.503 | 0.475 | 0.25 | 0.205 | 0.513 | -0.062 |
| A3SS | 3454 | ENSG00000011243 | AKAP8L | chr19 | - | 15397149 | 15397320 | 15397149 | 15397203 | 15397519 | 15397625 | 68 | 39 | 73 | 58 | 42 | 1 | 0 | 0 | 1 | 0 | 152 | 65 | 49 | 68 | 131 | 8 | 0 | 1 | 0 | 0 | 266 | 149 | 0.333387 | 1 | 0.974 | 1 | 1 | 0.97 | 1 | 0.914 | 1 | 0.965 | 1 | 1 | 0.013 |
| A3SS | 3821 | ENSG00000155229 | MMS19 | chr10 | - | 97467504 | 97467603 | 97467504 | 97467583 | 97468251 | 97468406 | 12 | 6 | 18 | 26 | 15 | 36 | 10 | 24 | 31 | 17 | 41 | 31 | 14 | 22 | 5 | 46 | 26 | 20 | 4 | 26 | 169 | 149 | 0.333630 | 1 | 0.227 | 0.346 | 0.398 | 0.425 | 0.438 | 0.44 | 0.512 | 0.382 | 0.829 | 0.145 | -0.095 |
| A3SS | 7086 | ENSG00000162664 | ZNF326 | chr1 | + | 90007344 | 90007750 | 90007611 | 90007750 | 90005132 | 90005244 | 92 | 63 | 90 | 107 | 91 | 2 | 2 | 2 | 2 | 0 | 35 | 94 | 66 | 8 | 6 | 3 | 2 | 2 | 2 | 0 | 298 | 149 | 0.333899 | 1 | 0.958 | 0.94 | 0.957 | 0.964 | 1 | 0.854 | 0.959 | 0.943 | 0.667 | 1 | 0.079 |
| A3SS | 11743 | ENSG00000133943 | DGLUCY | chr14 | + | 91188909 | 91189170 | 91188987 | 91189170 | 91181185 | 91181389 | 127 | 92 | 103 | 144 | 111 | 1 | 6 | 4 | 0 | 0 | 178 | 96 | 122 | 144 | 288 | 0 | 0 | 0 | 2 | 0 | 227 | 149 | 0.335055 | 1 | 0.988 | 0.91 | 0.944 | 1 | 1 | 1 | 1 | 1 | 0.979 | 1 | -0.027 |
| A3SS | 298 | ENSG00000239382 | ALKBH6 | chr19 | - | 36010737 | 36011077 | 36010737 | 36011045 | 36011403 | 36011464 | 11 | 6 | 10 | 6 | 15 | 36 | 13 | 32 | 18 | 34 | 5 | 4 | 5 | 0 | 45 | 32 | 61 | 21 | 10 | 15 | 181 | 149 | 0.335541 | 1 | 0.201 | 0.275 | 0.205 | 0.215 | 0.266 | 0.114 | 0.051 | 0.164 | 0 | 0.712 | 0.024 |
| A3SS | 11123 | ENSG00000100288 | CHKB | chr22 | - | 50579726 | 50579830 | 50579726 | 50579802 | 50579973 | 50580082 | 102 | 53 | 82 | 91 | 92 | 15 | 9 | 10 | 18 | 14 | 116 | 77 | 87 | 75 | 216 | 14 | 17 | 13 | 23 | 38 | 177 | 149 | 0.335792 | 1 | 0.851 | 0.832 | 0.873 | 0.81 | 0.847 | 0.875 | 0.792 | 0.849 | 0.733 | 0.827 | 0.027 |
| A3SS | 1309 | ENSG00000006025 | OSBPL7 | chr17 | - | 47808863 | 47808990 | 47808863 | 47808946 | 47809075 | 47809115 | 61 | 35 | 33 | 71 | 12 | 3 | 0 | 0 | 3 | 2 | 27 | 97 | 35 | 38 | 101 | 0 | 0 | 3 | 0 | 0 | 193 | 149 | 0.336409 | 1 | 0.94 | 1 | 1 | 0.948 | 0.822 | 1 | 1 | 0.9 | 1 | 1 | -0.038 |
| [illegible] | | | | | | | | | | | | | | | | | | | | | | | | | | | | | | | | | | | | | | | | | | | | | | |

| | | | | | | | | | | | | | | | | | | | | | | | | | | | | | | | | | | | | | | | | | | | | | | |
|---|---|---|---|---|---|---|---|---|---|---|---|---|---|---|---|---|---|---|---|---|---|---|---|---|---|---|---|---|---|---|---|---|---|---|---|---|---|---|---|---|---|---|---|---|---|---|
| A3SS | 8505 | ENSG00000184922 | FMNL1 | chr17 | + | 45245618 | 45245733 | 45245631 | 45245733 | 45245252 | 45245416 | 9 | 5 | 4 | 6 | 16 | 418 | 146 | 323 | 306 | 229 | 2 | 8 | 3 | 7 | 25 | 398 | 367 | 219 | 313 | 669 | 162 | 149 | 0.420431 | 1 | 0.019 | 0.031 | 0.011 | 0.018 | 0.06 | 0.005 | 0.02 | 0.012 | 0.02 | 0.033 | 0.01 |
| A3SS | 9584 | ENSG00000130544 | ZNF557 | chr19 | + | 7075654 | 7075743 | 7075675 | 7075743 | 7074995 | 7075105 | 18 | 13 | 21 | 30 | 5 | 12 | 12 | 5 | 5 | 8 | 24 | 26 | 29 | 13 | 30 | 10 | 17 | 2 | 8 | 10 | 170 | 149 | 0.420671 | 1 | 0.568 | 0.487 | 0.786 | 0.84 | 0.354 | 0.678 | 0.573 | 0.927 | 0.588 | 0.724 | -0.091 |
| A3SS | 637 | ENSG00000089639 | GMIP | chr19 | - | 19635414 | 19635721 | 19635414 | 19635569 | 19636706 | 19636796 | 41 | 14 | 23 | 39 | 33 | 11 | 2 | 3 | 3 | 3 | 55 | 39 | 36 | 55 | 112 | 0 | 12 | 5 | 11 | 2 | 298 | 149 | 0.420837 | 1 | 0.651 | 0.778 | 0.793 | 0.867 | 0.846 | 1 | 0.619 | 0.783 | 0.714 | 0.966 | -0.029 |
| A3SS | 1088 | ENSG00000141506 | PIK3R5 | chr17 | - | 8886476 | 8886605 | 8886476 | 8886602 | 8887095 | 8887221 | 40 | 7 | 16 | 18 | 6 | 17 | 5 | 19 | 8 | 2 | 20 | 39 | 18 | 22 | 34 | 27 | 14 | 28 | 5 | 0 | 152 | 149 | 0.421146 | 1 | 0.698 | 0.578 | 0.452 | 0.688 | 0.746 | 0.421 | 0.732 | 0.387 | 0.812 | 1 | -0.038 |
| A3SS | 151 | ENSG00000100239 | PPP6R2 | chr22 | + | 50419345 | 50419462 | 50419348 | 50419462 | 50418866 | 50418979 | 14 | 13 | 12 | 5 | 2 | 128 | 106 | 119 | 139 | 109 | 15 | 3 | 13 | 0 | 29 | 215 | 121 | 108 | 131 | 374 | 152 | 149 | 0.421150 | 1 | 0.097 | 0.107 | 0.09 | 0.034 | 0.018 | 0.064 | 0.024 | 0.106 | 0 | 0.071 | 0.016 |
| A3SS | 10790 | ENSG00000120699 | EXOSC8 | chr13 | + | 37008056 | 37008177 | 37008061 | 37008177 | 37006974 | 37007071 | 69 | 50 | 96 | 71 | 168 | 6 | 6 | 7 | 2 | 3 | 41 | 28 | 61 | 18 | 5 | 0 | 2 | 12 | 1 | 0 | 154 | 149 | 0.421536 | 1 | 0.918 | 0.89 | 0.93 | 0.972 | 0.982 | 1 | 0.931 | 0.831 | 0.946 | 1 | -0.003 |
| A3SS | 2769 | ENSG00000075399 | VPS9D1 | chr16 | - | 89712046 | 89712099 | 89712046 | 89712093 | 89712459 | 89712522 | 22 | 11 | 11 | 28 | 5 | 0 | 1 | 0 | 0 | 0 | 14 | 36 | 18 | 23 | 30 | 0 | 0 | 0 | 0 | 7 | 155 | 149 | 0.422158 | 1 | 1 | 0.914 | 1 | 1 | 1 | 1 | 1 | 1 | 1 | 0.805 | 0.022 |
| A3SS | 8078 | ENSG00000123240 | OPTN | chr10 | + | 13109096 | 13109288 | 13109111 | 13109288 | 13108961 | 13109015 | 21 | 14 | 12 | 10 | 27 | 43 | 14 | 29 | 31 | 12 | 21 | 22 | 10 | 20 | 7 | 18 | 34 | 36 | 11 | 2 | 164 | 149 | 0.423430 | 1 | 0.307 | 0.476 | 0.273 | 0.227 | 0.672 | 0.515 | 0.37 | 0.202 | 0.623 | 0.761 | -0.103 |
| A3SS | 3619 | ENSG00000125089 | SH3TC1 | chr4 | + | 8232407 | 8233513 | 8233362 | 8233513 | 8231975 | 8232156 | 14 | 18 | 29 | 19 | 10 | 22 | 14 | 17 | 18 | 1 | 67 | 57 | 68 | 42 | 97 | 37 | 31 | 22 | 24 | 70 | 298 | 149 | 0.423672 | 1 | 0.241 | 0.391 | 0.46 | 0.345 | 0.833 | 0.475 | 0.479 | 0.607 | 0.467 | 0.409 | -0.033 |
| A3SS | 11879 | ENSG00000154146 | NRGN | chr11 | + | 124745961 | 124745982 | 124745964 | 124745982 | 124745502 | 124745729 | 320 | 282 | 93 | 492 | 318 | 363 | 346 | 89 | 629 | 394 | 1038 | 667 | 251 | 485 | 6078 | 1052 | 696 | 277 | 555 | 7741 | 152 | 149 | 0.424034 | 1 | 0.464 | 0.444 | 0.506 | 0.434 | 0.442 | 0.492 | 0.484 | 0.47 | 0.461 | 0.435 | -0.01 |
| A3SS | 1162 | ENSG00000108479 | GALK1 | chr17 | - | 75763013 | 75763181 | 75763013 | 75763149 | 75763319 | 75763439 | 0 | 0 | 0 | 5 | 0 | 14 | 7 | 4 | 6 | 2 | 0 | 0 | 0 | 0 | 0 | 4 | 14 | 1 | 6 | 25 | 181 | 149 | 0.424899 | 1 | 0 | 0 | 0 | 0.407 | 0 | 0 | 0 | 0 | 0 | 0 | 0.081 |
| A3SS | 3493 | ENSG00000128694 | OSGEPL1 | chr2 | - | 189746659 | 189747331 | 189746659 | 189747168 | 189750549 | 189750656 | 22 | 30 | 23 | 20 | 49 | 10 | 3 | 5 | 0 | 16 | 8 | 17 | 16 | 2 | 4 | 0 | 6 | 3 | 1 | 0 | 298 | 149 | 0.424934 | 1 | 0.524 | 0.833 | 0.697 | 1 | 0.605 | 1 | 0.586 | 0.727 | 0.5 | 1 | -0.031 |
| A3SS | 7740 | ENSG00000115525 | ST3GAL5 | chr2 | - | 85852859 | 85853091 | 85852859 | 85853000 | 85861180 | 85861292 | 15 | 4 | 11 | 16 | 20 | 9 | 3 | 9 | 7 | 9 | 10 | 35 | 16 | 21 | 1 | 15 | 7 | 4 | 13 | 17 | 240 | 149 | 0.425340 | 1 | 0.509 | 0.453 | 0.431 | 0.587 | 0.58 | 0.293 | 0.756 | 0.713 | 0.501 | 0.035 | 0.052 |
| A3SS | 8955 | ENSG00000162419 | GMEB1 | chr1 | + | 28690073 | 28690186 | 28690103 | 28690186 | 28683582 | 28683740 | 18 | 17 | 31 | 24 | 41 | 40 | 22 | 45 | 44 | 31 | 55 | 22 | 15 | 40 | 54 | 74 | 44 | 26 | 46 | 18 | 179 | 149 | 0.425354 | 1 | 0.273 | 0.391 | 0.364 | 0.312 | 0.524 | 0.382 | 0.294 | 0.324 | 0.42 | 0.714 | -0.054 |
| A3SS | 4514 | ENSG00000187514 | PTMA | chr2 | + | 231711886 | 231711983 | 231711889 | 231711983 | 231711347 | 231711419 | 183 | 120 | 178 | 192 | 160 | 1492 | 994 | 1784 | 1749 | 1713 | 401 | 233 | 167 | 268 | 383 | 3155 | 1702 | 1805 | 2414 | 3460 | 152 | 149 | 0.426085 | 1 | 0.107 | 0.106 | 0.089 | 0.097 | 0.084 | 0.111 | 0.118 | 0.083 | 0.098 | 0.098 | -0.005 |
| A3SS | 13055 | ENSG00000179295 | PTPN11 | chr12 | + | 112486462 | 112486629 | 112486474 | 112486629 | 112482073 | 112482205 | 29 | 21 | 32 | 22 | 35 | 142 | 126 | 124 | 160 | 210 | 0 | 26 | 34 | 20 | 24 | 25 | 111 | 113 | 168 | 18 | 161 | 149 | 0.426533 | 1 | 0.159 | 0.134 | 0.193 | 0.113 | 0.134 | 0 | 0.178 | 0.218 | 0.099 | 0.552 | -0.063 |
| A3SS | 10674 | ENSG00000248323 | LUCAT1 | chr5 | - | 91312294 | 91312481 | 91312294 | 91312413 | 91313077 | 91313229 | 9 | 9 | 8 | 16 | 9 | 2 | 1 | 0 | 5 | 4 | 46 | 14 | 45 | 40 | 19 | 6 | 3 | 17 | 12 | 4 | 217 | 149 | 0.426941 | 1 | 0.755 | 0.861 | 1 | 0.687 | 0.607 | 0.84 | 0.762 | 0.645 | 0.696 | 0.765 | 0.04 |
| A3SS | 13120 | ENSG00000110906 | KCTD10 | chr12 | - | 109457991 | 109458184 | 109457991 | 109458078 | 109460635 | 109460805 | 10 | 8 | 7 | 13 | 13 | 22 | 19 | 15 | 38 | 13 | 21 | 33 | 11 | 8 | 21 | 54 | 21 | 15 | 34 | 34 | 255 | 149 | 0.427305 | 1 | 0.21 | 0.197 | 0.214 | 0.167 | 0.369 | 0.185 | 0.479 | 0.3 | 0.121 | 0.265 | -0.039 |
| A3SS | 5543 | ENSG00000286053 | ASDURF | chr2 | + | 189665356 | 189665451 | 189665375 | 189665451 | 189663900 | 189663954 | 1 | 7 | 4 | 6 | 5 | 219 | 210 | 237 | 209 | 390 | 0 | 6 | 2 | 0 | 0 | 49 | 121 | 144 | 76 | 9 | 168 | 149 | 0.427598 | 1 | 0.004 | 0.029 | 0.015 | 0.025 | 0.011 | 0 | 0.042 | 0.012 | 0 | 0 | 0.006 |
| A3SS | 2511 | ENSG00000197555 | SIPA1L1 | chr14 | + | 71738240 | 71738325 | 71738243 | 71738325 | 71735276 | 71735391 | 23 | 7 | 29 | 19 | 21 | 18 | 7 | 12 | 17 | 9 | 44 | 42 | 25 | 28 | 6 | 13 | 47 | 21 | 9 | 37 | 152 | 149 | 0.428915 | 1 | 0.556 | 0.495 | 0.703 | 0.523 | 0.696 | 0.768 | 0.467 | 0.539 | 0.753 | 0.137 | 0.062 |
| A3SS | 4758 | ENSG00000099330 | OCEL1 | chr19 | + | 17228430 | 17229215 | 17228802 | 17229215 | 17228255 | 17228309 | 10 | 4 | 9 | 12 | 21 | 20 | 23 | 23 | 22 | 51 | 3 | 13 | 16 | 17 | 0 | 41 | 26 | 5 | 30 | 63 | 298 | 149 | 0.429141 | 1 | 0.2 | 0.08 | 0.164 | 0.214 | 0.171 | 0.035 | 0.2 | 0.615 | 0.221 | 0 | -0.048 |
| A3SS | 10776 | ENSG00000101752 | MIB1 | chr18 | + | 21791334 | 21791557 | 21791373 | 21791557 | 21779480 | 21779685 | 15 | 7 | 18 | 16 | 6 | 148 | 112 | 104 | 144 | 182 | 21 | 6 | 3 | 0 | 6 | 68 | 93 | 42 | 52 | 37 | 188 | 149 | 0.429143 | 1 | 0.074 | 0.047 | 0.121 | 0.081 | 0.025 | 0.197 | 0.049 | 0.054 | 0 | 0.114 | -0.013 |
| A3SS | 7145 | ENSG00000011021 | CLCN6 | chr1 | + | 11827088 | 11827221 | 11827114 | 11827221 | 11826155 | 11826214 | 16 | 13 | 19 | 11 | 15 | 0 | 0 | 6 | 2 | 0 | 63 | 35 | 26 | 30 | 27 | 1 | 1 | 1 | 5 | 0 | 175 | 149 | 0.429262 | 1 | 1 | 1 | 0.729 | 0.824 | 1 | 0.982 | 0.968 | 0.957 | 0.836 | 1 | -0.038 |
| A3SS | 2201 | ENSG00000182511 | FES | chr15 | + | 90893099 | 90893194 | 90893157 | 90893194 | 90892706 | 90892825 | 83 | 21 | 49 | 50 | 28 | 0 | 0 | 0 | 0 | 0 | 61 | 87 | 29 | 66 | 119 | 0 | 0 | 0 | 0 | 9 | 207 | 149 | 0.429679 | 1 | 1 | 1 | 1 | 1 | 1 | 1 | 1 | 1 | 1 | 0.905 | 0.019 |
| A3SS | 2142 | ENSG00000103197 | TSC2 | chr16 | + | 2055979 | 2056244 | 2056195 | 2056244 | 2055401 | 2055519 | 8 | 2 | 3 | 0 | 1 | 60 | 25 | 34 | 41 | 66 | 0 | 1 | 1 | 4 | 8 | 25 | 35 | 13 | 14 | 60 | 298 | 149 | 0.430157 | 1 | 0.062 | 0.038 | 0.042 | 0 | 0.008 | 0 | 0.014 | 0.037 | 0.125 | 0.062 | -0.018 |
| A3SS | 13168 | ENSG00000132964 | CDK8 | chr13 | + | 26401465 | 26401624 | 26401468 | 26401624 | 26401268 | 26401347 | 43 | 48 | 28 | 33 | 67 | 14 | 5 | 9 | 8 | 15 | 40 | 26 | 33 | 30 | 19 | 16 | 7 | 16 | 9 | 0 | 152 | 149 | 0.430680 | 1 | 0.751 | 0.904 | 0.753 | 0.802 | 0.814 | 0.71 | 0.785 | 0.669 | 0.766 | 1 | 0.019 |
| A3SS | 9169 | ENSG00000175575 | PAAF1 | chr11 | + | 73900269 | 73900420 | 73900310 | 73900420 | 73899145 | 73899244 | 87 | 77 | 85 | 54 | 46 | 4 | 1 | 20 | 8 | 1 | 227 | 121 | 26 | 56 | 179 | 31 | 8 | 29 | 1 | 8 | 190 | 149 | 0.431025 | 1 | 0.945 | 0.984 | 0.769 | 0.841 | 0.973 | 0.852 | 0.922 | 0.413 | 0.978 | 0.946 | 0.08 |
| A3SS | 4021 | ENSG00000131042 | LILRB2 | chr19 | - | 54277549 | 54277600 | 54277549 | 54277597 | 54277888 | 54277939 | 23 | 16 | 11 | 35 | 12 | 29 | 8 | 10 | 20 | 13 | 88 | 53 | 17 | 48 | 101 | 50 | 17 | 14 | 51 | 78 | 152 | 149 | 0.431217 | 1 | 0.437 | 0.662 | 0.519 | 0.632 | 0.475 | 0.633 | 0.753 | 0.543 | 0.48 | 0.559 | -0.049 |
| A3SS | 4240 | ENSG00000211456 | SACM1L | chr3 | + | 45713136 | 45713196 | 45713140 | 45713196 | 45709497 | 45709647 | 102 | 85 | 73 | 75 | 126 | 1 | 1 | 3 | 3 | 4 | 18 | 19 | 80 | 33 | 12 | 0 | 0 | 1 | 5 | 0 | 153 | 149 | 0.431985 | 1 | 0.99 | 0.988 | 0.96 | 0.961 | 0.968 | 1 | 1 | 0.987 | 0.865 | 1 | 0.003 |
| A3SS | 12275 | ENSG00000247708 | STX18-AS1 | chr4 | + | 4649369 | 4649542 | 4649372 | 4649542 | 4648976 | 4649109 | 20 | 3 | 2 | 11 | 9 | 8 | 3 | 4 | 4 | 1 | 10 | 4 | 6 | 11 | 0 | 0 | 6 | 2 | 7 | 10 | 152 | 149 | 0.432420 | 1 | 0.71 | 0.495 | 0.329 | 0.729 | 0.898 | 1 | 0.395 | 0.746 | 0.606 | 0 | 0.083 |
| A3SS | 2632 | ENSG00000092108 | SCFD1 | chr14 | + | 30638106 | 30638247 | 30638183 | 30638247 | 30633946 | 30634037 | 122 | 98 | 87 | 101 | 210 | 2 | 0 | 0 | 7 | 2 | 25 | 54 | 53 | 38 | 26 | 0 | 0 | 2 | 0 | 0 | 226 | 149 | 0.432519 | 1 | 0.976 | 1 | 1 | 0.905 | 0.986 | 1 | 1 | 0.946 | 1 | 1 | -0.016 |
| A3SS | 2728 | ENSG00000100722 | ZC3H14 | chr14 | + | 88577984 | 88578140 | 88577992 | 88578140 | 88575839 | 88575940 | 187 | 129 | 192 | 177 | 464 | 2 | 0 | 3 | 2 | 1 | 118 | 152 | 78 | 100 | 57 | 7 | 0 | 5 | 0 | 0 | 157 | 149 | 0.432819 | 1 | 0.989 | 1 | 0.984 | 0.988 | 0.998 | 0.941 | 1 | 0.937 | 1 | 1 | 0.016 |
| A3SS | 9015 | ENSG00000111679 | PTPN6 | chr12 | + | 6951911 | 6952177 | 6952016 | 6952177 | 6951608 | 6951731 | 220 | 124 | 181 | 151 | 173 | 7 | 3 | 2 | 8 | 26 | 251 | 228 | 97 | 170 | 389 | 27 | 7 | 8 | 2 | 4 | 254 | 149 | 0.432973 | 1 | 0.949 | 0.96 | 0.982 | 0.917 | 0.796 | 0.845 | 0.95 | 0.877 | 0.98 | 0.983 | -0.006 |
| A3SS | 9455 | ENSG00000242802 | AP5Z1 | chr7 | + | 4787595 | 4787776 | 4787633 | 4787776 | 4786249 | 4786428 | 1 | 0 | 3 | 1 | 2 | 24 | 7 | 11 | 16 | 22 | 0 | 2 | 1 | 0 | 8 | 28 | 19 | 23 | 6 | 30 | 187 | 149 | 0.433106 | 1 | 0.032 | 0 | 0.179 | 0.047 | 0.068 | 0 | 0.077 | 0.033 | 0 | 0.175 | 0.008 |
| A3SS | 12626 | ENSG00000107959 | PITRM1 | chr10 | - | 3165412 | 3165523 | 3165412 | 3165483 | 3166228 | 3166380 | 41 | 30 | 29 | 35 | 61 | 7 | 6 | 9 | 12 | 1 | 40 | 20 | 33 | 39 | 64 | 0 | 19 | 6 | 9 | 19 | 189 | 149 | 0.433452 | 1 | 0.822 | 0.798 | 0.718 | 0.697 | 0.98 | 1 | 0.454 | 0.813 | 0.774 | 0.726 | 0.05 |
| A3SS | 8457 | ENSG00000132530 | XAF1 | chr17 | + | 6759255 | 6759718 | 6759661 | 6759718 | 6758088 | 6758224 | 34 | 38 | 23 | 36 | 20 | 75 | 76 | 38 | 88 | 83 | 77 | 24 | 13 | 40 | 16 | 59 | 70 | 44 | 184 | 9 | 298 | 149 | 0.433515 | 1 | 0.185 | 0.2 | 0.232 | 0.17 | 0.108 | 0.395 | 0.146 | 0.129 | 0.098 | 0.471 | -0.069 |
| A3SS | 900 | ENSG00000170471 | RALGAPB | chr20 | + | 38497352 | 38497516 | 38497479 | 38497516 | 38492929 | 38493132 | 259 | 182 | 197 | 189 | 260 | 2 | 0 | 2 | 2 | 11 | 149 | 212 | 207 | 206 | 73 | 0 | 6 | 6 | 6 | 0 | 276 | 149 | 0.433766 | 1 | 0.986 | 1 | 0.982 | 0.981 | 0.927 | 1 | 0.95 | 0.949 | 0.949 | 1 | 0.006 |
| A3SS | 3184 | ENSG00000111912 | NCOA7 | chr6 | + | 125888938 | 125888981 | 125888971 | 125888981 | 125885158 | 125885343 | 78 | 92 | 114 | 106 | 173 | 31 | 31 | 12 | 13 | 51 | 24 | 34 | 41 | 36 | 8 | 11 | 21 | 25 | 3 | 0 | 182 | 149 | 0.434174 | 1 | 0.673 | 0.708 | 0.886 | 0.87 | 0.735 | 0.641 | 0.57 | 0.573 | 0.908 | 1 | 0.036 |
| A3SS | 3548 | ENSG00000108039 | XPNPEP1 | chr10 | - | 109873366 | 109873577 | 109873366 | 109873427 | 109875527 | 109875599 | 9 | 9 | 14 | 9 | 1 | 126 | 73 | 64 | 136 | 108 | 17 | 12 | 18 | 14 | 8 | 253 | 147 | 100 | 143 | 439 | 298 | 149 | 0.434745 | 1 | 0.034 | 0.058 | 0.099 | 0.032 | 0.005 | 0.033 | 0.039 | 0.083 | 0.047 | 0.009 | 0.003 |
| A3SS | 5451 | ENSG00000177728 | TMEM94 | chr17 | + | 75498443 | 75498538 | 75498455 | 75498538 | 75498174 | 75498323 | 23 | 12 | 15 | 19 | 1 | 0 | 0 | 4 | 0 | 1 | 25 | 43 | 12 | 15 | 64 | 0 | 0 | 0 | 0 | 0 | 161 | 149 | 0.434746 | 1 | 1 | 1 | 0.776 | 1 | 0.481 | 1 | 1 | 1 | 1 | 1 | -0.149 |
| A3SS | 10271 | ENSG00000111581 | NUP107 | chr12 | + | 68696818 | 68696922 | 68696850 | 68696922 | 68691967 | 68692112 | 112 | 55 | 84 | 112 | 123 | 2 | 10 | 2 | 4 | 5 | 23 | 72 | 19 | 15 | 15 | 0 | 0 | 2 | 0 | 6 | 181 | 149 | 0.434850 | 1 | 0.979 | 0.819 | 0.972 | 0.958 | 0.953 | 1 | 1 | 0.887 | 1 | 0.673 | 0.024 |
| A3SS | 2176 | ENSG00000137871 | ZNF280D | chr15 | - | 56707081 | 56707150 | 56707081 | 56707143 | 56707262 | 56707306 | 105 | 98 | 93 | 114 | 125 | 77 | 45 | 42 | 70 | 63 | 20 | 78 | 32 | 40 | 11 | 8 | 27 | 11 | 12 | 18 | 156 | 149 | 0.435444 | 1 | 0.566 | 0.675 | 0.679 | 0.609 | 0.655 | 0.705 | 0.734 | 0.735 | 0.761 | 0.369 | -0.024 |
| A3SS | 2095 | ENSG00000140983 | RHOT2 | chr16 | + | 668352 | 668411 | 668357 | 668411 | 668150 | 668236 | 58 | 15 | 42 | 63 | 8 | 6 | 0 | 7 | 2 | 2 | 16 | 53 | 52 | 31 | 97 | 12 | 8 | 4 | 2 | 0 | 154 | 149 | 0.436236 | 1 | 0.903 | 1 | 0.853 | 0.968 | 0.795 | 0.563 | 0.865 | 0.926 | 0.937 | 1 | 0.046 |
| A3SS | 5890 | ENSG00000138386 | NAB1 | chr2 | + | 190685475 | 190685638 | 190685478 | 190685638 | 190683737 | 190683827 | 85 | 54 | 53 | 90 | 115 | 12 | 4 | 5 | 11 | 24 | 42 | 55 | 54 | 40 | 40 | 35 | 6 | 16 | 3 | 0 | 152 | 149 | 0.436428 | 1 | 0.874 | 0.93 | 0.912 | 0.889 | 0.824 | 0.541 | 0.9 | 0.768 | 0.929 | 1 | 0.058 |
| A3SS | 8568 | ENSG00000198848 | CES1 | chr16 | - | 55810926 | 55811010 | 55810926 | 55811007 | 55812902 | 55813043 | 86 | 22 | 52 | 21 | 90 | 21 | 1 | 5 | 0 | 17 | 333 | 637 | 112 | 548 | 67 | 110 | 36 | 7 | 97 | 31 | 152 | 149 | 0.437046 | 1 | 0.801 | 0.956 | 0.911 | 1 | 0.838 | 0.748 | 0.945 | 0.94 | 0.847 | 0.679 | 0.069 |
| A3SS | 4170 | ENSG00000100003 | SEC14L2 | chr22 | + | 30409106 | 30409282 | 30409186 | 30409282 | 30407414 | 30407603 | 6 | 2 | 2 | 2 | 1 | 6 | 1 | 17 | 5 | 5 | 0 | 13 | 9 | 0 | 80 | 7 | 8 | 39 | 1 | 59 | 229 | 149 | 0.437188 | 1 | 0.394 | 0.565 | 0.071 | 0.207 | 0.115 | 0 | 0.514 | 0.131 | 0 | 0.469 | 0.048 |
| A3SS | 11426 | ENSG00000136811 | ODF2 | chr9 | + | 128460512 | 128460666 | 128460549 | 128460666 | 128459566 | 128459667 | 72 | 32 | 31 | 67 | 29 | 5 | 4 | 6 | 11 | 0 | 47 | 56 | 44 | 70 | 116 | 0 | 6 | 3 | 16 | 23 | 186 | 149 | 0.437347 | 1 | 0.92 | 0.865 | 0.805 | 0.83 | 1 | 1 | 0.882 | 0.922 | 0.778 | 0.802 | 0.007 |
| A3SS | 6240 | ENSG00000111331 | OAS3 | chr12 | + | 112965743 | 112966029 | 112965896 | 112966029 | 112964234 | 112964408 | 85 | 83 | 54 | 181 | 51 | 4 | 5 | 0 | 1 | 9 | 555 | 188 | 87 | 508 | 115 | 34 | 4 | 0 | 0 | 4 | 298 | 149 | 0.437768 | 1 | 0.914 | 0.892 | 1 | 0.989 | 0.739 | 0.891 | 0.959 | 1 | 1 | 0.935 | -0.05 |
| A3SS | 13311 | ENSG00000120690 | ELF1 | chr13 | - | 40949805 | 40949973 | 40949805 | 40949901 | 40951328 | 40951436 | 1069 | 915 | 1095 | 1216 | 2114 | 4 | 12 | 10 | 21 | 17 | 608 | 1793 | 2522 | 1238 | 309 | 13 | 5 | 56 | 4 | 0 | 221 | 149 | 0.439291 | 1 | 0.994 | 0.981 | 0.987 | 0.975 | 0.988 | 0.969 | 0.996 | 0.968 | 0.995 | 1 | -0.001 |
| A3SS | 11464 | ENSG00000159069 | FBXW5 | chr9 | - | 136942768 | 136943003 | 136942768 | 136942943 | 136943348 | 136943506 | 11 | 0 | 7 | 2 | 0 | 86 | 30 | 68 | 75 | 34 | 0 | 1 | 3 | 3 | 11 | 67 | 80 | 30 | 18 | 169 | 209 | 149 | 0.440572 | 1 | 0.084 | 0 | 0.068 | 0.019 | 0 | 0 | 0.009 | 0.067 | 0.106 | 0.044 | -0.011 |

[illegible]

| | | | | | | | | | | | | | | | | | | | | | | | | |
|---|---|---|---|---|---|---|---|---|---|---|---|---|---|---|---|---|---|---|---|---|---|---|---|---|
| A3SS | 5819 | ENSG00000160058 | BSDC1 | chr1 | - | 32383829 | 32383997 | 32383829 | 32383939 | 32386778 | 32386895 | 138 | 104 | 85 | [illegible] | [illegible] | [illegible] | [illegible] | [illegible] | [illegible] | [illegible] | [illegible] | [illegible] | [illegible] |
| A3SS | 9719 | ENSG00000119616 | FCF1 | chr14 | + | 74715950 | 74716099 | 74715993 | 74716099 | 74713484 | 74713552 | 66 | 45 | 61 | 71 | 88 | 0 | 0 | 0 | 0 | 0 | 56 | 46 | 31 |
| A3SS | 331 | ENSG00000028277 | POU2F2 | chr19 | - | 42117249 | 42117432 | 42117249 | 42117366 | 42122125 | 42122182 | 5 | 3 | 3 | 3 | 9 | 57 | 38 | 34 | 41 | 82 | 2 | 0 | 17 |
| A3SS | 9890 | ENSG00000126883 | NUP214 | chr9 | + | 131232161 | 131232308 | 131232283 | 131232308 | 131230629 | 131230769 | 10 | 3 | 6 | 3 | 2 | 64 | 26 | 42 | 47 | 60 | 3 | 9 | 8 |
| A3SS | 10853 | ENSG00000152601 | MBNL1 | chr3 | + | 152447619 | 152447773 | 152447724 | 152447773 | 152445281 | 152445539 | 1406 | 1072 | 1050 | 1695 | 2222 | 18 | 21 | 19 | 29 | 31 | 491 | 1082 | 831 |
| A3SS | 8524 | ENSG00000144228 | SPOPL | chr2 | + | 138550902 | 138551054 | 138550973 | 138551054 | 138550482 | 138550604 | 270 | 224 | 161 | 234 | 241 | 53 | 36 | 36 | 51 | 51 | 39 | 137 | 185 |
| A3SS | 12227 | ENSG00000159063 | ALG8 | chr11 | - | 78106806 | 78106946 | 78106806 | 78106912 | 78109441 | 78109581 | 35 | 21 | 62 | 39 | 36 | 1 | 2 | 2 | 3 | 3 | 31 | 28 | 13 |
| A3SS | 6806 | ENSG00000169554 | ZEB2 | chr2 | - | 144398300 | 144400345 | 144398300 | 144400270 | 144401198 | 144401307 | 23 | 25 | 11 | 17 | 15 | 272 | 167 | 114 | 226 | 222 | 0 | 26 | 39 |
| A3SS | 2407 | ENSG00000128881 | TTBK2 | chr15 | - | 42872610 | 42872758 | 42872610 | 42872696 | 42878548 | 42878684 | 37 | 29 | 29 | 39 | 14 | 0 | 0 | 6 | 0 | 0 | 20 | 26 | 47 |
| A3SS | 8892 | ENSG00000119535 | CSF3R | chr1 | - | 36472516 | 36473622 | 36472516 | 36472686 | 36473718 | 36473887 | 32 | 18 | 29 | 27 | 11 | 0 | 0 | 0 | 0 | 0 | 42 | 31 | 55 |
| A3SS | 460 | ENSG00000104960 | PTOV1 | chr19 | + | 49860055 | 49860183 | 49860093 | 49860183 | 49858548 | 49858653 | 47 | 38 | 61 | 72 | 54 | 0 | 0 | 0 | 0 | 0 | 101 | 53 | 40 |
| A3SS | 11816 | ENSG00000173327 | MAP3K11 | chr11 | - | 65607640 | 65607816 | 65607640 | 65607697 | 65607921 | 65608070 | 77 | 27 | 30 | 73 | 14 | 0 | 0 | 3 | 0 | 3 | 72 | 58 | 15 |
| A3SS | 12736 | ENSG00000130640 | TUBGCP2 | chr10 | - | 133299984 | 133300547 | 133299984 | 133300100 | 133302791 | 133302980 | 39 | 31 | 27 | 39 | 99 | 0 | 3 | 0 | 1 | 0 | 38 | 28 | 57 |
| A3SS | 403 | ENSG00000090554 | FLT3LG | chr19 | + | 49475673 | 49475801 | 49475690 | 49475801 | 49474602 | 49474672 | 9 | 1 | 8 | 2 | 5 | 25 | 16 | 35 | 33 | 11 | 7 | 5 | 6 |
| A3SS | 266 | ENSG00000079432 | CIC | chr19 | + | 42294574 | 42294735 | 42294603 | 42294735 | 42294172 | 42294304 | 3 | 0 | 2 | 1 | 1 | 59 | 28 | 30 | 36 | 32 | 4 | 0 | 0 |
| A3SS | 3844 | ENSG00000102445 | RUBCNL | chr13 | - | 46371940 | 46372597 | 46371940 | 46372210 | 46377889 | 46378005 | 40 | 38 | 37 | 63 | 61 | 1 | 0 | 0 | 2 | 0 | 19 | 112 | 43 |
| A3SS | 10462 | ENSG00000167548 | KMT2D | chr12 | - | 49022279 | 49022411 | 49022279 | 49022353 | 49022589 | 49022875 | 3 | 6 | 1 | 3 | 2 | 184 | 104 | 160 | 205 | 148 | 0 | 5 | 5 |
| A3SS | 898 | ENSG00000080189 | SLC35C2 | chr20 | - | 46357622 | 46357773 | 46357622 | 46357702 | 46358386 | 46358763 | 47 | 39 | 76 | 51 | 37 | 4 | 0 | 7 | 0 | 0 | 63 | 44 | 39 |
| A3SS | 436 | ENSG00000161265 | U2AF1L4 | chr19 | - | 35743154 | 35743345 | 35743154 | 35743205 | 35743808 | 35743904 | 5 | 7 | 5 | 7 | 2 | 0 | 0 | 0 | 0 | 0 | 10 | 18 | 10 |
| A3SS | 10491 | ENSG00000177981 | ASB8 | chr12 | - | 48150086 | 48150170 | 48150086 | 48150140 | 48151200 | 48151305 | 10 | 1 | 0 | 2 | 3 | 3 | 1 | 2 | 1 | 2 | 6 | 3 | 5 |
| A3SS | 7218 | ENSG00000110200 | ANAPC15 | chr11 | - | 72109586 | 72109940 | 72109586 | 72109928 | 72110087 | 72110225 | 3 | 0 | 3 | 0 | 2 | 71 | 25 | 41 | 43 | 124 | 0 | 0 | 3 |
| A3SS | 4827 | ENSG00000215908 | CROCCP2 | chr1 | - | 16619844 | 16620038 | 16619844 | 16620024 | 16625774 | 16626005 | 5 | 1 | 7 | 9 | 0 | 19 | 7 | 4 | 21 | 5 | 0 | 5 | 2 |
| A3SS | 10600 | ENSG00000215021 | PHB2 | chr12 | - | 6968410 | 6968708 | 6968410 | 6968595 | 6969497 | 6969577 | 34 | 24 | 30 | 45 | 44 | 117 | 91 | 112 | 93 | 152 | 37 | 20 | 36 |
| A3SS | 11573 | ENSG00000060237 | WNK1 | chr12 | + | 878211 | 878361 | 878214 | 878361 | 871264 | 871348 | 39 | 14 | 37 | 46 | 26 | 27 | 14 | 21 | 14 | 35 | 51 | 62 | 36 |
| A3SS | 2253 | ENSG00000104093 | DMXL2 | chr15 | - | 51476588 | 51476719 | 51476588 | 51476716 | 51478270 | 51478347 | 123 | 119 | 55 | 86 | 127 | 38 | 54 | 25 | 38 | 94 | 9 | 50 | 33 |
| A3SS | 502 | ENSG00000076928 | ARHGEF1 | chr19 | + | 41903655 | 41903784 | 41903706 | 41903784 | 41903306 | 41903407 | 19 | 13 | 10 | 29 | 12 | 447 | 262 | 471 | 459 | 233 | 0 | 19 | 18 |
| A3SS | 10343 | ENSG00000174080 | CTSF | chr11 | - | 66564087 | 66564238 | 66564087 | 66564146 | 66564557 | 66564648 | 1 | 4 | 6 | 8 | 5 | 14 | 5 | 13 | 21 | 8 | 20 | 6 | 8 |
| A3SS | 354 | ENSG00000099804 | CDC34 | chr19 | + | 541338 | 542087 | 541345 | 542087 | 537012 | 537147 | 89 | 33 | 49 | 71 | 40 | 3 | 0 | 1 | 0 | 3 | 71 | 46 | 32 |
| A3SS | 7193 | ENSG00000001497 | LAS1L | chrX | - | 65522430 | 65522926 | 65522430 | 65522493 | 65523559 | 65523707 | 6 | 3 | 4 | 6 | 1 | 0 | 3 | 0 | 0 | 0 | 19 | 4 | 8 |
| A3SS | 997 | ENSG00000134444 | RELCH | chr18 | - | 62268868 | 62268948 | 62268880 | 62268948 | 62266700 | 62266749 | 59 | 31 | 56 | 68 | 64 | 3 | 1 | 2 | 3 | 0 | 13 | 28 | 21 |
| A3SS | 10344 | ENSG00000174080 | CTSF | chr11 | - | 66564087 | 66564241 | 66564087 | 66564146 | 66564557 | 66564648 | 1 | 4 | 6 | 8 | 5 | 14 | 5 | 13 | 21 | 8 | 20 | 6 | 8 |
| A3SS | 8828 | ENSG00000118495 | PLAGL1 | chr6 | - | 143966157 | 143966285 | 143966157 | 143966212 | 143968906 | 143968978 | 9 | 26 | 10 | 19 | 20 | 4 | 4 | 0 | 0 | 2 | 14 | 5 | 2 |
| A3SS | 10255 | ENSG00000169891 | REPS2 | chrX | + | 17025058 | 17025185 | 17025061 | 17025185 | 17022122 | 17022271 | 6 | 2 | 0 | 6 | 2 | 3 | 7 | 3 | 12 | 19 | 5 | 1 | 12 |
| A3SS | 67 | ENSG00000100139 | MICALL1 | chr22 | + | 37937079 | 37937194 | 37937084 | 37937194 | 37933038 | 37933112 | 68 | 45 | 54 | 60 | 57 | 5 | 1 | 2 | 0 | 0 | 93 | 139 | 68 |
| A3SS | 4215 | ENSG00000013441 | CLK1 | chr2 | - | 200859679 | 200859747 | 200859679 | 200859746 | 200861237 | 200861466 | 2 | 1 | 2 | 3 | 2 | 255 | 146 | 253 | 372 | 476 | 0 | 15 | 0 |
| A3SS | 5283 | ENSG00000127054 | INTS11 | chr1 | - | 1315519 | 1315614 | 1315519 | 1315552 | 1319295 | 1319524 | 66 | 29 | 34 | 71 | 52 | 1 | 2 | 1 | 3 | 0 | 67 | 71 | 34 |
| A3SS | 10314 | ENSG00000230590 | FTX | chrX | - | 74274341 | 74274410 | 74274341 | 74274371 | 74280393 | 74280494 | 19 | 22 | 19 | 35 | 44 | 5 | 0 | 1 | 3 | 0 | 30 | 24 | 28 |
| A3SS | 5730 | ENSG00000115446 | UNC50 | chr2 | + | 98618145 | 98618515 | 98618167 | 98618515 | 98616431 | 98616533 | 3 | 2 | 2 | 4 | 2 | 62 | 42 | 42 | 48 | 75 | 0 | 0 | 6 |
| A3SS | 8374 | ENSG00000204387 | SNHG32 | chr6 | + | 31836298 | 31836517 | 31836423 | 31836517 | 31835410 | 31835451 | 11 | 7 | 15 | 18 | 11 | 73 | 32 | 97 | 37 | 44 | 8 | 20 | 6 |
| A3SS | 1010 | ENSG00000104884 | ERCC2 | chr19 | - | 45368629 | 45368746 | 45368629 | 45368743 | 45368929 | 45369147 | 0 | 0 | 3 | 2 | 0 | 27 | 14 | 33 | 16 | 24 | 0 | 0 | 0 |
| A3SS | 6480 | ENSG00000163564 | PYHIN1 | chr1 | + | 158938396 | 158938542 | 158938423 | 158938542 | 158936890 | 158937175 | 438 | 231 | 221 | 252 | 525 | 63 | 26 | 43 | 52 | 78 | 15 | 141 | 77 |
| A3SS | 2970 | ENSG00000148218 | ALAD | chr9 | - | 113392118 | 113392335 | 113392118 | 113392169 | 113393446 | 113393634 | 3 | 5 | 4 | 4 | 0 | 8 | 15 | 18 | 21 | 10 | 21 | 2 | 0 |
| A3SS | 4598 | ENSG00000172830 | SSH3 | chr11 | + | 67310065 | 67310339 | 67310163 | 67310339 | 67309767 | 67309968 | 13 | 2 | 14 | 4 | 0 | 1 | 4 | 1 | 0 | 1 | 36 | 16 | 0 |
| A3SS | 12957 | ENSG00000204482 | LST1 | chr6 | + | 31588517 | 31588909 | 31588562 | 31588909 | 31587199 | 31587318 | 447 | 239 | 177 | 388 | 530 | 181 | 84 | 59 | 117 | 172 | 1125 | 255 | 245 |
| A3SS | 3847 | ENSG00000002586 | CD99 | chrX | + | 2738199 | 2738256 | 2738202 | 2738256 | 2733356 | 2733374 | 22 | 12 | 16 | 10 | 11 | 0 | 5 | 0 | 0 | 0 | 12 | 29 | 18 |
| A3SS | 7031 | ENSG00000126215 | XRCC3 | chr14 | - | 103711465 | 103711661 | 103711465 | 103711567 | 103712874 | 103712968 | 7 | 0 | 4 | 4 | 5 | 4 | 1 | 7 | 6 | 5 | 3 | 0 | 4 |
| A3SS | 832 | ENSG00000171552 | BCL2L1 | chr20 | - | 31721654 | 31722348 | 31721654 | 31722330 | 31723734 | 31723978 | 21 | 14 | 15 | 19 | 15 | 3 | 3 | 3 | 3 | 3 | 51 | 24 | 23 |
| A3SS | 9376 | ENSG00000197283 | SYNGAP1 | chr6 | + | 33440552 | 33440965 | 33440728 | 33440965 | 33438774 | 33438919 | 63 | 30 | 89 | 68 | 214 | 11 | 3 | 6 | 4 | 11 | 47 | 42 | 50 |
| A3SS | 9692 | ENSG00000134291 | TMEM106C | chr12 | + | 47964144 | 47964423 | 47964208 | 47964423 | 47963595 | 47963633 | 4 | 1 | 6 | 1 | 3 | 8 | 13 | 8 | 11 | 7 | 25 | 2 | 5 |
| A3SS | 12106 | ENSG00000103544 | VPS35L | chr16 | + | 19573118 | 19573241 | 19573123 | 19573241 | 19569423 | 19569591 | 88 | 29 | 52 | 48 | 66 | 3 | 2 | 0 | 1 | 1 | 100 | 66 | 46 |
| A3SS | 5907 | ENSG00000055609 | KMT2C | chr7 | - | 152156204 | 152156346 | 152156204 | 152156271 | 152157760 | 152157922 | 174 | 183 | 196 | 258 | 365 | 0 | 2 | 4 | 4 | 8 | 47 | 142 | 171 |
| A3SS | 4584 | ENSG00000140262 | TCF12 | chr15 | + | 57273029 | 57273262 | 57273032 | 57273262 | 57263111 | 57263274 | 143 | 106 | 88 | 98 | 169 | 14 | 10 | 17 | 19 | 12 | 39 | 65 | 90 |
| A3SS | 13324 | ENSG00000081189 | MEF2C | chr5 | - | 88718273 | 88722925 | 88718273 | 88722829 | 88728492 | 88728628 | 100 | 105 | 145 | 156 | 326 | 94 | 93 | 58 | 130 | 222 | 140 | 190 | 101 |
| A3SS | 3437 | ENSG00000134644 | PUM1 | chr1 | - | 30965981 | 30966278 | 30965981 | 30966197 | 30967166 | 30967310 | 170 | 61 | 120 | 128 | 69 | 12 | 5 | 26 | 15 | 3 | 54 | 113 | 72 |
| A3SS | 3639 | ENSG00000137802 | MAPKBP1 | chr15 | + | 41815258 | 41815799 | 41815623 | 41815799 | 41814549 | 41814739 | 28 | 9 | 15 | 25 | 15 | 4 | 0 | 1 | 0 | 3 | 55 | 21 | 15 |
| A3SS | 670 | ENSG00000064490 | RFXANK | chr19 | + | 19196962 | 19197046 | 19196965 | 19197046 | 19193938 | 19194133 | 20 | 15 | 30 | 36 | 36 | 38 | 30 | 49 | 42 | 61 | 83 | 51 | 17 |
| A3SS | 5731 | ENSG00000142330 | CAPN10 | chr2 | + | 240595933 | 240596521 | 240596318 | 240596521 | 240595023 | 240595304 | 23 | 2 | 11 | 19 | 7 | 20 | 3 | 12 | 7 | 4 | 10 | 9 | 10 |
| A3SS | 11908 | ENSG00000137500 | CCDC90B | chr11 | - | 83274638 | 83274763 | 83274638 | 83274740 | 83280140 | 83280260 | 3 | 0 | 0 | 1 | 10 | 33 | 20 | 12 | 31 | 25 | 10 | 1 | 1 |
| A3SS | 11400 | ENSG00000089053 | ANAPC5 | chr12 | - | 121318276 | 121318424 | 121318276 | 121318404 | 121318500 | 121318608 | 146 | 102 | 169 | 167 | 410 | 10 | 8 | 28 | 20 | 12 | 184 | 176 | 104 |
| A3SS | 5761 | ENSG00000144283 | PKP4 | chr2 | + | 158634069 | 158634289 | 158634072 | 158634289 | 158631752 | 158631941 | 5 | 10 | 13 | 12 | 13 | 2 | 6 | 5 | 2 | 2 | 10 | 17 | 2 |
| A3SS | 9142 | ENSG00000130023 | ERMARD | chr6 | + | 169753863 | 169754032 | 169753943 | 169754032 | 169753316 | 169753372 | 43 | 29 | 44 | 36 | 57 | 0 | 0 | 0 | 0 | 0 | 13 | 18 | 41 |
| A3SS | 6281 | ENSG00000074706 | IPCEF1 | chr6 | - | 154247448 | 154247488 | 154247448 | 154247485 | 154265911 | 154265964 | 15 | 21 | 17 | 19 | 30 | 3 | 16 | 9 | 8 | 7 | 36 | 30 | 24 |
| A3SS | 8186 | ENSG00000084623 | EIF3I | chr1 | + | 32226148 | 32226320 | 32226170 | 32226320 | 32224409 | 32224475 | 6 | 5 | 5 | 8 | 0 | 168 | 96 | 189 | 173 | 102 | 6 | 4 | 3 |
| A3SS | 10588 | ENSG00000104921 | FCER2 | chr19 | - | 7698355 | 7698854 | 7698355 | 7698409 | 7699738 | 7699845 | 147 | 49 | 125 | 91 | 104 | 1 | 1 | 4 | 2 | 0 | 29 | 97 | 16 |
| A3SS | 11973 | ENSG00000168067 | MAP4K2 | chr11 | - | 64802562 | 64802653 | 64802562 | 64802627 | 64802884 | 64802942 | 31 | 20 | 22 | 51 | 11 | 1 | 0 | 0 | 0 | 0 | 39 | 25 | 25 |
| A3SS | 9777 | ENSG00000074800 | ENO1 | chr1 | - | 8874823 | 8874917 | 8874823 | 8874914 | 8878579 | 8879092 | 711 | 324 | 673 | 708 | 552 | 3 | 1 | 4 | 4 | 0 | 1706 | 1381 | 622 |
| A3SS | 8412 | ENSG00000239779 | WBP1 | chr2 | + | 74459437 | 74459745 | 74459642 | 74459745 | 74458449 | 74458537 | 11 | 7 | 25 | 24 | 17 | 4 | 1 | 2 | 1 | 0 | 9 | 11 | 23 |
| A3SS | 3324 | ENSG00000113742 | CPEB4 | chr5 | + | 173949879 | 173950078 | 173949959 | 173950078 | 173949507 | 173949597 | 18 | 18 | 9 | 22 | 37 | 63 | 51 | 47 | 65 | 108 | 6 | 27 | 30 |
| A3SS | 9027 | ENSG00000197345 | MRPL21 | chr11 | - | 68896514 | 68896838 | 68896514 | 68896678 | 68897926 | 68898012 | 4 | 4 | 0 | 2 | 4 | 42 | 15 | 53 | 37 | 94 | 2 | 5 | 0 |
| A3SS | 4712 | ENSG00000092098 | RNF31 | chr14 | + | 24157831 | 24158011 | 24157897 | 24158011 | 24157519 | 24157638 | 3 | 0 | 4 | 2 | 1 | 37 | 24 | 34 | 30 | 32 | 0 | 0 | 6 |
| A3SS | 11724 | ENSG00000083799 | CYLD | chr16 | + | 50749548 | 50750202 | 50749575 | 50750202 | 50742761 | 50742841 | 27 | 21 | 26 | 29 | 20 | 156 | 136 | 192 | 150 | 117 | 6 | 38 | 33 |
| A3SS | 12831 | ENSG00000284770 | TBCE | chr1 | + | 235436385 | 235436608 | 235436543 | 235436608 | 235435744 | 235435840 | 72 | 58 | 58 | 47 | 157 | 0 | 3 | 0 | 2 | 1 | 57 | 64 | 61 |
| A3SS | 1797 | ENSG00000121281 | ADCY7 | chr16 | + | 50287881 | 50288350 | 50287911 | 50288350 | 50266550 | 50266587 | 7 | 2 | 3 | 1 | 0 | 16 | 9 | 9 | 9 | 13 | 0 | 0 | 0 |
| A3SS | 7658 | ENSG00000163904 | SENP2 | chr3 | + | 185598957 | 185599024 | 185598991 | 185599024 | 185598411 | 185598545 | 45 | 34 | 60 | 77 | 75 | 2 | 1 | 2 | 1 | 3 | 58 | 81 | 46 |
| [illegible] | | | | | | | | | | | | | | | | | | | | | | | | |

| | | | | | | | | | | | | | | | | | | | | | | | |
|---|---|---|---|---|---|---|---|---|---|---|---|---|---|---|---|---|---|---|---|---|---|---|---|
| A3SS | 1230 | ENSG00000108622 | ICAM2 | chr17 | - | 64005106 | 64005373 | 64005106 | 64005137 | 64006630 | 64006735 | 276 | 135 | 210 | 301 | 341 | 0 | 3 | 1 | 6 | 7 | 249 | 163 |
| A3SS | 3846 | ENSG00000002586 | CD99 | chrX | + | 2738199 | 2738256 | 2738202 | 2738256 | 2726259 | 2726373 | 1269 | 597 | 799 | 1115 | 1090 | 102 | 55 | 74 | 88 | 105 | 2064 | 1880 |
| A3SS | 10702 | ENSG00000046651 | OFD1 | chrX | + | 13752703 | 13753441 | 13753367 | 13753441 | 13751248 | 13751368 | 99 | 114 | 98 | 122 | 110 | 96 | 92 | 103 | 86 | 175 | 11 | 42 |
| A3SS | 5820 | ENSG00000160058 | BSDC1 | chr1 | - | 32383829 | 32384048 | 32383829 | 32383939 | 32386778 | 32386895 | 61 | 58 | 63 | 96 | 72 | 6 | 3 | 9 | 9 | 3 | 95 | 63 |
| A3SS | 1643 | ENSG00000142512 | SIGLEC10 | chr19 | - | 51416665 | 51416950 | 51416665 | 51416806 | 51417081 | 51417465 | 66 | 89 | 37 | 100 | 85 | 6 | 3 | 0 | 3 | 0 | 70 | 104 |
| A3SS | 3959 | ENSG00000141562 | NARF | chr17 | + | 82459991 | 82460072 | 82459999 | 82460072 | 82458710 | 82458830 | 45 | 34 | 32 | 36 | 15 | 0 | 0 | 0 | 0 | 5 | 90 | 51 |
| A3SS | 3494 | ENSG00000247774 | PCED1B-AS1 | chr12 | - | 47205897 | 47206145 | 47205897 | 47206069 | 47206510 | 47206588 | 352 | 225 | 374 | 352 | 861 | 73 | 40 | 177 | 102 | 244 | 197 | 158 |
| A3SS | 10937 | ENSG00000103266 | STUB1 | chr16 | + | 682019 | 682076 | 682058 | 682076 | 681792 | 681880 | 162 | 63 | 102 | 134 | 160 | 1 | 0 | 7 | 0 | 2 | 116 | 174 |
| A3SS | 11199 | ENSG00000104529 | EEF1D | chr8 | - | 143592714 | 143592942 | 143592714 | 143592893 | 143593874 | 143594071 | 29 | 24 | 27 | 20 | 9 | 0 | 0 | 0 | 0 | 4 | 6 | 19 |
| A3SS | 8401 | ENSG00000131236 | CAP1 | chr1 | + | 40059333 | 40059458 | 40059336 | 40059458 | 40041428 | 40041493 | 58 | 54 | 36 | 95 | 39 | 30 | 25 | 25 | 44 | 34 | 128 | 148 |
| A3SS | 983 | ENSG00000124155 | PIGT | chr20 | + | 45419142 | 45419395 | 45419294 | 45419395 | 45416516 | 45416694 | 10 | 3 | 19 | 11 | 7 | 7 | 3 | 10 | 14 | 4 | 7 | 43 |
| A3SS | 10659 | ENSG00000131269 | ABCB7 | chrX | - | 75114753 | 75114834 | 75114753 | 75114831 | 75156104 | 75156282 | 32 | 16 | 18 | 23 | 8 | 18 | 8 | 17 | 26 | 9 | 16 | 16 |
| A3SS | 9888 | ENSG00000126883 | NUP214 | chr9 | + | 131146128 | 131146304 | 131146164 | 131146304 | 131144279 | 131144754 | 45 | 15 | 23 | 21 | 29 | 2 | 2 | 0 | 0 | 2 | 82 | 16 |
| A3SS | 6883 | ENSG00000072110 | ACTN1 | chr14 | - | 68877081 | 68878523 | 68877081 | 68877240 | 68878988 | 68879069 | 13 | 12 | 29 | 19 | 4 | 116 | 141 | 146 | 127 | 149 | 12 | 29 |
| A3SS | 4388 | ENSG00000176261 | ZBTB8OS | chr1 | - | 32633950 | 32634110 | 32633950 | 32634072 | 32634767 | 32634792 | 24 | 13 | 12 | 12 | 37 | 68 | 40 | 70 | 43 | 82 | 0 | 23 |
| A3SS | 9913 | ENSG00000115274 | INO80B | chr2 | + | 74456102 | 74456272 | 74456147 | 74456272 | 74455817 | 74455936 | 134 | 115 | 115 | 197 | 166 | 3 | 2 | 2 | 4 | 1 | 220 | 143 |
| A3SS | 10229 | ENSG00000029363 | BCLAF1 | chr6 | - | 136275842 | 136276508 | 136275842 | 136275989 | 136277864 | 136278770 | 762 | 645 | 691 | 956 | 1323 | 34 | 12 | 22 | 46 | 26 | 60 | 386 |
| A3SS | 12964 | ENSG00000196498 | NCOR2 | chr12 | - | 124332318 | 124332467 | 124332318 | 124332464 | 124333129 | 124333279 | 37 | 47 | 37 | 49 | 34 | 7 | 0 | 12 | 21 | 16 | 162 | 71 |
| A3SS | 6388 | ENSG00000112182 | BACH2 | chr6 | - | 90206568 | 90206681 | 90206568 | 90206676 | 90296479 | 90296742 | 21 | 26 | 26 | 20 | 26 | 0 | 1 | 0 | 4 | 2 | 3 | 20 |
| A3SS | 1257 | ENSG00000129219 | PLD2 | chr17 | + | 4818001 | 4818385 | 4818296 | 4818385 | 4817145 | 4817259 | 13 | 10 | 14 | 16 | 10 | 0 | 0 | 4 | 0 | 0 | 46 | 11 |
| A3SS | 5253 | ENSG00000247828 | TMEM161B-AS1 | chr5 | + | 88287435 | 88287622 | 88287438 | 88287622 | 88285743 | 88285851 | 34 | 28 | 16 | 9 | 46 | 1 | 1 | 1 | 2 | 2 | 2 | 8 |
| A3SS | 3704 | ENSG00000164483 | SAMD3 | chr6 | - | 130184437 | 130184623 | 130184437 | 130184620 | 130209494 | 130209608 | 249 | 107 | 141 | 248 | 204 | 57 | 29 | 29 | 66 | 53 | 20 | 115 |
| A3SS | 4881 | ENSG00000132549 | VPS13B | chr8 | + | 99431520 | 99431664 | 99431536 | 99431664 | 99391556 | 99391704 | 12 | 9 | 7 | 9 | 11 | 110 | 94 | 104 | 67 | 81 | 0 | 16 |
| A3SS | 10241 | ENSG00000196998 | WDR45 | chrX | - | 49076644 | 49076825 | 49076644 | 49076753 | 49077642 | 49077747 | 8 | 1 | 10 | 10 | 0 | 7 | 1 | 1 | 2 | 2 | 45 | 0 |
| A3SS | 9590 | ENSG00000070669 | ASNS | chr7 | - | 97868907 | 97869179 | 97868907 | 97869105 | 97869777 | 97869813 | 37 | 17 | 22 | 44 | 31 | 1 | 0 | 2 | 1 | 2 | 26 | 17 |
| A3SS | 12021 | ENSG00000205531 | NAP1L4 | chr11 | - | 2989128 | 2989271 | 2989128 | 2989197 | 2991793 | 2992102 | 21 | 5 | 26 | 29 | 35 | 0 | 0 | 2 | 2 | 2 | 14 | 72 |
| A3SS | 5566 | ENSG00000108443 | RPS6KB1 | chr17 | + | 59936214 | 59936541 | 59936463 | 59936541 | 59935192 | 59935300 | 106 | 96 | 93 | 104 | 169 | 0 | 0 | 0 | 0 | 8 | 53 | 92 |
| A3SS | 9125 | ENSG00000105393 | BABAM1 | chr19 | + | 17268793 | 17269091 | 17268801 | 17269091 | 17267442 | 17267527 | 66 | 29 | 63 | 59 | 30 | 0 | 1 | 2 | 0 | 0 | 74 | 62 |
| A3SS | 1741 | ENSG00000179583 | CIITA | chr16 | + | 10906498 | 10909187 | 10909028 | 10909187 | 10904743 | 10904812 | 87 | 61 | 56 | 70 | 46 | 0 | 1 | 0 | 1 | 5 | 224 | 90 |
| A3SS | 4016 | ENSG00000163161 | ERCC3 | chr2 | - | 127292609 | 127292846 | 127292609 | 127292704 | 127293512 | 127293718 | 160 | 89 | 115 | 163 | 119 | 0 | 2 | 0 | 7 | 0 | 59 | 158 |
| A3SS | 4730 | ENSG00000182481 | KPNA2 | chr17 | + | 68037079 | 68037207 | 68037109 | 68037207 | 68035734 | 68035840 | 1 | 1 | 0 | 0 | 0 | 35 | 13 | 42 | 44 | 11 | 0 | 2 |
| A3SS | 8870 | ENSG00000182732 | RGS6 | chr14 | + | 72540040 | 72540094 | 72540044 | 72540094 | 72536185 | 72536275 | 4 | 10 | 2 | 16 | 11 | 2 | 0 | 0 | 1 | 0 | 16 | 5 |
| A3SS | 7284 | ENSG00000198898 | CAPZA2 | chr7 | + | 116910212 | 116910311 | 116910232 | 116910311 | 116906262 | 116906342 | 10 | 23 | 7 | 39 | 47 | 580 | 508 | 330 | 706 | 1242 | 0 | 14 |
| A3SS | 12535 | ENSG00000175197 | DDIT3 | chr12 | - | 57517268 | 57517438 | 57517268 | 57517424 | 57517705 | 57517753 | 21 | 10 | 10 | 16 | 21 | 4 | 0 | 0 | 1 | 1 | 26 | 15 |
| A3SS | 1182 | ENSG00000182173 | TSEN54 | chr17 | + | 75521354 | 75521510 | 75521408 | 75521510 | 75518994 | 75519047 | 8 | 1 | 6 | 2 | 6 | 49 | 17 | 49 | 48 | 18 | 0 | 6 |
| A3SS | 404 | ENSG00000090554 | FLT3LG | chr19 | + | 49476116 | 49476198 | 49476144 | 49476198 | 49475690 | 49475801 | 6 | 2 | 3 | 5 | 1 | 44 | 21 | 88 | 53 | 23 | 0 | 0 |
| A3SS | 11255 | ENSG00000010610 | CD4 | chr12 | + | 6815730 | 6816403 | 6816055 | 6816403 | 6814758 | 6814992 | 8 | 0 | 7 | 11 | 1 | 117 | 55 | 92 | 107 | 96 | 17 | 2 |
| A3SS | 6635 | ENSG00000159023 | EPB41 | chr1 | + | 29053103 | 29053312 | 29053106 | 29053312 | 29039253 | 29039426 | 246 | 168 | 301 | 224 | 273 | 46 | 27 | 36 | 60 | 39 | 292 | 246 |
| A3SS | 240 | ENSG00000127616 | SMARCA4 | chr19 | + | 11041297 | 11041560 | 11041306 | 11041560 | 11039457 | 11039553 | 1 | 2 | 0 | 1 | 0 | 14 | 7 | 8 | 12 | 9 | 0 | 5 |
| A3SS | 9717 | ENSG00000119616 | FCF1 | chr14 | + | 74715820 | 74716099 | 74715950 | 74716099 | 74713484 | 74713552 | 38 | 20 | 45 | 32 | 64 | 1 | 1 | 1 | 4 | 0 | 25 | 60 |
| A3SS | 4284 | ENSG00000068650 | ATP11A | chr13 | + | 112881815 | 112887168 | 112881875 | 112887168 | 112875775 | 112875941 | 22 | 10 | 22 | 12 | 32 | 10 | 2 | 6 | 6 | 7 | 56 | 14 |
| A3SS | 7685 | ENSG00000116922 | C1orf109 | chr1 | - | 37683318 | 37683441 | 37683318 | 37683437 | 37686272 | 37686452 | 4 | 3 | 3 | 4 | 8 | 20 | 10 | 21 | 20 | 20 | 0 | 2 |
| A3SS | 875 | ENSG00000203880 | PCMTD2 | chr20 | + | 64267886 | 64268010 | 64267967 | 64268010 | 64265257 | 64265429 | 234 | 175 | 238 | 209 | 416 | 20 | 6 | 18 | 23 | 21 | 81 | 138 |
| A3SS | 12616 | ENSG00000176986 | SEC24C | chr10 | + | 73770279 | 73770798 | 73770708 | 73770798 | 73769835 | 73770015 | 322 | 199 | 243 | 292 | 318 | 3 | 0 | 3 | 2 | 7 | 416 | 434 |
| A3SS | 4061 | ENSG00000205726 | ITSN1 | chr21 | + | 33772038 | 33772323 | 33772060 | 33772323 | 33767712 | 33767828 | 0 | 0 | 0 | 1 | 0 | 10 | 22 | 2 | 19 | 19 | 0 | 0 |
| A3SS | 663 | ENSG00000105705 | SUGP1 | chr19 | - | 19303723 | 19304043 | 19303723 | 19303912 | 19305848 | 19306076 | 17 | 13 | 17 | 13 | 5 | 1 | 3 | 9 | 2 | 2 | 5 | 8 |
| A3SS | 7081 | ENSG00000050748 | MAPK9 | chr5 | - | 180233593 | 180236531 | 180233593 | 180236526 | 180238331 | 180238403 | 10 | 20 | 12 | 33 | 62 | 131 | 102 | 101 | 89 | 241 | 5 | 2 |
| A3SS | 9528 | ENSG00000173200 | PARP15 | chr3 | + | 122617014 | 122617164 | 122617085 | 122617164 | 122615778 | 122615857 | 278 | 191 | 160 | 280 | 236 | 2 | 4 | 9 | 4 | 13 | 41 | 182 |
| A3SS | 5991 | ENSG00000213672 | NCKIPSD | chr3 | - | 48682044 | 48682156 | 48682044 | 48682135 | 48682347 | 48682552 | 4 | 5 | 2 | 13 | 6 | 3 | 2 | 4 | 6 | 7 | 19 | 11 |
| A3SS | 9591 | ENSG00000088038 | CNOT3 | chr19 | + | 54142925 | 54143003 | 54142928 | 54143003 | 54137761 | 54137993 | 3 | 1 | 3 | 8 | 3 | 9 | 20 | 30 | 25 | 7 | 0 | 3 |
| A3SS | 1844 | ENSG00000141002 | TCF25 | chr16 | + | 89904117 | 89905096 | 89904937 | 89905096 | 89900634 | 89900794 | 77 | 16 | 70 | 53 | 23 | 0 | 0 | 0 | 0 | 0 | 56 | 79 |
| A3SS | 12756 | ENSG00000132305 | IMMT | chr2 | - | 86171207 | 86171345 | 86171207 | 86171312 | 86173649 | 86173761 | 100 | 53 | 50 | 92 | 145 | 77 | 34 | 77 | 72 | 110 | 204 | 105 |
| A3SS | 5440 | ENSG00000121940 | CLCC1 | chr1 | - | 108950308 | 108950448 | 108950308 | 108950438 | 108962308 | 108962469 | 31 | 26 | 23 | 14 | 24 | 2 | 1 | 0 | 2 | 1 | 9 | 14 |
| A3SS | 1516 | ENSG00000011260 | UTP18 | chr17 | + | 51288028 | 51288203 | 51288123 | 51288203 | 51285244 | 51285368 | 179 | 131 | 122 | 139 | 330 | 2 | 1 | 1 | 0 | 6 | 72 | 103 |
| A3SS | 6096 | ENSG00000138095 | LRPPRC | chr2 | - | 43948121 | 43948230 | 43948121 | 43948199 | 43948411 | 43948518 | 6 | 1 | 2 | 3 | 6 | 96 | 68 | 88 | 71 | 144 | 0 | 0 |
| A3SS | 125 | ENSG00000100031 | GGT1 | chr22 | + | 24610249 | 24610531 | 24610357 | 24610531 | 24609961 | 24610033 | 5 | 5 | 7 | 12 | 11 | 1 | 3 | 3 | 0 | 1 | 22 | 16 |
| A3SS | 1389 | ENSG00000141574 | SECTM1 | chr17 | - | 82324581 | 82324890 | 82324581 | 82324783 | 82327146 | 82327292 | 70 | 68 | 52 | 112 | 43 | 2 | 0 | 0 | 5 | 0 | 187 | 106 |
| A3SS | 4148 | ENSG00000163932 | PRKCD | chr3 | + | 53183342 | 53183581 | 53183451 | 53183581 | 53183120 | 53183206 | 9 | 2 | 5 | 0 | 2 | 408 | 342 | 249 | 376 | 480 | 43 | 0 |
| A3SS | 1941 | ENSG00000213658 | LAT | chr16 | + | 28986354 | 28986446 | 28986384 | 28986446 | 28986134 | 28986216 | 13 | 4 | 8 | 16 | 13 | 25 | 8 | 26 | 16 | 12 | 10 | 33 |
| A3SS | 10561 | ENSG00000172534 | HCFC1 | chrX | - | 153952513 | 153953090 | 153952513 | 153952958 | 153953606 | 153953770 | 16 | 6 | 13 | 17 | 14 | 53 | 22 | 68 | 48 | 31 | 18 | 21 |
| A3SS | 11211 | ENSG00000104522 | GFUS | chr8 | - | 143616105 | 143616440 | 143616105 | 143616220 | 143616566 | 143616723 | 0 | 0 | 2 | 2 | 6 | 77 | 25 | 74 | 45 | 50 | 0 | 0 |
| A3SS | 11603 | ENSG00000148288 | GBGT1 | chr9 | - | 133156014 | 133156102 | 133156014 | 133156065 | 133161466 | 133161532 | 3 | 3 | 0 | 2 | 0 | 9 | 6 | 6 | 6 | 8 | 0 | 1 |
| A3SS | 10054 | ENSG00000223482 | NUTM2A-AS1 | chr10 | - | 87330891 | 87331096 | 87330891 | 87330965 | 87336785 | 87336889 | 54 | 77 | 16 | 64 | 73 | 0 | 0 | 0 | 2 | 0 | 89 | 106 |
| A3SS | 4554 | ENSG00000092931 | MFSD11 | chr17 | + | 76741968 | 76742048 | 76741993 | 76742048 | 76740956 | 76741064 | 81 | 52 | 39 | 45 | 40 | 4 | 0 | 4 | 2 | 1 | 73 | 37 |
| A3SS | 5760 | ENSG00000144283 | PKP4 | chr2 | + | 158624954 | 158625427 | 158625124 | 158625427 | 158621230 | 158621421 | 14 | 20 | 26 | 28 | 53 | 0 | 1 | 0 | 0 | 1 | 12 | 39 |
| A3SS | 714 | ENSG00000160633 | SAFB | chr19 | + | 5667346 | 5667450 | 5667352 | 5667450 | 5667045 | 5667164 | 50 | 50 | 71 | 54 | 13 | 18 | 20 | 20 | 18 | 21 | 77 | 46 |
| A3SS | 7180 | ENSG00000142405 | NLRP12 | chr19 | - | 53807494 | 53807668 | 53807494 | 53807665 | 53809586 | 53811288 | 7 | 9 | 2 | 5 | 4 | 47 | 12 | 17 | 23 | 6 | 11 | 12 |
| A3SS | 10606 | ENSG00000071889 | FAM3A | chrX | - | 154508473 | 154508811 | 154508473 | 154508597 | 154511847 | 154511871 | 8 | 6 | 3 | 3 | 3 | 50 | 14 | 51 | 11 | 7 | 0 | 3 |
| A3SS | 1832 | ENSG00000125149 | C16orf70 | chr16 | + | 67132445 | 67132911 | 67132816 | 67132911 | 67131285 | 67131329 | 75 | 34 | 37 | 45 | 82 | 0 | 0 | 0 | 1 | 0 | 131 | 44 |
| A3SS | 6638 | ENSG00000150455 | TIRAP | chr11 | + | 126291272 | 126291569 | 126291424 | 126291569 | 126290802 | 126290961 | 3 | 4 | 7 | 5 | 4 | 1 | 1 | 0 | 3 | 0 | 40 | 5 |
| A3SS | 8402 | ENSG00000131236 | CAP1 | chr1 | + | 40060066 | 40060170 | 40060069 | 40060170 | 40059333 | 40059458 | 1930 | 1253 | 1102 | 1585 | 1546 | 171 | 101 | 86 | 123 | 107 | 3715 | 2167 |
| A3SS | 11494 | ENSG00000070061 | ELP1 | chr9 | - | 108911009 | 108911211 | 108911009 | 108911180 | 108912263 | 108912494 | 1 | 0 | 6 | 4 | 0 | 65 | 37 | 52 | 69 | 58 | 0 | 4 |
| A3SS | 8380 | ENSG00000226281 | NA | chr6 | - | 6710871 | 6711059 | 6710871 | 6710981 | 6711696 | 6711858 | 4 | 0 | 0 | 0 | 0 | 1 | 1 | 1 | 5 | 11 | 3 | 1 |
| A3SS | 8805 | ENSG00000125503 | PPP1R12C | chr19 | - | 55095444 | 55095603 | 55095444 | 55095600 | 55095866 | 55095940 | 27 | 11 | 18 | 27 | 14 | 16 | 16 | 8 | 18 | 6 | 72 | 46 |
| A3SS | 7436 | ENSG00000142327 | RNPEPL1 | chr2 | + | 240576534 | 240576765 | 240576567 | 240576765 | 240575501 | 240575610 | 104 | 56 | 75 | 88 | 53 | 10 | 2 | 6 | 8 | 8 | 19 | 56 |
| A3SS | 6132 | ENSG00000198408 | OGA | chr10 | - | 101792838 | 101792943 | 101792838 | 101792901 | 101797979 | 101798154 | 142 | 78 | 120 | 113 | 149 | 1 | 0 | 0 | 0 | 0 | 81 | 79 |
| A3SS | 2150 | ENSG00000103197 | TSC2 | chr16 | + | 2086731 | 2086871 | 2086766 | 2086871 | 2086192 | 2086379 | 84 | 24 | 50 | 48 | 56 | 2 | 1 | 2 | 2 | 0 | 56 | 100 |
| A3SS | 3260 | ENSG00000140564 | FURIN | chr15 | + | 90877461 | 90877615 | 90877526 | 90877615 | 90877134 | 90877211 | 3 | 3 | 5 | 6 | 0 | 233 | 58 | 135 | 159 | 72 | 0 | 14 |
| A3SS | 3464 | ENSG00000011243 | AKAP8L | chr19 | - | 15397713 | 15400358 | 15397713 | 15397855 | 15400793 | 15400864 | 102 | 74 | 84 | 100 | 75 | 0 | 0 | 0 | 0 | 0 | 172 | 149 |
| A3SS | 12242 | ENSG00000172183 | ISG20 | chr15 | + | 88652109 | 88652310 | 88652169 | 88652310 | 88639342 | 88639594 | 296 | 134 | 313 | 199 | 88 | 17 | 4 | 12 | 11 | 0 | 87 | 396 |
| A3SS | 5118 | ENSG00000244682 | FCGR2C | chr1 | + | 161589561 | 161589819 | 161589564 | 161589819 | 161588421 | 161588442 | 333 | 125 | 51 | 151 | 121 | 29 | 12 | 9 | 17 | 8 | 121 | 86 |

| | | | | | | | | | | | | | | | | | | | | | | | | |
|---|---|---|---|---|---|---|---|---|---|---|---|---|---|---|---|---|---|---|---|---|---|---|---|---|
| A3SS | 8024 | ENSG00000125875 | TBC1D20 | chr20 | - | 440247 | 440389 | 440247 | 440317 | 441587 | 441689 | 158 | 80 | 122 | 197 | 173 | 2 | 0 | 0 | 5 | 6 | 190 | 156 | 75 |
| A3SS | 12737 | ENSG00000130640 | TUBGCP2 | chr10 | - | 133299984 | 133300547 | 133299984 | 133300113 | 133302791 | 133302980 | 31 | 28 | 14 | 23 | 74 | 107 | 40 | 90 | 92 | 101 | 13 | 29 | 37 |
| A3SS | 2718 | ENSG00000203485 | INF2 | chr14 | + | 104713444 | 104713606 | 104713448 | 104713606 | 104713206 | 104713309 | 36 | 27 | 39 | 60 | 6 | 1 | 0 | 1 | 1 | 2 | 19 | 15 | 23 |
| A3SS | 7472 | ENSG00000065485 | PDIA5 | chr3 | + | 123146098 | 123146259 | 123146117 | 123146259 | 123145521 | 123145592 | 17 | 12 | 14 | 13 | 37 | 0 | 1 | 0 | 2 | 0 | 69 | 22 | 34 |
| A3SS | 1690 | ENSG00000125124 | BBS2 | chr16 | - | 56509956 | 56510088 | 56509956 | 56510034 | 56510858 | 56510921 | 3 | 0 | 1 | 2 | 7 | 97 | 78 | 112 | 100 | 123 | 0 | 0 | 2 |
| A3SS | 13372 | ENSG00000275066 | SYNRG | chr17 | - | 37596222 | 37596344 | 37596222 | 37596341 | 37600362 | 37600403 | 103 | 32 | 90 | 94 | 49 | 12 | 10 | 22 | 31 | 0 | 18 | 64 | 34 |
| A3SS | 398 | ENSG00000142546 | NOSIP | chr19 | - | 49556548 | 49556736 | 49556548 | 49556657 | 49556874 | 49556993 | 266 | 83 | 297 | 204 | 147 | 0 | 2 | 4 | 0 | 7 | 67 | 88 | 61 |
| A3SS | 12536 | ENSG00000175197 | DDIT3 | chr12 | - | 57517268 | 57517542 | 57517268 | 57517424 | 57517705 | 57517753 | 28 | 15 | 34 | 32 | 11 | 4 | 0 | 0 | 1 | 1 | 58 | 89 | 90 |
| A3SS | 6101 | ENSG00000186716 | BCR | chr22 | + | 23309942 | 23310433 | 23310323 | 23310433 | 23309423 | 23309483 | 3 | 0 | 7 | 5 | 4 | 117 | 102 | 111 | 170 | 177 | 7 | 0 | 4 |
| A3SS | 7517 | ENSG00000160818 | GPATCH4 | chr1 | - | 156596080 | 156596248 | 156596080 | 156596131 | 156596386 | 156596478 | 19 | 6 | 12 | 7 | 42 | 17 | 5 | 15 | 19 | 16 | 35 | 22 | 13 |
| A3SS | 6418 | ENSG00000028839 | TBPL1 | chr6 | + | 133984375 | 133984479 | 133984448 | 133984479 | 133982816 | 133982880 | 245 | 123 | 164 | 171 | 443 | 3 | 2 | 5 | 2 | 1 | 95 | 205 | 176 |
| A3SS | 3658 | ENSG00000077232 | DNAJC10 | chr2 | + | 182732482 | 182732542 | 182732498 | 182732542 | 182731029 | 182731107 | 10 | 7 | 7 | 4 | 12 | 174 | 100 | 129 | 164 | 220 | 0 | 20 | 0 |
| A3SS | 3535 | ENSG00000131503 | ANKHD1 | chr5 | + | 140438493 | 140438617 | 140438502 | 140438617 | 140436103 | 140436257 | 15 | 16 | 12 | 18 | 12 | 41 | 25 | 27 | 49 | 38 | 9 | 10 | 17 |
| A3SS | 11824 | ENSG00000153827 | TRIP12 | chr2 | - | 229830755 | 229830839 | 229830755 | 229830821 | 229836847 | 229836984 | 151 | 129 | 133 | 164 | 102 | 1 | 3 | 1 | 1 | 2 | 125 | 184 | 108 |
| A3SS | 7830 | ENSG00000131748 | STARD3 | chr17 | + | 39657972 | 39658522 | 39658404 | 39658522 | 39657774 | 39657852 | 104 | 35 | 84 | 91 | 75 | 0 | 1 | 8 | 0 | 0 | 166 | 153 | 104 |
| A3SS | 2111 | ENSG00000167965 | MLST8 | chr16 | + | 2206030 | 2206214 | 2206060 | 2206214 | 2205453 | 2205512 | 2 | 2 | 2 | 5 | 0 | 0 | 1 | 3 | 4 | 1 | 0 | 3 | 3 |
| A3SS | 11823 | ENSG00000153827 | TRIP12 | chr2 | - | 229805729 | 229805883 | 229805729 | 229805802 | 229807707 | 229807864 | 233 | 140 | 173 | 235 | 275 | 36 | 13 | 30 | 27 | 33 | 112 | 171 | 150 |
| A3SS | 3367 | ENSG00000172785 | CBWD1 | chr9 | - | 121960 | 122136 | 121960 | 122090 | 123216 | 123282 | 1 | 9 | 5 | 7 | 2 | 40 | 98 | 42 | 50 | 132 | 11 | 9 | 0 |
| A3SS | 9872 | ENSG00000085514 | PILRA | chr7 | + | 100399784 | 100400096 | 100399822 | 100400096 | 100399580 | 100399612 | 113 | 56 | 63 | 60 | 74 | 4 | 4 | 1 | 0 | 0 | 165 | 88 | 86 |
| A3SS | 10237 | ENSG00000196998 | WDR45 | chrX | - | 49076644 | 49076753 | 49076644 | 49076750 | 49077642 | 49077747 | 12 | 1 | 2 | 4 | 2 | 50 | 15 | 40 | 35 | 49 | 8 | 6 | 5 |
| A3SS | 4985 | ENSG00000140307 | GTF2A2 | chr15 | - | 59650668 | 59650773 | 59650668 | 59650764 | 59652205 | 59652326 | 192 | 126 | 134 | 158 | 139 | 1 | 1 | 0 | 6 | 6 | 121 | 147 | 119 |
| A3SS | 888 | ENSG00000197586 | ENTPD6 | chr20 | + | 25207075 | 25207397 | 25207078 | 25207397 | 25206521 | 25206590 | 33 | 19 | 30 | 40 | 28 | 29 | 12 | 11 | 26 | 20 | 82 | 45 | 11 |
| A3SS | 12510 | ENSG00000095574 | IKZF5 | chr10 | - | 122998492 | 122998671 | 122998492 | 122998658 | 123007025 | 123007176 | 138 | 64 | 82 | 105 | 86 | 14 | 10 | 13 | 15 | 9 | 16 | 94 | 120 |
| A3SS | 8986 | ENSG00000155506 | LARP1 | chr5 | + | 154799872 | 154800042 | 154799899 | 154800042 | 154799590 | 154799759 | 326 | 208 | 411 | 333 | 399 | 2 | 2 | 1 | 4 | 0 | 426 | 500 | 352 |
| A3SS | 9819 | ENSG00000168137 | SETD5 | chr3 | + | 9445044 | 9445300 | 9445047 | 9445300 | 9443307 | 9443417 | 10 | 3 | 9 | 5 | 11 | 132 | 94 | 134 | 139 | 254 | 17 | 7 | 5 |
| A3SS | 13061 | ENSG00000111271 | ACAD10 | chr12 | + | 111749010 | 111749203 | 111749172 | 111749203 | 111748316 | 111748475 | 5 | 2 | 5 | 3 | 4 | 10 | 5 | 24 | 29 | 12 | 0 | 8 | 0 |
| A3SS | 7724 | ENSG00000128815 | WDFY4 | chr10 | + | 48982508 | 48982956 | 48982689 | 48982956 | 48981366 | 48981478 | 141 | 93 | 81 | 131 | 170 | 1 | 0 | 2 | 2 | 0 | 307 | 174 | 73 |
| A3SS | 5185 | ENSG00000162819 | BROX | chr1 | + | 222724091 | 222724164 | 222724099 | 222724164 | 222722418 | 222722514 | 188 | 134 | 113 | 172 | 261 | 1 | 3 | 4 | 0 | 0 | 56 | 131 | 107 |
| A3SS | 9557 | ENSG00000158560 | DYNC1I1 | chr7 | + | 95813195 | 95813337 | 95813246 | 95813337 | 95810391 | 95810506 | 0 | 0 | 0 | 0 | 0 | 1 | 3 | 2 | 9 | 1 | 0 | 0 | 0 |
| A3SS | 6615 | ENSG00000198019 | FCGR1B | chr1 | + | 121088653 | 121088908 | 121088682 | 121088908 | 121087820 | 121087841 | 14 | 11 | 4 | 2 | 19 | 0 | 0 | 0 | 0 | 0 | 89 | 4 | 0 |
| A3SS | 12203 | ENSG00000166483 | WEE1 | chr11 | + | 9586448 | 9586619 | 9586497 | 9586619 | 9585441 | 9585527 | 125 | 73 | 93 | 118 | 243 | 4 | 1 | 0 | 1 | 3 | 192 | 233 | 156 |
| A3SS | 3527 | ENSG00000181929 | PRKAG1 | chr12 | - | 49005742 | 49005881 | 49005742 | 49005852 | 49013061 | 49013110 | 1 | 0 | 3 | 1 | 3 | 38 | 24 | 23 | 24 | 45 | 0 | 0 | 4 |
| A3SS | 11442 | ENSG00000197070 | ARRDC1 | chr9 | + | 137614031 | 137614214 | 137614139 | 137614214 | 137613614 | 137613769 | 147 | 56 | 84 | 104 | 51 | 0 | 0 | 3 | 0 | 0 | 39 | 108 | 41 |
| A3SS | 3356 | ENSG00000082515 | MRPL22 | chr5 | + | 154950802 | 154950938 | 154950820 | 154950938 | 154941216 | 154941265 | 2 | 0 | 1 | 1 | 0 | 67 | 30 | 44 | 56 | 34 | 0 | 0 | 1 |
| A3SS | 6849 | ENSG00000138642 | HERC6 | chr4 | + | 88398107 | 88398209 | 88398141 | 88398209 | 88396850 | 88396987 | 4 | 5 | 3 | 4 | 2 | 112 | 89 | 100 | 113 | 98 | 0 | 9 | 0 |
| A3SS | 275 | ENSG00000105701 | FKBP8 | chr19 | - | 18538215 | 18538436 | 18538215 | 18538433 | 18539370 | 18539459 | 194 | 61 | 131 | 192 | 77 | 69 | 22 | 53 | 49 | 18 | 112 | 209 | 97 |
| A3SS | 2999 | ENSG00000137413 | TAF8 | chr6 | + | 42068464 | 42068607 | 42068503 | 42068607 | 42066311 | 42066459 | 113 | 50 | 87 | 94 | 71 | 0 | 7 | 0 | 0 | 0 | 100 | 101 | 72 |
| A3SS | 7742 | ENSG00000115525 | ST3GAL5 | chr2 | - | 85852859 | 85853099 | 85852859 | 85853000 | 85861180 | 85861292 | 15 | 5 | 12 | 20 | 20 | 9 | 3 | 9 | 7 | 9 | 10 | 42 | 21 |
| A3SS | 7826 | ENSG00000166986 | MARS1 | chr12 | + | 57489423 | 57489558 | 57489454 | 57489558 | 57489266 | 57489345 | 121 | 61 | 111 | 124 | 67 | 5 | 2 | 0 | 1 | 0 | 187 | 119 | 51 |
| A3SS | 1651 | ENSG00000168488 | ATXN2L | chr16 | + | 28832186 | 28832399 | 28832204 | 28832399 | 28830961 | 28831072 | 10 | 2 | 8 | 13 | 8 | 94 | 23 | 100 | 98 | 83 | 0 | 10 | 5 |
| A3SS | 4854 | ENSG00000137074 | APTX | chr9 | - | 32988082 | 32988129 | 32988082 | 32988124 | 32989758 | 32989895 | 15 | 11 | 18 | 16 | 13 | 12 | 11 | 5 | 8 | 11 | 16 | 13 | 29 |
| A3SS | 6961 | ENSG00000124181 | PLCG1 | chr20 | + | 41174123 | 41174311 | 41174126 | 41174311 | 41173922 | 41174011 | 79 | 38 | 100 | 97 | 158 | 24 | 15 | 30 | 43 | 32 | 56 | 70 | 41 |
| A3SS | 5348 | ENSG00000116001 | TIA1 | chr2 | - | 70214348 | 70214494 | 70214348 | 70214491 | 70215370 | 70215494 | 189 | 158 | 142 | 173 | 253 | 25 | 15 | 38 | 35 | 46 | 4 | 117 | 70 |
| A3SS | 8995 | ENSG00000166436 | TRIM66 | chr11 | - | 8621031 | 8621321 | 8621031 | 8621237 | 8621644 | 8621819 | 15 | 19 | 13 | 20 | 21 | 4 | 9 | 9 | 11 | 13 | 11 | 13 | 22 |
| A3SS | 160 | ENSG00000100150 | DEPDC5 | chr22 | + | 31836998 | 31837155 | 31837093 | 31837155 | 31833914 | 31833980 | 35 | 27 | 26 | 43 | 29 | 4 | 1 | 2 | 6 | 0 | 80 | 53 | 28 |
| A3SS | 7171 | ENSG00000147274 | RBMX | chrX | - | 136877914 | 136878086 | 136877914 | 136877964 | 136879016 | 136879123 | 611 | 427 | 672 | 620 | 1007 | 0 | 0 | 1 | 2 | 22 | 669 | 634 | 639 |
| A3SS | 9398 | ENSG00000135317 | SNX14 | chr6 | - | 85572136 | 85572242 | 85572136 | 85572215 | 85572297 | 85572374 | 7 | 2 | 0 | 1 | 5 | 154 | 96 | 116 | 172 | 217 | 0 | 0 | 8 |
| A3SS | 3228 | ENSG00000166888 | STAT6 | chr12 | - | 57106467 | 57106339 | 57105467 | 57105599 | 57106527 | 57106580 | 710 | 411 | 476 | 557 | 507 | 0 | 0 | 0 | 0 | 0 | 1202 | 658 | 395 |
| A3SS | 3163 | ENSG00000198612 | COPS8 | chr2 | + | 237093662 | 237094197 | 237094089 | 237094197 | 237088604 | 237088653 | 19 | 8 | 8 | 9 | 12 | 4 | 2 | 2 | 14 | 4 | 0 | 9 | 2 |
| A3SS | 9972 | ENSG00000091140 | DLD | chr7 | + | 107904862 | 107905058 | 107904957 | 107905058 | 107903477 | 107903547 | 11 | 24 | 26 | 22 | 21 | 94 | 46 | 87 | 97 | 71 | 0 | 20 | 19 |
| A3SS | 7913 | ENSG00000147533 | GOLGA7 | chr8 | + | 41507053 | 41507121 | 41507058 | 41507121 | 41505910 | 41506012 | 45 | 16 | 39 | 55 | 46 | 285 | 147 | 269 | 292 | 444 | 24 | 32 | 41 |
| A3SS | 2748 | ENSG00000129566 | TEP1 | chr14 | - | 20369343 | 20369576 | 20369343 | 20369554 | 20369673 | 20369779 | 100 | 86 | 74 | 95 | 205 | 1 | 1 | 2 | 3 | 0 | 298 | 106 | 79 |
| A3SS | 9752 | ENSG00000106617 | PRKAG2 | chr7 | - | 151595344 | 151595454 | 151595344 | 151595451 | 151632068 | 151632138 | 26 | 20 | 22 | 31 | 9 | 13 | 4 | 8 | 12 | 16 | 3 | 23 | 4 |
| A3SS | 4214 | ENSG00000013441 | CLK1 | chr2 | - | 200856890 | 200857884 | 200856890 | 200856985 | 200857972 | 200858089 | 658 | 451 | 571 | 691 | 876 | 6 | 5 | 4 | 12 | 1 | 120 | 553 | 516 |
| A3SS | 1539 | ENSG00000126246 | IGFLR1 | chr19 | - | 35739709 | 35740088 | 35739709 | 35739769 | 35741023 | 35741223 | 82 | 13 | 31 | 54 | 38 | 7 | 4 | 13 | 7 | 2 | 55 | 46 | 26 |
| A3SS | 10803 | ENSG00000182196 | ARL6IP4 | chr12 | + | 122981570 | 122981879 | 122981594 | 122981879 | 122981128 | 122981266 | 16 | 5 | 10 | 15 | 7 | 79 | 39 | 62 | 64 | 78 | 0 | 15 | 15 |
| A3SS | 1488 | ENSG00000182087 | TMEM259 | chr19 | - | 1011579 | 1011663 | 1011579 | 1011612 | 1011740 | 1011798 | 114 | 28 | 86 | 93 | 83 | 21 | 5 | 13 | 49 | 18 | 50 | 93 | 88 |
| A3SS | 9907 | ENSG00000140750 | ARHGAP17 | chr16 | - | 24968660 | 24968772 | 24968660 | 24968768 | 24970506 | 24970580 | 188 | 132 | 111 | 153 | 143 | 0 | 5 | 5 | 3 | 0 | 269 | 174 | 90 |
| A3SS | 8094 | ENSG00000183495 | EP400 | chr12 | + | 131979693 | 131979793 | 131979696 | 131979793 | 131960584 | 131961954 | 30 | 31 | 43 | 51 | 60 | 45 | 35 | 63 | 42 | 61 | 36 | 72 | 26 |
| A3SS | 3137 | ENSG00000173692 | PSMD1 | chr2 | + | 231163587 | 231163727 | 231163634 | 231163727 | 231161339 | 231161509 | 3 | 2 | 0 | 6 | 0 | 96 | 65 | 99 | 121 | 119 | 0 | 2 | 2 |
| A3SS | 2473 | ENSG00000154237 | LRRK1 | chr15 | + | 101024799 | 101024967 | 101024802 | 101024967 | 101022382 | 101022597 | 0 | 4 | 3 | 5 | 6 | 7 | 14 | 25 | 13 | 5 | 6 | 1 | 3 |
| A3SS | 8891 | ENSG00000119535 | CSF3R | chr1 | - | 36471432 | 36472139 | 36471432 | 36471646 | 36472237 | 36472391 | 121 | 45 | 63 | 76 | 54 | 0 | 0 | 0 | 1 | 0 | 134 | 103 | 141 |
| A3SS | 1819 | ENSG00000101624 | CEP76 | chr18 | - | 12698978 | 12699203 | 12698978 | 12699165 | 12699829 | 12699900 | 18 | 14 | 29 | 21 | 47 | 14 | 5 | 4 | 14 | 16 | 0 | 13 | 6 |
| A3SS | 1490 | ENSG00000185813 | PCYT2 | chr17 | - | 81909321 | 81909321 | 81908875 | 81909037 | 81909513 | 81909602 | 12 | 7 | 8 | 4 | 0 | 8 | 11 | 9 | 13 | 1 | 3 | 16 | 2 |
| A3SS | 2598 | ENSG00000133997 | MED6 | chr14 | - | 70596610 | 70596742 | 70596610 | 70596702 | 70597617 | 70597777 | 1 | 1 | 2 | 8 | 0 | 72 | 69 | 104 | 109 | 142 | 0 | 0 | 0 |
| A3SS | 8899 | ENSG00000144233 | AMMECR1L | chr2 | - | 127870761 | 127870957 | 127870761 | 127870928 | 127871248 | 127871359 | 0 | 0 | 3 | 1 | 0 | 20 | 15 | 10 | 13 | 20 | 0 | 4 | 1 |
| A3SS | 2313 | ENSG00000170776 | AKAP13 | chr15 | + | 85710578 | 85710645 | 85710581 | 85710645 | 85708018 | 85708086 | 702 | 388 | 496 | 573 | 800 | 44 | 27 | 34 | 14 | 48 | 308 | 704 | 465 |
| A3SS | 4916 | ENSG00000117054 | ACADM | chr1 | + | 75742167 | 75745914 | 75745834 | 75745914 | 75739979 | 75740110 | 18 | 19 | 23 | 24 | 22 | 1 | 0 | 0 | 0 | 4 | 13 | 17 | 20 |
| A3SS | 11452 | ENSG00000132740 | IGHMBP2 | chr11 | + | 68929747 | 68930434 | 68930322 | 68930434 | 68929182 | 68929357 | 18 | 16 | 11 | 11 | 18 | 0 | 0 | 0 | 0 | 0 | 56 | 49 | 31 |
| A3SS | 6795 | ENSG00000117139 | KDM5B | chr1 | - | 202727444 | 202729474 | 202727444 | 202729173 | 202729706 | 202730027 | 6 | 3 | 5 | 3 | 3 | 43 | 27 | 26 | 25 | 86 | 2 | 0 | 15 |
| A3SS | 9683 | ENSG00000145191 | EIF2B5 | chr3 | + | 184140056 | 184140157 | 184140079 | 184140157 | 184138165 | 184138246 | 0 | 3 | 4 | 0 | 3 | 106 | 52 | 105 | 107 | 136 | 0 | 0 | 2 |
| A3SS | 6912 | ENSG00000168610 | STAT3 | chr17 | - | 42348388 | 42348520 | 42348388 | 42348517 | 42388278 | 42388442 | 19 | 8 | 17 | 14 | 5 | 28 | 12 | 17 | 29 | 9 | 31 | 23 | 13 |
| A3SS | 8150 | ENSG00000131051 | RBM39 | chr20 | - | 35709223 | 35709309 | 35709223 | 35709274 | 35713018 | 35713096 | 12 | 20 | 17 | 10 | 16 | 320 | 200 | 221 | 296 | 405 | 0 | 8 | 13 |
| A3SS | 4977 | ENSG00000158793 | NIT1 | chr1 | + | 161119492 | 161119612 | 161119508 | 161119612 | 161119133 | 161119388 | 2 | 0 | 2 | 1 | 0 | 30 | 16 | 14 | 19 | 14 | 0 | 0 | 1 |

| | | | | | | | | | | | | | | |
|---|---|---|---|---|---|---|---|---|---|---|---|---|---|---|
| A3SS | 1624 | ENSG00000108559 | NUP88 | chr17 | - | 5386983 | 5387110 | 5386983 | 5387069 | 5387385 | 5387466 | 288 | 175 | 175 |
| A3SS | 7729 | ENSG00000115525 | ST3GAL5 | chr2 | - | 85846376 | 85846563 | 85846376 | 85846424 | 85847860 | 85848204 | 168 | 138 | 183 |
| A3SS | 7527 | ENSG00000237298 | TTN-AS1 | chr2 | + | 178583523 | 178583921 | 178583712 | 178583921 | 178542221 | 178542361 | 9 | 15 | 11 |
| A3SS | 9610 | ENSG00000171777 | RASGRP4 | chr19 | - | 38419859 | 38420013 | 38419859 | 38419971 | 38420130 | 38420262 | 40 | 4 | 32 |
| A3SS | 13103 | ENSG00000248810 | LINC02432 | chr4 | - | 141324801 | 141324912 | 141324801 | 141324909 | 141332158 | 141332617 | 29 | 55 | 14 |
| A3SS | 11316 | ENSG00000056586 | RC3H2 | chr9 | - | 122865348 | 122865657 | 122865348 | 122865541 | 122877470 | 122877583 | 207 | 168 | 198 |
| A3SS | 7694 | ENSG00000112031 | MTRF1L | chr6 | - | 152995135 | 152995319 | 152995135 | 152995294 | 152998549 | 152998629 | 58 | 56 | 45 |
| A3SS | 8610 | ENSG00000115762 | PLEKHB2 | chr2 | + | 131140166 | 131140275 | 131140169 | 131140275 | 131132901 | 131132991 | 245 | 162 | 214 |
| A3SS | 9538 | ENSG00000196923 | PDLIM7 | chr5 | - | 177491806 | 177491995 | 177491806 | 177491925 | 177492404 | 177492435 | 0 | 0 | 0 |
| A3SS | 2250 | ENSG00000128739 | SNRPN | chr15 | + | 24974386 | 24974456 | 24974397 | 24974456 | 24967931 | 24968082 | 41 | 35 | 34 |
| A3SS | 6601 | ENSG00000203705 | TATDN3 | chr1 | + | 212812226 | 212812328 | 212812250 | 212812328 | 212807735 | 212807848 | 8 | 7 | 4 |
| A3SS | 7219 | ENSG00000110200 | ANAPC15 | chr11 | - | 72111156 | 72111286 | 72111156 | 72111281 | 72111411 | 72111496 | 40 | 17 | 29 |
| A3SS | 2186 | ENSG00000179361 | ARID3B | chr15 | + | 74593134 | 74593236 | 74593137 | 74593236 | 74591559 | 74591814 | 5 | 0 | 4 |
| A3SS | 12095 | ENSG00000110274 | CEP164 | chr11 | + | 117394919 | 117395191 | 117395122 | 117395191 | 117394349 | 117394493 | 21 | 15 | 24 |
| A3SS | 10639 | ENSG00000147140 | NONO | chrX | + | 71297835 | 71298508 | 71298468 | 71298508 | 71297376 | 71297461 | 304 | 204 | 353 |
| A3SS | 1865 | ENSG00000197912 | SPG7 | chr16 | + | 89523482 | 89524247 | 89524005 | 89524247 | 89512947 | 89513037 | 20 | 4 | 13 |
| A3SS | 7666 | ENSG00000106992 | AK1 | chr9 | - | 127868320 | 127868512 | 127868320 | 127868475 | 127871822 | 127871939 | 78 | 39 | 55 |
| A3SS | 11755 | ENSG00000197563 | PIGN | chr18 | - | 62148213 | 62148338 | 62148213 | 62148253 | 62157686 | 62157808 | 24 | 24 | 25 |
| A3SS | 11131 | ENSG00000158473 | CD1D | chr1 | + | 158181454 | 158181721 | 158181489 | 158181721 | 158180894 | 158181162 | 30 | 6 | 21 |
| A3SS | 11226 | ENSG00000167325 | RRM1 | chr11 | + | 4121578 | 4121765 | 4121603 | 4121765 | 4119844 | 4119928 | 1 | 5 | 1 |
| A3SS | 9874 | ENSG00000147535 | PLPP5 | chr8 | - | 38268370 | 38268461 | 38268370 | 38268407 | 38268881 | 38268990 | 38 | 17 | 58 |
| A3SS | 2846 | ENSG00000103811 | CTSH | chr15 | - | 78937317 | 78937496 | 78937317 | 78937423 | 78939139 | 78939171 | 1 | 2 | 3 |
| A3SS | 10805 | ENSG00000182196 | ARL6IP4 | chr12 | + | 122982468 | 122982909 | 122982619 | 122982909 | 122981956 | 122982074 | 305 | 214 | 366 |
| A3SS | 2355 | ENSG00000140577 | CRTC3 | chr15 | + | 90641931 | 90645345 | 90641934 | 90645345 | 90641096 | 90641199 | 41 | 16 | 39 |
| A3SS | 12245 | ENSG00000167986 | DDB1 | chr11 | - | 61300808 | 61300936 | 61300808 | 61300932 | 61302256 | 61302359 | 1 | 0 | 0 |
| A3SS | 12378 | ENSG00000133816 | MICAL2 | chr11 | + | 12276258 | 12276715 | 12276619 | 12276715 | 12276079 | 12276166 | 4 | 11 | 1 |
| A3SS | 7229 | ENSG00000082701 | GSK3B | chr3 | - | 119912703 | 119912810 | 119912703 | 119912750 | 119916043 | 119916174 | 307 | 221 | 260 |
| A3SS | 12471 | ENSG00000196531 | NACA | chr12 | - | 56714601 | 56714703 | 56714601 | 56714675 | 56724451 | 56724523 | 122 | 60 | 89 |

| | | | | | | | | | | | | | | | | | | | | | | | | | | | | | | | | | | | | | | | | | | | | | | |
|---|---|---|---|---|---|---|---|---|---|---|---|---|---|---|---|---|---|---|---|---|---|---|---|---|---|---|---|---|---|---|---|---|---|---|---|---|---|---|---|---|---|---|---|---|---|---|
| A3SS | 7769 | ENSG00000196418 | ZNF124 | chr1 | - | 247159686 | 247159867 | 247159686 | 247159813 | 247171847 | 247172005 | 4 | 3 | 6 | 5 | 17 | 6 | 3 | 6 | 13 | 7 | 0 | 2 | 7 | 0 | 17 | 6 | 0 | 5 | 5 | 0 | 203 | 149 | 0.858075 | 1 | 0.329 | 0.423 | 0.423 | 0.22 | 0.641 | 0 | 1 | 0.507 | 0 | 1 | -0.094 |
| A3SS | 11079 | ENSG00000173064 | HECTD4 | chr12 | - | 112252423 | 112252558 | 112252423 | 112252528 | 112254042 | 112254162 | 7 | 1 | 3 | 4 | 4 | 77 | 48 | 51 | 62 | 92 | 0 | 13 | 3 | 0 | 11 | 125 | 87 | 51 | 52 | 121 | 179 | 149 | 0.858151 | 1 | 0.07 | 0.017 | 0.047 | 0.051 | 0.035 | 0 | 0.111 | 0.047 | 0 | 0.07 | -0.002 |
| A3SS | 7212 | ENSG00000164924 | YWHAZ | chr8 | - | 100924134 | 100924322 | 100924134 | 100924298 | 100924915 | 100925039 | 18 | 22 | 10 | 28 | 61 | 3595 | 2890 | 2724 | 3938 | 7072 | 29 | 44 | 43 | 28 | 23 | 3442 | 3280 | 3095 | 3660 | 3088 | 173 | 149 | 0.858377 | 1 | 0.004 | 0.007 | 0.003 | 0.006 | 0.007 | 0.007 | 0.011 | 0.012 | 0.007 | 0.006 | -0.003 |
| A3SS | 8559 | ENSG00000115282 | TTC31 | chr2 | + | 74490024 | 74490473 | 74490243 | 74490473 | 74483321 | 74483563 | 21 | 8 | 15 | 16 | 14 | 2 | 0 | 2 | 1 | 5 | 19 | 15 | 17 | 16 | 36 | 0 | 0 | 1 | 1 | 18 | 298 | 149 | 0.858619 | 1 | 0.84 | 1 | 0.789 | 0.889 | 0.583 | 1 | 1 | 0.895 | 0.889 | 0.5 | -0.037 |
| A3SS | 6269 | ENSG00000154222 | CC2D1B | chr1 | - | 52357780 | 52357916 | 52357780 | 52357898 | 52358330 | 52358461 | 12 | 4 | 16 | 11 | 5 | 32 | 8 | 23 | 15 | 12 | 4 | 0 | 5 | 9 | 4 | 0 | 11 | 9 | 8 | 18 | 167 | 149 | 0.859111 | 1 | 0.251 | 0.308 | 0.383 | 0.396 | 0.271 | 1 | 0 | 0.331 | 0.501 | 0.165 | -0.078 |
| A3SS | 534 | ENSG00000205744 | DENND1C | chr19 | - | 6478936 | 6479071 | 6478936 | 6479056 | 6479668 | 6479718 | 0 | 0 | 0 | 2 | 0 | 37 | 30 | 39 | 38 | 30 | 0 | 0 | 0 | 0 | 7 | 71 | 66 | 36 | 57 | 59 | 164 | 149 | 0.859402 | 1 | 0 | 0 | 0 | 0.046 | 0 | 1 | 0 | 0 | 0 | 0.097 | -0.01 |
| A3SS | 7967 | ENSG00000204642 | HLA-F | chr6 | + | 29725446 | 29725563 | 29725451 | 29725563 | 29725030 | 29725306 | 1494 | 819 | 1010 | 1179 | 905 | 38 | 11 | 9 | 14 | 19 | 1930 | 1558 | 1073 | 1383 | 2419 | 29 | 10 | 28 | 60 | 4 | 154 | 149 | 0.860083 | 1 | 0.974 | 0.986 | 0.991 | 0.988 | 0.979 | 0.985 | 0.993 | 0.974 | 0.957 | 0.998 | 0.002 |
| A3SS | 2621 | ENSG00000197249 | SERPINA1 | chr14 | - | 94388559 | 94388663 | 94388559 | 94388660 | 94388818 | 94389010 | 43 | 22 | 14 | 29 | 16 | 29 | 16 | 8 | 32 | 0 | 21 | 69 | 33 | 41 | 99 | 19 | 48 | 16 | 19 | 46 | 152 | 149 | 0.860207 | 1 | 0.592 | 0.574 | 0.632 | 0.47 | 1 | 0.52 | 0.585 | 0.669 | 0.679 | 0.678 | 0.027 |
| A3SS | 11785 | ENSG00000198561 | CTNND1 | chr11 | + | 57811398 | 57814373 | 57814310 | 57814373 | 57810108 | 57810223 | 90 | 157 | 67 | 112 | 169 | 0 | 2 | 0 | 0 | 0 | 322 | 60 | 71 | 129 | 180 | 0 | 1 | 4 | 2 | 0 | 298 | 149 | 0.860815 | 1 | 1 | 0.975 | 1 | 1 | 1 | 1 | 0.968 | 0.899 | 0.97 | 1 | 0.028 |
| A3SS | 1588 | ENSG00000214719 | NA | chr17 | + | 30610888 | 30610942 | 30610893 | 30610942 | 30608429 | 30608562 | 85 | 49 | 83 | 77 | 61 | 3 | 0 | 0 | 0 | 0 | 88 | 58 | 67 | 61 | 144 | 0 | 5 | 1 | 1 | 0 | 154 | 149 | 0.861066 | 1 | 0.965 | 1 | 1 | 1 | 1 | 1 | 0.918 | 0.985 | 0.983 | 1 | 0.016 |
| A3SS | 11707 | ENSG00000166199 | ALKBH3 | chr11 | + | 43886605 | 43886653 | 43886608 | 43886653 | 43883982 | 43884017 | 92 | 38 | 70 | 64 | 88 | 6 | 1 | 0 | 0 | 6 | 21 | 104 | 65 | 61 | 119 | 11 | 3 | 0 | 0 | 0 | 152 | 149 | 0.861626 | 1 | 0.938 | 0.974 | 1 | 1 | 0.935 | 0.652 | 0.971 | 1 | 1 | 1 | 0.045 |
| A3SS | 3229 | ENSG00000113282 | CLINT1 | chr5 | - | 157789362 | 157789567 | 157789362 | 157789513 | 157790599 | 157790651 | 51 | 50 | 56 | 83 | 124 | 7 | 0 | 4 | 6 | 8 | 13 | 57 | 68 | 36 | 30 | 0 | 1 | 13 | 4 | 0 | 203 | 149 | 0.861706 | 1 | 0.842 | 1 | 0.911 | 0.91 | 0.919 | 1 | 0.977 | 0.793 | 0.869 | 1 | -0.011 |
| A3SS | 6592 | ENSG00000029534 | ANK1 | chr8 | - | 41672353 | 41672912 | 41672353 | 41672426 | 41684543 | 41684690 | 11 | 11 | 5 | 9 | 14 | 0 | 6 | 1 | 1 | 3 | 35 | 31 | 16 | 4 | 306 | 9 | 2 | 4 | 0 | 46 | 298 | 149 | 0.861889 | 1 | 1 | 0.478 | 0.714 | 0.818 | 0.7 | 0.66 | 0.886 | 0.667 | 1 | 0.769 | -0.054 |
| A3SS | 2028 | ENSG00000074755 | ZZEF1 | chr17 | - | 4082436 | 4082507 | 4082436 | 4082504 | 4085669 | 4085803 | 64 | 30 | 64 | 78 | 62 | 1 | 2 | 5 | 1 | 0 | 108 | 82 | 54 | 55 | 133 | 0 | 4 | 0 | 0 | 6 | 152 | 149 | 0.862026 | 1 | 0.984 | 0.936 | 0.926 | 0.987 | 1 | 1 | 0.953 | 1 | 1 | 0.956 | -0.015 |
| A3SS | 3045 | ENSG00000137221 | TJAP1 | chr6 | + | 43498974 | 43499100 | 43498977 | 43499100 | 43497880 | 43497977 | 28 | 9 | 9 | 8 | 13 | 22 | 7 | 16 | 13 | 14 | 1 | 5 | 9 | 30 | 32 | 12 | 15 | 11 | 3 | 30 | 152 | 149 | 0.862121 | 1 | 0.555 | 0.558 | 0.355 | 0.376 | 0.477 | 0.076 | 0.246 | 0.445 | 0.907 | 0.511 | 0.027 |
| A3SS | 10804 | ENSG00000182196 | ARL6IP4 | chr12 | + | 122981570 | 122981879 | 122981594 | 122981879 | 122981128 | 122981299 | 15 | 9 | 6 | 11 | 2 | 42 | 31 | 43 | 55 | 30 | 0 | 15 | 14 | 12 | 13 | 55 | 40 | 22 | 59 | 72 | 173 | 149 | 0.862395 | 1 | 0.235 | 0.2 | 0.107 | 0.147 | 0.054 | 0 | 0.244 | 0.354 | 0.149 | 0.135 | -0.028 |
| A3SS | 1139 | ENSG00000109118 | PHF12 | chr17 | - | 28923908 | 28927063 | 28923908 | 28924302 | 28950064 | 28950246 | 298 | 157 | 224 | 273 | 191 | 0 | 0 | 0 | 0 | 0 | 310 | 290 | 392 | 285 | 409 | 0 | 0 | 0 | 0 | 10 | 298 | 149 | 0.863241 | 1 | 1 | 1 | 1 | 1 | 1 | 1 | 1 | 1 | 1 | 0.953 | 0.009 |
| A3SS | 6285 | ENSG00000134330 | IAH1 | chr2 | + | 9478221 | 9478370 | 9478253 | 9478370 | 9475986 | 9476039 | 83 | 44 | 62 | 58 | 120 | 2 | 0 | 2 | 5 | 0 | 122 | 114 | 52 | 86 | 68 | 0 | 0 | 1 | 0 | 2 | 181 | 149 | 0.863418 | 1 | 0.972 | 1 | 0.962 | 0.905 | 1 | 1 | 1 | 0.977 | 1 | 0.966 | -0.021 |
| A3SS | 5602 | ENSG00000154814 | OXNAD1 | chr3 | + | 16270928 | 16271071 | 16270944 | 16271071 | 16269125 | 16269275 | 4 | 3 | 2 | 5 | 6 | 21 | 6 | 13 | 16 | 16 | 6 | 0 | 1 | 3 | 3 | 14 | 30 | 2 | 4 | 0 | 165 | 149 | 0.863420 | 1 | 0.147 | 0.311 | 0.122 | 0.22 | 0.253 | 0.279 | 0 | 0.311 | 0.404 | 1 | -0.188 |
| A3SS | 10281 | ENSG00000182220 | ATP6AP2 | chrX | + | 40598651 | 40598734 | 40598680 | 40598734 | 40597526 | 40597664 | 4 | 0 | 3 | 3 | 3 | 389 | 268 | 278 | 313 | 550 | 0 | 0 | 6 | 10 | 0 | 163 | 330 | 296 | 294 | 117 | 178 | 149 | 0.864228 | 1 | 0.009 | 0 | 0.009 | 0.008 | 0.005 | 0 | 0 | 0.017 | 0.028 | 0 | -0.003 |
| A3SS | 9335 | ENSG00000155906 | RMND1 | chr6 | - | 151430137 | 151430203 | 151430137 | 151430177 | 151433154 | 151433230 | 2 | 2 | 2 | 0 | 2 | 39 | 37 | 30 | 37 | 43 | 0 | 5 | 2 | 1 | 0 | 27 | 48 | 22 | 54 | 2 | 175 | 149 | 0.864574 | 1 | 0.042 | 0.044 | 0.054 | 0 | 0.038 | 0 | 0.081 | 0.072 | 0.016 | 0 | 0.002 |
| A3SS | 7211 | ENSG00000079246 | XRCC5 | chr2 | + | 216117740 | 216117794 | 216117745 | 216117794 | 216116658 | 216116842 | 5 | 7 | 2 | 4 | 2 | 1265 | 992 | 1216 | 1279 | 1230 | 0 | 4 | 8 | 0 | 10 | 1371 | 1231 | 977 | 1409 | 196 | 154 | 149 | 0.865158 | 1 | 0.004 | 0.007 | 0.002 | 0.003 | 0.002 | 0 | 0.003 | 0.008 | 0 | 0.047 | -0.008 |
| A3SS | 8775 | ENSG00000115053 | NCL | chr2 | - | 231456009 | 231456159 | 231456009 | 231456136 | 231456630 | 231456764 | 3 | 1 | 3 | 2 | 16 | 601 | 357 | 600 | 525 | 550 | 0 | 1 | 2 | 0 | 0 | 315 | 279 | 463 | 425 | 102 | 172 | 149 | 0.865164 | 1 | 0.004 | 0.002 | 0.004 | 0.003 | 0.025 | 0 | 0.003 | 0.004 | 0 | 0 | 0.006 |
| A3SS | 296 | ENSG00000126247 | CAPNS1 | chr19 | + | 36146195 | 36146312 | 36146246 | 36146312 | 36145975 | 36146054 | 1618 | 705 | 919 | 1152 | 1385 | 0 | 1 | 0 | 0 | 19 | 2993 | 1769 | 1027 | 1403 | 4523 | 3 | 0 | 2 | 0 | 16 | 200 | 149 | 0.865252 | 1 | 1 | 0.998 | 1 | 1 | 0.982 | 0.999 | 1 | 0.997 | 1 | 0.995 | -0.002 |
| A3SS | 6094 | ENSG00000138095 | LRPPRC | chr2 | - | 43918024 | 43918151 | 43918024 | 43918133 | 43918255 | 43918398 | 3 | 1 | 1 | 14 | 2 | 493 | 317 | 350 | 357 | 715 | 0 | 0 | 1 | 0 | 0 | 62 | 247 | 296 | 96 | 86 | 167 | 149 | 0.865282 | 1 | 0.005 | 0.003 | 0.003 | 0.034 | 0.002 | 0 | 0 | 0.003 | 0 | 0 | 0.009 |
| A3SS | 3911 | ENSG00000124783 | SSR1 | chr6 | - | 7301309 | 7301572 | 7301309 | 7301368 | 7303549 | 7303637 | 1202 | 762 | 947 | 985 | 1705 | 9 | 6 | 5 | 2 | 13 | 329 | 860 | 887 | 608 | 177 | 0 | 9 | 3 | 1 | 0 | 298 | 149 | 0.867574 | 1 | 0.985 | 0.984 | 0.99 | 0.996 | 0.985 | 1 | 0.979 | 0.993 | 0.997 | 1 | -0.006 |
| A3SS | 11191 | ENSG00000104518 | GSDMD | chr8 | + | 143562661 | 143563062 | 143562669 | 143563062 | 143562447 | 143562521 | 110 | 32 | 93 | 73 | 12 | 1 | 0 | 1 | 0 | 0 | 120 | 124 | 32 | 64 | 118 | 4 | 4 | 0 | 0 | 0 | 157 | 149 | 0.867602 | 1 | 0.991 | 1 | 0.989 | 1 | 1 | 0.966 | 0.967 | 1 | 1 | 1 | 0.009 |
| A3SS | 5127 | ENSG00000168710 | AHCYL1 | chr1 | + | 110012361 | 110012462 | 110012386 | 110012462 | 110011213 | 110011357 | 162 | 73 | 75 | 116 | 71 | 3 | 0 | 0 | 2 | 0 | 231 | 136 | 83 | 140 | 267 | 0 | 12 | 0 | 0 | 0 | 174 | 149 | 0.868073 | 1 | 0.979 | 1 | 1 | 0.98 | 1 | 1 | 0.907 | 1 | 1 | 1 | 0.01 |
| A3SS | 12211 | ENSG00000142102 | PGGHG | chr11 | + | 293593 | 293727 | 293669 | 293727 | 293365 | 293502 | 111 | 52 | 133 | 105 | 48 | 2 | 2 | 5 | 8 | 11 | 16 | 74 | 76 | 61 | 89 | 21 | 5 | 3 | 0 | 0 | 225 | 149 | 0.868674 | 1 | 0.974 | 0.945 | 0.946 | 0.897 | 0.743 | 0.335 | 0.907 | 0.944 | 1 | 1 | 0.064 |
| A3SS | 2534 | ENSG00000092148 | HECTD1 | chr14 | - | 31128611 | 31129448 | 31128611 | 31129307 | 31133237 | 31133414 | 117 | 81 | 86 | 119 | 97 | 61 | 28 | 34 | 50 | 73 | 32 | 80 | 72 | 54 | 2 | 18 | 41 | 26 | 7 | 10 | 290 | 149 | 0.868921 | 1 | 0.496 | 0.598 | 0.565 | 0.55 | 0.406 | 0.477 | 0.501 | 0.587 | 0.799 | 0.093 | 0.032 |
| A3SS | 2282 | ENSG00000140521 | POLG | chr15 | - | 89326911 | 89327079 | 89326911 | 89327063 | 89327166 | 89327349 | 2 | 2 | 3 | 0 | 2 | 49 | 24 | 46 | 52 | 30 | 0 | 3 | 0 | 8 | 0 | 59 | 23 | 25 | 14 | 53 | 165 | 149 | 0.868951 | 1 | 0.036 | 0.07 | 0.056 | 0 | 0.057 | 0 | 0.105 | 0 | 0.34 | 0 | -0.045 |
| A3SS | 1541 | ENSG00000159210 | SNF8 | chr17 | - | 48931642 | 48931717 | 48931642 | 48931714 | 48933204 | 48933346 | 113 | 67 | 63 | 78 | 126 | 0 | 0 | 4 | 2 | 1 | 248 | 76 | 87 | 114 | 196 | 0 | 0 | 1 | 2 | 9 | 152 | 149 | 0.870106 | 1 | 1 | 1 | 0.939 | 0.975 | 0.992 | 1 | 1 | 0.988 | 0.982 | 0.955 | -0.004 |
| A3SS | 7722 | ENSG00000173597 | SULT1B1 | chr4 | - | 69755069 | 69755261 | 69755069 | 69755201 | 69760207 | 69760250 | 66 | 85 | 51 | 60 | 86 | 21 | 30 | 16 | 26 | 52 | 2 | 24 | 29 | 20 | 5 | 15 | 20 | 35 | 0 | 0 | 209 | 149 | 0.870780 | 1 | 0.691 | 0.669 | 0.694 | 0.622 | 0.541 | 0.087 | 0.461 | 0.371 | 1 | 1 | 0.06 |
| A3SS | 3082 | ENSG00000141279 | NPEPPS | chr17 | + | 47619277 | 47619784 | 47619736 | 47619784 | 47619008 | 47619164 | 16 | 36 | 24 | 24 | 17 | 72 | 52 | 90 | 64 | 105 | 34 | 12 | 27 | 15 | 9 | 91 | 52 | 22 | 66 | 68 | 298 | 149 | 0.870988 | 1 | 0.1 | 0.257 | 0.118 | 0.158 | 0.075 | 0.157 | 0.103 | 0.38 | 0.102 | 0.062 | -0.019 |
| A3SS | 3497 | ENSG00000247774 | PCED1B-AS1 | chr12 | - | 47209280 | 47209482 | 47209280 | 47209458 | 47210639 | 47210761 | 29 | 22 | 46 | 43 | 92 | 122 | 65 | 217 | 115 | 275 | 0 | 9 | 18 | 8 | 9 | 14 | 68 | 22 | 7 | 23 | 173 | 149 | 0.871456 | 1 | 0.17 | 0.226 | 0.154 | 0.244 | 0.224 | 0 | 0.102 | 0.413 | 0.496 | 0.252 | -0.049 |
| A3SS | 9084 | ENSG00000114388 | NPRL2 | chr3 | - | 50349385 | 50349494 | 50349385 | 50349490 | 50349664 | 50349828 | 38 | 16 | 42 | 29 | 35 | 8 | 3 | 2 | 6 | 3 | 18 | 27 | 6 | 21 | 51 | 0 | 0 | 1 | 4 | 15 | 153 | 149 | 0.871564 | 1 | 0.822 | 0.839 | 0.953 | 0.825 | 0.919 | 1 | 1 | 0.854 | 0.836 | 0.768 | -0.02 |
| A3SS | 6352 | ENSG00000173039 | RELA | chr11 | - | 65661686 | 65661868 | 65661686 | 65661835 | 65661936 | 65662088 | 2 | 3 | 5 | 1 | 19 | 30 | 9 | 42 | 36 | 31 | 19 | 8 | 4 | 0 | 9 | 62 | 34 | 42 | 54 | 50 | 182 | 149 | 0.871724 | 1 | 0.052 | 0.214 | 0.089 | 0.022 | 0.334 | 0.201 | 0.162 | 0.072 | 0 | 0.128 | 0.03 |
| A3SS | 5486 | ENSG00000160789 | LMNA | chr1 | + | 156135860 | 156136121 | 156135900 | 156136121 | 156135186 | 156135312 | 0 | 0 | 0 | 2 | 0 | 26 | 21 | 21 | 46 | 28 | 0 | 7 | 0 | 2 | 0 | 102 | 60 | 52 | 51 | 437 | 189 | 149 | 0.872253 | 1 | 0 | 0 | 0 | 0.033 | 0 | 0 | 0.084 | 0 | 0.03 | 0 | -0.016 |
| A3SS | 12635 | ENSG00000107862 | GBF1 | chr10 | + | 102359266 | 102359435 | 102359269 | 102359435 | 102358505 | 102358729 | 16 | 6 | 10 | 9 | 12 | 112 | 48 | 103 | 61 | 59 | 14 | 18 | 10 | 16 | 29 | 200 | 72 | 73 | 142 | 164 | 152 | 149 | 0.872929 | 1 | 0.123 | 0.109 | 0.087 | 0.126 | 0.166 | 0.064 | 0.197 | 0.118 | 0.099 | 0.148 | -0.003 |
| A3SS | 12300 | ENSG00000166340 | TPP1 | chr11 | - | 6616974 | 6617428 | 6616974 | 6617153 | 6617625 | 6617776 | 290 | 139 | 180 | 245 | 253 | 3 | 0 | 3 | 0 | 3 | 294 | 309 | 156 | 301 | 915 | 3 | 0 | 1 | 0 | 0 | 298 | 149 | 0.872931 | 1 | 0.98 | 1 | 0.968 | 1 | 0.977 | 0.98 | 1 | 0.987 | 1 | 1 | -0.008 |
| A3SS | 7370 | ENSG00000168216 | LMBRD1 | chr6 | - | 69700764 | 69700869 | 69700764 | 69700802 | 69701442 | 69701545 | 373 | 315 | 288 | 339 | 505 | 16 | 14 | 15 | 32 | 11 | 118 | 201 | 176 | 186 | 75 | 9 | 11 | 14 | 5 | 2 | 216 | 149 | 0.873170 | 1 | 0.941 | 0.939 | 0.93 | 0.88 | 0.969 | 0.9 | 0.926 | 0.897 | 0.962 | 0.963 | 0.002 |
| A3SS | 727 | ENSG00000099817 | POLR2E | chr19 | - | 1089471 | 1089573 | 1089471 | 1089551 | 1089883 | 1089962 | 2 | 3 | 5 | 2 | 0 | 80 | 31 | 32 | 44 | 21 | 0 | 0 | 9 | 4 | 2 | 36 | 80 | 55 | 36 | 152 | 171 | 149 | 0.873203 | 1 | 0.021 | 0.078 | 0.12 | 0.038 | 0 | 0 | 0 | 0.125 | 0.088 | 0.011 | 0.007 |
| A3SS | 1934 | ENSG00000089280 | FUS | chr16 | + | 31183857 | 31184002 | 31183860 | 31184002 | 31182512 | 31182664 | 160 | 87 | 155 | 192 | 56 | 88 | 45 | 72 | 76 | 42 | 153 | 192 | 145 | 176 | 236 | 55 | 119 | 45 | 125 | 125 | 152 | 149 | 0.873877 | 1 | 0.641 | 0.655 | 0.678 | 0.712 | 0.567 | 0.732 | 0.613 | 0.76 | 0.58 | 0.649 | -0.016 |
| A3SS | 3449 | ENSG00000113648 | MACROH2A1 | chr5 | - | 135360496 | 135360780 | 135360496 | 135360607 | 135369405 | 135369603 | 14 | 20 | 18 | 12 | 20 | 336 | 246 | 218 | 246 | 242 | 22 | 36 | 54 | 94 | 0 | 868 | 472 | 451 | 689 | 725 | 298 | 149 | 0.874404 | 1 | 0.02 | 0.039 | 0.04 | 0.024 | 0.04 | 0.013 | 0.037 | 0.056 | 0.064 | 0 | -0.001 |
| A3SS | 3130 | ENSG00000145730 | PAM | chr5 | + | 103028181 | 103028238 | 103028184 | 103028238 | 103019789 | 103019843 | 28 | 14 | 14 | 12 | 22 | 7 | 10 | 6 | 9 | 12 | 30 | 10 | 12 | 22 | 14 | 10 | 10 | 3 | 10 | 8 | 152 | 149 | 0.874600 | 1 | 0.797 | 0.578 | 0.696 | 0.567 | 0.642 | 0.746 | 0.495 | 0.797 | 0.683 | 0.632 | -0.015 |
| A3SS | 12281 | ENSG00000054654 | SYNE2 | chr14 | + | 64221533 | 64221704 | 64221575 | 64221704 | 64220436 | 64220637 | 29 | 7 | 10 | 11 | 6 | 87 | 34 | 46 | 46 | 58 | 0 | 9 | 15 | 3 | 0 | 14 | 30 | 17 | 4 | 25 | 191 | 149 | 0.874766 | 1 | 0.206 | 0.138 | 0.145 | 0.157 | 0.075 | 0 | 0.19 | 0.408 | 0.369 | 0 | -0.049 |
| A3SS | 10915 | ENSG00000177565 | TBL1XR1 | chr3 | - | 177026372 | 177026474 | 177026372 | 177026470 | 177032970 | 177033136 | 325 | 244 | 319 | 312 | 375 | 1 | 3 | 1 | 6 | 3 | 60 | 177 | 188 | 123 | 15 | 6 | 3 | 0 | 0 | 0 | 153 | 149 | 0.874858 | 1 | 0.997 | 0.988 | 0.997 | 0.981 | 0.992 | 0.907 | 0.983 | 1 | 1 | 1 | 0.013 |
| A3SS | 13177 | ENSG00000111596 | CNOT2 | chr12 | + | 70294070 | 70294294 | 70294236 | 70294294 | 70278131 | 70278274 | 12 | 25 | 11 | 18 | 6 | 18 | 9 | 10 | 14 | 26 | 0 | 37 | 14 | 20 | 2 | 6 | 29 | 11 | 10 | 0 | 298 | 149 | 0.875080 | 1 | 0.25 | 0.581 | 0.355 | 0.391 | 0.103 | 0 | 0.389 | 0.389 | 0.5 | 1 | -0.12 |
| A3SS | 1224 | ENSG00000275700 | AATF | chr17 | + | 37056600 | 37056871 | 37056605 | 37056871 | 37031613 | 37031685 | 170 | 109 | 129 | 194 | 408 | 1 | 0 | 2 | 1 | 5 | 267 | 265 | 147 | 174 | 163 | 11 | 3 | 0 | 0 | 0 | 154 | 149 | 0.875171 | 1 | 0.994 | 1 | 0.984 | 0.995 | 0.987 | 0.959 | 0.988 | 1 | 1 | 1 | 0.003 |
| A3SS | 6600 | ENSG00000203705 | TATDN3 | chr1 | + | 212812226 | 212812328 | 212812247 | 212812328 | 212807735 | 212807848 | 4 | 7 | 4 | 6 | 9 | 34 | 14 | 16 | 16 | 41 | 18 | 13 | 2 | 0 | 0 | 24 | 14 | 13 | 23 | 7 | 170 | 149 | 0.875532 | 1 | 0.093 | 0.305 | 0.18 | 0.247 | 0.161 | 0.397 | 0.449 | 0.119 | 0 | 0 | 0.004 |
| A3SS | 10530 | ENSG00000005302 | MSL3 | chrX | + | 11766960 | 11768682 | 11768574 | 11768682 | 11765466 | 11765729 | 78 | 72 | 70 | 50 | 111 | 1 | 2 | 0 | 4 | 0 | 29 | 58 | 82 | 63 | 21 | 0 | 0 | 0 | 0 | 0 | 298 | 149 | 0.876558 | 1 | 0.975 | 0.947 | 1 | 0.862 | 1 | 1 | 1 | 1 | 1 | 1 | -0.043 |
| A3SS | 12308 | ENSG00000130830 | MPP1 | chrX | - | 154786203 | 154786400 | 154786203 | 154786340 | 154789953 | 154790022 | 461 | 436 | 244 | 737 | 613 | 0 | 0 | 0 | 2 | 0 | 1719 | 750 | 572 | 958 | 5914 | 5 | 0 | 12 | 0 | 11 | 209 | 149 | 0.876564 | 1 | 1 | 1 | 1 | 0.996 | 1 | 0.996 | 1 | 0.971 | 1 | 0.997 | 0.006 |
| A3SS | 834 | ENSG00000125991 | ERGIC3 | chr20 | + | 35542785 | 35542941 | 35542821 | 35542941 | 35542512 | 35542600 | 4 | 0 | 1 | 2 | 3 | 282 | 143 | 297 | 249 | 230 | 0 | 3 | 1 | 10 | 0 | 297 | 218 | 210 | 224 | 354 | 185 | 149 | 0.876640 | 1 | 0.011 | 0 | 0.003 | 0.006 | 0.01 | 0 | 0.011 | 0.004 | 0.035 | 0 | -0.004 |
| A3SS | 682 | ENSG00000105397 | TYK2 | chr19 | - | 10379614 | 10379779 | 10379614 | 10379689 | 10380379 | 10380572 | 116 | 66 | 91 | 177 | 65 | 0 | 1 | 0 | 0 | 0 | 258 | 219 | 68 | 237 | 262 | 0 | 5 | 0 | 5 | 0 | 239 | 149 | 0.876825 | 1 | 1 | 0.976 | 1 | 1 | 1 | 1 | 0.965 | 1 | 0.967 | 1 | 0.009 |
| A3SS | 8090 | ENSG00000116809 | ZBTB17 | chr1 | - | 15946153 | 15946316 | 15946153 | 15946294 | 15946934 | 15947123 | 0 | 1 | 1 | 0 | 0 | 42 | 22 | 23 | 29 | 23 | 2 | 0 | 4 | 1 | 0 | 81 | 32 | 38 | 46 | 118 | 171 | 149 | 0.877035 | 1 | 0 | 0.038 | 0.037 | 0 | 0 | 0.021 | 0 | 0.084 | 0.019 | 0 | -0.01 |
| A3SS | 12152 | ENSG00000172663 | TMEM134 | chr11 | - | 67467500 | 67467590 | 67467500 | 67467563 | 67468027 | 67468092 | 9 | 6 | 11 | 8 | 9 | 4 | 0 | 3 | 3 | 3 | 20 | 15 | 6 | 0 | 24 | 0 | 4 | 4 | 1 | 9 | 176 | 149 | 0.877084 | 1 | 0.656 | 1 | 0.756 | 0.693 | 0.717 | 1 | 0.76 | 0.559 | 0 | 0.693 | 0.162 |
| A3SS | 5203 | ENSG00000117569 | PTBP2 | chr1 | + | 96813272 | 96813447 | 96813275 | 96813447 | 96813028 | 96813106 | 38 | 38 | 32 | 39 | 22 | 34 | 39 | 35 | 39 | 41 | 0 | 24 | 24 | 27 | 7 | 8 | 4 | 23 | 16 | 13 | 152 | 149 | 0.877244 | 1 | 0.523 | 0.489 | 0.473 | 0.495 | 0.345 | 0 | 0.855 | 0.506 | 0.623 | 0.345 | -0.001 |
| A3SS | 10399 | ENSG00000269556 | TMEM185A | chrX | - | 149608626 | 149608834 | 149608626 | 149608829 | 149611286 | 149611463 | 25 | 13 | 15 | 27 | 15 | 3 | 0 | 1 | 5 | 0 | 27 | 2 | 8 | 13 | 143 | 0 | 0 | 0 | 0 | 14 | 154 | 149 | 0.877345 | 1 | 0.89 | 1 | 0.936 | 0.839 | 1 | 1 | 1 | 1 | 1 | 0.908 | -0.049 |
| A3SS | 6772 | ENSG00000235162 | C12orf75 | chr12 | + | 105367471 | 105367509 | 105367476 | 105367509 | 105366616 | 105366696 | 503 | 253 | 265 | 402 | 547 | 1 | 0 | 5 | 1 | 5 | 371 | 357 | 248 | 257 | 2193 | 0 | 3 | 12 | 0 | 2 | 154 | 149 | 0.878192 | 1 | 0.998 | 1 | 0.981 | 0.997 | 0.991 | 1 | 0.991 | 0.952 | 1 | 0.999 | 0.005 |
| A3SS | 8213 | ENSG00000172493 | AFF1 | chr4 | + | 87134470 | 87134694 | 87134473 | 87134694 | 87132270 | 87132408 | 135 | 113 | 123 | 108 | 344 | 3 | 3 | 5 | 2 | 14 | 110 | 103 | 123 | 112 | 79 | 0 | 0 | 3 | 12 | 5 | 152 | 149 | 0.878494 | 1 | 0.978 | 0.974 | 0.96 | 0.981 | 0.96 | 1 | 1 | 0.976 | 0.901 | 0.939 | 0.007 |
| A3SS | 10531 | ENSG00000005302 | MSL3 | chrX | + | 11768572 | 11768682 | 11768574 | 11768682 | 11765466 | 11765729 | 58 | 69 | 97 | 69 | 63 | 1 | 2 | 0 | 4 | 0 | 134 | 101 | 49 | 62 | 49 | 0 | 0 | 0 | 0 | 0 | 151 | 149 | 0.879038 | 1 | 0.983 | 0.971 | 1 | 0.945 | 1 | 1 | 1 | 1 | 1 | 1 | -0.02 |
| A3SS | 7172 | ENSG00000147274 | RBMX | chrX | - | 136879016 | 136879158 | 136879016 | 136879123 | 136879318 | 136879453 | 12 | 11 | 9 | 11 | 10 | 363 | 295 | 326 | 403 | 359 | 14 | 4 | 17 | 15 | 4 | 480 | 416 | 377 | 299 | 308 | 184 | 149 | 0.879418 | 1 | 0.026 | 0.029 | 0.022 | 0.022 | 0.022 | 0.023 | 0.008 | 0.035 | 0.039 | 0.01 | 0.001 |
| A3SS | 4847 | ENSG00000137074 | APTX | chr9 | - | 32987543 | 32987846 | 32987543 | 32987731 | 32988082 | 32988124 | 22 | 24 | 19 | 26 | 45 | 14 | 4 | 14 | 3 | 13 | 33 | 45 | 51 | 24 | 6 | 8 | 6 | 3 | 22 | 9 | 264 | 149 | 0.881163 | 1 | 0.47 | 0.772 | 0.434 | 0.83 | 0.661 | 0.7 | 0.809 | 0.906 | 0.381 | 0.273 | 0.02 |
| A3SS | 4828 | ENSG00000215908 | CROCCP2 | chr1 | - | 16619844 | 16620245 | 16619844 | 16620024 | 16625774 | 16626005 | 12 | 1 | 35 | 51 | 10 | 19 | 7 | 4 | 21 | 5 | 0 | 16 | 19 | 21 | 52 | 5 | 31 | 10 | 4 | 4 | 298 | 149 | 0.881406 | 1 | 0.24 | 0.067 | 0.814 | 0.548 | 0.5 | 0 | 0.205 | 0.487 | 0.724 | 0.867 | -0.023 |
| A3SS | 2280 | ENSG00000140521 | POLG | chr15 | - | 89323403 | 89323552 | 89323403 | 89323511 | 89323814 | 89323901 | 10 | 5 | 7 | 12 | 4 | 136 | 96 | 136 | 143 | 88 | 12 | 9 | 0 | 3 | 7 | 135 | 62 | 77 | 76 | 170 | 190 | 149 | 0.881550 | 1 | 0.055 | 0.039 | 0.039 | 0.062 | 0.034 | 0.065 | 0.102 | 0 | 0.03 | 0.031 | 0 |
| A3SS | 1738 | ENSG00000179583 | CIITA | chr16 | + | 10898921 | 10899002 | 10898924 | 10899002 | 10898669 | 10898732 | 138 | 99 | 85 | 120 | 74 | 88 | 54 | 50 | 95 | 76 | 230 | 81 | 21 | 133 | 299 | 128 | 90 | 13 | 87 | 185 | 152 | 149 | 0.881579 | 1 | 0.606 | 0.642 | 0.625 | 0.553 | 0.488 | 0.638 | 0.469 | 0.613 | 0.6 | 0.613 | -0.004 |
| A3SS | 8468 | ENSG00000148248 | SURF4 | chr9 | - | 133367258 | 133367543 | 133367258 | 133367445 | 133375921 | 133376166 | 5 | 2 | 3 | 11 | 2 | 31 | 14 | 26 | 15 | 1 | 14 | 13 | 7 | 0 | 9 | 18 | 54 | 30 | 19 | 67 | 247 | 149 | 0.881716 | 1 | 0.089 | 0.079 | 0.065 | 0.307 | 0.547 | 0.319 | 0.127 | 0.123 | 0 | 0.075 | 0.089 |
| A3SS | 11020 | ENSG00000172269 | DPAGT1 | chr11 | - | 119097463 | 119098043 | 119097463 | 119097551 | 119098402 | 119098487 | 31 | 9 | 17 | 27 | 21 | 3 | 1 | 2 | 2 | 0 | 8 | 22 | 25 | 14 | 36 | 9 | 1 | 2 | 0 | 0 | 298 | 149 | 0.882000 | 1 | 0.838 | 0.818 | 0.81 | 0.871 | 1 | 0.308 | 0.917 | 0.862 | 1 | 1 | 0.05 |
| A3SS | 10544 | ENSG00000156531 | PHF6 | chrX | + | 134413487 | 134413657 | 134413490 | 134413657 | 134393908 | 134393952 | 22 | 20 | 25 | 23 | 30 | 87 | 99 | 85 | 93 | 158 | 0 | 31 | 17 | 5 | 0 | 16 | 42 | 62 | 8 | 17 | 152 | 149 | 0.882051 | 1 | 0.199 | 0.165 | 0.224 | 0.195 | 0.157 | 0 | 0.42 | 0.212 | 0.38 | 0 | -0.014 |
| A3SS | 8664 | ENSG00000110931 | CAMKK2 | chr12 | - | 121269527 | 121269627 | 121269527 | 121269581 | 121270897 | 121270945 | 4 | 0 | 0 | 8 | 10 | 48 | 49 | 42 | 33 | 56 | 31 | 1 | 0 | 13 | 0 | 96 | 51 | 28 | 108 | 186 | 195 | 149 | 0.882460 | 1 | 0.06 | 0 | 0 | 0.156 | 0.12 | 0.198 | 0.015 | 0 | 0.084 | 0 | 0.008 |
| A3SS | 6952 | ENSG00000116688 | MFN2 | chr1 | + | 11981969 | 11982114 | 11981973 | 11982114 | 11980214 | 11980484 | 53 | 29 | 27 | 28 | 40 | 13 | 7 | 7 | 2 | 2 | 39 | 35 | 27 | 49 | 115 | 19 | 12 | 6 | 6 | 0 | 153 | 149 | 0.882512 | 1 | 0.799 | 0.801 | 0.79 | 0.932 | 0.951 | 0.667 | 0.74 | 0.814 | 0.888 | 1 | 0.033 |
| A3SS | 11505 | ENSG00000233901 | LINC01503 | chr9 | + | 129341735 | 129342043 | 129341831 | 129342043 | 129338791 | 129338917 | 16 | 15 | 28 | 5 | 42 | 1 | 3 | 0 | 1 | 0 | 25 | 17 | 6 | 16 | 90 | 0 | 0 | 0 | 4 | 0 | 245 | 149 | 0.882758 | 1 | 0.907 | 0.753 | 1 | 0.753 | 1 | 1 | 1 | 1 | 0.709 | 1 | -0.059 |
| A3SS | 13396 | ENSG00000127022 | CANX | chr5 | + | 179709872 | 179710065 | 179709918 | 179710065 | 179708977 | 179709059 | 2180 | 1412 | 1864 | 1875 | 3833 | 6 | 2 | 11 | 3 | 6 | 1026 | 1346 | 1473 | 1764 | 533 | 5 | 1 | 5 | 2 | 13 | 195 | 149 | 0.882983 | 1 | 0.996 | 0.998 | 0.992 | 0.998 | 0.998 | 0.994 | 0.999 | 0.996 | 0.999 | 0.969 | 0.005 |
| A3SS | 7119 | ENSG00000143398 | PIP5K1A | chr1 | + | 151224244 | 151224279 | 151224247 | 151224279 | 151198552 | 151199081 | 22 | 12 | 30 | 12 | 41 | 52 | 40 | 60 | 38 | 84 | 29 | 4 | 15 | 16 | 28 | 88 | 46 | 30 | 25 | 34 | 152 | 149 | 0.883057 | 1 | 0.293 | 0.227 | 0.329 | 0.236 | 0.324 | 0.244 | 0.079 | 0.329 | 0.386 | 0.447 | -0.015 |
| A3SS | 9463 | ENSG00000167380 | ZNF226 | chr19 | + | 44165587 | 44165807 | 44165756 | 44165807 | 44165100 | 44165142 | 18 | 14 | 21 | 38 | 21 | 28 | 19 | 20 | 19 | 10 | 31 | 11 | 16 | 8 | 13 | 23 | 13 | 8 | 19 | 7 | 298 | 149 | 0.884285 | 1 | 0.243 | 0.269 | 0.344 | 0.5 | 0.512 | 0.403 | 0.297 | 0.5 | 0.174 | 0.481 | 0.003 |
| A3SS | 2604 | ENSG00000105669 | COPE | chr19 | - | 18899872 | 18900006 | 18899872 | 18899947 | 18900380 | 18900449 | 0 | 0 | 2 | 0 | 1 | 125 | 37 | 103 | 94 | 94 | 10 | 0 | 0 | 0 | 0 | 159 | 182 | 96 | 133 | 227 | 208 | 149 | 0.884731 | 1 | 0 | 0 | 0.014 | 0 | 0.008 | 0.043 | 0 | 0 | 0 | 0 | -0.004 |
| A3SS | 9533 | ENSG00000136250 | AOAH | chr7 | - | 36514471 | 36514676 | 36514471 | 36514590 | 36522038 | 36522115 | 36 | 29 | 20 | 30 | 11 | 12 | 7 | 5 | 6 | 6 | 3 | 51 | 35 | 31 | 5 | 17 | 12 | 4 | 4 | 0 | 235 | 149 | 0.885114 | 1 | 0.655 | 0.724 | 0.717 | 0.76 | 0.538 | 0.101 | 0.729 | 0.847 | 0.831 | 1 | -0.023 |
| A3SS | 5950 | ENSG00000095787 | WAC | chr10 | + | 28616172 | 28616362 | 28616175 | 28616362 | 28614566 | 28614685 | 322 | 192 | 320 | 283 | 423 | 114 | 66 | 93 | 132 | 120 | 41 | 248 | 225 | 183 | 31 | 26 | 68 | 120 | 43 | 7 | 152 | 149 | 0.885606 | 1 | 0.735 | 0.74 | 0.771 | 0.678 | 0.776 | 0.607 | 0.781 | 0.648 | 0.807 | 0.813 | 0.009 |
| A3SS | 9682 | ENSG00000145191 | EIF2B5 | chr3 | + | 184138143 | 184138246 | 184138165 | 184138246 | 184137897 | 184138075 | 2 | 0 | 0 | 0 | 0 | 112 | 53 | 128 | 100 | 103 | 0 | 0 | 1 | 8 | 0 | 111 | 75 | 69 | 102 | 123 | 171 | 149 | 0.885746 | 1 | 0.015 | 0 | 0 | 0 | 0 | 0 | 0 | 0.012 | 0.064 | 0 | -0.012 |
| A3SS | 9950 | ENSG00000155096 | AZIN1 | chr8 | - | 102849998 | 102850165 | 102849998 | 102850144 | 102858012 | 102858150 | 7 | 2 | 8 | 4 | 5 | 86 | 62 | 67 | 92 | 77 | 4 | 4 | 7 | 10 | 2 | 53 | 93 | 102 | 118 | 54 | 170 | 149 | 0.885753 | 1 | 0.067 | 0.027 | 0.095 | 0.037 | 0.054 | 0.062 | 0.036 | 0.057 | 0.069 | 0.031 | 0.005 |
| A3SS | 7812 | ENSG00000198520 | ARMH1 | chr1 | + | 44724319 | 44724668 | 44724336 | 44724668 | 44724121 | 44724244 | 34 | 16 | 34 | 27 | 27 | 2 | 1 | 7 | 5 | 3 | 21 | 17 | 10 | 54 | 52 | 0 | 7 | 5 | 10 | 0 | 166 | 149 | 0.886904 | 1 | 0.938 | 0.935 | 0.813 | 0.829 | 0.89 | 1 | 0.686 | 0.642 | 0.829 | 1 | 0.05 |
| A3SS | 11906 | ENSG00000137500 | CCDC90B | chr11 | - | 83274638 | 83274763 | 83274638 | 83274740 | 83278725 | 83278829 | 3 | 0 | 0 | 1 | 8 | 131 | 95 | 88 | 112 | 275 | 0 | 1 | 1 | 3 | 0 | 88 | 107 | 76 | 118 | 72 | 172 | 149 | 0.887816 | 1 | 0.019 | 0 | 0 | 0.008 | 0.025 | 0 | 0.008 | 0.011 | 0.022 | 0 | 0.002 |
| A3SS | 7867 | ENSG00000032444 | PNPLA6 | chr19 | + | 7553871 | 7554015 | 7553874 | 7554015 | 7551361 | 7551437 | 12 | 9 | 7 | 12 | 7 | 26 | 14 | 12 | 8 | 6 | 25 | 14 | 4 | 6 | 20 | 27 | 18 | 3 | 18 | 21 | 152 | 149 | 0.888086 | 1 | 0.311 | 0.387 | 0.364 | 0.595 | 0.534 | 0.476 | 0.433 | 0.567 | 0.246 | 0.483 | -0.003 |
| A3SS | 8737 | ENSG00000021574 | SPAST | chr2 | + | 32147194 | 32147258 | 32147217 | 32147258 | 32144936 | 32145007 | 2 | 0 | 1 | 0 | 6 | 112 | 84 | 73 | 99 | 129 | 0 | 0 | 0 | 0 | 0 | 8 | 104 | 50 | 60 | 6 | 172 | 149 | 0.888437 | 1 | 0.015 | 0 | 0.012 | 0 | 0.039 | 0 | 0 | 0 | 0 | 0 | 0.013 |
| A3SS | 602 | ENSG00000104859 | CLASRP | chr19 | + | 45069943 | 45070104 | 45070021 | 45070104 | 45069201 | 45069248 | 11 | 1 | 5 | 14 | 7 | 103 | 56 | 100 | 79 | 73 | 22 | 3 | 5 | 11 | 26 | 254 | 95 | 92 | 157 | 200 | 227 | 149 | 0.888895 | 1 | 0.066 | 0.012 | 0.032 | 0.104 | 0.059 | 0.054 | 0.02 | 0.034 | 0.044 | 0.079 | 0.008 |
| A3SS | 5799 | ENSG00000108654 | DDX5 | chr17 | - | 64506075 | 64506262 | 64506075 | 64506259 | 64506754 | 64507562 | 178 | 90 | 142 | 171 | 59 | 13 | 7 | 16 | 9 | 21 | 22 | 91 | 88 | 92 | 16 | 18 | 10 | 4 | 5 | 0 | 152 | 149 | 0.889124 | 1 | 0.931 | 0.926 | 0.897 | 0.949 | 0.734 | 0.545 | 0.899 | 0.956 | 0.947 | 1 | 0.018 |
| A3SS | 5210 | ENSG00000177989 | ODF3B | chr22 | - | 50531190 | 50531341 | 50531190 | 50531295 | 50531569 | 50531773 | 13 | 5 | 12 | 5 | 8 | 12 | 3 | 3 | 0 | 0 | 10 | 8 | 2 | 23 | 40 | 0 | 3 | 2 | 11 | 5 | 195 | 149 | 0.889209 | 1 | 0.453 | 0.56 | 0.753 | 1 | 1 | 1 | 0.671 | 0.433 | 0.615 | 0.859 | 0.038 |
| A3SS | 10277 | ENSG00000178761 | FAM219B | chr15 | - | 74902603 | 74902786 | 74902603 | 74902783 | 74904663 | 74904712 | 67 | 46 | 62 | 69 | 81 | 15 | 5 | 16 | 16 | 24 | 36 | 79 | 29 | 29 | 42 | 13 | 13 | 9 | 17 | 1 | 152 | 149 | 0.889737 | 1 | 0.814 | 0.9 | 0.792 | 0.809 | 0.768 | 0.731 | 0.856 | 0.76 | 0.626 | 0.976 | 0.027 |
| A3SS | 6330 | ENSG00000139636 | LMBR1L | chr12 | - | 49106960 | 49107062 | 49106960 | 49107045 | 49110483 | 49110847 | 0 | 0 | 0 | 0 | 1 | 49 | 22 | 59 | 38 | 62 | 7 | 0 | 1 | 0 | 0 | 45 | 75 | 29 | 44 | 73 | 166 | 149 | 0.889872 | 1 | 0 | 0 | 0 | 0 | 0.014 | 0.123 | 0 | 0.03 | 0 | 0 | -0.028 |
| A3SS | 2685 | ENSG00000100911 | PSME2 | chr14 | - | 24144399 | 24145155 | 24144399 | 24144468 | 24145242 | 24145273 | 456 | 353 | 479 | 481 | 668 | 2 | 0 | 2 | 0 | 0 | 1213 | 466 | 279 | 660 | 793 | 5 | 9 | 2 | 0 | 0 | 298 | 149 | 0.889956 | 1 | 0.991 | 1 | 0.992 | 1 | 1 | 0.992 | 0.963 | 0.986 | 1 | 1 | 0.008 |
| A3SS | 3756 | ENSG00000125386 | FAM193A | chr4 | + | 2662837 | 2662991 | 2662843 | 2662991 | 2659811 | 2660054 | 120 | 72 | 90 | 121 | 84 | 0 | 0 | 3 | 0 | 1 | 106 | 125 | 110 | 139 | 303 | 0 | 8 | 0 | 2 | 1 | 155 | 149 | 0.891533 | 1 | 1 | 1 | 0.966 | 1 | 0.988 | 1 | 0.938 | 1 | 0.985 | 0.997 | 0.007 |
| A3SS | 93 | ENSG00000242259 | C22orf39 | chr22 | - | 19440885 | 19444390 | 19440885 | 19441694 | 19447377 | 19447545 | 34 | 18 | 22 | 38 | 50 | 5 | 3 | 3 | 5 | 0 | 62 | 24 | 33 | 42 | 44 | 6 | 2 | 0 | 4 | 7 | 298 | 149 | 0.891745 | 1 | 0.773 | 0.75 | 0.786 | 0.792 | 1 | 0.838 | 0.857 | 1 | 0.84 | 0.759 | -0.039 |
| A3SS | 2740 | ENSG00000185215 | TNFAIP2 | chr14 | + | 103132701 | 103132872 | 103132749 | 103132872 | 103131638 | 103131762 | 0 | 1 | 1 | 2 | 0 | 175 | 83 | 90 | 125 | 62 | 0 | 8 | 1 | 1 | 4 | 251 | 88 | 58 | 146 | 426 | 197 | 149 | 0.892420 | 1 | 0 | 0.009 | 0.008 | 0.012 | 0 | 0 | 0.064 | 0.013 | 0.005 | 0.007 | -0.012 |
| A3SS | 5070 | ENSG00000170581 | STAT2 | chr12 | - | 56346441 | 56346652 | 56346441 | 56346624 | 56346818 | 56346955 | 4 | 0 | 4 | 7 | 5 | 182 | 123 | 195 | 238 | 181 | 10 | 6 | 3 | 11 | 6 | 880 | 281 | 130 | 349 | 391 | 177 | 149 | 0.892917 | 1 | 0.018 | 0 | 0.017 | 0.024 | 0.023 | 0.009 | 0.018 | 0.019 | 0.026 | 0.013 | -0.001 |
| A3SS | 9265 | ENSG00000135486 | HNRNPA1 | chr12 | + | 54283062 | 54283234 | 54283078 | 54283234 | 54282799 | 54282874 | 28 | 18 | 37 | 45 | 24 | 83 | 96 | 96 | 84 | 59 | 31 | 48 | 17 | 58 | 2 | 95 | 33 | 80 | 92 | 63 | 165 | 149 | 0.893045 | 1 | 0.234 | 0.145 | 0.258 | 0.326 | 0.269 | 0.228 | 0.568 | 0.161 | 0.363 | 0.028 | -0.023 |
| A3SS | 7493 | ENSG00000070814 | TCOF1 | chr5 | + | 150392704 | 150392790 | 150392707 | 150392790 | 150391956 | 150392176 | 33 | 16 | 38 | 34 | 37 | 12 | 5 | 8 | 10 | 4 | 23 | 16 | 14 | 23 | 26 | 14 | 0 | 7 | 0 | 14 | 152 | 149 | 0.894077 | 1 | 0.729 | 0.758 | 0.823 | 0.769 | 0.901 | 0.617 | 1 | 0.662 | 1 | 0.645 | 0.011 |
| A3SS | 10598 | ENSG00000215021 | PHB2 | chr12 | - | 6968410 | 6968595 | 6968410 | 6968587 | 6969497 | 6969577 | 121 | 96 | 120 | 99 | 170 | 0 | 0 | 0 | 5 | 0 | 181 | 89 | 80 | 116 | 224 | 6 | 0 | 3 | 1 | 0 | 157 | 149 | 0.894473 | 1 | 1 | 1 | 1 | 0.949 | 1 | 0.966 | 1 | 0.962 | 0.991 | 1 | 0.006 |
| A3SS | 760 | ENSG00000064601 | CTSA | chr20 | + | 45892272 | 45892484 | 45892397 | 45892484 | 45891915 | 45892027 | 231 | 98 | 114 | 184 | 143 | 2 | 1 | 1 | 0 | 2 | 383 | 534 | 149 | 355 | 2266 | 1 | 2 | 5 | 6 | 1 | 274 | 149 | 0.894542 | 1 | 0.984 | 0.982 | 0.984 | 1 | 0.975 | 0.995 | 0.993 | 0.942 | 0.97 | 0.999 | 0.005 |
| A3SS | 5068 | ENSG00000197746 | PSAP | chr10 | - | 71821875 | 71822016 | 71821875 | 71822007 | 71825836 | 71825893 | 54 | 30 | 48 | 49 | 10 | 5535 | 3954 | 3417 | 4706 | 4778 | 42 | 24 | 16 | 18 | 87 | 19901 | 7602 | 4741 | 8865 | 20191 | 158 | 149 | 0.894804 | 1 | 0.009 | 0.007 | 0.013 | 0.01 | 0.002 | 0.002 | 0.003 | 0.003 | 0.002 | 0.004 | 0.005 |
| A3SS | 11254 | ENSG00000080503 | SMARCA2 | chr9 | + | 2182140 | 2182242 | 2182145 | 2182242 | 2181570 | 2181676 | 255 | 259 | 284 | 303 | 396 | 1 | 0 | 2 | 6 | 3 | 340 | 256 | 237 | 201 | 153 | 0 | 4 | 0 | 0 | 9 | 154 | 149 | 0.895494 | 1 | 0.996 | 1 | 0.993 | 0.98 | 0.992 | 1 | 0.984 | 1 | 1 | 0.943 | 0.007 |
| A3SS | 8708 | ENSG00000145349 | CAMK2D | chr4 | - | 113509637 | 113509675 | 113509637 | 113509672 | 113513327 | 113513370 | 98 | 66 | 82 | 96 | 176 | 13 | 10 | 15 | 15 | 21 | 28 | 64 | 98 | 32 | 16 | 12 | 0 | 13 | 16 | 0 | 152 | 149 | 0.896025 | 1 | 0.881 | 0.866 | 0.843 | 0.863 | 0.891 | 0.696 | 1 | 0.881 | 0.662 | 1 | 0.021 |
| A3SS | 3998 | ENSG00000108591 | DRG2 | chr17 | + | 18099632 | 18100435 | 18100362 | 18100435 | 18099016 | 18099077 | 46 | 20 | 46 | 66 | 49 | 3 | 0 | 1 | 1 | 0 | 101 | 114 | 30 | 41 | 78 | 0 | 0 | 0 | 0 | 0 | 298 | 149 | 0.896596 | 1 | 0.885 | 1 | 0.958 | 0.971 | 1 | 1 | 1 | 1 | 1 | 1 | -0.037 |
| A3SS | 8804 | ENSG00000125503 | PPP1R12C | chr19 | - | 55092782 | 55092868 | 55092782 | 55092865 | 55093015 | 55093076 | 25 | 15 | 17 | 23 | 13 | 41 | 15 | 47 | 20 | 11 | 21 | 22 | 23 | 40 | 39 | 31 | 49 | 29 | 24 | 79 | 152 | 149 | 0.896885 | 1 | 0.374 | 0.495 | 0.262 | 0.53 | 0.537 | 0.399 | 0.306 | 0.437 | 0.62 | 0.326 | 0.022 |
| A3SS | 9589 | ENSG00000070669 | ASNS | chr7 | - | 97854579 | 97854754 | 97854579 | 97854680 | 97855352 | 97855459 | 8 | 14 | 11 | 15 | 15 | 47 | 43 | 70 | 75 | 77 | 17 | 5 | 8 | 0 | 2 | 40 | 35 | 38 | 16 | 18 | 223 | 149 | 0.897356 | 1 | 0.102 | 0.179 | 0.095 | 0.118 | 0.115 | 0.221 | 0.087 | 0.123 | 0 | 0.069 | 0.022 |
| A3SS | 4046 | ENSG00000258315 | C17orf49 | chr17 | + | 7016909 | 7017014 | 7016949 | 7017014 | 7016494 | 7016685 | 204 | 124 | 153 | 147 | 240 | 9 | 0 | 2 | 0 | 5 | 293 | 180 | 102 | 168 | 268 | 12 | 0 | 0 | 0 | 5 | 189 | 149 | 0.897771 | 1 | 0.947 | 1 | 0.984 | 1 | 0.974 | 0.951 | 1 | 1 | 1 | 0.977 | -0.005 |
| A3SS | 1762 | ENSG00000103549 | RNF40 | chr16 | + | 30769185 | 30769600 | 30769474 | 30769600 | 30768837 | 30768987 | 225 | 130 | 163 | 183 | 163 | 0 | 0 | 0 | 0 | 0 | 353 | 269 | 189 | 307 | 458 | 0 | 8 | 0 | 2 | 0 | 298 | 149 | 0.898165 | 1 | 1 | 1 | 1 | 1 | 1 | 1 | 0.944 | 1 | 0.987 | 1 | 0.014 |
| A3SS | 6040 | ENSG00000143337 | TOR1AIP1 | chr1 | + | 179889309 | 179889369 | 179889312 | 179889369 | 179884691 | 179884769 | 110 | 0 | 55 | 53 | 0 | 9 | 90 | 34 | 55 | 108 | 76 | 55 | 2 | 86 | 0 | 99 | 40 | 97 | 56 | 17 | 152 | 149 | 0.898274 | 1 | 0.923 | 0 | 0.613 | 0.486 | 0 | 0.429 | 0.574 | 0.02 | 0.601 | 0 | 0.08 |
| A3SS | 6446 | ENSG00000144136 | SLC20A1 | chr2 | + | 112658950 | 112659094 | 112658973 | 112659094 | 112657121 | 112657241 | 34 | 23 | 28 | 34 | 52 | 4 | 2 | 6 | 3 | 21 | 54 | 58 | 56 | 24 | 40 | 16 | 13 | 5 | 7 | 4 | 172 | 149 | 0.898393 | 1 | 0.88 | 0.909 | 0.802 | 0.908 | 0.682 | 0.745 | 0.794 | 0.907 | 0.748 | 0.897 | 0.018 |
| A3SS | 6409 | ENSG00000132781 | MUTYH | chr1 | - | 45333412 | 45333564 | 45333412 | 45333497 | 45334390 | 45334511 | 17 | 19 | 18 | 16 | 9 | 13 | 3 | 4 | 7 | 4 | 24 | 13 | 13 | 22 | 56 | 0 | 6 | 22 | 4 | 28 | 216 | 149 | 0.898501 | 1 | 0.474 | 0.814 | 0.756 | 0.612 | 0.608 | 1 | 0.599 | 0.29 | 0.791 | 0.58 | 0.001 |
| A3SS | 5621 | ENSG00000163874 | ZC3H12A | chr1 | + | 37482349 | 37482540 | 37482433 | 37482540 | 37481600 | 37481835 | 6 | 7 | 2 | 8 | 12 | 57 | 9 | 37 | 80 | 54 | 23 | 10 | 7 | 7 | 41 | 104 | 119 | 92 | 148 | 130 | 233 | 149 | 0.899572 | 1 | 0.063 | 0.332 | 0.033 | 0.06 | 0.124 | 0.124 | 0.051 | 0.046 | 0.029 | 0.168 | 0.039 |
| A3SS | 1822 | ENSG00000159720 | ATP6V0D1 | chr16 | - | 67438970 | 67439147 | 67438970 | 67439095 | 67439273 | 67439351 | 235 | 123 | 119 | 224 | 186 | 5 | 1 | 1 | 0 | 0 | 500 | 329 | 231 | 433 | 748 | 0 | 8 | 4 | 4 | 0 | 201 | 149 | 0.899753 | 1 | 0.972 | 0.989 | 0.989 | 1 | 1 | 1 | 0.968 | 0.977 | 0.988 | 1 | 0.003 |
| A3SS | 1101 | ENSG00000173065 | FAM222B | chr17 | - | 28755979 | 28759876 | 28755979 | 28759764 | 28766585 | 28766707 | 16 | 25 | 24 | 21 | 30 | 2 | 0 | 8 | 3 | 7 | 15 | 24 | 17 | 20 | 61 | 11 | 2 | 0 | 0 | 43 | 261 | 149 | 0.900865 | 1 | 0.82 | 1 | 0.631 | 0.8 | 0.71 | 0.438 | 0.873 | 1 | 1 | 0.447 | 0.041 |
| A3SS | 8099 | ENSG00000059804 | SLC2A3 | chr12 | - | 7932986 | 7933179 | 7932986 | 7933147 | 7933809 | 7933902 | 7 | 1 | 3 | 1 | 11 | 211 | 146 | 188 | 278 | 215 | 0 | 0 | 19 | 17 | 13 | 521 | 462 | 504 | 574 | 497 | 181 | 149 | 0.900891 | 1 | 0.027 | 0.006 | 0.013 | 0.003 | 0.04 | 0 | 0 | 0.03 | 0.024 | 0.021 | 0.003 |
| A3SS | 12078 | ENSG00000172932 | ANKRD13D | chr11 | + | 67290591 | 67291521 | 67291475 | 67291521 | 67290321 | 67290446 | 79 | 64 | 50 | 76 | 36 | 44 | 24 | 69 | 57 | 6 | 109 | 74 | 47 | 69 | 191 | 67 | 59 | 17 | 36 | 81 | 298 | 149 | 0.901125 | 1 | 0.473 | 0.571 | 0.266 | 0.4 | 0.75 | 0.449 | 0.385 | 0.58 | 0.489 | 0.541 | 0.003 |
| A3SS | 12959 | ENSG00000204482 | LST1 | chr6 | + | 31588517 | 31588909 | 31588562 | 31588909 | 31587640 | 31587733 | 262 | 189 | 118 | 223 | 319 | 28 | 36 | 34 | 29 | 49 | 631 | 162 | 159 | 336 | 524 | 64 | 42 | 51 | 41 | 57 | 194 | 149 | 0.902409 | 1 | 0.878 | 0.801 | 0.727 | 0.855 | 0.833 | 0.883 | 0.748 | 0.705 | 0.863 | 0.876 | 0.004 |
| A3SS | 1727 | ENSG00000273611 | ZNHIT3 | chr17 | + | 36493925 | 36494006 | 36493959 | 36494006 | 36492812 | 36492899 | 134 | 83 | 116 | 123 | 184 | 0 | 0 | 0 | 0 | 2 | 192 | 107 | 127 | 96 | 125 | 0 | 2 | 3 | 4 | 0 | 183 | 149 | 0.902529 | 1 | 1 | 1 | 1 | 1 | 0.987 | 1 | 0.978 | 0.972 | 0.951 | 1 | 0.017 |
| A3SS | 7562 | ENSG00000028528 | SNX1 | chr15 | + | 64129872 | 64130029 | 64129915 | 64130029 | 64127730 | 64127806 | 7 | 3 | 3 | 7 | 10 | 350 | 273 | 272 | 385 | 468 | 1 | 5 | 6 | 9 | 0 | 622 | 316 | 251 | 394 | 610 | 192 | 149 | 0.902649 | 1 | 0.015 | 0.008 | 0.008 | 0.014 | 0.016 | 0.001 | 0.012 | 0.018 | 0.017 | 0 | 0.003 |
| A3SS | 11831 | ENSG00000134627 | PIWIL4 | chr11 | + | 94608184 | 94608686 | 94608582 | 94608686 | 94607438 | 94607639 | 6 | 5 | 1 | 5 | 0 | 11 | 0 | 5 | 4 | 7 | 4 | 4 | 3 | 21 | 0 | 16 | 7 | 0 | 19 | 12 | 298 | 149 | 0.902941 | 1 | 0.214 | 1 | 0.091 | 0.385 | 0 | 0.111 | 0.222 | 1 | 0.356 | 0 | 0 |
| A3SS | 11093 | ENSG00000185728 | YTHDF3 | chr8 | + | 63175330 | 63175416 | 63175336 | 63175416 | 63169386 | 63169411 | 137 | 111 | 124 | 118 | 188 | 1 | 0 | 0 | 4 | 5 | 58 | 85 | 130 | 51 | 40 | 0 | 0 | 1 | 1 | 0 | 155 | 149 | 0.904122 | 1 | 0.992 | 1 | 1 | 0.966 | 0.973 | 1 | 1 | 0.992 | 0.98 | 1 | -0.008 |
| A3SS | 9801 | ENSG00000012124 | CD22 | chr19 | + | 35332546 | 35332924 | 35332552 | 35332924 | 35329211 | 35329230 | 6 | 6 | 6 | 21 | 23 | 10 | 2 | 4 | 6 | 5 | 8 | 17 | 0 | 10 | 19 | 0 | 18 | 2 | 4 | 5 | 155 | 149 | 0.904234 | 1 | 0.366 | 0.743 | 0.59 | 0.771 | 0.816 | 1 | 0.476 | 0 | 0.706 | 0.785 | 0.064 |
| A3SS | 5734 | ENSG00000169683 | LRRC45 | chr17 | + | 82029018 | 82029185 | 82029092 | 82029185 | 82028612 | 82028683 | 7 | 2 | 6 | 5 | 2 | 17 | 9 | 9 | 10 | 0 | 10 | 7 | 4 | 0 | 33 | 16 | 14 | 11 | 6 | 31 | 223 | 149 | 0.904425 | 1 | 0.216 | 0.129 | 0.308 | 0.25 | 1 | 0.295 | 0.25 | 0.195 | 0 | 0.416 | 0.149 |
| A3SS | 6457 | ENSG00000066427 | ATXN3 | chr14 | - | 92093745 | 92093850 | 92093745 | 92093831 | 92096092 | 92096137 | 1 | 0 | 0 | 4 | 2 | 76 | 57 | 38 | 74 | 43 | 1 | 0 | 0 | 0 | 0 | 12 | 71 | 31 | 19 | 9 | 168 | 149 | 0.904509 | 1 | 0.012 | 0 | 0 | 0.046 | 0.04 | 0.069 | 0 | 0 | 0 | 0 | 0.006 |
| A3SS | 11251 | ENSG00000143437 | ARNT | chr1 | - | 150829903 | 150829980 | 150829903 | 150829965 | 150831817 | 150831903 | 104 | 69 | 95 | 156 | 129 | 6 | 7 | 17 | 3 | 17 | 72 | 112 | 81 | 109 | 96 | 13 | 15 | 9 | 6 | 0 | 164 | 149 | 0.904961 | 1 | 0.94 | 0.9 | 0.835 | 0.979 | 0.873 | 0.834 | 0.872 | 0.891 | 0.943 | 1 | -0.003 |
| A3SS | 1947 | ENSG00000160014 | CALM3 | chr19 | + | 46608481 | 46608981 | 46608923 | 46608981 | 46608196 | 46608327 | 637 | 465 | 470 | 669 | 933 | 13 | 9 | 8 | 14 | 14 | 1335 | 695 | 480 | 663 | 3656 | 34 | 10 | 16 | 5 | 56 | 298 | 149 | 0.905761 | 1 | 0.961 | 0.963 | 0.967 | 0.96 | 0.971 | 0.952 | 0.972 | 0.938 | 0.985 | 0.97 | 0.001 |
| A3SS | 8337 | ENSG00000189046 | ALKBH2 | chr12 | - | 109092506 | 109092937 | 109092506 | 109092787 | 109093246 | 109093472 | 50 | 47 | 78 | 77 | 62 | 6 | 4 | 6 | 8 | 0 | 52 | 30 | 45 | 26 | 76 | 6 | 2 | 3 | 0 | 9 | 298 | 149 | 0.905831 | 1 | 0.806 | 0.855 | 0.867 | 0.828 | 1 | 0.812 | 0.882 | 0.882 | 1 | 0.809 | -0.006 |
| A3SS | 12927 | ENSG00000091039 | OSBPL8 | chr12 | - | 76450850 | 76450988 | 76450850 | 76450979 | 76487509 | 76487618 | 92 | 55 | 69 | 65 | 100 | 73 | 55 | 62 | 90 | 90 | 21 | 25 | 40 | 14 | 1 | 44 | 19 | 17 | 4 | 7 | 158 | 149 | 0.906330 | 1 | 0.543 | 0.485 | 0.512 | 0.405 | 0.512 | 0.31 | 0.554 | 0.689 | 0.767 | 0.119 | 0.004 |
| A3SS | 4544 | ENSG00000076108 | BAZ2A | chr12 | - | 56606246 | 56606395 | 56606246 | 56606312 | 56606632 | 56606733 | 14 | 28 | 31 | 21 | 28 | 142 | 144 | 143 | 144 | 272 | 125 | 43 | 46 | 57 | 53 | 718 | 338 | 347 | 460 | 335 | 232 | 149 | 0.906406 | 1 | 0.06 | 0.111 | 0.122 | 0.086 | 0.062 | 0.101 | 0.076 | 0.078 | 0.074 | 0.092 | 0.004 |
| A3SS | 13202 | ENSG00000050426 | LETMD1 | chr12 | + | 51053777 | 51053860 | 51053811 | 51053860 | 51049033 | 51049185 | 24 | 10 | 18 | 20 | 37 | 9 | 8 | 14 | 5 | 9 | 17 | 52 | 26 | 27 | 18 | 30 | 26 | 6 | 0 | 12 | 183 | 149 | 0.906494 | 1 | 0.685 | 0.504 | 0.511 | 0.765 | 0.77 | 0.316 | 0.62 | 0.779 | 1 | 0.55 | -0.006 |
| A3SS | 11634 | ENSG00000148356 | LRSAM1 | chr9 | + | 127489363 | 127489518 | 127489443 | 127489518 | 127487675 | 127487763 | 0 | 0 | 0 | 0 | 0 | 31 | 20 | 29 | 18 | 8 | 0 | 0 | 0 | 0 | 8 | 82 | 39 | 26 | 46 | 87 | 229 | 149 | 0.906611 | 1 | 0 | 0 | 0 | 0 | 0 | 0 | 0 | 0 | 0 | 0.056 | -0.011 |
| A3SS | 6067 | ENSG00000111602 | TIMELESS | chr12 | - | 56432368 | 56432524 | 56432368 | 56432521 | 56433025 | 56433127 | 25 | 10 | 27 | 32 | 8 | 2 | 1 | 2 | 2 | 5 | 25 | 20 | 21 | 45 | 43 | 6 | 3 | 5 | 1 | 0 | 152 | 149 | 0.906948 | 1 | 0.925 | 0.907 | 0.93 | 0.94 | 0.611 | 0.803 | 0.867 | 0.805 | 0.978 | 1 | -0.028 |
| A3SS | 2880 | ENSG00000112640 | PPP2R5D | chr6 | + | 43010633 | 43010736 | 43010663 | 43010736 | 43010467 | 43010569 | 0 | 1 | 0 | 0 | 4 | 54 | 33 | 77 | 81 | 93 | 9 | 0 | 0 | 0 | 0 | 112 | 79 | 76 | 93 | 88 | 179 | 149 | 0.907698 | 1 | 0 | 0.025 | 0 | 0 | 0.035 | 0.063 | 0 | 0 | 0 | 0 | -0.001 |
| A3SS | 8185 | ENSG00000084623 | EIF3I | chr1 | + | 32226104 | 32226320 | 32226170 | 32226320 | 32224409 | 32224475 | 7 | 9 | 8 | 10 | 1 | 168 | 96 | 189 | 173 | 102 | 17 | 4 | 3 | 16 | 7 | 189 | 191 | 125 | 124 | 328 | 215 | 149 | 0.908038 | 1 | 0.028 | 0.061 | 0.028 | 0.039 | 0.007 | 0.059 | 0.014 | 0.016 | 0.082 | 0.015 | -0.005 |
| A3SS | 5674 | ENSG00000162739 | SLAMF6 | chr1 | - | 160489087 | 160489170 | 160489087 | 160489167 | 160490197 | 160490236 | 141 | 56 | 84 | 112 | 161 | 115 | 49 | 70 | 72 | 112 | 46 | 144 | 22 | 42 | 61 | 45 | 84 | 38 | 29 | 29 | 152 | 149 | 0.908368 | 1 | 0.546 | 0.528 | 0.541 | 0.604 | 0.585 | 0.501 | 0.627 | 0.362 | 0.587 | 0.673 | 0.011 |
| A3SS | 9998 | ENSG00000160856 | FCRL3 | chr1 | - | 157697139 | 157697424 | 157697139 | 157697329 | 157697658 | 157697919 | 101 | 24 | 54 | 50 | 77 | 4 | 0 | 0 | 5 | 12 | 9 | 107 | 17 | 33 | 48 | 7 | 3 | 4 | 0 | 0 | 244 | 149 | 0.908722 | 1 | 0.939 | 1 | 1 | 0.859 | 0.797 | 0.44 | 0.956 | 0.722 | 1 | 1 | 0.095 |
| A3SS | 8188 | ENSG00000084623 | EIF3I | chr1 | + | 32231114 | 32231602 | 32231259 | 32231602 | 32230926 | 32231019 | 199 | 125 | 252 | 250 | 207 | 0 | 0 | 0 | 0 | 0 | 278 | 252 | 174 | 196 | 429 | 0 | 2 | 0 | 0 | 11 | 294 | 149 | 0.908851 | 1 | 1 | 1 | 1 | 1 | 1 | 1 | 0.985 | 1 | 1 | 0.952 | 0.013 |
| A3SS | 6721 | ENSG00000143851 | PTPN7 | chr1 | - | 202158117 | 202158320 | 202158117 | 202158248 | 202159280 | 202159454 | 9 | 6 | 11 | 19 | 4 | 2 | 1 | 1 | 3 | 3 | 11 | 32 | 25 | 15 | 2 | 0 | 11 | 1 | 0 | 10 | 221 | 149 | 0.910198 | 1 | 0.752 | 0.802 | 0.881 | 0.81 | 0.473 | 1 | 0.662 | 0.944 | 1 | 0.119 | -0.001 |
| A3SS | 9473 | ENSG00000105963 | ADAP1 | chr7 | - | 900532 | 900616 | 900532 | 900613 | 904125 | 904272 | 96 | 39 | 32 | 64 | 56 | 0 | 0 | 0 | 3 | 0 | 199 | 82 | 44 | 139 | 133 | 0 | 0 | 0 | 1 | 8 | 152 | 149 | 0.910526 | 1 | 1 | 1 | 1 | 0.954 | 1 | 1 | 1 | 1 | 0.993 | 0.942 | 0.004 |
| A3SS | 8507 | ENSG00000143390 | RFX5 | chr1 | - | 151345927 | 151345961 | 151345927 | 151345957 | 151346204 | 151346333 | 294 | 192 | 258 | 244 | 385 | 10 | 0 | 0 | 0 | 0 | 363 | 331 | 162 | 327 | 417 | 6 | 10 | 5 | 1 | 0 | 153 | 149 | 0.910678 | 1 | 0.966 | 1 | 1 | 1 | 1 | 0.983 | 0.97 | 0.969 | 0.997 | 1 | 0.009 |
| A3SS | 4086 | ENSG00000108509 | CAMTA2 | chr17 | - | 4982088 | 4982227 | 4982088 | 4982160 | 4982756 | 4982892 | 14 | 18 | 11 | 22 | 27 | 48 | 45 | 35 | 55 | 56 | 26 | 12 | 12 | 40 | 31 | 92 | 46 | 13 | 51 | 134 | 216 | 149 | 0.910995 | 1 | 0.167 | 0.216 | 0.178 | 0.216 | 0.25 | 0.163 | 0.153 | 0.389 | 0.351 | 0.138 | -0.033 |
| A3SS | 13356 | ENSG00000198176 | TFDP1 | chr13 | + | 113637619 | 113637896 | 113637817 | 113637896 | 113636533 | 113636700 | 11 | 5 | 5 | 9 | 18 | 23 | 18 | 44 | 25 | 51 | 9 | 8 | 2 | 16 | 6 | 13 | 12 | 22 | 55 | 35 | 298 | 149 | 0.911589 | 1 | 0.193 | 0.122 | 0.054 | 0.153 | 0.15 | 0.257 | 0.25 | 0.043 | 0.127 | 0.079 | -0.017 |
| A3SS | 13413 | ENSG00000134899 | ERCC5 | chr13 | + | 102873258 | 102873343 | 102873260 | 102873343 | 102872197 | 102872398 | 267 | 226 | 262 | 243 | 301 | 7 | 5 | 6 | 5 | 4 | 11 | 171 | 301 | 77 | 7 | 0 | 3 | 5 | 7 | 0 | 151 | 149 | 0.912039 | 1 | 0.974 | 0.978 | 0.977 | 0.98 | 0.987 | 1 | 0.983 | 0.983 | 0.916 | 1 | 0.003 |
| A3SS | 10132 | ENSG00000122643 | NT5C3A | chr7 | - | 33029628 | 33029714 | 33029628 | 33029709 | 33035933 | 33035988 | 15 | 5 | 12 | 15 | 35 | 2 | 3 | 3 | 9 | 12 | 2 | 8 | 11 | 10 | 3 | 0 | 17 | 0 | 2 | 5 | 154 | 149 | 0.912479 | 1 | 0.879 | 0.617 | 0.795 | 0.617 | 0.738 | 1 | 0.313 | 1 | 0.829 | 0.367 | 0.027 |
| A3SS | 8220 | ENSG00000143373 | ZNF687 | chr1 | + | 151290370 | 151290573 | 151290431 | 151290573 | 151290121 | 151290234 | 0 | 2 | 1 | 0 | 0 | 18 | 11 | 17 | 15 | 13 | 0 | 1 | 1 | 0 | 5 | 23 | 29 | 25 | 29 | 21 | 210 | 149 | 0.913685 | 1 | 0 | 0.114 | 0.04 | 0 | 0 | 0 | 0.024 | 0.028 | 0 | 0.145 | -0.009 |
| A3SS | 10318 | ENSG00000068323 | TFE3 | chrX | - | 49039106 | 49039410 | 49039106 | 49039345 | 49040454 | 49040568 | 30 | 10 | 12 | 22 | 18 | 9 | 1 | 2 | 8 | 2 | 73 | 23 | 40 | 36 | 54 | 18 | 1 | 5 | 18 | 8 | 214 | 149 | 0.913913 | 1 | 0.699 | 0.874 | 0.807 | 0.657 | 0.862 | 0.738 | 0.941 | 0.848 | 0.582 | 0.825 | -0.007 |
| A3SS | 10401 | ENSG00000157881 | PANK4 | chr1 | - | 2510677 | 2510782 | 2510677 | 2510748 | 2511337 | 2511387 | 22 | 5 | 25 | 7 | 11 | 8 | 6 | 22 | 22 | 14 | 16 | 5 | 9 | 5 | 22 | 11 | 5 | 21 | 3 | 19 | 183 | 149 | 0.914454 | 1 | 0.691 | 0.404 | 0.481 | 0.206 | 0.39 | 0.542 | 0.449 | 0.259 | 0.576 | 0.485 | -0.028 |
| A3SS | 7282 | ENSG00000141458 | NPC1 | chr18 | - | 23556242 | 23556664 | 23556242 | 23556613 | 23557116 | 23557190 | 21 | 5 | 0 | 9 | 8 | 72 | 23 | 32 | 36 | 16 | 5 | 11 | 8 | 5 | 5 | 28 | 60 | 48 | 25 | 3 | 200 | 149 | 0.914585 | 1 | 0.179 | 0.139 | 0 | 0.157 | 0.271 | 0.117 | 0.12 | 0.11 | 0.13 | 0.554 | -0.057 |
| A3SS | 10762 | ENSG00000143569 | UBAP2L | chr1 | + | 154268756 | 154268954 | 154268759 | 154268954 | 154266500 | 154266568 | 66 | 32 | 63 | 69 | 57 | 27 | 9 | 30 | 21 | 36 | 114 | 34 | 48 | 66 | 128 | 53 | 26 | 2 | 42 | 54 | 152 | 149 | 0.914784 | 1 | 0.706 | 0.777 | 0.673 | 0.763 | 0.608 | 0.678 | 0.562 | 0.959 | 0.606 | 0.699 | 0.005 |
| A3SS | 1270 | ENSG00000125450 | NUP85 | chr17 | + | 75226050 | 75226157 | 75226055 | 75226157 | 75225697 | 75225829 | 34 | 9 | 24 | 32 | 35 | 0 | 4 | 1 | 3 | 0 | 29 | 48 | 20 | 32 | 45 | 0 | 1 | 1 | 6 | 0 | 154 | 149 | 0.915350 | 1 | 1 | 0.685 | 0.959 | 0.912 | 1 | 1 | 0.979 | 0.951 | 0.838 | 1 | -0.042 |
| A3SS | 4078 | ENSG00000198931 | APRT | chr16 | - | 88809468 | 88809840 | 88809468 | 88809706 | 88810069 | 88810148 | 252 | 73 | 230 | 230 | 104 | 29 | 22 | 37 | 25 | 35 | 132 | 160 | 121 | 99 | 308 | 0 | 59 | 17 | 33 | 75 | 283 | 149 | 0.917129 | 1 | 0.821 | 0.636 | 0.766 | 0.829 | 0.61 | 1 | 0.588 | 0.789 | 0.612 | 0.684 | -0.002 |
| A3SS | 9570 | ENSG00000072609 | CHFR | chr12 | - | 132869618 | 132869798 | 132869618 | 132869675 | 132870723 | 132870783 | 154 | 130 | 154 | 172 | 231 | 3 | 8 | 12 | 4 | 9 | 208 | 218 | 96 | 140 | 231 | 11 | 9 | 2 | 6 | 9 | 272 | 149 | 0.917228 | 1 | 0.966 | 0.899 | 0.875 | 0.959 | 0.934 | 0.912 | 0.93 | 0.963 | 0.927 | 0.934 | -0.007 |
| A3SS | 8714 | ENSG00000130038 | CRACR2A | chr12 | - | 3638123 | 3638457 | 3638123 | 3638454 | 3641731 | 3641838 | 37 | 19 | 63 | 43 | 49 | 16 | 9 | 7 | 2 | 7 | 66 | 33 | 30 | 31 | 10 | 11 | 4 | 6 | 11 | 1 | 152 | 149 | 0.917509 | 1 | 0.694 | 0.674 | 0.898 | 0.955 | 0.873 | 0.855 | 0.89 | 0.831 | 0.734 | 0.907 | -0.025 |
| A3SS | 1118 | ENSG00000161939 | RNASEK-C17orf49 | chr17 | + | 7017256 | 7017520 | 7017260 | 7017520 | 7016494 | 7016685 | 25 | 4 | 11 | 14 | 48 | 9 | 0 | 2 | 1 | 11 | 23 | 7 | 13 | 5 | 12 | 1 | 13 | 3 | 1 | 0 | 153 | 149 | 0.918135 | 1 | 0.73 | 1 | 0.843 | 0.932 | 0.81 | 0.957 | 0.344 | 0.808 | 0.83 | 1 | 0.075 |
| A3SS | 4822 | ENSG00000197386 | HTT | chr4 | + | 3207280 | 3207357 | 3207285 | 3207357 | 3206806 | 3206983 | 107 | 48 | 74 | 122 | 112 | 10 | 2 | 5 | 6 | 9 | 143 | 123 | 83 | 166 | 317 | 17 | 1 | 12 | 11 | 8 | 154 | 149 | 0.918661 | 1 | 0.912 | 0.959 | 0.935 | 0.952 | 0.923 | 0.891 | 0.992 | 0.87 | 0.936 | 0.975 | 0.003 |
| A3SS | 2242 | ENSG00000168970 | JMJD7-PLA2G4B | chr15 | + | 41840024 | 41840230 | 41840157 | 41840230 | 41836474 | 41836551 | 48 | 13 | 64 | 65 | 28 | 1 | 4 | 1 | 7 | 4 | 71 | 39 | 18 | 57 | 99 | 12 | 4 | 0 | 3 | 4 | 282 | 149 | 0.919215 | 1 | 0.962 | 0.632 | 0.971 | 0.831 | 0.787 | 0.758 | 0.837 | 1 | 0.909 | 0.929 | -0.05 |
| A3SS | 6410 | ENSG00000132781 | MUTYH | chr1 | - | 45333412 | 45333594 | 45333412 | 45333497 | 45334390 | 45334511 | 21 | 22 | 25 | 19 | 12 | 13 | 3 | 4 | 7 | 4 | 14 | 21 | 13 | 38 | 49 | 0 | 6 | 22 | 4 | 28 | 246 | 149 | 0.920521 | 1 | 0.495 | 0.816 | 0.791 | 0.622 | 0.645 | 1 | 0.679 | 0.264 | 0.852 | 0.515 | 0.012 |
| A3SS | 12987 | ENSG00000198563 | DDX39B | chr6 | - | 31530778 | 31530949 | 31530778 | 31530926 | 31531052 | 31531197 | 4 | 4 | 3 | 8 | 0 | 831 | 570 | 831 | 773 | 942 | 17 | 8 | 10 | 15 | 17 | 1342 | 820 | 714 | 979 | 1407 | 172 | 149 | 0.921241 | 1 | 0.004 | 0.006 | 0.003 | 0.009 | 0 | 0.011 | 0.008 | 0.012 | 0.013 | 0.01 | -0.006 |
| A3SS | 479 | ENSG00000076662 | ICAM3 | chr19 | - | 10338681 | 10338948 | 10338681 | 10338863 | 10339538 | 10339624 | 364 | 128 | 213 | 219 | 134 | 2 | 0 | 0 | 3 | 1 | 186 | 361 | 204 | 221 | 428 | 0 | 0 | 2 | 8 | 0 | 234 | 149 | 0.921363 | 1 | 0.991 | 1 | 1 | 0.979 | 0.988 | 1 | 1 | 0.985 | 0.946 | 1 | 0.005 |
| A3SS | 9881 | ENSG00000101082 | SLA2 | chr20 | - | 36614304 | 36614437 | 36614304 | 36614387 | 36615224 | 36615374 | 32 | 25 | 28 | 36 | 20 | 8 | 4 | 5 | 11 | 5 | 22 | 80 | 50 | 34 | 200 | 11 | 15 | 9 | 7 | 38 | 199 | 149 | 0.921669 | 1 | 0.75 | 0.824 | 0.807 | 0.71 | 0.75 | 0.6 | 0.8 | 0.806 | 0.784 | 0.798 | 0.011 |
| A3SS | 683 | ENSG00000105401 | CDC37 | chr19 | - | 10395434 | 10395696 | 10395434 | 10395543 | 10395927 | 10396203 | 12 | 8 | 5 | 4 | 11 | 298 | 223 | 262 | 335 | 191 | 0 | 5 | 6 | 34 | 20 | 536 | 263 | 255 | 388 | 741 | 298 | 149 | 0.921982 | 1 | 0.02 | 0.018 | 0.009 | 0.006 | 0.028 | 0 | 0.009 | 0.012 | 0.042 | 0.013 | 0.001 |
| A3SS | 3969 | ENSG00000154144 | TBRG1 | chr11 | + | 124626812 | 124627050 | 124626903 | 124627050 | 124625670 | 124625903 | 62 | 69 | 57 | 89 | 36 | 20 | 17 | 21 | 19 | 67 | 40 | 56 | 88 | 20 | 40 | 18 | 12 | 25 | 19 | 25 | 240 | 149 | 0.922922 | 1 | 0.658 | 0.716 | 0.628 | 0.744 | 0.25 | 0.58 | 0.743 | 0.686 | 0.395 | 0.498 | 0.019 |
| A3SS | 726 | ENSG00000125912 | NCLN | chr19 | + | 3206261 | 3206425 | 3206264 | 3206425 | 3206151 | 3206190 | 25 | 24 | 41 | 44 | 13 | 49 | 10 | 23 | 20 | 32 | 71 | 42 | 12 | 43 | 101 | 49 | 42 | 30 | 23 | 62 | 152 | 149 | 0.924829 | 1 | 0.333 | 0.702 | 0.636 | 0.683 | 0.285 | 0.587 | 0.495 | 0.282 | 0.647 | 0.615 | 0.003 |
| A3SS | 10650 | ENSG00000163682 | RPL9 | chr4 | - | 39454863 | 39454944 | 39454863 | 39454929 | 39456405 | 39456538 | 3827 | 2173 | 1998 | 1609 | 4426 | 954 | 734 | 1209 | 1001 | 3357 | 1028 | 808 | 1573 | 1413 | 2414 | 892 | 634 | 563 | 422 | 881 | 164 | 149 | 0.925051 | 1 | 0.785 | 0.729 | 0.6 | 0.594 | 0.545 | 0.511 | 0.537 | 0.717 | 0.753 | 0.713 | 0.004 |
| A3SS | 2538 | ENSG00000126822 | PLEKHG3 | chr14 | + | 64731360 | 64731806 | 64731713 | 64731806 | 64731037 | 64731169 | 194 | 60 | 89 | 138 | 115 | 2 | 1 | 1 | 3 | 4 | 234 | 44 | 51 | 109 | 103 | 9 | 0 | 1 | 2 | 0 | 298 | 149 | 0.925235 | 1 | 0.98 | 0.968 | 0.978 | 0.958 | 0.935 | 0.929 | 1 | 0.962 | 0.965 | 1 | -0.007 |
| A3SS | 734 | ENSG00000087460 | GNAS | chr20 | + | 58903527 | 58903585 | 58903530 | 58903585 | 58898940 | 58898985 | 29 | 20 | 32 | 40 | 22 | 244 | 168 | 192 | 262 | 176 | 57 | 23 | 28 | 48 | 196 | 347 | 317 | 213 | 238 | 1594 | 152 | 149 | 0.926324 | 1 | 0.104 | 0.105 | 0.14 | 0.13 | 0.109 | 0.139 | 0.066 | 0.114 | 0.165 | 0.108 | -0.001 |
| A3SS | 7906 | ENSG00000166801 | FAM111A | chr11 | + | 59148796 | 59148953 | 59148835 | 59148953 | 59143082 | 59143304 | 29 | 25 | 21 | 24 | 82 | 22 | 21 | 25 | 20 | 22 | 0 | 16 | 24 | 36 | 3 | 14 | 11 | 10 | 12 | 0 | 188 | 149 | 0.927229 | 1 | 0.511 | 0.485 | 0.4 | 0.487 | 0.747 | 0 | 0.535 | 0.655 | 0.704 | 1 | -0.053 |
| A3SS | 12068 | ENSG00000087365 | SF3B2 | chr11 | + | 66067945 | 66068045 | 66067948 | 66068045 | 66063627 | 66063729 | 203 | 203 | 234 | 264 | 345 | 1 | 1 | 3 | 2 | 6 | 1026 | 320 | 285 | 476 | 827 | 0 | 0 | 1 | 3 | 17 | 152 | 149 | 0.927448 | 1 | 0.995 | 0.995 | 0.987 | 0.992 | 0.983 | 1 | 1 | 0.996 | 0.994 | 0.979 | -0.003 |
| A3SS | 12608 | ENSG00000196233 | LCOR | chr10 | + | 96944101 | 96944245 | 96944112 | 96944245 | 96907667 | 96907747 | 0 | 3 | 2 | 5 | 6 | 178 | 147 | 145 | 189 | 470 | 0 | 6 | 1 | 0 | 8 | 190 | 192 | 178 | 144 | 128 | 160 | 149 | 0.927568 | 1 | 0 | 0.019 | 0.013 | 0.024 | 0.012 | 0 | 0.028 | 0.005 | 0 | 0.055 | -0.004 |
| A3SS | 10609 | ENSG00000185485 | SDHAP1 | chr3 | - | 195965428 | 195965577 | 195965428 | 195965542 | 195967513 | 195967644 | 1 | 2 | 3 | 0 | 7 | 29 | 43 | 27 | 25 | 23 | 0 | 2 | 4 | 3 | 2 | 24 | 32 | 34 | 7 | 17 | 184 | 149 | 0.928161 | 1 | 0.027 | 0.036 | 0.083 | 0 | 0.198 | 0 | 0.048 | 0.087 | 0.258 | 0.087 | -0.027 |
| A3SS | 5978 | ENSG00000163867 | ZMYM6 | chr1 | - | 35030546 | 35030756 | 35030546 | 35030713 | 35031266 | 35031337 | 5 | 10 | 3 | 5 | 1 | 4 | 4 | 5 | 3 | 4 | 5 | 5 | 8 | 2 | 0 | 0 | 9 | 2 | 2 | 10 | 192 | 149 | 0.928301 | 1 | 0.492 | 0.66 | 0.318 | 0.564 | 0.162 | 1 | 0.301 | 0.756 | 0.437 | 0 | -0.06 |
| A3SS | 2089 | ENSG00000102858 | MGRN1 | chr16 | + | 4665098 | 4665151 | 4665101 | 4665151 | 4664708 | 4664775 | 19 | 7 | 19 | 11 | 12 | 113 | 60 | 103 | 118 | 174 | 14 | 19 | 17 | 28 | 23 | 102 | 209 | 95 | 212 | 263 | 152 | 149 | 0.928499 | 1 | 0.142 | 0.103 | 0.153 | 0.084 | 0.063 | 0.119 | 0.082 | 0.149 | 0.115 | 0.079 | 0 |
| A3SS | 1787 | ENSG00000166548 | TK2 | chr16 | - | 66513730 | 66513862 | 66513730 | 66513811 | 66517135 | 66517215 | 0 | 0 | 0 | 0 | 3 | 120 | 82 | 99 | 118 | 175 | 0 | 8 | 2 | 4 | 0 | 139 | 109 | 63 | 89 | 210 | 200 | 149 | 0.928579 | 1 | 0 | 0 | 0 | 0 | 0.013 | 0 | 0.052 | 0.023 | 0.032 | 0 | -0.019 |
| A3SS | 11791 | ENSG00000266173 | STRADA | chr17 | - | 63704483 | 63704908 | 63704483 | 63704582 | 63706634 | 63706739 | 4 | 6 | 3 | 0 | 1 | 157 | 81 | 143 | 145 | 270 | 0 | 1 | 2 | 3 | 10 | 181 | 109 | 68 | 119 | 137 | 298 | 149 | 0.930099 | 1 | 0.013 | 0.036 | 0.01 | 0 | 0.002 | 0 | 0.005 | 0.014 | 0.012 | 0.035 | -0.001 |
| A3SS | 1468 | ENSG00000062716 | VMP1 | chr17 | + | 59764970 | 59765138 | 59765051 | 59765138 | 59738836 | 59738947 | 755 | 462 | 477 | 555 | 1251 | 4 | 11 | 3 | 5 | 4 | 645 | 522 | 590 | 591 | 291 | 3 | 11 | 2 | 4 | 0 | 230 | 149 | 0.931685 | 1 | 0.992 | 0.965 | 0.99 | 0.986 | 0.995 | 0.993 | 0.968 | 0.995 | 0.99 | 1 | -0.004 |
| A3SS | 170 | ENSG00000100242 | SUN2 | chr22 | - | 38739341 | 38739426 | 38739341 | 38739423 | 38739721 | 38739943 | 960 | 314 | 748 | 688 | 610 | 12 | 3 | 11 | 5 | 7 | 400 | 578 | 353 | 288 | 850 | 4 | 0 | 2 | 0 | 8 | 152 | 149 | 0.932548 | 1 | 0.987 | 0.99 | 0.985 | 0.993 | 0.988 | 0.99 | 1 | 0.994 | 1 | 0.99 | -0.006 |
| A3SS | 7662 | ENSG00000196642 | RABL6 | chr9 | + | 136832261 | 136832370 | 136832264 | 136832370 | 136831720 | 136831861 | 5 | 0 | 2 | 6 | 0 | 68 | 23 | 46 | 56 | 51 | 0 | 7 | 3 | 3 | 0 | 48 | 85 | 18 | 40 | 67 | 152 | 149 | 0.933050 | 1 | 0.067 | 0 | 0.041 | 0.095 | 0 | 0 | 0.075 | 0.14 | 0.068 | 0 | -0.016 |
| A3SS | 549 | ENSG00000186818 | LILRB4 | chr19 | + | 54667634 | 54667793 | 54667637 | 54667793 | 54666696 | 54666749 | 22 | 34 | 7 | 19 | 11 | 21 | 13 | 9 | 7 | 15 | 49 | 87 | 34 | 15 | 63 | 59 | 41 | 18 | 22 | 36 | 152 | 149 | 0.933242 | 1 | 0.507 | 0.719 | 0.433 | 0.727 | 0.418 | 0.449 | 0.675 | 0.649 | 0.401 | 0.632 | 0 |
| A3SS | 13028 | ENSG00000166012 | TAF1D | chr11 | - | 93733349 | 93733505 | 93733349 | 93733419 | 93734624 | 93734660 | 38 | 20 | 24 | 21 | 28 | 7 | 14 | 13 | 16 | 8 | 13 | 15 | 20 | 10 | 17 | 7 | 8 | 5 | 0 | 26 | 235 | 149 | 0.934344 | 1 | 0.775 | 0.475 | 0.539 | 0.454 | 0.689 | 0.541 | 0.543 | 0.717 | 1 | 0.293 | -0.032 |
| A3SS | 11294 | ENSG00000105879 | CBLL1 | chr7 | + | 107753410 | 107753511 | 107753413 | 107753511 | 107748879 | 107749047 | 148 | 112 | 132 | 197 | 158 | 17 | 26 | 26 | 26 | 55 | 154 | 200 | 190 | 133 | 133 | 60 | 22 | 15 | 39 | 27 | 152 | 149 | 0.935456 | 1 | 0.895 | 0.809 | 0.833 | 0.881 | 0.738 | 0.716 | 0.899 | 0.925 | 0.77 | 0.828 | 0.004 |
| A3SS | 9687 | ENSG00000145191 | EIF2B5 | chr3 | + | 184142236 | 184142378 | 184142297 | 184142378 | 184141924 | 184142070 | 110 | 81 | 122 | 131 | 84 | 1 | 0 | 3 | 0 | 0 | 294 | 215 | 143 | 212 | 189 | 0 | 6 | 0 | 0 | 4 | 210 | 149 | 0.936994 | 1 | 0.987 | 1 | 0.967 | 1 | 1 | 1 | 0.962 | 1 | 1 | 0.971 | 0.004 |
| A3SS | 6411 | ENSG00000132781 | MUTYH | chr1 | - | 45333412 | 45333603 | 45333412 | 45333497 | 45334390 | 45334511 | 20 | 18 | 25 | 17 | 12 | 13 | 3 | 4 | 7 | 4 | 14 | 23 | 13 | 36 | 45 | 0 | 6 | 22 | 4 | 28 | 255 | 149 | 0.937706 | 1 | 0.473 | 0.778 | 0.785 | 0.587 | 0.637 | 1 | 0.691 | 0.257 | 0.84 | 0.484 | -0.002 |
| A3SS | 677 | ENSG00000127507 | ADGRE2 | chr19 | - | 14755653 | 14755877 | 14755653 | 14755844 | 14756237 | 14756345 | 80 | 55 | 44 | 67 | 80 | 1 | 2 | 1 | 6 | 21 | 200 | 79 | 92 | 98 | 85 | 0 | 1 | 16 | 18 | 8 | 182 | 149 | 0.938211 | 1 | 0.985 | 0.957 | 0.973 | 0.901 | 0.757 | 1 | 0.985 | 0.825 | 0.817 | 0.897 | 0.01 |
| A3SS | 6126 | ENSG00000132463 | GRSF1 | chr4 | - | 70825295 | 70826245 | 70825295 | 70825431 | 70827851 | 70828036 | 230 | 161 | 199 | 235 | 492 | 0 | 1 | 3 | 6 | 0 | 182 | 173 | 107 | 159 | 186 | 0 | 5 | 0 | 0 | 0 | 298 | 149 | 0.939151 | 1 | 1 | 0.988 | 0.971 | 0.951 | 1 | 1 | 0.945 | 1 | 1 | 1 | -0.007 |
| A3SS | 1237 | ENSG00000108622 | ICAM2 | chr17 | - | 64006630 | 64006735 | 64006630 | 64006671 | 64020522 | 64020625 | 142 | 87 | 130 | 145 | 111 | 4 | 3 | 5 | 6 | 0 | 135 | 64 | 40 | 73 | 156 | 0 | 11 | 1 | 0 | 5 | 213 | 149 | 0.939159 | 1 | 0.961 | 0.953 | 0.948 | 0.944 | 1 | 1 | 0.803 | 0.965 | 1 | 0.956 | 0.016 |
| A3SS | 10701 | ENSG00000046651 | OFD1 | chrX | + | 13752703 | 13753441 | 13753367 | 13753441 | 13749426 | 13749533 | 73 | 69 | 71 | 66 | 77 | 81 | 60 | 63 | 61 | 92 | 11 | 29 | 53 | 20 | 16 | 18 | 30 | 44 | 32 | 4 | 298 | 149 | 0.939182 | 1 | 0.311 | 0.365 | 0.36 | 0.351 | 0.295 | 0.234 | 0.326 | 0.376 | 0.238 | 0.667 | -0.032 |
| A3SS | 12800 | ENSG00000144579 | CTDSP1 | chr2 | + | 218403010 | 218403127 | 218403034 | 218403127 | 218402348 | 218402405 | 2 | 4 | 1 | 8 | 0 | 340 | 113 | 258 | 248 | 138 | 0 | 2 | 5 | 2 | 0 | 157 | 215 | 113 | 239 | 394 | 173 | 149 | 0.939405 | 1 | 0.005 | 0.03 | 0.003 | 0.027 | 0 | 0 | 0.008 | 0.037 | 0.007 | 0 | 0.003 |
| A3SS | 5570 | ENSG00000160271 | RALGDS | chr9 | - | 133109625 | 133109721 | 133109625 | 133109685 | 133110295 | 133110489 | 46 | 18 | 24 | 29 | 24 | 49 | 14 | 31 | 31 | 25 | 27 | 22 | 12 | 27 | 87 | 34 | 37 | 19 | 26 | 54 | 185 | 149 | 0.939852 | 1 | 0.431 | 0.509 | 0.384 | 0.43 | 0.436 | 0.39 | 0.324 | 0.337 | 0.455 | 0.565 | 0.024 |
| A3SS | 8456 | ENSG00000081059 | TCF7 | chr5 | + | 134146200 | 134148229 | 134146223 | 134148229 | 134143591 | 134143640 | 77 | 84 | 189 | 129 | 210 | 171 | 132 | 335 | 223 | 344 | 24 | 20 | 25 | 51 | 59 | 59 | 69 | 60 | 63 | 46 | 172 | 149 | 0.940021 | 1 | 0.281 | 0.355 | 0.328 | 0.334 | 0.346 | 0.261 | 0.201 | 0.265 | 0.412 | 0.526 | -0.004 |
| A3SS | 2539 | ENSG00000159110 | IFNAR2 | chr21 | + | 33241839 | 33241977 | 33241885 | 33241977 | 33230408 | 33230557 | 36 | 30 | 19 | 31 | 53 | 2 | 0 | 0 | 4 | 0 | 33 | 27 | 21 | 21 | 16 | 0 | 0 | 0 | 0 | 0 | 195 | 149 | 0.940576 | 1 | 0.932 | 1 | 1 | 0.856 | 1 | 1 | 1 | 1 | 1 | 1 | -0.042 |
| A3SS | 8558 | ENSG00000115282 | TTC31 | chr2 | + | 74490024 | 74490473 | 74490243 | 74490473 | 74483321 | 74483410 | 99 | 56 | 100 | 81 | 92 | 8 | 2 | 9 | 10 | 15 | 114 | 99 | 46 | 86 | 171 | 9 | 6 | 7 | 28 | 4 | 298 | 149 | 0.940889 | 1 | 0.861 | 0.933 | 0.847 | 0.802 | 0.754 | 0.864 | 0.892 | 0.767 | 0.606 | 0.955 | 0.023 |
| A3SS | 4467 | ENSG00000169714 | CNBP | chr3 | - | 129171078 | 129171280 | 129171078 | 129171277 | 129171445 | 129171538 | 383 | 222 | 377 | 338 | 357 | 678 | 353 | 620 | 631 | 448 | 442 | 278 | 412 | 338 | 353 | 695 | 577 | 491 | 534 | 617 | 152 | 149 | 0.942315 | 1 | 0.356 | 0.381 | 0.373 | 0.344 | 0.439 | 0.384 | 0.321 | 0.451 | 0.383 | 0.359 | -0.001 |
| A3SS | 2466 | ENSG00000140320 | BAHD1 | chr15 | + | 40461911 | 40462294 | 40461914 | 40462294 | 40458450 | 40459896 | 20 | 7 | 21 | 26 | 2 | 5 | 3 | 10 | 6 | 4 | 12 | 27 | 7 | 4 | 4 | 17 | 0 | 3 | 0 | 24 | 152 | 149 | 0.942422 | 1 | 0.797 | 0.696 | 0.673 | 0.809 | 0.329 | 0.409 | 1 | 0.696 | 1 | 0.14 | 0.012 |
| A3SS | 5423 | ENSG00000125637 | PSD4 | chr2 | + | 113197746 | 113197913 | 113197749 | 113197913 | 113197563 | 113197634 | 54 | 11 | 41 | 34 | 24 | 36 | 18 | 37 | 44 | 30 | 54 | 51 | 15 | 51 | 41 | 40 | 73 | 19 | 42 | 45 | 152 | 149 | 0.942563 | 1 | 0.595 | 0.375 | 0.521 | 0.431 | 0.44 | 0.57 | 0.406 | 0.436 | 0.543 | 0.472 | -0.013 |
| A3SS | 2160 | ENSG00000213614 | HEXA | chr15 | - | 72356524 | 72356710 | 72356524 | 72356617 | 72362375 | 72362466 | 10 | 7 | 16 | 14 | 13 | 1 | 0 | 2 | 5 | 0 | 14 | 25 | 1 | 30 | 28 | 0 | 3 | 4 | 12 | 0 | 242 | 149 | 0.942867 | 1 | 0.86 | 1 | 0.831 | 0.633 | 1 | 1 | 0.837 | 0.133 | 0.606 | 1 | 0.15 |
| A3SS | 2241 | ENSG00000168970 | JMJD7-PLA2G4B | chr15 | + | 41837067 | 41840230 | 41840157 | 41840230 | 41836474 | 41836551 | 48 | 14 | 65 | 67 | 28 | 1 | 4 | 1 | 7 | 4 | 71 | 38 | 19 | 58 | 106 | 12 | 4 | 0 | 3 | 4 | 298 | 149 | 0.943641 | 1 | 0.96 | 0.636 | 0.97 | 0.827 | 0.778 | 0.747 | 0.826 | 1 | 0.906 | 0.93 | -0.048 |
| A3SS | 7824 | ENSG00000162191 | UBXN1 | chr11 | - | 62677517 | 62677634 | 62677517 | 62677563 | 62677780 | 62677832 | 257 | 125 | 268 | 200 | 201 | 7 | 6 | 2 | 4 | 4 | 320 | 291 | 118 | 181 | 388 | 12 | 6 | 6 | 2 | 1 | 220 | 149 | 0.944007 | 1 | 0.961 | 0.934 | 0.989 | 0.971 | 0.971 | 0.948 | 0.97 | 0.93 | 0.984 | 0.996 | 0 |
| A3SS | 9720 | ENSG00000119616 | FCF1 | chr14 | + | 74715820 | 74716099 | 74715950 | 74716099 | 74714871 | 74714943 | 51 | 35 | 72 | 44 | 112 | 189 | 145 | 171 | 231 | 182 | 42 | 71 | 96 | 35 | 8 | 267 | 100 | 101 | 162 | 64 | 279 | 149 | 0.944351 | 1 | 0.126 | 0.114 | 0.184 | 0.092 | 0.247 | 0.077 | 0.275 | 0.337 | 0.103 | 0.063 | -0.018 |
| A3SS | 12892 | ENSG00000145901 | TNIP1 | chr5 | - | 151032286 | 151032400 | 151032286 | 151032383 | 151035001 | 151035067 | 5 | 3 | 2 | 5 | 3 | 9 | 6 | 10 | 17 | 4 | 12 | 10 | 0 | 0 | 13 | 19 | 11 | 14 | 46 | 2 | 166 | 149 | 0.944809 | 1 | 0.333 | 0.31 | 0.152 | 0.209 | 0.402 | 0.362 | 0.449 | 0 | 0 | 0.854 | -0.052 |
| A3SS | 1085 | ENSG00000108840 | HDAC5 | chr17 | - | 44110728 | 44110803 | 44110728 | 44110800 | 44117493 | 44117704 | 0 | 3 | 3 | 5 | 1 | 20 | 1 | 2 | 8 | 3 | 0 | 6 | 6 | 2 | 3 | 7 | 4 | 10 | 0 | 28 | 152 | 149 | 0.944943 | 1 | 0 | 0.746 | 0.595 | 0.38 | 0.246 | 0 | 0.595 | 0.37 | 1 | 0.095 | -0.019 |
| A3SS | 11776 | ENSG00000133884 | DPF2 | chr11 | + | 65343744 | 65343837 | 65343754 | 65343837 | 65341398 | 65341562 | 136 | 63 | 140 | 131 | 225 | 1 | 0 | 0 | 3 | 2 | 149 | 195 | 105 | 95 | 126 | 9 | 0 | 0 | 0 | 0 | 159 | 149 | 0.947017 | 1 | 0.992 | 1 | 1 | 0.976 | 0.991 | 0.939 | 1 | 1 | 1 | 1 | 0.004 |
| A3SS | 12479 | ENSG00000100109 | TFIP11 | chr22 | - | 26510616 | 26510692 | 26510616 | 26510667 | 26512098 | 26512175 | 23 | 30 | 15 | 33 | 36 | 0 | 0 | 3 | 2 | 5 | 47 | 52 | 30 | 90 | 52 | 4 | 2 | 1 | 8 | 0 | 174 | 149 | 0.948014 | 1 | 1 | 1 | 0.811 | 0.934 | 0.86 | 0.91 | 0.957 | 0.963 | 0.906 | 1 | -0.026 |
| A3SS | 4359 | ENSG00000156976 | EIF4A2 | chr3 | + | 186784428 | 186784477 | 186784431 | 186784477 | 186783576 | 186783639 | 348 | 215 | 378 | 366 | 272 | 1015 | 620 | 943 | 1077 | 848 | 97 | 166 | 184 | 114 | 16 | 188 | 499 | 486 | 309 | 106 | 152 | 149 | 0.950322 | 1 | 0.252 | 0.254 | 0.282 | 0.25 | 0.239 | 0.336 | 0.246 | 0.271 | 0.266 | 0.129 | 0.006 |
| A3SS | 1383 | ENSG00000132507 | EIF5A | chr17 | + | 7309596 | 7309800 | 7309614 | 7309800 | 7306998 | 7307115 | 6 | 2 | 7 | 0 | 3 | 8 | 1 | 13 | 12 | 27 | 2 | 5 | 5 | 2 | 0 | 3 | 24 | 3 | 6 | 7 | 167 | 149 | 0.951310 | 1 | 0.401 | 0.641 | 0.325 | 0 | 0.09 | 0.373 | 0.157 | 0.598 | 0.229 | 0 | 0.02 |
| A3SS | 8630 | ENSG00000102125 | TAZ | chrX | + | 154419704 | 154420094 | 154420031 | 154420094 | 154419542 | 154419623 | 36 | 9 | 56 | 30 | 36 | 13 | 6 | 18 | 10 | 3 | 16 | 54 | 12 | 68 | 61 | 6 | 28 | 3 | 3 | 39 | 298 | 149 | 0.951327 | 1 | 0.581 | 0.429 | 0.609 | 0.6 | 0.857 | 0.571 | 0.491 | 0.667 | 0.919 | 0.439 | -0.002 |
| A3SS | 13420 | ENSG00000071564 | TCF3 | chr19 | - | 1619315 | 1619474 | 1619315 | 1619471 | 1619779 | 1619853 | 30 | 12 | 24 | 28 | 17 | 1 | 8 | 5 | 10 | 2 | 31 | 32 | 8 | 13 | 64 | 12 | 5 | 6 | 7 | 0 | 152 | 149 | 0.951680 | 1 | 0.967 | 0.595 | 0.825 | 0.733 | 0.893 | 0.717 | 0.863 | 0.567 | 0.645 | 1 | 0.044 |
| A3SS | 9020 | ENSG00000111679 | PTPN6 | chr12 | + | 6960662 | 6960945 | 6960805 | 6960945 | 6960343 | 6960435 | 20 | 22 | 15 | 25 | 4 | 1196 | 850 | 949 | 1115 | 857 | 7 | 29 | 22 | 34 | 44 | 3425 | 1951 | 1004 | 2180 | 3456 | 292 | 149 | 0.951704 | 1 | 0.008 | 0.013 | 0.008 | 0.011 | 0.002 | 0.001 | 0.008 | 0.011 | 0.008 | 0.006 | 0.002 |
| A3SS | 8096 | ENSG00000183495 | EP400 | chr12 | + | 132066773 | 132066969 | 132066794 | 132066969 | 132064667 | 132064886 | 13 | 2 | 9 | 8 | 4 | 5 | 0 | 3 | 6 | 0 | 14 | 3 | 12 | 3 | 0 | 0 | 4 | 0 | 2 | 3 | 170 | 149 | 0.951963 | 1 | 0.695 | 1 | 0.724 | 0.539 | 1 | 1 | 0.397 | 1 | 0.568 | 0 | 0.199 |
| A3SS | 6866 | ENSG00000156983 | BRPF1 | chr3 | + | 9743577 | 9743901 | 9743580 | 9743901 | 9742943 | 9743253 | 26 | 24 | 15 | 6 | 11 | 26 | 12 | 28 | 33 | 23 | 28 | 28 | 26 | 37 | 43 | 78 | 46 | 33 | 41 | 56 | 152 | 149 | 0.952617 | 1 | 0.495 | 0.662 | 0.344 | 0.151 | 0.319 | 0.26 | 0.374 | 0.436 | 0.469 | 0.429 | 0.001 |
| A3SS | 3442 | ENSG00000130803 | ZNF317 | chr19 | + | 9157833 | 9158075 | 9157979 | 9158075 | 9157267 | 9157394 | 6 | 5 | 11 | 15 | 6 | 45 | 26 | 44 | 28 | 44 | 20 | 4 | 12 | 25 | 6 | 45 | 52 | 92 | 44 | 63 | 295 | 149 | 0.953250 | 1 | 0.063 | 0.089 | 0.112 | 0.213 | 0.064 | 0.183 | 0.037 | 0.062 | 0.223 | 0.046 | -0.002 |
| A3SS | 7245 | ENSG00000089127 | OAS1 | chr12 | + | 112911035 | 112911235 | 112911097 | 112911235 | 112908535 | 112908824 | 86 | 37 | 36 | 74 | 29 | 22 | 13 | 9 | 23 | 2 | 149 | 138 | 42 | 207 | 61 | 57 | 37 | 22 | 14 | 13 | 211 | 149 | 0.953922 | 1 | 0.734 | 0.668 | 0.739 | 0.694 | 0.911 | 0.649 | 0.725 | 0.574 | 0.913 | 0.768 | 0.023 |

| | | | | | | | | | | | | | | |
|---|---|---|---|---|---|---|---|---|---|---|---|---|---|---|
| A3SS | 9083 | ENSG00000114388 | NPRL2 | chr3 | - | 50348873 | 50349089 | 50348873 | 50349010 | 50349385 | 50349494 | 11 | 5 | 6 |
| A3SS | 2158 | ENSG00000213614 | HEXA | chr15 | - | 72345445 | 72346128 | 72345445 | 72345550 | 72346234 | 72346325 | 37 | 14 | 34 |
| A3SS | 5842 | ENSG00000213593 | TMX2 | chr11 | + | 57737785 | 57738026 | 57737835 | 57738026 | 57737607 | 57737668 | 4 | 4 | 7 |
| A3SS | 1904 | ENSG00000214078 | CPNE1 | chr20 | - | 35658929 | 35659014 | 35658929 | 35658982 | 35664759 | 35664900 | 372 | 226 | 400 |
| A3SS | 324 | ENSG00000196684 | HSH2D | chr19 | + | 16153031 | 16153208 | 16153042 | 16153208 | 16152551 | 16152641 | 16 | 6 | 7 |
| A3SS | 1529 | ENSG00000168259 | DNAJC7 | chr17 | - | 41994869 | 41994983 | 41994869 | 41994944 | 41996310 | 41996424 | 5 | 5 | 8 |
| A3SS | 661 | ENSG00000105705 | SUGP1 | chr19 | - | 19303723 | 19303912 | 19303723 | 19303847 | 19305848 | 19306076 | 20 | 10 | 30 |
| A3SS | 1359 | ENSG00000092929 | UNC13D | chr17 | - | 75840224 | 75840367 | 75840224 | 75840329 | 75840506 | 75840576 | 22 | 14 | 7 |
| A3SS | 12621 | ENSG00000107959 | PITRM1 | chr10 | - | 3143388 | 3143731 | 3143388 | 3143501 | 3144291 | 3144366 | 14 | 9 | 29 |
| A3SS | 2838 | ENSG00000119599 | DCAF4 | chr14 | + | 72954160 | 72954262 | 72954163 | 72954262 | 72951797 | 72951877 | 5 | 0 | 0 |
| A3SS | 9204 | ENSG00000204420 | MPIG6B | chr6 | + | 31724745 | 31724844 | 31724764 | 31724844 | 31724586 | 31724627 | 47 | 17 | 11 |
| A3SS | 2461 | ENSG00000166200 | COPS2 | chr15 | - | 49137347 | 49137458 | 49137347 | 49137437 | 49139527 | 49139653 | 196 | 179 | 175 |
| A3SS | 4768 | ENSG00000075826 | SEC31B | chr10 | - | 100507907 | 100508051 | 100507907 | 100508048 | 100509006 | 100509102 | 10 | 9 | 22 |
| A3SS | 8151 | ENSG00000131051 | RBM39 | chr20 | - | 35709223 | 35709274 | 35709223 | 35709256 | 35714184 | 35714389 | 204 | 148 | 175 |
| A3SS | 12310 | ENSG00000173992 | CCS | chr11 | + | 66605153 | 66605416 | 66605338 | 66605416 | 66600488 | 66600549 | 32 | 13 | 21 |
| A3SS | 3586 | ENSG00000214655 | ZSWIM8 | chr10 | + | 73796752 | 73797014 | 73796773 | 73797014 | 73795538 | 73795663 | 1 | 0 | 0 |
| A3SS | 10259 | ENSG00000112561 | TFEB | chr6 | - | 41686089 | 41686356 | 41686089 | 41686237 | 41687093 | 41687169 | 3 | 4 | 0 |
| A3SS | 10303 | ENSG00000169598 | DFFB | chr1 | + | 3865642 | 3866000 | 3865811 | 3866000 | 3858717 | 3858844 | 19 | 6 | 3 |
| A3SS | 4150 | ENSG00000278259 | MYO19 | chr17 | - | 36503949 | 36505177 | 36503949 | 36504020 | 36505296 | 36505404 | 4 | 1 | 6 |
| A3SS | 3193 | ENSG00000165819 | METTL3 | chr14 | - | 21501727 | 21501903 | 21501727 | 21501824 | 21503172 | 21503577 | 101 | 96 | 138 |
| A3SS | 3131 | ENSG00000145730 | PAM | chr5 | + | 103028181 | 103028238 | 103028184 | 103028238 | 103025130 | 103025334 | 23 | 7 | 6 |
| A3SS | 10250 | ENSG00000196696 | PDXDC2P-NPIPB14P | chr16 | - | 70038904 | 70039100 | 70038904 | 70038985 | 70042350 | 70042416 | 58 | 62 | 41 |
| A3SS | 1827 | ENSG00000135723 | FHOD1 | chr16 | - | 67231636 | 67231838 | 67231636 | 67231819 | 67232038 | 67232194 | 2 | 0 | 4 |
| A3SS | 4054 | ENSG00000003056 | M6PR | chr12 | - | 8943404 | 8943584 | 8943404 | 8943535 | 8943800 | 8943910 | 8 | 4 | 13 |
| A3SS | 10635 | ENSG00000147140 | NONO | chrX | + | 71297360 | 71298508 | 71297376 | 71298508 | 71296850 | 71297047 | 4 | 3 | 3 |
| A3SS | 7886 | ENSG00000101336 | HCK | chr20 | + | 32073715 | 32073818 | 32073718 | 32073818 | 32073318 | 32073361 | 282 | 171 | 157 |
| A3SS | 11009 | ENSG00000160932 | LY6E | chr8 | + | 143021313 | 143021433 | 143021348 | 143021433 | 143020882 | 143020991 | 525 | 342 | 410 |
| A3SS | 6364 | ENSG00000215788 | TNFRSF25 | chr1 | - | 6462628 | 6462666 | 6462628 | 6462663 | 6462862 | 6462970 | 15 | 10 | 24 |
| A3SS | 12171 | ENSG00000149187 | CELF1 | chr11 | - | 47476845 | 47476959 | 47476845 | 47476956 | 47477296 | 47477425 | 23 | 11 | 15 |

| | | | | | | | | | | | | | | | | | | | | | | | | | | | | | | | | | | | | | | | | | | | | | | |
|---|---|---|---|---|---|---|---|---|---|---|---|---|---|---|---|---|---|---|---|---|---|---|---|---|---|---|---|---|---|---|---|---|---|---|---|---|---|---|---|---|---|---|---|---|---|---|
| A3SS | 1242 | ENSG00000108788 | MLX | chr17 | + | 42569981 | 42570183 | 42570096 | 42570183 | 42569506 | 42569606 | 399 | 264 | 350 | 353 | 429 | 0 | 1 | 0 | 0 | 2 | 621 | 553 | 281 | 290 | 622 | 0 | 0 | 0 | 0 | 0 | 264 | 149 | 1.000000 | 1 | 1 | 0.993 | 1 | 1 | 0.992 | 1 | 1 | 1 | 1 | 1 | -0.003 |
| A3SS | 2157 | ENSG00000130703 | OSBPL2 | chr20 | + | 62259980 | 62260125 | 62260016 | 62260125 | 62256056 | 62256221 | 171 | 156 | 153 | 160 | 261 | 4 | 1 | 0 | 2 | 1 | 335 | 197 | 253 | 373 | 306 | 0 | 1 | 2 | 0 | 0 | 185 | 149 | 1.000000 | 1 | 0.972 | 0.992 | 1 | 0.985 | 0.995 | 1 | 0.994 | 0.99 | 1 | 1 | -0.008 |
| A3SS | 345 | ENSG00000129353 | SLC44A2 | chr19 | + | 10637643 | 10637747 | 10637649 | 10637747 | 10636661 | 10636756 | 212 | 98 | 167 | 179 | 156 | 0 | 0 | 3 | 0 | 0 | 153 | 241 | 89 | 155 | 730 | 0 | 0 | 0 | 0 | 0 | 155 | 149 | 1.000000 | 1 | 1 | 1 | 0.982 | 1 | 1 | 1 | 1 | 1 | 1 | 1 | -0.004 |
| A3SS | 1721 | ENSG00000086504 | MRPL28 | chr16 | - | 368500 | 369220 | 368500 | 369211 | 369930 | 370225 | 135 | 43 | 72 | 107 | 69 | 1 | 0 | 0 | 0 | 0 | 219 | 238 | 126 | 192 | 201 | 0 | 0 | 0 | 0 | 0 | 158 | 149 | 1.000000 | 1 | 0.992 | 1 | 1 | 1 | 1 | 1 | 1 | 1 | 1 | 1 | -0.002 |
| A3SS | 4471 | ENSG00000169714 | CNBP | chr3 | - | 129171445 | 129171538 | 129171445 | 129171508 | 129171654 | 129171771 | 629 | 354 | 579 | 596 | 473 | 1 | 2 | 3 | 0 | 0 | 791 | 587 | 615 | 585 | 595 | 0 | 6 | 0 | 0 | 0 | 179 | 149 | 1.000000 | 1 | 0.998 | 0.993 | 0.994 | 1 | 1 | 1 | 0.988 | 1 | 1 | 1 | -0.001 |
| A3SS | 8342 | ENSG00000172890 | NADSYN1 | chr11 | + | 71481922 | 71482025 | 71482000 | 71482025 | 71481355 | 71481404 | 290 | 203 | 323 | 252 | 343 | 1 | 3 | 1 | 5 | 0 | 501 | 376 | 269 | 277 | 527 | 0 | 0 | 3 | 0 | 0 | 227 | 149 | 1.000000 | 1 | 0.995 | 0.978 | 0.995 | 0.971 | 1 | 1 | 1 | 0.983 | 1 | 1 | -0.009 |
| A3SS | 4329 | ENSG00000247596 | TWF2 | chr3 | - | 52229919 | 52230995 | 52229919 | 52230070 | 52231126 | 52231231 | 222 | 112 | 152 | 190 | 173 | 0 | 0 | 0 | 0 | 0 | 316 | 289 | 92 | 249 | 442 | 0 | 0 | 0 | 6 | 0 | 298 | 149 | 1.000000 | 1 | 1 | 1 | 1 | 1 | 1 | 1 | 1 | 1 | 0.954 | 1 | 0.009 |
| A3SS | 5846 | ENSG00000100280 | AP1B1 | chr22 | - | 29334264 | 29334410 | 29334264 | 29334350 | 29338989 | 29339133 | 131 | 68 | 68 | 98 | 85 | 2 | 0 | 2 | 0 | 0 | 59 | 247 | 93 | 93 | 494 | 0 | 4 | 0 | 0 | 0 | 209 | 149 | 1.000000 | 1 | 0.979 | 1 | 0.96 | 1 | 1 | 1 | 0.978 | 1 | 1 | 1 | -0.008 |
| A3SS | 13151 | ENSG00000123349 | PFDN5 | chr12 | + | 53298044 | 53298150 | 53298048 | 53298150 | 53297849 | 53297924 | 1971 | 1111 | 1817 | 1601 | 1859 | 9 | 6 | 2 | 6 | 13 | 2537 | 1446 | 1070 | 1322 | 2508 | 0 | 3 | 5 | 4 | 13 | 153 | 149 | 1.000000 | 1 | 0.995 | 0.994 | 0.999 | 0.996 | 0.993 | 1 | 0.998 | 0.995 | 0.997 | 0.995 | -0.002 |
| A3SS | 4885 | ENSG00000116871 | MAP7D1 | chr1 | + | 36178418 | 36178596 | 36178470 | 36178596 | 36177872 | 36178201 | 145 | 90 | 106 | 99 | 50 | 1 | 0 | 0 | 1 | 0 | 180 | 160 | 125 | 187 | 359 | 0 | 0 | 0 | 0 | 0 | 201 | 149 | 1.000000 | 1 | 0.991 | 1 | 1 | 0.987 | 1 | 1 | 1 | 1 | 1 | 1 | -0.004 |
| A3SS | 11074 | ENSG00000079459 | FDFT1 | chr8 | + | 11809666 | 11809850 | 11809791 | 11809850 | 11808793 | 11808891 | 318 | 153 | 260 | 270 | 256 | 0 | 0 | 0 | 2 | 0 | 375 | 418 | 271 | 266 | 448 | 0 | 0 | 0 | 0 | 0 | 274 | 149 | 1.000000 | 1 | 1 | 1 | 1 | 0.987 | 1 | 1 | 1 | 1 | 1 | 1 | -0.003 |
| A3SS | 4942 | ENSG00000150753 | CCT5 | chr5 | + | 10254673 | 10254838 | 10254765 | 10254838 | 10254144 | 10254205 | 662 | 376 | 443 | 507 | 530 | 0 | 1 | 4 | 0 | 0 | 1031 | 455 | 334 | 830 | 384 | 0 | 0 | 2 | 1 | 0 | 241 | 149 | 1.000000 | 1 | 1 | 0.996 | 0.986 | 1 | 1 | 1 | 1 | 0.99 | 0.998 | 1 | -0.001 |
| A3SS | 1094 | ENSG00000161955 | TNFSF13 | chr17 | + | 7559845 | 7559893 | 7559850 | 7559893 | 7559623 | 7559702 | 256 | 174 | 98 | 158 | 114 | 0 | 1 | 0 | 0 | 2 | 411 | 176 | 93 | 195 | 439 | 0 | 0 | 0 | 0 | 0 | 154 | 149 | 1.000000 | 1 | 1 | 0.994 | 1 | 1 | 0.982 | 1 | 1 | 1 | 1 | 1 | -0.005 |
| A3SS | 1277 | ENSG00000185359 | HGS | chr17 | + | 81688703 | 81688827 | 81688751 | 81688827 | 81687002 | 81687095 | 156 | 112 | 178 | 166 | 177 | 0 | 0 | 0 | 0 | 1 | 327 | 223 | 139 | 168 | 438 | 0 | 0 | 0 | 0 | 0 | 197 | 149 | 1.000000 | 1 | 1 | 1 | 1 | 1 | 0.993 | 1 | 1 | 1 | 1 | 1 | -0.001 |
| A3SS | 5009 | ENSG00000143420 | ENSA | chr1 | - | 150625641 | 150625808 | 150625641 | 150625802 | 150627466 | 150627592 | 472 | 278 | 417 | 407 | 795 | 0 | 0 | 0 | 0 | 0 | 760 | 491 | 391 | 574 | 658 | 0 | 0 | 1 | 1 | 0 | 155 | 149 | 1.000000 | 1 | 1 | 1 | 1 | 1 | 1 | 1 | 1 | 0.997 | 0.998 | 1 | 0.001 |
| A3SS | 6907 | ENSG00000168610 | STAT3 | chr17 | - | 42316788 | 42317224 | 42316788 | 42316901 | 42322281 | 42322494 | 246 | 149 | 145 | 247 | 342 | 0 | 0 | 0 | 0 | 0 | 336 | 332 | 262 | 341 | 358 | 0 | 0 | 0 | 2 | 0 | 298 | 149 | 1.000000 | 1 | 1 | 1 | 1 | 1 | 1 | 1 | 1 | 1 | 0.988 | 1 | 0.002 |
| A3SS | 6260 | ENSG00000241878 | PISD | chr22 | - | 31620552 | 31620713 | 31620552 | 31620599 | 31620995 | 31621142 | 177 | 88 | 138 | 121 | 106 | 2 | 1 | 0 | 0 | 3 | 294 | 176 | 170 | 185 | 303 | 0 | 1 | 1 | 6 | 0 | 263 | 149 | 1.000000 | 1 | 0.98 | 0.98 | 1 | 1 | 0.952 | 1 | 0.99 | 0.99 | 0.946 | 1 | -0.003 |
| A3SS | 1215 | ENSG00000185624 | P4HB | chr17 | - | 81855452 | 81855579 | 81855452 | 81855505 | 81859180 | 81859387 | 173 | 88 | 180 | 162 | 83 | 0 | 0 | 0 | 3 | 0 | 201 | 179 | 197 | 243 | 362 | 0 | 0 | 0 | 0 | 0 | 223 | 149 | 1.000000 | 1 | 1 | 1 | 1 | 0.973 | 1 | 1 | 1 | 1 | 1 | 1 | -0.005 |
| A3SS | 1943 | ENSG00000213658 | LAT | chr16 | + | 28986539 | 28986714 | 28986656 | 28986714 | 28986381 | 28986446 | 176 | 125 | 182 | 241 | 78 | 0 | 3 | 1 | 2 | 0 | 251 | 292 | 143 | 160 | 452 | 0 | 7 | 0 | 0 | 0 | 266 | 149 | 1.000000 | 1 | 1 | 0.959 | 0.99 | 0.985 | 1 | 1 | 0.959 | 1 | 1 | 1 | -0.005 |
| A3SS | 3021 | ENSG00000139433 | GLTP | chr12 | - | 109855618 | 109855769 | 109855618 | 109855738 | 109857525 | 109857659 | 92 | 54 | 76 | 117 | 75 | 2 | 1 | 1 | 0 | 0 | 279 | 124 | 82 | 158 | 266 | 0 | 0 | 0 | 0 | 0 | 180 | 149 | 1.000000 | 1 | 0.974 | 0.978 | 0.984 | 1 | 1 | 1 | 1 | 1 | 1 | 1 | -0.013 |
| A3SS | 3764 | ENSG00000205937 | RNPS1 | chr16 | - | 2263095 | 2263287 | 2263095 | 2263161 | 2264572 | 2264760 | 458 | 254 | 470 | 469 | 652 | 0 | 0 | 1 | 0 | 0 | 619 | 573 | 376 | 483 | 745 | 6 | 0 | 0 | 0 | 0 | 275 | 149 | 1.000000 | 1 | 1 | 1 | 0.996 | 1 | 1 | 0.982 | 1 | 1 | 1 | 1 | 0.003 |
| A3SS | 4959 | ENSG00000158864 | NDUFS2 | chr1 | + | 161212350 | 161213475 | 161213379 | 161213475 | 161209482 | 161210710 | 176 | 120 | 159 | 207 | 170 | 1 | 2 | 1 | 3 | 0 | 346 | 227 | 189 | 248 | 312 | 0 | 0 | 0 | 5 | 0 | 298 | 149 | 1.000000 | 1 | 0.989 | 0.968 | 0.988 | 0.972 | 1 | 1 | 1 | 1 | 0.961 | 1 | -0.009 |
| A3SS | 2803 | ENSG00000089737 | DDX24 | chr14 | - | 94054995 | 94057897 | 94054995 | 94055184 | 94060097 | 94060613 | 280 | 163 | 385 | 328 | 286 | 0 | 0 | 0 | 0 | 0 | 273 | 471 | 469 | 469 | 248 | 0 | 2 | 0 | 0 | 0 | 298 | 149 | 1.000000 | 1 | 1 | 1 | 1 | 1 | 1 | 1 | 0.992 | 1 | 1 | 1 | 0.002 |
| A3SS | 10210 | ENSG00000213719 | CLIC1 | chr6 | - | 31736261 | 31736412 | 31736261 | 31736350 | 31736544 | 31736572 | 77 | 19 | 57 | 68 | 56 | 3 | 0 | 0 | 0 | 0 | 173 | 43 | 117 | 174 | 257 | 0 | 0 | 0 | 0 | 0 | 211 | 149 | 1.000000 | 1 | 0.948 | 1 | 1 | 1 | 1 | 1 | 1 | 1 | 1 | 1 | -0.01 |
| A3SS | 10760 | ENSG00000117676 | RPS6KA1 | chr1 | + | 26555150 | 26555221 | 26555183 | 26555221 | 26554595 | 26554738 | 280 | 135 | 158 | 184 | 246 | 1 | 2 | 0 | 3 | 0 | 294 | 307 | 171 | 226 | 486 | 0 | 1 | 1 | 2 | 0 | 182 | 149 | 1.000000 | 1 | 0.996 | 0.982 | 1 | 0.98 | 1 | 1 | 0.996 | 0.993 | 0.989 | 1 | -0.004 |
| A3SS | 5048 | ENSG00000125347 | IRF1 | chr5 | - | 132487925 | 132488052 | 132487925 | 132488022 | 132489391 | 132489483 | 96 | 45 | 90 | 100 | 63 | 0 | 0 | 0 | 0 | 0 | 293 | 124 | 59 | 193 | 204 | 0 | 3 | 0 | 0 | 0 | 179 | 149 | 1.000000 | 1 | 1 | 1 | 1 | 1 | 1 | 1 | 0.972 | 1 | 1 | 1 | 0.006 |
| A3SS | 13131 | ENSG00000168010 | ATG16L2 | chr11 | + | 72826517 | 72827293 | 72827187 | 72827293 | 72826172 | 72826243 | 138 | 62 | 82 | 92 | 80 | 1 | 0 | 0 | 0 | 0 | 280 | 89 | 91 | 205 | 262 | 0 | 0 | 0 | 0 | 0 | 298 | 149 | 1.000000 | 1 | 0.986 | 1 | 1 | 1 | 1 | 1 | 1 | 1 | 1 | 1 | -0.003 |
| A3SS | 792 | ENSG00000124214 | STAU1 | chr20 | - | 49123091 | 49123235 | 49123091 | 49123229 | 49124374 | 49124587 | 72 | 77 | 85 | 123 | 112 | 0 | 0 | 0 | 0 | 0 | 266 | 152 | 154 | 194 | 171 | 0 | 0 | 1 | 4 | 0 | 155 | 149 | 1.000000 | 1 | 1 | 1 | 1 | 1 | 1 | 1 | 1 | 0.993 | 0.979 | 1 | 0.006 |
| A3SS | 3245 | ENSG00000135074 | ADAM19 | chr5 | - | 157477303 | 157481002 | 157477303 | 157477735 | 157481790 | 157481943 | 173 | 79 | 237 | 182 | 263 | 0 | 0 | 6 | 0 | 0 | 254 | 395 | 230 | 195 | 326 | 0 | 1 | 0 | 0 | 0 | 298 | 149 | 1.000000 | 1 | 1 | 1 | 0.952 | 1 | 1 | 1 | 0.995 | 1 | 1 | 1 | -0.009 |
| A3SS | 5778 | ENSG00000058600 | POLR3E | chr16 | + | 22308924 | 22309040 | 22308974 | 22309040 | 22308147 | 22308225 | 118 | 66 | 115 | 108 | 124 | 2 | 0 | 0 | 1 | 1 | 140 | 174 | 157 | 198 | 326 | 0 | 3 | 0 | 3 | 0 | 199 | 149 | 1.000000 | 1 | 0.978 | 1 | 1 | 0.988 | 0.989 | 1 | 0.977 | 1 | 0.98 | 1 | 0 |
| A3SS | 8975 | ENSG00000167792 | NDUFV1 | chr11 | + | 67611402 | 67611569 | 67611521 | 67611569 | 67610994 | 67611207 | 268 | 163 | 261 | 202 | 280 | 0 | 0 | 2 | 3 | 0 | 398 | 368 | 243 | 220 | 404 | 0 | 1 | 0 | 3 | 0 | 268 | 149 | 1.000000 | 1 | 1 | 1 | 0.986 | 0.974 | 1 | 1 | 0.995 | 1 | 0.976 | 1 | -0.002 |
| A3SS | 4980 | ENSG00000186517 | ARHGAP30 | chr1 | - | 161052625 | 161053385 | 161052625 | 161052797 | 161054365 | 161054473 | 352 | 159 | 282 | 298 | 141 | 0 | 1 | 0 | 1 | 3 | 414 | 346 | 195 | 241 | 493 | 0 | 0 | 0 | 4 | 0 | 298 | 149 | 1.000000 | 1 | 1 | 0.988 | 1 | 0.993 | 0.959 | 1 | 1 | 1 | 0.968 | 1 | -0.006 |
| A3SS | 11313 | ENSG00000132334 | PTPRE | chr10 | + | 128049529 | 128049666 | 128049534 | 128049666 | 128047763 | 128047837 | 228 | 179 | 131 | 197 | 109 | 0 | 0 | 0 | 0 | 0 | 352 | 162 | 241 | 394 | 150 | 0 | 1 | 0 | 0 | 0 | 154 | 149 | 1.000000 | 1 | 1 | 1 | 1 | 1 | 1 | 1 | 0.994 | 1 | 1 | 1 | 0.001 |
| A3SS | 12197 | ENSG00000172977 | KAT5 | chr11 | + | 65714494 | 65714743 | 65714659 | 65714743 | 65713773 | 65713848 | 186 | 74 | 208 | 163 | 130 | 0 | 2 | 0 | 1 | 0 | 254 | 231 | 189 | 226 | 312 | 3 | 0 | 0 | 0 | 0 | 298 | 149 | 1.000000 | 1 | 1 | 0.949 | 1 | 0.988 | 1 | 0.977 | 1 | 1 | 1 | 1 | -0.008 |
| A3SS | 5994 | ENSG00000110619 | CARS1 | chr11 | - | 3001969 | 3002600 | 3001969 | 3002053 | 3005365 | 3005433 | 38 | 55 | 45 | 68 | 69 | 2 | 0 | 0 | 0 | 0 | 246 | 104 | 87 | 99 | 184 | 0 | 0 | 0 | 0 | 0 | 298 | 149 | 1.000000 | 1 | 0.905 | 1 | 1 | 1 | 1 | 1 | 1 | 1 | 1 | 1 | -0.019 |
| A3SS | 2800 | ENSG00000100764 | PSMC1 | chr14 | + | 90263661 | 90263847 | 90263800 | 90263847 | 90263317 | 90263442 | 228 | 126 | 179 | 197 | 260 | 0 | 0 | 0 | 0 | 1 | 484 | 198 | 254 | 327 | 170 | 0 | 6 | 0 | 1 | 0 | 288 | 149 | 1.000000 | 1 | 1 | 1 | 1 | 1 | 0.993 | 1 | 0.945 | 1 | 0.994 | 1 | 0.011 |
| A3SS | 2244 | ENSG00000128739 | SNRPN | chr15 | + | 24974310 | 24974456 | 24974397 | 24974456 | 24962113 | 24962209 | 177 | 86 | 146 | 173 | 130 | 0 | 0 | 0 | 1 | 0 | 135 | 186 | 109 | 136 | 586 | 0 | 2 | 0 | 0 | 0 | 236 | 149 | 1.000000 | 1 | 1 | 1 | 1 | 0.991 | 1 | 1 | 0.983 | 1 | 1 | 1 | 0.002 |
| A3SS | 797 | ENSG00000101265 | RASSF2 | chr20 | - | 4787632 | 4787754 | 4787632 | 4787701 | 4789595 | 4789697 | 242 | 149 | 152 | 186 | 196 | 0 | 0 | 0 | 0 | 0 | 308 | 185 | 219 | 286 | 366 | 0 | 0 | 2 | 0 | 0 | 202 | 149 | 1.000000 | 1 | 1 | 1 | 1 | 1 | 1 | 1 | 1 | 0.988 | 1 | 1 | 0.002 |
| A3SS | 2943 | ENSG00000135596 | MICAL1 | chr6 | - | 109452510 | 109452615 | 109452510 | 109452531 | 109453262 | 109453367 | 154 | 88 | 151 | 225 | 92 | 0 | 0 | 0 | 0 | 0 | 304 | 145 | 123 | 229 | 348 | 0 | 1 | 0 | 0 | 0 | 233 | 149 | 1.000000 | 1 | 1 | 1 | 1 | 1 | 1 | 1 | 0.989 | 1 | 1 | 1 | 0.002 |
| A3SS | 4081 | ENSG00000065882 | TBC1D1 | chr4 | + | 38124961 | 38125131 | 38125110 | 38125131 | 38115709 | 38115954 | 150 | 106 | 119 | 144 | 249 | 0 | 0 | 0 | 0 | 0 | 341 | 177 | 182 | 210 | 295 | 0 | 0 | 1 | 0 | 0 | 298 | 149 | 1.000000 | 1 | 1 | 1 | 1 | 1 | 1 | 1 | 1 | 0.989 | 1 | 1 | 0.002 |
| A3SS | 1166 | ENSG00000184640 | SEPTIN9 | chr17 | + | 77487423 | 77487552 | 77487481 | 77487552 | 77482143 | 77482335 | 1100 | 566 | 995 | 968 | 843 | 0 | 0 | 1 | 0 | 0 | 1091 | 976 | 583 | 678 | 1580 | 0 | 0 | 0 | 0 | 0 | 207 | 149 | 1.000000 | 1 | 1 | 1 | 0.999 | 1 | 1 | 1 | 1 | 1 | 1 | 1 | 0 |
| A3SS | 9323 | ENSG00000164116 | GUCY1A1 | chr4 | + | 155710541 | 155711251 | 155710681 | 155711251 | 155708235 | 155708294 | 30 | 99 | 19 | 160 | 123 | 0 | 1 | 0 | 0 | 0 | 103 | 83 | 52 | 83 | 546 | 0 | 0 | 0 | 1 | 0 | 289 | 149 | 1.000000 | 1 | 1 | 0.981 | 1 | 1 | 1 | 1 | 1 | 1 | 0.977 | 1 | 0.001 |
| A3SS | 440 | ENSG00000161265 | U2AF1L4 | chr19 | - | 35744322 | 35745187 | 35744322 | 35744421 | 35745347 | 35745442 | 64 | 49 | 50 | 62 | 77 | 0 | 0 | 0 | 2 | 0 | 231 | 136 | 101 | 111 | 158 | 0 | 0 | 0 | 0 | 0 | 298 | 149 | 1.000000 | 1 | 1 | 1 | 1 | 0.939 | 1 | 1 | 1 | 1 | 1 | 1 | -0.012 |
| A3SS | 10638 | ENSG00000147140 | NONO | chrX | + | 71297835 | 71297938 | 71297839 | 71297938 | 71297376 | 71297461 | 233 | 154 | 260 | 253 | 222 | 1 | 0 | 0 | 0 | 0 | 498 | 290 | 192 | 235 | 342 | 0 | 1 | 1 | 0 | 5 | 153 | 149 | 1.000000 | 1 | 0.996 | 1 | 1 | 1 | 1 | 1 | 0.996 | 0.995 | 1 | 0.985 | 0.004 |
| A3SS | 8795 | ENSG00000186635 | ARAP1 | chr11 | - | 72712437 | 72713243 | 72712437 | 72712568 | 72714151 | 72714321 | 137 | 84 | 97 | 140 | 33 | 0 | 0 | 0 | 0 | 0 | 218 | 133 | 112 | 168 | 293 | 0 | 1 | 2 | 2 | 0 | 298 | 149 | 1.000000 | 1 | 1 | 1 | 1 | 1 | 1 | 1 | 0.985 | 0.966 | 0.977 | 1 | 0.014 |
| A3SS | 174 | ENSG00000100055 | CYTH4 | chr22 | + | 37311455 | 37312174 | 37312019 | 37312174 | 37310987 | 37311064 | 160 | 84 | 138 | 160 | 84 | 0 | 0 | 0 | 1 | 0 | 251 | 226 | 159 | 165 | 257 | 0 | 0 | 0 | 0 | 0 | 298 | 149 | 1.000000 | 1 | 1 | 1 | 1 | 0.988 | 1 | 1 | 1 | 1 | 1 | 1 | -0.002 |
| A3SS | 2365 | ENSG00000175826 | CTDNEP1 | chr17 | - | 7247063 | 7247182 | 7247063 | 7247177 | 7247276 | 7247343 | 221 | 101 | 197 | 193 | 172 | 3 | 0 | 2 | 2 | 0 | 241 | 263 | 129 | 259 | 423 | 0 | 0 | 2 | 0 | 2 | 154 | 149 | 1.000000 | 1 | 0.986 | 1 | 0.99 | 0.989 | 1 | 1 | 1 | 0.984 | 1 | 0.995 | -0.003 |
| A3SS | 4377 | ENSG00000009790 | TRAF3IP3 | chr1 | + | 209759981 | 209760384 | 209760171 | 209760384 | 209759033 | 209759134 | 767 | 540 | 680 | 506 | 368 | 2 | 1 | 3 | 0 | 0 | 809 | 940 | 574 | 337 | 631 | 0 | 4 | 1 | 3 | 8 | 298 | 149 | 1.000000 | 1 | 0.995 | 0.996 | 0.991 | 1 | 1 | 1 | 0.992 | 0.997 | 0.983 | 0.975 | 0.007 |
| A3SS | 9796 | ENSG00000012124 | CD22 | chr19 | + | 35332018 | 35332924 | 35332652 | 35332924 | 35329186 | 35329230 | 103 | 61 | 93 | 119 | 166 | 1 | 0 | 0 | 1 | 0 | 304 | 278 | 31 | 149 | 210 | 0 | 0 | 3 | 0 | 0 | 298 | 149 | 1.000000 | 1 | 0.981 | 1 | 1 | 0.983 | 1 | 1 | 1 | 0.838 | 1 | 1 | 0.025 |
| A3SS | 8660 | ENSG00000163516 | ANKZF1 | chr2 | + | 219234132 | 219235875 | 219235707 | 219235875 | 219233285 | 219233433 | 120 | 100 | 144 | 115 | 91 | 0 | 2 | 0 | 0 | 0 | 177 | 243 | 196 | 177 | 206 | 0 | 0 | 1 | 4 | 0 | 298 | 149 | 1.000000 | 1 | 1 | 0.962 | 1 | 1 | 1 | 1 | 1 | 0.99 | 0.957 | 1 | 0.003 |
| A3SS | 6767 | ENSG00000158796 | DEDD | chr1 | - | 161124137 | 161124526 | 161124137 | 161124210 | 161130814 | 161130847 | 121 | 58 | 86 | 132 | 87 | 2 | 0 | 0 | 3 | 0 | 240 | 173 | 109 | 162 | 226 | 0 | 0 | 0 | 0 | 0 | 298 | 149 | 1.000000 | 1 | 0.968 | 1 | 1 | 0.957 | 1 | 1 | 1 | 1 | 1 | 1 | -0.015 |
| A3SS | 3996 | ENSG00000162430 | SELENON | chr1 | + | 25809025 | 25809150 | 25809094 | 25809150 | 25808579 | 25808789 | 47 | 17 | 49 | 45 | 17 | 0 | 0 | 1 | 0 | 0 | 119 | 90 | 46 | 74 | 266 | 0 | 0 | 0 | 2 | 0 | 218 | 149 | 1.000000 | 1 | 1 | 1 | 0.971 | 1 | 1 | 1 | 1 | 1 | 0.962 | 1 | 0.002 |
| A3SS | 2166 | ENSG00000103876 | FAH | chr15 | + | 80172148 | 80172248 | 80172161 | 80172248 | 80168263 | 80168316 | 12 | 12 | 11 | 29 | 32 | 1 | 0 | 0 | 0 | 0 | 42 | 32 | 36 | 41 | 362 | 0 | 0 | 1 | 0 | 0 | 162 | 149 | 1.000000 | 1 | 0.917 | 1 | 1 | 1 | 1 | 1 | 1 | 0.971 | 1 | 1 | -0.011 |
| A3SS | 2606 | ENSG00000155465 | SLC7A7 | chr14 | - | 22776194 | 22776318 | 22776194 | 22776232 | 22778792 | 22778937 | 214 | 98 | 119 | 204 | 123 | 0 | 0 | 0 | 0 | 1 | 327 | 220 | 84 | 138 | 406 | 0 | 0 | 0 | 0 | 0 | 235 | 149 | 1.000000 | 1 | 1 | 1 | 1 | 1 | 0.987 | 1 | 1 | 1 | 1 | 1 | -0.003 |
| A3SS | 1358 | ENSG00000092929 | UNC13D | chr17 | - | 75836571 | 75836696 | 75836571 | 75836593 | 75836800 | 75836918 | 188 | 63 | 132 | 182 | 125 | 1 | 0 | 0 | 3 | 0 | 148 | 242 | 109 | 130 | 477 | 0 | 5 | 0 | 0 | 1 | 252 | 149 | 1.000000 | 1 | 0.991 | 1 | 1 | 0.973 | 1 | 1 | 0.966 | 1 | 1 | 0.996 | 0 |
| A3SS | 9094 | ENSG00000169223 | LMAN2 | chr5 | - | 177337705 | 177337785 | 177337705 | 177337764 | 177338487 | 177338605 | 227 | 104 | 202 | 197 | 153 | 0 | 1 | 0 | 0 | 3 | 232 | 319 | 191 | 204 | 352 | 0 | 0 | 0 | 0 | 0 | 170 | 149 | 1.000000 | 1 | 1 | 0.989 | 1 | 1 | 0.978 | 1 | 1 | 1 | 1 | 1 | -0.007 |
| A3SS | 10213 | ENSG00000124222 | STX16 | chr20 | + | 58671153 | 58671297 | 58671163 | 58671297 | 58670511 | 58670603 | 270 | 191 | 226 | 247 | 157 | 0 | 2 | 0 | 0 | 1 | 433 | 255 | 247 | 282 | 287 | 0 | 0 | 0 | 0 | 0 | 159 | 149 | 1.000000 | 1 | 1 | 0.989 | 1 | 1 | 0.993 | 1 | 1 | 1 | 1 | 1 | -0.004 |
| A3SS | 12908 | ENSG00000012822 | CALCOCO1 | chr12 | - | 53723592 | 53723783 | 53723592 | 53723762 | 53724644 | 53724747 | 103 | 65 | 116 | 191 | 146 | 1 | 2 | 0 | 0 | 0 | 215 | 213 | 112 | 92 | 395 | 0 | 0 | 0 | 3 | 8 | 170 | 149 | 1.000000 | 1 | 0.989 | 0.966 | 1 | 1 | 1 | 1 | 1 | 1 | 0.964 | 0.977 | 0.003 |
| A3SS | 1740 | ENSG00000179583 | CIITA | chr16 | + | 10906498 | 10909187 | 10908982 | 10909187 | 10904743 | 10904812 | 90 | 68 | 63 | 65 | 49 | 0 | 0 | 0 | 0 | 0 | 275 | 97 | 30 | 103 | 235 | 0 | 0 | 0 | 0 | 1 | 298 | 149 | 1.000000 | 1 | 1 | 1 | 1 | 1 | 1 | 1 | 1 | 1 | 1 | 0.992 | 0.002 |
| A3SS | 10833 | ENSG00000147416 | ATP6V1B2 | chr8 | + | 20209432 | 20209531 | 20209440 | 20209531 | 20204483 | 20204539 | 567 | 348 | 292 | 360 | 340 | 0 | 0 | 0 | 0 | 0 | 614 | 309 | 297 | 403 | 686 | 1 | 0 | 0 | 0 | 0 | 157 | 149 | 1.000000 | 1 | 1 | 1 | 1 | 1 | 1 | 0.998 | 1 | 1 | 1 | 1 | 0 |
| A3SS | 4927 | ENSG00000004399 | PLXND1 | chr3 | - | 129555213 | 129556428 | 129555213 | 129555543 | 129556616 | 129556691 | 222 | 164 | 146 | 189 | 122 | 0 | 0 | 0 | 0 | 2 | 516 | 171 | 74 | 167 | 316 | 0 | 0 | 0 | 0 | 0 | 298 | 149 | 1.000000 | 1 | 1 | 1 | 1 | 1 | 0.968 | 1 | 1 | 1 | 1 | 1 | -0.006 |
| A3SS | 1072 | ENSG00000154889 | MPPE1 | chr18 | - | 11896983 | 11897356 | 11896983 | 11897131 | 11906202 | 11906309 | 143 | 112 | 128 | 160 | 217 | 0 | 0 | 0 | 0 | 0 | 262 | 232 | 180 | 181 | 306 | 0 | 0 | 0 | 0 | 7 | 298 | 149 | 1.000000 | 1 | 1 | 1 | 1 | 1 | 1 | 1 | 1 | 1 | 1 | 0.956 | 0.009 |
| A3SS | 6979 | ENSG00000117410 | ATP6V0B | chr1 | + | 43977025 | 43978295 | 43977980 | 43978295 | 43976772 | 43976824 | 202 | 49 | 78 | 103 | 78 | 1 | 0 | 0 | 0 | 0 | 198 | 156 | 107 | 139 | 309 | 0 | 0 | 0 | 0 | 0 | 298 | 149 | 1.000000 | 1 | 0.99 | 1 | 1 | 1 | 1 | 1 | 1 | 1 | 1 | 1 | -0.002 |
| A3SS | 6286 | ENSG00000126012 | KDM5C | chrX | - | 53198489 | 53198888 | 53198489 | 53198637 | 53198976 | 53199158 | 112 | 65 | 112 | 104 | 53 | 2 | 0 | 1 | 0 | 0 | 265 | 111 | 114 | 188 | 165 | 0 | 0 | 0 | 0 | 0 | 298 | 149 | 1.000000 | 1 | 0.966 | 1 | 0.982 | 1 | 1 | 1 | 1 | 1 | 1 | 1 | -0.01 |
| A3SS | 1609 | ENSG00000177731 | FLII | chr17 | - | 18254520 | 18254682 | 18254520 | 18254619 | 18254768 | 18254854 | 200 | 103 | 167 | 183 | 163 | 0 | 2 | 1 | 0 | 0 | 177 | 241 | 142 | 204 | 447 | 0 | 0 | 0 | 0 | 0 | 212 | 149 | 1.000000 | 1 | 1 | 0.973 | 0.992 | 1 | 1 | 1 | 1 | 1 | 1 | 1 | -0.007 |
| A3SS | 368 | ENSG00000104774 | MAN2B1 | chr19 | - | 12655693 | 12655879 | 12655693 | 12655862 | 12656570 | 12656687 | 287 | 177 | 190 | 209 | 222 | 1 | 0 | 0 | 0 | 0 | 405 | 264 | 151 | 220 | 439 | 0 | 0 | 0 | 0 | 0 | 166 | 149 | 1.000000 | 1 | 0.996 | 1 | 1 | 1 | 1 | 1 | 1 | 1 | 1 | 1 | -0.001 |
| A3SS | 3533 | ENSG00000131503 | ANKHD1 | chr5 | + | 140438460 | 140438617 | 140438493 | 140438617 | 140436103 | 140436257 | 423 | 355 | 351 | 384 | 497 | 3 | 4 | 3 | 3 | 1 | 571 | 454 | 611 | 464 | 303 | 0 | 0 | 7 | 7 | 0 | 182 | 149 | 1.000000 | 1 | 0.991 | 0.986 | 0.99 | 0.991 | 0.998 | 1 | 1 | 0.986 | 0.982 | 1 | -0.002 |
| A3SS | 5995 | ENSG00000110619 | CARS1 | chr11 | - | 3001969 | 3002600 | 3001969 | 3002174 | 3005365 | 3005433 | 30 | 44 | 33 | 47 | 58 | 1 | 0 | 0 | 0 | 0 | 223 | 46 | 81 | 99 | 152 | 0 | 0 | 0 | 0 | 0 | 298 | 149 | 1.000000 | 1 | 0.938 | 1 | 1 | 1 | 1 | 1 | 1 | 1 | 1 | 1 | -0.012 |
| A3SS | 6876 | ENSG00000117713 | ARID1A | chr1 | + | 26774328 | 26775220 | 26774979 | 26775220 | 26773801 | 26773898 | 154 | 102 | 171 | 129 | 101 | 0 | 0 | 2 | 0 | 0 | 210 | 219 | 146 | 178 | 292 | 0 | 0 | 0 | 0 | 0 | 298 | 149 | 1.000000 | 1 | 1 | 1 | 0.977 | 1 | 1 | 1 | 1 | 1 | 1 | 1 | -0.005 |
| A3SS | 4944 | ENSG00000197948 | FCHSD1 | chr5 | - | 141646086 | 141646722 | 141646086 | 141646191 | 141647134 | 141647230 | 95 | 66 | 109 | 117 | 107 | 0 | 0 | 0 | 2 | 0 | 230 | 193 | 106 | 117 | 231 | 0 | 0 | 0 | 0 | 0 | 298 | 149 | 1.000000 | 1 | 1 | 1 | 1 | 0.967 | 1 | 1 | 1 | 1 | 1 | 1 | -0.007 |
| A3SS | 12649 | ENSG00000107862 | GBF1 | chr10 | + | 102368217 | 102368454 | 102368340 | 102368454 | 102367477 | 102367560 | 231 | 129 | 204 | 176 | 241 | 1 | 0 | 0 | 0 | 0 | 379 | 242 | 163 | 246 | 327 | 0 | 0 | 0 | 0 | 0 | 272 | 149 | 1.000000 | 1 | 0.992 | 1 | 1 | 1 | 1 | 1 | 1 | 1 | 1 | 1 | -0.002 |
| A3SS | 485 | ENSG00000013275 | PSMC4 | chr19 | + | 39980069 | 39980454 | 39980285 | 39980454 | 39979816 | 39979984 | 139 | 70 | 112 | 140 | 66 | 0 | 0 | 0 | 0 | 0 | 281 | 155 | 91 | 139 | 235 | 0 | 0 | 2 | 0 | 0 | 298 | 149 | 1.000000 | 1 | 1 | 1 | 1 | 1 | 1 | 1 | 1 | 0.958 | 1 | 1 | 0.008 |
| A3SS | 11421 | ENSG00000148337 | CIZ1 | chr9 | - | 128178708 | 128179415 | 128178708 | 128178904 | 128180414 | 128180523 | 96 | 53 | 114 | 100 | 43 | 0 | 0 | 1 | 0 | 0 | 124 | 135 | 92 | 182 | 241 | 0 | 0 | 0 | 0 | 0 | 298 | 149 | 1.000000 | 1 | 1 | 1 | 0.983 | 1 | 1 | 1 | 1 | 1 | 1 | 1 | -0.003 |
| A3SS | 7217 | ENSG00000135899 | SP110 | chr2 | - | 230202578 | 230202728 | 230202578 | 230202722 | 230207990 | 230208059 | 79 | 95 | 72 | 120 | 97 | 1 | 0 | 1 | 0 | 0 | 261 | 112 | 113 | 219 | 122 | 0 | 0 | 0 | 0 | 0 | 155 | 149 | 1.000000 | 1 | 0.987 | 1 | 0.986 | 1 | 1 | 1 | 1 | 1 | 1 | 1 | -0.005 |
| A3SS | 8544 | ENSG00000108883 | EFTUD2 | chr17 | - | 44883092 | 44883158 | 44883092 | 44883128 | 44883648 | 44883724 | 284 | 175 | 165 | 247 | 436 | 3 | 1 | 0 | 0 | 0 | 377 | 300 | 208 | 267 | 516 | 0 | 0 | 0 | 4 | 2 | 179 | 149 | 1.000000 | 1 | 0.987 | 0.993 | 1 | 1 | 1 | 1 | 1 | 1 | 0.982 | 0.995 | 0.001 |
| A3SS | 366 | ENSG00000130816 | DNMT1 | chr19 | - | 10180777 | 10180885 | 10180777 | 10180836 | 10182040 | 10182077 | 202 | 143 | 141 | 190 | 307 | 0 | 0 | 0 | 0 | 0 | 426 | 186 | 114 | 299 | 317 | 0 | 0 | 0 | 2 | 2 | 198 | 149 | 1.000000 | 1 | 1 | 1 | 1 | 1 | 1 | 1 | 1 | 1 | 0.991 | 0.992 | 0.003 |
| A3SS | 4044 | ENSG00000100412 | ACO2 | chr22 | + | 41522829 | 41523278 | 41523204 | 41523278 | 41520170 | 41520276 | 202 | 117 | 210 | 190 | 247 | 0 | 0 | 1 | 0 | 0 | 371 | 328 | 144 | 202 | 278 | 0 | 2 | 0 | 0 | 0 | 298 | 149 | 1.000000 | 1 | 1 | 1 | 0.991 | 1 | 1 | 1 | 0.988 | 1 | 1 | 1 | 0.001 |
| A3SS | 13241 | ENSG00000101160 | CTSZ | chr20 | - | 59006321 | 59006485 | 59006321 | 59006450 | 59006985 | 59007230 | 50 | 24 | 31 | 57 | 27 | 2 | 1 | 1 | 0 | 0 | 156 | 117 | 24 | 52 | 197 | 4 | 9 | 0 | 5 | 7 | 184 | 149 | 1.000000 | 1 | 0.953 | 0.951 | 0.962 | 1 | 1 | 0.969 | 0.913 | 1 | 0.894 | 0.958 | 0.026 |
| A3SS | 7736 | ENSG00000115525 | ST3GAL5 | chr2 | - | 85847860 | 85848381 | 85847860 | 85847984 | 85861180 | 85861292 | 94 | 97 | 106 | 151 | 121 | 0 | 0 | 0 | 0 | 0 | 358 | 189 | 105 | 126 | 146 | 6 | 0 | 0 | 0 | 0 | 298 | 149 | 1.000000 | 1 | 1 | 1 | 1 | 1 | 1 | 0.968 | 1 | 1 | 1 | 1 | 0.006 |
| A3SS | 10559 | ENSG00000102032 | RENBP | chrX | - | 153942854 | 153943079 | 153942854 | 153943034 | 153943545 | 153943718 | 75 | 32 | 40 | 65 | 33 | 2 | 0 | 0 | 3 | 0 | 174 | 77 | 26 | 116 | 204 | 0 | 0 | 0 | 0 | 0 | 194 | 149 | 1.000000 | 1 | 0.966 | 1 | 1 | 0.943 | 1 | 1 | 1 | 1 | 1 | 1 | -0.018 |
| A3SS | 9311 | ENSG00000073050 | XRCC1 | chr19 | - | 43552796 | 43553091 | 43552796 | 43552843 | 43553400 | 43553512 | 81 | 59 | 78 | 64 | 127 | 1 | 0 | 0 | 1 | 0 | 203 | 108 | 63 | 166 | 220 | 0 | 0 | 0 | 0 | 0 | 298 | 149 | 1.000000 | 1 | 0.976 | 1 | 1 | 0.97 | 1 | 1 | 1 | 1 | 1 | 1 | -0.011 |
| A3SS | 5549 | ENSG00000142208 | AKT1 | chr14 | - | 104770744 | 104772452 | 104770744 | 104770847 | 104772877 | 104773092 | 196 | 64 | 150 | 196 | 126 | 0 | 0 | 1 | 0 | 0 | 246 | 245 | 137 | 129 | 326 | 0 | 0 | 0 | 0 | 0 | 298 | 149 | 1.000000 | 1 | 1 | 1 | 0.987 | 1 | 1 | 1 | 1 | 1 | 1 | 1 | -0.003 |
| A3SS | 10763 | ENSG00000102119 | EMD | chrX | + | 154380233 | 154380367 | 154380276 | 154380367 | 154379941 | 154380019 | 103 | 63 | 67 | 83 | 76 | 0 | 0 | 2 | 1 | 0 | 116 | 146 | 74 | 137 | 268 | 0 | 2 | 0 | 0 | 4 | 192 | 149 | 1.000000 | 1 | 1 | 1 | 0.963 | 0.985 | 1 | 1 | 0.983 | 1 | 1 | 0.981 | -0.003 |
| A3SS | 4617 | ENSG00000143554 | SLC27A3 | chr1 | + | 153778686 | 153778885 | 153778845 | 153778885 | 153778462 | 153778553 | 88 | 45 | 58 | 83 | 96 | 2 | 0 | 0 | 0 | 0 | 285 | 127 | 60 | 96 | 148 | 0 | 0 | 0 | 0 | 0 | 298 | 149 | 1.000000 | 1 | 0.957 | 1 | 1 | 1 | 1 | 1 | 1 | 1 | 1 | 1 | -0.009 |
| A3SS | 3309 | ENSG00000143774 | GUK1 | chr1 | + | 228147614 | 228147699 | 228147620 | 228147699 | 228147405 | 228147544 | 187 | 119 | 155 | 179 | 93 | 1 | 0 | 1 | 0 | 2 | 172 | 201 | 182 | 168 | 355 | 0 | 0 | 1 | 0 | 0 | 155 | 149 | 1.000000 | 1 | 0.994 | 1 | 0.993 | 1 | 0.978 | 1 | 1 | 0.994 | 1 | 1 | -0.006 |
| A3SS | 10105 | ENSG00000127824 | TUBA4A | chr2 | - | 219251193 | 219251713 | 219251193 | 219251323 | 219252007 | 219252230 | 50 | 46 | 49 | 86 | 73 | 0 | 0 | 1 | 3 | 0 | 116 | 124 | 90 | 51 | 268 | 0 | 0 | 0 | 0 | 12 | 298 | 149 | 1.000000 | 1 | 1 | 1 | 0.961 | 0.935 | 1 | 1 | 1 | 1 | 1 | 0.918 | -0.004 |
| A3SS | 8020 | ENSG00000166164 | BRD7 | chr16 | - | 50354424 | 50354482 | 50354424 | 50354479 | 50354792 | 50354922 | 183 | 166 | 166 | 205 | 238 | 3 | 1 | 0 | 0 | 5 | 320 | 154 | 368 | 322 | 138 | 0 | 2 | 1 | 1 | 0 | 152 | 149 | 1.000000 | 1 | 0.984 | 0.994 | 1 | 1 | 0.979 | 1 | 0.987 | 0.997 | 0.997 | 1 | -0.005 |
| A3SS | 4343 | ENSG00000188313 | PLSCR1 | chr3 | - | 146521543 | 146522053 | 146521543 | 146521705 | 146525604 | 146525647 | 231 | 232 | 151 | 279 | 118 | 2 | 0 | 0 | 2 | 0 | 272 | 265 | 229 | 506 | 81 | 0 | 3 | 0 | 4 | 0 | 298 | 149 | 1.000000 | 1 | 0.983 | 1 | 1 | 0.986 | 1 | 1 | 0.978 | 1 | 0.984 | 1 | 0.001 |
| A3SS | 6220 | ENSG00000120709 | FAM53C | chr5 | + | 138344824 | 138345609 | 138345395 | 138345609 | 138341808 | 138341866 | 97 | 44 | 60 | 95 | 68 | 1 | 0 | 0 | 2 | 0 | 166 | 138 | 155 | 96 | 150 | 0 | 0 | 0 | 0 | 0 | 298 | 149 | 1.000000 | 1 | 0.98 | 1 | 1 | 0.96 | 1 | 1 | 1 | 1 | 1 | 1 | -0.012 |
| A3SS | 4465 | ENSG00000204427 | ABHD16A | chr6 | - | 31688040 | 31688305 | 31688040 | 31688103 | 31688722 | 31688786 | 58 | 38 | 59 | 77 | 9 | 0 | 0 | 0 | 0 | 0 | 104 | 96 | 55 | 64 | 261 | 0 | 0 | 0 | 4 | 0 | 298 | 149 | 1.000000 | 1 | 1 | 1 | 1 | 1 | 1 | 1 | 1 | 1 | 0.889 | 1 | 0.022 |
| A3SS | 11302 | ENSG00000077721 | UBE2A | chrX | + | 119582587 | 119582676 | 119582596 | 119582676 | 119581506 | 119581596 | 194 | 119 | 142 | 173 | 266 | 0 | 1 | 0 | 0 | 0 | 363 | 202 | 154 | 248 | 263 | 0 | 0 | 0 | 0 | 0 | 158 | 149 | 1.000000 | 1 | 1 | 0.991 | 1 | 1 | 1 | 1 | 1 | 1 | 1 | 1 | -0.002 |
| A3SS | 12003 | ENSG00000149269 | PAK1 | chr11 | - | 77322016 | 77323360 | 77322016 | 77323154 | 77332729 | 77332867 | 239 | 226 | 198 | 249 | 423 | 0 | 1 | 0 | 0 | 0 | 516 | 176 | 201 | 310 | 467 | 0 | 0 | 0 | 4 | 0 | 298 | 149 | 1.000000 | 1 | 1 | 0.991 | 1 | 1 | 1 | 1 | 1 | 1 | 0.975 | 1 | 0.003 |
| A3SS | 12648 | ENSG00000107862 | GBF1 | chr10 | + | 102368217 | 102368454 | 102368297 | 102368454 | 102367477 | 102367560 | 194 | 104 | 160 | 154 | 200 | 0 | 0 | 0 | 0 | 0 | 331 | 208 | 143 | 213 | 252 | 0 | 8 | 0 | 0 | 0 | 229 | 149 | 1.000000 | 1 | 1 | 1 | 1 | 1 | 1 | 1 | 0.944 | 1 | 1 | 1 | 0.011 |
| A3SS | 10037 | ENSG00000065000 | AP3D1 | chr19 | - | 2109085 | 2109207 | 2109085 | 2109125 | 2109872 | 2109958 | 149 | 84 | 170 | 181 | 89 | 0 | 2 | 0 | 0 | 0 | 223 | 194 | 95 | 160 | 335 | 0 | 0 | 1 | 6 | 0 | 231 | 149 | 1.000000 | 1 | 1 | 0.964 | 1 | 1 | 1 | 1 | 1 | 0.984 | 0.945 | 1 | 0.007 |
| A3SS | 1674 | ENSG00000140829 | DHX38 | chr16 | + | 72110955 | 72111077 | 72111042 | 72111077 | 72109414 | 72109510 | 165 | 76 | 173 | 211 | 62 | 0 | 0 | 2 | 1 | 0 | 268 | 206 | 158 | 140 | 248 | 0 | 0 | 0 | 0 | 5 | 236 | 149 | 1.000000 | 1 | 1 | 1 | 0.982 | 0.993 | 1 | 1 | 1 | 1 | 1 | 0.969 | 0.001 |
| A3SS | 8023 | ENSG00000125875 | TBC1D20 | chr20 | - | 439107 | 439295 | 439107 | 439220 | 440247 | 440389 | 166 | 84 | 135 | 196 | 161 | 1 | 0 | 0 | 0 | 0 | 218 | 221 | 88 | 164 | 382 | 0 | 0 | 0 | 0 | 0 | 224 | 149 | 1.000000 | 1 | 0.991 | 1 | 1 | 1 | 1 | 1 | 1 | 1 | 1 | 1 | -0.002 |
| A3SS | 1465 | ENSG00000141582 | CBX4 | chr17 | - | 79835645 | 79835712 | 79835645 | 79835696 | 79837836 | 79837902 | 100 | 58 | 82 | 87 | 83 | 0 | 0 | 0 | 0 | 0 | 118 | 162 | 89 | 167 | 203 | 0 | 0 | 0 | 0 | 4 | 165 | 149 | 1.000000 | 1 | 1 | 1 | 1 | 1 | 1 | 1 | 1 | 1 | 1 | 0.979 | 0.004 |
| A3SS | 3030 | ENSG00000079616 | KIF22 | chr16 | + | 29798592 | 29799184 | 29798974 | 29799184 | 29798373 | 29798501 | 100 | 47 | 112 | 83 | 75 | 0 | 0 | 0 | 0 | 0 | 236 | 95 | 103 | 120 | 191 | 0 | 0 | 4 | 0 | 0 | 298 | 149 | 1.000000 | 1 | 1 | 1 | 1 | 1 | 1 | 1 | 1 | 0.928 | 1 | 1 | 0.014 |
| A3SS | 12153 | ENSG00000162236 | STX5 | chr11 | - | 62831018 | 62831262 | 62831018 | 62831110 | 62831953 | 62832042 | 123 | 68 | 127 | 138 | 63 | 1 | 0 | 1 | 0 | 0 | 204 | 201 | 117 | 178 | 142 | 0 | 1 | 3 | 0 | 0 | 298 | 149 | 1.000000 | 1 | 0.984 | 1 | 0.984 | 1 | 1 | 1 | 0.99 | 0.951 | 1 | 1 | 0.005 |
| A3SS | 1607 | ENSG00000177731 | FLII | chr17 | - | 18244835 | 18245419 | 18244835 | 18245272 | 18245554 | 18245660 | 189 | 111 | 129 | 173 | 192 | 0 | 0 | 0 | 1 | 0 | 251 | 203 | 121 | 156 | 385 | 0 | 0 | 0 | 0 | 0 | 296 | 149 | 1.000000 | 1 | 1 | 1 | 1 | 0.989 | 1 | 1 | 1 | 1 | 1 | 1 | -0.002 |
| A3SS | 2687 | ENSG00000100911 | PSME2 | chr14 | - | 24145709 | 24145896 | 24145709 | 24145752 | 24146207 | 24146240 | 152 | 119 | 126 | 176 | 216 | 0 | 0 | 1 | 0 | 0 | 382 | 175 | 70 | 208 | 275 | 0 | 0 | 0 | 2 | 0 | 293 | 149 | 1.000000 | 1 | 1 | 1 | 0.985 | 1 | 1 | 1 | 1 | 1 | 0.981 | 1 | 0.001 |
| A3SS | 4040 | ENSG00000100412 | ACO2 | chr22 | + | 41517526 | 41517631 | 41517596 | 41517631 | 41515766 | 41515917 | 171 | 102 | 133 | 148 | 133 | 0 | 0 | 3 | 0 | 0 | 156 | 345 | 112 | 119 | 274 | 0 | 2 | 0 | 2 | 0 | 219 | 149 | 1.000000 | 1 | 1 | 1 | 0.968 | 1 | 1 | 1 | 0.992 | 1 | 0.976 | 1 | 0 |
| A3SS | 6823 | ENSG00000179912 | R3HDM2 | chr12 | - | 57310263 | 57310463 | 57310263 | 57310460 | 57395748 | 57395818 | 164 | 111 | 167 | 166 | 127 | 1 | 0 | 0 | 0 | 0 | 282 | 173 | 162 | 215 | 220 | 0 | 0 | 0 | 0 | 0 | 152 | 149 | 1.000000 | 1 | 0.994 | 1 | 1 | 1 | 1 | 1 | 1 | 1 | 1 | 1 | -0.001 |
| A3SS | 9795 | ENSG00000012124 | CD22 | chr19 | + | 35332018 | 35332924 | 35332593 | 35332924 | 35329186 | 35329230 | 80 | 52 | 65 | 102 | 165 | 0 | 0 | 1 | 0 | 0 | 225 | 233 | 15 | 139 | 165 | 0 | 0 | 0 | 0 | 0 | 298 | 149 | 1.000000 | 1 | 1 | 1 | 0.97 | 1 | 1 | 1 | 1 | 1 | 1 | 1 | -0.006 |
| A3SS | 7735 | ENSG00000115525 | ST3GAL5 | chr2 | - | 85847860 | 85848204 | 85847860 | 85847984 | 85861180 | 85861292 | 178 | 195 | 189 | 234 | 309 | 0 | 0 | 0 | 0 | 0 | 571 | 232 | 152 | 200 | 263 | 6 | 0 | 0 | 0 | 0 | 298 | 149 | 1.000000 | 1 | 1 | 1 | 1 | 1 | 1 | 0.979 | 1 | 1 | 1 | 1 | 0.004 |
| A3SS | 12639 | ENSG00000107862 | GBF1 | chr10 | + | 102363709 | 102363798 | 102363760 | 102363798 | 102363255 | 102363396 | 167 | 115 | 101 | 176 | 157 | 0 | 0 | 0 | 0 | 0 | 253 | 172 | 97 | 196 | 311 | 7 | 0 | 0 | 3 | 0 | 200 | 149 | 1.000000 | 1 | 1 | 1 | 1 | 1 | 1 | 0.964 | 1 | 1 | 0.98 | 1 | 0.011 |
| A3SS | 4318 | ENSG00000172053 | QARS1 | chr3 | - | 49104323 | 49104471 | 49104323 | 49104465 | 49104616 | 49104752 | 60 | 10 | 49 | 55 | 3 | 0 | 0 | 0 | 1 | 0 | 117 | 64 | 64 | 70 | 174 | 0 | 0 | 0 | 0 | 0 | 155 | 149 | 1.000000 | 1 | 1 | 1 | 1 | 0.981 | 1 | 1 | 1 | 1 | 1 | 1 | -0.004 |
| A3SS | 1419 | ENSG00000179094 | PER1 | chr17 | - | 8144750 | 8144993 | 8144750 | 8144847 | 8145957 | 8146137 | 141 | 54 | 116 | 67 | 66 | 3 | 0 | 1 | 0 | 0 | 81 | 150 | 352 | 79 | 92 | 0 | 0 | 4 | 0 | 0 | 295 | 149 | 1.000000 | 1 | 0.96 | 1 | 0.983 | 1 | 1 | 1 | 1 | 0.978 | 1 | 1 | -0.007 |
| A3SS | 5094 | ENSG00000116977 | LGALS8 | chr1 | + | 236538878 | 236539089 | 236538885 | 236539089 | 236537496 | 236537585 | 339 | 305 | 284 | 342 | 270 | 0 | 1 | 0 | 0 | 4 | 282 | 717 | 378 | 343 | 130 | 0 | 0 | 0 | 0 | 0 | 156 | 149 | 1.000000 | 1 | 1 | 0.997 | 1 | 1 | 0.985 | 1 | 1 | 1 | 1 | 1 | -0.004 |
| A3SS | 4600 | ENSG00000051382 | PIK3CB | chr3 | - | 138742557 | 138742781 | 138742557 | 138742755 | 138755753 | 138755979 | 104 | 138 | 137 | 156 | 226 | 0 | 1 | 2 | 1 | 0 | 137 | 210 | 169 | 281 | 270 | 0 | 0 | 0 | 0 | 5 | 175 | 149 | 1.000000 | 1 | 1 | 0.992 | 0.983 | 0.993 | 1 | 1 | 1 | 1 | 1 | 0.979 | -0.002 |
| A3SS | 10809 | ENSG00000142046 | TMEM91 | chr19 | + | 41383560 | 41384083 | 41383885 | 41384083 | 41382771 | 41382921 | 24 | 15 | 17 | 34 | 7 | 1 | 1 | 0 | 1 | 0 | 66 | 41 | 43 | 24 | 229 | 18 | 3 | 5 | 0 | 24 | 298 | 149 | 1.000000 | 1 | 0.923 | 0.882 | 1 | 0.944 | 1 | 0.647 | 0.872 | 0.811 | 1 | 0.827 | 0.118 |
| A3SS | 412 | ENSG00000104872 | PIH1D1 | chr19 | - | 49447337 | 49447467 | 49447337 | 49447417 | 49447826 | 49447908 | 89 | 58 | 94 | 93 | 78 | 1 | 0 | 3 | 2 | 0 | 142 | 169 | 88 | 113 | 205 | 0 | 0 | 0 | 4 | 0 | 199 | 149 | 1.000000 | 1 | 0.985 | 1 | 0.959 | 0.972 | 1 | 1 | 1 | 1 | 0.955 | 1 | -0.008 |
| A3SS | 7668 | ENSG00000110446 | SLC15A3 | chr11 | - | 60942034 | 60942145 | 60942034 | 60942139 | 60943688 | 60943836 | 78 | 41 | 59 | 48 | 31 | 0 | 0 | 0 | 0 | 0 | 143 | 127 | 54 | 98 | 140 | 0 | 0 | 0 | 1 | 0 | 155 | 149 | 1.000000 | 1 | 1 | 1 | 1 | 1 | 1 | 1 | 1 | 1 | 0.989 | 1 | 0.002 |
| A3SS | 413 | ENSG00000104872 | PIH1D1 | chr19 | - | 49447337 | 49447467 | 49447337 | 49447420 | 49447826 | 49447908 | 89 | 58 | 93 | 93 | 78 | 1 | 2 | 2 | 0 | 0 | 142 | 169 | 88 | 111 | 205 | 0 | 0 | 0 | 0 | 0 | 196 | 149 | 1.000000 | 1 | 0.985 | 0.957 | 0.972 | 1 | 1 | 1 | 1 | 1 | 1 | 1 | -0.017 |
| A3SS | 11695 | ENSG00000141551 | CSNK1D | chr17 | - | 82252433 | 82252604 | 82252433 | 82252527 | 82265685 | 82265796 | 76 | 49 | 71 | 91 | 124 | 0 | 0 | 0 | 0 | 0 | 191 | 88 | 160 | 149 | 126 | 0 | 0 | 2 | 0 | 0 | 226 | 149 | 1.000000 | 1 | 1 | 1 | 1 | 1 | 1 | 1 | 1 | 0.981 | 1 | 1 | 0.004 |
| A3SS | 1455 | ENSG00000087157 | PGS1 | chr17 | + | 78400676 | 78400855 | 78400734 | 78400855 | 78399347 | 78399537 | 54 | 15 | 49 | 50 | 23 | 0 | 0 | 1 | 0 | 0 | 120 | 79 | 91 | 88 | 111 | 0 | 0 | 0 | 0 | 5 | 207 | 149 | 1.000000 | 1 | 1 | 1 | 0.972 | 1 | 1 | 1 | 1 | 1 | 1 | 0.941 | 0.006 |
| A3SS | 9798 | ENSG00000012124 | CD22 | chr19 | + | 35332924 | 35332924 | 35332652 | 35332924 | 35329186 | 35329230 | 58 | 47 | 63 | 78 | 134 | 1 | 0 | 0 | 1 | 0 | 202 | 195 | 19 | 116 | 146 | 0 | 0 | 3 | 0 | 0 | 255 | 149 | 1.000000 | 1 | 0.971 | 1 | 1 | 0.979 | 1 | 1 | 1 | 0.787 | 1 | 1 | 0.033 |
| A3SS | 4996 | ENSG00000132716 | DCAF8 | chr1 | - | 160261284 | 160261358 | 160261284 | 160261325 | 160262116 | 160262451 | 94 | 77 | 70 | 90 | 75 | 2 | 0 | 0 | 0 | 0 | 192 | 135 | 101 | 132 | 141 | 0 | 0 | 0 | 0 | 0 | 182 | 149 | 1.000000 | 1 | 0.975 | 1 | 1 | 1 | 1 | 1 | 1 | 1 | 1 | 1 | -0.005 |
| A3SS | 8187 | ENSG00000084623 | EIF3I | chr1 | + | 32226170 | 32226530 | 32226402 | 32226530 | 32224409 | 32224475 | 184 | 110 | 200 | 189 | 107 | 0 | 0 | 1 | 2 | 0 | 206 | 215 | 135 | 149 | 380 | 0 | 0 | 1 | 0 | 6 | 298 | 149 | 1.000000 | 1 | 1 | 1 | 0.99 | 0.979 | 1 | 1 | 1 | 0.985 | 1 | 0.969 | 0.003 |
| A3SS | 9035 | ENSG00000137959 | IFI44L | chr1 | + | 78641434 | 78641609 | 78641504 | 78641609 | 78641020 | 78641121 | 67 | 190 | 58 | 224 | 86 | 0 | 0 | 3 | 1 | 0 | 95 | 374 | 130 | 295 | 22 | 0 | 0 | 0 | 1 | 0 | 219 | 149 | 1.000000 | 1 | 1 | 1 | 0.929 | 0.993 | 1 | 1 | 1 | 1 | 0.995 | 1 | -0.015 |
| A3SS | 5119 | ENSG00000055332 | EIF2AK2 | chr2 | - | 37148856 | 37149023 | 37148856 | 37148939 | 37157366 | 37157522 | 38 | 86 | 50 | 83 | 71 | 0 | 0 | 0 | 0 | 0 | 105 | 124 | 105 | 255 | 30 | 0 | 0 | 2 | 0 | 0 | 233 | 149 | 1.000000 | 1 | 1 | 1 | 1 | 1 | 1 | 1 | 1 | 0.971 | 1 | 1 | 0.006 |
| A3SS | 13047 | ENSG00000123329 | ARHGAP9 | chr12 | - | 57479090 | 57479424 | 57479090 | 57479165 | 57479729 | 57479866 | 124 | 57 | 65 | 145 | 29 | 3 | 1 | 0 | 0 | 0 | 195 | 95 | 115 | 110 | 195 | 0 | 0 | 0 | 0 | 0 | 298 | 149 | 1.000000 | 1 | 0.954 | 0.966 | 1 | 1 | 1 | 1 | 1 | 1 | 1 | 1 | -0.016 |
| A3SS | 5818 | ENSG00000160058 | BSDC1 | chr1 | - | 32383829 | 32384048 | 32383829 | 32383906 | 32386778 | 32386895 | 91 | 95 | 96 | 136 | 108 | 0 | 2 | 1 | 0 | 0 | 158 | 119 | 141 | 152 | 246 | 0 | 0 | 1 | 0 | 5 | 291 | 149 | 1.000000 | 1 | 1 | 0.961 | 0.98 | 1 | 1 | 1 | 1 | 0.986 | 1 | 0.962 | -0.001 |
| A3SS | 13048 | ENSG00000123329 | ARHGAP9 | chr12 | - | 57479090 | 57479424 | 57479090 | 57479165 | 57480777 | 57480834 | 105 | 55 | 59 | 135 | 23 | 0 | 0 | 0 | 1 | 0 | 188 | 90 | 102 | 108 | 177 | 0 | 0 | 0 | 0 | 0 | 298 | 149 | 1.000000 | 1 | 1 | 1 | 1 | 0.985 | 1 | 1 | 1 | 1 | 1 | 1 | -0.003 |
| A3SS | 10808 | ENSG00000142046 | TMEM91 | chr19 | + | 41383560 | 41384083 | 41383873 | 41384083 | 41382771 | 41382921 | 25 | 20 | 17 | 38 | 14 | 0 | 0 | 0 | 2 | 0 | 66 | 46 | 40 | 26 | 222 | 0 | 0 | 1 | 0 | 0 | 298 | 149 | 1.000000 | 1 | 1 | 1 | 1 | 0.905 | 1 | 1 | 1 | 0.952 | 1 | 1 | -0.009 |
| A3SS | 11970 | ENSG00000168067 | MAP4K2 | chr11 | - | 64797103 | 64797192 | 64797103 | 64797168 | 64797274 | 64797380 | 43 | 30 | 52 | 55 | 50 | 3 | 0 | 1 | 1 | 0 | 75 | 77 | 25 | 74 | 263 | 0 | 2 | 1 | 4 | 4 | 173 | 149 | 1.000000 | 1 | 0.925 | 1 | 0.978 | 0.979 | 1 | 1 | 0.971 | 0.956 | 0.941 | 0.983 | 0.006 |
| A3SS | 11451 | ENSG00000132740 | IGHMBP2 | chr11 | + | 68929733 | 68930434 | 68930357 | 68930434 | 68929182 | 68929357 | 30 | 23 | 18 | 21 | 11 | 0 | 2 | 0 | 1 | 0 | 83 | 61 | 55 | 83 | 102 | 0 | 0 | 0 | 0 | 0 | 298 | 149 | 1.000000 | 1 | 1 | 0.852 | 1 | 0.913 | 1 | 1 | 1 | 1 | 1 | 1 | -0.047 |
| A3SS | 12735 | ENSG00000130640 | TUBGCP2 | chr10 | - | 133299984 | 133300113 | 133299984 | 133300100 | 133302791 | 133302980 | 115 | 44 | 103 | 109 | 132 | 0 | 3 | 0 | 1 | 0 | 198 | 107 | 117 | 99 | 262 | 0 | 0 | 3 | 4 | 0 | 162 | 149 | 1.000000 | 1 | 1 | 0.931 | 1 | 0.99 | 1 | 1 | 1 | 0.973 | 0.958 | 1 | -0.002 |
| A3SS | 11453 | ENSG00000132740 | IGHMBP2 | chr11 | + | 68929747 | 68930434 | 68930357 | 68930434 | 68929182 | 68929357 | 26 | 20 | 18 | 21 | 11 | 0 | 2 | 0 | 1 | 0 | 83 | 60 | 49 | 81 | 102 | 0 | 0 | 0 | 0 | 0 | 298 | 149 | 1.000000 | 1 | 1 | 0.833 | 1 | 0.913 | 1 | 1 | 1 | 1 | 1 | 1 | -0.051 |
| A3SS | 1489 | ENSG00000265148 | TSPOAP1-AS1 | chr17 | + | 58351734 | 58351914 | 58351743 | 58351914 | 58337447 | 58337679 | 60 | 45 | 63 | 84 | 85 | 0 | 1 | 0 | 1 | 0 | 101 | 72 | 40 | 74 | 329 | 0 | 0 | 0 | 0 | 0 | 158 | 149 | 1.000000 | 1 | 1 | 1 | 0.983 | 1 | 1 | 1 | 1 | 1 | 1 | 1 | -0.003 |
| A3SS | 12564 | ENSG00000177239 | MAN1B1 | chr9 | + | 137088868 | 137089005 | 137088952 | 137089005 | 137088074 | 137088183 | 250 | 158 | 231 | 138 | 198 | 0 | 0 | 0 | 0 | 0 | 325 | 272 | 185 | 138 | 334 | 0 | 0 | 2 | 0 | 0 | 233 | 149 | 1.000000 | 1 | 1 | 1 | 1 | 1 | 1 | 1 | 1 | 0.983 | 1 | 1 | 0.003 |
| A3SS | 9611 | ENSG00000106245 | BUD31 | chr7 | + | 99410033 | 99410169 | 99410036 | 99410169 | 99409084 | 99409245 | 22 | 11 | 19 | 35 | 22 | 0 | 0 | 0 | 2 | 0 | 106 | 43 | 62 | 74 | 100 | 0 | 0 | 2 | 0 | 4 | 152 | 149 | 1.000000 | 1 | 1 | 1 | 1 | 0.945 | 1 | 1 | 1 | 0.968 | 1 | 0.961 | 0.003 |
| A3SS | 3467 | ENSG00000011243 | AKAP8L | chr19 | - | 15401149 | 15403715 | 15401149 | 15401603 | 15404009 | 15404042 | 171 | 134 | 226 | 237 | 257 | 0 | 0 | 0 | 0 | 0 | 299 | 242 | 175 | 212 | 370 | 0 | 3 | 0 | 0 | 0 | 298 | 149 | 1.000000 | 1 | 1 | 1 | 1 | 1 | 1 | 1 | 0.976 | 1 | 1 | 1 | 0.005 |
| A3SS | 10640 | ENSG00000147140 | NONO | chrX | + | 71298468 | 71298508 | 71298477 | 71298508 | 71297835 | 71297938 | 282 | 148 | 279 | 294 | 267 | 0 | 0 | 0 | 0 | 0 | 396 | 287 | 224 | 249 | 386 | 0 | 0 | 0 | 0 | 0 | 158 | 149 | 1.000000 | 1 | 1 | 0.979 | 1 | 1 | 1 | 1 | 1 | 1 | 1 | 1 | -0.004 |
| A3SS | 7734 | ENSG00000115525 | ST3GAL5 | chr2 | - | 85847860 | 85848084 | 85847860 | 85847984 | 85861180 | 85861292 | 56 | 56 | 77 | 90 | 62 | 0 | 3 | 0 | 0 | 0 | 285 | 113 | 70 | 62 | 83 | 6 | 0 | 0 | 0 | 0 | 249 | 149 | 1.000000 | 1 | 1 | 1 | 1 | 1 | 1 | 0.966 | 1 | 1 | 1 | 1 | 0.007 |
| A3SS | 8689 | ENSG00000165280 | VCP | chr9 | - | 35057375 | 35057530 | 35057375 | 35057527 | 35059063 | 35059219 | 324 | 180 | 297 | 270 | 304 | 0 | 0 | 0 | 0 | 0 | 314 | 364 | 218 | 263 | 487 | 0 | 0 | 1 | 0 | 3 | 152 | 149 | 1.000000 | 1 | 1 | 1 | 1 | 1 | 1 | 1 | 1 | 0.995 | 1 | 0.994 | 0.002 |
| A3SS | 7335 | ENSG00000103365 | GGA2 | chr16 | - | 23493359 | 23494378 | 23493359 | 23493458 | 23495693 | 23495778 | 76 | 40 | 64 | 74 | 49 | 0 | 1 | 0 | 0 | 0 | 133 | 127 | 67 | 85 | 160 | 0 | 0 | 0 | 0 | 0 | 298 | 149 | 1.000000 | 1 | 1 | 0.952 | 1 | 1 | 1 | 1 | 1 | 1 | 1 | 1 | -0.01 |
| A3SS | 5845 | ENSG00000213593 | TMX2 | chr11 | + | 57738663 | 57739039 | 57738973 | 57739039 | 57737607 | 57737668 | 37 | 35 | 33 | 56 | 42 | 0 | 0 | 0 | 0 | 0 | 108 | 71 | 74 | 134 | 85 | 0 | 0 | 0 | 2 | 0 | 298 | 149 | 1.000000 | 1 | 1 | 1 | 1 | 1 | 1 | 1 | 1 | 1 | 0.971 | 1 | 0.006 |
| A3SS | 723 | ENSG00000104885 | DOT1L | chr19 | + | 2191011 | 2191240 | 2191083 | 2191240 | 2189731 | 2189795 | 65 | 38 | 58 | 89 | 41 | 0 | 0 | 0 | 1 | 1 | 152 | 76 | 49 | 151 | 130 | 0 | 0 | 0 | 0 | 0 | 221 | 149 | 1.000000 | 1 | 1 | 1 | 1 | 0.984 | 0.965 | 1 | 1 | 1 | 1 | 1 | -0.01 |
| A3SS | 13196 | ENSG00000177575 | CD163 | chr12 | - | 7479859 | 7481256 | 7479859 | 7479913 | 7482965 | 7483004 | 35 | 19 | 10 | 27 | 34 | 0 | 1 | 0 | 0 | 0 | 83 | 130 | 92 | 72 | 14 | 0 | 0 | 0 | 0 | 0 | 298 | 149 | 1.000000 | 1 | 1 | 0.905 | 1 | 1 | 1 | 1 | 1 | 1 | 1 | 1 | -0.019 |
| A3SS | 8956 | ENSG00000166508 | MCM7 | chr7 | - | 100099022 | 100099753 | 100099022 | 100099403 | 100100013 | 100100093 | 109 | 61 | 152 | 152 | 159 | 0 | 2 | 0 | 0 | 0 | 224 | 133 | 113 | 191 | 238 | 0 | 0 | 0 | 0 | 0 | 298 | 149 | 1.000000 | 1 | 1 | 0.938 | 1 | 1 | 1 | 1 | 1 | 1 | 1 | 1 | -0.012 |
| A3SS | 9799 | ENSG00000012124 | CD22 | chr19 | + | 35332593 | 35332924 | 35332652 | 35332924 | 35329186 | 35329230 | 38 | 30 | 41 | 42 | 43 | 1 | 0 | 0 | 1 | 0 | 133 | 141 | 19 | 74 | 92 | 0 | 0 | 3 | 0 | 0 | 208 | 149 | 1.000000 | 1 | 0.965 | 1 | 1 | 0.968 | 1 | 1 | 1 | 0.819 | 1 | 1 | 0.023 |
| A3SS | 11141 | ENSG00000197140 | ADAM32 | chr8 | + | 39151376 | 39151548 | 39151499 | 39151548 | 39149790 | 39149867 | 17 | 14 | 4 | 4 | 13 | 0 | 0 | 0 | 0 | 0 | 107 | 13 | 5 | 11 | 180 | 0 | 0 | 0 | 1 | 2 | 272 | 149 | 1.000000 | 1 | 1 | 1 | 1 | 1 | 1 | 1 | 1 | 1 | 0.858 | 0.98 | 0.032 |
| A3SS | 7449 | ENSG00000149532 | CPSF7 | chr11 | - | 61416104 | 61416519 | 61416104 | 61416162 | 61419948 | 61420094 | 197 | 150 | 203 | 210 | 332 | 0 | 0 | 0 | 0 | 1 | 395 | 211 | 138 | 243 | 367 | 0 | 0 | 0 | 0 | 6 | 298 | 149 | 1.000000 | 1 | 1 | 1 | 1 | 1 | 0.994 | 1 | 1 | 1 | 1 | 0.968 | 0.005 |
| A3SS | 988 | ENSG00000124155 | PIGT | chr20 | + | 45419503 | 45419590 | 45419507 | 45419590 | 45419294 | 45419395 | 117 | 40 | 92 | 77 | 75 | 3 | 0 | 1 | 0 | 0 | 89 | 189 | 111 | 107 | 166 | 0 | 3 | 5 | 0 | 0 | 153 | 149 | 1.000000 | 1 | 0.974 | 1 | 0.989 | 1 | 1 | 1 | 0.984 | 0.956 | 1 | 1 | 0.005 |
| A3SS | 6033 | ENSG00000169398 | PTK2 | chr8 | - | 140890542 | 140890769 | 140890542 | 140890603 | 140925660 | 140925749 | 46 | 85 | 72 | 111 | 142 | 0 | 0 | 0 | 0 | 0 | 141 | 106 | 70 | 94 | 303 | 0 | 0 | 0 | 0 | 5 | 298 | 149 | 1.000000 | 1 | 1 | 1 | 1 | 1 | 1 | 1 | 1 | 1 | 1 | 0.968 | 0.006 |
| A3SS | 6827 | ENSG00000100092 | SH3BP1 | chr22 | + | 37642894 | 37643006 | 37642898 | 37643006 | 37642538 | 37642615 | 127 | 52 | 83 | 125 | 66 | 0 | 0 | 0 | 2 | 0 | 171 | 114 | 67 | 145 | 212 | 0 | 0 | 2 | 0 | 3 | 153 | 149 | 1.000000 | 1 | 1 | 1 | 1 | 0.984 | 1 | 1 | 1 | 0.97 | 1 | 0.986 | 0.006 |
| A3SS | 287 | ENSG00000170889 | RPS9 | chr19 | + | 54206275 | 54206724 | 54206552 | 54206724 | 54201486 | 54201609 | 2412 | 1324 | 3199 | 2824 | 2831 | 0 | 0 | 0 | 0 | 0 | 2664 | 2809 | 1950 | 1909 | 3513 | 0 | 0 | 0 | 1 | 10 | 298 | 149 | 1.000000 | 1 | 1 | 1 | 1 | 1 | 1 | 1 | 1 | 1 | 0.999 | 0.994 | 0.001 |
| A3SS | 9469 | ENSG00000089820 | ARHGAP4 | chrX | - | 153913408 | 153913600 | 153913408 | 153913516 | 153913777 | 153913879 | 274 | 170 | 298 | 280 | 160 | 0 | 1 | 1 | 0 | 0 | 281 | 220 | 154 | 318 | 463 | 0 | 0 | 0 | 3 | 0 | 233 | 149 | 1.000000 | 1 | 1 | 0.991 | 0.995 | 1 | 1 | 1 | 1 | 1 | 0.985 | 1 | 0 |
| A3SS | 11100 | ENSG00000160948 | VPS28 | chr8 | - | 144424214 | 144424819 | 144424214 | 144424268 | 144424945 | 144425772 | 213 | 106 | 136 | 175 | 156 | 1 | 0 | 0 | 0 | 0 | 186 | 244 | 126 | 202 | 278 | 0 | 0 | 0 | 0 | 0 | 298 | 149 | 1.000000 | 1 | 0.991 | 1 | 1 | 1 | 1 | 1 | 1 | 1 | 1 | 1 | -0.002 |
| A3SS | 10066 | ENSG00000181444 | ZNF467 | chr7 | - | 149764181 | 149766239 | 149764181 | 149764678 | 149770439 | 149770556 | 31 | 13 | 28 | 38 | 14 | 2 | 0 | 1 | 0 | 0 | 68 | 91 | 24 | 51 | 138 | 0 | 2 | 0 | 0 | 0 | 298 | 149 | 1.000000 | 1 | 0.886 | 1 | 0.933 | 1 | 1 | 1 | 0.958 | 1 | 1 | 1 | -0.028 |
| A3SS | 1008 | ENSG00000130813 | SHFL | chr19 | + | 10089897 | 10090047 | 10089928 | 10090047 | 10089656 | 10089695 | 93 | 44 | 90 | 92 | 106 | 1 | 1 | 0 | 2 | 1 | 162 | 84 | 78 | 179 | 170 | 0 | 2 | 0 | 0 | 0 | 180 | 149 | 1.000000 | 1 | 0.987 | 0.973 | 1 | 0.974 | 0.989 | 1 | 0.972 | 1 | 1 | 1 | -0.01 |
| A3SS | 9739 | ENSG00000164808 | SPIDR | chr8 | + | 47700402 | 47700490 | 47700407 | 47700490 | 47673800 | 47673941 | 108 | 67 | 77 | 111 | 100 | 0 | 0 | 0 | 0 | 0 | 212 | 148 | 88 | 169 | 93 | 0 | 0 | 0 | 2 | 0 | 154 | 149 | 1.000000 | 1 | 1 | 1 | 1 | 1 | 1 | 1 | 1 | 1 | 0.988 | 1 | 0.002 |
| A3SS | 3316 | ENSG00000103496 | STX4 | chr16 | + | 31034225 | 31034536 | 31034461 | 31034536 | 31034012 | 31034114 | 98 | 45 | 72 | 78 | 16 | 0 | 0 | 0 | 0 | 0 | 127 | 66 | 94 | 118 | 147 | 0 | 0 | 2 | 0 | 0 | 298 | 149 | 1.000000 | 1 | 1 | 1 | 1 | 1 | 1 | 1 | 1 | 0.959 | 1 | 1 | 0.008 |
| A3SS | 1835 | ENSG00000172775 | PSME3IP1 | chr16 | - | 57173727 | 57173869 | 57173727 | 57173796 | 57178501 | 57178852 | 252 | 163 | 200 | 189 | 303 | 2 | 0 | 0 | 0 | 0 | 338 | 327 | 242 | 254 | 188 | 0 | 0 | 0 | 0 | 0 | 222 | 149 | 1.000000 | 1 | 0.988 | 1 | 1 | 1 | 1 | 1 | 1 | 1 | 1 | 1 | -0.002 |
| A3SS | 5296 | ENSG00000175203 | DCTN2 | chr12 | - | 57532014 | 57532643 | 57532014 | 57532106 | 57532732 | 57532810 | 90 | 48 | 79 | 79 | 88 | 0 | 0 | 0 | 1 | 0 | 163 | 110 | 77 | 55 | 220 | 0 | 0 | 0 | 0 | 0 | 298 | 149 | 1.000000 | 1 | 1 | 1 | 1 | 0.975 | 1 | 1 | 1 | 1 | 1 | 1 | -0.005 |
| A3SS | 11796 | ENSG00000259719 | LINC02284 | chr14 | + | 56317746 | 56318086 | 56317891 | 56318086 | 56317455 | 56317615 | 24 | 24 | 6 | 63 | 49 | 2 | 0 | 0 | 0 | 0 | 118 | 75 | 61 | 82 | 71 | 0 | 0 | 4 | 0 | 0 | 294 | 149 | 1.000000 | 1 | 0.859 | 1 | 1 | 1 | 1 | 1 | 1 | 0.885 | 1 | 1 | -0.005 |
| A3SS | 4813 | ENSG00000184014 | DENND5A | chr11 | - | 9141939 | 9144278 | 9141939 | 9142108 | 9144994 | 9145113 | 117 | 108 | 102 | 98 | 121 | 0 | 0 | 0 | 0 | 0 | 219 | 156 | 101 | 193 | 118 | 0 | 0 | 0 | 3 | 0 | 298 | 149 | 1.000000 | 1 | 1 | 1 | 1 | 1 | 1 | 1 | 1 | 1 | 0.97 | 1 | 0.006 |
| A3SS | 12096 | ENSG00000149136 | SSRP1 | chr11 | - | 57330290 | 57330927 | 57330290 | 57330429 | 57331667 | 57331889 | 111 | 66 | 134 | 118 | 124 | 0 | 0 | 0 | 1 | 0 | 239 | 115 | 113 | 168 | 158 | 0 | 0 | 0 | 0 | 0 | 298 | 149 | 1.000000 | 1 | 1 | 1 | 1 | 0.983 | 1 | 1 | 1 | 1 | 1 | 1 | -0.003 |
| A3SS | 344 | ENSG00000129351 | ILF3 | chr19 | + | 10684415 | 10685765 | 10684858 | 10685765 | 10683916 | 10683970 | 221 | 133 | 226 | 227 | 277 | 1 | 1 | 0 | 4 | 2 | 262 | 312 | 241 | 244 | 264 | 0 | 1 | 1 | 0 | 0 | 298 | 149 | 1.000000 | 1 | 0.991 | 0.985 | 1 | 0.966 | 0.986 | 1 | 0.994 | 0.992 | 1 | 1 | -0.012 |
| A3SS | 4466 | ENSG00000204427 | ABHD16A | chr6 | - | 31688722 | 31689119 | 31688722 | 31688786 | 31689580 | 31689704 | 49 | 20 | 42 | 40 | 21 | 0 | 0 | 0 | 1 | 0 | 65 | 53 | 58 | 56 | 179 | 0 | 0 | 0 | 4 | 0 | 298 | 149 | 1.000000 | 1 | 1 | 1 | 1 | 0.952 | 1 | 1 | 1 | 1 | 0.875 | 1 | 0.015 |
| A3SS | 11301 | ENSG00000077721 | UBE2A | chrX | + | 119582587 | 119582676 | 119582596 | 119582676 | 119575374 | 119575400 | 26 | 43 | 27 | 43 | 42 | 0 | 0 | 0 | 0 | 0 | 129 | 78 | 49 | 84 | 78 | 0 | 0 | 2 | 0 | 0 | 158 | 149 | 1.000000 | 1 | 1 | 1 | 1 | 1 | 1 | 1 | 1 | 0.959 | 1 | 1 | 0.008 |
| A3SS | 6243 | ENSG00000116337 | AMPD2 | chr1 | + | 109627773 | 109627903 | 109627777 | 109627903 | 109627428 | 109627518 | 114 | 61 | 73 | 98 | 64 | 1 | 0 | 0 | 0 | 0 | 182 | 101 | 98 | 106 | 158 | 0 | 0 | 1 | 0 | 5 | 153 | 149 | 1.000000 | 1 | 0.991 | 1 | 1 | 1 | 1 | 1 | 1 | 0.99 | 1 | 0.969 | 0.006 |
| A3SS | 12636 | ENSG00000107862 | GBF1 | chr10 | + | 102360183 | 102360395 | 102360189 | 102360395 | 102359269 | 102359435 | 62 | 31 | 79 | 53 | 61 | 2 | 0 | 0 | 1 | 0 | 103 | 66 | 64 | 102 | 184 | 0 | 0 | 0 | 0 | 0 | 155 | 149 | 1.000000 | 1 | 0.968 | 1 | 1 | 0.981 | 1 | 1 | 1 | 1 | 1 | 1 | -0.01 |
| A3SS | 8397 | ENSG00000087087 | SRRT | chr7 | + | 100888256 | 100888383 | 100888277 | 100888383 | 100888041 | 100888143 | 127 | 45 | 108 | 124 | 73 | 3 | 0 | 1 | 1 | 0 | 108 | 105 | 105 | 189 | 200 | 0 | 2 | 0 | 1 | 0 | 170 | 149 | 1.000000 | 1 | 0.974 | 1 | 0.99 | 0.991 | 1 | 1 | 0.979 | 1 | 0.994 | 1 | -0.004 |
| A3SS | 8126 | ENSG00000225190 | PLEKHM1 | chr17 | - | 45439476 | 45439634 | 45439476 | 45439577 | 45440162 | 45440226 | 158 | 56 | 73 | 127 | 129 | 2 | 0 | 1 | 0 | 2 | 115 | 250 | 103 | 145 | 156 | 0 | 0 | 2 | 0 | 0 | 206 | 149 | 1.000000 | 1 | 0.983 | 1 | 0.981 | 1 | 0.979 | 1 | 1 | 0.974 | 1 | 1 | -0.006 |
| A3SS | 4083 | ENSG00000108509 | CAMTA2 | chr17 | - | 4981677 | 4981831 | 4981677 | 4981742 | 4982088 | 4982227 | 65 | 44 | 63 | 71 | 68 | 0 | 0 | 0 | 1 | 0 | 97 | 101 | 62 | 82 | 193 | 0 | 0 | 0 | 0 | 0 | 238 | 149 | 1.000000 | 1 | 1 | 1 | 1 | 0.978 | 1 | 1 | 1 | 1 | 1 | 1 | -0.004 |
| A3SS | 3312 | ENSG00000124357 | NAGK | chr2 | + | 71075554 | 71075642 | 71075561 | 71075642 | 71073481 | 71073594 | 140 | 81 | 87 | 129 | 113 | 0 | 0 | 0 | 2 | 0 | 116 | 152 | 52 | 144 | 310 | 0 | 0 | 0 | 4 | 0 | 156 | 149 | 1.000000 | 1 | 1 | 1 | 1 | 0.984 | 1 | 1 | 1 | 1 | 0.972 | 1 | 0.002 |
| A3SS | 3588 | ENSG00000214655 | ZSWIM8 | chr10 | + | 73800010 | 73800194 | 73800119 | 73800194 | 73799376 | 73799490 | 58 | 24 | 48 | 59 | 24 | 0 | 0 | 0 | 0 | 0 | 101 | 85 | 55 | 85 | 111 | 0 | 0 | 3 | 0 | 0 | 258 | 149 | 1.000000 | 1 | 1 | 1 | 1 | 1 | 1 | 1 | 1 | 0.914 | 1 | 1 | 0.017 |
| A3SS | 12047 | ENSG00000149091 | DGKZ | chr11 | + | 46378456 | 46379111 | 46378990 | 46379111 | 46378197 | 46378229 | 102 | 46 | 127 | 113 | 54 | 0 | 0 | 0 | 0 | 0 | 129 | 145 | 80 | 83 | 228 | 0 | 0 | 0 | 2 | 0 | 298 | 149 | 1.000000 | 1 | 1 | 1 | 1 | 1 | 1 | 1 | 1 | 1 | 0.954 | 1 | 0.009 |
| A3SS | 4937 | ENSG00000121775 | TMEM39B | chr1 | + | 32075602 | 32075822 | 32075666 | 32075822 | 32074950 | 32075077 | 35 | 9 | 21 | 30 | 40 | 0 | 0 | 1 | 0 | 0 | 51 | 101 | 31 | 56 | 118 | 0 | 0 | 0 | 0 | 0 | 213 | 149 | 1.000000 | 1 | 1 | 1 | 0.936 | 1 | 1 | 1 | 1 | 1 | 1 | 1 | -0.013 |
| A3SS | 13176 | ENSG00000089693 | MLF2 | chr12 | - | 6750176 | 6750766 | 6750176 | 6750305 | 6751640 | 6751676 | 108 | 39 | 76 | 117 | 54 | 0 | 0 | 0 | 0 | 0 | 139 | 77 | 73 | 109 | 218 | 0 | 0 | 2 | 0 | 0 | 298 | 149 | 1.000000 | 1 | 1 | 1 | 1 | 1 | 1 | 1 | 1 | 0.948 | 1 | 1 | 0.01 |
| A3SS | 1884 | ENSG00000072864 | NDE1 | chr16 | + | 15724190 | 15727000 | 15724274 | 15727000 | 15696708 | 15696860 | 49 | 30 | 63 | 38 | 31 | 1 | 0 | 1 | 1 | 0 | 57 | 77 | 40 | 83 | 174 | 0 | 0 | 0 | 0 | 0 | 233 | 149 | 1.000000 | 1 | 0.969 | 1 | 0.976 | 0.96 | 1 | 1 | 1 | 1 | 1 | 1 | -0.019 |
| A3SS | 12289 | ENSG00000184428 | TOP1MT | chr8 | - | 143325345 | 143325533 | 143325345 | 143325478 | 143326221 | 143326344 | 60 | 66 | 74 | 85 | 121 | 1 | 0 | 0 | 2 | 1 | 235 | 40 | 53 | 81 | 217 | 0 | 0 | 0 | 0 | 6 | 204 | 149 | 1.000000 | 1 | 0.978 | 1 | 1 | 0.969 | 0.989 | 1 | 1 | 1 | 1 | 0.964 | -0.006 |
| A3SS | 13232 | ENSG00000139613 | SMARCC2 | chr12 | - | 56185029 | 56185111 | 56185029 | 56185092 | 56186154 | 56186240 | 146 | 134 | 185 | 241 | 211 | 0 | 0 | 1 | 1 | 5 | 282 | 210 | 113 | 214 | 317 | 0 | 0 | 1 | 0 | 7 | 168 | 149 | 1.000000 | 1 | 1 | 1 | 0.994 | 0.995 | 0.974 | 1 | 1 | 0.99 | 1 | 0.976 | -0.001 |
| A3SS | 1427 | ENSG00000187838 | PLSCR3 | chr17 | - | 7390633 | 7390795 | 7390633 | 7390693 | 7392790 | 7392952 | 218 | 115 | 191 | 228 | 83 | 1 | 4 | 1 | 1 | 1 | 231 | 207 | 135 | 200 | 280 | 0 | 0 | 0 | 0 | 4 | 251 | 149 | 1.000000 | 1 | 0.992 | 0.945 | 0.991 | 0.993 | 0.98 | 1 | 1 | 1 | 1 | 0.977 | -0.015 |
| A3SS | 7674 | ENSG00000092529 | CAPN3 | chr15 | + | 42410427 | 42410496 | 42410431 | 42410496 | 42409930 | 42409995 | 24 | 14 | 27 | 37 | 31 | 4 | 0 | 0 | 0 | 0 | 50 | 81 | 31 | 170 | 16 | 0 | 0 | 0 | 1 | 0 | 153 | 149 | 1.000000 | 1 | 0.854 | 1 | 1 | 1 | 1 | 1 | 1 | 1 | 0.994 | 1 | -0.028 |
| A3SS | 8751 | ENSG00000157637 | SLC38A10 | chr17 | - | 81246884 | 81247061 | 81246884 | 81247058 | 81251492 | 81251612 | 112 | 48 | 42 | 62 | 24 | 0 | 0 | 1 | 1 | 2 | 72 | 96 | 66 | 111 | 158 | 0 | 0 | 0 | 0 | 7 | 152 | 149 | 1.000000 | 1 | 1 | 1 | 0.976 | 0.984 | 0.922 | 1 | 1 | 1 | 1 | 0.957 | -0.015 |
| A3SS | 5636 | ENSG00000119402 | FBXW2 | chr9 | - | 120787768 | 120788278 | 120787768 | 120788186 | 120792886 | 120793257 | 130 | 98 | 116 | 126 | 130 | 0 | 0 | 0 | 0 | 5 | 104 | 217 | 198 | 192 | 103 | 6 | 0 | 0 | 0 | 0 | 241 | 149 | 1.000000 | 1 | 1 | 1 | 1 | 1 | 0.941 | 0.915 | 1 | 1 | 1 | 1 | 0.005 |
| A3SS | 728 | ENSG00000099817 | POLR2E | chr19 | - | 1090086 | 1090145 | 1090086 | 1090141 | 1090907 | 1090988 | 91 | 40 | 53 | 65 | 46 | 0 | 0 | 1 | 3 | 0 | 85 | 86 | 40 | 59 | 238 | 0 | 0 | 0 | 2 | 6 | 153 | 149 | 1.000000 | 1 | 1 | 1 | 0.981 | 0.955 | 1 | 1 | 1 | 1 | 0.966 | 0.975 | -0.001 |
| A3SS | 9321 | ENSG00000164116 | GUCY1A1 | chr4 | + | 155696755 | 155697122 | 155696934 | 155697122 | 155667345 | 155667419 | 15 | 39 | 11 | 45 | 19 | 0 | 0 | 0 | 2 | 0 | 51 | 16 | 14 | 42 | 218 | 0 | 0 | 0 | 0 | 0 | 298 | 149 | 1.000000 | 1 | 1 | 1 | 1 | 0.918 | 1 | 1 | 1 | 1 | 1 | 1 | -0.016 |
| A3SS | 8783 | ENSG00000130414 | NDUFA10 | chr2 | - | 240011616 | 240011696 | 240011616 | 240011660 | 240014738 | 240014860 | 114 | 70 | 172 | 116 | 165 | 0 | 0 | 0 | 0 | 5 | 263 | 180 | 131 | 129 | 145 | 0 | 0 | 0 | 0 | 0 | 185 | 149 | 1.000000 | 1 | 1 | 1 | 1 | 1 | 0.964 | 1 | 1 | 1 | 1 | 1 | -0.007 |
| A3SS | 13195 | ENSG00000177575 | CD163 | chr12 | - | 7479859 | 7479996 | 7479859 | 7479913 | 7482965 | 7483004 | 25 | 16 | 7 | 24 | 32 | 0 | 1 | 0 | 0 | 0 | 87 | 107 | 60 | 53 | 7 | 0 | 0 | 0 | 0 | 0 | 232 | 149 | 1.000000 | 1 | 1 | 0.911 | 1 | 1 | 1 | 1 | 1 | 1 | 1 | 1 | -0.018 |
| A3SS | 917 | ENSG00000101310 | SEC23B | chr20 | + | 18524432 | 18524669 | 18524486 | 18524669 | 18515649 | 18515736 | 215 | 119 | 164 | 170 | 146 | 0 | 0 | 2 | 0 | 0 | 203 | 181 | 219 | 189 | 230 | 0 | 0 | 0 | 0 | 0 | 203 | 149 | 1.000000 | 1 | 1 | 1 | 1 | 0.984 | 1 | 1 | 1 | 1 | 1 | 1 | -0.003 |
| A3SS | 8802 | ENSG00000126254 | RBM42 | chr19 | + | 35637157 | 35637352 | 35637249 | 35637352 | 35634255 | 35634373 | 119 | 63 | 122 | 125 | 123 | 0 | 4 | 0 | 1 | 0 | 165 | 190 | 75 | 144 | 183 | 0 | 0 | 4 | 0 | 0 | 241 | 149 | 1.000000 | 1 | 1 | 0.907 | 1 | 0.987 | 1 | 1 | 1 | 0.921 | 1 | 1 | -0.005 |
| A3SS | 4814 | ENSG00000184014 | DENND5A | chr11 | - | 9144096 | 9147151 | 9144096 | 9145113 | 9150080 | 9150209 | 165 | 115 | 155 | 138 | 376 | 0 | 0 | 0 | 0 | 0 | 413 | 138 | 166 | 260 | 177 | 0 | 0 | 0 | 5 | 0 | 298 | 149 | 1.000000 | 1 | 1 | 1 | 1 | 1 | 1 | 1 | 1 | 1 | 0.963 | 1 | 0.007 |
| A3SS | 9136 | ENSG00000168000 | BSCL2 | chr11 | - | 62691074 | 62691421 | 62691074 | 62691141 | 62692375 | 62692473 | 46 | 21 | 29 | 41 | 46 | 0 | 0 | 1 | 0 | 0 | 100 | 59 | 17 | 38 | 172 | 0 | 0 | 0 | 0 | 0 | 298 | 149 | 1.000000 | 1 | 1 | 1 | 0.935 | 1 | 1 | 1 | 1 | 1 | 1 | 1 | -0.013 |
| A3SS | 11097 | ENSG00000185122 | HSF1 | chr8 | + | 144313845 | 144314054 | 144313984 | 144314054 | 144313510 | 144313616 | 56 | 19 | 51 | 48 | 26 | 0 | 0 | 0 | 4 | 0 | 107 | 88 | 53 | 85 | 68 | 0 | 0 | 0 | 0 | 0 | 288 | 149 | 1.000000 | 1 | 1 | 1 | 1 | 0.861 | 1 | 1 | 1 | 1 | 1 | 1 | -0.028 |
| A3SS | 4824 | ENSG00000197386 | HTT | chr4 | + | 3238446 | 3238609 | 3238528 | 3238609 | 3236148 | 3236254 | 129 | 85 | 96 | 137 | 141 | 0 | 0 | 1 | 0 | 0 | 93 | 218 | 67 | 75 | 336 | 0 | 0 | 0 | 0 | 0 | 231 | 149 | 1.000000 | 1 | 1 | 1 | 0.984 | 1 | 1 | 1 | 1 | 1 | 1 | 1 | -0.003 |
| A3SS | 5297 | ENSG00000175203 | DCTN2 | chr12 | - | 57532732 | 57532810 | 57532732 | 57532807 | 57532983 | 57533303 | 129 | 87 | 96 | 82 | 113 | 1 | 0 | 0 | 0 | 0 | 142 | 94 | 111 | 67 | 294 | 0 | 0 | 0 | 0 | 0 | 152 | 149 | 1.000000 | 1 | 0.992 | 1 | 1 | 1 | 1 | 1 | 1 | 1 | 1 | 1 | -0.002 |
| A3SS | 9322 | ENSG00000164116 | GUCY1A1 | chr4 | + | 155696805 | 155697122 | 155696934 | 155697122 | 155667345 | 155667419 | 13 | 33 | 9 | 41 | 17 | 0 | 0 | 0 | 2 | 0 | 51 | 16 | 14 | 37 | 195 | 0 | 0 | 0 | 0 | 0 | 278 | 149 | 1.000000 | 1 | 1 | 1 | 1 | 0.917 | 1 | 1 | 1 | 1 | 1 | 1 | -0.017 |
| A3SS | 11341 | ENSG00000158850 | B4GALT3 | chr1 | - | 161173604 | 161174049 | 161173604 | 161173727 | 161174992 | 161175228 | 60 | 37 | 86 | 86 | 51 | 0 | 0 | 0 | 1 | 0 | 128 | 104 | 85 | 102 | 101 | 0 | 0 | 0 | 0 | 0 | 298 | 149 | 1.000000 | 1 | 1 | 1 | 1 | 0.977 | 1 | 1 | 1 | 1 | 1 | 1 | -0.005 |
| A3SS | 4189 | ENSG00000179344 | HLA-DQB1 | chr6 | - | 32661966 | 32662248 | 32661966 | 32662085 | 32664797 | 32665067 | 46 | 256 | 684 | 45 | 341 | 0 | 1 | 2 | 0 | 0 | 522 | 58 | 111 | 265 | 616 | 0 | 0 | 0 | 0 | 0 | 298 | 149 | 1.000000 | 1 | 1 | 0.992 | 0.994 | 1 | 1 | 1 | 1 | 1 | 1 | 1 | -0.003 |
| A3SS | 9988 | ENSG00000106348 | IMPDH1 | chr7 | - | 128409288 | 128409484 | 128409288 | 128409352 | 128409755 | 128409982 | 40 | 16 | 27 | 33 | 6 | 1 | 0 | 0 | 0 | 0 | 74 | 19 | 63 | 57 | 108 | 0 | 0 | 0 | 0 | 0 | 281 | 149 | 1.000000 | 1 | 0.955 | 1 | 1 | 1 | 1 | 1 | 1 | 1 | 1 | 1 | -0.009 |
| A3SS | 8942 | ENSG00000124356 | STAMBP | chr2 | + | 73845166 | 73845262 | 73845175 | 73845262 | 73844812 | 73844888 | 122 | 92 | 88 | 137 | 214 | 0 | 0 | 0 | 1 | 2 | 232 | 137 | 125 | 171 | 185 | 0 | 5 | 0 | 0 | 0 | 158 | 149 | 1.000000 | 1 | 1 | 1 | 1 | 0.992 | 0.99 | 1 | 0.963 | 1 | 1 | 1 | 0.004 |
| A3SS | 4039 | ENSG00000100412 | ACO2 | chr22 | + | 41517526 | 41517631 | 41517596 | 41517631 | 41515376 | 41515535 | 79 | 47 | 54 | 68 | 65 | 0 | 0 | 0 | 0 | 0 | 84 | 172 | 60 | 54 | 139 | 0 | 0 | 2 | 0 | 0 | 219 | 149 | 1.000000 | 1 | 1 | 1 | 1 | 1 | 1 | 1 | 1 | 0.953 | 1 | 1 | 0.009 |
| A3SS | 12705 | ENSG00000151651 | ADAM8 | chr10 | - | 133273253 | 133273838 | 133273253 | 133273443 | 133273950 | 133274029 | 172 | 45 | 103 | 108 | 20 | 0 | 0 | 0 | 0 | 0 | 182 | 129 | 87 | 73 | 172 | 0 | 2 | 0 | 0 | 0 | 298 | 149 | 1.000000 | 1 | 1 | 1 | 1 | 1 | 1 | 1 | 0.97 | 1 | 1 | 1 | 0.006 |
| A3SS | 1331 | ENSG00000007202 | KIAA0100 | chr17 | - | 28640559 | 28640683 | 28640559 | 28640627 | 28642274 | 28642481 | 78 | 41 | 65 | 73 | 120 | 0 | 1 | 0 | 0 | 0 | 155 | 107 | 65 | 81 | 162 | 0 | 0 | 0 | 0 | 0 | 205 | 149 | 1.000000 | 1 | 1 | 0.968 | 1 | 1 | 1 | 1 | 1 | 1 | 1 | 1 | -0.006 |
| A3SS | 9986 | ENSG00000106348 | IMPDH1 | chr7 | - | 128400332 | 128400539 | 128400332 | 128400536 | 128400816 | 128400891 | 60 | 38 | 30 | 53 | 50 | 3 | 0 | 0 | 0 | 0 | 104 | 79 | 58 | 64 | 118 | 0 | 0 | 0 | 0 | 0 | 152 | 149 | 1.000000 | 1 | 0.951 | 1 | 1 | 1 | 1 | 1 | 1 | 1 | 1 | 1 | -0.01 |
| A3SS | 4995 | ENSG00000132716 | DCAF8 | chr1 | - | 160261284 | 160261350 | 160261284 | 160261325 | 160262116 | 160262451 | 46 | 42 | 36 | 38 | 44 | 2 | 0 | 0 | 0 | 0 | 126 | 67 | 58 | 77 | 69 | 0 | 0 | 0 | 0 | 0 | 174 | 149 | 1.000000 | 1 | 0.952 | 1 | 1 | 1 | 1 | 1 | 1 | 1 | 1 | 1 | -0.01 |
| A3SS | 8666 | ENSG00000161202 | DVL3 | chr3 | + | 184166445 | 184166673 | 184166605 | 184166673 | 184166125 | 184166265 | 122 | 49 | 82 | 110 | 109 | 0 | 0 | 0 | 0 | 0 | 148 | 161 | 74 | 121 | 159 | 0 | 0 | 0 | 0 | 6 | 298 | 149 | 1.000000 | 1 | 1 | 1 | 1 | 1 | 1 | 1 | 1 | 1 | 1 | 0.93 | 0.014 |
| A3SS | 6591 | ENSG00000029534 | ANK1 | chr8 | - | 41672353 | 41672529 | 41672353 | 41672426 | 41684543 | 41684690 | 5 | 2 | 4 | 8 | 3 | 0 | 6 | 1 | 1 | 3 | 35 | 22 | 6 | 0 | 149 | 9 | 2 | 4 | 0 | 46 | 252 | 149 | 1.000000 | 1 | 1 | 0.165 | 0.703 | 0.825 | 0.372 | 0.697 | 0.867 | 0.47 | NA | 0.657 | -0.06 |
| A3SS | 10592 | ENSG00000102103 | PQBP1 | chrX | + | 48902232 | 48903084 | 48902927 | 48903084 | 48901929 | 48902042 | 185 | 115 | 125 | 165 | 148 | 0 | 0 | 0 | 0 | 0 | 265 | 140 | 157 | 131 | 235 | 1 | 0 | 0 | 0 | 0 | 298 | 149 | 1.000000 | 1 | 1 | 1 | 1 | 1 | 1 | 0.993 | 1 | 1 | 1 | 1 | 0.001 |
| A3SS | 8667 | ENSG00000161202 | DVL3 | chr3 | + | 184166825 | 184166975 | 184166876 | 184166975 | 184166605 | 184166673 | 106 | 56 | 88 | 97 | 126 | 0 | 0 | 0 | 0 | 3 | 162 | 135 | 102 | 91 | 173 | 0 | 2 | 0 | 4 | 0 | 200 | 149 | 1.000000 | 1 | 1 | 1 | 1 | 1 | 0.969 | 1 | 0.981 | 1 | 0.944 | 1 | 0.009 |
| A3SS | 7726 | ENSG00000187726 | DNAJB13 | chr11 | + | 73968215 | 73968458 | 73968344 | 73968458 | 73966137 | 73966251 | 55 | 25 | 33 | 24 | 6 | 0 | 0 | 0 | 1 | 2 | 90 | 63 | 42 | 46 | 91 | 0 | 0 | 0 | 0 | 0 | 278 | 149 | 1.000000 | 1 | 1 | 1 | 1 | 0.928 | 0.617 | 1 | 1 | 1 | 1 | 1 | -0.091 |
| A3SS | 11852 | ENSG00000142186 | SCYL1 | chr11 | + | 65536585 | 65536750 | 65536590 | 65536750 | 65536258 | 65536334 | 149 | 83 | 134 | 134 | 266 | 0 | 0 | 0 | 0 | 1 | 216 | 182 | 152 | 144 | 260 | 0 | 0 | 0 | 1 | 0 | 154 | 149 | 1.000000 | 1 | 1 | 1 | 1 | 1 | 0.996 | 1 | 1 | 1 | 0.993 | 1 | 0.001 |
| A3SS | 4992 | ENSG00000148290 | SURF1 | chr9 | - | 133352693 | 133352766 | 133352693 | 133352730 | 133353748 | 133353940 | 184 | 74 | 147 | 132 | 152 | 1 | 0 | 0 | 1 | 0 | 191 | 123 | 84 | 112 | 367 | 0 | 6 | 2 | 0 | 0 | 185 | 149 | 1.000000 | 1 | 0.993 | 1 | 1 | 0.991 | 1 | 1 | 0.943 | 0.971 | 1 | 1 | 0.014 |
| A3SS | 10614 | ENSG00000126759 | CFP | chrX | - | 47628101 | 47629674 | 47628101 | 47628277 | 47629768 | 47629895 | 172 | 65 | 86 | 82 | 52 | 1 | 0 | 2 | 0 | 0 | 110 | 85 | 44 | 142 | 264 | 0 | 0 | 2 | 3 | 0 | 298 | 149 | 1.000000 | 1 | 0.989 | 1 | 0.956 | 1 | 1 | 1 | 1 | 0.917 | 0.959 | 1 | 0.014 |
| A3SS | 6801 | ENSG00000170271 | FAXDC2 | chr5 | - | 154834842 | 154835119 | 154834842 | 154834934 | 154838130 | 154838178 | 22 | 9 | 1 | 16 | 5 | 1 | 0 | 0 | 0 | 1 | 10 | 30 | 17 | 2 | 181 | 0 | 0 | 0 | 0 | 0 | 298 | 149 | 1.000000 | 1 | 0.917 | 1 | 1 | 1 | 0.714 | 1 | 1 | 1 | 1 | 1 | -0.074 |

| | | | | | | | | | | | | | | | | | | | | | | | | | | | | | | | | | | | | | | | | | | | | | | |
|---|---|---|---|---|---|---|---|---|---|---|---|---|---|---|---|---|---|---|---|---|---|---|---|---|---|---|---|---|---|---|---|---|---|---|---|---|---|---|---|---|---|---|---|---|---|---|
| A3SS | 11343 | ENSG00000158850 | B4GALT3 | chr1 | - | 161175807 | 161176095 | 161175807 | 161175893 | 161176433 | 161176579 | 37 | 21 | 42 | 34 | 29 | 0 | 0 | 0 | 1 | 0 | 120 | 64 | 25 | 50 | 91 | 0 | 0 | 1 | 0 | 0 | 298 | 149 | 1.000000 | 1 | 1 | 1 | 1 | 0.944 | 1 | 1 | 1 | 0.926 | 1 | 1 | 0.004 |
| A3SS | 2197 | ENSG00000182511 | FES | chr15 | + | 90885036 | 90885258 | 90885041 | 90885258 | 90884468 | 90884544 | 27 | 6 | 10 | 18 | 1 | 1 | 0 | 0 | 1 | 0 | 77 | 60 | 15 | 46 | 50 | 0 | 6 | 0 | 0 | 0 | 154 | 149 | 1.000000 | 1 | 0.963 | 1 | 1 | 0.946 | 1 | 1 | 0.906 | 1 | 1 | 1 | 0.001 |
| A3SS | 3889 | ENSG00000163110 | PDLIM5 | chr4 | + | 94575615 | 94576034 | 94575942 | 94576034 | 94573350 | 94573393 | 135 | 102 | 95 | 140 | 224 | 0 | 0 | 0 | 0 | 0 | 174 | 89 | 93 | 189 | 337 | 0 | 0 | 0 | 0 | 2 | 298 | 149 | 1.000000 | 1 | 1 | 1 | 1 | 1 | 1 | 1 | 1 | 1 | 1 | 0.988 | 0.002 |
| A3SS | 8943 | ENSG00000124356 | STAMBP | chr2 | + | 73847386 | 73849487 | 73849362 | 73849487 | 73844812 | 73845262 | 158 | 111 | 106 | 162 | 263 | 0 | 0 | 0 | 1 | 0 | 260 | 172 | 175 | 183 | 195 | 0 | 0 | 0 | 0 | 0 | 298 | 149 | 1.000000 | 1 | 1 | 1 | 1 | 0.988 | 1 | 1 | 1 | 1 | 1 | 1 | -0.002 |
| A3SS | 2899 | ENSG00000143379 | SETDB1 | chr1 | + | 150962973 | 150963741 | 150963529 | 150963741 | 150962586 | 150962719 | 155 | 92 | 106 | 121 | 134 | 0 | 0 | 0 | 2 | 0 | 236 | 161 | 112 | 98 | 186 | 0 | 3 | 1 | 3 | 0 | 298 | 149 | 1.000000 | 1 | 1 | 1 | 1 | 0.968 | 1 | 1 | 0.964 | 0.982 | 0.942 | 1 | 0.016 |
| A3SS | 5180 | ENSG00000239969 | NA | chr7 | + | 102365973 | 102366076 | 102366017 | 102366076 | 102364297 | 102364411 | 24 | 7 | 15 | 21 | 11 | 0 | 1 | 0 | 0 | 0 | 72 | 51 | 29 | 51 | 59 | 0 | 0 | 0 | 0 | 2 | 193 | 149 | 1.000000 | 1 | 1 | 0.844 | 1 | 1 | 1 | 1 | 1 | 1 | 1 | 0.958 | -0.023 |
| A3SS | 4958 | ENSG00000158864 | NDUFS2 | chr1 | + | 161209856 | 161210188 | 161210110 | 161210188 | 161209482 | 161209595 | 199 | 108 | 232 | 186 | 252 | 0 | 0 | 0 | 1 | 0 | 193 | 247 | 170 | 198 | 353 | 0 | 0 | 0 | 0 | 0 | 298 | 149 | 1.000000 | 1 | 1 | 1 | 1 | 0.989 | 1 | 1 | 1 | 1 | 1 | 1 | -0.002 |
| A3SS | 4886 | ENSG00000116871 | MAP7D1 | chr1 | + | 36178445 | 36178596 | 36178470 | 36178596 | 36177872 | 36178201 | 42 | 26 | 41 | 46 | 17 | 1 | 0 | 0 | 1 | 0 | 58 | 60 | 30 | 85 | 122 | 0 | 0 | 0 | 0 | 0 | 174 | 149 | 1.000000 | 1 | 0.973 | 1 | 1 | 0.975 | 1 | 1 | 1 | 1 | 1 | 1 | -0.01 |
| A3SS | 13363 | ENSG00000185989 | RASA3 | chr13 | - | 114027388 | 114027461 | 114027388 | 114027456 | 114027846 | 114027927 | 303 | 133 | 323 | 266 | 338 | 0 | 0 | 1 | 0 | 0 | 210 | 278 | 190 | 222 | 646 | 0 | 0 | 0 | 0 | 0 | 154 | 149 | 1.000000 | 1 | 1 | 1 | 0.997 | 1 | 1 | 1 | 1 | 1 | 1 | 1 | -0.001 |
| A3SS | 8872 | ENSG00000204843 | DCTN1 | chr2 | - | 74366243 | 74366620 | 74366243 | 74366375 | 74366782 | 74366932 | 110 | 86 | 115 | 95 | 117 | 0 | 0 | 3 | 2 | 0 | 122 | 176 | 75 | 100 | 232 | 0 | 0 | 0 | 1 | 3 | 298 | 149 | 1.000000 | 1 | 1 | 1 | 0.95 | 0.96 | 1 | 1 | 1 | 1 | 0.98 | 0.975 | -0.009 |
| A3SS | 1871 | ENSG00000197912 | SPG7 | chr16 | + | 89541152 | 89542279 | 89541201 | 89542279 | 89540943 | 89541027 | 27 | 35 | 31 | 32 | 15 | 0 | 2 | 0 | 0 | 0 | 78 | 46 | 37 | 67 | 93 | 0 | 0 | 1 | 0 | 2 | 198 | 149 | 1.000000 | 1 | 1 | 0.929 | 1 | 1 | 1 | 1 | 1 | 0.965 | 1 | 0.972 | -0.002 |
| A3SS | 4121 | ENSG00000161013 | MGAT4B | chr5 | - | 179801553 | 179801694 | 179801553 | 179801692 | 179801783 | 179801969 | 53 | 14 | 28 | 21 | 24 | 1 | 1 | 0 | 2 | 0 | 50 | 41 | 8 | 16 | 205 | 0 | 0 | 0 | 0 | 0 | 151 | 149 | 1.000000 | 1 | 0.981 | 0.932 | 1 | 0.912 | 1 | 1 | 1 | 1 | 1 | 1 | -0.035 |
| A3SS | 4809 | ENSG00000196187 | TMEM63A | chr1 | - | 225865896 | 225866050 | 225865896 | 225865967 | 225871075 | 225871113 | 30 | 18 | 25 | 36 | 12 | 0 | 0 | 0 | 0 | 0 | 70 | 35 | 16 | 98 | 80 | 4 | 0 | 0 | 0 | 0 | 232 | 149 | 1.000000 | 1 | 1 | 1 | 1 | 1 | 1 | 0.918 | 1 | 1 | 1 | 1 | 0.016 |
| A3SS | 6797 | ENSG00000117139 | KDM5B | chr1 | - | 202773117 | 202773288 | 202773117 | 202773284 | 202774612 | 202774735 | 99 | 95 | 76 | 84 | 74 | 1 | 0 | 0 | 0 | 0 | 134 | 95 | 147 | 179 | 50 | 0 | 0 | 0 | 0 | 0 | 153 | 149 | 1.000000 | 1 | 0.99 | 1 | 1 | 1 | 1 | 1 | 1 | 1 | 1 | 1 | -0.002 |
| A3SS | 10438 | ENSG00000140332 | TLE3 | chr15 | - | 70054437 | 70054685 | 70054437 | 70054680 | 70055048 | 70055298 | 57 | 29 | 49 | 65 | 14 | 0 | 0 | 0 | 0 | 0 | 41 | 78 | 88 | 57 | 125 | 0 | 0 | 1 | 0 | 0 | 154 | 149 | 1.000000 | 1 | 1 | 1 | 1 | 1 | 1 | 1 | 1 | 0.988 | 1 | 1 | 0.002 |
| A3SS | 2711 | ENSG00000203485 | INF2 | chr14 | + | 104708435 | 104708587 | 104708531 | 104708587 | 104707252 | 104708002 | 39 | 24 | 58 | 83 | 38 | 0 | 0 | 0 | 0 | 0 | 95 | 35 | 36 | 45 | 206 | 0 | 0 | 0 | 0 | 6 | 245 | 149 | 1.000000 | 1 | 1 | 1 | 1 | 1 | 1 | 1 | 1 | 1 | 1 | 0.954 | 0.009 |
| A3SS | 1773 | ENSG00000140995 | DEF8 | chr16 | + | 89957510 | 89957660 | 89957537 | 89957660 | 89955168 | 89955266 | 60 | 27 | 49 | 43 | 17 | 0 | 0 | 0 | 0 | 0 | 79 | 63 | 56 | 68 | 104 | 0 | 0 | 0 | 3 | 0 | 176 | 149 | 1.000000 | 1 | 1 | 1 | 1 | 1 | 1 | 1 | 1 | 1 | 0.95 | 1 | 0.01 |
| A3SS | 6895 | ENSG00000119725 | ZNF410 | chr14 | + | 73896315 | 73896534 | 73896347 | 73896534 | 73894358 | 73894409 | 47 | 24 | 34 | 38 | 28 | 0 | 0 | 0 | 0 | 0 | 100 | 53 | 76 | 53 | 62 | 0 | 0 | 1 | 0 | 0 | 181 | 149 | 1.000000 | 1 | 1 | 1 | 1 | 1 | 1 | 1 | 1 | 0.984 | 1 | 1 | 0.003 |
| A3SS | 6256 | ENSG00000171793 | CTPS1 | chr1 | + | 40999829 | 41001117 | 41000972 | 41001117 | 40997393 | 40997526 | 1 | 5 | 12 | 26 | 15 | 0 | 1 | 1 | 0 | 0 | 77 | 41 | 43 | 50 | 20 | 0 | 0 | 0 | 2 | 0 | 298 | 149 | 1.000000 | 1 | 1 | 0.714 | 0.857 | 1 | 1 | 1 | 1 | 1 | 0.926 | 1 | -0.071 |
| A3SS | 2287 | ENSG00000102901 | CENPT | chr16 | - | 67831753 | 67831890 | 67831753 | 67831840 | 67832011 | 67832108 | 97 | 71 | 72 | 101 | 107 | 0 | 0 | 1 | 2 | 0 | 78 | 113 | 67 | 90 | 272 | 0 | 0 | 0 | 0 | 0 | 199 | 149 | 1.000000 | 1 | 1 | 1 | 0.982 | 0.974 | 1 | 1 | 1 | 1 | 1 | 1 | -0.009 |
| A3SS | 5422 | ENSG00000125637 | PSD4 | chr2 | + | 113185876 | 113186255 | 113185960 | 113186255 | 113185364 | 113185440 | 236 | 164 | 243 | 357 | 283 | 0 | 0 | 2 | 0 | 0 | 311 | 365 | 86 | 225 | 467 | 0 | 0 | 0 | 0 | 0 | 233 | 149 | 1.000000 | 1 | 1 | 1 | 0.987 | 1 | 1 | 1 | 1 | 1 | 1 | 1 | -0.003 |
| A3SS | 11342 | ENSG00000158850 | B4GALT3 | chr1 | - | 161175807 | 161176074 | 161175807 | 161175893 | 161176433 | 161176579 | 49 | 24 | 53 | 49 | 37 | 0 | 0 | 0 | 1 | 0 | 116 | 67 | 31 | 44 | 125 | 0 | 0 | 1 | 0 | 0 | 298 | 149 | 1.000000 | 1 | 1 | 1 | 1 | 0.961 | 1 | 1 | 1 | 0.939 | 1 | 1 | 0.004 |
| A3SS | 10425 | ENSG00000285171 | NA | chrX | - | 71105999 | 71106426 | 71105999 | 71106137 | 71106531 | 71106590 | 30 | 24 | 38 | 45 | 26 | 0 | 0 | 0 | 0 | 0 | 89 | 53 | 89 | 46 | 57 | 0 | 2 | 0 | 0 | 0 | 298 | 149 | 1.000000 | 1 | 1 | 1 | 1 | 1 | 1 | 1 | 0.93 | 1 | 1 | 1 | 0.014 |
| A3SS | 3108 | ENSG00000181991 | MRPS11 | chr15 | + | 88475109 | 88475239 | 88475171 | 88475239 | 88472626 | 88472725 | 92 | 46 | 63 | 67 | 84 | 0 | 1 | 0 | 0 | 0 | 142 | 84 | 54 | 87 | 155 | 0 | 1 | 0 | 0 | 0 | 211 | 149 | 1.000000 | 1 | 1 | 0.97 | 1 | 1 | 1 | 1 | 0.983 | 1 | 1 | 1 | -0.003 |
| A3SS | 495 | ENSG00000108106 | UBE2S | chr19 | - | 55404287 | 55404478 | 55404287 | 55404401 | 55406814 | 55406962 | 60 | 13 | 60 | 86 | 40 | 0 | 0 | 0 | 1 | 0 | 62 | 85 | 47 | 121 | 112 | 0 | 0 | 0 | 0 | 0 | 226 | 149 | 1.000000 | 1 | 1 | 1 | 1 | 0.983 | 1 | 1 | 1 | 1 | 1 | 1 | -0.003 |
| A3SS | 9470 | ENSG00000089820 | ARHGAP4 | chrX | - | 153913408 | 153913600 | 153913408 | 153913516 | 153918831 | 153919053 | 117 | 81 | 145 | 129 | 48 | 0 | 1 | 0 | 0 | 3 | 151 | 119 | 89 | 138 | 190 | 0 | 0 | 0 | 3 | 0 | 233 | 149 | 1.000000 | 1 | 1 | 0.981 | 1 | 1 | 0.911 | 1 | 1 | 1 | 0.967 | 1 | -0.015 |
| A3SS | 7663 | ENSG00000196642 | RABL6 | chr9 | + | 136839221 | 136839486 | 136839313 | 136839486 | 136838908 | 136839121 | 136 | 60 | 74 | 96 | 42 | 2 | 0 | 0 | 0 | 1 | 102 | 91 | 65 | 117 | 198 | 0 | 0 | 0 | 1 | 0 | 241 | 149 | 1.000000 | 1 | 0.977 | 1 | 1 | 1 | 0.963 | 1 | 1 | 1 | 0.986 | 1 | -0.009 |
| A3SS | 4092 | ENSG00000138031 | ADCY3 | chr2 | - | 24827901 | 24828161 | 24827901 | 24828023 | 24830708 | 24830825 | 23 | 19 | 30 | 47 | 24 | 0 | 0 | 0 | 0 | 0 | 40 | 47 | 25 | 74 | 122 | 0 | 1 | 0 | 0 | 0 | 287 | 149 | 1.000000 | 1 | 1 | 1 | 1 | 1 | 1 | 1 | 0.961 | 1 | 1 | 1 | 0.008 |
| A3SS | 7624 | ENSG00000061273 | HDAC7 | chr12 | - | 47797383 | 47797499 | 47797383 | 47797427 | 47798107 | 47798219 | 63 | 17 | 38 | 34 | 34 | 0 | 3 | 0 | 1 | 0 | 75 | 77 | 33 | 50 | 114 | 1 | 0 | 3 | 0 | 0 | 221 | 149 | 1.000000 | 1 | 1 | 0.793 | 1 | 0.958 | 1 | 0.981 | 1 | 0.881 | 1 | 1 | -0.022 |
| A3SS | 8348 | ENSG00000197858 | GPAA1 | chr8 | + | 144084133 | 144084330 | 144084285 | 144084330 | 144083938 | 144084040 | 115 | 37 | 78 | 73 | 58 | 5 | 0 | 0 | 0 | 0 | 113 | 97 | 59 | 84 | 170 | 0 | 0 | 0 | 0 | 0 | 298 | 149 | 1.000000 | 1 | 0.92 | 1 | 1 | 1 | 1 | 1 | 1 | 1 | 1 | 1 | -0.016 |
| A3SS | 2705 | ENSG00000099797 | TECR | chr19 | + | 14563799 | 14563903 | 14563834 | 14563903 | 14563657 | 14563702 | 200 | 82 | 259 | 208 | 161 | 0 | 0 | 0 | 0 | 0 | 129 | 206 | 212 | 158 | 367 | 5 | 0 | 0 | 0 | 0 | 184 | 149 | 1.000000 | 1 | 1 | 1 | 1 | 1 | 1 | 0.954 | 1 | 1 | 1 | 1 | 0.009 |
| A3SS | 4472 | ENSG00000169714 | CNBP | chr3 | - | 129171445 | 129171538 | 129171445 | 129171508 | 129183775 | 129183910 | 219 | 132 | 225 | 200 | 212 | 0 | 1 | 0 | 0 | 0 | 308 | 201 | 229 | 198 | 213 | 0 | 0 | 0 | 0 | 0 | 179 | 149 | 1.000000 | 1 | 1 | 0.991 | 1 | 1 | 1 | 1 | 1 | 1 | 1 | 1 | -0.002 |
| A3SS | 5817 | ENSG00000160058 | BSDC1 | chr1 | - | 32383829 | 32383997 | 32383829 | 32383906 | 32386778 | 32386895 | 168 | 141 | 118 | 171 | 211 | 0 | 2 | 1 | 0 | 0 | 150 | 137 | 152 | 183 | 347 | 0 | 0 | 1 | 0 | 5 | 240 | 149 | 1.000000 | 1 | 1 | 0.978 | 0.987 | 1 | 1 | 1 | 1 | 0.99 | 1 | 0.977 | 0 |
| A3SS | 1691 | ENSG00000140650 | PMM2 | chr16 | + | 8802080 | 8802389 | 8802260 | 8802389 | 8801798 | 8801910 | 17 | 14 | 10 | 15 | 40 | 1 | 0 | 0 | 0 | 0 | 156 | 18 | 45 | 16 | 20 | 0 | 0 | 0 | 0 | 0 | 298 | 149 | 1.000000 | 1 | 0.895 | 1 | 1 | 1 | 1 | 1 | 1 | 1 | 1 | 1 | -0.021 |
| A3SS | 10645 | ENSG00000169057 | MECP2 | chrX | - | 154056940 | 154057254 | 154056940 | 154057179 | 154057742 | 154057842 | 1 | 7 | 8 | 14 | 4 | 0 | 0 | 2 | 1 | 0 | 22 | 39 | 60 | 70 | 0 | 0 | 5 | 1 | 0 | 0 | 224 | 149 | 1.000000 | 1 | 1 | 1 | 0.727 | 0.903 | 1 | 1 | 0.838 | 0.976 | 1 | NA | -0.028 |
| A3SS | 1248 | ENSG00000132361 | CLUH | chr17 | - | 2700677 | 2700825 | 2700677 | 2700782 | 2701139 | 2701265 | 43 | 46 | 41 | 75 | 22 | 0 | 0 | 1 | 1 | 0 | 135 | 67 | 60 | 35 | 87 | 0 | 0 | 0 | 0 | 0 | 192 | 149 | 1.000000 | 1 | 1 | 1 | 0.97 | 0.983 | 1 | 1 | 1 | 1 | 1 | 1 | -0.009 |
| A3SS | 4710 | ENSG00000092098 | RNF31 | chr14 | + | 24149405 | 24149583 | 24149476 | 24149583 | 24148800 | 24148876 | 66 | 26 | 61 | 71 | 66 | 0 | 0 | 0 | 2 | 0 | 80 | 136 | 34 | 89 | 108 | 0 | 0 | 1 | 3 | 0 | 220 | 149 | 1.000000 | 1 | 1 | 1 | 1 | 0.96 | 1 | 1 | 1 | 0.958 | 0.953 | 1 | 0.01 |
| A3SS | 833 | ENSG00000171552 | BCL2L1 | chr20 | - | 31721654 | 31722348 | 31721654 | 31722330 | 31723863 | 31723989 | 21 | 14 | 16 | 21 | 13 | 9 | 1 | 14 | 1 | 1 | 68 | 28 | 23 | 44 | 78 | 0 | 2 | 0 | 3 | 0 | 167 | 149 | 1.000000 | 1 | 0.676 | 0.926 | 0.505 | 0.949 | 0.921 | 1 | 0.926 | 1 | 0.929 | 1 | -0.176 |
| A3SS | 8973 | ENSG00000167792 | NDUFV1 | chr11 | + | 67610380 | 67611207 | 67610994 | 67611207 | 67609451 | 67609635 | 58 | 31 | 63 | 37 | 30 | 0 | 2 | 0 | 0 | 0 | 62 | 83 | 50 | 34 | 146 | 0 | 0 | 0 | 0 | 0 | 298 | 149 | 1.000000 | 1 | 1 | 0.886 | 1 | 1 | 1 | 1 | 1 | 1 | 1 | 1 | -0.023 |
| A3SS | 1696 | ENSG00000140650 | PMM2 | chr16 | + | 8811637 | 8811713 | 8811641 | 8811713 | 8811078 | 8811178 | 74 | 64 | 71 | 96 | 115 | 0 | 0 | 1 | 0 | 2 | 125 | 126 | 93 | 95 | 137 | 0 | 0 | 0 | 3 | 0 | 153 | 149 | 1.000000 | 1 | 1 | 1 | 0.986 | 1 | 0.982 | 1 | 1 | 1 | 0.969 | 1 | 0 |
| A3SS | 7327 | ENSG00000159348 | CYB5R1 | chr1 | - | 202965886 | 202966600 | 202965886 | 202965993 | 202966748 | 202966898 | 60 | 27 | 21 | 31 | 52 | 2 | 0 | 2 | 0 | 0 | 35 | 83 | 41 | 39 | 148 | 0 | 0 | 0 | 0 | 0 | 298 | 149 | 1.000000 | 1 | 0.938 | 1 | 0.84 | 1 | 1 | 1 | 1 | 1 | 1 | 1 | -0.044 |
| A3SS | 12607 | ENSG00000172661 | WASHC2C | chr10 | + | 45769448 | 45769618 | 45769454 | 45769618 | 45765678 | 45765810 | 184 | 137 | 126 | 137 | 223 | 4 | 0 | 0 | 0 | 1 | 291 | 122 | 166 | 320 | 63 | 0 | 0 | 2 | 0 | 0 | 155 | 149 | 1.000000 | 1 | 0.978 | 1 | 1 | 1 | 0.995 | 1 | 1 | 0.988 | 1 | 1 | -0.003 |
| A3SS | 13096 | ENSG00000135124 | P2RX4 | chr12 | + | 121232610 | 121233092 | 121232996 | 121233092 | 121228962 | 121229099 | 5 | 10 | 3 | 13 | 4 | 0 | 2 | 0 | 0 | 0 | 63 | 3 | 6 | 32 | 85 | 0 | 0 | 0 | 0 | 0 | 298 | 149 | 1.000000 | 1 | 1 | 0.714 | 1 | 1 | 1 | 1 | 1 | 1 | 1 | 1 | -0.057 |
| A3SS | 6885 | ENSG00000072110 | ACTN1 | chr14 | - | 68877081 | 68878523 | 68877081 | 68877240 | 68879961 | 68880108 | 17 | 14 | 30 | 28 | 5 | 0 | 0 | 0 | 2 | 0 | 43 | 30 | 34 | 89 | 52 | 0 | 0 | 0 | 0 | 0 | 298 | 149 | 1.000000 | 1 | 1 | 1 | 1 | 0.875 | 1 | 1 | 1 | 1 | 1 | 1 | -0.025 |
| A3SS | 2515 | ENSG00000226777 | FAM30A | chr14 | + | 105921062 | 105921258 | 105921149 | 105921258 | 105918437 | 105918514 | 22 | 6 | 18 | 12 | 15 | 0 | 0 | 0 | 0 | 0 | 43 | 55 | 25 | 25 | 79 | 0 | 0 | 0 | 4 | 0 | 236 | 149 | 1.000000 | 1 | 1 | 1 | 1 | 1 | 1 | 1 | 1 | 1 | 0.798 | 1 | 0.04 |
| A3SS | 5803 | ENSG00000062485 | CS | chr12 | - | 56275995 | 56276195 | 56275995 | 56276143 | 56282419 | 56282608 | 295 | 182 | 184 | 235 | 369 | 1 | 0 | 1 | 2 | 6 | 414 | 169 | 183 | 280 | 371 | 0 | 4 | 2 | 2 | 0 | 201 | 149 | 1.000000 | 1 | 0.995 | 1 | 0.993 | 0.989 | 0.979 | 1 | 0.969 | 0.985 | 0.99 | 1 | 0.002 |
| A3SS | 6679 | ENSG00000172531 | PPP1CA | chr11 | - | 67398721 | 67399163 | 67398721 | 67398856 | 67399560 | 67399665 | 307 | 109 | 246 | 274 | 340 | 0 | 0 | 1 | 0 | 0 | 294 | 290 | 173 | 235 | 435 | 0 | 0 | 0 | 0 | 0 | 298 | 149 | 1.000000 | 1 | 1 | 1 | 0.992 | 1 | 1 | 1 | 1 | 1 | 1 | 1 | -0.002 |
| A3SS | 1003 | ENSG00000099956 | SMARCB1 | chr22 | + | 23800943 | 23803422 | 23803294 | 23803422 | 23793558 | 23793688 | 78 | 57 | 112 | 93 | 240 | 0 | 0 | 0 | 0 | 0 | 236 | 165 | 49 | 114 | 166 | 0 | 3 | 0 | 0 | 0 | 298 | 149 | 1.000000 | 1 | 1 | 1 | 1 | 1 | 1 | 1 | 0.965 | 1 | 1 | 1 | 0.007 |
| A3SS | 6832 | ENSG00000073578 | SDHA | chr5 | + | 256333 | 256679 | 256350 | 256679 | 254392 | 254506 | 255 | 137 | 225 | 221 | 203 | 0 | 1 | 0 | 0 | 0 | 135 | 327 | 187 | 193 | 348 | 0 | 0 | 0 | 0 | 0 | 166 | 149 | 1.000000 | 1 | 1 | 0.992 | 1 | 1 | 1 | 1 | 1 | 1 | 1 | 1 | -0.002 |
| A3SS | 1441 | ENSG00000004142 | POLDIP2 | chr17 | - | 28354487 | 28354585 | 28354487 | 28354531 | 28355794 | 28355876 | 103 | 38 | 101 | 87 | 218 | 0 | 0 | 0 | 0 | 0 | 140 | 129 | 85 | 94 | 248 | 0 | 2 | 1 | 0 | 0 | 203 | 149 | 1.000000 | 1 | 1 | 1 | 1 | 1 | 1 | 1 | 0.979 | 0.984 | 1 | 1 | 0.007 |
| A3SS | 11454 | ENSG00000132740 | IGHMBP2 | chr11 | + | 68930322 | 68930434 | 68930357 | 68930434 | 68929182 | 68929357 | 10 | 10 | 12 | 11 | 2 | 0 | 2 | 0 | 1 | 0 | 45 | 14 | 25 | 45 | 63 | 0 | 0 | 0 | 0 | 0 | 184 | 149 | 1.000000 | 1 | 1 | 0.802 | 1 | 0.899 | 1 | 1 | 1 | 1 | 1 | 1 | -0.06 |
| A3SS | 1111 | ENSG00000004975 | DVL2 | chr17 | - | 7227654 | 7227783 | 7227654 | 7227711 | 7227976 | 7228044 | 59 | 37 | 28 | 64 | 36 | 0 | 0 | 0 | 1 | 0 | 56 | 68 | 57 | 52 | 137 | 0 | 0 | 0 | 0 | 0 | 221 | 149 | 1.000000 | 1 | 1 | 1 | 1 | 0.977 | 1 | 1 | 1 | 1 | 1 | 1 | -0.005 |
| A3SS | 2749 | ENSG00000129566 | TEP1 | chr14 | - | 20371488 | 20371632 | 20371488 | 20371607 | 20372732 | 20372857 | 82 | 28 | 53 | 71 | 68 | 0 | 1 | 0 | 0 | 0 | 128 | 79 | 22 | 53 | 165 | 0 | 0 | 0 | 3 | 0 | 174 | 149 | 1.000000 | 1 | 1 | 0.96 | 1 | 1 | 1 | 1 | 1 | 1 | 0.938 | 1 | 0.004 |
| A3SS | 3824 | ENSG00000155229 | MMS19 | chr10 | - | 97477346 | 97477416 | 97477346 | 97477404 | 97477854 | 97477929 | 156 | 87 | 142 | 131 | 125 | 1 | 0 | 2 | 2 | 3 | 201 | 126 | 99 | 112 | 247 | 0 | 0 | 1 | 0 | 0 | 161 | 149 | 1.000000 | 1 | 0.993 | 1 | 0.985 | 0.984 | 0.975 | 1 | 1 | 0.989 | 1 | 1 | -0.01 |
| A3SS | 5816 | ENSG00000160058 | BSDC1 | chr1 | - | 32383829 | 32383939 | 32383829 | 32383906 | 32386778 | 32386895 | 36 | 40 | 42 | 49 | 39 | 0 | 2 | 1 | 0 | 0 | 75 | 59 | 68 | 49 | 98 | 0 | 0 | 1 | 0 | 5 | 182 | 149 | 1.000000 | 1 | 1 | 0.942 | 0.972 | 1 | 1 | 1 | 1 | 0.982 | 1 | 0.941 | -0.002 |
| A3SS | 9502 | ENSG00000158874 | APOA2 | chr1 | - | 161222917 | 161223266 | 161222917 | 161223050 | 161223349 | 161223425 | 0 | 14 | 7 | 14 | 11 | 0 | 0 | 0 | 0 | 0 | 52 | 13 | 32 | 75 | 17 | 0 | 0 | 1 | 0 | 0 | 298 | 149 | 1.000000 | 1 | NA | 1 | 1 | 1 | 1 | 1 | 1 | 0.941 | 1 | 1 | 0.012 |
| A3SS | 2096 | ENSG00000140983 | RHOT2 | chr16 | + | 668487 | 668699 | 668655 | 668699 | 668352 | 668411 | 80 | 25 | 68 | 82 | 14 | 1 | 0 | 0 | 0 | 0 | 40 | 92 | 92 | 31 | 155 | 0 | 4 | 0 | 0 | 0 | 298 | 149 | 1.000000 | 1 | 0.976 | 1 | 1 | 1 | 1 | 1 | 0.92 | 1 | 1 | 1 | 0.011 |
| A3SS | 4371 | ENSG00000145029 | NICN1 | chr3 | - | 49426251 | 49426428 | 49426251 | 49426407 | 49429107 | 49429326 | 19 | 18 | 30 | 21 | 22 | 0 | 0 | 0 | 1 | 0 | 79 | 34 | 24 | 64 | 48 | 0 | 0 | 0 | 0 | 0 | 170 | 149 | 1.000000 | 1 | 1 | 1 | 1 | 0.948 | 1 | 1 | 1 | 1 | 1 | 1 | -0.01 |
| A3SS | 7770 | ENSG00000006652 | IFRD1 | chr7 | + | 112458860 | 112459018 | 112458916 | 112459018 | 112456913 | 112457038 | 122 | 112 | 102 | 120 | 145 | 0 | 0 | 0 | 0 | 7 | 126 | 146 | 148 | 158 | 162 | 0 | 0 | 2 | 0 | 0 | 205 | 149 | 1.000000 | 1 | 1 | 1 | 1 | 1 | 0.938 | 1 | 1 | 0.982 | 1 | 1 | -0.009 |
| A3SS | 730 | ENSG00000064932 | SBNO2 | chr19 | - | 1117322 | 1117499 | 1117322 | 1117469 | 1119010 | 1119164 | 72 | 17 | 33 | 35 | 11 | 0 | 0 | 1 | 0 | 0 | 79 | 85 | 31 | 30 | 82 | 0 | 1 | 0 | 0 | 0 | 179 | 149 | 1.000000 | 1 | 1 | 1 | 0.965 | 1 | 1 | 1 | 0.986 | 1 | 1 | 1 | -0.004 |
| A3SS | 12963 | ENSG00000123384 | LRP1 | chr12 | + | 57148980 | 57150059 | 57149679 | 57150059 | 57145226 | 57145490 | 2 | 4 | 4 | 3 | 0 | 2 | 0 | 0 | 0 | 0 | 57 | 3 | 8 | 36 | 47 | 0 | 0 | 0 | 0 | 0 | 298 | 149 | 1.000000 | 1 | 0.333 | 1 | 1 | 1 | NA | 1 | 1 | 1 | 1 | 1 | -0.167 |
| A3SS | 5491 | ENSG00000099622 | CIRBP | chr19 | + | 1270927 | 1271467 | 1271328 | 1271467 | 1269331 | 1269410 | 70 | 46 | 50 | 58 | 56 | 0 | 2 | 0 | 0 | 0 | 50 | 100 | 60 | 68 | 140 | 0 | 0 | 0 | 0 | 0 | 298 | 149 | 1.000000 | 1 | 1 | 0.92 | 1 | 1 | 1 | 1 | 1 | 1 | 1 | 1 | -0.016 |
| A3SS | 2517 | ENSG00000226777 | FAM30A | chr14 | + | 105922198 | 105922344 | 105922202 | 105922344 | 105921149 | 105921258 | 26 | 4 | 16 | 23 | 5 | 0 | 0 | 1 | 0 | 0 | 51 | 39 | 23 | 35 | 64 | 0 | 0 | 1 | 0 | 0 | 153 | 149 | 1.000000 | 1 | 1 | 1 | 0.94 | 1 | 1 | 1 | 1 | 1 | 1 | 1 | -0.012 |
| A3SS | 2080 | ENSG00000140694 | PARN | chr16 | - | 14604145 | 14604226 | 14604145 | 14604223 | 14606483 | 14606526 | 198 | 162 | 198 | 179 | 311 | 0 | 0 | 6 | 2 | 0 | 321 | 260 | 259 | 265 | 81 | 0 | 0 | 1 | 0 | 0 | 152 | 149 | 1.000000 | 1 | 1 | 1 | 0.97 | 0.989 | 1 | 1 | 1 | 0.996 | 1 | 1 | -0.007 |
| A3SS | 9951 | ENSG00000141480 | ARRB2 | chr17 | + | 4715150 | 4715336 | 4715199 | 4715336 | 4715012 | 4715043 | 13 | 9 | 5 | 14 | 0 | 0 | 0 | 1 | 0 | 0 | 47 | 39 | 13 | 29 | 50 | 0 | 2 | 0 | 0 | 0 | 198 | 149 | 1.000000 | 1 | 1 | 1 | 0.79 | 1 | NA | 1 | 0.936 | 1 | 1 | 1 | -0.04 |
| A3SS | 10482 | ENSG00000177646 | ACAD9 | chr3 | + | 128884652 | 128884746 | 128884674 | 128884746 | 128879619 | 128879841 | 38 | 42 | 29 | 36 | 72 | 1 | 0 | 0 | 0 | 0 | 103 | 67 | 35 | 67 | 82 | 0 | 0 | 0 | 3 | 0 | 171 | 149 | 1.000000 | 1 | 0.971 | 1 | 1 | 1 | 1 | 1 | 1 | 1 | 0.951 | 1 | 0.004 |
| A3SS | 10676 | ENSG00000248323 | LUCAT1 | chr5 | - | 91310626 | 91313229 | 91310626 | 91311032 | 91313637 | 91313777 | 16 | 6 | 5 | 14 | 3 | 0 | 0 | 0 | 0 | 0 | 42 | 38 | 79 | 22 | 0 | 0 | 0 | 3 | 0 | 0 | 298 | 149 | 1.000000 | 1 | 1 | 1 | 1 | 1 | 1 | 1 | 1 | 0.929 | 1 | NA | 0.018 |
| A3SS | 3946 | ENSG00000203875 | SNHG5 | chr6 | - | 85677423 | 85678032 | 85677423 | 85677875 | 85678135 | 85678188 | 243 | 397 | 317 | 246 | 343 | 0 | 0 | 0 | 2 | 0 | 97 | 304 | 386 | 355 | 540 | 0 | 0 | 1 | 0 | 0 | 298 | 149 | 1.000000 | 1 | 1 | 1 | 1 | 0.984 | 1 | 1 | 1 | 0.995 | 1 | 1 | -0.002 |
| A3SS | 12460 | ENSG00000176155 | CCDC57 | chr17 | - | 82193755 | 82194139 | 82193755 | 82193830 | 82195262 | 82195364 | 22 | 25 | 33 | 42 | 8 | 0 | 0 | 0 | 0 | 0 | 71 | 42 | 27 | 47 | 79 | 0 | 0 | 2 | 0 | 0 | 298 | 149 | 1.000000 | 1 | 1 | 1 | 1 | 1 | 1 | 1 | 1 | 0.871 | 1 | 1 | 0.026 |
| A3SS | 13418 | ENSG00000071564 | TCF3 | chr19 | - | 1615284 | 1615520 | 1615284 | 1615515 | 1615685 | 1615821 | 20 | 8 | 23 | 14 | 4 | 0 | 0 | 0 | 0 | 0 | 50 | 51 | 22 | 28 | 53 | 4 | 0 | 0 | 0 | 0 | 154 | 149 | 1.000000 | 1 | 1 | 1 | 1 | 1 | 1 | 0.924 | 1 | 1 | 1 | 1 | 0.015 |
| A3SS | 7191 | ENSG00000001497 | LAS1L | chrX | - | 65520701 | 65522285 | 65520701 | 65520817 | 65523559 | 65523707 | 11 | 9 | 14 | 24 | 9 | 1 | 0 | 0 | 0 | 0 | 92 | 22 | 9 | 40 | 38 | 0 | 0 | 0 | 0 | 0 | 298 | 149 | 1.000000 | 1 | 0.846 | 1 | 1 | 1 | 1 | 1 | 1 | 1 | 1 | 1 | -0.031 |
| A3SS | 10793 | ENSG00000168066 | SF1 | chr11 | - | 64767191 | 64767251 | 64767191 | 64767246 | 64767566 | 64767844 | 29 | 28 | 33 | 47 | 6 | 0 | 0 | 0 | 0 | 0 | 50 | 46 | 37 | 86 | 58 | 0 | 3 | 0 | 0 | 0 | 154 | 149 | 1.000000 | 1 | 1 | 1 | 1 | 1 | 1 | 1 | 0.937 | 1 | 1 | 1 | 0.013 |
| A3SS | 5120 | ENSG00000163795 | ZNF513 | chr2 | - | 27378466 | 27379054 | 27378466 | 27378615 | 27380092 | 27380444 | 62 | 30 | 56 | 51 | 37 | 0 | 0 | 0 | 0 | 0 | 93 | 84 | 45 | 59 | 89 | 0 | 0 | 3 | 0 | 0 | 298 | 149 | 1.000000 | 1 | 1 | 1 | 1 | 1 | 1 | 1 | 1 | 0.882 | 1 | 1 | 0.024 |
| A3SS | 5976 | ENSG00000115155 | OTOF | chr2 | - | 26472518 | 26472649 | 26472518 | 26472589 | 26473131 | 26473294 | 0 | 0 | 4 | 2 | 0 | 6 | 0 | 0 | 1 | 4 | 50 | 28 | 0 | 62 | 0 | 191 | 165 | 18 | 273 | 22 | 209 | 149 | 1.000000 | 1 | 0 | NA | 1 | 0.588 | 0 | 0.157 | 0.108 | 0 | 0.139 | 0 | 0.316 |
| A3SS | 2563 | ENSG00000197102 | DYNC1H1 | chr14 | + | 102050434 | 102056443 | 102050471 | 102056443 | 102050070 | 102050198 | 183 | 117 | 176 | 205 | 98 | 0 | 0 | 0 | 3 | 0 | 229 | 185 | 102 | 207 | 189 | 0 | 4 | 0 | 4 | 2 | 186 | 149 | 1.000000 | 1 | 1 | 1 | 1 | 0.982 | 1 | 1 | 0.974 | 1 | 0.976 | 0.987 | 0.009 |
| A3SS | 276 | ENSG00000105607 | GCDH | chr19 | + | 12891486 | 12891974 | 12891830 | 12891974 | 12891270 | 12891395 | 45 | 25 | 50 | 46 | 12 | 1 | 0 | 0 | 0 | 0 | 78 | 61 | 28 | 45 | 98 | 0 | 0 | 0 | 0 | 0 | 298 | 149 | 1.000000 | 1 | 0.957 | 1 | 1 | 1 | 1 | 1 | 1 | 1 | 1 | 1 | -0.009 |
| A3SS | 2203 | ENSG00000140545 | MFGE8 | chr15 | - | 88901550 | 88901735 | 88901550 | 88901671 | 88905756 | 88905901 | 23 | 14 | 39 | 32 | 9 | 0 | 1 | 0 | 0 | 0 | 22 | 42 | 99 | 40 | 45 | 0 | 0 | 0 | 0 | 0 | 213 | 149 | 1.000000 | 1 | 1 | 0.907 | 1 | 1 | 1 | 1 | 1 | 1 | 1 | 1 | -0.019 |
| A3SS | 11989 | ENSG00000099783 | HNRNPM | chr19 | + | 8465323 | 8465515 | 8465479 | 8465515 | 8463592 | 8463686 | 843 | 606 | 711 | 823 | 1080 | 0 | 0 | 1 | 0 | 3 | 1087 | 836 | 611 | 907 | 753 | 0 | 1 | 2 | 0 | 0 | 298 | 149 | 1.000000 | 1 | 1 | 1 | 0.997 | 1 | 0.994 | 1 | 0.998 | 0.993 | 1 | 1 | 0 |
| A3SS | 1842 | ENSG00000141002 | TCF25 | chr16 | + | 89895037 | 89895137 | 89895082 | 89895137 | 89892192 | 89892275 | 67 | 27 | 28 | 44 | 17 | 0 | 0 | 0 | 1 | 0 | 90 | 65 | 27 | 36 | 95 | 0 | 0 | 0 | 0 | 0 | 194 | 149 | 1.000000 | 1 | 1 | 1 | 1 | 0.971 | 1 | 1 | 1 | 1 | 1 | 1 | -0.006 |
| A3SS | 6551 | ENSG00000127463 | EMC1 | chr1 | - | 19220763 | 19220848 | 19220763 | 19220810 | 19222623 | 19222834 | 50 | 23 | 35 | 38 | 45 | 1 | 0 | 0 | 2 | 1 | 74 | 68 | 22 | 63 | 94 | 0 | 0 | 2 | 0 | 0 | 187 | 149 | 1.000000 | 1 | 0.976 | 1 | 1 | 0.938 | 0.973 | 1 | 1 | 0.898 | 1 | 1 | -0.002 |
| A3SS | 6327 | ENSG00000139636 | LMBR1L | chr12 | - | 49100387 | 49101323 | 49100387 | 49100454 | 49101471 | 49101549 | 23 | 20 | 28 | 28 | 12 | 0 | 0 | 0 | 0 | 0 | 60 | 57 | 21 | 40 | 63 | 0 | 0 | 1 | 0 | 0 | 298 | 149 | 1.000000 | 1 | 1 | 1 | 1 | 1 | 1 | 1 | 1 | 0.913 | 1 | 1 | 0.017 |
| A3SS | 9513 | ENSG00000152457 | DCLRE1C | chr10 | - | 14935462 | 14935564 | 14935462 | 14935523 | 14936537 | 14936593 | 163 | 111 | 130 | 124 | 104 | 0 | 0 | 0 | 0 | 0 | 204 | 170 | 158 | 136 | 94 | 0 | 0 | 0 | 0 | 7 | 190 | 149 | 1.000000 | 1 | 1 | 1 | 1 | 1 | 1 | 1 | 1 | 1 | 1 | 0.913 | 0.017 |
| A3SS | 6167 | ENSG00000238113 | LINC01410 | chr9 | + | 62809767 | 62811589 | 62811399 | 62811589 | 62803983 | 62804140 | 15 | 15 | 6 | 21 | 16 | 0 | 0 | 0 | 0 | 0 | 15 | 22 | 12 | 96 | 58 | 0 | 0 | 0 | 0 | 3 | 298 | 149 | 1.000000 | 1 | 1 | 1 | 1 | 1 | 1 | 1 | 1 | 1 | 1 | 0.906 | 0.019 |
| A3SS | 1299 | ENSG00000185359 | HGS | chr17 | + | 81700694 | 81700814 | 81700799 | 81700814 | 81700466 | 81700600 | 23 | 1 | 16 | 35 | 7 | 0 | 0 | 1 | 0 | 0 | 22 | 77 | 6 | 41 | 65 | 0 | 0 | 0 | 0 | 0 | 254 | 149 | 1.000000 | 1 | 1 | 1 | 0.904 | 1 | 1 | 1 | 1 | 1 | 1 | 1 | -0.019 |
| A3SS | 7328 | ENSG00000159348 | CYB5R1 | chr1 | - | 202965886 | 202966600 | 202965886 | 202966109 | 202966748 | 202966898 | 66 | 44 | 33 | 42 | 59 | 0 | 0 | 1 | 1 | 0 | 43 | 64 | 49 | 46 | 171 | 0 | 0 | 1 | 0 | 0 | 298 | 149 | 1.000000 | 1 | 1 | 1 | 0.943 | 0.955 | 1 | 1 | 1 | 0.961 | 1 | 1 | -0.013 |
| A3SS | 11007 | ENSG00000160932 | LY6E | chr8 | + | 143020882 | 143020991 | 143020887 | 143020991 | 143018832 | 143019119 | 12 | 8 | 12 | 19 | 7 | 1 | 1 | 0 | 1 | 0 | 48 | 40 | 26 | 53 | 20 | 0 | 4 | 0 | 5 | 0 | 154 | 149 | 1.000000 | 1 | 0.921 | 0.886 | 1 | 0.948 | 1 | 1 | 0.906 | 1 | 0.911 | 1 | -0.012 |
| A3SS | 12895 | ENSG00000135720 | DYNC1LI2 | chr16 | - | 66742437 | 66742668 | 66742437 | 66742554 | 66749196 | 66749313 | 161 | 160 | 176 | 167 | 130 | 0 | 0 | 1 | 0 | 0 | 182 | 188 | 236 | 210 | 106 | 0 | 0 | 0 | 0 | 0 | 263 | 149 | 1.000000 | 1 | 1 | 1 | 0.99 | 1 | 1 | 1 | 1 | 1 | 1 | 1 | -0.002 |
| A3SS | 4612 | ENSG00000161204 | ABCF3 | chr3 | + | 184187663 | 184187983 | 184187860 | 184187983 | 184186795 | 184186875 | 18 | 26 | 20 | 18 | 6 | 0 | 0 | 0 | 0 | 2 | 37 | 71 | 27 | 29 | 51 | 0 | 0 | 0 | 0 | 0 | 298 | 149 | 1.000000 | 1 | 1 | 1 | 1 | 1 | 0.6 | 1 | 1 | 1 | 1 | 1 | -0.08 |
| A3SS | 370 | ENSG00000142541 | RPL13A | chr19 | + | 49490126 | 49490297 | 49490231 | 49490297 | 49489849 | 49489922 | 31 | 22 | 20 | 27 | 18 | 12021 | 4703 | 13109 | 9474 | 13052 | 78 | 50 | 31 | 30 | 56 | 5818 | 7716 | 5279 | 4104 | 9979 | 254 | 149 | 1.000000 | 1 | 0.002 | 0.003 | 0.001 | 0.002 | 0.001 | 0.008 | 0.004 | 0.003 | 0.004 | 0.003 | -0.003 |
| A3SS | 1271 | ENSG00000125450 | NUP85 | chr17 | + | 75234636 | 75235201 | 75235099 | 75235201 | 75233057 | 75233158 | 28 | 29 | 52 | 45 | 45 | 0 | 0 | 0 | 0 | 0 | 83 | 39 | 50 | 66 | 88 | 0 | 1 | 0 | 0 | 0 | 298 | 149 | 1.000000 | 1 | 1 | 1 | 1 | 1 | 1 | 1 | 1 | 0.962 | 1 | 1 | 0.008 |
| A3SS | 8983 | ENSG00000123600 | METTL8 | chr2 | - | 171330558 | 171330698 | 171330558 | 171330616 | 171331764 | 171331867 | 36 | 31 | 36 | 47 | 16 | 1 | 0 | 0 | 0 | 0 | 73 | 58 | 51 | 31 | 80 | 0 | 0 | 0 | 3 | 0 | 231 | 149 | 1.000000 | 1 | 0.959 | 1 | 1 | 1 | 1 | 1 | 1 | 1 | 0.87 | 1 | 0.018 |
| A3SS | 10268 | ENSG00000102145 | GATA1 | chrX | + | 48793792 | 48794311 | 48794133 | 48794311 | 48793171 | 48793297 | 8 | 9 | 2 | 12 | 2 | 2 | 0 | 0 | 0 | 0 | 61 | 9 | 8 | 15 | 66 | 0 | 0 | 0 | 0 | 0 | 298 | 149 | 1.000000 | 1 | 0.667 | 1 | 1 | 1 | 1 | 1 | 1 | 1 | 1 | 1 | -0.067 |
| A3SS | 6142 | ENSG00000159079 | CFAP298 | chr21 | - | 32601666 | 32603292 | 32601666 | 32601973 | 32604124 | 32604283 | 168 | 110 | 172 | 194 | 126 | 1 | 0 | 0 | 0 | 0 | 252 | 181 | 102 | 147 | 214 | 0 | 0 | 0 | 0 | 0 | 298 | 149 | 1.000000 | 1 | 0.988 | 1 | 1 | 1 | 1 | 1 | 1 | 1 | 1 | 1 | -0.002 |
| A3SS | 4327 | ENSG00000221978 | CCNL2 | chr1 | - | 1390765 | 1391575 | 1390765 | 1390865 | 1392789 | 1392803 | 201 | 160 | 267 | 202 | 180 | 0 | 0 | 2 | 0 | 0 | 293 | 194 | 187 | 223 | 239 | 0 | 1 | 0 | 0 | 0 | 298 | 149 | 1.000000 | 1 | 1 | 1 | 0.985 | 1 | 1 | 1 | 0.99 | 1 | 1 | 1 | -0.001 |
| A3SS | 10761 | ENSG00000143569 | UBAP2L | chr1 | + | 154258875 | 154259030 | 154258976 | 154259030 | 154257344 | 154257465 | 30 | 46 | 50 | 39 | 84 | 1 | 0 | 3 | 1 | 0 | 70 | 74 | 102 | 103 | 26 | 0 | 7 | 0 | 3 | 0 | 250 | 149 | 1.000000 | 1 | 0.947 | 1 | 0.909 | 0.959 | 1 | 1 | 0.863 | 1 | 0.953 | 1 | 0 |
| A3SS | 5759 | ENSG00000144283 | PKP4 | chr2 | + | 158624877 | 158625427 | 158625124 | 158625427 | 158621230 | 158621421 | 44 | 55 | 70 | 73 | 159 | 0 | 1 | 0 | 0 | 1 | 31 | 69 | 80 | 150 | 197 | 0 | 0 | 1 | 0 | 5 | 298 | 149 | 1.000000 | 1 | 1 | 0.965 | 1 | 1 | 0.988 | 1 | 1 | 0.976 | 1 | 0.952 | 0.005 |
| A3SS | 6115 | ENSG00000106610 | STAG3L4 | chr7 | + | 67308576 | 67309139 | 67308934 | 67309139 | 67302658 | 67302885 | 28 | 10 | 27 | 26 | 28 | 0 | 0 | 0 | 2 | 0 | 55 | 42 | 36 | 25 | 86 | 0 | 0 | 0 | 0 | 0 | 298 | 149 | 1.000000 | 1 | 1 | 1 | 1 | 0.867 | 1 | 1 | 1 | 1 | 1 | 1 | -0.027 |
| A3SS | 11300 | ENSG00000077721 | UBE2A | chrX | + | 119574900 | 119574981 | 119574907 | 119574981 | 119574562 | 119574755 | 29 | 22 | 32 | 28 | 9 | 1 | 0 | 0 | 0 | 0 | 54 | 56 | 36 | 30 | 69 | 0 | 0 | 0 | 0 | 0 | 156 | 149 | 1.000000 | 1 | 0.965 | 1 | 1 | 1 | 1 | 1 | 1 | 1 | 1 | 1 | -0.007 |
| A3SS | 9948 | ENSG00000155096 | AZIN1 | chr8 | - | 102843550 | 102843747 | 102843550 | 102843729 | 102849998 | 102850165 | 121 | 82 | 142 | 153 | 139 | 0 | 0 | 0 | 1 | 4 | 111 | 143 | 184 | 226 | 97 | 0 | 4 | 4 | 1 | 0 | 167 | 149 | 1.000000 | 1 | 1 | 1 | 1 | 0.993 | 0.969 | 1 | 0.97 | 0.976 | 0.995 | 1 | 0.004 |
| A3SS | 9715 | ENSG00000146918 | NCAPG2 | chr7 | - | 158675476 | 158675656 | 158675476 | 158675616 | 158679959 | 158680085 | 17 | 7 | 21 | 22 | 26 | 0 | 2 | 1 | 0 | 0 | 51 | 25 | 50 | 90 | 0 | 0 | 0 | 1 | 9 | 0 | 189 | 149 | 1.000000 | 1 | 1 | 0.734 | 0.943 | 1 | 1 | 1 | 1 | 0.975 | 0.887 | NA | -0.03 |
| A3SS | 5993 | ENSG00000110619 | CARS1 | chr11 | - | 3001969 | 3002174 | 3001969 | 3002053 | 3005365 | 3005433 | 24 | 37 | 25 | 38 | 36 | 2 | 0 | 0 | 0 | 0 | 64 | 71 | 40 | 48 | 60 | 0 | 0 | 0 | 0 | 0 | 270 | 149 | 1.000000 | 1 | 0.869 | 1 | 1 | 1 | 1 | 1 | 1 | 1 | 1 | 1 | -0.026 |
| A3SS | 4023 | ENSG00000189308 | LIN54 | chr4 | - | 82970326 | 82970469 | 82970326 | 82970465 | 82978882 | 82979006 | 54 | 62 | 65 | 60 | 91 | 0 | 0 | 1 | 0 | 0 | 96 | 89 | 128 | 82 | 60 | 0 | 0 | 3 | 4 | 0 | 153 | 149 | 1.000000 | 1 | 1 | 1 | 0.984 | 1 | 1 | 1 | 1 | 0.976 | 0.952 | 1 | 0.011 |
| A3SS | 8985 | ENSG00000123600 | METTL8 | chr2 | - | 171330558 | 171330698 | 171330558 | 171330616 | 171337452 | 171337502 | 36 | 30 | 31 | 40 | 16 | 0 | 1 | 0 | 0 | 0 | 73 | 54 | 47 | 26 | 74 | 0 | 0 | 0 | 0 | 0 | 231 | 149 | 1.000000 | 1 | 1 | 0.951 | 1 | 1 | 1 | 1 | 1 | 1 | 1 | 1 | -0.01 |
| A3SS | 12694 | ENSG00000138185 | ENTPD1 | chr10 | + | 95844475 | 95844635 | 95844554 | 95844635 | 95842343 | 95842494 | 131 | 82 | 79 | 92 | 135 | 0 | 3 | 3 | 1 | 0 | 104 | 138 | 114 | 202 | 82 | 0 | 3 | 4 | 1 | 0 | 228 | 149 | 1.000000 | 1 | 1 | 0.947 | 0.945 | 0.984 | 1 | 1 | 0.968 | 0.949 | 0.992 | 1 | -0.007 |
| A3SS | 3884 | ENSG00000120708 | TGFBI | chr5 | + | 136046334 | 136047015 | 136046850 | 136047015 | 136044057 | 136044122 | 484 | 262 | 273 | 299 | 525 | 0 | 2 | 0 | 0 | 0 | 484 | 162 | 203 | 346 | 769 | 0 | 0 | 0 | 0 | 0 | 298 | 149 | 1.000000 | 1 | 1 | 0.985 | 1 | 1 | 1 | 1 | 1 | 1 | 1 | 1 | -0.003 |
| A3SS | 11802 | ENSG00000089159 | PXN | chr12 | - | 120216272 | 120217116 | 120216272 | 120216581 | 120221622 | 120221758 | 24 | 19 | 7 | 7 | 11 | 0 | 0 | 1 | 0 | 3 | 16 | 39 | 33 | 42 | 58 | 0 | 0 | 0 | 0 | 0 | 298 | 149 | 1.000000 | 1 | 1 | 1 | 0.778 | 1 | 0.647 | 1 | 1 | 1 | 1 | 1 | -0.115 |
| A3SS | 707 | ENSG00000105355 | PLIN3 | chr19 | - | 4852015 | 4852301 | 4852015 | 4852265 | 4859589 | 4859672 | 33 | 10 | 28 | 25 | 8 | 0 | 0 | 0 | 1 | 0 | 67 | 37 | 27 | 41 | 52 | 0 | 0 | 0 | 0 | 0 | 185 | 149 | 1.000000 | 1 | 1 | 1 | 1 | 0.953 | 1 | 1 | 1 | 1 | 1 | 1 | -0.009 |
| A3SS | 7225 | ENSG00000175634 | RPS6KB2 | chr11 | + | 67433994 | 67434275 | 67434197 | 67434275 | 67433339 | 67433447 | 70 | 42 | 73 | 73 | 48 | 0 | 0 | 0 | 1 | 0 | 107 | 103 | 49 | 59 | 107 | 0 | 0 | 0 | 0 | 0 | 298 | 149 | 1.000000 | 1 | 1 | 1 | 1 | 0.973 | 1 | 1 | 1 | 1 | 1 | 1 | -0.005 |
| A3SS | 12540 | ENSG00000175197 | DDIT3 | chr12 | - | 57517268 | 57517595 | 57517268 | 57517542 | 57517705 | 57517753 | 9 | 9 | 11 | 18 | 29 | 0 | 0 | 0 | 1 | 0 | 45 | 37 | 32 | 10 | 71 | 0 | 0 | 1 | 0 | 0 | 202 | 149 | 1.000000 | 1 | 1 | 1 | 1 | 1 | 1 | 1 | 1 | 0.959 | 1 | 1 | 0.008 |
| A3SS | 484 | ENSG00000013275 | PSMC4 | chr19 | + | 39972368 | 39972555 | 39972461 | 39972555 | 39972145 | 39972244 | 194 | 82 | 138 | 159 | 168 | 2 | 0 | 0 | 1 | 0 | 162 | 209 | 116 | 140 | 233 | 0 | 2 | 2 | 4 | 0 | 242 | 149 | 1.000000 | 1 | 0.984 | 1 | 1 | 0.99 | 1 | 1 | 0.985 | 0.973 | 0.956 | 1 | 0.012 |
| A3SS | 6504 | ENSG00000166441 | RPL27A | chr11 | + | 8684685 | 8684892 | 8684717 | 8684892 | 8684005 | 8684081 | 51 | 31 | 40 | 71 | 77 | 8935 | 4893 | 9922 | 8633 | 4356 | 92 | 52 | 37 | 53 | 154 | 7636 | 7009 | 6101 | 5366 | 9677 | 181 | 149 | 1.000000 | 1 | 0.005 | 0.005 | 0.003 | 0.007 | 0.014 | 0.01 | 0.006 | 0.005 | 0.008 | 0.013 | -0.002 |
| A3SS | 373 | ENSG00000011304 | PTBP1 | chr19 | + | 804531 | 804702 | 804540 | 804702 | 804291 | 804438 | 143 | 66 | 120 | 169 | 92 | 2 | 0 | 0 | 0 | 0 | 94 | 171 | 89 | 144 | 209 | 0 | 0 | 0 | 0 | 0 | 158 | 149 | 1.000000 | 1 | 0.985 | 1 | 1 | 1 | 1 | 1 | 1 | 1 | 1 | 1 | -0.003 |
| A3SS | 550 | ENSG00000129347 | KRI1 | chr19 | - | 10562728 | 10562837 | 10562728 | 10562821 | 10564928 | 10565034 | 49 | 32 | 40 | 31 | 12 | 0 | 0 | 0 | 0 | 1 | 55 | 46 | 30 | 30 | 119 | 0 | 0 | 0 | 0 | 0 | 165 | 149 | 1.000000 | 1 | 1 | 1 | 1 | 1 | 0.916 | 1 | 1 | 1 | 1 | 1 | -0.017 |
| A3SS | 8273 | ENSG00000106636 | YKT6 | chr7 | + | 44208406 | 44211124 | 44208794 | 44211124 | 44208132 | 44208198 | 21 | 12 | 13 | 20 | 35 | 1 | 0 | 0 | 0 | 0 | 56 | 42 | 24 | 55 | 40 | 0 | 0 | 0 | 0 | 0 | 298 | 149 | 1.000000 | 1 | 0.913 | 1 | 1 | 1 | 1 | 1 | 1 | 1 | 1 | 1 | -0.017 |
| A3SS | 5569 | ENSG00000114626 | ABTB1 | chr3 | + | 127677464 | 127677843 | 127677675 | 127677843 | 127677167 | 127677286 | 62 | 13 | 71 | 43 | 47 | 0 | 0 | 1 | 0 | 0 | 81 | 80 | 28 | 49 | 114 | 0 | 0 | 2 | 0 | 0 | 298 | 149 | 1.000000 | 1 | 1 | 1 | 0.973 | 1 | 1 | 1 | 1 | 0.875 | 1 | 1 | 0.02 |
| A3SS | 654 | ENSG00000130475 | FCHO1 | chr19 | + | 17770424 | 17770577 | 17770463 | 17770577 | 17766668 | 17766810 | 116 | 65 | 116 | 120 | 84 | 0 | 1 | 1 | 1 | 1 | 46 | 136 | 98 | 159 | 177 | 0 | 0 | 0 | 0 | 0 | 188 | 149 | 1.000000 | 1 | 1 | 0.981 | 0.989 | 0.99 | 0.985 | 1 | 1 | 1 | 1 | 1 | -0.011 |
| A3SS | 1312 | ENSG00000136490 | LIMD2 | chr17 | - | 63698798 | 63698938 | 63698798 | 63698818 | 63699027 | 63699069 | 371 | 101 | 346 | 262 | 84 | 5 | 0 | 1 | 3 | 0 | 218 | 313 | 185 | 158 | 405 | 0 | 0 | 0 | 0 | 0 | 269 | 149 | 1.000000 | 1 | 0.976 | 1 | 0.995 | 0.98 | 1 | 1 | 1 | 1 | 1 | 1 | -0.01 |
| A3SS | 6841 | ENSG00000151694 | ADAM17 | chr2 | - | 9521202 | 9523338 | 9521202 | 9521321 | 9526110 | 9526264 | 132 | 109 | 91 | 96 | 162 | 0 | 0 | 0 | 0 | 0 | 130 | 153 | 166 | 209 | 47 | 0 | 0 | 1 | 0 | 0 | 298 | 149 | 1.000000 | 1 | 1 | 1 | 1 | 1 | 1 | 1 | 1 | 0.988 | 1 | 1 | 0.002 |
| A3SS | 6146 | ENSG00000134453 | RBM17 | chr10 | + | 6106140 | 6106238 | 6106143 | 6106238 | 6104930 | 6105097 | 148 | 99 | 109 | 204 | 170 | 1 | 0 | 0 | 0 | 0 | 172 | 126 | 203 | 250 | 94 | 0 | 0 | 0 | 5 | 0 | 152 | 149 | 1.000000 | 1 | 0.993 | 1 | 1 | 1 | 1 | 1 | 1 | 1 | 0.98 | 1 | 0.003 |
| A3SS | 4308 | ENSG00000178035 | IMPDH2 | chr3 | - | 49028757 | 49028806 | 49028757 | 49028765 | 49029252 | 49029416 | 108 | 42 | 116 | 98 | 88 | 0 | 0 | 0 | 0 | 0 | 125 | 102 | 101 | 77 | 162 | 4 | 2 | 0 | 0 | 0 | 190 | 149 | 1.000000 | 1 | 1 | 1 | 1 | 1 | 1 | 0.961 | 0.976 | 1 | 1 | 1 | 0.013 |
| A3SS | 5421 | ENSG00000125637 | PSD4 | chr2 | + | 113185661 | 113186255 | 113185960 | 113186255 | 113185364 | 113185440 | 85 | 45 | 77 | 89 | 64 | 0 | 0 | 2 | 0 | 0 | 138 | 101 | 37 | 98 | 100 | 0 | 0 | 0 | 0 | 0 | 298 | 149 | 1.000000 | 1 | 1 | 1 | 0.951 | 1 | 1 | 1 | 1 | 1 | 1 | 1 | -0.01 |
| A3SS | 11711 | ENSG00000170909 | OSCAR | chr19 | - | 54096861 | 54097164 | 54096861 | 54096991 | 54099747 | 54099780 | 50 | 33 | 30 | 45 | 26 | 0 | 0 | 0 | 0 | 2 | 63 | 65 | 23 | 73 | 73 | 0 | 5 | 0 | 0 | 0 | 298 | 149 | 1.000000 | 1 | 1 | 1 | 1 | 1 | 0.867 | 1 | 0.867 | 1 | 1 | 1 | 0 |
| A3SS | 1397 | ENSG00000129667 | RHBDF2 | chr17 | - | 76487711 | 76487909 | 76487711 | 76487851 | 76500935 | 76501371 | 35 | 20 | 10 | 6 | 8 | 0 | 0 | 0 | 1 | 0 | 37 | 64 | 7 | 24 | 59 | 0 | 1 | 0 | 0 | 0 | 207 | 149 | 1.000000 | 1 | 1 | 1 | 1 | 0.812 | 1 | 1 | 0.979 | 1 | 1 | 1 | -0.033 |
| A3SS | 6120 | ENSG00000127564 | PKMYT1 | chr16 | - | 2973137 | 2973445 | 2973137 | 2973215 | 2973999 | 2974157 | 12 | 3 | 5 | 9 | 0 | 2 | 0 | 1 | 0 | 0 | 40 | 21 | 10 | 13 | 56 | 8 | 0 | 0 | 0 | 0 | 298 | 149 | 1.000000 | 1 | 0.75 | 1 | 0.714 | 1 | NA | 0.714 | 1 | 1 | 1 | 1 | -0.077 |
| A3SS | 7516 | ENSG00000143303 | RRNAD1 | chr1 | + | 156732280 | 156732473 | 156732365 | 156732473 | 156731990 | 156732115 | 11 | 14 | 13 | 22 | 12 | 1 | 0 | 0 | 0 | 0 | 53 | 18 | 7 | 33 | 71 | 0 | 0 | 0 | 0 | 0 | 234 | 149 | 1.000000 | 1 | 0.875 | 1 | 1 | 1 | 1 | 1 | 1 | 1 | 1 | 1 | -0.025 |
| A3SS | 10960 | ENSG00000095485 | CWF19L1 | chr10 | - | 100243697 | 100243777 | 100243697 | 100243772 | 100245798 | 100245913 | 116 | 50 | 68 | 70 | 87 | 0 | 0 | 1 | 2 | 0 | 129 | 87 | 72 | 97 | 116 | 0 | 0 | 0 | 1 | 0 | 154 | 149 | 1.000000 | 1 | 1 | 1 | 0.985 | 0.971 | 1 | 1 | 1 | 1 | 0.989 | 1 | -0.007 |
| A3SS | 10048 | ENSG00000123066 | MED13L | chr12 | - | 115980749 | 115982603 | 115980749 | 115980938 | 115983116 | 115983540 | 106 | 58 | 90 | 96 | 70 | 0 | 0 | 0 | 0 | 1 | 103 | 137 | 155 | 120 | 15 | 0 | 0 | 0 | 0 | 0 | 298 | 149 | 1.000000 | 1 | 1 | 1 | 1 | 1 | 0.972 | 1 | 1 | 1 | 1 | 1 | -0.006 |
| A3SS | 1837 | ENSG00000172775 | PSME3IP1 | chr16 | - | 57173727 | 57173869 | 57173727 | 57173796 | 57185824 | 57186040 | 218 | 133 | 170 | 170 | 281 | 0 | 0 | 0 | 2 | 0 | 269 | 268 | 198 | 219 | 128 | 0 | 0 | 0 | 0 | 0 | 222 | 149 | 1.000000 | 1 | 1 | 1 | 1 | 0.983 | 1 | 1 | 1 | 1 | 1 | 1 | -0.003 |
| A3SS | 3315 | ENSG00000103496 | STX4 | chr16 | + | 31034203 | 31034536 | 31034461 | 31034536 | 31034012 | 31034114 | 29 | 17 | 14 | 25 | 7 | 0 | 0 | 0 | 0 | 0 | 29 | 29 | 47 | 49 | 48 | 0 | 0 | 2 | 0 | 0 | 298 | 149 | 1.000000 | 1 | 1 | 1 | 1 | 1 | 1 | 1 | 1 | 0.922 | 1 | 1 | 0.016 |
| A3SS | 7187 | ENSG00000001497 | LAS1L | chrX | - | 65517986 | 65522285 | 65517986 | 65518073 | 65522430 | 65522493 | 30 | 32 | 61 | 37 | 61 | 0 | 1 | 0 | 0 | 0 | 84 | 67 | 30 | 58 | 91 | 0 | 0 | 0 | 0 | 0 | 298 | 149 | 1.000000 | 1 | 1 | 0.941 | 1 | 1 | 1 | 1 | 1 | 1 | 1 | 1 | -0.012 |
| A3SS | 7188 | ENSG00000001497 | LAS1L | chrX | - | 65517986 | 65522313 | 65517986 | 65518073 | 65522430 | 65522493 | 30 | 32 | 61 | 37 | 61 | 0 | 1 | 0 | 0 | 0 | 84 | 67 | 30 | 58 | 91 | 0 | 0 | 0 | 0 | 0 | 298 | 149 | 1.000000 | 1 | 1 | 0.941 | 1 | 1 | 1 | 1 | 1 | 1 | 1 | 1 | -0.012 |
| A3SS | 6315 | ENSG00000196369 | SRGAP2B | chr1 | - | 144905858 | 144906074 | 144905858 | 144906039 | 144914691 | 144914754 | 33 | 20 | 17 | 37 | 14 | 0 | 1 | 0 | 2 | 0 | 30 | 58 | 67 | 54 | 21 | 6 | 2 | 0 | 7 | 28 | 184 | 149 | 1.000000 | 1 | 1 | 0.942 | 1 | 0.937 | 1 | 0.802 | 0.959 | 1 | 0.862 | 0.378 | 0.176 |
| A3SS | 9013 | ENSG00000111679 | PTPN6 | chr12 | + | 6951504 | 6951731 | 6951608 | 6951731 | 6951261 | 6951291 | 11 | 7 | 20 | 19 | 4 | 0 | 0 | 0 | 1 | 0 | 14 | 46 | 21 | 49 | 39 | 0 | 0 | 0 | 0 | 0 | 253 | 149 | 1.000000 | 1 | 1 | 1 | 1 | 0.918 | 1 | 1 | 1 | 1 | 1 | 1 | -0.016 |
| A3SS | 4974 | ENSG00000103507 | BCKDK | chr16 | + | 31110076 | 31110124 | 31110103 | 31110124 | 31109672 | 31109783 | 36 | 9 | 16 | 36 | 16 | 0 | 0 | 0 | 0 | 1 | 64 | 47 | 14 | 31 | 65 | 0 | 0 | 0 | 0 | 0 | 176 | 149 | 1.000000 | 1 | 1 | 1 | 1 | 1 | 0.931 | 1 | 1 | 1 | 1 | 1 | -0.014 |
| A3SS | 8499 | ENSG00000177479 | ARIH2 | chr3 | + | 48944981 | 48945181 | 48945135 | 48945181 | 48922747 | 48922811 | 4 | 6 | 17 | 5 | 4 | 1 | 0 | 4 | 0 | 0 | 34 | 23 | 13 | 36 | 37 | 0 | 1 | 1 | 0 | 2 | 298 | 149 | 1.000000 | 1 | 0.667 | 1 | 0.68 | 1 | 1 | 1 | 0.92 | 0.867 | 1 | 0.902 | -0.068 |
| A3SS | 10813 | ENSG00000142046 | TMEM91 | chr19 | + | 41383862 | 41384083 | 41383873 | 41384083 | 41382771 | 41382921 | 9 | 4 | 2 | 12 | 4 | 0 | 0 | 0 | 2 | 0 | 22 | 8 | 14 | 7 | 86 | 0 | 0 | 1 | 0 | 0 | 160 | 149 | 1.000000 | 1 | 1 | 1 | 1 | 0.848 | 1 | 1 | 1 | 0.929 | 1 | 1 | -0.016 |
| A3SS | 4997 | ENSG00000132716 | DCAF8 | chr1 | - | 160261284 | 160261358 | 160261284 | 160261350 | 160262116 | 160262451 | 48 | 36 | 35 | 52 | 31 | 0 | 1 | 1 | 0 | 0 | 66 | 72 | 43 | 55 | 72 | 0 | 4 | 0 | 0 | 0 | 157 | 149 | 1.000000 | 1 | 1 | 0.972 | 0.971 | 1 | 1 | 1 | 0.945 | 1 | 1 | 1 | 0 |
| A3SS | 9290 | ENSG00000166278 | C2 | chr6 | + | 31945177 | 31945672 | 31945461 | 31945672 | 31944979 | 31945029 | 21 | 5 | 4 | 6 | 9 | 2 | 0 | 0 | 0 | 0 | 34 | 34 | 14 | 49 | 20 | 0 | 0 | 2 | 0 | 0 | 298 | 149 | 1.000000 | 1 | 0.84 | 1 | 1 | 1 | 1 | 1 | 1 | 0.778 | 1 | 1 | 0.012 |
| A3SS | 4709 | ENSG00000092098 | RNF31 | chr14 | + | 24149346 | 24149583 | 24149476 | 24149583 | 24148800 | 24148876 | 29 | 17 | 32 | 42 | 33 | 0 | 0 | 0 | 2 | 0 | 80 | 54 | 21 | 47 | 57 | 0 | 0 | 1 | 3 | 0 | 279 | 149 | 1.000000 | 1 | 1 | 1 | 1 | 0.918 | 1 | 1 | 1 | 0.918 | 0.893 | 1 | 0.021 |
| A3SS | 9032 | ENSG00000137959 | IFI44L | chr1 | + | 78635336 | 78635489 | 78635341 | 78635489 | 78628950 | 78628999 | 0 | 9 | 7 | 21 | 7 | 0 | 0 | 0 | 0 | 0 | 2 | 96 | 13 | 38 | 0 | 0 | 3 | 0 | 0 | 0 | 154 | 149 | 1.000000 | 1 | NA | 1 | 1 | 1 | 1 | 1 | 0.969 | 1 | 1 | NA | 0.008 |
| A3SS | 6235 | ENSG00000111331 | OAS3 | chr12 | + | 112941569 | 112941852 | 112941573 | 112941852 | 112938523 | 112938707 | 22 | 24 | 22 | 37 | 9 | 0 | 0 | 0 | 0 | 1 | 47 | 24 | 8 | 102 | 37 | 0 | 0 | 0 | 0 | 0 | 153 | 149 | 1.000000 | 1 | 1 | 1 | 1 | 1 | 0.898 | 1 | 1 | 1 | 1 | 1 | -0.02 |
| A3SS | 4453 | ENSG00000114867 | EIF4G1 | chr3 | + | 184317716 | 184317816 | 184317749 | 184317816 | 184317320 | 184317497 | 217 | 75 | 107 | 152 | 102 | 1 | 0 | 2 | 1 | 0 | 224 | 132 | 69 | 112 | 220 | 0 | 4 | 1 | 2 | 0 | 182 | 149 | 1.000000 | 1 | 0.994 | 1 | 0.978 | 0.992 | 1 | 1 | 0.964 | 0.983 | 0.979 | 1 | 0.008 |
| A3SS | 546 | ENSG00000186818 | LILRB4 | chr19 | + | 54663753 | 54664038 | 54663833 | 54664038 | 54663531 | 54663567 | 34 | 17 | 20 | 28 | 7 | 0 | 0 | 0 | 0 | 0 | 34 | 31 | 28 | 37 | 80 | 0 | 0 | 4 | 0 | 0 | 229 | 149 | 1.000000 | 1 | 1 | 1 | 1 | 1 | 1 | 1 | 1 | 0.82 | 1 | 1 | 0.036 |
| A3SS | 664 | ENSG00000105705 | SUGP1 | chr19 | - | 19316421 | 19316593 | 19316421 | 19316522 | 19320462 | 19320509 | 79 | 53 | 70 | 86 | 258 | 1 | 0 | 0 | 0 | 2 | 162 | 136 | 67 | 124 | 160 | 0 | 0 | 2 | 0 | 0 | 220 | 149 | 1.000000 | 1 | 0.982 | 1 | 1 | 1 | 0.989 | 1 | 1 | 0.958 | 1 | 1 | 0.003 |
| A3SS | 9800 | ENSG00000012124 | CD22 | chr19 | + | 35336035 | 35336341 | 35336148 | 35336341 | 35329186 | 35329230 | 68 | 41 | 44 | 85 | 76 | 1 | 0 | 1 | 0 | 0 | 65 | 148 | 24 | 60 | 120 | 0 | 1 | 0 | 0 | 5 | 262 | 149 | 1.000000 | 1 | 0.975 | 1 | 0.962 | 1 | 1 | 1 | 0.988 | 1 | 1 | 0.932 | 0.003 |
| A3SS | 12683 | ENSG00000168209 | DDIT4 | chr10 | + | 72274156 | 72274421 | 72274224 | 72274421 | 72273923 | 72274060 | 96 | 27 | 61 | 73 | 3 | 0 | 0 | 0 | 1 | 0 | 13 | 94 | 159 | 23 | 73 | 0 | 0 | 0 | 0 | 0 | 217 | 149 | 1.000000 | 1 | 1 | 1 | 1 | 0.98 | 1 | 1 | 1 | 1 | 1 | 1 | -0.004 |
| A3SS | 2888 | ENSG00000162711 | NLRP3 | chr1 | + | 247434102 | 247434273 | 247434162 | 247434273 | 247429584 | 247429755 | 16 | 14 | 11 | 12 | 13 | 0 | 0 | 1 | 0 | 0 | 40 | 25 | 29 | 25 | 49 | 0 | 0 | 1 | 1 | 0 | 209 | 149 | 1.000000 | 1 | 1 | 1 | 0.887 | 1 | 1 | 1 | 1 | 0.954 | 0.947 | 1 | -0.003 |
| A3SS | 1228 | ENSG00000136478 | TEX2 | chr17 | - | 64154841 | 64154967 | 64154841 | 64154961 | 64160800 | 64160933 | 27 | 36 | 37 | 31 | 62 | 2 | 0 | 0 | 0 | 2 | 47 | 68 | 24 | 67 | 89 | 0 | 0 | 2 | 0 | 3 | 155 | 149 | 1.000000 | 1 | 0.928 | 1 | 1 | 1 | 0.968 | 1 | 1 | 0.92 | 1 | 0.966 | 0.002 |
| A3SS | 10523 | ENSG00000106976 | DNM1 | chr9 | + | 128247923 | 128248753 | 128248582 | 128248753 | 128247374 | 128247486 | 2 | 0 | 6 | 4 | 0 | 0 | 0 | 0 | 2 | 0 | 30 | 11 | 9 | 7 | 56 | 3 | 0 | 1 | 0 | 0 | 298 | 149 | 1.000000 | 1 | 1 | NA | 1 | 0.5 | NA | 0.833 | 1 | 0.818 | 1 | 1 | -0.097 |
| A3SS | 6168 | ENSG00000238113 | LINC01410 | chr9 | + | 62809767 | 62811589 | 62811510 | 62811589 | 62803983 | 62804140 | 24 | 23 | 10 | 26 | 32 | 0 | 0 | 0 | 2 | 0 | 15 | 26 | 11 | 93 | 71 | 0 | 0 | 0 | 0 | 0 | 298 | 149 | 1.000000 | 1 | 1 | 1 | 1 | 0.867 | 1 | 1 | 1 | 1 | 1 | 1 | -0.027 |
| A3SS | 7388 | ENSG00000135605 | TEC | chr4 | - | 48171367 | 48171449 | 48171367 | 48171408 | 48176081 | 48176186 | 43 | 29 | 25 | 47 | 78 | 0 | 0 | 0 | 0 | 0 | 86 | 59 | 46 | 71 | 60 | 0 | 1 | 0 | 4 | 0 | 190 | 149 | 1.000000 | 1 | 1 | 1 | 1 | 1 | 1 | 1 | 0.979 | 1 | 0.933 | 1 | 0.018 |
| A3SS | 2698 | ENSG00000100889 | PCK2 | chr14 | + | 24103509 | 24104125 | 24103841 | 24104125 | 24103159 | 24103255 | 76 | 23 | 46 | 39 | 17 | 1 | 0 | 1 | 1 | 0 | 84 | 83 | 16 | 33 | 85 | 0 | 0 | 1 | 0 | 15 | 298 | 149 | 1.000000 | 1 | 0.974 | 1 | 0.958 | 0.951 | 1 | 1 | 1 | 0.889 | 1 | 0.739 | 0.051 |
| A3SS | 12031 | ENSG00000157600 | TMEM164 | chrX | + | 110003500 | 110004164 | 110003882 | 110004164 | 110003094 | 110003156 | 18 | 4 | 6 | 8 | 9 | 0 | 0 | 0 | 0 | 0 | 42 | 21 | 0 | 9 | 72 | 0 | 3 | 0 | 0 | 2 | 298 | 149 | 1.000000 | 1 | 1 | 1 | 1 | 1 | 1 | 1 | 0.778 | NA | 1 | 0.947 | 0.069 |
| A3SS | 12073 | ENSG00000101158 | NELFCD | chr20 | + | 58993623 | 58994239 | 58994109 | 58994239 | 58993448 | 58993544 | 63 | 52 | 67 | 70 | 70 | 2 | 0 | 0 | 0 | 0 | 122 | 94 | 51 | 64 | 89 | 0 | 0 | 0 | 0 | 0 | 298 | 149 | 1.000000 | 1 | 0.94 | 1 | 1 | 1 | 1 | 1 | 1 | 1 | 1 | 1 | -0.012 |
| A3SS | 2099 | ENSG00000140983 | RHOT2 | chr16 | + | 668655 | 668699 | 668658 | 668699 | 668487 | 668569 | 79 | 41 | 57 | 70 | 30 | 0 | 0 | 0 | 2 | 0 | 51 | 67 | 58 | 62 | 137 | 0 | 0 | 0 | 0 | 0 | 152 | 149 | 1.000000 | 1 | 1 | 1 | 1 | 0.972 | 1 | 1 | 1 | 1 | 1 | 1 | -0.006 |
| A3SS | 5443 | ENSG00000079308 | TNS1 | chr2 | - | 217813214 | 217813307 | 217813214 | 217813304 | 217813684 | 217813816 | 1 | 2 | 0 | 10 | 8 | 0 | 1 | 0 | 0 | 2 | 63 | 2 | 1 | 25 | 27 | 0 | 0 | 1 | 1 | 0 | 152 | 149 | 1.000000 | 1 | 1 | 0.662 | NA | 1 | 0.797 | 1 | 1 | 0.495 | 0.961 | 1 | -0.026 |
| A3SS | 8725 | ENSG00000169247 | SH3TC2 | chr5 | - | 149006880 | 149007077 | 149006880 | 149007069 | 149008850 | 149009001 | 5 | 7 | 2 | 7 | 7 | 0 | 0 | 0 | 2 | 0 | 44 | 19 | 7 | 6 | 49 | 0 | 0 | 1 | 0 | 0 | 157 | 149 | 1.000000 | 1 | 1 | 1 | 1 | 0.769 | 1 | 1 | 1 | 0.869 | 1 | 1 | -0.02 |
| A3SS | 2887 | ENSG00000162711 | NLRP3 | chr1 | + | 247434102 | 247434273 | 247434162 | 247434273 | 247423846 | 247425599 | 18 | 11 | 8 | 16 | 11 | 1 | 0 | 0 | 1 | 0 | 41 | 25 | 31 | 22 | 41 | 0 | 0 | 0 | 0 | 0 | 209 | 149 | 1.000000 | 1 | 0.928 | 1 | 1 | 0.919 | 1 | 1 | 1 | 1 | 1 | 1 | -0.031 |
| A3SS | 11404 | ENSG00000184465 | WDR27 | chr6 | - | 169667135 | 169668776 | 169667135 | 169667187 | 169670568 | 169670693 | 53 | 70 | 70 | 74 | 29 | 0 | 0 | 2 | 0 | 2 | 107 | 96 | 58 | 64 | 66 | 0 | 0 | 0 | 0 | 0 | 298 | 149 | 1.000000 | 1 | 1 | 1 | 0.946 | 1 | 0.879 | 1 | 1 | 1 | 1 | 1 | -0.035 |
| A3SS | 4864 | ENSG00000134684 | YARS1 | chr1 | - | 32786361 | 32787075 | 32786361 | 32786447 | 32797762 | 32797843 | 26 | 16 | 8 | 20 | 8 | 0 | 1 | 0 | 0 | 0 | 26 | 50 | 56 | 41 | 0 | 0 | 0 | 0 | 0 | 0 | 298 | 149 | 1.000000 | 1 | 1 | 0.889 | 1 | 1 | 1 | 1 | 1 | 1 | 1 | NA | -0.022 |
| A3SS | 359 | ENSG00000167604 | NFKBID | chr19 | - | 35896725 | 35897058 | 35896725 | 35896831 | 35897650 | 35897856 | 17 | 9 | 18 | 21 | 9 | 0 | 0 | 0 | 1 | 0 | 25 | 19 | 13 | 55 | 57 | 0 | 0 | 0 | 0 | 0 | 298 | 149 | 1.000000 | 1 | 1 | 1 | 1 | 0.913 | 1 | 1 | 1 | 1 | 1 | 1 | -0.017 |
| A3SS | 3617 | ENSG00000125089 | SH3TC1 | chr4 | + | 8232407 | 8233065 | 8232706 | 8233065 | 8231975 | 8232156 | 14 | 6 | 9 | 9 | 0 | 0 | 1 | 0 | 2 | 0 | 6 | 20 | 40 | 14 | 52 | 0 | 0 | 0 | 0 | 0 | 298 | 149 | 1.000000 | 1 | 1 | 0.75 | 1 | 0.692 | NA | 1 | 1 | 1 | 1 | 1 | -0.139 |
| A3SS | 10251 | ENSG00000196696 | PDXDC2P-NPIPB14P | chr16 | - | 70038904 | 70039100 | 70038904 | 70038985 | 70042929 | 70043003 | 23 | 23 | 20 | 20 | 27 | 0 | 0 | 0 | 0 | 0 | 29 | 57 | 32 | 50 | 39 | 0 | 0 | 0 | 2 | 0 | 264 | 149 | 1.000000 | 1 | 1 | 1 | 1 | 1 | 1 | 1 | 1 | 1 | 0.934 | 1 | 0.013 |
| A3SS | 6263 | ENSG00000162298 | SYVN1 | chr11 | - | 65132247 | 65132351 | 65132247 | 65132308 | 65132731 | 65132780 | 117 | 25 | 81 | 89 | 62 | 0 | 0 | 3 | 0 | 0 | 67 | 115 | 45 | 46 | 194 | 0 | 0 | 0 | 0 | 0 | 192 | 149 | 1.000000 | 1 | 1 | 1 | 1 | 0.958 | 1 | 1 | 1 | 1 | 1 | 1 | -0.008 |
| A3SS | 532 | ENSG00000125734 | GPR108 | chr19 | - | 6731198 | 6731282 | 6731198 | 6731234 | 6731472 | 6731522 | 110 | 39 | 70 | 96 | 52 | 0 | 0 | 0 | 0 | 0 | 59 | 121 | 59 | 101 | 120 | 0 | 0 | 1 | 1 | 0 | 197 | 149 | 1.000000 | 1 | 1 | 1 | 1 | 1 | 1 | 1 | 1 | 0.978 | 0.987 | 1 | 0.007 |
| A3SS | 10087 | ENSG00000067182 | TNFRSF1A | chr12 | - | 6333366 | 6333516 | 6333366 | 6333451 | 6334090 | 6334244 | 125 | 64 | 73 | 87 | 125 | 0 | 0 | 0 | 0 | 0 | 130 | 134 | 57 | 89 | 156 | 1 | 0 | 0 | 0 | 0 | 214 | 149 | 1.000000 | 1 | 1 | 1 | 1 | 1 | 1 | 0.989 | 1 | 1 | 1 | 1 | 0.002 |
| A3SS | 9077 | ENSG00000128284 | APOL3 | chr22 | - | 36149045 | 36149371 | 36149045 | 36149185 | 36160668 | 36160929 | 101 | 67 | 125 | 97 | 48 | 1 | 0 | 0 | 0 | 0 | 184 | 37 | 74 | 100 | 134 | 0 | 3 | 0 | 0 | 0 | 298 | 149 | 1.000000 | 1 | 0.981 | 1 | 1 | 1 | 1 | 1 | 0.86 | 1 | 1 | 1 | 0.024 |
| A3SS | 1193 | ENSG00000100348 | TXN2 | chr22 | - | 36480574 | 36480837 | 36480574 | 36480748 | 36481142 | 36481340 | 17 | 6 | 24 | 21 | 8 | 0 | 0 | 0 | 0 | 0 | 37 | 32 | 41 | 12 | 45 | 0 | 0 | 0 | 2 | 0 | 238 | 149 | 1.000000 | 1 | 1 | 1 | 1 | 1 | 1 | 1 | 1 | 1 | 0.79 | 1 | 0.042 |
| A3SS | 2081 | ENSG00000140694 | PARN | chr16 | - | 14627105 | 14627187 | 14627105 | 14627136 | 14627268 | 14627336 | 399 | 335 | 286 | 392 | 365 | 5 | 2 | 0 | 0 | 0 | 484 | 491 | 316 | 377 | 199 | 0 | 2 | 3 | 0 | 0 | 200 | 149 | 1.000000 | 1 | 0.983 | 0.992 | 1 | 1 | 1 | 1 | 0.995 | 0.987 | 1 | 1 | -0.001 |
| A3SS | 12565 | ENSG00000177239 | MAN1B1 | chr9 | + | 137096236 | 137096391 | 137096239 | 137096391 | 137088868 | 137089005 | 147 | 74 | 122 | 75 | 146 | 0 | 0 | 0 | 0 | 2 | 124 | 153 | 105 | 76 | 196 | 0 | 0 | 1 | 0 | 0 | 152 | 149 | 1.000000 | 1 | 1 | 1 | 1 | 1 | 0.986 | 1 | 1 | 0.99 | 1 | 1 | -0.001 |
| A3SS | 2906 | ENSG00000175061 | SNHG29 | chr17 | + | 16439580 | 16440253 | 16440110 | 16440253 | 16439326 | 16439414 | 58 | 53 | 78 | 60 | 30 | 2 | 0 | 1 | 0 | 0 | 73 | 66 | 67 | 61 | 101 | 0 | 0 | 2 | 0 | 0 | 298 | 149 | 1.000000 | 1 | 0.935 | 1 | 1 | 0.968 | 1 | 1 | 1 | 0.944 | 1 | 1 | -0.008 |
| A3SS | 306 | ENSG00000132024 | CC2D1A | chr19 | + | 13927174 | 13928030 | 13927892 | 13928030 | 13926977 | 13927077 | 27 | 21 | 32 | 34 | 23 | 0 | 0 | 0 | 0 | 0 | 26 | 59 | 25 | 47 | 69 | 0 | 0 | 0 | 0 | 2 | 298 | 149 | 1.000000 | 1 | 1 | 1 | 1 | 1 | 1 | 1 | 1 | 1 | 1 | 0.945 | 0.011 |
| A3SS | 6225 | ENSG00000178927 | CYBC1 | chr17 | - | 82446622 | 82446696 | 82446622 | 82446692 | 82447579 | 82450109 | 96 | 26 | 68 | 67 | 27 | 0 | 0 | 0 | 1 | 0 | 111 | 86 | 30 | 60 | 85 | 0 | 0 | 0 | 0 | 0 | 153 | 149 | 1.000000 | 1 | 1 | 1 | 1 | 0.985 | 1 | 1 | 1 | 1 | 1 | 1 | -0.003 |
| A3SS | 4161 | ENSG00000137502 | RAB30 | chr11 | - | 82994038 | 82994122 | 82994038 | 82994106 | 82997223 | 82997324 | 46 | 48 | 85 | 82 | 97 | 0 | 0 | 0 | 0 | 0 | 108 | 100 | 150 | 78 | 10 | 0 | 3 | 0 | 0 | 0 | 165 | 149 | 1.000000 | 1 | 1 | 1 | 1 | 1 | 1 | 1 | 0.968 | 1 | 1 | 1 | 0.006 |
| A3SS | 4101 | ENSG00000185000 | DGAT1 | chr8 | - | 144318094 | 144318362 | 144318094 | 144318169 | 144318460 | 144318566 | 40 | 22 | 19 | 51 | 30 | 2 | 0 | 0 | 0 | 0 | 27 | 23 | 42 | 38 | 119 | 0 | 0 | 0 | 0 | 0 | 298 | 149 | 1.000000 | 1 | 0.909 | 1 | 1 | 1 | 1 | 1 | 1 | 1 | 1 | 1 | -0.018 |
| A3SS | 6512 | ENSG00000114841 | DNAH1 | chr3 | + | 52399544 | 52399779 | 52399620 | 52399779 | 52398849 | 52399201 | 29 | 22 | 29 | 21 | 44 | 0 | 0 | 2 | 0 | 0 | 55 | 33 | 26 | 25 | 93 | 0 | 0 | 0 | 2 | 0 | 225 | 149 | 1.000000 | 1 | 1 | 1 | 0.906 | 1 | 1 | 1 | 1 | 1 | 0.892 | 1 | 0.003 |
| A3SS | 481 | ENSG00000161847 | RAVER1 | chr19 | - | 10328641 | 10329111 | 10328641 | 10329052 | 10330459 | 10330526 | 41 | 14 | 46 | 30 | 40 | 0 | 0 | 0 | 0 | 0 | 66 | 42 | 16 | 36 | 98 | 0 | 1 | 0 | 0 | 0 | 208 | 149 | 1.000000 | 1 | 1 | 1 | 1 | 1 | 1 | 1 | 0.968 | 1 | 1 | 1 | 0.006 |
| A3SS | 196 | ENSG00000070371 | CLTCL1 | chr22 | - | 19196265 | 19196656 | 19196265 | 19196415 | 19199733 | 19199841 | 4 | 2 | 5 | 0 | 4 | 0 | 0 | 0 | 0 | 0 | 10 | 0 | 0 | 47 | 45 | 0 | 0 | 0 | 5 | 8 | 298 | 149 | 1.000000 | 1 | 1 | 1 | 1 | NA | 1 | 1 | NA | NA | 0.825 | 0.738 | 0.146 |
| A3SS | 11708 | ENSG00000166199 | ALKBH3 | chr11 | + | 43919037 | 43919136 | 43919041 | 43919136 | 43901515 | 43901725 | 33 | 28 | 35 | 23 | 12 | 0 | 3 | 0 | 0 | 0 | 83 | 56 | 28 | 21 | 29 | 0 | 0 | 0 | 0 | 0 | 153 | 149 | 1.000000 | 1 | 1 | 0.901 | 1 | 1 | 1 | 1 | 1 | 1 | 1 | 1 | -0.02 |
| A3SS | 1979 | ENSG00000122390 | NAA60 | chr16 | + | 3484690 | 3485083 | 3484761 | 3485083 | 3482501 | 3482598 | 55 | 18 | 20 | 34 | 37 | 1 | 0 | 1 | 0 | 0 | 48 | 48 | 28 | 36 | 90 | 0 | 0 | 0 | 0 | 0 | 220 | 149 | 1.000000 | 1 | 0.974 | 1 | 0.931 | 1 | 1 | 1 | 1 | 1 | 1 | 1 | -0.019 |
| A3SS | 9138 | ENSG00000168000 | BSCL2 | chr11 | - | 62692662 | 62692797 | 62692662 | 62692715 | 62694567 | 62694711 | 41 | 24 | 20 | 33 | 43 | 1 | 0 | 0 | 0 | 0 | 36 | 47 | 15 | 14 | 135 | 0 | 0 | 0 | 0 | 0 | 231 | 149 | 1.000000 | 1 | 0.964 | 1 | 1 | 1 | 1 | 1 | 1 | 1 | 1 | 1 | -0.007 |
| A3SS | 2311 | ENSG00000184056 | VPS33B | chr15 | - | 91006371 | 91007046 | 91006371 | 91006445 | 91007468 | 91007573 | 55 | 53 | 40 | 38 | 32 | 0 | 0 | 0 | 0 | 0 | 97 | 51 | 63 | 35 | 58 | 0 | 0 | 2 | 0 | 0 | 298 | 149 | 1.000000 | 1 | 1 | 1 | 1 | 1 | 1 | 1 | 1 | 0.94 | 1 | 1 | 0.012 |
| A3SS | 9602 | ENSG00000168014 | C2CD3 | chr11 | - | 74037477 | 74037698 | 74037477 | 74037652 | 74042053 | 74042218 | 36 | 20 | 29 | 33 | 36 | 0 | 0 | 0 | 0 | 1 | 40 | 50 | 65 | 62 | 22 | 0 | 0 | 0 | 0 | 0 | 195 | 149 | 1.000000 | 1 | 1 | 1 | 1 | 1 | 0.965 | 1 | 1 | 1 | 1 | 1 | -0.007 |
| A3SS | 249 | ENSG00000099624 | ATP5F1D | chr19 | + | 1244314 | 1244440 | 1244318 | 1244440 | 1244096 | 1244185 | 91 | 29 | 85 | 66 | 29 | 0 | 0 | 1 | 0 | 0 | 54 | 72 | 71 | 60 | 128 | 0 | 0 | 0 | 0 | 0 | 153 | 149 | 1.000000 | 1 | 1 | 1 | 0.988 | 1 | 1 | 1 | 1 | 1 | 1 | 1 | -0.002 |
| A3SS | 12661 | ENSG00000107862 | GBF1 | chr10 | + | 102380260 | 102380362 | 102380332 | 102380362 | 102379852 | 102379954 | 38 | 22 | 26 | 25 | 20 | 0 | 0 | 0 | 0 | 0 | 53 | 52 | 24 | 32 | 55 | 0 | 5 | 0 | 0 | 0 | 221 | 149 | 1.000000 | 1 | 1 | 1 | 1 | 1 | 1 | 1 | 0.875 | 1 | 1 | 1 | 0.025 |
| A3SS | 7030 | ENSG00000126215 | XRCC3 | chr14 | - | 103711032 | 103711245 | 103711032 | 103711180 | 103711465 | 103711567 | 18 | 8 | 30 | 26 | 23 | 0 | 2 | 0 | 0 | 0 | 23 | 26 | 37 | 45 | 58 | 0 | 0 | 0 | 0 | 0 | 214 | 149 | 1.000000 | 1 | 1 | 0.736 | 1 | 1 | 1 | 1 | 1 | 1 | 1 | 1 | -0.053 |
| A3SS | 11432 | ENSG00000159884 | CCDC107 | chr9 | + | 35660745 | 35660858 | 35660765 | 35660858 | 35660556 | 35660647 | 90 | 46 | 56 | 79 | 23 | 0 | 1 | 0 | 4 | 0 | 39 | 91 | 37 | 63 | 148 | 0 | 2 | 0 | 0 | 0 | 169 | 149 | 1.000000 | 1 | 1 | 0.976 | 1 | 0.946 | 1 | 1 | 0.976 | 1 | 1 | 1 | -0.011 |
| A3SS | 3313 | ENSG00000124357 | NAGK | chr2 | + | 71077557 | 71077636 | 71077569 | 71077636 | 71076603 | 71076701 | 175 | 81 | 95 | 134 | 103 | 1 | 0 | 1 | 0 | 1 | 142 | 106 | 34 | 120 | 270 | 0 | 0 | 0 | 0 | 5 | 161 | 149 | 1.000000 | 1 | 0.994 | 1 | 0.989 | 1 | 0.99 | 1 | 1 | 1 | 1 | 0.98 | -0.001 |
| A3SS | 4526 | ENSG00000156709 | AIFM1 | chrX | - | 130147751 | 130147876 | 130147751 | 130147870 | 130149468 | 130149568 | 74 | 48 | 45 | 51 | 88 | 0 | 0 | 0 | 0 | 0 | 96 | 56 | 51 | 84 | 103 | 0 | 0 | 0 | 0 | 2 | 155 | 149 | 1.000000 | 1 | 1 | 1 | 1 | 1 | 1 | 1 | 1 | 1 | 1 | 0.98 | 0.004 |
| A3SS | 10483 | ENSG00000177646 | ACAD9 | chr3 | + | 128884652 | 128884746 | 128884656 | 128884746 | 128879628 | 128880017 | 10 | 12 | 12 | 14 | 14 | 0 | 0 | 0 | 0 | 1 | 27 | 42 | 18 | 28 | 30 | 0 | 0 | 0 | 0 | 0 | 153 | 149 | 1.000000 | 1 | 1 | 1 | 1 | 1 | 0.932 | 1 | 1 | 1 | 1 | 1 | -0.014 |
| A3SS | 11557 | ENSG00000266714 | MYO15B | chr17 | + | 75625526 | 75625660 | 75625530 | 75625660 | 75625122 | 75625238 | 27 | 13 | 11 | 16 | 8 | 0 | 0 | 0 | 0 | 0 | 29 | 18 | 10 | 56 | 45 | 1 | 0 | 0 | 2 | 0 | 153 | 149 | 1.000000 | 1 | 1 | 1 | 1 | 1 | 1 | 0.966 | 1 | 1 | 0.965 | 1 | 0.014 |
| A3SS | 6262 | ENSG00000162298 | SYVN1 | chr11 | - | 65130001 | 65130175 | 65130001 | 65130172 | 65130250 | 65130379 | 4 | 2 | 4 | 8 | 0 | 2 | 1 | 4 | 3 | 0 | 19 | 7 | 4 | 0 | 71 | 29 | 9 | 3 | 8 | 5 | 152 | 149 | 1.000000 | 1 | 0.662 | 0.662 | 0.495 | 0.723 | NA | 0.391 | 0.433 | 0.567 | 0 | 0.933 | 0.171 |
| A3SS | 5883 | ENSG00000185864 | NPIPB4 | chr16 | - | 21857258 | 21857416 | 21857258 | 21857345 | 21857579 | 21857756 | 8 | 7 | 6 | 7 | 6 | 0 | 1 | 0 | 0 | 0 | 30 | 43 | 19 | 14 | 10 | 0 | 0 | 0 | 0 | 0 | 220 | 149 | 1.000000 | 1 | 1 | 0.826 | 1 | 1 | 1 | 1 | 1 | 1 | 1 | 1 | -0.035 |
| A3SS | 8976 | ENSG00000167792 | NDUFV1 | chr11 | + | 67611476 | 67611569 | 67611521 | 67611569 | 67610994 | 67611207 | 39 | 29 | 31 | 23 | 29 | 0 | 0 | 2 | 3 | 0 | 86 | 42 | 38 | 25 | 42 | 0 | 1 | 0 | 3 | 0 | 194 | 149 | 1.000000 | 1 | 1 | 1 | 0.923 | 0.855 | 1 | 1 | 0.97 | 1 | 0.865 | 1 | -0.011 |
| A3SS | 12659 | ENSG00000107862 | GBF1 | chr10 | + | 102380248 | 102380362 | 102380332 | 102380362 | 102379852 | 102379954 | 91 | 40 | 49 | 48 | 45 | 0 | 0 | 0 | 0 | 0 | 62 | 71 | 35 | 62 | 125 | 0 | 5 | 0 | 0 | 0 | 233 | 149 | 1.000000 | 1 | 1 | 1 | 1 | 1 | 1 | 1 | 0.901 | 1 | 1 | 1 | 0.02 |
| A3SS | 920 | ENSG00000105576 | TNPO2 | chr19 | - | 12701597 | 12701672 | 12701597 | 12701644 | 12701751 | 12701851 | 31 | 26 | 26 | 32 | 30 | 0 | 1 | 0 | 2 | 0 | 91 | 38 | 21 | 25 | 52 | 13 | 0 | 0 | 3 | 5 | 177 | 149 | 1.000000 | 1 | 1 | 0.956 | 1 | 0.931 | 1 | 0.855 | 1 | 1 | 0.875 | 0.897 | 0.052 |
| A3SS | 7348 | ENSG00000115204 | MPV17 | chr2 | - | 27317077 | 27317234 | 27317077 | 27317173 | 27322447 | 27322522 | 6 | 5 | 5 | 13 | 0 | 0 | 2 | 0 | 0 | 0 | 37 | 35 | 11 | 5 | 22 | 0 | 0 | 0 | 0 | 0 | 210 | 149 | 1.000000 | 1 | 1 | 0.639 | 1 | 1 | NA | 1 | 1 | 1 | 1 | 1 | -0.09 |
| A3SS | 4539 | ENSG00000169727 | GPS1 | chr17 | + | 82056839 | 82057470 | 82057052 | 82057470 | 82056628 | 82056766 | 77 | 37 | 76 | 87 | 60 | 1 | 0 | 0 | 0 | 0 | 131 | 92 | 47 | 50 | 98 | 0 | 0 | 0 | 0 | 0 | 298 | 149 | 1.000000 | 1 | 0.975 | 1 | 1 | 1 | 1 | 1 | 1 | 1 | 1 | 1 | -0.005 |
| A3SS | 12727 | ENSG00000130640 | TUBGCP2 | chr10 | - | 133283881 | 133285213 | 133283881 | 133284002 | 133285455 | 133285628 | 92 | 41 | 67 | 61 | 49 | 0 | 0 | 0 | 0 | 0 | 46 | 60 | 52 | 83 | 150 | 0 | 0 | 1 | 0 | 0 | 298 | 149 | 1.000000 | 1 | 1 | 1 | 1 | 1 | 1 | 1 | 1 | 0.963 | 1 | 1 | 0.007 |
| A3SS | 648 | ENSG00000130479 | MAP1S | chr19 | + | 17720935 | 17721037 | 17720985 | 17721037 | 17719479 | 17719620 | 44 | 18 | 14 | 36 | 20 | 0 | 0 | 0 | 0 | 0 | 53 | 47 | 28 | 36 | 49 | 0 | 0 | 1 | 0 | 0 | 199 | 149 | 1.000000 | 1 | 1 | 1 | 1 | 1 | 1 | 1 | 1 | 0.954 | 1 | 1 | 0.009 |
| A3SS | 11390 | ENSG00000168003 | SLC3A2 | chr11 | + | 62888330 | 62888860 | 62888651 | 62888860 | 62885464 | 62885608 | 63 | 16 | 37 | 63 | 39 | 0 | 0 | 0 | 0 | 0 | 67 | 45 | 46 | 42 | 99 | 0 | 0 | 0 | 2 | 0 | 298 | 149 | 1.000000 | 1 | 1 | 1 | 1 | 1 | 1 | 1 | 1 | 1 | 0.913 | 1 | 0.017 |
| A3SS | 11406 | ENSG00000184465 | WDR27 | chr6 | - | 169667135 | 169668776 | 169667135 | 169667187 | 169672254 | 169672396 | 50 | 69 | 64 | 66 | 30 | 0 | 0 | 0 | 0 | 1 | 97 | 79 | 58 | 59 | 66 | 0 | 0 | 0 | 0 | 0 | 298 | 149 | 1.000000 | 1 | 1 | 1 | 1 | 1 | 0.938 | 1 | 1 | 1 | 1 | 1 | -0.012 |
| A3SS | 12294 | ENSG00000179532 | DNHD1 | chr11 | + | 6497770 | 6498961 | 6498722 | 6498961 | 6497584 | 6497667 | 5 | 7 | 11 | 2 | 2 | 0 | 1 | 0 | 1 | 0 | 30 | 20 | 18 | 20 | 18 | 0 | 0 | 0 | 0 | 0 | 298 | 149 | 1.000000 | 1 | 1 | 0.778 | 1 | 0.5 | 1 | 1 | 1 | 1 | 1 | 1 | -0.144 |
| A3SS | 4491 | ENSG00000134802 | SLC43A3 | chr11 | - | 57425988 | 57426389 | 57425988 | 57426316 | 57426587 | 57426652 | 14 | 3 | 11 | 12 | 5 | 0 | 0 | 2 | 1 | 0 | 44 | 16 | 6 | 34 | 24 | 0 | 0 | 0 | 0 | 0 | 222 | 149 | 1.000000 | 1 | 1 | 1 | 0.787 | 0.89 | 1 | 1 | 1 | 1 | 1 | 1 | -0.065 |
| A3SS | 2383 | ENSG00000140400 | MAN2C1 | chr15 | - | 75359058 | 75359425 | 75359058 | 75359153 | 75359619 | 75359775 | 100 | 37 | 106 | 70 | 47 | 0 | 0 | 0 | 2 | 0 | 81 | 75 | 91 | 46 | 146 | 0 | 0 | 1 | 0 | 0 | 298 | 149 | 1.000000 | 1 | 1 | 1 | 1 | 0.946 | 1 | 1 | 1 | 0.978 | 1 | 1 | -0.006 |
| A3SS | 9649 | ENSG00000183735 | TBK1 | chr12 | + | 64457750 | 64458018 | 64457907 | 64458018 | 64455839 | 64455957 | 23 | 12 | 3 | 10 | 13 | 0 | 1 | 0 | 0 | 0 | 20 | 15 | 41 | 49 | 15 | 0 | 2 | 0 | 0 | 0 | 298 | 149 | 1.000000 | 1 | 1 | 0.857 | 1 | 1 | 1 | 1 | 0.789 | 1 | 1 | 1 | 0.014 |
| A3SS | 13198 | ENSG00000073060 | SCARB1 | chr12 | - | 124782682 | 124786503 | 124782682 | 124782811 | 124787405 | 124787457 | 19 | 7 | 17 | 12 | 14 | 0 | 0 | 0 | 1 | 0 | 47 | 20 | 1 | 22 | 57 | 0 | 0 | 0 | 0 | 0 | 298 | 149 | 1.000000 | 1 | 1 | 1 | 1 | 0.857 | 1 | 1 | 1 | 1 | 1 | 1 | -0.029 |
| A3SS | 5496 | ENSG00000162747 | FCGR3B | chr1 | - | 161629777 | 161630035 | 161629777 | 161630032 | 161630367 | 161630388 | 1 | 4 | 14 | 14 | 13 | 0 | 0 | 3 | 3 | 0 | 3 | 0 | 94 | 25 | 2 | 0 | 0 | 4 | 6 | 0 | 152 | 149 | 1.000000 | 1 | 1 | 1 | 0.821 | 0.821 | 1 | 1 | NA | 0.958 | 0.803 | 1 | -0.012 |
| A3SS | 13245 | ENSG00000110911 | SLC11A2 | chr12 | - | 51008475 | 51008624 | 51008475 | 51008544 | 51009116 | 51009270 | 17 | 28 | 34 | 21 | 32 | 2 | 0 | 0 | 0 | 0 | 62 | 18 | 30 | 63 | 35 | 0 | 0 | 0 | 0 | 0 | 229 | 149 | 1.000000 | 1 | 0.847 | 1 | 1 | 1 | 1 | 1 | 1 | 1 | 1 | 1 | -0.031 |

| | | | | | | | | | | | | | |
|---|---|---|---|---|---|---|---|---|---|---|---|---|---|
| A3SS | 10811 | ENSG00000142046 | TMEM91 | chr19 | + | 41383714 | 41384083 | 41383873 | 41384083 | 41382771 | 41382921 | 23 | [illegible] |
| A3SS | 486 | ENSG00000071655 | MBD3 | chr19 | - | 1578334 | 1578538 | 1578334 | 1578534 | 1581091 | 1581269 | 86 | [illegible] |
| A3SS | 4613 | ENSG00000161204 | ABCF3 | chr3 | + | 184189387 | 184189757 | 184189556 | 184189757 | 184189254 | 184189277 | 63 | [illegible] |
| A3SS | 1429 | ENSG00000187838 | PLSCR3 | chr17 | - | 7392790 | 7392952 | 7392790 | 7392880 | 7393466 | 7393509 | 21 | [illegible] |
| A3SS | 7313 | ENSG00000101638 | ST8SIA5 | chr18 | - | 46688774 | 46688919 | 46688774 | 46688867 | 46692168 | 46692255 | 1 | [illegible] |
| A3SS | 1692 | ENSG00000140650 | PMM2 | chr16 | + | 8802080 | 8802389 | 8802289 | 8802389 | 8801798 | 8801910 | 18 | [illegible] |
| A3SS | 720 | ENSG00000031823 | RANBP3 | chr19 | - | 5957917 | 5957973 | 5957917 | 5957935 | 5978060 | 5978140 | 74 | [illegible] |
| A3SS | 2765 | ENSG00000119705 | SLIRP | chr14 | + | 77717495 | 77717598 | 77717501 | 77717598 | 77715771 | 77715879 | 60 | [illegible] |
| A3SS | 374 | ENSG00000011304 | PTBP1 | chr19 | + | 805491 | 805569 | 805512 | 805569 | 805012 | 805187 | 75 | [illegible] |
| A3SS | 10370 | ENSG00000130734 | ATG4D | chr19 | + | 10546838 | 10547253 | 10547188 | 10547253 | 10544956 | 10545130 | 25 | [illegible] |
| A3SS | 5635 | ENSG00000101940 | WDR13 | chrX | + | 48599586 | 48599717 | 48599598 | 48599717 | 48599352 | 48599462 | 18 | [illegible] |
| A3SS | 4305 | ENSG00000164258 | NDUFS4 | chr5 | + | 53604723 | 53604895 | 53604806 | 53604895 | 53603451 | 53603530 | 17 | [illegible] |
| A3SS | 1402 | ENSG00000154957 | ZNF18 | chr17 | - | 11984112 | 11984197 | 11984112 | 11984191 | 11990461 | 11990550 | 27 | [illegible] |
| A3SS | 4450 | ENSG00000114867 | EIF4G1 | chr3 | + | 184317716 | 184317816 | 184317746 | 184317816 | 184315488 | 184315545 | 56 | [illegible] |
| A3SS | 1071 | ENSG00000154889 | MPPE1 | chr18 | - | 11893467 | 11893576 | 11893467 | 11893566 | 11895184 | 11895277 | 26 | [illegible] |
| A3SS | 2281 | ENSG00000140521 | POLG | chr15 | - | 89324106 | 89324227 | 89324106 | 89324187 | 89325489 | 89325686 | 26 | [illegible] |
| A3SS | 5945 | ENSG00000095787 | WAC | chr10 | + | 28583398 | 28583505 | 28583405 | 28583505 | 28535909 | 28536059 | 39 | [illegible] |
| A3SS | 2275 | ENSG00000140521 | POLG | chr15 | - | 89317375 | 89317536 | 89317375 | 89317476 | 89318540 | 89318749 | 146 | [illegible] |
| A3SS | 4771 | ENSG00000163681 | SLMAP | chr3 | + | 57861948 | 57862086 | 57861953 | 57862086 | 57860698 | 57860839 | 240 | [illegible] |
| A3SS | 8750 | ENSG00000157637 | SLC38A10 | chr17 | - | 81246884 | 81247061 | 81246884 | 81247058 | 81250069 | 81250093 | 10 | [illegible] |
| A3SS | 10487 | ENSG00000177646 | ACAD9 | chr3 | + | 128908184 | 128909099 | 128908972 | 128909099 | 128906120 | 128906249 | 26 | [illegible] |
| A3SS | 4981 | ENSG00000174547 | MRPL11 | chr11 | - | 66437349 | 66437443 | 66437349 | 66437435 | 66438163 | 66438259 | 73 | [illegible] |
| A3SS | 1771 | ENSG00000140995 | DEF8 | chr16 | + | 89957472 | 89957660 | 89957537 | 89957660 | 89955168 | 89955266 | 35 | [illegible] |
| A3SS | 2198 | ENSG00000182511 | FES | chr15 | + | 90885036 | 90885258 | 90885041 | 90885258 | 90884503 | 90884639 | 23 | [illegible] |
| A3SS | 2052 | ENSG00000038532 | CLEC16A | chr16 | + | 11039753 | 11042363 | 11042253 | 11042363 | 11024820 | 11024921 | 23 | [illegible] |
| A3SS | 903 | ENSG00000062598 | ELMO2 | chr20 | - | 46370442 | 46370525 | 46370442 | 46370513 | 46371351 | 46371459 | 148 | [illegible] |
| A3SS | 7273 | ENSG00000115594 | IL1R1 | chr2 | + | 102175477 | 102175645 | 102175570 | 102175645 | 102174586 | 102174730 | 11 | [illegible] |
| A3SS | 2309 | ENSG00000137812 | KNL1 | chr15 | + | 40629272 | 40629371 | 40629323 | 40629371 | 40628610 | 40628678 | 16 | [illegible] |
| [illegible] | | | | | | | | | | | | | |

| | | | | | | | | | | | | | | | | | | | | | | | | | | | | | | | | | | | | | | | | | | | | | | | | |
|---|---|---|---|---|---|---|---|---|---|---|---|---|---|---|---|---|---|---|---|---|---|---|---|---|---|---|---|---|---|---|---|---|---|---|---|---|---|---|---|---|---|---|---|---|---|---|---|---|
| A3SS | 12527 | ENSG00000248538 | NA | chr8 | + | 9202380 | 9202858 | 9202498 | 9202858 | 9198923 | 9198991 | 0 | 0 | 0 | 0 | 0 | 1 | 0 | 0 | 0 | 0 | 0 | 7 | 0 | 0 | 0 | 0 | 35 | 0 | 0 | 0 | 0 | 37 | 267 | 149 | 1.000000 | 1 | NA | NA | NA | | 1 | 0 | NA | NA | NA | NA | 0.346 | 0.154 |
| A3SS | 379 | ENSG00000104980 | TIMM44 | chr19 | - | 7934088 | 7934241 | 7934088 | 7934238 | 7938026 | 7938197 | 0 | 1 | 0 | 1 | 0 | 7 | 0 | 0 | 0 | 0 | 1 | 0 | 4 | 4 | 4 | 7 | 23 | 7 | 0 | 2 | 2 | 0 | 152 | 149 | 1.000000 | 1 | NA | 1 | NA | 0.873 | NA | 0.359 | 1 | 0.662 | 0.774 | 1 | 0.177 |
| A3SS | 12040 | ENSG00000165905 | LARGE2 | chr11 | + | 45926038 | 45926347 | 45926221 | 45926347 | 45924784 | 45924889 | 5 | 1 | 10 | 1 | 10 | 6 | 0 | 0 | 0 | 0 | 0 | 0 | 15 | 1 | 2 | 0 | 38 | 0 | 8 | 0 | 0 | 0 | 298 | 149 | 1.000000 | 1 | 1 | 1 | 1 | 1 | NA | 1 | 0.059 | 1 | NA | 1 | 0.235 |
| A3SS | 12103 | ENSG00000115414 | FN1 | chr2 | - | 215371908 | 215372375 | 215371908 | 215372300 | 215373321 | 215373411 | 2 | 34 | 3 | 34 | 3 | 4 | 0 | 0 | 2 | 1 | 0 | 0 | 26 | 0 | 17 | 34 | 0 | 11 | 6 | 1 | 13 | 5 | 224 | 149 | 1.000000 | 1 | 1 | 0.919 | 0.666 | 1 | NA | 0.611 | 0 | 0.919 | 0.635 | 0 | 0.463 |
| A3SS | 5357 | ENSG00000162772 | ATF3 | chr1 | + | 212615147 | 212615261 | 212615151 | 212615261 | 212615021 | 212615060 | 0 | 0 | 0 | 0 | 0 | 0 | 0 | 1 | 0 | 0 | 0 | 0 | 21 | 2 | 2 | 2 | 6 | 0 | 0 | 0 | 0 | 0 | 153 | 149 | 1.000000 | 1 | 0 | NA | NA | NA | NA | 1 | 1 | 1 | 1 | 1 | -1 |
| A3SS | 6613 | ENSG00000198019 | FCGR1B | chr1 | + | 121088650 | 121088908 | 121088653 | 121088908 | 121087820 | 121087841 | 2 | 1 | 1 | 1 | 1 | 0 | 5 | 14 | 6 | 4 | 2 | 16 | 6 | 30 | 0 | 6 | 0 | 81 | 0 | 0 | 11 | 41 | 152 | 149 | 1.000000 | 1 | 0.123 | 0.14 | 0.197 | 0 | 0.234 | 0.068 | 1 | NA | 0.348 | 0 | -0.215 |
| A3SS | 2392 | ENSG00000103942 | HOMER2 | chr15 | - | 82859028 | 82859168 | 82859028 | 82859135 | 82864166 | 82864259 | 0 | 2 | 0 | 2 | 0 | 2 | 0 | 19 | 21 | 16 | 29 | 29 | 15 | 0 | 0 | 1 | 21 | 66 | 59 | 26 | 48 | 1058 | 182 | 149 | 1.000000 | 1 | 0 | 0.072 | 0 | 0.053 | 0 | 0.157 | 0 | 0 | 0.017 | 0.016 | -0.013 |
| A3SS | 2284 | ENSG00000102901 | CENPT | chr16 | - | 67828473 | 67828578 | 67828473 | 67828559 | 67828666 | 67828843 | 76 | 29 | 49 | 29 | 49 | 61 | 96 | 1 | 0 | 1 | 1 | 0 | 82 | 43 | 43 | 66 | 110 | 0 | 0 | 0 | 0 | 6 | 168 | 149 | 1.000000 | 1 | 0.985 | 1 | 0.978 | 0.982 | 1 | 1 | 1 | 1 | 1 | 0.942 | 0.001 |
| A3SS | 10826 | ENSG00000104689 | TNFRSF10A | chr8 | - | 23202647 | 23202761 | 23202647 | 23202659 | 23212115 | 23212212 | 51 | 45 | 61 | 45 | 61 | 86 | 118 | 0 | 0 | 0 | 0 | 0 | 116 | 65 | 44 | 87 | 82 | 0 | 0 | 3 | 0 | 0 | 251 | 149 | 1.000000 | 1 | 1 | 1 | 1 | 1 | 1 | 1 | 1 | 0.897 | 1 | 1 | 0.021 |
| A3SS | 11588 | ENSG00000196507 | TCEAL3 | chrX | + | 103609037 | 103609927 | 103609042 | 103609927 | 103608603 | 103608673 | 12 | 3 | 6 | 3 | 6 | 7 | 9 | 0 | 0 | 0 | 0 | 0 | 8 | 17 | 4 | 6 | 35 | 0 | 0 | 0 | 1 | 0 | 154 | 149 | 1.000000 | 1 | 1 | 1 | 1 | 1 | 1 | 1 | 1 | 1 | 0.853 | 1 | 0.029 |
| A3SS | 5135 | ENSG00000123473 | STIL | chr1 | - | 47299904 | 47300152 | 47299904 | 47299968 | 47301560 | 47301748 | 5 | 6 | 3 | 6 | 3 | 5 | 19 | 0 | 0 | 0 | 0 | 0 | 9 | 9 | 17 | 36 | 0 | 0 | 2 | 0 | 0 | 0 | 298 | 149 | 1.000000 | 1 | 1 | 1 | 1 | 1 | 1 | 1 | 0.692 | 1 | 1 | NA | 0.077 |
| A3SS | 7070 | ENSG00000104970 | KIR3DX1 | chr19 | + | 54543264 | 54543428 | 54543375 | 54543428 | 54542523 | 54542646 | 2 | 0 | 0 | 0 | 0 | 0 | 1 | 3 | 0 | 3 | 2 | 0 | 14 | 8 | 4 | 0 | 10 | 9 | 12 | 18 | 1 | 15 | 260 | 149 | 1.000000 | 1 | 0.276 | NA | 0 | 0 | 1 | 0.471 | 0.276 | 0.113 | 0 | 0.276 | 0.092 |
| A3SS | 5292 | ENSG00000171163 | ZNF692 | chr1 | - | 248857827 | 248858017 | 248857827 | 248857859 | 248858130 | 248858321 | 7 | 1 | 9 | 1 | 9 | 5 | 0 | 13 | 2 | 5 | 3 | 2 | 16 | 7 | 0 | 24 | 7 | 0 | 5 | 0 | 1 | 6 | 298 | 149 | 1.000000 | 1 | 0.212 | 0.2 | 0.474 | 0.455 | 0 | 1 | 0.412 | NA | 0.923 | 0.368 | -0.408 |
| A3SS | 5293 | ENSG00000171163 | ZNF692 | chr1 | - | 248857827 | 248858034 | 248857827 | 248857859 | 248858130 | 248858321 | 7 | 1 | 9 | 1 | 9 | 5 | 0 | 13 | 2 | 5 | 3 | 2 | 16 | 7 | 0 | 24 | 7 | 0 | 5 | 0 | 1 | 6 | 298 | 149 | 1.000000 | 1 | 0.212 | 0.2 | 0.474 | 0.455 | 0 | 1 | 0.412 | NA | 0.923 | 0.368 | -0.408 |
| A3SS | 8404 | ENSG00000141013 | GAS8 | chr16 | + | 90031175 | 90031496 | 90031298 | 90031496 | 90027635 | 90027722 | 3 | 4 | 2 | 4 | 2 | 0 | 0 | 1 | 0 | 1 | 2 | 0 | 20 | 4 | 3 | 3 | 11 | 0 | 6 | 6 | 0 | 0 | 272 | 149 | 1.000000 | 1 | 0.622 | 1 | 0.523 | 0 | NA | 1 | 0.268 | 0.215 | 1 | 1 | -0.16 |
| A3SS | 826 | ENSG00000101473 | ACOT8 | chr20 | - | 45853922 | 45853984 | 45853922 | 45853979 | 45855158 | 45855292 | 5 | 0 | 0 | 0 | 0 | 1 | 4 | 0 | 0 | 1 | 0 | 3 | 12 | 19 | 4 | 3 | 4 | 10 | 3 | 0 | 0 | 8 | 154 | 149 | 1.000000 | 1 | 1 | NA | 0 | 1 | 0.563 | 0.537 | 0.86 | 1 | 1 | 0.326 | -0.104 |
| A3SS | 11140 | ENSG00000204685 | STARD7-AS1 | chr2 | + | 96239712 | 96240688 | 96240443 | 96240688 | 96237904 | 96238099 | 34 | 15 | 20 | 15 | 20 | 21 | 23 | 0 | 0 | 2 | 0 | 0 | 31 | 32 | 29 | 36 | 17 | 0 | 0 | 0 | 0 | 0 | 298 | 149 | 1.000000 | 1 | 1 | 1 | 0.833 | 1 | 1 | 1 | 1 | 1 | 1 | 1 | -0.033 |
| A3SS | 8800 | ENSG00000126254 | RBM42 | chr19 | + | 35633076 | 35633252 | 35633100 | 35633252 | 35632935 | 35633001 | 17 | 5 | 13 | 5 | 13 | 9 | 8 | 0 | 0 | 0 | 1 | 0 | 1 | 22 | 13 | 24 | 24 | 0 | 0 | 0 | 0 | 0 | 173 | 149 | 1.000000 | 1 | 1 | 1 | 1 | 0.886 | 1 | 1 | 1 | 1 | 1 | 1 | -0.023 |
| A3SS | 2931 | ENSG00000118520 | ARG1 | chr6 | + | 131579086 | 131579285 | 131579110 | 131579285 | 131576662 | 131576735 | 0 | 0 | 0 | 0 | 0 | 1 | 0 | 3 | 1 | 2 | 22 | 0 | 0 | 1 | 14 | 18 | 0 | 6 | 22 | 169 | 420 | 32 | 173 | 149 | 1.000000 | 1 | 0 | 0 | 0 | 0.038 | NA | 0 | 0.038 | 0.067 | 0.036 | 0 | -0.019 |
| A3SS | 9797 | ENSG00000012124 | CD22 | chr19 | + | 35332546 | 35332924 | 35332593 | 35332924 | 35329186 | 35329230 | 20 | 17 | 23 | 17 | 23 | 36 | 91 | 0 | 0 | 1 | 0 | 0 | 69 | 54 | 0 | 42 | 54 | 0 | 0 | 0 | 0 | 0 | 196 | 149 | 1.000000 | 1 | 1 | 1 | 0.946 | 1 | 1 | 1 | 1 | NA | 1 | 1 | -0.011 |
| A3SS | 2298 | ENSG00000140199 | SLC12A6 | chr15 | - | 34238231 | 34238401 | 34238231 | 34238331 | 34240660 | 34240829 | 46 | 56 | 58 | 56 | 58 | 94 | 37 | 0 | 0 | 0 | 1 | 0 | 74 | 35 | 95 | 90 | 29 | 0 | 0 | 0 | 1 | 0 | 219 | 149 | 1.000000 | 1 | 1 | 1 | 1 | 0.985 | 1 | 1 | 1 | 1 | 0.984 | 1 | 0 |
| A3SS | 6118 | ENSG00000142534 | RPS11 | chr19 | + | 49497172 | 49497325 | 49497193 | 49497325 | 49496433 | 49496471 | 22 | 24 | 43 | 24 | 43 | 40 | 37 | 5811 | 3569 | 5947 | 5437 | 5281 | 55 | 27 | 47 | 27 | 42 | 8143 | 5583 | 4651 | 4859 | 9897 | 170 | 149 | 1.000000 | 1 | 0.003 | 0.006 | 0.006 | 0.006 | 0.006 | 0.006 | 0.004 | 0.009 | 0.005 | 0.004 | 0 |
| A3SS | 5500 | ENSG00000145916 | RMND5B | chr5 | + | 178142582 | 178142728 | 178142632 | 178142728 | 178138107 | 178138258 | 16 | 7 | 16 | 7 | 16 | 8 | 1 | 0 | 0 | 1 | 0 | 0 | 18 | 15 | 9 | 9 | 29 | 0 | 0 | 0 | 1 | 0 | 199 | 149 | 1.000000 | 1 | 1 | 1 | 0.923 | 1 | 1 | 1 | 1 | 1 | 0.871 | 1 | 0.01 |
| A3SS | 3637 | ENSG00000144659 | SLC25A38 | chr3 | + | 39394409 | 39394576 | 39394449 | 39394576 | 39393680 | 39393776 | 20 | 16 | 18 | 16 | 18 | 16 | 9 | 0 | 0 | 0 | 0 | 0 | 14 | 30 | 17 | 19 | 31 | 0 | 0 | 1 | 0 | 0 | 189 | 149 | 1.000000 | 1 | 1 | 1 | 1 | 1 | 1 | 1 | 1 | 0.931 | 1 | 1 | 0.014 |
| A3SS | 10673 | ENSG00000248323 | LUCAT1 | chr5 | - | 91311645 | 91312413 | 91311645 | 91311737 | 91313077 | 91313229 | 5 | 4 | 1 | 4 | 1 | 15 | 10 | 0 | 0 | 0 | 0 | 0 | 6 | 3 | 28 | 22 | 8 | 0 | 0 | 2 | 0 | 0 | 298 | 149 | 1.000000 | 1 | 1 | 1 | 1 | 1 | 1 | 1 | 1 | 0.875 | 1 | 1 | 0.025 |
| [illegible] | | | | | | | | | | | | | | | | | | | | | | | | | | | | | | | | | | | | | | | | | | | | | | | | |

| | | | | | | | | | | | | | | | |
|---|---|---|---|---|---|---|---|---|---|---|---|---|---|---|---|
| A3SS | 7061 | ENSG00000131165 | CHMP1A | chr16 | - | 89647202 | 89648997 | 89647202 | 89648478 | 89649350 | 89649497 | 4 | 2 | 0 | [illegible] |
| A3SS | 8518 | ENSG00000177674 | AGTRAP | chr1 | + | 11745929 | 11746223 | 11745987 | 11746223 | 11745802 | 11745837 | 5 | 0 | 8 | [illegible] |
| A3SS | 13006 | ENSG00000135297 | MTO1 | chr6 | + | 73479731 | 73480126 | 73479935 | 73480126 | 73473364 | 73473654 | 95 | 85 | 71 | [illegible] |
| A3SS | 11134 | ENSG00000164733 | CTSB | chr8 | - | 11849045 | 11849215 | 11849045 | 11849164 | 11850865 | 11850980 | 5 | 0 | 0 | [illegible] |
| A3SS | 684 | ENSG00000167468 | GPX4 | chr19 | + | 1106180 | 1106266 | 1106241 | 1106266 | 1105657 | 1105809 | 10 | 4 | 3 | [illegible] |
| A3SS | 6193 | ENSG00000107551 | RASSF4 | chr10 | + | 44989616 | 44989721 | 44989669 | 44989721 | 44989273 | 44989375 | 8 | 2 | 4 | [illegible] |
| A3SS | 9229 | ENSG00000138071 | ACTR2 | chr2 | + | 65241971 | 65242063 | 65242048 | 65242063 | 65239851 | 65239962 | 36 | 23 | 17 | [illegible] |
| A3SS | 6727 | ENSG00000143851 | PTPN7 | chr1 | - | 202159280 | 202159454 | 202159280 | 202159416 | 202161386 | 202161588 | 8 | 1 | 5 | [illegible] |
| A3SS | 10686 | ENSG00000102312 | PORCN | chrX | + | 48509783 | 48509956 | 48509892 | 48509956 | 48509423 | 48509707 | 6 | 1 | 6 | [illegible] |
| A3SS | 2492 | ENSG00000206190 | ATP10A | chr15 | - | 25687702 | 25687996 | 25687702 | 25687828 | 25701915 | 25704020 | 2 | 9 | 4 | [illegible] |
| A3SS | 10468 | ENSG00000163697 | APBB2 | chr4 | - | 40944864 | 40945073 | 40944864 | 40945070 | 41013582 | 41014385 | 0 | 0 | 2 | [illegible] |
| A3SS | 9898 | ENSG00000235621 | LINC00494 | chr20 | + | 48365872 | 48367162 | 48366473 | 48367162 | 48364888 | 48365197 | 2 | 0 | 0 | [illegible] |
| A3SS | 1415 | ENSG00000178999 | AURKB | chr17 | - | 8206749 | 8206934 | 8206749 | 8206888 | 8207317 | 8207367 | 0 | 0 | 2 | [illegible] |
| A3SS | 4149 | ENSG00000278259 | MYO19 | chr17 | - | 36501851 | 36501961 | 36501851 | 36501901 | 36503096 | 36503200 | 1 | 0 | 8 | [illegible] |
| A3SS | 12812 | ENSG00000228784 | LINC00954 | chr2 | + | 19870401 | 19870546 | 19870424 | 19870546 | 19869495 | 19869613 | 0 | 1 | 0 | [illegible] |
| A3SS | 12004 | ENSG00000149269 | PAK1 | chr11 | - | 77392330 | 77392541 | 77392330 | 77392538 | 77408153 | 77408232 | 5 | 0 | 1 | [illegible] |
| A3SS | 3581 | ENSG00000083312 | TNPO1 | chr5 | + | 72851204 | 72851319 | 72851237 | 72851319 | 72848384 | 72848498 | 4 | 2 | 0 | [illegible] |
| A3SS | 7837 | ENSG00000160753 | RUSC1 | chr1 | + | 155326609 | 155327132 | 155326897 | 155327132 | 155325863 | 155325910 | 16 | 0 | 8 | [illegible] |
| A3SS | 1129 | ENSG00000136492 | BRIP1 | chr17 | - | 61776400 | 61776562 | 61776400 | 61776502 | 61778672 | 61778778 | 3 | 1 | 4 | [illegible] |
| A3SS | 7378 | ENSG00000168679 | SLC16A4 | chr1 | - | 110362850 | 110363913 | 110362850 | 110363893 | 110375457 | 110375551 | 0 | 3 | 0 | [illegible] |
| A3SS | 1188 | ENSG00000196704 | AMZ2 | chr17 | + | 68248639 | 68248705 | 68248643 | 68248705 | 68248034 | 68248058 | 2 | 2 | 1 | [illegible] |
| A3SS | 7470 | ENSG00000163534 | FCRL1 | chr1 | - | 157801914 | 157802193 | 157801914 | 157802162 | 157802614 | 157802664 | 20 | 6 | 7 | [illegible] |
| A3SS | 3863 | ENSG00000251600 | NA | chr4 | + | 143979849 | 143979952 | 143979854 | 143979952 | 143937866 | 143937923 | 3 | 2 | 0 | [illegible] |
| A3SS | 3086 | ENSG00000137070 | IL11RA | chr9 | + | 34660803 | 34660936 | 34660853 | 34660936 | 34660503 | 34660600 | 11 | 3 | 8 | [illegible] |
| A3SS | 6542 | ENSG00000274020 | LINC01138 | chr1 | - | 148435145 | 148435354 | 148435145 | 148435232 | 148436443 | 148436564 | 32 | 11 | 6 | [illegible] |
| A3SS | 11220 | ENSG00000147789 | ZNF7 | chr8 | + | 144829477 | 144829604 | 144829522 | 144829604 | 144829042 | 144829090 | 5 | 2 | 12 | [illegible] |
| A3SS | 7813 | ENSG00000198520 | ARMH1 | chr1 | + | 44724319 | 44724668 | 44724538 | 44724668 | 44724121 | 44724244 | 41 | 27 | 42 | [illegible] |
| A3SS | 12161 | ENSG00000173153 | ESRRA | chr11 | + | 64314740 | 64314911 | 64314743 | 64314911 | 64314238 | 64314367 | 22 | 6 | 20 | [illegible] |
| A3SS | 11287 | ENSG00000147162 | OGT | chrX | + | 71544566 | 71548023 | 71547906 | 71548023 | 71533103 | 71533336 | 246 | 183 | 235 | [illegible] |
| A3SS | 1181 | ENSG00000182173 | TSEN54 | chr17 | + | 75518994 | 75519047 | 75519002 | 75519047 | 75517556 | 75517655 | 52 | 19 | 48 | [illegible] |
| A3SS | 583 | ENSG00000204936 | CD177 | chr19 | + | 43355514 | 43355783 | 43355660 | 43355783 | 43354206 | 43354392 | 0 | 0 | 2 | [illegible] |
| A3SS | 7227 | ENSG00000082701 | GSK3B | chr3 | - | 119880244 | 119880510 | 119880244 | 119880498 | 119905754 | 119905852 | 0 | 1 | 0 | [illegible] |
| A3SS | 472 | ENSG00000105492 | SIGLEC6 | chr19 | - | 51531159 | 51531519 | 51531159 | 51531411 | 51531581 | 51531803 | 9 | 6 | 5 | [illegible] |
| A3SS | 3117 | ENSG00000250903 | GMDS-DT | chr6 | + | 2398865 | 2399037 | 2398882 | 2399037 | 2398576 | 2398778 | 0 | 8 | 0 | [illegible] |
| A3SS | 2377 | ENSG00000140326 | CDAN1 | chr15 | - | 42735874 | 42736081 | 42735874 | 42736078 | 42736301 | 42736780 | 3 | 2 | 2 | [illegible] |
| A3SS | 7428 | ENSG00000155367 | PPM1J | chr1 | - | 112709993 | 112710310 | 112709993 | 112710072 | 112710459 | 112710611 | 2 | 0 | 5 | [illegible] |
| A3SS | 9753 | ENSG00000106617 | PRKAG2 | chr7 | - | 151595344 | 151595454 | 151595344 | 151595451 | 151675419 | 151675637 | 0 | 0 | 1 | [illegible] |
| A3SS | 4460 | ENSG00000135083 | CCNJL | chr5 | - | 160280524 | 160280948 | 160280524 | 160280738 | 160311857 | 160311972 | 0 | 0 | 0 | [illegible] |
| A3SS | 1880 | ENSG00000197912 | SPG7 | chr16 | + | 89555877 | 89556085 | 89555920 | 89556085 | 89554485 | 89554563 | 1 | 0 | 10 | [illegible] |
| A3SS | 10687 | ENSG00000102312 | PORCN | chrX | + | 48511294 | 48511487 | 48511365 | 48511487 | 48509783 | 48509956 | 5 | 4 | 9 | [illegible] |
| A3SS | 3301 | ENSG00000027847 | B4GALT7 | chr5 | + | 177607282 | 177607527 | 177607301 | 177607527 | 177604178 | 177604541 | 2 | 0 | 0 | [illegible] |
| A3SS | 7032 | ENSG00000126215 | XRCC3 | chr14 | - | 103711465 | 103711661 | 103711465 | 103711567 | 103715423 | 103715484 | 5 | 0 | 4 | [illegible] |
| A3SS | 8603 | ENSG00000225889 | NA | chr2 | + | 64206971 | 64207045 | 64206974 | 64207045 | 64201747 | 64201894 | 0 | 1 | 1 | [illegible] |
| A3SS | 10201 | ENSG00000146707 | POMZP3 | chr7 | - | 76611453 | 76611627 | 76611453 | 76611591 | 76611721 | 76611813 | 0 | 3 | 1 | [illegible] |
| A3SS | 13174 | ENSG00000122966 | CIT | chr12 | - | 119718698 | 119718861 | 119718698 | 119718816 | 119720477 | 119720585 | 0 | 0 | 0 | [illegible] |
| A3SS | 2700 | ENSG00000099797 | TECR | chr19 | + | 14563140 | 14563257 | 14563173 | 14563257 | 14562524 | 14562575 | 0 | 0 | 0 | [illegible] |
| A3SS | 8073 | ENSG00000143537 | ADAM15 | chr1 | + | 155055701 | 155055852 | 155055786 | 155055852 | 155054313 | 155054506 | 1 | 0 | 1 | [illegible] |
| A3SS | 1695 | ENSG00000140650 | PMM2 | chr16 | + | 8802260 | 8802389 | 8802289 | 8802389 | 8802080 | 8802145 | 5 | 2 | 1 | [illegible] |
| A3SS | 798 | ENSG00000226648 | PLCG1-AS1 | chr20 | - | 41100657 | 41101289 | 41100657 | 41100792 | 41133486 | 41133645 | 4 | 5 | 4 | [illegible] |
| A3SS | 277 | ENSG00000105607 | GCDH | chr19 | + | 12896204 | 12896421 | 12896239 | 12896421 | 12895991 | 12896121 | 16 | 16 | 21 | [illegible] |
| A3SS | 10707 | ENSG00000147133 | TAF1 | chrX | + | 71392870 | 71392994 | 71392874 | 71392994 | 71392568 | 71392718 | 3 | 0 | 2 | [illegible] |
| A3SS | 5467 | ENSG00000181666 | ZNF875 | chr19 | + | 37347189 | 37347316 | 37347231 | 37347316 | 37324207 | 37324265 | 3 | 5 | 10 | [illegible] |
| A3SS | 10532 | ENSG00000005302 | MSL3 | chrX | + | 11768572 | 11768682 | 11768621 | 11768682 | 11765537 | 11765729 | 133 | 108 | 133 | [illegible] |
| A3SS | 2416 | ENSG00000140365 | COMMD4 | chr15 | + | 75338061 | 75338420 | 75338354 | 75338420 | 75336062 | 75336092 | 24 | 10 | 23 | [illegible] |
| A3SS | 7629 | ENSG00000186652 | PRG2 | chr11 | - | 57388576 | 57389317 | 57388576 | 57388708 | 57389886 | 57389956 | 2 | 3 | 2 | [illegible] |
| A3SS | 10071 | ENSG00000105889 | STEAP1B | chr7 | - | 22493323 | 22493836 | 22493323 | 22493779 | 22494771 | 22494886 | 1 | 1 | 5 | [illegible] |
| A3SS | 195 | ENSG00000274602 | PI4KAP1 | chr22 | - | 18547245 | 18547421 | 18547245 | 18547373 | 18548010 | 18548118 | 1 | 0 | 4 | [illegible] |
| A3SS | 10992 | ENSG00000140525 | FANCI | chr15 | + | 89306006 | 89306194 | 89306009 | 89306194 | 89305604 | 89305698 | 16 | 5 | 11 | [illegible] |
| A3SS | 4527 | ENSG00000156709 | AIFM1 | chrX | - | 130147751 | 130147935 | 130147751 | 130147870 | 130149468 | 130149568 | 2 | 0 | 2 | [illegible] |
| A3SS | 12221 | ENSG00000221926 | TRIM16 | chr17 | - | 15677175 | 15677280 | 15677175 | 15677277 | 15680864 | 15680953 | 2 | 0 | 0 | [illegible] |
| A3SS | 12158 | ENSG00000105808 | RASA4 | chr7 | - | 102596011 | 102596129 | 102596011 | 102596107 | 102599859 | 102599954 | 0 | 0 | 0 | [illegible] |
| A3SS | 4619 | ENSG00000175556 | LONRF3 | chrX | + | 119012879 | 119013201 | 119013038 | 119013201 | 119011814 | 119011973 | 3 | 8 | 3 | [illegible] |
| A3SS | 11598 | ENSG00000213930 | GALT | chr9 | + | 34648047 | 34648171 | 34648091 | 34648171 | 34647831 | 34647961 | 10 | 4 | 8 | [illegible] |
| A3SS | 4337 | ENSG00000243989 | ACY1 | chr3 | + | 51986944 | 51987061 | 51986987 | 51987061 | 51986604 | 51986661 | 0 | 0 | 0 | [illegible] |
| A3SS | 8104 | ENSG00000155366 | RHOC | chr1 | - | 112705099 | 112705201 | 112705099 | 112705123 | 112707077 | 112707102 | 12 | 1 | 11 | [illegible] |
| A3SS | 1703 | ENSG00000033011 | ALG1 | chr16 | + | 5074878 | 5075536 | 5075397 | 5075536 | 5073152 | 5073256 | 2 | 2 | 3 | [illegible] |
| A3SS | 5564 | ENSG00000077782 | FGFR1 | chr8 | - | 38421796 | 38423175 | 38421796 | 38421941 | 38424508 | 38424699 | 3 | 0 | 0 | [illegible] |
| A3SS | 8715 | ENSG00000130038 | CRACR2A | chr12 | - | 3638123 | 3638457 | 3638123 | 3638454 | 3648541 | 3648613 | 1 | 3 | 0 | [illegible] |
| A3SS | 8089 | ENSG00000116809 | ZBTB17 | chr1 | - | 15942330 | 15942441 | 15942330 | 15942420 | 15942528 | 15942738 | 1 | 0 | 0 | [illegible] |
| A3SS | 1881 | ENSG00000197912 | SPG7 | chr16 | + | 89555877 | 89556085 | 89555958 | 89556085 | 89554485 | 89554563 | 1 | 0 | 15 | [illegible] |
| A3SS | 8009 | ENSG00000149499 | EML3 | chr11 | - | 62610878 | 62610992 | 62610878 | 62610954 | 62611086 | 62611344 | 19 | 16 | 24 | [illegible] |
| A3SS | 9990 | ENSG00000160993 | ALKBH4 | chr7 | - | 102459603 | 102459801 | 102459603 | 102459710 | 102464713 | 102464863 | 7 | 4 | 4 | [illegible] |
| A3SS | 6723 | ENSG00000143851 | PTPN7 | chr1 | - | 202159280 | 202159454 | 202159280 | 202159416 | 202160544 | 202160608 | 61 | 15 | 45 | [illegible] |
| A3SS | 2393 | ENSG00000103942 | HOMER2 | chr15 | - | 82859028 | 82859186 | 82859028 | 82859135 | 82864166 | 82864259 | 0 | 2 | 0 | [illegible] |
| A3SS | 9579 | ENSG00000255036 | SUGT1P4-STRA6LP-CCDC180 | chr9 | + | 97362195 | 97362441 | 97362362 | 97362441 | 97361725 | 97361898 | 7 | 16 | 5 | [illegible] |
| A3SS | 3348 | ENSG00000214717 | ZBED1 | chrX | - | 2486413 | 2490772 | 2486413 | 2490769 | 2500816 | 2500967 | 3 | 2 | 6 | [illegible] |
| A3SS | 4953 | ENSG00000116514 | RNF19B | chr1 | - | 32948221 | 32948366 | 32948221 | 32948363 | 32964050 | 32964685 | 1 | 0 | 3 | [illegible] |
| A3SS | 8208 | ENSG00000066923 | STAG3 | chr7 | + | 100211434 | 100211539 | 100211437 | 100211539 | 100211010 | 100211185 | 1 | 0 | 0 | [illegible] |
| A3SS | 2356 | ENSG00000140534 | TICRR | chr15 | + | 89584285 | 89584527 | 89584288 | 89584527 | 89582685 | 89582965 | 0 | 0 | 4 | [illegible] |
| A3SS | 1665 | ENSG00000102878 | HSF4 | chr16 | + | 67166557 | 67166622 | 67166587 | 67166622 | 67166310 | 67166369 | 2 | 4 | 0 | [illegible] |
| A3SS | 4193 | ENSG00000164045 | CDC25A | chr3 | - | 48182928 | 48183030 | 48182928 | 48183027 | 48183799 | 48183836 | 1 | 0 | 4 | [illegible] |
| A3SS | 2154 | ENSG00000172349 | IL16 | chr15 | + | 81291923 | 81293037 | 81292555 | 81293037 | 81290452 | 81290540 | 0 | 0 | 0 | [illegible] |
| A3SS | 494 | ENSG00000063241 | ISOC2 | chr19 | - | 55455259 | 55455378 | 55455259 | 55455330 | 55456348 | 55456489 | 0 | 0 | 0 | [illegible] |
| A3SS | 53 | ENSG00000206140 | TMEM191C | chr22 | + | 21468314 | 21468431 | 21468353 | 21468431 | 21467893 | 21468017 | 0 | 0 | 1 | [illegible] |
| A3SS | 5129 | ENSG00000164776 | PHKG1 | chr7 | - | 56083649 | 56083724 | 56083649 | 56083715 | 56086969 | 56087024 | 0 | 0 | 0 | [illegible] |
| A3SS | 8403 | ENSG00000141013 | GAS8 | chr16 | + | 90031175 | 90031496 | 90031189 | 90031496 | 90027635 | 90027722 | 0 | 3 | 1 | [illegible] |
| A3SS | 11274 | ENSG00000106772 | PRUNE2 | chr9 | - | 76644738 | 76644912 | 76644738 | 76644909 | 76652482 | 76652683 | 0 | 5 | 0 | [illegible] |
| A3SS | 11189 | ENSG00000104518 | GSDMD | chr8 | + | 143558361 | 143558451 | 143558377 | 143558451 | 143555933 | 143556030 | 6 | 4 | 8 | [illegible] |
| A3SS | 11470 | ENSG00000107331 | ABCA2 | chr9 | - | 137022940 | 137023840 | 137022940 | 137023052 | 137024142 | 137024236 | 4 | 8 | 2 | [illegible] |
| A3SS | 7713 | ENSG00000049167 | ERCC8 | chr5 | - | 60924290 | 60924367 | 60924290 | 60924356 | 60928863 | 60928959 | 5 | 1 | 2 | [illegible] |
| A3SS | 12703 | ENSG00000151651 | ADAM8 | chr10 | - | 133267928 | 133268118 | 133267928 | 133268066 | 133269444 | 133269529 | 12 | 2 | 5 | [illegible] |
| A3SS | 6867 | ENSG00000160345 | C9orf116 | chr9 | - | 135495180 | 135495615 | 135495180 | 135495569 | 135498584 | 135498663 | 0 | 1 | 0 | [illegible] |
| A3SS | 3477 | ENSG00000123836 | PFKFB2 | chr1 | + | 207062546 | 207062716 | 207062619 | 207062716 | 207061952 | 207062078 | 2 | 4 | 4 | [illegible] |
| A3SS | 51 | ENSG00000187860 | CCDC157 | chr22 | + | 30370325 | 30371727 | 30371649 | 30371727 | 30369431 | 30369603 | 5 | 0 | 5 | [illegible] |
| A3SS | 254 | ENSG00000131351 | HAUS8 | chr19 | - | 17059996 | 17060092 | 17059996 | 17060089 | 17069030 | 17069086 | 4 | 5 | 2 | [illegible] |
| A3SS | 12681 | ENSG00000148634 | HERC4 | chr10 | - | 67992605 | 67995292 | 67992605 | 67992682 | 67996045 | 67996138 | 9 | 7 | 5 | [illegible] |
| A3SS | 8994 | ENSG00000144134 | RABL2A | chr2 | + | 113641352 | 113641450 | 113641355 | 113641450 | 113640893 | 113641005 | 5 | 1 | 5 | [illegible] |
| A3SS | 2757 | ENSG00000100526 | CDKN3 | chr14 | + | 54417847 | 54417951 | 54417850 | 54417951 | 54415898 | 54415930 | 20 | 5 | 6 | [illegible] |
| A3SS | 9713 | ENSG00000146918 | NCAPG2 | chr7 | - | 158654594 | 158654789 | 158654594 | 158654694 | 158655117 | 158655258 | 0 | 0 | 0 | [illegible] |
| A3SS | 7980 | ENSG00000178038 | ALS2CL | chr3 | - | 46671487 | 46671768 | 46671487 | 46671584 | 46671883 | 46672033 | 2 | 2 | 0 | [illegible] |
| A3SS | 4111 | ENSG00000155508 | CNOT8 | chr5 | + | 154863206 | 154863395 | 154863211 | 154863395 | 154858653 | 154858816 | 29 | 31 | 24 | [illegible] |
| A3SS | 10629 | ENSG00000196924 | FLNA | chrX | - | 154353000 | 154353221 | 154353000 | 154353204 | 154353295 | 154353457 | 4 | 2 | 0 | [illegible] |
| A3SS | 2050 | ENSG00000270049 | ATP6V0D1-DT | chr16 | + | 67492024 | 67492121 | 67492030 | 67492121 | 67484109 | 67484328 | 0 | 0 | 0 | [illegible] |
| A3SS | 1233 | ENSG00000108622 | ICAM2 | chr17 | - | 64006630 | 64006735 | 64006630 | 64006731 | 64007740 | 64007822 | 12 | 3 | 15 | [illegible] |
| A3SS | 8729 | ENSG00000169247 | SH3TC2 | chr5 | - | 149047861 | 149048019 | 149047861 | 149047989 | 149052141 | 149052240 | 0 | 2 | 0 | [illegible] |
| A3SS | 13112 | ENSG00000130396 | AFDN | chr6 | + | 167962432 | 167962567 | 167962438 | 167962567 | 167951185 | 167952187 | 3 | 5 | 2 | [illegible] |
| A3SS | 5256 | ENSG00000274372 | LINC02804 | chr1 | + | 148024503 | 148024719 | 148024529 | 148024719 | 148021913 | 148021998 | 0 | 2 | 0 | [illegible] |
| A3SS | 4268 | ENSG00000157601 | MX1 | chr21 | + | 41421837 | 41422095 | 41421974 | 41422095 | 41420609 | 41420817 | 0 | 0 | 0 | [illegible] |
| A3SS | 7130 | ENSG00000198929 | NOS1AP | chr1 | + | 162300632 | 162300706 | 162300647 | 162300706 | 162287343 | 162287436 | 0 | 0 | 2 | [illegible] |
| A3SS | 6758 | ENSG00000105609 | LILRB5 | chr19 | - | 54252870 | 54254068 | 54252870 | 54252987 | 54254364 | 54254415 | 0 | 0 | 0 | [illegible] |
| A3SS | 12925 | ENSG00000238105 | GOLGA2P5 | chr12 | - | 100166330 | 100166663 | 100166330 | 100166636 | 100167053 | 100167141 | 0 | 0 | 1 | [illegible] |
| A3SS | 6376 | ENSG00000181790 | ADGRB1 | chr8 | + | 142478186 | 142478360 | 142478195 | 142478360 | 142477384 | 142477549 | 0 | 0 | 0 | [illegible] |
| A3SS | 12175 | ENSG00000149150 | SLC43A1 | chr11 | - | 57501151 | 57501374 | 57501151 | 57501329 | 57513957 | 57514124 | 0 | 0 | 0 | [illegible] |
| A3SS | 1171 | ENSG00000120093 | HOXB3 | chr17 | - | 48552026 | 48552632 | 48552026 | 48552360 | 48555530 | 48555618 | 1 | 0 | 1 | [illegible] |
| A3SS | 7121 | ENSG00000143398 | PIP5K1A | chr1 | + | 151224244 | 151224279 | 151224247 | 151224279 | 151213516 | 151213547 | 2 | 0 | 2 | [illegible] |
| A3SS | 118 | ENSG00000100099 | HPS4 | chr22 | - | 26470718 | 26470922 | 26470718 | 26470813 | 26471212 | 26471388 | 11 | 4 | 7 | [illegible] |
| A3SS | 7120 | ENSG00000143398 | PIP5K1A | chr1 | + | 151224244 | 151224279 | 151224247 | 151224279 | 151198724 | 151199187 | 1 | 0 | 2 | [illegible] |
| A3SS | 1172 | ENSG00000120093 | HOXB3 | chr17 | - | 48552026 | 48552632 | 48552026 | 48552487 | 48555530 | 48555618 | 7 | 0 | 0 | [illegible] |
| A3SS | 1187 | ENSG00000182173 | TSEN54 | chr17 | + | 75523249 | 75523335 | 75523274 | 75523335 | 75521704 | 75522521 | 9 | 6 | 10 | [illegible] |
| A3SS | 9515 | ENSG00000105255 | FSD1 | chr19 | + | 4318345 | 4318505 | 4318383 | 4318505 | 4317181 | 4317280 | 4 | 2 | 4 | [illegible] |
| A3SS | 9401 | ENSG00000168661 | ZNF30 | chr19 | + | 34933624 | 34933723 | 34933627 | 34933723 | 34931842 | 34931993 | 5 | 2 | 3 | [illegible] |
| A3SS | 6496 | ENSG00000143224 | PPOX | chr1 | + | 161168431 | 161168576 | 161168444 | 161168576 | 161167994 | 161168127 | 4 | 0 | 11 | [illegible] |
| A3SS | 11043 | ENSG00000249859 | PVT1 | chr8 | + | 128096517 | 128096654 | 128096520 | 128096654 | 128082752 | 128082848 | 0 | 7 | 5 | [illegible] |
| A3SS | 712 | ENSG00000141994 | DUS3L | chr19 | - | 5786748 | 5786919 | 5786748 | 5786845 | 5787060 | 5787171 | 1 | 1 | 2 | [illegible] |
| A3SS | 9628 | ENSG00000197363 | ZNF517 | chr8 | + | 144804124 | 144804238 | 144804145 | 144804238 | 144803640 | 144803767 | 5 | 2 | 4 | [illegible] |
| A3SS | 4376 | ENSG00000241404 | EGFL8 | chr6 | + | 32167086 | 32167257 | 32167146 | 32167257 | 32166909 | 32167005 | 0 | 1 | 7 | [illegible] |
| A3SS | 10649 | ENSG00000166707 | ZCCHC18 | chrX | + | 104113433 | 104115842 | 104115251 | 104115842 | 104113194 | 104113246 | 12 | 13 | 19 | [illegible] |
| A3SS | 8087 | ENSG00000168890 | TMEM150A | chr2 | - | 85601020 | 85601482 | 85601020 | 85601107 | 85601883 | 85602064 | 8 | 3 | 15 | [illegible] |
| A3SS | 4490 | ENSG00000134802 | SLC43A3 | chr11 | - | 57425988 | 57426352 | 57425988 | 57426316 | 57426587 | 57426652 | 23 | 15 | 15 | [illegible] |
| A3SS | 7814 | ENSG00000198520 | ARMH1 | chr1 | + | 44724336 | 44724668 | 44724538 | 44724668 | 44724121 | 44724244 | 10 | 17 | 18 | [illegible] |
| A3SS | 4354 | ENSG00000096063 | SRPK1 | chr6 | - | 35890894 | 35891041 | 35890894 | 35891013 | 35904878 | 35904974 | 1 | 0 | 4 | [illegible] |
| A3SS | 11197 | ENSG00000104529 | EEF1D | chr8 | - | 143586728 | 143586852 | 143586728 | 143586849 | 143588990 | 143590460 | 17 | 10 | 31 | [illegible] |
| A3SS | 11941 | ENSG00000168060 | NAALADL1 | chr11 | - | 65044817 | 65045457 | 65044817 | 65045250 | 65045821 | 65045914 | 21 | 5 | 16 | [illegible] |
| A3SS | 11311 | ENSG00000110455 | ACCS | chr11 | + | 44077844 | 44078784 | 44078683 | 44078784 | 44077278 | 44077376 | 62 | 16 | 28 | [illegible] |
| A3SS | 1502 | ENSG00000006282 | SPATA20 | chr17 | + | 50550207 | 50550312 | 50550229 | 50550312 | 50549984 | 50550115 | 20 | 2 | 15 | [illegible] |
| A3SS | 13149 | ENSG00000123349 | PFDN5 | chr12 | + | 53298018 | 53298150 | 53298044 | 53298150 | 53297849 | 53297924 | 43 | 20 | 30 | [illegible] |
| A3SS | 5140 | ENSG00000137166 | FOXP4 | chr6 | + | 41587298 | 41587512 | 41587301 | 41587512 | 41587008 | 41587156 | 6 | 2 | 4 | [illegible] |
| A3SS | 1659 | ENSG00000168488 | ATXN2L | chr16 | + | 28836727 | 28837237 | 28836750 | 28837237 | 28836325 | 28836490 | 7 | 5 | 7 | [illegible] |
| A3SS | 4655 | ENSG00000174243 | DDX23 | chr12 | - | 48840012 | 48840151 | 48840012 | 48840106 | 48845573 | 48845782 | 4 | 5 | 5 | [illegible] |
| A3SS | 10954 | ENSG00000110074 | FOXRED1 | chr11 | + | 126273315 | 126273454 | 126273338 | 126273454 | 126272968 | 126273079 | 1 | 0 | 0 | [illegible] |
| A3SS | 1301 | ENSG00000185359 | HGS | chr17 | + | 81701214 | 81701422 | 81701243 | 81701422 | 81701044 | 81701131 | 0 | 0 | 3 | [illegible] |
| A3SS | 4954 | ENSG00000117245 | KIF17 | chr1 | - | 20670420 | 20670491 | 20670420 | 20670488 | 20671937 | 20672196 | 0 | 0 | 0 | [illegible] |
| A3SS | 64 | ENSG00000099957 | P2RX6 | chr22 | + | 21017920 | 21018060 | 21017988 | 21018060 | 21015941 | 21016092 | 0 | 0 | 0 | [illegible] |
| A3SS | 12277 | ENSG00000230583 | GTF2IRD1P1 | chr7 | - | 66823997 | 66824182 | 66823997 | 66824072 | 66825295 | 66825361 | 0 | 0 | 1 | [illegible] |
| A3SS | 11618 | ENSG00000165699 | TSC1 | chr9 | - | 132909806 | 132910692 | 132909806 | 132910689 | 132911452 | 132911568 | 0 | 0 | 0 | [illegible] |
| A3SS | 11016 | ENSG00000171045 | TSNARE1 | chr8 | - | 142315002 | 142315092 | 142315002 | 142315089 | 142330900 | 142330970 | 2 | 0 | 0 | [illegible] |
| A3SS | 816 | ENSG00000185513 | L3MBTL1 | chr20 | + | 43540021 | 43540308 | 43540150 | 43540308 | 43536408 | 43536458 | 1 | 0 | 3 | [illegible] |
| A3SS | 2169 | ENSG00000213471 | TTLL13P | chr15 | + | 90263739 | 90264058 | 90263946 | 90264058 | 90262965 | 90263145 | 1 | 1 | 0 | [illegible] |
| A3SS | 2170 | ENSG00000213471 | TTLL13P | chr15 | + | 90263754 | 90264058 | 90263946 | 90264058 | 90262965 | 90263145 | 1 | 1 | 0 | [illegible] |
| A3SS | 12224 | ENSG00000221926 | TRIM16 | chr17 | - | 15683035 | 15683371 | 15683035 | 15683157 | 15684195 | 15684311 | 10 | 5 | 4 | [illegible] |
| A3SS | 2094 | ENSG00000140983 | RHOT2 | chr16 | + | 668352 | 668411 | 668357 | 668411 | 668104 | 668154 | 3 | 0 | 0 | [illegible] |
| A3SS | 2317 | ENSG00000067369 | TP53BP1 | chr15 | - | 43446765 | 43446914 | 43446765 | 43446910 | 43447365 | 43447485 | 6 | 1 | 5 | [illegible] |
| A3SS | 13240 | ENSG00000101160 | CTSZ | chr20 | - | 58992761 | 58992841 | 58992761 | 58992821 | 58997602 | 58997753 | 3 | 3 | 2 | [illegible] |
| A3SS | 6874 | ENSG00000117713 | ARID1A | chr1 | + | 26772811 | 26772987 | 26772814 | 26772987 | 26771118 | 26771326 | 6 | 5 | 3 | [illegible] |
| A3SS | 12209 | ENSG00000142102 | PGGHG | chr11 | + | 290657 | 291113 | 290677 | 291113 | 290389 | 290600 | 4 | 0 | 1 | [illegible] |
| A3SS | 5622 | ENSG00000204435 | CSNK2B | chr6 | + | 31668512 | 31668654 | 31668538 | 31668654 | 31667867 | 31667970 | 2 | 0 | 0 | [illegible] |
| A3SS | 13230 | ENSG00000123411 | IKZF4 | chr12 | + | 56018122 | 56018177 | 56018125 | 56018177 | 56011557 | 56011737 | 0 | 0 | 0 | [illegible] |
| A3SS | 9993 | ENSG00000181090 | EHMT1 | chr9 | + | 137743370 | 137743528 | 137743391 | 137743528 | 137716625 | 137717182 | 23 | 4 | 9 | [illegible] |
| A3SS | 4454 | ENSG00000114867 | EIF4G1 | chr3 | + | 184317746 | 184317816 | 184317749 | 184317816 | 184317320 | 184317497 | 15 | 5 | 14 | [illegible] |
| A3SS | 10815 | ENSG00000142046 | TMEM91 | chr19 | + | 41383873 | 41384083 | 41383885 | 41384083 | 41382771 | 41382921 | 0 | 0 | 0 | [illegible] |
| A3SS | 7797 | ENSG00000167595 | PROSER3 | chr19 | + | 35762252 | 35762356 | 35762255 | 35762356 | 35762018 | 35762146 | 5 | 1 | 2 | [illegible] |
| A3SS | 906 | ENSG00000174365 | SNHG11 | chr20 | + | 38448661 | 38448730 | 38448665 | 38448730 | 38447465 | 38447623 | 4 | 1 | 8 | [illegible] |
| A3SS | 5727 | ENSG00000178809 | TRIM73 | chr7 | + | 75405181 | 75405605 | 75405184 | 75405605 | 75404838 | 75405069 | 0 | 2 | 3 | [illegible] |
| A3SS | 8818 | ENSG00000163482 | STK36 | chr2 | + | 218696526 | 218696601 | 218696589 | 218696601 | 218694524 | 218694635 | 2 | 0 | 8 | [illegible] |
| A3SS | 4894 | ENSG00000198925 | ATG9A | chr2 | - | 219228433 | 219228485 | 219228433 | 219228482 | 219229534 | 219229639 | 2 | 1 | 3 | [illegible] |
| A3SS | 4222 | ENSG00000225697 | SLC26A6 | chr3 | - | 48632244 | 48632396 | 48632244 | 48632337 | 48632973 | 48633084 | 20 | 5 | 5 | [illegible] |
| A3SS | 13388 | ENSG00000173581 | CCDC106 | chr19 | + | 55648757 | 55649077 | 55648945 | 55649077 | 55648232 | 55648277 | 3 | 1 | 1 | [illegible] |
| A3SS | 8369 | ENSG00000189223 | PAX8-AS1 | chr2 | + | 113235791 | 113236613 | 113236512 | 113236613 | 113235539 | 113235570 | 1 | 0 | 0 | [illegible] |
| A3SS | 1084 | ENSG00000073605 | GSDMB | chr17 | - | 39917081 | 39917330 | 39917081 | 39917321 | 39918533 | 39918635 | 0 | 0 | 0 | [illegible] |
| A3SS | 11410 | ENSG00000144524 | COPS7B | chr2 | + | 231801139 | 231801207 | 231801154 | 231801207 | 231798858 | 231798964 | 2 | 1 | 2 | [illegible] |
| A3SS | 8861 | ENSG00000228203 | GRASLND | chr2 | - | 6915822 | 6916044 | 6915822 | 6915967 | 6916815 | 6917224 | 0 | 1 | 1 | [illegible] |
| A3SS | 1169 | ENSG00000120093 | HOXB3 | chr17 | - | 48552026 | 48552632 | 48552026 | 48552090 | 48555530 | 48555618 | 2 | 2 | 0 | [illegible] |
| A3SS | 31 | ENSG00000142173 | COL6A2 | chr21 | + | 46128902 | 46129808 | 46129195 | 46129808 | 46126502 | 46126541 | 3 | 3 | 0 | [illegible] |
| A3SS | 4252 | ENSG00000150995 | ITPR1 | chr3 | + | 4727122 | 4727173 | 4727125 | 4727173 | 4725545 | 4725581 | 1 | 0 | 1 | [illegible] |
| A3SS | 13185 | ENSG00000198040 | ZNF84 | chr12 | + | 133047929 | 133048081 | 133047954 | 133048081 | 133041277 | 133041482 | 6 | 3 | 7 | [illegible] |
| A3SS | 8681 | ENSG00000155749 | FLACC1 | chr2 | - | 201289695 | 201289859 | 201289695 | 201289854 | 201299237 | 201299300 | 0 | 2 | 0 | [illegible] |
| A3SS | 3339 | ENSG00000125846 | ZNF133 | chr20 | + | 18305667 | 18305807 | 18305680 | 18305807 | 18305007 | 18305178 | 0 | 1 | 2 | [illegible] |
| A3SS | 11659 | ENSG00000130714 | POMT1 | chr9 | + | 131519083 | 131519486 | 131519393 | 131519486 | 131518836 | 131518957 | 0 | 0 | 3 | [illegible] |
| A3SS | 8002 | ENSG00000169379 | ARL13B | chr3 | + | 93996547 | 93996684 | 93996562 | 93996684 | 93995873 | 93995944 | 2 | 0 | 3 | [illegible] |
| A3SS | 4088 | ENSG00000099949 | LZTR1 | chr22 | + | 20995745 | 20995872 | 20995762 | 20995872 | 20995367 | 20995484 | 9 | 8 | 4 | [illegible] |
| A3SS | 6790 | ENSG00000090097 | PCBP4 | chr3 | - | 51961159 | 51961304 | 51961159 | 51961270 | 51962809 | 51963256 | 3 | 1 | 3 | [illegible] |
| A3SS | 4766 | ENSG00000075826 | SEC31B | chr10 | - | 100488857 | 100489398 | 100488857 | 100488974 | 100489702 | 100489761 | 15 | 9 | 23 | [illegible] |
| A3SS | 2625 | ENSG00000171469 | ZNF561 | chr19 | - | 9617044 | 9617226 | 9617044 | 9617171 | 9619431 | 9619582 | 13 | 5 | 10 | [illegible] |
| A3SS | 5698 | ENSG00000204599 | TRIM39 | chr6 | + | 30328888 | 30329041 | 30328908 | 30329041 | 30326478 | 30326887 | 16 | 10 | 10 | [illegible] |
| A3SS | 2267 | ENSG00000114062 | UBE3A | chr15 | - | 25407066 | 25407262 | 25407066 | 25407212 | 25408619 | 25408684 | 26 | 9 | 21 | [illegible] |
| A3SS | 2930 | ENSG00000077713 | SLC25A43 | chrX | + | 119451849 | 119452143 | 119452008 | 119452143 | 119426181 | 119426354 | 7 | 6 | 2 | [illegible] |
| A3SS | 6128 | ENSG00000158773 | USF1 | chr1 | - | 161040201 | 161040330 | 161040201 | 161040323 | 161040575 | 161040670 | 115 | 77 | 88 | [illegible] |
| A3SS | 552 | ENSG00000188493 | C19orf54 | chr19 | - | 40743904 | 40743981 | 40743904 | 40743975 | 40744558 | 40744683 | 7 | 0 | 1 | [illegible] |
| A3SS | 2124 | ENSG00000168096 | ANKS3 | chr16 | - | 4726658 | 4726854 | 4726658 | 4726776 | 4726978 | 4727177 | 1 | 1 | 7 | [illegible] |
| A3SS | 10167 | ENSG00000188878 | FBF1 | chr17 | - | 75923185 | 75923641 | 75923185 | 75923638 | 75925346 | 75925446 | 9 | 4 | 6 | [illegible] |
| A3SS | 12172 | ENSG00000149187 | CELF1 | chr11 | - | 47476845 | 47476959 | 47476845 | 47476956 | 47478876 | 47478940 | 1 | 1 | 2 | [illegible] |
| A3SS | 13060 | ENSG00000111206 | FOXM1 | chr12 | - | 2872095 | 2872247 | 2872095 | 2872244 | 2873976 | 2874525 | 2 | 0 | 6 | [illegible] |
| A3SS | 4183 | ENSG00000242086 | MUC20-OT1 | chr3 | + | 195717913 | 195718196 | 195718023 | 195718196 | 195708746 | 195708841 | 3 | 0 | 5 | [illegible] |
| A3SS | 7457 | ENSG00000183963 | SMTN | chr22 | + | 31082909 | 31083002 | 31082915 | 31083002 | 31081317 | 31081456 | 0 | 0 | 0 | [illegible] |
| A3SS | 2873 | ENSG00000078319 | PMS2P1 | chr7 | - | 100330908 | 100331043 | 100330908 | 100330961 | 100332441 | 100332581 | 0 | 0 | 0 | [illegible] |
| A3SS | 9300 | ENSG00000269313 | MAGIX | chrX | + | 49166073 | 49166632 | 49166088 | 49166632 | 49165189 | 49165361 | 0 | 0 | 0 | [illegible] |
| A3SS | 3037 | ENSG00000138778 | CENPE | chr4 | - | 103138350 | 103138452 | 103138350 | 103138449 | 103140255 | 103140414 | 0 | 4 | 0 | [illegible] |
| A3SS | 9044 | ENSG00000112787 | FBRSL1 | chr12 | + | 132508152 | 132508350 | 132508155 | 132508350 | 132507251 | 132507388 | 4 | 3 | 4 | [illegible] |
| A3SS | 9600 | ENSG00000168014 | C2CD3 | chr11 | - | 74037477 | 74037652 | 74037477 | 74037623 | 74042053 | 74042218 | 9 | 2 | 2 | [illegible] |
| A3SS | 3115 | ENSG00000155324 | GRAMD2B | chr5 | + | 126484398 | 126484524 | 126484401 | 126484524 | 126483462 | 126483574 | 0 | 1 | 0 | [illegible] |
| A3SS | 3993 | ENSG00000020577 | SAMD4A | chr14 | + | 54784339 | 54784620 | 54784536 | 54784620 | 54776413 | 54776540 | 0 | 1 | 2 | [illegible] |
| A3SS | 12420 | ENSG00000011478 | QPCTL | chr19 | + | 45695400 | 45695718 | 45695632 | 45695718 | 45693412 | 45693556 | 9 | 2 | 3 | [illegible] |
| A3SS | 13012 | ENSG00000135297 | MTO1 | chr6 | + | 73482028 | 73482244 | 73482039 | 73482244 | 73481000 | 73481116 | 4 | 3 | 4 | [illegible] |
| A3SS | 11834 | ENSG00000135424 | ITGA7 | chr12 | - | 55696898 | 55697068 | 55696898 | 55697050 | 55697215 | 55697277 | 2 | 0 | 2 | [illegible] |
| A3SS | 3510 | ENSG00000136235 | GPNMB | chr7 | + | 23266480 | 23266615 | 23266516 | 23266615 | 23260455 | 23260773 | 0 | 0 | 0 | [illegible] |
| A3SS | 13110 | ENSG00000130396 | AFDN | chr6 | + | 167872213 | 167872377 | 167872216 | 167872377 | 167870382 | 167870498 | 13 | 8 | 7 | [illegible] |
| A3SS | 8386 | ENSG00000101104 | PABPC1L | chr20 | + | 44938044 | 44938191 | 44938060 | 44938191 | 44936636 | 44936730 | 1 | 0 | 7 | [illegible] |
| A3SS | 6478 | ENSG00000143801 | PSEN2 | chr1 | + | 226898354 | 226899555 | 226899451 | 226899555 | 226897719 | 226897829 | 3 | 5 | 2 | [illegible] |
| A3SS | 7920 | ENSG00000148737 | TCF7L2 | chr10 | + | 113089396 | 113089540 | 113089399 | 113089540 | 113040024 | 113040126 | 0 | 0 | 0 | [illegible] |

| | | | | | | | | | | | | | | | | | | | | | | | | | | | | | | | | | | | | | | | | | | | | | | |
|---|---|---|---|---|---|---|---|---|---|---|---|---|---|---|---|---|---|---|---|---|---|---|---|---|---|---|---|---|---|---|---|---|---|---|---|---|---|---|---|---|---|---|---|---|---|---|
| A3SS | 8184 | ENSG00000084623 | EIF3I | chr1 | + | 32226104 | 32226320 | 32226148 | 32226320 | 32224409 | 32224475 | 2 | 4 | 4 | 2 | 1 | 1 | 0 | 1 | 0 | 0 | 11 | 0 | 0 | 6 | 5 | 0 | 0 | 0 | 0 | 0 | 193 | 149 | 1.000000 | 1 | 0.607 | 1 | 0.755 | 1 | 1 | 1 | NA | NA | 1 | 1 | -0.128 |
| A3SS | 13205 | ENSG00000050426 | LETMD1 | chr12 | + | 51056143 | 51056245 | 51056149 | 51056245 | 51053777 | 51053860 | 3 | 0 | 3 | 2 | 5 | 0 | 0 | 0 | 1 | 0 | 8 | 8 | 1 | 3 | 2 | 0 | 0 | 0 | 0 | 0 | 155 | 149 | 1.000000 | 1 | 1 | NA | 1 | 0.658 | 1 | 1 | 1 | 1 | 1 | 1 | -0.086 |
| A3SS | 9553 | ENSG00000164970 | FAM219A | chr9 | - | 34398183 | 34401122 | 34398183 | 34401119 | 34401665 | 34401720 | 10 | 9 | 15 | 15 | 16 | 11 | 2 | 3 | 4 | 0 | 0 | 7 | 20 | 17 | 30 | 0 | 3 | 6 | 0 | 0 | 152 | 149 | 1.000000 | 1 | 0.471 | 0.815 | 0.831 | 0.786 | 1 | NA | 0.696 | 0.766 | 1 | 1 | -0.085 |
| A3SS | 10219 | ENSG00000234129 | NA | chrX | - | 11055935 | 11056041 | 11055935 | 11056036 | 11106900 | 11106990 | 1 | 2 | 1 | 2 | 0 | 0 | 0 | 0 | 1 | 0 | 3 | 5 | 1 | 0 | 6 | 0 | 0 | 0 | 0 | 0 | 154 | 149 | 1.000000 | 1 | 1 | 1 | 1 | 0.659 | NA | 1 | 1 | 1 | NA | 1 | -0.085 |
| A3SS | 9662 | ENSG00000183735 | TBK1 | chr12 | + | 64496201 | 64496406 | 64496372 | 64496406 | 64495698 | 64495775 | 9 | 7 | 7 | 8 | 12 | 0 | 0 | 2 | 0 | 0 | 0 | 10 | 34 | 8 | 0 | 0 | 0 | 0 | 0 | 0 | 298 | 149 | 1.000000 | 1 | 1 | 1 | 0.636 | 1 | 1 | NA | 1 | 1 | 1 | NA | -0.073 |
| A3SS | 3530 | ENSG00000168487 | BMP1 | chr8 | + | 22176923 | 22177139 | 22176960 | 22177139 | 22176532 | 22176650 | 2 | 3 | 3 | 5 | 4 | 4 | 2 | 0 | 6 | 0 | 10 | 3 | 11 | 2 | 0 | 0 | 10 | 6 | 0 | 0 | 186 | 149 | 1.000000 | 1 | 0.286 | 0.546 | 1 | 0.4 | 1 | 1 | 0.194 | 0.595 | 1 | NA | -0.051 |
| A3SS | 9866 | ENSG00000121716 | PILRB | chr7 | + | 100353372 | 100354012 | 100353894 | 100354012 | 100353209 | 100353270 | 78 | 31 | 58 | 70 | 56 | 2 | 2 | 0 | 1 | 0 | 25 | 85 | 44 | 45 | 103 | 0 | 0 | 0 | 0 | 0 | 298 | 149 | 1.000000 | 1 | 0.951 | 0.886 | 1 | 0.972 | 1 | 1 | 1 | 1 | 1 | 1 | -0.038 |
| A3SS | 8842 | ENSG00000169679 | BUB1 | chr2 | - | 110641641 | 110641844 | 110641641 | 110641803 | 110642118 | 110642234 | 0 | 0 | 1 | 1 | 0 | 14 | 10 | 12 | 25 | 6 | 0 | 0 | 11 | 0 | 0 | 0 | 7 | 32 | 7 | 2 | 190 | 149 | 1.000000 | 1 | 0 | 0 | 0.061 | 0.03 | 0 | NA | 0 | 0.212 | 0 | 0 | -0.035 |
| A3SS | 11082 | ENSG00000104763 | ASAH1 | chr8 | - | 18061376 | 18062423 | 18061376 | 18061458 | 18067219 | 18067298 | 20 | 25 | 17 | 12 | 23 | 2 | 0 | 0 | 0 | 0 | 9 | 38 | 47 | 12 | 0 | 0 | 0 | 0 | 0 | 0 | 298 | 149 | 1.000000 | 1 | 0.833 | 1 | 1 | 1 | 1 | 1 | 1 | 1 | 1 | NA | -0.033 |
| A3SS | 6651 | ENSG00000124574 | ABCC10 | chr6 | + | 43442969 | 43444010 | 43443932 | 43444010 | 43441861 | 43441960 | 78 | 43 | 74 | 55 | 30 | 0 | 3 | 0 | 0 | 0 | 65 | 42 | 35 | 62 | 85 | 0 | 0 | 0 | 0 | 0 | 298 | 149 | 1.000000 | 1 | 1 | 0.878 | 1 | 1 | 1 | 1 | 1 | 1 | 1 | 1 | -0.024 |
| A3SS | 6690 | ENSG00000145020 | AMT | chr3 | - | 49416821 | 49417367 | 49416821 | 49417309 | 49417614 | 49417718 | 19 | 2 | 13 | 9 | 6 | 1 | 0 | 0 | 0 | 0 | 10 | 3 | 17 | 3 | 25 | 0 | 0 | 0 | 0 | 0 | 207 | 149 | 1.000000 | 1 | 0.932 | 1 | 1 | 1 | 1 | 1 | 1 | 1 | 1 | 1 | -0.014 |
| A3SS | 10284 | ENSG00000182220 | ATP6AP2 | chrX | + | 40599606 | 40599741 | 40599648 | 40599741 | 40598680 | 40598734 | 151 | 76 | 86 | 93 | 114 | 0 | 0 | 0 | 0 | 4 | 117 | 112 | 95 | 125 | 80 | 0 | 0 | 0 | 0 | 0 | 191 | 149 | 1.000000 | 1 | 1 | 1 | 1 | 1 | 0.957 | 1 | 1 | 1 | 1 | 1 | -0.009 |
| A3SS | 6164 | ENSG00000084754 | HADHA | chr2 | - | 26214442 | 26214576 | 26214442 | 26214561 | 26215052 | 26215175 | 1 | 0 | 0 | 1 | 0 | 355 | 233 | 330 | 307 | 359 | 0 | 3 | 0 | 8 | 0 | 696 | 383 | 183 | 316 | 655 | 164 | 149 | 1.000000 | 1 | 0.003 | 0 | 0 | 0.003 | 0 | 0 | 0.007 | 0 | 0.022 | 0 | -0.005 |
| A3SS | 12542 | ENSG00000175029 | CTBP2 | chr10 | - | 124994468 | 124994683 | 124994468 | 124994606 | 125038996 | 125039155 | 48 | 21 | 32 | 54 | 21 | 0 | 0 | 0 | 1 | 0 | 40 | 38 | 20 | 41 | 46 | 0 | 0 | 0 | 0 | 0 | 226 | 149 | 1.000000 | 1 | 1 | 1 | 1 | 0.973 | 1 | 1 | 1 | 1 | 1 | 1 | -0.005 |
| A3SS | 36 | ENSG00000241837 | ATP5PO | chr21 | - | 33903455 | 33903656 | 33903455 | 33903639 | 33903934 | 33904021 | 3 | 2 | 4 | 13 | 0 | 994 | 536 | 966 | 800 | 635 | 7 | 0 | 8 | 6 | 10 | 1203 | 855 | 702 | 800 | 1101 | 166 | 149 | 1.000000 | 1 | 0.003 | 0.003 | 0.004 | 0.014 | 0 | 0.005 | 0 | 0.01 | 0.007 | 0.008 | -0.001 |
| A3SS | 9483 | ENSG00000109339 | MAPK10 | chr4 | - | 86015268 | 86017375 | 86015268 | 86017370 | 86029196 | 86029274 | 0 | 0 | 0 | 1 | 0 | 1 | 0 | 0 | 0 | 1 | 10 | 0 | 0 | 0 | 0 | 0 | 5 | 0 | 0 | 2 | 154 | 149 | 1.000000 | 1 | 0 | NA | NA | 1 | 0 | 1 | 0 | NA | NA | 0 | 0 |
| A3SS | 2519 | ENSG00000100429 | HDAC10 | chr22 | - | 50248224 | 50248472 | 50248224 | 50248292 | 50248661 | 50248751 | 13 | 0 | 17 | 20 | 7 | 0 | 0 | 0 | 0 | 0 | 30 | 11 | 1 | 17 | 7 | 0 | 0 | 0 | 1 | 0 | 298 | 149 | 1.000000 | 1 | 1 | NA | 1 | 1 | 1 | 1 | 1 | 1 | 0.895 | 1 | 0.021 |
| A3SS | 7931 | ENSG00000136715 | SAP130 | chr2 | - | 127986784 | 127986962 | 127986784 | 127986959 | 127989563 | 127989866 | 33 | 17 | 26 | 31 | 25 | 0 | 2 | 1 | 1 | 0 | 15 | 51 | 26 | 45 | 4 | 0 | 10 | 3 | 1 | 0 | 152 | 149 | 1.000000 | 1 | 1 | 0.893 | 0.962 | 0.968 | 1 | 1 | 0.833 | 0.895 | 0.978 | 1 | 0.023 |
| A3SS | 4235 | ENSG00000213585 | VDAC1 | chr5 | - | 133992945 | 133993018 | 133992945 | 133993015 | 134004894 | 134004975 | 3 | 4 | 7 | 5 | 14 | 1 | 0 | 0 | 2 | 0 | 21 | 5 | 1 | 15 | 0 | 0 | 2 | 1 | 1 | 0 | 152 | 149 | 1.000000 | 1 | 0.746 | 1 | 1 | 0.71 | 1 | 1 | 0.71 | 0.495 | 0.936 | NA | 0.106 |
| A3SS | 8407 | ENSG00000065802 | ASB1 | chr2 | + | 238433548 | 238433695 | 238433553 | 238433695 | 238433312 | 238433447 | 2 | 0 | 0 | 0 | 0 | 0 | 0 | 0 | 0 | 0 | 0 | 4 | 2 | 5 | 0 | 0 | 2 | 0 | 0 | 0 | 154 | 149 | 1.000000 | 1 | 1 | NA | NA | NA | NA | NA | 0.659 | 1 | 1 | NA | 0.114 |

[illegible]

| | | | | | | | | | | | | | | | |
|---|---|---|---|---|---|---|---|---|---|---|---|---|---|---|---|
| A3SS | 6831 | ENSG00000073578 | SDHA | chr5 | + | 254357 | 254506 | 254392 | 254506 | 240357 | 240476 | 2 | 2 | 1 | [illegible] |
| A3SS | 10988 | ENSG00000078487 | ZCWPW1 | chr7 | - | 100415974 | 100416097 | 100415974 | 100416094 | 100416304 | 100416456 | 5 | 12 | 10 | [illegible] |
| A3SS | 1279 | ENSG00000185359 | HGS | chr17 | + | 81692384 | 81692553 | 81692422 | 81692553 | 81691446 | 81691571 | 0 | 0 | 1 | [illegible] |
| A3SS | 6416 | ENSG00000132781 | MUTYH | chr1 | - | 45333412 | 45333603 | 45333412 | 45333564 | 45334390 | 45334511 | 6 | 4 | 8 | [illegible] |
| A3SS | 5442 | ENSG00000121940 | CLCC1 | chr1 | - | 108962308 | 108962469 | 108962308 | 108962464 | 108963342 | 108963484 | 2 | 0 | 0 | [illegible] |
| A3SS | 7416 | ENSG00000169682 | SPNS1 | chr16 | + | 28975183 | 28975392 | 28975187 | 28975392 | 28974771 | 28974863 | 0 | 0 | 1 | [illegible] |
| A3SS | 7068 | ENSG00000104970 | KIR3DX1 | chr19 | + | 54536883 | 54537162 | 54536901 | 54537162 | 54533731 | 54534010 | 3 | 0 | 0 | [illegible] |
| A3SS | 9095 | ENSG00000055957 | ITIH1 | chr3 | + | 52787985 | 52788066 | 52788028 | 52788066 | 52787591 | 52787612 | 0 | 0 | 1 | [illegible] |
| A3SS | 7960 | ENSG00000246985 | SOCS2-AS1 | chr12 | - | 93545070 | 93545398 | 93545070 | 93545358 | 93565612 | 93565744 | 0 | 3 | 0 | [illegible] |
| A3SS | 1970 | ENSG00000099377 | HSD3B7 | chr16 | + | 30985459 | 30985824 | 30985652 | 30985824 | 30985206 | 30985297 | 0 | 0 | 0 | [illegible] |
| A3SS | 10838 | ENSG00000197467 | COL13A1 | chr10 | + | 69880413 | 69880553 | 69880502 | 69880553 | 69878038 | 69878065 | 0 | 0 | 0 | [illegible] |
| A3SS | 11932 | ENSG00000126705 | AHDC1 | chr1 | - | 27603396 | 27603494 | 27603396 | 27603491 | 27604159 | 27604431 | 0 | 0 | 0 | [illegible] |
| A3SS | 444 | ENSG00000196724 | ZNF418 | chr19 | - | 57930427 | 57930557 | 57930427 | 57930554 | 57933816 | 57933902 | 0 | 0 | 3 | [illegible] |
| A3SS | 11455 | ENSG00000132740 | IGHMBP2 | chr11 | + | 68933264 | 68933481 | 68933298 | 68933481 | 68930322 | 68930434 | 0 | 0 | 0 | [illegible] |
| A3SS | 1170 | ENSG00000120093 | HOXB3 | chr17 | - | 48552026 | 48552487 | 48552026 | 48552360 | 48555530 | 48555618 | 0 | 1 | 1 | [illegible] |
| A3SS | 3986 | ENSG00000178104 | PDE4DIP | chr1 | + | 148866410 | 148866532 | 148866413 | 148866532 | 148863249 | 148863305 | 0 | 0 | 0 | [illegible] |
| A3SS | 7303 | ENSG00000158571 | PFKFB1 | chrX | - | 54960823 | 54960917 | 54960823 | 54960851 | 54963256 | 54963382 | 2 | 0 | 0 | [illegible] |
| A3SS | 8335 | ENSG00000235703 | LINC00894 | chrX | + | 149945783 | 149947347 | 149945857 | 149947347 | 149944017 | 149944100 | 0 | 0 | 0 | [illegible] |
| A3SS | 9532 | ENSG00000164744 | SUN3 | chr7 | - | 47987150 | 47987449 | 47987150 | 47987360 | 47988787 | 47988880 | 0 | 0 | 0 | [illegible] |
| A3SS | 13307 | ENSG00000111452 | ADGRD1 | chr12 | + | 131137643 | 131138229 | 131138136 | 131138229 | 131136972 | 131137014 | 0 | 0 | 2 | [illegible] |
| A3SS | 138 | ENSG00000100124 | ANKRD54 | chr22 | - | 37833683 | 37833755 | 37833683 | 37833750 | 37833916 | 37834728 | 1 | 0 | 0 | [illegible] |
| A3SS | 1767 | ENSG00000140995 | DEF8 | chr16 | + | 89949416 | 89949513 | 89949419 | 89949513 | 89948754 | 89948814 | 2 | 0 | 0 | [illegible] |
| A3SS | 11335 | ENSG00000157657 | ZNF618 | chr9 | + | 114032644 | 114032728 | 114032647 | 114032728 | 114028732 | 114028972 | 0 | 0 | 0 | [illegible] |
| A3SS | 8988 | ENSG00000120896 | SORBS3 | chr8 | + | 22554862 | 22554980 | 22554907 | 22554980 | 22554451 | 22554608 | 4 | 0 | 0 | [illegible] |
| A3SS | 7551 | ENSG00000078808 | SDF4 | chr1 | - | 1218768 | 1219005 | 1218768 | 1218943 | 1223243 | 1223357 | 11 | 0 | 9 | [illegible] |
| A3SS | 8873 | ENSG00000143436 | MRPL9 | chr1 | - | 151761961 | 151762025 | 151761961 | 151762010 | 151762104 | 151762155 | 0 | 2 | 1 | [illegible] |
| A3SS | 5640 | ENSG00000026508 | CD44 | chr11 | + | 35201667 | 35201787 | 35201670 | 35201787 | 35201081 | 35201195 | 0 | 0 | 4 | [illegible] |
| A3SS | 4873 | ENSG00000065328 | MCM10 | chr10 | + | 13172624 | 13172765 | 13172627 | 13172765 | 13172375 | 13172480 | 0 | 0 | 0 | [illegible] |
| A3SS | 8662 | ENSG00000115486 | GGCX | chr2 | - | 85558439 | 85558662 | 85558439 | 85558605 | 85558916 | 85559075 | 0 | 0 | 0 | [illegible] |
| A3SS | 7359 | ENSG00000075702 | WDR62 | chr19 | + | 36102446 | 36102851 | 36102736 | 36102851 | 36102013 | 36102151 | 0 | 0 | 0 | [illegible] |
| A3SS | 6395 | ENSG00000143353 | LYPLAL1 | chr1 | + | 219210514 | 219210647 | 219210531 | 219210647 | 219179399 | 219179426 | 2 | 2 | 0 | [illegible] |
| A3SS | 9896 | ENSG00000130429 | ARPC1B | chr7 | + | 99377337 | 99377439 | 99377357 | 99377439 | 99374729 | 99374781 | 8 | 1 | 0 | [illegible] |
| A3SS | 7094 | ENSG00000075151 | EIF4G3 | chr1 | - | 21001198 | 21001312 | 21001198 | 21001307 | 21002712 | 21002808 | 5 | 1 | 1 | [illegible] |
| A3SS | 1669 | ENSG00000102878 | HSF4 | chr16 | + | 67169630 | 67169941 | 67169635 | 67169941 | 67169278 | 67169348 | 5 | 0 | 4 | [illegible] |
| A3SS | 1537 | ENSG00000126246 | IGFLR1 | chr19 | - | 35739709 | 35740088 | 35739709 | 35740086 | 35740379 | 35740564 | 13 | 8 | 11 | [illegible] |
| A3SS | 3860 | ENSG00000215424 | MCM3AP-AS1 | chr21 | + | 46240833 | 46242999 | 46240836 | 46242999 | 46236223 | 46236326 | 9 | 1 | 5 | [illegible] |
| A3SS | 12123 | ENSG00000070047 | PHRF1 | chr11 | + | 581961 | 582081 | 581964 | 582081 | 581491 | 581597 | 14 | 7 | 6 | [illegible] |
| A3SS | 9058 | ENSG00000116539 | ASH1L | chr1 | - | 155347655 | 155347904 | 155347655 | 155347901 | 155352705 | 155352858 | 1 | 1 | 2 | [illegible] |
| A3SS | 8822 | ENSG00000118495 | PLAGL1 | chr6 | - | 143960468 | 143960580 | 143960468 | 143960556 | 143964786 | 143964818 | 0 | 0 | 0 | [illegible] |
| A3SS | 10192 | ENSG00000087237 | CETP | chr16 | + | 56971216 | 56971381 | 56971320 | 56971381 | 56971032 | 56971102 | 0 | 0 | 0 | [illegible] |
| A3SS | 3171 | ENSG00000106123 | EPHB6 | chr7 | + | 142865959 | 142866605 | 142866480 | 142866605 | 142865474 | 142865630 | 6 | 1 | 3 | [illegible] |
| A3SS | 9668 | ENSG00000158636 | EMSY | chr11 | + | 76542167 | 76542367 | 76542215 | 76542367 | 76528267 | 76528466 | 2 | 1 | 5 | [illegible] |
| A3SS | 992 | ENSG00000124155 | PIGT | chr20 | + | 45429444 | 45429670 | 45429480 | 45429670 | 45427421 | 45427545 | 2 | 1 | 0 | [illegible] |
| A3SS | 5352 | ENSG00000160752 | FDPS | chr1 | + | 155310042 | 155310205 | 155310063 | 155310205 | 155309788 | 155309965 | 17 | 0 | 12 | [illegible] |
| A3SS | 6447 | ENSG00000081014 | AP4E1 | chr15 | + | 50915424 | 50915571 | 50915447 | 50915571 | 50912077 | 50912149 | 1 | 1 | 4 | [illegible] |
| A3SS | 77 | ENSG00000248751 | NA | chr22 | - | 30286015 | 30286142 | 30286015 | 30286098 | 30286262 | 30286376 | 0 | 0 | 3 | [illegible] |
| A3SS | 13175 | ENSG00000122966 | CIT | chr12 | - | 119734163 | 119734406 | 119734163 | 119734357 | 119735159 | 119735357 | 0 | 0 | 0 | [illegible] |
| A3SS | 3892 | ENSG00000157800 | SLC37A3 | chr7 | - | 140352061 | 140352181 | 140352061 | 140352146 | 140355667 | 140355764 | 1 | 2 | 0 | [illegible] |
| A3SS | 7582 | ENSG00000171503 | ETFDH | chr4 | + | 158703689 | 158706371 | 158706188 | 158706371 | 158703422 | 158703591 | 2 | 8 | 5 | [illegible] |
| A3SS | 8018 | ENSG00000164919 | COX6C | chr8 | - | 99891907 | 99892052 | 99891907 | 99892047 | 99893442 | 99893622 | 10 | 9 | 23 | [illegible] |
| A3SS | 5113 | ENSG00000073792 | IGF2BP2 | chr3 | - | 185692698 | 185692780 | 185692698 | 185692762 | 185696611 | 185696663 | 0 | 0 | 0 | [illegible] |
| A3SS | 10929 | ENSG00000094914 | AAAS | chr12 | - | 53308724 | 53308851 | 53308724 | 53308815 | 53308959 | 53309020 | 0 | 0 | 0 | [illegible] |
| A3SS | 11047 | ENSG00000070081 | NUCB2 | chr11 | + | 17311184 | 17311283 | 17311192 | 17311283 | 17310824 | 17311010 | 0 | 0 | 1 | [illegible] |
| A3SS | 12646 | ENSG00000107862 | GBF1 | chr10 | + | 102366356 | 102366506 | 102366382 | 102366506 | 102365396 | 102365599 | 1 | 0 | 0 | [illegible] |
| A3SS | 955 | ENSG00000141644 | MBD1 | chr18 | - | 50268846 | 50270085 | 50268846 | 50269818 | 50272823 | 50272955 | 94 | 66 | 73 | [illegible] |
| A3SS | 1113 | ENSG00000004975 | DVL2 | chr17 | - | 7230045 | 7230155 | 7230045 | 7230137 | 7230284 | 7230430 | 43 | 21 | 40 | [illegible] |
| A3SS | 5125 | ENSG00000144591 | GMPPA | chr2 | + | 219505694 | 219505761 | 219505714 | 219505761 | 219505457 | 219505555 | 2 | 0 | 1 | [illegible] |
| A3SS | 3186 | ENSG00000197375 | SLC22A5 | chr5 | + | 132390689 | 132390904 | 132390775 | 132390904 | 132388920 | 132389021 | 13 | 10 | 16 | [illegible] |
| A3SS | 11395 | ENSG00000116580 | GON4L | chr1 | - | 155751959 | 155752590 | 155751959 | 155752587 | 155754374 | 155754488 | 3 | 0 | 0 | [illegible] |
| A3SS | 7519 | ENSG00000160818 | GPATCH4 | chr1 | - | 156598967 | 156599104 | 156598967 | 156599030 | 156601408 | 156601470 | 17 | 9 | 20 | [illegible] |
| A3SS | 8528 | ENSG00000115163 | CENPA | chr2 | + | 26792501 | 26792833 | 26792755 | 26792833 | 26792130 | 26792240 | 0 | 0 | 0 | [illegible] |
| A3SS | 9526 | ENSG00000281103 | TRG-AS1 | chr7 | + | 38345972 | 38346127 | 38346020 | 38346127 | 38341614 | 38341799 | 5 | 1 | 0 | [illegible] |
| A3SS | 10363 | ENSG00000142528 | ZNF473 | chr19 | + | 50030891 | 50031091 | 50031013 | 50031091 | 50025892 | 50026122 | 9 | 0 | 4 | [illegible] |
| A3SS | 7254 | ENSG00000117262 | GPR89A | chr1 | + | 145646183 | 145646272 | 145646203 | 145646272 | 145645646 | 145645743 | 1 | 2 | 1 | [illegible] |
| A3SS | 8530 | ENSG00000160766 | GBAP1 | chr1 | - | 155215053 | 155215400 | 155215053 | 155215162 | 155215562 | 155215651 | 4 | 0 | 1 | [illegible] |
| A3SS | 961 | ENSG00000233527 | ZNF529-AS1 | chr19 | + | 36594400 | 36594708 | 36594441 | 36594708 | 36594155 | 36594299 | 0 | 2 | 0 | [illegible] |
| A3SS | 2292 | ENSG00000102901 | CENPT | chr16 | - | 67835171 | 67835300 | 67835171 | 67835295 | 67835537 | 67835658 | 1 | 5 | 1 | [illegible] |
| A3SS | 1790 | ENSG00000188603 | CLN3 | chr16 | - | 28479643 | 28479765 | 28479643 | 28479741 | 28482104 | 28482198 | 4 | 2 | 1 | [illegible] |
| A3SS | 12071 | ENSG00000174791 | RIN1 | chr11 | - | 66333918 | 66334224 | 66333918 | 66334038 | 66334513 | 66335251 | 2 | 0 | 4 | [illegible] |
| A3SS | 6279 | ENSG00000025708 | TYMP | chr22 | - | 50529495 | 50529754 | 50529495 | 50529747 | 50529903 | 50530012 | 3 | 3 | 1 | [illegible] |
| A3SS | 443 | ENSG00000196724 | ZNF418 | chr19 | - | 57930427 | 57930557 | 57930427 | 57930554 | 57932429 | 57932646 | 0 | 1 | 0 | [illegible] |
| A3SS | 9221 | ENSG00000112139 | MDGA1 | chr6 | - | 37647172 | 37647324 | 37647172 | 37647272 | 37648981 | 37649266 | 0 | 0 | 2 | [illegible] |
| A3SS | 5473 | ENSG00000111325 | OGFOD2 | chr12 | + | 122976600 | 122976767 | 122976653 | 122976767 | 122975810 | 122975867 | 2 | 1 | 3 | [illegible] |
| A3SS | 6023 | ENSG00000188004 | SNHG28 | chr1 | - | 159837916 | 159840881 | 159837916 | 159840866 | 159841125 | 159841235 | 0 | 0 | 0 | [illegible] |
| A3SS | 10802 | ENSG00000132793 | LPIN3 | chr20 | + | 41349088 | 41349172 | 41349091 | 41349172 | 41348618 | 41348887 | 0 | 0 | 0 | [illegible] |
| A3SS | 2860 | ENSG00000139974 | SLC38A6 | chr14 | + | 61078809 | 61078927 | 61078829 | 61078927 | 61052348 | 61052985 | 0 | 0 | 0 | [illegible] |
| A3SS | 12798 | ENSG00000079974 | RABL2B | chr22 | - | 50782187 | 50782351 | 50782187 | 50782347 | 50783500 | 50783662 | 0 | 0 | 0 | [illegible] |
| A3SS | 2000 | ENSG00000103121 | CMC2 | chr16 | - | 80998284 | 80998354 | 80998284 | 80998351 | 81001247 | 81001334 | 0 | 0 | 0 | [illegible] |
| A3SS | 6633 | ENSG00000237877 | LINC01473 | chr2 | - | 186083153 | 186083224 | 186083153 | 186083221 | 186215018 | 186215090 | 1 | 0 | 2 | [illegible] |
| A3SS | 6741 | ENSG00000231890 | DARS1-AS1 | chr2 | + | 136003244 | 136003367 | 136003269 | 136003367 | 136000412 | 136000479 | 0 | 0 | 0 | [illegible] |
| A3SS | 4408 | ENSG00000009830 | POMT2 | chr14 | - | 77278728 | 77278961 | 77278728 | 77278869 | 77279822 | 77279928 | 0 | 0 | 0 | [illegible] |
| A3SS | 1814 | ENSG00000226686 | LINC01535 | chr19 | + | 37257011 | 37257394 | 37257230 | 37257394 | 37256726 | 37256814 | 0 | 0 | 0 | [illegible] |
| A3SS | 7506 | ENSG00000147099 | HDAC8 | chrX | - | 72462003 | 72462102 | 72462003 | 72462098 | 72462361 | 72462442 | 0 | 0 | 0 | [illegible] |
| A3SS | 9487 | ENSG00000109339 | MAPK10 | chr4 | - | 86098523 | 86099285 | 86098523 | 86098595 | 86101051 | 86101217 | 0 | 0 | 0 | [illegible] |
| A3SS | 9154 | ENSG00000244355 | LY6G6D | chr6 | + | 31717459 | 31717804 | 31717580 | 31717804 | 31715501 | 31715624 | 0 | 0 | 0 | [illegible] |
| A3SS | 1668 | ENSG00000102878 | HSF4 | chr16 | + | 67167719 | 67167947 | 67167823 | 67167947 | 67167474 | 67167599 | 1 | 0 | 2 | [illegible] |
| A3SS | 2510 | ENSG00000197555 | SIPA1L1 | chr14 | + | 71738240 | 71738325 | 71738243 | 71738325 | 71733665 | 71733812 | 0 | 1 | 0 | [illegible] |
| A3SS | 11042 | ENSG00000249859 | PVT1 | chr8 | + | 128096517 | 128096654 | 128096520 | 128096654 | 128080302 | 128080430 | 0 | 0 | 0 | [illegible] |
| A3SS | 6577 | ENSG00000048707 | VPS13D | chr1 | + | 12341779 | 12341885 | 12341782 | 12341885 | 12333225 | 12333366 | 1 | 0 | 0 | [illegible] |
| A3SS | 2863 | ENSG00000204536 | CCHCR1 | chr6 | - | 31157022 | 31157155 | 31157022 | 31157089 | 31157384 | 31157500 | 0 | 0 | 0 | [illegible] |
| A3SS | 11934 | ENSG00000239523 | MYLK-AS1 | chr3 | + | 123629490 | 123629681 | 123629493 | 123629681 | 123585646 | 123585794 | 0 | 0 | 0 | [illegible] |
| A3SS | 8379 | ENSG00000244026 | FAM86DP | chr3 | - | 75426449 | 75426621 | 75426449 | 75426599 | 75430831 | 75430914 | 1 | 0 | 2 | [illegible] |
| A3SS | 12558 | ENSG00000198825 | INPP5F | chr10 | + | 119757891 | 119757977 | 119757894 | 119757977 | 119751075 | 119751156 | 0 | 0 | 0 | [illegible] |
| A3SS | 9792 | ENSG00000012124 | CD22 | chr19 | + | 35331905 | 35332074 | 35331926 | 35332074 | 35329186 | 35329230 | 2 | 3 | 3 | [illegible] |
| A3SS | 9010 | ENSG00000111011 | RSRC2 | chr12 | - | 122518838 | 122519051 | 122518838 | 122519029 | 122521384 | 122521428 | 36 | 41 | 42 | [illegible] |
| A3SS | 10564 | ENSG00000184634 | MED12 | chrX | + | 71137557 | 71137635 | 71137560 | 71137635 | 71137186 | 71137383 | 31 | 4 | 17 | [illegible] |
| A3SS | 4631 | ENSG00000262481 | TMEM256-PLSCR3 | chr17 | - | 7394103 | 7394271 | 7394103 | 7394267 | 7394907 | 7395090 | 0 | 1 | 1 | [illegible] |
| A3SS | 12776 | ENSG00000226711 | FAM66C | chr12 | + | 8194712 | 8194796 | 8194728 | 8194796 | 8194202 | 8194481 | 0 | 0 | 0 | [illegible] |
| A3SS | 12552 | ENSG00000235823 | OLMALINC | chr10 | + | 100388033 | 100388360 | 100388036 | 100388360 | 100381215 | 100381448 | 0 | 0 | 0 | [illegible] |
| A3SS | 11130 | ENSG00000158473 | CD1D | chr1 | + | 158181454 | 158181721 | 158181489 | 158181721 | 158180894 | 158181050 | 17 | 0 | 14 | [illegible] |
| A3SS | 4550 | ENSG00000113645 | WWC1 | chr5 | + | 168464710 | 168464962 | 168464728 | 168464962 | 168460649 | 168460742 | 0 | 1 | 1 | [illegible] |
| A3SS | 6609 | ENSG00000204054 | LINC00963 | chr9 | + | 129493415 | 129493595 | 129493418 | 129493595 | 129490804 | 129491128 | 0 | 0 | 0 | [illegible] |
| A3SS | 13058 | ENSG00000111206 | FOXM1 | chr12 | - | 2864682 | 2864755 | 2864682 | 2864752 | 2865354 | 2865399 | 0 | 0 | 0 | [illegible] |
| A3SS | 12170 | ENSG00000165915 | SLC39A13 | chr11 | + | 47415038 | 47415159 | 47415059 | 47415159 | 47414776 | 47414909 | 1 | 2 | 2 | [illegible] |
| A3SS | 6768 | ENSG00000158796 | DEDD | chr1 | - | 161130814 | 161130847 | 161130814 | 161130843 | 161131005 | 161131182 | 8 | 2 | 1 | [illegible] |
| A3SS | 12377 | ENSG00000133816 | MICAL2 | chr11 | + | 12275982 | 12276166 | 12275985 | 12276166 | 12259794 | 12259897 | 0 | 2 | 1 | [illegible] |
| A3SS | 10627 | ENSG00000012048 | BRCA1 | chr17 | - | 43076487 | 43076614 | 43076487 | 43076611 | 43082403 | 43082575 | 9 | 11 | 7 | [illegible] |
| A3SS | 5232 | ENSG00000117640 | MTFR1L | chr1 | + | 25823620 | 25823748 | 25823643 | 25823748 | 25823237 | 25823323 | 0 | 3 | 0 | [illegible] |
| A3SS | 1560 | ENSG00000276070 | CCL4L2 | chr17 | + | 36212292 | 36212878 | 36212333 | 36212878 | 36211775 | 36211890 | 11 | 2 | 1 | [illegible] |
| A3SS | 5196 | ENSG00000137411 | VARS2 | chr6 | + | 30922381 | 30922554 | 30922449 | 30922554 | 30922115 | 30922241 | 0 | 0 | 0 | [illegible] |
| A3SS | 7015 | ENSG00000122678 | POLM | chr7 | - | 44074130 | 44074233 | 44074130 | 44074220 | 44074397 | 44074530 | 11 | 2 | 8 | [illegible] |
| A3SS | 828 | ENSG00000125900 | SIRPD | chr20 | - | 1537154 | 1537313 | 1537154 | 1537310 | 1551690 | 1552038 | 0 | 0 | 1 | [illegible] |
| A3SS | 11661 | ENSG00000130714 | POMT1 | chr9 | + | 131521979 | 131522224 | 131522046 | 131522224 | 131521345 | 131521472 | 0 | 0 | 0 | [illegible] |
| A3SS | 2536 | ENSG00000198513 | ATL1 | chr14 | + | 50589183 | 50591075 | 50590940 | 50591075 | 50587830 | 50588078 | 2 | 0 | 0 | [illegible] |
| A3SS | 1263 | ENSG00000213339 | QTRT1 | chr19 | + | 10707465 | 10707615 | 10707499 | 10707615 | 10707301 | 10707380 | 2 | 0 | 0 | [illegible] |
| A3SS | 12586 | ENSG00000165702 | GFI1B | chr9 | + | 132989645 | 132989907 | 132989675 | 132989907 | 132989060 | 132989198 | 0 | 0 | 0 | [illegible] |
| A3SS | 9729 | ENSG00000272333 | KMT2B | chr19 | + | 35723402 | 35723502 | 35723446 | 35723502 | 35722994 | 35723274 | 0 | 0 | 1 | [illegible] |
| A3SS | 12292 | ENSG00000149292 | TTC12 | chr11 | + | 113334947 | 113335037 | 113334965 | 113335037 | 113329919 | 113329979 | 2 | 0 | 0 | [illegible] |
| A3SS | 7829 | ENSG00000131748 | STARD3 | chr17 | + | 39656909 | 39657085 | 39657007 | 39657085 | 39653480 | 39653750 | 1 | 0 | 0 | [illegible] |
| A3SS | 10474 | ENSG00000008294 | SPAG9 | chr17 | - | 50998443 | 50998632 | 50998443 | 50998617 | 50999660 | 50999717 | 0 | 0 | 0 | [illegible] |
| A3SS | 390 | ENSG00000183207 | RUVBL2 | chr19 | + | 49004206 | 49004418 | 49004276 | 49004418 | 49003278 | 49003334 | 1 | 0 | 1 | [illegible] |
| A3SS | 3343 | ENSG00000132780 | NASP | chr1 | + | 45604916 | 45605016 | 45604935 | 45605016 | 45602254 | 45602365 | 0 | 0 | 0 | [illegible] |
| A3SS | 6656 | ENSG00000131828 | PDHA1 | chrX | + | 19351259 | 19351407 | 19351280 | 19351407 | 19349936 | 19350110 | 0 | 1 | 0 | [illegible] |
| A3SS | 4209 | ENSG00000068745 | IP6K2 | chr3 | - | 48695089 | 48695421 | 48695089 | 48695418 | 48711366 | 48711459 | 0 | 0 | 0 | [illegible] |
| A3SS | 9942 | ENSG00000115677 | HDLBP | chr2 | - | 241235489 | 241235629 | 241235489 | 241235594 | 241236614 | 241236769 | 1 | 3 | 3 | [illegible] |
| A3SS | 9728 | ENSG00000272333 | KMT2B | chr19 | + | 35719783 | 35721804 | 35719849 | 35721804 | 35719468 | 35719541 | 23 | 17 | 31 | [illegible] |
| A3SS | 7272 | ENSG00000132906 | CASP9 | chr1 | - | 15506898 | 15507133 | 15506898 | 15507075 | 15507872 | 15507907 | 0 | 1 | 0 | [illegible] |
| A3SS | 4775 | ENSG00000130270 | ATP8B3 | chr19 | - | 1806096 | 1806508 | 1806096 | 1806169 | 1806627 | 1806689 | 2 | 1 | 1 | [illegible] |
| A3SS | 6733 | ENSG00000065526 | SPEN | chr1 | + | 15909320 | 15909481 | 15909323 | 15909481 | 15873501 | 15874395 | 6 | 2 | 3 | [illegible] |
| A3SS | 5660 | ENSG00000177873 | ZNF619 | chr3 | + | 40482199 | 40482349 | 40482229 | 40482349 | 40481862 | 40482016 | 0 | 1 | 1 | [illegible] |
| A3SS | 1778 | ENSG00000007038 | PRSS21 | chr16 | + | 2818676 | 2818969 | 2818682 | 2818969 | 2817800 | 2817966 | 4 | 1 | 0 | [illegible] |
| A3SS | 3574 | ENSG00000139625 | MAP3K12 | chr12 | - | 53486946 | 53487155 | 53486946 | 53487145 | 53487254 | 53487428 | 2 | 0 | 0 | [illegible] |
| A3SS | 3690 | ENSG00000197647 | ZNF433 | chr19 | - | 12018165 | 12018301 | 12018165 | 12018292 | 12030139 | 12030198 | 0 | 0 | 0 | [illegible] |
| A3SS | 9735 | ENSG00000168255 | POLR2J3 | chr7 | - | 102543524 | 102543697 | 102543524 | 102543661 | 102544705 | 102544776 | 0 | 2 | 0 | [illegible] |
| A3SS | 3758 | ENSG00000267364 | NA | chr17 | + | 35321734 | 35321865 | 35321736 | 35321865 | 35316355 | 35316409 | 4 | 1 | 1 | [illegible] |
| A3SS | 1816 | ENSG00000196557 | CACNA1H | chr16 | + | 1211945 | 1212138 | 1212007 | 1212138 | 1211715 | 1211805 | 0 | 0 | 0 | [illegible] |
| A3SS | 75 | ENSG00000133460 | SLC2A11 | chr22 | + | 23883873 | 23884024 | 23883948 | 23884024 | 23882758 | 23882869 | 0 | 0 | 0 | [illegible] |
| A3SS | 9539 | ENSG00000196923 | PDLIM7 | chr5 | - | 177491806 | 177491995 | 177491806 | 177491976 | 177492404 | 177492435 | 0 | 0 | 0 | [illegible] |
| A3SS | 10012 | ENSG00000105976 | MET | chr7 | + | 116759336 | 116759490 | 116759390 | 116759490 | 116758458 | 116758620 | 0 | 0 | 0 | [illegible] |
| A3SS | 10842 | ENSG00000186918 | ZNF395 | chr8 | - | 28375261 | 28375382 | 28375261 | 28375378 | 28385079 | 28385397 | 0 | 0 | 0 | [illegible] |
| A3SS | 11889 | ENSG00000166250 | CLMP | chr11 | - | 123074701 | 123074850 | 123074701 | 123074843 | 123083084 | 123083207 | 0 | 0 | 0 | [illegible] |
| A3SS | 7800 | ENSG00000130768 | SMPDL3B | chr1 | + | 27948993 | 27949162 | 27949064 | 27949162 | 27945231 | 27945445 | 0 | 0 | 3 | [illegible] |
| A3SS | 4978 | ENSG00000162755 | KLHDC9 | chr1 | + | 161099578 | 161099796 | 161099597 | 161099796 | 161099345 | 161099505 | 0 | 0 | 0 | [illegible] |
| A3SS | 4715 | ENSG00000006704 | GTF2IRD1 | chr7 | + | 74515344 | 74515596 | 74515440 | 74515596 | 74512829 | 74512971 | 0 | 0 | 0 | [illegible] |
| A3SS | 6655 | ENSG00000174885 | NLRP6 | chr11 | + | 282701 | 282797 | 282704 | 282797 | 280083 | 281839 | 0 | 0 | 0 | [illegible] |
| A3SS | 1564 | ENSG00000276070 | CCL4L2 | chr17 | + | 36212312 | 36212878 | 36212333 | 36212878 | 36211775 | 36211890 | 0 | 0 | 0 | [illegible] |
| A3SS | 12819 | ENSG00000139044 | B4GALNT3 | chr12 | + | 557890 | 558088 | 558015 | 558088 | 557607 | 557761 | 2 | 0 | 0 | [illegible] |
| A3SS | 9098 | ENSG00000197548 | ATG7 | chr3 | + | 11288645 | 11288740 | 11288648 | 11288740 | 11282192 | 11282438 | 4 | 0 | 0 | [illegible] |
| A3SS | 10849 | ENSG00000053108 | FSTL4 | chr5 | - | 133220747 | 133220993 | 133220747 | 133220866 | 133224189 | 133224216 | 0 | 0 | 0 | [illegible] |
| A3SS | 9007 | ENSG00000111011 | RSRC2 | chr12 | - | 122508217 | 122508450 | 122508217 | 122508447 | 122515104 | 122515227 | 0 | 0 | 0 | [illegible] |
| A3SS | 1550 | ENSG00000172057 | ORMDL3 | chr17 | - | 39924029 | 39924225 | 39924029 | 39924220 | 39925622 | 39926841 | 0 | 0 | 0 | [illegible] |
| A3SS | 3868 | ENSG00000078814 | MYH7B | chr20 | + | 35002158 | 35002437 | 35002176 | 35002437 | 35001947 | 35002085 | 0 | 0 | 0 | [illegible] |
| A3SS | 1806 | ENSG00000169951 | ZNF764 | chr16 | - | 30553763 | 30556110 | 30553763 | 30556107 | 30557732 | 30557846 | 1 | 0 | 3 | [illegible] |
| A3SS | 1168 | ENSG00000120093 | HOXB3 | chr17 | - | 48552026 | 48552487 | 48552026 | 48552090 | 48555530 | 48555618 | 1 | 3 | 0 | [illegible] |
| A3SS | 10607 | ENSG00000071889 | FAM3A | chrX | - | 154508473 | 154508811 | 154508473 | 154508652 | 154511847 | 154511871 | 7 | 3 | 2 | [illegible] |
| A3SS | 3655 | ENSG00000073331 | ALPK1 | chr4 | + | 112423725 | 112424003 | 112423943 | 112424003 | 112411826 | 112412025 | 2 | 8 | 8 | [illegible] |
| A3SS | 1247 | ENSG00000132361 | CLUH | chr17 | - | 2696433 | 2696538 | 2696433 | 2696535 | 2696718 | 2696942 | 5 | 6 | 6 | [illegible] |
| A3SS | 5979 | ENSG00000163867 | ZMYM6 | chr1 | - | 35030546 | 35030756 | 35030546 | 35030713 | 35031814 | 35031945 | 4 | 10 | 3 | [illegible] |
| A3SS | 1398 | ENSG00000176170 | SPHK1 | chr17 | + | 76385942 | 76386137 | 76385984 | 76386137 | 76385450 | 76385654 | 0 | 0 | 0 | [illegible] |
| A3SS | 1720 | ENSG00000242612 | DECR2 | chr16 | + | 410208 | 410367 | 410242 | 410367 | 407424 | 407560 | 0 | 0 | 0 | [illegible] |
| A3SS | 2044 | ENSG00000118557 | PMFBP1 | chr16 | - | 72119850 | 72120149 | 72119850 | 72120089 | 72122913 | 72122988 | 0 | 0 | 0 | [illegible] |
| A3SS | 2998 | ENSG00000204622 | HLA-J | chr6 | + | 30009037 | 30009177 | 30009144 | 30009177 | 30008584 | 30008701 | 1 | 1 | 0 | [illegible] |
| A3SS | 1244 | ENSG00000172123 | SLFN12 | chr17 | - | 35430470 | 35430613 | 35430470 | 35430592 | 35432187 | 35432257 | 2 | 1 | 1 | [illegible] |
| A3SS | 6406 | ENSG00000132781 | MUTYH | chr1 | - | 45331660 | 45331872 | 45331660 | 45331849 | 45332022 | 45332086 | 0 | 0 | 3 | [illegible] |
| A3SS | 11056 | ENSG00000171714 | ANO5 | chr11 | + | 22211263 | 22211314 | 22211266 | 22211314 | 22203803 | 22203850 | 3 | 0 | 1 | [illegible] |
| A3SS | 7811 | ENSG00000167291 | TBC1D16 | chr17 | - | 79950410 | 79950594 | 79950410 | 79950578 | 79951449 | 79951597 | 0 | 1 | 0 | [illegible] |
| A3SS | 6366 | ENSG00000215788 | TNFRSF25 | chr1 | - | 6462628 | 6462666 | 6462628 | 6462663 | 6464374 | 6464453 | 0 | 4 | 3 | [illegible] |
| A3SS | 3 | ENSG00000159267 | HLCS | chr21 | - | 36756541 | 36756980 | 36756541 | 36756755 | 36759726 | 36759841 | 8 | 13 | 8 | [illegible] |
| A3SS | 11894 | ENSG00000149548 | CCDC15 | chr11 | + | 125038887 | 125039069 | 125038920 | 125039069 | 125038430 | 125038604 | 0 | 0 | 0 | [illegible] |
| A3SS | 810 | ENSG00000185513 | L3MBTL1 | chr20 | + | 43529202 | 43529368 | 43529263 | 43529368 | 43528656 | 43528745 | 7 | 1 | 0 | [illegible] |
| A3SS | 7961 | ENSG00000246985 | SOCS2-AS1 | chr12 | - | 93545070 | 93545398 | 93545070 | 93545358 | 93566743 | 93566970 | 0 | 3 | 0 | [illegible] |
| A3SS | 8960 | ENSG00000081052 | COL4A4 | chr2 | - | 227144361 | 227144833 | 227144361 | 227144558 | 227147412 | 227147572 | 0 | 0 | 0 | [illegible] |
| A3SS | 7752 | ENSG00000196387 | ZNF140 | chr12 | + | 133083462 | 133083561 | 133083465 | 133083561 | 133083102 | 133083229 | 0 | 1 | 1 | [illegible] |
| A3SS | 10296 | ENSG00000184867 | ARMCX2 | chrX | - | 101658460 | 101658569 | 101658460 | 101658525 | 101659075 | 101659136 | 0 | 1 | 7 | [illegible] |
| A3SS | 148 | ENSG00000186951 | PPARA | chr22 | + | 46198341 | 46198591 | 46198344 | 46198591 | 46151887 | 46151970 | 0 | 0 | 0 | [illegible] |
| A3SS | 12150 | ENSG00000172663 | TMEM134 | chr11 | - | 67465055 | 67465267 | 67465055 | 67465100 | 67467311 | 67467388 | 11 | 6 | 6 | [illegible] |
| A3SS | 9876 | ENSG00000151208 | DLG5 | chr10 | - | 77805664 | 77805928 | 77805664 | 77805861 | 77806757 | 77806928 | 0 | 0 | 0 | [illegible] |
| A3SS | 13171 | ENSG00000122966 | CIT | chr12 | - | 119690150 | 119690481 | 119690150 | 119690454 | 119697658 | 119697838 | 0 | 0 | 0 | [illegible] |
| A3SS | 2174 | ENSG00000140463 | BBS4 | chr15 | + | 72722788 | 72722847 | 72722793 | 72722847 | 72716777 | 72716850 | 2 | 6 | 2 | [illegible] |
| A3SS | 8174 | ENSG00000226259 | GTF2H2B | chr5 | + | 70421734 | 70421771 | 70421736 | 70421771 | 70421279 | 70421357 | 0 | 0 | 0 | [illegible] |
| A3SS | 11629 | ENSG00000148356 | LRSAM1 | chr9 | + | 127487494 | 127487763 | 127487536 | 127487763 | 127485735 | 127485835 | 2 | 0 | 2 | [illegible] |
| A3SS | 2480 | ENSG00000127586 | CHTF18 | chr16 | + | 788927 | 789125 | 788930 | 789125 | 788624 | 788775 | 0 | 0 | 1 | [illegible] |
| A3SS | 7610 | ENSG00000213199 | ASIC3 | chr7 | + | 151052553 | 151052754 | 151052646 | 151052754 | 151052415 | 151052474 | 1 | 0 | 2 | [illegible] |
| A3SS | 2856 | ENSG00000116874 | WARS2 | chr1 | - | 119031215 | 119033388 | 119031215 | 119033359 | 119034094 | 119034213 | 1 | 0 | 2 | [illegible] |
| A3SS | 4623 | ENSG00000188687 | SLC4A5 | chr2 | - | 74224839 | 74225011 | 74224839 | 74224995 | 74226956 | 74227130 | 0 | 0 | 1 | [illegible] |
| A3SS | 7084 | ENSG00000188372 | ZP3 | chr7 | + | 76440438 | 76440611 | 76440474 | 76440611 | 76440249 | 76440341 | 0 | 0 | 0 | [illegible] |
| A3SS | 262 | ENSG00000104936 | DMPK | chr19 | - | 45777341 | 45777590 | 45777341 | 45777559 | 45777666 | 45777873 | 7 | 2 | 10 | [illegible] |
| A3SS | 1045 | ENSG00000090659 | CD209 | chr19 | - | 7744106 | 7744219 | 7744106 | 7744201 | 7744940 | 7745092 | 3 | 0 | 5 | [illegible] |
| A3SS | 3419 | ENSG00000171206 | TRIM8 | chr10 | + | 102656244 | 102656385 | 102656247 | 102656385 | 102656105 | 102656137 | 3 | 4 | 2 | [illegible] |
| A3SS | 11040 | ENSG00000249859 | PVT1 | chr8 | + | 128096517 | 128096654 | 128096520 | 128096654 | 128070159 | 128070272 | 12 | 8 | 14 | [illegible] |
| A3SS | 3458 | ENSG00000011243 | AKAP8L | chr19 | - | 15398629 | 15398766 | 15398629 | 15398703 | 15399301 | 15399410 | 13 | 12 | 14 | [illegible] |
| A3SS | 11487 | ENSG00000188649 | CC2D2B | chr10 | + | 95927236 | 95927332 | 95927248 | 95927332 | 95924778 | 95924844 | 5 | 6 | 1 | [illegible] |
| A3SS | 10178 | ENSG00000121380 | BCL2L14 | chr12 | + | 12094510 | 12094930 | 12094663 | 12094930 | 12090778 | 12090849 | 0 | 0 | 0 | [illegible] |
| A3SS | 13034 | ENSG00000120662 | MTRF1 | chr13 | - | 41257672 | 41257797 | 41257672 | 41257796 | 41260492 | 41260915 | 1 | 2 | 1 | [illegible] |
| A3SS | 13229 | ENSG00000204852 | TCTN1 | chr12 | + | 110647195 | 110647336 | 110647210 | 110647336 | 110644966 | 110645129 | 2 | 1 | 1 | [illegible] |
| A3SS | 4091 | ENSG00000138031 | ADCY3 | chr2 | - | 24826044 | 24826129 | 24826044 | 24826126 | 24827545 | 24827608 | 0 | 0 | 0 | [illegible] |
| A3SS | 5534 | ENSG00000108950 | FAM20A | chr17 | - | 68540848 | 68540983 | 68540848 | 68540958 | 68541984 | 68542165 | 0 | 1 | 0 | [illegible] |
| A3SS | 12168 | ENSG00000174483 | BBS1 | chr11 | + | 66519616 | 66519748 | 66519620 | 66519748 | 66515860 | 66515933 | 20 | 7 | 6 | [illegible] |
| A3SS | 12283 | ENSG00000054654 | SYNE2 | chr14 | + | 64224982 | 64225045 | 64224998 | 64225045 | 64224460 | 64224547 | 0 | 0 | 1 | [illegible] |
| A3SS | 7835 | ENSG00000160753 | RUSC1 | chr1 | + | 155326579 | 155327132 | 155326609 | 155327132 | 155325863 | 155325910 | 10 | 4 | 11 | [illegible] |
| A3SS | 9757 | ENSG00000068885 | IFT80 | chr3 | - | 160377429 | 160377565 | 160377429 | 160377540 | 160381502 | 160381724 | 0 | 0 | 1 | [illegible] |
| A3SS | 6242 | ENSG00000116337 | AMPD2 | chr1 | + | 109627412 | 109627518 | 109627428 | 109627518 | 109627174 | 109627316 | 0 | 0 | 0 | [illegible] |
| A3SS | 7418 | ENSG00000169682 | SPNS1 | chr16 | + | 28981454 | 28981615 | 28981469 | 28981615 | 28979404 | 28979471 | 0 | 0 | 0 | [illegible] |
| A3SS | 8039 | ENSG00000225138 | SLC9A3-AS1 | chr5 | + | 479781 | 480216 | 479865 | 480216 | 477805 | 477849 | 1 | 1 | 1 | [illegible] |
| A3SS | 9479 | ENSG00000127540 | UQCR11 | chr19 | - | 1598048 | 1598326 | 1598048 | 1598215 | 1599411 | 1599560 | 0 | 0 | 0 | [illegible] |
| A3SS | 3849 | ENSG00000185033 | SEMA4B | chr15 | + | 90227537 | 90227642 | 90227556 | 90227642 | 90225660 | 90225827 | 0 | 0 | 1 | [illegible] |
| A3SS | 10485 | ENSG00000177646 | ACAD9 | chr3 | + | 128904357 | 128904505 | 128904385 | 128904505 | 128904061 | 128904132 | 0 | 2 | 1 | [illegible] |
| A3SS | 4821 | ENSG00000197386 | HTT | chr4 | + | 3142744 | 3142886 | 3142765 | 3142886 | 3140509 | 3140656 | 1 | 0 | 0 | [illegible] |
| A3SS | 12259 | ENSG00000167770 | OTUB1 | chr11 | + | 63996853 | 63996941 | 63996856 | 63996941 | 63996529 | 63996648 | 0 | 3 | 0 | [illegible] |

| | | | | | | | | | | | | | | |
|---|---|---|---|---|---|---|---|---|---|---|---|---|---|---|
| A3SS | 2858 | ENSG00000139974 | SLC38A6 | chr14 | + | 61078809 | 61078927 | 61078829 | 61078927 | 61046066 | 61046167 | 0 | 0 | [illegible] |
| A3SS | 11163 | ENSG00000239672 | NME1 | chr17 | + | 51161097 | 51161272 | 51161159 | 51161272 | 51159979 | 51160081 | 3 | 1 | [illegible] |
| A3SS | 483 | ENSG00000013275 | PSMC4 | chr19 | + | 39972368 | 39972555 | 39972461 | 39972555 | 39971169 | 39971238 | 94 | 57 | [illegible] |
| A3SS | 10478 | ENSG00000108679 | LGALS3BP | chr17 | - | 78972969 | 78973343 | 78972969 | 78973222 | 78974687 | 78974819 | 0 | 3 | [illegible] |
| A3SS | 12533 | ENSG00000165752 | STK32C | chr10 | - | 132227976 | 132228189 | 132227976 | 132228128 | 132245899 | 132245955 | 0 | 3 | [illegible] |
| A3SS | 2857 | ENSG00000139974 | SLC38A6 | chr14 | + | 61078809 | 61078927 | 61078829 | 61078927 | 61030444 | 61030523 | 0 | 2 | [illegible] |
| A3SS | 3374 | ENSG00000105223 | PLD3 | chr19 | + | 40366383 | 40366510 | 40366418 | 40366510 | 40365552 | 40365585 | 3 | 2 | [illegible] |
| A3SS | 6415 | ENSG00000132781 | MUTYH | chr1 | - | 45333412 | 45333594 | 45333412 | 45333564 | 45334390 | 45334511 | 7 | 8 | [illegible] |
| A3SS | 6687 | ENSG00000135953 | MFSD9 | chr2 | - | 102723699 | 102723909 | 102723699 | 102723878 | 102731024 | 102731068 | 4 | 1 | [illegible] |
| A3SS | 651 | ENSG00000064545 | TMEM161A | chr19 | - | 19133129 | 19133306 | 19133129 | 19133210 | 19134783 | 19134887 | 2 | 1 | [illegible] |
| A3SS | 5218 | ENSG00000071127 | WDR1 | chr4 | - | 10083021 | 10083178 | 10083021 | 10083176 | 10084442 | 10084530 | 1 | 1 | [illegible] |
| A3SS | 1316 | ENSG00000008283 | CYB561 | chr17 | - | 63437345 | 63437560 | 63437345 | 63437484 | 63440191 | 63440211 | 2 | 1 | [illegible] |
| A3SS | 1972 | ENSG00000196118 | CCDC189 | chr16 | - | 30759341 | 30759557 | 30759341 | 30759458 | 30759653 | 30759724 | 1 | 0 | [illegible] |
| A3SS | 1623 | ENSG00000004660 | CAMKK1 | chr17 | - | 3881626 | 3882564 | 3881626 | 3881648 | 3883428 | 3883480 | 0 | 0 | [illegible] |
| A3SS | 4457 | ENSG00000114867 | EIF4G1 | chr3 | + | 184323404 | 184323593 | 184323407 | 184323593 | 184323082 | 184323218 | 1 | 0 | [illegible] |
| A3SS | 2007 | ENSG00000168367 | LINC00917 | chr16 | - | 86337360 | 86337608 | 86337360 | 86337592 | 86337681 | 86338661 | 0 | 0 | [illegible] |
| A3SS | 4191 | ENSG00000253438 | PCAT1 | chr8 | + | 127339279 | 127339435 | 127339280 | 127339435 | 127322204 | 127322226 | 0 | 0 | [illegible] |
| A3SS | 4334 | ENSG00000165621 | OXGR1 | chr13 | - | 96985712 | 96987833 | 96985712 | 96987775 | 96989757 | 96989809 | 0 | 0 | [illegible] |
| A3SS | 4573 | ENSG00000163958 | ZDHHC19 | chr3 | - | 196198788 | 196198874 | 196198788 | 196198869 | 196199345 | 196199652 | 0 | 0 | [illegible] |
| A3SS | 5346 | ENSG00000159214 | CCDC24 | chr1 | + | 43993886 | 43993964 | 43993889 | 43993964 | 43992522 | 43992639 | 0 | 0 | [illegible] |
| A3SS | 6759 | ENSG00000105609 | LILRB5 | chr19 | - | 54254017 | 54254068 | 54254017 | 54254065 | 54254364 | 54254415 | 0 | 0 | [illegible] |
| A3SS | 8534 | ENSG00000114923 | SLC4A3 | chr2 | + | 219630071 | 219630352 | 219630152 | 219630352 | 219629579 | 219629695 | 0 | 0 | [illegible] |
| A3SS | 12712 | ENSG00000171798 | KNDC1 | chr10 | + | 133168210 | 133168312 | 133168253 | 133168312 | 133167380 | 133167579 | 0 | 0 | [illegible] |
| A3SS | 8684 | ENSG00000236790 | LINC00299 | chr2 | - | 8383094 | 8383222 | 8383094 | 8383190 | 8391952 | 8392085 | 0 | 2 | [illegible] |
| A3SS | 2359 | ENSG00000103852 | TTC23 | chr15 | - | 99139316 | 99139467 | 99139316 | 99139399 | 99156147 | 99156297 | 0 | 1 | [illegible] |
| A3SS | 10168 | ENSG00000187260 | WDR86 | chr7 | - | 151385087 | 151385287 | 151385087 | 151385223 | 151395775 | 151396196 | 3 | 0 | [illegible] |
| A3SS | 103 | ENSG00000093009 | CDC45 | chr22 | + | 19516780 | 19516893 | 19516816 | 19516893 | 19516526 | 19516645 | 0 | 0 | [illegible] |
| A3SS | 104 | ENSG00000093009 | CDC45 | chr22 | + | 19516813 | 19516893 | 19516816 | 19516893 | 19516526 | 19516645 | 0 | 0 | [illegible] |
| A3SS | 510 | ENSG00000232677 | LINC00665 | chr19 | - | 36322129 | 36322225 | 36322129 | 36322222 | 36330252 | 36330492 | 0 | 0 | [illegible] |
| [illegible] | | | | | | | | | | | | | | |

| | | | | | | | | | | | |
|---|---|---|---|---|---|---|---|---|---|---|---|
| A3SS | 1549 | ENSG00000172057 | ORMDL3 | chr17 | - | 39924029 | 39924225 | 39924029 | 39924220 | 39925168 | 39925371 | [illegible] |
| A3SS | 6468 | ENSG00000115602 | IL1RL1 | chr2 | + | 102343035 | 102343177 | 102343077 | 102343177 | 102342222 | 102342294 | [illegible] |
| A3SS | 7745 | ENSG00000115525 | ST3GAL5 | chr2 | - | 85852859 | 85853099 | 85852859 | 85853097 | 85861180 | 85861292 | [illegible] |
| A3SS | 86 | ENSG00000182944 | EWSR1 | chr22 | + | 29291561 | 29291599 | 29291564 | 29291599 | 29286922 | 29287134 | [illegible] |
| A3SS | 9975 | ENSG00000160183 | TMPRSS3 | chr21 | - | 42376540 | 42376686 | 42376540 | 42376683 | 42380116 | 42380212 | [illegible] |
| A3SS | 3383 | ENSG00000250802 | ZBED3-AS1 | chr5 | + | 77100235 | 77100391 | 77100325 | 77100391 | 77098654 | 77098747 | [illegible] |
| A3SS | 4635 | ENSG00000187862 | TTC24 | chr1 | + | 156583796 | 156583895 | 156583799 | 156583895 | 156583041 | 156583170 | [illegible] |
| A3SS | 5474 | ENSG00000111325 | OGFOD2 | chr12 | + | 122976583 | 122976767 | 122976653 | 122976767 | 122976252 | 122976401 | [illegible] |
| A3SS | 7805 | ENSG00000231826 | LINC01819 | chr2 | - | 43032733 | 43033526 | 43032733 | 43032902 | 43036695 | 43036765 | [illegible] |
| A3SS | 5839 | ENSG00000106714 | CNTNAP3 | chr9 | - | 39132931 | 39133135 | 39132931 | 39133132 | 39140518 | 39140638 | [illegible] |
| A3SS | 8179 | ENSG00000237037 | NDUFA6-DT | chr22 | + | 42124476 | 42124562 | 42124519 | 42124562 | 42124046 | 42124130 | [illegible] |
| A3SS | 10887 | ENSG00000172954 | LCLAT1 | chr2 | + | 30476401 | 30476635 | 30476576 | 30476635 | 30459551 | 30459722 | [illegible] |
| A3SS | 5659 | ENSG00000189045 | ANKDD1B | chr5 | + | 75635648 | 75635882 | 75635783 | 75635882 | 75634897 | 75634996 | [illegible] |
| A3SS | 7963 | ENSG00000048342 | CC2D2A | chr4 | + | 15563309 | 15563522 | 15563354 | 15563522 | 15560530 | 15560622 | [illegible] |
| A3SS | 220 | ENSG00000270011 | ZNF559-ZNF177 | chr19 | + | 9380667 | 9382617 | 9381147 | 9382617 | 9380056 | 9380139 | [illegible] |
| A3SS | 4288 | ENSG00000068650 | ATP11A | chr13 | + | 112881812 | 112887168 | 112881815 | 112887168 | 112880545 | 112880649 | [illegible] |
| A3SS | 13009 | ENSG00000135297 | MTO1 | chr6 | + | 73480292 | 73480367 | 73480317 | 73480367 | 73479935 | 73480126 | [illegible] |
| A3SS | 8336 | ENSG00000235703 | LINC00894 | chrX | + | 150016333 | 150016500 | 150016336 | 150016500 | 150015950 | 150016140 | [illegible] |
| A3SS | 242 | ENSG00000127616 | SMARCA4 | chr19 | + | 11058254 | 11058363 | 11058257 | 11058363 | 11056216 | 11056382 | [illegible] |
| A3SS | 504 | ENSG00000076928 | ARHGEF1 | chr19 | + | 41928830 | 41928986 | 41928853 | 41928986 | 41928589 | 41928722 | [illegible] |
| A3SS | 525 | ENSG00000173480 | ZNF417 | chr19 | - | 57912059 | 57912189 | 57912059 | 57912186 | 57915337 | 57915531 | [illegible] |
| A3SS | 7082 | ENSG00000050748 | MAPK9 | chr5 | - | 180233593 | 180236531 | 180233593 | 180236526 | 180239923 | 180239987 | [illegible] |
| A3SS | 9774 | ENSG00000183742 | MACC1 | chr7 | - | 20170713 | 20170781 | 20170713 | 20170778 | 20184988 | 20185078 | [illegible] |
| A3SS | 11136 | ENSG00000164733 | CTSB | chr8 | - | 11849045 | 11849228 | 11849045 | 11849215 | 11850865 | 11850980 | [illegible] |
| A3SS | 6051 | ENSG00000197530 | MIB2 | chr1 | + | 1623388 | 1623482 | 1623430 | 1623482 | 1616507 | 1616614 | [illegible] |
| A3SS | 6189 | ENSG00000169744 | LDB2 | chr4 | - | 16508534 | 16508686 | 16508534 | 16508683 | 16511980 | 16512104 | [illegible] |
| A3SS | 10593 | ENSG00000067221 | STOML1 | chr15 | - | 73984658 | 73984871 | 73984658 | 73984868 | 73985317 | 73985513 | [illegible] |
| A3SS | 3602 | ENSG00000188818 | ZDHHC11 | chr5 | - | 840494 | 840650 | 840494 | 840597 | 843599 | 843724 | [illegible] |
| A3SS | 9327 | ENSG00000168538 | TRAPPC11 | chr4 | + | 183706786 | 183706940 | 183706806 | 183706940 | 183704978 | 183705070 | [illegible] |
| A3SS | 9073 | ENSG00000163512 | AZI2 | chr3 | - | 28341511 | 28341647 | 28341511 | 28341596 | 28348160 | 28348477 | [illegible] |
| A3SS | 6848 | ENSG00000138642 | HERC6 | chr4 | + | 88398107 | 88398209 | 88398141 | 88398209 | 88396014 | 88396142 | [illegible] |
| A3SS | 9402 | ENSG00000112578 | BYSL | chr6 | + | 41931703 | 41931830 | 41931727 | 41931830 | 41931395 | 41931556 | [illegible] |
| A3SS | 4004 | ENSG00000184271 | POU6F1 | chr12 | - | 51198549 | 51198775 | 51198549 | 51198686 | 51199746 | 51199868 | [illegible] |
| A3SS | 7226 | ENSG00000082701 | GSK3B | chr3 | - | 119840796 | 119840935 | 119840796 | 119840862 | 119843254 | 119843353 | [illegible] |
| A3SS | 7858 | ENSG00000283602 | NA | chr5 | - | 140571198 | 140571357 | 140571198 | 140571318 | 140571778 | 140571848 | [illegible] |
| A3SS | 11641 | ENSG00000171097 | KYAT1 | chr9 | - | 128845352 | 128845517 | 128845352 | 128845411 | 128846679 | 128846859 | [illegible] |
| A3SS | 1705 | ENSG00000033011 | ALG1 | chr16 | + | 5075371 | 5075536 | 5075397 | 5075536 | 5073152 | 5073256 | [illegible] |
| A3SS | 11259 | ENSG00000163378 | EOGT | chr3 | - | 68988505 | 68989017 | 68988505 | 68988577 | 68998010 | 68998114 | [illegible] |
| A3SS | 7560 | ENSG00000070495 | JMJD6 | chr17 | - | 76721262 | 76721356 | 76721262 | 76721351 | 76721797 | 76721933 | [illegible] |
| A3SS | 6306 | ENSG00000181847 | TIGIT | chr3 | + | 114293306 | 114294122 | 114293999 | 114294122 | 114291058 | 114293119 | [illegible] |
| A3SS | 5482 | ENSG00000113119 | TMCO6 | chr5 | + | 140642906 | 140643041 | 140642924 | 140643041 | 140642585 | 140642671 | [illegible] |
| A3SS | 3249 | ENSG00000164398 | ACSL6 | chr5 | - | 131971549 | 131971664 | 131971549 | 131971645 | 131972723 | 131972858 | [illegible] |
| A3SS | 10170 | ENSG00000055130 | CUL1 | chr7 | + | 148766560 | 148766723 | 148766564 | 148766723 | 148759547 | 148759638 | [illegible] |
| A3SS | 12475 | ENSG00000155254 | MARVELD1 | chr10 | + | 97715728 | 97715985 | 97715859 | 97715985 | 97715320 | 97715405 | [illegible] |
| A3SS | 7096 | ENSG00000075151 | EIF4G3 | chr1 | - | 21001198 | 21001324 | 21001198 | 21001312 | 21002712 | 21002808 | [illegible] |
| A3SS | 5489 | ENSG00000137274 | BPHL | chr6 | + | 3138040 | 3140509 | 3140385 | 3140509 | 3137361 | 3137493 | [illegible] |
| A3SS | 3806 | ENSG00000145907 | G3BP1 | chr5 | + | 151803861 | 151804141 | 151803884 | 151804141 | 151800759 | 151800869 | [illegible] |
| A3SS | 11615 | ENSG00000165699 | TSC1 | chr9 | - | 132902604 | 132902787 | 132902604 | 132902783 | 132903650 | 132903817 | [illegible] |
| A3SS | 3633 | ENSG00000160613 | PCSK7 | chr11 | - | 117208896 | 117209053 | 117208896 | 117209046 | 117218465 | 117218568 | [illegible] |
| A3SS | 12372 | ENSG00000162241 | SLC25A45 | chr11 | - | 65380131 | 65380234 | 65380131 | 65380175 | 65381874 | 65381969 | [illegible] |
| A3SS | 3917 | ENSG00000185684 | EP400P1 | chr12 | + | 132104374 | 132105304 | 132104566 | 132105304 | 132103996 | 132104170 | [illegible] |
| A3SS | 4436 | ENSG00000114742 | WDR48 | chr3 | + | 39065790 | 39065889 | 39065810 | 39065889 | 39063049 | 39063190 | [illegible] |
| A3SS | 5225 | ENSG00000084072 | PPIE | chr1 | + | 39741894 | 39743297 | 39743215 | 39743297 | 39741365 | 39741409 | [illegible] |
| A3SS | 9996 | ENSG00000181090 | EHMT1 | chr9 | + | 137814250 | 137814508 | 137814430 | 137814508 | 137813385 | 137813530 | [illegible] |
| A3SS | 8732 | ENSG00000077380 | DYNC1I2 | chr2 | + | 171707232 | 171707377 | 171707286 | 171707377 | 171692776 | 171692894 | [illegible] |
| A3SS | 11264 | ENSG00000156990 | RPUSD3 | chr3 | - | 9840183 | 9840335 | 9840183 | 9840307 | 9840531 | 9840616 | [illegible] |
| A3SS | 13401 | ENSG00000127022 | CANX | chr5 | + | 179726659 | 179726759 | 179726679 | 179726759 | 179724656 | 179724783 | [illegible] |
| A3SS | 10095 | ENSG00000061938 | TNK2 | chr3 | - | 195870113 | 195870404 | 195870113 | 195870205 | 195872275 | 195872470 | [illegible] |
| A3SS | 778 | ENSG00000173418 | NAA20 | chr20 | + | 20025668 | 20025767 | 20025676 | 20025767 | 20022455 | 20022480 | [illegible] |
| A3SS | 2286 | ENSG00000102901 | CENPT | chr16 | - | 67828473 | 67828605 | 67828473 | 67828578 | 67828666 | 67828843 | [illegible] |
| A3SS | 6245 | ENSG00000116337 | AMPD2 | chr1 | + | 109628626 | 109628806 | 109628642 | 109628806 | 109628363 | 109628495 | [illegible] |
| A3SS | 6793 | ENSG00000143442 | POGZ | chr1 | - | 151411624 | 151411798 | 151411624 | 151411771 | 151412295 | 151412396 | [illegible] |
| A3SS | 10837 | ENSG00000147471 | PLPBP | chr8 | + | 37775308 | 37775481 | 37775338 | 37775481 | 37772754 | 37772889 | [illegible] |
| A3SS | 3995 | ENSG00000164039 | BDH2 | chr4 | - | 103092599 | 103092718 | 103092599 | 103092696 | 103095202 | 103095281 | [illegible] |
| A3SS | 7540 | ENSG00000177542 | SLC25A22 | chr11 | - | 794457 | 794588 | 794457 | 794513 | 794775 | 794901 | [illegible] |
| A3SS | 11491 | ENSG00000070061 | ELP1 | chr9 | - | 108893942 | 108894088 | 108893942 | 108894066 | 108896495 | 108896644 | [illegible] |
| A3SS | 748 | ENSG00000258366 | RTEL1 | chr20 | + | 63662395 | 63662627 | 63662545 | 63662627 | 63661849 | 63661943 | [illegible] |
| A3SS | 380 | ENSG00000104980 | TIMM44 | chr19 | - | 7935064 | 7935253 | 7935064 | 7935145 | 7938026 | 7938197 | [illegible] |
| A3SS | 749 | ENSG00000258366 | RTEL1 | chr20 | + | 63662473 | 63662627 | 63662545 | 63662627 | 63661849 | 63661943 | [illegible] |
| A3SS | 1915 | ENSG00000090861 | AARS1 | chr16 | - | 70268270 | 70268395 | 70268270 | 70268379 | 70269617 | 70269763 | [illegible] |
| A3SS | 4627 | ENSG00000165272 | AQP3 | chr9 | - | 33443320 | 33443480 | 33443320 | 33443458 | 33443765 | 33443892 | [illegible] |
| A3SS | 9744 | ENSG00000091106 | NLRC4 | chr2 | - | 32241032 | 32241150 | 32241032 | 32241125 | 32249606 | 32251601 | [illegible] |
| A3SS | 13368 | ENSG00000085662 | AKR1B1 | chr7 | - | 134450785 | 134450957 | 134450785 | 134450902 | 134451585 | 134451753 | [illegible] |
| A3SS | 272 | ENSG00000006015 | REX1BD | chr19 | + | 18592088 | 18592336 | 18592107 | 18592336 | 18590853 | 18590933 | [illegible] |
| A3SS | 4065 | ENSG00000105443 | CYTH2 | chr19 | + | 48478065 | 48478145 | 48478068 | 48478145 | 48474837 | 48474949 | [illegible] |
| A3SS | 1912 | ENSG00000090861 | AARS1 | chr16 | - | 70258032 | 70258244 | 70258032 | 70258217 | 70258979 | 70259186 | [illegible] |
| A3SS | 4265 | ENSG00000169696 | ASPSCR1 | chr17 | + | 82012155 | 82012283 | 82012230 | 82012283 | 82011542 | 82011605 | [illegible] |
| A3SS | 7489 | ENSG00000146409 | SLC18B1 | chr6 | - | 132770889 | 132770959 | 132770889 | 132770939 | 132771035 | 132771129 | [illegible] |
| A3SS | 12584 | ENSG00000103024 | NME3 | chr16 | - | 1771069 | 1771191 | 1771069 | 1771171 | 1771279 | 1771410 | [illegible] |
| A3SS | 7509 | ENSG00000147099 | HDAC8 | chrX | - | 72462003 | 72462102 | 72462003 | 72462098 | 72464558 | 72464731 | [illegible] |
| A3SS | 2772 | ENSG00000213983 | AP1G2 | chr14 | - | 23563715 | 23564221 | 23563715 | 23563856 | 23564332 | 23564388 | [illegible] |
| A3SS | 5384 | ENSG00000169231 | THBS3 | chr1 | - | 155201956 | 155202047 | 155201956 | 155202034 | 155202260 | 155202401 | [illegible] |
| A3SS | 11650 | ENSG00000130714 | POMT1 | chr9 | + | 131507350 | 131507514 | 131507367 | 131507514 | 131506402 | 131506453 | [illegible] |
| A3SS | 2768 | ENSG00000100605 | ITPK1 | chr14 | - | 92962750 | 92962867 | 92962750 | 92962849 | 92993879 | 92993997 | [illegible] |
| A3SS | 10342 | ENSG00000174080 | CTSF | chr11 | - | 66563463 | 66564024 | 66563463 | 66564007 | 66564087 | 66564146 | [illegible] |
| A3SS | 6074 | ENSG00000157193 | LRP8 | chr1 | - | 53266472 | 53266699 | 53266472 | 53266647 | 53271027 | 53271153 | [illegible] |
| A3SS | 8936 | ENSG00000215559 | ANKRD20A11P | chr21 | - | 13945038 | 13945143 | 13945038 | 13945109 | 13951066 | 13951139 | [illegible] |
| A3SS | 13228 | ENSG00000204852 | TCTN1 | chr12 | + | 110644879 | 110645129 | 110644972 | 110645129 | 110641541 | 110641627 | [illegible] |
| A3SS | 2489 | ENSG00000166140 | ZFYVE19 | chr15 | + | 40813322 | 40813417 | 40813337 | 40813417 | 40810648 | 40810757 | [illegible] |
| A3SS | 2877 | ENSG00000112640 | PPP2R5D | chr6 | + | 43006445 | 43006679 | 43006462 | 43006679 | 42989610 | 42989688 | [illegible] |
| A3SS | 9156 | ENSG00000204410 | MSH5 | chr6 | + | 31758532 | 31758620 | 31758547 | 31758620 | 31758164 | 31758293 | [illegible] |
| A3SS | 7365 | ENSG00000143842 | SOX13 | chr1 | + | 204116503 | 204116679 | 204116506 | 204116679 | 204114518 | 204114605 | [illegible] |
| A3SS | 12077 | ENSG00000221968 | FADS3 | chr11 | - | 61875850 | 61876003 | 61875850 | 61875976 | 61876110 | 61876190 | [illegible] |
| A3SS | 8573 | ENSG00000104218 | CSPP1 | chr8 | + | 67172768 | 67172973 | 67172777 | 67172973 | 67172415 | 67172555 | [illegible] |
| A3SS | 5413 | ENSG00000141577 | CEP131 | chr17 | - | 81191192 | 81191350 | 81191192 | 81191335 | 81192317 | 81192392 | [illegible] |
| A3SS | 11192 | ENSG00000204839 | MROH6 | chr8 | - | 143570242 | 143570417 | 143570242 | 143570380 | 143570472 | 143570657 | [illegible] |
| A3SS | 4752 | ENSG00000125730 | C3 | chr19 | - | 6684387 | 6684478 | 6684387 | 6684439 | 6684559 | 6684650 | [illegible] |
| A3SS | 1999 | ENSG00000130304 | SLC27A1 | chr19 | + | 17487099 | 17487335 | 17487173 | 17487335 | 17486562 | 17486957 | [illegible] |
| A3SS | 12771 | ENSG00000142920 | AZIN2 | chr1 | + | 33098006 | 33098179 | 33098066 | 33098179 | 33096706 | 33096869 | [illegible] |
| A3SS | 571 | ENSG00000176531 | PHLDB3 | chr19 | - | 43495494 | 43495620 | 43495494 | 43495605 | 43497117 | 43497279 | [illegible] |
| A3SS | 12385 | ENSG00000150433 | TMEM218 | chr11 | - | 125102131 | 125102351 | 125102131 | 125102317 | 125102636 | 125102809 | [illegible] |
| A3SS | 7178 | ENSG00000226822 | LINC02785 | chr1 | + | 108072528 | 108072697 | 108072548 | 108072697 | 108071483 | 108071637 | [illegible] |
| A3SS | 5167 | ENSG00000013573 | DDX11 | chr12 | + | 31100559 | 31100707 | 31100634 | 31100707 | 31097884 | 31097997 | [illegible] |
| A3SS | 7484 | ENSG00000196440 | ARMCX4 | chrX | + | 101488007 | 101488099 | 101488043 | 101488099 | 101487607 | 101487691 | [illegible] |
| A3SS | 813 | ENSG00000185513 | L3MBTL1 | chr20 | + | 43536081 | 43536294 | 43536096 | 43536294 | 43535836 | 43535936 | [illegible] |
| A3SS | 11983 | ENSG00000242474 | NA | chr7 | - | 148531 | 148714 | 148531 | 148618 | 149328 | 149466 | [illegible] |
| A3SS | 4097 | ENSG00000008311 | AASS | chr7 | - | 122092842 | 122092982 | 122092842 | 122092951 | 122093047 | 122093158 | [illegible] |
| A3SS | 5014 | ENSG00000204580 | DDR1 | chr6 | + | 30897360 | 30897597 | 30897378 | 30897597 | 30897013 | 30897141 | [illegible] |
| A3SS | 1579 | ENSG00000133265 | HSPBP1 | chr19 | - | 55279398 | 55279705 | 55279398 | 55279702 | 55280034 | 55280108 | [illegible] |
| A3SS | 454 | ENSG00000178078 | STAP2 | chr19 | - | 4324454 | 4324664 | 4324454 | 4324529 | 4325215 | 4325308 | [illegible] |
| A3SS | 5964 | ENSG00000164136 | IL15 | chr4 | + | 141719365 | 141719476 | 141719377 | 141719476 | 141688821 | 141689035 | [illegible] |
| A3SS | 10608 | ENSG00000071889 | FAM3A | chrX | - | 154512313 | 154512431 | 154512313 | 154512411 | 154512822 | 154512936 | [illegible] |
| A3SS | 12402 | ENSG00000196155 | PLEKHG4 | chr16 | + | 67288475 | 67288604 | 67288488 | 67288604 | 67288166 | 67288400 | [illegible] |
| A3SS | 10528 | ENSG00000005302 | MSL3 | chrX | + | 11765469 | 11765729 | 11765521 | 11765729 | 11763779 | 11763938 | [illegible] |
| A3SS | 5168 | ENSG00000013573 | DDX11 | chr12 | + | 31100998 | 31101130 | 31101026 | 31101130 | 31100634 | 31100707 | [illegible] |
| A3SS | 8753 | ENSG00000144362 | PHOSPHO2 | chr2 | + | 169697864 | 169697982 | 169697892 | 169697982 | 169697360 | 169697531 | [illegible] |
| A3SS | 1321 | ENSG00000076604 | TRAF4 | chr17 | + | 28747821 | 28747947 | 28747842 | 28747947 | 28747212 | 28747264 | [illegible] |
| A3SS | 5509 | ENSG00000132561 | MATN2 | chr8 | + | 98003639 | 98003783 | 98003660 | 98003783 | 97994479 | 97994602 | [illegible] |
| A3SS | 7291 | ENSG00000120341 | SEC16B | chr1 | - | 177960790 | 177960942 | 177960790 | 177960939 | 177961589 | 177961734 | [illegible] |
| A3SS | 7864 | ENSG00000163126 | ANKRD23 | chr2 | - | 96839726 | 96839828 | 96839726 | 96839825 | 96839996 | 96840095 | [illegible] |
| A3SS | 11762 | ENSG00000025434 | NR1H3 | chr11 | + | 47259785 | 47259979 | 47259790 | 47259979 | 47259179 | 47259259 | [illegible] |
| A3SS | 12162 | ENSG00000100116 | GCAT | chr22 | + | 37813276 | 37813609 | 37813462 | 37813609 | 37812886 | 37812988 | [illegible] |
| A3SS | 4850 | ENSG00000137074 | APTX | chr9 | - | 32987543 | 32987846 | 32987543 | 32987842 | 32988082 | 32988540 | [illegible] |
| A3SS | 12108 | ENSG00000137760 | ALKBH8 | chr11 | - | 107553846 | 107553978 | 107553846 | 107553973 | 107560764 | 107560899 | [illegible] |
| A3SS | 6430 | ENSG00000227372 | TP73-AS1 | chr1 | - | 3742803 | 3742960 | 3742803 | 3742957 | 3743060 | 3743190 | [illegible] |
| A3SS | 11276 | ENSG00000228340 | MIR646HG | chr20 | + | 60100758 | 60101372 | 60101091 | 60101372 | 60100170 | 60100219 | [illegible] |
| A3SS | 6675 | ENSG00000237188 | NA | chr1 | + | 147264421 | 147264584 | 147264428 | 147264584 | 147256530 | 147256562 | [illegible] |
| A3SS | 718 | ENSG00000031823 | RANBP3 | chr19 | - | 5950716 | 5950875 | 5950716 | 5950872 | 5951392 | 5951596 | [illegible] |
| A3SS | 7956 | ENSG00000119777 | TMEM214 | chr2 | + | 27037789 | 27037892 | 27037814 | 27037892 | 27037560 | 27037702 | [illegible] |
| A3SS | 919 | ENSG00000089057 | SLC23A2 | chr20 | - | 4970792 | 4970919 | 4970792 | 4970915 | 5001405 | 5001510 | [illegible] |
| A3SS | 10678 | ENSG00000248323 | LUCAT1 | chr5 | - | 91313637 | 91313781 | 91313637 | 91313777 | 91314164 | 91314402 | [illegible] |
| A3SS | 1370 | ENSG00000183048 | SLC25A10 | chr17 | + | 81717693 | 81717861 | 81717783 | 81717861 | 81717398 | 81717491 | [illegible] |
| A3SS | 2656 | ENSG00000151332 | MBIP | chr14 | - | 36305088 | 36305305 | 36305088 | 36305304 | 36308091 | 36308189 | [illegible] |
| A3SS | 4285 | ENSG00000068650 | ATP11A | chr13 | + | 112881812 | 112887168 | 112881815 | 112887168 | 112878216 | 112878303 | [illegible] |
| A3SS | 6583 | ENSG00000162572 | SCNN1D | chr1 | + | 1282241 | 1282315 | 1282275 | 1282315 | 1281225 | 1281297 | [illegible] |
| A3SS | 9826 | ENSG00000169083 | AR | chrX | + | 67685940 | 67686126 | 67686009 | 67686126 | 67643255 | 67643407 | [illegible] |
| A3SS | 12412 | ENSG00000233673 | ANAPC1P1 | chr2 | + | 86894791 | 86894882 | 86894844 | 86894882 | 86893280 | 86893375 | [illegible] |
| A3SS | 611 | ENSG00000105048 | TNNT1 | chr19 | - | 55141856 | 55142057 | 55141856 | 55141920 | 55145543 | 55145565 | [illegible] |
| A3SS | 4716 | ENSG00000006704 | GTF2IRD1 | chr7 | + | 74555392 | 74555494 | 74555437 | 74555494 | 74555173 | 74555223 | [illegible] |
| A3SS | 2218 | ENSG00000137804 | NUSAP1 | chr15 | + | 41350990 | 41351129 | 41351032 | 41351129 | 41349097 | 41349241 | [illegible] |
| A3SS | 1847 | ENSG00000205336 | ADGRG1 | chr16 | + | 57641586 | 57641708 | 57641590 | 57641708 | 57628689 | 57628802 | [illegible] |
| A3SS | 5090 | ENSG00000131697 | NPHP4 | chr1 | - | 5905631 | 5905783 | 5905631 | 5905780 | 5927648 | 5927787 | [illegible] |
| A3SS | 6143 | ENSG00000182983 | ZNF662 | chr3 | + | 42908792 | 42908909 | 42908795 | 42908909 | 42908021 | 42908148 | [illegible] |
| A3SS | 9484 | ENSG00000109339 | MAPK10 | chr4 | - | 86029196 | 86029287 | 86029196 | 86029274 | 86031367 | 86031431 | [illegible] |
| A3SS | 2622 | ENSG00000197249 | SERPINA1 | chr14 | - | 94388559 | 94388663 | 94388559 | 94388660 | 94390419 | 94390692 | [illegible] |
| A3SS | 8937 | ENSG00000054148 | PHPT1 | chr9 | + | 136850502 | 136850560 | 136850505 | 136850560 | 136850012 | 136850137 | [illegible] |
| A3SS | 235 | ENSG00000127616 | SMARCA4 | chr19 | + | 10984120 | 10984373 | 10984123 | 10984373 | 10980776 | 10980852 | [illegible] |
| A3SS | 512 | ENSG00000232677 | LINC00665 | chr19 | - | 36322129 | 36322384 | 36322129 | 36322225 | 36330252 | 36330492 | [illegible] |
| A3SS | 1226 | ENSG00000125319 | HROB | chr17 | + | 44154552 | 44154664 | 44154555 | 44154664 | 44152636 | 44152777 | [illegible] |
| A3SS | 1243 | ENSG00000121101 | TEX14 | chr17 | - | 58616174 | 58616323 | 58616174 | 58616305 | 58617537 | 58617619 | [illegible] |
| A3SS | 7765 | ENSG00000143036 | SLC44A3 | chr1 | + | 94867330 | 94867417 | 94867333 | 94867417 | 94864742 | 94864899 | [illegible] |
| A3SS | 7876 | ENSG00000064225 | ST3GAL6 | chr3 | + | 98801421 | 98801617 | 98801548 | 98801617 | 98793674 | 98793792 | [illegible] |
| A3SS | 8250 | ENSG00000151338 | MIPOL1 | chr14 | + | 37247798 | 37247907 | 37247828 | 37247907 | 37247102 | 37247240 | [illegible] |
| A3SS | 8310 | ENSG00000181381 | DDX60L | chr4 | - | 168391539 | 168391644 | 168391539 | 168391640 | 168394464 | 168394617 | [illegible] |
| A3SS | 10832 | ENSG00000147465 | STAR | chr8 | - | 38148199 | 38148359 | 38148199 | 38148327 | 38148640 | 38148754 | [illegible] |
| A3SS | 6631 | ENSG00000237877 | LINC01473 | chr2 | - | 186080647 | 186080777 | 186080647 | 186080774 | 186081329 | 186081416 | [illegible] |
| A3SS | 112 | ENSG00000075275 | CELSR1 | chr22 | - | 46369691 | 46370237 | 46369691 | 46369804 | 46372882 | 46373057 | [illegible] |
| A3SS | 1408 | ENSG00000184381 | PLA2G6 | chr22 | - | 38114433 | 38114637 | 38114433 | 38114570 | 38115526 | 38115681 | [illegible] |
| A3SS | 3610 | ENSG00000083838 | ZNF446 | chr19 | + | 58479928 | 58480019 | 58479929 | 58480019 | 58478086 | 58478181 | [illegible] |
| A3SS | 5517 | ENSG00000196628 | TCF4 | chr18 | - | 55350873 | 55351003 | 55350873 | 55351000 | 55461018 | 55461115 | [illegible] |
| A3SS | 6316 | ENSG00000171574 | ZNF584 | chr19 | + | 58415792 | 58415940 | 58415818 | 58415940 | 58415523 | 58415646 | [illegible] |
| A3SS | 8276 | ENSG00000259425 | NA | chr15 | - | 22773695 | 22773867 | 22773695 | 22773725 | 22774540 | 22774669 | [illegible] |
| A3SS | 11212 | ENSG00000231527 | FAM27C | chr9 | + | 61920281 | 61920435 | 61920302 | 61920435 | 61917467 | 61917607 | [illegible] |
| A3SS | 12926 | ENSG00000151135 | TMEM263 | chr12 | + | 106959269 | 106959385 | 106959287 | 106959385 | 106957081 | 106957149 | [illegible] |
| A3SS | 9299 | ENSG00000269313 | MAGIX | chrX | + | 49164862 | 49165090 | 49164907 | 49165090 | 49164706 | 49164789 | [illegible] |
| A3SS | 10624 | ENSG00000148143 | ZNF462 | chr9 | + | 106932262 | 106932549 | 106932445 | 106932549 | 106930524 | 106930689 | [illegible] |
| A3SS | 8604 | ENSG00000178460 | MCMDC2 | chr8 | + | 66874325 | 66874586 | 66874394 | 66874586 | 66874052 | 66874234 | [illegible] |
| A3SS | 7122 | ENSG00000133619 | KRBA1 | chr7 | + | 149719532 | 149719672 | 149719545 | 149719672 | 149715010 | 149715066 | [illegible] |
| A3SS | 6460 | ENSG00000067606 | PRKCZ | chr1 | + | 2145337 | 2145511 | 2145388 | 2145511 | 2144209 | 2144341 | [illegible] |
| A3SS | 8685 | ENSG00000075391 | RASAL2 | chr1 | + | 178456699 | 178456899 | 178456720 | 178456899 | 178454446 | 178454648 | [illegible] |
| A3SS | 2691 | ENSG00000100908 | EMC9 | chr14 | - | 24139544 | 24139816 | 24139544 | 24139789 | 24140888 | 24140965 | [illegible] |
| A3SS | 10801 | ENSG00000255052 | FAM66D | chr8 | + | 12145931 | 12146015 | 12145947 | 12146015 | 12145638 | 12145700 | [illegible] |
| A3SS | 7939 | ENSG00000163006 | CCDC138 | chr2 | + | 108791554 | 108791802 | 108791674 | 108791802 | 108788851 | 108788966 | [illegible] |
| A3SS | 5108 | ENSG00000112294 | ALDH5A1 | chr6 | + | 24515124 | 24515310 | 24515166 | 24515310 | 24504868 | 24504985 | [illegible] |
| A3SS | 10562 | ENSG00000126953 | TIMM8A | chrX | - | 101345660 | 101348401 | 101345660 | 101346660 | 101348532 | 101348742 | [illegible] |
| A3SS | 4409 | ENSG00000009830 | POMT2 | chr14 | - | 77279536 | 77279622 | 77279536 | 77279586 | 77279822 | 77279928 | [illegible] |
| A3SS | 667 | ENSG00000130299 | GTPBP3 | chr19 | + | 17337898 | 17338255 | 17338007 | 17338255 | 17337558 | 17337664 | [illegible] |
| A3SS | 6851 | ENSG00000280670 | CCDC163 | chr1 | - | 45495417 | 45495545 | 45495417 | 45495473 | 45496555 | 45496623 | [illegible] |
| A3SS | 4757 | ENSG00000099330 | OCEL1 | chr19 | + | 17226974 | 17227199 | 17226993 | 17227199 | 17226692 | 17226869 | [illegible] |
| A3SS | 9158 | ENSG00000204410 | MSH5 | chr6 | + | 31761184 | 31761262 | 31761187 | 31761262 | 31760689 | 31760839 | [illegible] |
| A3SS | 11869 | ENSG00000150687 | PRSS23 | chr11 | + | 86823333 | 86823593 | 86823376 | 86823593 | 86800543 | 86800651 | [illegible] |
| A3SS | 6293 | ENSG00000197165 | SULT1A2 | chr16 | - | 28592262 | 28592481 | 28592262 | 28592443 | 28593251 | 28593346 | [illegible] |
| A3SS | 607 | ENSG00000161298 | ZNF382 | chr19 | + | 36609901 | 36610053 | 36609926 | 36610053 | 36607551 | 36607622 | [illegible] |
| A3SS | 1060 | ENSG00000101639 | CEP192 | chr18 | + | 13116373 | 13116503 | 13116376 | 13116503 | 13104983 | 13105079 | [illegible] |
| A3SS | 4911 | ENSG00000229180 | NA | chr7 | - | 66576754 | 66576932 | 66576754 | 66576929 | 66588511 | 66588580 | [illegible] |
| A3SS | 7995 | ENSG00000268362 | NA | chr19 | + | 24049404 | 24049548 | 24049409 | 24049548 | 24048936 | 24049014 | [illegible] |
| A3SS | 9818 | ENSG00000168137 | SETD5 | chr3 | + | 9434276 | 9434485 | 9434333 | 9434485 | 9434044 | 9434181 | [illegible] |
| A3SS | 6426 | ENSG00000227372 | TP73-AS1 | chr1 | - | 3737211 | 3738513 | 3737211 | 3737854 | 3740232 | 3740387 | [illegible] |
| A3SS | 4551 | ENSG00000113645 | WWC1 | chr5 | + | 168467839 | 168467964 | 168467842 | 168467964 | 168464710 | 168464962 | [illegible] |
| A3SS | 12797 | ENSG00000079974 | RABL2B | chr22 | - | 50769424 | 50769552 | 50769424 | 50769549 | 50775771 | 50775851 | [illegible] |
| A3SS | 4424 | ENSG00000109065 | NAT9 | chr17 | - | 74773865 | 74774055 | 74773865 | 74773939 | 74775621 | 74775707 | [illegible] |
| A3SS | 3999 | ENSG00000108591 | DRG2 | chr17 | + | 18099632 | 18100435 | 18100362 | 18100435 | 18099016 | 18099093 | [illegible] |
| A3SS | 6493 | ENSG00000143224 | PPOX | chr1 | + | 161167354 | 161167486 | 161167370 | 161167486 | 161167099 | 161167234 | [illegible] |
| A3SS | 11701 | ENSG00000148343 | MIGA2 | chr9 | + | 129063172 | 129063316 | 129063243 | 129063316 | 129061230 | 129061346 | [illegible] |
| A3SS | 522 | ENSG00000232677 | LINC00665 | chr19 | - | 36324923 | 36327275 | 36324923 | 36327225 | 36330252 | 36330492 | [illegible] |
| A3SS | 5554 | ENSG00000133067 | LGR6 | chr1 | + | 202306764 | 202306939 | 202306867 | 202306939 | 202305683 | 202305749 | [illegible] |
| A3SS | 6683 | ENSG00000111358 | GTF2H3 | chr12 | + | 123652523 | 123652561 | 123652531 | 123652561 | 123650993 | 123651056 | [illegible] |
| A3SS | 2438 | ENSG00000067141 | NEO1 | chr15 | + | 73270009 | 73270233 | 73270057 | 73270233 | 73266315 | 73266411 | [illegible] |
| A3SS | 11540 | ENSG00000128973 | CLN6 | chr15 | - | 68211674 | 68211921 | 68211674 | 68211863 | 68214289 | 68214388 | [illegible] |
| A3SS | 8932 | ENSG00000198721 | ECI2 | chr6 | - | 4130371 | 4130642 | 4130371 | 4130560 | 4130766 | 4130865 | [illegible] |
| A3SS | 6428 | ENSG00000227372 | TP73-AS1 | chr1 | - | 3742803 | 3742960 | 3742803 | 3742911 | 3743060 | 3743190 | [illegible] |
| A3SS | 7882 | ENSG00000185499 | MUC1 | chr1 | - | 155192182 | 155192310 | 155192182 | 155192283 | 155192785 | 155192867 | [illegible] |
| A3SS | 4243 | ENSG00000078618 | NRDC | chr1 | - | 51792376 | 51792425 | 51792376 | 51792424 | 51794471 | 51794610 | [illegible] |
| A3SS | 3992 | ENSG00000148154 | UGCG | chr9 | + | 111922816 | 111922951 | 111922848 | 111922951 | 111914604 | 111914746 | [illegible] |
| A3SS | 8770 | ENSG00000079785 | DDX1 | chr2 | + | 15617221 | 15617342 | 15617243 | 15617342 | 15613223 | 15613284 | [illegible] |
| A3SS | 12816 | ENSG00000087502 | ERGIC2 | chr12 | - | 29345442 | 29345557 | 29345442 | 29345540 | 29349078 | 29349177 | [illegible] |
| A3SS | 12962 | ENSG00000130177 | CDC16 | chr13 | + | 114239309 | 114239490 | 114239349 | 114239490 | 114238989 | 114239028 | [illegible] |
| A3SS | 8177 | ENSG00000170445 | HARS1 | chr5 | - | 140677326 | 140677457 | 140677326 | 140677420 | 140677654 | 140677753 | [illegible] |
| A3SS | 5690 | ENSG00000171611 | PTCRA | chr6 | + | 42925215 | 42925835 | 42925260 | 42925835 | 42924228 | 42924273 | [illegible] |

| | | | | | | | | | | | | | | |
|---|---|---|---|---|---|---|---|---|---|---|---|---|---|---|
| A3SS | 1254 | ENSG00000141503 | MINK1 | chr17 | + | 4881113 | 4881257 | 4881131 | 4881257 | 4880983 | 4881040 | 3 | 1 | 1 |
| A3SS | 11057 | ENSG00000134184 | GSTM1 | chr1 | + | 109688116 | 109688245 | 109688169 | 109688245 | 109687844 | 109687909 | 0 | 0 | 0 |
| A3SS | 12223 | ENSG00000221926 | TRIM16 | chr17 | - | 15682853 | 15682964 | 15682853 | 15682951 | 15683035 | 15683157 | 0 | 0 | 3 |
| A3SS | 10350 | ENSG00000174080 | CTSF | chr11 | - | 66564886 | 66565171 | 66564886 | 66565006 | 66565670 | 66565751 | 0 | 0 | 1 |
| A3SS | 1229 | ENSG00000136478 | TEX2 | chr17 | - | 64188167 | 64188436 | 64188167 | 64188415 | 64193558 | 64193889 | 2 | 0 | 0 |
| A3SS | 2714 | ENSG00000203485 | INF2 | chr14 | + | 104710918 | 104711007 | 104710936 | 104711007 | 104710087 | 104710188 | 2 | 0 | 0 |
| A3SS | 11468 | ENSG00000107331 | ABCA2 | chr9 | - | 137012484 | 137012637 | 137012484 | 137012590 | 137012711 | 137012925 | 1 | 0 | 0 |
| A3SS | 10754 | ENSG00000008838 | MED24 | chr17 | - | 40019784 | 40020010 | 40019784 | 40019933 | 40020272 | 40020353 | 0 | 1 | 1 |
| A3SS | 4180 | ENSG00000126970 | ZC4H2 | chrX | - | 64919041 | 64919236 | 64919041 | 64919204 | 64920080 | 64920253 | 1 | 0 | 1 |
| A3SS | 1886 | ENSG00000162063 | CCNF | chr16 | + | 2449110 | 2449462 | 2449281 | 2449462 | 2448854 | 2448978 | 2 | 0 | 0 |
| A3SS | 7424 | ENSG00000103168 | TAF1C | chr16 | - | 84183409 | 84183531 | 84183409 | 84183507 | 84183696 | 84183778 | 0 | 1 | 1 |
| A3SS | 12447 | ENSG00000151150 | ANK3 | chr10 | - | 60200128 | 60200263 | 60200128 | 60200227 | 60203001 | 60203100 | 0 | 0 | 0 |
| A3SS | 11896 | ENSG00000129255 | MPDU1 | chr17 | + | 7585694 | 7585797 | 7585731 | 7585797 | 7583859 | 7583965 | 2 | 2 | 0 |
| A3SS | 4495 | ENSG00000178802 | MPI | chr15 | + | 74890437 | 74890654 | 74890526 | 74890654 | 74890041 | 74890089 | 1 | 1 | 4 |
| A3SS | 6482 | ENSG00000155545 | MIER3 | chr5 | - | 56946925 | 56947086 | 56946925 | 56947071 | 56950627 | 56950652 | 2 | 0 | 0 |
| A3SS | 12405 | ENSG00000095794 | CREM | chr10 | + | 35178871 | 35178986 | 35178888 | 35178986 | 35148367 | 35148491 | 2 | 0 | 0 |
| A3SS | 1807 | ENSG00000007516 | BAIAP3 | chr16 | + | 1339112 | 1339244 | 1339163 | 1339244 | 1338901 | 1338989 | 0 | 0 | 0 |
| A3SS | 7051 | ENSG00000106462 | EZH2 | chr7 | - | 148817221 | 148817412 | 148817221 | 148817391 | 148817876 | 148818117 | 0 | 0 | 0 |
| A3SS | 1416 | ENSG00000178999 | AURKB | chr17 | - | 8207175 | 8207370 | 8207175 | 8207367 | 8207570 | 8207625 | 0 | 0 | 2 |
| A3SS | 5391 | ENSG00000116353 | MECR | chr1 | - | 29195940 | 29196042 | 29195940 | 29196013 | 29196197 | 29196258 | 1 | 0 | 0 |
| A3SS | 6476 | ENSG00000143801 | PSEN2 | chr1 | + | 226885460 | 226885679 | 226885537 | 226885679 | 226883704 | 226883919 | 0 | 0 | 1 |
| A3SS | 4144 | ENSG00000185278 | ZBTB37 | chr1 | + | 173870196 | 173871148 | 173870199 | 173871148 | 173868393 | 173868405 | 0 | 3 | 3 |
| A3SS | 9289 | ENSG00000166278 | C2 | chr6 | + | 31942914 | 31943099 | 31942958 | 31943099 | 31939230 | 31939320 | 0 | 0 | 0 |
| A3SS | 11060 | ENSG00000164953 | TMEM67 | chr8 | + | 93780857 | 93780982 | 93780873 | 93780982 | 93780592 | 93780747 | 1 | 0 | 0 |
| A3SS | 4336 | ENSG00000243989 | ACY1 | chr3 | + | 51985333 | 51985465 | 51985360 | 51985465 | 51985206 | 51985271 | 0 | 0 | 0 |
| A3SS | 5536 | ENSG00000138002 | IFT172 | chr2 | - | 27447514 | 27447657 | 27447514 | 27447634 | 27447811 | 27447922 | 2 | 0 | 1 |
| A3SS | 10055 | ENSG00000105926 | MPP6 | chr7 | + | 24623662 | 24623784 | 24623665 | 24623784 | 24620051 | 24620220 | 5 | 0 | 0 |
| A3SS | 3169 | ENSG00000106123 | EPHB6 | chr7 | + | 142861999 | 142862133 | 142862016 | 142862133 | 142861051 | 142861186 | 1 | 0 | 0 |
| A3SS | 11098 | ENSG00000182325 | FBXL6 | chr8 | - | 144356299 | 144356549 | 144356299 | 144356531 | 144356599 | 144356713 | 2 | 0 | 0 |
| A3SS | 6432 | ENSG00000227372 | TP73-AS1 | chr1 | - | 3742803 | 3742983 | 3742803 | 3742960 | 3743060 | 3743190 | 0 | 3 | 0 |
| A3SS | 12401 | ENSG00000095066 | HOOK2 | chr19 | - | 12772612 | 12772702 | 12772612 | 12772680 | 12772779 | 12772912 | 1 | 2 | 0 |
| A3SS | 9168 | ENSG00000175575 | PAAF1 | chr11 | + | 73887030 | 73887099 | 73887039 | 73887099 | 73878778 | 73878819 | 0 | 0 | 1 |
| A3SS | 4172 | ENSG00000163702 | IL17RC | chr3 | + | 9917863 | 9918075 | 9917922 | 9918075 | 9917712 | 9917734 | 2 | 0 | 0 |
| A3SS | 9865 | ENSG00000213420 | GPC2 | chr7 | - | 100171778 | 100171959 | 100171778 | 100171925 | 100172086 | 100172217 | 0 | 0 | 2 |
| A3SS | 6289 | ENSG00000157870 | PRXL2B | chr1 | + | 2588335 | 2588453 | 2588389 | 2588453 | 2587740 | 2587792 | 0 | 0 | 1 |
| A3SS | 6710 | ENSG00000158552 | ZFAND2B | chr2 | + | 219206966 | 219207042 | 219206969 | 219207042 | 219206783 | 219206845 | 0 | 0 | 3 |
| A3SS | 12810 | ENSG00000151491 | EPS8 | chr12 | - | 15682892 | 15684302 | 15682892 | 15682972 | 15789160 | 15789388 | 0 | 1 | 1 |
| A3SS | 665 | ENSG00000074855 | ANO8 | chr19 | - | 17328161 | 17329541 | 17328161 | 17328983 | 17329756 | 17329831 | 0 | 2 | 0 |
| A3SS | 11814 | ENSG00000173599 | PC | chr11 | - | 66853238 | 66853451 | 66853238 | 66853383 | 66863773 | 66863956 | 0 | 0 | 1 |
| A3SS | 1895 | ENSG00000103066 | PLA2G15 | chr16 | + | 68255705 | 68255990 | 68255765 | 68255990 | 68255281 | 68255380 | 1 | 0 | 1 |
| A3SS | 7984 | ENSG00000186073 | CDIN1 | chr15 | + | 36709837 | 36709961 | 36709855 | 36709961 | 36709222 | 36709288 | 0 | 1 | 0 |
| A3SS | 8131 | ENSG00000161813 | LARP4 | chr12 | + | 50427761 | 50427909 | 50427764 | 50427909 | 50402674 | 50402813 | 0 | 3 | 1 |
| A3SS | 9850 | ENSG00000179818 | PCBP1-AS1 | chr2 | - | 70053730 | 70053815 | 70053730 | 70053792 | 70055713 | 70055910 | 2 | 0 | 1 |
| A3SS | 2963 | ENSG00000127990 | SGCE | chr7 | - | 94623324 | 94624249 | 94623324 | 94623397 | 94628201 | 94628359 | 1 | 1 | 0 |
| A3SS | 4534 | ENSG00000169727 | GPS1 | chr17 | + | 82052255 | 82052459 | 82052258 | 82052459 | 82051896 | 82051964 | 1 | 1 | 2 |
| A3SS | 5158 | ENSG00000122483 | CCDC18 | chr1 | + | 93214742 | 93214966 | 93214745 | 93214966 | 93210801 | 93210926 | 1 | 0 | 0 |
| A3SS | 5007 | ENSG00000143452 | HORMAD1 | chr1 | - | 150717137 | 150717307 | 150717137 | 150717282 | 150719472 | 150719538 | 0 | 0 | 0 |
| A3SS | 386 | ENSG00000079435 | LIPE | chr19 | - | 42402606 | 42403055 | 42402606 | 42403031 | 42405384 | 42405561 | 0 | 0 | 2 |
| A3SS | 4177 | ENSG00000163702 | IL17RC | chr3 | + | 9929848 | 9929897 | 9929851 | 9929897 | 9928579 | 9928630 | 0 | 0 | 0 |
| A3SS | 9240 | ENSG00000251364 | NA | chr11 | - | 7457795 | 7457959 | 7457795 | 7457943 | 7461238 | 7461336 | 2 | 0 | 0 |
| A3SS | 6280 | ENSG00000198939 | ZFP2 | chr5 | + | 178912978 | 178913071 | 178912981 | 178913071 | 178912583 | 178912719 | 0 | 0 | 0 |
| A3SS | 1197 | ENSG00000141252 | VPS53 | chr17 | - | 623532 | 623674 | 623532 | 623669 | 627225 | 627316 | 3 | 1 | 1 |
| A3SS | 234 | ENSG00000127616 | SMARCA4 | chr19 | + | 10984120 | 10984373 | 10984123 | 10984373 | 10961320 | 10961729 | 0 | 0 | 0 |
| A3SS | 5762 | ENSG00000121933 | TMIGD3 | chr1 | - | 111488676 | 111489024 | 111488676 | 111488867 | 111489466 | 111489690 | 3 | 2 | 0 |
| A3SS | 6431 | ENSG00000227372 | TP73-AS1 | chr1 | - | 3742803 | 3742983 | 3742803 | 3742957 | 3743060 | 3743190 | 0 | 3 | 0 |
| A3SS | 2613 | ENSG00000129467 | ADCY4 | chr14 | - | 24330167 | 24330327 | 24330167 | 24330295 | 24331017 | 24331129 | 0 | 8 | 10 |
| A3SS | 3247 | ENSG00000113231 | PDE8B | chr5 | + | 77331345 | 77331459 | 77331401 | 77331459 | 77328997 | 77329057 | 0 | 0 | 0 |
| A3SS | 3299 | ENSG00000170074 | FAM153A | chr5 | - | 177747768 | 177747817 | 177747768 | 177747793 | 177748640 | 177748699 | 0 | 2 | 0 |
| A3SS | 11361 | ENSG00000158467 | AHCYL2 | chr7 | + | 129379637 | 129379749 | 129379640 | 129379749 | 129377537 | 129377623 | 3 | 0 | 2 |
| A3SS | 1866 | ENSG00000197912 | SPG7 | chr16 | + | 89528631 | 89529524 | 89529134 | 89529524 | 89526328 | 89526468 | 0 | 2 | 1 |
| A3SS | 11277 | ENSG00000228340 | MIR646HG | chr20 | + | 60100758 | 60101372 | 60101094 | 60101372 | 60100170 | 60100219 | 5 | 4 | 3 |
| A3SS | 11438 | ENSG00000130653 | PNPLA7 | chr9 | - | 137540641 | 137540855 | 137540641 | 137540722 | 137542641 | 137542801 | 0 | 0 | 0 |
| A3SS | 5201 | ENSG00000117569 | PTBP2 | chr1 | + | 96804784 | 96804939 | 96804799 | 96804939 | 96785058 | 96785254 | 1 | 2 | 2 |
| A3SS | 6020 | ENSG00000188004 | SNHG28 | chr1 | - | 159838364 | 159838655 | 159838364 | 159838451 | 159840768 | 159840866 | 2 | 4 | 2 |
| A3SS | 9063 | ENSG00000135525 | MAP7 | chr6 | - | 136372500 | 136372649 | 136372500 | 136372625 | 136377754 | 136377868 | 0 | 0 | 2 |
| A3SS | 10207 | ENSG00000219545 | UMAD1 | chr7 | + | 7673318 | 7673453 | 7673321 | 7673453 | 7640756 | 7640828 | 0 | 2 | 0 |
| A3SS | 1777 | ENSG00000102981 | PARD6A | chr16 | + | 67661892 | 67662774 | 67661895 | 67662774 | 67661454 | 67661677 | 0 | 0 | 0 |
| A3SS | 7461 | ENSG00000183963 | SMTN | chr22 | + | 31096997 | 31097338 | 31097268 | 31097338 | 31095533 | 31095609 | 0 | 0 | 2 |
| A3SS | 11334 | ENSG00000157657 | ZNF618 | chr9 | + | 114032644 | 114032728 | 114032647 | 114032728 | 114016694 | 114016784 | 2 | 0 | 0 |
| A3SS | 10291 | ENSG00000204001 | LCN8 | chr9 | - | 136755411 | 136755649 | 136755411 | 136755516 | 136756521 | 136756592 | 0 | 1 | 0 |
| A3SS | 11215 | ENSG00000147789 | ZNF7 | chr8 | + | 144829426 | 144829604 | 144829444 | 144829604 | 144829042 | 144829090 | 2 | 0 | 7 |
| A3SS | 1733 | ENSG00000125122 | LRRC29 | chr16 | - | 67210047 | 67211024 | 67210047 | 67210322 | 67223825 | 67223927 | 1 | 0 | 0 |
| A3SS | 1734 | ENSG00000125122 | LRRC29 | chr16 | - | 67210047 | 67211073 | 67210047 | 67210322 | 67223825 | 67223927 | 1 | 0 | 0 |
| A3SS | 1732 | ENSG00000125122 | LRRC29 | chr16 | - | 67210047 | 67210348 | 67210047 | 67210322 | 67223825 | 67223927 | 0 | 0 | 0 |
| A3SS | 4226 | ENSG00000254154 | CRYZL2P-SEC16B | chr1 | - | 177932697 | 177932809 | 177932697 | 177932806 | 177933213 | 177933312 | 0 | 0 | 0 |
| A3SS | 9852 | ENSG00000179818 | PCBP1-AS1 | chr2 | - | 70053730 | 70053815 | 70053730 | 70053792 | 70083537 | 70083687 | 2 | 0 | 1 |
| A3SS | 12633 | ENSG00000107862 | GBF1 | chr10 | + | 102314807 | 102314860 | 102314810 | 102314860 | 102260049 | 102260116 | 0 | 2 | 0 |
| A3SS | 6584 | ENSG00000162572 | SCNN1D | chr1 | + | 1282241 | 1282315 | 1282275 | 1282315 | 1281410 | 1281610 | 0 | 1 | 2 |
| A3SS | 10885 | ENSG00000183077 | AFMID | chr17 | + | 78197134 | 78197560 | 78197145 | 78197560 | 78190969 | 78191060 | 0 | 0 | 1 |
| A3SS | 1238 | ENSG00000259384 | GH1 | chr17 | - | 63918016 | 63918136 | 63918016 | 63918091 | 63918345 | 63918506 | 0 | 0 | 0 |
| A3SS | 11327 | ENSG00000236816 | ANKRD20A7P | chr9 | + | 42877732 | 42877817 | 42877735 | 42877817 | 42873720 | 42873790 | 2 | 0 | 0 |
| A3SS | 4430 | ENSG00000140451 | PIF1 | chr15 | - | 64819846 | 64820019 | 64819846 | 64819986 | 64820981 | 64821088 | 1 | 1 | 0 |
| A3SS | 11992 | ENSG00000177697 | CD151 | chr11 | + | 834457 | 834591 | 834529 | 834591 | 832985 | 833026 | 0 | 1 | 1 |
| A3SS | 9846 | ENSG00000179818 | PCBP1-AS1 | chr2 | - | 70053730 | 70053815 | 70053730 | 70053792 | 70055655 | 70055910 | 2 | 0 | 1 |
| A3SS | 11005 | ENSG00000254402 | LRRC24 | chr8 | - | 144524109 | 144524278 | 144524109 | 144524269 | 144524440 | 144524719 | 1 | 0 | 0 |
| A3SS | 24 | ENSG00000265590 | CFAP298-TCP10L | chr21 | - | 32573283 | 32574124 | 32573283 | 32573368 | 32574575 | 32574676 | 2 | 0 | 0 |
| A3SS | 886 | ENSG00000197586 | ENTPD6 | chr20 | + | 25207075 | 25207397 | 25207078 | 25207397 | 25196153 | 25196254 | 0 | 0 | 0 |
| A3SS | 958 | ENSG00000141644 | MBD1 | chr18 | - | 50274185 | 50274353 | 50274185 | 50274350 | 50275128 | 50275245 | 1 | 0 | 1 |
| A3SS | 1600 | ENSG00000274736 | CCL23 | chr17 | - | 36013743 | 36013909 | 36013743 | 36013858 | 36014333 | 36014393 | 1 | 0 | 0 |
| A3SS | 3717 | ENSG00000272690 | LINC02018 | chr3 | + | 75439753 | 75439846 | 75439784 | 75439846 | 75437249 | 75437275 | 0 | 0 | 0 |
| A3SS | 5342 | ENSG00000159214 | CCDC24 | chr1 | + | 43993803 | 43993964 | 43993886 | 43993964 | 43992522 | 43992639 | 0 | 1 | 1 |
| A3SS | 5365 | ENSG00000104381 | GDAP1 | chr8 | + | 74363371 | 74363449 | 74363383 | 74363449 | 74362938 | 74363053 | 0 | 0 | 2 |
| A3SS | 11608 | ENSG00000113356 | POLR3G | chr5 | + | 90485524 | 90485684 | 90485534 | 90485684 | 90471747 | 90471926 | 0 | 0 | 1 |
| A3SS | 11456 | ENSG00000132740 | IGHMBP2 | chr11 | + | 68934900 | 68934994 | 68934966 | 68934994 | 68934463 | 68934558 | 4 | 0 | 0 |
| A3SS | 3142 | ENSG00000043143 | JADE2 | chr5 | + | 134578493 | 134579317 | 134578496 | 134579317 | 134566115 | 134566580 | 3 | 0 | 0 |
| A3SS | 5105 | ENSG00000162613 | FUBP1 | chr1 | - | 77947452 | 77947552 | 77947452 | 77947547 | 77949154 | 77949300 | 3 | 0 | 1 |
| A3SS | 4718 | ENSG00000174564 | IL20RB | chr3 | + | 136980330 | 136980592 | 136980465 | 136980592 | 136946229 | 136946295 | 3 | 0 | 2 |
| A3SS | 11384 | ENSG00000168003 | SLC3A2 | chr11 | + | 62876783 | 62876879 | 62876786 | 62876879 | 62871576 | 62871669 | 1 | 0 | 4 |
| A3SS | 12165 | ENSG00000254986 | DPP3 | chr11 | + | 66482192 | 66482470 | 66482195 | 66482470 | 66480012 | 66480465 | 4 | 1 | 2 |
| A3SS | 7992 | ENSG00000196912 | ANKRD36B | chr2 | - | 97507563 | 97508065 | 97507563 | 97507657 | 97509580 | 97509733 | 0 | 1 | 0 |
| A3SS | 10305 | ENSG00000169598 | DFFB | chr1 | + | 3865761 | 3866000 | 3865811 | 3866000 | 3858717 | 3858866 | 3 | 0 | 2 |
| A3SS | 7095 | ENSG00000075151 | EIF4G3 | chr1 | - | 21001198 | 21001324 | 21001198 | 21001307 | 21002712 | 21002808 | 2 | 3 | 5 |
| A3SS | 12314 | ENSG00000149582 | TMEM25 | chr11 | + | 118532096 | 118532461 | 118532149 | 118532461 | 118531774 | 118531871 | 0 | 0 | 0 |
| A3SS | 4902 | ENSG00000085831 | TTC39A | chr1 | - | 51309260 | 51309430 | 51309260 | 51309325 | 51311253 | 51311321 | 4 | 0 | 0 |
| A3SS | 2235 | ENSG00000166783 | MARF1 | chr16 | - | 15622933 | 15623126 | 15622933 | 15623123 | 15631380 | 15631498 | 2 | 0 | 0 |
| A3SS | 12944 | ENSG00000184226 | PCDH9 | chr13 | - | 66631209 | 66631411 | 66631209 | 66631285 | 66903503 | 66903605 | 0 | 0 | 2 |
| A3SS | 3862 | ENSG00000251600 | NA | chr4 | + | 143979849 | 143979952 | 143979854 | 143979952 | 143865970 | 143866076 | 1 | 1 | 0 |
| A3SS | 8060 | ENSG00000224078 | SNHG14 | chr15 | + | 25101822 | 25103970 | 25103858 | 25103970 | 25100778 | 25100890 | 7 | 7 | 0 |
| A3SS | 5174 | ENSG00000188747 | NOXA1 | chr9 | + | 137434202 | 137434406 | 137434223 | 137434406 | 137433964 | 137434079 | 0 | 0 | 4 |
| A3SS | 6429 | ENSG00000227372 | TP73-AS1 | chr1 | - | 3742803 | 3742983 | 3742803 | 3742911 | 3743060 | 3743190 | 1 | 4 | 1 |
| A3SS | 463 | ENSG00000105655 | ISYNA1 | chr19 | - | 18436329 | 18437105 | 18436329 | 18436479 | 18437598 | 18437760 | 4 | 0 | 5 |
| A3SS | 5056 | ENSG00000242114 | MTFP1 | chr22 | + | 30427170 | 30427403 | 30427239 | 30427403 | 30426716 | 30426844 | 7 | 1 | 7 |
| A3SS | 9360 | ENSG00000161800 | RACGAP1 | chr12 | - | 50016630 | 50016719 | 50016630 | 50016715 | 50018515 | 50018589 | 0 | 3 | 0 |
| A3SS | 13224 | ENSG00000204852 | TCTN1 | chr12 | + | 110627929 | 110628224 | 110628051 | 110628224 | 110626361 | 110626492 | 0 | 0 | 0 |
| A3SS | 6367 | ENSG00000215788 | TNFRSF25 | chr1 | - | 6462862 | 6462970 | 6462862 | 6462956 | 6464374 | 6464453 | 0 | 1 | 11 |
| A3SS | 6755 | ENSG00000138614 | INTS14 | chr15 | - | 65605128 | 65605236 | 65605128 | 65605232 | 65607301 | 65607442 | 3 | 8 | 7 |
| A3SS | 3685 | ENSG00000131591 | C1orf159 | chr1 | - | 1090352 | 1090587 | 1090352 | 1090428 | 1091471 | 1091565 | 1 | 2 | 0 |
| A3SS | 5139 | ENSG00000137166 | FOXP4 | chr6 | + | 41585430 | 41585517 | 41585436 | 41585517 | 41584768 | 41584891 | 1 | 5 | 4 |
| A3SS | 5089 | ENSG00000131697 | NPHP4 | chr1 | - | 5905631 | 5905783 | 5905631 | 5905780 | 5907114 | 5907222 | 3 | 3 | 1 |
| A3SS | 1595 | ENSG00000166582 | CENPV | chr17 | - | 16348615 | 16349333 | 16348615 | 16348685 | 16349930 | 16350029 | 3 | 5 | 6 |
| A3SS | 11585 | ENSG00000165282 | PIGO | chr9 | - | 35091239 | 35092767 | 35091239 | 35091291 | 35093420 | 35093580 | 11 | 4 | 15 |
| A3SS | 11902 | ENSG00000149547 | EI24 | chr11 | + | 125575250 | 125575408 | 125575308 | 125575408 | 125572457 | 125572569 | 38 | 15 | 25 |
| A3SS | 12157 | ENSG00000110321 | EIF4G2 | chr11 | - | 10805906 | 10806047 | 10805906 | 10806010 | 10807254 | 10807381 | 94 | 79 | 96 |
| A3SS | 11176 | ENSG00000133794 | ARNTL | chr11 | + | 13378321 | 13378445 | 13378336 | 13378445 | 13376639 | 13376728 | 14 | 11 | 8 |
| A3SS | 12974 | ENSG00000257433 | NA | chr12 | + | 47719599 | 47719699 | 47719604 | 47719699 | 47719146 | 47719272 | 0 | 0 | 5 |
| A3SS | 2628 | ENSG00000092108 | SCFD1 | chr14 | + | 30633929 | 30634037 | 30633946 | 30634037 | 30630476 | 30630565 | 0 | 1 | 0 |
| A3SS | 427 | ENSG00000063177 | RPL18 | chr19 | - | 48616078 | 48616220 | 48616078 | 48616202 | 48616725 | 48616824 | 6 | 6 | 20 |
| A3SS | 1283 | ENSG00000185359 | HGS | chr17 | + | 81694796 | 81694853 | 81694814 | 81694853 | 81693869 | 81693965 | 2 | 0 | 0 |
| A3SS | 6974 | ENSG00000149474 | KAT14 | chr20 | + | 18181693 | 18181846 | 18181709 | 18181846 | 18162376 | 18162945 | 0 | 0 | 0 |
| A3SS | 8176 | ENSG00000170445 | HARS1 | chr5 | - | 140676988 | 140677142 | 140676988 | 140677116 | 140677326 | 140677420 | 0 | 0 | 2 |
| A3SS | 473 | ENSG00000248098 | BCKDHA | chr19 | + | 41422922 | 41423169 | 41422997 | 41423169 | 41422628 | 41422770 | 1 | 1 | 3 |
| A3SS | 498 | ENSG00000142453 | CARM1 | chr19 | + | 10919832 | 10919966 | 10919876 | 10919966 | 10919594 | 10919680 | 0 | 0 | 2 |
| A3SS | 2421 | ENSG00000156958 | GALK2 | chr15 | + | 49217172 | 49217313 | 49217189 | 49217313 | 49201161 | 49201250 | 1 | 0 | 0 |
| A3SS | 8875 | ENSG00000239665 | NA | chr10 | + | 13648619 | 13648768 | 13648625 | 13648768 | 13648084 | 13648230 | 4 | 0 | 0 |
| A3SS | 10914 | ENSG00000165806 | CASP7 | chr10 | + | 113721000 | 113721168 | 113721031 | 113721168 | 113697493 | 113697603 | 0 | 2 | 0 |
| A3SS | 3519 | ENSG00000069020 | MAST4 | chr5 | + | 67102506 | 67102611 | 67102535 | 67102611 | 67100434 | 67100592 | 0 | 1 | 2 |
| A3SS | 7222 | ENSG00000175634 | RPS6KB2 | chr11 | + | 67428912 | 67429022 | 67428981 | 67429022 | 67428493 | 67428623 | 1 | 0 | 3 |
| A3SS | 469 | ENSG00000105497 | ZNF175 | chr19 | + | 51581367 | 51581517 | 51581390 | 51581517 | 51573327 | 51573401 | 1 | 0 | 0 |
| A3SS | 9220 | ENSG00000105671 | DDX49 | chr19 | + | 18924852 | 18924979 | 18924881 | 18924979 | 18924622 | 18924699 | 1 | 1 | 1 |
| A3SS | 6902 | ENSG00000177225 | GATD1 | chr11 | - | 771332 | 771467 | 771332 | 771426 | 772426 | 772521 | 1 | 0 | 0 |
| A3SS | 7795 | ENSG00000084774 | CAD | chr2 | + | 27236727 | 27236830 | 27236748 | 27236830 | 27236283 | 27236523 | 0 | 0 | 2 |
| A3SS | 109 | ENSG00000183506 | PI4KAP2 | chr22 | - | 21477996 | 21478161 | 21477996 | 21478101 | 21478584 | 21478694 | 0 | 0 | 1 |
| A3SS | 5263 | ENSG00000168389 | MFSD2A | chr1 | + | 39958661 | 39958825 | 39958700 | 39958825 | 39957086 | 39957221 | 1 | 0 | 1 |
| A3SS | 8641 | ENSG00000160767 | FAM189B | chr1 | - | 155250275 | 155250929 | 155250275 | 155250821 | 155251094 | 155251164 | 0 | 2 | 0 |
| A3SS | 3266 | ENSG00000197603 | CPLANE1 | chr5 | - | 37125243 | 37125440 | 37125243 | 37125409 | 37138719 | 37138848 | 0 | 0 | 0 |
| A3SS | 12230 | ENSG00000159063 | ALG8 | chr11 | - | 78127357 | 78127439 | 78127357 | 78127436 | 78139493 | 78139622 | 0 | 0 | 0 |
| A3SS | 4274 | ENSG00000088387 | DOCK9 | chr13 | - | 98825871 | 98825975 | 98825871 | 98825940 | 98826829 | 98826887 | 0 | 1 | 0 |
| A3SS | 605 | ENSG00000197619 | ZNF615 | chr19 | - | 52004416 | 52004849 | 52004416 | 52004723 | 52007292 | 52007330 | 2 | 2 | 2 |
| A3SS | 9416 | ENSG00000160785 | SLC25A44 | chr1 | + | 156207861 | 156208013 | 156207885 | 156208013 | 156199834 | 156200472 | 0 | 1 | 3 |
| A3SS | 11231 | ENSG00000108666 | C17orf75 | chr17 | - | 32339812 | 32339962 | 32339812 | 32339938 | 32341203 | 32341284 | 0 | 2 | 1 |
| A3SS | 1315 | ENSG00000159640 | ACE | chr17 | + | 63488569 | 63488791 | 63488647 | 63488791 | 63486985 | 63487073 | 0 | 0 | 0 |
| A3SS | 10389 | ENSG00000204271 | SPIN3 | chrX | - | 56978579 | 56978762 | 56978579 | 56978660 | 56984265 | 56984478 | 1 | 1 | 0 |
| A3SS | 10388 | ENSG00000204271 | SPIN3 | chrX | - | 56978579 | 56978751 | 56978579 | 56978660 | 56984265 | 56984478 | 1 | 1 | 0 |
| A3SS | 4819 | ENSG00000095906 | NUBP2 | chr16 | + | 1786737 | 1786955 | 1786756 | 1786955 | 1786536 | 1786655 | 0 | 0 | 0 |
| A3SS | 10387 | ENSG00000204271 | SPIN3 | chrX | - | 56978579 | 56978747 | 56978579 | 56978660 | 56984265 | 56984478 | 1 | 1 | 0 |
| A3SS | 6494 | ENSG00000143224 | PPOX | chr1 | + | 161168412 | 161168576 | 161168431 | 161168576 | 161167994 | 161168127 | 0 | 0 | 1 |
| A3SS | 2883 | ENSG00000004809 | SLC22A16 | chr6 | - | 110424686 | 110425382 | 110424686 | 110425085 | 110431170 | 110431270 | 9 | 0 | 0 |
| A3SS | 3340 | ENSG00000125846 | ZNF133 | chr20 | + | 18306270 | 18306393 | 18306297 | 18306393 | 18305667 | 18305807 | 0 | 0 | 2 |
| A3SS | 1478 | ENSG00000108384 | RAD51C | chr17 | + | 58732480 | 58732544 | 58732483 | 58732544 | 58724039 | 58724100 | 0 | 3 | 0 |
| A3SS | 474 | ENSG00000187266 | EPOR | chr19 | - | 11381055 | 11381313 | 11381055 | 11381209 | 11381691 | 11381849 | 0 | 0 | 0 |
| A3SS | 12164 | ENSG00000174516 | PELI3 | chr11 | + | 66475571 | 66477337 | 66475597 | 66477337 | 66473736 | 66473925 | 1 | 0 | 0 |
| A3SS | 2776 | ENSG00000092068 | SLC7A8 | chr14 | - | 23128018 | 23128364 | 23128018 | 23128196 | 23129649 | 23129799 | 0 | 0 | 0 |
| A3SS | 4826 | ENSG00000215908 | CROCCP2 | chr1 | - | 16619844 | 16620245 | 16619844 | 16620024 | 16621945 | 16622115 | 12 | 1 | 35 |
| A3SS | 3531 | ENSG00000168487 | BMP1 | chr8 | + | 22196180 | 22196840 | 22196679 | 22196840 | 22195461 | 22195587 | 1 | 0 | 0 |
| A3SS | 11706 | ENSG00000188352 | FOCAD | chr9 | + | 20976321 | 20976548 | 20976419 | 20976548 | 20957501 | 20957617 | 1 | 7 | 2 |
| A3SS | 5519 | ENSG00000198794 | SCAMP5 | chr15 | + | 75018393 | 75018535 | 75018417 | 75018535 | 75017869 | 75017971 | 0 | 0 | 2 |
| A3SS | 12149 | ENSG00000172663 | TMEM134 | chr11 | - | 67465055 | 67465267 | 67465055 | 67465100 | 67467164 | 67467388 | 11 | 6 | 6 |
| A3SS | 1480 | ENSG00000267123 | SCAT1 | chr17 | - | 78617392 | 78619044 | 78617392 | 78617626 | 78624191 | 78624315 | 1 | 0 | 0 |
| A3SS | 1133 | ENSG00000161513 | FDXR | chr17 | - | 74866783 | 74866879 | 74866783 | 74866876 | 74872035 | 74872107 | 0 | 0 | 1 |
| A3SS | 5030 | ENSG00000117266 | CDK18 | chr1 | + | 205530185 | 205530349 | 205530258 | 205530349 | 205529520 | 205529563 | 0 | 0 | 3 |
| A3SS | 12381 | ENSG00000167535 | CACNB3 | chr12 | + | 48826244 | 48826518 | 48826419 | 48826518 | 48825659 | 48825769 | 5 | 1 | 4 |
| A3SS | 4798 | ENSG00000004838 | ZMYND10 | chr3 | - | 50342396 | 50342655 | 50342396 | 50342569 | 50342917 | 50343018 | 0 | 0 | 0 |
| A3SS | 470 | ENSG00000105492 | SIGLEC6 | chr19 | - | 51530436 | 51530517 | 51530436 | 51530484 | 51530680 | 51530959 | 0 | 3 | 0 |
| A3SS | 7609 | ENSG00000213199 | ASIC3 | chr7 | + | 151052553 | 151052754 | 151052573 | 151052754 | 151052415 | 151052474 | 0 | 0 | 0 |
| A3SS | 5805 | ENSG00000158488 | CD1E | chr1 | + | 158356727 | 158356896 | 158356763 | 158356896 | 158356497 | 158356591 | 2 | 1 | 1 |
| A3SS | 9577 | ENSG00000255036 | SUGT1P4-STRA6LP-CCDC180 | chr9 | + | 97349090 | 97349291 | 97349110 | 97349291 | 97347313 | 97347666 | 0 | 1 | 0 |
| A3SS | 11150 | ENSG00000139546 | TARBP2 | chr12 | + | 53503712 | 53503808 | 53503729 | 53503808 | 53503026 | 53503212 | 0 | 0 | 1 |
| A3SS | 9488 | ENSG00000109339 | MAPK10 | chr4 | - | 86151990 | 86152196 | 86151990 | 86152144 | 86152462 | 86152629 | 3 | 0 | 0 |
| A3SS | 11054 | ENSG00000171714 | ANO5 | chr11 | + | 22211236 | 22211314 | 22211263 | 22211314 | 22203803 | 22203850 | 0 | 0 | 1 |
| A3SS | 13303 | ENSG00000136104 | RNASEH2B | chr13 | + | 50960144 | 50960171 | 50960148 | 50960171 | 50949462 | 50949505 | 3 | 0 | 2 |
| A3SS | 5849 | ENSG00000152527 | PLEKHH2 | chr2 | + | 43740815 | 43741043 | 43740945 | 43741043 | 43738340 | 43738520 | 0 | 0 | 3 |
| A3SS | 7349 | ENSG00000075702 | WDR62 | chr19 | + | 36066207 | 36066427 | 36066256 | 36066427 | 36065957 | 36066015 | 0 | 0 | 2 |
| A3SS | 12697 | ENSG00000184988 | TMEM106A | chr17 | + | 43212204 | 43212394 | 43212226 | 43212394 | 43211874 | 43211922 | 0 | 1 | 0 |
| A3SS | 2518 | ENSG00000226777 | FAM30A | chr14 | + | 105922198 | 105922344 | 105922202 | 105922344 | 105921487 | 105921562 | 1 | 0 | 1 |
| A3SS | 11230 | ENSG00000108666 | C17orf75 | chr17 | - | 32339812 | 32339962 | 32339812 | 32339924 | 32341203 | 32341284 | 0 | 2 | 1 |
| A3SS | 11139 | ENSG00000164733 | CTSB | chr8 | - | 11864375 | 11864483 | 11864375 | 11864463 | 11868000 | 11868112 | 1 | 0 | 1 |
| A3SS | 10936 | ENSG00000120963 | ZNF706 | chr8 | - | 101202332 | 101202452 | 101202332 | 101202447 | 101205434 | 101205732 | 0 | 0 | 1 |
| A3SS | 526 | ENSG00000173480 | ZNF417 | chr19 | - | 57912059 | 57912189 | 57912059 | 57912186 | 57915564 | 57915665 | 2 | 0 | 0 |
| A3SS | 433 | ENSG00000161265 | U2AF1L4 | chr19 | - | 35743154 | 35743208 | 35743154 | 35743205 | 35743295 | 35743345 | 1 | 1 | 0 |
| A3SS | 5111 | ENSG00000225855 | RUSC1-AS1 | chr1 | - | 155319802 | 155319953 | 155319802 | 155319932 | 155321134 | 155321544 | 0 | 1 | 0 |
| A3SS | 11137 | ENSG00000164733 | CTSB | chr8 | - | 11864375 | 11864483 | 11864375 | 11864463 | 11867253 | 11867530 | 1 | 0 | 1 |
| A3SS | 11214 | ENSG00000147789 | ZNF7 | chr8 | + | 144837390 | 144837507 | 144837393 | 144837507 | 144827559 | 144827609 | 0 | 0 | 2 |
| A3SS | 887 | ENSG00000197586 | ENTPD6 | chr20 | + | 25207075 | 25207397 | 25207078 | 25207397 | 25197104 | 25197226 | 0 | 0 | 0 |
| A3SS | 3599 | ENSG00000188818 | ZDHHC11 | chr5 | - | 823388 | 824093 | 823388 | 823809 | 825163 | 825251 | 0 | 0 | 3 |
| A3SS | 5169 | ENSG00000013573 | DDX11 | chr12 | + | 31100998 | 31101130 | 31101044 | 31101130 | 31100634 | 31100707 | 0 | 0 | 0 |
| A3SS | 8574 | ENSG00000163016 | ALMS1P1 | chr2 | + | 73671029 | 73671150 | 73671079 | 73671150 | 73644918 | 73645154 | 1 | 0 | 0 |
| A3SS | 9000 | ENSG00000226025 | LGALS17A | chr19 | + | 39683486 | 39685418 | 39685301 | 39685418 | 39681997 | 39682208 | 0 | 0 | 3 |
| A3SS | 9827 | ENSG00000179698 | WDR97 | chr8 | + | 144110863 | 144110996 | 144110891 | 144110996 | 144110648 | 144110739 | 2 | 0 | 0 |
| A3SS | 10144 | ENSG00000244198 | ARHGEF35-AS1 | chr7 | + | 144239039 | 144239388 | 144239274 | 144239388 | 144207692 | 144207804 | 1 | 0 | 2 |
| A3SS | 10625 | ENSG00000148143 | ZNF462 | chr9 | + | 106932265 | 106932549 | 106932445 | 106932549 | 106930524 | 106930689 | 0 | 0 | 2 |
| A3SS | 8011 | ENSG00000105948 | TTC26 | chr7 | + | 139146983 | 139147290 | 139147144 | 139147290 | 139142255 | 139142305 | 0 | 0 | 2 |
| A3SS | 13318 | ENSG00000227354 | RBM26-AS1 | chr13 | + | 79415110 | 79415364 | 79415261 | 79415364 | 79412681 | 79412907 | 5 | 0 | 4 |
| A3SS | 2328 | ENSG00000092439 | TRPM7 | chr15 | - | 50583088 | 50583159 | 50583088 | 50583156 | 50589591 | 50589656 | 2 | 1 | 1 |
| A3SS | 6098 | ENSG00000138095 | LRPPRC | chr2 | - | 43948411 | 43948548 | 43948411 | 43948515 | 43949601 | 43949659 | 0 | 3 | 3 |
| A3SS | 13310 | ENSG00000187535 | IFT140 | chr16 | - | 1525886 | 1526796 | 1525886 | 1526077 | 1557934 | 1558134 | 0 | 0 | 4 |
| A3SS | 477 | ENSG00000171817 | ZNF540 | chr19 | + | 37601006 | 37601105 | 37601009 | 37601105 | 37599625 | 37599752 | 1 | 2 | 0 |
| A3SS | 7026 | ENSG00000185100 | ADSS1 | chr14 | + | 104739749 | 104739816 | 104739766 | 104739816 | 104739327 | 104739378 | 0 | 2 | 7 |
| A3SS | 147 | ENSG00000100095 | SEZ6L | chr22 | + | 26375574 | 26375689 | 26375577 | 26375689 | 26373450 | 26373483 | 12 | 3 | 0 |
| A3SS | 2784 | ENSG00000246084 | LINC02325 | chr14 | + | 97463560 | 97464159 | 97463631 | 97464159 | 97461803 | 97462332 | 4 | 0 | 1 |

| | | | | | | | | | | | | |
|---|---|---|---|---|---|---|---|---|---|---|---|---|
| A3SS | 11439 | ENSG00000197070 | ARRDC1 | chr9 | + | 137613412 | 137613510 | 137613422 | 137613510 | 137612895 | 137613006 | [illegible] |
| A3SS | 10096 | ENSG00000061938 | TNK2 | chr3 | - | 195870113 | 195870404 | 195870113 | 195870378 | 195872275 | 195872470 | [illegible] |
| A3SS | 7557 | ENSG00000198160 | MIER1 | chr1 | + | 66928885 | 66928963 | 66928889 | 66928963 | 66926141 | 66926242 | [illegible] |
| A3SS | 1580 | ENSG00000276234 | TADA2A | chr17 | + | 37440504 | 37440662 | 37440508 | 37440662 | 37426949 | 37427009 | [illegible] |
| A3SS | 8210 | ENSG00000172493 | AFF1 | chr4 | + | 87084119 | 87084184 | 87084122 | 87084184 | 87049601 | 87049762 | [illegible] |
| A3SS | 8372 | ENSG00000204387 | SNHG32 | chr6 | + | 31836294 | 31836517 | 31836298 | 31836517 | 31835410 | 31835451 | [illegible] |
| [illegible] | [illegible] | [illegible] | [illegible] | [illegible] | [illegible] | [illegible] | [illegible] | [illegible] | [illegible] | [illegible] | [illegible] | [illegible] |

| | | | | | | | | | | | | | | |
|---|---|---|---|---|---|---|---|---|---|---|---|---|---|---|
| A3SS | 9130 | ENSG00000143502 | SUSD4 | chr1 | - | 223229196 | 223229394 | 223229196 | 223229388 | 223264629 | 223264818 | 0 | 0 | 2 |
| A3SS | 1636 | ENSG00000167377 | ZNF23 | chr16 | - | 71454041 | 71454218 | 71454041 | 71454168 | 71456763 | 71456828 | 2 | 1 | 1 |
| A3SS | 4795 | ENSG00000174500 | GCSAM | chr3 | - | 112130444 | 112130519 | 112130444 | 112130513 | 112132636 | 112132910 | 3 | 0 | 1 |
| A3SS | 10453 | ENSG00000117143 | UAP1 | chr1 | + | 162592731 | 162592782 | 162592734 | 162592782 | 162590322 | 162590511 | 3 | 2 | 0 |
| A3SS | 11977 | ENSG00000176371 | ZSCAN2 | chr15 | + | 84622577 | 84622714 | 84622645 | 84622714 | 84620601 | 84620772 | 6 | 7 | 8 |
| A3SS | 6985 | ENSG00000215883 | CYB5RL | chr1 | - | 54187651 | 54187882 | 54187651 | 54187739 | 54190737 | 54190896 | 2 | 2 | 0 |
| A3SS | 1018 | ENSG00000133313 | CNDP2 | chr18 | + | 74516227 | 74519096 | 74518948 | 74519096 | 74512447 | 74512532 | 5 | 9 | 5 |
| A3SS | 11205 | ENSG00000104529 | EEF1D | chr8 | - | 143590607 | 143590738 | 143590607 | 143590735 | 143597347 | 143597415 | 3 | 4 | 3 |
| A3SS | 7108 | ENSG00000134255 | CEPT1 | chr1 | + | 111147641 | 111148053 | 111147646 | 111148053 | 111143481 | 111143545 | 6 | 4 | 2 |
| A3SS | 9954 | ENSG00000106608 | URGCP | chr7 | - | 43887789 | 43887816 | 43887789 | 43887811 | 43926131 | 43926411 | 2 | 1 | 1 |
| A3SS | 11984 | ENSG00000005801 | ZNF195 | chr11 | - | 3361742 | 3368855 | 3361742 | 3361889 | 3370974 | 3371070 | 6 | 3 | 7 |
| A3SS | 515 | ENSG00000232677 | LINC00665 | chr19 | - | 36324923 | 36327225 | 36324923 | 36325887 | 36330252 | 36330492 | 8 | 0 | 3 |
| A3SS | 6155 | ENSG00000000457 | SCYL3 | chr1 | - | 169868927 | 169869128 | 169868927 | 169869039 | 169870254 | 169870357 | 1 | 4 | 15 |
| A3SS | 7451 | ENSG00000149532 | CPSF7 | chr11 | - | 61421053 | 61421130 | 61421053 | 61421127 | 61421389 | 61421608 | 6 | 4 | 0 |
| A3SS | 9869 | ENSG00000121716 | PILRB | chr7 | + | 100358219 | 100358366 | 100358263 | 100358366 | 100356749 | 100356938 | 13 | 2 | 24 |
| A3SS | 9364 | ENSG00000232940 | HCG25 | chr6 | + | 33250608 | 33250711 | 33250615 | 33250711 | 33249789 | 33249889 | 1 | 2 | 5 |
| A3SS | 6639 | ENSG00000099840 | IZUMO4 | chr19 | + | 2097928 | 2098334 | 2098286 | 2098334 | 2097423 | 2097495 | 10 | 5 | 5 |
| A3SS | 5905 | ENSG00000055609 | KMT2C | chr7 | - | 152156204 | 152156274 | 152156204 | 152156271 | 152157760 | 152157922 | 21 | 11 | 8 |
| A3SS | 6221 | ENSG00000157429 | ZNF19 | chr16 | - | 71478878 | 71479100 | 71478878 | 71479005 | 71482081 | 71482143 | 4 | 0 | 1 |
| A3SS | 8063 | ENSG00000224078 | SNHG14 | chr15 | + | 25115265 | 25115656 | 25115298 | 25115656 | 25111764 | 25111894 | 1 | 1 | 4 |

[illegible]

| | | | | | | | | | | | | | | |
|---|---|---|---|---|---|---|---|---|---|---|---|---|---|---|
| A3SS | 8437 | ENSG00000124920 | MYRF | chr11 | + | 61785712 | 61785874 | 61785799 | 61785874 | 61784279 | 61784385 | 0 | 0 | 2 |
| A3SS | 3488 | ENSG00000176293 | ZNF135 | chr19 | + | 58066704 | 58069755 | 58066740 | 58069755 | 58063445 | 58063541 | 3 | 0 | 2 |
| A3SS | 5669 | ENSG00000167384 | ZNF180 | chr19 | - | 44497283 | 44497377 | 44497283 | 44497372 | 44500274 | 44500522 | 1 | 3 | 1 |
| A3SS | 12220 | ENSG00000221926 | TRIM16 | chr17 | - | 15677175 | 15677280 | 15677175 | 15677277 | 15677574 | 15677721 | 4 | 0 | 11 |
| A3SS | 13265 | ENSG00000256660 | CLEC12B | chr12 | + | 10014522 | 10014741 | 10014530 | 10014741 | 10012784 | 10012883 | 9 | 0 | 2 |
| A3SS | 2254 | ENSG00000104093 | DMXL2 | chr15 | - | 51476588 | 51476719 | 51476588 | 51476716 | 51479947 | 51480139 | 3 | 5 | 2 |
| A3SS | 12559 | ENSG00000198825 | INPP5F | chr10 | + | 119807867 | 119808060 | 119807931 | 119808060 | 119805383 | 119805461 | 2 | 3 | 3 |
| A3SS | 10661 | ENSG00000160959 | LRRC14 | chr8 | + | 144519614 | 144520054 | 144519700 | 144520054 | 144517991 | 144518041 | 9 | 0 | 6 |
| A3SS | 2338 | ENSG00000168781 | PPIP5K1 | chr15 | - | 43581226 | 43581429 | 43581226 | 43581388 | 43581852 | 43581954 | 8 | 6 | 9 |
| A3SS | 1551 | ENSG00000172057 | ORMDL3 | chr17 | - | 39924029 | 39924225 | 39924029 | 39924220 | 39926816 | 39926846 | 1 | 5 | 4 |
| A3SS | 3503 | ENSG00000001084 | GCLC | chr6 | - | 53516108 | 53516240 | 53516108 | 53516217 | 53520777 | 53520960 | 6 | 3 | 10 |
| A3SS | 11015 | ENSG00000171045 | TSNARE1 | chr8 | - | 142315002 | 142315092 | 142315002 | 142315089 | 142318543 | 142318634 | 17 | 3 | 7 |
| A3SS | 9412 | ENSG00000170485 | NPAS2 | chr2 | + | 100995399 | 100996055 | 100995795 | 100996055 | 100993346 | 100993527 | 4 | 3 | 8 |
| A3SS | 12560 | ENSG00000198825 | INPP5F | chr10 | + | 119811756 | 119811955 | 119811809 | 119811955 | 119805383 | 119805461 | 6 | 7 | 13 |
| A3SS | 10817 | ENSG00000139197 | PEX5 | chr12 | + | 7203431 | 7203551 | 7203455 | 7203551 | 7201750 | 7201841 | 5 | 2 | 6 |
| A3SS | 1992 | ENSG00000123815 | COQ8B | chr19 | - | 40703540 | 40703855 | 40703540 | 40703622 | 40710058 | 40710136 | 4 | 3 | 6 |
| A3SS | 1102 | ENSG00000167543 | TP53I13 | chr17 | + | 29571831 | 29572057 | 29571856 | 29572057 | 29571590 | 29571719 | 5 | 3 | 0 |
| A3SS | 3485 | ENSG00000106624 | AEBP1 | chr7 | + | 44112141 | 44112321 | 44112205 | 44112321 | 44111853 | 44112050 | 13 | 6 | 18 |
| A3SS | 10634 | ENSG00000147140 | NONO | chrX | + | 71291778 | 71291972 | 71291784 | 71291972 | 71290643 | 71290791 | 238 | 148 | 261 |
| A3SS | 5023 | ENSG00000080822 | CLDND1 | chr3 | - | 98521132 | 98521442 | 98521132 | 98521437 | 98521652 | 98521696 | 10 | 8 | 11 |
| A3SS | 893 | ENSG00000124224 | PPP4R1L | chr20 | - | 58232687 | 58232922 | 58232687 | 58232829 | 58235239 | 58235374 | 9 | 17 | 13 |
| A3SS | 3897 | ENSG00000069869 | NEDD4 | chr15 | - | 55951375 | 55951439 | 55951375 | 55951414 | 55951510 | 55951589 | 5 | 0 | 0 |
| A3SS | 8296 | ENSG00000152359 | POC5 | chr5 | - | 75702604 | 75702832 | 75702604 | 75702810 | 75705703 | 75705787 | 3 | 3 | 5 |
| A3SS | 7336 | ENSG00000107819 | SFXN3 | chr10 | + | 101036666 | 101036808 | 101036722 | 101036808 | 101036485 | 101036561 | 3 | 0 | 1 |
| A3SS | 2651 | ENSG00000075413 | MARK3 | chr14 | + | 103490951 | 103491107 | 103490977 | 103491107 | 103480386 | 103480490 | 4 | 15 | 1 |
| A3SS | 2827 | ENSG00000119711 | ALDH6A1 | chr14 | - | 74071194 | 74071660 | 74071194 | 74071497 | 74071895 | 74071974 | 5 | 0 | 2 |
| A3SS | 1635 | ENSG00000167377 | ZNF23 | chr16 | - | 71454041 | 71454218 | 71454041 | 71454168 | 71456071 | 71456121 | 2 | 1 | 6 |
| A3SS | 8527 | ENSG00000109805 | NCAPG | chr4 | + | 17823646 | 17823770 | 17823671 | 17823770 | 17822982 | 17823123 | 12 | 24 | 23 |
| A3SS | 6628 | ENSG00000175573 | C11orf68 | chr11 | - | 65916809 | 65918218 | 65916809 | 65918215 | 65918909 | 65919062 | 3 | 1 | 5 |
| A3SS | 12804 | ENSG00000110888 | CAPRIN2 | chr12 | - | 30714958 | 30715144 | 30714958 | 30715141 | 30716507 | 30716676 | 5 | 9 | 3 |
| A3SS | 8798 | ENSG00000126254 | RBM42 | chr19 | + | 35633097 | 35633252 | 35633100 | 35633252 | 35631330 | 35631405 | 16 | 0 | 10 |
| A3SS | 9002 | ENSG00000241316 | SUCLG2-AS1 | chr3 | + | 67670828 | 67671004 | 67670881 | 67671004 | 67669815 | 67669909 | 1 | 0 | 2 |
| A3SS | 7940 | ENSG00000163006 | CCDC138 | chr2 | + | 108791554 | 108791802 | 108791679 | 108791802 | 108788851 | 108788966 | 2 | 1 | 4 |
| A3SS | 754 | ENSG00000203896 | LIME1 | chr20 | + | 63737972 | 63738060 | 63738007 | 63738060 | 63737820 | 63737902 | 23 | 7 | 10 |
| A3SS | 430 | ENSG00000090661 | CERS4 | chr19 | + | 8251043 | 8251249 | 8251075 | 8251249 | 8210705 | 8210862 | 11 | 6 | 4 |
| A3SS | 12500 | ENSG00000166321 | NUDT13 | chr10 | + | 73126634 | 73126827 | 73126672 | 73126827 | 73125397 | 73125509 | 5 | 3 | 1 |
| A3SS | 9216 | ENSG00000096070 | BRPF3 | chr6 | + | 36225266 | 36225364 | 36225270 | 36225364 | 36217916 | 36218010 | 5 | 2 | 0 |
| A3SS | 10417 | ENSG00000103037 | SETD6 | chr16 | + | 58518019 | 58518231 | 58518050 | 58518231 | 58516477 | 58516672 | 5 | 1 | 5 |
| A3SS | 13380 | ENSG00000102710 | SUPT20H | chr13 | - | 37045246 | 37045376 | 37045246 | 37045373 | 37047345 | 37047408 | 14 | 2 | 4 |
| A3SS | 13243 | ENSG00000127311 | HELB | chr12 | + | 66324906 | 66325126 | 66324982 | 66325126 | 66323982 | 66324211 | 5 | 15 | 9 |
| A3SS | 8533 | ENSG00000105136 | ZNF419 | chr19 | + | 57491467 | 57491597 | 57491470 | 57491597 | 57490146 | 57490185 | 0 | 8 | 5 |
| A3SS | 4915 | ENSG00000117054 | ACADM | chr1 | + | 75742167 | 75745914 | 75745805 | 75745914 | 75739979 | 75740110 | 6 | 11 | 14 |
| A3SS | 9667 | ENSG00000158636 | EMSY | chr11 | + | 76459932 | 76460085 | 76459935 | 76460085 | 76458182 | 76458358 | 9 | 6 | 5 |
| A3SS | 2774 | ENSG00000213983 | AP1G2 | chr14 | - | 23567110 | 23567319 | 23567110 | 23567315 | 23567450 | 23567524 | 2 | 5 | 12 |
| A3SS | 8385 | ENSG00000101104 | PABPC1L | chr20 | + | 44933056 | 44933185 | 44933071 | 44933185 | 44932681 | 44932713 | 4 | 2 | 3 |
| A3SS | 8673 | ENSG00000197562 | RAB40C | chr16 | + | 625898 | 626121 | 625955 | 626121 | 625431 | 625509 | 16 | 6 | 18 |
| A3SS | 3323 | ENSG00000113742 | CPEB4 | chr5 | + | 173942883 | 173943049 | 173943025 | 173943049 | 173932449 | 173932500 | 24 | 63 | 19 |
| A3SS | 11059 | ENSG00000164953 | TMEM67 | chr8 | + | 93758482 | 93758576 | 93758486 | 93758576 | 93755777 | 93755866 | 3 | 5 | 6 |
| A3SS | 8132 | ENSG00000161813 | LARP4 | chr12 | + | 50427761 | 50427909 | 50427764 | 50427909 | 50420787 | 50420887 | 3 | 9 | 1 |
| A3SS | 3295 | ENSG00000172081 | MOB3A | chr19 | - | 2085174 | 2085328 | 2085174 | 2085325 | 2095240 | 2095323 | 31 | 14 | 17 |
| A3SS | 2322 | ENSG00000166411 | IDH3A | chr15 | + | 78157547 | 78157631 | 78157572 | 78157631 | 78156907 | 78157162 | 24 | 37 | 40 |
| A3SS | 11642 | ENSG00000095319 | NUP188 | chr9 | + | 128958009 | 128958054 | 128958013 | 128958054 | 128956951 | 128957032 | 93 | 49 | 84 |
| A3SS | 6002 | ENSG00000163479 | SSR2 | chr1 | - | 156014960 | 156015073 | 156014960 | 156015069 | 156018269 | 156018368 | 5 | 1 | 6 |
| A3SS | 7175 | ENSG00000160917 | CPSF4 | chr7 | + | 99454040 | 99454136 | 99454043 | 99454136 | 99452367 | 99452440 | 16 | 5 | 12 |
| A3SS | 10721 | ENSG00000215301 | DDX3X | chrX | + | 41346493 | 41346622 | 41346504 | 41346622 | 41346228 | 41346410 | 1 | 5 | 0 |
| A3SS | 13409 | ENSG00000134899 | ERCC5 | chr13 | + | 102852117 | 102852293 | 102852124 | 102852293 | 102849117 | 102849194 | 13 | 23 | 18 |
| A3SS | 2791 | ENSG00000100902 | PSMA6 | chr14 | + | 35312862 | 35313059 | 35312880 | 35313059 | 35310739 | 35310895 | 0 | 11 | 0 |
| A3SS | 5877 | ENSG00000115415 | STAT1 | chr2 | - | 190983641 | 190983758 | 190983641 | 190983740 | 190984309 | 190984393 | 3 | 0 | 3 |
| A3SS | 3435 | ENSG00000184584 | STING1 | chr5 | - | 139480789 | 139481704 | 139480789 | 139480898 | 139482209 | 139482302 | 147 | 67 | 103 |
| A3SS | 9133 | ENSG00000157450 | RNF111 | chr15 | + | 59084101 | 59084254 | 59084128 | 59084254 | 59080935 | 59081284 | 0 | 1 | 5 |
| A3SS | 7993 | ENSG00000064651 | SLC12A2 | chr5 | + | 128184722 | 128184856 | 128184788 | 128184856 | 128184365 | 128184501 | 1 | 3 | 2 |
| A3SS | 1000 | ENSG00000153391 | INO80C | chr18 | - | 35487385 | 35487823 | 35487385 | 35487439 | 35489331 | 35489385 | 10 | 8 | 1 |
| A3SS | 1933 | ENSG00000089280 | FUS | chr16 | + | 31183857 | 31184002 | 31183860 | 31184002 | 31182397 | 31182422 | 1 | 6 | 10 |
| A3SS | 4818 | ENSG00000095906 | NUBP2 | chr16 | + | 1786536 | 1787831 | 1787676 | 1787831 | 1782931 | 1783036 | 1 | 3 | 14 |
| A3SS | 6297 | ENSG00000227124 | ZNF717 | chr3 | - | 75741609 | 75741865 | 75741609 | 75741736 | 75783305 | 75783364 | 1 | 4 | 5 |
| A3SS | 4066 | ENSG00000017260 | ATP2C1 | chr3 | + | 130992930 | 130993001 | 130992950 | 130993001 | 130975331 | 130975488 | 8 | 11 | 6 |
| A3SS | 2992 | ENSG00000154309 | DISP1 | chr1 | + | 222991854 | 222992110 | 222992012 | 222992110 | 222991519 | 222991647 | 4 | 4 | 0 |
| A3SS | 2572 | ENSG00000100949 | RABGGTA | chr14 | - | 24265537 | 24265763 | 24265537 | 24265755 | 24266429 | 24266517 | 5 | 3 | 5 |
| A3SS | 4765 | ENSG00000075826 | SEC31B | chr10 | - | 100488857 | 100488977 | 100488857 | 100488974 | 100489251 | 100489398 | 2 | 0 | 10 |
| A3SS | 3473 | ENSG00000082397 | EPB41L3 | chr18 | - | 5398020 | 5401043 | 5398020 | 5398143 | 5406776 | 5406968 | 1 | 4 | 4 |
| A3SS | 5510 | ENSG00000132561 | MATN2 | chr8 | + | 98033041 | 98033176 | 98033098 | 98033176 | 98032245 | 98032317 | 6 | 0 | 2 |
| A3SS | 10935 | ENSG00000120963 | ZNF706 | chr8 | - | 101202332 | 101202452 | 101202332 | 101202447 | 101204649 | 101204779 | 2 | 5 | 1 |
| A3SS | 4476 | ENSG00000178149 | DALRD3 | chr3 | - | 49018022 | 49018318 | 49018022 | 49018213 | 49020845 | 49021251 | 12 | 0 | 0 |
| A3SS | 4007 | ENSG00000064886 | CHI3L2 | chr1 | + | 111227485 | 111227769 | 111227658 | 111227769 | 111225943 | 111226028 | 8 | 3 | 11 |
| A3SS | 8855 | ENSG00000113407 | TARS1 | chr5 | + | 33453288 | 33453412 | 33453293 | 33453412 | 33445323 | 33445404 | 8 | 0 | 4 |
| A3SS | 2440 | ENSG00000248508 | SRP14-AS1 | chr15 | + | 40064295 | 40064601 | 40064460 | 40064601 | 40045960 | 40046069 | 1 | 6 | 4 |
| A3SS | 7982 | ENSG00000178038 | ALS2CL | chr3 | - | 46681499 | 46681598 | 46681499 | 46681573 | 46682028 | 46682094 | 8 | 10 | 9 |
| A3SS | 11723 | ENSG00000083799 | CYLD | chr16 | + | 50749548 | 50750202 | 50749575 | 50750202 | 50742761 | 50742822 | 14 | 18 | 12 |
| A3SS | 8853 | ENSG00000146083 | RNF44 | chr5 | - | 176532003 | 176532193 | 176532003 | 176532132 | 176532365 | 176532516 | 19 | 5 | 7 |
| A3SS | 11360 | ENSG00000170264 | FAM161A | chr2 | - | 61839420 | 61840581 | 61839420 | 61840525 | 61842121 | 61842360 | 0 | 5 | 3 |
| A3SS | 8382 | ENSG00000101104 | PABPC1L | chr20 | + | 44933056 | 44933185 | 44933071 | 44933185 | 44930517 | 44930726 | 4 | 2 | 3 |
| A3SS | 7912 | ENSG00000147533 | GOLGA7 | chr8 | + | 41490832 | 41490965 | 41490869 | 41490965 | 41490611 | 41490689 | 6 | 7 | 14 |
| A3SS | 5461 | ENSG00000122335 | SERAC1 | chr6 | - | 158150452 | 158150589 | 158150452 | 158150563 | 158155314 | 158155351 | 17 | 11 | 10 |
| A3SS | 12561 | ENSG00000198825 | INPP5F | chr10 | + | 119807867 | 119808060 | 119807931 | 119808060 | 119806359 | 119806480 | 2 | 4 | 3 |
| A3SS | 5257 | ENSG00000159674 | SPON2 | chr4 | - | 1172543 | 1172684 | 1172543 | 1172679 | 1179447 | 1179541 | 12 | 0 | 2 |
| A3SS | 10422 | ENSG00000147586 | MRPS28 | chr8 | - | 80002998 | 80003180 | 80002998 | 80003120 | 80028694 | 80028796 | 43 | 35 | 40 |
| A3SS | 959 | ENSG00000141644 | MBD1 | chr18 | - | 50277089 | 50277204 | 50277089 | 50277196 | 50279882 | 50280017 | 39 | 17 | 29 |
| A3SS | 3648 | ENSG00000070756 | PABPC1 | chr8 | - | 100709458 | 100709731 | 100709458 | 100709673 | 100712361 | 100712457 | 2555 | 1947 | 2612 |
| A3SS | 8455 | ENSG00000180902 | D2HGDH | chr2 | + | 241767709 | 241768811 | 241767712 | 241768811 | 241755044 | 241756014 | 10 | 8 | 14 |
| A3SS | 8815 | ENSG00000163481 | RNF25 | chr2 | - | 218665915 | 218666084 | 218665915 | 218666059 | 218666158 | 218666230 | 3 | 0 | 7 |
| A3SS | 742 | ENSG00000101363 | MANBAL | chr20 | + | 37301170 | 37301413 | 37301207 | 37301413 | 37298762 | 37298879 | 3 | 1 | 5 |
| A3SS | 6451 | ENSG00000081014 | AP4E1 | chr15 | + | 50949823 | 50949938 | 50949825 | 50949938 | 50948019 | 50948159 | 2 | 5 | 2 |
| A3SS | 9598 | ENSG00000168014 | C2CD3 | chr11 | - | 74012717 | 74013525 | 74012717 | 74012773 | 74028286 | 74028398 | 17 | 34 | 16 |
| A3SS | 11522 | ENSG00000155636 | RBM45 | chr2 | + | 178121173 | 178121359 | 178121179 | 178121359 | 178120286 | 178120409 | 3 | 0 | 7 |
| A3SS | 10205 | ENSG00000128944 | KNSTRN | chr15 | + | 40386361 | 40386494 | 40386404 | 40386494 | 40383227 | 40383322 | 20 | 4 | 2 |
| A3SS | 263 | ENSG00000104936 | DMPK | chr19 | - | 45777666 | 45777873 | 45777666 | 45777723 | 45778126 | 45778220 | 21 | 12 | 16 |
| A3SS | 11147 | ENSG00000139546 | TARBP2 | chr12 | + | 53502014 | 53502184 | 53502019 | 53502184 | 53501618 | 53501708 | 1 | 4 | 9 |
| A3SS | 3133 | ENSG00000131504 | DIAPH1 | chr5 | - | 141534334 | 141534442 | 141534334 | 141534433 | 141571925 | 141572040 | 7 | 6 | 2 |
| A3SS | 6477 | ENSG00000143801 | PSEN2 | chr1 | + | 226891742 | 226891844 | 226891745 | 226891844 | 226891277 | 226891361 | 10 | 1 | 9 |
| A3SS | 3118 | ENSG00000250903 | GMDS-DT | chr6 | + | 2412486 | 2413548 | 2412779 | 2413548 | 2398865 | 2399037 | 7 | 3 | 9 |
| A3SS | 1717 | ENSG00000261611 | NA | chr16 | - | 71462209 | 71463148 | 71462209 | 71462370 | 71463545 | 71463648 | 3 | 1 | 1 |
| A3SS | 3580 | ENSG00000158290 | CUL4B | chrX | - | 120557528 | 120557579 | 120557528 | 120557560 | 120557923 | 120558039 | 3 | 6 | 0 |
| A3SS | 11473 | ENSG00000169519 | METTL15 | chr11 | + | 28296752 | 28296931 | 28296796 | 28296931 | 28211061 | 28211198 | 2 | 4 | 4 |
| A3SS | 6543 | ENSG00000274020 | LINC01138 | chr1 | - | 148435145 | 148435354 | 148435145 | 148435232 | 148460730 | 148461093 | 30 | 12 | 3 |
| A3SS | 12005 | ENSG00000149269 | PAK1 | chr11 | - | 77392330 | 77392541 | 77392330 | 77392538 | 77473551 | 77474094 | 4 | 3 | 7 |
| A3SS | 6630 | ENSG00000237877 | LINC01473 | chr2 | - | 186038330 | 186038435 | 186038330 | 186038432 | 186039499 | 186039555 | 0 | 3 | 5 |
| A3SS | 7954 | ENSG00000115109 | EPB41L5 | chr2 | + | 120167876 | 120168007 | 120167879 | 120168007 | 120167465 | 120167507 | 4 | 3 | 5 |
| A3SS | 13043 | ENSG00000111752 | PHC1 | chr12 | + | 8922632 | 8922788 | 8922656 | 8922788 | 8920984 | 8921065 | 3 | 2 | 3 |
| A3SS | 5537 | ENSG00000138002 | IFT172 | chr2 | - | 27449293 | 27449562 | 27449293 | 27449380 | 27449690 | 27449800 | 4 | 4 | 8 |
| A3SS | 7088 | ENSG00000114812 | VIPR1 | chr3 | + | 42527392 | 42527496 | 42527395 | 42527496 | 42525886 | 42525993 | 20 | 15 | 25 |
| A3SS | 8308 | ENSG00000176978 | DPP7 | chr9 | - | 137114242 | 137114382 | 137114242 | 137114378 | 137114462 | 137114576 | 26 | 14 | 16 |
| A3SS | 117 | ENSG00000100099 | HPS4 | chr22 | - | 26466225 | 26468623 | 26466225 | 26466262 | 26470718 | 26470813 | 53 | 32 | 50 |
| A3SS | 37 | ENSG00000237945 | LINC00649 | chr21 | + | 33969236 | 33969598 | 33969240 | 33969598 | 33945809 | 33945895 | 21 | 9 | 24 |
| A3SS | 11660 | ENSG00000130714 | POMT1 | chr9 | + | 131519388 | 131519486 | 131519393 | 131519486 | 131518836 | 131518957 | 33 | 20 | 33 |
| A3SS | 8236 | ENSG00000146414 | SHPRH | chr6 | - | 145945380 | 145945656 | 145945380 | 145945637 | 145946232 | 145946341 | 4 | 2 | 1 |
| A3SS | 490 | ENSG00000115286 | NDUFS7 | chr19 | + | 1388524 | 1388938 | 1388832 | 1388938 | 1387810 | 1387847 | 42 | 12 | 22 |
| A3SS | 2868 | ENSG00000197498 | RPF2 | chr6 | + | 110988987 | 110989065 | 110989027 | 110989065 | 110985005 | 110985138 | 0 | 1 | 2 |
| A3SS | 12444 | ENSG00000151150 | ANK3 | chr10 | - | 60059697 | 60060033 | 60059697 | 60060006 | 60063110 | 60063254 | 1 | 0 | 1 |
| A3SS | 12414 | ENSG00000233968 | NA | chr10 | - | 19722329 | 19722588 | 19722329 | 19722422 | 19724926 | 19724981 | 7 | 6 | 0 |
| A3SS | 9205 | ENSG00000112782 | CLIC5 | chr6 | - | 45914227 | 45914547 | 45914227 | 45914409 | 45941546 | 45941653 | 0 | 1 | 0 |
| A3SS | 9088 | ENSG00000114388 | NPRL2 | chr3 | - | 50349664 | 50349833 | 50349664 | 50349828 | 50349932 | 50350766 | 4 | 6 | 8 |
| A3SS | 12592 | ENSG00000269609 | C10orf95-AS1 | chr10 | + | 102451398 | 102452187 | 102452089 | 102452187 | 102450186 | 102450330 | 3 | 2 | 9 |
| A3SS | 11391 | ENSG00000156042 | CFAP70 | chr10 | - | 73298906 | 73299101 | 73298906 | 73299097 | 73299604 | 73299665 | 6 | 2 | 3 |
| A3SS | 1059 | ENSG00000101639 | CEP192 | chr18 | + | 13116373 | 13116503 | 13116376 | 13116503 | 13103508 | 13103588 | 0 | 2 | 12 |
| A3SS | 2427 | ENSG00000072071 | ADGRL1 | chr19 | - | 14152132 | 14152222 | 14152132 | 14152150 | 14152308 | 14152437 | 3 | 5 | 3 |
| A3SS | 7485 | ENSG00000196440 | ARMCX4 | chrX | + | 101509358 | 101509427 | 101509363 | 101509427 | 101504936 | 101505355 | 3 | 3 | 5 |
| A3SS | 5988 | ENSG00000144357 | UBR3 | chr2 | + | 170073403 | 170073607 | 170073427 | 170073607 | 170061317 | 170061443 | 5 | 1 | 4 |
| A3SS | 12257 | ENSG00000099910 | KLHL22 | chr22 | - | 20488984 | 20489244 | 20488984 | 20489212 | 20495759 | 20495795 | 21 | 2 | 11 |
| A3SS | 9347 | ENSG00000170919 | TPT1-AS1 | chr13 | + | 45344328 | 45344446 | 45344378 | 45344446 | 45341512 | 45341614 | 6 | 1 | 5 |
| A3SS | 780 | ENSG00000101247 | NDUFAF5 | chr20 | + | 13793159 | 13793227 | 13793179 | 13793227 | 13788588 | 13788652 | 7 | 13 | 3 |
| A3SS | 4404 | ENSG00000180376 | CCDC66 | chr3 | + | 56563651 | 56564125 | 56563683 | 56564125 | 56559568 | 56559594 | 10 | 6 | 9 |
| A3SS | 5055 | ENSG00000242114 | MTFP1 | chr22 | + | 30427121 | 30427403 | 30427239 | 30427403 | 30426716 | 30426844 | 8 | 0 | 8 |
| A3SS | 3651 | ENSG00000197020 | ZNF100 | chr19 | - | 21744016 | 21744115 | 21744016 | 21744112 | 21744940 | 21745067 | 7 | 8 | 6 |
| A3SS | 2269 | ENSG00000114062 | UBE3A | chr15 | - | 25407066 | 25407212 | 25407066 | 25407137 | 25409087 | 25409207 | 23 | 13 | 7 |
| A3SS | 3120 | ENSG00000131844 | MCCC2 | chr5 | + | 71643770 | 71643895 | 71643818 | 71643895 | 71636103 | 71636167 | 11 | 4 | 9 |
| A3SS | 1372 | ENSG00000072134 | EPN2 | chr17 | + | 19281954 | 19282077 | 19281978 | 19282077 | 19237496 | 19237531 | 8 | 4 | 4 |
| A3SS | 8368 | ENSG00000091972 | CD200 | chr3 | + | 112347557 | 112347830 | 112347562 | 112347830 | 112344961 | 112345288 | 10 | 4 | 16 |
| A3SS | 2236 | ENSG00000166783 | MARF1 | chr16 | - | 15625371 | 15625800 | 15625371 | 15625797 | 15631380 | 15631498 | 3 | 3 | 4 |
| A3SS | 3478 | ENSG00000203739 | PRDX6-AS1 | chr1 | - | 173441918 | 173442097 | 173441918 | 173442048 | 173442418 | 173442596 | 2 | 7 | 3 |
| A3SS | 3318 | ENSG00000104731 | KLHDC4 | chr16 | - | 87755193 | 87755292 | 87755193 | 87755254 | 87756398 | 87756477 | 107 | 52 | 97 |
| A3SS | 6966 | ENSG00000185379 | RAD51D | chr17 | - | 35106987 | 35107122 | 35106987 | 35107061 | 35107365 | 35107447 | 19 | 9 | 7 |
| A3SS | 13373 | ENSG00000055147 | FAM114A2 | chr5 | - | 154034743 | 154034967 | 154034743 | 154034946 | 154038856 | 154038910 | 14 | 7 | 10 |
| A3SS | 7466 | ENSG00000163534 | FCRL1 | chr1 | - | 157797100 | 157797132 | 157797100 | 157797129 | 157797680 | 157797820 | 5 | 9 | 17 |
| A3SS | 4239 | ENSG00000197808 | ZNF461 | chr19 | - | 36658298 | 36658446 | 36658298 | 36658425 | 36664697 | 36664786 | 6 | 2 | 0 |
| A3SS | 11037 | ENSG00000249859 | PVT1 | chr8 | + | 128096517 | 128096654 | 128096520 | 128096654 | 127989161 | 127989291 | 4 | 0 | 5 |
| A3SS | 6789 | ENSG00000090097 | PCBP4 | chr3 | - | 51960525 | 51960642 | 51960525 | 51960637 | 51960865 | 51960898 | 22 | 3 | 8 |
| A3SS | 2350 | ENSG00000186654 | PRR5 | chr22 | + | 44731678 | 44731821 | 44731734 | 44731821 | 44726576 | 44726634 | 5 | 3 | 7 |
| A3SS | 12593 | ENSG00000156374 | PCGF6 | chr10 | - | 103347237 | 103347319 | 103347237 | 103347297 | 103347394 | 103347450 | 2 | 1 | 2 |
| A3SS | 8434 | ENSG00000144895 | EIF2A | chr3 | + | 150568012 | 150568046 | 150568016 | 150568046 | 150567901 | 150567929 | 0 | 3 | 1 |
| A3SS | 5854 | ENSG00000093183 | SEC22C | chr3 | - | 42547968 | 42553448 | 42547968 | 42548657 | 42557577 | 42557696 | 16 | 13 | 18 |
| A3SS | 7981 | ENSG00000178038 | ALS2CL | chr3 | - | 46675617 | 46675976 | 46675617 | 46675686 | 46676244 | 46676402 | 0 | 1 | 11 |
| A3SS | 10183 | ENSG00000155428 | TRIM74 | chr7 | - | 72959484 | 72959908 | 72959484 | 72959905 | 72960020 | 72960251 | 1 | 2 | 8 |
| A3SS | 1445 | ENSG00000171916 | LGALS9C | chr17 | + | 18491944 | 18491989 | 18491947 | 18491989 | 18491272 | 18491323 | 5 | 2 | 5 |
| A3SS | 7210 | ENSG00000127989 | MTERF1 | chr7 | - | 91876877 | 91876978 | 91876877 | 91876926 | 91880054 | 91880113 | 5 | 8 | 0 |
| A3SS | 3032 | ENSG00000125459 | MSTO1 | chr1 | + | 155611426 | 155611603 | 155611548 | 155611603 | 155611215 | 155611291 | 5 | 5 | 5 |
| A3SS | 7199 | ENSG00000114354 | TFG | chr3 | + | 100744832 | 100744931 | 100744837 | 100744931 | 100736575 | 100736704 | 11 | 4 | 9 |
| A3SS | 5460 | ENSG00000122335 | SERAC1 | chr6 | - | 158119028 | 158119170 | 158119028 | 158119120 | 158120424 | 158120575 | 14 | 15 | 10 |
| A3SS | 4561 | ENSG00000188001 | TPRG1 | chr3 | + | 189150655 | 189151136 | 189150903 | 189151136 | 189147583 | 189147647 | 22 | 13 | 10 |
| A3SS | 4614 | ENSG00000120949 | TNFRSF8 | chr1 | + | 12138228 | 12138436 | 12138231 | 12138436 | 12135587 | 12135613 | 19 | 4 | 3 |
| A3SS | 4278 | ENSG00000184470 | TXNRD2 | chr22 | - | 19877039 | 19877234 | 19877039 | 19877212 | 19878089 | 19878187 | 29 | 9 | 18 |
| A3SS | 7510 | ENSG00000147099 | HDAC8 | chrX | - | 72490928 | 72491006 | 72490928 | 72491000 | 72495155 | 72495268 | 15 | 14 | 13 |
| A3SS | 7627 | ENSG00000234127 | TRIM26 | chr6 | - | 30201034 | 30201138 | 30201034 | 30201083 | 30213304 | 30213427 | 124 | 70 | 74 |
| A3SS | 6275 | ENSG00000132300 | PTCD3 | chr2 | + | 86110977 | 86111158 | 86111112 | 86111158 | 86108499 | 86108536 | 10 | 7 | 6 |
| A3SS | 12970 | ENSG00000089157 | RPLP0 | chr12 | - | 120198553 | 120198770 | 120198553 | 120198739 | 120198894 | 120199000 | 10 | 14 | 22 |
| A3SS | 6688 | ENSG00000135953 | MFSD9 | chr2 | - | 102732321 | 102732409 | 102732321 | 102732392 | 102736645 | 102736878 | 7 | 8 | 7 |
| A3SS | 11328 | ENSG00000182919 | C11orf54 | chr11 | + | 93747296 | 93747448 | 93747301 | 93747448 | 93741962 | 93742066 | 5 | 5 | 0 |
| A3SS | 12518 | ENSG00000138161 | CUZD1 | chr10 | - | 122836830 | 122837048 | 122836830 | 122836899 | 122837403 | 122837554 | 4 | 1 | 4 |
| A3SS | 10729 | ENSG00000234665 | LERFS | chr9 | - | 62867696 | 62868233 | 62867696 | 62867876 | 62869189 | 62869304 | 2 | 0 | 0 |
| A3SS | 8722 | ENSG00000164114 | MAP9 | chr4 | - | 155368585 | 155368812 | 155368585 | 155368809 | 155373135 | 155373456 | 1 | 3 | 4 |
| A3SS | 13227 | ENSG00000204852 | TCTN1 | chr12 | + | 110644966 | 110645129 | 110644991 | 110645129 | 110641023 | 110641149 | 1 | 7 | 10 |
| A3SS | 13184 | ENSG00000198040 | ZNF84 | chr12 | + | 133041277 | 133041482 | 133041284 | 133041482 | 133038874 | 133039012 | 4 | 10 | 7 |
| A3SS | 5673 | ENSG00000245146 | MALINC1 | chr5 | - | 140106295 | 140106490 | 140106295 | 140106461 | 140108472 | 140108603 | 2 | 1 | 5 |
| A3SS | 2347 | ENSG00000166965 | RCCD1 | chr15 | + | 90960327 | 90960498 | 90960333 | 90960498 | 90959899 | 90959998 | 16 | 11 | 10 |
| A3SS | 2238 | ENSG00000168970 | JMJD7-PLA2G4B | chr15 | + | 41834739 | 41835223 | 41834969 | 41835223 | 41828115 | 41828188 | 27 | 10 | 19 |
| A3SS | 11229 | ENSG00000108666 | C17orf75 | chr17 | - | 32339812 | 32339938 | 32339812 | 32339924 | 32341203 | 32341284 | 11 | 6 | 10 |
| A3SS | 1199 | ENSG00000120071 | KANSL1 | chr17 | - | 46032046 | 46032299 | 46032046 | 46032167 | 46033079 | 46033192 | 239 | 100 | 214 |
| A3SS | 12415 | ENSG00000233968 | NA | chr10 | - | 19722329 | 19722588 | 19722329 | 19722422 | 19728404 | 19728550 | 14 | 3 | 0 |
| A3SS | 6459 | ENSG00000066427 | ATXN3 | chr14 | - | 92096092 | 92096137 | 92096092 | 92096134 | 92106528 | 92106574 | 11 | 6 | 7 |
| A3SS | 1114 | ENSG00000004975 | DVL2 | chr17 | - | 7230284 | 7230797 | 7230284 | 7230430 | 7234068 | 7234517 | 32 | 20 | 42 |
| A3SS | 4928 | ENSG00000004399 | PLXND1 | chr3 | - | 129585951 | 129586081 | 129585951 | 129586048 | 129586171 | 129586272 | 17 | 5 | 9 |
| A3SS | 1710 | ENSG00000175595 | ERCC4 | chr16 | + | 13924529 | 13926756 | 13926560 | 13926756 | 13922030 | 13922211 | 7 | 6 | 5 |
| A3SS | 8144 | ENSG00000162227 | TAF6L | chr11 | + | 62776383 | 62776470 | 62776388 | 62776470 | 62775770 | 62775930 | 22 | 5 | 9 |
| A3SS | 6156 | ENSG00000000457 | SCYL3 | chr1 | - | 169868927 | 169869128 | 169868927 | 169869039 | 169873695 | 169873752 | 1 | 4 | 14 |
| A3SS | 12382 | ENSG00000167535 | CACNB3 | chr12 | + | 48826366 | 48826518 | 48826419 | 48826518 | 48825659 | 48825769 | 16 | 3 | 4 |
| A3SS | 3890 | ENSG00000166004 | CEP295 | chr11 | + | 93729610 | 93729781 | 93729613 | 93729781 | 93729433 | 93729530 | 2 | 2 | 2 |
| A3SS | 2441 | ENSG00000248508 | SRP14-AS1 | chr15 | + | 40064295 | 40064601 | 40064460 | 40064601 | 40060096 | 40060263 | 1 | 6 | 4 |
| A3SS | 2630 | ENSG00000092108 | SCFD1 | chr14 | + | 30634794 | 30635034 | 30634921 | 30635034 | 30633946 | 30634037 | 3 | 7 | 0 |
| A3SS | 11757 | ENSG00000142731 | PLK4 | chr4 | + | 127883261 | 127883357 | 127883265 | 127883357 | 127881830 | 127881926 | 4 | 3 | 8 |
| A3SS | 6166 | ENSG00000120334 | CENPL | chr1 | - | 173807266 | 173807518 | 173807266 | 173807310 | 173811131 | 173811306 | 12 | 10 | 18 |
| A3SS | 3141 | ENSG00000154134 | ROBO3 | chr11 | + | 124878583 | 124879341 | 124879189 | 124879341 | 124878297 | 124878436 | 15 | 5 | 9 |
| A3SS | 4339 | ENSG00000123737 | EXOSC9 | chr4 | + | 121813754 | 121814047 | 121813865 | 121814047 | 121811582 | 121811671 | 17 | 3 | 7 |
| A3SS | 2240 | ENSG00000168970 | JMJD7-PLA2G4B | chr15 | + | 41837067 | 41840230 | 41840024 | 41840230 | 41836474 | 41836551 | 35 | 15 | 55 |
| A3SS | 11113 | ENSG00000180900 | SCRIB | chr8 | - | 143792219 | 143792405 | 143792219 | 143792401 | 143792484 | 143792635 | 23 | 5 | 8 |
| A3SS | 12255 | ENSG00000162194 | LBHD1 | chr11 | - | 62669881 | 62670165 | 62669881 | 62670041 | 62671239 | 62671540 | 15 | 14 | 16 |
| A3SS | 4387 | ENSG00000176261 | ZBTB8OS | chr1 | - | 32633950 | 32634110 | 32633950 | 32634072 | 32634701 | 32634792 | 16 | 10 | 5 |
| A3SS | 11457 | ENSG00000132740 | IGHMBP2 | chr11 | + | 68935298 | 68935422 | 68935330 | 68935422 | 68934463 | 68934558 | 12 | 6 | 17 |
| A3SS | 10979 | ENSG00000246366 | LACTB2-AS1 | chr8 | + | 70652510 | 70652752 | 70652534 | 70652752 | 70651812 | 70651894 | 2 | 5 | 1 |
| A3SS | 6564 | ENSG00000007402 | CACNA2D2 | chr3 | - | 50364887 | 50364976 | 50364887 | 50364970 | 50365074 | 50365184 | 9 | 6 | 2 |
| A3SS | 218 | ENSG00000213793 | ZNF888 | chr19 | - | 52917858 | 52917997 | 52917858 | 52917931 | 52918818 | 52918937 | 7 | 3 | 7 |
| A3SS | 9699 | ENSG00000001631 | KRIT1 | chr7 | - | 92245426 | 92245595 | 92245426 | 92245574 | 92245789 | 92245920 | 12 | 13 | 11 |
| A3SS | 3459 | ENSG00000011243 | AKAP8L | chr19 | - | 15398629 | 15398766 | 15398629 | 15398731 | 15399301 | 15399410 | 13 | 10 | 13 |
| A3SS | 5886 | ENSG00000177311 | ZBTB38 | chr3 | + | 141438242 | 141438365 | 141438267 | 141438365 | 141403926 | 141404031 | 11 | 4 | 8 |
| A3SS | 11593 | ENSG00000213930 | GALT | chr9 | + | 34647540 | 34648171 | 34648047 | 34648171 | 34647088 | 34647258 | 18 | 8 | 18 |
| A3SS | 2042 | ENSG00000102977 | ACD | chr16 | - | 67657761 | 67657853 | 67657761 | 67657814 | 67657985 | 67658362 | 26 | 4 | 25 |
| A3SS | 9375 | ENSG00000112511 | PHF1 | chr6 | + | 33415589 | 33415670 | 33415592 | 33415670 | 33415260 | 33415329 | 13 | 12 | 22 |
| A3SS | 3319 | ENSG00000104731 | KLHDC4 | chr16 | - | 87755193 | 87755292 | 87755193 | 87755254 | 87756421 | 87756477 | 53 | 23 | 45 |
| A3SS | 12260 | ENSG00000167770 | OTUB1 | chr11 | + | 63997049 | 63997244 | 63997070 | 63997244 | 63996529 | 63996648 | 42 | 20 | 20 |
| A3SS | 3047 | ENSG00000109775 | UFSP2 | chr4 | - | 185405779 | 185405959 | 185405779 | 185405856 | 185407935 | 185408060 | 11 | 9 | 8 |
| A3SS | 4689 | ENSG00000147996 | CBWD5 | chr9 | + | 65732946 | 65733122 | 65732992 | 65733122 | 65731801 | 65731867 | 10 | 4 | 3 |

| | | | | | | | | | | | | | | | | | | | | | | | | | | | | | | | | | | | | | | | | | | | | | | |
|---|---|---|---|---|---|---|---|---|---|---|---|---|---|---|---|---|---|---|---|---|---|---|---|---|---|---|---|---|---|---|---|---|---|---|---|---|---|---|---|---|---|---|---|---|---|---|
| A3SS | 8121 | ENSG00000246090 | NA | chr4 | + | 99154544 | 99157579 | 99157490 | 99157579 | 99133554 | 99133805 | 0 | 3 | 4 | 3 | 11 | 1 | 0 | 1 | 5 | 2 | 0 | 0 | 1 | 0 | 0 | 0 | 1 | 0 | 0 | 0 | 298 | 149 | 1.000000 | 1 | 0 | 1 | 0.667 | 0.231 | 0.733 | NA | 0 | 1 | NA | NA | 0.026 |
| A3SS | 9379 | ENSG00000167930 | FAM234A | chr16 | + | 254380 | 254681 | 254387 | 254681 | 249548 | 249654 | 54 | 23 | 13 | 18 | 5 | 2 | 0 | 0 | 0 | 0 | 16 | 12 | 5 | 22 | 38 | 4 | 0 | 0 | 0 | 0 | 156 | 149 | 1.000000 | 1 | 0.963 | 1 | 1 | 1 | 1 | 0.793 | 1 | 1 | 1 | 1 | 0.034 |
| A3SS | 5184 | ENSG00000162819 | BROX | chr1 | + | 222724091 | 222724164 | 222724099 | 222724164 | 222719262 | 222719359 | 8 | 7 | 3 | 9 | 15 | 0 | 0 | 0 | 0 | 0 | 10 | 4 | 6 | 2 | 0 | 0 | 0 | 2 | 0 | 0 | 157 | 149 | 1.000000 | 1 | 1 | 1 | 1 | 1 | 1 | 1 | 1 | 0.74 | 1 | NA | 0.065 |
| A3SS | 11362 | ENSG00000203780 | FANK1 | chr10 | + | 126009053 | 126009592 | 126009372 | 126009592 | 126008406 | 126008550 | 4 | 4 | 2 | 8 | 3 | 0 | 0 | 0 | 0 | 0 | 0 | 0 | 1 | 0 | 0 | 0 | 0 | 0 | 0 | 2 | 298 | 149 | 1.000000 | 1 | 1 | 1 | 1 | 1 | 1 | NA | NA | 1 | NA | 0 | 0.5 |
| A3SS | 7408 | ENSG00000149231 | CCDC82 | chr11 | - | 96387529 | 96387653 | 96387529 | 96387613 | 96389843 | 96389905 | 10 | 5 | 7 | 3 | 3 | 13 | 4 | 9 | 19 | 0 | 0 | 0 | 4 | 3 | 0 | 0 | 0 | 1 | 0 | 0 | 189 | 149 | 1.000000 | 1 | 0.378 | 0.496 | 0.38 | 0.111 | 1 | NA | NA | 0.759 | 1 | NA | -0.406 |
| A3SS | 11405 | ENSG00000184465 | WDR27 | chr6 | - | 169667981 | 169668185 | 169667981 | 169668095 | 169670568 | 169670693 | 8 | 8 | 7 | 12 | 5 | 1 | 0 | 3 | 6 | 1 | 0 | 10 | 6 | 3 | 0 | 0 | 0 | 0 | 0 | 0 | 239 | 149 | 1.000000 | 1 | 0.833 | 1 | 0.593 | 0.555 | 0.757 | NA | 1 | 1 | 1 | NA | -0.252 |
| A3SS | 406 | ENSG00000090554 | FLT3LG | chr19 | + | 49485963 | 49486231 | 49486014 | 49486231 | 49476422 | 49476566 | 8 | 4 | 9 | 25 | 8 | 1 | 2 | 4 | 1 | 0 | 0 | 6 | 3 | 4 | 20 | 0 | 0 | 0 | 0 | 0 | 200 | 149 | 1.000000 | 1 | 0.856 | 0.598 | 0.626 | 0.949 | 1 | NA | 1 | 1 | 1 | 1 | -0.194 |
| A3SS | 6317 | ENSG00000161618 | ALDH16A1 | chr19 | + | 49461618 | 49462036 | 49461883 | 49462036 | 49460821 | 49460899 | 18 | 15 | 13 | 20 | 4 | 4 | 0 | 0 | 0 | 2 | 0 | 10 | 1 | 7 | 31 | 0 | 0 | 0 | 0 | 0 | 298 | 149 | 1.000000 | 1 | 0.692 | 1 | 1 | 1 | 0.5 | NA | 1 | 1 | 1 | 1 | -0.162 |
| A3SS | 8615 | ENSG00000095059 | DHPS | chr19 | - | 12676016 | 12676105 | 12676016 | 12676078 | 12677290 | 12677396 | 12 | 13 | 4 | 13 | 11 | 0 | 0 | 2 | 0 | 0 | 9 | 5 | 5 | 10 | 3 | 0 | 0 | 0 | 1 | 0 | 176 | 149 | 1.000000 | 1 | 1 | 1 | 0.629 | 1 | 1 | 1 | 1 | 1 | 0.894 | 1 | -0.053 |
| A3SS | 7434 | ENSG00000159363 | ATP13A2 | chr1 | - | 17004331 | 17004411 | 17004331 | 17004408 | 17004691 | 17004821 | 12 | 4 | 14 | 5 | 2 | 3 | 2 | 0 | 3 | 3 | 6 | 4 | 0 | 1 | 5 | 5 | 0 | 0 | 0 | 8 | 152 | 149 | 1.000000 | 1 | 0.797 | 0.662 | 1 | 0.62 | 0.395 | 0.541 | 1 | NA | 1 | 0.38 | -0.035 |
| A3SS | 10172 | ENSG00000160408 | ST6GALNAC6 | chr9 | - | 127885321 | 127886788 | 127885321 | 127886784 | 127887483 | 127887591 | 52 | 4 | 42 | 41 | 16 | 1 | 1 | 3 | 0 | 0 | 0 | 32 | 27 | 17 | 58 | 0 | 2 | 2 | 0 | 0 | 153 | 149 | 1.000000 | 1 | 0.981 | 0.796 | 0.932 | 1 | 1 | NA | 0.94 | 0.929 | 1 | 1 | -0.025 |
| A3SS | 9886 | ENSG00000126883 | NUP214 | chr9 | + | 131144567 | 131144754 | 131144576 | 131144754 | 131140548 | 131140710 | 5 | 11 | 13 | 8 | 30 | 0 | 0 | 1 | 0 | 0 | 4 | 2 | 11 | 8 | 21 | 0 | 0 | 0 | 0 | 0 | 158 | 149 | 1.000000 | 1 | 1 | 1 | 0.925 | 1 | 1 | 1 | 1 | 1 | 1 | 1 | -0.015 |
| A3SS | 4829 | ENSG00000215908 | CROCCP2 | chr1 | - | 16619844 | 16620245 | 16619844 | 16620038 | 16625774 | 16626005 | 7 | 0 | 30 | 44 | 10 | 0 | 0 | 2 | 2 | 0 | 0 | 12 | 17 | 15 | 26 | 0 | 1 | 0 | 0 | 0 | 298 | 149 | 1.000000 | 1 | 1 | NA | 0.882 | 0.917 | 1 | NA | 0.857 | 1 | 1 | 1 | -0.014 |
| A3SS | 3667 | ENSG00000166913 | YWHAB | chr20 | + | 44887544 | 44887742 | 44887647 | 44887742 | 44886878 | 44886978 | 33 | 28 | 35 | 35 | 38 | 0 | 1 | 1 | 1 | 0 | 7 | 44 | 51 | 31 | 15 | 0 | 4 | 0 | 0 | 0 | 252 | 149 | 1.000000 | 1 | 1 | 0.943 | 0.954 | 0.954 | 1 | 1 | 0.867 | 1 | 1 | 1 | -0.003 |
| A3SS | 2661 | ENSG00000089916 | GPATCH2L | chr14 | + | 76173545 | 76176690 | 76176622 | 76176690 | 76154353 | 76155025 | 8 | 22 | 16 | 15 | 14 | 0 | 0 | 0 | 0 | 1 | 8 | 3 | 40 | 3 | 0 | 0 | 0 | 2 | 0 | 0 | 298 | 149 | 1.000000 | 1 | 1 | 1 | 1 | 1 | 0.875 | 1 | 1 | 0.909 | 1 | NA | -0.002 |
| A3SS | 6927 | ENSG00000166938 | DIS3L | chr15 | + | 66306823 | 66306952 | 66306896 | 66306952 | 66294987 | 66295141 | 114 | 76 | 103 | 97 | 59 | 1 | 1 | 1 | 0 | 1 | 51 | 98 | 88 | 119 | 72 | 0 | 5 | 0 | 0 | 0 | 222 | 149 | 1.000000 | 1 | 0.987 | 0.981 | 0.986 | 1 | 0.975 | 1 | 0.929 | 1 | 1 | 1 | 0 |
| A3SS | 1343 | ENSG00000109083 | IFT20 | chr17 | - | 28328325 | 28328733 | 28328325 | 28328657 | 28329172 | 28329276 | 89 | 30 | 66 | 71 | 152 | 0 | 0 | 1 | 0 | 0 | 54 | 68 | 84 | 83 | 98 | 0 | 2 | 0 | 0 | 0 | 225 | 149 | 1.000000 | 1 | 1 | 1 | 0.978 | 1 | 1 | 1 | 0.957 | 1 | 1 | 1 | 0.004 |
| A3SS | 10789 | ENSG00000120699 | EXOSC8 | chr13 | + | 37004476 | 37004561 | 37004515 | 37004561 | 37002933 | 37003007 | 12 | 10 | 7 | 1 | 4 | 65 | 51 | 75 | 90 | 101 | 0 | 5 | 8 | 0 | 0 | 13 | 28 | 36 | 3 | 0 | 188 | 149 | 1.000000 | 1 | 0.128 | 0.135 | 0.069 | 0.009 | 0.03 | 0 | 0.124 | 0.15 | 0 | NA | 0.006 |
| A3SS | 2255 | ENSG00000137842 | TMEM62 | chr15 | + | 43149028 | 43149151 | 43149107 | 43149151 | 43148754 | 43148879 | 48 | 39 | 30 | 58 | 95 | 0 | 2 | 1 | 1 | 0 | 32 | 57 | 59 | 48 | 53 | 0 | 3 | 0 | 6 | 5 | 228 | 149 | 1.000000 | 1 | 1 | 0.927 | 0.951 | 0.974 | 1 | 1 | 0.925 | 1 | 0.839 | 0.874 | 0.043 |
| A3SS | 1980 | ENSG00000140987 | ZSCAN32 | chr16 | - | 3390009 | 3390183 | 3390009 | 3390133 | 3390422 | 3390517 | 5 | 0 | 6 | 4 | 7 | 3 | 5 | 11 | 9 | 5 | 0 | 0 | 1 | 0 | 0 | 0 | 1 | 7 | 0 | 2 | 199 | 149 | 1.000000 | 1 | 0.555 | 0 | 0.29 | 0.25 | 0.512 | NA | 0 | 0.097 | NA | 0 | 0.289 |
| A3SS | 8378 | ENSG00000115290 | GRB14 | chr2 | - | 164508454 | 164508603 | 164508454 | 164508550 | 164508741 | 164508852 | 0 | 4 | 1 | 8 | 15 | 0 | 2 | 1 | 5 | 4 | 0 | 0 | 7 | 0 | 0 | 0 | 3 | 1 | 5 | 0 | 202 | 149 | 1.000000 | 1 | NA | 0.596 | 0.425 | 0.541 | 0.734 | NA | 0 | 0.838 | 0 | NA | 0.295 |
| A3SS | 8878 | ENSG00000123607 | TTC21B | chr2 | - | 165876164 | 165879967 | 165876164 | 165876232 | 165880678 | 165880799 | 6 | 5 | 4 | 8 | 9 | 24 | 11 | 26 | 23 | 22 | 0 | 1 | 6 | 3 | 0 | 0 | 5 | 5 | 1 | 0 | 298 | 149 | 1.000000 | 1 | 0.111 | 0.185 | 0.071 | 0.148 | 0.17 | NA | 0.091 | 0.375 | 0.6 | NA | -0.218 |
| A3SS | 1353 | ENSG00000153944 | MSI2 | chr17 | + | 57674917 | 57675126 | 57674971 | 57675126 | 57652098 | 57652161 | 9 | 4 | 15 | 17 | 0 | 22 | 8 | 16 | 29 | 9 | 0 | 6 | 9 | 2 | 6 | 0 | 2 | 13 | 14 | 8 | 203 | 149 | 1.000000 | 1 | 0.231 | 0.268 | 0.408 | 0.301 | 0 | NA | 0.688 | 0.337 | 0.095 | 0.355 | -0.127 |
| A3SS | 6448 | ENSG00000081014 | AP4E1 | chr15 | + | 50934535 | 50934697 | 50934623 | 50934697 | 50930804 | 50930971 | 6 | 12 | 6 | 2 | 7 | 40 | 36 | 46 | 42 | 83 | 10 | 0 | 1 | 0 | 0 | 2 | 54 | 35 | 20 | 0 | 237 | 149 | 1.000000 | 1 | 0.086 | 0.173 | 0.076 | 0.029 | 0.05 | 0.759 | 0 | 0.018 | 0 | NA | -0.111 |
| A3SS | 11921 | ENSG00000139697 | SBNO1 | chr12 | - | 123345257 | 123345570 | 123345257 | 123345567 | 123350309 | 123350441 | 21 | 7 | 7 | 8 | 22 | 2 | 4 | 0 | 0 | 0 | 9 | 11 | 9 | 14 | 0 | 0 | 0 | 1 | 0 | 0 | 152 | 149 | 1.000000 | 1 | 0.911 | 0.632 | 1 | 1 | 1 | 1 | 1 | 0.898 | 1 | NA | -0.066 |
| A3SS | 9258 | ENSG00000136044 | APPL2 | chr12 | - | 105207970 | 105208047 | 105207970 | 105208001 | 105208157 | 105208199 | 9 | 2 | 3 | 8 | 13 | 0 | 0 | 1 | 0 | 0 | 3 | 0 | 9 | 1 | 0 | 0 | 0 | 0 | 0 | 0 | 195 | 149 | 1.000000 | 1 | 1 | 1 | 0.696 | 1 | 1 | 1 | NA | 1 | 1 | NA | -0.061 |
| A3SS | 7578 | ENSG00000171503 | ETFDH | chr4 | + | 158680466 | 158681614 | 158681570 | 158681614 | 158680123 | 158680182 | 18 | 5 | 6 | 7 | 40 | 0 | 0 | 1 | 0 | 0 | 0 | 15 | 25 | 7 | 7 | 0 | 0 | 0 | 0 | 0 | 298 | 149 | 1.000000 | 1 | 1 | 1 | 0.75 | 1 | 1 | NA | 1 | 1 | 1 | 1 | -0.05 |
| A3SS | 3425 | ENSG00000155329 | ZCCHC10 | chr5 | - | 133000131 | 133000201 | 133000131 | 133000173 | 133006758 | 133006920 | 7 | 1 | 5 | 10 | 7 | 65 | 46 | 40 | 43 | 100 | 0 | 3 | 5 | 0 | 0 | 10 | 29 | 8 | 5 | 0 | 177 | 149 | 1.000000 | 1 | 0.083 | 0.018 | 0.095 | 0.164 | 0.056 | 0 | 0.08 | 0.345 | 0 | NA | -0.023 |
| A3SS | 2285 | ENSG00000102901 | CENPT | chr16 | - | 67828473 | 67828605 | 67828473 | 67828559 | 67828666 | 67828843 | 18 | 6 | 8 | 12 | 23 | 1 | 0 | 1 | 1 | 0 | 0 | 8 | 9 | 3 | 25 | 0 | 0 | 0 | 0 | 6 | 195 | 149 | 1.000000 | 1 | 0.932 | 1 | 0.859 | 0.902 | 1 | NA | 1 | 1 | 1 | 0.761 | -0.002 |
| A3SS | 5161 | ENSG00000124406 | ATP8A1 | chr4 | - | 42455296 | 42455433 | 42455296 | 42455419 | 42455524 | 42455599 | 3 | 8 | 10 | 2 | 3 | 79 | 85 | 100 | 93 | 150 | 0 | 0 | 2 | 2 | 0 | 17 | 73 | 36 | 12 | 0 | 163 | 149 | 1.000000 | 1 | 0.034 | 0.079 | 0.084 | 0.019 | 0.018 | 0 | 0 | 0.048 | 0.132 | NA | 0.002 |
| A3SS | 11961 | ENSG00000160226 | CFAP410 | chr21 | - | 44330822 | 44330919 | 44330822 | 44330916 | 44331842 | 44332014 | 8 | 3 | 10 | 9 | 0 | 4 | 3 | 5 | 11 | 8 | 0 | 0 | 5 | 3 | 0 | 0 | 4 | 3 | 0 | 4 | 152 | 149 | 1.000000 | 1 | 0.662 | 0.495 | 0.662 | 0.445 | 0 | NA | 0 | 0.62 | 1 | 0 | 0.048 |
| A3SS | 2673 | ENSG00000100567 | PSMA3 | chr14 | + | 58260958 | 58261020 | 58260968 | 58261020 | 58257924 | 58257998 | 12 | 6 | 13 | 7 | 6 | 2 | 5 | 3 | 0 | 1 | 0 | 6 | 16 | 0 | 0 | 0 | 0 | 0 | 5 | 0 | 159 | 149 | 1.000000 | 1 | 0.849 | 0.529 | 0.802 | 1 | 0.849 | NA | 1 | 1 | 0 | NA | 0.139 |
| A3SS | 12375 | ENSG00000162241 | SLC25A45 | chr11 | - | 65380533 | 65380632 | 65380533 | 65380622 | 65381914 | 65382373 | 0 | 7 | 1 | 13 | 1 | 4 | 0 | 0 | 0 | 6 | 0 | 0 | 0 | 0 | 0 | 0 | 0 | 1 | 0 | 0 | 159 | 149 | 1.000000 | 1 | 0 | 1 | 1 | 1 | 0.135 | NA | NA | 0 | NA | NA | 0.627 |
| A3SS | 10313 | ENSG00000230590 | FTX | chrX | - | 74202650 | 74203953 | 74202650 | 74203909 | 74274341 | 74274410 | 10 | 14 | 20 | 9 | 1 | 8 | 5 | 5 | 11 | 15 | 0 | 24 | 6 | 1 | 0 | 0 | 1 | 0 | 0 | 6 | 193 | 149 | 1.000000 | 1 | 0.491 | 0.684 | 0.755 | 0.387 | 0.049 | NA | 0.949 | 1 | 1 | 0 | -0.264 |
| A3SS | 5651 | ENSG00000183137 | CEP57L1 | chr6 | + | 109150117 | 109150239 | 109150165 | 109150239 | 109146757 | 109146937 | 16 | 12 | 15 | 24 | 27 | 0 | 2 | 6 | 5 | 3 | 0 | 28 | 18 | 13 | 12 | 0 | 0 | 0 | 2 | 0 | 197 | 149 | 1.000000 | 1 | 1 | 0.819 | 0.654 | 0.784 | 0.872 | NA | 1 | 1 | 0.831 | 1 | -0.132 |
| A3SS | 9829 | ENSG00000142303 | ADAMTS10 | chr19 | - | 8589451 | 8589585 | 8589451 | 8589574 | 8589888 | 8589991 | 11 | 6 | 10 | 6 | 1 | 1 | 2 | 3 | 0 | 0 | 0 | 0 | 4 | 1 | 6 | 0 | 0 | 0 | 0 | 0 | 160 | 149 | 1.000000 | 1 | 0.911 | 0.736 | 0.756 | 1 | 1 | NA | NA | 1 | 1 | 1 | -0.119 |
| A3SS | 11038 | ENSG00000249859 | PVT1 | chr8 | + | 128096517 | 128096654 | 128096520 | 128096654 | 128010317 | 128010444 | 5 | 5 | 12 | 12 | 15 | 3 | 0 | 0 | 0 | 2 | 10 | 4 | 9 | 3 | 0 | 0 | 0 | 0 | 0 | 0 | 152 | 149 | 1.000000 | 1 | 0.62 | 1 | 1 | 1 | 0.88 | 1 | 1 | 1 | 1 | NA | -0.1 |
| A3SS | 10151 | ENSG00000196305 | IARS1 | chr9 | - | 92253361 | 92253453 | 92253361 | 92253446 | 92255534 | 92255604 | 9 | 8 | 9 | 3 | 9 | 0 | 0 | 0 | 0 | 5 | 1 | 3 | 6 | 5 | 0 | 0 | 0 | 0 | 0 | 0 | 156 | 149 | 1.000000 | 1 | 1 | 1 | 1 | 1 | 0.632 | 1 | 1 | 1 | 1 | NA | -0.074 |
| A3SS | 8422 | ENSG00000223960 | CHROMR | chr2 | + | 178430694 | 178430889 | 178430769 | 178430889 | 178426365 | 178426680 | 22 | 18 | 17 | 17 | 22 | 6 | 4 | 3 | 1 | 8 | 38 | 8 | 7 | 20 | 0 | 0 | 7 | 0 | 2 | 0 | 224 | 149 | 1.000000 | 1 | 0.709 | 0.75 | 0.79 | 0.919 | 0.647 | 1 | 0.432 | 1 | 0.869 | NA | -0.062 |
| A3SS | 3122 | ENSG00000131844 | MCCC2 | chr5 | + | 71643770 | 71643895 | 71643818 | 71643895 | 71641002 | 71641083 | 11 | 4 | 9 | 5 | 23 | 0 | 0 | 3 | 0 | 0 | 0 | 15 | 4 | 5 | 5 | 0 | 0 | 0 | 0 | 0 | 197 | 149 | 1.000000 | 1 | 1 | 1 | 0.694 | 1 | 1 | NA | 1 | 1 | 1 | 1 | -0.061 |
| A3SS | 6571 | ENSG00000135541 | AHI1 | chr6 | - | 135301407 | 135302841 | 135301407 | 135301539 | 135318518 | 135318616 | 6 | 14 | 1 | 7 | 4 | 1 | 0 | 0 | 0 | 0 | 0 | 6 | 3 | 0 | 0 | 0 | 0 | 0 | 0 | 0 | 298 | 149 | 1.000000 | 1 | 0.75 | 1 | 1 | 1 | 1 | NA | 1 | 1 | NA | NA | -0.05 |
| A3SS | 9714 | ENSG00000146918 | NCAPG2 | chr7 | - | 158655117 | 158655455 | 158655117 | 158655258 | 158656259 | 158656433 | 14 | 14 | 22 | 18 | 23 | 1 | 0 | 0 | 0 | 0 | 9 | 11 | 18 | 30 | 0 | 0 | 0 | 0 | 0 | 0 | 298 | 149 | 1.000000 | 1 | 0.875 | 1 | 1 | 1 | 1 | 1 | 1 | 1 | 1 | NA | -0.025 |
| A3SS | 4389 | ENSG00000176261 | ZBTB8OS | chr1 | - | 32633950 | 32634110 | 32633950 | 32634072 | 32650401 | 32650525 | 13 | 11 | 7 | 7 | 20 | 1 | 0 | 0 | 0 | 0 | 0 | 2 | 3 | 13 | 17 | 0 | 0 | 0 | 0 | 0 | 187 | 149 | 1.000000 | 1 | 0.912 | 1 | 1 | 1 | 1 | NA | 1 | 1 | 1 | 1 | -0.018 |
| A3SS | 11094 | ENSG00000185728 | YTHDF3 | chr8 | + | 63175330 | 63175416 | 63175336 | 63175416 | 63172652 | 63172798 | 15 | 10 | 15 | 20 | 26 | 0 | 0 | 0 | 0 | 2 | 0 | 21 | 13 | 28 | 1 | 0 | 0 | 0 | 0 | 0 | 155 | 149 | 1.000000 | 1 | 1 | 1 | 1 | 1 | 0.926 | NA | 1 | 1 | 1 | 1 | -0.015 |
| A3SS | 13106 | ENSG00000090686 | USP48 | chr1 | - | 21729703 | 21729832 | 21729703 | 21729829 | 21736429 | 21736625 | 17 | 8 | 13 | 12 | 24 | 1 | 2 | 2 | 0 | 1 | 0 | 14 | 14 | 18 | 5 | 0 | 6 | 1 | 0 | 0 | 152 | 149 | 1.000000 | 1 | 0.943 | 0.797 | 0.864 | 1 | 0.959 | NA | 0.696 | 0.932 | 1 | 1 | 0.006 |
| A3SS | 9195 | ENSG00000135678 | CPM | chr12 | - | 68885791 | 68885889 | 68885791 | 68885885 | 68932677 | 68932840 | 18 | 4 | 4 | 14 | 15 | 0 | 0 | 0 | 1 | 0 | 0 | 1 | 11 | 14 | 6 | 0 | 0 | 1 | 0 | 0 | 153 | 149 | 1.000000 | 1 | 1 | 1 | 1 | 0.932 | 1 | NA | 1 | 0.915 | 1 | 1 | 0.008 |
| A3SS | 8123 | ENSG00000165861 | ZFYVE1 | chr14 | - | 72977926 | 72978044 | 72977926 | 72978002 | 72978136 | 72978234 | 33 | 17 | 22 | 5 | 17 | 0 | 0 | 1 | 0 | 1 | 14 | 28 | 5 | 12 | 12 | 0 | 0 | 0 | 0 | 2 | 191 | 149 | 1.000000 | 1 | 1 | 1 | 0.945 | 1 | 0.93 | 1 | 1 | 1 | 1 | 0.824 | 0.01 |
| A3SS | 1471 | ENSG00000169738 | DCXR | chr17 | - | 82036190 | 82036448 | 82036190 | 82036308 | 82036543 | 82036643 | 74 | 43 | 69 | 52 | 33 | 0 | 0 | 0 | 1 | 0 | 70 | 28 | 41 | 33 | 76 | 0 | 1 | 1 | 0 | 0 | 289 | 149 | 1.000000 | 1 | 1 | 1 | 1 | 0.964 | 1 | 1 | 0.935 | 0.955 | 1 | 1 | 0.015 |
| A3SS | 3071 | ENSG00000205133 | TRIQK | chr8 | - | 92916928 | 92917010 | 92916928 | 92917007 | 92921502 | 92921660 | 7 | 6 | 4 | 13 | 12 | 0 | 3 | 2 | 4 | 10 | 0 | 10 | 1 | 3 | 5 | 0 | 0 | 4 | 4 | 0 | 152 | 149 | 1.000000 | 1 | 1 | 0.662 | 0.662 | 0.761 | 0.541 | NA | 1 | 0.197 | 0.424 | 1 | 0.07 |
| A3SS | 10653 | ENSG00000116754 | SRSF11 | chr1 | + | 70249930 | 70250047 | 70249951 | 70250047 | 70247116 | 70247149 | 2 | 6 | 12 | 4 | 7 | 19 | 17 | 35 | 21 | 9 | 6 | 0 | 2 | 0 | 0 | 6 | 8 | 17 | 2 | 0 | 170 | 149 | 1.000000 | 1 | 0.084 | 0.236 | 0.231 | 0.143 | 0.405 | 0.467 | 0 | 0.093 | 0 | NA | 0.08 |
| A3SS | 12386 | ENSG00000150433 | TMEM218 | chr11 | - | 125101200 | 125101303 | 125101200 | 125101256 | 125102733 | 125102809 | 14 | 9 | 12 | 19 | 18 | 0 | 0 | 0 | 0 | 0 | 7 | 13 | 3 | 18 | 8 | 0 | 0 | 1 | 3 | 0 | 196 | 149 | 1.000000 | 1 | 1 | 1 | 1 | 1 | 1 | 1 | 1 | 0.695 | 0.82 | 1 | 0.097 |
| A3SS | 8723 | ENSG00000164114 | MAP9 | chr4 | - | 155375775 | 155375914 | 155375775 | 155375870 | 155376770 | 155376813 | 3 | 4 | 4 | 9 | 18 | 0 | 0 | 0 | 0 | 0 | 0 | 2 | 4 | 9 | 0 | 0 | 1 | 0 | 0 | 0 | 193 | 149 | 1.000000 | 1 | 1 | 1 | 1 | 1 | 1 | NA | 0.607 | 1 | 1 | NA | 0.131 |
| A3SS | 8367 | ENSG00000107643 | MAPK8 | chr10 | + | 48434878 | 48439357 | 48434883 | 48439357 | 48431192 | 48431270 | 8 | 5 | 5 | 9 | 14 | 1 | 1 | 2 | 1 | 0 | 0 | 6 | 2 | 10 | 0 | 0 | 4 | 1 | 1 | 0 | 154 | 149 | 1.000000 | 1 | 0.886 | 0.829 | 0.708 | 0.897 | 1 | NA | 0.592 | 0.659 | 0.906 | NA | 0.145 |
| A3SS | 5143 | ENSG00000064687 | ABCA7 | chr19 | + | 1056906 | 1057429 | 1057313 | 1057429 | 1056329 | 1056499 | 36 | 15 | 46 | 48 | 28 | 1 | 1 | 0 | 2 | 0 | 32 | 25 | 33 | 26 | 34 | 0 | 9 | 5 | 8 | 0 | 298 | 149 | 1.000000 | 1 | 0.947 | 0.882 | 1 | 0.923 | 1 | 1 | 0.581 | 0.767 | 0.619 | 1 | 0.157 |
| A3SS | 5613 | ENSG00000072210 | ALDH3A2 | chr17 | + | 19648934 | 19649124 | 19648962 | 19649124 | 19648173 | 19648265 | 1 | 7 | 6 | 6 | 7 | 0 | 0 | 0 | 0 | 0 | 0 | 1 | 1 | 1 | 1 | 0 | 0 | 0 | 0 | 4 | 177 | 149 | 1.000000 | 1 | 1 | 1 | 1 | 1 | 1 | NA | 1 | 1 | 1 | 0.174 | 0.207 |
| A3SS | 8504 | ENSG00000138036 | DYNC2LI1 | chr2 | + | 43795886 | 43795958 | 43795889 | 43795958 | 43794456 | 43794643 | 2 | 4 | 4 | 4 | 15 | 3 | 2 | 5 | 4 | 20 | 0 | 0 | 0 | 6 | 0 | 0 | 2 | 7 | 4 | 0 | 152 | 149 | 1.000000 | 1 | 0.395 | 0.662 | 0.44 | 0.495 | 0.424 | NA | 0 | 0 | 0.595 | NA | 0.285 |
| A3SS | 7018 | ENSG00000122678 | POLM | chr7 | - | 44074397 | 44074530 | 44074397 | 44074522 | 44078755 | 44078811 | 6 | 5 | 4 | 5 | 10 | 0 | 0 | 0 | 0 | 0 | 0 | 6 | 1 | 0 | 0 | 6 | 0 | 0 | 0 | 0 | 157 | 149 | 1.000000 | 1 | 1 | 1 | 1 | 1 | 1 | 0 | 1 | 1 | NA | NA | 0.333 |
| A3SS | 1061 | ENSG00000101639 | CEP192 | chr18 | + | 13116373 | 13116503 | 13116376 | 13116503 | 13113585 | 13113705 | 1 | 5 | 13 | 7 | 2 | 1 | 0 | 3 | 0 | 3 | 0 | 1 | 4 | 0 | 0 | 0 | 4 | 2 | 3 | 26 | 152 | 149 | 1.000000 | 1 | 0.495 | 1 | 0.809 | 1 | 0.395 | NA | 0.197 | 0.662 | 0 | 0 | 0.525 |
| A3SS | 11129 | ENSG00000104408 | EIF3E | chr8 | - | 108239957 | 108240098 | 108239957 | 108240075 | 108241885 | 108241913 | 6 | 5 | 8 | 16 | 4 | 2 | 3 | 4 | 1 | 3 | 0 | 0 | 8 | 7 | 0 | 0 | 0 | 0 | 0 | 0 | 172 | 149 | 1.000000 | 1 | 0.722 | 0.591 | 0.634 | 0.933 | 0.536 | NA | NA | 1 | 1 | NA | -0.317 |
| A3SS | 2756 | ENSG00000197045 | GMFB | chr14 | - | 54480873 | 54480959 | 54480873 | 54480956 | 54481408 | 54481458 | 7 | 15 | 9 | 2 | 6 | 45 | 45 | 44 | 37 | 65 | 0 | 7 | 0 | 8 | 0 | 0 | 2 | 10 | 6 | 0 | 152 | 149 | 1.000000 | 1 | 0.132 | 0.246 | 0.167 | 0.05 | 0.083 | NA | 0.774 | 0 | 0.567 | NA | -0.311 |
| A3SS | 1531 | ENSG00000108848 | LUC7L3 | chr17 | + | 50738136 | 50738199 | 50738140 | 50738199 | 50736959 | 50737026 | 9 | 10 | 0 | 2 | 7 | 0 | 0 | 0 | 2 | 0 | 0 | 2 | 0 | 2 | 0 | 0 | 0 | 0 | 0 | 0 | 153 | 149 | 1.000000 | 1 | 1 | 1 | NA | 0.493 | 1 | NA | 1 | NA | 1 | NA | -0.127 |
| A3SS | 6169 | ENSG00000238113 | LINC01410 | chr9 | + | 62811399 | 62811589 | 62811510 | 62811589 | 62803983 | 62804140 | 18 | 18 | 4 | 7 | 17 | 0 | 0 | 0 | 2 | 0 | 0 | 8 | 3 | 13 | 16 | 0 | 0 | 0 | 0 | 0 | 260 | 149 | 1.000000 | 1 | 1 | 1 | 1 | 0.667 | 1 | NA | 1 | 1 | 1 | 1 | -0.067 |
| A3SS | 8876 | ENSG00000239665 | NA | chr10 | + | 13654183 | 13654427 | 13654352 | 13654427 | 13652407 | 13652489 | 9 | 9 | 10 | 11 | 2 | 0 | 1 | 0 | 0 | 0 | 0 | 8 | 0 | 9 | 0 | 0 | 0 | 0 | 0 | 0 | 298 | 149 | 1.000000 | 1 | 1 | 0.818 | 1 | 1 | 1 | NA | 1 | NA | 1 | NA | -0.036 |
| A3SS | 12383 | ENSG00000150433 | TMEM218 | chr11 | - | 125101200 | 125101303 | 125101200 | 125101256 | 125102131 | 125102317 | 12 | 10 | 12 | 21 | 20 | 0 | 0 | 1 | 0 | 1 | 10 | 2 | 5 | 5 | 29 | 0 | 0 | 0 | 0 | 0 | 196 | 149 | 1.000000 | 1 | 1 | 1 | 0.901 | 1 | 0.938 | 1 | 1 | 1 | 1 | 1 | -0.032 |
| A3SS | 8877 | ENSG00000239665 | NA | chr10 | + | 13654183 | 13654427 | 13654352 | 13654427 | 13652845 | 13652882 | 9 | 9 | 10 | 11 | 2 | 0 | 0 | 0 | 1 | 0 | 0 | 8 | 0 | 9 | 0 | 0 | 0 | 0 | 0 | 0 | 298 | 149 | 1.000000 | 1 | 1 | 1 | 1 | 0.846 | 1 | NA | 1 | NA | 1 | NA | -0.031 |
| A3SS | 4151 | ENSG00000163703 | CRELD1 | chr3 | + | 9938014 | 9938106 | 9938018 | 9938106 | 9937561 | 9937672 | 26 | 14 | 44 | 29 | 18 | 1 | 0 | 0 | 2 | 0 | 25 | 23 | 17 | 14 | 28 | 0 | 0 | 0 | 0 | 0 | 153 | 149 | 1.000000 | 1 | 0.962 | 1 | 1 | 0.934 | 1 | 1 | 1 | 1 | 1 | 1 | -0.021 |
| A3SS | 5190 | ENSG00000143498 | TAF1A | chr1 | - | 222579758 | 222579890 | 222579758 | 222579872 | 222584127 | 222584297 | 5 | 3 | 5 | 7 | 12 | 16 | 17 | 13 | 16 | 25 | 8 | 0 | 0 | 0 | 0 | 0 | 2 | 2 | 10 | 0 | 167 | 149 | 1.000000 | 1 | 0.218 | 0.136 | 0.255 | 0.281 | 0.3 | 1 | 0 | 0 | 0 | NA | -0.012 |
| A3SS | 11750 | ENSG00000132017 | DCAF15 | chr19 | + | 13959581 | 13959673 | 13959583 | 13959673 | 13959044 | 13959479 | 48 | 25 | 42 | 21 | 46 | 3 | 0 | 0 | 0 | 0 | 18 | 38 | 26 | 31 | 45 | 0 | 0 | 0 | 0 | 0 | 151 | 149 | 1.000000 | 1 | 0.94 | 1 | 1 | 1 | 1 | 1 | 1 | 1 | 1 | 1 | -0.012 |
| A3SS | 12318 | ENSG00000149582 | TMEM25 | chr11 | + | 118534265 | 118534355 | 118534268 | 118534355 | 118534028 | 118534129 | 5 | 3 | 18 | 3 | 9 | 0 | 0 | 1 | 0 | 0 | 0 | 3 | 11 | 0 | 0 | 0 | 0 | 0 | 0 | 0 | 152 | 149 | 1.000000 | 1 | 1 | 1 | 0.946 | 1 | 1 | NA | 1 | 1 | NA | NA | -0.011 |
| A3SS | 7803 | ENSG00000115966 | ATF2 | chr2 | - | 175074504 | 175074867 | 175074504 | 175074835 | 175080659 | 175080765 | 2 | 4 | 15 | 2 | 2 | 76 | 43 | 90 | 92 | 126 | 0 | 1 | 0 | 0 | 0 | 0 | 38 | 22 | 5 | 0 | 181 | 149 | 1.000000 | 1 | 0.021 | 0.071 | 0.121 | 0.018 | 0.013 | NA | 0.021 | 0 | 0 | NA | 0.042 |
| A3SS | 7052 | ENSG00000108523 | RNF167 | chr17 | + | 4940872 | 4940993 | 4940922 | 4940993 | 4940486 | 4940726 | 21 | 8 | 20 | 7 | 22 | 11 | 2 | 6 | 4 | 13 | 0 | 24 | 9 | 21 | 0 | 9 | 16 | 2 | 0 | 0 | 199 | 149 | 1.000000 | 1 | 0.588 | 0.75 | 0.714 | 0.567 | 0.559 | 0 | 0.529 | 0.771 | 1 | NA | 0.061 |
| A3SS | 3566 | ENSG00000113593 | PPWD1 | chr5 | + | 65569631 | 65569732 | 65569636 | 65569732 | 65568917 | 65568994 | 9 | 2 | 5 | 12 | 5 | 0 | 0 | 0 | 0 | 0 | 0 | 3 | 1 | 5 | 0 | 0 | 0 | 0 | 2 | 0 | 154 | 149 | 1.000000 | 1 | 1 | 1 | 1 | 1 | 1 | NA | 1 | 1 | 0.708 | NA | 0.097 |
| A3SS | 215 | ENSG00000197937 | ZNF347 | chr19 | - | 53148680 | 53148812 | 53148680 | 53148809 | 53149240 | 53149367 | 3 | 6 | 0 | 16 | 2 | 9 | 9 | 8 | 14 | 4 | 0 | 0 | 3 | 0 | 0 | 0 | 3 | 1 | 5 | 8 | 152 | 149 | 1.000000 | 1 | 0.246 | 0.395 | 0 | 0.528 | 0.329 | NA | 0 | 0.746 | 0 | 0 | 0.113 |
| A3SS | 7201 | ENSG00000114354 | TFG | chr3 | + | 100747360 | 100747454 | 100747413 | 100747454 | 100744832 | 100744931 | 2 | 4 | 14 | 8 | 3 | 0 | 0 | 0 | 1 | 0 | 0 | 2 | 0 | 5 | 0 | 0 | 0 | 0 | 2 | 0 | 202 | 149 | 1.000000 | 1 | 1 | 1 | 1 | 0.855 | 1 | NA | 1 | NA | 0.648 | NA | 0.147 |
| A3SS | 5889 | ENSG00000138386 | NAB1 | chr2 | + | 190685475 | 190685638 | 190685478 | 190685638 | 190673100 | 190673152 | 7 | 3 | 1 | 3 | 12 | 0 | 2 | 0 | 0 | 1 | 0 | 0 | 0 | 0 | 2 | 0 | 0 | 4 | 0 | 0 | 152 | 149 | 1.000000 | 1 | 1 | 0.595 | 1 | 1 | 0.922 | NA | NA | 0 | NA | 1 | 0.403 |
| A3SS | 8206 | ENSG00000285417 | NA | chr1 | + | 206051741 | 206052001 | 206051852 | 206052001 | 206035252 | 206035299 | 17 | 5 | 7 | 15 | 12 | 5 | 9 | 4 | 0 | 0 | 1 | 14 | 1 | 15 | 0 | 0 | 0 | 0 | 0 | 0 | 260 | 149 | 1.000000 | 1 | 0.661 | 0.241 | 0.501 | 1 | 1 | 1 | 1 | 1 | 1 | NA | -0.319 |
| A3SS | 5392 | ENSG00000116353 | MECR | chr1 | - | 29216587 | 29216812 | 29216587 | 29216685 | 29220833 | 29220921 | 11 | 4 | 8 | 11 | 15 | 0 | 1 | 1 | 0 | 3 | 0 | 10 | 0 | 9 | 5 | 0 | 0 | 0 | 0 | 0 | 276 | 149 | 1.000000 | 1 | 1 | 0.683 | 0.812 | 1 | 0.73 | NA | 1 | NA | 1 | 1 | -0.155 |
| A3SS | 4398 | ENSG00000105982 | RNF32 | chr7 | + | 156676294 | 156677130 | 156676418 | 156677130 | 156675695 | 156675863 | 18 | 11 | 19 | 33 | 23 | 1 | 0 | 0 | 0 | 3 | 7 | 27 | 30 | 11 | 4 | 0 | 0 | 0 | 0 | 0 | 273 | 149 | 1.000000 | 1 | 0.908 | 1 | 1 | 1 | 0.807 | 1 | 1 | 1 | 1 | 1 | -0.057 |
| A3SS | 8572 | ENSG00000104218 | CSPP1 | chr8 | + | 67158990 | 67159137 | 67158995 | 67159137 | 67158446 | 67158596 | 24 | 19 | 26 | 25 | 11 | 0 | 0 | 1 | 0 | 0 | 31 | 20 | 18 | 11 | 0 | 0 | 0 | 0 | 0 | 0 | 154 | 149 | 1.000000 | 1 | 1 | 1 | 0.962 | 1 | 1 | 1 | 1 | 1 | 1 | NA | -0.008 |
| A3SS | 11068 | ENSG00000100023 | PPIL2 | chr22 | + | 21695393 | 21695496 | 21695397 | 21695496 | 21694936 | 21695070 | 41 | 25 | 20 | 21 | 6 | 1 | 0 | 0 | 0 | 0 | 38 | 11 | 7 | 8 | 24 | 0 | 0 | 0 | 0 | 0 | 153 | 149 | 1.000000 | 1 | 0.976 | 1 | 1 | 1 | 1 | 1 | 1 | 1 | 1 | 1 | -0.005 |
| A3SS | 6569 | ENSG00000135541 | AHI1 | chr6 | - | 135300499 | 135302817 | 135300499 | 135300558 | 135318518 | 135318616 | 6 | 7 | 6 | 4 | 5 | 3 | 1 | 0 | 11 | 2 | 3 | 0 | 0 | 0 | 0 | 0 | 0 | 0 | 0 | 5 | 298 | 149 | 1.000000 | 1 | 0.5 | 0.778 | 1 | 0.154 | 0.556 | 1 | NA | NA | NA | 0 | 0.098 |
| A3SS | 2662 | ENSG00000089916 | GPATCH2L | chr14 | + | 76173545 | 76176690 | 76176622 | 76176690 | 76166662 | 76166727 | 9 | 22 | 16 | 15 | 14 | 0 | 0 | 0 | 0 | 0 | 8 | 2 | 40 | 1 | 0 | 0 | 0 | 0 | 3 | 0 | 298 | 149 | 1.000000 | 1 | 1 | 1 | 1 | 1 | 1 | 1 | 1 | 1 | 0.143 | NA | 0.214 |
| A3SS | 7802 | ENSG00000115966 | ATF2 | chr2 | - | 175074504 | 175074867 | 175074504 | 175074835 | 175074999 | 175075087 | 2 | 4 | 15 | 2 | 2 | 4 | 7 | 4 | 3 | 11 | 0 | 0 | 0 | 0 | 0 | 0 | 2 | 1 | 0 | 0 | 181 | 149 | 1.000000 | 1 | 0.292 | 0.32 | 0.755 | 0.354 | 0.13 | NA | 0 | 0 | NA | NA | 0.37 |
| A3SS | 1457 | ENSG00000181045 | SLC26A11 | chr17 | + | 80220915 | 80221115 | 80220918 | 80221115 | 80220437 | 80220496 | 6 | 3 | 6 | 7 | 3 | 0 | 1 | 0 | 0 | 0 | 0 | 0 | 0 | 0 | 0 | 0 | 6 | 0 | 0 | 0 | 152 | 149 | 1.000000 | 1 | 1 | 0.746 | 1 | 1 | 1 | NA | 0 | NA | NA | NA | 0.949 |
| A3SS | 11062 | ENSG00000164953 | TMEM67 | chr8 | + | 93785732 | 93786346 | 93786267 | 93786346 | 93785221 | 93785378 | 1 | 8 | 13 | 13 | 0 | 1 | 0 | 0 | 0 | 0 | 0 | 3 | 4 | 2 | 0 | 0 | 0 | 0 | 0 | 0 | 298 | 149 | 1.000000 | 1 | 0.333 | 1 | 1 | 1 | NA | NA | 1 | 1 | 1 | NA | -0.167 |
| A3SS | 2766 | ENSG00000100601 | ALKBH1 | chr14 | - | 77675655 | 77675849 | 77675655 | 77675794 | 77679879 | 77679970 | 21 | 10 | 16 | 14 | 13 | 3 | 0 | 0 | 3 | 4 | 0 | 19 | 16 | 13 | 0 | 0 | 0 | 2 | 0 | 0 | 204 | 149 | 1.000000 | 1 | 0.836 | 1 | 1 | 0.773 | 0.704 | NA | 1 | 0.854 | 1 | NA | -0.089 |
| A3SS | 8420 | ENSG00000223960 | CHROMR | chr2 | + | 178430694 | 178430889 | 178430769 | 178430889 | 178426365 | 178426505 | 22 | 25 | 16 | 22 | 25 | 5 | 4 | 8 | 10 | 7 | 38 | 16 | 11 | 19 | 0 | 3 | 4 | 2 | 13 | 0 | 224 | 149 | 1.000000 | 1 | 0.745 | 0.806 | 0.571 | 0.594 | 0.704 | 0.894 | 0.727 | 0.785 | 0.493 | NA | -0.041 |
| A3SS | 7655 | ENSG00000076242 | MLH1 | chr3 | + | 36996618 | 36996709 | 36996623 | 36996709 | 36993784 | 36993890 | 15 | 9 | 20 | 14 | 11 | 0 | 1 | 0 | 0 | 0 | 10 | 8 | 7 | 17 | 1 | 0 | 0 | 0 | 0 | 0 | 154 | 149 | 1.000000 | 1 | 1 | 0.897 | 1 | 1 | 1 | 1 | 1 | 1 | 1 | 1 | -0.021 |
| A3SS | 8612 | ENSG00000095059 | DHPS | chr19 | - | 12676016 | 12676105 | 12676016 | 12676078 | 12677107 | 12677211 | 10 | 15 | 5 | 13 | 11 | 0 | 0 | 0 | 0 | 1 | 2 | 5 | 5 | 13 | 3 | 0 | 0 | 0 | 0 | 0 | 176 | 149 | 1.000000 | 1 | 1 | 1 | 1 | 1 | 0.903 | 1 | 1 | 1 | 1 | 1 | -0.019 |
| A3SS | 5901 | ENSG00000059691 | GATB | chr4 | - | 151701328 | 151701518 | 151701328 | 151701456 | 151703850 | 151703895 | 39 | 22 | 37 | 30 | 60 | 0 | 0 | 1 | 1 | 0 | 25 | 34 | 8 | 45 | 50 | 0 | 0 | 0 | 0 | 0 | 211 | 149 | 1.000000 | 1 | 1 | 1 | 0.963 | 0.955 | 1 | 1 | 1 | 1 | 1 | 1 | -0.016 |
| A3SS | 1221 | ENSG00000184009 | ACTG1 | chr17 | - | 81511187 | 81511803 | 81511187 | 81511626 | 81511902 | 81512142 | 19 | 7 | 20 | 22 | 37 | 2380 | 1039 | 2005 | 1787 | 3117 | 3 | 8 | 27 | 21 | 20 | 1822 | 1453 | 938 | 1665 | 4361 | 298 | 149 | 1.000000 | 1 | 0.004 | 0.003 | 0.005 | 0.006 | 0.006 | 0.001 | 0.003 | 0.014 | 0.006 | 0.002 | 0 |
| A3SS | 12472 | ENSG00000196531 | NACA | chr12 | - | 56714601 | 56714703 | 56714601 | 56714687 | 56724451 | 56724523 | 16 | 4 | 8 | 5 | 14 | 2345 | 1245 | 2323 | 1903 | 1361 | 0 | 16 | 3 | 2 | 0 | 266 | 890 | 899 | 545 | 181 | 165 | 149 | 1.000000 | 1 | 0.006 | 0.003 | 0.003 | 0.002 | 0.009 | 0 | 0.016 | 0.003 | 0.003 | 0 | 0 |
| A3SS | 781 | ENSG00000101247 | NDUFAF5 | chr20 | + | 13801485 | 13801683 | 13801507 | 13801683 | 13798460 | 13798500 | 26 | 16 | 19 | 66 | 41 | 2 | 0 | 0 | 2 | 2 | 11 | 21 | 54 | 56 | 0 | 0 | 2 | 6 | 0 | 0 | 171 | 149 | 1.000000 | 1 | 0.919 | 1 | 1 | 0.966 | 0.947 | 1 | 0.901 | 0.887 | 1 | NA | 0.019 |
| A3SS | 5916 | ENSG00000055609 | KMT2C | chr7 | - | 152156204 | 152156346 | 152156204 | 152156343 | 152158862 | 152159072 | 11 | 2 | 5 | 10 | 8 | 0 | 0 | 0 | 1 | 0 | 0 | 4 | 6 | 0 | 0 | 0 | 0 | 2 | 0 | 0 | 152 | 149 | 1.000000 | 1 | 1 | 1 | 1 | 0.907 | 1 | NA | 1 | 0.746 | NA | NA | 0.108 |
| A3SS | 12043 | ENSG00000065717 | TLE2 | chr19 | - | 3019282 | 3019466 | 3019282 | 3019463 | 3019698 | 3019773 | 8 | 11 | 13 | 1 | 5 | 0 | 4 | 6 | 5 | 12 | 0 | 1 | 10 | 0 | 0 | 0 | 0 | 0 | 0 | 0 | 152 | 149 | 1.000000 | 1 | 1 | 0.729 | 0.68 | 0.164 | 0.29 | NA | 1 | 1 | NA | NA | -0.427 |
| A3SS | 9966 | ENSG00000258733 | LINC02328 | chr14 | + | 86128664 | 86128773 | 86128714 | 86128773 | 86127949 | 86128116 | 7 | 13 | 7 | 7 | 8 | 0 | 2 | 0 | 2 | 1 | 0 | 8 | 2 | 5 | 0 | 0 | 0 | 0 | 0 | 0 | 199 | 149 | 1.000000 | 1 | 1 | 0.83 | 1 | 0.724 | 0.857 | NA | 1 | 1 | 1 | NA | -0.118 |
| A3SS | 3127 | ENSG00000172262 | ZNF131 | chr5 | + | 43122038 | 43122177 | 43122043 | 43122177 | 43120919 | 43121123 | 7 | 5 | 8 | 14 | 18 | 0 | 1 | 0 | 1 | 2 | 14 | 4 | 3 | 4 | 0 | 0 | 0 | 0 | 0 | 0 | 154 | 149 | 1.000000 | 1 | 1 | 0.829 | 1 | 0.931 | 0.897 | 1 | 1 | 1 | 1 | NA | -0.069 |
| A3SS | 4449 | ENSG00000159147 | DONSON | chr21 | - | 33585977 | 33586181 | 33585977 | 33586060 | 33587521 | 33587602 | 8 | 15 | 26 | 18 | 23 | 0 | 0 | 4 | 0 | 0 | 2 | 19 | 29 | 13 | 0 | 0 | 0 | 0 | 0 | 0 | 270 | 149 | 1.000000 | 1 | 1 | 1 | 0.782 | 1 | 1 | 1 | 1 | 1 | 1 | NA | -0.044 |
| A3SS | 772 | ENSG00000101224 | CDC25B | chr20 | + | 3797621 | 3797749 | 3797663 | 3797749 | 3795738 | 3795879 | 32 | 24 | 38 | 29 | 15 | 0 | 3 | 0 | 0 | 0 | 30 | 22 | 21 | 22 | 16 | 0 | 0 | 0 | 0 | 0 | 191 | 149 | 1.000000 | 1 | 1 | 0.862 | 1 | 1 | 1 | 1 | 1 | 1 | 1 | 1 | -0.028 |
| A3SS | 11744 | ENSG00000167333 | TRIM68 | chr11 | - | 4602151 | 4602412 | 4602151 | 4602406 | 4603244 | 4603340 | 21 | 17 | 22 | 19 | 22 | 0 | 0 | 0 | 1 | 0 | 26 | 22 | 8 | 5 | 13 | 0 | 0 | 0 | 0 | 0 | 155 | 149 | 1.000000 | 1 | 1 | 1 | 1 | 0.948 | 1 | 1 | 1 | 1 | 1 | 1 | -0.01 |
| A3SS | 7778 | ENSG00000106105 | GARS1 | chr7 | + | 30601058 | 30601200 | 30601128 | 30601200 | 30599946 | 30600049 | 199 | 100 | 169 | 175 | 202 | 0 | 0 | 1 | 0 | 0 | 236 | 149 | 151 | 141 | 141 | 0 | 7 | 0 | 0 | 0 | 219 | 149 | 1.000000 | 1 | 1 | 1 | 0.991 | 1 | 1 | 1 | 0.935 | 1 | 1 | 1 | 0.011 |
| A3SS | 2650 | ENSG00000075413 | MARK3 | chr14 | + | 103489052 | 103491107 | 103490977 | 103491107 | 103480386 | 103480490 | 16 | 30 | 14 | 12 | 16 | 1 | 0 | 0 | 0 | 0 | 0 | 20 | 23 | 16 | 2 | 0 | 2 | 0 | 0 | 0 | 298 | 149 | 1.000000 | 1 | 0.889 | 1 | 1 | 1 | 1 | NA | 0.833 | 1 | 1 | 1 | 0.02 |
| A3SS | 12268 | ENSG00000235505 | CASP4LP | chr11 | - | 104901548 | 104901659 | 104901548 | 104901656 | 104903390 | 104903551 | 12 | 4 | 11 | 4 | 10 | 14 | 10 | 5 | 17 | 24 | 0 | 11 | 3 | 0 | 0 | 0 | 0 | 5 | 5 | 5 | 152 | 149 | 1.000000 | 1 | 0.457 | 0.282 | 0.683 | 0.187 | 0.29 | NA | 1 | 0.37 | 0 | 0 | 0.037 |
| A3SS | 10270 | ENSG00000111581 | NUP107 | chr12 | + | 68696818 | 68696922 | 68696850 | 68696922 | 68690630 | 68690746 | 9 | 8 | 5 | 23 | 19 | 0 | 0 | 0 | 0 | 0 | 16 | 12 | 4 | 5 | 0 | 0 | 0 | 1 | 0 | 0 | 181 | 149 | 1.000000 | 1 | 1 | 1 | 1 | 1 | 1 | 1 | 1 | 0.767 | 1 | NA | 0.058 |
| A3SS | 10156 | ENSG00000184863 | RBM33 | chr7 | + | 155711172 | 155711455 | 155711202 | 155711455 | 155707293 | 155707380 | 11 | 6 | 9 | 11 | 10 | 0 | 1 | 2 | 0 | 0 | 4 | 2 | 12 | 2 | 0 | 0 | 0 | 0 | 2 | 0 | 179 | 149 | 1.000000 | 1 | 1 | 0.833 | 0.789 | 1 | 1 | 1 | 1 | 1 | 0.454 | NA | 0.061 |
| A3SS | 9476 | ENSG00000071189 | SNX13 | chr7 | - | 17821508 | 17821681 | 17821508 | 17821648 | 17826021 | 17826091 | 4 | 8 | 4 | 7 | 7 | 47 | 55 | 38 | 51 | 54 | 0 | 0 | 3 | 0 | 0 | 0 | 21 | 25 | 10 | 9 | 182 | 149 | 1.000000 | 1 | 0.065 | 0.106 | 0.079 | 0.101 | 0.096 | NA | 0 | 0.089 | 0 | 0 | 0.067 |
| A3SS | 12384 | ENSG00000150433 | TMEM218 | chr11 | - | 125101200 | 125101303 | 125101200 | 125101256 | 125102636 | 125102809 | 9 | 8 | 9 | 14 | 10 | 0 | 0 | 0 | 1 | 0 | 7 | 2 | 1 | 4 | 9 | 0 | 2 | 0 | 0 | 0 | 196 | 149 | 1.000000 | 1 | 1 | 1 | 1 | 0.914 | 1 | 1 | 0.432 | 1 | 1 | 1 | 0.096 |
| A3SS | 5928 | ENSG00000141027 | NCOR1 | chr17 | - | 16065484 | 16065694 | 16065484 | 16065691 | 16070164 | 16070525 | 22 | 11 | 13 | 8 | 6 | 0 | 0 | 0 | 1 | 0 | 9 | 7 | 5 | 12 | 0 | 9 | 0 | 0 | 0 | 0 | 152 | 149 | 1.000000 | 1 | 1 | 1 | 1 | 0.887 | 1 | 0.495 | 1 | 1 | 1 | NA | 0.104 |
| A3SS | 2262 | ENSG00000114062 | UBE3A | chr15 | - | 25407066 | 25407262 | 25407066 | 25407212 | 25408571 | 25408684 | 10 | 5 | 12 | 5 | 17 | 0 | 0 | 0 | 1 | 0 | 0 | 7 | 1 | 7 | 7 | 0 | 0 | 0 | 0 | 7 | 199 | 149 | 1.000000 | 1 | 1 | 1 | 1 | 1 | 1 | NA | 1 | 1 | 1 | 0.428 | 0.143 |
| A3SS | 283 | ENSG00000105699 | LSR | chr19 | + | 35266835 | 35266967 | 35266838 | 35266967 | 35266678 | 35266738 | 16 | 3 | 12 | 8 | 3 | 3 | 2 | 2 | 2 | 4 | 0 | 0 | 15 | 0 | 0 | 0 | 0 | 0 | 0 | 11 | 152 | 149 | 1.000000 | 1 | 0.839 | 0.595 | 0.855 | 0.797 | 0.424 | NA | NA | 1 | NA | 0 | 0.202 |
| A3SS | 1784 | ENSG00000260565 | ERVK13-1 | chr16 | - | 2663083 | 2663430 | 2663083 | 2663407 | 2669410 | 2669561 | 13 | 4 | 6 | 3 | 12 | 4 | 8 | 6 | 15 | 0 | 0 | 2 | 9 | 0 | 0 | 16 | 11 | 1 | 10 | 0 | 172 | 149 | 1.000000 | 1 | 0.738 | 0.302 | 0.464 | 0.148 | 1 | 0 | 0.136 | 0.886 | 0 | NA | 0.275 |
| A3SS | 11595 | ENSG00000213930 | GALT | chr9 | + | 34648047 | 34648171 | 34648114 | 34648171 | 34647088 | 34647258 | 12 | 5 | 12 | 9 | 1 | 3 | 3 | 2 | 1 | 2 | 0 | 0 | 7 | 5 | 0 | 0 | 1 | 1 | 5 | 11 | 216 | 149 | 1.000000 | 1 | 0.734 | 0.535 | 0.805 | 0.861 | 0.256 | NA | 0 | 0.828 | 0.408 | 0 | 0.329 |
| A3SS | 5922 | ENSG00000055609 | KMT2C | chr7 | - | 152207299 | 152207428 | 152207299 | 152207425 | 152229922 | 152230027 | 8 | 13 | 6 | 10 | 4 | 0 | 0 | 0 | 1 | 2 | 2 | 8 | 2 | 1 | 0 | 0 | 0 | 0 | 0 | 0 | 152 | 149 | 1.000000 | 1 | 1 | 1 | 1 | 0.907 | 0.662 | 1 | 1 | 1 | 1 | NA | -0.086 |
| A3SS | 8989 | ENSG00000120896 | SORBS3 | chr8 | + | 22565825 | 22566484 | 22566344 | 22566484 | 22565267 | 22565354 | 19 | 16 | 10 | 11 | 13 | 0 | 0 | 2 | 0 | 0 | 7 | 18 | 10 | 0 | 6 | 0 | 0 | 0 | 0 | 0 | 298 | 149 | 1.000000 | 1 | 1 | 1 | 0.714 | 1 | 1 | 1 | 1 | 1 | NA | 1 | -0.057 |
| A3SS | 12219 | ENSG00000166261 | ZNF202 | chr11 | - | 123740116 | 123740232 | 123740116 | 123740192 | 123740416 | 123740536 | 18 | 4 | 12 | 11 | 5 | 0 | 0 | 1 | 2 | 0 | 0 | 14 | 2 | 0 | 6 | 0 | 0 | 0 | 0 | 0 | 189 | 149 | 1.000000 | 1 | 1 | 1 | 0.904 | 0.813 | 1 | NA | 1 | 1 | NA | 1 | -0.057 |
| A3SS | 9400 | ENSG00000172508 | CARNS1 | chr11 | + | 67416175 | 67417677 | 67417406 | 67417677 | 67415677 | 67415763 | 17 | 3 | 4 | 5 | 7 | 2 | 0 | 0 | 0 | 0 | 0 | 1 | 1 | 0 | 6 | 0 | 0 | 0 | 0 | 0 | 298 | 149 | 1.000000 | 1 | 0.81 | 1 | 1 | 1 | 1 | NA | 1 | 1 | NA | 1 | -0.038 |
| A3SS | 256 | ENSG00000160111 | CPAMD8 | chr19 | - | 16892950 | 16894437 | 16892950 | 16893339 | 16896175 | 16896326 | 17 | 3 | 14 | 8 | 9 | 1 | 0 | 0 | 0 | 0 | 11 | 0 | 5 | 1 | 6 | 0 | 0 | 0 | 0 | 0 | 298 | 149 | 1.000000 | 1 | 0.895 | 1 | 1 | 1 | 1 | 1 | NA | 1 | 1 | 1 | -0.021 |
| A3SS | 3823 | ENSG00000155229 | MMS19 | chr10 | - | 97476834 | 97476963 | 97476834 | 97476919 | 97477346 | 97477416 | 42 | 24 | 30 | 45 | 31 | 1 | 0 | 0 | 0 | 0 | 47 | 32 | 10 | 26 | 29 | 0 | 0 | 0 | 0 | 0 | 193 | 149 | 1.000000 | 1 | 0.97 | 1 | 1 | 1 | 1 | 1 | 1 | 1 | 1 | 1 | -0.006 |
| A3SS | 1232 | ENSG00000108622 | ICAM2 | chr17 | - | 64006630 | 64006735 | 64006630 | 64006671 | 64007740 | 64007822 | 51 | 40 | 58 | 66 | 59 | 0 | 0 | 0 | 2 | 2 | 96 | 37 | 9 | 22 | 82 | 0 | 3 | 0 | 0 | 0 | 213 | 149 | 1.000000 | 1 | 1 | 1 | 1 | 0.958 | 0.954 | 1 | 0.896 | 1 | 1 | 1 | 0.003 |
| A3SS | 7411 | ENSG00000185163 | DDX51 | chr12 | - | 132140830 | 132141420 | 132140830 | 132141020 | 132141497 | 132141606 | 22 | 11 | 11 | 17 | 9 | 0 | 0 | 0 | 0 | 0 | 0 | 27 | 10 | 4 | 1 | 0 | 0 | 2 | 0 | 0 | 298 | 149 | 1.000000 | 1 | 1 | 1 | 1 | 1 | 1 | NA | 1 | 0.714 | 1 | 1 | 0.072 |
| A3SS | 4194 | ENSG00000113971 | NPHP3 | chr3 | - | 132683398 | 132683544 | 132683398 | 132683524 | 132684553 | 132684794 | 12 | 1 | 7 | 3 | 7 | 46 | 58 | 32 | 31 | 76 | 0 | 0 | 2 | 0 | 0 | 0 | 31 | 29 | 4 | 0 | 169 | 149 | 1.000000 | 1 | 0.187 | 0.015 | 0.162 | 0.079 | 0.075 | NA | 0 | 0.057 | 0 | NA | 0.085 |
| A3SS | 4154 | ENSG00000163703 | CRELD1 | chr3 | + | 9943674 | 9944095 | 9943901 | 9944095 | 9943076 | 9943515 | 7 | 8 | 16 | 15 | 17 | 0 | 0 | 0 | 1 | 0 | 0 | 6 | 4 | 0 | 25 | 0 | 2 | 0 | 0 | 0 | 298 | 149 | 1.000000 | 1 | 1 | 1 | 1 | 0.882 | 1 | NA | 0.6 | 1 | NA | 1 | 0.11 |
| A3SS | 7986 | ENSG00000104728 | ARHGEF10 | chr8 | + | 1864369 | 1864436 | 1864372 | 1864436 | 1859896 | 1860184 | 3 | 1 | 31 | 0 | 0 | 2 | 1 | 11 | 1 | 1 | 0 | 2 | 0 | 4 | 1 | 0 | 2 | 3 | 5 | 13 | 152 | 149 | 1.000000 | 1 | 0.595 | 0.495 | 0.734 | 0 | 0 | NA | 0.495 | 0 | 0.44 | 0.07 | 0.114 |
| A3SS | 4603 | ENSG00000071794 | HLTF | chr3 | - | 149041489 | 149041680 | 149041489 | 149041668 | 149042165 | 149042290 | 5 | 3 | 4 | 1 | 15 | 22 | 18 | 23 | 24 | 58 | 0 | 0 | 0 | 0 | 0 | 0 | 11 | 14 | 0 | 0 | 161 | 149 | 1.000000 | 1 | 0.174 | 0.134 | 0.139 | 0.037 | 0.193 | NA | 0 | 0 | NA | NA | 0.135 |
| A3SS | 476 | ENSG00000198182 | ZNF607 | chr19 | - | 37707913 | 37708012 | 37707913 | 37708009 | 37709695 | 37709822 | 7 | 3 | 2 | 6 | 15 | 6 | 11 | 13 | 9 | 8 | 0 | 5 | 0 | 0 | 0 | 3 | 4 | 2 | 7 | 0 | 152 | 149 | 1.000000 | 1 | 0.534 | 0.211 | 0.131 | 0.395 | 0.648 | 0 | 0.551 | 0 | 0 | NA | 0.246 |
| A3SS | 140 | ENSG00000100124 | ANKRD54 | chr22 | - | 37838499 | 37838598 | 37838499 | 37838593 | 37840186 | 37840234 | 8 | 2 | 7 | 4 | 11 | 1 | 0 | 2 | 2 | 0 | 0 | 0 | 0 | 0 | 4 | 0 | 0 | 0 | 6 | 9 | 154 | 149 | 1.000000 | 1 | 0.886 | 1 | 0.772 | 0.659 | 1 | NA | NA | NA | 0 | 0.301 | 0.713 |
| A3SS | 11892 | ENSG00000245498 | NA | chr11 | + | 124805266 | 124805504 | 124805319 | 124805504 | 124804730 | 124804848 | 10 | 5 | 14 | 4 | 0 | 1 | 4 | 0 | 4 | 15 | 0 | 4 | 0 | 0 | 0 | 0 | 0 | 0 | 0 | 0 | 202 | 149 | 1.000000 | 1 | 0.881 | 0.48 | 1 | 0.425 | 0 | NA | 1 | NA | NA | NA | -0.443 |
| A3SS | 4333 | ENSG00000023330 | ALAS1 | chr3 | + | 52199209 | 52199440 | 52199220 | 52199440 | 52198671 | 52198848 | 14 | 4 | 10 | 10 | 10 | 2 | 0 | 1 | 1 | 1 | 0 | 4 | 3 | 4 | 8 | 0 | 0 | 0 | 0 | 0 | 160 | 149 | 1.000000 | 1 | 0.867 | 1 | 0.903 | 0.903 | 0.903 | NA | 1 | 1 | 1 | 1 | -0.085 |
| A3SS | 4714 | ENSG00000138459 | SLC35A5 | chr3 | + | 112581178 | 112581326 | 112581244 | 112581326 | 112580545 | 112580706 | 17 | 9 | 10 | 8 | 10 | 0 | 0 | 0 | 0 | 2 | 0 | 15 | 1 | 4 | 5 | 0 | 0 | 0 | 0 | 0 | 215 | 149 | 1.000000 | 1 | 1 | 1 | 1 | 1 | 0.776 | NA | 1 | 1 | 1 | 1 | -0.045 |
| A3SS | 6528 | ENSG00000143771 | CNIH4 | chr1 | + | 224375794 | 224379452 | 224379068 | 224379452 | 224365878 | 224365991 | 36 | 10 | 30 | 18 | 15 | 0 | 0 | 0 | 0 | 2 | 15 | 9 | 11 | 11 | 34 | 0 | 1 | 0 | 0 | 0 | 298 | 149 | 1.000000 | 1 | 1 | 1 | 1 | 1 | 0.789 | 1 | 0.818 | 1 | 1 | 1 | -0.006 |
| A3SS | 5798 | ENSG00000108654 | DDX5 | chr17 | - | 64504676 | 64504880 | 64504676 | 64504842 | 64506075 | 64506289 | 18 | 17 | 15 | 17 | 14 | 2226 | 1415 | 1827 | 2634 | 966 | 1 | 15 | 25 | 11 | 0 | 816 | 1766 | 1591 | 1315 | 484 | 187 | 149 | 1.000000 | 1 | 0.006 | 0.009 | 0.006 | 0.005 | 0.011 | 0.001 | 0.007 | 0.012 | 0.007 | 0 | 0.002 |
| A3SS | 7911 | ENSG00000147533 | GOLGA7 | chr8 | + | 41490832 | 41490965 | 41490869 | 41490965 | 41490561 | 41490617 | 29 | 21 | 33 | 41 | 19 | 0 | 0 | 0 | 0 | 0 | 18 | 29 | 29 | 21 | 17 | 0 | 1 | 0 | 0 | 0 | 186 | 149 | 1.000000 | 1 | 1 | 1 | 1 | 1 | 1 | 1 | 0.959 | 1 | 1 | 1 | 0.008 |
| A3SS | 2271 | ENSG00000114062 | UBE3A | chr15 | - | 25407066 | 25407262 | 25407066 | 25407212 | 25409087 | 25409207 | 10 | 5 | 13 | 7 | 23 | 0 | 1 | 0 | 0 | 0 | 0 | 14 | 1 | 7 | 7 | 0 | 0 | 0 | 2 | 0 | 199 | 149 | 1.000000 | 1 | 1 | 0.789 | 1 | 1 | 1 | NA | 1 | 1 | 0.724 | 1 | 0.027 |
| A3SS | 10203 | ENSG00000122545 | SEPTIN7 | chr7 | + | 35831352 | 35831496 | 35831424 | 35831496 | 35801218 | 35801270 | 9 | 5 | 13 | 7 | 8 | 0 | 1 | 3 | 3 | 0 | 0 | 4 | 9 | 0 | 0 | 0 | 6 | 0 | 0 | 0 | 221 | 149 | 1.000000 | 1 | 1 | 0.771 | 0.745 | 0.611 | 1 | NA | 0.31 | 1 | NA | NA | 0.17 |
| A3SS | 5893 | ENSG00000251136 | NA | chr8 | - | 89722014 | 89722156 | 89722014 | 89722081 | 89724628 | 89724966 | 5 | 1 | 1 | 11 | 21 | 0 | 0 | 0 | 1 | 2 | 0 | 8 | 2 | 0 | 0 | 0 | 0 | 0 | 1 | 0 | 224 | 149 | 1.000000 | 1 | 1 | 1 | 1 | 0.88 | 0.875 | NA | 1 | 1 | 0 | NA | 0.284 |
| A3SS | 12363 | ENSG00000187240 | DYNC2H1 | chr11 | + | 103192096 | 103192264 | 103192117 | 103192264 | 103191516 | 103191619 | 11 | 6 | 5 | 7 | 0 | 8 | 2 | 2 | 4 | 6 | 0 | 0 | 0 | 0 | 0 | 0 | 0 | 2 | 0 | 0 | 170 | 149 | 1.000000 | 1 | 0.547 | 0.724 | 0.687 | 0.605 | 0 | NA | NA | 0 | NA | NA | 0.513 |
| A3SS | 13300 | ENSG00000136104 | RNASEH2B | chr13 | + | 50949996 | 50950318 | 50950070 | 50950318 | 50949462 | 50949505 | 6 | 4 | 9 | 16 | 5 | 6 | 3 | 9 | 4 | 9 | 0 | 7 | 2 | 1 | 0 | 0 | 3 | 10 | 0 | 0 | 223 | 149 | 1.000000 | 1 | 0.401 | 0.471 | 0.401 | 0.728 | 0.271 | NA | 0.609 | 0.118 | 1 | NA | -0.121 |
| A3SS | 3346 | ENSG00000255330 | NA | chr6 | - | 127453846 | 127454101 | 127453846 | 127454087 | 127473079 | 127473391 | 7 | 14 | 12 | 2 | 13 | 0 | 3 | 1 | 0 | 4 | 0 | 5 | 13 | 0 | 0 | 0 | 0 | 0 | 0 | 0 | 163 | 149 | 1.000000 | 1 | 1 | 0.81 | 0.916 | 1 | 0.748 | NA | 1 | 1 | NA | NA | -0.105 |
| A3SS | 10315 | ENSG00000230590 | FTX | chrX | - | 74274341 | 74274457 | 74274341 | 74274371 | 74280393 | 74280494 | 13 | 16 | 21 | 17 | 35 | 5 | 0 | 1 | 0 | 0 | 22 | 18 | 18 | 14 | 0 | 0 | 0 | 0 | 0 | 0 | 235 | 149 | 1.000000 | 1 | 0.622 | 1 | 0.93 | 1 | 1 | 1 | 1 | 1 | 1 | NA | -0.09 |
| A3SS | 7932 | ENSG00000093144 | ECHDC1 | chr6 | - | 127315933 | 127316081 | 127315933 | 127315974 | 127316449 | 127316502 | 9 | 8 | 6 | 3 | 16 | 5 | 3 | 7 | 4 | 15 | 2 | 0 | 10 | 0 | 0 | 0 | 2 | 6 | 0 | 0 | 256 | 149 | 1.000000 | 1 | 0.512 | 0.608 | 0.333 | 0.304 | 0.383 | 1 | 0 | 0.492 | NA | NA | -0.069 |
| A3SS | 11118 | ENSG00000120533 | ENY2 | chr8 | + | 109340443 | 109340563 | 109340490 | 109340563 | 109339319 | 109339390 | 12 | 5 | 8 | 16 | 7 | 8 | 3 | 10 | 12 | 13 | 0 | 5 | 3 | 8 | 2 | 0 | 10 | 7 | 6 | 0 | 196 | 149 | 1.000000 | 1 | 0.533 | 0.559 | 0.378 | 0.503 | 0.29 | NA | 0.275 | 0.246 | 0.503 | 1 | -0.053 |
| A3SS | 6872 | ENSG00000138641 | HERC3 | chr4 | + | 88597949 | 88598087 | 88597968 | 88598087 | 88595556 | 88595614 | 30 | 24 | 33 | 53 | 16 | 4 | 0 | 2 | 2 | 2 | 40 | 37 | 28 | 21 | 0 | 4 | 0 | 1 | 0 | 0 | 168 | 149 | 1.000000 | 1 | 0.869 | 1 | 0.936 | 0.959 | 0.876 | 0.899 | 1 | 0.961 | 1 | NA | -0.037 |
| A3SS | 6821 | ENSG00000204220 | PFDN6 | chr6 | + | 33289769 | 33289920 | 33289774 | 33289920 | 33289596 | 33289628 | 12 | 2 | 6 | 17 | 24 | 4 | 0 | 0 | 0 | 0 | 6 | 18 | 0 | 7 | 0 | 0 | 0 | 0 | 1 | 0 | 154 | 149 | 1.000000 | 1 | 0.744 | 1 | 1 | 1 | 1 | 1 | 1 | NA | 0.871 | NA | -0.008 |
| A3SS | 4125 | ENSG00000171608 | PIK3CD | chr1 | + | 9716958 | 9717108 | 9717063 | 9717108 | 9716439 | 9716619 | 158 | 50 | 108 | 154 | 93 | 2 | 0 | 2 | 0 | 0 | 116 | 124 | 59 | 103 | 131 | 0 | 0 | 2 | 0 | 0 | 254 | 149 | 1.000000 | 1 | 0.979 | 1 | 0.969 | 1 | 1 | 1 | 1 | 0.945 | 1 | 1 | 0.001 |
| A3SS | 4153 | ENSG00000163703 | CRELD1 | chr3 | + | 9943380 | 9943515 | 9943481 | 9943515 | 9943076 | 9943253 | 47 | 15 | 28 | 41 | 43 | 1 | 0 | 0 | 0 | 0 | 26 | 20 | 13 | 27 | 58 | 0 | 0 | 0 | 2 | 0 | 250 | 149 | 1.000000 | 1 | 0.966 | 1 | 1 | 1 | 1 | 1 | 1 | 1 | 0.889 | 1 | 0.015 |
| A3SS | 8194 | ENSG00000129292 | PHF20L1 | chr8 | + | 132804580 | 132804740 | 132804611 | 132804740 | 132803818 | 132804032 | 8 | 8 | 2 | 8 | 15 | 40 | 27 | 17 | 33 | 46 | 0 | 3 | 8 | 0 | 0 | 0 | 36 | 19 | 3 | 7 | 180 | 149 | 1.000000 | 1 | 0.142 | 0.197 | 0.089 | 0.167 | 0.213 | NA | 0.065 | 0.258 | 0 | 0 | 0.081 |
| A3SS | 10477 | ENSG00000163389 | POGLUT1 | chr3 | + | 119479907 | 119480172 | 119480050 | 119480172 | 119478333 | 119478474 | 12 | 13 | 10 | 19 | 10 | 7 | 4 | 1 | 3 | 1 | 0 | 11 | 2 | 20 | 0 | 0 | 0 | 1 | 0 | 0 | 292 | 149 | 1.000000 | 1 | 0.467 | 0.624 | 0.836 | 0.764 | 0.836 | NA | 1 | 0.505 | 1 | NA | -0.13 |
| A3SS | 2823 | ENSG00000119723 | COQ6 | chr14 | + | 73958146 | 73959078 | 73958970 | 73959078 | 73955804 | 73955928 | 37 | 21 | 34 | 30 | 46 | 0 | 0 | 1 | 1 | 0 | 16 | 35 | 39 | 17 | 30 | 0 | 0 | 0 | 0 | 0 | 298 | 149 | 1.000000 | 1 | 1 | 1 | 0.944 | 0.938 | 1 | 1 | 1 | 1 | 1 | 1 | -0.024 |
| A3SS | 8992 | ENSG00000144134 | RABL2A | chr2 | + | 113641352 | 113641450 | 113641355 | 113641450 | 113634152 | 113634232 | 5 | 4 | 6 | 16 | 8 | 0 | 0 | 0 | 1 | 0 | 0 | 2 | 1 | 5 | 0 | 0 | 0 | 0 | 0 | 0 | 152 | 149 | 1.000000 | 1 | 1 | 1 | 1 | 0.94 | 1 | NA | 1 | 1 | 1 | NA | -0.012 |
| A3SS | 12216 | ENSG00000185101 | ANO9 | chr11 | - | 431693 | 431906 | 431693 | 431767 | 431998 | 432054 | 16 | 19 | 34 | 11 | 10 | 0 | 0 | 1 | 0 | 0 | 1 | 7 | 7 | 9 | 35 | 0 | 0 | 0 | 0 | 0 | 288 | 149 | 1.000000 | 1 | 1 | 1 | 0.946 | 1 | 1 | 1 | 1 | 1 | 1 | 1 | -0.011 |
| A3SS | 3965 | ENSG00000115289 | PCGF1 | chr2 | - | 74505551 | 74505638 | 74505551 | 74505612 | 74505736 | 74505770 | 76 | 37 | 36 | 46 | 112 | 0 | 0 | 0 | 0 | 0 | 85 | 26 | 38 | 28 | 99 | 0 | 2 | 0 | 0 | 0 | 175 | 149 | 1.000000 | 1 | 1 | 1 | 1 | 1 | 1 | 1 | 0.917 | 1 | 1 | 1 | 0.017 |
| A3SS | 10991 | ENSG00000140525 | FANCI | chr15 | + | 89294914 | 89295094 | 89294956 | 89295094 | 89293832 | 89293997 | 24 | 12 | 24 | 19 | 22 | 0 | 2 | 5 | 2 | 0 | 0 | 9 | 28 | 33 | 0 | 0 | 9 | 0 | 0 | 0 | 191 | 149 | 1.000000 | 1 | 1 | 0.824 | 0.789 | 0.881 | 1 | NA | 0.438 | 1 | 1 | NA | 0.086 |
| A3SS | 5667 | ENSG00000143493 | INTS7 | chr1 | - | 211968512 | 211968707 | 211968512 | 211968603 | 211978271 | 211978511 | 17 | 10 | 4 | 18 | 9 | 0 | 0 | 0 | 0 | 0 | 0 | 10 | 9 | 5 | 3 | 0 | 0 | 0 | 5 | 2 | 253 | 149 | 1.000000 | 1 | 1 | 1 | 1 | 1 | 1 | NA | 1 | 1 | 0.371 | 0.469 | 0.29 |
| A3SS | 6054 | ENSG00000197530 | MIB2 | chr1 | + | 1629033 | 1629311 | 1629132 | 1629311 | 1628488 | 1628722 | 17 | 3 | 10 | 4 | 0 | 1 | 2 | 8 | 2 | 0 | 0 | 0 | 0 | 2 | 1 | 0 | 0 | 1 | 1 | 8 | 248 | 149 | 1.000000 | 1 | 0.911 | 0.474 | 0.429 | 0.546 | NA | NA | NA | 0 | 0.546 | 0.07 | 0.385 |
| A3SS | 11063 | ENSG00000164953 | TMEM67 | chr8 | + | 93786222 | 93786346 | 93786267 | 93786346 | 93785221 | 93785378 | 1 | 4 | 10 | 13 | 11 | 1 | 0 | 0 | 0 | 0 | 0 | 1 | 5 | 1 | 0 | 0 | 0 | 0 | 0 | 0 | 194 | 149 | 1.000000 | 1 | 0.434 | 1 | 1 | 1 | 1 | NA | 1 | 1 | 1 | NA | -0.113 |
| A3SS | 6799 | ENSG00000189266 | PNRC2 | chr1 | + | 23960928 | 23961134 | 23960932 | 23961134 | 23959108 | 23959256 | 20 | 10 | 14 | 19 | 12 | 1 | 6 | 2 | 0 | 0 | 0 | 16 | 22 | 5 | 0 | 0 | 0 | 0 | 0 | 0 | 153 | 149 | 1.000000 | 1 | 0.951 | 0.619 | 0.872 | 1 | 1 | NA | 1 | 1 | 1 | NA | -0.112 |
| A3SS | 2535 | ENSG00000100473 | COCH | chr14 | + | 30884909 | 30885041 | 30884984 | 30885041 | 30884552 | 30884656 | 8 | 6 | 9 | 5 | 8 | 0 | 0 | 0 | 0 | 3 | 0 | 0 | 4 | 0 | 0 | 0 | 0 | 0 | 0 | 0 | 224 | 149 | 1.000000 | 1 | 1 | 1 | 1 | 1 | 0.639 | NA | NA | 1 | NA | NA | -0.072 |
| A3SS | 2697 | ENSG00000100889 | PCK2 | chr14 | + | 24098202 | 24098678 | 24098474 | 24098678 | 24096891 | 24097137 | 30 | 20 | 25 | 27 | 8 | 0 | 0 | 1 | 0 | 0 | 30 | 14 | 6 | 19 | 9 | 0 | 0 | 0 | 0 | 0 | 298 | 149 | 1.000000 | 1 | 1 | 1 | 0.926 | 1 | 1 | 1 | 1 | 1 | 1 | 1 | -0.015 |
| A3SS | 6575 | ENSG00000135541 | AHI1 | chr6 | - | 135447007 | 135447160 | 135447007 | 135447124 | 135448289 | 135448475 | 7 | 15 | 15 | 18 | 6 | 0 | 0 | 1 | 0 | 0 | 0 | 11 | 5 | 4 | 9 | 0 | 0 | 0 | 0 | 0 | 185 | 149 | 1.000000 | 1 | 1 | 1 | 0.924 | 1 | 1 | NA | 1 | 1 | 1 | 1 | -0.015 |
| A3SS | 7577 | ENSG00000171503 | ETFDH | chr4 | + | 158680466 | 158680607 | 158680527 | 158680607 | 158680123 | 158680182 | 5 | 13 | 19 | 15 | 26 | 0 | 2 | 0 | 0 | 0 | 12 | 18 | 11 | 5 | 0 | 0 | 0 | 1 | 0 | 0 | 210 | 149 | 1.000000 | 1 | 1 | 0.822 | 1 | 1 | 1 | 1 | 1 | 0.886 | 1 | NA | -0.007 |
| A3SS | 5303 | ENSG00000140403 | DNAJA4 | chr15 | + | 78275497 | 78275728 | 78275610 | 78275728 | 78274196 | 78274424 | 53 | 54 | 41 | 109 | 65 | 0 | 0 | 2 | 0 | 2 | 97 | 68 | 40 | 53 | 32 | 0 | 0 | 2 | 1 | 0 | 262 | 149 | 1.000000 | 1 | 1 | 1 | 0.921 | 1 | 0.949 | 1 | 1 | 0.919 | 0.968 | 1 | -0.003 |
| A3SS | 7425 | ENSG00000103168 | TAF1C | chr16 | - | 84184850 | 84185060 | 84184850 | 84184989 | 84186900 | 84187045 | 28 | 6 | 14 | 22 | 7 | 14 | 0 | 15 | 1 | 4 | 0 | 16 | 13 | 8 | 8 | 0 | 7 | 6 | 5 | 0 | 220 | 149 | 1.000000 | 1 | 0.575 | 1 | 0.387 | 0.937 | 0.542 | NA | 0.608 | 0.595 | 0.52 | 1 | 0.007 |
| A3SS | 1443 | ENSG00000154874 | CCDC144B | chr17 | - | 18607896 | 18607970 | 18607896 | 18607967 | 18610043 | 18610292 | 3 | 26 | 4 | 24 | 4 | 2 | 3 | 0 | 2 | 1 | 0 | 7 | 5 | 8 | 9 | 0 | 2 | 4 | 0 | 0 | 152 | 149 | 1.000000 | 1 | 0.595 | 0.895 | 1 | 0.922 | 0.797 | NA | 0.774 | 0.551 | 1 | 1 | 0.011 |
| A3SS | 7463 | ENSG00000135924 | DNAJB2 | chr2 | + | 219280577 | 219281771 | 219281717 | 219281771 | 219279797 | 219279898 | 28 | 27 | 40 | 52 | 23 | 0 | 0 | 0 | 0 | 0 | 27 | 25 | 34 | 22 | 30 | 0 | 2 | 0 | 0 | 0 | 298 | 149 | 1.000000 | 1 | 1 | 1 | 1 | 1 | 1 | 1 | 0.862 | 1 | 1 | 1 | 0.028 |
| A3SS | 13390 | ENSG00000134900 | TPP2 | chr13 | + | 102651338 | 102651397 | 102651358 | 102651397 | 102649407 | 102649486 | 10 | 8 | 3 | 7 | 11 | 36 | 38 | 49 | 55 | 69 | 0 | 0 | 7 | 0 | 0 | 16 | 27 | 48 | 20 | 0 | 169 | 149 | 1.000000 | 1 | 0.197 | 0.157 | 0.051 | 0.101 | 0.123 | 0 | 0 | 0.114 | 0 | NA | 0.097 |
| A3SS | 13302 | ENSG00000136104 | RNASEH2B | chr13 | + | 50959473 | 50960171 | 50960148 | 50960171 | 50949462 | 50949505 | 8 | 8 | 8 | 8 | 3 | 0 | 0 | 0 | 2 | 6 | 0 | 0 | 3 | 0 | 0 | 0 | 2 | 0 | 3 | 7 | 298 | 149 | 1.000000 | 1 | 1 | 1 | 1 | 0.667 | 0.2 | NA | 0 | 1 | 0 | 0 | 0.523 |
| A3SS | 4072 | ENSG00000137269 | LRRC1 | chr6 | + | 53897284 | 53897359 | 53897288 | 53897359 | 53896828 | 53896892 | 8 | 2 | 5 | 15 | 4 | 0 | 0 | 2 | 0 | 0 | 0 | 0 | 0 | 1 | 0 | 0 | 0 | 0 | 0 | 0 | 153 | 149 | 1.000000 | 1 | 1 | 1 | 0.709 | 1 | 1 | NA | NA | NA | 1 | NA | -0.058 |
| A3SS | 6570 | ENSG00000135541 | AHI1 | chr6 | - | 135301407 | 135302817 | 135301407 | 135301539 | 135318518 | 135318616 | 7 | 14 | 4 | 8 | 9 | 1 | 0 | 0 | 0 | 0 | 0 | 6 | 3 | 0 | 0 | 0 | 0 | 0 | 0 | 0 | 298 | 149 | 1.000000 | 1 | 0.778 | 1 | 1 | 1 | 1 | NA | 1 | 1 | NA | NA | -0.044 |
| A3SS | 13015 | ENSG00000198270 | TMEM116 | chr12 | - | 111933885 | 111936830 | 111933885 | 111934030 | 111938160 | 111938210 | 13 | 16 | 35 | 29 | 14 | 0 | 0 | 1 | 0 | 0 | 15 | 33 | 18 | 5 | 3 | 0 | 0 | 0 | 0 | 0 | 298 | 149 | 1.000000 | 1 | 1 | 1 | 0.946 | 1 | 1 | 1 | 1 | 1 | 1 | 1 | -0.011 |
| A3SS | 6086 | ENSG00000177030 | DEAF1 | chr11 | - | 687910 | 688057 | 687910 | 688049 | 688330 | 688460 | 84 | 36 | 56 | 41 | 39 | 1 | 0 | 2 | 2 | 0 | 50 | 39 | 24 | 42 | 68 | 0 | 1 | 1 | 0 | 0 | 157 | 149 | 1.000000 | 1 | 0.988 | 1 | 0.964 | 0.951 | 1 | 1 | 0.974 | 0.958 | 1 | 1 | -0.006 |
| A3SS | 13308 | ENSG00000122034 | GTF3A | chr13 | + | 27430532 | 27430621 | 27430537 | 27430621 | 27429869 | 27429966 | 167 | 91 | 144 | 176 | 296 | 0 | 0 | 1 | 3 | 7 | 204 | 144 | 141 | 117 | 235 | 0 | 0 | 2 | 0 | 4 | 154 | 149 | 1.000000 | 1 | 1 | 1 | 0.993 | 0.983 | 0.976 | 1 | 1 | 0.986 | 1 | 0.983 | -0.003 |
| A3SS | 7281 | ENSG00000141458 | NPC1 | chr18 | - | 23533354 | 23534559 | 23533354 | 23533517 | 23535468 | 23535700 | 144 | 52 | 61 | 98 | 33 | 0 | 0 | 0 | 1 | 0 | 67 | 80 | 85 | 62 | 61 | 0 | 0 | 1 | 0 | 0 | 298 | 149 | 1.000000 | 1 | 1 | 1 | 1 | 0.98 | 1 | 1 | 1 | 0.977 | 1 | 1 | 0.001 |
| A3SS | 2406 | ENSG00000128881 | TTBK2 | chr15 | - | 42872610 | 42872758 | 42872610 | 42872677 | 42878548 | 42878684 | 41 | 37 | 31 | 46 | 18 | 0 | 0 | 0 | 0 | 0 | 26 | 39 | 48 | 13 | 14 | 0 | 0 | 0 | 1 | 0 | 230 | 149 | 1.000000 | 1 | 1 | 1 | 1 | 1 | 1 | 1 | 1 | 1 | 0.894 | 1 | 0.021 |
| A3SS | 13301 | ENSG00000136104 | RNASEH2B | chr13 | + | 50959473 | 50960171 | 50960144 | 50960171 | 50949462 | 50949505 | 9 | 8 | 8 | 8 | 3 | 3 | 0 | 2 | 3 | 6 | 0 | 0 | 3 | 0 | 0 | 0 | 11 | 0 | 0 | 0 | 298 | 149 | 1.000000 | 1 | 0.6 | 1 | 0.667 | 0.571 | 0.2 | NA | 0 | 1 | NA | NA | 0.108 |
| A3SS | 567 | ENSG00000167613 | LAIR1 | chr19 | - | 54360915 | 54361209 | 54360915 | 54361206 | 54370261 | 54370558 | 7 | 7 | 13 | 13 | 4 | 2 | 2 | 1 | 4 | 0 | 0 | 1 | 0 | 7 | 2 | 0 | 0 | 0 | 0 | 0 | 152 | 149 | 1.000000 | 1 | 0.774 | 0.774 | 0.927 | 0.761 | 1 | NA | 1 | NA | 1 | 1 | -0.153 |
| A3SS | 13032 | ENSG00000135968 | GCC2 | chr2 | + | 108486434 | 108486648 | 108486510 | 108486648 | 108485830 | 108485908 | 6 | 6 | 12 | 4 | 20 | 136 | 118 | 161 | 191 | 297 | 0 | 0 | 12 | 2 | 0 | 0 | 10 | 40 | 6 | 0 | 225 | 149 | 1.000000 | 1 | 0.028 | 0.033 | 0.047 | 0.014 | 0.043 | NA | 0 | 0.166 | 0.181 | NA | -0.083 |
| A3SS | 1298 | ENSG00000185359 | HGS | chr17 | + | 81696823 | 81696998 | 81696829 | 81696998 | 81696606 | 81696747 | 24 | 6 | 15 | 23 | 11 | 0 | 0 | 0 | 3 | 0 | 13 | 23 | 2 | 7 | 0 | 0 | 0 | 0 | 0 | 0 | 155 | 149 | 1.000000 | 1 | 1 | 1 | 1 | 0.881 | 1 | 1 | 1 | 1 | 1 | NA | -0.024 |
| A3SS | 11169 | ENSG00000143748 | NVL | chr1 | - | 224317877 | 224317930 | 224317877 | 224317925 | 224326390 | 224326464 | 93 | 62 | 66 | 79 | 126 | 2 | 0 | 4 | 1 | 0 | 81 | 85 | 89 | 65 | 72 | 0 | 0 | 0 | 0 | 0 | 154 | 149 | 1.000000 | 1 | 0.978 | 1 | 0.941 | 0.987 | 1 | 1 | 1 | 1 | 1 | 1 | -0.019 |
| A3SS | 11202 | ENSG00000104529 | EEF1D | chr8 | - | 143581053 | 143581328 | 143581053 | 143581154 | 143597347 | 143597400 | 89 | 34 | 67 | 68 | 60 | 0 | 0 | 0 | 2 | 0 | 44 | 48 | 59 | 65 | 68 | 0 | 0 | 0 | 0 | 0 | 298 | 149 | 1.000000 | 1 | 1 | 1 | 1 | 0.944 | 1 | 1 | 1 | 1 | 1 | 1 | -0.011 |
| A3SS | 5459 | ENSG00000122335 | SERAC1 | chr6 | - | 158116184 | 158117821 | 158116184 | 158116282 | 158119028 | 158119170 | 18 | 17 | 8 | 13 | 15 | 0 | 4 | 3 | 0 | 0 | 10 | 13 | 8 | 6 | 0 | 0 | 0 | 0 | 6 | 0 | 298 | 149 | 1.000000 | 1 | 1 | 0.68 | 0.571 | 1 | 1 | 1 | 1 | 1 | 0.333 | NA | 0.017 |
| A3SS | 3650 | ENSG00000070756 | PABPC1 | chr8 | - | 100712651 | 100713181 | 100712651 | 100712789 | 100717772 | 100717888 | 9 | 73 | 19 | 69 | 57 | 0 | 0 | 0 | 0 | 0 | 13 | 31 | 47 | 88 | 14 | 0 | 0 | 3 | 1 | 0 | 298 | 149 | 1.000000 | 1 | 1 | 1 | 1 | 1 | 1 | 1 | 1 | 0.887 | 0.978 | 1 | 0.027 |
| A3SS | 3221 | ENSG00000153113 | CAST | chr5 | + | 96726753 | 96726859 | 96726793 | 96726859 | 96722638 | 96722718 | 19 | 29 | 11 | 6 | 15 | 0 | 0 | 0 | 0 | 0 | 0 | 18 | 19 | 9 | 0 | 0 | 0 | 4 | 1 | 0 | 189 | 149 | 1.000000 | 1 | 1 | 1 | 1 | 1 | 1 | NA | 1 | 0.789 | 0.876 | NA | 0.112 |
| A3SS | 11458 | ENSG00000173611 | SCAI | chr9 | - | 124971670 | 124971844 | 124971670 | 124971762 | 124975246 | 124975389 | 10 | 12 | 8 | 4 | 17 | 1 | 0 | 4 | 2 | 0 | 0 | 3 | 9 | 2 | 2 | 0 | 0 | 0 | 0 | 0 | 231 | 149 | 1.000000 | 1 | 0.866 | 1 | 0.563 | 0.563 | 1 | NA | 1 | 1 | 1 | 1 | -0.202 |
| A3SS | 1098 | ENSG00000166398 | GARRE1 | chr19 | + | 34349015 | 34349153 | 34349119 | 34349153 | 34347876 | 34348042 | 24 | 21 | 38 | 44 | 24 | 3 | 0 | 0 | 0 | 0 | 28 | 21 | 16 | 14 | 37 | 0 | 0 | 0 | 0 | 0 | 253 | 149 | 1.000000 | 1 | 0.825 | 1 | 1 | 1 | 1 | 1 | 1 | 1 | 1 | 1 | -0.035 |
| A3SS | 8509 | ENSG00000184708 | EIF4ENIF1 | chr22 | - | 31449347 | 31449531 | 31449347 | 31449408 | 31450288 | 31450360 | 122 | 70 | 98 | 118 | 145 | 0 | 1 | 0 | 0 | 2 | 69 | 119 | 66 | 106 | 158 | 0 | 0 | 0 | 0 | 0 | 272 | 149 | 1.000000 | 1 | 1 | 0.975 | 1 | 1 | 0.975 | 1 | 1 | 1 | 1 | 1 | -0.01 |
| A3SS | 11089 | ENSG00000104763 | ASAH1 | chr8 | - | 18069791 | 18069878 | 18069791 | 18069872 | 18071299 | 18071390 | 578 | 479 | 350 | 448 | 304 | 1 | 0 | 0 | 0 | 0 | 250 | 546 | 487 | 483 | 358 | 0 | 4 | 0 | 0 | 0 | 155 | 149 | 1.000000 | 1 | 0.998 | 1 | 1 | 1 | 1 | 1 | 0.992 | 1 | 1 | 1 | 0.001 |
| A3SS | 10148 | ENSG00000196305 | IARS1 | chr9 | - | 92249857 | 92249991 | 92249857 | 92249941 | 92250186 | 92250289 | 7 | 8 | 9 | 7 | 18 | 64 | 79 | 111 | 64 | 113 | 0 | 8 | 6 | 0 | 0 | 20 | 70 | 79 | 72 | 0 | 199 | 149 | 1.000000 | 1 | 0.076 | 0.07 | 0.057 | 0.076 | 0.107 | 0 | 0.079 | 0.054 | 0 | NA | 0.044 |
| A3SS | 392 | ENSG00000008382 | MPND | chr19 | + | 4357252 | 4357421 | 4357339 | 4357421 | 4355096 | 4355173 | 19 | 11 | 22 | 18 | 31 | 0 | 0 | 2 | 0 | 2 | 13 | 15 | 16 | 7 | 15 | 0 | 0 | 2 | 2 | 0 | 236 | 149 | 1.000000 | 1 | 1 | 1 | 0.874 | 1 | 0.907 | 1 | 1 | 0.835 | 0.688 | 1 | 0.052 |
| A3SS | 12404 | ENSG00000165309 | ARMC3 | chr10 | + | 23008814 | 23008931 | 23008835 | 23008931 | 23008275 | 23008374 | 1 | 10 | 2 | 10 | 14 | 1 | 0 | 0 | 2 | 8 | 0 | 1 | 1 | 0 | 0 | 4 | 0 | 0 | 0 | 0 | 170 | 149 | 1.000000 | 1 | 0.467 | 1 | 1 | 0.814 | 0.605 | 0 | 1 | 1 | NA | NA | 0.111 |
| A3SS | 10053 | ENSG00000223482 | NUTM2A-AS1 | chr10 | - | 87238666 | 87238872 | 87238666 | 87238849 | 87246114 | 87246212 | 4 | 2 | 0 | 10 | 19 | 3 | 4 | 0 | 4 | 3 | 0 | 0 | 0 | 0 | 0 | 0 | 7 | 4 | 1 | 0 | 172 | 149 | 1.000000 | 1 | 0.536 | 0.302 | NA | 0.684 | 0.846 | NA | 0 | 0 | 0 | NA | 0.592 |
| A3SS | 5093 | ENSG00000122644 | ARL4A | chr7 | + | 12688165 | 12689176 | 12688168 | 12689176 | 12686907 | 12687043 | 9 | 8 | 22 | 26 | 19 | 10 | 12 | 17 | 3 | 10 | 10 | 7 | 20 | 11 | 0 | 8 | 6 | 18 | 2 | 0 | 152 | 149 | 1.000000 | 1 | 0.469 | 0.395 | 0.559 | 0.895 | 0.651 | 0.551 | 0.534 | 0.521 | 0.844 | NA | -0.019 |
| A3SS | 8180 | ENSG00000064703 | DDX20 | chr1 | + | 111760473 | 111760588 | 111760477 | 111760588 | 111759399 | 111759568 | 45 | 30 | 32 | 32 | 30 | 0 | 0 | 1 | 0 | 1 | 10 | 31 | 40 | 39 | 13 | 0 | 0 | 0 | 0 | 0 | 153 | 149 | 1.000000 | 1 | 1 | 1 | 1 | 1 | 0.967 | 1 | 1 | 1 | 1 | 1 | -0.007 |
| A3SS | 9257 | ENSG00000136044 | APPL2 | chr12 | - | 105207970 | 105208029 | 105207970 | 105208001 | 105208157 | 105208199 | 53 | 25 | 45 | 60 | 42 | 0 | 0 | 1 | 0 | 0 | 37 | 33 | 38 | 48 | 33 | 0 | 0 | 0 | 0 | 0 | 177 | 149 | 1.000000 | 1 | 1 | 1 | 0.974 | 1 | 1 | 1 | 1 | 1 | 1 | 1 | -0.005 |
| A3SS | 6659 | ENSG00000083544 | TDRD3 | chr13 | + | 60509541 | 60509919 | 60509762 | 60509919 | 60494434 | 60494575 | 10 | 8 | 5 | 5 | 13 | 70 | 65 | 70 | 76 | 103 | 0 | 5 | 0 | 0 | 0 | 10 | 43 | 30 | 16 | 0 | 298 | 149 | 1.000000 | 1 | 0.067 | 0.058 | 0.034 | 0.032 | 0.059 | 0 | 0.055 | 0 | 0 | NA | 0.036 |
| A3SS | 7899 | ENSG00000160352 | ZNF714 | chr19 | + | 21098811 | 21098910 | 21098814 | 21098910 | 21098184 | 21098311 | 12 | 2 | 9 | 4 | 17 | 11 | 4 | 8 | 6 | 9 | 0 | 6 | 1 | 1 | 0 | 0 | 7 | 11 | 5 | 0 | 152 | 149 | 1.000000 | 1 | 0.517 | 0.329 | 0.524 | 0.395 | 0.649 | NA | 0.457 | 0.082 | 0.164 | NA | 0.248 |
| A3SS | 12752 | ENSG00000148672 | GLUD1 | chr10 | - | 87060160 | 87060825 | 87060160 | 87060241 | 87062655 | 87062835 | 20 | 4 | 4 | 7 | 15 | 0 | 0 | 0 | 0 | 0 | 0 | 0 | 13 | 1 | 0 | 0 | 0 | 0 | 3 | 0 | 298 | 149 | 1.000000 | 1 | 1 | 1 | 1 | 1 | 1 | NA | NA | 1 | 0.143 | NA | 0.428 |
| A3SS | 12850 | ENSG00000119801 | YPEL5 | chr2 | + | 30155814 | 30155910 | 30155849 | 30155910 | 30148244 | 30148541 | 10 | 12 | 14 | 14 | 16 | 2 | 0 | 1 | 8 | 10 | 0 | 14 | 10 | 6 | 0 | 13 | 14 | 10 | 7 | 0 | 184 | 149 | 1.000000 | 1 | 0.802 | 1 | 0.919 | 0.586 | 0.564 | 0 | 0.447 | 0.447 | 0.41 | NA | 0.448 |
| A3SS | 5749 | ENSG00000049246 | PER3 | chr1 | + | 7793956 | 7794008 | 7793959 | 7794008 | 7788044 | 7788246 | 6 | 3 | 6 | 14 | 7 | 2 | 1 | 1 | 1 | 3 | 0 | 0 | 0 | 0 | 0 | 0 | 2 | 0 | 0 | 0 | 152 | 149 | 1.000000 | 1 | 0.746 | 0.746 | 0.855 | 0.932 | 0.696 | NA | 0 | NA | NA | NA | 0.795 |
| A3SS | 13044 | ENSG00000111752 | PHC1 | chr12 | + | 8922632 | 8922788 | 8922656 | 8922788 | 8921600 | 8921750 | 9 | 5 | 20 | 1 | 21 | 2 | 2 | 1 | 0 | 3 | 4 | 5 | 0 | 0 | 10 | 0 | 1 | 0 | 0 | 0 | 173 | 149 | 1.000000 | 1 | 0.795 | 0.683 | 0.945 | 1 | 0.858 | 1 | 0.812 | NA | NA | 1 | -0.081 |
| A3SS | 12466 | ENSG00000156650 | KAT6B | chr10 | + | 74975398 | 74977437 | 74977315 | 74977437 | 74970019 | 74970101 | 16 | 23 | 11 | 28 | 4 | 1 | 1 | 0 | 0 | 2 | 7 | 14 | 16 | 7 | 1 | 0 | 0 | 1 | 1 | 0 | 298 | 149 | 1.000000 | 1 | 0.889 | 0.92 | 1 | 1 | 0.5 | 1 | 1 | 0.889 | 0.778 | 1 | -0.072 |
| A3SS | 7524 | ENSG00000187741 | FANCA | chr16 | - | 89738881 | 89738974 | 89738881 | 89738926 | 89739132 | 89739289 | 53 | 7 | 27 | 39 | 7 | 0 | 2 | 0 | 0 | 0 | 7 | 27 | 12 | 21 | 29 | 0 | 0 | 0 | 0 | 0 | 197 | 149 | 1.000000 | 1 | 1 | 0.726 | 1 | 1 | 1 | 1 | 1 | 1 | 1 | 1 | -0.055 |
| A3SS | 6456 | ENSG00000066427 | ATXN3 | chr14 | - | 92083125 | 92083258 | 92083125 | 92083235 | 92093251 | 92093318 | 14 | 13 | 15 | 19 | 56 | 4 | 0 | 0 | 0 | 0 | 0 | 24 | 9 | 41 | 6 | 0 | 0 | 0 | 0 | 0 | 172 | 149 | 1.000000 | 1 | 0.752 | 1 | 1 | 1 | 1 | NA | 1 | 1 | 1 | 1 | -0.05 |
| A3SS | 8058 | ENSG00000224078 | SNHG14 | chr15 | + | 25095626 | 25095828 | 25095718 | 25095828 | 25093833 | 25093974 | 33 | 24 | 12 | 18 | 16 | 2 | 0 | 0 | 0 | 0 | 16 | 9 | 14 | 27 | 0 | 0 | 0 | 0 | 0 | 0 | 241 | 149 | 1.000000 | 1 | 0.911 | 1 | 1 | 1 | 1 | 1 | 1 | 1 | 1 | NA | -0.018 |
| A3SS | 11584 | ENSG00000165282 | PIGO | chr9 | - | 35091239 | 35092767 | 35091239 | 35091291 | 35093029 | 35093209 | 40 | 14 | 29 | 29 | 29 | 0 | 0 | 1 | 0 | 0 | 6 | 24 | 29 | 9 | 36 | 0 | 0 | 0 | 0 | 0 | 298 | 149 | 1.000000 | 1 | 1 | 1 | 0.935 | 1 | 1 | 1 | 1 | 1 | 1 | 1 | -0.013 |
| A3SS | 9552 | ENSG00000223705 | NSUN5P1 | chr7 | + | 75415980 | 75416191 | 75416102 | 75416191 | 75415591 | 75415770 | 13 | 10 | 29 | 24 | 18 | 0 | 0 | 0 | 2 | 0 | 0 | 21 | 8 | 9 | 19 | 0 | 0 | 0 | 0 | 1 | 271 | 149 | 1.000000 | 1 | 1 | 1 | 1 | 0.868 | 1 | NA | 1 | 1 | 1 | 0.913 | -0.005 |
| A3SS | 1870 | ENSG00000197912 | SPG7 | chr16 | + | 89531903 | 89532066 | 89531924 | 89532066 | 89530682 | 89530808 | 33 | 27 | 34 | 45 | 37 | 0 | 0 | 0 | 0 | 0 | 6 | 30 | 19 | 42 | 42 | 0 | 0 | 1 | 0 | 0 | 170 | 149 | 1.000000 | 1 | 1 | 1 | 1 | 1 | 1 | 1 | 1 | 0.943 | 1 | 1 | 0.011 |
| A3SS | 12446 | ENSG00000151150 | ANK3 | chr10 | - | 60134270 | 60134373 | 60134270 | 60134358 | 60138963 | 60139087 | 15 | 4 | 29 | 9 | 18 | 0 | 0 | 0 | 0 | 0 | 0 | 14 | 7 | 13 | 4 | 0 | 0 | 3 | 0 | 0 | 164 | 149 | 1.000000 | 1 | 1 | 1 | 1 | 1 | 1 | NA | 1 | 0.679 | 1 | 1 | 0.08 |
| A3SS | 6982 | ENSG00000243406 | MRPS31P5 | chr13 | - | 52169986 | 52170099 | 52169986 | 52170056 | 52171157 | 52171248 | 7 | 9 | 8 | 15 | 10 | 13 | 14 | 24 | 7 | 23 | 0 | 3 | 9 | 0 | 0 | 11 | 21 | 12 | 5 | 0 | 192 | 149 | 1.000000 | 1 | 0.295 | 0.333 | 0.206 | 0.624 | 0.252 | 0 | 0.1 | 0.368 | 0 | NA | 0.225 |
| A3SS | 10994 | ENSG00000156787 | TBC1D31 | chr8 | + | 123129065 | 123129218 | 123129069 | 123129218 | 123126507 | 123126687 | 15 | 3 | 7 | 20 | 5 | 8 | 10 | 14 | 11 | 7 | 0 | 6 | 2 | 0 | 4 | 0 | 0 | 4 | 0 | 0 | 153 | 149 | 1.000000 | 1 | 0.646 | 0.226 | 0.327 | 0.639 | 0.41 | NA | 1 | 0.327 | NA | 1 | -0.326 |

| | | | | | | | | | | | | | | | |
|---|---|---|---|---|---|---|---|---|---|---|---|---|---|---|---|
| A3SS | 4530 | ENSG00000101811 | CSTF2 | chrX | + | 100826633 | 100826757 | 100826684 | 100826757 | 100824119 | 100824257 | 40 | 20 | 18 | [illegible] |
| A3SS | 8037 | ENSG00000179348 | GATA2 | chr3 | - | 128481818 | 128481944 | 128481818 | 128481902 | 128483859 | 128484005 | 30 | 23 | 40 | [illegible] |
| A3SS | 10109 | ENSG00000106628 | POLD2 | chr7 | - | 44115294 | 44115396 | 44115294 | 44115354 | 44115765 | 44115893 | 71 | 39 | 89 | [illegible] |
| A3SS | 9617 | ENSG00000158604 | TMED4 | chr7 | - | 44581722 | 44581823 | 44581722 | 44581806 | 44582046 | 44582287 | 120 | 22 | 94 | [illegible] |
| A3SS | 2597 | ENSG00000133997 | MED6 | chr14 | - | 70596610 | 70596742 | 70596610 | 70596698 | 70597617 | 70597777 | 8 | 7 | 14 | [illegible] |
| A3SS | 11403 | ENSG00000184465 | WDR27 | chr6 | - | 169663562 | 169663878 | 169663562 | 169663649 | 169664165 | 169664286 | 2 | 14 | 11 | [illegible] |
| A3SS | 4934 | ENSG00000196172 | ZNF681 | chr19 | - | 23754822 | 23754918 | 23754822 | 23754897 | 23755424 | 23755551 | 3 | 1 | 13 | [illegible] |
| A3SS | 12137 | ENSG00000132199 | ENOSF1 | chr18 | - | 690548 | 690745 | 690548 | 690631 | 693881 | 693908 | 13 | 1 | 20 | [illegible] |
| A3SS | 2528 | ENSG00000140043 | PTGR2 | chr14 | + | 73876997 | 73880176 | 73880054 | 73880176 | 73874022 | 73874214 | 10 | 9 | 19 | [illegible] |
| A3SS | 6273 | ENSG00000137628 | DDX60 | chr4 | - | 168216293 | 168217680 | 168216293 | 168217032 | 168220654 | 168220717 | 6 | 9 | 5 | [illegible] |
| A3SS | 11779 | ENSG00000198561 | CTNND1 | chr11 | + | 57791384 | 57791673 | 57791391 | 57791673 | 57789036 | 57789155 | 29 | 27 | 17 | [illegible] |
| A3SS | 1186 | ENSG00000182173 | TSEN54 | chr17 | + | 75521704 | 75523335 | 75523274 | 75523335 | 75521408 | 75521510 | 97 | 48 | 88 | [illegible] |
| A3SS | 12169 | ENSG00000206561 | COLQ | chr3 | - | 15488205 | 15488307 | 15488205 | 15488294 | 15489524 | 15489637 | 22 | 4 | 10 | [illegible] |
| A3SS | 8474 | ENSG00000112367 | FIG4 | chr6 | + | 109735131 | 109735298 | 109735155 | 109735298 | 109732636 | 109732687 | 14 | 8 | 7 | [illegible] |
| A3SS | 5875 | ENSG00000115415 | STAT1 | chr2 | - | 190975309 | 190975434 | 190975309 | 190975388 | 190975811 | 190975887 | 28 | 17 | 19 | [illegible] |
| A3SS | 4951 | ENSG00000141293 | SKAP1 | chr17 | - | 48162468 | 48162569 | 48162468 | 48162566 | 48180053 | 48180248 | 10 | 3 | 5 | [illegible] |
| A3SS | 13402 | ENSG00000127022 | CANX | chr5 | + | 179726659 | 179726759 | 179726682 | 179726759 | 179724656 | 179724783 | 6 | 4 | 9 | [illegible] |
| A3SS | 6372 | ENSG00000215788 | TNFRSF25 | chr1 | - | 6462862 | 6463147 | 6462862 | 6463127 | 6464374 | 6464453 | 11 | 3 | 12 | [illegible] |
| A3SS | 1304 | ENSG00000256525 | POLG2 | chr17 | - | 64482918 | 64482999 | 64482918 | 64482974 | 64483389 | 64483506 | 11 | 10 | 11 | [illegible] |
| A3SS | 6732 | ENSG00000065526 | SPEN | chr1 | + | 15909320 | 15909481 | 15909323 | 15909481 | 15872815 | 15873136 | 20 | 4 | 9 | [illegible] |
| A3SS | 5202 | ENSG00000117569 | PTBP2 | chr1 | + | 96804799 | 96806452 | 96806418 | 96806452 | 96785058 | 96785254 | 9 | 4 | 11 | [illegible] |
| A3SS | 9208 | ENSG00000078269 | SYNJ2 | chr6 | + | 158061991 | 158062164 | 158062054 | 158062164 | 158059256 | 158059353 | 34 | 19 | 32 | [illegible] |
| A3SS | 10741 | ENSG00000198954 | KIFBP | chr10 | + | 69005045 | 69005125 | 69005096 | 69005125 | 69000423 | 69000522 | 19 | 10 | 7 | [illegible] |
| A3SS | 3840 | ENSG00000102445 | RUBCNL | chr13 | - | 46368041 | 46368249 | 46368041 | 46368224 | 46368732 | 46368815 | 21 | 13 | 15 | [illegible] |
| A3SS | 3569 | ENSG00000105875 | WDR91 | chr7 | - | 135193230 | 135193399 | 135193230 | 135193346 | 135193577 | 135193672 | 51 | 43 | 51 | [illegible] |
| A3SS | 1231 | ENSG00000108622 | ICAM2 | chr17 | - | 64006630 | 64006731 | 64006630 | 64006671 | 64007740 | 64007822 | 39 | 37 | 43 | [illegible] |
| A3SS | 4647 | ENSG00000090263 | MRPS33 | chr7 | - | 141010418 | 141010660 | 141010418 | 141010617 | 141014550 | 141014679 | 42 | 26 | 53 | [illegible] |
| A3SS | 5832 | ENSG00000173230 | GOLGB1 | chr3 | - | 121719645 | 121719768 | 121719645 | 121719753 | 121722261 | 121722378 | 16 | 10 | 22 | [illegible] |
| A3SS | 11102 | ENSG00000185803 | SLC52A2 | chr8 | + | 144359622 | 144360493 | 144360114 | 144360493 | 144359183 | 144359423 | 34 | 4 | 34 | [illegible] |
| A3SS | 11208 | ENSG00000104529 | EEF1D | chr8 | - | 143592646 | 143592942 | 143592646 | 143592893 | 143597347 | 143597415 | 30 | 24 | 27 | [illegible] |
| A3SS | 8471 | ENSG00000112367 | FIG4 | chr6 | + | 109715077 | 109715176 | 109715079 | 109715176 | 109691264 | 109691501 | 54 | 36 | 49 | [illegible] |
| A3SS | 11270 | ENSG00000132386 | SERPINF1 | chr17 | + | 1770833 | 1771184 | 1771028 | 1771184 | 1769851 | 1770050 | 13 | 17 | 23 | [illegible] |
| A3SS | 391 | ENSG00000008382 | MPND | chr19 | + | 4357252 | 4357421 | 4357339 | 4357421 | 4354323 | 4354420 | 11 | 7 | 12 | [illegible] |
| A3SS | 10731 | ENSG00000068366 | ACSL4 | chrX | - | 109683135 | 109683375 | 109683135 | 109683372 | 109696143 | 109696196 | 6 | 8 | 15 | [illegible] |
| A3SS | 4883 | ENSG00000132549 | VPS13B | chr8 | + | 99870784 | 99871697 | 99871447 | 99871697 | 99868288 | 99868465 | 22 | 28 | 36 | [illegible] |
| A3SS | 7994 | ENSG00000268362 | NA | chr19 | + | 24049404 | 24049548 | 24049409 | 24049548 | 24046183 | 24046279 | 20 | 22 | 13 | [illegible] |
| A3SS | 7114 | ENSG00000228956 | SATB1-AS1 | chr3 | + | 18527784 | 18528139 | 18527950 | 18528139 | 18527173 | 18527326 | 7 | 7 | 14 | [illegible] |
| A3SS | 4565 | ENSG00000074935 | TUBE1 | chr6 | - | 112075936 | 112076171 | 112075936 | 112076112 | 112076321 | 112076509 | 3 | 9 | 10 | [illegible] |
| A3SS | 9103 | ENSG00000163939 | PBRM1 | chr3 | - | 52679573 | 52679726 | 52679573 | 52679723 | 52682091 | 52682233 | 19 | 9 | 25 | [illegible] |
| A3SS | 10138 | ENSG00000122786 | CALD1 | chr7 | + | 134958208 | 134958290 | 134958211 | 134958290 | 134958068 | 134958112 | 17 | 12 | 0 | [illegible] |
| A3SS | 8909 | ENSG00000168280 | KIF5C | chr2 | + | 148998399 | 148998509 | 148998402 | 148998509 | 148997263 | 148997340 | 6 | 21 | 27 | [illegible] |
| A3SS | 8565 | ENSG00000073737 | DHRS9 | chr2 | + | 169069452 | 169069717 | 169069561 | 169069717 | 169067034 | 169067267 | 12 | 9 | 5 | [illegible] |
| A3SS | 12136 | ENSG00000132199 | ENOSF1 | chr18 | - | 690548 | 690745 | 690548 | 690631 | 691203 | 691276 | 13 | 1 | 20 | [illegible] |
| A3SS | 3839 | ENSG00000102445 | RUBCNL | chr13 | - | 46350112 | 46350351 | 46350112 | 46350279 | 46356431 | 46356496 | 47 | 29 | 37 | [illegible] |
| A3SS | 1571 | ENSG00000276070 | CCL4L2 | chr17 | + | 36212363 | 36212878 | 36212481 | 36212878 | 36211775 | 36211890 | 35 | 10 | 2 | [illegible] |
| A3SS | 1466 | ENSG00000108592 | FTSJ3 | chr17 | - | 63820838 | 63821115 | 63820838 | 63820938 | 63821353 | 63821643 | 83 | 81 | 81 | [illegible] |
| A3SS | 11396 | ENSG00000116580 | GON4L | chr1 | - | 155757890 | 155758034 | 155757890 | 155757967 | 155760443 | 155760641 | 88 | 66 | 80 | [illegible] |
| A3SS | 6386 | ENSG00000163617 | CCDC191 | chr3 | - | 114042702 | 114042846 | 114042702 | 114042841 | 114046590 | 114046732 | 14 | 16 | 6 | [illegible] |
| A3SS | 1562 | ENSG00000276070 | CCL4L2 | chr17 | + | 36212292 | 36212878 | 36212481 | 36212878 | 36211775 | 36211890 | 57 | 14 | 4 | [illegible] |
| A3SS | 3009 | ENSG00000085382 | HACE1 | chr6 | - | 104777012 | 104777110 | 104777012 | 104777097 | 104777205 | 104777317 | 14 | 9 | 10 | [illegible] |
| A3SS | 1305 | ENSG00000256525 | POLG2 | chr17 | - | 64482918 | 64482999 | 64482918 | 64482974 | 64483963 | 64485868 | 21 | 10 | 12 | [illegible] |
| A3SS | 7779 | ENSG00000106105 | GARS1 | chr7 | + | 30615895 | 30616058 | 30615897 | 30616058 | 30612095 | 30612245 | 284 | 208 | 257 | [illegible] |
| A3SS | 8906 | ENSG00000114956 | DGUOK | chr2 | + | 73950584 | 73950732 | 73950600 | 73950732 | 73946718 | 73946906 | 60 | 26 | 60 | [illegible] |
| A3SS | 10216 | ENSG00000107897 | ACBD5 | chr10 | - | 27219722 | 27219857 | 27219722 | 27219803 | 27223206 | 27223239 | 49 | 58 | 56 | [illegible] |
| A3SS | 7579 | ENSG00000171503 | ETFDH | chr4 | + | 158685100 | 158685219 | 158685151 | 158685219 | 158684591 | 158684673 | 76 | 68 | 47 | [illegible] |
| A3SS | 6296 | ENSG00000060339 | CCAR1 | chr10 | + | 68742375 | 68742569 | 68742427 | 68742569 | 68736875 | 68737048 | 37 | 36 | 60 | [illegible] |
| A3SS | 1989 | ENSG00000123815 | COQ8B | chr19 | - | 40703540 | 40703855 | 40703540 | 40703622 | 40705095 | 40705447 | 25 | 7 | 23 | [illegible] |
| A3SS | 12403 | ENSG00000165309 | ARMC3 | chr10 | + | 22961883 | 22962078 | 22961938 | 22962078 | 22959398 | 22959574 | 8 | 11 | 1 | [illegible] |
| A3SS | 6346 | ENSG00000165046 | LETM2 | chr8 | + | 38392541 | 38392995 | 38392612 | 38392995 | 38387949 | 38388030 | 19 | 2 | 5 | [illegible] |
| A3SS | 6091 | ENSG00000138095 | LRPPRC | chr2 | - | 43890222 | 43890362 | 43890222 | 43890274 | 43894544 | 43894629 | 9 | 13 | 22 | [illegible] |
| A3SS | 6294 | ENSG00000131115 | ZNF227 | chr19 | + | 44228445 | 44228572 | 44228486 | 44228572 | 44217790 | 44217852 | 24 | 18 | 17 | [illegible] |
| A3SS | 9499 | ENSG00000117009 | KMO | chr1 | + | 241590011 | 241590113 | 241590050 | 241590113 | 241588747 | 241588830 | 15 | 12 | 9 | [illegible] |
| A3SS | 2270 | ENSG00000114062 | UBE3A | chr15 | - | 25407066 | 25407262 | 25407066 | 25407137 | 25409087 | 25409207 | 33 | 17 | 20 | [illegible] |
| A3SS | 949 | ENSG00000088876 | ZNF343 | chr20 | - | 2492698 | 2492825 | 2492698 | 2492785 | 2493518 | 2493577 | 19 | 21 | 20 | [illegible] |
| A3SS | 1905 | ENSG00000214078 | CPNE1 | chr20 | - | 35658929 | 35659014 | 35658929 | 35658982 | 35664878 | 35664932 | 180 | 140 | 205 | [illegible] |
| A3SS | 12803 | ENSG00000110888 | CAPRIN2 | chr12 | - | 30714958 | 30715144 | 30714958 | 30715081 | 30716507 | 30716676 | 23 | 29 | 18 | [illegible] |
| A3SS | 5393 | ENSG00000116353 | MECR | chr1 | - | 29216587 | 29216812 | 29216587 | 29216685 | 29220837 | 29220921 | 18 | 11 | 13 | [illegible] |
| A3SS | 3814 | ENSG00000106692 | FKTN | chr9 | + | 105619905 | 105620061 | 105619933 | 105620061 | 105617958 | 105618092 | 7 | 14 | 10 | [illegible] |
| A3SS | 8138 | ENSG00000102081 | FMR1 | chrX | + | 147944904 | 147945051 | 147944943 | 147945051 | 147940575 | 147940662 | 12 | 9 | 8 | [illegible] |
| A3SS | 11520 | ENSG00000184939 | ZFP90 | chr16 | + | 68558374 | 68558568 | 68558472 | 68558568 | 68557997 | 68558124 | 41 | 25 | 19 | [illegible] |
| A3SS | 6308 | ENSG00000164306 | PRIMPOL | chr4 | + | 184657081 | 184657320 | 184657084 | 184657320 | 184652022 | 184652100 | 14 | 20 | 14 | [illegible] |
| A3SS | 1566 | ENSG00000276070 | CCL4L2 | chr17 | + | 36212312 | 36212878 | 36212481 | 36212878 | 36211775 | 36211890 | 48 | 12 | 3 | [illegible] |
| A3SS | 6533 | ENSG00000123636 | BAZ2B | chr2 | - | 159448241 | 159448409 | 159448241 | 159448403 | 159478574 | 159478721 | 28 | 14 | 26 | [illegible] |
| A3SS | 6534 | ENSG00000153187 | HNRNPU | chr1 | - | 244847001 | 244847101 | 244847001 | 244847053 | 244848787 | 244848856 | 8 | 13 | 8 | [illegible] |
| A3SS | 966 | ENSG00000176225 | RTTN | chr18 | - | 70003803 | 70004236 | 70003803 | 70004168 | 70005197 | 70005267 | 18 | 20 | 15 | [illegible] |
| A3SS | 1556 | ENSG00000276070 | CCL4L2 | chr17 | + | 36212292 | 36212878 | 36212481 | 36212878 | 36211062 | 36211217 | 48 | 12 | 3 | [illegible] |
| A3SS | 377 | ENSG00000196268 | ZNF493 | chr19 | + | 21405757 | 21405856 | 21405760 | 21405856 | 21405128 | 21405255 | 15 | 12 | 19 | [illegible] |
| A3SS | 3269 | ENSG00000197603 | CPLANE1 | chr5 | - | 37206196 | 37206425 | 37206196 | 37206296 | 37213558 | 37213622 | 36 | 9 | 23 | [illegible] |
| A3SS | 4427 | ENSG00000157036 | EXOG | chr3 | + | 38501354 | 38501494 | 38501415 | 38501494 | 38500089 | 38500164 | 27 | 31 | 21 | [illegible] |
| A3SS | 10539 | ENSG00000244754 | N4BP2L2 | chr13 | - | 32442387 | 32444124 | 32442387 | 32442548 | 32517856 | 32518003 | 9 | 18 | 34 | [illegible] |
| A3SS | 5826 | ENSG00000170248 | PDCD6IP | chr3 | + | 33828837 | 33828969 | 33828852 | 33828969 | 33825186 | 33825340 | 24 | 18 | 17 | [illegible] |
| A3SS | 6931 | ENSG00000132635 | PCED1A | chr20 | - | 2838843 | 2839082 | 2838843 | 2838929 | 2839191 | 2839271 | 59 | 32 | 67 | [illegible] |
| A3SS | 4782 | ENSG00000168615 | ADAM9 | chr8 | + | 39090046 | 39090188 | 39090135 | 39090188 | 39082640 | 39082721 | 37 | 26 | 17 | [illegible] |
| A3SS | 3390 | ENSG00000101166 | PRELID3B | chr20 | - | 59037190 | 59037299 | 59037190 | 59037280 | 59038465 | 59038634 | 26 | 1 | 11 | [illegible] |
| A3SS | 11260 | ENSG00000216937 | CCDC7 | chr10 | + | 32728886 | 32728997 | 32728931 | 32728997 | 32726733 | 32726832 | 6 | 31 | 7 | [illegible] |
| A3SS | 3812 | ENSG00000106692 | FKTN | chr9 | + | 105615277 | 105615407 | 105615280 | 105615407 | 105607818 | 105607951 | 8 | 17 | 17 | [illegible] |
| A3SS | 2462 | ENSG00000166200 | COPS2 | chr15 | - | 49137347 | 49137458 | 49137347 | 49137437 | 49144226 | 49144304 | 9 | 6 | 15 | [illegible] |
| A3SS | 10420 | ENSG00000103037 | SETD6 | chr16 | + | 58518050 | 58518231 | 58518180 | 58518231 | 58516807 | 58516928 | 61 | 63 | 80 | [illegible] |
| A3SS | 10371 | ENSG00000158201 | ABHD3 | chr18 | - | 21703583 | 21703747 | 21703583 | 21703677 | 21704503 | 21704774 | 92 | 43 | 77 | [illegible] |
| A3SS | 8907 | ENSG00000114956 | DGUOK | chr2 | + | 73950584 | 73950732 | 73950626 | 73950732 | 73946718 | 73946906 | 82 | 39 | 74 | [illegible] |
| A3SS | 13004 | ENSG00000135297 | MTO1 | chr6 | + | 73466208 | 73466606 | 73466488 | 73466606 | 73461748 | 73462071 | 59 | 34 | 65 | [illegible] |
| A3SS | 11586 | ENSG00000165282 | PIGO | chr9 | - | 35091239 | 35094359 | 35091239 | 35092767 | 35095054 | 35095566 | 29 | 18 | 56 | [illegible] |
| A3SS | 1012 | ENSG00000150477 | KIAA1328 | chr18 | + | 36834202 | 36834355 | 36834319 | 36834355 | 36832673 | 36833388 | 5 | 22 | 10 | [illegible] |
| A3SS | 10017 | ENSG00000177189 | RPS6KA3 | chrX | - | 20240568 | 20240669 | 20240568 | 20240605 | 20241592 | 20241686 | 16 | 14 | 19 | [illegible] |
| A3SS | 1062 | ENSG00000101639 | CEP192 | chr18 | + | 13116373 | 13116503 | 13116376 | 13116503 | 13114129 | 13114251 | 16 | 9 | 17 | [illegible] |
| A3SS | 8181 | ENSG00000064703 | DDX20 | chr1 | + | 111760973 | 111761284 | 111761225 | 111761284 | 111760705 | 111760848 | 9 | 12 | 7 | [illegible] |
| A3SS | 5828 | ENSG00000170248 | PDCD6IP | chr3 | + | 33828837 | 33828969 | 33828852 | 33828969 | 33827621 | 33827739 | 23 | 16 | 16 | [illegible] |
| A3SS | 1582 | ENSG00000108599 | AKAP10 | chr17 | - | 19958013 | 19958571 | 19958013 | 19958567 | 19962839 | 19963022 | 34 | 20 | 18 | [illegible] |
| A3SS | 5382 | ENSG00000133256 | PDE6B | chr4 | + | 670045 | 670782 | 670048 | 670782 | 667855 | 668006 | 14 | 10 | 16 | [illegible] |
| A3SS | 10912 | ENSG00000148516 | ZEB1 | chr10 | + | 31495775 | 31495838 | 31495778 | 31495838 | 31461036 | 31461237 | 8 | 15 | 20 | [illegible] |
| A3SS | 3521 | ENSG00000013375 | PGM3 | chr6 | - | 83188613 | 83188798 | 83188613 | 83188794 | 83190808 | 83191014 | 25 | 20 | 33 | [illegible] |
| A3SS | 1569 | ENSG00000276070 | CCL4L2 | chr17 | + | 36212333 | 36212878 | 36212481 | 36212878 | 36211775 | 36211890 | 53 | 12 | 3 | [illegible] |
| A3SS | 1572 | ENSG00000276070 | CCL4L2 | chr17 | + | 36212363 | 36212878 | 36212496 | 36212878 | 36211775 | 36211890 | 42 | 10 | 2 | [illegible] |
| A3SS | 12361 | ENSG00000187240 | DYNC2H1 | chr11 | + | 103135701 | 103135948 | 103135866 | 103135948 | 103135494 | 103135634 | 16 | 19 | 5 | [illegible] |
| A3SS | 2107 | ENSG00000196678 | ERI2 | chr16 | - | 20803432 | 20803516 | 20803432 | 20803511 | 20803602 | 20803670 | 20 | 9 | 4 | [illegible] |
| A3SS | 4381 | ENSG00000009790 | TRAF3IP3 | chr1 | + | 209781344 | 209781458 | 209781349 | 209781458 | 209779314 | 209779374 | 26 | 19 | 42 | [illegible] |
| A3SS | 10932 | ENSG00000104325 | DECR1 | chr8 | + | 90017123 | 90017326 | 90017202 | 90017326 | 90005447 | 90005845 | 135 | 72 | 88 | [illegible] |
| A3SS | 2755 | ENSG00000197045 | GMFB | chr14 | - | 54480873 | 54480962 | 54480873 | 54480941 | 54481325 | 54481458 | 11 | 10 | 21 | [illegible] |
| A3SS | 169 | ENSG00000100307 | CBX7 | chr22 | - | 39134400 | 39134752 | 39134400 | 39134473 | 39141370 | 39141436 | 55 | 33 | 50 | [illegible] |
| A3SS | 7792 | ENSG00000173113 | TRMT112 | chr11 | - | 64317057 | 64317147 | 64317057 | 64317132 | 64317263 | 64317365 | 315 | 136 | 291 | [illegible] |
| A3SS | 7341 | ENSG00000168421 | RHOH | chr4 | + | 40197100 | 40197300 | 40197120 | 40197300 | 40193662 | 40193812 | 148 | 88 | 193 | [illegible] |
| A3SS | 3935 | ENSG00000146350 | TBC1D32 | chr6 | - | 121115171 | 121115747 | 121115171 | 121115241 | 121126377 | 121126461 | 10 | 13 | 7 | [illegible] |
| A3SS | 1563 | ENSG00000276070 | CCL4L2 | chr17 | + | 36212292 | 36212878 | 36212496 | 36212878 | 36211775 | 36211890 | 61 | 14 | 4 | [illegible] |
| A3SS | 2990 | ENSG00000154309 | DISP1 | chr1 | + | 222836922 | 222837384 | 222837218 | 222837384 | 222835118 | 222835215 | 29 | 8 | 25 | [illegible] |
| A3SS | 8570 | ENSG00000104218 | CSPP1 | chr8 | + | 67093542 | 67093641 | 67093547 | 67093641 | 67086006 | 67086110 | 17 | 13 | 15 | [illegible] |
| A3SS | 1706 | ENSG00000033011 | ALG1 | chr16 | + | 5075387 | 5075536 | 5075397 | 5075536 | 5073152 | 5073256 | 48 | 18 | 35 | [illegible] |
| A3SS | 10005 | ENSG00000113013 | HSPA9 | chr5 | - | 138574067 | 138574126 | 138574067 | 138574121 | 138575237 | 138575417 | 87 | 44 | 61 | [illegible] |
| A3SS | 284 | ENSG00000189144 | ZNF573 | chr19 | - | 37771563 | 37771696 | 37771563 | 37771690 | 37772937 | 37772982 | 16 | 8 | 11 | [illegible] |
| A3SS | 11272 | ENSG00000180776 | ZDHHC20 | chr13 | - | 21381433 | 21381549 | 21381433 | 21381546 | 21381900 | 21381994 | 37 | 32 | 22 | [illegible] |
| A3SS | 7115 | ENSG00000228956 | SATB1-AS1 | chr3 | + | 18527784 | 18528139 | 18528075 | 18528139 | 18527173 | 18527326 | 6 | 3 | 15 | [illegible] |
| A3SS | 8926 | ENSG00000240344 | PPIL3 | chr2 | - | 200885261 | 200885367 | 200885261 | 200885343 | 200885697 | 200885772 | 26 | 10 | 11 | [illegible] |
| A3SS | 3156 | ENSG00000133112 | TPT1 | chr13 | - | 45333470 | 45337553 | 45333470 | 45337388 | 45338659 | 45338776 | 89 | 65 | 132 | [illegible] |
| A3SS | 9999 | ENSG00000160856 | FCRL3 | chr1 | - | 157697139 | 157697919 | 157697139 | 157697424 | 157698383 | 157698629 | 288 | 99 | 131 | [illegible] |
| A3SS | 8175 | ENSG00000180488 | MIGA1 | chr1 | + | 77873000 | 77873120 | 77873003 | 77873120 | 77866337 | 77866391 | 8 | 4 | 18 | [illegible] |
| A3SS | 6995 | ENSG00000225470 | JPX | chrX | + | 73998767 | 74000496 | 73998953 | 74000496 | 73994839 | 73994961 | 12 | 3 | 14 | [illegible] |
| A3SS | 4296 | ENSG00000114127 | XRN1 | chr3 | - | 142332374 | 142332534 | 142332374 | 142332531 | 142335447 | 142335509 | 9 | 4 | 3 | [illegible] |
| A3SS | 3173 | ENSG00000106123 | EPHB6 | chr7 | + | 142870529 | 142870685 | 142870590 | 142870685 | 142870213 | 142870407 | 24 | 13 | 34 | [illegible] |
| A3SS | 9275 | ENSG00000135486 | HNRNPA1 | chr12 | + | 54284239 | 54284317 | 54284257 | 54284317 | 54283811 | 54283967 | 24 | 30 | 34 | [illegible] |
| A3SS | 2026 | ENSG00000005194 | CIAPIN1 | chr16 | - | 57439181 | 57439334 | 57439181 | 57439295 | 57440771 | 57440983 | 96 | 63 | 103 | [illegible] |
| A3SS | 10351 | ENSG00000174080 | CTSF | chr11 | - | 66565670 | 66565751 | 66565670 | 66565749 | 66565830 | 66565927 | 32 | 5 | 37 | [illegible] |
| A3SS | 5750 | ENSG00000049246 | PER3 | chr1 | + | 7829833 | 7830161 | 7829836 | 7830161 | 7827117 | 7827815 | 19 | 10 | 3 | [illegible] |
| A3SS | 9212 | ENSG00000107890 | ANKRD26 | chr10 | - | 27046352 | 27046523 | 27046352 | 27046520 | 27048800 | 27048979 | 3 | 11 | 19 | [illegible] |
| A3SS | 9029 | ENSG00000067248 | DHX29 | chr5 | - | 55273292 | 55273426 | 55273292 | 55273377 | 55274613 | 55274731 | 12 | 14 | 6 | [illegible] |
| A3SS | 5977 | ENSG00000163867 | ZMYM6 | chr1 | - | 34986164 | 34990311 | 34986164 | 34988935 | 34992233 | 34992387 | 15 | 14 | 11 | [illegible] |
| A3SS | 1567 | ENSG00000276070 | CCL4L2 | chr17 | + | 36212312 | 36212878 | 36212496 | 36212878 | 36211775 | 36211890 | 52 | 12 | 3 | [illegible] |
| A3SS | 12772 | ENSG00000066651 | TRMT11 | chr6 | + | 125993756 | 125993822 | 125993792 | 125993822 | 125986552 | 125986773 | 26 | 6 | 20 | [illegible] |
| A3SS | 6043 | ENSG00000113597 | TRAPPC13 | chr5 | + | 65664233 | 65664384 | 65664236 | 65664384 | 65662049 | 65662150 | 19 | 14 | 21 | [illegible] |
| A3SS | 5177 | ENSG00000118200 | CAMSAP2 | chr1 | + | 200847209 | 200847292 | 200847257 | 200847292 | 200844781 | 200844869 | 4 | 10 | 6 | [illegible] |
| A3SS | 9973 | ENSG00000091140 | DLD | chr7 | + | 107904862 | 107905058 | 107904957 | 107905058 | 107904662 | 107904746 | 11 | 24 | 26 | [illegible] |
| A3SS | 1028 | ENSG00000141469 | SLC14A1 | chr18 | + | 45749777 | 45752520 | 45752177 | 45752520 | 45748375 | 45748425 | 45 | 19 | 27 | [illegible] |
| A3SS | 12206 | ENSG00000073910 | FRY | chr13 | + | 32267169 | 32267359 | 32267242 | 32267359 | 32265449 | 32265616 | 29 | 9 | 8 | [illegible] |
| A3SS | 1726 | ENSG00000273611 | ZNHIT3 | chr17 | + | 36492812 | 36492899 | 36492817 | 36492899 | 36486934 | 36486966 | 104 | 46 | 111 | [illegible] |
| A3SS | 3751 | ENSG00000108961 | RANGRF | chr17 | + | 8289257 | 8290092 | 8289812 | 8290092 | 8288955 | 8289072 | 69 | 19 | 52 | [illegible] |
| A3SS | 10736 | ENSG00000068366 | ACSL4 | chrX | - | 109683135 | 109683666 | 109683135 | 109683552 | 109696143 | 109696196 | 21 | 22 | 10 | [illegible] |
| A3SS | 16 | ENSG00000154734 | ADAMTS1 | chr21 | - | 26839874 | 26840562 | 26839874 | 26840061 | 26840997 | 26841165 | 21 | 6 | 11 | [illegible] |
| A3SS | 8962 | ENSG00000146285 | SCML4 | chr6 | - | 107744948 | 107746889 | 107744948 | 107745143 | 107749683 | 107749813 | 25 | 22 | 38 | [illegible] |
| A3SS | 4783 | ENSG00000168615 | ADAM9 | chr8 | + | 39090046 | 39090188 | 39090135 | 39090188 | 39082967 | 39083073 | 87 | 81 | 57 | [illegible] |
| A3SS | 12438 | ENSG00000090621 | PABPC4 | chr1 | - | 39562322 | 39563922 | 39562322 | 39562416 | 39564470 | 39564542 | 54 | 21 | 58 | [illegible] |
| A3SS | 2437 | ENSG00000067141 | NEO1 | chr15 | + | 73236286 | 73236506 | 73236346 | 73236506 | 73178306 | 73178427 | 18 | 12 | 16 | [illegible] |
| A3SS | 11066 | ENSG00000136870 | ZNF189 | chr9 | + | 101399883 | 101400010 | 101399925 | 101400010 | 101399394 | 101399530 | 13 | 5 | 18 | [illegible] |
| A3SS | 1570 | ENSG00000276070 | CCL4L2 | chr17 | + | 36212333 | 36212878 | 36212496 | 36212878 | 36211775 | 36211890 | 57 | 12 | 3 | [illegible] |
| A3SS | 12825 | ENSG00000064102 | INTS13 | chr12 | - | 26914407 | 26914578 | 26914407 | 26914532 | 26917351 | 26917441 | 10 | 8 | 15 | [illegible] |
| A3SS | 12442 | ENSG00000151150 | ANK3 | chr10 | - | 60042671 | 60042759 | 60042671 | 60042756 | 60055657 | 60056036 | 12 | 8 | 32 | [illegible] |
| A3SS | 10503 | ENSG00000163605 | PPP4R2 | chr3 | + | 73064002 | 73064146 | 73064064 | 73064146 | 73061022 | 73061060 | 45 | 45 | 39 | [illegible] |
| A3SS | 2109 | ENSG00000162104 | ADCY9 | chr16 | - | 3992145 | 3992363 | 3992145 | 3992270 | 3993405 | 3993510 | 112 | 18 | 31 | [illegible] |
| A3SS | 6053 | ENSG00000197530 | MIB2 | chr1 | + | 1625545 | 1626754 | 1626649 | 1626754 | 1625285 | 1625428 | 30 | 7 | 28 | [illegible] |
| A3SS | 11052 | ENSG00000156802 | ATAD2 | chr8 | - | 123349284 | 123349444 | 123349284 | 123349421 | 123356388 | 123356477 | 18 | 8 | 38 | [illegible] |
| A3SS | 13099 | ENSG00000127328 | RAB3IP | chr12 | + | 69812777 | 69813033 | 69812963 | 69813033 | 69800208 | 69800332 | 9 | 11 | 35 | [illegible] |
| A3SS | 6027 | ENSG00000233184 | NA | chr1 | + | 101080388 | 101083574 | 101083460 | 101083574 | 101075394 | 101075716 | 8 | 15 | 22 | [illegible] |
| A3SS | 9107 | ENSG00000178904 | DPY19L3 | chr19 | + | 32467473 | 32468813 | 32468730 | 32468813 | 32464727 | 32464784 | 23 | 19 | 36 | [illegible] |
| A3SS | 12617 | ENSG00000138160 | KIF11 | chr10 | + | 92637386 | 92637545 | 92637390 | 92637545 | 92637183 | 92637309 | 13 | 17 | 17 | [illegible] |
| A3SS | 7590 | ENSG00000136891 | TEX10 | chr9 | - | 100303631 | 100303842 | 100303631 | 100303785 | 100308499 | 100308681 | 46 | 14 | 19 | [illegible] |
| A3SS | 1236 | ENSG00000108622 | ICAM2 | chr17 | - | 64006630 | 64006735 | 64006630 | 64006731 | 64020405 | 64020634 | 23 | 8 | 9 | [illegible] |
| A3SS | 8349 | ENSG00000197858 | GPAA1 | chr8 | + | 144084721 | 144085479 | 144085288 | 144085479 | 144084412 | 144084609 | 45 | 8 | 36 | [illegible] |
| A3SS | 6299 | ENSG00000003393 | ALS2 | chr2 | - | 201704453 | 201704603 | 201704453 | 201704583 | 201705138 | 201705200 | 10 | 13 | 14 | [illegible] |
| A3SS | 7718 | ENSG00000049167 | ERCC8 | chr5 | - | 60924290 | 60924508 | 60924290 | 60924429 | 60928863 | 60928959 | 12 | 10 | 15 | [illegible] |
| A3SS | 9067 | ENSG00000165792 | METTL17 | chr14 | + | 20996784 | 20997030 | 20996811 | 20997030 | 20996526 | 20996711 | 14 | 8 | 38 | [illegible] |
| A3SS | 2143 | ENSG00000103197 | TSC2 | chr16 | + | 2056643 | 2056769 | 2056655 | 2056769 | 2056195 | 2056244 | 57 | 24 | 32 | [illegible] |
| A3SS | 11071 | ENSG00000214706 | IFRD2 | chr3 | - | 50289254 | 50289360 | 50289254 | 50289356 | 50289446 | 50289628 | 24 | 10 | 30 | [illegible] |
| A3SS | 4924 | ENSG00000198818 | SFT2D1 | chr6 | - | 166322856 | 166324595 | 166322856 | 166322886 | 166328275 | 166328357 | 21 | 23 | 18 | [illegible] |
| A3SS | 1902 | ENSG00000214078 | CPNE1 | chr20 | - | 35630438 | 35631034 | 35630438 | 35630490 | 35631113 | 35631173 | 188 | 145 | 255 | [illegible] |
| A3SS | 773 | ENSG00000101181 | MTG2 | chr20 | + | 62200682 | 62203568 | 62203083 | 62203568 | 62199118 | 62199257 | 33 | 21 | 36 | [illegible] |
| A3SS | 4297 | ENSG00000114346 | ECT2 | chr3 | + | 172755294 | 172755374 | 172755297 | 172755374 | 172754508 | 172754660 | 22 | 16 | 20 | [illegible] |
| A3SS | 13352 | ENSG00000137822 | TUBGCP4 | chr15 | + | 43401715 | 43403799 | 43403682 | 43403799 | 43400043 | 43400221 | 32 | 59 | 39 | [illegible] |
| A3SS | 7534 | ENSG00000130713 | EXOSC2 | chr9 | + | 130702133 | 130702310 | 130702157 | 130702310 | 130700866 | 130700935 | 69 | 29 | 103 | [illegible] |
| A3SS | 8415 | ENSG00000237765 | FAM200B | chr4 | + | 15686931 | 15687145 | 15687000 | 15687145 | 15686235 | 15686462 | 15 | 9 | 13 | [illegible] |
| A3SS | 12802 | ENSG00000110888 | CAPRIN2 | chr12 | - | 30714958 | 30715141 | 30714958 | 30715081 | 30716507 | 30716676 | 39 | 28 | 21 | [illegible] |
| A3SS | 3962 | ENSG00000248049 | UBA6-AS1 | chr4 | + | 67730073 | 67730170 | 67730079 | 67730170 | 67728539 | 67728686 | 23 | 4 | 10 | [illegible] |
| A3SS | 2835 | ENSG00000050130 | JKAMP | chr14 | + | 59486712 | 59486804 | 59486730 | 59486804 | 59484989 | 59485116 | 24 | 16 | 20 | [illegible] |
| A3SS | 7942 | ENSG00000163006 | CCDC138 | chr2 | + | 108791674 | 108791802 | 108791679 | 108791802 | 108788851 | 108788966 | 8 | 12 | 13 | [illegible] |
| A3SS | 13406 | ENSG00000236778 | INTS6-AS1 | chr13 | + | 51454002 | 51454329 | 51454149 | 51454329 | 51453382 | 51453418 | 21 | 12 | 28 | [illegible] |
| A3SS | 8643 | ENSG00000119912 | IDE | chr10 | - | 92479276 | 92480916 | 92479276 | 92479421 | 92483254 | 92483337 | 13 | 14 | 25 | [illegible] |
| A3SS | 9604 | ENSG00000168014 | C2CD3 | chr11 | - | 74106370 | 74106493 | 74106370 | 74106489 | 74109033 | 74109152 | 29 | 24 | 20 | [illegible] |
| A3SS | 6030 | ENSG00000233184 | NA | chr1 | + | 101080886 | 101083574 | 101083460 | 101083574 | 101077362 | 101077508 | 11 | 15 | 26 | [illegible] |
| A3SS | 3716 | ENSG00000165521 | EML5 | chr14 | - | 88688270 | 88688473 | 88688270 | 88688359 | 88694306 | 88694407 | 15 | 23 | 16 | [illegible] |
| A3SS | 9021 | ENSG00000168785 | TSPAN5 | chr4 | - | 98478684 | 98478810 | 98478684 | 98478746 | 98482004 | 98482175 | 51 | 18 | 39 | [illegible] |
| A3SS | 584 | ENSG00000022556 | NLRP2 | chr19 | + | 54981616 | 54981682 | 54981625 | 54981682 | 54977751 | 54977823 | 31 | 10 | 6 | [illegible] |
| A3SS | 2581 | ENSG00000100749 | VRK1 | chr14 | + | 96846094 | 96846164 | 96846102 | 96846164 | 96837761 | 96837817 | 322 | 202 | 237 | [illegible] |
| A3SS | 10080 | ENSG00000105835 | NAMPT | chr7 | - | 106272529 | 106272658 | 106272529 | 106272655 | 106274945 | 106275049 | 606 | 540 | 404 | [illegible] |
| A3SS | 7255 | ENSG00000116667 | C1orf21 | chr1 | + | 184507587 | 184507682 | 184507597 | 184507682 | 184477385 | 184477603 | 273 | 85 | 57 | [illegible] |
| A3SS | 2619 | ENSG00000129484 | PARP2 | chr14 | + | 20354808 | 20354947 | 20354834 | 20354947 | 20354084 | 20354247 | 33 | 25 | 29 | [illegible] |
| A3SS | 13316 | ENSG00000136153 | LMO7 | chr13 | + | 75835151 | 75835339 | 75835232 | 75835339 | 75834225 | 75834387 | 10 | 10 | 22 | [illegible] |
| A3SS | 7959 | ENSG00000135945 | REV1 | chr2 | - | 99429839 | 99429948 | 99429839 | 99429945 | 99434331 | 99434448 | 22 | 15 | 16 | [illegible] |
| A3SS | 6310 | ENSG00000164306 | PRIMPOL | chr4 | + | 184685408 | 184685498 | 184685411 | 184685498 | 184682247 | 184682336 | 8 | 21 | 18 | [illegible] |
| A3SS | 13057 | ENSG00000090612 | ZNF268 | chr12 | + | 133191488 | 133191615 | 133191533 | 133191615 | 133187871 | 133188072 | 21 | 28 | 17 | [illegible] |
| A3SS | 10969 | ENSG00000120992 | LYPLA1 | chr8 | - | 54051011 | 54051188 | 54051011 | 54051146 | 54055059 | 54055133 | 40 | 32 | 52 | [illegible] |
| A3SS | 7338 | ENSG00000214941 | ZSWIM7 | chr17 | - | 15976559 | 15978163 | 15976559 | 15977092 | 15987265 | 15987368 | 36 | 26 | 22 | [illegible] |
| A3SS | 7116 | ENSG00000228956 | SATB1-AS1 | chr3 | + | 18527950 | 18528139 | 18528075 | 18528139 | 18527173 | 18527326 | 8 | 8 | 18 | [illegible] |
| A3SS | 7539 | ENSG00000138398 | PPIG | chr2 | + | 169631765 | 169631933 | 169631774 | 169631933 | 169631598 | 169631647 | 6 | 13 | 10 | [illegible] |
| A3SS | 10734 | ENSG00000068366 | ACSL4 | chrX | - | 109683135 | 109683552 | 109683135 | 109683375 | 109696143 | 109696196 | 28 | 14 | 11 | [illegible] |
| A3SS | 10540 | ENSG00000244754 | N4BP2L2 | chr13 | - | 32442387 | 32444126 | 32442387 | 32442548 | 32517856 | 32518003 | 10 | 21 | 39 | [illegible] |
| A3SS | 3697 | ENSG00000179088 | C12orf42 | chr12 | - | 103267852 | 103268962 | 103267852 | 103267886 | 103269806 | 103269853 | 20 | 6 | 15 | [illegible] |
| A3SS | 6031 | ENSG00000233184 | NA | chr1 | + | 101080886 | 101083574 | 101083460 | 101083574 | 101080388 | 101080613 | 15 | 15 | 22 | [illegible] |
| A3SS | 3508 | ENSG00000047621 | C12orf4 | chr12 | - | 4536016 | 4536219 | 4536016 | 4536211 | 4538408 | 4538508 | 15 | 14 | 14 | [illegible] |
| A3SS | 7632 | ENSG00000163382 | NAXE | chr1 | + | 156592100 | 156592475 | 156592364 | 156592475 | 156591775 | 156591986 | 65 | 26 | 67 | [illegible] |
| A3SS | 10732 | ENSG00000068366 | ACSL4 | chrX | - | 109683135 | 109683552 | 109683135 | 109683372 | 109696143 | 109696196 | 26 | 16 | 13 | [illegible] |
| A3SS | 5643 | ENSG00000157978 | LDLRAP1 | chr1 | + | 25565172 | 25565207 | 25565175 | 25565207 | 25564590 | 25564761 | 23 | 16 | 43 | [illegible] |
| A3SS | 5670 | ENSG00000134744 | TUT4 | chr1 | - | 52436754 | 52436975 | 52436754 | 52436963 | 52438219 | 52438335 | 14 | 8 | 20 | [illegible] |
| A3SS | 10419 | ENSG00000103037 | SETD6 | chr16 | + | 58518019 | 58518231 | 58518180 | 58518231 | 58516807 | 58516928 | 58 | 65 | 69 | [illegible] |

| | | | | | | | | | | | | | | | | |
|---|---|---|---|---|---|---|---|---|---|---|---|---|---|---|---|---|
| A3SS | 9868 | ENSG00000121716 | PILRB | chr7 | + | 100358219 | 100358366 | 100358263 | 100358366 | 100356749 | 100356883 | 70 | 25 | 71 | 47 | 48 |
| A3SS | 3355 | ENSG00000114107 | CEP70 | chr3 | - | 138572858 | 138572984 | 138572858 | 138572932 | 138591853 | 138591956 | 14 | 14 | 9 | 11 | 34 |
| A3SS | 11160 | ENSG00000134539 | KLRD1 | chr12 | + | 10313406 | 10313513 | 10313409 | 10313513 | 10311463 | 10311619 | 30 | 24 | 13 | 15 | 41 |
| A3SS | 12461 | ENSG00000107443 | CCNJ | chr10 | + | 96057087 | 96057247 | 96057120 | 96057247 | 96056700 | 96057000 | 22 | 10 | 27 | 15 | 49 |
| A3SS | 2829 | ENSG00000205659 | LIN52 | chr14 | + | 74091231 | 74091306 | 74091237 | 74091306 | 74084962 | 74084993 | 17 | 10 | 22 | 22 | 29 |
| A3SS | 10735 | ENSG00000068366 | ACSL4 | chrX | - | 109683135 | 109683666 | 109683135 | 109683375 | 109696143 | 109696196 | 33 | 14 | 12 | 14 | 24 |
| A3SS | 8632 | ENSG00000064201 | TSPAN32 | chr11 | + | 2316575 | 2316667 | 2316631 | 2316667 | 2316228 | 2316312 | 32 | 10 | 12 | 9 | 26 |
| A3SS | 872 | ENSG00000080839 | RBL1 | chr20 | - | 37066992 | 37067121 | 37066992 | 37067021 | 37067232 | 37067297 | 184 | 134 | 161 | 203 | 218 |
| A3SS | 967 | ENSG00000176225 | RTTN | chr18 | - | 70005197 | 70006484 | 70005197 | 70005267 | 70017406 | 70017674 | 22 | 20 | 23 | 12 | 47 |
| A3SS | 845 | ENSG00000125885 | MCM8 | chr20 | + | 5967829 | 5968025 | 5967877 | 5968025 | 5967435 | 5967587 | 34 | 34 | 22 | 37 | 27 |
| A3SS | 8406 | ENSG00000224660 | SH3BP5-AS1 | chr3 | + | 15258545 | 15260582 | 15258633 | 15260582 | 15258108 | 15258183 | 32 | 27 | 32 | 27 | 36 |
| A3SS | 3603 | ENSG00000159199 | ATP5MC1 | chr17 | + | 48895155 | 48895334 | 48895182 | 48895334 | 48894371 | 48894449 | 55 | 22 | 65 | 42 | 25 |
| A3SS | 10733 | ENSG00000068366 | ACSL4 | chrX | - | 109683135 | 109683666 | 109683135 | 109683372 | 109696143 | 109696196 | 31 | 16 | 14 | 14 | 24 |
| A3SS | 12462 | ENSG00000107443 | CCNJ | chr10 | + | 96057087 | 96057247 | 96057123 | 96057247 | 96056700 | 96057000 | 22 | 10 | 27 | 16 | 49 |
| A3SS | 11754 | ENSG00000197563 | PIGN | chr18 | - | 62148213 | 62148338 | 62148213 | 62148253 | 62157128 | 62157227 | 24 | 24 | 28 | 24 | 76 |
| A3SS | 2429 | ENSG00000140474 | ULK3 | chr15 | - | 74841404 | 74841509 | 74841404 | 74841505 | 74842074 | 74842195 | 44 | 29 | 54 | 53 | 89 |
| A3SS | 2991 | ENSG00000154309 | DISP1 | chr1 | + | 222836995 | 222837380 | 222837218 | 222837380 | 222835118 | 222835215 | 50 | 33 | 67 | 37 | 24 |
| A3SS | 4582 | ENSG00000164073 | MFSD8 | chr4 | - | 127957500 | 127957592 | 127957500 | 127957590 | 127965071 | 127965164 | 32 | 19 | 42 | 20 | 32 |
| A3SS | 4402 | ENSG00000128513 | POT1 | chr7 | - | 124928814 | 124928999 | 124928814 | 124928996 | 124929793 | 124929825 | 21 | 14 | 23 | 16 | 20 |
| A3SS | 5285 | ENSG00000127054 | INTS11 | chr1 | - | 1315519 | 1315618 | 1315519 | 1315614 | 1319295 | 1319524 | 89 | 40 | 62 | 79 | 74 |
| A3SS | 8616 | ENSG00000095059 | DHPS | chr19 | - | 12676016 | 12676142 | 12676016 | 12676078 | 12677290 | 12677396 | 47 | 31 | 36 | 35 | 42 |
| A3SS | 6930 | ENSG00000132635 | PCED1A | chr20 | - | 2836038 | 2836314 | 2836038 | 2836110 | 2838231 | 2838478 | 84 | 33 | 76 | 80 | 41 |
| A3SS | 6712 | ENSG00000188529 | SRSF10 | chr1 | - | 23974973 | 23975077 | 23974973 | 23975051 | 23977910 | 23978273 | 33 | 20 | 29 | 18 | 51 |
| A3SS | 7869 | ENSG00000143387 | CTSK | chr1 | - | 150799543 | 150799709 | 150799543 | 150799705 | 150804020 | 150804239 | 38 | 32 | 25 | 23 | 34 |
| A3SS | 7756 | ENSG00000114120 | SLC25A36 | chr3 | + | 140966933 | 140967036 | 140966999 | 140967036 | 140966347 | 140966434 | 16 | 15 | 23 | 22 | 20 |
| A3SS | 6130 | ENSG00000198408 | OGA | chr10 | - | 101792838 | 101792943 | 101792838 | 101792901 | 101793685 | 101793744 | 47 | 29 | 42 | 33 | 62 |
| A3SS | 3370 | ENSG00000204120 | GIGYF2 | chr2 | + | 232760479 | 232760591 | 232760483 | 232760591 | 232756222 | 232756334 | 173 | 129 | 108 | 156 | 135 |
| A3SS | 10359 | ENSG00000174080 | CTSF | chr11 | - | 66566290 | 66566404 | 66566290 | 66566401 | 66567245 | 66567321 | 29 | 7 | 51 | 21 | 67 |
| A3SS | 1273 | ENSG00000161381 | PLXDC1 | chr17 | - | 39079103 | 39079528 | 39079103 | 39079164 | 39087606 | 39087702 | 8 | 3 | 29 | 36 | 11 |
| A3SS | 3158 | ENSG00000175857 | GAPT | chr5 | + | 58494144 | 58497090 | 58496722 | 58497090 | 58493618 | 58493683 | 24 | 10 | 16 | 26 | 36 |
| A3SS | 10084 | ENSG00000105835 | NAMPT | chr7 | - | 106274945 | 106275049 | 106274945 | 106275043 | 106277083 | 106277179 | 29 | 16 | 21 | 40 | 116 |
| A3SS | 1467 | ENSG00000062716 | VMP1 | chr17 | + | 59764970 | 59765138 | 59765051 | 59765138 | 59735337 | 59735473 | 183 | 126 | 125 | 163 | 207 |
| A3SS | 11127 | ENSG00000104408 | EIF3E | chr8 | - | 108234997 | 108235102 | 108234997 | 108235097 | 108236162 | 108236203 | 18 | 11 | 23 | 10 | 26 |
| A3SS | 11250 | ENSG00000143437 | ARNT | chr1 | - | 150816258 | 150816406 | 150816258 | 150816400 | 150816787 | 150816890 | 92 | 58 | 80 | 79 | 220 |
| A3SS | 3631 | ENSG00000128059 | PPAT | chr4 | - | 56406494 | 56406701 | 56406494 | 56406618 | 56407649 | 56407716 | 17 | 13 | 27 | 22 | 32 |
| A3SS | 3541 | ENSG00000167766 | ZNF83 | chr19 | - | 52616717 | 52616841 | 52616717 | 52616815 | 52618935 | 52619062 | 24 | 16 | 10 | 15 | 21 |
| A3SS | 6028 | ENSG00000233184 | NA | chr1 | + | 101080388 | 101083574 | 101080886 | 101083574 | 101077362 | 101077508 | 5 | 22 | 23 | 13 | 35 |
| A3SS | 2659 | ENSG00000100564 | PIGH | chr14 | - | 67592634 | 67592718 | 67592634 | 67592715 | 67593742 | 67593952 | 37 | 11 | 30 | 37 | 44 |
| A3SS | 4643 | ENSG00000164463 | CREBRF | chr5 | + | 173080584 | 173080784 | 173080588 | 173080784 | 173056351 | 173056479 | 26 | 28 | 40 | 28 | 13 |
| A3SS | 9690 | ENSG00000145191 | EIF2B5 | chr3 | + | 184144098 | 184144224 | 184144127 | 184144224 | 184143051 | 184143565 | 97 | 55 | 85 | 67 | 102 |
| A3SS | 11166 | ENSG00000176623 | RMDN1 | chr8 | - | 86474819 | 86474953 | 86474819 | 86474914 | 86478922 | 86479010 | 14 | 30 | 7 | 18 | 42 |
| A3SS | 11459 | ENSG00000173611 | SCAI | chr9 | - | 124971670 | 124971844 | 124971670 | 124971762 | 124976113 | 124976186 | 32 | 18 | 19 | 10 | 36 |
| A3SS | 6523 | ENSG00000113812 | ACTR8 | chr3 | - | 53879938 | 53880109 | 53879938 | 53880072 | 53881978 | 53882152 | 44 | 35 | 32 | 44 | 35 |
| A3SS | 7929 | ENSG00000167220 | HDHD2 | chr18 | - | 47134495 | 47134704 | 47134495 | 47134677 | 47136301 | 47136449 | 40 | 33 | 20 | 29 | 30 |
| A3SS | 1634 | ENSG00000167377 | ZNF23 | chr16 | - | 71447596 | 71449885 | 71447596 | 71449882 | 71453242 | 71453350 | 15 | 23 | 28 | 18 | 20 |
| A3SS | 12562 | ENSG00000198825 | INPP5F | chr10 | + | 119811756 | 119811955 | 119811809 | 119811955 | 119810099 | 119810217 | 22 | 23 | 30 | 31 | 26 |
| A3SS | 5441 | ENSG00000121940 | CLCC1 | chr1 | - | 108950308 | 108950448 | 108950308 | 108950355 | 108962308 | 108963453 | 72 | 44 | 53 | 45 | 62 |
| A3SS | 6753 | ENSG00000138614 | INTS14 | chr15 | - | 65593427 | 65593572 | 65593427 | 65593569 | 65595732 | 65595825 | 61 | 20 | 55 | 32 | 53 |
| A3SS | 969 | ENSG00000176225 | RTTN | chr18 | - | 70017406 | 70017674 | 70017406 | 70017643 | 70020614 | 70020817 | 51 | 22 | 34 | 31 | 49 |
| A3SS | 11158 | ENSG00000134539 | KLRD1 | chr12 | + | 10313406 | 10313513 | 10313409 | 10313513 | 10309625 | 10309688 | 33 | 25 | 14 | 17 | 44 |
| A3SS | 7715 | ENSG00000049167 | ERCC8 | chr5 | - | 60924290 | 60924508 | 60924290 | 60924356 | 60928863 | 60928959 | 15 | 24 | 18 | 21 | 25 |
| A3SS | 1602 | ENSG00000172171 | TEFM | chr17 | - | 30900412 | 30900562 | 30900412 | 30900521 | 30904065 | 30904529 | 11 | 25 | 24 | 18 | 41 |
| A3SS | 7773 | ENSG00000197841 | ZNF181 | chr19 | + | 34739522 | 34739621 | 34739525 | 34739621 | 34739147 | 34739268 | 27 | 10 | 19 | 14 | 35 |
| A3SS | 10481 | ENSG00000161791 | FMNL3 | chr12 | - | 49647702 | 49647804 | 49647702 | 49647801 | 49648192 | 49648353 | 66 | 56 | 71 | 110 | 55 |
| A3SS | 1820 | ENSG00000101624 | CEP76 | chr18 | - | 12699829 | 12699905 | 12699829 | 12699900 | 12700957 | 12701113 | 23 | 14 | 19 | 19 | 47 |
| A3SS | 1585 | ENSG00000108599 | AKAP10 | chr17 | - | 19962839 | 19963022 | 19962839 | 19962991 | 19968413 | 19968461 | 34 | 21 | 23 | 23 | 35 |
| A3SS | 2215 | ENSG00000062524 | LTK | chr15 | - | 41507094 | 41507657 | 41507094 | 41507290 | 41508068 | 41508221 | 29 | 19 | 24 | 21 | 14 |
| A3SS | 11498 | ENSG00000070061 | ELP1 | chr9 | - | 108917546 | 108919349 | 108917546 | 108917670 | 108926522 | 108926603 | 33 | 16 | 24 | 43 | 15 |
| A3SS | 6065 | ENSG00000138821 | SLC39A8 | chr4 | - | 102315667 | 102315830 | 102315667 | 102315742 | 102344443 | 102344915 | 53 | 27 | 56 | 54 | 47 |
| A3SS | 6087 | ENSG00000143543 | JTB | chr1 | - | 153976692 | 153976922 | 153976692 | 153976775 | 153977338 | 153977394 | 60 | 32 | 48 | 56 | 47 |
| A3SS | 3542 | ENSG00000167766 | ZNF83 | chr19 | - | 52616717 | 52616875 | 52616717 | 52616815 | 52618935 | 52619062 | 28 | 19 | 17 | 13 | 16 |
| A3SS | 2828 | ENSG00000081177 | EXD2 | chr14 | + | 69234699 | 69235031 | 69234773 | 69235031 | 69230471 | 69230598 | 22 | 13 | 17 | 24 | 18 |
| A3SS | 9234 | ENSG00000134324 | LPIN1 | chr2 | + | 11804422 | 11804571 | 11804427 | 11804571 | 11802906 | 11803033 | 96 | 69 | 104 | 112 | 57 |
| A3SS | 10285 | ENSG00000182220 | ATP6AP2 | chrX | + | 40600761 | 40600881 | 40600768 | 40600881 | 40599591 | 40599741 | 722 | 463 | 421 | 486 | 780 |
| A3SS | 6070 | ENSG00000120798 | NR2C1 | chr12 | - | 95028386 | 95028524 | 95028386 | 95028509 | 95049067 | 95049233 | 14 | 8 | 8 | 15 | 45 |
| A3SS | 9627 | ENSG00000198839 | ZNF277 | chr7 | + | 112286872 | 112287074 | 112286876 | 112287074 | 112206694 | 112206807 | 26 | 6 | 29 | 18 | 19 |
| A3SS | 12788 | ENSG00000150045 | KLRF1 | chr12 | + | 9842301 | 9842433 | 9842320 | 9842433 | 9833302 | 9833452 | 45 | 18 | 8 | 15 | 3 |
| A3SS | 635 | ENSG00000089335 | ZNF302 | chr19 | + | 34683991 | 34686396 | 34684251 | 34686396 | 34683154 | 34683238 | 20 | 22 | 19 | 25 | 23 |
| A3SS | 634 | ENSG00000089335 | ZNF302 | chr19 | + | 34683151 | 34683238 | 34683154 | 34683238 | 34682776 | 34682897 | 9 | 25 | 20 | 17 | 26 |
| A3SS | 12138 | ENSG00000132199 | ENOSF1 | chr18 | - | 690548 | 691276 | 690548 | 690631 | 693881 | 693908 | 26 | 12 | 42 | 29 | 31 |
| A3SS | 6588 | ENSG00000231551 | PDE4DIPP6 | chr1 | + | 148425349 | 148425664 | 148425505 | 148425664 | 148424626 | 148424638 | 90 | 56 | 42 | 60 | 32 |
| A3SS | 1594 | ENSG00000006125 | AP2B1 | chr17 | + | 35650529 | 35650789 | 35650602 | 35650789 | 35641876 | 35641975 | 488 | 314 | 349 | 417 | 462 |
| A3SS | 2852 | ENSG00000197776 | KLHDC1 | chr14 | + | 49710262 | 49710381 | 49710288 | 49710381 | 49709708 | 49709826 | 70 | 45 | 51 | 76 | 78 |
| A3SS | 1115 | ENSG00000132514 | CLEC10A | chr17 | - | 7075984 | 7076152 | 7075984 | 7076071 | 7076732 | 7076804 | 16 | 38 | 14 | 26 | 33 |
| A3SS | 1573 | ENSG00000276070 | CCL4L2 | chr17 | + | 36212481 | 36212878 | 36212496 | 36212878 | 36211775 | 36211890 | 84 | 2 | 4 | 15 | 15 |
| A3SS | 3659 | ENSG00000077232 | DNAJC10 | chr2 | + | 182739544 | 182740388 | 182740298 | 182740388 | 182731029 | 182731107 | 11 | 13 | 27 | 8 | 40 |
| A3SS | 5349 | ENSG00000116001 | TIA1 | chr2 | - | 70214348 | 70214494 | 70214348 | 70214491 | 70216207 | 70216292 | 30 | 25 | 17 | 22 | 25 |
| A3SS | 9583 | ENSG00000164659 | ELAPOR2 | chr7 | - | 86912940 | 86913204 | 86912940 | 86913065 | 86914722 | 86914860 | 34 | 8 | 39 | 16 | 35 |
| A3SS | 6369 | ENSG00000215788 | TNFRSF25 | chr1 | - | 6462862 | 6463147 | 6462862 | 6462956 | 6464374 | 6464453 | 32 | 13 | 52 | 33 | 48 |
| A3SS | 6714 | ENSG00000188529 | SRSF10 | chr1 | - | 23974973 | 23975080 | 23974973 | 23975077 | 23977910 | 23978273 | 40 | 43 | 62 | 62 | 114 |
| A3SS | 8614 | ENSG00000095059 | DHPS | chr19 | - | 12676016 | 12676142 | 12676016 | 12676105 | 12677107 | 12677211 | 90 | 62 | 96 | 81 | 78 |
| A3SS | 6754 | ENSG00000138614 | INTS14 | chr15 | - | 65605128 | 65605236 | 65605128 | 65605232 | 65607158 | 65607442 | 49 | 27 | 48 | 52 | 43 |
| A3SS | 3810 | ENSG00000174652 | ZNF266 | chr19 | - | 9433667 | 9433786 | 9433667 | 9433783 | 9434074 | 9434235 | 18 | 18 | 19 | 17 | 30 |
| A3SS | 12179 | ENSG00000110075 | PPP6R3 | chr11 | + | 68587944 | 68588024 | 68587977 | 68588024 | 68575957 | 68576043 | 79 | 49 | 42 | 74 | 52 |
| A3SS | 10375 | ENSG00000125676 | THOC2 | chrX | - | 123619400 | 123619517 | 123619400 | 123619436 | 123620092 | 123620205 | 20 | 25 | 12 | 24 | 38 |
| A3SS | 6574 | ENSG00000135541 | AHI1 | chr6 | - | 135442581 | 135442714 | 135442581 | 135442671 | 135447007 | 135447160 | 30 | 36 | 18 | 29 | 25 |
| A3SS | 8261 | ENSG00000082269 | FAM135A | chr6 | + | 70524342 | 70526698 | 70525863 | 70526698 | 70523966 | 70524121 | 24 | 25 | 12 | 20 | 24 |
| A3SS | 6029 | ENSG00000233184 | NA | chr1 | + | 101080388 | 101083574 | 101083460 | 101083574 | 101077362 | 101077508 | 10 | 23 | 31 | 14 | 34 |
| A3SS | 3200 | ENSG00000145604 | SKP2 | chr5 | + | 36163644 | 36166662 | 36166518 | 36166662 | 36152770 | 36153042 | 36 | 15 | 30 | 22 | 29 |
| A3SS | 10325 | ENSG00000177853 | ZNF518A | chr10 | + | 96133555 | 96133648 | 96133582 | 96133648 | 96132585 | 96132670 | 23 | 20 | 15 | 27 | 30 |
| A3SS | 11752 | ENSG00000197563 | PIGN | chr18 | - | 62082672 | 62082746 | 62082672 | 62082706 | 62084530 | 62084606 | 51 | 32 | 37 | 40 | 63 |
| A3SS | 12305 | ENSG00000187147 | RNF220 | chr1 | + | 44649882 | 44651723 | 44650703 | 44651723 | 44649660 | 44649769 | 286 | 131 | 266 | 273 | 359 |
| A3SS | 9551 | ENSG00000223705 | NSUN5P1 | chr7 | + | 75412788 | 75415023 | 75414884 | 75415023 | 75411654 | 75411759 | 94 | 51 | 111 | 90 | 114 |
| A3SS | 7580 | ENSG00000171503 | ETFDH | chr4 | + | 158697558 | 158699130 | 158698986 | 158699130 | 158695496 | 158695643 | 36 | 49 | 34 | 32 | 32 |
| A3SS | 3191 | ENSG00000113240 | CLK4 | chr5 | - | 178619837 | 178620673 | 178619837 | 178619896 | 178622392 | 178622459 | 50 | 23 | 44 | 53 | 24 |
| A3SS | 3362 | ENSG00000170144 | HNRNPA3 | chr2 | + | 177216675 | 177216940 | 177216859 | 177216940 | 177216502 | 177216592 | 839 | 965 | 762 | 889 | 778 |
| A3SS | 973 | ENSG00000176225 | RTTN | chr18 | - | 70168854 | 70169067 | 70168854 | 70168973 | 70176674 | 70176845 | 58 | 45 | 27 | 50 | 38 |
| A3SS | 8136 | ENSG00000102081 | FMR1 | chrX | + | 147944868 | 147945051 | 147944904 | 147945051 | 147940575 | 147940662 | 30 | 37 | 35 | 42 | 23 |
| A3SS | 9885 | ENSG00000126883 | NUP214 | chr9 | + | 131144279 | 131144754 | 131144576 | 131144754 | 131140548 | 131140710 | 78 | 48 | 70 | 86 | 157 |
| A3SS | 2731 | ENSG00000100722 | ZC3H14 | chr14 | + | 88609678 | 88609803 | 88609726 | 88609803 | 88609266 | 88609403 | 16 | 19 | 25 | 27 | 79 |
| A3SS | 8721 | ENSG00000164114 | MAP9 | chr4 | - | 155360167 | 155360415 | 155360167 | 155360343 | 155362047 | 155362141 | 13 | 16 | 20 | 31 | 26 |
| A3SS | 6971 | ENSG00000213366 | GSTM2 | chr1 | + | 109671286 | 109671382 | 109671313 | 109671382 | 109670174 | 109670234 | 22 | 29 | 57 | 33 | 12 |
| A3SS | 11399 | ENSG00000089053 | ANAPC5 | chr12 | - | 121318276 | 121318424 | 121318276 | 121318351 | 121318500 | 121318608 | 203 | 130 | 211 | 242 | 506 |
| A3SS | 10073 | ENSG00000105835 | NAMPT | chr7 | - | 106269990 | 106270176 | 106269990 | 106270172 | 106272529 | 106272658 | 13 | 15 | 8 | 56 | 42 |
| A3SS | 1517 | ENSG00000073584 | SMARCE1 | chr17 | - | 40627760 | 40629568 | 40627760 | 40628993 | 40631591 | 40631693 | 55 | 57 | 30 | 42 | 36 |
| A3SS | 5571 | ENSG00000237943 | PRKCQ-AS1 | chr10 | + | 6615369 | 6615637 | 6615518 | 6615637 | 6613630 | 6613689 | 22 | 23 | 41 | 20 | 21 |
| A3SS | 9466 | ENSG00000167380 | ZNF226 | chr19 | + | 44172087 | 44172214 | 44172091 | 44172214 | 44170034 | 44170095 | 55 | 49 | 68 | 51 | 37 |
| A3SS | 7231 | ENSG00000082701 | GSK3B | chr3 | - | 120002045 | 120002239 | 120002045 | 120002236 | 120093346 | 120094447 | 183 | 122 | 142 | 183 | 189 |
| A3SS | 11324 | ENSG00000127084 | FGD3 | chr9 | + | 93032768 | 93034681 | 93034540 | 93034681 | 93029873 | 93029996 | 245 | 139 | 187 | 235 | 117 |
| A3SS | 8885 | ENSG00000123607 | TTC21B | chr2 | - | 165890837 | 165894824 | 165890837 | 165890988 | 165898685 | 165898767 | 22 | 16 | 15 | 33 | 25 |
| A3SS | 7970 | ENSG00000121579 | NAA50 | chr3 | - | 113723958 | 113724095 | 113723958 | 113724092 | 113745941 | 113746249 | 52 | 27 | 30 | 29 | 32 |
| A3SS | 8886 | ENSG00000123607 | TTC21B | chr2 | - | 165936920 | 165937067 | 165936920 | 165936991 | 165941026 | 165941184 | 47 | 23 | 31 | 31 | 38 |
| A3SS | 7587 | ENSG00000035115 | SH3YL1 | chr2 | - | 260084 | 261130 | 260084 | 260702 | 262630 | 262786 | 12 | 16 | 26 | 21 | 35 |
| A3SS | 10376 | ENSG00000125676 | THOC2 | chrX | - | 123619400 | 123619517 | 123619400 | 123619436 | 123620187 | 123620670 | 20 | 25 | 12 | 27 | 38 |
| A3SS | 3976 | ENSG00000177054 | ZDHHC13 | chr11 | + | 19166301 | 19170568 | 19170410 | 19170568 | 19165051 | 19165145 | 30 | 18 | 29 | 30 | 35 |
| A3SS | 11986 | ENSG00000070087 | PFN2 | chr3 | - | 149964903 | 149966586 | 149964903 | 149966264 | 149968357 | 149968550 | 12 | 25 | 16 | 19 | 38 |
| A3SS | 8583 | ENSG00000129680 | MAP7D3 | chrX | - | 136216813 | 136218493 | 136216813 | 136217258 | 136219397 | 136219495 | 30 | 22 | 19 | 25 | 54 |
| A3SS | 6368 | ENSG00000215788 | TNFRSF25 | chr1 | - | 6462862 | 6463127 | 6462862 | 6462956 | 6464374 | 6464453 | 37 | 16 | 57 | 38 | 49 |
| A3SS | 9908 | ENSG00000105865 | DUS4L | chr7 | + | 107576365 | 107576592 | 107576408 | 107576592 | 107575187 | 107575310 | 27 | 23 | 23 | 34 | 45 |
| A3SS | 8554 | ENSG00000143971 | ETAA1 | chr2 | + | 67400343 | 67402974 | 67402861 | 67402974 | 67399549 | 67399626 | 12 | 4 | 16 | 12 | 62 |
| A3SS | 3660 | ENSG00000077232 | DNAJC10 | chr2 | + | 182739544 | 182740388 | 182740298 | 182740388 | 182732498 | 182732542 | 12 | 14 | 28 | 10 | 39 |
| A3SS | 12828 | ENSG00000123200 | ZC3H13 | chr13 | - | 46045390 | 46045516 | 46045390 | 46045513 | 46052403 | 46052746 | 38 | 24 | 34 | 26 | 37 |
| A3SS | 1464 | ENSG00000265118 | NA | chr17 | - | 31313978 | 31314110 | 31313978 | 31314044 | 31314370 | 31314465 | 78 | 61 | 43 | 59 | 79 |
| A3SS | 6196 | ENSG00000136068 | FLNB | chr3 | + | 58130740 | 58132931 | 58132807 | 58132931 | 58126601 | 58126762 | 86 | 51 | 71 | 94 | 100 |
| A3SS | 8918 | ENSG00000115970 | THADA | chr2 | - | 43551788 | 43551925 | 43551788 | 43551922 | 43552203 | 43552339 | 56 | 27 | 37 | 50 | 38 |
| A3SS | 1544 | ENSG00000187607 | ZNF286A | chr17 | + | 15715999 | 15720787 | 15716058 | 15720787 | 15708154 | 15708247 | 22 | 23 | 31 | 33 | 57 |
| A3SS | 8061 | ENSG00000224078 | SNHG14 | chr15 | + | 25108997 | 25109403 | 25109197 | 25109403 | 25107909 | 25108038 | 35 | 15 | 29 | 24 | 56 |
| A3SS | 7987 | ENSG00000104728 | ARHGEF10 | chr8 | + | 1898432 | 1898525 | 1898435 | 1898525 | 1896332 | 1896449 | 4 | 4 | 110 | 4 | 1 |
| A3SS | 5572 | ENSG00000237943 | PRKCQ-AS1 | chr10 | + | 6615369 | 6615637 | 6615518 | 6615637 | 6613933 | 6613987 | 23 | 24 | 42 | 26 | 23 |
| A3SS | 10877 | ENSG00000178234 | GALNT11 | chr7 | + | 152094189 | 152094522 | 152094319 | 152094522 | 152025673 | 152025884 | 88 | 38 | 53 | 59 | 89 |
| A3SS | 10032 | ENSG00000104133 | SPG11 | chr15 | - | 44600466 | 44600632 | 44600466 | 44600498 | 44608443 | 44608605 | 89 | 79 | 75 | 89 | 81 |
| A3SS | 5729 | ENSG00000115446 | UNC50 | chr2 | + | 98609642 | 98610039 | 98609755 | 98610039 | 98608843 | 98609011 | 64 | 32 | 78 | 32 | 91 |
| A3SS | 13167 | ENSG00000134283 | PPHLN1 | chr12 | + | 42445990 | 42446290 | 42446065 | 42446290 | 42442640 | 42442741 | 20 | 15 | 21 | 25 | 44 |
| A3SS | 12591 | ENSG00000028116 | VRK2 | chr2 | + | 58154753 | 58154802 | 58154758 | 58154802 | 58146315 | 58146474 | 27 | 18 | 16 | 16 | 46 |
| A3SS | 6682 | ENSG00000111358 | GTF2H3 | chr12 | + | 123639263 | 123639343 | 123639293 | 123639343 | 123633828 | 123633872 | 27 | 22 | 40 | 22 | 84 |
| A3SS | 9141 | ENSG00000130023 | ERMARD | chr6 | + | 169753863 | 169754032 | 169753943 | 169754032 | 169753037 | 169753181 | 39 | 30 | 44 | 37 | 59 |
| A3SS | 10007 | ENSG00000000971 | CFH | chr1 | + | 196740618 | 196742051 | 196741874 | 196742051 | 196737474 | 196737660 | 26 | 13 | 20 | 32 | 25 |
| A3SS | 10581 | ENSG00000123562 | MORF4L2 | chrX | - | 103678498 | 103678651 | 103678498 | 103678620 | 103684680 | 103684729 | 28 | 29 | 34 | 24 | 91 |
| A3SS | 8219 | ENSG00000121542 | SEC22A | chr3 | + | 123223558 | 123223722 | 123223643 | 123223722 | 123209198 | 123209399 | 86 | 111 | 83 | 85 | 97 |
| A3SS | 3273 | ENSG00000124177 | CHD6 | chr20 | - | 41437273 | 41437334 | 41437273 | 41437332 | 41439999 | 41440129 | 88 | 63 | 118 | 85 | 82 |
| A3SS | 9389 | ENSG00000135317 | SNX14 | chr6 | - | 85507967 | 85508313 | 85507967 | 85508059 | 85513799 | 85513895 | 29 | 15 | 23 | 40 | 25 |
| A3SS | 2816 | ENSG00000033170 | FUT8 | chr14 | + | 65733230 | 65733381 | 65733277 | 65733381 | 65724146 | 65724323 | 67 | 64 | 80 | 77 | 249 |
| A3SS | 2582 | ENSG00000100749 | VRK1 | chr14 | + | 96847256 | 96847344 | 96847261 | 96847344 | 96846094 | 96846164 | 327 | 216 | 240 | 281 | 250 |
| A3SS | 1057 | ENSG00000101639 | CEP192 | chr18 | + | 13057584 | 13059312 | 13059081 | 13059312 | 13055779 | 13056698 | 26 | 11 | 15 | 35 | 41 |
| A3SS | 2637 | ENSG00000139990 | DCAF5 | chr14 | - | 69118138 | 69118275 | 69118138 | 69118194 | 69119193 | 69119230 | 104 | 88 | 125 | 107 | 129 |
| A3SS | 6589 | ENSG00000231551 | PDE4DIPP6 | chr1 | + | 148425349 | 148425664 | 148425505 | 148425664 | 148424626 | 148424678 | 95 | 57 | 48 | 63 | 45 |
| A3SS | 9468 | ENSG00000086827 | ZW10 | chr11 | - | 113736619 | 113736822 | 113736619 | 113736773 | 113737571 | 113737703 | 42 | 25 | 28 | 23 | 62 |
| A3SS | 4145 | ENSG00000113811 | SELENOK | chr3 | - | 53885825 | 53885912 | 53885825 | 53885904 | 53886850 | 53886934 | 376 | 280 | 369 | 422 | 781 |
| A3SS | 2019 | ENSG00000158805 | ZNF276 | chr16 | + | 89723259 | 89723709 | 89723278 | 89723709 | 89723136 | 89723183 | 83 | 32 | 77 | 71 | 13 |
| A3SS | 3250 | ENSG00000164398 | ACSL6 | chr5 | - | 131973265 | 131973400 | 131973265 | 131973358 | 131974892 | 131976240 | 28 | 25 | 47 | 26 | 58 |
| A3SS | 7309 | ENSG00000143156 | NME7 | chr1 | - | 169324392 | 169324500 | 169324392 | 169324469 | 169367707 | 169367946 | 46 | 45 | 32 | 33 | 37 |
| A3SS | 12006 | ENSG00000136527 | TRA2B | chr3 | - | 185918364 | 185918438 | 185918364 | 185918435 | 185919436 | 185919496 | 116 | 74 | 119 | 145 | 151 |
| A3SS | 7659 | ENSG00000205423 | CNEP1R1 | chr16 | + | 50034101 | 50034156 | 50034106 | 50034156 | 50033396 | 50033506 | 36 | 24 | 39 | 28 | 42 |
| A3SS | 9614 | ENSG00000105854 | PON2 | chr7 | - | 95411652 | 95411779 | 95411652 | 95411743 | 95412311 | 95412477 | 41 | 34 | 18 | 33 | 45 |
| A3SS | 5691 | ENSG00000133731 | IMPA1 | chr8 | - | 81676233 | 81676279 | 81676233 | 81676265 | 81679125 | 81679230 | 39 | 16 | 36 | 20 | 57 |
| A3SS | 285 | ENSG00000189144 | ZNF573 | chr19 | - | 37771563 | 37771696 | 37771563 | 37771690 | 37773660 | 37773751 | 52 | 36 | 60 | 37 | 36 |
| A3SS | 3759 | ENSG00000138796 | HADH | chr4 | + | 108027687 | 108028069 | 108027938 | 108028069 | 108023473 | 108023563 | 72 | 71 | 85 | 95 | 69 |
| A3SS | 11904 | ENSG00000149547 | EI24 | chr11 | + | 125575262 | 125575408 | 125575308 | 125575408 | 125572457 | 125572569 | 161 | 64 | 110 | 128 | 121 |
| A3SS | 13212 | ENSG00000166598 | HSP90B1 | chr12 | + | 103932278 | 103932418 | 103932280 | 103932418 | 103931520 | 103931623 | 25 | 22 | 39 | 17 | 63 |
| A3SS | 2752 | ENSG00000166428 | PLD4 | chr14 | + | 104927672 | 104927866 | 104927678 | 104927866 | 104927140 | 104927230 | 33 | 17 | 28 | 52 | 17 |
| A3SS | 13088 | ENSG00000187109 | NAP1L1 | chr12 | - | 76056032 | 76056161 | 76056032 | 76056158 | 76060137 | 76060279 | 92 | 82 | 81 | 77 | 102 |
| A3SS | 11012 | ENSG00000035928 | RFC1 | chr4 | - | 39306591 | 39306704 | 39306591 | 39306701 | 39308635 | 39309032 | 37 | 21 | 32 | 28 | 29 |
| A3SS | 12805 | ENSG00000110888 | CAPRIN2 | chr12 | - | 30719078 | 30720915 | 30719078 | 30719225 | 30723258 | 30723314 | 26 | 22 | 23 | 37 | 47 |
| A3SS | 12672 | ENSG00000150093 | ITGB1 | chr10 | - | 32920244 | 32922346 | 32920244 | 32920385 | 32923584 | 32923740 | 107 | 83 | 49 | 104 | 27 |
| A3SS | 6623 | ENSG00000169764 | UGP2 | chr2 | + | 63885770 | 63885886 | 63885788 | 63885886 | 63885588 | 63885677 | 33 | 29 | 25 | 26 | 49 |
| A3SS | 1249 | ENSG00000078687 | TNRC6C | chr17 | + | 78064721 | 78064937 | 78064730 | 78064937 | 78051066 | 78051448 | 69 | 25 | 60 | 59 | 24 |
| A3SS | 8613 | ENSG00000095059 | DHPS | chr19 | - | 12676016 | 12676142 | 12676016 | 12676078 | 12677107 | 12677211 | 98 | 75 | 100 | 94 | 89 |
| A3SS | 6812 | ENSG00000112303 | VNN2 | chr6 | - | 132751144 | 132751514 | 132751144 | 132751278 | 132752460 | 132752749 | 244 | 127 | 188 | 175 | 530 |
| A3SS | 3443 | ENSG00000112992 | NNT | chr5 | + | 43619031 | 43619119 | 43619086 | 43619119 | 43615847 | 43616065 | 133 | 75 | 90 | 112 | 161 |
| A3SS | 7124 | ENSG00000173744 | AGFG1 | chr2 | + | 227534844 | 227535025 | 227534907 | 227535025 | 227533548 | 227533758 | 112 | 61 | 70 | 103 | 150 |
| A3SS | 8762 | ENSG00000197635 | DPP4 | chr2 | - | 162047402 | 162047501 | 162047402 | 162047498 | 162073398 | 162073486 | 26 | 13 | 49 | 25 | 47 |
| A3SS | 9949 | ENSG00000155096 | AZIN1 | chr8 | - | 102843550 | 102843747 | 102843550 | 102843729 | 102858012 | 102858150 | 228 | 144 | 226 | 247 | 239 |
| A3SS | 7635 | ENSG00000163382 | NAXE | chr1 | + | 156593881 | 156594090 | 156593944 | 156594090 | 156593407 | 156593555 | 110 | 49 | 91 | 90 | 170 |
| A3SS | 1272 | ENSG00000161381 | PLXDC1 | chr17 | - | 39079103 | 39079528 | 39079103 | 39079164 | 39083458 | 39083540 | 13 | 5 | 44 | 63 | 16 |
| A3SS | 12362 | ENSG00000187240 | DYNC2H1 | chr11 | + | 103135719 | 103135948 | 103135866 | 103135948 | 103135494 | 103135634 | 32 | 26 | 15 | 39 | 15 |
| A3SS | 4947 | ENSG00000113810 | SMC4 | chr3 | + | 160404327 | 160404504 | 160404377 | 160404504 | 160401989 | 160402093 | 101 | 105 | 85 | 104 | 109 |
| A3SS | 3159 | ENSG00000175857 | GAPT | chr5 | + | 58494246 | 58497090 | 58496722 | 58497090 | 58493618 | 58493683 | 27 | 18 | 22 | 25 | 47 |
| A3SS | 2638 | ENSG00000139990 | DCAF5 | chr14 | - | 69118138 | 69118278 | 69118138 | 69118194 | 69119193 | 69119230 | 224 | 139 | 231 | 187 | 262 |
| A3SS | 11948 | ENSG00000149273 | RPS3 | chr11 | + | 75402303 | 75402446 | 75402351 | 75402446 | 75401639 | 75401733 | 44 | 19 | 54 | 41 | 18 |
| A3SS | 7311 | ENSG00000135336 | ORC3 | chr6 | + | 87656902 | 87656982 | 87656905 | 87656982 | 87653115 | 87653249 | 19 | 22 | 23 | 18 | 73 |
| A3SS | 13040 | ENSG00000176083 | ZNF683 | chr1 | - | 26367592 | 26367797 | 26367592 | 26367647 | 26368457 | 26368582 | 27 | 12 | 28 | 56 | 56 |
| A3SS | 9648 | ENSG00000183735 | TBK1 | chr12 | + | 64455839 | 64455957 | 64455916 | 64455957 | 64452130 | 64452187 | 143 | 119 | 75 | 106 | 10 |
| A3SS | 9763 | ENSG00000150768 | DLAT | chr11 | + | 112043465 | 112043533 | 112043471 | 112043533 | 112039243 | 112039397 | 71 | 41 | 73 | 62 | 49 |
| A3SS | 5665 | ENSG00000143493 | INTS7 | chr1 | - | 211946606 | 211946705 | 211946606 | 211946645 | 211952568 | 211952701 | 37 | 14 | 19 | 30 | 66 |
| A3SS | 1175 | ENSG00000129197 | RPAIN | chr17 | + | 5425970 | 5426299 | 5426235 | 5426299 | 5422768 | 5422829 | 70 | 34 | 103 | 83 | 90 |
| A3SS | 8155 | ENSG00000155561 | NUP205 | chr7 | + | 135614158 | 135614273 | 135614170 | 135614273 | 135607246 | 135607371 | 54 | 31 | 50 | 44 | 41 |
| A3SS | 3395 | ENSG00000104320 | NBN | chr8 | - | 89980733 | 89980893 | 89980733 | 89980852 | 89982721 | 89982855 | 15 | 11 | 31 | 30 | 62 |
| A3SS | 9334 | ENSG00000155906 | RMND1 | chr6 | - | 151421244 | 151422605 | 151421244 | 151421321 | 151423524 | 151423631 | 69 | 41 | 38 | 50 | 69 |
| A3SS | 10641 | ENSG00000128654 | MTX2 | chr2 | + | 176329300 | 176329426 | 176329369 | 176329426 | 176328873 | 176328912 | 49 | 33 | 49 | 45 | 71 |
| A3SS | 3394 | ENSG00000104320 | NBN | chr8 | - | 89980733 | 89980893 | 89980733 | 89980852 | 89981952 | 89982002 | 15 | 10 | 32 | 31 | 61 |
| A3SS | 6172 | ENSG00000214114 | MYCBP | chr1 | - | 38866879 | 38867009 | 38866879 | 38866999 | 38867561 | 38867610 | 56 | 55 | 44 | 68 | 101 |
| A3SS | 10898 | ENSG00000174197 | MGA | chr15 | + | 41739905 | 41740203 | 41740052 | 41740203 | 41736180 | 41736368 | 37 | 16 | 31 | 41 | 45 |
| A3SS | 8065 | ENSG00000224078 | SNHG14 | chr15 | + | 25118267 | 25118382 | 25118270 | 25118382 | 25117409 | 25117515 | 41 | 42 | 19 | 23 | 67 |
| A3SS | 6998 | ENSG00000225470 | JPX | chrX | + | 73998953 | 73999669 | 73999354 | 73999669 | 73994839 | 73994961 | 34 | 43 | 40 | 30 | 64 |
| A3SS | 5516 | ENSG00000196628 | TCF4 | chr18 | - | 55350873 | 55351003 | 55350873 | 55351000 | 55403453 | 55403518 | 40 | 42 | 35 | 48 | 85 |
| A3SS | 8038 | ENSG00000146463 | ZMYM4 | chr1 | + | 35396551 | 35396670 | 35396554 | 35396670 | 35393594 | 35393739 | 43 | 33 | 51 | 33 | 63 |
| A3SS | 6481 | ENSG00000155545 | MIER3 | chr5 | - | 56928766 | 56928861 | 56928766 | 56928858 | 56930663 | 56930745 | 28 | 26 | 34 | 39 | 39 |
| A3SS | 7583 | ENSG00000197056 | ZMYM1 | chr1 | + | 35111771 | 35111912 | 35111775 | 35111912 | 35110293 | 35110447 | 31 | 21 | 20 | 27 | 65 |
| A3SS | 9410 | ENSG00000131018 | SYNE1 | chr6 | - | 152308488 | 152308632 | 152308488 | 152308628 | 152309834 | 152310017 | 92 | 39 | 47 | 79 | 35 |
| A3SS | 5436 | ENSG00000121940 | CLCC1 | chr1 | - | 108941404 | 108941498 | 108941404 | 108941494 | 108943474 | 108943615 | 30 | 31 | 53 | 46 | 98 |
| A3SS | 6341 | ENSG00000087095 | NLK | chr17 | + | 28194581 | 28196381 | 28196179 | 28196381 | 28192119 | 28192213 | 80 | 103 | 94 | 92 | 189 |
| A3SS | 6756 | ENSG00000040275 | SPDL1 | chr5 | + | 169596560 | 169596701 | 169596584 | 169596701 | 169594570 | 169594681 | 17 | 21 | 28 | 45 | 29 |
| A3SS | 429 | ENSG00000167775 | CD320 | chr19 | - | 8303854 | 8304088 | 8303854 | 8303973 | 8305030 | 8305156 | 63 | 20 | 33 | 27 | 42 |
| A3SS | 12942 | ENSG00000111142 | METAP2 | chr12 | + | 95476070 | 95476178 | 95476142 | 95476178 | 95474151 | 95474330 | 67 | 61 | 66 | 75 | 64 |
| A3SS | 3496 | ENSG00000247774 | PCED1B-AS1 | chr12 | - | 47206510 | 47206588 | 47206510 | 47206585 | 47210639 | 47210761 | 31 | 11 | 50 | 35 | 84 |
| A3SS | 364 | ENSG00000130816 | DNMT1 | chr19 | - | 10139675 | 10140328 | 10139675 | 10139817 | 10140780 | 10140909 | 94 | 41 | 105 | 102 | 138 |
| A3SS | 9824 | ENSG00000006634 | DBF4 | chr7 | + | 87900220 | 87900349 | 87900281 | 87900349 | 87897293 | 87897339 | 6 | 4 | 52 | 69 | 68 |
| A3SS | 5672 | ENSG00000134744 | TUT4 | chr1 | - | 52436754 | 52436978 | 52436754 | 52436975 | 52438219 | 52438335 | 49 | 30 | 32 | 42 | 68 |
| A3SS | 1532 | ENSG00000108848 | LUC7L3 | chr17 | + | 50738136 | 50738199 | 50738140 | 50738199 | 50737286 | 50737479 | 49 | 22 | 40 | 37 | 40 |

| | | | | | | | | | | | | | | | |
|---|---|---|---|---|---|---|---|---|---|---|---|---|---|---|---|
| A3SS | 9906 | ENSG00000157764 | BRAF | chr7 | - | 140834608 | 140834872 | 140834608 | 140834703 | 140850110 | 140850212 | 192 | 138 | 181 | [illegible] |
| A3SS | 10078 | ENSG00000105835 | NAMPT | chr7 | - | 106269990 | 106270332 | 106269990 | 106270292 | 106272529 | 106272658 | 43 | 52 | 21 | [illegible] |
| A3SS | 12544 | ENSG00000107938 | EDRF1 | chr10 | + | 125737917 | 125737989 | 125737922 | 125737989 | 125735643 | 125735904 | 35 | 28 | 44 | [illegible] |
| A3SS | 4989 | ENSG00000091483 | FH | chr1 | - | 241508602 | 241508785 | 241508602 | 241508760 | 241511966 | 241512143 | 96 | 55 | 115 | [illegible] |
| A3SS | 4052 | ENSG00000134852 | CLOCK | chr4 | - | 55438281 | 55438537 | 55438281 | 55438407 | 55443686 | 55443896 | 42 | 25 | 26 | [illegible] |
| A3SS | 6807 | ENSG00000169554 | ZEB2 | chr2 | - | 144424795 | 144424867 | 144424795 | 144424864 | 144429768 | 144430026 | 369 | 297 | 198 | [illegible] |
| A3SS | 10899 | ENSG00000174197 | MGA | chr15 | + | 41739905 | 41740203 | 41740052 | 41740203 | 41736180 | 41736698 | 43 | 19 | 35 | [illegible] |
| A3SS | 9101 | ENSG00000163939 | PBRM1 | chr3 | - | 52589069 | 52589255 | 52589069 | 52589252 | 52603520 | 52603732 | 60 | 41 | 71 | [illegible] |
| A3SS | 6815 | ENSG00000112303 | VNN2 | chr6 | - | 132751144 | 132751518 | 132751144 | 132751514 | 132752672 | 132752749 | 45 | 15 | 38 | [illegible] |
| A3SS | 8137 | ENSG00000102081 | FMR1 | chrX | + | 147944868 | 147945051 | 147944943 | 147945051 | 147940575 | 147940662 | 41 | 46 | 41 | [illegible] |
| A3SS | 1185 | ENSG00000182173 | TSEN54 | chr17 | + | 75521704 | 75522333 | 75522006 | 75522333 | 75521408 | 75521510 | 128 | 53 | 102 | [illegible] |
| A3SS | 708 | ENSG00000007264 | MATK | chr19 | - | 3778508 | 3778595 | 3778508 | 3778592 | 3778991 | 3779187 | 93 | 21 | 54 | [illegible] |
| A3SS | 3110 | ENSG00000167881 | SRP68 | chr17 | - | 76070377 | 76070444 | 76070377 | 76070436 | 76072307 | 76072507 | 118 | 103 | 131 | [illegible] |
| A3SS | 12773 | ENSG00000066651 | TRMT11 | chr6 | + | 126010924 | 126011417 | 126011252 | 126011417 | 126008391 | 126008472 | 22 | 21 | 30 | [illegible] |
| A3SS | 5671 | ENSG00000134744 | TUT4 | chr1 | - | 52436754 | 52436978 | 52436754 | 52436963 | 52438219 | 52438335 | 54 | 31 | 35 | [illegible] |
| A3SS | 13005 | ENSG00000135297 | MTO1 | chr6 | + | 73466488 | 73466606 | 73466492 | 73466606 | 73466208 | 73466408 | 143 | 117 | 148 | [illegible] |
| A3SS | 5591 | ENSG00000115808 | STRN | chr2 | - | 36883931 | 36884075 | 36883931 | 36884065 | 36886715 | 36886826 | 94 | 67 | 69 | [illegible] |
| A3SS | 4100 | ENSG00000163104 | SMARCAD1 | chr4 | + | 94278922 | 94279050 | 94278928 | 94279050 | 94278615 | 94278733 | 39 | 19 | 26 | [illegible] |
| A3SS | 6323 | ENSG00000137575 | SDCBP | chr8 | + | 58565018 | 58565084 | 58565021 | 58565084 | 58561747 | 58561795 | 39 | 21 | 37 | [illegible] |
| A3SS | 8354 | ENSG00000163577 | EIF5A2 | chr3 | - | 170894291 | 170894423 | 170894291 | 170894383 | 170906988 | 170907093 | 81 | 36 | 57 | [illegible] |
| [illegible] | | | | | | | | | | | | | | | |

| | | | | | | | | | | | | | | |
|---|---|---|---|---|---|---|---|---|---|---|---|---|---|---|
| A3SS | 12200 | ENSG00000172543 | CTSW | chr11 | + | 65883214 | 65883741 | 65883507 | 65883741 | 65883068 | 65883133 | 616 | 102 | 249 |
| A3SS | 12205 | ENSG00000013725 | CD6 | chr11 | + | 61015712 | 61015835 | 61015732 | 61015835 | 61013918 | 61014014 | 224 | 117 | 221 |
| A3SS | 9758 | ENSG00000068885 | IFT80 | chr3 | - | 160381502 | 160381724 | 160381502 | 160381572 | 160384563 | 160384646 | 77 | 62 | 71 |
| A3SS | 2808 | ENSG00000070367 | EXOC5 | chr14 | - | 57246710 | 57246858 | 57246710 | 57246827 | 57247617 | 57247712 | 123 | 110 | 72 |
| A3SS | 2464 | ENSG00000137878 | GCOM1 | chr15 | + | 57708713 | 57709358 | 57709184 | 57709358 | 57706491 | 57706955 | 185 | 78 | 182 |
| A3SS | 6069 | ENSG00000120798 | NR2C1 | chr12 | - | 95028386 | 95028524 | 95028386 | 95028509 | 95031348 | 95031488 | 81 | 42 | 66 |
| A3SS | 5743 | ENSG00000204475 | NCR3 | chr6 | - | 31589781 | 31590126 | 31589781 | 31589899 | 31592856 | 31592959 | 111 | 23 | 78 |
| A3SS | 9759 | ENSG00000068885 | IFT80 | chr3 | - | 160381502 | 160381726 | 160381502 | 160381572 | 160384563 | 160384646 | 78 | 62 | 73 |
| A3SS | 3175 | ENSG00000152942 | RAD17 | chr5 | + | 69410492 | 69410550 | 69410513 | 69410550 | 69400048 | 69400169 | 83 | 64 | 80 |
| A3SS | 12467 | ENSG00000156650 | KAT6B | chr10 | + | 74975398 | 74977437 | 74977315 | 74977437 | 74972506 | 74972639 | 109 | 87 | 82 |
| A3SS | 7978 | ENSG00000115355 | CCDC88A | chr2 | - | 55328293 | 55328435 | 55328293 | 55328432 | 55332565 | 55332693 | 39 | 131 | 46 |
| A3SS | 4399 | ENSG00000128513 | POT1 | chr7 | - | 124822385 | 124824074 | 124822385 | 124822551 | 124825251 | 124825357 | 57 | 35 | 42 |
| A3SS | 5990 | ENSG00000163743 | RCHY1 | chr4 | - | 75490580 | 75490701 | 75490580 | 75490663 | 75491610 | 75491637 | 150 | 81 | 111 |
| A3SS | 1521 | ENSG00000073584 | SMARCE1 | chr17 | - | 40635930 | 40636102 | 40635930 | 40636098 | 40636394 | 40636526 | 207 | 124 | 202 |
| A3SS | 8497 | ENSG00000144566 | RAB5A | chr3 | + | 19975600 | 19975752 | 19975642 | 19975752 | 19950805 | 19951061 | 340 | 271 | 273 |
| A3SS | 12849 | ENSG00000133704 | IPO8 | chr12 | - | 30690495 | 30690577 | 30690495 | 30690548 | 30695563 | 30695869 | 104 | 78 | 115 |
| A3SS | 9193 | ENSG00000135316 | SYNCRIP | chr6 | - | 85608334 | 85612923 | 85608334 | 85612920 | 85614980 | 85615347 | 127 | 73 | 106 |
| A3SS | 7437 | ENSG00000076770 | MBNL3 | chrX | - | 132386660 | 132386811 | 132386660 | 132386706 | 132390846 | 132391083 | 145 | 88 | 105 |
| A3SS | 11850 | ENSG00000152404 | CWF19L2 | chr11 | - | 107353523 | 107353736 | 107353523 | 107353678 | 107390073 | 107390211 | 90 | 66 | 60 |
| A3SS | 12933 | ENSG00000028203 | VEZT | chr12 | + | 95274741 | 95274889 | 95274745 | 95274889 | 95270038 | 95270188 | 83 | 60 | 72 |
| A3SS | 5018 | ENSG00000164253 | WDR41 | chr5 | - | 77451290 | 77451353 | 77451290 | 77451346 | 77453816 | 77453928 | 176 | 72 | 72 |
| A3SS | 9650 | ENSG00000183735 | TBK1 | chr12 | + | 64460188 | 64460329 | 64460246 | 64460329 | 64455839 | 64455957 | 218 | 188 | 127 |
| A3SS | 8262 | ENSG00000082269 | FAM135A | chr6 | + | 70524930 | 70526698 | 70525863 | 70526698 | 70523966 | 70524121 | 79 | 74 | 56 |
| A3SS | 1520 | ENSG00000073584 | SMARCE1 | chr17 | - | 40631591 | 40632367 | 40631591 | 40631693 | 40635930 | 40636102 | 244 | 162 | 233 |
| A3SS | 3567 | ENSG00000113593 | PPWD1 | chr5 | + | 65579423 | 65579613 | 65579484 | 65579613 | 65576878 | 65577069 | 81 | 109 | 69 |
| A3SS | 2734 | ENSG00000100722 | ZC3H14 | chr14 | + | 88609714 | 88609803 | 88609726 | 88609803 | 88609266 | 88609403 | 65 | 59 | 56 |
| A3SS | 8931 | ENSG00000198721 | ECI2 | chr6 | - | 4125249 | 4126237 | 4125249 | 4125370 | 4127761 | 4127831 | 141 | 130 | 136 |
| A3SS | 3049 | ENSG00000109775 | UFSP2 | chr4 | - | 185415709 | 185415867 | 185415709 | 185415815 | 185418440 | 185418507 | 117 | 114 | 113 |
| A3SS | 536 | ENSG00000105193 | RPS16 | chr19 | - | 39435606 | 39435708 | 39435606 | 39435705 | 39435847 | 39435949 | 563 | 225 | 704 |
| A3SS | 13323 | ENSG00000148444 | COMMD3 | chr10 | + | 22318352 | 22318713 | 22318652 | 22318713 | 22318104 | 22318168 | 79 | 64 | 76 |
| A3SS | 9663 | ENSG00000183735 | TBK1 | chr12 | + | 64496366 | 64496406 | 64496372 | 64496406 | 64495698 | 64495775 | 92 | 73 | 75 |
| A3SS | 13371 | ENSG00000275066 | SYNRG | chr17 | - | 37568790 | 37568924 | 37568790 | 37568920 | 37570636 | 37570885 | 135 | 83 | 171 |
| A3SS | 6472 | ENSG00000036549 | ZZZ3 | chr1 | - | 77581775 | 77581891 | 77581775 | 77581888 | 77581978 | 77582126 | 63 | 59 | 70 |
| A3SS | 5222 | ENSG00000015171 | ZMYND11 | chr10 | + | 240055 | 240111 | 240058 | 240111 | 239437 | 239525 | 108 | 50 | 89 |
| A3SS | 13370 | ENSG00000275066 | SYNRG | chr17 | - | 37541971 | 37542565 | 37541971 | 37542430 | 37561194 | 37561257 | 167 | 109 | 176 |
| A3SS | 7002 | ENSG00000225470 | JPX | chrX | + | 73998953 | 74000205 | 73999608 | 74000205 | 73998767 | 73998844 | 82 | 90 | 70 |
| A3SS | 11492 | ENSG00000070061 | ELP1 | chr9 | - | 108897147 | 108897285 | 108897147 | 108897283 | 108898501 | 108898581 | 86 | 54 | 108 |
| A3SS | 9655 | ENSG00000183735 | TBK1 | chr12 | + | 64485925 | 64486017 | 64485957 | 64486017 | 64485454 | 64485513 | 193 | 131 | 122 |
| A3SS | 10171 | ENSG00000055130 | CUL1 | chr7 | + | 148766560 | 148766723 | 148766564 | 148766723 | 148760332 | 148760496 | 209 | 217 | 167 |
| A3SS | 3136 | ENSG00000173692 | PSMD1 | chr2 | + | 231072188 | 231072415 | 231072332 | 231072415 | 231070024 | 231070168 | 242 | 145 | 189 |
| A3SS | 10374 | ENSG00000125676 | THOC2 | chrX | - | 123610917 | 123610963 | 123610917 | 123610957 | 123611439 | 123611516 | 76 | 67 | 73 |
| A3SS | 7977 | ENSG00000115355 | CCDC88A | chr2 | - | 55296257 | 55296523 | 55296257 | 55296520 | 55299838 | 55299919 | 87 | 83 | 55 |
| A3SS | 3069 | ENSG00000205133 | TRIQK | chr8 | - | 92885319 | 92886735 | 92885319 | 92886667 | 92891988 | 92892074 | 106 | 60 | 62 |
| A3SS | 10843 | ENSG00000145868 | FBXO38 | chr5 | + | 148398998 | 148399132 | 148399014 | 148399132 | 148394711 | 148394904 | 122 | 110 | 98 |
| A3SS | 4277 | ENSG00000088387 | DOCK9 | chr13 | - | 98886531 | 98886624 | 98886531 | 98886610 | 98888157 | 98888223 | 87 | 69 | 124 |
| A3SS | 12199 | ENSG00000172543 | CTSW | chr11 | + | 65883214 | 65883424 | 65883265 | 65883424 | 65883068 | 65883133 | 840 | 152 | 335 |
| A3SS | 4587 | ENSG00000111642 | CHD4 | chr12 | - | 6578028 | 6578137 | 6578028 | 6578135 | 6578408 | 6578546 | 447 | 247 | 385 |
| A3SS | 8707 | ENSG00000145349 | CAMK2D | chr4 | - | 113500462 | 113500511 | 113500462 | 113500508 | 113509637 | 113509672 | 126 | 86 | 108 |
| A3SS | 9664 | ENSG00000183735 | TBK1 | chr12 | + | 64497162 | 64497259 | 64497166 | 64497259 | 64496948 | 64497050 | 126 | 92 | 99 |
| A3SS | 10161 | ENSG00000234741 | GAS5 | chr1 | - | 173864256 | 173864704 | 173864256 | 173864304 | 173865470 | 173865547 | 76 | 67 | 96 |
| A3SS | 1518 | ENSG00000073584 | SMARCE1 | chr17 | - | 40630713 | 40631693 | 40630713 | 40630924 | 40632194 | 40632367 | 327 | 296 | 320 |
| A3SS | 10033 | ENSG00000104133 | SPG11 | chr15 | - | 44622219 | 44622799 | 44622219 | 44622347 | 44626330 | 44626507 | 267 | 280 | 224 |
| A3SS | 9393 | ENSG00000135317 | SNX14 | chr6 | - | 85543181 | 85543760 | 85543181 | 85543306 | 85547111 | 85547226 | 161 | 94 | 91 |
| A3SS | 9653 | ENSG00000183735 | TBK1 | chr12 | + | 64481841 | 64484499 | 64484302 | 64484499 | 64480011 | 64480122 | 142 | 103 | 77 |
| A3SS | 10703 | ENSG00000046651 | OFD1 | chrX | + | 13753367 | 13755242 | 13755150 | 13755242 | 13751248 | 13751368 | 124 | 161 | 152 |
| A3SS | 3144 | ENSG00000154153 | RETREG1 | chr5 | - | 16478033 | 16478098 | 16478033 | 16478095 | 16478849 | 16478987 | 62 | 49 | 105 |
| A3SS | 11490 | ENSG00000070061 | ELP1 | chr9 | - | 108874894 | 108874970 | 108874894 | 108874963 | 108877994 | 108878149 | 112 | 67 | 98 |
| A3SS | 620 | ENSG00000130726 | TRIM28 | chr19 | + | 58547791 | 58547906 | 58547801 | 58547906 | 58547596 | 58547713 | 411 | 239 | 418 |
| A3SS | 8777 | ENSG00000115053 | NCL | chr2 | - | 231457000 | 231457800 | 231457000 | 231457124 | 231459000 | 231459125 | 232 | 213 | 242 |
| A3SS | 10022 | ENSG00000105819 | PMPCB | chr7 | + | 103303841 | 103304040 | 103303985 | 103304040 | 103300177 | 103300307 | 386 | 281 | 439 |
| A3SS | 11315 | ENSG00000056586 | RC3H2 | chr9 | - | 122854530 | 122854615 | 122854530 | 122854581 | 122855183 | 122855397 | 133 | 115 | 132 |
| A3SS | 5918 | ENSG00000055609 | KMT2C | chr7 | - | 152157760 | 152157925 | 152157760 | 152157854 | 152158862 | 152159072 | 107 | 97 | 81 |
| A3SS | 12350 | ENSG00000149311 | ATM | chr11 | + | 108253830 | 108253996 | 108253854 | 108253996 | 108252820 | 108252912 | 69 | 60 | 88 |
| A3SS | 11111 | ENSG00000121022 | COPS5 | chr8 | - | 67043081 | 67045960 | 67043081 | 67043317 | 67051229 | 67051341 | 117 | 99 | 125 |
| A3SS | 9612 | ENSG00000146802 | TMEM168 | chr7 | - | 112772780 | 112773055 | 112772780 | 112772818 | 112775175 | 112775318 | 94 | 98 | 112 |
| A3SS | 10743 | ENSG00000147050 | KDM6A | chrX | + | 45082575 | 45082789 | 45082714 | 45082789 | 45079145 | 45079351 | 213 | 221 | 185 |
| A3SS | 9652 | ENSG00000183735 | TBK1 | chr12 | + | 64481841 | 64482021 | 64481851 | 64482021 | 64480011 | 64480122 | 138 | 78 | 71 |
| A3SS | 3981 | ENSG00000005700 | IBTK | chr6 | - | 82196297 | 82196446 | 82196297 | 82196401 | 82200140 | 82200253 | 110 | 72 | 86 |
| A3SS | 7874 | ENSG00000117697 | NSL1 | chr1 | - | 212726152 | 212738686 | 212726152 | 212727179 | 212739533 | 212739601 | 134 | 103 | 136 |
| A3SS | 3872 | ENSG00000182150 | ERCC6L2 | chr9 | + | 95970575 | 95973088 | 95971932 | 95973088 | 95966561 | 95966714 | 103 | 83 | 85 |
| A3SS | 3699 | ENSG00000122008 | POLK | chr5 | + | 75593877 | 75594049 | 75593965 | 75594049 | 75587025 | 75587058 | 116 | 125 | 69 |
| A3SS | 1689 | ENSG00000125124 | BBS2 | chr16 | - | 56506119 | 56506224 | 56506119 | 56506220 | 56509956 | 56510034 | 133 | 104 | 136 |
| A3SS | 7695 | ENSG00000039319 | ZFYVE16 | chr5 | + | 80448025 | 80448404 | 80448282 | 80448404 | 80445262 | 80445405 | 96 | 87 | 78 |
| A3SS | 10283 | ENSG00000182220 | ATP6AP2 | chrX | + | 40599591 | 40599741 | 40599648 | 40599741 | 40598680 | 40598734 | 544 | 354 | 348 |
| A3SS | 755 | ENSG00000101347 | SAMHD1 | chr20 | - | 36911217 | 36912552 | 36911217 | 36911333 | 36916721 | 36916830 | 1940 | 1687 | 1438 |
| A3SS | 6813 | ENSG00000112303 | VNN2 | chr6 | - | 132751144 | 132751518 | 132751144 | 132751278 | 132752460 | 132752749 | 536 | 242 | 451 |
| A3SS | 4658 | ENSG00000168827 | GFM1 | chr3 | + | 158653309 | 158653467 | 158653313 | 158653467 | 158652197 | 158652246 | 182 | 160 | 147 |
| A3SS | 6617 | ENSG00000154027 | AK5 | chr1 | + | 77293792 | 77293960 | 77293864 | 77293960 | 77286940 | 77287127 | 57 | 44 | 96 |
| A3SS | 9269 | ENSG00000135486 | HNRNPA1 | chr12 | + | 54283815 | 54283967 | 54283836 | 54283967 | 54282799 | 54282878 | 404 | 332 | 438 |
| A3SS | 4244 | ENSG00000189079 | ARID2 | chr12 | + | 45839328 | 45839496 | 45839385 | 45839496 | 45837497 | 45837707 | 182 | 131 | 149 |
| A3SS | 11013 | ENSG00000035928 | RFC1 | chr4 | - | 39320382 | 39320669 | 39320382 | 39320660 | 39321286 | 39321374 | 108 | 99 | 138 |
| A3SS | 6266 | ENSG00000165359 | INTS6L | chrX | + | 135569336 | 135573033 | 135572814 | 135573033 | 135556167 | 135556300 | 130 | 155 | 161 |
| A3SS | 2584 | ENSG00000100749 | VRK1 | chr14 | + | 96860556 | 96860735 | 96860646 | 96860735 | 96856527 | 96856586 | 256 | 170 | 192 |
| A3SS | 12348 | ENSG00000149311 | ATM | chr11 | + | 108227594 | 108227696 | 108227598 | 108227696 | 108224783 | 108224871 | 137 | 93 | 103 |
| A3SS | 9656 | ENSG00000183735 | TBK1 | chr12 | + | 64495482 | 64495604 | 64495490 | 64495604 | 64490040 | 64490119 | 175 | 121 | 92 |
| A3SS | 3979 | ENSG00000005700 | IBTK | chr6 | - | 82194478 | 82194642 | 82194478 | 82194595 | 82196297 | 82196446 | 117 | 109 | 109 |
| A3SS | 5001 | ENSG00000182866 | LCK | chr1 | + | 32279670 | 32279994 | 32279840 | 32279994 | 32276606 | 32276786 | 581 | 239 | 576 |
| A3SS | 7407 | ENSG00000149231 | CCDC82 | chr11 | - | 96383961 | 96386293 | 96383961 | 96384761 | 96387529 | 96387613 | 149 | 127 | 115 |
| A3SS | 9329 | ENSG00000112739 | PRPF4B | chr6 | + | 4061654 | 4062299 | 4062212 | 4062299 | 4061014 | 4061147 | 153 | 117 | 109 |
| A3SS | 13164 | ENSG00000134283 | PPHLN1 | chr12 | + | 42374862 | 42375074 | 42374955 | 42375074 | 42355160 | 42355222 | 317 | 286 | 332 |
| A3SS | 4639 | ENSG00000109861 | CTSC | chr11 | - | 88296132 | 88300645 | 88296132 | 88296264 | 88309162 | 88309318 | 745 | 459 | 408 |
| A3SS | 9651 | ENSG00000183735 | TBK1 | chr12 | + | 64466900 | 64467082 | 64466978 | 64467082 | 64464333 | 64464463 | 237 | 144 | 155 |
| A3SS | 4307 | ENSG00000178035 | IMPDH2 | chr3 | - | 49026518 | 49026609 | 49026518 | 49026597 | 49026686 | 49026886 | 185 | 95 | 238 |
| A3SS | 9411 | ENSG00000131018 | SYNE1 | chr6 | - | 152471596 | 152471765 | 152471596 | 152471670 | 152472300 | 152472413 | 644 | 369 | 280 |
| A3SS | 10150 | ENSG00000196305 | IARS1 | chr9 | - | 92250712 | 92250834 | 92250712 | 92250759 | 92251807 | 92251885 | 151 | 91 | 208 |
| A3SS | 12189 | ENSG00000076053 | RBM7 | chr11 | + | 114402827 | 114402915 | 114402831 | 114402915 | 114401697 | 114401860 | 171 | 130 | 203 |
| A3SS | 2532 | ENSG00000079134 | THOC1 | chr18 | - | 225088 | 225403 | 225088 | 225139 | 226800 | 226901 | 110 | 109 | 115 |
| A3SS | 3217 | ENSG00000086189 | DIMT1 | chr5 | - | 62390875 | 62392234 | 62390875 | 62390982 | 62392925 | 62392990 | 259 | 265 | 271 |
| A3SS | 11044 | ENSG00000170633 | RNF34 | chr12 | + | 121417503 | 121417911 | 121417860 | 121417911 | 121416158 | 121416377 | 378 | 212 | 239 |
| A3SS | 8779 | ENSG00000115053 | NCL | chr2 | - | 231458265 | 231458620 | 231458265 | 231458344 | 231459000 | 231459125 | 410 | 403 | 363 |
| A3SS | 4393 | ENSG00000155975 | VPS37A | chr8 | + | 17274732 | 17274958 | 17274775 | 17274958 | 17268855 | 17268956 | 202 | 151 | 91 |
| A3SS | 3947 | ENSG00000171566 | PLRG1 | chr4 | - | 154537968 | 154538108 | 154537968 | 154538082 | 154539104 | 154539213 | 120 | 103 | 138 |
| A3SS | 9384 | ENSG00000168036 | CTNNB1 | chr3 | + | 41239133 | 41239900 | 41239659 | 41239900 | 41238015 | 41238076 | 334 | 198 | 338 |
| A3SS | 38 | ENSG00000156261 | CCT8 | chr21 | - | 29066715 | 29067071 | 29066715 | 29066777 | 29067555 | 29067705 | 184 | 141 | 192 |
| A3SS | 4364 | ENSG00000164327 | RICTOR | chr5 | - | 38945490 | 38945724 | 38945490 | 38945610 | 38947263 | 38947441 | 136 | 96 | 79 |
| A3SS | 8837 | ENSG00000197223 | C1D | chr2 | - | 68045987 | 68046410 | 68045987 | 68046043 | 68047172 | 68047319 | 307 | 173 | 228 |
| A3SS | 8834 | ENSG00000135426 | TESPA1 | chr12 | - | 54962430 | 54963242 | 54962430 | 54963238 | 54963741 | 54963950 | 159 | 159 | 221 |
| A3SS | 621 | ENSG00000130726 | TRIM28 | chr19 | + | 58548861 | 58548910 | 58548885 | 58548910 | 58548739 | 58548776 | 260 | 105 | 200 |
| A3SS | 8027 | ENSG00000119820 | YIPF4 | chr2 | + | 32301381 | 32301495 | 32301442 | 32301495 | 32298233 | 32298311 | 167 | 138 | 177 |
| A3SS | 2636 | ENSG00000092108 | SCFD1 | chr14 | + | 30700187 | 30700258 | 30700190 | 30700258 | 30694772 | 30694869 | 126 | 72 | 94 |
| A3SS | 13322 | ENSG00000148444 | COMMD3 | chr10 | + | 22318271 | 22318713 | 22318652 | 22318713 | 22318104 | 22318168 | 196 | 161 | 191 |
| A3SS | 5944 | ENSG00000095787 | WAC | chr10 | + | 28583398 | 28583505 | 28583405 | 28583505 | 28535561 | 28535757 | 464 | 359 | 444 |
| A3SS | 2533 | ENSG00000079134 | THOC1 | chr18 | - | 247848 | 247957 | 247848 | 247908 | 252538 | 252612 | 156 | 169 | 132 |
| A3SS | 10526 | ENSG00000005302 | MSL3 | chrX | + | 11765466 | 11765729 | 11765469 | 11765729 | 11763779 | 11763938 | 204 | 160 | 223 |
| A3SS | 10651 | ENSG00000116754 | SRSF11 | chr1 | + | 70228421 | 70228555 | 70228464 | 70228555 | 70222765 | 70222798 | 135 | 115 | 199 |
| A3SS | 10527 | ENSG00000005302 | MSL3 | chrX | + | 11765466 | 11765729 | 11765521 | 11765729 | 11763779 | 11763938 | 212 | 164 | 236 |
| A3SS | 2064 | ENSG00000180096 | SEPTIN1 | chr16 | - | 30381126 | 30381473 | 30381126 | 30381244 | 30381759 | 30381883 | 386 | 195 | 364 |
| A3SS | 7148 | ENSG00000197323 | TRIM33 | chr1 | - | 114397607 | 114397990 | 114397607 | 114397860 | 114399456 | 114399609 | 176 | 188 | 169 |
| A3SS | 5506 | ENSG00000110172 | CHORDC1 | chr11 | - | 90210535 | 90214175 | 90210535 | 90210594 | 90215173 | 90215230 | 168 | 148 | 139 |
| A3SS | 6267 | ENSG00000165359 | INTS6L | chrX | + | 135581527 | 135582510 | 135582240 | 135582510 | 135581049 | 135581143 | 159 | 143 | 162 |
| A3SS | 401 | ENSG00000142546 | NOSIP | chr19 | - | 49557089 | 49558978 | 49557089 | 49557249 | 49559933 | 49560039 | 305 | 143 | 337 |
| A3SS | 8556 | ENSG00000143971 | ETAA1 | chr2 | + | 67403224 | 67405335 | 67405235 | 67405335 | 67402861 | 67402974 | 111 | 76 | 104 |
| A3SS | 1725 | ENSG00000187555 | USP7 | chr16 | - | 8904434 | 8904565 | 8904434 | 8904524 | 8905186 | 8905331 | 445 | 293 | 409 |
| A3SS | 13410 | ENSG00000134899 | ERCC5 | chr13 | + | 102853756 | 102854374 | 102854287 | 102854374 | 102852117 | 102852293 | 279 | 241 | 298 |
| A3SS | 5638 | ENSG00000142676 | RPL11 | chr1 | + | 23694659 | 23695908 | 23695797 | 23695908 | 23693806 | 23693913 | 3813 | 2504 | 4100 |
| A3SS | 5369 | ENSG00000163946 | TASOR | chr3 | - | 56633043 | 56633966 | 56633043 | 56633843 | 56639985 | 56640130 | 139 | 167 | 127 |
| A3SS | 10204 | ENSG00000122545 | SEPTIN7 | chr7 | + | 35832797 | 35832900 | 35832803 | 35832900 | 35831491 | 35831496 | 182 | 129 | 130 |
| A3SS | 7774 | ENSG00000164134 | NAA15 | chr4 | + | 139384831 | 139384978 | 139384834 | 139384978 | 139378755 | 139378854 | 151 | 114 | 120 |
| A3SS | 11195 | ENSG00000104529 | EEF1D | chr8 | - | 143581053 | 143581328 | 143581053 | 143581154 | 143586218 | 143586290 | 1196 | 478 | 1254 |
| A3SS | 3048 | ENSG00000109775 | UFSP2 | chr4 | - | 185408270 | 185408435 | 185408270 | 185408390 | 185413725 | 185413872 | 199 | 178 | 207 |
| A3SS | 9451 | ENSG00000168944 | CEP120 | chr5 | - | 123389923 | 123391337 | 123389923 | 123390140 | 123393299 | 123393497 | 225 | 157 | 141 |
| A3SS | 12439 | ENSG00000090621 | PABPC4 | chr1 | - | 39565105 | 39565378 | 39565105 | 39565366 | 39567750 | 39567846 | 461 | 318 | 462 |
| A3SS | 6933 | ENSG00000135241 | PNPLA8 | chr7 | - | 108514435 | 108515574 | 108514435 | 108515447 | 108518189 | 108518301 | 161 | 167 | 153 |
| A3SS | 10745 | ENSG00000147050 | KDM6A | chrX | + | 45083459 | 45083608 | 45083518 | 45083608 | 45082714 | 45082789 | 319 | 328 | 256 |
| A3SS | 9974 | ENSG00000091140 | DLD | chr7 | + | 107906266 | 107906368 | 107906270 | 107906368 | 107905360 | 107905504 | 228 | 190 | 202 |
| A3SS | 10079 | ENSG00000105835 | NAMPT | chr7 | - | 106269153 | 106272658 | 106269153 | 106269312 | 106274945 | 106275049 | 712 | 777 | 438 |
| A3SS | 9391 | ENSG00000135317 | SNX14 | chr6 | - | 85526125 | 85526237 | 85526125 | 85526225 | 85528261 | 85528362 | 195 | 120 | 129 |
| A3SS | 13219 | ENSG00000188906 | LRRK2 | chr12 | + | 40367005 | 40367077 | 40367009 | 40367077 | 40364841 | 40365050 | 189 | 176 | 101 |
| A3SS | 10146 | ENSG00000196305 | IARS1 | chr9 | - | 92223345 | 92223489 | 92223345 | 92223429 | 92229000 | 92229126 | 171 | 105 | 183 |
| A3SS | 10993 | ENSG00000156787 | TBC1D31 | chr8 | + | 123109473 | 123109620 | 123109527 | 123109620 | 123109316 | 123109396 | 181 | 85 | 128 |
| A3SS | 6100 | ENSG00000138095 | LRPPRC | chr2 | - | 43979825 | 43979948 | 43979825 | 43979939 | 43982237 | 43982434 | 202 | 157 | 216 |
| A3SS | 4379 | ENSG00000009790 | TRAF3IP3 | chr1 | + | 209779314 | 209779655 | 209779430 | 209779655 | 209775598 | 209775736 | 308 | 230 | 294 |
| A3SS | 13395 | ENSG00000127022 | CANX | chr5 | + | 179709872 | 179710065 | 179709918 | 179710065 | 179708238 | 179708380 | 307 | 237 | 288 |
| A3SS | 12992 | ENSG00000132341 | RAN | chr12 | + | 130874545 | 130874733 | 130874550 | 130874733 | 130873002 | 130873128 | 559 | 362 | 619 |
| A3SS | 7405 | ENSG00000149231 | CCDC82 | chr11 | - | 96358992 | 96359178 | 96358992 | 96359097 | 96364979 | 96365150 | 141 | 109 | 140 |
| A3SS | 2223 | ENSG00000086666 | ZFAND6 | chr15 | + | 80121711 | 80121820 | 80121747 | 80121820 | 80120327 | 80120498 | 551 | 316 | 396 |
| A3SS | 4375 | ENSG00000175455 | CCDC14 | chr3 | - | 123913724 | 123915718 | 123913724 | 123915132 | 123931101 | 123931234 | 169 | 161 | 160 |
| A3SS | 5104 | ENSG00000162613 | FUBP1 | chr1 | - | 77944054 | 77948774 | 77944054 | 77944269 | 77949154 | 77949300 | 346 | 304 | 344 |
| A3SS | 6399 | ENSG00000104231 | ZFAND1 | chr8 | - | 81702968 | 81703124 | 81702968 | 81703103 | 81713917 | 81714039 | 158 | 186 | 163 |
| A3SS | 3113 | ENSG00000127603 | MACF1 | chr1 | + | 39484600 | 39486987 | 39485537 | 39486987 | 39479797 | 39480009 | 316 | 177 | 258 |
| A3SS | 12723 | ENSG00000099246 | RAB18 | chr10 | + | 27533927 | 27533994 | 27533940 | 27533994 | 27533734 | 27533853 | 354 | 274 | 233 |
| A3SS | 3416 | ENSG00000205302 | SNX2 | chr5 | + | 122799691 | 122799855 | 122799694 | 122799855 | 122795265 | 122795383 | 252 | 160 | 162 |
| A3SS | 6594 | ENSG00000153283 | CD96 | chr3 | + | 111637195 | 111637261 | 111637198 | 111637261 | 111624332 | 111624404 | 139 | 132 | 189 |
| A3SS | 6652 | ENSG00000144381 | HSPD1 | chr2 | - | 197489001 | 197490296 | 197489001 | 197489247 | 197493323 | 197493492 | 222 | 160 | 263 |
| A3SS | 4559 | ENSG00000158411 | MITD1 | chr2 | - | 99169262 | 99169610 | 99169262 | 99169470 | 99170536 | 99170652 | 407 | 384 | 286 |
| A3SS | 13031 | ENSG00000135968 | GCC2 | chr2 | + | 108481696 | 108481816 | 108481709 | 108481816 | 108475751 | 108475850 | 139 | 105 | 125 |
| A3SS | 974 | ENSG00000134758 | RNF138 | chr18 | + | 32126692 | 32126800 | 32126726 | 32126800 | 32124733 | 32124845 | 470 | 407 | 439 |
| A3SS | 6817 | ENSG00000112303 | VNN2 | chr6 | - | 132752460 | 132752749 | 132752460 | 132752577 | 132755842 | 132756035 | 505 | 133 | 382 |
| A3SS | 779 | ENSG00000101126 | ADNP | chr20 | - | 50889047 | 50894512 | 50889047 | 50889957 | 50902016 | 50902109 | 256 | 231 | 283 |
| A3SS | 3700 | ENSG00000122008 | POLK | chr5 | + | 75593877 | 75594049 | 75593965 | 75594049 | 75590343 | 75590440 | 188 | 223 | 129 |
| A3SS | 4365 | ENSG00000164327 | RICTOR | chr5 | - | 38978582 | 38978650 | 38978582 | 38978648 | 38981866 | 38982036 | 218 | 175 | 208 |
| A3SS | 4644 | ENSG00000164463 | CREBRF | chr5 | + | 173108623 | 173110711 | 173110521 | 173110711 | 173090314 | 173091401 | 251 | 187 | 201 |
| A3SS | 6845 | ENSG00000177409 | SAMD9L | chr7 | - | 93145384 | 93146040 | 93145384 | 93145532 | 93147087 | 93147146 | 189 | 175 | 121 |
| A3SS | 4868 | ENSG00000162777 | DENND2D | chr1 | - | 111195915 | 111197989 | 111195915 | 111196056 | 111198629 | 111198742 | 315 | 190 | 269 |
| A3SS | 3791 | ENSG00000171490 | RSL1D1 | chr16 | - | 11841694 | 11841820 | 11841694 | 11841817 | 11841906 | 11842000 | 490 | 489 | 576 |
| A3SS | 2850 | ENSG00000126773 | PCNX4 | chr14 | + | 60118328 | 60118692 | 60118376 | 60118692 | 60115940 | 60116060 | 138 | 117 | 152 |
| A3SS | 5465 | ENSG00000116044 | NFE2L2 | chr2 | - | 177232391 | 177232583 | 177232391 | 177232562 | 177233249 | 177233339 | 392 | 330 | 324 |
| A3SS | 8555 | ENSG00000143971 | ETAA1 | chr2 | + | 67403224 | 67405335 | 67404996 | 67405335 | 67402861 | 67402974 | 198 | 133 | 133 |
| A3SS | 3680 | ENSG00000008513 | ST3GAL1 | chr8 | - | 133475718 | 133476600 | 133475718 | 133476074 | 133499134 | 133499189 | 279 | 228 | 370 |
| A3SS | 6093 | ENSG00000138095 | LRPPRC | chr2 | - | 43896633 | 43896708 | 43896633 | 43896655 | 43899218 | 43899334 | 161 | 123 | 180 |
| A3SS | 5719 | ENSG00000116030 | SUMO1 | chr2 | - | 202210734 | 202210806 | 202210734 | 202210783 | 202214356 | 202214434 | 774 | 585 | 536 |
| A3SS | 4935 | ENSG00000172175 | MALT1 | chr18 | + | 58734306 | 58734566 | 58734450 | 58734566 | 58733396 | 58733574 | 233 | 218 | 232 |
| A3SS | 13299 | ENSG00000136104 | RNASEH2B | chr13 | + | 50947986 | 50949505 | 50949462 | 50949505 | 50945426 | 50945532 | 306 | 229 | 265 |
| A3SS | 4648 | ENSG00000041357 | PSMA4 | chr15 | + | 78542176 | 78542645 | 78542482 | 78542645 | 78541904 | 78541930 | 702 | 454 | 646 |
| A3SS | 2671 | ENSG00000100567 | PSMA3 | chr14 | + | 58260947 | 58261020 | 58260958 | 58261020 | 58257924 | 58257998 | 312 | 215 | 289 |
| A3SS | 4363 | ENSG00000164327 | RICTOR | chr5 | - | 38945490 | 38945724 | 38945490 | 38945610 | 38946467 | 38946552 | 205 | 144 | 139 |
| A3SS | 8129 | ENSG00000091527 | CDV3 | chr3 | + | 133586562 | 133586722 | 133586568 | 133586722 | 133584001 | 133584150 | 732 | 512 | 684 |
| A3SS | 9390 | ENSG00000135317 | SNX14 | chr6 | - | 85517755 | 85517875 | 85517755 | 85517871 | 85518007 | 85518048 | 249 | 122 | 132 |
| A3SS | 8925 | ENSG00000240344 | PPIL3 | chr2 | - | 200882341 | 200882435 | 200882341 | 200882382 | 200885697 | 200885772 | 308 | 198 | 302 |
| A3SS | 10016 | ENSG00000177189 | RPS6KA3 | chrX | - | 20172745 | 20172871 | 20172745 | 20172868 | 20175163 | 20175288 | 247 | 264 | 252 |
| A3SS | 7113 | ENSG00000228956 | SATB1-AS1 | chr3 | + | 18526386 | 18526540 | 18526425 | 18526540 | 18465982 | 18466039 | 127 | 92 | 181 |
| A3SS | 2672 | ENSG00000100567 | PSMA3 | chr14 | + | 58260947 | 58261020 | 58260968 | 58261020 | 58257924 | 58257998 | 324 | 221 | 302 |
| A3SS | 4241 | ENSG00000211456 | SACM1L | chr3 | + | 45735234 | 45735373 | 45735245 | 45735373 | 45732052 | 45732151 | 256 | 172 | 211 |
| A3SS | 6844 | ENSG00000177409 | SAMD9L | chr7 | - | 93145384 | 93146040 | 93145384 | 93145532 | 93146882 | 93147146 | 224 | 237 | 143 |
| A3SS | 1859 | ENSG00000167526 | RPL13 | chr16 | + | 89561226 | 89561751 | 89561595 | 89561751 | 89560939 | 89561063 | 2543 | 1014 | 2189 |
| A3SS | 3705 | ENSG00000164483 | SAMD3 | chr6 | - | 130209494 | 130209608 | 130209494 | 130209602 | 130214336 | 130214526 | 366 | 205 | 261 |
| A3SS | 13298 | ENSG00000136104 | RNASEH2B | chr13 | + | 50943320 | 50943394 | 50943329 | 50943394 | 50934884 | 50934999 | 522 | 355 | 512 |
| A3SS | 8224 | ENSG00000127870 | RNF6 | chr13 | - | 26218510 | 26218606 | 26218510 | 26218588 | 26219456 | 26219667 | 292 | 170 | 188 |
| A3SS | 12815 | ENSG00000087502 | ERGIC2 | chr12 | - | 29341733 | 29341816 | 29341733 | 29341761 | 29343119 | 29343282 | 248 | 159 | 198 |
| A3SS | 12351 | ENSG00000103479 | RBL2 | chr16 | + | 53465437 | 53467169 | 53467057 | 53467169 | 53462551 | 53462655 | 293 | 380 | 303 |
| A3SS | 3706 | ENSG00000164483 | SAMD3 | chr6 | - | 130214336 | 130214526 | 130214336 | 130214524 | 130215194 | 130215290 | 352 | 190 | 216 |
| A3SS | 3707 | ENSG00000164483 | SAMD3 | chr6 | - | 130214336 | 130214526 | 130214336 | 130214520 | 130215194 | 130215294 | 359 | 195 | 222 |
| A3SS | 2742 | ENSG00000092201 | SUPT16H | chr14 | - | 21371873 | 21372044 | 21371873 | 21371970 | 21373337 | 21373430 | 391 | 284 | 369 |
| A3SS | 2737 | ENSG00000100697 | DICER1 | chr14 | - | 95116452 | 95116695 | 95116452 | 95116580 | 95117621 | 95117754 | 322 | 336 | 258 |
| A3SS | 5946 | ENSG00000095787 | WAC | chr10 | + | 28590719 | 28590832 | 28590721 | 28590832 | 28589735 | 28589851 | 1208 | 913 | 944 |
| A3SS | 5360 | ENSG00000048740 | CELF2 | chr10 | + | 11328925 | 11334358 | 11330455 | 11334358 | 11325835 | 11325979 | 871 | 496 | 727 |
| A3SS | 5615 | ENSG00000134419 | RPS15A | chr16 | - | 18788062 | 18788159 | 18788062 | 18788105 | 18788980 | 18789118 | 556 | 245 | 595 |
| A3SS | 4292 | ENSG00000114127 | XRN1 | chr3 | - | 142332374 | 142332534 | 142332374 | 142332424 | 142332864 | 142332890 | 212 | 201 | 235 |
| A3SS | 5808 | ENSG00000162434 | JAK1 | chr1 | - | 64841244 | 64841339 | 64841244 | 64841333 | 64841450 | 64841601 | 811 | 537 | 617 |
| A3SS | 6678 | ENSG00000135837 | CEP350 | chr1 | + | 180110996 | 180113380 | 180112436 | 180113380 | 180098862 | 180098985 | 294 | 240 | 226 |
| A3SS | 6095 | ENSG00000138095 | LRPPRC | chr2 | - | 43943686 | 43943894 | 43943686 | 43943801 | 43945331 | 43945417 | 198 | 170 | 200 |
| A3SS | 2205 | ENSG00000185658 | BRWD1 | chr21 | - | 39264459 | 39264685 | 39264459 | 39264564 | 39264890 | 39265019 | 342 | 317 | 324 |
| A3SS | 10076 | ENSG00000105835 | NAMPT | chr7 | - | 106269990 | 106270292 | 106269990 | 106270176 | 106272529 | 106272658 | 201 | 390 | 104 |
| A3SS | 3579 | ENSG00000158290 | CUL4B | chrX | - | 120544480 | 120544643 | 120544480 | 120544550 | 120545443 | 120545517 | 304 | 223 | 301 |
| A3SS | 3382 | ENSG00000158987 | RAPGEF6 | chr5 | - | 131498442 | 131498607 | 131498442 | 131498518 | 131504625 | 131504778 | 227 | 264 | 345 |
| A3SS | 9711 | ENSG00000147894 | C9orf72 | chr9 | - | 27558490 | 27560299 | 27558490 | 27558607 | 27561584 | 27561649 | 212 | 170 | 145 |
| A3SS | 7852 | ENSG00000115464 | USP34 | chr2 | - | 61214059 | 61214694 | 61214059 | 61214278 | 61220309 | 61220457 | 346 | 224 | 271 |
| A3SS | 12817 | ENSG00000087502 | ERGIC2 | chr12 | - | 29370113 | 29370222 | 29370113 | 29370168 | 29371527 | 29371670 | 265 | 249 | 262 |
| A3SS | 13389 | ENSG00000134900 | TPP2 | chr13 | + | 102644908 | 102646390 | 102646293 | 102646390 | 102644556 | 102644673 | 328 | 265 | 333 |

| | | | | | | | | | | | | | | | | | | | | | | | | | | | | | | | | | | | | | | | | | | | | | | |
|---|---|---|---|---|---|---|---|---|---|---|---|---|---|---|---|---|---|---|---|---|---|---|---|---|---|---|---|---|---|---|---|---|---|---|---|---|---|---|---|---|---|---|---|---|---|---|
| A3SS | 11088 | ENSG00000104763 | ASAH1 | chr8 | - | 18064456 | 18064531 | 18064456 | 18064474 | 18067219 | 18067298 | 1154 | 941 | 720 | 1001 | 703 | 3 | 2 | 3 | 0 | 1 | 437 | 838 | 850 | 915 | 358 | 0 | 0 | 1 | 0 | 0 | 206 | 149 | 1.000000 | 1 | 0.996 | 0.997 | 0.994 | 1 | 0.998 | 1 | 1 | 0.998 | 1 | 1 | -0.003 |
| A3SS | 11087 | ENSG00000104763 | ASAH1 | chr8 | - | 18064456 | 18064531 | 18064456 | 18064471 | 18067219 | 18067298 | 1157 | 946 | 720 | 1003 | 707 | 2 | 0 | 4 | 0 | 2 | 437 | 842 | 858 | 915 | 358 | 0 | 7 | 0 | 0 | 0 | 209 | 149 | 1.000000 | 1 | 0.998 | 1 | 0.992 | 1 | 0.996 | 1 | 0.988 | 1 | 1 | 1 | 0 |
| A3SS | 10074 | ENSG00000105835 | NAMPT | chr7 | - | 106269990 | 106270292 | 106269990 | 106270172 | 106272529 | 106272658 | 206 | 398 | 107 | 293 | 456 | 3 | 1 | 0 | 9 | 4 | 14 | 111 | 190 | 21 | 0 | 0 | 5 | 1 | 0 | 0 | 269 | 149 | 1.000000 | 1 | 0.974 | 0.995 | 1 | 0.947 | 0.984 | 1 | 0.925 | 0.991 | 1 | NA | 0.001 |
| A3SS | 2633 | ENSG00000092108 | SCFD1 | chr14 | + | 30638124 | 30638247 | 30638183 | 30638247 | 30633946 | 30634037 | 284 | 273 | 237 | 289 | 525 | 2 | 0 | 0 | 7 | 2 | 46 | 146 | 144 | 99 | 44 | 0 | 0 | 2 | 0 | 0 | 208 | 149 | 1.000000 | 1 | 0.99 | 1 | 1 | 0.967 | 0.995 | 1 | 1 | 0.981 | 1 | 1 | -0.006 |
| A3SS | 8771 | ENSG00000079785 | DDX1 | chr2 | + | 15623435 | 15623582 | 15623468 | 15623582 | 15621064 | 15621116 | 335 | 241 | 310 | 330 | 471 | 0 | 2 | 3 | 0 | 0 | 33 | 232 | 188 | 82 | 11 | 0 | 0 | 5 | 0 | 0 | 182 | 149 | 1.000000 | 1 | 1 | 0.99 | 0.988 | 1 | 1 | 1 | 1 | 0.969 | 1 | 1 | 0.002 |
| A3SS | 8045 | ENSG00000088205 | DDX18 | chr2 | + | 117824939 | 117825101 | 117824996 | 117825101 | 117824568 | 117824708 | 268 | 190 | 243 | 200 | 499 | 4 | 2 | 4 | 3 | 9 | 11 | 79 | 126 | 43 | 0 | 0 | 0 | 8 | 5 | 0 | 206 | 149 | 1.000000 | 1 | 0.98 | 0.986 | 0.978 | 0.98 | 0.976 | 1 | 1 | 0.919 | 0.862 | NA | 0.035 |
| A3SS | 3529 | ENSG00000134480 | CCNH | chr5 | - | 87407975 | 87408186 | 87407975 | 87408006 | 87409289 | 87409363 | 378 | 310 | 416 | 387 | 657 | 0 | 3 | 0 | 4 | 5 | 144 | 268 | 408 | 134 | 52 | 0 | 0 | 0 | 0 | 0 | 298 | 149 | 1.000000 | 1 | 1 | 0.981 | 1 | 0.98 | 0.985 | 1 | 1 | 1 | 1 | 1 | -0.011 |
| A3SS | 13217 | ENSG00000188906 | LRRK2 | chr12 | + | 40237968 | 40238103 | 40238046 | 40238103 | 40235625 | 40235714 | 486 | 460 | 233 | 283 | 506 | 1 | 0 | 0 | 0 | 0 | 127 | 270 | 267 | 87 | 58 | 0 | 0 | 1 | 0 | 0 | 227 | 149 | 1.000000 | 1 | 0.997 | 1 | 1 | 1 | 1 | 1 | 1 | 0.994 | 1 | 1 | 0.001 |
| A3SS | 5801 | ENSG00000163714 | U2SURP | chr3 | + | 143028482 | 143028646 | 143028486 | 143028646 | 143028339 | 143028406 | 368 | 300 | 361 | 393 | 423 | 0 | 0 | 4 | 1 | 0 | 51 | 267 | 299 | 51 | 12 | 0 | 4 | 0 | 0 | 0 | 153 | 149 | 1.000000 | 1 | 1 | 1 | 0.989 | 0.997 | 1 | 1 | 0.985 | 1 | 1 | 1 | 0 |
| A3SS | 10077 | ENSG00000105835 | NAMPT | chr7 | - | 106269990 | 106270332 | 106269990 | 106270176 | 106272529 | 106272658 | 230 | 432 | 121 | 292 | 464 | 8 | 7 | 5 | 41 | 28 | 14 | 124 | 198 | 21 | 0 | 0 | 13 | 13 | 3 | 0 | 298 | 149 | 1.000000 | 1 | 0.935 | 0.969 | 0.924 | 0.781 | 0.892 | 1 | 0.827 | 0.884 | 0.778 | NA | 0.028 |
| A3SS | 3647 | ENSG00000070756 | PABPC1 | chr8 | - | 100704925 | 100705056 | 100704925 | 100705029 | 100705588 | 100705673 | 4999 | 3299 | 4742 | 4800 | 5734 | 0 | 1 | 3 | 3 | 0 | 5553 | 3215 | 3983 | 3332 | 6303 | 0 | 1 | 1 | 0 | 0 | 176 | 149 | 1.000000 | 1 | 1 | 1 | 0.999 | 0.999 | 1 | 1 | 1 | 1 | 1 | 1 | 0 |
| A3SS | 10075 | ENSG00000105835 | NAMPT | chr7 | - | 106269990 | 106270332 | 106269990 | 106270172 | 106272529 | 106272658 | 231 | 426 | 122 | 305 | 477 | 3 | 1 | 0 | 9 | 4 | 14 | 124 | 202 | 21 | 0 | 0 | 5 | 1 | 0 | 0 | 298 | 149 | 1.000000 | 1 | 0.975 | 0.995 | 1 | 0.944 | 0.984 | 1 | 0.925 | 0.99 | 1 | NA | 0.001 |
| A3SS | 12156 | ENSG00000110321 | EIF4G2 | chr11 | - | 10803479 | 10804050 | 10803479 | 10803590 | 10804136 | 10804203 | 918 | 649 | 667 | 879 | 1064 | 0 | 0 | 0 | 4 | 0 | 636 | 631 | 622 | 738 | 320 | 0 | 0 | 0 | 0 | 0 | 298 | 149 | 1.000000 | 1 | 1 | 1 | 1 | 0.991 | 1 | 1 | 1 | 1 | 1 | 1 | -0.002 |
| A3SS | 4638 | ENSG00000174953 | DHX36 | chr3 | - | 154295283 | 154295339 | 154295283 | 154295297 | 154299837 | 154299925 | 427 | 322 | 393 | 342 | 413 | 1 | 0 | 1 | 0 | 3 | 92 | 222 | 246 | 72 | 19 | 0 | 0 | 0 | 0 | 0 | 191 | 149 | 1.000000 | 1 | 0.997 | 1 | 0.997 | 1 | 0.991 | 1 | 1 | 1 | 1 | 1 | -0.003 |
| A3SS | 2451 | ENSG00000125107 | CNOT1 | chr16 | - | 58555250 | 58555537 | 58555250 | 58555429 | 58555783 | 58555908 | 677 | 545 | 605 | 738 | 862 | 0 | 0 | 1 | 0 | 0 | 291 | 693 | 479 | 620 | 85 | 0 | 0 | 0 | 0 | 0 | 257 | 149 | 1.000000 | 1 | 1 | 1 | 0.997 | 1 | 1 | 1 | 1 | 1 | 1 | 1 | -0.001 |
| A3SS | 13399 | ENSG00000127022 | CANX | chr5 | + | 179723659 | 179723779 | 179723663 | 179723779 | 179722803 | 179723019 | 1218 | 780 | 1055 | 1063 | 1110 | 0 | 0 | 0 | 0 | 1 | 588 | 807 | 1081 | 1244 | 240 | 0 | 0 | 0 | 0 | 0 | 153 | 149 | 1.000000 | 1 | 1 | 1 | 1 | 1 | 0.999 | 1 | 1 | 1 | 1 | 1 | 0 |
| A3SS | 3922 | ENSG00000100650 | SRSF5 | chr14 | + | 69768603 | 69768674 | 69768612 | 69768674 | 69768137 | 69768282 | 1972 | 1405 | 2051 | 2386 | 2446 | 1 | 0 | 0 | 0 | 9 | 1870 | 1991 | 1788 | 2240 | 1093 | 0 | 7 | 0 | 1 | 0 | 158 | 149 | 1.000000 | 1 | 0.999 | 1 | 1 | 1 | 0.996 | 1 | 0.996 | 1 | 1 | 1 | 0 |
| A3SS | 5021 | ENSG00000080822 | CLDND1 | chr3 | - | 98515482 | 98516879 | 98515482 | 98515832 | 98518884 | 98518995 | 256 | 181 | 329 | 349 | 545 | 0 | 0 | 0 | 0 | 2 | 35 | 96 | 158 | 42 | 22 | 0 | 0 | 0 | 0 | 0 | 298 | 149 | 1.000000 | 1 | 1 | 1 | 1 | 1 | 0.993 | 1 | 1 | 1 | 1 | 1 | -0.001 |
| A3SS | 12952 | ENSG00000127314 | RAP1B | chr12 | + | 68648698 | 68648781 | 68648705 | 68648781 | 68610907 | 68611043 | 693 | 535 | 404 | 670 | 577 | 4 | 0 | 1 | 5 | 4 | 179 | 549 | 422 | 304 | 117 | 0 | 7 | 1 | 0 | 0 | 156 | 149 | 1.000000 | 1 | 0.994 | 1 | 0.997 | 0.992 | 0.993 | 1 | 0.987 | 0.998 | 1 | 1 | -0.002 |
| A3SS | 10228 | ENSG00000029363 | BCLAF1 | chr6 | - | 136271994 | 136272079 | 136271994 | 136272076 | 136273081 | 136273187 | 347 | 301 | 355 | 466 | 457 | 2 | 0 | 5 | 3 | 0 | 59 | 175 | 304 | 70 | 4 | 0 | 0 | 0 | 0 | 0 | 152 | 149 | 1.000000 | 1 | 0.994 | 1 | 0.986 | 0.993 | 1 | 1 | 1 | 1 | 1 | 1 | -0.005 |
| A3SS | 2910 | ENSG00000175061 | SNHG29 | chr17 | + | 16441073 | 16442018 | 16441356 | 16442018 | 16440184 | 16440253 | 778 | 395 | 926 | 682 | 739 | 0 | 0 | 1 | 0 | 0 | 414 | 332 | 460 | 459 | 539 | 0 | 0 | 0 | 0 | 0 | 298 | 149 | 1.000000 | 1 | 1 | 1 | 0.998 | 1 | 1 | 1 | 1 | 1 | 1 | 1 | 0 |
| A3SS | 10067 | ENSG00000114503 | NCBP2 | chr3 | - | 196937509 | 196937648 | 196937509 | 196937631 | 196939250 | 196939432 | 573 | 473 | 495 | 474 | 696 | 1 | 1 | 1 | 5 | 0 | 265 | 279 | 383 | 323 | 114 | 0 | 0 | 0 | 0 | 0 | 166 | 149 | 1.000000 | 1 | 0.998 | 0.998 | 0.998 | 0.988 | 1 | 1 | 1 | 1 | 1 | 1 | -0.004 |
| A3SS | 8272 | ENSG00000163728 | TTC14 | chr3 | + | 180603123 | 180606372 | 180606252 | 180606372 | 180602890 | 180603015 | 332 | 224 | 320 | 351 | 545 | 0 | 0 | 0 | 0 | 0 | 74 | 157 | 116 | 49 | 21 | 0 | 0 | 0 | 2 | 0 | 298 | 149 | 1.000000 | 1 | 1 | 1 | 1 | 1 | 1 | 1 | 1 | 1 | 0.925 | 1 | 0.015 |
| A3SS | 2570 | ENSG00000100796 | PPP4R3A | chr14 | - | 91462048 | 91462239 | 91462048 | 91462221 | 91462734 | 91462877 | 468 | 503 | 408 | 444 | 822 | 3 | 3 | 1 | 3 | 7 | 162 | 312 | 431 | 323 | 55 | 0 | 5 | 4 | 5 | 0 | 167 | 149 | 1.000000 | 1 | 0.993 | 0.993 | 0.997 | 0.992 | 0.991 | 1 | 0.982 | 0.99 | 0.983 | 1 | 0.002 |
| A3SS | 3804 | ENSG00000145907 | G3BP1 | chr5 | + | 151799888 | 151800346 | 151800217 | 151800346 | 151799211 | 151799313 | 579 | 396 | 528 | 585 | 734 | 0 | 0 | 2 | 1 | 0 | 188 | 437 | 379 | 275 | 144 | 4 | 0 | 1 | 1 | 0 | 298 | 149 | 1.000000 | 1 | 1 | 1 | 0.992 | 0.997 | 1 | 0.959 | 1 | 0.995 | 0.993 | 1 | 0.008 |
| A3SS | 2996 | ENSG00000126777 | KTN1 | chr14 | + | 55634525 | 55634658 | 55634529 | 55634658 | 55633234 | 55633341 | 269 | 280 | 263 | 318 | 364 | 5 | 12 | 1 | 15 | 3 | 10 | 40 | 40 | 4 | 0 | 0 | 1 | 1 | 0 | 0 | 153 | 149 | 1.000000 | 1 | 0.981 | 0.958 | 0.996 | 0.954 | 0.992 | 1 | 0.975 | 0.975 | 1 | NA | -0.011 |
| A3SS | 10916 | ENSG00000177565 | TBL1XR1 | chr3 | - | 177038312 | 177038434 | 177038312 | 177038428 | 177046128 | 177046189 | 410 | 356 | 444 | 429 | 768 | 1 | 0 | 0 | 0 | 0 | 158 | 320 | 266 | 126 | 114 | 0 | 0 | 0 | 0 | 0 | 155 | 149 | 1.000000 | 1 | 0.997 | 1 | 1 | 1 | 1 | 1 | 1 | 1 | 1 | 1 | -0.001 |
| A3SS | 3803 | ENSG00000145907 | G3BP1 | chr5 | + | 151799888 | 151800000 | 151799898 | 151800000 | 151799211 | 151799313 | 577 | 389 | 532 | 585 | 749 | 4 | 0 | 1 | 0 | 3 | 194 | 434 | 352 | 264 | 137 | 0 | 0 | 0 | 0 | 0 | 159 | 149 | 1.000000 | 1 | 0.993 | 1 | 0.998 | 1 | 0.996 | 1 | 1 | 1 | 1 | 1 | -0.003 |
| A3SS | 4649 | ENSG00000041357 | PSMA4 | chr15 | + | 78542482 | 78542645 | 78542575 | 78542645 | 78542176 | 78542219 | 1051 | 691 | 959 | 1204 | 1016 | 3 | 11 | 1 | 3 | 5 | 901 | 649 | 605 | 1055 | 250 | 9 | 5 | 0 | 3 | 0 | 242 | 149 | 1.000000 | 1 | 0.995 | 0.975 | 0.998 | 0.996 | 0.992 | 0.984 | 0.988 | 1 | 0.995 | 1 | -0.002 |
| A3SS | 7482 | ENSG00000115875 | SRSF7 | chr2 | - | 38748897 | 38749346 | 38748897 | 38749173 | 38749528 | 38749705 | 656 | 554 | 723 | 774 | 1029 | 0 | 0 | 0 | 0 | 0 | 275 | 607 | 716 | 494 | 181 | 0 | 0 | 0 | 8 | 0 | 298 | 149 | 1.000000 | 1 | 1 | 1 | 1 | 1 | 1 | 1 | 1 | 1 | 0.969 | 1 | 0.006 |
| A3SS | 1647 | ENSG00000008517 | IL32 | chr16 | + | 3067376 | 3067613 | 3067442 | 3067613 | 3065783 | 3065826 | 1221 | 250 | 869 | 836 | 732 | 1 | 0 | 0 | 1 | 2 | 444 | 644 | 381 | 233 | 729 | 0 | 0 | 0 | 0 | 0 | 215 | 149 | 1.000000 | 1 | 0.999 | 1 | 1 | 0.998 | 0.996 | 1 | 1 | 1 | 1 | 1 | -0.001 |
| A3SS | 2911 | ENSG00000175061 | SNHG29 | chr17 | + | 16441073 | 16442018 | 16441367 | 16442018 | 16440184 | 16440253 | 813 | 407 | 967 | 711 | 859 | 84 | 33 | 116 | 59 | 109 | 420 | 336 | 474 | 473 | 577 | 60 | 41 | 30 | 45 | 65 | 298 | 149 | 1.000000 | 1 | 0.829 | 0.86 | 0.807 | 0.858 | 0.798 | 0.778 | 0.804 | 0.888 | 0.84 | 0.816 | 0.005 |
| A3SS | 9328 | ENSG00000112739 | PRPF4B | chr6 | + | 4060412 | 4061147 | 4061014 | 4061147 | 4058727 | 4058814 | 384 | 313 | 351 | 442 | 642 | 0 | 0 | 1 | 0 | 0 | 18 | 208 | 303 | 69 | 13 | 0 | 0 | 0 | 0 | 0 | 298 | 149 | 1.000000 | 1 | 1 | 1 | 0.994 | 1 | 1 | 1 | 1 | 1 | 1 | 1 | -0.001 |
| A3SS | 3912 | ENSG00000124783 | SSR1 | chr6 | - | 7303549 | 7303637 | 7303549 | 7303633 | 7309916 | 7310029 | 725 | 528 | 600 | 631 | 1167 | 1 | 3 | 0 | 1 | 3 | 307 | 540 | 638 | 474 | 123 | 0 | 4 | 0 | 0 | 0 | 153 | 149 | 1.000000 | 1 | 0.999 | 0.994 | 1 | 0.998 | 0.997 | 1 | 0.992 | 1 | 1 | 1 | -0.001 |
| A3SS | 10917 | ENSG00000171988 | JMJD1C | chr10 | - | 63192937 | 63193151 | 63192937 | 63193097 | 63193344 | 63193472 | 341 | 273 | 261 | 275 | 575 | 2 | 1 | 0 | 1 | 0 | 4 | 66 | 52 | 18 | 4 | 0 | 0 | 0 | 0 | 0 | 203 | 149 | 1.000000 | 1 | 0.992 | 0.995 | 1 | 0.995 | 1 | 1 | 1 | 1 | 1 | 1 | -0.004 |
| A3SS | 10015 | ENSG00000177189 | RPS6KA3 | chrX | - | 20164898 | 20165060 | 20164898 | 20164961 | 20167588 | 20167747 | 501 | 389 | 482 | 478 | 702 | 0 | 0 | 1 | 0 | 0 | 149 | 235 | 287 | 172 | 84 | 0 | 0 | 0 | 0 | 0 | 248 | 149 | 1.000000 | 1 | 1 | 1 | 0.997 | 1 | 1 | 1 | 1 | 1 | 1 | 1 | -0.001 |
| A3SS | 9674 | ENSG00000026950 | BTN3A1 | chr6 | + | 26405371 | 26405648 | 26405503 | 26405648 | 26402249 | 26402412 | 678 | 359 | 464 | 383 | 1002 | 1 | 0 | 0 | 0 | 0 | 211 | 442 | 181 | 205 | 210 | 0 | 1 | 0 | 0 | 0 | 281 | 149 | 1.000000 | 1 | 0.997 | 1 | 1 | 1 | 1 | 1 | 0.996 | 1 | 1 | 1 | 0 |
| A3SS | 5948 | ENSG00000095787 | WAC | chr10 | + | 28610698 | 28610821 | 28610710 | 28610821 | 28608185 | 28608431 | 759 | 619 | 691 | 805 | 1281 | 3 | 1 | 3 | 3 | 6 | 274 | 700 | 854 | 604 | 77 | 0 | 0 | 0 | 2 | 0 | 161 | 149 | 1.000000 | 1 | 0.996 | 0.998 | 0.995 | 0.996 | 0.995 | 1 | 1 | 1 | 0.996 | 1 | -0.003 |
| A3SS | 5501 | ENSG00000163565 | IFI16 | chr1 | + | 159016532 | 159016700 | 159016617 | 159016700 | 159015672 | 159015987 | 671 | 468 | 403 | 561 | 728 | 0 | 0 | 1 | 1 | 0 | 134 | 526 | 346 | 155 | 21 | 0 | 0 | 1 | 0 | 0 | 234 | 149 | 1.000000 | 1 | 1 | 1 | 0.996 | 0.997 | 1 | 1 | 1 | 0.995 | 1 | 1 | 0 |
| A3SS | 12955 | ENSG00000127314 | RAP1B | chr12 | + | 68654111 | 68654252 | 68654124 | 68654252 | 68652018 | 68652051 | 1891 | 1806 | 1058 | 2063 | 1811 | 0 | 0 | 0 | 0 | 0 | 1223 | 1884 | 1605 | 1343 | 900 | 0 | 0 | 1 | 0 | 0 | 162 | 149 | 1.000000 | 1 | 1 | 1 | 1 | 1 | 1 | 1 | 1 | 0.999 | 1 | 1 | 0 |
| A3SS | 10852 | ENSG00000152601 | MBNL1 | chr3 | + | 152447619 | 152447773 | 152447724 | 152447773 | 152268946 | 152269092 | 611 | 447 | 504 | 763 | 883 | 0 | 3 | 0 | 0 | 0 | 168 | 447 | 395 | 204 | 202 | 0 | 0 | 0 | 0 | 0 | 254 | 149 | 1.000000 | 1 | 1 | 0.989 | 1 | 1 | 1 | 1 | 1 | 1 | 1 | 1 | -0.002 |
| A3SS | 12674 | ENSG00000150093 | ITGB1 | chr10 | - | 32932514 | 32932600 | 32932514 | 32932590 | 32935491 | 32935558 | 889 | 580 | 395 | 918 | 1021 | 0 | 1 | 0 | 0 | 0 | 210 | 612 | 542 | 296 | 302 | 0 | 0 | 0 | 0 | 0 | 159 | 149 | 1.000000 | 1 | 1 | 0.998 | 1 | 1 | 1 | 1 | 1 | 1 | 1 | 1 | 0 |
| A3SS | 10062 | ENSG00000122566 | HNRNPA2B1 | chr7 | - | 26193574 | 26193694 | 26193574 | 26193598 | 26195846 | 26195909 | 1342 | 1290 | 1096 | 1480 | 1531 | 3 | 0 | 1 | 1 | 0 | 570 | 1200 | 1552 | 1266 | 302 | 0 | 0 | 0 | 1 | 0 | 245 | 149 | 1.000000 | 1 | 0.996 | 1 | 0.999 | 0.999 | 1 | 1 | 1 | 1 | 0.999 | 1 | -0.001 |
| A3SS | 3379 | ENSG00000204628 | RACK1 | chr5 | - | 181241491 | 181241639 | 181241491 | 181241563 | 181242169 | 181242345 | 1916 | 925 | 2201 | 1562 | 2260 | 2 | 2 | 1 | 0 | 9 | 1511 | 1192 | 977 | 1059 | 2268 | 0 | 3 | 2 | 0 | 0 | 225 | 149 | 1.000000 | 1 | 0.998 | 0.997 | 0.999 | 1 | 0.994 | 1 | 0.996 | 0.997 | 1 | 1 | -0.001 |
| A3SS | 10854 | ENSG00000152601 | MBNL1 | chr3 | + | 152447619 | 152447773 | 152447724 | 152447773 | 152446703 | 152446757 | 645 | 480 | 539 | 800 | 942 | 0 | 1 | 0 | 4 | 2 | 193 | 483 | 420 | 240 | 211 | 0 | 0 | 0 | 0 | 0 | 254 | 149 | 1.000000 | 1 | 1 | 0.996 | 1 | 0.992 | 0.996 | 1 | 1 | 1 | 1 | 1 | -0.003 |
| A3SS | 12865 | ENSG00000082898 | XPO1 | chr2 | - | 61483936 | 61485962 | 61483936 | 61484105 | 61488164 | 61488271 | 408 | 315 | 387 | 432 | 935 | 5 | 2 | 3 | 1 | 3 | 95 | 140 | 183 | 134 | 64 | 0 | 0 | 6 | 0 | 0 | 298 | 149 | 1.000000 | 1 | 0.976 | 0.987 | 0.985 | 0.995 | 0.994 | 1 | 1 | 0.938 | 1 | 1 | 0 |
| A3SS | 3246 | ENSG00000113263 | ITK | chr5 | + | 157240061 | 157241720 | 157241645 | 157241720 | 157238108 | 157238191 | 535 | 328 | 684 | 470 | 1314 | 0 | 1 | 2 | 0 | 0 | 200 | 312 | 580 | 224 | 134 | 0 | 0 | 3 | 0 | 0 | 298 | 149 | 1.000000 | 1 | 1 | 0.994 | 0.994 | 1 | 1 | 1 | 1 | 0.99 | 1 | 1 | 0 |
| A3SS | 6326 | ENSG00000137575 | SDCBP | chr8 | + | 58578032 | 58578208 | 58578047 | 58578208 | 58575902 | 58576061 | 708 | 483 | 472 | 505 | 972 | 2 | 0 | 0 | 0 | 0 | 70 | 315 | 631 | 141 | 43 | 0 | 0 | 0 | 0 | 0 | 164 | 149 | 1.000000 | 1 | 0.997 | 1 | 1 | 1 | 1 | 1 | 1 | 1 | 1 | 1 | -0.001 |
| A3SS | 12968 | ENSG00000089157 | RPLP0 | chr12 | - | 120198553 | 120198770 | 120198553 | 120198631 | 120198853 | 120199000 | 897 | 331 | 1143 | 850 | 1301 | 5 | 3 | 7 | 0 | 0 | 347 | 592 | 535 | 279 | 802 | 0 | 0 | 7 | 0 | 0 | 288 | 149 | 1.000000 | 1 | 0.989 | 0.983 | 0.988 | 1 | 1 | 1 | 1 | 0.975 | 1 | 1 | -0.003 |
| A3SS | 7395 | ENSG00000118816 | CCNI | chr4 | - | 77055961 | 77056102 | 77055961 | 77056100 | 77056248 | 77056485 | 1143 | 648 | 888 | 1017 | 3072 | 0 | 0 | 0 | 0 | 0 | 1272 | 761 | 669 | 856 | 1241 | 0 | 0 | 0 | 2 | 0 | 151 | 149 | 1.000000 | 1 | 1 | 1 | 1 | 1 | 1 | 1 | 1 | 1 | 0.998 | 1 | 0 |
| A3SS | 5452 | ENSG00000119048 | UBE2B | chr5 | + | 134388324 | 134388413 | 134388351 | 134388413 | 134380718 | 134380808 | 1084 | 710 | 790 | 823 | 1927 | 0 | 1 | 0 | 0 | 0 | 515 | 616 | 1036 | 763 | 364 | 0 | 0 | 2 | 0 | 0 | 176 | 149 | 1.000000 | 1 | 1 | 0.998 | 1 | 1 | 1 | 1 | 1 | 0.998 | 1 | 1 | 0 |
| A3SS | 10720 | ENSG00000215301 | DDX3X | chrX | + | 41345179 | 41345548 | 41345403 | 41345548 | 41344238 | 41344399 | 762 | 528 | 594 | 817 | 735 | 0 | 0 | 2 | 0 | 0 | 54 | 427 | 489 | 332 | 39 | 0 | 0 | 0 | 0 | 0 | 298 | 149 | 1.000000 | 1 | 1 | 1 | 0.993 | 1 | 1 | 1 | 1 | 1 | 1 | 1 | 0 |
| A3SS | 12867 | ENSG00000082898 | XPO1 | chr2 | - | 61492034 | 61492198 | 61492034 | 61492196 | 61492324 | 61492481 | 492 | 367 | 439 | 500 | 983 | 1 | 0 | 1 | 0 | 0 | 58 | 187 | 193 | 131 | 73 | 0 | 0 | 0 | 0 | 0 | 151 | 149 | 1.000000 | 1 | 0.998 | 1 | 0.998 | 1 | 1 | 1 | 1 | 1 | 1 | 1 | -0.001 |
| A3SS | 10162 | ENSG00000234741 | GAS5 | chr1 | - | 173864256 | 173865282 | 173864256 | 173864304 | 173865470 | 173865547 | 457 | 364 | 798 | 577 | 1053 | 1 | 0 | 1 | 0 | 5 | 216 | 118 | 396 | 156 | 219 | 0 | 0 | 0 | 0 | 0 | 298 | 149 | 1.000000 | 1 | 0.996 | 1 | 0.997 | 1 | 0.991 | 1 | 1 | 1 | 1 | 1 | -0.003 |
| A3SS | 12396 | ENSG00000124795 | DEK | chr6 | - | 18236451 | 18236600 | 18236451 | 18236596 | 18237380 | 18237516 | 623 | 479 | 413 | 686 | 882 | 0 | 4 | 2 | 3 | 14 | 185 | 293 | 220 | 117 | 104 | 0 | 0 | 1 | 0 | 0 | 153 | 149 | 1.000000 | 1 | 1 | 0.991 | 0.995 | 0.996 | 0.984 | 1 | 1 | 0.995 | 1 | 1 | -0.006 |
| A3SS | 10164 | ENSG00000234741 | GAS5 | chr1 | - | 173865856 | 173866206 | 173865856 | 173865894 | 173866527 | 173866567 | 872 | 485 | 1026 | 757 | 964 | 0 | 0 | 1 | 1 | 3 | 282 | 411 | 496 | 337 | 328 | 2 | 0 | 1 | 0 | 0 | 298 | 149 | 1.000000 | 1 | 1 | 1 | 0.998 | 0.997 | 0.994 | 0.986 | 1 | 0.996 | 1 | 1 | 0.001 |
| A3SS | 11413 | ENSG00000140988 | RPS2 | chr16 | - | 1962060 | 1962656 | 1962060 | 1962270 | 1962735 | 1962909 | 2738 | 1082 | 3092 | 2490 | 3238 | 1 | 0 | 0 | 2 | 0 | 1955 | 2322 | 1725 | 1176 | 3210 | 0 | 0 | 1 | 0 | 0 | 298 | 149 | 1.000000 | 1 | 0.999 | 1 | 1 | 0.998 | 1 | 1 | 1 | 0.999 | 1 | 1 | 0 |
| A3SS | 6535 | ENSG00000153187 | HNRNPU | chr1 | - | 244858728 | 244858841 | 244858728 | 244858835 | 244859274 | 244859374 | 1225 | 1034 | 1070 | 1225 | 2213 | 0 | 0 | 0 | 0 | 0 | 877 | 1021 | 1054 | 1229 | 333 | 0 | 0 | 1 | 0 | 0 | 155 | 149 | 1.000000 | 1 | 1 | 1 | 1 | 1 | 1 | 1 | 1 | 0.999 | 1 | 1 | 0 |
| A3SS | 8781 | ENSG00000115053 | NCL | chr2 | - | 231457000 | 231459125 | 231457000 | 231457800 | 231460151 | 231460293 | 1488 | 1122 | 1396 | 1444 | 1955 | 1 | 0 | 0 | 0 | 0 | 1003 | 1229 | 1306 | 1369 | 197 | 0 | 0 | 0 | 0 | 0 | 298 | 149 | 1.000000 | 1 | 0.999 | 1 | 1 | 1 | 1 | 1 | 1 | 1 | 1 | 1 | 0 |
| A3SS | 5809 | ENSG00000162434 | JAK1 | chr1 | - | 64845512 | 64846736 | 64845512 | 64845640 | 64847531 | 64847675 | 825 | 480 | 739 | 711 | 1375 | 0 | 0 | 1 | 5 | 0 | 371 | 515 | 517 | 311 | 107 | 0 | 0 | 2 | 4 | 0 | 298 | 149 | 1.000000 | 1 | 1 | 1 | 0.997 | 0.986 | 1 | 1 | 1 | 0.992 | 0.975 | 1 | 0.003 |
| A3SS | 6624 | ENSG00000169764 | UGP2 | chr2 | + | 63886340 | 63886538 | 63886479 | 63886538 | 63885588 | 63885886 | 651 | 494 | 589 | 644 | 1107 | 1 | 0 | 0 | 0 | 0 | 79 | 284 | 445 | 294 | 25 | 0 | 0 | 0 | 0 | 0 | 288 | 149 | 1.000000 | 1 | 0.997 | 1 | 1 | 1 | 1 | 1 | 1 | 1 | 1 | 1 | -0.001 |
| A3SS | 8776 | ENSG00000115053 | NCL | chr2 | - | 231457000 | 231457800 | 231457000 | 231457124 | 231458265 | 231458389 | 1749 | 1540 | 1742 | 1901 | 2344 | 0 | 0 | 0 | 0 | 0 | 1473 | 1526 | 1658 | 1846 | 387 | 0 | 2 | 0 | 0 | 0 | 298 | 149 | 1.000000 | 1 | 1 | 1 | 1 | 1 | 1 | 1 | 0.997 | 1 | 1 | 1 | 0.001 |
| A3SS | 10716 | ENSG00000215301 | DDX3X | chrX | + | 41343215 | 41343351 | 41343220 | 41343351 | 41342736 | 41342836 | 845 | 620 | 709 | 1015 | 1150 | 0 | 0 | 1 | 0 | 0 | 256 | 426 | 717 | 433 | 108 | 0 | 0 | 1 | 0 | 0 | 154 | 149 | 1.000000 | 1 | 1 | 1 | 0.999 | 1 | 1 | 1 | 1 | 0.999 | 1 | 1 | 0 |
| A3SS | 10227 | ENSG00000029363 | BCLAF1 | chr6 | - | 136268161 | 136268339 | 136268161 | 136268273 | 136269140 | 136269303 | 557 | 621 | 558 | 699 | 760 | 1 | 2 | 0 | 0 | 0 | 79 | 244 | 316 | 100 | 42 | 0 | 0 | 0 | 0 | 0 | 215 | 149 | 1.000000 | 1 | 0.997 | 0.995 | 1 | 1 | 1 | 1 | 1 | 1 | 1 | 1 | -0.002 |
| A3SS | 5603 | ENSG00000154814 | OXNAD1 | chr3 | + | 16303407 | 16303592 | 16303410 | 16303592 | 16302639 | 16302748 | 407 | 359 | 636 | 503 | 1684 | 3 | 2 | 2 | 2 | 6 | 249 | 240 | 331 | 179 | 126 | 0 | 0 | 0 | 0 | 0 | 152 | 149 | 1.000000 | 1 | 0.993 | 0.994 | 0.997 | 0.996 | 0.996 | 1 | 1 | 1 | 1 | 1 | -0.005 |
| A3SS | 2814 | ENSG00000198668 | CALM1 | chr14 | + | 90404378 | 90404514 | 90404398 | 90404514 | 90403861 | 90403968 | 3206 | 2057 | 2766 | 3486 | 5357 | 0 | 0 | 0 | 0 | 0 | 3143 | 2655 | 2615 | 3267 | 2681 | 6 | 0 | 0 | 0 | 0 | 169 | 149 | 1.000000 | 1 | 1 | 1 | 1 | 1 | 1 | 0.998 | 1 | 1 | 1 | 1 | 0 |
| A3SS | 9283 | ENSG00000132424 | PNISR | chr6 | - | 99412550 | 99412739 | 99412550 | 99412734 | 99414571 | 99414690 | 590 | 600 | 657 | 652 | 1263 | 0 | 1 | 5 | 3 | 2 | 268 | 316 | 255 | 204 | 149 | 0 | 0 | 0 | 0 | 0 | 154 | 149 | 1.000000 | 1 | 1 | 0.998 | 0.992 | 0.995 | 0.998 | 1 | 1 | 1 | 1 | 1 | -0.003 |
| A3SS | 8808 | ENSG00000169045 | HNRNPH1 | chr5 | - | 179620891 | 179621397 | 179620891 | 179621035 | 179623036 | 179623164 | 1320 | 876 | 1169 | 1289 | 1049 | 0 | 0 | 1 | 1 | 0 | 294 | 895 | 939 | 655 | 241 | 0 | 0 | 0 | 0 | 0 | 298 | 149 | 1.000000 | 1 | 1 | 1 | 0.998 | 0.998 | 1 | 1 | 1 | 1 | 1 | 1 | -0.001 |
| A3SS | 7682 | ENSG00000179639 | FCER1A | chr1 | + | 159303927 | 159304182 | 159304026 | 159304182 | 159302853 | 159302874 | 470 | 587 | 545 | 555 | 820 | 0 | 2 | 0 | 0 | 0 | 80 | 66 | 22 | 69 | 24 | 0 | 0 | 0 | 0 | 0 | 248 | 149 | 1.000000 | 1 | 1 | 0.994 | 1 | 1 | 1 | 1 | 1 | 1 | 1 | 1 | -0.001 |
| A3SS | 10160 | ENSG00000234741 | GAS5 | chr1 | - | 173864483 | 173864704 | 173864483 | 173864506 | 173865228 | 173865282 | 795 | 469 | 990 | 750 | 1337 | 0 | 0 | 0 | 0 | 0 | 264 | 305 | 453 | 261 | 301 | 0 | 0 | 0 | 1 | 0 | 298 | 149 | 1.000000 | 1 | 1 | 1 | 1 | 1 | 1 | 1 | 1 | 1 | 0.992 | 1 | 0.002 |
| A3SS | 10159 | ENSG00000234741 | GAS5 | chr1 | - | 173864256 | 173864704 | 173864256 | 173864304 | 173865228 | 173865282 | 832 | 490 | 1010 | 769 | 1366 | 4 | 0 | 0 | 2 | 0 | 264 | 311 | 454 | 270 | 301 | 0 | 0 | 0 | 0 | 0 | 298 | 149 | 1.000000 | 1 | 0.99 | 1 | 1 | 0.995 | 1 | 1 | 1 | 1 | 1 | 1 | -0.003 |
| A3SS | 10767 | ENSG00000229807 | XIST | chrX | - | 73831065 | 73831274 | 73831065 | 73831206 | 73841381 | 73841473 | 580 | 472 | 616 | 553 | 890 | 0 | 0 | 1 | 0 | 0 | 87 | 1 | 0 | 97 | 0 | 0 | 0 | 0 | 0 | 0 | 217 | 149 | 1.000000 | 1 | 1 | 1 | 0.998 | 1 | 1 | 1 | 1 | NA | 1 | NA | 0 |
| A3SS | 12673 | ENSG00000150093 | ITGB1 | chr10 | - | 32929821 | 32930044 | 32929821 | 32930032 | 32932514 | 32932600 | 1297 | 925 | 645 | 1418 | 1805 | 4 | 0 | 1 | 2 | 1 | 314 | 892 | 882 | 484 | 409 | 0 | 0 | 4 | 0 | 0 | 161 | 149 | 1.000000 | 1 | 0.997 | 1 | 0.998 | 0.998 | 0.999 | 1 | 1 | 0.995 | 1 | 1 | -0.001 |
| A3SS | 13215 | ENSG00000166598 | HSP90B1 | chr12 | + | 103941831 | 103942796 | 103942526 | 103942796 | 103941628 | 103941706 | 1186 | 702 | 1376 | 1158 | 1164 | 0 | 0 | 0 | 2 | 0 | 87 | 740 | 1415 | 206 | 25 | 0 | 2 | 0 | 0 | 0 | 298 | 149 | 1.000000 | 1 | 1 | 1 | 1 | 0.997 | 1 | 1 | 0.995 | 1 | 1 | 1 | 0 |
| A3SS | 10060 | ENSG00000122566 | HNRNPA2B1 | chr7 | - | 26190992 | 26192577 | 26190992 | 26192338 | 26193250 | 26193373 | 1480 | 1693 | 1292 | 1612 | 1151 | 5 | 0 | 0 | 0 | 0 | 435 | 904 | 1493 | 1054 | 200 | 0 | 5 | 4 | 7 | 0 | 298 | 149 | 1.000000 | 1 | 0.993 | 1 | 1 | 1 | 1 | 1 | 0.989 | 0.995 | 0.987 | 1 | 0.004 |
| A3SS | 2367 | ENSG00000137845 | ADAM10 | chr15 | - | 58640776 | 58640960 | 58640776 | 58640830 | 58643885 | 58643978 | 1108 | 1016 | 732 | 1095 | 1395 | 0 | 0 | 1 | 0 | 1 | 378 | 651 | 727 | 212 | 180 | 0 | 0 | 0 | 0 | 0 | 279 | 149 | 1.000000 | 1 | 1 | 1 | 0.997 | 1 | 0.999 | 1 | 1 | 1 | 1 | 1 | -0.001 |
| A3SS | 5789 | ENSG00000108654 | DDX5 | chr17 | - | 64500548 | 64500882 | 64500548 | 64500773 | 64500986 | 64501143 | 1622 | 1028 | 1570 | 1801 | 1306 | 2 | 0 | 0 | 0 | 0 | 226 | 1391 | 1482 | 875 | 85 | 0 | 0 | 0 | 0 | 0 | 258 | 149 | 1.000000 | 1 | 0.998 | 1 | 1 | 1 | 1 | 1 | 1 | 1 | 1 | 1 | 0 |
| A3SS | 10059 | ENSG00000122566 | HNRNPA2B1 | chr7 | - | 26190992 | 26192338 | 26190992 | 26191650 | 26192494 | 26192577 | 1476 | 1297 | 1120 | 1449 | 1355 | 1 | 1 | 4 | 2 | 2 | 361 | 849 | 1021 | 1009 | 182 | 0 | 1 | 1 | 2 | 0 | 298 | 149 | 1.000000 | 1 | 0.999 | 0.998 | 0.993 | 0.997 | 0.997 | 1 | 0.998 | 0.998 | 0.996 | 1 | -0.002 |
| A3SS | 3380 | ENSG00000204628 | RACK1 | chr5 | - | 181241491 | 181241639 | 181241491 | 181241563 | 181242173 | 181242345 | 3936 | 1845 | 4257 | 3201 | 3855 | 3 | 0 | 0 | 5 | 1 | 3012 | 2456 | 1923 | 1991 | 4434 | 0 | 0 | 0 | 0 | 0 | 225 | 149 | 1.000000 | 1 | 0.999 | 1 | 1 | 0.998 | 1 | 1 | 1 | 1 | 1 | 1 | -0.001 |
| A3SS | 2448 | ENSG00000174444 | RPL4 | chr15 | - | 66502611 | 66503164 | 66502611 | 66502871 | 66503357 | 66503529 | 2029 | 933 | 2244 | 1733 | 2521 | 0 | 2 | 1 | 1 | 0 | 1061 | 1177 | 1079 | 861 | 1986 | 0 | 4 | 0 | 1 | 0 | 298 | 149 | 1.000000 | 1 | 1 | 0.996 | 0.999 | 0.999 | 1 | 1 | 0.993 | 1 | 0.998 | 1 | 0.001 |
| A3SS | 6622 | ENSG00000169764 | UGP2 | chr2 | + | 63885588 | 63885886 | 63885770 | 63885886 | 63883959 | 63884093 | 1027 | 791 | 839 | 1051 | 1674 | 2 | 2 | 2 | 0 | 0 | 187 | 535 | 741 | 521 | 64 | 0 | 0 | 6 | 0 | 0 | 298 | 149 | 1.000000 | 1 | 0.996 | 0.995 | 0.995 | 1 | 1 | 1 | 1 | 0.984 | 1 | 1 | 0 |
| A3SS | 4797 | ENSG00000015479 | MATR3 | chr5 | + | 139317595 | 139317721 | 139317606 | 139317721 | 139317052 | 139317105 | 1085 | 821 | 966 | 1239 | 980 | 1 | 0 | 1 | 1 | 0 | 131 | 567 | 652 | 277 | 54 | 0 | 0 | 0 | 0 | 0 | 160 | 149 | 1.000000 | 1 | 0.999 | 1 | 0.999 | 0.999 | 1 | 1 | 1 | 1 | 1 | 1 | -0.001 |
| A3SS | 8778 | ENSG00000115053 | NCL | chr2 | - | 231458265 | 231458389 | 231458265 | 231458344 | 231459000 | 231459125 | 2208 | 1926 | 2115 | 2342 | 3351 | 1 | 1 | 1 | 3 | 3 | 1910 | 1757 | 1990 | 2302 | 389 | 0 | 2 | 1 | 2 | 0 | 194 | 149 | 1.000000 | 1 | 0.999 | 0.999 | 0.999 | 0.998 | 0.999 | 1 | 0.999 | 0.999 | 0.999 | 1 | -0.001 |
| A3SS | 10766 | ENSG00000229807 | XIST | chrX | - | 73831065 | 73831274 | 73831065 | 73831206 | 73837439 | 73837503 | 739 | 672 | 802 | 717 | 1074 | 1 | 5 | 2 | 2 | 1 | 153 | 1 | 0 | 150 | 0 | 0 | 0 | 0 | 0 | 0 | 217 | 149 | 1.000000 | 1 | 0.998 | 0.989 | 0.996 | 0.996 | 0.999 | 1 | 1 | NA | 1 | NA | -0.004 |
| A3SS | 12155 | ENSG00000110321 | EIF4G2 | chr11 | - | 10799551 | 10800348 | 10799551 | 10799756 | 10800431 | 10800647 | 1447 | 1050 | 1202 | 1483 | 2254 | 0 | 0 | 0 | 0 | 2 | 609 | 993 | 896 | 959 | 209 | 0 | 0 | 0 | 0 | 0 | 298 | 149 | 1.000000 | 1 | 1 | 1 | 1 | 1 | 0.998 | 1 | 1 | 1 | 1 | 1 | 0 |
| A3SS | 10764 | ENSG00000229807 | XIST | chrX | - | 73826114 | 73827984 | 73826114 | 73827928 | 73829067 | 73829231 | 946 | 795 | 862 | 706 | 1057 | 90 | 89 | 93 | 83 | 130 | 286 | 0 | 0 | 271 | 0 | 57 | 0 | 0 | 9 | 0 | 205 | 149 | 1.000000 | 1 | 0.884 | 0.867 | 0.871 | 0.861 | 0.855 | 0.785 | NA | NA | 0.956 | NA | -0.003 |
| A3SS | 11414 | ENSG00000140988 | RPS2 | chr16 | - | 1962496 | 1962909 | 1962496 | 1962656 | 1963148 | 1963256 | 7077 | 4083 | 8977 | 7245 | 8650 | 0 | 2 | 1 | 0 | 5 | 8731 | 7289 | 4213 | 3091 | 8886 | 0 | 2 | 1 | 1 | 12 | 298 | 149 | 1.000000 | 1 | 1 | 0.999 | 1 | 1 | 0.999 | 1 | 0.999 | 1 | 0.999 | 0.997 | 0.001 |
| A3SS | 11181 | ENSG00000156738 | MS4A1 | chr11 | + | 60462184 | 60462533 | 60462358 | 60462533 | 60461071 | 60461160 | 861 | 609 | 769 | 876 | 1577 | 4 | 0 | 0 | 0 | 0 | 142 | 470 | 59 | 45 | 87 | 0 | 0 | 0 | 0 | 0 | 298 | 149 | 1.000000 | 1 | 0.991 | 1 | 1 | 1 | 1 | 1 | 1 | 1 | 1 | 1 | -0.002 |
| A3SS | 8866 | ENSG00000110700 | RPS13 | chr11 | - | 17075096 | 17075197 | 17075096 | 17075163 | 17075453 | 17075623 | 2667 | 1185 | 2963 | 2064 | 2623 | 11 | 11 | 13 | 10 | 14 | 1710 | 1374 | 1180 | 1104 | 2161 | 7 | 9 | 5 | 5 | 9 | 183 | 149 | 1.000000 | 1 | 0.995 | 0.989 | 0.995 | 0.994 | 0.993 | 0.995 | 0.992 | 0.995 | 0.994 | 0.995 | -0.001 |
| A3SS | 5793 | ENSG00000108654 | DDX5 | chr17 | - | 64500548 | 64501891 | 64500548 | 64500773 | 64502009 | 64502069 | 2025 | 1257 | 1910 | 2188 | 1761 | 3200 | 2405 | 2808 | 3422 | 4374 | 249 | 1673 | 1853 | 1091 | 99 | 1232 | 2199 | 1885 | 1764 | 165 | 298 | 149 | 1.000000 | 1 | 0.24 | 0.207 | 0.254 | 0.242 | 0.168 | 0.092 | 0.276 | 0.33 | 0.236 | 0.231 | -0.011 |
| A3SS | 3166 | ENSG00000116459 | ATP5PB | chr1 | + | 111456085 | 111456249 | 111456226 | 111456249 | 111454210 | 111454356 | 1845 | 1158 | 1455 | 1595 | 2443 | 1 | 1 | 6 | 3 | 0 | 739 | 1105 | 1121 | 945 | 386 | 0 | 1 | 2 | 1 | 1 | 290 | 149 | 1.000000 | 1 | 0.999 | 0.998 | 0.992 | 0.996 | 1 | 1 | 0.998 | 0.997 | 0.998 | 0.995 | -0.001 |
| A3SS | 6282 | ENSG00000156482 | RPL30 | chr8 | - | 98042644 | 98042775 | 98042644 | 98042718 | 98044826 | 98045088 | 2470 | 1402 | 3288 | 2581 | 1777 | 0 | 0 | 1 | 1 | 0 | 1604 | 1401 | 1282 | 1159 | 1853 | 0 | 0 | 0 | 5 | 5 | 206 | 149 | 1.000000 | 1 | 1 | 1 | 1 | 0.999 | 1 | 1 | 1 | 1 | 0.994 | 0.996 | 0.002 |
| A3SS | 4216 | ENSG00000013441 | CLK1 | chr2 | - | 200859679 | 200861466 | 200859679 | 200859746 | 200861701 | 200861862 | 1374 | 1271 | 1353 | 1535 | 2015 | 1 | 0 | 2 | 0 | 0 | 431 | 1015 | 852 | 700 | 231 | 0 | 0 | 0 | 0 | 0 | 298 | 149 | 1.000000 | 1 | 0.999 | 1 | 0.997 | 1 | 1 | 1 | 1 | 1 | 1 | 1 | -0.001 |
| A3SS | 3852 | ENSG00000170571 | EMB | chr5 | - | 50411196 | 50411383 | 50411196 | 50411221 | 50428143 | 50428227 | 2056 | 1022 | 1473 | 1757 | 3594 | 1 | 2 | 11 | 6 | 19 | 591 | 1779 | 1515 | 1019 | 594 | 0 | 4 | 4 | 6 | 0 | 298 | 149 | 1.000000 | 1 | 0.999 | 0.996 | 0.985 | 0.993 | 0.99 | 1 | 0.996 | 0.995 | 0.988 | 1 | -0.003 |
| A3SS | 5810 | ENSG00000162434 | JAK1 | chr1 | - | 64864786 | 64864972 | 64864786 | 64864954 | 64866865 | 64867208 | 1977 | 1303 | 1732 | 1815 | 3865 | 2 | 2 | 2 | 2 | 10 | 1303 | 1486 | 1619 | 1204 | 592 | 2 | 5 | 0 | 4 | 0 | 167 | 149 | 1.000000 | 1 | 0.999 | 0.998 | 0.999 | 0.999 | 0.997 | 0.998 | 0.996 | 1 | 0.996 | 1 | 0 |
| A3SS | 10728 | ENSG00000215301 | DDX3X | chrX | + | 41347639 | 41347915 | 41347859 | 41347915 | 41347311 | 41347451 | 1232 | 845 | 1088 | 1313 | 1893 | 1 | 0 | 0 | 1 | 2 | 140 | 459 | 779 | 417 | 49 | 0 | 0 | 5 | 0 | 0 | 298 | 149 | 1.000000 | 1 | 0.998 | 1 | 1 | 0.998 | 0.998 | 1 | 1 | 0.987 | 1 | 1 | 0.001 |
| A3SS | 9051 | ENSG00000188846 | RPL14 | chr3 | + | 40461406 | 40461661 | 40461607 | 40461661 | 40458641 | 40458736 | 3353 | 1730 | 3985 | 3526 | 2988 | 0 | 0 | 0 | 0 | 8 | 2685 | 1963 | 2030 | 1777 | 2421 | 0 | 0 | 0 | 0 | 0 | 298 | 149 | 1.000000 | 1 | 1 | 1 | 1 | 1 | 0.995 | 1 | 1 | 1 | 1 | 1 | -0.001 |
| A3SS | 6506 | ENSG00000166441 | RPL27A | chr11 | + | 8684717 | 8684892 | 8684822 | 8684892 | 8684005 | 8684081 | 12640 | 7521 | 14504 | 12723 | 8696 | 15 | 12 | 34 | 21 | 1 | 11193 | 9279 | 8285 | 8178 | 14404 | 17 | 9 | 19 | 18 | 36 | 254 | 149 | 1.000000 | 1 | 0.998 | 0.997 | 0.996 | 0.997 | 1 | 0.997 | 0.998 | 0.996 | 0.996 | 0.996 | 0.001 |
| A3SS | 12154 | ENSG00000110321 | EIF4G2 | chr11 | - | 10799551 | 10799756 | 10799551 | 10799609 | 10800089 | 10800348 | 2996 | 2455 | 2201 | 3135 | 5342 | 0 | 0 | 0 | 0 | 2 | 2648 | 2393 | 2506 | 2944 | 838 | 0 | 0 | 0 | 0 | 0 | 296 | 149 | 1.000000 | 1 | 1 | 1 | 1 | 1 | 0.999 | 1 | 1 | 1 | 1 | 1 | 0 |
| A3SS | 5606 | ENSG00000115524 | SF3B1 | chr2 | - | 197392971 | 197393188 | 197392971 | 197393186 | 197396055 | 197396328 | 1496 | 984 | 1344 | 1447 | 1334 | 2 | 2 | 3 | 1 | 0 | 154 | 598 | 666 | 223 | 97 | 0 | 0 | 0 | 2 | 0 | 151 | 149 | 1.000000 | 1 | 0.999 | 0.998 | 0.998 | 0.999 | 1 | 1 | 1 | 1 | 0.991 | 1 | 0.001 |
| A3SS | 12967 | ENSG00000089157 | RPLP0 | chr12 | - | 120198553 | 120198739 | 120198553 | 120198631 | 120198853 | 120199000 | 2480 | 938 | 2967 | 2340 | 4273 | 5 | 3 | 7 | 0 | 0 | 1368 | 1678 | 1403 | 944 | 2252 | 0 | 0 | 7 | 0 | 0 | 257 | 149 | 1.000000 | 1 | 0.997 | 0.995 | 0.996 | 1 | 1 | 1 | 1 | 0.991 | 1 | 1 | -0.001 |
| A3SS | 7840 | ENSG00000181163 | NPM1 | chr5 | + | 171400152 | 171400925 | 171400838 | 171400925 | 171392913 | 171392978 | 1313 | 935 | 1732 | 1553 | 2989 | 0 | 0 | 0 | 0 | 0 | 316 | 862 | 939 | 654 | 111 | 0 | 3 | 0 | 0 | 0 | 298 | 149 | 1.000000 | 1 | 1 | 1 | 1 | 1 | 1 | 1 | 0.993 | 1 | 1 | 1 | 0.001 |
| A3SS | 4360 | ENSG00000156976 | EIF4A2 | chr3 | + | 186784563 | 186786051 | 186785882 | 186786051 | 186784431 | 186784477 | 2224 | 1377 | 2187 | 2375 | 1937 | 0 | 0 | 0 | 1 | 0 | 451 | 1396 | 1247 | 963 | 203 | 0 | 0 | 0 | 0 | 0 | 298 | 149 | 1.000000 | 1 | 1 | 1 | 1 | 0.999 | 1 | 1 | 1 | 1 | 1 | 1 | 0 |
| A3SS | 8172 | ENSG00000133872 | SARAF | chr8 | - | 30065987 | 30066139 | 30065987 | 30066067 | 30066776 | 30066918 | 2339 | 1375 | 2789 | 2770 | 3278 | 9 | 9 | 11 | 14 | 18 | 935 | 1202 | 2158 | 1112 | 868 | 0 | 7 | 8 | 2 | 1 | 221 | 149 | 1.000000 | 1 | 0.994 | 0.99 | 0.994 | 0.993 | 0.992 | 1 | 0.991 | 0.995 | 0.997 | 0.998 | -0.004 |
| A3SS | 10765 | ENSG00000229807 | XIST | chrX | - | 73831065 | 73831274 | 73831065 | 73831206 | 73833237 | 73833374 | 1443 | 1428 | 1410 | 1316 | 1731 | 18 | 16 | 23 | 18 | 33 | 296 | 2 | 0 | 340 | 0 | 23 | 0 | 0 | 3 | 0 | 217 | 149 | 1.000000 | 1 | 0.982 | 0.984 | 0.977 | 0.98 | 0.973 | 0.898 | 1 | NA | 0.987 | NA | 0.018 |
| A3SS | 1265 | ENSG00000108298 | RPL19 | chr17 | + | 39202316 | 39203109 | 39202988 | 39203109 | 39201212 | 39201319 | 4765 | 2605 | 5837 | 4447 | 7004 | 3 | 0 | 0 | 0 | 0 | 3347 | 3780 | 3053 | 2473 | 4908 | 0 | 0 | 0 | 0 | 0 | 298 | 149 | 1.000000 | 1 | 0.999 | 1 | 1 | 1 | 1 | 1 | 1 | 1 | 1 | 1 | 0 |
| A3SS | 7628 | ENSG00000009307 | CSDE1 | chr1 | - | 114739691 | 114739890 | 114739691 | 114739725 | 114749820 | 114750207 | 2455 | 1670 | 2411 | 2458 | 5004 | 0 | 0 | 0 | 0 | 0 | 394 | 1877 | 2165 | 1321 | 353 | 0 | 0 | 2 | 0 | 0 | 298 | 149 | 1.000000 | 1 | 1 | 1 | 1 | 1 | 1 | 1 | 1 | 0.998 | 1 | 1 | 0 |
| A3SS | 10063 | ENSG00000122566 | HNRNPA2B1 | chr7 | - | 26196556 | 26196658 | 26196556 | 26196653 | 26196806 | 26197017 | 5172 | 5380 | 4163 | 5490 | 6432 | 3 | 1 | 5 | 2 | 0 | 3066 | 3697 | 5005 | 5348 | 1179 | 32 | 1 | 2 | 2 | 0 | 154 | 149 | 1.000000 | 1 | 0.999 | 1 | 0.999 | 1 | 1 | 0.989 | 1 | 1 | 1 | 1 | 0.002 |
| A3SS | 8035 | ENSG00000148303 | RPL7A | chr9 | + | 133350239 | 133350727 | 133350596 | 133350727 | 133349911 | 133350052 | 8719 | 4714 | 10201 | 8381 | 10445 | 0 | 0 | 0 | 0 | 0 | 7671 | 6131 | 5520 | 4333 | 9597 | 4 | 3 | 0 | 7 | 0 | 298 | 149 | 1.000000 | 1 | 1 | 1 | 1 | 1 | 1 | 0.999 | 0.999 | 1 | 0.997 | 1 | 0.001 |
| A3SS | 9045 | ENSG00000147604 | RPL7 | chr8 | - | 73292688 | 73292797 | 73292688 | 73292795 | 73293598 | 73293633 | 2653 | 1647 | 2883 | 2327 | 3554 | 2 | 0 | 0 | 1 | 0 | 457 | 1080 | 1272 | 761 | 119 | 0 | 0 | 0 | 0 | 0 | 151 | 149 | 1.000000 | 1 | 0.999 | 1 | 1 | 1 | 1 | 1 | 1 | 1 | 1 | 1 | 0 |
| A3SS | 5608 | ENSG00000115524 | SF3B1 | chr2 | - | 197418508 | 197420542 | 197418508 | 197418588 | 197421028 | 197421133 | 2779 | 2189 | 2854 | 2991 | 3477 | 0 | 0 | 0 | 0 | 0 | 505 | 1761 | 1523 | 812 | 307 | 0 | 0 | 0 | 1 | 0 | 298 | 149 | 1.000000 | 1 | 1 | 1 | 1 | 1 | 1 | 1 | 1 | 1 | 0.998 | 1 | 0 |
| A3SS | 452 | ENSG00000115268 | RPS15 | chr19 | + | 1440018 | 1440253 | 1440025 | 1440253 | 1438806 | 1438892 | 5921 | 2514 | 6117 | 4575 | 6196 | 11 | 3 | 17 | 12 | 3 | 2394 | 4459 | 2612 | 2018 | 4329 | 10 | 6 | 7 | 9 | 12 | 156 | 149 | 1.000000 | 1 | 0.998 | 0.999 | 0.997 | 0.997 | 0.999 | 0.996 | 0.999 | 0.997 | 0.995 | 0.997 | 0.001 |
| A3SS | 5797 | ENSG00000108654 | DDX5 | chr17 | - | 64502161 | 64502549 | 64502161 | 64502223 | 64502925 | 64503098 | 5273 | 4947 | 4727 | 6792 | 8309 | 0 | 1 | 0 | 2 | 0 | 3337 | 5291 | 6255 | 4537 | 928 | 0 | 0 | 7 | 0 | 0 | 298 | 149 | 1.000000 | 1 | 1 | 1 | 1 | 0.999 | 1 | 1 | 1 | 0.998 | 1 | 1 | 0 |
| A3SS | 7838 | ENSG00000181163 | NPM1 | chr5 | + | 171391304 | 171391799 | 171391705 | 171391799 | 171390050 | 171390130 | 3961 | 3143 | 5135 | 4586 | 6068 | 0 | 0 | 3 | 0 | 0 | 1661 | 3387 | 4469 | 2937 | 452 | 0 | 0 | 0 | 0 | 0 | 298 | 149 | 1.000000 | 1 | 1 | 1 | 0.999 | 1 | 1 | 1 | 1 | 1 | 1 | 1 | 0 |
| A3SS | 428 | ENSG00000063177 | RPL18 | chr19 | - | 48617315 | 48617423 | 48617315 | 48617415 | 48617790 | 48617877 | 5308 | 2412 | 6169 | 4676 | 7198 | 7 | 2 | 9 | 4 | 3 | 3201 | 3217 | 2519 | 1903 | 4301 | 1 | 3 | 2 | 3 | 6 | 157 | 149 | 1.000000 | 1 | 0.999 | 0.999 | 0.998 | 0.999 | 1 | 1 | 0.999 | 0.999 | 0.998 | 0.999 | 0 |
| A3SS | 8034 | ENSG00000148303 | RPL7A | chr9 | + | 133348921 | 133349042 | 133348936 | 133349042 | 133348217 | 133348246 | 5245 | 3013 | 6864 | 5682 | 11702 | 0 | 0 | 0 | 0 | 0 | 4333 | 4584 | 3844 | 2765 | 5659 | 0 | 0 | 0 | 0 | 1 | 164 | 149 | 1.000000 | 1 | 1 | 1 | 1 | 1 | 1 | 1 | 1 | 1 | 1 | 1 | 0 |
| A3SS | 7839 | ENSG00000181163 | NPM1 | chr5 | + | 171392709 | 171392978 | 171392913 | 171392978 | 171391705 | 171391799 | 3849 | 3181 | 5121 | 4580 | 7692 | 0 | 0 | 0 | 0 | 0 | 1756 | 3295 | 4444 | 3089 | 445 | 0 | 2 | 0 | 0 | 0 | 298 | 149 | 1.000000 | 1 | 1 | 1 | 1 | 1 | 1 | 1 | 0.999 | 1 | 1 | 1 | 0 |
| A3SS | 11947 | ENSG00000149273 | RPS3 | chr11 | + | 75400693 | 75400824 | 75400694 | 75400824 | 75399556 | 75399577 | 5120 | 2490 | 5567 | 4717 | 6508 | 2 | 2 | 0 | 4 | 2 | 1750 | 2983 | 2861 | 1930 | 3356 | 0 | 0 | 2 | 0 | 0 | 150 | 149 | 1.000000 | 1 | 1 | 0.999 | 1 | 0.999 | 1 | 1 | 1 | 0.999 | 1 | 1 | 0 |
| A3SS | 11950 | ENSG00000149273 | RPS3 | chr11 | + | 75404019 | 75404207 | 75404053 | 75404207 | 75402351 | 75402446 | 4498 | 2024 | 5469 | 4368 | 8037 | 3 | 0 | 0 | 2 | 3 | 1877 | 2735 | 2625 | 1802 | 3665 | 0 | 1 | 1 | 0 | 0 | 183 | 149 | 1.000000 | 1 | 0.999 | 1 | 1 | 0.999 | 1 | 1 | 1 | 1 | 1 | 1 | 0 |
| A3SS | 11949 | ENSG00000149273 | RPS3 | chr11 | + | 75404671 | 75404868 | 75404759 | 75404868 | 75401639 | 75401733 | 4985 | 1936 | 5254 | 4101 | 5844 | 1 | 0 | 0 | 0 | 0 | 1133 | 2035 | 1692 | 1233 | 2676 | 0 | 0 | 0 | 0 | 0 | 237 | 149 | 1.000000 | 1 | 1 | 1 | 1 | 1 | 1 | 1 | 1 | 1 | 1 | 1 | 0 |
| A3SS | 10115 | ENSG00000171863 | RPS7 | chr2 | + | 3576486 | 3576630 | 3576558 | 3576630 | 3575816 | 3575888 | 9976 | 5549 | 10668 | 9306 | 14930 | 5 | 7 | 12 | 5 | 11 | 8183 | 7604 | 6309 | 5025 | 7928 | 8 | 31 | 4 | 2 | 0 | 221 | 149 | 1.000000 | 1 | 0.999 | 0.998 | 0.998 | 0.999 | 0.999 | 0.999 | 0.994 | 0.999 | 0.999 | 1 | 0 |
| A3SS | 6283 | ENSG00000156482 | RPL30 | chr8 | - | 98042644 | 98042775 | 98042644 | 98042718 | 98044942 | 98045088 | 9033 | 4660 | 10383 | 8025 | 8943 | 12 | 5 | 16 | 12 | 10 | 5653 | 4802 | 4278 | 3919 | 6538 | 13 | 3 | 10 | 3 | 9 | 206 | 149 | 1.000000 | 1 | 0.998 | 0.999 | 0.998 | 0.998 | 0.998 | 0.997 | 0.999 | 0.997 | 0.999 | 0.998 | 0 |
| A3SS | 11126 | ENSG00000104408 | EIF3E | chr8 | - | 108234997 | 108235102 | 108234997 | 108235097 | 108236160 | 108236203 | 4724 | 3311 | 4818 | 4084 | 4177 | 2 | 0 | 4 | 0 | 3 | 681 | 1529 | 1571 | 580 | 831 | 0 | 0 | 3 | 0 | 0 | 154 | 149 | 1.000000 | 1 | 1 | 1 | 0.999 | 1 | 0.999 | 1 | 1 | 0.998 | 1 | 1 | 0 |
| A3SS | 6008 | ENSG00000143947 | RPS27A | chr2 | + | 55234118 | 55234962 | 55234830 | 55234962 | 55233362 | 55233417 | 5531 | 2713 | 6511 | 5049 | 6277 | 1 | 0 | 3 | 1 | 1 | 1472 | 1707 | 1705 | 1024 | 1291 | 0 | 4 | 1 | 2 | 9 | 298 | 149 | 1.000000 | 1 | 1 | 1 | 0.999 | 1 | 1 | 1 | 0.995 | 0.999 | 0.996 | 0.986 | 0.005 |
| A3SS | 371 | ENSG00000142541 | RPL13A | chr19 | + | 49490231 | 49490576 | 49490474 | 49490576 | 49489849 | 49489922 | 12057 | 4728 | 13139 | 9517 | 13088 | 1 | 0 | 4 | 0 | 0 | 5874 | 7754 | 5314 | 4146 | 10049 | 0 | 0 | 0 | 0 | 0 | 298 | 149 | 1.000000 | 1 | 1 | 1 | 0.999 | 1 | 1 | 1 | 1 | 1 | 1 | 1 | 0 |
| A3SS | 372 | ENSG00000142541 | RPL13A | chr19 | + | 49490474 | 49490576 | 49490479 | 49490576 | 49490231 | 49490297 | 12002 | 4837 | 12603 | 9321 | 14678 | 5 | 8 | 1 | 6 | 1 | 6250 | 7585 | 5202 | 4225 | 10270 | 0 | 6 | 2 | 0 | 0 | 154 | 149 | 1.000000 | 1 | 1 | 0.998 | 1 | 0.999 | 1 | 1 | 0.999 | 1 | 1 | 1 | 0 |
| A3SS | 11809 | ENSG00000251562 | MALAT1 | chr11 | + | 65505503 | 65505662 | 65505590 | 65505662 | 65504518 | 65505019 | 10518 | 7910 | 10217 | 11337 | 4310 | 0 | 6 | 2 | 2 | 0 | 2032 | 8795 | 7670 | 2074 | 1803 | 0 | 7 | 0 | 0 | 0 | 236 | 149 | 1.000000 | 1 | 1 | 0.999 | 1 | 1 | 1 | 1 | 0.999 | 1 | 1 | 1 | 0 |
| A3SS | 5614 | ENSG00000134419 | RPS15A | chr16 | - | 18788062 | 18788142 | 18788062 | 18788105 | 18788980 | 18789118 | 14071 | 6201 | 15403 | 12606 | 14480 | 0 | 1 | 0 | 3 | 0 | 7210 | 7917 | 6723 | 5998 | 11366 | 0 | 0 | 0 | 8 | 0 | 186 | 149 | 1.000000 | 1 | 1 | 1 | 1 | 1 | 1 | 1 | 1 | 1 | 0.998 | 1 | 0 |
| A3SS | 3016 | ENSG00000118181 | RPS25 | chr11 | - | 119017361 | 119017545 | 119017361 | 119017433 | 119017957 | 119018053 | 11654 | 6297 | 13801 | 11215 | 19012 | 10 | 2 | 10 | 4 | 13 | 6084 | 6658 | 5937 | 5006 | 6259 | 0 | 7 | 6 | 0 | 7 | 261 | 149 | 1.000000 | 1 | 0.998 | 0.999 | 0.999 | 0.999 | 0.999 | 1 | 0.998 | 0.998 | 1 | 0.998 | 0 |
| A3SS | 8790 | ENSG00000145425 | RPS3A | chr4 | + | 151104176 | 151104286 | 151104184 | 151104286 | 151102870 | 151103079 | 14543 | 8477 | 13661 | 10951 | 29524 | 1 | 0 | 1 | 1 | 6 | 1947 | 5195 | 5607 | 3526 | 1407 | 0 | 0 | 0 | 0 | 0 | 157 | 149 | 1.000000 | 1 | 1 | 1 | 1 | 1 | 1 | 1 | 1 | 1 | 1 | 1 | 0 |
| A3SS | 9183 | ENSG00000156508 | EEF1A1 | chr6 | - | 73517514 | 73518264 | 73517514 | 73517934 | 73518353 | 73518610 | 33102 | 17190 | 37632 | 29265 | 40127 | 0 | 0 | 0 | 0 | 2 | 19232 | 14703 | 14653 | 12322 | 23295 | 0 | 0 | 0 | 0 | 0 | 298 | 149 | 1.000000 | 1 | 1 | 1 | 1 | 1 | 1 | 1 | 1 | 1 | 1 | 1 | 0 |
| A3SS | 9185 | ENSG00000156508 | EEF1A1 | chr6 | - | 73518931 | 73519516 | 73518931 | 73519228 | 73519882 | 73520056 | 57319 | 40364 | 64542 | 55003 | 98993 | 2 | 1 | 3 | 2 | 2 | 60850 | 36915 | 37337 | 35860 | 62164 | 0 | 1 | 0 | 8 | 0 | 298 | 149 | 1.000000 | 1 | 1 | 1 | 1 | 1 | 1 | 1 | 1 | 1 | 1 | 1 | 0 |
| A3SS | 9186 | ENSG00000156508 | EEF1A1 | chr6 | - | 73519336 | 73519516 | 73519336 | 73519507 | 73519882 | 73520056 | 58636 | 41239 | 65858 | 56188 | 101954 | 7 | 1 | 8 | 0 | 3 | 61933 | 37624 | 37949 | 36430 | 63388 | 17 | 5 | 2 | 0 | 0 | 158 | 149 | 1.000000 | 1 | 1 | 1 | 1 | 1 | 1 | 1 | 1 | 1 | 1 | 1 | 0 |
| A3SS | 9184 | ENSG00000156508 | EEF1A1 | chr6 | - | 73518029 | 73518610 | 73518029 | 73518264 | 73518697 | 73518848 | 47073 | 27233 | 53328 | 44229 | 59031 | 0 | 1 | 0 | 0 | 0 | 32816 | 23523 | 22226 | 19143 | 39101 | 0 | 0 | 0 | 0 | 0 | 298 | 149 | 1.000000 | 1 | 1 | 1 | 1 | 1 | 1 | 1 | 1 | 1 | 1 | 1 | 0 |
| A3SS | 11965 | ENSG00000144354 | CDCA7 | chr2 | + | 173359342 | 173359491 | 173359386 | 173359491 | 173358711 | 173358837 | 0 | 1 | 1 | 0 | 0 | 0 | 0 | 0 | 0 | 0 | 0 | 0 | 0 | 0 | 0 | 0 | 0 | 1 | 0 | 0 | 193 | 149 | 1.000000 | 1 | NA | 1 | 1 | NA | NA | NA | NA | 0 | NA | NA | 1 |

U12_A3SS_filtered

| Splice_type | ID | Gene_ID | Gene_symbol | Chromosome | Strand | LongExonStart_0base | LongExonEnd | ShortES | ShortEE | FlankingES | FlankingEE | Inclusion_counts_HC SRR13214146 | SRR13214147 | SRR13214148 | SRR13214149 | SRR13214150 | Skipping_counts_HC SRR13214146 | SRR13214147 | SRR13214148 | SRR13214149 | SRR13214150 | Inclusion counts_SLE SRR13214141 | SRR13214242 | SRR13214143 | SRR13214144 | SRR13214145 | Skipping_counts_SLE SRR13214141 | SRR13214242 | SRR13214143 | SRR13214144 | SRR13214145 | IncFormLen | SkipFormLen | p value | FDR | Inclusion_level(normalized) SRR13214146 | SRR13214147 | SRR13214148 | SRR13214149 | SRR13214150 | SRR13214141 | SRR13214242 | SRR13214143 | SRR13214144 | SRR13214145 | Inclevel_diff |
|---|---|---|---|---|---|---|---|---|---|---|---|---|---|---|---|---|---|---|---|---|---|---|---|---|---|---|---|---|---|---|---|---|---|---|---|---|---|---|---|---|---|---|---|---|---|---|
| A3SS | 12515 | ENSG00000120438 | TCP1 | chr6 | - | 159787742 | 159787902 | 159787742 | 159787871 | 159788057 | 159788143 | 12 | 8 | 13 | 32 | 18 | 171 | 91 | 142 | 148 | 196 | 20 | 33 | 45 | 24 | 7 | 79 | 56 | 63 | 99 | 13 | 180 | 149 | 6.66349E-11 | 1.62993E-08 | 0.055 | 0.068 | 0.07 | 0.152 | 0.071 | 0.173 | 0.328 | 0.372 | 0.167 | 0.308 | -0.186 |
| A3SS | 9126 | ENSG00000105393 | BABAM1 | chr19 | + | 17275776 | 17275825 | 17275788 | 17275825 | 17274106 | 17274185 | 3 | 1 | 1 | 1 | 5 | 0 | 0 | 0 | 0 | 0 | 8 | 0 | 2 | 3 | 10 | 11 | 2 | 2 | 0 | 8 | 161 | 149 | 1.29408E-10 | 2.90162E-08 | 1 | 1 | 1 | 1 | 1 | 0.402 | 0 | 0.481 | 1 | 0.536 | 0.516 |
| A3SS | 11969 | ENSG00000068831 | RASGRP2 | chr11 | - | 64742009 | 64742112 | 64742009 | 64742032 | 64742793 | 64742937 | 213 | 82 | 206 | 186 | 99 | 9 | 4 | 5 | 7 | 0 | 164 | 182 | 116 | 83 | 209 | 43 | 11 | 5 | 14 | 37 | 229 | 149 | 7.01685E-10 | 1.34857E-07 | 0.939 | 0.93 | 0.964 | 0.945 | 1 | 0.713 | 0.915 | 0.938 | 0.794 | 0.786 | 0.126 |
| A3SS | 12251 | ENSG00000167986 | DDB1 | chr11 | - | 61329957 | 61330183 | 61329957 | 61330074 | 61331542 | 61331691 | 26 | 24 | 23 | 23 | 12 | 221 | 88 | 165 | 131 | 288 | 115 | 32 | 40 | 75 | 36 | 151 | 161 | 101 | 171 | 210 | 258 | 149 | 7.82081E-10 | 1.46813E-07 | 0.064 | 0.136 | 0.075 | 0.092 | 0.023 | 0.305 | 0.103 | 0.186 | 0.202 | 0.09 | -0.099 |
| A3SS | 3582 | ENSG00000083312 | TNPO1 | chr5 | + | 72851204 | 72851319 | 72851243 | 72851319 | 72848384 | 72848498 | 5 | 2 | 0 | 1 | 3 | 138 | 66 | 83 | 99 | 88 | 0 | 13 | 13 | 4 | 3 | 59 | 62 | 23 | 19 | 16 | 188 | 149 | 3.20013E-09 | 4.23466E-07 | 0.028 | 0.023 | 0 | 0.008 | 0.026 | 0 | 0.142 | 0.309 | 0.143 | 0.129 | -0.128 |
| A3SS | 5415 | ENSG00000139842 | CUL4A | chr13 | + | 113233152 | 113233339 | 113233176 | 113233339 | 113229445 | 113229519 | 3 | 3 | 1 | 2 | 0 | 120 | 75 | 71 | 102 | 84 | 14 | 5 | 5 | 16 | 0 | 123 | 78 | 27 | 83 | 32 | 173 | 149 | 1.24005E-08 | 1.21512E-06 | 0.021 | 0.033 | 0.012 | 0.017 | 0 | 0.089 | 0.052 | 0.138 | 0.142 | 0 | -0.068 |
| A3SS | 11936 | ENSG00000135249 | RINT1 | chr7 | + | 105536564 | 105536749 | 105536657 | 105536749 | 105532823 | 105532869 | 104 | 68 | 76 | 102 | 56 | 1 | 3 | 0 | 6 | 0 | 133 | 56 | 105 | 108 | 90 | 16 | 12 | 5 | 10 | 13 | 242 | 149 | 3.51742E-08 | 2.9887E-06 | 0.985 | 0.933 | 1 | 0.913 | 1 | 0.837 | 0.742 | 0.928 | 0.869 | 0.81 | 0.129 |
| A3SS | 6001 | ENSG00000163479 | SSR2 | chr1 | - | 156011809 | 156012565 | 156011809 | 156011887 | 156014960 | 156015069 | 13 | 5 | 5 | 8 | 2 | 525 | 219 | 516 | 421 | 536 | 35 | 1 | 12 | 12 | 32 | 301 | 415 | 296 | 272 | 598 | 298 | 149 | 1.02302E-07 | 7.24371E-06 | 0.012 | 0.011 | 0.005 | 0.009 | 0.002 | 0.055 | 0.001 | 0.02 | 0.022 | 0.026 | -0.017 |
| A3SS | 4586 | ENSG00000111642 | CHD4 | chr12 | - | 6578028 | 6578137 | 6578028 | 6578117 | 6578408 | 6578546 | 468 | 261 | 419 | 433 | 654 | 1 | 1 | 4 | 2 | 16 | 489 | 326 | 304 | 286 | 455 | 16 | 9 | 27 | 5 | 5 | 169 | 149 | 4.5489E-07 | 2.49787E-05 | 0.998 | 0.996 | 0.989 | 0.995 | 0.973 | 0.964 | 0.97 | 0.908 | 0.981 | 0.988 | 0.028 |
| A3SS | 2977 | ENSG00000136986 | DERL1 | chr8 | - | 123019194 | 123019305 | 123019194 | 123019245 | 123021446 | 123021499 | 450 | 298 | 261 | 323 | 446 | 11 | 9 | 22 | 24 | 25 | 460 | 340 | 412 | 352 | 354 | 33 | 71 | 55 | 46 | 12 | 209 | 149 | 1.70174E-06 | 7.63137E-05 | 0.967 | 0.959 | 0.894 | 0.906 | 0.927 | 0.909 | 0.773 | 0.842 | 0.845 | 0.955 | 0.066 |
| A3SS | 7140 | ENSG00000124571 | XPO5 | chr6 | - | 43525838 | 43525954 | 43525838 | 43525921 | 43526684 | 43526747 | 0 | 1 | 2 | 0 | 0 | 31 | 23 | 28 | 24 | 28 | 6 | 0 | 1 | 3 | 10 | 35 | 41 | 12 | 22 | 15 | 182 | 149 | 3.69989E-06 | 0.000140214 | 0 | 0.034 | 0.055 | 0 | 0 | 0.123 | 0 | 0.064 | 0.1 | 0.353 | -0.11 |
| A3SS | 8391 | ENSG00000143799 | PARP1 | chr1 | - | 226392198 | 226392498 | 226392198 | 226392314 | 226402213 | 226402379 | 16 | 19 | 17 | 41 | 11 | 237 | 139 | 178 | 264 | 129 | 54 | 49 | 39 | 41 | 29 | 263 | 199 | 153 | 164 | 285 | 298 | 149 | 8.57994E-06 | 0.000283841 | 0.033 | 0.064 | 0.046 | 0.072 | 0.041 | 0.093 | 0.11 | 0.113 | 0.111 | 0.048 | -0.044 |
| A3SS | 12422 | ENSG00000143702 | CEP170 | chr1 | - | 243156220 | 243156455 | 243156220 | 243156347 | 243164283 | 243166116 | 104 | 108 | 95 | 126 | 208 | 10 | 7 | 9 | 0 | 13 | 8 | 64 | 69 | 8 | 9 | 0 | 3 | 0 | 0 | 0 | 257 | 149 | 9.27209E-06 | 0.000304868 | 0.858 | 0.899 | 0.86 | 1 | 0.903 | 1 | 0.925 | 1 | 1 | 1 | -0.081 |
| A3SS | 1889 | ENSG00000103043 | VAC14 | chr16 | - | 70697138 | 70697257 | 70697138 | 70697173 | 70698636 | 70698811 | 55 | 30 | 58 | 54 | 65 | 2 | 0 | 0 | 3 | 0 | 70 | 42 | 30 | 36 | 50 | 9 | 0 | 7 | 6 | 0 | 233 | 149 | 9.95357E-06 | 0.000321381 | 0.946 | 1 | 1 | 0.92 | 1 | 0.833 | 1 | 0.733 | 0.793 | 1 | 0.101 |
| A3SS | 4416 | ENSG00000041802 | LSG1 | chr3 | - | 194648680 | 194648804 | 194648680 | 194648753 | 194650880 | 194651024 | 99 | 71 | 71 | 100 | 127 | 1 | 2 | 3 | 0 | 0 | 78 | 54 | 40 | 52 | 40 | 14 | 3 | 1 | 2 | 0 | 200 | 149 | 1.04988E-05 | 0.000337634 | 0.987 | 0.964 | 0.946 | 1 | 1 | 0.806 | 0.931 | 0.968 | 0.951 | 1 | 0.048 |
| A3SS | 11911 | ENSG00000205758 | CRYZL1 | chr21 | - | 33591161 | 33591211 | 33591161 | 33591207 | 33595730 | 33595836 | 10 | 2 | 6 | 4 | 9 | 31 | 25 | 31 | 21 | 43 | 0 | 3 | 2 | 0 | 0 | 15 | 24 | 9 | 27 | 9 | 153 | 149 | 1.89626E-05 | 0.000558637 | 0.239 | 0.072 | 0.159 | 0.156 | 0.169 | 0 | 0.109 | 0.178 | 0 | 0 | 0.102 |
| A3SS | 3952 | ENSG00000141452 | RMC1 | chr18 | + | 23530386 | 23530612 | 23530510 | 23530612 | 23530228 | 23530297 | 267 | 119 | 131 | 174 | 125 | 9 | 3 | 6 | 6 | 0 | 271 | 249 | 169 | 217 | 206 | 0 | 0 | 1 | 0 | 6 | 273 | 149 | 3.31107E-05 | 0.000851176 | 0.942 | 0.956 | 0.923 | 0.941 | 1 | 1 | 1 | 0.989 | 1 | 0.949 | -0.035 |
| A3SS | 2449 | ENSG00000137841 | PLCB2 | chr15 | - | 40291264 | 40291924 | 40291264 | 40291487 | 40292063 | 40292158 | 201 | 111 | 118 | 150 | 78 | 3 | 0 | 0 | 3 | 0 | 203 | 205 | 110 | 166 | 315 | 17 | 7 | 4 | 3 | 0 | 298 | 149 | 4.13009E-05 | 0.000998145 | 0.971 | 1 | 1 | 0.962 | 1 | 0.857 | 0.936 | 0.932 | 0.965 | 1 | 0.049 |
| A3SS | 7076 | ENSG00000185946 | RNPC3 | chr1 | + | 103543295 | 103543447 | 103543298 | 103543447 | 103541349 | 103541475 | 147 | 192 | 106 | 162 | 213 | 35 | 61 | 31 | 46 | 72 | 76 | 75 | 109 | 61 | 16 | 8 | 19 | 10 | 11 | 3 | 152 | 149 | 5.74704E-05 | 0.001297763 | 0.805 | 0.755 | 0.77 | 0.775 | 0.744 | 0.903 | 0.795 | 0.914 | 0.845 | 0.839 | -0.089 |
| A3SS | 4893 | ENSG00000198925 | ATG9A | chr2 | - | 219227921 | 219228079 | 219227921 | 219228053 | 219228433 | 219228485 | 2 | 0 | 2 | 1 | 0 | 12 | 6 | 12 | 7 | 2 | 6 | 12 | 2 | 3 | 6 | 0 | 16 | 4 | 7 | 32 | 175 | 149 | 6.42599E-05 | 0.001401907 | 0.124 | 0 | 0.124 | 0.108 | 0 | 1 | 0.39 | 0.299 | 0.267 | 0.138 | -0.348 |
| A3SS | 5689 | ENSG00000088035 | ALG6 | chr1 | + | 63415817 | 63415957 | 63415872 | 63415957 | 63414060 | 63414146 | 2 | 6 | 2 | 4 | 11 | 53 | 29 | 35 | 48 | 76 | 0 | 0 | 0 | 2 | 0 | 5 | 22 | 17 | 9 | 10 | 204 | 149 | 9.51448E-05 | 0.001969253 | 0.027 | 0.131 | 0.04 | 0.057 | 0.096 | 0 | 0 | 0 | 0.14 | 0 | 0.042 |
| A3SS | 6565 | ENSG00000007402 | CACNA2D2 | chr3 | - | 50366577 | 50366628 | 50366577 | 50366625 | 50366830 | 50366919 | 8 | 1 | 3 | 9 | 2 | 11 | 0 | 4 | 3 | 3 | 0 | 1 | 0 | 0 | 5 | 8 | 0 | 4 | 1 | 9 | 152 | 149 | 0.00010419 | 0.002118444 | 0.416 | 1 | 0.424 | 0.746 | 0.395 | 0 | 1 | 0 | 0 | 0.353 | 0.326 |

[illegible]

| | | | | | | | | | | | | | | | | | | | | | | | | | |
|---|---|---|---|---|---|---|---|---|---|---|---|---|---|---|---|---|---|---|---|---|---|---|---|---|---|
| A3SS | 5765 | ENSG00000163608 | NEPRO | chr3 | - | 113011044 | 113011406 | 113011044 | 113011165 | 113012733 | 113012811 | 178 | 165 | 192 | 145 | 294 | 10 | 9 | 15 | 11 | 13 | 10 | 77 | 189 | 55 |
| A3SS | 2574 | ENSG00000100949 | RABGGTA | chr14 | - | 24270836 | 24270998 | 24270836 | 24270947 | 24271112 | 24271169 | 1 | 2 | 5 | 2 | 0 | 20 | 4 | 7 | 19 | 1 | 0 | 11 | 3 | 0 |
| A3SS | 12517 | ENSG00000175470 | PPP2R2D | chr10 | + | 131943967 | 131944145 | 131944094 | 131944145 | 131940581 | 131940694 | 255 | 201 | 186 | 154 | 395 | 9 | 6 | 6 | 8 | 14 | 197 | 171 | 130 | 196 |
| A3SS | 7677 | ENSG00000092529 | CAPN3 | chr15 | + | 42411746 | 42412285 | 42412085 | 42412285 | 42411286 | 42411345 | 41 | 29 | 29 | 65 | 58 | 0 | 0 | 0 | 3 | 2 | 74 | 50 | 33 | 164 |
| A3SS | 7686 | ENSG00000116922 | C1orf109 | chr1 | - | 37683318 | 37683476 | 37683318 | 37683437 | 37686272 | 37686452 | 1 | 0 | 0 | 0 | 4 | 20 | 10 | 21 | 20 | 20 | 0 | 0 | 0 | 1 |
| A3SS | 9366 | ENSG00000223501 | VPS52 | chr6 | - | 33269013 | 33269218 | 33269013 | 33269189 | 33269489 | 33269557 | 21 | 0 | 15 | 18 | 3 | 163 | 101 | 153 | 148 | 99 | 29 | 12 | 12 | 9 |
| A3SS | 3570 | ENSG00000105875 | WDR91 | chr7 | - | 135194933 | 135195084 | 135194933 | 135195002 | 135196143 | 135196337 | 34 | 10 | 61 | 32 | 28 | 1 | 0 | 4 | 1 | 0 | 27 | 34 | 52 | 40 |
| A3SS | 11968 | ENSG00000068831 | RASGRP2 | chr11 | - | 64740367 | 64740449 | 64740367 | 64740446 | 64740947 | 64741079 | 44 | 10 | 29 | 44 | 26 | 0 | 0 | 3 | 4 | 0 | 16 | 18 | 14 | 5 |
| A3SS | 5767 | ENSG00000163608 | NEPRO | chr3 | - | 113013271 | 113013403 | 113013271 | 113013361 | 113013952 | 113014032 | 31 | 28 | 37 | 29 | 35 | 0 | 0 | 0 | 2 | 3 | 13 | 20 | 28 | 5 |
| A3SS | 11115 | ENSG00000164941 | INTS8 | chr8 | + | 94838462 | 94838618 | 94838493 | 94838618 | 94836523 | 94836631 | 51 | 28 | 40 | 41 | 100 | 7 | 3 | 9 | 8 | 31 | 13 | 56 | 30 | 5 |
| A3SS | 11073 | ENSG00000174744 | BRMS1 | chr11 | - | 66340773 | 66341046 | 66340773 | 66340870 | 66341205 | 66341333 | 166 | 118 | 127 | 144 | 151 | 0 | 0 | 0 | 0 | 0 | 187 | 187 | 140 | 152 |
| A3SS | 7211 | ENSG00000079246 | XRCC5 | chr2 | + | 216117740 | 216117794 | 216117745 | 216117794 | 216116658 | 216116842 | 5 | 7 | 2 | 4 | 2 | 1265 | 992 | 1216 | 1279 | 1230 | 0 | 4 | 8 | 0 |
| A3SS | 296 | ENSG00000126247 | CAPNS1 | chr19 | + | 36146195 | 36146312 | 36146246 | 36146312 | 36145975 | 36146054 | 1618 | 705 | 919 | 1152 | 1385 | 0 | 1 | 0 | 0 | 19 | 2993 | 1769 | 1027 | 1403 |
| A3SS | 7812 | ENSG00000198520 | ARMH1 | chr1 | + | 44724319 | 44724668 | 44724336 | 44724668 | 44724121 | 44724244 | 34 | 16 | 34 | 27 | 27 | 2 | 1 | 7 | 5 | 3 | 21 | 17 | 10 | 54 |
| A3SS | 6330 | ENSG00000139636 | LMBR1L | chr12 | - | 49106960 | 49107062 | 49106960 | 49107045 | 49110483 | 49110847 | 0 | 0 | 0 | 0 | 1 | 49 | 22 | 59 | 38 | 62 | 7 | 0 | 1 | 0 |
| A3SS | 10598 | ENSG00000215021 | PHB2 | chr12 | - | 6968410 | 6968595 | 6968410 | 6968587 | 6969497 | 6969577 | 121 | 96 | 120 | 99 | 170 | 0 | 0 | 0 | 5 | 0 | 181 | 89 | 80 | 116 |
| A3SS | 3998 | ENSG00000108591 | DRG2 | chr17 | + | 18099632 | 18100435 | 18100362 | 18100435 | 18099016 | 18099077 | 46 | 20 | 46 | 66 | 49 | 3 | 0 | 1 | 1 | 0 | 101 | 114 | 30 | 41 |
| A3SS | 8185 | ENSG00000084623 | EIF3I | chr1 | + | 32226104 | 32226320 | 32226170 | 32226320 | 32224409 | 32224475 | 7 | 9 | 8 | 10 | 1 | 168 | 96 | 189 | 173 | 102 | 17 | 4 | 3 | 16 |
| A3SS | 8188 | ENSG00000084623 | EIF3I | chr1 | + | 32231114 | 32231602 | 32231259 | 32231602 | 32230926 | 32231019 | 199 | 125 | 252 | 250 | 207 | 0 | 0 | 0 | 0 | 0 | 278 | 252 | 174 | 196 |
| A3SS | 8507 | ENSG00000143390 | RFX5 | chr1 | - | 151345927 | 151345961 | 151345927 | 151345957 | 151346204 | 151346333 | 294 | 192 | 258 | 244 | 385 | 10 | 0 | 0 | 0 | 0 | 363 | 331 | 162 | 327 |
| A3SS | 13413 | ENSG00000134899 | ERCC5 | chr13 | + | 102873258 | 102873343 | 102873260 | 102873343 | 102872197 | 102872398 | 267 | 226 | 262 | 243 | 301 | 7 | 5 | 6 | 5 | 4 | 11 | 171 | 301 | 77 |
| A3SS | 4822 | ENSG00000197386 | HTT | chr4 | + | 3207280 | 3207357 | 3207285 | 3207357 | 3206806 | 3206983 | 107 | 48 | 74 | 122 | 112 | 10 | 2 | 5 | 6 | 9 | 143 | 123 | 83 | 166 |
| A3SS | 2838 | ENSG00000119599 | DCAF4 | chr14 | + | 72954160 | 72954262 | 72954163 | 72954262 | 72951797 | 72951877 | 5 | 0 | 0 | 5 | 4 | 6 | 1 | 3 | 6 | 2 | 0 | 3 | 3 | 2 |
| A3SS | 12246 | ENSG00000167986 | DDB1 | chr11 | - | 61302256 | 61302751 | 61302256 | 61302359 | 61303045 | 61303155 | 243 | 147 | 241 | 234 | 391 | 0 | 0 | 0 | 2 | 0 | 268 | 215 | 103 | 132 |
| A3SS | 12798 | ENSG00000079974 | RABL2B | chr22 | - | 50782187 | 50782351 | 50782187 | 50782347 | 50783500 | 50783662 | 0 | 0 | 0 | 0 | 0 | 0 | 1 | 1 | 1 | 0 | 0 | 1 | 2 | 0 |
| A3SS | 2359 | ENSG00000103852 | TTC23 | chr15 | - | 99139316 | 99139467 | 99139316 | 99139399 | 99156147 | 99156297 | 0 | 1 | 0 | 0 | 0 | 0 | 1 | 0 | 0 | 0 | 0 | 2 | 0 | 0 |
| A3SS | 103 | ENSG00000093009 | CDC45 | chr22 | + | 19516780 | 19516893 | 19516816 | 19516893 | 19516526 | 19516645 | 0 | 0 | 0 | 0 | 0 | 1 | 0 | 2 | 0 | 0 | 0 | 0 | 0 | 1 |
| A3SS | 104 | ENSG00000093009 | CDC45 | chr22 | + | 19516813 | 19516893 | 19516816 | 19516893 | 19516526 | 19516645 | 0 | 0 | 0 | 0 | 0 | 1 | 0 | 2 | 0 | 0 | 0 | 0 | 0 | 1 |
| A3SS | 9487 | ENSG00000109339 | MAPK10 | chr4 | - | 86098523 | 86099285 | 86098523 | 86098595 | 86101051 | 86101217 | 0 | 0 | 0 | 0 | 0 | 0 | 1 | 0 | 2 | 4 | 0 | 3 | 0 | 0 |
| A3SS | 4588 | ENSG00000111642 | CHD4 | chr12 | - | 6580581 | 6580699 | 6580581 | 6580679 | 6581043 | 6581173 | 0 | 1 | 0 | 1 | 1 | 0 | 4 | 2 | 0 | 3 | 0 | 7 | 2 | 7 |
| A3SS | 9600 | ENSG00000168014 | C2CD3 | chr11 | - | 74037477 | 74037652 | 74037477 | 74037623 | 74042053 | 74042218 | 9 | 2 | 2 | 2 | 2 | 0 | 0 | 2 | 4 | 3 | 0 | 22 | 0 | 4 |
| A3SS | 11913 | ENSG00000205758 | CRYZL1 | chr21 | - | 33591161 | 33591228 | 33591161 | 33591211 | 33595730 | 33595836 | 2 | 3 | 2 | 1 | 3 | 10 | 2 | 5 | 4 | 9 | 0 | 4 | 0 | 8 |
| A3SS | 5988 | ENSG00000144357 | UBR3 | chr2 | + | 170073403 | 170073607 | 170073427 | 170073607 | 170061317 | 170061443 | 5 | 1 | 4 | 0 | 12 | 69 | 48 | 43 | 51 | 94 | 0 | 1 | 3 | 2 |
| A3SS | 7082 | ENSG00000050748 | MAPK9 | chr5 | - | 180233593 | 180236531 | 180233593 | 180236526 | 180239923 | 180239987 | 0 | 1 | 1 | 0 | 0 | 0 | 0 | 0 | 0 | 2 | 0 | 0 | 1 | 0 |
| A3SS | 7953 | ENSG00000206530 | CFAP44 | chr3 | - | 113400544 | 113400672 | 113400544 | 113400644 | 113401239 | 113401287 | 0 | 0 | 0 | 0 | 0 | 16 | 27 | 34 | 28 | 17 | 0 | 0 | 0 | 1 |
| A3SS | 13147 | ENSG00000123349 | PFDN5 | chr12 | + | 53296429 | 53296551 | 53296453 | 53296551 | 53295838 | 53295941 | 0 | 3 | 7 | 2 | 0 | 0 | 6 | 1 | 0 | 3 | 0 | 3 | 0 | 4 |
| A3SS | 2756 | ENSG00000197045 | GMFB | chr14 | - | 54480873 | 54480959 | 54480873 | 54480956 | 54481408 | 54481458 | 7 | 15 | 9 | 2 | 6 | 45 | 45 | 44 | 37 | 65 | 0 | 7 | 0 | 8 |
| A3SS | 1817 | ENSG00000196557 | CACNA1H | chr16 | + | 1218969 | 1219130 | 1219002 | 1219130 | 1218209 | 1218651 | 0 | 0 | 3 | 0 | 0 | 0 | 0 | 1 | 0 | 0 | 0 | 2 | 0 | 1 |
| A3SS | 3581 | ENSG00000083312 | TNPO1 | chr5 | + | 72851204 | 72851319 | 72851237 | 72851319 | 72848384 | 72848498 | 4 | 2 | 0 | 1 | 3 | 18 | 0 | 7 | 13 | 5 | 0 | 11 | 9 | 3 |
| A3SS | 3580 | ENSG00000158290 | CUL4B | chrX | - | 120557528 | 120557579 | 120557528 | 120557560 | 120557923 | 120558039 | 3 | 6 | 0 | 3 | 4 | 0 | 0 | 4 | 2 | 0 | 0 | 1 | 0 | 0 |
| A3SS | 11405 | ENSG00000184465 | WDR27 | chr6 | - | 169667981 | 169668185 | 169667981 | 169668095 | 169670568 | 169670693 | 8 | 8 | 7 | 12 | 5 | 1 | 0 | 3 | 6 | 1 | 0 | 10 | 6 | 3 |
| A3SS | 3602 | ENSG00000188818 | ZDHHC11 | chr5 | - | 840494 | 840650 | 840494 | 840597 | 843599 | 843724 | 0 | 4 | 1 | 2 | 0 | 2 | 0 | 0 | 0 | 0 | 0 | 0 | 0 | 0 |
| A3SS | 3935 | ENSG00000146350 | TBC1D32 | chr6 | - | 121115171 | 121115747 | 121115171 | 121115241 | 121126377 | 121126461 | 10 | 13 | 7 | 17 | 11 | 7 | 7 | 2 | 2 | 8 | 0 | 3 | 2 | 0 |
| A3SS | 6564 | ENSG00000007402 | CACNA2D2 | chr3 | - | 50364887 | 50364976 | 50364887 | 50364970 | 50365074 | 50365184 | 9 | 6 | 2 | 5 | 6 | 24 | 6 | 4 | 3 | 9 | 6 | 0 | 2 | 0 |
| A3SS | 7214 | ENSG00000162604 | TM2D1 | chr1 | - | 61686791 | 61686921 | 61686791 | 61686895 | 61694696 | 61694770 | 5 | 2 | 5 | 8 | 0 | 0 | 0 | 0 | 0 | 1 | 8 | 5 | 2 | 0 |
| A3SS | 1910 | ENSG00000205084 | TMEM231 | chr16 | - | 75545825 | 75546077 | 75545825 | 75545954 | 75555803 | 75555973 | 0 | 1 | 0 | 0 | 0 | 0 | 0 | 5 | 0 | 0 | 0 | 4 | 2 | 0 |
| A3SS | 8990 | ENSG00000144134 | RABL2A | chr2 | + | 113628549 | 113628713 | 113628553 | 113628713 | 113627243 | 113627400 | 0 | 0 | 2 | 0 | 0 | 5 | 0 | 0 | 0 | 0 | 7 | 0 | 0 | 2 |
| A3SS | 9559 | ENSG00000173715 | C11orf80 | chr11 | + | 66803994 | 66804184 | 66803997 | 66804184 | 66801030 | 66801102 | 16 | 3 | 6 | 16 | 5 | 4 | 4 | 5 | 3 | 5 | 0 | 19 | 7 | 7 |
| A3SS | 2575 | ENSG00000100949 | RABGGTA | chr14 | - | 24271112 | 24271248 | 24271112 | 24271169 | 24271486 | 24271549 | 8 | 2 | 1 | 8 | 0 | 9 | 0 | 3 | 2 | 0 | 11 | 6 | 8 | 1 |
| A3SS | 2755 | ENSG00000197045 | GMFB | chr14 | - | 54480873 | 54480962 | 54480873 | 54480941 | 54481325 | 54481458 | 11 | 10 | 21 | 10 | 20 | 0 | 1 | 0 | 5 | 4 | 0 | 12 | 8 | 0 |
| A3SS | 1418 | ENSG00000179029 | TMEM107 | chr17 | - | 8175756 | 8175875 | 8175756 | 8175857 | 8175958 | 8176026 | 1 | 1 | 2 | 0 | 5 | 6 | 4 | 3 | 7 | 1 | 0 | 2 | 1 | 0 |
| A3SS | 8184 | ENSG00000084623 | EIF3I | chr1 | + | 32226104 | 32226320 | 32226148 | 32226320 | 32224409 | 32224475 | 2 | 4 | 4 | 2 | 1 | 1 | 0 | 1 | 0 | 0 | 11 | 0 | 0 | 6 |
| A3SS | 7648 | ENSG00000174371 | EXO1 | chr1 | + | 241858505 | 241858718 | 241858629 | 241858718 | 241857344 | 241857482 | 6 | 1 | 12 | 2 | 4 | 0 | 0 | 0 | 0 | 3 | 0 | 6 | 17 | 25 |

[illegible]

| | | | | | | | | | | | | | | | | | | | | | | | | | | | | | | | | | | | | | | | | | | | | | | |
|---|---|---|---|---|---|---|---|---|---|---|---|---|---|---|---|---|---|---|---|---|---|---|---|---|---|---|---|---|---|---|---|---|---|---|---|---|---|---|---|---|---|---|---|---|---|---|
| A3SS | 7254 | ENSG00000117262 | GPR89A | chr1 | + | 145646183 | 145646272 | 145646203 | 145646272 | 145645646 | 145645743 | 1 | 2 | 1 | 0 | 2 | 5 | 1 | 2 | 1 | 3 | 0 | 0 | 1 | 7 | 2 | 1 | 0 | 6 | 8 | 16 | 169 | 149 | 1 | 1 | 0.15 | 0.638 | 0.306 | 0 | 0.37 | 0 | NA | 0.128 | 0.435 | 0.099 | 0.127 |
| A3SS | 8367 | ENSG00000107643 | MAPK8 | chr10 | + | 48434878 | 48439357 | 48434883 | 48439357 | 48431192 | 48431270 | 8 | 5 | 5 | 9 | 14 | 1 | 1 | 2 | 1 | 0 | 0 | 6 | 2 | 10 | 0 | 0 | 4 | 1 | 1 | 0 | 154 | 149 | 1 | 1 | 0.886 | 0.829 | 0.708 | 0.897 | 1 | NA | 0.592 | 0.659 | 0.906 | NA | 0.145 |
| A3SS | 10374 | ENSG00000125676 | THOC2 | chrX | - | 123610917 | 123610963 | 123610917 | 123610957 | 123611439 | 123611516 | 76 | 67 | 73 | 73 | 119 | 29 | 22 | 12 | 18 | 5 | 0 | 19 | 12 | 3 | 0 | 0 | 15 | 5 | 1 | 0 | 155 | 149 | 1 | 1 | 0.716 | 0.745 | 0.854 | 0.796 | 0.958 | NA | 0.549 | 0.698 | 0.743 | NA | 0.15 |
| A3SS | 6834 | ENSG00000140992 | PDPK1 | chr16 | + | 2581294 | 2581363 | 2581299 | 2581363 | 2577424 | 2577500 | 4 | 4 | 3 | 6 | 0 | 0 | 0 | 0 | 0 | 5 | 0 | 9 | 5 | 4 | 0 | 0 | 0 | 0 | 3 | 7 | 154 | 149 | 1 | 1 | 1 | 1 | 1 | 1 | 0 | NA | 1 | 1 | 0.563 | 0 | 0.159 |
| A3SS | 2572 | ENSG00000100949 | RABGGTA | chr14 | - | 24265537 | 24265763 | 24265537 | 24265755 | 24266429 | 24266517 | 5 | 3 | 5 | 13 | 2 | 0 | 0 | 0 | 0 | 0 | 3 | 2 | 7 | 3 | 0 | 0 | 4 | 0 | 0 | 0 | 157 | 149 | 1 | 1 | 1 | 1 | 1 | 1 | 1 | 1 | 0.322 | 1 | 1 | NA | 0.169 |
| A3SS | 4894 | ENSG00000198925 | ATG9A | chr2 | - | 219228433 | 219228485 | 219228433 | 219228482 | 219229534 | 219229639 | 2 | 1 | 3 | 1 | 0 | 0 | 0 | 1 | 2 | 0 | 0 | 0 | 8 | 1 | 9 | 0 | 2 | 0 | 2 | 0 | 152 | 149 | 1 | 1 | 1 | 1 | 0.746 | 0.329 | NA | NA | 0 | 1 | 0.329 | 1 | 0.187 |
| A3SS | 651 | ENSG00000064545 | TMEM161A | chr19 | - | 19133129 | 19133306 | 19133129 | 19133210 | 19134783 | 19134887 | 2 | 1 | 2 | 0 | 0 | 3 | 0 | 2 | 8 | 0 | 0 | 3 | 0 | 4 | 0 | 0 | 6 | 0 | 6 | 4 | 245 | 149 | 1 | 1 | 0.288 | 1 | 0.378 | 0 | NA | NA | 0.233 | NA | 0.288 | 0 | 0.243 |
| A3SS | 10471 | ENSG00000100802 | C14orf93 | chr14 | - | 23000958 | 23001060 | 23000958 | 23001031 | 23010100 | 23010143 | 1 | 0 | 0 | 7 | 1 | 1 | 0 | 0 | 0 | 2 | 0 | 0 | 2 | 0 | 0 | 0 | 2 | 0 | 0 | 13 | 178 | 149 | 1 | 1 | 0.456 | NA | NA | 1 | 0.295 | NA | 0 | 1 | NA | 0 | 0.25 |
| A3SS | 5218 | ENSG00000071127 | WDR1 | chr4 | - | 10083021 | 10083178 | 10083021 | 10083176 | 10084442 | 10084530 | 1 | 1 | 1 | 3 | 2 | 0 | 0 | 0 | 0 | 0 | 2 | 0 | 2 | 0 | 6 | 0 | 0 | 6 | 0 | 0 | 151 | 149 | 1 | 1 | 1 | 1 | 1 | 1 | 1 | 1 | NA | 0.248 | NA | 1 | 0.251 |
| A3SS | 1816 | ENSG00000196557 | CACNA1H | chr16 | + | 1211945 | 1212138 | 1212007 | 1212138 | 1211715 | 1211805 | 0 | 0 | 0 | 2 | 0 | 0 | 0 | 0 | 0 | 0 | 0 | 0 | 4 | 1 | 0 | 0 | 0 | 0 | 1 | 0 | 211 | 149 | 1 | 1 | NA | NA | NA | 1 | NA | NA | NA | 1 | 0.414 | NA | 0.293 |
| A3SS | 10472 | ENSG00000100802 | C14orf93 | chr14 | - | 23000958 | 23001060 | 23000958 | 23001037 | 23010100 | 23010143 | 1 | 0 | 0 | 7 | 1 | 1 | 1 | 3 | 1 | 3 | 0 | 0 | 0 | 0 | 0 | 6 | 0 | 1 | 2 | 0 | 172 | 149 | 1 | 1 | 0.464 | 0 | 0 | 0.858 | 0.224 | 0 | NA | 0 | 0 | NA | 0.309 |
| A3SS | 3133 | ENSG00000131504 | DIAPH1 | chr5 | - | 141534334 | 141534442 | 141534334 | 141534433 | 141571925 | 141572040 | 7 | 6 | 2 | 5 | 13 | 0 | 0 | 0 | 0 | 0 | 1 | 0 | 3 | 0 | 15 | 2 | 0 | 1 | 0 | 0 | 158 | 149 | 1 | 1 | 1 | 1 | 1 | 1 | 1 | 0.32 | NA | 0.739 | NA | 1 | 0.314 |
| A3SS | 9484 | ENSG00000109339 | MAPK10 | chr4 | - | 86029196 | 86029287 | 86029196 | 86029274 | 86031367 | 86031431 | 0 | 0 | 0 | 1 | 0 | 1 | 0 | 0 | 0 | 1 | 0 | 0 | 0 | 0 | 0 | 12 | 8 | 0 | 0 | 2 | 162 | 149 | 1 | 1 | 0 | NA | NA | 1 | 0 | 0 | 0 | NA | NA | 0 | 0.333 |
| A3SS | 1012 | ENSG00000150477 | KIAA1328 | chr18 | + | 36834202 | 36834355 | 36834319 | 36834355 | 36832673 | 36833388 | 5 | 22 | 10 | 21 | 10 | 0 | 0 | 0 | 0 | 2 | 0 | 14 | 5 | 0 | 0 | 0 | 0 | 1 | 2 | 0 | 266 | 149 | 1 | 1 | 1 | 1 | 1 | 1 | 0.737 | NA | 1 | 0.737 | 0 | NA | 0.368 |
| A3SS | 9488 | ENSG00000109339 | MAPK10 | chr4 | - | 86151990 | 86152196 | 86151990 | 86152144 | 86152462 | 86152629 | 3 | 0 | 0 | 1 | 0 | 0 | 0 | 0 | 1 | 0 | 0 | 1 | 0 | 0 | 0 | 0 | 2 | 0 | 0 | 0 | 201 | 149 | 1 | 1 | 1 | NA | NA | 0.426 | NA | NA | 0.27 | NA | NA | NA | 0.443 |
| A3SS | 5510 | ENSG00000132561 | MATN2 | chr8 | + | 98033041 | 98033176 | 98033098 | 98033176 | 98032245 | 98032317 | 6 | 0 | 2 | 3 | 3 | 0 | 0 | 1 | 0 | 0 | 0 | 0 | 0 | 1 | 0 | 0 | 0 | 0 | 1 | 0 | 206 | 149 | 1 | 1 | 1 | NA | 0.591 | 1 | 1 | NA | NA | NA | 0.42 | NA | 0.478 |
| A3SS | 11403 | ENSG00000184465 | WDR27 | chr6 | - | 169663562 | 169663878 | 169663562 | 169663649 | 169664165 | 169664286 | 2 | 14 | 11 | 8 | 5 | 0 | 3 | 0 | 3 | 2 | 0 | 0 | 2 | 0 | 0 | 11 | 0 | 1 | 0 | 0 | 298 | 149 | 1 | 1 | 1 | 0.7 | 1 | 0.571 | 0.556 | 0 | NA | 0.5 | NA | NA | 0.515 |
| A3SS | 11230 | ENSG00000108666 | C17orf75 | chr17 | - | 32339812 | 32339962 | 32339812 | 32339924 | 32341203 | 32341284 | 0 | 2 | 1 | 0 | 0 | 1 | 1 | 0 | 0 | 0 | 0 | 0 | 0 | 0 | 0 | 0 | 0 | 0 | 1 | 0 | 187 | 149 | 1 | 1 | 0 | 0.614 | 1 | NA | NA | NA | NA | NA | 0 | NA | 0.538 |
| A3SS | 11922 | ENSG00000110318 | CEP126 | chr11 | + | 101955791 | 101956943 | 101956261 | 101956943 | 101948030 | 101948142 | 0 | 5 | 1 | 0 | 4 | 1 | 0 | 0 | 1 | 0 | 0 | 0 | 0 | 0 | 0 | 0 | 2 | 0 | 0 | 0 | 298 | 149 | 1 | 1 | 0 | 1 | 1 | 0 | 1 | NA | 0 | NA | NA | NA | 0.6 |
| A3SS | 4208 | ENSG00000068745 | IP6K2 | chr3 | - | 48695089 | 48695421 | 48695089 | 48695418 | 48708119 | 48708237 | 0 | 0 | 1 | 3 | 0 | 3 | 0 | 0 | 0 | 0 | 0 | 0 | 0 | 0 | 0 | 0 | 2 | 0 | 0 | 0 | 152 | 149 | 1 | 1 | 0 | NA | 1 | 1 | NA | NA | 0 | NA | NA | NA | 0.667 |
| A3SS | 7894 | ENSG00000151067 | CACNA1C | chr12 | + | 2674483 | 2674642 | 2674540 | 2674642 | 2668932 | 2669035 | 2 | 0 | 3 | 0 | 0 | 0 | 0 | 0 | 3 | 0 | 0 | 0 | 0 | 0 | 0 | 0 | 0 | 5 | 0 | 0 | 206 | 149 | 1 | 1 | 1 | NA | 1 | 0 | NA | NA | NA | 0 | NA | NA | 0.667 |
| A3SS | 13226 | ENSG00000204852 | TCTN1 | chr12 | + | 110647195 | 110647336 | 110647210 | 110647336 | 110640382 | 110640517 | 2 | 1 | 3 | 0 | 0 | 2 | 0 | 1 | 0 | 0 | 0 | 0 | 0 | 0 | 0 | 0 | 2 | 0 | 0 | 0 | 164 | 149 | 1 | 1 | 0.476 | 1 | 0.732 | NA | NA | NA | 0 | NA | NA | NA | 0.736 |
| A3SS | 2113 | ENSG00000167965 | MLST8 | chr16 | + | 2206478 | 2206659 | 2206483 | 2206659 | 2206357 | 2206409 | 2 | 0 | 0 | 2 | 0 | 0 | 0 | 0 | 0 | 0 | 0 | 0 | 0 | 0 | 0 | 0 | 0 | 0 | 0 | 3 | 154 | 149 | 1 | 1 | 1 | NA | NA | 1 | NA | NA | NA | NA | NA | 0 | 1 |
| A3SS | 3599 | ENSG00000188818 | ZDHHC11 | chr5 | - | 823388 | 824093 | 823388 | 823809 | 825163 | 825251 | 0 | 0 | 3 | 0 | 0 | 0 | 0 | 0 | 0 | 0 | 0 | 0 | 0 | 0 | 0 | 13 | 0 | 0 | 0 | 0 | 298 | 149 | 1 | 1 | NA | NA | 1 | NA | NA | 0 | NA | NA | NA | NA | 1 |

| Splicing_event | ID | Gene_ID | Gene_Symbol | Chromosome | Strand | ExonStart_0base | ExonEnd | UpstreamES | UpstreamEE | DownstreamES | DownstreamEE | SRR13214146 | SRR13214147 | SRR13214148 | SRR13214149 | SRR13214150 | SRR13214146 | SRR13214147 | SRR13214148 | SRR13214149 | SRR13214150 | SRR13214141 | SRR13214242 | SRR13214143 | SRR13214144 | SRR13214145 | SRR13214141 | SRR13214242 | SRR13214143 | SRR13214144 | SRR13214145 | IncFormLen | SkipFormLen | p value | FDR | SRR13214146 | SRR13214147 | SRR13214148 | SRR13214149 | SRR13214150 | SRR13214141 | SRR13214242 | SRR13214143 | SRR13214144 | SRR13214145 | Inclevel_diff |
|---|---|---|---|---|---|---|---|---|---|---|---|---|---|---|---|---|---|---|---|---|---|---|---|---|---|---|---|---|---|---|---|---|---|---|---|---|---|---|---|---|---|---|---|---|---|---|
| SE | 2382 | ENSG00000170889 | RPS9 | chr19 | + | 54206275 | 54206462 | 54201486 | 54201609 | 54207397 | 54207647 | 4621 | 2517 | 6056 | 5302 | 5268 | 6 | 14 | 31 | 35 | 18 | 4697 | 5121 | 3531 | 3394 | 6387 | 56 | 93 | 29 | 69 | 68 | 298 | 149 | 0 | 0 | 0.997 | 0.989 | 0.99 | 0.987 | 0.993 | 0.977 | 0.965 | 0.984 | 0.961 | 0.979 | 0.018 |
| SE | 2384 | ENSG00000170889 | RPS9 | chr19 | + | 54206275 | 54206724 | 54201486 | 54201609 | 54207397 | 54207644 | 2448 | 1346 | 3226 | 2837 | 2837 | 6 | 14 | 31 | 35 | 18 | 2769 | 2959 | 2005 | 1952 | 3488 | 56 | 93 | 29 | 69 | 68 | 298 | 149 | 0 | 0 | 0.995 | 0.98 | 0.981 | 0.976 | 0.987 | 0.961 | 0.941 | 0.972 | 0.934 | 0.962 | 0.03 |
| SE | 2386 | ENSG00000170889 | RPS9 | chr19 | + | 54206552 | 54206724 | 54206275 | 54206462 | 54207397 | 54207522 | 303 | 174 | 299 | 234 | 289 | 2265 | 1226 | 2895 | 2512 | 2463 | 797 | 672 | 410 | 465 | 728 | 2132 | 2377 | 1630 | 1587 | 3063 | 298 | 149 | 0 | 0 | 0.063 | 0.066 | 0.049 | 0.045 | 0.055 | 0.157 | 0.124 | 0.112 | 0.128 | 0.106 | -0.07 |
| SE | 2872 | ENSG00000104852 | SNRNP70 | chr19 | + | 49102113 | 49102185 | 49101389 | 49101471 | 49104633 | 49104735 | 132 | 83 | 154 | 141 | 243 | 165 | 149 | 134 | 194 | 129 | 122 | 53 | 47 | 73 | 148 | 662 | 227 | 212 | 282 | 568 | 221 | 149 | 0 | 0 | 0.35 | 0.273 | 0.437 | 0.329 | 0.559 | 0.111 | 0.136 | 0.13 | 0.149 | 0.149 | 0.255 |
| SE | 3861 | ENSG00000157540 | DYRK1A | chr21 | + | 37430268 | 37430400 | 37420298 | 37420384 | 37472683 | 37472829 | 23 | 6 | 10 | 21 | 27 | 182 | 120 | 170 | 202 | 263 | 43 | 104 | 119 | 49 | 10 | 157 | 163 | 142 | 119 | 52 | 281 | 149 | 0 | 0 | 0.063 | 0.026 | 0.03 | 0.052 | 0.052 | 0.127 | 0.253 | 0.308 | 0.179 | 0.093 | -0.147 |
| SE | 7693 | ENSG00000170004 | CHD3 | chr17 | + | 7902936 | 7903061 | 7902183 | 7902727 | 7903271 | 7903503 | 447 | 286 | 590 | 611 | 417 | 4 | 3 | 3 | 10 | 0 | 686 | 429 | 349 | 465 | 804 | 38 | 31 | 20 | 45 | 20 | 274 | 149 | 0 | 0 | 0.984 | 0.981 | 0.991 | 0.971 | 1 | 0.908 | 0.883 | 0.905 | 0.849 | 0.956 | 0.085 |
| SE | 7844 | ENSG00000108819 | PPP1R9B | chr17 | - | 50139428 | 50139581 | 50139262 | 50139316 | 50140092 | 50140228 | 1669 | 1207 | 1380 | 1710 | 1044 | 12 | 15 | 11 | 19 | 8 | 4374 | 2156 | 1213 | 2156 | 4401 | 156 | 40 | 63 | 68 | 158 | 298 | 149 | 0 | 0 | 0.986 | 0.976 | 0.984 | 0.978 | 0.985 | 0.933 | 0.964 | 0.906 | 0.941 | 0.933 | 0.046 |
| SE | 8645 | ENSG00000261371 | PECAM1 | chr17 | - | 64354932 | 64355040 | 64353490 | 64353518 | 64356110 | 64356398 | 926 | 685 | 619 | 916 | 1206 | 4 | 0 | 9 | 1 | 11 | 1543 | 780 | 482 | 1035 | 1033 | 71 | 16 | 21 | 45 | 63 | 257 | 149 | 0 | 0 | 0.993 | 1 | 0.976 | 0.998 | 0.985 | 0.926 | 0.966 | 0.93 | 0.93 | 0.905 | 0.059 |
| SE | 9423 | ENSG00000174231 | PRPF8 | chr17 | - | 1677564 | 1677694 | 1676975 | 1677172 | 1678517 | 1678652 | 858 | 432 | 820 | 767 | 773 | 3 | 0 | 1 | 3 | 3 | 908 | 767 | 478 | 604 | 1055 | 18 | 39 | 15 | 31 | 12 | 279 | 149 | 0 | 0 | 0.993 | 1 | 0.998 | 0.993 | 0.993 | 0.964 | 0.913 | 0.945 | 0.912 | 0.979 | 0.053 |
| SE | 9463 | ENSG00000277957 | SENP3-EIF4A1 | chr17 | + | 7574545 | 7574678 | 7574259 | 7574308 | 7575118 | 7575258 | 2156 | 1155 | 1761 | 1923 | 1705 | 10 | 3 | 3 | 17 | 9 | 3129 | 2061 | 1610 | 2052 | 2982 | 37 | 16 | 51 | 87 | 51 | 282 | 149 | 0 | 0 | 0.991 | 0.995 | 0.997 | 0.984 | 0.99 | 0.978 | 0.986 | 0.943 | 0.926 | 0.969 | 0.031 |
| SE | 9644 | ENSG00000173821 | RNF213 | chr17 | + | 80385038 | 80385171 | 80383676 | 80383928 | 80385537 | 80385621 | 1455 | 1001 | 1082 | 1512 | 1625 | 0 | 2 | 2 | 4 | 0 | 2713 | 1418 | 988 | 1921 | 1419 | 56 | 3 | 18 | 61 | 10 | 282 | 149 | 0 | 0 | 1 | 0.996 | 0.997 | 0.995 | 1 | 0.962 | 0.996 | 0.967 | 0.943 | 0.987 | 0.027 |
| SE | 10311 | ENSG00000108848 | LUC7L3 | chr17 | + | 50736959 | 50737026 | 50719564 | 50719831 | 50740305 | 50740345 | 541 | 403 | 538 | 506 | 392 | 4 | 1 | 9 | 6 | 0 | 95 | 113 | 96 | 44 | 67 | 15 | 32 | 8 | 21 | 15 | 216 | 149 | 0 | 0 | 0.989 | 0.996 | 0.976 | 0.983 | 1 | 0.814 | 0.709 | 0.892 | 0.591 | 0.755 | 0.237 |
| SE | 13323 | ENSG00000177663 | IL17RA | chr22 | + | 17100354 | 17100481 | 17098774 | 17098887 | 17101995 | 17102043 | 618 | 476 | 381 | 613 | 541 | 14 | 2 | 3 | 1 | 18 | 1462 | 1004 | 401 | 969 | 1188 | 140 | 30 | 21 | 75 | 61 | 276 | 149 | 0 | 0 | 0.96 | 0.992 | 0.986 | 0.997 | 0.942 | 0.849 | 0.948 | 0.912 | 0.875 | 0.913 | 0.076 |
| SE | 14143 | ENSG00000005339 | CREBBP | chr16 | - | 3738558 | 3738672 | 3736649 | 3736815 | 3740398 | 3740549 | 271 | 198 | 247 | 299 | 357 | 11 | 13 | 16 | 21 | 14 | 481 | 292 | 274 | 300 | 402 | 137 | 65 | 131 | 172 | 90 | 263 | 149 | 0 | 0 | 0.933 | 0.896 | 0.897 | 0.89 | 0.935 | 0.665 | 0.718 | 0.542 | 0.497 | 0.717 | 0.282 |
| SE | 14144 | ENSG00000005339 | CREBBP | chr16 | - | 3739577 | 3739724 | 3736649 | 3736815 | 3740398 | 3740549 | 138 | 71 | 138 | 143 | 155 | 11 | 13 | 16 | 21 | 14 | 171 | 159 | 90 | 135 | 179 | 137 | 65 | 131 | 172 | 90 | 296 | 149 | 0 | 0 | 0.863 | 0.733 | 0.813 | 0.774 | 0.848 | 0.386 | 0.552 | 0.257 | 0.283 | 0.5 | 0.411 |
| SE | 15465 | ENSG00000140199 | SLC12A6 | chr15 | - | 34257641 | 34257788 | 34256228 | 34256283 | 34258812 | 34258944 | 131 | 109 | 202 | 158 | 237 | 4 | 6 | 5 | 7 | 14 | 120 | 71 | 75 | 80 | 99 | 35 | 15 | 32 | 25 | 18 | 296 | 149 | 0 | 0 | 0.943 | 0.901 | 0.953 | 0.919 | 0.895 | 0.633 | 0.704 | 0.541 | 0.617 | 0.735 | 0.276 |
| SE | 15668 | ENSG00000092439 | TRPM7 | chr15 | - | 50634381 | 50634556 | 50632868 | 50632992 | 50637421 | 50637593 | 344 | 239 | 294 | 295 | 284 | 1 | 0 | 3 | 1 | 1 | 67 | 191 | 153 | 127 | 77 | 0 | 25 | 12 | 6 | 24 | 298 | 149 | 0 | 0 | 0.994 | 1 | 0.98 | 0.993 | 0.993 | 1 | 0.793 | 0.864 | 0.914 | 0.616 | 0.155 |
| SE | 15982 | ENSG00000185787 | MORF4L1 | chr15 | + | 78894057 | 78894230 | 78893538 | 78893627 | 78894819 | 78894904 | 3106 | 2147 | 2709 | 3337 | 5568 | 3 | 9 | 6 | 17 | 5 | 3611 | 2813 | 2983 | 3259 | 1697 | 19 | 57 | 18 | 64 | 38 | 298 | 149 | 0 | 0 | 0.998 | 0.992 | 0.996 | 0.99 | 0.998 | 0.99 | 0.961 | 0.988 | 0.962 | 0.957 | 0.023 |
| SE | 17128 | ENSG00000159128 | IFNGR2 | chr21 | + | 33432176 | 33432336 | 33426883 | 33427032 | 33432713 | 33432871 | 1958 | 1357 | 1102 | 1139 | 3210 | 6 | 1 | 1 | 9 | 24 | 3550 | 2626 | 1773 | 2467 | 3971 | 38 | 39 | 34 | 90 | 116 | 298 | 149 | 0 | 0 | 0.994 | 0.999 | 0.998 | 0.984 | 0.985 | 0.979 | 0.971 | 0.963 | 0.932 | 0.945 | 0.034 |
| SE | 17790 | ENSG00000100485 | SOS2 | chr14 | - | 50158564 | 50158646 | 50155891 | 50157121 | 50159430 | 50160086 | 447 | 343 | 294 | 379 | 567 | 8 | 8 | 4 | 5 | 19 | 37 | 179 | 365 | 177 | 8 | 5 | 33 | 74 | 16 | 0 | 231 | 149 | 0 | 0 | 0.973 | 0.965 | 0.979 | 0.98 | 0.951 | 0.827 | 0.778 | 0.761 | 0.877 | 1 | 0.121 |
| SE | 19144 | ENSG00000089902 | RCOR1 | chr14 | + | 102721319 | 102721377 | 102721006 | 102721084 | 102722186 | 102722416 | 453 | 323 | 260 | 338 | 419 | 3 | 2 | 4 | 0 | 3 | 935 | 403 | 442 | 915 | 559 | 42 | 17 | 4 | 73 | 29 | 207 | 149 | 0 | 0 | 0.991 | 0.991 | 0.979 | 1 | 0.99 | 0.941 | 0.945 | 0.988 | 0.9 | 0.933 | 0.049 |

| | | | | | | | | | | | | | | | | | | | | | | | | | | | | | | | | | | | | | | | | | | | | | | |
|---|---|---|---|---|---|---|---|---|---|---|---|---|---|---|---|---|---|---|---|---|---|---|---|---|---|---|---|---|---|---|---|---|---|---|---|---|---|---|---|---|---|---|---|---|---|---|
| SE | 85161 | ENSG00000136504 | KAT7 | chr17 | + | 49809118 | 49809208 | 49805362 | 49805445 | 49811475 | 49811574 | 547 | 297 | 484 | 573 | 745 | 61 | 45 | 42 | 53 | 99 | 920 | 693 | 550 | 682 | 612 | 266 | 134 | 144 | 241 | 124 | 239 | 149 | 9.21E-15 | 3.10E-12 | 0.848 | 0.804 | 0.878 | 0.871 | 0.824 | 0.683 | 0.763 | 0.704 | 0.638 | 0.755 | 0.136 |
| SE | 103268 | ENSG00000172661 | WASHC2C | chr10 | + | 45785508 | 45785631 | 45784820 | 45784901 | 45788870 | 45789491 | 130 | 87 | 93 | 114 | 211 | 0 | 2 | 1 | 1 | 0 | 63 | 65 | 61 | 118 | 16 | 0 | 7 | 18 | 17 | 4 | 272 | 149 | 1.15E-14 | 3.86E-12 | 1 | 0.96 | 0.981 | 0.984 | 1 | 1 | 0.836 | 0.65 | 0.792 | 0.687 | 0.192 |
| SE | 102687 | ENSG00000107938 | EDRF1 | chr10 | + | 125729002 | 125729104 | 125725681 | 125725838 | 125729357 | 125729479 | 72 | 44 | 76 | 61 | 66 | 47 | 35 | 43 | 41 | 54 | 19 | 3 | 8 | 5 | 0 | 21 | 48 | 51 | 32 | 13 | 251 | 149 | 1.19E-14 | 3.96E-12 | 0.476 | 0.427 | 0.512 | 0.469 | 0.42 | 0.349 | 0.036 | 0.085 | 0.085 | 0 | 0.35 |
| SE | 70425 | ENSG00000132323 | ILKAP | chr2 | - | 238183652 | 238183740 | 238182064 | 238182186 | 238184019 | 238184096 | 216 | 108 | 185 | 158 | 225 | 5 | 5 | 7 | 23 | 9 | 222 | 137 | 138 | 123 | 147 | 101 | 48 | 77 | 127 | 18 | 237 | 149 | 1.29E-14 | 4.27E-12 | 0.964 | 0.931 | 0.943 | 0.812 | 0.94 | 0.58 | 0.642 | 0.53 | 0.378 | 0.837 | 0.325 |
| SE | 92186 | ENSG00000122692 | SMU1 | chr9 | - | 33071739 | 33071892 | 33068823 | 33068934 | 33073595 | 33073806 | 473 | 327 | 416 | 503 | 768 | 18 | 14 | 14 | 19 | 20 | 579 | 460 | 295 | 594 | 259 | 47 | 60 | 67 | 84 | 15 | 298 | 149 | 1.32E-14 | 4.36E-12 | 0.929 | 0.921 | 0.937 | 0.93 | 0.95 | 0.86 | 0.793 | 0.688 | 0.78 | 0.896 | 0.13 |
| SE | 53039 | ENSG00000103657 | HERC1 | chr15 | - | 63632708 | 63632811 | 63630465 | 63630635 | 63633847 | 63633876 | 224 | 182 | 231 | 209 | 319 | 1 | 6 | 2 | 5 | 0 | 116 | 210 | 188 | 129 | 37 | 6 | 20 | 16 | 13 | 10 | 252 | 149 | 1.59E-14 | 5.22E-12 | 0.993 | 0.947 | 0.986 | 0.961 | 1 | 0.92 | 0.861 | 0.874 | 0.854 | 0.686 | 0.138 |
| SE | 69559 | ENSG00000177479 | ARIH2 | chr3 | + | 48961611 | 48961679 | 48927461 | 48927813 | 48964918 | 48964982 | 212 | 98 | 187 | 252 | 347 | 1 | 0 | 1 | 2 | 2 | 244 | 187 | 142 | 136 | 307 | 0 | 8 | 15 | 11 | 29 | 217 | 149 | 1.64E-14 | 5.38E-12 | 0.993 | 1 | 0.992 | 0.989 | 0.992 | 1 | 0.941 | 0.867 | 0.895 | 0.879 | 0.077 |
| SE | 75221 | ENSG00000100503 | NIN | chr14 | - | 50759993 | 50760359 | 50754741 | 50754867 | 50761789 | 50761867 | 156 | 143 | 150 | 137 | 213 | 12 | 8 | 13 | 12 | 14 | 42 | 98 | 168 | 69 | 12 | 8 | 28 | 64 | 30 | 4 | 298 | 149 | 1.84E-14 | 6.01E-12 | 0.867 | 0.899 | 0.852 | 0.851 | 0.884 | 0.724 | 0.636 | 0.568 | 0.535 | 0.6 | 0.258 |
| SE | 75219 | ENSG00000100503 | NIN | chr14 | - | 50759856 | 50760359 | 50754741 | 50754867 | 50761789 | 50761911 | 224 | 222 | 275 | 223 | 344 | 12 | 8 | 13 | 12 | 14 | 52 | 204 | 237 | 97 | 12 | 8 | 28 | 64 | 30 | 4 | 298 | 149 | 1.89E-14 | 6.13E-12 | 0.903 | 0.933 | 0.914 | 0.903 | 0.925 | 0.765 | 0.785 | 0.649 | 0.618 | 0.6 | 0.232 |
| SE | 79232 | ENSG00000116679 | IVNS1ABP | chr1 | - | 185309002 | 185309172 | 185308799 | 185308875 | 185309382 | 185309511 | 252 | 181 | 264 | 280 | 504 | 1 | 0 | 0 | 2 | 0 | 84 | 102 | 254 | 95 | 44 | 12 | 0 | 16 | 10 | 0 | 298 | 149 | 1.93E-14 | 6.25E-12 | 0.992 | 1 | 1 | 0.986 | 1 | 0.778 | 1 | 0.888 | 0.826 | 1 | 0.097 |
| SE | 31794 | ENSG00000204463 | BAG6 | chr6 | - | 31647590 | 31647826 | 31646393 | 31646523 | 31648676 | 31648751 | 522 | 225 | 514 | 469 | 397 | 2 | 4 | 3 | 3 | 0 | 743 | 533 | 306 | 460 | 804 | 4 | 29 | 24 | 14 | 29 | 298 | 149 | 1.95E-14 | 6.30E-12 | 0.992 | 0.966 | 0.988 | 0.987 | 1 | 0.989 | 0.902 | 0.864 | 0.943 | 0.933 | 0.06 |
| SE | 73547 | ENSG00000115232 | ITGA4 | chr2 | + | 181478756 | 181478824 | 181475158 | 181475288 | 181481597 | 181481683 | 348 | 251 | 339 | 341 | 409 | 2 | 4 | 0 | 1 | 1 | 93 | 229 | 190 | 97 | 89 | 32 | 1 | 9 | 1 | 16 | 217 | 149 | 2.16E-14 | 6.95E-12 | 0.992 | 0.977 | 1 | 0.996 | 0.996 | 0.666 | 0.994 | 0.935 | 0.985 | 0.793 | 0.118 |
| SE | 106635 | ENSG00000135968 | GCC2 | chr2 | + | 108469650 | 108472116 | 108468979 | 108469084 | 108475534 | 108475635 | 159 | 194 | 171 | 232 | 214 | 0 | 2 | 2 | 0 | 0 | 6 | 39 | 38 | 18 | 16 | 12 | 0 | 16 | 4 | 4 | 298 | 149 | 2.19E-14 | 6.99E-12 | 1 | 0.98 | 0.977 | 1 | 1 | 0.2 | 1 | 0.543 | 0.692 | 0.667 | 0.371 |
| SE | 92175 | ENSG00000153786 | ZDHHC7 | chr16 | - | 84988762 | 84988873 | 84981869 | 84981994 | 84990303 | 84990635 | 87 | 58 | 60 | 38 | 40 | 107 | 63 | 82 | 68 | 49 | 46 | 39 | 13 | 58 | 83 | 205 | 89 | 92 | 161 | 335 | 260 | 149 | 2.61E-14 | 8.30E-12 | 0.318 | 0.345 | 0.295 | 0.243 | 0.319 | 0.114 | 0.201 | 0.075 | 0.171 | 0.124 | 0.167 |
| SE | 19018 | ENSG00000198604 | BAZ1A | chr14 | - | 34862043 | 34862322 | 34826012 | 34826156 | 34874491 | 34874662 | 327 | 302 | 285 | 343 | 740 | 7 | 0 | 3 | 4 | 9 | 98 | 271 | 196 | 126 | 123 | 12 | 19 | 16 | 18 | 0 | 298 | 149 | 2.81E-14 | 8.90E-12 | 0.959 | 1 | 0.979 | 0.977 | 0.976 | 0.803 | 0.877 | 0.86 | 0.778 | 1 | 0.115 |
| SE | 24059 | ENSG00000172081 | MOB3A | chr19 | - | 2076810 | 2077013 | 2071035 | 2073424 | 2085174 | 2085328 | 628 | 272 | 416 | 511 | 324 | 9 | 3 | 6 | 10 | 1 | 862 | 449 | 336 | 440 | 798 | 16 | 24 | 16 | 45 | 60 | 298 | 149 | 2.85E-14 | 8.97E-12 | 0.972 | 0.978 | 0.972 | 0.962 | 0.994 | 0.964 | 0.903 | 0.913 | 0.83 | 0.869 | 0.08 |
| SE | 97849 | ENSG00000109971 | HSPA8 | chr11 | - | 123060592 | 123060798 | 123059813 | 123060268 | 123061119 | 123061258 | 4038 | 2699 | 3013 | 3575 | 3566 | 22 | 10 | 18 | 26 | 25 | 2775 | 1202 | 1411 | 1248 | 1772 | 20 | 38 | 42 | 35 | 11 | 298 | 149 | 2.85E-14 | 8.97E-12 | 0.989 | 0.993 | 0.988 | 0.986 | 0.986 | 0.986 | 0.941 | 0.944 | 0.947 | 0.988 | 0.027 |
| SE | 11474 | ENSG00000187555 | USP7 | chr16 | - | 8921156 | 8921295 | 8920358 | 8920447 | 8923214 | 8923413 | 823 | 668 | 776 | 869 | 904 | 9 | 3 | 3 | 5 | 4 | 1094 | 859 | 967 | 918 | 419 | 49 | 33 | 11 | 15 | 8 | 288 | 149 | 2.92E-14 | 9.14E-12 | 0.979 | 0.991 | 0.993 | 0.989 | 0.992 | 0.92 | 0.931 | 0.978 | 0.969 | 0.964 | 0.036 |
| SE | 18344 | ENSG00000257621 | PSMA3-AS1 | chr14 | - | 58288717 | 58288813 | 58274013 | 58274135 | 58291634 | 58291707 | 41 | 18 | 26 | 47 | 26 | 13 | 12 | 24 | 19 | 39 | 149 | 41 | 69 | 90 | 35 | 7 | 7 | 4 | 7 | 3 | 245 | 149 | 3.10E-14 | 9.62E-12 | 0.657 | 0.477 | 0.397 | 0.601 | 0.288 | 0.928 | 0.781 | 0.913 | 0.887 | 0.876 | -0.393 |
| SE | 104596 | ENSG00000069493 | CLEC2D | chr12 | + | 9693824 | 9693948 | 9693033 | 9693115 | 9694759 | 9695106 | 107 | 99 | 123 | 140 | 132 | 170 | 152 | 149 | 158 | 277 | 0 | 58 | 39 | 18 | 0 | 41 | 245 | 96 | 72 | 76 | 273 | 149 | 3.10E-14 | 9.62E-12 | 0.256 | 0.262 | 0.311 | 0.326 | 0.206 | 0 | 0.114 | 0.181 | 0.12 | 0 | 0.189 |
| SE | 10141 | ENSG00000011260 | UTP18 | chr17 | + | 51277129 | 51277304 | 51275865 | 51275991 | 51280004 | 51280105 | 144 | 127 | 159 | 152 | 227 | 6 | 3 | 22 | 23 | 26 | 62 | 52 | 81 | 83 | 36 | 24 | 30 | 24 | 53 | 15 | 298 | 149 | 3.41E-14 | 1.05E-11 | 0.923 | 0.955 | 0.783 | 0.768 | 0.814 | 0.564 | 0.464 | 0.628 | 0.439 | 0.545 | 0.321 |
| SE | 8744 | ENSG00000007202 | KIAA0100 | chr17 | - | 28615663 | 28615798 | 28614445 | 28615246 | 28616095 | 28616197 | 1423 | 1137 | 1105 | 1427 | 1650 | 7 | 4 | 2 | 4 | 4 | 4960 | 1902 | 1350 | 2093 | 3529 | 92 | 8 | 22 | 61 | 30 | 284 | 149 | 3.93E-14 | 1.21E-11 | 0.991 | 0.993 | 0.997 | 0.995 | 0.995 | 0.966 | 0.992 | 0.97 | 0.947 | 0.984 | 0.022 |

| | | | | | | | | | | | | | | | | | | | | | | | | | | | | | | | | | | | | | | | | | | | | | | |
|---|---|---|---|---|---|---|---|---|---|---|---|---|---|---|---|---|---|---|---|---|---|---|---|---|---|---|---|---|---|---|---|---|---|---|---|---|---|---|---|---|---|---|---|---|---|---|
| SE | 103574 | ENSG00000150093 | ITGB1 | chr10 | - | 32932514 | 32932590 | 32929821 | 32930044 | 32935491 | 32935558 | 610 | 451 | 326 | 720 | 826 | 9 | 15 | 6 | 6 | 11 | 184 | 430 | 466 | 253 | 172 | 9 | 21 | 35 | 17 | 5 | 225 | 149 | 2.47E-11 | 4.16E-09 | 0.978 | 0.952 | 0.973 | 0.988 | 0.98 | 0.931 | 0.931 | 0.898 | 0.908 | 0.958 | 0.049 |
| SE | 30736 | ENSG00000249437 | NAIP | chr5 | - | 71011711 | 71011786 | 71011274 | 71011374 | 71012347 | 71012918 | 12 | 3 | 3 | 6 | 4 | 239 | 239 | 121 | 227 | 278 | 35 | 49 | 44 | 6 | 16 | 395 | 428 | 226 | 499 | 175 | 224 | 149 | 2.53E-11 | 4.25E-09 | 0.032 | 0.008 | 0.016 | 0.017 | 0.009 | 0.056 | 0.071 | 0.115 | 0.008 | 0.057 | -0.045 |
| SE | 100591 | ENSG00000102317 | RBM3 | chrX | + | 48575560 | 48575667 | 48574951 | 48575283 | 48576313 | 48576419 | 1476 | 902 | 1765 | 1389 | 1355 | 15 | 5 | 17 | 22 | 18 | 3272 | 2021 | 2180 | 2000 | 2727 | 57 | 43 | 59 | 110 | 55 | 256 | 149 | 2.62E-11 | 4.39E-09 | 0.983 | 0.991 | 0.984 | 0.974 | 0.978 | 0.971 | 0.965 | 0.956 | 0.914 | 0.967 | 0.027 |
| SE | 33682 | ENSG00000176261 | ZBTB8OS | chr1 | - | 32633950 | 32634110 | 32627507 | 32627544 | 32634767 | 32634792 | 12 | 4 | 7 | 6 | 17 | 4 | 4 | 5 | 0 | 12 | 10 | 21 | 12 | 20 | 4 | 0 | 0 | 0 | 1 | 0 | 298 | 149 | 2.83E-11 | 4.72E-09 | 0.6 | 0.333 | 0.412 | 1 | 0.415 | 1 | 1 | 1 | 0.909 | 1 | -0.43 |
| SE | 98330 | ENSG00000149503 | INCENP | chr11 | + | 62129781 | 62130590 | 62128769 | 62128883 | 62137831 | 62137883 | 74 | 49 | 53 | 65 | 98 | 0 | 0 | 0 | 0 | 0 | 68 | 77 | 68 | 68 | 98 | 0 | 2 | 8 | 10 | 12 | 298 | 149 | 2.83E-11 | 4.72E-09 | 1 | 1 | 1 | 1 | 1 | 1 | 0.951 | 0.81 | 0.773 | 0.803 | 0.133 |
| SE | 88437 | ENSG00000179715 | PCED1B | chr12 | + | 47223923 | 47224039 | 47223544 | 47223672 | 47235006 | 47235238 | 21 | 10 | 21 | 19 | 38 | 5 | 8 | 1 | 8 | 2 | 12 | 18 | 36 | 45 | 18 | 0 | 0 | 0 | 0 | 0 | 265 | 149 | 3.16E-11 | 5.25E-09 | 0.703 | 0.413 | 0.922 | 0.572 | 0.914 | 1 | 1 | 1 | 1 | 1 | -0.295 |
| SE | 35817 | ENSG00000123130 | ACOT9 | chrX | - | 23734340 | 23734367 | 23733171 | 23733217 | 23735918 | 23736016 | 76 | 46 | 26 | 61 | 41 | 2 | 0 | 0 | 0 | 1 | 60 | 38 | 80 | 86 | 89 | 13 | 2 | 2 | 9 | 31 | 176 | 149 | 3.30E-11 | 5.47E-09 | 0.97 | 1 | 1 | 1 | 0.972 | 0.796 | 0.941 | 0.971 | 0.89 | 0.709 | 0.127 |
| SE | 54311 | ENSG00000143442 | POGZ | chr1 | - | 151429602 | 151429711 | 151428122 | 151428413 | 151430665 | 151430841 | 254 | 168 | 262 | 197 | 354 | 2 | 2 | 3 | 5 | 2 | 259 | 247 | 173 | 134 | 236 | 0 | 16 | 13 | 22 | 16 | 258 | 149 | 3.31E-11 | 5.47E-09 | 0.987 | 0.98 | 0.981 | 0.958 | 0.99 | 1 | 0.899 | 0.885 | 0.779 | 0.895 | 0.088 |
| SE | 95355 | ENSG00000115459 | ELMOD3 | chr2 | + | 85355554 | 85355716 | 85354725 | 85354829 | 85356966 | 85357252 | 3 | 7 | 8 | 13 | 6 | 1 | 4 | 2 | 7 | 3 | 12 | 10 | 14 | 12 | 5 | 1 | 0 | 0 | 0 | 0 | 298 | 149 | 3.34E-11 | 5.51E-09 | 0.6 | 0.467 | 0.667 | 0.481 | 0.5 | 0.857 | 1 | 1 | 1 | 1 | -0.428 |
| SE | 8634 | ENSG00000261371 | PECAM1 | chr17 | - | 64341633 | 64341690 | 64329699 | 64329722 | 64348259 | 64348322 | 1318 | 1033 | 789 | 1136 | 2555 | 31 | 15 | 6 | 19 | 41 | 2574 | 1191 | 726 | 1303 | 1627 | 73 | 28 | 34 | 82 | 58 | 206 | 149 | 3.35E-11 | 5.52E-09 | 0.969 | 0.98 | 0.99 | 0.977 | 0.978 | 0.962 | 0.969 | 0.939 | 0.92 | 0.953 | 0.03 |
| SE | 104024 | ENSG00000151923 | TIAL1 | chr10 | - | 119576846 | 119576905 | 119576610 | 119576750 | 119577079 | 119577214 | 63 | 60 | 69 | 73 | 146 | 379 | 256 | 342 | 407 | 597 | 3 | 75 | 34 | 35 | 6 | 344 | 331 | 314 | 292 | 341 | 208 | 149 | 3.46E-11 | 5.69E-09 | 0.106 | 0.144 | 0.126 | 0.114 | 0.149 | 0.006 | 0.14 | 0.072 | 0.079 | 0.012 | 0.066 |
| SE | 25317 | ENSG00000134644 | PUM1 | chr1 | - | 31005852 | 31006031 | 30995053 | 30995220 | 31006993 | 31007102 | 532 | 489 | 601 | 695 | 811 | 4 | 2 | 5 | 13 | 4 | 1064 | 767 | 671 | 797 | 563 | 32 | 14 | 18 | 26 | 31 | 298 | 149 | 3.59E-11 | 5.89E-09 | 0.985 | 0.992 | 0.984 | 0.964 | 0.99 | 0.943 | 0.965 | 0.949 | 0.939 | 0.901 | 0.044 |
| SE | 61780 | ENSG00000173041 | ZNF680 | chr7 | - | 64544305 | 64544432 | 64543706 | 64543802 | 64562924 | 64563041 | 28 | 14 | 11 | 19 | 41 | 4 | 6 | 4 | 2 | 6 | 39 | 11 | 7 | 1 | 16 | 0 | 0 | 0 | 0 | 0 | 276 | 149 | 3.63E-11 | 5.95E-09 | 0.791 | 0.557 | 0.598 | 0.837 | 0.787 | 1 | 1 | 1 | 1 | 1 | -0.286 |
| SE | 94180 | ENSG00000127080 | IPPK | chr9 | - | 92649452 | 92649574 | 92648058 | 92648148 | 92652572 | 92652639 | 28 | 19 | 29 | 28 | 51 | 0 | 0 | 2 | 0 | 1 | 39 | 3 | 23 | 19 | 49 | 9 | 4 | 10 | 2 | 14 | 271 | 149 | 3.68E-11 | 6.01E-09 | 1 | 1 | 0.889 | 1 | 0.966 | 0.704 | 0.292 | 0.558 | 0.839 | 0.658 | 0.361 |
| SE | 45231 | ENSG00000025800 | KPNA6 | chr1 | + | 32159399 | 32159531 | 32158266 | 32158361 | 32160614 | 32160703 | 219 | 197 | 218 | 289 | 435 | 0 | 1 | 0 | 0 | 0 | 326 | 168 | 204 | 233 | 340 | 29 | 7 | 0 | 2 | 6 | 281 | 149 | 3.79E-11 | 6.18E-09 | 1 | 0.991 | 1 | 1 | 1 | 0.856 | 0.927 | 1 | 0.984 | 0.968 | 0.051 |
| SE | 40701 | ENSG00000117448 | AKR1A1 | chr1 | + | 45568484 | 45568684 | 45567981 | 45568177 | 45568926 | 45568999 | 138 | 49 | 108 | 148 | 160 | 24 | 9 | 19 | 19 | 9 | 140 | 54 | 62 | 115 | 145 | 60 | 31 | 20 | 42 | 75 | 298 | 149 | 4.23E-11 | 6.89E-09 | 0.742 | 0.731 | 0.74 | 0.796 | 0.899 | 0.538 | 0.466 | 0.608 | 0.578 | 0.492 | 0.245 |
| SE | 73549 | ENSG00000115232 | ITGA4 | chr2 | + | 181480136 | 181480266 | 181475158 | 181475288 | 181481597 | 181481683 | 1174 | 1072 | 1175 | 1288 | 1719 | 2 | 4 | 0 | 1 | 1 | 582 | 835 | 924 | 433 | 416 | 32 | 1 | 9 | 1 | 16 | 279 | 149 | 4.42E-11 | 7.18E-09 | 0.997 | 0.993 | 1 | 0.999 | 0.999 | 0.907 | 0.998 | 0.982 | 0.996 | 0.933 | 0.034 |
| SE | 29480 | ENSG00000173020 | GRK2 | chr11 | + | 67283706 | 67283773 | 67283127 | 67283228 | 67283853 | 67283949 | 337 | 152 | 319 | 336 | 158 | 1 | 1 | 3 | 5 | 0 | 281 | 429 | 280 | 336 | 720 | 0 | 23 | 9 | 23 | 30 | 216 | 149 | 4.44E-11 | 7.19E-09 | 0.996 | 0.991 | 0.987 | 0.979 | 1 | 1 | 0.928 | 0.955 | 0.91 | 0.943 | 0.043 |
| SE | 79156 | ENSG00000145191 | EIF2B5 | chr3 | + | 184138846 | 184138939 | 184138165 | 184138246 | 184140079 | 184140157 | 0 | 3 | 11 | 1 | 5 | 106 | 52 | 105 | 107 | 136 | 16 | 7 | 13 | 16 | 30 | 132 | 80 | 58 | 92 | 94 | 242 | 149 | 4.53E-11 | 7.33E-09 | 0 | 0.034 | 0.061 | 0.006 | 0.022 | 0.069 | 0.051 | 0.121 | 0.097 | 0.164 | -0.076 |
| SE | 13857 | ENSG00000102893 | PHKB | chr16 | + | 47663676 | 47663734 | 47661718 | 47661800 | 47665941 | 47666032 | 70 | 81 | 69 | 123 | 95 | 0 | 0 | 2 | 0 | 0 | 33 | 98 | 73 | 74 | 49 | 0 | 8 | 6 | 10 | 7 | 207 | 149 | 4.54E-11 | 7.33E-09 | 1 | 1 | 0.961 | 1 | 1 | 1 | 0.898 | 0.898 | 0.842 | 0.834 | 0.098 |
| SE | 39095 | ENSG00000109320 | NFKB1 | chr4 | + | 102533844 | 102533885 | 102529835 | 102529914 | 102566986 | 102567135 | 28 | 13 | 18 | 20 | 35 | 4 | 1 | 1 | 5 | 0 | 27 | 11 | 15 | 31 | 51 | 23 | 9 | 50 | 29 | 20 | 190 | 149 | 4.80E-11 | 7.73E-09 | 0.846 | 0.911 | 0.934 | 0.758 | 1 | 0.479 | 0.489 | 0.19 | 0.456 | 0.667 | 0.434 |
| SE | 64967 | ENSG00000182973 | CNOT10 | chr3 | + | 32713226 | 32713369 | 32708669 | 32708820 | 32716224 | 32716311 | 243 | 175 | 194 | 225 | 458 | 0 | 2 | 0 | 3 | 0 | 235 | 115 | 121 | 158 | 65 | 29 | 4 | 1 | 4 | 0 | 292 | 149 | 4.94E-11 | 7.95E-09 | 1 | 0.978 | 1 | 0.975 | 1 | 0.805 | 0.936 | 0.984 | 0.953 | 1 | 0.055 |
| SE | 32741 | ENSG00000163349 | HIPK1 | chr1 | + | 113957123 | 113957286 | 113956626 | 113956811 | 113958065 | 113958291 | 584 | 436 | 470 | 576 | 1295 | 6 | 8 | 4 | 6 | 8 | 582 | 571 | 480 | 577 | 254 | 0 | 16 | 23 | 50 | 6 | 298 | 149 | 5.22E-11 | 8.38E-09 | 0.98 | 0.965 | 0.983 | 0.98 | 0.988 | 1 | 0.947 | 0.913 | 0.852 | 0.955 | 0.046 |
| SE | 16071 | ENSG00000128908 | INO80 | chr15 | - | 41027595 | 41027736 | 41020899 | 41021125 | 41044903 | 41045075 | 170 | 143 | 171 | 158 | 121 | 1 | 8 | 3 | 3 | 9 | 229 | 234 | 202 | 170 | 112 | 68 | 18 | 18 | 23 | 19 | 290 | 149 | 5.24E-11 | 8.39E-09 | 0.989 | 0.902 | 0.967 | 0.964 | 0.874 | 0.634 | 0.87 | 0.852 | 0.792 | 0.752 | 0.159 |
| SE | 18446 | ENSG00000100911 | PSME2 | chr14 | - | 24146207 | 24146240 | 24145133 | 24145772 | 24146533 | 24146578 | 314 | 211 | 297 | 300 | 207 | 2 | 1 | 1 | 3 | 0 | 699 | 439 | 190 | 424 | 634 | 49 | 10 | 8 | 13 | 9 | 182 | 149 | 5.31E-11 | 8.49E-09 | 0.992 | 0.994 | 0.996 | 0.988 | 1 | 0.921 | 0.973 | 0.951 | 0.964 | 0.983 | 0.036 |
| SE | 34077 | ENSG00000114867 | EIF4G1 | chr3 | + | 184317749 | 184317816 | 184314618 | 184314674 | 184319688 | 184319728 | 44 | 27 | 33 | 55 | 53 | 0 | 0 | 0 | 1 | 0 | 54 | 31 | 52 | 55 | 70 | 10 | 0 | 6 | 11 | 7 | 216 | 149 | 5.37E-11 | 8.57E-09 | 1 | 1 | 1 | 0.974 | 1 | 0.788 | 1 | 0.857 | 0.775 | 0.873 | 0.136 |
| SE | 11938 | ENSG00000005844 | ITGAL | chr16 | + | 30479330 | 30479461 | 30479089 | 30479208 | 30481438 | 30481584 | 1773 | 771 | 992 | 1177 | 1269 | 10 | 1 | 5 | 19 | 9 | 1987 | 1702 | 953 | 1348 | 2682 | 51 | 26 | 18 | 42 | 45 | 280 | 149 | 5.61E-11 | 8.91E-09 | 0.99 | 0.998 | 0.991 | 0.971 | 0.987 | 0.954 | 0.972 | 0.966 | 0.945 | 0.969 | 0.026 |
| SE | 109160 | ENSG00000197442 | MAP3K5 | chr6 | - | 136622847 | 136622981 | 136614158 | 136614286 | 136637324 | 136637406 | 330 | 213 | 184 | 215 | 343 | 1 | 0 | 2 | 0 | 1 | 142 | 203 | 184 | 146 | 148 | 10 | 8 | 6 | 4 | 7 | 283 | 149 | 5.62E-11 | 8.91E-09 | 0.994 | 1 | 0.98 | 1 | 0.994 | 0.882 | 0.93 | 0.942 | 0.951 | 0.918 | 0.069 |
| SE | 80295 | ENSG00000168137 | SETD5 | chr3 | + | 9443307 | 9443417 | 9442127 | 9442245 | 9445047 | 9445300 | 221 | 173 | 228 | 229 | 450 | 6 | 1 | 0 | 2 | 1 | 284 | 278 | 205 | 229 | 130 | 7 | 13 | 11 | 2 | 25 | 259 | 149 | 5.62E-11 | 8.91E-09 | 0.955 | 0.99 | 1 | 0.985 | 0.996 | 0.959 | 0.925 | 0.915 | 0.985 | 0.749 | 0.079 |
| SE | 96070 | ENSG00000132109 | TRIM21 | chr11 | - | 4386156 | 4386257 | 4384896 | 4385853 | 4386967 | 4386990 | 313 | 193 | 170 | 241 | 261 | 0 | 1 | 2 | 4 | 5 | 303 | 237 | 171 | 207 | 323 | 23 | 5 | 5 | 22 | 13 | 250 | 149 | 5.94E-11 | 9.39E-09 | 1 | 0.991 | 0.981 | 0.973 | 0.969 | 0.887 | 0.966 | 0.953 | 0.849 | 0.937 | 0.064 |
| SE | 34075 | ENSG00000114867 | EIF4G1 | chr3 | + | 184317746 | 184317816 | 184314653 | 184314674 | 184319688 | 184319801 | 55 | 30 | 43 | 68 | 57 | 0 | 0 | 0 | 1 | 0 | 64 | 40 | 59 | 65 | 78 | 10 | 0 | 6 | 11 | 7 | 219 | 149 | 6.24E-11 | 9.84E-09 | 1 | 1 | 1 | 0.979 | 1 | 0.813 | 1 | 0.87 | 0.801 | 0.883 | 0.122 |
| SE | 6930 | ENSG00000175354 | PTPN2 | chr18 | - | 12840713 | 12840962 | 12836790 | 12836891 | 12859163 | 12859254 | 8 | 11 | 9 | 10 | 14 | 183 | 119 | 84 | 142 | 121 | 24 | 65 | 30 | 32 | 0 | 79 | 61 | 113 | 100 | 3 | 298 | 149 | 6.30E-11 | 9.93E-09 | 0.021 | 0.044 | 0.051 | 0.034 | 0.055 | 0.132 | 0.348 | 0.117 | 0.138 | 0 | -0.106 |
| SE | 75941 | ENSG00000196730 | DAPK1 | chr9 | + | 87637942 | 87638081 | 87604953 | 87605175 | 87639353 | 87639483 | 201 | 204 | 85 | 204 | 193 | 4 | 0 | 0 | 3 | 0 | 494 | 136 | 109 | 120 | 430 | 19 | 7 | 18 | 16 | 4 | 288 | 149 | 6.41E-11 | 1.01E-08 | 0.963 | 1 | 1 | 0.972 | 1 | 0.931 | 0.91 | 0.758 | 0.795 | 0.982 | 0.112 |
| SE | 65306 | ENSG00000169621 | APLF | chr2 | + | 68531347 | 68531429 | 68526060 | 68526242 | 68537871 | 68538227 | 0 | 2 | 2 | 0 | 6 | 132 | 109 | 110 | 200 | 470 | 12 | 12 | 6 | 27 | 8 | 247 | 139 | 116 | 148 | 331 | 231 | 149 | 6.44E-11 | 1.01E-08 | 0 | 0.012 | 0.012 | 0 | 0.008 | 0.03 | 0.053 | 0.032 | 0.105 | 0.015 | -0.041 |
| SE | 7851 | ENSG00000108819 | PPP1R9B | chr17 | - | 50140092 | 50140359 | 50139459 | 50139581 | 50141268 | 50141373 | 581 | 372 | 492 | 591 | 377 | 10 | 0 | 4 | 4 | 3 | 1253 | 732 | 386 | 595 | 1349 | 54 | 10 | 9 | 31 | 34 | 298 | 149 | 6.75E-11 | 1.06E-08 | 0.967 | 1 | 0.984 | 0.987 | 0.984 | 0.921 | 0.973 | 0.955 | 0.906 | 0.952 | 0.043 |
| SE | 35058 | ENSG00000156030 | MIDEAS | chr14 | - | 73721296 | 73721509 | 73715121 | 73719501 | 73725271 | 73725360 | 93 | 69 | 92 | 105 | 70 | 0 | 1 | 1 | 1 | 3 | 114 | 61 | 54 | 61 | 99 | 0 | 9 | 5 | 16 | 27 | 298 | 149 | 7.06E-11 | 1.10E-08 | 1 | 0.972 | 0.979 | 0.981 | 0.921 | 1 | 0.772 | 0.844 | 0.656 | 0.647 | 0.187 |
| SE | 21581 | ENSG00000075568 | TMEM131 | chr2 | - | 97794929 | 97795115 | 97793289 | 97793553 | 97796217 | 97796404 | 227 | 155 | 189 | 194 | 352 | 15 | 6 | 15 | 10 | 6 | 50 | 177 | 92 | 153 | 29 | 12 | 34 | 50 | 15 | 7 | 298 | 149 | 7.18E-11 | 1.12E-08 | 0.883 | 0.928 | 0.863 | 0.907 | 0.967 | 0.676 | 0.722 | 0.479 | 0.836 | 0.674 | 0.232 |
| SE | 109330 | ENSG00000123104 | ITPR2 | chr12 | - | 26682573 | 26682673 | 26681873 | 26682034 | 26686480 | 26686632 | 209 | 200 | 188 | 150 | 202 | 2 | 5 | 2 | 4 | 1 | 98 | 121 | 128 | 67 | 54 | 8 | 17 | 9 | 16 | 0 | 249 | 149 | 7.27E-11 | 1.13E-08 | 0.984 | 0.96 | 0.983 | 0.957 | 0.992 | 0.88 | 0.81 | 0.895 | 0.715 | 1 | 0.115 |
| SE | 30592 | ENSG00000152784 | PRDM8 | chr4 | + | 80196012 | 80196124 | 80194112 | 80194239 | 80196351 | 80196419 | 8 | 8 | 8 | 6 | 12 | 9 | 5 | 2 | 4 | 4 | 17 | 14 | 18 | 4 | 23 | 0 | 1 | 2 | 0 | 0 | 261 | 149 | 7.27E-11 | 1.13E-08 | 0.337 | 0.477 | 0.695 | 0.461 | 0.631 | 1 | 0.889 | 0.837 | 1 | 1 | -0.425 |
| SE | 29606 | ENSG00000203875 | SNHG5 | chr6 | - | 85677790 | 85677875 | 85677423 | 85677492 | 85677953 | 85678032 | 130 | 327 | 174 | 110 | 288 | 32 | 22 | 29 | 20 | 98 | 82 | 196 | 290 | 295 | 479 | 0 | 16 | 16 | 0 | 40 | 234 | 149 | 7.38E-11 | 1.14E-08 | 0.721 | 0.904 | 0.793 | 0.778 | 0.652 | 1 | 0.886 | 0.92 | 1 | 0.884 | -0.168 |
| SE | 105990 | ENSG00000122965 | RBM19 | chr12 | - | 113959211 | 113959404 | 113957781 | 113958050 | 113959864 | 113959903 | 183 | 186 | 192 | 243 | 236 | 1 | 0 | 0 | 6 | 0 | 504 | 394 | 282 | 360 | 422 | 0 | 31 | 4 | 11 | 44 | 298 | 149 | 7.44E-11 | 1.15E-08 | 0.989 | 1 | 1 | 0.953 | 1 | 1 | 0.864 | 0.972 | 0.942 | 0.827 | 0.067 |
| SE | 87264 | ENSG00000147133 | TAF1 | chrX | + | 71388236 | 71388378 | 71387260 | 71387461 | 71388737 | 71388868 | 164 | 133 | 131 | 129 | 148 | 4 | 0 | 0 | 0 | 0 | 100 | 117 | 145 | 111 | 61 | 14 | 3 | 6 | 14 | 0 | 291 | 149 | 7.56E-11 | 1.17E-08 | 0.955 | 1 | 1 | 1 | 1 | 0.785 | 0.952 | 0.925 | 0.802 | 1 | 0.098 |
| SE | 92616 | ENSG00000083223 | TUT7 | chr9 | - | 86318957 | 86319058 | 86317218 | 86317276 | 86319583 | 86319644 | 417 | 436 | 266 | 306 | 530 | 20 | 50 | 23 | 26 | 43 | 30 | 170 | 187 | 46 | 29 | 10 | 47 | 80 | 9 | 4 | 250 | 149 | 7.85E-11 | 1.21E-08 | 0.926 | 0.839 | 0.873 | 0.875 | 0.88 | 0.641 | 0.683 | 0.582 | 0.753 | 0.812 | 0.184 |
| SE | 94583 | ENSG00000198771 | RCSD1 | chr1 | + | 167697098 | 167697842 | 167690048 | 167690120 | 167704663 | 167706248 | 390 | 228 | 369 | 300 | 238 | 0 | 0 | 0 | 0 | 0 | 383 | 443 | 177 | 292 | 490 | 30 | 9 | 2 | 5 | 2 | 298 | 149 | 7.89E-11 | 1.21E-08 | 1 | 1 | 1 | 1 | 1 | 0.865 | 0.961 | 0.978 | 0.967 | 0.992 | 0.047 |
| SE | 12932 | ENSG00000197818 | SLC9A8 | chr20 | + | 49849578 | 49849680 | 49845035 | 49845119 | 49855437 | 49855581 | 33 | 31 | 39 | 41 | 31 | 0 | 0 | 0 | 0 | 0 | 35 | 51 | 64 | 65 | 32 | 1 | 11 | 0 | 12 | 16 | 251 | 149 | 7.99E-11 | 1.23E-08 | 1 | 1 | 1 | 1 | 1 | 0.954 | 0.733 | 1 | 0.763 | 0.543 | 0.201 |
| SE | 4371 | ENSG00000118620 | ZNF430 | chr19 | + | 21022788 | 21022881 | 21020633 | 21020803 | 21033455 | 21033582 | 90 | 76 | 91 | 89 | 80 | 2 | 2 | 4 | 2 | 3 | 46 | 73 | 95 | 49 | 19 | 8 | 24 | 11 | 7 | 6 | 242 | 149 | 8.18E-11 | 1.25E-08 | 0.965 | 0.959 | 0.933 | 0.965 | 0.943 | 0.78 | 0.652 | 0.842 | 0.812 | 0.661 | 0.204 |
| SE | 49996 | ENSG00000241878 | PISD | chr22 | - | 31625762 | 31626016 | 31623663 | 31623849 | 31638365 | 31638405 | 37 | 7 | 26 | 19 | 28 | 0 | 0 | 0 | 0 | 0 | 22 | 13 | 9 | 39 | 19 | 0 | 5 | 7 | 2 | 21 | 298 | 149 | 8.38E-11 | 1.28E-08 | 1 | 1 | 1 | 1 | 1 | 1 | 0.565 | 0.391 | 0.907 | 0.311 | 0.365 |
| SE | 100368 | ENSG00000089597 | GANAB | chr11 | - | 62639358 | 62639467 | 62638982 | 62639110 | 62639626 | 62639731 | 718 | 369 | 641 | 638 | 485 | 5 | 7 | 9 | 16 | 0 | 689 | 613 | 435 | 451 | 938 | 41 | 12 | 39 | 19 | 18 | 258 | 149 | 8.48E-11 | 1.29E-08 | 0.988 | 0.968 | 0.976 | 0.958 | 1 | 0.907 | 0.967 | 0.866 | 0.932 | 0.968 | 0.05 |
| SE | 1074 | ENSG00000100330 | MTMR3 | chr22 | + | 30007113 | 30007319 | 30002879 | 30002993 | 30007900 | 30007975 | 220 | 102 | 152 | 172 | 256 | 1 | 3 | 2 | 1 | 2 | 331 | 302 | 285 | 369 | 336 | 16 | 11 | 19 | 35 | 6 | 298 | 149 | 8.62E-11 | 1.31E-08 | 0.991 | 0.944 | 0.974 | 0.989 | 0.985 | 0.912 | 0.932 | 0.882 | 0.841 | 0.966 | 0.07 |
| SE | 43469 | ENSG00000072786 | STK10 | chr5 | - | 172093411 | 172093960 | 172090231 | 172090362 | 172105655 | 172105737 | 414 | 216 | 365 | 416 | 277 | 0 | 3 | 0 | 1 | 0 | 390 | 462 | 275 | 326 | 692 | 33 | 5 | 5 | 1 | 15 | 298 | 149 | 8.82E-11 | 1.34E-08 | 1 | 0.973 | 1 | 0.995 | 1 | 0.855 | 0.979 | 0.965 | 0.994 | 0.958 | 0.043 |
| SE | 16006 | ENSG00000068305 | MEF2A | chr15 | + | 99671519 | 99671657 | 99671322 | 99671454 | 99674392 | 99674612 | 40 | 61 | 42 | 63 | 109 | 77 | 77 | 96 | 75 | 242 | 11 | 35 | 23 | 26 | 17 | 72 | 137 | 139 | 109 | 52 | 287 | 149 | 8.89E-11 | 1.35E-08 | 0.212 | 0.291 | 0.185 | 0.304 | 0.19 | 0.073 | 0.117 | 0.079 | 0.11 | 0.145 | 0.132 |
| SE | 41117 | ENSG00000115523 | GNLY | chr2 | + | 85694752 | 85694923 | 85694327 | 85694470 | 85695319 | 85695423 | 127 | 15 | 30 | 72 | 83 | 2496 | 480 | 702 | 845 | 838 | 2 | 19 | 25 | 7 | 10 | 645 | 335 | 714 | 334 | 509 | 298 | 149 | 8.98E-11 | 1.36E-08 | 0.025 | 0.015 | 0.021 | 0.041 | 0.047 | 0.002 | 0.028 | 0.017 | 0.01 | 0.01 | 0.016 |
| SE | 76466 | ENSG00000146476 | ARMT1 | chr6 | + | 151454547 | 151454653 | 151452465 | 151452586 | 151458327 | 151458572 | 117 | 61 | 63 | 49 | 140 | 2 | 5 | 3 | 5 | 0 | 0 | 36 | 13 | 5 | 2 | 5 | 8 | 5 | 4 | 7 | 255 | 149 | 9.38E-11 | 1.42E-08 | 0.972 | 0.877 | 0.925 | 0.851 | 1 | 0 | 0.724 | 0.603 | 0.422 | 0.143 | 0.547 |
| SE | 66654 | ENSG00000084623 | EIF3I | chr1 | + | 32226170 | 32226530 | 32224409 | 32224475 | 32228498 | 32228609 | 371 | 219 | 397 | 351 | 427 | 5 | 4 | 6 | 14 | 16 | 452 | 459 | 260 | 287 | 650 | 35 | 37 | 23 | 18 | 44 | 298 | 149 | 9.43E-11 | 1.42E-08 | 0.974 | 0.965 | 0.971 | 0.926 | 0.93 | 0.866 | 0.861 | 0.85 | 0.889 | 0.881 | 0.084 |
| SE | 27489 | ENSG00000104517 | UBR5 | chr8 | - | 102278825 | 102278939 | 102276971 | 102277192 | 102279035 | 102279208 | 371 | 335 | 332 | 435 | 388 | 5 | 12 | 0 | 9 | 3 | 55 | 361 | 349 | 178 | 21 | 53 | 30 | 65 | 23 | 0 | 263 | 149 | 9.61E-11 | 1.45E-08 | 0.977 | 0.941 | 1 | 0.965 | 0.987 | 0.37 | 0.872 | 0.753 | 0.814 | 1 | 0.212 |
| SE | 69231 | ENSG00000115484 | CCT4 | chr2 | - | 61877392 | 61877514 | 61876919 | 61877052 | 61878868 | 61879011 | 755 | 446 | 679 | 680 | 1201 | 3 | 4 | 4 | 11 | 15 | 460 | 439 | 378 | 344 | 225 | 23 | 20 | 7 | 11 | 9 | 271 | 149 | 9.63E-11 | 1.45E-08 | 0.993 | 0.984 | 0.989 | 0.971 | 0.978 | 0.917 | 0.923 | 0.967 | 0.945 | 0.932 | 0.046 |
| SE | 39351 | ENSG00000158864 | NDUFS2 | chr1 | + | 161210110 | 161210188 | 161209482 | 161209931 | 161210590 | 161210696 | 114 | 59 | 114 | 85 | 127 | 2 | 0 | 3 | 0 | 0 | 81 | 141 | 73 | 99 | 145 | 16 | 6 | 7 | 5 | 11 | 227 | 149 | 1.01E-10 | 1.52E-08 | 0.974 | 1 | 0.961 | 1 | 1 | 0.769 | 0.939 | 0.873 | 0.929 | 0.896 | 0.106 |
| SE | 40618 | ENSG00000213390 | ARHGAP19 | chr10 | - | 97256317 | 97256404 | 97243967 | 97244159 | 97259401 | 97259628 | 40 | 24 | 15 | 36 | 59 | 0 | 1 | 0 | 1 | 0 | 53 | 28 | 35 | 29 | 5 | 17 | 2 | 5 | 12 | 0 | 236 | 149 | 1.01E-10 | 1.52E-08 | 1 | 0.938 | 1 | 0.958 | 1 | 0.663 | 0.898 | 0.815 | 0.604 | 1 | 0.183 |
| SE | 25435 | ENSG00000143801 | PSEN2 | chr1 | + | 226883704 | 226883919 | 226880439 | 226882048 | 226885537 | 226885568 | 21 | 10 | 30 | 27 | 19 | 0 | 0 | 0 | 2 | 0 | 39 | 42 | 11 | 28 | 22 | 10 | 9 | 1 | 8 | 4 | 298 | 149 | 1.10E-10 | 1.64E-08 | 1 | 1 | 1 | 0.871 | 1 | 0.661 | 0.7 | 0.846 | 0.636 | 0.733 | 0.259 |
| SE | 19935 | ENSG00000092140 | G2E3 | chr14 | + | 30605504 | 30605812 | 30601998 | 30602131 | 30607887 | 30608002 | 125 | 116 | 87 | 102 | 156 | 11 | 9 | 13 | 9 | 19 | 5 | 39 | 47 | 15 | 1 | 0 | 0 | 0 | 0 | 0 | 298 | 149 | 1.13E-10 | 1.68E-08 | 0.85 | 0.866 | 0.77 | 0.85 | 0.804 | 1 | 1 | 1 | 1 | 1 | -0.172 |
| SE | 15641 | ENSG00000166411 | IDH3A | chr15 | + | 78164991 | 78165076 | 78163715 | 78163780 | 78166149 | 78166200 | 133 | 91 | 119 | 131 | 131 | 2 | 1 | 5 | 1 | 3 | 106 | 80 | 119 | 110 | 19 | 13 | 14 | 8 | 27 | 0 | 234 | 149 | 1.16E-10 | 1.73E-08 | 0.977 | 0.983 | 0.938 | 0.988 | 0.965 | 0.839 | 0.784 | 0.905 | 0.722 | 1 | 0.12 |
| SE | 22094 | ENSG00000111859 | NEDD9 | chr6 | - | 11305065 | 11305159 | 11213280 | 11213727 | 11305991 | 11306155 | 49 | 32 | 29 | 48 | 32 | 229 | 138 | 136 | 151 | 147 | 140 | 109 | 101 | 131 | 65 | 295 | 214 | 189 | 212 | 192 | 243 | 149 | 1.18E-10 | 1.75E-08 | 0.116 | 0.124 | 0.116 | 0.163 | 0.118 | 0.225 | 0.238 | 0.247 | 0.275 | 0.172 | -0.104 |
| SE | 33196 | ENSG00000163930 | BAP1 | chr3 | - | 52405109 | 52405294 | 52404452 | 52404586 | 52405764 | 52405912 | 202 | 102 | 207 | 232 | 222 | 0 | 0 | 1 | 2 | 3 | 291 | 306 | 125 | 181 | 309 | 18 | 6 | 14 | 9 | 7 | 298 | 149 | 1.18E-10 | 1.75E-08 | 1 | 1 | 0.99 | 0.983 | 0.974 | 0.89 | 0.962 | 0.817 | 0.91 | 0.957 | 0.082 |
| SE | 25886 | ENSG00000140350 | ANP32A | chr15 | - | 68787467 | 68787495 | 68784396 | 68784595 | 68787769 | 68787919 | 485 | 409 | 505 | 506 | 701 | 32 | 32 | 22 | 30 | 38 | 1157 | 725 | 500 | 826 | 439 | 116 | 67 | 172 | 121 | 89 | 232 | 149 | 1.20E-10 | 1.77E-08 | 0.907 | 0.891 | 0.936 | 0.915 | 0.922 | 0.865 | 0.874 | 0.651 | 0.814 | 0.76 | 0.121 |
| SE | 1199 | ENSG00000100227 | POLDIP3 | chr22 | - | 42601969 | 42602107 | 42599697 | 42599793 | 42602769 | 42603160 | 231 | 130 | 174 | 173 | 351 | 29 | 17 | 27 | 31 | 20 | 313 | 200 | 98 | 123 | 229 | 67 | 67 | 75 | 63 | 155 | 287 | 149 | 1.20E-10 | 1.77E-08 | 0.805 | 0.799 | 0.77 | 0.743 | 0.901 | 0.708 | 0.608 | 0.404 | 0.503 | 0.434 | 0.272 |
| SE | 99115 | ENSG00000130764 | LRRC47 | chr1 | - | 3783769 | 3783911 | 3782660 | 3782763 | 3783995 | 3784047 | 2 | 1 | 3 | 2 | 0 | 62 | 48 | 76 | 73 | 101 | 6 | 16 | 8 | 7 | 15 | 67 | 78 | 39 | 70 | 87 | 291 | 149 | 1.21E-10 | 1.78E-08 | 0.016 | 0.011 | 0.02 | 0.014 | 0 | 0.044 | 0.095 | 0.095 | 0.049 | 0.081 | -0.061 |
| SE | 104684 | ENSG00000100354 | TNRC6B | chr22 | + | 40281118 | 40281289 | 40279994 | 40280143 | 40285644 | 40285770 | 8 | 6 | 19 | 14 | 45 | 202 | 129 | 238 | 256 | 467 | 21 | 57 | 20 | 56 | 18 | 160 | 266 | 221 | 141 | 119 | 298 | 149 | 1.24E-10 | 1.82E-08 | 0.019 | 0.023 | 0.038 | 0.027 | 0.046 | 0.062 | 0.097 | 0.043 | 0.166 | 0.07 | -0.057 |
| SE | 81292 | ENSG00000155096 | AZIN1 | chr8 | - | 102849998 | 102850165 | 102843550 | 102843729 | 102858012 | 102858150 | 6 | 2 | 5 | 3 | 7 | 1 | 1 | 1 | 2 | 7 | 4 | 8 | 11 | 3 | 2 | 0 | 0 | 0 | 0 | 0 | 298 | 149 | 1.27E-10 | 1.86E-08 | 0.75 | 0.5 | 0.714 | 0.429 | 0.333 | 1 | 1 | 1 | 1 | 1 | -0.455 |
| SE | 11050 | ENSG00000185722 | ANKFY1 | chr17 | - | 4189381 | 4189479 | 4184817 | 4185046 | 4194977 | 4195177 | 119 | 86 | 87 | 128 | 112 | 1 | 0 | 1 | 0 | 0 | 160 | 99 | 77 | 147 | 147 | 10 | 5 | 7 | 12 | 4 | 247 | 149 | 1.29E-10 | 1.89E-08 | 0.986 | 1 | 0.981 | 1 | 1 | 0.906 | 0.923 | 0.869 | 0.881 | 0.957 | 0.086 |
| SE | 56835 | ENSG00000143398 | PIP5K1A | chr1 | + | 151242122 | 151242269 | 151239954 | 151240039 | 151242437 | 151242567 | 112 | 56 | 119 | 144 | 121 | 0 | 4 | 0 | 0 | 0 | 65 | 117 | 64 | 81 | 77 | 6 | 11 | 4 | 3 | 8 | 296 | 149 | 1.33E-10 | 1.94E-08 | 1 | 0.876 | 1 | 1 | 1 | 0.845 | 0.843 | 0.89 | 0.931 | 0.829 | 0.108 |
| SE | 64130 | ENSG00000163126 | ANKRD23 | chr2 | - | 96840826 | 96840912 | 96840415 | 96840514 | 96842059 | 96842117 | 4 | 2 | 1 | 4 | 5 | 0 | 0 | 1 | 0 | 1 | 0 | 8 | 1 | 0 | 0 | 6 | 0 | 2 | 6 | 3 | 235 | 149 | 1.37E-10 | 2.00E-08 | 1 | 1 | 0.388 | 1 | 0.76 | 0 | 1 | 0.241 | 0 | 0 | 0.581 |
| SE | 8747 | ENSG00000007202 | KIAA0100 | chr17 | - | 28616888 | 28616999 | 28616617 | 28616753 | 28617231 | 28617318 | 153 | 74 | 150 | 161 | 117 | 0 | 0 | 0 | 2 | 0 | 257 | 174 | 73 | 131 | 314 | 8 | 10 | 5 | 8 | 14 | 260 | 149 | 1.39E-10 | 2.03E-08 | 1 | 1 | 1 | 0.979 | 1 | 0.948 | 0.909 | 0.893 | 0.904 | 0.928 | 0.079 |
| SE | 26530 | ENSG00000138303 | ASCC1 | chr10 | - | 72198448 | 72198665 | 72196810 | 72196989 | 72203426 | 72203524 | 2 | 0 | 2 | 0 | 0 | 37 | 24 | 68 | 56 | 51 | 8 | 8 | 6 | 0 | 12 | 30 | 46 | 21 | 17 | 10 | 298 | 149 | 1.41E-10 | 2.05E-08 | 0.026 | 0 | 0.014 | 0 | 0 | 0.118 | 0.08 | 0.125 | 0 | 0.375 | -0.132 |
| SE | 5593 | ENSG00000126001 | CEP250 | chr20 | + | 35500048 | 35500169 | 35498594 | 35498716 | 35501844 | 35501966 | 28 | 27 | 40 | 32 | 27 | 0 | 0 | 5 | 4 | 0 | 12 | 8 | 11 | 26 | 6 | 7 | 8 | 3 | 13 | 14 | 270 | 149 | 1.41E-10 | 2.05E-08 | 1 | 1 | 0.815 | 0.815 | 1 | 0.486 | 0.356 | 0.669 | 0.525 | 0.191 | 0.481 |
| SE | 97733 | ENSG00000095139 | ARCN1 | chr11 | + | 118581245 | 118581509 | 118572408 | 118572550 | 118583178 | 118583358 | 392 | 309 | 490 | 367 | 329 | 3 | 2 | 12 | 14 | 0 | 344 | 265 | 257 | 271 | 154 | 35 | 8 | 9 | 37 | 4 | 298 | 149 | 1.43E-10 | 2.07E-08 | 0.985 | 0.987 | 0.953 | 0.929 | 1 | 0.831 | 0.943 | 0.935 | 0.786 | 0.951 | 0.082 |
| SE | 97660 | ENSG00000147457 | CHMP7 | chr8 | + | 23258329 | 23258449 | 23258032 | 23258081 | 23258731 | 23258830 | 224 | 125 | 286 | 240 | 314 | 2 | 1 | 1 | 1 | 1 | 108 | 193 | 92 | 76 | 144 | 4 | 6 | 10 | 8 | 4 | 269 | 149 | 1.44E-10 | 2.09E-08 | 0.984 | 0.986 | 0.994 | 0.993 | 0.994 | 0.937 | 0.947 | 0.836 | 0.84 | 0.952 | 0.088 |
| SE | 15807 | ENSG00000169375 | SIN3A | chr15 | - | 75400729 | 75400940 | 75400039 | 75400156 | 75401851 | 75401970 | 252 | 156 | 223 | 313 | 236 | 0 | 1 | 2 | 4 | 6 | 111 | 189 | 219 | 128 | 112 | 9 | 8 | 3 | 10 | 22 | 298 | 149 | 1.46E-10 | 2.10E-08 | 1 | 0.987 | 0.982 | 0.975 | 0.952 | 0.86 | 0.922 | 0.973 | 0.865 | 0.718 | 0.112 |
| SE | 95883 | ENSG00000148396 | SEC16A | chr9 | - | 136463678 | 136463740 | 136461176 | 136461274 | 136464419 | 136464562 | 57 | 27 | 41 | 29 | 42 | 0 | 3 | 0 | 1 | 0 | 32 | 27 | 23 | 19 | 36 | 0 | 5 | 24 | 7 | 15 | 211 | 149 | 1.46E-10 | 2.11E-08 | 1 | 0.864 | 1 | 0.953 | 1 | 1 | 0.792 | 0.404 | 0.657 | 0.629 | 0.267 |
| SE | 30218 | ENSG00000132376 | INPP5K | chr17 | - | 1509682 | 1509799 | 1508976 | 1509353 | 1513452 | 1513512 | 200 | 83 | 132 | 138 | 273 | 2 | 1 | 4 | 3 | 7 | 132 | 168 | 106 | 124 | 238 | 16 | 10 | 12 | 17 | 17 | 266 | 149 | 1.51E-10 | 2.18E-08 | 0.982 | 0.979 | 0.949 | 0.963 | 0.956 | 0.822 | 0.904 | 0.832 | 0.803 | 0.887 | 0.116 |
| SE | 2450 | ENSG00000105127 | AKAP8 | chr19 | - | 15369066 | 15369224 | 15368234 | 15368322 | 15370145 | 15370179 | 33 | 27 | 45 | 16 | 32 | 56 | 51 | 73 | 57 | 59 | 59 | 4 | 26 | 41 | 18 | 250 | 77 | 149 | 146 | 143 | 298 | 149 | 1.52E-10 | 2.18E-08 | 0.228 | 0.209 | 0.236 | 0.123 | 0.213 | 0.106 | 0.025 | 0.08 | 0.123 | 0.059 | 0.123 |
| SE | 1335 | ENSG00000093000 | NUP50 | chr22 | + | 45175893 | 45176080 | 45171599 | 45171683 | 45178237 | 45178430 | 548 | 428 | 527 | 543 | 1077 | 20 | 22 | 18 | 12 | 26 | 330 | 512 | 277 | 401 | 168 | 24 | 29 | 43 | 33 | 14 | 298 | 149 | 1.54E-10 | 2.21E-08 | 0.932 | 0.907 | 0.936 | 0.958 | 0.954 | 0.873 | 0.898 | 0.763 | 0.859 | 0.857 | 0.087 |
| SE | 58335 | ENSG00000115207 | GTF3C2 | chr2 | - | 27350313 | 27350632 | 27343330 | 27343578 | 27356738 | 27356999 | 3 | 1 | 1 | 1 | 0 | 23 | 20 | 33 | 25 | 14 | 5 | 15 | 9 | 6 | 12 | 35 | 16 | 13 | 22 | 20 | 298 | 149 | 1.65E-10 | 2.36E-08 | 0.061 | 0.024 | 0.015 | 0.02 | 0 | 0.067 | 0.319 | 0.257 | 0.12 | 0.231 | -0.175 |
| SE | 8850 | ENSG00000136436 | CALCOCO2 | chr17 | + | 48852505 | 48852628 | 48851558 | 48851628 | 48852925 | 48853012 | 502 | 500 | 422 | 635 | 947 | 4 | 1 | 5 | 4 | 0 | 802 | 520 | 594 | 770 | 181 | 20 | 2 | 16 | 31 | 1 | 272 | 149 | 1.69E-10 | 2.41E-08 | 0.986 | 0.996 | 0.979 | 0.989 | 1 | 0.956 | 0.993 | 0.953 | 0.932 | 0.99 | 0.025 |
| SE | 83511 | ENSG00000196313 | POM121 | chr7 | + | 72928384 | 72928465 | 72926801 | 72926963 | 72929939 | 72930111 | 320 | 211 | 268 | 323 | 523 | 1 | 1 | 1 | 3 | 4 | 519 | 415 | 291 | 382 | 487 | 17 | 4 | 7 | 55 | 0 | 230 | 149 | 1.69E-10 | 2.41E-08 | 0.995 | 0.993 | 0.994 | 0.986 | 0.988 | 0.952 | 0.985 | 0.964 | 0.818 | 1 | 0.047 |
| SE | 67573 | ENSG00000076003 | MCM6 | chr2 | - | 135866131 | 135866277 | 135865012 | 135865163 | 135866562 | 135866728 | 197 | 153 | 203 | 249 | 165 | 6 | 0 | 1 | 2 | 0 | 343 | 267 | 302 | 352 | 213 | 12 | 11 | 25 | 29 | 0 | 295 | 149 | 1.71E-10 | 2.43E-08 | 0.943 | 1 | 0.99 | 0.984 | 1 | 0.935 | 0.925 | 0.859 | 0.86 | 1 | 0.068 |
| SE | 12283 | ENSG00000103194 | USP10 | chr16 | + | 84772540 | 84772685 | 84768192 | 84768358 | 84775159 | 84775225 | 429 | 370 | 417 | 403 | 622 | 2 | 3 | 4 | 10 | 5 | 418 | 370 | 248 | 375 | 301 | 16 | 17 | 19 | 2 | 21 | 294 | 149 | 1.72E-10 | 2.45E-08 | 0.991 | 0.984 | 0.981 | 0.953 | 0.984 | 0.93 | 0.917 | 0.869 | 0.99 | 0.879 | 0.062 |
| SE | 86204 | ENSG00000172534 | HCFC1 | chrX | - | 153950812 | 153950998 | 153950242 | 153950543 | 153951349 | 153951487 | 169 | 85 | 131 | 148 | 172 | 0 | 0 | 0 | 1 | 0 | 162 | 182 | 155 | 146 | 257 | 10 | 3 | 0 | 25 | 6 | 298 | 149 | 1.76E-10 | 2.49E-08 | 1 | 1 | 1 | 0.987 | 1 | 0.89 | 0.968 | 1 | 0.745 | 0.955 | 0.086 |
| SE | 34925 | ENSG00000076108 | BAZ2A | chr12 | - | 56605829 | 56606063 | 56605072 | 56605327 | 56606246 | 56606312 | 213 | 148 | 191 | 200 | 334 | 0 | 0 | 0 | 1 | 4 | 622 | 333 | 307 | 301 | 352 | 18 | 11 | 14 | 14 | 8 | 298 | 149 | 1.83E-10 | 2.60E-08 | 1 | 1 | 1 | 0.99 | 0.977 | 0.945 | 0.938 | 0.916 | 0.915 | 0.957 | 0.059 |
| SE | 105443 | ENSG00000145901 | TNIP1 | chr5 | - | 151080072 | 151080359 | 151064959 | 151065131 | 151080879 | 151081015 | 4 | 2 | 2 | 0 | 1 | 48 | 12 | 23 | 32 | 10 | 44 | 6 | 9 | 21 | 29 | 54 | 44 | 14 | 30 | 96 | 298 | 149 | 1.86E-10 | 2.63E-08 | 0.04 | 0.077 | 0.042 | 0 | 0.048 | 0.289 | 0.064 | 0.243 | 0.259 | 0.131 | -0.156 |
| SE | 59153 | ENSG00000171492 | LRRC8D | chr1 | + | 89929807 | 89929878 | 89843637 | 89843782 | 89933066 | 89933160 | 8 | 2 | 7 | 13 | 1 | 45 | 31 | 57 | 71 | 25 | 0 | 0 | 0 | 0 | 0 | 63 | 50 | 49 | 52 | 45 | 220 | 149 | 1.93E-10 | 2.72E-08 | 0.107 | 0.042 | 0.077 | 0.11 | 0.026 | 0 | 0 | 0 | 0 | 0 | 0.072 |
| SE | 105486 | ENSG00000152256 | PDK1 | chr2 | + | 172564502 | 172564687 | 172562740 | 172562800 | 172564977 | 172565073 | 71 | 89 | 142 | 117 | 292 | 1 | 0 | 5 | 2 | 3 | 40 | 198 | 107 | 63 | 130 | 1 | 15 | 13 | 7 | 15 | 298 | 149 | 1.98E-10 | 2.78E-08 | 0.973 | 1 | 0.934 | 0.967 | 0.98 | 0.952 | 0.868 | 0.805 | 0.818 | 0.812 | 0.12 |
| SE | 81249 | ENSG00000115677 | HDLBP | chr2 | - | 241238648 | 241238787 | 241236614 | 241236769 | 241239601 | 241239736 | 477 | 308 | 436 | 494 | 362 | 3 | 2 | 4 | 6 | 7 | 1051 | 605 | 497 | 642 | 769 | 15 | 39 | 21 | 55 | 7 | 288 | 149 | 1.98E-10 | 2.79E-08 | 0.988 | 0.988 | 0.983 | 0.977 | 0.964 | 0.973 | 0.889 | 0.924 | 0.858 | 0.983 | 0.055 |
| SE | 94582 | ENSG00000198771 | RCSD1 | chr1 | + | 167694098 | 167694302 | 167690048 | 167690120 | 167704663 | 167706248 | 657 | 321 | 556 | 439 | 507 | 0 | 0 | 0 | 0 | 0 | 549 | 537 | 263 | 276 | 630 | 30 | 9 | 2 | 5 | 2 | 298 | 149 | 2.06E-10 | 2.89E-08 | 1 | 1 | 1 | 1 | 1 | 0.901 | 0.968 | 0.985 | 0.965 | 0.994 | 0.037 |
| SE | 65556 | ENSG00000153201 | RANBP2 | chr2 | + | 108762100 | 108762195 | 108758412 | 108758548 | 108763236 | 108768388 | 260 | 178 | 269 | 244 | 459 | 1 | 5 | 9 | 5 | 7 | 0 | 91 | 49 | 21 | 2 | 5 | 10 | 9 | 9 | 8 | 244 | 149 | 2.09E-10 | 2.93E-08 | 0.994 | 0.956 | 0.948 | 0.968 | 0.976 | 0 | 0.847 | 0.769 | 0.588 | 0.132 | 0.501 |
| SE | 45444 | ENSG00000170791 | CHCHD7 | chr8 | + | 56216679 | 56216754 | 56216388 | 56216531 | 56217330 | 56217412 | 12 | 7 | 20 | 18 | 26 | 99 | 85 | 112 | 91 | 274 | 22 | 12 | 24 | 19 | 15 | 32 | 45 | 39 | 59 | 27 | 224 | 149 | 2.16E-10 | 3.03E-08 | 0.075 | 0.052 | 0.106 | 0.116 | 0.059 | 0.314 | 0.151 | 0.29 | 0.176 | 0.27 | -0.159 |
| SE | 49392 | ENSG00000243943 | ZNF512 | chr2 | + | 27599969 | 27600053 | 27599582 | 27599678 | 27600690 | 27600728 | 180 | 148 | 246 | 182 | 290 | 10 | 2 | 3 | 5 | 0 | 205 | 167 | 139 | 123 | 51 | 10 | 10 | 26 | 13 | 4 | 233 | 149 | 2.21E-10 | 3.08E-08 | 0.92 | 0.979 | 0.981 | 0.959 | 1 | 0.929 | 0.914 | 0.774 | 0.858 | 0.891 | 0.095 |
| SE | 80399 | ENSG00000197694 | SPTAN1 | chr9 | + | 128592982 | 128593042 | 128591476 | 128591625 | 128594174 | 128594373 | 586 | 346 | 560 | 584 | 642 | 16 | 9 | 21 | 13 | 5 | 365 | 608 | 387 | 483 | 646 | 56 | 55 | 15 | 17 | 25 | 209 | 149 | 2.21E-10 | 3.08E-08 | 0.963 | 0.965 | 0.95 | 0.97 | 0.989 | 0.823 | 0.887 | 0.948 | 0.953 | 0.949 | 0.055 |
| SE | 78196 | ENSG00000124486 | USP9X | chrX | + | 41150920 | 41151057 | 41148368 | 41148575 | 41152947 | 41153081 | 368 | 263 | 292 | 324 | 615 | 4 | 2 | 3 | 4 | 2 | 55 | 257 | 275 | 149 | 15 | 24 | 8 | 20 | 6 | 0 | 286 | 149 | 2.22E-10 | 3.09E-08 | 0.98 | 0.986 | 0.981 | 0.977 | 0.994 | 0.544 | 0.944 | 0.878 | 0.928 | 1 | 0.125 |
| SE | 67470 | ENSG00000162923 | WDR26 | chr1 | - | 224413252 | 224413335 | 224411426 | 224411565 | 224418259 | 224418416 | 20 | 20 | 16 | 21 | 18 | 656 | 484 | 471 | 726 | 895 | 39 | 54 | 67 | 21 | 45 | 615 | 716 | 905 | 629 | 396 | 232 | 149 | 2.25E-10 | 3.12E-08 | 0.019 | 0.026 | 0.021 | 0.018 | 0.013 | 0.039 | 0.046 | 0.045 | 0.021 | 0.068 | -0.024 |
| SE | 12190 | ENSG00000141002 | TCF25 | chr16 | + | 89883350 | 89883512 | 89873569 | 89873859 | 89884581 | 89884613 | 437 | 277 | 447 | 420 | 276 | 0 | 0 | 0 | 0 | 0 | 986 | 675 | 425 | 570 | 1148 | 2 | 17 | 5 | 16 | 33 | 298 | 149 | 2.33E-10 | 3.23E-08 | 1 | 1 | 1 | 1 | 1 | 0.996 | 0.952 | 0.977 | 0.947 | 0.946 | 0.036 |
| SE | 78879 | ENSG00000196792 | STRN3 | chr14 | - | 30929200 | 30929311 | 30912006 | 30912182 | 30935162 | 30935304 | 28 | 37 | 26 | 39 | 53 | 3 | 4 | 0 | 4 | 12 | 85 | 58 | 51 | 56 | 35 | 0 | 0 | 1 | 0 | 0 | 260 | 149 | 2.41E-10 | 3.33E-08 | 0.842 | 0.841 | 1 | 0.848 | 0.717 | 1 | 1 | 0.967 | 1 | 1 | -0.144 |
| SE | 53067 | ENSG00000103657 | HERC1 | chr15 | - | 63713352 | 63713665 | 63706779 | 63706831 | 63716301 | 63716473 | 60 | 68 | 89 | 75 | 71 | 0 | 0 | 1 | 0 | 0 | 39 | 72 | 62 | 81 | 54 | 0 | 20 | 6 | 0 | 8 | 298 | 149 | 2.46E-10 | 3.40E-08 | 1 | 1 | 0.978 | 1 | 1 | 1 | 0.643 | 0.838 | 1 | 0.771 | 0.145 |
| SE | 95282 | ENSG00000172716 | SLFN11 | chr17 | - | 35366946 | 35367043 | 35363438 | 35363826 | 35367602 | 35367700 | 71 | 49 | 41 | 52 | 49 | 0 | 1 | 1 | 9 | 0 | 44 | 48 | 45 | 35 | 28 | 10 | 12 | 13 | 9 | 10 | 246 | 149 | 2.50E-10 | 3.45E-08 | 1 | 0.967 | 0.961 | 0.778 | 1 | 0.727 | 0.708 | 0.677 | 0.702 | 0.629 | 0.253 |
| SE | 43317 | ENSG00000009413 | REV3L | chr6 | - | 111472012 | 111472140 | 111416282 | 111416472 | 111482749 | 111482919 | 56 | 51 | 49 | 50 | 124 | 11 | 6 | 11 | 10 | 3 | 19 | 33 | 21 | 12 | 4 | 0 | 0 | 0 | 0 | 0 | 277 | 149 | 2.57E-10 | 3.54E-08 | 0.733 | 0.821 | 0.706 | 0.729 | 0.957 | 1 | 1 | 1 | 1 | 1 | -0.211 |
| SE | 80647 | ENSG00000166997 | CNPY4 | chr7 | + | 100122480 | 100122577 | 100122051 | 100122385 | 100122783 | 100122906 | 82 | 49 | 88 | 86 | 67 | 0 | 0 | 0 | 0 | 0 | 59 | 59 | 26 | 44 | 40 | 7 | 9 | 2 | 0 | 4 | 246 | 149 | 2.59E-10 | 3.56E-08 | 1 | 1 | 1 | 1 | 1 | 0.836 | 0.799 | 0.887 | 1 | 0.858 | 0.124 |
| SE | 28192 | ENSG00000164190 | NIPBL | chr5 | + | 36995621 | 36995804 | 36984675 | 36986301 | 37000816 | 37000888 | 419 | 388 | 286 | 382 | 686 | 9 | 4 | 4 | 5 | 0 | 11 | 93 | 156 | 27 | 0 | 6 | 7 | 11 | 6 | 7 | 298 | 149 | 2.62E-10 | 3.59E-08 | 0.959 | 0.98 | 0.973 | 0.974 | 1 | 0.478 | 0.869 | 0.876 | 0.692 | 0 | 0.394 |
| SE | 5965 | ENSG00000170471 | RALGAPB | chr20 | + | 38548688 | 38548795 | 38546242 | 38546430 | 38551070 | 38551223 | 222 | 210 | 177 | 202 | 256 | 5 | 3 | 3 | 4 | 4 | 219 | 197 | 135 | 140 | 124 | 12 | 23 | 33 | 8 | 3 | 256 | 149 | 2.66E-10 | 3.64E-08 | 0.963 | 0.976 | 0.972 | 0.967 | 0.974 | 0.914 | 0.833 | 0.704 | 0.911 | 0.96 | 0.106 |
| SE | 101464 | ENSG00000148481 | MINDY3 | chr10 | - | 15837541 | 15837633 | 15837203 | 15837318 | 15838227 | 15838279 | 30 | 21 | 34 | 18 | 35 | 129 | 42 | 66 | 95 | 98 | 0 | 7 | 26 | 21 | 9 | 168 | 79 | 86 | 115 | 217 | 241 | 149 | 2.70E-10 | 3.69E-08 | 0.126 | 0.236 | 0.242 | 0.105 | 0.181 | 0 | 0.052 | 0.157 | 0.101 | 0.025 | 0.111 |
| SE | 2277 | ENSG00000104824 | HNRNPL | chr19 | - | 38841612 | 38841682 | 38840487 | 38840559 | 38843841 | 38843914 | 55 | 36 | 51 | 67 | 100 | 829 | 461 | 790 | 831 | 1718 | 21 | 37 | 37 | 61 | 26 | 1058 | 1208 | 971 | 889 | 1433 | 219 | 149 | 2.73E-10 | 3.73E-08 | 0.043 | 0.05 | 0.042 | 0.052 | 0.038 | 0.013 | 0.02 | 0.025 | 0.045 | 0.012 | 0.022 |
| SE | 18365 | ENSG00000100888 | CHD8 | chr14 | - | 21391462 | 21391642 | 21390946 | 21391063 | 21391832 | 21391946 | 399 | 227 | 298 | 311 | 544 | 5 | 3 | 5 | 17 | 22 | 267 | 240 | 163 | 219 | 101 | 28 | 47 | 76 | 76 | 1 | 298 | 149 | 2.75E-10 | 3.74E-08 | 0.976 | 0.974 | 0.968 | 0.901 | 0.925 | 0.827 | 0.719 | 0.517 | 0.59 | 0.981 | 0.222 |
| SE | 35109 | ENSG00000092931 | MFSD11 | chr17 | + | 76743397 | 76743456 | 76742176 | 76742273 | 76744321 | 76744466 | 59 | 33 | 46 | 54 | 75 | 0 | 0 | 1 | 1 | 0 | 96 | 37 | 42 | 64 | 42 | 42 | 13 | 3 | 2 | 0 | 208 | 149 | 2.75E-10 | 3.74E-08 | 1 | 1 | 0.971 | 0.975 | 1 | 0.621 | 0.671 | 0.909 | 0.958 | 1 | 0.157 |
| SE | 86451 | ENSG00000102103 | PQBP1 | chrX | + | 48902218 | 48902517 | 48901929 | 48902042 | 48902731 | 48902795 | 85 | 42 | 81 | 59 | 96 | 3 | 2 | 1 | 2 | 0 | 70 | 61 | 42 | 37 | 94 | 37 | 5 | 14 | 3 | 12 | 298 | 149 | 2.78E-10 | 3.77E-08 | 0.934 | 0.913 | 0.976 | 0.937 | 1 | 0.486 | 0.859 | 0.6 | 0.86 | 0.797 | 0.232 |
| SE | 34823 | ENSG00000115827 | DCAF17 | chr2 | + | 171447385 | 171447503 | 171443522 | 171443613 | 171448680 | 171448817 | 13 | 6 | 8 | 6 | 17 | 53 | 46 | 56 | 36 | 75 | 0 | 0 | 2 | 0 | 0 | 40 | 34 | 30 | 27 | 9 | 267 | 149 | 2.80E-10 | 3.79E-08 | 0.12 | 0.068 | 0.074 | 0.085 | 0.112 | 0 | 0 | 0.036 | 0 | 0 | 0.085 |
| SE | 5383 | ENSG00000078747 | ITCH | chr20 | + | 34392736 | 34392914 | 34369393 | 34369470 | 34393790 | 34393881 | 6 | 1 | 5 | 3 | 1 | 199 | 92 | 170 | 226 | 285 | 11 | 14 | 34 | 5 | 0 | 34 | 314 | 192 | 158 | 12 | 298 | 149 | 2.86E-10 | 3.87E-08 | 0.015 | 0.005 | 0.014 | 0.007 | 0.002 | 0.139 | 0.022 | 0.081 | 0.016 | 0 | -0.043 |
| SE | 8760 | ENSG00000007202 | KIAA0100 | chr17 | - | 28641888 | 28642078 | 28640559 | 28640683 | 28642274 | 28642335 | 417 | 272 | 310 | 379 | 463 | 1 | 2 | 2 | 1 | 1 | 956 | 369 | 275 | 481 | 652 | 24 | 19 | 10 | 12 | 3 | 298 | 149 | 2.98E-10 | 4.02E-08 | 0.995 | 0.986 | 0.987 | 0.995 | 0.996 | 0.952 | 0.907 | 0.932 | 0.952 | 0.991 | 0.045 |
| SE | 41984 | ENSG00000172113 | NME6 | chr3 | - | 48295074 | 48295235 | 48290721 | 48294803 | 48296726 | 48296829 | 35 | 17 | 19 | 25 | 32 | 0 | 0 | 0 | 0 | 1 | 52 | 31 | 12 | 14 | 20 | 14 | 3 | 0 | 4 | 5 | 298 | 149 | 3.06E-10 | 4.13E-08 | 1 | 1 | 1 | 1 | 0.941 | 0.65 | 0.838 | 1 | 0.636 | 0.667 | 0.23 |
| SE | 52818 | ENSG00000169764 | UGP2 | chr2 | + | 63857549 | 63857718 | 63856305 | 63856433 | 63857828 | 63857936 | 40 | 31 | 49 | 28 | 43 | 348 | 258 | 303 | 395 | 315 | 28 | 54 | 64 | 38 | 28 | 120 | 249 | 195 | 174 | 71 | 298 | 149 | 3.12E-10 | 4.20E-08 | 0.054 | 0.057 | 0.075 | 0.034 | 0.064 | 0.104 | 0.098 | 0.141 | 0.098 | 0.165 | -0.064 |
| SE | 49283 | ENSG00000120733 | KDM3B | chr5 | + | 138386288 | 138386621 | 138381515 | 138381590 | 138391012 | 138392261 | 92 | 49 | 97 | 89 | 85 | 3 | 3 | 1 | 6 | 6 | 93 | 130 | 88 | 105 | 113 | 48 | 12 | 20 | 24 | 30 | 298 | 149 | 3.13E-10 | 4.20E-08 | 0.939 | 0.891 | 0.98 | 0.881 | 0.876 | 0.492 | 0.844 | 0.688 | 0.686 | 0.653 | 0.241 |
| SE | 7849 | ENSG00000108819 | PPP1R9B | chr17 | - | 50140092 | 50140228 | 50139428 | 50139581 | 50141268 | 50141373 | 1140 | 730 | 876 | 1169 | 590 | 10 | 0 | 4 | 4 | 3 | 2141 | 1398 | 712 | 1236 | 2603 | 54 | 10 | 9 | 31 | 34 | 285 | 149 | 3.19E-10 | 4.28E-08 | 0.983 | 1 | 0.991 | 0.993 | 0.99 | 0.954 | 0.987 | 0.976 | 0.954 | 0.976 | 0.022 |
| SE | 70695 | ENSG00000114268 | PFKFB4 | chr3 | - | 48543579 | 48543646 | 48539410 | 48539771 | 48549863 | 48549960 | 53 | 24 | 35 | 51 | 41 | 0 | 0 | 0 | 0 | 0 | 157 | 52 | 42 | 67 | 118 | 3 | 11 | 2 | 17 | 7 | 216 | 149 | 3.19E-10 | 4.28E-08 | 1 | 1 | 1 | 1 | 1 | 0.973 | 0.765 | 0.935 | 0.731 | 0.921 | 0.135 |
| SE | 52208 | ENSG00000127463 | EMC1 | chr1 | - | 19231260 | 19231422 | 19230843 | 19230963 | 19232623 | 19232773 | 144 | 54 | 103 | 88 | 149 | 0 | 0 | 0 | 2 | 10 | 147 | 70 | 61 | 99 | 92 | 23 | 15 | 1 | 11 | 17 | 298 | 149 | 3.26E-10 | 4.35E-08 | 1 | 1 | 1 | 0.957 | 0.882 | 0.762 | 0.7 | 0.968 | 0.818 | 0.73 | 0.172 |
| SE | 94730 | ENSG00000106948 | AKNA | chr9 | - | 114356862 | 114356969 | 114355924 | 114356136 | 114357920 | 114358167 | 475 | 173 | 298 | 396 | 229 | 1 | 1 | 1 | 0 | 2 | 352 | 609 | 341 | 225 | 624 | 19 | 15 | 12 | 6 | 8 | 256 | 149 | 3.28E-10 | 4.37E-08 | 0.996 | 0.99 | 0.994 | 1 | 0.985 | 0.915 | 0.959 | 0.943 | 0.956 | 0.978 | 0.043 |
| SE | 32854 | ENSG00000175105 | ZNF654 | chr3 | + | 88126133 | 88126269 | 88113114 | 88113196 | 88129686 | 88129826 | 25 | 8 | 21 | 20 | 22 | 16 | 2 | 7 | 7 | 7 | 10 | 19 | 10 | 3 | 10 | 0 | 0 | 2 | 0 | 0 | 285 | 149 | 3.33E-10 | 4.44E-08 | 0.45 | 0.677 | 0.611 | 0.599 | 0.622 | 1 | 1 | 0.723 | 1 | 1 | -0.353 |
| SE | 101651 | ENSG00000128245 | YWHAH | chr22 | + | 31945556 | 31945678 | 31944746 | 31944820 | 31956138 | 31956604 | 0 | 14 | 0 | 29 | 0 | 134 | 106 | 65 | 148 | 61 | 0 | 0 | 0 | 0 | 0 | 268 | 249 | 160 | 244 | 1704 | 271 | 149 | 3.34E-10 | 4.44E-08 | 0 | 0.068 | 0 | 0.097 | 0 | 0 | 0 | 0 | 0 | 0 | 0.033 |
| SE | 96757 | ENSG00000183431 | SF3A3 | chr1 | - | 37978987 | 37979055 | 37978719 | 37978827 | 37979464 | 37979533 | 321 | 243 | 284 | 299 | 472 | 2 | 4 | 2 | 1 | 2 | 407 | 321 | 251 | 287 | 294 | 25 | 3 | 1 | 37 | 0 | 217 | 149 | 3.35E-10 | 4.44E-08 | 0.991 | 0.977 | 0.99 | 0.995 | 0.994 | 0.918 | 0.987 | 0.994 | 0.842 | 1 | 0.041 |
| SE | 71294 | ENSG00000064012 | CASP8 | chr2 | + | 201272897 | 201272942 | 201272637 | 201272776 | 201274888 | 201274953 | 937 | 456 | 573 | 708 | 919 | 100 | 78 | 79 | 93 | 120 | 728 | 834 | 1290 | 637 | 227 | 62 | 64 | 44 | 53 | 26 | 194 | 149 | 3.52E-10 | 4.67E-08 | 0.878 | 0.818 | 0.848 | 0.854 | 0.855 | 0.9 | 0.909 | 0.957 | 0.902 | 0.87 | -0.057 |
| SE | 61854 | ENSG00000143393 | PI4KB | chr1 | - | 151310210 | 151310255 | 151307573 | 151307801 | 151326171 | 151326373 | 7 | 4 | 12 | 5 | 14 | 0 | 1 | 0 | 0 | 1 | 0 | 12 | 4 | 0 | 11 | 10 | 5 | 10 | 7 | 0 | 194 | 149 | 3.57E-10 | 4.71E-08 | 1 | 0.754 | 1 | 1 | 0.915 | 0 | 0.648 | 0.235 | 0 | 1 | 0.557 |
| SE | 47999 | ENSG00000110046 | ATG2A | chr11 | - | 64908990 | 64909150 | 64907747 | 64907890 | 64909270 | 64909367 | 125 | 66 | 70 | 67 | 65 | 0 | 0 | 0 | 1 | 0 | 61 | 124 | 168 | 89 | 127 | 10 | 3 | 14 | 10 | 0 | 298 | 149 | 3.57E-10 | 4.71E-08 | 1 | 1 | 1 | 0.971 | 1 | 0.753 | 0.954 | 0.857 | 0.817 | 1 | 0.118 |
| SE | 47684 | ENSG00000162869 | PPP1R21 | chr2 | + | 48471278 | 48471367 | 48471086 | 48471188 | 48474682 | 48474819 | 139 | 118 | 136 | 148 | 302 | 0 | 2 | 0 | 1 | 0 | 55 | 104 | 74 | 97 | 31 | 11 | 5 | 8 | 2 | 0 | 238 | 149 | 3.58E-10 | 4.71E-08 | 1 | 0.974 | 1 | 0.989 | 1 | 0.758 | 0.929 | 0.853 | 0.968 | 1 | 0.091 |
| SE | 51357 | ENSG00000066427 | ATXN3 | chr14 | - | 92083416 | 92083503 | 92083125 | 92083258 | 92088729 | 92088817 | 0 | 3 | 1 | 6 | 3 | 145 | 102 | 98 | 126 | 158 | 0 | 8 | 15 | 11 | 6 | 26 | 98 | 77 | 66 | 18 | 236 | 149 | 3.64E-10 | 4.79E-08 | 0 | 0.018 | 0.006 | 0.029 | 0.012 | 0 | 0.049 | 0.11 | 0.095 | 0.174 | -0.073 |
| SE | 3746 | ENSG00000187116 | LILRA5 | chr19 | - | 54312334 | 54312370 | 54311863 | 54312148 | 54312536 | 54312621 | 150 | 57 | 61 | 91 | 40 | 0 | 2 | 0 | 1 | 0 | 262 | 395 | 86 | 157 | 359 | 16 | 23 | 18 | 22 | 2 | 185 | 149 | 3.68E-10 | 4.83E-08 | 1 | 0.958 | 1 | 0.987 | 1 | 0.93 | 0.933 | 0.794 | 0.852 | 0.993 | 0.089 |
| SE | 12030 | ENSG00000176715 | ACSF3 | chr16 | + | 89112091 | 89112246 | 89102603 | 89102759 | 89133135 | 89133262 | 67 | 14 | 46 | 50 | 42 | 0 | 0 | 0 | 0 | 0 | 35 | 41 | 26 | 29 | 66 | 16 | 3 | 2 | 6 | 0 | 298 | 149 | 3.68E-10 | 4.83E-08 | 1 | 1 | 1 | 1 | 1 | 0.522 | 0.872 | 0.867 | 0.707 | 1 | 0.206 |
| SE | 39732 | ENSG00000132716 | DCAF8 | chr1 | - | 160261284 | 160261358 | 160243959 | 160244034 | 160262448 | 160262531 | 105 | 119 | 78 | 109 | 114 | 11 | 20 | 15 | 24 | 0 | 246 | 122 | 143 | 167 | 174 | 0 | 0 | 1 | 5 | 15 | 223 | 149 | 3.82E-10 | 5.00E-08 | 0.864 | 0.799 | 0.777 | 0.752 | 1 | 1 | 1 | 0.99 | 0.957 | 0.886 | -0.128 |
| SE | 64367 | ENSG00000163219 | ARHGAP25 | chr2 | + | 68813288 | 68813419 | 68807272 | 68807480 | 68816288 | 68816362 | 220 | 58 | 187 | 179 | 161 | 8 | 4 | 11 | 9 | 2 | 49 | 172 | 107 | 95 | 130 | 10 | 12 | 26 | 24 | 20 | 280 | 149 | 3.87E-10 | 5.06E-08 | 0.936 | 0.885 | 0.9 | 0.914 | 0.977 | 0.723 | 0.884 | 0.687 | 0.678 | 0.776 | 0.173 |
| SE | 48910 | ENSG00000204165 | CXorf65 | chrX | - | 71104768 | 71104837 | 71104297 | 71104404 | 71105999 | 71106137 | 17 | 19 | 30 | 22 | 45 | 2 | 0 | 3 | 0 | 2 | 43 | 30 | 48 | 28 | 28 | 14 | 6 | 14 | 24 | 23 | 218 | 149 | 3.95E-10 | 5.14E-08 | 0.853 | 1 | 0.872 | 1 | 0.939 | 0.677 | 0.774 | 0.701 | 0.444 | 0.454 | 0.323 |
| SE | 61526 | ENSG00000138757 | G3BP2 | chr4 | - | 75656923 | 75656962 | 75655767 | 75655870 | 75657556 | 75657730 | 28 | 17 | 40 | 36 | 113 | 2 | 0 | 0 | 4 | 4 | 0 | 38 | 45 | 30 | 0 | 5 | 29 | 10 | 34 | 2 | 188 | 149 | 3.95E-10 | 5.14E-08 | 0.917 | 1 | 1 | 0.877 | 0.957 | 0 | 0.509 | 0.781 | 0.412 | 0 | 0.61 |
| SE | 61860 | ENSG00000143393 | PI4KB | chr1 | - | 151324744 | 151324917 | 151307573 | 151307801 | 151326171 | 151326373 | 48 | 23 | 21 | 40 | 50 | 0 | 1 | 0 | 0 | 1 | 44 | 43 | 17 | 17 | 63 | 10 | 5 | 10 | 7 | 0 | 298 | 149 | 4.00E-10 | 5.20E-08 | 1 | 0.92 | 1 | 1 | 0.962 | 0.688 | 0.811 | 0.459 | 0.548 | 1 | 0.275 |
| SE | 51151 | ENSG00000165458 | INPPL1 | chr11 | + | 72233440 | 72233522 | 72233074 | 72233163 | 72233654 | 72233744 | 224 | 109 | 164 | 186 | 151 | 0 | 0 | 3 | 0 | 0 | 411 | 380 | 144 | 239 | 668 | 13 | 24 | 7 | 9 | 15 | 231 | 149 | 4.03E-10 | 5.24E-08 | 1 | 1 | 0.972 | 1 | 1 | 0.953 | 0.911 | 0.93 | 0.945 | 0.966 | 0.053 |
| SE | 74149 | ENSG00000111679 | PTPN6 | chr12 | + | 6960662 | 6960945 | 6960343 | 6960435 | 6961125 | 6961316 | 978 | 753 | 748 | 867 | 1278 | 3 | 0 | 2 | 6 | 0 | 1736 | 1283 | 601 | 1285 | 1902 | 48 | 4 | 7 | 31 | 10 | 298 | 149 | 4.20E-10 | 5.45E-08 | 0.994 | 1 | 0.995 | 0.986 | 1 | 0.948 | 0.994 | 0.977 | 0.954 | 0.99 | 0.022 |
| SE | 85157 | ENSG00000136504 | KAT7 | chr17 | + | 49805362 | 49805445 | 49798318 | 49798558 | 49809118 | 49809208 | 406 | 205 | 349 | 402 | 582 | 8 | 2 | 12 | 21 | 12 | 605 | 502 | 420 | 547 | 512 | 60 | 26 | 31 | 54 | 22 | 232 | 149 | 4.34E-10 | 5.61E-08 | 0.97 | 0.985 | 0.949 | 0.925 | 0.969 | 0.866 | 0.925 | 0.897 | 0.867 | 0.937 | 0.061 |
| SE | 35100 | ENSG00000234857 | HNRNPUL2-BSCL2 | chr11 | - | 62715863 | 62715937 | 62715499 | 62715607 | 62716988 | 62717189 | 660 | 475 | 693 | 654 | 890 | 23 | 24 | 37 | 22 | 30 | 1032 | 618 | 630 | 887 | 866 | 110 | 55 | 67 | 64 | 45 | 223 | 149 | 4.49E-10 | 5.80E-08 | 0.95 | 0.93 | 0.926 | 0.952 | 0.952 | 0.862 | 0.882 | 0.863 | 0.903 | 0.928 | 0.054 |
| SE | 24003 | ENSG00000112983 | BRD8 | chr5 | - | 138159554 | 138159599 | 138156881 | 138157267 | 138160068 | 138160129 | 7 | 10 | 18 | 18 | 23 | 89 | 64 | 93 | 95 | 167 | 0 | 6 | 2 | 2 | 0 | 130 | 77 | 69 | 58 | 4 | 194 | 149 | 4.56E-10 | 5.88E-08 | 0.057 | 0.107 | 0.129 | 0.127 | 0.096 | 0 | 0.056 | 0.022 | 0.026 | 0 | 0.082 |
| SE | 23632 | ENSG00000135074 | ADAM19 | chr5 | - | 157519838 | 157520031 | 157513423 | 157513505 | 157530806 | 157530883 | 126 | 71 | 148 | 122 | 149 | 1 | 1 | 2 | 4 | 1 | 258 | 275 | 160 | 174 | 260 | 19 | 26 | 11 | 6 | 20 | 298 | 149 | 4.65E-10 | 5.99E-08 | 0.984 | 0.973 | 0.974 | 0.938 | 0.987 | 0.872 | 0.841 | 0.879 | 0.935 | 0.867 | 0.092 |
| SE | 55019 | ENSG00000152223 | EPG5 | chr18 | - | 45887750 | 45887907 | 45884616 | 45884811 | 45889797 | 45889940 | 90 | 39 | 63 | 69 | 43 | 4 | 0 | 1 | 0 | 1 | 31 | 35 | 43 | 50 | 9 | 6 | 4 | 6 | 11 | 4 | 298 | 149 | 4.67E-10 | 6.01E-08 | 0.918 | 1 | 0.969 | 1 | 0.956 | 0.721 | 0.814 | 0.782 | 0.694 | 0.529 | 0.261 |
| SE | 34073 | ENSG00000114867 | EIF4G1 | chr3 | + | 184317716 | 184317816 | 184314550 | 184314674 | 184319688 | 184319801 | 114 | 44 | 65 | 99 | 69 | 0 | 0 | 0 | 1 | 0 | 106 | 63 | 92 | 100 | 145 | 10 | 0 | 6 | 11 | 7 | 249 | 149 | 4.70E-10 | 6.03E-08 | 1 | 1 | 1 | 0.983 | 1 | 0.864 | 1 | 0.902 | 0.845 | 0.925 | 0.089 |
| SE | 100030 | ENSG00000101972 | STAG2 | chrX | + | 124026564 | 124026687 | 124025839 | 124025918 | 124030960 | 124031125 | 36 | 28 | 34 | 58 | 49 | 234 | 215 | 223 | 271 | 453 | 30 | 12 | 25 | 12 | 29 | 21 | 71 | 42 | 10 | 20 | 272 | 149 | 4.70E-10 | 6.03E-08 | 0.078 | 0.067 | 0.077 | 0.105 | 0.056 | 0.439 | 0.085 | 0.246 | 0.397 | 0.443 | -0.245 |
| SE | 98205 | ENSG00000149273 | RPS3 | chr11 | + | 75402303 | 75402446 | 75401639 | 75401733 | 75404019 | 75404207 | 2009 | 948 | 2419 | 2020 | 3617 | 19 | 17 | 23 | 17 | 60 | 902 | 1291 | 1337 | 899 | 1630 | 40 | 29 | 26 | 38 | 35 | 292 | 149 | 4.73E-10 | 6.05E-08 | 0.982 | 0.966 | 0.982 | 0.984 | 0.969 | 0.92 | 0.958 | 0.963 | 0.924 | 0.96 | 0.032 |
| SE | 5502 | ENSG00000101294 | HM13 | chr20 | + | 31544946 | 31545035 | 31538178 | 31538261 | 31550063 | 31550121 | 83 | 27 | 44 | 57 | 62 | 0 | 0 | 0 | 1 | 0 | 165 | 70 | 100 | 111 | 131 | 13 | 11 | 8 | 0 | 12 | 238 | 149 | 4.78E-10 | 6.11E-08 | 1 | 1 | 1 | 0.973 | 1 | 0.888 | 0.799 | 0.887 | 1 | 0.872 | 0.105 |
| SE | 41749 | ENSG00000143222 | UFC1 | chr1 | + | 161157253 | 161157317 | 161156949 | 161157017 | 161157616 | 161157693 | 566 | 231 | 430 | 395 | 605 | 2 | 8 | 8 | 7 | 10 | 451 | 389 | 308 | 406 | 273 | 44 | 20 | 6 | 16 | 11 | 213 | 149 | 4.85E-10 | 6.18E-08 | 0.995 | 0.953 | 0.974 | 0.975 | 0.977 | 0.878 | 0.932 | 0.973 | 0.947 | 0.946 | 0.04 |
| SE | 8719 | ENSG00000070366 | SMG6 | chr17 | - | 2292870 | 2292977 | 2292551 | 2292630 | 2297242 | 2297353 | 108 | 95 | 99 | 80 | 125 | 0 | 0 | 2 | 0 | 3 | 263 | 124 | 140 | 227 | 282 | 0 | 16 | 7 | 12 | 37 | 256 | 149 | 4.89E-10 | 6.22E-08 | 1 | 1 | 0.966 | 1 | 0.96 | 1 | 0.819 | 0.921 | 0.917 | 0.816 | 0.091 |
| SE | 37157 | ENSG00000151726 | ACSL1 | chr4 | - | 184773080 | 184773154 | 184770398 | 184770476 | 184773662 | 184773714 | 192 | 142 | 111 | 138 | 294 | 0 | 0 | 4 | 0 | 0 | 302 | 398 | 697 | 435 | 154 | 0 | 10 | 28 | 43 | 0 | 223 | 149 | 4.89E-10 | 6.22E-08 | 1 | 1 | 0.949 | 1 | 1 | 1 | 0.964 | 0.943 | 0.871 | 1 | 0.034 |
| SE | 35955 | ENSG00000109861 | CTSC | chr11 | - | 88328111 | 88328196 | 88312387 | 88312554 | 88334936 | 88335082 | 134 | 68 | 76 | 93 | 97 | 364 | 218 | 189 | 226 | 213 | 74 | 94 | 36 | 35 | 129 | 574 | 325 | 181 | 317 | 430 | 234 | 149 | 4.90E-10 | 6.22E-08 | 0.19 | 0.166 | 0.204 | 0.208 | 0.225 | 0.076 | 0.156 | 0.112 | 0.066 | 0.16 | 0.085 |
| SE | 7468 | ENSG00000073605 | GSDMB | chr17 | - | 39906110 | 39906271 | 39905846 | 39905985 | 39909755 | 39909924 | 57 | 46 | 37 | 78 | 24 | 1 | 0 | 0 | 0 | 0 | 50 | 34 | 22 | 50 | 59 | 8 | 2 | 5 | 10 | 0 | 298 | 149 | 4.91E-10 | 6.23E-08 | 0.966 | 1 | 1 | 1 | 1 | 0.758 | 0.895 | 0.688 | 0.714 | 1 | 0.182 |
| SE | 49606 | ENSG00000076685 | NT5C2 | chr10 | - | 103139405 | 103139479 | 103106588 | 103106706 | 103174857 | 103174982 | 220 | 183 | 178 | 258 | 396 | 9 | 4 | 8 | 6 | 5 | 431 | 207 | 161 | 353 | 155 | 20 | 23 | 28 | 19 | 22 | 223 | 149 | 5.11E-10 | 6.46E-08 | 0.942 | 0.968 | 0.937 | 0.966 | 0.981 | 0.935 | 0.857 | 0.793 | 0.925 | 0.825 | 0.092 |
| SE | 31663 | ENSG00000131389 | SLC6A6 | chr3 | + | 14478465 | 14478568 | 14477204 | 14477342 | 14479084 | 14479185 | 159 | 91 | 133 | 185 | 174 | 0 | 0 | 0 | 0 | 6 | 342 | 219 | 164 | 144 | 353 | 29 | 0 | 8 | 4 | 23 | 252 | 149 | 5.21E-10 | 6.57E-08 | 1 | 1 | 1 | 1 | 0.945 | 0.875 | 1 | 0.924 | 0.955 | 0.901 | 0.058 |
| SE | 35191 | ENSG00000101596 | SMCHD1 | chr18 | + | 2750042 | 2750122 | 2747521 | 2747647 | 2750349 | 2750507 | 609 | 602 | 600 | 831 | 1022 | 9 | 17 | 15 | 34 | 10 | 157 | 355 | 437 | 106 | 25 | 5 | 30 | 34 | 12 | 5 | 229 | 149 | 5.30E-10 | 6.68E-08 | 0.978 | 0.958 | 0.963 | 0.941 | 0.985 | 0.953 | 0.885 | 0.893 | 0.852 | 0.765 | 0.095 |
| SE | 15463 | ENSG00000140199 | SLC12A6 | chr15 | - | 34242101 | 34242221 | 34241062 | 34241337 | 34243973 | 34244072 | 180 | 126 | 199 | 171 | 311 | 4 | 3 | 1 | 1 | 0 | 261 | 106 | 152 | 124 | 91 | 0 | 6 | 39 | 26 | 0 | 269 | 149 | 5.48E-10 | 6.90E-08 | 0.961 | 0.959 | 0.991 | 0.99 | 1 | 1 | 0.907 | 0.683 | 0.725 | 1 | 0.117 |
| SE | 95037 | ENSG00000060237 | WNK1 | chr12 | + | 882942 | 883059 | 881910 | 882073 | 883394 | 883419 | 1258 | 1172 | 898 | 1161 | 1208 | 8 | 2 | 5 | 9 | 3 | 1675 | 1268 | 1471 | 1687 | 727 | 24 | 26 | 25 | 19 | 19 | 266 | 149 | 5.60E-10 | 7.04E-08 | 0.989 | 0.997 | 0.99 | 0.986 | 0.996 | 0.975 | 0.965 | 0.971 | 0.98 | 0.955 | 0.022 |
| SE | 81161 | ENSG00000176142 | TMEM39A | chr3 | - | 119458017 | 119458083 | 119452508 | 119452530 | 119461961 | 119462148 | 15 | 5 | 10 | 14 | 15 | 0 | 0 | 2 | 0 | 0 | 7 | 3 | 13 | 10 | 13 | 0 | 13 | 7 | 9 | 7 | 215 | 149 | 5.83E-10 | 7.31E-08 | 1 | 1 | 0.776 | 1 | 1 | 1 | 0.138 | 0.563 | 0.435 | 0.563 | 0.415 |
| SE | 99904 | ENSG00000073910 | FRY | chr13 | + | 32179674 | 32179799 | 32178843 | 32179033 | 32182976 | 32183034 | 120 | 128 | 79 | 105 | 135 | 0 | 1 | 0 | 1 | 0 | 124 | 91 | 97 | 130 | 25 | 22 | 0 | 7 | 5 | 0 | 274 | 149 | 5.84E-10 | 7.32E-08 | 1 | 0.986 | 1 | 0.983 | 1 | 0.754 | 1 | 0.883 | 0.934 | 1 | 0.08 |
| SE | 70231 | ENSG00000160695 | VPS11 | chr11 | + | 119069441 | 119069577 | 119067820 | 119068202 | 119070233 | 119070397 | 156 | 89 | 159 | 155 | 189 | 2 | 0 | 2 | 0 | 1 | 265 | 173 | 153 | 219 | 215 | 17 | 0 | 3 | 22 | 10 | 285 | 149 | 5.96E-10 | 7.45E-08 | 0.976 | 1 | 0.977 | 1 | 0.99 | 0.891 | 1 | 0.964 | 0.839 | 0.918 | 0.066 |
| SE | 60131 | ENSG00000162373 | BEND5 | chr1 | - | 48742622 | 48742771 | 48736238 | 48736452 | 48758899 | 48759284 | 7 | 5 | 25 | 13 | 44 | 4 | 7 | 6 | 1 | 6 | 12 | 25 | 5 | 12 | 24 | 0 | 0 | 0 | 1 | 0 | 298 | 149 | 6.02E-10 | 7.51E-08 | 0.467 | 0.263 | 0.676 | 0.867 | 0.786 | 1 | 1 | 1 | 0.857 | 1 | -0.36 |
| SE | 58383 | ENSG00000232956 | SNHG15 | chr7 | - | 44984356 | 44984426 | 44983022 | 44984046 | 44986020 | 44986097 | 33 | 13 | 52 | 38 | 21 | 0 | 0 | 0 | 0 | 0 | 47 | 25 | 39 | 44 | 50 | 33 | 0 | 5 | 3 | 5 | 219 | 149 | 6.05E-10 | 7.54E-08 | 1 | 1 | 1 | 1 | 1 | 0.492 | 1 | 0.841 | 0.909 | 0.872 | 0.177 |
| SE | 81507 | ENSG00000092820 | EZR | chr6 | - | 158787107 | 158787203 | 158785308 | 158785583 | 158789287 | 158789371 | 1176 | 787 | 1019 | 1066 | 1956 | 5 | 12 | 8 | 13 | 8 | 1402 | 1381 | 1451 | 1026 | 1458 | 40 | 50 | 23 | 16 | 17 | 245 | 149 | 6.06E-10 | 7.55E-08 | 0.993 | 0.976 | 0.987 | 0.98 | 0.993 | 0.955 | 0.944 | 0.975 | 0.975 | 0.981 | 0.02 |
| SE | 5297 | ENSG00000101473 | ACOT8 | chr20 | - | 45853922 | 45853984 | 45848449 | 45848675 | 45855158 | 45855292 | 4 | 0 | 0 | 1 | 0 | 36 | 21 | 27 | 37 | 25 | 10 | 16 | 2 | 3 | 17 | 41 | 22 | 25 | 17 | 61 | 211 | 149 | 6.27E-10 | 7.80E-08 | 0.073 | 0 | 0 | 0.019 | 0 | 0.147 | 0.339 | 0.053 | 0.111 | 0.164 | -0.144 |
| SE | 103545 | ENSG00000107862 | GBF1 | chr10 | + | 102369860 | 102369984 | 102369710 | 102369775 | 102370173 | 102370245 | 112 | 69 | 76 | 137 | 101 | 6 | 3 | 0 | 4 | 3 | 62 | 104 | 35 | 73 | 196 | 22 | 21 | 12 | 38 | 5 | 273 | 149 | 6.34E-10 | 7.86E-08 | 0.911 | 0.926 | 1 | 0.949 | 0.948 | 0.606 | 0.73 | 0.614 | 0.512 | 0.955 | 0.263 |
| SE | 71551 | ENSG00000129691 | ASH2L | chr8 | + | 38116649 | 38116725 | 38114757 | 38115000 | 38119269 | 38119295 | 455 | 337 | 384 | 403 | 575 | 5 | 5 | 2 | 3 | 3 | 817 | 393 | 321 | 631 | 768 | 34 | 10 | 17 | 26 | 7 | 225 | 149 | 6.46E-10 | 8.00E-08 | 0.984 | 0.978 | 0.992 | 0.989 | 0.992 | 0.941 | 0.963 | 0.926 | 0.941 | 0.986 | 0.036 |
| SE | 24394 | ENSG00000169220 | RGS14 | chr5 | + | 177371474 | 177371589 | 177371349 | 177371396 | 177371872 | 177372166 | 279 | 128 | 181 | 228 | 198 | 16 | 14 | 20 | 17 | 5 | 252 | 226 | 90 | 170 | 255 | 44 | 42 | 19 | 34 | 48 | 264 | 149 | 6.57E-10 | 8.13E-08 | 0.908 | 0.838 | 0.836 | 0.883 | 0.957 | 0.764 | 0.752 | 0.728 | 0.738 | 0.75 | 0.138 |
| SE | 43468 | ENSG00000072786 | STK10 | chr5 | - | 172093411 | 172093960 | 172090231 | 172090362 | 172096425 | 172096560 | 895 | 381 | 653 | 707 | 625 | 1 | 0 | 1 | 1 | 0 | 592 | 790 | 441 | 517 | 1032 | 0 | 14 | 3 | 8 | 35 | 298 | 149 | 6.60E-10 | 8.15E-08 | 0.998 | 1 | 0.997 | 0.997 | 1 | 1 | 0.966 | 0.987 | 0.97 | 0.936 | 0.027 |
| SE | 76200 | ENSG00000112237 | CCNC | chr6 | - | 99550217 | 99550309 | 99549507 | 99549575 | 99550992 | 99551028 | 193 | 121 | 174 | 164 | 319 | 1 | 0 | 2 | 1 | 0 | 44 | 86 | 111 | 36 | 4 | 8 | 6 | 9 | 0 | 0 | 241 | 149 | 6.61E-10 | 8.15E-08 | 0.992 | 1 | 0.982 | 0.99 | 1 | 0.773 | 0.899 | 0.884 | 1 | 1 | 0.082 |
| SE | 51320 | ENSG00000144136 | SLC20A1 | chr2 | + | 112647323 | 112647464 | 112646562 | 112647162 | 112647652 | 112647738 | 250 | 125 | 180 | 166 | 126 | 5 | 2 | 1 | 7 | 0 | 80 | 154 | 105 | 121 | 115 | 0 | 11 | 19 | 22 | 10 | 290 | 149 | 6.67E-10 | 8.22E-08 | 0.963 | 0.97 | 0.989 | 0.924 | 1 | 1 | 0.878 | 0.74 | 0.739 | 0.855 | 0.127 |
| SE | 105043 | ENSG00000133316 | WDR74 | chr11 | - | 62833617 | 62833674 | 62832910 | 62833131 | 62833791 | 62833937 | 62 | 53 | 67 | 63 | 86 | 3 | 1 | 4 | 5 | 3 | 84 | 87 | 53 | 57 | 54 | 20 | 30 | 5 | 12 | 38 | 206 | 149 | 6.69E-10 | 8.23E-08 | 0.937 | 0.975 | 0.924 | 0.901 | 0.954 | 0.752 | 0.677 | 0.885 | 0.775 | 0.507 | 0.219 |
| SE | 95312 | ENSG00000125484 | GTF3C4 | chr9 | + | 132683562 | 132683693 | 132677976 | 132679803 | 132687238 | 132687327 | 73 | 45 | 80 | 68 | 135 | 0 | 0 | 1 | 0 | 4 | 51 | 61 | 27 | 48 | 97 | 5 | 7 | 9 | 14 | 0 | 280 | 149 | 6.86E-10 | 8.41E-08 | 1 | 1 | 0.977 | 1 | 0.947 | 0.844 | 0.823 | 0.615 | 0.646 | 1 | 0.199 |
| SE | 45920 | ENSG00000160058 | BSDC1 | chr1 | - | 32369229 | 32369338 | 32368446 | 32368550 | 32376261 | 32376741 | 2 | 0 | 3 | 3 | 0 | 88 | 42 | 65 | 74 | 34 | 23 | 3 | 0 | 4 | 31 | 71 | 40 | 51 | 43 | 152 | 258 | 149 | 6.93E-10 | 8.49E-08 | 0.013 | 0 | 0.026 | 0.023 | 0 | 0.158 | 0.042 | 0 | 0.051 | 0.105 | -0.059 |
| SE | 49426 | ENSG00000136068 | FLNB | chr3 | + | 58097814 | 58097977 | 58096140 | 58096218 | 58098710 | 58098908 | 188 | 107 | 191 | 250 | 236 | 0 | 0 | 1 | 1 | 0 | 377 | 179 | 184 | 81 | 215 | 0 | 6 | 8 | 16 | 8 | 298 | 149 | 6.95E-10 | 8.51E-08 | 1 | 1 | 0.99 | 0.992 | 1 | 1 | 0.937 | 0.92 | 0.717 | 0.931 | 0.095 |
| SE | 43470 | ENSG00000072786 | STK10 | chr5 | - | 172096425 | 172096560 | 172090231 | 172090362 | 172105655 | 172105737 | 162 | 70 | 121 | 144 | 98 | 0 | 3 | 0 | 1 | 0 | 79 | 167 | 123 | 100 | 163 | 33 | 5 | 5 | 1 | 15 | 284 | 149 | 7.15E-10 | 8.74E-08 | 1 | 0.924 | 1 | 0.987 | 1 | 0.557 | 0.946 | 0.928 | 0.981 | 0.851 | 0.13 |
| SE | 92059 | ENSG00000136573 | BLK | chr8 | + | 11543223 | 11543347 | 11494386 | 11494591 | 11546051 | 11546103 | 246 | 159 | 256 | 278 | 383 | 17 | 6 | 10 | 7 | 17 | 312 | 439 | 128 | 107 | 464 | 20 | 75 | 45 | 22 | 46 | 273 | 149 | 7.43E-10 | 9.04E-08 | 0.888 | 0.935 | 0.933 | 0.956 | 0.925 | 0.895 | 0.762 | 0.608 | 0.726 | 0.846 | 0.16 |
| SE | 4457 | ENSG00000181896 | ZNF101 | chr19 | + | 19678725 | 19678786 | 19677863 | 19677990 | 19679180 | 19679523 | 131 | 91 | 241 | 174 | 297 | 5 | 3 | 1 | 6 | 5 | 79 | 143 | 93 | 83 | 61 | 27 | 28 | 5 | 1 | 6 | 210 | 149 | 7.43E-10 | 9.04E-08 | 0.949 | 0.956 | 0.994 | 0.954 | 0.977 | 0.675 | 0.784 | 0.93 | 0.983 | 0.878 | 0.116 |
| SE | 23587 | ENSG00000169567 | HINT1 | chr5 | - | 131162063 | 131162477 | 131159291 | 131159611 | 131162571 | 131162676 | 51 | 24 | 63 | 37 | 111 | 380 | 179 | 430 | 380 | 991 | 52 | 47 | 41 | 64 | 55 | 257 | 213 | 145 | 147 | 282 | 298 | 149 | 7.44E-10 | 9.04E-08 | 0.063 | 0.063 | 0.068 | 0.046 | 0.053 | 0.092 | 0.099 | 0.124 | 0.179 | 0.089 | -0.058 |
| SE | 81944 | ENSG00000051825 | MPHOSPH9 | chr12 | - | 123162114 | 123162218 | 123161135 | 123161383 | 123163949 | 123163961 | 68 | 47 | 84 | 67 | 107 | 1 | 3 | 2 | 5 | 3 | 65 | 44 | 55 | 28 | 6 | 2 | 20 | 30 | 17 | 3 | 253 | 149 | 7.64E-10 | 9.28E-08 | 0.976 | 0.902 | 0.961 | 0.888 | 0.955 | 0.95 | 0.564 | 0.519 | 0.492 | 0.541 | 0.323 |
| SE | 39811 | ENSG00000144711 | IQSEC1 | chr3 | - | 12902772 | 12902822 | 12897042 | 12901522 | 12908348 | 12908525 | 83 | 58 | 79 | 71 | 65 | 380 | 200 | 318 | 303 | 384 | 27 | 47 | 39 | 75 | 110 | 578 | 497 | 339 | 409 | 673 | 199 | 149 | 7.76E-10 | 9.41E-08 | 0.141 | 0.178 | 0.157 | 0.149 | 0.112 | 0.034 | 0.066 | 0.079 | 0.121 | 0.109 | 0.066 |
| SE | 80168 | ENSG00000012124 | CD22 | chr19 | + | 35338167 | 35338431 | 35329186 | 35329230 | 35340880 | 35341138 | 150 | 98 | 142 | 163 | 275 | 5 | 1 | 5 | 8 | 5 | 310 | 405 | 59 | 116 | 386 | 19 | 36 | 18 | 17 | 53 | 298 | 149 | 7.90E-10 | 9.56E-08 | 0.938 | 0.98 | 0.934 | 0.911 | 0.965 | 0.891 | 0.849 | 0.621 | 0.773 | 0.785 | 0.162 |
| SE | 9241 | ENSG00000178971 | CTC1 | chr17 | - | 8229890 | 8229968 | 8229301 | 8229446 | 8230293 | 8230397 | 73 | 49 | 68 | 92 | 82 | 0 | 0 | 1 | 1 | 1 | 43 | 42 | 21 | 74 | 109 | 0 | 1 | 2 | 10 | 19 | 227 | 149 | 7.98E-10 | 9.64E-08 | 1 | 1 | 0.978 | 0.984 | 0.982 | 1 | 0.965 | 0.873 | 0.829 | 0.79 | 0.097 |
| SE | 85069 | ENSG00000147459 | DOCK5 | chr8 | + | 25368570 | 25368725 | 25366869 | 25366970 | 25369555 | 25369641 | 140 | 130 | 69 | 155 | 153 | 4 | 1 | 1 | 0 | 5 | 159 | 146 | 216 | 137 | 56 | 20 | 5 | 15 | 24 | 3 | 298 | 149 | 8.00E-10 | 9.64E-08 | 0.946 | 0.985 | 0.972 | 1 | 0.939 | 0.799 | 0.936 | 0.878 | 0.741 | 0.903 | 0.117 |
| SE | 42270 | ENSG00000160752 | FDPS | chr1 | + | 155309788 | 155309965 | 155308865 | 155308965 | 155310042 | 155310205 | 12 | 0 | 14 | 9 | 0 | 31 | 20 | 33 | 25 | 16 | 0 | 10 | 0 | 0 | 0 | 174 | 46 | 25 | 23 | 60 | 298 | 149 | 8.01E-10 | 9.64E-08 | 0.162 | 0 | 0.175 | 0.153 | 0 | 0 | 0.098 | 0 | 0 | 0 | 0.078 |
| SE | 63814 | ENSG00000181163 | NPM1 | chr5 | + | 171406396 | 171407161 | 171405301 | 171405403 | 171407699 | 171407774 | 171 | 162 | 197 | 182 | 354 | 1680 | 1039 | 1950 | 1516 | 2515 | 79 | 145 | 173 | 109 | 13 | 250 | 788 | 1017 | 454 | 32 | 298 | 149 | 8.15E-10 | 9.79E-08 | 0.048 | 0.072 | 0.048 | 0.057 | 0.066 | 0.136 | 0.084 | 0.078 | 0.107 | 0.169 | -0.057 |
| SE | 103102 | ENSG00000095585 | BLNK | chr10 | - | 96215320 | 96215389 | 96209837 | 96209907 | 96216652 | 96216734 | 20 | 36 | 34 | 43 | 68 | 6 | 6 | 5 | 1 | 6 | 15 | 64 | 19 | 33 | 40 | 0 | 0 | 0 | 0 | 0 | 218 | 149 | 8.16E-10 | 9.79E-08 | 0.695 | 0.804 | 0.823 | 0.967 | 0.886 | 1 | 1 | 1 | 1 | 1 | -0.165 |
| SE | 46216 | ENSG00000114439 | BBX | chr3 | + | 107772627 | 107773636 | 107755597 | 107755678 | 107774718 | 107774857 | 274 | 237 | 248 | 327 | 352 | 3 | 0 | 1 | 3 | 1 | 44 | 303 | 212 | 67 | 6 | 6 | 7 | 17 | 3 | 9 | 298 | 149 | 8.18E-10 | 9.80E-08 | 0.979 | 1 | 0.992 | 0.982 | 0.994 | 0.786 | 0.956 | 0.862 | 0.918 | 0.25 | 0.235 |
| SE | 89357 | ENSG00000115419 | GLS | chr2 | + | 190920214 | 190920248 | 190913192 | 190913364 | 190921023 | 190921056 | 15 | 12 | 13 | 15 | 12 | 28 | 19 | 40 | 27 | 49 | 39 | 24 | 16 | 14 | 34 | 0 | 0 | 5 | 5 | 41 | 183 | 149 | 8.47E-10 | 1.01E-07 | 0.304 | 0.34 | 0.209 | 0.311 | 0.166 | 1 | 1 | 0.723 | 0.695 | 0.403 | -0.498 |
| SE | 18710 | ENSG00000100678 | SLC8A3 | chr14 | - | 70060835 | 70060939 | 70051989 | 70052114 | 70166638 | 70168484 | 9 | 13 | 10 | 22 | 11 | 0 | 2 | 0 | 0 | 0 | 4 | 26 | 15 | 18 | 100 | 7 | 10 | 3 | 6 | 21 | 253 | 149 | 8.60E-10 | 1.03E-07 | 1 | 0.793 | 1 | 1 | 1 | 0.252 | 0.605 | 0.746 | 0.639 | 0.737 | 0.363 |
| SE | 68478 | ENSG00000242372 | EIF6 | chr20 | - | 35280653 | 35280829 | 35279941 | 35280118 | 35284175 | 35284261 | 185 | 74 | 151 | 173 | 168 | 6 | 6 | 2 | 3 | 4 | 145 | 169 | 66 | 107 | 239 | 29 | 20 | 10 | 14 | 12 | 298 | 149 | 9.07E-10 | 1.08E-07 | 0.939 | 0.86 | 0.974 | 0.966 | 0.955 | 0.714 | 0.809 | 0.767 | 0.793 | 0.909 | 0.14 |
| SE | 66659 | ENSG00000084623 | EIF3I | chr1 | + | 32229134 | 32229208 | 32228726 | 32228816 | 32230926 | 32231019 | 215 | 130 | 239 | 227 | 203 | 0 | 0 | 0 | 6 | 3 | 217 | 148 | 156 | 173 | 430 | 13 | 5 | 5 | 11 | 21 | 223 | 149 | 9.11E-10 | 1.09E-07 | 1 | 1 | 1 | 0.962 | 0.978 | 0.918 | 0.952 | 0.954 | 0.913 | 0.932 | 0.054 |
| SE | 81895 | ENSG00000177189 | RPS6KA3 | chrX | - | 20240568 | 20240605 | 20234757 | 20234814 | 20266563 | 20266606 | 0 | 2 | 3 | 0 | 0 | 55 | 32 | 57 | 70 | 32 | 10 | 8 | 12 | 0 | 0 | 22 | 38 | 50 | 8 | 3 | 186 | 149 | 9.13E-10 | 1.09E-07 | 0 | 0.048 | 0.04 | 0 | 0 | 0.267 | 0.144 | 0.161 | 0 | 0 | -0.097 |
| SE | 71568 | ENSG00000166197 | NOLC1 | chr10 | + | 102157448 | 102157555 | 102157018 | 102157074 | 102158048 | 102158214 | 76 | 59 | 79 | 56 | 156 | 7 | 1 | 0 | 0 | 0 | 55 | 53 | 42 | 52 | 28 | 14 | 2 | 3 | 12 | 6 | 256 | 149 | 9.17E-10 | 1.09E-07 | 0.863 | 0.972 | 1 | 1 | 1 | 0.696 | 0.939 | 0.891 | 0.716 | 0.731 | 0.172 |
| SE | 59492 | ENSG00000148660 | CAMK2G | chr10 | - | 73824039 | 73824084 | 73821681 | 73821730 | 73825278 | 73825347 | 148 | 111 | 176 | 167 | 182 | 0 | 0 | 2 | 2 | 0 | 249 | 146 | 118 | 199 | 241 | 0 | 3 | 1 | 18 | 27 | 194 | 149 | 9.69E-10 | 1.15E-07 | 1 | 1 | 0.985 | 0.985 | 1 | 1 | 0.974 | 0.989 | 0.895 | 0.873 | 0.048 |
| SE | 66833 | ENSG00000031003 | FAM13B | chr5 | - | 138018954 | 138019146 | 138018301 | 138018514 | 138019880 | 138019977 | 148 | 70 | 146 | 133 | 190 | 3 | 4 | 2 | 4 | 4 | 70 | 115 | 118 | 60 | 59 | 16 | 7 | 19 | 8 | 5 | 298 | 149 | 9.83E-10 | 1.16E-07 | 0.961 | 0.897 | 0.973 | 0.943 | 0.96 | 0.686 | 0.891 | 0.756 | 0.789 | 0.855 | 0.151 |
| SE | 93731 | ENSG00000107262 | BAG1 | chr9 | - | 33262104 | 33262246 | 33261086 | 33261169 | 33262701 | 33262830 | 66 | 35 | 50 | 99 | 67 | 229 | 88 | 156 | 201 | 227 | 313 | 164 | 101 | 211 | 144 | 168 | 207 | 116 | 239 | 267 | 291 | 149 | 9.96E-10 | 1.18E-07 | 0.129 | 0.169 | 0.141 | 0.201 | 0.131 | 0.488 | 0.289 | 0.308 | 0.311 | 0.216 | -0.168 |
| SE | 25287 | ENSG00000184584 | STING1 | chr5 | - | 139479103 | 139479204 | 139478461 | 139478508 | 139480789 | 139480898 | 0 | 1 | 4 | 1 | 1 | 126 | 56 | 107 | 75 | 87 | 0 | 17 | 8 | 7 | 35 | 214 | 101 | 75 | 99 | 212 | 250 | 149 | 9.97E-10 | 1.18E-07 | 0 | 0.011 | 0.022 | 0.008 | 0.007 | 0 | 0.091 | 0.06 | 0.04 | 0.09 | -0.047 |
| SE | 70562 | ENSG00000022840 | RNF10 | chr12 | + | 120560725 | 120560886 | 120557543 | 120557682 | 120562944 | 120563070 | 425 | 173 | 306 | 327 | 515 | 5 | 0 | 0 | 0 | 0 | 540 | 450 | 518 | 538 | 924 | 8 | 15 | 4 | 30 | 10 | 298 | 149 | 1.01E-09 | 1.19E-07 | 0.977 | 1 | 1 | 1 | 1 | 0.971 | 0.938 | 0.985 | 0.9 | 0.979 | 0.041 |
| SE | 43593 | ENSG00000058673 | ZC3H11A | chr1 | + | 203801574 | 203803016 | 203795622 | 203795794 | 203816925 | 203817124 | 299 | 225 | 305 | 302 | 482 | 9 | 3 | 6 | 6 | 7 | 20 | 185 | 239 | 113 | 21 | 6 | 25 | 25 | 11 | 0 | 298 | 149 | 1.01E-09 | 1.20E-07 | 0.943 | 0.974 | 0.962 | 0.962 | 0.972 | 0.625 | 0.787 | 0.827 | 0.837 | 1 | 0.147 |
| SE | 74556 | ENSG00000089818 | NECAP1 | chr12 | + | 8091768 | 8091850 | 8089935 | 8090299 | 8092675 | 8092784 | 102 | 80 | 95 | 99 | 251 | 13 | 5 | 9 | 13 | 38 | 85 | 106 | 116 | 118 | 73 | 28 | 39 | 40 | 37 | 18 | 231 | 149 | 1.05E-09 | 1.24E-07 | 0.835 | 0.912 | 0.872 | 0.831 | 0.81 | 0.662 | 0.637 | 0.652 | 0.673 | 0.723 | 0.183 |
| SE | 66354 | ENSG00000197037 | ZSCAN25 | chr7 | + | 99619560 | 99621574 | 99618524 | 99618651 | 99622548 | 99622640 | 47 | 39 | 60 | 68 | 44 | 0 | 0 | 0 | 0 | 1 | 19 | 61 | 52 | 39 | 25 | 4 | 12 | 1 | 1 | 5 | 298 | 149 | 1.06E-09 | 1.24E-07 | 1 | 1 | 1 | 1 | 0.957 | 0.704 | 0.718 | 0.963 | 0.951 | 0.714 | 0.181 |
| SE | 49177 | ENSG00000066136 | NFYC | chr1 | + | 40749572 | 40749686 | 40747533 | 40747605 | 40753150 | 40753216 | 360 | 273 | 288 | 317 | 487 | 1 | 6 | 2 | 1 | 5 | 667 | 450 | 254 | 466 | 576 | 13 | 7 | 11 | 50 | 9 | 263 | 149 | 1.07E-09 | 1.26E-07 | 0.995 | 0.963 | 0.988 | 0.994 | 0.982 | 0.967 | 0.973 | 0.929 | 0.841 | 0.973 | 0.048 |
| SE | 74147 | ENSG00000111679 | PTPN6 | chr12 | + | 6957918 | 6958073 | 6957653 | 6957785 | 6959926 | 6959994 | 1266 | 711 | 979 | 1049 | 796 | 4 | 4 | 7 | 9 | 8 | 2035 | 1577 | 691 | 1443 | 2823 | 37 | 30 | 14 | 46 | 37 | 298 | 149 | 1.11E-09 | 1.30E-07 | 0.994 | 0.989 | 0.986 | 0.983 | 0.98 | 0.965 | 0.963 | 0.961 | 0.94 | 0.974 | 0.026 |
| SE | 104785 | ENSG00000130119 | GNL3L | chrX | + | 54544222 | 54544326 | 54543145 | 54543342 | 54548228 | 54548373 | 177 | 72 | 130 | 156 | 140 | 1 | 0 | 1 | 1 | 1 | 189 | 103 | 97 | 132 | 221 | 18 | 0 | 7 | 6 | 9 | 253 | 149 | 1.11E-09 | 1.30E-07 | 0.99 | 1 | 0.987 | 0.989 | 0.988 | 0.861 | 1 | 0.891 | 0.928 | 0.935 | 0.068 |
| SE | 40636 | ENSG00000134248 | LAMTOR5 | chr1 | - | 110407032 | 110407128 | 110406317 | 110406379 | 110407585 | 110407668 | 11 | 13 | 11 | 24 | 75 | 153 | 67 | 123 | 146 | 148 | 61 | 49 | 68 | 66 | 55 | 50 | 100 | 102 | 125 | 140 | 245 | 149 | 1.12E-09 | 1.31E-07 | 0.042 | 0.106 | 0.052 | 0.091 | 0.236 | 0.426 | 0.23 | 0.288 | 0.243 | 0.193 | -0.171 |
| SE | 59287 | ENSG00000119004 | CYP20A1 | chr2 | + | 203272464 | 203272748 | 203266513 | 203266681 | 203278572 | 203278688 | 35 | 31 | 39 | 48 | 86 | 17 | 13 | 16 | 14 | 35 | 26 | 18 | 22 | 21 | 6 | 37 | 16 | 30 | 42 | 6 | 252 | 149 | 1.13E-09 | 1.32E-07 | 0.549 | 0.585 | 0.59 | 0.67 | 0.592 | 0.294 | 0.399 | 0.302 | 0.228 | 0.372 | 0.278 |
| SE | 64317 | ENSG00000166949 | SMAD3 | chr15 | + | 67165252 | 67165384 | 67164894 | 67165088 | 67166778 | 67166838 | 156 | 93 | 169 | 141 | 104 | 0 | 3 | 0 | 7 | 0 | 355 | 270 | 129 | 252 | 447 | 31 | 16 | 4 | 13 | 35 | 281 | 149 | 1.17E-09 | 1.37E-07 | 1 | 0.943 | 1 | 0.914 | 1 | 0.859 | 0.899 | 0.945 | 0.911 | 0.871 | 0.074 |
| SE | 32171 | ENSG00000164053 | ATRIP | chr3 | + | 48457299 | 48457416 | 48454299 | 48454418 | 48459358 | 48459397 | 89 | 43 | 87 | 87 | 132 | 3 | 2 | 4 | 8 | 11 | 112 | 60 | 31 | 35 | 80 | 20 | 16 | 9 | 7 | 31 | 298 | 149 | 1.18E-09 | 1.38E-07 | 0.937 | 0.915 | 0.916 | 0.845 | 0.857 | 0.737 | 0.652 | 0.633 | 0.714 | 0.563 | 0.234 |
| SE | 8328 | ENSG00000159259 | CHAF1B | chr21 | + | 36397414 | 36397511 | 36394546 | 36394650 | 36399520 | 36399605 | 26 | 19 | 13 | 9 | 11 | 0 | 0 | 0 | 0 | 0 | 12 | 2 | 22 | 38 | 6 | 9 | 2 | 0 | 5 | 4 | 246 | 149 | 1.20E-09 | 1.39E-07 | 1 | 1 | 1 | 1 | 1 | 0.447 | 0.377 | 1 | 0.822 | 0.476 | 0.376 |
| SE | 36287 | ENSG00000075420 | FNDC3B | chr3 | + | 172247532 | 172247776 | 172226870 | 172226947 | 172285925 | 172285984 | 184 | 125 | 143 | 162 | 248 | 0 | 1 | 4 | 0 | 3 | 221 | 216 | 230 | 168 | 76 | 0 | 15 | 24 | 16 | 0 | 298 | 149 | 1.20E-09 | 1.39E-07 | 1 | 0.984 | 0.947 | 1 | 0.976 | 1 | 0.878 | 0.827 | 0.84 | 1 | 0.072 |
| SE | 69242 | ENSG00000249141 | NA | chr6 | - | 166934090 | 166934136 | 166858093 | 166858259 | 166938894 | 166939008 | 70 | 67 | 85 | 107 | 129 | 1 | 1 | 5 | 3 | 5 | 171 | 131 | 81 | 133 | 138 | 38 | 11 | 8 | 9 | 45 | 195 | 149 | 1.22E-09 | 1.41E-07 | 0.982 | 0.981 | 0.929 | 0.965 | 0.952 | 0.775 | 0.901 | 0.886 | 0.919 | 0.701 | 0.125 |
| SE | 84768 | ENSG00000158201 | ABHD3 | chr18 | - | 21683919 | 21683965 | 21663677 | 21664230 | 21702315 | 21702498 | 114 | 78 | 110 | 107 | 127 | 6 | 7 | 7 | 5 | 13 | 53 | 77 | 64 | 40 | 88 | 0 | 0 | 1 | 0 | 0 | 195 | 149 | 1.23E-09 | 1.42E-07 | 0.936 | 0.895 | 0.923 | 0.942 | 0.882 | 1 | 1 | 0.98 | 1 | 1 | -0.08 |
| SE | 39729 | ENSG00000132716 | DCAF8 | chr1 | - | 160261284 | 160261350 | 160243959 | 160244034 | 160262448 | 160262528 | 89 | 96 | 66 | 81 | 104 | 11 | 20 | 15 | 24 | 0 | 211 | 108 | 117 | 158 | 143 | 0 | 0 | 1 | 5 | 15 | 215 | 149 | 1.24E-09 | 1.43E-07 | 0.849 | 0.769 | 0.753 | 0.701 | 1 | 1 | 1 | 0.988 | 0.956 | 0.869 | -0.148 |
| SE | 79428 | ENSG00000163513 | TGFBR2 | chr3 | + | 30623198 | 30623273 | 30606600 | 30606977 | 30644746 | 30644915 | 156 | 75 | 163 | 156 | 253 | 130 | 62 | 125 | 110 | 188 | 126 | 190 | 138 | 141 | 186 | 36 | 79 | 34 | 48 | 92 | 224 | 149 | 1.28E-09 | 1.47E-07 | 0.444 | 0.446 | 0.464 | 0.485 | 0.472 | 0.7 | 0.615 | 0.73 | 0.661 | 0.574 | -0.194 |
| SE | 45568 | ENSG00000117335 | CD46 | chr1 | + | 207767778 | 207767823 | 207767012 | 207767195 | 207770320 | 207770362 | 534 | 341 | 460 | 508 | 775 | 60 | 58 | 61 | 64 | 161 | 154 | 341 | 308 | 308 | 129 | 31 | 8 | 7 | 14 | 9 | 194 | 149 | 1.30E-09 | 1.49E-07 | 0.872 | 0.819 | 0.853 | 0.859 | 0.787 | 0.792 | 0.97 | 0.971 | 0.944 | 0.917 | -0.081 |
| SE | 54088 | ENSG00000115935 | WIPF1 | chr2 | - | 174575203 | 174575380 | 174571168 | 174572446 | 174581309 | 174581439 | 797 | 475 | 649 | 721 | 722 | 2 | 6 | 5 | 8 | 1 | 537 | 613 | 494 | 537 | 1172 | 0 | 12 | 26 | 11 | 33 | 298 | 149 | 1.31E-09 | 1.50E-07 | 0.995 | 0.975 | 0.985 | 0.978 | 0.997 | 1 | 0.962 | 0.905 | 0.961 | 0.947 | 0.031 |
| SE | 94152 | ENSG00000090054 | SPTLC1 | chr9 | - | 92079433 | 92079566 | 92067965 | 92068098 | 92080015 | 92080088 | 33 | 18 | 30 | 17 | 11 | 229 | 171 | 179 | 256 | 163 | 66 | 40 | 59 | 36 | 47 | 213 | 181 | 191 | 254 | 66 | 282 | 149 | 1.32E-09 | 1.51E-07 | 0.071 | 0.053 | 0.081 | 0.034 | 0.034 | 0.141 | 0.105 | 0.14 | 0.07 | 0.273 | -0.091 |
| SE | 34166 | ENSG00000163660 | CCNL1 | chr3 | - | 157149286 | 157149385 | 157147552 | 157148589 | 157149484 | 157149596 | 876 | 925 | 925 | 1278 | 1144 | 42 | 81 | 48 | 101 | 91 | 116 | 593 | 747 | 202 | 104 | 25 | 120 | 89 | 59 | 8 | 248 | 149 | 1.36E-09 | 1.55E-07 | 0.926 | 0.873 | 0.92 | 0.884 | 0.883 | 0.736 | 0.748 | 0.835 | 0.673 | 0.886 | 0.122 |
| SE | 8133 | ENSG00000120071 | KANSL1 | chr17 | - | 46082440 | 46082542 | 46067548 | 46067667 | 46094559 | 46094701 | 357 | 207 | 285 | 325 | 460 | 6 | 3 | 8 | 8 | 11 | 327 | 319 | 263 | 334 | 304 | 15 | 31 | 19 | 26 | 15 | 251 | 149 | 1.37E-09 | 1.56E-07 | 0.972 | 0.976 | 0.955 | 0.96 | 0.961 | 0.928 | 0.859 | 0.892 | 0.884 | 0.923 | 0.068 |
| SE | 38686 | ENSG00000167258 | CDK12 | chr17 | + | 39515730 | 39515808 | 39511528 | 39511630 | 39517439 | 39517556 | 175 | 111 | 190 | 195 | 358 | 6 | 5 | 10 | 8 | 9 | 120 | 152 | 175 | 169 | 63 | 25 | 14 | 22 | 21 | 5 | 227 | 149 | 1.37E-09 | 1.56E-07 | 0.95 | 0.936 | 0.926 | 0.941 | 0.963 | 0.759 | 0.877 | 0.839 | 0.841 | 0.892 | 0.102 |
| SE | 54441 | ENSG00000169554 | ZEB2 | chr2 | - | 144517277 | 144517419 | 144426512 | 144430206 | 144519938 | 144520006 | 548 | 385 | 237 | 408 | 325 | 0 | 2 | 1 | 1 | 0 | 291 | 450 | 347 | 238 | 133 | 0 | 8 | 40 | 0 | 0 | 291 | 149 | 1.39E-09 | 1.58E-07 | 1 | 0.99 | 0.992 | 0.995 | 1 | 1 | 0.966 | 0.816 | 1 | 1 | 0.039 |
| SE | 109256 | ENSG00000275066 | SYNRG | chr17 | - | 37577379 | 37577613 | 37576340 | 37576418 | 37584647 | 37584759 | 283 | 178 | 288 | 243 | 543 | 72 | 45 | 70 | 68 | 200 | 148 | 294 | 118 | 160 | 142 | 10 | 44 | 19 | 19 | 0 | 298 | 149 | 1.41E-09 | 1.61E-07 | 0.663 | 0.664 | 0.673 | 0.641 | 0.576 | 0.881 | 0.77 | 0.756 | 0.808 | 1 | -0.2 |
| SE | 66747 | ENSG00000129292 | PHF20L1 | chr8 | + | 132788803 | 132788999 | 132777788 | 132777911 | 132794409 | 132794581 | 1 | 1 | 0 | 3 | 0 | 166 | 165 | 153 | 273 | 243 | 6 | 4 | 2 | 0 | 36 | 60 | 112 | 168 | 79 | 67 | 298 | 149 | 1.42E-09 | 1.61E-07 | 0.003 | 0.003 | 0 | 0.005 | 0 | 0.048 | 0.018 | 0.006 | 0 | 0.212 | -0.055 |
| SE | 59640 | ENSG00000163466 | ARPC2 | chr2 | + | 218234351 | 218234397 | 218228737 | 218228850 | 218245419 | 218245546 | 852 | 640 | 627 | 766 | 725 | 12 | 0 | 0 | 7 | 20 | 1480 | 818 | 663 | 1150 | 1279 | 21 | 14 | 18 | 47 | 62 | 195 | 149 | 1.46E-09 | 1.65E-07 | 0.982 | 1 | 1 | 0.988 | 0.965 | 0.982 | 0.978 | 0.966 | 0.949 | 0.94 | 0.024 |
| SE | 95882 | ENSG00000148396 | SEC16A | chr9 | - | 136463585 | 136463740 | 136461176 | 136461274 | 136464419 | 136464526 | 66 | 28 | 43 | 29 | 42 | 0 | 3 | 0 | 1 | 0 | 32 | 28 | 27 | 38 | 30 | 0 | 5 | 24 | 7 | 15 | 298 | 149 | 1.47E-09 | 1.66E-07 | 1 | 0.824 | 1 | 0.935 | 1 | 1 | 0.737 | 0.36 | 0.731 | 0.5 | 0.286 |
| SE | 3982 | ENSG00000126261 | UBA2 | chr19 | + | 34438644 | 34438766 | 34434867 | 34434968 | 34443843 | 34443911 | 738 | 419 | 658 | 679 | 1324 | 6 | 1 | 3 | 1 | 1 | 421 | 420 | 511 | 405 | 298 | 8 | 13 | 1 | 8 | 16 | 271 | 149 | 1.49E-09 | 1.69E-07 | 0.985 | 0.996 | 0.992 | 0.997 | 0.999 | 0.967 | 0.947 | 0.996 | 0.965 | 0.911 | 0.037 |
| SE | 37919 | ENSG00000184014 | DENND5A | chr11 | - | 9176771 | 9176902 | 9170626 | 9170777 | 9178131 | 9178366 | 12 | 17 | 24 | 21 | 25 | 205 | 194 | 192 | 190 | 331 | 106 | 54 | 131 | 87 | 9 | 370 | 152 | 131 | 251 | 312 | 280 | 149 | 1.50E-09 | 1.70E-07 | 0.03 | 0.045 | 0.062 | 0.056 | 0.039 | 0.132 | 0.159 | 0.347 | 0.156 | 0.015 | -0.115 |
| SE | 15355 | ENSG00000140521 | POLG | chr15 | - | 89317375 | 89317476 | 89316329 | 89316827 | 89318540 | 89318749 | 63 | 38 | 53 | 51 | 101 | 0 | 0 | 0 | 0 | 1 | 103 | 48 | 16 | 33 | 72 | 11 | 2 | 0 | 1 | 12 | 250 | 149 | 1.52E-09 | 1.71E-07 | 1 | 1 | 1 | 1 | 0.984 | 0.848 | 0.935 | 1 | 0.952 | 0.781 | 0.094 |
| SE | 1085 | ENSG00000100330 | MTMR3 | chr22 | + | 30022028 | 30022139 | 30019479 | 30020884 | 30022608 | 30022697 | 135 | 52 | 71 | 89 | 85 | 12 | 14 | 11 | 13 | 20 | 82 | 114 | 94 | 62 | 117 | 38 | 61 | 40 | 53 | 52 | 260 | 149 | 1.54E-09 | 1.73E-07 | 0.866 | 0.68 | 0.787 | 0.797 | 0.709 | 0.553 | 0.517 | 0.574 | 0.401 | 0.563 | 0.246 |
| SE | 50384 | ENSG00000058866 | DGKG | chr3 | - | 186265246 | 186265306 | 186261698 | 186261778 | 186267684 | 186267777 | 7 | 9 | 8 | 12 | 26 | 1 | 0 | 0 | 0 | 0 | 16 | 35 | 8 | 30 | 73 | 12 | 2 | 1 | 12 | 15 | 209 | 149 | 1.54E-09 | 1.74E-07 | 0.833 | 1 | 1 | 1 | 1 | 0.487 | 0.926 | 0.851 | 0.641 | 0.776 | 0.23 |
| SE | 13969 | ENSG00000080603 | SRCAP | chr16 | + | 30720712 | 30720978 | 30720161 | 30720331 | 30721188 | 30721476 | 129 | 65 | 112 | 117 | 172 | 0 | 1 | 2 | 6 | 0 | 229 | 107 | 92 | 148 | 299 | 7 | 4 | 5 | 16 | 47 | 298 | 149 | 1.57E-09 | 1.76E-07 | 1 | 0.97 | 0.966 | 0.907 | 1 | 0.942 | 0.93 | 0.902 | 0.822 | 0.761 | 0.097 |
| SE | 105670 | ENSG00000156502 | SUPV3L1 | chr10 | + | 69189266 | 69189435 | 69187641 | 69187756 | 69191654 | 69191766 | 107 | 64 | 101 | 107 | 102 | 1 | 1 | 1 | 0 | 0 | 75 | 118 | 108 | 128 | 56 | 11 | 12 | 10 | 1 | 0 | 298 | 149 | 1.58E-09 | 1.77E-07 | 0.982 | 0.97 | 0.981 | 1 | 1 | 0.773 | 0.831 | 0.844 | 0.985 | 1 | 0.1 |

| | | | | | | | | | | | | | | | | | | | | | | | | | | | | | | | | | | | | | | | | | | | | | | |
|---|---|---|---|---|---|---|---|---|---|---|---|---|---|---|---|---|---|---|---|---|---|---|---|---|---|---|---|---|---|---|---|---|---|---|---|---|---|---|---|---|---|---|---|---|---|---|
| SE | 53797 | ENSG00000111653 | ING4 | chr12 | - | 6655637 | 6655913 | 6652935 | 6653050 | 6656726 | 6656798 | 7 | 4 | 3 | 2 | 12 | 7 | 8 | 12 | 10 | 12 | 0 | 2 | 0 | 0 | 0 | 25 | 11 | 11 | 3 | 3 | 298 | 149 | 1.58E-09 | 1.77E-07 | 0.333 | 0.2 | 0.111 | 0.091 | 0.333 | 0 | 0.083 | 0 | 0 | 0 | 0.197 |
| SE | 37100 | ENSG00000127481 | UBR4 | chr1 | - | 19156270 | 19156423 | 19155440 | 19155668 | 19156766 | 19156925 | 149 | 87 | 166 | 160 | 111 | 1 | 0 | 1 | 0 | 1 | 342 | 254 | 182 | 186 | 296 | 0 | 11 | 11 | 23 | 7 | 298 | 149 | 1.58E-09 | 1.77E-07 | 0.987 | 1 | 0.988 | 1 | 0.982 | 1 | 0.92 | 0.892 | 0.802 | 0.955 | 0.078 |
| SE | 44390 | ENSG00000072210 | ALDH3A2 | chr17 | + | 19673150 | 19673275 | 19671720 | 19671956 | 19675495 | 19675499 | 10 | 2 | 10 | 11 | 12 | 55 | 40 | 59 | 62 | 142 | 0 | 2 | 0 | 1 | 0 | 79 | 72 | 24 | 39 | 49 | 274 | 149 | 1.61E-09 | 1.80E-07 | 0.09 | 0.026 | 0.084 | 0.088 | 0.044 | 0 | 0.015 | 0 | 0.014 | 0 | 0.061 |
| SE | 88244 | ENSG00000147416 | ATP6V1B2 | chr8 | + | 20214817 | 20214968 | 20212781 | 20212905 | 20216412 | 20216495 | 977 | 577 | 604 | 559 | 1128 | 10 | 9 | 12 | 7 | 16 | 1143 | 499 | 327 | 457 | 1435 | 51 | 28 | 13 | 44 | 7 | 298 | 149 | 1.62E-09 | 1.81E-07 | 0.98 | 0.97 | 0.962 | 0.976 | 0.972 | 0.918 | 0.899 | 0.926 | 0.839 | 0.99 | 0.058 |
| SE | 108048 | ENSG00000139613 | SMARCC2 | chr12 | - | 56185029 | 56185092 | 56184843 | 56184936 | 56186154 | 56186240 | 130 | 71 | 131 | 144 | 179 | 0 | 1 | 0 | 1 | 0 | 157 | 149 | 87 | 89 | 185 | 0 | 7 | 8 | 4 | 16 | 212 | 149 | 1.64E-09 | 1.82E-07 | 1 | 0.98 | 1 | 0.99 | 1 | 1 | 0.937 | 0.884 | 0.94 | 0.89 | 0.064 |
| SE | 33413 | ENSG00000096063 | SRPK1 | chr6 | - | 35870280 | 35870494 | 35869481 | 35869901 | 35870933 | 35870959 | 96 | 110 | 89 | 124 | 194 | 2 | 6 | 5 | 6 | 5 | 17 | 140 | 178 | 111 | 0 | 10 | 16 | 93 | 41 | 7 | 298 | 149 | 1.65E-09 | 1.83E-07 | 0.96 | 0.902 | 0.899 | 0.912 | 0.951 | 0.459 | 0.814 | 0.489 | 0.575 | 0 | 0.457 |
| SE | 71214 | ENSG00000079263 | SP140 | chr2 | + | 230247915 | 230248065 | 230245862 | 230245940 | 230248884 | 230248968 | 191 | 173 | 223 | 165 | 160 | 2 | 7 | 2 | 6 | 4 | 205 | 292 | 232 | 216 | 72 | 15 | 23 | 19 | 19 | 12 | 298 | 149 | 1.68E-09 | 1.87E-07 | 0.979 | 0.925 | 0.982 | 0.932 | 0.952 | 0.872 | 0.864 | 0.859 | 0.85 | 0.75 | 0.115 |
| SE | 15044 | ENSG00000166783 | MARF1 | chr16 | - | 15600640 | 15600701 | 15600427 | 15600553 | 15601990 | 15602203 | 219 | 148 | 234 | 235 | 302 | 8 | 11 | 11 | 8 | 3 | 135 | 133 | 206 | 186 | 30 | 26 | 19 | 25 | 24 | 0 | 210 | 149 | 1.69E-09 | 1.88E-07 | 0.951 | 0.905 | 0.938 | 0.954 | 0.986 | 0.787 | 0.832 | 0.854 | 0.846 | 1 | 0.083 |
| SE | 99704 | ENSG00000110075 | PPP6R3 | chr11 | + | 68567013 | 68567166 | 68564302 | 68564432 | 68569747 | 68569897 | 826 | 605 | 762 | 902 | 966 | 7 | 5 | 17 | 18 | 16 | 313 | 764 | 750 | 601 | 214 | 11 | 33 | 52 | 22 | 0 | 298 | 149 | 1.73E-09 | 1.91E-07 | 0.983 | 0.984 | 0.957 | 0.962 | 0.968 | 0.934 | 0.92 | 0.878 | 0.932 | 1 | 0.038 |
| SE | 66469 | ENSG00000131051 | RBM39 | chr20 | - | 35734185 | 35735081 | 35731940 | 35732135 | 35738967 | 35739017 | 53 | 36 | 31 | 52 | 32 | 679 | 659 | 609 | 826 | 869 | 57 | 56 | 75 | 49 | 4 | 276 | 482 | 893 | 401 | 87 | 298 | 149 | 1.78E-09 | 1.96E-07 | 0.038 | 0.027 | 0.025 | 0.031 | 0.018 | 0.094 | 0.055 | 0.04 | 0.058 | 0.022 | -0.026 |
| SE | 92304 | ENSG00000204130 | RUFY2 | chr10 | - | 68386058 | 68386128 | 68384050 | 68384152 | 68393137 | 68393203 | 95 | 93 | 98 | 80 | 139 | 1 | 0 | 1 | 0 | 0 | 26 | 68 | 57 | 66 | 10 | 13 | 4 | 3 | 5 | 0 | 219 | 149 | 1.78E-09 | 1.97E-07 | 0.985 | 1 | 0.985 | 1 | 1 | 0.576 | 0.92 | 0.928 | 0.9 | 1 | 0.129 |
| SE | 19091 | ENSG00000259316 | NA | chr15 | - | 64207508 | 64207594 | 64204440 | 64204589 | 64213889 | 64214124 | 23 | 31 | 35 | 36 | 30 | 0 | 0 | 0 | 1 | 0 | 44 | 36 | 36 | 30 | 18 | 5 | 0 | 5 | 7 | 7 | 235 | 149 | 1.79E-09 | 1.98E-07 | 1 | 1 | 1 | 0.958 | 1 | 0.848 | 1 | 0.82 | 0.731 | 0.62 | 0.188 |
| SE | 92028 | ENSG00000135315 | CEP162 | chr6 | - | 84219102 | 84219214 | 84215775 | 84215922 | 84221056 | 84221171 | 18 | 18 | 23 | 29 | 18 | 23 | 29 | 12 | 19 | 23 | 27 | 21 | 6 | 16 | 13 | 0 | 6 | 0 | 0 | 3 | 261 | 149 | 1.81E-09 | 1.99E-07 | 0.309 | 0.262 | 0.522 | 0.466 | 0.309 | 1 | 0.666 | 1 | 1 | 0.712 | -0.502 |
| SE | 100590 | ENSG00000102317 | RBM3 | chrX | + | 48575560 | 48575630 | 48575167 | 48575283 | 48576313 | 48576419 | 344 | 224 | 379 | 367 | 306 | 15 | 5 | 17 | 22 | 18 | 734 | 446 | 522 | 434 | 569 | 57 | 43 | 59 | 110 | 55 | 219 | 149 | 1.82E-09 | 2.00E-07 | 0.94 | 0.968 | 0.938 | 0.919 | 0.92 | 0.898 | 0.876 | 0.858 | 0.729 | 0.876 | 0.09 |
| SE | 42341 | ENSG00000070759 | TESK2 | chr1 | - | 45346978 | 45347062 | 45346692 | 45346779 | 45347608 | 45347693 | 14 | 8 | 22 | 15 | 26 | 0 | 1 | 0 | 1 | 0 | 25 | 28 | 16 | 20 | 43 | 10 | 9 | 2 | 3 | 7 | 233 | 149 | 1.82E-09 | 2.00E-07 | 1 | 0.836 | 1 | 0.906 | 1 | 0.615 | 0.665 | 0.836 | 0.81 | 0.797 | 0.204 |
| SE | 102078 | ENSG00000181192 | DHTKD1 | chr10 | + | 12106245 | 12106396 | 12101041 | 12101181 | 12107908 | 12108015 | 76 | 39 | 51 | 61 | 68 | 0 | 1 | 1 | 0 | 0 | 71 | 63 | 47 | 62 | 146 | 9 | 9 | 0 | 0 | 19 | 298 | 149 | 1.85E-09 | 2.03E-07 | 1 | 0.951 | 0.962 | 1 | 1 | 0.798 | 0.778 | 1 | 1 | 0.793 | 0.109 |
| SE | 75334 | ENSG00000196262 | PPIA | chr7 | + | 44799391 | 44799480 | 44796978 | 44799277 | 44801286 | 44801624 | 4704 | 2747 | 4069 | 4232 | 3163 | 10 | 6 | 3 | 18 | 5 | 6029 | 4466 | 3684 | 5026 | 8045 | 70 | 41 | 59 | 53 | 75 | 238 | 149 | 1.86E-09 | 2.04E-07 | 0.997 | 0.997 | 0.999 | 0.993 | 0.997 | 0.982 | 0.986 | 0.975 | 0.983 | 0.985 | 0.014 |
| SE | 76017 | ENSG00000213654 | GPSM3 | chr6 | - | 32192147 | 32192250 | 32191708 | 32191908 | 32192471 | 32192530 | 397 | 133 | 283 | 358 | 492 | 26 | 9 | 21 | 21 | 2 | 735 | 413 | 396 | 475 | 1266 | 0 | 7 | 10 | 8 | 24 | 252 | 149 | 1.88E-09 | 2.05E-07 | 0.9 | 0.897 | 0.888 | 0.91 | 0.993 | 1 | 0.972 | 0.959 | 0.972 | 0.969 | -0.057 |
| SE | 38434 | ENSG00000132549 | VPS13B | chr8 | + | 99507769 | 99507911 | 99502835 | 99502950 | 99511103 | 99511340 | 27 | 23 | 17 | 13 | 18 | 18 | 15 | 5 | 21 | 36 | 22 | 9 | 7 | 2 | 14 | 0 | 2 | 2 | 0 | 0 | 291 | 149 | 1.89E-09 | 2.06E-07 | 0.434 | 0.44 | 0.635 | 0.241 | 0.204 | 1 | 0.697 | 0.642 | 1 | 1 | -0.477 |
| SE | 22218 | ENSG00000163933 | RFT1 | chr3 | - | 53121698 | 53121800 | 53119883 | 53120021 | 53122373 | 53122563 | 79 | 58 | 125 | 89 | 88 | 0 | 1 | 2 | 3 | 0 | 119 | 136 | 74 | 108 | 159 | 9 | 5 | 28 | 2 | 21 | 251 | 149 | 1.93E-09 | 2.10E-07 | 1 | 0.972 | 0.974 | 0.946 | 1 | 0.887 | 0.942 | 0.611 | 0.97 | 0.818 | 0.133 |
| SE | 26020 | ENSG00000134480 | CCNH | chr5 | - | 87404843 | 87405007 | 87401701 | 87401772 | 87407975 | 87408186 | 484 | 443 | 591 | 535 | 718 | 5 | 10 | 11 | 11 | 24 | 203 | 319 | 737 | 265 | 82 | 0 | 0 | 3 | 0 | 0 | 298 | 149 | 1.93E-09 | 2.10E-07 | 0.98 | 0.957 | 0.964 | 0.961 | 0.937 | 1 | 1 | 0.992 | 1 | 1 | -0.039 |
| SE | 53115 | ENSG00000166352 | IFTAP | chr11 | + | 36636050 | 36636117 | 36610103 | 36610239 | 36648015 | 36648155 | 8 | 8 | 5 | 5 | 30 | 4 | 2 | 3 | 4 | 8 | 15 | 15 | 3 | 17 | 8 | 0 | 0 | 0 | 2 | 0 | 216 | 149 | 2.04E-09 | 2.22E-07 | 0.58 | 0.734 | 0.535 | 0.463 | 0.721 | 1 | 1 | 1 | 0.854 | 1 | -0.364 |
| SE | 85193 | ENSG00000140332 | TLE3 | chr15 | - | 70064453 | 70064470 | 70060529 | 70060649 | 70074502 | 70074607 | 23 | 34 | 21 | 32 | 25 | 1 | 0 | 0 | 0 | 0 | 71 | 23 | 50 | 52 | 64 | 6 | 2 | 7 | 19 | 3 | 166 | 149 | 2.06E-09 | 2.23E-07 | 0.954 | 1 | 1 | 1 | 1 | 0.914 | 0.912 | 0.865 | 0.711 | 0.95 | 0.12 |
| SE | 69218 | ENSG00000148248 | SURF4 | chr9 | - | 133364839 | 133365026 | 133361449 | 133363759 | 133365984 | 133366028 | 292 | 127 | 241 | 287 | 171 | 6 | 1 | 0 | 1 | 2 | 290 | 317 | 160 | 231 | 331 | 10 | 2 | 14 | 17 | 16 | 298 | 149 | 2.06E-09 | 2.24E-07 | 0.961 | 0.984 | 1 | 0.993 | 0.977 | 0.935 | 0.988 | 0.851 | 0.872 | 0.912 | 0.071 |
| SE | 48387 | ENSG00000134215 | VAV3 | chr1 | - | 107777669 | 107777770 | 107777230 | 107777296 | 107779433 | 107779492 | 0 | 2 | 1 | 2 | 0 | 89 | 76 | 78 | 107 | 108 | 5 | 7 | 7 | 16 | 4 | 34 | 95 | 137 | 91 | 11 | 250 | 149 | 2.08E-09 | 2.25E-07 | 0 | 0.015 | 0.008 | 0.011 | 0 | 0.081 | 0.042 | 0.03 | 0.095 | 0.178 | -0.078 |
| SE | 76872 | ENSG00000001167 | NFYA | chr6 | + | 41080810 | 41080897 | 41079028 | 41079164 | 41084045 | 41084192 | 36 | 4 | 31 | 30 | 7 | 91 | 81 | 93 | 103 | 91 | 1 | 0 | 2 | 8 | 8 | 101 | 66 | 87 | 100 | 27 | 236 | 149 | 2.10E-09 | 2.27E-07 | 0.2 | 0.03 | 0.174 | 0.155 | 0.046 | 0.006 | 0 | 0.014 | 0.048 | 0.158 | 0.076 |
| SE | 81252 | ENSG00000115677 | HDLBP | chr2 | - | 241255373 | 241255580 | 241255050 | 241255158 | 241256183 | 241256399 | 323 | 236 | 323 | 359 | 366 | 1 | 1 | 0 | 0 | 0 | 582 | 420 | 429 | 463 | 487 | 25 | 7 | 7 | 9 | 5 | 298 | 149 | 2.11E-09 | 2.28E-07 | 0.994 | 0.992 | 1 | 1 | 1 | 0.921 | 0.968 | 0.968 | 0.963 | 0.98 | 0.037 |
| SE | 80459 | ENSG00000125812 | GZF1 | chr20 | + | 23369583 | 23369741 | 23368761 | 23368929 | 23370090 | 23373045 | 20 | 17 | 30 | 38 | 33 | 3 | 4 | 3 | 5 | 2 | 28 | 33 | 43 | 39 | 13 | 0 | 0 | 0 | 0 | 0 | 298 | 149 | 2.12E-09 | 2.28E-07 | 0.769 | 0.68 | 0.833 | 0.792 | 0.892 | 1 | 1 | 1 | 1 | 1 | -0.207 |
| SE | 91268 | ENSG00000154114 | TBCEL | chr11 | + | 121055051 | 121055308 | 121047527 | 121047667 | 121058344 | 121058471 | 92 | 59 | 81 | 107 | 163 | 11 | 4 | 3 | 3 | 10 | 74 | 100 | 38 | 41 | 88 | 0 | 1 | 0 | 0 | 0 | 298 | 149 | 2.14E-09 | 2.31E-07 | 0.807 | 0.881 | 0.931 | 0.947 | 0.891 | 1 | 0.98 | 1 | 1 | 1 | -0.105 |
| SE | 23585 | ENSG00000169567 | HINT1 | chr5 | - | 131162028 | 131162477 | 131159295 | 131159611 | 131162571 | 131162676 | 55 | 24 | 62 | 38 | 102 | 380 | 179 | 430 | 380 | 991 | 47 | 47 | 40 | 65 | 55 | 257 | 213 | 145 | 147 | 282 | 298 | 149 | 2.15E-09 | 2.31E-07 | 0.067 | 0.063 | 0.067 | 0.048 | 0.049 | 0.084 | 0.099 | 0.121 | 0.181 | 0.089 | -0.056 |
| SE | 3508 | ENSG00000130755 | GMFG | chr19 | - | 39335434 | 39335531 | 39335260 | 39335310 | 39335973 | 39336005 | 1367 | 868 | 1397 | 1458 | 1962 | 11 | 8 | 11 | 5 | 9 | 1709 | 1727 | 1313 | 1685 | 2030 | 36 | 11 | 23 | 30 | 46 | 246 | 149 | 2.16E-09 | 2.32E-07 | 0.987 | 0.985 | 0.987 | 0.994 | 0.992 | 0.966 | 0.99 | 0.972 | 0.971 | 0.964 | 0.016 |
| SE | 60904 | ENSG00000144674 | GOLGA4 | chr3 | + | 37321730 | 37321886 | 37319062 | 37319194 | 37328415 | 37328537 | 208 | 207 | 136 | 189 | 198 | 0 | 0 | 0 | 0 | 0 | 30 | 61 | 80 | 11 | 15 | 3 | 11 | 2 | 2 | 0 | 298 | 149 | 2.22E-09 | 2.39E-07 | 1 | 1 | 1 | 1 | 1 | 0.833 | 0.735 | 0.952 | 0.733 | 1 | 0.149 |
| SE | 30696 | ENSG00000132466 | ANKRD17 | chr4 | - | 73147240 | 73147432 | 73146763 | 73146873 | 73148812 | 73149050 | 427 | 302 | 340 | 375 | 403 | 5 | 1 | 1 | 2 | 1 | 81 | 262 | 280 | 325 | 14 | 19 | 9 | 13 | 0 | 0 | 298 | 149 | 2.23E-09 | 2.39E-07 | 0.977 | 0.993 | 0.994 | 0.989 | 0.995 | 0.681 | 0.936 | 0.915 | 1 | 1 | 0.083 |

| | | | | | | | | | |
|---|---|---|---|---|---|---|---|---|---|
| SE | 20935 | ENSG00000135596 | MICAL1 | chr6 | - | 109447874 | 109447963 | 109447356 | 109447440 |
| SE | 10545 | ENSG00000177302 | TOP3A | chr17 | - | 18280535 | 18280658 | 18277674 | 18278357 |
| SE | 60934 | ENSG00000138078 | PREPL | chr2 | - | 44344519 | 44344586 | 44343744 | 44343951 |
| SE | 74822 | ENSG00000042088 | TDP1 | chr14 | + | 89984515 | 89984683 | 89980539 | 89980632 |
| SE | 8343 | ENSG00000196535 | MYO18A | chr17 | - | 29086437 | 29086577 | 29082315 | 29082438 |
| SE | 90899 | ENSG00000185728 | YTHDF3 | chr8 | + | 63175330 | 63175416 | 63169386 | 63169411 |
| SE | 33012 | ENSG00000114316 | USP4 | chr3 | - | 49311513 | 49311654 | 49305714 | 49305888 |
| SE | 74255 | ENSG00000137959 | IFI44L | chr1 | + | 78620877 | 78620971 | 78620448 | 78620571 |
| SE | 79855 | ENSG00000106617 | PRKAG2 | chr7 | - | 151574890 | 151574949 | 151572407 | 151572709 |
| SE | 45812 | ENSG00000108654 | DDX5 | chr17 | - | 64501056 | 64501143 | 64500548 | 64500773 |
| SE | 44602 | ENSG00000026508 | CD44 | chr11 | + | 35221653 | 35221732 | 35219315 | 35219387 |
| SE | 53008 | ENSG00000158941 | CCAR2 | chr8 | + | 22618348 | 22618495 | 22617695 | 22617778 |
| SE | 25439 | ENSG00000011243 | AKAP8L | chr19 | - | 15398629 | 15398731 | 15397713 | 15397855 |
| SE | 95246 | ENSG00000159921 | GNE | chr9 | - | 36222776 | 36222998 | 36219837 | 36220020 |
| SE | 49444 | ENSG00000115020 | PIKFYVE | chr2 | + | 208273583 | 208273733 | 208271510 | 208271691 |
| SE | 95594 | ENSG00000130714 | POMT1 | chr9 | + | 131506113 | 131506220 | 131504188 | 131504340 |
| SE | 97147 | ENSG00000153827 | TRIP12 | chr2 | - | 229771518 | 229771632 | 229769230 | 229769325 |
| SE | 48902 | ENSG00000158773 | USF1 | chr1 | - | 161041650 | 161041846 | 161041323 | 161041411 |
| SE | 430 | ENSG00000237484 | LINC01684 | chr21 | + | 24438252 | 24438443 | 24428728 | 24428904 |
| SE | 42837 | ENSG00000140575 | IQGAP1 | chr15 | + | 90482701 | 90482749 | 90482196 | 90482281 |
| SE | 39352 | ENSG00000158864 | NDUFS2 | chr1 | + | 161210110 | 161210389 | 161209856 | 161209931 |
| SE | 82822 | ENSG00000155849 | ELMO1 | chr7 | - | 36887559 | 36887672 | 36878009 | 36878117 |
| SE | 27774 | ENSG00000121064 | SCPEP1 | chr17 | + | 56998384 | 56998498 | 56996961 | 56997055 |
| SE | 73426 | ENSG00000177463 | NR2C2 | chr3 | + | 14962408 | 14962492 | 14947728 | 14947906 |
| SE | 99447 | ENSG00000285509 | TBCEL-TECTA | chr11 | + | 121053550 | 121053732 | 121047527 | 121047667 |
| SE | 37582 | ENSG00000082074 | FYB1 | chr5 | - | 39125997 | 39126135 | 39124252 | 39124278 |
| SE | 23966 | ENSG00000124201 | ZNFX1 | chr20 | - | 49263333 | 49263483 | 49260462 | 49260577 |
| SE | 101678 | ENSG00000136935 | GOLGA1 | chr9 | - | 124928187 | 124928285 | 124926708 | 124926741 |
| SE | 39531 | ENSG00000111707 | SUDS3 | chr12 | + | 118401982 | 118402004 | 118401758 | 118401820 |
| SE | 46525 | ENSG00000223745 | CCDC18-AS1 | chr1 | - | 93305387 | 93305470 | 93264639 | 93264772 |
| SE | 75953 | ENSG00000066379 | POLR1H | chr6 | + | 30061916 | 30062017 | 30061264 | 30061669 |
| SE | 91265 | ENSG00000154114 | TBCEL | chr11 | + | 121053629 | 121053732 | 121047527 | 121047667 |
| SE | 31223 | ENSG00000025293 | PHF20 | chr20 | + | 35870954 | 35871134 | 35869437 | 35869551 |
| SE | 86468 | ENSG00000215021 | PHB2 | chr12 | - | 6968410 | 6968708 | 6967891 | 6968021 |
| SE | 75056 | ENSG00000198223 | CSF2RA | chrX | + | 1282408 | 1282560 | 1274754 | 1274818 |
| SE | 52547 | ENSG00000153283 | CD96 | chr3 | + | 111577501 | 111577549 | 111567522 | 111567647 |
| SE | 79957 | ENSG00000042980 | ADAM28 | chr8 | + | 24313387 | 24313580 | 24311360 | 24311437 |
| SE | 33231 | ENSG00000023330 | ALAS1 | chr3 | + | 52205838 | 52206023 | 52204692 | 52204915 |
| SE | 89165 | ENSG00000075856 | SART3 | chr12 | - | 108535358 | 108535468 | 108531203 | 108531280 |
| SE | 3614 | ENSG00000104974 | LILRA1 | chr19 | + | 54595754 | 54595935 | 54595099 | 54595402 |
| SE | 86466 | ENSG00000215021 | PHB2 | chr12 | - | 6968410 | 6968587 | 6966925 | 6968021 |
| SE | 102484 | ENSG00000119979 | DENND10 | chr10 | + | 119110192 | 119110314 | 119107967 | 119108164 |
| SE | 20444 | ENSG00000066933 | MYO9A | chr15 | - | 72019038 | 72019095 | 72010349 | 72010447 |
| SE | 81090 | ENSG00000140750 | ARHGAP17 | chr16 | - | 24952288 | 24952370 | 24948981 | 24949484 |
| SE | 81727 | ENSG00000117748 | RPA2 | chr1 | - | 27897632 | 27897707 | 27897004 | 27897121 |
| SE | 21291 | ENSG00000150867 | PIP4K2A | chr10 | - | 22591628 | 22591781 | 22573310 | 22573457 |
| SE | 1337 | ENSG00000093000 | NUP50 | chr22 | + | 45178237 | 45178900 | 45171599 | 45171683 |
| SE | 5057 | ENSG00000177410 | ZFAS1 | chr20 | + | 49280902 | 49280964 | 49280321 | 49280570 |
| SE | 83940 | ENSG00000147421 | HMBOX1 | chr8 | + | 29009683 | 29009821 | 29009071 | 29009182 |
| SE | 21774 | ENSG00000112081 | SRSF3 | chr6 | + | 36599820 | 36600276 | 36598848 | 36598983 |
| SE | 2651 | ENSG00000129351 | ILF3 | chr19 | + | 10680345 | 10680443 | 10679815 | 10679927 |
| SE | 104042 | ENSG00000065809 | FAM107B | chr10 | - | 14553321 | 14553387 | 14530331 | 14530515 |
| SE | 95387 | ENSG00000198431 | TXNRD1 | chr12 | + | 104309758 | 104309950 | 104288930 | 104289040 |
| SE | 95472 | ENSG00000095319 | NUP188 | chr9 | + | 128969399 | 128969514 | 128968505 | 128968717 |
| SE | 61 | ENSG00000160285 | LSS | chr21 | - | 46219475 | 46219572 | 46216388 | 46216524 |
| SE | 35135 | ENSG00000288520 | NA | chr6 | - | 154247448 | 154247485 | 154246590 | 154246760 |
| SE | 47596 | ENSG00000008869 | HEATR5B | chr2 | - | 37072109 | 37072281 | 37070229 | 37070387 |
| SE | 83139 | ENSG00000240225 | ZNF542P | chr19 | + | 56374685 | 56374757 | 56373530 | 56373623 |
| SE | 16765 | ENSG00000125107 | CNOT1 | chr16 | - | 58558472 | 58558674 | 58556846 | 58556993 |
| SE | 71735 | ENSG00000179051 | RCC2 | chr1 | - | 17413536 | 17413717 | 17413072 | 17413178 |
| SE | 48424 | ENSG00000086015 | MAST2 | chr1 | + | 46028767 | 46028933 | 46027730 | 46027863 |
| SE | 106229 | ENSG00000089154 | GCN1 | chr12 | - | 120131184 | 120131333 | 120130645 | 120130753 |
| SE | 67491 | ENSG00000151553 | FAM160B1 | chr10 | + | 114835536 | 114835641 | 114833232 | 114833402 |
| SE | 43992 | ENSG00000136450 | SRSF1 | chr17 | - | 58005413 | 58005600 | 58004587 | 58005041 |
| SE | 105792 | ENSG00000124164 | VAPB | chr20 | + | 58440906 | 58441083 | 58418210 | 58418363 |
| SE | 87225 | ENSG00000086758 | HUWE1 | chrX | - | 53648211 | 53648310 | 53645310 | 53645463 |
| SE | 3613 | ENSG00000104974 | LILRA1 | chr19 | + | 54595638 | 54595935 | 54595099 | 54595402 |
| SE | 12988 | ENSG00000103351 | CLUAP1 | chr16 | + | 3508288 | 3508468 | 3506330 | 3506415 |
| SE | 29737 | ENSG00000287542 | NA | chr4 | + | 88681158 | 88681325 | 88680092 | 88680236 |
| SE | 76018 | ENSG00000213654 | GPSM3 | chr6 | - | 32192147 | 32192272 | 32191708 | 32191908 |
| SE | 55126 | ENSG00000119725 | ZNF410 | chr14 | + | 73894358 | 73894409 | 73893796 | 73893932 |
| SE | 65213 | ENSG00000143870 | PDIA6 | chr2 | - | 10784933 | 10785030 | 10783390 | 10784326 |
| SE | 35665 | ENSG00000010404 | IDS | chrX | - | 149503311 | 149503489 | 149500948 | 149501037 |
| SE | 20195 | ENSG00000137210 | TMEM14B | chr6 | + | 10755141 | 10755232 | 10749621 | 10749698 |
| SE | 88636 | ENSG00000178234 | GALNT11 | chr7 | + | 152079286 | 152079408 | 152025797 | 152025884 |
| SE | 100436 | ENSG00000132275 | RRP8 | chr11 | - | 6600925 | 6601602 | 6600668 | 6600775 |
| SE | 33424 | ENSG00000135870 | RC3H1 | chr1 | - | 173943441 | 173943588 | 173941264 | 173941380 |
| SE | 50606 | ENSG00000134574 | DDB2 | chr11 | + | 47234756 | 47234934 | 47234572 | 47234672 |
| SE | 17024 | ENSG00000102921 | N4BP1 | chr16 | - | 48551385 | 48551482 | 48548006 | 48548114 |
| SE | 81299 | ENSG00000141480 | ARRB2 | chr17 | + | 4715150 | 4715336 | 4715012 | 4715043 |
| SE | 13128 | ENSG00000118900 | UBN1 | chr16 | + | 4859024 | 4859159 | 4858567 | 4858663 |
| SE | 73581 | ENSG00000129472 | RAB2B | chr14 | - | 21468669 | 21468752 | 21468356 | 21468449 |
| SE | 68726 | ENSG00000196295 | GARS1-DT | chr7 | - | 30579230 | 30579314 | 30566421 | 30568989 |
| SE | 91187 | ENSG00000164941 | INTS8 | chr8 | + | 94874551 | 94874602 | 94873373 | 94873477 |
| SE | 15357 | ENSG00000140521 | POLG | chr15 | - | 89317375 | 89317536 | 89316304 | 89316827 |
| SE | 60550 | ENSG00000116984 | MTR | chr1 | + | 236897005 | 236897118 | 236891222 | 236895550 |
| SE | 78187 | ENSG00000136273 | HUS1 | chr7 | - | 47976214 | 47976421 | 47975612 | 47975687 |
| SE | 76695 | ENSG00000231925 | TAPBP | chr6 | - | 33303954 | 33303989 | 33299693 | 33301771 |
| SE | 89056 | ENSG00000104081 | BMF | chr15 | - | 40104179 | 40104340 | 40087889 | 40091888 |
| SE | 37843 | ENSG00000114126 | TFDP2 | chr3 | - | 142000251 | 142000354 | 141995019 | 141995141 |
| SE | 40617 | ENSG00000213390 | ARHGAP19 | chr10 | - | 97246271 | 97246337 | 97243967 | 97244159 |
| SE | 31600 | ENSG00000082805 | ERC1 | chr12 | + | 1204500 | 1204512 | 1189858 | 1190052 |
| SE | 52626 | ENSG00000197872 | CYRIA | chr2 | - | 16588049 | 16588129 | 16565645 | 16565767 |
| SE | 1552 | ENSG00000198089 | SFI1 | chr22 | + | 31497247 | 31497348 | 31496557 | 31496637 |
| SE | 35059 | ENSG00000156030 | MIDEAS | chr14 | - | 73722697 | 73722847 | 73715121 | 73719501 |
| SE | 7785 | ENSG00000278053 | DDX52 | chr17 | - | 37620872 | 37620948 | 37619767 | 37619839 |
| SE | 97304 | ENSG00000139626 | ITGB7 | chr12 | - | 53193139 | 53193363 | 53192329 | 53192538 |
| SE | 79582 | ENSG00000139641 | ESYT1 | chr12 | + | 56142284 | 56142425 | 56138926 | 56139013 |
| SE | 71205 | ENSG00000079263 | SP140 | chr2 | + | 230243730 | 230243811 | 230241403 | 230241487 |
| SE | 68321 | ENSG00000168016 | TRANK1 | chr3 | - | 36874125 | 36874296 | 36864318 | 36864480 |
| SE | 82997 | ENSG00000265681 | RPL17 | chr18 | - | 49491745 | 49491833 | 49491531 | 49491584 |
| SE | 77501 | ENSG00000106415 | GLCCI1 | chr7 | + | 8024714 | 8024814 | 8022482 | 8022569 |
| SE | 101921 | ENSG00000138326 | RPS24 | chr10 | + | 78040203 | 78040225 | 78037193 | 78037304 |
| SE | 22942 | ENSG00000133112 | TPT1 | chr13 | - | 45340711 | 45340785 | 45339993 | 45340184 |
| SE | 51435 | ENSG00000107745 | MICU1 | chr10 | - | 72532994 | 72533162 | 72508154 | 72508269 |
| SE | 14142 | ENSG00000005339 | CREBBP | chr16 | - | 3738558 | 3738672 | 3736649 | 3736815 |
| SE | 94602 | ENSG00000106829 | TLE4 | chr9 | + | 79649819 | 79649858 | 79627373 | 79627448 |
| SE | 51418 | ENSG00000143621 | ILF2 | chr1 | - | 153663018 | 153663133 | 153662704 | 153662795 |
| SE | 32286 | ENSG00000163964 | PIGX | chr3 | + | 196717808 | 196717954 | 196716857 | 196716921 |
| SE | 9164 | ENSG00000154957 | ZNF18 | chr17 | - | 11983296 | 11983404 | 11977451 | 11978744 |
| SE | 98893 | ENSG00000013523 | ANGEL1 | chr14 | - | 76806415 | 76806849 | 76803785 | 76803912 |
| SE | 49869 | ENSG00000151136 | BTBD11 | chr12 | + | 107649205 | 107649274 | 107642089 | 107642171 |
| SE | 59597 | ENSG00000085377 | PREP | chr6 | - | 105285485 | 105285580 | 105282450 | 105282582 |
| SE | 26449 | ENSG00000083312 | TNPO1 | chr5 | + | 72888077 | 72888303 | 72887069 | 72887222 |
| SE | 76373 | ENSG00000164674 | SYTL3 | chr6 | + | 158745479 | 158745658 | 158725502 | 158725637 |
| SE | 50186 | ENSG00000074706 | IPCEF1 | chr6 | - | 154247448 | 154247488 | 154246590 | 154246760 |
| SE | 16958 | ENSG00000172575 | RASGRP1 | chr15 | - | 38501374 | 38501435 | 38501142 | 38501287 |
| SE | 99430 | ENSG00000110321 | EIF4G2 | chr11 | - | 10806819 | 10806885 | 10805906 | 10806047 |
| SE | 108845 | ENSG00000081189 | MEF2C | chr5 | - | 88880923 | 88881022 | 88823734 | 88823927 |
| SE | 50724 | ENSG00000173039 | RELA | chr11 | - | 65658286 | 65658317 | 65655854 | 65655935 |
| SE | 15561 | ENSG00000184056 | VPS33B | chr15 | - | 91006651 | 91006729 | 91006371 | 91006445 |
| SE | 76837 | ENSG00000124788 | ATXN1 | chr6 | - | 16432945 | 16433096 | 16326393 | 16328470 |
| SE | 93521 | ENSG00000119487 | MAPKAP1 | chr9 | - | 125471082 | 125471237 | 125467971 | 125468109 |
| SE | 43452 | ENSG00000095261 | PSMD5 | chr9 | - | 120829098 | 120829208 | 120824493 | 120824685 |
| SE | 78351 | ENSG00000072609 | CHFR | chr12 | - | 132843010 | 132843083 | 132832355 | 132841596 |
| SE | 47648 | ENSG00000288701 | PRRC2B | chr9 | + | 131482370 | 131482562 | 131481725 | 131481808 |
| SE | 15830 | ENSG00000198901 | PRC1 | chr15 | - | 90981498 | 90981669 | 90980883 | 90981033 |
| SE | 66812 | ENSG00000137075 | RNF38 | chr9 | - | 36390466 | 36390616 | 36375933 | 36376127 |
| SE | 79688 | ENSG00000134905 | CARS2 | chr13 | - | 110687946 | 110688018 | 110683050 | 110683134 |
| SE | 103593 | ENSG00000136754 | ABI1 | chr10 | - | 26759061 | 26759238 | 26751597 | 26751783 |
| SE | 75946 | ENSG00000196730 | DAPK1 | chr9 | + | 87697006 | 87697204 | 87686550 | 87686739 |
| SE | 47634 | ENSG00000260537 | NA | chr16 | + | 70317495 | 70317588 | 70315968 | 70316104 |
| SE | 60383 | ENSG00000242247 | ARFGAP3 | chr22 | - | 42810812 | 42810944 | 42808766 | 42808890 |
| SE | 58252 | ENSG00000089127 | OAS1 | chr12 | + | 112911097 | 112911258 | 112908535 | 112908824 |
| SE | 71228 | ENSG00000079263 | SP140 | chr2 | + | 230255451 | 230255532 | 230248884 | 230248968 |
| SE | 60359 | ENSG00000172766 | NAA16 | chr13 | + | 41362730 | 41362873 | 41362125 | 41362159 |
| SE | 34088 | ENSG00000114867 | EIF4G1 | chr3 | + | 184325850 | 184325957 | 184325479 | 184325639 |
| SE | 101649 | ENSG00000128245 | YWHAH | chr22 | + | 31945499 | 31945678 | 31944521 | 31944820 |
| SE | 5063 | ENSG00000101247 | NDUFAF5 | chr20 | + | 13794837 | 13794941 | 13793159 | 13793227 |
| SE | 91145 | ENSG00000116852 | KIF21B | chr1 | - | 201004540 | 201004933 | 201004339 | 201004455 |
| SE | 89269 | ENSG00000104325 | DECR1 | chr8 | + | 90017202 | 90017326 | 90001445 | 90001561 |
| SE | 98244 | ENSG00000131979 | GCH1 | chr14 | - | 54844000 | 54844143 | 54842007 | 54843054 |
| SE | 95534 | ENSG00000187764 | SEMA4D | chr9 | - | 89388635 | 89388792 | 89387385 | 89387608 |
| SE | 86336 | ENSG00000135048 | CEMIP2 | chr9 | - | 71746200 | 71746341 | 71740062 | 71740232 |
| SE | 101355 | ENSG00000170525 | PFKFB3 | chr10 | + | 6217134 | 6217191 | 6216705 | 6216780 |
| SE | 53798 | ENSG00000111653 | ING4 | chr12 | - | 6655637 | 6655913 | 6653229 | 6653396 |
| SE | 55478 | ENSG00000143258 | USP21 | chr1 | + | 161162614 | 161162726 | 161162269 | 161162390 |
| SE | 101342 | ENSG00000124795 | DEK | chr6 | - | 18244507 | 18244603 | 18237380 | 18237516 |
| SE | 57004 | ENSG00000079999 | KEAP1 | chr19 | - | 10489647 | 10489853 | 10489191 | 10489368 |
| SE | 83079 | ENSG00000184863 | RBM33 | chr7 | + | 155684932 | 155685052 | 155680589 | 155680908 |
| SE | 96325 | ENSG00000135387 | CAPRIN1 | chr11 | + | 34097195 | 34097296 | 34096478 | 34096673 |
| SE | 40183 | ENSG00000114544 | SLC41A3 | chr3 | - | 126050897 | 126051050 | 126033606 | 126033678 |
| SE | 23376 | ENSG00000147044 | CASK | chrX | - | 41671427 | 41671530 | 41665276 | 41665452 |
| SE | 58247 | ENSG00000089127 | OAS1 | chr12 | + | 112911050 | 112911258 | 112908535 | 112908824 |
| SE | 50725 | ENSG00000173039 | RELA | chr11 | - | 65658286 | 65658499 | 65655854 | 65655935 |
| SE | 8647 | ENSG00000261371 | PECAM1 | chr17 | - | 64360139 | 64360415 | 64356110 | 64356398 |
| SE | 63719 | ENSG00000166986 | MARS1 | chr12 | + | 57489895 | 57489971 | 57489266 | 57489558 |
| SE | 50811 | ENSG00000117481 | NSUN4 | chr1 | + | 46346920 | 46347075 | 46344800 | 46344962 |
| SE | 47649 | ENSG00000288701 | PRRC2B | chr9 | + | 131482373 | 131482562 | 131481725 | 131481808 |
| SE | 102838 | ENSG00000177239 | MAN1B1 | chr9 | + | 137105607 | 137105916 | 137101483 | 137101672 |
| SE | 57104 | ENSG00000162734 | PEA15 | chr1 | + | 160208463 | 160208665 | 160207132 | 160207237 |
| SE | 84356 | ENSG00000230590 | FTX | chrX | - | 74261911 | 74262030 | 74202646 | 74203953 |
| SE | 103832 | ENSG00000152582 | SPEF2 | chr5 | + | 35644354 | 35644525 | 35641430 | 35641683 |
| SE | 79361 | ENSG00000280828 | NA | chr7 | + | 128577963 | 128578033 | 128574719 | 128574791 |
| SE | 49098 | ENSG00000072736 | NFATC3 | chr16 | + | 68214328 | 68214432 | 68190767 | 68191775 |
| SE | 61879 | ENSG00000076242 | MLH1 | chr3 | + | 37008813 | 37008905 | 37006990 | 37007063 |
| SE | 35968 | ENSG00000078070 | MCCC1 | chr3 | - | 183017265 | 183017337 | 183015217 | 183015566 |
| SE | 86023 | ENSG00000244754 | N4BP2L2 | chr13 | - | 32536932 | 32537027 | 32527407 | 32527532 |
| SE | 10114 | ENSG00000108433 | GOSR2 | chr17 | + | 46935028 | 46935169 | 46932066 | 46932199 |
| SE | 79037 | ENSG00000160218 | TRAPPC10 | chr21 | + | 44091922 | 44092049 | 44088038 | 44089933 |
| SE | 68979 | ENSG00000009954 | BAZ1B | chr7 | - | 73476867 | 73478569 | 73470344 | 73470483 |
| SE | 48853 | ENSG00000163563 | MNDA | chr1 | + | 158845586 | 158846003 | 158843954 | 158844122 |
| SE | 86759 | ENSG00000124275 | MTRR | chr5 | + | 7886614 | 7886703 | 7885700 | 7885854 |
| SE | 34064 | ENSG00000114867 | EIF4G1 | chr3 | + | 184317320 | 184317431 | 184316131 | 184316218 |
| SE | 97716 | ENSG00000149212 | SESN3 | chr11 | - | 95185255 | 95185492 | 95184134 | 95184594 |
| SE | 60543 | ENSG00000116984 | MTR | chr1 | + | 236861124 | 236861277 | 236859832 | 236859922 |
| SE | 3793 | ENSG00000090372 | STRN4 | chr19 | - | 46733815 | 46733875 | 46733038 | 46733236 |
| SE | 14500 | ENSG00000181827 | RFX7 | chr15 | - | 56179058 | 56179133 | 56144400 | 56144483 |
| SE | 17766 | ENSG00000133997 | MED6 | chr14 | - | 70592879 | 70593009 | 70591265 | 70591381 |
| SE | 65824 | ENSG00000138795 | LEF1 | chr4 | - | 108081585 | 108081669 | 108079491 | 108079614 |
| SE | 105487 | ENSG00000152256 | PDK1 | chr2 | + | 172564502 | 172564687 | 172564051 | 172564130 |
| SE | 34701 | ENSG00000187514 | PTMA | chr2 | + | 231711889 | 231711983 | 231711347 | 231711419 |
| SE | 20856 | ENSG00000255819 | KLRC4-KLRK1 | chr12 | - | 10405849 | 10405950 | 10391834 | 10391891 |
| SE | 14759 | ENSG00000103994 | ZNF106 | chr15 | - | 42417804 | 42417951 | 42412822 | 42417360 |
| SE | 92455 | ENSG00000163378 | EOGT | chr3 | - | 69009636 | 69009860 | 69008427 | 69008528 |
| SE | 100687 | ENSG00000166387 | PPFIBP2 | chr11 | + | 7650839 | 7650965 | 7649531 | 7649654 |
| SE | 39444 | ENSG00000055917 | PUM2 | chr2 | - | 20326260 | 20326385 | 20318536 | 20318645 |
| SE | 71347 | ENSG00000085760 | MTIF2 | chr2 | - | 55268577 | 55268739 | 55267568 | 55267635 |
| SE | 29608 | ENSG00000203875 | SNHG5 | chr6 | - | 85677790 | 85678032 | 85677423 | 85677492 |
| SE | 2816 | ENSG00000104980 | TIMM44 | chr19 | - | 7935064 | 7935145 | 7934088 | 7934241 |
| SE | 52111 | ENSG00000158006 | PAFAH2 | chr1 | - | 25989447 | 25989601 | 25988230 | 25988327 |
| SE | 49749 | ENSG00000111331 | OAS3 | chr12 | + | 112965743 | 112966029 | 112964234 | 112964408 |
| SE | 39350 | ENSG00000158864 | NDUFS2 | chr1 | + | 161210110 | 161210188 | 161209482 | 161209931 |
| SE | 73626 | ENSG00000119707 | RBM25 | chr14 | + | 73103191 | 73103478 | 73099666 | 73099750 |
| SE | 10115 | ENSG00000108433 | GOSR2 | chr17 | + | 46935028 | 46935225 | 46932066 | 46932199 |
| SE | 10634 | ENSG00000277791 | PSMB3 | chr17 | + | 38760507 | 38760608 | 38755882 | 38755990 |
| SE | 105632 | ENSG00000238105 | GOLGA2P5 | chr12 | - | 100168451 | 100168633 | 100167549 | 100167639 |
| SE | 25351 | ENSG00000112992 | NNT | chr5 | + | 43619031 | 43619119 | 43615847 | 43616065 |
| SE | 66246 | ENSG00000123552 | USP45 | chr6 | - | 99468099 | 99468170 | 99466671 | 99466763 |
| SE | 62452 | ENSG00000120265 | PCMT1 | chr6 | + | 149772554 | 149772684 | 149771161 | 149771266 |
| SE | 37581 | ENSG00000082074 | FYB1 | chr5 | - | 39125971 | 39126135 | 39124252 | 39124278 |
| SE | 8514 | ENSG00000125447 | GGA3 | chr17 | - | 75243062 | 75243258 | 75242883 | 75242911 |
| SE | 20335 | ENSG00000111913 | RIPOR2 | chr6 | - | 24929281 | 24929447 | 24875690 | 24875817 |
| SE | 22981 | ENSG00000116459 | ATP5PB | chr1 | + | 111454210 | 111454356 | 111449681 | 111449873 |
| SE | 42125 | ENSG00000198952 | SMG5 | chr1 | - | 156250857 | 156250996 | 156249223 | 156250670 |
| SE | 11156 | ENSG00000168488 | ATXN2L | chr16 | + | 28826861 | 28826986 | 28826239 | 28826390 |
| SE | 59509 | ENSG00000148660 | CAMK2G | chr10 | - | 73848525 | 73848609 | 73847987 | 73848082 |
| SE | 50209 | ENSG00000148187 | MRRF | chr9 | + | 122270952 | 122271075 | 122264903 | 122264938 |
| SE | 95648 | ENSG00000137073 | UBAP2 | chr9 | - | 33956078 | 33956146 | 33948373 | 33948587 |
| SE | 7845 | ENSG00000108819 | PPP1R9B | chr17 | - | 50139428 | 50139581 | 50139262 | 50139316 |
| SE | 46765 | ENSG00000117115 | PADI2 | chr1 | - | 17079263 | 17079415 | 17074855 | 17075823 |
| SE | 100593 | ENSG00000102317 | RBM3 | chrX | + | 48575560 | 48576083 | 48575167 | 48575283 |
| SE | 39404 | ENSG00000160712 | IL6R | chr1 | + | 154454487 | 154454581 | 154448124 | 154448171 |
| SE | 20109 | ENSG00000078319 | PMS2P1 | chr7 | - | 100325690 | 100325779 | 100320438 | 100321084 |
| SE | 30686 | ENSG00000132466 | ANKRD17 | chr4 | - | 73118687 | 73118850 | 73115820 | 73115916 |
| SE | 26009 | ENSG00000181929 | PRKAG1 | chr12 | - | 49016606 | 49016839 | 49013061 | 49013110 |
| SE | 39354 | ENSG00000158864 | NDUFS2 | chr1 | + | 161210303 | 161210389 | 161209856 | 161209931 |
| SE | 87581 | ENSG00000112159 | MDN1 | chr6 | - | 89743575 | 89743714 | 89743149 | 89743280 |
| SE | 49887 | ENSG00000168884 | TNIP2 | chr4 | - | 2744696 | 2744945 | 2744386 | 2744506 |
| SE | 29888 | ENSG00000141562 | NARF | chr17 | + | 82478570 | 82478667 | 82472563 | 82472698 |
| SE | 9165 | ENSG00000154957 | ZNF18 | chr17 | - | 11983296 | 11983407 | 11977438 | 11978744 |
| SE | 28405 | ENSG00000204842 | ATXN2 | chr12 | - | 111518248 | 111518427 | 111516153 | 111516363 |
| SE | 81971 | ENSG00000051825 | MPHOSPH9 | chr12 | - | 123227462 | 123227616 | 123226287 | 123226351 |
| SE | 102885 | ENSG00000154124 | OTULIN | chr5 | + | 14690038 | 14690308 | 14687520 | 14687646 |
| SE | 95877 | ENSG00000148396 | SEC16A | chr9 | - | 136451255 | 136451408 | 136448083 | 136448161 |
| SE | 82364 | ENSG00000204257 | HLA-DMA | chr6 | - | 32949610 | 32949787 | 32949270 | 32949399 |
| SE | 87339 | ENSG00000215301 | DDX3X | chrX | + | 41347311 | 41347451 | 41346858 | 41347012 |
| SE | 79340 | ENSG00000231721 | LINC-PINT | chr7 | - | 130967388 | 130967517 | 130945609 | 130945835 |
| SE | 58248 | ENSG00000089127 | OAS1 | chr12 | + | 112911055 | 112911235 | 112908535 | 112908824 |
| SE | 54963 | ENSG00000169991 | IFFO2 | chr1 | - | 18919677 | 18919773 | 18918361 | 18918502 |
| SE | 58243 | ENSG00000089127 | OAS1 | chr12 | + | 112911035 | 112911235 | 112908535 | 112908824 |
| SE | 15574 | ENSG00000170776 | AKAP13 | chr15 | + | 85658536 | 85658590 | 85655416 | 85655787 |
| SE | 73430 | ENSG00000177463 | NR2C2 | chr3 | + | 15032378 | 15032500 | 15030274 | 15030452 |

| | | | | | | | | | | | | | | | | | | | | | | | | | | | | | | | | | | | | | | | | | | | | | | |
|---|---|---|---|---|---|---|---|---|---|---|---|---|---|---|---|---|---|---|---|---|---|---|---|---|---|---|---|---|---|---|---|---|---|---|---|---|---|---|---|---|---|---|---|---|---|---|
| SE | 64284 | ENSG00000068878 | PSME4 | chr2 | - | 53899880 | 53900017 | 53898300 | 53898354 | 53901349 | 53901559 | 255 | 180 | 210 | 278 | 445 | 18 | 4 | 3 | 4 | 7 | 145 | 206 | 217 | 194 | 61 | 7 | 25 | 32 | 18 | 0 | 286 | 149 | 7.52E-08 | 5.05E-06 | 0.881 | 0.959 | 0.973 | 0.973 | 0.971 | 0.915 | 0.811 | 0.779 | 0.849 | 1 | 0.081 |
| SE | 102577 | ENSG00000120438 | TCP1 | chr6 | - | 159779894 | 159780087 | 159778984 | 159779261 | 159780442 | 159780566 | 342 | 288 | 450 | 442 | 336 | 1 | 1 | 0 | 1 | 0 | 301 | 298 | 469 | 352 | 108 | 22 | 7 | 3 | 3 | 0 | 298 | 149 | 7.68E-08 | 5.16E-06 | 0.994 | 0.993 | 1 | 0.995 | 1 | 0.872 | 0.955 | 0.987 | 0.983 | 1 | 0.037 |
| SE | 68295 | ENSG00000122085 | MTERF4 | chr2 | - | 241097242 | 241097427 | 241094276 | 241094432 | 241099395 | 241099426 | 42 | 43 | 74 | 49 | 66 | 0 | 0 | 0 | 0 | 0 | 52 | 42 | 20 | 36 | 41 | 14 | 0 | 1 | 4 | 0 | 298 | 149 | 7.76E-08 | 5.21E-06 | 1 | 1 | 1 | 1 | 1 | 0.65 | 1 | 0.909 | 0.818 | 1 | 0.125 |
| SE | 41642 | ENSG00000115947 | ORC4 | chr2 | - | 147981927 | 147982081 | 147975901 | 147975975 | 148020632 | 148020724 | 69 | 39 | 22 | 57 | 53 | 48 | 36 | 45 | 59 | 13 | 188 | 49 | 32 | 53 | 108 | 0 | 15 | 14 | 13 | 0 | 298 | 149 | 7.85E-08 | 5.26E-06 | 0.418 | 0.351 | 0.196 | 0.326 | 0.671 | 1 | 0.62 | 0.533 | 0.671 | 1 | -0.372 |
| SE | 56689 | ENSG00000019582 | CD74 | chr5 | - | 150403120 | 150403312 | 150400040 | 150402250 | 150404679 | 150404767 | 490 | 271 | 284 | 350 | 408 | 1 | 0 | 0 | 0 | 0 | 792 | 441 | 192 | 334 | 1231 | 33 | 0 | 0 | 12 | 7 | 298 | 149 | 7.90E-08 | 5.29E-06 | 0.996 | 1 | 1 | 1 | 1 | 0.923 | 1 | 1 | 0.933 | 0.989 | 0.03 |
| SE | 108350 | ENSG00000135439 | AGAP2 | chr12 | - | 57732844 | 57732979 | 57732402 | 57732512 | 57734025 | 57734173 | 175 | 81 | 115 | 147 | 93 | 4 | 7 | 1 | 8 | 3 | 82 | 199 | 51 | 45 | 124 | 5 | 16 | 19 | 12 | 27 | 284 | 149 | 7.95E-08 | 5.32E-06 | 0.958 | 0.859 | 0.984 | 0.906 | 0.942 | 0.896 | 0.867 | 0.585 | 0.663 | 0.707 | 0.186 |
| SE | 32710 | ENSG00000058056 | USP13 | chr3 | + | 179721405 | 179721589 | 179719939 | 179720034 | 179730188 | 179730260 | 17 | 8 | 19 | 31 | 27 | 0 | 0 | 0 | 0 | 0 | 34 | 6 | 19 | 10 | 4 | 20 | 0 | 1 | 5 | 0 | 298 | 149 | 8.02E-08 | 5.36E-06 | 1 | 1 | 1 | 1 | 1 | 0.459 | 1 | 0.905 | 0.5 | 1 | 0.227 |
| SE | 106423 | ENSG00000135297 | MTO1 | chr6 | + | 73480317 | 73480367 | 73479935 | 73480126 | 73480674 | 73480805 | 17 | 5 | 11 | 9 | 15 | 130 | 101 | 111 | 106 | 161 | 0 | 0 | 8 | 1 | 0 | 90 | 101 | 84 | 74 | 60 | 199 | 149 | 8.02E-08 | 5.36E-06 | 0.089 | 0.036 | 0.069 | 0.06 | 0.065 | 0 | 0 | 0.067 | 0.01 | 0 | 0.048 |
| SE | 54087 | ENSG00000115935 | WIPF1 | chr2 | - | 174575203 | 174575371 | 174572323 | 174572446 | 174581309 | 174581439 | 398 | 219 | 345 | 361 | 423 | 2 | 6 | 5 | 8 | 1 | 291 | 333 | 279 | 336 | 674 | 0 | 12 | 26 | 11 | 33 | 298 | 149 | 8.08E-08 | 5.40E-06 | 0.99 | 0.948 | 0.972 | 0.958 | 0.995 | 1 | 0.933 | 0.843 | 0.939 | 0.911 | 0.047 |
| SE | 59295 | ENSG00000119004 | CYP20A1 | chr2 | + | 203285611 | 203285732 | 203280058 | 203280113 | 203289764 | 203289876 | 180 | 97 | 177 | 219 | 331 | 0 | 0 | 0 | 2 | 5 | 153 | 124 | 102 | 149 | 114 | 10 | 3 | 2 | 11 | 3 | 270 | 149 | 8.11E-08 | 5.41E-06 | 1 | 1 | 1 | 0.984 | 0.973 | 0.894 | 0.958 | 0.966 | 0.882 | 0.954 | 0.061 |
| SE | 56530 | ENSG00000189339 | SLC35E2B | chr1 | - | 1671508 | 1671629 | 1668326 | 1668472 | 1675462 | 1675590 | 28 | 19 | 56 | 65 | 25 | 0 | 0 | 0 | 0 | 3 | 16 | 53 | 11 | 30 | 64 | 0 | 10 | 2 | 6 | 5 | 270 | 149 | 8.19E-08 | 5.46E-06 | 1 | 1 | 1 | 1 | 0.821 | 1 | 0.745 | 0.752 | 0.734 | 0.876 | 0.143 |
| SE | 3673 | ENSG00000167613 | LAIR1 | chr19 | - | 54359842 | 54360072 | 54356927 | 54356966 | 54361009 | 54361209 | 5 | 13 | 7 | 11 | 12 | 0 | 1 | 0 | 0 | 1 | 12 | 5 | 1 | 19 | 17 | 10 | 2 | 2 | 8 | 0 | 298 | 149 | 8.20E-08 | 5.46E-06 | 1 | 0.867 | 1 | 1 | 0.857 | 0.375 | 0.556 | 0.2 | 0.543 | 1 | 0.41 |
| SE | 73116 | ENSG00000144231 | POLR2D | chr2 | - | 127850589 | 127850685 | 127843552 | 127848185 | 127852924 | 127853105 | 78 | 41 | 82 | 67 | 181 | 7 | 6 | 2 | 2 | 7 | 99 | 68 | 77 | 42 | 50 | 0 | 0 | 0 | 0 | 0 | 245 | 149 | 8.24E-08 | 5.48E-06 | 0.871 | 0.806 | 0.961 | 0.953 | 0.94 | 1 | 1 | 1 | 1 | 1 | -0.094 |
| SE | 89164 | ENSG00000075856 | SART3 | chr12 | - | 108532221 | 108532334 | 108531203 | 108531280 | 108536513 | 108536572 | 58 | 38 | 63 | 46 | 53 | 0 | 1 | 0 | 0 | 0 | 68 | 38 | 76 | 47 | 33 | 0 | 7 | 4 | 11 | 0 | 262 | 149 | 8.24E-08 | 5.48E-06 | 1 | 0.956 | 1 | 1 | 1 | 1 | 0.755 | 0.915 | 0.708 | 1 | 0.116 |
| SE | 66348 | ENSG00000197037 | ZSCAN25 | chr7 | + | 99619560 | 99619993 | 99618524 | 99618651 | 99622548 | 99622640 | 13 | 12 | 6 | 17 | 13 | 0 | 0 | 0 | 0 | 1 | 5 | 50 | 22 | 34 | 14 | 4 | 12 | 1 | 1 | 5 | 298 | 149 | 8.26E-08 | 5.49E-06 | 1 | 1 | 1 | 1 | 0.867 | 0.385 | 0.676 | 0.917 | 0.944 | 0.583 | 0.272 |
| SE | 27787 | ENSG00000177058 | SLC38A9 | chr5 | - | 55652528 | 55652723 | 55649206 | 55649314 | 55656714 | 55656774 | 73 | 35 | 45 | 54 | 104 | 10 | 2 | 5 | 1 | 10 | 11 | 40 | 15 | 26 | 15 | 0 | 0 | 1 | 0 | 0 | 298 | 149 | 8.39E-08 | 5.57E-06 | 0.785 | 0.897 | 0.818 | 0.964 | 0.839 | 1 | 1 | 0.882 | 1 | 1 | -0.116 |
| SE | 23450 | ENSG00000166888 | STAT6 | chr12 | - | 57107604 | 57107743 | 57107230 | 57107314 | 57108162 | 57108283 | 626 | 306 | 470 | 584 | 353 | 96 | 60 | 81 | 62 | 48 | 1002 | 580 | 304 | 477 | 1206 | 268 | 109 | 100 | 178 | 302 | 288 | 149 | 8.41E-08 | 5.57E-06 | 0.771 | 0.725 | 0.75 | 0.83 | 0.792 | 0.659 | 0.734 | 0.611 | 0.581 | 0.674 | 0.122 |
| SE | 88708 | ENSG00000183077 | AFMID | chr17 | + | 78205945 | 78206050 | 78202702 | 78202751 | 78206910 | 78207052 | 16 | 12 | 14 | 22 | 42 | 3 | 3 | 1 | 1 | 6 | 20 | 5 | 17 | 12 | 47 | 0 | 0 | 0 | 0 | 0 | 254 | 149 | 8.41E-08 | 5.57E-06 | 0.758 | 0.701 | 0.891 | 0.928 | 0.804 | 1 | 1 | 1 | 1 | 1 | -0.184 |
| SE | 49754 | ENSG00000111331 | OAS3 | chr12 | + | 112967417 | 112967593 | 112964234 | 112964408 | 112967935 | 112968174 | 46 | 61 | 33 | 55 | 17 | 2 | 0 | 0 | 0 | 0 | 243 | 64 | 30 | 256 | 62 | 28 | 0 | 0 | 27 | 0 | 298 | 149 | 8.49E-08 | 5.62E-06 | 0.92 | 1 | 1 | 1 | 1 | 0.813 | 1 | 1 | 0.826 | 1 | 0.056 |
| SE | 73480 | ENSG00000215559 | ANKRD20A11P | chr21 | - | 13916746 | 13916958 | 13911717 | 13911763 | 13917554 | 13917836 | 0 | 5 | 9 | 36 | 8 | 8 | 1 | 1 | 2 | 2 | 18 | 36 | 13 | 9 | 3 | 0 | 0 | 0 | 0 | 0 | 298 | 149 | 8.55E-08 | 5.65E-06 | 0 | 0.714 | 0.818 | 0.9 | 0.667 | 1 | 1 | 1 | 1 | 1 | -0.38 |
| SE | 20898 | ENSG00000152969 | JAKMIP1 | chr4 | - | 6078938 | 6078998 | 6064879 | 6065008 | 6080171 | 6080312 | 73 | 20 | 19 | 19 | 46 | 8 | 11 | 2 | 2 | 5 | 53 | 80 | 32 | 23 | 40 | 28 | 47 | 24 | 25 | 39 | 209 | 149 | 8.56E-08 | 5.66E-06 | 0.867 | 0.565 | 0.871 | 0.871 | 0.868 | 0.574 | 0.548 | 0.487 | 0.396 | 0.422 | 0.323 |
| SE | 14833 | ENSG00000069943 | PIGB | chr15 | + | 55350698 | 55350803 | 55341737 | 55341802 | 55354797 | 55354978 | 48 | 24 | 30 | 28 | 134 | 9 | 2 | 5 | 9 | 9 | 23 | 19 | 28 | 26 | 31 | 0 | 2 | 1 | 0 | 0 | 254 | 149 | 8.61E-08 | 5.68E-06 | 0.758 | 0.876 | 0.779 | 0.646 | 0.897 | 1 | 0.848 | 0.943 | 1 | 1 | -0.167 |
| SE | 81973 | ENSG00000051825 | MPHOSPH9 | chr12 | - | 123227545 | 123227616 | 123226287 | 123226351 | 123230260 | 123230522 | 59 | 45 | 72 | 43 | 95 | 0 | 2 | 0 | 0 | 1 | 38 | 53 | 27 | 37 | 5 | 8 | 0 | 1 | 19 | 0 | 220 | 149 | 8.67E-08 | 5.72E-06 | 1 | 0.938 | 1 | 1 | 0.985 | 0.763 | 1 | 0.948 | 0.569 | 1 | 0.129 |
| SE | 34316 | ENSG00000108219 | TSPAN14 | chr10 | + | 80468546 | 80468687 | 80459301 | 80459507 | 80489216 | 80489314 | 94 | 38 | 78 | 76 | 40 | 13 | 3 | 3 | 0 | 6 | 98 | 52 | 36 | 46 | 108 | 0 | 3 | 0 | 0 | 0 | 290 | 149 | 8.70E-08 | 5.73E-06 | 0.788 | 0.867 | 0.93 | 1 | 0.774 | 1 | 0.899 | 1 | 1 | 1 | -0.108 |
| SE | 58250 | ENSG00000089127 | OAS1 | chr12 | + | 112911097 | 112911235 | 112908535 | 112908824 | 112916508 | 112916738 | 267 | 187 | 154 | 227 | 172 | 0 | 1 | 0 | 0 | 0 | 870 | 491 | 148 | 508 | 269 | 0 | 7 | 1 | 18 | 24 | 287 | 149 | 8.76E-08 | 5.77E-06 | 1 | 0.99 | 1 | 1 | 1 | 1 | 0.973 | 0.987 | 0.936 | 0.853 | 0.048 |
| SE | 9679 | ENSG00000055483 | USP36 | chr17 | - | 78814411 | 78814552 | 78813772 | 78813873 | 78818666 | 78818778 | 105 | 47 | 115 | 90 | 79 | 0 | 0 | 0 | 0 | 0 | 54 | 49 | 125 | 68 | 75 | 0 | 10 | 3 | 8 | 0 | 290 | 149 | 8.77E-08 | 5.77E-06 | 1 | 1 | 1 | 1 | 1 | 1 | 0.716 | 0.955 | 0.814 | 1 | 0.103 |
| SE | 48503 | ENSG00000138095 | LRPPRC | chr2 | - | 43900507 | 43900583 | 43899465 | 43899605 | 43901319 | 43901524 | 0 | 3 | 3 | 1 | 6 | 183 | 136 | 184 | 200 | 235 | 0 | 11 | 6 | 0 | 16 | 16 | 85 | 120 | 38 | 19 | 225 | 149 | 8.78E-08 | 5.77E-06 | 0 | 0.014 | 0.011 | 0.003 | 0.017 | 0 | 0.079 | 0.032 | 0 | 0.358 | -0.085 |
| SE | 103653 | ENSG00000120616 | EPC1 | chr10 | - | 32292495 | 32292644 | 32291162 | 32291322 | 32292987 | 32293194 | 602 | 421 | 511 | 587 | 1020 | 1 | 4 | 7 | 4 | 15 | 294 | 377 | 532 | 344 | 182 | 12 | 13 | 14 | 15 | 3 | 298 | 149 | 8.80E-08 | 5.78E-06 | 0.997 | 0.981 | 0.973 | 0.987 | 0.971 | 0.925 | 0.935 | 0.95 | 0.92 | 0.968 | 0.042 |
| SE | 34955 | ENSG00000147854 | UHRF2 | chr9 | + | 6477621 | 6477808 | 6475390 | 6475500 | 6481642 | 6481766 | 237 | 176 | 213 | 304 | 338 | 0 | 0 | 0 | 0 | 2 | 85 | 128 | 178 | 94 | 116 | 12 | 2 | 8 | 1 | 0 | 298 | 149 | 8.82E-08 | 5.79E-06 | 1 | 1 | 1 | 1 | 0.988 | 0.78 | 0.97 | 0.918 | 0.979 | 1 | 0.068 |
| SE | 69406 | ENSG00000164951 | PDP1 | chr8 | + | 93917910 | 93917945 | 93916933 | 93917079 | 93922015 | 93922289 | 8 | 4 | 6 | 9 | 3 | 74 | 20 | 45 | 40 | 44 | 0 | 0 | 2 | 3 | 0 | 41 | 64 | 65 | 40 | 27 | 184 | 149 | 8.85E-08 | 5.80E-06 | 0.08 | 0.139 | 0.097 | 0.154 | 0.052 | 0 | 0 | 0.024 | 0.057 | 0 | 0.088 |
| SE | 60775 | ENSG00000187741 | FANCA | chr16 | - | 89758576 | 89758705 | 89752137 | 89752222 | 89761948 | 89762022 | 41 | 28 | 19 | 17 | 28 | 0 | 3 | 2 | 4 | 8 | 60 | 26 | 16 | 35 | 52 | 0 | 0 | 2 | 0 | 0 | 278 | 149 | 8.85E-08 | 5.80E-06 | 1 | 0.833 | 0.836 | 0.695 | 0.652 | 1 | 1 | 0.811 | 1 | 1 | -0.159 |
| SE | 62090 | ENSG00000143486 | EIF2D | chr1 | - | 206599732 | 206599836 | 206599002 | 206599092 | 206600262 | 206600308 | 122 | 63 | 86 | 75 | 80 | 1 | 0 | 1 | 2 | 0 | 99 | 104 | 64 | 67 | 101 | 22 | 3 | 0 | 11 | 0 | 253 | 149 | 8.86E-08 | 5.80E-06 | 0.986 | 1 | 0.981 | 0.957 | 1 | 0.726 | 0.953 | 1 | 0.782 | 1 | 0.093 |
| SE | 59619 | ENSG00000111801 | BTN3A3 | chr6 | + | 26445703 | 26445985 | 26443956 | 26444304 | 26448727 | 26448754 | 107 | 46 | 81 | 85 | 137 | 8 | 0 | 11 | 14 | 10 | 52 | 96 | 42 | 38 | 115 | 45 | 23 | 37 | 16 | 24 | 298 | 149 | 8.89E-08 | 5.82E-06 | 0.87 | 1 | 0.786 | 0.752 | 0.873 | 0.366 | 0.676 | 0.362 | 0.543 | 0.706 | 0.326 |
| SE | 195 | ENSG00000160216 | AGPAT3 | chr21 | + | 43978045 | 43978121 | 43971387 | 43971490 | 43980988 | 43981187 | 100 | 110 | 109 | 116 | 89 | 1 | 0 | 1 | 0 | 0 | 364 | 87 | 105 | 130 | 287 | 11 | 6 | 5 | 10 | 7 | 225 | 149 | 8.99E-08 | 5.88E-06 | 0.985 | 1 | 0.986 | 1 | 1 | 0.956 | 0.906 | 0.933 | 0.896 | 0.964 | 0.063 |
| SE | 48941 | ENSG00000102805 | CLN5 | chr13 | + | 76999929 | 76999995 | 76995901 | 76996127 | 77000457 | 77002345 | 2 | 0 | 0 | 1 | 1 | 59 | 37 | 39 | 40 | 132 | 8 | 0 | 4 | 7 | 0 | 12 | 17 | 13 | 26 | 21 | 215 | 149 | 9.00E-08 | 5.88E-06 | 0.023 | 0 | 0 | 0.017 | 0.005 | 0.316 | 0 | 0.176 | 0.157 | 0 | -0.121 |
| SE | 81949 | ENSG00000051825 | MPHOSPH9 | chr12 | - | 123202163 | 123202319 | 123198246 | 123198334 | 123202623 | 123203084 | 168 | 125 | 219 | 186 | 363 | 11 | 5 | 5 | 4 | 2 | 31 | 120 | 54 | 31 | 5 | 16 | 18 | 6 | 5 | 0 | 298 | 149 | 9.01E-08 | 5.88E-06 | 0.884 | 0.926 | 0.956 | 0.959 | 0.989 | 0.492 | 0.769 | 0.818 | 0.756 | 1 | 0.176 |
| SE | 41374 | ENSG00000114796 | KLHL24 | chr3 | + | 183636220 | 183636311 | 183635622 | 183635793 | 183643479 | 183643542 | 14 | 21 | 20 | 30 | 23 | 39 | 30 | 43 | 40 | 41 | 0 | 2 | 13 | 0 | 0 | 9 | 40 | 32 | 21 | 5 | 240 | 149 | 9.07E-08 | 5.91E-06 | 0.182 | 0.303 | 0.224 | 0.318 | 0.258 | 0 | 0.03 | 0.201 | 0 | 0 | 0.211 |
| SE | 51552 | ENSG00000036549 | ZZZ3 | chr1 | - | 77641383 | 77641446 | 77639448 | 77639602 | 77641521 | 77641655 | 82 | 50 | 48 | 77 | 173 | 4 | 6 | 9 | 1 | 18 | 22 | 26 | 29 | 27 | 17 | 0 | 0 | 1 | 0 | 0 | 212 | 149 | 9.07E-08 | 5.91E-06 | 0.935 | 0.854 | 0.789 | 0.982 | 0.871 | 1 | 1 | 0.953 | 1 | 1 | -0.104 |
| SE | 63956 | ENSG00000115902 | SLC1A4 | chr2 | + | 65011311 | 65011545 | 65010596 | 65010763 | 65016439 | 65016554 | 0 | 0 | 2 | 1 | 0 | 43 | 23 | 42 | 53 | 28 | 0 | 13 | 7 | 5 | 8 | 22 | 22 | 70 | 92 | 15 | 298 | 149 | 9.12E-08 | 5.94E-06 | 0 | 0 | 0.023 | 0.009 | 0 | 0 | 0.228 | 0.048 | 0.026 | 0.211 | -0.096 |
| SE | 87334 | ENSG00000215301 | DDX3X | chrX | + | 41346858 | 41347012 | 41346228 | 41346622 | 41347311 | 41347451 | 1624 | 1111 | 1333 | 1641 | 2306 | 4 | 6 | 11 | 16 | 16 | 154 | 618 | 1130 | 557 | 57 | 24 | 18 | 5 | 8 | 0 | 298 | 149 | 9.14E-08 | 5.95E-06 | 0.995 | 0.989 | 0.984 | 0.981 | 0.986 | 0.762 | 0.945 | 0.991 | 0.972 | 1 | 0.053 |
| SE | 94865 | ENSG00000196597 | ZNF782 | chr9 | - | 96844889 | 96845016 | 96827079 | 96827181 | 96851946 | 96852005 | 30 | 26 | 25 | 31 | 39 | 5 | 5 | 1 | 0 | 7 | 26 | 35 | 33 | 7 | 16 | 0 | 0 | 1 | 0 | 0 | 276 | 149 | 9.26E-08 | 6.02E-06 | 0.764 | 0.737 | 0.931 | 1 | 0.75 | 1 | 1 | 0.947 | 1 | 1 | -0.153 |
| SE | 91065 | ENSG00000104164 | BLOC1S6 | chr15 | + | 45597879 | 45597891 | 45587351 | 45587525 | 45605427 | 45605514 | 5 | 3 | 2 | 4 | 0 | 7 | 6 | 2 | 14 | 6 | 0 | 0 | 0 | 2 | 0 | 25 | 14 | 17 | 6 | 22 | 161 | 149 | 9.40E-08 | 6.10E-06 | 0.398 | 0.316 | 0.481 | 0.209 | 0 | 0 | 0 | 0 | 0.236 | 0 | 0.234 |
| SE | 37688 | ENSG00000147912 | FBXO10 | chr9 | - | 37529123 | 37529260 | 37525101 | 37525172 | 37531908 | 37532058 | 19 | 14 | 27 | 18 | 11 | 1 | 0 | 1 | 0 | 0 | 27 | 68 | 22 | 25 | 36 | 11 | 5 | 1 | 11 | 2 | 286 | 149 | 9.42E-08 | 6.11E-06 | 0.908 | 1 | 0.934 | 1 | 1 | 0.561 | 0.876 | 0.92 | 0.542 | 0.904 | 0.208 |
| SE | 57768 | ENSG00000159596 | TMEM69 | chr1 | + | 45690973 | 45691110 | 45688180 | 45688275 | 45693203 | 45694436 | 254 | 126 | 168 | 211 | 294 | 16 | 13 | 13 | 11 | 9 | 147 | 219 | 130 | 154 | 132 | 8 | 4 | 0 | 0 | 0 | 286 | 149 | 9.50E-08 | 6.16E-06 | 0.892 | 0.835 | 0.871 | 0.909 | 0.945 | 0.905 | 0.966 | 1 | 1 | 1 | -0.084 |
| SE | 18453 | ENSG00000100897 | DCAF11 | chr14 | + | 24115381 | 24115392 | 24114776 | 24115293 | 24115542 | 24115726 | 20 | 10 | 18 | 15 | 22 | 0 | 1 | 0 | 2 | 0 | 36 | 3 | 3 | 22 | 47 | 8 | 0 | 1 | 9 | 10 | 160 | 149 | 9.55E-08 | 6.18E-06 | 1 | 0.903 | 1 | 0.875 | 1 | 0.807 | 1 | 0.736 | 0.695 | 0.814 | 0.145 |
| SE | 106345 | ENSG00000076513 | ANKRD13A | chr12 | + | 110018344 | 110018488 | 110016387 | 110016433 | 110019138 | 110019328 | 542 | 387 | 571 | 632 | 984 | 5 | 2 | 3 | 15 | 12 | 1403 | 743 | 651 | 668 | 901 | 32 | 31 | 9 | 52 | 10 | 293 | 149 | 9.61E-08 | 6.22E-06 | 0.982 | 0.99 | 0.99 | 0.955 | 0.977 | 0.957 | 0.924 | 0.974 | 0.867 | 0.979 | 0.039 |
| SE | 81633 | ENSG00000130856 | ZNF236 | chr18 | + | 76905154 | 76905415 | 76904379 | 76904521 | 76908319 | 76908573 | 97 | 51 | 63 | 75 | 113 | 0 | 0 | 0 | 0 | 0 | 68 | 51 | 49 | 36 | 44 | 0 | 5 | 5 | 7 | 0 | 298 | 149 | 9.64E-08 | 6.23E-06 | 1 | 1 | 1 | 1 | 1 | 1 | 0.836 | 0.831 | 0.72 | 1 | 0.123 |
| SE | 100443 | ENSG00000165916 | PSMC3 | chr11 | - | 47420630 | 47420727 | 47420263 | 47420409 | 47422573 | 47422722 | 154 | 70 | 149 | 113 | 67 | 0 | 0 | 0 | 0 | 1 | 76 | 127 | 81 | 82 | 186 | 9 | 1 | 7 | 8 | 0 | 246 | 149 | 9.67E-08 | 6.25E-06 | 1 | 1 | 1 | 1 | 0.976 | 0.836 | 0.987 | 0.875 | 0.861 | 1 | 0.083 |
| SE | 3723 | ENSG00000167555 | ZNF528 | chr19 | + | 52402428 | 52402545 | 52401946 | 52402028 | 52405906 | 52405931 | 5 | 13 | 7 | 7 | 5 | 22 | 23 | 21 | 43 | 55 | 0 | 0 | 4 | 0 | 0 | 6 | 17 | 16 | 35 | 34 | 266 | 149 | 9.68E-08 | 6.25E-06 | 0.113 | 0.24 | 0.157 | 0.084 | 0.048 | 0 | 0 | 0.123 | 0 | 0 | 0.104 |
| SE | 10944 | ENSG00000091592 | NLRP1 | chr17 | - | 5520880 | 5521012 | 5515472 | 5515517 | 5521523 | 5521786 | 127 | 78 | 123 | 118 | 106 | 0 | 0 | 0 | 2 | 0 | 243 | 131 | 117 | 86 | 221 | 12 | 2 | 0 | 3 | 15 | 281 | 149 | 9.71E-08 | 6.26E-06 | 1 | 1 | 1 | 0.969 | 1 | 0.915 | 0.972 | 1 | 0.938 | 0.887 | 0.051 |
| SE | 61488 | ENSG00000124588 | NQO2 | chr6 | + | 3002049 | 3002286 | 2999973 | 3000085 | 3003668 | 3003789 | 8 | 6 | 7 | 10 | 7 | 2 | 0 | 0 | 0 | 0 | 6 | 11 | 13 | 10 | 25 | 13 | 7 | 0 | 3 | 7 | 298 | 149 | 9.80E-08 | 6.32E-06 | 0.667 | 1 | 1 | 1 | 1 | 0.188 | 0.44 | 1 | 0.625 | 0.641 | 0.355 |
| SE | 50838 | ENSG00000048052 | HDAC9 | chr7 | + | 18395114 | 18395183 | 18325511 | 18325592 | 18496261 | 18496324 | 3 | 3 | 4 | 2 | 4 | 12 | 12 | 3 | 11 | 12 | 0 | 0 | 0 | 0 | 0 | 4 | 8 | 2 | 15 | 3 | 218 | 149 | 9.83E-08 | 6.33E-06 | 0.146 | 0.146 | 0.477 | 0.111 | 0.186 | 0 | 0 | 0 | 0 | 0 | 0.213 |
| SE | 3675 | ENSG00000167613 | LAIR1 | chr19 | - | 54360021 | 54360072 | 54356711 | 54356966 | 54361009 | 54361193 | 32 | 42 | 32 | 35 | 44 | 0 | 1 | 0 | 0 | 1 | 23 | 39 | 7 | 45 | 53 | 10 | 2 | 2 | 8 | 0 | 200 | 149 | 9.84E-08 | 6.33E-06 | 1 | 0.969 | 1 | 1 | 0.97 | 0.631 | 0.936 | 0.723 | 0.807 | 1 | 0.168 |
| SE | 64309 | ENSG00000101336 | HCK | chr20 | + | 32084390 | 32084543 | 32083893 | 32084043 | 32086627 | 32086807 | 463 | 334 | 257 | 333 | 377 | 0 | 10 | 4 | 4 | 0 | 977 | 519 | 188 | 688 | 1228 | 54 | 9 | 13 | 14 | 24 | 298 | 149 | 9.88E-08 | 6.35E-06 | 1 | 0.944 | 0.97 | 0.977 | 1 | 0.9 | 0.966 | 0.879 | 0.961 | 0.962 | 0.045 |
| SE | 56195 | ENSG00000143106 | PSMA5 | chr1 | - | 109411010 | 109411113 | 109409927 | 109410014 | 109411876 | 109411935 | 435 | 217 | 292 | 296 | 365 | 7 | 1 | 1 | 5 | 7 | 288 | 359 | 191 | 302 | 250 | 10 | 2 | 18 | 27 | 7 | 252 | 149 | 9.93E-08 | 6.37E-06 | 0.974 | 0.992 | 0.994 | 0.972 | 0.969 | 0.945 | 0.991 | 0.863 | 0.869 | 0.955 | 0.056 |
| SE | 3359 | ENSG00000130164 | LDLR | chr19 | + | 11105219 | 11105600 | 11102663 | 11102786 | 11106564 | 11106687 | 37 | 14 | 38 | 28 | 6 | 5 | 2 | 4 | 8 | 0 | 31 | 9 | 15 | 19 | 16 | 49 | 11 | 4 | 44 | 14 | 298 | 149 | 9.93E-08 | 6.37E-06 | 0.787 | 0.778 | 0.826 | 0.636 | 1 | 0.24 | 0.29 | 0.652 | 0.178 | 0.364 | 0.461 |
| SE | 29983 | ENSG00000143624 | INTS3 | chr1 | + | 153764117 | 153764221 | 153763831 | 153763886 | 153764689 | 153764734 | 127 | 57 | 83 | 122 | 94 | 0 | 0 | 2 | 1 | 3 | 144 | 90 | 43 | 87 | 292 | 16 | 5 | 9 | 8 | 2 | 253 | 149 | 9.95E-08 | 6.38E-06 | 1 | 1 | 0.961 | 0.986 | 0.949 | 0.841 | 0.914 | 0.738 | 0.865 | 0.989 | 0.11 |
| SE | 9410 | ENSG00000105053 | VRK3 | chr19 | - | 49997503 | 49997570 | 49995190 | 49995275 | 50000789 | 50000854 | 93 | 36 | 63 | 54 | 48 | 3 | 2 | 2 | 0 | 1 | 86 | 60 | 17 | 59 | 84 | 11 | 12 | 3 | 3 | 13 | 216 | 149 | 9.97E-08 | 6.39E-06 | 0.955 | 0.925 | 0.956 | 1 | 0.971 | 0.844 | 0.775 | 0.796 | 0.931 | 0.817 | 0.129 |
| SE | 108398 | ENSG00000110851 | PRDM4 | chr12 | - | 107740945 | 107741260 | 107739382 | 107739551 | 107742220 | 107742348 | 208 | 141 | 200 | 191 | 395 | 3 | 0 | 0 | 0 | 3 | 128 | 222 | 231 | 191 | 205 | 29 | 0 | 4 | 7 | 2 | 298 | 149 | 1.01E-07 | 6.44E-06 | 0.972 | 1 | 1 | 1 | 0.985 | 0.688 | 1 | 0.967 | 0.932 | 0.981 | 0.078 |

| | | | | | | | | | | | | | | | | | | | | | | | | | | | | | | | | | | | | | | | | | | | | | | |
|---|---|---|---|---|---|---|---|---|---|---|---|---|---|---|---|---|---|---|---|---|---|---|---|---|---|---|---|---|---|---|---|---|---|---|---|---|---|---|---|---|---|---|---|---|---|---|
| SE | 10318 | ENSG00000108848 | LUC7L3 | chr17 | + | 50738140 | 50738199 | 50737286 | 50737479 | 50740305 | 50740345 | 41 | 35 | 59 | 46 | 57 | 36 | 53 | 76 | 58 | 43 | 4 | 8 | 20 | 12 | 7 | 26 | 46 | 31 | 32 | 36 | 208 | 149 | 2.76E-07 | 1.55E-05 | 0.449 | 0.321 | 0.357 | 0.362 | 0.487 | 0.099 | 0.111 | 0.316 | 0.212 | 0.122 | 0.223 |
| SE | 81942 | ENSG00000051825 | MPHOSPH9 | chr12 | - | 123162114 | 123162192 | 123161135 | 123161383 | 123163949 | 123164090 | 51 | 35 | 54 | 41 | 63 | 1 | 3 | 2 | 5 | 3 | 57 | 37 | 44 | 27 | 3 | 2 | 20 | 30 | 17 | 3 | 227 | 149 | 2.76E-07 | 1.55E-05 | 0.971 | 0.884 | 0.947 | 0.843 | 0.932 | 0.949 | 0.548 | 0.49 | 0.51 | 0.396 | 0.337 |
| SE | 49289 | ENSG00000120733 | KDM3B | chr5 | + | 138420705 | 138420962 | 138418952 | 138419232 | 138424074 | 138424341 | 312 | 167 | 269 | 235 | 451 | 0 | 0 | 2 | 0 | 0 | 319 | 229 | 191 | 325 | 381 | 12 | 14 | 4 | 3 | 0 | 298 | 149 | 2.77E-07 | 1.55E-05 | 1 | 1 | 0.985 | 1 | 1 | 0.93 | 0.891 | 0.96 | 0.982 | 1 | 0.044 |
| SE | 75471 | ENSG00000197081 | IGF2R | chr6 | + | 160068844 | 160069014 | 160068248 | 160068385 | 160069867 | 160070058 | 5 | 5 | 11 | 6 | 6 | 297 | 100 | 165 | 190 | 153 | 8 | 19 | 28 | 18 | 15 | 189 | 161 | 175 | 149 | 184 | 298 | 149 | 2.78E-07 | 1.55E-05 | 0.008 | 0.024 | 0.032 | 0.016 | 0.019 | 0.021 | 0.056 | 0.074 | 0.057 | 0.039 | -0.03 |
| SE | 38484 | ENSG00000198793 | MTOR | chr1 | - | 11117003 | 11117086 | 11115395 | 11115468 | 11121245 | 11121368 | 99 | 78 | 80 | 112 | 84 | 5 | 6 | 5 | 3 | 1 | 87 | 84 | 46 | 59 | 52 | 30 | 12 | 6 | 9 | 4 | 232 | 149 | 2.78E-07 | 1.56E-05 | 0.927 | 0.893 | 0.911 | 0.96 | 0.982 | 0.651 | 0.818 | 0.831 | 0.808 | 0.893 | 0.134 |
| SE | 24680 | ENSG00000120705 | ETF1 | chr5 | - | 138517560 | 138517700 | 138513567 | 138513706 | 138518691 | 138518867 | 578 | 435 | 464 | 664 | 768 | 4 | 2 | 0 | 5 | 15 | 306 | 640 | 952 | 905 | 47 | 15 | 18 | 22 | 20 | 0 | 289 | 149 | 2.80E-07 | 1.56E-05 | 0.987 | 0.991 | 1 | 0.986 | 0.963 | 0.913 | 0.948 | 0.957 | 0.959 | 1 | 0.03 |
| SE | 17413 | ENSG00000126822 | PLEKHG3 | chr14 | + | 64730242 | 64730312 | 64728995 | 64729093 | 64730641 | 64730688 | 140 | 50 | 43 | 107 | 101 | 1 | 0 | 1 | 0 | 2 | 127 | 43 | 39 | 43 | 84 | 11 | 0 | 3 | 7 | 3 | 219 | 149 | 2.81E-07 | 1.57E-05 | 0.99 | 1 | 0.967 | 1 | 0.972 | 0.887 | 1 | 0.898 | 0.807 | 0.95 | 0.077 |
| SE | 19857 | ENSG00000165516 | KLHDC2 | chr14 | + | 49779747 | 49779806 | 49779594 | 49779675 | 49780212 | 49780322 | 124 | 85 | 123 | 125 | 336 | 1 | 2 | 0 | 1 | 1 | 136 | 105 | 54 | 67 | 46 | 18 | 0 | 3 | 3 | 0 | 208 | 149 | 2.83E-07 | 1.58E-05 | 0.989 | 0.968 | 1 | 0.989 | 0.996 | 0.844 | 1 | 0.928 | 0.941 | 1 | 0.046 |
| SE | 11376 | ENSG00000166747 | AP1G1 | chr16 | - | 71756018 | 71756159 | 71753832 | 71753887 | 71758807 | 71758921 | 293 | 215 | 251 | 263 | 284 | 4 | 1 | 0 | 3 | 0 | 67 | 246 | 406 | 175 | 39 | 0 | 8 | 11 | 13 | 0 | 290 | 149 | 2.85E-07 | 1.59E-05 | 0.974 | 0.991 | 1 | 0.978 | 1 | 1 | 0.94 | 0.95 | 0.874 | 1 | 0.036 |
| SE | 33369 | ENSG00000187742 | SECISBP2 | chr9 | + | 89357284 | 89357565 | 89350631 | 89350852 | 89357998 | 89358129 | 70 | 48 | 58 | 71 | 83 | 1 | 1 | 2 | 8 | 0 | 65 | 38 | 29 | 29 | 13 | 16 | 1 | 18 | 6 | 5 | 298 | 149 | 2.87E-07 | 1.60E-05 | 0.972 | 0.96 | 0.935 | 0.816 | 1 | 0.67 | 0.95 | 0.446 | 0.707 | 0.565 | 0.269 |
| SE | 73449 | ENSG00000143374 | TARS2 | chr1 | + | 150491597 | 150491662 | 150491393 | 150491511 | 150492410 | 150492489 | 55 | 26 | 53 | 57 | 64 | 0 | 1 | 5 | 0 | 0 | 48 | 58 | 25 | 27 | 74 | 16 | 4 | 8 | 4 | 1 | 214 | 149 | 2.88E-07 | 1.60E-05 | 1 | 0.948 | 0.881 | 1 | 1 | 0.676 | 0.91 | 0.685 | 0.825 | 0.981 | 0.15 |
| SE | 12773 | ENSG00000168802 | CHTF8 | chr16 | - | 69120106 | 69120208 | 69118121 | 69118525 | 69121052 | 69121170 | 19 | 7 | 19 | 13 | 11 | 3 | 1 | 1 | 3 | 4 | 17 | 14 | 8 | 12 | 27 | 0 | 0 | 0 | 0 | 0 | 251 | 149 | 2.89E-07 | 1.61E-05 | 0.79 | 0.806 | 0.919 | 0.72 | 0.62 | 1 | 1 | 1 | 1 | 1 | -0.229 |
| SE | 712 | ENSG00000182944 | EWSR1 | chr22 | + | 29288605 | 29291599 | 29286922 | 29287134 | 29292136 | 29292169 | 284 | 147 | 228 | 182 | 135 | 6 | 3 | 8 | 14 | 12 | 274 | 249 | 157 | 288 | 334 | 14 | 30 | 51 | 34 | 50 | 298 | 149 | 2.90E-07 | 1.61E-05 | 0.959 | 0.961 | 0.934 | 0.867 | 0.849 | 0.907 | 0.806 | 0.606 | 0.809 | 0.77 | 0.134 |
| SE | 15710 | ENSG00000140259 | MFAP1 | chr15 | - | 43817228 | 43817448 | 43814944 | 43815074 | 43824490 | 43824690 | 152 | 144 | 121 | 158 | 200 | 0 | 1 | 0 | 0 | 0 | 51 | 89 | 225 | 64 | 20 | 0 | 5 | 5 | 10 | 0 | 298 | 149 | 2.91E-07 | 1.62E-05 | 1 | 0.986 | 1 | 1 | 1 | 1 | 0.899 | 0.957 | 0.762 | 1 | 0.074 |
| SE | 95769 | ENSG00000125944 | HNRNPR | chr1 | - | 23318482 | 23318697 | 23313552 | 23313702 | 23321527 | 23321663 | 954 | 871 | 758 | 862 | 1794 | 7 | 4 | 5 | 5 | 8 | 443 | 857 | 1088 | 931 | 180 | 6 | 3 | 22 | 31 | 0 | 298 | 149 | 2.93E-07 | 1.62E-05 | 0.986 | 0.991 | 0.987 | 0.989 | 0.991 | 0.974 | 0.993 | 0.961 | 0.938 | 1 | 0.016 |
| SE | 35090 | ENSG00000169359 | SLC33A1 | chr3 | - | 155833856 | 155834041 | 155829687 | 155829903 | 155842431 | 155842619 | 89 | 52 | 98 | 64 | 185 | 7 | 3 | 4 | 2 | 26 | 58 | 74 | 27 | 42 | 91 | 0 | 3 | 2 | 0 | 0 | 298 | 149 | 2.94E-07 | 1.63E-05 | 0.864 | 0.897 | 0.925 | 0.941 | 0.781 | 1 | 0.925 | 0.871 | 1 | 1 | -0.078 |
| SE | 17664 | ENSG00000176681 | LRRC37A | chr17 | + | 46332551 | 46332656 | 46330449 | 46331981 | 46335544 | 46335594 | 26 | 18 | 66 | 69 | 9 | 0 | 0 | 2 | 5 | 0 | 86 | 162 | 31 | 86 | 3 | 16 | 29 | 11 | 13 | 0 | 254 | 149 | 2.94E-07 | 1.63E-05 | 1 | 1 | 0.951 | 0.89 | 1 | 0.759 | 0.766 | 0.623 | 0.795 | 1 | 0.18 |
| SE | 20834 | ENSG00000107929 | LARP4B | chr10 | - | 870012 | 870083 | 864122 | 864270 | 884446 | 884506 | 11 | 11 | 12 | 18 | 24 | 78 | 43 | 54 | 75 | 31 | 3 | 4 | 9 | 5 | 10 | 42 | 89 | 56 | 99 | 112 | 220 | 149 | 2.95E-07 | 1.63E-05 | 0.087 | 0.148 | 0.131 | 0.14 | 0.344 | 0.046 | 0.03 | 0.098 | 0.033 | 0.057 | 0.117 |
| SE | 57664 | ENSG00000078177 | N4BP2 | chr4 | + | 40097226 | 40097569 | 40073454 | 40073551 | 40102074 | 40103218 | 41 | 61 | 46 | 43 | 49 | 3 | 4 | 8 | 0 | 4 | 4 | 11 | 21 | 7 | 6 | 0 | 0 | 0 | 0 | 0 | 298 | 149 | 2.98E-07 | 1.65E-05 | 0.872 | 0.884 | 0.742 | 1 | 0.86 | 1 | 1 | 1 | 1 | 1 | -0.128 |
| SE | 1336 | ENSG00000093000 | NUP50 | chr22 | + | 45175893 | 45176080 | 45171599 | 45171683 | 45181285 | 45181367 | 254 | 181 | 222 | 221 | 457 | 1 | 3 | 2 | 1 | 8 | 151 | 196 | 127 | 190 | 80 | 13 | 0 | 42 | 10 | 0 | 298 | 149 | 3.00E-07 | 1.66E-05 | 0.992 | 0.968 | 0.982 | 0.991 | 0.966 | 0.853 | 1 | 0.602 | 0.905 | 1 | 0.108 |
| SE | 30149 | ENSG00000005059 | MCUB | chr4 | + | 109664289 | 109664394 | 109660194 | 109660365 | 109682581 | 109682742 | 922 | 915 | 1034 | 1068 | 1742 | 3 | 6 | 5 | 9 | 6 | 1768 | 586 | 815 | 767 | 1072 | 16 | 18 | 10 | 7 | 27 | 254 | 149 | 3.01E-07 | 1.66E-05 | 0.994 | 0.989 | 0.992 | 0.986 | 0.994 | 0.985 | 0.95 | 0.98 | 0.985 | 0.959 | 0.019 |
| SE | 100696 | ENSG00000107372 | ZFAND5 | chr9 | - | 72363469 | 72363606 | 72360627 | 72360787 | 72364695 | 72365208 | 226 | 188 | 166 | 186 | 207 | 49 | 18 | 33 | 33 | 12 | 223 | 232 | 165 | 248 | 183 | 70 | 62 | 61 | 77 | 64 | 286 | 149 | 3.01E-07 | 1.66E-05 | 0.706 | 0.845 | 0.724 | 0.746 | 0.9 | 0.624 | 0.661 | 0.585 | 0.627 | 0.598 | 0.165 |
| SE | 48286 | ENSG00000229474 | PATL2 | chr15 | - | 44667105 | 44667203 | 44666391 | 44666541 | 44668979 | 44669139 | 30 | 11 | 18 | 33 | 25 | 0 | 0 | 1 | 0 | 0 | 8 | 68 | 30 | 40 | 49 | 0 | 11 | 1 | 4 | 4 | 247 | 149 | 3.02E-07 | 1.66E-05 | 1 | 1 | 0.916 | 1 | 1 | 1 | 0.789 | 0.948 | 0.858 | 0.881 | 0.088 |
| SE | 86494 | ENSG00000185485 | SDHAP1 | chr3 | - | 195968218 | 195968385 | 195967878 | 195967990 | 195971322 | 195971444 | 12 | 11 | 7 | 11 | 9 | 44 | 51 | 64 | 72 | 77 | 13 | 0 | 4 | 0 | 0 | 191 | 38 | 43 | 20 | 101 | 298 | 149 | 3.03E-07 | 1.67E-05 | 0.12 | 0.097 | 0.052 | 0.071 | 0.055 | 0.033 | 0 | 0.044 | 0 | 0 | 0.064 |
| SE | 35591 | ENSG00000114331 | ACAP2 | chr3 | - | 195312946 | 195312967 | 195308785 | 195308837 | 195320700 | 195320813 | 17 | 2 | 0 | 7 | 28 | 207 | 187 | 160 | 257 | 366 | 0 | 0 | 1 | 2 | 0 | 128 | 105 | 159 | 118 | 49 | 170 | 149 | 3.04E-07 | 1.67E-05 | 0.067 | 0.009 | 0 | 0.023 | 0.063 | 0 | 0 | 0.005 | 0.015 | 0 | 0.028 |
| SE | 15137 | ENSG00000104093 | DMXL2 | chr15 | - | 51464674 | 51464876 | 51463075 | 51463496 | 51465565 | 51465651 | 259 | 285 | 182 | 191 | 271 | 2 | 14 | 6 | 14 | 10 | 16 | 139 | 178 | 14 | 4 | 0 | 1 | 1 | 0 | 0 | 298 | 149 | 3.06E-07 | 1.68E-05 | 0.985 | 0.911 | 0.938 | 0.872 | 0.931 | 1 | 0.986 | 0.989 | 1 | 1 | -0.068 |
| SE | 62784 | ENSG00000138646 | HERC5 | chr4 | + | 88462134 | 88462356 | 88460094 | 88460171 | 88463531 | 88463623 | 96 | 96 | 107 | 107 | 94 | 8 | 0 | 2 | 3 | 2 | 69 | 121 | 117 | 134 | 15 | 11 | 7 | 26 | 27 | 0 | 298 | 149 | 3.17E-07 | 1.74E-05 | 0.857 | 1 | 0.964 | 0.947 | 0.959 | 0.758 | 0.896 | 0.692 | 0.713 | 1 | 0.134 |

| | | | | | | | | | | | | | | |
|---|---|---|---|---|---|---|---|---|---|---|---|---|---|---|
| SE | 40382 | ENSG00000163719 | MTMR14 | chr3 | + | 9668718 | 9668794 | 9653620 | 9653769 | 9669431 | 9669492 | 100 | 41 | 70 |
| SE | 68190 | ENSG00000020922 | MRE11 | chr11 | - | 94476288 | 94476403 | 94471573 | 94471759 | 94478734 | 94478876 | 216 | 170 | 135 |
| SE | 1828 | ENSG00000070371 | CLTCL1 | chr22 | - | 19234508 | 19234706 | 19233421 | 19233622 | 19235695 | 19235869 | 11 | 3 | 7 |
| SE | 106154 | ENSG00000089022 | MAPKAPK5 | chr12 | + | 111870270 | 111870360 | 111868752 | 111868861 | 111871084 | 111871180 | 107 | 44 | 153 |
| SE | 101422 | ENSG00000158545 | ZC3H18 | chr16 | + | 88611267 | 88611536 | 88599790 | 88599948 | 88622196 | 88622388 | 56 | 26 | 41 |
| SE | 86335 | ENSG00000135048 | CEMIP2 | chr9 | - | 71745017 | 71745579 | 71740062 | 71740232 | 71750042 | 71750385 | 197 | 89 | 110 |
| SE | 54232 | ENSG00000143643 | TTC13 | chr1 | - | 230943805 | 230943898 | 230940439 | 230940556 | 230945388 | 230945454 | 186 | 128 | 117 |
| SE | 35391 | ENSG00000161267 | BDH1 | chr3 | - | 197546360 | 197546486 | 197533488 | 197533561 | 197555780 | 197555977 | 15 | 20 | 27 |
| SE | 92477 | ENSG00000090470 | PDCD7 | chr15 | - | 65119717 | 65119954 | 65119375 | 65119463 | 65129031 | 65129170 | 362 | 242 | 297 |
| SE | 14627 | ENSG00000140157 | NIPA2 | chr15 | + | 22845145 | 22845267 | 22838657 | 22838947 | 22851638 | 22851665 | 51 | 20 | 41 |
| SE | 15304 | ENSG00000114062 | UBE3A | chr15 | - | 25407066 | 25407212 | 25405460 | 25405502 | 25408571 | 25408684 | 21 | 12 | 11 |
| SE | 75113 | ENSG00000114423 | CBLB | chr3 | - | 105749621 | 105749765 | 105745916 | 105746038 | 105751461 | 105751618 | 16 | 11 | 2 |
| SE | 87646 | ENSG00000117676 | RPS6KA1 | chr1 | + | 26555536 | 26555625 | 26555150 | 26555221 | 26556653 | 26556718 | 311 | 151 | 184 |
| SE | 22716 | ENSG00000164329 | TENT2 | chr5 | + | 79649061 | 79649190 | 79648616 | 79648693 | 79656957 | 79657001 | 376 | 293 | 331 |
| SE | 16946 | ENSG00000172575 | RASGRP1 | chr15 | - | 38494381 | 38494767 | 38488102 | 38490688 | 38498793 | 38498946 | 277 | 178 | 273 |
| SE | 79549 | ENSG00000117868 | ESYT2 | chr7 | - | 158773340 | 158773396 | 158764676 | 158764853 | 158788003 | 158788093 | 181 | 129 | 191 |
| SE | 54119 | ENSG00000124126 | PREX1 | chr20 | - | 48627547 | 48627615 | 48624251 | 48625927 | 48627860 | 48627963 | 139 | 36 | 85 |
| SE | 46029 | ENSG00000173230 | GOLGB1 | chr3 | - | 121722261 | 121722378 | 121718387 | 121718501 | 121726912 | 121727041 | 83 | 78 | 52 |
| SE | 92656 | ENSG00000197724 | PHF2 | chr9 | + | 93659510 | 93659600 | 93654412 | 93654575 | 93660191 | 93660560 | 36 | 16 | 26 |
| SE | 18925 | ENSG00000100461 | RBM23 | chr14 | - | 22909482 | 22909595 | 22905584 | 22906368 | 22911327 | 22911403 | 223 | 120 | 127 |
| SE | 46570 | ENSG00000138386 | NAB1 | chr2 | + | 190683737 | 190683827 | 190673100 | 190673152 | 190685475 | 190685638 | 111 | 63 | 70 |
| SE | 38829 | ENSG00000122674 | CCZ1 | chr7 | + | 5905093 | 5905269 | 5902595 | 5902744 | 5910034 | 5910116 | 5 | 61 | 57 |
| SE | 70538 | ENSG00000073849 | ST6GAL1 | chr3 | + | 187038394 | 187038491 | 187025482 | 187025517 | 187038741 | 187038873 | 12 | 2 | 13 |
| SE | 66951 | ENSG00000172493 | AFF1 | chr4 | + | 87132270 | 87132408 | 87131792 | 87131864 | 87134470 | 87134694 | 282 | 214 | 240 |
| SE | 82497 | ENSG00000128563 | PRKRIP1 | chr7 | + | 102407433 | 102407498 | 102399547 | 102399648 | 102425013 | 102425446 | 16 | 15 | 15 |
| SE | 8231 | ENSG00000185624 | P4HB | chr17 | - | 81855452 | 81855688 | 81855141 | 81855279 | 81859180 | 81859387 | 283 | 143 | 274 |
| SE | 68767 | ENSG00000163872 | YEATS2 | chr3 | + | 183772304 | 183772563 | 183762096 | 183762279 | 183773632 | 183773794 | 29 | 22 | 32 |
| SE | 18692 | ENSG00000129566 | TEP1 | chr14 | - | 20403722 | 20403884 | 20403376 | 20403448 | 20405450 | 20405585 | 43 | 27 | 50 |
| SE | 3674 | ENSG00000167613 | LAIR1 | chr19 | - | 54360021 | 54360072 | 54356711 | 54356966 | 54360915 | 54360949 | 81 | 77 | 93 |
| SE | 85194 | ENSG00000140332 | TLE3 | chr15 | - | 70064453 | 70064470 | 70060529 | 70060649 | 70074532 | 70074607 | 23 | 34 | 22 |
| SE | 40577 | ENSG00000163875 | MEAF6 | chr1 | - | 37496546 | 37496604 | 37495873 | 37495918 | 37496706 | 37496736 | 14 | 4 | 13 |
| SE | 48862 | ENSG00000184677 | ZBTB40 | chr1 | + | 22501491 | 22501684 | 22491399 | 22491533 | 22502298 | 22502441 | 126 | 123 | 153 |

[illegible]

| | | | | | | | | | | | | | | | |
|---|---|---|---|---|---|---|---|---|---|---|---|---|---|---|---|
| SE | 4786 | ENSG00000175221 | MED16 | chr19 | - | 876973 | 877180 | 875243 | 875454 | 879936 | 880148 | 58 | 6 | 27 | [illegible] |
| SE | 1093 | ENSG00000100030 | MAPK1 | chr22 | - | 21788693 | 21788808 | 21788256 | 21788388 | 21799011 | 21799128 | 1024 | 447 | 756 | [illegible] |
| SE | 83326 | ENSG00000161010 | MRNIP | chr5 | - | 179851242 | 179851337 | 179847977 | 179848066 | 179853377 | 179853437 | 1 | 2 | 5 | [illegible] |
| SE | 109153 | ENSG00000152213 | ARL11 | chr13 | + | 49630033 | 49630099 | 49628522 | 49628615 | 49630429 | 49633872 | 2 | 0 | 1 | [illegible] |
| SE | 96072 | ENSG00000132109 | TRIM21 | chr11 | - | 4388299 | 4388530 | 4386967 | 4386990 | 4389653 | 4389749 | 364 | 213 | 210 | [illegible] |
| SE | 91271 | ENSG00000154114 | TBCEL | chr11 | + | 121055051 | 121055308 | 121053550 | 121053732 | 121058344 | 121058471 | 149 | 75 | 122 | [illegible] |
| SE | 101546 | ENSG00000088298 | EDEM2 | chr20 | - | 35118597 | 35118719 | 35115363 | 35115933 | 35123889 | 35124034 | 128 | 63 | 136 | [illegible] |
| SE | 84932 | ENSG00000179195 | ZNF664 | chr12 | + | 124004877 | 124004935 | 123973122 | 123974020 | 124011380 | 124011441 | 19 | 11 | 12 | [illegible] |
| SE | 44004 | ENSG00000114098 | ARMC8 | chr3 | + | 138223388 | 138223531 | 138221925 | 138221997 | 138223635 | 138223733 | 406 | 284 | 306 | [illegible] |
| SE | 19011 | ENSG00000198604 | BAZ1A | chr14 | - | 34786125 | 34786221 | 34785776 | 34786001 | 34792774 | 34792921 | 251 | 172 | 186 | [illegible] |
| SE | 44685 | ENSG00000185344 | ATP6V0A2 | chr12 | + | 123751109 | 123751229 | 123748574 | 123748785 | 123752282 | 123752402 | 79 | 68 | 61 | [illegible] |
| SE | 25342 | ENSG00000130803 | ZNF317 | chr19 | + | 9157833 | 9158075 | 9157267 | 9157394 | 9158825 | 9158908 | 49 | 44 | 29 | [illegible] |
| SE | 64678 | ENSG00000136715 | SAP130 | chr2 | - | 127945455 | 127945559 | 127942423 | 127942537 | 127949868 | 127949994 | 129 | 74 | 94 | [illegible] |
| SE | 78188 | ENSG00000136273 | HUS1 | chr7 | - | 47976729 | 47976837 | 47975612 | 47975687 | 47978416 | 47978534 | 159 | 126 | 192 | [illegible] |
| SE | 13367 | ENSG00000005194 | CIAPIN1 | chr16 | - | 57440771 | 57440983 | 57439181 | 57439295 | 57447341 | 57447368 | 40 | 27 | 41 | [illegible] |
| SE | 38742 | ENSG00000086062 | B4GALT1 | chr9 | - | 33115990 | 33116113 | 33113773 | 33113878 | 33135188 | 33135424 | 437 | 185 | 328 | [illegible] |
| SE | 39297 | ENSG00000141293 | SKAP1 | chr17 | - | 48184722 | 48184847 | 48182393 | 48182457 | 48187842 | 48187926 | 425 | 315 | 384 | [illegible] |
| SE | 109098 | ENSG00000196312 | MFSD14C | chr9 | - | 96933794 | 96933915 | 96918642 | 96918826 | 96942602 | 96942730 | 30 | 40 | 27 | [illegible] |
| SE | 43366 | ENSG00000090863 | GLG1 | chr16 | - | 74468945 | 74469063 | 74467755 | 74467848 | 74469984 | 74470073 | 746 | 531 | 776 | [illegible] |
| SE | 32776 | ENSG00000168297 | PXK | chr3 | + | 58410089 | 58410159 | 58409531 | 58409618 | 58424751 | 58425151 | 75 | 44 | 41 | [illegible] |
| SE | 107593 | ENSG00000111596 | CNOT2 | chr12 | + | 70285399 | 70285586 | 70278131 | 70278274 | 70310894 | 70311017 | 10 | 4 | 4 | [illegible] |
| SE | 54023 | ENSG00000027001 | MIPEP | chr13 | - | 23869291 | 23869448 | 23864140 | 23864189 | 23870012 | 23870195 | 43 | 33 | 39 | [illegible] |
| SE | 42077 | ENSG00000266094 | RASSF5 | chr1 | + | 206585179 | 206585295 | 206584386 | 206584684 | 206586825 | 206589267 | 550 | 337 | 562 | [illegible] |
| SE | 23616 | ENSG00000113558 | SKP1 | chr5 | - | 134173700 | 134173766 | 134167169 | 134167243 | 134173925 | 134174022 | 3 | 5 | 15 | [illegible] |
| SE | 11717 | ENSG00000100266 | PACSIN2 | chr22 | - | 42912020 | 42912157 | 42893456 | 42893613 | 43002309 | 43002440 | 130 | 92 | 86 | [illegible] |
| SE | 74902 | ENSG00000277147 | LINC00869 | chr1 | + | 149658151 | 149658254 | 149655730 | 149655908 | 149660853 | 149661089 | 8 | 0 | 1 | [illegible] |
| SE | 74626 | ENSG00000085998 | POMGNT1 | chr1 | - | 46193563 | 46193639 | 46193304 | 46193388 | 46193854 | 46193925 | 35 | 8 | 18 | [illegible] |
| SE | 68100 | ENSG00000135838 | NPL | chr1 | + | 182818812 | 182818859 | 182816713 | 182816806 | 182822114 | 182822199 | 6 | 2 | 6 | [illegible] |
| SE | 4700 | ENSG00000160633 | SAFB | chr19 | + | 5648988 | 5649499 | 5648015 | 5648043 | 5649925 | 5649975 | 314 | 183 | 262 | [illegible] |
| SE | 94597 | ENSG00000106829 | TLE4 | chr9 | + | 79627373 | 79627448 | 79612655 | 79612718 | 79652592 | 79652794 | 169 | 130 | 131 | [illegible] |
| SE | 108131 | ENSG00000155957 | TMBIM4 | chr12 | - | 66168914 | 66169353 | 66153339 | 66153448 | 66169854 | 66169980 | 2 | 1 | 1 | [illegible] |
| SE | 71886 | ENSG00000074054 | CLASP1 | chr2 | - | 121402594 | 121402663 | 121401867 | 121401870 | 121404370 | 121404434 | 37 | 15 | 29 | [illegible] |
| SE | 4964 | ENSG00000124160 | NCOA5 | chr20 | - | 46065028 | 46065228 | 46063359 | 46063680 | 46067054 | 46067181 | 305 | 201 | 262 | [illegible] |
| SE | 45614 | ENSG00000132781 | MUTYH | chr1 | - | 45329888 | 45330068 | 45329241 | 45329437 | 45330515 | 45330557 | 9 | 6 | 12 | [illegible] |
| SE | 20939 | ENSG00000135596 | MICAL1 | chr6 | - | 109448202 | 109448393 | 109447680 | 109447722 | 109448731 | 109448870 | 61 | 42 | 57 | [illegible] |
| SE | 85 | ENSG00000186866 | POFUT2 | chr21 | - | 45282348 | 45282459 | 45278102 | 45278169 | 45285677 | 45285928 | 23 | 4 | 25 | [illegible] |
| SE | 88605 | ENSG00000134014 | ELP3 | chr8 | + | 28132277 | 28132404 | 28129318 | 28129663 | 28137697 | 28137891 | 274 | 171 | 237 | [illegible] |
| SE | 78464 | ENSG00000006715 | VPS41 | chr7 | - | 38898090 | 38898129 | 38869145 | 38869253 | 38909153 | 38909174 | 78 | 49 | 56 | [illegible] |
| SE | 81697 | ENSG00000171681 | ATF7IP | chr12 | + | 14446987 | 14447053 | 14438129 | 14438267 | 14456560 | 14456634 | 541 | 414 | 423 | [illegible] |
| SE | 63751 | ENSG00000152061 | RABGAP1L | chr1 | + | 174957890 | 174958173 | 174957456 | 174957549 | 174969276 | 174969387 | 311 | 230 | 140 | [illegible] |
| SE | 29680 | ENSG00000137656 | BUD13 | chr11 | - | 116758268 | 116758407 | 116757765 | 116757950 | 116760734 | 116760952 | 51 | 65 | 61 | [illegible] |
| SE | 49475 | ENSG00000115020 | PIKFYVE | chr2 | + | 208304845 | 208305013 | 208304170 | 208304318 | 208312235 | 208312295 | 90 | 70 | 107 | [illegible] |
| SE | 13944 | ENSG00000103426 | CORO7-PAM16 | chr16 | - | 4387985 | 4388068 | 4365490 | 4365545 | 4395288 | 4395339 | 20 | 5 | 14 | [illegible] |
| SE | 80135 | ENSG00000012124 | CD22 | chr19 | + | 35332018 | 35332074 | 35329165 | 35329230 | 35332546 | 35332830 | 29 | 8 | 23 | [illegible] |
| SE | 70363 | ENSG00000258984 | UBE2F-SCLY | chr2 | + | 238016565 | 238016633 | 237994743 | 237994809 | 238030555 | 238030613 | 228 | 138 | 156 | [illegible] |
| SE | 83661 | ENSG00000029363 | BCLAF1 | chr6 | - | 136269140 | 136269303 | 136268161 | 136268273 | 136269436 | 136269612 | 92 | 87 | 75 | [illegible] |
| SE | 24419 | ENSG00000055163 | CYFIP2 | chr5 | + | 157327972 | 157328049 | 157325599 | 157326267 | 157330741 | 157330850 | 1210 | 534 | 1046 | [illegible] |
| SE | 89355 | ENSG00000115419 | GLS | chr2 | + | 190913192 | 190913364 | 190910262 | 190910321 | 190921023 | 190921056 | 98 | 71 | 111 | [illegible] |
| SE | 106855 | ENSG00000166908 | PIP4K2C | chr12 | + | 57597957 | 57598062 | 57595887 | 57596031 | 57599064 | 57599211 | 4 | 6 | 12 | [illegible] |
| SE | 67963 | ENSG00000141522 | ARHGDIA | chr17 | - | 81869055 | 81869075 | 81868630 | 81868943 | 81869172 | 81869236 | 60 | 23 | 52 | [illegible] |
| SE | 75410 | ENSG00000169826 | CSGALNACT2 | chr10 | + | 43163865 | 43164044 | 43158714 | 43158931 | 43167003 | 43167098 | 156 | 109 | 122 | [illegible] |
| SE | 10652 | ENSG00000273559 | CWC25 | chr17 | - | 38815624 | 38815741 | 38814860 | 38815097 | 38820900 | 38821073 | 4 | 6 | 3 | [illegible] |
| SE | 84091 | ENSG00000197892 | KIF13B | chr8 | - | 29134039 | 29134210 | 29132307 | 29132465 | 29140062 | 29140191 | 141 | 94 | 114 | [illegible] |
| SE | 19090 | ENSG00000259316 | NA | chr15 | - | 64204864 | 64204949 | 64204440 | 64204589 | 64213889 | 64214124 | 92 | 87 | 98 | [illegible] |
| SE | 75499 | ENSG00000120802 | TMPO | chr12 | + | 98544441 | 98544537 | 98537474 | 98537572 | 98544950 | 98545061 | 287 | 215 | 333 | [illegible] |
| SE | 91070 | ENSG00000104164 | BLOC1S6 | chr15 | + | 45603099 | 45603187 | 45587235 | 45587525 | 45605427 | 45605470 | 151 | 241 | 115 | [illegible] |
| SE | 21485 | ENSG00000018189 | RUFY3 | chr4 | + | 70800140 | 70800205 | 70799404 | 70799875 | 70802955 | 70802983 | 12 | 19 | 8 | [illegible] |
| SE | 80361 | ENSG00000102531 | FNDC3A | chr13 | + | 49201132 | 49201229 | 49198361 | 49198574 | 49201799 | 49201966 | 17 | 28 | 18 | [illegible] |
| SE | 49630 | ENSG00000178927 | CYBC1 | chr17 | - | 82449169 | 82449292 | 82447579 | 82447621 | 82450699 | 82450717 | 92 | 20 | 52 | [illegible] |
| SE | 23930 | ENSG00000174989 | FBXW8 | chr12 | + | 116945363 | 116945528 | 116928022 | 116928127 | 116964696 | 116964854 | 35 | 26 | 34 | [illegible] |
| SE | 13932 | ENSG00000131797 | CLUHP3 | chr16 | + | 31702534 | 31702681 | 31701662 | 31701858 | 31703155 | 31703367 | 48 | 37 | 107 | [illegible] |
| SE | 67928 | ENSG00000091490 | SEL1L3 | chr4 | - | 25776276 | 25776360 | 25767739 | 25767830 | 25782241 | 25782418 | 88 | 76 | 102 | [illegible] |
| SE | 41481 | ENSG00000099194 | SCD | chr10 | + | 100352365 | 100352496 | 100348063 | 100348346 | 100354426 | 100354632 | 27 | 15 | 19 | [illegible] |
| SE | 33002 | ENSG00000114316 | USP4 | chr3 | - | 49300466 | 49300691 | 49298551 | 49298635 | 49302383 | 49302542 | 154 | 92 | 155 | [illegible] |
| SE | 97741 | ENSG00000095139 | ARCN1 | chr11 | + | 118597706 | 118597911 | 118593589 | 118593698 | 118600624 | 118600993 | 563 | 324 | 581 | [illegible] |
| SE | 36434 | ENSG00000121858 | TNFSF10 | chr3 | - | 172514860 | 172514998 | 172509216 | 172509321 | 172523252 | 172523310 | 151 | 135 | 117 | [illegible] |
| SE | 99280 | ENSG00000110079 | MS4A4A | chr11 | + | 60302558 | 60302717 | 60301000 | 60301057 | 60306099 | 60306201 | 43 | 20 | 3 | [illegible] |
| SE | 71413 | ENSG00000144048 | DUSP11 | chr2 | - | 73767160 | 73767207 | 73766827 | 73766903 | 73769264 | 73769325 | 288 | 197 | 274 | [illegible] |
| SE | 65441 | ENSG00000166164 | BRD7 | chr16 | - | 50323586 | 50323698 | 50321981 | 50322038 | 50325747 | 50325883 | 383 | 357 | 310 | [illegible] |
| SE | 27326 | ENSG00000196284 | SUPT3H | chr6 | - | 44954494 | 44954607 | 44953309 | 44953417 | 44961752 | 44961828 | 35 | 41 | 50 | [illegible] |
| SE | 44592 | ENSG00000026508 | CD44 | chr11 | + | 35209964 | 35210054 | 35208104 | 35208206 | 35214851 | 35214914 | 43 | 24 | 29 | [illegible] |
| SE | 8889 | ENSG00000121058 | COIL | chr17 | - | 56949680 | 56949767 | 56949386 | 56949434 | 56949888 | 56950996 | 126 | 62 | 67 | [illegible] |
| SE | 106237 | ENSG00000089154 | GCN1 | chr12 | - | 120154969 | 120155040 | 120153743 | 120153909 | 120155240 | 120155430 | 144 | 69 | 156 | [illegible] |
| SE | 46463 | ENSG00000115415 | STAT1 | chr2 | - | 191010366 | 191010449 | 191009875 | 191010004 | 191013524 | 191013678 | 1 | 0 | 3 | [illegible] |
| SE | 104592 | ENSG00000069493 | CLEC2D | chr12 | + | 9693033 | 9693115 | 9692827 | 9692931 | 9694759 | 9695129 | 273 | 226 | 240 | [illegible] |
| SE | 88069 | ENSG00000126432 | PRDX5 | chr11 | + | 64320660 | 64320792 | 64319733 | 64319868 | 64320864 | 64320903 | 256 | 133 | 187 | [illegible] |
| SE | 20282 | ENSG00000118482 | PHF3 | chr6 | + | 63684128 | 63685911 | 63679999 | 63680161 | 63691736 | 63692043 | 678 | 716 | 627 | [illegible] |
| SE | 107433 | ENSG00000166037 | CEP57 | chr11 | + | 95822498 | 95822576 | 95821870 | 95821978 | 95827785 | 95828027 | 91 | 43 | 90 | [illegible] |
| SE | 49545 | ENSG00000120158 | RCL1 | chr9 | + | 4844524 | 4844681 | 4841201 | 4841357 | 4849446 | 4849550 | 38 | 14 | 68 | [illegible] |
| SE | 70578 | ENSG00000022840 | RNF10 | chr12 | + | 120575630 | 120575688 | 120566824 | 120566980 | 120576589 | 120576882 | 5 | 1 | 5 | [illegible] |
| SE | 66670 | ENSG00000005483 | KMT2E | chr7 | + | 105064027 | 105064116 | 105063350 | 105063580 | 105066726 | 105066807 | 12 | 6 | 3 | [illegible] |
| SE | 45180 | ENSG00000107099 | DOCK8 | chr9 | + | 433874 | 433975 | 432165 | 432324 | 434782 | 434975 | 1976 | 1666 | 1389 | [illegible] |
| SE | 103123 | ENSG00000139651 | ZNF740 | chr12 | + | 53185386 | 53185476 | 53181676 | 53181992 | 53185953 | 53186031 | 98 | 71 | 76 | [illegible] |
| SE | 59251 | ENSG00000168118 | RAB4A | chr1 | + | 229288728 | 229288843 | 229286484 | 229286566 | 229295847 | 229295910 | 202 | 128 | 111 | [illegible] |
| SE | 50388 | ENSG00000058866 | DGKG | chr3 | - | 186284659 | 186284709 | 186280669 | 186280744 | 186288709 | 186288880 | 22 | 20 | 4 | [illegible] |
| SE | 29657 | ENSG00000196083 | IL1RAP | chr3 | + | 190556129 | 190556216 | 190514084 | 190514219 | 190564288 | 190564353 | 26 | 8 | 12 | [illegible] |
| SE | 22558 | ENSG00000167881 | SRP68 | chr17 | - | 76061491 | 76061574 | 76061109 | 76061219 | 76063975 | 76064171 | 366 | 289 | 400 | [illegible] |
| SE | 62669 | ENSG00000115525 | ST3GAL5 | chr2 | - | 85847381 | 85847562 | 85846376 | 85846563 | 85847860 | 85847984 | 35 | 24 | 21 | [illegible] |
| SE | 69258 | ENSG00000168385 | SEPTIN2 | chr2 | + | 241317498 | 241317599 | 241315943 | 241315982 | 241324215 | 241324241 | 38 | 23 | 40 | [illegible] |
| SE | 29610 | ENSG00000203875 | SNHG5 | chr6 | - | 85677794 | 85677868 | 85677423 | 85677492 | 85677953 | 85678032 | 91 | 193 | 139 | [illegible] |
| SE | 76337 | ENSG00000136861 | CDK5RAP2 | chr9 | - | 120437294 | 120437527 | 120422692 | 120422741 | 120439398 | 120439972 | 98 | 61 | 93 | [illegible] |
| SE | 8516 | ENSG00000125447 | GGA3 | chr17 | - | 75244618 | 75244717 | 75243446 | 75243570 | 75246508 | 75246584 | 262 | 199 | 274 | [illegible] |
| SE | 10427 | ENSG00000126581 | BECN1 | chr17 | - | 42813947 | 42814008 | 42810307 | 42810928 | 42814523 | 42814673 | 71 | 43 | 37 | [illegible] |
| SE | 4546 | ENSG00000127507 | ADGRE2 | chr19 | - | 14756237 | 14756345 | 14752328 | 14752526 | 14764432 | 14764610 | 77 | 60 | 45 | [illegible] |
| SE | 41302 | ENSG00000269893 | SNHG8 | chr4 | + | 118278933 | 118279137 | 118278754 | 118278792 | 118279388 | 118279431 | 76 | 32 | 121 | [illegible] |
| SE | 62806 | ENSG00000101882 | NKAP | chrX | - | 119938729 | 119938810 | 119936611 | 119936682 | 119943219 | 119943604 | 69 | 47 | 45 | [illegible] |
| SE | 55889 | ENSG00000074201 | CLNS1A | chr11 | - | 77622499 | 77622673 | 77619605 | 77619695 | 77624962 | 77625070 | 713 | 552 | 754 | [illegible] |
| SE | 77082 | ENSG00000086232 | EIF2AK1 | chr7 | - | 6054545 | 6054704 | 6048806 | 6048844 | 6058965 | 6059056 | 32 | 13 | 25 | [illegible] |
| SE | 95842 | ENSG00000141551 | CSNK1D | chr17 | - | 82255428 | 82255577 | 82253015 | 82253244 | 82265685 | 82265796 | 302 | 167 | 221 | [illegible] |
| SE | 103255 | ENSG00000172661 | WASHC2C | chr10 | + | 45740072 | 45740246 | 45737982 | 45738045 | 45743389 | 45743483 | 169 | 184 | 166 | [illegible] |
| SE | 102576 | ENSG00000120438 | TCP1 | chr6 | - | 159779626 | 159779790 | 159778497 | 159779261 | 159780442 | 159780566 | 453 | 398 | 470 | [illegible] |
| SE | 17786 | ENSG00000100485 | SOS2 | chr14 | - | 50145169 | 50145332 | 50139941 | 50140059 | 50145476 | 50145596 | 284 | 251 | 172 | [illegible] |
| SE | 295 | ENSG00000160305 | DIP2A | chr21 | + | 46534584 | 46534687 | 46534003 | 46534113 | 46537223 | 46537288 | 137 | 40 | 72 | [illegible] |
| SE | 81721 | ENSG00000113013 | HSPA9 | chr5 | - | 138574067 | 138574169 | 138573762 | 138573850 | 138575237 | 138575401 | 67 | 35 | 26 | [illegible] |
| SE | 14409 | ENSG00000167202 | TBC1D2B | chr15 | - | 78024155 | 78024539 | 78017846 | 78017957 | 78030006 | 78030170 | 44 | 52 | 63 | [illegible] |
| SE | 37065 | ENSG00000069535 | MAOB | chrX | - | 43838867 | 43839005 | 43803299 | 43803404 | 43843669 | 43843764 | 3 | 13 | 2 | [illegible] |
| SE | 35308 | ENSG00000165288 | BRWD3 | chrX | - | 80709427 | 80709577 | 80707426 | 80707503 | 80716156 | 80716250 | 264 | 250 | 227 | [illegible] |
| SE | 73431 | ENSG00000177463 | NR2C2 | chr3 | + | 15034669 | 15034809 | 15030274 | 15030452 | 15037999 | 15038137 | 51 | 47 | 63 | [illegible] |
| SE | 23391 | ENSG00000153113 | CAST | chr5 | + | 96726793 | 96726859 | 96722638 | 96722698 | 96727488 | 96727530 | 49 | 29 | 33 | [illegible] |
| SE | 4869 | ENSG00000196562 | SULF2 | chr20 | - | 47663069 | 47663212 | 47661772 | 47661896 | 47663452 | 47663554 | 317 | 176 | 175 | [illegible] |
| SE | 74257 | ENSG00000137959 | IFI44L | chr1 | + | 78627905 | 78628393 | 78620447 | 78620571 | 78628950 | 78628999 | 331 | 919 | 244 | [illegible] |
| SE | 37626 | ENSG00000082074 | FYB1 | chr5 | - | 39212677 | 39213017 | 39211057 | 39211091 | 39250670 | 39250806 | 3 | 4 | 3 | [illegible] |
| SE | 23563 | ENSG00000113141 | IK | chr5 | + | 140655828 | 140655992 | 140654680 | 140654727 | 140657553 | 140657662 | 950 | 589 | 792 | [illegible] |
| SE | 11843 | ENSG00000217555 | CKLF | chr16 | + | 66558189 | 66558348 | 66552568 | 66552793 | 66565885 | 66566072 | 128 | 56 | 136 | [illegible] |
| SE | 61682 | ENSG00000187098 | MITF | chr3 | + | 69939097 | 69939181 | 69937821 | 69938049 | 69941235 | 69941331 | 10 | 9 | 8 | [illegible] |
| SE | 59648 | ENSG00000163466 | ARPC2 | chr2 | + | 218239390 | 218239484 | 218228737 | 218228850 | 218245419 | 218245457 | 2152 | 1726 | 1491 | [illegible] |
| SE | 42361 | ENSG00000048740 | CELF2 | chr10 | + | 11217424 | 11217507 | 11165483 | 11165682 | 11249152 | 11249201 | 1435 | 1090 | 1233 | [illegible] |
| SE | 66650 | ENSG00000084623 | EIF3I | chr1 | + | 32226170 | 32226283 | 32224409 | 32224475 | 32228498 | 32228609 | 152 | 78 | 167 | [illegible] |
| SE | 99351 | ENSG00000255717 | SNHG1 | chr11 | - | 62854017 | 62854064 | 62853551 | 62853833 | 62854518 | 62854551 | 108 | 46 | 161 | [illegible] |
| SE | 97685 | ENSG00000122218 | COPA | chr1 | - | 160299101 | 160299264 | 160297555 | 160297745 | 160305432 | 160305571 | 168 | 147 | 165 | [illegible] |
| SE | 25720 | ENSG00000247774 | PCED1B-AS1 | chr12 | - | 47210639 | 47210761 | 47206510 | 47206588 | 47216164 | 47216200 | 235 | 125 | 354 | [illegible] |
| SE | 4672 | ENSG00000141968 | VAV1 | chr19 | + | 6825302 | 6825406 | 6825052 | 6825121 | 6826611 | 6826711 | 430 | 232 | 308 | [illegible] |
| SE | 97502 | ENSG00000173120 | KDM2A | chr11 | + | 67215855 | 67215949 | 67215339 | 67215446 | 67217755 | 67217884 | 133 | 92 | 145 | [illegible] |
| SE | 101047 | ENSG00000174775 | HRAS | chr11 | - | 533276 | 533358 | 532261 | 532755 | 533452 | 533612 | 21 | 3 | 9 | [illegible] |
| SE | 55817 | ENSG00000106615 | RHEB | chr7 | - | 151498088 | 151498180 | 151490942 | 151491014 | 151519459 | 151519851 | 4 | 7 | 0 | [illegible] |
| SE | 55691 | ENSG00000135749 | PCNX2 | chr1 | - | 233025145 | 233025399 | 233016920 | 233017154 | 233054267 | 233054483 | 26 | 25 | 37 | [illegible] |
| SE | 27953 | ENSG00000128609 | NDUFA5 | chr7 | - | 123551367 | 123551576 | 123545610 | 123550586 | 123557403 | 123557448 | 15 | 11 | 5 | [illegible] |
| SE | 39264 | ENSG00000113810 | SMC4 | chr3 | + | 160417722 | 160417956 | 160416250 | 160416415 | 160419357 | 160419543 | 114 | 111 | 128 | [illegible] |
| SE | 28681 | ENSG00000185033 | SEMA4B | chr15 | + | 90225281 | 90225397 | 90224967 | 90225178 | 90225660 | 90225827 | 21 | 10 | 17 | [illegible] |
| SE | 85203 | ENSG00000140332 | TLE3 | chr15 | - | 70095577 | 70095641 | 70094531 | 70094576 | 70096160 | 70096261 | 117 | 45 | 72 | [illegible] |
| SE | 57402 | ENSG00000101040 | ZMYND8 | chr20 | - | 47291795 | 47291888 | 47290186 | 47290274 | 47294665 | 47294779 | 346 | 333 | 345 | [illegible] |
| SE | 52823 | ENSG00000169764 | UGP2 | chr2 | + | 63862840 | 63862928 | 63857828 | 63857936 | 63882465 | 63882601 | 30 | 11 | 23 | [illegible] |
| SE | 28407 | ENSG00000204842 | ATXN2 | chr12 | - | 111519878 | 111520076 | 111516153 | 111516363 | 111520881 | 111520945 | 138 | 189 | 144 | [illegible] |
| SE | 97197 | ENSG00000147316 | MCPH1 | chr8 | + | 6436047 | 6436162 | 6431498 | 6431586 | 6438952 | 6439096 | 81 | 85 | 84 | [illegible] |
| SE | 20339 | ENSG00000162711 | NLRP3 | chr1 | + | 247418228 | 247418290 | 247416155 | 247416193 | 247418756 | 247419077 | 46 | 24 | 25 | [illegible] |
| SE | 94173 | ENSG00000152795 | HNRNPDL | chr4 | - | 82425562 | 82425667 | 82424628 | 82424883 | 82426036 | 82426129 | 52 | 32 | 52 | [illegible] |
| SE | 32826 | ENSG00000088387 | DOCK9 | chr13 | - | 98845923 | 98846060 | 98837493 | 98837609 | 98848591 | 98848639 | 92 | 44 | 84 | [illegible] |
| SE | 11384 | ENSG00000166747 | AP1G1 | chr16 | - | 71772959 | 71773053 | 71771155 | 71771252 | 71773220 | 71773362 | 8 | 5 | 6 | [illegible] |
| SE | 36687 | ENSG00000214922 | HLA-F-AS1 | chr6 | - | 29747147 | 29747425 | 29745472 | 29745727 | 29748899 | 29749049 | 0 | 11 | 4 | [illegible] |
| SE | 70627 | ENSG00000076201 | PTPN23 | chr3 | + | 47404651 | 47404779 | 47396142 | 47396217 | 47405004 | 47405081 | 68 | 29 | 68 | [illegible] |
| SE | 79386 | ENSG00000054523 | KIF1B | chr1 | + | 10282399 | 10282533 | 10277985 | 10278128 | 10291081 | 10291161 | 80 | 70 | 52 | [illegible] |
| SE | 91303 | ENSG00000089234 | BRAP | chr12 | - | 111665638 | 111665787 | 111660599 | 111660675 | 111672660 | 111672774 | 129 | 89 | 111 | [illegible] |
| SE | 48903 | ENSG00000158773 | USF1 | chr1 | - | 161041734 | 161041846 | 161041323 | 161041411 | 161042115 | 161042217 | 149 | 81 | 140 | [illegible] |
| SE | 20562 | ENSG00000226032 | NA | chr6 | - | 159043588 | 159043655 | 159042191 | 159042244 | 159043977 | 159044031 | 1076 | 740 | 1254 | [illegible] |
| SE | 49335 | ENSG00000101452 | DHX35 | chr20 | + | 39003748 | 39003907 | 39002771 | 39002868 | 39006145 | 39006356 | 58 | 42 | 79 | [illegible] |
| SE | 56649 | ENSG00000155307 | SAMSN1 | chr21 | - | 14516891 | 14517041 | 14512443 | 14512573 | 14521149 | 14521221 | 520 | 374 | 329 | [illegible] |
| SE | 76903 | ENSG00000204590 | GNL1 | chr6 | - | 30555054 | 30555191 | 30554763 | 30554915 | 30555554 | 30555720 | 173 | 82 | 164 | [illegible] |
| SE | 21821 | ENSG00000174353 | STAG3L3 | chr7 | - | 72999470 | 72999550 | 72998492 | 72998790 | 72999760 | 72999859 | 85 | 76 | 124 | [illegible] |
| SE | 66914 | ENSG00000152520 | PAN3 | chr13 | + | 28220230 | 28220378 | 28177864 | 28177935 | 28256291 | 28256539 | 205 | 107 | 152 | [illegible] |
| SE | 2281 | ENSG00000180448 | ARHGAP45 | chr19 | + | 1073673 | 1073746 | 1073505 | 1073590 | 1073947 | 1074014 | 631 | 393 | 692 | [illegible] |
| SE | 96181 | ENSG00000110756 | HPS5 | chr11 | - | 18291441 | 18292019 | 18287892 | 18288013 | 18295019 | 18295169 | 40 | 32 | 41 | [illegible] |
| SE | 74263 | ENSG00000137959 | IFI44L | chr1 | + | 78629719 | 78629915 | 78625669 | 78628999 | 78629992 | 78630187 | 142 | 412 | 112 | [illegible] |
| SE | 3207 | ENSG00000071655 | MBD3 | chr19 | - | 1581091 | 1581269 | 1576639 | 1578538 | 1582621 | 1582712 | 216 | 69 | 170 | [illegible] |
| SE | 23784 | ENSG00000113716 | HMGXB3 | chr5 | + | 150036635 | 150036937 | 150032453 | 150032603 | 150040747 | 150040879 | 36 | 31 | 60 | [illegible] |
| SE | 50914 | ENSG00000123064 | DDX54 | chr12 | - | 113163131 | 113163274 | 113162717 | 113163045 | 113164066 | 113164093 | 109 | 67 | 95 | [illegible] |
| SE | 5989 | ENSG00000062598 | ELMO2 | chr20 | - | 46374535 | 46374640 | 46374331 | 46374440 | 46375235 | 46375370 | 407 | 247 | 283 | [illegible] |
| SE | 48063 | ENSG00000143337 | TOR1AIP1 | chr1 | + | 179887828 | 179887959 | 179884691 | 179884769 | 179889312 | 179889369 | 1 | 4 | 1 | [illegible] |
| SE | 75046 | ENSG00000130023 | ERMARD | chr6 | + | 169776454 | 169776673 | 169775877 | 169776065 | 169779181 | 169779295 | 30 | 18 | 50 | [illegible] |
| SE | 50477 | ENSG00000165671 | NSD1 | chr5 | + | 177282464 | 177282581 | 177280564 | 177280834 | 177283786 | 177283928 | 264 | 201 | 271 | [illegible] |
| SE | 44581 | ENSG00000026508 | CD44 | chr11 | + | 35206114 | 35206243 | 35204511 | 35204640 | 35208104 | 35208206 | 18 | 13 | 17 | [illegible] |
| SE | 93192 | ENSG00000139112 | GABARAPL1 | chr12 | + | 10214668 | 10214779 | 10213835 | 10214187 | 10218062 | 10218141 | 7 | 6 | 8 | [illegible] |
| SE | 19247 | ENSG00000140943 | MBTPS1 | chr16 | - | 84084982 | 84085134 | 84081746 | 84081908 | 84087357 | 84087460 | 195 | 103 | 139 | [illegible] |
| SE | 13324 | ENSG00000177663 | IL17RA | chr22 | + | 17103493 | 17103577 | 17102138 | 17102302 | 17104725 | 17104810 | 648 | 380 | 374 | [illegible] |
| SE | 93463 | ENSG00000106771 | TMEM245 | chr9 | - | 109038016 | 109038117 | 109036205 | 109036380 | 109050282 | 109050428 | 205 | 167 | 233 | [illegible] |
| SE | 48199 | ENSG00000196204 | RNF216P1 | chr7 | + | 4983670 | 4983773 | 4976300 | 4976436 | 4984909 | 4985075 | 114 | 78 | 121 | [illegible] |
| SE | 88080 | ENSG00000196663 | TECPR2 | chr14 | + | 102431795 | 102432128 | 102428249 | 102428382 | 102434234 | 102435211 | 36 | 27 | 47 | [illegible] |
| SE | 74082 | ENSG00000085721 | RRN3 | chr16 | - | 15086364 | 15086454 | 15086128 | 15086258 | 15091314 | 15091371 | 179 | 143 | 187 | [illegible] |
| SE | 55980 | ENSG00000121892 | PDS5A | chr4 | - | 39848850 | 39848970 | 39845817 | 39845880 | 39849519 | 39849652 | 402 | 346 | 337 | [illegible] |
| SE | 5591 | ENSG00000126001 | CEP250 | chr20 | + | 35496565 | 35496715 | 35494523 | 35494657 | 35497718 | 35498067 | 129 | 108 | 110 | [illegible] |
| SE | 49201 | ENSG00000258830 | NA | chr12 | - | 57269322 | 57269449 | 57268921 | 57269082 | 57269751 | 57269994 | 85 | 27 | 66 | [illegible] |
| SE | 88416 | ENSG00000168172 | HOOK3 | chr8 | + | 42968012 | 42968214 | 42966472 | 42966613 | 42973288 | 42973399 | 354 | 319 | 217 | [illegible] |
| SE | 51548 | ENSG00000036549 | ZZZ3 | chr1 | - | 77639448 | 77639602 | 77631849 | 77633405 | 77641383 | 77641446 | 200 | 136 | 161 | [illegible] |
| SE | 74827 | ENSG00000042088 | TDP1 | chr14 | + | 90033105 | 90033214 | 90019315 | 90019418 | 90043069 | 90043252 | 74 | 37 | 82 | [illegible] |
| SE | 14442 | ENSG00000130703 | OSBPL2 | chr20 | + | 62272124 | 62272259 | 62263615 | 62263691 | 62279156 | 62279205 | 138 | 88 | 137 | [illegible] |
| SE | 69606 | ENSG00000184922 | FMNL1 | chr17 | + | 45246504 | 45246604 | 45246209 | 45246330 | 45246866 | 45247318 | 13 | 21 | 13 | [illegible] |
| SE | 16221 | ENSG00000077238 | IL4R | chr16 | + | 27360765 | 27360815 | 27358915 | 27358994 | 27362251 | 27362296 | 191 | 99 | 153 | [illegible] |
| SE | 4433 | ENSG00000105705 | SUGP1 | chr19 | - | 19303723 | 19303847 | 19303347 | 19303448 | 19305848 | 19306076 | 87 | 82 | 80 | [illegible] |
| SE | 23574 | ENSG00000254996 | ANKHD1-EIF4EBP3 | chr5 | + | 140512827 | 140512923 | 140510018 | 140510181 | 140513362 | 140513479 | 275 | 195 | 226 | [illegible] |
| SE | 29620 | ENSG00000203875 | SNHG5 | chr6 | - | 85677953 | 85678032 | 85677008 | 85677492 | 85678135 | 85678188 | 97 | 81 | 116 | [illegible] |
| SE | 51359 | ENSG00000066427 | ATXN3 | chr14 | - | 92088729 | 92088817 | 92083125 | 92083258 | 92093251 | 92093318 | 259 | 171 | 175 | [illegible] |
| SE | 70586 | ENSG00000188092 | GPR89B | chr1 | + | 147968894 | 147968963 | 147966770 | 147968428 | 147969866 | 147969959 | 3 | 5 | 8 | [illegible] |
| SE | 43555 | ENSG00000110172 | CHORDC1 | chr11 | - | 90206747 | 90206810 | 90206201 | 90206272 | 90210535 | 90210594 | 73 | 66 | 63 | [illegible] |
| SE | 74191 | ENSG00000197345 | MRPL21 | chr11 | - | 68893402 | 68893455 | 68892491 | 68892993 | 68896514 | 68896678 | 127 | 51 | 125 | [illegible] |
| SE | 7412 | ENSG00000168461 | RAB31 | chr18 | + | 9775277 | 9775357 | 9708300 | 9708444 | 9814019 | 9814091 | 19 | 9 | 6 | [illegible] |
| SE | 40419 | ENSG00000078304 | PPP2R5C | chr14 | + | 101831700 | 101831804 | 101786017 | 101786183 | 101856685 | 101856885 | 12 | 11 | 4 | [illegible] |
| SE | 96221 | ENSG00000110768 | GTF2H1 | chr11 | + | 18351880 | 18351969 | 18347831 | 18347919 | 18352328 | 18352446 | 192 | 129 | 184 | [illegible] |
| SE | 85600 | ENSG00000163389 | POGLUT1 | chr3 | + | 119478333 | 119478474 | 119477312 | 119477448 | 119480050 | 119480172 | 10 | 4 | 1 | [illegible] |
| SE | 8250 | ENSG00000275700 | AATF | chr17 | + | 37031613 | 37031685 | 37020933 | 37021014 | 37056600 | 37056842 | 246 | 157 | 166 | [illegible] |
| SE | 76696 | ENSG00000231925 | TAPBP | chr6 | - | 33304127 | 33304217 | 33299693 | 33301771 | 33304296 | 33304638 | 93 | 36 | 111 | [illegible] |
| SE | 60513 | ENSG00000143382 | ADAMTSL4 | chr1 | + | 150559760 | 150559905 | 150559286 | 150559466 | 150560059 | 150560667 | 59 | 18 | 38 | [illegible] |
| SE | 26008 | ENSG00000181929 | PRKAG1 | chr12 | - | 49008425 | 49008599 | 49005742 | 49005852 | 49013061 | 49013110 | 3 | 0 | 5 | [illegible] |
| SE | 81437 | ENSG00000011275 | RNF216 | chr7 | - | 5766894 | 5766939 | 5761010 | 5761138 | 5781540 | 5781620 | 2 | 9 | 1 | [illegible] |
| SE | 63598 | ENSG00000152127 | MGAT5 | chr2 | + | 134253094 | 134253218 | 134119982 | 134120291 | 134254261 | 134254644 | 33 | 27 | 14 | [illegible] |
| SE | 52619 | ENSG00000197872 | CYRIA | chr2 | - | 16563988 | 16564094 | 16562004 | 16562141 | 16565645 | 16565767 | 280 | 198 | 158 | [illegible] |
| SE | 30137 | ENSG00000150961 | SEC24D | chr4 | - | 118816743 | 118816850 | 118815450 | 118815726 | 118817263 | 118817412 | 0 | 0 | 0 | [illegible] |
| SE | 66170 | ENSG00000088179 | PTPN4 | chr2 | + | 119926597 | 119926666 | 119920068 | 119920241 | 119932423 | 119932549 | 135 | 87 | 126 | [illegible] |
| SE | 29338 | ENSG00000121210 | TMEM131L | chr4 | + | 153591052 | 153591194 | 153588889 | 153589007 | 153592474 | 153592584 | 204 | 196 | 254 | [illegible] |
| SE | 28006 | ENSG00000137601 | NEK1 | chr4 | - | 169587558 | 169587613 | 169585348 | 169585549 | 169588648 | 169588735 | 77 | 55 | 59 | [illegible] |
| SE | 1619 | ENSG00000244625 | MIATNB | chr22 | + | 26711829 | 26712008 | 26693692 | 26693836 | 26717252 | 26717459 | 12 | 7 | 9 | [illegible] |
| SE | 86690 | ENSG00000147140 | NONO | chrX | + | 71284396 | 71284453 | 71283589 | 71283721 | 71290628 | 71290791 | 303 | 184 | 293 | [illegible] |
| SE | 44249 | ENSG00000115808 | STRN | chr2 | - | 36855211 | 36855352 | 36850999 | 36851107 | 36857855 | 36858023 | 178 | 139 | 182 | [illegible] |
| SE | 61516 | ENSG00000138757 | G3BP2 | chr4 | - | 75646337 | 75646456 | 75641848 | 75645702 | 75647028 | 75647157 | 1392 | 1220 | 1183 | [illegible] |
| SE | 2886 | ENSG00000167670 | CHAF1A | chr19 | + | 4422565 | 4422795 | 4418019 | 4418076 | 4423334 | 4423395 | 11 | 12 | 23 | [illegible] |
| SE | 94601 | ENSG00000106829 | TLE4 | chr9 | + | 79649819 | 79649858 | 79627373 | 79627448 | 79652589 | 79652798 | 6 | 4 | 10 | [illegible] |
| SE | 60139 | ENSG00000171848 | RRM2 | chr2 | + | 10127086 | 10127220 | 10126511 | 10126969 | 10128847 | 10128952 | 62 | 39 | 241 | [illegible] |
| SE | 21307 | ENSG00000129534 | MIS18BP1 | chr14 | - | 45232508 | 45232627 | 45231140 | 45231298 | 45232732 | 45232820 | 18 | 8 | 32 | [illegible] |
| SE | 48180 | ENSG00000117543 | DPH5 | chr1 | - | 101025308 | 101025466 | 101021640 | 101021765 | 101025639 | 101025761 | 42 | 32 | 68 | [illegible] |
| SE | 104268 | ENSG00000132305 | IMMT | chr2 | - | 86158591 | 86158721 | 86153559 | 86153574 | 86159535 | 86159671 | 480 | 305 | 382 | [illegible] |
| SE | 98826 | ENSG00000135365 | PHF21A | chr11 | - | 45945839 | 45946098 | 45938218 | 45938312 | 45948885 | 45948946 | 68 | 46 | 24 | [illegible] |
| SE | 29617 | ENSG00000203875 | SNHG5 | chr6 | - | 85677794 | 85677899 | 85677423 | 85677492 | 85677953 | 85678032 | 88 | 194 | 135 | [illegible] |
| SE | 68777 | ENSG00000100523 | DDHD1 | chr14 | - | 53096116 | 53096218 | 53093315 | 53093444 | 53103682 | 53103856 | 31 | 30 | 34 | [illegible] |
| SE | 46762 | ENSG00000108179 | PPIF | chr10 | + | 79352316 | 79352392 | 79351486 | 79351583 | 79353706 | 79355334 | 114 | 81 | 111 | [illegible] |
| SE | 14000 | ENSG00000047578 | KATNIP | chr16 | + | 27740040 | 27740920 | 27721557 | 27721695 | 27749583 | 27750306 | 77 | 44 | 57 | [illegible] |
| SE | 8091 | ENSG00000141252 | VPS53 | chr17 | - | 628087 | 628231 | 627173 | 627316 | 631549 | 631628 | 208 | 183 | 216 | [illegible] |
| SE | 7675 | ENSG00000219200 | RNASEK | chr17 | + | 7013318 | 7013516 | 7012460 | 7012761 | 7013665 | 7013742 | 19 | 11 | 16 | [illegible] |
| SE | 75041 | ENSG00000130023 | ERMARD | chr6 | + | 169766637 | 169766667 | 169762428 | 169762531 | 169768102 | 169768171 | 112 | 88 | 97 | [illegible] |
| SE | 50141 | ENSG00000132300 | PTCD3 | chr2 | + | 86118920 | 86119044 | 86117054 | 86117159 | 86121478 | 86121594 | 442 | 313 | 415 | [illegible] |
| SE | 88705 | ENSG00000183077 | AFMID | chr17 | + | 78205602 | 78205753 | 78202702 | 78202751 | 78205945 | 78206050 | 0 | 0 | 0 | [illegible] |
| SE | 62962 | ENSG00000117222 | RBBP5 | chr1 | - | 205094986 | 205095064 | 205086141 | 205088815 | 205096681 | 205096911 | 58 | 58 | 69 | [illegible] |
| SE | 71469 | ENSG00000130038 | CRACR2A | chr12 | - | 3644594 | 3644640 | 3641731 | 3641838 | 3648541 | 3648613 | 73 | 48 | 69 | [illegible] |
| SE | 71410 | ENSG00000145349 | CAMK2D | chr4 | - | 113547643 | 113547716 | 113537340 | 113537443 | 113548656 | 113548729 | 17 | 14 | 30 | [illegible] |
| SE | 75371 | ENSG00000124593 | NA | chr6 | + | 41789231 | 41789336 | 41786439 | 41787825 | 41789567 | 41789891 | 703 | 287 | 674 | [illegible] |
| SE | 33342 | ENSG00000188313 | PLSCR1 | chr3 | - | 146537995 | 146538044 | 146536539 | 146536565 | 146544466 | 146544547 | 7 | 7 | 3 | [illegible] |
| SE | 49291 | ENSG00000120733 | KDM3B | chr5 | + | 138425410 | 138425582 | 138424074 | 138424341 | 138426974 | 138427065 | 345 | 260 | 364 | [illegible] |
| SE | 58061 | ENSG00000082701 | GSK3B | chr3 | - | 119912703 | 119912810 | 119863418 | 119863605 | 119916043 | 119916174 | 171 | 107 | 126 | [illegible] |
| SE | 6195 | ENSG00000116017 | ARID3A | chr19 | + | 932417 | 932742 | 929261 | 929896 | 960091 | 960164 | 28 | 15 | 42 | [illegible] |
| SE | 49625 | ENSG00000178927 | CYBC1 | chr17 | - | 82447579 | 82447646 | 82446622 | 82446696 | 82449169 | 82449320 | 11 | 9 | 14 | [illegible] |
| SE | 30204 | ENSG00000111266 | DUSP16 | chr12 | - | 12554535 | 12554659 | 12521080 | 12521463 | 12562116 | 12562863 | 14 | 12 | 11 | [illegible] |
| SE | 52001 | ENSG00000071054 | MAP4K4 | chr2 | + | 101888795 | 101888935 | 101887777 | 101887937 | 101891165 | 101891249 | 415 | 228 | 367 | [illegible] |
| SE | 67294 | ENSG00000164117 | FBXO8 | chr4 | - | 174269053 | 174269167 | 174262763 | 174263100 | 174283409 | 174283663 | 0 | 0 | 1 | [illegible] |
| SE | 66743 | ENSG00000138468 | SENP7 | chr3 | - | 101417592 | 101417790 | 101372007 | 101372126 | 101458954 | 101459052 | 101 | 111 | 112 | [illegible] |
| SE | 53151 | ENSG00000124574 | ABCC10 | chr6 | + | 43441861 | 43441960 | 43437933 | 43438795 | 43442969 | 43443159 | 73 | 40 | 64 | [illegible] |
| SE | 75788 | ENSG00000137200 | CMTR1 | chr6 | + | 37436308 | 37436382 | 37435623 | 37435762 | 37443998 | 37444083 | 1 | 0 | 0 | [illegible] |

| | | | | | | | | | | | | | | |
|---|---|---|---|---|---|---|---|---|---|---|---|---|---|---|
| SE | 61100 | ENSG00000067225 | PKM | chr15 | - | 72221167 | 72221284 | 72218943 | 72219110 | 72231323 | 72231624 | 2 | 0 | 0 |
| SE | 43847 | ENSG00000142208 | AKT1 | chr14 | - | 104793126 | 104794261 | 104792597 | 104792722 | 104795010 | 104795044 | 53 | 22 | 31 |
| SE | 10643 | ENSG00000171475 | WIPF2 | chr17 | + | 40262524 | 40262641 | 40260534 | 40260667 | 40264489 | 40265146 | 100 | 45 | 75 |
| SE | 95772 | ENSG00000125944 | HNRNPR | chr1 | - | 23323555 | 23323732 | 23321527 | 23321663 | 23337753 | 23337861 | 658 | 476 | 624 |
| SE | 103556 | ENSG00000107862 | GBF1 | chr10 | + | 102381126 | 102381255 | 102379283 | 102380686 | 102382055 | 102382336 | 52 | 27 | 45 |
| SE | 4604 | ENSG00000105397 | TYK2 | chr19 | - | 10378213 | 10378426 | 10368294 | 10368418 | 10379614 | 10379689 | 226 | 124 | 204 |
| SE | 20360 | ENSG00000120925 | RNF170 | chr8 | - | 42865415 | 42865489 | 42861329 | 42861855 | 42870003 | 42870112 | 66 | 58 | 89 |
| SE | 28422 | ENSG00000174652 | ZNF266 | chr19 | - | 9418765 | 9418895 | 9418504 | 9418631 | 9419216 | 9419299 | 50 | 29 | 28 |
| SE | 46703 | ENSG00000171103 | TRMT61B | chr2 | - | 28865016 | 28865119 | 28861117 | 28861308 | 28869578 | 28870295 | 115 | 97 | 101 |
| SE | 48190 | ENSG00000006831 | ADIPOR2 | chr12 | + | 1777853 | 1778025 | 1772713 | 1772961 | 1783879 | 1784073 | 111 | 90 | 79 |
| SE | 6077 | ENSG00000089057 | SLC23A2 | chr20 | - | 4883641 | 4883823 | 4874575 | 4874696 | 4884752 | 4884823 | 133 | 64 | 96 |
| SE | 8759 | ENSG00000007202 | KIAA0100 | chr17 | - | 28636964 | 28637170 | 28634516 | 28635609 | 28637835 | 28638094 | 478 | 190 | 315 |
| SE | 74108 | ENSG00000255587 | RAB44 | chr6 | + | 36727576 | 36727691 | 36725861 | 36725943 | 36728699 | 36728801 | 35 | 16 | 12 |
| SE | 39339 | ENSG00000144559 | TAMM41 | chr3 | - | 11830721 | 11830874 | 11829755 | 11829864 | 11833002 | 11833151 | 1 | 0 | 2 |
| SE | 91990 | ENSG00000119397 | CNTRL | chr9 | + | 121140640 | 121140786 | 121138544 | 121138679 | 121142090 | 121142270 | 145 | 167 | 166 |
| SE | 18958 | ENSG00000015133 | CCDC88C | chr14 | - | 91285394 | 91285836 | 91283328 | 91283517 | 91289104 | 91289331 | 508 | 200 | 507 |
| SE | 8527 | ENSG00000125450 | NUP85 | chr17 | + | 75218184 | 75218306 | 75213075 | 75213119 | 75225102 | 75225237 | 27 | 21 | 28 |
| SE | 106416 | ENSG00000135297 | MTO1 | chr6 | + | 73473364 | 73473654 | 73466208 | 73466606 | 73479731 | 73479844 | 229 | 187 | 226 |
| SE | 54967 | ENSG00000135842 | NIBAN1 | chr1 | - | 184805945 | 184806056 | 184803584 | 184803692 | 184808073 | 184808235 | 227 | 181 | 209 |
| SE | 15311 | ENSG00000114062 | UBE3A | chr15 | - | 25407066 | 25409207 | 25405460 | 25405502 | 25411907 | 25411971 | 100 | 63 | 91 |
| SE | 99965 | ENSG00000154310 | TNIK | chr3 | - | 171161269 | 171161336 | 171157459 | 171157664 | 171167094 | 171167270 | 93 | 122 | 130 |
| SE | 9884 | ENSG00000265148 | TSPOAP1-AS1 | chr17 | + | 58339242 | 58339447 | 58337335 | 58337679 | 58351734 | 58351743 | 1 | 0 | 0 |
| SE | 17868 | ENSG00000092199 | HNRNPC | chr14 | - | 21263666 | 21263829 | 21263310 | 21263336 | 21269297 | 21269406 | 200 | 158 | 187 |
| SE | 2487 | ENSG00000141867 | BRD4 | chr19 | - | 15267415 | 15267551 | 15265353 | 15265643 | 15268904 | 15269042 | 1329 | 1073 | 1154 |
| SE | 59925 | ENSG00000121931 | LRIF1 | chr1 | - | 110951287 | 110952815 | 110949850 | 110950123 | 110963620 | 110963922 | 43 | 47 | 50 |
| SE | 16131 | ENSG00000140553 | UNC45A | chr15 | + | 90942436 | 90942605 | 90940305 | 90940473 | 90942911 | 90943082 | 111 | 45 | 105 |
| SE | 10263 | ENSG00000108651 | UTP6 | chr17 | - | 31878701 | 31878781 | 31878249 | 31878327 | 31880572 | 31880754 | 90 | 69 | 85 |
| SE | 98813 | ENSG00000149091 | DGKZ | chr11 | + | 46372616 | 46372677 | 46372427 | 46372510 | 46372770 | 46372884 | 185 | 59 | 185 |
| SE | 36878 | ENSG00000122779 | TRIM24 | chr7 | + | 138515211 | 138515359 | 138504289 | 138504408 | 138519188 | 138519321 | 152 | 181 | 124 |
| SE | 90976 | ENSG00000132669 | RIN2 | chr20 | + | 19935438 | 19935607 | 19935098 | 19935199 | 19956614 | 19956807 | 0 | 2 | 2 |
| SE | 41455 | ENSG00000137776 | SLTM | chr15 | - | 58913498 | 58913696 | 58912562 | 58912610 | 58916934 | 58916999 | 265 | 265 | 222 |
| SE | 71062 | ENSG00000115486 | GGCX | chr2 | - | 85560814 | 85560985 | 85558724 | 85559075 | 85561385 | 85561473 | 30 | 16 | 37 |
| SE | 19098 | ENSG00000169118 | CSNK1G1 | chr15 | - | 64251511 | 64251581 | 64216561 | 64216713 | 64259200 | 64259241 | 72 | 70 | 89 |

| | | | | | | | |
|---|---|---|---|---|---|---|---|
| SE | 94256 | ENSG00000138663 | COPS4 | chr4 | + | 83065059 | 83065155 |
| SE | 40101 | ENSG00000095321 | CRAT | chr9 | - | 129108407 | 129108478 |
| SE | 108132 | ENSG00000155957 | TMBIM4 | chr12 | - | 66169289 | 66169353 |
| SE | 35098 | ENSG00000168000 | BSCL2 | chr11 | - | 62692375 | 62692473 |
| SE | 108651 | ENSG00000136147 | PHF11 | chr13 | + | 49506634 | 49506756 |
| SE | 20618 | ENSG00000130779 | CLIP1 | chr12 | - | 122355114 | 122355312 |
| SE | 85684 | ENSG00000177981 | ASB8 | chr12 | - | 48150086 | 48150140 |
| SE | 33082 | ENSG00000164258 | NDUFS4 | chr5 | + | 53604723 | 53604895 |
| SE | 17339 | ENSG00000258289 | CHURC1 | chr14 | + | 64926006 | 64926080 |
| SE | 38980 | ENSG00000004399 | PLXND1 | chr3 | - | 129559619 | 129559783 |
| SE | 49432 | ENSG00000136068 | FLNB | chr3 | + | 58141857 | 58141896 |
| SE | 12506 | ENSG00000132600 | PRMT7 | chr16 | + | 68329065 | 68329174 |
| SE | 7258 | ENSG00000101745 | ANKRD12 | chr18 | + | 9204475 | 9204544 |
| SE | 32535 | ENSG00000197808 | ZNF461 | chr19 | - | 36643793 | 36643862 |
| SE | 18693 | ENSG00000129566 | TEP1 | chr14 | - | 20404610 | 20404772 |
| SE | 26088 | ENSG00000174695 | TMEM167A | chr5 | - | 83065007 | 83065117 |
| SE | 358 | ENSG00000160294 | MCM3AP | chr21 | - | 46270400 | 46270563 |
| SE | 107722 | ENSG00000177575 | CD163 | chr12 | - | 7483366 | 7483675 |
| SE | 105789 | ENSG00000124164 | VAPB | chr20 | + | 58438944 | 58439025 |
| SE | 66402 | ENSG00000161813 | LARP4 | chr12 | + | 50402674 | 50402813 |
| SE | 6205 | ENSG00000125844 | RRBP1 | chr20 | - | 17616731 | 17616839 |
| SE | 82980 | ENSG00000186088 | GSAP | chr7 | - | 77349350 | 77349404 |
| SE | 51465 | ENSG00000162517 | PEF1 | chr1 | - | 31634833 | 31634943 |
| SE | 35089 | ENSG00000284952 | NA | chr3 | - | 155833467 | 155833585 |
| SE | 707 | ENSG00000182944 | EWSR1 | chr22 | + | 29288605 | 29288755 |
| SE | 85699 | ENSG00000175463 | TBC1D10C | chr11 | + | 67405906 | 67406017 |
| SE | 103516 | ENSG00000107862 | GBF1 | chr10 | + | 102361021 | 102361120 |
| SE | 10149 | ENSG00000011260 | UTP18 | chr17 | + | 51296964 | 51297003 |
| SE | 39277 | ENSG00000128050 | PAICS | chr4 | + | 56453602 | 56453761 |
| SE | 18375 | ENSG00000100888 | CHD8 | chr14 | - | 21430800 | 21431858 |
| SE | 82527 | ENSG00000061938 | TNK2 | chr3 | - | 195896052 | 195896297 |
| SE | 87821 | ENSG00000126767 | ELK1 | chrX | - | 47637750 | 47638182 |
| SE | 86792 | ENSG00000169057 | MECP2 | chrX | - | 154039791 | 154039903 |
| SE | 97536 | ENSG00000149761 | NUDT22 | chr11 | + | 64229246 | 64229344 |
| SE | 89536 | ENSG00000110074 | FOXRED1 | chr11 | + | 126274930 | 126275021 |
| SE | 101442 | ENSG00000165997 | ARL5B | chr10 | + | 18672621 | 18672705 |
| SE | 76167 | ENSG00000164691 | TAGAP | chr6 | - | 159042077 | 159042244 |
| SE | 67497 | ENSG00000068654 | POLR1A | chr2 | - | 86044138 | 86044304 |
| SE | 62610 | ENSG00000015676 | NUDCD3 | chr7 | - | 44413190 | 44413283 |
| SE | 71542 | ENSG00000145216 | FIP1L1 | chr4 | + | 53414614 | 53414722 |
| SE | 96895 | ENSG00000100731 | PCNX1 | chr14 | + | 71009633 | 71009724 |
| SE | 44962 | ENSG00000117682 | DHDDS | chr1 | + | 26460036 | 26460144 |
| SE | 35903 | ENSG00000146232 | NFKBIE | chr6 | - | 44262559 | 44262662 |
| SE | 4552 | ENSG00000127507 | ADGRE2 | chr19 | - | 14766977 | 14767109 |
| SE | 108841 | ENSG00000081189 | MEF2C | chr5 | - | 88839378 | 88839623 |
| SE | 69025 | ENSG00000132405 | TBC1D14 | chr4 | + | 7025003 | 7025262 |
| SE | 24964 | ENSG00000150712 | MTMR12 | chr5 | - | 32233772 | 32233934 |
| SE | 95248 | ENSG00000159921 | GNE | chr9 | - | 36249191 | 36249397 |
| SE | 54711 | ENSG00000137965 | IFI44 | chr1 | + | 78660553 | 78660691 |
| SE | 93937 | ENSG00000144524 | COPS7B | chr2 | + | 231801139 | 231801207 |
| SE | 48489 | ENSG00000138095 | LRPPRC | chr2 | - | 43890222 | 43890274 |
| SE | 104488 | ENSG00000013561 | RNF14 | chr5 | + | 141980122 | 141980351 |
| SE | 19971 | ENSG00000136758 | YME1L1 | chr10 | - | 27147497 | 27147668 |
| SE | 77784 | ENSG00000152457 | DCLRE1C | chr10 | - | 14942113 | 14942240 |
| SE | 108304 | ENSG00000115310 | RTN4 | chr2 | - | 54973820 | 54973867 |
| SE | 100839 | ENSG00000149311 | ATM | chr11 | + | 108223535 | 108223686 |
| SE | 106104 | ENSG00000108515 | ENO3 | chr17 | + | 4952024 | 4952147 |
| SE | 53393 | ENSG00000135837 | CEP350 | chr1 | + | 180036925 | 180037089 |
| SE | 62064 | ENSG00000110446 | SLC15A3 | chr11 | - | 60937515 | 60937566 |
| SE | 54314 | ENSG00000143442 | POGZ | chr1 | - | 151440927 | 151441086 |
| SE | 52239 | ENSG00000166825 | ANPEP | chr15 | - | 89797574 | 89797722 |
| SE | 69907 | ENSG00000064995 | TAF11 | chr6 | - | 34879625 | 34879736 |
| SE | 82618 | ENSG00000106144 | CASP2 | chr7 | + | 143292616 | 143292698 |
| SE | 35157 | ENSG00000188001 | TPRG1 | chr3 | + | 189121661 | 189121861 |
| SE | 90505 | ENSG00000160299 | PCNT | chr21 | + | 46421969 | 46422124 |
| SE | 99505 | ENSG00000163131 | CTSS | chr1 | - | 150750005 | 150750171 |
| SE | 10960 | ENSG00000091592 | NLRP1 | chr17 | - | 5536850 | 5536940 |
| SE | 15426 | ENSG00000102901 | CENPT | chr16 | - | 67832227 | 67832350 |
| SE | 67218 | ENSG00000144118 | RALB | chr2 | + | 120285873 | 120286082 |
| SE | 9557 | ENSG00000100360 | IFT27 | chr22 | - | 36762903 | 36763013 |
| SE | 38502 | ENSG00000198793 | MTOR | chr1 | - | 11256932 | 11257165 |
| SE | 50203 | ENSG00000148187 | MRRF | chr9 | + | 122265461 | 122265545 |
| SE | 58332 | ENSG00000115207 | GTF3C2 | chr2 | - | 27335916 | 27336028 |
| SE | 97632 | ENSG00000198879 | SFMBT2 | chr10 | - | 7197547 | 7197687 |
| SE | 22149 | ENSG00000156639 | ZFAND3 | chr6 | + | 38082391 | 38082457 |
| SE | 28017 | ENSG00000138698 | RAP1GDS1 | chr4 | + | 98379016 | 98379163 |
| SE | 49462 | ENSG00000115020 | PIKFYVE | chr2 | + | 208285725 | 208285933 |
| SE | 22734 | ENSG00000050393 | MCUR1 | chr6 | - | 13791877 | 13791992 |
| SE | 103542 | ENSG00000107862 | GBF1 | chr10 | + | 102368738 | 102368832 |
| SE | 5732 | ENSG00000154930 | ACSS1 | chr20 | - | 25015137 | 25015230 |
| SE | 77256 | ENSG00000141664 | ZCCHC2 | chr18 | + | 62558686 | 62558770 |
| SE | 109383 | ENSG00000166226 | CCT2 | chr12 | + | 69597637 | 69597766 |
| SE | 100846 | ENSG00000149311 | ATM | chr11 | + | 108224783 | 108224871 |
| SE | 22197 | ENSG00000137177 | KIF13A | chr6 | - | 17794248 | 17794395 |
| SE | 46268 | ENSG00000107625 | DDX50 | chr10 | + | 68906225 | 68906337 |
| SE | 51937 | ENSG00000074603 | DPP8 | chr15 | - | 65500605 | 65500779 |
| SE | 15935 | ENSG00000168904 | LRRC28 | chr15 | + | 99317463 | 99317546 |
| SE | 71435 | ENSG00000115234 | SNX17 | chr2 | + | 27372622 | 27372740 |
| SE | 25441 | ENSG00000011243 | AKAP8L | chr19 | - | 15398629 | 15399410 |
| SE | 9478 | ENSG00000248871 | TNFSF12-TNFSF13 | chr17 | + | 7559623 | 7559702 |
| SE | 91467 | ENSG00000197140 | ADAM32 | chr8 | + | 39160896 | 39160965 |
| SE | 38226 | ENSG00000113108 | APBB3 | chr5 | - | 140562662 | 140562723 |
| SE | 81257 | ENSG00000115677 | HDLBP | chr2 | - | 241268828 | 241268953 |
| SE | 57148 | ENSG00000143553 | SNAPIN | chr1 | + | 153659137 | 153659184 |
| SE | 102220 | ENSG00000100109 | TFIP11 | chr22 | - | 26510616 | 26510667 |
| SE | 104689 | ENSG00000100354 | TNRC6B | chr22 | + | 40310816 | 40310993 |
| SE | 28294 | ENSG00000146426 | TIAM2 | chr6 | + | 154860737 | 154860998 |
| SE | 49607 | ENSG00000076685 | NT5C2 | chr10 | - | 103139405 | 103139479 |
| SE | 65040 | ENSG00000091409 | ITGA6 | chr2 | + | 172501771 | 172501901 |
| SE | 53004 | ENSG00000158941 | CCAR2 | chr8 | + | 22613016 | 22613136 |
| SE | 61161 | ENSG00000028528 | SNX1 | chr15 | + | 64118202 | 64118244 |
| SE | 13436 | ENSG00000103111 | MON1B | chr16 | + | 77193450 | 77193777 |
| SE | 60213 | ENSG00000163788 | SNRK | chr3 | + | 43289739 | 43289789 |
| SE | 46617 | ENSG00000141556 | TBCD | chr17 | + | 82900650 | 82900731 |
| SE | 25833 | ENSG00000250334 | LINC00989 | chr4 | + | 79513784 | 79513902 |
| SE | 57277 | ENSG00000143119 | CD53 | chr1 | + | 110894959 | 110895055 |
| SE | 16321 | ENSG00000140374 | ETFA | chr15 | - | 76295590 | 76295737 |
| SE | 5460 | ENSG00000101337 | TM9SF4 | chr20 | + | 32133012 | 32133126 |
| SE | 335 | ENSG00000141959 | PFKL | chr21 | + | 44313912 | 44314021 |
| SE | 41028 | ENSG00000064687 | ABCA7 | chr19 | + | 1056906 | 1057084 |
| SE | 73960 | ENSG00000166436 | TRIM66 | chr11 | - | 8620445 | 8620572 |
| SE | 89178 | ENSG00000075856 | SART3 | chr12 | - | 108544426 | 108544532 |
| SE | 10316 | ENSG00000108848 | LUC7L3 | chr17 | + | 50738136 | 50738199 |
| SE | 11226 | ENSG00000102984 | ZNF821 | chr16 | - | 71861775 | 71861942 |
| SE | 30219 | ENSG00000132376 | INPP5K | chr17 | - | 1509682 | 1509799 |
| SE | 105593 | ENSG00000197971 | MBP | chr18 | - | 77097250 | 77097474 |
| SE | 4595 | ENSG00000105397 | TYK2 | chr19 | - | 10352433 | 10352551 |
| SE | 106562 | ENSG00000111790 | FGFR1OP2 | chr12 | + | 26960514 | 26960628 |
| SE | 91030 | ENSG00000120910 | PPP3CC | chr8 | + | 22513292 | 22513432 |
| SE | 3775 | ENSG00000105486 | LIG1 | chr19 | - | 48162261 | 48162351 |
| SE | 13999 | ENSG00000047578 | KATNIP | chr16 | + | 27701588 | 27701695 |
| SE | 78880 | ENSG00000196792 | STRN3 | chr14 | - | 30929200 | 30929311 |
| SE | 75697 | ENSG00000078269 | SYNJ2 | chr6 | + | 158078163 | 158078281 |
| SE | 27731 | ENSG00000152818 | UTRN | chr6 | + | 144437564 | 144437746 |
| SE | 67805 | ENSG00000055044 | NOP58 | chr2 | + | 202277949 | 202278002 |
| SE | 59621 | ENSG00000111801 | BTN3A3 | chr6 | + | 26448616 | 26448637 |
| SE | 850 | ENSG00000242259 | C22orf39 | chr22 | - | 19377868 | 19378066 |
| SE | 14204 | ENSG00000103381 | CPPED1 | chr16 | - | 12704623 | 12705049 |
| SE | 100362 | ENSG00000089597 | GANAB | chr11 | - | 62629813 | 62629957 |
| SE | 48864 | ENSG00000184677 | ZBTB40 | chr1 | + | 22508000 | 22508137 |
| SE | 28402 | ENSG00000204842 | ATXN2 | chr12 | - | 111516153 | 111516399 |
| SE | 102999 | ENSG00000197535 | MYO5A | chr15 | - | 52407291 | 52407399 |
| SE | 26698 | ENSG00000136709 | WDR33 | chr2 | - | 127713582 | 127714021 |
| SE | 868 | ENSG00000128159 | TUBGCP6 | chr22 | - | 50220191 | 50221874 |
| SE | 71071 | ENSG00000134490 | TMEM241 | chr18 | - | 23370215 | 23370261 |
| SE | 15427 | ENSG00000102901 | CENPT | chr16 | - | 67832227 | 67832354 |
| SE | 57332 | ENSG00000181722 | ZBTB20 | chr3 | - | 114713975 | 114714015 |
| SE | 43871 | ENSG00000153898 | MCOLN2 | chr1 | - | 84971935 | 84972034 |
| SE | 12666 | ENSG00000196712 | NF1 | chr17 | + | 31357268 | 31357369 |
| SE | 8200 | ENSG00000131462 | TUBG1 | chr17 | + | 42612426 | 42612506 |
| SE | 31059 | ENSG00000065882 | TBC1D1 | chr4 | + | 38044361 | 38044490 |
| SE | 9249 | ENSG00000178971 | CTC1 | chr17 | - | 8238030 | 8238242 |
| SE | 56934 | ENSG00000118197 | DDX59 | chr1 | - | 200650424 | 200650676 |
| SE | 100996 | ENSG00000100600 | LGMN | chr14 | - | 92706482 | 92706653 |
| SE | 40153 | ENSG00000138382 | METTL5 | chr2 | - | 169815476 | 169815528 |
| SE | 57656 | ENSG00000144535 | DIS3L2 | chr2 | + | 232136471 | 232136719 |
| SE | 14021 | ENSG00000090905 | TNRC6A | chr16 | + | 24778343 | 24778468 |
| SE | 36866 | ENSG00000133422 | MORC2 | chr22 | - | 30932888 | 30933030 |
| SE | 71841 | ENSG00000183486 | MX2 | chr21 | + | 41390564 | 41390703 |
| SE | 19785 | ENSG00000126790 | L3HYPDH | chr14 | - | 59474438 | 59474526 |
| SE | 78326 | ENSG00000228409 | CCT6P1 | chr7 | + | 65759859 | 65759998 |
| SE | 81599 | ENSG00000253729 | PRKDC | chr8 | - | 47887546 | 47887705 |
| SE | 55254 | ENSG00000135905 | DOCK10 | chr2 | - | 224796315 | 224796426 |
| SE | 40878 | ENSG00000067829 | IDH3G | chrX | - | 153786800 | 153786947 |
| SE | 28680 | ENSG00000185033 | SEMA4B | chr15 | + | 90220981 | 90221093 |
| SE | 62754 | ENSG00000162961 | DPY30 | chr2 | - | 31891991 | 31892134 |
| SE | 77654 | ENSG00000214021 | TTLL3 | chr3 | + | 9817644 | 9820741 |
| SE | 16405 | ENSG00000156958 | GALK2 | chr15 | + | 49318110 | 49318192 |
| SE | 87384 | ENSG00000134982 | APC | chr5 | + | 112834950 | 112835165 |
| SE | 52636 | ENSG00000197943 | PLCG2 | chr16 | + | 81883268 | 81883341 |
| SE | 86270 | ENSG00000134516 | DOCK2 | chr5 | + | 170008496 | 170008597 |
| SE | 27161 | ENSG00000159202 | UBE2Z | chr17 | + | 48912833 | 48912965 |
| SE | 15877 | ENSG00000167491 | GATAD2A | chr19 | + | 19427695 | 19427992 |
| SE | 28496 | ENSG00000155229 | MMS19 | chr10 | - | 97480240 | 97480357 |
| SE | 103057 | ENSG00000099290 | WASHC2A | chr10 | + | 50127159 | 50127222 |
| SE | 30674 | ENSG00000132466 | ANKRD17 | chr4 | - | 73113791 | 73113908 |
| SE | 7683 | ENSG00000170004 | CHD3 | chr17 | + | 7893285 | 7893569 |
| SE | 38967 | ENSG00000116406 | EDEM3 | chr1 | - | 184723750 | 184723856 |
| SE | 37333 | ENSG00000004534 | RBM6 | chr3 | + | 50092042 | 50092208 |
| SE | 15938 | ENSG00000168904 | LRRC28 | chr15 | + | 99333922 | 99334129 |
| SE | 60592 | ENSG00000070961 | ATP2B1 | chr12 | - | 89616801 | 89617039 |
| SE | 50061 | ENSG00000102763 | VWA8 | chr13 | - | 41690165 | 41690275 |
| SE | 57137 | ENSG00000023902 | PLEKHO1 | chr1 | + | 150157384 | 150157486 |
| SE | 28576 | ENSG00000119414 | PPP6C | chr9 | - | 125158240 | 125158382 |
| SE | 870 | ENSG00000128159 | TUBGCP6 | chr22 | - | 50221274 | 50221874 |
| SE | 16755 | ENSG00000125107 | CNOT1 | chr16 | - | 58553781 | 58553860 |
| SE | 26697 | ENSG00000136709 | WDR33 | chr2 | - | 127709489 | 127709582 |
| SE | 23197 | ENSG00000165819 | METTL3 | chr14 | - | 21501727 | 21501824 |
| SE | 72253 | ENSG00000162885 | B3GALNT2 | chr1 | - | 235458602 | 235458786 |
| SE | 8673 | ENSG00000198909 | MAP3K3 | chr17 | + | 63634708 | 63634801 |
| SE | 84979 | ENSG00000170832 | USP32 | chr17 | - | 60211012 | 60211118 |
| SE | 76390 | ENSG00000047579 | DTNBP1 | chr6 | - | 15651312 | 15651363 |
| SE | 8613 | ENSG00000256525 | POLG2 | chr17 | - | 64481245 | 64481446 |
| SE | 87337 | ENSG00000215301 | DDX3X | chrX | + | 41346858 | 41347451 |
| SE | 107010 | ENSG00000002016 | RAD52 | chr12 | - | 926795 | 926946 |
| SE | 39792 | ENSG00000106459 | NRF1 | chr7 | + | 129710373 | 129710571 |
| SE | 85783 | ENSG00000148832 | PAOX | chr10 | + | 133383959 | 133384212 |
| SE | 58297 | ENSG00000117262 | GPR89A | chr1 | + | 145608347 | 145608675 |
| SE | 74342 | ENSG00000187118 | CMC1 | chr3 | + | 28292735 | 28292964 |
| SE | 29193 | ENSG00000196850 | PPTC7 | chr12 | - | 110538143 | 110538273 |
| SE | 39167 | ENSG00000165240 | ATP7A | chrX | + | 78011175 | 78011252 |
| SE | 6961 | ENSG00000105656 | ELL | chr19 | - | 18475813 | 18475933 |
| SE | 82496 | ENSG00000128563 | PRKRIP1 | chr7 | + | 102404597 | 102404683 |
| SE | 74560 | ENSG00000089818 | NECAP1 | chr12 | + | 8092675 | 8092784 |
| SE | 95405 | ENSG00000148356 | LRSAM1 | chr9 | + | 127451928 | 127452084 |
| SE | 91201 | ENSG00000116604 | MEF2D | chr1 | - | 156483238 | 156483418 |
| SE | 55802 | ENSG00000144218 | AFF3 | chr2 | - | 100011496 | 100011571 |
| SE | 10479 | ENSG00000186566 | GPATCH8 | chr17 | - | 44464471 | 44464544 |
| SE | 22091 | ENSG00000111859 | NEDD9 | chr6 | - | 11251563 | 11251726 |
| SE | 92659 | ENSG00000197724 | PHF2 | chr9 | + | 93675679 | 93675789 |
| SE | 91794 | ENSG00000133794 | ARNTL | chr11 | + | 13365499 | 13365592 |
| SE | 90398 | ENSG00000170633 | RNF34 | chr12 | + | 121417503 | 121417911 |
| SE | 27734 | ENSG00000152818 | UTRN | chr6 | + | 144447210 | 144447318 |
| SE | 18931 | ENSG00000100461 | RBM23 | chr14 | - | 22911327 | 22911403 |
| SE | 58035 | ENSG00000082701 | GSK3B | chr3 | - | 119876412 | 119876508 |
| SE | 61624 | ENSG00000234127 | TRIM26 | chr6 | - | 30201034 | 30201138 |
| SE | 29796 | ENSG00000141452 | RMC1 | chr18 | + | 23506969 | 23507054 |
| SE | 59250 | ENSG00000168118 | RAB4A | chr1 | + | 229288728 | 229288843 |
| SE | 59762 | ENSG00000163444 | TMEM183A | chr1 | + | 203007773 | 203007863 |
| SE | 22147 | ENSG00000156639 | ZFAND3 | chr6 | + | 37929958 | 37929999 |
| SE | 10421 | ENSG00000126581 | BECN1 | chr17 | - | 42811654 | 42811797 |
| SE | 36375 | ENSG00000155621 | C9orf85 | chr9 | + | 71918424 | 71918555 |
| SE | 72177 | ENSG00000059588 | TARBP1 | chr1 | - | 234430086 | 234430301 |
| SE | 103371 | ENSG00000176986 | SEC24C | chr10 | + | 73762097 | 73762166 |
| SE | 42213 | ENSG00000116266 | STXBP3 | chr1 | + | 108782421 | 108782517 |
| SE | 99586 | ENSG00000149187 | CELF1 | chr11 | - | 47477296 | 47477425 |
| SE | 67929 | ENSG00000091490 | SEL1L3 | chr4 | - | 25779075 | 25779203 |
| SE | 11163 | ENSG00000168488 | ATXN2L | chr16 | + | 28836325 | 28836490 |
| SE | 10105 | ENSG00000108433 | GOSR2 | chr17 | + | 46931098 | 46931207 |
| SE | 47502 | ENSG00000143363 | PRUNE1 | chr1 | + | 151027232 | 151027327 |
| SE | 78355 | ENSG00000072609 | CHFR | chr12 | - | 132862332 | 132862425 |
| SE | 76253 | ENSG00000211450 | SELENOH | chr11 | + | 57742116 | 57742227 |
| SE | 86165 | ENSG00000113845 | TIMMDC1 | chr3 | + | 119503953 | 119504021 |
| SE | 65116 | ENSG00000173960 | UBXN2A | chr2 | + | 23958300 | 23958355 |
| SE | 12039 | ENSG00000132603 | NIP7 | chr16 | + | 69341179 | 69341320 |
| SE | 17870 | ENSG00000092199 | HNRNPC | chr14 | - | 21268622 | 21268668 |
| SE | 40588 | ENSG00000085433 | WDR47 | chr1 | - | 108982608 | 108982779 |
| SE | 88323 | ENSG00000145868 | FBXO38 | chr5 | + | 148397618 | 148397676 |
| SE | 41026 | ENSG00000064687 | ABCA7 | chr19 | + | 1055082 | 1055351 |
| SE | 41753 | ENSG00000143222 | UFC1 | chr1 | + | 161158120 | 161158211 |
| SE | 24365 | ENSG00000132780 | NASP | chr1 | + | 45600384 | 45600457 |
| SE | 93948 | ENSG00000196116 | TDRD7 | chr9 | + | 97472292 | 97472495 |
| SE | 71856 | ENSG00000105639 | JAK3 | chr19 | - | 17839179 | 17839293 |

| | | | | | | | | | | | | | | | | | | | | | | | | | | | | | | | | | | | | | | | | | | | | | | |
|---|---|---|---|---|---|---|---|---|---|---|---|---|---|---|---|---|---|---|---|---|---|---|---|---|---|---|---|---|---|---|---|---|---|---|---|---|---|---|---|---|---|---|---|---|---|---|
| SE | 2748 | ENSG00000130816 | DNMT1 | chr19 | - | 10180349 | 10180569 | 10177291 | 10177367 | 10180777 | 10180836 | 275 | 212 | 235 | 308 | 651 | 0 | 0 | 1 | 0 | 0 | 521 | 233 | 162 | 411 | 527 | 11 | 18 | 0 | 0 | 3 | 298 | 149 | 1.44E-05 | 0.000440552 | 1 | 1 | 0.992 | 1 | 1 | 0.959 | 0.866 | 1 | 1 | 0.989 | 0.036 |
| SE | 75794 | ENSG00000137200 | CMTR1 | chr6 | + | 37461548 | 37461645 | 37459565 | 37459684 | 37471020 | 37471077 | 119 | 75 | 120 | 132 | 96 | 0 | 1 | 0 | 1 | 0 | 387 | 187 | 103 | 256 | 211 | 21 | 1 | 4 | 6 | 0 | 246 | 149 | 1.44E-05 | 0.000440552 | 1 | 0.978 | 1 | 0.988 | 1 | 0.918 | 0.991 | 0.94 | 0.963 | 1 | 0.031 |
| SE | 107048 | ENSG00000134287 | ARF3 | chr12 | - | 48956097 | 48956350 | 48941093 | 48941188 | 48957309 | 48957487 | 21 | 33 | 8 | 35 | 36 | 136 | 78 | 82 | 100 | 81 | 168 | 29 | 31 | 66 | 715 | 116 | 83 | 50 | 155 | 314 | 298 | 149 | 1.45E-05 | 0.000442966 | 0.072 | 0.175 | 0.047 | 0.149 | 0.182 | 0.42 | 0.149 | 0.237 | 0.176 | 0.532 | -0.178 |
| SE | 103661 | ENSG00000148634 | HERC4 | chr10 | - | 67939587 | 67939654 | 67936152 | 67936235 | 67940938 | 67941105 | 203 | 187 | 197 | 197 | 264 | 2 | 4 | 2 | 3 | 22 | 44 | 133 | 64 | 41 | 6 | 0 | 0 | 0 | 0 | 0 | 216 | 149 | 1.45E-05 | 0.000443564 | 0.986 | 0.97 | 0.985 | 0.978 | 0.892 | 1 | 1 | 1 | 1 | 1 | -0.038 |
| SE | 8765 | ENSG00000007202 | KIAA0100 | chr17 | - | 28643163 | 28643311 | 28642868 | 28642987 | 28643609 | 28643659 | 164 | 97 | 138 | 159 | 183 | 7 | 3 | 1 | 1 | 0 | 296 | 171 | 126 | 195 | 313 | 6 | 11 | 10 | 5 | 22 | 297 | 149 | 1.46E-05 | 0.000444579 | 0.922 | 0.942 | 0.986 | 0.988 | 1 | 0.961 | 0.886 | 0.863 | 0.951 | 0.877 | 0.06 |
| SE | 97997 | ENSG00000205758 | CRYZL1 | chr21 | - | 33597279 | 33597401 | 33595730 | 33595836 | 33599149 | 33599248 | 223 | 210 | 254 | 229 | 184 | 9 | 19 | 7 | 20 | 20 | 72 | 174 | 124 | 184 | 13 | 0 | 7 | 4 | 3 | 0 | 271 | 149 | 1.46E-05 | 0.00044562 | 0.932 | 0.859 | 0.952 | 0.863 | 0.835 | 1 | 0.932 | 0.945 | 0.971 | 1 | -0.081 |
| SE | 63576 | ENSG00000206562 | METTL6 | chr3 | - | 15414020 | 15414162 | 15409759 | 15411437 | 15415497 | 15415583 | 68 | 19 | 41 | 80 | 65 | 0 | 0 | 0 | 2 | 0 | 86 | 39 | 29 | 12 | 67 | 17 | 0 | 1 | 0 | 0 | 291 | 149 | 1.47E-05 | 0.000448635 | 1 | 1 | 1 | 0.953 | 1 | 0.721 | 1 | 0.937 | 1 | 1 | 0.059 |
| SE | 68618 | ENSG00000163902 | RPN1 | chr3 | - | 128629950 | 128630143 | 128626732 | 128626832 | 128631947 | 128632157 | 898 | 451 | 745 | 701 | 980 | 4 | 3 | 3 | 3 | 4 | 1113 | 773 | 678 | 553 | 1220 | 9 | 3 | 8 | 19 | 20 | 298 | 149 | 1.48E-05 | 0.000451861 | 0.991 | 0.987 | 0.992 | 0.992 | 0.992 | 0.984 | 0.992 | 0.977 | 0.936 | 0.968 | 0.019 |
| SE | 71614 | ENSG00000184182 | UBE2F | chr2 | + | 237987962 | 237987992 | 237973091 | 237973225 | 238025341 | 238025412 | 27 | 10 | 23 | 42 | 21 | 0 | 0 | 0 | 1 | 0 | 18 | 24 | 22 | 56 | 44 | 0 | 5 | 2 | 2 | 6 | 179 | 149 | 1.49E-05 | 0.000454857 | 1 | 1 | 1 | 0.972 | 1 | 1 | 0.8 | 0.902 | 0.959 | 0.859 | 0.09 |
| SE | 24519 | ENSG00000170144 | HNRNPA3 | chr2 | + | 177219407 | 177221976 | 177219246 | 177219314 | 177222059 | 177223919 | 227 | 231 | 220 | 251 | 257 | 58 | 49 | 64 | 56 | 38 | 101 | 228 | 268 | 196 | 30 | 14 | 15 | 49 | 10 | 0 | 298 | 149 | 1.50E-05 | 0.000455659 | 0.662 | 0.702 | 0.632 | 0.691 | 0.772 | 0.783 | 0.884 | 0.732 | 0.907 | 1 | -0.169 |
| SE | 85698 | ENSG00000175463 | TBC1D10C | chr11 | + | 67405902 | 67406017 | 67405594 | 67405701 | 67406626 | 67406661 | 275 | 68 | 228 | 185 | 81 | 2 | 0 | 3 | 0 | 0 | 36 | 195 | 125 | 89 | 221 | 0 | 9 | 0 | 3 | 11 | 264 | 149 | 1.50E-05 | 0.000457321 | 0.987 | 1 | 0.977 | 1 | 1 | 1 | 0.924 | 1 | 0.944 | 0.919 | 0.035 |
| SE | 69781 | ENSG00000065675 | PRKCQ | chr10 | - | 6441892 | 6442081 | 6430809 | 6430938 | 6456673 | 6456812 | 230 | 138 | 197 | 200 | 343 | 3 | 0 | 0 | 6 | 1 | 96 | 171 | 185 | 99 | 63 | 0 | 1 | 11 | 13 | 1 | 298 | 149 | 1.51E-05 | 0.000460308 | 0.975 | 1 | 1 | 0.943 | 0.994 | 1 | 0.988 | 0.894 | 0.792 | 0.969 | 0.054 |
| SE | 14930 | ENSG00000159216 | RUNX1 | chr21 | - | 34834409 | 34834601 | 34799300 | 34799462 | 34859473 | 34859578 | 180 | 139 | 192 | 153 | 393 | 24 | 8 | 30 | 10 | 24 | 432 | 306 | 207 | 370 | 306 | 11 | 7 | 13 | 24 | 11 | 298 | 149 | 1.52E-05 | 0.000460594 | 0.789 | 0.897 | 0.762 | 0.884 | 0.891 | 0.952 | 0.956 | 0.888 | 0.885 | 0.933 | -0.078 |
| SE | 95929 | ENSG00000166199 | ALKBH3 | chr11 | + | 43892040 | 43892129 | 43889724 | 43889828 | 43901515 | 43901714 | 148 | 63 | 120 | 98 | 164 | 5 | 1 | 5 | 12 | 7 | 249 | 130 | 65 | 110 | 139 | 3 | 0 | 1 | 5 | 0 | 238 | 149 | 1.52E-05 | 0.000460594 | 0.949 | 0.975 | 0.938 | 0.836 | 0.936 | 0.981 | 1 | 0.976 | 0.932 | 1 | -0.051 |
| SE | 92197 | ENSG00000099810 | MTAP | chr9 | + | 21861766 | 21861817 | 21859322 | 21859425 | 21861975 | 21862220 | 2 | 0 | 0 | 1 | 0 | 53 | 59 | 62 | 45 | 34 | 9 | 4 | 2 | 2 | 0 | 23 | 26 | 30 | 24 | 43 | 200 | 149 | 1.52E-05 | 0.000460594 | 0.027 | 0 | 0 | 0.016 | 0 | 0.226 | 0.103 | 0.047 | 0.058 | 0 | -0.078 |
| SE | 82323 | ENSG00000146833 | TRIM4 | chr7 | - | 99908581 | 99908812 | 99903575 | 99903598 | 99909564 | 99909660 | 257 | 192 | 230 | 236 | 367 | 2 | 1 | 4 | 1 | 2 | 141 | 221 | 121 | 108 | 146 | 0 | 5 | 8 | 6 | 6 | 298 | 149 | 1.52E-05 | 0.000460594 | 0.985 | 0.99 | 0.966 | 0.992 | 0.989 | 1 | 0.957 | 0.883 | 0.9 | 0.924 | 0.052 |
| SE | 9752 | ENSG00000062716 | VMP1 | chr17 | + | 59738836 | 59738947 | 59735337 | 59735473 | 59764970 | 59765068 | 476 | 300 | 319 | 362 | 960 | 10 | 4 | 4 | 5 | 10 | 392 | 320 | 374 | 354 | 202 | 5 | 8 | 24 | 23 | 0 | 260 | 149 | 1.52E-05 | 0.000460594 | 0.965 | 0.977 | 0.979 | 0.976 | 0.982 | 0.978 | 0.958 | 0.899 | 0.898 | 1 | 0.029 |
| SE | 8526 | ENSG00000125450 | NUP85 | chr17 | + | 75215753 | 75215823 | 75213075 | 75213119 | 75225102 | 75225237 | 17 | 14 | 21 | 17 | 28 | 2 | 0 | 0 | 1 | 0 | 15 | 36 | 19 | 32 | 36 | 12 | 1 | 4 | 4 | 0 | 219 | 149 | 1.52E-05 | 0.000460812 | 0.853 | 1 | 1 | 0.92 | 1 | 0.46 | 0.961 | 0.764 | 0.845 | 1 | 0.149 |
| SE | 34207 | ENSG00000102349 | KLF8 | chrX | + | 56270181 | 56270321 | 56265179 | 56269489 | 56284312 | 56285097 | 27 | 6 | 11 | 31 | 18 | 2 | 0 | 2 | 0 | 1 | 0 | 20 | 14 | 14 | 14 | 4 | 7 | 6 | 6 | 0 | 289 | 149 | 1.52E-05 | 0.000461328 | 0.874 | 1 | 0.739 | 1 | 0.903 | 0 | 0.596 | 0.546 | 0.546 | 1 | 0.366 |
| SE | 47643 | ENSG00000288701 | PRRC2B | chr9 | + | 131474453 | 131476535 | 131473507 | 131473724 | 131477743 | 131477949 | 339 | 176 | 263 | 344 | 247 | 13 | 14 | 17 | 22 | 1 | 331 | 290 | 188 | 220 | 340 | 10 | 42 | 22 | 22 | 58 | 298 | 149 | 1.52E-05 | 0.000461482 | 0.929 | 0.863 | 0.886 | 0.887 | 0.992 | 0.943 | 0.775 | 0.81 | 0.833 | 0.746 | 0.09 |
| SE | 61272 | ENSG00000168175 | MAPK1IP1L | chr14 | + | 55061679 | 55061701 | 55051646 | 55051803 | 55062617 | 55063325 | 102 | 55 | 102 | 117 | 63 | 2 | 0 | 2 | 3 | 3 | 198 | 164 | 174 | 179 | 136 | 20 | 4 | 12 | 16 | 6 | 171 | 149 | 1.53E-05 | 0.000464647 | 0.978 | 1 | 0.978 | 0.971 | 0.948 | 0.896 | 0.973 | 0.927 | 0.907 | 0.952 | 0.044 |
| SE | 78804 | ENSG00000135269 | TES | chr7 | + | 116251759 | 116251975 | 116250160 | 116250496 | 116252317 | 116252476 | 743 | 352 | 585 | 564 | 685 | 11 | 19 | 6 | 15 | 16 | 551 | 400 | 296 | 310 | 227 | 13 | 12 | 48 | 24 | 5 | 298 | 149 | 1.54E-05 | 0.000465285 | 0.971 | 0.903 | 0.98 | 0.949 | 0.955 | 0.955 | 0.943 | 0.755 | 0.866 | 0.958 | 0.056 |
| SE | 87005 | ENSG00000258890 | CEP95 | chr17 | + | 64527110 | 64527264 | 64526070 | 64526200 | 64529287 | 64529427 | 139 | 166 | 169 | 158 | 245 | 2 | 3 | 0 | 0 | 1 | 154 | 162 | 278 | 128 | 66 | 7 | 8 | 3 | 7 | 0 | 298 | 149 | 1.54E-05 | 0.000467093 | 0.972 | 0.965 | 1 | 1 | 0.992 | 0.917 | 0.91 | 0.979 | 0.901 | 1 | 0.044 |
| SE | 76254 | ENSG00000211450 | SELENOH | chr11 | + | 57742116 | 57742247 | 57741698 | 57741954 | 57742862 | 57743411 | 222 | 107 | 191 | 209 | 103 | 1 | 0 | 0 | 4 | 0 | 279 | 188 | 110 | 155 | 330 | 7 | 20 | 2 | 1 | 0 | 280 | 149 | 1.55E-05 | 0.00046944 | 0.992 | 1 | 1 | 0.965 | 1 | 0.955 | 0.833 | 0.967 | 0.988 | 1 | 0.043 |
| SE | 20409 | ENSG00000204623 | ZNRD1ASP | chr6 | - | 30060740 | 30060928 | 30058114 | 30058190 | 30061078 | 30061184 | 22 | 8 | 19 | 25 | 11 | 2 | 0 | 5 | 0 | 4 | 7 | 1 | 5 | 8 | 20 | 0 | 0 | 0 | 0 | 0 | 298 | 149 | 1.56E-05 | 0.000472057 | 0.846 | 1 | 0.655 | 1 | 0.579 | 1 | 1 | 1 | 1 | 1 | -0.184 |
| SE | 69324 | ENSG00000112367 | FIG4 | chr6 | + | 109788797 | 109788956 | 109786301 | 109786449 | 109789593 | 109789677 | 0 | 0 | 2 | 0 | 4 | 74 | 62 | 41 | 65 | 151 | 20 | 3 | 2 | 0 | 0 | 76 | 49 | 33 | 69 | 17 | 298 | 149 | 1.57E-05 | 0.000475807 | 0 | 0 | 0.024 | 0 | 0.013 | 0.116 | 0.03 | 0.029 | 0 | 0 | -0.028 |
| SE | 62603 | ENSG00000143622 | RIT1 | chr1 | - | 155904730 | 155904804 | 155897807 | 155900618 | 155910449 | 155910506 | 44 | 40 | 28 | 60 | 20 | 0 | 1 | 0 | 0 | 0 | 45 | 35 | 46 | 28 | 46 | 14 | 3 | 0 | 0 | 0 | 223 | 149 | 1.58E-05 | 0.000476692 | 1 | 0.964 | 1 | 1 | 1 | 0.682 | 0.886 | 1 | 1 | 1 | 0.079 |
| SE | 93066 | ENSG00000163820 | FYCO1 | chr3 | - | 45975238 | 45975345 | 45969674 | 45969765 | 45979704 | 45979830 | 59 | 34 | 69 | 70 | 90 | 0 | 0 | 0 | 0 | 0 | 58 | 62 | 71 | 35 | 93 | 30 | 0 | 0 | 3 | 0 | 256 | 149 | 1.58E-05 | 0.000477824 | 1 | 1 | 1 | 1 | 1 | 0.529 | 1 | 1 | 0.872 | 1 | 0.12 |
| SE | 52171 | ENSG00000142599 | RERE | chr1 | - | 8359763 | 8359986 | 8358195 | 8358916 | 8360111 | 8361490 | 76 | 50 | 87 | 109 | 55 | 4 | 2 | 0 | 2 | 0 | 116 | 144 | 35 | 108 | 237 | 15 | 1 | 5 | 10 | 15 | 298 | 149 | 1.58E-05 | 0.000478012 | 0.905 | 0.926 | 1 | 0.965 | 1 | 0.795 | 0.986 | 0.778 | 0.844 | 0.888 | 0.101 |
| SE | 75009 | ENSG00000168000 | BSCL2 | chr11 | - | 62692375 | 62692797 | 62691279 | 62691421 | 62694567 | 62694711 | 40 | 30 | 24 | 42 | 37 | 0 | 0 | 0 | 0 | 0 | 101 | 45 | 14 | 10 | 169 | 10 | 0 | 5 | 2 | 5 | 298 | 149 | 1.58E-05 | 0.000478012 | 1 | 1 | 1 | 1 | 1 | 0.835 | 1 | 1 | 0.714 | 0.944 | 0.101 |
| SE | 28505 | ENSG00000169062 | UPF3A | chr13 | + | 114291488 | 114291544 | 114286373 | 114286629 | 114291633 | 114291718 | 90 | 69 | 97 | 94 | 123 | 1 | 1 | 0 | 1 | 3 | 168 | 47 | 83 | 97 | 88 | 13 | 5 | 5 | 3 | 0 | 205 | 149 | 1.58E-05 | 0.000478265 | 0.985 | 0.98 | 1 | 0.986 | 0.968 | 0.904 | 0.872 | 0.923 | 0.959 | 1 | 0.052 |
| SE | 97489 | ENSG00000173120 | KDM2A | chr11 | + | 67131951 | 67132073 | 67121268 | 67121358 | 67149054 | 67149207 | 4 | 14 | 7 | 10 | 19 | 4 | 0 | 0 | 3 | 5 | 19 | 15 | 2 | 9 | 3 | 18 | 7 | 9 | 18 | 29 | 271 | 149 | 1.59E-05 | 0.000478538 | 0.355 | 1 | 1 | 0.647 | 0.676 | 0.367 | 0.541 | 0.109 | 0.216 | 0.054 | 0.478 |
| SE | 54956 | ENSG00000088448 | ANKRD10 | chr13 | - | 110900529 | 110900694 | 110893196 | 110893263 | 110906032 | 110906124 | 4 | 2 | 5 | 7 | 12 | 115 | 63 | 86 | 105 | 79 | 19 | 21 | 5 | 1 | 48 | 112 | 115 | 72 | 71 | 184 | 298 | 149 | 1.60E-05 | 0.000481976 | 0.017 | 0.016 | 0.028 | 0.032 | 0.071 | 0.078 | 0.084 | 0.034 | 0.007 | 0.115 | -0.031 |
| SE | 13017 | ENSG00000140987 | ZSCAN32 | chr16 | - | 3390009 | 3390183 | 3384083 | 3384941 | 3393648 | 3393814 | 16 | 6 | 15 | 12 | 20 | 1 | 1 | 1 | 3 | 4 | 1 | 3 | 3 | 2 | 2 | 0 | 0 | 0 | 0 | 0 | 298 | 149 | 1.61E-05 | 0.000485105 | 0.889 | 0.75 | 0.882 | 0.667 | 0.714 | 1 | 1 | 1 | 1 | 1 | -0.22 |
| SE | 9725 | ENSG00000171302 | CANT1 | chr17 | - | 78995017 | 78995221 | 78991715 | 78993920 | 78996991 | 78997644 | 142 | 52 | 115 | 122 | 142 | 1 | 0 | 0 | 0 | 0 | 170 | 108 | 122 | 84 | 147 | 0 | 0 | 1 | 4 | 15 | 298 | 149 | 1.61E-05 | 0.000485451 | 0.986 | 1 | 1 | 1 | 1 | 1 | 1 | 0.984 | 0.913 | 0.831 | 0.052 |
| SE | 66450 | ENSG00000131051 | RBM39 | chr20 | - | 35710282 | 35710420 | 35709223 | 35709256 | 35713018 | 35713096 | 22 | 15 | 19 | 10 | 19 | 411 | 230 | 339 | 448 | 459 | 0 | 2 | 7 | 2 | 0 | 36 | 188 | 400 | 175 | 34 | 287 | 149 | 1.61E-05 | 0.000485534 | 0.027 | 0.033 | 0.028 | 0.011 | 0.021 | 0 | 0.005 | 0.009 | 0.006 | 0 | 0.02 |
| SE | 107423 | ENSG00000166037 | CEP57 | chr11 | + | 95795515 | 95795559 | 95790512 | 95790743 | 95799231 | 95799388 | 8 | 4 | 10 | 11 | 26 | 83 | 56 | 76 | 60 | 40 | 0 | 6 | 2 | 0 | 0 | 15 | 52 | 38 | 16 | 29 | 193 | 149 | 1.61E-05 | 0.000486294 | 0.069 | 0.052 | 0.092 | 0.124 | 0.334 | 0 | 0.082 | 0.039 | 0 | 0 | 0.11 |
| SE | 109309 | ENSG00000139597 | N4BP2L1 | chr13 | - | 32407249 | 32407338 | 32404320 | 32404397 | 32407644 | 32407772 | 125 | 94 | 73 | 88 | 95 | 3 | 3 | 6 | 11 | 0 | 126 | 80 | 42 | 65 | 88 | 0 | 0 | 4 | 0 | 0 | 238 | 149 | 1.62E-05 | 0.000487156 | 0.963 | 0.951 | 0.884 | 0.834 | 1 | 1 | 1 | 0.868 | 1 | 1 | -0.047 |
| SE | 25343 | ENSG00000130803 | ZNF317 | chr19 | + | 9157979 | 9158075 | 9156611 | 9157394 | 9158825 | 9158908 | 92 | 70 | 69 | 67 | 80 | 10 | 10 | 21 | 22 | 10 | 107 | 117 | 134 | 86 | 113 | 109 | 66 | 37 | 59 | 41 | 245 | 149 | 1.62E-05 | 0.000487343 | 0.848 | 0.81 | 0.666 | 0.649 | 0.83 | 0.374 | 0.519 | 0.688 | 0.47 | 0.626 | 0.225 |

[illegible]

| | | | | | | | | | | | | | | | |
|---|---|---|---|---|---|---|---|---|---|---|---|---|---|---|---|
| SE | 80946 | ENSG00000100997 | ABHD12 | chr20 | - | 25338884 | 25339004 | 25323324 | 25323430 | 25339226 | 25339351 | 7 | 8 | 10 | [illegible] |
| SE | 13330 | ENSG00000103248 | MTHFSD | chr16 | - | 86548034 | 86548111 | 86546558 | 86546649 | 86548463 | 86548577 | 9 | 8 | 5 | [illegible] |
| SE | 64952 | ENSG00000163644 | PPM1K | chr4 | - | 88278143 | 88278642 | 88275144 | 88277243 | 88284405 | 88284561 | 83 | 56 | 84 | [illegible] |
| SE | 74887 | ENSG00000187037 | GPR141 | chr7 | + | 37685178 | 37685298 | 37683796 | 37683903 | 37740379 | 37740705 | 33 | 12 | 8 | [illegible] |
| SE | 90679 | ENSG00000182872 | RBM10 | chrX | + | 47179683 | 47179781 | 47179314 | 47179495 | 47179879 | 47180037 | 0 | 0 | 0 | [illegible] |
| SE | 46189 | ENSG00000185158 | LRRC37B | chr17 | + | 32017971 | 32018054 | 32007987 | 32008132 | 32020696 | 32020803 | 24 | 11 | 31 | [illegible] |
| SE | 30229 | ENSG00000132376 | INPP5K | chr17 | - | 1515426 | 1515585 | 1513871 | 1513979 | 1516455 | 1516661 | 2 | 1 | 0 | [illegible] |
| SE | 79935 | ENSG00000006530 | AGK | chr7 | + | 141615470 | 141615569 | 141614145 | 141614178 | 141621731 | 141621801 | 58 | 36 | 45 | [illegible] |
| SE | 52648 | ENSG00000197943 | PLCG2 | chr16 | + | 81934428 | 81934531 | 81928557 | 81928624 | 81936168 | 81936210 | 137 | 65 | 81 | [illegible] |
| SE | 46656 | ENSG00000134222 | PSRC1 | chr1 | - | 109280395 | 109280489 | 109279555 | 109280156 | 109280776 | 109280916 | 16 | 6 | 9 | [illegible] |
| SE | 10329 | ENSG00000168961 | LGALS9 | chr17 | + | 27643524 | 27643620 | 27642237 | 27642348 | 27645313 | 27645349 | 49 | 22 | 26 | [illegible] |
| SE | 76830 | ENSG00000255339 | NA | chr10 | - | 100526398 | 100526554 | 100516873 | 100516960 | 100526974 | 100527074 | 199 | 130 | 267 | [illegible] |
| SE | 4897 | ENSG00000124243 | BCAS4 | chr20 | + | 50841765 | 50841900 | 50830278 | 50830380 | 50876485 | 50877177 | 40 | 19 | 54 | [illegible] |
| SE | 77329 | ENSG00000214022 | REPIN1 | chr7 | + | 150369741 | 150369868 | 150368808 | 150368941 | 150371230 | 150372162 | 3 | 1 | 6 | [illegible] |
| SE | 98596 | ENSG00000149269 | PAK1 | chr11 | - | 77408153 | 77408232 | 77392330 | 77392541 | 77411773 | 77411933 | 2 | 0 | 0 | [illegible] |
| SE | 89866 | ENSG00000137492 | THAP12 | chr11 | - | 76364337 | 76364432 | 76360955 | 76361063 | 76365851 | 76365972 | 95 | 117 | 145 | [illegible] |
| SE | 56215 | ENSG00000106462 | EZH2 | chr7 | - | 148816683 | 148817391 | 148813958 | 148815546 | 148817876 | 148818117 | 15 | 9 | 37 | [illegible] |
| SE | 57993 | ENSG00000175634 | RPS6KB2 | chr11 | + | 67432355 | 67432492 | 67431367 | 67431515 | 67432599 | 67432657 | 11 | 3 | 16 | [illegible] |
| SE | 56216 | ENSG00000106462 | EZH2 | chr7 | - | 148817221 | 148817391 | 148815505 | 148815546 | 148817876 | 148818117 | 8 | 5 | 19 | [illegible] |
| SE | 38311 | ENSG00000138172 | CALHM2 | chr10 | - | 103451082 | 103451285 | 103450295 | 103450800 | 103452155 | 103452370 | 38 | 24 | 21 | [illegible] |
| SE | 1075 | ENSG00000100330 | MTMR3 | chr22 | + | 30007113 | 30007319 | 30002879 | 30002993 | 30009017 | 30009129 | 104 | 42 | 60 | [illegible] |
| SE | 35762 | ENSG00000162736 | NCSTN | chr1 | + | 160357040 | 160357253 | 160356599 | 160356754 | 160358148 | 160358949 | 322 | 208 | 260 | [illegible] |
| SE | 58265 | ENSG00000065413 | ANKRD44 | chr2 | - | 196998906 | 196999052 | 196998336 | 196998419 | 197000418 | 197000502 | 397 | 299 | 347 | [illegible] |
| SE | 212 | ENSG00000160208 | RRP1B | chr21 | + | 43687515 | 43688240 | 43686131 | 43686935 | 43690287 | 43690440 | 217 | 153 | 203 | [illegible] |
| SE | 81139 | ENSG00000198556 | ZNF789 | chr7 | + | 99478273 | 99478356 | 99476402 | 99476480 | 99479660 | 99479787 | 4 | 4 | 3 | [illegible] |
| SE | 102622 | ENSG00000135972 | MRPS9 | chr2 | + | 105079982 | 105080062 | 105071458 | 105071489 | 105088983 | 105089069 | 234 | 193 | 175 | [illegible] |
| SE | 83319 | ENSG00000161010 | MRNIP | chr5 | - | 179847977 | 179848066 | 179844151 | 179844227 | 179853196 | 179853276 | 30 | 28 | 26 | [illegible] |
| SE | 8981 | ENSG00000161526 | SAP30BP | chr17 | + | 75690670 | 75690772 | 75671815 | 75671863 | 75693439 | 75693482 | 4 | 3 | 4 | [illegible] |
| SE | 5002 | ENSG00000101224 | CDC25B | chr20 | + | 3802729 | 3802808 | 3802280 | 3802376 | 3802909 | 3802972 | 11 | 27 | 34 | [illegible] |
| SE | 7298 | ENSG00000154889 | MPPE1 | chr18 | - | 11886498 | 11886621 | 11885675 | 11885816 | 11886916 | 11887025 | 43 | 18 | 23 | [illegible] |
| SE | 84375 | ENSG00000230590 | FTX | chrX | - | 74284742 | 74284948 | 74281701 | 74281848 | 74293351 | 74293543 | 3 | 5 | 2 | [illegible] |
| SE | 102182 | ENSG00000267699 | NA | chr18 | + | 51051353 | 51051415 | 51049294 | 51049324 | 51054780 | 51054993 | 12 | 7 | 16 | [illegible] |
| [illegible] | | | | | | | | | | | | | | | |

| | | | | | | | | | | | | | | | | | | | | | | | | | | | | | | | | | | | | | | | | | | | | | | |
|---|---|---|---|---|---|---|---|---|---|---|---|---|---|---|---|---|---|---|---|---|---|---|---|---|---|---|---|---|---|---|---|---|---|---|---|---|---|---|---|---|---|---|---|---|---|---|
| SE | 61881 | ENSG00000076242 | MLH1 | chr3 | + | 37014431 | 37014544 | 37012010 | 37012099 | 37020309 | 37020463 | 85 | 81 | 91 | 59 | 153 | 14 | 7 | 18 | 9 | 13 | 97 | 45 | 65 | 78 | 43 | 13 | 30 | 30 | 40 | 11 | 262 | 149 | 4.47E-05 | 0.001154741 | 0.775 | 0.868 | 0.742 | 0.789 | 0.87 | 0.809 | 0.46 | 0.552 | 0.526 | 0.69 | 0.201 |
| SE | 21935 | ENSG00000153044 | CENPH | chr5 | + | 69197052 | 69197109 | 69195716 | 69195791 | 69202505 | 69202569 | 41 | 52 | 51 | 39 | 143 | 4 | 0 | 0 | 0 | 3 | 107 | 34 | 49 | 51 | 53 | 22 | 0 | 0 | 9 | 0 | 206 | 149 | 4.47E-05 | 0.00115629 | 0.881 | 1 | 1 | 1 | 0.972 | 0.779 | 1 | 1 | 0.804 | 1 | 0.054 |
| SE | 1525 | ENSG00000100242 | SUN2 | chr22 | - | 38750897 | 38751035 | 38750224 | 38750320 | 38751209 | 38751373 | 742 | 309 | 646 | 609 | 235 | 1 | 0 | 10 | 4 | 0 | 262 | 530 | 236 | 230 | 534 | 0 | 7 | 5 | 8 | 15 | 287 | 149 | 4.48E-05 | 0.00115629 | 0.997 | 1 | 0.971 | 0.988 | 1 | 1 | 0.975 | 0.961 | 0.937 | 0.949 | 0.027 |
| SE | 80765 | ENSG00000136280 | CCM2 | chr7 | + | 45064462 | 45064646 | 45038252 | 45038426 | 45068442 | 45068579 | 23 | 9 | 27 | 24 | 24 | 0 | 0 | 3 | 2 | 6 | 44 | 23 | 6 | 35 | 30 | 0 | 0 | 0 | 0 | 0 | 298 | 149 | 4.48E-05 | 0.00115732 | 1 | 1 | 0.818 | 0.857 | 0.667 | 1 | 1 | 1 | 1 | 1 | -0.132 |
| SE | 57031 | ENSG00000268043 | NBPF12 | chr1 | + | 146959858 | 146960006 | 146951347 | 146951489 | 146960108 | 146960318 | 3 | 7 | 17 | 8 | 5 | 0 | 0 | 0 | 0 | 0 | 12 | 14 | 10 | 21 | 10 | 0 | 0 | 0 | 5 | 7 | 297 | 149 | 4.48E-05 | 0.00115732 | 1 | 1 | 1 | 1 | 1 | 1 | 1 | 1 | 0.678 | 0.417 | 0.181 |
| SE | 62157 | ENSG00000117625 | RCOR3 | chr1 | + | 211291523 | 211291599 | 211289177 | 211289396 | 211295675 | 211295753 | 15 | 12 | 12 | 10 | 7 | 116 | 119 | 132 | 115 | 199 | 0 | 5 | 6 | 6 | 0 | 105 | 117 | 96 | 122 | 111 | 225 | 149 | 4.49E-05 | 0.001159815 | 0.079 | 0.063 | 0.057 | 0.054 | 0.023 | 0 | 0.028 | 0.04 | 0.032 | 0 | 0.035 |
| SE | 109398 | ENSG00000134900 | TPP2 | chr13 | + | 102629481 | 102629609 | 102627847 | 102627924 | 102630095 | 102630195 | 530 | 393 | 535 | 526 | 657 | 3 | 0 | 2 | 6 | 0 | 117 | 384 | 342 | 213 | 70 | 0 | 15 | 8 | 2 | 0 | 277 | 149 | 4.50E-05 | 0.001160629 | 0.99 | 1 | 0.993 | 0.979 | 1 | 1 | 0.932 | 0.958 | 0.983 | 1 | 0.018 |
| SE | 42773 | ENSG00000137462 | TLR2 | chr4 | + | 153687901 | 153688047 | 153684260 | 153684360 | 153702891 | 153706260 | 69 | 38 | 23 | 45 | 50 | 2 | 1 | 1 | 2 | 0 | 21 | 43 | 27 | 20 | 4 | 0 | 26 | 6 | 2 | 0 | 295 | 149 | 4.51E-05 | 0.001163104 | 0.946 | 0.95 | 0.921 | 0.919 | 1 | 1 | 0.455 | 0.694 | 0.835 | 1 | 0.15 |
| SE | 23785 | ENSG00000113716 | HMGXB3 | chr5 | + | 150037399 | 150037527 | 150032453 | 150032603 | 150040747 | 150040879 | 39 | 23 | 41 | 41 | 33 | 3 | 1 | 1 | 1 | 2 | 82 | 57 | 68 | 61 | 44 | 19 | 0 | 10 | 10 | 15 | 277 | 149 | 4.54E-05 | 0.001170127 | 0.875 | 0.925 | 0.957 | 0.957 | 0.899 | 0.699 | 1 | 0.785 | 0.766 | 0.612 | 0.15 |
| SE | 82837 | ENSG00000162129 | CLPB | chr11 | - | 72354307 | 72354389 | 72329784 | 72329804 | 72358879 | 72359008 | 3 | 0 | 1 | 0 | 0 | 35 | 14 | 30 | 32 | 19 | 33 | 5 | 0 | 1 | 3 | 78 | 45 | 40 | 88 | 102 | 231 | 149 | 4.56E-05 | 0.001175466 | 0.052 | 0 | 0.021 | 0 | 0 | 0.214 | 0.067 | 0 | 0.007 | 0.019 | -0.047 |
| SE | 48904 | ENSG00000158773 | USF1 | chr1 | - | 161041833 | 161041846 | 161041323 | 161041411 | 161042115 | 161042217 | 12 | 6 | 8 | 5 | 5 | 4 | 1 | 6 | 9 | 4 | 37 | 9 | 5 | 5 | 6 | 72 | 36 | 15 | 10 | 58 | 162 | 149 | 4.58E-05 | 0.001178501 | 0.734 | 0.847 | 0.551 | 0.338 | 0.535 | 0.321 | 0.187 | 0.235 | 0.315 | 0.087 | 0.372 |
| SE | 54515 | ENSG00000197111 | PCBP2 | chr12 | + | 53464759 | 53464852 | 53462480 | 53462567 | 53467220 | 53467285 | 645 | 346 | 579 | 523 | 1111 | 41 | 24 | 61 | 44 | 64 | 547 | 484 | 393 | 465 | 999 | 73 | 90 | 64 | 62 | 50 | 242 | 149 | 4.58E-05 | 0.001178501 | 0.906 | 0.899 | 0.854 | 0.88 | 0.914 | 0.822 | 0.768 | 0.791 | 0.822 | 0.925 | 0.065 |
| SE | 59889 | ENSG00000175054 | ATR | chr3 | - | 142540903 | 142541034 | 142537291 | 142538625 | 142542664 | 142542757 | 196 | 145 | 184 | 248 | 309 | 11 | 4 | 4 | 2 | 7 | 4 | 92 | 52 | 36 | 15 | 0 | 0 | 0 | 0 | 0 | 280 | 149 | 4.58E-05 | 0.001178685 | 0.905 | 0.951 | 0.961 | 0.985 | 0.959 | 1 | 1 | 1 | 1 | 1 | -0.048 |
| SE | 107451 | ENSG00000215039 | CD27-AS1 | chr12 | - | 6448639 | 6448737 | 6446960 | 6448150 | 6450340 | 6450980 | 18 | 13 | 8 | 14 | 5 | 0 | 0 | 1 | 0 | 0 | 11 | 11 | 9 | 13 | 74 | 12 | 0 | 0 | 9 | 0 | 247 | 149 | 4.58E-05 | 0.001179177 | 1 | 1 | 0.828 | 1 | 1 | 0.356 | 1 | 1 | 0.466 | 1 | 0.201 |
| SE | 1556 | ENSG00000198089 | SFI1 | chr22 | + | 31521234 | 31521320 | 31508254 | 31508376 | 31528689 | 31528866 | 1 | 0 | 5 | 5 | 0 | 104 | 96 | 101 | 90 | 100 | 0 | 4 | 3 | 9 | 22 | 52 | 75 | 86 | 97 | 213 | 235 | 149 | 4.60E-05 | 0.001184646 | 0.006 | 0 | 0.03 | 0.034 | 0 | 0 | 0.033 | 0.022 | 0.056 | 0.061 | -0.02 |
| SE | 85590 | ENSG00000104870 | FCGRT | chr19 | + | 49521694 | 49521813 | 49513881 | 49514486 | 49524506 | 49524568 | 4 | 1 | 2 | 5 | 0 | 109 | 48 | 56 | 92 | 81 | 20 | 2 | 1 | 0 | 17 | 74 | 57 | 39 | 67 | 156 | 268 | 149 | 4.63E-05 | 0.001190051 | 0.02 | 0.011 | 0.019 | 0.029 | 0 | 0.131 | 0.019 | 0.014 | 0 | 0.057 | -0.028 |
| SE | 49393 | ENSG00000243943 | ZNF512 | chr2 | + | 27599969 | 27600053 | 27599582 | 27599678 | 27601355 | 27601442 | 110 | 81 | 175 | 125 | 214 | 2 | 8 | 2 | 5 | 2 | 127 | 130 | 82 | 81 | 42 | 0 | 8 | 9 | 12 | 12 | 233 | 149 | 4.63E-05 | 0.001190051 | 0.972 | 0.866 | 0.982 | 0.941 | 0.986 | 1 | 0.912 | 0.854 | 0.812 | 0.691 | 0.096 |
| SE | 48780 | ENSG00000153922 | CHD1 | chr5 | - | 98863407 | 98863586 | 98859971 | 98860068 | 98868494 | 98868629 | 839 | 745 | 658 | 850 | 1438 | 18 | 14 | 17 | 13 | 40 | 51 | 272 | 444 | 51 | 64 | 0 | 1 | 7 | 0 | 0 | 298 | 149 | 4.64E-05 | 0.001192744 | 0.959 | 0.964 | 0.951 | 0.97 | 0.947 | 1 | 0.993 | 0.969 | 1 | 1 | -0.034 |
| SE | 98647 | ENSG00000107036 | RIC1 | chr9 | + | 5738449 | 5738538 | 5732387 | 5732479 | 5742868 | 5743013 | 169 | 137 | 140 | 193 | 228 | 13 | 6 | 4 | 7 | 4 | 127 | 131 | 131 | 123 | 65 | 15 | 19 | 15 | 4 | 5 | 238 | 149 | 4.64E-05 | 0.001193029 | 0.891 | 0.935 | 0.956 | 0.945 | 0.973 | 0.841 | 0.812 | 0.845 | 0.951 | 0.891 | 0.072 |
| SE | 82187 | ENSG00000146872 | TLK2 | chr17 | + | 62597219 | 62597258 | 62596584 | 62596674 | 62600650 | 62600767 | 11 | 12 | 16 | 9 | 29 | 202 | 134 | 128 | 156 | 225 | 25 | 22 | 21 | 50 | 2 | 189 | 94 | 167 | 129 | 27 | 188 | 149 | 4.65E-05 | 0.001193029 | 0.041 | 0.066 | 0.09 | 0.044 | 0.093 | 0.095 | 0.156 | 0.091 | 0.235 | 0.055 | -0.06 |
| SE | 49792 | ENSG00000116337 | AMPD2 | chr1 | + | 109629795 | 109629916 | 109629326 | 109629490 | 109630232 | 109630406 | 109 | 34 | 62 | 60 | 49 | 5 | 2 | 2 | 4 | 2 | 138 | 47 | 44 | 79 | 123 | 5 | 5 | 8 | 33 | 21 | 270 | 149 | 4.65E-05 | 0.001193029 | 0.923 | 0.904 | 0.945 | 0.892 | 0.931 | 0.938 | 0.838 | 0.752 | 0.569 | 0.764 | 0.147 |
| SE | 58665 | ENSG00000125633 | CCDC93 | chr2 | - | 118000820 | 118000932 | 117996263 | 117996362 | 118006721 | 118006816 | 156 | 144 | 139 | 181 | 200 | 0 | 1 | 2 | 8 | 0 | 147 | 161 | 256 | 188 | 70 | 0 | 5 | 9 | 16 | 6 | 261 | 149 | 4.69E-05 | 0.00120443 | 1 | 0.988 | 0.975 | 0.928 | 1 | 1 | 0.948 | 0.942 | 0.87 | 0.869 | 0.052 |
| [illegible] | | | | | | | | | | | | | | | | | | | | | | | | | | | | | | | | | | | | | | | | | | | | | | |

| | | | | | | | | | | | | | | |
|---|---|---|---|---|---|---|---|---|---|---|---|---|---|---|
| SE | 25148 | ENSG00000155329 | ZCCHC10 | chr5 | - | 133000131 | 133000173 | 132996984 | 132998850 | 133006758 | 133006920 | 79 | 57 | 47 |
| SE | 47314 | ENSG00000133106 | EPSTI1 | chr13 | - | 42895008 | 42895108 | 42886387 | 42888502 | 42900309 | 42900383 | 237 | 263 | 231 |
| SE | 70496 | ENSG00000170266 | GLB1 | chr3 | - | 33072543 | 33072713 | 33068818 | 33068970 | 33097010 | 33097043 | 75 | 44 | 58 |
| SE | 44663 | ENSG00000203667 | COX20 | chr1 | + | 244836484 | 244836520 | 244835615 | 244835756 | 244841943 | 244842058 | 2 | 2 | 2 |
| SE | 56889 | ENSG00000125686 | MED1 | chr17 | - | 39445410 | 39445569 | 39443549 | 39443628 | 39447797 | 39447904 | 34 | 18 | 34 |
| SE | 53881 | ENSG00000166523 | CLEC4E | chr12 | - | 8536089 | 8536205 | 8533304 | 8534809 | 8539216 | 8539306 | 227 | 132 | 130 |
| SE | 82243 | ENSG00000128604 | IRF5 | chr7 | + | 128943058 | 128943134 | 128942070 | 128942276 | 128945844 | 128946034 | 2 | 1 | 3 |
| SE | 23240 | ENSG00000112851 | ERBIN | chr5 | + | 66068876 | 66069020 | 66053405 | 66054951 | 66075023 | 66075230 | 24 | 38 | 19 |
| SE | 61627 | ENSG00000234127 | TRIM26 | chr6 | - | 30204655 | 30204765 | 30198665 | 30199153 | 30213304 | 30213365 | 42 | 13 | 25 |
| SE | 22188 | ENSG00000137177 | KIF13A | chr6 | - | 17773477 | 17773583 | 17763692 | 17764946 | 17775014 | 17775062 | 54 | 35 | 28 |
| SE | 54225 | ENSG00000143643 | TTC13 | chr1 | - | 230940439 | 230940556 | 230933778 | 230933861 | 230943805 | 230943898 | 105 | 93 | 66 |
| SE | 89105 | ENSG00000064115 | TM7SF3 | chr12 | - | 26976259 | 26976357 | 26975495 | 26975658 | 26979783 | 26979936 | 131 | 60 | 71 |
| SE | 26583 | ENSG00000006459 | KDM7A | chr7 | - | 140126636 | 140126823 | 140124620 | 140124783 | 140127441 | 140127583 | 226 | 153 | 192 |
| SE | 24318 | ENSG00000058799 | YIPF1 | chr1 | - | 53882942 | 53883031 | 53878641 | 53878722 | 53883112 | 53883276 | 12 | 3 | 12 |
| SE | 64623 | ENSG00000148737 | TCF7L2 | chr10 | + | 113158666 | 113158717 | 113158020 | 113158069 | 113160618 | 113160691 | 1 | 2 | 1 |
| SE | 53069 | ENSG00000103657 | HERC1 | chr15 | - | 63718573 | 63718694 | 63716301 | 63716473 | 63723181 | 63723355 | 312 | 319 | 306 |
| SE | 18679 | ENSG00000129566 | TEP1 | chr14 | - | 20381552 | 20381686 | 20381312 | 20381401 | 20381912 | 20382063 | 40 | 17 | 20 |
| SE | 25697 | ENSG00000247774 | PCED1B-AS1 | chr12 | - | 47206510 | 47206585 | 47205898 | 47206069 | 47216164 | 47216200 | 50 | 29 | 124 |
| SE | 107183 | ENSG00000198715 | GLMP | chr1 | - | 156293319 | 156293576 | 156292686 | 156293209 | 156294017 | 156294236 | 100 | 31 | 84 |
| SE | 77327 | ENSG00000214022 | REPIN1 | chr7 | + | 150369670 | 150369868 | 150368789 | 150368941 | 150371227 | 150371392 | 56 | 18 | 46 |
| SE | 1829 | ENSG00000070371 | CLTCL1 | chr22 | - | 19235695 | 19235869 | 19233421 | 19233622 | 19239274 | 19239388 | 8 | 3 | 3 |
| SE | 90765 | ENSG00000086102 | NFX1 | chr9 | + | 33354085 | 33354187 | 33352645 | 33352719 | 33354850 | 33354892 | 57 | 96 | 96 |

| | | | | | | | | | | | | | | | | | | | | | | | | | | | | | | | | | | | | | | | | | | | | | | |
|---|---|---|---|---|---|---|---|---|---|---|---|---|---|---|---|---|---|---|---|---|---|---|---|---|---|---|---|---|---|---|---|---|---|---|---|---|---|---|---|---|---|---|---|---|---|---|
| SE | 50189 | ENSG00000074706 | IPCEF1 | chr6 | - | 154306694 | 154306762 | 154289712 | 154289756 | 154356671 | 154356792 | 0 | 0 | 0 | 1 | 2 | 15 | 30 | 15 | 16 | 26 | 16 | 0 | 3 | 4 | 0 | 31 | 52 | 32 | 72 | 7 | 217 | 149 | 0.000103673 | 0.002322522 | 0 | 0 | 0 | 0.041 | 0.05 | 0.262 | 0 | 0.06 | 0.037 | 0 | -0.054 |
| SE | 28578 | ENSG00000119414 | PPP6C | chr9 | - | 125160840 | 125160906 | 125158240 | 125158382 | 125171084 | 125171180 | 553 | 336 | 464 | 433 | 583 | 16 | 5 | 4 | 23 | 8 | 379 | 374 | 387 | 462 | 191 | 35 | 21 | 6 | 13 | 11 | 215 | 149 | 0.000103713 | 0.002322736 | 0.96 | 0.979 | 0.988 | 0.929 | 0.981 | 0.882 | 0.925 | 0.978 | 0.961 | 0.923 | 0.034 |
| SE | 25997 | ENSG00000181929 | PRKAG1 | chr12 | - | 49003756 | 49003922 | 49003143 | 49003290 | 49004506 | 49004633 | 136 | 92 | 111 | 81 | 225 | 0 | 1 | 1 | 2 | 3 | 149 | 135 | 77 | 66 | 97 | 8 | 4 | 1 | 9 | 0 | 298 | 149 | 0.000103826 | 0.002324603 | 1 | 0.979 | 0.982 | 0.953 | 0.974 | 0.903 | 0.944 | 0.975 | 0.786 | 1 | 0.056 |
| SE | 10331 | ENSG00000168961 | LGALS9 | chr17 | + | 27645313 | 27645349 | 27643524 | 27643620 | 27645860 | 27645899 | 18 | 13 | 16 | 21 | 28 | 0 | 0 | 3 | 1 | 0 | 115 | 65 | 12 | 57 | 143 | 11 | 0 | 0 | 20 | 12 | 185 | 149 | 0.000103881 | 0.002325187 | 1 | 1 | 0.811 | 0.944 | 1 | 0.894 | 1 | 1 | 0.697 | 0.906 | 0.052 |
| SE | 99961 | ENSG00000154310 | TNIK | chr3 | - | 171108064 | 171108162 | 171101448 | 171101633 | 171110713 | 171110877 | 94 | 85 | 93 | 127 | 70 | 8 | 1 | 4 | 5 | 1 | 61 | 103 | 129 | 63 | 115 | 0 | 0 | 3 | 1 | 0 | 247 | 149 | 0.000104126 | 0.00233 | 0.876 | 0.981 | 0.933 | 0.939 | 0.977 | 1 | 1 | 0.963 | 0.974 | 1 | -0.046 |
| SE | 34044 | ENSG00000114867 | EIF4G1 | chr3 | + | 184315488 | 184315545 | 184314501 | 184314674 | 184315762 | 184315856 | 64 | 27 | 38 | 35 | 35 | 3 | 2 | 2 | 2 | 5 | 21 | 41 | 28 | 34 | 67 | 0 | 0 | 2 | 0 | 0 | 206 | 149 | 0.000104278 | 0.002332724 | 0.939 | 0.907 | 0.932 | 0.927 | 0.835 | 1 | 1 | 0.91 | 1 | 1 | -0.074 |
| SE | 67789 | ENSG00000123106 | CCDC91 | chr12 | + | 28255580 | 28255662 | 28225794 | 28225861 | 28257201 | 28257245 | 90 | 57 | 62 | 75 | 219 | 7 | 2 | 0 | 7 | 4 | 32 | 19 | 21 | 30 | 64 | 0 | 0 | 1 | 0 | 0 | 231 | 149 | 0.000104314 | 0.002332861 | 0.892 | 0.948 | 1 | 0.874 | 0.972 | 1 | 1 | 0.931 | 1 | 1 | -0.049 |
| SE | 67695 | ENSG00000181381 | DDX60L | chr4 | - | 168419301 | 168419397 | 168416681 | 168416797 | 168420260 | 168420380 | 83 | 77 | 63 | 123 | 127 | 12 | 8 | 11 | 12 | 4 | 0 | 70 | 33 | 30 | 0 | 9 | 15 | 23 | 3 | 4 | 245 | 149 | 0.000104515 | 0.0023367 | 0.808 | 0.854 | 0.777 | 0.862 | 0.951 | 0 | 0.739 | 0.466 | 0.859 | 0 | 0.438 |
| SE | 32245 | ENSG00000156931 | VPS8 | chr3 | + | 184893233 | 184893412 | 184886109 | 184886156 | 184894702 | 184894925 | 35 | 22 | 23 | 28 | 43 | 71 | 77 | 97 | 106 | 118 | 13 | 55 | 18 | 22 | 22 | 18 | 42 | 64 | 26 | 29 | 298 | 149 | 0.000105105 | 0.002349223 | 0.198 | 0.125 | 0.106 | 0.117 | 0.154 | 0.265 | 0.396 | 0.123 | 0.297 | 0.275 | -0.131 |
| SE | 16048 | ENSG00000048471 | SNX29 | chr16 | + | 12061527 | 12061646 | 12051846 | 12052222 | 12069056 | 12069132 | 172 | 103 | 159 | 127 | 209 | 3 | 0 | 6 | 2 | 7 | 519 | 397 | 219 | 224 | 397 | 3 | 31 | 17 | 14 | 27 | 268 | 149 | 0.000105329 | 0.002353158 | 0.97 | 1 | 0.936 | 0.972 | 0.943 | 0.99 | 0.877 | 0.877 | 0.899 | 0.891 | 0.057 |
| SE | 93758 | ENSG00000143409 | MINDY1 | chr1 | - | 151002164 | 151002706 | 151001724 | 151001782 | 151006311 | 151006902 | 79 | 78 | 86 | 149 | 123 | 6 | 5 | 4 | 0 | 2 | 308 | 162 | 73 | 168 | 977 | 26 | 3 | 7 | 10 | 101 | 298 | 149 | 0.000105342 | 0.002353158 | 0.868 | 0.886 | 0.915 | 1 | 0.969 | 0.856 | 0.964 | 0.839 | 0.894 | 0.829 | 0.051 |
| SE | 10264 | ENSG00000108651 | UTP6 | chr17 | - | 31884423 | 31884505 | 31880180 | 31880754 | 31885979 | 31886061 | 272 | 250 | 182 | 267 | 323 | 1 | 2 | 1 | 4 | 0 | 260 | 151 | 128 | 232 | 71 | 10 | 0 | 2 | 12 | 0 | 231 | 149 | 0.000105448 | 0.002353975 | 0.994 | 0.988 | 0.992 | 0.977 | 1 | 0.944 | 1 | 0.976 | 0.926 | 1 | 0.021 |
| SE | 40379 | ENSG00000163719 | MTMR14 | chr3 | + | 9662266 | 9662375 | 9653620 | 9653769 | 9668718 | 9668752 | 307 | 161 | 249 | 303 | 254 | 0 | 0 | 2 | 6 | 4 | 600 | 380 | 298 | 430 | 945 | 36 | 1 | 8 | 12 | 5 | 258 | 149 | 0.000105462 | 0.002353975 | 1 | 1 | 0.986 | 0.967 | 0.973 | 0.906 | 0.995 | 0.956 | 0.954 | 0.991 | 0.025 |
| SE | 107454 | ENSG00000215039 | CD27-AS1 | chr12 | - | 6448639 | 6448757 | 6447615 | 6448150 | 6451468 | 6451515 | 19 | 9 | 7 | 15 | 3 | 0 | 0 | 0 | 0 | 0 | 11 | 7 | 9 | 15 | 68 | 0 | 0 | 0 | 0 | 12 | 267 | 149 | 0.000105468 | 0.002353975 | 1 | 1 | 1 | 1 | 1 | 1 | 1 | 1 | 1 | 0.76 | 0.048 |
| SE | 90456 | ENSG00000070081 | NUCB2 | chr11 | + | 17291239 | 17291365 | 17282788 | 17282943 | 17295323 | 17295441 | 1 | 2 | 8 | 17 | 4 | 114 | 76 | 161 | 117 | 168 | 15 | 4 | 11 | 0 | 0 | 14 | 34 | 29 | 32 | 7 | 275 | 149 | 0.000105627 | 0.002356839 | 0.005 | 0.014 | 0.026 | 0.073 | 0.013 | 0.367 | 0.06 | 0.17 | 0 | 0 | -0.093 |
| SE | 88457 | ENSG00000152601 | MBNL1 | chr3 | + | 152299404 | 152300367 | 152268039 | 152269092 | 152414940 | 152415111 | 1331 | 1027 | 1414 | 1426 | 2371 | 127 | 114 | 86 | 249 | 183 | 861 | 1020 | 1025 | 699 | 858 | 150 | 125 | 104 | 154 | 289 | 298 | 149 | 0.000105784 | 0.002359675 | 0.84 | 0.818 | 0.892 | 0.741 | 0.866 | 0.742 | 0.803 | 0.831 | 0.694 | 0.597 | 0.098 |
| SE | 32324 | ENSG00000124570 | SERPINB6 | chr6 | - | 2968778 | 2968913 | 2959167 | 2959342 | 2971116 | 2971276 | 14 | 8 | 11 | 16 | 13 | 21 | 5 | 16 | 9 | 10 | 6 | 1 | 10 | 17 | 13 | 31 | 12 | 11 | 27 | 86 | 284 | 149 | 0.000106097 | 0.002365692 | 0.259 | 0.456 | 0.265 | 0.483 | 0.405 | 0.092 | 0.042 | 0.323 | 0.248 | 0.073 | 0.218 |
| SE | 99060 | ENSG00000149577 | SIDT2 | chr11 | + | 117186989 | 117187001 | 117186583 | 117186636 | 117187377 | 117187449 | 9 | 3 | 6 | 16 | 0 | 70 | 54 | 25 | 125 | 35 | 0 | 6 | 0 | 3 | 0 | 49 | 94 | 23 | 38 | 66 | 161 | 149 | 0.000106114 | 0.002365692 | 0.106 | 0.049 | 0.182 | 0.106 | 0 | 0 | 0.056 | 0 | 0.068 | 0 | 0.064 |
| SE | 36376 | ENSG00000155621 | C9orf85 | chr9 | + | 71941804 | 71941934 | 71911640 | 71911836 | 71947005 | 71947112 | 0 | 0 | 2 | 0 | 0 | 38 | 17 | 23 | 11 | 29 | 0 | 7 | 1 | 0 | 7 | 35 | 16 | 13 | 12 | 10 | 279 | 149 | 0.000106504 | 0.002373705 | 0 | 0 | 0.044 | 0 | 0 | 0 | 0.189 | 0.039 | 0 | 0.272 | -0.091 |
| SE | 26436 | ENSG00000158290 | CUL4B | chrX | - | 120544480 | 120544643 | 120544113 | 120544203 | 120546546 | 120546616 | 176 | 118 | 164 | 175 | 325 | 0 | 0 | 1 | 1 | 0 | 21 | 83 | 56 | 48 | 31 | 16 | 1 | 3 | 0 | 0 | 298 | 149 | 0.000106552 | 0.002374093 | 1 | 1 | 0.988 | 0.989 | 1 | 0.396 | 0.976 | 0.903 | 1 | 1 | 0.14 |
| SE | 69333 | ENSG00000168958 | MFF | chr2 | + | 227330594 | 227330846 | 227328677 | 227328789 | 227332418 | 227332499 | 220 | 107 | 206 | 158 | 295 | 1 | 0 | 2 | 1 | 2 | 279 | 150 | 159 | 156 | 161 | 0 | 3 | 9 | 6 | 5 | 298 | 149 | 0.000106623 | 0.002374999 | 0.991 | 1 | 0.981 | 0.988 | 0.987 | 1 | 0.962 | 0.898 | 0.929 | 0.942 | 0.043 |
| SE | 77603 | ENSG00000127947 | PTPN12 | chr7 | + | 77583554 | 77583650 | 77581426 | 77581503 | 77585542 | 77585571 | 902 | 760 | 424 | 895 | 788 | 4 | 5 | 6 | 5 | 2 | 559 | 741 | 754 | 620 | 717 | 0 | 2 | 9 | 9 | 33 | 245 | 149 | 0.00010684 | 0.002379159 | 0.993 | 0.989 | 0.977 | 0.991 | 0.996 | 1 | 0.996 | 0.981 | 0.977 | 0.93 | 0.012 |
| SE | 657 | ENSG00000100226 | GTPBP1 | chr22 | + | 38734149 | 38734232 | 38730611 | 38730659 | 38734323 | 38734397 | 6 | 3 | 2 | 6 | 2 | 2 | 0 | 0 | 0 | 0 | 8 | 24 | 13 | 4 | 9 | 8 | 7 | 0 | 0 | 0 | 232 | 149 | 0.000107044 | 0.002383027 | 0.658 | 1 | 1 | 1 | 1 | 0.391 | 0.688 | 1 | 1 | 1 | 0.116 |
| SE | 97622 | ENSG00000149313 | AASDHPPT | chr11 | + | 106090556 | 106090678 | 106079466 | 106079692 | 106091315 | 106091459 | 198 | 110 | 154 | 179 | 202 | 3 | 2 | 2 | 3 | 9 | 32 | 93 | 47 | 41 | 9 | 0 | 11 | 6 | 4 | 1 | 271 | 149 | 0.000107456 | 0.002391515 | 0.973 | 0.968 | 0.977 | 0.97 | 0.925 | 1 | 0.823 | 0.812 | 0.849 | 0.832 | 0.099 |
| SE | 8690 | ENSG00000198909 | MAP3K3 | chr17 | + | 63692241 | 63692419 | 63691732 | 63691862 | 63693548 | 63693874 | 296 | 170 | 196 | 213 | 180 | 1 | 1 | 2 | 3 | 0 | 341 | 278 | 206 | 243 | 394 | 0 | 0 | 7 | 16 | 9 | 298 | 149 | 0.000107569 | 0.002393349 | 0.993 | 0.988 | 0.98 | 0.973 | 1 | 1 | 1 | 0.936 | 0.884 | 0.956 | 0.032 |
| SE | 80939 | ENSG00000100997 | ABHD12 | chr20 | - | 25317047 | 25317078 | 25309445 | 25309575 | 25320198 | 25320318 | 31 | 20 | 32 | 29 | 24 | 0 | 1 | 0 | 0 | 0 | 30 | 42 | 23 | 16 | 42 | 0 | 2 | 2 | 9 | 1 | 180 | 149 | 0.000107719 | 0.002396008 | 1 | 0.943 | 1 | 1 | 1 | 1 | 0.946 | 0.905 | 0.595 | 0.972 | 0.105 |
| SE | 22817 | ENSG00000173692 | PSMD1 | chr2 | + | 231138735 | 231138850 | 231087116 | 231087181 | 231146239 | 231146356 | 156 | 101 | 121 | 141 | 228 | 2 | 0 | 2 | 0 | 0 | 89 | 73 | 106 | 127 | 18 | 6 | 4 | 2 | 4 | 0 | 264 | 149 | 0.000108105 | 0.002403867 | 0.978 | 1 | 0.972 | 1 | 1 | 0.893 | 0.912 | 0.968 | 0.947 | 1 | 0.046 |
| SE | 21007 | ENSG00000126214 | KLC1 | chr14 | + | 103662701 | 103662921 | 103657545 | 103657776 | 103669510 | 103669598 | 0 | 0 | 1 | 1 | 1 | 1 | 2 | 0 | 1 | 0 | 12 | 1 | 2 | 7 | 0 | 9 | 0 | 0 | 0 | 7 | 298 | 149 | 0.000108133 | 0.002403867 | 0 | 0 | 1 | 0.333 | 1 | 0.4 | 1 | 1 | 1 | 0 | -0.213 |
| SE | 33070 | ENSG00000088726 | TMEM40 | chr3 | - | 12743899 | 12743989 | 12742453 | 12742507 | 12748654 | 12748792 | 18 | 24 | 6 | 43 | 36 | 8 | 8 | 1 | 4 | 0 | 49 | 18 | 22 | 38 | 187 | 20 | 32 | 11 | 13 | 158 | 239 | 149 | 0.000108247 | 0.002405682 | 0.584 | 0.652 | 0.789 | 0.87 | 1 | 0.604 | 0.26 | 0.555 | 0.646 | 0.425 | 0.281 |
| SE | 95487 | ENSG00000108424 | KPNB1 | chr17 | + | 47673489 | 47673561 | 47673017 | 47673165 | 47674637 | 47674755 | 632 | 421 | 608 | 571 | 705 | 1 | 1 | 0 | 0 | 4 | 508 | 472 | 451 | 454 | 187 | 15 | 0 | 4 | 12 | 6 | 221 | 149 | 0.000108277 | 0.002405682 | 0.998 | 0.996 | 1 | 1 | 0.992 | 0.958 | 1 | 0.987 | 0.962 | 0.955 | 0.025 |
| SE | 21565 | ENSG00000085382 | HACE1 | chr6 | - | 104843222 | 104843298 | 104833041 | 104833173 | 104849141 | 104849246 | 51 | 46 | 24 | 42 | 51 | 2 | 2 | 0 | 1 | 10 | 21 | 33 | 11 | 15 | 34 | 0 | 2 | 0 | 0 | 0 | 225 | 149 | 0.000108448 | 0.002408813 | 0.944 | 0.938 | 1 | 0.965 | 0.772 | 1 | 0.916 | 1 | 1 | 1 | -0.059 |
| SE | 47979 | ENSG00000154710 | RABGEF1 | chr7 | + | 66768830 | 66769025 | 66712166 | 66712224 | 66771882 | 66772078 | 5 | 8 | 12 | 11 | 10 | 0 | 0 | 0 | 0 | 0 | 12 | 11 | 10 | 15 | 5 | 0 | 1 | 0 | 6 | 3 | 298 | 149 | 0.000109287 | 0.002426759 | 1 | 1 | 1 | 1 | 1 | 1 | 0.846 | 1 | 0.556 | 0.455 | 0.229 |
| SE | 29618 | ENSG00000203875 | SNHG5 | chr6 | - | 85677794 | 85678032 | 85677078 | 85677492 | 85678135 | 85678188 | 165 | 259 | 247 | 187 | 381 | 4 | 6 | 2 | 1 | 11 | 89 | 236 | 237 | 190 | 408 | 0 | 0 | 2 | 0 | 0 | 298 | 149 | 0.00010934 | 0.002427246 | 0.954 | 0.956 | 0.984 | 0.989 | 0.945 | 1 | 1 | 0.983 | 1 | 1 | -0.031 |
| SE | 3709 | ENSG00000181027 | FKRP | chr19 | + | 46748774 | 46748884 | 46748514 | 46748665 | 46755411 | 46755634 | 8 | 6 | 0 | 8 | 11 | 3 | 4 | 4 | 14 | 20 | 0 | 0 | 7 | 0 | 6 | 9 | 16 | 10 | 20 | 48 | 259 | 149 | 0.000109462 | 0.002429256 | 0.605 | 0.463 | 0 | 0.247 | 0.24 | 0 | 0 | 0.287 | 0 | 0.067 | 0.24 |
| SE | 81303 | ENSG00000141480 | ARRB2 | chr17 | + | 4716146 | 4716191 | 4715588 | 4716033 | 4716411 | 4716515 | 357 | 174 | 222 | 220 | 283 | 9 | 3 | 20 | 13 | 11 | 437 | 293 | 284 | 406 | 597 | 26 | 50 | 16 | 90 | 14 | 194 | 149 | 0.000109673 | 0.002433263 | 0.968 | 0.978 | 0.895 | 0.929 | 0.952 | 0.928 | 0.818 | 0.932 | 0.776 | 0.97 | 0.06 |
| SE | 75329 | ENSG00000196262 | PPIA | chr7 | + | 44798747 | 44798904 | 44796680 | 44796793 | 44799246 | 44799277 | 170 | 93 | 148 | 182 | 139 | 4756 | 2726 | 4058 | 4201 | 3007 | 131 | 95 | 112 | 80 | 435 | 6185 | 4505 | 3624 | 4987 | 7974 | 298 | 149 | 0.000109733 | 0.002433889 | 0.018 | 0.017 | 0.018 | 0.021 | 0.023 | 0.01 | 0.01 | 0.015 | 0.008 | 0.027 | 0.005 |
| SE | 33827 | ENSG00000041802 | LSG1 | chr3 | - | 194660741 | 194660886 | 194660072 | 194660133 | 194665556 | 194665643 | 1 | 0 | 2 | 4 | 28 | 123 | 69 | 97 | 133 | 189 | 0 | 0 | 3 | 0 | 0 | 217 | 70 | 78 | 170 | 92 | 294 | 149 | 0.000109937 | 0.00243772 | 0.004 | 0 | 0.01 | 0.015 | 0.07 | 0 | 0 | 0.019 | 0 | 0 | 0.016 |
| SE | 100287 | ENSG00000167986 | DDB1 | chr11 | - | 61323010 | 61323094 | 61322295 | 61322412 | 61323978 | 61324137 | 656 | 470 | 539 | 567 | 532 | 0 | 0 | 0 | 0 | 0 | 757 | 562 | 384 | 467 | 695 | 18 | 0 | 9 | 4 | 13 | 233 | 149 | 0.000110126 | 0.002441233 | 1 | 1 | 1 | 1 | 1 | 0.964 | 1 | 0.965 | 0.987 | 0.972 | 0.022 |
| SE | 68093 | ENSG00000135838 | NPL | chr1 | + | 182806144 | 182806232 | 182803697 | 182803771 | 182812155 | 182812190 | 174 | 135 | 128 | 141 | 197 | 16 | 1 | 8 | 5 | 9 | 294 | 272 | 243 | 361 | 174 | 0 | 8 | 5 | 6 | 5 | 237 | 149 | 0.000110236 | 0.002442972 | 0.872 | 0.988 | 0.91 | 0.947 | 0.932 | 1 | 0.955 | 0.968 | 0.974 | 0.956 | -0.041 |
| SE | 86693 | ENSG00000147140 | NONO | chrX | + | 71284983 | 71285097 | 71284396 | 71284453 | 71290628 | 71290791 | 12 | 6 | 10 | 7 | 7 | 305 | 192 | 311 | 251 | 239 | 23 | 16 | 14 | 17 | 0 | 204 | 185 | 137 | 150 | 331 | 263 | 149 | 0.000110388 | 0.002445663 | 0.022 | 0.017 | 0.018 | 0.016 | 0.016 | 0.06 | 0.047 | 0.055 | 0.06 | 0 | -0.027 |
| SE | 44380 | ENSG00000072210 | ALDH3A2 | chr17 | + | 19655280 | 19655387 | 19652546 | 19652632 | 19656365 | 19656574 | 2 | 5 | 4 | 10 | 13 | 52 | 29 | 42 | 70 | 75 | 0 | 2 | 1 | 9 | 0 | 107 | 22 | 39 | 68 | 63 | 256 | 149 | 0.000110823 | 0.002454595 | 0.022 | 0.091 | 0.053 | 0.077 | 0.092 | 0 | 0.05 | 0.015 | 0.072 | 0 | 0.04 |
| SE | 35478 | ENSG00000172830 | SSH3 | chr11 | + | 67307548 | 67307737 | 67307370 | 67307436 | 67307845 | 67307906 | 13 | 3 | 5 | 23 | 3 | 0 | 0 | 0 | 0 | 0 | 23 | 12 | 9 | 17 | 55 | 0 | 0 | 0 | 0 | 13 | 298 | 149 | 0.000110979 | 0.002457353 | 1 | 1 | 1 | 1 | 1 | 1 | 1 | 1 | 1 | 0.679 | 0.064 |
| SE | 41744 | ENSG00000158882 | TOMM40L | chr1 | + | 161227635 | 161227737 | 161226887 | 161226955 | 161227883 | 161227989 | 9 | 6 | 9 | 13 | 2 | 0 | 0 | 0 | 3 | 0 | 15 | 17 | 9 | 5 | 42 | 0 | 0 | 2 | 14 | 9 | 251 | 149 | 0.000111588 | 0.002470147 | 1 | 1 | 1 | 0.72 | 1 | 1 | 1 | 0.728 | 0.175 | 0.735 | 0.216 |
| SE | 93868 | ENSG00000116580 | GON4L | chr1 | - | 155831534 | 155831620 | 155826869 | 155827028 | 155853275 | 155853809 | 5 | 3 | 8 | 5 | 0 | 145 | 147 | 150 | 148 | 220 | 0 | 21 | 4 | 15 | 8 | 172 | 185 | 137 | 160 | 51 | 235 | 149 | 0.000112249 | 0.002484063 | 0.021 | 0.013 | 0.033 | 0.021 | 0 | 0 | 0.067 | 0.018 | 0.056 | 0.09 | -0.029 |
| SE | 106187 | ENSG00000089094 | KDM2B | chr12 | - | 121549459 | 121549638 | 121548876 | 121548983 | 121574546 | 121574593 | 61 | 30 | 77 | 57 | 35 | 0 | 0 | 0 | 0 | 0 | 58 | 76 | 57 | 41 | 90 | 0 | 6 | 1 | 0 | 7 | 298 | 149 | 0.000112685 | 0.002493019 | 1 | 1 | 1 | 1 | 1 | 1 | 0.864 | 0.966 | 1 | 0.865 | 0.061 |
| SE | 67375 | ENSG00000187688 | TRPV2 | chr17 | + | 16426721 | 16426877 | 16423468 | 16423767 | 16427448 | 16427547 | 54 | 28 | 34 | 30 | 18 | 0 | 1 | 0 | 0 | 0 | 16 | 39 | 11 | 12 | 20 | 0 | 0 | 0 | 4 | 8 | 298 | 149 | 0.000112817 | 0.002495222 | 1 | 0.933 | 1 | 1 | 1 | 1 | 1 | 1 | 0.6 | 0.556 | 0.155 |
| SE | 91082 | ENSG00000165102 | HGSNAT | chr8 | + | 43173712 | 43173743 | 43172309 | 43172386 | 43178073 | 43178234 | 76 | 33 | 50 | 64 | 42 | 0 | 1 | 1 | 2 | 0 | 95 | 22 | 17 | 51 | 143 | 0 | 6 | 0 | 3 | 12 | 180 | 149 | 0.000113215 | 0.002503325 | 1 | 0.965 | 0.976 | 0.964 | 1 | 1 | 0.752 | 1 | 0.934 | 0.908 | 0.062 |
| SE | 70211 | ENSG00000085365 | SCAMP1 | chr5 | + | 78421800 | 78421960 | 78416540 | 78416649 | 78449932 | 78450034 | 105 | 61 | 64 | 87 | 71 | 0 | 3 | 2 | 6 | 4 | 36 | 50 | 44 | 5 | 21 | 0 | 0 | 0 | 0 | 0 | 298 | 149 | 0.000113294 | 0.002504362 | 1 | 0.91 | 0.941 | 0.879 | 0.899 | 1 | 1 | 1 | 1 | 1 | -0.074 |
| SE | 71411 | ENSG00000145349 | CAMK2D | chr4 | - | 113548656 | 113548729 | 113547643 | 113547716 | 113552030 | 113552096 | 31 | 20 | 43 | 36 | 24 | 6 | 3 | 1 | 6 | 4 | 3 | 16 | 40 | 15 | 5 | 0 | 0 | 1 | 3 | 0 | 222 | 149 | 0.000113368 | 0.002505289 | 0.776 | 0.817 | 0.967 | 0.801 | 0.801 | 1 | 1 | 0.964 | 0.77 | 1 | -0.114 |
| SE | 99855 | ENSG00000110713 | NUP98 | chr11 | - | 3700609 | 3700839 | 3699081 | 3699348 | 3702462 | 3702892 | 336 | 244 | 251 | 262 | 522 | 10 | 7 | 5 | 9 | 4 | 453 | 393 | 390 | 388 | 250 | 43 | 9 | 15 | 18 | 5 | 298 | 149 | 0.000113419 | 0.002505717 | 0.944 | 0.946 | 0.962 | 0.936 | 0.985 | 0.84 | 0.956 | 0.929 | 0.915 | 0.962 | 0.034 |
| SE | 52094 | ENSG00000117419 | ERI3 | chr1 | - | 44339044 | 44339322 | 44319627 | 44319744 | 44352849 | 44352925 | 75 | 34 | 54 | 76 | 84 | 4 | 1 | 7 | 1 | 5 | 37 | 86 | 52 | 85 | 78 | 0 | 2 | 0 | 3 | 0 | 298 | 149 | 0.000113785 | 0.002511884 | 0.904 | 0.944 | 0.794 | 0.974 | 0.894 | 1 | 0.956 | 1 | 0.934 | 1 | -0.076 |
| SE | 20761 | ENSG00000026652 | AGPAT4 | chr6 | - | 161154148 | 161154310 | 161153345 | 161153499 | 161166247 | 161166417 | 20 | 4 | 5 | 21 | 13 | 1 | 2 | 0 | 0 | 0 | 23 | 4 | 14 | 2 | 37 | 0 | 1 | 6 | 3 | 3 | 298 | 149 | 0.000113789 | 0.002511884 | 0.909 | 0.5 | 1 | 1 | 1 | 1 | 0.667 | 0.538 | 0.25 | 0.86 | 0.219 |
| SE | 78660 | ENSG00000274272 | NA | chr7 | - | 100577272 | 100577389 | 100577085 | 100577154 | 100577663 | 100577740 | 47 | 31 | 49 | 39 | 19 | 0 | 0 | 0 | 0 | 0 | 19 | 57 | 18 | 17 | 55 | 0 | 12 | 1 | 0 | 0 | 266 | 149 | 0.000113794 | 0.002511884 | 1 | 1 | 1 | 1 | 1 | 1 | 0.727 | 0.91 | 1 | 1 | 0.073 |
| SE | 29490 | ENSG00000185684 | EP400P1 | chr12 | + | 132114457 | 132114545 | 132108571 | 132108682 | 132120419 | 132120601 | 14 | 3 | 6 | 10 | 1 | 0 | 0 | 0 | 1 | 0 | 13 | 8 | 1 | 7 | 11 | 14 | 3 | 0 | 0 | 0 | 237 | 149 | 0.000113947 | 0.002514545 | 1 | 1 | 1 | 0.863 | 1 | 0.369 | 0.626 | 1 | 1 | 1 | 0.174 |
| SE | 82288 | ENSG00000122566 | HNRNPA2B1 | chr7 | - | 26190992 | 26191650 | 26189940 | 26190460 | 26192494 | 26192577 | 263 | 295 | 282 | 278 | 496 | 14 | 4 | 15 | 2 | 15 | 153 | 182 | 187 | 173 | 50 | 9 | 26 | 25 | 5 | 0 | 298 | 149 | 0.000114217 | 0.002519746 | 0.904 | 0.974 | 0.904 | 0.986 | 0.943 | 0.895 | 0.778 | 0.789 | 0.945 | 1 | 0.061 |
| SE | 63594 | ENSG00000119596 | YLPM1 | chr14 | + | 74824255 | 74824307 | 74821056 | 74821137 | 74855216 | 74859435 | 32 | 35 | 36 | 25 | 32 | 19 | 14 | 16 | 22 | 19 | 80 | 14 | 32 | 36 | 26 | 4 | 8 | 10 | 0 | 7 | 201 | 149 | 0.000114279 | 0.002519746 | 0.555 | 0.65 | 0.625 | 0.457 | 0.555 | 0.937 | 0.565 | 0.703 | 1 | 0.734 | -0.219 |
| SE | 43466 | ENSG00000072786 | STK10 | chr5 | - | 172090231 | 172090362 | 172082960 | 172083084 | 172105655 | 172105737 | 575 | 380 | 428 | 528 | 492 | 0 | 0 | 2 | 0 | 0 | 638 | 596 | 428 | 504 | 705 | 20 | 0 | 4 | 15 | 0 | 280 | 149 | 0.000114279 | 0.002519746 | 1 | 1 | 0.991 | 1 | 1 | 0.944 | 1 | 0.983 | 0.947 | 1 | 0.023 |
| SE | 33073 | ENSG00000088726 | TMEM40 | chr3 | - | 12748654 | 12748792 | 12742453 | 12742507 | 12749759 | 12749840 | 14 | 17 | 2 | 17 | 6 | 6 | 13 | 4 | 29 | 24 | 86 | 48 | 33 | 25 | 344 | 42 | 8 | 9 | 11 | 166 | 287 | 149 | 0.000114685 | 0.002527972 | 0.548 | 0.404 | 0.206 | 0.233 | 0.115 | 0.515 | 0.757 | 0.656 | 0.541 | 0.518 | -0.296 |
| SE | 102064 | ENSG00000181192 | DHTKD1 | chr10 | + | 12087534 | 12087729 | 12084539 | 12084751 | 12088985 | 12089255 | 168 | 100 | 124 | 137 | 110 | 7 | 1 | 1 | 1 | 9 | 214 | 178 | 69 | 128 | 303 | 0 | 3 | 0 | 2 | 0 | 298 | 149 | 0.00011516 | 0.002537732 | 0.923 | 0.98 | 0.984 | 0.986 | 0.859 | 1 | 0.967 | 1 | 0.97 | 1 | -0.041 |
| SE | 74058 | ENSG00000111011 | RSRC2 | chr12 | - | 122520503 | 122520581 | 122518838 | 122519029 | 122521384 | 122521428 | 87 | 56 | 64 | 65 | 98 | 85 | 38 | 32 | 56 | 71 | 38 | 112 | 79 | 63 | 30 | 10 | 32 | 54 | 12 | 0 | 227 | 149 | 0.000115364 | 0.0025415 | 0.402 | 0.492 | 0.568 | 0.432 | 0.475 | 0.714 | 0.697 | 0.49 | 0.775 | 1 | -0.261 |
| SE | 3794 | ENSG00000090372 | STRN4 | chr19 | - | 46736822 | 46736901 | 46730731 | 46730873 | 46738163 | 46738172 | 57 | 33 | 44 | 54 | 59 | 4 | 0 | 0 | 1 | 6 | 133 | 88 | 87 | 75 | 165 | 0 | 21 | 10 | 21 | 20 | 228 | 149 | 0.000115422 | 0.0025415 | 0.903 | 1 | 1 | 0.972 | 0.865 | 1 | 0.733 | 0.85 | 0.7 | 0.844 | 0.123 |
| SE | 84008 | ENSG00000104331 | BPNT2 | chr8 | - | 56967109 | 56967218 | 56966190 | 56966352 | 56978049 | 56978145 | 8 | 5 | 0 | 8 | 3 | 108 | 39 | 68 | 96 | 55 | 0 | 0 | 1 | 0 | 0 | 81 | 64 | 118 | 100 | 54 | 258 | 149 | 0.000115428 | 0.0025415 | 0.041 | 0.069 | 0 | 0.046 | 0.031 | 0 | 0 | 0.005 | 0 | 0 | 0.036 |
| SE | 68497 | ENSG00000087087 | SRRT | chr7 | + | 100880695 | 100880769 | 100875572 | 100875712 | 100881284 | 100881413 | 11 | 5 | 19 | 18 | 11 | 68 | 57 | 62 | 89 | 81 | 12 | 11 | 12 | 10 | 8 | 137 | 128 | 79 | 141 | 193 | 223 | 149 | 0.000115524 | 0.0025429 | 0.098 | 0.055 | 0.17 | 0.119 | 0.083 | 0.055 | 0.054 | 0.092 | 0.045 | 0.027 | 0.05 |
| SE | 43887 | ENSG00000179335 | CLK3 | chr15 | + | 74622493 | 74622560 | 74620008 | 74622216 | 74624901 | 74625018 | 153 | 66 | 122 | 150 | 269 | 0 | 0 | 1 | 0 | 3 | 174 | 195 | 163 | 144 | 205 | 6 | 4 | 0 | 6 | 9 | 216 | 149 | 0.0001161 | 0.002554849 | 1 | 1 | 0.988 | 1 | 0.984 | 0.952 | 0.971 | 1 | 0.943 | 0.94 | 0.033 |
| SE | 57289 | ENSG00000143119 | CD53 | chr1 | + | 110897808 | 110897892 | 110896652 | 110896733 | 110899123 | 110899222 | 2025 | 912 | 1407 | 1760 | 2024 | 15 | 5 | 7 | 12 | 17 | 1992 | 1815 | 854 | 1227 | 1949 | 3 | 42 | 19 | 15 | 25 | 233 | 149 | 0.000116142 | 0.002555053 | 0.989 | 0.991 | 0.992 | 0.989 | 0.987 | 0.998 | 0.965 | 0.966 | 0.981 | 0.98 | 0.012 |
| SE | 23024 | ENSG00000183474 | GTF2H2C | chr5 | + | 69566588 | 69566625 | 69565368 | 69566208 | 69566966 | 69567053 | 9 | 10 | 28 | 22 | 18 | 2 | 1 | 0 | 3 | 0 | 11 | 1 | 5 | 4 | 0 | 0 | 13 | 0 | 4 | 10 | 186 | 149 | 0.000116186 | 0.0025553 | 0.783 | 0.889 | 1 | 0.855 | 1 | 1 | 0.058 | 1 | 0.445 | 0 | 0.405 |
| SE | 17433 | ENSG00000140563 | MCTP2 | chr15 | + | 94314726 | 94314864 | 94314281 | 94314344 | 94315528 | 94315637 | 22 | 4 | 5 | 28 | 8 | 88 | 31 | 30 | 81 | 65 | 0 | 17 | 12 | 0 | 0 | 44 | 86 | 74 | 76 | 25 | 287 | 149 | 0.000116899 | 0.002570278 | 0.115 | 0.063 | 0.08 | 0.152 | 0.06 | 0 | 0.093 | 0.078 | 0 | 0 | 0.06 |
| SE | 98286 | ENSG00000068831 | RASGRP2 | chr11 | - | 64740367 | 64740449 | 64740012 | 64740163 | 64740947 | 64741079 | 52 | 14 | 47 | 54 | 32 | 114 | 67 | 109 | 137 | 133 | 19 | 30 | 18 | 5 | 10 | 44 | 137 | 65 | 63 | 175 | 231 | 149 | 0.000117161 | 0.002575314 | 0.227 | 0.119 | 0.218 | 0.203 | 0.134 | 0.218 | 0.124 | 0.152 | 0.049 | 0.036 | 0.064 |
| SE | 60716 | ENSG00000106346 | USP42 | chr7 | + | 6142931 | 6143014 | 6140913 | 6140984 | 6144084 | 6144196 | 171 | 77 | 102 | 147 | 138 | 0 | 4 | 0 | 1 | 0 | 86 | 157 | 165 | 117 | 55 | 8 | 1 | 2 | 10 | 0 | 232 | 149 | 0.000117803 | 0.002588695 | 1 | 0.925 | 1 | 0.99 | 1 | 0.873 | 0.99 | 0.981 | 0.883 | 1 | 0.038 |
| SE | 53349 | ENSG00000131778 | CHD1L | chr1 | + | 147280025 | 147280191 | 147276103 | 147276257 | 147284350 | 147284499 | 200 | 139 | 185 | 275 | 222 | 7 | 7 | 1 | 6 | 22 | 478 | 231 | 217 | 326 | 1023 | 3 | 5 | 10 | 14 | 2 | 298 | 149 | 0.000117915 | 0.002590418 | 0.935 | 0.908 | 0.989 | 0.958 | 0.835 | 0.988 | 0.959 | 0.916 | 0.921 | 0.996 | -0.031 |
| SE | 85792 | ENSG00000258529 | NA | chr11 | - | 111840654 | 111840809 | 111838248 | 111838378 | 111844600 | 111844723 | 79 | 48 | 76 | 54 | 114 | 4 | 1 | 1 | 7 | 8 | 113 | 68 | 71 | 35 | 53 | 5 | 8 | 22 | 10 | 13 | 298 | 149 | 0.000118079 | 0.002593307 | 0.908 | 0.96 | 0.974 | 0.794 | 0.877 | 0.919 | 0.81 | 0.617 | 0.636 | 0.671 | 0.172 |
| SE | 107415 | ENSG00000157538 | VPS26C | chr21 | - | 37233361 | 37233442 | 37232376 | 37232451 | 37267237 | 37267329 | 51 | 20 | 41 | 42 | 92 | 1 | 1 | 0 | 1 | 6 | 86 | 37 | 54 | 35 | 93 | 16 | 7 | 6 | 0 | 8 | 230 | 149 | 0.00011855 | 0.002602872 | 0.971 | 0.928 | 1 | 0.965 | 0.909 | 0.777 | 0.774 | 0.854 | 1 | 0.883 | 0.097 |
| SE | 39368 | ENSG00000174437 | ATP2A2 | chr12 | + | 110292019 | 110292124 | 110282712 | 110282795 | 110296598 | 110296712 | 444 | 343 | 329 | 407 | 727 | 1 | 1 | 1 | 10 | 3 | 733 | 365 | 383 | 424 | 564 | 12 | 4 | 3 | 4 | 24 | 254 | 149 | 0.000118581 | 0.002602872 | 0.996 | 0.995 | 0.995 | 0.96 | 0.993 | 0.973 | 0.982 | 0.987 | 0.984 | 0.932 | 0.016 |
| SE | 55508 | ENSG00000124181 | PLCG1 | chr20 | + | 41174126 | 41174311 | 41173922 | 41174011 | 41174466 | 41174967 | 83 | 33 | 121 | 111 | 131 | 1 | 2 | 2 | 0 | 0 | 87 | 69 | 51 | 49 | 106 | 0 | 15 | 2 | 2 | 0 | 298 | 149 | 0.000119161 | 0.002614859 | 0.976 | 0.892 | 0.968 | 1 | 1 | 1 | 0.697 | 0.927 | 0.925 | 1 | 0.057 |
| SE | 104144 | ENSG00000023041 | ZDHHC6 | chr10 | - | 112440533 | 112440695 | 112438335 | 112438389 | 112442191 | 112442259 | 269 | 256 | 230 | 309 | 355 | 23 | 3 | 28 | 3 | 10 | 324 | 250 | 234 | 229 | 99 | 40 | 23 | 36 | 14 | 4 | 298 | 149 | 0.000119326 | 0.002617749 | 0.854 | 0.977 | 0.804 | 0.981 | 0.947 | 0.802 | 0.845 | 0.765 | 0.891 | 0.925 | 0.067 |
| SE | 59478 | ENSG00000160310 | PRMT2 | chr21 | + | 46636438 | 46636547 | 46635673 | 46635763 | 46636895 | 46636990 | 226 | 133 | 130 | 200 | 159 | 95 | 29 | 33 | 65 | 52 | 239 | 389 | 149 | 151 | 314 | 51 | 33 | 26 | 43 | 52 | 258 | 149 | 0.000119389 | 0.0026181 | 0.579 | 0.726 | 0.695 | 0.64 | 0.638 | 0.73 | 0.872 | 0.768 | 0.67 | 0.777 | -0.108 |
| SE | 76719 | ENSG00000112511 | PHF1 | chr6 | + | 33412697 | 33412793 | 33412507 | 33412589 | 33413195 | 33413294 | 162 | 95 | 218 | 189 | 172 | 0 | 0 | 1 | 1 | 0 | 283 | 203 | 288 | 204 | 405 | 13 | 0 | 3 | 6 | 7 | 245 | 149 | 0.000119409 | 0.0026181 | 1 | 1 | 0.993 | 0.991 | 1 | 0.93 | 1 | 0.983 | 0.954 | 0.972 | 0.029 |
| SE | 43998 | ENSG00000114098 | ARMC8 | chr3 | + | 138188400 | 138188530 | 138187317 | 138187599 | 138209816 | 138209893 | 10 | 12 | 10 | 22 | 15 | 20 | 1 | 23 | 34 | 28 | 28 | 28 | 18 | 23 | 25 | 0 | 6 | 38 | 1 | 0 | 279 | 149 | 0.000119474 | 0.002618799 | 0.211 | 0.865 | 0.188 | 0.257 | 0.222 | 1 | 0.714 | 0.202 | 0.925 | 1 | -0.42 |
| SE | 14956 | ENSG00000182718 | ANXA2 | chr15 | - | 60390212 | 60390317 | 60386027 | 60386086 | 60396372 | 60396427 | 5 | 9 | 6 | 2 | 15 | 0 | 1 | 0 | 0 | 0 | 28 | 8 | 4 | 20 | 17 | 2 | 2 | 2 | 1 | 3 | 254 | 149 | 0.000119511 | 0.002618863 | 1 | 0.841 | 1 | 1 | 1 | 0.891 | 0.701 | 0.54 | 0.921 | 0.769 | 0.204 |
| SE | 98178 | ENSG00000171634 | BPTF | chr17 | + | 67928958 | 67929120 | 67928354 | 67928601 | 67929335 | 67929487 | 42 | 23 | 35 | 57 | 74 | 309 | 226 | 202 | 242 | 372 | 50 | 35 | 32 | 22 | 52 | 68 | 247 | 167 | 77 | 66 | 298 | 149 | 0.000119891 | 0.002626474 | 0.064 | 0.048 | 0.08 | 0.105 | 0.09 | 0.269 | 0.066 | 0.087 | 0.125 | 0.283 | -0.089 |
| SE | 78294 | ENSG00000134882 | UBAC2 | chr13 | + | 99231433 | 99231513 | 99200891 | 99200939 | 99238426 | 99238554 | 0 | 0 | 0 | 0 | 0 | 24 | 8 | 19 | 24 | 7 | 0 | 6 | 0 | 0 | 7 | 17 | 16 | 12 | 9 | 14 | 229 | 149 | 0.000119973 | 0.002627527 | 0 | 0 | 0 | 0 | 0 | 0 | 0.196 | 0 | 0 | 0.245 | -0.088 |
| SE | 5041 | ENSG00000101126 | ADNP | chr20 | - | 50902016 | 50902109 | 50888915 | 50894512 | 50928650 | 50928735 | 122 | 116 | 157 | 138 | 299 | 0 | 0 | 0 | 0 | 0 | 84 | 118 | 96 | 96 | 56 | 20 | 0 | 0 | 0 | 0 | 242 | 149 | 0.000120012 | 0.002627641 | 1 | 1 | 1 | 1 | 1 | 0.721 | 1 | 1 | 1 | 1 | 0.056 |
| SE | 41010 | ENSG00000164091 | WDR82 | chr3 | - | 52260384 | 52260501 | 52259716 | 52259872 | 52261379 | 52261479 | 714 | 470 | 812 | 830 | 1008 | 2 | 1 | 3 | 1 | 0 | 666 | 534 | 503 | 428 | 679 | 22 | 4 | 2 | 2 | 10 | 266 | 149 | 0.000120286 | 0.002632905 | 0.995 | 0.996 | 0.993 | 0.998 | 1 | 0.944 | 0.987 | 0.993 | 0.992 | 0.974 | 0.018 |
| SE | 1197 | ENSG00000100227 | POLDIP3 | chr22 | - | 42597672 | 42597737 | 42596185 | 42596365 | 42599697 | 42599793 | 26 | 31 | 33 | 43 | 61 | 269 | 165 | 210 | 218 | 408 | 60 | 88 | 58 | 39 | 70 | 307 | 263 | 195 | 197 | 354 | 214 | 149 | 0.00012053 | 0.002637513 | 0.063 | 0.116 | 0.099 | 0.121 | 0.094 | 0.12 | 0.189 | 0.172 | 0.121 | 0.121 | -0.046 |
| SE | 12654 | ENSG00000196712 | NF1 | chr17 | + | 31330295 | 31330498 | 31327498 | 31327839 | 31334837 | 31335031 | 110 | 112 | 94 | 100 | 216 | 7 | 7 | 4 | 9 | 16 | 33 | 122 | 115 | 51 | 5 | 0 | 0 | 2 | 2 | 2 | 298 | 149 | 0.000120653 | 0.002639473 | 0.887 | 0.889 | 0.922 | 0.847 | 0.871 | 1 | 1 | 0.966 | 0.927 | 0.556 | -0.007 |
| SE | 24868 | ENSG00000185627 | PSMD13 | chr11 | + | 238997 | 239076 | 236965 | 237144 | 244040 | 244075 | 124 | 89 | 129 | 213 | 60 | 0 | 2 | 1 | 3 | 2 | 339 | 195 | 150 | 183 | 377 | 16 | 0 | 6 | 18 | 4 | 228 | 149 | 0.000120773 | 0.002641348 | 1 | 0.967 | 0.988 | 0.979 | 0.951 | 0.933 | 1 | 0.942 | 0.869 | 0.984 | 0.031 |
| SE | 28486 | ENSG00000155229 | MMS19 | chr10 | - | 97461495 | 97461622 | 97460906 | 97461007 | 97461827 | 97461896 | 34 | 20 | 39 | 36 | 55 | 4 | 2 | 4 | 5 | 1 | 27 | 14 | 14 | 20 | 19 | 0 | 0 | 1 | 2 | 0 | 276 | 149 | 0.00012093 | 0.002644045 | 0.821 | 0.844 | 0.84 | 0.795 | 0.967 | 1 | 1 | 0.983 | 0.844 | 1 | -0.092 |
| SE | 74683 | ENSG00000197548 | ATG7 | chr3 | + | 11282192 | 11282438 | 11280993 | 11281102 | 11298685 | 11298855 | 23 | 29 | 28 | 27 | 42 | 2 | 8 | 3 | 8 | 8 | 51 | 43 | 42 | 83 | 52 | 0 | 5 | 3 | 6 | 0 | 298 | 149 | 0.000121125 | 0.002647583 | 0.852 | 0.644 | 0.824 | 0.628 | 0.724 | 1 | 0.811 | 0.875 | 0.874 | 1 | -0.178 |
| SE | 96119 | ENSG00000166405 | RIC3 | chr11 | - | 8126658 | 8126807 | 8106055 | 8111137 | 8137377 | 8137471 | 19 | 21 | 54 | 23 | 44 | 0 | 0 | 2 | 0 | 0 | 9 | 27 | 34 | 5 | 5 | 0 | 8 | 3 | 0 | 0 | 298 | 149 | 0.00012154 | 0.002655897 | 1 | 1 | 0.931 | 1 | 1 | 1 | 0.628 | 0.85 | 1 | 1 | 0.091 |
| SE | 18425 | ENSG00000213928 | IRF9 | chr14 | + | 24162143 | 24162324 | 24161264 | 24161338 | 24162965 | 24163149 | 346 | 206 | 320 | 418 | 159 | 0 | 3 | 1 | 5 | 1 | 750 | 546 | 308 | 663 | 502 | 10 | 4 | 4 | 30 | 3 | 298 | 149 | 0.000121751 | 0.002659774 | 1 | 0.972 | 0.994 | 0.977 | 0.988 | 0.974 | 0.986 | 0.975 | 0.917 | 0.988 | 0.018 |
| SE | 94688 | ENSG00000244187 | TMEM141 | chr9 | + | 136791625 | 136791777 | 136791367 | 136791424 | 136791946 | 136792030 | 18 | 9 | 10 | 21 | 7 | 5 | 0 | 2 | 5 | 1 | 11 | 17 | 3 | 14 | 17 | 0 | 0 | 2 | 1 | 0 | 298 | 149 | 0.00012184 | 0.002660967 | 0.643 | | 0.714 | 0.677 | 0.778 | 1 | 1 | 0.429 | 0.875 | 1 | -0.098 |
| SE | 95809 | ENSG00000215126 | CBWD6 | chr9 | - | 41132227 | 41132357 | 41131305 | 41131840 | 41133504 | 41133570 | 61 | 54 | 65 | 79 | 72 | 2 | 6 | 4 | 2 | 0 | 10 | 42 | 77 | 49 | 10 | 5 | 7 | 14 | 8 | 0 | 279 | 149 | 0.000121905 | 0.002661651 | 0.942 | 0.828 | 0.897 | 0.955 | 1 | 0.516 | 0.762 | 0.746 | 0.766 | 1 | 0.166 |
| SE | 93577 | ENSG00000165525 | NEMF | chr14 | - | 49813987 | 49814050 | 49806020 | 49806133 | 49814753 | 49814807 | 274 | 240 | 225 | 257 | 293 | 5 | 11 | 1 | 7 | 12 | 26 | 139 | 229 | 84 | 34 | 0 | 4 | 0 | 0 | 0 | 212 | 149 | 0.000122873 | 0.002682035 | 0.975 | 0.939 | 0.994 | 0.963 | 0.945 | 1 | 0.961 | 1 | | 1 | -0.029 |
| SE | 86462 | ENSG00000215021 | PHB2 | chr12 | - | 6967170 | 6967238 | 6966423 | 6966500 | 6967891 | 6968021 | 173 | 90 | 157 | 148 | 380 | 0 | 2 | 1 | 0 | 1 | 225 | 90 | 101 | 126 | 179 | 16 | 0 | 6 | 0 | 0 | 217 | 149 | 0.000123455 | 0.00269398 | 1 | 0.969 | 0.991 | 1 | 0.996 | 0.906 | 1 | 1 | 0.935 | 1 | 0.023 |
| SE | 93936 | ENSG00000144524 | COPS7B | chr2 | + | 231794262 | 231794351 | 231791603 | 231791808 | 231796105 | 231796145 | 116 | 55 | 115 | 91 | 118 | 0 | 0 | 0 | 0 | 0 | 107 | 117 | 47 | 78 | 107 | 9 | 0 | 0 | 8 | 0 | 238 | 149 | 0.000123547 | 0.002694913 | 1 | 1 | 1 | 1 | 1 | 0.882 | 1 | 1 | 0.859 | 1 | 0.052 |
| SE | 98537 | ENSG00000130935 | NOL11 | chr17 | + | 67736665 | 67736754 | 67735899 | 67736023 | 67737070 | 67737145 | 395 | 217 | 380 | 298 | 630 | 16 | 16 | 13 | 31 | 20 | 9 | 80 | 161 | 74 | 11 | 22 | 45 | 14 | 20 | 0 | 238 | 149 | 0.000123566 | 0.002694913 | 0.939 | 0.895 | 0.948 | 0.858 | 0.952 | 0.204 | 0.527 | 0.878 | 0.698 | 1 | 0.257 |
| SE | 83914 | ENSG00000167778 | SPRYD3 | chr12 | - | 53074648 | 53074784 | 53073159 | 53073471 | 53075094 | 53075219 | 229 | 98 | 206 | 221 | 147 | 2 | 2 | 1 | 0 | 2 | 184 | 82 | 75 | 123 | 200 | 0 | 6 | 3 | 7 | 5 | 285 | 149 | 0.000123609 | 0.002695097 | 0.984 | 0.962 | 0.991 | 1 | 0.975 | 1 | 0.877 | 0.929 | 0.902 | 0.954 | 0.05 |
| SE | 48592 | ENSG00000131374 | TBC1D5 | chr3 | - | 17233684 | 17233750 | 17214206 | 17214370 | 17238162 | 17238419 | 8 | 7 | 8 | 8 | 17 | 53 | 15 | 40 | 58 | 83 | 0 | 4 | 0 | 0 | 6 | 37 | 19 | 27 | 53 | 20 | 215 | 149 | 0.000123808 | 0.002698326 | 0.095 | 0.244 | 0.122 | 0.087 | 0.124 | 0 | 0.127 | 0 | 0 | 0.172 | 0.075 |
| SE | 101256 | ENSG00000166224 | SGPL1 | chr10 | + | 70871836 | 70871986 | 70871047 | 70871146 | 70875401 | 70875548 | 83 | 21 | 56 | 54 | 85 | 0 | 0 | 1 | 0 | 0 | 101 | 36 | 52 | 42 | 63 | 0 | 0 | 0 | 5 | 9 | 298 | 149 | 0.000123838 | 0.002698326 | 1 | 1 | 0.966 | 1 | 1 | 1 | 1 | 1 | 0.808 | 0.778 | 0.076 |
| SE | 58311 | ENSG00000204261 | PSMB8-AS1 | chr6 | - | 32844809 | 32844870 | 32844130 | 32844274 | 32845417 | 32846492 | 71 | 17 | 30 | 71 | 19 | 83 | 51 | 66 | 61 | 74 | 35 | 32 | 20 | 9 | 40 | 158 | 98 | 42 | 70 | 249 | 210 | 149 | 0.000123861 | 0.002698326 | 0.378 | 0.191 | 0.244 | 0.452 | 0.154 | 0.136 | 0.188 | 0.253 | 0.084 | 0.102 | 0.131 |
| SE | 68346 | ENSG00000125630 | POLR1B | chr2 | + | 112547082 | 112547179 | 112542449 | 112542671 | 112547420 | 112547567 | 22 | 13 | 34 | 29 | 48 | 0 | 0 | 0 | 0 | 0 | 4 | 34 | 13 | 14 | 16 | 4 | 0 | 6 | 1 | 0 | 246 | 149 | 0.000124001 | 0.00270064 | 1 | 1 | 1 | 1 | 1 | 0.377 | 1 | 0.568 | 0.895 | 1 | 0.232 |
| SE | 1271 | ENSG00000186951 | PPARA | chr22 | + | 46175988 | 46176144 | 46151887 | 46151970 | 46176752 | 46176836 | 0 | 0 | 0 | 0 | 2 | 15 | 2 | 5 | 5 | 1 | 0 | 0 | 2 | 2 | 15 | 4 | 7 | 8 | 8 | 9 | 298 | 149 | 0.000124396 | 0.002707757 | 0 | 0 | 0 | 0 | 0.5 | 0 | 0 | 0.111 | 0.111 | 0.455 | -0.035 |
| SE | 16865 | ENSG00000138613 | APH1B | chr15 | + | 63302344 | 63302472 | 63286557 | 63286628 | 63305613 | 63305735 | 27 | 16 | 23 | 14 | 24 | 0 | 0 | 0 | 2 | 1 | 25 | 58 | 25 | 28 | 24 | 0 | 11 | 3 | 0 | 0 | 277 | 149 | 0.000124397 | 0.002707757 | 1 | 1 | 1 | 0.79 | 0.928 | 1 | 0.739 | 0.818 | 1 | 1 | 0.032 |
| SE | 57854 | ENSG00000173846 | PLK3 | chr1 | + | 44801621 | 44801751 | 44801035 | 44801152 | 44801844 | 44801932 | 36 | 18 | 54 | 61 | 35 | 0 | 0 | 0 | 0 | 0 | 62 | 63 | 130 | 71 | 63 | 16 | 0 | 2 | 0 | 0 | 279 | 149 | 0.000124709 | 0.00271378 | 1 | 1 | 1 | 1 | 1 | 0.674 | 1 | 0.972 | 1 | 1 | 0.071 |
| SE | 31937 | ENSG00000278259 | MYO19 | chr17 | - | 36533159 | 36533423 | 36532526 | 36532681 | 36533952 | 36534104 | 0 | 2 | 0 | 1 | 0 | 28 | 10 | 21 | 32 | 20 | 10 | 3 | 3 | 0 | 0 | 8 | 7 | 15 | 14 | 37 | 298 | 149 | 0.00012478 | 0.002714571 | 0 | 0.091 | 0 | 0.015 | 0 | 0.385 | 0.176 | 0.091 | 0 | 0 | -0.109 |
| SE | 94553 | ENSG00000132950 | ZMYM5 | chr13 | - | 19838699 | 19838985 | 19837464 | 19837821 | 19851354 | 19851448 | 79 | 114 | 85 | 115 | 315 | 4 | 1 | 1 | 3 | 4 | 38 | 96 | 92 | 79 | 28 | 9 | 3 | 1 | 7 | 4 | 298 | 149 | 0.000125268 | 0.002724433 | 0.908 | 0.983 | 0.977 | 0.95 | 0.975 | 0.679 | 0.941 | 0.979 | 0.849 | 0.778 | 0.113 |
| SE | 46150 | ENSG00000126070 | AGO3 | chr1 | + | 36005718 | 36005858 | 36004340 | 36004475 | 36008689 | 36008777 | 21 | 35 | 38 | 50 | 46 | 54 | 45 | 35 | 54 | 83 | 29 | 56 | 71 | 44 | 55 | 31 | 38 | 22 | 29 | 9 | 289 | 149 | 0.00012538 | 0.002726119 | 0.167 | 0.286 | 0.359 | 0.323 | 0.222 | 0.325 | 0.432 | 0.625 | 0.439 | 0.759 | -0.245 |
| SE | 97244 | ENSG00000073921 | PICALM | chr11 | - | 86022366 | 86022469 | 86014869 | 86014963 | 86031468 | 86031611 | 887 | 677 | 504 | 750 | 908 | 3 | 9 | 9 | 8 | 3 | 584 | 600 | 847 | 706 | 394 | 0 | 4 | 19 | 8 | 31 | 252 | 149 | 0.000125428 | 0.002726234 | 0.994 | 0.978 | 0.971 | 0.982 | 0.994 | 1 | 0.989 | 0.963 | 0.981 | 0.883 | 0.021 |
| SE | 56910 | ENSG00000175970 | UNC119B | chr12 | + | 120716627 | 120716739 | 120713273 | 120713387 | 120719919 | 120723640 | 45 | 20 | 29 | 38 | 74 | 0 | 0 | 0 | 0 | 0 | 40 | 44 | 15 | 15 | 67 | 0 | 0 | 0 | 12 | 5 | 261 | 149 | 0.000125455 | 0.002726234 | 1 | 1 | 1 | 1 | 1 | 1 | 1 | 1 | 0.416 | 0.884 | 0.14 |
| SE | 61330 | ENSG00000102786 | INTS6 | chr13 | - | 51423029 | 51423090 | 51421111 | 51421334 | 51430293 | 51430383 | 20 | 15 | 9 | 21 | 14 | 2 | 0 | 1 | 5 | 5 | 5 | 23 | 27 | 13 | 5 | 0 | 4 | 0 | 0 | 0 | 210 | 149 | 0.000125495 | 0.002726348 | 0.876 | 1 | 0.865 | 0.749 | 0.665 | 1 | 0.803 | 1 | 1 | 1 | -0.13 |
| SE | 43885 | ENSG00000179335 | CLK3 | chr15 | + | 74622119 | 74622560 | 74620008 | 74620225 | 74624901 | 74625018 | 137 | 75 | 118 | 135 | 213 | 3 | 0 | 1 | 9 | 3 | 160 | 177 | 178 | 207 | 236 | 10 | 22 | 10 | 1 | 16 | 298 | 149 | 0.000125628 | 0.002728484 | 0.958 | 1 | 0.983 | 0.882 | 0.973 | 0.889 | 0.801 | 0.899 | 0.99 | 0.881 | 0.067 |
| SE | 49448 | ENSG00000115020 | PIKFYVE | chr2 | + | 208276711 | 208276830 | 208271510 | 208271691 | 208285725 | 208285933 | 9 | 0 | 3 | 16 | 13 | 5 | 2 | 7 | 4 | 0 | 7 | 13 | 4 | 5 | 13 | 0 | 0 | 2 | 2 | 0 | 268 | 149 | 0.000125827 | 0.002732048 | 0.5 | 0 | 0.192 | 0.69 | 1 | 1 | 1 | 0.527 | 0.582 | 1 | -0.345 |
| SE | 13695 | ENSG00000077235 | GTF3C1 | chr16 | - | 27508539 | 27508655 | 27506846 | 27507156 | 27511748 | 27511901 | 197 | 96 | 141 | 170 | 226 | 0 | 0 | 2 | 1 | 0 | 226 | 220 | 188 | 131 | 268 | 13 | 2 | 6 | 3 | 0 | 265 | 149 | 0.000125955 | 0.002734056 | 1 | 1 | 0.975 | 0.99 | 1 | 0.907 | 0.984 | 0.946 | 0.961 | 1 | 0.033 |
| SE | 39493 | ENSG00000010322 | NISCH | chr3 | + | 52476446 | 52476599 | 52473733 | 52473829 | 52477573 | 52477642 | 186 | 69 | 200 | 162 | 180 | 1 | 1 | 0 | 1 | 4 | 201 | 286 | 127 | 211 | 284 | 0 | 0 | 2 | 0 | 38 | 298 | 149 | 0.00012646 | 0.002744258 | 0.989 | 0.972 | 1 | 0.988 | 0.957 | 1 | 1 | 0.969 | 1 | 0.789 | 0.03 |
| SE | 105612 | ENSG00000139323 | POC1B | chr12 | - | 89471613 | 89471729 | 89470360 | 89470494 | 89472167 | 89472275 | 279 | 199 | 223 | 311 | 618 | 0 | 2 | 1 | 8 | 2 | 128 | 181 | 131 | 238 | 177 | 13 | 5 | 0 | 2 | 5 | 265 | 149 | 0.000126632 | 0.002745897 | 1 | 0.982 | 0.992 | 0.956 | 0.994 | 0.847 | 0.953 | 1 | 0.985 | 0.952 | 0.037 |
| SE | 75553 | ENSG00000103363 | ELOB | chr16 | - | 2775450 | 2775625 | 2771940 | 2772102 | 2776992 | 2777127 | 424 | 163 | 436 | 334 | 191 | 1 | 2 | 3 | 0 | 1 | 473 | 518 | 206 | 396 | 474 | 17 | 0 | 0 | 4 | 14 | 298 | 149 | 0.000126635 | 0.002745897 | 0.995 | 0.976 | 0.986 | 1 | 0.99 | 0.933 | 1 | 1 | 0.98 | 0.944 | 0.018 |
| SE | 62917 | ENSG00000185420 | SMYD3 | chr1 | - | 245764040 | 245764149 | 245749341 | 245749664 | 245858495 | 245858670 | 58 | 46 | 21 | 55 | 67 | 1 | 0 | 2 | 0 | 0 | 53 | 39 | 18 | 26 | 81 | 16 | 0 | 1 | 0 | 0 | 258 | 149 | 0.00012664 | 0.002745897 | 0.971 | 1 | 0.858 | 1 | 1 | 0.657 | 1 | 0.912 | 1 | 1 | 0.052 |
| SE | 42508 | ENSG00000176444 | CLK2 | chr1 | - | 155268707 | 155268792 | 155268292 | 155268359 | 155269487 | 155269713 | 73 | 49 | 38 | 49 | 23 | 9 | 0 | 1 | 3 | 3 | 49 | 34 | 7 | 29 | 33 | 6 | 20 | 4 | 1 | 20 | 234 | 149 | 0.00012672 | 0.002746493 | 0.838 | 1 | 0.96 | 0.912 | 0.83 | 0.839 | 0.52 | 0.527 | 0.949 | 0.512 | 0.239 |
| SE | 74130 | ENSG00000093167 | LRRFIP2 | chr3 | - | 37121634 | 37121691 | 37121491 | 37121536 | 37129062 | 37129149 | 15 | 9 | 18 | 9 | 26 | 1 | 0 | 1 | 1 | 0 | 90 | 26 | 32 | 33 | 62 | 17 | 0 | 2 | 4 | 0 | 206 | 149 | 0.000126738 | 0.002746493 | 0.916 | 1 | 0.929 | 0.867 | 1 | 0.793 | 1 | 0.92 | 0.856 | 1 | 0.029 |
| SE | 48762 | ENSG00000186432 | KPNA4 | chr3 | - | 160564504 | 160564640 | 160536795 | 160536840 | 160565213 | 160565389 | 0 | 3 | 2 | 0 | 0 | 95 | 74 | 81 | 85 | 135 | 0 | 8 | 2 | 4 | 8 | 8 | 68 | 79 | 40 | 4 | 285 | 149 | 0.000126908 | 0.002749422 | 0 | 0.021 | 0.013 | 0 | 0 | 0 | 0.058 | 0.013 | 0.05 | 0.511 | -0.12 |
| SE | 22836 | ENSG00000107968 | MAP3K8 | chr10 | + | 30451637 | 30451744 | 30450257 | 30450519 | 30458083 | 30458236 | 379 | 228 | 178 | 289 | 334 | 2 | 0 | 3 | 2 | 0 | 385 | 321 | 277 | 339 | 113 | 19 | 0 | 6 | 5 | 0 | 256 | 149 | 0.000127051 | 0.002751743 | 0.991 | 1 | 0.972 | 0.988 | 1 | 0.922 | 1 | 0.964 | 0.975 | 1 | 0.018 |
| SE | 22813 | ENSG00000173692 | PSMD1 | chr2 | + | 231079535 | 231079614 | 231078658 | 231078747 | 231080140 | 231080314 | 275 | 213 | 248 | 259 | 367 | 0 | 2 | 2 | 0 | 4 | 200 | 224 | 186 | 296 | 42 | 0 | 9 | 6 | 10 | 0 | 228 | 149 | 0.000127602 | 0.002762255 | 1 | 0.986 | 0.988 | 1 | 0.984 | 1 | 0.942 | 0.953 | 0.951 | 1 | 0.022 |
| SE | 65358 | ENSG00000153214 | TMEM87B | chr2 | + | 112074911 | 112074962 | 112066935 | 112067067 | 112077191 | 112077282 | 246 | 156 | 188 | 227 | 386 | 3 | 4 | 1 | 3 | 7 | 134 | 204 | 87 | 112 | 98 | 29 | 2 | 1 | 4 | 0 | 200 | 149 | 0.000127606 | 0.002762255 | 0.984 | 0.967 | 0.993 | 0.983 | 0.976 | 0.775 | 0.987 | 0.985 | 0.954 | 1 | 0.04 |
| SE | 40073 | ENSG00000125347 | IRF1 | chr5 | - | 132486250 | 132486348 | 132485666 | 132485716 | 132486556 | 132486686 | 306 | 144 | 327 | 323 | 277 | 29 | 21 | 44 | 29 | 45 | 751 | 170 | 267 | 372 | 417 | 134 | 91 | 39 | 142 | 71 | 247 | 149 | 0.000127706 | 0.002763645 | 0.864 | 0.805 | 0.818 | 0.87 | 0.788 | 0.772 | 0.53 | 0.805 | 0.612 | 0.78 | 0.129 |
| SE | 22089 | ENSG00000111859 | NEDD9 | chr6 | - | 11249138 | 11249214 | 11213721 | 11213727 | 11305991 | 11306155 | 1 | 8 | 4 | 11 | 18 | 229 | 138 | 136 | 151 | 147 | 28 | 7 | 48 | 25 | 10 | 295 | 214 | 189 | 212 | 192 | 225 | 149 | 0.000128042 | 0.002770143 | 0.003 | 0.037 | 0.019 | 0.046 | 0.075 | 0.059 | 0.021 | 0.144 | 0.072 | 0.033 | -0.03 |
| SE | 62288 | ENSG00000112031 | MTRF1L | chr6 | - | 152995243 | 152995319 | 152994512 | 152994676 | 152998549 | 152998629 | 19 | 25 | 20 | 23 | 9 | 0 | 0 | 0 | 0 | 0 | 14 | 41 | 9 | 13 | 31 | 6 | 0 | 2 | 3 | 0 | 225 | 149 | 0.000128101 | 0.00277045 | 1 | 1 | 1 | 1 | 1 | 0.607 | 1 | 0.749 | 0.742 | 1 | 0.18 |
| SE | 58240 | ENSG00000089127 | OAS1 | chr12 | + | 112908535 | 112908824 | 112906652 | 112907219 | 112911097 | 112911235 | 310 | 242 | 183 | 317 | 302 | 1 | 1 | 3 | 6 | 6 | 1061 | 625 | 175 | 949 | 367 | 18 | 24 | 6 | 25 | 15 | 298 | 149 | 0.000128146 | 0.00277045 | 0.994 | 0.992 | 0.968 | 0.964 | 0.962 | 0.967 | 0.929 | 0.936 | 0.95 | 0.924 | 0.035 |
| SE | 75743 | ENSG00000144580 | CNOT9 | chr2 | + | 218584611 | 218584721 | 218582970 | 218583086 | 218587585 | 218587695 | 179 | 124 | 200 | 184 | 177 | 6 | 1 | 16 | 5 | 5 | 108 | 142 | 89 | 123 | 146 | 20 | 8 | 37 | 16 | 0 | 259 | 149 | 0.000128162 | 0.00277045 | 0.945 | 0.986 | 0.878 | 0.955 | 0.953 | 0.756 | 0.911 | 0.581 | 0.816 | 1 | 0.131 |
| SE | 31574 | ENSG00000116668 | SWT1 | chr1 | + | 185214506 | 185214655 | 185206624 | 185206763 | 185231576 | 185231708 | 42 | 24 | 17 | 21 | 96 | 0 | 0 | 0 | 0 | 0 | 3 | 9 | 17 | 4 | 47 | 0 | 0 | 0 | 0 | 12 | 298 | 149 | 0.000128287 | 0.002772389 | 1 | 1 | 1 | 1 | 1 | 1 | 1 | 1 | 1 | 0.662 | 0.068 |
| SE | 30136 | ENSG00000150961 | SEC24D | chr4 | - | 118766528 | 118766823 | 118764801 | 118764917 | 118768172 | 118768311 | 8 | 5 | 9 | 4 | 2 | 140 | 63 | 102 | 102 | 120 | 7 | 6 | 16 | 12 | 8 | 51 | 111 | 84 | 87 | 31 | 298 | 149 | 0.000128431 | 0.002774194 | 0.028 | 0.038 | 0.042 | 0.019 | 0.008 | 0.064 | 0.026 | 0.087 | 0.065 | 0.114 | -0.044 |
| SE | 53056 | ENSG00000103657 | HERC1 | chr15 | - | 63648068 | 63648199 | 63645482 | 63645682 | 63649724 | 63649925 | 516 | 396 | 468 | 571 | 359 | 0 | 0 | 5 | 5 | 2 | 226 | 360 | 395 | 387 | 114 | 0 | 19 | 9 | 1 | 0 | 280 | 149 | 0.000128441 | 0.002774194 | 1 | 1 | 0.98 | 0.984 | 0.99 | 1 | 0.91 | 0.959 | 0.995 | 1 | 0.018 |
| SE | 103818 | ENSG00000225484 | NUTM2B-AS1 | chr10 | - | 79808758 | 79808963 | 79806038 | 79806122 | 79814612 | 79814716 | 79 | 49 | 73 | 72 | 99 | 28 | 5 | 8 | 11 | 8 | 179 | 119 | 98 | 104 | 149 | 11 | 7 | 12 | 5 | 0 | 298 | 149 | 0.000128484 | 0.002774298 | 0.585 | 0.831 | 0.82 | 0.766 | 0.861 | 0.891 | 0.895 | 0.803 | 0.912 | 1 | -0.128 |
| SE | 34763 | ENSG00000112149 | CD83 | chr6 | + | 14131519 | 14131748 | 14117949 | 14118065 | 14133648 | 14133755 | 30 | 5 | 13 | 30 | 39 | 0 | 0 | 0 | 3 | 0 | 59 | 18 | 62 | 122 | 90 | 9 | 4 | 1 | 7 | 0 | 298 | 149 | 0.000128517 | 0.002774298 | 1 | 1 | 1 | 0.833 | 1 | 0.766 | 0.692 | 0.969 | 0.897 | 1 | 0.102 |
| SE | 15026 | ENSG00000091157 | WDR7 | chr18 | + | 56938532 | 56938682 | 56935787 | 56935905 | 56939310 | 56939393 | 152 | 103 | 163 | 153 | 239 | 0 | 0 | 0 | 0 | 0 | 18 | 114 | 102 | 99 | 79 | 0 | 0 | 0 | 0 | 24 | 298 | 149 | 0.000128687 | 0.002776829 | 1 | 1 | 1 | 1 | 1 | 1 | 1 | 1 | 1 | 0.622 | 0.076 |
| SE | 106155 | ENSG00000089022 | MAPKAPK5 | chr12 | + | 111882789 | 111882853 | 111880446 | 111880527 | 111883580 | 111883768 | 11 | 9 | 13 | 1 | 13 | 63 | 70 | 60 | 98 | 131 | 20 | 5 | 26 | 32 | 16 | 40 | 81 | 37 | 71 | 128 | 213 | 149 | 0.000128705 | 0.002776829 | 0.109 | 0.083 | 0.132 | 0.007 | 0.065 | 0.259 | 0.041 | 0.33 | 0.24 | 0.08 | -0.111 |
| SE | 104300 | ENSG00000135655 | USP15 | chr12 | + | 62349220 | 62349307 | 62325871 | 62325933 | 62355330 | 62355475 | 524 | 401 | 313 | 307 | 997 | 89 | 67 | 50 | 60 | 136 | 79 | 201 | 99 | 40 | 56 | 0 | 15 | 4 | 9 | 7 | 236 | 149 | 0.000129067 | 0.002783867 | 0.788 | 0.791 | 0.798 | 0.764 | 0.822 | 1 | 0.894 | 0.94 | 0.737 | 0.835 | -0.089 |
| SE | 25551 | ENSG00000157916 | RER1 | chr1 | + | 2397115 | 2397220 | 2394004 | 2395871 | 2400856 | 2400935 | 160 | 83 | 136 | 144 | 159 | 0 | 0 | 0 | 0 | 0 | 86 | 115 | 70 | 61 | 167 | 0 | 3 | 2 | 0 | 12 | 254 | 149 | 0.00012918 | 0.00278554 | 1 | 1 | 1 | 1 | 1 | 1 | 0.957 | 0.954 | 1 | 0.891 | 0.04 |
| SE | 2178 | ENSG00000079805 | DNM2 | chr19 | + | 10797379 | 10797518 | 10796060 | 10796199 | 10798485 | 10798572 | 236 | 160 | 315 | 299 | 510 | 376 | 230 | 315 | 399 | 408 | 892 | 580 | 526 | 575 | 829 | 902 | 478 | 238 | 366 | 654 | 288 | 149 | 0.000129226 | 0.002785758 | 0.245 | 0.265 | 0.341 | 0.279 | 0.393 | 0.338 | 0.386 | 0.533 | 0.448 | 0.396 | -0.116 |
| SE | 107356 | ENSG00000110921 | MVK | chr12 | + | 109579801 | 109579946 | 109575997 | 109576145 | 109586021 | 109586125 | 25 | 12 | 21 | 23 | 11 | 3 | 1 | 1 | 3 | 3 | 16 | 17 | 3 | 2 | 6 | 0 | 7 | 7 | 10 | 29 | 294 | 149 | 0.000130031 | 0.002802333 | 0.809 | 0.859 | 0.914 | 0.795 | 0.65 | 1 | 0.552 | 0.178 | 0.092 | 0.095 | 0.422 |
| SE | 3972 | ENSG00000126261 | UBA2 | chr19 | + | 34430092 | 34430186 | 34428351 | 34428570 | 34430575 | 34430659 | 13 | 8 | 15 | 9 | 11 | 59 | 26 | 67 | 59 | 42 | 15 | 0 | 5 | 1 | 0 | 45 | 51 | 62 | 51 | 34 | 243 | 149 | 0.000130118 | 0.002803446 | 0.119 | 0.159 | 0.121 | 0.086 | 0.138 | 0.17 | 0 | 0.047 | 0.012 | 0 | 0.079 |
| SE | 66856 | ENSG00000082781 | ITGB5 | chr3 | - | 124817620 | 124817710 | 124809021 | 124809156 | 124819738 | 124819834 | 98 | 135 | 35 | 207 | 122 | 2 | 5 | 3 | 7 | 7 | 507 | 219 | 145 | 236 | 1651 | 0 | 3 | 4 | 4 | 19 | 239 | 149 | 0.000130635 | 0.002813807 | 0.968 | 0.944 | 0.879 | 0.949 | 0.916 | 1 | 0.978 | 0.958 | 0.974 | 0.982 | -0.047 |
| SE | 7989 | ENSG00000126858 | RHOT1 | chr17 | + | 32172936 | 32173060 | 32171042 | 32171101 | 32173820 | 32173912 | 46 | 32 | 32 | 37 | 52 | 64 | 42 | 48 | 71 | 51 | 68 | 72 | 60 | 77 | 38 | 24 | 56 | 35 | 49 | 45 | 273 | 149 | 0.000131207 | 0.002825346 | 0.282 | 0.294 | 0.267 | 0.221 | 0.358 | 0.607 | 0.412 | 0.483 | 0.462 | 0.315 | -0.171 |
| SE | 57991 | ENSG00000175634 | RPS6KB2 | chr11 | + | 67431367 | 67431515 | 67429526 | 67429595 | 67432599 | 67432657 | 164 | 84 | 155 | 130 | 135 | 2 | 2 | 2 | 2 | 4 | 292 | 194 | 123 | 161 | 272 | 5 | 13 | 19 | 5 | 8 | 297 | 149 | 0.000131253 | 0.002825572 | 0.976 | 0.955 | 0.975 | 0.97 | 0.944 | 0.967 | 0.882 | 0.765 | 0.942 | 0.945 | 0.064 |
| SE | 8530 | ENSG00000125450 | NUP85 | chr17 | + | 75218184 | 75218306 | 75215753 | 75215823 | 75225102 | 75225237 | 65 | 35 | 58 | 61 | 55 | 1 | 1 | 0 | 4 | 0 | 58 | 46 | 67 | 26 | 49 | 0 | 4 | 1 | 8 | 8 | 271 | 149 | 0.000131707 | 0.002834565 | 0.973 | 0.951 | 1 | 0.893 | 1 | 1 | 0.863 | 0.974 | 0.641 | 0.771 | 0.114 |
| SE | 89871 | ENSG00000034677 | RNF19A | chr8 | - | 100259853 | 100259997 | 100257059 | 100259246 | 100264670 | 100264785 | 310 | 208 | 220 | 270 | 437 | 1 | 3 | 2 | 5 | 2 | 56 | 392 | 643 | 178 | 35 | 0 | 19 | 12 | 4 | 0 | 293 | 149 | 0.000131797 | 0.002835712 | 0.994 | 0.972 | 0.982 | 0.965 | 0.991 | 1 | 0.913 | 0.965 | 0.958 | 1 | 0.014 |
| SE | 94261 | ENSG00000157954 | WIPI2 | chr7 | + | 5193117 | 5193345 | 5190258 | 5190493 | 5199575 | 5199658 | 2 | 1 | 2 | 6 | 0 | 62 | 33 | 47 | 36 | 27 | 15 | 8 | 0 | 5 | 18 | 102 | 82 | 30 | 41 | 43 | 298 | 149 | 0.000132034 | 0.002840038 | 0.016 | 0.015 | 0.021 | 0.077 | 0 | 0.068 | 0.047 | 0 | 0.057 | 0.173 | -0.043 |
| SE | 77714 | ENSG00000136205 | TNS3 | chr7 | - | 47302949 | 47303584 | 47302185 | 47302272 | 47304831 | 47305003 | 29 | 10 | 14 | 18 | 21 | 0 | 0 | 0 | 0 | 0 | 105 | 62 | 19 | 61 | 142 | 17 | 0 | 0 | 0 | 0 | 298 | 149 | 0.000132176 | 0.0028423 | 1 | 1 | 1 | 1 | 1 | 0.755 | 1 | 1 | 1 | 1 | 0.049 |
| SE | 103577 | ENSG00000150093 | ITGB1 | chr10 | + | 32933350 | 32933583 | 32932514 | 32932600 | 32935491 | 32935558 | 47 | 32 | 9 | 37 | 39 | 844 | 536 | 369 | 868 | 892 | 16 | 66 | 24 | 16 | 26 | 185 | 564 | 527 | 292 | 282 | 298 | 149 | 0.00013301 | 0.002859457 | 0.027 | 0.029 | 0.012 | 0.021 | 0.021 | 0.041 | 0.055 | 0.022 | 0.027 | 0.044 | -0.016 |
| SE | 76138 | ENSG00000135486 | HNRNPA1 | chr12 | + | 54283078 | 54283234 | 54282572 | 54282665 | 54283811 | 54283967 | 44 | 20 | 27 | 36 | 22 | 1 | 2 | 3 | 1 | 1 | 31 | 18 | 22 | 33 | 14 | 7 | 18 | 4 | 8 | 0 | 298 | 149 | 0.000133678 | 0.002873035 | 0.957 | 0.833 | 0.818 | 0.947 | 0.917 | 0.689 | 0.333 | 0.733 | 0.673 | 1 | 0.209 |
| SE | 104687 | ENSG00000100354 | TNRC6B | chr22 | + | 40308511 | 40308649 | 40301149 | 40301333 | 40312504 | 40312651 | 55 | 40 | 69 | 93 | 74 | 3 | 0 | 3 | 5 | 0 | 42 | 159 | 94 | 70 | 16 | 0 | 13 | 14 | 11 | 20 | 287 | 149 | 0.000133906 | 0.002877147 | 0.905 | 1 | 0.923 | 0.906 | 1 | 1 | 0.864 | 0.777 | 0.768 | 0.293 | 0.206 |
| SE | 108270 | ENSG00000139132 | FGD4 | chr12 | + | 32556873 | 32556995 | 32485971 | 32486143 | 32564136 | 32564289 | 1 | 0 | 0 | 0 | 5 | 52 | 74 | 23 | 62 | 46 | 8 | 0 | 4 | 3 | 9 | 5 | 77 | 128 | 54 | 14 | 271 | 149 | 0.000134336 | 0.002885598 | 0.01 | 0 | 0 | 0 | 0.056 | 0.468 | 0 | 0.017 | 0.03 | 0.261 | -0.142 |
| SE | 67013 | ENSG00000143373 | ZNF687 | chr1 | + | 151289094 | 151289271 | 151288527 | 151288706 | 151289377 | 151289540 | 41 | 21 | 36 | 26 | 16 | 2 | 0 | 0 | 0 | 0 | 59 | 58 | 62 | 55 | 111 | 0 | 0 | 2 | 10 | 5 | 298 | 149 | 0.000134451 | 0.002887271 | 0.911 | 1 | 1 | 1 | 1 | 1 | 1 | 0.939 | 0.733 | 0.917 | 0.064 |
| SE | 12447 | ENSG00000103043 | VAC14 | chr16 | - | 70786214 | 70786365 | 70785701 | 70785869 | 70800796 | 70800982 | 62 | 33 | 59 | 54 | 41 | 2 | 5 | 3 | 2 | 1 | 35 | 52 | 52 | 55 | 94 | 0 | 0 | 2 | 0 | 0 | 298 | 149 | 0.000134756 | 0.00289302 | 0.939 | 0.767 | 0.908 | 0.931 | 0.953 | 1 | 1 | 0.929 | 1 | 1 | -0.086 |
| SE | 43886 | ENSG00000179335 | CLK3 | chr15 | + | 74622493 | 74622560 | 74620008 | 74620225 | 74624901 | 74625018 | 94 | 43 | 73 | 91 | 161 | 3 | 0 | 1 | 9 | 3 | 129 | 113 | 114 | 141 | 140 | 10 | 22 | 10 | 1 | 16 | 216 | 149 | 0.000135009 | 0.002897652 | 0.956 | 1 | 0.981 | 0.875 | 0.974 | 0.899 | 0.78 | 0.887 | 0.99 | 0.858 | 0.074 |
| SE | 46007 | ENSG00000170322 | NFRKB | chr11 | - | 129876720 | 129876895 | 129875356 | 129875463 | 129878308 | 129878355 | 35 | 16 | 54 | 38 | 26 | 0 | 0 | 0 | 0 | 4 | 83 | 26 | 42 | 40 | 59 | 8 | 2 | 0 | 0 | 9 | 298 | 149 | 0.000135156 | 0.002899769 | 1 | 1 | 1 | 1 | 0.765 | 0.838 | 0.867 | 1 | 1 | 0.766 | 0.059 |
| SE | 21216 | ENSG00000150403 | TMCO3 | chr13 | + | 113510237 | 113510424 | 113503495 | 113503588 | 113520615 | 113520661 | 108 | 88 | 103 | 121 | 87 | 5 | 2 | 2 | 6 | 8 | 231 | 80 | 66 | 148 | 214 | 0 | 5 | 0 | 1 | 3 | 298 | 149 | 0.000135181 | 0.002899769 | 0.915 | 0.957 | 0.963 | 0.91 | 0.845 | 1 | 0.889 | 1 | 0.987 | 0.973 | -0.052 |
| SE | 103900 | ENSG00000066084 | DIP2B | chr12 | + | 50683137 | 50683248 | 50680671 | 50680763 | 50685832 | 50685956 | 218 | 173 | 272 | 235 | 389 | 2 | 0 | 2 | 5 | 0 | 374 | 219 | 281 | 382 | 87 | 0 | 13 | 3 | 9 | 6 | 260 | 149 | 0.000135239 | 0.002900212 | 0.984 | 1 | 0.987 | 0.964 | 1 | 1 | 0.906 | 0.982 | 0.961 | 0.893 | 0.039 |
| SE | 5196 | ENSG00000124214 | STAU1 | chr20 | - | 49165996 | 49166285 | 49158938 | 49159146 | 49174194 | 49174269 | 6 | 7 | 15 | 6 | 6 | 3 | 4 | 6 | 5 | 2 | 7 | 11 | 5 | 30 | 1 | 22 | 26 | 13 | 30 | 6 | 298 | 149 | 0.000135446 | 0.002903847 | 0.5 | 0.467 | 0.556 | 0.375 | 0.6 | 0.137 | 0.175 | 0.161 | 0.333 | 0.077 | 0.323 |
| SE | 88784 | ENSG00000140632 | GLYR1 | chr16 | - | 4813736 | 4813955 | 4811975 | 4812248 | 4814536 | 4814629 | 46 | 45 | 62 | 82 | 32 | 2 | 0 | 1 | 0 | 0 | 93 | 81 | 33 | 81 | 106 | 16 | 0 | 2 | 1 | 0 | 298 | 149 | 0.000136138 | 0.002917888 | 0.92 | 1 | 0.969 | 1 | 1 | 0.744 | 1 | 0.892 | 0.976 | 1 | 0.055 |
| SE | 85835 | ENSG00000134287 | ARF3 | chr12 | - | 48939996 | 48940107 | 48939654 | 48939779 | 48940947 | 48941188 | 654 | 438 | 443 | 646 | 982 | 6 | 4 | 1 | 1 | 1 | 1419 | 822 | 436 | 816 | 2505 | 10 | 7 | 13 | 10 | 34 | 260 | 149 | 0.000136283 | 0.002920204 | 0.984 | 0.984 | 0.996 | 0.997 | 0.998 | 0.988 | 0.985 | 0.951 | 0.979 | 0.977 | 0.016 |
| SE | 98285 | ENSG00000068831 | RASGRP2 | chr11 | - | 64740367 | 64740446 | 64740125 | 64740163 | 64740947 | 64741079 | 10 | 4 | 21 | 14 | 6 | 114 | 67 | 109 | 137 | 133 | 3 | 12 | 4 | 0 | 0 | 44 | 137 | 65 | 63 | 175 | 228 | 149 | 0.000136395 | 0.002921802 | 0.054 | 0.038 | 0.112 | 0.063 | 0.029 | 0.043 | 0.054 | 0.039 | 0 | 0 | 0.032 |
| SE | 38039 | ENSG00000215908 | CROCCP2 | chr1 | - | 16621945 | 16622115 | 16619844 | 16620024 | 16625774 | 16626005 | 0 | 0 | 0 | 3 | 0 | 19 | 7 | 4 | 21 | 5 | 1 | 14 | 0 | 0 | 0 | 5 | 31 | 10 | 4 | 4 | 298 | 149 | 0.00013656 | 0.002923826 | 0 | 0 | 0 | 0.067 | 0 | 0.091 | 0.184 | 0 | 0 | 0 | -0.042 |
| SE | 88744 | ENSG00000205581 | HMGN1 | chr21 | - | 39345829 | 39345958 | 39345145 | 39345274 | 39347246 | 39347292 | 234 | 301 | 344 | 323 | 342 | 3 | 0 | 2 | 2 | 0 | 61 | 215 | 239 | 191 | 80 | 4 | 1 | 0 | 11 | 4 | 278 | 149 | 0.000136564 | 0.002923826 | 0.977 | 1 | 0.989 | 0.989 | 1 | 0.891 | 0.991 | 1 | 0.903 | 0.915 | 0.051 |
| SE | 12696 | ENSG00000155666 | KDM8 | chr16 | + | 27213584 | 27213751 | 27210092 | 27210621 | 27214875 | 27215008 | 21 | 8 | 37 | 12 | 52 | 5 | 0 | 7 | 1 | 0 | 54 | 12 | 21 | 26 | 56 | 0 | 0 | 4 | 0 | 0 | 298 | 149 | 0.000136813 | 0.002928351 | 0.677 | 1 | 0.725 | 0.857 | 1 | 1 | 1 | 0.724 | 1 | 1 | -0.093 |
| SE | 17785 | ENSG00000100485 | SOS2 | chr14 | - | 50138611 | 50138784 | 50134122 | 50134239 | 50139941 | 50140059 | 433 | 372 | 294 | 324 | 786 | 1 | 5 | 4 | 6 | 10 | 33 | 278 | 277 | 157 | 21 | 6 | 6 | 14 | 5 | 0 | 298 | 149 | 0.00013688 | 0.002928993 | 0.995 | 0.974 | 0.974 | 0.964 | 0.975 | 0.733 | 0.959 | 0.908 | 0.94 | 1 | 0.068 |
| SE | 89876 | ENSG00000034677 | RNF19A | chr8 | - | 100261541 | 100261755 | 100259853 | 100259997 | 100264033 | 100264195 | 408 | 319 | 259 | 268 | 292 | 5 | 5 | 3 | 20 | 0 | 67 | 285 | 578 | 108 | 55 | 6 | 12 | 34 | 7 | 0 | 298 | 149 | 0.000137309 | 0.00293736 | 0.976 | 0.97 | 0.977 | 0.87 | 1 | 0.848 | 0.922 | 0.895 | 0.885 | 1 | 0.049 |
| SE | 60758 | ENSG00000141524 | TMC6 | chr17 | - | 78120408 | 78120555 | 78119296 | 78119392 | 78120652 | 78120719 | 15 | 0 | 16 | 0 | 27 | 205 | 80 | 135 | 227 | 81 | 15 | 0 | 0 | 0 | 0 | 37 | 189 | 56 | 69 | 192 | 296 | 149 | 0.000137714 | 0.002945226 | 0.036 | 0 | 0.056 | 0 | 0.144 | 0.169 | 0 | 0 | 0 | 0 | 0.013 |
| SE | 104593 | ENSG00000069493 | CLEC2D | chr12 | + | 9693415 | 9693948 | 9692827 | 9692931 | 9694759 | 9695138 | 109 | 99 | 124 | 139 | 123 | 375 | 239 | 353 | 349 | 583 | 5 | 53 | 35 | 18 | 0 | 35 | 208 | 175 | 60 | 72 | 298 | 149 | 0.000138351 | 0.002958045 | 0.127 | 0.172 | 0.149 | 0.166 | 0.095 | 0.067 | 0.113 | 0.091 | 0.13 | 0 | 0.062 |
| SE | 46500 | ENSG00000109184 | DCUN1D4 | chr4 | + | 51887122 | 51887244 | 51886567 | 51886638 | 51891759 | 51891851 | 2 | 6 | 10 | 6 | 4 | 74 | 65 | 44 | 60 | 108 | 0 | 0 | 1 | 1 | 0 | 21 | 29 | 18 | 9 | 57 | 271 | 149 | 0.000138739 | 0.002965528 | 0.015 | 0.048 | 0.111 | 0.052 | 0.02 | 0 | 0 | 0.03 | 0.058 | 0 | 0.032 |
| SE | 76506 | ENSG00000153406 | NMRAL1 | chr16 | - | 4474092 | 4474166 | 4469226 | 4469465 | 4474553 | 4474625 | 80 | 45 | 71 | 56 | 16 | 0 | 0 | 0 | 0 | 0 | 166 | 109 | 39 | 58 | 183 | 20 | 0 | 0 | 0 | 0 | 223 | 149 | 0.000139013 | 0.002970563 | 1 | 1 | 1 | 1 | 1 | 0.847 | 1 | 1 | 1 | 1 | 0.031 |
| SE | 43996 | ENSG00000160271 | RALGDS | chr9 | - | 133104447 | 133104551 | 133104262 | 133104331 | 133105931 | 133106016 | 7 | 2 | 4 | 3 | 3 | 130 | 77 | 97 | 113 | 142 | 8 | 5 | 17 | 3 | 16 | 174 | 121 | 77 | 68 | 122 | 253 | 149 | 0.000139253 | 0.002974877 | 0.031 | 0.015 | 0.024 | 0.015 | 0.012 | 0.026 | 0.024 | 0.115 | 0.025 | 0.072 | -0.033 |
| SE | 27465 | ENSG00000008513 | ST3GAL1 | chr8 | - | 133499134 | 133499189 | 133475718 | 133476600 | 133545773 | 133545926 | 148 | 136 | 177 | 156 | 426 | 2 | 6 | 2 | 4 | 3 | 112 | 164 | 60 | 109 | 172 | 24 | 1 | 6 | 0 | 4 | 204 | 149 | 0.00013959 | 0.002981261 | 0.982 | 0.943 | 0.985 | 0.966 | 0.99 | 0.773 | 0.992 | 0.88 | 1 | 0.969 | 0.05 |
| SE | 76207 | ENSG00000146070 | PLA2G7 | chr6 | - | 46714459 | 46714553 | 46712268 | 46712337 | 46716383 | 46716528 | 122 | 72 | 79 | 51 | 99 | 4 | 12 | 9 | 2 | 3 | 321 | 18 | 16 | 130 | 16 | 9 | 0 | 2 | 0 | 0 | 243 | 149 | 0.000139839 | 0.002985764 | 0.949 | 0.786 | 0.843 | 0.94 | 0.953 | 0.956 | 1 | 0.831 | 1 | 1 | -0.063 |
| SE | 2362 | ENSG00000104880 | ARHGEF18 | chr19 | + | 7376642 | 7376757 | 7375719 | 7375870 | 7378393 | 7378451 | 338 | 189 | 374 | 270 | 251 | 0 | 0 | 2 | 3 | 6 | 244 | 422 | 163 | 153 | 487 | 4 | 7 | 4 | 11 | 8 | 264 | 149 | 0.000139915 | 0.002986113 | 1 | 1 | 0.991 | 0.981 | 0.959 | 0.972 | 0.971 | 0.958 | 0.887 | 0.972 | 0.034 |
| SE | 52649 | ENSG00000197943 | PLCG2 | chr16 | + | 81934428 | 81934531 | 81931260 | 81931654 | 81936168 | 81936210 | 382 | 161 | 230 | 265 | 289 | 5 | 0 | 0 | 4 | 0 | 279 | 427 | 221 | 245 | 316 | 0 | 5 | 6 | 20 | 2 | 252 | 149 | 0.000139957 | 0.002986113 | 0.978 | | 1 | 0.975 | 1 | 1 | 0.981 | 0.956 | 0.879 | 0.989 | 0.03 |
| SE | 33128 | ENSG00000172053 | QARS1 | chr3 | - | 49102418 | 49102448 | 49102204 | 49102265 | 49103344 | 49103706 | 16 | 12 | 17 | 16 | 41 | 0 | 0 | 0 | 0 | 0 | 38 | 12 | 10 | 9 | 40 | 7 | 0 | 0 | 0 | 5 | 179 | 149 | 0.000139969 | 0.002986113 | 1 | 1 | 1 | 1 | 1 | 0.819 | 1 | 1 | 1 | 0.869 | 0.062 |
| SE | 93759 | ENSG00000143409 | MINDY1 | chr1 | - | 151002164 | 151002968 | 151001724 | 151001782 | 151006311 | 151006404 | 80 | 65 | 84 | 133 | 126 | 6 | 5 | 4 | 0 | 2 | 308 | 158 | 77 | 167 | 850 | 26 | 3 | 7 | 10 | 101 | 298 | 149 | 0.000140398 | 0.002994441 | 0.87 | 0.867 | 0.913 | 1 | 0.969 | 0.856 | 0.963 | 0.846 | 0.893 | 0.808 | 0.051 |
| SE | 79038 | ENSG00000160218 | TRAPPC10 | chr21 | + | 44096604 | 44096713 | 44094062 | 44094233 | 44098356 | 44098534 | 19 | 8 | 15 | 15 | 9 | 164 | 72 | 148 | 168 | 195 | 38 | 11 | 11 | 31 | 8 | 53 | 142 | 124 | 78 | 71 | 258 | 149 | 0.000140568 | 0.002997256 | 0.063 | 0.06 | 0.055 | 0.049 | 0.026 | 0.293 | 0.043 | 0.049 | 0.187 | 0.061 | -0.076 |
| SE | 90506 | ENSG00000160299 | PCNT | chr21 | + | 46425830 | 46425971 | 46421969 | 46422124 | 46427621 | 46427795 | 57 | 28 | 37 | 50 | 64 | 0 | 2 | 0 | 0 | 2 | 32 | 39 | 30 | 27 | 52 | 0 | 1 | 6 | 0 | 6 | 290 | 149 | 0.000140755 | 0.003000414 | 1 | 0.878 | 1 | 1 | 0.943 | 1 | 0.952 | 0.72 | 1 | 0.817 | 0.066 |
| SE | 15970 | ENSG00000137845 | ADAM10 | chr15 | - | 58698279 | 58698404 | 58682251 | 58682314 | 58717576 | 58717727 | 30 | 12 | 15 | 17 | 17 | 779 | 737 | 515 | 728 | 782 | 0 | 27 | 28 | 2 | 8 | 161 | 379 | 383 | 152 | 94 | 274 | 149 | 0.000141465 | 0.00301473 | 0.021 | 0.009 | 0.016 | 0.013 | 0.012 | 0 | 0.037 | 0.038 | 0.007 | 0.044 | -0.011 |
| SE | 71628 | ENSG00000184182 | UBE2F | chr2 | + | 238025341 | 238025412 | 237994743 | 237994809 | 238030555 | 238030595 | 165 | 83 | 119 | 159 | 171 | 0 | 4 | 0 | 1 | 2 | 292 | 178 | 167 | 233 | 334 | 14 | 1 | 4 | 3 | 14 | 220 | 149 | 0.000141986 | 0.003025014 | 1 | 0.934 | 1 | 0.991 | 0.983 | 0.934 | 0.992 | 0.966 | 0.981 | 0.942 | 0.019 |
| SE | 48007 | ENSG00000010810 | FYN | chr6 | - | 111700103 | 111700268 | 111696276 | 111696456 | 111702884 | 111703034 | 152 | 69 | 129 | 99 | 195 | 130 | 76 | 140 | 110 | 226 | 81 | 121 | 100 | 41 | 56 | 66 | 244 | 248 | 91 | 127 | 298 | 149 | 0.00014238 | 0.003032579 | 0.369 | 0.312 | 0.315 | 0.31 | 0.301 | 0.38 | 0.199 | 0.168 | 0.184 | 0.181 | 0.099 |
| SE | 96768 | ENSG00000052841 | TTC17 | chr11 | + | 43397973 | 43398113 | 43397346 | 43397491 | 43399887 | 43400048 | 322 | 237 | 249 | 282 | 290 | 4 | 7 | 4 | 6 | 4 | 259 | 247 | 226 | 268 | 79 | 0 | 24 | 25 | 7 | 0 | 289 | 149 | 0.000142679 | 0.003038136 | 0.976 | 0.946 | 0.97 | 0.96 | 0.974 | 1 | 0.841 | 0.823 | 0.952 | 1 | 0.042 |
| SE | 79944 | ENSG00000158169 | FANCC | chr9 | - | 95186500 | 95186645 | 95172036 | 95172147 | 95240648 | 95240743 | 1 | 2 | 1 | 5 | 4 | 10 | 9 | 4 | 13 | 30 | 0 | 0 | 0 | 0 | 0 | 12 | 2 | 7 | 6 | 3 | 294 | 149 | 0.000142727 | 0.003038329 | 0.048 | 0.101 | 0.112 | 0.163 | 0.063 | 0 | 0 | 0 | 0 | 0 | 0.097 |
| SE | 31035 | ENSG00000172795 | DCP2 | chr5 | + | 113003941 | 113004077 | 113001566 | 113001674 | 113007937 | 113008005 | 484 | 406 | 404 | 584 | 571 | 37 | 22 | 24 | 46 | 57 | 196 | 335 | 296 | 221 | 180 | 49 | 62 | 70 | 18 | 9 | 285 | 149 | 0.000142812 | 0.003039312 | 0.872 | 0.906 | 0.898 | 0.869 | 0.84 | 0.677 | 0.739 | 0.689 | 0.865 | 0.913 | 0.1 |
| SE | 54415 | ENSG00000169554 | ZEB2 | chr2 | - | 144394392 | 144394480 | 144388661 | 144390028 | 144396411 | 144396592 | 5 | 12 | 2 | 9 | 12 | 245 | 169 | 107 | 174 | 195 | 0 | 5 | 2 | 0 | 0 | 73 | 188 | 259 | 100 | 14 | 237 | 149 | 0.000142989 | 0.003042256 | 0.013 | 0.043 | 0.012 | 0.031 | 0.037 | 0 | 0.016 | 0.005 | 0 | 0 | 0.023 |
| SE | 78199 | ENSG00000124486 | USP9X | chrX | + | 41165871 | 41166214 | 41152947 | 41153081 | 41167481 | 41167577 | 261 | 175 | 206 | 249 | 373 | 0 | 0 | 1 | 0 | 0 | 53 | 141 | 161 | 69 | 11 | 7 | 3 | 0 | 5 | 0 | 298 | 149 | 0.000143424 | 0.003050666 | 1 | 1 | 0.99 | 1 | 1 | 0.791 | 0.959 | 1 | 0.873 | 1 | 0.073 |
| SE | 16220 | ENSG00000077238 | IL4R | chr16 | + | 27355807 | 27355907 | 27352539 | 27352696 | 27358915 | 27358994 | 145 | 79 | 129 | 108 | 116 | 2 | 0 | 1 | 1 | 0 | 122 | 191 | 142 | 136 | 177 | 6 | 5 | 5 | 7 | 0 | 249 | 149 | 0.000143519 | 0.003051866 | 0.977 | 1 | 0.987 | 0.985 | 1 | 0.924 | 0.958 | 0.944 | 0.921 | 1 | 0.04 |
| SE | 69790 | ENSG00000184007 | PTP4A2 | chr1 | - | 31910037 | 31910112 | 31906420 | 31908960 | 31911695 | 31911826 | 2246 | 1550 | 1522 | 2132 | 3234 | 49 | 30 | 16 | 43 | 52 | 4535 | 2143 | 2305 | 2988 | 2330 | 104 | 57 | 50 | 76 | 92 | 224 | 149 | 0.000143822 | 0.003057487 | 0.968 | 0.972 | 0.984 | 0.971 | 0.976 | 0.967 | 0.962 | 0.968 | 0.963 | 0.944 | 0.013 |
| SE | 36418 | ENSG00000179262 | RAD23A | chr19 | + | 12948513 | 12948552 | 12948176 | 12948358 | 12948685 | 12948813 | 18 | 8 | 17 | 6 | 6 | 5 | 3 | 2 | 1 | 2 | 26 | 12 | 10 | 14 | 31 | 18 | 12 | 20 | 17 | 11 | 188 | 149 | 0.000144448 | 0.003069942 | 0.74 | 0.679 | 0.871 | 0.826 | 0.704 | 0.534 | 0.442 | 0.284 | 0.395 | 0.691 | 0.295 |
| SE | 53622 | ENSG00000143851 | PTPN7 | chr1 | - | 202158117 | 202158248 | 202157738 | 202157823 | 202159280 | 202159454 | 59 | 28 | 46 | 46 | 82 | 2 | 0 | 0 | 0 | 0 | 67 | 107 | 28 | 47 | 70 | 0 | 2 | 7 | 0 | 6 | 280 | 149 | 0.000144851 | 0.003077672 | 0.94 | 1 | 1 | 1 | 1 | 1 | 0.966 | 0.68 | 1 | 0.861 | 0.087 |
| SE | 39662 | ENSG00000181555 | SETD2 | chr3 | - | 47080800 | 47081083 | 47067069 | 47067118 | 47083719 | 47083946 | 61 | 30 | 30 | 65 | 113 | 191 | 163 | 163 | 184 | 336 | 16 | 40 | 59 | 24 | 10 | 143 | 239 | 291 | 138 | 35 | 298 | 149 | 0.000145032 | 0.003080692 | 0.138 | 0.084 | 0.084 | 0.15 | 0.144 | 0.053 | 0.077 | 0.092 | 0.08 | 0.125 | 0.035 |
| SE | 104148 | ENSG00000023041 | ZDHHC6 | chr10 | - | 112441728 | 112441812 | 112440533 | 112440695 | 112442191 | 112442351 | 3 | 0 | 5 | 0 | 0 | 94 | 70 | 86 | 108 | 152 | 5 | 11 | 0 | 0 | 9 | 125 | 73 | 74 | 66 | 17 | 233 | 149 | 0.000145268 | 0.003084859 | 0.02 | 0 | 0.036 | 0 | 0 | 0.025 | 0.088 | 0 | 0 | 0.253 | -0.062 |
| SE | 1319 | ENSG00000100376 | FAM118A | chr22 | + | 45332424 | 45332710 | 45323174 | 45323427 | 45335349 | 45335382 | 101 | 12 | 33 | 201 | 93 | 0 | 0 | 1 | 0 | 0 | 54 | 48 | 6 | 243 | 33 | 12 | 0 | 0 | 0 | 3 | 298 | 149 | 0.000146586 | 0.003112003 | 1 | 1 | 0.943 | 1 | 1 | 0.692 | 1 | 1 | 1 | 0.846 | 0.081 |
| SE | 73324 | ENSG00000115970 | THADA | chr2 | - | 43344121 | 43344237 | 43320445 | 43320540 | 43397970 | 43398139 | 47 | 43 | 68 | 45 | 41 | 0 | 0 | 0 | 0 | 0 | 9 | 20 | 41 | 35 | 3 | 0 | 0 | 17 | 0 | 0 | 265 | 149 | 0.00014675 | 0.003114657 | 1 | 1 | 1 | 1 | 1 | 1 | 1 | 0.576 | 1 | 1 | 0.085 |
| SE | 100553 | ENSG00000054654 | SYNE2 | chr14 | + | 64202837 | 64202963 | 64190070 | 64190237 | 64208757 | 64208945 | 251 | 99 | 143 | 204 | 128 | 0 | 2 | 0 | 3 | 2 | 24 | 66 | 80 | 52 | 15 | 5 | 0 | 2 | 7 | 0 | 275 | 149 | 0.000146823 | 0.003115355 | 1 | 0.964 | 1 | 0.974 | 0.972 | 0.722 | 1 | 0.956 | 0.801 | 1 | 0.086 |
| SE | 96721 | ENSG00000266173 | STRADA | chr17 | - | 63726637 | 63726695 | 63714005 | 63714108 | 63728357 | 63728413 | 8 | 6 | 3 | 17 | 18 | 15 | 2 | 13 | 15 | 12 | 0 | 0 | 1 | 5 | 15 | 23 | 10 | 3 | 19 | 42 | 207 | 149 | 0.000147833 | 0.003135704 | 0.277 | 0.683 | 0.142 | 0.449 | 0.519 | 0 | 0 | 0.194 | 0.159 | 0.205 | 0.302 |
| SE | 34924 | ENSG00000076108 | BAZ2A | chr12 | - | 56604584 | 56604799 | 56604216 | 56604291 | 56605072 | 56605327 | 212 | 107 | 161 | 183 | 97 | 3 | 0 | 1 | 2 | 0 | 295 | 222 | 204 | 247 | 349 | 10 | 4 | 5 | 3 | 10 | 298 | 149 | 0.000147862 | 0.003135704 | 0.972 | 1 | 0.988 | 0.979 | 1 | 0.937 | 0.965 | 0.953 | 0.976 | 0.946 | 0.032 |
| SE | 84972 | ENSG00000170832 | USP32 | chr17 | - | 60180544 | 60180637 | 60177326 | 60179428 | 60181323 | 60181748 | 125 | 87 | 98 | 100 | 90 | 1 | 0 | 0 | 0 | 0 | 33 | 132 | 162 | 137 | 2 | 0 | 4 | 2 | 11 | 0 | 242 | 149 | 0.000147934 | 0.003136373 | 0.987 | 1 | 1 | 1 | 1 | 1 | 0.953 | 0.98 | 0.885 | 1 | 0.034 |
| SE | 9056 | ENSG00000072818 | ACAP1 | chr17 | + | 7348994 | 7349167 | 7348305 | 7348475 | 7349944 | 7350054 | 305 | 147 | 280 | 250 | 143 | 4 | 2 | 5 | 7 | 0 | 176 | 235 | 209 | 192 | 347 | 45 | 6 | 3 | 8 | 5 | 298 | 149 | 0.000148408 | 0.003145582 | 0.974 | 0.974 | 0.966 | 0.947 | 1 | 0.662 | 0.951 | 0.972 | 0.923 | 0.972 | 0.076 |
| SE | 85721 | ENSG00000168994 | PXDC1 | chr6 | - | 3727550 | 3727662 | 3722613 | 3723736 | 3737078 | 3737196 | 1 | 16 | 4 | 12 | 28 | 0 | 0 | 0 | 0 | 0 | 12 | 4 | 6 | 2 | 38 | 0 | 0 | 6 | 0 | 6 | 261 | 149 | 0.000148722 | 0.003150599 | 1 | 1 | 1 | 1 | 1 | 1 | 1 | 0.363 | 1 | 0.783 | 0.171 |
| SE | 19364 | ENSG00000119720 | NRDE2 | chr14 | - | 90317703 | 90317820 | 90316586 | 90316811 | 90318004 | 90318113 | 5 | 6 | 6 | 2 | 8 | 11 | 5 | 13 | 8 | 18 | 94 | 36 | 17 | 23 | 10 | 9 | 12 | 6 | 16 | 23 | 266 | 149 | 0.000148745 | 0.003150599 | 0.203 | 0.402 | 0.205 | 0.123 | 0.199 | 0.854 | 0.627 | 0.613 | 0.446 | 0.196 | -0.321 |
| SE | 62437 | ENSG00000113761 | ZNF346 | chr5 | + | 177050750 | 177050936 | 177044388 | 177044533 | 177062057 | 177062151 | 47 | 33 | 69 | 66 | 65 | 3 | 6 | 11 | 10 | 8 | 90 | 66 | 63 | 88 | 210 | 13 | 3 | 0 | 2 | 5 | 298 | 149 | 0.000148766 | 0.003150599 | 0.887 | 0.733 | 0.758 | 0.767 | 0.802 | 0.776 | 0.917 | 1 | 0.957 | 0.955 | -0.132 |
| SE | 92160 | ENSG00000090612 | ZNF268 | chr12 | + | 133193427 | 133193533 | 133191720 | 133192003 | 133202143 | 133202310 | 11 | 20 | 7 | 10 | 26 | 13 | 20 | 15 | 17 | 14 | 0 | 0 | 8 | 0 | 0 | 5 | 11 | 6 | 1 | 4 | 255 | 149 | 0.000148819 | 0.003150875 | 0.331 | 0.369 | 0.214 | 0.256 | 0.52 | 0 | 0 | 0.438 | 0 | 0 | 0.25 |
| SE | 60774 | ENSG00000187741 | FANCA | chr16 | - | 89752137 | 89752222 | 89749729 | 89749902 | 89758576 | 89758705 | 34 | 23 | 17 | 18 | 25 | 1 | 1 | 0 | 0 | 0 | 66 | 23 | 8 | 20 | 39 | 0 | 2 | 0 | 2 | 9 | 234 | 149 | 0.000149092 | 0.00315581 | 0.956 | 0.936 | 1 | 1 | 1 | 1 | 0.88 | 1 | 0.864 | 0.734 | 0.083 |
| SE | 56396 | ENSG00000151838 | CCDC175 | chr14 | - | 59531771 | 59531910 | 59521573 | 59521676 | 59538022 | 59538154 | 2 | 3 | 1 | 4 | 19 | 1 | 1 | 0 | 2 | 10 | 9 | 2 | 2 | 2 | 28 | 0 | 2 | 0 | 0 | 2 | 288 | 149 | 0.000149151 | 0.003156187 | 0.509 | 0.608 | 1 | 0.509 | 0.496 | 1 | 0.341 | 1 | 1 | 0.879 | -0.22 |
| SE | 8627 | ENSG00000271605 | MILR1 | chr17 | + | 64457399 | 64457684 | 64452596 | 64452866 | 64465451 | 64465541 | 36 | 29 | 19 | 41 | 37 | 3 | 0 | 0 | 1 | 0 | 128 | 20 | 45 | 102 | 74 | 7 | 3 | 3 | 10 | 0 | 298 | 149 | 0.000149211 | 0.003156608 | 0.857 | 1 | 1 | 0.953 | 1 | 0.901 | 0.769 | 0.882 | 0.836 | 1 | 0.084 |
| SE | 44750 | ENSG00000198265 | HELZ | chr17 | - | 67195418 | 67195470 | 67190156 | 67190355 | 67201128 | 67201185 | 63 | 33 | 41 | 67 | 19 | 5 | 0 | 6 | 1 | 6 | 12 | 100 | 45 | 52 | 17 | 0 | 0 | 5 | 0 | 0 | 201 | 149 | 0.000149297 | 0.003157578 | 0.903 | 1 | 0.835 | 0.98 | 0.701 | 1 | 1 | 0.87 | 1 | 1 | -0.09 |
| SE | 12663 | ENSG00000196712 | NF1 | chr17 | + | 31349119 | 31349251 | 31343008 | 31343135 | 31350182 | 31350318 | 86 | 44 | 88 | 74 | 126 | 0 | 0 | 0 | 0 | 0 | 10 | 91 | 68 | 16 | 19 | 0 | 10 | 0 | 0 | 3 | 281 | 149 | 0.000149438 | 0.003159701 | 1 | 1 | 1 | 1 | 1 | 1 | 0.828 | 1 | 1 | 0.771 | 0.08 |
| SE | 31099 | ENSG00000099949 | LZTR1 | chr22 | + | 20995367 | 20995484 | 20994869 | 20995026 | 20995745 | 20995872 | 9 | 0 | 2 | 6 | 2 | 62 | 20 | 32 | 47 | 10 | 4 | 0 | 0 | 0 | 0 | 66 | 20 | 14 | 35 | 48 | 266 | 149 | 0.00015009 | 0.003172638 | 0.075 | 0 | 0.034 | 0.067 | 0.101 | 0.033 | 0 | 0 | 0 | 0 | 0.049 |
| SE | 17570 | ENSG00000197102 | DYNC1H1 | chr14 | + | 102034334 | 102034452 | 102033975 | 102034188 | 102036488 | 102036642 | 271 | 114 | 211 | 215 | 248 | 0 | 3 | 7 | 0 | 1 | 245 | 264 | 166 | 190 | 223 | 4 | 6 | 1 | 1 | 23 | 267 | 149 | 0.000150195 | 0.003174002 | 1 | 0.955 | 0.944 | 1 | 0.993 | 0.972 | 0.961 | 0.989 | 0.991 | 0.844 | 0.027 |
| SE | 10892 | ENSG00000129250 | KIF1C | chr17 | + | 5000219 | 5000352 | 4999849 | 4999970 | 5000771 | 5000848 | 46 | 14 | 31 | 35 | 2 | 0 | 0 | 0 | 0 | 0 | 93 | 71 | 23 | 25 | 89 | 10 | 4 | 0 | 0 | 0 | 282 | 149 | 0.000150277 | 0.003174868 | 1 | 1 | 1 | 1 | 1 | 0.831 | 0.904 | 1 | 1 | 1 | 0.053 |
| SE | 47266 | ENSG00000113360 | DROSHA | chr5 | - | 31435764 | 31435864 | 31429474 | 31429545 | 31437238 | 31437298 | 52 | 37 | 35 | 46 | 31 | 1 | 0 | 0 | 0 | 0 | 33 | 39 | 36 | 37 | 42 | 14 | 0 | 2 | 0 | 0 | 249 | 149 | 0.000150524 | 0.00317924 | 0.969 | 1 | 1 | 1 | 1 | 0.585 | 1 | 0.915 | 1 | 1 | 0.094 |
| SE | 23275 | ENSG00000145604 | SKP2 | chr5 | + | 36170343 | 36170442 | 36168312 | 36168447 | 36171602 | 36171733 | 98 | 58 | 74 | 85 | 69 | 3 | 0 | 2 | 5 | 1 | 51 | 47 | 52 | 81 | 16 | 0 | 9 | 7 | 5 | 6 | 248 | 149 | 0.000150823 | 0.003184684 | 0.952 | 1 | 0.957 | 0.911 | 0.976 | 1 | 0.758 | 0.817 | 0.907 | 0.616 | 0.14 |
| SE | 60223 | ENSG00000286129 | NA | chr22 | + | 22307532 | 22307591 | 22307105 | 22307245 | 22308696 | 22308819 | 5 | 3 | 25 | 8 | 14 | 1 | 0 | 2 | 2 | 9 | 43 | 2 | 10 | 26 | 21 | 3 | 0 | 2 | 0 | 0 | 208 | 149 | 0.00015127 | 0.003193274 | 0.782 | 1 | 0.9 | 0.741 | 0.527 | 0.911 | 1 | 0.782 | 1 | 1 | -0.149 |
| SE | 31034 | ENSG00000172795 | DCP2 | chr5 | + | 112993105 | 112993173 | 112992671 | 112992770 | 113001083 | 113001236 | 37 | 25 | 40 | 41 | 144 | 138 | 150 | 129 | 209 | 479 | 16 | 49 | 55 | 30 | 30 | 65 | 123 | 87 | 69 | 58 | 217 | 149 | 0.000152105 | 0.003209353 | 0.155 | 0.103 | 0.176 | 0.119 | 0.171 | 0.145 | 0.215 | 0.303 | 0.23 | 0.262 | -0.086 |
| SE | 95817 | ENSG00000215126 | CBWD6 | chr9 | - | 41170283 | 41170423 | 41166783 | 41166830 | 41171846 | 41171934 | 4 | 0 | 0 | 0 | 0 | 8 | 3 | 12 | 26 | 22 | 0 | 1 | 5 | 4 | 9 | 11 | 0 | 5 | 6 | 10 | 289 | 149 | 0.000152122 | 0.003209353 | 0.205 | 0 | 0 | 0 | | 0 | 1 | 0.34 | 0.256 | 0.317 | -0.342 |
| SE | 16692 | ENSG00000137841 | PLCB2 | chr15 | - | 40291848 | 40291924 | 40291264 | 40291487 | 40292063 | 40292158 | 191 | 117 | 78 | 142 | 83 | 3 | 0 | 0 | 3 | 0 | 160 | 243 | 127 | 178 | 300 | 17 | 7 | 4 | 3 | 0 | 225 | 149 | 0.000152155 | 0.003209353 | 0.977 | 1 | 1 | 0.969 | 1 | 0.862 | 0.958 | 0.955 | 0.975 | 1 | 0.039 |
| SE | 90913 | ENSG00000185122 | HSF1 | chr8 | + | 144311173 | 144311249 | 144309771 | 144309896 | 144311320 | 144311367 | 179 | 80 | 145 | 140 | 230 | 0 | 0 | 2 | 0 | 3 | 225 | 148 | 124 | 141 | 247 | 8 | 2 | 5 | 7 | 3 | 225 | 149 | 0.000152422 | 0.003214117 | 1 | 1 | 0.98 | 1 | 0.981 | 0.949 | 0.98 | 0.943 | 0.93 | 0.982 | 0.035 |
| SE | 65688 | ENSG00000186141 | POLR3C | chr1 | - | 145838055 | 145838206 | 145836493 | 145836574 | 145839889 | 145839991 | 113 | 79 | 88 | 111 | 151 | 2 | 0 | 2 | 2 | 2 | 223 | 113 | 138 | 103 | 134 | 7 | 5 | 6 | 2 | 14 | 298 | 149 | 0.000152498 | 0.003214851 | 0.966 | 1 | 0.957 | 0.965 | 0.974 | 0.941 | 0.919 | 0.92 | 1 | 0.827 | 0.051 |
| SE | 10567 | ENSG00000181481 | RNF135 | chr17 | + | 30987943 | 30988106 | 30984616 | 30984760 | 30997241 | 30997248 | 105 | 42 | 53 | 75 | 84 | 5 | 0 | 0 | 6 | 1 | 164 | 131 | 46 | 64 | 134 | 21 | 3 | 0 | 4 | 36 | 298 | 149 | 0.000152551 | 0.003215104 | 0.913 | 1 | 1 | 0.862 | 0.977 | 0.796 | 0.956 | 0.92 | 0.889 | 0.65 | 0.108 |
| SE | 67474 | ENSG00000162923 | WDR26 | chr1 | - | 224418259 | 224418416 | 224411426 | 224411565 | 224419517 | 224419615 | 1024 | 737 | 725 | 1097 | 1373 | 2 | 1 | 0 | 2 | 3 | 1003 | 1101 | 1408 | 986 | 660 | 27 | 7 | 16 | 5 | 0 | 298 | 149 | 0.000152682 | 0.003217007 | 0.996 | 0.997 | 1 | 0.996 | 0.996 | 0.949 | 0.987 | 0.978 | 0.99 | 1 | 0.016 |
| SE | 108081 | ENSG00000079332 | SAR1A | chr10 | - | 70162689 | 70162759 | 70161857 | 70161931 | 70170412 | 70170523 | 65 | 50 | 62 | 93 | 71 | 138 | 49 | 88 | 100 | 54 | 29 | 168 | 216 | 118 | 22 | 21 | 43 | 30 | 37 | 30 | 219 | 149 | 0.00015315 | 0.003225988 | 0.243 | 0.41 | 0.324 | 0.388 | 0.472 | 0.484 | 0.727 | 0.83 | 0.685 | 0.333 | -0.244 |
| SE | 53910 | ENSG00000127526 | SLC35E1 | chr19 | - | 16565036 | 16565132 | 16555151 | 16555397 | 16566531 | 16566657 | 0 | 0 | 0 | 3 | 4 | 46 | 36 | 45 | 51 | 42 | 0 | 8 | 9 | 0 | 14 | 39 | 36 | 27 | 42 | 167 | 245 | 149 | 0.000153295 | 0.003228189 | 0 | 0 | 0 | 0.035 | 0.055 | 0 | 0.119 | 0.169 | 0 | 0.049 | -0.049 |
| SE | 79606 | ENSG00000111667 | USP5 | chr12 | + | 6855400 | 6855526 | 6852147 | 6852290 | 6855754 | 6855821 | 176 | 101 | 175 | 184 | 106 | 2 | 0 | 1 | 7 | 0 | 143 | 193 | 101 | 146 | 196 | 20 | 2 | 4 | 5 | 2 | 275 | 149 | 0.000153416 | 0.00322965 | 0.979 | 1 | 0.99 | 0.934 | 1 | 0.795 | 0.981 | 0.932 | 0.941 | 0.982 | 0.054 |
| SE | 30659 | ENSG00000145734 | BDP1 | chr5 | + | 71549419 | 71549606 | 71548681 | 71548745 | 71553115 | 71553320 | 262 | 267 | 216 | 230 | 503 | 10 | 17 | 11 | 6 | 9 | 42 | 87 | 42 | 10 | 49 | 0 | 4 | 0 | 0 | 0 | 298 | 149 | 0.000153447 | 0.00322965 | 0.929 | 0.887 | 0.908 | 0.95 | 0.965 | 1 | 0.916 | 1 | 1 | 1 | -0.055 |
| SE | 83238 | ENSG00000106077 | ABHD11 | chr7 | - | 73737875 | 73738072 | 73737567 | 73737735 | 73738327 | 73738463 | 0 | 3 | 0 | 0 | 0 | 8 | 0 | 2 | 9 | 2 | 20 | 1 | 1 | 1 | 13 | 0 | 13 | 0 | 5 | 1 | 298 | 149 | 0.000153786 | 0.003235907 | 0 | 1 | 0 | 0 | 0 | 1 | 0.037 | 1 | 0.091 | 0.867 | -0.399 |
| SE | 101237 | ENSG00000144802 | NFKBIZ | chr3 | + | 101855394 | 101855458 | 101855061 | 101855208 | 101855732 | 101855902 | 327 | 245 | 317 | 452 | 393 | 3 | 1 | 5 | 1 | 1 | 532 | 370 | 407 | 363 | 279 | 7 | 19 | 3 | 11 | 0 | 213 | 149 | 0.000154513 | 0.003250336 | 0.987 | 0.994 | 0.978 | 0.997 | 0.996 | 0.982 | 0.932 | 0.99 | 0.958 | 1 | 0.018 |
| SE | 95015 | ENSG00000135047 | CTSL | chr9 | + | 87730380 | 87730498 | 87728584 | 87729735 | 87731007 | 87731157 | 108 | 143 | 41 | 193 | 142 | 0 | 6 | 0 | 2 | 0 | 489 | 187 | 107 | 296 | 303 | 7 | 4 | 5 | 25 | 5 | 267 | 149 | 0.000154647 | 0.003252291 | 1 | 0.93 | 1 | 0.982 | 1 | 0.975 | 0.963 | 0.923 | 0.869 | 0.971 | 0.042 |
| SE | 106095 | ENSG00000196498 | NCOR2 | chr12 | - | 124430614 | 124430787 | 124426621 | 124426800 | 124437929 | 124437996 | 145 | 77 | 134 | 173 | 96 | 0 | 0 | 0 | 1 | 0 | 404 | 153 | 150 | 122 | 280 | 0 | 0 | 0 | 12 | 11 | 298 | 149 | 0.00015473 | 0.003253157 | 1 | 1 | 1 | 0.989 | 1 | 1 | 1 | 1 | 0.836 | 0.927 | 0.045 |
| SE | 64712 | ENSG00000167895 | TMC8 | chr17 | + | 78131881 | 78132214 | 78131280 | 78131737 | 78132358 | 78132465 | 47 | 31 | 60 | 76 | 16 | 8 | 0 | 4 | 5 | 0 | 10 | 23 | 23 | 18 | 7 | 0 | 0 | 2 | 1 | 0 | 298 | 149 | 0.000155577 | 0.003270088 | 0.746 | 1 | 0.882 | 0.884 | 1 | 1 | 1 | 0.852 | 0.9 | 1 | -0.048 |
| SE | 93757 | ENSG00000143409 | MINDY1 | chr1 | - | 151002164 | 151002400 | 151001724 | 151001782 | 151006311 | 151006485 | 69 | 60 | 75 | 125 | 117 | 6 | 5 | 4 | 0 | 2 | 263 | 145 | 62 | 133 | 797 | 26 | 3 | 7 | 10 | 101 | 298 | 149 | 0.00015645 | 0.003287543 | 0.852 | 0.857 | 0.904 | 1 | 0.967 | 0.835 | 0.96 | 0.816 | 0.869 | 0.798 | 0.06 |
| SE | 46032 | ENSG00000158517 | NCF1 | chr7 | + | 74778306 | 74778399 | 74777268 | 74777347 | 74779081 | 74779157 | 2 | 2 | 0 | 0 | 7 | 252 | 195 | 156 | 176 | 148 | 8 | 0 | 0 | 0 | 36 | 454 | 20 | 83 | 316 | 149 | 242 | 149 | 0.000156581 | 0.003289412 | 0.005 | 0.006 | 0 | 0 | 0.028 | 0.011 | 0 | 0 | 0 | 0.129 | -0.02 |

| | | | | | | | | | | | | | | | |
|---|---|---|---|---|---|---|---|---|---|---|---|---|---|---|---|
| SE | 75419 | ENSG00000244482 | LILRA6 | chr19 | - | 54238485 | 54238613 | 54236591 | 54237193 | 54240273 | 54240576 | 31 | 18 | 5 | [illegible] |
| SE | 83512 | ENSG00000196313 | POM121 | chr7 | + | 72938589 | 72938681 | 72929939 | 72930111 | 72939335 | 72939409 | 281 | 196 | 268 | [illegible] |
| SE | 16708 | ENSG00000166233 | ARIH1 | chr15 | + | 72563393 | 72563500 | 72561482 | 72561549 | 72566562 | 72566605 | 253 | 144 | 217 | [illegible] |
| SE | 94097 | ENSG00000136811 | ODF2 | chr9 | + | 128469182 | 128469353 | 128460941 | 128461067 | 128471307 | 128471346 | 160 | 86 | 68 | [illegible] |
| SE | 70575 | ENSG00000022840 | RNF10 | chr12 | + | 120571190 | 120571291 | 120566824 | 120566980 | 120575630 | 120575688 | 283 | 161 | 231 | [illegible] |
| SE | 109327 | ENSG00000123104 | ITPR2 | chr12 | - | 26653975 | 26654126 | 26631865 | 26632059 | 26655707 | 26655852 | 276 | 298 | 294 | [illegible] |
| SE | 7772 | ENSG00000109118 | PHF12 | chr17 | - | 28911329 | 28911384 | 28911111 | 28911237 | 28912481 | 28913277 | 2 | 2 | 1 | [illegible] |
| SE | 32969 | ENSG00000114127 | XRN1 | chr3 | - | 142426743 | 142426841 | 142425428 | 142425538 | 142432660 | 142432893 | 739 | 658 | 650 | [illegible] |
| SE | 28242 | ENSG00000157326 | DHRS4 | chr14 | + | 23966282 | 23966417 | 23955034 | 23955212 | 23967210 | 23967266 | 28 | 16 | 31 | [illegible] |
| SE | 101421 | ENSG00000158545 | ZC3H18 | chr16 | + | 88608933 | 88609051 | 88599790 | 88599948 | 88622196 | 88622388 | 213 | 131 | 149 | [illegible] |
| SE | 60339 | ENSG00000070785 | EIF2B3 | chr1 | - | 44969977 | 44970044 | 44941505 | 44941665 | 44978314 | 44978460 | 7 | 6 | 3 | [illegible] |
| SE | 55822 | ENSG00000143641 | GALNT2 | chr1 | + | 230243305 | 230243427 | 230236352 | 230236420 | 230246062 | 230246150 | 99 | 52 | 57 | [illegible] |
| SE | 42057 | ENSG00000092871 | RFFL | chr17 | - | 35017522 | 35017606 | 35016369 | 35016580 | 35021370 | 35021781 | 108 | 49 | 105 | [illegible] |
| SE | 47264 | ENSG00000113360 | DROSHA | chr5 | - | 31431575 | 31431678 | 31429474 | 31429545 | 31435764 | 31435864 | 90 | 70 | 60 | [illegible] |
| SE | 52934 | ENSG00000169981 | ZNF35 | chr3 | + | 44652556 | 44652701 | 44650940 | 44651259 | 44658700 | 44658765 | 26 | 14 | 14 | [illegible] |
| SE | 107596 | ENSG00000111596 | CNOT2 | chr12 | + | 70294070 | 70294294 | 70278131 | 70278274 | 70310894 | 70311017 | 6 | 4 | 5 | [illegible] |
| SE | 105433 | ENSG00000145901 | TNIP1 | chr5 | - | 151033479 | 151033579 | 151032286 | 151032383 | 151035001 | 151035067 | 0 | 0 | 0 | [illegible] |
| SE | 107452 | ENSG00000215039 | CD27-AS1 | chr12 | - | 6448639 | 6448737 | 6446960 | 6448150 | 6451468 | 6451502 | 15 | 8 | 9 | [illegible] |
| SE | 102850 | ENSG00000107554 | DNMBP | chr10 | - | 99896266 | 99896397 | 99894945 | 99895050 | 99898085 | 99898285 | 35 | 26 | 21 | [illegible] |
| SE | 103131 | ENSG00000180198 | RCC1 | chr1 | + | 28516724 | 28516853 | 28508829 | 28508905 | 28529857 | 28529939 | 52 | 26 | 28 | [illegible] |
| SE | 48713 | ENSG00000143401 | ANP32E | chr1 | - | 150229071 | 150229237 | 150226607 | 150226795 | 150230570 | 150230693 | 1034 | 1007 | 1373 | [illegible] |
| SE | 75844 | ENSG00000124523 | SIRT5 | chr6 | + | 13599031 | 13599155 | 13596962 | 13597016 | 13600833 | 13600949 | 38 | 9 | 43 | [illegible] |
| SE | 65263 | ENSG00000151729 | SLC25A4 | chr4 | + | 185145354 | 185145899 | 185144763 | 185145250 | 185146813 | 185147241 | 14 | 4 | 18 | [illegible] |
| SE | 100613 | ENSG00000149292 | TTC12 | chr11 | + | 113352069 | 113352207 | 113351238 | 113351299 | 113359362 | 113359461 | 36 | 25 | 40 | [illegible] |
| SE | 90507 | ENSG00000160299 | PCNT | chr21 | + | 46442496 | 46442573 | 46440854 | 46441084 | 46445283 | 46445437 | 34 | 12 | 19 | [illegible] |
| SE | 81878 | ENSG00000177189 | RPS6KA3 | chrX | - | 20167588 | 20167747 | 20164898 | 20165060 | 20172745 | 20172864 | 265 | 211 | 258 | [illegible] |
| SE | 54098 | ENSG00000115935 | WIPF1 | chr2 | - | 174625820 | 174625960 | 174585522 | 174585611 | 174634517 | 174634605 | 18 | 16 | 22 | [illegible] |
| SE | 54319 | ENSG00000117139 | KDM5B | chr1 | - | 202758390 | 202758510 | 202756357 | 202756516 | 202760414 | 202760573 | 106 | 68 | 100 | [illegible] |
| SE | 76967 | ENSG00000131018 | SYNE1 | chr6 | - | 152455885 | 152456044 | 152455425 | 152455590 | 152458756 | 152458930 | 589 | 318 | 259 | [illegible] |
| SE | 5483 | ENSG00000130589 | HELZ2 | chr20 | - | 63562071 | 63562203 | 63561822 | 63561984 | 63562287 | 63562382 | 67 | 45 | 59 | [illegible] |
| SE | 14919 | ENSG00000086666 | ZFAND6 | chr15 | + | 80098415 | 80098578 | 80072563 | 80072647 | 80120327 | 80120405 | 103 | 62 | 69 | [illegible] |
| SE | 31945 | ENSG00000163703 | CRELD1 | chr3 | + | 9938018 | 9938106 | 9937561 | 9937672 | 9940849 | 9941026 | 28 | 10 | 32 | [illegible] |
| [illegible] | [illegible] | [illegible] | [illegible] | [illegible] | [illegible] | [illegible] | [illegible] | [illegible] | [illegible] | [illegible] | [illegible] | [illegible] | [illegible] | [illegible] | [illegible] |

| | | | | | | | | | | | | | | | | | | | | | | | | | | | | | | | | | | | | | | | | | | | | | | |
|---|---|---|---|---|---|---|---|---|---|---|---|---|---|---|---|---|---|---|---|---|---|---|---|---|---|---|---|---|---|---|---|---|---|---|---|---|---|---|---|---|---|---|---|---|---|---|
| SE | 12294 | ENSG00000185324 | CDK10 | chr16 | + | 89689251 | 89689324 | 89686688 | 89686797 | 89690552 | 89690557 | 12 | 21 | 39 | 25 | 19 | 4 | 3 | 6 | 3 | 1 | 5 | 25 | 23 | 7 | 76 | 0 | 2 | 1 | 2 | 0 | 222 | 149 | 0.000227327 | 0.00449352 | 0.668 | 0.825 | 0.814 | 0.848 | 0.927 | 1 | 0.893 | 0.939 | 0.701 | 1 | -0.09 |
| SE | 101650 | ENSG00000128245 | YWHAH | chr22 | + | 31945499 | 31945678 | 31944965 | 31945235 | 31956138 | 31956482 | 0 | 14 | 0 | 23 | 0 | 3 | 6 | 1 | 2 | 1 | 0 | 0 | 3 | 0 | 0 | 22 | 14 | 3 | 8 | 84 | 298 | 149 | 0.000227592 | 0.004497608 | 0 | 0.538 | 0 | 0.852 | 0 | 0 | 0 | 0.333 | 0 | 0 | 0.211 |
| SE | 106897 | ENSG00000205885 | C1RL-AS1 | chr12 | + | 7110297 | 7110572 | 7108570 | 7108947 | 7111024 | 7111606 | 22 | 7 | 12 | 13 | 11 | 0 | 0 | 0 | 2 | 0 | 38 | 24 | 15 | 43 | 18 | 0 | 6 | 6 | 0 | 0 | 298 | 149 | 0.000227655 | 0.00449772 | 1 | 1 | 1 | 0.765 | 1 | 1 | 0.667 | 0.556 | 1 | 1 | 0.108 |
| SE | 9480 | ENSG00000248871 | TNFSF12-TNFSF13 | chr17 | + | 7559845 | 7559893 | 7550942 | 7550978 | 7560048 | 7560167 | 62 | 35 | 25 | 49 | 16 | 1 | 0 | 0 | 0 | 0 | 149 | 80 | 39 | 65 | 198 | 0 | 2 | 0 | 0 | 19 | 197 | 149 | 0.000228009 | 0.00450323 | 0.979 | 1 | 1 | 1 | 1 | 1 | 0.968 | 1 | 1 | 0.887 | 0.025 |
| SE | 18483 | ENSG00000186648 | CARMIL3 | chr14 | + | 24056298 | 24056393 | 24055700 | 24055789 | 24056621 | 24056708 | 18 | 6 | 4 | 5 | 2 | 1 | 0 | 0 | 0 | 0 | 19 | 13 | 1 | 6 | 19 | 0 | 0 | 0 | 3 | 10 | 244 | 149 | 0.000228049 | 0.00450323 | 0.917 | 1 | 1 | 1 | 1 | 1 | 1 | 1 | 0.55 | 0.537 | 0.166 |
| SE | 40922 | ENSG00000151883 | PARP8 | chr5 | + | 50821212 | 50821338 | 50815431 | 50815524 | 50822334 | 50822400 | 894 | 621 | 679 | 830 | 1274 | 24 | 13 | 24 | 36 | 52 | 92 | 398 | 480 | 169 | 101 | 0 | 16 | 48 | 18 | 8 | 275 | 149 | 0.000228577 | 0.004512517 | 0.953 | 0.963 | 0.939 | 0.926 | 0.93 | 1 | 0.931 | 0.844 | 0.836 | 0.872 | 0.046 |
| SE | 104410 | ENSG00000184047 | DIABLO | chr12 | - | 122209696 | 122209858 | 122208315 | 122208577 | 122216487 | 122216584 | 25 | 32 | 36 | 14 | 34 | 179 | 110 | 126 | 112 | 276 | 128 | 57 | 86 | 85 | 22 | 366 | 170 | 86 | 146 | 364 | 298 | 149 | 0.000229769 | 0.004534902 | 0.065 | 0.127 | 0.125 | 0.059 | 0.058 | 0.149 | 0.144 | 0.333 | 0.225 | 0.029 | -0.089 |
| SE | 9267 | ENSG00000179094 | PER1 | chr17 | - | 8145957 | 8146137 | 8143265 | 8143876 | 8146371 | 8146488 | 31 | 12 | 21 | 15 | 13 | 0 | 0 | 0 | 2 | 0 | 7 | 21 | 71 | 8 | 2 | 0 | 2 | 5 | 0 | 6 | 298 | 149 | 0.000230577 | 0.004549706 | 1 | 1 | 1 | 0.789 | 1 | 1 | 0.84 | 0.877 | 1 | 0.143 | 0.186 |
| SE | 107612 | ENSG00000111596 | CNOT2 | chr12 | + | 70330286 | 70330469 | 70329422 | 70329570 | 70332766 | 70332806 | 282 | 196 | 234 | 209 | 339 | 17 | 7 | 13 | 12 | 17 | 63 | 224 | 193 | 187 | 40 | 0 | 19 | 48 | 47 | 5 | 298 | 149 | 0.000231262 | 0.004562073 | 0.892 | 0.933 | 0.9 | 0.897 | 0.909 | 1 | 0.855 | 0.668 | 0.665 | 0.8 | 0.109 |
| SE | 40590 | ENSG00000085433 | WDR47 | chr1 | - | 108982608 | 108982785 | 108981732 | 108981864 | 108983281 | 108983451 | 45 | 23 | 24 | 35 | 25 | 1 | 0 | 0 | 0 | 4 | 13 | 12 | 8 | 28 | 7 | 0 | 8 | 8 | 2 | 0 | 298 | 149 | 0.000231659 | 0.004568755 | 0.957 | 1 | 1 | 1 | 0.758 | 1 | 0.429 | 0.333 | 0.875 | 1 | 0.216 |
| SE | 50675 | ENSG00000142687 | KIAA0319L | chr1 | - | 35448172 | 35448332 | 35444160 | 35444303 | 35449866 | 35450005 | 102 | 65 | 63 | 90 | 85 | 3 | 0 | 4 | 4 | 4 | 163 | 117 | 104 | 127 | 109 | 0 | 0 | 0 | 0 | 4 | 298 | 149 | 0.000231756 | 0.004569519 | 0.944 | 1 | 0.887 | 0.918 | 0.914 | 1 | 1 | 1 | 1 | 0.932 | -0.054 |
| SE | 3844 | ENSG00000157540 | DYRK1A | chr21 | + | 37421773 | 37421866 | 37420298 | 37420384 | 37428866 | 37428908 | 16 | 6 | 6 | 24 | 19 | 1 | 0 | 1 | 1 | 0 | 25 | 10 | 20 | 4 | 21 | 0 | 3 | 3 | 10 | 0 | 242 | 149 | 0.000232006 | 0.004573297 | 0.908 | 1 | 0.787 | 0.937 | 1 | 1 | 0.672 | 0.804 | 0.198 | 1 | 0.192 |
| SE | 83539 | ENSG00000106526 | ACTR3C | chr7 | - | 150293311 | 150293419 | 150289449 | 150289593 | 150295251 | 150295296 | 21 | 28 | 26 | 26 | 16 | 0 | 0 | 0 | 0 | 0 | 53 | 40 | 7 | 10 | 10 | 0 | 16 | 0 | 0 | 0 | 257 | 149 | 0.000232148 | 0.004574944 | 1 | 1 | 1 | 1 | 1 | 1 | 0.592 | 1 | 1 | 1 | 0.082 |
| SE | 100355 | ENSG00000174842 | GLMN | chr1 | - | 92286489 | 92286592 | 92271464 | 92271652 | 92288913 | 92289151 | 59 | 41 | 44 | 44 | 164 | 4 | 0 | 2 | 4 | 5 | 3 | 32 | 17 | 27 | 15 | 0 | 0 | 0 | 0 | 0 | 252 | 149 | 0.000232853 | 0.004587686 | 0.897 | 1 | 0.929 | 0.867 | 0.951 | 1 | 1 | 1 | 1 | 1 | -0.071 |
| SE | 94931 | ENSG00000119403 | PHF19 | chr9 | - | 120861917 | 120862005 | 120861088 | 120861174 | 120862612 | 120863026 | 29 | 28 | 51 | 60 | 44 | 0 | 0 | 0 | 1 | 3 | 33 | 18 | 29 | 45 | 40 | 0 | 1 | 0 | 2 | 12 | 237 | 149 | 0.000233118 | 0.004591681 | 1 | 1 | 1 | 0.974 | 0.902 | 1 | 0.919 | 1 | 0.934 | 0.677 | 0.069 |
| SE | 88415 | ENSG00000168172 | HOOK3 | chr8 | + | 42968012 | 42968214 | 42964310 | 42964474 | 42973288 | 42973399 | 119 | 108 | 58 | 123 | 190 | 0 | 0 | 0 | 5 | 0 | 15 | 143 | 153 | 72 | 9 | 0 | 3 | 10 | 0 | 8 | 298 | 149 | 0.000233173 | 0.004591681 | 1 | 1 | 1 | 0.925 | 1 | 1 | 0.96 | 0.884 | 1 | 0.36 | 0.144 |
| SE | 68345 | ENSG00000125630 | POLR1B | chr2 | + | 112547011 | 112547179 | 112542225 | 112542671 | 112547420 | 112547461 | 64 | 30 | 69 | 58 | 54 | 0 | 0 | 0 | 0 | 0 | 32 | 46 | 20 | 31 | 19 | 4 | 0 | 6 | 1 | 0 | 298 | 149 | 0.000233958 | 0.004605971 | 1 | 1 | 1 | 1 | 1 | 0.8 | 1 | 0.625 | 0.939 | 1 | 0.127 |
| SE | 61290 | ENSG00000134058 | CDK7 | chr5 | + | 69259817 | 69259936 | 69258042 | 69258153 | 69262204 | 69262207 | 114 | 87 | 133 | 150 | 278 | 0 | 4 | 0 | 0 | 4 | 62 | 156 | 135 | 100 | 17 | 0 | 18 | 3 | 0 | 0 | 268 | 149 | 0.000234217 | 0.004609927 | 1 | 0.924 | 1 | 1 | 0.975 | 1 | 0.828 | 0.962 | 1 | 1 | 0.022 |
| SE | 7259 | ENSG00000101745 | ANKRD12 | chr18 | + | 9204475 | 9204544 | 9195550 | 9195698 | 9208656 | 9208803 | 304 | 357 | 247 | 348 | 410 | 402 | 294 | 234 | 378 | 778 | 287 | 231 | 199 | 275 | 253 | 215 | 217 | 125 | 120 | 205 | 218 | 149 | 0.000234571 | 0.004614689 | 0.341 | 0.454 | 0.419 | 0.386 | 0.265 | 0.477 | 0.421 | 0.521 | 0.61 | 0.458 | -0.124 |
| SE | 93756 | ENSG00000143409 | MINDY1 | chr1 | - | 151002164 | 151002323 | 151001724 | 151001782 | 151006311 | 151006857 | 72 | 62 | 75 | 132 | 117 | 6 | 5 | 4 | 0 | 2 | 289 | 150 | 67 | 138 | 888 | 26 | 3 | 7 | 10 | 101 | 298 | 149 | 0.000234626 | 0.004614689 | 0.857 | 0.861 | 0.904 | 1 | 0.967 | 0.848 | 0.962 | 0.827 | 0.873 | 0.815 | 0.053 |
| SE | 64715 | ENSG00000167895 | TMC8 | chr17 | + | 78137234 | 78137323 | 78134948 | 78135009 | 78138004 | 78138071 | 59 | 30 | 62 | 119 | 29 | 6 | 1 | 0 | 4 | 0 | 57 | 67 | 30 | 28 | 53 | 5 | 3 | 1 | 6 | 16 | 238 | 149 | 0.000234636 | 0.004614689 | 0.86 | 0.949 | 1 | 0.949 | 1 | 0.877 | 0.933 | 0.949 | 0.745 | 0.675 | 0.116 |
| SE | 61118 | ENSG00000150782 | IL18 | chr11 | - | 112153591 | 112153603 | 112150071 | 112150206 | 112154974 | 112155061 | 40 | 23 | 22 | 30 | 63 | 0 | 1 | 0 | 1 | 0 | 91 | 35 | 48 | 111 | 86 | 9 | 0 | 3 | 8 | 0 | 161 | 149 | 0.000234789 | 0.004616531 | 1 | 0.955 | 1 | 0.965 | 1 | 0.903 | 1 | 0.937 | 0.928 | 1 | 0.03 |
| SE | 106194 | ENSG00000135090 | TAOK3 | chr12 | - | 118212913 | 118212995 | 118201295 | 118201463 | 118214016 | 118214110 | 361 | 322 | 298 | 348 | 493 | 4 | 12 | 6 | 7 | 7 | 172 | 350 | 291 | 369 | 69 | 0 | 17 | 4 | 32 | 6 | 231 | 149 | 0.000235192 | 0.004623352 | 0.983 | 0.945 | 0.97 | 0.97 | 0.978 | 1 | 0.93 | 0.979 | 0.881 | 0.881 | 0.035 |
| SE | 71400 | ENSG00000122203 | KIAA1191 | chr5 | - | 176359810 | 176359918 | 176355570 | 176355749 | 176361601 | 176361735 | 21 | 8 | 19 | 48 | 9 | 3 | 0 | 0 | 1 | 0 | 13 | 27 | 6 | 14 | 76 | 5 | 3 | 2 | 1 | 1 | 257 | 149 | 0.000235652 | 0.004631167 | 0.802 | 1 | 1 | 0.965 | 1 | 0.601 | 0.839 | 0.635 | 0.89 | 0.978 | 0.165 |
| SE | 52243 | ENSG00000166825 | ANPEP | chr15 | - | 89814113 | 89814294 | 89805969 | 89806806 | 89814771 | 89814852 | 0 | 2 | 1 | 0 | 1 | 39 | 9 | 16 | 20 | 27 | 15 | 1 | 6 | 0 | 0 | 42 | 43 | 8 | 21 | 18 | 298 | 149 | 0.000235931 | 0.004635493 | 0 | 0.1 | 0.03 | 0 | 0.018 | 0.152 | 0.011 | 0.273 | 0 | 0 | -0.058 |
| SE | 94353 | ENSG00000148090 | AUH | chr9 | - | 91216058 | 91216106 | 91213814 | 91214425 | 91217276 | 91217327 | 42 | 28 | 34 | 31 | 45 | 1 | 6 | 4 | 4 | 0 | 0 | 28 | 11 | 16 | 13 | 15 | 7 | 11 | 0 | 8 | 197 | 149 | 0.000236371 | 0.004641937 | 0.969 | 0.779 | 0.865 | 0.854 | 1 | 0 | 0.752 | 0.431 | 1 | 0.551 | 0.347 |
| SE | 10650 | ENSG00000273559 | CWC25 | chr17 | - | 38812794 | 38812860 | 38810353 | 38810595 | 38814860 | 38815097 | 10 | 8 | 11 | 16 | 11 | 0 | 0 | 0 | 3 | 0 | 52 | 24 | 13 | 49 | 8 | 19 | 0 | 4 | 0 | 0 | 215 | 149 | 0.000236421 | 0.004641937 | 1 | 1 | 1 | 0.787 | 1 | 0.655 | 1 | 0.693 | 1 | 1 | 0.088 |
| SE | 94614 | ENSG00000106829 | TLE4 | chr9 | + | 79722958 | 79723035 | 79722450 | 79722601 | 79725036 | 79725233 | 177 | 82 | 121 | 130 | 264 | 1 | 1 | 1 | 1 | 0 | 172 | 151 | 153 | 156 | 212 | 3 | 2 | 4 | 14 | 0 | 226 | 149 | 0.000236437 | 0.004641937 | 0.992 | 0.982 | 0.988 | 0.988 | 1 | 0.974 | 0.98 | 0.962 | 0.88 | 1 | 0.031 |
| SE | 85065 | ENSG00000147459 | DOCK5 | chr8 | + | 25368191 | 25368250 | 25366869 | 25366970 | 25368570 | 25368687 | 147 | 99 | 90 | 118 | 184 | 1 | 5 | 1 | 9 | 8 | 152 | 95 | 148 | 170 | 79 | 0 | 0 | 7 | 0 | 0 | 208 | 149 | 0.000236928 | 0.004650413 | 0.991 | 0.934 | 0.985 | 0.904 | 0.943 | 1 | 1 | 0.938 | 1 | 1 | -0.036 |
| SE | 21172 | ENSG00000057663 | ATG5 | chr6 | - | 106308363 | 106308491 | 106293027 | 106293106 | 106316100 | 106316208 | 152 | 105 | 140 | 184 | 245 | 18 | 18 | 8 | 13 | 23 | 61 | 137 | 73 | 79 | 71 | 0 | 3 | 8 | 6 | 0 | 277 | 149 | 0.000237175 | 0.004654107 | 0.82 | 0.758 | 0.904 | 0.884 | 0.851 | 1 | 0.961 | 0.831 | 0.876 | 1 | -0.09 |
| SE | 106981 | ENSG00000111271 | ACAD10 | chr12 | + | 111736830 | 111737004 | 111727961 | 111728143 | 111744642 | 111745043 | 33 | 11 | 29 | 23 | 19 | 0 | 0 | 0 | 1 | 0 | 51 | 15 | 16 | 10 | 43 | 0 | 7 | 0 | 0 | 5 | 298 | 149 | 0.000237384 | 0.004657037 | 1 | 1 | 1 | 0.92 | 1 | 1 | 0.517 | 1 | 1 | 0.811 | 0.118 |
| SE | 61622 | ENSG00000234127 | TRIM26 | chr6 | - | 30201034 | 30201083 | 30198665 | 30199153 | 30213304 | 30213427 | 16 | 5 | 15 | 29 | 7 | 2 | 0 | 0 | 2 | 0 | 9 | 14 | 17 | 18 | 21 | 8 | 0 | 0 | 1 | 13 | 198 | 149 | 0.000237529 | 0.004658701 | 0.858 | 1 | 1 | 0.916 | 1 | 0.458 | 1 | 1 | 0.931 | 0.549 | 0.167 |
| SE | 67635 | ENSG00000162735 | PEX19 | chr1 | - | 160282416 | 160282502 | 160282038 | 160282200 | 160282943 | 160283109 | 216 | 181 | 214 | 201 | 275 | 6 | 6 | 16 | 3 | 4 | 513 | 350 | 213 | 251 | 292 | 33 | 16 | 15 | 36 | 6 | 235 | 149 | 0.000237625 | 0.004659417 | 0.958 | 0.95 | 0.895 | 0.977 | 0.978 | 0.908 | 0.933 | 0.9 | 0.816 | 0.969 | 0.046 |
| SE | 18948 | ENSG00000160256 | FAM207A | chr21 | + | 44973196 | 44973265 | 44967033 | 44967181 | 44976679 | 44976973 | 31 | 23 | 35 | 25 | 37 | 1 | 1 | 1 | 0 | 2 | 15 | 31 | 20 | 18 | 39 | 0 | 3 | 6 | 1 | 6 | 218 | 149 | 0.000237851 | 0.004662683 | 0.955 | 0.94 | 0.96 | 1 | 0.927 | 1 | 0.876 | 0.695 | 0.925 | 0.816 | 0.094 |
| SE | 101251 | ENSG00000166224 | SGPL1 | chr10 | + | 70871047 | 70871146 | 70869791 | 70869897 | 70873350 | 70873589 | 72 | 23 | 36 | 44 | 49 | 0 | 0 | 0 | 0 | 0 | 41 | 41 | 27 | 9 | 14 | 15 | 0 | 0 | 0 | 0 | 248 | 149 | 0.000238038 | 0.004665191 | 1 | 1 | 1 | 1 | 1 | 0.622 | 1 | 1 | 1 | 1 | 0.076 |
| SE | 25494 | ENSG00000082397 | EPB41L3 | chr18 | - | 5443837 | 5443880 | 5438034 | 5438110 | 5445139 | 5445244 | 139 | 171 | 57 | 133 | 160 | 1 | 0 | 0 | 0 | 0 | 312 | 152 | 94 | 180 | 47 | 15 | 4 | 2 | 0 | 2 | 192 | 149 | 0.000238502 | 0.004672782 | 0.991 | 1 | 1 | 1 | 1 | 0.942 | 0.967 | 0.973 | 1 | 0.948 | 0.032 |
| SE | 84928 | ENSG00000179195 | ZNF664 | chr12 | + | 123988042 | 123988138 | 123973122 | 123974020 | 124011380 | 124011441 | 68 | 38 | 76 | 65 | 64 | 5 | 4 | 3 | 7 | 0 | 16 | 34 | 31 | 39 | 53 | 0 | 3 | 0 | 1 | 0 | 245 | 149 | 0.000238598 | 0.004672782 | 0.892 | 0.852 | 0.939 | 0.85 | 1 | 1 | 0.873 | 1 | 0.96 | 1 | -0.06 |
| SE | 88785 | ENSG00000140632 | GLYR1 | chr16 | - | 4821379 | 4821453 | 4817597 | 4817697 | 4822874 | 4822931 | 63 | 34 | 54 | 91 | 53 | 3 | 0 | 0 | 1 | 2 | 134 | 44 | 30 | 80 | 134 | 5 | 0 | 3 | 22 | 0 | 223 | 149 | 0.000238605 | 0.004672782 | 0.933 | 1 | 1 | 0.984 | 0.947 | 0.947 | 1 | 0.87 | 0.708 | 1 | 0.068 |
| SE | 83670 | ENSG00000044446 | PHKA2 | chrX | - | 18902791 | 18902910 | 18901484 | 18901603 | 18905757 | 18905815 | 6 | 4 | 5 | 3 | 3 | 69 | 32 | 53 | 60 | 69 | 2 | 12 | 8 | 16 | 22 | 64 | 86 | 29 | 44 | 108 | 268 | 149 | 0.000238714 | 0.004673765 | 0.046 | 0.065 | 0.05 | 0.027 | 0.024 | 0.017 | 0.072 | 0.133 | 0.168 | 0.102 | -0.056 |
| SE | 99621 | ENSG00000109920 | FNBP4 | chr11 | - | 47722975 | 47723316 | 47719928 | 47720086 | 47724027 | 47724172 | 251 | 176 | 238 | 237 | 326 | 1 | 5 | 12 | 5 | 1 | 102 | 246 | 238 | 101 | 49 | 0 | 0 | 2 | 0 | 0 | 298 | 149 | 0.000238845 | 0.004675153 | 0.992 | 0.946 | 0.908 | 0.96 | 0.994 | 1 | 1 | 1 | 0.962 | 1 | -0.032 |
| SE | 3790 | ENSG00000090372 | STRN4 | chr19 | - | 46733038 | 46733236 | 46730731 | 46730873 | 46738163 | 46738172 | 105 | 52 | 64 | 77 | 51 | 4 | 0 | 0 | 1 | 6 | 169 | 141 | 89 | 132 | 341 | 0 | 21 | 10 | 21 | 20 | 298 | 149 | 0.000239195 | 0.004680829 | 0.929 | 1 | 1 | 0.975 | 0.81 | 1 | 0.77 | 0.817 | 0.759 | 0.895 | 0.095 |
| SE | 24151 | ENSG00000164574 | GALNT10 | chr5 | + | 154329571 | 154329738 | 154297940 | 154298079 | 154376276 | 154376367 | 168 | 69 | 113 | 175 | 178 | 4 | 0 | 1 | 2 | 11 | 222 | 253 | 162 | 149 | 364 | 0 | 0 | 0 | 0 | 4 | 298 | 149 | 0.00023958 | 0.004687204 | 0.955 | 1 | 0.983 | 0.978 | 0.89 | 1 | 1 | 1 | 1 | 0.978 | -0.034 |
| SE | 22136 | ENSG00000175727 | MLXIP | chr12 | + | 122138423 | 122138551 | 122138193 | 122138295 | 122138814 | 122138938 | 89 | 53 | 106 | 127 | 76 | 3 | 1 | 4 | 2 | 1 | 17 | 96 | 29 | 46 | 341 | 5 | 3 | 8 | 6 | 10 | 277 | 149 | 0.000239775 | 0.004689841 | 0.941 | 0.966 | 0.934 | 0.972 | 0.976 | 0.647 | 0.945 | 0.661 | 0.805 | 0.948 | 0.157 |
| SE | 36997 | ENSG00000233452 | STXBP5-AS1 | chr6 | - | 146842409 | 146842809 | 146841052 | 146841133 | 146844318 | 146844712 | 7 | 10 | 4 | 26 | 14 | 2 | 1 | 0 | 0 | 0 | 10 | 28 | 18 | 2 | 38 | 9 | 3 | 2 | 0 | 0 | 298 | 149 | 0.000239994 | 0.004692949 | 0.636 | 0.833 | 1 | 1 | 1 | 0.357 | 0.824 | 0.818 | 1 | 1 | 0.094 |
| SE | 95102 | ENSG00000148358 | GPR107 | chr9 | + | 130104419 | 130104550 | 130101105 | 130101223 | 130107495 | 130107539 | 113 | 67 | 93 | 97 | 116 | 1 | 0 | 8 | 2 | 11 | 181 | 94 | 93 | 126 | 120 | 24 | 19 | 9 | 7 | 9 | 280 | 149 | 0.000240457 | 0.004700431 | 0.984 | 1 | 0.861 | 0.963 | 0.849 | 0.801 | 0.725 | 0.846 | 0.905 | 0.876 | 0.101 |
| SE | 63341 | ENSG00000151092 | NGLY1 | chr3 | - | 25736003 | 25736149 | 25733871 | 25733982 | 25740402 | 25740450 | 145 | 107 | 112 | 157 | 155 | 0 | 0 | 1 | 0 | 3 | 31 | 134 | 75 | 83 | 18 | 0 | 3 | 5 | 6 | 0 | 295 | 149 | 0.000240497 | 0.004700431 | 1 | 1 | 0.983 | 1 | 0.963 | 1 | 0.958 | 0.883 | 0.875 | 1 | 0.046 |
| SE | 60459 | ENSG00000146409 | SLC18B1 | chr6 | - | 132772131 | 132772249 | 132771035 | 132771129 | 132772992 | 132773088 | 18 | 8 | 11 | 22 | 50 | 1 | 0 | 0 | 1 | 0 | 9 | 14 | 2 | 10 | 5 | 0 | 3 | 0 | 11 | 0 | 267 | 149 | 0.000240602 | 0.00470132 | 0.909 | 1 | 1 | 0.925 | 1 | 1 | 0.723 | 1 | 0.337 | 1 | 0.155 |
| SE | 7074 | ENSG00000175387 | SMAD2 | chr18 | - | 47870474 | 47870564 | 47869242 | 47869436 | 47896520 | 47896809 | 686 | 482 | 486 | 626 | 1327 | 35 | 21 | 33 | 23 | 39 | 407 | 482 | 443 | 395 | 564 | 8 | 12 | 15 | 12 | 8 | 239 | 149 | 0.000241142 | 0.004710698 | 0.924 | 0.935 | 0.902 | 0.944 | 0.955 | 0.969 | 0.962 | 0.948 | 0.954 | 0.978 | -0.03 |
| SE | 99026 | ENSG00000087053 | MTMR2 | chr11 | - | 95841616 | 95841709 | 95838093 | 95838207 | 95844952 | 95845159 | 118 | 54 | 55 | 90 | 196 | 3 | 0 | 0 | 0 | 4 | 24 | 54 | 60 | 29 | 35 | 0 | 3 | 10 | 2 | 0 | 242 | 149 | 0.000241204 | 0.004710725 | 0.96 | 1 | 1 | 1 | 0.968 | 1 | 0.917 | 0.787 | 0.899 | 1 | 0.065 |
| SE | 101172 | ENSG00000133816 | MICAL2 | chr11 | + | 12255642 | 12255750 | 12239435 | 12239585 | 12256784 | 12256909 | 0 | 3 | 6 | 0 | 1 | 85 | 49 | 63 | 100 | 61 | 1 | 0 | 2 | 8 | 25 | 201 | 56 | 30 | 65 | 102 | 257 | 149 | 0.000241615 | 0.004717585 | 0 | 0.034 | 0.052 | 0 | 0.009 | 0.003 | 0 | 0.037 | 0.067 | 0.124 | -0.027 |
| SE | 2628 | ENSG00000105723 | GSK3A | chr19 | - | 42232495 | 42232594 | 42232056 | 42232149 | 42233109 | 42233205 | 69 | 25 | 44 | 66 | 82 | 2 | 0 | 1 | 1 | 1 | 137 | 87 | 58 | 37 | 198 | 0 | 12 | 0 | 16 | 2 | 248 | 149 | 0.000241802 | 0.004720052 | 0.954 | 1 | 0.964 | 0.975 | 0.98 | 1 | 0.813 | 1 | 0.581 | 0.983 | 0.099 |
| SE | 22708 | ENSG00000164329 | TENT2 | chr5 | + | 79623251 | 79623489 | 79612434 | 79613075 | 79640850 | 79640965 | 180 | 136 | 123 | 148 | 287 | 1 | 0 | 4 | 1 | 3 | 42 | 93 | 54 | 31 | 35 | 16 | 5 | 0 | 0 | 0 | 298 | 149 | 0.000242401 | 0.004730563 | 0.989 | 1 | 0.939 | 0.987 | 0.98 | 0.568 | 0.903 | 1 | 1 | 1 | 0.085 |
| SE | 2280 | ENSG00000180448 | ARHGAP45 | chr19 | + | 1068413 | 1068744 | 1067166 | 1067495 | 1073148 | 1073223 | 356 | 174 | 273 | 316 | 174 | 3 | 0 | 2 | 4 | 2 | 169 | 253 | 114 | 194 | 420 | 0 | 3 | 4 | 4 | 19 | 298 | 149 | 0.000243186 | 0.0047447 | 0.983 | 1 | 0.986 | 0.975 | 0.978 | 1 | 0.977 | 0.934 | 0.96 | 0.917 | 0.027 |
| SE | 56551 | ENSG00000069702 | TGFBR3 | chr1 | - | 91786185 | 91786227 | 91758612 | 91758750 | 91797286 | 91797471 | 4 | 4 | 1 | 1 | 1 | 170 | 45 | 70 | 132 | 151 | 16 | 5 | 3 | 4 | 0 | 8 | 163 | 85 | 24 | 22 | 191 | 149 | 0.000243526 | 0.004750124 | 0.018 | 0.065 | 0.011 | 0.006 | 0.005 | 0.609 | 0.023 | 0.027 | 0.115 | 0 | -0.134 |
| SE | 6040 | ENSG00000196756 | SNHG17 | chr20 | - | 38434503 | 38434626 | 38430779 | 38431112 | 38435026 | 38435313 | 3 | 4 | 4 | 3 | 4 | 0 | 0 | 0 | 0 | 0 | 11 | 20 | 3 | 14 | 27 | 0 | 0 | 0 | 8 | 3 | 272 | 149 | 0.000243585 | 0.004750124 | 1 | 1 | 1 | 1 | 1 | 1 | 1 | 1 | 0.489 | 0.831 | 0.136 |
| SE | 8782 | ENSG00000109103 | UNC119 | chr17 | - | 28547998 | 28548101 | 28546707 | 28547849 | 28548591 | 28548656 | 155 | 78 | 98 | 126 | 183 | 0 | 0 | 2 | 0 | 1 | 184 | 102 | 94 | 100 | 238 | 7 | 3 | 7 | 3 | 0 | 252 | 149 | 0.000244076 | 0.004758506 | 1 | 1 | 0.967 | 1 | 0.991 | 0.94 | 0.953 | 0.888 | 0.952 | 1 | 0.045 |
| SE | 51529 | ENSG00000069248 | NUP133 | chr1 | - | 229499683 | 229499818 | 229498135 | 229498306 | 229500755 | 229500863 | 156 | 96 | 102 | 121 | 62 | 0 | 0 | 0 | 0 | 0 | 47 | 107 | 75 | 50 | 17 | 0 | 0 | 1 | 15 | 0 | 284 | 149 | 0.000244214 | 0.004760018 | 1 | 1 | 1 | 1 | 1 | 1 | 1 | 0.975 | 0.636 | 1 | 0.078 |
| SE | 58668 | ENSG00000141458 | NPC1 | chr18 | - | 23533354 | 23533517 | 23531441 | 23532284 | 23534445 | 23534559 | 232 | 68 | 122 | 137 | 60 | 1 | 0 | 0 | 0 | 0 | 85 | 89 | 95 | 41 | 112 | 11 | 0 | 0 | 4 | 0 | 298 | 149 | 0.000244351 | 0.004761428 | 0.991 | 1 | 1 | 1 | 1 | 0.794 | 1 | 1 | 0.837 | 1 | 0.072 |
| SE | 63241 | ENSG00000106105 | GARS1 | chr7 | + | 30608407 | 30608483 | 30603495 | 30603572 | 30609584 | 30609730 | 6 | 3 | 3 | 9 | 12 | 126 | 101 | 138 | 114 | 201 | 0 | 19 | 6 | 16 | 23 | 130 | 122 | 109 | 96 | 130 | 225 | 149 | 0.000244483 | 0.004761428 | 0.031 | 0.019 | 0.014 | 0.05 | 0.038 | 0 | 0.093 | 0.035 | 0.099 | 0.105 | -0.036 |
| SE | 73942 | ENSG00000144134 | RABL2A | chr2 | + | 113635050 | 113635130 | 113634152 | 113634232 | 113641352 | 113641450 | 18 | 3 | 7 | 1 | 8 | 5 | 3 | 6 | 16 | 6 | 5 | 17 | 4 | 10 | 5 | 0 | 2 | 1 | 5 | 0 | 229 | 149 | 0.000244513 | 0.004761428 | 0.701 | 0.394 | 0.432 | 0.039 | 0.465 | 1 | 0.847 | 0.722 | 0.565 | 1 | -0.421 |
| SE | 33359 | ENSG00000187742 | SECISBP2 | chr9 | + | 89325547 | 89325676 | 89319651 | 89319797 | 89325896 | 89326038 | 130 | 128 | 127 | 129 | 261 | 0 | 2 | 5 | 0 | 4 | 39 | 125 | 128 | 96 | 47 | 0 | 3 | 12 | 3 | 5 | 278 | 149 | 0.00024453 | 0.004761428 | 1 | 0.972 | 0.932 | 1 | 0.972 | 1 | 0.957 | 0.851 | 0.945 | 0.834 | 0.058 |
| SE | 34507 | ENSG00000116350 | SRSF4 | chr1 | - | 29159863 | 29160091 | 29159373 | 29159486 | 29160374 | 29160517 | 0 | 5 | 3 | 8 | 0 | 190 | 128 | 148 | 161 | 139 | 19 | 11 | 21 | 12 | 14 | 317 | 260 | 346 | 287 | 364 | 298 | 149 | 0.000244635 | 0.004762288 | 0 | 0.019 | 0.01 | 0.024 | 0 | 0.029 | 0.021 | 0.029 | 0.02 | 0.019 | -0.013 |
| SE | 26087 | ENSG00000174695 | TMEM167A | chr5 | - | 83065007 | 83065117 | 83052845 | 83057154 | 83077320 | 83077394 | 55 | 31 | 58 | 60 | 23 | 0 | 0 | 0 | 1 | 0 | 37 | 28 | 20 | 21 | 53 | 0 | 0 | 0 | 0 | 14 | 259 | 149 | 0.000244821 | 0.004764721 | 1 | 1 | 1 | 0.972 | 1 | 1 | 1 | 1 | 1 | 0.685 | 0.057 |
| SE | 54125 | ENSG00000124126 | PREX1 | chr20 | - | 48644408 | 48644497 | 48642406 | 48642489 | 48645850 | 48646057 | 767 | 356 | 486 | 698 | 354 | 1 | 0 | 5 | 1 | 0 | 750 | 911 | 815 | 635 | 735 | 4 | 8 | 5 | 26 | 9 | 238 | 149 | 0.000245139 | 0.004769732 | 0.998 | 1 | 0.984 | 0.998 | 1 | 0.992 | 0.986 | 0.99 | 0.939 | 0.981 | 0.018 |
| SE | 33773 | ENSG00000180376 | CCDC66 | chr3 | + | 56563683 | 56564125 | 56559568 | 56559594 | 56566593 | 56566739 | 404 | 378 | 388 | 367 | 655 | 0 | 0 | 0 | 6 | 2 | 100 | 127 | 198 | 136 | 63 | 16 | 2 | 0 | 0 | 0 | 298 | 149 | 0.000246124 | 0.004787018 | 1 | 1 | 1 | 0.968 | 0.994 | 0.758 | 0.969 | 1 | 1 | 1 | 0.047 |
| SE | 31748 | ENSG00000211897 | IGHG3 | chr14 | - | 105770292 | 105770337 | 105770104 | 105770149 | 105770480 | 105770525 | 20 | 5 | 12 | 30 | 38 | 0 | 0 | 0 | 0 | 20 | 44 | 207 | 108 | 92 | 23 | 0 | 0 | 0 | 0 | 4 | 194 | 149 | 0.00024615 | 0.004787018 | 1 | 1 | 1 | 1 | 0.593 | 1 | 1 | 1 | 1 | 0.815 | -0.044 |
| SE | 31505 | ENSG00000174227 | PIGG | chr4 | + | 501100 | 501170 | 500395 | 500601 | 505717 | 505927 | 25 | 20 | 28 | 33 | 45 | 96 | 83 | 131 | 116 | 118 | 119 | 34 | 88 | 93 | 98 | 232 | 124 | 76 | 153 | 247 | 219 | 149 | 0.000246406 | 0.004790821 | 0.151 | 0.141 | 0.127 | 0.162 | 0.206 | 0.259 | 0.157 | 0.441 | 0.293 | 0.213 | -0.115 |
| SE | 94695 | ENSG00000198382 | UVRAG | chr11 | + | 76012926 | 76013102 | 76008806 | 76008867 | 76016814 | 76016980 | 7 | 5 | 8 | 5 | 11 | 112 | 112 | 88 | 104 | 253 | 5 | 0 | 1 | 0 | 0 | 108 | 168 | 113 | 134 | 65 | 298 | 149 | 0.000246516 | 0.004791753 | 0.03 | 0.022 | 0.043 | 0.023 | 0.021 | 0.023 | 0 | 0.004 | 0 | 0 | 0.022 |
| SE | 94 | ENSG00000186866 | POFUT2 | chr21 | - | 45285268 | 45285328 | 45283351 | 45283527 | 45285677 | 45285928 | 0 | 0 | 4 | 0 | 0 | 11 | 11 | 21 | 14 | 19 | 0 | 19 | 0 | 6 | 0 | 11 | 22 | 11 | 6 | 30 | 209 | 149 | 0.000247297 | 0.004805678 | 0 | 0 | 0.12 | 0 | 0 | 0 | 0.381 | 0 | 0.416 | 0 | -0.135 |
| SE | 94336 | ENSG00000165802 | NSMF | chr9 | - | 137453730 | 137453820 | 137452735 | 137453180 | 137454390 | 137454432 | 60 | 35 | 45 | 55 | 40 | 6 | 2 | 2 | 6 | 0 | 45 | 53 | 43 | 34 | 112 | 1 | 1 | 0 | 0 | 3 | 239 | 149 | 0.000247355 | 0.004805678 | 0.862 | 0.916 | 0.933 | 0.851 | 1 | 0.966 | 0.971 | 1 | 1 | 0.959 | -0.067 |
| SE | 76918 | ENSG00000167483 | NIBAN3 | chr19 | + | 17543523 | 17543631 | 17543316 | 17543433 | 17546685 | 17546797 | 51 | 50 | 78 | 86 | 45 | 2 | 3 | 0 | 3 | 0 | 133 | 170 | 21 | 85 | 357 | 6 | 5 | 0 | 12 | 31 | 257 | 149 | 0.000247487 | 0.004807052 | 0.937 | 0.906 | 1 | 0.943 | 1 | 0.928 | 0.952 | 1 | 0.804 | 0.87 | 0.046 |
| SE | 75740 | ENSG00000144580 | CNOT9 | chr2 | + | 218580994 | 218581114 | 218580560 | 218580740 | 218582970 | 218583086 | 5 | 3 | 7 | 4 | 8 | 63 | 54 | 54 | 63 | 121 | 10 | 11 | 3 | 10 | 37 | 117 | 45 | 100 | 46 | 74 | 269 | 149 | 0.000247714 | 0.004810281 | 0.042 | 0.03 | 0.067 | 0.034 | 0.035 | 0.045 | 0.119 | 0.016 | 0.107 | 0.217 | -0.059 |
| SE | 38757 | ENSG00000148399 | DPH7 | chr9 | - | 137564892 | 137564958 | 137564433 | 137564606 | 137565084 | 137565154 | 57 | 34 | 30 | 57 | 19 | 1 | 1 | 6 | 1 | 1 | 20 | 12 | 17 | 46 | 66 | 0 | 3 | 1 | 12 | 20 | 215 | 149 | 0.000248162 | 0.004817779 | 0.975 | 0.959 | 0.776 | 0.975 | 0.929 | 1 | 0.735 | 0.922 | 0.727 | 0.696 | 0.107 |
| SE | 26533 | ENSG00000138303 | ASCC1 | chr10 | - | 72204421 | 72204505 | 72203426 | 72203524 | 72210731 | 72210831 | 4 | 0 | 2 | 3 | 0 | 95 | 58 | 109 | 87 | 88 | 0 | 11 | 12 | 4 | 6 | 88 | 181 | 115 | 66 | 81 | 233 | 149 | 0.000248254 | 0.004818371 | 0.026 | 0 | 0.012 | 0.022 | 0 | 0 | 0.037 | 0.063 | 0.037 | 0.045 | -0.024 |
| SE | 46887 | ENSG00000055609 | KMT2C | chr7 | - | 152157760 | 152157925 | 152156204 | 152156346 | 152158862 | 152159072 | 139 | 131 | 142 | 195 | 313 | 8 | 2 | 1 | 6 | 4 | 13 | 113 | 89 | 42 | 36 | 0 | 0 | 0 | 0 | 0 | 298 | 149 | 0.000248735 | 0.004826516 | 0.897 | 0.97 | 0.986 | 0.942 | 0.975 | 1 | 1 | 1 | 1 | 1 | -0.046 |
| SE | 17018 | ENSG00000102921 | N4BP1 | chr16 | - | 48546146 | 48546254 | 48538725 | 48543261 | 48553538 | 48553669 | 135 | 103 | 129 | 123 | 283 | 2 | 1 | 2 | 2 | 1 | 30 | 135 | 132 | 71 | 47 | 0 | 2 | 5 | 12 | 0 | 257 | 149 | 0.000249237 | 0.004835056 | 0.975 | 0.984 | 0.974 | 0.973 | 0.994 | 1 | 0.975 | 0.939 | 0.774 | 1 | 0.042 |
| SE | 40845 | ENSG00000117020 | AKT3 | chr1 | - | 243564284 | 243564387 | 243563719 | 243563848 | 243572925 | 243573048 | 45 | 32 | 30 | 22 | 54 | 202 | 125 | 153 | 180 | 162 | 0 | 12 | 10 | 0 | 13 | 86 | 83 | 60 | 48 | 37 | 252 | 149 | 0.000249385 | 0.004836732 | 0.116 | 0.131 | 0.104 | 0.067 | 0.165 | 0 | 0.079 | 0.09 | 0 | 0.172 | 0.048 |
| SE | 77328 | ENSG00000214022 | REPIN1 | chr7 | + | 150369741 | 150369868 | 150368808 | 150368941 | 150371227 | 150372242 | 41 | 15 | 27 | 25 | 19 | 11 | 3 | 6 | 8 | 1 | 37 | 25 | 8 | 16 | 23 | 0 | 5 | 1 | 1 | 0 | 276 | 149 | 0.000249612 | 0.004839537 | 0.668 | 0.73 | 0.708 | 0.628 | 0.911 | 1 | 0.73 | 0.812 | 0.896 | 1 | -0.159 |
| SE | 62289 | ENSG00000112031 | MTRF1L | chr6 | - | 152995262 | 152995319 | 152994512 | 152994676 | 152998549 | 152998629 | 9 | 11 | 6 | 17 | 6 | 0 | 0 | 0 | 0 | 0 | 14 | 25 | 7 | 1 | 23 | 6 | 0 | 2 | 3 | 0 | 206 | 149 | 0.000249654 | 0.004839537 | 1 | 1 | 1 | 1 | 1 | 0.628 | 1 | 0.717 | 0.194 | 1 | 0.292 |
| SE | 2999 | ENSG00000161265 | U2AF1L4 | chr19 | - | 35744625 | 35744738 | 35744069 | 35744421 | 35745124 | 35745176 | 36 | 19 | 35 | 30 | 61 | 2 | 2 | 1 | 0 | 0 | 34 | 24 | 30 | 7 | 77 | 0 | 10 | 3 | 12 | 0 | 262 | 149 | 0.000250082 | 0.00484663 | 0.911 | 0.844 | 0.952 | 1 | 1 | 1 | 0.577 | 0.85 | 0.249 | 1 | 0.206 |
| SE | 149 | ENSG00000183527 | PSMG1 | chr21 | - | 39179923 | 39179986 | 39178448 | 39178647 | 39180284 | 39180436 | 68 | 27 | 74 | 39 | 170 | 5 | 3 | 8 | 2 | 2 | 45 | 41 | 27 | 30 | 18 | 8 | 3 | 8 | 4 | 6 | 212 | 149 | 0.000250159 | 0.00484693 | 0.905 | 0.863 | 0.867 | 0.932 | 0.984 | 0.798 | 0.906 | 0.703 | 0.841 | 0.678 | 0.125 |
| SE | 74314 | ENSG00000114166 | KAT2B | chr3 | + | 20099861 | 20099954 | 20095262 | 20095408 | 20111595 | 20111787 | 284 | 268 | 195 | 242 | 295 | 0 | 0 | 0 | 0 | 0 | 129 | 175 | 191 | 163 | 88 | 0 | 7 | 0 | 0 | 13 | 242 | 149 | 0.00025092 | 0.00486048 | 1 | 1 | 1 | 1 | 1 | 1 | 0.939 | 1 | 1 | 0.806 | 0.051 |
| SE | 56318 | ENSG00000134186 | PRPF38B | chr1 | + | 108693589 | 108693642 | 108692326 | 108692867 | 108695701 | 108695770 | 32 | 15 | 39 | 26 | 9 | 74 | 30 | 46 | 86 | 22 | 34 | 24 | 36 | 8 | 10 | 0 | 15 | 24 | 9 | 13 | 202 | 149 | 0.000251031 | 0.004861425 | 0.242 | 0.269 | 0.385 | 0.182 | 0.232 | 1 | 0.541 | 0.525 | 0.396 | 0.362 | -0.303 |
| SE | 80711 | ENSG00000147535 | PLPP5 | chr8 | - | 38267896 | 38267960 | 38266979 | 38267391 | 38268370 | 38268407 | 49 | 25 | 73 | 32 | 68 | 0 | 0 | 0 | 0 | 0 | 41 | 92 | 51 | 65 | 75 | 0 | 0 | 0 | 0 | 17 | 213 | 149 | 0.000251356 | 0.004866504 | 1 | 1 | 1 | 1 | 1 | 1 | 1 | 1 | 1 | 0.755 | 0.049 |
| SE | 101786 | ENSG00000170759 | KIF5B | chr10 | - | 32039326 | 32039431 | 32038162 | 32038218 | 32040383 | 32040457 | 253 | 156 | 189 | 229 | 265 | 0 | 0 | 3 | 3 | 1 | 31 | 155 | 139 | 88 | 37 | 0 | 15 | 2 | 0 | 0 | 254 | 149 | 0.000251419 | 0.004866528 | 1 | 1 | 0.974 | 0.978 | 0.994 | 1 | 0.858 | 0.976 | 1 | 1 | 0.022 |
| SE | 96977 | ENSG00000136270 | TBRG4 | chr7 | - | 45104537 | 45104709 | 45103030 | 45103443 | 45105440 | 45105478 | 49 | 12 | 70 | 52 | 78 | 14 | 7 | 8 | 7 | 8 | 15 | 44 | 38 | 19 | 74 | 0 | 1 | 0 | 0 | 10 | 298 | 149 | 0.000251592 | 0.004868681 | 0.636 | 0.462 | 0.814 | 0.788 | 0.83 | 1 | 0.957 | 1 | 1 | 0.787 | -0.243 |
| SE | 87562 | ENSG00000112159 | MDN1 | chr6 | - | 89662791 | 89662967 | 89662086 | 89662239 | 89664486 | 89664628 | 95 | 93 | 136 | 142 | 174 | 9 | 4 | 9 | 13 | 13 | 39 | 44 | 78 | 59 | 19 | 11 | 19 | 29 | 10 | 0 | 298 | 149 | 0.000251859 | 0.004872638 | 0.841 | 0.921 | 0.883 | 0.845 | 0.87 | 0.639 | 0.537 | 0.574 | 0.747 | 1 | 0.173 |
| SE | 10878 | ENSG00000198920 | KIAA0753 | chr17 | - | 6600379 | 6600458 | 6599236 | 6599320 | 6606872 | 6606928 | 48 | 43 | 43 | 61 | 65 | 1 | 1 | 0 | 1 | 0 | 49 | 51 | 30 | 42 | 33 | 0 | 0 | 0 | 8 | 6 | 228 | 149 | 0.00025212 | 0.004876481 | 0.969 | 0.966 | 1 | 0.976 | 1 | 1 | 1 | 1 | 0.774 | 0.782 | 0.071 |
| SE | 46410 | ENSG00000168724 | DNAJC21 | chr5 | + | 34949542 | 34949677 | 34945721 | 34945803 | 34950169 | 34950257 | 24 | 6 | 27 | 19 | 33 | 90 | 83 | 83 | 102 | 87 | 0 | 4 | 13 | 13 | 0 | 14 | 62 | 78 | 95 | 9 | 284 | 149 | 0.000252298 | 0.004878717 | 0.123 | 0.037 | 0.146 | 0.089 | 0.166 | 0 | 0.033 | 0.08 | 0.067 | 0 | 0.076 |
| SE | 52093 | ENSG00000117419 | ERI3 | chr1 | - | 44339044 | 44339316 | 44319627 | 44319744 | 44352849 | 44352925 | 58 | 23 | 45 | 73 | 63 | 4 | 1 | 7 | 1 | 5 | 17 | 49 | 39 | 59 | 58 | 0 | 2 | 0 | 3 | 0 | 298 | 149 | 0.000252556 | 0.004881245 | 0.879 | 0.92 | 0.763 | 0.973 | 0.863 | 1 | 0.925 | 1 | 0.908 | 1 | -0.087 |
| SE | 72895 | ENSG00000168763 | CNNM3 | chr2 | + | 96828566 | 96828700 | 96828098 | 96828195 | 96828995 | 96829134 | 217 | 106 | 152 | 259 | 237 | 3 | 4 | 3 | 11 | 3 | 76 | 103 | 98 | 92 | 257 | 0 | 0 | 2 | 2 | 0 | 283 | 149 | 0.000252598 | 0.004881245 | 0.974 | 0.933 | 0.964 | 0.925 | 0.977 | 1 | 1 | 0.963 | 0.96 | 1 | -0.03 |
| SE | 66552 | ENSG00000133872 | SARAF | chr8 | - | 30073876 | 30074028 | 30069873 | 30070059 | 30082846 | 30083074 | 793 | 520 | 905 | 844 | 1436 | 32 | 27 | 47 | 34 | 44 | 344 | 440 | 648 | 261 | 252 | 12 | 45 | 45 | 15 | 19 | 298 | 149 | 0.000252616 | 0.004881245 | 0.925 | 0.906 | 0.906 | 0.925 | 0.942 | 0.935 | 0.83 | 0.878 | 0.897 | 0.869 | 0.039 |
| SE | 8315 | ENSG00000100038 | TOP3B | chr22 | - | 21975904 | 21975953 | 21975639 | 21975807 | 21982729 | 21982766 | 0 | 1 | 0 | 0 | 0 | 32 | 25 | 40 | 37 | 22 | 0 | 0 | 4 | 1 | 11 | 51 | 51 | 22 | 28 | 32 | 198 | 149 | 0.000253698 | 0.004900957 | 0 | 0.029 | 0 | 0 | 0 | 0 | 0 | 0.12 | 0.026 | 0.206 | -0.065 |
| SE | 86322 | ENSG00000135048 | CEMIP2 | chr9 | - | 71690091 | 71690246 | 71685742 | 71685846 | 71694508 | 71694607 | 201 | 94 | 114 | 188 | 121 | 0 | 1 | 1 | 2 | 0 | 131 | 200 | 219 | 169 | 34 | 0 | 9 | 10 | 1 | 0 | 298 | 149 | 0.000253971 | 0.004905002 | 1 | 0.979 | 0.983 | 0.979 | 1 | 1 | 0.917 | 0.916 | 0.988 | 1 | 0.024 |
| SE | 59756 | ENSG00000158711 | ELK4 | chr1 | - | 205603200 | 205603353 | 205600650 | 205600794 | 205616477 | 205616644 | 1 | 3 | 7 | 7 | 16 | 1 | 1 | 0 | 0 | 1 | 13 | 7 | 7 | 6 | 16 | 4 | 3 | 5 | 0 | 0 | 298 | 149 | 0.000254804 | 0.004919881 | 0.333 | 0.6 | 1 | 1 | 0.889 | 0.619 | 0.538 | 0.412 | 1 | 1 | 0.051 |
| SE | 53941 | ENSG00000040275 | SPDL1 | chr5 | + | 169590755 | 169590829 | 169588393 | 169588575 | 169591047 | 169591074 | 3 | 0 | 1 | 2 | 0 | 51 | 26 | 38 | 43 | 42 | 0 | 12 | 2 | 0 | 0 | 5 | 30 | 16 | 9 | 6 | 223 | 149 | 0.000255118 | 0.004924187 | 0.038 | 0 | 0.017 | 0.03 | 0 | 0 | 0.211 | 0.077 | 0 | 0 | -0.041 |
| SE | 50570 | ENSG00000139636 | LMBR1L | chr12 | - | 49102886 | 49102951 | 49102467 | 49102540 | 49103090 | 49103120 | 26 | 4 | 13 | 15 | 22 | 1 | 0 | 4 | 4 | 0 | 0 | 10 | 15 | 14 | 12 | 12 | 6 | 0 | 14 | 10 | 214 | 149 | 0.000255153 | 0.004924187 | 0.948 | 1 | 0.694 | 0.723 | 1 | 0 | 0.537 | 1 | 0.41 | 0.455 | 0.393 |
| SE | 105377 | ENSG00000114302 | PRKAR2A | chr3 | - | 48756378 | 48756444 | 48752175 | 48752317 | 48765003 | 48765078 | 124 | 93 | 100 | 104 | 164 | 1 | 0 | 0 | 1 | 2 | 281 | 111 | 127 | 148 | 173 | 14 | 0 | 1 | 0 | 11 | 215 | 149 | 0.000255669 | 0.004932945 | 1 | 0.97 | 1 | 0.986 | 0.983 | 0.933 | 1 | 0.989 | 1 | 0.916 | 0.02 |
| SE | 77307 | ENSG00000099968 | BCL2L13 | chr22 | + | 17702242 | 17702386 | 17683213 | 17683321 | 17726676 | 17728699 | 80 | 64 | 88 | 94 | 89 | 0 | 0 | 0 | 0 | 0 | 110 | 130 | 47 | 101 | 190 | 14 | 0 | 0 | 3 | 0 | 293 | 149 | 0.000255827 | 0.004934773 | 1 | 1 | 1 | 1 | 1 | 0.8 | 1 | 1 | 0.945 | 1 | 0.051 |
| SE | 99308 | ENSG00000101017 | CD40 | chr20 | + | 46123125 | 46123219 | 46122609 | 46122756 | 46128137 | 46128224 | 35 | 6 | 18 | 28 | 30 | 0 | 1 | 0 | 3 | 7 | 26 | 32 | 11 | 16 | 27 | 0 | 0 | 0 | 0 | 0 | 243 | 149 | 0.000256453 | 0.004945623 | 1 | 0.786 | 1 | 0.851 | 0.724 | 1 | 1 | 1 | 1 | 1 | -0.128 |
| SE | 10986 | ENSG00000083454 | P2RX5 | chr17 | - | 3689491 | 3689627 | 3688625 | 3688759 | 3690069 | 3690150 | 97 | 21 | 91 | 102 | 116 | 0 | 0 | 0 | 0 | 0 | 67 | 200 | 58 | 68 | 148 | 0 | 17 | 0 | 0 | 0 | 285 | 149 | 0.000256668 | 0.004948552 | 1 | 1 | 1 | 1 | 1 | 1 | 0.86 | 1 | 1 | 1 | 0.028 |
| SE | 7949 | ENSG00000108448 | TRIM16L | chr17 | + | 18723769 | 18723831 | 18722117 | 18722332 | 18727524 | 18727758 | 0 | 0 | 0 | 0 | 1 | 6 | 2 | 5 | 6 | 10 | 7 | 0 | 0 | 0 | 8 | 10 | 12 | 3 | 4 | 64 | 211 | 149 | 0.000256856 | 0.004950959 | 0 | 0 | 0 | 0 | 0.066 | 0.331 | 0 | 0 | 0 | 0.081 | -0.069 |
| SE | 58787 | ENSG00000077254 | USP33 | chr1 | - | 77715741 | 77715868 | 77713199 | 77713281 | 77717866 | 77718047 | 199 | 157 | 140 | 189 | 187 | 7 | 3 | 2 | 3 | 0 | 26 | 132 | 64 | 99 | 2 | 0 | 12 | 11 | 1 | 0 | 276 | 149 | 0.000257076 | 0.004953988 | 0.939 | 0.966 | 0.974 | 0.971 | 1 | 1 | 0.856 | 0.759 | 0.982 | 1 | 0.051 |
| SE | 74326 | ENSG00000151789 | ZNF385D | chr3 | - | 21510860 | 21511023 | 21436969 | 21437203 | 21563261 | 21563312 | 9 | 19 | 1 | 16 | 19 | 0 | 0 | 0 | 0 | 0 | 23 | 12 | 20 | 59 | 72 | 32 | 0 | 4 | 4 | 0 | 298 | 149 | 0.000257148 | 0.004954148 | 1 | 1 | 1 | 1 | 1 | 0.264 | 1 | 1 | 0.881 | 1 | 0.171 |
| SE | 40865 | ENSG00000167196 | FBXO22 | chr15 | + | 75905577 | 75905657 | 75904490 | 75904629 | 75913202 | 75913290 | 4 | 3 | 0 | 8 | 1 | 14 | 11 | 23 | 16 | 11 | 0 | 3 | 0 | 0 | 0 | 21 | 4 | 5 | 11 | 12 | 229 | 149 | 0.000257216 | 0.004954248 | 0.157 | 0.151 | 0 | 0.245 | 0.056 | 0 | 0.328 | 0 | 0 | 0 | 0.056 |
| SE | 74816 | ENSG00000042088 | TDP1 | chr14 | + | 89963224 | 89963673 | 89956577 | 89956800 | 89966146 | 89966190 | 29 | 20 | 42 | 36 | 47 | 0 | 0 | 0 | 4 | 0 | 19 | 41 | 18 | 47 | 54 | 0 | 15 | 5 | 0 | 0 | 298 | 149 | 0.000257469 | 0.004957884 | 1 | 1 | 1 | 1 | 0.855 | 1 | 0.577 | 0.643 | 1 | 1 | 0.127 |
| SE | 20481 | ENSG00000164080 | RAD54L2 | chr3 | + | 51645023 | 51645229 | 51643874 | 51643974 | 51645590 | 51645763 | 74 | 54 | 96 | 77 | 82 | 0 | 1 | 0 | 0 | 0 | 118 | 104 | 64 | 50 | 81 | 11 | 4 | 0 | 0 | 0 | 298 | 149 | 0.000257932 | 0.004965588 | 1 | 0.964 | 1 | 1 | 1 | 0.843 | 0.929 | 1 | 1 | 1 | 0.038 |
| SE | 49938 | ENSG00000133657 | ATP13A3 | chr3 | - | 194441310 | 194441461 | 194438855 | 194438972 | 194444724 | 194444786 | 268 | 134 | 175 | 205 | 469 | 2 | 0 | 2 | 4 | 0 | 50 | 111 | 168 | 38 | 19 | 0 | 5 | 10 | 0 | 0 | 298 | 149 | 0.000258182 | 0.004969169 | 0.985 | 1 | 0.978 | 0.962 | 1 | 1 | 0.917 | 0.894 | 1 | 1 | 0.023 |
| SE | 70905 | ENSG00000130706 | ADRM1 | chr20 | + | 62303567 | 62303781 | 62302895 | 62303049 | 62306196 | 62306320 | 44 | 19 | 20 | 43 | 9 | 0 | 3 | 0 | 0 | 0 | 49 | 76 | 44 | 32 | 79 | 0 | 3 | 0 | 4 | 8 | 298 | 149 | 0.000258329 | 0.004969998 | 1 | 0.76 | 1 | 1 | 1 | 1 | 0.927 | 1 | 0.8 | 0.832 | 0.04 |
| SE | 30339 | ENSG00000227345 | PARG | chr10 | - | 49935075 | 49935142 | 49933176 | 49934163 | 49941508 | 49941824 | 57 | 27 | 52 | 47 | 76 | 6 | 0 | 5 | 1 | 2 | 10 | 31 | 17 | 14 | 14 | 0 | 0 | 0 | 0 | 0 | 216 | 149 | 0.000258352 | 0.004969998 | 0.868 | 1 | 0.878 | 0.97 | 0.963 | 1 | 1 | 1 | 1 | 1 | -0.064 |
| SE | 87341 | ENSG00000215301 | DDX3X | chrX | + | 41347314 | 41347451 | 41346228 | 41346622 | 41347639 | 41347694 | 424 | 284 | 414 | 441 | 488 | 10 | 6 | 7 | 20 | 12 | 53 | 164 | 296 | 163 | 17 | 10 | 11 | 8 | 18 | 0 | 286 | 149 | 0.000258606 | 0.004973668 | 0.957 | 0.961 | 0.969 | 0.92 | 0.955 | 0.734 | 0.886 | 0.951 | 0.825 | 1 | 0.073 |
| SE | 15425 | ENSG00000102901 | CENPT | chr16 | - | 67832227 | 67832315 | 67832011 | 67832108 | 67832454 | 67832545 | 103 | 77 | 93 | 119 | 124 | 0 | 0 | 1 | 0 | 0 | 61 | 117 | 62 | 74 | 326 | 6 | 0 | 0 | 6 | 7 | 237 | 149 | 0.000258703 | 0.004974229 | 1 | 1 | 0.983 | 1 | 1 | 0.865 | 1 | 1 | 0.886 | 0.967 | 0.053 |
| SE | 28577 | ENSG00000119414 | PPP6C | chr9 | - | 125160840 | 125160906 | 125153905 | 125153985 | 125171084 | 125171180 | 230 | 133 | 176 | 178 | 182 | 9 | 8 | 4 | 4 | 6 | 154 | 117 | 181 | 194 | 74 | 22 | 0 | 13 | 9 | 19 | 215 | 149 | 0.000258762 | 0.004974229 | 0.947 | 0.92 | 0.968 | 0.969 | 0.955 | 0.829 | 1 | 0.906 | 0.937 | 0.73 | 0.071 |
| SE | 100323 | ENSG00000081237 | PTPRC | chr1 | + | 198696711 | 198696909 | 198692346 | 198692373 | 198699563 | 198699620 | 1556 | 776 | 1106 | 1299 | 2628 | 1003 | 892 | 987 | 1227 | 3005 | 270 | 1028 | 441 | 156 | 125 | 53 | 566 | 377 | 93 | 43 | 298 | 149 | 0.000259462 | 0.004986451 | 0.437 | 0.303 | 0.359 | 0.346 | 0.304 | 0.718 | 0.476 | 0.369 | 0.456 | 0.592 | -0.172 |
| SE | 92243 | ENSG00000114209 | PDCD10 | chr3 | - | 167725400 | 167725473 | 167720061 | 167720273 | 167734458 | 167734510 | 56 | 50 | 29 | 45 | 85 | 2 | 6 | 0 | 3 | 5 | 14 | 32 | 22 | 0 | 17 | 8 | 2 | 5 | 3 | 10 | 222 | 149 | 0.000259579 | 0.004987272 | 0.949 | 0.848 | 1 | 0.91 | 0.919 | 0.54 | 0.915 | 0.747 | 1 | 0.533 | 0.378 |
| SE | 92850 | ENSG00000130958 | SLC35D2 | chr9 | - | 96357410 | 96357551 | 96352037 | 96352109 | 96360153 | 96360221 | 5 | 0 | 0 | 3 | 3 | 64 | 90 | 85 | 104 | 120 | 0 | 8 | 6 | 0 | 48 | 183 | 72 | 78 | 143 | 583 | 290 | 149 | 0.000259632 | 0.004987272 | 0.039 | 0 | 0 | 0.01 | 0.013 | 0 | 0.054 | 0.038 | 0 | 0.041 | -0.014 |
| SE | 25700 | ENSG00000247774 | PCED1B-AS1 | chr12 | - | 47206510 | 47206588 | 47205897 | 47206069 | 47216164 | 47216200 | 63 | 45 | 144 | 89 | 187 | 12 | 2 | 13 | 9 | 13 | 37 | 34 | 47 | 27 | 6 | 0 | 0 | 4 | 2 | 0 | 227 | 149 | 0.00026006 | 0.004994204 | 0.775 | 0.937 | 0.879 | 0.867 | 0.904 | 1 | 1 | 0.885 | 0.899 | 1 | -0.084 |
| SE | 71223 | ENSG00000079263 | SP140 | chr2 | + | 230250980 | 230251061 | 230248884 | 230248968 | 230255451 | 230255532 | 21 | 25 | 19 | 26 | 46 | 37 | 39 | 37 | 38 | 78 | 20 | 22 | 14 | 22 | 1 | 45 | 58 | 76 | 63 | 22 | 230 | 149 | 0.00026012 | 0.004994204 | 0.269 | 0.293 | 0.25 | 0.307 | 0.276 | 0.224 | 0.197 | 0.107 | 0.184 | 0.029 | 0.131 |
| SE | 58661 | ENSG00000125633 | CCDC93 | chr2 | - | 117973907 | 117973994 | 117958364 | 117958481 | 117974849 | 117974900 | 124 | 55 | 87 | 131 | 126 | 0 | 1 | 0 | 3 | 0 | 56 | 109 | 80 | 70 | 18 | 0 | 0 | 3 | 13 | 0 | 236 | 149 | 0.00026053 | 0.005000844 | 1 | 0.972 | 1 | 0.965 | 1 | 1 | 1 | 0.944 | 0.773 | 1 | 0.044 |
| SE | 18003 | ENSG00000092108 | SCFD1 | chr14 | + | 30707816 | 30707876 | 30705822 | 30705885 | 30707989 | 30708065 | 1 | 5 | 8 | 4 | 12 | 92 | 71 | 73 | 108 | 104 | 0 | 2 | 1 | 0 | 0 | 15 | 41 | 27 | 20 | 34 | 209 | 149 | 0.000260653 | 0.005001979 | 0.008 | 0.048 | 0.072 | 0.026 | 0.076 | 0 | 0.034 | 0.026 | 0 | 0 | 0.034 |
| SE | 70756 | ENSG00000040933 | INPP4A | chr2 | + | 98545968 | 98546073 | 98543876 | 98544007 | 98552785 | 98552969 | 179 | 82 | 160 | 166 | 141 | 6 | 4 | 18 | 14 | 9 | 43 | 206 | 73 | 118 | 152 | 0 | 10 | 3 | 4 | 0 | 254 | 149 | 0.000260954 | 0.005006531 | 0.946 | 0.923 | 0.839 | 0.874 | 0.902 | 1 | 0.924 | 0.935 | 0.945 | 1 | -0.064 |
| SE | 4834 | ENSG00000132670 | PTPRA | chr20 | + | 2947981 | 2948024 | 2923206 | 2923285 | 2986764 | 2986849 | 16 | 15 | 8 | 20 | 5 | 0 | 0 | 0 | 0 | 0 | 39 | 26 | 17 | 28 | 59 | 0 | 0 | 0 | 0 | 14 | 192 | 149 | 0.000261113 | 0.005008354 | 1 | 1 | 1 | 1 | 1 | 1 | 1 | 1 | 1 | 0.766 | 0.047 |
| SE | 39780 | ENSG00000072274 | TFRC | chr3 | - | 196073929 | 196074125 | 196072002 | 196072152 | 196077063 | 196077122 | 116 | 80 | 142 | 140 | 78 | 5 | 1 | 1 | 0 | 0 | 78 | 209 | 187 | 102 | 27 | 0 | 10 | 3 | 12 | 0 | 298 | 149 | 0.000261258 | 0.005009903 | 0.921 | 0.976 | 0.986 | 1 | 1 | 1 | 0.913 | 0.969 | 0.81 | 1 | 0.038 |
| SE | 74328 | ENSG00000151789 | ZNF385D | chr3 | - | 21511621 | 21511757 | 21510860 | 21511023 | 21564573 | 21564684 | 0 | 0 | 0 | 0 | 1 | 4 | 14 | 1 | 15 | 15 | 0 | 20 | 0 | 8 | 0 | 16 | 10 | 17 | 40 | 42 | 285 | 149 | 0.000261801 | 0.005019086 | 0 | 0 | 0 | 0 | 0.034 | 0 | 0.511 | 0 | 0.095 | 0 | -0.114 |
| SE | 13779 | ENSG00000205629 | LCMT1 | chr16 | + | 25170713 | 25170805 | 25169111 | 25169213 | 25174936 | 25175034 | 67 | 34 | 64 | 64 | 87 | 1 | 0 | 3 | 2 | 0 | 82 | 96 | 32 | 56 | 87 | 0 | 8 | 5 | 9 | 0 | 241 | 149 | 0.000262168 | 0.005024898 | 0.976 | 1 | 0.93 | 0.952 | 1 | 1 | 0.881 | 0.798 | 0.794 | 1 | 0.077 |
| SE | 42483 | ENSG00000251503 | CENPS-CORT | chr1 | + | 10433841 | 10433965 | 10431257 | 10431401 | 10434656 | 10434690 | 25 | 13 | 13 | 19 | 33 | 1 | 0 | 0 | 0 | 0 | 13 | 14 | 21 | 3 | 35 | 18 | 3 | 0 | 0 | 0 | 273 | 149 | 0.00026231 | 0.00502639 | 0.932 | 1 | 1 | 1 | 1 | 0.283 | 0.718 | 1 | 1 | 1 | 0.186 |
| SE | 43888 | ENSG00000179335 | CLK3 | chr15 | + | 74625801 | 74625967 | 74624901 | 74625018 | 74627351 | 74627446 | 221 | 141 | 233 | 250 | 611 | 11 | 16 | 16 | 15 | 45 | 394 | 271 | 216 | 237 | 431 | 112 | 6 | 23 | 104 | 132 | 298 | 149 | 0.000262848 | 0.005035468 | 0.909 | 0.815 | 0.879 | 0.893 | 0.872 | 0.638 | 0.958 | 0.824 | 0.533 | 0.62 | 0.159 |
| SE | 48927 | ENSG00000198408 | OGA | chr10 | - | 101806044 | 101806143 | 101803734 | 101804019 | 101807729 | 101807901 | 798 | 610 | 699 | 824 | 1370 | 13 | 5 | 9 | 21 | 24 | 506 | 785 | 1053 | 524 | 336 | 0 | 13 | 7 | 5 | 0 | 248 | 149 | 0.00026292 | 0.005035615 | 0.974 | 0.987 | 0.979 | 0.959 | 0.972 | 1 | 0.973 | 0.989 | 0.984 | 1 | -0.015 |
| SE | 67341 | ENSG00000106636 | YKT6 | chr7 | + | 44206384 | 44206485 | 44204567 | 44204650 | 44207387 | 44207492 | 214 | 122 | 209 | 213 | 249 | 0 | 0 | 0 | 1 | 0 | 433 | 265 | 246 | 245 | 374 | 22 | 4 | 1 | 2 | 0 | 250 | 149 | 0.000263205 | 0.005039841 | 1 | 1 | 1 | 0.992 | 1 | 0.921 | 0.975 | 0.993 | 0.986 | 1 | 0.023 |
| SE | 88783 | ENSG00000140632 | GLYR1 | chr16 | - | 4813736 | 4813838 | 4812085 | 4812248 | 4814536 | 4814629 | 144 | 101 | 120 | 159 | 100 | 2 | 0 | 1 | 0 | 0 | 128 | 116 | 58 | 102 | 205 | 16 | 0 | 2 | 1 | 0 | 251 | 149 | 0.00026378 | 0.005049623 | 0.977 | 1 | 0.986 | 1 | 1 | 0.826 | 1 | 0.945 | 0.984 | 1 | 0.042 |
| SE | 106325 | ENSG00000125458 | NT5C | chr17 | - | 75130964 | 75131105 | 75130567 | 75130867 | 75131180 | 75131281 | 18 | 19 | 16 | 23 | 26 | 2 | 0 | 0 | 0 | 0 | 9 | 12 | 21 | 9 | 43 | 0 | 2 | 5 | 1 | 2 | 290 | 149 | 0.000265057 | 0.00507281 | 0.822 | 1 | 1 | 1 | 1 | 1 | 0.755 | 0.683 | 0.822 | 0.917 | 0.129 |
| SE | 71927 | ENSG00000157637 | SLC38A10 | chr17 | - | 81272515 | 81272627 | 81270917 | 81271024 | 81275968 | 81276151 | 195 | 75 | 146 | 152 | 94 | 2 | 2 | 0 | 1 | 0 | 329 | 345 | 150 | 257 | 398 | 7 | 4 | 9 | 4 | 10 | 261 | 149 | 0.000266407 | 0.005097404 | 0.982 | 0.955 | 1 | 0.989 | 1 | 0.964 | 0.98 | 0.905 | 0.973 | 0.958 | 0.029 |
| SE | 41013 | ENSG00000164091 | WDR82 | chr3 | - | 52261379 | 52261479 | 52260384 | 52260501 | 52266951 | 52267018 | 586 | 351 | 660 | 677 | 549 | 3 | 1 | 4 | 5 | 3 | 608 | 484 | 377 | 344 | 531 | 2 | 3 | 2 | 5 | 27 | 249 | 149 | 0.000266656 | 0.005100929 | 0.992 | 0.995 | 0.99 | 0.988 | 0.991 | 0.995 | 0.99 | 0.991 | 0.976 | 0.922 | 0.016 |
| SE | 40498 | ENSG00000285708 | NA | chr3 | - | 71335159 | 71335281 | 71299819 | 71299880 | 71359149 | 71359244 | 1 | 8 | 8 | 6 | 8 | 80 | 68 | 105 | 80 | 90 | 0 | 7 | 2 | 0 | 0 | 58 | 85 | 88 | 50 | 99 | 271 | 149 | 0.000268456 | 0.005134106 | 0.007 | 0.061 | 0.04 | 0.04 | 0.047 | 0 | 0.043 | 0.012 | 0 | 0 | 0.028 |
| SE | 9618 | ENSG00000173821 | RNF213 | chr17 | + | 80278763 | 80278910 | 80273240 | 80273404 | 80287814 | 80288363 | 58 | 33 | 33 | 46 | 50 | 253 | 119 | 175 | 219 | 225 | 210 | 64 | 42 | 143 | 100 | 636 | 214 | 161 | 351 | 317 | 296 | 149 | 0.000269897 | 0.005158742 | 0.103 | 0.122 | 0.087 | 0.096 | 0.101 | 0.143 | 0.131 | 0.116 | 0.17 | 0.137 | -0.038 |
| SE | 95045 | ENSG00000148334 | PTGES2 | chr9 | - | 128123701 | 128123851 | 128120758 | 128123134 | 128124491 | 128124550 | 119 | 80 | 130 | 91 | 50 | 0 | 0 | 0 | 0 | 4 | 151 | 167 | 54 | 168 | 175 | 6 | 0 | 0 | 6 | 10 | 298 | 149 | 0.000269958 | 0.005158742 | 1 | 1 | 1 | 1 | 0.862 | 0.926 | 1 | 1 | 0.933 | 0.897 | 0.021 |
| SE | 105113 | ENSG00000278311 | GGNBP2 | chr17 | + | 36567662 | 36567776 | 36560772 | 36560871 | 36577982 | 36578186 | 404 | 287 | 353 | 354 | 486 | 5 | 2 | 0 | 0 | 4 | 94 | 305 | 331 | 220 | 65 | 16 | 0 | 1 | 2 | 4 | 263 | 149 | 0.000270002 | 0.005158742 | 0.979 | 0.988 | 1 | 1 | 0.986 | 0.769 | 1 | 0.995 | 0.984 | 0.902 | 0.061 |
| SE | 79304 | ENSG00000001631 | KRIT1 | chr7 | - | 92244901 | 92245171 | 92242033 | 92242137 | 92245426 | 92245595 | 3 | 11 | 7 | 4 | 15 | 2 | 1 | 13 | 11 | 1 | 0 | 1 | 1 | 5 | 0 | 7 | 3 | 7 | 3 | 2 | 298 | 149 | 0.000270008 | 0.005158742 | 0.429 | 0.846 | 0.212 | 0.154 | 0.882 | 0 | 0.143 | 0.067 | 0.455 | 0 | 0.372 |
| SE | 5133 | ENSG00000196476 | C20orf96 | chr20 | - | 289558 | 289676 | 283962 | 284081 | 290258 | 290307 | 16 | 7 | 12 | 7 | 7 | 2 | 2 | 4 | 2 | 0 | 72 | 4 | 10 | 19 | 42 | 0 | 3 | 0 | 0 | 0 | 267 | 149 | 0.00027093 | 0.005175103 | 0.817 | 0.661 | 0.626 | 0.661 | 1 | 1 | 0.427 | 1 | 1 | 1 | -0.132 |
| SE | 12630 | ENSG00000090857 | PDPR | chr16 | + | 70131301 | 70131419 | 70128958 | 70129122 | 70132150 | 70132300 | 36 | 5 | 37 | 19 | 49 | 3 | 0 | 0 | 1 | 0 | 15 | 23 | 19 | 18 | 104 | 0 | 0 | 0 | 0 | 20 | 267 | 149 | 0.000271012 | 0.0051754 | 0.87 | 1 | 1 | 0.914 | 1 | 1 | 1 | 1 | 1 | 0.744 | 0.008 |
| SE | 4747 | ENSG00000031823 | RANBP3 | chr19 | - | 5957917 | 5957968 | 5951392 | 5951596 | 5978060 | 5978093 | 12 | 2 | 18 | 11 | 22 | 1 | 1 | 1 | 2 | 0 | 14 | 12 | 3 | 5 | 24 | 0 | 3 | 2 | 3 | 13 | 200 | 149 | 0.000271827 | 0.005189695 | 0.899 | 0.598 | 0.931 | 0.804 | 1 | 1 | 0.749 | 0.528 | 0.554 | 0.579 | 0.164 |
| SE | 101253 | ENSG00000166224 | SGPL1 | chr10 | + | 70871836 | 70871986 | 70869791 | 70869897 | 70873350 | 70873589 | 54 | 32 | 52 | 47 | 66 | 0 | 0 | 0 | 0 | 0 | 54 | 27 | 41 | 11 | 68 | 15 | 0 | 0 | 0 | 0 | 298 | 149 | 0.000271913 | 0.005190079 | 1 | 1 | 1 | 1 | 1 | 0.643 | 1 | 1 | 1 | 1 | 0.071 |
| SE | 70389 | ENSG00000125257 | ABCC4 | chr13 | - | 95163121 | 95163216 | 95161188 | 95161335 | 95163609 | 95163647 | 161 | 108 | 73 | 238 | 147 | 1 | 1 | 0 | 1 | 1 | 80 | 181 | 45 | 120 | 566 | 0 | 5 | 0 | 1 | 19 | 244 | 149 | 0.000272666 | 0.005203184 | 0.99 | 0.985 | 1 | 0.993 | 0.989 | 1 | 0.957 | 1 | 0.987 | 0.948 | 0.013 |
| SE | 17075 | ENSG00000273749 | CYFIP1 | chr15 | - | 22873490 | 22873729 | 22872636 | 22872972 | 22874549 | 22874644 | 104 | 59 | 79 | 71 | 54 | 0 | 0 | 0 | 0 | 0 | 346 | 83 | 52 | 168 | 313 | 10 | 0 | 6 | 5 | 0 | 298 | 149 | 0.000273376 | 0.005215463 | 1 | 1 | 1 | 1 | 1 | 0.945 | 1 | 0.812 | 0.944 | 1 | 0.06 |
| SE | 66553 | ENSG00000133872 | SARAF | chr8 | - | 30073876 | 30074051 | 30069641 | 30070059 | 30082846 | 30083124 | 794 | 522 | 913 | 852 | 1437 | 32 | 27 | 47 | 34 | 44 | 344 | 445 | 650 | 261 | 263 | 12 | 45 | 45 | 15 | 19 | 298 | 149 | 0.00027495 | 0.005244207 | 0.925 | 0.906 | 0.907 | 0.926 | 0.942 | 0.935 | 0.832 | 0.878 | 0.897 | 0.874 | 0.038 |
| SE | 99594 | ENSG00000149187 | CELF1 | chr11 | - | 47494330 | 47494443 | 47488971 | 47489024 | 47499452 | 47499604 | 6 | 0 | 3 | 8 | 11 | 40 | 24 | 29 | 39 | 53 | 22 | 0 | 2 | 18 | 10 | 19 | 8 | 13 | 23 | 9 | 262 | 149 | 0.000275057 | 0.005244965 | 0.079 | 0 | 0.056 | 0.104 | 0.106 | 0.397 | 0 | 0.08 | 0.308 | 0.387 | -0.165 |
| SE | 6296 | ENSG00000141644 | MBD1 | chr18 | - | 50274185 | 50274353 | 50273563 | 50273863 | 50274976 | 50275045 | 113 | 62 | 76 | 111 | 15 | 6 | 9 | 2 | 12 | 3 | 68 | 38 | 59 | 67 | 170 | 15 | 20 | 11 | 19 | 20 | 298 | 149 | 0.000275358 | 0.005249427 | 0.904 | 0.775 | 0.95 | 0.822 | 0.714 | 0.694 | 0.487 | 0.728 | 0.638 | 0.81 | 0.162 |
| SE | 76069 | ENSG00000137692 | DCUN1D5 | chr11 | - | 103082747 | 103082839 | 103066458 | 103066567 | 103083255 | 103083326 | 92 | 65 | 44 | 67 | 87 | 0 | 5 | 8 | 6 | 13 | 2 | 44 | 51 | 16 | 17 | 0 | 3 | 1 | 1 | 0 | 241 | 149 | 0.000275477 | 0.005250433 | 1 | 0.889 | 0.773 | 0.873 | 0.805 | 1 | 0.901 | 0.969 | 0.908 | 1 | -0.088 |
| SE | 14438 | ENSG00000130703 | OSBPL2 | chr20 | + | 62260016 | 62260125 | 62256056 | 62256221 | 62263615 | 62263691 | 117 | 73 | 106 | 104 | 125 | 4 | 0 | 2 | 3 | 8 | 180 | 104 | 95 | 100 | 114 | 9 | 0 | 9 | 13 | 20 | 258 | 149 | 0.000275673 | 0.005252885 | 0.944 | 1 | 0.968 | 0.952 | 0.9 | 0.92 | 1 | 0.859 | 0.816 | 0.767 | 0.08 |
| SE | 41732 | ENSG00000163319 | MRPS18C | chr4 | + | 83458345 | 83458429 | 83456908 | 83456958 | 83459739 | 83459797 | 231 | 138 | 179 | 156 | 262 | 21 | 4 | 8 | 8 | 11 | 149 | 154 | 103 | 220 | 115 | 22 | 13 | 17 | 23 | 6 | 233 | 149 | 0.000275844 | 0.005254871 | 0.876 | 0.957 | 0.935 | 0.926 | 0.938 | 0.812 | 0.883 | 0.795 | 0.859 | 0.925 | 0.072 |
| SE | 20000 | ENSG00000198663 | C6orf89 | chr6 | + | 36894503 | 36894603 | 36885951 | 36886028 | 36899425 | 36899633 | 26 | 19 | 26 | 21 | 20 | 5 | 1 | 0 | 6 | 5 | 57 | 20 | 10 | 37 | 41 | 0 | 2 | 2 | 2 | 0 | 249 | 149 | 0.000277338 | 0.005282036 | 0.757 | 0.919 | 1 | 0.677 | 0.705 | 1 | 0.857 | 0.749 | 0.917 | 1 | -0.093 |
| SE | 45007 | ENSG00000275131 | PDE4DIPP2 | chr1 | - | 120523893 | 120524126 | 120519467 | 120519821 | 120530295 | 120530384 | 36 | 29 | 33 | 27 | 33 | 0 | 0 | 0 | 2 | 0 | 22 | 44 | 35 | 24 | 12 | 8 | 0 | 0 | 4 | 0 | 298 | 149 | 0.000278416 | 0.005301277 | 1 | 1 | 1 | 0.871 | 1 | 0.579 | 1 | 1 | 0.75 | 1 | 0.108 |
| SE | 46247 | ENSG00000198301 | SDAD1 | chr4 | - | 75974075 | 75974133 | 75973316 | 75973391 | 75975743 | 75975844 | 326 | 255 | 249 | 241 | 468 | 6 | 24 | 20 | 12 | 18 | 76 | 153 | 78 | 39 | 49 | 8 | 20 | 18 | 7 | 0 | 207 | 149 | 0.000278756 | 0.005306456 | 0.975 | 0.884 | 0.9 | 0.935 | 0.949 | 0.872 | 0.846 | 0.757 | 0.8 | 1 | 0.074 |
| SE | 16968 | ENSG00000137767 | SQOR | chr15 | + | 45676100 | 45676310 | 45673606 | 45673801 | 45682477 | 45682661 | 459 | 265 | 357 | 380 | 663 | 3 | 4 | 0 | 5 | 2 | 889 | 635 | 293 | 711 | 881 | 16 | 4 | 2 | 27 | 5 | 298 | 149 | 0.000278975 | 0.005309344 | 0.987 | 0.971 | 1 | 0.974 | 0.994 | 0.965 | 0.988 | 0.987 | 0.929 | 0.989 | 0.014 |
| SE | 68584 | ENSG00000163684 | RPP14 | chr3 | + | 58310308 | 58310406 | 58306244 | 58306417 | 58310506 | 58310591 | 7 | 4 | 10 | 15 | 19 | 9 | 6 | 1 | 16 | 4 | 26 | 44 | 10 | 13 | 51 | 0 | 6 | 3 | 5 | 0 | 247 | 149 | 0.000279056 | 0.005309597 | 0.319 | 0.287 | 0.858 | 0.361 | 0.741 | 1 | 0.816 | 0.668 | 0.611 | 1 | -0.306 |
| SE | 22491 | ENSG00000165097 | KDM1B | chr6 | + | 18205536 | 18205664 | 18200438 | 18200576 | 18207397 | 18207529 | 61 | 88 | 41 | 101 | 97 | 1 | 1 | 3 | 0 | 3 | 72 | 68 | 51 | 113 | 19 | 0 | 9 | 2 | 14 | 0 | 277 | 149 | 0.000279298 | 0.0053129 | 0.97 | 0.979 | 0.88 | 1 | 0.946 | 1 | 0.803 | 0.932 | 0.813 | 1 | 0.045 |
| SE | 80937 | ENSG00000100997 | ABHD12 | chr20 | - | 25314924 | 25314970 | 25309445 | 25309575 | 25317047 | 25317074 | 59 | 21 | 45 | 35 | 11 | 1 | 2 | 0 | 0 | 0 | 50 | 46 | 52 | 42 | 50 | 6 | 6 | 2 | 2 | 0 | 195 | 149 | 0.000279594 | 0.005317239 | 0.978 | 0.889 | 1 | 1 | 1 | 0.864 | 0.854 | 0.952 | 0.941 | 1 | 0.051 |
| SE | 106324 | ENSG00000125458 | NT5C | chr17 | - | 75130964 | 75131081 | 75130752 | 75130867 | 75131180 | 75131281 | 12 | 6 | 10 | 16 | 12 | 2 | 0 | 0 | 0 | 0 | 5 | 0 | 11 | 2 | 23 | 0 | 2 | 5 | 1 | 2 | 266 | 149 | 0.000279795 | 0.005319765 | 0.771 | 1 | 1 | 1 | 1 | 1 | 0 | 0.552 | 0.528 | 0.866 | 0.365 |
| SE | 7496 | ENSG00000108840 | HDAC5 | chr17 | - | 44085021 | 44085155 | 44084554 | 44084675 | 44086571 | 44086737 | 81 | 52 | 66 | 76 | 78 | 3 | 2 | 4 | 2 | 2 | 70 | 51 | 27 | 39 | 122 | 4 | 3 | 5 | 3 | 21 | 283 | 149 | 0.000280017 | 0.005322702 | 0.934 | 0.932 | 0.897 | 0.952 | 0.954 | 0.902 | 0.9 | 0.74 | 0.873 | 0.754 | 0.1 |
| SE | 75937 | ENSG00000160185 | UBASH3A | chr21 | + | 42434831 | 42435509 | 42432102 | 42432202 | 42437487 | 42437580 | 39 | 32 | 52 | 61 | 50 | 0 | 0 | 0 | 12 | 10 | 39 | 65 | 17 | 18 | 91 | 0 | 0 | 0 | 0 | 5 | 298 | 149 | 0.000280412 | 0.005328919 | 1 | 1 | 1 | 0.718 | 0.714 | 1 | 1 | 1 | 1 | 0.901 | -0.094 |
| SE | 105923 | ENSG00000234608 | MAPKAPK5-AS1 | chr12 | - | 111841326 | 111841470 | 111839776 | 111840003 | 111841683 | 111841686 | 20 | 14 | 11 | 18 | 18 | 2 | 0 | 2 | 5 | 3 | 17 | 3 | 11 | 4 | 47 | 0 | 0 | 4 | 0 | 0 | 293 | 149 | 0.000280528 | 0.005329395 | 0.836 | 1 | 0.737 | 0.647 | 0.753 | 1 | 1 | 0.583 | 1 | 1 | -0.122 |
| SE | 47588 | ENSG00000008869 | HEATR5B | chr2 | - | 37020654 | 37020836 | 37019808 | 37019877 | 37027922 | 37028174 | 226 | 100 | 187 | 169 | 292 | 4 | 2 | 5 | 6 | 7 | 11 | 125 | 110 | 76 | 12 | 0 | 0 | 2 | 0 | 0 | 298 | 149 | 0.000280573 | 0.005329395 | 0.966 | 0.962 | 0.949 | 0.934 | 0.954 | 1 | 1 | 0.965 | 1 | 1 | -0.04 |
| SE | 28702 | ENSG00000112249 | ASCC3 | chr6 | - | 100864063 | 100864214 | 100848003 | 100848707 | 100867907 | 100868032 | 760 | 778 | 477 | 794 | 813 | 2 | 6 | 1 | 9 | 2 | 231 | 548 | 436 | 231 | 68 | 0 | 2 | 7 | 4 | 14 | 298 | 149 | 0.000280708 | 0.005330658 | 0.995 | 0.985 | 0.996 | 0.978 | 0.995 | 1 | 0.993 | 0.969 | 0.967 | 0.708 | 0.062 |
| SE | 89271 | ENSG00000104325 | DECR1 | chr8 | + | 90017301 | 90017326 | 90001462 | 90001561 | 90018908 | 90018966 | 57 | 32 | 49 | 34 | 13 | 25 | 5 | 15 | 19 | 1 | 84 | 56 | 16 | 55 | 16 | 0 | 4 | 11 | 4 | 0 | 174 | 149 | 0.000282049 | 0.005354823 | 0.661 | 0.846 | 0.737 | 0.605 | 0.918 | 1 | 0.923 | 0.555 | 0.922 | 1 | -0.127 |
| SE | 55155 | ENSG00000116586 | LAMTOR2 | chr1 | + | 156057977 | 156058067 | 156055084 | 156055425 | 156058314 | 156058506 | 240 | 228 | 292 | 226 | 105 | 1 | 4 | 0 | 1 | 0 | 769 | 399 | 205 | 396 | 826 | 20 | 6 | 5 | 15 | 0 | 239 | 149 | 0.000282177 | 0.005355035 | 0.993 | 0.973 | 1 | 0.993 | 1 | 0.96 | 0.976 | 0.962 | 0.943 | 1 | 0.024 |
| SE | 104932 | ENSG00000064102 | INTS13 | chr12 | - | 26916001 | 26916180 | 26914407 | 26914532 | 26917351 | 26917441 | 112 | 100 | 86 | 93 | 171 | 3 | 0 | 0 | 5 | 15 | 66 | 69 | 90 | 34 | 6 | 0 | 0 | 4 | 0 | 0 | 298 | 149 | 0.000282253 | 0.005355035 | 0.949 | 1 | 1 | 0.903 | 0.851 | 1 | 1 | 0.918 | 1 | 1 | -0.043 |
| SE | 40063 | ENSG00000125347 | IRF1 | chr5 | - | 132484361 | 132484497 | 132481608 | 132484075 | 132486556 | 132486686 | 145 | 87 | 168 | 127 | 120 | 10 | 0 | 6 | 5 | 2 | 179 | 74 | 70 | 289 | 153 | 0 | 2 | 2 | 4 | 0 | 285 | 149 | 0.000282299 | 0.005355035 | 0.883 | 1 | 0.936 | 0.93 | 0.969 | 1 | 0.951 | 0.948 | 0.974 | 1 | -0.031 |
| SE | 3507 | ENSG00000130755 | GMFG | chr19 | - | 39335434 | 39335531 | 39332903 | 39333126 | 39335973 | 39336005 | 236 | 112 | 192 | 222 | 144 | 9 | 3 | 5 | 6 | 8 | 304 | 243 | 198 | 186 | 305 | 35 | 2 | 6 | 18 | 34 | 246 | 149 | 0.000282333 | 0.005355035 | 0.941 | 0.958 | 0.959 | 0.957 | 0.916 | 0.84 | 0.987 | 0.952 | 0.862 | 0.845 | 0.049 |
| SE | 21183 | ENSG00000143252 | SDHC | chr1 | + | 161356676 | 161356840 | 161340593 | 161340655 | 161362328 | 161362433 | 503 | 214 | 470 | 469 | 576 | 13 | 3 | 14 | 20 | 11 | 426 | 516 | 232 | 334 | 569 | 26 | 31 | 17 | 21 | 9 | 298 | 149 | 0.000282411 | 0.005355208 | 0.951 | 0.973 | 0.944 | 0.921 | 0.963 | 0.891 | 0.893 | 0.872 | 0.888 | 0.969 | 0.048 |
| SE | 39996 | ENSG00000107566 | ERLIN1 | chr10 | - | 100164003 | 100164095 | 100156144 | 100156234 | 100167347 | 100167406 | 59 | 63 | 52 | 57 | 111 | 2 | 2 | 0 | 0 | 0 | 76 | 76 | 120 | 126 | 50 | 10 | 2 | 6 | 1 | 0 | 241 | 149 | 0.000282898 | 0.005362379 | 0.948 | 0.951 | 1 | 1 | 1 | 0.825 | 0.959 | 0.925 | 0.987 | 1 | 0.041 |
| SE | 86926 | ENSG00000122122 | SASH3 | chrX | + | 129788430 | 129788577 | 129787974 | 129788070 | 129790939 | 129791081 | 1139 | 497 | 694 | 820 | 967 | 2 | 0 | 2 | 6 | 0 | 967 | 962 | 496 | 796 | 983 | 5 | 0 | 8 | 14 | 20 | 296 | 149 | 0.000282926 | 0.005362379 | 0.997 | 1 | 0.994 | 0.986 | 1 | 0.99 | 1 | 0.969 | 0.966 | 0.961 | 0.018 |
| SE | 104819 | ENSG00000136404 | TM6SF1 | chr15 | + | 83126754 | 83126847 | 83124671 | 83124776 | 83127357 | 83127477 | 46 | 48 | 30 | 54 | 64 | 4 | 1 | 1 | 2 | 7 | 70 | 32 | 14 | 42 | 10 | 16 | 3 | 1 | 13 | 32 | 242 | 149 | 0.000283298 | 0.00536813 | 0.876 | 0.967 | 0.949 | 0.943 | 0.849 | 0.729 | 0.868 | 0.896 | 0.665 | 0.161 | 0.253 |
| SE | 9245 | ENSG00000178971 | CTC1 | chr17 | - | 8235830 | 8235959 | 8235052 | 8235285 | 8236057 | 8236342 | 76 | 60 | 55 | 106 | 53 | 1 | 1 | 0 | 8 | 0 | 14 | 43 | 41 | 47 | 37 | 0 | 10 | 1 | 4 | 7 | 278 | 149 | 0.000283429 | 0.005369303 | 0.976 | 0.97 | 1 | 0.877 | 1 | 1 | 0.697 | 0.956 | 0.863 | 0.739 | 0.114 |
| SE | 85158 | ENSG00000136504 | KAT7 | chr17 | + | 49805362 | 49805445 | 49798318 | 49798558 | 49811475 | 49811574 | 145 | 86 | 124 | 127 | 241 | 2 | 1 | 3 | 3 | 12 | 409 | 260 | 240 | 389 | 271 | 24 | 7 | 8 | 32 | 24 | 232 | 149 | 0.000283527 | 0.005369894 | 0.979 | 0.982 | 0.964 | 0.965 | 0.928 | 0.916 | 0.96 | 0.951 | 0.886 | 0.879 | 0.045 |
| SE | 10457 | ENSG00000141698 | NT5C3B | chr17 | - | 41828789 | 41828952 | 41827425 | 41827626 | 41830800 | 41830890 | 67 | 61 | 103 | 101 | 99 | 1 | 3 | 4 | 7 | 1 | 73 | 123 | 39 | 158 | 90 | 0 | 0 | 0 | 4 | 0 | 298 | 149 | 0.000283765 | 0.005373067 | 0.971 | 0.91 | 0.928 | 0.878 | 0.98 | 1 | 1 | 1 | 0.952 | 1 | -0.057 |
| SE | 48279 | ENSG00000113532 | ST8SIA4 | chr5 | - | 100856102 | 100856396 | 100806932 | 100812129 | 100886342 | 100886600 | 323 | 229 | 234 | 294 | 499 | 7 | 12 | 1 | 12 | 8 | 70 | 175 | 161 | 103 | 50 | 14 | 6 | 15 | 6 | 0 | 298 | 149 | 0.000283869 | 0.005373752 | 0.958 | 0.905 | 0.992 | 0.925 | 0.969 | 0.714 | 0.936 | 0.843 | 0.896 | 1 | 0.072 |
| SE | 24888 | ENSG00000113441 | LNPEP | chr5 | + | 97024520 | 97024682 | 97022299 | 97022484 | 97027732 | 97027814 | 216 | 160 | 203 | 262 | 317 | 11 | 1 | 1 | 4 | 1 | 5 | 140 | 133 | 54 | 24 | 0 | 2 | 17 | 1 | 6 | 298 | 149 | 0.000284023 | 0.005375351 | 0.908 | 0.988 | 0.99 | 0.97 | 0.994 | 1 | 0.972 | 0.796 | 0.964 | 0.667 | 0.09 |
| SE | 84319 | ENSG00000164062 | APEH | chr3 | + | 49676377 | 49676515 | 49676055 | 49676219 | 49676608 | 49676700 | 423 | 209 | 302 | 347 | 345 | 11 | 10 | 12 | 10 | 19 | 558 | 355 | 204 | 310 | 675 | 14 | 10 | 10 | 0 | 3 | 287 | 149 | 0.000284096 | 0.00537545 | 0.952 | 0.916 | 0.929 | 0.947 | 0.904 | 0.954 | 0.949 | 0.914 | 1 | 0.992 | -0.032 |
| SE | 43263 | ENSG00000139372 | TDG | chr12 | + | 103976917 | 103977060 | 103965871 | 103966060 | 103979830 | 103980072 | 97 | 52 | 95 | 103 | 70 | 1 | 1 | 0 | 1 | 0 | 34 | 89 | 88 | 90 | 23 | 5 | 3 | 5 | 1 | 0 | 292 | 149 | 0.000284499 | 0.005381769 | 0.98 | 0.964 | 1 | 0.981 | 1 | 0.776 | 0.938 | 0.9 | 0.979 | 1 | 0.066 |
| SE | 5902 | ENSG00000124228 | DDX27 | chr20 | + | 49231022 | 49231115 | 49230198 | 49230349 | 49233305 | 49233405 | 3 | 5 | 5 | 3 | 0 | 22 | 7 | 27 | 28 | 5 | 7 | 23 | 13 | 14 | 24 | 44 | 30 | 18 | 21 | 39 | 242 | 149 | 0.0002852 | 0.005393729 | 0.077 | 0.305 | 0.102 | 0.062 | 0 | 0.089 | 0.321 | 0.308 | 0.291 | 0.275 | -0.148 |
| SE | 105227 | ENSG00000102967 | DHODH | chr16 | + | 72023473 | 72023633 | 72023164 | 72023318 | 72024144 | 72027659 | 21 | 12 | 15 | 11 | 8 | 1 | 0 | 3 | 0 | 0 | 14 | 11 | 7 | 13 | 29 | 0 | 5 | 2 | 3 | 0 | 298 | 149 | 0.000286694 | 0.005420665 | 0.913 | 1 | 0.714 | 1 | 1 | 1 | 0.524 | 0.636 | 0.684 | 1 | 0.157 |
| SE | 91730 | ENSG00000176623 | RMDN1 | chr8 | - | 86494828 | 86494966 | 86488551 | 86488639 | 86506994 | 86507112 | 12 | 3 | 5 | 4 | 20 | 60 | 38 | 34 | 62 | 18 | 0 | 7 | 2 | 2 | 0 | 43 | 56 | 14 | 35 | 18 | 287 | 149 | 0.00028742 | 0.005433092 | 0.094 | 0.039 | 0.071 | 0.032 | 0.366 | 0 | 0.061 | 0.069 | 0.029 | 0 | 0.089 |
| SE | 96327 | ENSG00000135387 | CAPRIN1 | chr11 | + | 34097697 | 34097761 | 34096478 | 34096673 | 34099302 | 34099332 | 61 | 48 | 51 | 77 | 101 | 0 | 0 | 2 | 0 | 2 | 27 | 51 | 36 | 41 | 11 | 0 | 6 | 0 | 7 | 0 | 213 | 149 | 0.000287671 | 0.005436526 | 1 | 1 | 0.947 | 1 | 0.972 | 1 | 0.856 | 1 | 0.804 | 1 | 0.052 |
| SE | 55123 | ENSG00000119725 | ZNF410 | chr14 | + | 73892026 | 73892107 | 73886860 | 73886915 | 73893796 | 73893932 | 19 | 16 | 17 | 14 | 14 | 3 | 1 | 0 | 0 | 0 | 36 | 6 | 41 | 21 | 27 | 6 | 0 | 6 | 4 | 0 | 230 | 149 | 0.000287813 | 0.005437885 | 0.804 | 0.912 | 1 | 1 | 1 | 0.795 | 1 | 0.816 | 0.773 | 1 | 0.066 |
| SE | 67667 | ENSG00000163848 | ZNF148 | chr3 | - | 125314936 | 125315053 | 125313307 | 125313656 | 125323308 | 125323444 | 9 | 8 | 6 | 16 | 19 | 167 | 105 | 125 | 135 | 230 | 0 | 10 | 10 | 16 | 29 | 14 | 89 | 95 | 10 | 41 | 266 | 149 | 0.000288263 | 0.005445078 | 0.029 | 0.041 | 0.026 | 0.062 | 0.044 | 0 | 0.059 | 0.056 | 0.473 | 0.284 | -0.134 |
| SE | 69656 | ENSG00000143390 | RFX5 | chr1 | - | 151346421 | 151346606 | 151346204 | 151346333 | 151347210 | 151347292 | 41 | 21 | 39 | 26 | 30 | 0 | 0 | 0 | 0 | 0 | 47 | 30 | 36 | 28 | 67 | 4 | 0 | 0 | 0 | 8 | 298 | 149 | 0.000288363 | 0.005445658 | 1 | 1 | 1 | 1 | 1 | 0.855 | 1 | 1 | 1 | 0.807 | 0.068 |
| SE | 19966 | ENSG00000136758 | YME1L1 | chr10 | - | 27142386 | 27142485 | 27136275 | 27136385 | 27145427 | 27145590 | 533 | 375 | 478 | 524 | 719 | 10 | 13 | 12 | 8 | 5 | 141 | 317 | 404 | 378 | 75 | 7 | 9 | 22 | 16 | 0 | 248 | 149 | 0.000288779 | 0.005452194 | 0.97 | 0.945 | 0.96 | 0.975 | 0.989 | 0.924 | 0.955 | 0.917 | 0.934 | 1 | 0.022 |
| SE | 52647 | ENSG00000197943 | PLCG2 | chr16 | + | 81931496 | 81931654 | 81928557 | 81928624 | 81936168 | 81936378 | 153 | 76 | 99 | 116 | 145 | 1 | 1 | 1 | 0 | 0 | 139 | 179 | 140 | 176 | 171 | 6 | 4 | 5 | 5 | 0 | 298 | 149 | 0.000289668 | 0.005467666 | 0.987 | 0.974 | 0.98 | 1 | 1 | 0.921 | 0.957 | 0.933 | 0.946 | 1 | 0.037 |
| SE | 68094 | ENSG00000135838 | NPL | chr1 | + | 182806144 | 182806531 | 182803697 | 182803771 | 182812155 | 182812213 | 131 | 73 | 76 | 104 | 128 | 16 | 1 | 8 | 5 | 9 | 193 | 173 | 152 | 219 | 128 | 0 | 8 | 5 | 6 | 5 | 298 | 149 | 0.000289861 | 0.005469171 | 0.804 | 0.973 | 0.826 | 0.912 | 0.877 | 1 | 0.915 | 0.938 | 0.948 | 0.928 | -0.067 |
| SE | 105591 | ENSG00000197971 | MBP | chr18 | - | 77095367 | 77095550 | 77066297 | 77066385 | 77105210 | 77105286 | 4 | 2 | 2 | 7 | 11 | 210 | 143 | 164 | 201 | 282 | 8 | 2 | 11 | 3 | 18 | 182 | 120 | 123 | 91 | 121 | 298 | 149 | 0.000289888 | 0.005469171 | 0.009 | 0.007 | 0.006 | 0.017 | 0.019 | 0.022 | 0.008 | 0.043 | 0.016 | 0.069 | -0.02 |
| SE | 105228 | ENSG00000102967 | DHODH | chr16 | + | 72023479 | 72023633 | 72023164 | 72023318 | 72024144 | 72024512 | 15 | 10 | 11 | 9 | 8 | 1 | 0 | 3 | 0 | 0 | 9 | 11 | 7 | 12 | 20 | 0 | 5 | 2 | 3 | 0 | 298 | 149 | 0.000290097 | 0.005471064 | 0.882 | 1 | 0.647 | 1 | 1 | 1 | 0.524 | 0.636 | 0.667 | 1 | 0.14 |
| SE | 20709 | ENSG00000115760 | BIRC6 | chr2 | + | 32453807 | 32453954 | 32445520 | 32445668 | 32463193 | 32463381 | 33 | 18 | 30 | 27 | 67 | 0 | 0 | 8 | 4 | 0 | 16 | 22 | 23 | 9 | 33 | 0 | 0 | 1 | 0 | 0 | 296 | 149 | 0.000290128 | 0.005471064 | 1 | 1 | 0.654 | 0.773 | 1 | 1 | 1 | 0.92 | 1 | 1 | -0.099 |
| SE | 73481 | ENSG00000215559 | ANKRD20A11P | chr21 | - | 13916746 | 13916958 | 13915186 | 13915409 | 13917554 | 13917836 | 0 | 15 | 18 | 56 | 16 | 13 | 0 | 4 | 1 | 0 | 28 | 79 | 25 | 9 | 8 | 0 | 2 | 0 | 0 | 0 | 298 | 149 | 0.000290316 | 0.005473296 | 0 | 1 | 0.692 | 0.966 | 1 | 1 | 0.952 | 1 | 1 | 1 | -0.259 |
| SE | 61847 | ENSG00000162929 | KIAA1841 | chr2 | + | 61117356 | 61117385 | 61115978 | 61116070 | 61117466 | 61117540 | 32 | 37 | 19 | 36 | 18 | 2 | 0 | 0 | 0 | 0 | 61 | 21 | 15 | 10 | 41 | 8 | 4 | 0 | 1 | 0 | 178 | 149 | 0.000290812 | 0.005481317 | 0.931 | 1 | 1 | 1 | 1 | 0.865 | 0.815 | 1 | 0.893 | 1 | 0.072 |
| SE | 65454 | ENSG00000204344 | STK19 | chr6 | + | 31979413 | 31979553 | 31978902 | 31978998 | 31980450 | 31980548 | 24 | 3 | 14 | 30 | 13 | 0 | 1 | 0 | 0 | 0 | 32 | 34 | 30 | 26 | 13 | 18 | 0 | 0 | 2 | 0 | 289 | 149 | 0.000291426 | 0.00549157 | 1 | 0.607 | 1 | 1 | 1 | 0.478 | 1 | 1 | 0.87 | 1 | 0.052 |
| SE | 84374 | ENSG00000230590 | FTX | chrX | - | 74284742 | 74284948 | 74281701 | 74281848 | 74292426 | 74292620 | 6 | 7 | 2 | 1 | 1 | 17 | 28 | 33 | 32 | 19 | 0 | 0 | 4 | 0 | 0 | 80 | 17 | 14 | 23 | 36 | 298 | 149 | 0.000291809 | 0.005496853 | 0.15 | 0.111 | 0.029 | 0.015 | 0.026 | 0 | 0 | 0.125 | 0 | 0 | 0.041 |
| SE | 108405 | ENSG00000110851 | PRDM4 | chr12 | - | 107744542 | 107744682 | 107743196 | 107743282 | 107746274 | 107746424 | 97 | 42 | 69 | 78 | 79 | 3 | 2 | 4 | 7 | 2 | 40 | 72 | 85 | 67 | 43 | 17 | 13 | 2 | 9 | 6 | 289 | 149 | 0.000291847 | 0.005496853 | 0.943 | 0.915 | 0.899 | 0.852 | 0.953 | 0.548 | 0.741 | 0.956 | 0.793 | 0.787 | 0.147 |
| SE | 31606 | ENSG00000181789 | COPG1 | chr3 | + | 129250681 | 129250894 | 129249652 | 129249746 | 129252280 | 129252361 | 46 | 17 | 33 | 31 | 33 | 0 | 0 | 4 | 2 | 0 | 51 | 61 | 29 | 50 | 86 | 8 | 4 | 2 | 4 | 11 | 298 | 149 | 0.000292145 | 0.005501151 | 1 | 1 | 0.805 | 0.886 | 1 | 0.761 | 0.884 | 0.879 | 0.862 | 0.796 | 0.102 |
| SE | 17120 | ENSG00000159128 | IFNGR2 | chr21 | + | 33410840 | 33410892 | 33403465 | 33403616 | 33414887 | 33415020 | 2 | 0 | 1 | 0 | 0 | 10 | 5 | 6 | 11 | 2 | 0 | 19 | 4 | 2 | 0 | 37 | 28 | 20 | 19 | 25 | 201 | 149 | 0.000292932 | 0.005513985 | 0.129 | 0 | 0.11 | 0 | 0 | 0 | 0.335 | 0.129 | 0.072 | 0 | -0.059 |
| SE | 65539 | ENSG00000284292 | NA | chr7 | + | 99348851 | 99348959 | 99344292 | 99344515 | 99353908 | 99354121 | 101 | 91 | 78 | 103 | 70 | 4 | 2 | 3 | 3 | 1 | 155 | 127 | 92 | 77 | 98 | 16 | 5 | 4 | 12 | 6 | 257 | 149 | 0.000292967 | 0.005513985 | 0.936 | 0.963 | 0.938 | 0.952 | 0.976 | 0.849 | 0.936 | 0.93 | 0.788 | 0.904 | 0.072 |
| SE | 13773 | ENSG00000205629 | LCMT1 | chr16 | + | 25128474 | 25128566 | 25111734 | 25111996 | 25132401 | 25132523 | 73 | 29 | 54 | 68 | 53 | 0 | 0 | 0 | 0 | 0 | 69 | 96 | 47 | 55 | 107 | 19 | 0 | 0 | 0 | 0 | 241 | 149 | 0.000293471 | 0.005522135 | 1 | 1 | 1 | 1 | 1 | 0.692 | 1 | 1 | 1 | 1 | 0.062 |
| SE | 25372 | ENSG00000092020 | PPP2R3C | chr14 | - | 35095047 | 35095184 | 35091069 | 35091207 | 35096557 | 35096633 | 223 | 157 | 193 | 207 | 479 | 4 | 1 | 2 | 5 | 16 | 14 | 160 | 115 | 59 | 30 | 0 | 7 | 6 | 5 | 12 | 286 | 149 | 0.000293774 | 0.005526515 | 0.967 | 0.988 | 0.98 | 0.956 | 0.94 | 1 | 0.923 | 0.909 | 0.86 | 0.566 | 0.115 |
| SE | 31642 | ENSG00000131389 | SLC6A6 | chr3 | + | 14457949 | 14458082 | 14447581 | 14447816 | 14466515 | 14466650 | 101 | 52 | 78 | 122 | 48 | 0 | 0 | 0 | 0 | 0 | 204 | 101 | 129 | 181 | 249 | 8 | 3 | 6 | 3 | 0 | 282 | 149 | 0.000293942 | 0.005527704 | 1 | 1 | 1 | 1 | 1 | 0.931 | 0.947 | 0.919 | 0.97 | 1 | 0.047 |
| SE | 73825 | ENSG00000043462 | LCP2 | chr5 | - | 170275319 | 170275351 | 170273780 | 170274338 | 170275794 | 170275860 | 566 | 404 | 515 | 551 | 1312 | 2 | 2 | 1 | 9 | 0 | 857 | 617 | 478 | 610 | 665 | 8 | 1 | 4 | 38 | 5 | 181 | 149 | 0.000293979 | 0.005527704 | 0.996 | 0.994 | 0.998 | 0.981 | 1 | 0.989 | 0.998 | 0.99 | 0.93 | 0.991 | 0.014 |
| SE | 96207 | ENSG00000117751 | PPP1R8 | chr1 | + | 27841013 | 27841234 | 27838698 | 27838852 | 27843185 | 27843330 | 320 | 203 | 314 | 309 | 575 | 12 | 5 | 2 | 13 | 5 | 237 | 184 | 200 | 233 | 185 | 0 | 8 | 25 | 11 | 10 | 298 | 149 | 0.000294259 | 0.005531637 | 0.93 | 0.953 | 0.987 | 0.922 | 0.983 | 1 | 0.92 | 0.8 | 0.914 | 0.902 | 0.048 |
| SE | 52663 | ENSG00000203705 | TATDN3 | chr1 | + | 212812226 | 212812328 | 212807735 | 212807848 | 212815012 | 212815597 | 26 | 17 | 21 | 16 | 40 | 3 | 0 | 0 | 1 | 0 | 44 | 46 | 22 | 38 | 13 | 10 | 0 | 0 | 7 | 0 | 251 | 149 | 0.000294367 | 0.005532348 | 0.837 | 1 | 1 | 0.905 | 1 | 0.723 | 1 | 1 | 0.763 | 1 | 0.051 |
| SE | 104289 | ENSG00000096717 | SIRT1 | chr10 | + | 67906162 | 67906287 | 67891483 | 67891554 | 67906789 | 67906937 | 35 | 27 | 47 | 47 | 99 | 123 | 63 | 79 | 85 | 186 | 16 | 61 | 63 | 35 | 25 | 45 | 52 | 35 | 13 | 4 | 274 | 149 | 0.000294592 | 0.00553525 | 0.134 | 0.189 | 0.244 | 0.231 | 0.224 | 0.162 | 0.389 | 0.495 | 0.594 | 0.773 | -0.278 |
| SE | 29682 | ENSG00000137656 | BUD13 | chr11 | - | 116759073 | 116759179 | 116757765 | 116757950 | 116760734 | 116760952 | 82 | 34 | 41 | 40 | 34 | 12 | 2 | 0 | 6 | 0 | 41 | 35 | 38 | 54 | 4 | 26 | 33 | 4 | 4 | 7 | 255 | 149 | 0.000295099 | 0.005543447 | 0.8 | 0.909 | 1 | 0.796 | 1 | 0.48 | 0.383 | 0.847 | 0.887 | 0.25 | 0.332 |
| SE | 87251 | ENSG00000046651 | OFD1 | chrX | + | 13753367 | 13755242 | 13751248 | 13751368 | 13756577 | 13756767 | 139 | 170 | 138 | 144 | 290 | 2 | 0 | 0 | 1 | 5 | 54 | 121 | 171 | 67 | 21 | 11 | 6 | 1 | 0 | 0 | 298 | 149 | 0.000295542 | 0.005550432 | 0.972 | 1 | 1 | 0.986 | 0.967 | 0.711 | 0.91 | 0.988 | 1 | 1 | 0.063 |
| SE | 34066 | ENSG00000114867 | EIF4G1 | chr3 | + | 184317320 | 184317497 | 184314541 | 184314674 | 184317746 | 184317816 | 17 | 6 | 10 | 9 | 6 | 0 | 0 | 0 | 0 | 0 | 4 | 12 | 6 | 16 | 29 | 0 | 4 | 1 | 2 | 2 | 298 | 149 | 0.000296222 | 0.00556187 | 1 | 1 | 1 | 1 | 1 | 1 | 0.6 | 0.75 | 0.8 | 0.879 | 0.194 |
| SE | 106074 | ENSG00000196498 | NCOR2 | chr12 | - | 124340601 | 124340751 | 124340293 | 124340443 | 124341822 | 124342074 | 48 | 16 | 26 | 25 | 6 | 0 | 0 | 2 | 2 | 0 | 37 | 12 | 15 | 52 | 91 | 0 | 0 | 0 | 9 | 10 | 298 | 149 | 0.000296452 | 0.005564848 | 1 | 1 | 0.867 | 0.862 | 1 | 1 | 1 | 1 | 0.743 | 0.82 | 0.033 |
| SE | 49689 | ENSG00000263956 | NBPF11 | chr1 | - | 148124898 | 148125001 | 148123852 | 148124067 | 148126828 | 148127038 | 33 | 24 | 39 | 26 | 41 | 16 | 13 | 15 | 9 | 21 | 18 | 5 | 14 | 20 | 11 | 23 | 35 | 11 | 20 | 6 | 252 | 149 | 0.000296924 | 0.005571197 | 0.549 | 0.522 | 0.606 | 0.631 | 0.536 | 0.316 | 0.078 | 0.429 | 0.372 | 0.52 | 0.226 |
| SE | 80835 | ENSG00000151948 | GLT1D1 | chr12 | + | 128927104 | 128927154 | 128899235 | 128899287 | 128945325 | 128945369 | 15 | 16 | 10 | 13 | 13 | 1 | 7 | 0 | 8 | 5 | 45 | 27 | 27 | 22 | 22 | 58 | 27 | 11 | 34 | 30 | 199 | 149 | 0.000296951 | 0.005571197 | 0.918 | 0.631 | 1 | 0.549 | 0.661 | 0.367 | 0.428 | 0.648 | 0.326 | 0.354 | 0.327 |
| SE | 53438 | ENSG00000145020 | AMT | chr3 | - | 49417817 | 49417973 | 49416777 | 49417718 | 49418970 | 49419026 | 44 | 23 | 41 | 29 | 19 | 0 | 0 | 0 | 0 | 0 | 46 | 45 | 29 | 42 | 108 | 18 | 1 | 0 | 0 | 0 | 298 | 149 | 0.000297003 | 0.005571197 | 1 | 1 | 1 | 1 | 1 | 0.561 | 0.957 | 1 | 1 | 1 | 0.096 |
| SE | 2930 | ENSG00000102575 | ACP5 | chr19 | - | 11578584 | 11578657 | 11577592 | 11577679 | 11578843 | 11578954 | 10 | 3 | 10 | 4 | 4 | 4 | 0 | 0 | 4 | 0 | 0 | 5 | 4 | 3 | 3 | 6 | 4 | 2 | 4 | 4 | 222 | 149 | 0.000297166 | 0.005572912 | 0.627 | 1 | 1 | 0.402 | 1 | 0 | 0.456 | 0.573 | 0.335 | 0.335 | 0.466 |
| SE | 85336 | ENSG00000014257 | ACP3 | chr3 | + | 132337455 | 132337554 | 132332191 | 132332344 | 132342551 | 132342579 | 91 | 98 | 68 | 97 | 107 | 1 | 4 | 2 | 2 | 0 | 300 | 148 | 56 | 195 | 89 | 4 | 15 | 0 | 4 | 20 | 248 | 149 | 0.000297283 | 0.005573763 | 0.982 | 0.936 | 0.953 | 0.967 | 1 | 0.978 | 0.856 | 1 | 0.967 | 0.728 | 0.062 |
| SE | 13034 | ENSG00000006194 | ZNF263 | chr16 | + | 3285058 | 3285239 | 3283490 | 3284205 | 3285680 | 3285754 | 25 | 8 | 10 | 21 | 6 | 4 | 0 | 2 | 1 | 0 | 0 | 5 | 2 | 0 | 3 | 8 | 5 | 0 | 4 | 0 | 298 | 149 | 0.000297527 | 0.005577016 | 0.758 | 1 | 0.714 | 0.913 | 1 | 0 | 0.333 | 1 | 0 | 1 | 0.41 |
| SE | 103214 | ENSG00000157625 | TAB3 | chrX | - | 30845932 | 30846071 | 30842987 | 30843049 | 30846550 | 30846644 | 1 | 0 | 9 | 3 | 6 | 44 | 46 | 47 | 61 | 111 | 4 | 7 | 10 | 6 | 14 | 71 | 57 | 58 | 40 | 21 | 288 | 149 | 0.000298239 | 0.005588083 | 0.012 | 0 | 0.09 | 0.025 | 0.027 | 0.028 | 0.06 | 0.082 | 0.072 | 0.256 | -0.069 |
| SE | 58173 | ENSG00000172965 | MIR4435-2HG | chr2 | - | 111208755 | 111208936 | 111207773 | 111208661 | 111248147 | 111248241 | 7 | 4 | 3 | 14 | 17 | 0 | 2 | 0 | 1 | 3 | 26 | 8 | 5 | 19 | 10 | 10 | 13 | 5 | 4 | 5 | 298 | 149 | 0.00029826 | 0.005588083 | 1 | 0.5 | 1 | 0.875 | 0.739 | 0.565 | 0.235 | 0.333 | 0.704 | 0.5 | 0.355 |
| SE | 82111 | ENSG00000136048 | DRAM1 | chr12 | + | 101908185 | 101908363 | 101901290 | 101901433 | 101914173 | 101914232 | 54 | 33 | 31 | 46 | 77 | 0 | 0 | 0 | 2 | 0 | 60 | 77 | 64 | 122 | 113 | 0 | 5 | 0 | 9 | 2 | 298 | 149 | 0.000298495 | 0.005591142 | 1 | 1 | 1 | 0.92 | 1 | 1 | 0.885 | 1 | 0.871 | 0.966 | 0.04 |
| SE | 60311 | ENSG00000143418 | CERS2 | chr1 | - | 150968082 | 150968201 | 150967819 | 150967877 | 150968394 | 150968478 | 678 | 334 | 324 | 577 | 655 | 0 | 1 | 3 | 3 | 5 | 943 | 563 | 344 | 475 | 1689 | 0 | 8 | 7 | 36 | 0 | 268 | 149 | 0.000299244 | 0.005603821 | 1 | 0.995 | 0.984 | 0.991 | 0.986 | 1 | 0.975 | 0.965 | 0.88 | 1 | 0.027 |
| SE | 60926 | ENSG00000138398 | PPIG | chr2 | + | 169631598 | 169631647 | 169630849 | 169630987 | 169631765 | 169631933 | 3 | 2 | 8 | 13 | 17 | 69 | 39 | 64 | 48 | 160 | 0 | 5 | 2 | 0 | 0 | 19 | 39 | 64 | 23 | 11 | 198 | 149 | 0.000299497 | 0.005607228 | 0.032 | 0.037 | 0.086 | 0.169 | 0.074 | 0 | 0.088 | 0.023 | 0 | 0 | 0.057 |
| SE | 89669 | ENSG00000198851 | CD3E | chr11 | + | 118312617 | 118312866 | 118312152 | 118312170 | 118314447 | 118314494 | 2127 | 1281 | 2302 | 2339 | 3557 | 11 | 8 | 8 | 4 | 23 | 851 | 2026 | 990 | 833 | 1508 | 15 | 28 | 16 | 5 | 0 | 298 | 149 | 0.000299682 | 0.005613083 | 0.99 | 0.988 | 0.993 | 0.997 | 0.987 | 0.966 | 0.973 | 0.969 | 0.988 | 1 | 0.012 |
| SE | 103060 | ENSG00000099290 | WASHC2A | chr10 | + | 50127582 | 50127795 | 50125368 | 50125449 | 50129418 | 50129784 | 41 | 34 | 42 | 51 | 37 | 0 | 1 | 0 | 4 | 0 | 53 | 35 | 31 | 33 | 21 | 0 | 5 | 5 | 3 | 1 | 298 | 149 | 0.000300219 | 0.005618058 | 1 | 0.944 | 1 | 0.864 | 1 | 1 | 0.778 | 0.756 | 0.846 | 0.913 | 0.103 |
| SE | 63718 | ENSG00000166986 | MARS1 | chr12 | + | 57489895 | 57489971 | 57489266 | 57489345 | 57490206 | 57490216 | 31 | 11 | 28 | 20 | 24 | 3 | 1 | 1 | 0 | 0 | 51 | 24 | 16 | 12 | 64 | 0 | 1 | 4 | 18 | 8 | 225 | 149 | 0.000300325 | 0.005618702 | 0.872 | 0.879 | 0.949 | 1 | 1 | 1 | 0.941 | 0.726 | 0.306 | 0.841 | 0.177 |
| SE | 27514 | ENSG00000198189 | HSD17B11 | chr4 | - | 87374360 | 87374414 | 87372708 | 87372815 | 87374698 | 87374830 | 6 | 4 | 9 | 11 | 20 | 608 | 425 | 523 | 424 | 976 | 0 | 16 | 16 | 15 | 0 | 167 | 351 | 315 | 316 | 27 | 203 | 149 | 0.00030106 | 0.005631097 | 0.007 | 0.007 | 0.012 | 0.019 | 0.015 | 0 | 0.032 | 0.036 | 0.034 | 0 | -0.008 |
| SE | 62316 | ENSG00000185811 | IKZF1 | chr7 | + | 50368005 | 50368065 | 50327637 | 50327757 | 50376532 | 50377327 | 151 | 82 | 153 | 136 | 158 | 355 | 211 | 274 | 289 | 161 | 112 | 158 | 123 | 182 | 186 | 152 | 226 | 137 | 176 | 202 | 209 | 149 | 0.000301634 | 0.005639047 | 0.233 | 0.217 | 0.285 | 0.251 | 0.412 | 0.344 | 0.333 | 0.39 | 0.424 | 0.396 | -0.098 |
| SE | 61334 | ENSG00000124772 | CPNE5 | chr6 | - | 36752579 | 36752620 | 36748220 | 36748267 | 36753033 | 36753095 | 8 | 0 | 15 | 9 | 9 | 19 | 11 | 51 | 33 | 92 | 3 | 16 | 5 | 0 | 2 | 64 | 108 | 52 | 42 | 102 | 190 | 149 | 0.00030169 | 0.005639047 | 0.248 | 0 | 0.187 | 0.176 | 0.071 | 0.035 | 0.104 | 0.07 | 0 | 0.015 | 0.092 |
| SE | 72607 | ENSG00000115977 | AAK1 | chr2 | - | 69482710 | 69482812 | 69480859 | 69480961 | 69505568 | 69505673 | 5 | 7 | 13 | 4 | 2 | 0 | 5 | 6 | 4 | 3 | 37 | 17 | 15 | 16 | 2 | 0 | 0 | 1 | 13 | 6 | 251 | 149 | 0.000301732 | 0.005639047 | 1 | 0.454 | 0.563 | 0.372 | 0.284 | 1 | 1 | 1 | 0.905 | 0.165 | -0.279 |
| SE | 99255 | ENSG00000133318 | RTN3 | chr11 | + | 63756111 | 63756170 | 63753661 | 63753708 | 63758155 | 63758421 | 650 | 406 | 416 | 588 | 912 | 15 | 7 | 8 | 9 | 26 | 1257 | 774 | 682 | 1274 | 2148 | 17 | 22 | 1 | 13 | 11 | 208 | 149 | 0.000301773 | 0.005639047 | 0.969 | 0.976 | 0.974 | 0.979 | 0.962 | 0.981 | 0.962 | 0.998 | 0.986 | 0.993 | -0.012 |
| SE | 66034 | ENSG00000183495 | EP400 | chr12 | + | 131963531 | 131963642 | 131960584 | 131961954 | 131979693 | 131979793 | 3 | 4 | 5 | 2 | 4 | 30 | 29 | 43 | 49 | 58 | 0 | 0 | 2 | 0 | 0 | 36 | 71 | 26 | 61 | 58 | 260 | 149 | 0.000302732 | 0.005655612 | 0.054 | 0.073 | 0.062 | 0.023 | 0.038 | 0 | 0 | 0.042 | 0 | 0 | 0.042 |
| SE | 81415 | ENSG00000129103 | SUMF2 | chr7 | + | 56074585 | 56074736 | 56074113 | 56074218 | 56076833 | 56076889 | 247 | 133 | 270 | 246 | 462 | 14 | 3 | 11 | 5 | 21 | 196 | 329 | 111 | 126 | 347 | 23 | 25 | 16 | 4 | 20 | 298 | 149 | 0.00030316 | 0.005662263 | 0.898 | 0.957 | 0.925 | 0.961 | 0.917 | 0.81 | 0.868 | 0.776 | 0.94 | 0.897 | 0.073 |
| SE | 47737 | ENSG00000145687 | SSBP2 | chr5 | - | 81446867 | 81446922 | 81442650 | 81442723 | 81448789 | 81448825 | 49 | 43 | 58 | 58 | 92 | 2 | 2 | 1 | 5 | 9 | 21 | 37 | 13 | 16 | 37 | 0 | 3 | 0 | 0 | 0 | 204 | 149 | 0.000304596 | 0.005687737 | 0.947 | 0.94 | 0.977 | 0.894 | 0.882 | 1 | 0.9 | 1 | 1 | 1 | -0.052 |

| | | | | | | | | | | | | | | |
|---|---|---|---|---|---|---|---|---|---|---|---|---|---|---|
| SE | 44547 | ENSG00000166526 | ZNF3 | chr7 | - | 100079535 | 100079656 | 100077017 | 100077433 | 100081634 | 100081703 | 18 | 11 | 35 |
| SE | 108169 | ENSG00000075089 | ACTR6 | chr12 | + | 100210072 | 100210208 | 100207662 | 100207786 | 100210294 | 100210320 | 327 | 178 | 267 |
| SE | 76391 | ENSG00000047579 | DTNBP1 | chr6 | - | 15651312 | 15651408 | 15637743 | 15637804 | 15652086 | 15652140 | 61 | 37 | 56 |
| SE | 65519 | ENSG00000140386 | SCAPER | chr15 | - | 76862415 | 76862533 | 76857808 | 76857879 | 76883811 | 76883876 | 144 | 138 | 128 |
| SE | 8525 | ENSG00000125450 | NUP85 | chr17 | + | 75215753 | 75215823 | 75213075 | 75213119 | 75218184 | 75218306 | 55 | 27 | 51 |
| SE | 86844 | ENSG00000116754 | SRSF11 | chr1 | + | 70231094 | 70231203 | 70228421 | 70228555 | 70232267 | 70232377 | 8 | 13 | 32 |
| SE | 16504 | ENSG00000140367 | UBE2Q2 | chr15 | + | 75859877 | 75859982 | 75854385 | 75854487 | 75868950 | 75869010 | 326 | 256 | 252 |
| SE | 109646 | ENSG00000005810 | MYCBP2 | chr13 | - | 77233155 | 77233263 | 77225434 | 77225554 | 77243058 | 77243160 | 226 | 199 | 190 |
| SE | 3030 | ENSG00000007255 | TRAPPC6A | chr19 | - | 45165126 | 45165194 | 45164724 | 45164970 | 45178092 | 45178211 | 82 | 13 | 84 |
| SE | 23061 | ENSG00000111641 | NOP2 | chr12 | - | 6566100 | 6566336 | 6563890 | 6563946 | 6566528 | 6566559 | 197 | 146 | 227 |
| SE | 26480 | ENSG00000038382 | TRIO | chr5 | + | 14507121 | 14507260 | 14504392 | 14504593 | 14507879 | 14508676 | 57 | 39 | 24 |
| SE | 106777 | ENSG00000176083 | ZNF683 | chr1 | - | 26362115 | 26362175 | 26361633 | 26362022 | 26363025 | 26363154 | 1 | 0 | 3 |
| SE | 81967 | ENSG00000051825 | MPHOSPH9 | chr12 | - | 123226287 | 123226351 | 123223037 | 123223127 | 123230260 | 123230522 | 7 | 12 | 0 |
| SE | 382 | ENSG00000237945 | LINC00649 | chr21 | + | 33943033 | 33943145 | 33931162 | 33931742 | 33945809 | 33945895 | 0 | 0 | 2 |
| SE | 102327 | ENSG00000166171 | DPCD | chr10 | + | 101605139 | 101605212 | 101601202 | 101601336 | 101608834 | 101608937 | 0 | 1 | 0 |
| SE | 109023 | ENSG00000185896 | LAMP1 | chr13 | + | 113306484 | 113306606 | 113297238 | 113297495 | 113309642 | 113309862 | 301 | 165 | 183 |
| SE | 10068 | ENSG00000108406 | DHX40 | chr17 | + | 59570517 | 59570663 | 59566626 | 59566794 | 59573115 | 59573235 | 200 | 127 | 160 |
| SE | 32463 | ENSG00000131375 | CAPN7 | chr3 | + | 15245525 | 15245671 | 15242177 | 15242253 | 15246731 | 15246794 | 265 | 259 | 301 |
| SE | 83762 | ENSG00000196696 | PDXDC2P-NPIPB14P | chr16 | - | 69976810 | 69978003 | 69976296 | 69976732 | 69978076 | 69978112 | 22 | 22 | 9 |
| SE | 84934 | ENSG00000170734 | POLH | chr6 | + | 43582315 | 43582456 | 43576184 | 43576440 | 43583006 | 43583141 | 86 | 63 | 85 |
| SE | 40080 | ENSG00000125347 | IRF1 | chr5 | - | 132486794 | 132486840 | 132486556 | 132486686 | 132486953 | 132487130 | 503 | 285 | 506 |
| SE | 95751 | ENSG00000070476 | ZXDC | chr3 | - | 126459652 | 126459737 | 126441764 | 126441946 | 126461534 | 126462220 | 42 | 23 | 23 |
| SE | 76864 | ENSG00000112576 | CCND3 | chr6 | - | 41940951 | 41941161 | 41940493 | 41940585 | 41941451 | 41941808 | 0 | 0 | 0 |
| SE | 66455 | ENSG00000131051 | RBM39 | chr20 | - | 35713701 | 35713857 | 35713018 | 35713096 | 35714184 | 35714389 | 50 | 42 | 58 |
| SE | 14998 | ENSG00000157823 | AP3S2 | chr15 | - | 89888520 | 89888632 | 89878264 | 89878300 | 89889048 | 89889136 | 67 | 48 | 53 |
| SE | 48872 | ENSG00000143514 | TP53BP2 | chr1 | - | 223789007 | 223789174 | 223784114 | 223784314 | 223792388 | 223792522 | 138 | 90 | 118 |
| SE | 44401 | ENSG00000163346 | PBXIP1 | chr1 | - | 154951794 | 154951919 | 154951470 | 154951535 | 154953670 | 154953757 | 217 | 124 | 217 |

[illegible]

| | | | | | | | | | | | | | | | | | | | | | | | | | | | | | | | | | | | | | | | | | | | | | | |
|---|---|---|---|---|---|---|---|---|---|---|---|---|---|---|---|---|---|---|---|---|---|---|---|---|---|---|---|---|---|---|---|---|---|---|---|---|---|---|---|---|---|---|---|---|---|---|
| SE | 86779 | ENSG00000151689 | INPP1 | chr2 | + | 190366694 | 190366895 | 190362626 | 190362687 | 190369102 | 190369208 | 66 | 45 | 44 | 58 | 56 | 0 | 3 | 0 | 0 | 1 | 87 | 104 | 33 | 47 | 74 | 14 | 0 | 0 | 0 | 3 | 298 | 149 | 0.000431177 | 0.007625809 | 1 | 0.882 | 1 | 1 | 0.966 | 0.757 | 1 | 1 | 1 | 0.925 | 0.033 |
| SE | 57433 | ENSG00000159399 | HK2 | chr2 | + | 74873275 | 74873371 | 74872299 | 74872419 | 74873843 | 74873943 | 158 | 120 | 116 | 120 | 191 | 3 | 1 | 3 | 8 | 3 | 284 | 227 | 152 | 186 | 308 | 21 | 14 | 7 | 25 | 0 | 245 | 149 | 0.000431395 | 0.007627938 | 0.97 | 0.986 | 0.959 | 0.901 | 0.975 | 0.892 | 0.908 | 0.93 | 0.819 | 1 | 0.048 |
| SE | 77959 | ENSG00000173200 | PARP15 | chr3 | + | 122605935 | 122606055 | 122577627 | 122577853 | 122610493 | 122610730 | 277 | 203 | 159 | 277 | 135 | 3 | 0 | 0 | 1 | 6 | 96 | 279 | 162 | 102 | 123 | 0 | 13 | 4 | 7 | 0 | 269 | 149 | 0.000431951 | 0.007636047 | 0.981 | 1 | 1 | 0.994 | 0.926 | 1 | 0.922 | 0.957 | 0.89 | 1 | 0.026 |
| SE | 104828 | ENSG00000205356 | TECPR1 | chr7 | - | 98236775 | 98236921 | 98232826 | 98232972 | 98238508 | 98238610 | 68 | 49 | 89 | 72 | 112 | 0 | 0 | 0 | 0 | 0 | 93 | 82 | 63 | 51 | 106 | 0 | 16 | 0 | 0 | 0 | 295 | 149 | 0.000432523 | 0.00764442 | 1 | 1 | 1 | 1 | 1 | 1 | 0.721 | 1 | 1 | 1 | 0.056 |
| SE | 66554 | ENSG00000133872 | SARAF | chr8 | - | 30073876 | 30074055 | 30069641 | 30070059 | 30082846 | 30082948 | 939 | 588 | 1047 | 985 | 1501 | 32 | 27 | 47 | 34 | 44 | 367 | 520 | 759 | 321 | 289 | 12 | 45 | 45 | 15 | 19 | 298 | 149 | 0.000433005 | 0.007651018 | 0.936 | 0.916 | 0.918 | 0.935 | 0.945 | 0.939 | 0.852 | 0.894 | 0.915 | 0.884 | 0.033 |
| SE | 1042 | ENSG00000100099 | HPS4 | chr22 | - | 26458444 | 26458577 | 26450881 | 26453404 | 26463916 | 26464015 | 52 | 43 | 36 | 43 | 23 | 1 | 0 | 1 | 5 | 0 | 35 | 54 | 37 | 44 | 34 | 35 | 2 | 3 | 3 | 3 | 282 | 149 | 0.000433091 | 0.007651018 | 0.965 | 1 | 0.95 | 0.82 | 1 | 0.346 | 0.934 | 0.867 | 0.886 | 0.857 | 0.169 |
| SE | 27183 | ENSG00000156500 | PABIR3 | chrX | + | 134838726 | 134839603 | 134829225 | 134829282 | 134845204 | 134845248 | 5 | 1 | 9 | 6 | 17 | 1 | 17 | 2 | 5 | 4 | 0 | 0 | 0 | 3 | 10 | 16 | 1 | 2 | 11 | 18 | 298 | 149 | 0.000433463 | 0.007655467 | 0.714 | 0.029 | 0.692 | 0.375 | 0.68 | 0 | 0 | 0 | 0.12 | 0.217 | 0.431 |
| SE | 102810 | ENSG00000165832 | TRUB1 | chr10 | + | 114959725 | 114959807 | 114951093 | 114951149 | 114970367 | 114970440 | 66 | 45 | 58 | 69 | 90 | 4 | 2 | 8 | 4 | 1 | 7 | 46 | 45 | 35 | 31 | 0 | 4 | 0 | 0 | 0 | 231 | 149 | 0.000433539 | 0.007655467 | 0.914 | 0.936 | 0.824 | 0.918 | 0.983 | 1 | 0.881 | 1 | 1 | 1 | -0.061 |
| SE | 45301 | ENSG00000002822 | MAD1L1 | chr7 | - | 2225409 | 2225550 | 2222574 | 2222754 | 2229983 | 2230143 | 42 | 48 | 49 | 76 | 35 | 5 | 3 | 6 | 4 | 3 | 103 | 30 | 31 | 71 | 185 | 43 | 18 | 14 | 2 | 34 | 290 | 149 | 0.000433642 | 0.007655567 | 0.812 | 0.892 | 0.808 | 0.907 | 0.857 | 0.552 | 0.461 | 0.532 | 0.948 | 0.737 | 0.209 |
| SE | 58785 | ENSG00000077254 | USP33 | chr1 | - | 77714613 | 77714783 | 77713199 | 77713281 | 77715741 | 77715868 | 565 | 431 | 451 | 510 | 637 | 2 | 0 | 2 | 1 | 0 | 222 | 346 | 238 | 328 | 43 | 0 | 17 | 2 | 3 | 0 | 298 | 149 | 0.000435094 | 0.00767946 | 0.993 | 1 | 0.991 | 0.996 | 1 | 1 | 0.911 | 0.983 | 0.982 | 1 | 0.021 |
| SE | 64609 | ENSG00000100644 | HIF1A | chr14 | + | 61745690 | 61745817 | 61744704 | 61744813 | 61746933 | 61747771 | 803 | 590 | 623 | 722 | 647 | 24 | 8 | 5 | 11 | 5 | 93 | 455 | 1008 | 193 | 65 | 0 | 0 | 1 | 10 | 0 | 276 | 149 | 0.000435741 | 0.007689149 | 0.948 | 0.975 | 0.985 | 0.973 | 0.986 | 1 | 1 | 0.998 | 0.912 | 1 | -0.009 |
| SE | 22469 | ENSG00000066135 | KDM4A | chr1 | + | 43666953 | 43667091 | 43666451 | 43666555 | 43667771 | 43668019 | 217 | 174 | 190 | 159 | 149 | 0 | 0 | 0 | 0 | 0 | 278 | 248 | 234 | 218 | 298 | 11 | 0 | 0 | 5 | 7 | 287 | 149 | 0.000436059 | 0.007691652 | 1 | 1 | 1 | 1 | 1 | 0.929 | 1 | 1 | 0.958 | 0.957 | 0.031 |
| SE | 53624 | ENSG00000143851 | PTPN7 | chr1 | - | 202158117 | 202158301 | 202157738 | 202157823 | 202159280 | 202159341 | 117 | 48 | 81 | 93 | 98 | 2 | 0 | 0 | 0 | 0 | 78 | 143 | 50 | 86 | 98 | 0 | 2 | 7 | 0 | 6 | 298 | 149 | 0.000436417 | 0.007695872 | 0.967 | 1 | 1 | 1 | 1 | 1 | 0.973 | 0.781 | 1 | 0.891 | 0.064 |
| SE | 54860 | ENSG00000133056 | PIK3C2B | chr1 | - | 204489815 | 204489931 | 204468869 | 204469886 | 204494355 | 204494701 | 3 | 7 | 6 | 6 | 11 | 7 | 8 | 10 | 2 | 6 | 3 | 16 | 6 | 9 | 39 | 0 | 5 | 1 | 2 | 9 | 265 | 149 | 0.000436701 | 0.007699162 | 0.194 | 0.33 | 0.252 | 0.628 | 0.508 | 1 | 0.643 | 0.771 | 0.717 | 0.709 | -0.386 |
| SE | 87252 | ENSG00000046651 | OFD1 | chrX | + | 13755150 | 13755242 | 13751248 | 13751368 | 13756577 | 13756767 | 96 | 104 | 76 | 88 | 157 | 2 | 0 | 0 | 1 | 5 | 50 | 63 | 94 | 33 | 19 | 11 | 6 | 1 | 0 | 0 | 241 | 149 | 0.000437184 | 0.007705941 | 0.967 | 1 | 1 | 0.982 | 0.951 | 0.738 | 0.867 | 0.983 | 1 | 1 | 0.062 |
| SE | 31723 | ENSG00000145012 | LPP | chr3 | + | 188759262 | 188759500 | 188708266 | 188708393 | 188760112 | 188760282 | 0 | 9 | 0 | 2 | 10 | 63 | 34 | 50 | 81 | 37 | 0 | 0 | 0 | 0 | 4 | 80 | 66 | 86 | 50 | 93 | 298 | 149 | 0.00043758 | 0.007711173 | 0 | 0.117 | 0 | 0.012 | 0.119 | 0 | 0 | 0 | 0 | 0.021 | 0.045 |
| SE | 76206 | ENSG00000146070 | PLA2G7 | chr6 | - | 46709326 | 46709418 | 46707990 | 46708161 | 46710544 | 46710658 | 129 | 111 | 82 | 78 | 163 | 2 | 3 | 9 | 11 | 1 | 226 | 8 | 18 | 100 | 0 | 34 | 4 | 1 | 13 | 10 | 241 | 149 | 0.000437679 | 0.007711182 | 0.976 | 0.958 | 0.849 | 0.814 | 0.99 | 0.804 | 0.553 | 0.918 | 0.826 | 0 | 0.297 |
| SE | 66952 | ENSG00000172493 | AFF1 | chr4 | + | 87132270 | 87132408 | 87131792 | 87131864 | 87134473 | 87134694 | 156 | 106 | 123 | 110 | 324 | 1 | 0 | 1 | 0 | 0 | 164 | 110 | 118 | 97 | 74 | 6 | 0 | 8 | 2 | 0 | 287 | 149 | 0.000437847 | 0.007712413 | 0.988 | 1 | 0.985 | 1 | 1 | 0.934 | 1 | 0.884 | 0.962 | 1 | 0.039 |
| SE | 70185 | ENSG00000115282 | TTC31 | chr2 | + | 74490655 | 74490739 | 74490024 | 74490473 | 74491127 | 74491184 | 72 | 53 | 77 | 91 | 88 | 1 | 0 | 3 | 5 | 0 | 53 | 61 | 58 | 120 | 140 | 0 | 4 | 10 | 6 | 10 | 233 | 149 | 0.000437981 | 0.007713041 | 0.979 | 1 | 0.943 | 0.921 | 1 | 1 | 0.907 | 0.788 | 0.927 | 0.9 | 0.064 |
| SE | 105621 | ENSG00000139323 | POC1B | chr12 | - | 89491935 | 89492115 | 89472167 | 89472275 | 89525119 | 89525204 | 93 | 58 | 79 | 100 | 106 | 0 | 5 | 3 | 2 | 5 | 82 | 39 | 42 | 46 | 95 | 0 | 0 | 2 | 0 | 0 | 298 | 149 | 0.000438288 | 0.007716693 | 1 | 0.853 | 0.929 | 0.962 | 0.914 | 1 | 1 | 0.913 | 1 | 1 | -0.051 |
| SE | 72199 | ENSG00000145425 | RPS3A | chr4 | + | 151100974 | 151101162 | 151099627 | 151099714 | 151102870 | 151102932 | 10343 | 5877 | 10845 | 8496 | 25522 | 140 | 77 | 126 | 102 | 180 | 1317 | 4035 | 4378 | 2327 | 1031 | 42 | 39 | 48 | 40 | 31 | 298 | 149 | 0.000438394 | 0.007716693 | 0.974 | 0.974 | 0.977 | 0.977 | 0.986 | 0.94 | 0.981 | 0.979 | 0.967 | 0.943 | 0.016 |
| SE | 86737 | ENSG00000101868 | POLA1 | chrX | + | 24716883 | 24716971 | 24716361 | 24716454 | 24717289 | 24717491 | 24 | 13 | 27 | 33 | 21 | 3 | 1 | 0 | 2 | 0 | 0 | 7 | 10 | 14 | 6 | 5 | 0 | 8 | 8 | 0 | 237 | 149 | 0.000438484 | 0.007716693 | 0.834 | 0.891 | 1 | 0.912 | 1 | 0 | 1 | 0.44 | 0.524 | 1 | 0.335 |
| [illegible] | | | | | | | | | | | | | | | | | | | | | | | | | | | | | | | | | | | | | | | | | | | | | | |

| | | | | | | | | | | | | | | | | | | |
|---|---|---|---|---|---|---|---|---|---|---|---|---|---|---|---|---|---|---|
| SE | 37189 | ENSG00000033327 | GAB2 | chr11 | - | 78220318 | 78220444 | 78215292 | 78219415 | 78221676 | 78221779 | 109 | 51 | 80 | 77 | 107 | 1 | 0 |
| SE | 66922 | ENSG00000152520 | PAN3 | chr13 | + | 28271980 | 28272071 | 28270700 | 28270866 | 28277236 | 28277376 | 158 | 104 | 142 | 154 | 303 | 2 | 0 |
| SE | 70553 | ENSG00000073849 | ST6GAL1 | chr3 | + | 187064519 | 187064616 | 187051248 | 187051346 | 187072848 | 187072947 | 12 | 11 | 24 | 12 | 30 | 354 | 215 |
| SE | 45793 | ENSG00000118217 | ATF6 | chr1 | + | 161778243 | 161778316 | 161766341 | 161766442 | 161781911 | 161781999 | 41 | 32 | 41 | 40 | 75 | 2 | 0 |
| SE | 74327 | ENSG00000151789 | ZNF385D | chr3 | - | 21510860 | 21511023 | 21436969 | 21437203 | 21564573 | 21564684 | 12 | 32 | 2 | 31 | 32 | 0 | 3 |
| SE | 34853 | ENSG00000159433 | STARD9 | chr15 | + | 42637906 | 42637939 | 42634855 | 42634972 | 42638025 | 42638087 | 18 | 13 | 15 | 12 | 12 | 0 | 4 |
| SE | 1346 | ENSG00000100150 | DEPDC5 | chr22 | + | 31758526 | 31758633 | 31754861 | 31754979 | 31760655 | 31760702 | 26 | 14 | 36 | 27 | 50 | 0 | 0 |
| SE | 103020 | ENSG00000156374 | PCGF6 | chr10 | - | 103314185 | 103314272 | 103302795 | 103303961 | 103326533 | 103326632 | 58 | 56 | 46 | 60 | 100 | 2 | 0 |
| SE | 11162 | ENSG00000168488 | ATXN2L | chr16 | + | 28832816 | 28832887 | 28832495 | 28832567 | 28833058 | 28833354 | 375 | 242 | 370 | 389 | 483 | 0 | 2 |
| SE | 82363 | ENSG00000228649 | SNHG26 | chr7 | + | 22858570 | 22858830 | 22856972 | 22857016 | 22860546 | 22860638 | 6 | 4 | 11 | 9 | 11 | 0 | 0 |
| SE | 36014 | ENSG00000164463 | CREBRF | chr5 | + | 173080588 | 173080784 | 173056351 | 173056479 | 173086500 | 173086626 | 326 | 229 | 307 | 299 | 661 | 0 | 0 |
| SE | 9711 | ENSG00000141568 | FOXK2 | chr17 | + | 82584012 | 82584188 | 82571723 | 82571870 | 82585903 | 82586200 | 50 | 17 | 42 | 44 | 65 | 5 | 0 |
| SE | 22475 | ENSG00000188428 | BLOC1S5 | chr6 | - | 8041138 | 8041268 | 8026366 | 8026425 | 8062533 | 8062616 | 197 | 145 | 150 | 180 | 255 | 4 | 2 |
| SE | 65443 | ENSG00000166164 | BRD7 | chr16 | - | 50326283 | 50326391 | 50325747 | 50325883 | 50328668 | 50328744 | 184 | 192 | 187 | 247 | 181 | 10 | 11 |
| SE | 72445 | ENSG00000074582 | BCS1L | chr2 | + | 218662509 | 218662679 | 218662196 | 218662260 | 218662882 | 218663000 | 37 | 19 | 42 | 73 | 50 | 0 | 0 |
| SE | 74178 | ENSG00000116698 | SMG7 | chr1 | + | 183542075 | 183542364 | 183540983 | 183541103 | 183544352 | 183544497 | 191 | 169 | 200 | 279 | 303 | 0 | 0 |
| SE | 54124 | ENSG00000124126 | PREX1 | chr20 | - | 48642167 | 48642258 | 48639765 | 48639894 | 48642406 | 48642489 | 597 | 253 | 387 | 506 | 338 | 0 | 0 |
| SE | 16764 | ENSG00000125107 | CNOT1 | chr16 | - | 58558472 | 58558674 | 58556846 | 58556990 | 58560211 | 58560362 | 245 | 131 | 199 | 195 | 306 | 0 | 0 |
| SE | 53074 | ENSG00000103657 | HERC1 | chr15 | - | 63725291 | 63725513 | 63718782 | 63718897 | 63727646 | 63727838 | 83 | 64 | 64 | 99 | 69 | 0 | 0 |
| SE | 56244 | ENSG00000108523 | RNF167 | chr17 | + | 4942850 | 4942941 | 4942576 | 4942664 | 4944557 | 4944601 | 154 | 62 | 95 | 123 | 140 | 1 | 1 |
| SE | 82854 | ENSG00000146776 | ATXN7L1 | chr7 | - | 105620199 | 105620321 | 105613473 | 105614816 | 105624074 | 105624266 | 90 | 43 | 88 | 84 | 115 | 0 | 0 |
| SE | 10353 | ENSG00000131467 | PSME3 | chr17 | + | 42838092 | 42838205 | 42837648 | 42837697 | 42838730 | 42838798 | 380 | 227 | 305 | 360 | 537 | 1 | 1 |
| SE | 95660 | ENSG00000137073 | UBAP2 | chr9 | - | 33973182 | 33973237 | 33963725 | 33963791 | 33986759 | 33986837 | 32 | 28 | 54 | 31 | 80 | 1 | 6 |
| SE | 94966 | ENSG00000266714 | MYO15B | chr17 | + | 75594474 | 75594588 | 75592678 | 75592840 | 75594839 | 75594972 | 3 | 2 | 4 | 14 | 7 | 1 | 0 |
| SE | 18935 | ENSG00000100461 | RBM23 | chr14 | - | 22913723 | 22913821 | 22911327 | 22911403 | 22917478 | 22917556 | 1 | 1 | 6 | 3 | 8 | 0 | 0 |
| SE | 103447 | ENSG00000148840 | PPRC1 | chr10 | + | 102144249 | 102144307 | 102143044 | 102143098 | 102146671 | 102147392 | 19 | 11 | 27 | 13 | 23 | 3 | 0 |
| SE | 60650 | ENSG00000147099 | HDAC8 | chrX | - | 72462361 | 72462442 | 72460258 | 72462098 | 72464558 | 72464564 | 1 | 1 | 5 | 2 | 2 | 12 | 4 |
| SE | 65453 | ENSG00000204344 | STK19 | chr6 | + | 31972620 | 31972919 | 31972351 | 31972511 | 31978902 | 31978998 | 20 | 7 | 14 | 4 | 12 | 1 | 0 |
| SE | 58783 | ENSG00000077254 | USP33 | chr1 | - | 77701368 | 77701471 | 77697862 | 77697931 | 77711746 | 77711855 | 237 | 180 | 193 | 203 | 234 | 20 | 9 |
| SE | 91652 | ENSG00000105339 | DENND3 | chr8 | + | 141163332 | 141163429 | 141160631 | 141160787 | 141166189 | 141166389 | 32 | 18 | 32 | 21 | 32 | 2 | 0 |
| SE | 33329 | ENSG00000188313 | PLSCR1 | chr3 | - | 146521543 | 146522053 | 146517005 | 146517167 | 146525604 | 146525647 | 271 | 267 | 199 | 355 | 202 | 6 | 2 |
| SE | 87125 | ENSG00000102312 | PORCN | chrX | + | 48512834 | 48512852 | 48512588 | 48512719 | 48514239 | 48514365 | 2 | 0 | 0 | 0 | 0 | 6 | 2 |
| SE | 60483 | ENSG00000070814 | TCOF1 | chr5 | + | 150387901 | 150388088 | 150379531 | 150379732 | 150389886 | 150390023 | 30 | 19 | 35 | 61 | 8 | 0 | 0 |
| SE | 50602 | ENSG00000085871 | MGST2 | chr4 | + | 139695196 | 139695267 | 139678542 | 139678642 | 139703454 | 139703536 | 131 | 51 | 55 | 102 | 102 | 3 | 5 |
| SE | 38116 | ENSG00000132394 | EEFSEC | chr3 | + | 128246835 | 128247043 | 128153480 | 128153823 | 128262127 | 128262224 | 48 | 14 | 15 | 44 | 27 | 0 | 0 |
| SE | 60681 | ENSG00000248333 | CDK11B | chr1 | - | 1645125 | 1645262 | 1642346 | 1642508 | 1649498 | 1649637 | 12 | 3 | 23 | 31 | 64 | 0 | 3 |
| SE | 66657 | ENSG00000084623 | EIF3I | chr1 | + | 32226402 | 32226530 | 32226170 | 32226283 | 32228498 | 32228609 | 201 | 123 | 208 | 177 | 326 | 1 | 0 |
| SE | 60647 | ENSG00000147099 | HDAC8 | chrX | - | 72330815 | 72330876 | 72329567 | 72330076 | 72351732 | 72351838 | 4 | 4 | 1 | 5 | 0 | 18 | 9 |
| SE | 52627 | ENSG00000197872 | CYRIA | chr2 | - | 16622938 | 16623137 | 16588049 | 16588129 | 16623863 | 16624019 | 2 | 4 | 2 | 6 | 0 | 153 | 83 |
| SE | 29804 | ENSG00000141452 | RMC1 | chr18 | + | 23509192 | 23509279 | 23506969 | 23507054 | 23515855 | 23515886 | 128 | 97 | 85 | 102 | 139 | 0 | 1 |
| SE | 107140 | ENSG00000175582 | RAB6A | chr11 | - | 73718612 | 73718718 | 73716250 | 73716362 | 73718784 | 73718890 | 643 | 449 | 409 | 594 | 709 | 470 | 355 |
| SE | 25164 | ENSG00000164219 | PGGT1B | chr5 | - | 115221823 | 115222008 | 115216864 | 115216973 | 115230975 | 115231021 | 207 | 185 | 153 | 206 | 261 | 2 | 2 |
| SE | 10476 | ENSG00000186566 | GPATCH8 | chr17 | - | 44438670 | 44438715 | 44436477 | 44436545 | 44464471 | 44464544 | 32 | 29 | 29 | 28 | 63 | 169 | 91 |
| SE | 41232 | ENSG00000153815 | CMIP | chr16 | + | 81702621 | 81702669 | 81701659 | 81701800 | 81703938 | 81704085 | 122 | 72 | 90 | 96 | 47 | 2 | 1 |
| SE | 93266 | ENSG00000132334 | PTPRE | chr10 | + | 128060938 | 128061015 | 128056122 | 128056213 | 128061678 | 128061715 | 482 | 328 | 281 | 308 | 441 | 1 | 0 |
| SE | 21286 | ENSG00000150867 | PIP4K2A | chr10 | - | 22591628 | 22591781 | 22550658 | 22550772 | 22607926 | 22608023 | 1068 | 638 | 789 | 1098 | 1416 | 8 | 5 |
| SE | 14326 | ENSG00000103197 | TSC2 | chr16 | + | 2053341 | 2053452 | 2050399 | 2050486 | 2056195 | 2056244 | 62 | 40 | 50 | 39 | 45 | 4 | 1 |
| SE | 8725 | ENSG00000070366 | SMG6 | chr17 | - | 2297862 | 2298055 | 2297242 | 2297353 | 2298905 | 2300664 | 106 | 75 | 87 | 95 | 151 | 0 | 0 |
| SE | 18674 | ENSG00000129566 | TEP1 | chr14 | - | 20372732 | 20372857 | 20371488 | 20371632 | 20373010 | 20373100 | 145 | 44 | 82 | 119 | 88 | 26 | 10 |
| SE | 68627 | ENSG00000223960 | CHROMR | chr2 | + | 178426365 | 178426661 | 178413634 | 178414113 | 178430694 | 178430889 | 60 | 32 | 20 | 53 | 30 | 1 | 0 |
| SE | 73997 | ENSG00000096433 | ITPR3 | chr6 | + | 33671164 | 33671306 | 33670670 | 33670815 | 33672028 | 33672228 | 66 | 24 | 73 | 56 | 64 | 3 | 1 |
| SE | 55419 | ENSG00000070540 | WIPI1 | chr17 | - | 68429995 | 68430160 | 68427134 | 68427253 | 68433467 | 68433571 | 36 | 21 | 30 | 62 | 76 | 0 | 0 |
| SE | 23837 | ENSG00000172273 | HINFP | chr11 | + | 119132242 | 119132297 | 119131829 | 119131982 | 119132495 | 119132573 | 22 | 15 | 18 | 40 | 35 | 37 | 13 |
| SE | 85761 | ENSG00000075539 | FRYL | chr4 | - | 48553214 | 48553383 | 48551493 | 48551578 | 48556977 | 48557118 | 370 | 236 | 375 | 367 | 418 | 7 | 7 |
| SE | 54640 | ENSG00000140992 | PDPK1 | chr16 | + | 2595792 | 2595850 | 2586675 | 2586893 | 2597650 | 2597928 | 49 | 20 | 46 | 68 | 17 | 0 | 0 |
| SE | 91202 | ENSG00000116604 | MEF2D | chr1 | - | 156483238 | 156483430 | 156482436 | 156482640 | 156500485 | 156500753 | 281 | 117 | 256 | 293 | 201 | 0 | 0 |
| SE | 48442 | ENSG00000104643 | MTMR9 | chr8 | + | 11297899 | 11297951 | 11295193 | 11295302 | 11300022 | 11300148 | 8 | 1 | 4 | 3 | 0 | 29 | 21 |
| SE | 38900 | ENSG00000151466 | SCLT1 | chr4 | - | 128952342 | 128952532 | 128948495 | 128948570 | 128952768 | 128952840 | 1 | 0 | 0 | 0 | 0 | 96 | 109 |
| SE | 68593 | ENSG00000239779 | WBP1 | chr2 | + | 74458996 | 74459098 | 74458399 | 74458671 | 74459642 | 74459745 | 2 | 0 | 0 | 4 | 0 | 15 | 2 |
| SE | 82273 | ENSG00000223482 | NUTM2A-AS1 | chr10 | - | 87333825 | 87334814 | 87330891 | 87331096 | 87336785 | 87336829 | 8 | 10 | 6 | 15 | 21 | 9 | 10 |
| SE | 91836 | ENSG00000104365 | IKBKB | chr8 | + | 42319584 | 42319646 | 42319269 | 42319421 | 42320734 | 42320844 | 304 | 168 | 187 | 219 | 255 | 14 | 1 |
| SE | 41063 | ENSG00000160741 | CRTC2 | chr1 | - | 153951259 | 153951666 | 153949114 | 153949384 | 153952017 | 153952262 | 22 | 11 | 10 | 30 | 10 | 4 | 2 |
| SE | 103819 | ENSG00000225484 | NUTM2B-AS1 | chr10 | - | 79808762 | 79808832 | 79806038 | 79806122 | 79814612 | 79814716 | 44 | 29 | 43 | 73 | 43 | 28 | 5 |
| SE | 39859 | ENSG00000169239 | CA5B | chrX | + | 15766911 | 15767026 | 15764595 | 15764775 | 15772495 | 15772614 | 6 | 0 | 2 | 3 | 0 | 130 | 79 |
| SE | 35601 | ENSG00000114331 | ACAP2 | chr3 | - | 195410706 | 195411009 | 195392089 | 195392147 | 195442794 | 195442905 | 19 | 6 | 17 | 13 | 39 | 45 | 25 |
| SE | 80253 | ENSG00000164535 | DAGLB | chr7 | - | 6424751 | 6424835 | 6416840 | 6416921 | 6425987 | 6426114 | 25 | 15 | 22 | 24 | 30 | 0 | 0 |
| SE | 41492 | ENSG00000270629 | NBPF14 | chr1 | - | 148593597 | 148593700 | 148592551 | 148592766 | 148595542 | 148595717 | 160 | 113 | 50 | 88 | 108 | 23 | 28 |
| SE | 42726 | ENSG00000182473 | EXOC7 | chr17 | - | 76091142 | 76091235 | 76089174 | 76089320 | 76094413 | 76094581 | 17 | 2 | 13 | 2 | 7 | 52 | 29 |
| SE | 55696 | ENSG00000135749 | PCNX2 | chr1 | - | 233160282 | 233160433 | 233139713 | 233139855 | 233161270 | 233161363 | 60 | 35 | 106 | 63 | 79 | 2 | 0 |
| SE | 67515 | ENSG00000115159 | GPD2 | chr2 | + | 156495561 | 156495691 | 156476097 | 156476207 | 156496043 | 156496215 | 0 | 1 | 1 | 2 | 2 | 12 | 16 |
| SE | 14110 | ENSG00000102897 | LYRM1 | chr16 | + | 20902485 | 20902717 | 20900665 | 20900889 | 20915555 | 20915714 | 5 | 4 | 1 | 7 | 3 | 1 | 1 |
| SE | 9974 | ENSG00000170921 | TANC2 | chr17 | + | 63354782 | 63355390 | 63351249 | 63351416 | 63379717 | 63379826 | 60 | 53 | 43 | 71 | 56 | 5 | 3 |
| SE | 46101 | ENSG00000166912 | MTMR10 | chr15 | - | 30976818 | 30976955 | 30974930 | 30975003 | 30990776 | 30990837 | 478 | 474 | 395 | 488 | 681 | 13 | 8 |
| SE | 61758 | ENSG00000153029 | MR1 | chr1 | + | 181052321 | 181052510 | 181050010 | 181050286 | 181053572 | 181053677 | 40 | 11 | 17 | 14 | 37 | 70 | 61 |
| SE | 2185 | ENSG00000079805 | DNM2 | chr19 | + | 10808568 | 10808580 | 10805915 | 10805967 | 10812263 | 10812377 | 240 | 146 | 245 | 303 | 302 | 18 | 21 |
| SE | 30684 | ENSG00000132466 | ANKRD17 | chr4 | - | 73118687 | 73118850 | 73102375 | 73102547 | 73120161 | 73120337 | 80 | 59 | 96 | 84 | 79 | 1 | 0 |
| SE | 43894 | ENSG00000179335 | CLK3 | chr15 | + | 74627969 | 74628052 | 74627538 | 74627668 | 74628603 | 74628683 | 477 | 334 | 452 | 566 | 472 | 1 | 5 |
| SE | 101258 | ENSG00000166224 | SGPL1 | chr10 | + | 70873350 | 70873589 | 70871047 | 70871146 | 70875401 | 70875548 | 90 | 63 | 85 | 90 | 102 | 0 | 0 |
| SE | 99013 | ENSG00000110324 | IL10RA | chr11 | + | 117989441 | 117989620 | 117986736 | 117988502 | 117993240 | 117993410 | 1590 | 652 | 823 | 950 | 804 | 15 | 5 |
| SE | 8013 | ENSG00000129197 | RPAIN | chr17 | + | 5425970 | 5426299 | 5422768 | 5422829 | 5432541 | 5432876 | 47 | 23 | 58 | 52 | 61 | 10 | 10 |
| SE | 102221 | ENSG00000100109 | TFIP11 | chr22 | - | 26510616 | 26510667 | 26510063 | 26510281 | 26512098 | 26512175 | 36 | 39 | 31 | 47 | 75 | 0 | 0 |
| SE | 77671 | ENSG00000214021 | TTLL3 | chr3 | + | 9834426 | 9834503 | 9833103 | 9833273 | 9834680 | 9834907 | 3 | 1 | 3 | 2 | 0 | 5 | 0 |
| SE | 41030 | ENSG00000064687 | ABCA7 | chr19 | + | 1056906 | 1057084 | 1056329 | 1056499 | 1058617 | 1058747 | 31 | 7 | 38 | 42 | 21 | 0 | 0 |
| SE | 12081 | ENSG00000135723 | FHOD1 | chr16 | - | 67236410 | 67236434 | 67234356 | 67234472 | 67236556 | 67236616 | 3 | 0 | 0 | 3 | 0 | 16 | 12 |
| SE | 47484 | ENSG00000143363 | PRUNE1 | chr1 | + | 151018466 | 151018669 | 151017811 | 151017904 | 151024610 | 151024795 | 75 | 62 | 88 | 132 | 134 | 25 | 21 |
| SE | 42640 | ENSG00000008130 | NADK | chr1 | - | 1755373 | 1755476 | 1754543 | 1754698 | 1756257 | 1756327 | 237 | 92 | 154 | 151 | 183 | 19 | 6 |
| SE | 62369 | ENSG00000168826 | ZBTB49 | chr4 | + | 4313040 | 4313114 | 4302310 | 4303091 | 4320639 | 4320697 | 3 | 8 | 6 | 4 | 9 | 1 | 0 |
| SE | 67593 | ENSG00000008083 | JARID2 | chr6 | + | 15410223 | 15410365 | 15374116 | 15374252 | 15468541 | 15468718 | 189 | 99 | 132 | 202 | 198 | 0 | 2 |
| SE | 17023 | ENSG00000102921 | N4BP1 | chr16 | - | 48551385 | 48551482 | 48538725 | 48543261 | 48553538 | 48553669 | 76 | 49 | 58 | 68 | 122 | 2 | 1 |
| SE | 41032 | ENSG00000064687 | ABCA7 | chr19 | + | 1056906 | 1057429 | 1056329 | 1056499 | 1058617 | 1058747 | 31 | 7 | 40 | 40 | 21 | 0 | 0 |
| SE | 38551 | ENSG00000180098 | TRNAU1AP | chr1 | + | 28574588 | 28574718 | 28571866 | 28571900 | 28577499 | 28577775 | 0 | 2 | 0 | 0 | 2 | 35 | 13 |
| SE | 14814 | ENSG00000137804 | NUSAP1 | chr15 | + | 41375711 | 41375828 | 41371526 | 41371684 | 41377195 | 41377288 | 18 | 33 | 36 | 38 | 50 | 1 | 0 |
| SE | 76654 | ENSG00000161800 | RACGAP1 | chr12 | - | 50016630 | 50016715 | 50006433 | 50006636 | 50025327 | 50025436 | 20 | 15 | 16 | 33 | 19 | 0 | 0 |
| SE | 48770 | ENSG00000116161 | CACYBP | chr1 | + | 175000088 | 175000195 | 174999434 | 174999689 | 175004613 | 175004750 | 65 | 23 | 51 | 36 | 37 | 3 | 6 |
| SE | 19493 | ENSG00000033170 | FUT8 | chr14 | + | 65733230 | 65733381 | 65724146 | 65724323 | 65742092 | 65742936 | 91 | 101 | 128 | 103 | 358 | 4 | 0 |
| SE | 16780 | ENSG00000125107 | CNOT1 | chr16 | - | 58623281 | 58623409 | 58599262 | 58599511 | 58629727 | 58629885 | 4 | 5 | 15 | 19 | 4 | 75 | 43 |
| SE | 98566 | ENSG00000134955 | SLC37A2 | chr11 | + | 125084233 | 125084319 | 125083814 | 125083877 | 125084824 | 125084873 | 37 | 12 | 5 | 22 | 4 | 1 | 0 |
| SE | 83711 | ENSG00000196998 | WDR45 | chrX | - | 49075135 | 49075281 | 49074432 | 49074912 | 49075363 | 49075378 | 198 | 84 | 139 | 200 | 132 | 1 | 1 |
| SE | 18936 | ENSG00000100461 | RBM23 | chr14 | - | 22913723 | 22913821 | 22911327 | 22911403 | 22918998 | 22919149 | 1 | 0 | 6 | 5 | 8 | 183 | 86 |
| SE | 87763 | ENSG00000063601 | MTMR1 | chrX | + | 150755688 | 150755865 | 150750729 | 150750843 | 150762564 | 150762737 | 97 | 78 | 115 | 99 | 107 | 1 | 2 |
| SE | 14046 | ENSG00000140694 | PARN | chr16 | - | 14586317 | 14586361 | 14584748 | 14584791 | 14593300 | 14593378 | 193 | 157 | 200 | 207 | 444 | 2 | 12 |
| SE | 68877 | ENSG00000114450 | GNB4 | chr3 | - | 179424111 | 179424284 | 179420888 | 179420927 | 179426143 | 179426200 | 0 | 9 | 0 | 6 | 5 | 31 | 51 |
| SE | 71956 | ENSG00000160710 | ADAR | chr1 | - | 154586180 | 154586363 | 154585216 | 154585865 | 154588124 | 154588258 | 602 | 362 | 587 | 781 | 629 | 0 | 1 |
| SE | 9280 | ENSG00000063244 | U2AF2 | chr19 | + | 55658861 | 55658981 | 55655108 | 55655153 | 55659209 | 55659345 | 6 | 8 | 2 | 4 | 0 | 70 | 31 |
| SE | 72534 | ENSG00000135426 | TESPA1 | chr12 | - | 54966052 | 54966151 | 54963740 | 54963950 | 54966387 | 54966424 | 160 | 131 | 247 | 191 | 372 | 3 | 0 |
| SE | 33429 | ENSG00000135870 | RC3H1 | chr1 | - | 173972508 | 173972627 | 173970504 | 173970617 | 173978487 | 173978620 | 192 | 108 | 150 | 169 | 129 | 1 | 0 |
| SE | 11014 | ENSG00000108559 | NUP88 | chr17 | - | 5418014 | 5418116 | 5416512 | 5416682 | 5419353 | 5419661 | 7 | 4 | 8 | 11 | 2 | 140 | 35 |
| SE | 89138 | ENSG00000154589 | LY96 | chr8 | + | 74004795 | 74004885 | 73991391 | 73991554 | 74010000 | 74010129 | 55 | 31 | 38 | 75 | 86 | 9 | 5 |
| SE | 10332 | ENSG00000168961 | LGALS9 | chr17 | + | 27647269 | 27647432 | 27647029 | 27647118 | 27648835 | 27649541 | 599 | 442 | 434 | 519 | 374 | 12 | 9 |
| SE | 57006 | ENSG00000127124 | HIVEP3 | chr1 | - | 41512815 | 41513750 | 41506364 | 41511263 | 41518401 | 41518488 | 33 | 23 | 9 | 21 | 13 | 0 | 0 |
| SE | 42999 | ENSG00000065911 | MTHFD2 | chr2 | + | 74205704 | 74205889 | 74198609 | 74198742 | 74207703 | 74207826 | 147 | 82 | 154 | 138 | 82 | 5 | 2 |
| SE | 52340 | ENSG00000142197 | DOP1B | chr21 | + | 36200330 | 36200501 | 36164707 | 36164871 | 36208714 | 36208904 | 21 | 4 | 12 | 14 | 14 | 0 | 2 |
| SE | 82291 | ENSG00000122566 | HNRNPA2B1 | chr7 | - | 26191055 | 26191128 | 26189730 | 26190460 | 26192494 | 26192577 | 111 | 83 | 129 | 127 | 221 | 14 | 4 |
| SE | 44696 | ENSG00000159592 | GPBP1L1 | chr1 | - | 45685575 | 45685620 | 45660183 | 45661225 | 45686411 | 45686533 | 96 | 76 | 103 | 102 | 121 | 25 | 27 |
| SE | 43688 | ENSG00000163291 | PAQR3 | chr4 | - | 78922140 | 78922421 | 78917939 | 78920681 | 78923856 | 78923947 | 3 | 1 | 0 | 4 | 6 | 18 | 8 |
| SE | 40390 | ENSG00000163719 | MTMR14 | chr3 | + | 9677316 | 9677387 | 9672684 | 9672758 | 9683177 | 9683217 | 228 | 140 | 161 | 177 | 246 | 2 | 0 |
| SE | 79459 | ENSG00000166135 | HIF1AN | chr10 | + | 100540633 | 100540782 | 100536410 | 100536661 | 100544950 | 100545096 | 284 | 174 | 231 | 274 | 293 | 5 | 5 |
| SE | 55275 | ENSG00000260269 | NA | chr15 | - | 75600636 | 75600890 | 75598278 | 75598681 | 75601137 | 75601186 | 4 | 1 | 3 | 6 | 15 | 65 | 25 |
| SE | 5204 | ENSG00000124207 | CSE1L | chr20 | + | 49085282 | 49085386 | 49084025 | 49084162 | 49088008 | 49088106 | 188 | 198 | 168 | 162 | 673 | 9 | 7 |
| SE | 44715 | ENSG00000143772 | ITPKB | chr1 | - | 226641920 | 226642125 | 226639556 | 226639658 | 226648671 | 226648771 | 176 | 107 | 157 | 207 | 187 | 0 | 0 |
| SE | 75819 | ENSG00000153147 | SMARCA5 | chr4 | + | 143545924 | 143546047 | 143545469 | 143545583 | 143546775 | 143546908 | 418 | 341 | 342 | 403 | 554 | 6 | 6 |
| SE | 107826 | ENSG00000135457 | TFCP2 | chr12 | - | 51101934 | 51102025 | 51099654 | 51099779 | 51106524 | 51106613 | 48 | 32 | 52 | 56 | 76 | 0 | 0 |
| SE | 208 | ENSG00000160209 | PDXK | chr21 | + | 43752517 | 43752629 | 43750499 | 43750545 | 43753582 | 43753594 | 258 | 121 | 176 | 237 | 193 | 0 | 0 |
| SE | 85790 | ENSG00000258529 | NA | chr11 | - | 111838248 | 111838378 | 111837467 | 111837615 | 111844600 | 111844723 | 56 | 31 | 52 | 61 | 73 | 0 | 3 |
| SE | 13778 | ENSG00000205629 | LCMT1 | chr16 | + | 25170713 | 25170797 | 25169111 | 25169213 | 25174936 | 25175034 | 36 | 9 | 18 | 26 | 29 | 1 | 0 |
| SE | 10641 | ENSG00000171475 | WIPF2 | chr17 | + | 40260534 | 40260667 | 40256390 | 40256522 | 40262524 | 40262638 | 98 | 46 | 64 | 79 | 84 | 0 | 1 |
| SE | 81548 | ENSG00000106348 | IMPDH1 | chr7 | - | 128405766 | 128405865 | 128403705 | 128403754 | 128409755 | 128409982 | 52 | 20 | 30 | 30 | 6 | 4 | 0 |
| SE | 1567 | ENSG00000198089 | SFI1 | chr22 | + | 31546860 | 31546971 | 31508254 | 31508376 | 31556941 | 31557059 | 16 | 19 | 24 | 18 | 12 | 0 | 0 |
| SE | 98102 | ENSG00000171466 | ZNF562 | chr19 | - | 9656546 | 9656653 | 9652712 | 9653878 | 9658008 | 9658135 | 89 | 90 | 71 | 86 | 60 | 3 | 0 |
| SE | 22411 | ENSG00000151612 | ZNF827 | chr4 | - | 145870244 | 145870478 | 145849321 | 145849561 | 145885677 | 145885769 | 19 | 20 | 33 | 29 | 29 | 3 | 0 |
| SE | 26902 | ENSG00000160613 | PCSK7 | chr11 | - | 117207066 | 117207160 | 117206679 | 117206798 | 117207969 | 117208064 | 133 | 60 | 100 | 110 | 87 | 15 | 10 |
| SE | 89303 | ENSG00000166169 | POLL | chr10 | - | 101585315 | 101585478 | 101584601 | 101584919 | 101587245 | 101587382 | 10 | 10 | 4 | 11 | 8 | 2 | 1 |
| SE | 25149 | ENSG00000155329 | ZCCHC10 | chr5 | - | 133000131 | 133000201 | 132998407 | 132998850 | 133006758 | 133006920 | 21 | 12 | 11 | 13 | 35 | 8 | 4 |
| SE | 46903 | ENSG00000055609 | KMT2C | chr7 | - | 152224434 | 152224616 | 152224014 | 152224179 | 152229922 | 152230027 | 252 | 176 | 236 | 287 | 245 | 0 | 0 |
| SE | 103538 | ENSG00000107862 | GBF1 | chr10 | + | 102368217 | 102368454 | 102367477 | 102367560 | 102368738 | 102368832 | 185 | 108 | 179 | 146 | 175 | 0 | 0 |
| SE | 71969 | ENSG00000160710 | ADAR | chr1 | - | 154606005 | 154606376 | 154601040 | 154602626 | 154627854 | 154627991 | 2 | 0 | 0 | 1 | 0 | 6 | 10 |
| SE | 85895 | ENSG00000163421 | PROK2 | chr3 | - | 71774444 | 71774507 | 71771654 | 71772828 | 71781466 | 71781592 | 61 | 4 | 9 | 38 | 50 | 9 | 3 |
| SE | 37465 | ENSG00000163681 | SLMAP | chr3 | + | 57865241 | 57865292 | 57864806 | 57864857 | 57890040 | 57890100 | 29 | 31 | 32 | 31 | 52 | 68 | 26 |
| SE | 22596 | ENSG00000127603 | MACF1 | chr1 | + | 39331202 | 39336653 | 39327217 | 39327353 | 39337181 | 39337331 | 459 | 305 | 386 | 469 | 403 | 10 | 3 |
| SE | 97185 | ENSG00000106290 | TAF6 | chr7 | - | 100113615 | 100113769 | 100113348 | 100113405 | 100113867 | 100113954 | 107 | 39 | 77 | 81 | 61 | 1 | 2 |
| SE | 107781 | ENSG00000102780 | DGKH | chr13 | + | 42166514 | 42166674 | 42165330 | 42165433 | 42168439 | 42168548 | 20 | 14 | 22 | 26 | 15 | 0 | 1 |
| SE | 5411 | ENSG00000198053 | SIRPA | chr20 | + | 1924763 | 1924877 | 1915098 | 1915455 | 1927874 | 1927899 | 193 | 143 | 92 | 151 | 189 | 0 | 0 |
| SE | 78155 | ENSG00000254641 | NA | chr11 | - | 6608836 | 6608978 | 6608666 | 6608725 | 6609907 | 6610135 | 0 | 0 | 0 | 2 | 0 | 7 | 3 |
| SE | 44596 | ENSG00000026508 | CD44 | chr11 | + | 35211245 | 35211449 | 35208104 | 35208206 | 35214851 | 35214861 | 171 | 97 | 114 | 111 | 145 | 0 | 5 |
| SE | 16689 | ENSG00000137841 | PLCB2 | chr15 | - | 40290024 | 40290082 | 40289065 | 40289358 | 40290576 | 40290672 | 656 | 487 | 502 | 713 | 652 | 35 | 29 |
| SE | 38464 | ENSG00000147251 | DOCK11 | chrX | + | 118624538 | 118624655 | 118618549 | 118618728 | 118627503 | 118627579 | 384 | 295 | 320 | 456 | 601 | 6 | 0 |
| SE | 49191 | ENSG00000066136 | NFYC | chr1 | + | 40769355 | 40769415 | 40766595 | 40766703 | 40770708 | 40770940 | 58 | 19 | 34 | 56 | 25 | 0 | 0 |
| SE | 49782 | ENSG00000116337 | AMPD2 | chr1 | + | 109625302 | 109625433 | 109619830 | 109620278 | 109625661 | 109625792 | 60 | 19 | 22 | 38 | 14 | 1 | 1 |
| SE | 91261 | ENSG00000154114 | TBCEL | chr11 | + | 121053550 | 121053732 | 121047527 | 121047667 | 121055051 | 121055308 | 120 | 57 | 88 | 124 | 165 | 2 | 5 |
| SE | 59643 | ENSG00000163466 | ARPC2 | chr2 | + | 218236368 | 218236733 | 218234848 | 218235044 | 218238663 | 218238730 | 18 | 4 | 6 | 11 | 0 | 8 | 9 |
| SE | 5043 | ENSG00000101126 | ADNP | chr20 | - | 50903888 | 50904001 | 50888917 | 50894512 | 50928650 | 50928732 | 25 | 19 | 24 | 36 | 34 | 0 | 0 |
| SE | 3929 | ENSG00000105568 | PPP2R1A | chr19 | + | 52220188 | 52220249 | 52219690 | 52219864 | 52220978 | 52221133 | 150 | 65 | 134 | 169 | 215 | 0 | 0 |
| SE | 101240 | ENSG00000144802 | NFKBIZ | chr3 | + | 101857072 | 101857183 | 101855732 | 101855902 | 101857291 | 101857459 | 186 | 146 | 172 | 241 | 202 | 4 | 0 |
| SE | 5956 | ENSG00000170471 | RALGAPB | chr20 | + | 38524777 | 38524945 | 38517783 | 38518000 | 38525403 | 38525518 | 104 | 95 | 87 | 98 | 180 | 1 | 0 |
| SE | 57389 | ENSG00000101040 | ZMYND8 | chr20 | - | 47238895 | 47239138 | 47236325 | 47236516 | 47249286 | 47249439 | 28 | 18 | 31 | 28 | 25 | 1 | 0 |
| SE | 20111 | ENSG00000078319 | PMS2P1 | chr7 | - | 100328835 | 100329019 | 100320912 | 100321084 | 100330181 | 100330284 | 61 | 26 | 92 | 51 | 222 | 2 | 0 |
| SE | 81035 | ENSG00000168092 | PAFAH1B2 | chr11 | + | 117152440 | 117152528 | 117144283 | 117144418 | 117159933 | 117160023 | 192 | 162 | 176 | 216 | 194 | 2 | 1 |
| SE | 92364 | ENSG00000143437 | ARNT | chr1 | - | 150861246 | 150861302 | 150858348 | 150858460 | 150876542 | 150876708 | 0 | 3 | 0 | 0 | 0 | 8 | 7 |
| SE | 53391 | ENSG00000135837 | CEP350 | chr1 | + | 180006453 | 180006567 | 180003173 | 180003287 | 180011928 | 180012075 | 305 | 328 | 270 | 334 | 478 | 11 | 1 |
| SE | 74179 | ENSG00000116698 | SMG7 | chr1 | + | 183542075 | 183542502 | 183540983 | 183541103 | 183544352 | 183544497 | 203 | 154 | 189 | 269 | 303 | 0 | 0 |
| SE | 24526 | ENSG00000113719 | ERGIC1 | chr5 | + | 172915517 | 172915690 | 172914713 | 172914838 | 172924004 | 172924024 | 6 | 6 | 7 | 1 | 0 | 136 | 86 |
| SE | 67844 | ENSG00000018280 | SLC11A1 | chr2 | + | 218382959 | 218383070 | 218382301 | 218382375 | 218384242 | 218384365 | 48 | 27 | 21 | 15 | 26 | 6 | 1 |
| SE | 38313 | ENSG00000138172 | CALHM2 | chr10 | - | 103451878 | 103452036 | 103451082 | 103451182 | 103452155 | 103452336 | 7 | 0 | 7 | 4 | 0 | 60 | 39 |
| SE | 39233 | ENSG00000197948 | FCHSD1 | chr5 | - | 141642396 | 141642503 | 141641701 | 141641757 | 141644217 | 141644438 | 7 | 1 | 2 | 2 | 16 | 1 | 0 |
| SE | 32619 | ENSG00000078618 | NRDC | chr1 | - | 51837486 | 51837620 | 51836368 | 51836438 | 51840225 | 51840514 | 27 | 39 | 27 | 22 | 50 | 6 | 7 |
| SE | 11554 | ENSG00000125814 | NAPB | chr20 | - | 23394921 | 23394999 | 23390208 | 23390264 | 23402992 | 23403072 | 24 | 12 | 20 | 27 | 54 | 0 | 0 |
| SE | 32992 | ENSG00000114316 | USP4 | chr3 | - | 49278813 | 49278902 | 49277143 | 49278451 | 49283986 | 49284136 | 144 | 115 | 106 | 154 | 186 | 0 | 2 |
| SE | 49451 | ENSG00000115020 | PIKFYVE | chr2 | + | 208276711 | 208276830 | 208273583 | 208273733 | 208285725 | 208285933 | 73 | 53 | 53 | 80 | 71 | 8 | 4 |
| SE | 61331 | ENSG00000102786 | INTS6 | chr13 | - | 51451024 | 51451174 | 51430234 | 51430383 | 51451977 | 51452055 | 227 | 141 | 150 | 243 | 182 | 8 | 0 |
| SE | 53377 | ENSG00000135763 | URB2 | chr1 | + | 229645858 | 229645969 | 229643532 | 229643693 | 229647509 | 229647752 | 33 | 6 | 23 | 20 | 32 | 1 | 1 |
| SE | 1165 | ENSG00000100395 | L3MBTL2 | chr22 | + | 41229539 | 41229656 | 41227088 | 41227869 | 41230138 | 41230654 | 78 | 36 | 73 | 63 | 114 | 2 | 3 |
| SE | 105518 | ENSG00000084110 | HAL | chr12 | - | 95990392 | 95990532 | 95988192 | 95988240 | 95992679 | 95992805 | 38 | 20 | 29 | 48 | 49 | 1 | 0 |
| SE | 36282 | ENSG00000075420 | FNDC3B | chr3 | + | 172184244 | 172184383 | 172133470 | 172133546 | 172186711 | 172187218 | 0 | 0 | 1 | 0 | 0 | 4 | 4 |
| SE | 4331 | ENSG00000064607 | SUGP2 | chr19 | - | 18991339 | 18991553 | 18990892 | 18991149 | 18993647 | 18993740 | 10 | 2 | 4 | 4 | 14 | 4 | 3 |
| SE | 8423 | ENSG00000078687 | TNRC6C | chr17 | + | 78064730 | 78064937 | 78051066 | 78051448 | 78067756 | 78067910 | 69 | 23 | 72 | 43 | 31 | 2 | 1 |
| SE | 22453 | ENSG00000151835 | SACS | chr13 | - | 23358334 | 23358481 | 23353967 | 23356007 | 23365165 | 23365277 | 105 | 100 | 117 | 110 | 181 | 0 | 4 |
| SE | 51112 | ENSG00000132781 | MUTYH | chr1 | - | 45333412 | 45333561 | 45332762 | 45333324 | 45334390 | 45334511 | 49 | 32 | 46 | 32 | 18 | 0 | 0 |
| SE | 84980 | ENSG00000170832 | USP32 | chr17 | - | 60211375 | 60211514 | 60209369 | 60209543 | 60212023 | 60212098 | 191 | 190 | 136 | 158 | 206 | 0 | 0 |
| SE | 103924 | ENSG00000188234 | AGAP4 | chr10 | - | 45841652 | 45841687 | 45834015 | 45834116 | 45844325 | 45844394 | 24 | 16 | 42 | 32 | 54 | 0 | 1 |
| SE | 82744 | ENSG00000140396 | NCOA2 | chr8 | - | 70148272 | 70148483 | 70141178 | 70141399 | 70155970 | 70157240 | 158 | 152 | 165 | 177 | 326 | 7 | 7 |
| SE | 83676 | ENSG00000044446 | PHKA2 | chrX | - | 18943709 | 18943808 | 18941528 | 18941675 | 18945077 | 18945158 | 126 | 78 | 116 | 90 | 122 | 3 | 7 |
| SE | 63773 | ENSG00000131748 | STARD3 | chr17 | + | 39658721 | 39658820 | 39657972 | 39658522 | 39659050 | 39659106 | 112 | 40 | 97 | 92 | 90 | 9 | 1 |
| SE | 39102 | ENSG00000109320 | NFKB1 | chr4 | + | 102582552 | 102582634 | 102580534 | 102580639 | 102582865 | 102582957 | 1 | 0 | 3 | 0 | 5 | 100 | 72 |
| SE | 24475 | ENSG00000197321 | SVIL | chr10 | - | 29526960 | 29527056 | 29524471 | 29524715 | 29529704 | 29529844 | 8 | 13 | 3 | 1 | 7 | 63 | 43 |
| SE | 2179 | ENSG00000079805 | DNM2 | chr19 | + | 10798485 | 10798572 | 10795371 | 10795439 | 10802287 | 10802358 | 414 | 215 | 347 | 373 | 535 | 1 | 0 |
| SE | 99226 | ENSG00000086848 | ALG9 | chr11 | - | 111838248 | 111838399 | 111837467 | 111837615 | 111844600 | 111844723 | 67 | 37 | 61 | 65 | 82 | 0 | 3 |
| SE | 84165 | ENSG00000137642 | SORL1 | chr11 | + | 121470006 | 121470123 | 121452313 | 121452616 | 121478117 | 121478243 | 995 | 418 | 719 | 649 | 996 | 6 | 8 |
| SE | 42854 | ENSG00000152495 | CAMK4 | chr5 | + | 111376859 | 111376942 | 111374849 | 111374912 | 111394709 | 111394749 | 206 | 254 | 401 | 256 | 932 | 1 | 2 |
| SE | 43966 | ENSG00000108443 | RPS6KB1 | chr17 | + | 59893795 | 59893924 | 59893045 | 59893325 | 59910561 | 59910611 | 0 | 2 | 5 | 4 | 1 | 35 | 33 |
| SE | 78504 | ENSG00000112592 | TBP | chr6 | + | 170567225 | 170567505 | 170566917 | 170567009 | 170569611 | 170569779 | 6 | 4 | 3 | 3 | 7 | 39 | 34 |
| SE | 76113 | ENSG00000100220 | RTCB | chr22 | - | 32406661 | 32406761 | 32401561 | 32401903 | 32408174 | 32408242 | 295 | 231 | 245 | 281 | 376 | 0 | 0 |
| SE | 8989 | ENSG00000161526 | SAP30BP | chr17 | + | 75703807 | 75703859 | 75703310 | 75703371 | 75704755 | 75704768 | 324 | 206 | 218 | 197 | 498 | 0 | 0 |
| SE | 89648 | ENSG00000245164 | LINC00861 | chr8 | - | 125922335 | 125922547 | 125921059 | 125921226 | 125936239 | 125936333 | 193 | 191 | 261 | 167 | 355 | 6 | 4 |
| SE | 71145 | ENSG00000089123 | TASP1 | chr20 | - | 13600542 | 13600617 | 13587249 | 13587370 | 13623445 | 13623514 | 4 | 3 | 1 | 2 | 5 | 12 | 27 |
| SE | 29231 | ENSG00000163110 | PDLIM5 | chr4 | + | 94584985 | 94585003 | 94575942 | 94576034 | 94585564 | 94585737 | 1 | 0 | 1 | 4 | 0 | 52 | 25 |
| SE | 86328 | ENSG00000135048 | CEMIP2 | chr9 | - | 71734805 | 71734994 | 71732350 | 71732520 | 71745017 | 71745579 | 260 | 149 | 163 | 163 | 155 | 0 | 0 |
| SE | 35662 | ENSG00000010404 | IDS | chrX | - | 149498106 | 149498307 | 149496345 | 149496516 | 149500948 | 149501037 | 987 | 563 | 735 | 807 | 1242 | 10 | 2 |
| SE | 16790 | ENSG00000241839 | PLEKHO2 | chr15 | + | 64859893 | 64859998 | 64854920 | 64855037 | 64864898 | 64868000 | 180 | 82 | 113 | 135 | 90 | 3 | 2 |
| SE | 14463 | ENSG00000213614 | HEXA | chr15 | - | 72346526 | 72346710 | 72343455 | 72346325 | 72347695 | 72347758 | 419 | 247 | 286 | 280 | 382 | 5 | 2 |
| SE | 87799 | ENSG00000111237 | VPS29 | chr12 | - | 110492995 | 110493231 | 110491082 | 110492122 | 110496011 | 110496203 | 1304 | 874 | 1002 | 1122 | 1103 | 5 | 5 |
| SE | 13612 | ENSG00000038358 | EDC4 | chr16 | + | 67875737 | 67875837 | 67873051 | 67873343 | 67875944 | 67876101 | 1 | 2 | 3 | 0 | 0 | 25 | 9 |
| SE | 179 | ENSG00000182871 | COL18A1 | chr21 | + | 45494544 | 45494571 | 45493500 | 45493575 | 45494861 | 45494915 | 38 | 31 | 124 | 60 | 31 | 1 | 0 |
| SE | 31909 | ENSG00000163932 | PRKCD | chr3 | + | 53165103 | 53165215 | 53161520 | 53161575 | 53178403 | 53178537 | 38 | 24 | 13 | 45 | 11 | 2 | 0 |
| SE | 27291 | ENSG00000064726 | BTBD1 | chr15 | - | 83028615 | 83028728 | 83020674 | 83020762 | 83030135 | 83030328 | 21 | 51 | 42 | 49 | 51 | 184 | 140 |
| SE | 11818 | ENSG00000188603 | CLN3 | chr16 | - | 28488590 | 28488662 | 28487455 | 28487541 | 28489289 | 28489386 | 36 | 15 | 30 | 29 | 24 | 0 | 0 |
| SE | 108035 | ENSG00000139613 | SMARCC2 | chr12 | - | 56170143 | 56170208 | 56169775 | 56169911 | 56171270 | 56171432 | 396 | 267 | 414 | 431 | 318 | 1 | 0 |
| SE | 93689 | ENSG00000198780 | FAM169A | chr5 | - | 74838964 | 74839050 | 74813839 | 74814019 | 74840073 | 74840173 | 66 | 26 | 17 | 24 | 32 | 8 | 0 |
| SE | 97226 | ENSG00000073921 | PICALM | chr11 | - | 85974707 | 85974812 | 85957174 | 85959060 | 85981128 | 85981228 | 365 | 258 | 220 | 320 | 524 | 11 | 4 |
| SE | 6711 | ENSG00000130813 | SHFL | chr19 | + | 10089110 | 10089378 | 10087250 | 10087300 | 10089656 | 10089695 | 22 | 10 | 20 | 22 | 20 | 72 | 36 |
| SE | 43478 | ENSG00000145293 | ENOPH1 | chr4 | + | 82447919 | 82448021 | 82430647 | 82430884 | 82454721 | 82454854 | 33 | 6 | 28 | 14 | 12 | 0 | 0 |
| SE | 54972 | ENSG00000135842 | NIBAN1 | chr1 | - | 184823637 | 184823742 | 184823166 | 184823329 | 184831846 | 184831962 | 193 | 204 | 171 | 264 | 258 | 1 | 2 |
| SE | 40392 | ENSG00000163719 | MTMR14 | chr3 | + | 9677983 | 9678058 | 9677316 | 9677387 | 9683177 | 9683217 | 187 | 81 | 137 | 139 | 253 | 18 | 15 |
| SE | 108055 | ENSG00000257727 | CNPY2 | chr12 | - | 56310309 | 56310595 | 56309841 | 56310029 | 56310957 | 56311054 | 102 | 45 | 82 | 81 | 73 | 3 | 4 |
| SE | 35757 | ENSG00000162736 | NCSTN | chr1 | + | 160355862 | 160355958 | 160354117 | 160354290 | 160356259 | 160356347 | 97 | 56 | 93 | 83 | 208 | 3 | 1 |
| SE | 69003 | ENSG00000107521 | HPS1 | chr10 | - | 98431130 | 98431291 | 98430570 | 98430670 | 98433939 | 98434091 | 28 | 13 | 35 | 20 | 23 | 18 | 8 |
| SE | 95841 | ENSG00000141551 | CSNK1D | chr17 | - | 82255428 | 82255577 | 82252433 | 82252604 | 82265685 | 82265796 | 112 | 37 | 54 | 84 | 69 | 0 | 0 |
| SE | 80842 | ENSG00000164885 | CDK5 | chr7 | - | 151055752 | 151055848 | 151055531 | 151055606 | 151056579 | 151056636 | 46 | 28 | 38 | 54 | 69 | 2 | 4 |
| SE | 10879 | ENSG00000198920 | KIAA0753 | chr17 | - | 6611918 | 6612148 | 6609993 | 6610160 | 6620787 | 6620998 | 24 | 28 | 28 | 33 | 40 | 0 | 5 |
| SE | 57635 | ENSG00000142405 | NLRP12 | chr19 | - | 53805593 | 53805783 | 53805279 | 53805450 | 53807494 | 53808705 | 2 | 0 | 1 | 1 | 0 | 24 | 13 |
| SE | 95692 | ENSG00000132589 | FLOT2 | chr17 | - | 28884224 | 28884315 | 28882978 | 28883231 | 28897525 | 28897652 | 143 | 48 | 99 | 108 | 75 | 0 | 3 |
| SE | 11220 | ENSG00000140829 | DHX38 | chr16 | + | 72098926 | 72099045 | 72098644 | 72098792 | 72099203 | 72099280 | 218 | 119 | 176 | 223 | 165 | 0 | 0 |
| SE | 83616 | ENSG00000130066 | SAT1 | chrX | + | 23784293 | 23784403 | 23783799 | 23783883 | 23785327 | 23785429 | 293 | 210 | 177 | 434 | 282 | 428 | 291 |
| SE | 8456 | ENSG00000141503 | MINK1 | chr17 | + | 4890221 | 4890308 | 4889646 | 4889763 | 4890516 | 4890735 | 3 | 3 | 1 | 1 | 0 | 34 | 5 |
| SE | 11852 | ENSG00000121274 | TENT4B | chr16 | + | 50222305 | 50222434 | 50217555 | 50217663 | 50223173 | 50223387 | 274 | 185 | 238 | 249 | 281 | 4 | 4 |
| SE | 78354 | ENSG00000072609 | CHFR | chr12 | - | 132862332 | 132862425 | 132861466 | 132861634 | 132869618 | 132869834 | 1 | 2 | 1 | 3 | 7 | 84 | 81 |
| SE | 76859 | ENSG00000112578 | BYSL | chr6 | + | 41930131 | 41930270 | 41927373 | 41927536 | 41930634 | 41930768 | 23 | 10 | 18 | 14 | 10 | 1 | 4 |
| SE | 15368 | ENSG00000140521 | POLG | chr15 | - | 89321961 | 89322015 | 89319291 | 89319350 | 89322741 | 89322902 | 27 | 14 | 19 | 28 | 25 | 0 | 0 |
| SE | 14780 | ENSG00000242028 | HYPK | chr15 | + | 43801131 | 43801187 | 43800494 | 43800784 | 43801517 | 43801569 | 71 | 60 | 74 | 76 | 52 | 0 | 3 |
| SE | 24819 | ENSG00000164576 | SAP30L | chr5 | + | 154452455 | 154452499 | 154451090 | 154451213 | 154453401 | 154453485 | 34 | 21 | 48 | 66 | 60 | 143 | 83 |
| SE | 25965 | ENSG00000005156 | LIG3 | chr17 | + | 34985941 | 34986131 | 34983451 | 34983552 | 34989465 | 34989663 | 83 | 72 | 86 | 74 | 84 | 0 | 0 |
| SE | 83962 | ENSG00000068400 | GRIPAP1 | chrX | - | 48981794 | 48981872 | 48981595 | 48981691 | 48982978 | 48983092 | 143 | 90 | 98 | 127 | 65 | 3 | 0 |
| SE | 107517 | ENSG00000205352 | PRR13 | chr12 | + | 53443390 | 53443773 | 53442694 | 53442733 | 53446014 | 53446094 | 515 | 267 | 331 | 449 | 296 | 1 | 0 |
| SE | 1132 | ENSG00000099992 | TBC1D10A | chr22 | - | 30304530 | 30304651 | 30299443 | 30299551 | 30326672 | 30326901 | 90 | 51 | 86 | 74 | 25 | 11 | 1 |

| | | | | | | | | | |
|---|---|---|---|---|---|---|---|---|---|
| SE | 81551 | ENSG00000106348 | IMPDH1 | chr7 | - | 128409288 | 128409352 | 128405766 | 128405865 |
| SE | 92851 | ENSG00000130958 | SLC35D2 | chr9 | - | 96360153 | 96360221 | 96324090 | 96324169 |
| SE | 89106 | ENSG00000064115 | TM7SF3 | chr12 | - | 26976259 | 26976357 | 26975495 | 26975658 |
| SE | 26526 | ENSG00000138303 | ASCC1 | chr10 | - | 72132233 | 72132341 | 72128081 | 72128167 |
| SE | 101769 | ENSG00000165406 | MARCHF8 | chr10 | - | 45464238 | 45464327 | 45461082 | 45461411 |
| SE | 88914 | ENSG00000158669 | GPAT4 | chr8 | + | 41612785 | 41612960 | 41612179 | 41612273 |
| SE | 41904 | ENSG00000175203 | DCTN2 | chr12 | - | 57538252 | 57538347 | 57535748 | 57535845 |
| SE | 46870 | ENSG00000162894 | FCMR | chr1 | - | 206913861 | 206914094 | 206912928 | 206913042 |
| SE | 46779 | ENSG00000163798 | SLC4A1AP | chr2 | + | 27688699 | 27688767 | 27687933 | 27688020 |
| SE | 98099 | ENSG00000171466 | ZNF562 | chr19 | - | 9656546 | 9656653 | 9641806 | 9653881 |
| SE | 49350 | ENSG00000120942 | UBIAD1 | chr1 | + | 11293466 | 11293626 | 11285643 | 11285732 |
| SE | 72724 | ENSG00000172765 | TMCC1 | chr3 | - | 129867009 | 129867098 | 129848342 | 129848412 |
| SE | 78889 | ENSG00000198874 | TYW1 | chr7 | + | 67055834 | 67055887 | 67049948 | 67050066 |
| SE | 9498 | ENSG00000171928 | TVP23B | chr17 | + | 18790895 | 18791040 | 18789352 | 18789435 |
| SE | 8014 | ENSG00000129197 | RPAIN | chr17 | + | 5425970 | 5428211 | 5422768 | 5422829 |
| SE | 8187 | ENSG00000108799 | EZH1 | chr17 | - | 42724304 | 42724424 | 42720272 | 42720449 |
| SE | 25525 | ENSG00000131732 | ZCCHC9 | chr5 | + | 81304740 | 81305141 | 81301586 | 81301698 |
| SE | 54326 | ENSG00000117139 | KDM5B | chr1 | - | 202766925 | 202767051 | 202764048 | 202764145 |
| SE | 98270 | ENSG00000144354 | CDCA7 | chr2 | + | 173363999 | 173364161 | 173363817 | 173363895 |
| SE | 61751 | ENSG00000153029 | MR1 | chr1 | + | 181036996 | 181037112 | 181034009 | 181034074 |
| SE | 45891 | ENSG00000145740 | SLC30A5 | chr5 | + | 69128003 | 69128132 | 69123198 | 69123425 |
| SE | 49903 | ENSG00000108061 | SHOC2 | chr10 | + | 110985627 | 110985765 | 110919603 | 110919657 |
| SE | 50026 | ENSG00000165359 | INTS6L | chrX | + | 135570435 | 135570546 | 135569336 | 135569431 |
| SE | 4723 | ENSG00000031823 | RANBP3 | chr19 | - | 5924826 | 5924905 | 5921200 | 5921321 |
| SE | 86265 | ENSG00000134516 | DOCK2 | chr5 | + | 169761518 | 169761625 | 169747395 | 169747504 |
| SE | 8020 | ENSG00000129197 | RPAIN | chr17 | + | 5428070 | 5428211 | 5425970 | 5426082 |
| SE | 17511 | ENSG00000172590 | MRPL52 | chr14 | + | 22830963 | 22831037 | 22830186 | 22830254 |
| SE | 15939 | ENSG00000168904 | LRRC28 | chr15 | + | 99334017 | 99334129 | 99287813 | 99287951 |
| SE | 89436 | ENSG00000121879 | PIK3CA | chr3 | + | 179220985 | 179221157 | 179219948 | 179220052 |
| SE | 81987 | ENSG00000110917 | MLEC | chr12 | + | 120694823 | 120695000 | 120693993 | 120694269 |
| SE | 48551 | ENSG00000126091 | ST3GAL3 | chr1 | + | 43824730 | 43824892 | 43814890 | 43814933 |
| SE | 1507 | ENSG00000054611 | TBC1D22A | chr22 | + | 47037070 | 47037198 | 46997633 | 46997709 |
| SE | 24148 | ENSG00000143774 | GUK1 | chr1 | + | 228148370 | 228148456 | 228147465 | 228147699 |
| SE | 50903 | ENSG00000156515 | HK1 | chr10 | + | 69398594 | 69398828 | 69394949 | 69395105 |
| SE | 97057 | ENSG00000124469 | CEACAM8 | chr19 | - | 42593540 | 42593900 | 42589456 | 42589735 |
| SE | 99042 | ENSG00000087053 | MTMR2 | chr11 | - | 95887611 | 95887684 | 95882373 | 95882496 |
| SE | 18072 | ENSG00000075413 | MARK3 | chr14 | + | 103428386 | 103428440 | 103405075 | 103405267 |
| SE | 52440 | ENSG00000117308 | GALE | chr1 | - | 23798886 | 23799012 | 23798116 | 23798730 |
| SE | 20604 | ENSG00000198945 | L3MBTL3 | chr6 | + | 130133451 | 130133621 | 130120878 | 130120958 |
| SE | 39781 | ENSG00000072274 | TFRC | chr3 | - | 196075158 | 196075360 | 196072002 | 196072152 |
| SE | 25693 | ENSG00000247774 | PCED1B-AS1 | chr12 | - | 47206510 | 47206585 | 47205897 | 47206145 |
| SE | 12806 | ENSG00000102882 | MAPK3 | chr16 | - | 30117136 | 30117285 | 30116893 | 30117003 |
| SE | 15366 | ENSG00000140521 | POLG | chr15 | - | 89321124 | 89321260 | 89320765 | 89321012 |
| SE | 1539 | ENSG00000100055 | CYTH4 | chr22 | + | 37304171 | 37304286 | 37303253 | 37303402 |
| SE | 51114 | ENSG00000132781 | MUTYH | chr1 | - | 45333412 | 45333594 | 45333284 | 45333324 |
| SE | 63582 | ENSG00000206562 | METTL6 | chr3 | - | 15414729 | 15414914 | 15414020 | 15414162 |
| SE | 89894 | ENSG00000147576 | ADHFE1 | chr8 | + | 66442797 | 66442844 | 66440161 | 66440199 |
| SE | 26403 | ENSG00000138744 | NAAA | chr4 | - | 75936108 | 75936235 | 75925734 | 75925811 |
| SE | 74347 | ENSG00000187118 | CMC1 | chr3 | + | 28294433 | 28294526 | 28292735 | 28292964 |
| SE | 25301 | ENSG00000134644 | PUM1 | chr1 | - | 30945345 | 30945483 | 30941997 | 30942123 |
| SE | 8060 | ENSG00000141337 | ARSG | chr17 | + | 68395072 | 68395193 | 68385063 | 68385172 |
| SE | 31203 | ENSG00000025293 | PHF20 | chr20 | + | 35782327 | 35782609 | 35772021 | 35772079 |
| SE | 10393 | ENSG00000159210 | SNF8 | chr17 | - | 48931642 | 48931717 | 48929315 | 48930612 |
| SE | 100764 | ENSG00000173914 | RBM4B | chr11 | - | 66666271 | 66666493 | 66665547 | 66665578 |
| SE | 17259 | ENSG00000176438 | SYNE3 | chr14 | - | 95457176 | 95457338 | 95455376 | 95455724 |
| SE | 55448 | ENSG00000116688 | MFN2 | chr1 | + | 12007052 | 12007249 | 12006537 | 12006693 |
| SE | 14013 | ENSG00000090905 | TNRC6A | chr16 | + | 24730252 | 24730300 | 24729650 | 24729846 |
| SE | 27148 | ENSG00000178562 | CD28 | chr2 | + | 203729647 | 203729772 | 203726632 | 203726989 |
| SE | 4740 | ENSG00000031823 | RANBP3 | chr19 | - | 5951392 | 5951596 | 5941798 | 5941835 |
| SE | 106450 | ENSG00000135127 | BICDL1 | chr12 | + | 120091246 | 120091401 | 120089950 | 120090071 |
| SE | 64716 | ENSG00000167895 | TMC8 | chr17 | + | 78137234 | 78137358 | 78134869 | 78135009 |
| SE | 20903 | ENSG00000152969 | JAKMIP1 | chr4 | - | 6105472 | 6105967 | 6085419 | 6085629 |
| SE | 15602 | ENSG00000169032 | MAP2K1 | chr15 | + | 66484989 | 66485191 | 66481754 | 66481879 |
| SE | 19987 | ENSG00000112584 | FAM120B | chr6 | + | 170388286 | 170388493 | 170358225 | 170358318 |
| SE | 26182 | ENSG00000048140 | TSPAN17 | chr5 | + | 176656699 | 176656816 | 176656077 | 176656125 |
| SE | 34152 | ENSG00000181458 | TMEM45A | chr3 | + | 100556759 | 100556972 | 100555208 | 100555401 |
| SE | 46820 | ENSG00000183291 | SELENOF | chr1 | - | 86868052 | 86868102 | 86862444 | 86863605 |
| SE | 45695 | ENSG00000049541 | RFC2 | chr7 | - | 74239937 | 74240095 | 74238922 | 74238988 |
| SE | 90978 | ENSG00000132669 | RIN2 | chr20 | + | 19970837 | 19970929 | 19960699 | 19960811 |
| SE | 6572 | ENSG00000074657 | ZNF532 | chr18 | + | 58931645 | 58931755 | 58920329 | 58920633 |
| SE | 18881 | ENSG00000170956 | CEACAM3 | chr19 | + | 41807110 | 41807617 | 41797588 | 41797948 |
| SE | 48674 | ENSG00000131374 | TBC1D5 | chr3 | - | 17646960 | 17647056 | 17623848 | 17623913 |
| SE | 452 | ENSG00000159131 | GART | chr21 | - | 33505973 | 33506104 | 33505560 | 33505702 |
| SE | 64624 | ENSG00000148737 | TCF7L2 | chr10 | + | 113158666 | 113158717 | 113158020 | 113158069 |
| SE | 64383 | ENSG00000264538 | SUZ12P1 | chr17 | + | 30768833 | 30768867 | 30766502 | 30766588 |
| SE | 107355 | ENSG00000110921 | MVK | chr12 | + | 109579801 | 109579946 | 109575997 | 109576145 |
| SE | 71888 | ENSG00000074054 | CLASP1 | chr2 | - | 121409023 | 121409047 | 121407470 | 121407715 |
| SE | 14341 | ENSG00000007545 | CRAMP1 | chr16 | + | 1667334 | 1667400 | 1666421 | 1666600 |
| SE | 2782 | ENSG00000011304 | PTBP1 | chr19 | + | 799412 | 799443 | 797074 | 797505 |
| SE | 4474 | ENSG00000127527 | EPS15L1 | chr19 | - | 16361778 | 16361984 | 16355238 | 16355851 |
| SE | 36151 | ENSG00000131263 | RLIM | chrX | - | 74612581 | 74612643 | 74595808 | 74596000 |
| SE | 34659 | ENSG00000166501 | PRKCB | chr16 | + | 24172269 | 24172361 | 24154683 | 24154857 |
| SE | 30716 | ENSG00000215158 | NA | chr5 | - | 34239374 | 34239448 | 34218294 | 34218450 |
| SE | 13589 | ENSG00000167261 | DPEP2 | chr16 | - | 67991837 | 67991979 | 67991114 | 67991184 |
| SE | 41483 | ENSG00000270629 | NBPF14 | chr1 | - | 148579073 | 148579237 | 148577982 | 148578034 |
| SE | 64696 | ENSG00000093144 | ECHDC1 | chr6 | - | 127329821 | 127329882 | 127327001 | 127327144 |
| SE | 92891 | ENSG00000136874 | STX17 | chr9 | + | 99913989 | 99914151 | 99906676 | 99906706 |
| SE | 70436 | ENSG00000144306 | SCRN3 | chr2 | + | 174398274 | 174398442 | 174397335 | 174397404 |
| SE | 68083 | ENSG00000135838 | NPL | chr1 | + | 182792231 | 182792286 | 182789292 | 182789805 |
| SE | 62640 | ENSG00000128815 | WDFY4 | chr10 | + | 48777418 | 48777495 | 48776749 | 48776984 |
| SE | 82164 | ENSG00000087077 | TRIP6 | chr7 | + | 100867910 | 100867988 | 100867386 | 100867606 |
| SE | 16218 | ENSG00000077238 | IL4R | chr16 | + | 27346466 | 27346618 | 27344868 | 27345020 |
| SE | 4670 | ENSG00000141968 | VAV1 | chr19 | + | 6821790 | 6821859 | 6821621 | 6821680 |
| SE | 85661 | ENSG00000161791 | FMNL3 | chr12 | - | 49646638 | 49646753 | 49644930 | 49645903 |
| SE | 29298 | ENSG00000254870 | ATP6V1G2-DDX39B | chr6 | - | 31539146 | 31539274 | 31536499 | 31536683 |
| SE | 15328 | ENSG00000114062 | UBE3A | chr15 | - | 25415709 | 25415842 | 25411907 | 25411971 |
| SE | 32288 | ENSG00000163964 | PIGX | chr3 | + | 196721176 | 196721356 | 196717808 | 196717954 |
| SE | 47012 | ENSG00000122417 | ODF2L | chr1 | - | 86385457 | 86385590 | 86384675 | 86384801 |
| SE | 86719 | ENSG00000198814 | GK | chrX | + | 30674301 | 30674466 | 30668011 | 30668118 |
| SE | 79129 | ENSG00000026950 | BTN3A1 | chr6 | + | 26409893 | 26409914 | 26409532 | 26409733 |
| SE | 1747 | ENSG00000100325 | ASCC2 | chr22 | - | 29832244 | 29832342 | 29825621 | 29825780 |
| SE | 78356 | ENSG00000072609 | CHFR | chr12 | - | 132869618 | 132869798 | 132861466 | 132861634 |
| SE | 27789 | ENSG00000177058 | SLC38A9 | chr5 | - | 55664692 | 55664863 | 55656714 | 55656774 |
| SE | 9878 | ENSG00000204650 | LINC02210 | chr17 | + | 45627375 | 45627682 | 45621907 | 45622041 |
| SE | 77257 | ENSG00000141664 | ZCCHC2 | chr18 | + | 62558686 | 62558770 | 62556202 | 62556297 |
| SE | 69713 | ENSG00000177674 | AGTRAP | chr1 | + | 11746126 | 11746223 | 11745800 | 11745837 |
| SE | 45794 | ENSG00000118217 | ATF6 | chr1 | + | 161778243 | 161778320 | 161766297 | 161766442 |
| SE | 35189 | ENSG00000101596 | SMCHD1 | chr18 | + | 2740702 | 2740821 | 2739431 | 2739520 |
| SE | 57889 | ENSG00000162604 | TM2D1 | chr1 | - | 61723712 | 61723772 | 61709328 | 61709437 |
| SE | 59639 | ENSG00000163466 | ARPC2 | chr2 | + | 218234351 | 218234397 | 218228737 | 218228850 |
| SE | 7539 | ENSG00000167851 | CD300A | chr17 | + | 74469945 | 74470242 | 74466861 | 74466993 |
| SE | 39528 | ENSG00000186517 | ARHGAP30 | chr1 | - | 161059613 | 161059716 | 161054622 | 161054705 |
| SE | 18078 | ENSG00000075413 | MARK3 | chr14 | + | 103438024 | 103438333 | 103428386 | 103428440 |
| SE | 69319 | ENSG00000112367 | FIG4 | chr6 | + | 109760249 | 109760383 | 109743109 | 109743772 |
| SE | 70128 | ENSG00000105953 | OGDH | chr7 | + | 44645326 | 44645518 | 44624316 | 44624565 |
| SE | 12371 | ENSG00000072864 | NDE1 | chr16 | + | 15696708 | 15696860 | 15694164 | 15694256 |
| SE | 103646 | ENSG00000120616 | EPC1 | chr10 | - | 32273013 | 32273082 | 32272025 | 32272167 |
| SE | 26086 | ENSG00000174695 | TMEM167A | chr5 | - | 83061876 | 83061911 | 83052845 | 83057154 |
| SE | 51840 | ENSG00000087206 | UIMC1 | chr5 | - | 176968554 | 176969291 | 176958092 | 176958154 |
| SE | 8517 | ENSG00000125447 | GGA3 | chr17 | - | 75248839 | 75249008 | 75246711 | 75246796 |
| SE | 77068 | ENSG00000104312 | RIPK2 | chr8 | + | 89762828 | 89762982 | 89757815 | 89758233 |
| SE | 25943 | ENSG00000119522 | DENND1A | chr9 | - | 123557569 | 123557695 | 123457792 | 123457897 |
| SE | 83089 | ENSG00000184863 | RBM33 | chr7 | + | 155763811 | 155764018 | 155744960 | 155745602 |
| SE | 83380 | ENSG00000274523 | RCC1L | chr7 | - | 75066663 | 75066792 | 75064581 | 75064648 |
| SE | 51111 | ENSG00000132781 | MUTYH | chr1 | - | 45333412 | 45333497 | 45332676 | 45333324 |
| SE | 9361 | ENSG00000141030 | COPS3 | chr17 | - | 17276034 | 17276164 | 17270757 | 17271008 |
| SE | 13383 | ENSG00000074755 | ZZEF1 | chr17 | - | 4056215 | 4056345 | 4054056 | 4054195 |
| SE | 82256 | ENSG00000158457 | TSPAN33 | chr7 | + | 129167398 | 129167560 | 129166439 | 129166906 |
| SE | 63521 | ENSG00000115966 | ATF2 | chr2 | - | 175092542 | 175092616 | 175080659 | 175080765 |
| SE | 22591 | ENSG00000127603 | MACF1 | chr1 | + | 39315512 | 39315691 | 39310830 | 39311000 |
| SE | 7563 | ENSG00000161955 | TNFSF13 | chr17 | + | 7559845 | 7559893 | 7559623 | 7559702 |
| SE | 26735 | ENSG00000125089 | SH3TC1 | chr4 | + | 8215171 | 8215242 | 8214474 | 8214580 |
| SE | 72366 | ENSG00000169045 | HNRNPH1 | chr5 | - | 179615545 | 179615595 | 179614177 | 179614959 |
| SE | 68604 | ENSG00000237765 | FAM200B | chr4 | + | 15684849 | 15684959 | 15681795 | 15681968 |
| SE | 18350 | ENSG00000257621 | PSMA3-AS1 | chr14 | - | 58293065 | 58293138 | 58291634 | 58291707 |
| SE | 57930 | ENSG00000168439 | STIP1 | chr11 | + | 64194260 | 64194330 | 64193077 | 64193287 |
| SE | 44419 | ENSG00000056050 | HPF1 | chr4 | - | 169750535 | 169750725 | 169748743 | 169748842 |
| SE | 61487 | ENSG00000124588 | NQO2 | chr6 | + | 2989527 | 2989710 | 2988645 | 2988709 |
| SE | 82070 | ENSG00000086730 | LAT2 | chr7 | + | 74221636 | 74221692 | 74220703 | 74220734 |
| SE | 67595 | ENSG00000008083 | JARID2 | chr6 | + | 15452005 | 15452175 | 15374116 | 15374252 |
| SE | 90651 | ENSG00000100023 | PPIL2 | chr22 | + | 21670298 | 21670360 | 21669912 | 21669962 |
| SE | 105368 | ENSG00000110719 | TCIRG1 | chr11 | + | 68044131 | 68044344 | 68043788 | 68043907 |
| SE | 85131 | ENSG00000285171 | NA | chrX | - | 71107852 | 71107921 | 71106531 | 71106590 |
| SE | 86785 | ENSG00000102393 | GLA | chrX | - | 101400665 | 101400757 | 101398369 | 101398946 |
| SE | 94665 | ENSG00000165209 | STRBP | chr9 | - | 123160366 | 123160462 | 123159095 | 123159207 |
| SE | 11287 | ENSG00000125124 | BBS2 | chr16 | - | 56511158 | 56511284 | 56510858 | 56510921 |
| SE | 64323 | ENSG00000138363 | ATIC | chr2 | + | 215325179 | 215325329 | 215319664 | 215319731 |
| SE | 98077 | ENSG00000126705 | AHDC1 | chr1 | - | 27558304 | 27558530 | 27553073 | 27553185 |
| SE | 27455 | ENSG00000126522 | ASL | chr7 | + | 66087333 | 66087386 | 66086584 | 66086662 |
| SE | 75580 | ENSG00000204568 | MRPS18B | chr6 | + | 30622831 | 30622898 | 30619492 | 30619989 |
| SE | 34534 | ENSG00000186298 | PPP1CC | chr12 | - | 110734717 | 110734986 | 110731815 | 110731901 |
| SE | 77362 | ENSG00000171316 | CHD7 | chr8 | + | 60860903 | 60861125 | 60856444 | 60856888 |
| SE | 29245 | ENSG00000166004 | CEP295 | chr11 | + | 93702775 | 93702919 | 93702459 | 93702637 |
| SE | 35174 | ENSG00000101596 | SMCHD1 | chr18 | + | 2694526 | 2694693 | 2688627 | 2688747 |
| SE | 72313 | ENSG00000126254 | RBM42 | chr19 | + | 35632935 | 35633001 | 35631330 | 35631405 |
| SE | 23824 | ENSG00000160293 | VAV2 | chr9 | - | 133770377 | 133770501 | 133768441 | 133768596 |
| SE | 18812 | ENSG00000119689 | DLST | chr14 | + | 74894311 | 74894409 | 74893347 | 74893424 |
| SE | 37621 | ENSG00000082074 | FYB1 | chr5 | - | 39212677 | 39213017 | 39208863 | 39208964 |
| SE | 25308 | ENSG00000134644 | PUM1 | chr1 | - | 30980061 | 30980163 | 30974648 | 30974802 |
| SE | 108656 | ENSG00000136147 | PHF11 | chr13 | + | 49523174 | 49523241 | 49521928 | 49522107 |
| SE | 41555 | ENSG00000015171 | ZMYND11 | chr10 | + | 242020 | 242139 | 240892 | 240970 |
| SE | 20172 | ENSG00000146457 | WTAP | chr6 | + | 159742087 | 159742146 | 159738989 | 159739045 |
| SE | 88786 | ENSG00000140632 | GLYR1 | chr16 | - | 4821379 | 4821664 | 4817663 | 4817697 |
| SE | 52856 | ENSG00000160691 | SHC1 | chr1 | - | 154973292 | 154973508 | 154969377 | 154969448 |
| SE | 89903 | ENSG00000104375 | STK3 | chr8 | - | 98594860 | 98595045 | 98579758 | 98579789 |
| SE | 56833 | ENSG00000143398 | PIP5K1A | chr1 | + | 151238181 | 151238265 | 151236557 | 151236763 |
| SE | 100551 | ENSG00000054654 | SYNE2 | chr14 | + | 64202800 | 64202963 | 64190070 | 64190237 |
| SE | 102891 | ENSG00000107679 | PLEKHA1 | chr10 | + | 122406575 | 122406673 | 122397917 | 122397974 |
| SE | 44543 | ENSG00000166526 | ZNF3 | chr7 | - | 100077302 | 100077433 | 100075229 | 100075630 |
| SE | 107467 | ENSG00000100599 | RIN3 | chr14 | + | 92676474 | 92676606 | 92659160 | 92659469 |
| SE | 21 | ENSG00000155313 | USP25 | chr21 | + | 15811136 | 15811210 | 15805120 | 15805258 |
| SE | 50705 | ENSG00000113318 | MSH3 | chr5 | + | 80744505 | 80744615 | 80728850 | 80728965 |
| SE | 61164 | ENSG00000028528 | SNX1 | chr15 | + | 64123502 | 64123546 | 64118787 | 64118854 |
| SE | 57989 | ENSG00000175634 | RPS6KB2 | chr11 | + | 67428912 | 67429022 | 67428493 | 67428623 |
| SE | 4988 | ENSG00000000419 | DPM1 | chr20 | - | 50941104 | 50941209 | 50940864 | 50940933 |
| SE | 53254 | ENSG00000138688 | KIAA1109 | chr4 | + | 122275971 | 122276022 | 122274342 | 122274436 |
| SE | 73941 | ENSG00000144134 | RABL2A | chr2 | + | 113635050 | 113635130 | 113634152 | 113634232 |
| SE | 82950 | ENSG00000075142 | SRI | chr7 | - | 88214794 | 88214936 | 88210881 | 88210925 |
| SE | 59680 | ENSG00000152683 | SLC30A6 | chr2 | + | 32175318 | 32175361 | 32174062 | 32174147 |
| SE | 53812 | ENSG00000173575 | CHD2 | chr15 | + | 92944414 | 92944515 | 92942842 | 92943068 |
| SE | 69832 | ENSG00000163803 | PLB1 | chr2 | + | 28620599 | 28620643 | 28620264 | 28620332 |
| SE | 71692 | ENSG00000135976 | ANKRD36 | chr2 | + | 97142762 | 97142835 | 97142639 | 97142668 |
| SE | 11469 | ENSG00000187555 | USP7 | chr16 | - | 8916501 | 8916556 | 8915444 | 8915525 |
| SE | 47978 | ENSG00000154710 | RABGEF1 | chr7 | + | 66739995 | 66740792 | 66712166 | 66712224 |
| SE | 2541 | ENSG00000104814 | MAP4K1 | chr19 | - | 38614244 | 38614292 | 38614042 | 38614085 |
| SE | 52788 | ENSG00000087191 | PSMC5 | chr17 | + | 63827673 | 63827923 | 63827459 | 63827514 |
| SE | 31421 | ENSG00000155508 | CNOT8 | chr5 | + | 154863206 | 154863395 | 154858626 | 154858768 |
| SE | 49584 | ENSG00000076685 | NT5C2 | chr10 | - | 103090610 | 103090787 | 103087184 | 103089908 |
| SE | 95801 | ENSG00000167721 | TSR1 | chr17 | - | 2329342 | 2329475 | 2325303 | 2325420 |
| SE | 73111 | ENSG00000057252 | SOAT1 | chr1 | + | 179344946 | 179345076 | 179343589 | 179343635 |
| SE | 26073 | ENSG00000038427 | VCAN | chr5 | + | 83493803 | 83493931 | 83493545 | 83493720 |
| SE | 41697 | ENSG00000247828 | TMEM161B-AS1 | chr5 | + | 88283009 | 88283086 | 88282714 | 88282866 |
| SE | 70131 | ENSG00000105953 | OGDH | chr7 | + | 44647656 | 44647759 | 44624316 | 44624565 |
| SE | 83256 | ENSG00000137501 | SYTL2 | chr11 | - | 85727895 | 85727967 | 85720857 | 85720959 |
| SE | 12041 | ENSG00000213380 | COG8 | chr16 | - | 69330812 | 69331095 | 69326427 | 69329179 |
| SE | 45623 | ENSG00000132781 | MUTYH | chr1 | - | 45333412 | 45333603 | 45333284 | 45333324 |
| SE | 96560 | ENSG00000214413 | BBIP1 | chr10 | - | 110907679 | 110907836 | 110901537 | 110901612 |
| SE | 35193 | ENSG00000101596 | SMCHD1 | chr18 | + | 2760651 | 2760739 | 2752487 | 2752552 |
| SE | 9026 | ENSG00000182446 | NPLOC4 | chr17 | - | 81610209 | 81610258 | 81608727 | 81608822 |
| SE | 59660 | ENSG00000167671 | UBXN6 | chr19 | - | 4446048 | 4446197 | 4444998 | 4445623 |
| SE | 79652 | ENSG00000167081 | PBX3 | chr9 | + | 125915685 | 125915927 | 125748181 | 125748623 |
| SE | 32660 | ENSG00000150995 | ITPR1 | chr3 | + | 4702829 | 4702950 | 4699812 | 4699941 |
| SE | 78865 | ENSG00000158321 | AUTS2 | chr7 | + | 70775356 | 70775386 | 70771548 | 70771644 |
| SE | 91125 | ENSG00000180900 | SCRIB | chr8 | - | 143791665 | 143791740 | 143791388 | 143791440 |
| SE | 77106 | ENSG00000122512 | PMS2 | chr7 | - | 5999107 | 5999226 | 5997325 | 5997423 |
| SE | 76432 | ENSG00000112739 | PRPF4B | chr6 | + | 4062556 | 4062760 | 4062103 | 4062299 |
| SE | 75066 | ENSG00000198223 | CSF2RA | chrX | + | 1300490 | 1300626 | 1295426 | 1295456 |
| SE | 104043 | ENSG00000065809 | FAM107B | chr10 | - | 14553321 | 14553387 | 14530331 | 14530515 |
| SE | 6487 | ENSG00000101605 | MYOM1 | chr18 | - | 3085044 | 3085132 | 3083988 | 3084027 |
| SE | 68629 | ENSG00000223960 | CHROMR | chr2 | + | 178426365 | 178426680 | 178413676 | 178414113 |
| SE | 2175 | ENSG00000079805 | DNM2 | chr19 | + | 10796060 | 10796199 | 10795371 | 10795439 |
| SE | 25011 | ENSG00000150593 | PDCD4 | chr10 | + | 110876669 | 110876753 | 110875965 | 110876070 |
| SE | 58141 | ENSG00000117523 | PRRC2C | chr1 | + | 171589368 | 171589605 | 171588378 | 171588505 |
| SE | 79806 | ENSG00000136286 | MYO1G | chr7 | - | 44964944 | 44965089 | 44964414 | 44964519 |
| SE | 105065 | ENSG00000119801 | YPEL5 | chr2 | + | 30155849 | 30155910 | 30148244 | 30148541 |
| SE | 33425 | ENSG00000135870 | RC3H1 | chr1 | - | 173943441 | 173943615 | 173941264 | 173941380 |
| SE | 67703 | ENSG00000145332 | KLHL8 | chr4 | - | 87170446 | 87170615 | 87170078 | 87170238 |
| SE | 54816 | ENSG00000204138 | PHACTR4 | chr1 | + | 28432930 | 28433046 | 28407409 | 28407463 |
| SE | 18402 | ENSG00000260669 | NA | chr14 | - | 24210653 | 24210768 | 24210347 | 24210483 |
| SE | 108714 | ENSG00000198246 | SLC29A3 | chr10 | + | 71351561 | 71351788 | 71322755 | 71323054 |
| SE | 85491 | ENSG00000167548 | KMT2D | chr12 | - | 49034581 | 49034666 | 49034409 | 49034476 |
| SE | 105865 | ENSG00000090006 | LTBP4 | chr19 | + | 40622400 | 40622667 | 40619346 | 40619493 |
| SE | 17412 | ENSG00000126822 | PLEKHG3 | chr14 | + | 64730242 | 64730312 | 64727592 | 64727982 |
| SE | 24523 | ENSG00000186532 | SMYD4 | chr17 | - | 1814565 | 1814795 | 1812061 | 1812115 |
| SE | 94499 | ENSG00000109689 | STIM2 | chr4 | + | 27019399 | 27021068 | 27017710 | 27017984 |
| SE | 56818 | ENSG00000119950 | MXI1 | chr10 | + | 110244827 | 110244873 | 110228188 | 110228321 |
| SE | 88716 | ENSG00000104738 | MCM4 | chr8 | + | 47971340 | 47971468 | 47970510 | 47970876 |
| SE | 35096 | ENSG00000284952 | NA | chr3 | - | 155842431 | 155842997 | 155833856 | 155834041 |
| SE | 88467 | ENSG00000152601 | MBNL1 | chr3 | + | 152335110 | 152335230 | 152300197 | 152300367 |
| SE | 41972 | ENSG00000120008 | WDR11 | chr10 | + | 120890715 | 120890887 | 120889894 | 120890009 |
| SE | 87779 | ENSG00000165675 | ENOX2 | chrX | - | 130783546 | 130783690 | 130703119 | 130703254 |
| SE | 58851 | ENSG00000182552 | RWDD4 | chr4 | - | 183651217 | 183651323 | 183650983 | 183651131 |
| SE | 22501 | ENSG00000119812 | FAM98A | chr2 | - | 33588334 | 33588519 | 33587239 | 33587320 |

| | | | | | | | | | | | | | | | | | | | | | | | | | | | | | | | | | | | | | | | | | | | | | | |
|---|---|---|---|---|---|---|---|---|---|---|---|---|---|---|---|---|---|---|---|---|---|---|---|---|---|---|---|---|---|---|---|---|---|---|---|---|---|---|---|---|---|---|---|---|---|---|
| SE | 42368 | ENSG00000048740 | CELF2 | chr10 | + | 11270665 | 11270824 | 11257737 | 11257872 | 11275056 | 11275120 | 330 | 176 | 247 | 250 | 329 | 1 | 0 | 0 | 8 | 0 | 145 | 336 | 328 | 114 | 126 | 26 | 0 | 1 | 0 | 0 | 298 | 149 | 0.001075105 | 0.016418262 | 0.994 | 1 | 1 | 0.94 | 1 | 0.736 | 1 | 0.994 | 1 | 1 | 0.041 |
| SE | 58175 | ENSG00000172965 | MIR4435-2HG | chr2 | - | 111208755 | 111208977 | 111207775 | 111208661 | 111233627 | 111233739 | 8 | 4 | 5 | 13 | 17 | 0 | 1 | 1 | 7 | 2 | 26 | 15 | 7 | 15 | 4 | 5 | 0 | 0 | 0 | 0 | 298 | 149 | 0.001076227 | 0.016431631 | 1 | 0.667 | 0.714 | 0.481 | 0.81 | 0.722 | 1 | 1 | 1 | 1 | -0.21 |
| SE | 59368 | ENSG00000116127 | ALMS1 | chr2 | + | 73426452 | 73426553 | 73424429 | 73424902 | 73432197 | 73432291 | 117 | 101 | 98 | 89 | 188 | 4 | 1 | 6 | 6 | 0 | 50 | 56 | 72 | 31 | 28 | 0 | 0 | 1 | 0 | 0 | 250 | 149 | 0.0010764 | 0.016431631 | 0.946 | 0.984 | 0.907 | 0.898 | 1 | 1 | 1 | 0.977 | 1 | 1 | -0.048 |
| SE | 93928 | ENSG00000144524 | COPS7B | chr2 | + | 231790921 | 231791173 | 231788554 | 231788732 | 231791732 | 231791762 | 1 | 0 | 13 | 3 | 4 | 82 | 33 | 61 | 65 | 52 | 10 | 11 | 4 | 4 | 21 | 99 | 79 | 23 | 51 | 60 | 298 | 149 | 0.00107727 | 0.0164417 | 0.006 | 0 | 0.096 | 0.023 | 0.037 | 0.048 | 0.065 | 0.08 | 0.038 | 0.149 | -0.044 |
| SE | 39791 | ENSG00000106459 | NRF1 | chr7 | + | 129709074 | 129709233 | 129690405 | 129690546 | 129711474 | 129711576 | 97 | 31 | 78 | 88 | 76 | 0 | 0 | 1 | 0 | 0 | 173 | 117 | 91 | 155 | 79 | 0 | 12 | 4 | 0 | 0 | 298 | 149 | 0.001083033 | 0.016526431 | 1 | 1 | 0.975 | 1 | 1 | 1 | 0.83 | 0.919 | 1 | 1 | 0.045 |
| SE | 94767 | ENSG00000107077 | KDM4C | chr9 | + | 7105401 | 7105500 | 7103684 | 7103870 | 7128065 | 7128236 | 23 | 14 | 14 | 22 | 16 | 107 | 75 | 116 | 108 | 117 | 7 | 22 | 2 | 8 | 1 | 105 | 119 | 63 | 86 | 62 | 248 | 149 | 0.001083336 | 0.016527836 | 0.114 | 0.101 | 0.068 | 0.109 | 0.076 | 0.039 | 0.1 | 0.019 | 0.053 | 0.01 | 0.049 |
| SE | 104448 | ENSG00000149100 | EIF3M | chr11 | + | 32602078 | 32602114 | 32601761 | 32601822 | 32602278 | 32602377 | 9 | 5 | 4 | 15 | 15 | 937 | 490 | 746 | 651 | 1454 | 10 | 13 | 20 | 12 | 2 | 497 | 533 | 469 | 396 | 417 | 185 | 149 | 0.00108457 | 0.01654342 | 0.008 | 0.008 | 0.004 | 0.018 | 0.008 | 0.016 | 0.019 | 0.033 | 0.024 | 0.004 | -0.01 |
| SE | 6645 | ENSG00000100342 | APOL1 | chr22 | + | 36253837 | 36253991 | 36253077 | 36253219 | 36254897 | 36254999 | 12 | 1 | 1 | 3 | 0 | 4 | 3 | 1 | 3 | 1 | 10 | 5 | 7 | 4 | 13 | 0 | 0 | 1 | 0 | 10 | 298 | 149 | 0.001084925 | 0.016545636 | 0.6 | 0.143 | 0.333 | 0.333 | 0 | 1 | 1 | 0.778 | 1 | 0.394 | -0.553 |
| SE | 12031 | ENSG00000176715 | ACSF3 | chr16 | + | 89114338 | 89114487 | 89112091 | 89112246 | 89120800 | 89120913 | 185 | 107 | 172 | 150 | 139 | 16 | 9 | 24 | 30 | 22 | 325 | 253 | 135 | 228 | 277 | 75 | 44 | 49 | 32 | 99 | 298 | 149 | 0.001088398 | 0.016592669 | 0.853 | 0.856 | 0.782 | 0.714 | 0.76 | 0.684 | 0.742 | 0.579 | 0.781 | 0.583 | 0.119 |
| SE | 19921 | ENSG00000100852 | ARHGAP5 | chr14 | + | 32117139 | 32117287 | 32090501 | 32094386 | 32146262 | 32146340 | 153 | 146 | 169 | 180 | 309 | 7 | 4 | 0 | 6 | 3 | 19 | 52 | 23 | 30 | 46 | 0 | 0 | 0 | 0 | 0 | 297 | 149 | 0.001088433 | 0.016592669 | 0.916 | 0.948 | 1 | 0.938 | 0.981 | 1 | 1 | 1 | 1 | 1 | -0.043 |
| SE | 25009 | ENSG00000150593 | PDCD4 | chr10 | + | 110875965 | 110876070 | 110871794 | 110872018 | 110881274 | 110881535 | 52 | 30 | 61 | 58 | 45 | 0 | 0 | 0 | 0 | 0 | 30 | 26 | 24 | 12 | 12 | 0 | 0 | 6 | 0 | 4 | 254 | 149 | 0.001088817 | 0.016595293 | 1 | 1 | 1 | 1 | 1 | 1 | 1 | 0.701 | 1 | 0.638 | 0.132 |
| SE | 29526 | ENSG00000119242 | CCDC92 | chr12 | - | 123956283 | 123956444 | 123943346 | 123943493 | 123972528 | 123972616 | 2 | 3 | 0 | 1 | 5 | 1 | 3 | 3 | 3 | 2 | 0 | 0 | 0 | 0 | 7 | 4 | 1 | 4 | 2 | 58 | 298 | 149 | 0.001092835 | 0.016653297 | 0.5 | 0.333 | 0 | 0.143 | 0.556 | 0 | 0 | 0 | 0 | 0.057 | 0.295 |
| SE | 1625 | ENSG00000244625 | MIATNB | chr22 | + | 26717252 | 26717459 | 26693680 | 26693836 | 26726643 | 26726720 | 27 | 53 | 20 | 51 | 47 | 2 | 1 | 0 | 2 | 0 | 41 | 80 | 54 | 98 | 22 | 0 | 7 | 1 | 2 | 18 | 298 | 149 | 0.001093099 | 0.01665407 | 0.871 | 0.964 | 1 | 0.927 | 1 | 1 | 0.851 | 0.964 | 0.961 | 0.379 | 0.121 |
| SE | 36532 | ENSG00000167566 | NCKAP5L | chr12 | - | 49798349 | 49798463 | 49794764 | 49797394 | 49801847 | 49801967 | 34 | 32 | 31 | 46 | 43 | 1 | 1 | 0 | 0 | 0 | 33 | 58 | 15 | 10 | 25 | 1 | 5 | 0 | 5 | 0 | 263 | 149 | 0.001094144 | 0.016666757 | 0.951 | 0.948 | 1 | 1 | 1 | 0.949 | 0.868 | 1 | 0.531 | 1 | 0.11 |
| SE | 1388 | ENSG00000100150 | DEPDC5 | chr22 | + | 31895358 | 31895954 | 31893581 | 31893751 | 31897481 | 31897653 | 3 | 2 | 3 | 5 | 2 | 10 | 0 | 10 | 9 | 4 | 13 | 15 | 21 | 6 | 19 | 4 | 5 | 2 | 6 | 2 | 298 | 149 | 0.001094815 | 0.01667372 | 0.13 | 1 | 0.13 | 0.217 | 0.2 | 0.619 | 0.6 | 0.84 | 0.333 | 0.826 | -0.308 |
| SE | 22893 | ENSG00000140853 | NLRC5 | chr16 | + | 57023784 | 57023853 | 57020700 | 57021007 | 57025367 | 57027018 | 57 | 27 | 55 | 61 | 42 | 0 | 0 | 0 | 0 | 0 | 66 | 68 | 25 | 20 | 123 | 0 | 0 | 0 | 0 | 14 | 218 | 149 | 0.00109623 | 0.016687851 | 1 | 1 | 1 | 1 | 1 | 1 | 1 | 1 | 1 | 0.857 | 0.029 |
| SE | 71401 | ENSG00000122203 | KIAA1191 | chr5 | - | 176359810 | 176359918 | 176359461 | 176359567 | 176361601 | 176361735 | 54 | 40 | 61 | 84 | 33 | 2 | 0 | 2 | 2 | 0 | 24 | 76 | 37 | 68 | 163 | 0 | 9 | 1 | 2 | 10 | 257 | 149 | 0.001096303 | 0.016687851 | 0.94 | 1 | 0.946 | 0.961 | 1 | 1 | 0.83 | 0.955 | 0.952 | 0.904 | 0.041 |
| SE | 70757 | ENSG00000040933 | INPP4A | chr2 | + | 98546585 | 98546694 | 98543876 | 98544007 | 98552785 | 98552969 | 178 | 90 | 146 | 192 | 163 | 6 | 4 | 18 | 14 | 9 | 98 | 128 | 126 | 114 | 103 | 0 | 10 | 3 | 4 | 0 | 258 | 149 | 0.001096382 | 0.016687851 | 0.945 | 0.929 | 0.824 | 0.888 | 0.913 | 1 | 0.881 | 0.96 | 0.943 | 1 | -0.057 |
| SE | 4912 | ENSG00000258366 | RTEL1 | chr20 | + | 63659232 | 63659504 | 63658359 | 63658459 | 63661297 | 63661462 | 24 | 10 | 17 | 30 | 34 | 0 | 0 | 0 | 0 | 0 | 10 | 31 | 26 | 34 | 10 | 0 | 6 | 0 | 0 | 4 | 298 | 149 | 0.001097065 | 0.016694025 | 1 | 1 | 1 | 1 | 1 | 1 | 0.721 | 1 | 1 | 0.556 | 0.145 |
| SE | 50091 | ENSG00000167528 | ZNF641 | chr12 | - | 48350045 | 48350218 | 48347906 | 48348115 | 48350785 | 48350794 | 0 | 5 | 1 | 2 | 3 | 11 | 7 | 10 | 16 | 5 | 0 | 0 | 0 | 1 | 0 | 19 | 11 | 12 | 14 | 12 | 298 | 149 | 0.001097214 | 0.016694025 | 0 | 0.263 | 0.048 | 0.059 | 0.231 | 0 | 0 | 0 | 0.034 | 0 | 0.113 |
| SE | 18543 | ENSG00000100722 | ZC3H14 | chr14 | + | 88596733 | 88596808 | 88577984 | 88578140 | 88601923 | 88602083 | 57 | 42 | 78 | 68 | 168 | 3 | 3 | 6 | 5 | 5 | 67 | 53 | 37 | 19 | 34 | 0 | 2 | 3 | 0 | 0 | 224 | 149 | 0.001098365 | 0.016707416 | 0.927 | 0.903 | 0.896 | 0.9 | 0.957 | 1 | 0.946 | 0.891 | 1 | 1 | -0.051 |
| SE | 92396 | ENSG00000080503 | SMARCA2 | chr9 | + | 2191265 | 2191408 | 2186095 | 2186228 | 2192703 | 2192751 | 393 | 278 | 314 | 411 | 269 | 24 | 10 | 11 | 13 | 12 | 234 | 233 | 198 | 203 | 38 | 22 | 13 | 10 | 21 | 30 | 292 | 149 | 0.001098726 | 0.016707416 | 0.893 | 0.934 | 0.936 | 0.942 | 0.92 | 0.844 | 0.901 | 0.91 | 0.831 | 0.393 | 0.149 |
| SE | 67857 | ENSG00000018280 | SLC11A1 | chr2 | + | 218384318 | 218384365 | 218382959 | 218383102 | 218385146 | 218385266 | 58 | 20 | 20 | 29 | 30 | 0 | 0 | 0 | 1 | 0 | 46 | 82 | 57 | 58 | 128 | 12 | 0 | 0 | 4 | 0 | 196 | 149 | 0.001098734 | 0.016707416 | 1 | 1 | 1 | 0.957 | 1 | 0.745 | 1 | 1 | 0.917 | 1 | 0.059 |
| SE | 75092 | ENSG00000188554 | NBR1 | chr17 | + | 43202654 | 43202712 | 43201685 | 43201780 | 43203680 | 43203786 | 185 | 96 | 150 | 186 | 135 | 2 | 4 | 2 | 7 | 0 | 63 | 91 | 111 | 88 | 59 | 0 | 11 | 6 | 8 | 0 | 207 | 149 | 0.001099609 | 0.016717473 | 0.985 | 0.945 | 0.982 | 0.95 | 1 | 1 | 0.856 | 0.93 | 0.888 | 1 | 0.038 |
| SE | 19738 | ENSG00000100714 | MTHFD1 | chr14 | + | 64417887 | 64418024 | 64415637 | 64415739 | 64419813 | 64419925 | 71 | 55 | 103 | 111 | 81 | 0 | 0 | 0 | 3 | 0 | 211 | 102 | 123 | 192 | 211 | 5 | 10 | 3 | 2 | 0 | 286 | 149 | 0.001100736 | 0.01672591 | 1 | 1 | 1 | 0.951 | 1 | 0.956 | 0.842 | 0.955 | 0.98 | 1 | 0.044 |
| SE | 13344 | ENSG00000103091 | WDR59 | chr16 | - | 74909821 | 74909917 | 74909500 | 74909657 | 74912197 | 74912235 | 33 | 17 | 39 | 20 | 56 | 0 | 0 | 1 | 0 | 3 | 16 | 17 | 9 | 13 | 11 | 0 | 2 | 5 | 3 | 0 | 245 | 149 | 0.00110091 | 0.01672591 | 1 | 1 | 0.96 | 1 | 0.919 | 1 | 0.838 | 0.523 | 0.725 | 1 | 0.159 |
| SE | 42261 | ENSG00000116001 | TIA1 | chr2 | - | 70228343 | 70228462 | 70227734 | 70227791 | 70229058 | 70229091 | 24 | 15 | 29 | 21 | 46 | 6 | 6 | 5 | 2 | 16 | 9 | 16 | 24 | 20 | 14 | 3 | 1 | 0 | 3 | 0 | 268 | 149 | 0.001100992 | 0.01672591 | 0.69 | 0.582 | 0.763 | 0.854 | 0.615 | 0.625 | 0.899 | 1 | 0.788 | 1 | -0.162 |
| SE | 5107 | ENSG00000124198 | ARFGEF2 | chr20 | + | 48974765 | 48974874 | 48973144 | 48973284 | 48976015 | 48976199 | 121 | 94 | 123 | 121 | 140 | 13 | 12 | 14 | 11 | 23 | 87 | 91 | 74 | 58 | 55 | 5 | 28 | 32 | 17 | 19 | 258 | 149 | 0.001101018 | 0.01672591 | 0.843 | 0.819 | 0.835 | 0.864 | 0.779 | 0.909 | 0.652 | 0.572 | 0.663 | 0.626 | 0.144 |
| SE | 103541 | ENSG00000107862 | GBF1 | chr10 | + | 102368738 | 102368832 | 102367477 | 102367560 | 102369210 | 102369387 | 28 | 10 | 6 | 21 | 10 | 0 | 4 | 0 | 0 | 2 | 25 | 15 | 15 | 13 | 15 | 15 | 2 | 0 | 8 | 4 | 243 | 149 | 0.001101358 | 0.016727822 | 1 | 0.605 | 1 | 1 | 0.754 | 0.505 | 0.821 | 1 | 0.499 | 0.697 | 0.167 |
| SE | 106705 | ENSG00000141376 | BCAS3 | chr17 | + | 61353587 | 61354502 | 61084504 | 61084564 | 61368326 | 61368494 | 2 | 0 | 0 | 3 | 0 | 53 | 40 | 45 | 48 | 55 | 3 | 7 | 6 | 4 | 0 | 62 | 53 | 27 | 36 | 60 | 298 | 149 | 0.00110195 | 0.016733572 | 0.019 | 0 | 0 | 0.03 | 0 | 0.024 | 0.062 | 0.1 | 0.053 | 0 | -0.038 |
| SE | 28595 | ENSG00000113369 | ARRDC3 | chr5 | - | 91376620 | 91376768 | 91375510 | 91375613 | 91378693 | 91378775 | 316 | 215 | 392 | 330 | 772 | 56 | 29 | 51 | 43 | 94 | 105 | 269 | 299 | 165 | 43 | 0 | 24 | 35 | 17 | 0 | 297 | 149 | 0.001106829 | 0.016804393 | 0.739 | 0.788 | 0.794 | 0.794 | 0.805 | 1 | 0.849 | 0.811 | 0.83 | 1 | -0.114 |

[illegible]

| | | | | | | | | | | | | | | | | | | | | | | | | | | | | | | | | | | |
|---|---|---|---|---|---|---|---|---|---|---|---|---|---|---|---|---|---|---|---|---|---|---|---|---|---|---|---|---|---|---|---|---|---|---|
| SE | 57788 | ENSG00000093010 | COMT | chr22 | + | 19967163 | 19967621 | 19964167 | 19964299 | 19968535 | 19968954 | 3 | 0 | 2 | 3 | 0 | 120 | 60 | 60 | 87 | 37 | 0 | 16 | 1 | 3 | 11 | 120 | 91 | 60 | 74 | 153 | 298 | 149 | [illegible] |
| SE | 35429 | ENSG00000186001 | LRCH3 | chr3 | + | 197854391 | 197854445 | 197852327 | 197852620 | 197865422 | 197865471 | 78 | 38 | 83 | 39 | 85 | 4 | 2 | 3 | 12 | 9 | 19 | 40 | 12 | 27 | 39 | 0 | 0 | 1 | 0 | 5 | 203 | 149 | [illegible] |
| SE | 107224 | ENSG00000135108 | FBXO21 | chr12 | - | 117166897 | 117167077 | 117165484 | 117165617 | 117172470 | 117172607 | 82 | 34 | 73 | 70 | 88 | 5 | 0 | 2 | 0 | 0 | 36 | 28 | 54 | 84 | 23 | 3 | 1 | 17 | 0 | 0 | 298 | 149 | [illegible] |
| SE | 31872 | ENSG00000143669 | LYST | chr1 | - | 235753043 | 235753274 | 235746335 | 235746527 | 235755477 | 235755647 | 136 | 102 | 90 | 85 | 246 | 3 | 4 | 1 | 8 | 0 | 20 | 70 | 35 | 4 | 4 | 0 | 0 | 0 | 0 | 0 | 298 | 149 | [illegible] |
| SE | 90403 | ENSG00000170633 | RNF34 | chr12 | + | 121420241 | 121420334 | 121417503 | 121417911 | 121423385 | 121423517 | 88 | 42 | 57 | 82 | 60 | 3 | 0 | 2 | 3 | 3 | 23 | 56 | 31 | 40 | 31 | 8 | 6 | 3 | 4 | 0 | 242 | 149 | [illegible] |
| SE | 47444 | ENSG00000198178 | CLEC4C | chr12 | - | 7746330 | 7746423 | 7741420 | 7741531 | 7747317 | 7747339 | 70 | 57 | 72 | 113 | 57 | 5 | 7 | 7 | 14 | 2 | 35 | 4 | 13 | 20 | 18 | 3 | 0 | 1 | 1 | 0 | 242 | 149 | [illegible] |
| SE | 36869 | ENSG00000133422 | MORC2 | chr22 | - | 30941432 | 30941558 | 30940757 | 30940837 | 30941890 | 30942002 | 126 | 100 | 133 | 111 | 96 | 2 | 1 | 2 | 0 | 0 | 120 | 89 | 59 | 85 | 24 | 7 | 2 | 0 | 5 | 0 | 275 | 149 | [illegible] |
| SE | 105944 | ENSG00000127314 | RAP1B | chr12 | + | 68647992 | 68648095 | 68624670 | 68624873 | 68648698 | 68648757 | 1 | 0 | 0 | 3 | 3 | 34 | 31 | 14 | 49 | 40 | 13 | 0 | 5 | 0 | 5 | 0 | 61 | 36 | 29 | 3 | 252 | 149 | [illegible] |
| SE | 107652 | ENSG00000139718 | SETD1B | chr12 | + | 121808207 | 121808320 | 121805834 | 121806105 | 121809602 | 121810835 | 73 | 60 | 85 | 101 | 91 | 0 | 0 | 1 | 0 | 0 | 188 | 150 | 64 | 101 | 184 | 9 | 1 | 2 | 5 | 0 | 262 | 149 | [illegible] |
| SE | 25043 | ENSG00000164308 | ERAP2 | chr5 | + | 96892298 | 96892453 | 96889184 | 96889305 | 96895245 | 96895359 | 501 | 714 | 845 | 808 | 455 | 2 | 6 | 2 | 0 | 5 | 39 | 461 | 119 | 125 | 7 | 10 | 5 | 2 | 0 | 0 | 298 | 149 | [illegible] |
| SE | 72446 | ENSG00000074582 | BCS1L | chr2 | + | 218662514 | 218662679 | 218662196 | 218662260 | 218662882 | 218663000 | 24 | 14 | 34 | 52 | 28 | 0 | 0 | 0 | 0 | 0 | 32 | 20 | 17 | 3 | 42 | 13 | 0 | 0 | 0 | 0 | 298 | 149 | [illegible] |
| SE | 19979 | ENSG00000112584 | FAM120B | chr6 | + | 170348150 | 170348323 | 170330448 | 170330550 | 170388286 | 170388493 | 54 | 30 | 55 | 53 | 54 | 0 | 0 | 0 | 2 | 0 | 42 | 42 | 20 | 27 | 29 | 0 | 0 | 2 | 0 | 9 | 298 | 149 | [illegible] |
| SE | 33272 | ENSG00000257335 | MGAM | chr7 | + | 142096330 | 142096415 | 142095564 | 142095713 | 142097592 | 142097649 | 19 | 1 | 15 | 8 | 16 | 4 | 0 | 0 | 0 | 0 | 17 | 38 | 39 | 62 | 16 | 0 | 0 | 12 | 9 | 0 | 234 | 149 | [illegible] |
| SE | 55319 | ENSG00000164111 | ANXA5 | chr4 | - | 121678351 | 121678494 | 121677893 | 121677950 | 121681670 | 121681761 | 1100 | 1138 | 733 | 977 | 1256 | 13 | 17 | 9 | 3 | 1 | 6002 | 1500 | 1023 | 2210 | 3296 | 104 | 12 | 21 | 58 | 0 | 292 | 149 | [illegible] |
| SE | 5516 | ENSG00000101294 | HM13 | chr20 | + | 31566209 | 31566295 | 31561633 | 31561736 | 31569119 | 31569174 | 184 | 74 | 133 | 157 | 78 | 14 | 10 | 9 | 20 | 3 | 412 | 159 | 175 | 366 | 284 | 37 | 61 | 22 | 59 | 45 | 235 | 149 | [illegible] |
| SE | 19662 | ENSG00000119688 | ABCD4 | chr14 | - | 74290026 | 74290118 | 74289300 | 74289519 | 74290290 | 74290452 | 46 | 21 | 32 | 49 | 11 | 3 | 3 | 1 | 2 | 3 | 39 | 30 | 29 | 27 | 61 | 5 | 13 | 6 | 2 | 16 | 241 | 149 | [illegible] |
| SE | 52120 | ENSG00000274020 | LINC01138 | chr1 | - | 148436443 | 148436564 | 148435145 | 148435354 | 148459601 | 148459959 | 7 | 3 | 5 | 5 | 16 | 19 | 19 | 2 | 17 | 12 | 20 | 18 | 6 | 10 | 16 | 10 | 4 | 2 | 8 | 13 | 270 | 149 | [illegible] |
| SE | 95385 | ENSG00000198431 | TXNRD1 | chr12 | + | 104290992 | 104291104 | 104288930 | 104289040 | 104311289 | 104311412 | 2 | 0 | 2 | 0 | 0 | 104 | 76 | 76 | 94 | 82 | 8 | 12 | 1 | 0 | 0 | 80 | 77 | 60 | 59 | 62 | 261 | 149 | [illegible] |
| SE | 46713 | ENSG00000055070 | SZRD1 | chr1 | + | 16387054 | 16387125 | 16367234 | 16367308 | 16391374 | 16391424 | 3 | 2 | 0 | 0 | 0 | 46 | 39 | 48 | 56 | 50 | 17 | 0 | 0 | 6 | 0 | 36 | 93 | 40 | 65 | 111 | 220 | 149 | [illegible] |
| SE | 82562 | ENSG00000235944 | ZNF815P | chr7 | + | 5839948 | 5840075 | 5823159 | 5823526 | 5840689 | 5840856 | 27 | 21 | 48 | 33 | 6 | 5 | 4 | 1 | 0 | 0 | 28 | 47 | 22 | 41 | 41 | 0 | 0 | 0 | 0 | 0 | 276 | 149 | [illegible] |
| SE | 106594 | ENSG00000166012 | TAF1D | chr11 | - | 93736140 | 93736304 | 93734624 | 93734660 | 93737063 | 93737239 | 213 | 227 | 235 | 172 | 448 | 52 | 42 | 39 | 30 | 103 | 13 | 72 | 90 | 14 | 2 | 6 | 23 | 42 | 5 | 13 | 298 | 149 | [illegible] |
| SE | 55181 | ENSG00000143158 | MPC2 | chr1 | - | 167919978 | 167920090 | 167916674 | 167918359 | 167920546 | 167920631 | 155 | 84 | 125 | 168 | 229 | 1 | 0 | 0 | 0 | 0 | 164 | 92 | 151 | 241 | 336 | 0 | 7 | 2 | 11 | 0 | 261 | 149 | [illegible] |
| SE | 62191 | ENSG00000113569 | NUP155 | chr5 | - | 37307296 | 37307432 | 37305056 | 37305210 | 37309128 | 37309267 | 158 | 125 | 189 | 241 | 482 | 9 | 7 | 11 | 8 | 9 | 85 | 145 | 90 | 113 | 7 | 0 | 11 | 29 | 19 | 0 | 285 | 149 | [illegible] |
| SE | 99227 | ENSG00000086848 | ALG9 | chr11 | - | 111840654 | 111840809 | 111838248 | 111838399 | 111844600 | 111844723 | 65 | 34 | 51 | 34 | 76 | 3 | 0 | 3 | 1 | 0 | 131 | 50 | 45 | 23 | 23 | 0 | 2 | 10 | 5 | 3 | 298 | 149 | [illegible] |
| SE | 60893 | ENSG00000144674 | GOLGA4 | chr3 | + | 37273535 | 37273601 | 37251394 | 37251484 | 37281957 | 37281985 | 120 | 102 | 92 | 113 | 125 | 14 | 13 | 14 | 15 | 10 | 41 | 60 | 31 | 28 | 42 | 0 | 5 | 6 | 0 | 0 | 215 | 149 | [illegible] |
| SE | 82996 | ENSG00000265681 | RPL17 | chr18 | - | 49491735 | 49491833 | 49491531 | 49491584 | 49492457 | 49492508 | 22 | 17 | 31 | 33 | 24 | 474 | 254 | 551 | 423 | 418 | 26 | 25 | 21 | 32 | 6 | 126 | 296 | 260 | 215 | 224 | 247 | 149 | [illegible] |
| SE | 56584 | ENSG00000105700 | KXD1 | chr19 | + | 18565133 | 18565190 | 18564868 | 18565021 | 18567131 | 18567178 | 14 | 10 | 6 | 7 | 11 | 77 | 56 | 62 | 89 | 125 | 0 | 11 | 3 | 3 | 19 | 133 | 120 | 119 | 101 | 174 | 206 | 149 | [illegible] |
| SE | 71477 | ENSG00000130038 | CRACR2A | chr12 | - | 3680237 | 3680349 | 3673445 | 3673592 | 3696771 | 3697035 | 44 | 30 | 38 | 22 | 24 | 0 | 0 | 0 | 2 | 0 | 10 | 32 | 36 | 15 | 32 | 9 | 0 | 2 | 1 | 0 | 261 | 149 | [illegible] |
| SE | 30003 | ENSG00000185305 | ARL15 | chr5 | - | 54307887 | 54307946 | 54171783 | 54171928 | 54310294 | 54310415 | 0 | 4 | 0 | 1 | 0 | 5 | 13 | 3 | 19 | 2 | 12 | 10 | 0 | 1 | 0 | 23 | 18 | 1 | 2 | 44 | 208 | 149 | [illegible] |
| SE | 91400 | ENSG00000158473 | CD1D | chr1 | + | 158182877 | 158183156 | 158182031 | 158182310 | 158184128 | 158186427 | 131 | 94 | 55 | 82 | 34 | 0 | 0 | 2 | 2 | 5 | 296 | 147 | 37 | 90 | 211 | 12 | 4 | 0 | 17 | 11 | 298 | 149 | [illegible] |
| SE | 58056 | ENSG00000082701 | GSK3B | chr3 | - | 119905754 | 119905852 | 119863418 | 119863605 | 119916043 | 119916174 | 10 | 10 | 3 | 17 | 12 | 0 | 0 | 0 | 0 | 0 | 7 | 38 | 33 | 29 | 5 | 20 | 3 | 0 | 0 | 0 | 247 | 149 | [illegible] |

[illegible]

| | | | | | | | | | | | |
|---|---|---|---|---|---|---|---|---|---|---|---|
| SE | 106896 | ENSG00000205885 | C1RL-AS1 | chr12 | + | 7110063 | 7110147 | 7108672 | 7108947 | 7110297 | 7110572 |
| SE | 103588 | ENSG00000136754 | ABI1 | chr10 | - | 26755654 | 26755741 | 26751597 | 26751783 | 26759061 | 26759235 |
| SE | 26197 | ENSG00000059378 | PARP12 | chr7 | - | 140041643 | 140041839 | 140034234 | 140034331 | 140046883 | 140047007 |
| SE | 72469 | ENSG00000118495 | PLAGL1 | chr6 | - | 143947984 | 143948460 | 143940299 | 143942663 | 143960468 | 143960542 |
| SE | 97257 | ENSG00000134627 | PIWIL4 | chr11 | + | 94608184 | 94608686 | 94607438 | 94607639 | 94616492 | 94616563 |
| SE | 103379 | ENSG00000176986 | SEC24C | chr10 | + | 73766349 | 73766541 | 73766085 | 73766210 | 73766759 | 73766853 |
| SE | 107021 | ENSG00000159403 | C1R | chr12 | - | 7089586 | 7089820 | 7089357 | 7089489 | 7090055 | 7090248 |
| SE | 88920 | ENSG00000176731 | RBIS | chr8 | - | 85214920 | 85215037 | 85214047 | 85214631 | 85217385 | 85217502 |
| SE | 75780 | ENSG00000204231 | RXRB | chr6 | - | 33195362 | 33195466 | 33194944 | 33195050 | 33195569 | 33195702 |
| SE | 715 | ENSG00000182944 | EWSR1 | chr22 | + | 29291561 | 29291599 | 29286922 | 29287134 | 29292136 | 29292169 |
| SE | 66673 | ENSG00000005483 | KMT2E | chr7 | + | 105066726 | 105066807 | 105063350 | 105063580 | 105073618 | 105073677 |
| SE | 32443 | ENSG00000131370 | SH3BP5 | chr3 | - | 15273115 | 15273405 | 15269712 | 15269877 | 15304102 | 15304231 |
| SE | 6139 | ENSG00000131069 | ACSS2 | chr20 | + | 34926104 | 34926281 | 34925697 | 34925766 | 34926876 | 34926938 |
| SE | 62751 | ENSG00000162961 | DPY30 | chr2 | - | 31883385 | 31883462 | 31869847 | 31869952 | 31917925 | 31918037 |
| SE | 59671 | ENSG00000186283 | TOR3A | chr1 | + | 179085627 | 179085893 | 179082069 | 179082387 | 179087910 | 179088083 |
| SE | 96788 | ENSG00000259719 | LINC02284 | chr14 | + | 56317455 | 56317615 | 56310879 | 56311102 | 56317891 | 56318086 |
| SE | 17357 | ENSG00000133961 | NUMB | chr14 | - | 73366896 | 73366981 | 73355625 | 73355766 | 73409936 | 73410068 |
| SE | 66984 | ENSG00000135930 | EIF4E2 | chr2 | + | 232557883 | 232558018 | 232556415 | 232556530 | 232564246 | 232564351 |
| SE | 28234 | ENSG00000205937 | RNPS1 | chr16 | - | 2267174 | 2267237 | 2264572 | 2264760 | 2268054 | 2268114 |
| SE | 76111 | ENSG00000100220 | RTCB | chr22 | - | 32397940 | 32398100 | 32396073 | 32396249 | 32399602 | 32399759 |
| SE | 46466 | ENSG00000185864 | NPIPB4 | chr16 | - | 21839939 | 21840008 | 21839249 | 21839310 | 21843385 | 21843412 |
| SE | 76784 | ENSG00000135317 | SNX14 | chr6 | - | 85538837 | 85538864 | 85536791 | 85536924 | 85541984 | 85542043 |
| SE | 45861 | ENSG00000062485 | CS | chr12 | - | 56291630 | 56291793 | 56286594 | 56286645 | 56298608 | 56298691 |
| SE | 9364 | ENSG00000264187 | NA | chr17 | - | 17231793 | 17231882 | 17227989 | 17228161 | 17232787 | 17232901 |
| SE | 57880 | ENSG00000164924 | YWHAZ | chr8 | - | 100951928 | 100952308 | 100948613 | 100948900 | 100952849 | 100952883 |
| SE | 19051 | ENSG00000196405 | EVL | chr14 | + | 100135904 | 100135968 | 100132718 | 100132779 | 100141179 | 100141246 |
| SE | 68084 | ENSG00000135838 | NPL | chr1 | + | 182794355 | 182794439 | 182789292 | 182789805 | 182803697 | 182803771 |
| SE | 71528 | ENSG00000144504 | ANKMY1 | chr2 | - | 240529036 | 240529509 | 240525684 | 240525849 | 240552913 | 240553057 |
| SE | 18736 | ENSG00000080824 | HSP90AA1 | chr14 | - | 102082110 | 102082444 | 102080741 | 102081821 | 102083033 | 102083302 |
| SE | 61973 | ENSG00000205423 | CNEP1R1 | chr16 | + | 50025650 | 50025701 | 50024409 | 50025340 | 50026395 | 50026467 |
| SE | 6094 | ENSG00000171456 | ASXL1 | chr20 | + | 32367726 | 32367729 | 32366383 | 32366466 | 32369014 | 32369123 |
| SE | 82324 | ENSG00000146833 | TRIM4 | chr7 | - | 99916701 | 99916779 | 99909564 | 99909660 | 99919008 | 99919531 |
| SE | 101791 | ENSG00000204161 | TMEM273 | chr10 | - | 49165204 | 49165361 | 49160347 | 49161622 | 49165765 | 49165796 |
| SE | 12592 | ENSG00000090861 | AARS1 | chr16 | - | 70262365 | 70262524 | 70261043 | 70261157 | 70264957 | 70265102 |
| SE | 86343 | ENSG00000074966 | TXK | chr4 | - | 48073934 | 48074053 | 48071516 | 48071674 | 48076401 | 48076466 |
| SE | 2062 | ENSG00000099331 | MYO9B | chr19 | + | 17210332 | 17210380 | 17209585 | 17209709 | 17210714 | 17210848 |
| SE | 106698 | ENSG00000141376 | BCAS3 | chr17 | + | 60889694 | 60889771 | 60874661 | 60874738 | 60902619 | 60902703 |
| SE | 40940 | ENSG00000168710 | AHCYL1 | chr1 | + | 110020730 | 110020851 | 110019547 | 110019626 | 110021673 | 110022045 |
| SE | 32618 | ENSG00000078618 | NRDC | chr1 | - | 51836368 | 51836438 | 51836130 | 51836212 | 51840225 | 51840454 |
| SE | 68501 | ENSG00000087087 | SRRT | chr7 | + | 100884367 | 100884552 | 100884069 | 100884239 | 100884739 | 100884838 |
| SE | 58786 | ENSG00000077254 | USP33 | chr1 | - | 77714613 | 77714783 | 77713199 | 77713281 | 77717866 | 77718047 |
| SE | 74050 | ENSG00000111011 | RSRC2 | chr12 | - | 122513766 | 122513874 | 122511108 | 122511188 | 122514711 | 122514728 |
| SE | 5121 | ENSG00000125826 | RBCK1 | chr20 | + | 417219 | 417342 | 409883 | 410025 | 417525 | 417619 |
| SE | 14937 | ENSG00000159216 | RUNX1 | chr21 | - | 34886842 | 34887096 | 34880556 | 34880713 | 34892924 | 34892963 |
| SE | 22251 | ENSG00000141279 | NPEPPS | chr17 | + | 47619277 | 47619322 | 47619008 | 47619164 | 47619736 | 47619784 |
| SE | 104766 | ENSG00000011258 | MBTD1 | chr17 | - | 51209313 | 51209430 | 51206887 | 51207005 | 51217333 | 51217416 |
| SE | 15089 | ENSG00000128739 | SNRPN | chr15 | + | 24967028 | 24967240 | 24962113 | 24962209 | 24967931 | 24968082 |
| SE | 14437 | ENSG00000130703 | OSBPL2 | chr20 | + | 62259980 | 62260125 | 62256056 | 62256221 | 62263615 | 62263691 |
| SE | 50041 | ENSG00000154222 | CC2D1B | chr1 | - | 52357000 | 52357174 | 52356383 | 52356442 | 52357525 | 52357698 |
| SE | 17996 | ENSG00000092108 | SCFD1 | chr14 | + | 30643879 | 30644018 | 30643315 | 30643405 | 30649527 | 30649583 |
| SE | 96372 | ENSG00000184060 | ADAP2 | chr17 | + | 30926808 | 30926918 | 30922939 | 30923070 | 30931888 | 30931968 |
| SE | 84844 | ENSG00000094631 | HDAC6 | chrX | + | 48824165 | 48824294 | 48823671 | 48823785 | 48824543 | 48824860 |
| SE | 3582 | ENSG00000129347 | KRI1 | chr19 | - | 10558163 | 10558239 | 10557971 | 10558060 | 10559358 | 10559529 |
| SE | 79235 | ENSG00000196511 | TPK1 | chr7 | - | 144591422 | 144591569 | 144548490 | 144548602 | 144623165 | 144623261 |
| SE | 82176 | ENSG00000146872 | TLK2 | chr17 | + | 62552301 | 62552397 | 62536169 | 62536337 | 62560015 | 62560126 |
| SE | 66810 | ENSG00000137075 | RNF38 | chr9 | - | 36390466 | 36390616 | 36375933 | 36376127 | 36400553 | 36400923 |
| SE | 21863 | ENSG00000265354 | TIMM23 | chr10 | + | 45982845 | 45982930 | 45982522 | 45982616 | 45988736 | 45988847 |
| SE | 8947 | ENSG00000132478 | UNK | chr17 | + | 75808947 | 75809039 | 75784861 | 75784984 | 75809759 | 75809969 |
| SE | 15687 | ENSG00000174485 | DENND4A | chr15 | - | 65706090 | 65706224 | 65702872 | 65703008 | 65715477 | 65715623 |
| SE | 99309 | ENSG00000101017 | CD40 | chr20 | + | 46126639 | 46126701 | 46122609 | 46122756 | 46128137 | 46128224 |
| SE | 25722 | ENSG00000247774 | PCED1B-AS1 | chr12 | - | 47210639 | 47210761 | 47209280 | 47209458 | 47216164 | 47216200 |
| SE | 51065 | ENSG00000104231 | ZFAND1 | chr8 | - | 81708787 | 81708829 | 81702968 | 81703124 | 81713917 | 81714039 |
| SE | 62069 | ENSG00000110446 | SLC15A3 | chr11 | - | 60942034 | 60942139 | 60939479 | 60939638 | 60943688 | 60943836 |
| SE | 93457 | ENSG00000169925 | BRD3 | chr9 | - | 134036031 | 134036324 | 134034700 | 134034829 | 134041759 | 134041794 |
| SE | 23952 | ENSG00000133835 | HSD17B4 | chr5 | + | 119509429 | 119509494 | 119509140 | 119509244 | 119514980 | 119515046 |
| SE | 32421 | ENSG00000136040 | PLXNC1 | chr12 | + | 94297188 | 94297220 | 94294485 | 94294540 | 94297315 | 94297423 |
| SE | 54132 | ENSG00000124126 | PREX1 | chr20 | - | 48666282 | 48666355 | 48658135 | 48658228 | 48676192 | 48676268 |
| SE | 84130 | ENSG00000094841 | UPRT | chrX | + | 75299734 | 75299896 | 75297490 | 75297553 | 75300866 | 75300965 |
| SE | 1052 | ENSG00000100099 | HPS4 | chr22 | - | 26482745 | 26482873 | 26481834 | 26482240 | 26483673 | 26483836 |
| SE | 30198 | ENSG00000168397 | ATG4B | chr2 | + | 241655270 | 241655343 | 241654545 | 241654647 | 241666644 | 241666838 |
| SE | 10020 | ENSG00000006282 | SPATA20 | chr17 | + | 50549285 | 50549487 | 50548809 | 50549186 | 50549984 | 50550115 |
| SE | 28767 | ENSG00000170571 | EMB | chr5 | - | 50410876 | 50410965 | 50405724 | 50405852 | 50411196 | 50411221 |
| SE | 75100 | ENSG00000114423 | CBLB | chr3 | - | 105671020 | 105671046 | 105670232 | 105670352 | 105678430 | 105678571 |
| SE | 39734 | ENSG00000132581 | SDF2 | chr17 | - | 28651485 | 28651542 | 28648841 | 28649276 | 28655286 | 28655483 |
| SE | 57279 | ENSG00000143119 | CD53 | chr1 | + | 110896652 | 110896733 | 110892344 | 110892533 | 110899123 | 110899922 |
| SE | 99215 | ENSG00000149196 | HIKESHI | chr11 | + | 86326512 | 86326676 | 86306244 | 86306482 | 86337378 | 86337530 |
| SE | 48184 | ENSG00000006831 | ADIPOR2 | chr12 | + | 1702885 | 1702982 | 1691162 | 1691213 | 1754257 | 1754514 |
| SE | 100321 | ENSG00000081237 | PTPRC | chr1 | + | 198694017 | 198694140 | 198692346 | 198692373 | 198703297 | 198703372 |
| SE | 22875 | ENSG00000043143 | JADE2 | chr5 | + | 134576767 | 134577038 | 134573644 | 134573762 | 134578496 | 134579475 |
| SE | 90262 | ENSG00000249859 | PVT1 | chr8 | + | 127795893 | 127796071 | 127794530 | 127794734 | 127890588 | 127890998 |
| SE | 20560 | ENSG00000108588 | CCDC47 | chr17 | - | 63772776 | 63772863 | 63765911 | 63766194 | 63773411 | 63773570 |
| SE | 97587 | ENSG00000110721 | CHKA | chr11 | - | 68061492 | 68061830 | 68053514 | 68054047 | 68061952 | 68062034 |
| SE | 28215 | ENSG00000204084 | INPP5B | chr1 | - | 37869924 | 37870014 | 37868500 | 37868614 | 37872929 | 37873028 |
| SE | 104979 | ENSG00000111371 | SLC38A1 | chr12 | - | 46201097 | 46201198 | 46198624 | 46198743 | 46203009 | 46203089 |
| SE | 19383 | ENSG00000258875 | NA | chr14 | + | 91247489 | 91247659 | 91244354 | 91244499 | 91250764 | 91251228 |
| SE | 4576 | ENSG00000123146 | ADGRE5 | chr19 | + | 14407060 | 14407229 | 14406868 | 14406960 | 14407907 | 14408009 |
| SE | 69670 | ENSG00000184708 | EIF4ENIF1 | chr22 | - | 31463680 | 31463967 | 31462931 | 31463133 | 31468174 | 31468302 |
| SE | 30468 | ENSG00000196209 | SIRPB2 | chr20 | - | 1479699 | 1479794 | 1478503 | 1478607 | 1479925 | 1480065 |
| SE | 11624 | ENSG00000198736 | MSRB1 | chr16 | - | 1940777 | 1940892 | 1938228 | 1939143 | 1941256 | 1941405 |
| SE | 30629 | ENSG00000189308 | LIN54 | chr4 | - | 82935980 | 82936118 | 82930942 | 82931145 | 82936278 | 82936381 |
| SE | 34887 | ENSG00000127483 | HP1BP3 | chr1 | - | 20780344 | 20780540 | 20779811 | 20779911 | 20787194 | 20787307 |
| SE | 101187 | ENSG00000186174 | BCL9L | chr11 | - | 118909913 | 118910015 | 118908269 | 118908655 | 118918825 | 118918879 |
| SE | 9969 | ENSG00000266173 | STRADA | chr17 | - | 63726637 | 63726695 | 63714005 | 63714108 | 63728333 | 63728384 |
| SE | 57199 | ENSG00000084112 | SSH1 | chr12 | - | 108798999 | 108799200 | 108791918 | 108792829 | 108800779 | 108800926 |
| SE | 325 | ENSG00000182670 | TTC3 | chr21 | + | 37158130 | 37158203 | 37156654 | 37156906 | 37159698 | 37159745 |
| SE | 33394 | ENSG00000254087 | LYN | chr8 | + | 55941854 | 55941928 | 55879834 | 55880103 | 55946447 | 55946493 |
| SE | 25694 | ENSG00000247774 | PCED1B-AS1 | chr12 | - | 47206510 | 47206585 | 47205897 | 47206145 | 47216164 | 47216200 |
| SE | 59107 | ENSG00000214941 | ZSWIM7 | chr17 | - | 15986799 | 15986892 | 15981039 | 15981144 | 15987265 | 15987368 |
| SE | 15882 | ENSG00000167491 | GATAD2A | chr19 | + | 19436130 | 19436219 | 19405837 | 19406019 | 19465339 | 19465577 |
| SE | 14439 | ENSG00000130703 | OSBPL2 | chr20 | + | 62264628 | 62264718 | 62263615 | 62263691 | 62272124 | 62272259 |
| SE | 36099 | ENSG00000100578 | KIAA0586 | chr14 | + | 58448339 | 58448493 | 58443953 | 58444175 | 58450578 | 58450746 |
| SE | 63539 | ENSG00000119844 | AFTPH | chr2 | + | 64581189 | 64581273 | 64579485 | 64579546 | 64585421 | 64585545 |
| SE | 57878 | ENSG00000164924 | YWHAZ | chr8 | - | 100951928 | 100952125 | 100948595 | 100948900 | 100952849 | 100952983 |
| SE | 44103 | ENSG00000114982 | KANSL3 | chr2 | - | 96602752 | 96602862 | 96602115 | 96602338 | 96604249 | 96604380 |
| SE | 26101 | ENSG00000164181 | ELOVL7 | chr5 | - | 60771948 | 60772093 | 60767822 | 60767903 | 60787333 | 60787465 |
| SE | 27137 | ENSG00000197020 | ZNF100 | chr19 | - | 21744940 | 21745067 | 21722770 | 21727989 | 21765693 | 21765783 |
| SE | 20858 | ENSG00000255819 | KLRC4-KLRK1 | chr12 | - | 10409388 | 10409478 | 10408911 | 10409010 | 10410098 | 10410128 |
| SE | 56628 | ENSG00000075151 | EIF4G3 | chr1 | - | 21001198 | 21001312 | 20981047 | 20981227 | 21002712 | 21002808 |
| SE | 9843 | ENSG00000170836 | PPM1D | chr17 | + | 60633852 | 60633977 | 60623520 | 60623749 | 60647891 | 60648082 |
| SE | 91807 | ENSG00000156738 | MS4A1 | chr11 | + | 60461071 | 60461160 | 60455846 | 60455945 | 60462184 | 60462298 |
| SE | 54423 | ENSG00000169554 | ZEB2 | chr2 | - | 144398300 | 144400345 | 144396411 | 144396592 | 144401198 | 144401307 |
| SE | 20248 | ENSG00000124789 | NUP153 | chr6 | - | 17646066 | 17646154 | 17639938 | 17640064 | 17647806 | 17647905 |
| SE | 88322 | ENSG00000119523 | ALG2 | chr9 | - | 99220978 | 99221137 | 99216425 | 99218836 | 99221546 | 99221942 |
| SE | 95100 | ENSG00000148358 | GPR107 | chr9 | + | 130100628 | 130100702 | 130099456 | 130099532 | 130101105 | 130101223 |
| SE | 8937 | ENSG00000092929 | UNC13D | chr17 | - | 75840017 | 75840110 | 75839838 | 75839942 | 75840224 | 75840304 |
| SE | 75413 | ENSG00000169826 | CSGALNACT2 | chr10 | + | 43163865 | 43164044 | 43160493 | 43160595 | 43175950 | 43176032 |
| SE | 28231 | ENSG00000205937 | RNPS1 | chr16 | - | 2264572 | 2264760 | 2264175 | 2264331 | 2268054 | 2268091 |
| SE | 105902 | ENSG00000137955 | RABGGTB | chr1 | + | 75789158 | 75789356 | 75786700 | 75787604 | 75789951 | 75789996 |
| SE | 8429 | ENSG00000078687 | TNRC6C | chr17 | + | 78077184 | 78077334 | 78075135 | 78075278 | 78086502 | 78086586 |
| SE | 68760 | ENSG00000163872 | YEATS2 | chr3 | + | 183724418 | 183724531 | 183721890 | 183722136 | 183728689 | 183728851 |
| SE | 93354 | ENSG00000225783 | MIAT | chr22 | + | 26667184 | 26667310 | 26665456 | 26666937 | 26668157 | 26669655 |
| SE | 12527 | ENSG00000103066 | PLA2G15 | chr16 | + | 68255281 | 68255380 | 68254352 | 68255037 | 68255890 | 68255990 |
| SE | 897 | ENSG00000100084 | HIRA | chr22 | - | 19351357 | 19351446 | 19330697 | 19331556 | 19353355 | 19353519 |
| SE | 24985 | ENSG00000196189 | SEMA4A | chr1 | + | 156175085 | 156175243 | 156172825 | 156173006 | 156175555 | 156175647 |
| SE | 31851 | ENSG00000066777 | ARFGEF1 | chr8 | - | 67266875 | 67266984 | 67266005 | 67266207 | 67267342 | 67267442 |
| SE | 109643 | ENSG00000005810 | MYCBP2 | chr13 | - | 77185870 | 77186063 | 77185102 | 77185377 | 77188950 | 77189047 |
| SE | 27462 | ENSG00000008513 | ST3GAL1 | chr8 | - | 133475718 | 133476600 | 133465893 | 133466090 | 133499134 | 133499189 |
| SE | 36752 | ENSG00000173193 | PARP14 | chr3 | + | 122720254 | 122720388 | 122718358 | 122718958 | 122727811 | 122727986 |
| SE | 28658 | ENSG00000002586 | CD99 | chrX | + | 2717604 | 2717652 | 2714421 | 2714454 | 2719660 | 2719705 |
| SE | 79267 | ENSG00000133574 | GIMAP4 | chr7 | + | 150569887 | 150569959 | 150567389 | 150567437 | 150572128 | 150573953 |
| SE | 12941 | ENSG00000167397 | VKORC1 | chr16 | - | 31093311 | 31093482 | 31090841 | 31091342 | 31094556 | 31094760 |
| SE | 69491 | ENSG00000109743 | BST1 | chr4 | + | 15705514 | 15705641 | 15703064 | 15703332 | 15707510 | 15707646 |
| SE | 106540 | ENSG00000183955 | KMT5A | chr12 | + | 123403572 | 123403632 | 123396344 | 123396432 | 123407492 | 123407568 |
| SE | 68577 | ENSG00000140368 | PSTPIP1 | chr15 | + | 77032861 | 77032952 | 77031179 | 77031278 | 77035507 | 77035563 |
| SE | 40395 | ENSG00000163719 | MTMR14 | chr3 | + | 9689963 | 9690143 | 9688943 | 9689082 | 9701789 | 9702393 |
| SE | 44664 | ENSG00000203667 | COX20 | chr1 | + | 244841943 | 244842058 | 244835657 | 244835756 | 244842194 | 244842258 |
| SE | 18766 | ENSG00000170348 | TMED10 | chr14 | - | 75139073 | 75139171 | 75135759 | 75135886 | 75147663 | 75147737 |
| SE | 17040 | ENSG00000128829 | EIF2AK4 | chr15 | + | 40016501 | 40016672 | 40011280 | 40011346 | 40017107 | 40017242 |
| SE | 102751 | ENSG00000040341 | STAU2 | chr8 | - | 73697307 | 73697442 | 73688653 | 73688813 | 73709031 | 73709162 |
| SE | 36358 | ENSG00000173889 | PHC3 | chr3 | - | 170128339 | 170128387 | 170122590 | 170122744 | 170128683 | 170128807 |
| SE | 107882 | ENSG00000188906 | LRRK2 | chr12 | + | 40259479 | 40259604 | 40252964 | 40253016 | 40263788 | 40263901 |
| SE | 93883 | ENSG00000089053 | ANAPC5 | chr12 | - | 121331226 | 121331428 | 121330582 | 121330672 | 121335532 | 121335723 |
| SE | 81561 | ENSG00000167703 | SLC43A2 | chr17 | - | 1616561 | 1616630 | 1614978 | 1615034 | 1627714 | 1627911 |
| SE | 71219 | ENSG00000079263 | SP140 | chr2 | + | 230248884 | 230248968 | 230245862 | 230245940 | 230255451 | 230255532 |
| SE | 55663 | ENSG00000138119 | MYOF | chr10 | - | 93379862 | 93379987 | 93377322 | 93377429 | 93381218 | 93381396 |
| SE | 52759 | ENSG00000136682 | CBWD2 | chr2 | + | 113471032 | 113471089 | 113465128 | 113465173 | 113482176 | 113482228 |
| SE | 63262 | ENSG00000013810 | TACC3 | chr4 | + | 1723727 | 1723870 | 1723420 | 1723583 | 1727707 | 1728094 |
| SE | 82678 | ENSG00000141380 | SS18 | chr18 | - | 26082362 | 26082528 | 26078075 | 26078160 | 26087500 | 26087577 |
| SE | 66345 | ENSG00000197037 | ZSCAN25 | chr7 | + | 99619047 | 99619132 | 99618524 | 99618651 | 99619560 | 99619816 |
| SE | 30942 | ENSG00000017260 | ATP2C1 | chr3 | + | 130975331 | 130975488 | 130969291 | 130969396 | 130992950 | 130993001 |
| SE | 29128 | ENSG00000248672 | LY75-CD302 | chr2 | - | 159850361 | 159850467 | 159849979 | 159850140 | 159852200 | 159852340 |
| SE | 73563 | ENSG00000115998 | C2orf42 | chr2 | - | 70165091 | 70165192 | 70160624 | 70160787 | 70165527 | 70165635 |
| SE | 29323 | ENSG00000067082 | KLF6 | chr10 | - | 3780105 | 3780229 | 3775995 | 3779590 | 3781640 | 3782214 |
| SE | 28312 | ENSG00000123545 | NDUFAF4 | chr6 | - | 96896083 | 96896153 | 96891150 | 96891391 | 96896743 | 96896847 |
| SE | 7769 | ENSG00000109118 | PHF12 | chr17 | - | 28910711 | 28910833 | 28910054 | 28910369 | 28911111 | 28911237 |
| SE | 19860 | ENSG00000165516 | KLHDC2 | chr14 | + | 49780212 | 49780322 | 49779594 | 49779806 | 49780702 | 49780775 |
| SE | 12159 | ENSG00000172775 | PSME3IP1 | chr16 | - | 57178501 | 57178852 | 57175538 | 57175590 | 57185820 | 57186039 |
| SE | 3339 | ENSG00000012061 | ERCC1 | chr19 | - | 45413676 | 45413745 | 45407333 | 45409725 | 45416820 | 45416897 |
| SE | 89338 | ENSG00000165392 | WRN | chr8 | + | 31090464 | 31090532 | 31088889 | 31088965 | 31090833 | 31090942 |
| SE | 1763 | ENSG00000100077 | GRK3 | chr22 | + | 25663629 | 25663704 | 25604376 | 25604453 | 25667738 | 25667800 |
| SE | 92578 | ENSG00000137054 | POLR1E | chr9 | + | 37492281 | 37492375 | 37489314 | 37489400 | 37492656 | 37492715 |
| SE | 25705 | ENSG00000247774 | PCED1B-AS1 | chr12 | - | 47209280 | 47209458 | 47205897 | 47206145 | 47216164 | 47216249 |
| SE | 60713 | ENSG00000106346 | USP42 | chr7 | + | 6128137 | 6128230 | 6116794 | 6116939 | 6135840 | 6135951 |
| SE | 24316 | ENSG00000058799 | YIPF1 | chr1 | - | 53866757 | 53866924 | 53866194 | 53866382 | 53871371 | 53871488 |
| SE | 23547 | ENSG00000163635 | ATXN7 | chr3 | + | 63982185 | 63982445 | 63979914 | 63980167 | 63982938 | 63983021 |
| SE | 77979 | ENSG00000106635 | BCL7B | chr7 | - | 73539881 | 73540052 | 73537933 | 73538013 | 73543547 | 73543601 |
| SE | 35531 | ENSG00000175066 | GK5 | chr3 | - | 142186193 | 142186267 | 142185928 | 142185989 | 142186451 | 142186513 |
| SE | 69955 | ENSG00000108883 | EFTUD2 | chr17 | - | 44872445 | 44872570 | 44868286 | 44868350 | 44875933 | 44876100 |
| SE | 67644 | ENSG00000126768 | TIMM17B | chrX | - | 48895351 | 48895501 | 48895037 | 48895101 | 48896758 | 48896858 |
| SE | 83239 | ENSG00000106077 | ABHD11 | chr7 | - | 73737875 | 73738142 | 73737645 | 73737735 | 73738327 | 73738383 |
| SE | 24168 | ENSG00000124357 | NAGK | chr2 | + | 71071685 | 71071827 | 71068315 | 71068712 | 71072640 | 71072690 |
| SE | 96949 | ENSG00000062822 | POLD1 | chr19 | + | 50395024 | 50395150 | 50384341 | 50384390 | 50398850 | 50399053 |
| SE | 57212 | ENSG00000136231 | IGF2BP3 | chr7 | - | 23343717 | 23343853 | 23342063 | 23342189 | 23345939 | 23346062 |
| SE | 99331 | ENSG00000132199 | ENOSF1 | chr18 | - | 677742 | 677872 | 677344 | 677444 | 678695 | 678737 |
| SE | 90475 | ENSG00000151461 | UPF2 | chr10 | - | 12001675 | 12001825 | 11999905 | 12000009 | 12004529 | 12004727 |
| SE | 67936 | ENSG00000091490 | SEL1L3 | chr4 | - | 25830097 | 25830156 | 25821995 | 25822128 | 25847293 | 25847864 |
| SE | 27106 | ENSG00000105755 | ETHE1 | chr19 | - | 43526200 | 43526349 | 43511436 | 43511566 | 43526514 | 43526659 |
| SE | 8457 | ENSG00000141503 | MINK1 | chr17 | + | 4890950 | 4891124 | 4889646 | 4890735 | 4891455 | 4891716 |
| SE | 41938 | ENSG00000163374 | YY1AP1 | chr1 | - | 155674684 | 155674818 | 155672559 | 155672727 | 155675009 | 155675096 |
| SE | 1076 | ENSG00000100330 | MTMR3 | chr22 | + | 30007900 | 30008032 | 30002879 | 30002993 | 30009017 | 30009129 |
| SE | 38717 | ENSG00000228794 | LINC01128 | chr1 | + | 850180 | 850351 | 847653 | 847806 | 851926 | 852014 |
| SE | 91466 | ENSG00000197140 | ADAM32 | chr8 | + | 39149790 | 39149867 | 39147129 | 39147205 | 39151376 | 39151406 |
| SE | 70438 | ENSG00000144306 | SCRN3 | chr2 | + | 174398365 | 174398442 | 174397335 | 174397404 | 174399921 | 174400103 |
| SE | 31782 | ENSG00000204463 | BAG6 | chr6 | - | 31639499 | 31639646 | 31639027 | 31639226 | 31640198 | 31640306 |
| SE | 98100 | ENSG00000171466 | ZNF562 | chr19 | - | 9656546 | 9656653 | 9641806 | 9653881 | 9660719 | 9660874 |
| SE | 31515 | ENSG00000174227 | PIGG | chr4 | + | 508828 | 508970 | 507404 | 507593 | 515972 | 516185 |
| SE | 43520 | ENSG00000163565 | IFI16 | chr1 | + | 159053532 | 159053724 | 159051678 | 159052098 | 159054820 | 159055124 |
| SE | 103444 | ENSG00000148840 | PPRC1 | chr10 | + | 102138878 | 102138980 | 102138618 | 102138765 | 102139099 | 102142004 |
| SE | 70358 | ENSG00000184182 | UBE2F | chr2 | + | 237987962 | 237987992 | 237973091 | 237973225 | 238016565 | 238016633 |
| SE | 98616 | ENSG00000137710 | RDX | chr11 | - | 110263959 | 110264234 | 110258105 | 110258189 | 110272535 | 110272619 |
| SE | 82841 | ENSG00000162129 | CLPB | chr11 | - | 72408097 | 72408204 | 72402965 | 72403052 | 72430311 | 72430363 |
| SE | 8734 | ENSG00000182628 | SKA2 | chr17 | - | 59131280 | 59131367 | 59119298 | 59119495 | 59155130 | 59155174 |
| SE | 97645 | ENSG00000149823 | VPS51 | chr11 | + | 65107186 | 65107372 | 65096997 | 65097127 | 65107580 | 65107727 |
| SE | 11028 | ENSG00000072849 | DERL2 | chr17 | - | 5480386 | 5480543 | 5480053 | 5480144 | 5481100 | 5481142 |
| SE | 54647 | ENSG00000131791 | PRKAB2 | chr1 | - | 147167766 | 147167933 | 147166845 | 147166939 | 147171988 | 147172167 |
| SE | 40691 | ENSG00000076641 | PAG1 | chr8 | - | 81070111 | 81070170 | 81029995 | 81030089 | 81111590 | 81112068 |
| SE | 85780 | ENSG00000162065 | TBC1D24 | chr16 | + | 2499834 | 2499930 | 2499356 | 2499420 | 2500267 | 2500332 |
| SE | 51872 | ENSG00000138448 | ITGAV | chr2 | + | 186667149 | 186667230 | 186666703 | 186666783 | 186667670 | 186667776 |
| SE | 91509 | ENSG00000135679 | MDM2 | chr12 | + | 68815627 | 68815720 | 68813553 | 68813628 | 68816811 | 68816945 |
| SE | 67495 | ENSG00000068654 | POLR1A | chr2 | - | 86039997 | 86040200 | 86039324 | 86039462 | 86040391 | 86040519 |
| SE | 26693 | ENSG00000160179 | ABCG1 | chr21 | + | 42276894 | 42276945 | 42273302 | 42273435 | 42282273 | 42282419 |
| SE | 75068 | ENSG00000198223 | CSF2RA | chrX | + | 1303922 | 1304019 | 1295426 | 1295456 | 1305445 | 1305527 |
| SE | 108846 | ENSG00000081189 | MEF2C | chr5 | - | 88880923 | 88881022 | 88823900 | 88823930 | 88882954 | 88883227 |
| SE | 106693 | ENSG00000141376 | BCAS3 | chr17 | + | 60688019 | 60688195 | 60683981 | 60684036 | 60688931 | 60689046 |
| SE | 48647 | ENSG00000131374 | TBC1D5 | chr3 | - | 17586352 | 17586557 | 17518901 | 17518995 | 17623848 | 17623913 |
| SE | 70052 | ENSG00000151665 | PIGF | chr2 | - | 46588074 | 46588179 | 46581284 | 46581591 | 46591802 | 46591925 |
| SE | 92571 | ENSG00000148057 | IDNK | chr9 | + | 83628180 | 83628211 | 83623142 | 83623221 | 83628872 | 83628959 |
| SE | 85094 | ENSG00000139146 | SINHCAF | chr12 | - | 31296909 | 31297044 | 31295233 | 31295333 | 31298076 | 31298175 |
| SE | 3162 | ENSG00000213020 | ZNF611 | chr19 | - | 52714014 | 52714141 | 52702812 | 52706864 | 52715831 | 52715913 |
| SE | 41138 | ENSG00000124406 | ATP8A1 | chr4 | - | 42590810 | 42590884 | 42588259 | 42588329 | 42600477 | 42600518 |
| SE | 1256 | ENSG00000100083 | GGA1 | chr22 | + | 37624968 | 37625076 | 37623551 | 37623633 | 37629461 | 37629526 |
| SE | 80623 | ENSG00000258227 | CLEC5A | chr7 | - | 141935813 | 141935950 | 141931506 | 141931826 | 141946806 | 141946933 |
| SE | 46265 | ENSG00000162851 | TFB2M | chr1 | - | 246557380 | 246557534 | 246556572 | 246556721 | 246564345 | 246564434 |
| SE | 15180 | ENSG00000159459 | UBR1 | chr15 | - | 42983896 | 42983993 | 42977879 | 42977947 | 42984886 | 42984942 |
| SE | 87344 | ENSG00000215301 | DDX3X | chrX | + | 41347314 | 41347454 | 41346504 | 41346622 | 41347639 | 41349884 |
| SE | 15811 | ENSG00000169375 | SIN3A | chr15 | - | 75422646 | 75422823 | 75414204 | 75414311 | 75430186 | 75430408 |
| SE | 32131 | ENSG00000084463 | WBP11 | chr12 | - | 14796806 | 14797003 | 14794536 | 14794736 | 14799634 | 14799728 |
| SE | 22253 | ENSG00000141279 | NPEPPS | chr17 | + | 47619277 | 47619495 | 47619008 | 47619164 | 47619736 | 47619784 |
| SE | 12014 | ENSG00000170100 | ZNF778 | chr16 | + | 89222091 | 89222183 | 89220996 | 89221152 | 89223156 | 89223282 |
| SE | 86522 | ENSG00000085224 | ATRX | chrX | - | 77676225 | 77676298 | 77664644 | 77664778 | 77681519 | 77683451 |
| SE | 26649 | ENSG00000119729 | RHOQ | chr2 | + | 46543753 | 46543812 | 46542489 | 46543188 | 46576086 | 46576251 |
| SE | 11406 | ENSG00000040199 | PHLPP2 | chr16 | - | 71702597 | 71702731 | 71690518 | 71690709 | 71714511 | 71714801 |
| SE | 61598 | ENSG00000061273 | HDAC7 | chr12 | - | 47786584 | 47786703 | 47785450 | 47785885 | 47787711 | 47787809 |
| SE | 87539 | ENSG00000147050 | KDM6A | chrX | + | 45060021 | 45060156 | 45059246 | 45059466 | 45060608 | 45060764 |
| SE | 73845 | ENSG00000167792 | NDUFV1 | chr11 | + | 67611896 | 67611978 | 67611147 | 67611569 | 67612119 | 67612209 |
| SE | 92332 | ENSG00000137509 | PRCP | chr11 | - | 82849048 | 82849218 | 82843183 | 82843304 | 82849913 | 82850071 |
| SE | 1391 | ENSG00000100150 | DEPDC5 | chr22 | + | 31897481 | 31897653 | 31895358 | 31895954 | 31901741 | 31901802 |
| SE | 36918 | ENSG00000135622 | SEMA4F | chr2 | + | 74673456 | 74673576 | 74657564 | 74657624 | 74674497 | 74674676 |
| SE | 54321 | ENSG00000117139 | KDM5B | chr1 | - | 202762698 | 202762808 | 202760414 | 202760573 | 202764048 | 202764145 |
| SE | 11448 | ENSG00000187555 | USP7 | chr16 | - | 8896998 | 8897099 | 8895641 | 8895741 | 8898359 | 8898437 |
| SE | 39407 | ENSG00000240230 | COX19 | chr7 | - | 900532 | 900616 | 900101 | 900164 | 904125 | 904272 |
| SE | 16189 | ENSG00000196547 | MAN2A2 | chr15 | + | 90903230 | 90903734 | 90902925 | 90902975 | 90904189 | 90904339 |
| SE | 96107 | ENSG00000123360 | PDE1B | chr12 | + | 54576570 | 54576701 | 54575991 | 54576100 | 54577224 | 54577236 |

| | | | | | | | | | | | | | | |
|---|---|---|---|---|---|---|---|---|---|---|---|---|---|---|
| SE | 105201 | ENSG00000139624 | CERS5 | chr12 | - | 50144768 | 50144969 | 50143951 | 50144057 | 50167100 | 50167303 | 10 | 0 | 4 |
| SE | 66284 | ENSG00000149716 | LTO1 | chr11 | - | 69667478 | 69667587 | 69665648 | 69666409 | 69667894 | 69668012 | 44 | 19 | 39 |
| SE | 85683 | ENSG00000177646 | ACAD9 | chr3 | + | 128910740 | 128910813 | 128909343 | 128910149 | 128912506 | 128912705 | 52 | 44 | 66 |
| SE | 99531 | ENSG00000188690 | UROS | chr10 | - | 125796789 | 125796861 | 125796102 | 125796188 | 125798064 | 125798145 | 7 | 4 | 7 |
| SE | 102227 | ENSG00000100109 | TFIP11 | chr22 | - | 26511118 | 26511153 | 26510616 | 26510702 | 26512098 | 26512175 | 6 | 5 | 3 |
| SE | 12837 | ENSG00000288632 | NA | chr16 | - | 29554415 | 29554488 | 29553085 | 29553183 | 29554866 | 29554992 | 18 | 3 | 9 |
| SE | 49553 | ENSG00000070269 | TMEM260 | chr14 | + | 56603814 | 56603992 | 56585692 | 56585912 | 56605569 | 56605683 | 18 | 13 | 17 |
| SE | 22577 | ENSG00000137265 | IRF4 | chr6 | + | 398827 | 398935 | 397107 | 397252 | 401423 | 401777 | 94 | 32 | 217 |
| SE | 72715 | ENSG00000172765 | TMCC1 | chr3 | - | 129857249 | 129857405 | 129848342 | 129848412 | 129880308 | 129880559 | 26 | 15 | 10 |
| SE | 100734 | ENSG00000130830 | MPP1 | chrX | - | 154804696 | 154804827 | 154792141 | 154792285 | 154805271 | 154805456 | 0 | 3 | 0 |
| SE | 31647 | ENSG00000131389 | SLC6A6 | chr3 | + | 14468087 | 14468212 | 14467852 | 14467956 | 14479084 | 14479185 | 23 | 14 | 10 |
| SE | 578 | ENSG00000254413 | CHKB-CPT1B | chr22 | - | 50578548 | 50578659 | 50577774 | 50577934 | 50579035 | 50579255 | 58 | 35 | 14 |
| SE | 80076 | ENSG00000198198 | SZT2 | chr1 | + | 43419733 | 43419961 | 43416534 | 43416641 | 43420152 | 43420323 | 34 | 28 | 35 |
| SE | 46846 | ENSG00000155463 | OXA1L | chr14 | + | 22769796 | 22769934 | 22767957 | 22768171 | 22770192 | 22770625 | 207 | 135 | 208 |
| SE | 109352 | ENSG00000102710 | SUPT20H | chr13 | - | 37047345 | 37047408 | 37045246 | 37045373 | 37047534 | 37047601 | 11 | 8 | 12 |
| SE | 82044 | ENSG00000104133 | SPG11 | chr15 | - | 44610839 | 44610985 | 44608443 | 44608605 | 44613429 | 44613536 | 384 | 303 | 329 |
| SE | 101797 | ENSG00000090621 | PABPC4 | chr1 | - | 39563835 | 39563922 | 39563526 | 39563741 | 39564422 | 39564491 | 98 | 38 | 70 |
| SE | 74890 | ENSG00000187037 | GPR141 | chr7 | + | 37685459 | 37685583 | 37683765 | 37683903 | 37740379 | 37740705 | 67 | 29 | 26 |
| SE | 50707 | ENSG00000113318 | MSH3 | chr5 | + | 80761545 | 80761678 | 80744505 | 80744615 | 80767932 | 80768120 | 139 | 74 | 99 |
| SE | 54101 | ENSG00000115935 | WIPF1 | chr2 | - | 174629665 | 174629868 | 174585522 | 174585611 | 174634517 | 174634536 | 17 | 5 | 5 |
| SE | 15470 | ENSG00000140199 | SLC12A6 | chr15 | - | 34275344 | 34275389 | 34260925 | 34261020 | 34336409 | 34336601 | 258 | 203 | 227 |
| SE | 3628 | ENSG00000007047 | MARK4 | chr19 | + | 45263318 | 45263367 | 45262990 | 45263166 | 45264683 | 45264749 | 61 | 27 | 51 |
| SE | 80758 | ENSG00000136280 | CCM2 | chr7 | + | 45038252 | 45038426 | 45027025 | 45027797 | 45063917 | 45064001 | 178 | 116 | 185 |
| SE | 3446 | ENSG00000125734 | GPR108 | chr19 | - | 6735907 | 6735958 | 6735373 | 6735704 | 6736591 | 6736711 | 114 | 41 | 71 |
| SE | 71224 | ENSG00000079263 | SP140 | chr2 | + | 230253315 | 230253417 | 230245862 | 230245940 | 230255451 | 230255532 | 28 | 48 | 43 |
| SE | 96839 | ENSG00000149516 | MS4A3 | chr11 | + | 60062467 | 60062605 | 60061145 | 60061316 | 60064261 | 60064318 | 21 | 12 | 24 |
| SE | 20827 | ENSG00000107929 | LARP4B | chr10 | - | 825064 | 825316 | 820799 | 820845 | 825763 | 825870 | 386 | 218 | 319 |
| SE | 47317 | ENSG00000133106 | EPSTI1 | chr13 | - | 42919299 | 42919332 | 42917540 | 42917624 | 42926335 | 42926429 | 35 | 20 | 20 |
| SE | 48731 | ENSG00000076650 | GPATCH1 | chr19 | + | 33100172 | 33100274 | 33097754 | 33097902 | 33101494 | 33101574 | 6 | 0 | 0 |
| SE | 57321 | ENSG00000181722 | ZBTB20 | chr3 | - | 114582113 | 114582266 | 114500351 | 114500391 | 114693527 | 114693575 | 2 | 4 | 3 |
| SE | 93526 | ENSG00000119487 | MAPKAP1 | chr9 | - | 125506309 | 125506417 | 125467971 | 125468109 | 125543058 | 125543168 | 49 | 39 | 61 |
| SE | 41139 | ENSG00000124406 | ATP8A1 | chr4 | - | 42594288 | 42594362 | 42588259 | 42588329 | 42600477 | 42600518 | 126 | 80 | 129 |
| SE | 56199 | ENSG00000124942 | AHNAK | chr11 | - | 62491731 | 62491831 | 62433545 | 62433891 | 62535002 | 62535190 | 60 | 33 | 54 |
| SE | 106067 | ENSG00000123384 | LRP1 | chr12 | + | 57201033 | 57201153 | 57200698 | 57200815 | 57201496 | 57201619 | 261 | 110 | 154 |
| SE | 31318 | ENSG00000087266 | SH3BP2 | chr4 | + | 2825125 | 2825196 | 2824612 | 2824730 | 2827229 | 2827318 | 189 | 94 | 121 |
| SE | 41645 | ENSG00000119185 | ITGB1BP1 | chr2 | - | 9407448 | 9407598 | 9403474 | 9406905 | 9408112 | 9408205 | 356 | 152 | 254 |
| SE | 64560 | ENSG00000144036 | EXOC6B | chr2 | - | 72731154 | 72731245 | 72731006 | 72731052 | 72741303 | 72741469 | 66 | 59 | 61 |
| SE | 26004 | ENSG00000181929 | PRKAG1 | chr12 | - | 49005742 | 49005852 | 49005461 | 49005543 | 49013061 | 49013110 | 114 | 92 | 80 |
| SE | 106976 | ENSG00000111271 | ACAD10 | chr12 | + | 111729805 | 111729956 | 111727961 | 111728143 | 111736830 | 111737004 | 27 | 27 | 27 |
| SE | 37942 | ENSG00000138964 | PARVG | chr22 | + | 44181863 | 44181917 | 44181021 | 44181185 | 44183317 | 44183408 | 203 | 84 | 107 |
| SE | 35653 | ENSG00000161204 | ABCF3 | chr3 | + | 184187396 | 184187443 | 184186795 | 184186875 | 184187663 | 184187761 | 148 | 59 | 86 |
| SE | 76969 | ENSG00000131018 | SYNE1 | chr6 | - | 152463352 | 152463517 | 152462286 | 152462890 | 152465257 | 152465460 | 291 | 191 | 136 |
| SE | 108442 | ENSG00000067167 | TRAM1 | chr8 | - | 70598133 | 70598255 | 70597890 | 70598011 | 70600018 | 70600082 | 547 | 270 | 406 |
| SE | 17622 | ENSG00000092036 | HAUS4 | chr14 | - | 22950313 | 22950410 | 22947600 | 22947731 | 22952327 | 22952459 | 14 | 12 | 14 |
| SE | 12632 | ENSG00000090857 | PDPR | chr16 | + | 70136193 | 70136386 | 70132150 | 70132300 | 70138898 | 70139023 | 162 | 63 | 128 |
| SE | 42176 | ENSG00000255733 | IFNG-AS1 | chr12 | + | 68013530 | 68013648 | 68006089 | 68006146 | 68019861 | 68020331 | 1 | 1 | 2 |
| SE | 8434 | ENSG00000078687 | TNRC6C | chr17 | + | 78083046 | 78083166 | 78075135 | 78075278 | 78086502 | 78086586 | 61 | 32 | 116 |
| SE | 61493 | ENSG00000124588 | NQO2 | chr6 | + | 3004494 | 3004651 | 2993930 | 2994053 | 3006467 | 3006559 | 17 | 3 | 7 |
| SE | 30594 | ENSG00000152784 | PRDM8 | chr4 | + | 80200078 | 80200299 | 80197502 | 80197763 | 80201289 | 80201521 | 32 | 24 | 37 |
| SE | 67435 | ENSG00000115425 | PECR | chr2 | - | 216058894 | 216058976 | 216051448 | 216051545 | 216065311 | 216065477 | 51 | 7 | 49 |
| SE | 34015 | ENSG00000060971 | ACAA1 | chr3 | - | 38133951 | 38134009 | 38131925 | 38132005 | 38136591 | 38136624 | 91 | 79 | 79 |
| SE | 89851 | ENSG00000137770 | CTDSPL2 | chr15 | + | 44501961 | 44502048 | 44499726 | 44499813 | 44514597 | 44514660 | 15 | 6 | 16 |
| SE | 106221 | ENSG00000070214 | SLC44A1 | chr9 | + | 105356211 | 105356381 | 105348357 | 105348451 | 105358343 | 105358433 | 129 | 83 | 131 |
| SE | 71914 | ENSG00000143324 | XPR1 | chr1 | + | 180836521 | 180836716 | 180834873 | 180835045 | 180863707 | 180863874 | 102 | 66 | 79 |
| SE | 10827 | ENSG00000109046 | WSB1 | chr17 | + | 27301766 | 27301956 | 27294075 | 27294435 | 27303366 | 27303635 | 515 | 314 | 349 |
| SE | 109440 | ENSG00000121644 | DESI2 | chr1 | + | 244689248 | 244689342 | 244686596 | 244686669 | 244691883 | 244692020 | 62 | 42 | 39 |
| SE | 8132 | ENSG00000120071 | KANSL1 | chr17 | - | 46039701 | 46039884 | 46039026 | 46039215 | 46050532 | 46050694 | 344 | 129 | 268 |
| SE | 60075 | ENSG00000149532 | CPSF7 | chr11 | - | 61421053 | 61421130 | 61420469 | 61420573 | 61421389 | 61421608 | 13 | 9 | 3 |
| SE | 87758 | ENSG00000063601 | MTMR1 | chrX | + | 150736594 | 150736780 | 150732541 | 150732730 | 150737241 | 150737448 | 336 | 260 | 305 |
| SE | 86083 | ENSG00000169359 | SLC33A1 | chr3 | - | 155842431 | 155842619 | 155833467 | 155833585 | 155853222 | 155854250 | 148 | 153 | 147 |
| SE | 107364 | ENSG00000033030 | ZCCHC8 | chr12 | - | 122482634 | 122482695 | 122481944 | 122482087 | 122489385 | 122489463 | 75 | 60 | 64 |
| SE | 41615 | ENSG00000117640 | MTFR1L | chr1 | + | 25823345 | 25823439 | 25823018 | 25823128 | 25823643 | 25823748 | 2 | 4 | 1 |
| SE | 87434 | ENSG00000183354 | KIAA2026 | chr9 | - | 5933750 | 5933881 | 5929043 | 5929159 | 5944870 | 5944960 | 83 | 52 | 71 |
| SE | 105812 | ENSG00000111785 | RIC8B | chr12 | + | 106843851 | 106843947 | 106825725 | 106825820 | 106851449 | 106851594 | 46 | 38 | 48 |
| SE | 48894 | ENSG00000115107 | STEAP3 | chr2 | + | 119230619 | 119231034 | 119223833 | 119223888 | 119245488 | 119245988 | 3 | 1 | 1 |
| SE | 79980 | ENSG00000118058 | KMT2A | chr11 | + | 118482421 | 118482495 | 118481714 | 118482092 | 118484182 | 118484314 | 131 | 108 | 98 |
| SE | 92296 | ENSG00000157045 | NTAN1 | chr16 | - | 15047854 | 15047920 | 15047215 | 15047550 | 15055890 | 15055917 | 15 | 7 | 11 |
| SE | 51421 | ENSG00000143621 | ILF2 | chr1 | - | 153664042 | 153664130 | 153663018 | 153663276 | 153664395 | 153664474 | 873 | 613 | 850 |
| SE | 55933 | ENSG00000137338 | PGBD1 | chr6 | + | 28285550 | 28285707 | 28283755 | 28284209 | 28287079 | 28287168 | 10 | 2 | 3 |
| SE | 106201 | ENSG00000135090 | TAOK3 | chr12 | - | 118261496 | 118261673 | 118255503 | 118255655 | 118266654 | 118266759 | 74 | 81 | 44 |
| SE | 16519 | ENSG00000140367 | UBE2Q2 | chr15 | + | 75877960 | 75878021 | 75868950 | 75869010 | 75879097 | 75879188 | 116 | 110 | 84 |
| SE | 96708 | ENSG00000135362 | PRR5L | chr11 | + | 36451208 | 36451335 | 36446299 | 36446440 | 36462341 | 36464965 | 154 | 57 | 55 |
| SE | 72267 | ENSG00000186635 | ARAP1 | chr11 | - | 72687438 | 72687502 | 72685694 | 72686191 | 72687687 | 72687738 | 672 | 289 | 298 |

| | | | | | | | | | | | | | | | |
|---|---|---|---|---|---|---|---|---|---|---|---|---|---|---|---|
| SE | 107988 | ENSG00000204852 | TCTN1 | chr12 | + | 110634669 | 110634779 | 110632471 | 110632559 | 110636480 | 110636501 | 25 | 24 | 19 | [illegible] |
| SE | 29681 | ENSG00000137656 | BUD13 | chr11 | - | 116759073 | 116759179 | 116757145 | 116757227 | 116760734 | 116760952 | 78 | 30 | 38 | [illegible] |
| SE | 52390 | ENSG00000048707 | VPS13D | chr1 | + | 12358458 | 12358601 | 12356397 | 12356524 | 12362719 | 12362850 | 93 | 56 | 111 | [illegible] |
| SE | 27065 | ENSG00000062194 | GPBP1 | chr5 | + | 57221375 | 57221396 | 57214073 | 57214193 | 57230845 | 57230969 | 76 | 51 | 114 | [illegible] |
| SE | 103322 | ENSG00000134452 | FBH1 | chr10 | + | 5923620 | 5923696 | 5921447 | 5921569 | 5924310 | 5924508 | 166 | 118 | 206 | [illegible] |
| SE | 9106 | ENSG00000129657 | SEC14L1 | chr17 | + | 77213313 | 77213492 | 77209860 | 77212201 | 77213917 | 77214019 | 312 | 114 | 143 | [illegible] |
| SE | 106042 | ENSG00000204482 | LST1 | chr6 | + | 31587948 | 31587966 | 31587315 | 31587318 | 31588517 | 31588676 | 29 | 7 | 14 | [illegible] |
| SE | 79850 | ENSG00000131931 | THAP1 | chr8 | - | 42839185 | 42839381 | 42836673 | 42838336 | 42843023 | 42843201 | 79 | 82 | 74 | [illegible] |
| SE | 63577 | ENSG00000206562 | METTL6 | chr3 | - | 15414020 | 15414162 | 15409759 | 15411437 | 15415771 | 15415907 | 83 | 43 | 55 | [illegible] |
| SE | 89057 | ENSG00000104081 | BMF | chr15 | - | 40104242 | 40104340 | 40091786 | 40091888 | 40105794 | 40106086 | 15 | 2 | 4 | [illegible] |
| SE | 98294 | ENSG00000168067 | MAP4K2 | chr11 | - | 64798793 | 64798837 | 64797625 | 64797664 | 64799420 | 64799479 | 97 | 49 | 70 | [illegible] |
| SE | 51667 | ENSG00000136643 | RPS6KC1 | chr1 | + | 213077695 | 213077816 | 213071005 | 213071041 | 213104453 | 213104569 | 67 | 76 | 87 | [illegible] |
| SE | 36156 | ENSG00000041357 | PSMA4 | chr15 | + | 78542575 | 78542645 | 78542176 | 78542219 | 78544189 | 78544267 | 161 | 132 | 186 | [illegible] |
| SE | 100469 | ENSG00000235505 | CASP4LP | chr11 | - | 104906449 | 104907618 | 104905447 | 104905557 | 104908834 | 104909008 | 40 | 45 | 29 | [illegible] |
| SE | 77056 | ENSG00000125779 | PANK2 | chr20 | + | 3912457 | 3912634 | 3910576 | 3910830 | 3916926 | 3917050 | 517 | 405 | 493 | [illegible] |
| SE | 75935 | ENSG00000160185 | UBASH3A | chr21 | + | 42416441 | 42416611 | 42413023 | 42413222 | 42418400 | 42418609 | 75 | 45 | 73 | [illegible] |
| SE | 100118 | ENSG00000159063 | ALG8 | chr11 | - | 78102873 | 78102948 | 78100931 | 78101195 | 78103979 | 78104052 | 8 | 3 | 27 | [illegible] |
| SE | 8628 | ENSG00000271605 | MILR1 | chr17 | + | 64460821 | 64460932 | 64452596 | 64452866 | 64465451 | 64465541 | 60 | 64 | 31 | [illegible] |
| SE | 66341 | ENSG00000197037 | ZSCAN25 | chr7 | + | 99618134 | 99618209 | 99616965 | 99617016 | 99618524 | 99618651 | 2 | 0 | 3 | [illegible] |
| SE | 41745 | ENSG00000158882 | TOMM40L | chr1 | + | 161227635 | 161227737 | 161227257 | 161227350 | 161227883 | 161227989 | 29 | 13 | 22 | [illegible] |
| SE | 65023 | ENSG00000091409 | ITGA6 | chr2 | + | 172467480 | 172467560 | 172465538 | 172465663 | 172469124 | 172469380 | 129 | 80 | 192 | [illegible] |
| SE | 86209 | ENSG00000172534 | HCFC1 | chrX | - | 153958568 | 153958766 | 153958024 | 153958249 | 153959330 | 153959491 | 129 | 44 | 134 | [illegible] |
| SE | 80354 | ENSG00000102531 | FNDC3A | chr13 | + | 49175366 | 49175541 | 49168612 | 49168751 | 49178567 | 49178654 | 141 | 127 | 150 | [illegible] |
| SE | 105492 | ENSG00000152256 | PDK1 | chr2 | + | 172568740 | 172568817 | 172566855 | 172566933 | 172570725 | 172570812 | 56 | 79 | 97 | [illegible] |
| SE | 38419 | ENSG00000132549 | VPS13B | chr8 | + | 99156548 | 99156743 | 99147840 | 99148010 | 99170038 | 99170163 | 309 | 212 | 269 | [illegible] |
| SE | 104827 | ENSG00000205356 | TECPR1 | chr7 | - | 98233420 | 98233911 | 98232826 | 98232972 | 98238508 | 98238610 | 40 | 18 | 38 | [illegible] |
| SE | 67556 | ENSG00000150540 | HNMT | chr2 | + | 137967096 | 137967386 | 137964478 | 137964628 | 137970164 | 137970217 | 19 | 9 | 16 | [illegible] |
| SE | 105909 | ENSG00000111144 | LTA4H | chr12 | - | 96003837 | 96003981 | 96002989 | 96003064 | 96006313 | 96006409 | 652 | 384 | 410 | [illegible] |
| SE | 80815 | ENSG00000151948 | GLT1D1 | chr12 | + | 128899235 | 128899293 | 128888638 | 128888744 | 128914932 | 128914989 | 16 | 10 | 10 | [illegible] |
| SE | 104269 | ENSG00000132305 | IMMT | chr2 | - | 86161951 | 86162079 | 86159535 | 86159671 | 86166507 | 86166644 | 123 | 71 | 73 | [illegible] |
| SE | 82572 | ENSG00000127824 | TUBA4A | chr2 | - | 219252007 | 219252230 | 219251193 | 219251713 | 219253855 | 219253916 | 79 | 71 | 71 | [illegible] |
| SE | 51143 | ENSG00000165458 | INPPL1 | chr11 | + | 72228347 | 72228450 | 72227880 | 72228253 | 72228453 | 72228498 | 8 | 2 | 13 | [illegible] |
| SE | 29632 | ENSG00000110536 | PTPMT1 | chr11 | + | 47569699 | 47569891 | 47565457 | 47565986 | 47571470 | 47572360 | 81 | 53 | 131 | [illegible] |
| SE | 23858 | ENSG00000197603 | CPLANE1 | chr5 | - | 37243012 | 37243119 | 37239712 | 37239869 | 37244374 | 37244607 | 46 | 42 | 40 | [illegible] |
| SE | 17866 | ENSG00000092199 | HNRNPC | chr14 | - | 21263310 | 21263336 | 21233952 | 21234229 | 21269297 | 21269403 | 1457 | 774 | 1420 | [illegible] |
| SE | 74441 | ENSG00000138376 | BARD1 | chr2 | - | 214780967 | 214781509 | 214769231 | 214769312 | 214792296 | 214792445 | 47 | 38 | 30 | [illegible] |
| SE | 32546 | ENSG00000134602 | STK26 | chrX | + | 132069477 | 132069663 | 132068411 | 132068569 | 132071068 | 132071217 | 737 | 438 | 576 | [illegible] |
| SE | 50088 | ENSG00000167528 | ZNF641 | chr12 | - | 48345344 | 48345474 | 48344598 | 48344712 | 48347251 | 48347343 | 156 | 83 | 109 | [illegible] |
| SE | 9288 | ENSG00000170037 | CNTROB | chr17 | + | 7949081 | 7949157 | 7948486 | 7948619 | 7949384 | 7949578 | 12 | 16 | 8 | [illegible] |
| SE | 40806 | ENSG00000055332 | EIF2AK2 | chr2 | - | 37114730 | 37114859 | 37109193 | 37109295 | 37119958 | 37120139 | 347 | 460 | 254 | [illegible] |
| SE | 48654 | ENSG00000131374 | TBC1D5 | chr3 | - | 17586352 | 17586557 | 17576481 | 17576549 | 17623848 | 17623913 | 6 | 2 | 15 | [illegible] |
| SE | 100011 | ENSG00000166261 | ZNF202 | chr11 | - | 123727475 | 123727595 | 123723913 | 123726991 | 123728132 | 123728262 | 54 | 55 | 39 | [illegible] |
| SE | 46821 | ENSG00000183291 | SELENOF | chr1 | - | 86887155 | 86887196 | 86880661 | 86880725 | 86903280 | 86903352 | 23 | 6 | 9 | [illegible] |
| SE | 63894 | ENSG00000073614 | KDM5A | chr12 | - | 320994 | 321109 | 318105 | 318461 | 322416 | 322567 | 308 | 181 | 211 | [illegible] |
| SE | 91144 | ENSG00000116852 | KIF21B | chr1 | - | 201004339 | 201004455 | 201001406 | 201003781 | 201005307 | 201005442 | 123 | 64 | 158 | [illegible] |
| SE | 102690 | ENSG00000107938 | EDRF1 | chr10 | + | 125730287 | 125730399 | 125729357 | 125729479 | 125733403 | 125733551 | 141 | 111 | 119 | [illegible] |
| SE | 106336 | ENSG00000213923 | CSNK1E | chr22 | - | 38295386 | 38295447 | 38294341 | 38294534 | 38298785 | 38298934 | 7 | 2 | 0 | [illegible] |
| SE | 56622 | ENSG00000075151 | EIF4G3 | chr1 | - | 20997600 | 20997633 | 20981047 | 20981227 | 21002712 | 21002808 | 0 | 0 | 0 | [illegible] |
| SE | 68585 | ENSG00000163684 | RPP14 | chr3 | + | 58310308 | 58310406 | 58306261 | 58306651 | 58310506 | 58310591 | 14 | 5 | 6 | [illegible] |
| SE | 95880 | ENSG00000148396 | SEC16A | chr9 | - | 136462886 | 136463132 | 136461176 | 136461274 | 136463462 | 136463601 | 333 | 209 | 213 | [illegible] |
| SE | 60591 | ENSG00000070961 | ATP2B1 | chr12 | - | 89603711 | 89603925 | 89603042 | 89603254 | 89604154 | 89604346 | 490 | 472 | 339 | [illegible] |
| SE | 76845 | ENSG00000124788 | ATXN1 | chr6 | - | 16753232 | 16753347 | 16735453 | 16739918 | 16761297 | 16761370 | 36 | 9 | 17 | [illegible] |
| SE | 80268 | ENSG00000164535 | DAGLB | chr7 | - | 6434761 | 6435020 | 6432836 | 6432959 | 6436361 | 6436533 | 90 | 73 | 80 | [illegible] |
| SE | 60863 | ENSG00000237298 | TTN-AS1 | chr2 | + | 178591138 | 178591314 | 178542221 | 178542361 | 178597595 | 178597681 | 26 | 8 | 14 | [illegible] |
| SE | 80812 | ENSG00000151948 | GLT1D1 | chr12 | + | 128899235 | 128899287 | 128888638 | 128888744 | 128926358 | 128926454 | 23 | 29 | 26 | [illegible] |
| SE | 75516 | ENSG00000184281 | TSSC4 | chr11 | + | 2401838 | 2402147 | 2400762 | 2400798 | 2402293 | 2402450 | 9 | 2 | 4 | [illegible] |
| SE | 16269 | ENSG00000124299 | PEPD | chr19 | - | 33396089 | 33396155 | 33391294 | 33391479 | 33401720 | 33401869 | 0 | 0 | 0 | [illegible] |
| SE | 9135 | ENSG00000161542 | PRPSAP1 | chr17 | - | 76344670 | 76344737 | 76332262 | 76332435 | 76348528 | 76348581 | 115 | 52 | 65 | [illegible] |
| SE | 77651 | ENSG00000214021 | TTLL3 | chr3 | + | 9817644 | 9817759 | 9816073 | 9816202 | 9820545 | 9820741 | 18 | 14 | 13 | [illegible] |
| SE | 49965 | ENSG00000162976 | SLC66A3 | chr2 | + | 11172750 | 11172824 | 11171924 | 11172045 | 11174967 | 11175009 | 3 | 4 | 1 | [illegible] |
| SE | 56397 | ENSG00000151838 | CCDC175 | chr14 | - | 59531771 | 59531910 | 59525281 | 59525434 | 59538022 | 59538154 | 2 | 8 | 1 | [illegible] |
| SE | 74628 | ENSG00000085998 | POMGNT1 | chr1 | - | 46194552 | 46194651 | 46194273 | 46194401 | 46194843 | 46194961 | 84 | 28 | 57 | [illegible] |
| SE | 32968 | ENSG00000114127 | XRN1 | chr3 | - | 142426743 | 142426841 | 142423559 | 142423642 | 142432660 | 142432893 | 248 | 219 | 242 | [illegible] |
| SE | 5471 | ENSG00000088356 | PDRG1 | chr20 | - | 31950311 | 31950387 | 31948807 | 31948882 | 31951874 | 31952046 | 51 | 11 | 58 | [illegible] |
| SE | 57276 | ENSG00000143119 | CD53 | chr1 | + | 110894959 | 110895055 | 110894308 | 110894401 | 110897808 | 110897892 | 596 | 257 | 468 | [illegible] |
| SE | 31243 | ENSG00000121060 | TRIM25 | chr17 | - | 56912104 | 56912165 | 56908467 | 56908563 | 56913391 | 56914049 | 5 | 0 | 0 | [illegible] |
| SE | 16493 | ENSG00000140474 | ULK3 | chr15 | - | 74838265 | 74838344 | 74838151 | 74838192 | 74838439 | 74838504 | 45 | 23 | 50 | [illegible] |
| SE | 69684 | ENSG00000004897 | CDC27 | chr17 | - | 47141852 | 47142025 | 47138738 | 47138891 | 47142228 | 47142436 | 161 | 142 | 148 | [illegible] |
| SE | 38689 | ENSG00000167258 | CDK12 | chr17 | + | 39551037 | 39551142 | 39547820 | 39547830 | 39556285 | 39556413 | 24 | 18 | 19 | [illegible] |
| SE | 23311 | ENSG00000119862 | LGALSL | chr2 | + | 64455588 | 64455677 | 64455343 | 64455415 | 64456287 | 64456465 | 59 | 104 | 13 | [illegible] |
| SE | 46529 | ENSG00000223745 | CCDC18-AS1 | chr1 | - | 93309714 | 93309790 | 93305387 | 93305470 | 93324634 | 93324649 | 10 | 0 | 6 | [illegible] |
| SE | 35796 | ENSG00000173226 | IQCB1 | chr3 | - | 121795456 | 121795566 | 121790072 | 121790215 | 121799195 | 121799374 | 85 | 74 | 83 | [illegible] |
| SE | 54664 | ENSG00000163877 | SNIP1 | chr1 | - | 37548820 | 37548987 | 37540156 | 37540755 | 37552644 | 37552747 | 2 | 0 | 0 | [illegible] |
| SE | 20307 | ENSG00000111913 | RIPOR2 | chr6 | - | 24832255 | 24832391 | 24830508 | 24830670 | 24835702 | 24835871 | 613 | 327 | 614 | [illegible] |
| SE | 43479 | ENSG00000145293 | ENOPH1 | chr4 | + | 82451042 | 82451245 | 82447919 | 82448021 | 82454721 | 82454854 | 305 | 155 | 212 | [illegible] |
| SE | 12816 | ENSG00000149923 | PPP4C | chr16 | + | 30082745 | 30082847 | 30082483 | 30082534 | 30083393 | 30083567 | 226 | 106 | 178 | [illegible] |
| SE | 43072 | ENSG00000178917 | ZNF852 | chr3 | - | 44502964 | 44503056 | 44502541 | 44502668 | 44510552 | 44510636 | 37 | 34 | 42 | [illegible] |
| SE | 33563 | ENSG00000009790 | TRAF3IP3 | chr1 | + | 209760171 | 209760384 | 209759033 | 209759134 | 209762514 | 209762596 | 253 | 158 | 195 | [illegible] |
| SE | 79778 | ENSG00000164808 | SPIDR | chr8 | + | 47729411 | 47729465 | 47728932 | 47729047 | 47735306 | 47735374 | 48 | 35 | 29 | [illegible] |
| SE | 74635 | ENSG00000134440 | NARS1 | chr18 | - | 57607133 | 57607333 | 57606615 | 57606751 | 57607443 | 57607483 | 368 | 228 | 312 | [illegible] |
| SE | 13290 | ENSG00000102974 | CTCF | chr16 | + | 67610823 | 67611613 | 67570772 | 67571264 | 67616744 | 67616878 | 95 | 57 | 77 | [illegible] |
| SE | 35795 | ENSG00000173226 | IQCB1 | chr3 | - | 121795456 | 121795566 | 121790072 | 121790215 | 121797117 | 121797227 | 135 | 128 | 137 | [illegible] |
| SE | 108089 | ENSG00000116212 | LRRC42 | chr1 | + | 53947787 | 53947821 | 53946349 | 53946549 | 53951985 | 53952472 | 3 | 2 | 5 | [illegible] |
| SE | 51914 | ENSG00000074603 | DPP8 | chr15 | - | 65454262 | 65454415 | 65451959 | 65452102 | 65455653 | 65455691 | 93 | 57 | 64 | [illegible] |
| SE | 92486 | ENSG00000130749 | ZC3H4 | chr19 | - | 47069091 | 47069343 | 47064186 | 47067869 | 47071777 | 47072121 | 193 | 91 | 124 | [illegible] |
| SE | 7957 | ENSG00000184640 | SEPTIN9 | chr17 | + | 77451366 | 77451528 | 77450575 | 77450786 | 77482143 | 77482335 | 34 | 8 | 28 | [illegible] |
| SE | 34086 | ENSG00000114867 | EIF4G1 | chr3 | + | 184325850 | 184325951 | 184325298 | 184325639 | 184326526 | 184326629 | 554 | 364 | 517 | [illegible] |
| SE | 28003 | ENSG00000137601 | NEK1 | chr4 | - | 169555719 | 169555851 | 169537808 | 169537911 | 169555931 | 169556095 | 29 | 36 | 22 | [illegible] |
| SE | 58260 | ENSG00000089127 | OAS1 | chr12 | + | 112918596 | 112918617 | 112917546 | 112917700 | 112919486 | 112919909 | 7 | 4 | 1 | [illegible] |
| SE | 42118 | ENSG00000106400 | ZNHIT1 | chr7 | + | 101218318 | 101218485 | 101218165 | 101218217 | 101222603 | 101222774 | 4 | 0 | 1 | [illegible] |
| SE | 28506 | ENSG00000169062 | UPF3A | chr13 | + | 114291604 | 114291792 | 114286518 | 114286629 | 114298839 | 114299000 | 18 | 23 | 20 | [illegible] |
| SE | 34804 | ENSG00000136731 | UGGT1 | chr2 | + | 128103078 | 128103145 | 128097428 | 128097564 | 128103931 | 128104014 | 6 | 8 | 9 | [illegible] |
| SE | 49401 | ENSG00000085231 | AK6 | chr5 | - | 69366502 | 69366595 | 69355894 | 69355953 | 69369462 | 69369490 | 43 | 19 | 39 | [illegible] |
| SE | 82271 | ENSG00000223482 | NUTM2A-AS1 | chr10 | - | 87330891 | 87330965 | 87326629 | 87326672 | 87336785 | 87336889 | 7 | 6 | 3 | [illegible] |
| SE | 28221 | ENSG00000204084 | INPP5B | chr1 | - | 37885637 | 37885825 | 37882806 | 37882918 | 37886887 | 37887004 | 168 | 99 | 155 | [illegible] |
| SE | 105987 | ENSG00000122965 | RBM19 | chr12 | - | 113920610 | 113920690 | 113918391 | 113918447 | 113924696 | 113924757 | 115 | 60 | 123 | [illegible] |
| SE | 46606 | ENSG00000141556 | TBCD | chr17 | + | 82781588 | 82781721 | 82772451 | 82772507 | 82797756 | 82797802 | 53 | 32 | 34 | [illegible] |
| SE | 82368 | ENSG00000204257 | HLA-DMA | chr6 | - | 32952283 | 32952461 | 32950083 | 32950803 | 32952948 | 32953034 | 2 | 3 | 3 | [illegible] |
| SE | 25699 | ENSG00000247774 | PCED1B-AS1 | chr12 | - | 47206510 | 47206588 | 47205897 | 47206069 | 47210639 | 47210761 | 70 | 37 | 151 | [illegible] |
| SE | 59167 | ENSG00000115204 | MPV17 | chr2 | - | 27312993 | 27313154 | 27312679 | 27312772 | 27322447 | 27322522 | 40 | 27 | 22 | [illegible] |
| SE | 55339 | ENSG00000204469 | PRRC2A | chr6 | + | 31623731 | 31623909 | 31622689 | 31622901 | 31624260 | 31624360 | 285 | 193 | 278 | [illegible] |
| SE | 100300 | ENSG00000162144 | CYB561A3 | chr11 | - | 61357202 | 61357371 | 61356529 | 61356728 | 61357732 | 61357828 | 1 | 2 | 5 | [illegible] |
| SE | 90390 | ENSG00000249859 | PVT1 | chr8 | + | 128099369 | 128099573 | 128096517 | 128096654 | 128100979 | 128101252 | 1 | 0 | 1 | [illegible] |
| SE | 28047 | ENSG00000108961 | RANGRF | chr17 | + | 8289502 | 8289588 | 8289257 | 8289414 | 8289812 | 8290087 | 38 | 15 | 33 | [illegible] |
| SE | 14803 | ENSG00000137815 | RTF1 | chr15 | + | 41452900 | 41453048 | 41438320 | 41438431 | 41457671 | 41457876 | 143 | 89 | 126 | [illegible] |
| SE | 16930 | ENSG00000171606 | ZNF274 | chr19 | + | 58209960 | 58210073 | 58206719 | 58207202 | 58211559 | 58211686 | 28 | 13 | 21 | [illegible] |
| SE | 40420 | ENSG00000078304 | PPP2R5C | chr14 | + | 101831700 | 101831804 | 101809868 | 101810036 | 101836168 | 101836432 | 5 | 1 | 15 | [illegible] |
| SE | 18955 | ENSG00000015133 | CCDC88C | chr14 | - | 91281161 | 91281308 | 91279237 | 91279306 | 91281456 | 91281525 | 29 | 8 | 20 | [illegible] |
| SE | 22947 | ENSG00000170234 | PWWP2A | chr5 | - | 160080650 | 160080770 | 160075881 | 160078168 | 160089505 | 160089610 | 53 | 9 | 17 | [illegible] |
| SE | 21065 | ENSG00000196182 | STK40 | chr1 | - | 36361220 | 36361340 | 36358736 | 36358822 | 36367851 | 36368084 | 200 | 138 | 125 | [illegible] |
| SE | 39462 | ENSG00000213024 | NUP62 | chr19 | - | 49927693 | 49927815 | 49906824 | 49909884 | 49929325 | 49929429 | 105 | 77 | 98 | [illegible] |
| SE | 41598 | ENSG00000118308 | IRAG2 | chr12 | + | 25066364 | 25066512 | 25063719 | 25063816 | 25069349 | 25069431 | 290 | 236 | 260 | [illegible] |
| SE | 10042 | ENSG00000154217 | PITPNC1 | chr17 | + | 67687065 | 67687184 | 67675478 | 67675542 | 67692571 | 67693367 | 599 | 305 | 538 | [illegible] |
| SE | 13018 | ENSG00000140987 | ZSCAN32 | chr16 | - | 3390009 | 3390183 | 3384083 | 3384941 | 3397191 | 3397787 | 17 | 6 | 16 | [illegible] |
| SE | 107689 | ENSG00000128585 | MKLN1 | chr7 | + | 131445773 | 131445903 | 131443480 | 131443702 | 131463216 | 131463364 | 277 | 218 | 208 | [illegible] |
| SE | 86014 | ENSG00000244754 | N4BP2L2 | chr13 | - | 32527457 | 32527532 | 32522181 | 32522270 | 32535768 | 32537027 | 176 | 96 | 135 | [illegible] |
| SE | 46614 | ENSG00000141556 | TBCD | chr17 | + | 82895700 | 82895967 | 82893546 | 82893632 | 82900650 | 82900731 | 0 | 0 | 7 | [illegible] |
| SE | 65597 | ENSG00000112078 | KCTD20 | chr6 | + | 36451533 | 36451632 | 36443037 | 36443111 | 36470051 | 36470257 | 5 | 0 | 1 | [illegible] |
| SE | 47536 | ENSG00000121766 | ZCCHC17 | chr1 | + | 31348828 | 31348974 | 31346639 | 31346740 | 31364031 | 31364552 | 123 | 99 | 80 | [illegible] |
| SE | 11446 | ENSG00000187555 | USP7 | chr16 | - | 8894549 | 8894640 | 8892096 | 8894104 | 8894783 | 8894855 | 485 | 306 | 489 | [illegible] |
| SE | 61655 | ENSG00000009307 | CSDE1 | chr1 | - | 114737470 | 114737563 | 114736757 | 114736855 | 114737962 | 114738072 | 122 | 76 | 85 | [illegible] |
| SE | 38650 | ENSG00000073417 | PDE8A | chr15 | + | 85089337 | 85089416 | 85083555 | 85083644 | 85091043 | 85091181 | 46 | 31 | 37 | [illegible] |
| SE | 81963 | ENSG00000051825 | MPHOSPH9 | chr12 | - | 123223037 | 123223127 | 123221371 | 123221895 | 123227462 | 123227572 | 127 | 110 | 156 | [illegible] |
| SE | 101661 | ENSG00000112425 | EPM2A | chr6 | - | 145720900 | 145721155 | 145686121 | 145686296 | 145735197 | 145735470 | 5 | 2 | 5 | [illegible] |
| SE | 18676 | ENSG00000129566 | TEP1 | chr14 | - | 20378753 | 20378853 | 20378379 | 20378535 | 20378980 | 20379105 | 32 | 23 | 37 | [illegible] |
| SE | 89196 | ENSG00000147649 | MTDH | chr8 | + | 97719048 | 97719189 | 97713661 | 97713769 | 97722878 | 97723035 | 851 | 753 | 868 | [illegible] |
| SE | 95925 | ENSG00000188352 | FOCAD | chr9 | + | 20978338 | 20978454 | 20976321 | 20976548 | 20981425 | 20981461 | 21 | 12 | 36 | [illegible] |
| SE | 4902 | ENSG00000101363 | MANBAL | chr20 | + | 37298762 | 37298879 | 37296895 | 37297534 | 37301207 | 37301413 | 12 | 3 | 7 | [illegible] |
| SE | 19243 | ENSG00000140943 | MBTPS1 | chr16 | - | 84074596 | 84074741 | 84070587 | 84070776 | 84081746 | 84081908 | 222 | 127 | 228 | [illegible] |
| SE | 49284 | ENSG00000120733 | KDM3B | chr5 | + | 138391012 | 138392261 | 138381515 | 138381590 | 138393170 | 138393372 | 87 | 65 | 121 | [illegible] |
| SE | 19481 | ENSG00000033170 | FUT8 | chr14 | + | 65617089 | 65617191 | 65616210 | 65616373 | 65629491 | 65629606 | 3 | 11 | 2 | [illegible] |
| SE | 61910 | ENSG00000124067 | SLC12A4 | chr16 | - | 67950653 | 67951060 | 67950318 | 67950493 | 67951139 | 67951304 | 7 | 2 | 3 | [illegible] |
| SE | 40697 | ENSG00000117448 | AKR1A1 | chr1 | + | 45552435 | 45552563 | 45550542 | 45551155 | 45561878 | 45561878 | 42 | 33 | 34 | [illegible] |
| SE | 32507 | ENSG00000114857 | NKTR | chr3 | + | 42627199 | 42627227 | 42621004 | 42621516 | 42630545 | 42630575 | 32 | 33 | 41 | [illegible] |
| SE | 77834 | ENSG00000256043 | CTSO | chr4 | - | 155928335 | 155928424 | 155924383 | 155926070 | 155929541 | 155929705 | 88 | 51 | 54 | [illegible] |
| SE | 83523 | ENSG00000161203 | AP2M1 | chr3 | + | 184178856 | 184179191 | 184176950 | 184177067 | 184180168 | 184180212 | 454 | 231 | 293 | [illegible] |
| SE | 1018 | ENSG00000100335 | MIEF1 | chr22 | + | 39509421 | 39509571 | 39504202 | 39504534 | 39511287 | 39511438 | 9 | 4 | 8 | [illegible] |
| SE | 16927 | ENSG00000171606 | ZNF274 | chr19 | + | 58186946 | 58187042 | 58183920 | 58183998 | 58206719 | 58207089 | 35 | 31 | 44 | [illegible] |
| SE | 84079 | ENSG00000102316 | MAGED2 | chrX | + | 54812944 | 54813024 | 54812156 | 54812251 | 54813487 | 54813550 | 96 | 63 | 105 | [illegible] |
| SE | 48203 | ENSG00000196204 | RNF216P1 | chr7 | + | 4986566 | 4986631 | 4984909 | 4985075 | 4989062 | 4989177 | 0 | 2 | 3 | [illegible] |
| SE | 61028 | ENSG00000066027 | PPP2R5A | chr1 | + | 212306782 | 212306900 | 212285409 | 212286291 | 212329134 | 212329331 | 1 | 0 | 0 | [illegible] |
| SE | 106439 | ENSG00000135297 | MTO1 | chr6 | + | 73486597 | 73486761 | 73482448 | 73482620 | 73492233 | 73492352 | 8 | 6 | 10 | [illegible] |
| SE | 68576 | ENSG00000140368 | PSTPIP1 | chr15 | + | 77032297 | 77032428 | 77031179 | 77031278 | 77035507 | 77035563 | 73 | 17 | 86 | [illegible] |
| SE | 26761 | ENSG00000127804 | METTL16 | chr17 | - | 2473523 | 2473664 | 2464207 | 2464350 | 2477685 | 2477885 | 128 | 87 | 122 | [illegible] |
| SE | 74785 | ENSG00000011198 | ABHD5 | chr3 | + | 43717670 | 43717857 | 43714946 | 43715058 | 43718442 | 43718561 | 198 | 129 | 125 | [illegible] |
| SE | 25978 | ENSG00000137364 | TPMT | chr6 | - | 18132132 | 18132177 | 18128310 | 18130780 | 18133803 | 18133889 | 85 | 63 | 52 | [illegible] |
| SE | 11307 | ENSG00000140650 | PMM2 | chr16 | + | 8849368 | 8849655 | 8812990 | 8813106 | 8849848 | 8849993 | 3 | 12 | 4 | [illegible] |
| SE | 39096 | ENSG00000109320 | NFKB1 | chr4 | + | 102533847 | 102533885 | 102529835 | 102529914 | 102537857 | 102537899 | 99 | 78 | 89 | [illegible] |
| SE | 21202 | ENSG00000100219 | XBP1 | chr22 | - | 28796046 | 28796192 | 28794554 | 28795732 | 28797076 | 28797205 | 649 | 287 | 758 | [illegible] |
| SE | 5991 | ENSG00000062598 | ELMO2 | chr20 | - | 46386123 | 46386275 | 46383415 | 46383494 | 46389038 | 46389220 | 89 | 34 | 54 | [illegible] |
| SE | 91193 | ENSG00000288596 | C8orf44 | chr8 | + | 66677641 | 66677954 | 66667617 | 66667701 | 66679720 | 66681000 | 1 | 1 | 0 | [illegible] |
| SE | 9008 | ENSG00000182446 | NPLOC4 | chr17 | - | 81563920 | 81563979 | 81559090 | 81559416 | 81565504 | 81565607 | 9 | 9 | 2 | [illegible] |
| SE | 27778 | ENSG00000177058 | SLC38A9 | chr5 | - | 55627890 | 55627980 | 55625844 | 55626659 | 55645788 | 55645895 | 8 | 14 | 11 | [illegible] |
| SE | 26395 | ENSG00000138744 | NAAA | chr4 | - | 75914867 | 75914985 | 75913659 | 75914338 | 75918760 | 75918789 | 136 | 112 | 130 | [illegible] |
| SE | 58214 | ENSG00000172965 | MIR4435-2HG | chr2 | - | 111365560 | 111367575 | 111344060 | 111344237 | 111494884 | 111494933 | 123 | 57 | 59 | [illegible] |
| SE | 951 | ENSG00000100418 | DESI1 | chr22 | - | 41607261 | 41607331 | 41604043 | 41604153 | 41620751 | 41621043 | 33 | 8 | 18 | [illegible] |
| SE | 73343 | ENSG00000115970 | THADA | chr2 | - | 43577021 | 43577242 | 43574335 | 43575027 | 43578512 | 43578607 | 124 | 77 | 115 | [illegible] |
| SE | 39439 | ENSG00000055917 | PUM2 | chr2 | - | 20290651 | 20290790 | 20283342 | 20283486 | 20294375 | 20294518 | 121 | 69 | 98 | [illegible] |
| SE | 15829 | ENSG00000284946 | NA | chr15 | - | 90980883 | 90981669 | 90980241 | 90980389 | 90981747 | 90981981 | 18 | 18 | 23 | [illegible] |
| SE | 30167 | ENSG00000163811 | WDR43 | chr2 | + | 28941460 | 28941574 | 28937930 | 28937994 | 28942311 | 28942381 | 101 | 81 | 175 | [illegible] |
| SE | 6958 | ENSG00000105656 | ELL | chr19 | - | 18458204 | 18458329 | 18451551 | 18451648 | 18461577 | 18461852 | 57 | 9 | 26 | [illegible] |
| SE | 31575 | ENSG00000116668 | SWT1 | chr1 | + | 185221848 | 185222036 | 185206624 | 185206763 | 185231576 | 185231708 | 54 | 43 | 40 | [illegible] |
| SE | 77790 | ENSG00000152457 | DCLRE1C | chr10 | - | 14947524 | 14947609 | 14945434 | 14945565 | 14949035 | 14949087 | 12 | 6 | 3 | [illegible] |
| SE | 43884 | ENSG00000179335 | CLK3 | chr15 | + | 74622119 | 74622216 | 74619966 | 74620225 | 74624901 | 74625018 | 49 | 31 | 54 | [illegible] |
| SE | 14874 | ENSG00000128989 | ARPP19 | chr15 | - | 52568847 | 52568902 | 52557099 | 52557222 | 52569115 | 52569154 | 156 | 89 | 120 | [illegible] |
| SE | 81965 | ENSG00000051825 | MPHOSPH9 | chr12 | - | 123226287 | 123226351 | 123223037 | 123223127 | 123227462 | 123227616 | 16 | 31 | 0 | [illegible] |
| SE | 98959 | ENSG00000172932 | ANKRD13D | chr11 | + | 67299057 | 67299124 | 67291990 | 67292180 | 67299529 | 67299611 | 155 | 51 | 120 | [illegible] |
| SE | 15624 | ENSG00000072062 | PRKACA | chr19 | - | 14093627 | 14093792 | 14091687 | 14093237 | 14097360 | 14097483 | 546 | 203 | 285 | [illegible] |
| SE | 16150 | ENSG00000140400 | MAN2C1 | chr15 | - | 75363998 | 75364188 | 75362641 | 75362748 | 75364487 | 75364665 | 128 | 53 | 154 | [illegible] |
| SE | 106471 | ENSG00000198270 | TMEM116 | chr12 | - | 111943264 | 111943369 | 111938160 | 111938210 | 111991757 | 111991889 | 68 | 44 | 52 | [illegible] |
| SE | 12817 | ENSG00000149923 | PPP4C | chr16 | + | 30082745 | 30082963 | 30082483 | 30082534 | 30083393 | 30083567 | 124 | 52 | 102 | [illegible] |
| SE | 88192 | ENSG00000136279 | DBNL | chr7 | + | 44051829 | 44051942 | 44050224 | 44050280 | 44056756 | 44056903 | 67 | 27 | 72 | [illegible] |
| SE | 29897 | ENSG00000010165 | EEF1AKNMT | chr1 | + | 171792016 | 171792235 | 171790451 | 171790616 | 171794395 | 171794527 | 73 | 34 | 76 | [illegible] |
| SE | 81071 | ENSG00000157764 | BRAF | chr7 | - | 140753274 | 140754233 | 140749286 | 140749418 | 140776911 | 140777088 | 152 | 115 | 142 | [illegible] |
| SE | 83364 | ENSG00000087274 | ADD1 | chr4 | + | 2926641 | 2926675 | 2926013 | 2926112 | 2928170 | 2928523 | 1 | 3 | 5 | [illegible] |
| SE | 58656 | ENSG00000125633 | CCDC93 | chr2 | - | 117944023 | 117944086 | 117941188 | 117941297 | 117945528 | 117945582 | 157 | 139 | 154 | [illegible] |
| SE | 26108 | ENSG00000164181 | ELOVL7 | chr5 | - | 60799179 | 60799230 | 60787333 | 60787431 | 60844159 | 60844215 | 23 | 40 | 10 | [illegible] |
| SE | 41498 | ENSG00000126107 | HECTD3 | chr1 | - | 45009985 | 45010121 | 45009567 | 45009683 | 45010200 | 45010293 | 71 | 32 | 63 | [illegible] |
| SE | 4015 | ENSG00000104856 | RELB | chr19 | + | 45009813 | 45009822 | 45002948 | 45002996 | 45011935 | 45012229 | 6 | 6 | 6 | [illegible] |
| SE | 102372 | ENSG00000019995 | ZRANB1 | chr10 | + | 124983174 | 124983304 | 124974199 | 124974398 | 124984773 | 124988189 | 3 | 6 | 6 | [illegible] |
| SE | 44177 | ENSG00000027869 | SH2D2A | chr1 | - | 156809202 | 156809490 | 156807174 | 156807345 | 156809660 | 156809807 | 213 | 53 | 81 | [illegible] |
| SE | 39665 | ENSG00000181555 | SETD2 | chr3 | - | 47083719 | 47084382 | 47066991 | 47067118 | 47097954 | 47097961 | 191 | 163 | 163 | [illegible] |
| SE | 79338 | ENSG00000231721 | LINC-PINT | chr7 | - | 130959868 | 130959970 | 130945719 | 130945835 | 130984052 | 130984109 | 8 | 6 | 1 | [illegible] |
| SE | 51919 | ENSG00000074603 | DPP8 | chr15 | - | 65455711 | 65455841 | 65454262 | 65454415 | 65456224 | 65456371 | 12 | 4 | 13 | [illegible] |
| SE | 24142 | ENSG00000143774 | GUK1 | chr1 | + | 228141123 | 228141507 | 228140317 | 228140363 | 228145510 | 228145624 | 5 | 1 | 3 | [illegible] |
| SE | 9332 | ENSG00000108439 | PNPO | chr17 | + | 47945860 | 47945989 | 47944615 | 47945612 | 47946322 | 47946393 | 60 | 22 | 62 | [illegible] |
| SE | 11301 | ENSG00000140650 | PMM2 | chr16 | + | 8806315 | 8806407 | 8801798 | 8801910 | 8811078 | 8811111 | 53 | 41 | 66 | [illegible] |
| SE | 36699 | ENSG00000158062 | UBXN11 | chr1 | - | 26302812 | 26302918 | 26297961 | 26298062 | 26306591 | 26306609 | 16 | 33 | 15 | [illegible] |
| SE | 32241 | ENSG00000156931 | VPS8 | chr3 | + | 184868945 | 184869036 | 184868023 | 184868059 | 184869481 | 184869528 | 88 | 90 | 76 | [illegible] |
| SE | 51066 | ENSG00000104231 | ZFAND1 | chr8 | - | 81717248 | 81717288 | 81714986 | 81715114 | 81718181 | 81718224 | 132 | 107 | 169 | [illegible] |
| SE | 99110 | ENSG00000110274 | CEP164 | chr11 | + | 117408889 | 117409028 | 117407924 | 117408032 | 117409617 | 117409965 | 46 | 43 | 61 | [illegible] |
| SE | 104146 | ENSG00000023041 | ZDHHC6 | chr10 | - | 112440533 | 112441812 | 112438335 | 112438389 | 112442191 | 112442351 | 175 | 186 | 148 | [illegible] |
| SE | 62641 | ENSG00000128815 | WDFY4 | chr10 | + | 48779940 | 48780119 | 48778610 | 48778832 | 48786638 | 48786870 | 98 | 62 | 88 | [illegible] |
| SE | 53147 | ENSG00000124574 | ABCC10 | chr6 | + | 43430747 | 43430866 | 43427967 | 43428139 | 43432141 | 43432176 | 4 | 0 | 0 | [illegible] |
| SE | 25957 | ENSG00000139278 | GLIPR1 | chr12 | + | 75490405 | 75490518 | 75481833 | 75482079 | 75495576 | 75495662 | 1545 | 769 | 895 | [illegible] |
| SE | 24174 | ENSG00000124357 | NAGK | chr2 | + | 71075561 | 71075642 | 71073481 | 71073594 | 71076603 | 71076701 | 85 | 35 | 42 | [illegible] |
| SE | 70060 | ENSG00000151665 | PIGF | chr2 | - | 46614936 | 46615185 | 46613693 | 46613785 | 46616969 | 46617041 | 85 | 42 | 65 | [illegible] |
| SE | 41806 | ENSG00000136059 | VILL | chr3 | + | 38003167 | 38003313 | 38002395 | 38002575 | 38004254 | 38004399 | 24 | 11 | 39 | [illegible] |
| SE | 103928 | ENSG00000188234 | AGAP4 | chr10 | - | 45848842 | 45848912 | 45847124 | 45847312 | 45853654 | 45853875 | 4 | 2 | 3 | [illegible] |
| SE | 82702 | ENSG00000127914 | AKAP9 | chr7 | + | 91979435 | 91979574 | 91973710 | 91973968 | 91980288 | 91980333 | 13 | 6 | 13 | [illegible] |
| SE | 63272 | ENSG00000095002 | MSH2 | chr2 | + | 47414268 | 47414418 | 47412413 | 47412560 | 47416295 | 47416429 | 234 | 188 | 264 | [illegible] |
| SE | 80831 | ENSG00000151948 | GLT1D1 | chr12 | + | 128926358 | 128926454 | 128899235 | 128899287 | 128945325 | 128945369 | 7 | 17 | 8 | [illegible] |
| SE | 12626 | ENSG00000090857 | PDPR | chr16 | + | 70114869 | 70114930 | 70114371 | 70114440 | 70120460 | 70120719 | 3 | 1 | 3 | [illegible] |
| SE | 92357 | ENSG00000143437 | ARNT | chr1 | - | 150823193 | 150823345 | 150817919 | 150818030 | 150826542 | 150826617 | 312 | 246 | 227 | [illegible] |
| SE | 6407 | ENSG00000176225 | RTTN | chr18 | - | 70020614 | 70020817 | 70017406 | 70017674 | 70024721 | 70024848 | 58 | 21 | 38 | [illegible] |
| SE | 31958 | ENSG00000137502 | RAB30 | chr11 | - | 83034171 | 83034272 | 82997223 | 82997324 | 83071690 | 83071866 | 8 | 6 | 12 | [illegible] |
| SE | 3863 | ENSG00000157540 | DYRK1A | chr21 | + | 37430268 | 37430400 | 37421773 | 37421866 | 37472683 | 37472829 | 26 | 12 | 11 | [illegible] |
| SE | 12792 | ENSG00000102879 | CORO1A | chr16 | + | 30184225 | 30184343 | 30183608 | 30183725 | 30185208 | 30185407 | 5 | 7 | 8 | [illegible] |
| SE | 105383 | ENSG00000114302 | PRKAR2A | chr3 | - | 48782985 | 48783092 | 48765247 | 48765349 | 48790543 | 48790627 | 160 | 112 | 96 | [illegible] |
| SE | 27225 | ENSG00000163812 | ZDHHC3 | chr3 | - | 44929305 | 44929436 | 44915260 | 44926847 | 44932906 | 44932990 | 68 | 52 | 48 | [illegible] |
| SE | 8735 | ENSG00000073969 | NSF | chr17 | + | 46630410 | 46630450 | 46626612 | 46626712 | 46637375 | 46637542 | 0 | 0 | 1 | [illegible] |
| SE | 35719 | ENSG00000185507 | IRF7 | chr11 | - | 614173 | 614399 | 613950 | 614037 | 614475 | 614534 | 36 | 20 | 26 | [illegible] |
| SE | 72818 | ENSG00000163938 | GNL3 | chr3 | + | 52686768 | 52686827 | 52686004 | 52686105 | 52687245 | 52687383 | 75 | 32 | 55 | [illegible] |
| SE | 67612 | ENSG00000109466 | KLHL2 | chr4 | + | 165263196 | 165263359 | 165238777 | 165238899 | 165294358 | 165294468 | 98 | 65 | 86 | [illegible] |
| SE | 3617 | ENSG00000104812 | GYS1 | chr19 | - | 48974212 | 48974339 | 48970749 | 48971023 | 48974619 | 48974733 | 124 | 77 | 87 | [illegible] |
| SE | 95283 | ENSG00000172716 | SLFN11 | chr17 | - | 35366946 | 35367043 | 35363438 | 35363826 | 35373473 | 35373622 | 68 | 45 | 41 | [illegible] |
| SE | 106767 | ENSG00000078674 | PCM1 | chr8 | + | 17980590 | 17980755 | 17972328 | 17972687 | 17985446 | 17985619 | 382 | 387 | 290 | [illegible] |
| SE | 65763 | ENSG00000146463 | ZMYM4 | chr1 | + | 35397051 | 35397123 | 35396551 | 35396670 | 35397376 | 35397462 | 2 | 10 | 5 | [illegible] |
| SE | 7982 | ENSG00000126561 | STAT5A | chr17 | + | 42291971 | 42292061 | 42289865 | 42290022 | 42295618 | 42295793 | 170 | 106 | 179 | [illegible] |
| SE | 2279 | ENSG00000180448 | ARHGAP45 | chr19 | + | 1068332 | 1068744 | 1067326 | 1067495 | 1073148 | 1073292 | 117 | 37 | 94 | [illegible] |
| SE | 106492 | ENSG00000185482 | STAC3 | chr12 | - | 57244536 | 57244622 | 57244314 | 57244366 | 57244915 | 57244965 | 15 | 23 | 12 | [illegible] |
| SE | 60484 | ENSG00000070814 | TCOF1 | chr5 | + | 150387942 | 150388088 | 150379531 | 150379732 | 150389886 | 150390023 | 16 | 14 | 23 | [illegible] |
| SE | 81086 | ENSG00000140750 | ARHGAP17 | chr16 | - | 24929612 | 24929660 | 24927683 | 24927732 | 24930783 | 24931023 | 0 | 0 | 0 | [illegible] |
| SE | 87813 | ENSG00000111237 | VPS29 | chr12 | - | 110499534 | 110499546 | 110496011 | 110496203 | 110502048 | 110502099 | 93 | 57 | 79 | [illegible] |
| SE | 67624 | ENSG00000175215 | CTDSP2 | chr12 | - | 57826995 | 57827123 | 57826345 | 57826402 | 57827551 | 57827590 | 213 | 126 | 217 | [illegible] |
| SE | 17706 | ENSG00000100749 | VRK1 | chr14 | + | 96841517 | 96841626 | 96837761 | 96837817 | 96846094 | 96846164 | 2 | 1 | 7 | [illegible] |
| SE | 8750 | ENSG00000007202 | KIAA0100 | chr17 | - | 28628264 | 28628559 | 28624232 | 28624390 | 28631478 | 28631681 | 377 | 200 | 283 | [illegible] |
| SE | 72558 | ENSG00000068724 | TTC7A | chr2 | + | 47021861 | 47021979 | 47011330 | 47011435 | 47023407 | 47023465 | 91 | 22 | 50 | [illegible] |
| SE | 62835 | ENSG00000197451 | HNRNPAB | chr5 | + | 178205841 | 178206010 | 178204547 | 178205046 | 178206731 | 178206890 | 527 | 423 | 421 | [illegible] |
| SE | 10640 | ENSG00000171475 | WIPF2 | chr17 | + | 40260534 | 40260627 | 40256390 | 40256522 | 40262524 | 40262641 | 46 | 26 | 27 | [illegible] |
| SE | 82847 | ENSG00000181513 | ACBD4 | chr17 | + | 45137367 | 45137454 | 45137018 | 45137139 | 45137759 | 45137830 | 36 | 13 | 24 | [illegible] |
| SE | 85806 | ENSG00000163605 | PPP4R2 | chr3 | + | 73063672 | 73063747 | 73061022 | 73061060 | 73064002 | 73064146 | 433 | 267 | 325 | [illegible] |
| SE | 60538 | ENSG00000116984 | MTR | chr1 | + | 236850343 | 236850523 | 236838413 | 236838599 | 236859832 | 236859922 | 88 | 76 | 107 | [illegible] |

| | | | | | | | | | | | | | | |
|---|---|---|---|---|---|---|---|---|---|---|---|---|---|---|
| SE | 7478 | ENSG00000073605 | GSDMB | chr17 | - | 39908957 | 39909042 | 39906110 | 39906987 | 39909755 | 39909924 | 40 | 45 | 39 |
| SE | 42163 | ENSG00000197343 | ZNF655 | chr7 | + | 99563193 | 99563266 | 99560532 | 99560695 | 99563862 | 99564346 | 0 | 0 | 0 |
| SE | 37229 | ENSG00000284707 | NA | chr7 | - | 97872350 | 97872449 | 97869777 | 97869813 | 97928081 | 97928317 | 13 | 4 | 5 |
| SE | 31121 | ENSG00000198586 | TLK1 | chr2 | - | 171022871 | 171023129 | 171014850 | 171014948 | 171046173 | 171046362 | 19 | 21 | 13 |
| SE | 72133 | ENSG00000172936 | MYD88 | chr3 | + | 38139863 | 38139998 | 38138477 | 38139028 | 38140756 | 38140848 | 136 | 57 | 64 |
| SE | 106040 | ENSG00000204482 | LST1 | chr6 | + | 31587943 | 31587966 | 31587640 | 31587733 | 31588517 | 31588909 | 149 | 141 | 104 |
| SE | 94453 | ENSG00000122490 | SLC66A2 | chr18 | - | 79933968 | 79934022 | 79902419 | 79904183 | 79943328 | 79943462 | 2 | 5 | 2 |
| SE | 41681 | ENSG00000079277 | MKNK1 | chr1 | - | 46585873 | 46585974 | 46583227 | 46583329 | 46586233 | 46586259 | 10 | 2 | 8 |
| SE | 32494 | ENSG00000107140 | TESK1 | chr9 | + | 35607326 | 35607409 | 35606836 | 35606983 | 35607581 | 35607672 | 66 | 39 | 42 |
| SE | 90499 | ENSG00000160299 | PCNT | chr21 | + | 46351428 | 46351540 | 46349683 | 46349820 | 46353103 | 46353326 | 264 | 118 | 125 |
| SE | 79734 | ENSG00000168255 | POLR2J3 | chr7 | - | 102566581 | 102566736 | 102545302 | 102545420 | 102566996 | 102567083 | 27 | 10 | 21 |
| SE | 14116 | ENSG00000102897 | LYRM1 | chr16 | + | 20911182 | 20911264 | 20902485 | 20902717 | 20915555 | 20915714 | 2 | 9 | 4 |
| SE | 40058 | ENSG00000125347 | IRF1 | chr5 | - | 132484361 | 132484497 | 132472148 | 132472178 | 132486556 | 132486686 | 13 | 16 | 17 |
| SE | 23523 | ENSG00000100284 | TOM1 | chr22 | + | 35333403 | 35333497 | 35332980 | 35333014 | 35334327 | 35334448 | 175 | 60 | 120 |
| SE | 8967 | ENSG00000108395 | TRIM37 | chr17 | - | 59073093 | 59073141 | 59070822 | 59070947 | 59075646 | 59075714 | 0 | 1 | 3 |
| SE | 83088 | ENSG00000184863 | RBM33 | chr7 | + | 155763808 | 155764018 | 155744960 | 155745602 | 155766466 | 155766556 | 62 | 35 | 51 |
| SE | 61709 | ENSG00000198836 | OPA1 | chr3 | + | 193617177 | 193617285 | 193615673 | 193615770 | 193618868 | 193618936 | 193 | 183 | 229 |
| SE | 69506 | ENSG00000106799 | TGFBR1 | chr9 | + | 99146484 | 99146609 | 99144731 | 99144888 | 99147653 | 99147784 | 564 | 365 | 328 |
| SE | 94508 | ENSG00000070061 | ELP1 | chr9 | - | 108875632 | 108875735 | 108874894 | 108874970 | 108877994 | 108878149 | 2 | 0 | 4 |
| SE | 28008 | ENSG00000137601 | NEK1 | chr4 | - | 169599099 | 169599197 | 169590725 | 169590809 | 169602007 | 169602104 | 78 | 51 | 53 |
| SE | 30010 | ENSG00000248049 | UBA6-AS1 | chr4 | + | 67720421 | 67721051 | 67701260 | 67701484 | 67721476 | 67723217 | 15 | 14 | 21 |
| SE | 25832 | ENSG00000250334 | LINC00989 | chr4 | + | 79513784 | 79513902 | 79493151 | 79493585 | 79540132 | 79540780 | 61 | 72 | 28 |
| SE | 105061 | ENSG00000119801 | YPEL5 | chr2 | + | 30155814 | 30155910 | 30148244 | 30148541 | 30156627 | 30156787 | 13 | 17 | 16 |
| SE | 15975 | ENSG00000185787 | MORF4L1 | chr15 | + | 78884966 | 78885083 | 78880511 | 78880579 | 78886140 | 78886227 | 44 | 20 | 47 |
| SE | 69683 | ENSG00000004897 | CDC27 | chr17 | - | 47141852 | 47142022 | 47138738 | 47138891 | 47142228 | 47142381 | 190 | 135 | 173 |
| SE | 49101 | ENSG00000049249 | TNFRSF9 | chr1 | - | 7935012 | 7935143 | 7933126 | 7933296 | 7937689 | 7937756 | 28 | 10 | 2 |
| SE | 1701 | ENSG00000228315 | GUSBP11 | chr22 | - | 23686837 | 23686995 | 23683724 | 23683867 | 23690235 | 23695512 | 8 | 8 | 10 |
| SE | 52357 | ENSG00000142197 | DOP1B | chr21 | + | 36263747 | 36263814 | 36260676 | 36260732 | 36270012 | 36270157 | 10 | 17 | 17 |
| SE | 76921 | ENSG00000167483 | NIBAN3 | chr19 | + | 17546685 | 17546797 | 17543316 | 17543433 | 17549443 | 17549527 | 63 | 65 | 76 |
| SE | 69711 | ENSG00000177674 | AGTRAP | chr1 | + | 11745929 | 11746223 | 11745802 | 11745837 | 11747439 | 11747545 | 12 | 3 | 16 |
| SE | 17942 | ENSG00000197249 | SERPINA1 | chr14 | - | 94388818 | 94389010 | 94388559 | 94388660 | 94390419 | 94390635 | 34 | 17 | 8 |
| SE | 12136 | ENSG00000172775 | PSME3IP1 | chr16 | - | 57174619 | 57174688 | 57173727 | 57173869 | 57185820 | 57186022 | 30 | 14 | 23 |
| SE | 48472 | ENSG00000143543 | JTB | chr1 | - | 153976692 | 153976775 | 153975816 | 153975905 | 153976975 | 153977013 | 290 | 117 | 272 |

| | | | | | | | | | | | |
|---|---|---|---|---|---|---|---|---|---|---|---|
| SE | 40698 | ENSG00000117448 | AKR1A1 | chr1 | + | 45552435 | 45552563 | 45550542 | 45551155 | 45566568 | 45566688 |
| SE | 42662 | ENSG00000088854 | C20orf194 | chr20 | - | 3304439 | 3304543 | 3295493 | 3296949 | 3315032 | 3315120 |
| SE | 7075 | ENSG00000175387 | SMAD2 | chr18 | - | 47920071 | 47920169 | 47896520 | 47896809 | 47930360 | 47930599 |
| SE | 27227 | ENSG00000163812 | ZDHHC3 | chr3 | - | 44932906 | 44932990 | 44917408 | 44918143 | 44933117 | 44933199 |
| SE | 102752 | ENSG00000040341 | STAU2 | chr8 | - | 73697307 | 73697442 | 73688653 | 73688813 | 73738283 | 73738349 |
| SE | 25056 | ENSG00000064652 | SNX24 | chr5 | + | 122891026 | 122891125 | 122845604 | 122845693 | 122936733 | 122936817 |
| SE | 72447 | ENSG00000163481 | RNF25 | chr2 | - | 218665154 | 218665247 | 218664738 | 218664873 | 218665915 | 218666059 |
| SE | 44691 | ENSG00000159592 | GPBP1L1 | chr1 | - | 45634095 | 45634236 | 45630481 | 45630606 | 45640209 | 45640403 |
| SE | 1130 | ENSG00000099992 | TBC1D10A | chr22 | - | 30299443 | 30299551 | 30295736 | 30295843 | 30304530 | 30304630 |
| SE | 64031 | ENSG00000115464 | USP34 | chr2 | - | 61220309 | 61220457 | 61214059 | 61214694 | 61221501 | 61221606 |
| SE | 70400 | ENSG00000125257 | ABCC4 | chr13 | - | 95266111 | 95266264 | 95247642 | 95247753 | 95301240 | 95301302 |
| SE | 59919 | ENSG00000164144 | ARFIP1 | chr4 | + | 152870752 | 152870848 | 152863605 | 152863714 | 152872451 | 152872564 |
| SE | 83768 | ENSG00000196696 | PDXDC2P-NPIPB14P | chr16 | - | 69992910 | 69993248 | 69990596 | 69990683 | 69996068 | 69996226 |
| SE | 74997 | ENSG00000157450 | RNF111 | chr15 | + | 59084101 | 59084254 | 59080935 | 59081284 | 59085658 | 59085785 |
| SE | 1806 | ENSG00000099917 | MED15 | chr22 | + | 20554935 | 20555148 | 20553096 | 20553174 | 20564449 | 20564466 |
| SE | 53208 | ENSG00000083544 | TDRD3 | chr13 | + | 60483774 | 60483846 | 60467237 | 60467379 | 60485798 | 60485899 |
| SE | 21661 | ENSG00000130449 | ZSWIM6 | chr5 | + | 61490785 | 61490934 | 61472680 | 61473037 | 61494259 | 61494410 |
| SE | 35613 | ENSG00000167618 | LAIR2 | chr19 | + | 54509034 | 54509085 | 54507890 | 54508184 | 54510525 | 54510687 |
| SE | 21484 | ENSG00000018189 | RUFY3 | chr4 | + | 70800140 | 70800205 | 70794672 | 70794894 | 70802955 | 70802983 |
| SE | 46394 | ENSG00000178028 | DMAP1 | chr1 | + | 44214538 | 44214616 | 44214349 | 44214441 | 44214702 | 44214880 |
| SE | 14396 | ENSG00000167202 | TBC1D2B | chr15 | - | 78008996 | 78009114 | 78002481 | 78003490 | 78012822 | 78013317 |
| SE | 41123 | ENSG00000115523 | GNLY | chr2 | + | 85698397 | 85698447 | 85695957 | 85696056 | 85698563 | 85698747 |
| SE | 98116 | ENSG00000171466 | ZNF562 | chr19 | - | 9659378 | 9659467 | 9658008 | 9658135 | 9675014 | 9675052 |
| SE | 80532 | ENSG00000179818 | PCBP1-AS1 | chr2 | - | 70053730 | 70053792 | 70051044 | 70051305 | 70085546 | 70085573 |
| SE | 89372 | ENSG00000108946 | PRKAR1A | chr17 | + | 68514986 | 68515119 | 68511779 | 68512142 | 68515393 | 68515536 |
| SE | 83886 | ENSG00000168807 | SNTB2 | chr16 | + | 69235747 | 69235833 | 69187560 | 69187746 | 69245601 | 69245815 |
| SE | 23195 | ENSG00000165819 | METTL3 | chr14 | - | 21500494 | 21500682 | 21499763 | 21499802 | 21500912 | 21501129 |
| SE | 45497 | ENSG00000168894 | RNF181 | chr2 | + | 85596856 | 85596931 | 85596518 | 85596649 | 85597103 | 85597178 |
| SE | 41456 | ENSG00000137776 | SLTM | chr15 | - | 58916934 | 58916999 | 58901259 | 58901287 | 58932355 | 58932443 |
| SE | 53581 | ENSG00000188529 | SRSF10 | chr1 | - | 23971572 | 23971623 | 23966444 | 23967723 | 23971849 | 23972012 |
| SE | 50184 | ENSG00000185808 | PIGP | chr21 | - | 37072144 | 37072310 | 37069551 | 37069624 | 37072433 | 37072537 |
| SE | 10524 | ENSG00000125970 | RALY | chr20 | + | 34065043 | 34065160 | 34031521 | 34031604 | 34072065 | 34072330 |
| SE | 81623 | ENSG00000135093 | USP30 | chr12 | + | 109056681 | 109056791 | 109052575 | 109052761 | 109067523 | 109067627 |

[illegible]

| | | | | | | | | | | | | | | | | | | | | | | | | | | | | | | | | | | | | | | | | | | | | | | |
|---|---|---|---|---|---|---|---|---|---|---|---|---|---|---|---|---|---|---|---|---|---|---|---|---|---|---|---|---|---|---|---|---|---|---|---|---|---|---|---|---|---|---|---|---|---|---|
| SE | 21929 | ENSG00000112624 | BICRAL | chr6 | + | 42853637 | 42853738 | 42852091 | 42852197 | 42855855 | 42855917 | 160 | 86 | 177 | 155 | 106 | 3 | 3 | 4 | 1 | 1 | 49 | 120 | 83 | 96 | 38 | 15 | 3 | 0 | 4 | 0 | 250 | 149 | 0.005340556 | 0.061859929 | 0.969 | 0.945 | 0.963 | 0.989 | 0.984 | 0.661 | 0.96 | 1 | 0.935 | 1 | 0.059 |
| SE | 27067 | ENSG00000062194 | GPBP1 | chr5 | + | 57237112 | 57237172 | 57235965 | 57236032 | 57246299 | 57246484 | 62 | 40 | 78 | 52 | 101 | 143 | 129 | 146 | 123 | 381 | 3 | 43 | 33 | 5 | 18 | 0 | 51 | 53 | 13 | 0 | 209 | 149 | 0.005340926 | 0.061859929 | 0.236 | 0.181 | 0.276 | 0.232 | 0.159 | 1 | 0.375 | 0.307 | 0.215 | 1 | -0.363 |
| SE | 17768 | ENSG00000213463 | SYNJ2BP | chr14 | - | 70388469 | 70388606 | 70375675 | 70375771 | 70416899 | 70417090 | 89 | 82 | 111 | 144 | 179 | 3 | 1 | 10 | 1 | 0 | 121 | 93 | 54 | 63 | 42 | 0 | 0 | 3 | 0 | 0 | 286 | 149 | 0.005345315 | 0.061901608 | 0.939 | 0.977 | 0.853 | 0.987 | 1 | 1 | 1 | 0.904 | 1 | 1 | -0.03 |
| SE | 67647 | ENSG00000176978 | DPP7 | chr9 | - | 137111940 | 137112029 | 137111689 | 137111754 | 137112111 | 137112230 | 42 | 18 | 59 | 36 | 35 | 0 | 1 | 0 | 0 | 1 | 51 | 85 | 30 | 35 | 115 | 6 | 0 | 2 | 4 | 0 | 238 | 149 | 0.005350823 | 0.061956229 | 1 | 0.918 | 1 | 1 | 0.956 | 0.842 | 1 | 0.904 | 0.846 | 1 | 0.056 |
| SE | 33662 | ENSG00000176261 | ZBTB8OS | chr1 | - | 32632207 | 32632345 | 32631826 | 32631879 | 32633644 | 32633727 | 5 | 14 | 8 | 12 | 13 | 92 | 57 | 79 | 65 | 182 | 15 | 4 | 11 | 1 | 7 | 23 | 34 | 31 | 28 | 30 | 287 | 149 | 0.005353609 | 0.06197932 | 0.027 | 0.113 | 0.05 | 0.087 | 0.036 | 0.253 | 0.058 | 0.156 | 0.018 | 0.108 | -0.056 |
| SE | 93146 | ENSG00000077721 | UBE2A | chrX | + | 119581506 | 119581596 | 119575374 | 119575400 | 119582587 | 119582647 | 175 | 79 | 122 | 144 | 221 | 0 | 2 | 0 | 6 | 2 | 282 | 162 | 115 | 178 | 245 | 8 | 13 | 2 | 0 | 6 | 239 | 149 | 0.005357642 | 0.062007927 | 1 | 0.961 | 1 | 0.937 | 0.986 | 0.956 | 0.886 | 0.973 | 1 | 0.962 | 0.021 |
| SE | 29975 | ENSG00000143624 | INTS3 | chr1 | + | 153761569 | 153761676 | 153757636 | 153757763 | 153762727 | 153762847 | 85 | 66 | 63 | 95 | 45 | 0 | 2 | 0 | 1 | 0 | 164 | 118 | 77 | 154 | 182 | 10 | 0 | 7 | 0 | 0 | 256 | 149 | 0.005357664 | 0.062007927 | 1 | 0.951 | 1 | 0.982 | 1 | 0.905 | 1 | 0.865 | 1 | 1 | 0.033 |
| SE | 38021 | ENSG00000197386 | HTT | chr4 | + | 3223991 | 3224131 | 3223405 | 3223560 | 3225660 | 3225743 | 128 | 70 | 124 | 109 | 68 | 0 | 0 | 0 | 0 | 0 | 204 | 115 | 67 | 151 | 231 | 0 | 0 | 0 | 6 | 10 | 289 | 149 | 0.005358502 | 0.062008461 | 1 | 1 | 1 | 1 | 1 | 1 | 1 | 1 | 0.928 | 0.923 | 0.03 |
| SE | 7123 | ENSG00000168234 | TTC39C | chr18 | + | 24069156 | 24069271 | 24066011 | 24066140 | 24114553 | 24114647 | 171 | 114 | 194 | 175 | 335 | 3 | 4 | 2 | 4 | 4 | 120 | 168 | 125 | 136 | 84 | 7 | 4 | 15 | 1 | 0 | 264 | 149 | 0.005360155 | 0.062018423 | 0.97 | 0.941 | 0.982 | 0.961 | 0.979 | 0.906 | 0.96 | 0.825 | 0.987 | 1 | 0.031 |
| SE | 87542 | ENSG00000147050 | KDM6A | chrX | + | 45060608 | 45060764 | 45060021 | 45060156 | 45061323 | 45061419 | 61 | 37 | 28 | 35 | 68 | 17 | 14 | 20 | 36 | 40 | 32 | 36 | 48 | 34 | 31 | 24 | 0 | 7 | 21 | 0 | 298 | 149 | 0.005362946 | 0.062041546 | 0.642 | 0.569 | 0.412 | 0.327 | 0.459 | 0.4 | 1 | 0.774 | 0.447 | 1 | -0.242 |
| SE | 34514 | ENSG00000106733 | NMRK1 | chr9 | - | 75069741 | 75069813 | 75068995 | 75069102 | 75069894 | 75070042 | 268 | 179 | 371 | 283 | 243 | 0 | 24 | 32 | 19 | 25 | 125 | 93 | 160 | 75 | 58 | 0 | 2 | 21 | 0 | 0 | 221 | 149 | 0.005367589 | 0.062086093 | 1 | 0.834 | 0.887 | 0.909 | 0.868 | 1 | 0.969 | 0.837 | 1 | 1 | -0.062 |
| SE | 62758 | ENSG00000162961 | DPY30 | chr2 | - | 32023676 | 32023808 | 32023421 | 32023474 | 32029593 | 32029736 | 28 | 6 | 12 | 10 | 30 | 0 | 2 | 2 | 3 | 7 | 12 | 2 | 7 | 21 | 14 | 2 | 0 | 1 | 0 | 2 | 281 | 149 | 0.005370371 | 0.062109099 | 1 | 0.614 | 0.761 | 0.639 | 0.694 | 0.761 | 1 | 0.788 | 1 | 0.788 | -0.126 |
| SE | 11482 | ENSG00000273611 | ZNHIT3 | chr17 | + | 36486934 | 36486966 | 36486680 | 36486785 | 36492812 | 36492899 | 103 | 46 | 109 | 97 | 106 | 13 | 8 | 14 | 11 | 14 | 91 | 95 | 88 | 44 | 93 | 2 | 5 | 12 | 2 | 8 | 181 | 149 | 0.005372242 | 0.062121559 | 0.867 | 0.826 | 0.865 | 0.879 | 0.862 | 0.974 | 0.94 | 0.858 | 0.948 | 0.905 | -0.065 |
| SE | 33368 | ENSG00000187742 | SECISBP2 | chr9 | + | 89339863 | 89339953 | 89338457 | 89338580 | 89341346 | 89341479 | 155 | 121 | 131 | 149 | 187 | 14 | 23 | 23 | 14 | 14 | 100 | 145 | 137 | 130 | 40 | 15 | 41 | 42 | 32 | 2 | 239 | 149 | 0.005376546 | 0.062153723 | 0.873 | 0.766 | 0.78 | 0.869 | 0.893 | 0.806 | 0.688 | 0.67 | 0.717 | 0.926 | 0.075 |
| SE | 16332 | ENSG00000255769 | GOLGA2P10 | chr15 | - | 82477837 | 82477926 | 82477393 | 82477582 | 82513665 | 82513950 | 8 | 5 | 30 | 15 | 1 | 0 | 0 | 0 | 0 | 0 | 1 | 0 | 4 | 3 | 5 | 0 | 2 | 0 | 0 | 14 | 238 | 149 | 0.005376803 | 0.062153723 | 1 | 1 | 1 | 1 | 1 | 1 | 0 | 1 | 1 | 0.183 | 0.363 |
| SE | 89026 | ENSG00000177565 | TBL1XR1 | chr3 | - | 177110808 | 177110887 | 177098486 | 177098541 | 177197120 | 177197227 | 2 | 1 | 9 | 0 | 1 | 35 | 28 | 30 | 28 | 29 | 0 | 0 | 0 | 2 | 0 | 21 | 34 | 28 | 13 | 19 | 228 | 149 | 0.005377405 | 0.062153723 | 0.036 | 0.023 | 0.164 | 0 | 0.022 | 0 | 0 | 0 | 0.091 | 0 | 0.031 |
| SE | 107554 | ENSG00000111725 | PRKAB1 | chr12 | + | 119671494 | 119671668 | 119668149 | 119668403 | 119672300 | 119672427 | 3 | 1 | 3 | 3 | 11 | 22 | 12 | 18 | 18 | 10 | 20 | 3 | 13 | 8 | 15 | 17 | 6 | 20 | 28 | 27 | 298 | 149 | 0.005380057 | 0.062175193 | 0.064 | 0.04 | 0.077 | 0.077 | 0.355 | 0.37 | 0.2 | 0.245 | 0.125 | 0.217 | -0.109 |
| SE | 27408 | ENSG00000214367 | HAUS3 | chr4 | - | 2241519 | 2241781 | 2240846 | 2241093 | 2242050 | 2242121 | 36 | 25 | 29 | 25 | 21 | 18 | 16 | 12 | 18 | 5 | 5 | 14 | 29 | 29 | 7 | 0 | 17 | 6 | 0 | 0 | 298 | 149 | 0.005384862 | 0.062203183 | 0.5 | 0.439 | 0.547 | 0.41 | 0.677 | 1 | 0.292 | 0.707 | 1 | 1 | -0.285 |
| SE | 27409 | ENSG00000214367 | HAUS3 | chr4 | - | 2241519 | 2241790 | 2240037 | 2241093 | 2242050 | 2242121 | 36 | 25 | 29 | 25 | 21 | 18 | 16 | 12 | 18 | 5 | 5 | 14 | 29 | 29 | 7 | 0 | 17 | 6 | 0 | 0 | 298 | 149 | 0.005384862 | 0.062203183 | 0.5 | 0.439 | 0.547 | 0.41 | 0.677 | 1 | 0.292 | 0.707 | 1 | 1 | -0.285 |
| SE | 27410 | ENSG00000214367 | HAUS3 | chr4 | - | 2241519 | 2241837 | 2240915 | 2241093 | 2242050 | 2242128 | 36 | 25 | 29 | 25 | 21 | 18 | 16 | 12 | 18 | 5 | 5 | 14 | 29 | 29 | 7 | 0 | 17 | 6 | 0 | 0 | 298 | 149 | 0.005384862 | 0.062203183 | 0.5 | 0.439 | 0.547 | 0.41 | 0.677 | 1 | 0.292 | 0.707 | 1 | 1 | -0.285 |
| SE | 37902 | ENSG00000184014 | DENND5A | chr11 | - | 9143402 | 9143485 | 9138848 | 9142108 | 9144096 | 9144278 | 31 | 27 | 33 | 31 | 17 | 1 | 4 | 2 | 1 | 0 | 104 | 47 | 25 | 46 | 83 | 7 | 3 | 6 | 16 | 4 | 232 | 149 | 0.005386273 | 0.062205196 | 0.952 | 0.813 | 0.914 | 0.952 | 1 | 0.905 | 0.91 | 0.728 | 0.649 | 0.93 | 0.102 |
| SE | 92531 | ENSG00000216937 | CCDC7 | chr10 | + | 32623718 | 32623817 | 32584231 | 32584304 | 32675664 | 32675813 | 34 | 26 | 20 | 17 | 77 | 3 | 1 | 0 | 2 | 4 | 16 | 47 | 20 | 6 | 25 | 0 | 0 | 0 | 0 | 0 | 248 | 149 | 0.005386626 | 0.062205196 | 0.872 | 0.94 | 1 | 0.836 | 0.92 | 1 | 1 | 1 | 1 | 1 | -0.086 |
| SE | 9419 | ENSG00000174231 | PRPF8 | chr17 | - | 1651980 | 1652037 | 1651662 | 1651788 | 1653541 | 1653683 | 1 | 1 | 0 | 0 | 0 | 261 | 133 | 210 | 277 | 282 | 18 | 0 | 3 | 0 | 12 | 303 | 212 | 143 | 141 | 268 | 206 | 149 | 0.00539186 | 0.062256459 | 0.003 | 0.005 | 0 | 0 | 0 | 0.041 | 0 | 0.015 | 0 | 0.031 | -0.016 |
| SE | 101356 | ENSG00000170525 | PFKFB3 | chr10 | + | 6217134 | 6217191 | 6216705 | 6216780 | 6220657 | 6220865 | 4 | 4 | 3 | 2 | 7 | 0 | 0 | 0 | 0 | 0 | 20 | 19 | 39 | 24 | 3 | 18 | 0 | 0 | 0 | 0 | 206 | 149 | 0.005404733 | 0.062394194 | 1 | 1 | 1 | 1 | 1 | 0.446 | 1 | 1 | 1 | 1 | 0.111 |
| SE | 74955 | ENSG00000105393 | BABAM1 | chr19 | + | 17273903 | 17274024 | 17271596 | 17271655 | 17276494 | 17276555 | 157 | 82 | 114 | 99 | 192 | 6 | 2 | 1 | 3 | 5 | 167 | 116 | 98 | 134 | 188 | 17 | 0 | 1 | 5 | 18 | 270 | 149 | 0.005405382 | 0.062394194 | 0.935 | 0.958 | 0.984 | 0.948 | 0.955 | 0.844 | 1 | 0.982 | 0.937 | 0.852 | 0.033 |
| SE | 79506 | ENSG00000146918 | NCAPG2 | chr7 | - | 158650831 | 158650972 | 158646459 | 158646563 | 158652292 | 158652480 | 25 | 47 | 81 | 41 | 63 | 3 | 2 | 1 | 2 | 2 | 8 | 24 | 15 | 86 | 14 | 0 | 5 | 17 | 8 | 0 | 290 | 149 | 0.005408616 | 0.062422318 | 0.811 | 0.924 | 0.977 | 0.913 | 0.942 | 1 | 0.712 | 0.312 | 0.847 | 1 | 0.139 |
| SE | 80783 | ENSG00000136280 | CCM2 | chr7 | + | 45072725 | 45072783 | 45069825 | 45069961 | 45073459 | 45073519 | 146 | 49 | 109 | 113 | 67 | 3 | 2 | 0 | 6 | 4 | 87 | 114 | 39 | 108 | 127 | 0 | 0 | 1 | 2 | 0 | 207 | 149 | 0.005410539 | 0.062434507 | 0.972 | 0.946 | 1 | 0.931 | 0.923 | 1 | 1 | 0.966 | 0.975 | 1 | -0.034 |
| SE | 105191 | ENSG00000139624 | CERS5 | chr12 | - | 50132961 | 50134068 | 50129797 | 50130694 | 50134545 | 50134702 | 6 | 1 | 5 | 4 | 2 | 16 | 16 | 16 | 27 | 20 | 0 | 4 | 2 | 3 | 0 | 24 | 31 | 19 | 15 | 51 | 298 | 149 | 0.005411765 | 0.062434507 | 0.158 | 0.03 | 0.135 | 0.069 | 0.048 | 0 | 0.061 | 0.05 | 0.091 | 0 | 0.048 |
| SE | 84843 | ENSG00000094631 | HDAC6 | chrX | + | 48823921 | 48824068 | 48823671 | 48823785 | 48824543 | 48824860 | 67 | 31 | 47 | 91 | 26 | 0 | 1 | 0 | 0 | 0 | 81 | 68 | 46 | 60 | 151 | 13 | 0 | 0 | 0 | 0 | 296 | 149 | 0.005412064 | 0.062434507 | 1 | 0.94 | 1 | 1 | 1 | 0.758 | 1 | 1 | 1 | 1 | 0.036 |
| SE | 33382 | ENSG00000168591 | TMUB2 | chr17 | + | 44187906 | 44188009 | 44187271 | 44187743 | 44188934 | 44189299 | 104 | 28 | 59 | 90 | 70 | 18 | 11 | 14 | 23 | 27 | 124 | 100 | 93 | 107 | 146 | 8 | 14 | 9 | 23 | 34 | 252 | 149 | 0.005418067 | 0.062494544 | 0.774 | 0.601 | 0.714 | 0.698 | 0.605 | 0.902 | 0.809 | 0.859 | 0.733 | 0.717 | -0.126 |
| SE | 49586 | ENSG00000076685 | NT5C2 | chr10 | - | 103097290 | 103097374 | 103095938 | 103095980 | 103098930 | 103098984 | 284 | 232 | 264 | 360 | 524 | 0 | 0 | 0 | 3 | 0 | 679 | 305 | 353 | 484 | 236 | 0 | 14 | 4 | 5 | 7 | 233 | 149 | 0.005432608 | 0.062653039 | 1 | 1 | 1 | 0.987 | 1 | 1 | 0.933 | 0.983 | 0.984 | 0.956 | 0.026 |
| SE | 65365 | ENSG00000153214 | TMEM87B | chr2 | + | 112081334 | 112081518 | 112081056 | 112081118 | 112086004 | 112086104 | 327 | 304 | 217 | 412 | 418 | 6 | 11 | 7 | 18 | 16 | 285 | 239 | 135 | 182 | 177 | 2 | 6 | 11 | 16 | 41 | 298 | 149 | 0.005441235 | 0.062743293 | 0.965 | 0.933 | 0.939 | 0.92 | 0.929 | 0.986 | 0.952 | 0.86 | 0.85 | 0.683 | 0.071 |
| SE | 68205 | ENSG00000103707 | MTFMT | chr15 | - | 65004853 | 65004936 | 65001511 | 65003256 | 65006112 | 65006191 | 73 | 56 | 67 | 82 | 103 | 0 | 0 | 0 | 0 | 0 | 42 | 50 | 25 | 62 | 77 | 0 | 0 | 0 | 0 | 12 | 232 | 149 | 0.005443336 | 0.062758278 | 1 | 1 | 1 | 1 | 1 | 1 | 1 | 1 | 1 | 0.805 | 0.039 |
| SE | 69347 | ENSG00000168958 | MFF | chr2 | + | 227352513 | 227352573 | 227342744 | 227342819 | 227355676 | 227355761 | 12 | 4 | 15 | 16 | 32 | 27 | 15 | 29 | 13 | 44 | 11 | 9 | 18 | 23 | 3 | 75 | 67 | 30 | 36 | 34 | 209 | 149 | 0.00544826 | 0.062805797 | 0.241 | 0.16 | 0.269 | 0.467 | 0.341 | 0.095 | 0.087 | 0.3 | 0.313 | 0.059 | 0.125 |
| SE | 33326 | ENSG00000188313 | PLSCR1 | chr3 | - | 146521543 | 146521701 | 146517005 | 146517167 | 146525604 | 146525647 | 113 | 122 | 85 | 169 | 112 | 6 | 2 | 0 | 10 | 10 | 212 | 126 | 105 | 413 | 59 | 12 | 27 | 19 | 27 | 0 | 298 | 149 | 0.005451708 | 0.062836293 | 0.904 | 0.968 | 1 | 0.894 | 0.848 | 0.898 | 0.7 | 0.734 | 0.884 | 1 | 0.08 |
| SE | 39094 | ENSG00000109320 | NFKB1 | chr4 | + | 102533844 | 102533885 | 102529835 | 102529914 | 102537857 | 102537956 | 132 | 85 | 91 | 97 | 178 | 0 | 2 | 12 | 1 | 10 | 179 | 75 | 77 | 117 | 196 | 10 | 3 | 13 | 19 | 8 | 190 | 149 | 0.005454537 | 0.06285964 | 1 | 0.971 | 0.856 | 0.987 | 0.933 | 0.933 | 0.951 | 0.823 | 0.828 | 0.951 | 0.052 |
| SE | 99850 | ENSG00000110713 | NUP98 | chr11 | - | 3685972 | 3686194 | 3683199 | 3683441 | 3691346 | 3691489 | 133 | 81 | 112 | 97 | 70 | 2 | 0 | 1 | 2 | 11 | 68 | 93 | 62 | 121 | 118 | 0 | 5 | 0 | 1 | 0 | 298 | 149 | 0.005457033 | 0.062879159 | 0.971 | 1 | 0.982 | 0.96 | 0.761 | 1 | 0.903 | 1 | 0.984 | 1 | -0.043 |
| SE | 51971 | ENSG00000071054 | MAP4K4 | chr2 | + | 101835899 | 101835978 | 101834408 | 101834463 | 101839818 | 101839994 | 341 | 296 | 350 | 413 | 407 | 2 | 10 | 3 | 5 | 1 | 674 | 537 | 537 | 566 | 482 | 7 | 11 | 8 | 17 | 16 | 228 | 149 | 0.005464561 | 0.062956627 | 0.991 | 0.951 | 0.987 | 0.982 | 0.996 | 0.984 | 0.97 | 0.978 | 0.956 | 0.952 | 0.013 |
| SE | 81346 | ENSG00000173852 | DPY19L1 | chr7 | - | 34957983 | 34958070 | 34955307 | 34955367 | 34966893 | 34966971 | 52 | 42 | 17 | 38 | 41 | 0 | 2 | 1 | 7 | 2 | 33 | 33 | 46 | 31 | 18 | 0 | 2 | 11 | 9 | 7 | 236 | 149 | 0.005466651 | 0.062971439 | 1 | 0.93 | 0.915 | 0.774 | 0.928 | 1 | 0.912 | 0.725 | 0.685 | 0.619 | 0.121 |
| SE | 21720 | ENSG00000111860 | CEP85L | chr6 | - | 118491685 | 118491865 | 118483705 | 118483858 | 118511297 | 118511415 | 383 | 276 | 425 | 330 | 575 | 3 | 9 | 6 | 5 | 3 | 52 | 124 | 127 | 61 | 39 | 0 | 0 | 0 | 0 | 0 | 298 | 149 | 0.005470609 | 0.063007771 | 0.985 | 0.939 | 0.973 | 0.971 | 0.99 | 1 | 1 | 1 | 1 | 1 | -0.028 |
| SE | 5859 | ENSG00000101391 | CDK5RAP1 | chr20 | - | 33395012 | 33395116 | 33394031 | 33394066 | 33396760 | 33396936 | 57 | 28 | 56 | 59 | 35 | 0 | 0 | 1 | 1 | 4 | 118 | 50 | 62 | 87 | 65 | 18 | 2 | 1 | 0 | 0 | 253 | 149 | 0.00547347 | 0.063031447 | 1 | 1 | 0.971 | 0.972 | 0.837 | 0.794 | 0.936 | 0.973 | 1 | 1 | 0.015 |
| SE | 32289 | ENSG00000163964 | PIGX | chr3 | + | 196726654 | 196726771 | 196722414 | 196722556 | 196727922 | 196728037 | 9 | 3 | 6 | 2 | 3 | 36 | 21 | 18 | 33 | 51 | 10 | 11 | 5 | 6 | 4 | 15 | 22 | 10 | 32 | 28 | 266 | 149 | 0.005476787 | 0.063060371 | 0.123 | 0.074 | 0.157 | 0.033 | 0.032 | 0.272 | 0.219 | 0.219 | 0.095 | 0.074 | -0.092 |
| SE | 99020 | ENSG00000160593 | JAML | chr11 | - | 118205914 | 118205991 | 118203427 | 118203665 | 118210486 | 118210712 | 316 | 318 | 217 | 239 | 298 | 0 | 0 | 2 | 3 | 3 | 514 | 170 | 127 | 312 | 344 | 14 | 1 | 1 | 4 | 6 | 226 | 149 | 0.005480651 | 0.063089652 | 1 | 1 | 0.986 | 0.981 | 0.985 | 0.96 | 0.991 | 0.988 | 0.981 | 0.974 | 0.012 |
| SE | 68468 | ENSG00000143799 | PARP1 | chr1 | - | 226392916 | 226392980 | 226392257 | 226392314 | 226402213 | 226402379 | 5 | 0 | 1 | 4 | 1 | 237 | 139 | 178 | 264 | 129 | 9 | 14 | 2 | 7 | 2 | 263 | 199 | 153 | 164 | 285 | 213 | 149 | 0.005481335 | 0.063089652 | 0.015 | 0 | 0.004 | 0.01 | 0.005 | 0.023 | 0.047 | 0.009 | 0.029 | 0.005 | -0.016 |
| SE | 52676 | ENSG00000198860 | TSEN15 | chr1 | + | 184072156 | 184072194 | 184054727 | 184054863 | 184072826 | 184074212 | 12 | 2 | 13 | 15 | 11 | 27 | 10 | 29 | 30 | 60 | 0 | 6 | 1 | 1 | 4 | 20 | 24 | 10 | 12 | 8 | 187 | 149 | 0.005481749 | 0.063089652 | 0.262 | 0.137 | 0.263 | 0.285 | 0.127 | 0 | 0.166 | 0.074 | 0.062 | 0.285 | 0.097 |
| SE | 82046 | ENSG00000104133 | SPG11 | chr15 | - | 44610917 | 44610985 | 44608443 | 44608605 | 44613429 | 44613536 | 91 | 99 | 89 | 91 | 162 | 4 | 5 | 3 | 2 | 0 | 25 | 144 | 70 | 74 | 18 | 0 | 5 | 5 | 4 | 10 | 217 | 149 | 0.005482553 | 0.063089652 | 0.94 | 0.931 | 0.953 | 0.969 | 1 | 1 | 0.952 | 0.906 | 0.927 | 0.553 | 0.091 |
| SE | 85251 | ENSG00000197021 | EOLA2 | chrX | - | 149934376 | 149934573 | 149934004 | 149934137 | 149937432 | 149937480 | 6 | 1 | 7 | 8 | 0 | 37 | 13 | 17 | 24 | 10 | 0 | 7 | 1 | 0 | 0 | 12 | 16 | 10 | 14 | 33 | 298 | 149 | 0.005485337 | 0.063112414 | 0.075 | 0.037 | 0.171 | 0.143 | 0 | 0 | 0.179 | 0.048 | 0 | 0 | 0.04 |
| SE | 55266 | ENSG00000135905 | DOCK10 | chr2 | - | 224886062 | 224886185 | 224885670 | 224885805 | 224886458 | 224886526 | 316 | 276 | 364 | 408 | 488 | 10 | 8 | 5 | 8 | 14 | 118 | 191 | 221 | 85 | 76 | 0 | 4 | 23 | 16 | 0 | 272 | 149 | 0.005489974 | 0.063150042 | 0.945 | 0.95 | 0.976 | 0.965 | 0.95 | 1 | 0.963 | 0.84 | 0.744 | 1 | 0.048 |
| SE | 60759 | ENSG00000141524 | TMC6 | chr17 | - | 78120652 | 78120832 | 78119296 | 78119392 | 78121012 | 78121164 | 303 | 113 | 189 | 352 | 93 | 21 | 6 | 10 | 22 | 0 | 61 | 285 | 81 | 86 | 274 | 18 | 7 | 8 | 9 | 41 | 298 | 149 | 0.005490221 | 0.063150042 | 0.878 | 0.904 | 0.904 | 0.889 | 1 | 0.629 | 0.953 | 0.835 | 0.827 | 0.77 | 0.112 |
| SE | 81283 | ENSG00000155096 | AZIN1 | chr8 | - | 102833055 | 102833218 | 102829820 | 102829936 | 102834188 | 102834263 | 627 | 364 | 505 | 654 | 994 | 9 | 1 | 3 | 7 | 11 | 342 | 462 | 541 | 537 | 263 | 10 | 12 | 9 | 11 | 0 | 298 | 149 | 0.005492378 | 0.063161859 | 0.972 | 0.995 | 0.988 | 0.979 | 0.978 | 0.945 | 0.951 | 0.968 | 0.961 | 1 | 0.017 |
| SE | 23420 | ENSG00000148110 | MFSD14B | chr9 | + | 94440984 | 94441099 | 94438400 | 94438528 | 94444963 | 94445182 | 135 | 63 | 104 | 131 | 119 | 13 | 6 | 3 | 1 | 0 | 190 | 225 | 51 | 123 | 272 | 0 | 0 | 3 | 0 | 11 | 264 | 149 | 0.005492862 | 0.063161859 | 0.854 | 0.856 | 0.951 | 0.987 | 1 | 1 | 1 | 0.906 | 1 | 0.933 | -0.038 |
| SE | 14470 | ENSG00000213614 | HEXA | chr15 | - | 72353067 | 72353178 | 72351131 | 72351234 | 72355558 | 72355624 | 148 | 75 | 127 | 132 | 289 | 3 | 1 | 0 | 10 | 4 | 152 | 174 | 133 | 190 | 207 | 0 | 0 | 3 | 4 | 0 | 260 | 149 | 0.005497352 | 0.063204212 | 0.966 | 0.977 | 1 | 0.883 | 0.976 | 1 | 1 | 0.962 | 0.965 | 1 | -0.025 |
| SE | 24103 | ENSG00000143889 | HNRNPLL | chr2 | - | 38591529 | 38591648 | 38585643 | 38585881 | 38602437 | 38602600 | 106 | 65 | 91 | 72 | 68 | 2 | 0 | 2 | 2 | 0 | 59 | 79 | 69 | 110 | 68 | 0 | 3 | 1 | 3 | 9 | 268 | 149 | 0.005500768 | 0.063234201 | 0.967 | 1 | 0.962 | 0.952 | 1 | 1 | 0.936 | 0.975 | 0.953 | 0.808 | 0.042 |
| SE | 42452 | ENSG00000163946 | TASOR | chr3 | - | 56666259 | 56666384 | 56663540 | 56663572 | 56668396 | 56668489 | 418 | 205 | 279 | 366 | 540 | 25 | 36 | 16 | 39 | 59 | 43 | 188 | 141 | 81 | 84 | 16 | 20 | 19 | 10 | 25 | 274 | 149 | 0.005506777 | 0.063293983 | 0.901 | 0.756 | 0.905 | 0.836 | 0.833 | 0.594 | 0.836 | 0.801 | 0.815 | 0.646 | 0.108 |
| SE | 101959 | ENSG00000188404 | SELL | chr1 | - | 169704035 | 169704094 | 169703316 | 169703440 | 169704567 | 169704753 | 19 | 5 | 27 | 14 | 33 | 900 | 417 | 888 | 657 | 1498 | 0 | 9 | 26 | 1 | 0 | 823 | 934 | 636 | 861 | 411 | 208 | 149 | 0.005515962 | 0.063390243 | 0.015 | 0.009 | 0.021 | 0.015 | 0.016 | 0 | 0.007 | 0.028 | 0.001 | 0 | 0.008 |
| SE | 91791 | ENSG00000133794 | ARNTL | chr11 | + | 13355232 | 13355296 | 13354300 | 13354448 | 13356738 | 13356777 | 8 | 1 | 4 | 1 | 8 | 42 | 26 | 24 | 31 | 38 | 0 | 3 | 7 | 0 | 0 | 32 | 35 | 49 | 55 | 11 | 213 | 149 | 0.005520434 | 0.063432334 | 0.118 | 0.026 | 0.104 | 0.022 | 0.128 | 0 | 0.057 | 0.091 | 0 | 0 | 0.05 |
| SE | 65452 | ENSG00000204344 | STK19 | chr6 | + | 31972620 | 31972757 | 31971868 | 31972511 | 31978902 | 31978998 | 44 | 16 | 28 | 16 | 25 | 1 | 0 | 0 | 0 | 1 | 46 | 53 | 57 | 39 | 80 | 0 | 5 | 0 | 0 | 6 | 286 | 149 | 0.005527094 | 0.063499541 | 0.958 | 1 | 1 | 1 | 0.929 | 1 | 0.847 | 1 | 1 | 0.874 | 0.033 |
| SE | 29385 | ENSG00000170242 | USP47 | chr11 | + | 11880176 | 11880380 | 11841971 | 11842224 | 11884466 | 11884499 | 173 | 101 | 149 | 196 | 211 | 5 | 4 | 3 | 3 | 1 | 43 | 77 | 74 | 36 | 68 | 0 | 0 | 0 | 0 | 0 | 298 | 149 | 0.005529656 | 0.063519651 | 0.945 | 0.927 | 0.961 | 0.97 | 0.991 | 1 | 1 | 1 | 1 | 1 | -0.041 |
| SE | 27592 | ENSG00000132694 | ARHGEF11 | chr1 | - | 156979228 | 156979286 | 156978203 | 156978382 | 156984338 | 156984437 | 23 | 18 | 17 | 14 | 38 | 1 | 0 | 1 | 0 | 0 | 50 | 32 | 14 | 46 | 50 | 0 | 3 | 4 | 0 | 4 | 207 | 149 | 0.005532727 | 0.063545603 | 0.943 | 1 | 0.924 | 1 | 1 | 1 | 0.885 | 0.716 | 1 | 0.9 | 0.073 |
| SE | 15593 | ENSG00000170776 | AKAP13 | chr15 | + | 85741045 | 85741495 | 85736089 | 85736134 | 85743491 | 85743825 | 176 | 62 | 95 | 121 | 63 | 1 | 0 | 0 | 1 | 3 | 78 | 125 | 95 | 56 | 61 | 6 | 0 | 1 | 0 | 6 | 298 | 149 | 0.005537045 | 0.063585868 | 0.989 | 1 | 1 | 0.984 | 0.913 | 0.867 | 1 | 0.979 | 1 | 0.836 | 0.041 |
| SE | 66109 | ENSG00000000938 | FGR | chr1 | - | 27623041 | 27623144 | 27621558 | 27621657 | 27625088 | 27625151 | 573 | 300 | 247 | 320 | 347 | 1 | 1 | 3 | 0 | 2 | 922 | 679 | 324 | 647 | 876 | 0 | 3 | 0 | 10 | 29 | 252 | 149 | 0.005545178 | 0.063669935 | 0.997 | 0.994 | 0.98 | 1 | 0.99 | 1 | 0.993 | 1 | 0.975 | 0.947 | 0.009 |
| SE | 87940 | ENSG00000104361 | NIPAL2 | chr8 | - | 98194727 | 98194822 | 98189825 | 98193090 | 98195941 | 98196005 | 55 | 38 | 20 | 26 | 89 | 1 | 0 | 0 | 1 | 9 | 7 | 23 | 28 | 18 | 6 | 0 | 0 | 0 | 0 | 0 | 244 | 149 | 0.005552487 | 0.063744509 | 0.971 | 1 | 1 | 0.941 | 0.858 | 1 | 1 | 1 | 1 | 1 | -0.046 |
| SE | 80677 | ENSG00000085514 | PILRA | chr7 | + | 100389887 | 100390106 | 100374043 | 100374433 | 100397878 | 100397912 | 421 | 275 | 216 | 304 | 279 | 89 | 59 | 50 | 69 | 64 | 1426 | 515 | 322 | 713 | 1327 | 227 | 94 | 62 | 105 | 230 | 298 | 149 | 0.005560692 | 0.06382934 | 0.703 | 0.7 | 0.684 | 0.688 | 0.686 | 0.759 | 0.733 | 0.722 | 0.772 | 0.743 | -0.054 |
| SE | 1682 | ENSG00000185721 | DRG1 | chr22 | + | 31400619 | 31400743 | 31399603 | 31399725 | 31411011 | 31411081 | 44 | 22 | 60 | 57 | 61 | 0 | 0 | 0 | 0 | 0 | 36 | 72 | 48 | 90 | 78 | 15 | 0 | 0 | 0 | 0 | 273 | 149 | 0.005562756 | 0.06384368 | 1 | 1 | 1 | 1 | 1 | 0.567 | 1 | 1 | 1 | 1 | 0.087 |
| SE | 78759 | ENSG00000179406 | LINC00174 | chr7 | - | 66455658 | 66455853 | 66450070 | 66450236 | 66476258 | 66476327 | 10 | 2 | 6 | 3 | 8 | 3 | 3 | 6 | 1 | 2 | 0 | 16 | 0 | 3 | 2 | 7 | 5 | 3 | 3 | 13 | 298 | 149 | 0.005569144 | 0.063907627 | 0.625 | 0.25 | 0.333 | 0.6 | 0.667 | 0 | 0.615 | 0 | 0.333 | 0.071 | 0.291 |
| SE | 105546 | ENSG00000012822 | CALCOCO1 | chr12 | - | 53712774 | 53712846 | 53711118 | 53712121 | 53713099 | 53713206 | 3 | 1 | 2 | 7 | 3 | 89 | 54 | 57 | 54 | 131 | 1 | 9 | 0 | 16 | 19 | 116 | 87 | 47 | 67 | 219 | 221 | 149 | 0.005572104 | 0.063926575 | 0.022 | 0.012 | 0.023 | 0.08 | 0.015 | 0.006 | 0.065 | 0 | 0.139 | 0.055 | -0.023 |
| SE | 16004 | ENSG00000068305 | MEF2A | chr15 | + | 99671322 | 99671454 | 99645560 | 99645764 | 99671519 | 99671657 | 77 | 85 | 86 | 77 | 118 | 57 | 59 | 51 | 75 | 126 | 71 | 103 | 109 | 86 | 50 | 56 | 56 | 39 | 47 | 15 | 281 | 149 | 0.005572429 | 0.063926575 | 0.417 | 0.433 | 0.472 | 0.352 | 0.332 | 0.402 | 0.494 | 0.597 | 0.492 | 0.639 | -0.124 |
| SE | 31817 | ENSG00000112701 | SENP6 | chr6 | + | 75622750 | 75622858 | 75602223 | 75602576 | 75623899 | 75623960 | 15 | 15 | 16 | 31 | 26 | 3 | 5 | 3 | 1 | 2 | 4 | 7 | 7 | 0 | 10 | 0 | 0 | 2 | 1 | 0 | 257 | 149 | 0.005578872 | 0.063991121 | 0.744 | 0.635 | 0.756 | 0.947 | 0.883 | 1 | 1 | 0.67 | 0 | 1 | 0.059 |
| SE | 92912 | ENSG00000006607 | FARP2 | chr2 | + | 241475856 | 241475987 | 241468139 | 241468377 | 241483464 | 241483533 | 9 | 9 | 20 | 10 | 3 | 0 | 0 | 0 | 0 | 0 | 9 | 6 | 8 | 17 | 24 | 0 | 5 | 1 | 2 | 0 | 280 | 149 | 0.005583532 | 0.064035183 | 1 | 1 | 1 | 1 | 1 | 1 | 0.39 | 0.81 | 0.819 | 1 | 0.196 |
| SE | 81083 | ENSG00000140750 | ARHGAP17 | chr16 | - | 24927683 | 24927732 | 24919394 | 24920260 | 24929612 | 24929660 | 1 | 1 | 1 | 3 | 4 | 2 | 2 | 2 | 4 | 3 | 12 | 7 | 1 | 6 | 18 | 14 | 0 | 0 | 2 | 4 | 198 | 149 | 0.005587861 | 0.064066066 | 0.273 | 0.273 | 0.273 | 0.361 | 0.501 | 0.392 | 1 | 1 | 0.693 | 0.772 | -0.435 |
| SE | 56911 | ENSG00000175970 | UNC119B | chr12 | + | 120716869 | 120717042 | 120716627 | 120716739 | 120719919 | 120723640 | 157 | 80 | 145 | 125 | 157 | 7 | 2 | 2 | 5 | 0 | 205 | 79 | 55 | 106 | 266 | 24 | 4 | 3 | 0 | 7 | 298 | 149 | 0.005593486 | 0.064121169 | 0.918 | 0.952 | 0.973 | 0.926 | 1 | 0.81 | 0.908 | 0.902 | 1 | 0.95 | 0.04 |
| SE | 64078 | ENSG00000110066 | KMT5B | chr11 | - | 68180131 | 68180200 | 68175017 | 68175183 | 68185780 | 68185928 | 346 | 260 | 285 | 275 | 279 | 12 | 8 | 10 | 12 | 14 | 73 | 184 | 112 | 179 | 88 | 0 | 13 | 21 | 20 | 0 | 218 | 149 | 0.005599254 | 0.064177898 | 0.952 | 0.957 | 0.951 | 0.94 | 0.932 | 1 | 0.906 | 0.785 | 0.859 | 1 | 0.036 |
| SE | 58915 | ENSG00000135336 | ORC3 | chr6 | + | 87594526 | 87594613 | 87594352 | 87594407 | 87601783 | 87601881 | 22 | 24 | 34 | 16 | 38 | 77 | 47 | 97 | 88 | 94 | 0 | 11 | 18 | 4 | 0 | 25 | 67 | 40 | 18 | 18 | 236 | 149 | 0.005601993 | 0.06419989 | 0.153 | 0.244 | 0.181 | 0.103 | 0.203 | 0 | 0.094 | 0.221 | 0.123 | 0 | 0.089 |
| SE | 18617 | ENSG00000256053 | COA8 | chr14 | + | 103563214 | 103563379 | 103563052 | 103563124 | 103571622 | 103571820 | 3 | 0 | 6 | 0 | 4 | 11 | 4 | 25 | 8 | 10 | 12 | 8 | 5 | 12 | 3 | 26 | 30 | 7 | 3 | 6 | 298 | 149 | 0.005603148 | 0.064203732 | 0.12 | 0 | 0.107 | 0 | 0.167 | 0.188 | 0.118 | 0.263 | 0.667 | 0.2 | -0.208 |
| SE | 9621 | ENSG00000173821 | RNF213 | chr17 | + | 80306251 | 80306468 | 80298320 | 80298518 | 80307127 | 80307201 | 1178 | 681 | 864 | 1182 | 879 | 6 | 0 | 2 | 2 | 5 | 1892 | 1316 | 845 | 1374 | 1348 | 18 | 6 | 9 | 19 | 7 | 298 | 149 | 0.005610318 | 0.06427648 | 0.99 | 1 | 0.995 | 0.997 | 0.989 | 0.981 | 0.991 | 0.979 | 0.973 | 0.99 | 0.011 |
| SE | 50022 | ENSG00000157873 | TNFRSF14 | chr1 | + | 2561776 | 2561815 | 2560681 | 2560714 | 2562864 | 2562896 | 81 | 22 | 54 | 64 | 72 | 0 | 4 | 2 | 3 | 0 | 104 | 81 | 42 | 57 | 74 | 8 | 15 | 0 | 3 | 0 | 188 | 149 | 0.005634164 | 0.064540231 | 1 | 0.813 | 0.955 | 0.944 | 1 | 0.912 | 0.811 | 1 | 0.938 | 1 | 0.01 |
| SE | 41080 | ENSG00000092841 | MYL6 | chr12 | + | 56159622 | 56159730 | 56158683 | 56158711 | 56159974 | 56160148 | 1056 | 782 | 664 | 1090 | 874 | 62 | 62 | 51 | 94 | 48 | 2938 | 1249 | 754 | 1465 | 5374 | 261 | 146 | 120 | 264 | 176 | 257 | 149 | 0.005637763 | 0.064572019 | 0.908 | 0.88 | 0.883 | 0.871 | 0.913 | 0.867 | 0.832 | 0.785 | 0.763 | 0.947 | 0.052 |
| SE | 32185 | ENSG00000113971 | NPHP3 | chr3 | - | 132683398 | 132683524 | 132681281 | 132682818 | 132684553 | 132684685 | 100 | 106 | 65 | 75 | 155 | 0 | 0 | 2 | 0 | 0 | 23 | 53 | 67 | 4 | 12 | 0 | 4 | 1 | 5 | 0 | 275 | 149 | 0.005640728 | 0.064596523 | 1 | 1 | 0.946 | 1 | 1 | 1 | 0.878 | 0.973 | 0.302 | 1 | 0.159 |
| SE | 17382 | ENSG00000092148 | HECTD1 | chr14 | - | 31128611 | 31129307 | 31122913 | 31123047 | 31133237 | 31133414 | 61 | 28 | 34 | 50 | 73 | 0 | 0 | 0 | 1 | 1 | 18 | 41 | 28 | 7 | 10 | 0 | 0 | 11 | 0 | 0 | 298 | 149 | 0.00564281 | 0.06461092 | 1 | 1 | 1 | 0.962 | 0.973 | 1 | 1 | 0.56 | 1 | 1 | 0.075 |
| SE | 16826 | ENSG00000137878 | GCOM1 | chr15 | + | 57646136 | 57646669 | 57639439 | 57639545 | 57661449 | 57661533 | 6 | 5 | 0 | 7 | 8 | 14 | 6 | 3 | 18 | 15 | 0 | 25 | 1 | 2 | 8 | 16 | 47 | 9 | 10 | 169 | 298 | 149 | 0.005658267 | 0.064778427 | 0.176 | 0.294 | 0 | 0.163 | 0.211 | 0 | 0.21 | 0.053 | 0.091 | 0.023 | 0.093 |
| SE | 14153 | ENSG00000005339 | CREBBP | chr16 | - | 3770569 | 3770986 | 3769173 | 3769353 | 3773750 | 3773930 | 154 | 61 | 118 | 149 | 121 | 1 | 1 | 4 | 5 | 0 | 245 | 199 | 193 | 196 | 268 | 5 | 15 | 5 | 8 | 6 | 298 | 149 | 0.005663708 | 0.064831239 | 0.987 | 0.968 | 0.937 | 0.937 | 1 | 0.961 | 0.869 | 0.951 | 0.925 | 0.957 | 0.033 |
| SE | 14432 | ENSG00000130703 | OSBPL2 | chr20 | + | 62242320 | 62242434 | 62238563 | 62238597 | 62256056 | 62256221 | 0 | 0 | 3 | 2 | 0 | 90 | 67 | 65 | 83 | 37 | 8 | 4 | 5 | 6 | 2 | 127 | 76 | 94 | 127 | 89 | 263 | 149 | 0.005669677 | 0.064890073 | 0 | 0 | 0.025 | 0.013 | 0 | 0.034 | 0.029 | 0.029 | 0.026 | 0.013 | -0.019 |
| SE | 15290 | ENSG00000114062 | UBE3A | chr15 | - | 25354527 | 25354683 | 25352281 | 25354426 | 25355891 | 25356056 | 666 | 507 | 568 | 547 | 1413 | 6 | 13 | 6 | 22 | 23 | 188 | 452 | 359 | 263 | 36 | 8 | 4 | 28 | 8 | 0 | 298 | 149 | 0.005672389 | 0.064911631 | 0.982 | 0.951 | 0.979 | 0.926 | 0.968 | 0.922 | 0.983 | 0.865 | 0.943 | 1 | 0.019 |
| SE | 106398 | ENSG00000089041 | P2RX7 | chr12 | + | 121155235 | 121155374 | 121154784 | 121154953 | 121156078 | 121156147 | 0 | 5 | 3 | 0 | 1 | 64 | 56 | 52 | 65 | 123 | 23 | 0 | 1 | 6 | 0 | 222 | 68 | 53 | 92 | 75 | 288 | 149 | 0.005681035 | 0.065001068 | 0 | 0.044 | 0.029 | 0 | 0.004 | 0.051 | 0 | 0.01 | 0.033 | 0 | -0.003 |
| SE | 13940 | ENSG00000131797 | CLUHP3 | chr16 | + | 31706420 | 31706454 | 31705889 | 31706034 | 31706890 | 31707351 | 1 | 0 | 0 | 7 | 2 | 8 | 11 | 18 | 24 | 13 | 0 | 1 | 0 | 0 | 0 | 4 | 2 | 12 | 18 | 45 | 183 | 149 | 0.005682171 | 0.065004568 | 0.092 | 0 | 0 | 0.192 | 0.111 | 0 | 0.289 | 0 | 0 | 0 | 0.021 |
| SE | 51431 | ENSG00000107745 | MICU1 | chr10 | - | 72508154 | 72508269 | 72477169 | 72477256 | 72533745 | 72533789 | 217 | 186 | 136 | 196 | 504 | 3 | 0 | 2 | 1 | 0 | 333 | 191 | 118 | 271 | 850 | 7 | 2 | 5 | 10 | 2 | 264 | 149 | 0.005685388 | 0.065031868 | 0.976 | 1 | 0.975 | 0.991 | 1 | 0.964 | 0.982 | 0.93 | 0.939 | 0.996 | 0.026 |
| SE | 40640 | ENSG00000143322 | ABL2 | chr1 | - | 179117331 | 179117516 | 179114877 | 179115030 | 179118586 | 179118764 | 33 | 32 | 70 | 57 | 50 | 0 | 0 | 0 | 1 | 0 | 38 | 39 | 70 | 27 | 9 | 0 | 0 | 4 | 6 | 0 | 298 | 149 | 0.005695969 | 0.065143383 | 1 | 1 | 1 | 0.966 | 1 | 1 | 1 | 0.897 | 0.692 | 1 | 0.075 |
| SE | 92517 | ENSG00000216937 | CCDC7 | chr10 | + | 32574428 | 32574526 | 32571858 | 32571893 | 32584231 | 32584304 | 26 | 10 | 9 | 19 | 45 | 7 | 5 | 0 | 8 | 103 | 0 | 23 | 6 | 1 | 22 | 1 | 28 | 0 | 0 | 0 | 247 | 149 | 0.005702964 | 0.06521386 | 0.691 | 0.547 | 1 | 0.589 | 0.209 | 0 | 0.331 | 1 | 1 | 1 | -0.059 |
| SE | 85136 | ENSG00000285171 | NA | chrX | - | 71110503 | 71110688 | 71109227 | 71109390 | 71111424 | 71111571 | 32 | 12 | 30 | 29 | 24 | 12 | 2 | 5 | 5 | 0 | 31 | 66 | 118 | 81 | 79 | 5 | 9 | 1 | 5 | 0 | 298 | 149 | 0.005705265 | 0.065230638 | 0.571 | 0.75 | 0.75 | 0.744 | 1 | 0.756 | 0.786 | 0.983 | 0.89 | 1 | -0.12 |
| SE | 85687 | ENSG00000177981 | ASB8 | chr12 | - | 48151564 | 48151665 | 48151200 | 48151305 | 48153367 | 48153525 | 4 | 5 | 7 | 4 | 1 | 79 | 27 | 62 | 62 | 83 | 0 | 12 | 5 | 11 | 18 | 83 | 88 | 68 | 20 | 68 | 250 | 149 | 0.00571322 | 0.065312053 | 0.029 | 0.099 | 0.063 | 0.037 | 0.007 | 0 | 0.075 | 0.042 | 0.247 | 0.136 | -0.053 |
| SE | 106144 | ENSG00000089022 | MAPKAPK5 | chr12 | + | 111866155 | 111866231 | 111865249 | 111865323 | 111867571 | 111867669 | 287 | 166 | 287 | 282 | 335 | 2 | 2 | 4 | 9 | 7 | 494 | 227 | 200 | 236 | 230 | 0 | 1 | 0 | 0 | 0 | 225 | 149 | 0.005714175 | 0.065313438 | 0.99 | 0.982 | 0.979 | 0.954 | 0.969 | 1 | 0.993 | 1 | 1 | 1 | -0.024 |
| SE | 101974 | ENSG00000107771 | CCSER2 | chr10 | + | 84438511 | 84438707 | 84425730 | 84425893 | 84463932 | 84463945 | 180 | 115 | 223 | 197 | 528 | 10 | 2 | 1 | 5 | 9 | 60 | 72 | 147 | 18 | 63 | 0 | 0 | 5 | 0 | 0 | 298 | 149 | 0.005716248 | 0.065323648 | 0.9 | 0.966 | 0.991 | 0.952 | 0.967 | 1 | 1 | 0.936 | 1 | 1 | -0.032 |
| SE | 90105 | ENSG00000205038 | PKHD1L1 | chr8 | + | 109425084 | 109425232 | 109420517 | 109420690 | 109427001 | 109427156 | 31 | 40 | 3 | 56 | 35 | 1 | 0 | 0 | 0 | 0 | 41 | 53 | 20 | 58 | 212 | 0 | 0 | 2 | 0 | 10 | 297 | 149 | 0.005716739 | 0.065323648 | 0.94 | 1 | 1 | 1 | 1 | 1 | 0.93 | 1 | 1 | 0.914 | 0.019 |
| SE | 86167 | ENSG00000113845 | TIMMDC1 | chr3 | + | 119513640 | 119513719 | 119498566 | 119498927 | 119517204 | 119517315 | 50 | 54 | 59 | 74 | 114 | 0 | 0 | 0 | 0 | 0 | 155 | 57 | 22 | 101 | 116 | 0 | 0 | 1 | 8 | 5 | 228 | 149 | 0.00571797 | 0.065323648 | 1 | 1 | 1 | 1 | 1 | 1 | 1 | 0.935 | 0.892 | 0.938 | 0.047 |
| SE | 1333 | ENSG00000093000 | NUP50 | chr22 | + | 45170987 | 45171177 | 45168165 | 45168246 | 45171599 | 45171683 | 43 | 30 | 87 | 54 | 70 | 158 | 138 | 137 | 138 | 215 | 22 | 53 | 39 | 58 | 4 | 172 | 181 | 131 | 142 | 83 | 298 | 149 | 0.005718406 | 0.065323648 | 0.12 | 0.098 | 0.241 | 0.164 | 0.14 | 0.06 | 0.128 | 0.13 | 0.17 | 0.024 | 0.05 |
| SE | 31792 | ENSG00000204463 | BAG6 | chr6 | - | 31644524 | 31644629 | 31644081 | 31644194 | 31644945 | 31645069 | 13 | 1 | 9 | 11 | 5 | 1 | 0 | 0 | 0 | 2 | 8 | 8 | 7 | 5 | 4 | 0 | 0 | 5 | 4 | 0 | 254 | 149 | 0.005725524 | 0.0653886 | 0.884 | 1 | 1 | 1 | 0.595 | 1 | 1 | 0.451 | 0.423 | 1 | 0.121 |
| SE | 65710 | ENSG00000138434 | ITPRID2 | chr2 | + | 181918322 | 181918470 | 181915215 | 181916427 | 181918597 | 181918675 | 0 | 1 | 4 | 0 | 0 | 37 | 31 | 26 | 31 | 27 | 0 | 9 | 6 | 0 | 0 | 31 | 24 | 46 | 34 | 6 | 297 | 149 | 0.005725762 | 0.0653886 | 0 | 0.016 | 0.072 | 0 | 0 | 0 | 0.158 | 0.061 | 0 | 0 | -0.026 |
| SE | 39532 | ENSG00000111707 | SUDS3 | chr12 | - | 118404055 | 118404473 | 118403500 | 118403517 | 118411072 | 118411147 | 4 | 0 | 3 | 5 | 0 | 81 | 57 | 126 | 92 | 124 | 0 | 6 | 5 | 21 | 0 | 165 | 115 | 84 | 119 | 93 | 298 | 149 | 0.005727137 | 0.065394771 | 0.024 | 0 | 0.012 | 0.026 | 0 | 0 | 0.025 | 0.029 | 0.081 | 0 | -0.015 |
| SE | 88637 | ENSG00000178234 | GALNT11 | chr7 | + | 152079286 | 152079408 | 152027596 | 152027718 | 152094189 | 152094522 | 3 | 0 | 4 | 3 | 5 | 10 | 2 | 9 | 9 | 1 | 0 | 0 | 2 | 0 | 1 | 6 | 8 | 0 | 7 | 4 | 271 | 149 | 0.005728481 | 0.065400577 | 0.142 | 0 | 0.196 | 0.155 | 0.733 | 0 | 0 | 1 | 0 | 0.121 | 0.021 |
| SE | 71741 | ENSG00000179051 | RCC2 | chr1 | - | 17425540 | 17425684 | 17422704 | 17422836 | 17429105 | 17429199 | 321 | 146 | 236 | 284 | 136 | 0 | 0 | 0 | 1 | 1 | 682 | 437 | 172 | 377 | 742 | 8 | 0 | 2 | 12 | 7 | 293 | 149 | 0.005731872 | 0.065429754 | 1 | 1 | 1 | 0.993 | 0.986 | 0.977 | 1 | 0.978 | 0.941 | 0.982 | 0.02 |
| SE | 94653 | ENSG00000011454 | RABGAP1 | chr9 | + | 123074284 | 123074428 | 123073551 | 123073677 | 123076244 | 123076286 | 130 | 125 | 102 | 150 | 249 | 1 | 0 | 0 | 1 | 0 | 133 | 132 | 129 | 117 | 81 | 0 | 3 | 5 | 6 | 0 | 293 | 149 | 0.00573845 | 0.065495351 | 0.985 | 1 | 1 | 0.987 | 1 | 1 | 0.957 | 0.929 | 0.908 | 1 | 0.036 |
| SE | 43088 | ENSG00000239900 | ADSL | chr22 | + | 40354247 | 40354327 | 40349831 | 40350035 | 40358863 | 40358912 | 44 | 43 | 49 | 64 | 74 | 4 | 3 | 2 | 6 | 6 | 86 | 79 | 38 | 50 | 97 | 23 | 0 | 14 | 40 | 8 | 229 | 149 | 0.00574274 | 0.065534701 | 0.877 | 0.903 | 0.941 | 0.874 | 0.889 | 0.709 | 1 | 0.638 | 0.449 | 0.888 | 0.16 |
| SE | 54725 | ENSG00000177409 | SAMD9L | chr7 | - | 93145384 | 93145532 | 93144731 | 93144833 | 93148193 | 93148345 | 154 | 215 | 133 | 220 | 166 | 1 | 0 | 3 | 4 | 4 | 5 | 46 | 28 | 24 | 4 | 0 | 2 | 14 | 0 | 0 | 297 | 149 | 0.005745661 | 0.065558473 | 0.987 | 1 | 0.957 | 0.965 | 0.954 | 1 | 0.92 | 0.501 | 1 | 1 | 0.088 |
| SE | 74337 | ENSG00000187118 | CMC1 | chr3 | + | 28263290 | 28263380 | 28241583 | 28241812 | 28316332 | 28316423 | 8 | 17 | 20 | 20 | 16 | 3 | 0 | 0 | 0 | 6 | 13 | 3 | 6 | 10 | 50 | 0 | 0 | 1 | 0 | 0 | 239 | 149 | 0.005758691 | 0.065697575 | 0.624 | 1 | 1 | 1 | 0.624 | 1 | 1 | 0.789 | 1 | 1 | -0.108 |
| SE | 11328 | ENSG00000033011 | ALG1 | chr16 | + | 5079063 | 5079102 | 5078756 | 5078878 | 5079747 | 5079807 | 31 | 7 | 14 | 30 | 21 | 2 | 1 | 1 | 1 | 1 | 36 | 31 | 8 | 12 | 38 | 15 | 4 | 3 | 0 | 0 | 188 | 149 | 0.005765578 | 0.065766565 | 0.925 | 0.847 | 0.917 | 0.96 | 0.943 | 0.655 | 0.86 | 0.679 | 1 | 1 | 0.08 |
| SE | 64355 | ENSG00000163219 | ARHGAP25 | chr2 | + | 68757637 | 68757730 | 68734951 | 68735260 | 68775220 | 68775417 | 5 | 6 | 1 | 5 | 13 | 155 | 157 | 227 | 169 | 234 | 29 | 0 | 0 | 3 | 11 | 195 | 124 | 129 | 78 | 141 | 242 | 149 | 0.005769253 | 0.065789763 | 0.019 | 0.023 | 0.003 | 0.018 | 0.033 | 0.084 | 0 | 0 | 0.023 | 0.046 | -0.011 |
| SE | 44016 | ENSG00000114098 | ARMC8 | chr3 | + | 138241782 | 138241983 | 138238019 | 138239528 | 138245087 | 138245183 | 312 | 282 | 264 | 335 | 382 | 14 | 17 | 11 | 14 | 28 | 77 | 249 | 293 | 197 | 183 | 26 | 40 | 23 | 23 | 0 | 298 | 149 | 0.005769292 | 0.065789763 | 0.918 | 0.892 | 0.923 | 0.923 | 0.872 | 0.597 | 0.757 | 0.864 | 0.811 | 1 | 0.1 |
| SE | 104679 | ENSG00000100354 | TNRC6B | chr22 | + | 40270121 | 40270280 | 40264687 | 40267036 | 40273424 | 40273600 | 79 | 61 | 121 | 107 | 90 | 122 | 113 | 110 | 150 | 131 | 70 | 187 | 163 | 152 | 88 | 111 | 147 | 140 | 153 | 50 | 298 | 149 | 0.005771563 | 0.065806074 | 0.245 | 0.213 | 0.355 | 0.263 | 0.256 | 0.24 | 0.389 | 0.368 | 0.332 | 0.468 | -0.093 |
| SE | 99442 | ENSG00000109458 | GAB1 | chr4 | + | 143434087 | 143434168 | 143433490 | 143433716 | 143437998 | 143438041 | 1 | 0 | 1 | 3 | 6 | 20 | 11 | 23 | 5 | 27 | 0 | 0 | 0 | 0 | 0 | 26 | 27 | 15 | 2 | 5 | 230 | 149 | 0.005778215 | 0.065872322 | 0.031 | 0 | 0.027 | 0.28 | 0.126 | 0 | 0 | 0 | 0 | 0 | 0.093 |
| SE | 103261 | ENSG00000172661 | WASHC2C | chr10 | + | 45759314 | 45759401 | 45757009 | 45757139 | 45763387 | 45763489 | 133 | 100 | 131 | 138 | 245 | 26 | 28 | 11 | 11 | 54 | 168 | 145 | 105 | 166 | 53 | 96 | 27 | 24 | 78 | 9 | 236 | 149 | 0.005780234 | 0.065885751 | 0.764 | 0.693 | 0.883 | 0.888 | 0.741 | 0.525 | 0.772 | 0.734 | 0.573 | 0.788 | 0.115 |
| SE | 34524 | ENSG00000134802 | SLC43A3 | chr11 | - | 57409934 | 57410121 | 57409174 | 57409298 | 57414614 | 57414731 | 106 | 46 | 65 | 111 | 78 | 0 | 1 | 0 | 0 | 0 | 190 | 181 | 83 | 181 | 419 | 0 | 7 | 1 | 10 | 0 | 298 | 149 | 0.005791138 | 0.066000426 | 1 | 0.958 | 1 | 1 | 1 | 1 | 0.928 | 0.976 | 0.9 | 1 | 0.031 |
| SE | 86665 | ENSG00000196924 | FLNA | chrX | - | 154366023 | 154366224 | 154365348 | 154365486 | 154366307 | 154366334 | 3787 | 1477 | 1996 | 2963 | 1173 | 26 | 9 | 18 | 24 | 10 | 4988 | 3796 | 2182 | 2754 | 18200 | 69 | 33 | 33 | 60 | 109 | 298 | 149 | 0.005813653 | 0.066247385 | 0.986 | 0.988 | 0.982 | 0.984 | 0.983 | 0.973 | 0.983 | 0.971 | 0.958 | 0.988 | 0.01 |
| SE | 84820 | ENSG00000107185 | RGP1 | chr9 | + | 35751626 | 35751754 | 35751265 | 35751412 | 35752650 | 35752802 | 59 | 25 | 59 | 47 | 44 | 7 | 0 | 0 | 5 | 1 | 45 | 51 | 19 | 25 | 92 | 0 | 6 | 5 | 11 | 11 | 277 | 149 | 0.005816477 | 0.066269924 | 0.819 | 1 | 1 | 0.835 | 0.959 | 1 | 0.821 | 0.671 | 0.55 | 0.818 | 0.151 |
| SE | 54616 | ENSG00000073578 | SDHA | chr5 | + | 220239 | 220337 | 218326 | 218418 | 223481 | 223568 | 1 | 3 | 0 | 1 | 0 | 84 | 38 | 69 | 70 | 15 | 2 | 2 | 1 | 6 | 14 | 97 | 113 | 60 | 58 | 177 | 247 | 149 | 0.005819273 | 0.066292134 | 0.007 | 0.045 | 0 | 0.009 | 0 | 0.012 | 0.011 | 0.01 | 0.059 | 0.046 | -0.015 |
| SE | 81291 | ENSG00000155096 | AZIN1 | chr8 | - | 102849998 | 102850144 | 102843550 | 102843747 | 102858012 | 102858150 | 185 | 116 | 169 | 195 | 156 | 206 | 116 | 187 | 198 | 179 | 139 | 205 | 243 | 303 | 124 | 111 | 204 | 146 | 213 | 128 | 295 | 149 | 0.005823079 | 0.066325838 | 0.312 | 0.336 | 0.313 | 0.332 | 0.306 | 0.387 | 0.337 | 0.457 | 0.418 | 0.329 | -0.066 |
| SE | 15473 | ENSG00000140526 | ABHD2 | chr15 | + | 89103902 | 89104044 | 89088421 | 89088563 | 89113724 | 89113824 | 2 | 0 | 1 | 0 | 1 | 55 | 26 | 52 | 64 | 24 | 10 | 1 | 4 | 0 | 0 | 36 | 39 | 34 | 49 | 14 | 291 | 149 | 0.005828611 | 0.066379189 | 0.018 | 0 | 0.01 | 0 | 0.021 | 0.125 | 0.013 | 0.057 | 0 | 0 | -0.029 |
| SE | 30222 | ENSG00000132376 | INPP5K | chr17 | - | 1513452 | 1513561 | 1509177 | 1509353 | 1513871 | 1513979 | 38 | 22 | 19 | 21 | 26 | 1 | 2 | 3 | 1 | 1 | 47 | 45 | 21 | 24 | 81 | 3 | 8 | 0 | 18 | 11 | 258 | 149 | 0.005831736 | 0.066389008 | 0.956 | 0.864 | 0.785 | 0.924 | 0.938 | 0.9 | 0.765 | 1 | 0.435 | 0.81 | 0.111 |
| SE | 83436 | ENSG00000122545 | SEPTIN7 | chr7 | + | 35838563 | 35838657 | 35832797 | 35832900 | 35863551 | 35863658 | 14 | 13 | 20 | 15 | 39 | 701 | 524 | 473 | 693 | 857 | 0 | 15 | 8 | 12 | 0 | 55 | 275 | 119 | 38 | 27 | 243 | 149 | 0.005831878 | 0.066389008 | 0.012 | 0.015 | 0.025 | 0.013 | 0.027 | 0 | 0.032 | 0.04 | 0.162 | 0 | -0.028 |
| SE | 11986 | ENSG00000156860 | FBRS | chr16 | + | 30666511 | 30666541 | 30665637 | 30665706 | 30666918 | 30666990 | 77 | 40 | 57 | 71 | 66 | 1 | 0 | 1 | 3 | 2 | 27 | 89 | 80 | 79 | 224 | 19 | 1 | 3 | 5 | 0 | 179 | 149 | 0.005832017 | 0.066389008 | 0.985 | 1 | 0.979 | 0.952 | 0.965 | 0.542 | 0.987 | 0.957 | 0.929 | 1 | 0.093 |
| SE | 24518 | ENSG00000170144 | HNRNPA3 | chr2 | + | 177217704 | 177217845 | 177215977 | 177216940 | 177219036 | 177219156 | 342 | 334 | 322 | 349 | 370 | 15 | 26 | 21 | 54 | 23 | 212 | 300 | 388 | 328 | 84 | 19 | 27 | 80 | 38 | 8 | 290 | 149 | 0.005836253 | 0.066427567 | 0.921 | 0.868 | 0.887 | 0.769 | 0.892 | 0.851 | 0.851 | 0.714 | 0.816 | 0.844 | 0.052 |
| SE | 52352 | ENSG00000142197 | DOP1B | chr21 | + | 36239764 | 36239955 | 36232803 | 36233075 | 36245047 | 36246677 | 20 | 24 | 14 | 14 | 12 | 0 | 0 | 0 | 0 | 0 | 32 | 24 | 7 | 55 | 25 | 0 | 0 | 0 | 0 | 13 | 298 | 149 | 0.005850899 | 0.066577054 | 1 | 1 | 1 | 1 | 1 | 1 | 1 | 1 | 1 | 0.49 | 0.102 |
| SE | 49687 | ENSG00000263956 | NBPF11 | chr1 | - | 148117571 | 148117786 | 148116462 | 148116535 | 148118619 | 148118722 | 38 | 23 | 21 | 22 | 22 | 0 | 0 | 1 | 1 | 0 | 9 | 14 | 14 | 19 | 4 | 0 | 1 | 3 | 4 | 0 | 298 | 149 | 0.005851087 | 0.066577054 | 1 | 1 | 0.913 | 0.917 | 1 | 1 | 0.875 | 0.7 | 0.704 | 1 | 0.11 |
| SE | 67926 | ENSG00000091490 | SEL1L3 | chr4 | - | 25758940 | 25759068 | 25747432 | 25748564 | 25765325 | 25765360 | 84 | 48 | 95 | 72 | 164 | 0 | 0 | 0 | 0 | 0 | 28 | 103 | 46 | 45 | 148 | 21 | 1 | 0 | 0 | 0 | 277 | 149 | 0.005852082 | 0.066578695 | 1 | 1 | 1 | 1 | 1 | 0.418 | 0.982 | 1 | 1 | 1 | 0.12 |
| SE | 57992 | ENSG00000175634 | RPS6KB2 | chr11 | + | 67432352 | 67432492 | 67431367 | 67431515 | 67432599 | 67432657 | 13 | 4 | 5 | 9 | 14 | 84 | 44 | 71 | 67 | 57 | 12 | 12 | 7 | 3 | 6 | 138 | 96 | 53 | 125 | 157 | 289 | 149 | 0.005856109 | 0.066614839 | 0.074 | 0.045 | 0.035 | 0.065 | 0.112 | 0.043 | 0.061 | 0.064 | 0.012 | 0.019 | 0.026 |
| SE | 91122 | ENSG00000121022 | COPS5 | chr8 | - | 67060496 | 67060545 | 67059210 | 67059445 | 67061853 | 67062024 | 1 | 1 | 2 | 4 | 0 | 150 | 83 | 119 | 135 | 105 | 0 | 6 | 7 | 0 | 6 | 78 | 72 | 94 | 99 | 76 | 198 | 149 | 0.005863956 | 0.066694403 | 0.005 | 0.009 | 0.012 | 0.022 | 0 | 0 | 0.059 | 0.053 | 0 | 0.056 | -0.024 |
| SE | 26732 | ENSG00000125089 | SH3TC1 | chr4 | + | 8205166 | 8205366 | 8199289 | 8199405 | 8209664 | 8209822 | 19 | 6 | 11 | 1 | 8 | 0 | 0 | 0 | 0 | 0 | 7 | 14 | 8 | 2 | 13 | 0 | 0 | 0 | 2 | 9 | 298 | 149 | 0.005866951 | 0.066711138 | 1 | 1 | 1 | 1 | 1 | 1 | 1 | 1 | 0.333 | 0.419 | 0.25 |
| SE | 41118 | ENSG00000115523 | GNLY | chr2 | + | 85694752 | 85694996 | 85694292 | 85694470 | 85695319 | 85695423 | 342 | 56 | 69 | 134 | 185 | 2496 | 480 | 702 | 845 | 838 | 67 | 42 | 61 | 22 | 94 | 645 | 335 | 714 | 334 | 509 | 298 | 149 | 0.005867131 | 0.066711138 | 0.064 | 0.055 | 0.047 | 0.073 | 0.099 | 0.049 | 0.059 | 0.041 | 0.032 | 0.085 | 0.014 |
| SE | 57282 | ENSG00000143119 | CD53 | chr1 | + | 110896652 | 110896733 | 110894959 | 110895055 | 110897808 | 110897892 | 1889 | 802 | 1200 | 1523 | 1769 | 27 | 10 | 21 | 8 | 21 | 1792 | 1490 | 745 | 1068 | 1763 | 25 | 24 | 12 | 11 | 56 | 230 | 149 | 0.005873643 | 0.066766974 | 0.978 | 0.981 | 0.974 | 0.992 | 0.982 | 0.979 | 0.976 | 0.976 | 0.984 | 0.953 | 0.008 |
| SE | 52556 | ENSG00000153283 | CD96 | chr3 | + | 111600725 | 111600914 | 111598119 | 111598210 | 111606699 | 111606792 | 795 | 604 | 819 | 861 | 1825 | 9 | 1 | 1 | 10 | 30 | 109 | 308 | 358 | 152 | 87 | 0 | 0 | 3 | 0 | 0 | 298 | 149 | 0.005873747 | 0.066766974 | 0.978 | 0.997 | 0.998 | 0.977 | 0.968 | 1 | 1 | 0.984 | 1 | 1 | -0.013 |
| SE | 107284 | ENSG00000090686 | USP48 | chr1 | - | 21721649 | 21721764 | 21721035 | 21721166 | 21723897 | 21724095 | 199 | 202 | 238 | 252 | 308 | 4 | 4 | 1 | 6 | 16 | 56 | 200 | 182 | 107 | 97 | 5 | 6 | 9 | 16 | 0 | 264 | 149 | 0.005878113 | 0.066806899 | 0.966 | 0.966 | 0.993 | 0.96 | 0.916 | 0.863 | 0.95 | 0.919 | 0.791 | 1 | 0.056 |
| SE | 10107 | ENSG00000108433 | GOSR2 | chr17 | + | 46931101 | 46931207 | 46929519 | 46929584 | 46935028 | 46935169 | 3 | 2 | 4 | 5 | 0 | 1 | 0 | 2 | 0 | 1 | 12 | 7 | 6 | 0 | 16 | 10 | 7 | 0 | 2 | 3 | 255 | 149 | 0.005888686 | 0.066917356 | 0.637 | 1 | 0.539 | 1 | 0 | 0.412 | 0.369 | 1 | 0 | 0.757 | 0.128 |
| SE | 44130 | ENSG00000136717 | BIN1 | chr2 | - | 127050420 | 127050522 | 127048026 | 127048633 | 127059010 | 127059155 | 46 | 14 | 37 | 63 | 21 | 6 | 1 | 0 | 0 | 3 | 23 | 45 | 19 | 34 | 44 | 0 | 1 | 0 | 0 | 0 | 251 | 149 | 0.005889744 | 0.066919668 | 0.82 | 0.893 | 1 | 1 | 0.806 | 1 | 0.964 | 1 | 1 | 1 | -0.089 |
| SE | 42724 | ENSG00000182473 | EXOC7 | chr17 | - | 76090328 | 76090397 | 76089174 | 76089320 | 76094413 | 76094581 | 24 | 2 | 8 | 21 | 8 | 52 | 29 | 39 | 46 | 36 | 13 | 33 | 7 | 22 | 31 | 42 | 45 | 19 | 32 | 50 | 218 | 149 | 0.005895003 | 0.066969701 | 0.24 | 0.045 | 0.123 | 0.238 | 0.132 | 0.175 | 0.334 | 0.201 | 0.32 | 0.298 | -0.11 |
| SE | 7651 | ENSG00000170296 | GABARAP | chr17 | - | 7241600 | 7241679 | 7241270 | 7241460 | 7242240 | 7242447 | 3481 | 2253 | 2678 | 3090 | 4650 | 12 | 17 | 14 | 17 | 16 | 7632 | 4038 | 2614 | 3980 | 8619 | 62 | 9 | 29 | 76 | 58 | 228 | 149 | 0.005896393 | 0.066975766 | 0.995 | 0.989 | 0.992 | 0.992 | 0.995 | 0.988 | 0.997 | 0.983 | 0.972 | 0.99 | 0.007 |
| SE | 1280 | ENSG00000186951 | PPARA | chr22 | + | 46231791 | 46232239 | 46218262 | 46218401 | 46235132 | 46235403 | 27 | 14 | 24 | 23 | 16 | 0 | 3 | 0 | 1 | 1 | 24 | 23 | 14 | 27 | 17 | 0 | 4 | 0 | 9 | 0 | 298 | 149 | 0.005904053 | 0.067053049 | 1 | 0.7 | 1 | 0.92 | 0.889 | 1 | 0.742 | 1 | 0.6 | 1 | 0.033 |
| SE | 10776 | ENSG00000105556 | MIER2 | chr19 | - | 330263 | 330380 | 327863 | 327989 | 334399 | 334542 | 1 | 0 | 1 | 0 | 0 | 51 | 21 | 18 | 26 | 11 | 0 | 1 | 6 | 0 | 10 | 74 | 51 | 21 | 32 | 67 | 266 | 149 | 0.005921074 | 0.067236608 | 0.011 | 0 | 0.03 | 0 | 0 | 0 | 0.011 | 0.138 | 0 | 0.077 | -0.037 |
| SE | 25696 | ENSG00000247774 | PCED1B-AS1 | chr12 | - | 47206510 | 47206585 | 47205898 | 47206069 | 47210639 | 47210737 | 51 | 31 | 123 | 79 | 154 | 21 | 7 | 17 | 13 | 42 | 37 | 33 | 48 | 26 | 6 | 0 | 1 | 4 | 13 | 0 | 224 | 149 | 0.005925924 | 0.067281927 | 0.618 | 0.747 | 0.828 | 0.802 | 0.709 | 1 | 0.956 | 0.889 | 0.571 | 1 | -0.142 |
| SE | 9968 | ENSG00000266173 | STRADA | chr17 | - | 63723297 | 63723326 | 63714005 | 63714108 | 63726637 | 63726695 | 2 | 3 | 2 | 1 | 1 | 18 | 8 | 6 | 23 | 33 | 5 | 3 | 1 | 5 | 2 | 0 | 4 | 2 | 14 | 22 | 178 | 149 | 0.005948794 | 0.067531793 | 0.085 | 0.239 | 0.218 | 0.035 | 0.025 | 1 | 0.386 | 0.295 | 0.23 | 0.071 | -0.276 |
| SE | 4744 | ENSG00000031823 | RANBP3 | chr19 | - | 5957917 | 5957935 | 5941798 | 5941835 | 5978060 | 5978140 | 9 | 2 | 6 | 8 | 15 | 0 | 2 | 1 | 1 | 5 | 21 | 1 | 5 | 8 | 23 | 0 | 9 | 11 | 5 | 24 | 167 | 149 | 0.005952897 | 0.067568574 | 1 | 0.472 | 0.843 | 0.877 | 0.728 | 1 | 0.09 | 0.289 | 0.588 | 0.461 | 0.298 |
| SE | 28396 | ENSG00000204842 | ATXN2 | chr12 | - | 111485270 | 111485331 | 111470557 | 111470742 | 111485712 | 111485744 | 16 | 22 | 18 | 23 | 20 | 0 | 1 | 2 | 0 | 0 | 10 | 19 | 15 | 18 | 57 | 0 | 3 | 0 | 0 | 7 | 210 | 149 | 0.005956283 | 0.067597202 | 1 | 0.94 | 0.865 | 1 | 1 | 1 | 0.818 | 1 | 1 | 0.852 | 0.027 |
| SE | 107814 | ENSG00000050426 | LETMD1 | chr12 | + | 51056143 | 51056245 | 51049033 | 51049185 | 51056349 | 51056428 | 50 | 26 | 48 | 41 | 61 | 32 | 13 | 32 | 44 | 60 | 49 | 77 | 45 | 21 | 30 | 13 | 16 | 19 | 17 | 17 | 251 | 149 | 0.005958879 | 0.067604963 | 0.481 | 0.543 | 0.471 | 0.356 | 0.376 | 0.691 | 0.741 | 0.584 | 0.423 | 0.512 | -0.145 |
| SE | 1827 | ENSG00000070371 | CLTCL1 | chr22 | - | 19233165 | 19233318 | 19232475 | 19232598 | 19233421 | 19233622 | 6 | 4 | 5 | 1 | 12 | 0 | 0 | 0 | 0 | 0 | 57 | 12 | 3 | 85 | 42 | 4 | 0 | 0 | 6 | 0 | 298 | 149 | 0.005959146 | 0.067604963 | 1 | 1 | 1 | 1 | 1 | 0.877 | 1 | 1 | 0.876 | 1 | 0.049 |
| SE | 84369 | ENSG00000230590 | FTX | chrX | - | 74284742 | 74284944 | 74281701 | 74281848 | 74292168 | 74292359 | 3 | 5 | 2 | 1 | 1 | 15 | 19 | 12 | 18 | 29 | 0 | 0 | 0 | 0 | 0 | 36 | 5 | 15 | 7 | 36 | 298 | 149 | 0.005959969 | 0.067604963 | 0.091 | 0.116 | 0.077 | 0.027 | 0.017 | 0 | 0 | 0 | 0 | 0 | 0.066 |
| SE | 20840 | ENSG00000213809 | KLRK1 | chr12 | - | 10379446 | 10379482 | 10378553 | 10378705 | 10379699 | 10379792 | 1511 | 763 | 867 | 1184 | 1588 | 20 | 7 | 12 | 25 | 20 | 165 | 817 | 558 | 333 | 46 | 7 | 0 | 7 | 28 | 6 | 185 | 149 | 0.00596042 | 0.067604963 | 0.984 | 0.989 | 0.983 | 0.974 | 0.985 | 0.95 | 1 | 0.985 | 0.905 | 0.861 | 0.043 |
| SE | 26355 | ENSG00000048028 | USP28 | chr11 | - | 113831919 | 113831993 | 113830866 | 113830946 | 113833419 | 113833470 | 68 | 25 | 34 | 54 | 45 | 7 | 1 | 0 | 4 | 0 | 10 | 32 | 22 | 19 | 26 | 0 | 3 | 0 | 0 | 0 | 223 | 149 | 0.005975836 | 0.067770001 | 0.867 | 0.944 | 1 | 0.9 | 1 | 1 | 0.877 | 1 | 1 | 1 | -0.033 |
| SE | 72555 | ENSG00000068724 | TTC7A | chr2 | + | 46994356 | 46994514 | 46993449 | 46993528 | 46995135 | 46995199 | 280 | 138 | 167 | 234 | 173 | 3 | 2 | 2 | 0 | 2 | 426 | 286 | 180 | 215 | 414 | 9 | 16 | 1 | 3 | 0 | 298 | 149 | 0.005981186 | 0.067820847 | 0.979 | 0.972 | 0.977 | 1 | 0.977 | 0.959 | 0.899 | 0.989 | 0.973 | 1 | 0.017 |
| SE | 6053 | ENSG00000132792 | CTNNBL1 | chr20 | + | 37694926 | 37695033 | 37694031 | 37694152 | 37732878 | 37733067 | 2 | 0 | 4 | 8 | 3 | 26 | 47 | 46 | 72 | 39 | 0 | 5 | 0 | 5 | 0 | 134 | 48 | 57 | 85 | 106 | 256 | 149 | 0.005995699 | 0.06797557 | 0.043 | 0 | 0.048 | 0.061 | 0.043 | 0 | 0.057 | 0 | 0.033 | 0 | 0.021 |
| SE | 54233 | ENSG00000143643 | TTC13 | chr1 | - | 230945388 | 230945454 | 230933778 | 230933861 | 230954332 | 230954403 | 104 | 68 | 64 | 76 | 78 | 11 | 3 | 2 | 3 | 5 | 57 | 60 | 58 | 47 | 109 | 0 | 7 | 1 | 1 | 0 | 215 | 149 | 0.00600211 | 0.068038395 | 0.868 | 0.94 | 0.957 | 0.946 | 0.915 | 1 | 0.856 | 0.976 | 0.97 | 1 | -0.035 |
| SE | 39564 | ENSG00000144554 | FANCD2 | chr3 | + | 10064355 | 10064429 | 10063791 | 10063911 | 10064728 | 10064875 | 28 | 28 | 35 | 38 | 14 | 2 | 6 | 0 | 3 | 1 | 27 | 34 | 54 | 47 | 21 | 6 | 12 | 8 | 12 | 0 | 223 | 149 | 0.006005717 | 0.068069424 | 0.903 | 0.757 | 1 | 0.894 | 0.903 | 0.75 | 0.654 | 0.819 | 0.724 | 1 | 0.102 |
| SE | 25046 | ENSG00000164308 | ERAP2 | chr5 | + | 96900120 | 96900189 | 96896731 | 96896863 | 96901505 | 96901681 | 322 | 522 | 681 | 576 | 720 | 2 | 1 | 1 | 2 | 0 | 29 | 219 | 36 | 124 | 0 | 0 | 12 | 3 | 2 | 8 | 218 | 149 | 0.006012731 | 0.068139065 | 0.991 | 0.997 | 0.998 | 0.995 | 1 | 1 | 0.98 | 0.925 | 0.977 | 0 | 0.22 |
| SE | 82471 | ENSG00000119471 | HSDL2 | chr9 | + | 112438742 | 112438812 | 112438430 | 112438625 | 112441698 | 112441770 | 5 | 8 | 7 | 9 | 10 | 44 | 59 | 47 | 62 | 116 | 3 | 7 | 14 | 19 | 9 | 0 | 49 | 50 | 29 | 12 | 219 | 149 | 0.006016468 | 0.068169387 | 0.072 | 0.084 | 0.092 | 0.09 | 0.055 | 1 | 0.089 | 0.16 | 0.308 | 0.338 | -0.3 |
| SE | 39496 | ENSG00000010322 | NISCH | chr3 | + | 52480878 | 52481037 | 52480183 | 52480295 | 52484512 | 52484637 | 7 | 12 | 32 | 15 | 25 | 52 | 15 | 31 | 21 | 42 | 14 | 37 | 14 | 6 | 13 | 68 | 80 | 17 | 30 | 120 | 298 | 149 | 0.006018019 | 0.068169387 | 0.063 | 0.286 | 0.34 | 0.263 | 0.229 | 0.093 | 0.188 | 0.292 | 0.091 | 0.051 | 0.093 |
| SE | 39497 | ENSG00000010322 | NISCH | chr3 | + | 52480878 | 52481109 | 52480138 | 52480295 | 52484512 | 52484637 | 7 | 12 | 32 | 15 | 25 | 52 | 15 | 31 | 21 | 42 | 14 | 37 | 14 | 6 | 13 | 68 | 80 | 17 | 30 | 120 | 298 | 149 | 0.006018019 | 0.068169387 | 0.063 | 0.286 | 0.34 | 0.263 | 0.229 | 0.093 | 0.188 | 0.292 | 0.091 | 0.051 | 0.093 |
| SE | 23942 | ENSG00000133835 | HSD17B4 | chr5 | + | 119471635 | 119471700 | 119456314 | 119456368 | 119473907 | 119474015 | 10 | 25 | 12 | 24 | 14 | 111 | 75 | 80 | 77 | 99 | 35 | 35 | 35 | 25 | 3 | 100 | 93 | 45 | 114 | 25 | 214 | 149 | 0.006021512 | 0.068199092 | 0.059 | 0.188 | 0.095 | 0.178 | 0.09 | 0.196 | 0.208 | 0.351 | 0.132 | 0.077 | -0.071 |
| SE | 44245 | ENSG00000102606 | ARHGEF7 | chr13 | + | 111280271 | 111280350 | 111277586 | 111277673 | 111280537 | 111280677 | 189 | 167 | 196 | 212 | 311 | 2 | 2 | 9 | 8 | 1 | 195 | 179 | 162 | 126 | 110 | 16 | 3 | 9 | 6 | 0 | 228 | 149 | 0.006024672 | 0.068225009 | 0.984 | 0.982 | 0.934 | 0.945 | 0.995 | 0.888 | 0.975 | 0.922 | 0.932 | 1 | 0.025 |
| SE | 31645 | ENSG00000131389 | SLC6A6 | chr3 | + | 14468087 | 14468212 | 14467852 | 14467956 | 14472204 | 14472317 | 61 | 26 | 47 | 64 | 51 | 4 | 7 | 1 | 10 | 1 | 103 | 40 | 45 | 37 | 155 | 0 | 0 | 5 | 5 | 2 | 274 | 149 | 0.00604088 | 0.068390387 | 0.892 | 0.669 | 0.962 | 0.777 | 0.965 | 1 | 1 | 0.83 | 0.801 | 0.977 | -0.069 |
| SE | 76194 | ENSG00000112237 | CCNC | chr6 | - | 99544205 | 99544273 | 99543463 | 99543609 | 99545111 | 99545230 | 2 | 3 | 3 | 2 | 11 | 55 | 42 | 70 | 62 | 107 | 0 | 5 | 0 | 0 | 0 | 28 | 71 | 53 | 9 | 6 | 217 | 149 | 0.006041023 | 0.068390387 | 0.024 | 0.047 | 0.029 | 0.022 | 0.066 | 0 | 0.046 | 0 | 0 | 0 | 0.028 |
| SE | 72185 | ENSG00000164902 | PHAX | chr5 | + | 126617249 | 126617333 | 126608356 | 126608484 | 126624574 | 126625209 | 201 | 153 | 128 | 127 | 219 | 4 | 1 | 1 | 1 | 10 | 75 | 82 | 142 | 95 | 1 | 5 | 13 | 3 | 1 | 0 | 233 | 149 | 0.006042638 | 0.068398782 | 0.97 | 0.99 | 0.988 | 0.988 | 0.933 | 0.906 | 0.801 | 0.968 | 0.984 | 1 | 0.042 |
| SE | 75738 | ENSG00000144580 | CNOT9 | chr2 | + | 218580560 | 218580740 | 218568579 | 218568978 | 218582970 | 218583086 | 117 | 91 | 97 | 100 | 184 | 0 | 0 | 0 | 4 | 0 | 195 | 90 | 135 | 119 | 138 | 0 | 0 | 0 | 0 | 16 | 298 | 149 | 0.006049107 | 0.068462107 | 1 | 1 | 1 | 0.926 | 1 | 1 | 1 | 1 | 1 | 0.812 | 0.023 |
| SE | 20337 | ENSG00000111913 | RIPOR2 | chr6 | - | 25015757 | 25015835 | 25014951 | 25015583 | 25041850 | 25042170 | 11 | 6 | 9 | 26 | 25 | 2 | 7 | 4 | 4 | 4 | 29 | 137 | 19 | 49 | 91 | 0 | 3 | 4 | 12 | 7 | 227 | 149 | 0.006050222 | 0.068464834 | 0.783 | 0.36 | 0.596 | 0.81 | 0.804 | 1 | 0.968 | 0.757 | 0.728 | 0.895 | -0.199 |
| SE | 66616 | ENSG00000170445 | HARS1 | chr5 | - | 140677326 | 140677457 | 140676988 | 140677116 | 140677654 | 140677753 | 65 | 40 | 58 | 60 | 77 | 0 | 0 | 0 | 3 | 1 | 22 | 43 | 35 | 55 | 34 | 4 | 0 | 3 | 3 | 0 | 280 | 149 | 0.006056204 | 0.068522618 | 1 | 1 | 1 | 0.914 | 0.976 | 0.745 | 1 | 0.861 | 0.907 | 1 | 0.075 |
| SE | 1986 | ENSG00000083814 | ZNF671 | chr19 | - | 57726907 | 57727025 | 57723264 | 57723340 | 57727390 | 57727614 | 1 | 2 | 4 | 0 | 10 | 19 | 10 | 22 | 11 | 1 | 0 | 0 | 0 | 4 | 0 | 2 | 25 | 15 | 5 | 18 | 267 | 149 | 0.00606476 | 0.068609515 | 0.029 | 0.1 | 0.092 | 0 | 0.848 | 0 | 0 | 0 | 0.309 | 0 | 0.152 |
| SE | 18928 | ENSG00000100461 | RBM23 | chr14 | - | 22909482 | 22909595 | 22908332 | 22908380 | 22918998 | 22919138 | 36 | 28 | 23 | 40 | 37 | 0 | 1 | 0 | 0 | 0 | 71 | 36 | 38 | 58 | 63 | 0 | 0 | 4 | 7 | 0 | 262 | 149 | 0.00607248 | 0.068686927 | 1 | 0.941 | 1 | 1 | 1 | 1 | 1 | 0.844 | 0.825 | 1 | 0.054 |
| SE | 103663 | ENSG00000148634 | HERC4 | chr10 | - | 67959112 | 67959136 | 67956877 | 67956976 | 67966682 | 67966802 | 9 | 3 | 10 | 19 | 19 | 161 | 123 | 141 | 141 | 221 | 0 | 2 | 2 | 5 | 3 | 35 | 69 | 74 | 31 | 74 | 173 | 149 | 0.006075088 | 0.068706499 | 0.046 | 0.021 | 0.058 | 0.104 | 0.069 | 0 | 0.024 | 0.023 | 0.122 | 0.034 | 0.019 |
| SE | 48260 | ENSG00000111602 | TIMELESS | chr12 | - | 56422846 | 56422992 | 56422105 | 56422191 | 56423273 | 56423475 | 26 | 27 | 18 | 34 | 19 | 0 | 0 | 0 | 0 | 0 | 69 | 26 | 22 | 32 | 31 | 7 | 0 | 3 | 0 | 0 | 295 | 149 | 0.006084251 | 0.068800185 | 1 | 1 | 1 | 1 | 1 | 0.833 | 1 | 0.787 | 1 | 1 | 0.076 |
| SE | 72831 | ENSG00000129158 | SERGEF | chr11 | - | 17926737 | 17926846 | 17878207 | 17878244 | 17988596 | 17988755 | 0 | 0 | 1 | 0 | 0 | 36 | 20 | 26 | 26 | 35 | 0 | 0 | 0 | 0 | 14 | 47 | 19 | 29 | 11 | 45 | 258 | 149 | 0.006091906 | 0.068876802 | 0 | 0 | 0.022 | 0 | 0 | 0 | 0 | 0 | 0 | 0.152 | -0.026 |
| SE | 67621 | ENSG00000175215 | CTDSP2 | chr12 | - | 57826995 | 57827097 | 57826345 | 57826402 | 57827533 | 57827590 | 220 | 136 | 248 | 272 | 199 | 12 | 0 | 7 | 2 | 1 | 315 | 254 | 163 | 168 | 475 | 0 | 2 | 1 | 3 | 5 | 251 | 149 | 0.006095468 | 0.068899657 | 0.916 | 1 | 0.955 | 0.988 | 0.992 | 1 | 0.987 | 0.99 | 0.971 | 0.983 | -0.016 |
| SE | 689 | ENSG00000186575 | NF2 | chr22 | + | 29636750 | 29636876 | 29603555 | 29604112 | 29639089 | 29639212 | 183 | 97 | 148 | 205 | 121 | 0 | 3 | 1 | 0 | 0 | 324 | 219 | 147 | 127 | 314 | 8 | 11 | 0 | 1 | 0 | 275 | 149 | 0.006095687 | 0.068899657 | 1 | 0.946 | 0.988 | 1 | 1 | 0.956 | 0.915 | 1 | 0.986 | 1 | 0.015 |
| SE | 104758 | ENSG00000011258 | MBTD1 | chr17 | - | 51202021 | 51202077 | 51201591 | 51201696 | 51202700 | 51202935 | 96 | 83 | 72 | 131 | 164 | 9 | 8 | 5 | 8 | 14 | 53 | 84 | 83 | 53 | 51 | 7 | 19 | 14 | 18 | 0 | 205 | 149 | 0.006096726 | 0.068901457 | 0.886 | 0.883 | 0.913 | 0.922 | 0.895 | 0.846 | 0.763 | 0.812 | 0.682 | 1 | 0.079 |
| SE | 61421 | ENSG00000053900 | ANAPC4 | chr4 | + | 25417615 | 25417739 | 25416424 | 25416598 | 25418154 | 25418261 | 319 | 398 | 363 | 408 | 478 | 10 | 3 | 0 | 2 | 2 | 117 | 174 | 178 | 92 | 29 | 3 | 0 | 3 | 11 | 0 | 273 | 149 | 0.006100756 | 0.068937042 | 0.946 | 0.986 | 1 | 0.991 | 0.992 | 0.955 | 1 | 0.97 | 0.82 | 1 | 0.034 |
| SE | 82389 | ENSG00000008282 | SYPL1 | chr7 | - | 106092547 | 106092669 | 106091406 | 106091939 | 106092948 | 106093137 | 5 | 0 | 6 | 15 | 5 | 327 | 286 | 349 | 432 | 343 | 2 | 6 | 35 | 10 | 0 | 324 | 362 | 274 | 376 | 164 | 271 | 149 | 0.006103942 | 0.068963096 | 0.008 | 0 | 0.009 | 0.019 | 0.008 | 0.003 | 0.009 | 0.066 | 0.014 | 0 | -0.01 |
| SE | 81746 | ENSG00000176454 | LPCAT4 | chr15 | - | 34359588 | 34359745 | 34358632 | 34359302 | 34360110 | 34360209 | 76 | 37 | 73 | 67 | 111 | 0 | 0 | 0 | 0 | 0 | 69 | 57 | 56 | 62 | 108 | 11 | 0 | 1 | 0 | 0 | 298 | 149 | 0.006108136 | 0.069000521 | 1 | 1 | 1 | 1 | 1 | 0.758 | 1 | 0.966 | 1 | 1 | 0.055 |
| SE | 33150 | ENSG00000183726 | TMEM50A | chr1 | + | 25342960 | 25343073 | 25340473 | 25340579 | 25351625 | 25351693 | 743 | 469 | 437 | 647 | 971 | 25 | 9 | 3 | 26 | 17 | 985 | 603 | 572 | 639 | 1354 | 19 | 22 | 32 | 18 | 61 | 262 | 149 | 0.006120528 | 0.069130535 | 0.944 | 0.967 | 0.988 | 0.934 | 0.97 | 0.967 | 0.94 | 0.91 | 0.953 | 0.927 | 0.021 |
| SE | 89880 | ENSG00000034677 | RNF19A | chr8 | - | 100264033 | 100264195 | 100261541 | 100261755 | 100264670 | 100264785 | 223 | 168 | 169 | 143 | 136 | 24 | 16 | 18 | 23 | 7 | 37 | 135 | 302 | 52 | 85 | 8 | 71 | 134 | 50 | 0 | 298 | 149 | 0.00612364 | 0.069155704 | 0.823 | 0.84 | 0.824 | 0.757 | 0.907 | 0.698 | 0.487 | 0.53 | 0.342 | 1 | 0.219 |
| SE | 98832 | ENSG00000135365 | PHF21A | chr11 | - | 45971115 | 45971367 | 45969814 | 45969904 | 45979759 | 45979966 | 124 | 75 | 102 | 126 | 127 | 3 | 4 | 0 | 1 | 1 | 268 | 179 | 254 | 254 | 162 | 6 | 12 | 1 | 4 | 12 | 298 | 149 | 0.006124851 | 0.069159405 | 0.954 | 0.904 | 1 | 0.984 | 0.984 | 0.957 | 0.882 | 0.992 | 0.969 | 0.871 | 0.031 |
| SE | 100701 | ENSG00000109917 | ZPR1 | chr11 | - | 116784377 | 116784498 | 116783529 | 116783619 | 116784858 | 116784921 | 78 | 45 | 71 | 91 | 50 | 0 | 0 | 0 | 0 | 0 | 70 | 94 | 65 | 110 | 110 | 6 | 1 | 1 | 5 | 0 | 270 | 149 | 0.006126525 | 0.069168334 | 1 | 1 | 1 | 1 | 1 | 0.866 | 0.981 | 0.973 | 0.924 | 1 | 0.051 |
| SE | 7283 | ENSG00000101019 | UQCC1 | chr20 | - | 35374183 | 35374256 | 35366556 | 35366614 | 35394091 | 35394196 | 68 | 52 | 51 | 72 | 50 | 0 | 0 | 0 | 0 | 0 | 89 | 76 | 89 | 80 | 31 | 0 | 3 | 2 | 8 | 0 | 222 | 149 | 0.006130326 | 0.069201265 | 1 | 1 | 1 | 1 | 1 | 1 | 0.944 | 0.968 | 0.87 | 1 | 0.044 |
| SE | 50128 | ENSG00000132704 | FCRL2 | chr1 | - | 157769865 | 157770150 | 157768413 | 157768701 | 157770408 | 157770666 | 48 | 24 | 30 | 30 | 79 | 4 | 0 | 3 | 7 | 1 | 23 | 72 | 6 | 7 | 5 | 0 | 5 | 0 | 0 | 0 | 298 | 149 | 0.006132872 | 0.069220032 | 0.857 | 1 | 0.833 | 0.682 | 0.975 | 1 | 0.878 | 1 | 1 | 1 | -0.106 |
| SE | 14028 | ENSG00000090905 | TNRC6A | chr16 | + | 24805673 | 24805733 | 24805013 | 24805151 | 24806205 | 24806283 | 22 | 11 | 15 | 22 | 15 | 0 | 0 | 2 | 0 | 2 | 14 | 10 | 11 | 9 | 2 | 0 | 3 | 0 | 7 | 0 | 209 | 149 | 0.006138602 | 0.069274716 | 1 | 1 | 0.842 | 1 | 0.842 | 1 | 0.704 | 1 | 0.478 | 1 | 0.1 |
| SE | 68825 | ENSG00000082258 | CCNT2 | chr2 | + | 134945786 | 134945982 | 134942611 | 134942674 | 134946100 | 134946214 | 6 | 3 | 12 | 3 | 15 | 29 | 27 | 26 | 27 | 40 | 0 | 9 | 0 | 1 | 0 | 7 | 32 | 29 | 7 | 2 | 298 | 149 | 0.006146531 | 0.069354195 | 0.094 | 0.053 | 0.188 | 0.053 | 0.158 | 0 | 0.123 | 0 | 0.067 | 0 | 0.071 |
| SE | 45328 | ENSG00000144320 | LNPK | chr2 | - | 175994133 | 175994280 | 175993180 | 175993223 | 175995557 | 175995646 | 30 | 12 | 14 | 29 | 25 | 46 | 23 | 42 | 18 | 37 | 0 | 11 | 8 | 1 | 8 | 13 | 17 | 15 | 27 | 15 | 296 | 149 | 0.006147964 | 0.069360371 | 0.247 | 0.208 | 0.144 | 0.448 | 0.254 | 0 | 0.246 | 0.212 | 0.018 | 0.212 | 0.123 |
| SE | 46205 | ENSG00000114439 | BBX | chr3 | + | 107641790 | 107641864 | 107526326 | 107526398 | 107645835 | 107645909 | 2 | 7 | 9 | 9 | 5 | 111 | 78 | 89 | 83 | 64 | 12 | 14 | 19 | 0 | 0 | 30 | 81 | 69 | 41 | 40 | 223 | 149 | 0.006154341 | 0.069422309 | 0.012 | 0.057 | 0.063 | 0.068 | 0.05 | 0.211 | 0.104 | 0.155 | 0 | 0 | -0.044 |
| SE | 11053 | ENSG00000185722 | ANKFY1 | chr17 | - | 4235771 | 4235890 | 4216516 | 4217118 | 4242255 | 4242448 | 135 | 81 | 76 | 104 | 52 | 0 | 3 | 5 | 5 | 0 | 152 | 105 | 48 | 93 | 85 | 0 | 0 | 0 | 0 | 0 | 268 | 149 | 0.006156361 | 0.069435098 | 1 | 0.938 | 0.894 | 0.92 | 1 | 1 | 1 | 1 | 1 | 1 | -0.05 |
| SE | 98636 | ENSG00000149179 | C11orf49 | chr11 | + | 46991542 | 46991647 | 46987207 | 46987310 | 47052387 | 47052518 | 1 | 0 | 0 | 1 | 0 | 13 | 16 | 10 | 7 | 14 | 0 | 4 | 0 | 0 | 6 | 17 | 6 | 6 | 11 | 9 | 254 | 149 | 0.006157647 | 0.0694396 | 0.043 | 0 | 0 | 0.077 | 0 | 0 | 0.281 | 0 | 0 | 0.281 | -0.088 |
| SE | 183 | ENSG00000185917 | SETD4 | chr21 | - | 36048307 | 36048396 | 36045581 | 36046011 | 36053582 | 36053620 | 46 | 28 | 38 | 48 | 50 | 8 | 1 | 7 | 5 | 5 | 16 | 42 | 26 | 14 | 66 | 0 | 0 | 0 | 1 | 8 | 238 | 149 | 0.006160509 | 0.069461868 | 0.783 | 0.946 | 0.773 | 0.857 | 0.862 | 1 | 1 | 1 | 0.898 | 0.838 | -0.103 |
| SE | 47951 | ENSG00000169398 | PTK2 | chr8 | - | 140769574 | 140769592 | 140764233 | 140764290 | 140789473 | 140789526 | 4 | 1 | 7 | 2 | 6 | 25 | 33 | 15 | 47 | 41 | 0 | 2 | 4 | 3 | 0 | 58 | 45 | 18 | 37 | 22 | 167 | 149 | 0.006169532 | 0.069545375 | 0.125 | 0.026 | 0.294 | 0.037 | 0.115 | 0 | 0.038 | 0.165 | 0.067 | 0 | 0.065 |
| SE | 64429 | ENSG00000162813 | BPNT1 | chr1 | - | 220062756 | 220062954 | 220059685 | 220059791 | 220067301 | 220067393 | 132 | 67 | 115 | 88 | 117 | 5 | 0 | 2 | 0 | 7 | 103 | 111 | 54 | 65 | 111 | 0 | 0 | 0 | 4 | 0 | 298 | 149 | 0.006169692 | 0.069545375 | 0.93 | 1 | 0.966 | 1 | 0.893 | 1 | 1 | 1 | 0.89 | 1 | -0.02 |
| SE | 97490 | ENSG00000173120 | KDM2A | chr11 | + | 67131951 | 67132073 | 67121268 | 67121358 | 67180078 | 67180217 | 9 | 18 | 9 | 14 | 23 | 266 | 204 | 271 | 300 | 383 | 15 | 15 | 5 | 10 | 15 | 519 | 358 | 308 | 449 | 343 | 271 | 149 | 0.006170599 | 0.069545596 | 0.018 | 0.046 | 0.018 | 0.025 | 0.032 | 0.016 | 0.023 | 0.009 | 0.012 | 0.023 | 0.011 |
| SE | 68379 | ENSG00000136710 | CCDC115 | chr2 | - | 130339559 | 130339762 | 130338242 | 130339232 | 130340907 | 130341039 | 3 | 5 | 4 | 3 | 7 | 77 | 44 | 53 | 50 | 40 | 0 | 2 | 3 | 2 | 5 | 99 | 72 | 40 | 76 | 79 | 298 | 149 | 0.006176265 | 0.069599426 | 0.019 | 0.054 | 0.036 | 0.029 | 0.08 | 0 | 0.014 | 0.036 | 0.013 | 0.031 | 0.025 |
| SE | 75683 | ENSG00000139163 | ETNK1 | chr12 | + | 22668597 | 22668700 | 22661062 | 22661205 | 22671271 | 22671355 | 10 | 7 | 4 | 4 | 4 | 102 | 50 | 57 | 100 | 129 | 8 | 2 | 8 | 12 | 0 | 35 | 31 | 18 | 2 | 7 | 252 | 149 | 0.006180858 | 0.069641167 | 0.055 | 0.076 | 0.04 | 0.023 | 0.018 | 0.119 | 0.037 | 0.208 | 0.78 | 0 | -0.186 |
| SE | 18096 | ENSG00000075413 | MARK3 | chr14 | + | 103457141 | 103457212 | 103451917 | 103451983 | 103462404 | 103462461 | 500 | 444 | 394 | 431 | 648 | 7 | 2 | 2 | 4 | 1 | 854 | 524 | 391 | 652 | 424 | 5 | 6 | 9 | 10 | 11 | 220 | 149 | 0.006183178 | 0.069657277 | 0.98 | 0.993 | 0.993 | 0.986 | 0.998 | 0.991 | 0.983 | 0.967 | 0.978 | 0.963 | 0.014 |
| SE | 37075 | ENSG00000005812 | FBXL3 | chr13 | - | 77026166 | 77026861 | 77018599 | 77018722 | 77026826 | 77027195 | 67 | 36 | 47 | 49 | 29 | 2 | 1 | 1 | 6 | 0 | 9 | 19 | 22 | 12 | 9 | 0 | 0 | 1 | 0 | 0 | 298 | 149 | 0.006186999 | 0.069690301 | 0.944 | 0.947 | 0.959 | 0.803 | 1 | 1 | 1 | 0.917 | 1 | 1 | -0.053 |
| SE | 66042 | ENSG00000183495 | EP400 | chr12 | + | 131994866 | 131994956 | 131992172 | 131992230 | 132005076 | 132005184 | 139 | 90 | 135 | 114 | 112 | 4 | 1 | 0 | 4 | 4 | 272 | 119 | 58 | 140 | 144 | 14 | 10 | 1 | 0 | 9 | 239 | 149 | 0.006191308 | 0.06972881 | 0.956 | 0.982 | 1 | 0.947 | 0.946 | 0.924 | 0.881 | 0.973 | 1 | 0.909 | 0.029 |
| SE | 77630 | ENSG00000146592 | CREB5 | chr7 | + | 28818760 | 28818789 | 28818070 | 28818179 | 28819115 | 28819559 | 5 | 5 | 4 | 5 | 5 | 33 | 18 | 8 | 15 | 29 | 21 | 19 | 43 | 15 | 0 | 20 | 27 | 60 | 19 | 19 | 178 | 149 | 0.006192237 | 0.069729243 | 0.113 | 0.189 | 0.295 | 0.218 | 0.126 | 0.468 | 0.371 | 0.375 | 0.398 | 0 | -0.134 |
| SE | 28392 | ENSG00000204842 | ATXN2 | chr12 | - | 111482786 | 111482831 | 111470557 | 111470742 | 111485270 | 111485331 | 2 | 0 | 2 | 2 | 1 | 1 | 5 | 2 | 1 | 2 | 3 | 3 | 2 | 10 | 34 | 0 | 2 | 5 | 2 | 4 | 194 | 149 | 0.006193963 | 0.069738643 | 0.606 | 0 | 0.434 | 0.606 | 0.277 | 1 | 0.535 | 0.235 | 0.793 | 0.867 | -0.301 |
| SE | 72675 | ENSG00000113387 | SUB1 | chr5 | + | 32586491 | 32586501 | 32585587 | 32585625 | 32588511 | 32588584 | 2 | 2 | 5 | 5 | 0 | 463 | 201 | 331 | 295 | 246 | 0 | 7 | 14 | 4 | 8 | 134 | 444 | 376 | 250 | 166 | 159 | 149 | 0.006201624 | 0.069814861 | 0.004 | 0.009 | 0.014 | 0.016 | 0 | 0 | 0.015 | 0.034 | 0.015 | 0.043 | -0.013 |
| SE | 39205 | ENSG00000150753 | CCT5 | chr5 | + | 10254673 | 10254838 | 10254144 | 10254205 | 10255954 | 10256153 | 766 | 448 | 541 | 630 | 646 | 5 | 11 | 8 | 14 | 19 | 1208 | 543 | 352 | 926 | 479 | 57 | 17 | 19 | 15 | 0 | 298 | 149 | 0.006203008 | 0.069820402 | 0.987 | 0.953 | 0.971 | 0.957 | 0.944 | 0.914 | 0.941 | 0.903 | 0.969 | 1 | 0.017 |
| SE | 41482 | ENSG00000099194 | SCD | chr10 | + | 100356531 | 100356764 | 100354426 | 100354632 | 100360733 | 100364826 | 17 | 18 | 37 | 23 | 40 | 0 | 0 | 2 | 3 | 2 | 46 | 46 | 26 | 162 | 88 | 0 | 11 | 8 | 32 | 0 | 298 | 149 | 0.00621039 | 0.069893449 | 1 | 1 | 0.902 | 0.793 | 0.909 | 1 | 0.676 | 0.619 | 0.717 | 1 | 0.118 |
| SE | 98580 | ENSG00000025796 | SEC63 | chr6 | - | 107955964 | 107956106 | 107929414 | 107929514 | 107957885 | 107958042 | 0 | 4 | 4 | 15 | 3 | 105 | 80 | 119 | 87 | 105 | 0 | 0 | 3 | 6 | 0 | 72 | 102 | 160 | 128 | 19 | 291 | 149 | 0.006215496 | 0.069940857 | 0 | 0.025 | 0.017 | 0.081 | 0.014 | 0 | 0 | 0.01 | 0.023 | 0 | 0.021 |
| SE | 3214 | ENSG00000071626 | DAZAP1 | chr19 | + | 1425877 | 1425960 | 1422347 | 1422396 | 1428841 | 1428995 | 351 | 230 | 329 | 420 | 431 | 53 | 26 | 64 | 65 | 87 | 790 | 451 | 362 | 645 | 698 | 108 | 77 | 39 | 61 | 79 | 232 | 149 | 0.006232811 | 0.070125619 | 0.81 | 0.85 | 0.768 | 0.806 | 0.761 | 0.824 | 0.79 | 0.856 | 0.872 | 0.85 | -0.039 |
| SE | 13925 | ENSG00000171208 | NETO2 | chr16 | - | 47086225 | 47086339 | 47077702 | 47083801 | 47109482 | 47109711 | 32 | 22 | 20 | 23 | 32 | 4 | 1 | 2 | 24 | 15 | 52 | 15 | 18 | 35 | 29 | 0 | 3 | 6 | 2 | 0 | 263 | 149 | 0.006242796 | 0.070219143 | 0.819 | 0.926 | 0.85 | 0.352 | 0.547 | 1 | 0.739 | 0.63 | 0.908 | 1 | -0.157 |
| SE | 12388 | ENSG00000188559 | RALGAPA2 | chr20 | - | 20396694 | 20396734 | 20389529 | 20393253 | 20412026 | 20412113 | 23 | 17 | 23 | 20 | 41 | 5 | 6 | 0 | 4 | 9 | 18 | 24 | 19 | 45 | 39 | 14 | 15 | 17 | 10 | 8 | 189 | 149 | 0.006242917 | 0.070219143 | 0.784 | 0.691 | 1 | 0.798 | 0.782 | 0.503 | 0.558 | 0.468 | 0.78 | 0.794 | 0.19 |
| SE | 18828 | ENSG00000127663 | KDM4B | chr19 | + | 5071009 | 5071059 | 5047475 | 5047669 | 5077366 | 5077470 | 71 | 14 | 55 | 50 | 36 | 3 | 0 | 3 | 0 | 3 | 21 | 81 | 57 | 60 | 150 | 0 | 6 | 7 | 11 | 6 | 199 | 149 | 0.006244461 | 0.07022271 | 0.947 | 1 | 0.932 | 0.926 | 1 | 1 | 0.91 | 0.859 | 0.803 | 0.949 | 0.057 |
| SE | 28623 | ENSG00000102445 | RUBCNL | chr13 | - | 46368488 | 46368615 | 46368041 | 46368249 | 46368732 | 46368815 | 10 | 5 | 13 | 10 | 23 | 10 | 7 | 12 | 16 | 48 | 25 | 29 | 20 | 10 | 19 | 20 | 8 | 4 | 17 | 13 | 276 | 149 | 0.006245028 | 0.07022271 | 0.351 | 0.278 | 0.369 | 0.252 | 0.206 | 0.403 | 0.662 | 0.73 | 0.241 | 0.441 | -0.204 |
| SE | 97434 | ENSG00000168876 | ANKRD49 | chr11 | + | 94497901 | 94497964 | 94496603 | 94496951 | 94498070 | 94498214 | 34 | 18 | 25 | 30 | 55 | 22 | 26 | 26 | 23 | 33 | 4 | 21 | 21 | 9 | 0 | 20 | 19 | 20 | 16 | 12 | 212 | 149 | 0.006248376 | 0.070240584 | 0.521 | 0.327 | 0.403 | 0.478 | 0.539 | 0.123 | 0.437 | 0.425 | 0.283 | 0 | 0.2 |
| SE | 43137 | ENSG00000186470 | BTN3A2 | chr6 | + | 26373275 | 26373296 | 26370321 | 26370603 | 26373386 | 26373413 | 89 | 35 | 63 | 34 | 120 | 0 | 1 | 0 | 0 | 0 | 134 | 70 | 61 | 51 | 229 | 8 | 5 | 2 | 2 | 0 | 170 | 149 | 0.006248412 | 0.070240584 | 1 | 0.968 | 1 | 1 | 1 | 0.936 | 0.925 | 0.964 | 0.957 | 1 | 0.037 |
| SE | 13378 | ENSG00000074755 | ZZEF1 | chr17 | - | 4024918 | 4025118 | 4022415 | 4022828 | 4032125 | 4032258 | 155 | 99 | 148 | 154 | 163 | 2 | 0 | 6 | 5 | 2 | 287 | 232 | 154 | 271 | 271 | 0 | 0 | 4 | 1 | 0 | 298 | 149 | 0.006250118 | 0.070249674 | 0.975 | 1 | 0.925 | 0.939 | 0.976 | 1 | 1 | 0.951 | 0.993 | 1 | -0.026 |
| SE | 105013 | ENSG00000284770 | TBCE | chr1 | + | 235432945 | 235433098 | 235430704 | 235430804 | 235434203 | 235434280 | 2 | 7 | 7 | 3 | 3 | 19 | 14 | 15 | 19 | 55 | 0 | 0 | 2 | 5 | 0 | 34 | 12 | 11 | 8 | 21 | 298 | 149 | 0.006263446 | 0.070389373 | 0.05 | 0.2 | 0.189 | 0.073 | 0.027 | 0 | 0 | 0.083 | 0.238 | 0 | 0.044 |
| SE | 39447 | ENSG00000055917 | PUM2 | chr2 | - | 20331716 | 20331937 | 20327309 | 20327378 | 20350809 | 20350834 | 31 | 22 | 33 | 39 | 55 | 10 | 3 | 4 | 5 | 1 | 13 | 25 | 36 | 25 | 26 | 0 | 0 | 7 | 0 | 0 | 298 | 149 | 0.00626999 | 0.070452801 | 0.608 | 0.786 | 0.805 | 0.796 | 0.965 | 1 | 1 | 0.72 | 1 | 1 | -0.152 |
| SE | 86657 | ENSG00000196924 | FLNA | chrX | - | 154357250 | 154357274 | 154354824 | 154355072 | 154357433 | 154357623 | 1539 | 533 | 845 | 1144 | 549 | 1633 | 863 | 839 | 1500 | 784 | 1459 | 1507 | 758 | 900 | 3546 | 2952 | 1636 | 1027 | 1761 | 11256 | 173 | 149 | 0.006272161 | 0.070461235 | 0.448 | 0.347 | 0.465 | 0.396 | 0.376 | 0.299 | 0.442 | 0.389 | 0.306 | 0.213 | 0.077 |
| SE | 32296 | ENSG00000241553 | ARPC4 | chr3 | + | 9799878 | 9800067 | 9797658 | 9797777 | 9801660 | 9801756 | 15 | 17 | 12 | 9 | 28 | 1 | 0 | 1 | 3 | 3 | 19 | 24 | 9 | 22 | 13 | 0 | 0 | 0 | 0 | 0 | 298 | 149 | 0.00627254 | 0.070461235 | 0.882 | 1 | 0.857 | 0.6 | 0.824 | 1 | 1 | 1 | 1 | 1 | -0.167 |
| SE | 93929 | ENSG00000144524 | COPS7B | chr2 | + | 231790946 | 231791173 | 231788554 | 231788732 | 231791732 | 231791808 | 1 | 0 | 13 | 3 | 4 | 82 | 33 | 61 | 65 | 52 | 10 | 10 | 4 | 4 | 16 | 99 | 79 | 23 | 51 | 60 | 298 | 149 | 0.006277248 | 0.070493886 | 0.006 | 0 | 0.096 | 0.023 | 0.037 | 0.048 | 0.06 | 0.08 | 0.038 | 0.118 | -0.036 |
| SE | 93930 | ENSG00000144524 | COPS7B | chr2 | + | 231790950 | 231791173 | 231788554 | 231788732 | 231791732 | 231791786 | 1 | 0 | 13 | 3 | 4 | 82 | 33 | 61 | 65 | 52 | 10 | 10 | 4 | 4 | 16 | 99 | 79 | 23 | 51 | 60 | 298 | 149 | 0.006277248 | 0.070493886 | 0.006 | 0 | 0.096 | 0.023 | 0.037 | 0.048 | 0.06 | 0.08 | 0.038 | 0.118 | -0.036 |
| SE | 6247 | ENSG00000088812 | ATRN | chr20 | + | 3559392 | 3559483 | 3549169 | 3549338 | 3560661 | 3560905 | 263 | 203 | 241 | 212 | 270 | 0 | 2 | 3 | 4 | 1 | 394 | 277 | 181 | 200 | 411 | 23 | 4 | 1 | 0 | 0 | 240 | 149 | 0.006284278 | 0.070562715 | 1 | 0.984 | 0.98 | 0.971 | 0.994 | 0.914 | 0.977 | 0.991 | 1 | 1 | 0.009 |
| SE | 19448 | ENSG00000198668 | CALM1 | chr14 | + | 90398990 | 90399087 | 90397028 | 90397233 | 90400064 | 90400095 | 14 | 2 | 9 | 8 | 1 | 235 | 115 | 244 | 241 | 87 | 0 | 11 | 15 | 18 | 20 | 233 | 204 | 172 | 208 | 176 | 246 | 149 | 0.006287806 | 0.070592209 | 0.035 | 0.01 | 0.022 | 0.02 | 0.007 | 0 | 0.032 | 0.05 | 0.05 | 0.064 | -0.02 |
| SE | 39332 | ENSG00000230092 | NA | chr1 | - | 808573 | 808623 | 805798 | 805891 | 810066 | 810161 | 62 | 51 | 80 | 44 | 116 | 0 | 0 | 2 | 0 | 0 | 128 | 26 | 46 | 64 | 58 | 0 | 0 | 2 | 0 | 11 | 199 | 149 | 0.006305828 | 0.070784391 | 1 | 1 | 0.968 | 1 | 1 | 1 | 1 | 0.945 | 1 | 0.798 | 0.045 |
| SE | 38445 | ENSG00000132549 | VPS13B | chr8 | + | 99854409 | 99854519 | 99853450 | 99854256 | 99859303 | 99859480 | 1 | 0 | 1 | 2 | 0 | 16 | 7 | 14 | 7 | 7 | 0 | 4 | 4 | 0 | 6 | 1 | 15 | 14 | 4 | 0 | 259 | 149 | 0.006306751 | 0.070784603 | 0.035 | 0 | 0.039 | 0.141 | 0 | 0 | 0.133 | 0.141 | 0 | 1 | -0.212 |
| SE | 31556 | ENSG00000163913 | IFT122 | chr3 | + | 129458598 | 129458677 | 129451913 | 129451998 | 129461227 | 129461304 | 37 | 21 | 23 | 31 | 17 | 0 | 0 | 1 | 0 | 0 | 106 | 36 | 23 | 67 | 65 | 6 | 6 | 0 | 0 | 0 | 228 | 149 | 0.006307707 | 0.070785182 | 1 | 1 | 0.938 | 1 | 1 | 0.92 | 0.797 | 1 | 1 | 1 | 0.044 |
| SE | 103921 | ENSG00000184988 | TMEM106A | chr17 | + | 43216448 | 43216589 | 43215787 | 43215941 | 43216696 | 43216740 | 53 | 37 | 60 | 68 | 108 | 0 | 1 | 3 | 2 | 0 | 242 | 81 | 59 | 121 | 177 | 8 | 3 | 0 | 6 | 7 | 290 | 149 | 0.006311277 | 0.070806041 | 1 | 0.95 | 0.911 | 0.946 | 1 | 0.94 | 0.933 | 1 | 0.912 | 0.929 | 0.019 |
| SE | 26469 | ENSG00000214655 | ZSWIM8 | chr10 | + | 73791323 | 73791499 | 73790974 | 73791176 | 73791858 | 73792490 | 63 | 6 | 26 | 38 | 22 | 7 | 1 | 3 | 3 | 1 | 44 | 31 | 20 | 3 | 25 | 0 | 12 | 14 | 1 | 13 | 298 | 149 | 0.006311374 | 0.070806041 | 0.818 | 0.75 | 0.812 | 0.864 | 0.917 | 1 | 0.564 | 0.417 | 0.6 | 0.49 | 0.218 |
| SE | 42339 | ENSG00000070759 | TESK2 | chr1 | - | 45346692 | 45346779 | 45343882 | 45345558 | 45346978 | 45347062 | 6 | 6 | 14 | 2 | 8 | 0 | 0 | 0 | 0 | 0 | 24 | 33 | 8 | 36 | 63 | 14 | 0 | 0 | 0 | 0 | 236 | 149 | 0.006314952 | 0.070836028 | 1 | 1 | 1 | 1 | 1 | 0.52 | 1 | 1 | 1 | 1 | 0.096 |
| SE | 46559 | ENSG00000223745 | CCDC18-AS1 | chr1 | + | 93340457 | 93340510 | 93339648 | 93339679 | 93345723 | 93345799 | 67 | 62 | 44 | 50 | 32 | 0 | 1 | 0 | 0 | 0 | 21 | 30 | 42 | 27 | 8 | 0 | 7 | 0 | 3 | 0 | 202 | 149 | 0.006317165 | 0.070850702 | 1 | 0.979 | 1 | 1 | 1 | 1 | 0.76 | 1 | 0.869 | 1 | 0.07 |
| SE | 23579 | ENSG00000254996 | ANKHD1-EIF4EBP3 | chr5 | + | 140538085 | 140538234 | 140537337 | 140537589 | 140538918 | 140539083 | 182 | 116 | 201 | 183 | 271 | 0 | 0 | 1 | 2 | 1 | 73 | 147 | 144 | 149 | 83 | 0 | 7 | 0 | 7 | 0 | 298 | 149 | 0.006329023 | 0.070973529 | 1 | 1 | 0.99 | 0.979 | 0.993 | 1 | 0.913 | 1 | 0.914 | 1 | 0.027 |
| SE | 52904 | ENSG00000159023 | EPB41 | chr1 | + | 29058588 | 29058645 | 29053103 | 29053312 | 29060421 | 29060484 | 0 | 0 | 1 | 0 | 0 | 11 | 8 | 10 | 13 | 19 | 0 | 0 | 0 | 6 | 13 | 6 | 17 | 5 | 47 | 0 | 206 | 149 | 0.006332205 | 0.070999039 | 0 | 0 | 0.067 | 0 | 0 | 0 | 0 | 0 | 0.085 | 1 | -0.204 |
| SE | 80739 | ENSG00000086300 | SNX10 | chr7 | + | 26354056 | 26354184 | 26346419 | 26346466 | 26360974 | 26361061 | 6 | 16 | 3 | 12 | 15 | 144 | 95 | 82 | 97 | 120 | 7 | 10 | 9 | 11 | 0 | 15 | 48 | 64 | 27 | 19 | 277 | 149 | 0.006333714 | 0.07100579 | 0.022 | 0.083 | 0.019 | 0.062 | 0.063 | 0.201 | 0.101 | 0.07 | 0.18 | 0 | -0.061 |
| SE | 74693 | ENSG00000197548 | ATG7 | chr3 | + | 11331339 | 11331428 | 11315343 | 11315493 | 11340644 | 11340735 | 98 | 44 | 60 | 62 | 97 | 2 | 4 | 1 | 3 | 1 | 136 | 79 | 39 | 128 | 132 | 0 | 9 | 5 | 5 | 14 | 238 | 149 | 0.006340429 | 0.071070904 | 0.968 | 0.873 | 0.974 | 0.928 | 0.984 | 1 | 0.846 | 0.83 | 0.941 | 0.855 | 0.051 |
| SE | 5040 | ENSG00000101126 | ADNP | chr20 | - | 50902016 | 50902109 | 50888915 | 50894512 | 50903888 | 50904001 | 330 | 266 | 332 | 352 | 766 | 5 | 0 | 6 | 0 | 2 | 207 | 268 | 265 | 216 | 138 | 0 | 6 | 12 | 3 | 0 | 242 | 149 | 0.006347359 | 0.071138389 | 0.976 | 1 | 0.971 | 1 | 0.996 | 1 | 0.965 | 0.931 | 0.978 | 1 | 0.014 |
| SE | 67852 | ENSG00000018280 | SLC11A1 | chr2 | + | 218384242 | 218384365 | 218382202 | 218382375 | 218386634 | 218386741 | 5 | 1 | 2 | 8 | 2 | 2 | 2 | 0 | 3 | 0 | 12 | 33 | 27 | 17 | 11 | 0 | 0 | 0 | 0 | 3 | 272 | 149 | 0.006351404 | 0.071173545 | 0.578 | 0.215 | 1 | 0.594 | 1 | 1 | 1 | 1 | 1 | 0.668 | -0.256 |
| SE | 17884 | ENSG00000129467 | ADCY4 | chr14 | - | 24319741 | 24319888 | 24319328 | 24319436 | 24322065 | 24322224 | 4 | 1 | 5 | 6 | 4 | 0 | 0 | 0 | 0 | 0 | 17 | 22 | 22 | 33 | 10 | 0 | 14 | 0 | 0 | 0 | 296 | 149 | 0.006357363 | 0.071230127 | 1 | 1 | 1 | 1 | 1 | 1 | 0.442 | 1 | 1 | 1 | 0.112 |
| SE | 91798 | ENSG00000133794 | ARNTL | chr11 | + | 13378321 | 13378445 | 13376639 | 13376728 | 13381160 | 13381254 | 76 | 36 | 61 | 65 | 89 | 2 | 0 | 0 | 0 | 0 | 4 | 75 | 106 | 40 | 7 | 3 | 2 | 0 | 0 | 4 | 273 | 149 | 0.006358406 | 0.071231618 | 0.954 | 1 | 1 | 1 | 1 | 0.421 | 0.953 | 1 | 1 | 0.489 | 0.218 |
| SE | 36246 | ENSG00000168827 | GFM1 | chr3 | + | 158649040 | 158649157 | 158646742 | 158646947 | 158652095 | 158652246 | 125 | 76 | 93 | 130 | 120 | 4 | 3 | 5 | 4 | 8 | 61 | 78 | 35 | 23 | 0 | 4 | 0 | 12 | 7 | 7 | 266 | 149 | 0.006365096 | 0.071296364 | 0.946 | 0.934 | 0.912 | 0.948 | 0.894 | 0.895 | 1 | 0.62 | 0.648 | 0 | 0.294 |
| SE | 18058 | ENSG00000100450 | GZMH | chr14 | - | 24607148 | 24607406 | 24606479 | 24606746 | 24608264 | 24608412 | 984 | 211 | 232 | 491 | 468 | 7 | 7 | 2 | 4 | 1 | 492 | 924 | 361 | 463 | 740 | 35 | 6 | 2 | 2 | 5 | 298 | 149 | 0.006366087 | 0.071297177 | 0.986 | 0.938 | 0.983 | 0.984 | 0.996 | 0.875 | 0.987 | 0.989 | 0.991 | 0.987 | 0.012 |
| SE | 29199 | ENSG00000120708 | TGFBI | chr5 | + | 136048252 | 136048349 | 136047273 | 136047420 | 136049438 | 136049580 | 3 | 0 | 6 | 2 | 0 | 351 | 157 | 215 | 204 | 249 | 8 | 11 | 12 | 0 | 24 | 673 | 154 | 142 | 212 | 711 | 246 | 149 | 0.00636699 | 0.071297177 | 0.005 | 0 | 0.017 | 0.006 | 0 | 0.007 | 0.041 | 0.049 | 0 | 0.02 | -0.018 |

| | | | | | | | | | | | | | | |
|---|---|---|---|---|---|---|---|---|---|---|---|---|---|---|
| SE | 40237 | ENSG00000117408 | IPO13 | chr1 | + | 43957401 | 43957549 | 43956809 | 43956976 | 43957976 | 43958158 | 32 | 6 | 19 |
| SE | 42129 | ENSG00000198952 | SMG5 | chr1 | - | 156266506 | 156266678 | 156266136 | 156266380 | 156267469 | 156267588 | 85 | 36 | 113 |
| SE | 73313 | ENSG00000136699 | SMPD4 | chr2 | - | 130173513 | 130173656 | 130172786 | 130172895 | 130174913 | 130175000 | 30 | 12 | 24 |
| SE | 92191 | ENSG00000099810 | MTAP | chr9 | + | 21808806 | 21809000 | 21802661 | 21802781 | 21815432 | 21815519 | 2 | 0 | 0 |
| SE | 14465 | ENSG00000213614 | HEXA | chr15 | - | 72347685 | 72347758 | 72345269 | 72346325 | 72348047 | 72348134 | 52 | 40 | 46 |
| SE | 11665 | ENSG00000167207 | NOD2 | chr16 | + | 50699487 | 50699954 | 50693587 | 50693662 | 50707854 | 50707960 | 15 | 10 | 11 |
| SE | 1435 | ENSG00000075234 | TTC38 | chr22 | + | 46269141 | 46269289 | 46268513 | 46268591 | 46272334 | 46272416 | 14 | 1 | 4 |
| SE | 103438 | ENSG00000265479 | DTX2P1-UPK3BP1-PMS2P11 | chr7 | + | 77041976 | 77042063 | 77018908 | 77018997 | 77043594 | 77043775 | 40 | 18 | 8 |
| SE | 20 | ENSG00000155313 | USP25 | chr21 | + | 15808808 | 15808885 | 15805120 | 15805258 | 15811136 | 15811210 | 361 | 243 | 238 |
| SE | 49920 | ENSG00000144455 | SUMF1 | chr3 | - | 4420063 | 4420146 | 4418009 | 4418132 | 4449265 | 4449340 | 184 | 142 | 179 |
| SE | 42776 | ENSG00000137462 | TLR2 | chr4 | + | 153702226 | 153702510 | 153687901 | 153688047 | 153702891 | 153702962 | 4 | 11 | 4 |
| SE | 36131 | ENSG00000169221 | TBC1D10B | chr16 | - | 30359171 | 30359361 | 30358662 | 30358817 | 30359537 | 30359603 | 63 | 15 | 46 |
| SE | 105111 | ENSG00000278311 | GGNBP2 | chr17 | + | 36545618 | 36545817 | 36544918 | 36545097 | 36554819 | 36554900 | 100 | 62 | 107 |
| SE | 23728 | ENSG00000140564 | FURIN | chr15 | + | 90877461 | 90877615 | 90877134 | 90877211 | 90878131 | 90878304 | 65 | 17 | 40 |
| SE | 89318 | ENSG00000120963 | ZNF706 | chr8 | - | 101199194 | 101199239 | 101177996 | 101178387 | 101199989 | 101200093 | 34 | 28 | 35 |
| SE | 43447 | ENSG00000099622 | CIRBP | chr19 | + | 1273600 | 1273715 | 1271980 | 1272051 | 1274306 | 1274440 | 40 | 17 | 46 |
| SE | 49059 | ENSG00000010270 | STARD3NL | chr7 | + | 38198210 | 38198365 | 38178394 | 38178420 | 38207446 | 38207729 | 0 | 0 | 0 |
| SE | 68997 | ENSG00000138375 | SMARCAL1 | chr2 | + | 216478202 | 216478299 | 216477108 | 216477209 | 216482737 | 216483017 | 59 | 58 | 49 |
| SE | 57687 | ENSG00000082068 | WDR70 | chr5 | + | 37687909 | 37688007 | 37605063 | 37605238 | 37697654 | 37697907 | 0 | 4 | 0 |
| SE | 37450 | ENSG00000189180 | ZNF33A | chr10 | + | 38054374 | 38054511 | 38017290 | 38017386 | 38064087 | 38065088 | 47 | 54 | 64 |
| SE | 52764 | ENSG00000198019 | FCGR1B | chr1 | + | 121088650 | 121088908 | 121087820 | 121087841 | 121092995 | 121093250 | 15 | 6 | 5 |
| SE | 51924 | ENSG00000074603 | DPP8 | chr15 | - | 65474208 | 65474288 | 65467070 | 65467223 | 65478879 | 65479039 | 121 | 106 | 154 |
| SE | 8459 | ENSG00000141503 | MINK1 | chr17 | + | 4892401 | 4892512 | 4892148 | 4892234 | 4892655 | 4892768 | 101 | 28 | 55 |
| SE | 80604 | ENSG00000232859 | LYRM9 | chr17 | - | 27880120 | 27880183 | 27879050 | 27879490 | 27880273 | 27880366 | 5 | 1 | 2 |
| SE | 79996 | ENSG00000084070 | SMAP2 | chr1 | + | 40362299 | 40362436 | 40344849 | 40344910 | 40406735 | 40406869 | 126 | 92 | 34 |
| SE | 18697 | ENSG00000105122 | RASAL3 | chr19 | - | 15452044 | 15452108 | 15451623 | 15451938 | 15452657 | 15452815 | 173 | 59 | 133 |
| SE | 55418 | ENSG00000070540 | WIPI1 | chr17 | - | 68428828 | 68428936 | 68427134 | 68427253 | 68433467 | 68433571 | 34 | 25 | 16 |
| SE | 49950 | ENSG00000037474 | NSUN2 | chr5 | - | 6622015 | 6622100 | 6620105 | 6620298 | 6623213 | 6623285 | 370 | 309 | 345 |
| SE | 79726 | ENSG00000168255 | POLR2J3 | chr7 | - | 102543524 | 102543702 | 102541500 | 102541662 | 102544705 | 102544776 | 11 | 1 | 12 |
| SE | 99833 | ENSG00000013725 | CD6 | chr11 | + | 61011069 | 61011135 | 61008533 | 61008845 | 61013422 | 61013563 | 86 | 31 | 89 |
| SE | 13559 | ENSG00000141101 | NOB1 | chr16 | - | 69752240 | 69752371 | 69749558 | 69749630 | 69754593 | 69754726 | 90 | 46 | 126 |
| SE | 73569 | ENSG00000115998 | C2orf42 | chr2 | - | 70182666 | 70182935 | 70181162 | 70181997 | 70190972 | 70191019 | 98 | 58 | 88 |
| SE | 11392 | ENSG00000166747 | AP1G1 | chr16 | - | 71808504 | 71808677 | 71789278 | 71789482 | 71808762 | 71808813 | 5 | 4 | 1 |
| SE | 99848 | ENSG00000110713 | NUP98 | chr11 | - | 3679553 | 3679708 | 3676508 | 3676620 | 3683199 | 3683441 | 150 | 95 | 142 |
| SE | 4812 | ENSG00000087460 | GNAS | chr20 | + | 58898940 | 58898985 | 58893094 | 58895684 | 58903527 | 58903585 | 137 | 118 | 121 |
| SE | 56570 | ENSG00000253710 | ALG11 | chr13 | + | 52023712 | 52023779 | 52018912 | 52019143 | 52024005 | 52024937 | 8 | 15 | 3 |
| SE | 59814 | ENSG00000166377 | ATP9B | chr18 | + | 79306985 | 79307234 | 79303603 | 79303716 | 79329140 | 79329213 | 109 | 77 | 103 |
| SE | 3290 | ENSG00000142453 | CARM1 | chr19 | + | 10908549 | 10908668 | 10908038 | 10908145 | 10909102 | 10909207 | 5 | 0 | 6 |
| SE | 57315 | ENSG00000181722 | ZBTB20 | chr3 | - | 114388064 | 114388123 | 114380881 | 114380940 | 114389004 | 114389105 | 0 | 1 | 0 |
| SE | 95768 | ENSG00000125944 | HNRNPR | chr1 | - | 23318482 | 23318688 | 23313552 | 23313702 | 23321527 | 23321663 | 2120 | 1780 | 1802 |
| SE | 37527 | ENSG00000105281 | SLC1A5 | chr19 | - | 46784516 | 46784559 | 46784096 | 46784144 | 46787399 | 46788594 | 39 | 12 | 19 |
| SE | 35384 | ENSG00000140262 | TCF12 | chr15 | + | 57232290 | 57232430 | 57231151 | 57231257 | 57232711 | 57232856 | 260 | 186 | 258 |
| SE | 53227 | ENSG00000181852 | RNF41 | chr12 | - | 56207167 | 56207310 | 56205449 | 56206798 | 56207645 | 56207749 | 2 | 0 | 4 |
| SE | 62225 | ENSG00000119778 | ATAD2B | chr2 | - | 23883593 | 23883635 | 23880638 | 23880755 | 23884764 | 23884873 | 19 | 6 | 7 |
| SE | 35172 | ENSG00000101596 | SMCHD1 | chr18 | + | 2674014 | 2674145 | 2673280 | 2673363 | 2688393 | 2688508 | 1506 | 1457 | 1486 |
| SE | 56563 | ENSG00000279457 | WASH9P | chr1 | - | 187754 | 187890 | 187375 | 187577 | 188129 | 188266 | 64 | 16 | 10 |
| SE | 60712 | ENSG00000106346 | USP42 | chr7 | + | 6128137 | 6128230 | 6115322 | 6115523 | 6135840 | 6135951 | 1 | 0 | 1 |
| SE | 21605 | ENSG00000283352 | NA | chr6 | - | 57930103 | 57930291 | 57926425 | 57926510 | 57959027 | 57959111 | 15 | 17 | 19 |
| SE | 9307 | ENSG00000133030 | MPRIP | chr17 | + | 17164108 | 17167915 | 17161239 | 17161356 | 17171717 | 17171865 | 69 | 20 | 63 |
| SE | 4479 | ENSG00000127527 | EPS15L1 | chr19 | - | 16404587 | 16404749 | 16403732 | 16403930 | 16413772 | 16413845 | 273 | 116 | 186 |
| SE | 80303 | ENSG00000168137 | SETD5 | chr3 | + | 9454004 | 9454101 | 9453738 | 9453868 | 9464424 | 9464954 | 33 | 43 | 27 |
| SE | 10556 | ENSG00000177302 | TOP3A | chr17 | - | 18306890 | 18306966 | 18305111 | 18305220 | 18308350 | 18308424 | 108 | 74 | 55 |
| SE | 86851 | ENSG00000156467 | UQCRB | chr8 | - | 96231378 | 96231515 | 96230941 | 96231132 | 96231773 | 96231940 | 24 | 3 | 16 |
| SE | 51496 | ENSG00000108021 | TASOR2 | chr10 | + | 5709529 | 5709663 | 5684941 | 5685175 | 5712822 | 5712918 | 22 | 31 | 23 |
| SE | 98556 | ENSG00000051009 | FAM160A2 | chr11 | - | 6222810 | 6222897 | 6222441 | 6222609 | 6223079 | 6223238 | 80 | 27 | 31 |
| SE | 86408 | ENSG00000131381 | RBSN | chr3 | - | 15082366 | 15082608 | 15080731 | 15080802 | 15084734 | 15084896 | 84 | 33 | 54 |
| SE | 27213 | ENSG00000173085 | COQ2 | chr4 | - | 83267089 | 83267208 | 83262588 | 83262655 | 83267585 | 83267774 | 4 | 3 | 2 |
| SE | 75256 | ENSG00000175575 | PAAF1 | chr11 | + | 73900310 | 73900420 | 73899145 | 73899244 | 73909398 | 73909593 | 14 | 6 | 29 |
| SE | 84997 | ENSG00000101901 | ALG13 | chrX | + | 111689086 | 111689197 | 111684964 | 111685103 | 111708026 | 111708166 | 1 | 2 | 0 |
| SE | 18629 | ENSG00000256053 | COA8 | chr14 | + | 103579370 | 103579491 | 103574106 | 103574170 | 103587273 | 103587364 | 6 | 10 | 6 |
| SE | 36704 | ENSG00000158062 | UBXN11 | chr1 | - | 26303042 | 26303167 | 26302812 | 26302918 | 26306591 | 26306636 | 8 | 12 | 3 |
| SE | 67061 | ENSG00000127720 | METTL25 | chr12 | + | 82369448 | 82369560 | 82358893 | 82358916 | 82386802 | 82386967 | 33 | 10 | 21 |
| SE | 97657 | ENSG00000147457 | CHMP7 | chr8 | + | 23255246 | 23255432 | 23246658 | 23246994 | 23256459 | 23256563 | 52 | 27 | 48 |
| SE | 83615 | ENSG00000130066 | SAT1 | chrX | + | 23784293 | 23784403 | 23783657 | 23783709 | 23785327 | 23785429 | 113 | 95 | 76 |
| SE | 8950 | ENSG00000132478 | UNK | chr17 | + | 75818102 | 75818168 | 75817325 | 75817526 | 75818641 | 75818816 | 17 | 9 | 34 |
| SE | 52450 | ENSG00000072803 | FBXW11 | chr5 | - | 171872871 | 171872990 | 171870747 | 171870858 | 171876284 | 171876291 | 167 | 93 | 153 |
| SE | 79076 | ENSG00000158636 | EMSY | chr11 | + | 76547027 | 76547100 | 76545947 | 76546297 | 76549951 | 76550006 | 5 | 4 | 4 |
| SE | 59232 | ENSG00000117153 | KLHL12 | chr1 | - | 202893238 | 202893425 | 202891119 | 202892659 | 202894183 | 202894282 | 74 | 37 | 47 |
| SE | 35798 | ENSG00000173226 | IQCB1 | chr3 | - | 121797117 | 121797227 | 121790072 | 121790215 | 121799195 | 121799374 | 40 | 42 | 44 |
| SE | 39005 | ENSG00000003400 | CASP10 | chr2 | + | 201192983 | 201193119 | 201187679 | 201187799 | 201195841 | 201195948 | 239 | 181 | 205 |
| SE | 21175 | ENSG00000143252 | SDHC | chr1 | + | 161323613 | 161323670 | 161314380 | 161314425 | 161340593 | 161340655 | 16 | 6 | 16 |
| SE | 20460 | ENSG00000006744 | ELAC2 | chr17 | - | 13004988 | 13005110 | 13003478 | 13003574 | 13005752 | 13005825 | 64 | 21 | 45 |
| SE | 55693 | ENSG00000135749 | PCNX2 | chr1 | - | 233112844 | 233112966 | 233057231 | 233057290 | 233135012 | 233135190 | 12 | 11 | 29 |
| SE | 91768 | ENSG00000143748 | NVL | chr1 | - | 224326390 | 224326464 | 224317877 | 224317930 | 224330070 | 224330137 | 98 | 67 | 67 |
| SE | 58266 | ENSG00000065413 | ANKRD44 | chr2 | - | 197000418 | 197000502 | 196998906 | 196999052 | 197001752 | 197001840 | 415 | 277 | 412 |
| SE | 34990 | ENSG00000204104 | TRAF3IP1 | chr2 | + | 238333959 | 238334035 | 238328925 | 238329342 | 238347454 | 238347475 | 7 | 2 | 8 |
| SE | 87926 | ENSG00000173273 | TNKS | chr8 | + | 9772368 | 9772423 | 9770130 | 9770262 | 9776649 | 9777036 | 11 | 5 | 13 |
| SE | 94457 | ENSG00000122490 | SLC66A2 | chr18 | - | 79949518 | 79949636 | 79943328 | 79943462 | 79950723 | 79951025 | 0 | 2 | 2 |
| SE | 89305 | ENSG00000166169 | POLL | chr10 | - | 101585861 | 101586156 | 101583507 | 101583681 | 101587245 | 101587382 | 12 | 9 | 9 |
| SE | 106196 | ENSG00000135090 | TAOK3 | chr12 | - | 118239226 | 118239272 | 118238072 | 118238169 | 118243414 | 118243490 | 301 | 215 | 258 |
| SE | 106072 | ENSG00000196498 | NCOR2 | chr12 | - | 124335134 | 124335280 | 124334423 | 124334617 | 124335482 | 124335632 | 136 | 52 | 88 |
| SE | 57363 | ENSG00000116815 | CD58 | chr1 | - | 116519230 | 116519302 | 116514738 | 116514822 | 116521905 | 116521983 | 17 | 7 | 5 |
| SE | 54149 | ENSG00000266053 | NDUFV2-AS1 | chr18 | - | 9132222 | 9132398 | 9121264 | 9121456 | 9133390 | 9133510 | 9 | 3 | 23 |
| SE | 67608 | ENSG00000109466 | KLHL2 | chr4 | + | 165228806 | 165228913 | 165219933 | 165220059 | 165238777 | 165238831 | 92 | 62 | 85 |
| SE | 52283 | ENSG00000007402 | CACNA2D2 | chr3 | - | 50384195 | 50384337 | 50380994 | 50381126 | 50387567 | 50387612 | 106 | 43 | 35 |
| SE | 7311 | ENSG00000154889 | MPPE1 | chr18 | - | 11897525 | 11897620 | 11896982 | 11897356 | 11908200 | 11908308 | 27 | 1 | 7 |
| SE | 91057 | ENSG00000104164 | BLOC1S6 | chr15 | + | 45592134 | 45592276 | 45587354 | 45587525 | 45605427 | 45605514 | 53 | 48 | 47 |
| SE | 71505 | ENSG00000170325 | PRDM10 | chr11 | - | 129942425 | 129942629 | 129937597 | 129937670 | 129947144 | 129947292 | 28 | 21 | 25 |
| SE | 2245 | ENSG00000189068 | VSTM1 | chr19 | - | 54055505 | 54055799 | 54051409 | 54051448 | 54058305 | 54058590 | 15 | 7 | 4 |
| SE | 10799 | ENSG00000091542 | ALKBH5 | chr17 | + | 18206814 | 18206970 | 18194954 | 18195035 | 18208218 | 18209398 | 586 | 274 | 504 |
| SE | 15216 | ENSG00000101266 | CSNK2A1 | chr20 | - | 500030 | 500115 | 499832 | 499934 | 505117 | 505229 | 6 | 7 | 12 |
| SE | 9948 | ENSG00000109062 | SLC9A3R1 | chr17 | + | 74762011 | 74762173 | 74748627 | 74749287 | 74766935 | 74766976 | 137 | 79 | 111 |
| SE | 84065 | ENSG00000197157 | SND1 | chr7 | + | 128008033 | 128008144 | 127990946 | 127991056 | 128074501 | 128074690 | 3 | 0 | 2 |
| SE | 28566 | ENSG00000241258 | CRCP | chr7 | + | 66130743 | 66130842 | 66127703 | 66127740 | 66134279 | 66134374 | 138 | 77 | 118 |
| SE | 15831 | ENSG00000198901 | PRC1 | chr15 | - | 90984017 | 90984140 | 90981498 | 90981981 | 90984692 | 90984825 | 15 | 18 | 23 |
| SE | 98308 | ENSG00000140022 | STON2 | chr14 | - | 81324016 | 81324187 | 81276900 | 81278739 | 81370987 | 81371185 | 11 | 17 | 2 |
| SE | 42661 | ENSG00000088854 | C20orf194 | chr20 | - | 3298028 | 3298175 | 3295493 | 3296949 | 3315032 | 3315120 | 19 | 17 | 8 |
| SE | 49894 | ENSG00000141068 | KSR1 | chr17 | + | 27577491 | 27577639 | 27550567 | 27550708 | 27582645 | 27583002 | 31 | 16 | 46 |
| SE | 46872 | ENSG00000162894 | FCMR | chr1 | - | 206921340 | 206921433 | 206912972 | 206913042 | 206921817 | 206921997 | 18 | 5 | 19 |
| SE | 70380 | ENSG00000132330 | SCLY | chr2 | + | 238069296 | 238069477 | 238068064 | 238068165 | 238073782 | 238073847 | 48 | 31 | 29 |
| SE | 59908 | ENSG00000170606 | HSPA4 | chr5 | + | 133091192 | 133091374 | 133089561 | 133089695 | 133092699 | 133092789 | 323 | 220 | 266 |
| SE | 17680 | ENSG00000119682 | AREL1 | chr14 | - | 74669946 | 74670126 | 74669648 | 74669774 | 74670761 | 74670871 | 189 | 98 | 142 |
| SE | 89125 | ENSG00000064115 | TM7SF3 | chr12 | - | 26996741 | 26996862 | 26995234 | 26995408 | 27003235 | 27003390 | 50 | 12 | 45 |
| SE | 17370 | ENSG00000079134 | THOC1 | chr18 | - | 254272 | 254355 | 252538 | 252612 | 259179 | 259275 | 128 | 137 | 133 |
| SE | 102233 | ENSG00000100109 | TFIP11 | chr22 | - | 26511118 | 26511317 | 26510616 | 26510702 | 26512098 | 26512175 | 9 | 4 | 3 |
| SE | 46461 | ENSG00000115415 | STAT1 | chr2 | - | 190998216 | 190998308 | 190997855 | 190998007 | 190999625 | 190999704 | 2082 | 2603 | 2186 |
| SE | 77107 | ENSG00000122512 | PMS2 | chr7 | - | 5999107 | 5999275 | 5997325 | 5997423 | 6002452 | 6002636 | 41 | 24 | 44 |
| SE | 92673 | ENSG00000090060 | PAPOLA | chr14 | + | 96562818 | 96562893 | 96556174 | 96556413 | 96564954 | 96564988 | 213 | 189 | 159 |
| SE | 86548 | ENSG00000109111 | SUPT6H | chr17 | + | 28678075 | 28678192 | 28677714 | 28677816 | 28678544 | 28678634 | 193 | 133 | 254 |
| SE | 59166 | ENSG00000115204 | MPV17 | chr2 | - | 27312993 | 27313109 | 27312679 | 27312772 | 27322447 | 27322522 | 73 | 37 | 54 |
| SE | 5723 | ENSG00000158470 | B4GALT5 | chr20 | - | 49639677 | 49639800 | 49637340 | 49637442 | 49640477 | 49640665 | 271 | 157 | 159 |
| SE | 836 | ENSG00000198911 | SREBF2 | chr22 | + | 41900019 | 41900122 | 41898648 | 41898781 | 41900329 | 41900498 | 4 | 1 | 5 |
| SE | 82292 | ENSG00000122566 | HNRNPA2B1 | chr7 | - | 26191055 | 26191650 | 26189956 | 26190460 | 26192494 | 26192577 | 72 | 57 | 95 |
| SE | 46894 | ENSG00000055609 | KMT2C | chr7 | - | 152195906 | 152196011 | 152194397 | 152194568 | 152199278 | 152199459 | 326 | 291 | 298 |
| SE | 81877 | ENSG00000177189 | RPS6KA3 | chrX | - | 20167588 | 20167747 | 20164898 | 20165060 | 20169401 | 20169491 | 492 | 496 | 524 |
| SE | 54408 | ENSG00000141127 | PRPSAP2 | chr17 | + | 18867281 | 18867334 | 18865801 | 18865952 | 18872582 | 18872649 | 175 | 130 | 182 |
| SE | 18258 | ENSG00000171723 | GPHN | chr14 | + | 66709356 | 66709458 | 66681106 | 66681185 | 66776463 | 66776521 | 0 | 2 | 0 |
| SE | 23566 | ENSG00000204574 | ABCF1 | chr6 | + | 30582393 | 30582507 | 30580405 | 30580519 | 30583065 | 30583188 | 172 | 122 | 172 |
| SE | 44857 | ENSG00000143493 | INTS7 | chr1 | - | 212020720 | 212020754 | 212020121 | 212020268 | 212021082 | 212021212 | 5 | 2 | 0 |
| SE | 18479 | ENSG00000100889 | PCK2 | chr14 | + | 24103159 | 24103255 | 24102674 | 24102890 | 24103509 | 24103722 | 57 | 13 | 38 |
| SE | 59868 | ENSG00000103168 | TAF1C | chr16 | - | 84184850 | 84184899 | 84183696 | 84183778 | 84186900 | 84187049 | 18 | 9 | 16 |
| SE | 59068 | ENSG00000103365 | GGA2 | chr16 | - | 23493359 | 23493587 | 23491776 | 23491800 | 23494302 | 23494378 | 172 | 81 | 141 |
| SE | 60760 | ENSG00000141524 | TMC6 | chr17 | - | 78121012 | 78121164 | 78120176 | 78120832 | 78122604 | 78122748 | 102 | 33 | 55 |
| SE | 26268 | ENSG00000137449 | CPEB2 | chr4 | + | 15040463 | 15040487 | 15033160 | 15033211 | 15052413 | 15052584 | 13 | 8 | 18 |
| SE | 61941 | ENSG00000136932 | TRMO | chr9 | - | 97912872 | 97912975 | 97910282 | 97910616 | 97913400 | 97913558 | 28 | 30 | 24 |
| SE | 31565 | ENSG00000163913 | IFT122 | chr3 | + | 129492140 | 129492194 | 129488256 | 129488397 | 129495445 | 129495607 | 7 | 9 | 17 |
| SE | 17923 | ENSG00000197249 | SERPINA1 | chr14 | - | 94388559 | 94388660 | 94379463 | 94379611 | 94390456 | 94390662 | 62 | 33 | 21 |
| SE | 64307 | ENSG00000101336 | HCK | chr20 | + | 32073718 | 32073818 | 32073318 | 32073361 | 32074622 | 32074721 | 249 | 168 | 142 |
| SE | 50390 | ENSG00000143384 | MCL1 | chr1 | - | 150578243 | 150578491 | 150574175 | 150577491 | 150578842 | 150578963 | 3596 | 2736 | 3515 |
| SE | 7281 | ENSG00000101019 | UQCC1 | chr20 | - | 35366556 | 35366614 | 35344109 | 35347272 | 35374183 | 35374256 | 124 | 95 | 99 |
| SE | 24172 | ENSG00000124357 | NAGK | chr2 | + | 71075554 | 71075642 | 71072640 | 71073594 | 71076603 | 71076641 | 216 | 114 | 125 |
| SE | 101565 | ENSG00000088298 | EDEM2 | chr20 | - | 35144978 | 35145018 | 35142372 | 35142478 | 35147151 | 35147319 | 21 | 8 | 4 |
| SE | 20283 | ENSG00000118482 | PHF3 | chr6 | + | 63684128 | 63685911 | 63679999 | 63680161 | 63694580 | 63694690 | 426 | 434 | 390 |
| SE | 5432 | ENSG00000118705 | RPN2 | chr20 | + | 37238401 | 37238449 | 37236579 | 37236709 | 37241302 | 37241619 | 16 | 7 | 9 |
| SE | 96083 | ENSG00000159445 | THEM4 | chr1 | - | 151889982 | 151890214 | 151889213 | 151889373 | 151895007 | 151895194 | 21 | 37 | 51 |
| SE | 49595 | ENSG00000076685 | NT5C2 | chr10 | - | 103106588 | 103106706 | 103105705 | 103105801 | 103139405 | 103139479 | 383 | 313 | 310 |
| SE | 52905 | ENSG00000159023 | EPB41 | chr1 | + | 29058588 | 29058645 | 29053103 | 29053312 | 29064981 | 29065158 | 2 | 1 | 6 |
| SE | 69268 | ENSG00000168385 | SEPTIN2 | chr2 | + | 241320202 | 241320287 | 241317498 | 241317599 | 241324215 | 241324241 | 6 | 1 | 2 |
| SE | 23885 | ENSG00000008394 | MGST1 | chr12 | + | 16357604 | 16357699 | 16354230 | 16354378 | 16363794 | 16363806 | 234 | 110 | 90 |
| SE | 4675 | ENSG00000141968 | VAV1 | chr19 | + | 6828075 | 6828171 | 6826611 | 6826711 | 6828418 | 6828487 | 223 | 130 | 170 |
| SE | 78327 | ENSG00000228409 | CCT6P1 | chr7 | + | 65761645 | 65761748 | 65760423 | 65760556 | 65762946 | 65763020 | 14 | 4 | 10 |
| SE | 78146 | ENSG00000214783 | POLR2J4 | chr7 | - | 43969731 | 43969861 | 43966263 | 43966395 | 43972629 | 43972691 | 5 | 3 | 2 |
| SE | 42555 | ENSG00000169241 | SLC50A1 | chr1 | + | 155137578 | 155137722 | 155136827 | 155136951 | 155137978 | 155138098 | 50 | 22 | 46 |
| SE | 94637 | ENSG00000011454 | RABGAP1 | chr9 | + | 122971664 | 122971952 | 122957010 | 122957209 | 122984484 | 122984719 | 20 | 9 | 5 |
| SE | 31049 | ENSG00000103174 | NAGPA | chr16 | - | 5028296 | 5028398 | 5027979 | 5028185 | 5028879 | 5029008 | 2 | 5 | 3 |
| SE | 11719 | ENSG00000100266 | PACSIN2 | chr22 | - | 42993619 | 42993780 | 42912020 | 42912157 | 43015020 | 43015135 | 0 | 0 | 0 |
| SE | 34078 | ENSG00000114867 | EIF4G1 | chr3 | + | 184319688 | 184319801 | 184314494 | 184314674 | 184320629 | 184320722 | 136 | 65 | 91 |
| SE | 7141 | ENSG00000141425 | RPRD1A | chr18 | - | 36033707 | 36033837 | 36030990 | 36031097 | 36067253 | 36067556 | 138 | 101 | 114 |
| SE | 28270 | ENSG00000138443 | ABI2 | chr2 | + | 203376056 | 203376132 | 203366876 | 203367044 | 203380207 | 203380273 | 5 | 4 | 3 |
| SE | 19746 | ENSG00000100714 | MTHFD1 | chr14 | + | 64458213 | 64458307 | 64454722 | 64454875 | 64459758 | 64459996 | 57 | 48 | 79 |
| SE | 75746 | ENSG00000144580 | CNOT9 | chr2 | + | 218592615 | 218592707 | 218592303 | 218592402 | 218594107 | 218594526 | 218 | 126 | 202 |
| SE | 36919 | ENSG00000135622 | SEMA4F | chr2 | + | 74673456 | 74673576 | 74662731 | 74662825 | 74674497 | 74674676 | 10 | 5 | 15 |
| SE | 10368 | ENSG00000197381 | ADARB1 | chr21 | + | 45134714 | 45134832 | 45128401 | 45128573 | 45171609 | 45171684 | 23 | 6 | 25 |
| SE | 10932 | ENSG00000091592 | NLRP1 | chr17 | - | 5515472 | 5515517 | 5502291 | 5505977 | 5517745 | 5517887 | 47 | 46 | 71 |
| SE | 56664 | ENSG00000144749 | LRIG1 | chr3 | - | 66398969 | 66399041 | 66398111 | 66398183 | 66405197 | 66405278 | 40 | 27 | 43 |
| SE | 28395 | ENSG00000204842 | ATXN2 | chr12 | - | 111485264 | 111485331 | 111470557 | 111470742 | 111485712 | 111485793 | 18 | 24 | 20 |
| SE | 11431 | ENSG00000167522 | ANKRD11 | chr16 | - | 89291680 | 89291799 | 89291012 | 89291183 | 89305205 | 89305344 | 26 | 6 | 20 |
| SE | 107941 | ENSG00000177119 | ANO6 | chr12 | + | 45228167 | 45228273 | 45216119 | 45216391 | 45270461 | 45270524 | 3 | 4 | 4 |
| SE | 88018 | ENSG00000147439 | BIN3 | chr8 | - | 22636524 | 22636586 | 22630441 | 22630578 | 22636921 | 22636962 | 106 | 104 | 115 |
| SE | 103513 | ENSG00000107862 | GBF1 | chr10 | + | 102360183 | 102360395 | 102358505 | 102358729 | 102361021 | 102361120 | 74 | 37 | 67 |
| SE | 22340 | ENSG00000166091 | CMTM5 | chr14 | + | 23379030 | 23379123 | 23377036 | 23377377 | 23379298 | 23379383 | 16 | 20 | 4 |
| SE | 673 | ENSG00000133460 | SLC2A11 | chr22 | + | 23877384 | 23877869 | 23877041 | 23877171 | 23882458 | 23882646 | 1 | 0 | 1 |
| SE | 72215 | ENSG00000223546 | LINC00630 | chrX | + | 102825992 | 102826169 | 102816991 | 102817082 | 102826968 | 102827144 | 18 | 13 | 14 |
| SE | 13853 | ENSG00000102893 | PHKB | chr16 | + | 47562397 | 47562495 | 47547432 | 47547548 | 47580294 | 47580358 | 5 | 0 | 5 |
| SE | 101592 | ENSG00000172671 | ZFAND4 | chr10 | - | 45639815 | 45639963 | 45625950 | 45627105 | 45648293 | 45648534 | 67 | 53 | 67 |
| SE | 35628 | ENSG00000138246 | DNAJC13 | chr3 | + | 132446474 | 132446550 | 132434537 | 132434618 | 132447320 | 132447470 | 320 | 241 | 176 |
| SE | 35466 | ENSG00000161217 | PCYT1A | chr3 | - | 196283474 | 196283538 | 196270414 | 196270541 | 196287614 | 196287713 | 6 | 0 | 3 |
| SE | 49433 | ENSG00000136068 | FLNB | chr3 | + | 58141857 | 58141929 | 58138281 | 58138529 | 58142649 | 58142752 | 40 | 10 | 13 |
| SE | 25279 | ENSG00000163590 | PPM1L | chr3 | + | 160939655 | 160939755 | 160756230 | 160756707 | 160961735 | 160961910 | 8 | 4 | 18 |
| SE | 85791 | ENSG00000258529 | NA | chr11 | - | 111840654 | 111840809 | 111837467 | 111837615 | 111844600 | 111844723 | 46 | 25 | 38 |
| SE | 67769 | ENSG00000036257 | CUL3 | chr2 | - | 224511353 | 224511582 | 224506857 | 224507003 | 224513523 | 224513638 | 564 | 408 | 452 |
| SE | 1803 | ENSG00000099917 | MED15 | chr22 | + | 20552538 | 20552709 | 20551435 | 20551487 | 20553144 | 20553174 | 9 | 3 | 6 |
| SE | 10760 | ENSG00000236320 | SLFN14 | chr17 | - | 35559726 | 35559805 | 35557002 | 35558106 | 35560766 | 35560819 | 4 | 2 | 4 |
| SE | 80749 | ENSG00000004864 | SLC25A13 | chr7 | - | 96234801 | 96234917 | 96208837 | 96208977 | 96277195 | 96277338 | 20 | 37 | 84 |
| SE | 85010 | ENSG00000101901 | ALG13 | chrX | + | 111716952 | 111717004 | 111713224 | 111713297 | 111717845 | 111717927 | 4 | 5 | 11 |
| SE | 85704 | ENSG00000072778 | ACADVL | chr17 | + | 7220463 | 7220529 | 7219843 | 7220197 | 7220603 | 7220676 | 63 | 18 | 51 |
| SE | 33199 | ENSG00000163930 | BAP1 | chr3 | - | 52407173 | 52407316 | 52406252 | 52406907 | 52407957 | 52408047 | 68 | 32 | 50 |
| SE | 17682 | ENSG00000119682 | AREL1 | chr14 | - | 74669946 | 74670142 | 74669686 | 74669774 | 74670761 | 74670871 | 108 | 64 | 85 |
| SE | 68504 | ENSG00000087087 | SRRT | chr7 | + | 100888277 | 100888383 | 100888041 | 100888143 | 100888473 | 100888656 | 84 | 28 | 61 |
| SE | 80319 | ENSG00000241973 | PI4KA | chr22 | - | 20796145 | 20796314 | 20793192 | 20793243 | 20798583 | 20798687 | 243 | 132 | 231 |
| SE | 53363 | ENSG00000130939 | UBE4B | chr1 | + | 10121961 | 10122076 | 10119512 | 10119613 | 10126793 | 10126877 | 136 | 68 | 107 |
| SE | 96704 | ENSG00000135362 | PRR5L | chr11 | + | 36437384 | 36437476 | 36431852 | 36431910 | 36446299 | 36446440 | 381 | 97 | 84 |
| SE | 82860 | ENSG00000146776 | ATXN7L1 | chr7 | - | 105788603 | 105788708 | 105665065 | 105665288 | 105875811 | 105875880 | 94 | 46 | 95 |
| SE | 70907 | ENSG00000130706 | ADRM1 | chr20 | + | 62308367 | 62308470 | 62308020 | 62308178 | 62308654 | 62308860 | 150 | 51 | 92 |
| SE | 34517 | ENSG00000106733 | NMRK1 | chr9 | - | 75078325 | 75078582 | 75077489 | 75077580 | 75083086 | 75083150 | 81 | 57 | 154 |
| SE | 59280 | ENSG00000119004 | CYP20A1 | chr2 | + | 203246754 | 203246921 | 203245845 | 203245895 | 203251966 | 203252109 | 103 | 61 | 106 |
| SE | 93807 | ENSG00000136869 | TLR4 | chr9 | + | 117708198 | 117708318 | 117704191 | 117704565 | 117708562 | 117708729 | 65 | 19 | 22 |
| SE | 81671 | ENSG00000181090 | EHMT1 | chr9 | + | 137802856 | 137803010 | 137800879 | 137800984 | 137811460 | 137811491 | 2 | 6 | 8 |
| SE | 67622 | ENSG00000175215 | CTDSP2 | chr12 | - | 57826995 | 57827097 | 57826345 | 57826402 | 57827551 | 57827590 | 470 | 224 | 412 |
| SE | 1798 | ENSG00000254709 | IGLL5 | chr22 | + | 22893699 | 22893818 | 22887815 | 22888259 | 22895374 | 22896111 | 43 | 11 | 83 |
| SE | 77040 | ENSG00000160785 | SLC25A44 | chr1 | + | 156207885 | 156208013 | 156194103 | 156194247 | 156210239 | 156212795 | 26 | 23 | 30 |
| SE | 95766 | ENSG00000178177 | LCORL | chr4 | - | 17960264 | 17960311 | 17909093 | 17909345 | 17961902 | 17962032 | 9 | 5 | 4 |
| SE | 95423 | ENSG00000148356 | LRSAM1 | chr9 | + | 127495319 | 127495418 | 127492801 | 127492897 | 127495963 | 127496075 | 45 | 30 | 23 |
| SE | 15833 | ENSG00000189136 | UBE2Q2P1 | chr15 | - | 84539703 | 84539788 | 84538531 | 84538616 | 84542144 | 84542229 | 3 | 0 | 2 |
| SE | 89568 | ENSG00000095485 | CWF19L1 | chr10 | - | 100256261 | 100256476 | 100253420 | 100253539 | 100260217 | 100260319 | 82 | 46 | 94 |
| SE | 85805 | ENSG00000163605 | PPP4R2 | chr3 | + | 73047185 | 73047356 | 72998076 | 72998158 | 73059036 | 73059051 | 567 | 437 | 495 |
| SE | 96847 | ENSG00000083535 | PIBF1 | chr13 | + | 72783422 | 72783721 | 72782132 | 72782349 | 72792446 | 72792547 | 129 | 105 | 140 |
| SE | 92474 | ENSG00000100815 | TRIP11 | chr14 | - | 92011754 | 92011795 | 92010985 | 92011072 | 92014214 | 92014577 | 102 | 136 | 100 |
| SE | 95771 | ENSG00000125944 | HNRNPR | chr1 | - | 23323555 | 23323732 | 23321527 | 23321663 | 23333517 | 23333631 | 862 | 601 | 904 |
| SE | 40763 | ENSG00000114026 | OGG1 | chr3 | + | 9754703 | 9754885 | 9751769 | 9751949 | 9781512 | 9781600 | 4 | 3 | 7 |
| SE | 66362 | ENSG00000197037 | ZSCAN25 | chr7 | + | 99624056 | 99624180 | 99621060 | 99622640 | 99629190 | 99629468 | 76 | 51 | 82 |
| SE | 3421 | ENSG00000127452 | FBXL12 | chr19 | - | 9814137 | 9814342 | 9813230 | 9813274 | 9818544 | 9818617 | 0 | 2 | 0 |
| SE | 83015 | ENSG00000152484 | USP12 | chr13 | - | 27105730 | 27105944 | 27095600 | 27095830 | 27116515 | 27116596 | 209 | 216 | 159 |
| SE | 103719 | ENSG00000107863 | ARHGAP21 | chr10 | - | 24622732 | 24622762 | 24619472 | 24621369 | 24629995 | 24630050 | 1 | 7 | 7 |
| SE | 82941 | ENSG00000148606 | POLR3A | chr10 | - | 78025621 | 78025759 | 78024970 | 78025142 | 78026093 | 78026229 | 75 | 57 | 80 |
| SE | 103659 | ENSG00000185009 | AP3M1 | chr10 | - | 74138805 | 74138931 | 74138106 | 74138382 | 74150754 | 74150842 | 75 | 55 | 74 |
| SE | 12566 | ENSG00000214078 | CPNE1 | chr20 | - | 35658929 | 35658982 | 35632794 | 35632923 | 35664801 | 35664956 | 8 | 2 | 7 |
| SE | 71648 | ENSG00000163832 | ELP6 | chr3 | - | 47501649 | 47501851 | 47498265 | 47498432 | 47504329 | 47504448 | 79 | 45 | 73 |
| SE | 996 | ENSG00000243156 | MICAL3 | chr22 | - | 17817310 | 17819129 | 17816689 | 17816784 | 17821426 | 17821509 | 17 | 27 | 44 |
| SE | 106170 | ENSG00000120805 | ARL1 | chr12 | - | 101405843 | 101405981 | 101401061 | 101401173 | 101407641 | 101407746 | 101 | 46 | 79 |
| SE | 60350 | ENSG00000257267 | ZNF271P | chr18 | + | 35291009 | 35291232 | 35290281 | 35290391 | 35305975 | 35308549 | 178 | 162 | 92 |
| SE | 101644 | ENSG00000198258 | UBL5 | chr19 | + | 9828591 | 9828675 | 9828326 | 9828389 | 9828836 | 9828874 | 103 | 56 | 91 |
| SE | 10478 | ENSG00000186566 | GPATCH8 | chr17 | - | 44464471 | 44464544 | 44424348 | 44424492 | 44474828 | 44474903 | 35 | 23 | 31 |
| SE | 37352 | ENSG00000075826 | SEC31B | chr10 | - | 100501653 | 100501710 | 100500109 | 100500159 | 100502253 | 100502484 | 5 | 2 | 9 |
| SE | 39529 | ENSG00000186517 | ARHGAP30 | chr1 | - | 161059613 | 161059716 | 161056387 | 161056532 | 161069527 | 161069666 | 521 | 251 | 431 |
| SE | 8809 | ENSG00000181038 | METTL23 | chr17 | + | 76729689 | 76729794 | 76726829 | 76726879 | 76732977 | 76733215 | 25 | 10 | 30 |
| SE | 47741 | ENSG00000145687 | SSBP2 | chr5 | - | 81489249 | 81489309 | 81474495 | 81474562 | 81513627 | 81513717 | 85 | 72 | 87 |
| SE | 66900 | ENSG00000196504 | PRPF40A | chr2 | - | 152679055 | 152679109 | 152677450 | 152677475 | 152679351 | 152679472 | 340 | 323 | 287 |
| SE | 19449 | ENSG00000198668 | CALM1 | chr14 | + | 90398990 | 90399087 | 90398261 | 90398488 | 90400064 | 90400095 | 12 | 1 | 10 |
| SE | 19428 | ENSG00000151445 | VIPAS39 | chr14 | - | 77441065 | 77441093 | 77437807 | 77437881 | 77442559 | 77442662 | 85 | 32 | 71 |
| SE | 32791 | ENSG00000157601 | MX1 | chr21 | + | 41427205 | 41427304 | 41426141 | 41426263 | 41427754 | 41427866 | 38 | 57 | 39 |
| SE | 84884 | ENSG00000157881 | PANK4 | chr1 | - | 2515561 | 2515717 | 2514353 | 2514466 | 2518163 | 2518264 | 27 | 9 | 22 |
| SE | 55173 | ENSG00000054267 | ARID4B | chr1 | - | 235213768 | 235214026 | 235196030 | 235196115 | 235219792 | 235219968 | 199 | 233 | 203 |
| SE | 52447 | ENSG00000116954 | RRAGC | chr1 | - | 38851614 | 38851757 | 38845938 | 38846087 | 38855707 | 38855907 | 165 | 88 | 137 |
| SE | 26545 | ENSG00000117091 | CD48 | chr1 | - | 160681201 | 160681468 | 160678745 | 160679131 | 160684886 | 160685189 | 2724 | 1499 | 2409 |
| SE | 25761 | ENSG00000224032 | EPB41L4A-AS1 | chr5 | + | 112161702 | 112161775 | 112160525 | 112161295 | 112162150 | 112162311 | 104 | 89 | 96 |
| SE | 27838 | ENSG00000101493 | ZNF516 | chr18 | - | 76370527 | 76370595 | 76357681 | 76362557 | 76371466 | 76371571 | 27 | 22 | 27 |
| SE | 26100 | ENSG00000164181 | ELOVL7 | chr5 | - | 60771902 | 60772093 | 60767822 | 60767903 | 60787333 | 60787431 | 136 | 246 | 39 |
| SE | 95288 | ENSG00000172716 | SLFN11 | chr17 | - | 35366946 | 35367063 | 35362738 | 35363826 | 35373581 | 35373620 | 83 | 62 | 44 |
| SE | 32129 | ENSG00000084463 | WBP11 | chr12 | - | 14794970 | 14795104 | 14794536 | 14794736 | 14799634 | 14799728 | 209 | 126 | 215 |
| SE | 32424 | ENSG00000136040 | PLXNC1 | chr12 | + | 94298631 | 94298795 | 94297315 | 94297423 | 94303755 | 94303896 | 131 | 133 | 82 |
| SE | 103578 | ENSG00000150093 | ITGB1 | chr10 | - | 32933481 | 32933583 | 32932514 | 32932600 | 32935491 | 32935558 | 15 | 18 | 12 |
| SE | 21074 | ENSG00000196182 | STK40 | chr1 | - | 36372648 | 36372774 | 36367853 | 36368084 | 36385722 | 36385876 | 5 | 1 | 0 |

| | | | | | | | | | | | | | |
|---|---|---|---|---|---|---|---|---|---|---|---|---|---|
| SE | 42656 | ENSG00000088854 | C20orf194 | chr20 | - | 3270426 | 3270562 | 3264437 | 3264524 | 3278911 | 3278949 | 18 | 9 | 15 |
| SE | 103092 | ENSG00000108055 | SMC3 | chr10 | + | 110581922 | 110582098 | 110580903 | 110581021 | 110582561 | 110582642 | 312 | 332 | 283 |
| SE | 28621 | ENSG00000102445 | RUBCNL | chr13 | - | 46362537 | 46362598 | 46361440 | 46361573 | 46363114 | 46363213 | 76 | 64 | 84 |
| SE | 30325 | ENSG00000100298 | APOBEC3H | chr22 | + | 39102511 | 39102605 | 39101917 | 39102042 | 39103691 | 39104067 | 0 | 1 | 2 |
| SE | 939 | ENSG00000100138 | SNU13 | chr22 | - | 41681225 | 41681559 | 41680243 | 41680364 | 41688793 | 41688981 | 7 | 2 | 4 |
| SE | 57685 | ENSG00000082068 | WDR70 | chr5 | + | 37443238 | 37443372 | 37437921 | 37437981 | 37479833 | 37479987 | 290 | 224 | 239 |
| SE | 57285 | ENSG00000143119 | CD53 | chr1 | + | 110897808 | 110897892 | 110891391 | 110891471 | 110899123 | 110899922 | 623 | 299 | 527 |
| SE | 11401 | ENSG00000074356 | NCBP3 | chr17 | - | 3840099 | 3840205 | 3829242 | 3829368 | 3843085 | 3843151 | 159 | 103 | 76 |
| SE | 1276 | ENSG00000186951 | PPARA | chr22 | + | 46188722 | 46188822 | 46176752 | 46176836 | 46198341 | 46198591 | 2 | 3 | 5 |
| SE | 8428 | ENSG00000078687 | TNRC6C | chr17 | + | 78077184 | 78077334 | 78075135 | 78075278 | 78083046 | 78083166 | 82 | 40 | 85 |
| SE | 85074 | ENSG00000147459 | DOCK5 | chr8 | + | 25407982 | 25408154 | 25403557 | 25403724 | 25408801 | 25408940 | 129 | 93 | 57 |
| SE | 15976 | ENSG00000185787 | MORF4L1 | chr15 | + | 78884969 | 78885083 | 78880511 | 78880579 | 78886140 | 78886227 | 30 | 9 | 28 |
| SE | 65069 | ENSG00000115355 | CCDC88A | chr2 | - | 55312433 | 55312579 | 55309161 | 55309254 | 55315927 | 55315952 | 269 | 438 | 220 |
| SE | 107357 | ENSG00000110921 | MVK | chr12 | + | 109581394 | 109581550 | 109575997 | 109576145 | 109586021 | 109586096 | 29 | 7 | 24 |
| SE | 55343 | ENSG00000204469 | PRRC2A | chr6 | + | 31626771 | 31626862 | 31625753 | 31626162 | 31626981 | 31627074 | 372 | 230 | 347 |
| SE | 78837 | ENSG00000105854 | PON2 | chr7 | - | 95406986 | 95407068 | 95406118 | 95406247 | 95409900 | 95410101 | 67 | 49 | 19 |
| SE | 107747 | ENSG00000135677 | GNS | chr12 | - | 64756666 | 64756762 | 64752697 | 64752757 | 64759084 | 64759406 | 1 | 4 | 1 |
| SE | 47028 | ENSG00000175309 | PHYKPL | chr5 | - | 178223319 | 178223451 | 178222851 | 178222934 | 178224615 | 178224729 | 0 | 2 | 4 |
| SE | 106712 | ENSG00000141376 | BCAS3 | chr17 | + | 61380505 | 61380571 | 61368326 | 61368494 | 61388617 | 61388708 | 4 | 2 | 3 |
| SE | 80725 | ENSG00000163636 | PSMD6 | chr3 | - | 64013438 | 64013607 | 64010877 | 64010955 | 64018598 | 64018707 | 430 | 295 | 407 |
| SE | 73262 | ENSG00000114956 | DGUOK | chr2 | + | 73957124 | 73957240 | 73946718 | 73946906 | 73958145 | 73958245 | 82 | 44 | 52 |
| SE | 55789 | ENSG00000187231 | SESTD1 | chr2 | - | 179116667 | 179116790 | 179115064 | 179115256 | 179117531 | 179117613 | 79 | 74 | 77 |
| SE | 27150 | ENSG00000109534 | GAR1 | chr4 | + | 109816152 | 109816378 | 109815634 | 109815804 | 109817935 | 109818090 | 82 | 101 | 80 |
| SE | 86386 | ENSG00000123562 | MORF4L2 | chrX | - | 103684680 | 103684729 | 103678498 | 103678600 | 103685170 | 103685260 | 70 | 45 | 68 |
| SE | 52057 | ENSG00000178685 | PARP10 | chr8 | - | 143984034 | 143984104 | 143983166 | 143983811 | 143984209 | 143984431 | 87 | 31 | 113 |
| SE | 83945 | ENSG00000147421 | HMBOX1 | chr8 | + | 29049289 | 29049361 | 29048956 | 29049048 | 29051017 | 29051549 | 16 | 16 | 21 |
| SE | 42260 | ENSG00000116001 | TIA1 | chr2 | - | 70225327 | 70225393 | 70224553 | 70224629 | 70227734 | 70227822 | 19 | 15 | 16 |
| SE | 73517 | ENSG00000138380 | CARF | chr2 | + | 202941859 | 202941980 | 202924296 | 202924415 | 202952558 | 202952679 | 22 | 10 | 21 |
| SE | 38653 | ENSG00000073417 | PDE8A | chr15 | + | 85090717 | 85090915 | 85089337 | 85089416 | 85091043 | 85091181 | 6 | 3 | 3 |
| SE | 32820 | ENSG00000088387 | DOCK9 | chr13 | - | 98825871 | 98825940 | 98824397 | 98824504 | 98826829 | 98826887 | 5 | 9 | 25 |
| SE | 96241 | ENSG00000133943 | DGLUCY | chr14 | + | 91170002 | 91170186 | 91167224 | 91167378 | 91173288 | 91173403 | 46 | 33 | 48 |
| SE | 91281 | ENSG00000111530 | CAND1 | chr12 | + | 67292621 | 67292776 | 67281909 | 67282053 | 67295032 | 67295156 | 419 | 303 | 303 |
| SE | 78081 | ENSG00000196923 | PDLIM7 | chr5 | - | 177491806 | 177491976 | 177491009 | 177491146 | 177492404 | 177492435 | 18 | 2 | 12 |
| SE | 51970 | ENSG00000071054 | MAP4K4 | chr2 | + | 101831720 | 101831851 | 101829503 | 101829594 | 101834408 | 101834463 | 558 | 468 | 591 |
| SE | 69348 | ENSG00000168958 | MFF | chr2 | + | 227352513 | 227352573 | 227346724 | 227347384 | 227355676 | 227355761 | 18 | 13 | 38 |
| SE | 99092 | ENSG00000110274 | CEP164 | chr11 | + | 117362403 | 117362538 | 117361834 | 117361993 | 117371079 | 117371466 | 46 | 30 | 41 |
| SE | 5543 | ENSG00000101057 | MYBL2 | chr20 | + | 43681783 | 43681855 | 43673805 | 43673899 | 43682793 | 43682886 | 10 | 6 | 42 |
| SE | 58811 | ENSG00000140416 | TPM1 | chr15 | + | 63061712 | 63061788 | 63060868 | 63061273 | 63062214 | 63062277 | 4 | 6 | 1 |
| SE | 20272 | ENSG00000118482 | PHF3 | chr6 | + | 63679999 | 63680161 | 63635822 | 63636150 | 63684128 | 63684245 | 460 | 456 | 423 |
| SE | 52762 | ENSG00000198019 | FCGR1B | chr1 | + | 121087820 | 121087841 | 121087344 | 121087425 | 121092995 | 121093250 | 0 | 0 | 0 |
| SE | 49623 | ENSG00000178927 | CYBC1 | chr17 | - | 82447579 | 82447621 | 82446622 | 82446696 | 82449169 | 82449292 | 95 | 26 | 57 |
| SE | 63741 | ENSG00000152061 | RABGAP1L | chr1 | + | 174304985 | 174305127 | 174278612 | 174278779 | 174370978 | 174371072 | 376 | 262 | 211 |
| SE | 11170 | ENSG00000213918 | DNASE1 | chr16 | + | 3655848 | 3655966 | 3655372 | 3655520 | 3656101 | 3656185 | 1 | 0 | 2 |
| SE | 99946 | ENSG00000230724 | LINC01001 | chr11 | - | 134563 | 134796 | 133572 | 133778 | 134889 | 134947 | 22 | 13 | 29 |
| SE | 31564 | ENSG00000163913 | IFT122 | chr3 | + | 129466889 | 129467066 | 129464634 | 129464781 | 129469341 | 129469417 | 1 | 0 | 4 |
| SE | 94608 | ENSG00000106829 | TLE4 | chr9 | + | 79706746 | 79706899 | 79705888 | 79705942 | 79708117 | 79708162 | 322 | 188 | 252 |
| SE | 74764 | ENSG00000163939 | PBRM1 | chr3 | - | 52682091 | 52682233 | 52679573 | 52679726 | 52685748 | 52685850 | 21 | 10 | 22 |
| SE | 89281 | ENSG00000166169 | POLL | chr10 | - | 101582762 | 101582891 | 101580247 | 101580416 | 101587245 | 101587406 | 18 | 12 | 33 |
| SE | 10351 | ENSG00000131467 | PSME3 | chr17 | + | 42834514 | 42834577 | 42833410 | 42834376 | 42837648 | 42837697 | 52 | 8 | 34 |
| SE | 94334 | ENSG00000136936 | XPA | chr9 | - | 97689533 | 97689639 | 97687095 | 97687261 | 97693648 | 97693759 | 305 | 251 | 313 |
| SE | 10142 | ENSG00000011260 | UTP18 | chr17 | + | 51280004 | 51280105 | 51275867 | 51275991 | 51280388 | 51280479 | 37 | 11 | 60 |
| SE | 75022 | ENSG00000100814 | CCNB1IP1 | chr14 | - | 20315780 | 20315850 | 20313467 | 20313801 | 20316226 | 20316265 | 5 | 0 | 7 |
| SE | 35142 | ENSG00000107758 | PPP3CB | chr10 | - | 73439871 | 73439901 | 73436432 | 73438420 | 73444724 | 73444822 | 136 | 79 | 139 |
| SE | 89227 | ENSG00000164024 | METAP1 | chr4 | + | 99048732 | 99048876 | 99045178 | 99045310 | 99057752 | 99057818 | 316 | 219 | 271 |
| SE | 48456 | ENSG00000131238 | PPT1 | chr1 | - | 40076841 | 40076913 | 40072709 | 40074183 | 40078559 | 40078658 | 1522 | 813 | 870 |
| SE | 60442 | ENSG00000077157 | PPP1R12B | chr1 | + | 202434655 | 202434768 | 202431479 | 202431619 | 202437820 | 202438024 | 82 | 44 | 49 |
| SE | 2564 | ENSG00000214046 | SMIM7 | chr19 | - | 16659394 | 16659447 | 16646341 | 16647261 | 16659958 | 16660000 | 23 | 8 | 12 |
| SE | 56500 | ENSG00000176853 | FAM91A1 | chr8 | + | 123786494 | 123786610 | 123785628 | 123785741 | 123787260 | 123787355 | 214 | 128 | 140 |
| SE | 30987 | ENSG00000116237 | ICMT | chr1 | - | 6233473 | 6233643 | 6231901 | 6232119 | 6234885 | 6234974 | 59 | 19 | 28 |
| SE | 61664 | ENSG00000227232 | WASH7P | chr1 | - | 17232 | 17368 | 16857 | 17055 | 17605 | 17742 | 42 | 18 | 39 |
| SE | 15146 | ENSG00000104093 | DMXL2 | chr15 | - | 51495023 | 51495134 | 51491577 | 51491747 | 51498851 | 51500231 | 209 | 161 | 82 |
| SE | 44136 | ENSG00000136717 | BIN1 | chr2 | - | 127057472 | 127057601 | 127050801 | 127050912 | 127059010 | 127059155 | 7 | 1 | 3 |
| SE | 54559 | ENSG00000179912 | R3HDM2 | chr12 | - | 57296410 | 57296551 | 57295398 | 57295507 | 57297327 | 57297387 | 974 | 756 | 765 |
| SE | 69926 | ENSG00000169016 | E2F6 | chr2 | - | 11455341 | 11455474 | 11453581 | 11453798 | 11457178 | 11457233 | 17 | 3 | 4 |
| SE | 42519 | ENSG00000133256 | PDE6B | chr4 | + | 667855 | 667941 | 666530 | 666614 | 670045 | 670314 | 10 | 11 | 19 |
| SE | 8748 | ENSG00000007202 | KIAA0100 | chr17 | - | 28620508 | 28620630 | 28619859 | 28620015 | 28620987 | 28621170 | 438 | 271 | 351 |
| SE | 75253 | ENSG00000175575 | PAAF1 | chr11 | + | 73887039 | 73887099 | 73878778 | 73878819 | 73887353 | 73887457 | 0 | 1 | 1 |
| SE | 66463 | ENSG00000131051 | RBM39 | chr20 | - | 35729461 | 35729527 | 35729311 | 35729365 | 35731940 | 35732135 | 968 | 793 | 913 |
| SE | 41022 | ENSG00000064687 | ABCA7 | chr19 | + | 1041830 | 1041972 | 1041509 | 1041603 | 1042063 | 1042176 | 85 | 53 | 111 |
| SE | 26006 | ENSG00000181929 | PRKAG1 | chr12 | - | 49005742 | 49005881 | 49005461 | 49005543 | 49013061 | 49013110 | 77 | 68 | 60 |
| SE | 7673 | ENSG00000258315 | C17orf49 | chr17 | + | 7016909 | 7017014 | 7016494 | 7016685 | 7017256 | 7017375 | 180 | 97 | 125 |
| SE | 39180 | ENSG00000121775 | TMEM39B | chr1 | + | 32077163 | 32077318 | 32075602 | 32075822 | 32091674 | 32091744 | 7 | 1 | 5 |
| SE | 92510 | ENSG00000216937 | CCDC7 | chr10 | + | 32567669 | 32567891 | 32565557 | 32565620 | 32584231 | 32584304 | 315 | 238 | 166 |
| SE | 14163 | ENSG00000005339 | CREBBP | chr16 | - | 3791980 | 3792094 | 3782683 | 3782926 | 3793385 | 3793626 | 193 | 107 | 227 |
| SE | 91580 | ENSG00000070501 | POLB | chr8 | + | 42369848 | 42369988 | 42368935 | 42369335 | 42371562 | 42371765 | 174 | 147 | 154 |
| SE | 39469 | ENSG00000213024 | NUP62 | chr19 | - | 49927693 | 49927848 | 49908104 | 49909884 | 49929325 | 49929435 | 103 | 77 | 99 |
| SE | 44796 | ENSG00000054282 | SDCCAG8 | chr1 | + | 243293398 | 243293494 | 243293090 | 243293219 | 243304712 | 243304777 | 9 | 3 | 25 |
| SE | 54677 | ENSG00000070010 | UFD1 | chr22 | - | 19475067 | 19475100 | 19471686 | 19471808 | 19479082 | 19479172 | 46 | 22 | 29 |
| SE | 3183 | ENSG00000161847 | RAVER1 | chr19 | - | 10328641 | 10329052 | 10323505 | 10323566 | 10330459 | 10330526 | 13 | 5 | 15 |
| SE | 47405 | ENSG00000156804 | FBXO32 | chr8 | - | 123513197 | 123513382 | 123506391 | 123506574 | 123531897 | 123531990 | 62 | 34 | 45 |
| SE | 91757 | ENSG00000143748 | NVL | chr1 | - | 224296500 | 224296618 | 224294266 | 224294411 | 224300561 | 224300663 | 71 | 59 | 77 |
| SE | 7005 | ENSG00000101782 | RIOK3 | chr18 | + | 23467398 | 23467526 | 23466132 | 23466276 | 23473428 | 23473483 | 297 | 395 | 195 |
| SE | 98489 | ENSG00000254703 | SENCR | chr11 | - | 128692654 | 128693391 | 128691671 | 128692121 | 128695912 | 128696023 | 10 | 26 | 9 |

| | | | | | | | | | | | | | | | |
|---|---|---|---|---|---|---|---|---|---|---|---|---|---|---|---|
| SE | 24866 | ENSG00000155846 | PPARGC1B | chr5 | + | 149840040 | 149840116 | 149836262 | 149837073 | 149842255 | 149842377 | 10 | 6 | 22 | [illegible] |
| SE | 83556 | ENSG00000124222 | STX16 | chr20 | + | 58659622 | 58659634 | 58651271 | 58652087 | 58667200 | 58667318 | 3 | 3 | 2 | [illegible] |
| SE | 102222 | ENSG00000100109 | TFIP11 | chr22 | - | 26510616 | 26510692 | 26510063 | 26510281 | 26512098 | 26512175 | 57 | 67 | 41 | [illegible] |
| SE | 106401 | ENSG00000089041 | P2RX7 | chr12 | + | 121160901 | 121160974 | 121156078 | 121156147 | 121165356 | 121165437 | 20 | 22 | 20 | [illegible] |
| SE | 3708 | ENSG00000181027 | FKRP | chr19 | + | 46748514 | 46748884 | 46748026 | 46748088 | 46755411 | 46755449 | 15 | 9 | 5 | [illegible] |
| SE | 25167 | ENSG00000164219 | PGGT1B | chr5 | - | 115230975 | 115231021 | 115221823 | 115222008 | 115236389 | 115236522 | 213 | 135 | 126 | [illegible] |
| SE | 17965 | ENSG00000171469 | ZNF561 | chr19 | - | 9620915 | 9620963 | 9619431 | 9619582 | 9621161 | 9621222 | 4 | 3 | 9 | [illegible] |
| SE | 54589 | ENSG00000164048 | ZNF589 | chr3 | + | 48260812 | 48260939 | 48247624 | 48247677 | 48267914 | 48267981 | 92 | 79 | 97 | [illegible] |
| SE | 96935 | ENSG00000089159 | PXN | chr12 | - | 120222548 | 120222750 | 120221622 | 120221758 | 120222862 | 120222999 | 957 | 317 | 476 | [illegible] |
| SE | 34094 | ENSG00000114867 | EIF4G1 | chr3 | + | 184328908 | 184328990 | 184328615 | 184328756 | 184331265 | 184331364 | 293 | 132 | 231 | [illegible] |
| SE | 52789 | ENSG00000087191 | PSMC5 | chr17 | + | 63827828 | 63827923 | 63827445 | 63827514 | 63828137 | 63828209 | 4 | 2 | 2 | [illegible] |
| SE | 77648 | ENSG00000214021 | TTLL3 | chr3 | + | 9817644 | 9817759 | 9813247 | 9813345 | 9820545 | 9820741 | 21 | 18 | 22 | [illegible] |
| SE | 36670 | ENSG00000180530 | NRIP1 | chr21 | - | 15014343 | 15014466 | 14961234 | 14968526 | 15043494 | 15043574 | 77 | 62 | 70 | [illegible] |
| SE | 60779 | ENSG00000187741 | FANCA | chr16 | - | 89779868 | 89779957 | 89778942 | 89779003 | 89782858 | 89782918 | 2 | 10 | 11 | [illegible] |
| SE | 39449 | ENSG00000055917 | PUM2 | chr2 | - | 20332966 | 20333063 | 20327309 | 20327378 | 20350809 | 20350834 | 21 | 24 | 11 | [illegible] |
| SE | 50615 | ENSG00000143033 | MTF2 | chr1 | + | 93120548 | 93120672 | 93119332 | 93119401 | 93127231 | 93127299 | 136 | 105 | 107 | [illegible] |
| SE | 65661 | ENSG00000111729 | CLEC4A | chr12 | + | 8129263 | 8129362 | 8123878 | 8123960 | 8135584 | 8135736 | 287 | 183 | 158 | [illegible] |
| SE | 74612 | ENSG00000179222 | MAGED1 | chrX | + | 51894268 | 51894349 | 51803006 | 51803117 | 51895052 | 51895760 | 65 | 21 | 58 | [illegible] |
| SE | 96532 | ENSG00000143549 | TPM3 | chr1 | - | 154169304 | 154169383 | 154156612 | 154159048 | 154170399 | 154170469 | 24 | 16 | 23 | [illegible] |
| SE | 98602 | ENSG00000136527 | TRA2B | chr3 | - | 185923795 | 185924008 | 185922042 | 185922126 | 185925463 | 185925626 | 941 | 746 | 798 | [illegible] |
| SE | 28479 | ENSG00000104972 | LILRB1 | chr19 | + | 54635103 | 54635179 | 54634370 | 54634763 | 54635258 | 54635296 | 120 | 61 | 32 | [illegible] |
| SE | 26421 | ENSG00000165416 | SUGT1 | chr13 | + | 52658098 | 52658226 | 52657531 | 52657622 | 52658398 | 52658468 | 33 | 15 | 15 | [illegible] |
| SE | 97384 | ENSG00000113163 | CERT1 | chr5 | - | 75399309 | 75399387 | 75389591 | 75389687 | 75400204 | 75400297 | 12 | 21 | 14 | [illegible] |
| SE | 56950 | ENSG00000244005 | NFS1 | chr20 | - | 35675044 | 35675202 | 35674511 | 35674617 | 35680736 | 35680871 | 74 | 42 | 39 | [illegible] |
| SE | 13613 | ENSG00000038358 | EDC4 | chr16 | + | 67876487 | 67876599 | 67875944 | 67876101 | 67876872 | 67876972 | 98 | 65 | 110 | [illegible] |
| SE | 67268 | ENSG00000135632 | SMYD5 | chr2 | + | 73224865 | 73224960 | 73223946 | 73224003 | 73225630 | 73225701 | 33 | 14 | 56 | [illegible] |
| [illegible] | | | | | | | | | | | | | | | |

| | | | | | | | | | | | | | | | |
|---|---|---|---|---|---|---|---|---|---|---|---|---|---|---|---|
| SE | 17229 | ENSG00000197555 | SIPA1L1 | chr14 | + | 71413947 | 71414014 | 71400793 | 71400918 | 71512742 | 71512845 | 21 | 20 | 17 | [illegible] |
| SE | 60124 | ENSG00000122223 | CD244 | chr1 | - | 160841209 | 160841500 | 160838938 | 160839049 | 160841583 | 160841901 | 356 | 234 | 142 | [illegible] |
| SE | 94065 | ENSG00000167985 | SDHAF2 | chr11 | + | 61436760 | 61436960 | 61430041 | 61430182 | 61437624 | 61437848 | 15 | 16 | 6 | [illegible] |
| SE | 40046 | ENSG00000143190 | POU2F1 | chr1 | + | 167337969 | 167338258 | 167332469 | 167332535 | 167365466 | 167365567 | 5 | 0 | 6 | [illegible] |
| SE | 49959 | ENSG00000171793 | CTPS1 | chr1 | + | 41007448 | 41007545 | 41006050 | 41006094 | 41008658 | 41008714 | 75 | 31 | 68 | [illegible] |
| SE | 48857 | ENSG00000165661 | QSOX2 | chr9 | - | 136224860 | 136224909 | 136224006 | 136224112 | 136226773 | 136226874 | 29 | 14 | 49 | [illegible] |
| SE | 75517 | ENSG00000184281 | TSSC4 | chr11 | + | 2401975 | 2402147 | 2400627 | 2400798 | 2402293 | 2402450 | 7 | 2 | 3 | [illegible] |
| SE | 95697 | ENSG00000132589 | FLOT2 | chr17 | - | 28885856 | 28885947 | 28884224 | 28884315 | 28888944 | 28889026 | 11 | 0 | 16 | [illegible] |
| SE | 62419 | ENSG00000113761 | ZNF346 | chr5 | + | 177041777 | 177041870 | 177041125 | 177041229 | 177044391 | 177044533 | 41 | 22 | 27 | [illegible] |
| SE | 46671 | ENSG00000198740 | ZNF652 | chr17 | - | 49311926 | 49312042 | 49289205 | 49298924 | 49312697 | 49312845 | 157 | 100 | 146 | [illegible] |
| SE | 45694 | ENSG00000049541 | RFC2 | chr7 | - | 74233774 | 74233894 | 74232103 | 74232216 | 74235531 | 74235645 | 1 | 0 | 1 | [illegible] |
| SE | 4081 | ENSG00000011422 | PLAUR | chr19 | - | 43655438 | 43655573 | 43648578 | 43649143 | 43656478 | 43656573 | 223 | 167 | 152 | [illegible] |
| SE | 92518 | ENSG00000216937 | CCDC7 | chr10 | + | 32583033 | 32583307 | 32565557 | 32565620 | 32584231 | 32584304 | 208 | 167 | 123 | [illegible] |
| SE | 3165 | ENSG00000213020 | ZNF611 | chr19 | - | 52716110 | 52716233 | 52715831 | 52715913 | 52728729 | 52728786 | 8 | 3 | 2 | [illegible] |
| SE | 26803 | ENSG00000163281 | GNPDA2 | chr4 | - | 44717112 | 44717295 | 44710952 | 44711137 | 44726473 | 44726523 | 32 | 37 | 51 | [illegible] |
| SE | 26396 | ENSG00000138744 | NAAA | chr4 | - | 75914867 | 75914985 | 75913659 | 75914338 | 75919908 | 75919975 | 66 | 49 | 66 | [illegible] |
| SE | 87448 | ENSG00000123131 | PRDX4 | chrX | + | 23671393 | 23671433 | 23667498 | 23667811 | 23671528 | 23671646 | 0 | 0 | 0 | [illegible] |
| SE | 98843 | ENSG00000135365 | PHF21A | chr11 | - | 46117921 | 46118145 | 46092182 | 46092223 | 46118741 | 46118800 | 23 | 17 | 15 | [illegible] |
| SE | 86139 | ENSG00000197713 | RPE | chr2 | + | 210016203 | 210016257 | 210015972 | 210016112 | 210016506 | 210016597 | 22 | 11 | 19 | [illegible] |
| SE | 74995 | ENSG00000157450 | RNF111 | chr15 | + | 59004001 | 59004174 | 58987676 | 58988068 | 59030803 | 59031138 | 10 | 15 | 22 | [illegible] |
| SE | 47839 | ENSG00000267009 | NA | chr17 | + | 68414026 | 68414182 | 68413729 | 68413795 | 68420188 | 68420506 | 5 | 0 | 8 | [illegible] |
| SE | 15061 | ENSG00000166783 | MARF1 | chr16 | - | 15625371 | 15625800 | 15625015 | 15625173 | 15630331 | 15630504 | 284 | 188 | 238 | [illegible] |
| SE | 14838 | ENSG00000140479 | PCSK6 | chr15 | - | 101331650 | 101331689 | 101326376 | 101326479 | 101331851 | 101332031 | 0 | 7 | 0 | [illegible] |
| SE | 36297 | ENSG00000075420 | FNDC3B | chr3 | + | 172343010 | 172343116 | 172341112 | 172341231 | 172344085 | 172344258 | 107 | 56 | 48 | [illegible] |
| SE | 96313 | ENSG00000110422 | HIPK3 | chr11 | + | 33348165 | 33348228 | 33347851 | 33348013 | 33349146 | 33349287 | 73 | 60 | 34 | [illegible] |
| SE | 25023 | ENSG00000171467 | ZNF318 | chr6 | - | 43352376 | 43352476 | 43348323 | 43348625 | 43354663 | 43356145 | 204 | 207 | 186 | [illegible] |
| SE | 12895 | ENSG00000178952 | TUFM | chr16 | - | 28845313 | 28845480 | 28844950 | 28845055 | 28845911 | 28846106 | 301 | 177 | 333 | [illegible] |
| SE | 103942 | ENSG00000150347 | ARID5B | chr10 | + | 62000090 | 62000321 | 61940182 | 61940408 | 62050887 | 62051000 | 195 | 98 | 132 | [illegible] |
| SE | 72336 | ENSG00000123739 | PLA2G12A | chr4 | - | 109718682 | 109718759 | 109717547 | 109717713 | 109729601 | 109730070 | 35 | 38 | 37 | [illegible] |
| SE | 92465 | ENSG00000100815 | TRIP11 | chr14 | - | 91995351 | 91995515 | 91993808 | 91993912 | 91999239 | 91999379 | 148 | 135 | 119 | [illegible] |
| SE | 54501 | ENSG00000151414 | NEK7 | chr1 | + | 198297126 | 198297240 | 198292944 | 198293039 | 198319411 | 198319522 | 373 | 296 | 348 | [illegible] |
| SE | 95589 | ENSG00000130714 | POMT1 | chr9 | + | 131504453 | 131504649 | 131504188 | 131504340 | 131506113 | 131506220 | 0 | 2 | 0 | [illegible] |
| SE | 67215 | ENSG00000144118 | RALB | chr2 | + | 120280834 | 120280907 | 120278617 | 120278778 | 120285873 | 120286082 | 12 | 15 | 9 | [illegible] |
| SE | 102057 | ENSG00000138134 | STAMBPL1 | chr10 | + | 88916679 | 88916817 | 88914533 | 88914658 | 88921282 | 88921395 | 52 | 72 | 85 | [illegible] |
| SE | 40263 | ENSG00000170581 | STAT2 | chr12 | - | 56349162 | 56349261 | 56348751 | 56348851 | 56349425 | 56349473 | 76 | 32 | 36 | [illegible] |
| SE | 22903 | ENSG00000140853 | NLRC5 | chr16 | + | 57046551 | 57046641 | 57045296 | 57045492 | 57047544 | 57047628 | 155 | 53 | 106 | [illegible] |
| SE | 31789 | ENSG00000204463 | BAG6 | chr6 | - | 31644306 | 31644414 | 31644081 | 31644194 | 31644945 | 31645198 | 9 | 4 | 13 | [illegible] |
| SE | 55034 | ENSG00000152223 | EPG5 | chr18 | - | 45917678 | 45917819 | 45916437 | 45916582 | 45923267 | 45923387 | 32 | 19 | 40 | [illegible] |
| SE | 55090 | ENSG00000086200 | IPO11 | chr5 | + | 62601763 | 62601848 | 62591576 | 62591672 | 62627153 | 62627569 | 68 | 62 | 62 | [illegible] |
| SE | 14217 | ENSG00000156858 | PRR14 | chr16 | + | 30654628 | 30655214 | 30654108 | 30654339 | 30655350 | 30655420 | 53 | 39 | 46 | [illegible] |
| SE | 95516 | ENSG00000137145 | DENND4C | chr9 | + | 19336685 | 19336832 | 19336269 | 19336414 | 19340991 | 19341114 | 362 | 292 | 254 | [illegible] |
| SE | 42130 | ENSG00000198952 | SMG5 | chr1 | - | 156266540 | 156266678 | 156265780 | 156266380 | 156267469 | 156267678 | 174 | 92 | 197 | [illegible] |
| SE | 89326 | ENSG00000120963 | ZNF706 | chr8 | - | 101202332 | 101202452 | 101201715 | 101201743 | 101205434 | 101205732 | 46 | 32 | 55 | [illegible] |
| SE | 51499 | ENSG00000108021 | TASOR2 | chr10 | + | 5712393 | 5712539 | 5684941 | 5685175 | 5712822 | 5712918 | 101 | 41 | 50 | [illegible] |
| SE | 46738 | ENSG00000163125 | RPRD2 | chr1 | + | 150464526 | 150464727 | 150460059 | 150460317 | 150470560 | 150473556 | 344 | 225 | 322 | [illegible] |
| SE | 41733 | ENSG00000163319 | MRPS18C | chr4 | + | 83458345 | 83458429 | 83456908 | 83456958 | 83460972 | 83461032 | 100 | 53 | 64 | [illegible] |
| SE | 44242 | ENSG00000102606 | ARHGEF7 | chr13 | + | 111273813 | 111273952 | 111267547 | 111267670 | 111274730 | 111274790 | 124 | 113 | 163 | [illegible] |
| SE | 6152 | ENSG00000282826 | FRG1CP | chr20 | - | 28601602 | 28601756 | 28600391 | 28600461 | 28602608 | 28602773 | 14 | 14 | 7 | [illegible] |
| SE | 88525 | ENSG00000138378 | STAT4 | chr2 | - | 191054489 | 191054534 | 191041064 | 191041148 | 191058017 | 191058111 | 452 | 295 | 390 | [illegible] |
| SE | 13379 | ENSG00000074755 | ZZEF1 | chr17 | - | 4032827 | 4033002 | 4032125 | 4032258 | 4034014 | 4034292 | 211 | 117 | 178 | [illegible] |
| SE | 68947 | ENSG00000135926 | TMBIM1 | chr2 | - | 218285762 | 218285938 | 218281939 | 218282181 | 218292465 | 218292529 | 16 | 6 | 4 | [illegible] |
| SE | 44522 | ENSG00000119402 | FBXW2 | chr9 | - | 120776930 | 120777021 | 120776092 | 120776226 | 120778350 | 120778545 | 20 | 6 | 4 | [illegible] |
| SE | 6677 | ENSG00000099956 | SMARCB1 | chr22 | + | 23791755 | 23791867 | 23786930 | 23787262 | 23793558 | 23793688 | 153 | 113 | 140 | [illegible] |
| SE | 25269 | ENSG00000119685 | TTLL5 | chr14 | + | 75709178 | 75709239 | 75707622 | 75707707 | 75717860 | 75717962 | 2 | 4 | 2 | [illegible] |
| SE | 74809 | ENSG00000042088 | TDP1 | chr14 | + | 89956577 | 89956800 | 89956036 | 89956278 | 89963107 | 89963552 | 24 | 24 | 38 | [illegible] |
| SE | 22966 | ENSG00000198612 | COPS8 | chr2 | + | 237086731 | 237086796 | 237085881 | 237086042 | 237087126 | 237087197 | 0 | 3 | 2 | [illegible] |
| SE | 8247 | ENSG00000275700 | AATF | chr17 | + | 36953769 | 36953907 | 36951830 | 36953296 | 36986616 | 36986731 | 533 | 376 | 373 | [illegible] |
| SE | 39427 | ENSG00000137393 | RNF144B | chr6 | + | 18457154 | 18457359 | 18439683 | 18439744 | 18463290 | 18463380 | 107 | 117 | 58 | [illegible] |
| SE | 69363 | ENSG00000171720 | HDAC3 | chr5 | - | 141626193 | 141626283 | 141626012 | 141626071 | 141627892 | 141627936 | 204 | 100 | 140 | [illegible] |
| SE | 29676 | ENSG00000137656 | BUD13 | chr11 | - | 116757765 | 116757950 | 116757145 | 116757227 | 116760734 | 116760952 | 66 | 36 | 38 | [illegible] |
| SE | 104586 | ENSG00000069493 | CLEC2D | chr12 | + | 9680922 | 9681033 | 9669712 | 9669795 | 9687901 | 9688086 | 517 | 284 | 445 | [illegible] |
| SE | 59369 | ENSG00000116127 | ALMS1 | chr2 | + | 73426452 | 73426553 | 73424429 | 73424902 | 73447959 | 73454067 | 30 | 32 | 23 | [illegible] |
| SE | 52963 | ENSG00000150455 | TIRAP | chr11 | + | 126289664 | 126289817 | 126283117 | 126283153 | 126290461 | 126290585 | 1 | 1 | 7 | [illegible] |
| SE | 63665 | ENSG00000144451 | SPAG16 | chr2 | + | 213310058 | 213310177 | 213297261 | 213297357 | 213317218 | 213317356 | 67 | 50 | 114 | [illegible] |
| SE | 40264 | ENSG00000170581 | STAT2 | chr12 | - | 56349162 | 56349261 | 56348923 | 56349059 | 56349344 | 56349509 | 31 | 15 | 17 | [illegible] |
| SE | 32909 | ENSG00000075711 | DLG1 | chr3 | - | 197076585 | 197076685 | 197075836 | 197075870 | 197081050 | 197081117 | 50 | 31 | 39 | [illegible] |
| SE | 89722 | ENSG00000120899 | PTK2B | chr8 | + | 27312303 | 27312387 | 27311481 | 27311709 | 27313204 | 27313287 | 4 | 11 | 16 | [illegible] |
| SE | 68807 | ENSG00000279765 | NA | chr15 | + | 92901166 | 92901299 | 92892003 | 92892072 | 92924320 | 92924552 | 136 | 103 | 152 | [illegible] |
| SE | 40456 | ENSG00000184432 | COPB2 | chr3 | - | 139373665 | 139373808 | 139373212 | 139373412 | 139374488 | 139374588 | 658 | 531 | 552 | [illegible] |
| SE | 21321 | ENSG00000137055 | PLAA | chr9 | - | 26920226 | 26920384 | 26919309 | 26919529 | 26923177 | 26923233 | 148 | 103 | 129 | [illegible] |
| SE | 55476 | ENSG00000143258 | USP21 | chr1 | + | 161160328 | 161160506 | 161159520 | 161159678 | 161160619 | 161160992 | 9 | 13 | 6 | [illegible] |
| SE | 104707 | ENSG00000110888 | CAPRIN2 | chr12 | - | 30731342 | 30731510 | 30730238 | 30730282 | 30734967 | 30735206 | 155 | 117 | 137 | [illegible] |
| SE | 32667 | ENSG00000150995 | ITPR1 | chr3 | + | 4727122 | 4727173 | 4711756 | 4711868 | 4733087 | 4733220 | 9 | 5 | 3 | [illegible] |
| SE | 106006 | ENSG00000007392 | LUC7L | chr16 | - | 228332 | 228402 | 227241 | 227336 | 229278 | 229374 | 16 | 31 | 26 | [illegible] |
| SE | 44891 | ENSG00000134744 | TUT4 | chr1 | - | 52488908 | 52489035 | 52477707 | 52477882 | 52490731 | 52490801 | 80 | 75 | 81 | [illegible] |
| SE | 85242 | ENSG00000147905 | ZCCHC7 | chr9 | + | 37147361 | 37147445 | 37126311 | 37126942 | 37302187 | 37302231 | 12 | 8 | 13 | [illegible] |
| SE | 18637 | ENSG00000256500 | NA | chr14 | + | 103662701 | 103662927 | 103657545 | 103657776 | 103669510 | 103669598 | 105 | 80 | 109 | [illegible] |
| SE | 49597 | ENSG00000076685 | NT5C2 | chr10 | - | 103111692 | 103111805 | 103106588 | 103106706 | 103139405 | 103139479 | 5 | 5 | 10 | [illegible] |
| SE | 36598 | ENSG00000169180 | XPO6 | chr16 | - | 28126617 | 28126760 | 28125831 | 28125848 | 28132333 | 28132403 | 4 | 0 | 3 | [illegible] |
| SE | 80836 | ENSG00000151948 | GLT1D1 | chr12 | + | 128927104 | 128927154 | 128914932 | 128914989 | 128945325 | 128945369 | 16 | 16 | 13 | [illegible] |
| SE | 71895 | ENSG00000074054 | CLASP1 | chr2 | - | 121515664 | 121515762 | 121503166 | 121503234 | 121525844 | 121525920 | 208 | 133 | 160 | [illegible] |
| SE | 25730 | ENSG00000196302 | GUSBP15 | chr5 | - | 70522000 | 70522188 | 70518320 | 70518405 | 70555217 | 70555266 | 10 | 8 | 4 | [illegible] |
| SE | 71526 | ENSG00000144504 | ANKMY1 | chr2 | - | 240525684 | 240525849 | 240521937 | 240524381 | 240552913 | 240553057 | 58 | 26 | 56 | [illegible] |
| SE | 3494 | ENSG00000130755 | GMFG | chr19 | - | 39329547 | 39329626 | 39328999 | 39329073 | 39333076 | 39333126 | 395 | 304 | 434 | [illegible] |
| SE | 15614 | ENSG00000067369 | TP53BP1 | chr15 | - | 43432193 | 43432677 | 43428015 | 43428168 | 43438323 | 43438416 | 62 | 80 | 71 | [illegible] |
| SE | 57222 | ENSG00000223519 | KIF28P | chr1 | - | 246776062 | 246776253 | 246771836 | 246771991 | 246776539 | 246776684 | 3 | 9 | 9 | [illegible] |
| SE | 79685 | ENSG00000134905 | CARS2 | chr13 | - | 110651033 | 110651100 | 110646850 | 110647239 | 110663450 | 110663518 | 171 | 105 | 129 | [illegible] |
| SE | 55246 | ENSG00000158161 | EYA3 | chr1 | - | 28042570 | 28042650 | 28038838 | 28038905 | 28048382 | 28048426 | 90 | 78 | 83 | [illegible] |
| SE | 69881 | ENSG00000109171 | SLAIN2 | chr4 | + | 48394575 | 48394653 | 48383646 | 48383784 | 48420124 | 48420443 | 129 | 84 | 116 | [illegible] |
| SE | 50454 | ENSG00000171574 | ZNF584 | chr19 | + | 58415818 | 58415940 | 58415523 | 58415646 | 58416810 | 58417091 | 0 | 0 | 0 | [illegible] |
| SE | 23219 | ENSG00000038210 | PI4K2B | chr4 | + | 25252320 | 25252475 | 25234032 | 25234431 | 25255064 | 25255265 | 87 | 77 | 52 | [illegible] |
| SE | 17491 | ENSG00000100580 | TMED8 | chr14 | - | 77346348 | 77346478 | 77343696 | 77343823 | 77351672 | 77351751 | 29 | 33 | 40 | [illegible] |
| SE | 68316 | ENSG00000168016 | TRANK1 | chr3 | - | 36850153 | 36851417 | 36847199 | 36847346 | 36851718 | 36851856 | 7 | 7 | 11 | [illegible] |
| SE | 12346 | ENSG00000197912 | SPG7 | chr16 | + | 89548979 | 89549336 | 89548002 | 89548113 | 89550493 | 89550609 | 8 | 8 | 12 | [illegible] |
| SE | 65613 | ENSG00000112078 | KCTD20 | chr6 | + | 36479120 | 36479223 | 36470051 | 36470257 | 36481561 | 36481759 | 14 | 16 | 6 | [illegible] |
| SE | 105943 | ENSG00000127314 | RAP1B | chr12 | + | 68641037 | 68641153 | 68610912 | 68611043 | 68648698 | 68648781 | 15 | 22 | 6 | [illegible] |
| SE | 63839 | ENSG00000075292 | ZNF638 | chr2 | + | 71428530 | 71428651 | 71427349 | 71427414 | 71431326 | 71431428 | 112 | 76 | 77 | [illegible] |
| SE | 28409 | ENSG00000204842 | ATXN2 | chr12 | - | 111552905 | 111552977 | 111552279 | 111552430 | 111554157 | 111554217 | 158 | 132 | 89 | [illegible] |
| SE | 24084 | ENSG00000146282 | RARS2 | chr6 | - | 87524556 | 87524652 | 87521463 | 87521524 | 87529541 | 87529648 | 163 | 145 | 170 | [illegible] |
| SE | 4874 | ENSG00000196562 | SULF2 | chr20 | - | 47702518 | 47702670 | 47689547 | 47690295 | 47736702 | 47736942 | 501 | 286 | 320 | [illegible] |
| SE | 1295 | ENSG00000100239 | PPP6R2 | chr22 | + | 50418866 | 50418979 | 50416091 | 50416157 | 50419348 | 50419462 | 163 | 130 | 158 | [illegible] |
| SE | 20502 | ENSG00000143379 | SETDB1 | chr1 | + | 150962129 | 150962158 | 150960562 | 150961191 | 150962973 | 150963139 | 3 | 2 | 2 | [illegible] |
| SE | 101270 | ENSG00000134460 | IL2RA | chr10 | - | 6019869 | 6019941 | 6019427 | 6019499 | 6024243 | 6024354 | 15 | 12 | 14 | [illegible] |
| SE | 65663 | ENSG00000111729 | CLEC4A | chr12 | + | 8129263 | 8129362 | 8125560 | 8125677 | 8135584 | 8135736 | 415 | 287 | 244 | [illegible] |
| SE | 24362 | ENSG00000132780 | NASP | chr1 | + | 45591222 | 45591270 | 45583987 | 45584205 | 45602254 | 45602278 | 82 | 70 | 77 | [illegible] |
| SE | 13044 | ENSG00000263072 | ZNF213-AS1 | chr16 | - | 3132849 | 3132942 | 3131746 | 3132103 | 3134629 | 3134882 | 6 | 4 | 13 | [illegible] |
| SE | 8150 | ENSG00000171861 | MRM3 | chr17 | + | 783082 | 783327 | 782352 | 782692 | 787964 | 788132 | 38 | 35 | 62 | [illegible] |
| SE | 24547 | ENSG00000172785 | CBWD1 | chr9 | - | 152033 | 152078 | 134978 | 135030 | 154708 | 154795 | 19 | 22 | 9 | [illegible] |
| SE | 7556 | ENSG00000167851 | CD300A | chr17 | + | 74481725 | 74481833 | 74481288 | 74481326 | 74484000 | 74484167 | 322 | 123 | 161 | [illegible] |
| SE | 77669 | ENSG00000214021 | TTLL3 | chr3 | + | 9833103 | 9833273 | 9828959 | 9829395 | 9834680 | 9835096 | 24 | 2 | 11 | [illegible] |
| SE | 105786 | ENSG00000124164 | VAPB | chr20 | + | 58434601 | 58434705 | 58418210 | 58418363 | 58438944 | 58439025 | 132 | 146 | 117 | [illegible] |
| SE | 88787 | ENSG00000140632 | GLYR1 | chr16 | - | 4821546 | 4821597 | 4821203 | 4821453 | 4822874 | 4822931 | 255 | 178 | 234 | [illegible] |
| SE | 27004 | ENSG00000137802 | MAPKBP1 | chr15 | + | 41813620 | 41813781 | 41812515 | 41813101 | 41814549 | 41814739 | 64 | 24 | 39 | [illegible] |
| SE | 83702 | ENSG00000177324 | BEND2 | chrX | - | 18195295 | 18195442 | 18191000 | 18191108 | 18203500 | 18203915 | 83 | 100 | 12 | [illegible] |
| SE | 12098 | ENSG00000125149 | C16orf70 | chr16 | + | 67131285 | 67131329 | 67125958 | 67126042 | 67132445 | 67132525 | 81 | 35 | 36 | [illegible] |
| SE | 95477 | ENSG00000095319 | NUP188 | chr9 | + | 128994372 | 128994442 | 128993524 | 128993694 | 128994855 | 128994890 | 100 | 49 | 84 | [illegible] |
| SE | 25715 | ENSG00000247774 | PCED1B-AS1 | chr12 | - | 47210639 | 47210757 | 47205901 | 47206145 | 47216164 | 47216249 | 15 | 8 | 18 | [illegible] |
| SE | 98098 | ENSG00000171466 | ZNF562 | chr19 | - | 9656546 | 9656653 | 9641806 | 9653881 | 9658008 | 9658135 | 155 | 132 | 112 | [illegible] |
| SE | 64041 | ENSG00000115464 | USP34 | chr2 | - | 61284874 | 61284957 | 61283408 | 61283449 | 61288676 | 61288877 | 98 | 50 | 102 | [illegible] |
| SE | 37662 | ENSG00000168615 | ADAM9 | chr8 | + | 39025802 | 39025884 | 39023155 | 39023325 | 39026676 | 39026810 | 66 | 65 | 44 | [illegible] |
| SE | 18333 | ENSG00000032219 | ARID4A | chr14 | + | 58347019 | 58347117 | 58346410 | 58346505 | 58347646 | 58347878 | 185 | 171 | 158 | [illegible] |
| SE | 86702 | ENSG00000147140 | NONO | chrX | + | 71297839 | 71297938 | 71297376 | 71297461 | 71298468 | 71298508 | 151 | 83 | 139 | [illegible] |
| SE | 9694 | ENSG00000154265 | ABCA5 | chr17 | - | 69251746 | 69251866 | 69250471 | 69250621 | 69253572 | 69253667 | 46 | 50 | 47 | [illegible] |
| SE | 28494 | ENSG00000155229 | MMS19 | chr10 | - | 97476834 | 97476963 | 97476682 | 97476744 | 97477346 | 97477404 | 54 | 34 | 39 | [illegible] |
| SE | 39735 | ENSG00000132581 | SDF2 | chr17 | - | 28651485 | 28651577 | 28648742 | 28649137 | 28655286 | 28655483 | 10 | 9 | 7 | [illegible] |
| SE | 76687 | ENSG00000223501 | VPS52 | chr6 | - | 33269998 | 33270051 | 33269743 | 33269819 | 33270198 | 33270283 | 54 | 30 | 46 | [illegible] |
| SE | 49954 | ENSG00000171793 | CTPS1 | chr1 | + | 40995916 | 40996068 | 40991764 | 40991845 | 40997393 | 40997526 | 94 | 71 | 103 | [illegible] |
| SE | 33492 | ENSG00000173540 | GMPPB | chr3 | - | 49722863 | 49723114 | 49722595 | 49722754 | 49723253 | 49723302 | 14 | 7 | 15 | [illegible] |
| SE | 55895 | ENSG00000132510 | KDM6B | chr17 | + | 7839905 | 7840024 | 7834216 | 7834350 | 7844900 | 7845020 | 19 | 6 | 6 | [illegible] |
| SE | 94149 | ENSG00000090054 | SPTLC1 | chr9 | - | 92076797 | 92077035 | 92067965 | 92068098 | 92080015 | 92080076 | 0 | 1 | 11 | [illegible] |
| SE | 73346 | ENSG00000115970 | THADA | chr2 | - | 43592843 | 43592890 | 43592316 | 43592416 | 43595930 | 43596007 | 2 | 2 | 3 | [illegible] |
| SE | 14464 | ENSG00000213614 | HEXA | chr15 | - | 72346526 | 72346710 | 72343455 | 72346325 | 72348047 | 72348134 | 220 | 129 | 151 | [illegible] |
| SE | 23204 | ENSG00000082213 | C5orf22 | chr5 | + | 31548489 | 31548687 | 31545670 | 31545712 | 31551292 | 31551432 | 7 | 2 | 7 | [illegible] |
| SE | 88616 | ENSG00000169139 | UBE2V2 | chr8 | + | 48050259 | 48050508 | 48049852 | 48049978 | 48060681 | 48060863 | 2 | 0 | 2 | [illegible] |
| SE | 18878 | ENSG00000075399 | VPS9D1 | chr16 | - | 89709776 | 89709906 | 89709226 | 89709435 | 89710585 | 89711010 | 20 | 6 | 20 | [illegible] |
| SE | 64268 | ENSG00000145919 | BOD1 | chr5 | - | 173609237 | 173609434 | 173607144 | 173608292 | 173613130 | 173613255 | 68 | 35 | 88 | [illegible] |
| SE | 17354 | ENSG00000133961 | NUMB | chr14 | - | 73316389 | 73316422 | 73297210 | 73297285 | 73323129 | 73323204 | 14 | 5 | 2 | [illegible] |
| SE | 94777 | ENSG00000119314 | PTBP3 | chr9 | - | 112250928 | 112251103 | 112234819 | 112234897 | 112252677 | 112252788 | 751 | 476 | 571 | [illegible] |
| SE | 58182 | ENSG00000172965 | MIR4435-2HG | chr2 | - | 111233627 | 111233739 | 111207754 | 111208661 | 111248147 | 111248241 | 1 | 0 | 1 | [illegible] |
| SE | 25473 | ENSG00000082397 | EPB41L3 | chr18 | - | 5394676 | 5394793 | 5392380 | 5393478 | 5395066 | 5395147 | 125 | 164 | 54 | [illegible] |
| SE | 90911 | ENSG00000185122 | HSF1 | chr8 | + | 144292482 | 144292619 | 144291604 | 144291874 | 144308905 | 144309014 | 1 | 1 | 2 | [illegible] |
| SE | 65950 | ENSG00000123240 | OPTN | chr10 | + | 13108961 | 13109015 | 13100172 | 13100302 | 13109111 | 13109288 | 20 | 10 | 18 | [illegible] |
| SE | 327 | ENSG00000182670 | TTC3 | chr21 | + | 37165549 | 37166615 | 37164050 | 37164215 | 37167554 | 37167620 | 957 | 885 | 887 | [illegible] |
| SE | 102836 | ENSG00000177239 | MAN1B1 | chr9 | + | 137088267 | 137088398 | 137088074 | 137088183 | 137088868 | 137089005 | 11 | 9 | 6 | [illegible] |
| SE | 1690 | ENSG00000228315 | GUSBP11 | chr22 | - | 23652478 | 23652576 | 23638486 | 23638942 | 23659864 | 23660000 | 1 | 1 | 3 | [illegible] |
| SE | 81290 | ENSG00000155096 | AZIN1 | chr8 | - | 102849986 | 102850144 | 102843550 | 102843747 | 102858012 | 102858150 | 108 | 92 | 92 | [illegible] |
| SE | 64433 | ENSG00000162813 | BPNT1 | chr1 | - | 220079726 | 220079854 | 220073966 | 220074071 | 220089643 | 220089769 | 41 | 28 | 48 | [illegible] |
| SE | 20414 | ENSG00000125871 | MGME1 | chr20 | + | 17975683 | 17975903 | 17969800 | 17970370 | 17989938 | 17991122 | 20 | 7 | 18 | [illegible] |
| SE | 11219 | ENSG00000140829 | DHX38 | chr16 | + | 72098644 | 72098792 | 72097159 | 72097781 | 72098926 | 72099045 | 141 | 77 | 131 | [illegible] |
| SE | 39819 | ENSG00000144711 | IQSEC1 | chr3 | - | 12920429 | 12920596 | 12915593 | 12915733 | 12922119 | 12922242 | 415 | 206 | 319 | [illegible] |
| SE | 46197 | ENSG00000185158 | LRRC37B | chr17 | + | 32045699 | 32045818 | 32035564 | 32035639 | 32047760 | 32047901 | 59 | 71 | 76 | [illegible] |
| SE | 4195 | ENSG00000105186 | ANKRD27 | chr19 | - | 32658913 | 32659045 | 32649681 | 32649792 | 32675070 | 32675147 | 73 | 29 | 75 | [illegible] |
| SE | 304 | ENSG00000157557 | ETS2 | chr21 | + | 38806450 | 38806808 | 38805950 | 38806120 | 38810034 | 38810106 | 2 | 1 | 0 | [illegible] |
| SE | 54917 | ENSG00000010803 | SCMH1 | chr1 | - | 41152598 | 41152741 | 41151613 | 41151684 | 41160874 | 41160898 | 12 | 11 | 5 | [illegible] |
| SE | 40765 | ENSG00000114026 | OGG1 | chr3 | + | 9756470 | 9756621 | 9751769 | 9751949 | 9756766 | 9756816 | 35 | 13 | 15 | [illegible] |
| SE | 5940 | ENSG00000170471 | RALGAPB | chr20 | + | 38497352 | 38497516 | 38492929 | 38493132 | 38499446 | 38499633 | 247 | 177 | 195 | [illegible] |
| SE | 69401 | ENSG00000135636 | DYSF | chr2 | + | 71674196 | 71674296 | 71665161 | 71665304 | 71679056 | 71679235 | 39 | 45 | 30 | [illegible] |
| SE | 51917 | ENSG00000074603 | DPP8 | chr15 | - | 65454262 | 65454415 | 65451959 | 65452102 | 65463760 | 65463906 | 86 | 56 | 65 | [illegible] |
| SE | 42096 | ENSG00000026751 | SLAMF7 | chr1 | + | 160751344 | 160751448 | 160749820 | 160750093 | 160752185 | 160752248 | 155 | 87 | 120 | [illegible] |
| SE | 68943 | ENSG00000135926 | TMBIM1 | chr2 | - | 218285762 | 218285934 | 218281939 | 218282181 | 218292465 | 218292535 | 12 | 3 | 2 | [illegible] |
| SE | 5192 | ENSG00000124214 | STAU1 | chr20 | - | 49158436 | 49158497 | 49153932 | 49154071 | 49158938 | 49159146 | 2 | 2 | 5 | [illegible] |
| SE | 52965 | ENSG00000150455 | TIRAP | chr11 | + | 126289664 | 126289833 | 126283072 | 126283153 | 126290461 | 126290585 | 2 | 3 | 6 | [illegible] |
| SE | 32049 | ENSG00000081307 | UBA5 | chr3 | + | 132675247 | 132675383 | 132671776 | 132672177 | 132675604 | 132675680 | 159 | 121 | 167 | [illegible] |
| SE | 26405 | ENSG00000165494 | PCF11 | chr11 | + | 83161326 | 83161452 | 83156987 | 83157631 | 83163678 | 83163867 | 483 | 392 | 398 | [illegible] |
| SE | 38634 | ENSG00000115325 | DOK1 | chr2 | + | 74555893 | 74556078 | 74555205 | 74555453 | 74556307 | 74556621 | 60 | 32 | 23 | [illegible] |
| SE | 45297 | ENSG00000286192 | NA | chr7 | - | 2014501 | 2014642 | 1980452 | 1980541 | 2069193 | 2069338 | 25 | 9 | 9 | [illegible] |
| SE | 56458 | ENSG00000127337 | YEATS4 | chr12 | + | 69364176 | 69364299 | 69362787 | 69362907 | 69365632 | 69365699 | 5 | 3 | 1 | [illegible] |
| SE | 93839 | ENSG00000116580 | GON4L | chr1 | - | 155751766 | 155751869 | 155749658 | 155750733 | 155751959 | 155752587 | 53 | 53 | 65 | [illegible] |
| SE | 109363 | ENSG00000186184 | POLR1D | chr13 | + | 27650041 | 27650241 | 27648378 | 27648453 | 27665685 | 27667410 | 24 | 15 | 23 | [illegible] |
| SE | 77931 | ENSG00000095015 | MAP3K1 | chr5 | + | 56887377 | 56887520 | 56885931 | 56886063 | 56888225 | 56888357 | 679 | 467 | 514 | [illegible] |
| SE | 83975 | ENSG00000068400 | GRIPAP1 | chrX | - | 49000565 | 49000689 | 48999437 | 48999504 | 49000860 | 49000959 | 0 | 1 | 4 | [illegible] |
| SE | 11651 | ENSG00000153066 | TXNDC11 | chr16 | - | 11733981 | 11734079 | 11730644 | 11730774 | 11736016 | 11736233 | 111 | 67 | 153 | [illegible] |
| SE | 79242 | ENSG00000196511 | TPK1 | chr7 | - | 144681806 | 144681864 | 144648806 | 144648879 | 144682908 | 144682978 | 17 | 10 | 8 | [illegible] |
| SE | 35891 | ENSG00000197415 | VEPH1 | chr3 | - | 157493186 | 157493293 | 157470418 | 157470529 | 157495211 | 157495506 | 2 | 8 | 5 | [illegible] |
| SE | 57650 | ENSG00000144535 | DIS3L2 | chr2 | + | 232111196 | 232111255 | 232087486 | 232087721 | 232130618 | 232130719 | 3 | 3 | 4 | [illegible] |
| SE | 48477 | ENSG00000117461 | PIK3R3 | chr1 | - | 46066910 | 46067091 | 46066053 | 46066179 | 46077514 | 46077613 | 32 | 27 | 8 | [illegible] |
| SE | 74992 | ENSG00000189077 | TMEM120A | chr7 | - | 75987537 | 75987602 | 75986982 | 75987428 | 75987717 | 75987810 | 18 | 3 | 11 | [illegible] |
| SE | 108967 | ENSG00000111877 | MCM9 | chr6 | - | 118856370 | 118856545 | 118829047 | 118829250 | 118894026 | 118894356 | 30 | 20 | 27 | [illegible] |
| SE | 22252 | ENSG00000141279 | NPEPPS | chr17 | + | 47619277 | 47619372 | 47619008 | 47619164 | 47619736 | 47619784 | 2 | 4 | 5 | [illegible] |
| SE | 80866 | ENSG00000122557 | HERPUD2 | chr7 | - | 35670214 | 35670328 | 35667433 | 35667588 | 35694183 | 35694566 | 65 | 61 | 136 | [illegible] |
| SE | 96390 | ENSG00000197563 | PIGN | chr18 | - | 62123506 | 62123544 | 62114560 | 62114639 | 62138242 | 62138298 | 2 | 2 | 1 | [illegible] |
| SE | 27092 | ENSG00000156265 | MAP3K7CL | chr21 | + | 29109039 | 29109183 | 29092253 | 29092581 | 29130794 | 29130923 | 2 | 1 | 6 | [illegible] |
| SE | 17885 | ENSG00000129467 | ADCY4 | chr14 | - | 24319741 | 24319904 | 24319328 | 24319436 | 24322065 | 24322224 | 2 | 1 | 3 | [illegible] |
| SE | 46484 | ENSG00000065978 | YBX1 | chr1 | + | 42696198 | 42696288 | 42693489 | 42693523 | 42696641 | 42696944 | 1817 | 1390 | 1543 | [illegible] |
| SE | 100557 | ENSG00000054654 | SYNE2 | chr14 | + | 64215285 | 64215354 | 64214143 | 64214470 | 64216247 | 64216387 | 21 | 3 | 5 | [illegible] |
| SE | 25414 | ENSG00000020633 | RUNX3 | chr1 | - | 24907258 | 24907417 | 24899510 | 24902666 | 24919239 | 24919344 | 450 | 97 | 217 | [illegible] |
| SE | 33634 | ENSG00000114405 | C3orf14 | chr3 | + | 62319720 | 62319832 | 62319014 | 62319055 | 62320459 | 62320524 | 12 | 3 | 6 | [illegible] |
| SE | 94609 | ENSG00000106829 | TLE4 | chr9 | + | 79707122 | 79707218 | 79705888 | 79705942 | 79708117 | 79708250 | 5 | 2 | 0 | [illegible] |
| SE | 86862 | ENSG00000090316 | MAEA | chr4 | + | 1322380 | 1322503 | 1315396 | 1315600 | 1327626 | 1327703 | 159 | 49 | 98 | [illegible] |
| SE | 74694 | ENSG00000197548 | ATG7 | chr3 | + | 11332971 | 11333093 | 11315343 | 11315493 | 11340644 | 11340735 | 61 | 28 | 38 | [illegible] |
| SE | 80291 | ENSG00000168137 | SETD5 | chr3 | + | 9436067 | 9436140 | 9435727 | 9435906 | 9436849 | 9436888 | 2 | 6 | 4 | [illegible] |
| SE | 4280 | ENSG00000089639 | GMIP | chr19 | - | 19642536 | 19642619 | 19641975 | 19642051 | 19643510 | 19643601 | 22 | 7 | 19 | [illegible] |
| SE | 12563 | ENSG00000214078 | CPNE1 | chr20 | - | 35655201 | 35655344 | 35632794 | 35632923 | 35664759 | 35664886 | 8 | 4 | 13 | [illegible] |
| SE | 24551 | ENSG00000172785 | CBWD1 | chr9 | - | 160000 | 160140 | 156480 | 156527 | 162431 | 162469 | 0 | 1 | 0 | [illegible] |
| SE | 86037 | ENSG00000156313 | RPGR | chrX | - | 38287864 | 38288041 | 38276586 | 38276772 | 38290958 | 38291024 | 27 | 17 | 16 | [illegible] |
| SE | 48302 | ENSG00000229474 | PATL2 | chr15 | - | 44710095 | 44710214 | 44676474 | 44676565 | 44710870 | 44710929 | 30 | 26 | 43 | [illegible] |
| SE | 96371 | ENSG00000184060 | ADAP2 | chr17 | + | 30922939 | 30923070 | 30921679 | 30922108 | 30931888 | 30931968 | 16 | 5 | 3 | [illegible] |
| SE | 76240 | ENSG00000204351 | SKIV2L | chr6 | + | 31960219 | 31960337 | 31960009 | 31960119 | 31960665 | 31960740 | 50 | 40 | 50 | [illegible] |
| SE | 27949 | ENSG00000128609 | NDUFA5 | chr7 | - | 123546582 | 123546817 | 123545610 | 123545676 | 123550469 | 123550586 | 7 | 12 | 2 | [illegible] |
| SE | 66664 | ENSG00000005483 | KMT2E | chr7 | + | 105038384 | 105038525 | 105038125 | 105038199 | 105040838 | 105041023 | 12 | 4 | 11 | [illegible] |
| SE | 32737 | ENSG00000163349 | HIPK1 | chr1 | + | 113952765 | 113952889 | 113940381 | 113941459 | 113955562 | 113955649 | 256 | 115 | 166 | [illegible] |
| SE | 51759 | ENSG00000114841 | DNAH1 | chr3 | + | 52378601 | 52378780 | 52375954 | 52375993 | 52379904 | 52380135 | 36 | 38 | 38 | [illegible] |
| SE | 105526 | ENSG00000188342 | GTF2F2 | chr13 | + | 45220865 | 45220909 | 45207423 | 45207505 | 45252870 | 45252970 | 5 | 1 | 0 | [illegible] |
| SE | 7535 | ENSG00000169718 | DUS1L | chr17 | - | 82063467 | 82063518 | 82062860 | 82062973 | 82064125 | 82064201 | 90 | 42 | 93 | [illegible] |
| SE | 87006 | ENSG00000258890 | CEP95 | chr17 | + | 64529287 | 64529427 | 64526070 | 64526200 | 64530925 | 64531018 | 43 | 64 | 75 | [illegible] |
| SE | 77780 | ENSG00000152457 | DCLRE1C | chr10 | - | 14936537 | 14936593 | 14935462 | 14935564 | 14939809 | 14939869 | 137 | 93 | 107 | [illegible] |
| SE | 76098 | ENSG00000136044 | APPL2 | chr12 | - | 105197764 | 105197953 | 105195584 | 105195627 | 105203702 | 105203785 | 57 | 30 | 38 | [illegible] |
| SE | 13029 | ENSG00000140987 | ZSCAN32 | chr16 | - | 3393648 | 3393814 | 3390422 | 3390517 | 3397191 | 3397744 | 22 | 8 | 22 | [illegible] |
| SE | 48939 | ENSG00000102805 | CLN5 | chr13 | + | 76995456 | 76995513 | 76995062 | 76995228 | 76995901 | 76996099 | 0 | 2 | 2 | [illegible] |
| SE | 18884 | ENSG00000170956 | CEACAM3 | chr19 | + | 41810831 | 41810897 | 41810322 | 41810354 | 41811171 | 41811553 | 11 | 10 | 9 | [illegible] |
| SE | 81114 | ENSG00000106484 | MEST | chr7 | + | 130500788 | 130500890 | 130500461 | 130500532 | 130502643 | 130502720 | 29 | 14 | 23 | [illegible] |
| SE | 37377 | ENSG00000166927 | MS4A7 | chr11 | + | 60389389 | 60389596 | 60383128 | 60383288 | 60392684 | 60392780 | 222 | 184 | 93 | [illegible] |
| SE | 16705 | ENSG00000166233 | ARIH1 | chr15 | + | 72544819 | 72544964 | 72518066 | 72518134 | 72555270 | 72555363 | 261 | 170 | 272 | [illegible] |
| SE | 93797 | ENSG00000116701 | NCF2 | chr1 | - | 183573184 | 183573292 | 183570779 | 183570839 | 183577598 | 183577707 | 548 | 302 | 373 | [illegible] |
| SE | 31589 | ENSG00000183624 | HMCES | chr3 | + | 129290678 | 129290804 | 129288853 | 129288997 | 129298353 | 129298535 | 287 | 194 | 265 | [illegible] |
| SE | 69898 | ENSG00000142961 | MOB3C | chr1 | - | 46612903 | 46613371 | 46610001 | 46610204 | 46616710 | 46616884 | 70 | 34 | 31 | [illegible] |
| SE | 89295 | ENSG00000166169 | POLL | chr10 | - | 101584601 | 101584919 | 101583507 | 101583681 | 101587245 | 101587406 | 31 | 12 | 17 | [illegible] |
| SE | 31180 | ENSG00000163104 | SMARCAD1 | chr4 | + | 94281471 | 94281590 | 94280591 | 94280780 | 94283120 | 94283303 | 201 | 207 | 168 | [illegible] |
| SE | 23580 | ENSG00000254996 | ANKHD1-EIF4EBP3 | chr5 | + | 140538085 | 140538261 | 140537337 | 140537589 | 140538918 | 140539083 | 272 | 171 | 294 | [illegible] |
| SE | 35914 | ENSG00000187862 | TTC24 | chr1 | + | 156583799 | 156583895 | 156583337 | 156583450 | 156584876 | 156584974 | 7 | 16 | 15 | [illegible] |
| SE | 56592 | ENSG00000183386 | FHL3 | chr1 | - | 37997962 | 37998132 | 37997683 | 37997870 | 37998973 | 37999148 | 55 | 14 | 23 | [illegible] |
| SE | 87629 | ENSG00000140678 | ITGAX | chr16 | + | 31369980 | 31370098 | 31363164 | 31363374 | 31371083 | 31371214 | 3 | 3 | 2 | [illegible] |
| SE | 1082 | ENSG00000100330 | MTMR3 | chr22 | + | 30013355 | 30013541 | 30012367 | 30012563 | 30016527 | 30016698 | 322 | 126 | 165 | [illegible] |
| SE | 82294 | ENSG00000122566 | HNRNPA2B1 | chr7 | - | 26191421 | 26191650 | 26190992 | 26191128 | 26192494 | 26192577 | 16 | 9 | 19 | [illegible] |
| SE | 37837 | ENSG00000114126 | TFDP2 | chr3 | - | 141995019 | 141995141 | 141978519 | 141978682 | 142005440 | 142005469 | 83 | 41 | 59 | [illegible] |
| SE | 25956 | ENSG00000139278 | GLIPR1 | chr12 | + | 75487738 | 75487809 | 75481833 | 75482079 | 75490405 | 75490518 | 58 | 17 | 27 | [illegible] |
| SE | 38087 | ENSG00000160584 | SIK3 | chr11 | - | 116867945 | 116868089 | 116863667 | 116863818 | 116870330 | 116870401 | 119 | 51 | 111 | [illegible] |
| SE | 21385 | ENSG00000154309 | DISP1 | chr1 | + | 222827427 | 222827601 | 222823855 | 222823945 | 222835118 | 222835215 | 23 | 16 | 17 | [illegible] |
| SE | 33105 | ENSG00000178035 | IMPDH2 | chr3 | - | 49028247 | 49028322 | 49026686 | 49027916 | 49028430 | 49028532 | 115 | 78 | 135 | [illegible] |
| SE | 40193 | ENSG00000231607 | DLEU2 | chr13 | - | 50044650 | 50044768 | 50027089 | 50027406 | 50049583 | 50049662 | 35 | 20 | 17 | [illegible] |
| SE | 52198 | ENSG00000162511 | LAPTM5 | chr1 | - | 30742455 | 30742549 | 30741639 | 30741716 | 30757658 | 30757774 | 1598 | 483 | 1157 | [illegible] |
| SE | 82668 | ENSG00000141380 | SS18 | chr18 | - | 26068493 | 26068557 | 26057588 | 26057742 | 26078075 | 26078160 | 6 | 11 | 11 | [illegible] |
| SE | 92509 | ENSG00000216937 | CCDC7 | chr10 | + | 32567669 | 32567891 | 32565557 | 32565620 | 32571858 | 32571893 | 479 | 349 | 254 | [illegible] |

| | | | | | | | | | | | | | | | | | | | | | | | | | |
|---|---|---|---|---|---|---|---|---|---|---|---|---|---|---|---|---|---|---|---|---|---|---|---|---|---|
| SE | 55022 | ENSG00000152223 | EPG5 | chr18 | - | 45889797 | 45889940 | 45887750 | 45887907 | 45899403 | 45899566 | 92 | 47 | 73 | 68 | 48 | 0 | 0 | 0 | 1 | 0 | 52 | 57 | 51 | 42 |
| SE | 45917 | ENSG00000141076 | UTP4 | chr16 | + | 69153583 | 69153680 | 69150812 | 69150904 | 69155870 | 69155993 | 31 | 9 | 31 | 18 | 31 | 0 | 0 | 1 | 0 | 0 | 13 | 20 | 25 | 23 |
| SE | 95591 | ENSG00000130714 | POMT1 | chr9 | + | 131506113 | 131506220 | 131502894 | 131503073 | 131506402 | 131506453 | 23 | 5 | 13 | 6 | 18 | 1 | 0 | 0 | 4 | 0 | 8 | 23 | 11 | 2 |
| SE | 26397 | ENSG00000138744 | NAAA | chr4 | - | 75918760 | 75918789 | 75913659 | 75914338 | 75919908 | 75919975 | 32 | 4 | 20 | 30 | 22 | 5 | 2 | 3 | 3 | 6 | 23 | 5 | 5 | 15 |
| SE | 87335 | ENSG00000215301 | DDX3X | chrX | + | 41346858 | 41347012 | 41346228 | 41346622 | 41347314 | 41347451 | 1289 | 837 | 1126 | 1287 | 1856 | 1 | 0 | 3 | 2 | 0 | 106 | 562 | 944 | 443 |
| SE | 67773 | ENSG00000036257 | CUL3 | chr2 | - | 224556294 | 224556420 | 224535527 | 224535641 | 224557658 | 224557820 | 36 | 19 | 16 | 18 | 60 | 319 | 243 | 258 | 250 | 450 | 25 | 8 | 20 | 18 |
| SE | 28929 | ENSG00000144959 | NCEH1 | chr3 | - | 172647885 | 172648114 | 172645622 | 172645692 | 172710846 | 172711067 | 32 | 14 | 24 | 17 | 42 | 1 | 0 | 1 | 5 | 2 | 35 | 31 | 10 | 23 |
| SE | 84519 | ENSG00000198960 | ARMCX6 | chrX | - | 101617282 | 101617370 | 101615125 | 101616766 | 101617502 | 101617584 | 9 | 0 | 6 | 1 | 11 | 193 | 143 | 118 | 177 | 414 | 0 | 17 | 0 | 3 |
| SE | 8229 | ENSG00000185624 | P4HB | chr17 | - | 81855452 | 81855586 | 81855141 | 81855279 | 81859180 | 81859289 | 630 | 314 | 579 | 573 | 332 | 1 | 0 | 0 | 2 | 0 | 769 | 677 | 607 | 695 |
| SE | 106242 | ENSG00000089154 | GCN1 | chr12 | - | 120183568 | 120183677 | 120178851 | 120178950 | 120184111 | 120184243 | 140 | 78 | 168 | 149 | 205 | 0 | 0 | 0 | 2 | 0 | 195 | 154 | 113 | 98 |
| SE | 91357 | ENSG00000238083 | LRRC37A2 | chr17 | + | 46523083 | 46523155 | 46520211 | 46520283 | 46523803 | 46523884 | 49 | 21 | 53 | 56 | 75 | 8 | 0 | 1 | 2 | 2 | 61 | 134 | 36 | 57 |
| SE | 16722 | ENSG00000214013 | GANC | chr15 | + | 42280916 | 42281133 | 42278481 | 42278590 | 42287690 | 42287818 | 12 | 19 | 20 | 17 | 43 | 66 | 35 | 25 | 46 | 99 | 28 | 22 | 11 | 25 |
| SE | 49786 | ENSG00000116337 | AMPD2 | chr1 | + | 109625661 | 109625792 | 109619836 | 109620278 | 109626159 | 109626195 | 64 | 26 | 43 | 70 | 3 | 0 | 0 | 0 | 0 | 0 | 60 | 68 | 54 | 109 |
| SE | 35641 | ENSG00000138246 | DNAJC13 | chr3 | + | 132484362 | 132484488 | 132483374 | 132483577 | 132484587 | 132484672 | 11 | 4 | 3 | 10 | 32 | 97 | 62 | 83 | 112 | 214 | 0 | 8 | 7 | 13 |
| SE | 68571 | ENSG00000140368 | PSTPIP1 | chr15 | + | 77026115 | 77026172 | 77025497 | 77025604 | 77027851 | 77027914 | 5 | 2 | 2 | 1 | 8 | 213 | 134 | 189 | 199 | 130 | 11 | 10 | 11 | 14 |
| SE | 52465 | ENSG00000072803 | FBXW11 | chr5 | - | 171996889 | 171997051 | 171910571 | 171910797 | 172006457 | 172006649 | 39 | 13 | 17 | 23 | 23 | 11 | 10 | 9 | 14 | 6 | 54 | 11 | 27 | 9 |
| SE | 84957 | ENSG00000139436 | GIT2 | chr12 | - | 109951211 | 109951316 | 109939164 | 109939247 | 109953091 | 109953234 | 85 | 80 | 76 | 92 | 94 | 1 | 2 | 3 | 12 | 0 | 96 | 94 | 99 | 137 |
| SE | 56407 | ENSG00000151838 | CCDC175 | chr14 | - | 59545162 | 59545299 | 59543343 | 59543454 | 59551354 | 59551436 | 4 | 0 | 4 | 10 | 14 | 3 | 6 | 3 | 6 | 10 | 38 | 9 | 3 | 11 |
| SE | 49154 | ENSG00000077044 | DGKD | chr2 | + | 233438216 | 233438379 | 233437376 | 233437479 | 233441886 | 233441995 | 228 | 139 | 201 | 225 | 207 | 5 | 10 | 13 | 8 | 6 | 238 | 287 | 154 | 263 |
| SE | 100929 | ENSG00000133895 | MEN1 | chr11 | - | 64805887 | 64806013 | 64805634 | 64805770 | 64806231 | 64806368 | 7 | 1 | 3 | 3 | 0 | 25 | 14 | 20 | 30 | 16 | 11 | 0 | 0 | 6 |
| SE | 10896 | ENSG00000176974 | SHMT1 | chr17 | - | 18335558 | 18335675 | 18333165 | 18333288 | 18340042 | 18340255 | 10 | 6 | 17 | 8 | 5 | 0 | 0 | 1 | 0 | 3 | 0 | 18 | 13 | 8 |
| SE | 17742 | ENSG00000100934 | SEC23A | chr14 | - | 39074414 | 39074530 | 39067172 | 39067296 | 39075934 | 39076093 | 197 | 143 | 233 | 263 | 243 | 0 | 2 | 0 | 0 | 1 | 56 | 159 | 125 | 126 |
| SE | 62533 | ENSG00000004866 | ST7 | chr7 | + | 117134123 | 117134192 | 117131884 | 117131960 | 117136080 | 117136235 | 5 | 7 | 5 | 4 | 11 | 29 | 16 | 15 | 33 | 31 | 0 | 0 | 1 | 0 |
| SE | 30726 | ENSG00000125375 | DMAC2L | chr14 | + | 50323944 | 50324116 | 50322510 | 50322719 | 50325608 | 50325645 | 66 | 34 | 55 | 66 | 79 | 8 | 1 | 0 | 2 | 1 | 12 | 30 | 37 | 25 |
| SE | 71743 | ENSG00000179051 | RCC2 | chr1 | - | 17438229 | 17438522 | 17429105 | 17429199 | 17439544 | 17439677 | 105 | 70 | 55 | 110 | 26 | 0 | 0 | 0 | 1 | 0 | 272 | 150 | 42 | 117 |
| SE | 9420 | ENSG00000174231 | PRPF8 | chr17 | - | 1660992 | 1661162 | 1660697 | 1660827 | 1661270 | 1661329 | 894 | 481 | 869 | 702 | 982 | 1 | 0 | 4 | 1 | 0 | 933 | 782 | 509 | 695 |
| SE | 52766 | ENSG00000198019 | FCGR1B | chr1 | + | 121088682 | 121088908 | 121087820 | 121087841 | 121092995 | 121093250 | 13 | 5 | 4 | 13 | 6 | 0 | 0 | 0 | 1 | 3 | 57 | 7 | 10 | 55 |
| SE | 695 | ENSG00000186575 | NF2 | chr22 | + | 29661204 | 29661339 | 29658188 | 29658264 | 29664989 | 29665064 | 318 | 196 | 277 | 318 | 399 | 2 | 6 | 0 | 1 | 5 | 599 | 400 | 267 | 288 |
| SE | 58801 | ENSG00000077254 | USP33 | chr1 | - | 77745420 | 77745599 | 77741616 | 77741748 | 77759642 | 77759852 | 54 | 47 | 23 | 47 | 53 | 24 | 6 | 21 | 18 | 11 | 21 | 29 | 34 | 35 |
| SE | 18230 | ENSG00000119686 | FLVCR2 | chr14 | + | 75639351 | 75639462 | 75634909 | 75635013 | 75640954 | 75641060 | 29 | 33 | 27 | 22 | 25 | 13 | 2 | 1 | 6 | 17 | 58 | 72 | 24 | 21 |
| SE | 27610 | ENSG00000001629 | ANKIB1 | chr7 | + | 92347946 | 92348016 | 92344977 | 92345066 | 92350949 | 92351094 | 20 | 7 | 3 | 12 | 11 | 70 | 57 | 43 | 56 | 83 | 0 | 11 | 3 | 0 |
| SE | 82393 | ENSG00000105835 | NAMPT | chr7 | - | 106254116 | 106254504 | 106253016 | 106253151 | 106261587 | 106261707 | 254 | 238 | 202 | 385 | 690 | 4 | 0 | 0 | 4 | 0 | 64 | 234 | 1925 | 160 |
| SE | 18060 | ENSG00000100450 | GZMH | chr14 | - | 24607611 | 24607747 | 24606484 | 24606746 | 24608264 | 24608412 | 762 | 162 | 151 | 251 | 300 | 7 | 7 | 2 | 4 | 1 | 393 | 578 | 334 | 279 |
| SE | 7095 | ENSG00000134046 | MBD2 | chr18 | - | 54187713 | 54187837 | 54164522 | 54164700 | 54188873 | 54189011 | 12 | 11 | 18 | 22 | 32 | 1 | 1 | 2 | 3 | 3 | 18 | 12 | 4 | 17 |
| SE | 80671 | ENSG00000121716 | PILRB | chr7 | + | 100353894 | 100354012 | 100353372 | 100353483 | 100355142 | 100355240 | 104 | 46 | 91 | 105 | 97 | 7 | 6 | 2 | 6 | 0 | 35 | 121 | 58 | 85 |
| SE | 69527 | ENSG00000177479 | ARIH2 | chr3 | + | 48924717 | 48924839 | 48922747 | 48922811 | 48927461 | 48927813 | 62 | 26 | 44 | 45 | 41 | 69 | 42 | 36 | 70 | 36 | 66 | 110 | 45 | 43 |
| SE | 29708 | ENSG00000182621 | PLCB1 | chr20 | + | 8739260 | 8739360 | 8737027 | 8737192 | 8740343 | 8740448 | 75 | 62 | 46 | 54 | 74 | 3 | 0 | 0 | 1 | 0 | 31 | 41 | 49 | 13 |
| SE | 63516 | ENSG00000125962 | ARMCX5 | chrX | + | 102600919 | 102601032 | 102599521 | 102599859 | 102601463 | 102601509 | 7 | 6 | 10 | 16 | 33 | 1 | 5 | 5 | 1 | 0 | 16 | 3 | 0 | 4 |
| SE | 66834 | ENSG00000031003 | FAM13B | chr5 | - | 138018954 | 138019146 | 138018301 | 138018514 | 138021030 | 138021197 | 250 | 117 | 224 | 214 | 291 | 39 | 26 | 23 | 33 | 21 | 83 | 163 | 131 | 74 |
| SE | 87748 | ENSG00000063601 | MTMR1 | chrX | + | 150712341 | 150712365 | 150699194 | 150699300 | 150718624 | 150718700 | 65 | 45 | 93 | 84 | 109 | 3 | 0 | 1 | 4 | 0 | 74 | 76 | 45 | 34 |
| SE | 75682 | ENSG00000139163 | ETNK1 | chr12 | + | 22656431 | 22656745 | 22643762 | 22644022 | 22659013 | 22659154 | 4 | 2 | 1 | 4 | 6 | 117 | 71 | 69 | 116 | 167 | 0 | 3 | 1 | 11 |
| SE | 87535 | ENSG00000147050 | KDM6A | chrX | + | 45020609 | 45020730 | 45010960 | 45011019 | 45034930 | 45034985 | 274 | 184 | 190 | 200 | 268 | 20 | 12 | 9 | 11 | 8 | 171 | 140 | 140 | 174 |
| SE | 66828 | ENSG00000031003 | FAM13B | chr5 | - | 137987460 | 137987616 | 137985256 | 137985389 | 137988273 | 137988315 | 860 | 667 | 583 | 869 | 1100 | 6 | 9 | 2 | 12 | 3 | 634 | 672 | 514 | 547 |
| SE | 62500 | ENSG00000183943 | PRKX | chrX | - | 3611221 | 3611334 | 3608624 | 3608945 | 3612176 | 3612281 | 57 | 16 | 34 | 45 | 59 | 44 | 48 | 57 | 50 | 94 | 71 | 40 | 10 | 56 |
| SE | 82235 | ENSG00000123066 | MED13L | chr12 | - | 116236820 | 116236890 | 116111427 | 116111512 | 116237467 | 116237569 | 0 | 10 | 28 | 0 | 0 | 280 | 293 | 271 | 286 | 344 | 11 | 12 | 26 | 24 |
| SE | 87575 | ENSG00000112159 | MDN1 | chr6 | - | 89712035 | 89712256 | 89708495 | 89708628 | 89712574 | 89712786 | 54 | 24 | 57 | 48 | 42 | 0 | 0 | 0 | 0 | 0 | 28 | 32 | 41 | 37 |
| SE | 71543 | ENSG00000145216 | FIP1L1 | chr4 | + | 53440581 | 53440608 | 53428026 | 53428183 | 53442652 | 53442707 | 37 | 12 | 25 | 36 | 13 | 40 | 29 | 46 | 47 | 26 | 25 | 54 | 28 | 31 |
| SE | 41128 | ENSG00000124406 | ATP8A1 | chr4 | - | 42485495 | 42485668 | 42464892 | 42465076 | 42503449 | 42503514 | 150 | 111 | 184 | 150 | 179 | 5 | 4 | 0 | 5 | 0 | 39 | 52 | 76 | 55 |
| SE | 34811 | ENSG00000136731 | UGGT1 | chr2 | + | 128156391 | 128156415 | 128155488 | 128155587 | 128157251 | 128157346 | 21 | 7 | 20 | 11 | 28 | 165 | 119 | 147 | 149 | 233 | 16 | 38 | 5 | 28 |
| SE | 98676 | ENSG00000205531 | NAP1L4 | chr11 | - | 2949226 | 2949264 | 2944436 | 2945646 | 2951258 | 2951315 | 100 | 58 | 92 | 113 | 150 | 62 | 57 | 89 | 68 | 85 | 62 | 44 | 63 | 58 |
| SE | 9572 | ENSG00000213246 | SUPT4H1 | chr17 | - | 58347528 | 58347584 | 58347187 | 58347241 | 58351401 | 58351508 | 746 | 462 | 432 | 625 | 861 | 9 | 7 | 11 | 12 | 5 | 1623 | 860 | 608 | 921 |
| SE | 7897 | ENSG00000159842 | ABR | chr17 | - | 1011144 | 1011388 | 1010728 | 1010863 | 1011845 | 1012589 | 1 | 2 | 1 | 3 | 8 | 197 | 67 | 116 | 109 | 96 | 8 | 5 | 7 | 1 |
| SE | 102618 | ENSG00000196177 | ACADSB | chr10 | + | 123037746 | 123037847 | 123034355 | 123034515 | 123040465 | 123040657 | 116 | 59 | 77 | 92 | 89 | 2 | 2 | 2 | 9 | 1 | 13 | 100 | 66 | 30 |
| SE | 33021 | ENSG00000114316 | USP4 | chr3 | - | 49325718 | 49325845 | 49324893 | 49325039 | 49327685 | 49327816 | 707 | 427 | 593 | 678 | 1018 | 0 | 11 | 0 | 3 | 0 | 1342 | 725 | 574 | 899 |
| SE | 66395 | ENSG00000114023 | FAM162A | chr3 | + | 122401375 | 122401535 | 122384175 | 122384299 | 122402759 | 122402882 | 2 | 0 | 1 | 0 | 0 | 54 | 46 | 64 | 56 | 16 | 0 | 0 | 0 | 2 |
| SE | 12591 | ENSG00000090861 | AARS1 | chr16 | - | 70262345 | 70262484 | 70261043 | 70261157 | 70264957 | 70265102 | 83 | 34 | 71 | 43 | 58 | 0 | 2 | 11 | 5 | 0 | 61 | 51 | 23 | 47 |

[illegible]

| | | | | | | | | | | | | | | | | | | | | | | | | | | | | | | | | | | | | | | | | | | | | | | |
|---|---|---|---|---|---|---|---|---|---|---|---|---|---|---|---|---|---|---|---|---|---|---|---|---|---|---|---|---|---|---|---|---|---|---|---|---|---|---|---|---|---|---|---|---|---|---|
| SE | 43580 | ENSG00000058673 | ZC3H11A | chr1 | + | 203796292 | 203796496 | 203795653 | 203795794 | 203801574 | 203801628 | 29 | 44 | 59 | 55 | 60 | 33 | 13 | 34 | 21 | 45 | 10 | 73 | 65 | 25 | 13 | 0 | 10 | 7 | 11 | 11 | 298 | 149 | 0.015293473 | 0.146760356 | 0.305 | 0.629 | 0.465 | 0.567 | 0.4 | 1 | 0.785 | 0.823 | 0.532 | 0.371 | -0.229 |
| SE | 28032 | ENSG00000120519 | SLC10A7 | chr4 | - | 146442782 | 146442821 | 146325960 | 146325996 | 146503848 | 146503924 | 48 | 21 | 47 | 49 | 47 | 0 | 0 | 3 | 2 | 6 | 7 | 29 | 31 | 8 | 2 | 0 | 0 | 0 | 2 | 0 | 188 | 149 | 0.015314794 | 0.14694695 | 1 | 1 | 0.925 | 0.951 | 0.861 | 1 | 1 | 1 | 0.76 | 1 | -0.005 |
| SE | 1700 | ENSG00000228315 | GUSBP11 | chr22 | - | 23686837 | 23686995 | 23683724 | 23683867 | 23690231 | 23690293 | 4 | 6 | 8 | 22 | 14 | 15 | 26 | 20 | 16 | 15 | 48 | 9 | 6 | 20 | 21 | 17 | 10 | 15 | 8 | 20 | 298 | 149 | 0.015321183 | 0.146990242 | 0.118 | 0.103 | 0.167 | 0.407 | 0.318 | 0.585 | 0.31 | 0.167 | 0.556 | 0.344 | -0.17 |
| SE | 43149 | ENSG00000113161 | HMGCR | chr5 | + | 75359397 | 75359556 | 75359169 | 75359310 | 75359984 | 75360139 | 242 | 123 | 231 | 246 | 174 | 2 | 8 | 2 | 5 | 0 | 45 | 77 | 141 | 84 | 4 | 2 | 2 | 10 | 3 | 0 | 298 | 149 | 0.015365884 | 0.147401033 | 0.984 | 0.885 | 0.983 | 0.961 | 1 | 0.918 | 0.951 | 0.876 | 0.933 | 1 | 0.027 |
| SE | 83013 | ENSG00000152484 | USP12 | chr13 | - | 27071070 | 27071149 | 27066155 | 27069384 | 27075190 | 27075388 | 114 | 75 | 68 | 144 | 142 | 3 | 1 | 8 | 2 | 0 | 38 | 88 | 68 | 25 | 4 | 0 | 0 | 1 | 0 | 0 | 228 | 149 | 0.015379577 | 0.147514312 | 0.961 | 0.98 | 0.847 | 0.979 | 1 | 1 | 1 | 0.978 | 1 | 1 | -0.042 |
| SE | 48614 | ENSG00000131374 | TBC1D5 | chr3 | - | 17562018 | 17562142 | 17508473 | 17508605 | 17575202 | 17575310 | 33 | 10 | 11 | 31 | 13 | 0 | 0 | 0 | 0 | 0 | 18 | 23 | 21 | 27 | 28 | 11 | 0 | 0 | 0 | 0 | 273 | 149 | 0.015388614 | 0.147573596 | 1 | 1 | 1 | 1 | 1 | 0.472 | 1 | 1 | 1 | 1 | 0.106 |
| SE | 75217 | ENSG00000100503 | NIN | chr14 | - | 50752517 | 50752733 | 50747991 | 50748105 | 50754562 | 50754632 | 657 | 756 | 592 | 662 | 1199 | 4 | 4 | 7 | 10 | 10 | 158 | 514 | 745 | 308 | 18 | 0 | 1 | 18 | 13 | 0 | 298 | 149 | 0.015389527 | 0.147573596 | 0.988 | 0.99 | 0.977 | 0.971 | 0.984 | 1 | 0.996 | 0.954 | 0.922 | 1 | 0.008 |
| SE | 50574 | ENSG00000139636 | LMBR1L | chr12 | - | 49104745 | 49104885 | 49103700 | 49103813 | 49105923 | 49105957 | 23 | 32 | 47 | 25 | 67 | 0 | 0 | 0 | 0 | 0 | 36 | 21 | 14 | 6 | 57 | 0 | 0 | 1 | 0 | 8 | 289 | 149 | 0.015400172 | 0.147657586 | 1 | 1 | 1 | 1 | 1 | 1 | 1 | 0.878 | 1 | 0.786 | 0.067 |
| SE | 40396 | ENSG00000163719 | MTMR14 | chr3 | + | 9697710 | 9697866 | 9688943 | 9689082 | 9701789 | 9702216 | 23 | 11 | 27 | 21 | 9 | 8 | 9 | 8 | 8 | 1 | 47 | 24 | 25 | 19 | 45 | 20 | 16 | 15 | 29 | 61 | 298 | 149 | 0.015418989 | 0.147819903 | 0.59 | 0.379 | 0.628 | 0.568 | 0.818 | 0.54 | 0.429 | 0.455 | 0.247 | 0.269 | 0.209 |
| SE | 86405 | ENSG00000131381 | RBSN | chr3 | - | 15075605 | 15075710 | 15070072 | 15074930 | 15077061 | 15077164 | 58 | 20 | 57 | 66 | 55 | 0 | 0 | 1 | 0 | 0 | 112 | 56 | 65 | 120 | 96 | 0 | 0 | 0 | 0 | 13 | 254 | 149 | 0.015424514 | 0.147854773 | 1 | 1 | 0.971 | 1 | 1 | 1 | 1 | 1 | 1 | 0.812 | 0.032 |
| SE | 80821 | ENSG00000151948 | GLT1D1 | chr12 | + | 128914932 | 128914989 | 128888638 | 128888744 | 128926358 | 128926454 | 23 | 19 | 6 | 14 | 15 | 1 | 0 | 0 | 0 | 4 | 60 | 33 | 49 | 80 | 79 | 14 | 3 | 1 | 0 | 3 | 206 | 149 | 0.015439695 | 0.14798217 | 0.943 | 1 | 1 | 1 | 0.731 | 0.756 | 0.888 | 0.973 | 1 | 0.95 | 0.021 |
| SE | 42206 | ENSG00000132155 | RAF1 | chr3 | - | 12599112 | 12599225 | 12591707 | 12591792 | 12599690 | 12599808 | 5 | 2 | 0 | 4 | 11 | 138 | 87 | 130 | 124 | 185 | 16 | 5 | 5 | 13 | 14 | 201 | 122 | 169 | 238 | 207 | 262 | 149 | 0.015465516 | 0.148211513 | 0.02 | 0.013 | 0 | 0.018 | 0.033 | 0.043 | 0.023 | 0.017 | 0.03 | 0.037 | -0.013 |
| SE | 58579 | ENSG00000072858 | SIDT1 | chr3 | + | 113611007 | 113611144 | 113608418 | 113608536 | 113612085 | 113612194 | 54 | 22 | 57 | 58 | 60 | 2 | 0 | 2 | 1 | 0 | 14 | 48 | 23 | 28 | 41 | 0 | 0 | 4 | 0 | 7 | 286 | 149 | 0.015478712 | 0.148319817 | 0.934 | 1 | 0.937 | 0.968 | 1 | 1 | 1 | 0.75 | 1 | 0.753 | 0.067 |
| SE | 20124 | ENSG00000078319 | PMS2P1 | chr7 | - | 100335205 | 100335428 | 100332441 | 100332581 | 100335837 | 100335977 | 4 | 0 | 1 | 0 | 1 | 28 | 19 | 48 | 19 | 5 | 6 | 4 | 4 | 4 | 0 | 18 | 53 | 20 | 35 | 66 | 298 | 149 | 0.015484774 | 0.148342541 | 0.067 | 0 | 0.01 | 0 | 0.091 | 0.143 | 0.036 | 0.091 | 0.054 | 0 | -0.031 |
| SE | 51553 | ENSG00000143801 | PSEN2 | chr1 | + | 226871261 | 226871404 | 226870516 | 226870649 | 226875364 | 226875550 | 14 | 0 | 5 | 4 | 24 | 0 | 1 | 4 | 2 | 0 | 15 | 35 | 12 | 34 | 44 | 0 | 0 | 0 | 0 | 0 | 292 | 149 | 0.015484873 | 0.148342541 | 1 | 0 | 0.389 | 0.505 | 1 | 1 | 1 | 1 | 1 | 1 | -0.421 |
| SE | 83381 | ENSG00000162959 | MEMO1 | chr2 | - | 31893241 | 31893321 | 31892052 | 31892134 | 31917925 | 31918037 | 27 | 13 | 16 | 16 | 7 | 114 | 99 | 102 | 105 | 133 | 12 | 4 | 5 | 20 | 20 | 172 | 55 | 77 | 176 | 190 | 229 | 149 | 0.015494493 | 0.14841654 | 0.134 | 0.079 | 0.093 | 0.09 | 0.033 | 0.043 | 0.045 | 0.041 | 0.069 | 0.064 | 0.033 |
| SE | 95671 | ENSG00000137073 | UBAP2 | chr9 | - | 34032798 | 34032908 | 34017049 | 34017189 | 34048824 | 34048879 | 0 | 0 | 0 | 0 | 0 | 16 | 8 | 10 | 14 | 1 | 12 | 0 | 0 | 0 | 0 | 34 | 21 | 10 | 12 | 38 | 259 | 149 | 0.015499803 | 0.148446136 | 0 | 0 | 0 | 0 | 0 | 0.169 | 0 | 0 | 0 | 0 | -0.034 |
| SE | 19769 | ENSG00000100614 | PPM1A | chr14 | + | 60277312 | 60277365 | 60277004 | 60277091 | 60282683 | 60283089 | 11 | 13 | 9 | 5 | 4 | 12 | 16 | 9 | 14 | 10 | 10 | 36 | 26 | 19 | 20 | 10 | 14 | 25 | 22 | 7 | 202 | 149 | 0.015501374 | 0.148446136 | 0.403 | 0.375 | 0.425 | 0.209 | 0.228 | 0.425 | 0.655 | 0.434 | 0.389 | 0.678 | -0.188 |
| SE | 46655 | ENSG00000126749 | EMG1 | chr12 | + | 6983001 | 6983206 | 6975228 | 6975378 | 6987781 | 6987838 | 13 | 28 | 22 | 17 | 15 | 0 | 0 | 0 | 0 | 0 | 24 | 19 | 3 | 8 | 1 | 0 | 9 | 0 | 0 | 0 | 298 | 149 | 0.015513256 | 0.148541752 | 1 | 1 | 1 | 1 | 1 | 1 | 0.514 | 1 | 1 | 1 | 0.097 |
| SE | 80273 | ENSG00000164535 | DAGLB | chr7 | - | 6436361 | 6436533 | 6434761 | 6435020 | 6445952 | 6446104 | 44 | 30 | 32 | 44 | 31 | 2 | 6 | 1 | 0 | 0 | 41 | 51 | 30 | 32 | 46 | 0 | 0 | 0 | 0 | 0 | 298 | 149 | 0.015523473 | 0.148610757 | 0.917 | 0.714 | 0.941 | 1 | 1 | 1 | 1 | 1 | 1 | 1 | -0.086 |
| SE | 5464 | ENSG00000101337 | TM9SF4 | chr20 | + | 32136876 | 32136997 | 32136073 | 32136173 | 32141540 | 32141665 | 2 | 2 | 3 | 1 | 4 | 4 | 5 | 10 | 8 | 17 | 17 | 0 | 10 | 2 | 11 | 7 | 6 | 4 | 12 | 10 | 270 | 149 | 0.015524259 | 0.148610757 | 0.216 | 0.181 | 0.142 | 0.065 | 0.115 | 0.573 | 0 | 0.58 | 0.084 | 0.378 | -0.179 |
| SE | 88400 | ENSG00000145041 | DCAF1 | chr3 | - | 51466805 | 51466876 | 51463113 | 51463227 | 51470928 | 51471005 | 64 | 51 | 61 | 55 | 51 | 3 | 0 | 5 | 3 | 8 | 60 | 74 | 67 | 64 | 33 | 4 | 0 | 1 | 4 | 0 | 220 | 149 | 0.015530311 | 0.148650516 | 0.935 | 1 | 0.892 | 0.925 | 0.812 | 0.91 | 1 | 0.978 | 0.916 | 1 | -0.048 |
| SE | 20452 | ENSG00000148814 | LRRC27 | chr10 | + | 132348962 | 132349057 | 132347983 | 132348356 | 132351606 | 132351753 | 5 | 3 | 10 | 2 | 8 | 10 | 10 | 12 | 3 | 7 | 7 | 25 | 14 | 31 | 12 | 9 | 8 | 9 | 6 | 20 | 244 | 149 | 0.015544551 | 0.14876863 | 0.234 | 0.155 | 0.337 | 0.289 | 0.411 | 0.322 | 0.656 | 0.487 | 0.759 | 0.268 | -0.213 |
| SE | 83664 | ENSG00000029363 | BCLAF1 | chr6 | - | 136273962 | 136274189 | 136273081 | 136273187 | 136275531 | 136275701 | 31 | 26 | 40 | 43 | 29 | 521 | 452 | 575 | 671 | 728 | 0 | 22 | 7 | 0 | 0 | 37 | 242 | 369 | 68 | 21 | 298 | 149 | 0.015551618 | 0.148800988 | 0.029 | 0.028 | 0.034 | 0.031 | 0.02 | 0 | 0.043 | 0.009 | 0 | 0 | 0.018 |
| SE | 84953 | ENSG00000139436 | GIT2 | chr12 | - | 109951166 | 109951316 | 109939164 | 109939247 | 109953091 | 109953234 | 117 | 92 | 96 | 113 | 109 | 1 | 2 | 3 | 12 | 0 | 118 | 110 | 114 | 169 | 118 | 0 | 0 | 5 | 3 | 1 | 298 | 149 | 0.015551733 | 0.148800988 | 0.983 | 0.958 | 0.941 | 0.825 | 1 | 1 | 1 | 0.919 | 0.966 | 0.983 | -0.032 |
| SE | 33026 | ENSG00000114316 | USP4 | chr3 | - | 49335468 | 49335596 | 49327685 | 49327816 | 49339923 | 49340033 | 607 | 365 | 426 | 517 | 509 | 1 | 0 | 1 | 1 | 8 | 940 | 598 | 580 | 824 | 1193 | 7 | 16 | 3 | 9 | 7 | 277 | 149 | 0.01555518 | 0.148815787 | 0.997 | 1 | 0.996 | 0.996 | 0.972 | 0.986 | 0.953 | 0.99 | 0.98 | 0.989 | 0.013 |
| SE | 47407 | ENSG00000156804 | FBXO32 | chr8 | - | 123514239 | 123514333 | 123506391 | 123506574 | 123531897 | 123531990 | 46 | 26 | 66 | 39 | 81 | 7 | 0 | 4 | 2 | 14 | 47 | 53 | 22 | 27 | 8 | 0 | 0 | 5 | 3 | 0 | 243 | 149 | 0.015558415 | 0.148828548 | 0.801 | 1 | 0.91 | 0.923 | 0.78 | 1 | 1 | 0.73 | 0.847 | 1 | -0.033 |
| SE | 60769 | ENSG00000187741 | FANCA | chr16 | - | 89742799 | 89742938 | 89740803 | 89740866 | 89744958 | 89745062 | 51 | 23 | 29 | 39 | 29 | 0 | 0 | 0 | 10 | 0 | 34 | 62 | 31 | 38 | 120 | 0 | 0 | 2 | 0 | 0 | 288 | 149 | 0.015582349 | 0.149039283 | 1 | 1 | 1 | 0.669 | 1 | 1 | 1 | 0.889 | 1 | 1 | -0.044 |
| SE | 47318 | ENSG00000133106 | EPSTI1 | chr13 | - | 42953947 | 42954021 | 42926335 | 42926429 | 42963254 | 42963338 | 264 | 284 | 226 | 391 | 357 | 0 | 2 | 0 | 3 | 6 | 1492 | 862 | 760 | 1404 | 349 | 54 | 6 | 4 | 0 | 0 | 223 | 149 | 0.015607299 | 0.149259694 | 1 | 0.99 | 1 | 0.989 | 0.975 | 0.949 | 0.99 | 0.992 | 1 | 1 | 0.005 |
| SE | 10322 | ENSG00000108848 | LUC7L3 | chr17 | + | 50749218 | 50749344 | 50746541 | 50746702 | 50750500 | 50750550 | 66 | 71 | 65 | 108 | 70 | 338 | 396 | 442 | 440 | 659 | 4 | 30 | 20 | 0 | 15 | 32 | 225 | 172 | 85 | 52 | 275 | 149 | 0.015613374 | 0.149293047 | 0.096 | 0.089 | 0.074 | 0.117 | 0.054 | 0.063 | 0.067 | 0.059 | 0 | 0.135 | 0.021 |
| SE | 97335 | ENSG00000116747 | RO60 | chr1 | + | 193069033 | 193069181 | 193059618 | 193059776 | 193075819 | 193075890 | 73 | 24 | 46 | 76 | 48 | 19 | 6 | 10 | 6 | 6 | 4 | 31 | 14 | 18 | 7 | 0 | 0 | 23 | 0 | 0 | 297 | 149 | 0.015614601 | 0.149293047 | 0.658 | 0.667 | 0.698 | 0.864 | 0.801 | 1 | 1 | 0.234 | 1 | 1 | -0.109 |
| SE | 32783 | ENSG00000168297 | PXK | chr3 | + | 58423460 | 58423494 | 58412903 | 58412963 | 58424751 | 58426019 | 0 | 0 | 0 | 0 | 1 | 86 | 34 | 57 | 73 | 210 | 0 | 11 | 2 | 4 | 0 | 183 | 74 | 67 | 70 | 38 | 183 | 149 | 0.015618399 | 0.149311128 | 0 | 0 | 0 | 0 | 0.004 | 0 | 0.108 | 0.024 | 0.044 | 0 | -0.034 |
| SE | 34574 | ENSG00000104388 | RAB2A | chr8 | + | 60576233 | 60576297 | 60572045 | 60572113 | 60584207 | 60584290 | 5 | 1 | 6 | 10 | 7 | 202 | 175 | 164 | 203 | 334 | 22 | 16 | 19 | 0 | 0 | 345 | 172 | 219 | 308 | 211 | 213 | 149 | 0.015622777 | 0.149334752 | 0.017 | 0.004 | 0.025 | 0.033 | 0.014 | 0.043 | 0.061 | 0.057 | 0 | 0 | -0.014 |
| SE | 105934 | ENSG00000127314 | RAP1B | chr12 | + | 68624670 | 68624873 | 68610907 | 68611043 | 68648698 | 68648781 | 35 | 31 | 16 | 51 | 40 | 635 | 465 | 365 | 613 | 429 | 0 | 61 | 39 | 34 | 3 | 157 | 485 | 366 | 251 | 95 | 298 | 149 | 0.015643548 | 0.149515044 | 0.027 | 0.032 | 0.021 | 0.04 | 0.045 | 0 | 0.059 | 0.051 | 0.063 | 0.016 | -0.005 |
| SE | 33069 | ENSG00000088726 | TMEM40 | chr3 | - | 12742453 | 12742507 | 12738552 | 12738588 | 12749759 | 12749840 | 28 | 37 | 15 | 71 | 51 | 3 | 2 | 0 | 0 | 1 | 127 | 61 | 44 | 33 | 372 | 0 | 0 | 0 | 4 | 26 | 203 | 149 | 0.015646744 | 0.14951697 | 0.873 | 0.931 | 1 | 1 | 0.974 | 1 | 1 | 1 | 0.858 | 0.913 | 0.001 |
| SE | 21383 | ENSG00000154309 | DISP1 | chr1 | + | 222827427 | 222827601 | 222815046 | 222815078 | 222835118 | 222835215 | 14 | 11 | 10 | 10 | 9 | 2 | 0 | 10 | 2 | 0 | 18 | 34 | 8 | 19 | 5 | 0 | 0 | 0 | 0 | 7 | 298 | 149 | 0.015647569 | 0.14951697 | 0.778 | 1 | 0.333 | 0.714 | 1 | 1 | 1 | 1 | 1 | 0.263 | -0.088 |
| SE | 47838 | ENSG00000267009 | NA | chr17 | + | 68413910 | 68414182 | 68413739 | 68413795 | 68420188 | 68420506 | 3 | 0 | 6 | 3 | 3 | 1 | 1 | 1 | 1 | 0 | 16 | 4 | 2 | 2 | 7 | 0 | 3 | 0 | 4 | 3 | 298 | 149 | 0.015657335 | 0.14957406 | 0.6 | 0 | 0.75 | 0.6 | 1 | 1 | 0.4 | 1 | 0.2 | 0.538 | -0.038 |
| SE | 51509 | ENSG00000108021 | TASOR2 | chr10 | + | 5717658 | 5717750 | 5684837 | 5685175 | 5720543 | 5720611 | 48 | 18 | 28 | 32 | 24 | 6 | 2 | 1 | 2 | 0 | 32 | 13 | 12 | 6 | 20 | 0 | 0 | 4 | 0 | 0 | 241 | 149 | 0.015657364 | 0.14957406 | 0.832 | 0.848 | 0.945 | 0.908 | 1 | 1 | 1 | 0.65 | 1 | 1 | -0.023 |
| SE | 29419 | ENSG00000169762 | TAPT1 | chr4 | - | 16196657 | 16196723 | 16191360 | 16191523 | 16202461 | 16202582 | 5 | 6 | 3 | 9 | 1 | 28 | 21 | 14 | 17 | 56 | 0 | 0 | 1 | 6 | 0 | 4 | 42 | 8 | 8 | 12 | 215 | 149 | 0.015671509 | 0.149682432 | 0.11 | 0.165 | 0.129 | 0.268 | 0.012 | 0 | 0 | 0.08 | 0.342 | 0 | 0.052 |
| SE | 101156 | ENSG00000133816 | MICAL2 | chr11 | + | 12209496 | 12209598 | 12207724 | 12208139 | 12213254 | 12213410 | 220 | 146 | 132 | 223 | 184 | 15 | 7 | 3 | 13 | 13 | 441 | 110 | 85 | 135 | 320 | 35 | 6 | 19 | 21 | 13 | 251 | 149 | 0.015672532 | 0.149682432 | 0.897 | 0.925 | 0.963 | 0.911 | 0.894 | 0.882 | 0.916 | 0.726 | 0.792 | 0.936 | 0.068 |
| SE | 13913 | ENSG00000067365 | METTL22 | chr16 | + | 8644528 | 8644725 | 8642462 | 8642565 | 8646107 | 8646218 | 21 | 10 | 8 | 30 | 21 | 3 | 0 | 2 | 0 | 0 | 40 | 44 | 15 | 10 | 109 | 4 | 2 | 0 | 4 | 4 | 298 | 149 | 0.015677186 | 0.149708617 | 0.778 | 1 | 0.667 | 1 | 1 | 0.833 | 0.917 | 1 | 0.556 | 0.932 | 0.041 |
| SE | 64299 | ENSG00000131873 | CHSY1 | chr15 | - | 101187992 | 101188219 | 101177100 | 101178980 | 101235081 | 101235147 | 8 | 9 | 7 | 11 | 28 | 44 | 29 | 36 | 53 | 45 | 81 | 21 | 85 | 27 | 21 | 99 | 73 | 59 | 117 | 54 | 298 | 149 | 0.015680118 | 0.149709373 | 0.083 | 0.134 | 0.089 | 0.094 | 0.237 | 0.29 | 0.126 | 0.419 | 0.103 | 0.163 | -0.093 |
| SE | 102821 | ENSG00000148700 | ADD3 | chr10 | + | 110075617 | 110075709 | 110008230 | 110008299 | 110100624 | 110100848 | 0 | 0 | 1 | 3 | 4 | 162 | 84 | 138 | 176 | 211 | 0 | 12 | 7 | 0 | 0 | 23 | 145 | 124 | 67 | 24 | 241 | 149 | 0.015681289 | 0.149709373 | 0 | 0 | 0.004 | 0.01 | 0.012 | 0 | 0.049 | 0.034 | 0 | 0 | -0.011 |
| SE | 97756 | ENSG00000166900 | STX3 | chr11 | + | 59797282 | 59797396 | 59795371 | 59795482 | 59800854 | 59802708 | 77 | 69 | 81 | 65 | 113 | 2 | 4 | 2 | 1 | 6 | 138 | 56 | 121 | 190 | 77 | 5 | 5 | 6 | 10 | 13 | 263 | 149 | 0.015683001 | 0.149709373 | 0.956 | 0.907 | 0.958 | 0.974 | 0.914 | 0.94 | 0.864 | 0.92 | 0.915 | 0.77 | 0.06 |
| SE | 74589 | ENSG00000125703 | ATG4C | chr1 | + | 62829039 | 62829176 | 62821138 | 62821209 | 62834037 | 62834116 | 63 | 46 | 87 | 72 | 159 | 4 | 1 | 2 | 0 | 8 | 26 | 35 | 12 | 8 | 8 | 0 | 0 | 0 | 3 | 0 | 286 | 149 | 0.015693056 | 0.149787093 | 0.891 | 0.96 | 0.958 | 1 | 0.912 | 1 | 1 | 1 | 0.581 | 1 | 0.028 |
| SE | 73543 | ENSG00000115232 | ITGA4 | chr2 | + | 181475158 | 181475288 | 181474959 | 181475066 | 181478756 | 181478824 | 1148 | 960 | 1110 | 1199 | 1600 | 28 | 17 | 35 | 27 | 25 | 443 | 663 | 657 | 294 | 244 | 8 | 15 | 25 | 13 | 16 | 279 | 149 | 0.015695086 | 0.149788209 | 0.956 | 0.968 | 0.944 | 0.96 | 0.972 | 0.967 | 0.959 | 0.933 | 0.924 | 0.891 | 0.025 |
| SE | 89007 | ENSG00000177565 | TBL1XR1 | chr3 | - | 177079826 | 177079945 | 177064919 | 177065022 | 177098465 | 177098541 | 34 | 17 | 14 | 26 | 32 | 67 | 66 | 92 | 69 | 51 | 4 | 20 | 10 | 8 | 5 | 37 | 65 | 52 | 35 | 20 | 268 | 149 | 0.015702246 | 0.14981776 | 0.22 | 0.125 | 0.078 | 0.173 | 0.259 | 0.057 | 0.146 | 0.097 | 0.113 | 0.122 | 0.064 |
| SE | 49967 | ENSG00000162976 | SLC66A3 | chr2 | + | 11174967 | 11175009 | 11171924 | 11172045 | 11177736 | 11177957 | 93 | 54 | 67 | 61 | 96 | 23 | 18 | 13 | 20 | 25 | 23 | 62 | 28 | 54 | 67 | 9 | 4 | 1 | 8 | 14 | 191 | 149 | 0.015703032 | 0.14981776 | 0.759 | 0.701 | 0.801 | 0.704 | 0.75 | 0.666 | 0.924 | 0.956 | 0.84 | 0.789 | -0.092 |
| SE | 62843 | ENSG00000197451 | HNRNPAB | chr5 | + | 178210131 | 178210272 | 178209329 | 178209447 | 178210552 | 178211142 | 103 | 45 | 83 | 145 | 80 | 12 | 6 | 7 | 20 | 13 | 156 | 99 | 41 | 128 | 171 | 0 | 7 | 7 | 23 | 5 | 290 | 149 | 0.015703923 | 0.14981776 | 0.815 | 0.794 | 0.859 | 0.788 | 0.76 | 1 | 0.879 | 0.751 | 0.741 | 0.946 | -0.06 |
| SE | 106648 | ENSG00000139178 | C1RL | chr12 | - | 7108250 | 7108479 | 7101604 | 7102087 | 7109109 | 7109197 | 49 | 46 | 32 | 30 | 27 | 1 | 0 | 3 | 0 | 0 | 18 | 72 | 73 | 115 | 164 | 0 | 0 | 10 | 4 | 0 | 298 | 149 | 0.01571156 | 0.149872354 | 0.961 | 1 | 0.842 | 1 | 1 | 1 | 1 | 0.785 | 0.935 | 1 | 0.017 |
| SE | 2300 | ENSG00000142002 | DPP9 | chr19 | - | 4722194 | 4722304 | 4719850 | 4719941 | 4722498 | 4722551 | 7 | 0 | 4 | 6 | 0 | 36 | 2 | 18 | 11 | 2 | 36 | 3 | 19 | 3 | 14 | 25 | 31 | 15 | 7 | 43 | 259 | 149 | 0.015717047 | 0.149897641 | 0.101 | 0 | 0.113 | 0.239 | 0 | 0.453 | 0.053 | 0.422 | 0.198 | 0.158 | -0.166 |
| SE | 66758 | ENSG00000129292 | PHF20L1 | chr8 | + | 132799082 | 132799172 | 132798771 | 132798860 | 132803818 | 132804032 | 16 | 13 | 12 | 10 | 30 | 65 | 54 | 45 | 77 | 84 | 43 | 24 | 7 | 7 | 16 | 31 | 29 | 40 | 46 | 29 | 239 | 149 | 0.015719331 | 0.149897641 | 0.133 | 0.13 | 0.143 | 0.075 | 0.182 | 0.464 | 0.34 | 0.098 | 0.087 | 0.256 | -0.116 |
| SE | 15300 | ENSG00000114062 | UBE3A | chr15 | - | 25393745 | 25393858 | 25375464 | 25375763 | 25405460 | 25405502 | 9 | 9 | 4 | 8 | 17 | 150 | 116 | 140 | 151 | 160 | 0 | 16 | 12 | 4 | 13 | 39 | 113 | 123 | 57 | 41 | 262 | 149 | 0.015720563 | 0.149897641 | 0.033 | 0.042 | 0.016 | 0.029 | 0.057 | 0 | 0.075 | 0.053 | 0.038 | 0.153 | -0.028 |
| SE | 59806 | ENSG00000166377 | ATP9B | chr18 | + | 79209547 | 79209722 | 79206936 | 79207012 | 79213961 | 79214038 | 1 | 1 | 4 | 0 | 0 | 39 | 41 | 59 | 41 | 55 | 0 | 11 | 0 | 1 | 4 | 56 | 34 | 34 | 31 | 59 | 298 | 149 | 0.015721869 | 0.149897641 | 0.013 | 0.012 | 0.033 | 0 | 0 | 0 | 0.139 | 0 | 0.016 | 0.033 | -0.026 |
| SE | 80249 | ENSG00000164535 | DAGLB | chr7 | - | 6416258 | 6416754 | 6412986 | 6413034 | 6416840 | 6416921 | 12 | 9 | 27 | 12 | 8 | 2 | 1 | 2 | 1 | 5 | 7 | 27 | 7 | 12 | 0 | 0 | 0 | 0 | 0 | 4 | 298 | 149 | 0.015768535 | 0.150324271 | 0.75 | 0.818 | 0.871 | 0.857 | 0.444 | 1 | 1 | 1 | 1 | 0 | -0.052 |
| SE | 37079 | ENSG00000127481 | UBR4 | chr1 | - | 19077975 | 19078066 | 19076739 | 19076902 | 19081348 | 19081573 | 335 | 156 | 278 | 271 | 341 | 1 | 0 | 0 | 1 | 0 | 518 | 450 | 216 | 413 | 477 | 0 | 2 | 10 | 8 | 8 | 240 | 149 | 0.015779596 | 0.150411398 | 0.995 | 1 | 1 | 0.994 | 1 | 1 | 0.993 | 0.931 | 0.97 | 0.974 | 0.024 |
| [illegible] | | | | | | | | | | | | | | | | | | | | | | | | | | | | | | | | | | | | | | | | | | | | | | |

| | | | | | | | | | | | | | | | | | | | | | | | | | | | | | | | | | | | | | | | | | | | | | | |
|---|---|---|---|---|---|---|---|---|---|---|---|---|---|---|---|---|---|---|---|---|---|---|---|---|---|---|---|---|---|---|---|---|---|---|---|---|---|---|---|---|---|---|---|---|---|---|
| SE | 66304 | ENSG00000225190 | PLEKHM1 | chr17 | - | 45468208 | 45468593 | 45458168 | 45458439 | 45475099 | 45475726 | 179 | 80 | 120 | 177 | 187 | 26 | 3 | 21 | 14 | 7 | 224 | 218 | 185 | 216 | 234 | 25 | 15 | 33 | 46 | 43 | 298 | 149 | 0.017854131 | 0.166576148 | 0.775 | 0.93 | 0.741 | 0.863 | 0.93 | 0.818 | 0.879 | 0.737 | 0.701 | 0.731 | 0.075 |
| SE | 39777 | ENSG00000072274 | TFRC | chr3 | - | 196071395 | 196071498 | 196069454 | 196069568 | 196072002 | 196072152 | 189 | 134 | 232 | 194 | 158 | 4 | 6 | 5 | 5 | 7 | 99 | 244 | 334 | 160 | 86 | 0 | 17 | 9 | 13 | 9 | 252 | 149 | 0.017858817 | 0.16660002 | 0.965 | 0.93 | 0.965 | 0.958 | 0.93 | 1 | 0.895 | 0.956 | 0.879 | 0.85 | 0.034 |
| SE | 83335 | ENSG00000161010 | MRNIP | chr5 | - | 179853196 | 179853276 | 179847977 | 179848066 | 179853377 | 179853437 | 8 | 6 | 11 | 11 | 12 | 25 | 15 | 24 | 22 | 8 | 33 | 28 | 17 | 25 | 44 | 57 | 20 | 21 | 21 | 34 | 229 | 149 | 0.017870474 | 0.166688898 | 0.172 | 0.207 | 0.23 | 0.245 | 0.494 | 0.274 | 0.477 | 0.345 | 0.436 | 0.457 | -0.128 |
| SE | 70558 | ENSG00000022840 | RNF10 | chr12 | + | 120552498 | 120552698 | 120546404 | 120546601 | 120554717 | 120554808 | 500 | 167 | 301 | 412 | 781 | 2 | 9 | 2 | 2 | 1 | 665 | 447 | 585 | 473 | 883 | 2 | 16 | 5 | 17 | 0 | 298 | 149 | 0.017877918 | 0.166738473 | 0.992 | 0.903 | 0.987 | 0.99 | 0.997 | 0.994 | 0.933 | 0.983 | 0.933 | 1 | 0.005 |
| SE | 92134 | ENSG00000108666 | C17orf75 | chr17 | - | 32333416 | 32333520 | 32328440 | 32331978 | 32334468 | 32334605 | 69 | 68 | 88 | 81 | 90 | 0 | 0 | 0 | 0 | 1 | 63 | 25 | 49 | 28 | 10 | 0 | 5 | 0 | 0 | 4 | 253 | 149 | 0.017888457 | 0.166816891 | 1 | 1 | 1 | 1 | 0.981 | 1 | 0.746 | 1 | 1 | 0.596 | 0.128 |
| SE | 57879 | ENSG00000164924 | YWHAZ | chr8 | - | 100951928 | 100952125 | 100948595 | 100948900 | 100953268 | 100953388 | 803 | 672 | 628 | 1007 | 263 | 547 | 419 | 506 | 498 | 384 | 1063 | 907 | 841 | 1189 | 1099 | 189 | 673 | 662 | 850 | 92 | 298 | 149 | 0.017901113 | 0.166915038 | 0.423 | 0.445 | 0.383 | 0.503 | 0.255 | 0.738 | 0.403 | 0.388 | 0.412 | 0.857 | -0.158 |
| SE | 63861 | ENSG00000219626 | FAM228B | chr2 | + | 24137908 | 24138100 | 24135118 | 24135187 | 24139369 | 24139450 | 115 | 124 | 119 | 212 | 415 | 14 | 8 | 27 | 27 | 33 | 271 | 124 | 83 | 139 | 309 | 24 | 12 | 8 | 14 | 8 | 298 | 149 | 0.017917542 | 0.167048327 | 0.804 | 0.886 | 0.688 | 0.797 | 0.863 | 0.85 | 0.838 | 0.838 | 0.832 | 0.951 | -0.054 |
| SE | 34783 | ENSG00000075790 | BCAP29 | chr7 | + | 107613653 | 107613732 | 107613331 | 107613432 | 107618327 | 107618664 | 11 | 3 | 6 | 3 | 6 | 69 | 28 | 41 | 50 | 145 | 20 | 0 | 2 | 4 | 6 | 17 | 30 | 24 | 21 | 3 | 228 | 149 | 0.01793316 | 0.16717403 | 0.094 | 0.065 | 0.087 | 0.038 | 0.026 | 0.435 | 0 | 0.052 | 0.111 | 0.567 | -0.171 |
| SE | 57162 | ENSG00000011021 | CLCN6 | chr1 | + | 11807130 | 11807190 | 11806095 | 11806349 | 11815845 | 11815911 | 28 | 22 | 31 | 30 | 50 | 0 | 0 | 0 | 0 | 0 | 62 | 32 | 26 | 48 | 138 | 0 | 0 | 0 | 0 | 12 | 209 | 149 | 0.017964146 | 0.167440166 | 1 | 1 | 1 | 1 | 1 | 1 | 1 | 1 | 1 | 0.891 | 0.022 |
| SE | 107883 | ENSG00000188906 | LRRK2 | chr12 | + | 40259479 | 40259604 | 40257247 | 40257377 | 40263788 | 40263901 | 151 | 183 | 99 | 125 | 223 | 0 | 0 | 0 | 0 | 0 | 32 | 127 | 95 | 31 | 4 | 11 | 0 | 0 | 0 | 0 | 274 | 149 | 0.017965986 | 0.167440166 | 1 | 1 | 1 | 1 | 1 | 0.613 | 1 | 1 | 1 | 1 | 0.077 |
| SE | 94782 | ENSG00000119314 | PTBP3 | chr9 | - | 112275843 | 112276013 | 112268048 | 112268195 | 112297831 | 112297916 | 793 | 488 | 599 | 663 | 1193 | 37 | 20 | 20 | 30 | 49 | 265 | 570 | 626 | 421 | 319 | 8 | 19 | 17 | 19 | 2 | 298 | 149 | 0.017971697 | 0.167459289 | 0.915 | 0.924 | 0.937 | 0.917 | 0.924 | 0.943 | 0.937 | 0.948 | 0.917 | 0.988 | -0.023 |
| SE | 69802 | ENSG00000184007 | PTP4A2 | chr1 | - | 31915894 | 31915987 | 31911695 | 31911826 | 31918969 | 31919109 | 3627 | 2599 | 2501 | 3219 | 4400 | 174 | 120 | 76 | 164 | 234 | 7898 | 3799 | 3773 | 5134 | 3827 | 483 | 214 | 155 | 212 | 316 | 242 | 149 | 0.017974416 | 0.167459289 | 0.928 | 0.93 | 0.953 | 0.924 | 0.92 | 0.91 | 0.916 | 0.937 | 0.937 | 0.882 | 0.015 |
| SE | 86897 | ENSG00000022267 | FHL1 | chrX | + | 136208454 | 136208641 | 136207791 | 136207961 | 136209870 | 136209954 | 78 | 57 | 47 | 168 | 72 | 0 | 0 | 0 | 0 | 0 | 92 | 98 | 43 | 23 | 1776 | 0 | 0 | 3 | 0 | 23 | 298 | 149 | 0.017974454 | 0.167459289 | 1 | 1 | 1 | 1 | 1 | 1 | 1 | 0.878 | 1 | 0.975 | 0.029 |
| SE | 13146 | ENSG00000103121 | CMC2 | chr16 | - | 80980769 | 80980879 | 80975801 | 80976179 | 80981805 | 80981877 | 24 | 17 | 26 | 34 | 36 | 70 | 45 | 42 | 73 | 88 | 23 | 73 | 52 | 58 | 157 | 72 | 117 | 54 | 123 | 139 | 259 | 149 | 0.017982089 | 0.16751049 | 0.165 | 0.179 | 0.263 | 0.211 | 0.191 | 0.155 | 0.264 | 0.356 | 0.213 | 0.394 | -0.075 |
| SE | 37747 | ENSG00000186469 | GNG2 | chr14 | + | 51877616 | 51877657 | 51860641 | 51860790 | 51950649 | 51950765 | 652 | 199 | 304 | 377 | 274 | 20 | 3 | 4 | 6 | 9 | 87 | 283 | 314 | 189 | 84 | 0 | 7 | 3 | 3 | 0 | 190 | 149 | 0.01798685 | 0.167534906 | 0.962 | 0.981 | 0.983 | 0.98 | 0.96 | 1 | 0.969 | 0.988 | 0.98 | 1 | -0.014 |
| SE | 318 | ENSG00000182670 | TTC3 | chr21 | + | 37150819 | 37150884 | 37150077 | 37150170 | 37151892 | 37151958 | 354 | 377 | 418 | 344 | 844 | 8 | 12 | 8 | 14 | 23 | 73 | 190 | 249 | 45 | 49 | 0 | 0 | 6 | 0 | 5 | 214 | 149 | 0.017990623 | 0.167550116 | 0.969 | 0.956 | 0.973 | 0.945 | 0.962 | 1 | 1 | 0.967 | 1 | 0.872 | -0.007 |
| SE | 55124 | ENSG00000119725 | ZNF410 | chr14 | + | 73892026 | 73892208 | 73886616 | 73886915 | 73893796 | 73893932 | 201 | 95 | 159 | 145 | 214 | 3 | 1 | 0 | 0 | 0 | 261 | 213 | 143 | 186 | 157 | 6 | 0 | 6 | 4 | 0 | 298 | 149 | 0.018019543 | 0.16779949 | 0.971 | 0.979 | 1 | 1 | 1 | 0.956 | 1 | 0.923 | 0.959 | 1 | 0.022 |
| SE | 34053 | ENSG00000114867 | EIF4G1 | chr3 | + | 184315762 | 184315856 | 184315165 | 184315545 | 184317716 | 184317773 | 50 | 24 | 28 | 28 | 29 | 0 | 0 | 0 | 0 | 0 | 21 | 31 | 27 | 43 | 70 | 0 | 1 | 0 | 9 | 0 | 243 | 149 | 0.018022325 | 0.167805443 | 1 | 1 | 1 | 1 | 1 | 1 | 0.95 | 1 | 0.746 | 1 | 0.061 |
| SE | 6286 | ENSG00000141644 | MBD1 | chr18 | - | 50271468 | 50271540 | 50269556 | 50269811 | 50272676 | 50272738 | 49 | 28 | 35 | 40 | 53 | 4 | 8 | 12 | 12 | 14 | 67 | 29 | 60 | 41 | 64 | 3 | 8 | 3 | 0 | 22 | 221 | 149 | 0.018048491 | 0.168029093 | 0.892 | 0.702 | 0.663 | 0.692 | 0.718 | 0.938 | 0.71 | 0.931 | 1 | 0.662 | -0.115 |
| SE | 20946 | ENSG00000135596 | MICAL1 | chr6 | - | 109451599 | 109451700 | 109446695 | 109446772 | 109452245 | 109452401 | 42 | 35 | 42 | 40 | 45 | 2 | 0 | 0 | 1 | 0 | 63 | 37 | 23 | 119 | 128 | 0 | 0 | 0 | 0 | 12 | 250 | 149 | 0.018061635 | 0.168127115 | 0.926 | 1 | 1 | 0.96 | 1 | 1 | 1 | 1 | 1 | 0.864 | 0.004 |
| SE | 38010 | ENSG00000197386 | HTT | chr4 | + | 3154292 | 3154419 | 3148004 | 3148207 | 3157071 | 3157199 | 152 | 96 | 136 | 149 | 183 | 2 | 0 | 0 | 0 | 2 | 303 | 188 | 113 | 178 | 316 | 10 | 0 | 2 | 0 | 6 | 276 | 149 | 0.018065057 | 0.168127115 | 0.976 | 1 | 1 | 1 | 0.98 | 0.942 | 1 | 0.968 | 1 | 0.966 | 0.016 |
| SE | 102208 | ENSG00000179988 | PSTK | chr10 | + | 122982732 | 122983024 | 122980039 | 122980695 | 122983271 | 122983470 | 18 | 13 | 13 | 14 | 21 | 1 | 0 | 0 | 1 | 6 | 13 | 15 | 10 | 13 | 7 | 0 | 0 | 0 | 0 | 0 | 298 | 149 | 0.018065462 | 0.168127115 | 0.9 | 1 | 1 | 0.875 | 0.636 | 1 | 1 | 1 | 1 | 1 | -0.118 |
| SE | 58081 | ENSG00000223804 | NA | chr1 | - | 120340755 | 120340810 | 120267333 | 120267838 | 120341716 | 120341839 | 13 | 4 | 1 | 4 | 9 | 37 | 25 | 13 | 27 | 46 | 10 | 7 | 1 | 0 | 0 | 27 | 17 | 31 | 40 | 91 | 204 | 149 | 0.018072283 | 0.168170605 | 0.204 | 0.105 | 0.053 | 0.098 | 0.125 | 0.213 | 0.231 | 0.023 | 0 | 0 | 0.024 |
| SE | 4477 | ENSG00000127527 | EPS15L1 | chr19 | - | 16377121 | 16377254 | 16361578 | 16361984 | 16385128 | 16385211 | 30 | 19 | 16 | 28 | 42 | 8 | 2 | 3 | 3 | 5 | 60 | 53 | 39 | 46 | 109 | 18 | 12 | 13 | 19 | 21 | 282 | 149 | 0.018081373 | 0.168235201 | 0.665 | 0.834 | 0.738 | 0.831 | 0.816 | 0.638 | 0.7 | 0.613 | 0.561 | 0.733 | 0.128 |
| SE | 13492 | ENSG00000135698 | MPHOSPH6 | chr16 | - | 82149308 | 82149403 | 82148161 | 82148863 | 82151423 | 82151514 | 118 | 96 | 106 | 123 | 224 | 2 | 3 | 3 | 6 | 1 | 31 | 69 | 56 | 35 | 13 | 0 | 0 | 0 | 2 | 0 | 244 | 149 | 0.018087884 | 0.168258728 | 0.973 | 0.951 | 0.956 | 0.926 | 0.993 | 1 | 1 | 1 | 0.914 | 1 | -0.023 |
| SE | 85768 | ENSG00000075539 | FRYL | chr4 | - | 48684672 | 48684795 | 48634290 | 48634490 | 48710518 | 48710698 | 251 | 231 | 236 | 247 | 174 | 15 | 16 | 5 | 6 | 5 | 43 | 145 | 88 | 51 | 48 | 0 | 7 | 3 | 0 | 0 | 272 | 149 | 0.0180882 | 0.168258728 | 0.902 | 0.888 | 0.963 | 0.958 | 0.95 | 1 | 0.919 | 0.941 | 1 | 1 | -0.04 |
| SE | 80832 | ENSG00000151948 | GLT1D1 | chr12 | + | 128926358 | 128926454 | 128914899 | 128914989 | 128927104 | 128927154 | 96 | 98 | 42 | 90 | 93 | 1 | 0 | 3 | 3 | 2 | 469 | 176 | 151 | 387 | 325 | 11 | 1 | 14 | 21 | 3 | 245 | 149 | 0.018117353 | 0.168509893 | 0.983 | 1 | 0.895 | 0.948 | 0.966 | 0.963 | 0.991 | 0.868 | 0.918 | 0.985 | 0.013 |
| SE | 70409 | ENSG00000115652 | UXS1 | chr2 | - | 106098715 | 106098773 | 106096717 | 106096821 | 106101057 | 106101118 | 102 | 101 | 98 | 148 | 168 | 0 | 0 | 0 | 3 | 0 | 281 | 134 | 104 | 179 | 735 | 3 | 5 | 5 | 11 | 0 | 207 | 149 | 0.018125185 | 0.168562713 | 1 | 1 | 1 | 0.973 | 1 | 0.985 | 0.951 | 0.937 | 0.921 | 1 | 0.036 |
| SE | 26506 | ENSG00000140471 | LINS1 | chr15 | - | 100586223 | 100586332 | 100580443 | 100580945 | 100602116 | 100602157 | 11 | 4 | 9 | 6 | 17 | 9 | 2 | 9 | 12 | 11 | 9 | 5 | 2 | 0 | 4 | 0 | 1 | 0 | 18 | 0 | 258 | 149 | 0.018158883 | 0.168856044 | 0.414 | 0.536 | 0.366 | 0.224 | 0.472 | 1 | 0.743 | 1 | 0 | 1 | -0.346 |
| SE | 56361 | ENSG00000142875 | PRKACB | chr1 | + | 84184036 | 84184135 | 84182199 | 84182328 | 84185099 | 84185182 | 1255 | 826 | 1010 | 1210 | 1867 | 10 | 11 | 5 | 5 | 7 | 215 | 408 | 153 | 112 | 256 | 0 | 0 | 0 | 0 | 20 | 248 | 149 | 0.018175094 | 0.168986716 | 0.987 | 0.978 | 0.992 | 0.993 | 0.994 | 1 | 1 | 1 | 1 | 0.885 | 0.012 |
| SE | 101041 | ENSG00000118276 | B4GALT6 | chr18 | - | 31645354 | 31645479 | 31638643 | 31638760 | 31657975 | 31658089 | 61 | 32 | 28 | 44 | 43 | 1 | 0 | 0 | 2 | 3 | 50 | 10 | 10 | 20 | 38 | 13 | 0 | 0 | 0 | 0 | 274 | 149 | 0.018192374 | 0.169112257 | 0.971 | 1 | 1 | 0.923 | 0.886 | 0.677 | 1 | 1 | 1 | 1 | 0.021 |
| SE | 24651 | ENSG00000105223 | PLD3 | chr19 | + | 40365717 | 40365930 | 40348684 | 40348768 | 40366418 | 40366464 | 27 | 20 | 17 | 28 | 17 | 2 | 0 | 1 | 5 | 0 | 71 | 63 | 23 | 33 | 51 | 0 | 1 | 0 | 1 | 0 | 298 | 149 | 0.018192916 | 0.169112257 | 0.871 | 1 | 0.895 | 0.737 | 1 | 1 | 0.969 | 1 | 0.943 | 1 | -0.082 |
| [illegible] | | | | | | | | | | | | | | | | | | | | | | | | | | | | | | | | | | | | | | | | | | | | | | |

| | | | | | | | | | | | | | |
|---|---|---|---|---|---|---|---|---|---|---|---|---|---|
| SE | 1784 | ENSG00000100077 | GRK3 | chr22 | + | 25704108 | 25704209 | 25703509 | 25703576 | 25709897 | 25709964 | 195 | [illegible] |
| SE | 98705 | ENSG00000140299 | BNIP2 | chr15 | - | 59669275 | 59669362 | 59668754 | 59668990 | 59671182 | 59671314 | 586 | [illegible] |
| SE | 30706 | ENSG00000196954 | CASP4 | chr11 | - | 104951895 | 104952005 | 104949401 | 104949777 | 104954746 | 104955001 | 209 | [illegible] |
| SE | 76776 | ENSG00000135317 | SNX14 | chr6 | - | 85514069 | 85514234 | 85507967 | 85513895 | 85514505 | 85514629 | 462 | [illegible] |
| SE | 33672 | ENSG00000176261 | ZBTB8OS | chr1 | - | 32633644 | 32633727 | 32631824 | 32631879 | 32650432 | 32650525 | 47 | [illegible] |
| SE | 62319 | ENSG00000185811 | IKZF1 | chr7 | + | 50376532 | 50376793 | 50327378 | 50327757 | 50382539 | 50382707 | 608 | [illegible] |
| SE | 54176 | ENSG00000164300 | SERINC5 | chr5 | - | 80169334 | 80169546 | 80166382 | 80166478 | 80174953 | 80175047 | 158 | [illegible] |
| SE | 79764 | ENSG00000164808 | SPIDR | chr8 | + | 47701955 | 47702015 | 47701720 | 47701864 | 47712661 | 47712872 | 77 | [illegible] |
| SE | 45696 | ENSG00000049541 | RFC2 | chr7 | - | 74246661 | 74246763 | 74243145 | 74243246 | 74249011 | 74249118 | 90 | [illegible] |
| SE | 53078 | ENSG00000103657 | HERC1 | chr15 | - | 63746917 | 63747083 | 63734642 | 63734849 | 63747723 | 63747858 | 213 | [illegible] |
| SE | 60671 | ENSG00000147099 | HDAC8 | chrX | - | 72568753 | 72568884 | 72567581 | 72568030 | 72572056 | 72572104 | 85 | [illegible] |
| SE | 43809 | ENSG00000286053 | ASDURF | chr2 | + | 189665375 | 189665451 | 189663900 | 189663954 | 189666040 | 189666401 | 265 | [illegible] |
| SE | 60074 | ENSG00000149532 | CPSF7 | chr11 | - | 61421053 | 61421127 | 61420469 | 61420573 | 61421389 | 61421608 | 9 | [illegible] |
| SE | 74806 | ENSG00000042088 | TDP1 | chr14 | + | 89956577 | 89956800 | 89955901 | 89955970 | 89963107 | 89963552 | 32 | [illegible] |
| SE | 89484 | ENSG00000064313 | TAF2 | chr8 | - | 119796989 | 119797103 | 119795531 | 119795631 | 119797661 | 119797846 | 164 | [illegible] |
| SE | 69521 | ENSG00000177479 | ARIH2 | chr3 | + | 48922348 | 48922468 | 48918856 | 48918998 | 48922747 | 48922811 | 0 | [illegible] |
| SE | 23798 | ENSG00000153071 | DAB2 | chr5 | - | 39382617 | 39383271 | 39376649 | 39377282 | 39388798 | 39388852 | 6 | [illegible] |
| SE | 26342 | ENSG00000048028 | USP28 | chr11 | - | 113817657 | 113817837 | 113815173 | 113815382 | 113823604 | 113823700 | 112 | [illegible] |
| SE | 49416 | ENSG00000107551 | RASSF4 | chr10 | + | 44984021 | 44984113 | 44970164 | 44971848 | 44984812 | 44984970 | 54 | [illegible] |
| SE | 98207 | ENSG00000149273 | RPS3 | chr11 | + | 75402351 | 75402446 | 75401639 | 75401733 | 75404019 | 75404156 | 6863 | [illegible] |
| SE | 43318 | ENSG00000135597 | REPS1 | chr6 | - | 138907494 | 138907600 | 138903492 | 138905132 | 138908667 | 138908816 | 215 | [illegible] |
| SE | 25549 | ENSG00000157916 | RER1 | chr1 | + | 2395607 | 2395648 | 2391842 | 2391958 | 2395783 | 2395871 | 0 | [illegible] |
| SE | 109279 | ENSG00000085832 | EPS15 | chr1 | - | 51399031 | 51399165 | 51394380 | 51394447 | 51400917 | 51400953 | 535 | [illegible] |
| SE | 21876 | ENSG00000189091 | SF3B3 | chr16 | + | 70550649 | 70550678 | 70548369 | 70548442 | 70554445 | 70554597 | 18 | [illegible] |
| SE | 105693 | ENSG00000028203 | VEZT | chr12 | + | 95256549 | 95256622 | 95251939 | 95252071 | 95257149 | 95257181 | 55 | [illegible] |
| SE | 25856 | ENSG00000169118 | CSNK1G1 | chr15 | - | 64188374 | 64188485 | 64180347 | 64180454 | 64203081 | 64203189 | 8 | [illegible] |
| SE | 48528 | ENSG00000111481 | COPZ1 | chr12 | + | 54343224 | 54343316 | 54342205 | 54342287 | 54345459 | 54345475 | 822 | [illegible] |
| SE | 28919 | ENSG00000145495 | MARCHF6 | chr5 | + | 10417269 | 10417410 | 10415487 | 10415669 | 10423734 | 10423835 | 201 | [illegible] |
| SE | 32146 | ENSG00000198964 | SGMS1 | chr10 | - | 50466889 | 50466932 | 50460672 | 50460814 | 50519830 | 50519921 | 90 | [illegible] |
| SE | 49082 | ENSG00000188739 | RBM34 | chr1 | - | 235160510 | 235160647 | 235154880 | 235155112 | 235160892 | 235161067 | 285 | [illegible] |
| SE | 50921 | ENSG00000000460 | C1orf112 | chr1 | + | 169823407 | 169823472 | 169821678 | 169821759 | 169827050 | 169827199 | 8 | [illegible] |
| SE | 93086 | ENSG00000113924 | HGD | chr3 | - | 120638454 | 120638581 | 120633333 | 120633590 | 120641588 | 120641693 | 15 | [illegible] |
| [illegible] | | | | | | | | | | | | | |

| | | | | | | | | | |
|---|---|---|---|---|---|---|---|---|---|
| SE | 106554 | ENSG00000134532 | SOX5 | chr12 | - | 24213342 | 24213417 | 23895792 | 23896024 |
| SE | 48225 | ENSG00000007237 | GAS7 | chr17 | - | 9969676 | 9969762 | 9959201 | 9959255 |
| SE | 102702 | ENSG00000180229 | HERC2P3 | chr15 | - | 20476924 | 20477056 | 20466194 | 20466370 |
| SE | 93099 | ENSG00000105879 | CBLL1 | chr7 | + | 107752544 | 107752617 | 107748879 | 107749047 |
| SE | 7550 | ENSG00000167851 | CD300A | chr17 | + | 74477435 | 74477530 | 74474531 | 74474685 |
| SE | 88696 | ENSG00000183077 | AFMID | chr17 | + | 78204655 | 78204741 | 78202702 | 78202751 |
| SE | 41065 | ENSG00000160741 | CRTC2 | chr1 | - | 153953265 | 153953369 | 153952804 | 153952834 |
| SE | 38805 | ENSG00000072422 | RHOBTB1 | chr10 | - | 60910886 | 60910990 | 60892809 | 60892995 |
| SE | 75054 | ENSG00000198223 | CSF2RA | chrX | + | 1274754 | 1274818 | 1268813 | 1268879 |
| SE | 55676 | ENSG00000272886 | DCP1A | chr3 | - | 53342143 | 53342271 | 53319406 | 53319473 |
| SE | 7477 | ENSG00000073605 | GSDMB | chr17 | - | 39908957 | 39909042 | 39906110 | 39906271 |
| SE | 14230 | ENSG00000142192 | APP | chr21 | - | 25997359 | 25997416 | 25982343 | 25982477 |
| SE | 79021 | ENSG00000160218 | TRAPPC10 | chr21 | + | 44063052 | 44063168 | 44059102 | 44059214 |
| SE | 92505 | ENSG00000216937 | CCDC7 | chr10 | + | 32517944 | 32517975 | 32473966 | 32474023 |
| SE | 52761 | ENSG00000136682 | CBWD2 | chr2 | + | 113495119 | 113495249 | 113493928 | 113493994 |
| SE | 28042 | ENSG00000049656 | CLPTM1L | chr5 | - | 1330279 | 1330383 | 1325750 | 1325816 |
| SE | 67840 | ENSG00000122025 | FLT3 | chr13 | - | 28025283 | 28025415 | 28024860 | 28024943 |
| SE | 2111 | ENSG00000127616 | SMARCA4 | chr19 | + | 11033766 | 11033865 | 11033081 | 11033517 |
| SE | 65516 | ENSG00000140386 | SCAPER | chr15 | - | 76841733 | 76841931 | 76804532 | 76804633 |
| SE | 101558 | ENSG00000088298 | EDEM2 | chr20 | - | 35134737 | 35134949 | 35131641 | 35131783 |
| SE | 75964 | ENSG00000118873 | RAB3GAP2 | chr1 | - | 220170891 | 220171120 | 220167501 | 220167675 |
| SE | 81829 | ENSG00000187953 | PMS2CL | chr7 | + | 6740173 | 6740324 | 6737634 | 6737936 |
| SE | 46192 | ENSG00000185158 | LRRC37B | chr17 | + | 32027768 | 32027840 | 32024710 | 32024782 |
| SE | 8427 | ENSG00000078687 | TNRC6C | chr17 | + | 78077184 | 78077334 | 78075135 | 78075278 |
| SE | 53287 | ENSG00000066654 | THUMPD1 | chr16 | - | 20738087 | 20738269 | 20737707 | 20737956 |
| SE | 20515 | ENSG00000175061 | SNHG29 | chr17 | + | 16439326 | 16439703 | 16439036 | 16439060 |
| [illegible] | | | | | | | | | |

| | | | | | | | | | | | | | | | | | | | | | | | | | | | | | | | | | | | | | | | | | | | | | | |
|---|---|---|---|---|---|---|---|---|---|---|---|---|---|---|---|---|---|---|---|---|---|---|---|---|---|---|---|---|---|---|---|---|---|---|---|---|---|---|---|---|---|---|---|---|---|---|
| SE | 68998 | ENSG00000147454 | SLC25A37 | chr8 | + | 23543087 | 23543211 | 23529059 | 23529212 | 23566107 | 23566378 | 0 | 0 | 8 | 2 | 0 | 47 | 26 | 52 | 45 | 38 | 0 | 0 | 28 | 10 | 7 | 118 | 78 | 178 | 182 | 56 | 273 | 149 | 0.026610566 | 0.229182802 | 0 | 0 | 0.077 | 0.024 | 0 | 0 | 0 | 0.079 | 0.029 | 0.064 | -0.014 |
| SE | 102465 | ENSG00000108100 | CCNY | chr10 | + | 35516522 | 35516623 | 35501500 | 35501535 | 35525963 | 35525999 | 735 | 535 | 538 | 704 | 903 | 18 | 4 | 14 | 5 | 5 | 1856 | 574 | 555 | 1064 | 2108 | 40 | 13 | 27 | 21 | 14 | 250 | 149 | 0.026680497 | 0.229732499 | 0.961 | 0.988 | 0.958 | 0.988 | 0.991 | 0.965 | 0.963 | 0.925 | 0.968 | 0.989 | 0.015 |
| SE | 50040 | ENSG00000154222 | CC2D1B | chr1 | - | 52357000 | 52357126 | 52356383 | 52356442 | 52357525 | 52357698 | 78 | 43 | 109 | 77 | 53 | 1 | 0 | 0 | 0 | 0 | 54 | 48 | 63 | 54 | 65 | 3 | 0 | 1 | 6 | 0 | 275 | 149 | 0.026682863 | 0.229732499 | 0.977 | 1 | 1 | 1 | 1 | 0.907 | 1 | 0.972 | 0.83 | 1 | 0.054 |
| SE | 5939 | ENSG00000240849 | PEDS1 | chr20 | - | 50130855 | 50130947 | 50129545 | 50129690 | 50143501 | 50143621 | 33 | 15 | 43 | 54 | 27 | 0 | 0 | 0 | 0 | 1 | 58 | 49 | 15 | 29 | 183 | 0 | 1 | 5 | 2 | 3 | 241 | 149 | 0.026683194 | 0.229732499 | 1 | 1 | 1 | 1 | 0.943 | 1 | 0.968 | 0.65 | 0.9 | 0.974 | 0.09 |
| SE | 24333 | ENSG00000185404 | SP140L | chr2 | + | 230391714 | 230391807 | 230388558 | 230388633 | 230392086 | 230392229 | 8 | 6 | 4 | 6 | 7 | 14 | 6 | 21 | 8 | 24 | 0 | 28 | 0 | 4 | 0 | 20 | 34 | 34 | 13 | 8 | 242 | 149 | 0.026688545 | 0.229748468 | 0.26 | 0.381 | 0.105 | 0.316 | 0.152 | 0 | 0.336 | 0 | 0.159 | 0 | 0.144 |
| SE | 88733 | ENSG00000172954 | LCLAT1 | chr2 | + | 30525586 | 30525755 | 30459551 | 30459722 | 30533115 | 30533266 | 43 | 29 | 43 | 51 | 64 | 1 | 0 | 0 | 0 | 0 | 33 | 36 | 31 | 19 | 32 | 0 | 0 | 0 | 10 | 0 | 298 | 149 | 0.026690917 | 0.229748468 | 0.956 | 1 | 1 | 1 | 1 | 1 | 1 | 1 | 0.487 | 1 | 0.094 |
| SE | 46091 | ENSG00000166912 | MTMR10 | chr15 | - | 30942889 | 30943072 | 30938940 | 30942072 | 30947129 | 30947300 | 150 | 153 | 130 | 214 | 191 | 0 | 3 | 1 | 1 | 5 | 48 | 137 | 159 | 128 | 58 | 6 | 8 | 2 | 0 | 0 | 298 | 149 | 0.026695872 | 0.229765854 | 1 | 0.962 | 0.985 | 0.991 | 0.95 | 0.8 | 0.895 | 0.975 | 1 | 1 | 0.044 |
| SE | 25500 | ENSG00000134986 | NREP | chr5 | - | 111755769 | 111755926 | 111735429 | 111735507 | 111757135 | 111757179 | 21 | 1 | 20 | 23 | 27 | 1 | 1 | 1 | 2 | 4 | 32 | 16 | 17 | 45 | 64 | 0 | 3 | 0 | 2 | 0 | 298 | 149 | 0.026714087 | 0.229897355 | 0.913 | 0.333 | 0.909 | 0.852 | 0.771 | 1 | 0.727 | 1 | 0.918 | 1 | -0.173 |
| SE | 85243 | ENSG00000147905 | ZCCHC7 | chr9 | + | 37302187 | 37302231 | 37126311 | 37126942 | 37304187 | 37304313 | 276 | 237 | 268 | 186 | 808 | 2 | 6 | 3 | 2 | 17 | 87 | 247 | 153 | 54 | 93 | 0 | 3 | 0 | 2 | 0 | 193 | 149 | 0.026722348 | 0.229943176 | 0.991 | 0.968 | 0.986 | 0.986 | 0.973 | 1 | 0.985 | 1 | 0.954 | 1 | -0.007 |
| SE | 103708 | ENSG00000139517 | LNX2 | chr13 | - | 27562412 | 27562781 | 27556235 | 27556413 | 27567639 | 27567839 | 17 | 13 | 28 | 20 | 30 | 0 | 0 | 2 | 1 | 0 | 42 | 36 | 9 | 15 | 6 | 0 | 2 | 3 | 3 | 0 | 298 | 149 | 0.026729245 | 0.229977245 | 1 | 1 | 0.875 | 0.909 | 1 | 1 | 0.9 | 0.6 | 0.714 | 1 | 0.114 |
| SE | 84406 | ENSG00000188917 | TRMT2B | chrX | - | 101037916 | 101038051 | 101036973 | 101037073 | 101042232 | 101042312 | 40 | 19 | 30 | 35 | 70 | 0 | 0 | 0 | 0 | 0 | 15 | 50 | 25 | 20 | 49 | 11 | 0 | 0 | 0 | 0 | 284 | 149 | 0.026777664 | 0.230368533 | 1 | 1 | 1 | 1 | 1 | 0.417 | 1 | 1 | 1 | 1 | 0.117 |
| SE | 105473 | ENSG00000111145 | ELK3 | chr12 | + | 96198025 | 96198142 | 96194401 | 96194705 | 96223564 | 96223682 | 18 | 9 | 17 | 14 | 21 | 105 | 87 | 113 | 110 | 154 | 19 | 30 | 24 | 10 | 4 | 57 | 85 | 78 | 56 | 137 | 266 | 149 | 0.026786012 | 0.230415028 | 0.088 | 0.055 | 0.078 | 0.067 | 0.071 | 0.157 | 0.165 | 0.147 | 0.091 | 0.016 | -0.043 |
| SE | 18219 | ENSG00000119686 | FLVCR2 | chr14 | + | 75622078 | 75622220 | 75578619 | 75579641 | 75640954 | 75641060 | 8 | 6 | 8 | 5 | 3 | 2 | 0 | 0 | 5 | 0 | 9 | 10 | 4 | 7 | 0 | 6 | 5 | 3 | 0 | 2 | 291 | 149 | 0.026795023 | 0.230467221 | 0.672 | 1 | 1 | 0.339 | 1 | 0.434 | 0.506 | 0.406 | 1 | 0 | 0.333 |
| SE | 40271 | ENSG00000170581 | STAT2 | chr12 | - | 56350096 | 56350190 | 56349588 | 56349636 | 56350411 | 56350432 | 357 | 190 | 227 | 438 | 237 | 6 | 1 | 0 | 1 | 3 | 1301 | 571 | 227 | 914 | 722 | 2 | 1 | 6 | 17 | 20 | 243 | 149 | 0.026810356 | 0.230573777 | 0.973 | 0.991 | 1 | 0.996 | 0.98 | 0.997 | 0.997 | 0.959 | 0.971 | 0.957 | 0.012 |
| SE | 105787 | ENSG00000124164 | VAPB | chr20 | + | 58434601 | 58434705 | 58418210 | 58418363 | 58440906 | 58441083 | 34 | 41 | 35 | 55 | 56 | 1 | 0 | 2 | 2 | 7 | 52 | 39 | 33 | 41 | 106 | 9 | 6 | 2 | 2 | 8 | 253 | 149 | 0.026828326 | 0.230702974 | 0.952 | 1 | 0.912 | 0.942 | 0.825 | 0.773 | 0.793 | 0.907 | 0.924 | 0.886 | 0.07 |
| SE | 106223 | ENSG00000070214 | SLC44A1 | chr9 | + | 105366345 | 105366429 | 105361190 | 105361330 | 105374597 | 105374735 | 58 | 46 | 46 | 56 | 105 | 0 | 0 | 0 | 0 | 0 | 131 | 61 | 58 | 78 | 169 | 0 | 0 | 5 | 2 | 6 | 233 | 149 | 0.026842382 | 0.230798499 | 1 | 1 | 1 | 1 | 1 | 1 | 1 | 0.881 | 0.961 | 0.947 | 0.042 |
| SE | 67807 | ENSG00000055044 | NOP58 | chr2 | + | 202282350 | 202282472 | 202275112 | 202275189 | 202284344 | 202284378 | 25 | 8 | 29 | 26 | 29 | 0 | 0 | 1 | 4 | 3 | 23 | 13 | 43 | 23 | 6 | 0 | 0 | 0 | 0 | 0 | 271 | 149 | 0.026848244 | 0.230808802 | 1 | 1 | 0.941 | 0.781 | 0.842 | 1 | 1 | 1 | 1 | 1 | -0.087 |
| SE | 19903 | ENSG00000214900 | LINC01588 | chr14 | - | 50047466 | 50047589 | 50039391 | 50039469 | 50052773 | 50052904 | 15 | 4 | 8 | 16 | 24 | 4 | 3 | 3 | 7 | 5 | 27 | 9 | 0 | 11 | 19 | 0 | 0 | 8 | 2 | 0 | 272 | 149 | 0.026849476 | 0.230808802 | 0.673 | 0.422 | 0.594 | 0.556 | 0.724 | 1 | 1 | 0 | 0.751 | 1 | -0.156 |
| SE | 102097 | ENSG00000155629 | PIK3AP1 | chr10 | - | 96602279 | 96602398 | 96593314 | 96595634 | 96603978 | 96604049 | 312 | 210 | 158 | 242 | 275 | 8 | 3 | 3 | 10 | 2 | 596 | 357 | 163 | 543 | 153 | 12 | 24 | 9 | 5 | 10 | 268 | 149 | 0.026857388 | 0.230851472 | 0.956 | 0.975 | 0.967 | 0.931 | 0.987 | 0.965 | 0.892 | 0.91 | 0.984 | 0.895 | 0.034 |
| SE | 63353 | ENSG00000151092 | NGLY1 | chr3 | - | 25737333 | 25737455 | 25736003 | 25736149 | 25739576 | 25739799 | 208 | 158 | 162 | 219 | 239 | 3 | 3 | 0 | 0 | 0 | 63 | 141 | 113 | 134 | 15 | 0 | 10 | 2 | 0 | 0 | 271 | 149 | 0.026862615 | 0.230871054 | 0.974 | 0.967 | 1 | 1 | 1 | 1 | 0.886 | 0.969 | 1 | 1 | 0.017 |
| SE | 98873 | ENSG00000175294 | CATSPER1 | chr11 | - | 66020563 | 66020627 | 66020316 | 66020389 | 66020810 | 66020954 | 22 | 5 | 2 | 2 | 2 | 8 | 10 | 3 | 8 | 0 | 11 | 8 | 9 | 2 | 31 | 6 | 5 | 1 | 0 | 5 | 213 | 149 | 0.026868393 | 0.230895366 | 0.658 | 0.259 | 0.318 | 0.149 | 1 | 0.562 | 0.528 | 0.863 | 1 | 0.813 | -0.276 |
| SE | 6045 | ENSG00000196839 | ADA | chr20 | - | 44623006 | 44623078 | 44622828 | 44622930 | 44624201 | 44624329 | 63 | 29 | 32 | 61 | 39 | 2 | 0 | 5 | 3 | 0 | 13 | 56 | 34 | 14 | 40 | 0 | 0 | 2 | 1 | 0 | 221 | 149 | 0.026881635 | 0.230975456 | 0.955 | 1 | 0.812 | 0.932 | 1 | 1 | 1 | 0.92 | 0.904 | 1 | -0.025 |
| SE | 35920 | ENSG00000131323 | TRAF3 | chr14 | + | 102889559 | 102889634 | 102876357 | 102876525 | 102891324 | 102891417 | 19 | 20 | 36 | 30 | 19 | 2 | 0 | 3 | 1 | 6 | 28 | 23 | 19 | 36 | 37 | 0 | 2 | 12 | 10 | 11 | 224 | 149 | 0.026883613 | 0.230975456 | 0.863 | 1 | 0.889 | 0.952 | 0.678 | 1 | 0.884 | 0.513 | 0.705 | 0.691 | 0.118 |
| SE | 13996 | ENSG00000047578 | KATNIP | chr16 | + | 27699533 | 27699599 | 27698327 | 27698500 | 27701588 | 27701695 | 13 | 13 | 23 | 19 | 16 | 0 | 2 | 4 | 2 | 0 | 12 | 22 | 18 | 17 | 71 | 0 | 12 | 14 | 5 | 0 | 215 | 149 | 0.026906397 | 0.231145846 | 1 | 0.818 | 0.799 | 0.868 | 1 | 1 | 0.56 | 0.471 | 0.702 | 1 | 0.15 |
| SE | 11880 | ENSG00000103222 | ABCC1 | chr16 | + | 16009775 | 16009901 | 16007815 | 16007992 | 16014490 | 16014628 | 203 | 76 | 178 | 213 | 239 | 0 | 1 | 1 | 0 | 3 | 282 | 256 | 255 | 199 | 518 | 0 | 9 | 0 | 1 | 11 | 275 | 149 | 0.026918215 | 0.231222001 | 1 | 0.976 | 0.99 | 1 | 0.977 | 1 | 0.939 | 1 | 0.991 | 0.962 | 0.01 |
| SE | 54599 | ENSG00000151422 | FER | chr5 | + | 108768092 | 108768238 | 108747896 | 108748000 | 108798123 | 108798389 | 18 | 16 | 12 | 19 | 13 | 2 | 0 | 0 | 0 | 1 | 3 | 14 | 3 | 4 | 4 | 0 | 6 | 2 | 0 | 0 | 295 | 149 | 0.026933702 | 0.231329657 | 0.82 | 1 | 1 | 1 | 0.868 | 1 | 0.541 | 0.431 | 1 | 1 | 0.143 |
| SE | 8252 | ENSG00000108312 | UBTF | chr17 | - | 44208618 | 44209001 | 44207863 | 44207911 | 44209351 | 44209422 | 20 | 15 | 20 | 10 | 3 | 148 | 99 | 227 | 200 | 250 | 3 | 6 | 3 | 5 | 18 | 166 | 146 | 70 | 139 | 199 | 298 | 149 | 0.026940556 | 0.231363144 | 0.063 | 0.07 | 0.042 | 0.024 | 0.006 | 0.009 | 0.02 | 0.021 | 0.018 | 0.043 | 0.019 |
| SE | 61944 | ENSG00000136932 | TRMO | chr9 | - | 97916163 | 97916338 | 97913251 | 97913558 | 97922417 | 97922487 | 68 | 50 | 93 | 74 | 82 | 5 | 4 | 1 | 15 | 9 | 12 | 35 | 34 | 16 | 6 | 0 | 13 | 34 | 20 | 0 | 298 | 149 | 0.026961053 | 0.231513779 | 0.872 | 0.862 | 0.979 | 0.712 | 0.82 | 1 | 0.574 | 0.333 | 0.286 | 1 | 0.21 |
| SE | 28416 | ENSG00000174652 | ZNF266 | chr19 | - | 9415653 | 9415742 | 9412428 | 9414720 | 9418504 | 9418631 | 126 | 74 | 70 | 113 | 127 | 0 | 0 | 0 | 0 | 0 | 13 | 98 | 61 | 84 | 18 | 12 | 0 | 0 | 0 | 0 | 238 | 149 | 0.026985078 | 0.231694666 | 1 | 1 | 1 | 1 | 1 | 0.404 | 1 | 1 | 1 | 1 | 0.119 |
| SE | 53839 | ENSG00000173575 | CHD2 | chr15 | + | 92998148 | 92998261 | 92997252 | 92997403 | 92998498 | 92998621 | 94 | 57 | 74 | 59 | 106 | 219 | 194 | 233 | 277 | 398 | 70 | 43 | 72 | 42 | 44 | 160 | 270 | 372 | 287 | 135 | 262 | 149 | 0.026989305 | 0.231705553 | 0.196 | 0.143 | 0.153 | 0.108 | 0.132 | 0.199 | 0.083 | 0.099 | 0.077 | 0.156 | 0.024 |
| SE | 30560 | ENSG00000118503 | TNFAIP3 | chr6 | + | 137880070 | 137880252 | 137878431 | 137879351 | 137881034 | 137881160 | 987 | 544 | 1307 | 1248 | 2308 | 23 | 18 | 14 | 33 | 15 | 1120 | 1649 | 2763 | 1480 | 1114 | 6 | 48 | 24 | 51 | 49 | 298 | 149 | 0.026993589 | 0.231716919 | 0.955 | 0.938 | 0.979 | 0.95 | 0.987 | 0.989 | 0.945 | 0.983 | 0.936 | 0.919 | 0.007 |
| SE | 51671 | ENSG00000136643 | RPS6KC1 | chr1 | + | 213125765 | 213125843 | 213117316 | 213117410 | 213129526 | 213129889 | 13 | 13 | 4 | 15 | 17 | 84 | 67 | 67 | 73 | 69 | 13 | 4 | 10 | 6 | 0 | 74 | 82 | 82 | 35 | 35 | 227 | 149 | 0.027038836 | 0.232079885 | 0.092 | 0.113 | 0.038 | 0.119 | 0.139 | 0.103 | 0.031 | 0.074 | 0.101 | 0 | 0.038 |
| SE | 38408 | ENSG00000143458 | GABPB2 | chr1 | + | 151117391 | 151117516 | 151107036 | 151107222 | 151117956 | 151118671 | 248 | 236 | 213 | 280 | 342 | 1 | 0 | 0 | 6 | 7 | 445 | 199 | 204 | 336 | 290 | 17 | 4 | 8 | 0 | 0 | 274 | 149 | 0.027044468 | 0.232102777 | 0.993 | 1 | 1 | 0.962 | 0.964 | 0.934 | 0.964 | 0.933 | 1 | 1 | 0.018 |
| SE | 12765 | ENSG00000007541 | PIGQ | chr16 | + | 581198 | 581341 | 580857 | 580972 | 582247 | 582309 | 0 | 0 | 0 | 0 | 0 | 9 | 0 | 5 | 6 | 9 | 1 | 0 | 0 | 10 | 0 | 10 | 1 | 4 | 7 | 4 | 292 | 149 | 0.027076626 | 0.232346044 | 0 | 0 | 0 | 0 | 0 | 0 | 0 | 0 | 0.422 | 0 | -0.084 |
| SE | 100381 | ENSG00000133812 | SBF2 | chr11 | - | 9795830 | 9795957 | 9790555 | 9790683 | 9807999 | 9808185 | 53 | 44 | 46 | 41 | 48 | 3 | 1 | 0 | 2 | 7 | 22 | 83 | 48 | 60 | 8 | 7 | 19 | 3 | 2 | 0 | 276 | 149 | 0.027081406 | 0.232346044 | 0.905 | 0.96 | 1 | 0.917 | 0.787 | 0.629 | 0.702 | 0.896 | 0.942 | 1 | 0.08 |
| SE | 13860 | ENSG00000102893 | PHKB | chr16 | + | 47665941 | 47666032 | 47661718 | 47661800 | 47669214 | 47669417 | 55 | 60 | 66 | 82 | 82 | 1 | 0 | 2 | 4 | 4 | 47 | 59 | 56 | 60 | 35 | 0 | 0 | 2 | 0 | 0 | 240 | 149 | 0.027081716 | 0.232346044 | 0.972 | 1 | 0.953 | 0.927 | 0.927 | 1 | 1 | 0.946 | 1 | 1 | -0.033 |
| SE | 90205 | ENSG00000104324 | CPQ | chr8 | + | 96659283 | 96659323 | 96645374 | 96645402 | 96784863 | 96785330 | 6 | 0 | 0 | 0 | 0 | 57 | 17 | 20 | 55 | 13 | 0 | 0 | 0 | 7 | 11 | 74 | 54 | 29 | 18 | 90 | 189 | 149 | 0.02709011 | 0.23236757 | 0.077 | 0 | 0 | 0 | 0 | 0 | 0 | 0 | 0.235 | 0.088 | -0.049 |
| SE | 12389 | ENSG00000188559 | RALGAPA2 | chr20 | - | 20524405 | 20524543 | 20520916 | 20521100 | 20524829 | 20524898 | 151 | 79 | 110 | 107 | 150 | 1 | 1 | 0 | 1 | 0 | 109 | 126 | 114 | 203 | 78 | 0 | 5 | 7 | 1 | 0 | 287 | 149 | 0.02709016 | 0.23236757 | 0.987 | 0.976 | 1 | 0.982 | 1 | 1 | 0.929 | 0.894 | 0.991 | 1 | 0.026 |
| SE | 47112 | ENSG00000174839 | DENND6A | chr3 | - | 57661445 | 57661551 | 57660759 | 57660839 | 57663635 | 57663716 | 352 | 174 | 278 | 281 | 535 | 0 | 0 | 0 | 0 | 2 | 165 | 325 | 325 | 184 | 109 | 18 | 0 | 0 | 0 | 0 | 255 | 149 | 0.027101668 | 0.232426968 | 1 | 1 | 1 | 1 | 0.994 | 0.843 | 1 | 1 | 1 | 1 | 0.03 |
| SE | 42162 | ENSG00000197343 | ZNF655 | chr7 | + | 99563187 | 99563266 | 99562343 | 99562494 | 99563862 | 99564686 | 1 | 0 | 1 | 0 | 0 | 140 | 67 | 105 | 132 | 206 | 0 | 2 | 12 | 7 | 8 | 144 | 181 | 170 | 159 | 265 | 228 | 149 | 0.027103022 | 0.232426968 | 0.005 | 0 | 0.006 | 0 | 0 | 0 | 0.007 | 0.044 | 0.028 | 0.019 | -0.017 |
| SE | 100995 | ENSG00000100600 | LGMN | chr14 | - | 92706482 | 92706653 | 92703808 | 92704361 | 92709671 | 92709872 | 24 | 8 | 22 | 29 | 6 | 0 | 4 | 0 | 4 | 0 | 26 | 25 | 13 | 7 | 60 | 0 | 0 | 1 | 0 | 2 | 298 | 149 | 0.027114045 | 0.232496033 | 1 | 0.5 | 1 | 0.784 | 1 | 1 | 1 | 0.867 | 1 | 0.938 | -0.104 |
| SE | 64735 | ENSG00000042493 | CAPG | chr2 | - | 85398689 | 85398782 | 85398019 | 85398152 | 85399135 | 85399285 | 553 | 252 | 367 | 365 | 429 | 1 | 0 | 3 | 2 | 7 | 1425 | 678 | 305 | 913 | 1217 | 12 | 10 | 6 | 17 | 6 | 242 | 149 | 0.027121673 | 0.232535976 | 0.997 | 1 | 0.987 | 0.991 | 0.974 | 0.987 | 0.977 | 0.969 | 0.971 | 0.992 | 0.011 |
| SE | 52817 | ENSG00000169764 | UGP2 | chr2 | + | 63857549 | 63857642 | 63856305 | 63856433 | 63857828 | 63857914 | 6 | 3 | 4 | 1 | 14 | 348 | 258 | 303 | 395 | 315 | 5 | 11 | 10 | 3 | 0 | 120 | 249 | 195 | 174 | 71 | 242 | 149 | 0.027197459 | 0.233160218 | 0.011 | 0.007 | 0.008 | 0.002 | 0.027 | 0.025 | 0.026 | 0.031 | 0.011 | 0 | -0.008 |
| SE | 100264 | ENSG00000171940 | ZNF217 | chr20 | - | 53578333 | 53578450 | 53575726 | 53577280 | 53581460 | 53583097 | 498 | 294 | 402 | 372 | 635 | 19 | 16 | 10 | 11 | 9 | 107 | 154 | 277 | 114 | 44 | 0 | 14 | 17 | 0 | 12 | 266 | 149 | 0.027204122 | 0.233191803 | 0.936 | 0.911 | 0.957 | 0.95 | 0.975 | 1 | 0.86 | 0.901 | 1 | 0.673 | 0.059 |
| SE | 30387 | ENSG00000120137 | PANK3 | chr5 | - | 168563888 | 168564065 | 168561392 | 168561516 | 168566012 | 168566266 | 253 | 232 | 211 | 233 | 312 | 3 | 6 | 1 | 8 | 18 | 101 | 176 | 201 | 112 | 65 | 21 | 8 | 10 | 1 | 0 | 298 | 149 | 0.027221101 | 0.233311805 | 0.977 | 0.951 | 0.991 | 0.936 | 0.897 | 0.706 | 0.917 | 0.91 | 0.982 | 1 | 0.047 |
| SE | 87975 | ENSG00000168066 | SF1 | chr11 | - | 64777986 | 64778146 | 64776497 | 64776626 | 64778361 | 64778507 | 5 | 3 | 9 | 11 | 1 | 156 | 62 | 152 | 139 | 104 | 5 | 30 | 4 | 18 | 6 | 244 | 163 | 119 | 173 | 153 | 298 | 149 | 0.027224277 | 0.233313484 | 0.016 | 0.024 | 0.029 | 0.038 | 0.005 | 0.01 | 0.084 | 0.017 | 0.049 | 0.019 | -0.013 |
| SE | 31452 | ENSG00000136261 | BZW2 | chr7 | + | 16685904 | 16686040 | 16682779 | 16682845 | 16689796 | 16689881 | 85 | 53 | 111 | 106 | 186 | 4 | 11 | 9 | 7 | 13 | 80 | 89 | 72 | 89 | 85 | 14 | 0 | 2 | 4 | 0 | 285 | 149 | 0.027279137 | 0.233758052 | 0.917 | 0.716 | 0.866 | 0.888 | 0.882 | 0.749 | 1 | 0.95 | 0.921 | 1 | -0.07 |
| SE | 68734 | ENSG00000196295 | GARS1-DT | chr7 | - | 30584422 | 30584453 | 30579230 | 30579314 | 30584710 | 30584808 | 4 | 2 | 1 | 1 | 3 | 38 | 18 | 45 | 37 | 61 | 8 | 5 | 4 | 0 | 11 | 64 | 51 | 37 | 49 | 73 | 180 | 149 | 0.027298877 | 0.23390161 | 0.08 | 0.084 | 0.018 | 0.022 | 0.039 | 0.094 | 0.075 | 0.082 | 0 | 0.111 | -0.024 |
| SE | 78757 | ENSG00000179406 | LINC00174 | chr7 | - | 66439134 | 66439304 | 66429883 | 66430017 | 66450070 | 66450236 | 18 | 12 | 22 | 27 | 10 | 6 | 9 | 5 | 5 | 16 | 20 | 12 | 10 | 28 | 17 | 11 | 5 | 0 | 3 | 0 | 298 | 149 | 0.027310115 | 0.233972289 | 0.6 | 0.4 | 0.688 | 0.73 | 0.238 | 0.476 | 0.545 | 1 | 0.824 | 1 | -0.238 |
| SE | 65465 | ENSG00000130348 | QRSL1 | chr6 | + | 106640100 | 106640206 | 106629598 | 106629705 | 106640348 | 106640508 | 2 | 2 | 2 | 4 | 2 | 21 | 19 | 29 | 30 | 20 | 4 | 12 | 5 | 10 | 2 | 3 | 31 | 45 | 49 | 13 | 255 | 149 | 0.027333477 | 0.234146821 | 0.1 | 0.058 | 0.039 | 0.072 | 0.055 | 0.438 | 0.184 | 0.061 | 0.107 | 0.082 | -0.11 |
| SE | 39768 | ENSG00000132185 | FCRLA | chr1 | + | 161711207 | 161711474 | 161710759 | 161710912 | 161711933 | 161712218 | 82 | 35 | 44 | 78 | 63 | 1 | 0 | 3 | 4 | 0 | 61 | 89 | 18 | 20 | 173 | 7 | 3 | 0 | 0 | 8 | 298 | 149 | 0.027341276 | 0.234188004 | 0.976 | 1 | 0.88 | 0.907 | 1 | 0.813 | 0.937 | 1 | 1 | 0.915 | 0.02 |
| SE | 49258 | ENSG00000025156 | HSF2 | chr6 | + | 122422149 | 122422298 | 122420134 | 122420222 | 122422717 | 122422957 | 195 | 192 | 242 | 267 | 430 | 3 | 2 | 3 | 6 | 12 | 5 | 75 | 146 | 55 | 7 | 0 | 1 | 4 | 0 | 0 | 298 | 149 | 0.027345796 | 0.234201061 | 0.97 | 0.98 | 0.976 | 0.957 | 0.947 | 1 | 0.974 | 0.948 | 1 | 1 | -0.018 |
| SE | 62575 | ENSG00000107614 | TRDMT1 | chr10 | - | 17168840 | 17168917 | 17162165 | 17162237 | 17174550 | 17174660 | 67 | 35 | 39 | 49 | 177 | 2 | 3 | 6 | 1 | 4 | 8 | 19 | 4 | 8 | 3 | 0 | 0 | 3 | 0 | 0 | 226 | 149 | 0.027348783 | 0.234201061 | 0.957 | 0.885 | 0.811 | 0.97 | 0.967 | 1 | 1 | 0.468 | 1 | 1 | 0.024 |
| SE | 34874 | ENSG00000117597 | UTP25 | chr1 | + | 209843454 | 209843696 | 209842264 | 209842447 | 209851203 | 209857565 | 55 | 11 | 49 | 39 | 89 | 2 | 2 | 2 | 8 | 7 | 3 | 14 | 24 | 12 | 28 | 18 | 0 | 19 | 12 | 0 | 298 | 149 | 0.027362423 | 0.23427816 | 0.932 | 0.733 | 0.925 | 0.709 | 0.864 | 0.077 | 1 | 0.387 | 0.333 | 1 | 0.273 |
| SE | 11094 | ENSG00000135709 | KIAA0513 | chr16 | + | 85082563 | 85082593 | 85081314 | 85081392 | 85086643 | 85086724 | 0 | 0 | 0 | 1 | 0 | 111 | 78 | 76 | 93 | 112 | 10 | 0 | 1 | 4 | 7 | 275 | 84 | 43 | 91 | 253 | 179 | 149 | 0.027363771 | 0.23427816 | 0 | 0 | | 0.009 | 0 | 0.029 | | 0.019 | 0.035 | 0.023 | -0.019 |
| SE | 14979 | ENSG00000182718 | ANXA2 | chr15 | - | 60397257 | 60397338 | 60386027 | 60386086 | 60397942 | 60397964 | 117 | 73 | 68 | 88 | 44 | 298 | 135 | 160 | 185 | 87 | 314 | 192 | 78 | 186 | 292 | 521 | 257 | 158 | 344 | 622 | 230 | 149 | 0.027367218 | 0.234282056 | 0.203 | 0.259 | 0.216 | 0.236 | 0.247 | 0.281 | 0.326 | 0.242 | 0.259 | 0.233 | -0.036 |
| SE | 48352 | ENSG00000157193 | LRP8 | chr1 | - | 53257239 | 53257464 | 53255116 | 53255185 | 53258318 | 53258465 | 19 | 17 | 15 | 16 | 0 | 0 | 3 | 0 | 2 | 2 | 21 | 26 | 29 | 25 | 72 | 0 | 3 | 0 | 0 | 0 | 298 | 149 | 0.027390016 | 0.234431851 | 1 | 0.739 | 1 | 0.8 | 0 | 1 | 0.812 | 1 | 1 | 1 | -0.255 |
| SE | 59642 | ENSG00000163466 | ARPC2 | chr2 | + | 218234848 | 218235044 | 218234351 | 218234397 | 218245419 | 218245546 | 4 | 4 | 2 | 8 | 6 | 1 | 0 | 0 | 5 | 0 | 2 | 7 | 19 | 14 | 15 | 0 | 3 | 4 | 5 | 8 | 298 | 149 | 0.027390704 | 0.234431851 | 0.667 | 1 | 1 | 0.444 | 1 | 1 | 0.538 | 0.704 | 0.583 | 0.484 | 0.16 |
| SE | 21262 | ENSG00000155111 | CDK19 | chr6 | - | 110632029 | 110632161 | 110627001 | 110627145 | 110667433 | 110667574 | 62 | 53 | 71 | 48 | 103 | 2 | 2 | 0 | 0 | 7 | 10 | 20 | 49 | 34 | 16 | 0 | 3 | 0 | 0 | 0 | 281 | 149 | 0.027404438 | 0.234468085 | 0.943 | 0.934 | 1 | 1 | 0.886 | 1 | 0.779 | 1 | 1 | 1 | -0.003 |
| SE | 91197 | ENSG00000116604 | MEF2D | chr1 | - | 156475107 | 156475237 | 156468779 | 156469020 | 156477011 | 156477202 | 77 | 26 | 53 | 63 | 25 | 0 | 0 | 0 | 2 | 2 | 236 | 72 | 109 | 174 | 297 | 0 | 2 | 0 | 7 | 9 | 279 | 149 | 0.027404675 | 0.234468085 | 1 | 1 | 1 | 0.944 | 0.87 | 1 | 0.951 | 1 | 0.93 | 0.946 | -0.003 |
| SE | 51366 | ENSG00000066427 | ATXN3 | chr14 | - | 92093251 | 92093318 | 92088729 | 92088817 | 92096092 | 92096134 | 70 | 66 | 61 | 81 | 115 | 3 | 1 | 0 | 3 | 5 | 11 | 62 | 95 | 61 | 4 | 0 | 2 | 0 | 0 | 0 | 216 | 149 | 0.027405892 | 0.234468085 | 0.942 | 0.979 | 1 | 0.949 | 0.941 | 1 | 0.955 | 1 | 1 | 1 | -0.029 |
| SE | 105270 | ENSG00000082898 | XPO1 | chr2 | - | 61534557 | 61534678 | 61533771 | 61533903 | 61537561 | 61537587 | 8 | 8 | 18 | 5 | 7 | 334 | 259 | 307 | 321 | 240 | 0 | 1 | 1 | 4 | 2 | 55 | 197 | 184 | 153 | 33 | 270 | 149 | 0.027408285 | 0.234468085 | 0.013 | 0.017 | 0.031 | 0.009 | 0.016 | 0 | 0.003 | 0.003 | 0.014 | 0.032 | 0.007 |
| SE | 38281 | ENSG00000134684 | YARS1 | chr1 | - | 32810910 | 32811057 | 32806481 | 32806611 | 32817187 | 32817272 | 83 | 36 | 62 | 59 | 18 | 1 | 1 | 3 | 5 | 0 | 47 | 51 | 55 | 74 | 70 | 3 | 4 | 7 | 3 | 5 | 296 | 149 | 0.027412612 | 0.234468085 | 0.977 | 0.948 | 0.912 | 0.856 | 1 | 0.887 | 0.865 | 0.798 | 0.925 | 0.876 | 0.068 |
| SE | 67947 | ENSG00000091490 | SEL1L3 | chr4 | - | 25835196 | 25835323 | 25833447 | 25833569 | 25847293 | 25847864 | 234 | 184 | 361 | 260 | 319 | 3 | 6 | 9 | 12 | 0 | 120 | 494 | 273 | 167 | 452 | 17 | 24 | 4 | 7 | 2 | 276 | 149 | 0.027412906 | 0.234468085 | 0.977 | 0.943 | 0.956 | 0.921 | 1 | 0.792 | 0.917 | 0.974 | 0.928 | 0.992 | 0.039 |
| SE | 24339 | ENSG00000185404 | SP140L | chr2 | + | 230400954 | 230401063 | 230400126 | 230400242 | 230401365 | 230401443 | 53 | 44 | 70 | 73 | 50 | 0 | 0 | 0 | 0 | 0 | 103 | 47 | 47 | 47 | 36 | 0 | 0 | 5 | 0 | 5 | 258 | 149 | 0.027421947 | 0.234519796 | 1 | 1 | 1 | 1 | 1 | 1 | 1 | 0.844 | 1 | 0.806 | 0.07 |
| SE | 97093 | ENSG00000173327 | MAP3K11 | chr11 | - | 65605592 | 65605642 | 65599450 | 65599768 | 65605760 | 65605852 | 6 | 0 | 1 | 3 | 4 | 72 | 38 | 61 | 51 | 60 | 3 | 3 | 0 | 0 | 0 | 104 | 77 | 50 | 81 | 183 | 199 | 149 | 0.027442366 | 0.234668795 | 0.059 | 0 | 0.012 | 0.042 | 0.048 | 0.021 | 0.028 | 0 | 0 | 0 | 0.022 |
| SE | 30517 | ENSG00000163138 | PACRGL | chr4 | + | 20712787 | 20712922 | 20709682 | 20709773 | 20713431 | 20713516 | 31 | 22 | 21 | 19 | 58 | 0 | 1 | 2 | 0 | 0 | 26 | 14 | 6 | 4 | 2 | 0 | 0 | 2 | 0 | 5 | 284 | 149 | 0.027488339 | 0.235036259 | 1 | 0.92 | 0.846 | 1 | 1 | 1 | 1 | 0.611 | 1 | 0.173 | 0.196 |
| SE | 21352 | ENSG00000056972 | TRAF3IP2 | chr6 | - | 111595702 | 111595820 | 111591544 | 111592094 | 111605775 | 111605868 | 0 | 0 | 0 | 0 | 0 | 12 | 2 | 4 | 11 | 12 | 0 | 0 | 0 | 0 | 18 | 7 | 11 | 13 | 12 | 6 | 267 | 149 | 0.027492071 | 0.235042497 | 0 | 0 | 0 | 0 | 0 | 0 | 0 | 0 | 0 | 0.626 | -0.125 |
| SE | 79151 | ENSG00000104299 | INTS9 | chr8 | - | 28862039 | 28862212 | 28859435 | 28859563 | 28889873 | 28889934 | 2 | 2 | 5 | 14 | 15 | 36 | 11 | 10 | 26 | 36 | 0 | 1 | 4 | 0 | 18 | 14 | 38 | 20 | 31 | 26 | 298 | 149 | 0.027510888 | 0.235177689 | 0.027 | 0.083 | 0.2 | 0.212 | 0.172 | 0 | 0.013 | 0.091 | 0 | 0.257 | 0.067 |

| | | | | | | | | | | | | | | | | | | | | | | | | | |
|---|---|---|---|---|---|---|---|---|---|---|---|---|---|---|---|---|---|---|---|---|---|---|---|---|---|
| SE | 38874 | ENSG00000117054 | ACADM | chr1 | + | 75745834 | 75745914 | 75739979 | 75740110 | 75749418 | 75749559 | 28 | 18 | 22 | [illegible] | [illegible] | [illegible] | [illegible] | [illegible] | [illegible] | [illegible] | [illegible] | [illegible] | [illegible] | [illegible] |
| SE | 62237 | ENSG00000197043 | ANXA6 | chr5 | - | 151110626 | 151110644 | 151109752 | 151109846 | 151117126 | 151117180 | 1017 | 545 | 922 | [illegible] | [illegible] | [illegible] | [illegible] | [illegible] | [illegible] | [illegible] | [illegible] | [illegible] | [illegible] | [illegible] |
| SE | 30671 | ENSG00000132466 | ANKRD17 | chr4 | - | 73085248 | 73085446 | 73078641 | 73078890 | 73090666 | 73092300 | 287 | 179 | 235 | [illegible] | [illegible] | [illegible] | [illegible] | [illegible] | [illegible] | [illegible] | [illegible] | [illegible] | [illegible] | [illegible] |
| SE | 96448 | ENSG00000110514 | MADD | chr11 | + | 47286432 | 47286534 | 47284940 | 47285194 | 47289390 | 47289493 | 123 | 72 | 93 | [illegible] | [illegible] | [illegible] | [illegible] | [illegible] | [illegible] | [illegible] | [illegible] | [illegible] | [illegible] | [illegible] |
| SE | 75840 | ENSG00000124523 | SIRT5 | chr6 | + | 13597727 | 13597864 | 13596962 | 13597016 | 13599031 | 13599155 | 0 | 0 | 0 | [illegible] | [illegible] | [illegible] | [illegible] | [illegible] | [illegible] | [illegible] | [illegible] | [illegible] | [illegible] | [illegible] |
| SE | 76992 | ENSG00000137404 | NRM | chr6 | - | 30689275 | 30689452 | 30688046 | 30688942 | 30690046 | 30690243 | 30 | 5 | 13 | [illegible] | [illegible] | [illegible] | [illegible] | [illegible] | [illegible] | [illegible] | [illegible] | [illegible] | [illegible] | [illegible] |
| SE | 6133 | ENSG00000131069 | ACSS2 | chr20 | + | 34913392 | 34913496 | 34913092 | 34913187 | 34913752 | 34913810 | 116 | 72 | 66 | [illegible] | [illegible] | [illegible] | [illegible] | [illegible] | [illegible] | [illegible] | [illegible] | [illegible] | [illegible] | [illegible] |
| SE | 80306 | ENSG00000107816 | LZTS2 | chr10 | + | 101003506 | 101004166 | 100999475 | 101002946 | 101005457 | 101005715 | 25 | 16 | 30 | [illegible] | [illegible] | [illegible] | [illegible] | [illegible] | [illegible] | [illegible] | [illegible] | [illegible] | [illegible] | [illegible] |
| SE | 32677 | ENSG00000150995 | ITPR1 | chr3 | + | 4795064 | 4795187 | 4787946 | 4788139 | 4800424 | 4800600 | 200 | 145 | 186 | [illegible] | [illegible] | [illegible] | [illegible] | [illegible] | [illegible] | [illegible] | [illegible] | [illegible] | [illegible] | [illegible] |
| SE | 63237 | ENSG00000106105 | GARS1 | chr7 | + | 30595823 | 30595933 | 30594831 | 30595143 | 30598795 | 30598897 | 2 | 2 | 6 | [illegible] | [illegible] | [illegible] | [illegible] | [illegible] | [illegible] | [illegible] | [illegible] | [illegible] | [illegible] | [illegible] |
| SE | 60622 | ENSG00000128191 | DGCR8 | chr22 | + | 20092808 | 20092907 | 20091868 | 20091970 | 20094712 | 20094795 | 41 | 39 | 56 | [illegible] | [illegible] | [illegible] | [illegible] | [illegible] | [illegible] | [illegible] | [illegible] | [illegible] | [illegible] | [illegible] |
| SE | 92800 | ENSG00000165650 | PDZD8 | chr10 | - | 117340979 | 117341102 | 117318871 | 117318974 | 117374355 | 117375440 | 109 | 39 | 57 | 56 | 85 | 3 | 2 | 1 | 3 | 6 | 47 | 67 | 84 | 110 |
| SE | 96834 | ENSG00000149516 | MS4A3 | chr11 | + | 60061145 | 60061316 | 60056627 | 60056740 | 60064261 | 60064318 | 2 | 9 | 13 | 6 | 9 | 10 | 3 | 12 | 11 | 15 | 14 | 10 | 2 | 193 |
| SE | 31319 | ENSG00000087266 | SH3BP2 | chr4 | + | 2829492 | 2830147 | 2827605 | 2827674 | 2831570 | 2831679 | 220 | 109 | 141 | 167 | 139 | 6 | 1 | 0 | 7 | 0 | 370 | 250 | 179 | 314 |
| SE | 1586 | ENSG00000198089 | SFI1 | chr22 | + | 31580271 | 31580364 | 31578381 | 31578452 | 31583874 | 31583877 | 72 | 72 | 121 | 96 | 134 | 11 | 2 | 6 | 5 | 7 | 67 | 95 | 63 | 84 |
| SE | 52632 | ENSG00000197943 | PLCG2 | chr16 | + | 81854443 | 81854587 | 81785745 | 81786182 | 81859115 | 81859163 | 132 | 46 | 66 | 73 | 72 | 0 | 0 | 0 | 0 | 0 | 65 | 137 | 73 | 93 |
| SE | 7763 | ENSG00000109118 | PHF12 | chr17 | - | 28907589 | 28907672 | 28906855 | 28906994 | 28910225 | 28910269 | 112 | 60 | 83 | 104 | 96 | 3 | 0 | 0 | 2 | 7 | 115 | 104 | 116 | 120 |
| SE | 26239 | ENSG00000141084 | RANBP10 | chr16 | - | 67805427 | 67805539 | 67772033 | 67772086 | 67806301 | 67806518 | 70 | 42 | 82 | 94 | 115 | 0 | 0 | 0 | 0 | 0 | 86 | 62 | 18 | 66 |
| SE | 49286 | ENSG00000120733 | KDM3B | chr5 | + | 138415131 | 138415239 | 138399859 | 138400012 | 138417482 | 138417610 | 382 | 233 | 303 | 311 | 537 | 0 | 3 | 0 | 7 | 0 | 535 | 408 | 229 | 447 |
| SE | 62335 | ENSG00000185811 | IKZF1 | chr7 | + | 50387344 | 50387470 | 50382539 | 50382707 | 50391728 | 50391863 | 922 | 599 | 889 | 785 | 1183 | 58 | 31 | 37 | 28 | 37 | 702 | 804 | 495 | 455 |
| SE | 3664 | ENSG00000142227 | EMP3 | chr19 | + | 48326829 | 48326922 | 48325546 | 48325610 | 48327520 | 48327561 | 631 | 245 | 361 | 449 | 282 | 20 | 3 | 12 | 11 | 8 | 983 | 693 | 352 | 861 |
| SE | 3535 | ENSG00000105483 | CARD8 | chr19 | - | 48232452 | 48232493 | 48231659 | 48231810 | 48234402 | 48234543 | 249 | 167 | 131 | 304 | 424 | 30 | 21 | 51 | 46 | 90 | 108 | 76 | 35 | 82 |
| SE | 50059 | ENSG00000137628 | DDX60 | chr4 | - | 168308005 | 168308195 | 168306378 | 168306720 | 168310997 | 168311067 | 193 | 181 | 164 | 231 | 331 | 0 | 0 | 3 | 0 | 2 | 32 | 194 | 91 | 138 |
| SE | 78269 | ENSG00000173715 | C11orf80 | chr11 | + | 66821642 | 66821764 | 66815833 | 66816199 | 66828265 | 66828364 | 18 | 8 | 25 | 18 | 55 | 0 | 4 | 4 | 2 | 12 | 36 | 13 | 13 | 46 |
| SE | 98831 | ENSG00000135365 | PHF21A | chr11 | - | 45971115 | 45971367 | 45969814 | 45969901 | 45979759 | 45979966 | 72 | 24 | 46 | 48 | 30 | 0 | 0 | 0 | 0 | 1 | 51 | 86 | 109 | 89 |
| SE | 38871 | ENSG00000117054 | ACADM | chr1 | + | 75742167 | 75745914 | 75739979 | 75740110 | 75749418 | 75749559 | 39 | 30 | 34 | 57 | 47 | 3 | 0 | 0 | 0 | 0 | 14 | 42 | 27 | 24 |
| SE | 86202 | ENSG00000171311 | EXOSC1 | chr10 | - | 97445452 | 97445606 | 97443236 | 97443311 | 97445731 | 97445847 | 0 | 0 | 2 | 0 | 0 | 56 | 32 | 49 | 51 | 14 | 14 | 0 | 0 | 0 |
| SE | 32745 | ENSG00000163349 | HIPK1 | chr1 | + | 113971823 | 113971954 | 113968441 | 113968648 | 113973023 | 113973571 | 34 | 19 | 28 | 22 | 29 | 0 | 0 | 0 | 0 | 0 | 38 | 39 | 39 | 61 |
| SE | 35092 | ENSG00000169359 | SLC33A1 | chr3 | - | 155833856 | 155834041 | 155833467 | 155833585 | 155842431 | 155842619 | 121 | 89 | 125 | 93 | 243 | 1 | 0 | 4 | 2 | 10 | 67 | 103 | 32 | 68 |
| SE | 30533 | ENSG00000163161 | ERCC3 | chr2 | - | 127290223 | 127290273 | 127289336 | 127289824 | 127292609 | 127292653 | 50 | 25 | 51 | 40 | 26 | 0 | 1 | 0 | 2 | 0 | 33 | 63 | 43 | 39 |
| SE | 104549 | ENSG00000150045 | KLRF1 | chr12 | + | 9841811 | 9841951 | 9832315 | 9832414 | 9842320 | 9842433 | 1121 | 370 | 337 | 436 | 429 | 14 | 4 | 6 | 5 | 0 | 18 | 124 | 77 | 25 |
| SE | 25320 | ENSG00000134644 | PUM1 | chr1 | - | 31006993 | 31007102 | 30995053 | 30995220 | 31028795 | 31028864 | 128 | 74 | 117 | 138 | 119 | 15 | 10 | 5 | 19 | 20 | 225 | 137 | 116 | 134 |
| SE | 21592 | ENSG00000075568 | TMEM131 | chr2 | - | 97844144 | 97844261 | 97841814 | 97841937 | 97859303 | 97859427 | 320 | 219 | 252 | 297 | 499 | 10 | 15 | 23 | 14 | 33 | 134 | 260 | 149 | 180 |
| SE | 36879 | ENSG00000122779 | TRIM24 | chr7 | + | 138554697 | 138554864 | 138551062 | 138551180 | 138567480 | 138567654 | 70 | 35 | 67 | 55 | 122 | 0 | 0 | 0 | 0 | 0 | 1 | 89 | 38 | 70 |
| SE | 88465 | ENSG00000152601 | MBNL1 | chr3 | + | 152335110 | 152335230 | 152269102 | 152269407 | 152414940 | 152415109 | 36 | 34 | 27 | 34 | 79 | 7 | 6 | 2 | 11 | 3 | 6 | 12 | 9 | 8 |
| SE | 101147 | ENSG00000175224 | ATG13 | chr11 | + | 46665382 | 46665519 | 46659334 | 46659485 | 46672254 | 46672660 | 79 | 39 | 88 | 81 | 116 | 0 | 0 | 0 | 0 | 0 | 50 | 95 | 83 | 83 |
| SE | 10485 | ENSG00000186566 | GPATCH8 | chr17 | - | 44474828 | 44474903 | 44436477 | 44436545 | 44503325 | 44503389 | 54 | 33 | 70 | 44 | 68 | 0 | 0 | 3 | 0 | 0 | 60 | 86 | 109 | 109 |
| SE | 72651 | ENSG00000153094 | BCL2L11 | chr2 | + | 111150043 | 111150147 | 111123732 | 111124139 | 111164132 | 111164231 | 99 | 39 | 74 | 56 | 91 | 9 | 1 | 6 | 15 | 7 | 131 | 156 | 253 | 118 |
| SE | 19157 | ENSG00000183484 | GPR132 | chr14 | - | 105055386 | 105056166 | 105049394 | 105052102 | 105057166 | 105057280 | 17 | 26 | 17 | 24 | 8 | 23 | 14 | 11 | 16 | 20 | 11 | 43 | 34 | 8 |
| SE | 80148 | ENSG00000012124 | CD22 | chr19 | + | 35332577 | 35332924 | 35329165 | 35329230 | 35336035 | 35336134 | 61 | 42 | 53 | 66 | 100 | 0 | 0 | 0 | 1 | 3 | 91 | 113 | 24 | 52 |
| SE | 31273 | ENSG00000047365 | ARAP2 | chr4 | - | 36212395 | 36212487 | 36210389 | 36210743 | 36213242 | 36213319 | 1048 | 725 | 623 | 872 | 806 | 4 | 5 | 0 | 8 | 0 | 189 | 487 | 367 | 207 |
| SE | 8908 | ENSG00000257949 | TEN1 | chr17 | + | 75986186 | 75986284 | 75979239 | 75979511 | 75991465 | 75991540 | 41 | 27 | 42 | 51 | 16 | 0 | 0 | 0 | 0 | 0 | 95 | 44 | 34 | 24 |
| SE | 51949 | ENSG00000023171 | GRAMD1B | chr11 | + | 123594734 | 123594838 | 123594081 | 123594166 | 123595941 | 123596037 | 32 | 36 | 22 | 24 | 29 | 1 | 0 | 3 | 0 | 0 | 53 | 41 | 16 | 61 |
| SE | 54310 | ENSG00000143442 | POGZ | chr1 | - | 151429602 | 151429711 | 151423948 | 151424286 | 151430665 | 151430841 | 82 | 55 | 90 | 68 | 116 | 0 | 0 | 0 | 1 | 0 | 92 | 77 | 71 | 48 |
| SE | 37652 | ENSG00000136933 | RABEPK | chr9 | + | 125207563 | 125207721 | 125200604 | 125200906 | 125213369 | 125213522 | 49 | 11 | 28 | 24 | 58 | 0 | 0 | 0 | 0 | 0 | 46 | 10 | 10 | 12 |
| SE | 72466 | ENSG00000023516 | AKAP11 | chr13 | + | 42313046 | 42313130 | 42308453 | 42308609 | 42313893 | 42313940 | 65 | 27 | 23 | 31 | 49 | 114 | 107 | 157 | 191 | 179 | 0 | 5 | 10 | 6 |
| SE | 33513 | ENSG00000134072 | CAMK1 | chr3 | - | 9766124 | 9766213 | 9765758 | 9765890 | 9767666 | 9767781 | 3 | 0 | 1 | 9 | 2 | 30 | 26 | 10 | 26 | 14 | 16 | 6 | 0 | 4 |
| SE | 53413 | ENSG00000172531 | PPP1CA | chr11 | - | 67401067 | 67401199 | 67398939 | 67400919 | 67401727 | 67401788 | 466 | 183 | 269 | 377 | 208 | 10 | 1 | 6 | 6 | 0 | 194 | 416 | 191 | 236 |
| SE | 101502 | ENSG00000095794 | CREM | chr10 | + | 35188199 | 35188388 | 35179133 | 35179276 | 35206894 | 35207006 | 20 | 8 | 5 | 18 | 10 | 16 | 18 | 5 | 9 | 21 | 0 | 6 | 0 | 0 |
| SE | 33515 | ENSG00000175455 | CCDC14 | chr3 | - | 123931307 | 123931526 | 123931101 | 123931234 | 123933672 | 123933755 | 286 | 284 | 201 | 306 | 286 | 2 | 1 | 3 | 0 | 3 | 114 | 76 | 106 | 48 |
| SE | 19994 | ENSG00000180694 | TMEM64 | chr8 | - | 90631551 | 90631707 | 90621999 | 90625862 | 90645110 | 90646083 | 154 | 143 | 93 | 203 | 108 | 2 | 4 | 2 | 5 | 0 | 76 | 128 | 113 | 68 |
| SE | 23403 | ENSG00000153113 | CAST | chr5 | + | 96740744 | 96740783 | 96740037 | 96740118 | 96741265 | 96741317 | 558 | 379 | 334 | 201 | 477 | 0 | 0 | 0 | 244 | 367 | 566 | 478 | 484 | 259 |
| SE | 22761 | ENSG00000145730 | PAM | chr5 | + | 103017333 | 103017433 | 103009750 | 103009866 | 103019789 | 103019843 | 205 | 116 | 110 | 105 | 127 | 3 | 1 | 5 | 4 | 1 | 159 | 58 | 39 | 111 |
| SE | 77835 | ENSG00000256043 | CTSO | chr4 | - | 155928335 | 155928428 | 155924117 | 155926070 | 155929541 | 155929705 | 266 | 150 | 183 | 160 | 323 | 2 | 1 | 0 | 0 | 1 | 77 | 142 | 122 | 165 |
| SE | 11910 | ENSG00000103222 | ABCC1 | chr16 | + | 16124788 | 16124915 | 16121974 | 16122174 | 16125809 | 16125911 | 74 | 39 | 62 | 70 | 66 | 2 | 0 | 0 | 4 | 3 | 42 | 114 | 59 | 49 |
| SE | 16066 | ENSG00000128908 | INO80 | chr15 | - | 41016087 | 41016215 | 41005592 | 41005687 | 41020899 | 41021125 | 73 | 40 | 81 | 73 | 50 | 0 | 1 | 4 | 2 | 3 | 64 | 84 | 75 | 101 |
| SE | 29475 | ENSG00000077684 | JADE1 | chr4 | + | 128867855 | 128867973 | 128861703 | 128862225 | 128871354 | 128872334 | 102 | 94 | 129 | 142 | 84 | 5 | 1 | 1 | 5 | 4 | 26 | 47 | 51 | 62 |
| SE | 98540 | ENSG00000118096 | IFT46 | chr11 | - | 118557737 | 118557890 | 118556905 | 118557045 | 118559784 | 118559864 | 2 | 15 | 14 | 21 | 14 | 17 | 22 | 19 | 24 | 13 | 0 | 12 | 0 | 1 |
| SE | 82937 | ENSG00000148606 | POLR3A | chr10 | - | 77985902 | 77985985 | 77985169 | 77985340 | 77986072 | 77986159 | 30 | 15 | 18 | 33 | 40 | 1 | 0 | 0 | 0 | 0 | 39 | 36 | 21 | 28 |
| SE | 21792 | ENSG00000198625 | MDM4 | chr1 | + | 204544534 | 204544684 | 204542783 | 204542944 | 204546796 | 204546832 | 597 | 741 | 675 | 732 | 841 | 8 | 16 | 10 | 15 | 21 | 1050 | 1129 | 1126 | 1285 |
| SE | 52023 | ENSG00000135250 | SRPK2 | chr7 | - | 105126247 | 105126340 | 105125781 | 105125874 | 105126992 | 105127062 | 54 | 28 | 37 | 61 | 47 | 32 | 10 | 11 | 18 | 16 | 32 | 40 | 35 | 12 |
| SE | 63104 | ENSG00000138050 | THUMPD2 | chr2 | - | 39769709 | 39770119 | 39768900 | 39769093 | 39771504 | 39771640 | 29 | 35 | 32 | 32 | 36 | 0 | 0 | 0 | 2 | 0 | 8 | 17 | 50 | 19 |
| SE | 35148 | ENSG00000107758 | PPP3CB | chr10 | - | 73470686 | 73470781 | 73466957 | 73467678 | 73470886 | 73470964 | 413 | 304 | 294 | 328 | 527 | 2 | 0 | 1 | 1 | 0 | 278 | 272 | 215 | 209 |
| SE | 37262 | ENSG00000099330 | OCEL1 | chr19 | + | 17228802 | 17228915 | 17228253 | 17228309 | 17229008 | 17229215 | 14 | 13 | 16 | 12 | 32 | 6 | 0 | 2 | 0 | 0 | 41 | 14 | 4 | 21 |
| SE | 52888 | ENSG00000159023 | EPB41 | chr1 | + | 28993329 | 28993542 | 28987430 | 28987905 | 29011864 | 29011907 | 433 | 358 | 508 | 556 | 888 | 17 | 8 | 17 | 11 | 32 | 368 | 391 | 420 | 505 |
| SE | 63526 | ENSG00000115966 | ATF2 | chr2 | - | 175121443 | 175121540 | 175118250 | 175118369 | 175130137 | 175130205 | 288 | 227 | 249 | 232 | 621 | 98 | 81 | 71 | 83 | 125 | 126 | 184 | 124 | 95 |
| SE | 102056 | ENSG00000138134 | STAMBPL1 | chr10 | + | 88913100 | 88913458 | 88910915 | 88911011 | 88914533 | 88914658 | 75 | 78 | 56 | 81 | 118 | 2 | 8 | 4 | 7 | 12 | 13 | 38 | 18 | 39 |
| SE | 109178 | ENSG00000124496 | TRERF1 | chr6 | - | 42232680 | 42232892 | 42224930 | 42228669 | 42243247 | 42243361 | 45 | 23 | 40 | 53 | 27 | 0 | 2 | 2 | 2 | 3 | 40 | 69 | 100 | 65 |
| SE | 35840 | ENSG00000143612 | C1orf43 | chr1 | - | 154214456 | 154214574 | 154213892 | 154213946 | 154219835 | 154219937 | 265 | 109 | 254 | 226 | 305 | 4 | 2 | 5 | 4 | 3 | 230 | 282 | 90 | 191 |
| SE | 94748 | ENSG00000107077 | KDM4C | chr9 | + | 6805598 | 6805774 | 6792971 | 6793132 | 6814630 | 6814745 | 674 | 542 | 708 | 755 | 468 | 4 | 7 | 8 | 9 | 0 | 761 | 625 | 755 | 720 |
| SE | 73323 | ENSG00000115970 | THADA | chr2 | - | 43333154 | 43333263 | 43320445 | 43320540 | 43344121 | 43344237 | 2 | 2 | 8 | 3 | 7 | 24 | 15 | 34 | 23 | 19 | 9 | 6 | 4 | 6 |
| SE | 27593 | ENSG00000132694 | ARHGEF11 | chr1 | - | 156980436 | 156980486 | 156978203 | 156978382 | 156984338 | 156984437 | 23 | 12 | 11 | 20 | 24 | 1 | 0 | 1 | 0 | 0 | 76 | 55 | 21 | 60 |
| SE | 13836 | ENSG00000068903 | SIRT2 | chr19 | - | 38898378 | 38898425 | 38893818 | 38893867 | 38898793 | 38898918 | 11 | 0 | 0 | 5 | 2 | 0 | 2 | 2 | 0 | 0 | 5 | 1 | 2 | 12 |
| SE | 77642 | ENSG00000214021 | TTLL3 | chr3 | + | 9816073 | 9816202 | 9813247 | 9813345 | 9817644 | 9817658 | 45 | 30 | 42 | 59 | 55 | 23 | 20 | 28 | 27 | 36 | 87 | 73 | 72 | 67 |
| SE | 47742 | ENSG00000145687 | SSBP2 | chr5 | - | 81489249 | 81489309 | 81474495 | 81474562 | 81615472 | 81615557 | 18 | 10 | 26 | 26 | 36 | 0 | 1 | 0 | 0 | 8 | 43 | 12 | 18 | 8 |
| SE | 68843 | ENSG00000168769 | TET2 | chr4 | + | 105190359 | 105190505 | 105146785 | 105146979 | 105233896 | 105234272 | 106 | 74 | 70 | 146 | 70 | 26 | 22 | 21 | 21 | 30 | 32 | 97 | 80 | 21 |
| SE | 96392 | ENSG00000197563 | PIGN | chr18 | - | 62140419 | 62140479 | 62138982 | 62139075 | 62143305 | 62143346 | 53 | 22 | 48 | 40 | 105 | 2 | 2 | 0 | 2 | 6 | 40 | 69 | 26 | 35 |
| SE | 20932 | ENSG00000135596 | MICAL1 | chr6 | - | 109447680 | 109447722 | 109447356 | 109447440 | 109447874 | 109447963 | 31 | 26 | 44 | 54 | 26 | 0 | 0 | 0 | 0 | 0 | 56 | 53 | 36 | 66 |
| SE | 75735 | ENSG00000144580 | CNOT9 | chr2 | + | 218573720 | 218573791 | 218568930 | 218568978 | 218580560 | 218580740 | 7 | 0 | 1 | 4 | 1 | 54 | 37 | 43 | 37 | 63 | 0 | 12 | 2 | 10 |
| SE | 43752 | ENSG00000148019 | CEP78 | chr9 | + | 78262906 | 78262984 | 78254473 | 78254964 | 78264149 | 78264228 | 290 | 116 | 132 | 218 | 285 | 9 | 3 | 1 | 2 | 4 | 40 | 155 | 153 | 84 |
| SE | 56968 | ENSG00000130775 | THEMIS2 | chr1 | + | 27876587 | 27876728 | 27872542 | 27872665 | 27885294 | 27885451 | 112 | 57 | 78 | 63 | 13 | 2 | 1 | 2 | 2 | 0 | 147 | 86 | 67 | 83 |
| SE | 3293 | ENSG00000142453 | CARM1 | chr19 | + | 10921374 | 10921443 | 10921049 | 10921127 | 10921614 | 10921900 | 25 | 7 | 10 | 6 | 1 | 37 | 17 | 24 | 30 | 21 | 17 | 36 | 30 | 7 |
| SE | 93635 | ENSG00000107362 | ABHD17B | chr9 | - | 71870082 | 71870262 | 71865151 | 71867006 | 71874613 | 71875083 | 102 | 53 | 71 | 72 | 73 | 12 | 1 | 0 | 3 | 4 | 51 | 108 | 64 | 63 |
| SE | 78164 | ENSG00000223705 | NSUN5P1 | chr7 | + | 75414639 | 75414798 | 75412788 | 75412932 | 75414884 | 75415024 | 1 | 3 | 5 | 0 | 6 | 20 | 11 | 15 | 10 | 35 | 0 | 8 | 3 | 0 |
| SE | 54532 | ENSG00000161929 | SCIMP | chr17 | - | 5223353 | 5223456 | 5214924 | 5214998 | 5234734 | 5234823 | 134 | 80 | 63 | 66 | 85 | 0 | 0 | 0 | 0 | 0 | 121 | 106 | 19 | 46 |
| SE | 80768 | ENSG00000136280 | CCM2 | chr7 | + | 45068442 | 45068579 | 45027025 | 45027797 | 45069825 | 45069961 | 29 | 11 | 25 | 26 | 20 | 0 | 0 | 1 | 1 | 5 | 57 | 11 | 14 | 34 |
| SE | 105161 | ENSG00000174718 | RESF1 | chr12 | + | 31970188 | 31970356 | 31960776 | 31960871 | 31980877 | 31982979 | 2279 | 1832 | 1897 | 1959 | 4777 | 55 | 26 | 52 | 30 | 52 | 217 | 683 | 497 | 291 |
| SE | 73094 | ENSG00000198001 | IRAK4 | chr12 | + | 43763301 | 43763456 | 43760924 | 43760972 | 43768102 | 43768272 | 7 | 0 | 1 | 1 | 1 | 22 | 17 | 20 | 14 | 29 | 0 | 9 | 3 | 3 |
| SE | 35683 | ENSG00000143554 | SLC27A3 | chr1 | + | 153777760 | 153777885 | 153777061 | 153777220 | 153778160 | 153778355 | 98 | 61 | 55 | 120 | 37 | 3 | 2 | 0 | 4 | 2 | 174 | 150 | 27 | 166 |
| SE | 25779 | ENSG00000113391 | FAM172A | chr5 | - | 94053125 | 94053227 | 93964457 | 93964523 | 94111510 | 94111648 | 56 | 51 | 36 | 82 | 35 | 0 | 0 | 1 | 0 | 0 | 9 | 34 | 35 | 29 |
| SE | 102014 | ENSG00000148572 | NRBF2 | chr10 | + | 63146208 | 63146293 | 63133327 | 63133500 | 63152149 | 63152190 | 56 | 43 | 56 | 39 | 74 | 0 | 2 | 0 | 1 | 0 | 17 | 51 | 101 | 31 |
| SE | 98649 | ENSG00000107036 | RIC1 | chr9 | + | 5742868 | 5743013 | 5732387 | 5732479 | 5743688 | 5743737 | 83 | 66 | 82 | 62 | 144 | 0 | 0 | 0 | 2 | 7 | 48 | 112 | 84 | 62 |
| SE | 44513 | ENSG00000285215 | NA | chr1 | + | 120973107 | 120973246 | 120969118 | 120969354 | 120973346 | 120973561 | 12 | 6 | 4 | 9 | 10 | 0 | 5 | 0 | 0 | 2 | 7 | 13 | 10 | 14 |
| [illegible] | [illegible] | [illegible] | [illegible] | [illegible] | [illegible] | [illegible] | [illegible] | [illegible] | [illegible] | [illegible] | [illegible] | [illegible] | [illegible] | [illegible] | [illegible] | [illegible] | [illegible] | [illegible] | [illegible] | [illegible] | [illegible] | [illegible] | [illegible] | [illegible] | [illegible] |

| | | | | | | | | | | | | | | | | | | | | | | | | | | | | | | | | | | | | | | | | | | | | | | |
|---|---|---|---|---|---|---|---|---|---|---|---|---|---|---|---|---|---|---|---|---|---|---|---|---|---|---|---|---|---|---|---|---|---|---|---|---|---|---|---|---|---|---|---|---|---|---|
| SE | 8966 | ENSG00000108395 | TRIM37 | chr17 | - | 59064354 | 59064405 | 59062566 | 59062648 | 59070822 | 59070947 | 135 | 79 | 65 | 112 | 106 | 2 | 1 | 0 | 1 | 2 | 53 | 84 | 115 | 90 | 15 | 7 | 3 | 2 | 0 | 0 | 200 | 149 | 0.034233957 | 0.28043334 | 0.981 | 0.983 | 1 | 0.988 | 0.975 | 0.849 | 0.954 | 0.977 | 1 | 1 | 0.029 |
| SE | 71234 | ENSG00000079263 | SP140 | chr2 | + | 230292645 | 230292788 | 230290459 | 230290564 | 230294270 | 230294318 | 205 | 184 | 209 | 216 | 273 | 7 | 5 | 13 | 16 | 6 | 309 | 264 | 111 | 166 | 146 | 9 | 21 | 25 | 12 | 3 | 292 | 149 | 0.034240203 | 0.28045517 | 0.937 | 0.949 | 0.891 | 0.873 | 0.959 | 0.946 | 0.865 | 0.694 | 0.876 | 0.961 | 0.053 |
| SE | 95585 | ENSG00000130714 | POMT1 | chr9 | + | 131504188 | 131504649 | 131503009 | 131503073 | 131506113 | 131506220 | 11 | 8 | 8 | 17 | 4 | 1 | 0 | 0 | 0 | 3 | 22 | 4 | 5 | 20 | 33 | 0 | 13 | 3 | 0 | 0 | 298 | 149 | 0.034293659 | 0.28086363 | 0.846 | 1 | 1 | 1 | 0.4 | 1 | 0.133 | 0.455 | 1 | 1 | 0.132 |
| SE | 94921 | ENSG00000157693 | TMEM268 | chr9 | + | 114633767 | 114633878 | 114628100 | 114628250 | 114636989 | 114637070 | 59 | 45 | 53 | 63 | 48 | 3 | 2 | 3 | 2 | 5 | 76 | 66 | 69 | 93 | 113 | 6 | 0 | 0 | 4 | 0 | 260 | 149 | 0.034330418 | 0.281135279 | 0.919 | 0.928 | 0.91 | 0.948 | 0.846 | 0.879 | 1 | 1 | 0.93 | 1 | -0.052 |
| SE | 71113 | ENSG00000198363 | ASPH | chr8 | - | 61633682 | 61633727 | 61618977 | 61619019 | 61637946 | 61637986 | 53 | 30 | 31 | 41 | 83 | 4 | 6 | 3 | 7 | 10 | 99 | 48 | 67 | 32 | 55 | 0 | 18 | 3 | 1 | 0 | 194 | 149 | 0.034340631 | 0.281182364 | 0.911 | 0.793 | 0.888 | 0.818 | 0.864 | 1 | 0.672 | 0.945 | 0.961 | 1 | -0.061 |
| SE | 43883 | ENSG00000179335 | CLK3 | chr15 | + | 74622119 | 74622216 | 74619966 | 74620225 | 74622493 | 74622560 | 124 | 59 | 116 | 119 | 162 | 17 | 5 | 14 | 17 | 9 | 105 | 149 | 124 | 84 | 213 | 24 | 9 | 16 | 29 | 34 | 246 | 149 | 0.034344803 | 0.281182364 | 0.815 | 0.877 | 0.834 | 0.809 | 0.916 | 0.726 | 0.909 | 0.824 | 0.637 | 0.791 | 0.073 |
| SE | 84435 | ENSG00000188917 | TRMT2B | chrX | - | 101051250 | 101051366 | 101042041 | 101042312 | 101051816 | 101052054 | 22 | 13 | 21 | 14 | 15 | 2 | 4 | 11 | 3 | 2 | 8 | 14 | 18 | 32 | 21 | 0 | 4 | 3 | 0 | 3 | 265 | 149 | 0.034346941 | 0.281182364 | 0.861 | 0.646 | 0.518 | 0.724 | 0.808 | 1 | 0.663 | 0.771 | 1 | 0.797 | -0.135 |
| SE | 3555 | ENSG00000105483 | CARD8 | chr19 | - | 48249522 | 48249633 | 48242539 | 48242644 | 48249750 | 48249810 | 150 | 59 | 107 | 142 | 111 | 12 | 2 | 2 | 5 | 9 | 52 | 81 | 71 | 60 | 36 | 15 | 13 | 5 | 2 | 0 | 260 | 149 | 0.034371948 | 0.281334222 | 0.878 | 0.944 | 0.968 | 0.942 | 0.876 | 0.665 | 0.781 | 0.891 | 0.945 | 1 | 0.065 |
| SE | 6873 | ENSG00000263753 | LINC00667 | chr18 | + | 5240647 | 5241255 | 5238080 | 5239049 | 5241698 | 5241812 | 15 | 2 | 13 | 15 | 22 | 12 | 33 | 15 | 41 | 29 | 7 | 5 | 2 | 15 | 6 | 24 | 21 | 19 | 29 | 29 | 298 | 149 | 0.034372678 | 0.281334222 | 0.385 | 0.029 | 0.302 | 0.155 | 0.275 | 0.127 | 0.106 | 0.05 | 0.205 | 0.094 | 0.113 |
| SE | 72938 | ENSG00000048649 | RSF1 | chr11 | - | 77698486 | 77698693 | 77693506 | 77693611 | 77725544 | 77725680 | 45 | 34 | 35 | 63 | 35 | 0 | 0 | 0 | 0 | 0 | 0 | 34 | 48 | 3 | 7 | 9 | 0 | 0 | 0 | 0 | 298 | 149 | 0.03438707 | 0.281415153 | 1 | 1 | 1 | 1 | 1 | 0 | 1 | 1 | 1 | 1 | 0.2 |
| SE | 104725 | ENSG00000005189 | REXO5 | chr16 | + | 20845053 | 20845241 | 20844628 | 20844845 | 20846220 | 20846339 | 1 | 6 | 5 | 10 | 5 | 0 | 0 | 0 | 0 | 0 | 0 | 2 | 6 | 14 | 13 | 9 | 0 | 0 | 0 | 0 | 298 | 149 | 0.034389754 | 0.281415153 | 1 | 1 | 1 | 1 | 1 | 0 | 1 | 1 | 1 | 1 | 0.2 |
| SE | 81625 | ENSG00000135093 | USP30 | chr12 | + | 109057925 | 109058108 | 109056681 | 109056791 | 109067523 | 109067627 | 16 | 22 | 21 | 14 | 45 | 0 | 0 | 4 | 3 | 3 | 42 | 11 | 12 | 18 | 16 | 21 | 2 | 2 | 3 | 0 | 298 | 149 | 0.034394358 | 0.281423416 | 1 | 1 | 0.724 | 0.7 | 0.882 | 0.5 | 0.733 | 0.75 | 0.75 | 1 | 0.115 |
| SE | 48162 | ENSG00000117543 | DPH5 | chr1 | - | 101001466 | 101001587 | 100995109 | 100995149 | 101013709 | 101013818 | 163 | 117 | 166 | 181 | 324 | 1 | 2 | 4 | 3 | 3 | 102 | 75 | 114 | 57 | 48 | 15 | 0 | 1 | 0 | 0 | 270 | 149 | 0.03441576 | 0.281552204 | 0.989 | 0.97 | 0.958 | 0.971 | 0.983 | 0.79 | 1 | 0.984 | 1 | 1 | 0.019 |
| SE | 80144 | ENSG00000012124 | CD22 | chr19 | + | 35332552 | 35332924 | 35329211 | 35329230 | 35336035 | 35336042 | 71 | 42 | 56 | 72 | 102 | 0 | 0 | 0 | 1 | 3 | 82 | 131 | 26 | 56 | 142 | 0 | 6 | 0 | 2 | 3 | 298 | 149 | 0.03441729 | 0.281552204 | 1 | 1 | 1 | 0.973 | 0.944 | 1 | 0.916 | 1 | 0.933 | 0.959 | 0.022 |
| SE | 54449 | ENSG00000164305 | CASP3 | chr4 | - | 184648464 | 184648631 | 184638400 | 184638468 | 184649394 | 184649447 | 28 | 10 | 11 | 6 | 1 | 28 | 35 | 27 | 40 | 25 | 16 | 22 | 4 | 5 | 4 | 9 | 24 | 39 | 2 | 0 | 298 | 149 | 0.034424773 | 0.281583997 | 0.333 | 0.125 | 0.169 | 0.07 | 0.02 | 0.471 | 0.314 | 0.049 | 0.556 | 1 | -0.335 |
| SE | 101844 | ENSG00000151150 | ANK3 | chr10 | - | 60166590 | 60166653 | 60138963 | 60139087 | 60166823 | 60166896 | 10 | 2 | 13 | 6 | 8 | 4 | 3 | 13 | 8 | 9 | 0 | 20 | 26 | 6 | 12 | 7 | 1 | 3 | 1 | 3 | 212 | 149 | 0.034454534 | 0.281788768 | 0.637 | 0.319 | 0.413 | 0.345 | 0.385 | 0 | 0.934 | 0.859 | 0.808 | 0.738 | -0.248 |
| SE | 29432 | ENSG00000124783 | SSR1 | chr6 | - | 7295391 | 7295485 | 7281142 | 7289931 | 7298746 | 7298823 | 312 | 143 | 277 | 247 | 435 | 2 | 0 | 0 | 0 | 0 | 19 | 189 | 145 | 87 | 39 | 0 | 0 | 0 | 0 | 12 | 243 | 149 | 0.034457005 | 0.281788768 | 0.99 | 1 | 1 | 1 | 1 | 1 | 1 | 1 | 1 | 0.666 | 0.065 |
| SE | 6671 | ENSG00000141627 | DYM | chr18 | - | 49391384 | 49391451 | 49379745 | 49379758 | 49391592 | 49391645 | 0 | 0 | 1 | 2 | 4 | 79 | 58 | 57 | 89 | 98 | 0 | 12 | 2 | 0 | 0 | 56 | 68 | 46 | 48 | 42 | 216 | 149 | 0.034474016 | 0.281892433 | 0 | 0 | 0.012 | 0.015 | 0.027 | 0 | 0.109 | 0.029 | 0 | 0 | -0.017 |
| SE | 104742 | ENSG00000122694 | GLIPR2 | chr9 | + | 36150871 | 36150949 | 36148546 | 36148650 | 36162361 | 36163902 | 390 | 212 | 254 | 330 | 236 | 3 | 0 | 0 | 3 | 2 | 824 | 459 | 300 | 508 | 964 | 18 | 13 | 3 | 2 | 1 | 227 | 149 | 0.034476882 | 0.281892433 | 0.988 | 1 | 1 | 0.986 | 0.987 | 0.968 | 0.959 | 0.985 | 0.994 | 0.998 | 0.011 |
| SE | 106874 | ENSG00000005379 | TSPOAP1 | chr17 | - | 58304610 | 58304723 | 58304337 | 58304399 | 58305060 | 58305171 | 1 | 0 | 2 | 6 | 0 | 13 | 8 | 11 | 23 | 4 | 0 | 0 | 0 | 0 | 0 | 7 | 15 | 11 | 8 | 36 | 262 | 149 | 0.034545656 | 0.282425258 | 0.042 | 0 | 0.094 | 0.129 | 0 | 0 | 0 | 0 | 0 | 0 | 0.053 |
| SE | 70544 | ENSG00000073849 | ST6GAL1 | chr3 | + | 187038741 | 187038873 | 187021855 | 187022004 | 187051248 | 187051346 | 19 | 6 | 24 | 39 | 23 | 0 | 0 | 0 | 0 | 0 | 12 | 30 | 18 | 51 | 237 | 0 | 0 | 0 | 0 | 11 | 281 | 149 | 0.034572979 | 0.282616852 | 1 | 1 | 1 | 1 | 1 | 1 | 1 | 1 | 1 | 0.92 | 0.016 |
| SE | 68795 | ENSG00000204634 | TBC1D8 | chr2 | - | 101029490 | 101029776 | 101028302 | 101028432 | 101032267 | 101032385 | 54 | 24 | 20 | 53 | 15 | 9 | 0 | 0 | 0 | 0 | 28 | 54 | 30 | 57 | 23 | 0 | 0 | 0 | 0 | 0 | 298 | 149 | 0.034576311 | 0.282616852 | 0.75 | 1 | 1 | 1 | 1 | 1 | 1 | 1 | 1 | 1 | -0.05 |
| SE | 4682 | ENSG00000141968 | VAV1 | chr19 | + | 6852964 | 6853079 | 6850669 | 6850757 | 6853946 | 6854098 | 427 | 292 | 371 | 459 | 332 | 6 | 2 | 4 | 6 | 2 | 816 | 688 | 382 | 562 | 922 | 16 | 19 | 1 | 10 | 9 | 264 | 149 | 0.034580588 | 0.282622306 | 0.976 | 0.988 | 0.981 | 0.977 | 0.989 | 0.966 | 0.953 | 0.995 | 0.969 | 0.983 | 0.009 |
| SE | 44381 | ENSG00000072210 | ALDH3A2 | chr17 | + | 19656365 | 19656574 | 19652546 | 19652632 | 19661126 | 19661268 | 52 | 29 | 42 | 72 | 75 | 0 | 0 | 0 | 2 | 0 | 107 | 22 | 39 | 71 | 63 | 0 | 0 | 0 | 10 | 0 | 298 | 149 | 0.034585087 | 0.28262958 | 1 | 1 | 1 | 0.947 | 1 | 1 | 1 | 1 | 0.78 | 1 | 0.033 |
| SE | 71021 | ENSG00000163516 | ANKZF1 | chr2 | + | 219230227 | 219230350 | 219229805 | 219229900 | 219232259 | 219232362 | 74 | 38 | 61 | 72 | 24 | 0 | 0 | 0 | 1 | 0 | 13 | 51 | 39 | 40 | 60 | 0 | 0 | 0 | 5 | 4 | 272 | 149 | 0.034601004 | 0.282730141 | 1 | 1 | 1 | 0.975 | 1 | 1 | 1 | 1 | 0.814 | 0.892 | 0.054 |
| SE | 24951 | ENSG00000163297 | ANTXR2 | chr4 | - | 80071582 | 80071654 | 80069435 | 80069507 | 80079352 | 80079600 | 55 | 42 | 32 | 41 | 45 | 1 | 0 | 0 | 0 | 0 | 14 | 34 | 34 | 35 | 13 | 0 | 0 | 1 | 0 | 8 | 221 | 149 | 0.034649861 | 0.283099816 | 0.974 | 1 | 1 | 1 | 1 | 1 | 1 | 0.958 | 1 | 0.523 | 0.099 |
| SE | 108913 | ENSG00000162695 | SLC30A7 | chr1 | + | 100961827 | 100961918 | 100921705 | 100921841 | 100965768 | 100965918 | 160 | 98 | 108 | 107 | 242 | 5 | 1 | 2 | 2 | 9 | 66 | 57 | 40 | 38 | 28 | 0 | 0 | 5 | 0 | 0 | 240 | 149 | 0.034668045 | 0.28321883 | 0.952 | 0.984 | 0.971 | 0.971 | 0.943 | 1 | 1 | 0.832 | 1 | 1 | -0.002 |
| SE | 108634 | ENSG00000136104 | RNASEH2B | chr13 | + | 50953904 | 50953985 | 50947986 | 50949505 | 50956357 | 50956405 | 125 | 74 | 96 | 62 | 250 | 0 | 0 | 3 | 0 | 11 | 81 | 53 | 62 | 75 | 46 | 0 | 0 | 0 | 1 | 0 | 230 | 149 | 0.034672022 | 0.283221769 | 1 | 1 | 0.954 | 1 | 0.936 | 1 | 1 | 1 | 0.98 | 1 | -0.018 |
| SE | 76371 | ENSG00000164674 | SYTL3 | chr6 | + | 158718086 | 158718211 | 158708321 | 158708391 | 158725502 | 158725637 | 108 | 43 | 69 | 45 | 78 | 31 | 15 | 22 | 12 | 13 | 31 | 65 | 68 | 26 | 40 | 24 | 23 | 36 | 4 | 26 | 274 | 149 | 0.034685611 | 0.283303215 | 0.655 | 0.609 | 0.63 | 0.671 | 0.765 | 0.413 | 0.606 | 0.507 | 0.779 | 0.456 | 0.114 |
| SE | 88916 | ENSG00000158669 | GPAT4 | chr8 | - | 41612844 | 41612960 | 41612179 | 41612273 | 41614385 | 41614441 | 357 | 211 | 284 | 360 | 275 | 1 | 0 | 4 | 1 | 1 | 789 | 504 | 304 | 345 | 752 | 0 | 3 | 5 | 12 | 10 | 265 | 149 | 0.034699374 | 0.283386061 | 0.995 | 1 | 0.976 | 0.995 | 0.994 | 1 | 0.99 | 0.972 | 0.942 | 0.977 | 0.016 |
| SE | 50380 | ENSG00000058866 | DGKG | chr3 | - | 186260438 | 186260513 | 186257763 | 186257939 | 186261698 | 186261778 | 3 | 7 | 4 | 10 | 4 | 3 | 0 | 1 | 2 | 7 | 29 | 13 | 7 | 46 | 27 | 0 | 10 | 0 | 5 | 0 | 224 | 149 | 0.034722704 | 0.283539796 | 0.399 | 1 | 0.727 | 0.769 | 0.275 | 1 | 0.464 | 1 | 0.86 | 1 | -0.231 |
| SE | 84767 | ENSG00000158201 | ABHD3 | chr18 | - | 21683919 | 21683965 | 21659169 | 21659343 | 21702315 | 21702498 | 72 | 41 | 73 | 47 | 60 | 2 | 2 | 2 | 1 | 4 | 32 | 33 | 51 | 16 | 49 | 0 | 2 | 0 | 0 | 0 | 195 | 149 | 0.034725441 | 0.283539796 | 0.965 | 0.94 | 0.965 | 0.973 | 0.92 | 1 | 0.927 | 1 | 1 | 1 | -0.033 |
| SE | 61192 | ENSG00000198873 | GRK5 | chr10 | + | 119441998 | 119442088 | 119439730 | 119439768 | 119443543 | 119443752 | 170 | 113 | 108 | 178 | 146 | 3 | 2 | 2 | 0 | 0 | 232 | 145 | 145 | 133 | 624 | 0 | 2 | 1 | 8 | 11 | 239 | 149 | 0.034735595 | 0.283593135 | 0.972 | 0.972 | 0.971 | 1 | 1 | 1 | 0.978 | 0.989 | 0.912 | 0.973 | 0.013 |
| SE | 36605 | ENSG00000169180 | XPO6 | chr16 | - | 28152658 | 28152785 | 28135215 | 28135324 | 28156073 | 28156527 | 119 | 59 | 135 | 111 | 175 | 0 | 0 | 0 | 0 | 0 | 107 | 62 | 119 | 67 | 124 | 0 | 0 | 0 | 0 | 12 | 276 | 149 | 0.034784595 | 0.283963571 | 1 | 1 | 1 | 1 | 1 | 1 | 1 | 1 | 1 | 0.848 | 0.03 |
| SE | 20355 | ENSG00000162711 | NLRP3 | chr1 | + | 247435969 | 247436140 | 247434102 | 247434273 | 247443971 | 247444142 | 52 | 68 | 40 | 38 | 57 | 9 | 8 | 3 | 7 | 3 | 82 | 42 | 70 | 58 | 74 | 14 | 7 | 5 | 13 | 55 | 298 | 149 | 0.034793613 | 0.284007586 | 0.743 | 0.81 | 0.87 | 0.731 | 0.905 | 0.745 | 0.75 | 0.875 | 0.69 | 0.402 | 0.119 |
| SE | 56932 | ENSG00000118197 | DDX59 | chr1 | - | 200648438 | 200648567 | 200644036 | 200644517 | 200649073 | 200649149 | 147 | 68 | 135 | 148 | 236 | 9 | 9 | 9 | 4 | 11 | 86 | 62 | 99 | 73 | 33 | 10 | 4 | 9 | 14 | 1 | 278 | 149 | 0.034809569 | 0.284108204 | 0.897 | 0.802 | 0.889 | 0.952 | 0.92 | 0.822 | 0.893 | 0.855 | 0.736 | 0.946 | 0.042 |
| SE | 75759 | ENSG00000096070 | BRPF3 | chr6 | + | 36217916 | 36218010 | 36213879 | 36214386 | 36222167 | 36222265 | 89 | 36 | 67 | 52 | 60 | 0 | 0 | 0 | 0 | 0 | 60 | 102 | 40 | 94 | 126 | 0 | 12 | 0 | 0 | 0 | 243 | 149 | 0.034831714 | 0.284259316 | 1 | 1 | 1 | 1 | 1 | 1 | 0.839 | 1 | 1 | 1 | 0.032 |
| SE | 100521 | ENSG00000054654 | SYNE2 | chr14 | + | 64009965 | 64010116 | 64007042 | 64007222 | 64016472 | 64016631 | 605 | 464 | 328 | 420 | 543 | 7 | 6 | 3 | 3 | 12 | 83 | 237 | 227 | 37 | 51 | 0 | 11 | 8 | 1 | 0 | 298 | 149 | 0.034848394 | 0.284365798 | 0.977 | 0.975 | 0.982 | 0.986 | 0.958 | 1 | 0.915 | 0.934 | 0.949 | 1 | 0.016 |
| SE | 99622 | ENSG00000109920 | FNBP4 | chr11 | - | 47725736 | 47725836 | 47724471 | 47724778 | 47731373 | 47731561 | 138 | 140 | 145 | 144 | 698 | 188 | 150 | 197 | 212 | 803 | 88 | 178 | 180 | 84 | 9 | 71 | 196 | 122 | 75 | 17 | 249 | 149 | 0.034853095 | 0.284374524 | 0.305 | 0.358 | 0.306 | 0.289 | 0.342 | 0.426 | 0.352 | 0.469 | 0.401 | 0.241 | -0.058 |
| SE | 72694 | ENSG00000104047 | DTWD1 | chr15 | + | 49631000 | 49631058 | 49625112 | 49625431 | 49632158 | 49632302 | 20 | 18 | 29 | 19 | 58 | 50 | 58 | 51 | 70 | 99 | 41 | 22 | 10 | 12 | 9 | 8 | 33 | 16 | 15 | 26 | 207 | 149 | 0.03485691 | 0.284376014 | 0.224 | 0.183 | 0.29 | 0.163 | 0.297 | 0.787 | 0.324 | 0.31 | 0.365 | 0.199 | -0.166 |
| SE | 96606 | ENSG00000108854 | SMURF2 | chr17 | - | 64606601 | 64606640 | 64598381 | 64598490 | 64661828 | 64661928 | 80 | 44 | 54 | 55 | 36 | 3 | 2 | 4 | 2 | 0 | 31 | 50 | 36 | 39 | 8 | 0 | 0 | 1 | 0 | 0 | 188 | 149 | 0.034869115 | 0.284445954 | 0.955 | 0.946 | 0.915 | 0.956 | 1 | 1 | 1 | 0.966 | 1 | 1 | -0.039 |
| SE | 58132 | ENSG00000117523 | PRRC2C | chr1 | + | 171568246 | 171568339 | 171566591 | 171566843 | 171574926 | 171575128 | 272 | 202 | 212 | 224 | 302 | 4 | 8 | 8 | 10 | 11 | 97 | 120 | 185 | 57 | 45 | 0 | 12 | 19 | 3 | 0 | 242 | 149 | 0.034927141 | 0.284889617 | 0.977 | 0.94 | 0.942 | 0.932 | 0.944 | 1 | 0.86 | 0.857 | 0.921 | 1 | 0.019 |
| SE | 74786 | ENSG00000011198 | ABHD5 | chr3 | + | 43717765 | 43717857 | 43714946 | 43715058 | 43718442 | 43718606 | 43 | 47 | 23 | 41 | 54 | 6 | 6 | 7 | 7 | 18 | 32 | 33 | 30 | 15 | 30 | 0 | 1 | 5 | 11 | 0 | 241 | 149 | 0.034953169 | 0.285072221 | 0.816 | 0.829 | 0.67 | 0.784 | 0.65 | 1 | 0.953 | 0.788 | 0.457 | 1 | -0.09 |
| SE | 80591 | ENSG00000179818 | PCBP1-AS1 | chr2 | - | 70083537 | 70083687 | 70048698 | 70051305 | 70085546 | 70085573 | 37 | 19 | 27 | 23 | 43 | 15 | 8 | 8 | 13 | 15 | 12 | 72 | 7 | 64 | 20 | 5 | 6 | 4 | 8 | 6 | 298 | 149 | 0.03495938 | 0.285093177 | 0.552 | 0.543 | 0.628 | 0.469 | 0.589 | 0.545 | 0.857 | 0.467 | 0.8 | 0.625 | -0.103 |
| SE | 13677 | ENSG00000006007 | GDE1 | chr16 | - | 19507654 | 19507779 | 19503860 | 19505092 | 19510838 | 19510944 | 75 | 53 | 43 | 66 | 104 | 4 | 1 | 1 | 3 | 2 | 231 | 90 | 110 | 124 | 230 | 0 | 0 | 0 | 8 | 0 | 274 | 149 | 0.03499751 | 0.285374402 | 0.911 | 0.966 | 0.959 | 0.923 | 0.966 | 1 | 1 | 1 | 0.894 | 1 | -0.034 |
| SE | 60531 | ENSG00000116984 | MTR | chr1 | + | 236815603 | 236815663 | 236812737 | 236812844 | 236816448 | 236816543 | 174 | 124 | 146 | 179 | 182 | 0 | 0 | 0 | 0 | 5 | 167 | 160 | 142 | 140 | 137 | 1 | 2 | 1 | 9 | 3 | 209 | 149 | 0.035001495 | 0.285377178 | 1 | 1 | 1 | 1 | 0.963 | 0.992 | 0.983 | 0.99 | 0.917 | 0.97 | 0.022 |
| SE | 72402 | ENSG00000138835 | RGS3 | chr9 | + | 113582071 | 113582242 | 113536795 | 113536918 | 113583449 | 113583671 | 38 | 25 | 14 | 13 | 6 | 85 | 13 | 42 | 39 | 24 | 15 | 25 | 9 | 11 | 30 | 9 | 44 | 10 | 12 | 28 | 298 | 149 | 0.035010837 | 0.285423627 | 0.183 | 0.49 | 0.143 | 0.143 | 0.111 | 0.455 | 0.221 | 0.31 | 0.314 | 0.349 | -0.116 |
| SE | 99572 | ENSG00000206561 | COLQ | chr3 | - | 15488205 | 15488307 | 15479337 | 15479382 | 15489524 | 15489637 | 30 | 10 | 15 | 33 | 5 | 0 | 0 | 3 | 1 | 0 | 16 | 6 | 4 | 0 | 15 | 0 | 0 | 2 | 6 | 0 | 251 | 149 | 0.035043491 | 0.285660088 | 1 | 1 | 0.748 | 0.951 | 1 | 1 | 1 | 0.543 | 0 | 1 | 0.231 |
| SE | 35739 | ENSG00000162736 | NCSTN | chr1 | + | 160343703 | 160343888 | 160343392 | 160343481 | 160344721 | 160344826 | 7 | 5 | 9 | 11 | 5 | 164 | 111 | 104 | 138 | 124 | 92 | 30 | 25 | 23 | 10 | 593 | 392 | 218 | 381 | 662 | 298 | 149 | 0.035053748 | 0.285713955 | 0.021 | 0.022 | 0.041 | 0.038 | 0.02 | 0.072 | 0.037 | 0.054 | 0.029 | 0.007 | -0.011 |
| SE | 78103 | ENSG00000141258 | SGSM2 | chr17 | + | 2373330 | 2373513 | 2372952 | 2373081 | 2375491 | 2375875 | 23 | 12 | 32 | 40 | 14 | 0 | 0 | 2 | 0 | 0 | 23 | 57 | 36 | 41 | 50 | 0 | 1 | 0 | 0 | 8 | 298 | 149 | 0.035060674 | 0.285740657 | 1 | 1 | 0.889 | 1 | 1 | 1 | 0.966 | 1 | 1 | 0.758 | 0.033 |
| SE | 4855 | ENSG00000132670 | PTPRA | chr20 | + | 3022688 | 3022824 | 3017815 | 3017913 | 3024471 | 3024621 | 432 | 234 | 323 | 426 | 270 | 10 | 2 | 7 | 8 | 5 | 876 | 483 | 374 | 627 | 1023 | 16 | 32 | 16 | 5 | 22 | 285 | 149 | 0.035085385 | 0.285912293 | 0.958 | 0.984 | 0.96 | 0.965 | 0.966 | 0.966 | 0.888 | 0.924 | 0.985 | 0.96 | 0.022 |
| SE | 18455 | ENSG00000100897 | DCAF11 | chr14 | + | 24116971 | 24117044 | 24115325 | 24115749 | 24117265 | 24117330 | 165 | 64 | 127 | 108 | 166 | 1 | 6 | 13 | 9 | 6 | 165 | 218 | 93 | 130 | 231 | 28 | 4 | 14 | 11 | 16 | 222 | 149 | 0.035108512 | 0.286070983 | 0.991 | 0.877 | 0.868 | 0.89 | 0.949 | 0.798 | 0.973 | 0.817 | 0.888 | 0.906 | 0.039 |
| SE | 109339 | ENSG00000102710 | SUPT20H | chr13 | - | 37021447 | 37021602 | 37019341 | 37019397 | 37022010 | 37022080 | 349 | 235 | 318 | 386 | 339 | 0 | 0 | 2 | 0 | 5 | 181 | 317 | 294 | 214 | 77 | 5 | 3 | 0 | 0 | 8 | 298 | 149 | 0.035112975 | 0.286077575 | 1 | 1 | 0.988 | 1 | 0.971 | 0.948 | 0.981 | 1 | 1 | 0.828 | 0.04 |
| SE | 21735 | ENSG00000188994 | ZNF292 | chr6 | + | 87214997 | 87215142 | 87155620 | 87155759 | 87215902 | 87216057 | 11 | 3 | 3 | 2 | 8 | 73 | 30 | 51 | 42 | 31 | 0 | 9 | 6 | 5 | 2 | 7 | 10 | 27 | 13 | 18 | 294 | 149 | 0.035133418 | 0.28621435 | 0.071 | 0.048 | 0.029 | 0.024 | 0.116 | 0 | 0.313 | 0.101 | 0.163 | 0.053 | -0.068 |
| SE | 49724 | ENSG00000171735 | CAMTA1 | chr1 | + | 6886239 | 6886346 | 6871756 | 6871840 | 6887653 | 6887842 | 5 | 2 | 8 | 2 | 11 | 51 | 37 | 22 | 66 | 81 | 0 | 0 | 2 | 2 | 18 | 78 | 70 | 16 | 8 | 205 | 256 | 149 | 0.035190613 | 0.286650463 | 0.054 | 0.031 | 0.175 | 0.017 | 0.073 | 0 | 0 | 0.068 | 0.127 | 0.049 | 0.021 |
| SE | 83671 | ENSG00000044446 | PHKA2 | chrX | - | 18902817 | 18902910 | 18901484 | 18901603 | 18905757 | 18905859 | 1 | 3 | 2 | 3 | 3 | 69 | 32 | 53 | 60 | 69 | 8 | 12 | 0 | 1 | 5 | 64 | 86 | 29 | 44 | 108 | 242 | 149 | 0.035200243 | 0.286699074 | 0.009 | 0.055 | 0.023 | 0.03 | 0.026 | 0.071 | 0.079 | 0 | 0.014 | 0.028 | -0.01 |
| SE | 33871 | ENSG00000132153 | DHX30 | chr3 | + | 47815962 | 47816128 | 47810656 | 47810711 | 47818021 | 47818117 | 18 | 4 | 20 | 8 | 7 | 29 | 20 | 35 | 38 | 56 | 20 | 20 | 8 | 17 | 10 | 34 | 14 | 30 | 18 | 37 | 298 | 149 | 0.035212118 | 0.286765966 | 0.237 | 0.091 | 0.222 | 0.095 | 0.059 | 0.227 | 0.417 | 0.118 | 0.321 | 0.119 | -0.1 |
| SE | 11381 | ENSG00000166747 | AP1G1 | chr16 | - | 71767876 | 71767885 | 71765488 | 71765584 | 71769622 | 71769699 | 5 | 2 | 3 | 1 | 5 | 136 | 67 | 74 | 96 | 116 | 0 | 0 | 0 | 0 | 0 | 38 | 99 | 131 | 72 | 24 | 158 | 149 | 0.035238512 | 0.286936735 | 0.034 | 0.027 | 0.037 | 0.01 | 0.039 | 0 | 0 | 0 | 0 | 0 | 0.029 |
| SE | 23564 | ENSG00000204574 | ABCF1 | chr6 | + | 30577408 | 30577455 | 30571392 | 30571560 | 30577817 | 30577913 | 34 | 28 | 36 | 35 | 23 | 0 | 0 | 0 | 1 | 0 | 75 | 24 | 21 | 59 | 71 | 0 | 0 | 0 | 11 | 0 | 196 | 149 | 0.035240416 | 0.286936735 | 1 | 1 | 1 | 0.964 | 1 | 1 | 1 | 1 | 0.803 | 1 | 0.032 |
| SE | 85521 | ENSG00000139083 | ETV6 | chr12 | + | 11752449 | 11752579 | 11649673 | 11650160 | 11839139 | 11839304 | 191 | 147 | 195 | 219 | 325 | 1 | 5 | 3 | 12 | 6 | 459 | 265 | 222 | 261 | 358 | 26 | 0 | 8 | 4 | 24 | 279 | 149 | 0.03524951 | 0.286980935 | 0.99 | 0.94 | 0.972 | 0.907 | 0.967 | 0.904 | 1 | 0.937 | 0.972 | 0.888 | 0.015 |
| SE | 1301 | ENSG00000138942 | RNF185 | chr22 | + | 31187046 | 31187270 | 31160181 | 31160303 | 31192683 | 31192702 | 52 | 15 | 27 | 42 | 10 | 1 | 7 | 1 | 1 | 4 | 51 | 25 | 22 | 37 | 79 | 19 | 12 | 1 | 6 | 15 | 298 | 149 | 0.035274243 | 0.287125624 | 0.963 | 0.517 | 0.931 | 0.955 | 0.556 | 0.573 | 0.51 | 0.917 | 0.755 | 0.725 | 0.088 |
| SE | 636 | ENSG00000100129 | EIF3L | chr22 | + | 37874369 | 37874524 | 37870164 | 37870347 | 37875840 | 37876011 | 1429 | 806 | 1492 | 1106 | 2351 | 0 | 1 | 2 | 1 | 3 | 1260 | 696 | 755 | 549 | 1268 | 0 | 8 | 14 | 7 | 10 | 298 | 149 | 0.035274784 | 0.287125624 | 1 | 0.998 | 0.997 | 0.998 | 0.997 | 1 | 0.978 | 0.964 | 0.975 | 0.984 | 0.018 |
| SE | 53414 | ENSG00000172531 | PPP1CA | chr11 | - | 67401067 | 67401232 | 67400688 | 67400919 | 67401727 | 67401793 | 275 | 108 | 175 | 224 | 169 | 10 | 1 | 6 | 6 | 0 | 133 | 251 | 115 | 153 | 304 | 3 | 7 | 5 | 19 | 5 | 298 | 149 | 0.035278284 | 0.287125624 | 0.932 | 0.982 | 0.936 | 0.949 | 1 | 0.957 | 0.947 | 0.92 | 0.801 | 0.968 | 0.041 |
| SE | 31927 | ENSG00000278259 | MYO19 | chr17 | - | 36498265 | 36498559 | 36495635 | 36496406 | 36499074 | 36499160 | 34 | 21 | 29 | 52 | 34 | 2 | 1 | 3 | 3 | 1 | 39 | 40 | 17 | 20 | 54 | 0 | 0 | 1 | 4 | 0 | 298 | 149 | 0.035336715 | 0.287571292 | 0.895 | 0.913 | 0.829 | 0.897 | 0.944 | 1 | 1 | 0.895 | 0.714 | 1 | -0.026 |
| SE | 26998 | ENSG00000026297 | RNASET2 | chr6 | - | 166955483 | 166955627 | 166952487 | 166952548 | 166956096 | 166956182 | 16 | 10 | 9 | 14 | 7 | 311 | 196 | 217 | 281 | 114 | 15 | 18 | 5 | 5 | 19 | 156 | 143 | 87 | 145 | 269 | 293 | 149 | 0.03534438 | 0.28760378 | 0.025 | 0.025 | 0.021 | 0.025 | 0.03 | 0.047 | 0.06 | 0.028 | 0.017 | 0.035 | -0.012 |
| SE | 97437 | ENSG00000186197 | EDARADD | chr1 | + | 236468230 | 236468276 | 236427391 | 236427450 | 236482266 | 236482319 | 12 | 7 | 9 | 13 | 32 | 0 | 2 | 1 | 1 | 0 | 6 | 10 | 19 | 6 | 12 | 0 | 10 | 1 | 0 | 0 | 195 | 149 | 0.035354754 | 0.287627018 | 1 | 0.728 | 0.873 | 0.909 | 1 | 1 | 0.433 | 0.936 | 1 | 1 | 0.028 |
| SE | 37250 | ENSG00000178694 | NSUN3 | chr3 | + | 94063811 | 94063979 | 94063010 | 94063138 | 94064436 | 94064518 | 4 | 9 | 9 | 5 | 6 | 43 | 23 | 41 | 35 | 32 | 7 | 14 | 5 | 10 | 2 | 4 | 15 | 68 | 23 | 6 | 298 | 149 | 0.035355669 | 0.287627018 | 0.044 | 0.164 | 0.099 | 0.067 | 0.086 | 0.467 | 0.318 | 0.035 | 0.179 | 0.143 | -0.136 |
| SE | 37983 | ENSG00000114670 | NEK11 | chr3 | + | 131080422 | 131080588 | 131029597 | 131029878 | 131109802 | 131109921 | 20 | 12 | 3 | 10 | 6 | 1 | 0 | 0 | 0 | 2 | 18 | 4 | 2 | 4 | 17 | 0 | 0 | 0 | 0 | 10 | 298 | 149 | 0.035358257 | 0.287627018 | 0.909 | 1 | 1 | 1 | 0.6 | 1 | 1 | 1 | 1 | 0.459 | 0.01 |
| SE | 89859 | ENSG00000284341 | NA | chr9 | + | 136980182 | 136980284 | 136979945 | 136980062 | 136980832 | 136980855 | 255 | 39 | 31 | 91 | 71 | 0 | 0 | 0 | 0 | 0 | 15 | 54 | 31 | 12 | 4 | 0 | 0 | 0 | 10 | 0 | 251 | 149 | 0.035390728 | 0.287840281 | 1 | 1 | 1 | 1 | 1 | 1 | 1 | 1 | 0.416 | 1 | 0.117 |
| SE | 3889 | ENSG00000105383 | CD33 | chr19 | + | 51235156 | 51235253 | 51226308 | 51226356 | 51239517 | 51239942 | 48 | 20 | 4 | 27 | 12 | 3 | 0 | 1 | 4 | 0 | 51 | 46 | 14 | 20 | 49 | 0 | 0 | 0 | 0 | 0 | 246 | 149 | 0.035392594 | 0.287840281 | 0.906 | 1 | 0.708 | 0.803 | 1 | 1 | 1 | 1 | 1 | 1 | -0.117 |
| SE | 6015 | ENSG00000025772 | TOMM34 | chr20 | - | 44948729 | 44948877 | 44943452 | 44943579 | 44951832 | 44952002 | 70 | 26 | 49 | 37 | 108 | 7 | 3 | 5 | 9 | 22 | 39 | 77 | 21 | 35 | 65 | 0 | 4 | 2 | 9 | 5 | 297 | 149 | 0.035395502 | 0.287840281 | 0.834 | 0.813 | 0.831 | 0.673 | 0.711 | 1 | 0.906 | 0.84 | 0.661 | 0.867 | -0.082 |
| SE | 69447 | ENSG00000145782 | ATG12 | chr5 | - | 115840817 | 115840934 | 115837627 | 115837764 | 115841389 | 115841544 | 6 | 13 | 6 | 22 | 3 | 35 | 20 | 42 | 40 | 34 | 0 | 6 | 11 | 2 | 0 | 5 | 21 | 37 | 51 | 4 | 266 | 149 | 0.035419264 | 0.288003601 | 0.088 | 0.267 | 0.074 | 0.236 | 0.047 | 0 | 0.138 | 0.143 | 0.021 | 0 | 0.082 |
| SE | 14997 | ENSG00000157823 | AP3S2 | chr15 | - | 89878264 | 89878300 | 89877305 | 89877424 | 89888520 | 89888632 | 15 | 7 | 5 | 10 | 1 | 25 | 24 | 18 | 22 | 26 | 17 | 11 | 32 | 29 | 12 | 44 | 50 | 13 | 19 | 29 | 185 | 149 | 0.035459061 | 0.288297259 | 0.326 | 0.19 | 0.183 | 0.268 | 0.03 | 0.237 | 0.151 | 0.665 | 0.551 | 0.25 | -0.171 |
| SE | 58854 | ENSG00000182552 | RWDD4 | chr4 | - | 183651217 | 183651327 | 183650983 | 183651131 | 183658928 | 183659178 | 31 | 61 | 29 | 34 | 111 | 0 | 0 | 0 | 0 | 2 | 5 | 41 | 10 | 26 | 6 | 10 | 0 | 0 | 0 | 0 | 259 | 149 | 0.035474995 | 0.288396863 | 1 | 1 | 1 | 1 | 0.97 | 0.223 | 1 | 1 | 1 | 1 | 0.149 |
| SE | 29204 | ENSG00000120708 | TGFBI | chr5 | + | 136062662 | 136062687 | 136061189 | 136061579 | 136063185 | 136063665 | 798 | 415 | 498 | 499 | 659 | 1 | 0 | 7 | 4 | 0 | 1193 | 372 | 302 | 491 | 1467 | 34 | 0 | 0 | 18 | 0 | 174 | 149 | 0.035482285 | 0.288415357 | 0.999 | 1 | 0.984 | 0.991 | 1 | 0.968 | 1 | 1 | 0.959 | 1 | 0.009 |
| SE | 16009 | ENSG00000068305 | MEF2A | chr15 | + | 99703361 | 99703385 | 99690240 | 99690428 | 99706728 | 99706855 | 0 | 4 | 3 | 1 | 2 | 91 | 90 | 83 | 133 | 154 | 12 | 0 | 7 | 0 | 0 | 70 | 152 | 105 | 125 | 58 | 173 | 149 | 0.035484637 | 0.288415357 | 0 | 0.037 | 0.03 | 0.006 | 0.011 | 0.129 | 0 | 0.054 | 0 | 0 | -0.02 |
| SE | 34056 | ENSG00000114867 | EIF4G1 | chr3 | + | 184316131 | 184316218 | 184314494 | 184314674 | 184317716 | 184317816 | 0 | 0 | 1 | 0 | 2 | 7 | 2 | 6 | 6 | 0 | 0 | 0 | 4 | 6 | 4 | 13 | 11 | 15 | 11 | 12 | 236 | 149 | 0.035503007 | 0.288482348 | 0 | 0 | 0.095 | 0 | 1 | 0 | 0 | 0.144 | 0.256 | 0.174 | 0.104 |
| SE | 73884 | ENSG00000123600 | METTL8 | chr2 | - | 171330558 | 171330616 | 171326041 | 171326148 | 171337452 | 171337502 | 22 | 21 | 36 | 37 | 27 | 4 | 4 | 4 | 2 | 6 | 61 | 22 | 24 | 15 | 15 | 0 | 15 | 12 | 6 | 6 | 207 | 149 | 0.035503541 | 0.288482348 | 0.798 | 0.791 | 0.866 | 0.93 | 0.764 | 1 | 0.514 | 0.59 | 0.643 | 0.643 | 0.152 |
| SE | 49620 | ENSG00000178927 | CYBC1 | chr17 | - | 82446622 | 82446696 | 82445250 | 82445960 | 82450699 | 82450734 | 37 | 23 | 26 | 40 | 19 | 0 | 0 | 2 | 0 | 0 | 11 | 43 | 18 | 63 | 24 | 0 | 0 | 4 | 5 | 0 | 223 | 149 | 0.035503933 | 0.288482348 | 1 | 1 | 0.897 | 1 | 1 | 1 | 1 | 0.75 | 0.894 | 1 | 0.051 |
| SE | 30789 | ENSG00000197976 | AKAP17A | chrX | + | 1600137 | 1600207 | 1599191 | 1599432 | 1600658 | 1602502 | 81 | 125 | 33 | 68 | 66 | 52 | 88 | 82 | 74 | 62 | 120 | 50 | 68 | 157 | 84 | 187 | 108 | 107 | 131 | 197 | 219 | 149 | 0.035511531 | 0.288514148 | 0.515 | 0.491 | 0.215 | 0.385 | 0.42 | 0.304 | 0.24 | 0.302 | 0.449 | 0.225 | 0.101 |
| SE | 22156 | ENSG00000007944 | MYLIP | chr6 | + | 16141624 | 16141810 | 16130556 | 16130747 | 16143019 | 16143217 | 339 | 199 | 354 | 442 | 447 | 5 | 1 | 1 | 2 | 1 | 390 | 503 | 380 | 250 | 158 | 7 | 4 | 1 | 9 | 0 | 298 | 149 | 0.035530207 | 0.288635931 | 0.971 | 0.99 | 0.994 | 0.991 | 0.996 | 0.965 | 0.984 | 0.995 | 0.933 | 1 | 0.013 |
| SE | 87345 | ENSG00000215301 | DDX3X | chrX | + | 41347314 | 41347454 | 41346858 | 41347012 | 41347639 | 41349884 | 299 | 194 | 294 | 303 | 411 | 39 | 22 | 19 | 58 | 50 | 17 | 168 | 243 | 120 | 36 | 12 | 54 | 56 | 33 | 0 | 289 | 149 | 0.035541464 | 0.288696532 | 0.798 | 0.82 | 0.889 | 0.729 | 0.809 | 0.422 | 0.616 | 0.691 | 0.652 | 1 | 0.133 |
| SE | 41363 | ENSG00000144401 | METTL21A | chr2 | - | 207621805 | 207621917 | 207609114 | 207613443 | 207624228 | 207624404 | 45 | 21 | 36 | 78 | 21 | 0 | 0 | 1 | 1 | 0 | 29 | 24 | 10 | 60 | 61 | 0 | 0 | 11 | 0 | 0 | 261 | 149 | 0.035545041 | 0.288696532 | 1 | 1 | 0.954 | 0.978 | 1 | 1 | 1 | 0.342 | 1 | 1 | 0.118 |
| SE | 59198 | ENSG00000113615 | SEC24A | chr5 | + | 134666822 | 134666996 | 134661118 | 134661586 | 134671808 | 134671886 | 36 | 31 | 22 | 23 | 63 | 4 | 7 | 0 | 7 | 7 | 65 | 36 | 77 | 59 | 29 | 0 | 12 | 4 | 2 | 0 | 298 | 149 | 0.035593727 | 0.28906197 | 0.818 | 0.689 | 1 | 0.622 | 0.818 | 1 | 0.6 | 0.906 | 0.937 | 1 | -0.099 |
| SE | 44250 | ENSG00000115808 | STRN | chr2 | - | 36857855 | 36858023 | 36855211 | 36855352 | 36861131 | 36861253 | 148 | 93 | 142 | 204 | 196 | 0 | 0 | 0 | 0 | 0 | 8 | 220 | 157 | 72 | 10 | 10 | 0 | 0 | 0 | 0 | 298 | 149 | 0.03560249 | 0.289100055 | 1 | 1 | 1 | 1 | 1 | 0.286 | 1 | 1 | 1 | 1 | 0.143 |
| SE | 56174 | ENSG00000094975 | SUCO | chr1 | + | 172553259 | 172553370 | 172551511 | 172551626 | 172555868 | 172556023 | 114 | 82 | 98 | 131 | 233 | 15 | 7 | 8 | 1 | 13 | 33 | 104 | 76 | 51 | 109 | 0 | 14 | 2 | 0 | 0 | 260 | 149 | 0.035605802 | 0.289100055 | 0.813 | 0.87 | 0.875 | 0.987 | 0.911 | 1 | 0.81 | 0.956 | 1 | 1 | -0.062 |
| SE | 46412 | ENSG00000164010 | ERMAP | chr1 | + | 42817533 | 42817815 | 42817121 | 42817253 | 42825622 | 42825735 | 10 | 3 | 1 | 10 | 1 | 7 | 7 | 1 | 6 | 0 | 40 | 19 | 1 | 9 | 27 | 15 | 6 | 0 | 0 | 9 | 298 | 149 | 0.035615016 | 0.28914489 | 0.417 | 0.176 | 0.333 | 0.455 | 1 | 0.571 | 0.613 | 1 | 1 | 0.6 | -0.281 |
| SE | 91781 | ENSG00000133794 | ARNTL | chr11 | + | 13326412 | 13326467 | 13309986 | 13310059 | 13349944 | 13350070 | 26 | 3 | 14 | 20 | 24 | 30 | 23 | 24 | 24 | 19 | 6 | 25 | 31 | 19 | 16 | 8 | 18 | 23 | 9 | 18 | 204 | 149 | 0.035629524 | 0.289232681 | 0.388 | 0.087 | 0.299 | 0.378 | 0.48 | 0.354 | 0.504 | 0.496 | 0.607 | 0.394 | -0.145 |
| SE | 33701 | ENSG00000176261 | ZBTB8OS | chr1 | - | 32634767 | 32634792 | 32633950 | 32634072 | 32650432 | 32650535 | 66 | 35 | 65 | 44 | 82 | 0 | 0 | 3 | 7 | 2 | 53 | 54 | 58 | 29 | 60 | 0 | 4 | 12 | 0 | 5 | 174 | 149 | 0.03565063 | 0.28937401 | 1 | 1 | 0.949 | 0.843 | 0.972 | 1 | 0.92 | 0.805 | 1 | 0.911 | 0.026 |
| SE | 106044 | ENSG00000204482 | LST1 | chr6 | + | 31587948 | 31587966 | 31587640 | 31587733 | 31588517 | 31588676 | 24 | 9 | 22 | 20 | 37 | 36 | 49 | 23 | 30 | 39 | 17 | 25 | 5 | 12 | 47 | 66 | 34 | 28 | 74 | 84 | 167 | 149 | 0.035691908 | 0.289679036 | 0.373 | 0.141 | 0.46 | 0.373 | 0.458 | 0.187 | 0.396 | 0.137 | 0.126 | 0.333 | 0.125 |
| SE | 81645 | ENSG00000133195 | SLC39A11 | chr17 | - | 73091823 | 73092116 | 73088656 | 73088775 | 73092610 | 73092712 | 1 | 2 | 0 | 0 | 0 | 9 | 0 | 6 | 9 | 1 | 7 | 6 | 1 | 1 | 0 | 0 | 8 | 7 | 6 | 3 | 298 | 149 | 0.03570101 | 0.289722873 | 0.053 | 1 | 0 | 0 | 0 | 1 | 0.273 | 0.067 | 0.077 | 0 | -0.073 |
| SE | 13918 | ENSG00000069345 | DNAJA2 | chr16 | - | 46967512 | 46967646 | 46964610 | 46964807 | 46968083 | 46968164 | 773 | 564 | 697 | 927 | 1411 | 2 | 2 | 1 | 6 | 2 | 1250 | 818 | 798 | 1135 | 689 | 0 | 18 | 4 | 15 | 5 | 283 | 149 | 0.035708745 | 0.289755615 | 0.995 | 0.993 | 0.997 | 0.988 | 0.997 | 1 | 0.96 | 0.991 | 0.976 | 0.986 | 0.011 |
| SE | 91181 | ENSG00000164941 | INTS8 | chr8 | + | 94866157 | 94866191 | 94859510 | 94859632 | 94867139 | 94867196 | 34 | 39 | 28 | 43 | 55 | 1 | 5 | 2 | 2 | 0 | 6 | 4 | 18 | 10 | 1 | 0 | 0 | 0 | 0 | 0 | 183 | 149 | 0.035740723 | 0.289985042 | 0.965 | 0.864 | 0.919 | 0.946 | 1 | 1 | 1 | 1 | 1 | 1 | -0.061 |
| SE | 63342 | ENSG00000151092 | NGLY1 | chr3 | - | 25736003 | 25736149 | 25733871 | 25733982 | 25751097 | 25751263 | 145 | 108 | 114 | 158 | 153 | 7 | 5 | 9 | 7 | 10 | 31 | 132 | 78 | 83 | 18 | 0 | 0 | 2 | 10 | 0 | 295 | 149 | 0.035746017 | 0.28999794 | 0.913 | 0.916 | 0.865 | 0.919 | 0.885 | 1 | 1 | 0.952 | 0.807 | 1 | -0.052 |
| SE | 93261 | ENSG00000114686 | MRPL3 | chr3 | - | 131498178 | 131498277 | 131489980 | 131490080 | 131500429 | 131500484 | 475 | 412 | 462 | 476 | 764 | 4 | 0 | 4 | 2 | 2 | 552 | 354 | 349 | 344 | 203 | 2 | 4 | 16 | 2 | 0 | 248 | 149 | 0.035754013 | 0.290032763 | 0.986 | 1 | 0.986 | 0.993 | 0.996 | 0.994 | 0.982 | 0.929 | 0.99 | 1 | 0.013 |
| SE | 30192 | ENSG00000168397 | ATG4B | chr2 | + | 241653306 | 241653369 | 241651263 | 241651335 | 241653511 | 241653524 | 33 | 28 | 32 | 25 | 28 | 128 | 90 | 75 | 115 | 215 | 24 | 85 | 21 | 52 | 87 | 186 | 192 | 82 | 117 | 166 | 212 | 149 | 0.035770346 | 0.290135193 | 0.153 | 0.179 | 0.231 | 0.133 | 0.084 | 0.083 | 0.237 | 0.153 | 0.238 | 0.269 | -0.04 |
| SE | 33364 | ENSG00000187742 | SECISBP2 | chr9 | + | 89334521 | 89334730 | 89332907 | 89332986 | 89338457 | 89338580 | 162 | 133 | 159 | 166 | 295 | 0 | 0 | 0 | 0 | 0 | 98 | 189 | 121 | 123 | 23 | 0 | 6 | 5 | 1 | 0 | 298 | 149 | 0.035789659 | 0.290261773 | 1 | 1 | 1 | 1 | 1 | 1 | 0.94 | 0.924 | 0.984 | 1 | 0.03 |
| SE | 39476 | ENSG00000103507 | BCKDK | chr16 | + | 31110687 | 31110761 | 31110400 | 31110499 | 31111090 | 31111219 | 118 | 50 | 89 | 87 | 187 | 6 | 0 | 1 | 1 | 0 | 164 | 125 | 102 | 115 | 232 | 0 | 0 | 4 | 8 | 8 | 223 | 149 | 0.035797965 | 0.290299067 | 0.929 | 1 | 0.983 | 0.983 | 1 | 1 | 1 | 0.945 | 0.906 | 0.951 | 0.019 |
| SE | 89101 | ENSG00000064115 | TM7SF3 | chr12 | - | 26975495 | 26975658 | 26971578 | 26974227 | 26980565 | 26980646 | 155 | 85 | 100 | 123 | 155 | 0 | 0 | 0 | 1 | 0 | 143 | 150 | 102 | 165 | 233 | 0 | 4 | 0 | 0 | 10 | 298 | 149 | 0.035815424 | 0.290410568 | 1 | 1 | 1 | 0.984 | 1 | 1 | 0.949 | 1 | 1 | 0.921 | 0.023 |
| SE | 10906 | ENSG00000074370 | ATP2A3 | chr17 | - | 3928226 | 3928327 | 3925345 | 3925438 | 3928662 | 3928780 | 2 | 2 | 6 | 6 | 0 | 33 | 4 | 21 | 37 | 15 | 9 | 21 | 4 | 0 | 14 | 25 | 29 | 16 | 16 | 106 | 250 | 149 | 0.035895429 | 0.29101832 | 0.035 | 0.23 | 0.146 | 0.088 | 0 | 0.177 | 0.301 | 0.13 | 0 | 0.073 | -0.036 |
| SE | 79678 | ENSG00000134905 | CARS2 | chr13 | - | 110644064 | 110644182 | 110642314 | 110642521 | 110644384 | 110644483 | 78 | 83 | 81 | 83 | 98 | 94 | 59 | 69 | 79 | 100 | 152 | 126 | 107 | 153 | 131 | 105 | 79 | 19 | 69 | 184 | 267 | 149 | 0.03589781 | 0.29101832 | 0.317 | 0.44 | 0.396 | 0.37 | 0.354 | 0.447 | 0.471 | 0.759 | 0.553 | 0.284 | -0.127 |
| SE | 5648 | ENSG00000080839 | RBL1 | chr20 | - | 37067232 | 37067297 | 37066992 | 37067121 | 37067985 | 37068186 | 100 | 68 | 84 | 115 | 148 | 11 | 18 | 10 | 16 | 21 | 85 | 93 | 37 | 83 | 74 | 18 | 23 | 29 | 19 | 7 | 214 | 149 | 0.03591206 | 0.291103698 | 0.864 | 0.725 | 0.854 | 0.833 | 0.831 | 0.767 | 0.738 | 0.47 | 0.753 | 0.88 | 0.1 |
| SE | 87610 | ENSG00000102078 | SLC25A14 | chrX | + | 130346543 | 130346691 | 130345181 | 130345275 | 130349250 | 130349345 | 30 | 31 | 26 | 44 | 25 | 1 | 0 | 3 | 1 | 3 | 40 | 26 | 34 | 29 | 4 | 0 | 0 | 0 | 1 | 0 | 297 | 149 | 0.035926296 | 0.29118895 | 0.938 | 1 | 0.813 | 0.957 | 0.807 | 1 | 1 | 1 | 0.936 | 1 | -0.084 |
| SE | 57996 | ENSG00000175634 | RPS6KB2 | chr11 | + | 67433125 | 67433216 | 67432951 | 67433042 | 67433339 | 67433447 | 68 | 58 | 69 | 73 | 70 | 5 | 2 | 3 | 6 | 2 | 101 | 117 | 84 | 83 | 103 | 5 | 9 | 14 | 4 | 11 | 240 | 149 | 0.035943461 | 0.291297925 | 0.894 | 0.947 | 0.935 | 0.883 | 0.956 | 0.926 | 0.89 | 0.788 | 0.928 | 0.853 | 0.046 |
| SE | 87443 | ENSG00000183354 | KIAA2026 | chr9 | - | 5954015 | 5954095 | 5944870 | 5944960 | 5968018 | 5968081 | 298 | 294 | 253 | 286 | 602 | 6 | 6 | 8 | 18 | 13 | 55 | 305 | 237 | 117 | 20 | 11 | 10 | 16 | 2 | 0 | 229 | 149 | 0.035966192 | 0.291451971 | 0.97 | 0.97 | 0.954 | 0.912 | 0.968 | 0.765 | 0.952 | 0.906 | 0.974 | 1 | 0.035 |
| SE | 45847 | ENSG00000163714 | U2SURP | chr3 | + | 143054942 | 143055119 | 143053675 | 143053794 | 143056311 | 143056500 | 359 | 280 | 341 | 303 | 570 | 18 | 13 | 11 | 16 | 37 | 6 | 206 | 192 | 21 | 10 | 2 | 5 | 7 | 0 | 0 | 298 | 149 | 0.035989661 | 0.291586702 | 0.909 | 0.915 | 0.939 | 0.904 | 0.885 | 0.6 | 0.954 | 0.932 | 1 | 1 | 0.013 |
| SE | 55073 | ENSG00000104894 | CD37 | chr19 | + | 49339329 | 49339413 | 49338699 | 49338936 | 49339657 | 49340027 | 518 | 166 | 322 | 343 | 342 | 3 | 4 | 7 | 9 | 1 | 368 | 457 | 265 | 321 | 782 | 0 | 5 | 2 | 2 | 0 | 233 | 149 | 0.035990266 | 0.291586702 | 0.991 | 0.964 | 0.967 | 0.961 | 0.995 | 1 | 0.983 | 0.988 | 0.99 | 1 | -0.017 |
| SE | 81570 | ENSG00000132256 | TRIM5 | chr11 | - | 5667688 | 5667711 | 5665980 | 5666081 | 5678203 | 5678434 | 68 | 90 | 81 | 123 | 102 | 4 | 1 | 3 | 2 | 4 | 284 | 43 | 55 | 88 | 83 | 3 | 0 | 0 | 2 | 0 | 172 | 149 | 0.035999341 | 0.291613685 | 0.936 | 0.987 | 0.959 | 0.982 | 0.957 | 0.988 | 1 | 1 | 0.974 | 1 | -0.028 |
| SE | 97661 | ENSG00000147457 | CHMP7 | chr8 | + | 23259065 | 23259126 | 23258731 | 23258830 | 23260143 | 23260286 | 305 | 171 | 370 | 273 | 553 | 11 | 6 | 13 | 10 | 6 | 115 | 260 | 143 | 122 | 137 | 0 | 4 | 2 | 8 | 0 | 210 | 149 | 0.036001046 | 0.291613685 | 0.952 | 0.953 | 0.953 | 0.951 | 0.985 | 1 | 0.979 | 0.981 | 0.915 | 1 | -0.016 |
| SE | 60515 | ENSG00000116984 | MTR | chr1 | + | 236803427 | 236803642 | 236795259 | 236795737 | 236812737 | 236812844 | 51 | 15 | 43 | 53 | 17 | 0 | 1 | 0 | 0 | 0 | 23 | 46 | 25 | 22 | 24 | 0 | 0 | 9 | 0 | 0 | 298 | 149 | 0.036005636 | 0.291620697 | 1 | 0.882 | 1 | 1 | 1 | 1 | 1 | 0.581 | 1 | 1 | 0.06 |
| SE | 56118 | ENSG00000163001 | CFAP36 | chr2 | + | 55544219 | 55544369 | 55543937 | 55544074 | 55544906 | 55545079 | 111 | 94 | 158 | 120 | 204 | 3 | 0 | 0 | 1 | 4 | 88 | 113 | 180 | 65 | 16 | 15 | 0 | 0 | 0 | 0 | 298 | 149 | 0.036046441 | 0.291920989 | 0.949 | 1 | 1 | 0.984 | 0.962 | 0.746 | 1 | 1 | 1 | 1 | 0.03 |
| SE | 58852 | ENSG00000182552 | RWDD4 | chr4 | - | 183651217 | 183651323 | 183650983 | 183651131 | 183658928 | 183659156 | 31 | 59 | 29 | 34 | 107 | 0 | 0 | 0 | 0 | 2 | 5 | 41 | 10 | 26 | 6 | 10 | 0 | 0 | 0 | 0 | 255 | 149 | 0.036060568 | 0.292005198 | 1 | 1 | 1 | 1 | 0.969 | 0.226 | 1 | 1 | 1 | 1 | 0.149 |
| SE | 91036 | ENSG00000120910 | PPP3CC | chr8 | + | 22539468 | 22539498 | 22532920 | 22533018 | 22540614 | 22541056 | 97 | 51 | 117 | 128 | 232 | 30 | 13 | 23 | 31 | 55 | 90 | 176 | 96 | 71 | 81 | 7 | 24 | 25 | 12 | 20 | 179 | 149 | 0.036091627 | 0.292216558 | 0.729 | 0.766 | 0.809 | 0.775 | 0.778 | 0.915 | 0.859 | 0.762 | 0.831 | 0.771 | -0.056 |
| SE | 17109 | ENSG00000276550 | HERC2P2 | chr15 | + | 22542482 | 22542615 | 22541994 | 22542209 | 22542722 | 22542850 | 100 | 78 | 130 | 78 | 59 | 0 | 0 | 0 | 0 | 0 | 80 | 111 | 83 | 34 | 35 | 0 | 0 | 1 | 4 | 5 | 282 | 149 | 0.036094134 | 0.292216558 | 1 | 1 | 1 | 1 | 1 | 1 | 1 | 0.978 | 0.818 | 0.787 | 0.083 |
| SE | 106675 | ENSG00000120662 | MTRF1 | chr13 | - | 41257672 | 41257797 | 41254528 | 41254620 | 41260492 | 41260915 | 6 | 15 | 8 | 7 | 34 | 129 | 82 | 83 | 101 | 247 | 30 | 23 | 9 | 3 | 0 | 54 | 76 | 73 | 74 | 48 | 274 | 149 | 0.036147328 | 0.292616953 | 0.025 | 0.09 | 0.05 | 0.036 | 0.07 | 0.232 | 0.141 | 0.063 | 0.022 | 0 | -0.037 |
| SE | 36626 | ENSG00000169180 | XPO6 | chr16 | - | 28180940 | 28181031 | 28177219 | 28177332 | 28211365 | 28211965 | 444 | 300 | 517 | 461 | 975 | 4 | 7 | 2 | 13 | 13 | 552 | 456 | 733 | 404 | 545 | 10 | 16 | 18 | 6 | 10 | 240 | 149 | 0.036173788 | 0.29280088 | 0.986 | 0.964 | 0.994 | 0.957 | 0.979 | 0.972 | 0.947 | 0.962 | 0.977 | 0.971 | 0.01 |
| SE | 16246 | ENSG00000166557 | TMED3 | chr15 | + | 79314540 | 79314655 | 79313756 | 79314005 | 79321977 | 79322837 | 5 | 1 | 0 | 6 | 0 | 71 | 40 | 59 | 55 | 58 | 0 | 23 | 4 | 0 | 6 | 120 | 68 | 67 | 43 | 160 | 264 | 149 | 0.03618658 | 0.292874147 | 0.038 | 0.014 | 0 | 0.058 | 0 | 0 | 0.16 | 0.033 | 0 | 0.021 | -0.021 |
| SE | 78892 | ENSG00000173875 | ZNF791 | chr19 | + | 12612225 | 12612365 | 12610919 | 12611082 | 12623699 | 12623826 | 2 | 2 | 1 | 0 | 0 | 23 | 25 | 50 | 43 | 11 | 0 | 0 | 5 | 9 | 0 | 37 | 43 | 65 | 30 | 13 | 289 | 149 | 0.03621634 | 0.293084708 | 0.043 | 0.04 | 0.01 | 0 | 0 | 0 | 0 | 0.038 | 0.134 | 0 | -0.016 |
| SE | 16944 | ENSG00000154237 | LRRK1 | chr15 | + | 101066639 | 101066741 | 101065351 | 101066205 | 101068670 | 101069068 | 10 | 4 | 19 | 8 | 6 | 1 | 0 | 0 | 5 | 4 | 25 | 14 | 8 | 34 | 45 | 2 | 0 | 4 | 1 | 0 | 251 | 149 | 0.036246563 | 0.293298974 | 0.856 | 1 | 1 | 0.487 | 0.471 | 0.881 | 1 | 0.543 | 0.953 | 1 | -0.113 |
| SE | 57608 | ENSG00000101361 | NOP56 | chr20 | + | 2657372 | 2657488 | 2657080 | 2657218 | 2657928 | 2658393 | 4 | 1 | 6 | 1 | 2 | 32 | 16 | 16 | 37 | 17 | 13 | 2 | 8 | 0 | 10 | 12 | 33 | 22 | 26 | 46 | 265 | 149 | 0.036253308 | 0.293323239 | 0.066 | 0.034 | 0.174 | 0.015 | 0.062 | 0.379 | 0.033 | 0.17 | 0 | 0.109 | -0.068 |
| SE | 5276 | ENSG00000101417 | PXMP4 | chr20 | - | 33710554 | 33710753 | 33702757 | 33707969 | 33714673 | 33714736 | 27 | 12 | 36 | 38 | 29 | 0 | 1 | 4 | 2 | 1 | 7 | 33 | 17 | 6 | 42 | 0 | 0 | 0 | 0 | 0 | 298 | 149 | 0.036280782 | 0.293515204 | 1 | 0.857 | 0.818 | 0.905 | 0.935 | 1 | 1 | 1 | 1 | 1 | -0.097 |
| SE | 37233 | ENSG00000150457 | LATS2 | chr13 | - | 20979690 | 20979797 | 20973035 | 20975364 | 20981465 | 20981648 | 62 | 33 | 28 | 64 | 39 | 0 | 1 | 0 | 0 | 0 | 40 | 19 | 44 | 47 | 58 | 0 | 4 | 5 | 0 | 0 | 256 | 149 | 0.036292926 | 0.293583111 | 1 | 0.951 | 1 | 1 | 1 | 1 | 0.734 | 0.837 | 1 | 1 | 0.076 |
| SE | 25711 | ENSG00000247774 | PCED1B-AS1 | chr12 | - | 47209280 | 47209482 | 47205900 | 47206069 | 47210639 | 47210761 | 58 | 24 | 82 | 78 | 169 | 21 | 7 | 17 | 13 | 42 | 18 | 36 | 45 | 6 | 9 | 0 | 1 | 4 | 13 | 0 | 298 | 149 | 0.036320342 | 0.293774536 | 0.58 | 0.632 | 0.707 | 0.75 | 0.668 | 1 | 0.947 | 0.849 | 0.188 | 1 | -0.129 |
| SE | 96888 | ENSG00000100731 | PCNX1 | chr14 | + | 70995740 | 70995843 | 70976941 | 70978648 | 71009633 | 71009724 | 8 | 3 | 7 | 6 | 5 | 82 | 47 | 69 | 69 | 92 | 14 | 6 | 10 | 8 | 0 | 7 | 112 | 90 | 17 | 28 | 252 | 149 | 0.036346314 | 0.293954242 | 0.055 | 0.036 | 0.057 | 0.049 | 0.031 | 0.542 | 0.031 | 0.062 | 0.218 | 0 | -0.125 |
| SE | 69912 | ENSG00000064995 | TAF11 | chr6 | - | 34880288 | 34880376 | 34877777 | 34878720 | 34882931 | 34883080 | 60 | 41 | 36 | 56 | 61 | 0 | 3 | 7 | 4 | 6 | 98 | 84 | 101 | 57 | 62 | 10 | 26 | 12 | 13 | 0 | 237 | 149 | 0.036373461 | 0.294143412 | 1 | 0.896 | 0.764 | 0.898 | 0.865 | 0.86 | 0.67 | 0.841 | 0.734 | 1 | 0.064 |
| SE | 22478 | ENSG00000165097 | KDM1B | chr6 | + | 18162834 | 18162924 | 18161326 | 18161454 | 18166266 | 18166378 | 158 | 128 | 105 | 173 | 199 | 16 | 10 | 6 | 4 | 12 | 239 | 149 | 113 | 334 | 118 | 7 | 2 | 0 | 17 | 5 | 239 | 149 | 0.036380444 | 0.294169501 | 0.86 | 0.889 | 1 | 0.964 | 0.912 | 0.955 | 0.979 | 1 | 0.925 | 0.936 | -0.034 |
| SE | 92848 | ENSG00000130958 | SLC35D2 | chr9 | - | 96351102 | 96351171 | 96345298 | 96345401 | 96360153 | 96360192 | 28 | 24 | 38 | 37 | 41 | 0 | 3 | 4 | 0 | 2 | 60 | 43 | 40 | 16 | 100 | 0 | 1 | 0 | 0 | 0 | 218 | 149 | 0.036397208 | 0.294272241 | 1 | 0.845 | 0.867 | 1 | 0.933 | 1 | 0.967 | 1 | 1 | 1 | -0.064 |
| SE | 23778 | ENSG00000113716 | HMGXB3 | chr5 | + | 150024261 | 150024680 | 150018565 | 150018697 | 150026705 | 150026881 | 130 | 44 | 70 | 74 | 44 | 0 | 1 | 2 | 0 | 0 | 59 | 77 | 78 | 27 | 98 | 0 | 0 | 5 | 0 | 5 | 298 | 149 | 0.036400667 | 0.294272241 | 1 | 0.957 | 0.946 | 1 | 1 | 1 | 1 | 0.886 | 1 | 0.907 | 0.022 |
| SE | 84587 | ENSG00000078589 | P2RY10 | chrX | + | 78952192 | 78952335 | 78945420 | 78945495 | 78960507 | 78961080 | 87 | 71 | 165 | 151 | 342 | 49 | 44 | 78 | 75 | 162 | 80 | 122 | 89 | 65 | 30 | 16 | 56 | 40 | 19 | 13 | 292 | 149 | 0.036417939 | 0.294381475 | 0.475 | 0.452 | 0.519 | 0.507 | 0.519 | 0.718 | 0.526 | 0.532 | 0.636 | 0.541 | -0.096 |
| SE | 59839 | ENSG00000120913 | PDLIM2 | chr8 | + | 22589297 | 22589374 | 22585320 | 22585395 | 22589595 | 22589684 | 30 | 11 | 24 | 11 | 8 | 0 | 1 | 1 | 6 | 0 | 13 | 34 | 3 | 11 | 17 | 0 | 0 | 0 | 0 | 0 | 226 | 149 | 0.036438267 | 0.294487729 | 1 | 0.879 | 0.941 | 0.547 | 1 | 1 | 1 | 1 | 1 | 1 | -0.127 |
| SE | 24149 | ENSG00000143774 | GUK1 | chr1 | + | 228148370 | 228148539 | 228147620 | 228147699 | 228148664 | 228148808 | 92 | 41 | 52 | 73 | 47 | 4 | 2 | 3 | 3 | 0 | 76 | 128 | 73 | 104 | 146 | 15 | 0 | 1 | 10 | 11 | 298 | 149 | 0.036438606 | 0.294487729 | 0.92 | 0.911 | 0.897 | 0.924 | 1 | 0.717 | 1 | 0.973 | 0.839 | 0.869 | 0.051 |
| SE | 44113 | ENSG00000114982 | KANSL3 | chr2 | - | 96619671 | 96619762 | 96619358 | 96619544 | 96631311 | 96631482 | 84 | 48 | 66 | 57 | 66 | 2 | 1 | 0 | 6 | 1 | 81 | 73 | 46 | 75 | 190 | 0 | 3 | 7 | 3 | 12 | 240 | 149 | 0.036450617 | 0.29455439 | 0.963 | 0.968 | 1 | 0.855 | 0.976 | 1 | 0.938 | 0.803 | 0.939 | 0.908 | 0.035 |
| SE | 17544 | ENSG00000100813 | ACIN1 | chr14 | - | 23071124 | 23071186 | 23069475 | 23069617 | 23071426 | 23071538 | 2 | 4 | 4 | 8 | 2 | 4 | 2 | 1 | 2 | 0 | 33 | 8 | 5 | 2 | 21 | 26 | 8 | 9 | 16 | 14 | 211 | 149 | 0.036497828 | 0.294875032 | 0.261 | 0.585 | 0.739 | 0.739 | 1 | 0.473 | 0.414 | 0.282 | 0.081 | 0.514 | 0.312 |
| SE | 14364 | ENSG00000047346 | FAM214A | chr15 | - | 52600126 | 52600235 | 52593416 | 52593736 | 52601085 | 52601164 | 163 | 127 | 178 | 162 | 284 | 2 | 3 | 2 | 7 | 2 | 27 | 139 | 140 | 22 | 25 | 0 | 0 | 0 | 0 | 0 | 258 | 149 | 0.036523132 | 0.295049024 | 0.979 | 0.961 | 0.981 | 0.93 | 0.988 | 1 | 1 | 1 | 1 | 1 | -0.032 |
| SE | 14689 | ENSG00000182511 | FES | chr15 | + | 90889516 | 90889636 | 90889305 | 90889443 | 90889839 | 90889962 | 109 | 52 | 58 | 96 | 28 | 1 | 4 | 1 | 0 | 3 | 197 | 234 | 105 | 201 | 302 | 22 | 9 | 4 | 1 | 4 | 269 | 149 | 0.03654023 | 0.295151049 | 0.984 | 0.878 | 0.97 | 1 | 0.838 | 0.832 | 0.935 | 0.936 | 0.991 | 0.977 | 0 |
| SE | 103586 | ENSG00000150093 | ITGB1 | chr10 | - | 32953592 | 32953809 | 32935491 | 32935558 | 32958144 | 32958196 | 18 | 11 | 6 | 15 | 6 | 266 | 137 | 117 | 281 | 97 | 2 | 14 | 12 | 18 | 16 | 76 | 219 | 179 | 86 | 90 | 298 | 149 | 0.036543301 | 0.295151049 | 0.033 | 0.039 | 0.025 | 0.026 | 0.03 | 0.013 | 0.031 | 0.032 | 0.095 | 0.082 | -0.02 |
| SE | 6724 | ENSG00000186111 | PIP5K1C | chr19 | - | 3633436 | 3633520 | 3630182 | 3633169 | 3638883 | 3639016 | 18 | 5 | 7 | 13 | 2 | 27 | 3 | 14 | 10 | 4 | 30 | 15 | 7 | 27 | 42 | 13 | 5 | 1 | 14 | 75 | 233 | 149 | 0.036580178 | 0.295418422 | 0.299 | 0.516 | 0.242 | 0.454 | 0.242 | 0.596 | 0.657 | 0.817 | 0.552 | 0.264 | -0.227 |
| SE | 33116 | ENSG00000172053 | QARS1 | chr3 | - | 49097991 | 49098117 | 49095931 | 49096079 | 49098352 | 49098480 | 33 | 41 | 30 | 53 | 38 | 0 | 0 | 0 | 0 | 0 | 39 | 22 | 11 | 26 | 110 | 0 | 0 | 10 | 0 | 0 | 275 | 149 | 0.036593434 | 0.295466035 | 1 | 1 | 1 | 1 | 1 | 1 | 1 | 0.373 | 1 | 1 | 0.125 |
| SE | 67767 | ENSG00000036257 | CUL3 | chr2 | - | 224506857 | 224507003 | 224505748 | 224506132 | 224513523 | 224513638 | 295 | 207 | 228 | 260 | 355 | 0 | 0 | 0 | 0 | 1 | 101 | 125 | 292 | 111 | 38 | 0 | 5 | 5 | 3 | 0 | 295 | 149 | 0.036593621 | 0.295466035 | 1 | 1 | 1 | 1 | 0.994 | 1 | 0.927 | 0.967 | 0.949 | 1 | 0.03 |
| SE | 69454 | ENSG00000132912 | DCTN4 | chr5 | - | 150755530 | 150755567 | 150753478 | 150753657 | 150756416 | 150756487 | 3 | 2 | 5 | 4 | 19 | 101 | 80 | 78 | 84 | 87 | 0 | 13 | 30 | 29 | 0 | 86 | 122 | 118 | 100 | 90 | 186 | 149 | 0.03660126 | 0.295497245 | 0.023 | 0.02 | 0.049 | 0.037 | 0.149 | 0 | 0.079 | 0.169 | 0.189 | 0 | -0.032 |
| SE | 40833 | ENSG00000144591 | GMPPA | chr2 | + | 219505292 | 219505362 | 219504082 | 219504213 | 219505457 | 219505555 | 10 | 5 | 11 | 11 | 10 | 3 | 0 | 2 | 0 | 1 | 47 | 35 | 14 | 12 | 11 | 11 | 0 | 2 | 4 | 5 | 219 | 149 | 0.036613902 | 0.295568829 | 0.694 | 1 | 0.789 | 1 | 0.872 | 0.744 | 1 | 0.826 | 0.671 | 0.599 | 0.103 |
| SE | 44805 | ENSG00000054282 | SDCCAG8 | chr1 | + | 243415701 | 243415829 | 243378720 | 243378863 | 243426426 | 243426558 | 25 | 17 | 13 | 22 | 24 | 2 | 0 | 3 | 1 | 2 | 10 | 24 | 8 | 10 | 7 | 3 | 1 | 3 | 5 | 0 | 277 | 149 | 0.036622226 | 0.295605549 | 0.871 | 1 | 0.7 | 0.922 | 0.866 | 0.642 | 0.928 | 0.589 | 0.518 | 1 | 0.136 |
| SE | 11216 | ENSG00000140829 | DHX38 | chr16 | + | 72097676 | 72097781 | 72096821 | 72097009 | 72098644 | 72098792 | 112 | 74 | 105 | 120 | 163 | 7 | 2 | 3 | 6 | 2 | 278 | 145 | 132 | 184 | 303 | 0 | 15 | 2 | 11 | 32 | 254 | 149 | 0.036639515 | 0.295714612 | 0.904 | 0.956 | 0.954 | 0.921 | 0.98 | 1 | 0.85 | 0.975 | 0.908 | 0.847 | 0.027 |
| SE | 78125 | ENSG00000108590 | MED31 | chr17 | - | 6649981 | 6650078 | 6643310 | 6644659 | 6650355 | 6650433 | 108 | 43 | 57 | 91 | 125 | 12 | 4 | 10 | 6 | 2 | 58 | 56 | 34 | 74 | 44 | 4 | 7 | 3 | 11 | 16 | 246 | 149 | 0.036647212 | 0.295746247 | 0.845 | 0.867 | 0.775 | 0.902 | 0.974 | 0.898 | 0.829 | 0.873 | 0.803 | 0.625 | 0.067 |
| SE | 6153 | ENSG00000282826 | FRG1CP | chr20 | - | 28601656 | 28601756 | 28600391 | 28600461 | 28602608 | 28602761 | 21 | 24 | 10 | 24 | 27 | 36 | 60 | 67 | 67 | 58 | 16 | 11 | 8 | 16 | 56 | 20 | 24 | 36 | 25 | 21 | 249 | 149 | 0.036658594 | 0.295807612 | 0.259 | 0.193 | 0.082 | 0.177 | 0.218 | 0.324 | 0.215 | 0.117 | 0.277 | 0.615 | -0.124 |
| SE | 47526 | ENSG00000121766 | ZCCHC17 | chr1 | + | 31337174 | 31337275 | 31319108 | 31319166 | 31338956 | 31339048 | 294 | 190 | 186 | 293 | 247 | 0 | 0 | 2 | 3 | 0 | 414 | 292 | 198 | 342 | 805 | 10 | 0 | 4 | 0 | 12 | 250 | 149 | 0.036668716 | 0.2958588 | 1 | 1 | 0.982 | 0.983 | 1 | 0.961 | 1 | 0.967 | 1 | 0.976 | 0.012 |
| SE | 53435 | ENSG00000135953 | MFSD9 | chr2 | - | 102726844 | 102726974 | 102723699 | 102723878 | 102731024 | 102731068 | 20 | 4 | 12 | 3 | 2 | 0 | 0 | 1 | 3 | 0 | 5 | 23 | 3 | 9 | 11 | 0 | 0 | 4 | 9 | 0 | 279 | 149 | 0.036697526 | 0.296060745 | 1 | 1 | 0.865 | 0.348 | 1 | 1 | 1 | 0.286 | 0.348 | 1 | 0.116 |
| SE | 75907 | ENSG00000134324 | LPIN1 | chr2 | + | 11763022 | 11763286 | 11746633 | 11746671 | 11765532 | 11765733 | 7 | 1 | 2 | 1 | 2 | 4 | 1 | 5 | 7 | 1 | 0 | 3 | 3 | 2 | 0 | 7 | 15 | 5 | 3 | 6 | 298 | 149 | 0.036702759 | 0.296072449 | 0.467 | 0.333 | 0.167 | 0.067 | 0.5 | 0 | 0.091 | 0.231 | 0.25 | 0 | 0.192 |
| SE | 71623 | ENSG00000184182 | UBE2F | chr2 | + | 238016565 | 238016633 | 237973091 | 237973225 | 238025341 | 238025412 | 146 | 90 | 99 | 132 | 108 | 0 | 0 | 0 | 1 | 0 | 183 | 118 | 67 | 142 | 161 | 0 | 5 | 2 | 2 | 6 | 217 | 149 | 0.036739803 | 0.296333848 | 1 | 1 | 1 | 0.989 | 1 | 1 | 0.942 | 0.958 | 0.98 | 0.949 | 0.032 |
| SE | 6678 | ENSG00000099956 | SMARCB1 | chr22 | + | 23791755 | 23791894 | 23786965 | 23787262 | 23793558 | 23793688 | 139 | 71 | 109 | 103 | 145 | 6 | 0 | 0 | 2 | 2 | 136 | 230 | 91 | 134 | 191 | 1 | 0 | 0 | 0 | 32 | 288 | 149 | 0.036743487 | 0.296333848 | 0.923 | 1 | 1 | 0.964 | 0.974 | 0.986 | 1 | 1 | 1 | 0.755 | 0.024 |
| SE | 50119 | ENSG00000132704 | FCRL2 | chr1 | - | 157748552 | 157748618 | 157746870 | 157746899 | 157748874 | 157748960 | 73 | 62 | 65 | 88 | 94 | 6 | 0 | 2 | 4 | 0 | 58 | 114 | 46 | 20 | 48 | 0 | 0 | 2 | 2 | 0 | 215 | 149 | 0.036746517 | 0.296333848 | 0.894 | 1 | 0.957 | 0.938 | 1 | 1 | 1 | 0.941 | 0.874 | 1 | -0.005 |
| SE | 21577 | ENSG00000075568 | TMEM131 | chr2 | - | 97772296 | 97772424 | 97766477 | 97766602 | 97775842 | 97776018 | 145 | 121 | 136 | 173 | 170 | 3 | 2 | 1 | 0 | 0 | 125 | 145 | 82 | 104 | 62 | 0 | 4 | 3 | 5 | 0 | 277 | 149 | 0.036764357 | 0.296429371 | 0.963 | 0.97 | 0.987 | 1 | 1 | 1 | 0.951 | 0.936 | 0.918 | 1 | 0.023 |
| SE | 107828 | ENSG00000135457 | TFCP2 | chr12 | - | 51104154 | 51104203 | 51103669 | 51103763 | 51106524 | 51106613 | 251 | 231 | 277 | 286 | 752 | 5 | 9 | 10 | 15 | 17 | 340 | 280 | 127 | 306 | 162 | 17 | 15 | 3 | 25 | 0 | 198 | 149 | 0.036765935 | 0.296429371 | 0.974 | 0.951 | 0.954 | 0.935 | 0.971 | 0.938 | 0.934 | 0.97 | 0.902 | 1 | 0.008 |
| SE | 105597 | ENSG00000139323 | POC1B | chr12 | - | 89425160 | 89425379 | 89419717 | 89421257 | 89459637 | 89459718 | 132 | 116 | 110 | 155 | 439 | 19 | 10 | 9 | 11 | 15 | 43 | 119 | 98 | 72 | 30 | 0 | 3 | 11 | 3 | 0 | 298 | 149 | 0.036770563 | 0.296436159 | 0.776 | 0.853 | 0.859 | 0.876 | 0.936 | 1 | 0.952 | 0.817 | 0.923 | 1 | -0.078 |
| SE | 1764 | ENSG00000100077 | GRK3 | chr22 | + | 25663629 | 25663704 | 25644591 | 25644665 | 25667738 | 25667800 | 102 | 116 | 46 | 78 | 148 | 2 | 2 | 0 | 1 | 1 | 268 | 110 | 69 | 122 | 124 | 8 | 0 | 0 | 8 | 0 | 224 | 149 | 0.036789849 | 0.296561109 | 0.971 | 0.975 | 1 | 0.981 | 0.99 | 0.957 | 1 | 1 | 0.91 | 1 | 0.01 |
| SE | 106278 | ENSG00000251301 | LINC02384 | chr12 | - | 68442063 | 68442141 | 68433283 | 68433371 | 68451366 | 68451385 | 113 | 85 | 16 | 17 | 47 | 13 | 0 | 0 | 1 | 9 | 52 | 130 | 32 | 21 | 84 | 14 | 2 | 5 | 4 | 20 | 227 | 149 | 0.036808894 | 0.29666822 | 0.851 | 1 | 1 | 0.918 | 0.774 | 0.709 | 0.977 | 0.808 | 0.775 | 0.734 | 0.108 |
| SE | 99975 | ENSG00000185187 | SIGIRR | chr11 | - | 408694 | 408893 | 408072 | 408206 | 409859 | 410027 | 112 | 27 | 78 | 62 | 30 | 5 | 1 | 1 | 5 | 0 | 8 | 61 | 35 | 43 | 67 | 4 | 0 | 0 | 0 | 0 | 298 | 149 | 0.036811204 | 0.29666822 | 0.918 | 0.931 | 0.975 | 0.861 | 1 | 0.5 | 1 | 1 | 1 | 1 | 0.037 |
| SE | 12529 | ENSG00000103066 | PLA2G15 | chr16 | + | 68255705 | 68255990 | 68255281 | 68255380 | 68259145 | 68259252 | 21 | 8 | 20 | 12 | 30 | 0 | 0 | 0 | 0 | 0 | 9 | 22 | 9 | 32 | 29 | 0 | 0 | 2 | 0 | 6 | 298 | 149 | 0.036814504 | 0.29666822 | 1 | 1 | 1 | 1 | 1 | 1 | 1 | 0.692 | 1 | 0.707 | 0.12 |
| SE | 97186 | ENSG00000106290 | TAF6 | chr7 | - | 100113679 | 100113769 | 100113348 | 100113405 | 100113867 | 100113954 | 18 | 6 | 18 | 15 | 10 | 1 | 2 | 1 | 2 | 0 | 18 | 6 | 6 | 10 | 47 | 0 | 0 | 0 | 3 | 23 | 239 | 149 | 0.03681914 | 0.296675046 | 0.918 | 0.652 | 0.918 | 0.824 | 1 | 1 | 1 | 1 | 0.675 | 0.56 | 0.015 |
| SE | 48656 | ENSG00000131374 | TBC1D5 | chr3 | - | 17622647 | 17622765 | 17508473 | 17508605 | 17623848 | 17623913 | 63 | 40 | 38 | 58 | 126 | 88 | 86 | 50 | 75 | 125 | 45 | 33 | 40 | 55 | 7 | 60 | 99 | 51 | 101 | 29 | 267 | 149 | 0.036826304 | 0.296702237 | 0.285 | 0.206 | 0.298 | 0.301 | 0.36 | 0.295 | 0.157 | 0.304 | 0.233 | 0.119 | 0.068 |
| SE | 99556 | ENSG00000254986 | DPP3 | chr11 | + | 66504611 | 66504774 | 66497297 | 66497477 | 66509078 | 66509345 | 26 | 9 | 22 | 25 | 7 | 0 | 0 | 0 | 2 | 0 | 6 | 58 | 15 | 29 | 40 | 13 | 0 | 0 | 0 | 0 | 298 | 149 | 0.036844463 | 0.296817992 | 1 | 1 | 1 | 0.862 | 1 | 0.188 | 1 | 1 | 1 | 1 | 0.135 |
| SE | 21970 | ENSG00000012660 | ELOVL5 | chr6 | - | 53290400 | 53290535 | 53287861 | 53287942 | 53291775 | 53291963 | 4 | 3 | 7 | 8 | 4 | 1 | 2 | 0 | 0 | 3 | 3 | 20 | 16 | 11 | 9 | 0 | 0 | 0 | 0 | 0 | 284 | 149 | 0.036856408 | 0.296883678 | 0.677 | 0.44 | 1 | 1 | 0.412 | 1 | 1 | 1 | 1 | 1 | -0.294 |
| SE | 5548 | ENSG00000101052 | IFT52 | chr20 | + | 43605001 | 43605073 | 43604182 | 43604258 | 43613849 | 43613976 | 57 | 71 | 78 | 89 | 178 | 2 | 0 | 0 | 5 | 2 | 40 | 67 | 51 | 95 | 20 | 0 | 5 | 0 | 4 | 5 | 221 | 149 | 0.036894852 | 0.297162778 | 0.951 | 1 | 1 | 0.923 | 0.984 | 1 | 0.9 | 1 | 0.941 | 0.729 | 0.058 |
| SE | 27481 | ENSG00000104517 | UBR5 | chr8 | - | 102261145 | 102261259 | 102258984 | 102259129 | 102261914 | 102261986 | 255 | 164 | 211 | 205 | 309 | 0 | 0 | 2 | 0 | 0 | 0 | 196 | 174 | 60 | 24 | 9 | 1 | 1 | 0 | 0 | 263 | 149 | 0.036950098 | 0.29757713 | 1 | 1 | 0.984 | 1 | 1 | 0 | 0.991 | 0.99 | 1 | 1 | 0.201 |
| SE | 35969 | ENSG00000078070 | MCCC1 | chr3 | - | 183017265 | 183017337 | 183015217 | 183015566 | 183022416 | 183022554 | 26 | 12 | 22 | 21 | 56 | 0 | 0 | 0 | 0 | 0 | 24 | 38 | 17 | 16 | 2 | 0 | 0 | 0 | 9 | 0 | 221 | 149 | 0.036961061 | 0.297634806 | 1 | 1 | 1 | 1 | 1 | 1 | 1 | 1 | 0.545 | 1 | 0.091 |
| SE | 8963 | ENSG00000108395 | TRIM37 | chr17 | - | 59062566 | 59062648 | 59061031 | 59061108 | 59070822 | 59070947 | 55 | 32 | 22 | 48 | 42 | 0 | 0 | 2 | 0 | 0 | 38 | 36 | 40 | 44 | 17 | 5 | 1 | 0 | 3 | 0 | 231 | 149 | 0.036991312 | 0.297847775 | 1 | 1 | 0.876 | 1 | 1 | 0.831 | 0.959 | 1 | 0.904 | 1 | 0.036 |
| SE | 102668 | ENSG00000175029 | CTBP2 | chr10 | - | 124993201 | 124993329 | 124992694 | 124992812 | 124993854 | 124993985 | 228 | 75 | 119 | 250 | 86 | 1 | 0 | 0 | 3 | 0 | 210 | 118 | 122 | 157 | 150 | 0 | 4 | 0 | 0 | 10 | 277 | 149 | 0.037019942 | 0.298047646 | 0.992 | 1 | 1 | 0.978 | 1 | 1 | 0.941 | 1 | 1 | 0.89 | 0.028 |
| SE | 89590 | ENSG00000174799 | CEP135 | chr4 | + | 55974745 | 55974969 | 55971269 | 55971408 | 55980142 | 55980295 | 70 | 72 | 69 | 91 | 143 | 9 | 11 | 7 | 7 | 27 | 1 | 33 | 21 | 17 | 14 | 0 | 15 | 7 | 2 | 10 | 298 | 149 | 0.037076988 | 0.298476234 | 0.795 | 0.766 | 0.831 | 0.867 | 0.726 | 1 | 0.524 | 0.6 | 0.81 | 0.412 | 0.128 |
| SE | 69638 | ENSG00000120451 | SNX19 | chr11 | - | 130880621 | 130880806 | 130879623 | 130879711 | 130903254 | 130903384 | 102 | 74 | 107 | 92 | 184 | 30 | 1 | 0 | 0 | 0 | 79 | 127 | 46 | 72 | 85 | 32 | 5 | 10 | 0 | 0 | 298 | 149 | 0.037096052 | 0.298599004 | 0.63 | 0.974 | 1 | 1 | 1 | 0.552 | 0.927 | 0.697 | 1 | 1 | 0.086 |
| SE | 30947 | ENSG00000017260 | ATP2C1 | chr3 | + | 130980581 | 130980679 | 130975331 | 130975488 | 130992950 | 130993001 | 181 | 183 | 127 | 177 | 108 | 4 | 2 | 10 | 9 | 0 | 140 | 152 | 170 | 157 | 35 | 0 | 5 | 6 | 1 | 0 | 247 | 149 | 0.03711221 | 0.298698351 | 0.965 | 0.982 | 0.885 | 0.922 | 1 | 1 | 0.948 | 0.945 | 0.99 | 1 | -0.026 |
| SE | 100868 | ENSG00000149311 | ATM | chr11 | + | 108248932 | 108249102 | 108246963 | 108247127 | 108250700 | 108251072 | 1241 | 1097 | 1325 | 1423 | 3762 | 11 | 25 | 15 | 20 | 56 | 61 | 398 | 305 | 85 | 94 | 0 | 0 | 1 | 9 | 0 | 298 | 149 | 0.037125187 | 0.298772088 | 0.983 | 0.956 | 0.978 | 0.973 | 0.971 | 1 | 1 | 0.993 | 0.825 | 1 | 0.009 |
| SE | 93216 | ENSG00000113328 | CCNG1 | chr5 | + | 163444336 | 163444481 | 163443756 | 163443925 | 163445903 | 163446151 | 34 | 21 | 24 | 20 | 47 | 4 | 2 | 1 | 1 | 0 | 9 | 11 | 19 | 2 | 11 | 0 | 5 | 3 | 4 | 0 | 294 | 149 | 0.037139631 | 0.298857607 | 0.812 | 0.842 | 0.924 | 0.91 | 1 | 1 | 0.527 | 0.762 | 0.202 | 1 | 0.199 |
| SE | 92489 | ENSG00000130749 | ZC3H4 | chr19 | - | 47110894 | 47110956 | 47094388 | 47094608 | 47112147 | 47112247 | 4 | 5 | 17 | 9 | 6 | 22 | 15 | 23 | 19 | 17 | 14 | 19 | 13 | 23 | 29 | 28 | 37 | 3 | 29 | 36 | 211 | 149 | 0.037151276 | 0.298892902 | 0.114 | 0.191 | 0.343 | 0.251 | 0.2 | 0.261 | 0.266 | 0.754 | 0.359 | 0.363 | -0.181 |
| SE | 85527 | ENSG00000139083 | ETV6 | chr12 | + | 11884444 | 11884587 | 11869423 | 11869969 | 11890940 | 11895377 | 127 | 75 | 81 | 99 | 84 | 2 | 0 | 4 | 0 | 0 | 319 | 130 | 108 | 218 | 221 | 9 | 2 | 0 | 6 | 2 | 292 | 149 | 0.037151652 | 0.298892902 | 0.97 | 1 | 0.912 | 1 | 1 | 0.948 | 0.971 | 1 | 0.949 | 0.983 | 0.006 |
| SE | 93869 | ENSG00000116580 | GON4L | chr1 | - | 155832831 | 155832950 | 155826836 | 155827028 | 155853275 | 155853405 | 10 | 15 | 6 | 7 | 30 | 145 | 147 | 150 | 148 | 220 | 7 | 19 | 15 | 30 | 14 | 172 | 185 | 137 | 160 | 51 | 268 | 149 | 0.037179938 | 0.299089738 | 0.037 | 0.054 | 0.022 | 0.026 | 0.07 | 0.022 | 0.054 | 0.057 | 0.094 | 0.132 | -0.03 |
| SE | 37317 | ENSG00000004534 | RBM6 | chr3 | + | 49967197 | 49968748 | 49962575 | 49962685 | 49972058 | 49972148 | 221 | 186 | 185 | 165 | 200 | 4 | 0 | 5 | 2 | 1 | 160 | 209 | 187 | 197 | 90 | 0 | 10 | 2 | 4 | 5 | 298 | 149 | 0.037188732 | 0.299129746 | 0.965 | 1 | 0.949 | 0.976 | 0.99 | 1 | 0.913 | 0.979 | 0.961 | 0.9 | 0.025 |
| SE | 80424 | ENSG00000126746 | ZNF384 | chr12 | - | 6678662 | 6678710 | 6678126 | 6678460 | 6678945 | 6679183 | 72 | 41 | 61 | 72 | 69 | 3 | 4 | 6 | 11 | 2 | 119 | 72 | 36 | 71 | 137 | 8 | 1 | 1 | 5 | 4 | 197 | 149 | 0.037215277 | 0.299312516 | 0.948 | 0.886 | 0.885 | 0.832 | 0.963 | 0.918 | 0.982 | 0.965 | 0.915 | 0.963 | -0.046 |
| SE | 5386 | ENSG00000078747 | ITCH | chr20 | + | 34397207 | 34397321 | 34393790 | 34393881 | 34408650 | 34408792 | 6 | 4 | 17 | 12 | 20 | 321 | 185 | 259 | 337 | 532 | 0 | 34 | 22 | 7 | 0 | 111 | 437 | 281 | 249 | 28 | 263 | 149 | 0.037226463 | 0.299371727 | 0.01 | 0.012 | 0.036 | 0.02 | 0.021 | 0 | 0.042 | 0.042 | 0.016 | 0 | 0 |
| SE | 106263 | ENSG00000116198 | CEP104 | chr1 | - | 3828884 | 3828971 | 3826707 | 3826744 | 3829265 | 3829373 | 2 | 1 | 1 | 0 | 6 | 35 | 24 | 35 | 28 | 48 | 0 | 0 | 2 | 18 | 0 | 16 | 23 | 24 | 44 | 37 | 236 | 149 | 0.037261018 | 0.29961884 | 0.035 | 0.026 | 0.018 | 0 | 0.073 | 0 | 0 | 0.05 | 0.205 | 0 | -0.021 |
| SE | 60500 | ENSG00000115211 | EIF2B4 | chr2 | - | 27367745 | 27367822 | 27367456 | 27367559 | 27368024 | 27368113 | 82 | 48 | 64 | 69 | 101 | 2 | 0 | 1 | 0 | 1 | 29 | 73 | 43 | 43 | 73 | 6 | 0 | 0 | 4 | 0 | 226 | 149 | 0.037283077 | 0.299765432 | 0.964 | 1 | 0.977 | 1 | 0.985 | 0.761 | 1 | 1 | 0.876 | 1 | 0.058 |
| SE | 38373 | ENSG00000112365 | ZBTB24 | chr6 | - | 109467652 | 109467734 | 109462593 | 109466574 | 109481074 | 109482054 | 19 | 16 | 19 | 9 | 13 | 0 | 0 | 0 | 0 | 0 | 5 | 9 | 11 | 4 | 0 | 0 | 0 | 0 | 0 | 10 | 231 | 149 | 0.037295462 | 0.299834224 | 1 | 1 | 1 | 1 | 1 | 1 | 1 | 1 | 1 | 0 | 0.2 |
| SE | 29422 | ENSG00000169762 | TAPT1 | chr4 | - | 16202461 | 16202580 | 16191360 | 16191523 | 16226258 | 16226471 | 21 | 17 | 10 | 13 | 38 | 3 | 0 | 0 | 2 | 5 | 4 | 36 | 7 | 3 | 12 | 0 | 1 | 0 | 3 | 0 | 268 | 149 | 0.037323743 | 0.300030777 | 0.796 | 1 | 1 | 0.783 | 0.809 | 1 | 0.952 | 1 | 0.357 | 1 | 0.016 |
| SE | 54558 | ENSG00000179912 | R3HDM2 | chr12 | - | 57292571 | 57292667 | 57289008 | 57289040 | 57295398 | 57295507 | 534 | 532 | 423 | 644 | 700 | 38 | 49 | 27 | 35 | 76 | 1262 | 643 | 501 | 803 | 1068 | 161 | 44 | 55 | 51 | 120 | 245 | 149 | 0.037335779 | 0.300096721 | 0.895 | 0.868 | 0.905 | 0.918 | 0.849 | 0.827 | 0.899 | 0.847 | 0.905 | 0.844 | 0.023 |
| SE | 1035 | ENSG00000189306 | RRP7A | chr22 | - | 42518612 | 42518728 | 42518004 | 42518147 | 42519713 | 42519794 | 8 | 0 | 0 | 0 | 4 | 41 | 14 | 17 | 31 | 2 | 0 | 0 | 0 | 0 | 0 | 43 | 51 | 37 | 33 | 11 | 265 | 149 | 0.037340212 | 0.300101541 | 0.099 | 0 | 0 | 0 | 0.529 | 0 | 0 | 0 | 0 | 0 | 0.126 |
| SE | 15559 | ENSG00000184056 | VPS33B | chr15 | - | 91006371 | 91007046 | 91006000 | 91006059 | 91007468 | 91007573 | 60 | 49 | 51 | 40 | 55 | 0 | 0 | 0 | 0 | 0 | 104 | 63 | 81 | 43 | 61 | 0 | 4 | 0 | 6 | 0 | 298 | 149 | 0.037348524 | 0.300137536 | 1 | 1 | 1 | 1 | 1 | 1 | 0.887 | 1 | 0.782 | 1 | 0.066 |
| SE | 4922 | ENSG00000101246 | ARFRP1 | chr20 | - | 63701829 | 63701900 | 63699477 | 63700702 | 63702135 | 63702217 | 80 | 34 | 70 | 63 | 35 | 4 | 1 | 1 | 1 | 3 | 21 | 63 | 26 | 53 | 50 | 0 | 0 | 0 | 0 | 0 | 220 | 149 | 0.03739224 | 0.300458006 | 0.931 | 0.958 | 0.979 | 0.977 | 0.888 | 1 | 1 | 1 | 1 | 1 | -0.053 |
| SE | 10956 | ENSG00000091592 | NLRP1 | chr17 | - | 5533896 | 5533988 | 5532821 | 5532984 | 5536850 | 5536940 | 71 | 42 | 99 | 76 | 45 | 11 | 13 | 12 | 6 | 9 | 215 | 71 | 62 | 131 | 250 | 11 | 19 | 13 | 4 | 10 | 241 | 149 | 0.037399477 | 0.300477778 | 0.8 | 0.666 | 0.836 | 0.887 | 0.756 | 0.924 | 0.698 | 0.747 | 0.953 | 0.939 | -0.063 |
| SE | 72404 | ENSG00000138835 | RGS3 | chr9 | + | 113582071 | 113582242 | 113581558 | 113581658 | 113583449 | 113583671 | 39 | 25 | 15 | 12 | 6 | 1 | 0 | 0 | 0 | 8 | 15 | 23 | 9 | 11 | 30 | 0 | 0 | 0 | 1 | 0 | 298 | 149 | 0.037402376 | 0.300477778 | 0.951 | 1 | 1 | 1 | 0.273 | 1 | 1 | 1 | 0.846 | 1 | -0.124 |
| SE | 8678 | ENSG00000198909 | MAP3K3 | chr17 | + | 63668145 | 63668334 | 63666985 | 63667060 | 63681765 | 63681899 | 2 | 4 | 2 | 3 | 8 | 299 | 161 | 190 | 229 | 439 | 20 | 14 | 7 | 1 | 4 | 359 | 268 | 189 | 209 | 347 | 298 | 149 | 0.037433833 | 0.300699636 | 0.003 | 0.012 | 0.005 | 0.007 | 0.009 | 0.027 | 0.025 | 0.018 | 0.002 | 0.006 | -0.008 |
| SE | 26035 | ENSG00000131503 | ANKHD1 | chr5 | + | 140472238 | 140472295 | 140464666 | 140464776 | 140482579 | 140482667 | 335 | 224 | 241 | 311 | 238 | 67 | 55 | 67 | 69 | 37 | 96 | 211 | 347 | 249 | 146 | 25 | 55 | 57 | 24 | 10 | 206 | 149 | 0.037533714 | 0.301471041 | 0.783 | 0.747 | 0.722 | 0.765 | 0.823 | 0.735 | 0.735 | 0.815 | 0.882 | 0.913 | -0.048 |
| SE | 83631 | ENSG00000152082 | MZT2B | chr2 | + | 130182626 | 130182775 | 130182256 | 130182452 | 130190468 | 130190695 | 59 | 25 | 61 | 44 | 10 | 5 | 0 | 1 | 2 | 1 | 43 | 36 | 43 | 36 | 18 | 0 | 4 | 0 | 0 | 0 | 298 | 149 | 0.037544589 | 0.301527455 | 0.855 | 1 | 0.968 | 0.917 | 0.833 | 1 | 0.818 | 1 | 1 | 1 | -0.049 |
| SE | 83196 | ENSG00000183773 | AIFM3 | chr22 | + | 20977031 | 20977095 | 20976866 | 20976938 | 20977699 | 20977776 | 11 | 4 | 3 | 3 | 6 | 0 | 0 | 0 | 0 | 0 | 9 | 3 | 7 | 8 | 47 | 10 | 0 | 0 | 0 | 0 | 213 | 149 | 0.037553234 | 0.301548723 | 1 | 1 | 1 | 1 | 1 | 0.386 | 1 | 1 | 1 | 1 | 0.123 |
| SE | 50409 | ENSG00000164306 | PRIMPOL | chr4 | + | 184661773 | 184661903 | 184659339 | 184659437 | 184665916 | 184666064 | 54 | 63 | 66 | 62 | 133 | 6 | 1 | 2 | 3 | 1 | 63 | 68 | 21 | 22 | 21 | 0 | 3 | 2 | 0 | 0 | 279 | 149 | 0.03755494 | 0.301548723 | 0.828 | 0.971 | 0.946 | 0.917 | 0.986 | 1 | 0.924 | 0.849 | 1 | 1 | -0.025 |
| SE | 56613 | ENSG00000075151 | EIF4G3 | chr1 | - | 20950002 | 20950108 | 20941490 | 20942330 | 20969473 | 20969596 | 71 | 55 | 44 | 87 | 61 | 3 | 0 | 1 | 3 | 3 | 67 | 88 | 85 | 113 | 26 | 0 | 0 | 0 | 0 | 0 | 255 | 149 | 0.037564182 | 0.301586358 | 0.933 | 1 | 0.963 | 0.944 | 0.922 | 1 | 1 | 1 | 1 | 1 | -0.048 |
| SE | 41607 | ENSG00000118308 | IRAG2 | chr12 | + | 25104360 | 25104462 | 25104008 | 25104058 | 25106942 | 25107050 | 203 | 163 | 253 | 242 | 388 | 2 | 1 | 4 | 7 | 5 | 27 | 146 | 143 | 60 | 26 | 0 | 0 | 0 | 2 | 0 | 251 | 149 | 0.037567331 | 0.301586358 | 0.984 | 0.99 | 0.974 | 0.954 | 0.979 | 1 | 1 | 1 | 0.947 | 1 | -0.013 |
| SE | 14404 | ENSG00000167202 | TBC1D2B | chr15 | - | 78017846 | 78017957 | 78016545 | 78016739 | 78030006 | 78030170 | 77 | 45 | 68 | 74 | 74 | 0 | 3 | 0 | 4 | 0 | 172 | 77 | 41 | 79 | 171 | 0 | 0 | 1 | 1 | 16 | 260 | 149 | 0.037572905 | 0.301600185 | 1 | 0.896 | 1 | 0.914 | 1 | 1 | 1 | 0.959 | 0.978 | 0.86 | 0.003 |
| SE | 12141 | ENSG00000172775 | PSME3IP1 | chr16 | - | 57175538 | 57175590 | 57173727 | 57173869 | 57185820 | 57186032 | 12 | 19 | 8 | 21 | 15 | 88 | 55 | 63 | 72 | 57 | 5 | 47 | 55 | 39 | 25 | 114 | 77 | 40 | 67 | 99 | 201 | 149 | 0.037577231 | 0.301603983 | 0.092 | 0.204 | 0.086 | 0.178 | 0.163 | 0.031 | 0.312 | 0.505 | 0.301 | 0.158 | -0.117 |
| SE | 46405 | ENSG00000168724 | DNAJC21 | chr5 | + | 34936143 | 34936266 | 34935709 | 34935833 | 34937325 | 34937630 | 316 | 249 | 264 | 338 | 381 | 4 | 0 | 1 | 2 | 0 | 195 | 212 | 272 | 231 | 58 | 4 | 4 | 4 | 0 | 4 | 272 | 149 | 0.037586104 | 0.301644284 | 0.977 | 1 | 0.993 | 0.989 | 1 | 0.964 | 0.967 | 0.974 | 1 | 0.888 | 0.033 |
| SE | 24864 | ENSG00000155846 | PPARGC1B | chr5 | + | 149761431 | 149761612 | 149730302 | 149730420 | 149820432 | 149820606 | 0 | 0 | 0 | 0 | 0 | 2 | 2 | 2 | 2 | 2 | 25 | 0 | 0 | 0 | 0 | 0 | 2 | 1 | 6 | 1 | 298 | 149 | 0.037597082 | 0.301701461 | 0 | 0 | 0 | 0 | 0 | 1 | 0 | 0 | 0 | 0 | -0.2 |
| SE | 594 | ENSG00000100207 | TCF20 | chr22 | - | 42168608 | 42168736 | 42160012 | 42161358 | 42169846 | 42169896 | 30 | 17 | 21 | 32 | 12 | 79 | 55 | 90 | 84 | 189 | 1 | 27 | 18 | 11 | 7 | 89 | 82 | 71 | 78 | 72 | 277 | 149 | 0.037602728 | 0.30171584 | 0.17 | 0.143 | 0.112 | 0.17 | 0.033 | 0.006 | 0.15 | 0.12 | 0.071 | 0.05 | 0.046 |
| SE | 71953 | ENSG00000197930 | ERO1A | chr14 | - | 52683787 | 52683907 | 52682324 | 52682408 | 52695367 | 52695529 | 155 | 73 | 84 | 115 | 128 | 0 | 0 | 0 | 0 | 0 | 71 | 149 | 110 | 151 | 18 | 0 | 8 | 4 | 0 | 0 | 269 | 149 | 0.037611569 | 0.301755856 | 1 | 1 | 1 | 1 | 1 | 1 | 0.912 | 0.938 | 1 | 1 | 0.03 |
| SE | 64076 | ENSG00000110066 | KMT5B | chr11 | - | 68179469 | 68179537 | 68175017 | 68175183 | 68180131 | 68180200 | 21 | 17 | 19 | 9 | 22 | 272 | 195 | 183 | 209 | 197 | 13 | 19 | 16 | 6 | 0 | 50 | 122 | 71 | 120 | 33 | 217 | 149 | 0.037620503 | 0.301796606 | 0.05 | 0.056 | 0.067 | 0.029 | 0.071 | 0.151 | 0.097 | 0.134 | 0.033 | 0 | -0.028 |
| SE | 94046 | ENSG00000100296 | THOC5 | chr22 | - | 29550727 | 29550843 | 29549051 | 29549158 | 29553670 | 29553741 | 15 | 5 | 20 | 32 | 10 | 15 | 19 | 13 | 22 | 3 | 17 | 47 | 48 | 45 | 34 | 11 | 14 | 8 | 18 | 31 | 265 | 149 | 0.037689902 | 0.302322361 | 0.36 | 0.129 | 0.464 | 0.45 | 0.652 | 0.465 | 0.654 | 0.771 | 0.584 | 0.381 | -0.16 |
| SE | 107131 | ENSG00000111424 | VDR | chr12 | - | 47882693 | 47882774 | 47878967 | 47879115 | 47904954 | 47904994 | 10 | 2 | 5 | 8 | 2 | 0 | 0 | 0 | 1 | 0 | 27 | 2 | 2 | 7 | 20 | 0 | 4 | 2 | 0 | 0 | 230 | 149 | 0.037732658 | 0.302634312 | 1 | 1 | 1 | 0.838 | 1 | 1 | 0.245 | 0.393 | 1 | 1 | 0.24 |
| SE | 6485 | ENSG00000101608 | MYL12A | chr18 | + | 3249561 | 3249723 | 3249360 | 3249443 | 3253232 | 3253284 | 7 | 5 | 14 | 11 | 16 | 8 | 5 | 4 | 14 | 5 | 0 | 15 | 13 | 12 | 0 | 20 | 18 | 6 | 4 | 10 | 298 | 149 | 0.037763898 | 0.302853851 | 0.304 | 0.333 | 0.636 | 0.282 | 0.615 | 0 | 0.294 | 0.52 | 0.6 | 0 | 0.151 |
| SE | 12958 | ENSG00000103415 | HMOX2 | chr16 | + | 4483636 | 4483754 | 4476339 | 4476487 | 4505483 | 4505610 | 10 | 5 | 8 | 2 | 3 | 33 | 15 | 19 | 22 | 10 | 0 | 5 | 0 | 0 | 4 | 2 | 13 | 14 | 11 | 12 | 267 | 149 | 0.037838745 | 0.303423021 | 0.145 | 0.157 | 0.19 | 0.048 | 0.143 | 0 | 0.177 | 0 | 0 | 0.157 | 0.07 |
| SE | 67202 | ENSG00000124151 | NCOA3 | chr20 | + | 47623910 | 47624083 | 47622228 | 47622330 | 47625380 | 47625481 | 695 | 621 | 751 | 649 | 1000 | 2 | 1 | 9 | 8 | 4 | 786 | 957 | 1103 | 1240 | 432 | 0 | 2 | 11 | 8 | 25 | 298 | 149 | 0.03785118 | 0.303491654 | 0.994 | 0.997 | 0.977 | 0.976 | 0.992 | 1 | 0.996 | 0.98 | 0.987 | 0.896 | 0.015 |
| SE | 84421 | ENSG00000188917 | TRMT2B | chrX | - | 101042041 | 101042312 | 101036973 | 101037073 | 101051250 | 101051366 | 24 | 17 | 17 | 19 | 24 | 0 | 2 | 0 | 0 | 0 | 8 | 19 | 22 | 22 | 26 | 0 | 0 | 3 | 0 | 5 | 298 | 149 | 0.037892877 | 0.303794866 | 1 | 0.81 | 1 | 1 | 1 | 1 | 1 | 0.786 | 1 | 0.722 | 0.06 |
| SE | 86034 | ENSG00000156313 | RPGR | chrX | - | 38287093 | 38288041 | 38276586 | 38276772 | 38290958 | 38291024 | 46 | 20 | 27 | 31 | 39 | 5 | 4 | 4 | 2 | 4 | 5 | 25 | 30 | 8 | 0 | 0 | 0 | 1 | 0 | 4 | 298 | 149 | 0.037906343 | 0.303811227 | 0.821 | 0.714 | 0.771 | 0.886 | 0.83 | 1 | 1 | 0.938 | 1 | 0 | 0.017 |
| SE | 25499 | ENSG00000134986 | NREP | chr5 | - | 111755769 | 111755830 | 111734983 | 111735507 | 111757135 | 111757173 | 32 | 12 | 33 | 44 | 51 | 1 | 1 | 1 | 2 | 4 | 87 | 22 | 22 | 63 | 118 | 0 | 3 | 0 | 2 | 0 | 210 | 149 | 0.037907657 | 0.303811227 | 0.958 | 0.895 | 0.959 | 0.94 | 0.9 | 1 | 0.839 | 1 | 0.957 | 1 | -0.029 |
| SE | 42753 | ENSG00000134371 | CDC73 | chr1 | + | 193141849 | 193142066 | 193138084 | 193138173 | 193147866 | 193147965 | 450 | 380 | 339 | 454 | 661 | 1 | 0 | 0 | 1 | 2 | 137 | 306 | 347 | 228 | 214 | 13 | 2 | 2 | 0 | 0 | 298 | 149 | 0.037909113 | 0.303811227 | 0.996 | 1 | 1 | 0.996 | 0.994 | 0.84 | 0.987 | 0.989 | 1 | 1 | 0.034 |
| SE | 88262 | ENSG00000104756 | KCTD9 | chr8 | - | 25444291 | 25444335 | 25440542 | 25440673 | 25446128 | 25446250 | 69 | 63 | 76 | 58 | 55 | 2 | 0 | 0 | 1 | 3 | 28 | 52 | 31 | 26 | 0 | 0 | 5 | 0 | 0 | 4 | 193 | 149 | 0.037910438 | 0.303811227 | 0.964 | 1 | 1 | 0.978 | 0.934 | 1 | 0.889 | 1 | 1 | 0 | 0.197 |
| SE | 1709 | ENSG00000228315 | GUSBP11 | chr22 | - | 23700725 | 23700845 | 23694118 | 23695517 | 23705428 | 23705541 | 16 | 12 | 18 | 27 | 19 | 13 | 13 | 7 | 34 | 16 | 5 | 13 | 12 | 17 | 37 | 62 | 5 | 15 | 28 | 57 | 269 | 149 | 0.038010931 | 0.304571066 | 0.405 | 0.338 | 0.588 | 0.305 | 0.397 | 0.043 | 0.59 | 0.307 | 0.252 | 0.264 | 0.115 |
| SE | 74805 | ENSG00000094880 | CDC23 | chr5 | - | 138212990 | 138213063 | 138205870 | 138206684 | 138213151 | 138213215 | 78 | 47 | 94 | 72 | 38 | 2 | 0 | 5 | 3 | 0 | 41 | 61 | 80 | 41 | 7 | 7 | 0 | 0 | 9 | 0 | 222 | 149 | 0.038013033 | 0.304571066 | 0.963 | 1 | 0.927 | 0.942 | 1 | 0.797 | 1 | 1 | 0.754 | 1 | 0.056 |
| SE | 93775 | ENSG00000148484 | RSU1 | chr10 | - | 16816972 | 16817084 | 16782033 | 16782084 | 16817314 | 16817424 | 462 | 548 | 292 | 479 | 379 | 24 | 12 | 32 | 46 | 21 | 1337 | 714 | 397 | 1043 | 6444 | 46 | 49 | 40 | 51 | 212 | 261 | 149 | 0.038055863 | 0.30488303 | 0.917 | 0.963 | 0.839 | 0.856 | 0.912 | 0.943 | 0.893 | 0.85 | 0.921 | 0.946 | -0.013 |
| SE | 84173 | ENSG00000137642 | SORL1 | chr11 | + | 121514151 | 121514321 | 121496868 | 121497049 | 121520656 | 121520849 | 571 | 360 | 461 | 505 | 548 | 2 | 0 | 1 | 3 | 3 | 418 | 280 | 605 | 456 | 284 | 0 | 5 | 10 | 0 | 8 | 298 | 149 | 0.03806321 | 0.304910687 | 0.993 | 1 | 0.996 | 0.988 | 0.989 | 1 | 0.966 | 0.968 | 1 | 0.947 | 0.017 |
| SE | 43183 | ENSG00000169188 | APEX2 | chrX | + | 55002250 | 55002431 | 55001545 | 55001629 | 55002961 | 55003108 | 59 | 54 | 46 | 44 | 58 | 4 | 3 | 2 | 7 | 8 | 11 | 35 | 7 | 13 | 24 | 26 | 9 | 4 | 1 | 0 | 298 | 149 | 0.03807235 | 0.304952705 | 0.881 | 0.9 | 0.92 | 0.759 | 0.784 | 0.175 | 0.66 | 0.467 | 0.867 | 1 | 0.215 |
| SE | 59804 | ENSG00000166377 | ATP9B | chr18 | + | 79176812 | 79176907 | 79154501 | 79154555 | 79193182 | 79193263 | 161 | 155 | 206 | 163 | 172 | 24 | 30 | 32 | 15 | 20 | 206 | 199 | 159 | 173 | 264 | 10 | 27 | 12 | 19 | 34 | 244 | 149 | 0.038101516 | 0.305155107 | 0.804 | 0.759 | 0.797 | 0.869 | 0.84 | 0.926 | 0.818 | 0.89 | 0.848 | 0.826 | -0.048 |
| SE | 72610 | ENSG00000115977 | AAK1 | chr2 | - | 69495984 | 69496080 | 69481055 | 69482812 | 69505568 | 69505673 | 14 | 8 | 21 | 17 | 14 | 3 | 5 | 7 | 3 | 0 | 10 | 22 | 19 | 5 | 10 | 19 | 15 | 8 | 11 | 0 | 245 | 149 | 0.038149605 | 0.305508994 | 0.739 | 0.493 | 0.646 | 0.775 | 1 | 0.242 | 0.471 | 0.591 | 0.217 | 1 | 0.226 |
| SE | 40061 | ENSG00000125347 | IRF1 | chr5 | - | 132484361 | 132484497 | 132481608 | 132484075 | 132486250 | 132486373 | 134 | 75 | 158 | 114 | 115 | 4 | 3 | 0 | 11 | 0 | 141 | 58 | 67 | 222 | 129 | 7 | 0 | 8 | 0 | 20 | 285 | 149 | 0.038153993 | 0.305512888 | 0.946 | 0.929 | 1 | 0.844 | 1 | 0.913 | 1 | 0.814 | 1 | 0.771 | 0.044 |
| SE | 70577 | ENSG00000022840 | RNF10 | chr12 | + | 120574467 | 120574537 | 120571190 | 120571291 | 120575630 | 120575688 | 4 | 1 | 5 | 3 | 6 | 168 | 94 | 131 | 126 | 133 | 0 | 15 | 16 | 26 | 0 | 218 | 197 | 192 | 221 | 496 | 219 | 149 | 0.038209085 | 0.305922741 | 0.016 | 0.007 | 0.025 | 0.016 | 0.03 | 0 | 0.049 | 0.054 | 0.074 | 0 | -0.017 |
| SE | 79203 | ENSG00000242588 | NA | chr7 | + | 128579998 | 128580094 | 128578889 | 128578933 | 128591849 | 128592017 | 193 | 145 | 152 | 247 | 173 | 25 | 30 | 3 | 8 | 50 | 675 | 338 | 268 | 320 | 685 | 166 | 64 | 39 | 169 | 39 | 245 | 149 | 0.038231723 | 0.306072699 | 0.824 | 0.746 | 0.969 | 0.949 | 0.678 | 0.712 | 0.763 | 0.807 | 0.535 | 0.914 | 0.087 |
| SE | 19220 | ENSG00000100784 | RPS6KA5 | chr14 | - | 90939536 | 90939599 | 90923112 | 90923196 | 90943077 | 90943185 | 0 | 1 | 5 | 2 | 2 | 93 | 100 | 85 | 86 | 103 | 0 | 1 | 0 | 15 | 0 | 52 | 86 | 101 | 52 | 17 | 212 | 149 | 0.03825175 | 0.306196342 | 0 | 0.007 | 0.04 | 0.016 | 0.013 | 0 | 0.008 | 0 | 0.169 | 0 | -0.02 |
| SE | 25890 | ENSG00000161911 | TREML1 | chr6 | - | 41150260 | 41150313 | 41149602 | 41149918 | 41150818 | 41150907 | 14 | 24 | 12 | 27 | 36 | 52 | 49 | 10 | 142 | 257 | 41 | 23 | 19 | 9 | 449 | 215 | 208 | 78 | 121 | 2168 | 202 | 149 | 0.038254989 | 0.306196342 | 0.166 | 0.265 | 0.47 | 0.123 | 0.094 | 0.123 | 0.075 | 0.152 | 0.052 | 0.133 | 0.117 |
| SE | 16400 | ENSG00000156958 | GALK2 | chr15 | + | 49283565 | 49283718 | 49239220 | 49239367 | 49292326 | 49292537 | 126 | 74 | 77 | 85 | 136 | 4 | 6 | 1 | 4 | 1 | 73 | 116 | 96 | 71 | 117 | 0 | 8 | 0 | 0 | 0 | 298 | 149 | 0.038284871 | 0.306379863 | 0.94 | 0.86 | 0.975 | 0.914 | 0.986 | 1 | 0.879 | 1 | 1 | 1 | -0.041 |
| SE | 49748 | ENSG00000111331 | OAS3 | chr12 | + | 112965743 | 112966029 | 112964234 | 112964408 | 112967417 | 112967583 | 96 | 104 | 68 | 155 | 70 | 0 | 1 | 0 | 3 | 0 | 632 | 180 | 95 | 555 | 134 | 0 | 0 | 0 | 14 | 7 | 298 | 149 | 0.038285744 | 0.306379863 | 1 | 0.981 | 1 | 0.963 | 1 | 1 | 1 | 1 | 0.952 | 0.905 | 0.017 |
| SE | 3706 | ENSG00000181027 | FKRP | chr19 | + | 46748514 | 46748717 | 46748026 | 46748088 | 46755411 | 46755610 | 8 | 3 | 6 | 23 | 6 | 0 | 4 | 0 | 0 | 0 | 42 | 16 | 12 | 12 | 27 | 0 | 0 | 0 | 3 | 7 | 298 | 149 | 0.038305708 | 0.306488995 | 1 | 0.273 | 1 | 1 | 1 | 1 | 1 | 1 | 0.818 | 0.702 | -0.049 |
| SE | 92248 | ENSG00000114209 | PDCD10 | chr3 | - | 167734213 | 167734350 | 167720061 | 167720273 | 167734805 | 167734836 | 12 | 17 | 11 | 8 | 11 | 180 | 106 | 105 | 153 | 154 | 0 | 12 | 12 | 8 | 15 | 57 | 160 | 68 | 37 | 14 | 286 | 149 | 0.03830721 | 0.306488995 | 0.034 | 0.077 | 0.052 | 0.027 | 0.036 | 0 | 0.038 | 0.084 | 0.101 | 0.358 | -0.071 |
| SE | 90692 | ENSG00000110077 | MS4A6A | chr11 | - | 60179830 | 60179965 | 60178170 | 60178316 | 60181580 | 60181741 | 1336 | 912 | 760 | 1016 | 1740 | 55 | 67 | 44 | 90 | 100 | 4187 | 1326 | 924 | 1793 | 2252 | 260 | 78 | 81 | 180 | 169 | 284 | 149 | 0.038354302 | 0.306834418 | 0.927 | 0.877 | 0.901 | 0.856 | 0.901 | 0.894 | 0.899 | 0.857 | 0.839 | 0.875 | 0.02 |
| SE | 101169 | ENSG00000133816 | MICAL2 | chr11 | + | 12249183 | 12249246 | 12239435 | 12239585 | 12256784 | 12256968 | 49 | 14 | 29 | 28 | 34 | 85 | 49 | 63 | 100 | 61 | 100 | 16 | 29 | 49 | 91 | 201 | 56 | 30 | 65 | 102 | 212 | 149 | 0.038358595 | 0.306837406 | 0.288 | 0.167 | 0.244 | 0.164 | 0.281 | 0.259 | 0.167 | 0.405 | 0.346 | 0.385 | -0.084 |
| SE | 64957 | ENSG00000053371 | AKR7A2 | chr1 | - | 19307313 | 19307410 | 19307001 | 19307101 | 19308157 | 19308262 | 115 | 74 | 82 | 91 | 46 | 3 | 3 | 6 | 2 | 2 | 117 | 83 | 54 | 63 | 420 | 0 | 0 | 4 | 5 | 3 | 246 | 149 | 0.038362826 | 0.306839902 | 0.959 | 0.937 | 0.892 | 0.965 | 0.933 | 1 | 1 | 0.891 | 0.884 | 0.988 | -0.015 |
| SE | 8293 | ENSG00000108622 | ICAM2 | chr17 | - | 64007740 | 64007822 | 64006630 | 64006735 | 64020522 | 64020607 | 10 | 3 | 19 | 5 | 3 | 100 | 50 | 86 | 86 | 57 | 9 | 11 | 0 | 10 | 17 | 48 | 39 | 31 | 56 | 91 | 231 | 149 | 0.038390691 | 0.307031417 | 0.061 | 0.037 | 0.125 | 0.036 | 0.033 | 0.108 | 0.154 | 0 | 0.103 | 0.108 | -0.036 |
| SE | 59086 | ENSG00000282246 | NA | chr10 | + | 13630259 | 13630465 | 13616530 | 13616553 | 13644916 | 13645072 | 140 | 48 | 119 | 100 | 184 | 3 | 0 | 0 | 2 | 7 | 7 | 43 | 74 | 24 | 8 | 0 | 0 | 1 | 0 | 0 | 298 | 149 | 0.038398055 | 0.307058942 | 0.959 | 1 | 1 | 0.962 | 0.929 | 1 | 1 | 0.974 | 1 | 1 | -0.025 |

| | | | | | | | | |
|---|---|---|---|---|---|---|---|---|
| SE | 97636 | ENSG00000198879 | SFMBT2 | chr10 | - | 7276891 | 7276989 | [illegible] |
| SE | 18620 | ENSG00000256053 | COA8 | chr14 | + | 103571622 | 103571820 | [illegible] |
| SE | 24330 | ENSG00000185404 | SP140L | chr2 | + | 230389918 | 230390023 | [illegible] |
| SE | 31728 | ENSG00000047849 | MAP4 | chr3 | - | 47867245 | 47867338 | [illegible] |
| SE | 29876 | ENSG00000154743 | TSEN2 | chr3 | + | 12530854 | 12530941 | [illegible] |
| SE | 90731 | ENSG00000168404 | MLKL | chr16 | - | 74678898 | 74678980 | [illegible] |
| SE | 85863 | ENSG00000223959 | AFG3L1P | chr16 | + | 89988681 | 89988951 | [illegible] |
| SE | 49228 | ENSG00000142864 | SERBP1 | chr1 | - | 67424887 | 67424977 | [illegible] |
| SE | 45962 | ENSG00000117593 | DARS2 | chr1 | + | 173853794 | 173853905 | [illegible] |
| SE | 98822 | ENSG00000135365 | PHF21A | chr11 | - | 45935120 | 45935246 | [illegible] |
| SE | 74688 | ENSG00000197548 | ATG7 | chr3 | + | 11299361 | 11299416 | [illegible] |
| SE | 37286 | ENSG00000174738 | NR1D2 | chr3 | + | 23967812 | 23968023 | [illegible] |
| SE | 77394 | ENSG00000089820 | ARHGAP4 | chrX | - | 153919566 | 153919686 | [illegible] |
| SE | 55825 | ENSG00000143641 | GALNT2 | chr1 | + | 230255217 | 230255344 | [illegible] |
| SE | 73096 | ENSG00000198001 | IRAK4 | chr12 | + | 43771037 | 43771104 | [illegible] |
| SE | 88051 | ENSG00000239213 | NCK1-DT | chr3 | - | 136858517 | 136858603 | [illegible] |
| SE | 17924 | ENSG00000197249 | SERPINA1 | chr14 | - | 94388559 | 94388660 | [illegible] |
| SE | 18356 | ENSG00000257621 | PSMA3-AS1 | chr14 | - | 58293765 | 58295811 | [illegible] |
| SE | 71370 | ENSG00000122203 | KIAA1191 | chr5 | - | 176350612 | 176350737 | [illegible] |
| SE | 24104 | ENSG00000187522 | HSPA14 | chr10 | + | 14848789 | 14848895 | [illegible] |
| SE | 23167 | ENSG00000176783 | RUFY1 | chr5 | + | 179580946 | 179581012 | [illegible] |
| SE | 24314 | ENSG00000058799 | YIPF1 | chr1 | - | 53866199 | 53866382 | [illegible] |
| SE | 69099 | ENSG00000136536 | MARCHF7 | chr2 | + | 159764625 | 159764674 | [illegible] |
| SE | 71343 | ENSG00000085760 | MTIF2 | chr2 | - | 55254040 | 55254201 | [illegible] |
| SE | 41585 | ENSG00000118308 | IRAG2 | chr12 | + | 25062819 | 25062900 | [illegible] |
| SE | 95173 | ENSG00000213930 | GALT | chr9 | + | 34647491 | 34647705 | [illegible] |
| SE | 100793 | ENSG00000135272 | MDFIC | chr7 | + | 115014462 | 115014533 | [illegible] |
| SE | 22607 | ENSG00000127603 | MACF1 | chr1 | + | 39381952 | 39382152 | [illegible] |
| SE | 42805 | ENSG00000051341 | POLQ | chr3 | - | 121460049 | 121460234 | [illegible] |
| SE | 52781 | ENSG00000149476 | TKFC | chr11 | + | 61341822 | 61341912 | [illegible] |
| SE | 108219 | ENSG00000184752 | NDUFA12 | chr12 | - | 94994169 | 94994257 | [illegible] |
| SE | 89096 | ENSG00000160588 | MPZL3 | chr11 | - | 118237049 | 118237260 | [illegible] |
| SE | 83309 | ENSG00000182487 | NCF1B | chr7 | + | 73229566 | 73230260 | [illegible] |
| SE | 68850 | ENSG00000168769 | TET2 | chr4 | + | 105233896 | 105237351 | [illegible] |
| SE | 31624 | ENSG00000168026 | TTC21A | chr3 | + | 39130991 | 39131095 | [illegible] |
| SE | 83572 | ENSG00000124222 | STX16 | chr20 | + | 58671163 | 58671297 | [illegible] |
| SE | 11667 | ENSG00000167207 | NOD2 | chr16 | + | 50709943 | 50710016 | [illegible] |
| SE | 31365 | ENSG00000005436 | GCFC2 | chr2 | - | 75702312 | 75702423 | [illegible] |
| SE | 49460 | ENSG00000115020 | PIKFYVE | chr2 | + | 208277536 | 208277708 | [illegible] |
| SE | 72358 | ENSG00000163214 | DHX57 | chr2 | - | 38828339 | 38828436 | [illegible] |
| SE | 19567 | ENSG00000198718 | TOGARAM1 | chr14 | + | 45027298 | 45027474 | [illegible] |
| SE | 80392 | ENSG00000150787 | PTS | chr11 | + | 112230625 | 112230682 | [illegible] |
| SE | 36757 | ENSG00000163539 | CLASP2 | chr3 | - | 33501651 | 33501768 | [illegible] |
| SE | 103752 | ENSG00000047056 | WDR37 | chr10 | + | 1103598 | 1103836 | [illegible] |
| SE | 85961 | ENSG00000111203 | ITFG2 | chr12 | + | 2821542 | 2821596 | [illegible] |
| SE | 101488 | ENSG00000095794 | CREM | chr10 | + | 35148367 | 35148491 | [illegible] |
| SE | 82866 | ENSG00000161048 | NAPEPLD | chr7 | - | 103115059 | 103115174 | [illegible] |
| SE | 71023 | ENSG00000163516 | ANKZF1 | chr2 | + | 219230227 | 219230405 | [illegible] |
| SE | 6491 | ENSG00000101654 | RNMT | chr18 | + | 13730626 | 13730755 | [illegible] |
| SE | 2772 | ENSG00000104774 | MAN2B1 | chr19 | - | 12658427 | 12658510 | [illegible] |
| SE | 92570 | ENSG00000148057 | IDNK | chr9 | + | 83628006 | 83628211 | [illegible] |
| SE | 39050 | ENSG00000118985 | ELL2 | chr5 | - | 95932631 | 95932713 | [illegible] |
| SE | 80813 | ENSG00000151948 | GLT1D1 | chr12 | + | 128899235 | 128899287 | [illegible] |
| SE | 11816 | ENSG00000188603 | CLN3 | chr16 | - | 28487661 | 28487741 | [illegible] |
| SE | 23932 | ENSG00000174989 | FBXW8 | chr12 | + | 116949617 | 116949706 | [illegible] |
| SE | 89783 | ENSG00000104490 | NCALD | chr8 | - | 102029009 | 102029096 | [illegible] |
| SE | 33583 | ENSG00000009790 | TRAF3IP3 | chr1 | + | 209778110 | 209778173 | [illegible] |
| SE | 100183 | ENSG00000078124 | ACER3 | chr11 | + | 76998762 | 76998821 | [illegible] |
| SE | 6921 | ENSG00000175354 | PTPN2 | chr18 | - | 12819213 | 12819282 | [illegible] |
| SE | 56614 | ENSG00000075151 | EIF4G3 | chr1 | - | 20950002 | 20950111 | [illegible] |
| SE | 95410 | ENSG00000148356 | LRSAM1 | chr9 | + | 127467739 | 127467830 | [illegible] |
| SE | 99951 | ENSG00000142102 | PGGHG | chr11 | + | 292885 | 292983 | [illegible] |
| SE | 50852 | ENSG00000048052 | HDAC9 | chr7 | + | 18654762 | 18654869 | [illegible] |
| SE | 40535 | ENSG00000008128 | CDK11A | chr1 | - | 1722587 | 1722634 | [illegible] |
| SE | 46494 | ENSG00000109184 | DCUN1D4 | chr4 | + | 51848204 | 51848367 | [illegible] |
| SE | 94160 | ENSG00000198000 | NOL8 | chr9 | - | 92311145 | 92311259 | [illegible] |
| SE | 12353 | ENSG00000197912 | SPG7 | chr16 | + | 89555877 | 89556085 | [illegible] |
| SE | 82492 | ENSG00000128563 | PRKRIP1 | chr7 | + | 102395976 | 102396537 | [illegible] |
| SE | 4133 | ENSG00000167578 | RAB4B | chr19 | + | 40783777 | 40783840 | [illegible] |
| SE | 54529 | ENSG00000161929 | SCIMP | chr17 | - | 5221286 | 5221350 | [illegible] |
| SE | 40262 | ENSG00000170581 | STAT2 | chr12 | - | 56349162 | 56349261 | [illegible] |
| SE | 38310 | ENSG00000138172 | CALHM2 | chr10 | - | 103451082 | 103451285 | [illegible] |
| SE | 76998 | ENSG00000149657 | LSM14B | chr20 | + | 62126225 | 62126439 | [illegible] |
| SE | 81522 | ENSG00000071462 | BUD23 | chr7 | + | 73690915 | 73691012 | [illegible] |
| SE | 68594 | ENSG00000239779 | WBP1 | chr2 | + | 74459437 | 74459552 | [illegible] |
| SE | 89976 | ENSG00000153317 | ASAP1 | chr8 | - | 130091972 | 130092143 | [illegible] |
| SE | 85216 | ENSG00000099204 | ABLIM1 | chr10 | - | 114453378 | 114453483 | [illegible] |
| SE | 68081 | ENSG00000175489 | LRRC25 | chr19 | - | 18396184 | 18396972 | [illegible] |
| SE | 93471 | ENSG00000106771 | TMEM245 | chr9 | - | 109090921 | 109091155 | [illegible] |
| SE | 44721 | ENSG00000135999 | EPC2 | chr2 | + | 148761781 | 148761930 | [illegible] |
| SE | 95931 | ENSG00000166199 | ALKBH3 | chr11 | + | 43897194 | 43897354 | [illegible] |
| SE | 997 | ENSG00000243156 | MICAL3 | chr22 | - | 17822946 | 17823060 | [illegible] |
| SE | 15169 | ENSG00000137842 | TMEM62 | chr15 | + | 43178606 | 43178711 | [illegible] |
| SE | 62068 | ENSG00000110446 | SLC15A3 | chr11 | - | 60941121 | 60942145 | [illegible] |
| SE | 38654 | ENSG00000073417 | PDE8A | chr15 | + | 85091043 | 85091181 | [illegible] |
| SE | 30289 | ENSG00000148154 | UGCG | chr9 | + | 111915758 | 111915836 | [illegible] |
| SE | 74251 | ENSG00000137478 | FCHSD2 | chr11 | - | 73015808 | 73015885 | [illegible] |
| SE | 73679 | ENSG00000151687 | ANKAR | chr2 | + | 189730494 | 189730601 | [illegible] |
| SE | 92346 | ENSG00000027697 | IFNGR1 | chr6 | - | 137215266 | 137215377 | [illegible] |
| SE | 54342 | ENSG00000049618 | ARID1B | chr6 | + | 156871613 | 156871652 | [illegible] |
| SE | 19967 | ENSG00000136758 | YME1L1 | chr10 | - | 27142386 | 27142485 | [illegible] |
| SE | 52067 | ENSG00000043514 | TRIT1 | chr1 | - | 39847547 | 39847660 | [illegible] |
| SE | 54331 | ENSG00000117139 | KDM5B | chr1 | - | 202776104 | 202776220 | [illegible] |
| SE | 31626 | ENSG00000168026 | TTC21A | chr3 | + | 39136898 | 39137060 | [illegible] |
| SE | 2792 | ENSG00000011304 | PTBP1 | chr19 | + | 806407 | 806556 | [illegible] |
| SE | 9290 | ENSG00000170037 | CNTROB | chr17 | + | 7949084 | 7949157 | [illegible] |
| SE | 90915 | ENSG00000185122 | HSF1 | chr8 | + | 144311962 | 144312244 | [illegible] |
| SE | 12059 | ENSG00000159720 | ATP6V0D1 | chr16 | - | 67453543 | 67453715 | [illegible] |
| SE | 60145 | ENSG00000236810 | ELOA-AS1 | chr1 | - | 23771701 | 23771883 | [illegible] |
| SE | 103051 | ENSG00000099290 | WASHC2A | chr10 | + | 50126056 | 50126179 | [illegible] |
| SE | 55312 | ENSG00000166938 | DIS3L | chr15 | + | 66311723 | 66311900 | [illegible] |
| SE | 28052 | ENSG00000114738 | MAPKAPK3 | chr3 | + | 50644408 | 50644532 | [illegible] |
| SE | 1017 | ENSG00000100335 | MIEF1 | chr22 | + | 39509337 | 39509571 | [illegible] |
| SE | 107829 | ENSG00000135457 | TFCP2 | chr12 | - | 51109120 | 51109273 | [illegible] |
| SE | 40716 | ENSG00000135932 | CAB39 | chr2 | + | 230810262 | 230810322 | [illegible] |
| SE | 85312 | ENSG00000147394 | ZNF185 | chrX | + | 152920327 | 152920411 | [illegible] |
| SE | 74340 | ENSG00000187118 | CMC1 | chr3 | + | 28292735 | 28292964 | [illegible] |
| SE | 80073 | ENSG00000198198 | SZT2 | chr1 | + | 43419733 | 43419944 | [illegible] |
| SE | 41721 | ENSG00000116497 | S100PBP | chr1 | + | 32825312 | 32825485 | [illegible] |
| SE | 27779 | ENSG00000177058 | SLC38A9 | chr5 | - | 55633694 | 55635657 | [illegible] |
| SE | 36488 | ENSG00000164211 | STARD4 | chr5 | - | 111501961 | 111502088 | [illegible] |
| SE | 1689 | ENSG00000228315 | GUSBP11 | chr22 | - | 23640307 | 23640423 | [illegible] |
| SE | 77300 | ENSG00000099968 | BCL2L13 | chr22 | + | 17688985 | 17689142 | [illegible] |
| SE | 81567 | ENSG00000132256 | TRIM5 | chr11 | - | 5665980 | 5666081 | [illegible] |
| SE | 74815 | ENSG00000042088 | TDP1 | chr14 | + | 89963224 | 89963673 | [illegible] |
| SE | 108239 | ENSG00000118454 | ANKRD13C | chr1 | - | 70306223 | 70306290 | [illegible] |
| SE | 20187 | ENSG00000137210 | TMEM14B | chr6 | + | 10751132 | 10751234 | [illegible] |
| SE | 104845 | ENSG00000011105 | TSPAN9 | chr12 | + | 3281733 | 3281817 | [illegible] |
| SE | 98250 | ENSG00000173208 | ABCD2 | chr12 | - | 39586151 | 39586297 | [illegible] |
| SE | 77455 | ENSG00000106100 | NOD1 | chr7 | - | 30428503 | 30428609 | [illegible] |
| SE | 46333 | ENSG00000095564 | BTAF1 | chr10 | + | 91966636 | 91966757 | [illegible] |
| SE | 66607 | ENSG00000115687 | PASK | chr2 | - | 241108466 | 241108807 | [illegible] |
| SE | 93373 | ENSG00000119509 | INVS | chr9 | + | 100215400 | 100215488 | [illegible] |
| SE | 54950 | ENSG00000088448 | ANKRD10 | chr13 | - | 110893027 | 110893263 | [illegible] |
| SE | 17146 | ENSG00000011052 | NME1-NME2 | chr17 | + | 51154224 | 51154444 | [illegible] |
| SE | 46473 | ENSG00000185864 | NPIPB4 | chr16 | - | 21857658 | 21857756 | [illegible] |
| SE | 28056 | ENSG00000114738 | MAPKAPK3 | chr3 | + | 50647122 | 50647203 | [illegible] |
| SE | 83537 | ENSG00000171115 | GIMAP8 | chr7 | + | 150474011 | 150474638 | [illegible] |
| SE | 40487 | ENSG00000285708 | NA | chr3 | - | 71015548 | 71015653 | [illegible] |
| SE | 57275 | ENSG00000143119 | CD53 | chr1 | + | 110894959 | 110895055 | [illegible] |
| SE | 33416 | ENSG00000096063 | SRPK1 | chr6 | - | 35904878 | 35904974 | [illegible] |
| SE | 11312 | ENSG00000122257 | RBBP6 | chr16 | + | 24567791 | 24567893 | [illegible] |
| SE | 92672 | ENSG00000090060 | PAPOLA | chr14 | + | 96560648 | 96560711 | [illegible] |
| SE | 84327 | ENSG00000008256 | CYTH3 | chr7 | - | 6170829 | 6170978 | [illegible] |
| SE | 38628 | ENSG00000115325 | DOK1 | chr2 | + | 74555153 | 74555453 | [illegible] |
| SE | 33200 | ENSG00000163930 | BAP1 | chr3 | - | 52407398 | 52407460 | [illegible] |
| SE | 28398 | ENSG00000204842 | ATXN2 | chr12 | - | 111510384 | 111510444 | [illegible] |
| SE | 96310 | ENSG00000110422 | HIPK3 | chr11 | + | 33337074 | 33337194 | [illegible] |
| SE | 95586 | ENSG00000130714 | POMT1 | chr9 | + | 131504188 | 131504649 | [illegible] |
| SE | 90217 | ENSG00000132294 | EFR3A | chr8 | + | 131996405 | 131996497 | [illegible] |
| SE | 43194 | ENSG00000171853 | TRAPPC12 | chr2 | + | 3443778 | 3443891 | [illegible] |
| SE | 1173 | ENSG00000196236 | XPNPEP3 | chr22 | + | 40870047 | 40870139 | [illegible] |
| SE | 106038 | ENSG00000204482 | LST1 | chr6 | + | 31587943 | 31587966 | [illegible] |
| SE | 34855 | ENSG00000159433 | STARD9 | chr15 | + | 42663280 | 42663490 | [illegible] |
| SE | 6870 | ENSG00000263753 | LINC00667 | chr18 | + | 5240647 | 5240819 | [illegible] |
| SE | 3688 | ENSG00000167613 | LAIR1 | chr19 | - | 54364294 | 54364486 | [illegible] |
| SE | 49270 | ENSG00000176148 | TCP11L1 | chr11 | + | 33057114 | 33057235 | [illegible] |
| SE | 99994 | ENSG00000122359 | ANXA11 | chr10 | - | 80176106 | 80176155 | [illegible] |
| SE | 4068 | ENSG00000105771 | SMG9 | chr19 | - | 43734388 | 43734495 | [illegible] |
| SE | 83325 | ENSG00000161010 | MRNIP | chr5 | - | 179851242 | 179851337 | [illegible] |
| SE | 66165 | ENSG00000088179 | PTPN4 | chr2 | + | 119920068 | 119920241 | [illegible] |
| SE | 25357 | ENSG00000112992 | NNT | chr5 | + | 43647923 | 43648440 | [illegible] |
| SE | 63100 | ENSG00000138050 | THUMPD2 | chr2 | - | 39757247 | 39757368 | [illegible] |
| SE | 28098 | ENSG00000084676 | NCOA1 | chr2 | + | 24768078 | 24768135 | [illegible] |
| SE | 77062 | ENSG00000125779 | PANK2 | chr20 | + | 3918670 | 3918796 | [illegible] |
| SE | 75067 | ENSG00000198223 | CSF2RA | chrX | + | 1303260 | 1303362 | [illegible] |
| SE | 8454 | ENSG00000141503 | MINK1 | chr17 | + | 4859212 | 4859328 | [illegible] |
| SE | 59949 | ENSG00000117280 | RAB29 | chr1 | - | 205771471 | 205771653 | [illegible] |
| SE | 43808 | ENSG00000286053 | ASDURF | chr2 | + | 189665356 | 189665451 | [illegible] |
| SE | 57639 | ENSG00000142405 | NLRP12 | chr19 | - | 53807494 | 53807668 | [illegible] |
| SE | 97439 | ENSG00000023445 | BIRC3 | chr11 | + | 102319876 | 102320006 | [illegible] |
| SE | 53594 | ENSG00000115756 | HPCAL1 | chr2 | + | 10424524 | 10424613 | [illegible] |
| SE | 100740 | ENSG00000143756 | FBXO28 | chr1 | + | 224153141 | 224153337 | [illegible] |
| SE | 40044 | ENSG00000143190 | POU2F1 | chr1 | + | 167319013 | 167319056 | [illegible] |
| SE | 88346 | ENSG00000188938 | FAM120AOS | chr9 | - | 93448368 | 93448489 | [illegible] |
| SE | 79403 | ENSG00000133624 | ZNF767P | chr7 | - | 149552800 | 149553013 | [illegible] |
| SE | 57327 | ENSG00000181722 | ZBTB20 | chr3 | - | 114710267 | 114710313 | [illegible] |
| SE | 101922 | ENSG00000138326 | RPS24 | chr10 | + | 78040614 | 78040747 | [illegible] |
| SE | 100520 | ENSG00000054654 | SYNE2 | chr14 | + | 64009965 | 64010116 | [illegible] |
| SE | 102797 | ENSG00000198825 | INPP5F | chr10 | + | 119806359 | 119806480 | [illegible] |
| SE | 1291 | ENSG00000100239 | PPP6R2 | chr22 | + | 50391866 | 50392015 | [illegible] |
| SE | 71009 | ENSG00000173867 | NA | chr15 | - | 88464702 | 88464873 | [illegible] |
| SE | 8494 | ENSG00000125454 | SLC25A19 | chr17 | - | 75278151 | 75278335 | [illegible] |
| SE | 87967 | ENSG00000168066 | SF1 | chr11 | - | 64766899 | 64767079 | [illegible] |
| SE | 28179 | ENSG00000170448 | NFXL1 | chr4 | - | 47851855 | 47851942 | [illegible] |
| SE | 19938 | ENSG00000112651 | MRPL2 | chr6 | - | 43055544 | 43055618 | [illegible] |
| SE | 26435 | ENSG00000158290 | CUL4B | chrX | - | 120544480 | 120544643 | [illegible] |
| SE | 104538 | ENSG00000075303 | SLC25A40 | chr7 | - | 87836729 | 87836810 | [illegible] |
| SE | 50607 | ENSG00000134574 | DDB2 | chr11 | + | 47235269 | 47235412 | [illegible] |
| SE | 48667 | ENSG00000131374 | TBC1D5 | chr3 | - | 17623848 | 17623913 | [illegible] |
| SE | 77000 | ENSG00000149657 | LSM14B | chr20 | + | 62126303 | 62126439 | [illegible] |
| SE | 9310 | ENSG00000133030 | MPRIP | chr17 | + | 17175292 | 17175412 | [illegible] |
| SE | 57885 | ENSG00000162604 | TM2D1 | chr1 | - | 61686791 | 61686921 | [illegible] |
| SE | 802 | ENSG00000100297 | MCM5 | chr22 | + | 35401665 | 35401814 | [illegible] |
| SE | 71355 | ENSG00000014641 | MDH1 | chr2 | + | 63594487 | 63594586 | [illegible] |
| SE | 14461 | ENSG00000140548 | ZNF710 | chr15 | + | 90074115 | 90074290 | [illegible] |
| SE | 70670 | ENSG00000115762 | PLEKHB2 | chr2 | + | 131132901 | 131132991 | [illegible] |
| SE | 59367 | ENSG00000116127 | ALMS1 | chr2 | + | 73408621 | 73408747 | [illegible] |
| SE | 89684 | ENSG00000120992 | LYPLA1 | chr8 | - | 54062253 | 54062324 | [illegible] |
| SE | 65012 | ENSG00000167107 | ACSF2 | chr17 | + | 50473893 | 50474004 | [illegible] |
| SE | 32426 | ENSG00000136040 | PLXNC1 | chr12 | + | 94300909 | 94301057 | [illegible] |
| SE | 80423 | ENSG00000126746 | ZNF384 | chr12 | - | 6678662 | 6678710 | [illegible] |
| SE | 108191 | ENSG00000121749 | TBC1D15 | chr12 | + | 71907862 | 71907989 | [illegible] |
| SE | 8313 | ENSG00000100038 | TOP3B | chr22 | - | 21970979 | 21971061 | [illegible] |
| SE | 104884 | ENSG00000109685 | NSD2 | chr4 | + | 1954536 | 1954685 | [illegible] |
| SE | 97557 | ENSG00000109452 | INPP4B | chr4 | - | 142425206 | 142425263 | [illegible] |
| SE | 29035 | ENSG00000182150 | ERCC6L2 | chr9 | + | 95966561 | 95966714 | [illegible] |
| SE | 64697 | ENSG00000093144 | ECHDC1 | chr6 | - | 127330808 | 127331030 | [illegible] |
| SE | 17353 | ENSG00000133961 | NUMB | chr14 | - | 73316389 | 73316422 | [illegible] |
| SE | 1693 | ENSG00000228315 | GUSBP11 | chr22 | - | 23659864 | 23660000 | [illegible] |
| SE | 100301 | ENSG00000162144 | CYB561A3 | chr11 | - | 61357732 | 61357828 | [illegible] |
| SE | 59511 | ENSG00000148660 | CAMK2G | chr10 | - | 73852253 | 73852319 | [illegible] |
| SE | 61789 | ENSG00000104343 | UBE2W | chr8 | - | 73870268 | 73870301 | [illegible] |
| SE | 9708 | ENSG00000141568 | FOXK2 | chr17 | + | 82582740 | 82582934 | [illegible] |
| SE | 26815 | ENSG00000138760 | SCARB2 | chr4 | - | 76179516 | 76179705 | [illegible] |
| SE | 76683 | ENSG00000223501 | VPS52 | chr6 | - | 33267681 | 33267739 | [illegible] |
| SE | 41651 | ENSG00000119185 | ITGB1BP1 | chr2 | - | 9414177 | 9414256 | [illegible] |
| SE | 58262 | ENSG00000089127 | OAS1 | chr12 | + | 112919486 | 112919615 | [illegible] |
| SE | 67733 | ENSG00000135912 | TTLL4 | chr2 | + | 218753114 | 218753185 | [illegible] |
| SE | 52862 | ENSG00000116138 | DNAJC16 | chr1 | + | 15562141 | 15562325 | [illegible] |
| SE | 65840 | ENSG00000204713 | TRIM27 | chr6 | - | 28908807 | 28908840 | [illegible] |
| SE | 24171 | ENSG00000124357 | NAGK | chr2 | + | 71075554 | 71075610 | [illegible] |
| SE | 18454 | ENSG00000100897 | DCAF11 | chr14 | + | 24116916 | 24117044 | [illegible] |
| SE | 32079 | ENSG00000242086 | MUC20-OT1 | chr3 | + | 195689338 | 195689438 | [illegible] |
| SE | 49866 | ENSG00000151136 | BTBD11 | chr12 | + | 107617300 | 107617436 | [illegible] |
| SE | 39820 | ENSG00000144711 | IQSEC1 | chr3 | - | 12941570 | 12941865 | [illegible] |
| SE | 103357 | ENSG00000221817 | PPP3CB-AS1 | chr10 | + | 73504804 | 73504911 | [illegible] |
| SE | 5117 | ENSG00000125826 | RBCK1 | chr20 | + | 409829 | 410025 | [illegible] |
| SE | 71436 | ENSG00000115234 | SNX17 | chr2 | + | 27373039 | 27373113 | [illegible] |
| SE | 55544 | ENSG00000117010 | ZNF684 | chr1 | + | 40532732 | 40532808 | [illegible] |
| SE | 67267 | ENSG00000135632 | SMYD5 | chr2 | + | 73220660 | 73220782 | [illegible] |
| SE | 46093 | ENSG00000166912 | MTMR10 | chr15 | - | 30951967 | 30952038 | [illegible] |
| SE | 88968 | ENSG00000148516 | ZEB1 | chr10 | + | 31387123 | 31387266 | [illegible] |
| SE | 99624 | ENSG00000109920 | FNBP4 | chr11 | - | 47743952 | 47744163 | [illegible] |
| SE | 98669 | ENSG00000130592 | LSP1 | chr11 | + | 1881431 | 1881596 | [illegible] |
| SE | 32217 | ENSG00000114650 | SCAP | chr3 | - | 47428512 | 47428670 | [illegible] |
| SE | 10945 | ENSG00000091592 | NLRP1 | chr17 | - | 5520880 | 5521012 | [illegible] |
| SE | 60947 | ENSG00000117155 | SSX2IP | chr1 | - | 84664416 | 84664552 | [illegible] |
| SE | 8050 | ENSG00000196704 | AMZ2 | chr17 | + | 68249010 | 68249096 | [illegible] |
| SE | 26133 | ENSG00000167766 | ZNF83 | chr19 | - | 52616717 | 52616875 | [illegible] |
| SE | 20727 | ENSG00000115760 | BIRC6 | chr2 | + | 32487646 | 32487801 | [illegible] |
| SE | 76318 | ENSG00000085491 | SLC25A24 | chr1 | - | 108156700 | 108156885 | [illegible] |
| SE | 106575 | ENSG00000103356 | EARS2 | chr16 | - | 23553043 | 23553123 | [illegible] |

| | | | | | | | | | | | | | | |
|---|---|---|---|---|---|---|---|---|---|---|---|---|---|---|
| SE | 61467 | ENSG00000096080 | MRPS18A | chr6 | - | 43675201 | 43675271 | 43671201 | 43671906 | 43675493 | 43675617 | 76 | 45 | 75 |
| SE | 36378 | ENSG00000155621 | C9orf85 | chr9 | + | 71947005 | 71947112 | 71911544 | 71911836 | 71971504 | 71971618 | 82 | 48 | 59 |
| SE | 65797 | ENSG00000116521 | SCAMP3 | chr1 | - | 155256673 | 155256791 | 155255978 | 155256419 | 155257284 | 155257386 | 88 | 24 | 108 |
| SE | 38652 | ENSG00000073417 | PDE8A | chr15 | + | 85090717 | 85090915 | 85083555 | 85083644 | 85091043 | 85091181 | 7 | 3 | 3 |
| SE | 48470 | ENSG00000259494 | MRPL46 | chr15 | - | 88464702 | 88464876 | 88459477 | 88459863 | 88465586 | 88465773 | 65 | 38 | 75 |
| SE | 81348 | ENSG00000173852 | DPY19L1 | chr7 | - | 35013567 | 35013705 | 35011329 | 35011450 | 35017881 | 35017969 | 142 | 71 | 46 |
| SE | 68881 | ENSG00000031081 | ARHGAP31 | chr3 | + | 119368371 | 119368516 | 119365315 | 119365418 | 119380903 | 119380986 | 114 | 67 | 72 |
| SE | 56533 | ENSG00000103489 | XYLT1 | chr16 | - | 17127665 | 17127861 | 17101768 | 17109017 | 17134472 | 17134735 | 35 | 15 | 33 |
| SE | 35389 | ENSG00000161267 | BDH1 | chr3 | - | 197538305 | 197538952 | 197533488 | 197533561 | 197546360 | 197546486 | 5 | 3 | 19 |
| SE | 100787 | ENSG00000162408 | NOL9 | chr1 | - | 6545044 | 6545180 | 6533279 | 6533441 | 6549570 | 6549698 | 81 | 44 | 65 |
| SE | 43148 | ENSG00000113161 | HMGCR | chr5 | + | 75359397 | 75359511 | 75359169 | 75359310 | 75359984 | 75360139 | 76 | 46 | 82 |
| SE | 6582 | ENSG00000074657 | ZNF532 | chr18 | + | 58948066 | 58948229 | 58939444 | 58939621 | 58953517 | 58953799 | 85 | 61 | 50 |
| SE | 60796 | ENSG00000160714 | UBE2Q1 | chr1 | - | 154552735 | 154552820 | 154552381 | 154552464 | 154553031 | 154553172 | 361 | 201 | 354 |
| SE | 11221 | ENSG00000140829 | DHX38 | chr16 | + | 72101499 | 72101612 | 72101081 | 72101193 | 72103073 | 72103211 | 261 | 133 | 188 |
| SE | 37920 | ENSG00000184014 | DENND5A | chr11 | - | 9193493 | 9193681 | 9180766 | 9181084 | 9203659 | 9204317 | 173 | 91 | 155 |
| SE | 82953 | ENSG00000127954 | STEAP4 | chr7 | - | 88290915 | 88291014 | 88283813 | 88284271 | 88306791 | 88306880 | 5 | 0 | 4 |
| SE | 67513 | ENSG00000148426 | PROSER2 | chr10 | + | 11862911 | 11863042 | 11851996 | 11852215 | 11866530 | 11866776 | 2 | 1 | 4 |
| SE | 83571 | ENSG00000124222 | STX16 | chr20 | + | 58671153 | 58671297 | 58670511 | 58670603 | 58673630 | 58673711 | 349 | 260 | 332 |
| SE | 71681 | ENSG00000135976 | ANKRD36 | chr2 | + | 97122886 | 97122993 | 97118343 | 97118517 | 97127066 | 97127134 | 32 | 36 | 43 |
| SE | 62401 | ENSG00000122912 | SLC25A16 | chr10 | - | 68496433 | 68496512 | 68493448 | 68493570 | 68503631 | 68503695 | 5 | 8 | 0 |
| SE | 22057 | ENSG00000205133 | TRIQK | chr8 | - | 92921502 | 92921660 | 92916928 | 92917010 | 92954405 | 92954564 | 18 | 13 | 14 |
| SE | 4603 | ENSG00000105397 | TYK2 | chr19 | - | 10369756 | 10369877 | 10368328 | 10368418 | 10378213 | 10378426 | 5 | 0 | 8 |
| SE | 57884 | ENSG00000162604 | TM2D1 | chr1 | - | 61686791 | 61686895 | 61683496 | 61683546 | 61694696 | 61694770 | 10 | 3 | 15 |
| SE | 106106 | ENSG00000089157 | RPLP0 | chr12 | - | 120198553 | 120198631 | 120197321 | 120197462 | 120198853 | 120199000 | 1271 | 528 | 1489 |
| SE | 65655 | ENSG00000143924 | EML4 | chr2 | + | 42317426 | 42317524 | 42304483 | 42304551 | 42325466 | 42325554 | 95 | 62 | 122 |
| SE | 43135 | ENSG00000186470 | BTN3A2 | chr6 | + | 26372896 | 26373097 | 26370321 | 26370603 | 26373275 | 26373296 | 691 | 467 | 536 |
| SE | 9731 | ENSG00000171302 | CANT1 | chr17 | - | 79000099 | 79000270 | 78997392 | 78997963 | 79009663 | 79009764 | 8 | 4 | 4 |
| [illegible] | [illegible] | [illegible] | [illegible] | [illegible] | [illegible] | [illegible] | [illegible] | [illegible] | [illegible] | [illegible] | [illegible] | [illegible] | [illegible] | [illegible] |

| | | | | | | | | | | | |
|---|---|---|---|---|---|---|---|---|---|---|---|
| SE | 12538 | ENSG00000103148 | NPRL3 | chr16 | - | 119125 | 119255 | 112621 | 112775 | 130521 | 130591 |
| SE | 57674 | ENSG00000057757 | PITHD1 | chr1 | + | 23779437 | 23779481 | 23778417 | 23778713 | 23779863 | 23779941 |
| SE | 10480 | ENSG00000186566 | GPATCH8 | chr17 | - | 44464471 | 44464544 | 44436477 | 44436545 | 44503325 | 44503391 |
| SE | 56540 | ENSG00000069702 | TGFBR3 | chr1 | - | 91716235 | 91716394 | 91712242 | 91712542 | 91716567 | 91716708 |
| SE | 52765 | ENSG00000198019 | FCGR1B | chr1 | + | 121088653 | 121088908 | 121087820 | 121087841 | 121092995 | 121093250 |
| SE | 13980 | ENSG00000103522 | IL21R | chr16 | + | 27434346 | 27434449 | 27430055 | 27430120 | 27437487 | 27437687 |
| SE | 83409 | ENSG00000146707 | POMZP3 | chr7 | - | 76618182 | 76618300 | 76611721 | 76611813 | 76625521 | 76625683 |
| SE | 105770 | ENSG00000125651 | GTF2F1 | chr19 | - | 6381358 | 6381615 | 6381121 | 6381195 | 6381696 | 6381775 |
| SE | 45418 | ENSG00000142330 | CAPN10 | chr2 | + | 240595933 | 240596521 | 240595023 | 240595304 | 240597887 | 240598087 |
| SE | 77157 | ENSG00000136295 | TTYH3 | chr7 | + | 2659939 | 2660040 | 2658939 | 2659015 | 2661667 | 2661842 |
| SE | 43907 | ENSG00000112118 | MCM3 | chr6 | - | 52283293 | 52283406 | 52282652 | 52282861 | 52284596 | 52284742 |
| SE | 99304 | ENSG00000119699 | TGFB3 | chr14 | - | 75960922 | 75961076 | 75958096 | 75959345 | 75963315 | 75963487 |
| SE | 53574 | ENSG00000158552 | ZFAND2B | chr2 | + | 219207647 | 219207774 | 219207326 | 219207421 | 219207886 | 219208003 |
| SE | 17545 | ENSG00000100813 | ACIN1 | chr14 | - | 23071124 | 23071186 | 23069475 | 23069617 | 23078150 | 23078266 |
| SE | 20905 | ENSG00000153975 | ZUP1 | chr6 | - | 116651571 | 116651737 | 116647458 | 116647610 | 116652003 | 116652192 |
| SE | 5534 | ENSG00000244687 | UBE2V1 | chr20 | - | 50096671 | 50096820 | 50083877 | 50084254 | 50113106 | 50113125 |
| SE | 59333 | ENSG00000213064 | SFT2D2 | chr1 | + | 168231513 | 168231600 | 168225949 | 168226142 | 168235100 | 168235182 |
| SE | 71117 | ENSG00000198363 | ASPH | chr8 | - | 61643944 | 61644001 | 61643385 | 61643433 | 61644599 | 61644632 |
| SE | 19933 | ENSG00000092140 | G2E3 | chr14 | + | 30598482 | 30598599 | 30597419 | 30597526 | 30601769 | 30601894 |
| SE | 71120 | ENSG00000148297 | MED22 | chr9 | - | 133346539 | 133346700 | 133345171 | 133345252 | 133347921 | 133348117 |
| SE | 96743 | ENSG00000185238 | PRMT3 | chr11 | + | 20393790 | 20393854 | 20392896 | 20392999 | 20395802 | 20395962 |
| SE | 6761 | ENSG00000133313 | CNDP2 | chr18 | + | 74501211 | 74501472 | 74499936 | 74500033 | 74505848 | 74505942 |
| SE | 3005 | ENSG00000267120 | NA | chr19 | - | 35739709 | 35740088 | 35739251 | 35739626 | 35741023 | 35741228 |
| SE | 91089 | ENSG00000165102 | HGSNAT | chr8 | + | 43197839 | 43197952 | 43197671 | 43197742 | 43199387 | 43199858 |
| SE | 106326 | ENSG00000125458 | NT5C | chr17 | - | 75131044 | 75131105 | 75130752 | 75130867 | 75131180 | 75131258 |
| SE | 24977 | ENSG00000150712 | MTMR12 | chr5 | - | 32276681 | 32276742 | 32273979 | 32274122 | 32312757 | 32312938 |
| SE | 78015 | ENSG00000125977 | EIF2S2 | chr20 | - | 34093674 | 34093731 | 34090516 | 34090602 | 34096656 | 34096805 |
| SE | 53887 | ENSG00000166523 | CLEC4E | chr12 | - | 8537114 | 8537266 | 8536089 | 8536205 | 8539216 | 8539255 |
| SE | 50276 | ENSG00000131115 | ZNF227 | chr19 | + | 44217790 | 44217852 | 44213068 | 44213244 | 44228445 | 44228572 |
| SE | 50110 | ENSG00000163322 | ABRAXAS1 | chr4 | - | 83472221 | 83472288 | 83470202 | 83470396 | 83476642 | 83476679 |
| SE | 40476 | ENSG00000164088 | PPM1M | chr3 | + | 52248636 | 52248700 | 52248152 | 52248453 | 52248955 | 52249063 |
| SE | 104022 | ENSG00000165983 | PTER | chr10 | + | 16505019 | 16505160 | 16486351 | 16486617 | 16511045 | 16513737 |

| | | | | | | | | | | | | | | | | | | | | | | | | | | | | | | | | | | | | | | | | | | | | | | |
|---|---|---|---|---|---|---|---|---|---|---|---|---|---|---|---|---|---|---|---|---|---|---|---|---|---|---|---|---|---|---|---|---|---|---|---|---|---|---|---|---|---|---|---|---|---|---|
| SE | 31862 | ENSG00000143669 | LYST | chr1 | - | 235728075 | 235728131 | 235724027 | 235724180 | 235729595 | 235729657 | 283 | 345 | 210 | 252 | 376 | 0 | 2 | 3 | 3 | 0 | 30 | 121 | 88 | 13 | 69 | 0 | 5 | 4 | 2 | 0 | 205 | 149 | 0.053357689 | 0.398216915 | 1 | 0.992 | 0.981 | 0.984 | 1 | 1 | 0.946 | 0.941 | 0.825 | 1 | 0.049 |
| SE | 63630 | ENSG00000198520 | ARMH1 | chr1 | + | 44724761 | 44724839 | 44724319 | 44724392 | 44725032 | 44725217 | 4 | 3 | 7 | 8 | 1 | 4 | 0 | 1 | 4 | 4 | 21 | 0 | 0 | 13 | 0 | 0 | 1 | 2 | 12 | 10 | 227 | 149 | 0.053365872 | 0.398240025 | 0.396 | 1 | 0.821 | 0.568 | 0.141 | 1 | 0 | 0 | 0.416 | 0 | 0.302 |
| SE | 59435 | ENSG00000112339 | HBS1L | chr6 | - | 135037810 | 135038026 | 135002620 | 135002842 | 135039572 | 135039767 | 34 | 21 | 32 | 33 | 63 | 97 | 70 | 107 | 92 | 215 | 0 | 17 | 29 | 6 | 0 | 19 | 55 | 90 | 46 | 3 | 298 | 149 | 0.05338397 | 0.398327793 | 0.149 | 0.13 | 0.13 | 0.152 | 0.128 | 0 | 0.134 | 0.139 | 0.061 | 0 | 0.071 |
| SE | 37848 | ENSG00000114126 | TFDP2 | chr3 | - | 142005440 | 142005544 | 141994269 | 141995141 | 142054114 | 142054149 | 215 | 78 | 119 | 122 | 151 | 2 | 0 | 0 | 1 | 4 | 136 | 109 | 80 | 64 | 93 | 2 | 4 | 0 | 6 | 0 | 253 | 149 | 0.053388442 | 0.398327793 | 0.984 | 1 | 1 | 0.986 | 0.957 | 0.976 | 0.941 | 1 | 0.863 | 1 | 0.029 |
| SE | 104487 | ENSG00000013561 | RNF14 | chr5 | + | 141978302 | 141978830 | 141974803 | 141974955 | 141983379 | 141983552 | 37 | 31 | 30 | 41 | 62 | 3 | 0 | 0 | 0 | 6 | 17 | 29 | 21 | 27 | 49 | 0 | 0 | 3 | 0 | 0 | 298 | 149 | 0.053392896 | 0.398327793 | 0.86 | 1 | 1 | 1 | 0.838 | 1 | 1 | 0.778 | 1 | 1 | -0.016 |
| SE | 94722 | ENSG00000106948 | AKNA | chr9 | - | 114342008 | 114342125 | 114334155 | 114337306 | 114345862 | 114346009 | 39 | 21 | 39 | 34 | 28 | 0 | 4 | 3 | 0 | 2 | 103 | 168 | 150 | 89 | 177 | 0 | 0 | 1 | 8 | 0 | 266 | 149 | 0.053405596 | 0.398384579 | 1 | 0.746 | 0.879 | 1 | 0.887 | 1 | 1 | 0.988 | 0.862 | 1 | -0.068 |
| SE | 26813 | ENSG00000138760 | SCARB2 | chr4 | - | 76179516 | 76179695 | 76176436 | 76176528 | 76180953 | 76181101 | 214 | 180 | 154 | 237 | 371 | 5 | 1 | 0 | 6 | 5 | 370 | 226 | 215 | 370 | 136 | 8 | 3 | 6 | 13 | 2 | 298 | 149 | 0.053440855 | 0.398571653 | 0.955 | 0.989 | 1 | 0.952 | 0.974 | 0.959 | 0.974 | 0.947 | 0.934 | 0.971 | 0.017 |
| SE | 26816 | ENSG00000138760 | SCARB2 | chr4 | - | 76179516 | 76179771 | 76176436 | 76176528 | 76180953 | 76181101 | 214 | 180 | 154 | 237 | 371 | 5 | 1 | 0 | 6 | 5 | 370 | 226 | 215 | 370 | 136 | 8 | 3 | 6 | 13 | 2 | 298 | 149 | 0.053440855 | 0.398571653 | 0.955 | 0.989 | 1 | 0.952 | 0.974 | 0.959 | 0.974 | 0.947 | 0.934 | 0.971 | 0.017 |
| SE | 50214 | ENSG00000148187 | MRRF | chr9 | + | 122286038 | 122286166 | 122285168 | 122285287 | 122291748 | 122291798 | 15 | 14 | 29 | 19 | 9 | 77 | 68 | 60 | 64 | 96 | 17 | 25 | 11 | 11 | 0 | 87 | 104 | 76 | 83 | 40 | 277 | 149 | 0.053498229 | 0.398947363 | 0.095 | 0.1 | 0.206 | 0.138 | 0.048 | 0.095 | 0.114 | 0.072 | 0.067 | 0 | 0.048 |
| SE | 38473 | ENSG00000147251 | DOCK11 | chrX | + | 118643456 | 118643594 | 118641189 | 118641305 | 118648944 | 118649127 | 519 | 361 | 425 | 410 | 719 | 29 | 13 | 15 | 27 | 2 | 22 | 308 | 217 | 52 | 33 | 0 | 24 | 9 | 7 | 1 | 287 | 149 | 0.053504887 | 0.398947363 | 0.903 | 0.935 | 0.936 | 0.887 | 0.995 | 1 | 0.869 | 0.926 | 0.794 | 0.945 | 0.024 |
| SE | 38825 | ENSG00000229180 | NA | chr7 | - | 66588511 | 66588580 | 66576754 | 66576932 | 66592256 | 66592386 | 2 | 1 | 7 | 1 | 1 | 8 | 1 | 7 | 8 | 5 | 0 | 0 | 0 | 1 | 6 | 18 | 7 | 5 | 12 | 0 | 218 | 149 | 0.053506517 | 0.398947363 | 0.146 | 0.406 | 0.406 | 0.079 | 0.12 | 0 | 0 | 0 | 0.054 | 1 | 0.021 |
| SE | 70266 | ENSG00000104218 | CSPP1 | chr8 | + | 67103036 | 67103135 | 67095292 | 67095732 | 67105904 | 67105975 | 98 | 91 | 86 | 108 | 116 | 4 | 0 | 1 | 0 | 7 | 59 | 51 | 51 | 43 | 34 | 0 | 0 | 0 | 0 | 0 | 248 | 149 | 0.053516503 | 0.398983821 | 0.936 | 1 | 0.981 | 1 | 0.909 | 1 | 1 | 1 | 1 | 1 | -0.035 |
| SE | 61233 | ENSG00000171503 | ETFDH | chr4 | + | 158679289 | 158679677 | 158672275 | 158672490 | 158680466 | 158680607 | 2 | 5 | 9 | 2 | 3 | 16 | 8 | 9 | 7 | 7 | 3 | 2 | 14 | 5 | 5 | 5 | 8 | 0 | 2 | 7 | 298 | 149 | 0.053663381 | 0.400040757 | 0.059 | 0.238 | 0.333 | 0.125 | 0.176 | 0.231 | 0.111 | 1 | 0.556 | 0.263 | -0.246 |
| SE | 9129 | ENSG00000163597 | SNHG16 | chr17 | + | 76558373 | 76559043 | 76557777 | 76557857 | 76561287 | 76561398 | 81 | 18 | 82 | 73 | 123 | 81 | 48 | 84 | 64 | 56 | 24 | 47 | 59 | 55 | 42 | 82 | 49 | 98 | 72 | 43 | 298 | 149 | 0.053679169 | 0.400091155 | 0.333 | 0.158 | 0.328 | 0.363 | 0.523 | 0.128 | 0.324 | 0.231 | 0.276 | 0.328 | 0.084 |
| SE | 9574 | ENSG00000108604 | SMARCD2 | chr17 | - | 63835411 | 63835567 | 63834704 | 63834800 | 63836921 | 63837044 | 284 | 142 | 285 | 293 | 301 | 2 | 0 | 0 | 1 | 2 | 391 | 219 | 162 | 214 | 410 | 0 | 0 | 4 | 4 | 10 | 298 | 149 | 0.053680362 | 0.400091155 | 0.986 | 1 | 0.993 | 0.987 | 0.987 | 1 | 1 | 0.953 | 0.964 | 0.953 | 0.017 |
| SE | 102962 | ENSG00000028116 | VRK2 | chr2 | + | 58084088 | 58084138 | 58048826 | 58048967 | 58088340 | 58088446 | 37 | 45 | 37 | 34 | 30 | 0 | 0 | 0 | 3 | 0 | 13 | 14 | 16 | 1 | 0 | 0 | 0 | 0 | 0 | 8 | 199 | 149 | 0.053707917 | 0.400258431 | 1 | 1 | 1 | 0.895 | 1 | 1 | 1 | 1 | 1 | 0 | 0.179 |
| SE | 64432 | ENSG00000162813 | BPNT1 | chr1 | - | 220072849 | 220072957 | 220069383 | 220069432 | 220073966 | 220074067 | 124 | 71 | 150 | 106 | 157 | 9 | 6 | 6 | 5 | 15 | 76 | 105 | 48 | 114 | 130 | 6 | 15 | 11 | 6 | 12 | 257 | 149 | 0.053715897 | 0.400274471 | 0.889 | 0.873 | 0.935 | 0.925 | 0.859 | 0.88 | 0.802 | 0.717 | 0.917 | 0.863 | 0.06 |
| SE | 27096 | ENSG00000156265 | MAP3K7CL | chr21 | + | 29130794 | 29130923 | 29122054 | 29122213 | 29133305 | 29133414 | 6 | 2 | 6 | 3 | 27 | 0 | 0 | 0 | 0 | 0 | 7 | 14 | 4 | 5 | 8 | 8 | 0 | 0 | 0 | 0 | 278 | 149 | 0.053720294 | 0.400274471 | 1 | 1 | 1 | 1 | 1 | 0.319 | 1 | 1 | 1 | 1 | 0.136 |
| SE | 15131 | ENSG00000104093 | DMXL2 | chr15 | - | 51451644 | 51451697 | 51450128 | 51450346 | 51453549 | 51453641 | 159 | 179 | 96 | 139 | 130 | 6 | 5 | 4 | 10 | 3 | 15 | 107 | 70 | 16 | 30 | 0 | 0 | 15 | 0 | 9 | 202 | 149 | 0.053771936 | 0.400621138 | 0.951 | 0.964 | 0.947 | 0.911 | 0.97 | 1 | 1 | 0.775 | 1 | 0.711 | 0.051 |
| SE | 9390 | ENSG00000121067 | SPOP | chr17 | - | 49675894 | 49676016 | 49668026 | 49668078 | 49677932 | 49678000 | 2 | 2 | 3 | 8 | 0 | 12 | 9 | 5 | 15 | 6 | 0 | 0 | 0 | 0 | 7 | 1 | 7 | 3 | 4 | 22 | 271 | 149 | 0.053804873 | 0.400828383 | 0.084 | 0.109 | 0.248 | 0.227 | 0 | 0 | 0 | 0 | 0 | 0.149 | 0.104 |
| SE | 9088 | ENSG00000141574 | SECTM1 | chr17 | - | 82322877 | 82322905 | 82321886 | 82322370 | 82324581 | 82324890 | 9 | 3 | 2 | 6 | 0 | 0 | 2 | 0 | 3 | 2 | 26 | 4 | 2 | 14 | 13 | 0 | 0 | 3 | 1 | 0 | 177 | 149 | 0.05381512 | 0.40086658 | 1 | 0.558 | 1 | 0.627 | 0 | 1 | 1 | 0.359 | 0.922 | 1 | -0.219 |
| SE | 106713 | ENSG00000141376 | BCAS3 | chr17 | + | 61380505 | 61380571 | 61368326 | 61368494 | 61391976 | 61392182 | 4 | 3 | 4 | 3 | 13 | 26 | 16 | 23 | 18 | 19 | 16 | 9 | 3 | 2 | 12 | 7 | 22 | 8 | 9 | 30 | 215 | 149 | 0.053834848 | 0.40097539 | 0.096 | 0.115 | 0.108 | 0.104 | 0.322 | 0.613 | 0.221 | 0.206 | 0.133 | 0.217 | -0.129 |
| SE | 16872 | ENSG00000166734 | GOLM2 | chr15 | + | 44402886 | 44403054 | 44380805 | 44380976 | 44413335 | 44415729 | 313 | 233 | 238 | 290 | 383 | 24 | 19 | 28 | 30 | 32 | 83 | 226 | 245 | 181 | 70 | 0 | 14 | 19 | 19 | 3 | 298 | 149 | 0.053865547 | 0.401165877 | 0.867 | 0.86 | 0.81 | 0.829 | 0.857 | 1 | 0.89 | 0.866 | 0.826 | 0.921 | -0.056 |
| SE | 89246 | ENSG00000137574 | TGS1 | chr8 | + | 55786237 | 55787060 | 55785718 | 55785891 | 55790181 | 55790299 | 153 | 114 | 144 | 200 | 452 | 0 | 1 | 0 | 1 | 9 | 42 | 109 | 182 | 129 | 15 | 4 | 0 | 2 | 0 | 6 | 298 | 149 | 0.053891443 | 0.401289059 | 1 | 0.983 | 1 | 0.99 | 0.962 | 0.84 | 1 | 0.978 | 1 | 0.556 | 0.112 |
| SE | 58076 | ENSG00000082701 | GSK3B | chr3 | - | 120000551 | 120000723 | 119947267 | 119947351 | 120002045 | 120002239 | 11 | 5 | 16 | 11 | 7 | 72 | 73 | 64 | 75 | 104 | 0 | 11 | 6 | 1 | 0 | 32 | 53 | 39 | 54 | 29 | 298 | 149 | 0.053892337 | 0.401289059 | 0.071 | 0.033 | 0.111 | 0.068 | 0.033 | 0 | 0.094 | 0.071 | 0.009 | 0 | 0.028 |
| SE | 88494 | ENSG00000152601 | MBNL1 | chr3 | + | 152456266 | 152456361 | 152447619 | 152447773 | 152459270 | 152459327 | 210 | 189 | 195 | 290 | 306 | 3 | 3 | 2 | 3 | 9 | 51 | 130 | 145 | 104 | 105 | 0 | 5 | 1 | 0 | 0 | 244 | 149 | 0.053903415 | 0.401333374 | 0.977 | 0.975 | 0.983 | 0.983 | 0.954 | 1 | 0.941 | 0.989 | 1 | 1 | -0.012 |
| SE | 24315 | ENSG00000058799 | YIPF1 | chr1 | - | 53866757 | 53866924 | 53860055 | 53860153 | 53871371 | 53871488 | 44 | 35 | 34 | 28 | 64 | 0 | 0 | 0 | 1 | 0 | 67 | 49 | 23 | 33 | 71 | 1 | 1 | 7 | 0 | 0 | 298 | 149 | 0.053931377 | 0.401503382 | 1 | 1 | 1 | 0.933 | 1 | 0.971 | 0.961 | 0.622 | 1 | 1 | 0.076 |
| SE | 54095 | ENSG00000115935 | WIPF1 | chr2 | - | 174597600 | 174597858 | 174585522 | 174585611 | 174625820 | 174625960 | 631 | 364 | 476 | 688 | 1024 | 2 | 6 | 8 | 4 | 15 | 930 | 662 | 846 | 756 | 1814 | 8 | 13 | 10 | 1 | 0 | 298 | 149 | 0.053942055 | 0.401544698 | 0.994 | 0.968 | 0.967 | 0.989 | 0.972 | 0.983 | 0.962 | 0.977 | 0.997 | 1 | -0.006 |
| SE | 92066 | ENSG00000165934 | CPSF2 | chr14 | + | 92158982 | 92159282 | 92157658 | 92157884 | 92161111 | 92161246 | 430 | 425 | 416 | 465 | 800 | 2 | 0 | 0 | 6 | 9 | 94 | 425 | 299 | 275 | 17 | 6 | 9 | 2 | 0 | 0 | 298 | 149 | 0.053951001 | 0.401573116 | 0.991 | 1 | 1 | 0.975 | 0.978 | 0.887 | 0.959 | 0.987 | 1 | 1 | 0.022 |
| SE | 40077 | ENSG00000125347 | IRF1 | chr5 | - | 132486250 | 132486373 | 132484361 | 132484497 | 132486556 | 132486639 | 616 | 379 | 601 | 788 | 682 | 14 | 16 | 17 | 22 | 5 | 2008 | 414 | 846 | 995 | 1249 | 46 | 17 | 11 | 76 | 24 | 272 | 149 | 0.053963544 | 0.401628289 | 0.96 | 0.928 | 0.951 | 0.952 | 0.987 | 0.96 | 0.93 | 0.977 | 0.878 | 0.966 | 0.013 |
| SE | 43085 | ENSG00000239900 | ADSL | chr22 | + | 40353072 | 40353117 | 40349831 | 40350035 | 40358863 | 40359035 | 24 | 11 | 20 | 7 | 19 | 4 | 3 | 2 | 6 | 6 | 25 | 12 | 2 | 19 | 31 | 23 | 0 | 14 | 40 | 8 | 194 | 149 | 0.053991081 | 0.401795043 | 0.822 | 0.738 | 0.885 | 0.473 | 0.709 | 0.455 | 1 | 0.099 | 0.267 | 0.749 | 0.211 |
| SE | 83822 | ENSG00000169891 | REPS2 | chrX | + | 17025058 | 17025185 | 17022122 | 17022271 | 17029525 | 17029623 | 13 | 9 | 7 | 14 | 6 | 0 | 0 | 0 | 0 | 0 | 29 | 3 | 18 | 9 | 33 | 0 | 6 | 1 | 0 | 0 | 276 | 149 | 0.054022035 | 0.401987191 | 1 | 1 | 1 | 1 | 1 | 1 | 0.213 | 0.907 | 1 | 1 | 0.176 |
| SE | 78150 | ENSG00000242073 | NA | chr7 | - | 75484197 | 75484256 | 75475061 | 75475188 | 75485863 | 75486108 | 8 | 7 | 9 | 4 | 3 | 2 | 3 | 4 | 3 | 5 | 20 | 25 | 14 | 28 | 15 | 6 | 4 | 9 | 2 | 2 | 208 | 149 | 0.054044787 | 0.402093923 | 0.741 | 0.626 | 0.617 | 0.489 | 0.301 | 0.705 | 0.817 | 0.527 | 0.909 | 0.843 | -0.205 |
| SE | 61250 | ENSG00000171503 | ETFDH | chr4 | + | 158697558 | 158697699 | 158694500 | 158695643 | 158698986 | 158699130 | 68 | 76 | 76 | 61 | 80 | 0 | 0 | 0 | 0 | 0 | 21 | 85 | 50 | 43 | 0 | 0 | 1 | 0 | 0 | 6 | 290 | 149 | 0.054046649 | 0.402093923 | 1 | 1 | 1 | 1 | 1 | 1 | 0.978 | 1 | 1 | 0 | 0.204 |
| SE | 13914 | ENSG00000067365 | METTL22 | chr16 | + | 8644556 | 8644725 | 8642462 | 8642565 | 8646107 | 8646191 | 37 | 21 | 11 | 54 | 39 | 3 | 0 | 2 | 0 | 0 | 70 | 63 | 20 | 28 | 208 | 4 | 2 | 0 | 4 | 4 | 298 | 149 | 0.054092148 | 0.402394184 | 0.86 | 1 | 0.733 | 1 | 1 | 0.897 | 0.94 | 1 | 0.778 | 0.963 | 0.003 |
| SE | 38222 | ENSG00000113108 | APBB3 | chr5 | - | 140561017 | 140561101 | 140560312 | 140560504 | 140561364 | 140561449 | 8 | 5 | 5 | 6 | 2 | 0 | 0 | 0 | 0 | 0 | 10 | 8 | 2 | 20 | 6 | 0 | 0 | 0 | 0 | 8 | 233 | 149 | 0.054103521 | 0.402440555 | 1 | 1 | 1 | 1 | 1 | 1 | 1 | 1 | 1 | 0.324 | 0.135 |
| SE | 11429 | ENSG00000167522 | ANKRD11 | chr16 | - | 89291680 | 89291777 | 89291012 | 89291183 | 89305205 | 89305344 | 15 | 10 | 19 | 10 | 18 | 46 | 37 | 49 | 44 | 43 | 8 | 6 | 21 | 14 | 22 | 85 | 27 | 71 | 51 | 87 | 246 | 149 | 0.054130844 | 0.402605547 | 0.165 | 0.141 | 0.19 | 0.121 | 0.202 | 0.054 | 0.119 | 0.152 | 0.143 | 0.133 | 0.044 |
| SE | 61603 | ENSG00000061273 | HDAC7 | chr12 | - | 47792537 | 47792669 | 47791870 | 47792004 | 47793368 | 47793542 | 1 | 0 | 5 | 2 | 1 | 23 | 6 | 10 | 14 | 7 | 0 | 0 | 9 | 0 | 11 | 9 | 12 | 15 | 3 | 8 | 281 | 149 | 0.054152143 | 0.402692954 | 0.023 | 0 | 0.21 | 0.07 | 0.07 | 0 | 0 | 0.241 | 0 | 0.422 | -0.058 |
| SE | 99566 | ENSG00000206561 | COLQ | chr3 | - | 15455898 | 15456019 | 15453828 | 15453931 | 15456459 | 15456579 | 63 | 20 | 25 | 45 | 7 | 0 | 0 | 0 | 0 | 0 | 13 | 9 | 9 | 10 | 9 | 8 | 0 | 0 | 0 | 0 | 270 | 149 | 0.054152883 | 0.402692954 | 1 | 1 | 1 | 1 | 1 | 0.473 | 1 | 1 | 1 | 1 | 0.105 |
| SE | 31397 | ENSG00000123933 | MXD4 | chr4 | - | 2258884 | 2259057 | 2257981 | 2258011 | 2261724 | 2261824 | 5 | 2 | 0 | 0 | 0 | 115 | 24 | 25 | 45 | 12 | 5 | 6 | 0 | 7 | 1 | 64 | 73 | 48 | 29 | 134 | 298 | 149 | 0.05417121 | 0.402790983 | 0.021 | 0.04 | 0 | 0 | 0 | 0.038 | 0.039 | 0 | 0.108 | 0.004 | -0.026 |
| SE | 102737 | ENSG00000040341 | STAU2 | chr8 | - | 73561410 | 73561539 | 73551895 | 73552319 | 73582769 | 73582830 | 6 | 7 | 2 | 3 | 9 | 99 | 68 | 75 | 91 | 189 | 0 | 4 | 6 | 16 | 0 | 22 | 70 | 62 | 39 | 63 | 278 | 149 | 0.054257526 | 0.40339448 | 0.031 | 0.052 | 0.014 | 0.017 | 0.025 | 0 | 0.03 | 0.049 | 0.18 | 0 | -0.024 |
| SE | 75191 | ENSG00000147419 | CCDC25 | chr8 | - | 27748494 | 27748598 | 27748071 | 27748279 | 27752511 | 27752587 | 152 | 106 | 167 | 155 | 241 | 12 | 6 | 11 | 16 | 40 | 120 | 138 | 143 | 152 | 20 | 5 | 22 | 36 | 18 | 16 | 253 | 149 | 0.05438371 | 0.40429424 | 0.882 | 0.912 | 0.899 | 0.851 | 0.78 | 0.934 | 0.787 | 0.701 | 0.833 | 0.424 | 0.129 |
| SE | 90309 | ENSG00000249859 | PVT1 | chr8 | + | 127998199 | 127998359 | 127989161 | 127989291 | 128009589 | 128009718 | 8 | 2 | 5 | 1 | 0 | 24 | 4 | 23 | 15 | 50 | 3 | 8 | 7 | 4 | 11 | 22 | 38 | 15 | 31 | 30 | 298 | 149 | 0.054388907 | 0.404294491 | 0.143 | 0.2 | 0.098 | 0.032 | 0 | 0.064 | 0.095 | 0.189 | 0.061 | 0.155 | -0.018 |
| SE | 91226 | ENSG00000245910 | SNHG6 | chr8 | - | 66922613 | 66922725 | 66922323 | 66922392 | 66925436 | 66925541 | 549 | 532 | 571 | 485 | 812 | 5 | 1 | 4 | 1 | 4 | 705 | 369 | 727 | 573 | 804 | 12 | 3 | 12 | 6 | 0 | 261 | 149 | 0.05445801 | 0.404753549 | 0.984 | 0.997 | 0.988 | 0.996 | 0.991 | 0.971 | 0.986 | 0.972 | 0.982 | 1 | 0.009 |
| SE | 95172 | ENSG00000213930 | GALT | chr9 | + | 34647491 | 34647567 | 34646667 | 34647258 | 34647831 | 34647961 | 34 | 15 | 59 | 30 | 16 | 3 | 0 | 4 | 0 | 0 | 28 | 27 | 13 | 27 | 11 | 10 | 1 | 0 | 0 | 0 | 225 | 149 | 0.054461002 | 0.404753549 | 0.882 | 1 | 0.907 | 1 | 1 | 0.65 | 0.947 | 1 | 1 | 1 | 0.038 |
| SE | 59628 | ENSG00000138279 | ANXA7 | chr10 | - | 73396043 | 73396109 | 73388311 | 73388414 | 73396518 | 73396583 | 1 | 1 | 4 | 5 | 4 | 137 | 114 | 158 | 128 | 115 | 14 | 4 | 17 | 11 | 0 | 345 | 174 | 206 | 222 | 423 | 215 | 149 | 0.054469355 | 0.404777201 | 0.005 | 0.006 | 0.017 | 0.026 | 0.024 | 0.027 | 0.016 | 0.054 | 0.033 | 0 | -0.01 |
| SE | 11648 | ENSG00000153066 | TXNDC11 | chr16 | - | 11730644 | 11730774 | 11721576 | 11721670 | 11736016 | 11736233 | 145 | 73 | 193 | 158 | 241 | 0 | 0 | 1 | 4 | 4 | 144 | 277 | 281 | 214 | 289 | 0 | 0 | 5 | 9 | 8 | 279 | 149 | 0.054496522 | 0.404940656 | 1 | 1 | 0.99 | 0.955 | 0.97 | 1 | 1 | 0.968 | 0.927 | 0.951 | 0.014 |
| SE | 7154 | ENSG00000101639 | CEP192 | chr18 | + | 13037236 | 13037301 | 13030464 | 13030608 | 13040829 | 13040956 | 83 | 39 | 65 | 73 | 141 | 11 | 3 | 9 | 8 | 21 | 36 | 40 | 50 | 46 | 30 | 3 | 6 | 0 | 8 | 0 | 214 | 149 | 0.05451896 | 0.405050419 | 0.84 | 0.901 | 0.834 | 0.864 | 0.824 | 0.893 | 0.823 | 1 | 0.8 | 1 | -0.051 |
| SE | 48652 | ENSG00000131374 | TBC1D5 | chr3 | - | 17586352 | 17586557 | 17562018 | 17562142 | 17742480 | 17742612 | 23 | 16 | 3 | 18 | 18 | 1 | 1 | 3 | 2 | 0 | 8 | 6 | 9 | 17 | 16 | 0 | 0 | 1 | 1 | 0 | 298 | 149 | 0.05452164 | 0.405050419 | 0.92 | 0.889 | 0.333 | 0.818 | 1 | 1 | 1 | 0.818 | 0.895 | 1 | -0.151 |
| SE | 93853 | ENSG00000116580 | GON4L | chr1 | - | 155763311 | 155763564 | 155762189 | 155762374 | 155764999 | 155765350 | 196 | 171 | 219 | 214 | 299 | 0 | 0 | 0 | 0 | 0 | 129 | 198 | 157 | 150 | 83 | 13 | 0 | 0 | 0 | 0 | 298 | 149 | 0.054589045 | 0.405512701 | 1 | 1 | 1 | 1 | 1 | 0.832 | 1 | 1 | 1 | 1 | 0.034 |
| SE | 7069 | ENSG00000041353 | RAB27B | chr18 | + | 54877566 | 54877738 | 54828476 | 54828700 | 54879368 | 54879454 | 196 | 339 | 104 | 493 | 339 | 4 | 2 | 0 | 6 | 0 | 262 | 187 | 176 | 221 | 248 | 12 | 2 | 2 | 0 | 2 | 298 | 149 | 0.054611852 | 0.405643639 | 0.961 | 0.988 | 1 | 0.976 | 1 | 0.916 | 0.979 | 0.978 | 1 | 0.984 | 0.014 |
| SE | 47583 | ENSG00000008869 | HEATR5B | chr2 | - | 37002305 | 37002572 | 37000585 | 37000813 | 37003541 | 37003686 | 291 | 230 | 208 | 231 | 315 | 10 | 2 | 2 | 1 | 2 | 64 | 330 | 153 | 57 | 58 | 0 | 1 | 2 | 1 | 3 | 298 | 149 | 0.054629004 | 0.405723514 | 0.936 | 0.983 | 0.981 | 0.991 | 0.987 | 1 | 0.994 | 0.962 | 1 | 1 | -0.016 |
| SE | 94498 | ENSG00000109689 | STIM2 | chr4 | + | 27007982 | 27008006 | 27007532 | 27007700 | 27008427 | 27008528 | 14 | 11 | 27 | 34 | 63 | 121 | 97 | 127 | 109 | 223 | 19 | 26 | 39 | 16 | 16 | 114 | 99 | 89 | 55 | 31 | 173 | 149 | 0.05463297 | 0.405723514 | 0.091 | 0.089 | 0.155 | 0.212 | 0.196 | 0.126 | 0.184 | 0.274 | 0.2 | 0.308 | -0.07 |
| SE | 52920 | ENSG00000159023 | EPB41 | chr1 | + | 29097806 | 29097935 | 29064956 | 29065158 | 29109335 | 29109346 | 1112 | 758 | 1149 | 1231 | 2711 | 31 | 19 | 25 | 17 | 48 | 1303 | 1083 | 1115 | 913 | 1264 | 47 | 26 | 24 | 22 | 32 | 278 | 149 | 0.054638327 | 0.405724814 | 0.951 | 0.955 | 0.961 | 0.975 | 0.968 | 0.937 | 0.957 | 0.961 | 0.957 | 0.955 | 0.009 |
| SE | 32329 | ENSG00000068745 | IP6K2 | chr3 | - | 48693269 | 48693338 | 48693058 | 48693179 | 48695089 | 48695421 | 7 | 6 | 0 | 1 | 10 | 142 | 77 | 92 | 123 | 136 | 0 | 0 | 1 | 16 | 0 | 226 | 110 | 161 | 121 | 132 | 218 | 149 | 0.054648067 | 0.405758655 | 0.033 | 0.051 | 0 | 0.006 | 0.048 | 0 | 0 | 0.004 | 0.083 | 0 | 0.01 |
| SE | 9885 | ENSG00000265148 | TSPOAP1-AS1 | chr17 | + | 58346279 | 58346343 | 58337307 | 58337679 | 58351734 | 58351914 | 7 | 15 | 18 | 17 | 25 | 58 | 44 | 54 | 80 | 82 | 50 | 10 | 1 | 8 | 24 | 101 | 69 | 30 | 52 | 297 | 213 | 149 | 0.054683202 | 0.405980706 | 0.078 | 0.193 | 0.189 | 0.129 | 0.176 | 0.257 | 0.092 | 0.023 | 0.097 | 0.054 | 0.048 |
| SE | 47905 | ENSG00000233184 | NA | chr1 | + | 101072659 | 101072830 | 101025868 | 101026061 | 101075394 | 101075716 | 37 | 12 | 50 | 27 | 28 | 1 | 1 | 6 | 0 | 0 | 10 | 28 | 16 | 24 | 2 | 0 | 2 | 0 | 0 | 0 | 298 | 149 | 0.054688343 | 0.405980706 | 0.949 | 0.857 | 0.806 | 1 | 1 | 1 | 0.875 | 1 | 1 | 1 | -0.053 |
| SE | 478 | ENSG00000142207 | URB1 | chr21 | - | 32355448 | 32355565 | 32354858 | 32354997 | 32357536 | 32357656 | 49 | 25 | 58 | 74 | 114 | 0 | 0 | 0 | 0 | 0 | 78 | 54 | 28 | 25 | 99 | 10 | 0 | 0 | 0 | 0 | 266 | 149 | 0.054701367 | 0.406026157 | 1 | 1 | 1 | 1 | 1 | 0.814 | 1 | 1 | 1 | 1 | 0.037 |
| SE | 108577 | ENSG00000136169 | SETDB2 | chr13 | + | 49485629 | 49485723 | 49483463 | 49483563 | 49488289 | 49488630 | 156 | 181 | 181 | 185 | 301 | 4 | 6 | 1 | 5 | 2 | 71 | 119 | 129 | 54 | 12 | 1 | 0 | 4 | 0 | 0 | 243 | 149 | 0.054704837 | 0.406026157 | 0.96 | 0.949 | 0.991 | 0.958 | 0.989 | 0.978 | 1 | 0.952 | 1 | 1 | -0.017 |
| SE | 28310 | ENSG00000001036 | FUCA2 | chr6 | - | 143503912 | 143504252 | 143501931 | 143502122 | 143507236 | 143507424 | 87 | 80 | 26 | 67 | 64 | 1 | 1 | 3 | 4 | 0 | 146 | 63 | 69 | 153 | 126 | 24 | 5 | 0 | 0 | 0 | 298 | 149 | 0.054719556 | 0.406096908 | 0.978 | 0.976 | 0.812 | 0.893 | 1 | 0.753 | 0.863 | 1 | 1 | 1 | 0.009 |
| SE | 67283 | ENSG00000109084 | TMEM97 | chr17 | + | 28325502 | 28325647 | 28319199 | 28319365 | 28326533 | 28326875 | 8 | 1 | 15 | 9 | 10 | 1 | 0 | 0 | 0 | 0 | 0 | 7 | 5 | 19 | 5 | 6 | 0 | 0 | 0 | 0 | 294 | 149 | 0.054742184 | 0.406211172 | 0.802 | 1 | 1 | 1 | 1 | 0 | 1 | 1 | 1 | 1 | 0.16 |
| SE | 96971 | ENSG00000183597 | TANGO2 | chr22 | + | 20061529 | 20061683 | 20055942 | 20056013 | 20063337 | 20063411 | 91 | 32 | 49 | 73 | 118 | 1 | 0 | 0 | 0 | 9 | 106 | 89 | 25 | 64 | 349 | 0 | 1 | 2 | 10 | 15 | 298 | 149 | 0.054745329 | 0.406211172 | 0.978 | 1 | 1 | 1 | 0.868 | 1 | 0.978 | 0.862 | 0.762 | 0.921 | 0.065 |
| SE | 23294 | ENSG00000164182 | NDUFAF2 | chr5 | + | 61045523 | 61045604 | 60945182 | 60945382 | 61073124 | 61073214 | 0 | 0 | 7 | 2 | 0 | 18 | 13 | 5 | 23 | 6 | 0 | 0 | 0 | 0 | 0 | 42 | 10 | 3 | 20 | 7 | 230 | 149 | 0.054758809 | 0.406272688 | 0 | 0 | 0.476 | 0.053 | 0 | 0 | 0 | 0 | 0 | 0 | 0.106 |
| SE | 83545 | ENSG00000101846 | STS | chrX | + | 7190879 | 7191008 | 7147711 | 7148083 | 7253195 | 7253336 | 19 | 3 | 16 | 16 | 27 | 0 | 0 | 0 | 0 | 8 | 18 | 23 | 21 | 26 | 15 | 0 | 0 | 0 | 0 | 0 | 278 | 149 | 0.054792059 | 0.406480869 | 1 | 1 | 1 | 1 | 0.644 | 1 | 1 | 1 | 1 | 1 | -0.071 |
| SE | 36369 | ENSG00000173889 | PHC3 | chr3 | - | 170149085 | 170149244 | 170136365 | 170136665 | 170171372 | 170171450 | 48 | 33 | 38 | 44 | 27 | 36 | 9 | 19 | 12 | 19 | 13 | 87 | 36 | 12 | 26 | 0 | 14 | 29 | 10 | 0 | 298 | 149 | 0.054841531 | 0.406809336 | 0.4 | 0.647 | 0.5 | 0.647 | 0.415 | 1 | 0.757 | 0.383 | 0.375 | 1 | -0.181 |
| SE | 16683 | ENSG00000140464 | PML | chr15 | + | 74033155 | 74033422 | 74024856 | 74024927 | 74034477 | 74034530 | 12 | 13 | 7 | 20 | 6 | 0 | 4 | 3 | 0 | 0 | 43 | 22 | 10 | 17 | 41 | 0 | 0 | 0 | 2 | 0 | 298 | 149 | 0.054877969 | 0.407041065 | 1 | 0.619 | 0.538 | 1 | 1 | 1 | 1 | 1 | 0.81 | 1 | -0.131 |
| SE | 8380 | ENSG00000172123 | SLFN12 | chr17 | - | 35430470 | 35430613 | 35422820 | 35423068 | 35432904 | 35433145 | 11 | 2 | 8 | 20 | 6 | 19 | 25 | 28 | 32 | 17 | 0 | 8 | 6 | 5 | 4 | 19 | 33 | 36 | 14 | 5 | 292 | 149 | 0.054932437 | 0.407406465 | 0.228 | 0.039 | 0.127 | 0.242 | 0.153 | 0 | 0.11 | 0.078 | 0.154 | 0.29 | 0.031 |
| SE | 44911 | ENSG00000162889 | MAPKAPK2 | chr1 | + | 206730687 | 206730763 | 206729971 | 206730098 | 206731137 | 206731262 | 487 | 232 | 352 | 438 | 657 | 0 | 0 | 1 | 1 | 0 | 883 | 537 | 661 | 741 | 1649 | 41 | 0 | 0 | 2 | 4 | 225 | 149 | 0.054944513 | 0.407457436 | 1 | 1 | 0.996 | 0.997 | 1 | 0.934 | 1 | 1 | 0.996 | 0.996 | 0.013 |
| SE | 58595 | ENSG00000186854 | TRABD2A | chr2 | - | 84841860 | 84842007 | 84839066 | 84839323 | 84870224 | 84870785 | 135 | 119 | 328 | 219 | 197 | 1 | 0 | 8 | 4 | 4 | 29 | 39 | 48 | 48 | 48 | 0 | 2 | 2 | 0 | 8 | 296 | 149 | 0.054963259 | 0.407557849 | 0.985 | 1 | 0.954 | 0.965 | 0.961 | 1 | 0.908 | 0.924 | 1 | 0.751 | 0.056 |
| SE | 32214 | ENSG00000114650 | SCAP | chr3 | - | 47423932 | 47424045 | 47422441 | 47422536 | 47435007 | 47435137 | 59 | 17 | 38 | 55 | 24 | 6 | 0 | 2 | 1 | 3 | 30 | 14 | 7 | 13 | 55 | 2 | 11 | 1 | 4 | 0 | 262 | 149 | 0.054980077 | 0.407643947 | 0.848 | 1 | 0.915 | 0.969 | 0.82 | 0.895 | 0.42 | 0.799 | 0.649 | 1 | 0.158 |
| SE | 5995 | ENSG00000062598 | ELMO2 | chr20 | - | 46394404 | 46394503 | 46394048 | 46394089 | 46398696 | 46398771 | 98 | 37 | 49 | 61 | 51 | 0 | 0 | 0 | 0 | 0 | 109 | 98 | 68 | 75 | 131 | 0 | 0 | 0 | 1 | 11 | 248 | 149 | 0.054986426 | 0.407652418 | 1 | 1 | 1 | 1 | 1 | 1 | 1 | 1 | 0.978 | 0.877 | 0.029 |
| SE | 76288 | ENSG00000124508 | BTN2A2 | chr6 | + | 26385014 | 26385362 | 26383107 | 26383181 | 26388012 | 26388289 | 25 | 20 | 18 | 25 | 34 | 5 | 0 | 0 | 1 | 7 | 13 | 65 | 11 | 37 | 27 | 4 | 3 | 0 | 0 | 0 | 298 | 149 | 0.055105611 | 0.408495134 | 0.714 | 1 | 1 | 0.926 | 0.708 | 0.619 | 0.915 | 1 | 1 | 1 | -0.037 |
| SE | 20461 | ENSG00000006744 | ELAC2 | chr17 | - | 13010612 | 13010671 | 13005920 | 13005979 | 13011662 | 13011670 | 153 | 89 | 127 | 113 | 205 | 0 | 0 | 0 | 0 | 0 | 121 | 109 | 74 | 124 | 82 | 7 | 0 | 0 | 6 | 0 | 208 | 149 | 0.055110531 | 0.408495134 | 1 | 1 | 1 | 1 | 1 | 0.925 | 1 | 1 | 0.937 | 1 | 0.028 |
| SE | 60217 | ENSG00000163788 | SNRK | chr3 | + | 43303097 | 43303792 | 43286537 | 43286675 | 43332168 | 43332310 | 162 | 86 | 194 | 149 | 261 | 2 | 0 | 1 | 0 | 3 | 62 | 78 | 130 | 78 | 7 | 0 | 1 | 2 | 0 | 6 | 298 | 149 | 0.055116366 | 0.408499717 | 0.976 | 1 | 0.99 | 1 | 0.978 | 1 | 0.975 | 0.97 | 1 | 0.368 | 0.126 |
| SE | 108168 | ENSG00000075089 | ACTR6 | chr12 | + | 100210072 | 100210208 | 100205675 | 100205757 | 100210294 | 100210320 | 132 | 83 | 123 | 120 | 229 | 0 | 0 | 0 | 0 | 0 | 0 | 60 | 60 | 24 | 16 | 5 | 0 | 0 | 1 | 0 | 285 | 149 | 0.055145927 | 0.408680124 | 1 | 1 | 1 | 1 | 1 | 0 | 1 | 1 | 0.926 | 1 | 0.215 |
| SE | 87438 | ENSG00000183354 | KIAA2026 | chr9 | - | 5944870 | 5944960 | 5929043 | 5929159 | 5968018 | 5968081 | 13 | 26 | 18 | 34 | 32 | 6 | 0 | 2 | 4 | 5 | 19 | 39 | 37 | 20 | 8 | 0 | 0 | 2 | 6 | 0 | 239 | 149 | 0.0551796 | 0.408890965 | 0.575 | 1 | 0.849 | 0.841 | 0.8 | 1 | 1 | 0.92 | 0.675 | 1 | -0.106 |
| SE | 34122 | ENSG00000115129 | TP53I3 | chr2 | - | 24079443 | 24079640 | 24077432 | 24077761 | 24080818 | 24081031 | 22 | 10 | 16 | 22 | 13 | 2 | 3 | 4 | 5 | 7 | 73 | 73 | 43 | 93 | 166 | 4 | 11 | 11 | 18 | 5 | 298 | 149 | 0.055191812 | 0.40892702 | 0.846 | 0.625 | 0.667 | 0.688 | 0.481 | 0.901 | 0.768 | 0.662 | 0.721 | 0.943 | -0.138 |
| SE | 74802 | ENSG00000146063 | TRIM41 | chr5 | + | 181233412 | 181233435 | 181232658 | 181232889 | 181233635 | 181233763 | 72 | 32 | 53 | 39 | 36 | 5 | 2 | 5 | 1 | 1 | 26 | 47 | 23 | 37 | 18 | 6 | 0 | 4 | 1 | 8 | 172 | 149 | 0.055194911 | 0.40892702 | 0.926 | 0.933 | 0.902 | 0.971 | 0.969 | 0.79 | 1 | 0.833 | 0.97 | 0.661 | 0.089 |
| SE | 34651 | ENSG00000166501 | PRKCB | chr16 | + | 23988507 | 23988590 | 23837374 | 23837406 | 24032135 | 24032247 | 247 | 118 | 210 | 263 | 184 | 1 | 2 | 0 | 0 | 0 | 401 | 349 | 269 | 376 | 801 | 7 | 0 | 0 | 0 | 21 | 232 | 149 | 0.055211642 | 0.409012274 | 0.994 | 0.974 | 1 | 1 | 1 | 0.974 | 1 | 1 | 1 | 0.961 | 0.007 |
| SE | 93384 | ENSG00000065923 | SLC9A7 | chrX | - | 46682335 | 46682535 | 46679677 | 46679755 | 46758704 | 46759055 | 26 | 26 | 17 | 11 | 20 | 0 | 0 | 0 | 0 | 0 | 21 | 50 | 16 | 21 | 20 | 0 | 3 | 5 | 0 | 0 | 298 | 149 | 0.055230067 | 0.409110056 | 1 | 1 | 1 | 1 | 1 | 1 | 0.893 | 0.615 | 1 | 1 | 0.098 |
| SE | 9301 | ENSG00000133030 | MPRIP | chr17 | + | 17142626 | 17142765 | 17137915 | 17138429 | 17146035 | 17146092 | 34 | 17 | 53 | 57 | 30 | 0 | 0 | 0 | 0 | 0 | 35 | 74 | 29 | 26 | 83 | 0 | 2 | 0 | 2 | 5 | 288 | 149 | 0.055271546 | 0.409378578 | 1 | 1 | 1 | 1 | 1 | 1 | 0.95 | 1 | 0.871 | 0.896 | 0.057 |
| SE | 45147 | ENSG00000107099 | DOCK8 | chr9 | + | 276993 | 277086 | 273083 | 273100 | 286460 | 286636 | 18 | 2 | 12 | 18 | 17 | 84 | 56 | 95 | 91 | 56 | 5 | 1 | 11 | 14 | 5 | 47 | 124 | 72 | 34 | 53 | 242 | 149 | 0.055341096 | 0.40985494 | 0.117 | 0.022 | 0.072 | 0.109 | 0.157 | 0.061 | 0.005 | 0.086 | 0.202 | 0.055 | 0.014 |
| SE | 63643 | ENSG00000112855 | HARS2 | chr5 | + | 140693490 | 140693508 | 140691457 | 140691756 | 140693590 | 140693665 | 0 | 2 | 1 | 0 | 2 | 28 | 14 | 31 | 22 | 10 | 2 | 0 | 8 | 0 | 0 | 5 | 9 | 19 | 19 | 36 | 167 | 149 | 0.055385953 | 0.410148357 | 0 | 0.113 | 0.028 | 0 | 0.151 | 0.263 | 0 | 0.273 | 0 | 0 | -0.049 |
| SE | 78424 | ENSG00000161405 | IKZF3 | chr17 | - | 39791415 | 39791583 | 39788257 | 39788374 | 39792672 | 39792933 | 692 | 403 | 624 | 691 | 667 | 8 | 2 | 2 | 2 | 2 | 517 | 962 | 488 | 460 | 379 | 0 | 3 | 0 | 25 | 0 | 298 | 149 | 0.055405012 | 0.410250696 | 0.977 | 0.99 | 0.994 | 0.994 | 0.994 | 1 | 0.994 | 1 | 0.902 | 1 | 0.011 |
| SE | 45773 | ENSG00000284770 | TBCE | chr1 | + | 235419472 | 235419561 | 235414432 | 235414618 | 235427139 | 235427147 | 43 | 29 | 32 | 35 | 85 | 3 | 2 | 0 | 1 | 3 | 64 | 50 | 28 | 33 | 51 | 0 | 0 | 0 | 0 | 0 | 238 | 149 | 0.055477358 | 0.410747538 | 0.9 | 0.901 | 1 | 0.956 | 0.947 | 1 | 1 | 1 | 1 | 1 | -0.059 |
| SE | 49968 | ENSG00000167208 | SNX20 | chr16 | - | 50677396 | 50677535 | 50675769 | 50675921 | 50681189 | 50681206 | 153 | 117 | 140 | 128 | 64 | 2 | 0 | 0 | 0 | 0 | 180 | 219 | 107 | 137 | 204 | 0 | 4 | 10 | 0 | 0 | 288 | 149 | 0.055528635 | 0.411070141 | 0.975 | 1 | 1 | 1 | 1 | 1 | 0.966 | 0.847 | 1 | 1 | 0.032 |
| SE | 35848 | ENSG00000143612 | C1orf43 | chr1 | - | 154219835 | 154219937 | 154214456 | 154214574 | 154220341 | 154220533 | 188 | 100 | 186 | 172 | 116 | 48 | 28 | 38 | 48 | 68 | 208 | 195 | 35 | 95 | 308 | 21 | 113 | 97 | 65 | 171 | 251 | 149 | 0.055531431 | 0.411070141 | 0.699 | 0.679 | 0.744 | 0.68 | 0.503 | 0.855 | 0.506 | 0.176 | 0.465 | 0.517 | 0.157 |
| SE | 33216 | ENSG00000221978 | CCNL2 | chr1 | - | 1392789 | 1392803 | 1390765 | 1390865 | 1393395 | 1393460 | 2 | 2 | 7 | 3 | 2 | 63 | 32 | 44 | 45 | 62 | 0 | 4 | 3 | 1 | 0 | 76 | 64 | 40 | 19 | 31 | 163 | 149 | 0.055563133 | 0.411265939 | 0.028 | 0.054 | 0.127 | 0.057 | 0.029 | 0 | 0.054 | 0.064 | 0.046 | 0 | 0.026 |
| SE | 51395 | ENSG00000067606 | PRKCZ | chr1 | + | 2145337 | 2145511 | 2144209 | 2144341 | 2146026 | 2146108 | 0 | 1 | 2 | 3 | 0 | 32 | 17 | 36 | 31 | 12 | 0 | 4 | 10 | 0 | 0 | 21 | 40 | 22 | 26 | 42 | 298 | 149 | 0.05562244 | 0.411665993 | 0 | 0.029 | 0.027 | 0.046 | 0 | 0 | 0.048 | 0.185 | 0 | 0 | -0.026 |
| SE | 73968 | ENSG00000166436 | TRIM66 | chr11 | - | 8645740 | 8645887 | 8643008 | 8643126 | 8646446 | 8646561 | 25 | 16 | 21 | 28 | 27 | 0 | 0 | 0 | 0 | 0 | 13 | 7 | 15 | 43 | 7 | 0 | 6 | 1 | 0 | 0 | 296 | 149 | 0.05567859 | 0.412042614 | 1 | 1 | 1 | 1 | 1 | 1 | 0.37 | 0.883 | 1 | 1 | 0.149 |
| SE | 61880 | ENSG00000076242 | MLH1 | chr3 | + | 37014431 | 37014544 | 37012010 | 37012099 | 37017505 | 37017599 | 220 | 230 | 249 | 181 | 507 | 2 | 4 | 2 | 5 | 10 | 267 | 118 | 154 | 232 | 73 | 7 | 2 | 10 | 7 | 0 | 262 | 149 | 0.055701988 | 0.412176812 | 0.984 | 0.97 | 0.986 | 0.954 | 0.966 | 0.956 | 0.971 | 0.898 | 0.95 | 1 | 0.017 |
| SE | 105258 | ENSG00000082898 | XPO1 | chr2 | - | 61501995 | 61502040 | 61499712 | 61499894 | 61502248 | 61502310 | 378 | 248 | 328 | 349 | 381 | 0 | 4 | 1 | 4 | 2 | 46 | 188 | 123 | 141 | 63 | 0 | 0 | 1 | 0 | 13 | 194 | 149 | 0.055709448 | 0.412192984 | 1 | 0.979 | 0.996 | 0.985 | 0.993 | 1 | 1 | 0.99 | 1 | 0.788 | 0.035 |
| SE | 69223 | ENSG00000204272 | NBDY | chrX | + | 56811019 | 56811179 | 56732062 | 56732199 | 56817319 | 56817569 | 6 | 1 | 11 | 10 | 1 | 231 | 116 | 170 | 206 | 190 | 5 | 6 | 2 | 2 | 33 | 189 | 178 | 129 | 133 | 326 | 298 | 149 | 0.05571901 | 0.412192984 | 0.013 | 0.004 | 0.031 | 0.024 | 0.003 | 0.013 | 0.017 | 0.008 | 0.007 | 0.048 | -0.004 |
| SE | 20311 | ENSG00000111913 | RIPOR2 | chr6 | - | 24835702 | 24835871 | 24832255 | 24832391 | 24842861 | 24843554 | 356 | 177 | 347 | 328 | 502 | 3 | 2 | 1 | 2 | 2 | 149 | 171 | 115 | 111 | 64 | 0 | 0 | 3 | 9 | 0 | 298 | 149 | 0.055719967 | 0.412192984 | 0.983 | 0.978 | 0.994 | 0.988 | 0.992 | 1 | 1 | 0.95 | 0.86 | 1 | 0.025 |
| SE | 101556 | ENSG00000088298 | EDEM2 | chr20 | - | 35134737 | 35134949 | 35126250 | 35126375 | 35137879 | 35138005 | 36 | 30 | 37 | 36 | 64 | 5 | 4 | 2 | 8 | 7 | 29 | 44 | 47 | 45 | 60 | 23 | 17 | 3 | 4 | 16 | 298 | 149 | 0.055792149 | 0.412687965 | 0.783 | 0.789 | 0.902 | 0.692 | 0.821 | 0.387 | 0.564 | 0.887 | 0.849 | 0.652 | 0.13 |
| SE | 4615 | ENSG00000167468 | GPX4 | chr19 | + | 1106180 | 1106266 | 1105657 | 1105809 | 1106399 | 1106443 | 77 | 35 | 37 | 58 | 86 | 11 | 1 | 4 | 10 | 1 | 39 | 44 | 24 | 46 | 216 | 21 | 2 | 6 | 12 | 11 | 235 | 149 | 0.055853602 | 0.4131035 | 0.816 | 0.957 | 0.854 | 0.786 | 0.982 | 0.541 | 0.933 | 0.717 | 0.708 | 0.926 | 0.114 |
| SE | 52159 | ENSG00000163393 | SLC22A15 | chr1 | + | 115992030 | 115992243 | 115976512 | 115976714 | 116019581 | 116019714 | 21 | 8 | 2 | 13 | 13 | 0 | 1 | 0 | 1 | 0 | 9 | 8 | 3 | 12 | 10 | 0 | 1 | 6 | 0 | 0 | 298 | 149 | 0.055882791 | 0.413280343 | 1 | 0.8 | 1 | 0.867 | 1 | 1 | 0.8 | 0.2 | 1 | 1 | 0.133 |
| SE | 98115 | ENSG00000171466 | ZNF562 | chr19 | - | 9659378 | 9659467 | 9658008 | 9658135 | 9660719 | 9660821 | 53 | 33 | 64 | 49 | 57 | 3 | 1 | 0 | 3 | 3 | 131 | 90 | 62 | 64 | 27 | 0 | 5 | 10 | 3 | 6 | 238 | 149 | 0.055914413 | 0.413412199 | 0.917 | 0.954 | 1 | 0.911 | 0.922 | 1 | 0.918 | 0.795 | 0.93 | 0.738 | 0.065 |
| SE | 80901 | ENSG00000126883 | NUP214 | chr9 | + | 131201646 | 131201717 | 131197215 | 131199015 | 131222777 | 131222930 | 11 | 7 | 8 | 19 | 6 | 0 | 0 | 0 | 0 | 0 | 6 | 19 | 6 | 12 | 7 | 8 | 0 | 0 | 0 | 0 | 220 | 149 | 0.055915615 | 0.413412199 | 1 | 1 | 1 | 1 | 1 | 0.337 | 1 | 1 | 1 | 1 | 0.133 |
| SE | 49 | ENSG00000159267 | HLCS | chr21 | - | 36756541 | 36756755 | 36748625 | 36754417 | 36759726 | 36759841 | 54 | 11 | 39 | 49 | 76 | 0 | 0 | 0 | 0 | 0 | 5 | 38 | 28 | 20 | 16 | 5 | 0 | 2 | 0 | 0 | 298 | 149 | 0.05591646 | 0.413412199 | 1 | 1 | 1 | 1 | 1 | 0.333 | 1 | 0.875 | 1 | 1 | 0.158 |
| SE | 28549 | ENSG00000105202 | FBL | chr19 | - | 39840655 | 39840787 | 39840413 | 39840515 | 39846290 | 39846329 | 42 | 12 | 53 | 23 | 6 | 4 | 1 | 2 | 2 | 0 | 20 | 2 | 9 | 33 | 27 | 0 | 4 | 3 | 0 | 12 | 281 | 149 | 0.055941411 | 0.413557618 | 0.848 | 0.864 | 0.934 | 0.859 | 1 | 1 | 0.21 | 0.614 | 1 | 0.544 | 0.227 |
| SE | 90223 | ENSG00000172269 | DPAGT1 | chr11 | - | 119097463 | 119097551 | 119096527 | 119097297 | 119098402 | 119098487 | 35 | 24 | 49 | 49 | 54 | 5 | 0 | 3 | 2 | 1 | 43 | 27 | 19 | 13 | 49 | 0 | 0 | 5 | 3 | 9 | 237 | 149 | 0.055952381 | 0.413599659 | 0.815 | 1 | 0.911 | 0.939 | 0.971 | 1 | 1 | 0.705 | 0.731 | 0.774 | 0.085 |
| SE | 37787 | ENSG00000015479 | MATR3 | chr5 | + | 139307238 | 139308327 | 139293777 | 139294028 | 139314674 | 139314736 | 541 | 400 | 460 | 588 | 781 | 9 | 3 | 6 | 8 | 1 | 23 | 248 | 274 | 80 | 23 | 0 | 1 | 0 | 0 | 0 | 298 | 149 | 0.055989925 | 0.413838116 | 0.968 | 0.985 | 0.975 | 0.974 | 0.997 | 1 | 0.992 | 1 | 1 | 1 | -0.019 |
| SE | 61354 | ENSG00000189319 | FAM53B | chr10 | - | 124706635 | 124706887 | 124696157 | 124696212 | 124744012 | 124744269 | 314 | 95 | 142 | 174 | 153 | 3 | 0 | 0 | 0 | 0 | 247 | 415 | 164 | 174 | 412 | 0 | 6 | 4 | 1 | 7 | 298 | 149 | 0.055997588 | 0.413849029 | 0.981 | 1 | 1 | 1 | 1 | 1 | 0.972 | 0.953 | 0.989 | 0.967 | 0.02 |
| SE | 73594 | ENSG00000124356 | STAMBP | chr2 | + | 73830844 | 73831050 | 73828973 | 73828996 | 73844812 | 73844888 | 14 | 8 | 10 | 13 | 18 | 3 | 4 | 1 | 2 | 3 | 42 | 21 | 17 | 9 | 22 | 6 | 1 | 1 | 2 | 0 | 298 | 149 | 0.056001973 | 0.413849029 | 0.7 | 0.5 | 0.833 | 0.765 | 0.75 | 0.778 | 0.913 | 0.895 | 0.692 | 1 | -0.146 |
| SE | 93028 | ENSG00000147162 | OGT | chrX | + | 71555189 | 71555385 | 71554512 | 71554592 | 71555953 | 71556094 | 1310 | 1131 | 1357 | 1475 | 1397 | 356 | 305 | 282 | 267 | 206 | 578 | 1203 | 1382 | 1102 | 396 | 176 | 165 | 250 | 180 | 26 | 298 | 149 | 0.056019061 | 0.41393624 | 0.648 | 0.65 | 0.706 | 0.734 | 0.772 | 0.622 | 0.785 | 0.734 | 0.754 | 0.884 | -0.054 |
| SE | 54444 | ENSG00000169554 | ZEB2 | chr2 | - | 144518292 | 144518446 | 144517277 | 144517419 | 144519938 | 144520361 | 13 | 2 | 3 | 7 | 7 | 233 | 168 | 104 | 205 | 119 | 0 | 5 | 14 | 16 | 10 | 145 | 212 | 180 | 117 | 95 | 298 | 149 | 0.056035178 | 0.414016257 | 0.027 | 0.006 | 0.014 | 0.017 | 0.029 | 0 | 0.012 | 0.037 | 0.064 | 0.05 | -0.014 |
| SE | 72288 | ENSG00000006062 | MAP3K14 | chr17 | - | 45286430 | 45287045 | 45284811 | 45284949 | 45289235 | 45289305 | 41 | 18 | 26 | 29 | 27 | 0 | 0 | 0 | 0 | 0 | 55 | 54 | 26 | 47 | 36 | 7 | 0 | 2 | 0 | 0 | 298 | 149 | 0.056068874 | 0.414210549 | 1 | 1 | 1 | 1 | 1 | 0.797 | 1 | 0.867 | 1 | 1 | 0.067 |
| SE | 68607 | ENSG00000237765 | FAM200B | chr4 | + | 15684885 | 15684959 | 15681822 | 15681901 | 15686235 | 15686606 | 8 | 6 | 7 | 7 | 10 | 22 | 16 | 26 | 28 | 25 | 0 | 4 | 11 | 4 | 7 | 40 | 9 | 19 | 33 | 46 | 223 | 149 | 0.056072055 | 0.414210549 | 0.195 | 0.2 | 0.152 | 0.143 | 0.211 | 0 | 0.229 | 0.279 | 0.075 | 0.092 | 0.045 |
| SE | 95576 | ENSG00000130714 | POMT1 | chr9 | + | 131504188 | 131504340 | 131502788 | 131503073 | 131506402 | 131506453 | 11 | 7 | 8 | 18 | 4 | 1 | 0 | 0 | 4 | 0 | 44 | 2 | 12 | 25 | 31 | 0 | 13 | 6 | 0 | 0 | 298 | 149 | 0.056164787 | 0.414856429 | 0.846 | 1 | 1 | 0.692 | 1 | 1 | 0.071 | 0.5 | 1 | 1 | 0.193 |
| SE | 17327 | ENSG00000258461 | NA | chr15 | + | 42386166 | 42386285 | 42384482 | 42384552 | 42387752 | 42387886 | 44 | 41 | 44 | 38 | 59 | 0 | 0 | 0 | 0 | 0 | 88 | 85 | 40 | 214 | 94 | 12 | 0 | 0 | 0 | 0 | 268 | 149 | 0.056239843 | 0.415371636 | 1 | 1 | 1 | 1 | 1 | 0.803 | 1 | 1 | 1 | 1 | 0.039 |
| SE | 11893 | ENSG00000103222 | ABCC1 | chr16 | + | 16068155 | 16068302 | 16056091 | 16056295 | 16071641 | 16071729 | 131 | 60 | 112 | 116 | 126 | 1 | 4 | 5 | 0 | 2 | 123 | 154 | 138 | 120 | 206 | 0 | 0 | 0 | 7 | 0 | 296 | 149 | 0.056252752 | 0.415427795 | 0.985 | 0.883 | 0.919 | 1 | 0.969 | 1 | 1 | 1 | 0.896 | 1 | -0.028 |
| SE | 100930 | ENSG00000133895 | MEN1 | chr11 | - | 64807010 | 64807098 | 64805634 | 64806368 | 64807178 | 64807219 | 17 | 18 | 33 | 44 | 23 | 0 | 0 | 0 | 0 | 0 | 15 | 24 | 20 | 18 | 90 | 9 | 0 | 0 | 0 | 0 | 237 | 149 | 0.056283192 | 0.41557951 | 1 | 1 | 1 | 1 | 1 | 0.512 | 1 | 1 | 1 | 1 | 0.098 |
| SE | 46324 | ENSG00000095564 | BTAF1 | chr10 | + | 91953736 | 91953873 | 91951402 | 91951566 | 91956527 | 91956657 | 470 | 486 | 509 | 568 | 448 | 7 | 3 | 1 | 1 | 1 | 221 | 413 | 403 | 202 | 94 | 0 | 8 | 10 | 0 | 0 | 286 | 149 | 0.056283911 | 0.41557951 | 0.972 | 0.988 | 0.996 | 0.997 | 0.996 | 1 | 0.964 | 0.955 | 1 | 1 | 0.006 |
| SE | 93124 | ENSG00000079387 | SENP1 | chr12 | - | 48089162 | 48089293 | 48088800 | 48088960 | 48096342 | 48096427 | 30 | 14 | 29 | 17 | 29 | 60 | 55 | 55 | 62 | 103 | 0 | 18 | 12 | 3 | 16 | 27 | 53 | 51 | 50 | 19 | 280 | 149 | 0.056291999 | 0.415600031 | 0.21 | 0.119 | 0.219 | 0.127 | 0.13 | 0 | 0.153 | 0.111 | 0.031 | 0.309 | 0.04 |
| SE | 56990 | ENSG00000131725 | WDR44 | chrX | + | 118387339 | 118387414 | 118378418 | 118378452 | 118394054 | 118394185 | 20 | 11 | 17 | 20 | 25 | 0 | 0 | 0 | 0 | 0 | 5 | 14 | 16 | 4 | 22 | 0 | 0 | 0 | 8 | 0 | 224 | 149 | 0.056336993 | 0.415893002 | 1 | 1 | 1 | 1 | 1 | 1 | 1 | 1 | 0.25 | 1 | 0.15 |
| SE | 54555 | ENSG00000179912 | R3HDM2 | chr12 | - | 57289008 | 57289040 | 57283823 | 57284056 | 57295398 | 57295507 | 53 | 53 | 35 | 40 | 88 | 18 | 25 | 23 | 27 | 46 | 156 | 48 | 59 | 43 | 125 | 42 | 14 | 20 | 41 | 16 | 181 | 149 | 0.056361426 | 0.416034145 | 0.708 | 0.636 | 0.556 | 0.549 | 0.612 | 0.754 | 0.738 | 0.708 | 0.463 | 0.865 | -0.093 |
| SE | 3340 | ENSG00000012061 | ERCC1 | chr19 | - | 45413962 | 45414034 | 45413433 | 45413745 | 45414860 | 45414960 | 118 | 71 | 119 | 115 | 169 | 11 | 6 | 7 | 18 | 13 | 279 | 156 | 84 | 120 | 332 | 6 | 8 | 10 | 17 | 15 | 221 | 149 | 0.056402255 | 0.416296282 | 0.879 | 0.889 | 0.92 | 0.812 | 0.898 | 0.969 | 0.929 | 0.85 | 0.826 | 0.937 | -0.023 |
| SE | 56846 | ENSG00000133619 | KRBA1 | chr7 | + | 149722782 | 149722956 | 149722421 | 149722553 | 149723733 | 149723889 | 10 | 8 | 13 | 19 | 4 | 0 | 0 | 0 | 0 | 0 | 14 | 3 | 4 | 0 | 19 | 0 | 0 | 0 | 6 | 0 | 298 | 149 | 0.056426346 | 0.416434838 | 1 | 1 | 1 | 1 | 1 | 1 | 1 | 1 | 0 | 1 | 0.2 |
| SE | 98259 | ENSG00000168071 | CCDC88B | chr11 | + | 64354003 | 64354028 | 64353714 | 64353813 | 64355193 | 64355400 | 18 | 8 | 5 | 3 | 0 | 8 | 0 | 0 | 5 | 2 | 10 | 17 | 0 | 9 | 32 | 16 | 14 | 7 | 13 | 23 | 174 | 149 | 0.056435054 | 0.416459845 | 0.658 | 1 | 1 | 0.339 | 0 | 0.349 | 0.51 | 0 | 0.372 | 0.544 | 0.244 |
| SE | 79122 | ENSG00000160219 | GAB3 | chrX | - | 154680131 | 154680248 | 154675248 | 154678294 | 154695916 | 154696019 | 247 | 127 | 122 | 174 | 185 | 6 | 3 | 0 | 5 | 9 | 299 | 197 | 166 | 153 | 210 | 0 | 4 | 7 | 3 | 35 | 266 | 149 | 0.056461949 | 0.416619051 | 0.958 | 0.96 | 1 | 0.951 | 0.92 | 1 | 0.965 | 0.93 | 0.966 | 0.771 | 0.031 |
| SE | 99247 | ENSG00000133318 | RTN3 | chr11 | + | 63718701 | 63721032 | 63704850 | 63704907 | 63749990 | 63750198 | 1 | 3 | 3 | 4 | 10 | 0 | 0 | 0 | 0 | 0 | 0 | 2 | 2 | 6 | 2 | 6 | 0 | 0 | 0 | 0 | 298 | 149 | 0.056475332 | 0.416678534 | 1 | 1 | 1 | 1 | 1 | 0 | 1 | 1 | 1 | 1 | 0.2 |
| SE | 61317 | ENSG00000102786 | INTS6 | chr13 | - | 51389318 | 51389444 | 51387385 | 51387540 | 51395299 | 51395483 | 209 | 127 | 159 | 186 | 216 | 2 | 0 | 4 | 0 | 1 | 38 | 132 | 99 | 111 | 68 | 0 | 8 | 0 | 3 | 0 | 275 | 149 | 0.056492 | 0.416762241 | 0.983 | 1 | 0.956 | 1 | 0.992 | 1 | 0.899 | 1 | 0.952 | 1 | 0.016 |
| SE | 23822 | ENSG00000115661 | STK16 | chr2 | + | 219248192 | 219248314 | 219247414 | 219247535 | 219248420 | 219248702 | 33 | 20 | 19 | 62 | 45 | 0 | 0 | 0 | 1 | 0 | 33 | 73 | 76 | 19 | 86 | 0 | 10 | 0 | 0 | 0 | 271 | 149 | 0.056515038 | 0.416892915 | 1 | 1 | 1 | 0.972 | 1 | 1 | 0.801 | 1 | 1 | 1 | 0.034 |
| SE | 59089 | ENSG00000180182 | MED14 | chrX | - | 40666719 | 40666851 | 40664306 | 40664489 | 40671860 | 40671972 | 54 | 46 | 43 | 64 | 69 | 3 | 4 | 1 | 3 | 11 | 32 | 85 | 54 | 46 | 21 | 0 | 10 | 1 | 0 | 0 | 281 | 149 | 0.056539559 | 0.41703451 | 0.905 | 0.859 | 0.958 | 0.919 | 0.769 | 1 | 0.818 | 0.966 | 1 | 1 | -0.075 |
| SE | 66334 | ENSG00000143207 | COP1 | chr1 | - | 176163814 | 176163891 | 176149005 | 176149074 | 176175909 | 176176007 | 147 | 126 | 97 | 115 | 297 | 8 | 8 | 6 | 2 | 20 | 100 | 105 | 140 | 104 | 94 | 3 | 8 | 4 | 5 | 0 | 226 | 149 | 0.05654957 | 0.41706906 | 0.924 | 0.912 | 0.914 | 0.974 | 0.907 | 0.956 | 0.896 | 0.958 | 0.932 | 1 | -0.022 |
| SE | 20849 | ENSG00000255819 | KLRC4-KLRK1 | chr12 | - | 10391834 | 10391891 | 10389942 | 10390027 | 10407635 | 10407766 | 49 | 15 | 16 | 43 | 43 | 7 | 1 | 1 | 7 | 5 | 29 | 32 | 27 | 25 | 6 | 0 | 5 | 1 | 3 | 0 | 206 | 149 | 0.056568015 | 0.417165798 | 0.835 | 0.916 | 0.92 | 0.816 | 0.862 | 1 | 0.822 | 0.951 | 0.858 | 1 | -0.056 |
| SE | 32914 | ENSG00000075711 | DLG1 | chr3 | - | 197104902 | 197105005 | 197090911 | 197091026 | 197115926 | 197116083 | 291 | 243 | 247 | 343 | 489 | 0 | 0 | 0 | 0 | 0 | 57 | 159 | 302 | 172 | 149 | 0 | 3 | 9 | 3 | 0 | 252 | 149 | 0.05659816 | 0.417348796 | 1 | 1 | 1 | 1 | 1 | 1 | 0.969 | 0.952 | 0.971 | 1 | 0.022 |
| SE | 80848 | ENSG00000164896 | FASTK | chr7 | - | 151079499 | 151079922 | 151078841 | 151079021 | 151080684 | 151080790 | 31 | 13 | 35 | 25 | 16 | 6 | 2 | 0 | 0 | 0 | 11 | 12 | 10 | 23 | 27 | 0 | 0 | 0 | 0 | 0 | 298 | 149 | 0.056666712 | 0.417814944 | 0.721 | 0.765 | 1 | 1 | 1 | 1 | 1 | 1 | 1 | 1 | -0.103 |
| SE | 97498 | ENSG00000173120 | KDM2A | chr11 | + | 67215339 | 67215446 | 67181845 | 67181892 | 67217730 | 67217884 | 8 | 9 | 6 | 15 | 1 | 0 | 0 | 0 | 0 | 0 | 46 | 31 | 37 | 111 | 50 | 10 | 0 | 0 | 0 | 0 | 256 | 149 | 0.056678779 | 0.417864567 | 1 | 1 | 1 | 1 | 1 | 0.728 | 1 | 1 | 1 | 1 | 0.054 |
| SE | 49473 | ENSG00000115020 | PIKFYVE | chr2 | + | 208302241 | 208302353 | 208300936 | 208301094 | 208304170 | 208304318 | 230 | 134 | 180 | 209 | 332 | 2 | 3 | 1 | 12 | 1 | 95 | 218 | 148 | 88 | 66 | 0 | 0 | 3 | 4 | 0 | 261 | 149 | 0.056696235 | 0.417953901 | 0.985 | 0.962 | 0.99 | 0.909 | 0.995 | 1 | 1 | 0.966 | 0.926 | 1 | -0.01 |
| SE | 97552 | ENSG00000109452 | INPP4B | chr4 | - | 142314711 | 142314762 | 142305457 | 142305537 | 142402937 | 142403054 | 276 | 226 | 348 | 261 | 349 | 3 | 5 | 0 | 9 | 0 | 76 | 225 | 189 | 100 | 181 | 14 | 0 | 0 | 0 | 5 | 200 | 149 | 0.056704129 | 0.417972745 | 0.986 | 0.971 | 1 | 0.956 | 1 | 0.802 | 1 | 1 | 1 | 0.964 | 0.029 |
| SE | 70792 | ENSG00000064201 | TSPAN32 | chr11 | + | 2316631 | 2316667 | 2316228 | 2316312 | 2317343 | 2317525 | 1 | 2 | 2 | 4 | 3 | 0 | 0 | 0 | 0 | 0 | 5 | 6 | 4 | 0 | 15 | 0 | 0 | 0 | 8 | 0 | 185 | 149 | 0.056743376 | 0.418222666 | 1 | 1 | 1 | 1 | 1 | 1 | 1 | 1 | 0 | 1 | 0.2 |
| SE | 37654 | ENSG00000136933 | RABEPK | chr9 | + | 125213369 | 125213522 | 125207563 | 125207721 | 125220538 | 125220658 | 112 | 20 | 55 | 66 | 130 | 3 | 1 | 4 | 6 | 0 | 120 | 41 | 33 | 35 | 95 | 0 | 10 | 5 | 0 | 8 | 298 | 149 | 0.056779785 | 0.418451622 | 0.949 | 0.909 | 0.873 | 0.846 | 1 | 1 | 0.672 | 0.767 | 1 | 0.856 | 0.056 |
| SE | 76268 | ENSG00000065427 | KARS1 | chr16 | - | 75640183 | 75640349 | 75635911 | 75636098 | 75641563 | 75641723 | 297 | 175 | 264 | 273 | 313 | 1 | 2 | 3 | 2 | 0 | 323 | 249 | 225 | 263 | 330 | 5 | 1 | 7 | 1 | 4 | 298 | 149 | 0.056835753 | 0.418824671 | 0.993 | 0.978 | 0.978 | 0.986 | 1 | 0.97 | 0.992 | 0.941 | 0.992 | 0.976 | 0.013 |
| SE | 58571 | ENSG00000072858 | SIDT1 | chr3 | + | 113576921 | 113576967 | 113567539 | 113567710 | 113580607 | 113580709 | 66 | 17 | 54 | 66 | 55 | 0 | 1 | 2 | 4 | 3 | 10 | 61 | 22 | 23 | 55 | 0 | 2 | 1 | 0 | 0 | 195 | 149 | 0.056846803 | 0.418857977 | 1 | 0.929 | 0.954 | 0.927 | 0.933 | 1 | 0.959 | 0.944 | 1 | 1 | -0.032 |
| SE | 30368 | ENSG00000145348 | TBCK | chr4 | - | 106164391 | 106164461 | 106116318 | 106116378 | 106171094 | 106171240 | 14 | 4 | 5 | 5 | 12 | 84 | 53 | 48 | 57 | 90 | 0 | 7 | 8 | 0 | 0 | 17 | 66 | 24 | 54 | 32 | 219 | 149 | 0.056850972 | 0.418857977 | 0.102 | 0.049 | 0.066 | 0.056 | 0.083 | 0 | 0.067 | 0.185 | 0 | 0 | 0.021 |
| SE | 2114 | ENSG00000127616 | SMARCA4 | chr19 | + | 11039457 | 11039553 | 11034913 | 11035132 | 11041306 | 11041560 | 22 | 15 | 14 | 16 | 17 | 113 | 55 | 120 | 72 | 70 | 8 | 13 | 9 | 12 | 29 | 103 | 104 | 50 | 76 | 178 | 245 | 149 | 0.056870378 | 0.418961524 | 0.106 | 0.142 | 0.066 | 0.119 | 0.129 | 0.045 | 0.071 | 0.099 | 0.088 | 0.09 | 0.034 |
| SE | 63950 | ENSG00000275052 | PPP4R3B | chr2 | - | 55578246 | 55578342 | 55577314 | 55577356 | 55579678 | 55579781 | 319 | 232 | 291 | 277 | 919 | 152 | 116 | 135 | 144 | 406 | 7 | 186 | 133 | 64 | 23 | 4 | 85 | 39 | 14 | 9 | 245 | 149 | 0.056930207 | 0.419362827 | 0.561 | 0.549 | 0.567 | 0.539 | 0.579 | 0.516 | 0.571 | 0.675 | 0.735 | 0.608 | -0.062 |
| SE | 1691 | ENSG00000228315 | GUSBP11 | chr22 | - | 23652478 | 23652576 | 23640307 | 23640423 | 23652860 | 23652886 | 11 | 14 | 15 | 19 | 10 | 1 | 0 | 0 | 0 | 0 | 3 | 11 | 5 | 23 | 21 | 0 | 0 | 0 | 0 | 8 | 247 | 149 | 0.056962714 | 0.419562808 | 0.869 | 1 | 1 | 1 | 1 | 1 | 1 | 1 | 1 | 0.613 | 0.051 |
| SE | 19506 | ENSG00000100479 | POLE2 | chr14 | - | 49654783 | 49654838 | 49654154 | 49654214 | 49655004 | 49655094 | 5 | 4 | 8 | 11 | 5 | 4 | 2 | 1 | 0 | 18 | 26 | 8 | 6 | 12 | 5 | 4 | 0 | 0 | 7 | 0 | 204 | 149 | 0.057008595 | 0.419861249 | 0.477 | 0.594 | 0.854 | 1 | 0.169 | 0.826 | 1 | 1 | 0.556 | 1 | -0.258 |
| SE | 39438 | ENSG00000055917 | PUM2 | chr2 | - | 20278582 | 20278819 | 20263192 | 20263460 | 20282946 | 20283231 | 171 | 120 | 116 | 164 | 230 | 140 | 78 | 90 | 98 | 143 | 40 | 70 | 124 | 101 | 36 | 5 | 89 | 62 | 44 | 5 | 298 | 149 | 0.057032953 | 0.420001139 | 0.379 | 0.435 | 0.392 | 0.456 | 0.446 | 0.8 | 0.282 | 0.5 | 0.534 | 0.783 | -0.158 |
| SE | 38107 | ENSG00000110871 | COQ5 | chr12 | - | 120523884 | 120524002 | 120522213 | 120522363 | 120526456 | 120526546 | 0 | 0 | 0 | 3 | 6 | 7 | 11 | 12 | 8 | 17 | 0 | 0 | 0 | 0 | 0 | 2 | 15 | 5 | 25 | 22 | 267 | 149 | 0.05706944 | 0.420230309 | 0 | 0 | 0 | 0.173 | 0.165 | 0 | 0 | 0 | 0 | 0 | 0.068 |
| SE | 49140 | ENSG00000163528 | CHCHD4 | chr3 | - | 14121916 | 14122086 | 14116425 | 14116524 | 14124654 | 14124749 | 2 | 1 | 3 | 2 | 0 | 15 | 2 | 14 | 10 | 13 | 20 | 1 | 0 | 3 | 0 | 21 | 18 | 4 | 7 | 14 | 298 | 149 | 0.057154093 | 0.420814073 | 0.062 | 0.2 | 0.097 | 0.091 | 0 | 0.323 | 0.027 | 0 | 0.176 | 0 | -0.015 |
| SE | 1112 | ENSG00000100014 | SPECC1L | chr22 | + | 24321404 | 24322918 | 24321287 | 24321323 | 24324219 | 24324427 | 98 | 101 | 86 | 108 | 100 | 17 | 9 | 23 | 13 | 10 | 83 | 89 | 55 | 114 | 33 | 0 | 6 | 6 | 7 | 12 | 298 | 149 | 0.057175405 | 0.420906657 | 0.742 | 0.849 | 0.652 | 0.806 | 0.833 | 1 | 0.881 | 0.821 | 0.891 | 0.579 | -0.058 |
| SE | 57342 | ENSG00000181722 | ZBTB20 | chr3 | - | 114874038 | 114874108 | 114801100 | 114801174 | 114900303 | 114900342 | 5 | 6 | 9 | 7 | 17 | 114 | 109 | 130 | 110 | 162 | 6 | 3 | 0 | 0 | 0 | 6 | 68 | 37 | 70 | 23 | 219 | 149 | 0.057177419 | 0.420906657 | 0.029 | 0.036 | 0.045 | 0.041 | 0.067 | 0.405 | 0.029 | 0 | 0 | 0 | -0.043 |
| SE | 106853 | ENSG00000166908 | PIP4K2C | chr12 | + | 57595887 | 57596031 | 57594024 | 57594122 | 57597957 | 57598062 | 3 | 4 | 8 | 5 | 6 | 0 | 0 | 0 | 0 | 0 | 10 | 0 | 2 | 3 | 2 | 0 | 6 | 0 | 0 | 0 | 293 | 149 | 0.057194714 | 0.420994391 | 1 | 1 | 1 | 1 | 1 | 1 | 0 | 1 | 1 | 1 | 0.2 |
| SE | 42549 | ENSG00000169241 | SLC50A1 | chr1 | + | 155136827 | 155136951 | 155136298 | 155136396 | 155137560 | 155137596 | 21 | 3 | 12 | 20 | 20 | 0 | 0 | 0 | 0 | 0 | 13 | 23 | 10 | 3 | 68 | 5 | 0 | 0 | 2 | 0 | 273 | 149 | 0.057217248 | 0.421120667 | 1 | 1 | 1 | 1 | 1 | 0.587 | 1 | 1 | 0.45 | 1 | 0.193 |
| SE | 105815 | ENSG00000111785 | RIC8B | chr12 | + | 106851449 | 106851594 | 106843851 | 106843947 | 106860267 | 106860316 | 123 | 75 | 133 | 148 | 189 | 4 | 9 | 5 | 4 | 4 | 90 | 60 | 45 | 68 | 134 | 0 | 7 | 0 | 0 | 3 | 294 | 149 | 0.057234403 | 0.421207332 | 0.94 | 0.809 | 0.931 | 0.949 | 0.96 | 1 | 0.813 | 1 | 1 | 0.958 | -0.036 |
| SE | 63651 | ENSG00000112855 | HARS2 | chr5 | + | 140696102 | 140696201 | 140695737 | 140695845 | 140696520 | 140696614 | 81 | 44 | 88 | 83 | 86 | 1 | 1 | 3 | 2 | 11 | 67 | 55 | 41 | 33 | 35 | 0 | 0 | 2 | 5 | 0 | 248 | 149 | 0.057256149 | 0.42132777 | 0.98 | 0.964 | 0.946 | 0.961 | 0.824 | 1 | 1 | 0.925 | 0.799 | 1 | -0.01 |
| SE | 95291 | ENSG00000172716 | SLFN11 | chr17 | - | 35367540 | 35367700 | 35366946 | 35367043 | 35373473 | 35373620 | 10 | 5 | 7 | 17 | 0 | 5 | 4 | 4 | 3 | 2 | 22 | 11 | 3 | 5 | 6 | 0 | 0 | 6 | 3 | 1 | 298 | 149 | 0.057277228 | 0.421443268 | 0.5 | 0.385 | 0.467 | 0.739 | 0 | 1 | 1 | 0.2 | 0.455 | 0.75 | -0.263 |
| SE | 97955 | ENSG00000137500 | CCDC90B | chr11 | - | 83274638 | 83274740 | 83273792 | 83273992 | 83278725 | 83278829 | 356 | 269 | 245 | 280 | 571 | 2 | 1 | 1 | 6 | 0 | 365 | 287 | 207 | 263 | 303 | 0 | 9 | 1 | 1 | 9 | 251 | 149 | 0.057290627 | 0.421502247 | 0.991 | 0.994 | 0.993 | 0.965 | 1 | 1 | 0.95 | 0.992 | 0.994 | 0.952 | 0.011 |
| SE | 935 | ENSG00000100209 | HSCB | chr22 | + | 28745863 | 28746008 | 28744614 | 28744709 | 28751240 | 28751288 | 31 | 16 | 25 | 28 | 42 | 2 | 0 | 0 | 0 | 0 | 12 | 9 | 11 | 7 | 13 | 0 | 0 | 5 | 2 | 0 | 294 | 149 | 0.057296934 | 0.421509045 | 0.887 | 1 | 1 | 1 | 1 | 1 | 1 | 0.527 | 0.639 | 1 | 0.144 |
| SE | 82050 | ENSG00000104133 | SPG11 | chr15 | - | 44620124 | 44620403 | 44615362 | 44615566 | 44621758 | 44621934 | 310 | 279 | 245 | 272 | 452 | 7 | 2 | 2 | 2 | 7 | 63 | 319 | 282 | 219 | 49 | 0 | 1 | 3 | 3 | 0 | 298 | 149 | 0.057331793 | 0.421725861 | 0.957 | 0.986 | 0.984 | 0.986 | 0.97 | 1 | 0.994 | 0.979 | 0.973 | 1 | -0.013 |
| SE | 12957 | ENSG00000103415 | HMOX2 | chr16 | + | 4483636 | 4483738 | 4476411 | 4476487 | 4505483 | 4505610 | 5 | 1 | 1 | 1 | 1 | 33 | 15 | 19 | 22 | 10 | 0 | 0 | 0 | 0 | 0 | 2 | 13 | 14 | 11 | 12 | 251 | 149 | 0.05733787 | 0.421730939 | 0.083 | 0.038 | 0.03 | 0.026 | 0.056 | 0 | 0 | 0 | 0 | 0 | 0.047 |
| SE | 50001 | ENSG00000241878 | PISD | chr22 | - | 31628852 | 31628981 | 31625877 | 31626016 | 31638365 | 31638445 | 3 | 0 | 1 | 1 | 2 | 1 | 2 | 3 | 8 | 1 | 0 | 0 | 0 | 0 | 0 | 10 | 1 | 2 | 11 | 10 | 278 | 149 | 0.057388796 | 0.422065862 | 0.617 | 0 | 0.152 | 0.063 | 0.517 | 0 | 0 | 0 | 0 | 0 | 0.27 |
| SE | 29097 | ENSG00000109762 | SNX25 | chr4 | + | 185332594 | 185332713 | 185323527 | 185323800 | 185339378 | 185339510 | 68 | 38 | 59 | 85 | 139 | 5 | 0 | 0 | 1 | 0 | 26 | 108 | 57 | 29 | 4 | 0 | 2 | 2 | 0 | 6 | 268 | 149 | 0.057436648 | 0.422378118 | 0.883 | 1 | 1 | 0.979 | 1 | 1 | 0.968 | 0.941 | 1 | 0.27 | 0.137 |
| SE | 99615 | ENSG00000109919 | MTCH2 | chr11 | - | 47625673 | 47625741 | 47622700 | 47622776 | 47627079 | 47627127 | 144 | 79 | 105 | 105 | 189 | 5 | 1 | 2 | 18 | 8 | 238 | 184 | 84 | 126 | 170 | 23 | 15 | 5 | 9 | 13 | 217 | 149 | 0.05750041 | 0.422807298 | 0.952 | 0.982 | 0.973 | 0.8 | 0.942 | 0.877 | 0.894 | 0.92 | 0.906 | 0.9 | 0.03 |
| SE | 12528 | ENSG00000103066 | PLA2G15 | chr16 | + | 68255281 | 68255380 | 68254352 | 68255037 | 68259145 | 68259174 | 12 | 5 | 1 | 13 | 8 | 1 | 0 | 0 | 0 | 0 | 0 | 21 | 3 | 17 | 31 | 5 | 0 | 0 | 1 | 0 | 248 | 149 | 0.057543009 | 0.423080802 | 0.878 | 1 | 1 | 1 | 1 | 0 | 1 | 1 | 0.911 | 1 | 0.193 |
| SE | 35924 | ENSG00000131323 | TRAF3 | chr14 | + | 102891324 | 102891417 | 102876357 | 102876525 | 102897260 | 102897401 | 71 | 30 | 62 | 50 | 53 | 1 | 1 | 2 | 2 | 1 | 15 | 48 | 42 | 65 | 76 | 0 | 7 | 1 | 6 | 0 | 242 | 149 | 0.057556884 | 0.423124427 | 0.978 | 0.949 | 0.95 | 0.939 | 0.97 | 1 | 0.809 | 0.963 | 0.87 | 1 | 0.029 |
| SE | 18887 | ENSG00000213983 | AP1G2 | chr14 | - | 23561961 | 23562066 | 23561511 | 23561635 | 23562287 | 23562415 | 122 | 73 | 122 | 110 | 91 | 1 | 0 | 2 | 1 | 0 | 112 | 114 | 58 | 56 | 86 | 0 | 0 | 1 | 8 | 2 | 254 | 149 | 0.057559751 | 0.423124427 | 0.986 | 1 | 0.973 | 0.985 | 1 | 1 | 1 | 0.971 | 0.804 | 0.962 | 0.041 |
| SE | 8522 | ENSG00000125450 | NUP85 | chr17 | + | 75208536 | 75208620 | 75205719 | 75205794 | 75209822 | 75209985 | 40 | 26 | 23 | 45 | 49 | 4 | 0 | 0 | 1 | 0 | 36 | 37 | 17 | 33 | 37 | 0 | 2 | 2 | 0 | 5 | 233 | 149 | 0.057570796 | 0.423165889 | 0.865 | 1 | 1 | 0.936 | 1 | 1 | 0.922 | 0.845 | 1 | 0.826 | 0.042 |
| SE | 89029 | ENSG00000177565 | TBL1XR1 | chr3 | - | 177131362 | 177131449 | 177098465 | 177098541 | 177197120 | 177197163 | 3 | 3 | 2 | 1 | 3 | 35 | 28 | 30 | 28 | 29 | 2 | 3 | 3 | 16 | 0 | 21 | 34 | 28 | 13 | 19 | 236 | 149 | 0.057579843 | 0.423192004 | 0.051 | 0.063 | 0.04 | 0.022 | 0.061 | 0.057 | 0.053 | 0.063 | 0.437 | 0 | -0.075 |
| SE | 92290 | ENSG00000125743 | SNRPD2 | chr19 | - | 45689232 | 45689347 | 45688386 | 45688566 | 45691886 | 45691941 | 1 | 0 | 0 | 1 | 6 | 170 | 71 | 188 | 129 | 176 | 0 | 0 | 2 | 0 | 18 | 120 | 107 | 53 | 73 | 117 | 264 | 149 | 0.057585159 | 0.423192004 | 0.003 | 0 | 0 | 0.004 | 0.019 | 0 | 0 | 0.021 | 0 | 0.08 | -0.015 |
| SE | 61200 | ENSG00000131408 | NR1H2 | chr19 | + | 50376718 | 50376826 | 50376234 | 50376576 | 50377586 | 50377648 | 144 | 78 | 104 | 104 | 170 | 6 | 0 | 2 | 0 | 0 | 224 | 128 | 142 | 168 | 253 | 12 | 0 | 2 | 6 | 0 | 257 | 149 | 0.057617432 | 0.423389439 | 0.933 | 1 | 0.968 | 1 | 1 | 0.915 | 1 | 0.976 | 0.942 | 1 | 0.014 |
| SE | 108639 | ENSG00000136104 | RNASEH2B | chr13 | + | 50956357 | 50956421 | 50953904 | 50953985 | 50969931 | 50970407 | 35 | 22 | 32 | 20 | 42 | 2 | 1 | 3 | 1 | 0 | 31 | 15 | 21 | 34 | 16 | 0 | 0 | 3 | 13 | 0 | 213 | 149 | 0.057629861 | 0.423441029 | 0.924 | 0.939 | 0.882 | 0.933 | 1 | 1 | 1 | 0.83 | 0.647 | 1 | 0.04 |
| SE | 95511 | ENSG00000137145 | DENND4C | chr9 | + | 19324361 | 19324507 | 19316620 | 19316839 | 19325938 | 19325974 | 306 | 296 | 277 | 312 | 458 | 0 | 0 | 0 | 0 | 0 | 57 | 209 | 145 | 63 | 29 | 0 | 0 | 4 | 7 | 0 | 295 | 149 | 0.05764089 | 0.423458883 | 1 | 1 | 1 | 1 | 1 | 1 | 1 | 0.948 | 0.82 | 1 | 0.046 |
| SE | 80232 | ENSG00000198961 | PJA2 | chr5 | - | 109390647 | 109390758 | 109383395 | 109383520 | 109409841 | 109409974 | 6 | 2 | 6 | 9 | 2 | 166 | 100 | 131 | 131 | 62 | 0 | 16 | 9 | 0 | 0 | 38 | 87 | 114 | 33 | 33 | 270 | 149 | 0.057643108 | 0.423458883 | 0.02 | 0.011 | 0.025 | 0.037 | 0.017 | 0 | 0.092 | 0.042 | 0 | 0 | -0.005 |
| SE | 85969 | ENSG00000111203 | ITFG2 | chr12 | + | 2836744 | 2836956 | 2824089 | 2824252 | 2840841 | 2840995 | 6 | 7 | 12 | 5 | 9 | 0 | 0 | 0 | 0 | 0 | 10 | 4 | 4 | 6 | 8 | 8 | 0 | 0 | 0 | 0 | 298 | 149 | 0.057653183 | 0.423481541 | 1 | 1 | 1 | 1 | 1 | 0.385 | 1 | 1 | 1 | 1 | 0.123 |
| SE | 24624 | ENSG00000204120 | GIGYF2 | chr2 | + | 232847347 | 232847571 | 232845731 | 232845886 | 232856792 | 232858608 | 149 | 91 | 103 | 126 | 239 | 0 | 0 | 0 | 0 | 0 | 28 | 75 | 84 | 71 | 10 | 0 | 0 | 0 | 0 | 8 | 298 | 149 | 0.05765701 | 0.423481541 | 1 | 1 | 1 | 1 | 1 | 1 | 1 | 1 | 1 | 0.385 | 0.123 |
| SE | 65648 | ENSG00000143924 | EML4 | chr2 | + | 42264918 | 42264951 | 42264705 | 42264731 | 42280849 | 42280973 | 5 | 6 | 9 | 6 | 1 | 198 | 176 | 198 | 168 | 189 | 1 | 1 | 13 | 9 | 7 | 56 | 126 | 184 | 148 | 93 | 182 | 149 | 0.057726667 | 0.423921882 | 0.02 | 0.027 | 0.036 | 0.028 | 0.004 | 0.014 | 0.006 | 0.055 | 0.047 | 0.058 | -0.013 |
| SE | 93565 | ENSG00000095383 | TBC1D2 | chr9 | - | 98251784 | 98251926 | 98243993 | 98244129 | 98255172 | 98255580 | 87 | 41 | 78 | 52 | 51 | 5 | 4 | 1 | 2 | 4 | 175 | 188 | 111 | 193 | 285 | 0 | 3 | 11 | 6 | 0 | 291 | 149 | 0.057727791 | 0.423921882 | 0.899 | 0.84 | 0.976 | 0.93 | 0.867 | 1 | 0.97 | 0.838 | 0.943 | 1 | -0.048 |
| SE | 61499 | ENSG00000124588 | NQO2 | chr6 | + | 3015529 | 3015643 | 3012543 | 3012674 | 3016883 | 3016985 | 110 | 30 | 84 | 112 | 104 | 8 | 2 | 2 | 8 | 12 | 152 | 181 | 56 | 271 | 290 | 16 | 23 | 3 | 68 | 9 | 263 | 149 | 0.057743455 | 0.42394808 | 0.886 | 0.895 | 0.96 | 0.888 | 0.831 | 0.843 | 0.817 | 0.914 | 0.693 | 0.948 | 0.049 |
| SE | 25695 | ENSG00000247774 | PCED1B-AS1 | chr12 | - | 47206510 | 47206585 | 47205898 | 47206069 | 47209280 | 47209458 | 85 | 33 | 156 | 104 | 197 | 37 | 9 | 52 | 45 | 103 | 82 | 61 | 59 | 29 | 32 | 18 | 28 | 34 | 4 | 0 | 224 | 149 | 0.057746258 | 0.42394808 | 0.604 | 0.709 | 0.666 | 0.606 | 0.56 | 0.752 | 0.592 | 0.536 | 0.828 | 1 | -0.113 |
| SE | 66757 | ENSG00000129292 | PHF20L1 | chr8 | + | 132799082 | 132799172 | 132794732 | 132794817 | 132803818 | 132804032 | 7 | 9 | 8 | 6 | 17 | 1 | 0 | 2 | 0 | 0 | 14 | 18 | 9 | 2 | 8 | 0 | 0 | 1 | 9 | 0 | 239 | 149 | 0.057747602 | 0.42394808 | 0.814 | 1 | 0.714 | 1 | 1 | 1 | 1 | 0.849 | 0.122 | 1 | 0.111 |
| SE | 71876 | ENSG00000115514 | TXNDC9 | chr2 | - | 99321954 | 99322209 | 99318981 | 99319799 | 99327534 | 99327653 | 486 | 316 | 418 | 403 | 509 | 1 | 1 | 0 | 0 | 0 | 89 | 334 | 276 | 131 | 37 | 6 | 5 | 3 | 0 | 0 | 298 | 149 | 0.057780675 | 0.424099129 | 0.996 | 0.994 | 1 | 1 | 1 | 0.881 | 0.971 | 0.979 | 1 | 1 | 0.032 |
| SE | 71772 | ENSG00000021574 | SPAST | chr2 | + | 32144936 | 32145007 | 32143335 | 32143415 | 32147217 | 32147258 | 148 | 108 | 100 | 135 | 177 | 5 | 7 | 2 | 5 | 4 | 8 | 111 | 71 | 78 | 6 | 0 | 4 | 0 | 3 | 0 | 220 | 149 | 0.057784227 | 0.424099129 | 0.952 | 0.913 | 0.971 | 0.948 | 0.968 | 1 | 0.949 | 1 | 0.946 | 1 | -0.029 |
| SE | 63182 | ENSG00000175662 | TOM1L2 | chr17 | - | 17882704 | 17882863 | 17879626 | 17879743 | 17898595 | 17898674 | 19 | 16 | 14 | 26 | 13 | 0 | 0 | 0 | 0 | 0 | 46 | 30 | 11 | 19 | 52 | 0 | 0 | 2 | 0 | 6 | 298 | 149 | 0.057787095 | 0.424099129 | 1 | 1 | 1 | 1 | 1 | 1 | 1 | 0.733 | 1 | 0.812 | 0.091 |
| SE | 79040 | ENSG00000160218 | TRAPPC10 | chr21 | + | 44102777 | 44102972 | 44096579 | 44098534 | 44103292 | 44104733 | 67 | 21 | 48 | 52 | 59 | 0 | 0 | 0 | 1 | 0 | 23 | 75 | 47 | 40 | 11 | 0 | 0 | 0 | 0 | 8 | 298 | 149 | 0.057789844 | 0.424099129 | 1 | 1 | 1 | 0.963 | 1 | 1 | 1 | 1 | 1 | 0.407 | 0.111 |
| SE | 11111 | ENSG00000142512 | SIGLEC10 | chr19 | - | 51414823 | 51415108 | 51414421 | 51414515 | 51415180 | 51415438 | 35 | 24 | 18 | 36 | 4 | 22 | 7 | 11 | 24 | 17 | 49 | 38 | 4 | 7 | 16 | 15 | 32 | 18 | 41 | 45 | 298 | 149 | 0.057795715 | 0.424102466 | 0.443 | 0.632 | 0.45 | 0.429 | 0.105 | 0.62 | 0.373 | 0.1 | 0.079 | 0.151 | 0.147 |
| SE | 17939 | ENSG00000197249 | SERPINA1 | chr14 | - | 94388800 | 94389010 | 94388559 | 94388663 | 94390456 | 94390635 | 160 | 72 | 43 | 88 | 73 | 98 | 36 | 34 | 38 | 38 | 108 | 181 | 63 | 104 | 319 | 97 | 145 | 75 | 61 | 168 | 298 | 149 | 0.057862034 | 0.424549324 | 0.449 | 0.5 | 0.387 | 0.537 | 0.49 | 0.358 | 0.384 | 0.296 | 0.46 | 0.487 | 0.076 |
| SE | 60533 | ENSG00000116984 | MTR | chr1 | + | 236825337 | 236825399 | 236816448 | 236816543 | 236826828 | 236826896 | 52 | 29 | 28 | 26 | 45 | 7 | 2 | 0 | 2 | 0 | 27 | 44 | 24 | 59 | 20 | 20 | 3 | 4 | 1 | 0 | 211 | 149 | 0.057868057 | 0.424553727 | 0.84 | 0.911 | 1 | 0.902 | 1 | 0.488 | 0.912 | 0.809 | 0.977 | 1 | 0.093 |
| SE | 102937 | ENSG00000084073 | ZMPSTE24 | chr1 | + | 40259349 | 40259532 | 40258235 | 40258394 | 40260838 | 40260985 | 1 | 0 | 1 | 1 | 4 | 55 | 32 | 31 | 51 | 18 | 2 | 2 | 2 | 0 | 6 | 10 | 45 | 34 | 35 | 12 | 298 | 149 | 0.057952153 | 0.425130867 | 0.009 | 0 | 0.016 | 0.01 | 0.1 | 0.091 | 0.022 | 0.029 | 0 | 0.2 | -0.041 |
| SE | 106495 | ENSG00000185482 | STAC3 | chr12 | - | 57246803 | 57246901 | 57245144 | 57245211 | 57248125 | 57248198 | 67 | 74 | 42 | 80 | 116 | 1 | 1 | 0 | 0 | 0 | 543 | 226 | 76 | 345 | 249 | 0 | 0 | 0 | 8 | 12 | 247 | 149 | 0.057963195 | 0.425145373 | 0.976 | 0.978 | 1 | 1 | 1 | 1 | 1 | 1 | 0.963 | 0.926 | 0.013 |
| SE | 65620 | ENSG00000112078 | KCTD20 | chr6 | + | 36479590 | 36479711 | 36474788 | 36475062 | 36481561 | 36481759 | 201 | 206 | 139 | 240 | 330 | 6 | 4 | 3 | 5 | 17 | 430 | 224 | 227 | 310 | 199 | 0 | 3 | 8 | 7 | 8 | 270 | 149 | 0.05796499 | 0.425145373 | 0.949 | 0.966 | 0.962 | 0.964 | 0.915 | 1 | 0.976 | 0.94 | 0.961 | 0.932 | -0.011 |
| SE | 4567 | ENSG00000123146 | ADGRE5 | chr19 | + | 14397076 | 14397223 | 14396249 | 14396473 | 14397657 | 14397804 | 278 | 141 | 134 | 167 | 197 | 138 | 70 | 114 | 111 | 98 | 300 | 190 | 221 | 227 | 170 | 183 | 78 | 71 | 107 | 83 | 296 | 149 | 0.058010026 | 0.425435835 | 0.503 | 0.503 | 0.372 | 0.431 | 0.503 | 0.452 | 0.551 | 0.61 | 0.516 | 0.508 | -0.065 |
| SE | 36365 | ENSG00000173889 | PHC3 | chr3 | - | 170145422 | 170145521 | 170136365 | 170136665 | 170171372 | 170171450 | 154 | 97 | 147 | 195 | 302 | 36 | 9 | 19 | 12 | 19 | 109 | 136 | 122 | 106 | 224 | 0 | 14 | 29 | 10 | 0 | 248 | 149 | 0.05812471 | 0.426236986 | 0.72 | 0.866 | 0.823 | 0.907 | 0.905 | 1 | 0.854 | 0.717 | 0.864 | 1 | -0.043 |
| SE | 31566 | ENSG00000163913 | IFT122 | chr3 | + | 129495445 | 129495607 | 129492140 | 129492194 | 129499901 | 129500068 | 21 | 9 | 22 | 16 | 19 | 0 | 0 | 0 | 0 | 8 | 65 | 32 | 19 | 27 | 34 | 0 | 0 | 0 | 0 | 0 | 298 | 149 | 0.058130408 | 0.426238849 | 1 | 1 | 1 | 1 | 0.543 | 1 | 1 | 1 | 1 | 1 | -0.091 |
| SE | 22140 | ENSG00000065060 | UHRF1BP1 | chr6 | + | 34836155 | 34836348 | 34835293 | 34835464 | 34855616 | 34855703 | 50 | 34 | 32 | 46 | 90 | 1 | 3 | 0 | 1 | 0 | 57 | 89 | 55 | 149 | 78 | 0 | 0 | 0 | 4 | 8 | 298 | 149 | 0.058143427 | 0.426294385 | 0.962 | 0.85 | 1 | 0.958 | 1 | 1 | 1 | 1 | 0.949 | 0.83 | -0.002 |
| SE | 88491 | ENSG00000152601 | MBNL1 | chr3 | + | 152455541 | 152455577 | 152447619 | 152447773 | 152459270 | 152459327 | 207 | 119 | 126 | 203 | 232 | 3 | 3 | 2 | 3 | 9 | 25 | 145 | 122 | 57 | 23 | 0 | 5 | 1 | 0 | 0 | 185 | 149 | 0.058221362 | 0.426825824 | 0.982 | 0.97 | 0.981 | 0.982 | 0.954 | 1 | 0.959 | 0.99 | 1 | 1 | -0.016 |
| SE | 36939 | ENSG00000092098 | RNF31 | chr14 | + | 24151298 | 24151379 | 24150597 | 24150888 | 24151484 | 24151670 | 51 | 21 | 40 | 33 | 18 | 0 | 0 | 0 | 0 | 0 | 63 | 30 | 17 | 20 | 26 | 0 | 9 | 0 | 0 | 0 | 230 | 149 | 0.05824095 | 0.426929449 | 1 | 1 | 1 | 1 | 1 | 1 | 0.683 | 1 | 1 | 1 | 0.063 |
| SE | 95853 | ENSG00000107263 | RAPGEF1 | chr9 | - | 131603960 | 131604053 | 131602060 | 131602149 | 131621795 | 131621838 | 21 | 1 | 0 | 0 | 9 | 179 | 120 | 187 | 160 | 102 | 3 | 5 | 19 | 2 | 0 | 326 | 300 | 194 | 135 | 277 | 242 | 149 | 0.058284594 | 0.427209376 | 0.067 | 0.005 | 0 | 0 | 0.052 | 0.006 | 0.01 | 0.057 | 0.009 | 0 | 0.008 |
| SE | 55973 | ENSG00000136161 | RCBTB2 | chr13 | - | 48524658 | 48524757 | 48515585 | 48515741 | 48533027 | 48533080 | 8 | 3 | 4 | 6 | 10 | 15 | 1 | 6 | 11 | 1 | 0 | 12 | 6 | 0 | 0 | 17 | 0 | 5 | 2 | 2 | 248 | 149 | 0.058344112 | 0.42757406 | 0.243 | 0.643 | 0.286 | 0.247 | 0.857 | 0 | 1 | 0.419 | 0 | 0 | 0.171 |
| SE | 60688 | ENSG00000118961 | LDAH | chr2 | - | 20790254 | 20790398 | 20774809 | 20774979 | 20801291 | 20801465 | 38 | 26 | 44 | 45 | 65 | 0 | 0 | 0 | 0 | 0 | 6 | 47 | 30 | 14 | 3 | 0 | 0 | 0 | 0 | 7 | 293 | 149 | 0.05834527 | 0.42757406 | 1 | 1 | 1 | 1 | 1 | 1 | 1 | 1 | 1 | 0.179 | 0.164 |
| SE | 101866 | ENSG00000026103 | FAS | chr10 | + | 89008888 | 89008997 | 89007699 | 89007837 | 89010538 | 89010600 | 168 | 122 | 132 | 130 | 259 | 5 | 1 | 4 | 0 | 6 | 1 | 88 | 106 | 46 | 16 | 5 | 0 | 1 | 2 | 4 | 258 | 149 | 0.058365361 | 0.427681265 | 0.951 | 0.986 | 0.95 | 1 | 0.961 | 0.104 | 1 | 0.984 | 0.93 | 0.698 | 0.226 |
| SE | 9204 | ENSG00000141576 | RNF157 | chr17 | - | 76155251 | 76155317 | 76154279 | 76154328 | 76155661 | 76155734 | 11 | 13 | 51 | 11 | 7 | 0 | 0 | 0 | 0 | 1 | 26 | 11 | 7 | 12 | 3 | 8 | 0 | 0 | 0 | 0 | 215 | 149 | 0.058377985 | 0.427733738 | 1 | 1 | 1 | 1 | 0.829 | 0.693 | 1 | 1 | 1 | 1 | 0.027 |
| SE | 30634 | ENSG00000189308 | LIN54 | chr4 | - | 82970326 | 82970469 | 82946257 | 82946474 | 82984160 | 82984876 | 50 | 33 | 38 | 61 | 78 | 5 | 9 | 3 | 8 | 11 | 29 | 31 | 24 | 41 | 17 | 0 | 2 | 7 | 3 | 0 | 292 | 149 | 0.058406972 | 0.427906083 | 0.836 | 0.652 | 0.866 | 0.796 | 0.783 | 1 | 0.888 | 0.636 | 0.875 | 1 | -0.093 |
| SE | 60607 | ENSG00000147121 | KRBOX4 | chrX | + | 46471141 | 46471185 | 46463197 | 46463308 | 46472749 | 46474639 | 5 | 5 | 3 | 8 | 11 | 22 | 7 | 5 | 12 | 19 | 0 | 2 | 1 | 1 | 17 | 15 | 15 | 12 | 2 | 16 | 193 | 149 | 0.058423866 | 0.427989806 | 0.149 | 0.355 | 0.317 | 0.34 | 0.309 | 0 | 0.093 | 0.06 | 0.279 | 0.451 | 0.117 |
| SE | 96567 | ENSG00000138660 | AP1AR | chr4 | + | 112262987 | 112263086 | 112260765 | 112260862 | 112265008 | 112265067 | 112 | 101 | 98 | 143 | 226 | 14 | 21 | 12 | 16 | 16 | 42 | 62 | 17 | 11 | 14 | 0 | 5 | 7 | 0 | 0 | 248 | 149 | 0.05843763 | 0.42805058 | 0.828 | 0.743 | 0.831 | 0.843 | 0.895 | 1 | 0.882 | 0.593 | 1 | 1 | -0.067 |
| SE | 107081 | ENSG00000010626 | LRRC23 | chr12 | + | 6912729 | 6913027 | 6909889 | 6910026 | 6913890 | 6914194 | 3 | 12 | 6 | 12 | 11 | 2 | 0 | 2 | 1 | 0 | 5 | 3 | 5 | 9 | 15 | 0 | 8 | 0 | 0 | 17 | 298 | 149 | 0.058480044 | 0.428318191 | 0.429 | 1 | 0.6 | 0.857 | 1 | 1 | 0.158 | 1 | 1 | 0.306 | 0.084 |
| SE | 97166 | ENSG00000153827 | TRIP12 | chr2 | - | 229829192 | 229829288 | 229818363 | 229818512 | 229836847 | 229836984 | 140 | 119 | 107 | 135 | 75 | 0 | 0 | 5 | 1 | 0 | 68 | 119 | 90 | 144 | 70 | 6 | 3 | 0 | 3 | 0 | 245 | 149 | 0.058485105 | 0.428318191 | 1 | 1 | 0.929 | 0.988 | 1 | 0.873 | 0.96 | 1 | 0.967 | 1 | 0.023 |
| SE | 68938 | ENSG00000135926 | TMBIM1 | chr2 | - | 218284038 | 218284136 | 218281939 | 218282181 | 218285762 | 218285938 | 2 | 0 | 0 | 3 | 0 | 11 | 3 | 2 | 12 | 3 | 12 | 0 | 0 | 0 | 8 | 51 | 3 | 9 | 11 | 118 | 247 | 149 | 0.058493019 | 0.428336081 | 0.099 | 0 | 0 | 0.131 | 0 | 0.124 | 0 | 0 | 0 | 0.039 | 0.013 |
| SE | 32277 | ENSG00000114745 | GORASP1 | chr3 | - | 39100787 | 39100877 | 39100304 | 39100503 | 39101015 | 39101102 | 9 | 10 | 14 | 5 | 13 | 1 | 0 | 0 | 1 | 0 | 37 | 19 | 13 | 21 | 39 | 5 | 3 | 0 | 0 | 0 | 239 | 149 | 0.058515187 | 0.428458346 | 0.849 | 1 | 1 | 0.757 | 1 | 0.822 | 0.798 | 1 | 1 | 1 | -0.003 |
| SE | 15693 | ENSG00000174485 | DENND4A | chr15 | - | 65741714 | 65741784 | 65738705 | 65738875 | 65752378 | 65752628 | 230 | 217 | 170 | 267 | 339 | 2 | 0 | 6 | 0 | 0 | 327 | 334 | 251 | 215 | 219 | 5 | 11 | 2 | 0 | 3 | 219 | 149 | 0.058536549 | 0.428574682 | 0.987 | 1 | 0.951 | 1 | 1 | 0.978 | 0.954 | 0.988 | 1 | 0.98 | 0.008 |
| SE | 106602 | ENSG00000259431 | THTPA | chr14 | + | 23556742 | 23557034 | 23556270 | 23556343 | 23558694 | 23558898 | 2 | 2 | 10 | 11 | 2 | 6 | 2 | 6 | 13 | 0 | 0 | 4 | 1 | 0 | 10 | 8 | 13 | 2 | 4 | 11 | 298 | 149 | 0.058587099 | 0.428904675 | 0.143 | 0.333 | 0.455 | 0.297 | 1 | 0 | 0.133 | 0.2 | 0 | 0.312 | 0.317 |
| SE | 86087 | ENSG00000169359 | SLC33A1 | chr3 | - | 155845012 | 155845076 | 155842431 | 155842619 | 155853107 | 155854456 | 0 | 2 | 0 | 1 | 3 | 3 | 11 | 9 | 3 | 4 | 0 | 0 | 0 | 8 | 7 | 3 | 8 | 12 | 14 | 7 | 213 | 149 | 0.058612154 | 0.429047982 | 0 | 0.113 | 0 | 0.189 | 0.344 | 0 | 0 | 0 | 0.286 | 0.412 | -0.01 |
| SE | 24017 | ENSG00000112983 | BRD8 | chr5 | - | 138166517 | 138166727 | 138165827 | 138166108 | 138167933 | 138168078 | 40 | 13 | 35 | 25 | 40 | 9 | 6 | 6 | 5 | 12 | 18 | 3 | 12 | 4 | 13 | 10 | 14 | 10 | 3 | 0 | 298 | 149 | 0.058634003 | 0.429167793 | 0.69 | 0.52 | 0.745 | 0.714 | 0.625 | 0.474 | 0.097 | 0.375 | 0.4 | 1 | 0.19 |
| SE | 41918 | ENSG00000143110 | C1orf162 | chr1 | + | 111477333 | 111477428 | 111476797 | 111477078 | 111477650 | 111477775 | 56 | 14 | 10 | 54 | 90 | 2 | 0 | 2 | 0 | 4 | 255 | 43 | 54 | 118 | 126 | 12 | 3 | 0 | 6 | 8 | 244 | 149 | 0.058639632 | 0.429168875 | 0.945 | 1 | 0.753 | 1 | 0.932 | 0.928 | 0.897 | 1 | 0.923 | 0.906 | -0.005 |
| SE | 12032 | ENSG00000176715 | ACSF3 | chr16 | + | 89116989 | 89117086 | 89114338 | 89114487 | 89120800 | 89120913 | 4 | 4 | 9 | 6 | 4 | 95 | 58 | 85 | 69 | 54 | 23 | 4 | 3 | 15 | 22 | 166 | 136 | 69 | 115 | 131 | 246 | 149 | 0.058704904 | 0.42958434 | 0.025 | 0.04 | 0.06 | 0.05 | 0.043 | 0.077 | 0.018 | 0.026 | 0.073 | 0.092 | -0.014 |
| SE | 24442 | ENSG00000114107 | CEP70 | chr3 | - | 138572858 | 138572932 | 138571265 | 138571356 | 138591853 | 138591956 | 54 | 38 | 65 | 54 | 71 | 0 | 1 | 4 | 1 | 3 | 24 | 33 | 34 | 15 | 27 | 0 | 0 | 2 | 0 | 0 | 223 | 149 | 0.058708089 | 0.42958434 | 1 | 0.962 | 0.916 | 0.973 | 0.941 | 1 | 1 | 0.919 | 1 | 1 | -0.025 |
| SE | 106503 | ENSG00000111666 | CHPT1 | chr12 | + | 101714883 | 101715076 | 101714503 | 101714645 | 101716727 | 101716812 | 0 | 1 | 2 | 6 | 5 | 61 | 35 | 43 | 38 | 32 | 4 | 0 | 4 | 4 | 0 | 133 | 96 | 54 | 110 | 124 | 298 | 149 | 0.058712859 | 0.42958434 | 0 | 0.014 | 0.023 | 0.073 | 0.072 | 0.015 | 0 | 0.036 | 0.018 | 0 | 0.023 |
| SE | 11235 | ENSG00000102984 | ZNF821 | chr16 | - | 71867892 | 71868037 | 71864753 | 71865048 | 71879906 | 71880023 | 20 | 16 | 32 | 22 | 21 | 3 | 0 | 1 | 3 | 0 | 26 | 46 | 17 | 26 | 43 | 0 | 6 | 4 | 0 | 6 | 294 | 149 | 0.058757873 | 0.429872797 | 0.772 | 1 | 0.942 | 0.788 | 1 | 1 | 0.795 | 0.683 | 1 | 0.784 | 0.048 |
| SE | 73617 | ENSG00000119707 | RBM25 | chr14 | + | 73064971 | 73065093 | 73058532 | 73058705 | 73071626 | 73071747 | 3 | 2 | 19 | 7 | 0 | 31 | 30 | 42 | 29 | 14 | 0 | 1 | 11 | 24 | 24 | 21 | 20 | 22 | 17 | 8 | 271 | 149 | 0.058763264 | 0.429872797 | 0.051 | 0.035 | 0.199 | 0.117 | 0 | 0 | 0.027 | 0.216 | 0.437 | 0.623 | -0.18 |
| SE | 70791 | ENSG00000064201 | TSPAN32 | chr11 | + | 2316575 | 2316667 | 2316228 | 2316312 | 2317343 | 2317525 | 28 | 5 | 11 | 11 | 26 | 0 | 0 | 0 | 0 | 0 | 7 | 7 | 7 | 7 | 15 | 0 | 0 | 0 | 8 | 0 | 241 | 149 | 0.058832516 | 0.430311275 | 1 | 1 | 1 | 1 | 1 | 1 | 1 | 1 | 0.351 | 1 | 0.13 |
| SE | 58172 | ENSG00000172965 | MIR4435-2HG | chr2 | - | 111208755 | 111208936 | 111207773 | 111208661 | 111239801 | 111239966 | 6 | 6 | 3 | 13 | 17 | 0 | 1 | 0 | 0 | 0 | 26 | 9 | 5 | 15 | 2 | 0 | 0 | 0 | 0 | 7 | 298 | 149 | 0.058834196 | 0.430311275 | 1 | 0.75 | 1 | 1 | 1 | 1 | 1 | 1 | 1 | 0.125 | 0.125 |
| SE | 108142 | ENSG00000127311 | HELB | chr12 | + | 66315241 | 66315383 | 66309705 | 66310608 | 66318637 | 66318792 | 159 | 99 | 175 | 127 | 314 | 4 | 1 | 5 | 3 | 8 | 5 | 88 | 86 | 24 | 2 | 0 | 4 | 8 | 2 | 0 | 291 | 149 | 0.058853092 | 0.430409278 | 0.953 | 0.981 | 0.947 | 0.956 | 0.953 | 1 | 0.918 | 0.846 | 0.86 | 1 | 0.033 |
| SE | 65373 | ENSG00000087008 | ACOX3 | chr4 | - | 8373560 | 8373628 | 8370907 | 8370994 | 8374977 | 8375152 | 11 | 4 | 5 | 3 | 6 | 3 | 2 | 0 | 5 | 4 | 2 | 9 | 7 | 11 | 35 | 0 | 0 | 0 | 8 | 5 | 217 | 149 | 0.0588817 | 0.430578271 | 0.716 | 0.579 | 1 | 0.34 | 0.452 | 1 | 1 | 1 | 0.486 | 0.828 | -0.245 |
| SE | 30285 | ENSG00000114480 | GBE1 | chr3 | - | 81646391 | 81646504 | 81642570 | 81642990 | 81648855 | 81648991 | 25 | 17 | 19 | 15 | 21 | 2 | 0 | 1 | 5 | 0 | 17 | 4 | 10 | 8 | 18 | 0 | 2 | 1 | 0 | 0 | 262 | 149 | 0.058919016 | 0.430810913 | 0.877 | 1 | 0.915 | 0.63 | 1 | 1 | 0.532 | 0.85 | 1 | 1 | 0.008 |
| SE | 70615 | ENSG00000225889 | NA | chr2 | + | 64229170 | 64229288 | 64228540 | 64228629 | 64229496 | 64229640 | 9 | 2 | 11 | 2 | 2 | 3 | 2 | 3 | 6 | 0 | 10 | 7 | 9 | 4 | 1 | 18 | 5 | 10 | 17 | 4 | 267 | 149 | 0.058931442 | 0.430861536 | 0.626 | 0.358 | 0.672 | 0.157 | 1 | 0.237 | 0.439 | 0.334 | 0.116 | 0.122 | 0.313 |
| SE | 84449 | ENSG00000189376 | C8orf76 | chr8 | - | 123226299 | 123226632 | 123219966 | 123220297 | 123237797 | 123237941 | 12 | 12 | 14 | 26 | 40 | 1 | 3 | 1 | 6 | 1 | 11 | 11 | 17 | 4 | 10 | 4 | 3 | 4 | 0 | 4 | 282 | 149 | 0.058965667 | 0.431071509 | 0.864 | 0.679 | 0.881 | 0.696 | 0.955 | 0.592 | 0.66 | 0.692 | 1 | 0.569 | 0.112 |
| SE | 87630 | ENSG00000140678 | ITGAX | chr16 | + | 31371083 | 31371214 | 31363164 | 31363374 | 31371629 | 31371784 | 89 | 42 | 75 | 49 | 27 | 2 | 2 | 4 | 0 | 0 | 95 | 105 | 48 | 67 | 121 | 0 | 2 | 1 | 9 | 8 | 280 | 149 | 0.058986362 | 0.431182544 | 0.959 | 0.918 | 0.909 | 1 | 1 | 1 | 0.965 | 0.962 | 0.798 | 0.889 | 0.034 |
| SE | 77668 | ENSG00000214021 | TTLL3 | chr3 | + | 9833103 | 9833245 | 9828678 | 9829395 | 9834680 | 9834907 | 22 | 2 | 8 | 9 | 2 | 3 | 7 | 9 | 9 | 4 | 16 | 13 | 19 | 13 | 35 | 2 | 0 | 6 | 13 | 32 | 291 | 149 | 0.05899402 | 0.431198263 | 0.79 | 0.128 | 0.313 | 0.339 | 0.204 | 0.804 | 1 | 0.619 | 0.339 | 0.359 | -0.269 |
| SE | 28097 | ENSG00000084676 | NCOA1 | chr2 | + | 24768078 | 24768135 | 24762686 | 24762776 | 24768220 | 24770702 | 191 | 106 | 111 | 148 | 207 | 75 | 46 | 34 | 94 | 69 | 118 | 91 | 209 | 146 | 58 | 64 | 50 | 43 | 34 | 0 | 206 | 149 | 0.059129867 | 0.432150848 | 0.648 | 0.625 | 0.703 | 0.532 | 0.685 | 0.571 | 0.568 | 0.779 | 0.756 | 1 | -0.096 |
| SE | 53785 | ENSG00000185800 | DMWD | chr19 | - | 45784640 | 45784715 | 45782946 | 45784290 | 45785593 | 45786871 | 10 | 0 | 2 | 3 | 6 | 18 | 13 | 5 | 14 | 2 | 0 | 8 | 7 | 3 | 33 | 10 | 7 | 0 | 0 | 39 | 224 | 149 | 0.059166817 | 0.432376446 | 0.27 | 0 | 0.21 | 0.125 | 0.666 | 0 | 0.432 | 1 | 1 | 0.36 | -0.304 |
| SE | 61387 | ENSG00000151893 | CACUL1 | chr10 | - | 118729294 | 118729397 | 118707491 | 118707587 | 118730283 | 118730410 | 199 | 177 | 178 | 138 | 325 | 1 | 3 | 0 | 4 | 0 | 127 | 133 | 195 | 166 | 100 | 0 | 0 | 5 | 1 | 8 | 252 | 149 | 0.059175429 | 0.432376446 | 0.992 | 0.972 | 1 | 0.953 | 1 | 1 | 1 | 0.958 | 0.99 | 0.881 | 0.018 |
| SE | 86242 | ENSG00000134516 | DOCK2 | chr5 | + | 169669297 | 169669328 | 169654402 | 169654486 | 169671077 | 169671174 | 61 | 20 | 42 | 28 | 6 | 0 | 0 | 0 | 1 | 7 | 14 | 63 | 21 | 61 | 45 | 0 | 0 | 6 | 9 | 4 | 190 | 149 | 0.059177301 | 0.432376446 | 1 | 1 | 1 | 0.956 | 0.402 | 1 | 1 | 0.733 | 0.842 | 0.898 | -0.023 |
| SE | 51981 | ENSG00000071054 | MAP4K4 | chr2 | + | 101860824 | 101860986 | 101859642 | 101859864 | 101864929 | 101865036 | 35 | 27 | 28 | 39 | 40 | 42 | 38 | 32 | 50 | 24 | 12 | 27 | 36 | 36 | 59 | 40 | 67 | 62 | 82 | 33 | 298 | 149 | 0.059216795 | 0.432624635 | 0.294 | 0.262 | 0.304 | 0.281 | 0.455 | 0.13 | 0.168 | 0.225 | 0.18 | 0.472 | 0.084 |
| SE | 8918 | ENSG00000161533 | ACOX1 | chr17 | - | 75960214 | 75960375 | 75957420 | 75957566 | 75978533 | 75978693 | 121 | 61 | 78 | 69 | 98 | 3 | 3 | 1 | 0 | 1 | 138 | 136 | 83 | 70 | 138 | 0 | 8 | 10 | 0 | 1 | 298 | 149 | 0.059257703 | 0.432883103 | 0.953 | 0.91 | 0.975 | 1 | 0.98 | 1 | 0.895 | 0.806 | 1 | 0.986 | 0.026 |
| SE | 25503 | ENSG00000103540 | CCP110 | chr16 | + | 19525718 | 19525861 | 19523810 | 19524088 | 19527866 | 19528022 | 16 | 17 | 9 | 17 | 29 | 11 | 10 | 8 | 5 | 14 | 3 | 11 | 10 | 8 | 9 | 0 | 4 | 4 | 4 | 0 | 292 | 149 | 0.059339328 | 0.433438943 | 0.426 | 0.465 | 0.365 | 0.634 | 0.514 | 1 | 0.584 | 0.561 | 0.505 | 1 | -0.249 |
| SE | 94888 | ENSG00000119446 | RBM18 | chr9 | - | 122243743 | 122243868 | 122241669 | 122242043 | 122245255 | 122245341 | 2 | 0 | 7 | 5 | 8 | 108 | 57 | 56 | 74 | 117 | 18 | 2 | 9 | 2 | 0 | 5 | 64 | 77 | 24 | 21 | 274 | 149 | 0.059356751 | 0.433525761 | 0.01 | 0 | 0.064 | 0.035 | 0.036 | 0.662 | 0.017 | 0.06 | 0.043 | 0 | -0.127 |
| SE | 73310 | ENSG00000136699 | SMPD4 | chr2 | - | 130173278 | 130173354 | 130172786 | 130172895 | 130174913 | 130175000 | 30 | 11 | 26 | 21 | 65 | 0 | 0 | 0 | 0 | 0 | 43 | 40 | 39 | 20 | 56 | 0 | 0 | 1 | 2 | 6 | 225 | 149 | 0.059380688 | 0.433660141 | 1 | 1 | 1 | 1 | 1 | 1 | 1 | 0.963 | 0.869 | 0.861 | 0.061 |
| SE | 99200 | ENSG00000148925 | BTBD10 | chr11 | - | 13459665 | 13459736 | 13421746 | 13421838 | 13463091 | 13463252 | 1 | 0 | 0 | 1 | 0 | 31 | 19 | 16 | 16 | 14 | 12 | 0 | 0 | 0 | 0 | 0 | 22 | 9 | 3 | 10 | 220 | 149 | 0.059394976 | 0.433724029 | 0.021 | 0 | 0 | 0.041 | 0 | 1 | 0 | 0 | 0 | 0 | -0.188 |
| SE | 37186 | ENSG00000114446 | IFT57 | chr3 | - | 108165430 | 108165493 | 108163662 | 108163729 | 108167792 | 108167864 | 5 | 12 | 11 | 13 | 21 | 0 | 0 | 0 | 0 | 0 | 5 | 23 | 13 | 3 | 7 | 8 | 0 | 0 | 0 | 0 | 212 | 149 | 0.059417875 | 0.433850786 | 1 | 1 | 1 | 1 | 1 | 0.305 | 1 | 1 | 1 | 1 | 0.139 |

| | | | | | | | | | | | | | | | | | | | | | | | | | |
|---|---|---|---|---|---|---|---|---|---|---|---|---|---|---|---|---|---|---|---|---|---|---|---|---|---|
| SE | 57494 | ENSG00000134313 | KIDINS220 | chr2 | - | 8749370 | 8749421 | 8746456 | 8748000 | 8750111 | 8750335 | 6 | 1 | 10 | 16 | 13 | 101 | 57 | 70 | 98 | 159 | 0 | 18 | 12 | 1 |
| SE | 35128 | ENSG00000288520 | NA | chr6 | - | 154167919 | 154168113 | 154159827 | 154160040 | 154199667 | 154200040 | 374 | 271 | 461 | 364 | 955 | 0 | 0 | 1 | 2 | 7 | 59 | 175 | 266 | 180 |
| SE | 65654 | ENSG00000143924 | EML4 | chr2 | + | 42315961 | 42316050 | 42304483 | 42304551 | 42325466 | 42325554 | 80 | 71 | 104 | 103 | 131 | 3 | 1 | 8 | 0 | 0 | 27 | 40 | 84 | 53 |
| SE | 45862 | ENSG00000062485 | CS | chr12 | - | 56291630 | 56291793 | 56286594 | 56286645 | 56300159 | 56300391 | 3 | 1 | 5 | 3 | 5 | 83 | 38 | 52 | 69 | 45 | 21 | 0 | 6 | 2 |
| SE | 106796 | ENSG00000173442 | EHBP1L1 | chr11 | + | 65581538 | 65582016 | 65581210 | 65581373 | 65584240 | 65584398 | 293 | 212 | 219 | 265 | 186 | 6 | 0 | 1 | 1 | 9 | 620 | 315 | 195 | 390 |
| SE | 2430 | ENSG00000132024 | CC2D1A | chr19 | + | 13919817 | 13919951 | 13919129 | 13919202 | 13920556 | 13920668 | 21 | 12 | 11 | 27 | 8 | 0 | 0 | 0 | 0 | 0 | 16 | 51 | 21 | 29 |
| SE | 18583 | ENSG00000184990 | SIVA1 | chr14 | + | 104756603 | 104756760 | 104755629 | 104755824 | 104766789 | 104766897 | 30 | 15 | 34 | 51 | 14 | 1 | 0 | 0 | 0 | 0 | 40 | 43 | 23 | 35 |
| SE | 73586 | ENSG00000147443 | DOK2 | chr8 | - | 21911900 | 21911988 | 21910672 | 21910857 | 21912228 | 21912510 | 226 | 106 | 130 | 181 | 121 | 34 | 9 | 12 | 31 | 12 | 202 | 154 | 55 | 111 |
| SE | 15998 | ENSG00000068305 | MEF2A | chr15 | + | 99620890 | 99621080 | 99598429 | 99598511 | 99632977 | 99633173 | 7 | 12 | 7 | 6 | 9 | 105 | 158 | 100 | 139 | 79 | 0 | 5 | 3 | 10 |
| SE | 1802 | ENSG00000099917 | MED15 | chr22 | + | 20551435 | 20551487 | 20537116 | 20537204 | 20564449 | 20564500 | 34 | 16 | 28 | 14 | 12 | 0 | 0 | 0 | 1 | 0 | 41 | 33 | 18 | 32 |
| SE | 66419 | ENSG00000161813 | LARP4 | chr12 | + | 50461134 | 50461347 | 50454313 | 50454417 | 50462581 | 50462630 | 82 | 67 | 103 | 94 | 96 | 3 | 4 | 0 | 4 | 5 | 27 | 73 | 65 | 26 |
| SE | 71308 | ENSG00000163754 | GYG1 | chr3 | + | 148996301 | 148996427 | 148994141 | 148994277 | 148996741 | 148996847 | 35 | 17 | 38 | 41 | 26 | 0 | 0 | 0 | 1 | 0 | 32 | 78 | 34 | 108 |
| SE | 85707 | ENSG00000072778 | ACADVL | chr17 | + | 7220923 | 7221058 | 7220757 | 7220830 | 7221537 | 7221655 | 284 | 144 | 187 | 275 | 249 | 1 | 0 | 5 | 0 | 0 | 275 | 323 | 189 | 227 |
| SE | 89572 | ENSG00000095485 | CWF19L1 | chr10 | - | 100260217 | 100260319 | 100256261 | 100256476 | 100260965 | 100261028 | 116 | 85 | 128 | 87 | 216 | 3 | 4 | 5 | 1 | 0 | 142 | 110 | 104 | 153 |
| SE | 14395 | ENSG00000167202 | TBC1D2B | chr15 | - | 77998502 | 77998900 | 77998243 | 77998355 | 78001618 | 78001666 | 11 | 12 | 7 | 17 | 13 | 55 | 38 | 68 | 65 | 93 | 45 | 2 | 7 | 6 |
| SE | 1603 | ENSG00000198089 | SFI1 | chr22 | + | 31604868 | 31604945 | 31589446 | 31589577 | 31606327 | 31606430 | 8 | 8 | 11 | 8 | 3 | 0 | 4 | 3 | 0 | 0 | 22 | 8 | 4 | 2 |
| SE | 50300 | ENSG00000100591 | AHSA1 | chr14 | + | 77462641 | 77462759 | 77462159 | 77462242 | 77464597 | 77464686 | 328 | 193 | 292 | 372 | 594 | 5 | 4 | 3 | 1 | 9 | 562 | 216 | 192 | 231 |
| SE | 82853 | ENSG00000146776 | ATXN7L1 | chr7 | - | 105617791 | 105618097 | 105613473 | 105614816 | 105620199 | 105620321 | 2 | 5 | 5 | 2 | 8 | 55 | 28 | 56 | 50 | 78 | 20 | 0 | 0 | 0 |
| SE | 29997 | ENSG00000185305 | ARL15 | chr5 | - | 54171783 | 54171928 | 54113132 | 54113410 | 54310431 | 54310573 | 35 | 34 | 29 | 43 | 11 | 0 | 0 | 0 | 0 | 8 | 17 | 57 | 36 | 20 |
| SE | 31690 | ENSG00000174776 | WDR49 | chr3 | - | 167575917 | 167576151 | 167566829 | 167566894 | 167602126 | 167602275 | 14 | 12 | 10 | 15 | 18 | 1 | 0 | 0 | 0 | 1 | 5 | 9 | 3 | 6 |
| SE | 12019 | ENSG00000101624 | CEP76 | chr18 | - | 12691358 | 12691487 | 12684295 | 12686450 | 12695253 | 12695351 | 23 | 21 | 28 | 29 | 50 | 0 | 0 | 1 | 0 | 0 | 0 | 24 | 11 | 38 |
| SE | 49459 | ENSG00000115020 | PIKFYVE | chr2 | + | 208277536 | 208277708 | 208276711 | 208276830 | 208285725 | 208285800 | 185 | 128 | 149 | 156 | 211 | 9 | 0 | 5 | 20 | 18 | 115 | 124 | 79 | 101 |
| SE | 108158 | ENSG00000075089 | ACTR6 | chr12 | + | 100204939 | 100205057 | 100199864 | 100200919 | 100207662 | 100207786 | 71 | 27 | 82 | 65 | 58 | 0 | 2 | 0 | 0 | 3 | 7 | 36 | 21 | 9 |
| SE | 47887 | ENSG00000147324 | MFHAS1 | chr8 | - | 8790323 | 8790434 | 8785758 | 8786055 | 8797364 | 8797491 | 6 | 4 | 16 | 6 | 4 | 22 | 29 | 23 | 27 | 13 | 6 | 3 | 2 | 9 |
| SE | 72100 | ENSG00000122484 | RPAP2 | chr1 | + | 92380723 | 92380873 | 92345845 | 92345914 | 92387010 | 92387233 | 123 | 58 | 130 | 147 | 391 | 1 | 6 | 5 | 0 | 2 | 22 | 49 | 33 | 44 |
| SE | 39730 | ENSG00000132716 | DCAF8 | chr1 | - | 160261284 | 160261358 | 160243959 | 160244034 | 160262116 | 160262425 | 129 | 126 | 96 | 130 | 127 | 22 | 8 | 12 | 16 | 13 | 273 | 168 | 156 | 202 |
| SE | 7390 | ENSG00000101557 | USP14 | chr18 | + | 178932 | 179037 | 166786 | 166819 | 180235 | 180339 | 154 | 74 | 117 | 103 | 143 | 6 | 3 | 5 | 5 | 0 | 58 | 36 | 135 | 94 |
| SE | 32799 | ENSG00000157601 | MX1 | chr21 | + | 41439693 | 41439848 | 41437014 | 41437152 | 41440886 | 41441025 | 502 | 709 | 511 | 1006 | 768 | 2 | 0 | 0 | 4 | 4 | 3956 | 1920 | 806 | 3197 |
| SE | 57334 | ENSG00000181722 | ZBTB20 | chr3 | - | 114743711 | 114743779 | 114693527 | 114693575 | 114801100 | 114801177 | 1 | 2 | 0 | 1 | 0 | 80 | 61 | 85 | 70 | 140 | 10 | 0 | 0 | 0 |
| SE | 14359 | ENSG00000175265 | GOLGA8A | chr15 | - | 34407523 | 34407735 | 34406673 | 34406870 | 34435382 | 34435471 | 75 | 59 | 97 | 75 | 81 | 0 | 0 | 0 | 0 | 0 | 65 | 4 | 35 | 37 |
| SE | 87746 | ENSG00000171100 | MTM1 | chrX | + | 150663432 | 150663609 | 150660370 | 150660484 | 150671430 | 150672874 | 85 | 51 | 94 | 86 | 239 | 0 | 0 | 0 | 0 | 0 | 4 | 75 | 40 | 42 |
| SE | 105868 | ENSG00000090006 | LTBP4 | chr19 | + | 40623603 | 40623732 | 40619346 | 40619493 | 40625856 | 40626006 | 7 | 1 | 12 | 13 | 7 | 0 | 0 | 0 | 0 | 0 | 14 | 1 | 6 | 2 |
| SE | 50098 | ENSG00000163322 | ABRAXAS1 | chr4 | - | 83463493 | 83463608 | 83459516 | 83462902 | 83467453 | 83467538 | 149 | 165 | 115 | 132 | 210 | 1 | 3 | 2 | 0 | 8 | 35 | 38 | 50 | 12 |
| SE | 9340 | ENSG00000167280 | ENGASE | chr17 | + | 79079488 | 79079637 | 79077662 | 79077864 | 79080206 | 79080364 | 91 | 53 | 79 | 100 | 52 | 9 | 2 | 16 | 5 | 4 | 52 | 69 | 44 | 70 |
| SE | 20807 | ENSG00000181036 | FCRL6 | chr1 | + | 159814220 | 159814292 | 159810093 | 159810216 | 159815535 | 159815802 | 33 | 14 | 10 | 18 | 27 | 0 | 0 | 0 | 0 | 0 | 32 | 26 | 9 | 4 |
| SE | 32878 | ENSG00000068650 | ATP11A | chr13 | + | 112858144 | 112858290 | 112857817 | 112857920 | 112860286 | 112860414 | 70 | 50 | 40 | 83 | 81 | 0 | 0 | 0 | 0 | 0 | 91 | 68 | 45 | 141 |
| SE | 19180 | ENSG00000140105 | WARS1 | chr14 | - | 100374135 | 100374179 | 100369086 | 100369258 | 100375282 | 100375347 | 0 | 0 | 11 | 0 | 12 | 110 | 121 | 112 | 101 | 71 | 0 | 0 | 0 | 0 |
| SE | 67485 | ENSG00000120159 | CAAP1 | chr9 | - | 26869925 | 26869977 | 26861065 | 26861139 | 26884809 | 26884885 | 3 | 5 | 0 | 9 | 4 | 50 | 65 | 41 | 56 | 91 | 0 | 6 | 11 | 2 |
| SE | 70853 | ENSG00000243970 | PPIEL | chr1 | - | 39537190 | 39537343 | 39529054 | 39529164 | 39541719 | 39541841 | 1 | 0 | 1 | 0 | 8 | 15 | 18 | 13 | 35 | 2 | 0 | 0 | 0 | 0 |
| SE | 31087 | ENSG00000108509 | CAMTA2 | chr17 | - | 4982088 | 4982160 | 4981677 | 4981831 | 4982756 | 4982892 | 54 | 46 | 46 | 57 | 61 | 6 | 9 | 10 | 6 | 1 | 98 | 61 | 23 | 63 |
| SE | 59494 | ENSG00000148660 | CAMK2G | chr10 | - | 73824039 | 73824084 | 73821681 | 73821730 | 73837467 | 73837511 | 65 | 46 | 62 | 75 | 92 | 0 | 0 | 6 | 2 | 4 | 142 | 63 | 62 | 50 |
| SE | 58304 | ENSG00000117262 | GPR89A | chr1 | + | 145646203 | 145646272 | 145644085 | 145645743 | 145647174 | 145647267 | 5 | 2 | 4 | 1 | 3 | 2 | 0 | 0 | 2 | 0 | 14 | 0 | 6 | 0 |
| SE | 100255 | ENSG00000166471 | TMEM41B | chr11 | - | 9286454 | 9286593 | 9280653 | 9283593 | 9287701 | 9287806 | 125 | 100 | 181 | 131 | 214 | 0 | 1 | 2 | 4 | 4 | 87 | 121 | 139 | 93 |
| SE | 101787 | ENSG00000170759 | KIF5B | chr10 | - | 32039326 | 32039431 | 32038777 | 32038826 | 32040383 | 32040457 | 770 | 551 | 536 | 701 | 1046 | 0 | 2 | 3 | 2 | 0 | 81 | 404 | 373 | 167 |
| SE | 73691 | ENSG00000151687 | ANKAR | chr2 | + | 189738564 | 189738682 | 189728659 | 189728821 | 189741341 | 189741451 | 34 | 27 | 13 | 13 | 46 | 5 | 5 | 7 | 7 | 19 | 26 | 18 | 17 | 24 |
| SE | 52154 | ENSG00000162367 | TAL1 | chr1 | - | 47223383 | 47223619 | 47218687 | 47220174 | 47224003 | 47224076 | 0 | 3 | 0 | 3 | 9 | 15 | 30 | 1 | 42 | 12 | 0 | 7 | 3 | 4 |
| SE | 61839 | ENSG00000170871 | KIAA0232 | chr4 | + | 6876659 | 6876757 | 6871573 | 6871682 | 6880786 | 6884164 | 241 | 203 | 160 | 215 | 291 | 3 | 5 | 8 | 5 | 4 | 53 | 157 | 247 | 179 |
| SE | 13165 | ENSG00000103121 | CMC2 | chr16 | - | 80998284 | 80998351 | 80997313 | 80997429 | 81006733 | 81006839 | 23 | 12 | 1 | 11 | 12 | 36 | 13 | 24 | 43 | 1 | 0 | 20 | 8 | 11 |
| SE | 66740 | ENSG00000138468 | SENP7 | chr3 | - | 101367829 | 101368011 | 101366429 | 101366769 | 101372007 | 101372126 | 364 | 304 | 313 | 257 | 738 | 5 | 13 | 2 | 2 | 8 | 16 | 127 | 71 | 26 |
| SE | 19132 | ENSG00000090061 | CCNK | chr14 | + | 99491945 | 99492320 | 99481424 | 99481479 | 99492625 | 99492874 | 9 | 11 | 6 | 6 | 11 | 75 | 40 | 85 | 74 | 49 | 7 | 31 | 28 | 15 |
| SE | 28110 | ENSG00000125386 | FAM193A | chr4 | + | 2708145 | 2708268 | 2699679 | 2700544 | 2716022 | 2716104 | 0 | 0 | 0 | 0 | 4 | 62 | 46 | 50 | 51 | 54 | 9 | 1 | 3 | 4 |
| SE | 105249 | ENSG00000082898 | XPO1 | chr2 | - | 61490641 | 61490776 | 61488587 | 61488771 | 61492034 | 61492196 | 874 | 563 | 768 | 881 | 1460 | 10 | 3 | 3 | 9 | 18 | 96 | 340 | 308 | 208 |
| SE | 22601 | ENSG00000127603 | MACF1 | chr1 | + | 39346976 | 39347210 | 39340803 | 39340953 | 39349477 | 39349627 | 541 | 291 | 364 | 578 | 477 | 1 | 0 | 1 | 2 | 0 | 62 | 597 | 410 | 99 |
| SE | 14020 | ENSG00000090905 | TNRC6A | chr16 | + | 24776932 | 24777358 | 24758338 | 24758360 | 24789231 | 24791817 | 63 | 52 | 69 | 66 | 37 | 0 | 3 | 0 | 5 | 1 | 38 | 47 | 60 | 51 |
| SE | 27337 | ENSG00000157869 | RAB28 | chr4 | - | 13460698 | 13460828 | 13381490 | 13381594 | 13474317 | 13474322 | 165 | 103 | 129 | 169 | 182 | 5 | 8 | 4 | 2 | 4 | 91 | 101 | 95 | 70 |
| SE | 45821 | ENSG00000108654 | DDX5 | chr17 | - | 64506075 | 64506262 | 64504676 | 64504842 | 64506754 | 64507314 | 2316 | 1459 | 1896 | 2715 | 1015 | 6 | 6 | 8 | 18 | 12 | 824 | 1821 | 1623 | 1349 |
| SE | 96326 | ENSG00000135387 | CAPRIN1 | chr11 | + | 34097195 | 34097296 | 34096478 | 34096673 | 34099302 | 34099332 | 177 | 121 | 164 | 151 | 139 | 0 | 0 | 2 | 0 | 2 | 171 | 134 | 132 | 152 |
| SE | 80772 | ENSG00000136280 | CCM2 | chr7 | + | 45068442 | 45068579 | 45064462 | 45064646 | 45072725 | 45072783 | 22 | 8 | 27 | 19 | 24 | 9 | 6 | 11 | 8 | 14 | 45 | 20 | 6 | 35 |
| SE | 8944 | ENSG00000132478 | UNK | chr17 | + | 75792312 | 75792478 | 75784848 | 75784984 | 75809759 | 75809969 | 0 | 0 | 6 | 0 | 1 | 15 | 4 | 17 | 9 | 0 | 0 | 8 | 7 | 0 |
| SE | 93474 | ENSG00000136826 | KLF4 | chr9 | - | 107487294 | 107488267 | 107487027 | 107487192 | 107488929 | 107489050 | 58 | 23 | 29 | 91 | 30 | 2 | 0 | 1 | 5 | 0 | 136 | 35 | 21 | 101 |
| SE | 48657 | ENSG00000131374 | TBC1D5 | chr3 | - | 17622647 | 17622765 | 17518901 | 17518995 | 17623848 | 17623913 | 43 | 23 | 31 | 40 | 90 | 13 | 9 | 5 | 9 | 9 | 21 | 21 | 26 | 48 |
| SE | 68105 | ENSG00000135838 | NPL | chr1 | + | 182822114 | 182822199 | 182818812 | 182818859 | 182825780 | 182825820 | 76 | 41 | 59 | 70 | 103 | 16 | 3 | 6 | 10 | 5 | 87 | 99 | 80 | 163 |
| SE | 55594 | ENSG00000137275 | RIPK1 | chr6 | + | 3089580 | 3089657 | 3083084 | 3085408 | 3104224 | 3104315 | 141 | 129 | 126 | 167 | 143 | 3 | 0 | 0 | 3 | 0 | 75 | 134 | 170 | 134 |
| SE | 94593 | ENSG00000106829 | TLE4 | chr9 | + | 79627373 | 79627448 | 79576132 | 79576177 | 79652589 | 79652629 | 10 | 8 | 10 | 7 | 13 | 1 | 0 | 5 | 3 | 0 | 1 | 9 | 8 | 0 |
| SE | 57727 | ENSG00000114354 | TFG | chr3 | + | 100710970 | 100711068 | 100709461 | 100709721 | 100713642 | 100713869 | 2 | 7 | 1 | 2 | 0 | 26 | 12 | 22 | 14 | 9 | 5 | 0 | 4 | 7 |
| SE | 105963 | ENSG00000102225 | CDK16 | chrX | + | 47222260 | 47222380 | 47218690 | 47219105 | 47223551 | 47223759 | 0 | 2 | 0 | 1 | 11 | 12 | 6 | 9 | 10 | 0 | 23 | 12 | 11 | 5 |
| SE | 75124 | ENSG00000112343 | TRIM38 | chr6 | + | 25969324 | 25969420 | 25963048 | 25963282 | 25971868 | 25972099 | 214 | 94 | 177 | 181 | 128 | 2 | 1 | 1 | 0 | 1 | 109 | 195 | 168 | 217 |
| SE | 44812 | ENSG00000177873 | ZNF619 | chr3 | + | 40482199 | 40482349 | 40481862 | 40482016 | 40482587 | 40482704 | 0 | 0 | 0 | 0 | 1 | 10 | 6 | 17 | 11 | 1 | 16 | 0 | 0 | 0 |
| SE | 9373 | ENSG00000121067 | SPOP | chr17 | - | 49618503 | 49618563 | 49611279 | 49611457 | 49618980 | 49619108 | 4 | 2 | 3 | 4 | 1 | 114 | 86 | 142 | 157 | 102 | 0 | 5 | 7 | 8 |
| SE | 30541 | ENSG00000132182 | NUP210 | chr3 | - | 13371833 | 13372032 | 13365945 | 13366091 | 13375503 | 13375641 | 277 | 109 | 204 | 216 | 251 | 0 | 0 | 0 | 0 | 0 | 179 | 261 | 93 | 123 |
| SE | 99952 | ENSG00000142102 | PGGHG | chr11 | + | 292885 | 292997 | 292545 | 292677 | 293067 | 293235 | 37 | 9 | 51 | 54 | 28 | 0 | 0 | 0 | 0 | 0 | 14 | 30 | 42 | 33 |
| SE | 41992 | ENSG00000172113 | NME6 | chr3 | - | 48296726 | 48296829 | 48294196 | 48294803 | 48298426 | 48298523 | 29 | 12 | 27 | 17 | 8 | 0 | 0 | 0 | 0 | 0 | 37 | 11 | 2 | 14 |
| SE | 28243 | ENSG00000157326 | DHRS4 | chr14 | + | 23966282 | 23966417 | 23959901 | 23960003 | 23967210 | 23967266 | 28 | 16 | 31 | 25 | 25 | 5 | 0 | 4 | 1 | 3 | 14 | 34 | 9 | 42 |

[illegible]

| | | | | | | | | | | | | | |
|---|---|---|---|---|---|---|---|---|---|---|---|---|---|
| [illegible] | | | | | | | | | | | | | |
| SE | 106488 | ENSG00000176915 | ANKLE2 | chr12 | - | 132741808 | 132741891 | 132741418 | 132741485 | 132743153 | 132743276 | 7 | 4 | 12 |
| SE | 109017 | ENSG00000137822 | TUBGCP4 | chr15 | + | 43403682 | 43403842 | 43401817 | 43401850 | 43404412 | 43404552 | 5 | 16 | 9 |
| SE | 61194 | ENSG00000185619 | PCGF3 | chr4 | + | 732443 | 732498 | 730969 | 731110 | 733671 | 733789 | 55 | 50 | 45 |
| SE | 42090 | ENSG00000026751 | SLAMF7 | chr1 | + | 160750303 | 160750423 | 160749820 | 160750093 | 160751344 | 160751448 | 267 | 149 | 213 |
| SE | 52821 | ENSG00000169764 | UGP2 | chr2 | + | 63859179 | 63859391 | 63857828 | 63857936 | 63862840 | 63862928 | 2 | 3 | 3 |
| SE | 95262 | ENSG00000165219 | GAPVD1 | chr9 | + | 125295457 | 125295574 | 125268935 | 125268984 | 125298889 | 125299106 | 12 | 9 | 13 |
| SE | 72291 | ENSG00000006062 | MAP3K14 | chr17 | - | 45312284 | 45312437 | 45290489 | 45290765 | 45316959 | 45317029 | 0 | 0 | 0 |
| SE | 57920 | ENSG00000110200 | ANAPC15 | chr11 | - | 72111156 | 72111281 | 72110543 | 72110603 | 72111411 | 72111496 | 62 | 30 | 20 |
| SE | 37597 | ENSG00000082074 | FYB1 | chr5 | - | 39208863 | 39208964 | 39202955 | 39202987 | 39274402 | 39274528 | 19 | 4 | 2 |
| SE | 103184 | ENSG00000011376 | LARS2 | chr3 | + | 45519692 | 45519790 | 45517962 | 45518072 | 45520218 | 45520296 | 2 | 1 | 4 |
| SE | 31860 | ENSG00000143669 | LYST | chr1 | - | 235702746 | 235702977 | 235697082 | 235697272 | 235709090 | 235709308 | 197 | 240 | 140 |
| SE | 77820 | ENSG00000134250 | NOTCH2 | chr1 | - | 120097179 | 120097279 | 120029905 | 120029987 | 120100653 | 120100779 | 9 | 7 | 12 |
| SE | 58160 | ENSG00000160688 | FLAD1 | chr1 | + | 154983841 | 154984066 | 154983358 | 154983397 | 154988104 | 154988433 | 91 | 39 | 55 |
| SE | 35970 | ENSG00000078070 | MCCC1 | chr3 | - | 183020129 | 183020237 | 183015217 | 183015566 | 183022416 | 183022554 | 60 | 34 | 39 |
| SE | 107274 | ENSG00000090686 | USP48 | chr1 | - | 21703511 | 21703618 | 21701497 | 21701602 | 21704261 | 21704392 | 342 | 303 | 262 |
| SE | 38318 | ENSG00000143742 | SRP9 | chr1 | + | 225786861 | 225786985 | 225783299 | 225783368 | 225789239 | 225790466 | 99 | 45 | 103 |
| SE | 49617 | ENSG00000178927 | CYBC1 | chr17 | - | 82446622 | 82446692 | 82445863 | 82445960 | 82450699 | 82450728 | 35 | 22 | 24 |
| SE | 66390 | ENSG00000128708 | HAT1 | chr2 | + | 171946707 | 171946783 | 171925536 | 171925641 | 171965337 | 171965517 | 59 | 29 | 54 |
| SE | 33039 | ENSG00000183323 | CCDC125 | chr5 | - | 69320236 | 69320345 | 69313984 | 69314043 | 69320464 | 69320580 | 72 | 41 | 47 |
| SE | 37212 | ENSG00000153551 | CMTM7 | chr3 | + | 32449453 | 32449552 | 32441839 | 32442013 | 32452391 | 32452473 | 81 | 52 | 106 |
| SE | 72833 | ENSG00000129158 | SERGEF | chr11 | - | 17959469 | 17959636 | 17878207 | 17878244 | 17992930 | 17992993 | 41 | 21 | 27 |
| SE | 94551 | ENSG00000132950 | ZMYM5 | chr13 | - | 19838699 | 19838985 | 19835476 | 19835689 | 19851354 | 19851448 | 59 | 80 | 60 |
| SE | 45838 | ENSG00000163714 | U2SURP | chr3 | + | 143023974 | 143024018 | 143022852 | 143023064 | 143027148 | 143027253 | 435 | 317 | 406 |
| SE | 46147 | ENSG00000126070 | AGO3 | chr1 | + | 35982620 | 35982712 | 35973374 | 35973511 | 36004340 | 36004475 | 21 | 12 | 14 |
| SE | 62428 | ENSG00000113761 | ZNF346 | chr5 | + | 177044391 | 177044533 | 177041777 | 177041870 | 177062057 | 177062151 | 5 | 7 | 12 |
| SE | 45412 | ENSG00000070190 | DAPP1 | chr4 | + | 99866556 | 99866618 | 99866033 | 99866121 | 99868116 | 99870154 | 6 | 25 | 6 |
| SE | 19757 | ENSG00000100614 | PPM1A | chr14 | + | 60268292 | 60268435 | 60250430 | 60250481 | 60282683 | 60283266 | 7 | 6 | 10 |
| SE | 86736 | ENSG00000101868 | POLA1 | chrX | + | 24703250 | 24703347 | 24699424 | 24699549 | 24704388 | 24704469 | 41 | 31 | 30 |
| SE | 71959 | ENSG00000160710 | ADAR | chr1 | - | 154589368 | 154589462 | 154588124 | 154588673 | 154589756 | 154589928 | 533 | 408 | 518 |
| SE | 103095 | ENSG00000108055 | SMC3 | chr10 | + | 110582561 | 110582642 | 110581922 | 110582098 | 110583383 | 110583548 | 366 | 424 | 369 |
| SE | 49414 | ENSG00000107551 | RASSF4 | chr10 | + | 44982520 | 44982663 | 44970164 | 44971848 | 44984021 | 44984113 | 244 | 117 | 130 |
| SE | 17378 | ENSG00000092148 | HECTD1 | chr14 | - | 31113876 | 31114059 | 31113276 | 31113484 | 31114285 | 31114340 | 301 | 187 | 232 |
| SE | 55919 | ENSG00000122678 | POLM | chr7 | - | 44074130 | 44074220 | 44073782 | 44074025 | 44074397 | 44074530 | 5 | 7 | 5 |
| SE | 22873 | ENSG00000043143 | JADE2 | chr5 | + | 134576767 | 134576896 | 134573644 | 134573762 | 134578493 | 134578818 | 200 | 113 | 160 |
| SE | 74679 | ENSG00000197548 | ATG7 | chr3 | + | 11281036 | 11281102 | 11272308 | 11272430 | 11298685 | 11298855 | 2 | 3 | 2 |
| SE | 68745 | ENSG00000196141 | SPATS2L | chr2 | + | 200308886 | 200309090 | 200306375 | 200306601 | 200329430 | 200329480 | 0 | 0 | 0 |
| [illegible] | | | | | | | | | | | | | |

| | | | | | | | | | | | | | | | | | | | | | | | | | | | | | | | | | | | | | | | | | | | | | | |
|---|---|---|---|---|---|---|---|---|---|---|---|---|---|---|---|---|---|---|---|---|---|---|---|---|---|---|---|---|---|---|---|---|---|---|---|---|---|---|---|---|---|---|---|---|---|---|
| SE | 74808 | ENSG00000042088 | TDP1 | chr14 | + | 89956577 | 89956800 | 89955901 | 89955970 | 89966146 | 89966190 | 9 | 5 | 4 | 1 | 5 | 2 | 0 | 0 | 0 | 0 | 4 | 17 | 19 | 8 | 5 | 0 | 8 | 0 | 0 | 0 | 298 | 149 | 0.07150997 | 0.500352509 | 0.692 | 1 | 1 | 1 | 1 | 1 | 0.515 | 1 | 1 | 1 | 0.035 |
| SE | 699 | ENSG00000186575 | NF2 | chr22 | + | 29683019 | 29683079 | 29681438 | 29681601 | 29694751 | 29694811 | 1 | 0 | 1 | 2 | 6 | 66 | 17 | 45 | 57 | 25 | 9 | 6 | 2 | 0 | 0 | 15 | 51 | 19 | 64 | 68 | 209 | 149 | 0.071569762 | 0.500726128 | 0.011 | 0 | 0.016 | 0.024 | 0.146 | 0.3 | 0.077 | 0.07 | 0 | 0 | -0.05 |
| SE | 12943 | ENSG00000255439 | NA | chr16 | - | 31091091 | 31091342 | 31087805 | 31087826 | 31093311 | 31093421 | 40 | 12 | 25 | 27 | 7 | 1 | 0 | 1 | 0 | 5 | 40 | 45 | 13 | 30 | 99 | 0 | 0 | 0 | 0 | 0 | 298 | 149 | 0.071615641 | 0.501002343 | 0.952 | 1 | 0.926 | 1 | 0.412 | 1 | 1 | 1 | 1 | 1 | -0.142 |
| SE | 16517 | ENSG00000140367 | UBE2Q2 | chr15 | + | 75876186 | 75876271 | 75873427 | 75873568 | 75879097 | 75879188 | 195 | 129 | 145 | 162 | 136 | 8 | 10 | 6 | 12 | 11 | 72 | 104 | 90 | 46 | 61 | 14 | 6 | 12 | 0 | 7 | 234 | 149 | 0.071652954 | 0.501218589 | 0.939 | 0.891 | 0.939 | 0.896 | 0.887 | 0.766 | 0.917 | 0.827 | 1 | 0.847 | 0.039 |
| SE | 21578 | ENSG00000075568 | TMEM131 | chr2 | - | 97775842 | 97776018 | 97766477 | 97766602 | 97792385 | 97792984 | 102 | 60 | 72 | 57 | 68 | 0 | 0 | 0 | 0 | 0 | 37 | 89 | 34 | 87 | 23 | 0 | 8 | 1 | 0 | 0 | 298 | 149 | 0.071663178 | 0.501245329 | 1 | 1 | 1 | 1 | 1 | 1 | 0.848 | 0.944 | 1 | 1 | 0.042 |
| SE | 48678 | ENSG00000038002 | AGA | chr4 | - | 177436297 | 177436351 | 177434383 | 177434489 | 177437404 | 177437519 | 43 | 23 | 36 | 45 | 44 | 1 | 4 | 1 | 3 | 0 | 27 | 36 | 19 | 33 | 19 | 0 | 0 | 2 | 1 | 0 | 203 | 149 | 0.071722256 | 0.501613731 | 0.969 | 0.808 | 0.964 | 0.917 | 1 | 1 | 1 | 0.875 | 0.96 | 1 | -0.035 |
| SE | 67145 | ENSG00000089060 | SLC8B1 | chr12 | - | 113320330 | 113320498 | 113318963 | 113319071 | 113320580 | 113320686 | 29 | 9 | 26 | 24 | 21 | 0 | 0 | 0 | 0 | 8 | 39 | 3 | 9 | 10 | 20 | 0 | 0 | 0 | 0 | 0 | 298 | 149 | 0.071757343 | 0.50181021 | 1 | 1 | 1 | 1 | 0.568 | 1 | 1 | 1 | 1 | 1 | -0.086 |
| SE | 101953 | ENSG00000122033 | MTIF3 | chr13 | - | 27449851 | 27449918 | 27445087 | 27445156 | 27450508 | 27450564 | 3 | 2 | 9 | 0 | 3 | 48 | 37 | 47 | 56 | 55 | 0 | 5 | 2 | 0 | 0 | 65 | 24 | 20 | 29 | 9 | 216 | 149 | 0.071767132 | 0.50181021 | 0.041 | 0.036 | 0.117 | 0 | 0.036 | 0 | 0.126 | 0.065 | 0 | 0 | 0.008 |
| SE | 3407 | ENSG00000204514 | ZNF814 | chr19 | - | 57885882 | 57885969 | 57876915 | 57877042 | 57888766 | 57888958 | 1 | 0 | 0 | 0 | 0 | 4 | 12 | 9 | 6 | 6 | 0 | 0 | 0 | 0 | 10 | 6 | 4 | 1 | 4 | 0 | 236 | 149 | 0.071769576 | 0.50181021 | 0.136 | 0 | 0 | 0 | 0 | 0 | 0 | 0 | 0 | 1 | -0.173 |
| SE | 57550 | ENSG00000147274 | RBMX | chrX | - | 136877915 | 136878086 | 136876599 | 136876655 | 136879016 | 136879158 | 368 | 288 | 383 | 410 | 522 | 7 | 4 | 6 | 1 | 5 | 486 | 402 | 423 | 303 | 307 | 12 | 1 | 0 | 0 | 0 | 298 | 149 | 0.071786766 | 0.501885581 | 0.963 | 0.973 | 0.97 | 0.995 | 0.981 | 0.953 | 0.995 | 1 | 1 | 1 | -0.013 |
| SE | 47626 | ENSG00000110619 | CARS1 | chr11 | - | 3028995 | 3029084 | 3026675 | 3026797 | 3029302 | 3029443 | 114 | 39 | 97 | 74 | 85 | 0 | 2 | 0 | 0 | 0 | 90 | 93 | 77 | 84 | 173 | 8 | 0 | 4 | 0 | 0 | 238 | 149 | 0.07182457 | 0.502105045 | 1 | 0.924 | 1 | 1 | 1 | 0.876 | 1 | 0.923 | 1 | 1 | 0.025 |
| SE | 25592 | ENSG00000105341 | DMAC2 | chr19 | - | 41436391 | 41436472 | 41433439 | 41433673 | 41438217 | 41438414 | 63 | 16 | 31 | 34 | 30 | 10 | 5 | 12 | 2 | 16 | 27 | 40 | 7 | 41 | 162 | 11 | 25 | 14 | 12 | 40 | 230 | 149 | 0.071835079 | 0.502133677 | 0.803 | 0.675 | 0.626 | 0.917 | 0.548 | 0.614 | 0.509 | 0.245 | 0.689 | 0.724 | 0.158 |
| SE | 99099 | ENSG00000110274 | CEP164 | chr11 | + | 117382795 | 117382942 | 117381700 | 117381868 | 117390776 | 117390908 | 4 | 2 | 3 | 5 | 3 | 0 | 1 | 0 | 4 | 0 | 18 | 4 | 8 | 6 | 25 | 10 | 1 | 1 | 0 | 0 | 296 | 149 | 0.071855898 | 0.502234365 | 1 | 0.502 | 1 | 0.386 | 1 | 0.475 | 0.668 | 0.801 | 1 | 1 | -0.011 |
| SE | 100561 | ENSG00000054654 | SYNE2 | chr14 | + | 64221575 | 64221704 | 64220436 | 64220637 | 64223188 | 64223380 | 255 | 101 | 131 | 138 | 120 | 0 | 0 | 0 | 2 | 1 | 57 | 81 | 73 | 15 | 39 | 1 | 4 | 0 | 2 | 2 | 278 | 149 | 0.071880429 | 0.502360984 | 1 | 1 | 1 | 0.974 | 0.985 | 0.968 | 0.916 | 1 | 0.801 | 0.913 | 0.072 |
| SE | 91098 | ENSG00000185900 | POMK | chr8 | + | 43102504 | 43102600 | 43097523 | 43097615 | 43103527 | 43103830 | 35 | 10 | 64 | 43 | 30 | 5 | 0 | 5 | 0 | 1 | 4 | 11 | 4 | 31 | 4 | 0 | 2 | 2 | 0 | 0 | 245 | 149 | 0.071897796 | 0.502437505 | 0.81 | 1 | 0.886 | 1 | 0.948 | 1 | 0.77 | 0.549 | 1 | 1 | 0.065 |
| SE | 19248 | ENSG00000140943 | MBTPS1 | chr16 | - | 84087357 | 84087460 | 84081746 | 84081908 | 84090874 | 84090920 | 85 | 77 | 95 | 84 | 137 | 3 | 1 | 0 | 1 | 0 | 289 | 142 | 161 | 202 | 135 | 0 | 0 | 16 | 0 | 0 | 252 | 149 | 0.071928698 | 0.502608597 | 0.944 | 0.979 | 1 | 0.98 | 1 | 1 | 1 | 0.856 | 1 | 1 | 0.009 |
| SE | 9691 | ENSG00000214176 | PLEKHM1P1 | chr17 | - | 64824624 | 64824872 | 64821766 | 64822393 | 64829160 | 64829208 | 107 | 49 | 64 | 46 | 33 | 2 | 6 | 9 | 13 | 7 | 183 | 68 | 45 | 150 | 154 | 32 | 19 | 28 | 37 | 9 | 298 | 149 | 0.071968225 | 0.502839919 | 0.964 | 0.803 | 0.78 | 0.639 | 0.702 | 0.741 | 0.642 | 0.446 | 0.67 | 0.895 | 0.099 |
| SE | 51974 | ENSG00000071054 | MAP4K4 | chr2 | + | 101855976 | 101856138 | 101844100 | 101844311 | 101858995 | 101859082 | 296 | 243 | 303 | 363 | 544 | 1 | 0 | 1 | 1 | 5 | 605 | 385 | 510 | 656 | 490 | 2 | 0 | 7 | 14 | 3 | 298 | 149 | 0.071977225 | 0.502857928 | 0.993 | 1 | 0.993 | 0.995 | 0.982 | 0.993 | 1 | 0.973 | 0.959 | 0.988 | 0.01 |
| SE | 28079 | ENSG00000084676 | NCOA1 | chr2 | + | 24584475 | 24584560 | 24564294 | 24564430 | 24643965 | 24644122 | 287 | 210 | 206 | 288 | 411 | 11 | 5 | 5 | 9 | 6 | 325 | 365 | 370 | 208 | 204 | 0 | 9 | 1 | 2 | 12 | 234 | 149 | 0.07205963 | 0.503384011 | 0.943 | 0.964 | 0.963 | 0.953 | 0.978 | 1 | 0.963 | 0.996 | 0.985 | 0.915 | -0.012 |
| SE | 59715 | ENSG00000183530 | PRR14L | chr22 | - | 31738386 | 31738911 | 31725537 | 31725610 | 31750071 | 31750140 | 129 | 77 | 72 | 108 | 122 | 2 | 0 | 2 | 0 | 7 | 16 | 120 | 40 | 30 | 20 | 0 | 0 | 0 | 0 | 0 | 298 | 149 | 0.072065385 | 0.503384011 | 0.97 | 1 | 0.947 | 1 | 0.897 | 1 | 1 | 1 | 1 | 1 | -0.037 |
| SE | 25067 | ENSG00000171206 | TRIM8 | chr10 | + | 102656746 | 102656920 | 102656269 | 102656385 | 102657922 | 102658315 | 241 | 152 | 171 | 189 | 191 | 0 | 0 | 0 | 0 | 4 | 220 | 210 | 137 | 127 | 348 | 2 | 7 | 1 | 0 | 5 | 298 | 149 | 0.072077708 | 0.503425179 | 1 | 1 | 1 | 1 | 0.96 | 0.982 | 0.938 | 0.986 | 1 | 0.972 | 0.016 |
| SE | 71188 | ENSG00000057657 | PRDM1 | chr6 | + | 106095614 | 106095734 | 106088200 | 106088449 | 106099299 | 106099552 | 92 | 75 | 160 | 94 | 139 | 0 | 0 | 0 | 0 | 0 | 86 | 296 | 415 | 117 | 84 | 14 | 0 | 0 | 0 | 0 | 269 | 149 | 0.072095298 | 0.503503118 | 1 | 1 | 1 | 1 | 1 | 0.773 | 1 | 1 | 1 | 1 | 0.045 |
| SE | 11452 | ENSG00000187555 | USP7 | chr16 | - | 8899120 | 8899188 | 8898530 | 8898639 | 8900989 | 8901057 | 180 | 123 | 127 | 190 | 119 | 2 | 3 | 1 | 0 | 1 | 53 | 171 | 161 | 206 | 64 | 6 | 3 | 2 | 3 | 0 | 217 | 149 | 0.072108714 | 0.503551899 | 0.984 | 0.966 | 0.989 | 1 | 0.988 | 0.858 | 0.975 | 0.982 | 0.979 | 1 | 0.027 |
| SE | 5305 | ENSG00000125900 | SIRPD | chr20 | - | 1537154 | 1537313 | 1534250 | 1534441 | 1551690 | 1552038 | 4 | 3 | 3 | 3 | 4 | 2 | 0 | 0 | 4 | 0 | 41 | 23 | 17 | 21 | 19 | 0 | 3 | 0 | 0 | 0 | 298 | 149 | 0.072206564 | 0.504190235 | 0.5 | 1 | 1 | 0.273 | 1 | 1 | 0.793 | 1 | 1 | 1 | -0.204 |
| SE | 35495 | ENSG00000051382 | PIK3CB | chr3 | - | 138688874 | 138688974 | 138684624 | 138684803 | 138690999 | 138691143 | 174 | 154 | 170 | 226 | 241 | 6 | 7 | 1 | 5 | 5 | 63 | 228 | 222 | 248 | 229 | 12 | 6 | 1 | 14 | 5 | 249 | 149 | 0.072217565 | 0.504222088 | 0.946 | 0.929 | 0.99 | 0.964 | 0.966 | 0.759 | 0.958 | 0.993 | 0.914 | 0.965 | 0.041 |
| SE | 104708 | ENSG00000110888 | CAPRIN2 | chr12 | - | 30733628 | 30733711 | 30731342 | 30731510 | 30734967 | 30735014 | 4 | 8 | 12 | 14 | 33 | 84 | 63 | 87 | 80 | 101 | 0 | 5 | 5 | 1 | 16 | 70 | 63 | 43 | 50 | 22 | 232 | 149 | 0.072238842 | 0.50431155 | 0.03 | 0.075 | 0.081 | 0.101 | 0.173 | 0 | 0.048 | 0.069 | 0.013 | 0.318 | 0.002 |
| SE | 100248 | ENSG00000084234 | APLP2 | chr11 | + | 130137255 | 130137291 | 130135562 | 130135715 | 130140397 | 130140483 | 190 | 95 | 114 | 122 | 109 | 281 | 195 | 167 | 251 | 219 | 661 | 435 | 235 | 432 | 469 | 1253 | 453 | 289 | 793 | 700 | 185 | 149 | 0.072243261 | 0.50431155 | 0.353 | 0.282 | 0.355 | 0.281 | 0.286 | 0.298 | 0.436 | 0.396 | 0.305 | 0.35 | -0.046 |
| SE | 50672 | ENSG00000142687 | KIAA0319L | chr1 | - | 35444160 | 35444303 | 35442905 | 35443028 | 35451627 | 35451776 | 76 | 36 | 47 | 59 | 58 | 0 | 0 | 0 | 0 | 0 | 87 | 63 | 48 | 61 | 76 | 0 | 0 | 0 | 0 | 10 | 292 | 149 | 0.072357182 | 0.505061771 | 1 | 1 | 1 | 1 | 1 | 1 | 1 | 1 | 1 | 0.795 | 0.041 |
| SE | 15329 | ENSG00000114062 | UBE3A | chr15 | - | 25415714 | 25415842 | 25411907 | 25411971 | 25420655 | 25420733 | 6 | 5 | 25 | 11 | 19 | 0 | 0 | 0 | 0 | 1 | 29 | 19 | 13 | 8 | 5 | 6 | 0 | 1 | 0 | 0 | 277 | 149 | 0.072406564 | 0.505361411 | 1 | 1 | 1 | 1 | 0.911 | 0.722 | 1 | 0.875 | 1 | 1 | 0.063 |
| SE | 42513 | ENSG00000172071 | EIF2AK3 | chr2 | - | 88593912 | 88594108 | 88593271 | 88593405 | 88595468 | 88595663 | 9 | 5 | 9 | 13 | 13 | 82 | 53 | 70 | 70 | 76 | 8 | 11 | 17 | 3 | 6 | 15 | 47 | 120 | 19 | 0 | 298 | 149 | 0.072426994 | 0.505458945 | 0.052 | 0.045 | 0.06 | 0.085 | 0.079 | 0.211 | 0.105 | 0.066 | 0.073 | 1 | -0.227 |
| SE | 57423 | ENSG00000042445 | RETSAT | chr2 | - | 85343638 | 85343798 | 85341954 | 85343381 | 85344238 | 85344348 | 18 | 12 | 11 | 8 | 14 | 0 | 0 | 0 | 0 | 0 | 2 | 18 | 24 | 11 | 52 | 5 | 0 | 1 | 0 | 0 | 298 | 149 | 0.072439391 | 0.505500407 | 1 | 1 | 1 | 1 | 1 | 0.167 | 1 | 0.923 | 1 | 1 | 0.182 |
| SE | 105336 | ENSG00000172819 | RARG | chr12 | - | 53214445 | 53214606 | 53213939 | 53214235 | 53215292 | 53215352 | 86 | 63 | 80 | 108 | 67 | 0 | 0 | 0 | 0 | 0 | 90 | 68 | 14 | 58 | 165 | 0 | 0 | 8 | 0 | 0 | 298 | 149 | 0.072451844 | 0.505532494 | 1 | 1 | 1 | 1 | 1 | 1 | 1 | 0.467 | 1 | 1 | 0.107 |
| SE | 41342 | ENSG00000145287 | PLAC8 | chr4 | - | 83107803 | 83107950 | 83104895 | 83105020 | 83114665 | 83114715 | 1750 | 953 | 1369 | 1693 | 1319 | 89 | 70 | 127 | 72 | 188 | 2108 | 1263 | 747 | 1771 | 1347 | 177 | 166 | 110 | 157 | 60 | 296 | 149 | 0.072466756 | 0.505532494 | 0.908 | 0.873 | 0.844 | 0.922 | 0.779 | 0.857 | 0.793 | 0.774 | 0.85 | 0.919 | 0.027 |
| [illegible] | | | | | | | | | | | | | | | | | | | | | | | | | | | | | | | | | | | | | | | | | | | | | | |

| | | | | | | | | | | | | | | | | | | | | | | | | | |
|---|---|---|---|---|---|---|---|---|---|---|---|---|---|---|---|---|---|---|---|---|---|---|---|---|---|
| SE | 94363 | ENSG00000021776 | AQR | chr15 | - | 34927034 | 34927138 | 34920331 | 34920434 | 34930257 | 34930371 | 122 | 90 | 116 | 106 | 256 | 1 | 1 | 0 | 0 | 0 | 31 | 79 | 67 | 53 |
| SE | 17233 | ENSG00000197555 | SIPA1L1 | chr14 | + | 71497853 | 71497889 | 71418129 | 71418196 | 71512742 | 71512845 | 0 | 0 | 0 | 0 | 0 | 12 | 5 | 7 | 5 | 15 | 8 | 0 | 0 | 0 |
| SE | 40172 | ENSG00000115306 | SPTBN1 | chr2 | + | 54621399 | 54621512 | 54618077 | 54618193 | 54622299 | 54622487 | 618 | 452 | 1024 | 825 | 1164 | 4 | 2 | 6 | 8 | 0 | 1058 | 687 | 525 | 823 |
| SE | 96826 | ENSG00000152270 | PDE3B | chr11 | + | 14786436 | 14786532 | 14771936 | 14771987 | 14789105 | 14789242 | 108 | 94 | 182 | 136 | 251 | 3 | 0 | 6 | 0 | 5 | 17 | 71 | 100 | 34 |
| SE | 86240 | ENSG00000184634 | MED12 | chrX | + | 71137186 | 71137392 | 71136878 | 71137029 | 71137557 | 71137635 | 46 | 14 | 41 | 44 | 30 | 0 | 0 | 0 | 2 | 0 | 1 | 10 | 6 | 34 |
| SE | 83867 | ENSG00000130741 | EIF2S3 | chrX | + | 24073090 | 24073263 | 24071557 | 24071727 | 24076721 | 24078810 | 2782 | 2319 | 2892 | 2617 | 3633 | 22 | 5 | 26 | 15 | 22 | 2002 | 1079 | 1773 | 2399 |
| SE | 2717 | ENSG00000011600 | TYROBP | chr19 | - | 35907729 | 35907762 | 35907293 | 35907580 | 35908174 | 35908295 | 341 | 140 | 184 | 268 | 64 | 66 | 15 | 25 | 42 | 20 | 759 | 417 | 232 | 409 |
| SE | 63230 | ENSG00000068784 | SRBD1 | chr2 | - | 45551124 | 45551282 | 45547521 | 45547612 | 45553622 | 45553730 | 321 | 158 | 194 | 170 | 348 | 2 | 5 | 1 | 3 | 5 | 143 | 160 | 101 | 98 |
| SE | 705 | ENSG00000182944 | EWSR1 | chr22 | + | 29282389 | 29282557 | 29275628 | 29278216 | 29286922 | 29287106 | 766 | 376 | 703 | 641 | 831 | 14 | 5 | 10 | 12 | 12 | 832 | 824 | 522 | 979 |
| SE | 48136 | ENSG00000226688 | ENTPD1-AS1 | chr10 | - | 95904829 | 95904935 | 95872113 | 95876660 | 95907662 | 95907842 | 7 | 14 | 15 | 9 | 12 | 0 | 0 | 0 | 0 | 0 | 6 | 2 | 3 | 1 |
| SE | 74653 | ENSG00000114383 | TUSC2 | chr3 | - | 50327027 | 50327224 | 50326355 | 50326477 | 50327953 | 50328195 | 0 | 0 | 1 | 0 | 0 | 13 | 7 | 11 | 11 | 3 | 2 | 7 | 0 | 0 |
| SE | 106852 | ENSG00000166908 | PIP4K2C | chr12 | + | 57595125 | 57595222 | 57594024 | 57594122 | 57595887 | 57596031 | 66 | 28 | 52 | 48 | 74 | 0 | 0 | 0 | 0 | 0 | 29 | 85 | 68 | 47 |
| SE | 95741 | ENSG00000164978 | NUDT2 | chr9 | + | 34338712 | 34338847 | 34329505 | 34329599 | 34339023 | 34339166 | 50 | 48 | 68 | 99 | 75 | 1 | 0 | 1 | 1 | 0 | 105 | 36 | 49 | 64 |
| SE | 100720 | ENSG00000167286 | CD3D | chr11 | - | 118340374 | 118340593 | 118339450 | 118339494 | 118342552 | 118342631 | 1087 | 566 | 979 | 912 | 1241 | 6 | 2 | 6 | 8 | 5 | 338 | 932 | 454 | 368 |
| SE | 66437 | ENSG00000162227 | TAF6L | chr11 | + | 62781893 | 62781968 | 62778868 | 62778963 | 62782112 | 62782333 | 15 | 9 | 17 | 10 | 20 | 0 | 0 | 0 | 0 | 0 | 8 | 30 | 2 | 14 |
| SE | 6096 | ENSG00000171456 | ASXL1 | chr20 | + | 32371777 | 32371874 | 32369014 | 32369123 | 32428127 | 32428248 | 28 | 6 | 16 | 10 | 16 | 58 | 35 | 65 | 66 | 40 | 18 | 17 | 15 | 13 |
| SE | 72635 | ENSG00000153094 | BCL2L11 | chr2 | + | 111123732 | 111123869 | 111120913 | 111121188 | 111150043 | 111150147 | 85 | 22 | 57 | 52 | 29 | 4 | 6 | 0 | 2 | 0 | 43 | 117 | 100 | 129 |
| SE | 2290 | ENSG00000142002 | DPP9 | chr19 | - | 4683189 | 4683324 | 4682695 | 4682838 | 4683476 | 4683629 | 30 | 7 | 19 | 38 | 4 | 51 | 21 | 32 | 71 | 33 | 2 | 27 | 17 | 9 |
| SE | 106404 | ENSG00000089041 | P2RX7 | chr12 | + | 121162423 | 121162520 | 121156078 | 121156147 | 121165356 | 121165437 | 38 | 34 | 27 | 40 | 61 | 0 | 0 | 0 | 0 | 0 | 123 | 90 | 63 | 87 |
| SE | 69830 | ENSG00000163803 | PLB1 | chr2 | + | 28605852 | 28605948 | 28603965 | 28604047 | 28606495 | 28606567 | 7 | 1 | 1 | 7 | 1 | 0 | 1 | 0 | 0 | 0 | 7 | 3 | 0 | 9 |
| SE | 76402 | ENSG00000081377 | CDC14B | chr9 | - | 96509672 | 96509789 | 96500107 | 96503789 | 96515648 | 96515792 | 1 | 6 | 1 | 16 | 4 | 3 | 3 | 1 | 4 | 4 | 7 | 10 | 2 | 8 |
| SE | 6016 | ENSG00000130684 | ZNF337 | chr20 | - | 25685566 | 25685662 | 25673194 | 25677037 | 25685995 | 25686122 | 56 | 53 | 52 | 68 | 29 | 6 | 5 | 8 | 3 | 0 | 38 | 34 | 65 | 38 |
| SE | 17333 | ENSG00000092529 | CAPN3 | chr15 | + | 42408210 | 42408324 | 42403740 | 42403777 | 42409302 | 42409380 | 22 | 27 | 18 | 19 | 17 | 8 | 0 | 7 | 7 | 2 | 24 | 56 | 19 | 120 |
| SE | 95889 | ENSG00000080298 | RFX3 | chr9 | - | 3277339 | 3277461 | 3275499 | 3275612 | 3288130 | 3288250 | 66 | 30 | 60 | 63 | 92 | 2 | 1 | 4 | 1 | 0 | 26 | 22 | 60 | 23 |
| SE | 65608 | ENSG00000112078 | KCTD20 | chr6 | + | 36474788 | 36475062 | 36470051 | 36470257 | 36479590 | 36479711 | 139 | 106 | 70 | 159 | 217 | 3 | 3 | 1 | 2 | 11 | 175 | 117 | 81 | 126 |
| SE | 86382 | ENSG00000123562 | MORF4L2 | chrX | - | 103678498 | 103678620 | 103676825 | 103677051 | 103685170 | 103685260 | 285 | 253 | 316 | 335 | 509 | 15 | 5 | 22 | 15 | 19 | 415 | 194 | 306 | 247 |
| SE | 58256 | ENSG00000089127 | OAS1 | chr12 | + | 112917546 | 112917700 | 112913636 | 112916738 | 112931877 | 112933219 | 170 | 199 | 111 | 177 | 201 | 1 | 0 | 2 | 0 | 0 | 725 | 366 | 102 | 484 |
| SE | 72887 | ENSG00000121716 | PILRB | chr7 | + | 100353372 | 100354012 | 100353209 | 100353270 | 100355142 | 100355240 | 126 | 53 | 102 | 131 | 111 | 2 | 2 | 10 | 5 | 4 | 42 | 124 | 56 | 106 |
| SE | 46830 | ENSG00000155330 | C16orf87 | chr16 | - | 46824385 | 46824482 | 46796606 | 46803070 | 46831083 | 46831162 | 46 | 20 | 42 | 64 | 49 | 0 | 0 | 0 | 0 | 0 | 20 | 45 | 58 | 20 |
| SE | 85123 | ENSG00000285171 | NA | chrX | - | 71104768 | 71104837 | 71104297 | 71104404 | 71108276 | 71108346 | 6 | 6 | 15 | 9 | 29 | 0 | 0 | 0 | 0 | 0 | 7 | 17 | 23 | 11 |
| SE | 33232 | ENSG00000023330 | ALAS1 | chr3 | + | 52208082 | 52208247 | 52206571 | 52206751 | 52211282 | 52211551 | 117 | 61 | 106 | 116 | 142 | 3 | 1 | 0 | 1 | 2 | 232 | 104 | 85 | 280 |
| SE | 59249 | ENSG00000168118 | RAB4A | chr1 | + | 229286485 | 229286566 | 229271074 | 229271370 | 229295847 | 229295910 | 11 | 3 | 7 | 16 | 2 | 6 | 6 | 4 | 1 | 2 | 0 | 11 | 6 | 9 |
| SE | 16388 | ENSG00000156958 | GALK2 | chr15 | + | 49161739 | 49161861 | 49155758 | 49156016 | 49201161 | 49201250 | 9 | 3 | 3 | 14 | 2 | 21 | 4 | 12 | 12 | 22 | 0 | 5 | 3 | 5 |
| SE | 95803 | ENSG00000167721 | TSR1 | chr17 | - | 2330946 | 2331109 | 2330514 | 2330625 | 2332168 | 2332359 | 164 | 128 | 183 | 168 | 300 | 0 | 0 | 1 | 0 | 0 | 156 | 136 | 146 | 119 |
| SE | 87863 | ENSG00000156504 | PABIR2 | chrX | - | 134772142 | 134772283 | 134771095 | 134771354 | 134781820 | 134781917 | 73 | 38 | 49 | 71 | 61 | 75 | 40 | 41 | 59 | 51 | 33 | 36 | 34 | 34 |
| SE | 55305 | ENSG00000065457 | ADAT1 | chr16 | - | 75618585 | 75618698 | 75617141 | 75617272 | 75620265 | 75620334 | 30 | 25 | 30 | 22 | 99 | 9 | 8 | 11 | 10 | 1 | 3 | 30 | 20 | 15 |
| SE | 24729 | ENSG00000158987 | RAPGEF6 | chr5 | - | 131440145 | 131440228 | 131439580 | 131439715 | 131442348 | 131442537 | 6 | 2 | 10 | 5 | 1 | 127 | 100 | 183 | 126 | 172 | 3 | 0 | 1 | 0 |
| SE | 96425 | ENSG00000159314 | ARHGAP27 | chr17 | - | 45404268 | 45404334 | 45404028 | 45404096 | 45404444 | 45404528 | 109 | 55 | 53 | 70 | 40 | 33 | 8 | 20 | 27 | 9 | 62 | 106 | 61 | 73 |
| SE | 28936 | ENSG00000112079 | STK38 | chr6 | - | 36507499 | 36507602 | 36506582 | 36506644 | 36515337 | 36515492 | 739 | 488 | 549 | 571 | 854 | 5 | 3 | 1 | 3 | 2 | 353 | 559 | 244 | 252 |
| SE | 67709 | ENSG00000145332 | KLHL8 | chr4 | - | 87185250 | 87185571 | 87183202 | 87183389 | 87195323 | 87195690 | 28 | 22 | 15 | 17 | 40 | 18 | 11 | 7 | 15 | 20 | 5 | 12 | 11 | 20 |
| SE | 11814 | ENSG00000188603 | CLN3 | chr16 | - | 28487661 | 28487741 | 28486577 | 28487541 | 28488590 | 28488604 | 78 | 23 | 40 | 49 | 51 | 0 | 3 | 6 | 1 | 0 | 67 | 96 | 46 | 23 |
| SE | 47006 | ENSG00000122417 | ODF2L | chr1 | - | 86368635 | 86368722 | 86360425 | 86360536 | 86371017 | 86371153 | 1 | 1 | 7 | 1 | 11 | 33 | 56 | 60 | 45 | 200 | 0 | 3 | 1 | 0 |
| SE | 101806 | ENSG00000090621 | PABPC4 | chr1 | - | 39575758 | 39576223 | 39572392 | 39572586 | 39576551 | 39576790 | 179 | 41 | 147 | 167 | 55 | 0 | 0 | 3 | 1 | 1 | 178 | 117 | 109 | 98 |
| SE | 62410 | ENSG00000033867 | SLC4A7 | chr3 | - | 27421621 | 27421779 | 27420699 | 27420787 | 27424036 | 27424152 | 122 | 59 | 80 | 90 | 140 | 3 | 3 | 2 | 4 | 1 | 48 | 49 | 44 | 6 |
| SE | 30800 | ENSG00000100412 | ACO2 | chr22 | + | 41499725 | 41499862 | 41469105 | 41469182 | 41507790 | 41508049 | 192 | 79 | 143 | 136 | 184 | 1 | 1 | 4 | 1 | 0 | 162 | 187 | 86 | 171 |
| SE | 101213 | ENSG00000150433 | TMEM218 | chr11 | - | 125102131 | 125102351 | 125101200 | 125101303 | 125102636 | 125102809 | 5 | 3 | 5 | 9 | 11 | 1 | 1 | 1 | 2 | 1 | 3 | 2 | 4 | 1 |
| SE | 87193 | ENSG00000086758 | HUWE1 | chrX | - | 53558654 | 53558809 | 53557381 | 53557427 | 53558970 | 53559060 | 279 | 154 | 230 | 284 | 373 | 0 | 0 | 0 | 0 | 0 | 348 | 272 | 234 | 305 |
| SE | 57479 | ENSG00000154845 | PPP4R1 | chr18 | - | 9583116 | 9583275 | 9577063 | 9577191 | 9584514 | 9584580 | 209 | 100 | 170 | 183 | 173 | 3 | 1 | 2 | 0 | 0 | 111 | 223 | 177 | 132 |
| SE | 98563 | ENSG00000134955 | SLC37A2 | chr11 | + | 125079111 | 125079247 | 125077449 | 125077528 | 125079683 | 125079760 | 58 | 28 | 37 | 89 | 48 | 0 | 0 | 0 | 0 | 0 | 118 | 57 | 26 | 20 |
| SE | 21061 | ENSG00000196182 | STK40 | chr1 | - | 36358736 | 36358822 | 36358238 | 36358382 | 36361220 | 36361340 | 220 | 157 | 148 | 237 | 167 | 1 | 1 | 1 | 3 | 8 | 371 | 189 | 106 | 222 |
| SE | 41751 | ENSG00000143222 | UFC1 | chr1 | + | 161157616 | 161157693 | 161156949 | 161157017 | 161158411 | 161158682 | 20 | 12 | 12 | 16 | 39 | 0 | 0 | 0 | 2 | 0 | 79 | 51 | 37 | 71 |
| SE | 106493 | ENSG00000185482 | STAC3 | chr12 | - | 57245144 | 57245211 | 57244756 | 57244965 | 57246803 | 57246901 | 44 | 52 | 29 | 55 | 91 | 0 | 0 | 4 | 1 | 4 | 283 | 129 | 56 | 204 |
| SE | 53932 | ENSG00000138614 | INTS14 | chr15 | - | 65607301 | 65607442 | 65605128 | 65605236 | 65610795 | 65611121 | 6 | 6 | 8 | 13 | 3 | 0 | 2 | 6 | 0 | 0 | 4 | 16 | 5 | 2 |
| SE | 73589 | ENSG00000147443 | DOK2 | chr8 | - | 21912228 | 21912510 | 21911900 | 21911988 | 21913538 | 21913676 | 155 | 107 | 106 | 168 | 105 | 6 | 0 | 1 | 3 | 2 | 168 | 122 | 63 | 105 |
| SE | 26307 | ENSG00000105875 | WDR91 | chr7 | - | 135188432 | 135188545 | 135186971 | 135187169 | 135189343 | 135189452 | 26 | 17 | 35 | 44 | 16 | 5 | 0 | 2 | 2 | 0 | 25 | 44 | 45 | 28 |
| SE | 48501 | ENSG00000138095 | LRPPRC | chr2 | - | 43896633 | 43896708 | 43894544 | 43894629 | 43899218 | 43899334 | 218 | 157 | 213 | 183 | 470 | 23 | 10 | 3 | 24 | 19 | 6 | 80 | 98 | 7 |
| SE | 74124 | ENSG00000093167 | LRRFIP2 | chr3 | - | 37112914 | 37112980 | 37096615 | 37096660 | 37121491 | 37121536 | 0 | 0 | 1 | 1 | 0 | 12 | 12 | 14 | 5 | 15 | 5 | 0 | 3 | 0 |
| SE | 43785 | ENSG00000138002 | IFT172 | chr2 | - | 27476640 | 27476726 | 27471232 | 27472362 | 27477216 | 27477320 | 25 | 12 | 18 | 22 | 10 | 0 | 0 | 0 | 0 | 0 | 14 | 16 | 8 | 12 |
| SE | 80255 | ENSG00000164535 | DAGLB | chr7 | - | 6424751 | 6424835 | 6421726 | 6421804 | 6430479 | 6430607 | 27 | 13 | 19 | 29 | 21 | 0 | 0 | 0 | 0 | 0 | 6 | 16 | 8 | 8 |
| [illegible] | | | | | | | | | | | | | | | | | | | | | | | | | |

| | | | | | | | | | | | | | | | | | | | | | | | | | | | | | | | | | | | | | | | | | | | | | | |
|---|---|---|---|---|---|---|---|---|---|---|---|---|---|---|---|---|---|---|---|---|---|---|---|---|---|---|---|---|---|---|---|---|---|---|---|---|---|---|---|---|---|---|---|---|---|---|
| SE | 17445 | ENSG00000140563 | MCTP2 | chr15 | + | 94440175 | 94440298 | 94401899 | 94402019 | 94442918 | 94442960 | 123 | 94 | 105 | 152 | 143 | 53 | 2 | 2 | 9 | 3 | 65 | 134 | 127 | 83 | 46 | 6 | 15 | 28 | 10 | 0 | 272 | 149 | 0.085803552 | 0.576357138 | 0.56 | 0.963 | 0.966 | 0.902 | 0.963 | 0.856 | 0.83 | 0.713 | 0.82 | 1 | 0.027 |
| SE | 57755 | ENSG00000118007 | STAG1 | chr3 | - | 136518247 | 136518415 | 136502627 | 136502779 | 136521212 | 136521417 | 10 | 20 | 12 | 22 | 5 | 258 | 237 | 226 | 278 | 545 | 5 | 6 | 20 | 8 | 2 | 60 | 204 | 216 | 74 | 17 | 298 | 149 | 0.085810197 | 0.576357138 | 0.019 | 0.04 | 0.026 | 0.038 | 0.005 | 0.04 | 0.014 | 0.044 | 0.051 | 0.056 | -0.015 |
| SE | 105969 | ENSG00000102225 | CDK16 | chrX | + | 47225771 | 47225866 | 47224987 | 47225102 | 47225964 | 47226027 | 115 | 83 | 120 | 126 | 151 | 6 | 2 | 4 | 1 | 1 | 188 | 140 | 144 | 143 | 355 | 17 | 3 | 2 | 11 | 0 | 244 | 149 | 0.085873633 | 0.576733742 | 0.921 | 0.962 | 0.948 | 0.987 | 0.989 | 0.871 | 0.966 | 0.978 | 0.888 | 1 | 0.021 |
| SE | 30633 | ENSG00000189308 | LIN54 | chr4 | - | 82970326 | 82970469 | 82946257 | 82946474 | 82978882 | 82979006 | 104 | 95 | 103 | 117 | 169 | 6 | 3 | 4 | 1 | 3 | 125 | 119 | 152 | 123 | 77 | 6 | 0 | 0 | 4 | 0 | 292 | 149 | 0.08601764 | 0.577629083 | 0.898 | 0.942 | 0.929 | 0.984 | 0.966 | 0.914 | 1 | 1 | 0.94 | 1 | -0.027 |
| SE | 4896 | ENSG00000124243 | BCAS4 | chr20 | + | 50840536 | 50840930 | 50830278 | 50830380 | 50841765 | 50841900 | 0 | 0 | 1 | 0 | 10 | 15 | 11 | 34 | 18 | 39 | 0 | 0 | 0 | 0 | 0 | 17 | 27 | 14 | 14 | 27 | 298 | 149 | 0.086021701 | 0.577629083 | 0 | 0 | 0.014 | 0 | 0.114 | 0 | 0 | 0 | 0 | 0 | 0.026 |
| SE | 32753 | ENSG00000136521 | NDUFB5 | chr3 | + | 179614970 | 179615059 | 179604689 | 179604939 | 179615982 | 179616049 | 295 | 134 | 216 | 264 | 295 | 3 | 1 | 0 | 3 | 2 | 235 | 234 | 213 | 301 | 309 | 5 | 5 | 0 | 5 | 4 | 238 | 149 | 0.086059407 | 0.577832719 | 0.984 | 0.988 | 1 | 0.982 | 0.989 | 0.967 | 0.967 | 1 | 0.974 | 0.98 | 0.011 |
| SE | 65768 | ENSG00000146463 | ZMYM4 | chr1 | + | 35413971 | 35414083 | 35405372 | 35405468 | 35415465 | 35415714 | 74 | 46 | 82 | 74 | 118 | 1 | 0 | 0 | 0 | 0 | 8 | 29 | 34 | 29 | 32 | 7 | 0 | 0 | 0 | 0 | 261 | 149 | 0.086067003 | 0.577834169 | 0.977 | 1 | 1 | 1 | 1 | 0.395 | 1 | 1 | 1 | 1 | 0.116 |
| SE | 38931 | ENSG00000067066 | SP100 | chr2 | + | 230449560 | 230449710 | 230449087 | 230449150 | 230461261 | 230461414 | 307 | 249 | 277 | 308 | 684 | 12 | 6 | 7 | 12 | 2 | 182 | 372 | 342 | 312 | 217 | 0 | 8 | 8 | 5 | 0 | 298 | 149 | 0.086078652 | 0.577862824 | 0.927 | 0.954 | 0.952 | 0.928 | 0.994 | 1 | 0.959 | 0.955 | 0.969 | 1 | -0.026 |
| SE | 60659 | ENSG00000147099 | HDAC8 | chrX | - | 72494195 | 72494381 | 72490928 | 72491006 | 72495155 | 72495268 | 3 | 8 | 10 | 11 | 5 | 15 | 13 | 11 | 8 | 15 | 0 | 14 | 0 | 4 | 2 | 7 | 21 | 9 | 8 | 7 | 298 | 149 | 0.086146749 | 0.578270393 | 0.091 | 0.235 | 0.312 | 0.407 | 0.143 | 0 | 0.25 | 0 | 0.2 | 0.125 | 0.123 |
| SE | 4066 | ENSG00000099326 | MZF1 | chr19 | - | 58572549 | 58572719 | 58571199 | 58571429 | 58573054 | 58573311 | 0 | 0 | 9 | 2 | 0 | 5 | 2 | 13 | 4 | 4 | 33 | 4 | 2 | 0 | 21 | 8 | 16 | 0 | 1 | 24 | 298 | 149 | 0.086178948 | 0.578436939 | 0 | 0 | 0.257 | 0.2 | 0 | 0.673 | 0.111 | 1 | 0 | 0.304 | -0.326 |
| SE | 83109 | ENSG00000146909 | NOM1 | chr7 | + | 156958605 | 156958856 | 156954111 | 156954298 | 156959850 | 156960118 | 4 | 2 | 7 | 6 | 5 | 42 | 20 | 30 | 41 | 49 | 10 | 0 | 0 | 4 | 0 | 81 | 32 | 16 | 29 | 18 | 298 | 149 | 0.0861916 | 0.57847227 | 0.045 | 0.048 | 0.104 | 0.068 | 0.049 | 0.058 | 0 | 0 | 0.065 | 0 | 0.038 |
| SE | 37485 | ENSG00000163681 | SLMAP | chr3 | + | 57908167 | 57908227 | 57907883 | 57908006 | 57909075 | 57909150 | 0 | 2 | 1 | 0 | 0 | 88 | 93 | 90 | 93 | 160 | 0 | 0 | 0 | 0 | 8 | 4 | 48 | 53 | 33 | 0 | 209 | 149 | 0.086199861 | 0.57847813 | 0 | 0.015 | 0.008 | 0 | 0 | 0 | 0 | 0 | 0 | 1 | -0.195 |
| SE | 6330 | ENSG00000166347 | CYB5A | chr18 | - | 74256829 | 74256853 | 74255740 | 74255775 | 74260914 | 74260944 | 0 | 4 | 0 | 2 | 3 | 27 | 12 | 26 | 31 | 33 | 0 | 0 | 0 | 0 | 0 | 21 | 30 | 5 | 7 | 29 | 173 | 149 | 0.086276293 | 0.578941433 | 0 | 0.223 | 0 | 0.053 | 0.073 | 0 | 0 | 0 | 0 | 0 | 0.07 |
| SE | 104438 | ENSG00000149100 | EIF3M | chr11 | + | 32587045 | 32587144 | 32583837 | 32583929 | 32588593 | 32588732 | 535 | 401 | 529 | 438 | 1352 | 4 | 1 | 5 | 3 | 3 | 391 | 398 | 331 | 235 | 252 | 11 | 0 | 4 | 3 | 3 | 248 | 149 | 0.086330438 | 0.579255116 | 0.988 | 0.996 | 0.985 | 0.989 | 0.996 | 0.955 | 1 | 0.98 | 0.979 | 0.981 | 0.012 |
| SE | 40474 | ENSG00000164088 | PPM1M | chr3 | + | 52248334 | 52248453 | 52248152 | 52248237 | 52248636 | 52248700 | 385 | 178 | 298 | 247 | 378 | 0 | 0 | 3 | 5 | 9 | 298 | 443 | 214 | 244 | 518 | 0 | 9 | 2 | 4 | 16 | 268 | 149 | 0.086368564 | 0.579461274 | 1 | 1 | 0.982 | 0.965 | 0.959 | 1 | 0.965 | 0.983 | 0.971 | 0.947 | 0.008 |
| SE | 31821 | ENSG00000112701 | SENP6 | chr6 | + | 75640686 | 75640704 | 75634706 | 75634811 | 75647730 | 75647801 | 98 | 74 | 61 | 51 | 140 | 28 | 34 | 24 | 36 | 36 | 19 | 29 | 27 | 32 | 24 | 6 | 0 | 12 | 10 | 7 | 167 | 149 | 0.086406438 | 0.579665708 | 0.757 | 0.66 | 0.694 | 0.558 | 0.776 | 0.739 | 1 | 0.667 | 0.741 | 0.754 | -0.091 |
| SE | 102262 | ENSG00000178440 | TIMM23B-AGAP6 | chr10 | + | 49981188 | 49981281 | 49972834 | 49973142 | 49987164 | 49987234 | 10 | 7 | 6 | 5 | 1 | 10 | 4 | 4 | 10 | 6 | 0 | 3 | 2 | 3 | 3 | 11 | 6 | 2 | 3 | 4 | 242 | 149 | 0.086448629 | 0.579899064 | 0.381 | 0.519 | 0.48 | 0.235 | 0.093 | 0 | 0.235 | 0.381 | 0.381 | 0.316 | 0.079 |
| SE | 785 | ENSG00000138867 | GUCD1 | chr22 | - | 24543841 | 24544083 | 24540422 | 24543100 | 24546913 | 24547005 | 172 | 82 | 109 | 129 | 86 | 0 | 0 | 0 | 0 | 0 | 171 | 208 | 108 | 128 | 323 | 8 | 0 | 0 | 0 | 6 | 298 | 149 | 0.086475969 | 0.580032765 | 1 | 1 | 1 | 1 | 1 | 0.914 | 1 | 1 | 1 | 0.964 | 0.024 |
| SE | 23927 | ENSG00000124177 | CHD6 | chr20 | - | 41564010 | 41564151 | 41551304 | 41551360 | 41618339 | 41618375 | 2 | 6 | 3 | 3 | 5 | 94 | 58 | 86 | 93 | 76 | 0 | 7 | 2 | 0 | 0 | 36 | 116 | 133 | 63 | 21 | 290 | 149 | 0.086508378 | 0.580200443 | 0.011 | 0.05 | 0.018 | 0.016 | 0.033 | 0 | 0.03 | 0.008 | 0 | 0 | 0.018 |
| SE | 95815 | ENSG00000215126 | CBWD6 | chr9 | - | 41165000 | 41165087 | 41162332 | 41162377 | 41166783 | 41166830 | 20 | 20 | 15 | 46 | 72 | 0 | 1 | 3 | 2 | 3 | 14 | 14 | 30 | 20 | 4 | 0 | 4 | 0 | 0 | 0 | 236 | 149 | 0.0866079 | 0.580818169 | 1 | 0.927 | 0.759 | 0.936 | 0.938 | 1 | 0.688 | 1 | 1 | 1 | -0.026 |
| SE | 76500 | ENSG00000153406 | NMRAL1 | chr16 | - | 4466526 | 4466685 | 4466152 | 4466402 | 4469503 | 4469685 | 2 | 1 | 0 | 0 | 0 | 2 | 2 | 3 | 2 | 7 | 0 | 3 | 6 | 0 | 0 | 18 | 11 | 7 | 3 | 5 | 298 | 149 | 0.086634856 | 0.580918257 | 0.333 | 0.2 | 0 | 0 | 0 | 0 | 0.12 | 0.3 | 0 | 0 | 0.023 |
| SE | 31506 | ENSG00000174227 | PIGG | chr4 | + | 501100 | 501170 | 500395 | 500601 | 507404 | 507593 | 10 | 11 | 10 | 16 | 17 | 1 | 0 | 0 | 0 | 1 | 11 | 13 | 28 | 31 | 37 | 9 | 0 | 0 | 0 | 0 | 219 | 149 | 0.086637663 | 0.580918257 | 0.872 | 1 | 1 | 1 | 0.92 | 0.454 | 1 | 1 | 1 | 1 | 0.068 |
| SE | 66924 | ENSG00000152520 | PAN3 | chr13 | + | 28281314 | 28281379 | 28280411 | 28280541 | 28287983 | 28288122 | 257 | 193 | 217 | 220 | 419 | 0 | 0 | 0 | 0 | 0 | 90 | 184 | 219 | 125 | 31 | 0 | 0 | 0 | 0 | 11 | 214 | 149 | 0.086664529 | 0.581048637 | 1 | 1 | 1 | 1 | 1 | 1 | 1 | 1 | 1 | 0.662 | 0.068 |
| SE | 65240 | ENSG00000169379 | ARL13B | chr3 | + | 93996562 | 93996684 | 93995873 | 93995944 | 94003658 | 94003908 | 5 | 1 | 4 | 0 | 4 | 6 | 3 | 6 | 5 | 1 | 0 | 2 | 1 | 3 | 4 | 5 | 0 | 0 | 5 | 0 | 271 | 149 | 0.086672161 | 0.581050048 | 0.314 | 0.155 | 0.268 | 0 | 0.687 | 0 | 1 | 1 | 0.248 | 1 | -0.365 |
| SE | 73687 | ENSG00000151687 | ANKAR | chr2 | + | 189737682 | 189737841 | 189728659 | 189728821 | 189738564 | 189738682 | 45 | 56 | 38 | 30 | 99 | 4 | 2 | 0 | 2 | 4 | 79 | 33 | 42 | 34 | 96 | 0 | 3 | 1 | 0 | 3 | 298 | 149 | 0.086707548 | 0.581237512 | 0.849 | 0.933 | 1 | 0.882 | 0.925 | 1 | 0.846 | 0.955 | 1 | 0.941 | -0.031 |
| SE | 13439 | ENSG00000167508 | MVD | chr16 | - | 88657145 | 88657247 | 88656104 | 88656304 | 88657435 | 88657582 | 8 | 0 | 2 | 0 | 0 | 24 | 4 | 6 | 13 | 9 | 6 | 0 | 2 | 0 | 14 | 0 | 3 | 1 | 7 | 13 | 251 | 149 | 0.086741647 | 0.581416315 | 0.165 | 0 | 0.165 | 0 | 0 | 1 | 0 | 0.543 | 0 | 0.39 | -0.321 |
| SE | 61840 | ENSG00000148773 | MKI67 | chr10 | - | 128114927 | 128116007 | 128113426 | 128113602 | 128116490 | 128116536 | 21 | 9 | 16 | 24 | 19 | 1 | 0 | 12 | 14 | 9 | 48 | 49 | 69 | 61 | 5 | 2 | 7 | 14 | 8 | 0 | 298 | 149 | 0.086777279 | 0.58160536 | 0.913 | 1 | 0.4 | 0.462 | 0.514 | 0.923 | 0.778 | 0.711 | 0.792 | 1 | -0.183 |
| SE | 79402 | ENSG00000065970 | FOXJ2 | chr12 | + | 8050349 | 8050440 | 8049361 | 8049571 | 8050521 | 8050620 | 6 | 0 | 1 | 1 | 5 | 54 | 23 | 39 | 52 | 77 | 1 | 0 | 4 | 0 | 5 | 178 | 43 | 60 | 62 | 164 | 240 | 149 | 0.086817418 | 0.58182458 | 0.065 | 0 | 0.016 | 0.012 | 0.039 | 0.003 | 0 | 0.04 | 0 | 0.019 | 0.014 |
| SE | 69109 | ENSG00000105374 | NKG7 | chr19 | - | 51372160 | 51372202 | 51371939 | 51372074 | 51372378 | 51372410 | 248 | 40 | 78 | 105 | 18 | 85 | 14 | 32 | 37 | 7 | 58 | 160 | 77 | 72 | 160 | 32 | 106 | 49 | 31 | 40 | 191 | 149 | 0.086836122 | 0.581900119 | 0.695 | 0.69 | 0.655 | 0.689 | 0.667 | 0.586 | 0.541 | 0.551 | 0.644 | 0.757 | 0.063 |
| SE | 12129 | ENSG00000172775 | PSME3IP1 | chr16 | - | 57174168 | 57174286 | 57173727 | 57173869 | 57185820 | 57186055 | 14 | 7 | 17 | 15 | 7 | 88 | 55 | 63 | 72 | 57 | 0 | 12 | 10 | 17 | 9 | 114 | 77 | 40 | 67 | 99 | 267 | 149 | 0.086844663 | 0.581907553 | 0.082 | 0.066 | 0.131 | 0.104 | 0.064 | 0 | 0.08 | 0.122 | 0.124 | 0.048 | 0.015 |
| SE | 5105 | ENSG00000124198 | ARFGEF2 | chr20 | + | 48973144 | 48973284 | 48972325 | 48972425 | 48974765 | 48974874 | 159 | 159 | 163 | 164 | 188 | 0 | 0 | 0 | 0 | 0 | 77 | 120 | 77 | 62 | 69 | 9 | 0 | 1 | 0 | 0 | 289 | 149 | 0.086954918 | 0.582596463 | 1 | 1 | 1 | 1 | 1 | 0.815 | 1 | 0.975 | 1 | 1 | 0.042 |
| SE | 41125 | ENSG00000124406 | ATP8A1 | chr4 | - | 42423616 | 42423705 | 42422806 | 42422899 | 42443564 | 42443672 | 175 | 96 | 164 | 180 | 158 | 4 | 1 | 0 | 0 | 0 | 7 | 98 | 75 | 32 | 27 | 7 | 0 | 0 | 0 | 0 | 238 | 149 | 0.086974157 | 0.582675505 | 0.965 | 0.984 | 1 | 1 | 1 | 0.385 | 1 | 1 | 1 | 1 | 0.113 |
| SE | 79677 | ENSG00000186591 | UBE2H | chr7 | - | 129948958 | 129949102 | 129880894 | 129880971 | 129952502 | 129952722 | 0 | 0 | 0 | 0 | 0 | 46 | 28 | 58 | 38 | 63 | 18 | 0 | 0 | 0 | 0 | 37 | 116 | 102 | 68 | 120 | 293 | 149 | 0.08700765 | 0.582850012 | 0 | 0 | 0 | 0 | 0 | 0.198 | 0 | 0 | 0 | 0 | -0.04 |
| SE | 19307 | ENSG00000156414 | TDRD9 | chr14 | + | 103986216 | 103986320 | 103975388 | 103975553 | 103991159 | 103991224 | 15 | 28 | 7 | 41 | 12 | 1 | 2 | 1 | 3 | 0 | 14 | 33 | 24 | 17 | 11 | 13 | 3 | 0 | 0 | 2 | 253 | 149 | 0.087110442 | 0.583468728 | 0.898 | 0.892 | 0.805 | 0.889 | 1 | 0.388 | 0.866 | 1 | 1 | 0.764 | 0.093 |
| SE | 89985 | ENSG00000153317 | ASAP1 | chr8 | - | 130187235 | 130187285 | 130180750 | 130180880 | 130188108 | 130188183 | 297 | 361 | 252 | 310 | 420 | 2 | 1 | 1 | 1 | 0 | 1437 | 701 | 523 | 1258 | 1148 | 4 | 7 | 2 | 32 | 11 | 199 | 149 | 0.087114916 | 0.583468728 | 0.991 | 0.996 | 0.995 | 0.996 | 1 | 0.996 | 0.987 | 0.995 | 0.967 | 0.987 | 0.009 |
| SE | 21042 | ENSG00000135506 | OS9 | chr12 | + | 57695961 | 57696038 | 57695779 | 57695843 | 57715759 | 57715970 | 109 | 96 | 88 | 103 | 135 | 0 | 0 | 2 | 0 | 0 | 211 | 171 | 95 | 156 | 299 | 0 | 1 | 5 | 10 | 0 | 226 | 149 | 0.087136761 | 0.58356512 | 1 | 1 | 0.967 | 1 | 1 | 1 | 0.991 | 0.926 | 0.911 | 1 | 0.028 |
| SE | 6291 | ENSG00000141644 | MBD1 | chr18 | - | 50273333 | 50273426 | 50272823 | 50272955 | 50273563 | 50273863 | 59 | 37 | 38 | 65 | 14 | 4 | 8 | 21 | 6 | 9 | 87 | 39 | 34 | 13 | 92 | 23 | 32 | 12 | 17 | 16 | 242 | 149 | 0.087150395 | 0.583606513 | 0.901 | 0.74 | 0.527 | 0.87 | 0.489 | 0.7 | 0.429 | 0.636 | 0.32 | 0.78 | 0.132 |
| SE | 63430 | ENSG00000167595 | PROSER3 | chr19 | + | 35764853 | 35764936 | 35762252 | 35762356 | 35766767 | 35766955 | 5 | 1 | 3 | 3 | 4 | 0 | 0 | 0 | 0 | 0 | 7 | 12 | 4 | 6 | 7 | 7 | 0 | 0 | 0 | 0 | 232 | 149 | 0.087188738 | 0.583813343 | 1 | 1 | 1 | 1 | 1 | 0.391 | 1 | 1 | 1 | 1 | 0.122 |
| SE | 90123 | ENSG00000160932 | LY6E | chr8 | + | 143020261 | 143020377 | 143018516 | 143018586 | 143020887 | 143020991 | 1 | 2 | 2 | 3 | 0 | 40 | 27 | 35 | 40 | 4 | 22 | 3 | 1 | 11 | 0 | 183 | 167 | 46 | 107 | 67 | 265 | 149 | 0.087199403 | 0.583834829 | 0.014 | 0.04 | 0.031 | 0.04 | 0 | 0.063 | 0.01 | 0.012 | 0.055 | 0 | -0.003 |

| | | | | | | | | | |
|---|---|---|---|---|---|---|---|---|---|
| SE | 99730 | ENSG00000173264 | GPR137 | chr11 | + | 64286509 | 64286881 | 64285056 | 64285380 |
| SE | 93569 | ENSG00000165525 | NEMF | chr14 | - | 49796232 | 49796328 | 49795790 | 49795944 |
| SE | 107572 | ENSG00000120837 | NFYB | chr12 | - | 104126113 | 104126244 | 104123225 | 104123423 |
| SE | 17928 | ENSG00000197249 | SERPINA1 | chr14 | - | 94388559 | 94388660 | 94382591 | 94383241 |
| SE | 101683 | ENSG00000180398 | MCFD2 | chr2 | - | 46907809 | 46907969 | 46901869 | 46905594 |
| SE | 69282 | ENSG00000168385 | SEPTIN2 | chr2 | + | 241324408 | 241324662 | 241324215 | 241324241 |
| SE | 98332 | ENSG00000149503 | INCENP | chr11 | + | 62141499 | 62141511 | 62140912 | 62141044 |
| SE | 81056 | ENSG00000188243 | COMMD6 | chr13 | - | 75530310 | 75530405 | 75526435 | 75526639 |
| SE | 34039 | ENSG00000159147 | DONSON | chr21 | - | 33582164 | 33582246 | 33581950 | 33582055 |
| SE | 3553 | ENSG00000105483 | CARD8 | chr19 | - | 48249522 | 48249633 | 48240961 | 48241063 |
| SE | 83798 | ENSG00000165168 | CYBB | chrX | + | 37793664 | 37793810 | 37791974 | 37792059 |
| SE | 39127 | ENSG00000134352 | IL6ST | chr5 | - | 55951463 | 55951604 | 55947492 | 55947589 |
| SE | 6594 | ENSG00000134444 | RELCH | chr18 | + | 62264728 | 62264852 | 62263988 | 62264145 |
| SE | 6691 | ENSG00000177426 | TGIF1 | chr18 | + | 3450324 | 3450505 | 3450035 | 3450070 |
| SE | 23408 | ENSG00000153113 | CAST | chr5 | + | 96746341 | 96746425 | 96741493 | 96742756 |
| SE | 92382 | ENSG00000080503 | SMARCA2 | chr9 | + | 2115821 | 2116049 | 2110253 | 2110417 |
| SE | 40201 | ENSG00000231607 | DLEU2 | chr13 | - | 50071515 | 50071629 | 50070581 | 50070641 |
| SE | 51508 | ENSG00000108021 | TASOR2 | chr10 | + | 5714134 | 5714207 | 5712822 | 5712918 |
| SE | 33580 | ENSG00000009790 | TRAF3IP3 | chr1 | + | 209778110 | 209778173 | 209777351 | 209777487 |
| SE | 43314 | ENSG00000009413 | REV3L | chr6 | - | 111381324 | 111381444 | 111379981 | 111380219 |
| SE | 85643 | ENSG00000172922 | RNASEH2C | chr11 | - | 65719680 | 65719809 | 65714004 | 65715317 |
| SE | 49415 | ENSG00000107551 | RASSF4 | chr10 | + | 44982520 | 44982663 | 44970164 | 44971848 |
| SE | 95437 | ENSG00000119383 | PTPA | chr9 | + | 129123051 | 129123138 | 129120512 | 129120610 |
| SE | 71852 | ENSG00000185414 | MRPL30 | chr2 | + | 99186918 | 99187018 | 99186176 | 99186254 |
| SE | 97265 | ENSG00000156110 | ADK | chr10 | + | 74525255 | 74525426 | 74398470 | 74398579 |
| SE | 201 | ENSG00000160209 | PDXK | chr21 | + | 43732138 | 43732647 | 43719222 | 43719381 |
| SE | 17940 | ENSG00000197249 | SERPINA1 | chr14 | - | 94388818 | 94389010 | 94382591 | 94383241 |
| SE | 41868 | ENSG00000171163 | ZNF692 | chr1 | - | 248856287 | 248856422 | 248855557 | 248855946 |
| SE | 88399 | ENSG00000145041 | DCAF1 | chr3 | - | 51466802 | 51466876 | 51463113 | 51463227 |
| SE | 108658 | ENSG00000136147 | PHF11 | chr13 | + | 49525772 | 49525937 | 49524084 | 49524216 |
| SE | 50175 | ENSG00000025708 | TYMP | chr22 | - | 50529495 | 50529731 | 50527587 | 50527717 |
| SE | 19434 | ENSG00000151445 | VIPAS39 | chr14 | - | 77457094 | 77457365 | 77454009 | 77454102 |
| SE | 44527 | ENSG00000076321 | KLHL20 | chr1 | + | 173715522 | 173715654 | 173714940 | 173715078 |
| SE | 83766 | ENSG00000196696 | PDXDC2P-NPIPB14P | chr16 | - | 69990596 | 69990683 | 69986380 | 69986509 |
| SE | 54615 | ENSG00000188897 | NA | chr16 | - | 11487261 | 11487386 | 11484878 | 11485073 |
| SE | 71059 | ENSG00000115486 | GGCX | chr2 | - | 85552415 | 85552567 | 85551811 | 85551981 |
| SE | 61662 | ENSG00000009307 | CSDE1 | chr1 | - | 114749973 | 114750207 | 114739752 | 114739890 |
| SE | 43184 | ENSG00000169188 | APEX2 | chrX | + | 55002250 | 55002431 | 55001545 | 55001629 |
| SE | 70602 | ENSG00000164548 | TRA2A | chr7 | - | 23522131 | 23522432 | 23521706 | 23521840 |
| SE | 96809 | ENSG00000166889 | PATL1 | chr11 | - | 59655955 | 59656045 | 59655522 | 59655740 |
| SE | 86498 | ENSG00000185485 | SDHAP1 | chr3 | - | 195984491 | 195984737 | 195984111 | 195984276 |
| SE | 90486 | ENSG00000033800 | PIAS1 | chr15 | + | 68183629 | 68183667 | 68181211 | 68181354 |
| SE | 104337 | ENSG00000155256 | ZFYVE27 | chr10 | + | 97748268 | 97748364 | 97738476 | 97738649 |
| SE | 61043 | ENSG00000066027 | PPP2R5A | chr1 | + | 212349188 | 212349242 | 212345802 | 212345933 |
| SE | 47650 | ENSG00000288701 | PRRC2B | chr9 | + | 131482709 | 131482907 | 131481725 | 131481808 |
| SE | 29886 | ENSG00000141562 | NARF | chr17 | + | 82472563 | 82472698 | 82468498 | 82468896 |
| SE | 62340 | ENSG00000185811 | IKZF1 | chr7 | + | 50391728 | 50391863 | 50382539 | 50382707 |
| SE | 35691 | ENSG00000175556 | LONRF3 | chrX | + | 118986924 | 118987080 | 118982820 | 118982943 |
| SE | 26662 | ENSG00000215845 | TSTD1 | chr1 | - | 161038550 | 161038673 | 161037630 | 161038075 |
| SE | 5959 | ENSG00000170471 | RALGAPB | chr20 | + | 38535073 | 38535207 | 38532729 | 38532859 |
| SE | 53285 | ENSG00000116213 | WRAP73 | chr1 | - | 3635159 | 3635294 | 3634996 | 3635074 |
| SE | 12565 | ENSG00000214078 | CPNE1 | chr20 | - | 35658929 | 35658982 | 35632794 | 35632923 |
| SE | 24816 | ENSG00000101166 | PRELID3B | chr20 | - | 59037190 | 59037299 | 59036689 | 59036760 |
| SE | 64375 | ENSG00000264538 | SUZ12P1 | chr17 | + | 30743300 | 30743369 | 30731705 | 30732123 |
| SE | 99176 | ENSG00000139131 | YARS2 | chr12 | - | 32749936 | 32750107 | 32746543 | 32747363 |
| SE | 86499 | ENSG00000185485 | SDHAP1 | chr3 | - | 195984593 | 195984737 | 195984111 | 195984276 |
| SE | 100161 | ENSG00000078124 | ACER3 | chr11 | + | 76868119 | 76868267 | 76860923 | 76861079 |
| SE | 27693 | ENSG00000186431 | FCAR | chr19 | + | 54875329 | 54875365 | 54874234 | 54874323 |
| SE | 2291 | ENSG00000142002 | DPP9 | chr19 | - | 4695377 | 4695555 | 4694316 | 4694823 |
| SE | 19603 | ENSG00000255185 | PDXDC2P | chr16 | - | 70014103 | 70014167 | 70011840 | 70011929 |
| SE | 70842 | ENSG00000243970 | PPIEL | chr1 | - | 39523561 | 39523697 | 39522279 | 39522505 |
| SE | 45075 | ENSG00000007341 | ST7L | chr1 | - | 112616812 | 112616895 | 112610458 | 112611003 |
| SE | 3549 | ENSG00000105483 | CARD8 | chr19 | - | 48246074 | 48246381 | 48242539 | 48242644 |
| SE | 29596 | ENSG00000203875 | SNHG5 | chr6 | - | 85677092 | 85677191 | 85664959 | 85666147 |
| SE | 15201 | ENSG00000159459 | UBR1 | chr15 | - | 43085983 | 43086240 | 43082637 | 43082716 |
| SE | 36667 | ENSG00000180530 | NRIP1 | chr21 | - | 14991200 | 14991306 | 14968154 | 14968526 |
| SE | 48676 | ENSG00000131374 | TBC1D5 | chr3 | - | 17662989 | 17663110 | 17623848 | 17623913 |
| SE | 50783 | ENSG00000103351 | CLUAP1 | chr16 | + | 3519902 | 3520036 | 3515375 | 3515591 |
| SE | 78020 | ENSG00000136250 | AOAH | chr7 | - | 36514471 | 36514590 | 36513001 | 36513380 |

| | | | | | | | | | | | | | | |
|---|---|---|---|---|---|---|---|---|---|---|---|---|---|---|
| SE | 81600 | ENSG00000253729 | PRKDC | chr8 | - | 47893138 | 47893387 | 47890256 | 47890480 | 47897160 | 47897294 | 98 | 51 | 104 |
| SE | 32437 | ENSG00000131370 | SH3BP5 | chr3 | - | 15256852 | 15257113 | 15254352 | 15256303 | 15258830 | 15258883 | 284 | 125 | 147 |
| SE | 108175 | ENSG00000121749 | TBC1D15 | chr12 | + | 71841100 | 71841231 | 71839706 | 71839811 | 71872069 | 71872168 | 27 | 7 | 11 |
| SE | 34504 | ENSG00000116350 | SRSF4 | chr1 | - | 29150102 | 29150192 | 29147742 | 29149226 | 29154695 | 29154910 | 277 | 152 | 236 |
| SE | 101772 | ENSG00000012779 | ALOX5 | chr10 | + | 45395854 | 45395936 | 45382482 | 45382681 | 45412190 | 45412313 | 525 | 357 | 304 |
| SE | 94796 | ENSG00000165275 | TRMT10B | chr9 | + | 37762576 | 37762685 | 37761902 | 37762117 | 37768075 | 37768228 | 26 | 14 | 46 |
| SE | 67609 | ENSG00000109466 | KLHL2 | chr4 | + | 165228806 | 165228913 | 165219933 | 165220059 | 165263196 | 165263359 | 53 | 36 | 40 |
| SE | 9246 | ENSG00000178971 | CTC1 | chr17 | - | 8235830 | 8235959 | 8235052 | 8235285 | 8236082 | 8236342 | 43 | 29 | 21 |
| SE | 33454 | ENSG00000156976 | EIF4A2 | chr3 | + | 186788309 | 186788416 | 186787802 | 186787882 | 186789124 | 186789881 | 361 | 311 | 412 |
| SE | 29090 | ENSG00000163625 | WDFY3 | chr4 | - | 84932269 | 84932363 | 84896910 | 84897010 | 84966208 | 84966391 | 9 | 7 | 3 |
| SE | 100187 | ENSG00000078124 | ACER3 | chr11 | + | 77011329 | 77011375 | 76998762 | 76998821 | 77016674 | 77016779 | 1 | 3 | 1 |
| SE | 29336 | ENSG00000121210 | TMEM131L | chr4 | + | 153585457 | 153585611 | 153584834 | 153584931 | 153586208 | 153586379 | 377 | 344 | 428 |
| SE | 6693 | ENSG00000078043 | PIAS2 | chr18 | - | 46844733 | 46844839 | 46844053 | 46844127 | 46846706 | 46846841 | 84 | 93 | 88 |
| SE | 107064 | ENSG00000136521 | NDUFB5 | chr3 | + | 179615982 | 179616049 | 179604808 | 179604939 | 179616982 | 179617044 | 93 | 49 | 60 |
| SE | 23619 | ENSG00000113558 | SKP1 | chr5 | - | 134174461 | 134174514 | 134173925 | 134174022 | 134176854 | 134176950 | 2 | 2 | 12 |
| SE | 93154 | ENSG00000197969 | VPS13A | chr9 | + | 77283354 | 77283471 | 77282120 | 77282274 | 77283546 | 77283650 | 56 | 52 | 34 |
| SE | 61406 | ENSG00000185630 | PBX1 | chr1 | + | 164821536 | 164821626 | 164811989 | 164812149 | 164846583 | 164846955 | 5 | 22 | 12 |
| SE | 54604 | ENSG00000151422 | FER | chr5 | + | 108832769 | 108832943 | 108798113 | 108798389 | 108835707 | 108835807 | 109 | 127 | 78 |
| SE | 62318 | ENSG00000185811 | IKZF1 | chr7 | + | 50368005 | 50368065 | 50327637 | 50327757 | 50387344 | 50387470 | 48 | 31 | 59 |
| SE | 43350 | ENSG00000075188 | NUP37 | chr12 | - | 102112107 | 102112232 | 102101031 | 102101104 | 102118362 | 102118583 | 95 | 49 | 113 |
| SE | 21855 | ENSG00000068796 | KIF2A | chr5 | + | 62365242 | 62365353 | 62363177 | 62363899 | 62373686 | 62373837 | 263 | 278 | 209 |
| SE | 105175 | ENSG00000117151 | CTBS | chr1 | - | 84569930 | 84570139 | 84565840 | 84566012 | 84570581 | 84570720 | 195 | 124 | 160 |
| SE | 9411 | ENSG00000105053 | VRK3 | chr19 | - | 50009235 | 50009289 | 50007568 | 50007826 | 50016023 | 50016163 | 18 | 2 | 5 |
| SE | 77873 | ENSG00000198815 | FOXJ3 | chr1 | - | 42205761 | 42205863 | 42199101 | 42199230 | 42227882 | 42227966 | 216 | 155 | 232 |
| SE | 81343 | ENSG00000135164 | DMTF1 | chr7 | + | 87190944 | 87191027 | 87185995 | 87188301 | 87193197 | 87193301 | 238 | 171 | 229 |
| SE | 62987 | ENSG00000137497 | NUMA1 | chr11 | - | 72008091 | 72008145 | 72007188 | 72007435 | 72008687 | 72008845 | 8 | 0 | 11 |
| SE | 36966 | ENSG00000153914 | SREK1 | chr5 | + | 66156064 | 66158932 | 66153462 | 66153596 | 66159218 | 66159334 | 59 | 53 | 47 |
| SE | 42621 | ENSG00000126088 | UROD | chr1 | + | 45013291 | 45013354 | 45013135 | 45013215 | 45013593 | 45013703 | 83 | 53 | 87 |
| SE | 104484 | ENSG00000013561 | RNF14 | chr5 | + | 141978302 | 141978830 | 141973582 | 141973742 | 141980122 | 141980351 | 74 | 47 | 70 |
| SE | 103669 | ENSG00000148634 | HERC4 | chr10 | - | 67990210 | 67990400 | 67988662 | 67988835 | 67990903 | 67991015 | 334 | 231 | 249 |
| SE | 79760 | ENSG00000164808 | SPIDR | chr8 | + | 47440322 | 47440542 | 47407860 | 47407961 | 47595810 | 47595984 | 158 | 106 | 137 |
| SE | 49171 | ENSG00000066136 | NFYC | chr1 | + | 40735600 | 40735750 | 40691837 | 40691867 | 40738835 | 40738948 | 4 | 4 | 1 |
| SE | 107494 | ENSG00000123349 | PFDN5 | chr12 | + | 53295838 | 53295941 | 53295290 | 53295639 | 53298044 | 53298150 | 669 | 407 | 530 |

| | | | | | | | | | | | | | | | | | |
|---|---|---|---|---|---|---|---|---|---|---|---|---|---|---|---|---|---|
| SE | 56106 | ENSG00000165689 | ENTR1 | chr9 | - | 136408998 | 136409067 | 136407825 | 136407938 | 136410089 | 136410239 | 16 | 6 | 11 | [illegible] | [illegible] | [illegible] |
| SE | 29831 | ENSG00000129187 | DCTD | chr4 | - | 182916418 | 182916539 | 182915460 | 182915575 | 182917287 | 182917389 | 6 | 1 | 3 | [illegible] | [illegible] | [illegible] |
| SE | 82008 | ENSG00000104133 | SPG11 | chr15 | - | 44570524 | 44570658 | 44567423 | 44567592 | 44572682 | 44572820 | 107 | 61 | 75 | [illegible] | [illegible] | [illegible] |
| SE | 17713 | ENSG00000100749 | VRK1 | chr14 | + | 96848359 | 96848479 | 96847256 | 96847344 | 96852830 | 96852939 | 0 | 0 | 1 | [illegible] | [illegible] | [illegible] |
| SE | 73692 | ENSG00000151687 | ANKAR | chr2 | + | 189738564 | 189738682 | 189733106 | 189733229 | 189741341 | 189741451 | 32 | 31 | 15 | 16 | 48 | 5 |
| SE | 52990 | ENSG00000077585 | GPR137B | chr1 | + | 236179878 | 236180028 | 236168700 | 236168755 | 236205125 | 236205198 | 4 | 1 | 1 | 1 | 6 | 0 |
| SE | 80358 | ENSG00000102531 | FNDC3A | chr13 | + | 49191014 | 49191120 | 49188514 | 49188633 | 49191208 | 49191384 | 152 | 104 | 134 | 164 | 155 | 5 |
| SE | 80712 | ENSG00000147535 | PLPP5 | chr8 | - | 38268370 | 38268407 | 38267896 | 38267960 | 38268881 | 38268990 | 6 | 4 | 10 | 9 | 8 | 0 |
| SE | 51510 | ENSG00000108021 | TASOR2 | chr10 | + | 5717658 | 5717750 | 5712822 | 5712918 | 5720543 | 5720611 | 54 | 32 | 31 | 35 | 24 | 14 |
| SE | 105683 | ENSG00000028203 | VEZT | chr12 | + | 95242163 | 95242264 | 95217779 | 95217886 | 95251939 | 95252071 | 3 | 0 | 0 | 4 | 2 | 49 |
| SE | 62726 | ENSG00000151779 | NBAS | chr2 | - | 15536417 | 15536551 | 15534542 | 15534641 | 15539222 | 15539356 | 147 | 105 | 133 | 128 | 250 | 1 |
| SE | 108251 | ENSG00000121073 | SLC35B1 | chr17 | - | 49706203 | 49706334 | 49705865 | 49705896 | 49706964 | 49707068 | 54 | 33 | 84 | 71 | 80 | 2 |
| SE | 90017 | ENSG00000155926 | SLA | chr8 | - | 133044983 | 133045115 | 133039997 | 133040130 | 133047829 | 133047933 | 369 | 154 | 283 | 234 | 200 | 7 |
| SE | 6690 | ENSG00000177426 | TGIF1 | chr18 | + | 3450321 | 3450505 | 3449881 | 3450070 | 3456353 | 3456580 | 30 | 13 | 28 | 32 | 31 | 10 |
| SE | 16792 | ENSG00000241839 | PLEKHO2 | chr15 | + | 64861476 | 64861575 | 64859893 | 64859998 | 64864898 | 64868000 | 286 | 180 | 191 | 294 | 342 | 2 |
| SE | 39789 | ENSG00000106459 | NRF1 | chr7 | + | 129671428 | 129671543 | 129657342 | 129657574 | 129677631 | 129677758 | 173 | 93 | 189 | 147 | 136 | 0 |
| SE | 69446 | ENSG00000145782 | ATG12 | chr5 | - | 115840496 | 115840612 | 115837627 | 115837764 | 115841389 | 115841837 | 13 | 10 | 23 | 34 | 3 | 35 |
| SE | 10236 | ENSG00000108960 | MMD | chr17 | - | 55407745 | 55407820 | 55403766 | 55403868 | 55411256 | 55411417 | 288 | 391 | 112 | 511 | 300 | 0 |
| SE | 68602 | ENSG00000237765 | FAM200B | chr4 | + | 15684849 | 15684959 | 15681755 | 15681784 | 15686235 | 15686396 | 19 | 7 | 11 | 11 | 22 | 71 |
| SE | 24364 | ENSG00000132780 | NASP | chr1 | + | 45594693 | 45594773 | 45591222 | 45591270 | 45602254 | 45602365 | 2 | 1 | 9 | 15 | 2 | 59 |
| SE | 20038 | ENSG00000139974 | SLC38A6 | chr14 | + | 61052348 | 61052985 | 61052040 | 61052135 | 61078829 | 61078927 | 15 | 15 | 31 | 6 | 22 | 8 |
| SE | 76116 | ENSG00000065029 | ZNF76 | chr6 | + | 35293750 | 35293915 | 35292880 | 35293044 | 35294455 | 35294552 | 35 | 18 | 40 | 42 | 41 | 18 |
| SE | 68012 | ENSG00000137414 | FAM8A1 | chr6 | + | 17605873 | 17606013 | 17604905 | 17605029 | 17608194 | 17611715 | 206 | 106 | 159 | 206 | 243 | 1 |
| SE | 87902 | ENSG00000119328 | ABITRAM | chr9 | + | 108936307 | 108936437 | 108935637 | 108935689 | 108950506 | 108950744 | 20 | 4 | 16 | 25 | 15 | 0 |
| SE | 31864 | ENSG00000143669 | LYST | chr1 | - | 235747169 | 235747331 | 235746467 | 235746527 | 235751209 | 235751362 | 2 | 6 | 4 | 4 | 8 | 194 |
| SE | 53464 | ENSG00000249915 | PDCD6 | chr5 | + | 306601 | 306754 | 304176 | 304221 | 311292 | 311402 | 187 | 85 | 108 | 138 | 90 | 0 |
| SE | 63983 | ENSG00000049089 | COL9A2 | chr1 | - | 40307702 | 40307756 | 40305714 | 40305768 | 40308191 | 40308245 | 3 | 1 | 2 | 4 | 2 | 0 |
| SE | 29795 | ENSG00000141452 | RMC1 | chr18 | + | 23506969 | 23507054 | 23504370 | 23504447 | 23507984 | 23508041 | 163 | 126 | 149 | 171 | 150 | 1 |
| SE | 14219 | ENSG00000167965 | MLST8 | chr16 | + | 2207034 | 2207110 | 2206357 | 2206659 | 2207192 | 2207250 | 33 | 12 | 27 | 18 | 24 | 1 |
| SE | 60818 | ENSG00000237298 | TTN-AS1 | chr2 | + | 178537360 | 178537447 | 178535731 | 178535828 | 178537603 | 178537691 | 5 | 4 | 2 | 8 | 2 | 71 |
| SE | 79445 | ENSG00000147894 | C9orf72 | chr9 | - | 27560226 | 27560299 | 27556455 | 27556796 | 27561584 | 27561649 | 25 | 18 | 13 | 14 | 83 | 1 |
| SE | 34792 | ENSG00000136152 | COG3 | chr13 | + | 45480124 | 45480290 | 45479004 | 45479066 | 45481229 | 45481304 | 176 | 98 | 105 | 87 | 222 | 2 |
| SE | 39999 | ENSG00000107566 | ERLIN1 | chr10 | - | 100167347 | 100167406 | 100164003 | 100164095 | 100174207 | 100174281 | 83 | 61 | 72 | 72 | 167 | 1 |
| SE | 32567 | ENSG00000113851 | CRBN | chr3 | - | 3152455 | 3152587 | 3148991 | 3151045 | 3153423 | 3153488 | 453 | 338 | 374 | 408 | 457 | 46 |
| SE | 64374 | ENSG00000264538 | SUZ12P1 | chr17 | + | 30743300 | 30743369 | 30709298 | 30709811 | 30759490 | 30759539 | 39 | 44 | 33 | 46 | 92 | 3 |
| SE | 99758 | ENSG00000131626 | PPFIA1 | chr11 | + | 70382087 | 70382158 | 70378029 | 70378195 | 70383002 | 70384387 | 56 | 77 | 56 | 52 | 128 | 11 |
| SE | 22202 | ENSG00000137177 | KIF13A | chr6 | - | 17808767 | 17808930 | 17805474 | 17805615 | 17817019 | 17817200 | 47 | 69 | 63 | 51 | 37 | 0 |
| SE | 87980 | ENSG00000198373 | WWP2 | chr16 | + | 69783031 | 69783152 | 69776188 | 69776241 | 69786995 | 69787080 | 14 | 2 | 6 | 11 | 0 | 35 |
| SE | 71644 | ENSG00000113916 | BCL6 | chr3 | - | 187733532 | 187733703 | 187731708 | 187731930 | 187734868 | 187734882 | 102 | 83 | 99 | 131 | 129 | 0 |
| SE | 8451 | ENSG00000108784 | NAGLU | chr17 | + | 42537397 | 42537545 | 42536240 | 42536655 | 42538338 | 42538485 | 13 | 5 | 15 | 11 | 9 | 7 |
| SE | 32780 | ENSG00000168297 | PXK | chr3 | + | 58420535 | 58420568 | 58410089 | 58410159 | 58424751 | 58425175 | 0 | 2 | 1 | 0 | 5 | 19 |
| SE | 71458 | ENSG00000123983 | ACSL3 | chr2 | + | 222928861 | 222928936 | 222924455 | 222924595 | 222930620 | 222930812 | 36 | 52 | 46 | 78 | 102 | 2 |
| SE | 35350 | ENSG00000164073 | MFSD8 | chr4 | - | 127921859 | 127921963 | 127917932 | 127921771 | 127930682 | 127930817 | 56 | 28 | 55 | 44 | 70 | 42 |
| SE | 79214 | ENSG00000242588 | NA | chr7 | + | 128609486 | 128609579 | 128591849 | 128592017 | 128615859 | 128615970 | 79 | 57 | 54 | 76 | 68 | 4 |
| SE | 87647 | ENSG00000117676 | RPS6KA1 | chr1 | + | 26555536 | 26555625 | 26555150 | 26555221 | 26556997 | 26557036 | 112 | 57 | 60 | 71 | 66 | 0 |
| SE | 84417 | ENSG00000188917 | TRMT2B | chrX | - | 101041316 | 101041371 | 101037916 | 101038051 | 101042232 | 101042312 | 43 | 16 | 30 | 35 | 41 | 0 |
| SE | 20668 | ENSG00000196505 | GDAP2 | chr1 | - | 117886576 | 117886653 | 117882903 | 117883627 | 117896832 | 117896989 | 48 | 32 | 23 | 29 | 57 | 1 |
| SE | 60640 | ENSG00000135829 | DHX9 | chr1 | + | 182881519 | 182881647 | 182881263 | 182881425 | 182883138 | 182883216 | 576 | 444 | 542 | 591 | 1174 | 0 |
| SE | 41870 | ENSG00000171163 | ZNF692 | chr1 | - | 248856372 | 248856422 | 248855557 | 248855946 | 248856513 | 248856562 | 17 | 6 | 15 | 27 | 29 | 2 |
| SE | 60936 | ENSG00000138078 | PREPL | chr2 | - | 44359496 | 44359750 | 44346267 | 44346390 | 44361379 | 44361806 | 68 | 76 | 65 | 39 | 169 | 8 |
| SE | 42032 | ENSG00000117281 | CD160 | chr1 | + | 145730743 | 145731070 | 145728243 | 145728400 | 145735996 | 145736134 | 297 | 102 | 133 | 221 | 106 | 27 |
| SE | 52141 | ENSG00000143157 | POGK | chr1 | + | 166846611 | 166846738 | 166840954 | 166841088 | 166848937 | 166849208 | 37 | 31 | 47 | 41 | 43 | 15 |
| SE | 1185 | ENSG00000100351 | GRAP2 | chr22 | + | 39955818 | 39955910 | 39946816 | 39947184 | 39960054 | 39960174 | 826 | 840 | 541 | 1544 | 1255 | 38 |
| SE | 107332 | ENSG00000151148 | UBE3B | chr12 | + | 109481636 | 109481742 | 109477401 | 109477764 | 109483530 | 109483712 | 147 | 113 | 121 | 180 | 120 | 5 |
| SE | 83270 | ENSG00000146373 | RNF217 | chr6 | + | 125048151 | 125048259 | 125046606 | 125046661 | 125057941 | 125058106 | 0 | 0 | 0 | 0 | 4 | 4 |
| SE | 107896 | ENSG00000188906 | LRRK2 | chr12 | + | 40316277 | 40316403 | 40315211 | 40315300 | 40319987 | 40320175 | 6 | 6 | 4 | 8 | 12 | 124 |
| SE | 36632 | ENSG00000180964 | TCEAL8 | chrX | - | 103254604 | 103254677 | 103252994 | 103254017 | 103255077 | 103255155 | 142 | 116 | 95 | 153 | 111 | 28 |
| SE | 85822 | ENSG00000138594 | TMOD3 | chr15 | + | 51834993 | 51835058 | 51829627 | 51829836 | 51862810 | 51862983 | 3 | 2 | 6 | 5 | 7 | 120 |
| SE | 6532 | ENSG00000124155 | PIGT | chr20 | + | 45419294 | 45419395 | 45416474 | 45416694 | 45419503 | 45419590 | 95 | 37 | 86 | 74 | 73 | 0 |
| SE | 48775 | ENSG00000058804 | NDC1 | chr1 | - | 53809694 | 53809746 | 53807655 | 53807791 | 53818970 | 53819079 | 84 | 45 | 66 | 76 | 171 | 1 |
| SE | 69633 | ENSG00000197622 | CDC42SE1 | chr1 | - | 151056655 | 151056786 | 151055015 | 151055126 | 151059478 | 151059773 | 88 | 63 | 87 | 75 | 36 | 12 |
| SE | 14812 | ENSG00000137804 | NUSAP1 | chr15 | + | 41371526 | 41371684 | 41365401 | 41365589 | 41375711 | 41375751 | 21 | 22 | 52 | 55 | 59 | 1 |
| SE | 15256 | ENSG00000089195 | TRMT6 | chr20 | - | 5944804 | 5944914 | 5944161 | 5944253 | 5946405 | 5946533 | 153 | 94 | 124 | 110 | 186 | 0 |
| SE | 35195 | ENSG00000101596 | SMCHD1 | chr18 | + | 2763875 | 2763969 | 2763636 | 2763789 | 2769693 | 2769820 | 0 | 2 | 6 | 6 | 6 | 415 |
| SE | 35919 | ENSG00000131323 | TRAF3 | chr14 | + | 102886188 | 102886269 | 102876357 | 102876525 | 102897260 | 102897307 | 27 | 19 | 34 | 30 | 61 | 1 |
| SE | 57928 | ENSG00000110200 | ANAPC15 | chr11 | - | 72111441 | 72111496 | 72111156 | 72111286 | 72112649 | 72112774 | 27 | 11 | 7 | 15 | 23 | 3 |
| SE | 99843 | ENSG00000013725 | CD6 | chr11 | + | 61015712 | 61015835 | 61013422 | 61013563 | 61017478 | 61017550 | 187 | 140 | 274 | 253 | 333 | 15 |
| SE | 102846 | ENSG00000023191 | RNH1 | chr11 | - | 504823 | 504996 | 502061 | 502249 | 507112 | 507195 | 49 | 31 | 42 | 58 | 20 | 13 |
| SE | 18238 | ENSG00000174837 | ADGRE1 | chr19 | + | 6924677 | 6924872 | 6921712 | 6921883 | 6926365 | 6926601 | 60 | 54 | 31 | 46 | 107 | 3 |
| SE | 49676 | ENSG00000256269 | HMBS | chr11 | + | 119088254 | 119088308 | 119084865 | 119085066 | 119088634 | 119088707 | 19 | 14 | 19 | 26 | 13 | 0 |
| SE | 67067 | ENSG00000225205 | NA | chr2 | - | 172512333 | 172512518 | 172495374 | 172495516 | 172556088 | 172556529 | 11 | 9 | 11 | 24 | 33 | 3 |
| SE | 95871 | ENSG00000148396 | SEC16A | chr9 | - | 136445051 | 136445111 | 136443822 | 136443900 | 136445644 | 136445719 | 16 | 9 | 3 | 14 | 7 | 42 |
| SE | 40686 | ENSG00000076641 | PAG1 | chr8 | - | 80980434 | 80980494 | 80967809 | 80976906 | 80984775 | 80985377 | 226 | 293 | 318 | 286 | 526 | 0 |
| SE | 91155 | ENSG00000183049 | CAMK1D | chr10 | + | 12816249 | 12816328 | 12791157 | 12791233 | 12824464 | 12824552 | 32 | 31 | 33 | 31 | 21 | 0 |
| SE | 90169 | ENSG00000035928 | RFC1 | chr4 | - | 39323339 | 39323417 | 39321286 | 39321374 | 39326562 | 39326640 | 190 | 171 | 197 | 231 | 357 | 19 |
| SE | 99236 | ENSG00000184743 | ATL3 | chr11 | - | 63643356 | 63643495 | 63636206 | 63636334 | 63644168 | 63644261 | 537 | 464 | 386 | 566 | 997 | 1 |
| SE | 59135 | ENSG00000168421 | RHOH | chr4 | + | 40197120 | 40197300 | 40191052 | 40191338 | 40242713 | 40242834 | 290 | 161 | 347 | 224 | 588 | 2 |
| SE | 12184 | ENSG00000260528 | FAM157C | chr16 | + | 90111371 | 90111485 | 90106528 | 90106705 | 90116287 | 90116483 | 11 | 7 | 25 | 5 | 6 | 0 |
| SE | 46310 | ENSG00000093183 | SEC22C | chr3 | - | 42563764 | 42563880 | 42563522 | 42563686 | 42568864 | 42569073 | 13 | 7 | 14 | 9 | 7 | 24 |
| SE | 107225 | ENSG00000135108 | FBXO21 | chr12 | - | 117166897 | 117167077 | 117165484 | 117165617 | 117174204 | 117174341 | 29 | 14 | 32 | 25 | 43 | 2 |
| SE | 35266 | ENSG00000103064 | SLC7A6 | chr16 | + | 68273817 | 68273904 | 68266592 | 68266721 | 68274690 | 68274811 | 32 | 35 | 77 | 43 | 22 | 34 |
| SE | 51669 | ENSG00000136643 | RPS6KC1 | chr1 | + | 213117316 | 213117410 | 213104453 | 213104569 | 213129526 | 213129889 | 137 | 127 | 117 | 132 | 123 | 2 |
| SE | 21888 | ENSG00000189091 | SF3B3 | chr16 | + | 70565064 | 70565270 | 70563728 | 70564050 | 70567410 | 70567536 | 183 | 106 | 154 | 191 | 219 | 5 |
| SE | 16781 | ENSG00000125107 | CNOT1 | chr16 | - | 58624652 | 58624794 | 58623281 | 58623409 | 58629727 | 58629815 | 10 | 6 | 25 | 11 | 12 | 2 |
| SE | 82640 | ENSG00000103145 | HCFC1R1 | chr16 | - | 3023473 | 3023530 | 3023232 | 3023361 | 3023846 | 3023942 | 28 | 12 | 11 | 3 | 3 | 1 |
| SE | 37903 | ENSG00000184014 | DENND5A | chr11 | - | 9143402 | 9143485 | 9142721 | 9142845 | 9144096 | 9144278 | 177 | 124 | 167 | 160 | 159 | 10 |
| SE | 7924 | ENSG00000108953 | YWHAE | chr17 | - | 1361929 | 1362008 | 1354210 | 1354347 | 1364858 | 1365058 | 269 | 251 | 200 | 267 | 359 | 6 |
| SE | 62324 | ENSG00000185811 | IKZF1 | chr7 | + | 50382539 | 50382707 | 50327637 | 50327757 | 50391728 | 50391863 | 185 | 103 | 173 | 117 | 116 | 5 |
| SE | 62918 | ENSG00000185420 | SMYD3 | chr1 | - | 245814341 | 245814415 | 245764040 | 245764149 | 245858495 | 245858670 | 0 | 5 | 2 | 3 | 0 | 32 |
| SE | 75072 | ENSG00000198223 | CSF2RA | chrX | + | 1305445 | 1305527 | 1303922 | 1304019 | 1309401 | 1309638 | 117 | 97 | 88 | 85 | 115 | 0 |
| SE | 90296 | ENSG00000249859 | PVT1 | chr8 | + | 127969415 | 127969537 | 127939507 | 127939676 | 127989161 | 127989291 | 1 | 0 | 2 | 0 | 0 | 6 |
| SE | 66447 | ENSG00000131051 | RBM39 | chr20 | - | 35707939 | 35707992 | 35707119 | 35707201 | 35709223 | 35709256 | 1 | 9 | 0 | 1 | 11 | 721 |
| SE | 96352 | ENSG00000149428 | HYOU1 | chr11 | - | 119051818 | 119051951 | 119051437 | 119051625 | 119052089 | 119052172 | 209 | 96 | 230 | 204 | 152 | 0 |
| SE | 69634 | ENSG00000197622 | CDC42SE1 | chr1 | - | 151056655 | 151056786 | 151055582 | 151055993 | 151059478 | 151059773 | 155 | 111 | 159 | 132 | 161 | 632 |
| SE | 58251 | ENSG00000089127 | OAS1 | chr12 | + | 112911097 | 112911258 | 112906961 | 112907219 | 112916508 | 112916738 | 6 | 2 | 4 | 8 | 7 | 2 |
| SE | 48016 | ENSG00000010810 | FYN | chr6 | - | 111740980 | 111741093 | 111719804 | 111720062 | 111759844 | 111759925 | 67 | 18 | 29 | 37 | 49 | 97 |
| SE | 64152 | ENSG00000032444 | PNPLA6 | chr19 | + | 7554208 | 7554272 | 7553871 | 7554015 | 7554554 | 7554723 | 45 | 37 | 29 | 54 | 13 | 0 |
| SE | 1772 | ENSG00000100077 | GRK3 | chr22 | + | 25674436 | 25674528 | 25667738 | 25667800 | 25678815 | 25678915 | 107 | 146 | 69 | 139 | 156 | 8 |
| SE | 54104 | ENSG00000115935 | WIPF1 | chr2 | - | 174629665 | 174629868 | 174625623 | 174625960 | 174634517 | 174634536 | 14 | 13 | 10 | 14 | 13 | 16 |
| SE | 96230 | ENSG00000133943 | DGLUCY | chr14 | + | 91145220 | 91145313 | 91114685 | 91114994 | 91157638 | 91157690 | 17 | 11 | 18 | 32 | 22 | 9 |
| SE | 82646 | ENSG00000131373 | HACL1 | chr3 | - | 15585247 | 15585342 | 15582876 | 15582989 | 15586524 | 15586602 | 61 | 23 | 46 | 63 | 81 | 1 |
| SE | 65806 | ENSG00000116521 | SCAMP3 | chr1 | - | 155261656 | 155261734 | 155260536 | 155260659 | 155262085 | 155262360 | 82 | 18 | 53 | 85 | 20 | 10 |
| SE | 22128 | ENSG00000237296 | SMG1P1 | chr16 | + | 22454701 | 22454865 | 22437400 | 22437492 | 22459354 | 22459491 | 187 | 91 | 144 | 232 | 61 | 9 |
| SE | 77387 | ENSG00000089820 | ARHGAP4 | chrX | - | 153911128 | 153911189 | 153910921 | 153910999 | 153912699 | 153912737 | 330 | 133 | 280 | 226 | 238 | 2 |
| SE | 93902 | ENSG00000184465 | WDR27 | chr6 | - | 169662303 | 169662424 | 169660662 | 169660766 | 169672254 | 169672396 | 9 | 1 | 16 | 13 | 14 | 0 |
| SE | 107994 | ENSG00000204852 | TCTN1 | chr12 | + | 110641023 | 110641149 | 110640382 | 110640517 | 110644966 | 110645057 | 5 | 12 | 7 | 6 | 16 | 0 |
| SE | 96403 | ENSG00000197563 | PIGN | chr18 | - | 62162252 | 62162329 | 62161132 | 62161385 | 62163530 | 62163656 | 11 | 15 | 22 | 32 | 16 | 17 |
| SE | 83158 | ENSG00000055130 | CUL1 | chr7 | + | 148767618 | 148767749 | 148766560 | 148766723 | 148783782 | 148783890 | 441 | 326 | 357 | 410 | 495 | 0 |
| SE | 86157 | ENSG00000113845 | TIMMDC1 | chr3 | + | 119500694 | 119500860 | 119498546 | 119498927 | 119503531 | 119503620 | 149 | 98 | 111 | 114 | 153 | 1 |
| SE | 32768 | ENSG00000168297 | PXK | chr3 | + | 58390581 | 58390659 | 58382513 | 58382700 | 58394997 | 58395102 | 41 | 31 | 30 | 42 | 98 | 3 |
| SE | 82328 | ENSG00000198742 | SMURF1 | chr7 | - | 99049562 | 99049709 | 99047683 | 99047882 | 99051356 | 99051441 | 86 | 56 | 96 | 90 | 147 | 0 |
| SE | 87923 | ENSG00000173273 | TNKS | chr8 | + | 9751608 | 9751846 | 9748023 | 9748212 | 9752543 | 9752626 | 104 | 55 | 111 | 70 | 98 | 0 |
| SE | 30724 | ENSG00000125375 | DMAC2L | chr14 | + | 50314590 | 50314626 | 50312323 | 50312389 | 50321482 | 50321594 | 2 | 2 | 2 | 2 | 0 | 2 |
| SE | 96564 | ENSG00000214413 | BBIP1 | chr10 | - | 110918120 | 110918213 | 110901537 | 110901612 | 110919120 | 110919157 | 60 | 47 | 91 | 70 | 79 | 12 |
| SE | 93017 | ENSG00000147162 | OGT | chrX | + | 71544566 | 71544635 | 71537828 | 71538072 | 71547906 | 71548023 | 371 | 245 | 345 | 361 | 192 | 7 |
| SE | 7180 | ENSG00000101639 | CEP192 | chr18 | + | 13088753 | 13088904 | 13087530 | 13087646 | 13089455 | 13089565 | 13 | 9 | 3 | 10 | 5 | 120 |
| SE | 47478 | ENSG00000183023 | SLC8A1 | chr2 | - | 40446414 | 40446648 | 40428472 | 40430301 | 40451903 | 40452008 | 1 | 3 | 2 | 7 | 0 | 5 |
| SE | 60939 | ENSG00000138078 | PREPL | chr2 | - | 44359496 | 44360038 | 44346267 | 44346390 | 44361379 | 44361494 | 68 | 75 | 65 | 42 | 166 | 8 |
| SE | 78144 | ENSG00000214765 | SEPTIN7P2 | chr7 | - | 45756908 | 45756992 | 45751528 | 45751646 | 45758042 | 45758177 | 58 | 34 | 76 | 74 | 75 | 2 |
| SE | 40285 | ENSG00000271383 | NBPF19 | chr1 | + | 149478879 | 149479094 | 149475759 | 149476107 | 149480141 | 149480214 | 15 | 22 | 16 | 24 | 23 | 0 |
| SE | 101686 | ENSG00000180398 | MCFD2 | chr2 | - | 46909022 | 46909177 | 46901873 | 46905594 | 46915722 | 46915761 | 12 | 10 | 6 | 16 | 9 | 0 |
| SE | 101207 | ENSG00000150433 | TMEM218 | chr11 | - | 125102131 | 125102317 | 125101200 | 125101303 | 125102636 | 125102809 | 4 | 9 | 5 | 9 | 13 | 1 |
| SE | 76294 | ENSG00000124508 | BTN2A2 | chr6 | + | 26385142 | 26385362 | 26383791 | 26383915 | 26388012 | 26388159 | 23 | 20 | 23 | 30 | 30 | 0 |
| SE | 38831 | ENSG00000122674 | CCZ1 | chr7 | + | 5912842 | 5912954 | 5910034 | 5910116 | 5918866 | 5918900 | 29 | 13 | 29 | 21 | 104 | 0 |
| SE | 104582 | ENSG00000256751 | PLBD1-AS1 | chr12 | + | 14619151 | 14619373 | 14614521 | 14614570 | 14619737 | 14619803 | 17 | 3 | 5 | 16 | 28 | 0 |
| SE | 33016 | ENSG00000114316 | USP4 | chr3 | - | 49324893 | 49325039 | 49324696 | 49324763 | 49325718 | 49325845 | 427 | 267 | 383 | 423 | 563 | 0 |
| SE | 38204 | ENSG00000186501 | TMEM222 | chr1 | + | 27330984 | 27331137 | 27330719 | 27330804 | 27332069 | 27332101 | 20 | 2 | 9 | 16 | 16 | 34 |
| SE | 19576 | ENSG00000165355 | FBXO33 | chr14 | - | 39402400 | 39402511 | 39401175 | 39401861 | 39431563 | 39432500 | 180 | 87 | 169 | 182 | 117 | 3 |
| SE | 40522 | ENSG00000285708 | NA | chr3 | - | 71710411 | 71710484 | 71699613 | 71699708 | 71728475 | 71728623 | 76 | 94 | 57 | 91 | 111 | 0 |
| SE | 58647 | ENSG00000134759 | ELP2 | chr18 | + | 36146249 | 36146381 | 36145947 | 36146048 | 36154849 | 36154999 | 138 | 77 | 141 | 139 | 271 | 0 |
| SE | 12321 | ENSG00000197912 | SPG7 | chr16 | + | 89529476 | 89529579 | 89526328 | 89526468 | 89530682 | 89530808 | 66 | 57 | 94 | 59 | 49 | 1 |
| SE | 78357 | ENSG00000072609 | CHFR | chr12 | - | 132869618 | 132869834 | 132861466 | 132861634 | 132870723 | 132870783 | 110 | 84 | 105 | 121 | 137 | 36 |
| SE | 10126 | ENSG00000108433 | GOSR2 | chr17 | + | 46963696 | 46965039 | 46938598 | 46938704 | 46966533 | 46966851 | 2 | 3 | 6 | 5 | 10 | 0 |
| SE | 2002 | ENSG00000198131 | ZNF544 | chr19 | + | 58246745 | 58246794 | 58246300 | 58246427 | 58260850 | 58260965 | 13 | 11 | 18 | 7 | 12 | 0 |
| SE | 82173 | ENSG00000146872 | TLK2 | chr17 | + | 62524235 | 62524331 | 62523133 | 62523177 | 62536169 | 62536337 | 105 | 92 | 121 | 123 | 148 | 24 |
| SE | 68192 | ENSG00000020922 | MRE11 | chr11 | - | 94479673 | 94479761 | 94478734 | 94478876 | 94485923 | 94486084 | 177 | 142 | 119 | 98 | 131 | 4 |
| SE | 72324 | ENSG00000197299 | BLM | chr15 | + | 90782821 | 90782928 | 90769437 | 90769586 | 90784920 | 90785081 | 36 | 23 | 41 | 70 | 72 | 0 |
| SE | 16521 | ENSG00000140367 | UBE2Q2 | chr15 | + | 75877960 | 75878021 | 75873427 | 75873568 | 75879097 | 75879188 | 121 | 117 | 103 | 114 | 119 | 8 |
| SE | 44012 | ENSG00000114098 | ARMC8 | chr3 | + | 138237308 | 138237384 | 138235033 | 138235114 | 138239467 | 138239528 | 31 | 28 | 23 | 53 | 58 | 2 |
| SE | 3367 | ENSG00000130164 | LDLR | chr19 | + | 11120369 | 11120522 | 11120091 | 11120233 | 11123173 | 11123344 | 190 | 62 | 171 | 143 | 211 | 0 |
| SE | 69897 | ENSG00000142961 | MOB3C | chr1 | - | 46610001 | 46610204 | 46607714 | 46609684 | 46612903 | 46613371 | 91 | 29 | 51 | 62 | 39 | 0 |
| SE | 3988 | ENSG00000126261 | UBA2 | chr19 | + | 34458768 | 34458924 | 34454443 | 34454556 | 34460469 | 34460566 | 365 | 248 | 358 | 382 | 592 | 10 |
| SE | 96151 | ENSG00000115084 | SLC35F5 | chr2 | - | 113717752 | 113717850 | 113706708 | 113715195 | 113719153 | 113719280 | 63 | 64 | 63 | 68 | 74 | 5 |
| SE | 51880 | ENSG00000285791 | NA | chr8 | + | 66456816 | 66456895 | 66454058 | 66454157 | 66457069 | 66457166 | 32 | 17 | 25 | 31 | 16 | 0 |
| SE | 14910 | ENSG00000086666 | ZFAND6 | chr15 | + | 80073265 | 80073330 | 80059713 | 80059809 | 80120327 | 80120498 | 1 | 0 | 0 | 0 | 1 | 47 |
| SE | 49147 | ENSG00000160072 | ATAD3B | chr1 | + | 1489691 | 1489811 | 1489203 | 1489274 | 1490256 | 1490424 | 3 | 2 | 2 | 3 | 2 | 7 |
| SE | 49312 | ENSG00000087589 | CASS4 | chr20 | + | 56451818 | 56453129 | 56450598 | 56450679 | 56458339 | 56458780 | 16 | 23 | 41 | 30 | 30 | 3 |
| SE | 94132 | ENSG00000139644 | TMBIM6 | chr12 | + | 49743416 | 49743565 | 49741956 | 49742115 | 49752463 | 49752549 | 5 | 1 | 0 | 7 | 0 | 26 |
| SE | 101114 | ENSG00000162241 | SLC25A45 | chr11 | - | 65380533 | 65380632 | 65380131 | 65380175 | 65381914 | 65382373 | 7 | 16 | 0 | 6 | 13 | 13 |
| SE | 78534 | ENSG00000164054 | SHISA5 | chr3 | - | 48503094 | 48503193 | 48501136 | 48501293 | 48504018 | 48504140 | 4 | 0 | 2 | 3 | 0 | 15 |
| SE | 6192 | ENSG00000125772 | GPCPD1 | chr20 | - | 5586193 | 5586269 | 5584280 | 5584322 | 5593326 | 5593411 | 207 | 165 | 178 | 158 | 376 | 75 |
| SE | 52174 | ENSG00000142599 | RERE | chr1 | - | 8423166 | 8423247 | 8422726 | 8422807 | 8424014 | 8424234 | 5 | 3 | 6 | 10 | 4 | 8 |
| SE | 81594 | ENSG00000253729 | PRKDC | chr8 | - | 47855221 | 47855373 | 47854082 | 47854214 | 47857155 | 47857299 | 225 | 159 | 233 | 267 | 301 | 3 |
| SE | 87277 | ENSG00000147133 | TAF1 | chrX | + | 71454169 | 71454237 | 71424153 | 71424238 | 71454740 | 71454857 | 152 | 148 | 132 | 142 | 307 | 1 |
| SE | 6483 | ENSG00000101608 | MYL12A | chr18 | + | 3249360 | 3249443 | 3247812 | 3247909 | 3253232 | 3253428 | 8 | 4 | 3 | 10 | 9 | 473 |
| SE | 18393 | ENSG00000136305 | CIDEB | chr14 | - | 24307817 | 24307920 | 24307370 | 24307515 | 24308478 | 24308830 | 132 | 45 | 101 | 102 | 241 | 14 |
| SE | 38949 | ENSG00000067066 | SP100 | chr2 | + | 230498460 | 230498535 | 230494415 | 230494460 | 230503065 | 230503110 | 207 | 238 | 190 | 209 | 347 | 6 |
| SE | 76635 | ENSG00000135503 | ACVR1B | chr12 | + | 51980968 | 51981199 | 51976326 | 51976575 | 51986817 | 51986942 | 32 | 18 | 29 | 37 | 3 | 0 |
| SE | 80622 | ENSG00000258227 | CLEC5A | chr7 | - | 141935813 | 141935950 | 141931506 | 141931826 | 141945340 | 141945400 | 22 | 12 | 12 | 6 | 17 | 0 |
| SE | 57922 | ENSG00000110200 | ANAPC15 | chr11 | - | 72111156 | 72111286 | 72110543 | 72110603 | 72111411 | 72111443 | 82 | 32 | 44 | 38 | 68 | 0 |
| SE | 17025 | ENSG00000102921 | N4BP1 | chr16 | - | 48553538 | 48553669 | 48538725 | 48543261 | 48560753 | 48562444 | 110 | 68 | 92 | 98 | 163 | 2 |
| SE | 70657 | ENSG00000168214 | RBPJ | chr4 | + | 26414048 | 26414155 | 26406174 | 26406270 | 26415474 | 26415640 | 4 | 17 | 1 | 5 | 4 | 98 |
| SE | 95439 | ENSG00000119383 | PTPA | chr9 | + | 129127963 | 129128068 | 129123051 | 129123138 | 129128984 | 129129110 | 32 | 7 | 19 | 19 | 59 | 58 |
| SE | 21031 | ENSG00000126214 | KLC1 | chr14 | + | 103693552 | 103693634 | 103692358 | 103692425 | 103700654 | 103700727 | 26 | 9 | 17 | 10 | 15 | 2 |
| SE | 312 | ENSG00000182670 | TTC3 | chr21 | + | 37094004 | 37094090 | 37091292 | 37091413 | 37095349 | 37095444 | 359 | 308 | 344 | 377 | 538 | 3 |
| SE | 106053 | ENSG00000130177 | CDC16 | chr13 | + | 114243256 | 114243348 | 114242120 | 114242280 | 114243855 | 114243989 | 206 | 145 | 136 | 144 | 281 | 0 |
| SE | 106604 | ENSG00000259431 | THTPA | chr14 | + | 23556742 | 23557304 | 23556006 | 23556343 | 23558694 | 23559574 | 22 | 24 | 30 | 16 | 2 | 6 |
| SE | 20520 | ENSG00000175061 | SNHG29 | chr17 | + | 16439659 | 16439703 | 16439037 | 16439060 | 16440184 | 16440253 | 53 | 24 | 59 | 56 | 55 | 8 |
| SE | 24444 | ENSG00000114107 | CEP70 | chr3 | - | 138572858 | 138572984 | 138571265 | 138571356 | 138591853 | 138591956 | 58 | 42 | 52 | 56 | 83 | 0 |
| SE | 105315 | ENSG00000240505 | TNFRSF13B | chr17 | - | 16952445 | 16952583 | 16948737 | 16948983 | 16972014 | 16972078 | 14 | 10 | 22 | 23 | 11 | 1 |
| SE | 36479 | ENSG00000124596 | OARD1 | chr6 | - | 41070075 | 41070134 | 41066787 | 41067437 | 41071131 | 41071217 | 79 | 60 | 75 | 62 | 59 | 4 |
| SE | 13148 | ENSG00000103121 | CMC2 | chr16 | - | 80980769 | 80980879 | 80975801 | 80976179 | 81006733 | 81006839 | 4 | 2 | 6 | 10 | 13 | 0 |
| SE | 65304 | ENSG00000169621 | APLF | chr2 | + | 68513547 | 68513680 | 68502730 | 68502903 | 68526060 | 68526242 | 52 | 64 | 53 | 63 | 136 | 0 |
| SE | 39517 | ENSG00000118260 | CREB1 | chr2 | + | 207577109 | 207577224 | 207575271 | 207575454 | 207577504 | 207577655 | 5 | 25 | 4 | 12 | 27 | 101 |
| SE | 81150 | ENSG00000110665 | C11orf21 | chr11 | - | 2300519 | 2300613 | 2299427 | 2299707 | 2301755 | 2301918 | 62 | 10 | 22 | 42 | 11 | 11 |
| SE | 31348 | ENSG00000157426 | AASDH | chr4 | - | 56384069 | 56384341 | 56381758 | 56382597 | 56387361 | 56387459 | 47 | 34 | 69 | 43 | 76 | 5 |
| SE | 12458 | ENSG00000149932 | TMEM219 | chr16 | + | 29972041 | 29972140 | 29968024 | 29968254 | 29972949 | 29973048 | 13 | 7 | 3 | 6 | 8 | 2 |
| SE | 41677 | ENSG00000079277 | MKNK1 | chr1 | - | 46580529 | 46580627 | 46573733 | 46576654 | 46583227 | 46583329 | 201 | 135 | 152 | 126 | 194 | 1 |
| SE | 102995 | ENSG00000197535 | MYO5A | chr15 | - | 52348817 | 52348826 | 52346360 | 52346461 | 52351253 | 52351481 | 74 | 83 | 71 | 73 | 121 | 7 |
| SE | 17069 | ENSG00000166140 | ZFYVE19 | chr15 | + | 40809851 | 40810216 | 40808120 | 40809458 | 40810648 | 40810757 | 79 | 67 | 56 | 125 | 54 | 0 |
| SE | 91642 | ENSG00000105339 | DENND3 | chr8 | + | 141150833 | 141150953 | 141136508 | 141136791 | 141151618 | 141151837 | 45 | 23 | 12 | 38 | 54 | 0 |
| SE | 97537 | ENSG00000149761 | NUDT22 | chr11 | + | 64229477 | 64229571 | 64226289 | 64227132 | 64229849 | 64229983 | 39 | 13 | 21 | 30 | 16 | 0 |
| SE | 56128 | ENSG00000204406 | MBD5 | chr2 | + | 148233244 | 148233395 | 148178699 | 148178793 | 148342213 | 148342336 | 63 | 79 | 104 | 70 | 116 | 0 |
| SE | 44247 | ENSG00000102606 | ARHGEF7 | chr13 | + | 111288353 | 111288443 | 111286146 | 111286240 | 111300747 | 111300847 | 204 | 138 | 178 | 190 | 167 | 1 |
| SE | 49155 | ENSG00000077044 | DGKD | chr2 | + | 233446711 | 233446796 | 233445622 | 233445762 | 233448086 | 233448181 | 133 | 70 | 107 | 134 | 93 | 4 |
| SE | 94143 | ENSG00000090054 | SPTLC1 | chr9 | - | 92046274 | 92046334 | 92045998 | 92046053 | 92047171 | 92047268 | 3 | 10 | 5 | 5 | 3 | 226 |
| SE | 999 | ENSG00000243156 | MICAL3 | chr22 | - | 17827643 | 17827781 | 17822946 | 17823060 | 17831853 | 17832107 | 2 | 0 | 1 | 4 | 2 | 10 |
| SE | 97238 | ENSG00000073921 | PICALM | chr11 | - | 86000642 | 86000779 | 85996825 | 85996929 | 86003365 | 86003451 | 548 | 301 | 311 | 419 | 653 | 1 |
| SE | 100622 | ENSG00000132254 | ARFIP2 | chr11 | - | 6480322 | 6480423 | 6479971 | 6480068 | 6481230 | 6481312 | 52 | 27 | 49 | 50 | 31 | 2 |
| SE | 87259 | ENSG00000046651 | OFD1 | chrX | + | 13767126 | 13767284 | 13763744 | 13763855 | 13768053 | 13768224 | 110 | 94 | 119 | 92 | 161 | 1 |
| SE | 80698 | ENSG00000175471 | MCTP1 | chr5 | - | 94917895 | 94917973 | 94912805 | 94912976 | 94923961 | 94924021 | 82 | 88 | 49 | 96 | 151 | 41 |
| SE | 86207 | ENSG00000172534 | HCFC1 | chrX | - | 153956190 | 153956411 | 153954065 | 153955542 | 153956624 | 153956763 | 138 | 58 | 114 | 107 | 88 | 0 |
| SE | 52379 | ENSG00000048707 | VPS13D | chr1 | + | 12304505 | 12304728 | 12299201 | 12299384 | 12308430 | 12308641 | 163 | 104 | 139 | 120 | 126 | 0 |
| SE | 20011 | ENSG00000112679 | DUSP22 | chr6 | + | 335113 | 335163 | 311879 | 311962 | 345853 | 345912 | 85 | 60 | 110 | 135 | 134 | 87 |
| SE | 93933 | ENSG00000144524 | COPS7B | chr2 | + | 231791732 | 231791808 | 231788554 | 231788732 | 231796105 | 231796308 | 36 | 13 | 29 | 40 | 33 | 1 |
| SE | 8000 | ENSG00000126858 | RHOT1 | chr17 | + | 32209349 | 32209445 | 32208106 | 32208309 | 32211115 | 32211238 | 32 | 16 | 31 | 28 | 49 | 10 |
| SE | 36703 | ENSG00000158062 | UBXN11 | chr1 | - | 26303042 | 26303167 | 26302812 | 26302918 | 26306287 | 26306576 | 7 | 10 | 3 | 13 | 4 | 0 |
| SE | 83048 | ENSG00000196305 | IARS1 | chr9 | - | 92242613 | 92242687 | 92242153 | 92242330 | 92243215 | 92243311 | 1 | 7 | 2 | 2 | 0 | 132 |
| SE | 17867 | ENSG00000092199 | HNRNPC | chr14 | - | 21263310 | 21263582 | 21234030 | 21234229 | 21269297 | 21269425 | 506 | 233 | 455 | 513 | 249 | 424 |
| SE | 93221 | ENSG00000166507 | NDST2 | chr10 | - | 73806656 | 73806811 | 73805899 | 73806028 | 73807107 | 73807195 | 54 | 16 | 54 | 41 | 56 | 0 |
| SE | 104124 | ENSG00000130783 | CCDC62 | chr12 | + | 122781163 | 122781330 | 122777490 | 122777683 | 122785718 | 122785820 | 8 | 4 | 10 | 16 | 30 | 0 |
| SE | 3868 | ENSG00000157540 | DYRK1A | chr21 | + | 37456129 | 37456308 | 37430268 | 37430400 | 37472683 | 37472880 | 3 | 2 | 2 | 8 | 0 | 24 |
| SE | 109615 | ENSG00000005810 | MYCBP2 | chr13 | - | 77088831 | 77089031 | 77087483 | 77087633 | 77090105 | 77090263 | 469 | 440 | 421 | 431 | 989 | 14 |
| SE | 101471 | ENSG00000165309 | ARMC3 | chr10 | + | 22946143 | 22946261 | 22931995 | 22932044 | 22955806 | 22955932 | 8 | 39 | 9 | 50 | 83 | 1 |
| SE | 95722 | ENSG00000185920 | PTCH1 | chr9 | - | 95476758 | 95476857 | 95476033 | 95476159 | 95478054 | 95478186 | 17 | 9 | 7 | 24 | 28 | 0 |
| SE | 95306 | ENSG00000148400 | NOTCH1 | chr9 | - | 136515980 | 136516094 | 136515482 | 136515716 | 136517751 | 136517937 | 38 | 13 | 35 | 36 | 11 | 0 |
| SE | 54053 | ENSG00000158796 | DEDD | chr1 | - | 161123074 | 161123221 | 161120973 | 161122523 | 161123838 | 161123946 | 159 | 94 | 134 | 149 | 142 | 1 |
| SE | 39727 | ENSG00000132716 | DCAF8 | chr1 | - | 160261284 | 160261350 | 160243959 | 160244034 | 160262116 | 160262451 | 89 | 96 | 67 | 81 | 104 | 22 |
| SE | 40767 | ENSG00000114026 | OGG1 | chr3 | + | 9756470 | 9756621 | 9754703 | 9754885 | 9756766 | 9756816 | 49 | 19 | 24 | 30 | 15 | 0 |
| SE | 106125 | ENSG00000187778 | MCRS1 | chr12 | - | 49566076 | 49566215 | 49565528 | 49565667 | 49568047 | 49568100 | 108 | 37 | 67 | 74 | 60 | 1 |
| SE | 25443 | ENSG00000011243 | AKAP8L | chr19 | - | 15398629 | 15400358 | 15397713 | 15397855 | 15400793 | 15400864 | 56 | 44 | 47 | 52 | 44 | 0 |
| SE | 90286 | ENSG00000249859 | PVT1 | chr8 | + | 127902455 | 127902539 | 127890195 | 127890998 | 127939507 | 127940454 | 1 | 0 | 1 | 0 | 0 | 8 |
| SE | 93752 | ENSG00000143409 | MINDY1 | chr1 | - | 150998081 | 150998273 | 150997623 | 150997779 | 150999368 | 150999511 | 134 | 121 | 90 | 187 | 133 | 1 |
| SE | 54641 | ENSG00000140992 | PDPK1 | chr16 | + | 2597122 | 2597275 | 2586675 | 2586893 | 2597650 | 2597928 | 155 | 96 | 137 | 134 | 195 | 0 |
| SE | 43128 | ENSG00000186470 | BTN3A2 | chr6 | + | 26367989 | 26368267 | 26365257 | 26365352 | 26368564 | 26368765 | 125 | 104 | 102 | 68 | 168 | 0 |
| SE | 8720 | ENSG00000070366 | SMG6 | chr17 | - | 2292870 | 2292977 | 2292551 | 2292630 | 2298905 | 2300664 | 37 | 26 | 32 | 17 | 26 | 1 |
| SE | 51774 | ENSG00000042429 | MED17 | chr11 | + | 93797534 | 93797719 | 93794907 | 93796540 | 93801834 | 93801942 | 87 | 64 | 101 | 83 | 112 | 12 |
| SE | 24942 | ENSG00000072364 | AFF4 | chr5 | - | 132927120 | 132927207 | 132904367 | 132904404 | 132932177 | 132932222 | 169 | 134 | 181 | 177 | 67 | 2 |
| SE | 31258 | ENSG00000047365 | ARAP2 | chr4 | - | 36080215 | 36080279 | 36073688 | 36073823 | 36082250 | 36082286 | 355 | 247 | 199 | 280 | 356 | 5 |
| SE | 108957 | ENSG00000139496 | NUP58 | chr13 | + | 25319325 | 25319350 | 25315356 | 25315467 | 25320529 | 25320595 | 112 | 58 | 82 | 42 | 124 | 2 |
| SE | 94667 | ENSG00000165209 | STRBP | chr9 | - | 123168188 | 123168258 | 123160976 | 123161068 | 123169901 | 123170046 | 2 | 3 | 5 | 3 | 6 | 78 |
| SE | 83880 | ENSG00000157168 | NRG1 | chr8 | + | 32742007 | 32742075 | 32727948 | 32728078 | 32742674 | 32743224 | 28 | 12 | 10 | 6 | 31 | 17 |
| SE | 57978 | ENSG00000160746 | ANO10 | chr3 | - | 43580352 | 43580472 | 43565652 | 43565727 | 43598531 | 43598666 | 25 | 23 | 27 | 21 | 33 | 0 |
| SE | 55315 | ENSG00000166938 | DIS3L | chr15 | + | 66323492 | 66323585 | 66322686 | 66322934 | 66325830 | 66325988 | 92 | 46 | 90 | 100 | 150 | 13 |
| SE | 98847 | ENSG00000077458 | FAM76B | chr11 | - | 95779077 | 95779135 | 95778821 | 95778957 | 95779606 | 95779687 | 70 | 82 | 70 | 64 | 70 | 125 |
| SE | 88750 | ENSG00000205581 | HMGN1 | chr21 | - | 39347246 | 39347292 | 39345145 | 39345274 | 39347378 | 39347483 | 7 | 4 | 4 | 3 | 1 | 11 |
| SE | 31652 | ENSG00000131389 | SLC6A6 | chr3 | + | 14472204 | 14472317 | 14468087 | 14468212 | 14479084 | 14479185 | 34 | 8 | 30 | 27 | 22 | 0 |
| SE | 51124 | ENSG00000233355 | CHRM3-AS2 | chr1 | - | 239707614 | 239707760 | 239703891 | 239705175 | 239714176 | 239714264 | 38 | 52 | 103 | 41 | 78 | 2 |
| SE | 4593 | ENSG00000105397 | TYK2 | chr19 | - | 10351051 | 10351162 | 10350528 | 10350968 | 10352433 | 10352551 | 260 | 129 | 196 | 243 | 221 | 1 |
| SE | 53726 | ENSG00000151503 | NCAPD3 | chr11 | - | 134157927 | 134158067 | 134157017 | 134157095 | 134158328 | 134158495 | 38 | 30 | 43 | 32 | 32 | 1 |
| SE | 103902 | ENSG00000066084 | DIP2B | chr12 | + | 50691048 | 50691151 | 50686572 | 50686682 | 50695266 | 50695360 | 122 | 87 | 119 | 124 | 131 | 10 |
| SE | 48455 | ENSG00000131238 | PPT1 | chr1 | - | 40076729 | 40076913 | 40072718 | 40074183 | 40078559 | 40078658 | 589 | 329 | 333 | 388 | 641 | 48 |
| SE | 10097 | ENSG00000108423 | TUBD1 | chr17 | - | 59890830 | 59891041 | 59886082 | 59886230 | 59892696 | 59892715 | 38 | 34 | 57 | 30 | 57 | 11 |

| | | | | | | | | | | | | | | | | | | | | | | | | | | | | | | | | | | | | | | | | | | | | | | |
|---|---|---|---|---|---|---|---|---|---|---|---|---|---|---|---|---|---|---|---|---|---|---|---|---|---|---|---|---|---|---|---|---|---|---|---|---|---|---|---|---|---|---|---|---|---|---|
| SE | 80078 | ENSG00000198198 | SZT2 | chr1 | + | 43422082 | 43422225 | 43421173 | 43421303 | 43422479 | 43422632 | 53 | 24 | 44 | 88 | 14 | 1 | 0 | 0 | 0 | 0 | 52 | 71 | 45 | 55 | 107 | 0 | 6 | 0 | 3 | 0 | 292 | 149 | 0.122077532 | 0.759276788 | 0.964 | 1 | 1 | 1 | 1 | 1 | 0.858 | 1 | 0.903 | 1 | 0.041 |
| SE | 63390 | ENSG00000118263 | KLF7 | chr2 | - | 207148981 | 207149200 | 207124352 | 207124404 | 207165466 | 207165706 | 11 | 11 | 8 | 7 | 19 | 31 | 25 | 53 | 48 | 71 | 26 | 9 | 23 | 49 | 0 | 45 | 49 | 76 | 59 | 27 | 298 | 149 | 0.122082041 | 0.759276788 | 0.151 | 0.18 | 0.07 | 0.068 | 0.118 | 0.224 | 0.084 | 0.131 | 0.293 | 0 | -0.029 |
| SE | 100175 | ENSG00000078124 | ACER3 | chr11 | + | 76994155 | 76994226 | 76990538 | 76990574 | 76997205 | 76997273 | 5 | 6 | 0 | 9 | 6 | 1 | 9 | 4 | 6 | 5 | 29 | 8 | 6 | 9 | 5 | 15 | 0 | 9 | 0 | 9 | 220 | 149 | 0.122128812 | 0.759507345 | 0.772 | 0.311 | 0 | 0.504 | 0.448 | 0.567 | 1 | 0.311 | 1 | 0.273 | -0.223 |
| SE | 54859 | ENSG00000133056 | PIK3C2B | chr1 | - | 204464449 | 204464604 | 204464011 | 204464132 | 204465218 | 204465319 | 51 | 33 | 69 | 55 | 71 | 0 | 0 | 0 | 0 | 0 | 65 | 114 | 25 | 14 | 154 | 0 | 0 | 0 | 7 | 0 | 298 | 149 | 0.122212987 | 0.759970455 | 1 | 1 | 1 | 1 | 1 | 1 | 1 | 1 | 0.5 | 1 | 0.1 |
| SE | 72608 | ENSG00000115977 | AAK1 | chr2 | - | 69495984 | 69496080 | 69457996 | 69466551 | 69505568 | 69505673 | 17 | 15 | 34 | 25 | 22 | 98 | 67 | 71 | 94 | 143 | 18 | 16 | 30 | 16 | 10 | 22 | 67 | 92 | 47 | 44 | 245 | 149 | 0.122227718 | 0.760001703 | 0.095 | 0.12 | 0.226 | 0.139 | 0.086 | 0.332 | 0.127 | 0.165 | 0.172 | 0.121 | -0.05 |
| SE | 33559 | ENSG00000009790 | TRAF3IP3 | chr1 | + | 209759981 | 209760324 | 209759033 | 209759134 | 209762514 | 209762662 | 408 | 303 | 360 | 285 | 198 | 0 | 1 | 0 | 0 | 0 | 436 | 476 | 279 | 193 | 356 | 19 | 0 | 0 | 0 | 0 | 298 | 149 | 0.122313215 | 0.76047292 | 1 | 0.993 | 1 | 1 | 1 | 0.92 | 1 | 1 | 1 | 1 | 0.015 |
| SE | 47475 | ENSG00000183023 | SLC8A1 | chr2 | - | 40177751 | 40177855 | 40174824 | 40174842 | 40428472 | 40430280 | 110 | 155 | 51 | 82 | 171 | 0 | 15 | 4 | 7 | 6 | 203 | 116 | 136 | 145 | 33 | 0 | 8 | 3 | 12 | 0 | 253 | 149 | 0.122342544 | 0.760594873 | 1 | 0.859 | 0.882 | 0.873 | 0.944 | 1 | 0.895 | 0.964 | 0.877 | 1 | -0.036 |
| SE | 7416 | ENSG00000168461 | RAB31 | chr18 | + | 9792153 | 9792235 | 9775277 | 9775357 | 9814019 | 9814091 | 283 | 254 | 165 | 368 | 256 | 4 | 2 | 2 | 2 | 0 | 646 | 513 | 272 | 685 | 821 | 0 | 4 | 5 | 5 | 19 | 231 | 149 | 0.122452074 | 0.761215371 | 0.979 | 0.988 | 0.982 | 0.992 | 1 | 1 | 0.988 | 0.972 | 0.989 | 0.965 | 0.005 |
| SE | 64430 | ENSG00000162813 | BPNT1 | chr1 | - | 220066072 | 220066117 | 220062756 | 220062954 | 220067301 | 220067393 | 5 | 4 | 3 | 0 | 4 | 96 | 46 | 80 | 71 | 93 | 0 | 15 | 5 | 3 | 0 | 62 | 74 | 37 | 56 | 98 | 194 | 149 | 0.12246335 | 0.761216201 | 0.038 | 0.063 | 0.028 | 0 | 0.032 | 0 | 0.135 | 0.094 | 0.04 | 0 | -0.022 |
| SE | 65524 | ENSG00000196652 | ZKSCAN5 | chr7 | + | 99519826 | 99519909 | 99512452 | 99512591 | 99525812 | 99526418 | 6 | 1 | 7 | 8 | 0 | 0 | 3 | 0 | 1 | 1 | 21 | 1 | 14 | 13 | 13 | 0 | 0 | 1 | 0 | 0 | 232 | 149 | 0.122471652 | 0.761216201 | 1 | 0.176 | 1 | 0.837 | 0 | 1 | 1 | 0.9 | 1 | 1 | -0.377 |
| SE | 31726 | ENSG00000047849 | MAP4 | chr3 | - | 47867245 | 47867338 | 47857430 | 47857512 | 47869213 | 47869327 | 109 | 67 | 72 | 108 | 40 | 5 | 4 | 4 | 11 | 1 | 140 | 122 | 65 | 82 | 99 | 0 | 2 | 9 | 8 | 0 | 242 | 149 | 0.122485483 | 0.761241734 | 0.931 | 0.912 | 0.917 | 0.858 | 0.961 | 1 | 0.974 | 0.816 | 0.863 | 1 | -0.015 |
| SE | 24836 | ENSG00000104320 | NBN | chr8 | - | 89946139 | 89946295 | 89943252 | 89943366 | 89947823 | 89947892 | 404 | 364 | 308 | 328 | 800 | 1 | 5 | 1 | 8 | 7 | 6 | 178 | 109 | 71 | 63 | 0 | 0 | 0 | 0 | 0 | 298 | 149 | 0.122603522 | 0.761856525 | 0.995 | 0.973 | 0.994 | 0.953 | 0.983 | 1 | 1 | 1 | 1 | 1 | -0.02 |
| SE | 90242 | ENSG00000066855 | MTFR1 | chr8 | + | 65682327 | 65682451 | 65669872 | 65670018 | 65693643 | 65693759 | 22 | 24 | 8 | 21 | 21 | 1 | 0 | 2 | 1 | 0 | 6 | 3 | 13 | 19 | 6 | 0 | 0 | 0 | 0 | 7 | 273 | 149 | 0.122603865 | 0.761856525 | 0.923 | 1 | 0.686 | 0.92 | 1 | 1 | 1 | 1 | 1 | 0.319 | 0.042 |
| SE | 20738 | ENSG00000115760 | BIRC6 | chr2 | + | 32590805 | 32590978 | 32575155 | 32575366 | 32593914 | 32594060 | 29 | 15 | 16 | 28 | 51 | 159 | 132 | 104 | 147 | 211 | 33 | 5 | 3 | 11 | 40 | 8 | 94 | 135 | 30 | 15 | 298 | 149 | 0.12263442 | 0.761985919 | 0.084 | 0.054 | 0.071 | 0.087 | 0.108 | 0.673 | 0.026 | 0.011 | 0.155 | 0.571 | -0.206 |
| SE | 106037 | ENSG00000204482 | LST1 | chr6 | + | 31587943 | 31587966 | 31587053 | 31587318 | 31588517 | 31588909 | 132 | 56 | 50 | 125 | 174 | 221 | 99 | 82 | 195 | 250 | 249 | 90 | 26 | 98 | 333 | 560 | 127 | 114 | 255 | 390 | 172 | 149 | 0.122660094 | 0.762084965 | 0.341 | 0.329 | 0.346 | 0.357 | 0.376 | 0.278 | 0.38 | 0.165 | 0.25 | 0.425 | 0.05 |
| SE | 10303 | ENSG00000168259 | DNAJC7 | chr17 | - | 42003358 | 42003557 | 42000481 | 42000570 | 42017339 | 42017425 | 7 | 3 | 5 | 6 | 4 | 76 | 48 | 53 | 35 | 16 | 16 | 14 | 8 | 4 | 6 | 62 | 49 | 64 | 66 | 68 | 298 | 149 | 0.122731835 | 0.762470184 | 0.044 | 0.03 | 0.045 | 0.079 | 0.111 | 0.114 | 0.125 | 0.059 | 0.029 | 0.042 | -0.012 |
| SE | 45704 | ENSG00000014164 | ZC3H3 | chr8 | - | 143465716 | 143465848 | 143440935 | 143441120 | 143468208 | 143468278 | 34 | 17 | 38 | 24 | 16 | 0 | 0 | 0 | 0 | 0 | 23 | 50 | 13 | 22 | 51 | 0 | 0 | 0 | 8 | 0 | 281 | 149 | 0.122782869 | 0.762726714 | 1 | 1 | 1 | 1 | 1 | 1 | 1 | 1 | 0.593 | 1 | 0.081 |
| SE | 106349 | ENSG00000076513 | ANKRD13A | chr12 | + | 110024045 | 110024112 | 110019138 | 110019328 | 110025741 | 110025823 | 1025 | 795 | 967 | 1154 | 1958 | 7 | 4 | 2 | 4 | 15 | 2994 | 1309 | 1270 | 1490 | 2344 | 25 | 23 | 16 | 16 | 0 | 216 | 149 | 0.122841001 | 0.763027291 | 0.99 | 0.993 | 0.997 | 0.995 | 0.989 | 0.988 | 0.975 | 0.982 | 0.985 | 1 | 0.007 |
| SE | 33118 | ENSG00000172053 | QARS1 | chr3 | - | 49098191 | 49098258 | 49095931 | 49096079 | 49098352 | 49098480 | 77 | 34 | 65 | 66 | 52 | 0 | 0 | 0 | 0 | 0 | 81 | 53 | 49 | 37 | 76 | 0 | 0 | 10 | 0 | 0 | 216 | 149 | 0.122924807 | 0.763487285 | 1 | 1 | 1 | 1 | 1 | 1 | 1 | 0.772 | 1 | 1 | 0.046 |
| SE | 61219 | ENSG00000162402 | USP24 | chr1 | - | 55141615 | 55141731 | 55138943 | 55139010 | 55142741 | 55142795 | 450 | 287 | 356 | 304 | 525 | 0 | 0 | 0 | 1 | 1 | 78 | 251 | 168 | 96 | 30 | 0 | 10 | 2 | 0 | 0 | 265 | 149 | 0.122978393 | 0.76375952 | 1 | 1 | 1 | 0.994 | 0.997 | 1 | 0.934 | 0.979 | 1 | 1 | 0.016 |
| SE | 5910 | ENSG00000089050 | RBBP9 | chr20 | - | 18495837 | 18495880 | 18493957 | 18494063 | 18497068 | 18497225 | 29 | 15 | 30 | 38 | 14 | 2 | 0 | 4 | 1 | 0 | 44 | 35 | 20 | 31 | 25 | 0 | 14 | 0 | 0 | 0 | 192 | 149 | 0.123007321 | 0.763878585 | 0.918 | 1 | 0.853 | 0.967 | 1 | 1 | 0.66 | 1 | 1 | 1 | 0.016 |
| SE | 31361 | ENSG00000005436 | GCFC2 | chr2 | - | 75691976 | 75692100 | 75690487 | 75690719 | 75694240 | 75694427 | 178 | 129 | 138 | 134 | 381 | 7 | 5 | 4 | 14 | 1 | 19 | 84 | 56 | 18 | 35 | 7 | 0 | 0 | 0 | 0 | 273 | 149 | 0.123033739 | 0.763954378 | 0.933 | 0.934 | 0.95 | 0.839 | 0.995 | 0.597 | 1 | 1 | 1 | 1 | 0.011 |
| SE | 25349 | ENSG00000112992 | NNT | chr5 | + | 43612907 | 43613481 | 43609142 | 43609346 | 43615847 | 43616065 | 260 | 170 | 201 | 249 | 224 | 7 | 4 | 6 | 4 | 2 | 231 | 121 | 118 | 125 | 270 | 0 | 5 | 7 | 0 | 0 | 298 | 149 | 0.12303904 | 0.763954378 | 0.949 | 0.955 | 0.944 | 0.969 | 0.982 | 1 | 0.924 | 0.894 | 1 | 1 | -0.004 |
| SE | 102551 | ENSG00000203791 | EEF1AKMT2 | chr10 | - | 124789038 | 124789157 | 124774674 | 124774782 | 124790272 | 124790338 | 58 | 55 | 60 | 36 | 55 | 1 | 0 | 2 | 0 | 2 | 67 | 31 | 30 | 50 | 31 | 0 | 1 | 0 | 4 | 4 | 268 | 149 | 0.123118637 | 0.764387976 | 0.97 | 1 | 0.943 | 1 | 0.939 | 1 | 0.945 | 1 | 0.874 | 0.812 | 0.044 |
| SE | 107559 | ENSG00000122966 | CIT | chr12 | - | 119734163 | 119734357 | 119730494 | 119730630 | 119735159 | 119735357 | 3 | 3 | 19 | 20 | 6 | 0 | 0 | 0 | 0 | 0 | 50 | 16 | 33 | 42 | 1 | 0 | 0 | 8 | 0 | 0 | 298 | 149 | 0.123134385 | 0.764425133 | 1 | 1 | 1 | 1 | 1 | 1 | 1 | 0.673 | 1 | 1 | 0.065 |
| SE | 66306 | ENSG00000225190 | PLEKHM1 | chr17 | - | 45475887 | 45476073 | 45475596 | 45475726 | 45477899 | 45478147 | 11 | 2 | 1 | 4 | 0 | 36 | 11 | 46 | 36 | 11 | 0 | 3 | 3 | 0 | 6 | 26 | 33 | 15 | 62 | 32 | 298 | 149 | 0.123151788 | 0.764472562 | 0.133 | 0.083 | 0.011 | 0.053 | 0 | 0 | 0.043 | 0.091 | 0 | 0.086 | 0.012 |
| SE | 21020 | ENSG00000126214 | KLC1 | chr14 | + | 103687080 | 103687211 | 103679383 | 103679518 | 103693552 | 103693688 | 5 | 10 | 9 | 16 | 18 | 1 | 2 | 2 | 0 | 0 | 23 | 1 | 6 | 10 | 6 | 6 | 0 | 0 | 2 | 0 | 280 | 149 | 0.123206174 | 0.764740639 | 0.727 | 0.727 | 0.705 | 1 | 1 | 0.671 | 1 | 1 | 0.727 | 1 | -0.048 |
| SE | 51122 | ENSG00000233355 | CHRM3-AS2 | chr1 | - | 239707533 | 239707760 | 239703891 | 239705715 | 239714176 | 239714264 | 38 | 52 | 103 | 40 | 82 | 2 | 0 | 0 | 5 | 3 | 8 | 17 | 19 | 6 | 3 | 0 | 0 | 4 | 0 | 0 | 298 | 149 | 0.123214509 | 0.764740639 | 0.905 | 1 | 1 | 0.8 | 0.932 | 1 | 1 | 0.704 | 1 | 1 | -0.013 |
| SE | 57757 | ENSG00000118007 | STAG1 | chr3 | - | 136568764 | 136568861 | 136542118 | 136542195 | 136604308 | 136604473 | 160 | 149 | 136 | 209 | 241 | 2 | 0 | 1 | 4 | 0 | 54 | 133 | 133 | 56 | 8 | 0 | 6 | 4 | 0 | 0 | 246 | 149 | 0.12322473 | 0.764743459 | 0.98 | 1 | 0.988 | 0.969 | 1 | 1 | 0.931 | 0.953 | 1 | 1 | 0.011 |
| SE | 76282 | ENSG00000124508 | BTN2A2 | chr6 | + | 26383791 | 26383915 | 26383101 | 26383181 | 26385014 | 26385362 | 39 | 22 | 28 | 55 | 11 | 2 | 0 | 0 | 0 | 5 | 48 | 35 | 35 | 11 | 32 | 0 | 0 | 0 | 0 | 0 | 273 | 149 | 0.123263145 | 0.764921235 | 0.914 | 1 | 1 | 1 | 0.546 | 1 | 1 | 1 | 1 | 1 | -0.108 |
| SE | 85296 | ENSG00000117143 | UAP1 | chr1 | + | 162576776 | 162576981 | 162566011 | 162566348 | 162579427 | 162579603 | 197 | 68 | 142 | 126 | 100 | 3 | 1 | 0 | 0 | 2 | 39 | 113 | 201 | 111 | 8 | 0 | 6 | 0 | 4 | 0 | 298 | 149 | 0.12329333 | 0.765047915 | 0.97 | 0.971 | 1 | 1 | 0.962 | 1 | 0.904 | 1 | 0.933 | 1 | 0.013 |
| SE | 105460 | ENSG00000139318 | DUSP6 | chr12 | - | 89350587 | 89351025 | 89347234 | 89349561 | 89351639 | 89352501 | 743 | 500 | 372 | 643 | 995 | 11 | 10 | 11 | 9 | 6 | 1683 | 204 | 160 | 1372 | 509 | 18 | 2 | 0 | 21 | 0 | 298 | 149 | 0.123324724 | 0.765182073 | 0.971 | 0.962 | 0.944 | 0.973 | 0.988 | 0.979 | 0.981 | 1 | 0.97 | 1 | -0.018 |
| [illegible] | [illegible] | [illegible] | [illegible] | [illegible] | [illegible] | [illegible] | [illegible] | [illegible] | [illegible] | [illegible] | [illegible] | [illegible] | [illegible] | [illegible] | [illegible] | [illegible] | [illegible] | [illegible] | [illegible] | [illegible] | [illegible] | [illegible] | [illegible] | [illegible] | [illegible] | [illegible] | [illegible] | [illegible] | [illegible] | [illegible] | [illegible] | [illegible] | [illegible] | [illegible] | [illegible] | [illegible] | [illegible] | [illegible] | [illegible] | [illegible] | [illegible] | [illegible] | [illegible] | [illegible] | [illegible] | [illegible] |

| | | | | | | | | | | | | | | |
|---|---|---|---|---|---|---|---|---|---|---|---|---|---|---|
| SE | 70618 | ENSG00000225889 | NA | chr2 | + | 64248323 | 64248538 | 64229496 | 64229640 | 64251118 | 64251126 | 0 | 0 | 0 |
| SE | 37298 | ENSG00000151576 | QTRT2 | chr3 | + | 114082962 | 114083003 | 114082768 | 114082794 | 114085672 | 114086486 | 13 | 12 | 10 |
| SE | 72864 | ENSG00000080345 | RIF1 | chr2 | + | [illegible] | [illegible] | [illegible] | [illegible] | [illegible] | [illegible] | 363 | 289 | 301 |

[illegible]

| | | | | | | | | | | | | | | | | | | | | | | | | | | | | | | | | | | | | | | | | | | | | | | |
|---|---|---|---|---|---|---|---|---|---|---|---|---|---|---|---|---|---|---|---|---|---|---|---|---|---|---|---|---|---|---|---|---|---|---|---|---|---|---|---|---|---|---|---|---|---|---|
| SE | 106476 | ENSG00000198270 | TMEM116 | chr12 | - | 111965594 | 111965759 | 111943264 | 111943369 | 111991757 | 111991889 | 7 | 5 | 9 | 10 | 6 | 22 | 18 | 19 | 12 | 30 | 0 | 7 | 1 | 0 | 9 | 19 | 5 | 16 | 15 | 11 | 298 | 149 | 0.143877512 | 0.862956461 | 0.137 | 0.122 | 0.191 | 0.294 | 0.091 | 0 | 0.412 | 0.03 | 0 | 0.29 | 0.021 |
| SE | 4413 | ENSG00000105662 | CRTC1 | chr19 | + | 18765403 | 18765528 | 18760007 | 18760228 | 18775640 | 18775821 | 10 | 14 | 6 | 3 | 7 | 0 | 0 | 0 | 0 | 0 | 8 | 2 | 4 | 5 | 20 | 0 | 0 | 2 | 0 | 4 | 274 | 149 | 0.143946259 | 0.863302667 | 1 | 1 | 1 | 1 | 1 | 1 | 1 | 0.521 | 1 | 0.731 | 0.15 |
| SE | 11874 | ENSG00000121281 | ADCY7 | chr16 | + | 50307049 | 50307147 | 50305776 | 50305849 | 50308666 | 50308792 | 75 | 18 | 31 | 53 | 26 | 0 | 0 | 0 | 0 | 5 | 18 | 59 | 35 | 34 | 41 | 15 | 0 | 0 | 0 | 0 | 247 | 149 | 0.143984756 | 0.863467406 | 1 | 1 | 1 | 1 | 0.758 | 0.42 | 1 | 1 | 1 | 1 | 0.068 |
| SE | 94234 | ENSG00000213722 | DDAH2 | chr6 | - | 31728864 | 31729221 | 31728645 | 31728745 | 31730050 | 31730263 | 30 | 19 | 29 | 36 | 30 | 1 | 0 | 0 | 0 | 0 | 47 | 32 | 38 | 43 | 63 | 0 | 0 | 0 | 0 | 8 | 298 | 149 | 0.14401573 | 0.863587009 | 0.938 | 1 | 1 | 1 | 1 | 1 | 1 | 1 | 1 | 0.797 | 0.028 |
| SE | 75815 | ENSG00000153147 | SMARCA5 | chr4 | + | 143527867 | 143528023 | 143526280 | 143526460 | 143528582 | 143528714 | 980 | 925 | 812 | 947 | 1082 | 5 | 3 | 4 | 7 | 2 | 372 | 556 | 604 | 378 | 165 | 0 | 8 | 5 | 8 | 0 | 298 | 149 | 0.144026913 | 0.86358793 | 0.99 | 0.994 | 0.99 | 0.985 | 0.996 | 1 | 0.972 | 0.984 | 0.959 | 1 | 0.008 |
| SE | 33354 | ENSG00000226149 | NA | chr6 | - | 129546394 | 129546628 | 129540446 | 129540542 | 129558057 | 129558166 | 0 | 2 | 0 | 6 | 4 | 5 | 13 | 2 | 6 | 3 | 8 | 0 | 0 | 1 | 3 | 5 | 3 | 3 | 0 | 45 | 298 | 149 | 0.144060409 | 0.863722626 | 0 | 0.071 | 0 | 0.333 | 0.4 | 0.444 | 0 | 0 | 1 | 0.032 | -0.134 |
| SE | 11057 | ENSG00000172530 | BANP | chr16 | + | 87952180 | 87952273 | 87951444 | 87951515 | 87975047 | 87975185 | 6 | 0 | 1 | 1 | 1 | 37 | 25 | 17 | 46 | 18 | 0 | 0 | 1 | 0 | 2 | 2 | 34 | 23 | 25 | 28 | 242 | 149 | 0.144103893 | 0.863917182 | 0.091 | 0 | 0.035 | 0.013 | 0.033 | 0 | 0 | 0.026 | 0 | 0.042 | 0.021 |
| SE | 4911 | ENSG00000258366 | RTEL1 | chr20 | + | 63659232 | 63659504 | 63657809 | 63658466 | 63661297 | 63661462 | 32 | 11 | 28 | 36 | 34 | 0 | 0 | 0 | 0 | 0 | 34 | 43 | 42 | 40 | 19 | 0 | 0 | 0 | 0 | 7 | 298 | 149 | 0.144141143 | 0.864074337 | 1 | 1 | 1 | 1 | 1 | 1 | 1 | 1 | 1 | 0.576 | 0.085 |
| SE | 20908 | ENSG00000153975 | ZUP1 | chr6 | - | 116660232 | 116660369 | 116658802 | 116658924 | 116660735 | 116660846 | 3 | 1 | 6 | 10 | 10 | 32 | 40 | 47 | 29 | 91 | 0 | 9 | 4 | 0 | 0 | 21 | 24 | 72 | 13 | 14 | 286 | 149 | 0.14416446 | 0.864147953 | 0.047 | 0.013 | 0.062 | 0.152 | 0.054 | 0 | 0.163 | 0.028 | 0 | 0 | 0.027 |
| SE | 13981 | ENSG00000103522 | IL21R | chr16 | + | 27442961 | 27443116 | 27437487 | 27437687 | 27444541 | 27444719 | 25 | 21 | 25 | 31 | 35 | 0 | 0 | 5 | 2 | 0 | 16 | 40 | 36 | 15 | 16 | 0 | 0 | 0 | 0 | 0 | 298 | 149 | 0.14418048 | 0.864177822 | 1 | 1 | 0.714 | 0.886 | 1 | 1 | 1 | 1 | 1 | 1 | -0.08 |
| SE | 43102 | ENSG00000177728 | TMEM94 | chr17 | + | 75492238 | 75492337 | 75491698 | 75491900 | 75492473 | 75492742 | 0 | 2 | 3 | 1 | 5 | 21 | 14 | 8 | 4 | 1 | 7 | 8 | 0 | 1 | 23 | 4 | 14 | 18 | 7 | 19 | 248 | 149 | 0.144206756 | 0.864269147 | 0 | 0.079 | 0.184 | 0.131 | 0.75 | 0.513 | 0.256 | 0 | 0.079 | 0.421 | -0.025 |
| SE | 56912 | ENSG00000174446 | SNAPC5 | chr15 | - | 66494902 | 66495419 | 66493467 | 66494552 | 66497641 | 66497780 | 25 | 20 | 31 | 35 | 21 | 4 | 3 | 14 | 12 | 1 | 11 | 38 | 14 | 37 | 23 | 33 | 6 | 6 | 8 | 6 | 298 | 149 | 0.144236429 | 0.864380821 | 0.758 | 0.769 | 0.525 | 0.593 | 0.913 | 0.143 | 0.76 | 0.538 | 0.698 | 0.657 | 0.152 |
| SE | 43551 | ENSG00000105323 | HNRNPUL1 | chr19 | + | 41305831 | 41305867 | 41303971 | 41304261 | 41306448 | 41307029 | 135 | 66 | 151 | 113 | 109 | 5 | 1 | 0 | 8 | 6 | 111 | 132 | 102 | 109 | 159 | 0 | 2 | 11 | 12 | 6 | 185 | 149 | 0.144250998 | 0.864401968 | 0.956 | 0.982 | 1 | 0.919 | 0.936 | 1 | 0.982 | 0.882 | 0.88 | 0.955 | 0.019 |
| SE | 748 | ENSG00000099991 | CABIN1 | chr22 | + | 24176110 | 24176275 | 24171712 | 24171995 | 24177503 | 24177817 | 98 | 50 | 108 | 117 | 43 | 0 | 0 | 0 | 2 | 8 | 65 | 147 | 63 | 94 | 181 | 12 | 0 | 2 | 0 | 7 | 298 | 149 | 0.144282872 | 0.864526804 | 1 | 1 | 1 | 0.967 | 0.729 | 0.73 | 1 | 0.94 | 1 | 0.928 | 0.02 |
| SE | 12869 | ENSG00000103472 | RRN3P2 | chr16 | + | 29096130 | 29096261 | 29094237 | 29094330 | 29097131 | 29097316 | 83 | 71 | 55 | 47 | 50 | 1 | 0 | 0 | 0 | 0 | 144 | 78 | 82 | 111 | 40 | 0 | 6 | 0 | 4 | 0 | 280 | 149 | 0.144477563 | 0.865568053 | 0.978 | 1 | 1 | 1 | 1 | 1 | 0.874 | 1 | 0.937 | 1 | 0.033 |
| SE | 5264 | ENSG00000185513 | L3MBTL1 | chr20 | + | 43515291 | 43515415 | 43514953 | 43515159 | 43515918 | 43516177 | 8 | 13 | 7 | 8 | 8 | 0 | 0 | 1 | 0 | 0 | 22 | 3 | 4 | 21 | 25 | 0 | 0 | 5 | 0 | 0 | 273 | 149 | 0.144478759 | 0.865568053 | 1 | 1 | 0.793 | 1 | 1 | 1 | 1 | 0.304 | 1 | 1 | 0.098 |
| SE | 98937 | ENSG00000175115 | PACS1 | chr11 | + | 66242911 | 66243031 | 66241226 | 66241653 | 66243164 | 66243234 | 397 | 159 | 301 | 294 | 305 | 0 | 1 | 1 | 2 | 10 | 320 | 354 | 290 | 343 | 862 | 0 | 4 | 3 | 14 | 8 | 269 | 149 | 0.144548553 | 0.865915297 | 1 | 0.989 | 0.994 | 0.988 | 0.944 | 1 | 0.98 | 0.982 | 0.931 | 0.984 | 0.008 |
| SE | 35265 | ENSG00000103064 | SLC7A6 | chr16 | + | 68273794 | 68273904 | 68266601 | 68266721 | 68274690 | 68275095 | 17 | 16 | 44 | 26 | 19 | 34 | 45 | 60 | 51 | 39 | 6 | 15 | 3 | 20 | 13 | 72 | 10 | 43 | 29 | 22 | 259 | 149 | 0.144558839 | 0.865915297 | 0.223 | 0.17 | 0.297 | 0.227 | 0.219 | 0.046 | 0.463 | 0.039 | 0.284 | 0.254 | 0.01 |
| SE | 102671 | ENSG00000175029 | CTBP2 | chr10 | - | 125038996 | 125039155 | 125003337 | 125003492 | 125110989 | 125111093 | 169 | 67 | 84 | 168 | 77 | 3 | 0 | 3 | 4 | 1 | 54 | 45 | 38 | 88 | 74 | 0 | 3 | 0 | 2 | 0 | 298 | 149 | 0.144587429 | 0.866020298 | 0.966 | 1 | 0.933 | 0.955 | 0.975 | 1 | 0.882 | 1 | 0.957 | 1 | -0.002 |
| SE | 30708 | ENSG00000196954 | CASP4 | chr11 | - | 104951895 | 104952063 | 104949542 | 104949777 | 104954746 | 104955001 | 27 | 17 | 13 | 9 | 22 | 13 | 10 | 4 | 18 | 12 | 3 | 20 | 24 | 7 | 11 | 19 | 26 | 60 | 22 | 0 | 298 | 149 | 0.144715725 | 0.866722435 | 0.509 | 0.459 | 0.619 | 0.2 | 0.478 | 0.073 | 0.278 | 0.167 | 0.137 | 1 | 0.122 |
| SE | 44387 | ENSG00000072210 | ALDH3A2 | chr17 | + | 19671720 | 19671956 | 19663332 | 19665047 | 19675495 | 19675499 | 117 | 82 | 113 | 132 | 245 | 0 | 1 | 0 | 1 | 5 | 149 | 116 | 49 | 92 | 128 | 0 | 0 | 4 | 7 | 0 | 298 | 149 | 0.144785969 | 0.86707681 | 1 | 0.976 | 1 | 0.985 | 0.961 | 1 | 1 | 0.86 | 0.868 | 1 | 0.039 |
| SE | 15100 | ENSG00000128739 | SNRPN | chr15 | + | 24974386 | 24974456 | 24967931 | 24968082 | 24975357 | 24975509 | 340 | 264 | 310 | 374 | 478 | 1 | 1 | 3 | 8 | 10 | 254 | 297 | 205 | 151 | 980 | 5 | 1 | 0 | 0 | 7 | 219 | 149 | 0.144861903 | 0.86740399 | 0.996 | 0.994 | 0.986 | 0.97 | 0.97 | 0.972 | 0.995 | 1 | 1 | 0.99 | -0.008 |
| [illegible] | | | | | | | | | | | | | | | | | | | | | | | | | | | | | | | | | | | | | | | | | | | | | | |

| | | | | | | | | | | | | | | | | | | | | | | | | | | | | | | | | | | | | | | | | | | | | | | |
|---|---|---|---|---|---|---|---|---|---|---|---|---|---|---|---|---|---|---|---|---|---|---|---|---|---|---|---|---|---|---|---|---|---|---|---|---|---|---|---|---|---|---|---|---|---|---|
| SE | 72842 | ENSG00000129158 | SERGEF | chr11 | - | 17995795 | 17995909 | 17992930 | 17992993 | 17999493 | 17999572 | 131 | 61 | 85 | 89 | 106 | 0 | 0 | 0 | 0 | 0 | 56 | 125 | 49 | 36 | 84 | 0 | 0 | 9 | 0 | 0 | 263 | 149 | 0.154101193 | 0.908040125 | 1 | 1 | 1 | 1 | 1 | 1 | 1 | 0.755 | 1 | 1 | 0.049 |
| SE | 66903 | ENSG00000196504 | PRPF40A | chr2 | - | 152714549 | 152714629 | 152694062 | 152694622 | 152715994 | 152716130 | 6 | 2 | 0 | 3 | 4 | 78 | 42 | 87 | 74 | 35 | 0 | 16 | 0 | 0 | 0 | 19 | 39 | 39 | 27 | 13 | 229 | 149 | 0.154110913 | 0.908040125 | 0.048 | 0.03 | 0 | 0.026 | 0.069 | 0 | 0.211 | 0 | 0 | 0 | -0.008 |
| SE | 95368 | ENSG00000115459 | ELMOD3 | chr2 | + | 85363096 | 85363166 | 85362185 | 85362260 | 85368685 | 85368754 | 67 | 39 | 33 | 46 | 33 | 2 | 4 | 0 | 5 | 7 | 113 | 48 | 35 | 39 | 41 | 0 | 7 | 6 | 0 | 0 | 219 | 149 | 0.154119562 | 0.908040125 | 0.958 | 0.869 | 1 | 0.862 | 0.762 | 1 | 0.823 | 0.799 | 1 | 1 | -0.034 |
| SE | 92860 | ENSG00000147872 | PLIN2 | chr9 | - | 19119649 | 19119831 | 19118320 | 19118455 | 19120879 | 19121165 | 286 | 167 | 190 | 178 | 353 | 2 | 2 | 0 | 1 | 0 | 214 | 392 | 241 | 181 | 307 | 4 | 12 | 0 | 0 | 0 | 298 | 149 | 0.154203826 | 0.908458034 | 0.986 | 0.977 | 1 | 0.989 | 1 | 0.964 | 0.942 | 1 | 1 | 1 | 0.009 |
| SE | 72535 | ENSG00000135426 | TESPA1 | chr12 | - | 54981773 | 54981945 | 54974399 | 54974607 | 54983993 | 54984166 | 10 | 21 | 35 | 20 | 60 | 7 | 10 | 19 | 11 | 54 | 0 | 12 | 2 | 8 | 0 | 16 | 1 | 0 | 7 | 4 | 298 | 149 | 0.154213698 | 0.908458034 | 0.417 | 0.512 | 0.479 | 0.476 | 0.357 | 0 | 0.857 | 1 | 0.364 | 0 | 0.004 |
| SE | 21423 | ENSG00000164180 | TMEM161B | chr5 | - | 88205753 | 88205954 | 88197668 | 88203075 | 88206438 | 88206499 | 87 | 37 | 72 | 69 | 181 | 0 | 0 | 3 | 0 | 8 | 11 | 27 | 12 | 7 | 2 | 0 | 0 | 0 | 3 | 0 | 298 | 149 | 0.154239795 | 0.90847815 | 1 | 1 | 0.923 | 1 | 0.919 | 1 | 1 | 1 | 0.538 | 1 | 0.061 |
| SE | 16849 | ENSG00000151575 | TEX9 | chr15 | + | 56391242 | 56391418 | 56389317 | 56389400 | 56412301 | 56412436 | 21 | 18 | 11 | 16 | 25 | 0 | 0 | 6 | 0 | 0 | 12 | 5 | 13 | 14 | 26 | 0 | 0 | 0 | 0 | 0 | 298 | 149 | 0.154240319 | 0.90847815 | 1 | 1 | 0.478 | 1 | 1 | 1 | 1 | 1 | 1 | 1 | -0.104 |
| SE | 34317 | ENSG00000108219 | TSPAN14 | chr10 | + | 80468546 | 80468687 | 80463087 | 80463127 | 80489216 | 80489314 | 96 | 34 | 82 | 78 | 45 | 7 | 1 | 1 | 0 | 0 | 98 | 61 | 36 | 44 | 108 | 0 | 0 | 3 | 0 | 10 | 290 | 149 | 0.154385582 | 0.909265347 | 0.876 | 0.946 | 0.977 | 1 | 1 | 1 | 1 | 0.86 | 1 | 0.847 | 0.018 |
| SE | 7043 | ENSG00000152229 | PSTPIP2 | chr18 | - | 46039946 | 46040047 | 46015902 | 46015937 | 46072155 | 46072257 | 32 | 14 | 7 | 14 | 27 | 0 | 0 | 0 | 0 | 0 | 15 | 54 | 29 | 66 | 53 | 7 | 0 | 0 | 0 | 0 | 250 | 149 | 0.154436884 | 0.909466727 | 1 | 1 | 1 | 1 | 1 | 0.561 | 1 | 1 | 1 | 1 | 0.088 |
| SE | 34068 | ENSG00000114867 | EIF4G1 | chr3 | + | 184317320 | 184317497 | 184315163 | 184315545 | 184317716 | 184317773 | 150 | 59 | 74 | 100 | 85 | 0 | 0 | 0 | 0 | 0 | 154 | 104 | 44 | 74 | 147 | 0 | 1 | 0 | 9 | 0 | 298 | 149 | 0.154443006 | 0.909466727 | 1 | 1 | 1 | 1 | 1 | 1 | 0.981 | 1 | 0.804 | 1 | 0.043 |
| SE | 81565 | ENSG00000167703 | SLC43A2 | chr17 | - | 1627714 | 1627920 | 1614978 | 1615034 | 1628793 | 1628834 | 16 | 1 | 12 | 8 | 3 | 0 | 0 | 0 | 0 | 0 | 6 | 8 | 8 | 5 | 61 | 0 | 0 | 0 | 5 | 0 | 298 | 149 | 0.154456224 | 0.909476159 | 1 | 1 | 1 | 1 | 1 | 1 | 1 | 1 | 0.333 | 1 | 0.133 |
| SE | 106663 | ENSG00000139350 | NEDD1 | chr12 | + | 96917620 | 96917737 | 96907603 | 96907624 | 96919984 | 96920118 | 61 | 34 | 74 | 64 | 73 | 0 | 0 | 0 | 0 | 0 | 7 | 37 | 20 | 4 | 25 | 0 | 0 | 0 | 5 | 0 | 266 | 149 | 0.154472359 | 0.90950277 | 1 | 1 | 1 | 1 | 1 | 1 | 1 | 1 | 0.309 | 1 | 0.138 |
| SE | 57384 | ENSG00000101040 | ZMYND8 | chr20 | - | 47236325 | 47236516 | 47229725 | 47229806 | 47246007 | 47246517 | 40 | 20 | 38 | 45 | 46 | 0 | 0 | 0 | 0 | 0 | 2 | 71 | 25 | 45 | 43 | 4 | 0 | 0 | 1 | 0 | 298 | 149 | 0.154508711 | 0.909648399 | 1 | 1 | 1 | 1 | 1 | 0.2 | 1 | 0.926 | 1 | 1 | 0.175 |
| SE | 109217 | ENSG00000173473 | SMARCC1 | chr3 | - | 47662333 | 47662592 | 47661293 | 47661455 | 47670657 | 47670717 | 238 | 162 | 242 | 221 | 283 | 3 | 1 | 1 | 4 | 5 | 167 | 249 | 222 | 195 | 24 | 4 | 8 | 6 | 0 | 0 | 298 | 149 | 0.154594038 | 0.910082317 | 0.975 | 0.988 | 0.992 | 0.965 | 0.966 | 0.954 | 0.94 | 0.949 | 1 | 1 | 0.009 |
| SE | 72884 | ENSG00000272752 | STAG3L5P-PVRIG2P-PILRB | chr7 | + | 100352760 | 100352914 | 100352563 | 100352667 | 100352996 | 100353123 | 23 | 4 | 13 | 6 | 4 | 0 | 0 | 0 | 0 | 0 | 14 | 6 | 1 | 3 | 5 | 0 | 0 | 0 | 0 | 5 | 298 | 149 | 0.154612158 | 0.910120557 | 1 | 1 | 1 | 1 | 1 | 1 | 1 | 1 | 1 | 0.333 | 0.133 |
| SE | 66848 | ENSG00000198598 | MMP17 | chr12 | + | 131845296 | 131845449 | 131844869 | 131845200 | 131849938 | 131850059 | 22 | 6 | 5 | 18 | 2 | 0 | 0 | 0 | 0 | 0 | 3 | 5 | 5 | 10 | 22 | 0 | 5 | 0 | 0 | 0 | 298 | 149 | 0.154664304 | 0.910359072 | 1 | 1 | 1 | 1 | 1 | 1 | 0.333 | 1 | 1 | 1 | 0.133 |
| SE | 27062 | ENSG00000062194 | GPBP1 | chr5 | + | 57214073 | 57214193 | 57175447 | 57176400 | 57230845 | 57230969 | 485 | 383 | 425 | 493 | 955 | 5 | 7 | 7 | 10 | 18 | 26 | 284 | 308 | 74 | 25 | 0 | 6 | 9 | 4 | 3 | 269 | 149 | 0.154711926 | 0.910570923 | 0.982 | 0.968 | 0.971 | 0.965 | 0.967 | 1 | 0.963 | 0.95 | 0.911 | 0.822 | 0.041 |
| SE | 50100 | ENSG00000163322 | ABRAXAS1 | chr4 | - | 83469031 | 83469151 | 83467453 | 83467538 | 83470202 | 83470333 | 135 | 137 | 118 | 133 | 196 | 4 | 2 | 2 | 2 | 2 | 22 | 47 | 59 | 13 | 38 | 0 | 0 | 0 | 0 | 0 | 269 | 149 | 0.154735021 | 0.910638397 | 0.949 | 0.974 | 0.97 | 0.974 | 0.982 | 1 | 1 | 1 | 1 | 1 | -0.03 |
| SE | 46425 | ENSG00000162341 | TPCN2 | chr11 | + | 69079833 | 69079883 | 69078891 | 69079020 | 69081399 | 69081499 | 8 | 8 | 9 | 15 | 4 | 2 | 3 | 0 | 0 | 1 | 22 | 16 | 9 | 14 | 35 | 0 | 1 | 0 | 0 | 20 | 199 | 149 | 0.154752257 | 0.910671382 | 0.75 | 0.666 | 1 | 1 | 0.75 | 1 | 0.923 | 1 | 1 | 0.567 | -0.065 |
| SE | 75153 | ENSG00000059758 | CDK17 | chr12 | - | 96282508 | 96282599 | 96280423 | 96280885 | 96283602 | 96283645 | 275 | 183 | 249 | 333 | 385 | 0 | 0 | 0 | 1 | 0 | 54 | 275 | 358 | 138 | 26 | 0 | 3 | 3 | 8 | 0 | 240 | 149 | 0.154780615 | 0.910769807 | 1 | 1 | 1 | 0.995 | 1 | 1 | 0.983 | 0.987 | 0.915 | 1 | 0.022 |
| SE | 98 | ENSG00000160255 | ITGB2 | chr21 | - | 44889977 | 44890222 | 44888692 | 44888895 | 44893403 | 44893544 | 3 | 6 | 5 | 4 | 1 | 0 | 0 | 0 | 0 | 0 | 21 | 10 | 5 | 9 | 23 | 0 | 3 | 5 | 0 | 0 | 298 | 149 | 0.154861526 | 0.91117743 | 1 | 1 | 1 | 1 | 1 | 1 | 1 | 0.333 | 1 | 1 | 0.133 |
| SE | 26198 | ENSG00000059378 | PARP12 | chr7 | - | 140041643 | 140041839 | 140034234 | 140034331 | 140054661 | 140054763 | 6 | 2 | 8 | 5 | 1 | 1 | 0 | 0 | 0 | 0 | 25 | 5 | 2 | 12 | 14 | 0 | 1 | 0 | 5 | 0 | 298 | 149 | 0.155048374 | 0.91220826 | 0.75 | 1 | 1 | 1 | 1 | 1 | 0.714 | 1 | 0.545 | 1 | 0.098 |
| SE | 1278 | ENSG00000186951 | PPARA | chr22 | + | 46219811 | 46220014 | 46218262 | 46218401 | 46231791 | 46232239 | 45 | 33 | 18 | 44 | 39 | 4 | 3 | 1 | 1 | 4 | 19 | 10 | 14 | 26 | 34 | 2 | 0 | 4 | 0 | 17 | 298 | 149 | 0.155162164 | 0.912809137 | 0.849 | 0.846 | 0.9 | 0.957 | 0.83 | 0.826 | 1 | 0.636 | 1 | 0.5 | 0.084 |
| SE | 80145 | ENSG00000012124 | CD22 | chr19 | + | 35332552 | 35332924 | 35329211 | 35329230 | 35336148 | 35336341 | 11 | 3 | 5 | 7 | 11 | 1 | 0 | 1 | 0 | 0 | 10 | 26 | 7 | 10 | 8 | 0 | 1 | 0 | 0 | 5 | 298 | 149 | 0.155206492 | 0.913001316 | 0.846 | 1 | 0.714 | 1 | 1 | 1 | 0.929 | 1 | 1 | 0.444 | 0.037 |
| SE | 55351 | ENSG00000136732 | GYPC | chr2 | + | 126690254 | 126690311 | 126656157 | 126656312 | 126693863 | 126693947 | 126 | 49 | 103 | 101 | 58 | 18 | 4 | 5 | 6 | 1 | 93 | 193 | 112 | 110 | 177 | 11 | 5 | 22 | 10 | 21 | 206 | 149 | 0.155323968 | 0.913623726 | 0.835 | 0.899 | 0.937 | 0.924 | 0.977 | 0.859 | 0.965 | 0.786 | 0.888 | 0.859 | 0.043 |
| SE | 6320 | ENSG00000060069 | CTDP1 | chr18 | + | 79736354 | 79736521 | 79728906 | 79729069 | 79753651 | 79754503 | 96 | 46 | 146 | 74 | 86 | 18 | 13 | 6 | 7 | 8 | 98 | 86 | 43 | 46 | 144 | 0 | 23 | 12 | 3 | 0 | 298 | 149 | 0.15539532 | 0.913938444 | 0.727 | 0.639 | 0.924 | 0.841 | 0.843 | 1 | 0.652 | 0.642 | 0.885 | 1 | -0.041 |
| SE | 42664 | ENSG00000088854 | C20orf194 | chr20 | - | 3316722 | 3316793 | 3315734 | 3315785 | 3322216 | 3322262 | 11 | 11 | 5 | 11 | 6 | 1 | 0 | 0 | 0 | 0 | 17 | 4 | 10 | 8 | 13 | 0 | 2 | 4 | 0 | 0 | 220 | 149 | 0.155400819 | 0.913938444 | 0.882 | 1 | 1 | 1 | 1 | 1 | 0.575 | 0.629 | 1 | 1 | 0.136 |
| SE | 68429 | ENSG00000101104 | PABPC1L | chr20 | + | 44934500 | 44934563 | 44933056 | 44933185 | 44935390 | 44935497 | 0 | 0 | 0 | 0 | 0 | 7 | 4 | 8 | 16 | 3 | 0 | 8 | 0 | 0 | 0 | 4 | 23 | 4 | 6 | 9 | 212 | 149 | 0.155436434 | 0.914079242 | 0 | 0 | 0 | 0 | 0 | 0 | 0.196 | 0 | 0 | 0 | -0.039 |
| SE | 7887 | ENSG00000141580 | WDR45B | chr17 | - | 82629530 | 82629628 | 82627203 | 82627291 | 82630920 | 82631022 | 11 | 0 | 6 | 2 | 2 | 107 | 79 | 97 | 102 | 240 | 12 | 0 | 1 | 0 | 1 | 210 | 95 | 96 | 120 | 225 | 247 | 149 | 0.155452282 | 0.914103782 | 0.058 | 0 | 0.036 | 0.012 | 0.005 | 0.033 | 0 | 0.006 | 0 | 0.003 | 0.014 |
| SE | 92804 | ENSG00000131669 | NINJ1 | chr9 | - | 93131484 | 93131541 | 93126409 | 93126638 | 93134142 | 93134219 | 5 | 0 | 1 | 3 | 0 | 53 | 12 | 30 | 27 | 3 | 0 | 0 | 1 | 0 | 0 | 109 | 32 | 62 | 101 | 103 | 206 | 149 | 0.155500969 | 0.914257397 | 0.064 | 0 | 0.024 | 0.074 | 0 | 0 | 0 | 0.012 | 0 | 0 | 0.03 |
| SE | 52607 | ENSG00000115548 | KDM3A | chr2 | + | 86460778 | 86460831 | 86456982 | 86457071 | 86464052 | 86464216 | 19 | 5 | 34 | 18 | 41 | 120 | 93 | 149 | 126 | 227 | 20 | 3 | 11 | 12 | 0 | 63 | 142 | 102 | 65 | 4 | 202 | 149 | 0.155502907 | 0.914257397 | 0.105 | 0.038 | 0.144 | 0.095 | 0.118 | 0.19 | 0.015 | 0.074 | 0.12 | 0 | 0.02 |
| SE | 24536 | ENSG00000172785 | CBWD1 | chr9 | - | 121960 | 122090 | 121037 | 121573 | 123216 | 123282 | 101 | 263 | 111 | 125 | 293 | 7 | 33 | 6 | 6 | 39 | 209 | 245 | 137 | 44 | 49 | 4 | 30 | 46 | 5 | 16 | 279 | 149 | 0.155513436 | 0.914257397 | 0.885 | 0.81 | 0.908 | 0.918 | 0.8 | 0.965 | 0.813 | 0.614 | 0.825 | 0.621 | 0.097 |
| SE | 7143 | ENSG00000141425 | RPRD1A | chr18 | - | 36040806 | 36040865 | 36033707 | 36033837 | 36067253 | 36067530 | 25 | 14 | 35 | 16 | 10 | 51 | 23 | 50 | 31 | 9 | 16 | 2 | 12 | 6 | 7 | 32 | 30 | 18 | 6 | 16 | 208 | 149 | 0.155548584 | 0.914395367 | 0.26 | 0.304 | 0.334 | 0.27 | 0.443 | 0.264 | 0.046 | 0.323 | 0.417 | 0.239 | 0.064 |
| SE | 59175 | ENSG00000080819 | CPOX | chr3 | - | 98589661 | 98589756 | 98588712 | 98588854 | 98590631 | 98590657 | 6 | 3 | 1 | 4 | 6 | 58 | 31 | 53 | 60 | 100 | 0 | 6 | 3 | 1 | 0 | 37 | 44 | 21 | 55 | 54 | 244 | 149 | 0.155601464 | 0.914637555 | 0.059 | 0.056 | 0.011 | 0.039 | 0.035 | 0 | 0.077 | 0.08 | 0.011 | 0 | 0.006 |
| SE | 102994 | ENSG00000197535 | MYO5A | chr15 | - | 52343116 | 52343197 | 52340195 | 52340394 | 52346360 | 52346461 | 81 | 98 | 71 | 82 | 83 | 12 | 22 | 22 | 22 | 10 | 50 | 79 | 106 | 68 | 0 | 0 | 37 | 94 | 23 | 6 | 230 | 149 | 0.155628445 | 0.914727479 | 0.814 | 0.743 | 0.676 | 0.707 | 0.843 | 1 | 0.58 | 0.422 | 0.657 | 0 | 0.225 |
| SE | 54493 | ENSG00000151414 | NEK7 | chr1 | + | 198220998 | 198221110 | 198156993 | 198157276 | 198232552 | 198232637 | 6 | 19 | 10 | 16 | 44 | 81 | 33 | 44 | 59 | 8 | 27 | 9 | 2 | 8 | 20 | 0 | 29 | 17 | 10 | 10 | 261 | 149 | 0.155643035 | 0.914744562 | 0.041 | 0.247 | 0.115 | 0.134 | 0.758 | 1 | 0.151 | 0.063 | 0.314 | 0.533 | -0.153 |
| SE | 99086 | ENSG00000110274 | CEP164 | chr11 | + | 117335604 | 117335680 | 117327928 | 117328208 | 117338565 | 117338668 | 33 | 21 | 29 | 41 | 15 | 14 | 22 | 16 | 20 | 7 | 48 | 35 | 21 | 28 | 34 | 27 | 59 | 22 | 13 | 24 | 225 | 149 | 0.15568705 | 0.91493457 | 0.61 | 0.387 | 0.546 | 0.576 | 0.587 | 0.541 | 0.282 | 0.387 | 0.588 | 0.484 | 0.085 |
| SE | 93283 | ENSG00000056586 | RC3H2 | chr9 | - | 122865623 | 122865657 | 122859916 | 122860131 | 122877470 | 122877583 | 17 | 17 | 27 | 31 | 68 | 3 | 0 | 6 | 7 | 2 | 2 | 23 | 20 | 3 | 4 | 0 | 11 | 5 | 1 | 0 | 183 | 149 | 0.15579717 | 0.915480175 | 0.822 | 1 | 0.786 | 0.783 | 0.965 | 1 | 0.63 | 0.765 | 0.71 | 1 | 0.05 |
| SE | 59172 | ENSG00000115204 | MPV17 | chr2 | - | 27317077 | 27317234 | 27312993 | 27313109 | 27322447 | 27322522 | 3 | 2 | 1 | 11 | 5 | 34 | 10 | 32 | 13 | 10 | 46 | 9 | 4 | 1 | 6 | 25 | 30 | 16 | 11 | 44 | 298 | 149 | 0.155803276 | 0.915480175 | 0.042 | 0.091 | 0.015 | 0.297 | 0.2 | 0.479 | 0.13 | 0.111 | 0.043 | 0.064 | -0.036 |
| SE | 41734 | ENSG00000163319 | MRPS18C | chr4 | + | 83459739 | 83459797 | 83456908 | 83456958 | 83460972 | 83461032 | 57 | 20 | 16 | 21 | 48 | 0 | 2 | 0 | 0 | 7 | 32 | 35 | 24 | 47 | 16 | 0 | 0 | 2 | 18 | 0 | 207 | 149 | 0.155827565 | 0.915554182 | 1 | 0.878 | 1 | 1 | 0.832 | 1 | 1 | 0.896 | 0.653 | 1 | 0.032 |
| SE | 26251 | ENSG00000111224 | PARP11 | chr12 | - | 3863835 | 3863957 | 3829889 | 3830018 | 3873211 | 3873355 | 37 | 26 | 47 | 43 | 46 | 32 | 21 | 26 | 16 | 24 | 22 | 41 | 39 | 33 | 9 | 12 | 9 | 20 | 19 | 1 | 271 | 149 | 0.155931066 | 0.916093553 | 0.389 | 0.405 | 0.498 | 0.596 | 0.513 | 0.502 | 0.715 | 0.517 | 0.488 | 0.832 | -0.131 |
| SE | 94100 | ENSG00000136811 | ODF2 | chr9 | + | 128471307 | 128471468 | 128460941 | 128461067 | 128472912 | 128473042 | 72 | 56 | 69 | 104 | 129 | 0 | 0 | 0 | 0 | 0 | 153 | 61 | 87 | 120 | 192 | 11 | 0 | 0 | 0 | 0 | 298 | 149 | 0.155972432 | 0.916267825 | 1 | 1 | 1 | 1 | 1 | 0.874 | 1 | 1 | 1 | 1 | 0.025 |
| SE | 101564 | ENSG00000088298 | EDEM2 | chr20 | - | 35144978 | 35145018 | 35142372 | 35142478 | 35146824 | 35146935 | 59 | 17 | 41 | 25 | 25 | 0 | 0 | 0 | 4 | 0 | 73 | 81 | 34 | 49 | 61 | 0 | 0 | 5 | 5 | 0 | 189 | 149 | 0.156135112 | 0.917145284 | 1 | 1 | 1 | 0.831 | 1 | 1 | 1 | 0.843 | 0.885 | 1 | 0.021 |
| SE | 31788 | ENSG00000204463 | BAG6 | chr6 | - | 31644306 | 31644414 | 31644081 | 31644194 | 31644524 | 31644602 | 67 | 36 | 85 | 68 | 70 | 70 | 23 | 33 | 52 | 43 | 60 | 105 | 35 | 71 | 89 | 30 | 46 | 19 | 42 | 50 | 257 | 149 | 0.156145226 | 0.917145284 | 0.357 | 0.476 | 0.599 | 0.431 | 0.486 | 0.537 | 0.57 | 0.516 | 0.495 | 0.508 | -0.055 |

[illegible]

| | | | | | | | | | | | | | | | | | | | | | | | | | | | | | | | | | | | | | | | | | | | | | | |
|---|---|---|---|---|---|---|---|---|---|---|---|---|---|---|---|---|---|---|---|---|---|---|---|---|---|---|---|---|---|---|---|---|---|---|---|---|---|---|---|---|---|---|---|---|---|---|
| SE | 8412 | ENSG00000173757 | STAT5B | chr17 | - | 42227528 | 42227685 | 42224778 | 42224868 | 42231999 | 42232137 | 410 | 229 | 404 | 325 | 368 | 4 | 1 | 10 | 1 | 14 | 369 | 329 | 303 | 357 | 607 | 6 | 2 | 10 | 1 | 4 | 298 | 149 | 0.1659074 | 0.960784089 | 0.981 | 0.991 | 0.953 | 0.994 | 0.929 | 0.969 | 0.988 | 0.938 | 0.994 | 0.987 | -0.006 |
| SE | 21713 | ENSG00000178385 | PLEKHM3 | chr2 | - | 207861104 | 207861262 | 207821287 | 207828496 | 207908513 | 207908577 | 45 | 12 | 23 | 45 | 37 | 1 | 0 | 0 | 0 | 0 | 30 | 24 | 33 | 44 | 8 | 6 | 0 | 1 | 0 | 0 | 298 | 149 | 0.165928804 | 0.960784089 | 0.957 | 1 | 1 | 1 | 1 | 0.714 | 1 | 0.943 | 1 | 1 | 0.06 |
| SE | 54325 | ENSG00000117139 | KDM5B | chr1 | - | 202766396 | 202766504 | 202764048 | 202764145 | 202766925 | 202767060 | 16 | 1 | 10 | 0 | 5 | 20 | 18 | 18 | 22 | 15 | 5 | 0 | 12 | 6 | 0 | 31 | 16 | 21 | 21 | 30 | 257 | 149 | 0.165930858 | 0.960784089 | 0.317 | 0.031 | 0.244 | 0 | 0.162 | 0.086 | 0 | 0.249 | 0.142 | 0 | 0.055 |
| SE | 98108 | ENSG00000171466 | ZNF562 | chr19 | - | 9658008 | 9658135 | 9656546 | 9656653 | 9659378 | 9659467 | 108 | 99 | 93 | 106 | 116 | 17 | 3 | 2 | 4 | 1 | 217 | 140 | 118 | 131 | 92 | 0 | 5 | 11 | 0 | 6 | 276 | 149 | 0.166008159 | 0.9611606 | 0.774 | 0.947 | 0.962 | 0.935 | 0.984 | 1 | 0.938 | 0.853 | 1 | 0.892 | -0.016 |
| SE | 107610 | ENSG00000111596 | CNOT2 | chr12 | + | 70329422 | 70329570 | 70319297 | 70319364 | 70332766 | 70332846 | 232 | 187 | 224 | 218 | 337 | 0 | 0 | 7 | 3 | 5 | 49 | 243 | 204 | 176 | 45 | 0 | 0 | 0 | 1 | 0 | 297 | 149 | 0.166040938 | 0.961279304 | 1 | 1 | 0.941 | 0.973 | 0.971 | 1 | 1 | 1 | 0.989 | 1 | -0.021 |
| SE | 108250 | ENSG00000121073 | SLC35B1 | chr17 | - | 49706203 | 49706309 | 49705865 | 49705896 | 49706964 | 49707068 | 42 | 22 | 49 | 42 | 62 | 2 | 2 | 5 | 10 | 7 | 64 | 37 | 60 | 57 | 70 | 13 | 7 | 10 | 0 | 29 | 255 | 149 | 0.166121087 | 0.961672211 | 0.925 | 0.865 | 0.851 | 0.71 | 0.838 | 0.742 | 0.755 | 0.778 | 1 | 0.585 | 0.066 |
| SE | 60521 | ENSG00000116984 | MTR | chr1 | + | 236806143 | 236806233 | 236803427 | 236803642 | 236812737 | 236812844 | 59 | 41 | 69 | 92 | 141 | 0 | 0 | 2 | 8 | 0 | 38 | 52 | 15 | 39 | 44 | 0 | 2 | 1 | 0 | 0 | 239 | 149 | 0.166156346 | 0.961805211 | 1 | 1 | 0.956 | 0.878 | 1 | 1 | 0.942 | 0.903 | 1 | 1 | -0.002 |
| SE | 99788 | ENSG00000172977 | KAT5 | chr11 | + | 65713347 | 65713503 | 65712921 | 65713058 | 65713592 | 65713664 | 44 | 17 | 61 | 52 | 24 | 40 | 22 | 55 | 57 | 56 | 41 | 25 | 40 | 41 | 23 | 50 | 96 | 25 | 50 | 76 | 298 | 149 | 0.166277027 | 0.962353527 | 0.355 | 0.279 | 0.357 | 0.313 | 0.176 | 0.291 | 0.115 | 0.444 | 0.291 | 0.131 | 0.042 |
| SE | 99996 | ENSG00000122359 | ANXA11 | chr10 | - | 80180940 | 80181136 | 80176106 | 80176155 | 80205342 | 80205572 | 16 | 10 | 7 | 15 | 22 | 25 | 5 | 15 | 13 | 10 | 16 | 5 | 6 | 20 | 31 | 26 | 15 | 16 | 33 | 19 | 298 | 149 | 0.166287908 | 0.962353527 | 0.242 | 0.5 | 0.189 | 0.366 | 0.524 | 0.235 | 0.143 | 0.158 | 0.233 | 0.449 | 0.121 |
| SE | 6940 | ENSG00000100983 | GSS | chr20 | - | 34935575 | 34935642 | 34931938 | 34932133 | 34936762 | 34936840 | 69 | 37 | 63 | 74 | 66 | 0 | 0 | 8 | 1 | 0 | 92 | 70 | 39 | 81 | 70 | 3 | 0 | 0 | 0 | 9 | 216 | 149 | 0.166287944 | 0.962353527 | 1 | 1 | 0.845 | 0.981 | 1 | 0.955 | 1 | 1 | 1 | 0.843 | 0.006 |
| SE | 38316 | ENSG00000143742 | SRP9 | chr1 | + | 225786861 | 225786985 | 225783299 | 225783368 | 225787970 | 225788266 | 56 | 24 | 59 | 45 | 46 | 11 | 6 | 11 | 9 | 4 | 19 | 84 | 38 | 28 | 20 | 7 | 0 | 6 | 2 | 5 | 273 | 149 | 0.166308981 | 0.962404139 | 0.735 | 0.686 | 0.745 | 0.732 | 0.863 | 0.597 | 1 | 0.776 | 0.884 | 0.686 | -0.036 |
| SE | 95777 | ENSG00000125944 | HNRNPR | chr1 | - | 23340851 | 23341020 | 23338489 | 23338608 | 23344210 | 23344322 | 148 | 169 | 176 | 173 | 217 | 118 | 87 | 69 | 118 | 61 | 144 | 211 | 267 | 350 | 82 | 77 | 102 | 83 | 72 | 43 | 298 | 149 | 0.166340671 | 0.962451951 | 0.385 | 0.493 | 0.561 | 0.423 | 0.64 | 0.483 | 0.508 | 0.617 | 0.709 | 0.488 | -0.061 |
| SE | 8445 | ENSG00000141699 | RETREG3 | chr17 | - | 42596532 | 42596627 | 42592055 | 42592162 | 42609085 | 42609324 | 14 | 7 | 4 | 4 | 3 | 103 | 18 | 64 | 66 | 30 | 0 | 3 | 11 | 2 | 0 | 3 | 64 | 30 | 21 | 84 | 244 | 149 | 0.166341828 | 0.962451951 | 0.077 | 0.192 | 0.037 | 0.036 | 0.058 | 0 | 0.028 | 0.183 | 0.055 | 0 | 0.027 |
| SE | 36435 | ENSG00000121858 | TNFSF10 | chr3 | - | 172514860 | 172514998 | 172511444 | 172511659 | 172523252 | 172523310 | 436 | 414 | 287 | 360 | 520 | 0 | 1 | 0 | 0 | 0 | 314 | 235 | 189 | 307 | 114 | 0 | 8 | 0 | 7 | 0 | 287 | 149 | 0.166365878 | 0.962512664 | 1 | 0.995 | 1 | 1 | 1 | 1 | 0.938 | 1 | 0.958 | 1 | 0.02 |
| SE | 59507 | ENSG00000148660 | CAMK2G | chr10 | - | 73848525 | 73848609 | 73842457 | 73842541 | 73849012 | 73849115 | 12 | 6 | 18 | 7 | 15 | 0 | 0 | 1 | 0 | 0 | 6 | 16 | 7 | 12 | 33 | 0 | 0 | 2 | 5 | 0 | 233 | 149 | 0.166376908 | 0.962512664 | 1 | 1 | 0.92 | 1 | 1 | 1 | 1 | 0.691 | 0.605 | 1 | 0.125 |
| SE | 76913 | ENSG00000167483 | NIBAN3 | chr19 | + | 17530754 | 17530816 | 17527290 | 17527395 | 17532262 | 17532388 | 31 | 20 | 24 | 24 | 7 | 15 | 3 | 17 | 26 | 6 | 0 | 43 | 4 | 19 | 81 | 38 | 5 | 2 | 17 | 116 | 211 | 149 | 0.166448606 | 0.962847563 | 0.593 | 0.825 | 0.499 | 0.395 | 0.452 | 0 | 0.859 | 0.585 | 0.441 | 0.33 | 0.11 |
| SE | 10774 | ENSG00000108578 | BLMH | chr17 | - | 30289372 | 30289482 | 30287805 | 30287947 | 30291310 | 30291508 | 159 | 106 | 138 | 120 | 136 | 0 | 0 | 1 | 0 | 0 | 145 | 132 | 136 | 83 | 163 | 4 | 0 | 6 | 2 | 0 | 259 | 149 | 0.166459393 | 0.962847563 | 1 | 1 | 0.988 | 1 | 1 | 0.954 | 1 | 0.929 | 0.96 | 1 | 0.029 |
| SE | 7413 | ENSG00000168461 | RAB31 | chr18 | + | 9775277 | 9775357 | 9708300 | 9708444 | 9845581 | 9845691 | 15 | 7 | 4 | 11 | 2 | 0 | 0 | 0 | 0 | 0 | 23 | 23 | 3 | 40 | 41 | 8 | 0 | 0 | 0 | 0 | 229 | 149 | 0.166491075 | 0.962959679 | 1 | 1 | 1 | 1 | 1 | 0.652 | 1 | 1 | 1 | 1 | 0.07 |
| SE | 74690 | ENSG00000197548 | ATG7 | chr3 | + | 11313303 | 11313420 | 11308983 | 11309061 | 11315343 | 11315482 | 124 | 78 | 97 | 100 | 143 | 1 | 0 | 0 | 0 | 2 | 101 | 151 | 52 | 193 | 194 | 0 | 2 | 0 | 0 | 10 | 266 | 149 | 0.166535791 | 0.963137748 | 0.986 | 1 | 1 | 1 | 0.976 | 1 | 0.977 | 1 | 1 | 0.916 | 0.014 |
| SE | 72046 | ENSG00000115053 | NCL | chr2 | - | 231457285 | 231457357 | 231457000 | 231457124 | 231457642 | 231457800 | 7 | 7 | 5 | 8 | 12 | 666 | 459 | 600 | 604 | 816 | 6 | 11 | 13 | 10 | 0 | 430 | 343 | 450 | 413 | 134 | 221 | 149 | 0.166546464 | 0.963137748 | 0.007 | 0.01 | 0.006 | 0.009 | 0.01 | 0.009 | 0.021 | 0.019 | 0.016 | 0 | -0.005 |
| SE | 53985 | ENSG00000188641 | DPYD | chr1 | - | 97720824 | 97720926 | 97699350 | 97699547 | 97721509 | 97721654 | 4 | 2 | 2 | 5 | 12 | 179 | 141 | 165 | 133 | 151 | 0 | 0 | 3 | 9 | 0 | 107 | 157 | 76 | 108 | 81 | 251 | 149 | 0.166563282 | 0.963163867 | 0.013 | 0.008 | 0.007 | 0.022 | 0.045 | 0 | 0 | 0.023 | 0.047 | 0 | 0.005 |
| SE | 12928 | ENSG00000197818 | SLC9A8 | chr20 | + | 49839540 | 49839599 | 49823060 | 49823141 | 49845035 | 49845119 | 64 | 47 | 64 | 57 | 47 | 1 | 0 | 3 | 0 | 0 | 122 | 125 | 140 | 182 | 98 | 4 | 0 | 1 | 3 | 5 | 208 | 149 | 0.166636517 | 0.963510095 | 0.979 | 1 | 0.939 | 1 | 1 | 0.956 | 1 | 0.99 | 0.978 | 0.934 | 0.012 |
| SE | 42091 | ENSG00000026751 | SLAMF7 | chr1 | + | 160750303 | 160750423 | 160749820 | 160750093 | 160752185 | 160752248 | 166 | 87 | 148 | 146 | 117 | 15 | 8 | 10 | 11 | 9 | 156 | 182 | 172 | 120 | 137 | 11 | 16 | 10 | 14 | 30 | 269 | 149 | 0.166647769 | 0.963510095 | 0.86 | 0.858 | 0.891 | 0.88 | 0.878 | 0.887 | 0.863 | 0.905 | 0.826 | 0.717 | 0.034 |
| SE | 63971 | ENSG00000109332 | UBE2D3 | chr4 | - | 102827591 | 102827741 | 102826484 | 102826636 | 102827943 | 102828029 | 16 | 3 | 7 | 12 | 14 | 31 | 23 | 25 | 20 | 18 | 12 | 4 | 10 | 7 | 0 | 6 | 28 | 40 | 43 | 7 | 298 | 149 | 0.166712308 | 0.96381207 | 0.205 | 0.061 | 0.123 | 0.231 | 0.28 | 0.5 | 0.067 | 0.111 | 0.075 | 0 | 0.029 |
| SE | 55071 | ENSG00000104894 | CD37 | chr19 | + | 49335885 | 49336029 | 49335713 | 49335782 | 49336908 | 49337033 | 5 | 2 | 3 | 0 | 5 | 68 | 39 | 76 | 54 | 60 | 0 | 1 | 2 | 4 | 5 | 135 | 119 | 41 | 73 | 296 | 293 | 149 | 0.166769614 | 0.964072185 | 0.036 | 0.025 | 0.02 | 0 | 0.041 | 0 | 0.004 | 0.024 | 0.027 | 0.009 | 0.012 |
| SE | 97225 | ENSG00000073921 | PICALM | chr11 | - | 85960674 | 85960777 | 85958754 | 85959060 | 85974707 | 85974812 | 19 | 5 | 9 | 17 | 11 | 438 | 309 | 284 | 392 | 626 | 10 | 18 | 10 | 11 | 0 | 170 | 338 | 332 | 299 | 34 | 252 | 149 | 0.166832212 | 0.964320318 | 0.025 | 0.009 | 0.018 | 0.025 | 0.01 | 0.034 | 0.031 | 0.017 | 0.021 | 0 | -0.003 |
| SE | 91698 | ENSG00000285130 | NA | chr9 | + | 69064937 | 69065035 | 69053139 | 69053260 | 69099900 | 69100045 | 33 | 20 | 37 | 24 | 22 | 0 | 0 | 0 | 0 | 0 | 56 | 28 | 13 | 27 | 11 | 0 | 8 | 0 | 0 | 0 | 247 | 149 | 0.16683717 | 0.964320318 | 1 | 1 | 1 | 1 | 1 | 1 | 0.679 | 1 | 1 | 1 | 0.064 |
| SE | 28500 | ENSG00000169062 | UPF3A | chr13 | + | 114282836 | 114282943 | 114281924 | 114282127 | 114286518 | 114286629 | 62 | 42 | 71 | 72 | 67 | 6 | 0 | 0 | 3 | 0 | 64 | 66 | 56 | 79 | 68 | 0 | 0 | 0 | 0 | 5 | 256 | 149 | 0.166891138 | 0.96456105 | 0.857 | 1 | 1 | 0.933 | 1 | 1 | 1 | 1 | 1 | 0.888 | -0.02 |
| SE | 55829 | ENSG00000126709 | IFI6 | chr1 | - | 27669220 | 27669346 | 27668457 | 27668535 | 27672122 | 27672178 | 24 | 24 | 26 | 51 | 21 | 0 | 0 | 2 | 0 | 0 | 192 | 228 | 49 | 292 | 78 | 0 | 4 | 2 | 7 | 0 | 275 | 149 | 0.166925637 | 0.964689228 | 1 | 1 | 0.876 | 1 | 1 | 1 | 0.969 | 0.93 | 0.958 | 1 | 0.004 |
| SE | 5016 | ENSG00000088833 | NSFL1C | chr20 | - | 1463334 | 1463441 | 1458199 | 1458274 | 1464328 | 1464426 | 14 | 5 | 2 | 5 | 0 | 124 | 94 | 93 | 130 | 106 | 0 | 10 | 1 | 15 | 0 | 267 | 139 | 114 | 149 | 203 | 256 | 149 | 0.166969444 | 0.964841082 | 0.062 | 0.03 | 0.012 | 0.022 | 0 | 0 | 0.04 | 0.005 | 0.055 | 0 | 0.005 |
| SE | 3097 | ENSG00000167600 | CYP2S1 | chr19 | + | 41198461 | 41198622 | 41197778 | 41197928 | 41198708 | 41198888 | 18 | 17 | 2 | 13 | 6 | 0 | 0 | 0 | 6 | 0 | 24 | 8 | 6 | 23 | 14 | 0 | 0 | 0 | 0 | 0 | 298 | 149 | 0.166976559 | 0.964841082 | 1 | 1 | 1 | 0.52 | 1 | 1 | 1 | 1 | 1 | 1 | -0.096 |
| SE | 16783 | ENSG00000125107 | CNOT1 | chr16 | - | 58624672 | 58624794 | 58623281 | 58623409 | 58629727 | 58629825 | 5 | 5 | 17 | 8 | 12 | 2 | 4 | 12 | 15 | 2 | 0 | 20 | 8 | 6 | 0 | 32 | 4 | 4 | 1 | 7 | 271 | 149 | 0.167004222 | 0.964849751 | 0.579 | 0.407 | 0.438 | 0.227 | 0.767 | 0 | 0.733 | 0.524 | 0.767 | 0 | 0.079 |
| SE | 2670 | ENSG00000129353 | SLC44A2 | chr19 | + | 10635162 | 10635515 | 10634755 | 10634887 | 10636322 | 10636496 | 42 | 22 | 28 | 28 | 62 | 10 | 2 | 2 | 11 | 7 | 16 | 31 | 24 | 31 | 104 | 23 | 29 | 6 | 4 | 5 | 298 | 149 | 0.167016556 | 0.964849751 | 0.677 | 0.846 | 0.875 | 0.56 | 0.816 | 0.258 | 0.348 | 0.667 | 0.795 | 0.912 | 0.159 |

[illegible]

| | | | | | | | | | | | |
|---|---|---|---|---|---|---|---|---|---|---|---|
| SE | 75097 | ENSG00000198948 | MFAP3L | chr4 | - | 170005579 | 170006010 | 169986601 | 169992309 | 170026233 | 170026395 |
| SE | 44093 | ENSG00000182923 | CEP63 | chr3 | + | 134558141 | 134558347 | 134551925 | 134552012 | 134561376 | 134561749 |
| SE | 24219 | ENSG00000113742 | CPEB4 | chr5 | + | 173943025 | 173943049 | 173932449 | 173932500 | 173944966 | 173945140 |
| SE | 43968 | ENSG00000108443 | RPS6KB1 | chr17 | + | 59914634 | 59914703 | 59912683 | 59912804 | 59926434 | 59926582 |
| SE | 74173 | ENSG00000116698 | SMG7 | chr1 | + | 183512836 | 183512868 | 183472215 | 183472649 | 183515873 | 183515894 |
| SE | 98799 | ENSG00000121671 | CRY2 | chr11 | + | 45862059 | 45862148 | 45860847 | 45861032 | 45867611 | 45867752 |
| SE | 57753 | ENSG00000118007 | STAG1 | chr3 | - | 136472412 | 136472492 | 136464880 | 136464988 | 136473538 | 136473603 |
| SE | 15202 | ENSG00000159459 | UBR1 | chr15 | - | 43091960 | 43092080 | 43085983 | 43086240 | 43105941 | 43106107 |
| SE | 2620 | ENSG00000028277 | POU2F2 | chr19 | - | 42110700 | 42110756 | 42099715 | 42099821 | 42117249 | 42117366 |
| SE | 7501 | ENSG00000108840 | HDAC5 | chr17 | - | 44117493 | 44117704 | 44110728 | 44110800 | 44123503 | 44123618 |
| SE | 75417 | ENSG00000169826 | CSGALNACT2 | chr10 | + | 43167003 | 43167098 | 43163865 | 43164044 | 43175950 | 43176032 |
| SE | 109414 | ENSG00000134900 | TPP2 | chr13 | + | 102651358 | 102651397 | 102649407 | 102649486 | 102657055 | 102657207 |
| SE | 56979 | ENSG00000171603 | CLSTN1 | chr1 | - | 9746888 | 9747012 | 9744394 | 9744643 | 9749460 | 9749646 |
| SE | 78372 | ENSG00000176102 | CSTF3 | chr11 | - | 33156707 | 33156803 | 33141884 | 33141986 | 33161298 | 33161494 |
| SE | 73397 | ENSG00000240344 | PPIL3 | chr2 | - | 200885261 | 200885367 | 200882341 | 200882435 | 200885697 | 200885772 |
| SE | 78770 | ENSG00000179406 | LINC00174 | chr7 | - | 66485094 | 66485153 | 66481540 | 66481599 | 66489937 | 66490107 |
| SE | 92312 | ENSG00000188921 | HACD4 | chr9 | - | 21008020 | 21008146 | 20999508 | 21007119 | 21011588 | 21011695 |
| SE | 732 | ENSG00000099991 | CABIN1 | chr22 | + | 24050824 | 24050974 | 24049090 | 24049220 | 24054872 | 24055159 |
| SE | 68176 | ENSG00000020922 | MRE11 | chr11 | - | 94445809 | 94445890 | 94437176 | 94437235 | 94447218 | 94447438 |
| SE | 100531 | ENSG00000054654 | SYNE2 | chr14 | + | 64075944 | 64076100 | 64073967 | 64074136 | 64078465 | 64078606 |
| SE | 14772 | ENSG00000103994 | ZNF106 | chr15 | - | 42449770 | 42451954 | 42448071 | 42448705 | 42466052 | 42466114 |
| SE | 31426 | ENSG00000155508 | CNOT8 | chr5 | + | 154863211 | 154863395 | 154858653 | 154858816 | 154865191 | 154865344 |

[illegible]

| | | | | | | | | | | |
|---|---|---|---|---|---|---|---|---|---|---|
| SE | 34667 | ENSG00000176473 | WDR25 | chr14 | + | 100380909 | 100381746 | 100376484 | 100376495 | 100468020 |
| SE | 25347 | ENSG00000112992 | NNT | chr5 | + | 43612907 | 43613137 | 43609142 | 43609346 | 43615847 |
| SE | 80074 | ENSG00000198198 | SZT2 | chr1 | + | 43419733 | 43419961 | 43404403 | 43404550 | 43420152 |
| SE | 19907 | ENSG00000100890 | PRORP | chr14 | + | 35126734 | 35126782 | 35122721 | 35122910 | 35127478 |
| SE | 85720 | ENSG00000104356 | POP1 | chr8 | + | 98148814 | 98149006 | 98146567 | 98146683 | 98150484 |
| SE | 41145 | ENSG00000009780 | FAM76A | chr1 | + | 27749067 | 27749154 | 27744653 | 27744811 | 27755194 |
| SE | 18631 | ENSG00000256053 | COA8 | chr14 | + | 103581568 | 103581656 | 103574106 | 103574170 | 103587273 |
| SE | 101944 | ENSG00000122033 | MTIF3 | chr13 | - | 27445087 | 27445137 | 27441016 | 27441077 | 27450508 |
| SE | 53262 | ENSG00000138688 | KIAA1109 | chr4 | + | 122344381 | 122344561 | 122343409 | 122343610 | 122346622 |
| SE | 16478 | ENSG00000247240 | UBL7-AS1 | chr15 | + | 74480620 | 74480914 | 74479059 | 74479278 | 74481050 |
| SE | 94644 | ENSG00000011454 | RABGAP1 | chr9 | + | 123070349 | 123070424 | 123065347 | 123065461 | 123076244 |
| SE | 10407 | ENSG00000282851 | BISPR | chr19 | + | 17413096 | 17413305 | 17406076 | 17406296 | 17414390 |
| SE | 48300 | ENSG00000229474 | PATL2 | chr15 | - | 44675485 | 44675691 | 44674292 | 44674411 | 44676474 |
| SE | 31309 | ENSG00000013288 | MAN2B2 | chr4 | + | 6610879 | 6610990 | 6609797 | 6610050 | 6611085 |
| SE | 95313 | ENSG00000165699 | TSC1 | chr9 | - | 132900714 | 132900837 | 132897183 | 132897610 | 132901588 |
| SE | 66651 | ENSG00000084623 | EIF3I | chr1 | + | 32226170 | 32226320 | 32222439 | 32222630 | 32228498 |
| SE | 87514 | ENSG00000239306 | RBM14 | chr11 | + | 66623900 | 66623968 | 66616629 | 66617057 | 66624213 |
| SE | 11256 | ENSG00000182149 | IST1 | chr16 | + | 71923287 | 71923380 | 71922473 | 71922680 | 71924768 |
| SE | 47787 | ENSG00000154358 | OBSCN | chr1 | + | 228299258 | 228299522 | 228298453 | 228298717 | 228299881 |
| SE | 38309 | ENSG00000138172 | CALHM2 | chr10 | - | 103451082 | 103451182 | 103449386 | 103450099 | 103452155 |
| SE | 57777 | ENSG00000182831 | C16orf72 | chr16 | + | 9092920 | 9093001 | 9091643 | 9092270 | 9102983 |
| SE | 27900 | ENSG00000138495 | COX17 | chr3 | - | 119674672 | 119674788 | 119666918 | 119666982 | 119675144 |
| SE | 57271 | ENSG00000143119 | CD53 | chr1 | + | 110894959 | 110895055 | 110891391 | 110891471 | 110899123 |
| SE | 48767 | ENSG00000142534 | RPS11 | chr19 | + | 49497193 | 49497282 | 49496453 | 49496471 | 49497519 |
| SE | 43127 | ENSG00000186470 | BTN3A2 | chr6 | + | 26367989 | 26368050 | 26365158 | 26365352 | 26368564 |
| SE | 61141 | ENSG00000070495 | JMJD6 | chr17 | - | 76721262 | 76721351 | 76720359 | 76720498 | 76721797 |

| | | | | | | | | | | | | | | | |
|---|---|---|---|---|---|---|---|---|---|---|---|---|---|---|---|
| SE | 60489 | ENSG00000070814 | TCOF1 | chr5 | + | 150391543 | 150391795 | 150389886 | 150390023 | 150391956 | 150394916 | 32 | 6 | 27 | [illegible] |
| SE | 21984 | ENSG00000120318 | ARAP3 | chr5 | - | 141655361 | 141655400 | 141653400 | 141654435 | 141655620 | 141655758 | 8 | 9 | 7 | [illegible] |
| SE | 52910 | ENSG00000159023 | EPB41 | chr1 | + | 29060421 | 29060484 | 29053103 | 29053312 | 29064981 | 29065158 | 17 | 12 | 12 | [illegible] |
| SE | 22240 | ENSG00000141279 | NPEPPS | chr17 | + | 47586147 | 47586244 | 47585499 | 47585745 | 47586364 | 47586398 | 98 | 60 | 82 | [illegible] |
| SE | 88638 | ENSG00000178234 | GALNT11 | chr7 | + | 152100797 | 152100921 | 152093990 | 152094522 | 152103111 | 152103172 | 171 | 110 | 135 | [illegible] |
| SE | 102070 | ENSG00000181192 | DHTKD1 | chr10 | + | 12091512 | 12091684 | 12088985 | 12089255 | 12094072 | 12094271 | 85 | 52 | 76 | [illegible] |
| SE | 72432 | ENSG00000047932 | GOPC | chr6 | - | 117577447 | 117577471 | 117575176 | 117575352 | 117578899 | 117579064 | 14 | 4 | 10 | [illegible] |
| SE | 68946 | ENSG00000135926 | TMBIM1 | chr2 | - | 218285762 | 218285938 | 218281939 | 218282181 | 218289965 | 218289994 | 11 | 3 | 2 | [illegible] |
| SE | 1177 | ENSG00000100105 | PATZ1 | chr22 | - | 31328786 | 31328859 | 31325808 | 31327309 | 31335691 | 31335863 | 10 | 9 | 21 | [illegible] |
| SE | 105811 | ENSG00000111785 | RIC8B | chr12 | + | 106842588 | 106842817 | 106825725 | 106825820 | 106843851 | 106843947 | 109 | 103 | 94 | [illegible] |
| SE | 43054 | ENSG00000182287 | AP1S2 | chrX | - | 15833443 | 15833524 | 15827028 | 15827372 | 15845378 | 15845516 | 12 | 8 | 5 | [illegible] |
| SE | 17049 | ENSG00000128829 | EIF2AK4 | chr15 | + | 40032756 | 40032801 | 40030358 | 40030456 | 40034325 | 40034444 | 15 | 11 | 7 | [illegible] |
| SE | 17747 | ENSG00000006432 | MAP3K9 | chr14 | - | 70733756 | 70733798 | 70732538 | 70733342 | 70734385 | 70734498 | 2 | 2 | 5 | [illegible] |
| SE | 65680 | ENSG00000179104 | TMTC2 | chr12 | + | 82857009 | 82857580 | 82686879 | 82687669 | 82895817 | 82896646 | 21 | 18 | 18 | [illegible] |
| SE | 5068 | ENSG00000101247 | NDUFAF5 | chr20 | + | 13814417 | 13814515 | 13808841 | 13808902 | 13816874 | 13816957 | 16 | 10 | 8 | [illegible] |
| SE | 30237 | ENSG00000005700 | IBTK | chr6 | - | 82196297 | 82196446 | 82194478 | 82194642 | 82200140 | 82200253 | 140 | 102 | 104 | [illegible] |
| SE | 104768 | ENSG00000011258 | MBTD1 | chr17 | - | 51259142 | 51259206 | 51225007 | 51225209 | 51260119 | 51260163 | 9 | 13 | 9 | [illegible] |
| SE | 84207 | ENSG00000068354 | TBC1D25 | chrX | + | 48544868 | 48545023 | 48541284 | 48541442 | 48558896 | 48559024 | 82 | 19 | 47 | [illegible] |
| SE | 84388 | ENSG00000136143 | SUCLA2 | chr13 | - | 47994799 | 47994896 | 47988881 | 47988981 | 47996842 | 47997023 | 4 | 6 | 8 | [illegible] |
| SE | 37617 | ENSG00000082074 | FYB1 | chr5 | - | 39212677 | 39213017 | 39202773 | 39202987 | 39219442 | 39219563 | 34 | 28 | 28 | [illegible] |
| SE | 28542 | ENSG00000170088 | TMEM192 | chr4 | - | 165088467 | 165088602 | 165085585 | 165085688 | 165102949 | 165103096 | 64 | 38 | 64 | [illegible] |
| SE | 19574 | ENSG00000165355 | FBXO33 | chr14 | - | 39401175 | 39401861 | 39397668 | 39399787 | 39431563 | 39432500 | 97 | 79 | 90 | [illegible] |
| SE | 107358 | ENSG00000110921 | MVK | chr12 | + | 109581394 | 109581550 | 109579801 | 109579946 | 109586021 | 109586096 | 49 | 16 | 38 | [illegible] |
| SE | 70054 | ENSG00000151665 | PIGF | chr2 | - | 46591802 | 46591925 | 46581284 | 46581591 | 46592474 | 46592583 | 16 | 10 | 9 | [illegible] |
| SE | 99217 | ENSG00000149196 | HIKESHI | chr11 | + | 86345093 | 86345198 | 86344602 | 86344721 | 86345583 | 86345663 | 5 | 2 | 4 | [illegible] |
| SE | 72401 | ENSG00000138835 | RGS3 | chr9 | + | 113582071 | 113582242 | 113529220 | 113529264 | 113583449 | 113583671 | 38 | 27 | 14 | [illegible] |
| SE | 65937 | ENSG00000143537 | ADAM15 | chr1 | + | 155061414 | 155061489 | 155060762 | 155060832 | 155061903 | 155061975 | 11 | 7 | 8 | [illegible] |
| SE | 81059 | ENSG00000133597 | ADCK2 | chr7 | + | 140681041 | 140681137 | 140679154 | 140679283 | 140686989 | 140687153 | 54 | 32 | 48 | [illegible] |
| SE | 40625 | ENSG00000213390 | ARHGAP19 | chr10 | - | 97265859 | 97266125 | 97264825 | 97264906 | 97292571 | 97292623 | 81 | 47 | 79 | [illegible] |
| SE | 38401 | ENSG00000143458 | GABPB2 | chr1 | + | 151090405 | 151090573 | 151088157 | 151088297 | 151097851 | 151097915 | 32 | 45 | 34 | [illegible] |
| SE | 34309 | ENSG00000108219 | TSPAN14 | chr10 | + | 80463087 | 80463127 | 80454309 | 80454371 | 80468546 | 80468687 | 1 | 1 | 4 | [illegible] |
| SE | 24676 | ENSG00000120705 | ETF1 | chr5 | - | 138512763 | 138512954 | 138511103 | 138511604 | 138513567 | 138513706 | 655 | 509 | 497 | [illegible] |
| SE | 30231 | ENSG00000132376 | INPP5K | chr17 | - | 1515887 | 1516118 | 1513871 | 1513979 | 1516455 | 1516612 | 14 | 3 | 4 | [illegible] |
| [illegible] | | | | | | | | | | | | | | | |

| | | | | | | | | | | | | | | | | | | | | | | | | | | | | | | | | | | | | | | | | | | | | | | |
|---|---|---|---|---|---|---|---|---|---|---|---|---|---|---|---|---|---|---|---|---|---|---|---|---|---|---|---|---|---|---|---|---|---|---|---|---|---|---|---|---|---|---|---|---|---|---|
| SE | 76370 | ENSG00000164674 | SYTL3 | chr6 | + | 158713799 | 158713878 | 158708321 | 158708391 | 158725502 | 158725637 | 50 | 13 | 36 | 33 | 37 | 31 | 15 | 22 | 12 | 13 | 4 | 39 | 33 | 17 | 41 | 24 | 23 | 36 | 4 | 26 | 228 | 149 | 0.218489113 | 1 | 0.513 | 0.362 | 0.517 | 0.642 | 0.65 | 0.098 | 0.526 | 0.375 | 0.735 | 0.508 | 0.088 |
| SE | 66214 | ENSG00000137817 | PARP6 | chr15 | - | 72253949 | 72254070 | 72253436 | 72253504 | 72254454 | 72254481 | 8 | 15 | 11 | 9 | 21 | 101 | 71 | 124 | 114 | 261 | 5 | 20 | 11 | 15 | 0 | 114 | 99 | 142 | 117 | 217 | 270 | 149 | 0.218497672 | 1 | 0.042 | 0.104 | 0.047 | 0.042 | 0.043 | 0.024 | 0.1 | 0.041 | 0.066 | 0 | 0.009 |
| SE | 11150 | ENSG00000103404 | USP31 | chr16 | - | 23079945 | 23080171 | 23073721 | 23073880 | 23084859 | 23084989 | 9 | 2 | 7 | 1 | 2 | 0 | 0 | 0 | 0 | 0 | 3 | 5 | 0 | 9 | 11 | 0 | 0 | 3 | 0 | 0 | 298 | 149 | 0.218592102 | 1 | 1 | 1 | 1 | 1 | 1 | 1 | 1 | 0 | 1 | 1 | 0.2 |
| SE | 83211 | ENSG00000150637 | CD226 | chr18 | - | 69873143 | 69873246 | 69867356 | 69867411 | 69895700 | 69896045 | 528 | 520 | 340 | 968 | 804 | 0 | 10 | 14 | 33 | 13 | 462 | 340 | 365 | 389 | 526 | 11 | 4 | 18 | 4 | 0 | 252 | 149 | 0.21860513 | 1 | 1 | 0.968 | 0.935 | 0.945 | 0.973 | 0.961 | 0.98 | 0.923 | 0.983 | 1 | -0.005 |
| SE | 9626 | ENSG00000173821 | RNF213 | chr17 | + | 80349769 | 80349906 | 80344677 | 80348286 | 80350300 | 80350396 | 798 | 400 | 678 | 728 | 444 | 1 | 2 | 2 | 5 | 0 | 1394 | 1054 | 637 | 1215 | 932 | 14 | 0 | 0 | 13 | 13 | 286 | 149 | 0.21863852 | 1 | 0.998 | 0.99 | 0.994 | 0.987 | 1 | 0.981 | 1 | 1 | 0.98 | 0.974 | 0.007 |
| SE | 25485 | ENSG00000082397 | EPB41L3 | chr18 | - | 5400968 | 5401043 | 5397057 | 5397426 | 5406776 | 5406968 | 0 | 4 | 0 | 5 | 5 | 22 | 24 | 11 | 20 | 41 | 4 | 0 | 0 | 6 | 0 | 32 | 31 | 12 | 46 | 6 | 224 | 149 | 0.218675767 | 1 | 0 | 0.1 | 0 | 0.143 | 0.075 | 0.077 | 0 | 0 | 0.08 | 0 | 0.032 |
| SE | 2216 | ENSG00000221983 | UBA52 | chr19 | + | 18572350 | 18572607 | 18571874 | 18571909 | 18573292 | 18573403 | 21 | 17 | 23 | 13 | 38 | 229 | 79 | 193 | 187 | 87 | 28 | 25 | 5 | 19 | 46 | 83 | 196 | 100 | 115 | 159 | 298 | 149 | 0.218764979 | 1 | 0.044 | 0.097 | 0.056 | 0.034 | 0.179 | 0.144 | 0.06 | 0.024 | 0.076 | 0.126 | -0.004 |
| SE | 94924 | ENSG00000148655 | LRMDA | chr10 | + | 76346429 | 76346553 | 76324400 | 76324485 | 76557208 | 76557306 | 2 | 0 | 4 | 1 | 4 | 3 | 14 | 5 | 7 | 19 | 0 | 0 | 6 | 14 | 16 | 35 | 7 | 13 | 6 | 41 | 273 | 149 | 0.218774957 | 1 | 0.267 | 0 | 0.304 | 0.072 | 0.103 | 0 | 0 | 0.201 | 0.56 | 0.176 | -0.038 |
| SE | 106647 | ENSG00000139178 | C1RL | chr12 | - | 7101897 | 7102087 | 7099900 | 7100026 | 7108250 | 7108299 | 39 | 30 | 42 | 24 | 27 | 0 | 0 | 0 | 0 | 0 | 46 | 39 | 68 | 99 | 131 | 0 | 0 | 0 | 0 | 9 | 298 | 149 | 0.218790675 | 1 | 1 | 1 | 1 | 1 | 1 | 1 | 1 | 1 | 1 | 0.879 | 0.024 |
| SE | 12991 | ENSG00000122390 | NAA60 | chr16 | + | 3447481 | 3447620 | 3443693 | 3443837 | 3448470 | 3448540 | 5 | 6 | 3 | 0 | 3 | 8 | 1 | 7 | 11 | 10 | 15 | 8 | 5 | 11 | 18 | 23 | 18 | 2 | 16 | 21 | 288 | 149 | 0.218933188 | 1 | 0.244 | 0.756 | 0.181 | 0 | 0.134 | 0.252 | 0.187 | 0.564 | 0.262 | 0.307 | -0.051 |
| SE | 697 | ENSG00000186575 | NF2 | chr22 | + | 29681438 | 29681601 | 29678195 | 29678323 | 29694751 | 29694945 | 141 | 49 | 130 | 142 | 81 | 3 | 2 | 2 | 3 | 0 | 181 | 158 | 81 | 123 | 199 | 16 | 0 | 0 | 4 | 0 | 298 | 149 | 0.218979004 | 1 | 0.959 | 0.925 | 0.97 | 0.959 | 1 | 0.85 | 1 | 1 | 0.939 | 1 | 0.005 |
| SE | 25982 | ENSG00000137364 | TPMT | chr6 | - | 18147822 | 18147915 | 18139664 | 18139717 | 18148987 | 18149171 | 35 | 45 | 15 | 43 | 82 | 0 | 0 | 1 | 0 | 0 | 70 | 43 | 22 | 73 | 43 | 0 | 4 | 4 | 0 | 0 | 242 | 149 | 0.219097067 | 1 | 1 | 1 | 0.902 | 1 | 1 | 1 | 0.869 | 0.772 | 1 | 1 | 0.052 |
| SE | 11801 | ENSG00000166780 | BMERB1 | chr16 | + | 15532986 | 15533135 | 15515304 | 15515428 | 15567982 | 15568056 | 0 | 0 | 0 | 1 | 0 | 4 | 1 | 3 | 2 | 3 | 0 | 0 | 6 | 0 | 0 | 5 | 12 | 3 | 3 | 21 | 298 | 149 | 0.219114116 | 1 | 0 | 0 | 0 | 0.2 | 0 | 0 | 0 | 0.5 | 0 | 0 | -0.06 |
| SE | 22092 | ENSG00000111859 | NEDD9 | chr6 | - | 11251563 | 11251726 | 11248932 | 11249214 | 11305065 | 11305347 | 6 | 3 | 3 | 3 | 8 | 0 | 3 | 0 | 1 | 2 | 7 | 5 | 14 | 7 | 10 | 7 | 0 | 20 | 0 | 0 | 298 | 149 | 0.219272257 | 1 | 1 | 0.333 | 1 | 0.6 | 0.667 | 0.333 | 1 | 0.259 | 1 | 1 | 0.002 |
| SE | 54143 | ENSG00000142765 | SYTL1 | chr1 | + | 27349651 | 27349765 | 27349397 | 27349498 | 27350388 | 27350485 | 66 | 17 | 40 | 40 | 32 | 2 | 0 | 1 | 3 | 0 | 18 | 38 | 11 | 25 | 38 | 0 | 1 | 0 | 0 | 8 | 263 | 149 | 0.219275624 | 1 | 0.949 | 1 | 0.958 | 0.883 | 1 | 1 | 0.956 | 1 | 1 | 0.729 | 0.021 |
| SE | 45448 | ENSG00000100991 | TRPC4AP | chr20 | - | 35021189 | 35021332 | 35016007 | 35016139 | 35035122 | 35035308 | 333 | 166 | 218 | 252 | 235 | 14 | 9 | 6 | 9 | 4 | 495 | 285 | 207 | 313 | 543 | 0 | 1 | 11 | 13 | 22 | 292 | 149 | 0.219311027 | 1 | 0.924 | 0.904 | 0.949 | 0.935 | 0.968 | 1 | 0.993 | 0.906 | 0.925 | 0.926 | -0.014 |
| SE | 69224 | ENSG00000115484 | CCT4 | chr2 | - | 61869439 | 61869553 | 61868084 | 61868706 | 61872081 | 61872316 | 207 | 100 | 217 | 181 | 432 | 1 | 7 | 0 | 4 | 7 | 109 | 85 | 67 | 55 | 36 | 0 | 5 | 0 | 8 | 0 | 263 | 149 | 0.219416742 | 1 | 0.992 | 0.89 | 1 | 0.962 | 0.972 | 1 | 0.906 | 1 | 0.796 | 1 | 0.023 |
| SE | 61630 | ENSG00000234127 | TRIM26 | chr6 | - | 30204655 | 30204765 | 30201034 | 30201138 | 30213304 | 30213365 | 125 | 49 | 82 | 73 | 96 | 20 | 8 | 8 | 14 | 12 | 60 | 62 | 83 | 92 | 82 | 15 | 29 | 10 | 6 | 18 | 259 | 149 | 0.219615079 | 1 | 0.782 | 0.779 | 0.855 | 0.75 | 0.822 | 0.697 | 0.552 | 0.827 | 0.898 | 0.724 | 0.058 |
| SE | 84773 | ENSG00000125676 | THOC2 | chrX | - | 123623643 | 123623709 | 123623104 | 123623283 | 123623786 | 123623971 | 15 | 19 | 13 | 25 | 33 | 82 | 90 | 98 | 138 | 214 | 0 | 23 | 10 | 0 | 0 | 1 | 53 | 34 | 1 | 6 | 215 | 149 | 0.219708444 | 1 | 0.113 | 0.128 | 0.084 | 0.112 | 0.097 | 0 | 0.231 | 0.169 | 0 | 0 | 0.027 |
| SE | 12904 | ENSG00000261832 | NA | chr16 | - | 28491481 | 28491513 | 28489289 | 28489386 | 28491713 | 28491835 | 22 | 11 | 11 | 19 | 20 | 2 | 0 | 6 | 4 | 1 | 14 | 34 | 18 | 15 | 36 | 9 | 3 | 4 | 1 | 10 | 181 | 149 | 0.219712263 | 1 | 0.901 | 1 | 0.601 | 0.796 | 0.943 | 0.562 | 0.903 | 0.787 | 0.925 | 0.748 | 0.063 |
| SE | 108993 | ENSG00000160961 | ZNF333 | chr19 | + | 14716111 | 14716238 | 14714737 | 14715470 | 14716993 | 14717089 | 61 | 55 | 92 | 79 | 31 | 0 | 0 | 2 | 0 | 0 | 76 | 70 | 61 | 31 | 72 | 0 | 1 | 0 | 7 | 0 | 276 | 149 | 0.219729438 | 1 | 1 | 1 | 0.961 | 1 | 1 | 1 | 0.974 | 1 | 0.705 | 1 | 0.056 |
| SE | 5456 | ENSG00000100294 | MCAT | chr22 | - | 43137080 | 43137298 | 43132208 | 43133486 | 43141161 | 43141249 | 13 | 12 | 11 | 19 | 2 | 0 | 1 | 7 | 6 | 4 | 0 | 1 | 8 | 6 | 21 | 12 | 2 | 0 | 1 | 9 | 298 | 149 | 0.219864617 | 1 | 1 | 0.857 | 0.44 | 0.613 | 0.2 | 0 | 0.2 | 1 | 0.75 | 0.538 | 0.124 |
| SE | 100537 | ENSG00000054654 | SYNE2 | chr14 | + | 64113340 | 64113571 | 64107490 | 64107607 | 64119426 | 64119609 | 319 | 176 | 208 | 246 | 160 | 1 | 1 | 0 | 0 | 5 | 62 | 78 | 98 | 34 | 50 | 0 | 0 | 0 | 0 | 9 | 298 | 149 | 0.219919575 | 1 | 0.994 | 0.989 | 1 | 1 | 0.941 | 1 | 1 | 1 | 1 | 0.735 | 0.038 |
| SE | 83828 | ENSG00000169891 | REPS2 | chrX | + | 17074113 | 17074159 | 17069939 | 17069993 | 17103717 | 17103779 | 4 | 4 | 3 | 3 | 25 | 12 | 30 | 6 | 22 | 42 | 13 | 6 | 10 | 6 | 4 | 55 | 16 | 21 | 26 | 85 | 195 | 149 | 0.219935721 | 1 | 0.203 | 0.092 | 0.276 | 0.094 | 0.313 | 0.153 | 0.223 | 0.267 | 0.15 | 0.035 | 0.03 |
| SE | 12828 | ENSG00000169592 | INO80E | chr16 | + | 30001413 | 30001530 | 30000928 | 30001040 | 30005220 | 30005791 | 132 | 66 | 120 | 113 | 224 | 7 | 6 | 14 | 3 | 14 | 78 | 93 | 37 | 68 | 100 | 25 | 1 | 5 | 12 | 1 | 266 | 149 | 0.219980681 | 1 | 0.914 | 0.86 | 0.828 | 0.955 | 0.9 | 0.636 | 0.981 | 0.806 | 0.76 | 0.982 | 0.058 |
| SE | 55326 | ENSG00000132635 | PCED1A | chr20 | - | 2836038 | 2836110 | 2835313 | 2835709 | 2838231 | 2838478 | 44 | 21 | 34 | 52 | 68 | 0 | 1 | 5 | 2 | 2 | 50 | 33 | 46 | 37 | 64 | 0 | 0 | 0 | 0 | 7 | 221 | 149 | 0.219997613 | 1 | 1 | 0.934 | 0.821 | 0.946 | 0.958 | 1 | 1 | 1 | 1 | 0.86 | -0.04 |
| SE | 3356 | ENSG00000167565 | SERTAD3 | chr19 | - | 40442322 | 40442515 | 40441805 | 40442086 | 40444214 | 40444275 | 4 | 2 | 2 | 0 | 2 | 17 | 1 | 11 | 5 | 10 | 3 | 3 | 5 | 5 | 6 | 12 | 9 | 7 | 18 | 29 | 298 | 149 | 0.220186853 | 1 | 0.105 | 0.5 | 0.083 | 0 | 0.091 | 0.111 | 0.143 | 0.263 | 0.122 | 0.094 | 0.009 |
| SE | 55786 | ENSG00000012174 | MBTPS2 | chrX | + | 21869497 | 21869678 | 21853375 | 21853503 | 21878041 | 21878136 | 23 | 11 | 33 | 18 | 54 | 1 | 0 | 0 | 0 | 0 | 22 | 13 | 14 | 13 | 4 | 0 | 0 | 6 | 0 | 0 | 298 | 149 | 0.220217863 | 1 | 0.92 | 1 | 1 | 1 | 1 | 1 | 1 | 0.538 | 1 | 1 | 0.076 |
| SE | 47114 | ENSG00000174839 | DENND6A | chr3 | - | 57672255 | 57672298 | 57665960 | 57666235 | 57672399 | 57672438 | 71 | 44 | 71 | 60 | 96 | 0 | 0 | 0 | 0 | 10 | 32 | 72 | 104 | 61 | 59 | 0 | 0 | 0 | 0 | 0 | 192 | 149 | 0.220240355 | 1 | 1 | 1 | 1 | 1 | 0.882 | 1 | 1 | 1 | 1 | 1 | -0.024 |
| SE | 64185 | ENSG00000048991 | R3HDM1 | chr2 | + | 135641535 | 135641790 | 135638895 | 135639122 | 135645378 | 135645527 | 9 | 5 | 8 | 12 | 0 | 41 | 42 | 29 | 40 | 65 | 14 | 0 | 1 | 7 | 0 | 44 | 53 | 30 | 19 | 42 | 298 | 149 | 0.220276816 | 1 | 0.099 | 0.056 | 0.121 | 0.13 | 0 | 0.137 | 0 | 0.016 | 0.156 | 0 | 0.019 |
| SE | 106088 | ENSG00000196498 | NCOR2 | chr12 | - | 124385744 | 124385887 | 124348173 | 124348314 | 124398118 | 124398181 | 103 | 72 | 71 | 90 | 113 | 0 | 0 | 0 | 0 | 0 | 430 | 143 | 104 | 130 | 297 | 9 | 0 | 0 | 0 | 5 | 292 | 149 | 0.220313356 | 1 | 1 | 1 | 1 | 1 | 1 | 0.961 | 1 | 1 | 1 | 0.968 | 0.014 |
| SE | 64179 | ENSG00000048991 | R3HDM1 | chr2 | + | 135616151 | 135616193 | 135604805 | 135605016 | 135616667 | 135616757 | 18 | 6 | 15 | 3 | 14 | 21 | 24 | 28 | 46 | 67 | 5 | 15 | 11 | 5 | 9 | 24 | 16 | 20 | 13 | 23 | 191 | 149 | 0.220342862 | 1 | 0.401 | 0.163 | 0.295 | 0.048 | 0.14 | 0.14 | 0.422 | 0.3 | 0.231 | 0.234 | -0.056 |
| SE | 47322 | ENSG00000133106 | EPSTI1 | chr13 | - | 42970611 | 42970670 | 42969093 | 42969177 | 42991977 | 42992165 | 125 | 168 | 128 | 202 | 93 | 2 | 1 | 4 | 12 | 0 | 768 | 348 | 309 | 726 | 158 | 2 | 1 | 3 | 21 | 7 | 208 | 149 | 0.220349822 | 1 | 0.978 | 0.992 | 0.958 | 0.923 | 1 | 0.996 | 0.996 | 0.987 | 0.961 | 0.942 | -0.006 |
| SE | 105754 | ENSG00000061987 | MON2 | chr12 | + | 62493914 | 62494042 | 62484169 | 62484233 | 62498918 | 62499048 | 78 | 71 | 60 | 78 | 55 | 0 | 0 | 0 | 1 | 4 | 37 | 50 | 63 | 41 | 47 | 0 | 0 | 8 | 0 | 0 | 277 | 149 | 0.220352211 | 1 | 1 | 1 | 1 | 0.977 | 0.881 | 1 | 1 | 0.809 | 1 | 1 | 0.01 |
| SE | 50170 | ENSG00000025708 | TYMP | chr22 | - | 50528511 | 50528610 | 50527196 | 50527717 | 50529135 | 50529338 | 488 | 444 | 258 | 372 | 165 | 1 | 1 | 1 | 1 | 0 | 1735 | 708 | 220 | 1176 | 1240 | 0 | 9 | 7 | 15 | 10 | 248 | 149 | 0.220499853 | 1 | 0.997 | 0.996 | 0.994 | 0.996 | 1 | 1 | 0.979 | 0.95 | 0.979 | 0.987 | 0.018 |
| SE | 107892 | ENSG00000188906 | LRRK2 | chr12 | + | 40305784 | 40305966 | 40303947 | 40304134 | 40308466 | 40308696 | 407 | 410 | 210 | 263 | 578 | 34 | 15 | 9 | 7 | 22 | 4 | 231 | 201 | 22 | 10 | 0 | 7 | 12 | 0 | 0 | 298 | 149 | 0.220509443 | 1 | 0.857 | 0.932 | 0.921 | 0.949 | 0.929 | 1 | 0.943 | 0.893 | 1 | 1 | -0.05 |
| SE | 66174 | ENSG00000088179 | PTPN4 | chr2 | + | 119955156 | 119955323 | 119951972 | 119952129 | 119956843 | 119956934 | 787 | 351 | 457 | 497 | 783 | 7 | 0 | 1 | 4 | 13 | 33 | 264 | 183 | 30 | 8 | 0 | 0 | 1 | 0 | 0 | 298 | 149 | 0.220616706 | 1 | 0.983 | 1 | 0.996 | 0.984 | 0.968 | 1 | 1 | 0.989 | 1 | 1 | -0.012 |
| SE | 2857 | ENSG00000176619 | LMNB2 | chr19 | - | 2431782 | 2431902 | 2431547 | 2431658 | 2432415 | 2432523 | 75 | 44 | 86 | 95 | 70 | 0 | 0 | 0 | 0 | 0 | 135 | 47 | 54 | 126 | 127 | 9 | 0 | 0 | 1 | 0 | 269 | 149 | 0.2206195 | 1 | 1 | 1 | 1 | 1 | 1 | 0.893 | 1 | 1 | 0.986 | 1 | 0.024 |
| SE | 33726 | ENSG00000196455 | PIK3R4 | chr3 | - | 130728462 | 130728684 | 130723413 | 130723587 | 130730307 | 130730442 | 134 | 80 | 109 | 129 | 381 | 4 | 0 | 0 | 2 | 1 | 20 | 47 | 49 | 26 | 9 | 0 | 4 | 3 | 0 | 0 | 298 | 149 | 0.220631216 | 1 | 0.944 | 1 | 1 | 0.97 | 0.995 | 1 | 0.855 | 0.891 | 1 | 1 | 0.033 |
| SE | 30063 | ENSG00000138814 | PPP3CA | chr4 | - | 101029165 | 101029195 | 101023408 | 101026061 | 101032266 | 101032364 | 6 | 3 | 7 | 7 | 14 | 136 | 74 | 100 | 102 | 151 | 0 | 3 | 4 | 23 | 0 | 19 | 84 | 62 | 77 | 25 | 179 | 149 | 0.220632873 | 1 | 0.035 | 0.033 | 0.055 | 0.054 | 0.072 | 0 | 0.029 | 0.051 | 0.199 | 0 | -0.006 |
| SE | 103386 | ENSG00000122958 | VPS26A | chr10 | + | 69155811 | 69155887 | 69132897 | 69133047 | 69157006 | 69157163 | 327 | 298 | 242 | 367 | 387 | 11 | 9 | 7 | 8 | 6 | 23 | 201 | 149 | 59 | 54 | 0 | 1 | 1 | 1 | 8 | 225 | 149 | 0.220831339 | 1 | 0.952 | 0.956 | 0.958 | 0.968 | 0.977 | 1 | 0.993 | 0.99 | 0.975 | 0.817 | 0.007 |
| SE | 28077 | ENSG00000084676 | NCOA1 | chr2 | + | 24564294 | 24564430 | 24556014 | 24556055 | 24584475 | 24584484 | 234 | 173 | 165 | 231 | 306 | 1 | 5 | 2 | 5 | 0 | 280 | 291 | 284 | 204 | 224 | 0 | 1 | 2 | 0 | 19 | 285 | 149 | 0.220875766 | 1 | 0.992 | 0.948 | 0.977 | 0.96 | 1 | 1 | 0.993 | 0.987 | 1 | 0.86 | 0.007 |
| SE | 18932 | ENSG00000100461 | RBM23 | chr14 | - | 22911327 | 22911403 | 22908332 | 22908380 | 22918998 | 22919110 | 108 | 47 | 64 | 82 | 78 | 0 | 1 | 0 | 0 | 0 | 108 | 86 | 71 | 120 | 126 | 0 | 0 | 4 | 7 | 0 | 225 | 149 | 0.220969473 | 1 | 1 | 0.969 | 1 | 1 | 1 | 1 | 1 | 0.922 | 0.919 | 1 | 0.026 |
| SE | 104867 | ENSG00000109685 | NSD2 | chr4 | + | 1912090 | 1912194 | 1904215 | 1904378 | 1918140 | 1918623 | 10 | 6 | 9 | 10 | 25 | 1 | 1 | 1 | 0 | 0 | 25 | 2 | 8 | 44 | 10 | 0 | 3 | 0 | 0 | 2 | 253 | 149 | 0.220996206 | 1 | 0.855 | 0.779 | 0.841 | 1 | 1 | 1 | 0.282 | 1 | 1 | 0.746 | 0.089 |
| SE | 79134 | ENSG00000169660 | HEXD | chr17 | + | 82428029 | 82428127 | 82419748 | 82419883 | 82428557 | 82428645 | 3 | 0 | 1 | 0 | 4 | 4 | 2 | 4 | 3 | 9 | 9 | 0 | 3 | 0 | 0 | 0 | 2 | 1 | 3 | 12 | 247 | 149 | 0.221069849 | 1 | 0.311 | 0 | 0.131 | 0 | 0.211 | 1 | 0 | 0.644 | 0 | 0 | -0.198 |
| SE | 1380 | ENSG00000100150 | DEPDC5 | chr22 | + | 31838684 | 31838721 | 31837093 | 31837155 | 31843094 | 31843212 | 15 | 9 | 8 | 13 | 2 | 0 | 0 | 5 | 1 | 0 | 23 | 29 | 5 | 14 | 20 | 0 | 0 | 0 | 0 | 0 | 186 | 149 | 0.221113907 | 1 | 1 | 1 | 0.562 | 0.912 | 1 | 1 | 1 | 1 | 1 | 1 | -0.105 |
| SE | 51128 | ENSG00000028839 | TBPL1 | chr6 | + | 133953184 | 133953425 | 133952285 | 133952730 | 133980081 | 133980260 | 40 | 24 | 22 | 50 | 20 | 2 | 0 | 2 | 0 | 0 | 15 | 57 | 18 | 37 | 28 | 8 | 0 | 0 | 0 | 0 | 298 | 149 | 0.221125738 | 1 | 0.909 | 1 | 0.846 | 1 | 1 | 0.484 | 1 | 1 | 1 | 1 | 0.054 |
| SE | 105629 | ENSG00000238105 | GOLGA2P5 | chr12 | - | 100157348 | 100157445 | 100157164 | 100157256 | 100157527 | 100157739 | 0 | 8 | 2 | 1 | 0 | 5 | 5 | 2 | 7 | 2 | 0 | 0 | 1 | 3 | 3 | 7 | 11 | 13 | 7 | 0 | 246 | 149 | 0.221202935 | 1 | 0 | 0.492 | 0.377 | 0.08 | 0 | 0 | 0 | 0.045 | 0.206 | 1 | -0.06 |
| SE | 93121 | ENSG00000079387 | SENP1 | chr12 | - | 48048000 | 48048080 | 48046977 | 48047062 | 48048928 | 48049132 | 38 | 40 | 51 | 35 | 99 | 1 | 1 | 0 | 0 | 3 | 7 | 66 | 56 | 18 | 10 | 0 | 0 | 2 | 6 | 0 | 229 | 149 | 0.221241695 | 1 | 0.961 | 0.963 | 1 | 1 | 0.955 | 1 | 1 | 0.948 | 0.661 | 1 | 0.054 |
| SE | 85818 | ENSG00000118922 | KLF12 | chr13 | - | 73943980 | 73944070 | 73845826 | 73846373 | 73994989 | 73995053 | 510 | 344 | 406 | 378 | 627 | 2 | 8 | 6 | 2 | 7 | 190 | 222 | 252 | 85 | 146 | 0 | 1 | 7 | 0 | 0 | 239 | 149 | 0.221293147 | 1 | 0.994 | 0.964 | 0.977 | 0.992 | 0.982 | 1 | 0.993 | 0.957 | 1 | 1 | -0.008 |
| SE | 51841 | ENSG00000087206 | UIMC1 | chr5 | - | 176968554 | 176969291 | 176958092 | 176958154 | 176970741 | 176970866 | 55 | 39 | 44 | 45 | 93 | 8 | 3 | 5 | 6 | 20 | 80 | 50 | 32 | 29 | 66 | 3 | 10 | 22 | 4 | 25 | 298 | 149 | 0.221311561 | 1 | 0.775 | 0.867 | 0.815 | 0.789 | 0.699 | 0.93 | 0.714 | 0.421 | 0.784 | 0.569 | 0.105 |
| SE | 51837 | ENSG00000087206 | UIMC1 | chr5 | - | 176919169 | 176919251 | 176911310 | 176911389 | 176943334 | 176943488 | 3 | 4 | 4 | 1 | 6 | 97 | 85 | 56 | 118 | 101 | 0 | 10 | 17 | 4 | 6 | 124 | 123 | 167 | 188 | 124 | 231 | 149 | 0.221324993 | 1 | 0.02 | 0.029 | 0.044 | 0.005 | 0.037 | 0 | 0.05 | 0.062 | 0.014 | 0.03 | -0.004 |
| SE | 48855 | ENSG00000165661 | QSOX2 | chr9 | - | 136223762 | 136223853 | 136219029 | 136219164 | 136224006 | 136224112 | 24 | 7 | 21 | 22 | 14 | 0 | 0 | 2 | 0 | 1 | 47 | 20 | 13 | 10 | 25 | 0 | 0 | 0 | 0 | 7 | 240 | 149 | 0.221328748 | 1 | 1 | 1 | 0.867 | 1 | 0.897 | 1 | 1 | 1 | 1 | 0.689 | 0.015 |
| SE | 17877 | ENSG00000187097 | ENTPD5 | chr14 | - | 73975935 | 73976015 | 73974923 | 73974985 | 73976323 | 73976412 | 31 | 44 | 30 | 49 | 23 | 0 | 2 | 0 | 3 | 1 | 57 | 32 | 36 | 51 | 125 | 5 | 5 | 0 | 3 | 0 | 229 | 149 | 0.221339007 | 1 | 1 | 0.935 | 1 | 0.914 | 0.937 | 0.881 | 0.806 | 1 | 0.917 | 1 | 0.036 |
| SE | 90426 | ENSG00000123908 | AGO2 | chr8 | - | 140585118 | 140585311 | 140572811 | 140572932 | 140635484 | 140635546 | 49 | 36 | 38 | 25 | 44 | 0 | 0 | 0 | 0 | 0 | 30 | 41 | 44 | 25 | 48 | 6 | 0 | 0 | 1 | 0 | 298 | 149 | 0.221344744 | 1 | 1 | 1 | 1 | 1 | 1 | 0.714 | 1 | 1 | 0.926 | 1 | 0.072 |
| SE | 52460 | ENSG00000072803 | FBXW11 | chr5 | - | 171957596 | 171957698 | 171910571 | 171910797 | 171996889 | 171997051 | 0 | 0 | 0 | 0 | 0 | 17 | 8 | 7 | 10 | 12 | 0 | 0 | 4 | 3 | 0 | 24 | 2 | 13 | 4 | 9 | 251 | 149 | 0.221354232 | 1 | 0 | 0 | 0 | 0 | 0 | 0 | 0 | 0.154 | 0.308 | 0 | -0.092 |
| SE | 51236 | ENSG00000146376 | ARHGAP18 | chr6 | - | 129605876 | 129605959 | 129600641 | 129600848 | 129611532 | 129611610 | 64 | 98 | 21 | 144 | 100 | 1 | 1 | 3 | 5 | 0 | 28 | 82 | 79 | 28 | 25 | 0 | 2 | 1 | 0 | 0 | 232 | 149 | 0.221378578 | 1 | 0.976 | 0.984 | 0.818 | 0.949 | 1 | 1 | 0.963 | 0.981 | 1 | 1 | -0.043 |
| SE | 94235 | ENSG00000213722 | DDAH2 | chr6 | - | 31728864 | 31729225 | 31728180 | 31728745 | 31730050 | 31730245 | 35 | 21 | 37 | 44 | 34 | 1 | 0 | 0 | 0 | 0 | 49 | 32 | 48 | 52 | 86 | 0 | 0 | 0 | 0 | 8 | 298 | 149 | 0.221449681 | 1 | 0.946 | 1 | 1 | 1 | 1 | 1 | 1 | 1 | 1 | 0.843 | 0.021 |
| SE | 74232 | ENSG00000137478 | FCHSD2 | chr11 | - | 72841453 | 72841583 | 72840876 | 72840959 | 72842620 | 72842841 | 234 | 146 | 193 | 206 | 293 | 1 | 0 | 0 | 5 | 2 | 185 | 183 | 223 | 200 | 145 | 0 | 5 | 7 | 1 | 0 | 279 | 149 | 0.221467723 | 1 | 0.992 | 1 | 1 | 0.957 | 0.987 | 1 | 0.951 | 0.944 | 0.991 | 1 | 0.01 |
| SE | 69168 | ENSG00000132530 | XAF1 | chr17 | + | 6759661 | 6759718 | 6758088 | 6758224 | 6760405 | 6760479 | 91 | 102 | 42 | 106 | 99 | 2 | 1 | 1 | 0 | 7 | 54 | 78 | 49 | 202 | 9 | 0 | 0 | 0 | 7 | 0 | 206 | 149 | 0.221468315 | 1 | 0.971 | 0.987 | 0.968 | 1 | 0.911 | 1 | 1 | 1 | 0.954 | 1 | -0.023 |
| SE | 57621 | ENSG00000142405 | NLRP12 | chr19 | - | 53798242 | 53798413 | 53795858 | 53796029 | 53801226 | 53801397 | 59 | 19 | 32 | 40 | 39 | 7 | 1 | 2 | 6 | 0 | 78 | 95 | 22 | 37 | 86 | 17 | 5 | 4 | 1 | 10 | 298 | 149 | 0.22154196 | 1 | 0.808 | 0.905 | 0.889 | 0.769 | 1 | 0.696 | 0.905 | 0.733 | 0.949 | 0.811 | 0.055 |
| SE | 108420 | ENSG00000197111 | PCBP2 | chr12 | + | 53467804 | 53467843 | 53467220 | 53467293 | 53468779 | 53468832 | 279 | 130 | 309 | 230 | 207 | 14 | 2 | 6 | 9 | 26 | 209 | 249 | 165 | 173 | 326 | 13 | 3 | 9 | 5 | 13 | 188 | 149 | 0.221545108 | 1 | 0.94 | 0.981 | 0.976 | 0.953 | 0.863 | 0.927 | 0.985 | 0.936 | 0.965 | 0.952 | -0.01 |
| SE | 19284 | ENSG00000226174 | TEX22 | chr14 | + | 105445417 | 105445511 | 105438671 | 105438739 | 105450057 | 105450106 | 36 | 11 | 25 | 26 | 17 | 1 | 3 | 1 | 2 | 3 | 48 | 43 | 12 | 11 | 44 | 0 | 6 | 2 | 0 | 0 | 243 | 149 | 0.221573271 | 1 | 0.957 | 0.692 | 0.939 | 0.889 | 0.777 | 1 | 0.815 | 0.786 | 1 | 1 | -0.069 |
| SE | 70614 | ENSG00000225889 | NA | chr2 | + | 64229170 | 64229288 | 64228452 | 64228522 | 64229496 | 64229640 | 9 | 2 | 11 | 2 | 2 | 9 | 10 | 10 | 4 | 1 | 10 | 11 | 9 | 4 | 1 | 0 | 10 | 12 | 13 | 0 | 267 | 149 | 0.221577716 | 1 | 0.358 | 0.1 | 0.38 | 0.218 | 0.527 | 1 | 0.38 | 0.295 | 0.147 | 1 | -0.248 |
| SE | 22683 | ENSG00000131844 | MCCC2 | chr5 | + | 71626639 | 71626753 | 71604355 | 71604468 | 71632120 | 71632185 | 44 | 15 | 37 | 22 | 40 | 10 | 11 | 15 | 17 | 26 | 62 | 26 | 28 | 30 | 46 | 25 | 5 | 18 | 3 | 23 | 263 | 149 | 0.221750268 | 1 | 0.714 | 0.436 | 0.583 | 0.423 | 0.466 | 0.584 | 0.747 | 0.468 | 0.85 | 0.531 | -0.112 |
| SE | 98638 | ENSG00000149179 | C11orf49 | chr11 | + | 47023338 | 47023394 | 46987207 | 46987310 | 47052387 | 47052518 | 0 | 2 | 0 | 6 | 0 | 13 | 16 | 10 | 7 | 14 | 7 | 11 | 0 | 0 | 0 | 17 | 6 | 6 | 11 | 9 | 205 | 149 | 0.221780304 | 1 | 0 | 0.083 | 0 | 0.384 | 0 | 0.23 | 0.571 | 0 | 0 | 0 | -0.067 |
| SE | 71079 | ENSG00000134490 | TMEM241 | chr18 | - | 23399565 | 23399647 | 23384989 | 23385067 | 23421366 | 23421444 | 9 | 6 | 4 | 8 | 7 | 2 | 0 | 0 | 0 | 0 | 10 | 26 | 6 | 13 | 4 | 6 | 0 | 0 | 0 | 0 | 231 | 149 | 0.221797278 | 1 | 0.744 | 1 | 1 | 1 | 1 | 0.518 | 1 | 1 | 1 | 1 | 0.045 |
| SE | 80278 | ENSG00000168137 | SETD5 | chr3 | + | 9429568 | 9429667 | 9428822 | 9429009 | 9429826 | 9430235 | 0 | 2 | 2 | 3 | 1 | 24 | 21 | 22 | 33 | 26 | 0 | 0 | 4 | 0 | 0 | 16 | 32 | 30 | 29 | 16 | 248 | 149 | 0.221808231 | 1 | 0 | 0.054 | 0.052 | 0.052 | 0.023 | 0 | 0 | 0.074 | 0 | 0 | 0.021 |
| SE | 50016 | ENSG00000162298 | SYVN1 | chr11 | - | 65132731 | 65132780 | 65132247 | 65132351 | 65132921 | 65133074 | 100 | 28 | 74 | 78 | 69 | 0 | 0 | 0 | 0 | 3 | 30 | 105 | 39 | 26 | 173 | 0 | 5 | 1 | 4 | 0 | 198 | 149 | 0.221818548 | 1 | 1 | 1 | 1 | 1 | 0.945 | 1 | 0.94 | 0.967 | 0.83 | 1 | 0.042 |
| SE | 106069 | ENSG00000196498 | NCOR2 | chr12 | - | 124327408 | 124327633 | 124326190 | 124326370 | 124330844 | 124330898 | 83 | 34 | 54 | 77 | 39 | 47 | 14 | 29 | 31 | 48 | 87 | 98 | 40 | 54 | 114 | 40 | 73 | 39 | 39 | 111 | 298 | 149 | 0.221903246 | 1 | 0.469 | 0.548 | 0.482 | 0.554 | 0.289 | 0.521 | 0.402 | 0.339 | 0.409 | 0.339 | 0.066 |
| SE | 101134 | ENSG00000175224 | ATG13 | chr11 | + | 46630044 | 46630100 | 46618259 | 46618375 | 46644278 | 46644360 | 36 | 20 | 30 | 32 | 24 | 10 | 4 | 12 | 6 | 14 | 39 | 30 | 8 | 14 | 56 | 9 | 10 | 0 | 4 | 10 | 205 | 149 | 0.222009826 | 1 | 0.723 | 0.784 | 0.645 | 0.795 | 0.555 | 0.759 | 0.686 | 1 | 0.718 | 0.803 | -0.093 |
| SE | 1330 | ENSG00000093000 | NUP50 | chr22 | + | 45166308 | 45166430 | 45163986 | 45164296 | 45168167 | 45168246 | 1 | 4 | 10 | 6 | 4 | 76 | 61 | 83 | 74 | 47 | 0 | 20 | 4 | 17 | 0 | 104 | 110 | 77 | 68 | 59 | 271 | 149 | 0.222053793 | 1 | 0.007 | 0.035 | 0.062 | 0.043 | 0.045 | 0 | 0.091 | 0.028 | 0.121 | 0 | -0.01 |
| SE | 41299 | ENSG00000058668 | ATP2B4 | chr1 | + | 203723880 | 203723988 | 203722477 | 203722689 | 203727394 | 203727571 | 658 | 253 | 416 | 515 | 655 | 8 | 5 | 1 | 10 | 11 | 295 | 327 | 296 | 334 | 223 | 7 | 3 | 3 | 9 | 8 | 257 | 149 | 0.222116402 | 1 | 0.979 | 0.967 | 0.996 | 0.968 | 0.972 | 0.961 | 0.984 | 0.983 | 0.956 | 0.942 | 0.011 |
| SE | 80954 | ENSG00000133393 | CEP20 | chr16 | - | 15882948 | 15883020 | 15879803 | 15879888 | 15884007 | 15884205 | 6 | 0 | 5 | 2 | 11 | 156 | 113 | 148 | 133 | 161 | 0 | 0 | 2 | 0 | 10 | 109 | 72 | 86 | 134 | 62 | 221 | 149 | 0.222159396 | 1 | 0.025 | 0 | 0.022 | 0.01 | 0.044 | 0 | 0 | 0.015 | 0 | 0.098 | -0.002 |
| SE | 27907 | ENSG00000118564 | FBXL5 | chr4 | - | 15627884 | 15628033 | 15626872 | 15626955 | 15630665 | 15630791 | 517 | 332 | 386 | 489 | 635 | 2 | 6 | 2 | 1 | 8 | 232 | 435 | 418 | 383 | 79 | 0 | 5 | 12 | 3 | 0 | 298 | 149 | 0.222259866 | 1 | 0.992 | 0.965 | 0.99 | 0.996 | 0.975 | 1 | 0.978 | 0.946 | 0.985 | 1 | 0.002 |
| SE | 82856 | ENSG00000146776 | ATXN7L1 | chr7 | - | 105639486 | 105639569 | 105638352 | 105638609 | 105642837 | 105643121 | 93 | 72 | 114 | 112 | 165 | 14 | 11 | 14 | 15 | 11 | 166 | 139 | 142 | 138 | 227 | 48 | 11 | 17 | 38 | 15 | 232 | 149 | 0.222278685 | 1 | 0.81 | 0.808 | 0.839 | 0.827 | 0.906 | 0.69 | 0.89 | 0.843 | 0.7 | 0.907 | 0.032 |
| SE | 28484 | ENSG00000155229 | MMS19 | chr10 | - | 97459361 | 97459505 | 97459222 | 97459282 | 97459658 | 97459741 | 40 | 23 | 32 | 31 | 33 | 0 | 0 | 1 | 3 | 3 | 63 | 22 | 11 | 40 | 24 | 0 | 0 | 0 | 0 | 0 | 293 | 149 | 0.222292589 | 1 | 1 | 1 | 0.942 | 0.84 | 0.848 | 1 | 1 | 1 | 1 | 1 | -0.074 |
| SE | 75156 | ENSG00000059758 | CDK17 | chr12 | - | 96297269 | 96297332 | 96294998 | 96295122 | 96297626 | 96297721 | 308 | 181 | 257 | 226 | 453 | 13 | 13 | 15 | 14 | 23 | 27 | 126 | 164 | 57 | 33 | 0 | 0 | 35 | 13 | 0 | 212 | 149 | 0.222308625 | 1 | 0.943 | 0.907 | 0.923 | 0.919 | 0.933 | 1 | 1 | 0.767 | 0.755 | 1 | 0.021 |
| SE | 20477 | ENSG00000164080 | RAD54L2 | chr3 | + | 51633576 | 51633759 | 51630704 | 51630931 | 51633901 | 51634035 | 64 | 25 | 60 | 42 | 58 | 0 | 0 | 0 | 0 | 0 | 8 | 83 | 33 | 79 | 61 | 0 | 0 | 0 | 8 | 0 | 298 | 149 | 0.222390176 | 1 | 1 | 1 | 1 | 1 | 1 | 1 | 1 | 1 | 0.832 | 1 | 0.034 |
| SE | 82703 | ENSG00000127914 | AKAP9 | chr7 | + | 91980872 | 91980958 | 91980288 | 91980333 | 91992157 | 91992211 | 18 | 13 | 18 | 22 | 21 | 166 | 174 | 126 | 179 | 224 | 4 | 14 | 1 | 6 | 0 | 30 | 61 | 15 | 6 | 4 | 235 | 149 | 0.222398194 | 1 | 0.064 | 0.045 | 0.083 | 0.072 | 0.056 | 0.078 | 0.127 | 0.041 | 0.388 | 0 | -0.063 |
| SE | 77308 | ENSG00000099968 | BCL2L13 | chr22 | + | 17702242 | 17702386 | 17688985 | 17689142 | 17726676 | 17728699 | 94 | 70 | 97 | 105 | 98 | 10 | 6 | 7 | 3 | 4 | 96 | 138 | 58 | 114 | 191 | 6 | 14 | 0 | 7 | 0 | 293 | 149 | 0.222438722 | 1 | 0.827 | 0.856 | 0.876 | 0.947 | 0.926 | 0.891 | 0.834 | 1 | 0.892 | 1 | -0.037 |
| SE | 97621 | ENSG00000166002 | SMCO4 | chr11 | - | 93499275 | 93499348 | 93478471 | 93479269 | 93537845 | 93537915 | 42 | 37 | 39 | 44 | 50 | 0 | 0 | 0 | 0 | 0 | 107 | 42 | 21 | 30 | 91 | 0 | 0 | 0 | 0 | 9 | 222 | 149 | 0.222454376 | 1 | 1 | 1 | 1 | 1 | 1 | 1 | 1 | 1 | 1 | 0.872 | 0.026 |
| SE | 16549 | ENSG00000137764 | MAP2K5 | chr15 | + | 67580753 | 67580823 | 67563282 | 67563350 | 67585889 | 67585930 | 101 | 94 | 126 | 116 | 162 | 6 | 5 | 11 | 7 | 6 | 164 | 156 | 61 | 82 | 228 | 17 | 16 | 9 | 5 | 5 | 219 | 149 | 0.222526992 | 1 | 0.92 | 0.927 | 0.886 | 0.919 | 0.948 | 0.868 | 0.869 | 0.822 | 0.918 | 0.969 | 0.031 |
| SE | 74156 | ENSG00000168785 | TSPAN5 | chr4 | - | 98486737 | 98486884 | 98482004 | 98482175 | 98507677 | 98507728 | 48 | 20 | 29 | 45 | 29 | 29 | 13 | 10 | 21 | 11 | 21 | 11 | 4 | 42 | 18 | 11 | 8 | 9 | 17 | 14 | 296 | 149 | 0.222553438 | 1 | 0.454 | 0.436 | 0.593 | 0.519 | 0.57 | 0.49 | 0.409 | 0.183 | 0.554 | 0.393 | 0.109 |
| SE | 88874 | ENSG00000183255 | PTTG1IP | chr21 | - | 44861162 | 44861271 | 44856192 | 44856364 | 44865394 | 44865447 | 365 | 314 | 224 | 306 | 320 | 3 | 3 | 3 | 7 | 5 | 900 | 410 | 201 | 608 | 1366 | 14 | 11 | 6 | 0 | 27 | 258 | 149 | 0.222605638 | 1 | 0.986 | 0.984 | 0.977 | 0.962 | 0.974 | 0.974 | 0.956 | 0.951 | 1 | 0.967 | 0.007 |
| SE | 7585 | ENSG00000132388 | UBE2G1 | chr17 | - | 4289229 | 4289408 | 4282797 | 4282921 | 4296716 | 4296814 | 527 | 315 | 553 | 514 | 678 | 0 | 2 | 0 | 3 | 5 | 546 | 570 | 559 | 484 | 400 | 10 | 3 | 1 | 9 | 0 | 298 | 149 | 0.222743394 | 1 | 1 | 0.987 | 1 | 0.988 | 0.985 | 0.965 | 0.99 | 0.996 | 0.964 | 1 | 0.009 |
| SE | 52592 | ENSG00000163607 | GTPBP8 | chr3 | + | 112995136 | 112995265 | 112993074 | 112993124 | 112996891 | 112996991 | 89 | 37 | 76 | 66 | 63 | 16 | 11 | 18 | 20 | 18 | 100 | 37 | 39 | 27 | 20 | 0 | 17 | 3 | 15 | 5 | 278 | 149 | 0.222852156 | 1 | 0.749 | 0.643 | 0.694 | 0.639 | 0.652 | 1 | 0.538 | 0.874 | 0.491 | 0.682 | -0.042 |
| SE | 76572 | ENSG00000170919 | TPT1-AS1 | chr13 | + | 45369958 | 45370182 | 45341404 | 45341614 | 45377060 | 45377154 | 6 | 6 | 6 | 1 | 9 | 0 | 0 | 0 | 1 | 4 | 20 | 6 | 6 | 9 | 5 | 0 | 0 | 0 | 0 | 0 | 298 | 149 | 0.222895215 | 1 | 1 | 1 | 1 | 0.333 | 0.529 | 1 | 1 | 1 | 1 | 1 | -0.228 |
| SE | 52099 | ENSG00000158006 | PAFAH2 | chr1 | - | 25972557 | 25972712 | 25959766 | 25962083 | 25974479 | 25974650 | 134 | 71 | 96 | 106 | 150 | 4 | 4 | 5 | 4 | 9 | 81 | 83 | 72 | 47 | 39 | 0 | 3 | 18 | 0 | 9 | 298 | 149 | 0.222903563 | 1 | 0.944 | 0.899 | 0.906 | 0.93 | 0.893 | 1 | 0.933 | 0.667 | 1 | 0.684 | 0.058 |
| SE | 34527 | ENSG00000134802 | SLC43A3 | chr11 | - | 57425540 | 57425709 | 57423981 | 57424028 | 57425988 | 57426316 | 37 | 20 | 35 | 34 | 54 | 1 | 0 | 0 | 2 | 0 | 122 | 77 | 34 | 79 | 91 | 0 | 0 | 0 | 9 | 0 | 298 | 149 | 0.222943707 | 1 | 0.949 | 1 | 1 | 0.895 | 1 | 1 | 1 | 1 | 0.814 | 1 | 0.006 |
| SE | 99108 | ENSG00000110274 | CEP164 | chr11 | + | 117408525 | 117408755 | 117407924 | 117408032 | 117408889 | 117409028 | 4 | 0 | 4 | 1 | 6 | 26 | 31 | 49 | 31 | 84 | 4 | 3 | 4 | 0 | 0 | 72 | 36 | 24 | 30 | 98 | 298 | 149 | 0.222973108 | 1 | 0.071 | 0 | 0.039 | 0.016 | 0.034 | 0.027 | 0.04 | 0.077 | 0 | 0 | 0.003 |
| SE | 86258 | ENSG00000134516 | DOCK2 | chr5 | + | 169708168 | 169708267 | 169702302 | 169702427 | 169711934 | 169712007 | 1126 | 819 | 902 | 1130 | 1334 | 1 | 8 | 0 | 5 | 40 | 1771 | 1425 | 1240 | 1337 | 870 | 21 | 15 | 0 | 9 | 6 | 248 | 149 | 0.223110077 | 1 | 0.999 | 0.984 | 1 | 0.993 | 0.952 | 0.981 | 0.983 | 1 | 0.989 | 0.989 | -0.003 |
| SE | 75102 | ENSG00000114423 | CBLB | chr3 | - | 105671023 | 105671046 | 105670232 | 105670352 | 105678430 | 105678571 | 1 | 6 | 5 | 7 | 4 | 96 | 50 | 44 | 77 | 98 | 0 | 7 | 5 | 6 | 0 | 35 | 33 | 58 | 19 | 12 | 172 | 149 | 0.223146137 | 1 | 0.009 | 0.094 | 0.09 | 0.073 | 0.034 | 0 | 0.155 | 0.069 | 0.215 | 0 | -0.028 |
| SE | 17935 | ENSG00000197249 | SERPINA1 | chr14 | - | 94388559 | 94388663 | 94382591 | 94383241 | 94390456 | 94390650 | 422 | 224 | 164 | 221 | 174 | 1096 | 525 | 550 | 643 | 1047 | 587 | 602 | 330 | 252 | 982 | 3262 | 1930 | 690 | 1662 | 2598 | 253 | 149 | 0.22321459 | 1 | 0.185 | 0.201 | 0.149 | 0.168 | 0.089 | 0.096 | 0.155 | 0.22 | 0.082 | 0.182 | 0.011 |
| SE | 63992 | ENSG00000068912 | ERLEC1 | chr2 | + | 53801712 | 53801842 | 53799046 | 53799081 | 53809213 | 53809273 | 27 | 16 | 26 | 34 | 84 | 1 | 1 | 0 | 0 | 13 | 4 | 26 | 33 | 17 | 18 | 8 | 0 | 1 | 0 | 0 | 279 | 149 | 0.223303984 | 1 | 0.935 | 0.895 | 1 | 1 | 0.775 | 0.211 | 1 | 0.946 | 1 | 1 | 0.09 |
| SE | 22740 | ENSG00000050393 | MCUR1 | chr6 | - | 13801287 | 13801389 | 13800340 | 13800382 | 13802242 | 13802346 | 160 | 183 | 85 | 365 | 349 | 1 | 1 | 4 | 0 | 4 | 552 | 259 | 220 | 430 | 1924 | 24 | 0 | 3 | 1 | 5 | 251 | 149 | 0.223378984 | 1 | 0.99 | 0.991 | 0.927 | 1 | 0.981 | 0.932 | 1 | 0.978 | 0.996 | 0.996 | -0.003 |
| SE | 101257 | ENSG00000166224 | SGPL1 | chr10 | + | 70873350 | 70873589 | 70869791 | 70869897 | 70875401 | 70875548 | 90 | 62 | 85 | 90 | 100 | 0 | 0 | 0 | 1 | 0 | 104 | 71 | 65 | 58 | 129 | 0 | 7 | 0 | 2 | 0 | 298 | 149 | 0.223424818 | 1 | 1 | 1 | 1 | 0.978 | 1 | 1 | 0.835 | 1 | 0.935 | 1 | 0.042 |
| SE | 21823 | ENSG00000174353 | STAG3L3 | chr7 | - | 73000357 | 73000481 | 72999760 | 72999859 | 73003020 | 73003137 | 6 | 1 | 15 | 12 | 10 | 90 | 84 | 167 | 163 | 159 | 17 | 4 | 9 | 2 | 11 | 134 | 126 | 139 | 226 | 266 | 273 | 149 | 0.223428448 | 1 | 0.035 | 0.006 | 0.047 | 0.039 | 0.033 | 0.065 | 0.017 | 0.034 | 0.005 | 0.022 | 0.003 |
| SE | 65161 | ENSG00000196912 | ANKRD36B | chr2 | - | 97576378 | 97576446 | 97560831 | 97560860 | 97580461 | 97580568 | 16 | 14 | 15 | 25 | 31 | 27 | 13 | 19 | 22 | 23 | 8 | 9 | 9 | 4 | 27 | 4 | 5 | 13 | 6 | 8 | 217 | 149 | 0.223435396 | 1 | 0.289 | 0.425 | 0.352 | 0.438 | 0.481 | 0.579 | 0.553 | 0.322 | 0.314 | 0.699 | -0.096 |
| SE | 15383 | ENSG00000138600 | SPPL2A | chr15 | - | 50726072 | 50726126 | 50725220 | 50725323 | 50726320 | 50726377 | 22 | 15 | 14 | 10 | 42 | 39 | 25 | 22 | 37 | 36 | 41 | 22 | 16 | 22 | 0 | 17 | 14 | 15 | 13 | 16 | 203 | 149 | 0.223477531 | 1 | 0.293 | 0.306 | 0.318 | 0.166 | 0.461 | 0.639 | 0.536 | 0.439 | 0.554 | 0 | -0.125 |
| SE | 33181 | ENSG00000145545 | SRD5A1 | chr5 | + | 6644807 | 6645309 | 6633426 | 6633869 | 6651841 | 6652008 | 0 | 2 | 0 | 2 | 2 | 16 | 7 | 12 | 17 | 5 | 0 | 15 | 0 | 3 | 0 | 92 | 35 | 5 | 18 | 63 | 298 | 149 | 0.223605823 | 1 | 0 | 0.125 | 0 | 0.056 | 0.167 | 0 | 0.176 | 0 | 0.077 | 0 | 0.019 |
| SE | 44201 | ENSG00000237094 | NA | chr1 | - | 498046 | 498305 | 497239 | 497299 | 498398 | 498456 | 1 | 0 | 2 | 2 | 0 | 10 | 8 | 9 | 6 | 16 | 7 | 0 | 0 | 6 | 1 | 58 | 39 | 5 | 55 | 70 | 298 | 149 | 0.223627102 | 1 | 0.048 | 0 | 0.1 | 0.143 | 0 | 0.057 | 0 | 0 | 0.052 | 0.007 | 0.035 |
| SE | 56744 | ENSG00000117616 | RSRP1 | chr1 | - | 25244453 | 25244498 | 25243549 | 25244224 | 25245149 | 25245301 | 116 | 103 | 126 | 133 | 111 | 31 | 23 | 10 | 41 | 40 | 264 | 48 | 65 | 88 | 139 | 36 | 15 | 3 | 45 | 12 | 194 | 149 | 0.223717488 | 1 | 0.742 | 0.775 | 0.906 | 0.714 | 0.681 | 0.849 | 0.711 | 0.943 | 0.6 | 0.899 | -0.037 |
| SE | 100784 | ENSG00000162408 | NOL9 | chr1 | - | 6541829 | 6541927 | 6533279 | 6533441 | 6544825 | 6544922 | 228 | 91 | 163 | 149 | 319 | 0 | 3 | 2 | 2 | 3 | 126 | 155 | 79 | 137 | 120 | 0 | 3 | 6 | 0 | 3 | 247 | 149 | 0.223736551 | 1 | 1 | 0.948 | 0.98 | 0.978 | 0.985 | 1 | 0.969 | 0.888 | 1 | 0.96 | 0.015 |
| SE | 56275 | ENSG00000248429 | FAM198B-AS1 | chr4 | + | 158198787 | 158198870 | 158179660 | 158179735 | 158199087 | 158199486 | 23 | 46 | 12 | 32 | 17 | 2 | 0 | 0 | 3 | 5 | 32 | 48 | 18 | 38 | 15 | 9 | 17 | 0 | 0 | 0 | 232 | 149 | 0.223880905 | 1 | 0.881 | 1 | 1 | 0.873 | 0.686 | 0.695 | 0.645 | 1 | 1 | 1 | 0.02 |
| SE | 67242 | ENSG00000123338 | NCKAP1L | chr12 | + | 54518595 | 54518732 | 54517805 | 54517938 | 54518913 | 54518972 | 458 | 322 | 347 | 377 | 1116 | 0 | 0 | 0 | 3 | 2 | 576 | 530 | 264 | 343 | 590 | 0 | 10 | 0 | 0 | 12 | 286 | 149 | 0.223942254 | 1 | 1 | 1 | 1 | 0.985 | 0.997 | 1 | 0.965 | 1 | 1 | 0.962 | 0.011 |
| SE | 49152 | ENSG00000077044 | DGKD | chr2 | + | 233434768 | 233434901 | 233434379 | 233434484 | 233435817 | 233435894 | 376 | 220 | 267 | 347 | 296 | 4 | 8 | 8 | 8 | 9 | 377 | 420 | 260 | 390 | 889 | 0 | 17 | 0 | 30 | 26 | 282 | 149 | 0.223950276 | 1 | 0.98 | 0.936 | 0.946 | 0.958 | 0.946 | 1 | 0.929 | 1 | 0.873 | 0.948 | 0.003 |
| SE | 107715 | ENSG00000177575 | CD163 | chr12 | - | 7481160 | 7481256 | 7479859 | 7479996 | 7482965 | 7483004 | 29 | 9 | 7 | 24 | 11 | 0 | 0 | 0 | 2 | 0 | 101 | 101 | 69 | 37 | 22 | 6 | 3 | 0 | 0 | 0 | 245 | 149 | 0.22403603 | 1 | 1 | 1 | 1 | 0.879 | 1 | 0.911 | 0.953 | 1 | 1 | 1 | 0.003 |
| SE | 37905 | ENSG00000184014 | DENND5A | chr11 | - | 9145669 | 9145815 | 9144096 | 9145113 | 9147029 | 9147151 | 201 | 141 | 184 | 159 | 274 | 0 | 0 | 1 | 1 | 2 | 386 | 174 | 154 | 304 | 246 | 0 | 2 | 2 | 11 | 0 | 295 | 149 | 0.224120895 | 1 | 1 | 1 | 0.989 | 0.988 | 0.986 | 1 | 0.978 | 0.975 | 0.933 | 1 | 0.015 |
| SE | 17769 | ENSG00000133983 | COX16 | chr14 | - | 70342657 | 70342729 | 70329173 | 70329236 | 70359518 | 70359683 | 110 | 59 | 78 | 69 | 186 | 17 | 2 | 3 | 11 | 3 | 107 | 86 | 30 | 50 | 56 | 0 | 12 | 5 | 11 | 6 | 221 | 149 | 0.224225986 | 1 | 0.814 | 0.952 | 0.946 | 0.809 | 0.977 | 1 | 0.829 | 0.802 | 0.754 | 0.863 | 0.05 |
| SE | 15379 | ENSG00000138600 | SPPL2A | chr15 | - | 50719939 | 50720100 | 50702265 | 50707874 | 50722123 | 50722201 | 299 | 165 | 164 | 177 | 321 | 1 | 1 | 1 | 0 | 5 | 105 | 140 | 139 | 142 | 26 | 0 | 0 | 9 | 1 | 0 | 298 | 149 | 0.224323658 | 1 | 0.993 | 0.988 | 0.988 | 1 | 0.97 | 1 | 1 | 0.885 | 0.986 | 1 | 0.014 |
| SE | 19457 | ENSG00000125746 | EML2 | chr19 | - | 45617629 | 45617697 | 45616764 | 45616853 | 45619059 | 45619090 | 69 | 27 | 35 | 48 | 14 | 4 | 3 | 1 | 0 | 0 | 40 | 43 | 22 | 22 | 62 | 0 | 0 | 0 | 3 | 0 | 217 | 149 | 0.224347249 | 1 | 0.922 | 0.861 | 0.96 | 1 | 1 | 1 | 1 | 1 | 0.834 | 1 | -0.018 |
| SE | 76085 | ENSG00000112697 | TMEM30A | chr6 | - | 75267640 | 75267748 | 75265230 | 75265338 | 75284401 | 75284763 | 506 | 278 | 423 | 439 | 578 | 8 | 4 | 1 | 4 | 6 | 56 | 292 | 376 | 135 | 112 | 0 | 0 | 2 | 0 | 0 | 257 | 149 | 0.224350475 | 1 | 0.973 | 0.976 | 0.996 | 0.985 | 0.982 | 1 | 1 | 0.991 | 1 | 1 | -0.016 |
| SE | 19762 | ENSG00000100614 | PPM1A | chr14 | + | 60277004 | 60277091 | 60249659 | 60249677 | 60282683 | 60283537 | 10 | 8 | 5 | 12 | 10 | 9 | 6 | 9 | 15 | 1 | 0 | 12 | 21 | 6 | 7 | 10 | 13 | 21 | 5 | 8 | 236 | 149 | 0.224461806 | 1 | 0.412 | 0.457 | 0.26 | 0.336 | 0.863 | 0 | 0.368 | 0.387 | 0.431 | 0.356 | 0.157 |
| SE | 16665 | ENSG00000180304 | OAZ2 | chr15 | - | 64691433 | 64691591 | 64690166 | 64690549 | 64703028 | 64703194 | 124 | 62 | 65 | 95 | 90 | 6 | 1 | 4 | 2 | 1 | 136 | 83 | 79 | 104 | 249 | 0 | 7 | 4 | 1 | 0 | 298 | 149 | 0.224535434 | 1 | 0.912 | 0.969 | 0.89 | 0.96 | 0.978 | 1 | 0.856 | 0.908 | 0.981 | 1 | -0.007 |
| SE | 13943 | ENSG00000103426 | CORO7-PAM16 | chr16 | - | 4361748 | 4361803 | 4361360 | 4361469 | 4361984 | 4362160 | 5 | 3 | 8 | 2 | 1 | 60 | 10 | 29 | 30 | 31 | 15 | 9 | 3 | 6 | 7 | 44 | 67 | 26 | 12 | 125 | 204 | 149 | 0.224617432 | 1 | 0.057 | 0.18 | 0.168 | 0.046 | 0.023 | 0.199 | 0.089 | 0.078 | 0.268 | 0.039 | -0.04 |
| SE | 17927 | ENSG00000197249 | SERPINA1 | chr14 | - | 94388559 | 94388660 | 94382591 | 94383241 | 94390419 | 94390692 | 328 | 189 | 133 | 185 | 136 | 13 | 12 | 1 | 11 | 6 | 496 | 452 | 260 | 203 | 832 | 10 | 29 | 3 | 19 | 8 | 250 | 149 | 0.224643199 | 1 | 0.938 | 0.904 | 0.988 | 0.909 | 0.931 | 0.967 | 0.903 | 0.981 | 0.864 | 0.984 | -0.006 |
| SE | 13108 | ENSG00000167264 | DUS2 | chr16 | + | 68025380 | 68025494 | 68023264 | 68023351 | 68038005 | 68038149 | 24 | 19 | 29 | 34 | 23 | 0 | 6 | 3 | 8 | 0 | 63 | 68 | 61 | 32 | 61 | 3 | 0 | 11 | 5 | 0 | 263 | 149 | 0.224702214 | 1 | 1 | 0.642 | 0.846 | 0.707 | 1 | 0.922 | 1 | 0.759 | 0.784 | 1 | -0.054 |
| SE | 21883 | ENSG00000189091 | SF3B3 | chr16 | + | 70560468 | 70560591 | 70556885 | 70557029 | 70563875 | 70564050 | 109 | 46 | 110 | 116 | 112 | 0 | 0 | 2 | 2 | 11 | 77 | 115 | 74 | 82 | 158 | 3 | 5 | 3 | 6 | 5 | 272 | 149 | 0.224758108 | 1 | 1 | 1 | 0.968 | 0.969 | 0.848 | 0.934 | 0.926 | 0.931 | 0.882 | 0.945 | 0.033 |
| SE | 71512 | ENSG00000184787 | UBE2G2 | chr21 | - | 44779143 | 44779216 | 44777298 | 44777417 | 44787919 | 44787965 | 3 | 3 | 5 | 5 | 1 | 102 | 107 | 105 | 103 | 102 | 0 | 0 | 4 | 3 | 0 | 146 | 108 | 98 | 78 | 118 | 222 | 149 | 0.224761473 | 1 | 0.019 | 0.018 | 0.031 | 0.032 | 0.007 | 0 | 0 | 0.027 | 0.025 | 0 | 0.011 |
| SE | 50545 | ENSG00000001461 | NIPAL3 | chr1 | + | 24419290 | 24419640 | 24415801 | 24415904 | 24440171 | 24440240 | 69 | 42 | 98 | 61 | 80 | 0 | 1 | 0 | 4 | 4 | 24 | 51 | 37 | 37 | 34 | 0 | 0 | 2 | 0 | 3 | 298 | 149 | 0.224799891 | 1 | 1 | 0.955 | 1 | 0.884 | 0.909 | 1 | 1 | 0.902 | 1 | 0.85 | -0.001 |
| SE | 64421 | ENSG00000143919 | CAMKMT | chr2 | + | 44549554 | 44549728 | 44390240 | 44390305 | 44704282 | 44704343 | 1 | 5 | 13 | 5 | 2 | 15 | 12 | 28 | 28 | 36 | 0 | 6 | 4 | 9 | 4 | 6 | 17 | 11 | 2 | 15 | 298 | 149 | 0.22497154 | 1 | 0.032 | 0.172 | 0.188 | 0.082 | 0.027 | 0 | 0.15 | 0.154 | 0.692 | 0.118 | -0.123 |
| SE | 4523 | ENSG00000131355 | ADGRE3 | chr19 | - | 14641418 | 14641616 | 14638104 | 14638340 | 14644107 | 14644275 | 35 | 27 | 35 | 60 | 58 | 4 | 0 | 0 | 0 | 0 | 9 | 6 | 23 | 5 | 31 | 0 | 0 | 2 | 4 | 0 | 298 | 149 | 0.225059108 | 1 | 0.814 | 1 | 1 | 1 | 1 | 1 | 1 | 0.852 | 0.385 | 1 | 0.115 |
| SE | 108377 | ENSG00000111554 | MDM1 | chr12 | - | 68316580 | 68316610 | 68316077 | 68316253 | 68321346 | 68321446 | 79 | 52 | 40 | 95 | 106 | 1 | 1 | 2 | 12 | 15 | 3 | 87 | 38 | 54 | 6 | 0 | 10 | 14 | 3 | 0 | 179 | 149 | 0.225123166 | 1 | 0.985 | 0.977 | 0.943 | 0.868 | 0.855 | 1 | 0.879 | 0.693 | 0.937 | 1 | 0.024 |
| SE | 13790 | ENSG00000161999 | JMJD8 | chr16 | - | 682957 | 683087 | 681698 | 682874 | 683166 | 683234 | 43 | 14 | 26 | 35 | 14 | 1 | 0 | 3 | 4 | 2 | 35 | 31 | 12 | 25 | 68 | 0 | 3 | 6 | 9 | 0 | 279 | 149 | 0.22543574 | 1 | 0.958 | 1 | 0.822 | 0.824 | 0.789 | 1 | 0.847 | 0.516 | 0.597 | 1 | 0.087 |
| SE | 14118 | ENSG00000102897 | LYRM1 | chr16 | + | 20915555 | 20915714 | 20900234 | 20900535 | 20920121 | 20920214 | 158 | 113 | 144 | 138 | 214 | 1 | 3 | 2 | 0 | 7 | 189 | 159 | 130 | 135 | 70 | 0 | 7 | 0 | 0 | 1 | 298 | 149 | 0.225437368 | 1 | 0.988 | 0.95 | 0.973 | 1 | 0.939 | 1 | 0.919 | 1 | 1 | 0.972 | -0.008 |
| SE | 82064 | ENSG00000086730 | LAT2 | chr7 | + | 74214821 | 74215099 | 74210005 | 74210088 | 74215946 | 74216069 | 42 | 19 | 28 | 26 | 21 | 78 | 28 | 21 | 47 | 18 | 52 | 51 | 11 | 70 | 67 | 121 | 59 | 48 | 76 | 121 | 298 | 149 | 0.22550202 | 1 | 0.212 | 0.253 | 0.4 | 0.217 | 0.368 | 0.177 | 0.302 | 0.103 | 0.315 | 0.217 | 0.067 |
| SE | 105480 | ENSG00000152256 | PDK1 | chr2 | + | 172562740 | 172562800 | 172562219 | 172562291 | 172564977 | 172565073 | 5 | 10 | 15 | 9 | 11 | 0 | 7 | 2 | 6 | 12 | 12 | 30 | 20 | 19 | 15 | 0 | 7 | 6 | 7 | 4 | 209 | 149 | 0.225590029 | 1 | 1 | 0.505 | 0.842 | 0.517 | 0.395 | 1 | 0.753 | 0.704 | 0.659 | 0.728 | -0.117 |
| SE | 39380 | ENSG00000220804 | LINC01881 | chr2 | + | 242116182 | 242116232 | 242094886 | 242095027 | 242118964 | 242119057 | 44 | 56 | 46 | 46 | 107 | 16 | 0 | 7 | 20 | 4 | 49 | 31 | 57 | 51 | 9 | 48 | 0 | 2 | 21 | 9 | 199 | 149 | 0.225600054 | 1 | 0.673 | 1 | 0.831 | 0.633 | 0.952 | 0.433 | 1 | 0.955 | 0.645 | 0.428 | 0.126 |
| SE | 9336 | ENSG00000108774 | RAB5C | chr17 | - | 42131606 | 42131822 | 42130336 | 42130590 | 42154892 | 42154988 | 1 | 0 | 5 | 1 | 1 | 104 | 43 | 89 | 119 | 28 | 0 | 7 | 4 | 6 | 12 | 223 | 160 | 86 | 118 | 301 | 298 | 149 | 0.22561203 | 1 | 0.005 | 0 | 0.027 | 0.004 | 0.018 | 0 | 0.021 | 0.023 | 0.025 | 0.02 | -0.007 |
| SE | 66763 | ENSG00000129292 | PHF20L1 | chr8 | + | 132825034 | 132825157 | 132824003 | 132824060 | 132825263 | 132825337 | 20 | 25 | 13 | 25 | 24 | 18 | 33 | 27 | 42 | 42 | 16 | 26 | 24 | 10 | 0 | 6 | 11 | 22 | 10 | 9 | 272 | 149 | 0.225705305 | 1 | 0.378 | 0.293 | 0.209 | 0.246 | 0.238 | 0.594 | 0.564 | 0.374 | 0.354 | 0 | -0.104 |
| SE | 67864 | ENSG00000018280 | SLC11A1 | chr2 | + | 218385411 | 218385485 | 218382168 | 218382375 | 218386634 | 218386667 | 13 | 6 | 3 | 12 | 10 | 2 | 2 | 0 | 3 | 0 | 27 | 21 | 29 | 9 | 34 | 0 | 0 | 0 | 0 | 3 | 223 | 149 | 0.225712324 | 1 | 0.813 | 0.667 | 1 | 0.728 | 1 | 1 | 1 | 1 | 1 | 0.883 | -0.135 |
| SE | 10439 | ENSG00000126581 | BECN1 | chr17 | - | 42818543 | 42818680 | 42818220 | 42818415 | 42819547 | 42819609 | 187 | 123 | 141 | 173 | 226 | 0 | 0 | 0 | 0 | 0 | 401 | 252 | 211 | 304 | 364 | 2 | 3 | 0 | 4 | 8 | 286 | 149 | 0.22578141 | 1 | 1 | 1 | 1 | 1 | 1 | 0.991 | 0.978 | 1 | 0.975 | 0.96 | 0.019 |
| SE | 66544 | ENSG00000168438 | CDC40 | chr6 | + | 110201557 | 110201687 | 110193181 | 110193268 | 110207505 | 110207589 | 513 | 425 | 350 | 427 | 659 | 9 | 1 | 1 | 14 | 7 | 331 | 448 | 737 | 450 | 93 | 0 | 12 | 20 | 6 | 0 | 279 | 149 | 0.225785366 | 1 | 0.968 | 0.996 | 0.995 | 0.942 | 0.98 | 1 | 0.952 | 0.952 | 0.976 | 1 | 0 |
| SE | 41750 | ENSG00000143222 | UFC1 | chr1 | + | 161157253 | 161157317 | 161156949 | 161157017 | 161158411 | 161158682 | 111 | 42 | 79 | 99 | 132 | 0 | 1 | 0 | 2 | 0 | 141 | 71 | 69 | 91 | 37 | 0 | 0 | 10 | 0 | 0 | 213 | 149 | 0.225825987 | 1 | 1 | 1 | 1 | 0.972 | 1 | 1 | 1 | 0.828 | 1 | 1 | 0.029 |
| SE | 42279 | ENSG00000160752 | FDPS | chr1 | + | 155319856 | 155319928 | 155319610 | 155319688 | 155320408 | 155320665 | 92 | 44 | 66 | 59 | 93 | 1 | 0 | 5 | 5 | 7 | 104 | 57 | 59 | 88 | 113 | 0 | 0 | 2 | 12 | 0 | 221 | 149 | 0.22583292 | 1 | 0.984 | 1 | 0.899 | 0.888 | 0.9 | 1 | 1 | 0.952 | 0.832 | 1 | -0.023 |
| SE | 56473 | ENSG00000162664 | ZNF326 | chr1 | + | 90007344 | 90007750 | 90005132 | 90005244 | 90010087 | 90010286 | 48 | 54 | 50 | 62 | 62 | 0 | 1 | 0 | 1 | 6 | 5 | 46 | 89 | 9 | 3 | 0 | 0 | 0 | 0 | 0 | 298 | 149 | 0.225866406 | 1 | 1 | 0.964 | 1 | 0.969 | 0.838 | 1 | 1 | 1 | 1 | 1 | -0.046 |
| SE | 10578 | ENSG00000108306 | FBXL20 | chr17 | - | 39265396 | 39265453 | 39263875 | 39264387 | 39268826 | 39268871 | 145 | 87 | 94 | 141 | 179 | 0 | 0 | 1 | 0 | 0 | 148 | 124 | 144 | 132 | 20 | 5 | 0 | 7 | 0 | 0 | 206 | 149 | 0.225921632 | 1 | 1 | 1 | 0.986 | 1 | 1 | 0.955 | 1 | 1 | 0.932 | 1 | 0.02 |
| SE | 10195 | ENSG00000178852 | EFCAB13 | chr17 | + | 47328792 | 47329038 | 47328268 | 47328383 | 47335195 | 47335356 | 0 | 0 | 4 | 7 | 8 | 23 | 76 | 66 | 77 | 176 | 0 | 6 | 5 | 4 | 0 | 48 | 52 | 15 | 20 | 41 | 298 | 149 | 0.225987943 | 1 | 0 | 0 | 0.029 | 0.043 | 0.022 | 0 | 0.055 | 0.143 | 0.091 | 0 | -0.039 |
| SE | 15310 | ENSG00000114062 | UBE3A | chr15 | - | 25407066 | 25407262 | 25405460 | 25405502 | 25409087 | 25409207 | 22 | 12 | 12 | 16 | 32 | 80 | 69 | 73 | 78 | 103 | 43 | 33 | 3 | 17 | 12 | 16 | 71 | 124 | 40 | 59 | 298 | 149 | 0.226025565 | 1 | 0.121 | 0.08 | 0.076 | 0.093 | 0.134 | 0.573 | 0.189 | 0.012 | 0.175 | 0.092 | -0.107 |
| SE | 66615 | ENSG00000170445 | HARS1 | chr5 | - | 140677326 | 140677420 | 140676988 | 140677116 | 140677654 | 140677753 | 154 | 79 | 133 | 139 | 178 | 0 | 0 | 0 | 3 | 1 | 75 | 139 | 75 | 87 | 91 | 4 | 0 | 3 | 3 | 0 | 243 | 149 | 0.226105822 | 1 | 1 | 1 | 1 | 0.966 | 0.991 | 0.92 | 1 | 0.939 | 0.947 | 1 | 0.03 |
| SE | 85205 | ENSG00000233382 | NKAPP1 | chrX | - | 120240226 | 120240355 | 120178487 | 120178556 | 120244763 | 120245233 | 17 | 4 | 11 | 10 | 0 | 0 | 0 | 0 | 0 | 3 | 6 | 2 | 6 | 6 | 2 | 0 | 0 | 0 | 0 | 0 | 278 | 149 | 0.226218052 | 1 | 1 | 1 | 1 | 1 | 0 | 1 | 1 | 1 | 1 | 1 | -0.2 |
| SE | 34195 | ENSG00000154767 | XPC | chr3 | - | 14168256 | 14168380 | 14167168 | 14167253 | 14170437 | 14170460 | 57 | 38 | 75 | 84 | 84 | 6 | 6 | 1 | 0 | 8 | 82 | 44 | 84 | 63 | 76 | 9 | 0 | 2 | 4 | 0 | 273 | 149 | 0.226244611 | 1 | 0.838 | 0.776 | 0.976 | 1 | 0.851 | 0.833 | 1 | 0.958 | 0.896 | 1 | -0.049 |
| SE | 50464 | ENSG00000161618 | ALDH16A1 | chr19 | + | 49461883 | 49462036 | 49461618 | 49461800 | 49462569 | 49462755 | 49 | 11 | 9 | 30 | 2 | 2 | 0 | 1 | 2 | 0 | 4 | 23 | 10 | 19 | 51 | 0 | 0 | 0 | 0 | 8 | 298 | 149 | 0.226249928 | 1 | 0.925 | 1 | 0.818 | 0.882 | 1 | 1 | 1 | 1 | 1 | 0.761 | -0.027 |
| SE | 27606 | ENSG00000001629 | ANKIB1 | chr7 | + | 92325927 | 92326008 | 92319329 | 92319512 | 92327782 | 92327900 | 7 | 8 | 4 | 4 | 13 | 74 | 61 | 85 | 76 | 153 | 4 | 7 | 5 | 2 | 8 | 27 | 64 | 45 | 44 | 31 | 230 | 149 | 0.226287382 | 1 | 0.058 | 0.078 | 0.03 | 0.033 | 0.052 | 0.088 | 0.066 | 0.067 | 0.029 | 0.143 | -0.028 |
| SE | 70859 | ENSG00000243970 | PPIEL | chr1 | - | 39556133 | 39556201 | 39554348 | 39554392 | 39556598 | 39556660 | 3 | 3 | 1 | 4 | 1 | 5 | 8 | 13 | 6 | 4 | 0 | 4 | 6 | 2 | 0 | 13 | 12 | 6 | 16 | 31 | 217 | 149 | 0.226535797 | 1 | 0.292 | 0.205 | 0.05 | 0.314 | 0.147 | 0 | 0.186 | 0.407 | 0.079 | 0 | 0.067 |
| SE | 13062 | ENSG00000123815 | COQ8B | chr19 | - | 40695988 | 40696661 | 40692950 | 40693037 | 40700066 | 40700174 | 26 | 12 | 14 | 19 | 5 | 0 | 0 | 0 | 0 | 0 | 15 | 8 | 1 | 8 | 21 | 0 | 0 | 0 | 5 | 0 | 298 | 149 | 0.226547464 | 1 | 1 | 1 | 1 | 1 | 1 | 1 | 1 | 1 | 0.444 | 1 | 0.111 |
| SE | 32431 | ENSG00000182048 | TRPC2 | chr11 | + | 3635044 | 3635172 | 3634291 | 3634484 | 3635274 | 3635353 | 2 | 2 | 2 | 8 | 6 | 0 | 0 | 0 | 0 | 0 | 0 | 3 | 4 | 5 | 24 | 3 | 0 | 0 | 0 | 0 | 277 | 149 | 0.226622405 | 1 | 1 | 1 | 1 | 1 | 1 | 0 | 1 | 1 | 1 | 1 | 0.2 |
| SE | 104020 | ENSG00000165983 | PTER | chr10 | + | 16484336 | 16484816 | 16437009 | 16437047 | 16486351 | 16486617 | 86 | 45 | 97 | 67 | 99 | 2 | 1 | 0 | 0 | 6 | 23 | 31 | 12 | 14 | 29 | 0 | 0 | 0 | 0 | 0 | 298 | 149 | 0.226637622 | 1 | 0.956 | 0.957 | 1 | 1 | 0.892 | 1 | 1 | 1 | 1 | 1 | -0.039 |
| SE | 52739 | ENSG00000143970 | ASXL2 | chr2 | - | 25835537 | 25835540 | 25806228 | 25806337 | 25845480 | 25845563 | 113 | 118 | 117 | 146 | 175 | 0 | 3 | 4 | 1 | 5 | 46 | 110 | 81 | 100 | 55 | 0 | 10 | 1 | 3 | 0 | 152 | 149 | 0.226665994 | 1 | 1 | 0.975 | 0.966 | 0.993 | 0.972 | 1 | 0.915 | 0.988 | 0.97 | 1 | 0.007 |
| SE | 98417 | ENSG00000110002 | VWA5A | chr11 | + | 124136128 | 124136293 | 124134919 | 124135034 | 124136573 | 124136674 | 74 | 21 | 41 | 45 | 115 | 2 | 0 | 0 | 0 | 8 | 50 | 25 | 23 | 11 | 59 | 7 | 0 | 0 | 0 | 0 | 298 | 149 | 0.226723933 | 1 | 0.949 | 1 | 1 | 1 | 0.878 | 0.781 | 1 | 1 | 1 | 1 | 0.009 |
| SE | 22762 | ENSG00000145730 | PAM | chr5 | + | 103025130 | 103025334 | 103019789 | 103019843 | 103028181 | 103028238 | 92 | 38 | 38 | 51 | 49 | 21 | 14 | 13 | 7 | 21 | 61 | 37 | 30 | 44 | 15 | 30 | 9 | 8 | 17 | 11 | 298 | 149 | 0.22674039 | 1 | 0.687 | 0.576 | 0.594 | 0.785 | 0.538 | 0.504 | 0.673 | 0.652 | 0.564 | 0.405 | 0.076 |
| SE | 4206 | ENSG00000187994 | RINL | chr19 | - | 38871646 | 38871711 | 38871077 | 38871227 | 38871797 | 38871870 | 83 | 15 | 38 | 38 | 42 | 2 | 3 | 4 | 1 | 3 | 60 | 93 | 18 | 39 | 58 | 0 | 3 | 1 | 6 | 0 | 214 | 149 | 0.226759686 | 1 | 0.967 | 0.777 | 0.869 | 0.964 | 0.907 | 1 | 0.956 | 0.926 | 0.819 | 1 | -0.043 |
| SE | 103323 | ENSG00000134452 | FBH1 | chr10 | + | 5924750 | 5924801 | 5924310 | 5924508 | 5925366 | 5925492 | 5 | 5 | 1 | 2 | 4 | 65 | 46 | 81 | 77 | 74 | 0 | 14 | 2 | 0 | 9 | 118 | 54 | 52 | 44 | 80 | 200 | 149 | 0.226763481 | 1 | 0.054 | 0.075 | 0.009 | 0.019 | 0.039 | 0 | 0.162 | 0.028 | 0 | 0.077 | -0.014 |
| SE | 38693 | ENSG00000167258 | CDK12 | chr17 | + | 39556863 | 39557058 | 39556285 | 39556413 | 39564759 | 39564907 | 3 | 3 | 2 | 10 | 18 | 15 | 17 | 19 | 29 | 32 | 8 | 15 | 6 | 5 | 7 | 0 | 21 | 15 | 21 | 39 | 298 | 149 | 0.227004544 | 1 | 0.091 | 0.081 | 0.05 | 0.147 | 0.22 | 1 | 0.263 | 0.167 | 0.106 | 0.082 | -0.206 |
| SE | 32588 | ENSG00000072756 | TRNT1 | chr3 | + | 3140509 | 3140648 | 3137259 | 3137453 | 3144583 | 3144710 | 153 | 119 | 148 | 177 | 213 | 7 | 2 | 1 | 0 | 2 | 95 | 78 | 73 | 45 | 96 | 0 | 0 | 3 | 0 | 0 | 288 | 149 | 0.227024866 | 1 | 0.919 | 0.969 | 0.987 | 1 | 0.982 | 1 | 1 | 0.926 | 1 | 1 | -0.014 |
| SE | 85798 | ENSG00000255561 | FDXACB1 | chr11 | - | 111878555 | 111878712 | 111876807 | 111877011 | 111878960 | 111879165 | 36 | 12 | 11 | 13 | 13 | 3 | 0 | 9 | 6 | 0 | 21 | 28 | 26 | 27 | 6 | 14 | 0 | 0 | 0 | 2 | 298 | 149 | 0.227117569 | 1 | 0.857 | 1 | 0.379 | 0.52 | 1 | 0.429 | 1 | 1 | 1 | 0.6 | -0.055 |
| SE | 3641 | ENSG00000239998 | LILRA2 | chr19 | + | 54575255 | 54575542 | 54574730 | 54575033 | 54575806 | 54575836 | 295 | 145 | 125 | 278 | 141 | 1 | 3 | 1 | 0 | 0 | 434 | 285 | 209 | 341 | 502 | 0 | 0 | 1 | 9 | 8 | 298 | 149 | 0.227247206 | 1 | 0.993 | 0.96 | 0.984 | 1 | 1 | 1 | 1 | 0.991 | 0.95 | 0.969 | 0.005 |
| SE | 38564 | ENSG00000198925 | ATG9A | chr2 | - | 219227921 | 219228053 | 219227769 | 219227813 | 219228433 | 219228482 | 40 | 19 | 38 | 22 | 16 | 0 | 0 | 1 | 0 | 0 | 16 | 24 | 9 | 15 | 47 | 1 | 2 | 4 | 0 | 0 | 281 | 149 | 0.22732695 | 1 | 1 | 1 | 0.953 | 1 | 1 | 0.895 | 0.864 | 0.544 | 1 | 1 | 0.13 |
| SE | 13444 | ENSG00000131149 | GSE1 | chr16 | + | 85633913 | 85634132 | 85555966 | 85556363 | 85648551 | 85648751 | 24 | 17 | 20 | 14 | 30 | 31 | 14 | 12 | 17 | 27 | 43 | 18 | 17 | 17 | 84 | 40 | 55 | 27 | 28 | 55 | 298 | 149 | 0.227350288 | 1 | 0.279 | 0.378 | 0.455 | 0.292 | 0.357 | 0.35 | 0.141 | 0.239 | 0.233 | 0.433 | 0.073 |
| SE | 18024 | ENSG00000249624 | NA | chr21 | + | 33268393 | 33268517 | 33262792 | 33263637 | 33279751 | 33279918 | 11 | 3 | 1 | 3 | 5 | 0 | 0 | 0 | 0 | 0 | 0 | 17 | 2 | 1 | 18 | 3 | 0 | 0 | 0 | 0 | 273 | 149 | 0.227615888 | 1 | 1 | 1 | 1 | 1 | 1 | 0 | 1 | 1 | 1 | 1 | 0.2 |
| SE | 78624 | ENSG00000262246 | CORO7 | chr16 | - | 4415714 | 4415852 | 4413307 | 4413404 | 4416458 | 4416627 | 1 | 3 | 1 | 5 | 0 | 43 | 18 | 31 | 31 | 9 | 0 | 2 | 0 | 0 | 0 | 23 | 72 | 36 | 26 | 49 | 287 | 149 | 0.227618521 | 1 | 0.012 | 0.08 | 0.016 | 0.077 | 0 | 0 | 0.014 | 0 | 0 | 0 | 0.034 |
| SE | 69811 | ENSG00000184007 | PTP4A2 | chr1 | - | 31930932 | 31931142 | 31918999 | 31919658 | 31937986 | 31938042 | 11 | 8 | 4 | 12 | 6 | 365 | 201 | 248 | 325 | 59 | 33 | 17 | 2 | 18 | 0 | 427 | 326 | 260 | 370 | 215 | 298 | 149 | 0.227624702 | 1 | 0.015 | 0.02 | 0.008 | 0.018 | 0.048 | 0.037 | 0.025 | 0.004 | 0.024 | 0 | 0.004 |
| SE | 102756 | ENSG00000040341 | STAU2 | chr8 | - | 73709031 | 73709162 | 73697307 | 73697442 | 73738283 | 73738344 | 27 | 34 | 28 | 47 | 45 | 0 | 0 | 1 | 3 | 1 | 22 | 48 | 26 | 28 | 17 | 0 | 0 | 0 | 4 | 0 | 280 | 149 | 0.227656262 | 1 | 1 | 1 | 0.937 | 0.893 | 0.96 | 1 | 1 | 1 | 0.788 | 1 | 0 |
| SE | 20380 | ENSG00000213066 | CEP43 | chr6 | + | 167027866 | 167027993 | 167026546 | 167026615 | 167032602 | 167032642 | 4 | 0 | 4 | 7 | 6 | 33 | 45 | 48 | 23 | 75 | 0 | 3 | 1 | 0 | 4 | 4 | 40 | 25 | 5 | 15 | 276 | 149 | 0.227793418 | 1 | 0.061 | 0 | 0.043 | 0.141 | 0.041 | 0 | 0.039 | 0.021 | 0 | 0.126 | 0.02 |
| SE | 103736 | ENSG00000107937 | GTPBP4 | chr10 | + | 1005817 | 1005907 | 1000947 | 1001013 | 1007017 | 1007128 | 167 | 106 | 139 | 129 | 142 | 0 | 7 | 2 | 2 | 1 | 100 | 190 | 238 | 189 | 113 | 0 | 0 | 2 | 6 | 0 | 239 | 149 | 0.227795681 | 1 | 1 | 0.904 | 0.977 | 0.976 | 0.989 | 1 | 1 | 0.987 | 0.952 | 1 | -0.019 |
| SE | 69759 | ENSG00000196247 | ZNF107 | chr7 | + | 64679176 | 64679336 | 64666129 | 64666285 | 64691247 | 64691374 | 11 | 3 | 1 | 21 | 8 | 5 | 3 | 3 | 1 | 2 | 0 | 24 | 19 | 36 | 12 | 8 | 0 | 4 | 5 | 0 | 298 | 149 | 0.227802824 | 1 | 0.524 | 0.333 | 0.143 | 0.913 | 0.667 | 0 | 1 | 0.704 | 0.783 | 1 | -0.181 |
| SE | 28917 | ENSG00000145495 | MARCHF6 | chr5 | + | 10417269 | 10417404 | 10415472 | 10415669 | 10423734 | 10423824 | 300 | 144 | 320 | 269 | 207 | 3 | 1 | 0 | 0 | 2 | 108 | 265 | 215 | 167 | 206 | 8 | 0 | 5 | 0 | 0 | 284 | 149 | 0.227868112 | 1 | 0.981 | 0.987 | 1 | 1 | 0.982 | 0.876 | 1 | 0.958 | 1 | 1 | 0.023 |
| SE | 23014 | ENSG00000106123 | EPHB6 | chr7 | + | 142862016 | 142862133 | 142861051 | 142861186 | 142862755 | 142862833 | 31 | 15 | 59 | 20 | 108 | 5 | 9 | 16 | 18 | 17 | 59 | 33 | 11 | 22 | 62 | 31 | 14 | 2 | 9 | 41 | 266 | 149 | 0.227979496 | 1 | 0.776 | 0.483 | 0.674 | 0.384 | 0.781 | 0.516 | 0.569 | 0.755 | 0.578 | 0.459 | 0.044 |
| SE | 29803 | ENSG00000141452 | RMC1 | chr18 | + | 23509192 | 23509279 | 23504370 | 23504447 | 23515855 | 23515886 | 131 | 102 | 85 | 102 | 143 | 1 | 0 | 0 | 0 | 3 | 182 | 140 | 95 | 151 | 211 | 0 | 1 | 0 | 0 | 12 | 236 | 149 | 0.227992972 | 1 | 0.988 | 1 | 1 | 1 | 0.968 | 1 | 0.989 | 1 | 1 | 0.917 | 0.01 |
| SE | 5457 | ENSG00000101346 | POFUT1 | chr20 | + | 32210070 | 32210192 | 32207879 | 32208065 | 32215268 | 32215451 | 26 | 13 | 16 | 14 | 10 | 3 | 1 | 1 | 1 | 0 | 21 | 22 | 12 | 21 | 17 | 0 | 0 | 0 | 3 | 0 | 271 | 149 | 0.228058076 | 1 | 0.827 | 0.877 | 0.898 | 0.885 | 1 | 1 | 1 | 1 | 0.794 | 1 | -0.061 |
| SE | 29045 | ENSG00000182150 | ERCC6L2 | chr9 | + | 95997245 | 95997413 | 95978060 | 95978215 | 96004519 | 96004701 | 11 | 6 | 16 | 1 | 0 | 109 | 154 | 178 | 184 | 303 | 16 | 5 | 0 | 0 | 1 | 13 | 127 | 96 | 47 | 35 | 298 | 149 | 0.228361209 | 1 | 0.048 | 0.019 | 0.043 | 0.003 | 0 | 0.381 | 0.019 | 0 | 0 | 0.014 | -0.06 |
| SE | 63000 | ENSG00000137497 | NUMA1 | chr11 | - | 72068078 | 72068231 | 72035901 | 72035975 | 72069841 | 72069911 | 7 | 2 | 0 | 7 | 3 | 41 | 17 | 34 | 33 | 20 | 6 | 14 | 8 | 14 | 4 | 82 | 65 | 36 | 43 | 65 | 298 | 149 | 0.228363859 | 1 | 0.079 | 0.056 | 0 | 0.096 | 0.07 | 0.035 | 0.097 | 0.1 | 0.14 | 0.03 | -0.02 |
| SE | 64630 | ENSG00000148737 | TCF7L2 | chr10 | + | 113160618 | 113160691 | 113158666 | 113158717 | 113165554 | 113165972 | 4 | 4 | 2 | 2 | 6 | 3 | 8 | 0 | 7 | 0 | 23 | 0 | 6 | 12 | 21 | 40 | 5 | 8 | 13 | 16 | 222 | 149 | 0.228473098 | 1 | 0.472 | 0.251 | 1 | 0.161 | 1 | 0.278 | 0 | 0.335 | 0.383 | 0.468 | 0.284 |
| SE | 83998 | ENSG00000112715 | VEGFA | chr6 | + | 43781955 | 43782052 | 43778888 | 43778918 | 43784540 | 43784562 | 6 | 3 | 3 | 8 | 0 | 5 | 8 | 5 | 4 | 7 | 4 | 19 | 4 | 7 | 3 | 17 | 28 | 9 | 48 | 15 | 246 | 149 | 0.22852945 | 1 | 0.421 | 0.185 | 0.267 | 0.548 | 0 | 0.125 | 0.291 | 0.212 | 0.081 | 0.108 | 0.121 |
| SE | 95703 | ENSG00000067064 | IDI1 | chr10 | - | 1043998 | 1044171 | 1043300 | 1043393 | 1048863 | 1049119 | 76 | 33 | 44 | 58 | 78 | 40 | 10 | 28 | 43 | 9 | 42 | 40 | 94 | 47 | 5 | 5 | 20 | 32 | 11 | 1 | 298 | 149 | 0.228549986 | 1 | 0.487 | 0.623 | 0.44 | 0.403 | 0.812 | 0.808 | 0.5 | 0.595 | 0.681 | 0.714 | -0.107 |
| SE | 90395 | ENSG00000170633 | RNF34 | chr12 | + | 121404848 | 121405113 | 121402741 | 121402807 | 121416158 | 121416377 | 8 | 2 | 3 | 2 | 0 | 21 | 8 | 12 | 1 | 12 | 9 | 0 | 1 | 0 | 0 | 0 | 20 | 4 | 11 | 4 | 298 | 149 | 0.228628851 | 1 | 0.16 | 0.111 | 0.111 | 0.5 | 0 | 1 | 0 | 0.111 | 0 | 0 | -0.046 |
| SE | 14426 | ENSG00000129007 | CALML4 | chr15 | - | 68205120 | 68205151 | 68199540 | 68199681 | 68205244 | 68205332 | 11 | 32 | 7 | 8 | 17 | 0 | 0 | 0 | 8 | 0 | 19 | 20 | 14 | 27 | 19 | 0 | 0 | 0 | 0 | 4 | 180 | 149 | 0.228713675 | 1 | 1 | 1 | 1 | 0.453 | 1 | 1 | 1 | 1 | 1 | 0.797 | -0.069 |
| SE | 24522 | ENSG00000186532 | SMYD4 | chr17 | - | 1811970 | 1812115 | 1804345 | 1804715 | 1827860 | 1828006 | 74 | 50 | 84 | 95 | 97 | 3 | 0 | 5 | 1 | 6 | 109 | 56 | 19 | 80 | 82 | 12 | 0 | 4 | 5 | 0 | 294 | 149 | 0.228744897 | 1 | 0.926 | 1 | 0.895 | 0.98 | 0.891 | 0.822 | 1 | 0.707 | 0.89 | 1 | 0.055 |
| SE | 47234 | ENSG00000182134 | TDRKH | chr1 | - | 151774412 | 151774504 | 151770106 | 151770562 | 151774709 | 151774806 | 20 | 5 | 30 | 17 | 40 | 0 | 0 | 1 | 1 | 0 | 5 | 20 | 2 | 4 | 14 | 0 | 0 | 4 | 0 | 0 | 241 | 149 | 0.228853701 | 1 | 1 | 1 | 0.949 | 0.913 | 1 | 1 | 1 | 0.236 | 1 | 1 | 0.125 |
| SE | 18776 | ENSG00000160190 | SLC37A1 | chr21 | + | 42535471 | 42535550 | 42525775 | 42525857 | 42539511 | 42539647 | 18 | 8 | 9 | 11 | 2 | 0 | 0 | 0 | 0 | 0 | 23 | 6 | 15 | 6 | 45 | 0 | 5 | 0 | 0 | 0 | 228 | 149 | 0.228894748 | 1 | 1 | 1 | 1 | 1 | 1 | 1 | 0.44 | 1 | 1 | 1 | 0.112 |
| SE | 54970 | ENSG00000135842 | NIBAN1 | chr1 | - | 184818637 | 184818825 | 184808073 | 184808235 | 184823166 | 184823329 | 276 | 180 | 176 | 231 | 352 | 2 | 0 | 2 | 3 | 8 | 300 | 263 | 398 | 242 | 185 | 0 | 4 | 20 | 0 | 0 | 298 | 149 | 0.228931321 | 1 | 0.986 | 1 | 0.978 | 0.975 | 0.957 | 1 | 0.97 | 0.909 | 1 | 1 | 0.003 |
| SE | 34107 | ENSG00000129235 | TXNDC17 | chr17 | + | 6641752 | 6641834 | 6641015 | 6641152 | 6642248 | 6642324 | 23 | 6 | 11 | 32 | 34 | 0 | 0 | 2 | 2 | 1 | 9 | 31 | 20 | 24 | 13 | 0 | 1 | 3 | 0 | 3 | 231 | 149 | 0.228961113 | 1 | 1 | 1 | 0.78 | 0.912 | 0.956 | 1 | 0.952 | 0.811 | 1 | 0.737 | 0.03 |
| SE | 15522 | ENSG00000129003 | VPS13C | chr15 | - | 61884127 | 61884269 | 61882595 | 61882736 | 61890164 | 61890400 | 510 | 536 | 393 | 490 | 606 | 3 | 2 | 2 | 8 | 5 | 49 | 187 | 202 | 41 | 17 | 0 | 0 | 0 | 0 | 10 | 291 | 149 | 0.228968624 | 1 | 0.989 | 0.993 | 0.99 | 0.969 | 0.984 | 1 | 1 | 1 | 1 | 0.465 | 0.092 |
| SE | 102757 | ENSG00000040341 | STAU2 | chr8 | - | 73709031 | 73709177 | 73673106 | 73673242 | 73738283 | 73738349 | 0 | 4 | 3 | 0 | 5 | 2 | 2 | 0 | 5 | 0 | 0 | 3 | 0 | 0 | 7 | 13 | 0 | 3 | 2 | 0 | 295 | 149 | 0.228992403 | 1 | 0 | 0.503 | 1 | 0 | 1 | 0 | 1 | 0 | 0 | 1 | 0.101 |
| SE | 66912 | ENSG00000152520 | PAN3 | chr13 | + | 28197184 | 28197346 | 28177864 | 28177935 | 28220230 | 28220378 | 143 | 136 | 119 | 149 | 235 | 98 | 47 | 69 | 73 | 137 | 109 | 124 | 92 | 85 | 108 | 19 | 58 | 79 | 37 | 36 | 298 | 149 | 0.228996421 | 1 | 0.422 | 0.591 | 0.463 | 0.505 | 0.462 | 0.741 | 0.517 | 0.368 | 0.535 | 0.6 | -0.064 |
| SE | 86138 | ENSG00000197713 | RPE | chr2 | + | 210009656 | 210009736 | 210003437 | 210003468 | 210015972 | 210016112 | 77 | 65 | 67 | 68 | 26 | 2 | 1 | 0 | 4 | 2 | 69 | 61 | 32 | 69 | 15 | 0 | 0 | 0 | 0 | 0 | 229 | 149 | 0.2291772 | 1 | 0.962 | 0.977 | 1 | 0.917 | 0.894 | 1 | 1 | 1 | 1 | 1 | -0.05 |
| SE | 93328 | ENSG00000148331 | ASB6 | chr9 | - | 129638572 | 129638659 | 129634604 | 129638457 | 129639201 | 129639310 | 53 | 13 | 25 | 33 | 39 | 0 | 0 | 0 | 0 | 0 | 27 | 35 | 40 | 38 | 92 | 0 | 5 | 0 | 0 | 3 | 236 | 149 | 0.229273622 | 1 | 1 | 1 | 1 | 1 | 1 | 1 | 0.815 | 1 | 1 | 0.951 | 0.047 |
| SE | 38478 | ENSG00000147251 | DOCK11 | chrX | + | 118681693 | 118681794 | 118681057 | 118681248 | 118683078 | 118683217 | 643 | 463 | 370 | 622 | 883 | 3 | 3 | 3 | 2 | 2 | 24 | 284 | 282 | 23 | 18 | 7 | 2 | 1 | 0 | 0 | 250 | 149 | 0.229278239 | 1 | 0.992 | 0.989 | 0.987 | 0.995 | 0.996 | 0.671 | 0.988 | 0.994 | 1 | 1 | 0.061 |
| SE | 19234 | ENSG00000066739 | ATG2B | chr14 | - | 96333687 | 96333873 | 96332500 | 96332655 | 96334404 | 96334501 | 70 | 57 | 51 | 97 | 169 | 0 | 3 | 3 | 4 | 9 | 17 | 80 | 64 | 78 | 18 | 0 | 2 | 5 | 0 | 2 | 298 | 149 | 0.229293484 | 1 | 1 | 0.905 | 0.895 | 0.924 | 0.904 | 1 | 0.952 | 0.865 | 1 | 0.818 | -0.001 |
| SE | 3651 | ENSG00000142230 | SAE1 | chr19 | + | 47151995 | 47152053 | 47150201 | 47150375 | 47152897 | 47153040 | 1 | 14 | 6 | 6 | 2 | 300 | 211 | 262 | 299 | 403 | 17 | 9 | 0 | 20 | 0 | 438 | 352 | 222 | 267 | 532 | 207 | 149 | 0.229293499 | 1 | 0.002 | 0.046 | 0.016 | 0.014 | 0.004 | 0.027 | 0.018 | 0 | 0.051 | 0 | -0.003 |
| SE | 51909 | ENSG00000074603 | DPP8 | chr15 | - | 65451959 | 65452102 | 65450738 | 65451110 | 65454262 | 65454415 | 158 | 123 | 121 | 127 | 183 | 1 | 4 | 1 | 4 | 8 | 10 | 102 | 139 | 92 | 12 | 0 | 21 | 0 | 0 | 0 | 292 | 149 | 0.229379242 | 1 | 0.988 | 0.94 | 0.984 | 0.942 | 0.921 | 1 | 0.713 | 1 | 1 | 1 | 0.012 |
| SE | 90104 | ENSG00000205038 | PKHD1L1 | chr8 | + | 109413420 | 109413545 | 109412264 | 109412414 | 109419096 | 109419260 | 138 | 197 | 37 | 284 | 272 | 0 | 0 | 0 | 0 | 0 | 589 | 248 | 128 | 234 | 1016 | 7 | 6 | 0 | 0 | 9 | 274 | 149 | 0.229383985 | 1 | 1 | 1 | 1 | 1 | 1 | 0.979 | 0.957 | 1 | 1 | 0.984 | 0.016 |
| SE | 46643 | ENSG00000162614 | NEXN | chr1 | + | 77917565 | 77917757 | 77916054 | 77916133 | 77917959 | 77918038 | 2 | 3 | 2 | 2 | 6 | 16 | 15 | 4 | 15 | 10 | 41 | 2 | 2 | 10 | 42 | 97 | 11 | 52 | 60 | 26 | 298 | 149 | 0.229395969 | 1 | 0.059 | 0.091 | 0.2 | 0.062 | 0.231 | 0.174 | 0.083 | 0.019 | 0.077 | 0.447 | -0.031 |
| SE | 8066 | ENSG00000214087 | ARL16 | chr17 | - | 81682490 | 81682675 | 81682033 | 81682149 | 81683012 | 81683126 | 27 | 11 | 29 | 17 | 18 | 30 | 8 | 8 | 6 | 34 | 10 | 44 | 12 | 11 | 48 | 34 | 18 | 8 | 43 | 32 | 298 | 149 | 0.229446432 | 1 | 0.31 | 0.407 | 0.644 | 0.586 | 0.209 | 0.128 | 0.55 | 0.429 | 0.113 | 0.429 | 0.101 |
| SE | 21786 | ENSG00000198625 | MDM4 | chr1 | + | 204537429 | 204537497 | 204530683 | 204530817 | 204538208 | 204538308 | 23 | 25 | 32 | 34 | 53 | 0 | 0 | 1 | 1 | 0 | 20 | 48 | 31 | 68 | 9 | 0 | 3 | 0 | 4 | 0 | 217 | 149 | 0.229564881 | 1 | 1 | 1 | 0.956 | 0.959 | 1 | 1 | 0.917 | 1 | 0.921 | 1 | 0.015 |
| SE | 8609 | ENSG00000185359 | HGS | chr17 | + | 81701214 | 81701422 | 81701044 | 81701131 | 81701507 | 81701929 | 0 | 0 | 0 | 0 | 0 | 17 | 3 | 16 | 22 | 1 | 7 | 0 | 0 | 0 | 0 | 5 | 43 | 7 | 44 | 47 | 298 | 149 | 0.229631062 | 1 | 0 | 0 | 0 | 0 | 0 | 0.412 | 0 | 0 | 0 | 0 | -0.082 |
| SE | 93201 | ENSG00000137207 | YIPF3 | chr6 | - | 43515605 | 43515701 | 43513587 | 43513633 | 43515888 | 43516187 | 92 | 50 | 58 | 61 | 60 | 3 | 1 | 0 | 4 | 6 | 145 | 105 | 61 | 91 | 159 | 9 | 0 | 0 | 3 | 2 | 245 | 149 | 0.229725797 | 1 | 0.949 | 0.968 | 1 | 0.903 | 0.859 | 0.907 | 1 | 1 | 0.949 | 0.98 | -0.031 |
| SE | 49601 | ENSG00000076685 | NT5C2 | chr10 | - | 103111744 | 103111805 | 103106588 | 103106706 | 103139405 | 103139479 | 67 | 35 | 69 | 55 | 85 | 169 | 152 | 132 | 193 | 294 | 71 | 53 | 68 | 78 | 15 | 357 | 156 | 117 | 295 | 119 | 210 | 149 | 0.229742601 | 1 | 0.22 | 0.14 | 0.271 | 0.168 | 0.17 | 0.124 | 0.194 | 0.292 | 0.158 | 0.082 | 0.024 |
| SE | 76679 | ENSG00000223501 | VPS52 | chr6 | - | 33251517 | 33251636 | 33250271 | 33250987 | 33251859 | 33251971 | 82 | 46 | 86 | 57 | 111 | 0 | 0 | 0 | 6 | 0 | 116 | 65 | 70 | 82 | 90 | 10 | 0 | 0 | 0 | 0 | 268 | 149 | 0.229890326 | 1 | 1 | 1 | 1 | 0.841 | 1 | 0.866 | 1 | 1 | 1 | 1 | -0.005 |
| SE | 4793 | ENSG00000282508 | LINC01002 | chr19 | - | 204843 | 205076 | 203961 | 204065 | 205169 | 205227 | 6 | 8 | 7 | 4 | 0 | 14 | 13 | 22 | 2 | 23 | 9 | 8 | 31 | 5 | 13 | 3 | 41 | 14 | 38 | 30 | 298 | 149 | 0.229935258 | 1 | 0.176 | 0.235 | 0.137 | 0.5 | 0 | 0.6 | 0.089 | 0.525 | 0.062 | 0.178 | -0.081 |
| SE | 91129 | ENSG00000180900 | SCRIB | chr8 | - | 143807551 | 143807614 | 143806923 | 143807013 | 143808608 | 143809025 | 66 | 29 | 63 | 49 | 24 | 7 | 5 | 1 | 4 | 5 | 65 | 65 | 34 | 36 | 93 | 0 | 5 | 8 | 14 | 13 | 212 | 149 | 0.230040016 | 1 | 0.869 | 0.803 | 0.978 | 0.896 | 0.771 | 1 | 0.901 | 0.749 | 0.644 | 0.834 | 0.038 |
| SE | 81747 | ENSG00000176454 | LPCAT4 | chr15 | - | 34361399 | 34361532 | 34359588 | 34360209 | 34362195 | 34362321 | 43 | 39 | 37 | 52 | 51 | 11 | 7 | 9 | 4 | 14 | 19 | 39 | 26 | 34 | 79 | 55 | 2 | 2 | 15 | 34 | 282 | 149 | 0.230072196 | 1 | 0.674 | 0.746 | 0.685 | 0.873 | 0.658 | 0.154 | 0.912 | 0.873 | 0.545 | 0.551 | 0.12 |
| SE | 45172 | ENSG00000107099 | DOCK8 | chr9 | + | 420948 | 421078 | 420400 | 420583 | 422047 | 422135 | 1157 | 1053 | 820 | 1170 | 1174 | 0 | 1 | 0 | 4 | 6 | 3112 | 1588 | 1377 | 1930 | 858 | 0 | 0 | 18 | 19 | 14 | 279 | 149 | 0.230086583 | 1 | 1 | 0.998 | 1 | 0.994 | 0.991 | 1 | 1 | 0.976 | 0.982 | 0.97 | 0.011 |
| SE | 82523 | ENSG00000061938 | TNK2 | chr3 | - | 195885412 | 195885543 | 195884829 | 195885033 | 195886976 | 195887047 | 28 | 16 | 24 | 21 | 32 | 88 | 61 | 63 | 89 | 66 | 72 | 29 | 19 | 15 | 32 | 153 | 140 | 36 | 115 | 173 | 280 | 149 | 0.230215898 | 1 | 0.145 | 0.122 | 0.169 | 0.112 | 0.205 | 0.2 | 0.099 | 0.219 | 0.065 | 0.09 | 0.016 |
| SE | 54620 | ENSG00000073578 | SDHA | chr5 | + | 235143 | 235339 | 233476 | 233645 | 236427 | 236599 | 365 | 145 | 325 | 333 | 301 | 3 | 0 | 1 | 2 | 2 | 309 | 236 | 201 | 175 | 379 | 0 | 2 | 10 | 0 | 3 | 298 | 149 | 0.230365235 | 1 | 0.984 | 1 | 0.994 | 0.988 | 0.987 | 1 | 0.983 | 0.91 | 1 | 0.984 | 0.015 |
| SE | 104992 | ENSG00000111371 | SLC38A1 | chr12 | - | 46254813 | 46254936 | 46243199 | 46243314 | 46268525 | 46268997 | 92 | 50 | 105 | 91 | 148 | 94 | 80 | 55 | 65 | 46 | 41 | 73 | 39 | 30 | 30 | 10 | 77 | 60 | 22 | 0 | 272 | 149 | 0.230402028 | 1 | 0.349 | 0.255 | 0.511 | 0.434 | 0.638 | 0.692 | 0.342 | 0.263 | 0.517 | 1 | -0.125 |
| SE | 78328 | ENSG00000205413 | SAMD9 | chr7 | - | 93114794 | 93114895 | 93099517 | 93106105 | 93117862 | 93117979 | 1017 | 777 | 590 | 839 | 982 | 17 | 9 | 6 | 18 | 12 | 98 | 271 | 44 | 65 | 36 | 0 | 9 | 0 | 0 | 0 | 250 | 149 | 0.230424421 | 1 | 0.973 | 0.981 | 0.983 | 0.965 | 0.98 | 1 | 0.947 | 1 | 1 | 1 | -0.013 |
| SE | 80825 | ENSG00000151948 | GLT1D1 | chr12 | + | 128914932 | 128914989 | 128899235 | 128899287 | 128927104 | 128927154 | 5 | 2 | 5 | 2 | 1 | 0 | 0 | 0 | 0 | 0 | 11 | 5 | 18 | 18 | 12 | 6 | 0 | 0 | 0 | 0 | 206 | 149 | 0.230444663 | 1 | 1 | 1 | 1 | 1 | 1 | 0.57 | 1 | 1 | 1 | 1 | 0.086 |
| SE | 98640 | ENSG00000149179 | C11orf49 | chr11 | + | 47137315 | 47137560 | 47052387 | 47052518 | 47137644 | 47137729 | 2 | 2 | 0 | 2 | 3 | 8 | 6 | 11 | 17 | 28 | 0 | 0 | 4 | 9 | 2 | 8 | 12 | 11 | 19 | 24 | 298 | 149 | 0.230539569 | 1 | 0.111 | 0.143 | 0 | 0.056 | 0.051 | 0 | 0 | 0.154 | 0.191 | 0.04 | -0.005 |
| SE | 14555 | ENSG00000159322 | ADPGK | chr15 | - | 72760406 | 72760527 | 72755555 | 72755654 | 72771782 | 72771845 | 154 | 104 | 92 | 124 | 133 | 1 | 1 | 0 | 0 | 7 | 180 | 95 | 91 | 107 | 201 | 0 | 11 | 4 | 0 | 0 | 270 | 149 | 0.23059768 | 1 | 0.988 | 0.983 | 1 | 1 | 0.913 | 1 | 0.827 | 0.926 | 1 | 1 | 0.026 |
| SE | 101159 | ENSG00000133816 | MICAL2 | chr11 | + | 12236176 | 12236245 | 12227024 | 12227131 | 12239435 | 12239585 | 112 | 86 | 71 | 96 | 113 | 3 | 3 | 4 | 2 | 0 | 119 | 75 | 38 | 93 | 171 | 6 | 3 | 2 | 1 | 6 | 218 | 149 | 0.230602243 | 1 | 0.962 | 0.951 | 0.924 | 0.97 | 1 | 0.931 | 0.945 | 0.929 | 0.985 | 0.951 | 0.013 |
| SE | 79425 | ENSG00000170260 | ZNF212 | chr7 | + | 149251209 | 149251327 | 149250709 | 149250807 | 149252705 | 149252795 | 0 | 0 | 3 | 4 | 0 | 23 | 15 | 13 | 22 | 35 | 0 | 0 | 11 | 0 | 0 | 23 | 16 | 30 | 20 | 10 | 267 | 149 | 0.230684225 | 1 | 0 | 0 | 0.114 | 0.092 | 0 | 0 | 0 | 0.17 | 0 | 0 | 0.007 |
| SE | 9412 | ENSG00000105053 | VRK3 | chr19 | - | 50009235 | 50009385 | 50007568 | 50007826 | 50016023 | 50016163 | 154 | 76 | 106 | 103 | 175 | 23 | 18 | 8 | 22 | 23 | 184 | 163 | 85 | 142 | 253 | 15 | 15 | 31 | 14 | 17 | 298 | 149 | 0.230807227 | 1 | 0.77 | 0.679 | 0.869 | 0.701 | 0.792 | 0.86 | 0.845 | 0.578 | 0.835 | 0.882 | -0.038 |
| SE | 62665 | ENSG00000115525 | ST3GAL5 | chr2 | - | 85844395 | 85844554 | 85837119 | 85840392 | 85846376 | 85846424 | 253 | 210 | 244 | 228 | 479 | 3 | 2 | 4 | 8 | 11 | 231 | 187 | 177 | 262 | 216 | 5 | 6 | 1 | 4 | 0 | 298 | 149 | 0.230815026 | 1 | 0.977 | 0.981 | 0.968 | 0.934 | 0.956 | 0.959 | 0.94 | 0.989 | 0.97 | 1 | -0.008 |
| SE | 67313 | ENSG00000157570 | TSPAN18 | chr11 | + | 44909704 | 44909899 | 44906406 | 44906479 | 44917971 | 44918046 | 24 | 24 | 44 | 44 | 50 | 1 | 1 | 2 | 2 | 2 | 40 | 33 | 0 | 22 | 47 | 0 | 0 | 8 | 0 | 22 | 298 | 149 | 0.230942369 | 1 | 0.923 | 0.923 | 0.917 | 0.917 | 0.926 | 1 | 1 | 0 | 1 | 0.516 | 0.218 |
| SE | 78765 | ENSG00000179406 | LINC00174 | chr7 | - | 66476258 | 66476327 | 66450070 | 66450236 | 66479008 | 66479096 | 9 | 4 | 20 | 11 | 13 | 7 | 0 | 1 | 9 | 5 | 24 | 8 | 8 | 12 | 23 | 0 | 12 | 8 | 13 | 16 | 218 | 149 | 0.231070536 | 1 | 0.468 | 1 | 0.932 | 0.455 | 0.64 | 1 | 0.313 | 0.406 | 0.387 | 0.496 | 0.179 |
| SE | 78192 | ENSG00000124486 | USP9X | chrX | + | 41140655 | 41140771 | 41136803 | 41137022 | 41141292 | 41141431 | 102 | 68 | 78 | 89 | 131 | 0 | 0 | 0 | 1 | 0 | 0 | 74 | 70 | 35 | 6 | 3 | 0 | 0 | 0 | 0 | 265 | 149 | 0.231182828 | 1 | 1 | 1 | 1 | 0.98 | 1 | 0 | 1 | 1 | 1 | 1 | 0.196 |
| SE | 21430 | ENSG00000164180 | TMEM161B | chr5 | - | 88225768 | 88225866 | 88220562 | 88220719 | 88228444 | 88228528 | 48 | 45 | 70 | 65 | 79 | 4 | 2 | 6 | 9 | 15 | 21 | 36 | 14 | 13 | 11 | 0 | 3 | 0 | 0 | 8 | 247 | 149 | 0.231183679 | 1 | 0.879 | 0.931 | 0.876 | 0.813 | 0.761 | 1 | 0.879 | 1 | 1 | 0.453 | -0.014 |
| SE | 55963 | ENSG00000136161 | RCBTB2 | chr13 | - | 48515204 | 48515355 | 48512728 | 48512895 | 48521897 | 48521962 | 100 | 55 | 44 | 69 | 72 | 0 | 0 | 0 | 0 | 0 | 81 | 65 | 28 | 53 | 42 | 8 | 0 | 0 | 0 | 0 | 298 | 149 | 0.23139052 | 1 | 1 | 1 | 1 | 1 | 1 | 0.835 | 1 | 1 | 1 | 1 | 0.033 |
| SE | 56260 | ENSG00000143995 | MEIS1 | chr2 | + | 66439842 | 66439984 | 66439320 | 66439671 | 66440561 | 66440612 | 9 | 23 | 6 | 27 | 22 | 0 | 2 | 2 | 0 | 0 | 117 | 90 | 45 | 95 | 131 | 0 | 0 | 2 | 0 | 9 | 291 | 149 | 0.231460965 | 1 | 1 | 0.855 | 0.606 | 1 | 1 | 1 | 1 | 0.92 | 1 | 0.882 | -0.068 |
| SE | 44786 | ENSG00000134717 | BTF3L4 | chr1 | + | 52083339 | 52083541 | 52064824 | 52064938 | 52086111 | 52086171 | 354 | 250 | 283 | 304 | 636 | 27 | 16 | 31 | 39 | 33 | 277 | 271 | 168 | 292 | 105 | 18 | 17 | 21 | 29 | 24 | 298 | 149 | 0.231480912 | 1 | 0.868 | 0.887 | 0.82 | 0.796 | 0.906 | 0.885 | 0.889 | 0.8 | 0.834 | 0.686 | 0.037 |
| SE | 19197 | ENSG00000140105 | WARS1 | chr14 | - | 100375282 | 100375350 | 100369086 | 100369258 | 100376259 | 100376307 | 61 | 80 | 52 | 61 | 18 | 75 | 29 | 52 | 42 | 11 | 247 | 16 | 22 | 112 | 165 | 65 | 80 | 75 | 78 | 123 | 217 | 149 | 0.231502454 | 1 | 0.358 | 0.654 | 0.407 | 0.499 | 0.529 | 0.723 | 0.121 | 0.168 | 0.496 | 0.479 | 0.092 |
| SE | 86877 | ENSG00000125354 | SEPTIN6 | chrX | - | 119625334 | 119625379 | 119615723 | 119616742 | 119629317 | 119629508 | 280 | 248 | 416 | 380 | 1311 | 3 | 10 | 15 | 13 | 62 | 490 | 343 | 174 | 305 | 830 | 20 | 25 | 7 | 43 | 0 | 194 | 149 | 0.231563809 | 1 | 0.986 | 0.95 | 0.955 | 0.957 | 0.942 | 0.95 | 0.913 | 0.95 | 0.845 | 1 | 0.026 |
| SE | 2702 | ENSG00000105011 | ASF1B | chr19 | - | 14126121 | 14126237 | 14121286 | 14121708 | 14136347 | 14136553 | 10 | 12 | 14 | 14 | 12 | 0 | 0 | 0 | 0 | 0 | 9 | 23 | 11 | 23 | 5 | 3 | 3 | 0 | 0 | 0 | 265 | 149 | 0.23166963 | 1 | 1 | 1 | 1 | 1 | 1 | 0.628 | 0.812 | 1 | 1 | 1 | 0.112 |
| SE | 26234 | ENSG00000141084 | RANBP10 | chr16 | - | 67759572 | 67759732 | 67744287 | 67744455 | 67772033 | 67772086 | 0 | 0 | 7 | 6 | 2 | 41 | 32 | 63 | 78 | 108 | 0 | 4 | 0 | 0 | 4 | 87 | 36 | 10 | 47 | 79 | 298 | 149 | 0.231670509 | 1 | 0 | 0 | 0.053 | 0.037 | 0.009 | 0 | 0.053 | 0 | 0 | 0.025 | 0.004 |
| SE | 83585 | ENSG00000107897 | ACBD5 | chr10 | - | 27210921 | 27211081 | 27208245 | 27208445 | 27215534 | 27215641 | 206 | 176 | 143 | 153 | 197 | 3 | 1 | 3 | 0 | 5 | 85 | 152 | 197 | 116 | 13 | 0 | 1 | 2 | 2 | 0 | 298 | 149 | 0.231714131 | 1 | 0.972 | 0.989 | 0.96 | 1 | 0.952 | 1 | 0.987 | 0.98 | 0.967 | 1 | -0.012 |
| SE | 70051 | ENSG00000111644 | ACRBP | chr12 | - | 6646482 | 6646577 | 6645219 | 6645337 | 6646793 | 6647012 | 3 | 17 | 7 | 12 | 4 | 0 | 0 | 0 | 0 | 0 | 39 | 14 | 5 | 38 | 41 | 0 | 0 | 0 | 0 | 8 | 244 | 149 | 0.231740156 | 1 | 1 | 1 | 1 | 1 | 1 | 1 | 1 | 1 | 1 | 0.758 | 0.048 |
| SE | 103419 | ENSG00000107669 | ATE1 | chr10 | - | 121898840 | 121898969 | 121870005 | 121870038 | 121902390 | 121902620 | 62 | 64 | 93 | 74 | 80 | 3 | 2 | 0 | 4 | 0 | 39 | 65 | 39 | 70 | 108 | 0 | 0 | 0 | 0 | 0 | 278 | 149 | 0.231767805 | 1 | 0.917 | 0.945 | 1 | 0.908 | 1 | 1 | 1 | 1 | 1 | 1 | -0.046 |
| SE | 63346 | ENSG00000151092 | NGLY1 | chr3 | - | 25736308 | 25736400 | 25736003 | 25736149 | 25737333 | 25737455 | 2 | 9 | 15 | 19 | 14 | 126 | 104 | 105 | 141 | 120 | 0 | 10 | 12 | 9 | 14 | 42 | 110 | 63 | 98 | 12 | 241 | 149 | 0.231915972 | 1 | 0.01 | 0.051 | 0.081 | 0.077 | 0.067 | 0 | 0.053 | 0.105 | 0.054 | 0.419 | -0.069 |
| SE | 54915 | ENSG00000010803 | SCMH1 | chr1 | - | 41152598 | 41152741 | 41142877 | 41143112 | 41160874 | 41160898 | 0 | 1 | 0 | 1 | 0 | 3 | 4 | 15 | 1 | 5 | 0 | 0 | 0 | 2 | 3 | 8 | 11 | 22 | 0 | 9 | 292 | 149 | 0.231929355 | 1 | 0 | 0.113 | 0 | 0.338 | 0 | 0 | 0 | 0 | 1 | 0.145 | -0.139 |
| SE | 59479 | ENSG00000160310 | PRMT2 | chr21 | + | 46636895 | 46636990 | 46635594 | 46635763 | 46648457 | 46648619 | 56 | 17 | 14 | 46 | 22 | 0 | 0 | 0 | 0 | 0 | 14 | 14 | 26 | 33 | 41 | 0 | 6 | 0 | 0 | 0 | 244 | 149 | 0.231986168 | 1 | 1 | 1 | 1 | 1 | 1 | 1 | 0.588 | 1 | 1 | 1 | 0.082 |
| SE | 91645 | ENSG00000105339 | DENND3 | chr8 | + | 141160631 | 141160787 | 141155848 | 141155970 | 141163332 | 141163429 | 113 | 66 | 112 | 88 | 76 | 9 | 14 | 4 | 11 | 6 | 309 | 189 | 283 | 354 | 421 | 38 | 20 | 9 | 25 | 16 | 298 | 149 | 0.232002058 | 1 | 0.863 | 0.702 | 0.933 | 0.8 | 0.864 | 0.803 | 0.825 | 0.94 | 0.876 | 0.929 | -0.042 |

| | | | | | | | | | | | | |
|---|---|---|---|---|---|---|---|---|---|---|---|---|
| SE | 66613 | ENSG00000115687 | PASK | chr2 | - | 241145749 | 241145860 | 241142836 | 241143074 | 241149413 | 241149449 | [illegible] |
| SE | 38400 | ENSG00000143458 | GABPB2 | chr1 | + | 151090405 | 151090573 | 151088157 | 151088297 | 151093191 | 151093386 | [illegible] |
| SE | 62860 | ENSG00000112335 | SNX3 | chr6 | - | 108222233 | 108222320 | 108211512 | 108212254 | 108222949 | 108223045 | [illegible] |
| SE | 18318 | ENSG00000100575 | TIMM9 | chr14 | - | 58423827 | 58423849 | 58411906 | 58411971 | 58424007 | 58424095 | [illegible] |
| SE | 65940 | ENSG00000143537 | ADAM15 | chr1 | + | 155061417 | 155061489 | 155060762 | 155060832 | 155061903 | 155061975 | [illegible] |
| SE | 18068 | ENSG00000075413 | MARK3 | chr14 | + | 103398950 | 103399041 | 103385979 | 103386080 | 103405075 | 103405267 | [illegible] |
| SE | 44928 | ENSG00000182095 | TNRC18 | chr7 | - | 5370374 | 5371364 | 5362649 | 5362825 | 5374054 | 5374484 | [illegible] |
| SE | 54253 | ENSG00000011405 | PIK3C2A | chr11 | - | 17136481 | 17136625 | 17135110 | 17135159 | 17145667 | 17145731 | [illegible] |
| SE | 18575 | ENSG00000168411 | RFWD3 | chr16 | - | 74664623 | 74664745 | 74660779 | 74661451 | 74666785 | 74666832 | [illegible] |
| SE | 102550 | ENSG00000203791 | EEF1AKMT2 | chr10 | - | 124762299 | 124762558 | 124757830 | 124760503 | 124765391 | 124765608 | [illegible] |
| SE | 108186 | ENSG00000121749 | TBC1D15 | chr12 | + | 71880468 | 71880607 | 71872928 | 71873003 | 71893221 | 71893306 | [illegible] |
| SE | 60392 | ENSG00000174796 | THAP6 | chr4 | + | 75515432 | 75515532 | 75514437 | 75514520 | 75516771 | 75516789 | [illegible] |
| SE | 637 | ENSG00000100129 | EIF3L | chr22 | + | 37886764 | 37886845 | 37877673 | 37878171 | 37888425 | 37888479 | [illegible] |
| SE | 38655 | ENSG00000073417 | PDE8A | chr15 | + | 85091043 | 85091181 | 85089337 | 85089416 | 85097947 | 85098036 | [illegible] |
| SE | 58950 | ENSG00000131778 | CHD1L | chr1 | + | 147270931 | 147271113 | 147268781 | 147268878 | 147272170 | 147272281 | [illegible] |
| SE | 66315 | ENSG00000225190 | PLEKHM1 | chr17 | - | 45482436 | 45482525 | 45477073 | 45478147 | 45490651 | 45490721 | [illegible] |
| SE | 70589 | ENSG00000169689 | CENPX | chr17 | - | 82019509 | 82019694 | 82019292 | 82019381 | 82019857 | 82019909 | [illegible] |
| SE | 60490 | ENSG00000070814 | TCOF1 | chr5 | + | 150391543 | 150392176 | 150389946 | 150390023 | 150392704 | 150392891 | [illegible] |
| SE | 103264 | ENSG00000172661 | WASHC2C | chr10 | + | 45777272 | 45777425 | 45773255 | 45773358 | 45778952 | 45779135 | [illegible] |
| SE | 45516 | ENSG00000163960 | UBXN7 | chr3 | - | 196407245 | 196407393 | 196393553 | 196393619 | 196432326 | 196432397 | [illegible] |
| SE | 8917 | ENSG00000161533 | ACOX1 | chr17 | - | 75960214 | 75960375 | 75957420 | 75957566 | 75973624 | 75973785 | [illegible] |
| SE | 89 | ENSG00000186866 | POFUT2 | chr21 | - | 45282803 | 45282938 | 45282368 | 45282398 | 45283382 | 45283527 | [illegible] |
| SE | 59397 | ENSG00000139631 | CSAD | chr12 | - | 53161126 | 53161191 | 53160762 | 53160844 | 53161272 | 53161389 | [illegible] |
| SE | 18604 | ENSG00000166887 | VPS39 | chr15 | - | 42191124 | 42191167 | 42189113 | 42189208 | 42191495 | 42191560 | [illegible] |
| SE | 8830 | ENSG00000109083 | IFT20 | chr17 | - | 28332145 | 28332181 | 28331602 | 28331987 | 28335339 | 28335439 | [illegible] |
| SE | 70819 | ENSG00000169224 | GCSAML | chr1 | + | 247565930 | 247565959 | 247563589 | 247563639 | 247574142 | 247574215 | [illegible] |
| SE | 70155 | ENSG00000010219 | DYRK4 | chr12 | + | 4580823 | 4580977 | 4573329 | 4573356 | 4588936 | 4589017 | [illegible] |
| SE | 72245 | ENSG00000152348 | ATG10 | chr5 | + | 82252561 | 82252659 | 82178489 | 82178587 | 82276144 | 82276297 | [illegible] |
| SE | 23098 | ENSG00000111912 | NCOA7 | chr6 | + | 125796679 | 125796797 | 125786136 | 125786213 | 125815290 | 125815404 | [illegible] |
| SE | 34933 | ENSG00000076108 | BAZ2A | chr12 | - | 56615013 | 56615607 | 56613952 | 56614138 | 56617394 | 56617532 | [illegible] |
| SE | 81677 | ENSG00000160856 | FCRL3 | chr1 | - | 157697139 | 157697329 | 157696039 | 157696327 | 157697658 | 157697919 | [illegible] |
| SE | 50686 | ENSG00000142687 | KIAA0319L | chr1 | - | 35456012 | 35456241 | 35454361 | 35454485 | 35460304 | 35460437 | [illegible] |
| SE | 40198 | ENSG00000231607 | DLEU2 | chr13 | - | 50070581 | 50070641 | 50068096 | 50068163 | 50071515 | 50071629 | [illegible] |
| SE | 33690 | ENSG00000176261 | ZBTB8OS | chr1 | - | 32633950 | 32634110 | 32633644 | 32633727 | 32650432 | 32650903 | [illegible] |
| SE | 56566 | ENSG00000253710 | ALG11 | chr13 | + | 52015782 | 52017685 | 52012397 | 52012462 | 52018912 | 52019143 | [illegible] |
| SE | 12531 | ENSG00000103066 | PLA2G15 | chr16 | + | 68255890 | 68255990 | 68254918 | 68255037 | 68259145 | 68259174 | [illegible] |
| SE | 95835 | ENSG00000141551 | CSNK1D | chr17 | - | 82249697 | 82250179 | 82249430 | 82249602 | 82251378 | 82251414 | [illegible] |
| SE | 12549 | ENSG00000214078 | CPNE1 | chr20 | - | 35626562 | 35626803 | 35626043 | 35626381 | 35627279 | 35627413 | [illegible] |
| SE | 16511 | ENSG00000140367 | UBE2Q2 | chr15 | + | 75873427 | 75873568 | 75868950 | 75869010 | 75879097 | 75879188 | [illegible] |
| SE | 62995 | ENSG00000137497 | NUMA1 | chr11 | - | 72049418 | 72049539 | 72035901 | 72035975 | 72068104 | 72068231 | [illegible] |
| SE | 95873 | ENSG00000148396 | SEC16A | chr9 | - | 136445644 | 136445719 | 136440096 | 136441823 | 136446854 | 136446949 | [illegible] |
| SE | 13772 | ENSG00000205629 | LCMT1 | chr16 | + | 25125963 | 25126126 | 25111730 | 25111996 | 25128474 | 25128566 | [illegible] |
| SE | 27715 | ENSG00000164483 | SAMD3 | chr6 | - | 130221636 | 130221775 | 130216570 | 130216616 | 130222693 | 130222813 | [illegible] |
| SE | 78010 | ENSG00000173890 | GPR160 | chr3 | + | 170077122 | 170077923 | 170038914 | 170039043 | 170079773 | 170079897 | [illegible] |
| SE | 73166 | ENSG00000145391 | SETD7 | chr4 | - | 139546919 | 139547049 | 139532578 | 139533366 | 139556097 | 139556219 | [illegible] |
| SE | 43296 | ENSG00000134698 | AGO4 | chr1 | + | 35832436 | 35832570 | 35832056 | 35832185 | 35833989 | 35834174 | [illegible] |
| SE | 102184 | ENSG00000267699 | NA | chr18 | + | 51052634 | 51052725 | 51051353 | 51051415 | 51054780 | 51054993 | [illegible] |
| SE | 22373 | ENSG00000140455 | USP3 | chr15 | + | 63542054 | 63542192 | 63537137 | 63537156 | 63553714 | 63553798 | [illegible] |
| SE | 53733 | ENSG00000151503 | NCAPD3 | chr11 | - | 134223862 | 134224000 | 134220568 | 134220726 | 134225096 | 134225454 | [illegible] |
| SE | 75044 | ENSG00000130023 | ERMARD | chr6 | + | 169775269 | 169775346 | 169773043 | 169773402 | 169775939 | 169776065 | [illegible] |
| SE | 84095 | ENSG00000197892 | KIF13B | chr8 | - | 29155725 | 29155856 | 29150296 | 29150383 | 29160732 | 29160867 | [illegible] |
| SE | 57344 | ENSG00000181722 | ZBTB20 | chr3 | - | 114904326 | 114904368 | 114900303 | 114900342 | 114974365 | 114974416 | [illegible] |
| SE | 35339 | ENSG00000184203 | PPP1R2 | chr3 | - | 195523691 | 195523786 | 195519017 | 195519185 | 195524818 | 195524896 | [illegible] |
| SE | 8029 | ENSG00000073350 | LLGL2 | chr17 | + | 75574452 | 75574495 | 75574110 | 75574260 | 75574609 | 75574668 | [illegible] |
| SE | 96438 | ENSG00000110514 | MADD | chr11 | + | 47273826 | 47273976 | 47269692 | 47269906 | 47274562 | 47274726 | [illegible] |
| SE | 64929 | ENSG00000204642 | HLA-F | chr6 | + | 29726010 | 29726043 | 29725446 | 29725563 | 29738121 | 29738188 | [illegible] |
| SE | 69004 | ENSG00000107521 | HPS1 | chr10 | - | 98431130 | 98431291 | 98430570 | 98430670 | 98433982 | 98434091 | [illegible] |
| SE | 5017 | ENSG00000088833 | NSFL1C | chr20 | - | 1464328 | 1464426 | 1458029 | 1458274 | 1466719 | 1466824 | [illegible] |
| SE | 75330 | ENSG00000196262 | PPIA | chr7 | + | 44798747 | 44798904 | 44797135 | 44797621 | 44799246 | 44799277 | [illegible] |
| SE | 46873 | ENSG00000162894 | FCMR | chr1 | - | 206921340 | 206921433 | 206913800 | 206914094 | 206921817 | 206921997 | [illegible] |
| SE | 54705 | ENSG00000137965 | IFI44 | chr1 | + | 78650185 | 78650652 | 78649830 | 78649905 | 78654242 | 78654279 | [illegible] |
| SE | 59645 | ENSG00000163466 | ARPC2 | chr2 | + | 218238663 | 218238850 | 218228737 | 218228850 | 218245419 | 218245457 | [illegible] |
| SE | 32887 | ENSG00000068650 | ATP11A | chr13 | + | 112878216 | 112878303 | 112875775 | 112875941 | 112881815 | 112887168 | [illegible] |
| SE | 2417 | ENSG00000239382 | ALKBH6 | chr19 | - | 36013513 | 36013588 | 36013343 | 36013422 | 36014174 | 36014227 | [illegible] |
| SE | 25241 | ENSG00000183621 | ZNF438 | chr10 | - | 30984368 | 30984426 | 30941574 | 30941651 | 30999337 | 30999517 | [illegible] |
| SE | 99134 | ENSG00000162341 | TPCN2 | chr11 | + | 69067502 | 69067605 | 69063894 | 69063967 | 69070429 | 69070495 | [illegible] |
| SE | 31942 | ENSG00000163703 | CRELD1 | chr3 | + | 9938014 | 9938106 | 9934834 | 9934917 | 9940849 | 9941016 | [illegible] |
| SE | 41814 | ENSG00000168389 | MFSD2A | chr1 | + | 39967627 | 39967916 | 39967085 | 39967169 | 39968333 | 39968477 | [illegible] |
| SE | 107793 | ENSG00000050426 | LETMD1 | chr12 | + | 51052091 | 51052207 | 51048342 | 51049185 | 51055834 | 51056021 | [illegible] |
| SE | 8420 | ENSG00000078687 | TNRC6C | chr17 | + | 78031515 | 78031842 | 78005055 | 78005079 | 78048844 | 78051448 | [illegible] |
| SE | 35793 | ENSG00000173226 | IQCB1 | chr3 | - | 121788283 | 121788432 | 121781742 | 121781874 | 121790072 | 121790215 | [illegible] |
| SE | 42971 | ENSG00000121940 | CLCC1 | chr1 | - | 108943474 | 108943615 | 108941404 | 108941498 | 108947610 | 108947718 | [illegible] |
| SE | 61388 | ENSG00000151893 | CACUL1 | chr10 | - | 118730283 | 118730410 | 118707491 | 118707587 | 118754395 | 118754477 | [illegible] |
| SE | 63003 | ENSG00000137497 | NUMA1 | chr11 | - | 72068104 | 72068231 | 72049418 | 72049539 | 72069841 | 72070063 | [illegible] |
| SE | 51295 | ENSG00000174780 | SRP72 | chr4 | + | 56478591 | 56478649 | 56476670 | 56476702 | 56483138 | 56483270 | [illegible] |
| SE | 34339 | ENSG00000157020 | SEC13 | chr3 | - | 10319138 | 10319316 | 10318049 | 10318094 | 10321049 | 10321084 | [illegible] |
| SE | 6726 | ENSG00000186111 | PIP5K1C | chr19 | - | 3637535 | 3637613 | 3633436 | 3633520 | 3638883 | 3639016 | [illegible] |
| SE | 28934 | ENSG00000112079 | STK38 | chr6 | - | 36499872 | 36499990 | 36497779 | 36497875 | 36506582 | 36506644 | [illegible] |
| SE | 83412 | ENSG00000175274 | TP53I11 | chr11 | - | 44936558 | 44936689 | 44935619 | 44935662 | 44936802 | 44936899 | [illegible] |
| SE | 64212 | ENSG00000117697 | NSL1 | chr1 | - | 212739533 | 212739601 | 212726152 | 212738686 | 212782371 | 212782426 | [illegible] |
| SE | 17543 | ENSG00000100813 | ACIN1 | chr14 | - | 23067313 | 23068671 | 23065965 | 23066008 | 23069475 | 23069525 | [illegible] |
| SE | 77081 | ENSG00000086232 | EIF2AK1 | chr7 | - | 6049911 | 6050045 | 6048806 | 6048844 | 6054545 | 6054704 | [illegible] |
| SE | 94238 | ENSG00000130653 | PNPLA7 | chr9 | - | 137547584 | 137547659 | 137547308 | 137547396 | 137550167 | 137550384 | [illegible] |
| SE | 86068 | ENSG00000164031 | DNAJB14 | chr4 | - | 99928541 | 99928617 | 99923039 | 99923185 | 99930449 | 99930534 | [illegible] |
| SE | 71887 | ENSG00000074054 | CLASP1 | chr2 | - | 121409023 | 121409047 | 121407470 | 121407712 | 121410865 | 121410969 | [illegible] |
| SE | 35596 | ENSG00000114331 | ACAP2 | chr3 | - | 195357690 | 195357784 | 195345258 | 195345317 | 195381008 | 195381062 | [illegible] |
| SE | 38183 | ENSG00000137074 | APTX | chr9 | - | 32988082 | 32988124 | 32987543 | 32987731 | 32989758 | 32989895 | [illegible] |
| SE | 23025 | ENSG00000183474 | GTF2H2C | chr5 | + | 69567726 | 69567777 | 69566966 | 69567053 | 69568152 | 69568207 | [illegible] |
| SE | 23305 | ENSG00000242110 | AMACR | chr5 | - | 33994077 | 33994120 | 33988083 | 33989502 | 33998640 | 33998827 | [illegible] |
| SE | 83999 | ENSG00000112715 | VEGFA | chr6 | + | 43781955 | 43782087 | 43778888 | 43778918 | 43784540 | 43784562 | [illegible] |
| SE | 96707 | ENSG00000135362 | PRR5L | chr11 | + | 36451208 | 36451335 | 36437384 | 36437476 | 36462341 | 36464965 | [illegible] |
| SE | 101744 | ENSG00000198056 | PRIM1 | chr12 | - | 56740011 | 56740128 | 56739293 | 56739363 | 56741434 | 56741576 | [illegible] |
| SE | 76575 | ENSG00000170919 | TPT1-AS1 | chr13 | + | 45370079 | 45370182 | 45341452 | 45341614 | 45377060 | 45377215 | [illegible] |
| SE | 78608 | ENSG00000083168 | KAT6A | chr8 | - | 42049619 | 42049707 | 42048377 | 42049302 | 42051900 | 42051987 | [illegible] |
| SE | 46175 | ENSG00000100280 | AP1B1 | chr22 | - | 29341500 | 29341760 | 29340655 | 29340857 | 29342284 | 29342383 | [illegible] |
| SE | 93846 | ENSG00000116580 | GON4L | chr1 | - | 155753203 | 155753414 | 155751959 | 155752587 | 155754374 | 155754488 | [illegible] |
| SE | 73864 | ENSG00000164609 | SLU7 | chr5 | - | 160418595 | 160418685 | 160415124 | 160415310 | 160419022 | 160419081 | [illegible] |
| SE | 84426 | ENSG00000188917 | TRMT2B | chrX | - | 101042041 | 101042312 | 101041316 | 101041371 | 101051816 | 101052054 | [illegible] |
| SE | 48150 | ENSG00000117543 | DPH5 | chr1 | - | 100995109 | 100995149 | 100990062 | 100990631 | 101001466 | 101001544 | [illegible] |
| SE | 56400 | ENSG00000151838 | CCDC175 | chr14 | - | 59540674 | 59540746 | 59538704 | 59538840 | 59551354 | 59551436 | [illegible] |
| SE | 21258 | ENSG00000155111 | CDK19 | chr6 | - | 110621103 | 110621370 | 110609977 | 110614666 | 110622087 | 110622166 | [illegible] |
| SE | 93747 | ENSG00000168003 | SLC3A2 | chr11 | + | 62888100 | 62888218 | 62885464 | 62885608 | 62888330 | 62888849 | [illegible] |
| SE | 106448 | ENSG00000135127 | BICDL1 | chr12 | + | 120074442 | 120074586 | 120072510 | 120072729 | 120080886 | 120081017 | [illegible] |
| SE | 45577 | ENSG00000117335 | CD46 | chr1 | + | 207790252 | 207790345 | 207785618 | 207785682 | 207793518 | 207793873 | [illegible] |
| SE | 90293 | ENSG00000249859 | PVT1 | chr8 | + | 127939507 | 127939676 | 127890586 | 127890998 | 127989161 | 127989291 | [illegible] |
| SE | 73304 | ENSG00000136699 | SMPD4 | chr2 | - | 130161185 | 130161272 | 130157250 | 130157396 | 130164373 | 130164445 | [illegible] |
| SE | 43986 | ENSG00000270617 | URGCP-MRPS24 | chr7 | - | 43881906 | 43881957 | 43881658 | 43881697 | 43887414 | 43887485 | [illegible] |
| SE | 20351 | ENSG00000162711 | NLRP3 | chr1 | + | 247434162 | 247434273 | 247423846 | 247425599 | 247435969 | 247436140 | [illegible] |
| SE | 104147 | ENSG00000023041 | ZDHHC6 | chr10 | - | 112441728 | 112441812 | 112438335 | 112438389 | 112442191 | 112442351 | [illegible] |
| SE | 17519 | ENSG00000125952 | MAX | chr14 | - | 65077327 | 65077428 | 65075123 | 65076663 | 65077912 | 65078036 | [illegible] |
| SE | 105259 | ENSG00000082898 | XPO1 | chr2 | - | 61502248 | 61502310 | 61499712 | 61499894 | 61522610 | 61522683 | [illegible] |
| SE | 95218 | ENSG00000148300 | REXO4 | chr9 | - | 133412298 | 133412492 | 133410984 | 133411073 | 133412777 | 133412921 | [illegible] |
| SE | 17850 | ENSG00000089775 | ZBTB25 | chr14 | - | 64487113 | 64487196 | 64449329 | 64449638 | 64490360 | 64490533 | [illegible] |
| SE | 1808 | ENSG00000099917 | MED15 | chr22 | + | 20575112 | 20575232 | 20568520 | 20568631 | 20582610 | 20582747 | [illegible] |
| SE | 69175 | ENSG00000132530 | XAF1 | chr17 | + | 6762154 | 6762240 | 6760405 | 6760601 | 6770642 | 6770969 | [illegible] |
| SE | 8755 | ENSG00000007202 | KIAA0100 | chr17 | - | 28633548 | 28633751 | 28633282 | 28633401 | 28633894 | 28634086 | [illegible] |
| SE | 47154 | ENSG00000138617 | PARP16 | chr15 | - | 65266561 | 65266768 | 65263148 | 65263320 | 65270934 | 65271072 | [illegible] |
| SE | 102213 | ENSG00000100109 | TFIP11 | chr22 | - | 26494138 | 26494304 | 26491224 | 26492368 | 26494796 | 26494939 | [illegible] |
| SE | 63681 | ENSG00000110934 | BIN2 | chr12 | - | 51295795 | 51295878 | 51291590 | 51292344 | 51299202 | 51299288 | [illegible] |
| SE | 68529 | ENSG00000131236 | CAP1 | chr1 | + | 40041428 | 40041493 | 40040731 | 40040801 | 40059333 | 40059458 | [illegible] |
| SE | 65517 | ENSG00000140386 | SCAPER | chr15 | - | 76855809 | 76855876 | 76841733 | 76841931 | 76857808 | 76857879 | [illegible] |
| SE | 74814 | ENSG00000042088 | TDP1 | chr14 | + | 89963224 | 89963673 | 89955940 | 89955970 | 89966146 | 89966190 | [illegible] |
| SE | 65282 | ENSG00000115295 | CLIP4 | chr2 | + | 29121373 | 29121521 | 29113995 | 29114312 | 29131257 | 29131397 | [illegible] |
| SE | 93200 | ENSG00000137207 | YIPF3 | chr6 | - | 43515594 | 43515701 | 43513448 | 43513633 | 43515888 | 43516046 | [illegible] |
| SE | 22065 | ENSG00000159579 | RSPRY1 | chr16 | + | 57207592 | 57207685 | 57204596 | 57205008 | 57208057 | 57208110 | [illegible] |
| SE | 553 | ENSG00000100221 | JOSD1 | chr22 | - | 38688934 | 38689129 | 38685542 | 38688001 | 38699802 | 38699970 | [illegible] |
| SE | 56430 | ENSG00000050748 | MAPK9 | chr5 | - | 180247662 | 180247728 | 180246544 | 180247510 | 180248972 | 180249138 | [illegible] |
| SE | 73969 | ENSG00000166436 | TRIM66 | chr11 | - | 8647969 | 8648080 | 8646446 | 8646561 | 8648415 | 8648548 | [illegible] |
| SE | 108198 | ENSG00000186908 | ZDHHC17 | chr12 | + | 76809042 | 76809120 | 76805316 | 76805439 | 76809712 | 76809750 | [illegible] |
| SE | 63155 | ENSG00000251474 | RPL32P3 | chr3 | - | 129397039 | 129397140 | 129393606 | 129393795 | 129399007 | 129399405 | [illegible] |
| SE | 2378 | ENSG00000170889 | RPS9 | chr19 | + | 54201486 | 54201609 | 54201132 | 54201281 | 54206275 | 54206462 | [illegible] |
| SE | 51979 | ENSG00000071054 | MAP4K4 | chr2 | + | 101859735 | 101859864 | 101855976 | 101856138 | 101864929 | 101865036 | [illegible] |
| SE | 17410 | ENSG00000126822 | PLEKHG3 | chr14 | + | 64728995 | 64729093 | 64727592 | 64727982 | 64730242 | 64730278 | [illegible] |
| SE | 20350 | ENSG00000162711 | NLRP3 | chr1 | + | 247434102 | 247434273 | 247429584 | 247429755 | 247443971 | 247444142 | [illegible] |
| SE | 57680 | ENSG00000167110 | GOLGA2 | chr9 | - | 128272784 | 128272865 | 128268419 | 128268524 | 128273849 | 128273972 | [illegible] |
| SE | 57975 | ENSG00000160746 | ANO10 | chr3 | - | 43574808 | 43574864 | 43565652 | 43565727 | 43580352 | 43580472 | [illegible] |
| SE | 20854 | ENSG00000255819 | KLRC4-KLRK1 | chr12 | - | 10405849 | 10405950 | 10388770 | 10388875 | 10407635 | 10407759 | [illegible] |
| SE | 53570 | ENSG00000145979 | TBC1D7 | chr6 | - | 13307599 | 13307745 | 13306397 | 13306527 | 13316570 | 13316708 | [illegible] |
| SE | 53191 | ENSG00000227671 | ZNF731P | chr1 | - | 247202015 | 247202095 | 247200066 | 247200193 | 247210690 | 247210856 | [illegible] |
| SE | 103699 | ENSG00000125485 | DDX31 | chr9 | - | 132618329 | 132618441 | 132612086 | 132612255 | 132630263 | 132630403 | [illegible] |
| SE | 12281 | ENSG00000103194 | USP10 | chr16 | + | 84759890 | 84759946 | 84744632 | 84745673 | 84760171 | 84760236 | [illegible] |
| SE | 53968 | ENSG00000188641 | DPYD | chr1 | - | 97082329 | 97082470 | 97077742 | 97079146 | 97098488 | 97098632 | [illegible] |
| SE | 92651 | ENSG00000165233 | CARD19 | chr9 | + | 93108990 | 93109092 | 93107673 | 93107827 | 93110567 | 93110721 | [illegible] |
| SE | 32561 | ENSG00000240682 | ISY1 | chr3 | - | 129144067 | 129144189 | 129140367 | 129140485 | 129145760 | 129145873 | [illegible] |
| SE | 94122 | ENSG00000139644 | TMBIM6 | chr12 | + | 49741956 | 49742115 | 49741607 | 49741768 | 49752463 | 49752549 | [illegible] |
| SE | 12173 | ENSG00000051108 | HERPUD1 | chr16 | + | 56939894 | 56940245 | 56936723 | 56936817 | 56942131 | 56942202 | [illegible] |
| SE | 7176 | ENSG00000101639 | CEP192 | chr18 | + | 13071038 | 13071212 | 13069737 | 13069856 | 13072754 | 13072845 | [illegible] |
| SE | 49115 | ENSG00000153363 | LINC00467 | chr1 | + | 211384441 | 211384534 | 211382799 | 211382925 | 211391615 | 211391697 | [illegible] |
| SE | 3639 | ENSG00000239998 | LILRA2 | chr19 | + | 54574075 | 54574111 | 54573697 | 54573912 | 54574300 | 54574544 | [illegible] |
| SE | 77428 | ENSG00000071189 | SNX13 | chr7 | - | 17803418 | 17803580 | 17801587 | 17801659 | 17814833 | 17814944 | [illegible] |
| SE | 53957 | ENSG00000138101 | DTNB | chr2 | - | 25455404 | 25455494 | 25451547 | 25451635 | 25482795 | 25482873 | [illegible] |
| SE | 30308 | ENSG00000135299 | ANKRD6 | chr6 | + | 89616873 | 89616956 | 89616558 | 89616657 | 89617953 | 89618031 | [illegible] |
| SE | 68796 | ENSG00000204634 | TBC1D8 | chr2 | - | 101032267 | 101032385 | 101029490 | 101029776 | 101033543 | 101033758 | [illegible] |
| SE | 63305 | ENSG00000138018 | SELENOI | chr2 | + | 26383298 | 26383347 | 26373366 | 26373629 | 26389004 | 26389067 | [illegible] |
| SE | 32501 | ENSG00000181061 | HIGD1A | chr3 | - | 42794156 | 42794275 | 42786027 | 42786162 | 42804435 | 42804490 | [illegible] |
| SE | 16203 | ENSG00000140598 | EFL1 | chr15 | - | 82238306 | 82238521 | 82230847 | 82230971 | 82240417 | 82240555 | [illegible] |
| SE | 100037 | ENSG00000101972 | STAG2 | chrX | + | 124090853 | 124090964 | 124090574 | 124090764 | 124094017 | 124094144 | [illegible] |
| SE | 106461 | ENSG00000198270 | TMEM116 | chr12 | - | 111936691 | 111936830 | 111933885 | 111934030 | 111937159 | 111937243 | [illegible] |
| SE | 34013 | ENSG00000060971 | ACAA1 | chr3 | - | 38131595 | 38131638 | 38129289 | 38129388 | 38131925 | 38131946 | [illegible] |
| SE | 53817 | ENSG00000173575 | CHD2 | chr15 | + | 92953356 | 92954336 | 92948951 | 92949076 | 92955422 | 92955512 | [illegible] |
| SE | 12725 | ENSG00000157106 | SMG1 | chr16 | - | 18829925 | 18830115 | 18829285 | 18829755 | 18832939 | 18833166 | [illegible] |
| SE | 14630 | ENSG00000140157 | NIPA2 | chr15 | + | 22845145 | 22845267 | 22839654 | 22839790 | 22851638 | 22851665 | [illegible] |
| SE | 19456 | ENSG00000125746 | EML2 | chr19 | - | 45616460 | 45616558 | 45615801 | 45615889 | 45616764 | 45616853 | [illegible] |
| SE | 7164 | ENSG00000101639 | CEP192 | chr18 | + | 13040829 | 13040956 | 13030464 | 13030608 | 13042203 | 13042334 | [illegible] |
| SE | 45120 | ENSG00000119203 | CPSF3 | chr2 | + | 9430751 | 9430880 | 9429922 | 9430020 | 9432510 | 9432688 | [illegible] |
| SE | 106707 | ENSG00000141376 | BCAS3 | chr17 | + | 61354361 | 61354502 | 61084466 | 61084564 | 61368326 | 61368494 | [illegible] |
| SE | 23257 | ENSG00000108262 | GIT1 | chr17 | - | 29578961 | 29578988 | 29578730 | 29578779 | 29581337 | 29581380 | [illegible] |
| SE | 65169 | ENSG00000064651 | SLC12A2 | chr5 | + | 128177104 | 128177152 | 128174540 | 128174666 | 128178566 | 128178689 | [illegible] |
| SE | 55171 | ENSG00000054267 | ARID4B | chr1 | - | 235176747 | 235176903 | 235175183 | 235175399 | 235177799 | 235177913 | [illegible] |
| SE | 66013 | ENSG00000009335 | UBE3C | chr7 | + | 157220688 | 157220776 | 157216866 | 157216971 | 157223253 | 157223351 | [illegible] |
| SE | 24470 | ENSG00000197321 | SVIL | chr10 | - | 29471137 | 29471243 | 29470275 | 29470483 | 29473837 | 29473989 | [illegible] |
| SE | 13856 | ENSG00000102893 | PHKB | chr16 | + | 47649099 | 47649204 | 47648532 | 47648616 | 47650543 | 47650626 | [illegible] |
| SE | 38943 | ENSG00000067066 | SP100 | chr2 | + | 230467119 | 230467215 | 230466300 | 230466354 | 230469042 | 230469096 | [illegible] |
| SE | 22766 | ENSG00000145730 | PAM | chr5 | + | 103025133 | 103025334 | 103019789 | 103019843 | 103028184 | 103028238 | [illegible] |
| SE | 84377 | ENSG00000230590 | FTX | chrX | - | 74285914 | 74286170 | 74281701 | 74281848 | 74292168 | 74292349 | [illegible] |
| SE | 99708 | ENSG00000110075 | PPP6R3 | chr11 | + | 68575957 | 68576043 | 68574108 | 68574224 | 68583042 | 68583129 | [illegible] |
| SE | 47730 | ENSG00000145687 | SSBP2 | chr5 | - | 81442652 | 81442723 | 81437429 | 81437458 | 81446867 | 81446922 | [illegible] |
| SE | 20848 | ENSG00000255819 | KLRC4-KLRK1 | chr12 | - | 10391834 | 10391891 | 10389942 | 10390027 | 10405849 | 10405950 | [illegible] |
| SE | 93984 | ENSG00000213625 | LEPROT | chr1 | + | 65421327 | 65421459 | 65420715 | 65420740 | 65425302 | 65425378 | [illegible] |
| SE | 67850 | ENSG00000018280 | SLC11A1 | chr2 | + | 218384242 | 218384365 | 218382202 | 218382375 | 218385146 | 218385266 | [illegible] |
| SE | 80335 | ENSG00000006634 | DBF4 | chr7 | + | 87904291 | 87904416 | 87900763 | 87900878 | 87907187 | 87907219 | [illegible] |
| SE | 63748 | ENSG00000152061 | RABGAP1L | chr1 | + | 174877391 | 174877605 | 174811831 | 174811960 | 174957456 | 174957549 | [illegible] |
| SE | 48149 | ENSG00000117543 | DPH5 | chr1 | - | 100992636 | 100992740 | 100990223 | 100990628 | 100995109 | 100995149 | [illegible] |
| SE | 30446 | ENSG00000055955 | ITIH4 | chr3 | - | 52819939 | 52819990 | 52819753 | 52819792 | 52820290 | 52820317 | [illegible] |
| SE | 83344 | ENSG00000087274 | ADD1 | chr4 | + | 2867971 | 2868129 | 2843944 | 2844024 | 2875895 | 2876110 | [illegible] |
| SE | 69574 | ENSG00000134262 | AP4B1 | chr1 | - | 113897839 | 113897943 | 113895756 | 113896038 | 113898717 | 113898763 | [illegible] |
| SE | 80371 | ENSG00000204371 | EHMT2 | chr6 | - | 31893361 | 31893662 | 31892825 | 31892910 | 31896262 | 31896504 | [illegible] |
| SE | 68762 | ENSG00000163872 | YEATS2 | chr3 | + | 183736717 | 183736829 | 183728689 | 183728851 | 183747671 | 183747716 | [illegible] |
| SE | 39181 | ENSG00000121775 | TMEM39B | chr1 | + | 32077163 | 32077318 | 32076762 | 32076846 | 32091674 | 32091744 | [illegible] |
| SE | 81152 | ENSG00000110665 | C11orf21 | chr11 | - | 2300683 | 2300768 | 2300519 | 2300613 | 2301755 | 2301913 | [illegible] |
| SE | 33588 | ENSG00000009790 | TRAF3IP3 | chr1 | + | 209779314 | 209779374 | 209777351 | 209777487 | 209782055 | 209782307 | [illegible] |
| SE | 106094 | ENSG00000196498 | NCOR2 | chr12 | - | 124429612 | 124429706 | 124426621 | 124426800 | 124430614 | 124430787 | [illegible] |
| SE | 48800 | ENSG00000064763 | FAR2 | chr12 | + | 29291415 | 29291494 | 29270448 | 29270638 | 29293299 | 29293475 | [illegible] |
| SE | 23004 | ENSG00000171617 | ENC1 | chr5 | - | 74637493 | 74637618 | 74636336 | 74636498 | 74640306 | 74641424 | [illegible] |
| SE | 13147 | ENSG00000103121 | CMC2 | chr16 | - | 80980769 | 80980879 | 80975801 | 80976179 | 80997313 | 80997429 | [illegible] |
| SE | 59160 | ENSG00000115204 | MPV17 | chr2 | - | 27312679 | 27312772 | 27312213 | 27312246 | 27312993 | 27313074 | [illegible] |
| SE | 21021 | ENSG00000126214 | KLC1 | chr14 | + | 103687080 | 103687211 | 103679383 | 103679518 | 103700654 | 103700727 | [illegible] |
| SE | 44406 | ENSG00000163346 | PBXIP1 | chr1 | - | 154953670 | 154953757 | 154951470 | 154951921 | 154956068 | 154956099 | [illegible] |
| SE | 32064 | ENSG00000242086 | MUC20-OT1 | chr3 | + | 195685817 | 195685939 | 195683749 | 195683865 | 195688532 | 195688797 | [illegible] |
| SE | 15160 | ENSG00000137842 | TMEM62 | chr15 | + | 43138573 | 43138619 | 43135511 | 43135649 | 43146492 | 43146521 | [illegible] |
| SE | 77253 | ENSG00000141664 | ZCCHC2 | chr18 | + | 62544801 | 62544873 | 62542497 | 62542574 | 62550347 | 62550460 | [illegible] |
| SE | 108659 | ENSG00000136147 | PHF11 | chr13 | + | 49527205 | 49527354 | 49526386 | 49526458 | 49528510 | 49528981 | [illegible] |
| SE | 76412 | ENSG00000168538 | TRAPPC11 | chr4 | + | 183667961 | 183668117 | 183667059 | 183667130 | 183674712 | 183674870 | [illegible] |
| SE | 29958 | ENSG00000136051 | WASHC4 | chr12 | + | 105156725 | 105156792 | 105152342 | 105152451 | 105157235 | 105157322 | [illegible] |
| SE | 59500 | ENSG00000148660 | CAMK2G | chr10 | - | 73828088 | 73828121 | 73824039 | 73824084 | 73837467 | 73837511 | [illegible] |
| SE | 80562 | ENSG00000179818 | PCBP1-AS1 | chr2 | - | 70059642 | 70059680 | 70048706 | 70051305 | 70083537 | 70083687 | [illegible] |
| SE | 12277 | ENSG00000103194 | USP10 | chr16 | + | 84740308 | 84740369 | 84733137 | 84733503 | 84758715 | 84758807 | [illegible] |
| SE | 97477 | ENSG00000166575 | TMEM135 | chr11 | + | 87157340 | 87157406 | 87091361 | 87091395 | 87236637 | 87236684 | [illegible] |
| SE | 74622 | ENSG00000100319 | ZMAT5 | chr22 | - | 29738329 | 29738441 | 29730955 | 29731354 | 29740649 | 29740730 | [illegible] |
| SE | 93748 | ENSG00000168003 | SLC3A2 | chr11 | + | 62888112 | 62888218 | 62885464 | 62885608 | 62888330 | 62888849 | [illegible] |
| SE | 88079 | ENSG00000196663 | TECPR2 | chr14 | + | 102431795 | 102432128 | 102414635 | 102414793 | 102434234 | 102435211 | [illegible] |
| SE | 81265 | ENSG00000123505 | AMD1 | chr6 | + | 110887504 | 110887591 | 110874783 | 110875215 | 110888856 | 110888983 | [illegible] |
| SE | 65203 | ENSG00000010244 | ZNF207 | chr17 | + | 32366664 | 32366757 | 32365329 | 32365487 | 32367771 | 32368014 | [illegible] |
| SE | 56732 | ENSG00000270231 | NBPF8 | chr1 | + | 120425746 | 120425845 | 120419849 | 120420118 | 120427705 | 120427847 | [illegible] |
| SE | 68747 | ENSG00000196141 | SPATS2L | chr2 | + | 200329430 | 200329480 | 200306150 | 200306601 | 200389222 | 200389283 | [illegible] |
| SE | 79363 | ENSG00000280828 | NA | chr7 | + | 128578889 | 128578933 | 128533651 | 128533730 | 128591849 | 128592017 | [illegible] |
| SE | 31014 | ENSG00000145247 | OCIAD2 | chr4 | - | 48894005 | 48894053 | 48892770 | 48892889 | 48897803 | 48897835 | [illegible] |
| SE | 58116 | ENSG00000109445 | ZNF330 | chr4 | + | 141230165 | 141230270 | 141229570 | 141229697 | 141231438 | 141231485 | [illegible] |
| SE | 101010 | ENSG00000261008 | LINC01572 | chr16 | - | 72290449 | 72290607 | 72285378 | 72285453 | 72291169 | 72291413 | [illegible] |
| SE | 58131 | ENSG00000117523 | PRRC2C | chr1 | + | 171568246 | 171568339 | 171566591 | 171566843 | 171571319 | 171571421 | [illegible] |
| SE | 50836 | ENSG00000048052 | HDAC9 | chr7 | + | 18325511 | 18325592 | 18290440 | 18290515 | 18496261 | 18496746 | [illegible] |
| SE | 26096 | ENSG00000285000 | NA | chr5 | + | 77048160 | 77048272 | 77046346 | 77046677 | 77048935 | 77048987 | [illegible] |
| SE | 68531 | ENSG00000131236 | CAP1 | chr1 | + | 40041428 | 40041493 | 40040732 | 40040783 | 40059333 | 40059458 | [illegible] |
| SE | 93229 | ENSG00000110455 | ACCS | chr11 | + | 44073446 | 44073517 | 44071255 | 44071315 | 44074611 | 44074681 | [illegible] |
| SE | 40064 | ENSG00000125347 | IRF1 | chr5 | - | 132484548 | 132484639 | 132483978 | 132484075 | 132485666 | 132485716 | [illegible] |
| SE | 13435 | ENSG00000103111 | MON1B | chr16 | + | 77193450 | 77193777 | 77191189 | 77191258 | 77194334 | 77194360 | [illegible] |
| SE | 95550 | ENSG00000196363 | WDR5 | chr9 | + | 134151982 | 134152029 | 134148287 | 134148343 | 134154465 | 134154541 | [illegible] |
| SE | 72241 | ENSG00000152348 | ATG10 | chr5 | + | 82058494 | 82058602 | 81987557 | 81987678 | 82164398 | 82164537 | [illegible] |
| SE | 76815 | ENSG00000135317 | SNX14 | chr6 | - | 85574257 | 85574378 | 85572297 | 85572374 | 85588862 | 85588950 | [illegible] |
| SE | 93673 | ENSG00000148153 | INIP | chr9 | - | 112716460 | 112716541 | 112694033 | 112694233 | 112717986 | 112718086 | [illegible] |
| SE | 37620 | ENSG00000082074 | FYB1 | chr5 | - | 39212677 | 39213017 | 39202773 | 39202987 | 39274402 | 39274528 | [illegible] |
| SE | 1123 | ENSG00000128340 | RAC2 | chr22 | - | 37231230 | 37231390 | 37226670 | 37226803 | 37241586 | 37241658 | [illegible] |
| SE | 40734 | ENSG00000073792 | IGF2BP2 | chr3 | - | 185649402 | 185649534 | 185647024 | 185647138 | 185652093 | 185652168 | [illegible] |

| | | | | | | | | | | | | | | | | |
|---|---|---|---|---|---|---|---|---|---|---|---|---|---|---|---|---|
| SE | 74175 | ENSG00000116698 | SMG7 | chr1 | + | 183527625 | 183527731 | 183526595 | 183526767 | 183527955 | 183528027 | 13 | 7 | 19 | [illegible] | [illegible] |
| SE | 14626 | ENSG00000140157 | NIPA2 | chr15 | + | 22839654 | 22839790 | 22838675 | 22839174 | 22851638 | 22851661 | 34 | 24 | 21 | [illegible] | [illegible] |
| SE | 76418 | ENSG00000168538 | TRAPPC11 | chr4 | + | 183684695 | 183684841 | 183684304 | 183684359 | 183685083 | 183685145 | 277 | 254 | 229 | [illegible] | [illegible] |
| SE | 52020 | ENSG00000135250 | SRPK2 | chr7 | - | 105125781 | 105125874 | 105117387 | 105118022 | 105126247 | 105126340 | 22 | 8 | 18 | [illegible] | [illegible] |
| SE | 76170 | ENSG00000132423 | COQ3 | chr6 | - | 99375939 | 99376182 | 99371427 | 99371587 | 99377385 | 99377485 | 37 | 17 | 20 | 26 | 48 |
| SE | 41024 | ENSG00000064687 | ABCA7 | chr19 | + | 1050920 | 1051052 | 1049265 | 1049437 | 1051154 | 1051294 | 41 | 20 | 37 | 28 | 27 |
| SE | 44402 | ENSG00000163346 | PBXIP1 | chr1 | - | 154951794 | 154951919 | 154951470 | 154951535 | 154956068 | 154956121 | 218 | 114 | 208 | 238 | 227 |
| SE | 86244 | ENSG00000134516 | DOCK2 | chr5 | + | 169670541 | 169670597 | 169654402 | 169654486 | 169671077 | 169671174 | 252 | 182 | 159 | 173 | 207 |
| SE | 60487 | ENSG00000070814 | TCOF1 | chr5 | + | 150391543 | 150391657 | 150389886 | 150390023 | 150391956 | 150392176 | 82 | 14 | 46 | 61 | 30 |
| SE | 75060 | ENSG00000198223 | CSF2RA | chrX | + | 1285777 | 1285920 | 1282677 | 1282779 | 1288518 | 1288642 | 250 | 224 | 150 | 231 | 400 |
| SE | 41602 | ENSG00000118308 | IRAG2 | chr12 | + | 25069349 | 25069431 | 25066364 | 25066512 | 25079243 | 25079290 | 201 | 187 | 186 | 224 | 278 |
| SE | 653 | ENSG00000100226 | GTPBP1 | chr22 | + | 38708844 | 38708956 | 38705943 | 38706147 | 38715906 | 38715962 | 104 | 64 | 60 | 110 | 88 |
| SE | 77108 | ENSG00000122512 | PMS2 | chr7 | - | 6003689 | 6003792 | 6001965 | 6002636 | 6003971 | 6004058 | 91 | 66 | 102 | 117 | 205 |
| SE | 10287 | ENSG00000168259 | DNAJC7 | chr17 | - | 41981854 | 41982007 | 41977260 | 41977323 | 41983562 | 41983617 | 101 | 121 | 95 | 143 | 147 |
| SE | 62197 | ENSG00000113569 | NUP155 | chr5 | - | 37342548 | 37342646 | 37341089 | 37341242 | 37348504 | 37348596 | 124 | 75 | 105 | 80 | 205 |
| SE | 17280 | ENSG00000100429 | HDAC10 | chr22 | - | 50250426 | 50250523 | 50250062 | 50250160 | 50250770 | 50250905 | 39 | 17 | 14 | 15 | 13 |
| SE | 56750 | ENSG00000117616 | RSRP1 | chr1 | - | 25318062 | 25318271 | 25306895 | 25306949 | 25337039 | 25337401 | 14 | 8 | 3 | 9 | 21 |
| SE | 35961 | ENSG00000144395 | CCDC150 | chr2 | + | 196672344 | 196672437 | 196669832 | 196669876 | 196674240 | 196674348 | 1 | 5 | 1 | 1 | 7 |
| SE | 80391 | ENSG00000150787 | PTS | chr11 | + | 112230207 | 112230230 | 112228593 | 112228673 | 112230625 | 112230682 | 4 | 4 | 3 | 7 | 10 |
| SE | 31939 | ENSG00000163703 | CRELD1 | chr3 | + | 9937561 | 9937672 | 9934834 | 9934917 | 9940849 | 9941026 | 15 | 19 | 29 | 38 | 14 |
| SE | 59407 | ENSG00000139631 | CSAD | chr12 | - | 53172345 | 53172436 | 53171877 | 53171988 | 53173344 | 53173374 | 9 | 1 | 3 | 11 | 0 |
| SE | 87296 | ENSG00000083457 | ITGAE | chr17 | - | 3723287 | 3723383 | 3720306 | 3720402 | 3723687 | 3723744 | 21 | 24 | 20 | 31 | 19 |
| SE | 37623 | ENSG00000082074 | FYB1 | chr5 | - | 39212677 | 39213017 | 39208863 | 39208964 | 39270568 | 39270828 | 5 | 5 | 1 | 6 | 4 |
| SE | 106036 | ENSG00000204482 | LST1 | chr6 | + | 31587943 | 31587966 | 31587053 | 31587318 | 31588496 | 31588908 | 28 | 6 | 6 | 17 | 25 |
| SE | 28627 | ENSG00000102445 | RUBCNL | chr13 | - | 46371940 | 46372210 | 46368732 | 46368815 | 46377889 | 46378005 | 21 | 23 | 22 | 30 | 30 |
| SE | 26022 | ENSG00000134480 | CCNH | chr5 | - | 87409087 | 87409129 | 87408037 | 87408186 | 87409289 | 87409363 | 14 | 12 | 18 | 15 | 19 |
| SE | 20122 | ENSG00000078319 | PMS2P1 | chr7 | - | 100332441 | 100332581 | 100330462 | 100330549 | 100335837 | 100335860 | 102 | 53 | 139 | 98 | 150 |
| SE | 101363 | ENSG00000170525 | PFKFB3 | chr10 | + | 6226191 | 6226365 | 6224148 | 6224213 | 6232894 | 6232959 | 48 | 17 | 37 | 48 | 16 |
| SE | 56196 | ENSG00000143106 | PSMA5 | chr1 | - | 109413067 | 109413135 | 109412076 | 109412184 | 109415236 | 109415363 | 546 | 344 | 515 | 492 | 742 |
| SE | 109073 | ENSG00000198176 | TFDP1 | chr13 | + | 113636533 | 113636700 | 113635976 | 113636128 | 113640119 | 113640310 | 34 | 26 | 55 | 63 | 83 |
| SE | 65947 | ENSG00000123240 | OPTN | chr10 | + | 13108137 | 13108289 | 13100162 | 13100302 | 13108961 | 13109015 | 56 | 24 | 63 | 61 | 32 |
| SE | 49922 | ENSG00000144455 | SUMF1 | chr3 | - | 4449265 | 4449340 | 4420063 | 4420146 | 4452875 | 4453049 | 152 | 102 | 158 | 115 | 99 |
| SE | 65736 | ENSG00000138073 | PREB | chr2 | - | 27133531 | 27133654 | 27133116 | 27133337 | 27134286 | 27134615 | 52 | 25 | 24 | 45 | 21 |
| SE | 53122 | ENSG00000185436 | IFNLR1 | chr1 | - | 24159051 | 24159182 | 24154167 | 24157891 | 24159473 | 24159633 | 21 | 10 | 9 | 9 | 3 |
| SE | 14965 | ENSG00000182718 | ANXA2 | chr15 | - | 60396150 | 60396427 | 60386027 | 60386086 | 60397257 | 60397338 | 0 | 9 | 3 | 3 | 2 |
| SE | 42344 | ENSG00000070759 | TESK2 | chr1 | - | 45385911 | 45385960 | 45355302 | 45355449 | 45421724 | 45421846 | 16 | 4 | 11 | 14 | 11 |
| SE | 100357 | ENSG00000149541 | B3GAT3 | chr11 | - | 62616505 | 62616796 | 62615295 | 62615799 | 62616986 | 62617347 | 81 | 34 | 65 | 78 | 47 |
| SE | 100326 | ENSG00000081237 | PTPRC | chr1 | + | 198696711 | 198696909 | 198694017 | 198694140 | 198699563 | 198699620 | 964 | 420 | 611 | 675 | 1590 |
| SE | 30309 | ENSG00000135299 | ANKRD6 | chr6 | + | 89617953 | 89618031 | 89616397 | 89616657 | 89621921 | 89622026 | 12 | 8 | 8 | 6 | 3 |
| SE | 47720 | ENSG00000153560 | UBP1 | chr3 | - | 33408689 | 33408797 | 33402800 | 33402904 | 33409235 | 33409346 | 345 | 238 | 358 | 384 | 836 |
| SE | 29854 | ENSG00000154743 | TSEN2 | chr3 | + | 12501667 | 12501724 | 12496517 | 12496554 | 12503261 | 12503784 | 2 | 0 | 7 | 4 | 1 |
| SE | 85321 | ENSG00000147394 | ZNF185 | chrX | + | 152928574 | 152928661 | 152922172 | 152922256 | 152931671 | 152931776 | 33 | 42 | 11 | 72 | 23 |
| SE | 70604 | ENSG00000164548 | TRA2A | chr7 | - | 23531180 | 23531270 | 23521706 | 23521840 | 23531788 | 23532041 | 16 | 3 | 13 | 12 | 22 |
| SE | 70356 | ENSG00000184182 | UBE2F | chr2 | + | 237973091 | 237973225 | 237966826 | 237967132 | 237987962 | 237987992 | 323 | 151 | 207 | 273 | 160 |
| SE | 103229 | ENSG00000165732 | DDX21 | chr10 | + | 68959805 | 68960249 | 68956169 | 68956312 | 68962081 | 68962157 | 649 | 488 | 532 | 625 | 920 |
| SE | 46845 | ENSG00000155463 | OXA1L | chr14 | + | 22769790 | 22769934 | 22767957 | 22768171 | 22770192 | 22770278 | 609 | 352 | 573 | 545 | 454 |
| SE | 2815 | ENSG00000104980 | TIMM44 | chr19 | - | 7935064 | 7935145 | 7934088 | 7934238 | 7938026 | 7938197 | 109 | 57 | 78 | 67 | 73 |
| SE | 107791 | ENSG00000050426 | LETMD1 | chr12 | + | 51052091 | 51052207 | 51048342 | 51049185 | 51053777 | 51053860 | 102 | 56 | 102 | 68 | 224 |
| SE | 90106 | ENSG00000205038 | PKHD1L1 | chr8 | + | 109476507 | 109476667 | 109475117 | 109475269 | 109477224 | 109477396 | 16 | 26 | 4 | 53 | 39 |
| SE | 108341 | ENSG00000111300 | NAA25 | chr12 | - | 112097240 | 112097441 | 112093050 | 112093136 | 112108715 | 112108779 | 0 | 0 | 2 | 7 | 0 |
| SE | 108334 | ENSG00000111300 | NAA25 | chr12 | - | 112061180 | 112061388 | 112060269 | 112060359 | 112068879 | 112068992 | 32 | 11 | 25 | 39 | 50 |
| SE | 51396 | ENSG00000067606 | PRKCZ | chr1 | + | 2145388 | 2145511 | 2144209 | 2144341 | 2146026 | 2146108 | 2 | 0 | 2 | 0 | 0 |
| SE | 33072 | ENSG00000088726 | TMEM40 | chr3 | - | 12748654 | 12748792 | 12738552 | 12738588 | 12749759 | 12749840 | 7 | 9 | 1 | 13 | 6 |
| SE | 35659 | ENSG00000010404 | IDS | chrX | - | 149486924 | 149487098 | 149476987 | 149483218 | 149490313 | 149490440 | 688 | 389 | 557 | 601 | 533 |
| SE | 5604 | ENSG00000215440 | NPEPL1 | chr20 | + | 58705497 | 58705569 | 58701015 | 58701158 | 58707122 | 58707200 | 9 | 7 | 12 | 6 | 11 |
| SE | 92117 | ENSG00000066117 | SMARCD1 | chr12 | + | 50096849 | 50096972 | 50094436 | 50094572 | 50098713 | 50098815 | 188 | 162 | 222 | 209 | 259 |
| SE | 2613 | ENSG00000160570 | DEDD2 | chr19 | - | 42216679 | 42217045 | 42215132 | 42215252 | 42220073 | 42220123 | 49 | 15 | 34 | 36 | 11 |
| SE | 20076 | ENSG00000145088 | EAF2 | chr3 | + | 121836644 | 121836794 | 121835276 | 121835391 | 121844452 | 121844547 | 1 | 4 | 7 | 0 | 2 |
| SE | 50004 | ENSG00000241878 | PISD | chr22 | - | 31632182 | 31632291 | 31625762 | 31626016 | 31638365 | 31638445 | 4 | 0 | 2 | 0 | 0 |
| SE | 8254 | ENSG00000108312 | UBTF | chr17 | - | 44210791 | 44210947 | 44210317 | 44210473 | 44211038 | 44211152 | 418 | 231 | 497 | 425 | 356 |
| SE | 68634 | ENSG00000197714 | ZNF460 | chr19 | + | 57284550 | 57284677 | 57280484 | 57280836 | 57290698 | 57291045 | 127 | 61 | 103 | 115 | 42 |
| SE | 23881 | ENSG00000164338 | UTP15 | chr5 | + | 73572488 | 73572624 | 73570585 | 73570711 | 73576951 | 73577036 | 125 | 113 | 142 | 115 | 184 |
| SE | 29218 | ENSG00000164118 | CEP44 | chr4 | + | 174297965 | 174298062 | 174283910 | 174283943 | 174299071 | 174299210 | 84 | 38 | 53 | 45 | 16 |
| SE | 54052 | ENSG00000158796 | DEDD | chr1 | - | 161123074 | 161123174 | 161121743 | 161122523 | 161123838 | 161123946 | 82 | 36 | 64 | 65 | 78 |
| SE | 35303 | ENSG00000163961 | RNF168 | chr3 | - | 196488606 | 196488683 | 196487398 | 196487578 | 196502872 | 196503768 | 302 | 235 | 228 | 255 | 232 |
| SE | 8111 | ENSG00000141252 | VPS53 | chr17 | - | 695077 | 695225 | 661808 | 661895 | 697417 | 697484 | 9 | 2 | 0 | 1 | 2 |
| SE | 18868 | ENSG00000100605 | ITPK1 | chr14 | - | 92993879 | 92993997 | 92962750 | 92962849 | 93016675 | 93016801 | 210 | 101 | 180 | 181 | 272 |
| SE | 78631 | ENSG00000168014 | C2CD3 | chr11 | - | 74054606 | 74054671 | 74049336 | 74049542 | 74057405 | 74057544 | 43 | 11 | 40 | 33 | 44 |
| SE | 61576 | ENSG00000131876 | SNRPA1 | chr15 | - | 101286213 | 101286293 | 101285725 | 101285801 | 101286907 | 101287010 | 117 | 84 | 125 | 117 | 112 |
| SE | 66960 | ENSG00000152056 | AP1S3 | chr2 | - | 223775900 | 223776009 | 223765176 | 223765350 | 223777690 | 223777869 | 18 | 15 | 21 | 12 | 25 |
| SE | 50356 | ENSG00000003393 | ALS2 | chr2 | - | 201718076 | 201718210 | 201715671 | 201715839 | 201723042 | 201723120 | 56 | 24 | 40 | 81 | 72 |
| SE | 38460 | ENSG00000147251 | DOCK11 | chrX | + | 118590209 | 118590302 | 118588412 | 118588478 | 118593213 | 118593337 | 702 | 514 | 557 | 635 | 1157 |
| SE | 57204 | ENSG00000084112 | SSH1 | chr12 | - | 108823257 | 108823361 | 108818248 | 108818313 | 108852637 | 108852678 | 34 | 9 | 17 | 40 | 12 |
| SE | 84868 | ENSG00000164074 | ABHD18 | chr4 | + | 128017362 | 128017501 | 128011672 | 128011700 | 128020079 | 128020169 | 111 | 75 | 113 | 148 | 65 |
| SE | 61405 | ENSG00000185630 | PBX1 | chr1 | + | 164820071 | 164820184 | 164811989 | 164812149 | 164821536 | 164821626 | 1 | 18 | 5 | 33 | 13 |
| SE | 108180 | ENSG00000121749 | TBC1D15 | chr12 | + | 71854784 | 71854853 | 71839758 | 71839811 | 71872069 | 71872168 | 0 | 4 | 2 | 2 | 0 |
| SE | 22896 | ENSG00000140853 | NLRC5 | chr16 | + | 57028071 | 57028203 | 57025367 | 57027018 | 57028301 | 57028385 | 51 | 31 | 87 | 79 | 45 |
| SE | 42582 | ENSG00000198380 | GFPT1 | chr2 | - | 69370000 | 69370108 | 69363544 | 69363670 | 69374005 | 69374113 | 120 | 88 | 105 | 154 | 170 |
| SE | 45650 | ENSG00000144283 | PKP4 | chr2 | + | 158625124 | 158625427 | 158621230 | 158621421 | 158631752 | 158631932 | 7 | 8 | 14 | 17 | 11 |
| SE | 53900 | ENSG00000196510 | ANAPC7 | chr12 | - | 110377392 | 110377617 | 110375476 | 110376216 | 110381751 | 110381948 | 261 | 236 | 223 | 238 | 302 |
| SE | 48058 | ENSG00000266028 | SRGAP2 | chr1 | + | 206458370 | 206458409 | 206454877 | 206455024 | 206458622 | 206458947 | 1 | 3 | 0 | 3 | 0 |
| SE | 73148 | ENSG00000144233 | AMMECR1L | chr2 | - | 127870813 | 127870928 | 127866899 | 127866996 | 127871248 | 127871359 | 16 | 14 | 7 | 13 | 20 |
| SE | 70745 | ENSG00000102125 | TAZ | chrX | + | 154419704 | 154419746 | 154419542 | 154419623 | 154420031 | 154420094 | 16 | 6 | 28 | 26 | 31 |
| [illegible] | | | | | | | | | | | | | | | | |

| | | | | | | | | | | | | | | |
|---|---|---|---|---|---|---|---|---|---|---|---|---|---|---|
| SE | 105599 | ENSG00000139323 | POC1B | chr12 | - | 89459637 | 89459718 | 89425160 | 89425379 | 89466769 | 89466922 | 94 | 62 | 70 |
| SE | 29335 | ENSG00000121210 | TMEM131L | chr4 | + | 153583563 | 153583672 | 153583189 | 153583248 | 153584837 | 153584931 | 150 | 151 | 199 |
| SE | 61151 | ENSG00000111711 | GOLT1B | chr12 | + | 21507943 | 21508109 | 21507116 | 21507151 | 21508382 | 21508561 | 2 | 6 | 12 |
| SE | 57976 | ENSG00000160746 | ANO10 | chr3 | - | 43576691 | 43577261 | 43565652 | 43565727 | 43580352 | 43580472 | 11 | 7 | 4 |
| SE | 31596 | ENSG00000082805 | ERC1 | chr12 | + | 1112214 | 1112298 | 1110191 | 1110347 | 1115865 | 1116033 | 105 | 87 | 111 |
| SE | 12080 | ENSG00000135723 | FHOD1 | chr16 | - | 67231187 | 67231349 | 67230600 | 67230791 | 67231429 | 67231549 | 84 | 33 | 40 |
| SE | 9388 | ENSG00000121067 | SPOP | chr17 | - | 49675894 | 49676016 | 49622732 | 49622876 | 49677932 | 49678000 | 1 | 4 | 3 |
| SE | 29523 | ENSG00000119242 | CCDC92 | chr12 | - | 123944668 | 123944740 | 123944271 | 123944364 | 123972528 | 123972831 | 4 | 2 | 2 |
| SE | 89071 | ENSG00000171988 | JMJD1C | chr10 | - | 63198512 | 63198727 | 63197410 | 63197563 | 63206594 | 63208801 | 266 | 267 | 231 |
| SE | 52696 | ENSG00000160055 | TMEM234 | chr1 | - | 32216383 | 32216517 | 32215846 | 32216281 | 32216888 | 32216947 | 16 | 6 | 20 |
| SE | 81166 | ENSG00000089289 | IGBP1 | chrX | + | 70146632 | 70146828 | 70134522 | 70134816 | 70148760 | 70148840 | 786 | 564 | 657 |
| SE | 95174 | ENSG00000213930 | GALT | chr9 | + | 34647491 | 34647705 | 34647088 | 34647258 | 34648114 | 34648171 | 32 | 15 | 65 |
| SE | 98141 | ENSG00000162769 | FLVCR1 | chr1 | + | 212864392 | 212864519 | 212863630 | 212863869 | 212870053 | 212870166 | 27 | 3 | 27 |
| SE | 54768 | ENSG00000138642 | HERC6 | chr4 | + | 88417424 | 88417579 | 88413076 | 88413266 | 88423859 | 88423973 | 73 | 79 | 81 |
| SE | 46582 | ENSG00000058272 | PPP1R12A | chr12 | - | 79786373 | 79786478 | 79781814 | 79781862 | 79788647 | 79788752 | 1010 | 795 | 661 |
| SE | 35815 | ENSG00000123130 | ACOT9 | chrX | - | 23713134 | 23713208 | 23707716 | 23707944 | 23721880 | 23721984 | 190 | 249 | 146 |
| SE | 86709 | ENSG00000128654 | MTX2 | chr2 | + | 176329300 | 176329426 | 176328873 | 176328912 | 176330583 | 176330660 | 53 | 42 | 58 |
| SE | 68591 | ENSG00000239779 | WBP1 | chr2 | + | 74458831 | 74459117 | 74458471 | 74458671 | 74459642 | 74459745 | 5 | 2 | 4 |
| SE | 12153 | ENSG00000172775 | PSME3IP1 | chr16 | - | 57178501 | 57178852 | 57173727 | 57173869 | 57185820 | 57186039 | 52 | 41 | 47 |
| SE | 107468 | ENSG00000100599 | RIN3 | chr14 | + | 92676532 | 92676606 | 92659384 | 92659469 | 92684986 | 92685150 | 44 | 29 | 31 |
| SE | 48873 | ENSG00000143514 | TP53BP2 | chr1 | - | 223798214 | 223798677 | 223795814 | 223796590 | 223799898 | 223800047 | 96 | 48 | 78 |
| SE | 69323 | ENSG00000112367 | FIG4 | chr6 | + | 109788797 | 109788913 | 109786301 | 109786449 | 109789593 | 109789677 | 1 | 0 | 3 |
| SE | 92802 | ENSG00000131669 | NINJ1 | chr9 | - | 93126409 | 93126585 | 93124898 | 93125062 | 93134142 | 93134238 | 95 | 30 | 51 |
| SE | 13189 | ENSG00000103495 | MAZ | chr16 | + | 29809539 | 29809764 | 29808569 | 29808741 | 29810076 | 29811164 | 141 | 37 | 75 |
| SE | 3544 | ENSG00000105483 | CARD8 | chr19 | - | 48242539 | 48242644 | 48240961 | 48241063 | 48246074 | 48246381 | 13 | 7 | 11 |
| SE | 59170 | ENSG00000115204 | MPV17 | chr2 | - | 27317077 | 27317234 | 27312679 | 27312772 | 27322447 | 27322522 | 1 | 2 | 2 |
| SE | 25010 | ENSG00000150593 | PDCD4 | chr10 | + | 110876669 | 110876753 | 110875965 | 110876070 | 110881232 | 110881535 | 102 | 61 | 106 |
| SE | 56836 | ENSG00000143398 | PIP5K1A | chr1 | + | 151242437 | 151242567 | 151239954 | 151240039 | 151246919 | 151246965 | 22 | 28 | 24 |
| SE | 13157 | ENSG00000103121 | CMC2 | chr16 | - | 80991753 | 80991957 | 80981805 | 80981877 | 80997313 | 80997429 | 2 | 0 | 0 |
| SE | 37938 | ENSG00000138964 | PARVG | chr22 | + | 44181741 | 44181917 | 44172955 | 44173191 | 44183317 | 44183408 | 376 | 223 | 210 |
| SE | 15267 | ENSG00000035664 | DAPK2 | chr15 | - | 63924815 | 63924861 | 63912107 | 63912197 | 63925940 | 63926093 | 13 | 3 | 5 |
| SE | 92279 | ENSG00000132326 | PER2 | chr2 | - | 238275742 | 238275897 | 238273069 | 238273191 | 238277130 | 238277193 | 31 | 16 | 29 |
| SE | 96379 | ENSG00000110042 | DTX4 | chr11 | + | 59192097 | 59192250 | 59188734 | 59188796 | 59195207 | 59195369 | 6 | 4 | 8 |

[illegible]

| | | | | | | | | | | | | | | | | | | | | | | | | | | | | | | | | | | | | | | | | | | | | | | |
|---|---|---|---|---|---|---|---|---|---|---|---|---|---|---|---|---|---|---|---|---|---|---|---|---|---|---|---|---|---|---|---|---|---|---|---|---|---|---|---|---|---|---|---|---|---|---|
| SE | 47088 | ENSG00000095787 | WAC | chr10 | + | 28537583 | 28537815 | 28535909 | 28536059 | 28583398 | 28583505 | 2 | 3 | 1 | 5 | 0 | 8 | 12 | 25 | 10 | 10 | 2 | 0 | 1 | 20 | 0 | 4 | 12 | 48 | 29 | 4 | 298 | 149 | 0.27790214 | 1 | 0.111 | 0.111 | 0.02 | 0.2 | 0 | 0.2 | 0 | 0.01 | 0.256 | 0 | -0.005 |
| SE | 68024 | ENSG00000172890 | NADSYN1 | chr11 | + | 71477344 | 71477438 | 71474394 | 71474526 | 71478394 | 71478469 | 11 | 0 | 12 | 1 | 2 | 46 | 22 | 47 | 56 | 66 | 8 | 6 | 2 | 5 | 13 | 129 | 91 | 50 | 107 | 87 | 243 | 149 | 0.278154872 | 1 | 0.128 | 0 | 0.135 | 0.011 | 0.018 | 0.037 | 0.039 | 0.024 | 0.028 | 0.084 | 0.016 |
| SE | 30284 | ENSG00000114480 | GBE1 | chr3 | - | 81646391 | 81646482 | 81642780 | 81642990 | 81648855 | 81648991 | 74 | 51 | 48 | 43 | 91 | 2 | 0 | 1 | 5 | 0 | 37 | 34 | 34 | 66 | 54 | 0 | 2 | 1 | 0 | 0 | 240 | 149 | 0.278338568 | 1 | 0.958 | 1 | 0.968 | 0.842 | 1 | 1 | 0.913 | 0.955 | 1 | 1 | -0.02 |
| SE | 49600 | ENSG00000076685 | NT5C2 | chr10 | - | 103111744 | 103111805 | 103106588 | 103106706 | 103136248 | 103136372 | 22 | 16 | 26 | 19 | 36 | 7 | 1 | 4 | 4 | 6 | 30 | 9 | 28 | 38 | 4 | 0 | 0 | 17 | 4 | 0 | 210 | 149 | 0.278443853 | 1 | 0.69 | 0.919 | 0.822 | 0.771 | 0.81 | 1 | 1 | 0.539 | 0.871 | 1 | -0.08 |
| SE | 46597 | ENSG00000058272 | PPP1R12A | chr12 | - | 79872807 | 79872938 | 79845301 | 79845420 | 79934694 | 79934715 | 794 | 511 | 515 | 702 | 384 | 4 | 1 | 2 | 6 | 7 | 344 | 526 | 551 | 206 | 292 | 0 | 3 | 7 | 0 | 10 | 280 | 149 | 0.278487245 | 1 | 0.991 | 0.996 | 0.993 | 0.984 | 0.967 | 1 | 0.989 | 0.977 | 1 | 0.94 | 0.005 |
| SE | 94454 | ENSG00000122490 | SLC66A2 | chr18 | - | 79933968 | 79934022 | 79919183 | 79919400 | 79943328 | 79943462 | 12 | 5 | 8 | 8 | 1 | 9 | 4 | 4 | 8 | 19 | 11 | 18 | 5 | 16 | 33 | 0 | 20 | 9 | 17 | 26 | 203 | 149 | 0.278669811 | 1 | 0.495 | 0.478 | 0.595 | 0.423 | 0.037 | 1 | 0.398 | 0.29 | 0.409 | 0.482 | -0.11 |
| SE | 23917 | ENSG00000124177 | CHD6 | chr20 | - | 41499294 | 41499357 | 41496808 | 41498226 | 41512845 | 41512995 | 317 | 235 | 306 | 329 | 269 | 1 | 1 | 0 | 2 | 0 | 160 | 336 | 347 | 250 | 85 | 0 | 3 | 5 | 9 | 0 | 212 | 149 | 0.27867614 | 1 | 0.996 | 0.994 | 1 | 0.991 | 1 | 1 | 0.987 | 0.98 | 0.951 | 1 | 0.013 |
| SE | 26230 | ENSG00000141084 | RANBP10 | chr16 | - | 67734857 | 67735042 | 67731028 | 67731584 | 67744287 | 67744455 | 25 | 12 | 15 | 25 | 12 | 0 | 0 | 0 | 4 | 3 | 13 | 41 | 16 | 21 | 55 | 0 | 0 | 0 | 0 | 0 | 298 | 149 | 0.278678101 | 1 | 1 | 1 | 1 | 0.758 | 0.667 | 1 | 1 | 1 | 1 | 1 | -0.115 |
| SE | 106839 | ENSG00000123329 | ARHGAP9 | chr12 | - | 57473602 | 57473708 | 57472263 | 57472688 | 57474041 | 57474176 | 431 | 214 | 294 | 423 | 517 | 54 | 37 | 47 | 53 | 56 | 542 | 322 | 330 | 413 | 835 | 124 | 61 | 34 | 77 | 75 | 255 | 149 | 0.278824565 | 1 | 0.823 | 0.772 | 0.785 | 0.823 | 0.844 | 0.719 | 0.755 | 0.85 | 0.758 | 0.867 | 0.02 |
| SE | 81475 | ENSG00000091140 | DLD | chr7 | + | 107902324 | 107902393 | 107901737 | 107901817 | 107903477 | 107903547 | 262 | 177 | 235 | 314 | 171 | 0 | 2 | 4 | 4 | 5 | 20 | 169 | 147 | 58 | 57 | 0 | 0 | 10 | 2 | 0 | 218 | 149 | 0.278838672 | 1 | 1 | 0.984 | 0.976 | 0.982 | 0.959 | 1 | 1 | 0.909 | 0.952 | 1 | 0.008 |
| SE | 24945 | ENSG00000163297 | ANTXR2 | chr4 | - | 79907576 | 79907664 | 79905590 | 79907467 | 79977620 | 79977701 | 7 | 0 | 1 | 11 | 0 | 138 | 128 | 94 | 144 | 113 | 0 | 6 | 2 | 2 | 0 | 132 | 83 | 97 | 113 | 28 | 237 | 149 | 0.278847536 | 1 | 0.031 | 0 | 0.007 | 0.046 | 0 | 0 | 0.043 | 0.013 | 0.011 | 0 | 0.003 |
| SE | 59971 | ENSG00000076770 | MBNL3 | chrX | - | 132381358 | 132381432 | 132379203 | 132379677 | 132382177 | 132382272 | 44 | 42 | 34 | 30 | 70 | 39 | 25 | 24 | 41 | 27 | 0 | 36 | 11 | 9 | 6 | 1 | 20 | 23 | 5 | 9 | 223 | 149 | 0.278881761 | 1 | 0.43 | 0.529 | 0.486 | 0.328 | 0.634 | 0 | 0.546 | 0.242 | 0.546 | 0.308 | 0.153 |
| SE | 66467 | ENSG00000131051 | RBM39 | chr20 | - | 35734185 | 35734257 | 35731915 | 35732135 | 35738967 | 35739017 | 36 | 21 | 11 | 31 | 10 | 679 | 659 | 609 | 826 | 869 | 7 | 6 | 29 | 10 | 3 | 276 | 482 | 893 | 401 | 87 | 221 | 149 | 0.27897738 | 1 | 0.035 | 0.021 | 0.012 | 0.025 | 0.008 | 0.017 | 0.008 | 0.021 | 0.017 | 0.023 | 0.003 |
| SE | 35664 | ENSG00000010404 | IDS | chrX | - | 149502074 | 149502177 | 149500948 | 149501037 | 149503311 | 149503489 | 99 | 35 | 53 | 81 | 103 | 368 | 214 | 278 | 357 | 437 | 74 | 84 | 103 | 54 | 68 | 472 | 382 | 327 | 250 | 530 | 252 | 149 | 0.27901509 | 1 | 0.137 | 0.088 | 0.101 | 0.118 | 0.122 | 0.085 | 0.115 | 0.157 | 0.113 | 0.071 | 0.005 |
| SE | 50463 | ENSG00000161618 | ALDH16A1 | chr19 | + | 49461618 | 49461800 | 49460821 | 49460899 | 49461883 | 49462036 | 43 | 21 | 12 | 34 | 4 | 4 | 0 | 0 | 0 | 2 | 4 | 16 | 4 | 17 | 58 | 0 | 0 | 0 | 0 | 0 | 298 | 149 | 0.279062137 | 1 | 0.843 | 1 | 1 | 1 | 0.5 | 1 | 1 | 1 | 1 | 1 | -0.131 |
| SE | 81053 | ENSG00000188243 | COMMD6 | chr13 | - | 75527812 | 75527851 | 75526435 | 75526639 | 75530113 | 75530266 | 5 | 5 | 25 | 14 | 20 | 897 | 446 | 830 | 818 | 1468 | 14 | 9 | 0 | 11 | 10 | 454 | 420 | 347 | 380 | 523 | 188 | 149 | 0.279071257 | 1 | 0.004 | 0.009 | 0.023 | 0.013 | 0.011 | 0.024 | 0.017 | 0 | 0.022 | 0.015 | -0.004 |
| SE | 52968 | ENSG00000150455 | TIRAP | chr11 | + | 126290461 | 126290585 | 126289664 | 126289817 | 126290802 | 126290897 | 29 | 3 | 13 | 22 | 6 | 1 | 0 | 2 | 0 | 0 | 11 | 14 | 4 | 13 | 15 | 0 | 0 | 0 | 6 | 0 | 273 | 149 | 0.279099628 | 1 | 0.941 | 1 | 0.78 | 1 | 1 | 1 | 1 | 1 | 0.542 | 1 | 0.036 |
| SE | 4071 | ENSG00000105771 | SMG9 | chr19 | - | 43744771 | 43744884 | 43739672 | 43740218 | 43747441 | 43747539 | 70 | 42 | 96 | 73 | 63 | 4 | 0 | 1 | 5 | 0 | 103 | 60 | 20 | 75 | 120 | 5 | 0 | 0 | 7 | 1 | 262 | 149 | 0.279201743 | 1 | 0.909 | 1 | 0.982 | 0.893 | 1 | 0.921 | 1 | 1 | 0.859 | 0.986 | 0.004 |
| SE | 86427 | ENSG00000196092 | PAX5 | chr9 | - | 37002647 | 37002776 | 36966548 | 36966724 | 37006472 | 37006537 | 104 | 29 | 63 | 114 | 76 | 0 | 0 | 0 | 0 | 0 | 133 | 203 | 63 | 85 | 188 | 8 | 2 | 0 | 1 | 0 | 278 | 149 | 0.279302412 | 1 | 1 | 1 | 1 | 1 | 1 | 0.899 | 0.982 | 1 | 0.979 | 1 | 0.028 |
| SE | 51240 | ENSG00000146376 | ARHGAP18 | chr6 | - | 129634041 | 129634105 | 129618686 | 129618852 | 129638393 | 129638629 | 9 | 17 | 7 | 36 | 15 | 0 | 0 | 0 | 0 | 6 | 17 | 15 | 6 | 10 | 18 | 0 | 0 | 0 | 0 | 0 | 213 | 149 | 0.279367484 | 1 | 1 | 1 | 1 | 1 | 0.636 | 1 | 1 | 1 | 1 | 1 | -0.073 |
| SE | 107944 | ENSG00000177119 | ANO6 | chr12 | + | 45270461 | 45270524 | 45216084 | 45216391 | 45302013 | 45302093 | 48 | 50 | 29 | 69 | 69 | 163 | 150 | 118 | 248 | 185 | 50 | 67 | 71 | 90 | 71 | 285 | 155 | 166 | 257 | 155 | 212 | 149 | 0.279477704 | 1 | 0.171 | 0.19 | 0.147 | 0.164 | 0.208 | 0.11 | 0.233 | 0.231 | 0.198 | 0.244 | -0.027 |
| SE | 83458 | ENSG00000156831 | NSMCE2 | chr8 | + | 125102013 | 125102146 | 125091843 | 125091958 | 125102316 | 125102487 | 27 | 20 | 26 | 20 | 13 | 4 | 2 | 2 | 8 | 1 | 30 | 18 | 29 | 46 | 12 | 0 | 12 | 14 | 2 | 11 | 282 | 149 | 0.279536791 | 1 | 0.781 | 0.841 | 0.873 | 0.569 | 0.873 | 1 | 0.442 | 0.523 | 0.924 | 0.366 | 0.136 |
| SE | 80699 | ENSG00000175471 | MCTP1 | chr5 | - | 94917895 | 94917973 | 94912805 | 94912976 | 94931952 | 94931991 | 26 | 30 | 9 | 20 | 28 | 27 | 28 | 6 | 29 | 21 | 34 | 32 | 15 | 16 | 24 | 10 | 27 | 7 | 26 | 20 | 227 | 149 | 0.279551173 | 1 | 0.387 | 0.413 | 0.496 | 0.312 | 0.467 | 0.691 | 0.438 | 0.584 | 0.288 | 0.441 | -0.073 |
| SE | 104026 | ENSG00000151923 | TIAL1 | chr10 | - | 119580470 | 119580538 | 119579934 | 119580010 | 119581921 | 119582009 | 42 | 24 | 25 | 25 | 71 | 266 | 131 | 193 | 227 | 414 | 12 | 16 | 28 | 26 | 35 | 142 | 118 | 84 | 139 | 165 | 217 | 149 | 0.279580059 | 1 | 0.098 | 0.112 | 0.082 | 0.07 | 0.105 | 0.055 | 0.085 | 0.186 | 0.114 | 0.127 | -0.02 |
| SE | 70407 | ENSG00000115652 | UXS1 | chr2 | - | 106098715 | 106098773 | 106093307 | 106094157 | 106101057 | 106101118 | 33 | 31 | 35 | 35 | 39 | 1 | 0 | 2 | 2 | 5 | 86 | 79 | 25 | 27 | 241 | 0 | 0 | 0 | 0 | 12 | 207 | 149 | 0.279634555 | 1 | 0.96 | 1 | 0.926 | 0.926 | 0.849 | 1 | 1 | 1 | 1 | 0.935 | -0.055 |
| SE | 18595 | ENSG00000100664 | EIF5 | chr14 | + | 103337115 | 103337227 | 103336505 | 103336849 | 103338326 | 103338472 | 282 | 308 | 322 | 406 | 858 | 0 | 0 | 0 | 0 | 0 | 106 | 368 | 417 | 296 | 138 | 13 | 0 | 0 | 0 | 0 | 261 | 149 | 0.279650213 | 1 | 1 | 1 | 1 | 1 | 1 | 0.823 | 1 | 1 | 1 | 1 | 0.035 |
| SE | 33927 | ENSG00000160796 | NBEAL2 | chr3 | + | 46992474 | 46992555 | 46991839 | 46991946 | 46993936 | 46994020 | 57 | 16 | 26 | 18 | 27 | 29 | 14 | 23 | 45 | 27 | 42 | 37 | 27 | 33 | 98 | 103 | 47 | 17 | 63 | 92 | 230 | 149 | 0.279671264 | 1 | 0.56 | 0.425 | 0.423 | 0.206 | 0.393 | 0.209 | 0.338 | 0.507 | 0.253 | 0.408 | 0.058 |
| SE | 51782 | ENSG00000042429 | MED17 | chr11 | + | 93807517 | 93807635 | 93800660 | 93801972 | 93809716 | 93809858 | 148 | 91 | 107 | 102 | 178 | 1 | 2 | 0 | 4 | 0 | 15 | 71 | 86 | 60 | 22 | 7 | 0 | 0 | 0 | 0 | 267 | 149 | 0.27971752 | 1 | 0.988 | 0.962 | 1 | 0.934 | 1 | 0.545 | 1 | 1 | 1 | 1 | 0.068 |
| SE | 1063 | ENSG00000239900 | ADSL | chr22 | + | 40364879 | 40365056 | 40364275 | 40364365 | 40366435 | 40366489 | 186 | 131 | 202 | 160 | 253 | 19 | 7 | 10 | 7 | 39 | 137 | 88 | 98 | 148 | 225 | 37 | 1 | 20 | 34 | 4 | 298 | 149 | 0.279734276 | 1 | 0.83 | 0.903 | 0.91 | 0.92 | 0.764 | 0.649 | 0.978 | 0.71 | 0.685 | 0.966 | 0.068 |
| SE | 46353 | ENSG00000028310 | BRD9 | chr5 | - | 869244 | 869404 | 865413 | 865581 | 870472 | 870553 | 14 | 1 | 1 | 6 | 7 | 35 | 23 | 43 | 44 | 33 | 0 | 7 | 5 | 14 | 7 | 44 | 40 | 27 | 36 | 167 | 298 | 149 | 0.279770099 | 1 | 0.167 | 0.021 | 0.011 | 0.064 | 0.096 | 0 | 0.08 | 0.085 | 0.163 | 0.021 | 0.002 |
| SE | 4136 | ENSG00000269547 | NA | chr19 | - | 38917920 | 38918008 | 38917723 | 38917833 | 38918421 | 38918530 | 8 | 2 | 9 | 4 | 13 | 0 | 0 | 0 | 0 | 0 | 6 | 8 | 3 | 4 | 19 | 0 | 0 | 4 | 0 | 0 | 237 | 149 | 0.279859495 | 1 | 1 | 1 | 1 | 1 | 1 | 1 | 1 | 0.32 | 1 | 1 | 0.136 |
| SE | 44942 | ENSG00000118193 | KIF14 | chr1 | - | 200614317 | 200614405 | 200608829 | 200608928 | 200615354 | 200615609 | 9 | 4 | 4 | 2 | 15 | 0 | 0 | 0 | 0 | 0 | 19 | 8 | 2 | 3 | 1 | 0 | 0 | 0 | 4 | 0 | 237 | 149 | 0.279867545 | 1 | 1 | 1 | 1 | 1 | 1 | 1 | 1 | 1 | 0.32 | 1 | 0.136 |
| SE | 96236 | ENSG00000133943 | DGLUCY | chr14 | + | 91148573 | 91148852 | 91147780 | 91147858 | 91157638 | 91157690 | 0 | 0 | 0 | 1 | 0 | 46 | 34 | 33 | 38 | 53 | 0 | 0 | 1 | 7 | 7 | 83 | 70 | 19 | 48 | 110 | 298 | 149 | 0.279878661 | 1 | 0 | 0 | 0 | 0.013 | 0 | 0 | 0 | 0.026 | 0.068 | 0.031 | -0.022 |
| SE | 62986 | ENSG00000066697 | MSANTD3 | chr9 | + | 100445168 | 100445237 | 100441905 | 100442356 | 100450556 | 100450705 | 1 | 2 | 0 | 1 | 5 | 57 | 76 | 31 | 123 | 84 | 0 | 0 | 6 | 3 | 22 | 278 | 119 | 74 | 114 | 644 | 218 | 149 | 0.279896992 | 1 | 0.012 | 0.018 | 0 | 0.006 | 0.039 | 0 | 0 | 0.053 | 0.018 | 0.023 | -0.004 |
| SE | 8631 | ENSG00000261371 | PECAM1 | chr17 | - | 64329699 | 64329722 | 64319414 | 64323845 | 64341633 | 64341690 | 1259 | 965 | 804 | 1123 | 2667 | 32 | 17 | 9 | 24 | 63 | 2387 | 1124 | 642 | 1189 | 1522 | 65 | 10 | 10 | 43 | 41 | 172 | 149 | 0.280043488 | 1 | 0.971 | 0.98 | 0.987 | 0.976 | 0.973 | 0.97 | 0.99 | 0.982 | 0.96 | 0.97 | 0.003 |
| SE | 30578 | ENSG00000128039 | SRD5A3 | chr4 | + | 55367587 | 55367722 | 55364073 | 55364271 | 55369831 | 55370109 | 79 | 28 | 44 | 54 | 79 | 4 | 2 | 2 | 1 | 0 | 49 | 23 | 35 | 16 | 115 | 6 | 0 | 0 | 0 | 0 | 284 | 149 | 0.280086818 | 1 | 0.912 | 0.88 | 0.92 | 0.966 | 1 | 0.811 | 1 | 1 | 1 | 1 | -0.027 |
| SE | 75241 | ENSG00000065883 | CDK13 | chr7 | + | 39998043 | 39998134 | 39997493 | 39997664 | 39999360 | 39999386 | 7 | 11 | 5 | 5 | 15 | 329 | 255 | 295 | 363 | 275 | 23 | 1 | 2 | 8 | 17 | 286 | 346 | 201 | 355 | 226 | 240 | 149 | 0.2801542 | 1 | 0.013 | 0.026 | 0.01 | 0.008 | 0.033 | 0.048 | 0.002 | 0.006 | 0.014 | 0.045 | -0.005 |
| SE | 44676 | ENSG00000072501 | SMC1A | chrX | - | 53421894 | 53422040 | 53414980 | 53415169 | 53422491 | 53422728 | 10 | 1 | 10 | 4 | 4 | 85 | 47 | 67 | 74 | 62 | 18 | 4 | 1 | 8 | 6 | 107 | 48 | 36 | 114 | 21 | 295 | 149 | 0.280205344 | 1 | 0.056 | 0.011 | 0.07 | 0.027 | 0.032 | 0.078 | 0.04 | 0.014 | 0.034 | 0.126 | -0.019 |
| SE | 19176 | ENSG00000140105 | WARS1 | chr14 | - | 100373979 | 100374064 | 100369086 | 100369258 | 100376259 | 100376311 | 0 | 4 | 1 | 0 | 5 | 75 | 29 | 52 | 42 | 11 | 0 | 4 | 0 | 3 | 0 | 65 | 80 | 75 | 78 | 123 | 234 | 149 | 0.280261003 | 1 | 0 | 0.081 | 0.012 | 0 | 0.224 | 0 | 0.031 | 0 | 0.024 | 0 | 0.052 |
| SE | 55423 | ENSG00000116688 | MFN2 | chr1 | + | 11981809 | 11981853 | 11980298 | 11980484 | 11981969 | 11982114 | 6 | 4 | 1 | 3 | 9 | 50 | 27 | 27 | 22 | 33 | 0 | 0 | 9 | 4 | 7 | 22 | 29 | 21 | 42 | 96 | 193 | 149 | 0.280288736 | 1 | 0.085 | 0.103 | 0.028 | 0.095 | 0.174 | 0 | 0 | 0.249 | 0.068 | 0.053 | 0.023 |
| SE | 18876 | ENSG00000100721 | TCL1A | chr14 | - | 95712550 | 95712774 | 95712219 | 95712396 | 95713946 | 95714127 | 2 | 0 | 1 | 0 | 6 | 16 | 9 | 15 | 22 | 12 | 8 | 10 | 0 | 0 | 0 | 14 | 87 | 9 | 8 | 58 | 298 | 149 | 0.280288917 | 1 | 0.059 | 0 | 0.032 | 0 | 0.2 | 0.222 | 0.054 | 0 | 0 | 0 | 0.003 |
| SE | 86173 | ENSG00000073464 | CLCN4 | chrX | + | 10197938 | 10198061 | 10194910 | 10195098 | 10206357 | 10206564 | 23 | 5 | 8 | 28 | 2 | 0 | 0 | 2 | 1 | 0 | 12 | 18 | 5 | 13 | 122 | 0 | 3 | 1 | 1 | 2 | 272 | 149 | 0.280383216 | 1 | 1 | 1 | 0.687 | 0.939 | 1 | 1 | 0.767 | 0.733 | 0.877 | 0.971 | 0.056 |
| SE | 89233 | ENSG00000094914 | AAAS | chr12 | - | 53314750 | 53314849 | 53314297 | 53314441 | 53315093 | 53315140 | 64 | 42 | 69 | 85 | 98 | 0 | 6 | 0 | 1 | 10 | 32 | 67 | 40 | 87 | 80 | 20 | 9 | 1 | 0 | 0 | 248 | 149 | 0.28052697 | 1 | 1 | 0.808 | 1 | 0.981 | 0.855 | 0.49 | 0.817 | 0.96 | 1 | 1 | 0.075 |
| SE | 9712 | ENSG00000141568 | FOXK2 | chr17 | + | 82584012 | 82584188 | 82582740 | 82582934 | 82585903 | 82586200 | 107 | 57 | 105 | 115 | 129 | 0 | 0 | 0 | 2 | 0 | 138 | 104 | 31 | 60 | 103 | 0 | 5 | 0 | 0 | 4 | 298 | 149 | 0.280597431 | 1 | 1 | 1 | 1 | 0.966 | 1 | 1 | 0.912 | 1 | 1 | 0.928 | 0.025 |
| SE | 103134 | ENSG00000180198 | RCC1 | chr1 | + | 28530522 | 28530573 | 28529857 | 28529939 | 28531802 | 28531818 | 14 | 12 | 17 | 28 | 8 | 46 | 33 | 26 | 47 | 60 | 6 | 25 | 6 | 9 | 16 | 68 | 24 | 21 | 42 | 74 | 200 | 149 | 0.280604794 | 1 | 0.185 | 0.213 | 0.328 | 0.307 | 0.09 | 0.062 | 0.437 | 0.176 | 0.138 | 0.139 | 0.034 |
| SE | 90734 | ENSG00000168404 | MLKL | chr16 | - | 74678898 | 74678980 | 74676300 | 74676425 | 74685485 | 74685583 | 16 | 16 | 8 | 24 | 19 | 0 | 0 | 0 | 0 | 0 | 39 | 7 | 24 | 22 | 4 | 0 | 3 | 3 | 0 | 0 | 231 | 149 | 0.280658668 | 1 | 1 | 1 | 1 | 1 | 1 | 1 | 0.601 | 0.838 | 1 | 1 | 0.112 |
| SE | 12977 | ENSG00000126602 | TRAP1 | chr16 | - | 3671721 | 3671791 | 3665970 | 3666118 | 3674338 | 3674494 | 59 | 12 | 46 | 22 | 16 | 0 | 0 | 0 | 0 | 0 | 23 | 26 | 13 | 46 | 75 | 7 | 0 | 0 | 0 | 0 | 219 | 149 | 0.280701294 | 1 | 1 | 1 | 1 | 1 | 1 | 0.691 | 1 | 1 | 1 | 1 | 0.062 |
| SE | 107281 | ENSG00000090686 | USP48 | chr1 | - | 21706743 | 21706907 | 21706466 | 21706589 | 21715388 | 21715457 | 50 | 54 | 53 | 80 | 57 | 3 | 0 | 1 | 4 | 0 | 6 | 78 | 52 | 54 | 8 | 0 | 0 | 0 | 3 | 0 | 298 | 149 | 0.280754848 | 1 | 0.893 | 1 | 0.964 | 0.909 | 1 | 1 | 1 | 1 | 0.9 | 1 | -0.027 |
| SE | 41148 | ENSG00000013573 | DDX11 | chr12 | + | 31084582 | 31084669 | 31083812 | 31084061 | 31087937 | 31087983 | 3 | 1 | 3 | 3 | 10 | 0 | 0 | 0 | 4 | 0 | 21 | 14 | 7 | 37 | 49 | 0 | 0 | 0 | 0 | 0 | 236 | 149 | 0.280771875 | 1 | 1 | 1 | 1 | 0.321 | 1 | 1 | 1 | 1 | 1 | 1 | -0.136 |
| SE | 17956 | ENSG00000171469 | ZNF561 | chr19 | - | 9617048 | 9617171 | 9614020 | 9614103 | 9618090 | 9618179 | 67 | 45 | 47 | 55 | 55 | 7 | 6 | 9 | 4 | 10 | 19 | 66 | 68 | 42 | 15 | 0 | 2 | 12 | 10 | 0 | 272 | 149 | 0.281045035 | 1 | 0.84 | 0.804 | 0.741 | 0.883 | 0.751 | 1 | 0.948 | 0.756 | 0.697 | 1 | -0.076 |
| SE | 12482 | ENSG00000169896 | ITGAM | chr16 | + | 31321463 | 31321627 | 31321240 | 31321371 | 31324398 | 31324553 | 1207 | 402 | 505 | 685 | 671 | 3 | 0 | 0 | 0 | 0 | 1744 | 1160 | 577 | 1419 | 1974 | 2 | 7 | 10 | 11 | 16 | 298 | 149 | 0.281260865 | 1 | 0.995 | 1 | 1 | 1 | 1 | 0.998 | 0.988 | 0.966 | 0.985 | 0.984 | 0.015 |
| SE | 36494 | ENSG00000163626 | COX18 | chr4 | - | 73064777 | 73064899 | 73061812 | 73061920 | 73065249 | 73065413 | 41 | 33 | 45 | 45 | 29 | 0 | 1 | 0 | 2 | 2 | 14 | 26 | 33 | 19 | 23 | 10 | 0 | 0 | 0 | 0 | 271 | 149 | 0.281343619 | 1 | 1 | 0.948 | 1 | 0.925 | 0.889 | 0.435 | 1 | 1 | 1 | 1 | 0.065 |
| SE | 59073 | ENSG00000181409 | AATK | chr17 | - | 81123193 | 81123343 | 81120200 | 81122823 | 81124726 | 81124848 | 20 | 6 | 2 | 14 | 28 | 0 | 0 | 0 | 1 | 12 | 26 | 20 | 17 | 20 | 12 | 18 | 0 | 2 | 0 | 12 | 298 | 149 | 0.281413439 | 1 | 1 | 1 | 1 | 0.875 | 0.538 | 0.419 | 1 | 0.81 | 1 | 0.333 | 0.17 |
| SE | 88969 | ENSG00000148516 | ZEB1 | chr10 | + | 31387719 | 31387798 | 31387123 | 31387266 | 31461036 | 31461237 | 7 | 14 | 11 | 7 | 11 | 32 | 37 | 33 | 29 | 101 | 0 | 5 | 7 | 8 | 8 | 47 | 6 | 15 | 21 | 4 | 228 | 149 | 0.281423944 | 1 | 0.125 | 0.198 | 0.179 | 0.136 | 0.066 | 0 | 0.353 | 0.234 | 0.199 | 0.567 | -0.13 |
| SE | 64629 | ENSG00000148737 | TCF7L2 | chr10 | + | 113160618 | 113160691 | 113158020 | 113158069 | 113165554 | 113165972 | 11 | 9 | 1 | 5 | 16 | 15 | 14 | 2 | 9 | 2 | 31 | 2 | 3 | 26 | 15 | 16 | 12 | 1 | 2 | 9 | 222 | 149 | 0.281452188 | 1 | 0.33 | 0.301 | 0.251 | 0.272 | 0.843 | 0.565 | 0.101 | 0.668 | 0.897 | 0.528 | -0.152 |
| SE | 80417 | ENSG00000117906 | RCN2 | chr15 | + | 76941636 | 76941690 | 76935525 | 76935722 | 76943757 | 76943871 | 1 | 1 | 3 | 3 | 9 | 118 | 73 | 124 | 112 | 180 | 0 | 0 | 0 | 2 | 0 | 59 | 61 | 93 | 55 | 25 | 203 | 149 | 0.28146905 | 1 | 0.006 | 0.01 | 0.017 | 0.019 | 0.035 | 0 | 0 | 0 | 0.026 | 0 | 0.012 |
| SE | 13297 | ENSG00000102974 | CTCF | chr16 | + | 67621441 | 67621591 | 67614962 | 67616878 | 67626554 | 67626715 | 147 | 77 | 122 | 115 | 175 | 0 | 0 | 0 | 3 | 0 | 71 | 93 | 96 | 123 | 54 | 0 | 5 | 4 | 0 | 0 | 298 | 149 | 0.281499803 | 1 | 1 | 1 | 1 | 0.95 | 1 | 1 | 0.903 | 0.923 | 1 | 1 | 0.025 |
| SE | 28620 | ENSG00000102445 | RUBCNL | chr13 | - | 46362537 | 46362598 | 46359485 | 46359631 | 46363114 | 46363213 | 10 | 13 | 14 | 10 | 12 | 1 | 0 | 5 | 0 | 1 | 22 | 29 | 14 | 24 | 10 | 12 | 0 | 0 | 3 | 0 | 210 | 149 | 0.281527875 | 1 | 0.876 | 1 | 0.665 | 1 | 0.895 | 0.565 | 1 | 1 | 0.85 | 1 | 0.004 |
| SE | 22951 | ENSG00000170234 | PWWP2A | chr5 | - | 160089505 | 160089610 | 160077843 | 160078168 | 160093100 | 160093180 | 17 | 10 | 12 | 9 | 15 | 9 | 12 | 14 | 5 | 13 | 21 | 3 | 1 | 2 | 15 | 0 | 5 | 3 | 7 | 0 | 254 | 149 | 0.281535799 | 1 | 0.526 | 0.328 | 0.335 | 0.514 | 0.404 | 1 | 0.26 | 0.164 | 0.144 | 1 | -0.092 |
| SE | 22364 | ENSG00000140455 | USP3 | chr15 | + | 63529013 | 63529166 | 63504611 | 63504830 | 63532646 | 63532707 | 27 | 10 | 25 | 21 | 37 | 89 | 43 | 65 | 58 | 31 | 31 | 23 | 25 | 26 | 32 | 79 | 74 | 96 | 80 | 81 | 298 | 149 | 0.28158198 | 1 | 0.132 | 0.104 | 0.161 | 0.153 | 0.374 | 0.164 | 0.135 | 0.115 | 0.14 | 0.165 | 0.041 |
| SE | 100742 | ENSG00000173992 | CCS | chr11 | + | 66593641 | 66593714 | 66593152 | 66593300 | 66599115 | 66599253 | 49 | 21 | 46 | 39 | 9 | 0 | 0 | 0 | 0 | 0 | 20 | 24 | 11 | 36 | 44 | 0 | 0 | 0 | 0 | 7 | 222 | 149 | 0.281651958 | 1 | 1 | 1 | 1 | 1 | 1 | 1 | 1 | 1 | 1 | 0.808 | 0.038 |
| SE | 39956 | ENSG00000164253 | WDR41 | chr5 | - | 77453816 | 77453910 | 77451290 | 77451353 | 77459061 | 77459124 | 132 | 57 | 51 | 108 | 159 | 0 | 0 | 0 | 0 | 0 | 42 | 68 | 34 | 75 | 22 | 0 | 0 | 2 | 6 | 0 | 243 | 149 | 0.28171734 | 1 | 1 | 1 | 1 | 1 | 1 | 1 | 1 | 0.912 | 0.885 | 1 | 0.041 |
| SE | 80086 | ENSG00000198198 | SZT2 | chr1 | + | 43432061 | 43432180 | 43431715 | 43431901 | 43432271 | 43432439 | 3 | 1 | 4 | 8 | 1 | 21 | 8 | 17 | 11 | 5 | 0 | 8 | 1 | 10 | 8 | 47 | 35 | 17 | 21 | 64 | 268 | 149 | 0.281769312 | 1 | 0.074 | 0.065 | 0.116 | 0.288 | 0.1 | 0 | 0.113 | 0.032 | 0.209 | 0.065 | 0.045 |
| SE | 21516 | ENSG00000137413 | TAF8 | chr6 | + | 42066311 | 42066459 | 42057388 | 42057513 | 42068464 | 42068607 | 163 | 68 | 131 | 129 | 99 | 1 | 2 | 3 | 4 | 0 | 143 | 136 | 104 | 128 | 196 | 0 | 5 | 0 | 1 | 0 | 297 | 149 | 0.281802783 | 1 | 0.988 | 0.945 | 0.956 | 0.942 | 1 | 1 | 0.932 | 1 | 0.985 | 1 | -0.017 |
| SE | 46148 | ENSG00000126070 | AGO3 | chr1 | + | 35984469 | 35984548 | 35982620 | 35982712 | 36004340 | 36004475 | 11 | 6 | 7 | 8 | 8 | 3 | 5 | 1 | 6 | 1 | 0 | 4 | 7 | 0 | 6 | 5 | 0 | 0 | 2 | 0 | 228 | 149 | 0.281822604 | 1 | 0.706 | 0.44 | 0.821 | 0.466 | 0.839 | 0 | 1 | 1 | 0 | 1 | 0.054 |
| SE | 89776 | ENSG00000104490 | NCALD | chr8 | - | 101852571 | 101852670 | 101779923 | 101779979 | 101915808 | 101915858 | 6 | 2 | 0 | 1 | 1 | 10 | 5 | 9 | 9 | 0 | 0 | 1 | 0 | 4 | 0 | 26 | 5 | 10 | 1 | 6 | 248 | 149 | 0.281920563 | 1 | 0.265 | 0.194 | 0 | 0.063 | 1 | 0 | 0.107 | 0 | 0.706 | 0 | 0.142 |
| SE | 52266 | ENSG00000267787 | NA | chr18 | + | 57731413 | 57732023 | 57641558 | 57642052 | 57735150 | 57735592 | 14 | 17 | 12 | 15 | 26 | 1 | 3 | 3 | 6 | 13 | 25 | 20 | 10 | 8 | 20 | 0 | 1 | 9 | 0 | 16 | 298 | 149 | 0.282106021 | 1 | 0.875 | 0.739 | 0.667 | 0.556 | 0.5 | 1 | 0.909 | 0.357 | 1 | 0.385 | -0.063 |
| SE | 103163 | ENSG00000057608 | GDI2 | chr10 | - | 5794884 | 5795019 | 5785851 | 5786050 | 5796762 | 5796862 | 2121 | 1457 | 1652 | 1921 | 3412 | 34 | 17 | 14 | 14 | 39 | 1557 | 1417 | 931 | 1521 | 862 | 4 | 6 | 10 | 41 | 0 | 284 | 149 | 0.282111845 | 1 | 0.97 | 0.978 | 0.984 | 0.986 | 0.979 | 0.995 | 0.992 | 0.98 | 0.951 | 1 | -0.004 |
| SE | 14890 | ENSG00000166855 | CLPX | chr15 | - | 65156777 | 65156932 | 65155691 | 65155856 | 65157745 | 65157910 | 94 | 84 | 79 | 71 | 139 | 0 | 2 | 0 | 7 | 0 | 42 | 55 | 100 | 53 | 14 | 0 | 0 | 0 | 0 | 0 | 298 | 149 | 0.282206528 | 1 | 1 | 0.955 | 1 | 0.835 | 1 | 1 | 1 | 1 | 1 | 1 | -0.042 |
| SE | 102008 | ENSG00000176155 | CCDC57 | chr17 | - | 82179026 | 82179189 | 82178473 | 82178605 | 82183773 | 82183932 | 26 | 39 | 48 | 79 | 47 | 3 | 5 | 4 | 7 | 1 | 133 | 89 | 56 | 97 | 83 | 23 | 6 | 3 | 4 | 14 | 298 | 149 | 0.282213952 | 1 | 0.812 | 0.796 | 0.857 | 0.849 | 0.959 | 0.743 | 0.881 | 0.903 | 0.924 | 0.748 | 0.015 |
| SE | 45319 | ENSG00000116030 | SUMO1 | chr2 | - | 202220031 | 202220106 | 202214356 | 202214434 | 202238439 | 202238462 | 446 | 352 | 337 | 470 | 653 | 6 | 4 | 3 | 5 | 8 | 349 | 391 | 477 | 439 | 53 | 0 | 20 | 3 | 5 | 0 | 224 | 149 | 0.28232657 | 1 | 0.98 | 0.983 | 0.987 | 0.984 | 0.982 | 1 | 0.929 | 0.991 | 0.983 | 1 | 0.003 |
| SE | 9524 | ENSG00000083290 | ULK2 | chr17 | - | 19783696 | 19783905 | 19780985 | 19781104 | 19785936 | 19786005 | 11 | 10 | 14 | 17 | 15 | 0 | 0 | 0 | 0 | 0 | 1 | 6 | 4 | 3 | 20 | 0 | 0 | 4 | 0 | 0 | 298 | 149 | 0.282345256 | 1 | 1 | 1 | 1 | 1 | 1 | 1 | 1 | 0.333 | 1 | 1 | 0.133 |
| SE | 97630 | ENSG00000152558 | TMEM123 | chr11 | - | 102448235 | 102448375 | 102402098 | 102402206 | 102448811 | 102448868 | 6 | 2 | 1 | 13 | 27 | 477 | 413 | 598 | 528 | 949 | 9 | 1 | 8 | 11 | 0 | 92 | 238 | 323 | 216 | 58 | 289 | 149 | 0.28235212 | 1 | 0.006 | 0.002 | 0.001 | 0.013 | 0.014 | 0.048 | 0.002 | 0.013 | 0.026 | 0 | -0.011 |
| SE | 58331 | ENSG00000115207 | GTF3C2 | chr2 | - | 27335597 | 27335706 | 27333973 | 27333999 | 27335916 | 27336028 | 58 | 24 | 55 | 40 | 39 | 0 | 0 | 0 | 0 | 0 | 40 | 41 | 22 | 23 | 40 | 0 | 0 | 2 | 0 | 5 | 258 | 149 | 0.282389359 | 1 | 1 | 1 | 1 | 1 | 1 | 1 | 1 | 0.864 | 1 | 0.822 | 0.063 |
| SE | 98486 | ENSG00000151702 | FLI1 | chr11 | + | 128807179 | 128807239 | 128805365 | 128805431 | 128809156 | 128809204 | 850 | 587 | 719 | 867 | 1026 | 20 | 8 | 14 | 8 | 19 | 1050 | 1063 | 537 | 718 | 603 | 14 | 8 | 7 | 18 | 8 | 209 | 149 | 0.282523306 | 1 | 0.968 | 0.981 | 0.973 | 0.987 | 0.975 | 0.982 | 0.99 | 0.982 | 0.966 | 0.982 | -0.004 |
| SE | 63424 | ENSG00000122507 | BBS9 | chr7 | + | 33352858 | 33352873 | 33351026 | 33351323 | 33357854 | 33357995 | 103 | 64 | 93 | 105 | 240 | 3 | 4 | 2 | 1 | 4 | 22 | 28 | 31 | 51 | 29 | 0 | 2 | 1 | 2 | 0 | 164 | 149 | 0.282596324 | 1 | 0.969 | 0.936 | 0.977 | 0.99 | 0.982 | 1 | 0.927 | 0.966 | 0.959 | 1 | 0 |
| SE | 31308 | ENSG00000013288 | MAN2B2 | chr4 | + | 6609396 | 6609574 | 6609106 | 6609298 | 6609797 | 6610041 | 5 | 1 | 2 | 6 | 4 | 124 | 56 | 75 | 113 | 85 | 0 | 1 | 0 | 0 | 0 | 131 | 164 | 85 | 93 | 176 | 298 | 149 | 0.282689337 | 1 | 0.02 | 0.009 | 0.013 | 0.026 | 0.023 | 0 | 0.003 | 0 | 0 | 0 | 0.018 |
| SE | 92657 | ENSG00000197724 | PHF2 | chr9 | + | 93659510 | 93659600 | 93658144 | 93658236 | 93660191 | 93660560 | 98 | 38 | 88 | 97 | 125 | 0 | 2 | 0 | 1 | 1 | 118 | 95 | 111 | 141 | 222 | 0 | 0 | 4 | 4 | 3 | 239 | 149 | 0.282716469 | 1 | 1 | 0.922 | 1 | 0.984 | 0.987 | 1 | 1 | 0.945 | 0.956 | 0.979 | 0.003 |
| SE | 9811 | ENSG00000172794 | RAB37 | chr17 | + | 74740767 | 74740878 | 74737218 | 74737365 | 74742253 | 74742295 | 267 | 127 | 170 | 198 | 182 | 2 | 3 | 9 | 2 | 1 | 215 | 265 | 95 | 152 | 1026 | 0 | 4 | 0 | 0 | 16 | 260 | 149 | 0.282763781 | 1 | 0.987 | 0.96 | 0.915 | 0.983 | 0.991 | 1 | 0.974 | 1 | 1 | 0.974 | -0.022 |
| SE | 7282 | ENSG00000101019 | UQCC1 | chr20 | - | 35374183 | 35374256 | 35366556 | 35366614 | 35381917 | 35382025 | 148 | 129 | 108 | 127 | 128 | 2 | 3 | 4 | 2 | 1 | 137 | 129 | 165 | 170 | 90 | 0 | 0 | 0 | 3 | 0 | 222 | 149 | 0.282875482 | 1 | 0.98 | 0.967 | 0.948 | 0.977 | 0.988 | 1 | 1 | 1 | 0.974 | 1 | -0.023 |
| SE | 12255 | ENSG00000103313 | MEFV | chr16 | - | 3244472 | 3244588 | 3244135 | 3244286 | 3246524 | 3246547 | 423 | 243 | 219 | 214 | 606 | 1 | 0 | 0 | 1 | 0 | 789 | 332 | 224 | 379 | 637 | 13 | 0 | 2 | 5 | 3 | 265 | 149 | 0.282942984 | 1 | 0.996 | 1 | 1 | 0.992 | 1 | 0.972 | 1 | 0.984 | 0.977 | 0.992 | 0.013 |
| SE | 36573 | ENSG00000147996 | CBWD5 | chr9 | + | 65690555 | 65690601 | 65682765 | 65682857 | 65690966 | 65691026 | 3 | 0 | 1 | 0 | 9 | 77 | 61 | 56 | 47 | 127 | 0 | 2 | 3 | 0 | 0 | 12 | 32 | 29 | 23 | 16 | 195 | 149 | 0.282944913 | 1 | 0.029 | 0 | 0.013 | 0 | 0.051 | 0 | 0.046 | 0.073 | 0 | 0 | -0.005 |
| SE | 72848 | ENSG00000129158 | SERGEF | chr11 | - | 18010070 | 18010139 | 18007940 | 18008076 | 18012850 | 18012898 | 17 | 18 | 12 | 19 | 19 | 1 | 2 | 5 | 9 | 0 | 19 | 6 | 8 | 11 | 12 | 0 | 4 | 0 | 0 | 4 | 218 | 149 | 0.28300138 | 1 | 0.921 | 0.86 | 0.621 | 0.591 | 1 | 1 | 0.506 | 1 | 1 | 0.672 | -0.037 |
| SE | 49691 | ENSG00000263956 | NBPF11 | chr1 | - | 148127140 | 148127288 | 148126828 | 148127038 | 148135123 | 148135247 | 5 | 6 | 5 | 7 | 4 | 0 | 0 | 0 | 0 | 4 | 18 | 20 | 1 | 3 | 5 | 0 | 0 | 0 | 0 | 0 | 297 | 149 | 0.283083082 | 1 | 1 | 1 | 1 | 1 | 0.334 | 1 | 1 | 1 | 1 | 1 | -0.133 |
| SE | 7918 | ENSG00000108953 | YWHAE | chr17 | - | 1361091 | 1361298 | 1344274 | 1345499 | 1364858 | 1365058 | 2 | 1 | 3 | 5 | 12 | 0 | 0 | 0 | 1 | 0 | 7 | 1 | 8 | 9 | 2 | 0 | 3 | 0 | 1 | 0 | 298 | 149 | 0.283180345 | 1 | 1 | 1 | 1 | 0.714 | 1 | 1 | 0.143 | 1 | 0.818 | 1 | 0.151 |
| SE | 65931 | ENSG00000143537 | ADAM15 | chr1 | + | 155055701 | 155055852 | 155054313 | 155054506 | 155055931 | 155056000 | 13 | 9 | 7 | 10 | 2 | 0 | 1 | 0 | 0 | 0 | 18 | 20 | 14 | 20 | 10 | 0 | 4 | 0 | 2 | 0 | 298 | 149 | 0.283281075 | 1 | 1 | 0.818 | 1 | 1 | 1 | 1 | 0.714 | 1 | 0.833 | 1 | 0.054 |
| SE | 68297 | ENSG00000122085 | MTERF4 | chr2 | - | 241097322 | 241097427 | 241095129 | 241096438 | 241099395 | 241099426 | 169 | 141 | 235 | 233 | 205 | 0 | 2 | 3 | 7 | 3 | 155 | 169 | 160 | 244 | 133 | 8 | 0 | 4 | 0 | 7 | 298 | 149 | 0.283304641 | 1 | 1 | 0.972 | 0.975 | 0.943 | 0.972 | 0.906 | 1 | 0.952 | 1 | 0.905 | 0.02 |
| SE | 81937 | ENSG00000182165 | TP53TG1 | chr7 | - | 87345042 | 87345233 | 87341401 | 87341684 | 87345304 | 87345515 | 34 | 16 | 29 | 36 | 24 | 4 | 6 | 4 | 2 | 1 | 11 | 28 | 13 | 40 | 30 | 0 | 1 | 1 | 7 | 2 | 298 | 149 | 0.283421689 | 1 | 0.81 | 0.571 | 0.784 | 0.9 | 0.923 | 1 | 0.933 | 0.867 | 0.741 | 0.882 | -0.087 |
| SE | 42035 | ENSG00000117281 | CD160 | chr1 | - | 145735996 | 145736134 | 145730743 | 145731070 | 145738485 | 145738493 | 278 | 108 | 165 | 248 | 168 | 8 | 4 | 0 | 1 | 0 | 10 | 114 | 25 | 64 | 112 | 0 | 3 | 2 | 0 | 0 | 287 | 149 | 0.283434782 | 1 | 0.947 | 0.933 | 1 | 0.992 | 1 | 1 | 0.952 | 0.866 | 1 | 1 | 0.011 |
| SE | 44717 | ENSG00000143772 | ITPKB | chr1 | - | 226735526 | 226737663 | 226707189 | 226707915 | 226739040 | 226739282 | 326 | 196 | 370 | 352 | 269 | 0 | 0 | 0 | 0 | 1 | 132 | 240 | 241 | 186 | 221 | 0 | 9 | 2 | 2 | 0 | 298 | 149 | 0.28353446 | 1 | 1 | 1 | 1 | 1 | 0.993 | 1 | 0.93 | 0.984 | 0.979 | 1 | 0.02 |
| SE | 23530 | ENSG00000104497 | SNX16 | chr8 | - | 81829429 | 81829516 | 81823791 | 81823940 | 81839611 | 81839751 | 62 | 52 | 52 | 73 | 99 | 8 | 0 | 3 | 0 | 16 | 12 | 44 | 23 | 15 | 15 | 0 | 0 | 8 | 0 | 0 | 236 | 149 | 0.283552435 | 1 | 0.83 | 1 | 0.916 | 1 | 0.796 | 1 | 1 | 0.645 | 1 | 1 | -0.021 |
| SE | 25448 | ENSG00000011243 | AKAP8L | chr19 | - | 15400294 | 15400358 | 15397713 | 15397855 | 15400793 | 15400864 | 33 | 32 | 25 | 42 | 32 | 0 | 0 | 0 | 0 | 0 | 93 | 41 | 60 | 89 | 135 | 0 | 0 | 2 | 8 | 0 | 213 | 149 | 0.283586341 | 1 | 1 | 1 | 1 | 1 | 1 | 1 | 1 | 0.955 | 0.886 | 1 | 0.032 |
| SE | 9237 | ENSG00000178971 | CTC1 | chr17 | - | 8228726 | 8228892 | 8228502 | 8228629 | 8229301 | 8229446 | 61 | 37 | 52 | 49 | 21 | 18 | 11 | 11 | 24 | 16 | 64 | 35 | 22 | 45 | 42 | 9 | 11 | 1 | 12 | 31 | 298 | 149 | 0.283599215 | 1 | 0.629 | 0.627 | 0.703 | 0.505 | 0.396 | 0.78 | 0.614 | 0.917 | 0.652 | 0.404 | -0.101 |
| SE | 98002 | ENSG00000205758 | CRYZL1 | chr21 | - | 33631485 | 33631527 | 33624682 | 33624760 | 33641680 | 33641730 | 54 | 35 | 31 | 41 | 65 | 0 | 0 | 0 | 0 | 0 | 24 | 23 | 23 | 50 | 37 | 8 | 0 | 0 | 0 | 0 | 191 | 149 | 0.283784073 | 1 | 1 | 1 | 1 | 1 | 1 | 0.701 | 1 | 1 | 1 | 1 | 0.06 |
| SE | 63115 | ENSG00000115761 | NOL10 | chr2 | - | 10657741 | 10657891 | 10654480 | 10654547 | 10659171 | 10659250 | 132 | 96 | 123 | 162 | 338 | 3 | 8 | 2 | 2 | 2 | 105 | 131 | 105 | 119 | 55 | 3 | 4 | 9 | 1 | 0 | 298 | 149 | 0.283904442 | 1 | 0.957 | 0.857 | 0.969 | 0.976 | 0.988 | 0.946 | 0.942 | 0.854 | 0.983 | 1 | 0.004 |
| SE | 6278 | ENSG00000260861 | NA | chr20 | - | 1557580 | 1557744 | 1551959 | 1552049 | 1578337 | 1578694 | 3 | 1 | 4 | 3 | 3 | 0 | 1 | 1 | 0 | 0 | 14 | 2 | 2 | 3 | 6 | 0 | 2 | 2 | 0 | 0 | 298 | 149 | 0.283979526 | 1 | 1 | 0.333 | 0.667 | 1 | 1 | 1 | 0.333 | 0.333 | 1 | 1 | 0.067 |
| SE | 32551 | ENSG00000211456 | SACM1L | chr3 | + | 45706779 | 45706907 | 45705134 | 45705209 | 45709497 | 45709643 | 151 | 76 | 148 | 149 | 256 | 5 | 1 | 5 | 1 | 2 | 2 | 39 | 35 | 32 | 22 | 0 | 0 | 5 | 0 | 0 | 277 | 149 | 0.28408946 | 1 | 0.942 | 0.976 | 0.941 | 0.988 | 0.986 | 1 | 1 | 0.79 | 1 | 1 | 0.009 |
| SE | 88462 | ENSG00000152601 | MBNL1 | chr3 | + | 152335110 | 152335230 | 152244320 | 152244440 | 152414940 | 152415109 | 34 | 40 | 31 | 37 | 79 | 0 | 12 | 19 | 18 | 42 | 6 | 12 | 9 | 8 | 12 | 0 | 10 | 1 | 5 | 0 | 269 | 149 | 0.284122154 | 1 | 1 | 0.649 | 0.475 | 0.532 | 0.51 | 1 | 0.399 | 0.833 | 0.47 | 1 | -0.107 |
| SE | 96905 | ENSG00000100731 | PCNX1 | chr14 | + | 71045132 | 71045283 | 71036064 | 71036157 | 71046963 | 71047105 | 131 | 56 | 136 | 130 | 264 | 0 | 0 | 0 | 0 | 0 | 41 | 100 | 102 | 25 | 29 | 0 | 0 | 8 | 0 | 0 | 298 | 149 | 0.284181717 | 1 | 1 | 1 | 1 | 1 | 1 | 1 | 1 | 0.864 | 1 | 1 | 0.027 |
| SE | 71116 | ENSG00000198363 | ASPH | chr8 | - | 61638321 | 61638363 | 61637946 | 61638003 | 61642887 | 61642920 | 84 | 62 | 68 | 82 | 110 | 5 | 2 | 1 | 7 | 17 | 111 | 97 | 95 | 62 | 91 | 0 | 12 | 25 | 3 | 9 | 191 | 149 | 0.284206812 | 1 | 0.929 | 0.96 | 0.981 | 0.901 | 0.835 | 1 | 0.863 | 0.748 | 0.942 | 0.887 | 0.033 |
| SE | 107502 | ENSG00000123349 | PFDN5 | chr12 | + | 53296429 | 53296551 | 53295524 | 53295639 | 53297849 | 53297924 | 17 | 9 | 22 | 11 | 11 | 208 | 128 | 218 | 168 | 123 | 18 | 15 | 6 | 23 | 31 | 212 | 136 | 153 | 132 | 306 | 271 | 149 | 0.284406076 | 1 | 0.043 | 0.037 | 0.053 | 0.035 | 0.047 | 0.045 | 0.057 | 0.021 | 0.087 | 0.053 | -0.01 |
| SE | 82476 | ENSG00000160999 | SH2B2 | chr7 | + | 102300521 | 102301279 | 102286887 | 102287094 | 102306720 | 102306822 | 21 | 6 | 9 | 4 | 2 | 0 | 0 | 0 | 0 | 0 | 12 | 23 | 14 | 11 | 49 | 0 | 2 | 4 | 0 | 0 | 298 | 149 | 0.284712688 | 1 | 1 | 1 | 1 | 1 | 1 | 1 | 0.852 | 0.636 | 1 | 1 | 0.102 |
| SE | 50535 | ENSG00000173436 | MICOS10 | chr1 | + | 19600786 | 19600971 | 19596982 | 19597109 | 19622099 | 19622147 | 0 | 0 | 0 | 1 | 0 | 104 | 65 | 109 | 106 | 119 | 11 | 2 | 2 | 1 | 0 | 98 | 65 | 60 | 99 | 48 | 298 | 149 | 0.28474431 | 1 | 0 | 0 | 0 | 0.005 | 0 | 0.053 | 0.015 | 0.016 | 0.005 | 0 | -0.017 |
| SE | 86065 | ENSG00000164031 | DNAJB14 | chr4 | - | 99927716 | 99927838 | 99923039 | 99923185 | 99930449 | 99930589 | 1 | 3 | 1 | 5 | 20 | 231 | 173 | 159 | 204 | 285 | 0 | 0 | 0 | 2 | 0 | 48 | 58 | 47 | 16 | 8 | 271 | 149 | 0.284747589 | 1 | 0.002 | 0.009 | 0.003 | 0.013 | 0.037 | 0 | 0 | 0 | 0.064 | 0 | 0 |
| SE | 82663 | ENSG00000104447 | TRPS1 | chr8 | - | 115637233 | 115637330 | 115619912 | 115620060 | 115668544 | 115668975 | 5 | 4 | 1 | 1 | 5 | 43 | 27 | 29 | 43 | 30 | 7 | 3 | 4 | 3 | 2 | 13 | 31 | 24 | 25 | 46 | 246 | 149 | 0.284814755 | 1 | 0.066 | 0.082 | 0.02 | 0.014 | 0.092 | 0.246 | 0.055 | 0.092 | 0.068 | 0.026 | -0.043 |
| SE | 34529 | ENSG00000134802 | SLC43A3 | chr11 | - | 57426587 | 57426652 | 57425988 | 57426352 | 57427061 | 57427100 | 8 | 11 | 5 | 18 | 16 | 0 | 1 | 2 | 3 | 0 | 25 | 12 | 4 | 5 | 1 | 0 | 1 | 0 | 0 | 0 | 214 | 149 | 0.284893273 | 1 | 1 | 0.885 | 0.635 | 0.807 | 1 | 1 | 0.893 | 1 | 1 | 1 | -0.113 |
| SE | 52055 | ENSG00000197355 | UAP1L1 | chr9 | + | 137079255 | 137079449 | 137078975 | 137079148 | 137080001 | 137080142 | 40 | 27 | 28 | 47 | 13 | 0 | 0 | 0 | 0 | 0 | 117 | 87 | 41 | 94 | 113 | 0 | 0 | 0 | 0 | 9 | 298 | 149 | 0.285035554 | 1 | 1 | 1 | 1 | 1 | 1 | 1 | 1 | 1 | 1 | 0.863 | 0.027 |
| SE | 9236 | ENSG00000178971 | CTC1 | chr17 | - | 8228726 | 8228892 | 8228502 | 8228629 | 8229141 | 8229206 | 88 | 57 | 61 | 70 | 26 | 6 | 3 | 1 | 5 | 0 | 77 | 44 | 35 | 48 | 76 | 0 | 4 | 0 | 2 | 5 | 298 | 149 | 0.285263867 | 1 | 0.88 | 0.905 | 0.968 | 0.875 | 1 | 1 | 0.846 | 1 | 0.923 | 0.884 | -0.005 |
| SE | 95866 | ENSG00000107263 | RAPGEF1 | chr9 | - | 131650128 | 131650242 | 131638634 | 131638791 | 131650809 | 131650949 | 103 | 62 | 62 | 112 | 137 | 0 | 0 | 5 | 0 | 1 | 177 | 156 | 60 | 71 | 174 | 10 | 0 | 1 | 0 | 0 | 263 | 149 | 0.285320941 | 1 | 1 | 1 | 0.875 | 1 | 0.987 | 0.909 | 1 | 0.971 | 1 | 1 | -0.004 |
| SE | 72537 | ENSG00000135426 | TESPA1 | chr12 | - | 54983993 | 54984166 | 54974419 | 54974607 | 54984584 | 54984731 | 15 | 18 | 52 | 24 | 129 | 78 | 53 | 66 | 125 | 72 | 16 | 10 | 0 | 14 | 10 | 0 | 47 | 40 | 32 | 170 | 298 | 149 | 0.285504731 | 1 | 0.088 | 0.145 | 0.283 | 0.088 | 0.473 | 1 | 0.096 | 0 | 0.179 | 0.029 | -0.045 |
| SE | 103140 | ENSG00000120539 | MASTL | chr10 | + | 27167101 | 27167274 | 27165388 | 27165539 | 27169943 | 27171083 | 23 | 18 | 32 | 45 | 51 | 1 | 0 | 2 | 8 | 1 | 55 | 27 | 44 | 33 | 27 | 0 | 3 | 6 | 2 | 7 | 298 | 149 | 0.285749868 | 1 | 0.92 | 1 | 0.889 | 0.738 | 0.962 | 1 | 0.818 | 0.786 | 0.892 | 0.659 | 0.071 |
| SE | 16989 | ENSG00000248905 | FMN1 | chr15 | - | 33064956 | 33065074 | 33008013 | 33008075 | 33153047 | 33155045 | 2 | 55 | 19 | 21 | 25 | 1 | 2 | 0 | 1 | 0 | 47 | 21 | 47 | 53 | 53 | 0 | 1 | 1 | 0 | 6 | 267 | 149 | 0.285876511 | 1 | 0.527 | 0.939 | 1 | 0.921 | 1 | 1 | 0.921 | 0.963 | 1 | 0.831 | -0.066 |
| SE | 68700 | ENSG00000196295 | GARS1-DT | chr7 | - | 30551317 | 30551395 | 30550567 | 30550781 | 30561465 | 30561516 | 1 | 7 | 7 | 13 | 0 | 31 | 34 | 28 | 48 | 32 | 9 | 4 | 3 | 0 | 0 | 21 | 47 | 44 | 8 | 12 | 227 | 149 | 0.285968197 | 1 | 0.021 | 0.119 | 0.141 | 0.151 | 0 | 0.22 | 0.053 | 0.043 | 0 | 0 | 0.023 |
| SE | 97340 | ENSG00000116747 | RO60 | chr1 | + | 193075819 | 193076040 | 193059453 | 193059776 | 193076500 | 193076647 | 394 | 297 | 217 | 309 | 465 | 1 | 7 | 2 | 6 | 1 | 31 | 219 | 190 | 50 | 55 | 0 | 0 | 0 | 0 | 0 | 298 | 149 | 0.286225137 | 1 | 0.995 | 0.955 | 0.982 | 0.963 | 0.996 | 1 | 1 | 1 | 1 | 1 | -0.022 |
| SE | 14119 | ENSG00000102897 | LYRM1 | chr16 | + | 20915555 | 20915714 | 20900770 | 20900889 | 20920121 | 20920214 | 144 | 100 | 130 | 147 | 203 | 0 | 1 | 2 | 0 | 0 | 149 | 148 | 125 | 118 | 70 | 0 | 0 | 3 | 1 | 6 | 298 | 149 | 0.286303341 | 1 | 1 | 0.98 | 0.97 | 1 | 1 | 1 | 1 | 0.954 | 0.983 | 0.854 | 0.032 |
| SE | 58244 | ENSG00000089127 | OAS1 | chr12 | + | 112911050 | 112911235 | 112906652 | 112907219 | 112916508 | 112916738 | 295 | 196 | 166 | 239 | 184 | 2 | 2 | 4 | 2 | 7 | 874 | 523 | 138 | 574 | 265 | 30 | 4 | 4 | 4 | 2 | 298 | 149 | 0.286339752 | 1 | 0.987 | 0.98 | 0.954 | 0.984 | 0.929 | 0.936 | 0.985 | 0.945 | 0.986 | 0.985 | -0.001 |
| SE | 104767 | ENSG00000011258 | MBTD1 | chr17 | - | 51259142 | 51259206 | 51225007 | 51225209 | 51259834 | 51259941 | 12 | 14 | 12 | 29 | 27 | 3 | 5 | 5 | 4 | 4 | 0 | 22 | 20 | 6 | 12 | 8 | 4 | 5 | 5 | 0 | 213 | 149 | 0.286401846 | 1 | 0.737 | 0.662 | 0.627 | 0.835 | 0.825 | 0 | 0.794 | 0.737 | 0.456 | 1 | 0.14 |
| SE | 82548 | ENSG00000139737 | SLAIN1 | chr13 | + | 77743015 | 77743165 | 77719531 | 77719671 | 77744282 | 77744432 | 34 | 7 | 19 | 23 | 37 | 29 | 36 | 68 | 44 | 53 | 0 | 26 | 10 | 1 | 34 | 24 | 47 | 16 | 30 | 23 | 298 | 149 | 0.28648712 | 1 | 0.37 | 0.089 | 0.123 | 0.207 | 0.259 | 0 | 0.217 | 0.238 | 0.016 | 0.425 | 0.03 |
| SE | 81924 | ENSG00000105819 | PMPCB | chr7 | + | 103298567 | 103298708 | 103297434 | 103297558 | 103299442 | 103299529 | 317 | 232 | 337 | 355 | 332 | 0 | 0 | 0 | 2 | 0 | 260 | 281 | 358 | 257 | 60 | 0 | 0 | 3 | 2 | 8 | 290 | 149 | 0.286563389 | 1 | 1 | 1 | 1 | 0.989 | 1 | 1 | 1 | 0.984 | 0.985 | 0.794 | 0.045 |
| SE | 24634 | ENSG00000196950 | SLC39A10 | chr2 | + | 195706615 | 195706785 | 195683698 | 195683906 | 195708655 | 195708844 | 181 | 115 | 179 | 162 | 288 | 4 | 2 | 1 | 2 | 3 | 29 | 83 | 73 | 30 | 18 | 0 | 4 | 0 | 0 | 0 | 298 | 149 | 0.2866489 | 1 | 0.958 | 0.966 | 0.989 | 0.976 | 0.98 | 1 | 0.912 | 1 | 1 | 1 | -0.009 |
| SE | 48600 | ENSG00000131374 | TBC1D5 | chr3 | - | 17428449 | 17428519 | 17406417 | 17406526 | 17623848 | 17623913 | 37 | 21 | 27 | 41 | 83 | 0 | 0 | 0 | 0 | 4 | 23 | 37 | 17 | 33 | 15 | 0 | 0 | 0 | 7 | 0 | 219 | 149 | 0.286750435 | 1 | 1 | 1 | 1 | 1 | 0.934 | 1 | 1 | 1 | 0.762 | 1 | 0.034 |
| SE | 3471 | ENSG00000090924 | PLEKHG2 | chr19 | + | 39420948 | 39420996 | 39420750 | 39420852 | 39421074 | 39421113 | 23 | 16 | 29 | 21 | 34 | 6 | 2 | 3 | 0 | 0 | 57 | 26 | 31 | 20 | 6 | 16 | 0 | 6 | 0 | 0 | 197 | 149 | 0.286807659 | 1 | 0.744 | 0.858 | 0.88 | 1 | 1 | 0.729 | 1 | 0.796 | 1 | 1 | -0.009 |
| SE | 67707 | ENSG00000145332 | KLHL8 | chr4 | - | 87183202 | 87183389 | 87178476 | 87178620 | 87195323 | 87195690 | 118 | 82 | 38 | 82 | 117 | 13 | 11 | 2 | 1 | 0 | 31 | 91 | 70 | 48 | 19 | 0 | 7 | 22 | 3 | 0 | 298 | 149 | 0.286856618 | 1 | 0.819 | 0.788 | 0.905 | 0.976 | 1 | 1 | 0.867 | 0.614 | 0.889 | 1 | 0.024 |
| SE | 4524 | ENSG00000131355 | ADGRE3 | chr19 | - | 14654981 | 14655165 | 14651084 | 14651204 | 14658512 | 14658550 | 104 | 83 | 44 | 82 | 204 | 1 | 1 | 2 | 1 | 3 | 66 | 15 | 113 | 50 | 64 | 0 | 0 | 4 | 0 | 0 | 298 | 149 | 0.286975527 | 1 | 0.981 | 0.976 | 0.917 | 0.976 | 0.971 | 1 | 1 | 0.934 | 1 | 1 | -0.023 |
| SE | 31582 | ENSG00000129071 | MBD4 | chr3 | - | 129434062 | 129437308 | 129434061 | 129434136 | 129437719 | 129437950 | 296 | 249 | 260 | 362 | 504 | 2 | 5 | 5 | 4 | 1 | 170 | 290 | 248 | 257 | 88 | 0 | 0 | 3 | 2 | 0 | 298 | 149 | 0.287358 | 1 | 0.987 | 0.961 | 0.963 | 0.978 | 0.996 | 1 | 1 | 0.976 | 0.985 | 1 | -0.015 |
| SE | 15694 | ENSG00000174485 | DENND4A | chr15 | - | 65741714 | 65741784 | 65738705 | 65738875 | 65756139 | 65756472 | 99 | 86 | 65 | 105 | 126 | 0 | 0 | 5 | 2 | 0 | 99 | 132 | 93 | 65 | 30 | 0 | 10 | 0 | 0 | 0 | 219 | 149 | 0.28737889 | 1 | 1 | 1 | 0.898 | 0.973 | 1 | 1 | 0.9 | 1 | 1 | 1 | -0.006 |
| SE | 49573 | ENSG00000074964 | ARHGEF10L | chr1 | + | 17648553 | 17648675 | 17640201 | 17640302 | 17654635 | 17654665 | 23 | 24 | 20 | 15 | 35 | 0 | 0 | 0 | 1 | 1 | 56 | 17 | 4 | 27 | 39 | 0 | 0 | 2 | 4 | 0 | 271 | 149 | 0.287405357 | 1 | 1 | 1 | 1 | 0.892 | 0.951 | 1 | 1 | 0.524 | 0.788 | 1 | 0.106 |
| SE | 98711 | ENSG00000140299 | BNIP2 | chr15 | - | 59679591 | 59679768 | 59677812 | 59678087 | 59682407 | 59682502 | 437 | 453 | 340 | 440 | 684 | 0 | 0 | 0 | 0 | 0 | 155 | 355 | 445 | 296 | 179 | 14 | 0 | 0 | 0 | 0 | 298 | 149 | 0.287457782 | 1 | 1 | 1 | 1 | 1 | 1 | 0.847 | 1 | 1 | 1 | 1 | 0.031 |
| SE | 13598 | ENSG00000167261 | DPEP2 | chr16 | - | 67992949 | 67993257 | 67991642 | 67992193 | 67999374 | 67999484 | 13 | 2 | 4 | 1 | 5 | 14 | 13 | 15 | 26 | 32 | 0 | 4 | 8 | 3 | 0 | 9 | 5 | 14 | 8 | 55 | 298 | 149 | 0.287461435 | 1 | 0.317 | 0.071 | 0.118 | 0.019 | 0.072 | 0 | 0.286 | 0.222 | 0.158 | 0 | -0.014 |
| SE | 80700 | ENSG00000175471 | MCTP1 | chr5 | - | 94923961 | 94924021 | 94912805 | 94912976 | 94931952 | 94931991 | 57 | 80 | 25 | 82 | 108 | 27 | 28 | 6 | 29 | 21 | 72 | 35 | 64 | 27 | 9 | 10 | 27 | 7 | 26 | 20 | 209 | 149 | 0.287492963 | 1 | 0.601 | 0.671 | 0.748 | 0.668 | 0.786 | 0.837 | 0.48 | 0.867 | 0.425 | 0.243 | 0.124 |
| SE | 6282 | ENSG00000141644 | MBD1 | chr18 | - | 50271468 | 50271540 | 50268840 | 50269818 | 50272676 | 50272738 | 66 | 29 | 46 | 60 | 56 | 13 | 5 | 24 | 17 | 12 | 61 | 39 | 67 | 66 | 72 | 36 | 26 | 9 | 21 | 20 | 221 | 149 | 0.287524759 | 1 | 0.774 | 0.796 | 0.564 | 0.704 | 0.759 | 0.533 | 0.503 | 0.834 | 0.679 | 0.708 | 0.068 |
| SE | 36012 | ENSG00000258653 | NA | chr14 | + | 73923394 | 73923522 | 73920979 | 73921105 | 73931502 | 73932278 | 113 | 64 | 104 | 109 | 188 | 0 | 0 | 0 | 0 | 2 | 203 | 111 | 111 | 154 | 108 | 11 | 0 | 0 | 0 | 0 | 277 | 149 | 0.28759223 | 1 | 1 | 1 | 1 | 1 | 0.981 | 0.908 | 1 | 1 | 1 | 1 | 0.015 |
| SE | 31790 | ENSG00000204463 | BAG6 | chr6 | - | 31644524 | 31644602 | 31644081 | 31644194 | 31644945 | 31645198 | 116 | 40 | 65 | 84 | 55 | 1 | 0 | 0 | 0 | 2 | 57 | 68 | 30 | 86 | 135 | 0 | 0 | 5 | 4 | 0 | 227 | 149 | 0.287714371 | 1 | 0.987 | 1 | 1 | 1 | 0.948 | 1 | 1 | 0.798 | 0.934 | 1 | 0.041 |
| SE | 109255 | ENSG00000275066 | SYNRG | chr17 | - | 37577379 | 37577613 | 37576340 | 37576418 | 37579131 | 37579434 | 246 | 143 | 263 | 236 | 422 | 4 | 4 | 25 | 28 | 6 | 173 | 214 | 137 | 150 | 108 | 8 | 3 | 12 | 4 | 5 | 298 | 149 | 0.287894079 | 1 | 0.969 | 0.947 | 0.84 | 0.808 | 0.972 | 0.915 | 0.973 | 0.851 | 0.949 | 0.915 | -0.013 |
| SE | 32191 | ENSG00000113971 | NPHP3 | chr3 | - | 132684553 | 132684794 | 132682702 | 132682818 | 132686259 | 132686387 | 20 | 18 | 22 | 13 | 28 | 0 | 1 | 0 | 0 | 0 | 1 | 35 | 16 | 7 | 11 | 0 | 5 | 0 | 1 | 0 | 298 | 149 | 0.28796363 | 1 | 1 | 0.9 | 1 | 1 | 1 | 1 | 0.778 | 1 | 0.778 | 1 | 0.069 |
| SE | 40462 | ENSG00000184432 | COPB2 | chr3 | - | 139385003 | 139385200 | 139383297 | 139383435 | 139389547 | 139389560 | 16 | 6 | 29 | 10 | 7 | 146 | 52 | 93 | 96 | 87 | 0 | 15 | 21 | 8 | 0 | 32 | 59 | 23 | 72 | 6 | 298 | 149 | 0.288033712 | 1 | 0.052 | 0.055 | 0.135 | 0.05 | 0.039 | 0 | 0.113 | 0.313 | 0.053 | 0 | -0.03 |
| SE | 13291 | ENSG00000102974 | CTCF | chr16 | + | 67611950 | 67612121 | 67570772 | 67571264 | 67616744 | 67616878 | 361 | 259 | 336 | 345 | 649 | 0 | 0 | 0 | 0 | 0 | 225 | 252 | 193 | 204 | 104 | 0 | 2 | 0 | 0 | 10 | 298 | 149 | 0.288125446 | 1 | 1 | 1 | 1 | 1 | 1 | 1 | 0.984 | 1 | 1 | 0.839 | 0.035 |
| SE | 88925 | ENSG00000164823 | OSGIN2 | chr8 | + | 89914554 | 89914746 | 89909566 | 89909721 | 89921079 | 89921171 | 35 | 44 | 41 | 56 | 87 | 0 | 0 | 0 | 0 | 0 | 44 | 19 | 63 | 38 | 7 | 0 | 6 | 0 | 0 | 0 | 298 | 149 | 0.288246825 | 1 | 1 | 1 | 1 | 1 | 1 | 1 | 0.613 | 1 | 1 | 1 | 0.077 |
| SE | 46074 | ENSG00000068097 | HEATR6 | chr17 | - | 60072214 | 60072329 | 60070705 | 60070807 | 60078695 | 60078921 | 50 | 13 | 32 | 37 | 35 | 0 | 0 | 1 | 0 | 0 | 10 | 23 | 24 | 19 | 38 | 0 | 0 | 7 | 0 | 0 | 264 | 149 | 0.288279809 | 1 | 1 | 1 | 0.948 | 1 | 1 | 1 | 1 | 0.659 | 1 | 1 | 0.058 |
| SE | 107069 | ENSG00000136521 | NDUFB5 | chr3 | + | 179618414 | 179618521 | 179616974 | 179617044 | 179623919 | 179624091 | 436 | 278 | 381 | 444 | 771 | 0 | 0 | 1 | 0 | 1 | 535 | 366 | 308 | 430 | 364 | 18 | 3 | 0 | 0 | 0 | 256 | 149 | 0.288359771 | 1 | 1 | 1 | 0.996 | 1 | 0.998 | 0.945 | 0.986 | 1 | 1 | 1 | 0.013 |
| SE | 3657 | ENSG00000142230 | SAE1 | chr19 | + | 47197232 | 47197377 | 47169817 | 47169923 | 47203670 | 47203740 | 272 | 200 | 293 | 296 | 394 | 9 | 1 | 11 | 9 | 9 | 427 | 200 | 198 | 226 | 478 | 6 | 8 | 7 | 10 | 0 | 294 | 149 | 0.288388726 | 1 | 0.939 | 0.99 | 0.931 | 0.943 | 0.957 | 0.973 | 0.927 | 0.935 | 0.92 | 1 | 0.001 |
| SE | 43956 | ENSG00000115165 | CYTIP | chr2 | - | 157434369 | 157434424 | 157430859 | 157430962 | 157434697 | 157434747 | 1531 | 1066 | 1410 | 1520 | 1661 | 10 | 8 | 11 | 16 | 15 | 476 | 1081 | 1981 | 581 | 404 | 6 | 7 | 15 | 3 | 17 | 204 | 149 | 0.288552154 | 1 | 0.991 | 0.99 | 0.989 | 0.986 | 0.988 | 0.983 | 0.991 | 0.99 | 0.993 | 0.946 | 0.008 |
| SE | 70979 | ENSG00000127081 | ZNF484 | chr9 | - | 92862139 | 92862165 | 92856191 | 92856318 | 92875014 | 92875059 | 8 | 10 | 11 | 8 | 2 | 24 | 11 | 13 | 11 | 20 | 16 | 4 | 4 | 0 | 0 | 9 | 21 | 3 | 9 | 9 | 175 | 149 | 0.28862335 | 1 | 0.221 | 0.436 | 0.419 | 0.382 | 0.078 | 0.602 | 0.14 | 0.532 | 0 | 0 | 0.052 |
| SE | 22162 | ENSG00000214160 | ALG3 | chr3 | - | 184245197 | 184245358 | 184244600 | 184244721 | 184245467 | 184245615 | 19 | 9 | 13 | 16 | 20 | 0 | 0 | 0 | 0 | 0 | 19 | 27 | 10 | 20 | 21 | 0 | 7 | 0 | 0 | 0 | 298 | 149 | 0.288661333 | 1 | 1 | 1 | 1 | 1 | 1 | 1 | 0.659 | 1 | 1 | 1 | 0.068 |
| SE | 747 | ENSG00000099991 | CABIN1 | chr22 | + | 24168446 | 24168521 | 24166638 | 24167313 | 24171712 | 24171995 | 87 | 42 | 54 | 97 | 67 | 0 | 0 | 1 | 5 | 0 | 115 | 51 | 26 | 78 | 102 | 0 | 2 | 0 | 3 | 4 | 224 | 149 | 0.288672725 | 1 | 1 | 1 | 0.973 | 0.928 | 1 | 1 | 0.944 | 1 | 0.945 | 0.944 | 0.014 |
| SE | 63420 | ENSG00000122507 | BBS9 | chr7 | + | 33351026 | 33351323 | 33349067 | 33349170 | 33352858 | 33352873 | 87 | 57 | 71 | 86 | 172 | 1 | 3 | 3 | 1 | 1 | 18 | 23 | 19 | 40 | 17 | 0 | 0 | 0 | 0 | 0 | 298 | 149 | 0.288692624 | 1 | 0.978 | 0.905 | 0.922 | 0.977 | 0.989 | 1 | 1 | 1 | 1 | 1 | -0.046 |
| SE | 47987 | ENSG00000154710 | RABGEF1 | chr7 | + | 66783674 | 66783841 | 66775226 | 66775393 | 66797373 | 66797506 | 70 | 63 | 71 | 100 | 70 | 1 | 0 | 0 | 2 | 0 | 38 | 127 | 110 | 144 | 18 | 0 | 0 | 1 | 0 | 6 | 298 | 149 | 0.288703885 | 1 | 0.972 | 1 | 1 | 0.962 | 1 | 1 | 1 | 0.982 | 1 | 0.6 | 0.07 |
| SE | 79816 | ENSG00000136286 | MYO1G | chr7 | - | 44976563 | 44976657 | 44975475 | 44975649 | 44978866 | 44979069 | 538 | 299 | 376 | 482 | 286 | 18 | 4 | 9 | 13 | 1 | 1410 | 983 | 515 | 706 | 1462 | 12 | 6 | 15 | 23 | 29 | 243 | 149 | 0.288735775 | 1 | 0.948 | 0.979 | 0.962 | 0.958 | 0.994 | 0.986 | 0.99 | 0.955 | 0.95 | 0.969 | -0.002 |
| SE | 56598 | ENSG00000075151 | EIF4G3 | chr1 | - | 20810737 | 20810884 | 20806291 | 20807500 | 20813157 | 20813239 | 214 | 194 | 100 | 205 | 181 | 5 | 13 | 10 | 16 | 16 | 74 | 183 | 137 | 203 | 23 | 9 | 12 | 12 | 17 | 3 | 296 | 149 | 0.288752572 | 1 | 0.956 | 0.883 | 0.834 | 0.866 | 0.851 | 0.805 | 0.885 | 0.852 | 0.857 | 0.794 | 0.039 |
| SE | 21885 | ENSG00000189091 | SF3B3 | chr16 | + | 70561629 | 70561784 | 70556885 | 70557029 | 70563875 | 70564033 | 271 | 126 | 267 | 300 | 268 | 0 | 0 | 2 | 2 | 11 | 362 | 334 | 180 | 151 | 347 | 3 | 5 | 3 | 6 | 5 | 298 | 149 | 0.288785632 | 1 | 1 | 1 | 0.985 | 0.987 | 0.924 | 0.984 | 0.971 | 0.968 | 0.926 | 0.972 | 0.015 |
| SE | 1482 | ENSG00000100368 | CSF2RB | chr22 | + | 36930654 | 36930830 | 36930374 | 36930510 | 36932764 | 36932904 | 116 | 46 | 110 | 113 | 71 | 1 | 0 | 2 | 0 | 0 | 158 | 39 | 110 | 150 | 131 | 10 | 0 | 0 | 0 | 0 | 298 | 149 | 0.288795352 | 1 | 0.983 | 1 | 0.965 | 1 | 1 | 0.888 | 1 | 1 | 1 | 1 | 0.012 |
| SE | 41233 | ENSG00000153815 | CMIP | chr16 | + | 81705498 | 81705604 | 81703856 | 81704085 | 81707013 | 81707084 | 177 | 122 | 151 | 183 | 79 | 3 | 1 | 0 | 1 | 11 | 425 | 291 | 174 | 322 | 1071 | 0 | 1 | 5 | 13 | 9 | 255 | 149 | 0.288824134 | 1 | 0.972 | 0.986 | 1 | 0.991 | 0.808 | 1 | 0.994 | 0.953 | 0.935 | 0.986 | -0.022 |
| SE | 64388 | ENSG00000101577 | LPIN2 | chr18 | - | 2923774 | 2923861 | 2922046 | 2922199 | 2924397 | 2924546 | 372 | 186 | 328 | 263 | 318 | 2 | 0 | 2 | 3 | 0 | 137 | 181 | 178 | 155 | 156 | 0 | 6 | 7 | 0 | 0 | 236 | 149 | 0.288876221 | 1 | 0.992 | 1 | 0.99 | 0.982 | 1 | 1 | 0.95 | 0.941 | 1 | 1 | 0.015 |
| SE | 75446 | ENSG00000135049 | AGTPBP1 | chr9 | - | 85678334 | 85678398 | 85677435 | 85677582 | 85681267 | 85681335 | 598 | 401 | 413 | 484 | 660 | 17 | 5 | 5 | 9 | 13 | 235 | 409 | 446 | 285 | 208 | 5 | 4 | 11 | 5 | 0 | 213 | 149 | 0.288925985 | 1 | 0.961 | 0.982 | 0.983 | 0.974 | 0.973 | 0.97 | 0.986 | 0.966 | 0.976 | 1 | -0.005 |
| SE | 8646 | ENSG00000261371 | PECAM1 | chr17 | - | 64356110 | 64356398 | 64353490 | 64353518 | 64363148 | 64363397 | 13 | 9 | 12 | 10 | 27 | 0 | 0 | 0 | 2 | 0 | 137 | 46 | 32 | 67 | 120 | 0 | 0 | 0 | 0 | 9 | 298 | 149 | 0.289013738 | 1 | 1 | 1 | 1 | 0.714 | 1 | 1 | 1 | 1 | 1 | 0.87 | -0.031 |
| SE | 108800 | ENSG00000105983 | LMBR1 | chr7 | - | 156779654 | 156779732 | 156756392 | 156756465 | 156796388 | 156796492 | 4 | 3 | 3 | 3 | 10 | 0 | 4 | 0 | 0 | 0 | 11 | 18 | 4 | 2 | 16 | 0 | 0 | 0 | 0 | 0 | 227 | 149 | 0.289037709 | 1 | 1 | 0.33 | 1 | 1 | 1 | 1 | 1 | 1 | 1 | 1 | -0.134 |
| SE | 55327 | ENSG00000132635 | PCED1A | chr20 | - | 2836038 | 2836314 | 2835313 | 2835709 | 2838231 | 2838478 | 101 | 53 | 90 | 116 | 99 | 0 | 1 | 5 | 2 | 2 | 102 | 73 | 87 | 75 | 96 | 0 | 0 | 0 | 0 | 7 | 298 | 149 | 0.28912769 | 1 | 1 | 0.964 | 0.9 | 0.967 | 0.961 | 1 | 1 | 1 | 1 | 0.873 | -0.016 |
| SE | 48005 | ENSG00000010810 | FYN | chr6 | - | 111699514 | 111699670 | 111696276 | 111696456 | 111700103 | 111700268 | 524 | 318 | 353 | 480 | 656 | 69 | 31 | 58 | 52 | 90 | 444 | 399 | 191 | 320 | 594 | 35 | 69 | 52 | 18 | 33 | 298 | 149 | 0.289159611 | 1 | 0.792 | 0.837 | 0.753 | 0.822 | 0.785 | 0.864 | 0.743 | 0.647 | 0.899 | 0.9 | -0.013 |
| SE | 28914 | ENSG00000145495 | MARCHF6 | chr5 | + | 10411332 | 10411537 | 10410138 | 10410276 | 10414432 | 10414502 | 551 | 377 | 534 | 562 | 469 | 2 | 0 | 1 | 1 | 0 | 368 | 568 | 429 | 340 | 484 | 0 | 4 | 12 | 4 | 0 | 298 | 149 | 0.289244243 | 1 | 0.993 | 1 | 0.996 | 0.996 | 1 | 1 | 0.986 | 0.947 | 0.977 | 1 | 0.015 |
| SE | 87541 | ENSG00000147050 | KDM6A | chrX | + | 45060608 | 45060764 | 45059246 | 45059466 | 45061323 | 45061419 | 83 | 49 | 45 | 57 | 112 | 11 | 12 | 16 | 14 | 32 | 56 | 65 | 82 | 65 | 36 | 0 | 12 | 17 | 21 | 11 | 298 | 149 | 0.289263011 | 1 | 0.79 | 0.671 | 0.584 | 0.671 | 0.636 | 1 | 0.73 | 0.707 | 0.607 | 0.621 | -0.063 |
| SE | 51690 | ENSG00000198837 | DENND4B | chr1 | - | 153941868 | 153942113 | 153941373 | 153941440 | 153942877 | 153943130 | 34 | 9 | 19 | 24 | 12 | 0 | 0 | 0 | 0 | 0 | 64 | 40 | 13 | 32 | 79 | 0 | 1 | 0 | 6 | 0 | 298 | 149 | 0.289321569 | 1 | 1 | 1 | 1 | 1 | 1 | 1 | 0.952 | 1 | 0.727 | 1 | 0.064 |
| SE | 60202 | ENSG00000163534 | FCRL1 | chr1 | - | 157803844 | 157804111 | 157802376 | 157802664 | 157807101 | 157807122 | 76 | 60 | 77 | 91 | 72 | 4 | 0 | 0 | 0 | 2 | 48 | 249 | 49 | 45 | 72 | 0 | 10 | 0 | 0 | 0 | 298 | 149 | 0.289414051 | 1 | 0.905 | 1 | 1 | 1 | 0.947 | 1 | 0.926 | 1 | 1 | 1 | -0.015 |
| SE | 101470 | ENSG00000148481 | MINDY3 | chr10 | - | 15847863 | 15847943 | 15843211 | 15843272 | 15860205 | 15860498 | 184 | 128 | 171 | 195 | 118 | 4 | 2 | 2 | 1 | 0 | 125 | 128 | 126 | 75 | 129 | 0 | 5 | 6 | 0 | 0 | 229 | 149 | 0.289501509 | 1 | 0.968 | 0.977 | 0.982 | 0.992 | 1 | 1 | 0.943 | 0.932 | 1 | 1 | 0.009 |
| SE | 73768 | ENSG00000146285 | SCML4 | chr6 | - | 107772171 | 107772386 | 107749683 | 107749813 | 107824125 | 107824313 | 84 | 53 | 101 | 66 | 34 | 14 | 0 | 17 | 10 | 3 | 23 | 28 | 26 | 21 | 31 | 0 | 6 | 4 | 6 | 10 | 298 | 149 | 0.28964893 | 1 | 0.75 | 1 | 0.748 | 0.767 | 0.85 | 1 | 0.7 | 0.765 | 0.636 | 0.608 | 0.081 |
| SE | 104472 | ENSG00000228063 | LYPLAL1-DT | chr1 | - | 219156119 | 219156199 | 219149938 | 219150031 | 219173469 | 219173493 | 4 | 48 | 6 | 122 | 52 | 0 | 0 | 0 | 4 | 2 | 96 | 68 | 20 | 32 | 2 | 0 | 0 | 0 | 4 | 0 | 229 | 149 | 0.289731983 | 1 | 1 | 1 | 1 | 0.952 | 0.944 | 1 | 1 | 1 | 0.839 | 1 | 0.011 |
| SE | 105860 | ENSG00000090006 | LTBP4 | chr19 | + | 40616888 | 40617020 | 40614191 | 40614446 | 40619346 | 40619382 | 15 | 18 | 20 | 11 | 17 | 6 | 3 | 16 | 4 | 2 | 27 | 11 | 21 | 2 | 11 | 0 | 7 | 10 | 8 | 19 | 281 | 149 | 0.289891781 | 1 | 0.57 | 0.761 | 0.399 | 0.593 | 0.818 | 1 | 0.455 | 0.527 | 0.117 | 0.235 | 0.161 |
| SE | 74115 | ENSG00000093167 | LRRFIP2 | chr3 | - | 37091466 | 37091538 | 37083635 | 37083806 | 37094791 | 37094908 | 98 | 115 | 94 | 98 | 124 | 8 | 4 | 15 | 6 | 6 | 173 | 111 | 138 | 152 | 81 | 0 | 0 | 22 | 5 | 11 | 221 | 149 | 0.289992426 | 1 | 0.892 | 0.951 | 0.809 | 0.917 | 0.933 | 1 | 1 | 0.809 | 0.953 | 0.832 | -0.018 |
| SE | 52610 | ENSG00000164983 | TMEM65 | chr8 | - | 124327353 | 124327421 | 124323320 | 124323375 | 124371853 | 124372701 | 48 | 27 | 47 | 45 | 53 | 2 | 2 | 0 | 4 | 0 | 24 | 15 | 32 | 14 | 28 | 0 | 0 | 0 | 0 | 0 | 217 | 149 | 0.290015272 | 1 | 0.943 | 0.903 | 1 | 0.885 | 1 | 1 | 1 | 1 | 1 | 1 | -0.054 |
| SE | 72832 | ENSG00000129158 | SERGEF | chr11 | - | 17959469 | 17959636 | 17878207 | 17878244 | 17988596 | 17988755 | 96 | 42 | 70 | 73 | 105 | 36 | 20 | 26 | 26 | 35 | 47 | 55 | 63 | 27 | 145 | 47 | 19 | 29 | 11 | 45 | 298 | 149 | 0.290022356 | 1 | 0.571 | 0.512 | 0.574 | 0.584 | 0.6 | 0.333 | 0.591 | 0.521 | 0.551 | 0.617 | 0.046 |
| SE | 17697 | ENSG00000119682 | AREL1 | chr14 | - | 74692040 | 74692328 | 74685599 | 74685660 | 74712932 | 74713067 | 110 | 66 | 85 | 83 | 57 | 0 | 0 | 0 | 0 | 0 | 38 | 69 | 125 | 93 | 125 | 0 | 0 | 0 | 0 | 9 | 298 | 149 | 0.290059122 | 1 | 1 | 1 | 1 | 1 | 1 | 1 | 1 | 1 | 1 | 0.874 | 0.025 |
| SE | 36074 | ENSG00000155657 | TTN | chr2 | - | 178723475 | 178724143 | 178723417 | 178723696 | 178725367 | 178725649 | 4 | 31 | 18 | 34 | 19 | 2 | 1 | 0 | 3 | 0 | 7 | 15 | 12 | 8 | 9 | 0 | 0 | 0 | 0 | 0 | 298 | 149 | 0.290077644 | 1 | 0.5 | 0.939 | 1 | 0.85 | 1 | 1 | 1 | 1 | 1 | 1 | -0.142 |
| SE | 51843 | ENSG00000087206 | UIMC1 | chr5 | - | 176969052 | 176969291 | 176958092 | 176958154 | 176970741 | 176970866 | 13 | 15 | 6 | 12 | 20 | 8 | 3 | 5 | 6 | 20 | 34 | 18 | 17 | 27 | 48 | 3 | 10 | 22 | 4 | 25 | 298 | 149 | 0.290092624 | 1 | 0.448 | 0.714 | 0.375 | 0.5 | 0.333 | 0.85 | 0.474 | 0.279 | 0.771 | 0.49 | -0.099 |
| SE | 82974 | ENSG00000186088 | GSAP | chr7 | - | 77312100 | 77312202 | 77311840 | 77311940 | 77313487 | 77313549 | 130 | 100 | 87 | 76 | 124 | 2 | 0 | 0 | 1 | 1 | 22 | 51 | 27 | 7 | 16 | 7 | 0 | 0 | 0 | 0 | 251 | 149 | 0.290213989 | 1 | 0.975 | 1 | 0.981 | 1 | 0.987 | 0.651 | 1 | 1 | 1 | 1 | 0.058 |
| SE | 100186 | ENSG00000078124 | ACER3 | chr11 | + | 77011329 | 77011375 | 76998762 | 76998821 | 77015015 | 77015117 | 3 | 8 | 6 | 5 | 4 | 45 | 44 | 28 | 39 | 94 | 3 | 4 | 1 | 0 | 9 | 15 | 53 | 35 | 79 | 36 | 195 | 149 | 0.290233279 | 1 | 0.048 | 0.122 | 0.141 | 0.089 | 0.031 | 0.133 | 0.055 | 0.021 | 0 | 0.16 | 0.012 |
| SE | 68360 | ENSG00000114988 | LMAN2L | chr2 | - | 96733518 | 96733601 | 96711863 | 96712025 | 96734408 | 96734526 | 25 | 11 | 22 | 23 | 43 | 0 | 0 | 0 | 1 | 0 | 34 | 36 | 10 | 9 | 21 | 0 | 0 | 0 | 0 | 7 | 232 | 149 | 0.290237817 | 1 | 1 | 1 | 1 | 0.937 | 1 | 1 | 1 | 1 | 1 | 0.658 | 0.056 |
| SE | 74358 | ENSG00000112787 | FBRSL1 | chr12 | + | 132507251 | 132507388 | 132490150 | 132490861 | 132508152 | 132508350 | 2 | 3 | 1 | 3 | 3 | 26 | 9 | 16 | 11 | 6 | 6 | 2 | 0 | 6 | 5 | 10 | 2 | 6 | 8 | 62 | 286 | 149 | 0.290294866 | 1 | 0.039 | 0.148 | 0.032 | 0.124 | 0.207 | 0.238 | 0.343 | 0 | 0.281 | 0.04 | -0.07 |
| SE | 75568 | ENSG00000204569 | PPP1R10 | chr6 | - | 30607839 | 30607891 | 30606778 | 30606856 | 30608778 | 30608914 | 94 | 50 | 85 | 78 | 70 | 0 | 1 | 0 | 0 | 0 | 20 | 90 | 19 | 30 | 57 | 7 | 0 | 0 | 0 | 0 | 201 | 149 | 0.290297735 | 1 | 1 | 0.974 | 1 | 1 | 1 | 0.679 | 1 | 1 | 1 | 1 | 0.059 |
| SE | 106964 | ENSG00000111271 | ACAD10 | chr12 | + | 111710272 | 111710365 | 111709525 | 111709684 | 111712497 | 111712657 | 5 | 0 | 6 | 9 | 17 | 26 | 24 | 34 | 46 | 39 | 15 | 0 | 7 | 1 | 14 | 78 | 26 | 21 | 44 | 64 | 242 | 149 | 0.290368661 | 1 | 0.106 | 0 | 0.098 | 0.108 | 0.212 | 0.106 | 0 | 0.17 | 0.014 | 0.119 | 0.023 |
| SE | 78392 | ENSG00000169919 | GUSB | chr7 | - | 65974914 | 65975071 | 65974525 | 65974704 | 65976014 | 65976202 | 68 | 22 | 51 | 71 | 45 | 25 | 4 | 15 | 13 | 13 | 68 | 126 | 69 | 104 | 113 | 52 | 40 | 19 | 21 | 35 | 298 | 149 | 0.290377552 | 1 | 0.576 | 0.733 | 0.63 | 0.732 | 0.634 | 0.395 | 0.612 | 0.645 | 0.712 | 0.617 | 0.065 |
| SE | 6580 | ENSG00000074657 | ZNF532 | chr18 | + | 58946066 | 58948229 | 58918270 | 58920633 | 58953517 | 58953799 | 60 | 39 | 33 | 56 | 50 | 2 | 1 | 2 | 1 | 0 | 115 | 51 | 54 | 87 | 15 | 0 | 0 | 0 | 5 | 0 | 298 | 149 | 0.290390472 | 1 | 0.938 | 0.951 | 0.892 | 0.966 | 1 | 1 | 1 | 1 | 0.897 | 1 | -0.03 |
| SE | 13197 | ENSG00000140836 | ZFHX3 | chr16 | - | 72957426 | 72960194 | 72950468 | 72950965 | 73047751 | 73048128 | 24 | 19 | 21 | 33 | 5 | 1 | 0 | 0 | 0 | 0 | 50 | 12 | 10 | 11 | 27 | 0 | 0 | 0 | 0 | 7 | 298 | 149 | 0.290445666 | 1 | 0.923 | 1 | 1 | 1 | 1 | 1 | 1 | 1 | 1 | 0.659 | 0.053 |
| SE | 13855 | ENSG00000102893 | PHKB | chr16 | + | 47594136 | 47594214 | 47593499 | 47593557 | 47596372 | 47596531 | 324 | 301 | 237 | 369 | 434 | 7 | 4 | 1 | 4 | 4 | 275 | 199 | 258 | 251 | 212 | 0 | 5 | 1 | 2 | 1 | 227 | 149 | 0.290490328 | 1 | 0.968 | 0.98 | 0.994 | 0.984 | 0.986 | 1 | 0.963 | 0.994 | 0.988 | 0.993 | -0.005 |
| SE | 49092 | ENSG00000072736 | NFATC3 | chr16 | + | 68166842 | 68167015 | 68157868 | 68158068 | 68174373 | 68174514 | 765 | 410 | 627 | 698 | 1091 | 0 | 0 | 6 | 7 | 8 | 534 | 696 | 386 | 458 | 349 | 0 | 9 | 5 | 8 | 0 | 298 | 149 | 0.290605602 | 1 | 1 | 1 | 0.981 | 0.98 | 0.986 | 1 | 0.975 | 0.975 | 0.966 | 1 | 0.006 |
| SE | 46947 | ENSG00000141027 | NCOR1 | chr17 | - | 16151589 | 16151616 | 16149450 | 16149517 | 16151945 | 16151998 | 85 | 74 | 56 | 98 | 96 | 260 | 185 | 207 | 278 | 249 | 68 | 65 | 46 | 107 | 36 | 217 | 240 | 248 | 278 | 77 | 176 | 149 | 0.290643462 | 1 | 0.217 | 0.253 | 0.186 | 0.23 | 0.246 | 0.21 | 0.187 | 0.136 | 0.246 | 0.284 | 0.014 |
| SE | 60896 | ENSG00000144674 | GOLGA4 | chr3 | + | 37286013 | 37286061 | 37281957 | 37282272 | 37294978 | 37295077 | 31 | 27 | 20 | 19 | 29 | 2 | 3 | 1 | 3 | 1 | 3 | 12 | 6 | 0 | 12 | 0 | 0 | 0 | 3 | 0 | 197 | 149 | 0.290684441 | 1 | 0.921 | 0.872 | 0.938 | 0.827 | 0.956 | 1 | 1 | 1 | 0 | 1 | 0.103 |
| SE | 25455 | ENSG00000011243 | AKAP8L | chr19 | - | 15403657 | 15403715 | 15403474 | 15403555 | 15404009 | 15404042 | 61 | 40 | 96 | 99 | 62 | 4 | 0 | 3 | 0 | 1 | 74 | 97 | 77 | 67 | 119 | 0 | 0 | 5 | 3 | 5 | 207 | 149 | 0.290816162 | 1 | 0.917 | 1 | 0.958 | 1 | 0.978 | 1 | 1 | 0.917 | 0.941 | 0.945 | 0.01 |
| SE | 38562 | ENSG00000165801 | ARHGEF40 | chr14 | + | 21087319 | 21087463 | 21087000 | 21087105 | 21087967 | 21088098 | 97 | 25 | 36 | 58 | 68 | 3 | 3 | 4 | 4 | 0 | 52 | 49 | 49 | 82 | 141 | 6 | 4 | 4 | 6 | 4 | 293 | 149 | 0.290846445 | 1 | 0.943 | 0.809 | 0.821 | 0.881 | 1 | 0.815 | 0.862 | 0.862 | 0.874 | 0.947 | 0.019 |
| SE | 81329 | ENSG00000135164 | DMTF1 | chr7 | + | 87163238 | 87163332 | 87155454 | 87155497 | 87163494 | 87163539 | 5 | 0 | 1 | 5 | 6 | 0 | 1 | 0 | 0 | 0 | 0 | 2 | 2 | 5 | 3 | 3 | 0 | 0 | 0 | 0 | 243 | 149 | 0.290850519 | 1 | 1 | 0 | 1 | 1 | 1 | 0 | 1 | 1 | 1 | 1 | 0 |
| SE | 68833 | ENSG00000113456 | RAD1 | chr5 | - | 34913463 | 34913578 | 34911553 | 34911812 | 34914694 | 34914961 | 38 | 28 | 47 | 38 | 81 | 8 | 20 | 6 | 14 | 22 | 67 | 11 | 11 | 10 | 10 | 0 | 30 | 9 | 1 | 0 | 258 | 149 | 0.291111884 | 1 | 0.733 | 0.447 | 0.819 | 0.611 | 0.68 | 1 | 0.175 | 0.414 | 0.852 | 1 | -0.03 |
| SE | 82469 | ENSG00000119471 | HSDL2 | chr9 | + | 112438430 | 112438625 | 112418859 | 112418958 | 112441698 | 112441770 | 103 | 101 | 88 | 102 | 193 | 2 | 4 | 0 | 1 | 9 | 6 | 80 | 93 | 66 | 22 | 0 | 6 | 2 | 5 | 0 | 298 | 149 | 0.291116275 | 1 | 0.963 | 0.927 | 1 | 0.981 | 0.915 | 1 | 0.87 | 0.959 | 0.868 | 1 | 0.018 |
| SE | 102790 | ENSG00000198825 | INPP5F | chr10 | + | 119781634 | 119781771 | 119757891 | 119757977 | 119791516 | 119791645 | 8 | 17 | 16 | 19 | 16 | 0 | 0 | 0 | 0 | 0 | 7 | 11 | 9 | 4 | 17 | 0 | 0 | 0 | 4 | 0 | 286 | 149 | 0.291455114 | 1 | 1 | 1 | 1 | 1 | 1 | 1 | 1 | 1 | 0.343 | 1 | 0.131 |
| SE | 3525 | ENSG00000105483 | CARD8 | chr19 | - | 48206383 | 48206520 | 48203145 | 48204243 | 48215339 | 48215384 | 4 | 2 | 1 | 3 | 20 | 11 | 0 | 0 | 5 | 6 | 0 | 5 | 0 | 8 | 3 | 14 | 5 | 8 | 4 | 0 | 286 | 149 | 0.291501015 | 1 | 0.159 | 1 | 0.094 | 0.343 | 0.635 | 0 | 0.343 | 0 | 0.51 | 1 | 0.076 |
| SE | 72229 | ENSG00000154473 | BUB3 | chr10 | + | 123160406 | 123160565 | 123157728 | 123157880 | 123162235 | 123162413 | 638 | 437 | 587 | 672 | 827 | 3 | 4 | 4 | 9 | 15 | 280 | 429 | 360 | 271 | 265 | 0 | 5 | 14 | 7 | 0 | 298 | 149 | 0.291721246 | 1 | 0.991 | 0.982 | 0.987 | 0.974 | 0.965 | 1 | 0.977 | 0.928 | 0.951 | 1 | 0.009 |
| SE | 26168 | ENSG00000108039 | XPNPEP1 | chr10 | - | 109884066 | 109884148 | 109882431 | 109882642 | 109886245 | 109886341 | 135 | 106 | 118 | 157 | 219 | 0 | 0 | 1 | 3 | 0 | 237 | 235 | 106 | 152 | 437 | 0 | 0 | 5 | 4 | 7 | 231 | 149 | 0.291908682 | 1 | 1 | 1 | 0.987 | 0.971 | 1 | 1 | 1 | 0.932 | 0.961 | 0.976 | 0.018 |
| SE | 41443 | ENSG00000119285 | HEATR1 | chr1 | - | 236599482 | 236599624 | 236597877 | 236597979 | 236603159 | 236603376 | 108 | 55 | 141 | 104 | 142 | 0 | 0 | 3 | 0 | 5 | 18 | 85 | 80 | 54 | 18 | 0 | 0 | 0 | 0 | 4 | 291 | 149 | 0.292007657 | 1 | 1 | 1 | 0.96 | 1 | 0.936 | 1 | 1 | 1 | 1 | 0.697 | 0.04 |
| SE | 79721 | ENSG00000168255 | POLR2J3 | chr7 | - | 102541092 | 102541303 | 102540014 | 102540091 | 102566996 | 102567070 | 50 | 27 | 25 | 39 | 19 | 2 | 0 | 2 | 1 | 0 | 9 | 45 | 28 | 22 | 28 | 0 | 0 | 0 | 5 | 0 | 298 | 149 | 0.292185512 | 1 | 0.926 | 1 | 0.862 | 0.951 | 1 | 1 | 1 | 1 | 0.688 | 1 | 0.01 |
| SE | 89170 | ENSG00000075856 | SART3 | chr12 | - | 108538064 | 108538203 | 108537487 | 108537595 | 108538933 | 108539089 | 267 | 126 | 232 | 193 | 174 | 0 | 2 | 0 | 0 | 2 | 131 | 161 | 124 | 159 | 143 | 1 | 0 | 1 | 9 | 0 | 288 | 149 | 0.292323948 | 1 | 1 | 0.97 | 1 | 1 | 0.978 | 0.985 | 1 | 0.985 | 0.901 | 1 | 0.015 |
| SE | 32984 | ENSG00000146192 | FGD2 | chr6 | + | 37013961 | 37014100 | 37013024 | 37013765 | 37014645 | 37014704 | 267 | 202 | 131 | 256 | 199 | 6 | 25 | 10 | 8 | 1 | 631 | 367 | 112 | 242 | 549 | 63 | 5 | 4 | 16 | 25 | 288 | 149 | 0.292343177 | 1 | 0.958 | 0.807 | 0.871 | 0.943 | 0.99 | 0.838 | 0.974 | 0.935 | 0.887 | 0.919 | 0.003 |
| SE | 68636 | ENSG00000166295 | ANAPC16 | chr10 | + | 72220053 | 72220379 | 72215999 | 72216138 | 72230365 | 72230440 | 46 | 15 | 48 | 35 | 32 | 18 | 14 | 18 | 14 | 20 | 25 | 77 | 27 | 24 | 8 | 42 | 0 | 19 | 12 | 1 | 298 | 149 | 0.292454984 | 1 | 0.561 | 0.349 | 0.571 | 0.556 | 0.444 | 0.229 | 1 | 0.415 | 0.5 | 0.8 | -0.093 |
| SE | 83221 | ENSG00000150637 | CD226 | chr18 | - | 69947360 | 69947532 | 69895700 | 69896045 | 69961591 | 69961803 | 6 | 17 | 6 | 23 | 11 | 14 | 22 | 2 | 35 | 14 | 17 | 4 | 33 | 20 | 20 | 0 | 27 | 17 | 27 | 16 | 298 | 149 | 0.292509605 | 1 | 0.176 | 0.279 | 0.6 | 0.247 | 0.282 | 1 | 0.069 | 0.493 | 0.27 | 0.385 | -0.127 |
| SE | 30178 | ENSG00000114487 | MORC1 | chr3 | - | 109054727 | 109054882 | 109035339 | 109035468 | 109057342 | 109057486 | 10 | 35 | 14 | 72 | 66 | 0 | 0 | 0 | 0 | 0 | 128 | 10 | 18 | 38 | 93 | 0 | 5 | 0 | 0 | 0 | 298 | 149 | 0.292525313 | 1 | 1 | 1 | 1 | 1 | 1 | 1 | 0.5 | 1 | 1 | 1 | 0.1 |
| SE | 72150 | ENSG00000115904 | SOS1 | chr2 | - | 39058672 | 39058804 | 39056701 | 39056866 | 39067627 | 39067753 | 311 | 280 | 224 | 312 | 339 | 6 | 2 | 0 | 1 | 2 | 113 | 296 | 230 | 238 | 127 | 12 | 0 | 1 | 0 | 0 | 281 | 149 | 0.292526566 | 1 | 0.965 | 0.987 | 1 | 0.994 | 0.989 | 0.833 | 1 | 0.992 | 1 | 1 | 0.022 |
| SE | 33681 | ENSG00000176261 | ZBTB8OS | chr1 | - | 32633950 | 32634072 | 32633644 | 32633727 | 32650432 | 32650525 | 201 | 92 | 154 | 134 | 213 | 3 | 1 | 3 | 5 | 0 | 39 | 127 | 88 | 76 | 143 | 0 | 0 | 0 | 0 | 0 | 271 | 149 | 0.292582798 | 1 | 0.974 | 0.981 | 0.966 | 0.936 | 1 | 1 | 1 | 1 | 1 | 1 | -0.029 |
| SE | 73125 | ENSG00000145414 | NAF1 | chr4 | - | 163158224 | 163158327 | 163156963 | 163157454 | 163164216 | 163164391 | 5 | 2 | 4 | 2 | 0 | 3 | 4 | 0 | 17 | 3 | 0 | 2 | 1 | 7 | 0 | 2 | 6 | 9 | 14 | 4 | 252 | 149 | 0.29265036 | 1 | 0.496 | 0.228 | 1 | 0.065 | 0 | 0 | 0.165 | 0.062 | 0.228 | 0 | 0.267 |
| SE | 91157 | ENSG00000164941 | INTS8 | chr8 | + | 94824892 | 94825067 | 94823286 | 94823561 | 94827721 | 94827793 | 52 | 27 | 59 | 44 | 35 | 6 | 1 | 0 | 1 | 0 | 6 | 42 | 23 | 12 | 1 | 0 | 2 | 4 | 2 | 0 | 298 | 149 | 0.292713412 | 1 | 0.812 | 0.931 | 1 | 0.957 | 1 | 1 | 0.913 | 0.742 | 0.75 | 1 | 0.059 |
| SE | 100624 | ENSG00000283977 | NA | chr11 | + | 6481756 | 6481852 | 6481500 | 6481555 | 6482044 | 6482289 | 22 | 7 | 22 | 13 | 7 | 0 | 0 | 3 | 0 | 0 | 12 | 15 | 12 | 13 | 3 | 6 | 0 | 0 | 0 | 0 | 245 | 149 | 0.292713511 | 1 | 1 | 1 | 0.817 | 1 | 1 | 0.549 | 1 | 1 | 1 | 1 | 0.054 |
| SE | 20300 | ENSG00000164465 | DCBLD1 | chr6 | + | 117540667 | 117540815 | 117539254 | 117539379 | 117540917 | 117541025 | 34 | 4 | 2 | 16 | 17 | 2 | 2 | 0 | 1 | 1 | 30 | 2 | 8 | 10 | 17 | 0 | 0 | 3 | 0 | 0 | 297 | 149 | 0.292779791 | 1 | 0.895 | 0.501 | 1 | 0.889 | 0.895 | 1 | 1 | 0.572 | 1 | 1 | -0.078 |
| SE | 13571 | ENSG00000167261 | DPEP2 | chr16 | - | 67987861 | 67987987 | 67987393 | 67987760 | 67989322 | 67989398 | 150 | 123 | 131 | 183 | 254 | 10 | 7 | 14 | 7 | 36 | 201 | 98 | 111 | 83 | 241 | 6 | 4 | 13 | 24 | 41 | 285 | 149 | 0.292796807 | 1 | 0.887 | 0.902 | 0.83 | 0.932 | 0.787 | 0.946 | 0.928 | 0.817 | 0.644 | 0.754 | 0.05 |
| SE | 107687 | ENSG00000128585 | MKLN1 | chr7 | + | 131443480 | 131443702 | 131437784 | 131437997 | 131463216 | 131463364 | 134 | 157 | 88 | 129 | 139 | 9 | 0 | 0 | 0 | 0 | 29 | 151 | 121 | 65 | 6 | 0 | 0 | 4 | 3 | 3 | 298 | 149 | 0.292896916 | 1 | 0.882 | 1 | 1 | 1 | 1 | 1 | 1 | 0.938 | 0.915 | 0.5 | 0.106 |

| | | | | | | | | | | | | | | | |
|---|---|---|---|---|---|---|---|---|---|---|---|---|---|---|---|
| SE | 35154 | ENSG00000158411 | MITD1 | chr2 | - | 99171342 | 99171429 | 99170534 | 99170652 | 99171509 | 99171646 | 520 | 478 | 372 | [illegible] |
| SE | 37394 | ENSG00000010327 | STAB1 | chr3 | + | 52513129 | 52513241 | 52512827 | 52512958 | 52513716 | 52513794 | 27 | 15 | 29 | [illegible] |
| SE | 5948 | ENSG00000170471 | RALGAPB | chr20 | + | 38499446 | 38499633 | 38497352 | 38497516 | 38509076 | 38509208 | 113 | 72 | 120 | [illegible] |
| SE | 12054 | ENSG00000159720 | ATP6V0D1 | chr16 | - | 67444527 | 67444706 | 67443098 | 67443178 | 67453543 | 67453715 | 744 | 345 | 379 | [illegible] |
| SE | 41963 | ENSG00000134864 | GGACT | chr13 | - | 100533562 | 100533774 | 100530182 | 100532601 | 100583824 | 100583997 | 3 | 0 | 3 | [illegible] |
| SE | 92855 | ENSG00000130958 | SLC35D2 | chr9 | - | 96360153 | 96360221 | 96352037 | 96352109 | 96364463 | 96364550 | 86 | 108 | 102 | [illegible] |
| SE | 80980 | ENSG00000130429 | ARPC1B | chr7 | + | 99377333 | 99377439 | 99374706 | 99374781 | 99385701 | 99385778 | 6 | 0 | 5 | [illegible] |
| SE | 28639 | ENSG00000102445 | RUBCNL | chr13 | - | 46377889 | 46378005 | 46371940 | 46372597 | 46387133 | 46387230 | 4 | 10 | 8 | [illegible] |
| SE | 69847 | ENSG00000107581 | EIF3A | chr10 | - | 119050520 | 119050674 | 119037118 | 119037309 | 119051198 | 119051321 | 734 | 762 | 521 | [illegible] |
| SE | 40164 | ENSG00000185291 | IL3RA | chrX | + | 1345315 | 1345434 | 1341727 | 1341829 | 1348430 | 1348545 | 55 | 73 | 53 | [illegible] |
| SE | 17215 | ENSG00000023608 | SNAPC1 | chr14 | + | 61776094 | 61776253 | 61768635 | 61768740 | 61778071 | 61778140 | 80 | 61 | 70 | [illegible] |
| SE | 70286 | ENSG00000104218 | CSPP1 | chr8 | + | 67172768 | 67172973 | 67172415 | 67172555 | 67175295 | 67175436 | 3 | 2 | 2 | [illegible] |
| SE | 55857 | ENSG00000177200 | CHD9 | chr16 | + | 53223449 | 53223507 | 53222643 | 53222755 | 53226365 | 53226403 | 6 | 16 | 15 | [illegible] |
| SE | 29021 | ENSG00000123352 | SPATS2 | chr12 | + | 49519072 | 49519182 | 49514554 | 49514613 | 49522750 | 49522853 | 17 | 4 | 27 | [illegible] |
| SE | 93809 | ENSG00000136869 | TLR4 | chr9 | + | 117708562 | 117708729 | 117704402 | 117704565 | 117712388 | 117715775 | 200 | 79 | 80 | [illegible] |
| SE | 9230 | ENSG00000184381 | PLA2G6 | chr22 | - | 38147349 | 38147475 | 38145437 | 38145653 | 38169217 | 38169467 | 0 | 0 | 5 | [illegible] |
| SE | 62372 | ENSG00000168826 | ZBTB49 | chr4 | + | 4315635 | 4315718 | 4313040 | 4313114 | 4320639 | 4321398 | 21 | 5 | 4 | [illegible] |
| SE | 32372 | ENSG00000013441 | CLK1 | chr2 | - | 200857717 | 200857884 | 200856681 | 200856811 | 200857972 | 200858089 | 562 | 365 | 482 | [illegible] |
| SE | 4736 | ENSG00000031823 | RANBP3 | chr19 | - | 5950716 | 5950872 | 5941798 | 5941835 | 5951392 | 5951596 | 13 | 3 | 1 | [illegible] |
| SE | 60915 | ENSG00000138398 | PPIG | chr2 | + | 169592650 | 169592739 | 169584366 | 169584490 | 169603641 | 169603694 | 5 | 6 | 0 | [illegible] |
| SE | 76688 | ENSG00000237441 | RGL2 | chr6 | - | 33293603 | 33293699 | 33293412 | 33293524 | 33293794 | 33293916 | 68 | 33 | 34 | [illegible] |
| SE | 7641 | ENSG00000270647 | TAF15 | chr17 | + | 35833906 | 35833941 | 35824077 | 35824198 | 35834565 | 35834598 | 292 | 223 | 277 | [illegible] |
| SE | 30049 | ENSG00000155744 | FAM126B | chr2 | - | 201045495 | 201045593 | 201023958 | 201024039 | 201071453 | 201071579 | 45 | 40 | 40 | [illegible] |
| SE | 12332 | ENSG00000197912 | SPG7 | chr16 | + | 89540943 | 89541384 | 89532462 | 89532636 | 89544647 | 89544772 | 17 | 3 | 10 | [illegible] |
| SE | 40054 | ENSG00000204186 | ZDBF2 | chr2 | + | 206292013 | 206292116 | 206281800 | 206281909 | 206297245 | 206297373 | 25 | 14 | 47 | [illegible] |
| SE | 40435 | ENSG00000078304 | PPP2R5C | chr14 | + | 101890236 | 101890296 | 101883431 | 101883562 | 101892999 | 101893053 | 957 | 494 | 675 | [illegible] |
| SE | 98980 | ENSG00000076053 | RBM7 | chr11 | + | 114402827 | 114402915 | 114400612 | 114400767 | 114405705 | 114405799 | 48 | 23 | 35 | [illegible] |
| SE | 84302 | ENSG00000147202 | DIAPH2 | chrX | + | 96758153 | 96758258 | 96738585 | 96738762 | 96881578 | 96881718 | 110 | 129 | 71 | [illegible] |
| SE | 60491 | ENSG00000070814 | TCOF1 | chr5 | + | 150391956 | 150392176 | 150389886 | 150390023 | 150392704 | 150392790 | 41 | 30 | 58 | [illegible] |
| SE | 105017 | ENSG00000284770 | TBCE | chr1 | + | 235438768 | 235438988 | 235437321 | 235437474 | 235441813 | 235441882 | 33 | 27 | 29 | [illegible] |
| SE | 35987 | ENSG00000078070 | MCCC1 | chr3 | - | 183094558 | 183094605 | 183086692 | 183086788 | 183099351 | 183099495 | 14 | 2 | 19 | [illegible] |
| SE | 62128 | ENSG00000115364 | MRPL19 | chr2 | + | 75655063 | 75655302 | 75654735 | 75654917 | 75670578 | 75670753 | 140 | 68 | 102 | [illegible] |
| SE | 64306 | ENSG00000101336 | HCK | chr20 | + | 32073715 | 32073818 | 32073318 | 32073361 | 32074622 | 32074721 | 458 | 282 | 264 | [illegible] |
| SE | 36842 | ENSG00000168038 | ULK4 | chr3 | - | 41937228 | 41937340 | 41935861 | 41935940 | 41938097 | 41938197 | 11 | 6 | 10 | [illegible] |
| SE | 97786 | ENSG00000166452 | AKIP1 | chr11 | + | 8912452 | 8912533 | 8911191 | 8911671 | 8914825 | 8914930 | 24 | 14 | 5 | [illegible] |
| SE | 33693 | ENSG00000176261 | ZBTB8OS | chr1 | - | 32634701 | 32634792 | 32633950 | 32634110 | 32650432 | 32650525 | 14 | 5 | 4 | [illegible] |
| SE | 106135 | ENSG00000136021 | SCYL2 | chr12 | + | 100337386 | 100337506 | 100335590 | 100335906 | 100338527 | 100338633 | 381 | 306 | 261 | [illegible] |
| SE | 62988 | ENSG00000137497 | NUMA1 | chr11 | - | 72010785 | 72010854 | 72009267 | 72009387 | 72012894 | 72013312 | 153 | 112 | 150 | [illegible] |
| SE | 99541 | ENSG00000188690 | UROS | chr10 | - | 125816176 | 125816260 | 125812213 | 125812288 | 125816436 | 125816510 | 18 | 6 | 13 | [illegible] |
| SE | 83973 | ENSG00000068400 | GRIPAP1 | chrX | - | 48998153 | 48998180 | 48993427 | 48993578 | 48999237 | 48999299 | 49 | 33 | 29 | [illegible] |
| SE | 91270 | ENSG00000154114 | TBCEL | chr11 | + | 121055051 | 121055308 | 121053550 | 121053732 | 121057507 | 121057614 | 59 | 21 | 45 | [illegible] |
| SE | 24537 | ENSG00000172785 | CBWD1 | chr9 | - | 121960 | 122136 | 121502 | 121573 | 123216 | 123282 | 61 | 172 | 71 | [illegible] |
| SE | 12392 | ENSG00000188559 | RALGAPA2 | chr20 | - | 20535744 | 20535803 | 20531686 | 20531795 | 20546703 | 20546832 | 14 | 16 | 34 | [illegible] |
| SE | 81941 | ENSG00000051825 | MPHOSPH9 | chr12 | - | 123162114 | 123162192 | 123161135 | 123161383 | 123163013 | 123163036 | 61 | 52 | 60 | [illegible] |
| [illegible] | | | | | | | | | | | | | | | |

| | | | | | | | | | | | | | | | | | | | | | | | | | | | | | | | | | | | | | | | |
|---|---|---|---|---|---|---|---|---|---|---|---|---|---|---|---|---|---|---|---|---|---|---|---|---|---|---|---|---|---|---|---|---|---|---|---|---|---|---|---|
| [illegible] | [illegible] | [illegible] | [illegible] | [illegible] | [illegible] | [illegible] | [illegible] | [illegible] | [illegible] | [illegible] | [illegible] | [illegible] | [illegible] | [illegible] | [illegible] | [illegible] | [illegible] | [illegible] | [illegible] | [illegible] | [illegible] | [illegible] | [illegible] | [illegible] | [illegible] | [illegible] | [illegible] | [illegible] | [illegible] | [illegible] | [illegible] | [illegible] | [illegible] | [illegible] | [illegible] | [illegible] | [illegible] | [illegible] | [illegible] |

| | | | | | | | | | |
|---|---|---|---|---|---|---|---|---|---|
| SE | 38901 | ENSG00000151466 | SCLT1 | chr4 | - | 128965226 | 128965318 | 128959599 | 128959777 |
| SE | 26590 | ENSG00000143891 | GALM | chr2 | + | 38698365 | 38698436 | 38689812 | 38689894 |
| SE | 106769 | ENSG00000078674 | PCM1 | chr8 | + | 17983232 | 17983289 | 17972582 | 17972687 |
| SE | 47874 | ENSG00000146247 | PHIP | chr6 | - | 78970780 | 78970888 | 78969834 | 78969917 |
| SE | 53541 | ENSG00000138668 | HNRNPD | chr4 | - | 82371527 | 82371584 | 82359470 | 82359639 |
| SE | 35267 | ENSG00000103064 | SLC7A6 | chr16 | + | 68273821 | 68273904 | 68266601 | 68266721 |
| SE | 63119 | ENSG00000115761 | NOL10 | chr2 | - | 10675793 | 10675871 | 10673321 | 10673557 |
| SE | 59400 | ENSG00000139631 | CSAD | chr12 | - | 53171272 | 53171441 | 53170422 | 53170502 |
| SE | 43359 | ENSG00000234171 | RNASEH1-AS1 | chr2 | + | 3559403 | 3559729 | 3558627 | 3558998 |
| SE | 19207 | ENSG00000100902 | PSMA6 | chr14 | + | 35292952 | 35293277 | 35292442 | 35292552 |
| SE | 45087 | ENSG00000100181 | TPTEP1 | chr22 | + | 16622840 | 16622897 | 16614076 | 16614178 |
| SE | 30685 | ENSG00000132466 | ANKRD17 | chr4 | - | 73118687 | 73118850 | 73113791 | 73113908 |
| SE | 79997 | ENSG00000084070 | SMAP2 | chr1 | + | 40362299 | 40362436 | 40344849 | 40344910 |
| SE | 99002 | ENSG00000136444 | RSAD1 | chr17 | + | 50484441 | 50484545 | 50482090 | 50482456 |
| SE | 67125 | ENSG00000123136 | DDX39A | chr19 | - | 14410547 | 14410605 | 14409772 | 14410334 |
| SE | 35205 | ENSG00000102362 | SYTL4 | chrX | - | 100703092 | 100703185 | 100701927 | 100702107 |
| SE | 10756 | ENSG00000154760 | SLFN13 | chr17 | - | 35446716 | 35446753 | 35444614 | 35445693 |
| SE | 91250 | ENSG00000085719 | CPNE3 | chr8 | + | 86556101 | 86556338 | 86554850 | 86554984 |
| SE | 5176 | ENSG00000101191 | DIDO1 | chr20 | - | 62891986 | 62892076 | 62887213 | 62891155 |
| SE | 23577 | ENSG00000254996 | ANKHD1-EIF4EBP3 | chr5 | + | 140535361 | 140535538 | 140528183 | 140529796 |
| SE | 97482 | ENSG00000076984 | MAP2K7 | chr19 | + | 7905807 | 7905855 | 7903884 | 7904068 |
| SE | 5052 | ENSG00000101126 | ADNP | chr20 | - | 50918738 | 50918781 | 50904765 | 50904849 |
| SE | 107945 | ENSG00000177119 | ANO6 | chr12 | + | 45270461 | 45270524 | 45216084 | 45216391 |
| SE | 5815 | ENSG00000197586 | ENTPD6 | chr20 | + | 25207075 | 25207397 | 25195692 | 25195867 |
| SE | 63417 | ENSG00000122507 | BBS9 | chr7 | + | 33155637 | 33155702 | 33152700 | 33152851 |
| SE | 62414 | ENSG00000033867 | SLC4A7 | chr3 | - | 27431297 | 27431669 | 27424036 | 27424152 |
| SE | 92228 | ENSG00000119421 | NDUFA8 | chr9 | - | 122152244 | 122152408 | 122148111 | 122148277 |
| SE | 11666 | ENSG00000167207 | NOD2 | chr16 | + | 50707854 | 50707960 | 50699487 | 50699954 |
| SE | 54418 | ENSG00000169554 | ZEB2 | chr2 | - | 144396411 | 144396592 | 144388015 | 144388882 |
| SE | 1531 | ENSG00000100242 | SUN2 | chr22 | - | 38754641 | 38754706 | 38752506 | 38752665 |
| SE | 107895 | ENSG00000188906 | LRRK2 | chr12 | + | 40313971 | 40314173 | 40310430 | 40310649 |
| SE | 1290 | ENSG00000100239 | PPP6R2 | chr22 | + | 50388985 | 50389111 | 50372071 | 50372150 |
| [illegible] | | | | | | | | | |

| | | | | | | | | | | | | | | | | | | | | | | | | | | |
|---|---|---|---|---|---|---|---|---|---|---|---|---|---|---|---|---|---|---|---|---|---|---|---|---|---|---|
| SE | 18352 | ENSG00000257621 | PSMA3-AS1 | chr14 | - | 58293313 | 58293544 | 58291634 | 58291707 | 58293765 | 58295811 | 4 | 4 | 3 | 0 | 10 | 23 | 10 | 29 | 27 | 35 | 0 | 18 | 7 | 0 | 0 |
| SE | 61877 | ENSG00000076242 | MLH1 | chr3 | + | 37004400 | 37004474 | 37000954 | 37001053 | 37006990 | 37007063 | 251 | 255 | 307 | 304 | 302 | 1 | 0 | 0 | 0 | 0 | 388 | 316 | 310 | 343 | 83 |
| SE | 90641 | ENSG00000120885 | CLU | chr8 | - | 27613577 | 27613683 | 27610474 | 27610600 | 27614654 | 27614698 | 0 | 4 | 1 | 0 | 0 | 35 | 41 | 10 | 82 | 10 | 0 | 2 | 0 | 0 | 34 |
| SE | 42458 | ENSG00000163946 | TASOR | chr3 | - | 56670072 | 56670145 | 56669699 | 56669791 | 56671599 | 56671692 | 1206 | 1137 | 939 | 1076 | 1373 | 6 | 17 | 5 | 10 | 14 | 1066 | 980 | 809 | 468 | 682 |
| SE | 41946 | ENSG00000163374 | YY1AP1 | chr1 | - | 155680415 | 155680456 | 155679408 | 155679512 | 155688070 | 155688201 | 144 | 86 | 116 | 130 | 190 | 0 | 6 | 7 | 3 | 2 | 258 | 173 | 102 | 173 | 257 |
| SE | 39347 | ENSG00000158864 | NDUFS2 | chr1 | + | 161209192 | 161209313 | 161205784 | 161206597 | 161209482 | 161209595 | 241 | 140 | 291 | 299 | 264 | 0 | 3 | 0 | 0 | 0 | 650 | 398 | 317 | 389 | 611 |
| SE | 22948 | ENSG00000170234 | PWWP2A | chr5 | - | 160080650 | 160080770 | 160075881 | 160078168 | 160093100 | 160094065 | 60 | 16 | 28 | 46 | 71 | 9 | 12 | 14 | 5 | 13 | 24 | 11 | 14 | 16 | 3 |
| SE | 68703 | ENSG00000196295 | GARS1-DT | chr7 | - | 30551317 | 30551479 | 30548358 | 30550781 | 30561465 | 30562128 | 1 | 9 | 8 | 13 | 0 | 31 | 34 | 28 | 48 | 32 | 17 | 9 | 9 | 0 | 0 |
| SE | 85871 | ENSG00000041988 | THAP3 | chr1 | + | 6632390 | 6632495 | 6630287 | 6630353 | 6632795 | 6633121 | 44 | 6 | 30 | 29 | 31 | 0 | 0 | 0 | 0 | 0 | 39 | 39 | 18 | 15 | 67 |
| SE | 54522 | ENSG00000130997 | POLN | chr4 | - | 2176265 | 2176334 | 2174690 | 2174751 | 2179307 | 2179387 | 7 | 1 | 3 | 2 | 4 | 0 | 0 | 0 | 0 | 4 | 6 | 9 | 5 | 8 | 16 |
| SE | 94591 | ENSG00000106829 | TLE4 | chr9 | + | 79612655 | 79612718 | 79576132 | 79576177 | 79652589 | 79652703 | 72 | 69 | 79 | 70 | 102 | 1 | 0 | 5 | 3 | 0 | 84 | 78 | 83 | 83 | 69 |
| SE | 62126 | ENSG00000168495 | POLR3D | chr8 | + | 22248480 | 22248649 | 22248153 | 22248278 | 22250074 | 22250227 | 64 | 49 | 49 | 67 | 101 | 0 | 0 | 0 | 0 | 0 | 76 | 47 | 21 | 34 | 45 |
| SE | 91897 | ENSG00000104529 | EEF1D | chr8 | - | 143581285 | 143581328 | 143581053 | 143581154 | 143586728 | 143586852 | 46 | 21 | 75 | 56 | 71 | 0 | 0 | 0 | 4 | 3 | 44 | 24 | 49 | 30 | 141 |
| SE | 49452 | ENSG00000115020 | PIKFYVE | chr2 | + | 208276711 | 208276830 | 208273583 | 208273733 | 208288728 | 208288818 | 67 | 53 | 51 | 62 | 58 | 1 | 3 | 0 | 4 | 0 | 70 | 85 | 67 | 46 | 51 |
| SE | 12825 | ENSG00000169592 | INO80E | chr16 | + | 30001211 | 30001530 | 30000928 | 30001040 | 30005220 | 30005323 | 44 | 40 | 60 | 38 | 49 | 7 | 6 | 14 | 3 | 14 | 60 | 55 | 34 | 19 | 125 |
| SE | 14549 | ENSG00000159322 | ADPGK | chr15 | - | 72756250 | 72756447 | 72751368 | 72752892 | 72760406 | 72760527 | 198 | 83 | 108 | 150 | 187 | 2 | 1 | 1 | 3 | 9 | 265 | 126 | 43 | 141 | 133 |
| SE | 10571 | ENSG00000188895 | MSL1 | chr17 | + | 40131219 | 40131345 | 40129244 | 40129627 | 40131536 | 40131584 | 5 | 4 | 4 | 9 | 15 | 146 | 80 | 126 | 152 | 261 | 3 | 13 | 21 | 18 | 0 |
| SE | 64757 | ENSG00000144677 | CTDSPL | chr3 | + | 37957110 | 37957143 | 37947056 | 37947211 | 37964570 | 37964672 | 12 | 4 | 5 | 5 | 9 | 0 | 3 | 1 | 4 | 0 | 8 | 10 | 4 | 19 | 55 |
| SE | 50772 | ENSG00000165185 | KIAA1958 | chr9 | + | 112617870 | 112617954 | 112574056 | 112575251 | 112645649 | 112645822 | 3 | 1 | 0 | 4 | 0 | 5 | 5 | 6 | 4 | 1 | 0 | 0 | 2 | 2 | 3 |
| SE | 39860 | ENSG00000169239 | CA5B | chrX | + | 15772495 | 15772614 | 15764577 | 15764775 | 15774301 | 15774397 | 273 | 168 | 219 | 224 | 251 | 0 | 0 | 0 | 0 | 1 | 141 | 129 | 94 | 73 | 138 |
| SE | 51639 | ENSG00000132128 | LRRC41 | chr1 | - | 46280395 | 46280560 | 46280191 | 46280290 | 46281124 | 46281161 | 84 | 38 | 76 | 62 | 61 | 1 | 0 | 1 | 5 | 0 | 77 | 76 | 40 | 55 | 95 |
| SE | 16318 | ENSG00000140374 | ETFA | chr15 | - | 76292618 | 76292700 | 76292430 | 76292513 | 76295590 | 76295737 | 208 | 182 | 145 | 267 | 212 | 21 | 6 | 6 | 9 | 12 | 234 | 288 | 131 | 168 | 648 |
| SE | 22618 | ENSG00000127603 | MACF1 | chr1 | + | 39433047 | 39433155 | 39432534 | 39432654 | 39434413 | 39434632 | 724 | 534 | 575 | 781 | 910 | 4 | 8 | 9 | 1 | 0 | 178 | 674 | 664 | 323 | 60 |
| SE | 85224 | ENSG00000099204 | ABLIM1 | chr10 | - | 114473878 | 114473956 | 114472976 | 114473132 | 114487957 | 114487993 | 309 | 273 | 566 | 512 | 974 | 25 | 7 | 21 | 8 | 21 | 178 | 218 | 222 | 154 | 397 |
| SE | 107155 | ENSG00000087448 | KLHL42 | chr12 | + | 27797246 | 27797349 | 27791707 | 27791901 | 27797714 | 27798156 | 12 | 3 | 6 | 9 | 7 | 16 | 9 | 29 | 8 | 27 | 12 | 0 | 8 | 6 | 6 |
| SE | 21179 | ENSG00000143252 | SDHC | chr1 | + | 161328395 | 161328497 | 161323613 | 161323670 | 161340593 | 161340655 | 268 | 115 | 240 | 252 | 279 | 16 | 6 | 16 | 18 | 15 | 302 | 255 | 140 | 157 | 302 |
| SE | 23295 | ENSG00000164182 | NDUFAF2 | chr5 | + | 61045839 | 61045978 | 60945182 | 60945382 | 61073124 | 61073214 | 4 | 1 | 2 | 0 | 8 | 18 | 13 | 5 | 23 | 6 | 8 | 5 | 0 | 0 | 0 |
| SE | 99248 | ENSG00000133318 | RTN3 | chr11 | + | 63718980 | 63721032 | 63681495 | 63681778 | 63749990 | 63750198 | 1 | 3 | 3 | 4 | 10 | 151 | 82 | 107 | 170 | 88 | 0 | 2 | 2 | 11 | 2 |
| SE | 43237 | ENSG00000181666 | ZNF875 | chr19 | + | 37347189 | 37347316 | 37325153 | 37325274 | 37347776 | 37347810 | 14 | 15 | 25 | 21 | 38 | 0 | 0 | 0 | 0 | 0 | 40 | 35 | 23 | 12 | 12 |
| SE | 91479 | ENSG00000141569 | TRIM65 | chr17 | - | 75892010 | 75892185 | 75891812 | 75891878 | 75892266 | 75892303 | 46 | 35 | 28 | 49 | 18 | 0 | 0 | 0 | 0 | 0 | 11 | 13 | 7 | 19 | 66 |
| SE | 73106 | ENSG00000152642 | GPD1L | chr3 | + | 32138586 | 32138727 | 32128075 | 32128253 | 32140227 | 32140361 | 82 | 43 | 67 | 52 | 63 | 6 | 1 | 3 | 0 | 2 | 46 | 33 | 47 | 41 | 56 |
| SE | 20441 | ENSG00000066933 | MYO9A | chr15 | - | 71951701 | 71951896 | 71938851 | 71938927 | 71959900 | 71960096 | 25 | 21 | 29 | 37 | 61 | 16 | 9 | 3 | 7 | 31 | 6 | 13 | 12 | 7 | 1 |
| SE | 35363 | ENSG00000164073 | MFSD8 | chr4 | - | 127939852 | 127939997 | 127938782 | 127938838 | 127942044 | 127942158 | 80 | 62 | 70 | 73 | 160 | 3 | 3 | 4 | 5 | 0 | 32 | 38 | 27 | 26 | 14 |
| SE | 59100 | ENSG00000214941 | ZSWIM7 | chr17 | - | 15977799 | 15977913 | 15976559 | 15977092 | 15978043 | 15978163 | 6 | 9 | 10 | 7 | 8 | 35 | 13 | 4 | 6 | 17 | 7 | 1 | 4 | 8 | 28 |
| SE | 102675 | ENSG00000175029 | CTBP2 | chr10 | - | 125110989 | 125111093 | 125038996 | 125039155 | 125158634 | 125158716 | 69 | 22 | 27 | 65 | 39 | 0 | 0 | 0 | 3 | 0 | 2 | 9 | 15 | 15 | 24 |
| SE | 57771 | ENSG00000254469 | NA | chr11 | - | 71923308 | 71923475 | 71918884 | 71919026 | 71928364 | 71928421 | 16 | 20 | 26 | 12 | 8 | 2 | 1 | 3 | 0 | 0 | 19 | 43 | 6 | 11 | 20 |
| SE | 28582 | ENSG00000158481 | CD1C | chr1 | + | 158293211 | 158293302 | 158292595 | 158292874 | 158293454 | 158294774 | 90 | 84 | 79 | 106 | 145 | 24 | 25 | 44 | 33 | 58 | 147 | 1 | 12 | 25 | 50 |
| SE | 11098 | ENSG00000178188 | SH2B1 | chr16 | + | 28866011 | 28867033 | 28863667 | 28863754 | 28867373 | 28867432 | 11 | 4 | 7 | 14 | 19 | 2 | 1 | 0 | 1 | 0 | 21 | 11 | 2 | 8 | 11 |
| SE | 21593 | ENSG00000112234 | FBXL4 | chr6 | - | 98880552 | 98880624 | 98875414 | 98875727 | 98899267 | 98899481 | 68 | 45 | 67 | 115 | 76 | 2 | 2 | 4 | 2 | 2 | 11 | 23 | 35 | 38 | 48 |
| SE | 5499 | ENSG00000101294 | HM13 | chr20 | + | 31538178 | 31538554 | 31527483 | 31527582 | 31550063 | 31550121 | 77 | 33 | 41 | 67 | 47 | 1 | 0 | 0 | 1 | 0 | 114 | 77 | 62 | 114 | 172 |
| SE | 104906 | ENSG00000170456 | DENND5B | chr12 | - | 31413435 | 31413564 | 31409262 | 31409384 | 31415366 | 31415448 | 23 | 6 | 26 | 15 | 38 | 0 | 0 | 0 | 4 | 2 | 19 | 31 | 39 | 11 | 9 |
| SE | 105920 | ENSG00000234608 | MAPKAPK5-AS1 | chr12 | - | 111840332 | 111840477 | 111839767 | 111840003 | 111841326 | 111841470 | 43 | 13 | 46 | 37 | 74 | 10 | 4 | 5 | 9 | 8 | 61 | 37 | 23 | 27 | 7 |
| SE | 106996 | ENSG00000133858 | ZFC3H1 | chr12 | - | 71614833 | 71614938 | 71614534 | 71614700 | 71615205 | 71615316 | 240 | 284 | 200 | 200 | 359 | 1 | 8 | 2 | 2 | 4 | 25 | 93 | 91 | 19 | 8 |
| SE | 41224 | ENSG00000153815 | CMIP | chr16 | + | 81620875 | 81620926 | 81607563 | 81607692 | 81652202 | 81652364 | 295 | 190 | 255 | 321 | 382 | 3 | 0 | 0 | 0 | 4 | 775 | 379 | 270 | 569 | 1112 |
| SE | 38640 | ENSG00000156875 | MFSD14A | chr1 | + | 100068472 | 100068586 | 100067976 | 100068195 | 100077164 | 100077277 | 102 | 44 | 84 | 96 | 94 | 1 | 0 | 1 | 4 | 1 | 64 | 42 | 132 | 61 | 9 |
| SE | 40613 | ENSG00000162613 | FUBP1 | chr1 | - | 77969021 | 77969084 | 77968164 | 77968203 | 77969924 | 77970015 | 47 | 31 | 57 | 85 | 66 | 211 | 175 | 165 | 238 | 231 | 21 | 32 | 120 | 94 | 32 |
| SE | 106672 | ENSG00000120662 | MTRF1 | chr13 | - | 41240260 | 41240433 | 41233889 | 41234007 | 41252644 | 41252752 | 113 | 47 | 94 | 83 | 68 | 11 | 0 | 6 | 4 | 2 | 49 | 32 | 23 | 39 | 51 |
| SE | 98658 | ENSG00000107036 | RIC1 | chr9 | + | 5753535 | 5753646 | 5753199 | 5753238 | 5754840 | 5754930 | 257 | 186 | 204 | 226 | 230 | 4 | 4 | 10 | 10 | 8 | 181 | 219 | 198 | 265 | 58 |
| SE | 103502 | ENSG00000107862 | GBF1 | chr10 | + | 102332407 | 102332526 | 102260116 | 102260116 | 102344050 | 102344182 | 5 | 5 | 3 | 4 | 14 | 41 | 32 | 40 | 34 | 38 | 0 | 10 | 2 | 5 | 18 |
| SE | 87973 | ENSG00000168066 | SF1 | chr11 | - | 64773429 | 64773505 | 64770255 | 64770408 | 64776497 | 64776626 | 699 | 397 | 649 | 739 | 931 | 50 | 28 | 55 | 59 | 82 | 1276 | 788 | 610 | 996 | 1473 |
| SE | 14687 | ENSG00000182511 | FES | chr15 | + | 90885411 | 90885585 | 90885036 | 90885258 | 90886960 | 90887015 | 148 | 47 | 56 | 88 | 77 | 21 | 1 | 12 | 14 | 2 | 248 | 265 | 60 | 188 | 197 |
| SE | 31209 | ENSG00000025293 | PHF20 | chr20 | + | 35842572 | 35842744 | 35801490 | 35801605 | 35858301 | 35858381 | 113 | 61 | 79 | 80 | 70 | 0 | 1 | 1 | 0 | 0 | 105 | 52 | 168 | 70 | 44 |
| SE | 41255 | ENSG00000162819 | BROX | chr1 | + | 222715653 | 222715800 | 222713320 | 222713416 | 222718924 | 222719031 | 107 | 58 | 81 | 83 | 106 | 4 | 0 | 1 | 2 | 1 | 20 | 53 | 67 | 52 | 7 |
| SE | 100451 | ENSG00000103429 | BFAR | chr16 | + | 14648387 | 14648592 | 14644273 | 14644609 | 14655065 | 14655171 | 87 | 45 | 77 | 77 | 124 | 2 | 3 | 3 | 6 | 0 | 140 | 62 | 84 | 53 | 93 |
| SE | 100757 | ENSG00000106723 | SPIN1 | chr9 | + | 88462495 | 88462749 | 88448940 | 88448989 | 88468371 | 88468477 | 173 | 104 | 114 | 156 | 164 | 23 | 5 | 14 | 20 | 9 | 126 | 78 | 111 | 105 | 68 |
| SE | 10851 | ENSG00000029725 | RABEP1 | chr17 | + | 5356507 | 5356573 | 5354400 | 5354490 | 5361207 | 5361629 | 31 | 44 | 43 | 30 | 60 | 233 | 128 | 143 | 215 | 223 | 8 | 100 | 33 | 30 | 0 |
| SE | 106033 | ENSG00000204482 | LST1 | chr6 | + | 31587640 | 31587733 | 31587116 | 31587318 | 31588517 | 31588909 | 176 | 221 | 138 | 153 | 198 | 221 | 99 | 82 | 195 | 250 | 263 | 120 | 165 | 325 | 341 |
| SE | 44011 | ENSG00000114098 | ARMC8 | chr3 | + | 138237308 | 138237384 | 138235033 | 138235114 | 138237481 | 138237572 | 192 | 154 | 152 | 217 | 274 | 1 | 1 | 0 | 12 | 0 | 72 | 151 | 125 | 99 | 103 |
| SE | 17530 | ENSG00000087111 | PIGS | chr17 | - | 28561421 | 28561629 | 28560048 | 28560191 | 28563430 | 28563522 | 160 | 90 | 146 | 155 | 168 | 3 | 2 | 3 | 2 | 0 | 255 | 217 | 87 | 134 | 309 |
| SE | 69778 | ENSG00000065675 | PRKCQ | chr10 | - | 6430809 | 6430938 | 6427142 | 6428362 | 6441892 | 6442081 | 191 | 83 | 152 | 155 | 259 | 0 | 2 | 2 | 1 | 8 | 46 | 99 | 146 | 55 | 80 |
| SE | 24937 | ENSG00000072364 | AFF4 | chr5 | - | 132893029 | 132893118 | 132891816 | 132892404 | 132896322 | 132897240 | 284 | 245 | 304 | 282 | 393 | 3 | 1 | 7 | 6 | 7 | 88 | 342 | 437 | 245 | 44 |
| SE | 102042 | ENSG00000196151 | WDSUB1 | chr2 | - | 159256195 | 159256375 | 159248371 | 159248512 | 159275545 | 159275638 | 104 | 73 | 61 | 62 | 109 | 0 | 1 | 1 | 0 | 0 | 78 | 79 | 53 | 47 | 90 |
| SE | 83392 | ENSG00000174428 | GTF2IRD2B | chr7 | + | 75142481 | 75142665 | 75141102 | 75141186 | 75143949 | 75143978 | 16 | 7 | 19 | 20 | 23 | 1 | 1 | 1 | 0 | 2 | 36 | 20 | 14 | 17 | 12 |
| SE | 75635 | ENSG00000112146 | FBXO9 | chr6 | + | 53080967 | 53081098 | 53076485 | 53076543 | 53082503 | 53082618 | 138 | 83 | 127 | 123 | 108 | 3 | 2 | 1 | 2 | 6 | 132 | 178 | 83 | 169 | 125 |
| SE | 39458 | ENSG00000213024 | NUP62 | chr19 | - | 49918507 | 49918621 | 49909394 | 49909884 | 49927693 | 49927815 | 0 | 6 | 0 | 7 | 2 | 87 | 64 | 84 | 77 | 73 | 10 | 5 | 0 | 0 | 0 |
| SE | 35763 | ENSG00000198843 | SELENOT | chr3 | + | 150618708 | 150618803 | 150603327 | 150603499 | 150622384 | 150622495 | 2 | 0 | 2 | 0 | 0 | 75 | 34 | 54 | 86 | 49 | 0 | 0 | 8 | 3 | 0 |
| SE | 20346 | ENSG00000162711 | NLRP3 | chr1 | + | 247429584 | 247429755 | 247423846 | 247425599 | 247443971 | 247444142 | 20 | 13 | 17 | 16 | 8 | 3 | 3 | 3 | 1 | 5 | 8 | 15 | 11 | 39 | 36 |
| SE | 64690 | ENSG00000093144 | ECHDC1 | chr6 | - | 127315933 | 127315974 | 127314815 | 127314896 | 127327001 | 127327012 | 3 | 0 | 4 | 4 | 0 | 71 | 55 | 40 | 64 | 62 | 0 | 0 | 0 | 0 | 0 |
| SE | 78221 | ENSG00000183696 | UPP1 | chr7 | + | 48103290 | 48103411 | 48101823 | 48101982 | 48106872 | 48107082 | 162 | 72 | 77 | 98 | 58 | 4 | 5 | 5 | 2 | 2 | 289 | 91 | 96 | 135 | 311 |
| SE | 32085 | ENSG00000242086 | MUC20-OT1 | chr3 | + | 195696360 | 195696589 | 195689336 | 195689438 | 195699029 | 195699116 | 49 | 66 | 63 | 69 | 60 | 23 | 12 | 7 | 14 | 18 | 127 | 62 | 108 | 125 | 76 |
| SE | 31950 | ENSG00000163703 | CRELD1 | chr3 | + | 9943674 | 9944095 | 9943076 | 9943515 | 9944364 | 9944985 | 2 | 1 | 3 | 10 | 3 | 38 | 16 | 23 | 29 | 33 | 0 | 0 | 0 | 1 | 11 |
| SE | 95891 | ENSG00000080298 | RFX3 | chr9 | - | 3301545 | 3301620 | 3293076 | 3293258 | 3330258 | 3330517 | 36 | 25 | 40 | 33 | 32 | 15 | 7 | 11 | 20 | 16 | 37 | 9 | 29 | 4 | 41 |
| SE | 61584 | ENSG00000131876 | SNRPA1 | chr15 | - | 101291961 | 101292040 | 101287655 | 101287702 | 101295096 | 101295204 | 104 | 79 | 121 | 99 | 80 | 1 | 6 | 5 | 8 | 0 | 74 | 77 | 116 | 71 | 18 |
| SE | 57772 | ENSG00000254469 | NA | chr11 | - | 71923308 | 71923475 | 71918884 | 71919026 | 71928448 | 71928624 | 11 | 17 | 26 | 12 | 12 | 0 | 2 | 4 | 0 | 0 | 14 | 33 | 4 | 11 | 16 |
| SE | 50825 | ENSG00000215788 | TNFRSF25 | chr1 | - | 6462862 | 6462970 | 6462628 | 6462663 | 6464374 | 6464453 | 6 | 4 | 15 | 11 | 5 | 1 | 5 | 1 | 3 | 11 | 2 | 1 | 4 | 2 | 9 |
| SE | 34960 | ENSG00000147854 | UHRF2 | chr9 | + | 6492302 | 6492401 | 6486820 | 6486925 | 6493825 | 6493932 | 111 | 85 | 102 | 130 | 144 | 140 | 111 | 122 | 128 | 136 | 17 | 115 | 134 | 64 | 3 |
| SE | 82661 | ENSG00000104447 | TRPS1 | chr8 | - | 115623600 | 115623758 | 115619131 | 115620060 | 115668544 | 115668975 | 30 | 33 | 67 | 70 | 108 | 43 | 27 | 29 | 43 | 30 | 47 | 62 | 51 | 44 | 80 |
| SE | 63302 | ENSG00000138018 | SELENOI | chr2 | + | 26375039 | 26375148 | 26373366 | 26373629 | 26383298 | 26383347 | 20 | 3 | 15 | 13 | 16 | 6 | 4 | 6 | 4 | 20 | 0 | 11 | 11 | 13 | 20 |
| SE | 76642 | ENSG00000135503 | ACVR1B | chr12 | + | 51983998 | 51984166 | 51980968 | 51981199 | 51991862 | 51991993 | 34 | 25 | 28 | 42 | 61 | 0 | 1 | 1 | 2 | 1 | 114 | 52 | 56 | 64 | 116 |
| SE | 73336 | ENSG00000115970 | THADA | chr2 | - | 43556344 | 43556555 | 43552067 | 43552339 | 43560233 | 43560385 | 147 | 84 | 108 | 99 | 138 | 1 | 0 | 2 | 2 | 4 | 48 | 85 | 76 | 69 | 32 |

| | | | | | | | | | | | | | | |
|---|---|---|---|---|---|---|---|---|---|---|---|---|---|---|
| SE | 55808 | ENSG00000144218 | AFF3 | chr2 | - | 100112385 | 100112524 | 100008856 | 100008932 | 100129223 | 100129306 | 12 | 11 | 8 |
| SE | 18682 | ENSG00000129566 | TEP1 | chr14 | - | 20384599 | 20384713 | 20384037 | 20384232 | 20384984 | 20385109 | 20 | 4 | 17 |
| SE | 41150 | ENSG00000013573 | DDX11 | chr12 | + | 31084968 | 31085126 | 31084582 | 31084669 | 31087937 | 31087983 | 10 | 2 | 21 |
| SE | 11343 | ENSG00000175595 | ERCC4 | chr16 | + | 13934191 | 13934302 | 13932156 | 13932285 | 13935145 | 13935743 | 41 | 37 | 38 |
| SE | 98774 | ENSG00000157600 | TMEM164 | chrX | + | 110109079 | 110109146 | 110067346 | 110067396 | 110144797 | 110144876 | 71 | 37 | 45 |
| SE | 45775 | ENSG00000284770 | TBCE | chr1 | + | 235430704 | 235430804 | 235427139 | 235427239 | 235434203 | 235434233 | 34 | 17 | 26 |
| SE | 92833 | ENSG00000130958 | SLC35D2 | chr9 | - | 96343903 | 96343996 | 96324090 | 96324169 | 96360153 | 96360192 | 2 | 5 | 1 |
| SE | 57237 | ENSG00000143815 | LBR | chr1 | - | 225410290 | 225410818 | 225406663 | 225406832 | 225411336 | 225411440 | 180 | 92 | 147 |
| SE | 4639 | ENSG00000142347 | MYO1F | chr19 | - | 8545649 | 8545736 | 8544296 | 8544464 | 8548035 | 8548043 | 1089 | 467 | 634 |
| SE | 100978 | ENSG00000151348 | EXT2 | chr11 | + | 44126815 | 44126955 | 44124788 | 44124984 | 44130044 | 44130138 | 138 | 65 | 97 |
| SE | 43450 | ENSG00000162747 | FCGR3B | chr1 | - | 161626144 | 161626402 | 161623195 | 161624639 | 161629777 | 161630032 | 142 | 123 | 143 |
| SE | 72281 | ENSG00000131368 | MRPS25 | chr3 | - | 15053290 | 15053467 | 15049880 | 15049935 | 15059368 | 15059475 | 78 | 69 | 72 |
| SE | 37669 | ENSG00000168615 | ADAM9 | chr8 | + | 39071297 | 39071403 | 39055576 | 39055772 | 39077227 | 39077411 | 129 | 115 | 81 |
| SE | 70576 | ENSG00000022840 | RNF10 | chr12 | + | 120574467 | 120574537 | 120566952 | 120566980 | 120575630 | 120575688 | 2 | 1 | 1 |
| SE | 54909 | ENSG00000010803 | SCMH1 | chr1 | - | 41075218 | 41075451 | 41070594 | 41070721 | 41113282 | 41113526 | 34 | 28 | 37 |
| SE | 87114 | ENSG00000101935 | AMMECR1 | chrX | - | 110202445 | 110202536 | 110200953 | 110201050 | 110216517 | 110216632 | 63 | 32 | 71 |
| SE | 75228 | ENSG00000100503 | NIN | chr14 | - | 50766306 | 50766396 | 50763825 | 50763964 | 50766779 | 50766795 | 303 | 367 | 306 |
| SE | 77594 | ENSG00000127947 | PTPN12 | chr7 | + | 77563935 | 77564195 | 77537553 | 77537645 | 77571077 | 77571186 | 17 | 11 | 7 |
| SE | 46462 | ENSG00000115415 | STAT1 | chr2 | - | 191001073 | 191001163 | 190999625 | 190999704 | 191007562 | 191007661 | 1999 | 2517 | 2014 |
| SE | 90983 | ENSG00000132669 | RIN2 | chr20 | + | 19992167 | 19992299 | 19990005 | 19990311 | 19996678 | 19996842 | 160 | 156 | 60 |
| SE | 31700 | ENSG00000164284 | GRPEL2 | chr5 | + | 149349653 | 149349735 | 149348271 | 149348425 | 149350917 | 149354583 | 62 | 41 | 80 |
| SE | 59066 | ENSG00000103365 | GGA2 | chr16 | - | 23493359 | 23493458 | 23491676 | 23491800 | 23494302 | 23494378 | 279 | 110 | 244 |
| SE | 8749 | ENSG00000007202 | KIAA0100 | chr17 | - | 28621398 | 28621498 | 28620987 | 28621170 | 28623759 | 28623966 | 192 | 97 | 156 |
| SE | 63589 | ENSG00000206562 | METTL6 | chr3 | - | 15424954 | 15425089 | 15415771 | 15415942 | 15426286 | 15426635 | 106 | 61 | 69 |
| SE | 45427 | ENSG00000176542 | USF3 | chr3 | - | 113673347 | 113673376 | 113664312 | 113664409 | 113674831 | 113674896 | 20 | 11 | 16 |
| SE | 85552 | ENSG00000008294 | SPAG9 | chr17 | - | 50975862 | 50975901 | 50974770 | 50974947 | 50977107 | 50977221 | 8 | 12 | 14 |
| SE | 48276 | ENSG00000069424 | KCNAB2 | chr1 | + | 6091262 | 6091307 | 6090388 | 6090475 | 6094399 | 6094485 | 2 | 0 | 0 |
| SE | 75833 | ENSG00000124523 | SIRT5 | chr6 | + | 13591668 | 13591894 | 13584075 | 13584225 | 13595476 | 13595564 | 18 | 8 | 29 |
| SE | 35239 | ENSG00000119231 | SENP5 | chr3 | + | 196885150 | 196886694 | 196867855 | 196868073 | 196899665 | 196899771 | 210 | 253 | 226 |
| SE | 76675 | ENSG00000204227 | RING1 | chr6 | + | 33209914 | 33210130 | 33209559 | 33209786 | 33211157 | 33211547 | 124 | 76 | 132 |
| SE | 62528 | ENSG00000004866 | ST7 | chr7 | + | 117098612 | 117098815 | 116953502 | 116953691 | 117099761 | 117099844 | 3 | 3 | 0 |
| SE | 27076 | ENSG00000002549 | LAP3 | chr4 | + | 17597045 | 17597134 | 17588818 | 17588977 | 17598455 | 17598558 | 56 | 59 | 63 |
| SE | 11754 | ENSG00000260565 | ERVK13-1 | chr16 | - | 2672706 | 2672976 | 2671344 | 2672570 | 2673282 | 2673427 | 12 | 0 | 6 |
| SE | 106877 | ENSG00000005379 | TSPOAP1 | chr17 | - | 58308540 | 58309380 | 58307841 | 58307941 | 58309966 | 58310158 | 89 | 29 | 30 |
| SE | 106156 | ENSG00000089022 | MAPKAPK5 | chr12 | + | 111883580 | 111883768 | 111880446 | 111880527 | 111885915 | 111886036 | 130 | 112 | 156 |
| SE | 9354 | ENSG00000141030 | COPS3 | chr17 | - | 17260300 | 17260474 | 17248925 | 17249039 | 17261965 | 17262106 | 105 | 51 | 94 |
| SE | 108630 | ENSG00000136104 | RNASEH2B | chr13 | + | 50934884 | 50934999 | 50930682 | 50930759 | 50943320 | 50943352 | 799 | 562 | 772 |
| SE | 106431 | ENSG00000135297 | MTO1 | chr6 | + | 73482028 | 73482244 | 73481000 | 73481116 | 73482448 | 73482620 | 42 | 25 | 35 |
| SE | 109065 | ENSG00000198176 | TFDP1 | chr13 | + | 113585579 | 113585677 | 113584720 | 113584888 | 113585773 | 113585849 | 0 | 0 | 3 |
| SE | 46701 | ENSG00000171103 | TRMT61B | chr2 | - | 28861117 | 28861308 | 28852407 | 28852499 | 28869578 | 28870295 | 23 | 13 | 25 |
| SE | 39674 | ENSG00000181555 | SETD2 | chr3 | - | 47101457 | 47101555 | 47097954 | 47098081 | 47103345 | 47103423 | 469 | 429 | 429 |
| SE | 9930 | ENSG00000262879 | NA | chr17 | - | 47035043 | 47035126 | 47033277 | 47033381 | 47054117 | 47054213 | 0 | 2 | 0 |
| SE | 100247 | ENSG00000084234 | APLP2 | chr11 | + | 130135562 | 130135715 | 130133628 | 130133728 | 130140397 | 130140483 | 907 | 530 | 553 |
| SE | 71544 | ENSG00000145216 | FIP1L1 | chr4 | + | 53452919 | 53453133 | 53444047 | 53444103 | 53458652 | 53458790 | 184 | 117 | 149 |
| SE | 9800 | ENSG00000108384 | RAD51C | chr17 | + | 58706488 | 58706608 | 58703195 | 58703329 | 58709858 | 58709876 | 0 | 1 | 0 |
| SE | 21805 | ENSG00000126091 | ST3GAL3 | chr1 | + | 43792101 | 43792149 | 43736232 | 43736380 | 43814890 | 43814933 | 43 | 20 | 23 |
| SE | 74882 | ENSG00000162585 | FAAP20 | chr1 | - | 2192844 | 2192975 | 2189712 | 2189781 | 2193638 | 2193910 | 8 | 3 | 10 |
| SE | 61036 | ENSG00000066027 | PPP2R5A | chr1 | + | 212347346 | 212347406 | 212345802 | 212345933 | 212348388 | 212348497 | 311 | 175 | 205 |
| SE | 27245 | ENSG00000073331 | ALPK1 | chr4 | + | 112302338 | 112302485 | 112297368 | 112297469 | 112315800 | 112315852 | 2 | 9 | 10 |
| SE | 41659 | ENSG00000119185 | ITGB1BP1 | chr2 | - | 9419989 | 9420100 | 9418625 | 9418732 | 9422478 | 9422566 | 32 | 7 | 20 |
| SE | 97746 | ENSG00000146223 | RPL7L1 | chr6 | + | 42880860 | 42881082 | 42879638 | 42879951 | 42883450 | 42883614 | 16 | 14 | 19 |
| SE | 107531 | ENSG00000134283 | PPHLN1 | chr12 | + | 42332659 | 42332702 | 42326144 | 42326229 | 42335882 | 42335974 | 68 | 68 | 41 |
| SE | 58196 | ENSG00000172965 | MIR4435-2HG | chr2 | - | 111239801 | 111239996 | 111233627 | 111233739 | 111248147 | 111248241 | 4 | 1 | 6 |
| SE | 102975 | ENSG00000269609 | C10orf95-AS1 | chr10 | + | 102451398 | 102451555 | 102450186 | 102450330 | 102455336 | 102456267 | 4 | 2 | 15 |
| SE | 12774 | ENSG00000168802 | CHTF8 | chr16 | - | 69121052 | 69121170 | 69118009 | 69120649 | 69121435 | 69121492 | 183 | 111 | 92 |
| SE | 6648 | ENSG00000100342 | APOL1 | chr22 | + | 36259668 | 36259856 | 36257318 | 36257407 | 36261595 | 36261644 | 6 | 2 | 1 |
| SE | 84436 | ENSG00000188917 | TRMT2B | chrX | - | 101051250 | 101051733 | 101036973 | 101037073 | 101051816 | 101052111 | 21 | 16 | 15 |
| SE | 106032 | ENSG00000204482 | LST1 | chr6 | + | 31587640 | 31587733 | 31587116 | 31587318 | 31588496 | 31588810 | 144 | 177 | 119 |
| SE | 19867 | ENSG00000165501 | LRR1 | chr14 | + | 49602369 | 49602468 | 49598760 | 49599203 | 49614255 | 49614670 | 4 | 2 | 4 |
| SE | 94523 | ENSG00000070061 | ELP1 | chr9 | - | 108920690 | 108920783 | 108919252 | 108919349 | 108922841 | 108922927 | 2 | 0 | 1 |
| SE | 41766 | ENSG00000162521 | RBBP4 | chr1 | + | 32668738 | 32668854 | 32668224 | 32668398 | 32668971 | 32669088 | 655 | 412 | 545 |
| SE | 66320 | ENSG00000143207 | COP1 | chr1 | - | 176116623 | 176116681 | 176085775 | 176085890 | 176149005 | 176149074 | 97 | 86 | 56 |
| SE | 72885 | ENSG00000121716 | PILRB | chr7 | + | 100353209 | 100353270 | 100352563 | 100353123 | 100355142 | 100355240 | 13 | 6 | 22 |
| SE | 19181 | ENSG00000140105 | WARS1 | chr14 | - | 100374135 | 100374220 | 100361830 | 100361921 | 100375282 | 100375344 | 0 | 0 | 12 |
| SE | 4445 | ENSG00000105058 | FAM32A | chr19 | + | 16189387 | 16189511 | 16185623 | 16185765 | 16190519 | 16190573 | 0 | 2 | 2 |
| SE | 98289 | ENSG00000068831 | RASGRP2 | chr11 | - | 64744332 | 64744482 | 64742793 | 64742937 | 64744741 | 64744867 | 15 | 6 | 12 |
| SE | 9350 | ENSG00000141030 | COPS3 | chr17 | - | 17247479 | 17247560 | 17246615 | 17247151 | 17248925 | 17249039 | 567 | 505 | 476 |
| SE | 3137 | ENSG00000121417 | ZNF211 | chr19 | + | 57639911 | 57640067 | 57634628 | 57634755 | 57640703 | 57642279 | 12 | 22 | 19 |
| SE | 27226 | ENSG00000163812 | ZDHHC3 | chr3 | - | 44929305 | 44929436 | 44915260 | 44926847 | 44933117 | 44933199 | 187 | 129 | 99 |
| SE | 79375 | ENSG00000122482 | ZNF644 | chr1 | - | 90937484 | 90938090 | 90918051 | 90918154 | 90982309 | 90982370 | 110 | 86 | 119 |
| SE | 53960 | ENSG00000138101 | DTNB | chr2 | - | 25596085 | 25596240 | 25580720 | 25580826 | 25628170 | 25628384 | 32 | 30 | 26 |
| SE | 95566 | ENSG00000180104 | EXOC3 | chr5 | + | 459358 | 459459 | 457899 | 458025 | 461959 | 462070 | 142 | 69 | 121 |
| SE | 36756 | ENSG00000138231 | DBR1 | chr3 | - | 138171632 | 138171713 | 138170106 | 138170192 | 138173501 | 138173626 | 36 | 34 | 36 |
| SE | 60992 | ENSG00000109381 | ELF2 | chr4 | - | 139067683 | 139067770 | 139061864 | 139062057 | 139071865 | 139072003 | 134 | 103 | 132 |
| SE | 70418 | ENSG00000115652 | UXS1 | chr2 | - | 106164735 | 106164799 | 106163666 | 106163710 | 106166055 | 106166083 | 31 | 22 | 27 |
| SE | 44922 | ENSG00000182095 | TNRC18 | chr7 | - | 5329936 | 5329969 | 5325095 | 5325248 | 5332621 | 5332878 | 16 | 9 | 14 |
| SE | 93089 | ENSG00000113924 | HGD | chr3 | - | 120644318 | 120644443 | 120641588 | 120641693 | 120646266 | 120646335 | 1 | 1 | 2 |
| SE | 48623 | ENSG00000131374 | TBC1D5 | chr3 | - | 17562018 | 17562142 | 17518901 | 17518995 | 17623848 | 17623913 | 5 | 5 | 18 |
| SE | 72571 | ENSG00000068724 | TTC7A | chr2 | + | 47060768 | 47060971 | 47051745 | 47051880 | 47073701 | 47073923 | 95 | 29 | 70 |
| SE | 48285 | ENSG00000229474 | PATL2 | chr15 | - | 44667105 | 44667203 | 44666391 | 44666541 | 44668341 | 44668482 | 50 | 13 | 23 |
| SE | 26956 | ENSG00000172869 | DMXL1 | chr5 | + | 119081573 | 119081697 | 119071528 | 119071656 | 119097978 | 119098104 | 1 | 4 | 3 |
| SE | 48462 | ENSG00000177030 | DEAF1 | chr11 | - | 651161 | 651407 | 644232 | 644654 | 653961 | 654051 | 6 | 0 | 1 |
| SE | 90972 | ENSG00000132669 | RIN2 | chr20 | + | 19815222 | 19815298 | 19799621 | 19799747 | 19871081 | 19872284 | 8 | 0 | 0 |
| SE | 59603 | ENSG00000085377 | PREP | chr6 | - | 105333313 | 105333505 | 105328828 | 105329026 | 105352971 | 105353077 | 477 | 270 | 382 |
| SE | 101947 | ENSG00000122033 | MTIF3 | chr13 | - | 27445087 | 27445156 | 27441016 | 27441077 | 27450508 | 27450564 | 15 | 10 | 10 |
| SE | 4616 | ENSG00000167468 | GPX4 | chr19 | + | 1106241 | 1106266 | 1105657 | 1105809 | 1106399 | 1106459 | 822 | 513 | 632 |
| SE | 81124 | ENSG00000143226 | FCGR2A | chr1 | + | 161506336 | 161506591 | 161505986 | 161506007 | 161509819 | 161510074 | 408 | 230 | 233 |
| SE | 12547 | ENSG00000103148 | NPRL3 | chr16 | - | 130521 | 130591 | 119125 | 119255 | 138149 | 138334 | 46 | 11 | 15 |
| SE | 73133 | ENSG00000175198 | PCCA | chr13 | + | 100262728 | 100262831 | 100257594 | 100257673 | 100268688 | 100268783 | 61 | 68 | 44 |
| SE | 105464 | ENSG00000135720 | DYNC1LI2 | chr16 | - | 66727687 | 66727805 | 66725827 | 66725944 | 66729039 | 66729099 | 46 | 27 | 40 |
| SE | 109126 | ENSG00000126217 | MCF2L | chr13 | + | 113060592 | 113060712 | 113045270 | 113045361 | 113064303 | 113064311 | 7 | 12 | 23 |
| SE | 103982 | ENSG00000151651 | ADAM8 | chr10 | - | 133263687 | 133263765 | 133262419 | 133263233 | 133267351 | 133267417 | 117 | 26 | 67 |
| SE | 73590 | ENSG00000124356 | STAMBP | chr2 | + | 73829404 | 73829510 | 73828960 | 73828996 | 73830844 | 73831059 | 9 | 1 | 4 |
| SE | 100705 | ENSG00000187147 | RNF220 | chr1 | + | 44635544 | 44635588 | 44632254 | 44632385 | 44636029 | 44636162 | 166 | 93 | 121 |
| SE | 57028 | ENSG00000268043 | NBPF12 | chr1 | + | 146951347 | 146951489 | 146949313 | 146949422 | 146960108 | 146960318 | 19 | 6 | 6 |
| SE | 41805 | ENSG00000136059 | VILL | chr3 | + | 38003167 | 38003313 | 38001701 | 38001860 | 38004254 | 38004399 | 15 | 7 | 14 |
| SE | 56068 | ENSG00000160229 | ZNF66 | chr19 | + | 20792511 | 20792638 | 20776303 | 20776481 | 20793782 | 20793878 | 6 | 1 | 2 |
| SE | 64120 | ENSG00000135205 | CCDC146 | chr7 | + | 77273693 | 77273789 | 77262120 | 77262307 | 77274481 | 77274652 | 48 | 37 | 42 |
| SE | 52532 | ENSG00000029534 | ANK1 | chr8 | - | 41661800 | 41661941 | 41653219 | 41655753 | 41663658 | 41663742 | 15 | 22 | 15 |
| SE | 59443 | ENSG00000145246 | ATP10D | chr4 | + | 47515475 | 47515670 | 47512503 | 47512830 | 47523011 | 47523216 | 152 | 99 | 91 |
| SE | 67684 | ENSG00000181381 | DDX60L | chr4 | - | 168377727 | 168377843 | 168375376 | 168375524 | 168378353 | 168378475 | 30 | 89 | 76 |
| SE | 47631 | ENSG00000157349 | DDX19B | chr16 | + | 70312608 | 70312657 | 70299185 | 70299354 | 70314901 | 70314955 | 33 | 17 | 27 |
| SE | 54678 | ENSG00000070010 | UFD1 | chr22 | - | 19475469 | 19475602 | 19467402 | 19468003 | 19479082 | 19479172 | 29 | 6 | 23 |
| SE | 82170 | ENSG00000146872 | TLK2 | chr17 | + | 62508407 | 62508553 | 62481120 | 62481206 | 62520772 | 62520844 | 4 | 18 | 14 |
| SE | 39762 | ENSG00000132185 | FCRLA | chr1 | + | 161710439 | 161710492 | 161707238 | 161707343 | 161710759 | 161710863 | 1 | 0 | 0 |
| SE | 41940 | ENSG00000163374 | YY1AP1 | chr1 | - | 155675009 | 155675096 | 155670319 | 155670464 | 155676547 | 155676746 | 97 | 76 | 92 |
| SE | 82149 | ENSG00000065000 | AP3D1 | chr19 | - | 2127151 | 2127201 | 2123829 | 2123879 | 2129089 | 2129121 | 234 | 126 | 208 |
| SE | 49305 | ENSG00000120029 | ARMH3 | chr10 | - | 102023674 | 102023749 | 102013967 | 102014024 | 102025155 | 102025248 | 50 | 33 | 56 |
| SE | 17084 | ENSG00000206190 | ATP10A | chr15 | - | 25694818 | 25695146 | 25691714 | 25691791 | 25701915 | 25702100 | 51 | 26 | 56 |
| SE | 17900 | ENSG00000197249 | SERPINA1 | chr14 | - | 94379463 | 94379611 | 94376746 | 94378640 | 94382591 | 94383241 | 1273 | 743 | 568 |
| SE | 26913 | ENSG00000078967 | UBE2D4 | chr7 | + | 43942825 | 43942857 | 43938430 | 43938494 | 43942953 | 43943031 | 11 | 13 | 10 |
| SE | 65491 | ENSG00000125875 | TBC1D20 | chr20 | - | 447888 | 448074 | 445049 | 445130 | 462335 | 462361 | 88 | 54 | 84 |
| SE | 68852 | ENSG00000168769 | TET2 | chr4 | + | 105241338 | 105241429 | 105233896 | 105237351 | 105242833 | 105242927 | 221 | 203 | 135 |
| SE | 71057 | ENSG00000169914 | OTUD3 | chr1 | + | 19894367 | 19894480 | 19890384 | 19890533 | 19897539 | 19897662 | 37 | 51 | 37 |
| SE | 80986 | ENSG00000130429 | ARPC1B | chr7 | + | 99377357 | 99377455 | 99374729 | 99374781 | 99385701 | 99385778 | 12 | 4 | 10 |
| SE | 67347 | ENSG00000156475 | PPP2R2B | chr5 | - | 146592048 | 146592193 | 146590037 | 146590226 | 146592970 | 146593062 | 11 | 4 | 4 |
| SE | 13169 | ENSG00000103121 | CMC2 | chr16 | - | 81001247 | 81001334 | 80997313 | 80997429 | 81006733 | 81006858 | 3 | 1 | 3 |
| SE | 98166 | ENSG00000171634 | BPTF | chr17 | + | 67886127 | 67886316 | 67874919 | 67875020 | 67891843 | 67891962 | 137 | 96 | 108 |
| SE | 68006 | ENSG00000189046 | ALKBH2 | chr12 | - | 109090008 | 109090207 | 109088188 | 109088512 | 109092506 | 109092787 | 25 | 27 | 40 |
| SE | 88035 | ENSG00000156171 | DRAM2 | chr1 | - | 111122615 | 111122643 | 111120515 | 111120693 | 111124741 | 111124881 | 16 | 7 | 13 |
| SE | 19558 | ENSG00000198718 | TOGARAM1 | chr14 | + | 44995745 | 44995902 | 44962189 | 44964467 | 44999362 | 44999497 | 61 | 54 | 56 |
| SE | 102331 | ENSG00000166171 | DPCD | chr10 | + | 101608834 | 101608937 | 101605139 | 101605212 | 101609366 | 101609662 | 2 | 1 | 1 |
| SE | 71970 | ENSG00000160710 | ADAR | chr1 | - | 154625062 | 154625153 | 154601040 | 154602626 | 154627854 | 154627991 | 0 | 0 | 0 |
| SE | 65201 | ENSG00000010244 | ZNF207 | chr17 | + | 32361467 | 32361515 | 32360891 | 32360967 | 32362913 | 32362984 | 488 | 325 | 475 |
| SE | 98271 | ENSG00000144354 | CDCA7 | chr2 | + | 173366282 | 173366432 | 173365451 | 173365592 | 173367149 | 173367286 | 30 | 11 | 34 |
| SE | 6510 | ENSG00000124155 | PIGT | chr20 | + | 45416516 | 45416654 | 45416106 | 45416343 | 45419294 | 45419395 | 85 | 24 | 46 |
| SE | 103436 | ENSG00000265479 | DTX2P1-UPK3BP1-PMS2P11 | chr7 | + | 77039950 | 77040089 | 77015671 | 77015855 | 77041976 | 77042063 | 102 | 43 | 33 |
| SE | 42131 | ENSG00000198952 | SMG5 | chr1 | - | 156272319 | 156272398 | 156268289 | 156268415 | 156273360 | 156273450 | 172 | 129 | 145 |
| SE | 9387 | ENSG00000121067 | SPOP | chr17 | - | 49668026 | 49668078 | 49636758 | 49636809 | 49677932 | 49678003 | 9 | 6 | 9 |
| SE | 71266 | ENSG00000121741 | ZMYM2 | chr13 | + | 19970024 | 19970071 | 19959957 | 19960026 | 19993062 | 19993919 | 35 | 24 | 51 |
| SE | 39284 | ENSG00000130695 | CEP85 | chr1 | + | 26259616 | 26259802 | 26258142 | 26258260 | 26268482 | 26268635 | 76 | 69 | 78 |
| SE | 55958 | ENSG00000136161 | RCBTB2 | chr13 | - | 48499620 | 48499760 | 48496190 | 48496321 | 48501741 | 48501868 | 101 | 77 | 48 |
| SE | 19313 | ENSG00000156414 | TDRD9 | chr14 | + | 104022156 | 104022330 | 104015980 | 104016088 | 104024568 | 104024680 | 7 | 13 | 8 |
| SE | 9677 | ENSG00000055483 | USP36 | chr17 | - | 78813772 | 78813873 | 78812859 | 78813001 | 78818666 | 78818778 | 57 | 23 | 57 |
| SE | 99104 | ENSG00000110274 | CEP164 | chr11 | + | 117393003 | 117393126 | 117392495 | 117392627 | 117394349 | 117394493 | 96 | 55 | 105 |
| SE | 58298 | ENSG00000117262 | GPR89A | chr1 | + | 145612201 | 145612247 | 145608047 | 145608175 | 145616233 | 145616293 | 21 | 2 | 26 |
| SE | 2043 | ENSG00000099331 | MYO9B | chr19 | + | 17159394 | 17159484 | 17156908 | 17157038 | 17162349 | 17162466 | 431 | 200 | 319 |
| SE | 8120 | ENSG00000120071 | KANSL1 | chr17 | - | 46032046 | 46032167 | 46029984 | 46031703 | 46033079 | 46033192 | 27 | 9 | 33 |
| SE | 48148 | ENSG00000117543 | DPH5 | chr1 | - | 100992636 | 100992740 | 100989622 | 100990631 | 101001466 | 101001544 | 63 | 52 | 86 |
| SE | 10317 | ENSG00000108848 | LUC7L3 | chr17 | + | 50738140 | 50738199 | 50736959 | 50737026 | 50740305 | 50740345 | 40 | 35 | 59 |
| SE | 40660 | ENSG00000162994 | CLHC1 | chr2 | - | 55209403 | 55209516 | 55208625 | 55208710 | 55209629 | 55209652 | 7 | 26 | 4 |
| SE | 1527 | ENSG00000100242 | SUN2 | chr22 | - | 38754641 | 38754690 | 38752506 | 38752665 | 38755271 | 38755429 | 88 | 42 | 93 |
| SE | 104695 | ENSG00000211455 | STK38L | chr12 | + | 27315014 | 27315117 | 27314503 | 27314658 | 27315288 | 27315350 | 181 | 188 | 168 |
| SE | 3201 | ENSG00000128000 | ZNF780B | chr19 | - | 40055415 | 40055576 | 40050323 | 40050377 | 40056100 | 40056175 | 23 | 13 | 12 |
| SE | 52712 | ENSG00000204054 | LINC00963 | chr9 | + | 129493477 | 129493595 | 129488708 | 129489299 | 129503322 | 129503672 | 16 | 12 | 10 |
| SE | 71487 | ENSG00000155438 | NIFK | chr2 | - | 121735612 | 121735750 | 121730382 | 121731104 | 121736745 | 121736872 | 23 | 7 | 36 |
| SE | 59505 | ENSG00000148660 | CAMK2G | chr10 | - | 73847987 | 73848082 | 73847224 | 73847347 | 73848525 | 73848609 | 77 | 40 | 75 |
| SE | 34244 | ENSG00000136560 | TANK | chr2 | + | 161161284 | 161161412 | 161160348 | 161160486 | 161179613 | 161179761 | 8 | 9 | 13 |
| SE | 21069 | ENSG00000196182 | STK40 | chr1 | - | 36367847 | 36368084 | 36361220 | 36361340 | 36385722 | 36385884 | 32 | 8 | 10 |
| SE | 26741 | ENSG00000125089 | SH3TC1 | chr4 | + | 8232706 | 8233065 | 8231975 | 8232156 | 8233362 | 8233513 | 4 | 2 | 0 |
| SE | 64107 | ENSG00000283602 | NA | chr5 | - | 140591669 | 140591752 | 140590874 | 140590991 | 140592023 | 140592101 | 6 | 2 | 0 |
| SE | 91562 | ENSG00000070501 | POLB | chr8 | + | 42339011 | 42339069 | 42338481 | 42338685 | 42344952 | 42345019 | 12 | 8 | 22 |
| SE | 90895 | ENSG00000185728 | YTHDF3 | chr8 | + | 63172652 | 63172798 | 63169386 | 63169411 | 63175330 | 63175416 | 12 | 13 | 17 |
| SE | 1615 | ENSG00000244625 | MIATNB | chr22 | + | 26693680 | 26693836 | 26672766 | 26672987 | 26726643 | 26726794 | 8 | 24 | 11 |
| SE | 7861 | ENSG00000100425 | BRD1 | chr22 | - | 49787389 | 49787494 | 49777677 | 49777813 | 49794033 | 49794294 | 84 | 37 | 106 |
| SE | 57927 | ENSG00000110200 | ANAPC15 | chr11 | - | 72111411 | 72111496 | 72111156 | 72111286 | 72112649 | 72112754 | 69 | 30 | 35 |
| SE | 69315 | ENSG00000112367 | FIG4 | chr6 | + | 109741443 | 109741544 | 109735149 | 109735298 | 109743109 | 109743272 | 45 | 43 | 44 |
| SE | 52591 | ENSG00000163607 | GTPBP8 | chr3 | + | 112995134 | 112995265 | 112993025 | 112993124 | 112996891 | 112996991 | 164 | 87 | 144 |
| SE | 54011 | ENSG00000197989 | SNHG12 | chr1 | - | 28581093 | 28581229 | 28580334 | 28580646 | 28581589 | 28581623 | 46 | 16 | 57 |
| SE | 52129 | ENSG00000197312 | DDI2 | chr1 | + | 15656616 | 15656679 | 15651705 | 15651895 | 15659836 | 15661057 | 34 | 14 | 28 |
| SE | 41297 | ENSG00000058668 | ATP2B4 | chr1 | + | 203713164 | 203713252 | 203711959 | 203712139 | 203714170 | 203714277 | 390 | 161 | 268 |
| SE | 8799 | ENSG00000011143 | MKS1 | chr17 | - | 58214258 | 58214387 | 58212981 | 58213869 | 58216087 | 58216243 | 13 | 12 | 13 |
| SE | 73057 | ENSG00000115993 | TRAK2 | chr2 | - | 201394797 | 201394872 | 201392908 | 201393046 | 201395313 | 201395444 | 296 | 246 | 278 |
| SE | 40520 | ENSG00000285708 | NA | chr3 | - | 71710411 | 71710484 | 71693874 | 71693941 | 71728475 | 71728623 | 6 | 12 | 5 |
| SE | 16369 | ENSG00000140395 | WDR61 | chr15 | - | 78288842 | 78288917 | 78288338 | 78288382 | 78289618 | 78289712 | 3 | 2 | 4 |
| SE | 99773 | ENSG00000186104 | CYP2R1 | chr11 | - | 14880135 | 14880768 | 14879113 | 14879207 | 14885775 | 14885917 | 48 | 25 | 36 |
| SE | 16808 | ENSG00000137814 | HAUS2 | chr15 | + | 42559338 | 42559408 | 42558197 | 42558290 | 42563748 | 42563857 | 8 | 12 | 13 |
| SE | 65318 | ENSG00000285756 | NA | chrX | - | 3818656 | 3818703 | 3818441 | 3818500 | 3828505 | 3829392 | 7 | 7 | 10 |
| SE | 78156 | ENSG00000239713 | APOBEC3G | chr22 | + | 39078931 | 39079085 | 39077274 | 39077378 | 39080932 | 39081194 | 731 | 324 | 361 |
| SE | 84944 | ENSG00000139436 | GIT2 | chr12 | - | 109945259 | 109945349 | 109939164 | 109939247 | 109951211 | 109951316 | 22 | 19 | 21 |
| SE | 97953 | ENSG00000137500 | CCDC90B | chr11 | - | 83266866 | 83267038 | 83265864 | 83265979 | 83273646 | 83273700 | 4 | 1 | 9 |
| SE | 40546 | ENSG00000116977 | LGALS8 | chr1 | + | 236525423 | 236525545 | 236523969 | 236524061 | 236525967 | 236526115 | 1 | 1 | 2 |
| SE | 59399 | ENSG00000139631 | CSAD | chr12 | - | 53171176 | 53171441 | 53170422 | 53170502 | 53171881 | 53171988 | 20 | 10 | 11 |
| SE | 61905 | ENSG00000076242 | MLH1 | chr3 | + | 37048516 | 37048609 | 37046753 | 37047683 | 37048903 | 37049017 | 78 | 37 | 55 |
| SE | 10983 | ENSG00000083454 | P2RX5 | chr17 | - | 3688625 | 3688759 | 3688011 | 3688105 | 3689491 | 3689627 | 172 | 61 | 193 |
| SE | 16782 | ENSG00000125107 | CNOT1 | chr16 | - | 58624672 | 58624794 | 58599235 | 58599511 | 58629727 | 58629825 | 9 | 7 | 18 |
| SE | 73963 | ENSG00000166436 | TRIM66 | chr11 | - | 8621031 | 8621237 | 8619382 | 8619535 | 8621644 | 8621819 | 4 | 9 | 12 |
| SE | 71212 | ENSG00000079263 | SP140 | chr2 | + | 230247915 | 230248065 | 230244987 | 230245071 | 230248884 | 230248968 | 32 | 46 | 35 |
| SE | 89870 | ENSG00000034677 | RNF19A | chr8 | - | 100259853 | 100259997 | 100257059 | 100259246 | 100264033 | 100264080 | 311 | 211 | 217 |
| SE | 56283 | ENSG00000182183 | SHISAL2A | chr1 | + | 52642862 | 52643002 | 52633167 | 52633675 | 52667398 | 52667688 | 37 | 16 | 49 |
| SE | 103258 | ENSG00000172661 | WASHC2C | chr10 | + | 45751481 | 45751553 | 45750750 | 45750838 | 45752587 | 45752706 | 141 | 129 | 113 |
| SE | 36408 | ENSG00000116191 | RALGPS2 | chr1 | + | 178892229 | 178892307 | 178889641 | 178889696 | 178893918 | 178894024 | 35 | 23 | 17 |
| SE | 87467 | ENSG00000271147 | ARMCX5-GPRASP2 | chrX | + | 102639399 | 102639533 | 102605481 | 102605653 | 102713781 | 102713849 | 22 | 12 | 14 |
| SE | 17313 | ENSG00000100445 | SDR39U1 | chr14 | - | 24442177 | 24442260 | 24441673 | 24441795 | 24442345 | 24442448 | 88 | 56 | 96 |
| SE | 108971 | ENSG00000111877 | MCM9 | chr6 | - | 118932606 | 118932740 | 118931419 | 118931738 | 118934890 | 118935024 | 10 | 13 | 9 |
| SE | 63615 | ENSG00000138029 | HADHB | chr2 | + | 26284116 | 26284204 | 26283003 | 26283051 | 26284882 | 26284957 | 576 | 532 | 454 |
| SE | 69789 | ENSG00000065675 | PRKCQ | chr10 | - | 6507435 | 6507496 | 6498395 | 6498558 | 6510994 | 6511194 | 245 | 124 | 233 |
| SE | 56991 | ENSG00000131725 | WDR44 | chrX | + | 118392631 | 118393271 | 118378418 | 118378452 | 118394054 | 118394185 | 107 | 102 | 73 |
| SE | 65475 | ENSG00000227811 | INKA2-AS1 | chr1 | + | 111744374 | 111744529 | 111739578 | 111740077 | 111746059 | 111747765 | 26 | 12 | 1 |
| SE | 70636 | ENSG00000188647 | PTAR1 | chr9 | - | 69732138 | 69732352 | 69723325 | 69723630 | 69734149 | 69734254 | 186 | 161 | 188 |
| SE | 58070 | ENSG00000082701 | GSK3B | chr3 | - | 119947267 | 119947351 | 119863418 | 119863605 | 120002045 | 120002236 | 45 | 42 | 44 |
| SE | 103816 | ENSG00000225484 | NUTM2B-AS1 | chr10 | - | 79808758 | 79808832 | 79804484 | 79804527 | 79814612 | 79814716 | 24 | 27 | 30 |
| SE | 24323 | ENSG00000185404 | SP140L | chr2 | + | 230328756 | 230328831 | 230327183 | 230327301 | 230357804 | 230357967 | 114 | 85 | 106 |
| SE | 39079 | ENSG00000172175 | MALT1 | chr18 | + | 58737767 | 58737926 | 58735201 | 58735329 | 58741864 | 58741879 | 1 | 5 | 3 |
| SE | 11023 | ENSG00000213977 | TAX1BP3 | chr17 | - | 3664194 | 3664272 | 3662894 | 3663885 | 3664678 | 3664798 | 123 | 84 | 63 |
| SE | 33527 | ENSG00000082996 | RNF13 | chr3 | + | 149846010 | 149846140 | 149813214 | 149813353 | 149852515 | 149852596 | 424 | 236 | 275 |
| SE | 14422 | ENSG00000129007 | CALML4 | chr15 | - | 68199540 | 68199681 | 68197439 | 68197628 | 68205120 | 68205151 | 22 | 34 | 5 |
| SE | 107795 | ENSG00000050426 | LETMD1 | chr12 | + | 51052091 | 51052207 | 51048342 | 51049185 | 51056349 | 51056391 | 42 | 17 | 44 |
| SE | 96235 | ENSG00000133943 | DGLUCY | chr14 | + | 91147780 | 91147858 | 91145220 | 91145313 | 91157638 | 91157690 | 67 | 65 | 61 |
| SE | 57303 | ENSG00000015532 | XYLT2 | chr17 | + | 50355500 | 50355581 | 50354407 | 50354583 | 50355780 | 50355997 | 23 | 12 | 24 |
| SE | 101171 | ENSG00000133816 | MICAL2 | chr11 | + | 12249183 | 12249246 | 12243986 | 12244112 | 12256784 | 12256968 | 27 | 5 | 16 |
| SE | 43316 | ENSG00000009413 | REV3L | chr6 | - | 111460299 | 111460406 | 111416282 | 111416472 | 111472012 | 111472140 | 0 | 2 | 4 |
| SE | 32673 | ENSG00000150995 | ITPR1 | chr3 | + | 4777263 | 4777374 | 4775241 | 4775442 | 4779549 | 4779645 | 161 | 109 | 116 |
| SE | 78325 | ENSG00000228409 | CCT6P1 | chr7 | + | 65759174 | 65759271 | 65756468 | 65756574 | 65759859 | 65759998 | 21 | 21 | 16 |
| SE | 25713 | ENSG00000247774 | PCED1B-AS1 | chr12 | - | 47209280 | 47209482 | 47206510 | 47206585 | 47210639 | 47210761 | 74 | 27 | 91 |
| SE | 81827 | ENSG00000187953 | PMS2CL | chr7 | + | 6733398 | 6733527 | 6731896 | 6731955 | 6735304 | 6735389 | 12 | 7 | 25 |
| SE | 36197 | ENSG00000116731 | PRDM2 | chr1 | + | 13816426 | 13816570 | 13773077 | 13773188 | 13823158 | 13823419 | 30 | 8 | 31 |
| SE | 104874 | ENSG00000109685 | NSD2 | chr4 | + | 1939653 | 1939778 | 1935143 | 1935262 | 1951071 | 1951180 | 39 | 28 | 31 |
| SE | 70058 | ENSG00000151665 | PIGF | chr2 | - | 46613693 | 46613785 | 46612227 | 46612344 | 46616969 | 46617041 | 32 | 20 | 23 |
| SE | 2297 | ENSG00000142002 | DPP9 | chr19 | - | 4715528 | 4715726 | 4714080 | 4714337 | 4719850 | 4719941 | 1 | 0 | 3 |
| SE | 1071 | ENSG00000100330 | MTMR3 | chr22 | + | 29970975 | 29971062 | 29957035 | 29957088 | 29978441 | 29978531 | 226 | 99 | 130 |
| SE | 102261 | ENSG00000178440 | TIMM23B-AGAP6 | chr10 | + | 49976083 | 49976185 | 49973011 | 49973142 | 49987164 | 49987234 | 1 | 2 | 0 |
| SE | 79166 | ENSG00000145191 | EIF2B5 | chr3 | + | 184143441 | 184143565 | 184143051 | 184143142 | 184144098 | 184144224 | 180 | 103 | 146 |
| SE | 95784 | ENSG00000148120 | AOPEP | chr9 | + | 94923985 | 94924175 | 94800756 | 94801002 | 94955176 | 94955279 | 8 | 5 | 12 |
| SE | 18265 | ENSG00000171723 | GPHN | chr14 | + | 66842677 | 66842716 | 66824473 | 66824566 | 66879938 | 66880032 | 3 | 0 | 0 |
| SE | 86713 | ENSG00000128654 | MTX2 | chr2 | + | 176330583 | 176330660 | 176328873 | 176328912 | 176337492 | 176337630 | 9 | 15 | 5 |
| SE | 82103 | ENSG00000128891 | CCDC32 | chr15 | - | 40553594 | 40554127 | 40539238 | 40539355 | 40557215 | 40557372 | 59 | 51 | 60 |
| SE | 85166 | ENSG00000136504 | KAT7 | chr17 | + | 49817819 | 49818011 | 49815802 | 49815913 | 49821336 | 49821426 | 303 | 201 | 327 |
| SE | 82829 | ENSG00000155849 | ELMO1 | chr7 | - | 37393821 | 37393917 | 37342612 | 37342763 | 37448674 | 37449186 | 0 | 0 | 2 |
| SE | 107178 | ENSG00000063046 | EIF4B | chr12 | + | 53025226 | 53025345 | 53022492 | 53022627 | 53027781 | 53027919 | 18 | 7 | 5 |
| SE | 88239 | ENSG00000100918 | REC8 | chr14 | + | 24175542 | 24175624 | 24173290 | 24173411 | 24176821 | 24176884 | 19 | 28 | 38 |
| SE | 79526 | ENSG00000105778 | AVL9 | chr7 | + | 32580218 | 32580272 | 32575954 | 32576072 | 32580801 | 32580890 | 83 | 48 | 73 |
| SE | 97754 | ENSG00000166900 | STX3 | chr11 | + | 59795371 | 59795482 | 59793379 | 59793514 | 59797282 | 59797396 | 105 | 98 | 100 |
| SE | 3629 | ENSG00000007047 | MARK4 | chr19 | + | 45277922 | 45278042 | 45271471 | 45271708 | 45278515 | 45278615 | 93 | 69 | 81 |
| SE | 47533 | ENSG00000121766 | ZCCHC17 | chr1 | + | 31346639 | 31346740 | 31338956 | 31339048 | 31364031 | 31364953 | 102 | 55 | 56 |
| SE | 33925 | ENSG00000160796 | NBEAL2 | chr3 | + | 46991218 | 46991304 | 46989510 | 46989593 | 46991839 | 46991946 | 50 | 17 | 34 |
| SE | 37354 | ENSG00000075826 | SEC31B | chr10 | - | 100507424 | 100507567 | 100506320 | 100506420 | 100507907 | 100508048 | 44 | 43 | 54 |
| SE | 35114 | ENSG00000266338 | NBPF15 | chr1 | - | 144429699 | 144429863 | 144428605 | 144428657 | 144433772 | 144433824 | 113 | 60 | 83 |
| SE | 70950 | ENSG00000131781 | FMO5 | chr1 | - | 147201151 | 147201343 | 147190176 | 147190249 | 147208851 | 147209051 | 22 | 14 | 4 |
| SE | 70557 | ENSG00000022840 | RNF10 | chr12 | + | 120539367 | 120539468 | 120534587 | 120534968 | 120546404 | 120546601 | 16 | 12 | 18 |
| SE | 23443 | ENSG00000166888 | STAT6 | chr12 | - | 57099766 | 57099903 | 57098791 | 57099440 | 57099995 | 57100090 | 603 | 374 | 534 |
| SE | 71319 | ENSG00000165280 | VCP | chr9 | - | 35059063 | 35059219 | 35053927 | 35057530 | 35059442 | 35059801 | 315 | 170 | 292 |
| SE | 68427 | ENSG00000101104 | PABPC1L | chr20 | + | 44933056 | 44933185 | 44930459 | 44930726 | 44935390 | 44935497 | 6 | 5 | 8 |

| | | | | | | | | | | | |
|---|---|---|---|---|---|---|---|---|---|---|---|
| SE | 20607 | ENSG00000130779 | CLIP1 | chr12 | - | 122276438 | 122276444 | 122274037 | 122274162 | 122278153 | 122278203 |
| SE | 39025 | ENSG00000003400 | CASP10 | chr2 | + | 201205881 | 201205973 | 201203729 | 201203766 | 201208074 | 201208183 |
| SE | 12204 | ENSG00000141002 | TCF25 | chr16 | + | 89908965 | 89909083 | 89907242 | 89907322 | 89910590 | 89910663 |
| SE | 75030 | ENSG00000100814 | CCNB1IP1 | chr14 | - | 20329173 | 20329373 | 20326715 | 20326793 | 20333253 | 20333279 |
| SE | 79618 | ENSG00000162688 | AGL | chr1 | + | 99862256 | 99862423 | 99850974 | 99851124 | 99864385 | 99864589 |
| SE | 9385 | ENSG00000121067 | SPOP | chr17 | - | 49668026 | 49668078 | 49622732 | 49622876 | 49677932 | 49678003 |
| SE | 92842 | ENSG00000130958 | SLC35D2 | chr9 | - | 96345298 | 96345401 | 96343903 | 96343996 | 96351102 | 96351171 |
| SE | 103445 | ENSG00000148840 | PPRC1 | chr10 | + | 102143044 | 102143098 | 102139099 | 102142004 | 102144249 | 102144307 |
| SE | 32081 | ENSG00000242086 | MUC20-OT1 | chr3 | + | 195689545 | 195689608 | 195689338 | 195689438 | 195696360 | 195696589 |
| SE | 73211 | ENSG00000143499 | SMYD2 | chr1 | + | 214305186 | 214305250 | 214281101 | 214281427 | 214314761 | 214314872 |
| SE | 25674 | ENSG00000179152 | TCAIM | chr3 | + | 44357740 | 44357876 | 44354738 | 44354811 | 44361364 | 44361518 |
| SE | 10984 | ENSG00000083454 | P2RX5 | chr17 | - | 3688625 | 3688759 | 3688011 | 3688105 | 3690069 | 3690150 |
| SE | 101113 | ENSG00000162241 | SLC25A45 | chr11 | - | 65380533 | 65380632 | 65380131 | 65380175 | 65381874 | 65381969 |
| SE | 8949 | ENSG00000132478 | UNK | chr17 | + | 75815168 | 75815253 | 75813760 | 75813878 | 75816769 | 75816912 |
| SE | 9978 | ENSG00000133193 | FAM104A | chr17 | - | 73212677 | 73212740 | 73207352 | 73209768 | 73227164 | 73227264 |
| SE | 80740 | ENSG00000086300 | SNX10 | chr7 | + | 26357006 | 26357127 | 26346419 | 26346466 | 26360974 | 26361031 |
| SE | 17311 | ENSG00000100445 | SDR39U1 | chr14 | - | 24442108 | 24442260 | 24441677 | 24441795 | 24442345 | 24442375 |
| SE | 39973 | ENSG00000135537 | AFG1L | chr6 | + | 108355653 | 108355755 | 108346987 | 108347039 | 108356689 | 108356751 |
| SE | 17221 | ENSG00000126804 | ZBTB1 | chr14 | + | 64516599 | 64516751 | 64511248 | 64511384 | 64521486 | 64521606 |
| SE | 87105 | ENSG00000165591 | FAAH2 | chrX | + | 57431917 | 57432037 | 57380911 | 57381029 | 57446927 | 57447039 |
| SE | 104436 | ENSG00000149100 | EIF3M | chr11 | + | 32587011 | 32587144 | 32583797 | 32583929 | 32588593 | 32588681 |
| SE | 5497 | ENSG00000101294 | HM13 | chr20 | + | 31538178 | 31538359 | 31527483 | 31527582 | 31550063 | 31550471 |
| SE | 4924 | ENSG00000101246 | ARFRP1 | chr20 | - | 63706998 | 63707097 | 63706650 | 63706738 | 63707866 | 63707916 |
| SE | 16417 | ENSG00000185088 | RPS27L | chr15 | - | 63156786 | 63156973 | 63156412 | 63156521 | 63157399 | 63157477 |
| SE | 81054 | ENSG00000188243 | COMMD6 | chr13 | - | 75527812 | 75527851 | 75526435 | 75526639 | 75530310 | 75530408 |
| SE | 14904 | ENSG00000090487 | SPG21 | chr15 | - | 64983506 | 64983593 | 64980863 | 64981025 | 64989664 | 64989815 |
| SE | 19150 | ENSG00000257341 | NA | chr14 | + | 105488662 | 105488947 | 105488470 | 105488516 | 105490980 | 105491126 |
| SE | 7559 | ENSG00000161955 | TNFSF13 | chr17 | + | 7559623 | 7559702 | 7558475 | 7559297 | 7559845 | 7559893 |
| SE | 42965 | ENSG00000121940 | CLCC1 | chr1 | - | 108941404 | 108941498 | 108940044 | 108940142 | 108943474 | 108943615 |
| SE | 71437 | ENSG00000115234 | SNX17 | chr2 | + | 27373039 | 27373113 | 27372622 | 27372740 | 27373246 | 27373311 |
| SE | 42823 | ENSG00000140575 | IQGAP1 | chr15 | + | 90466268 | 90466436 | 90466000 | 90466091 | 90472839 | 90473010 |
| SE | 52222 | ENSG00000155363 | MOV10 | chr1 | + | 112686965 | 112687106 | 112674736 | 112675049 | 112688934 | 112689138 |
| SE | 27293 | ENSG00000064726 | BTBD1 | chr15 | - | 83050072 | 83050178 | 83041727 | 83041925 | 83056388 | 83056545 |
| SE | 5637 | ENSG00000080839 | RBL1 | chr20 | - | 37032664 | 37032876 | 37022649 | 37022826 | 37035241 | 37035508 |
| SE | 102693 | ENSG00000107938 | EDRF1 | chr10 | + | 125740462 | 125740651 | 125738294 | 125738445 | 125741000 | 125741201 |
| SE | 57946 | ENSG00000171791 | BCL2 | chr18 | - | 63280886 | 63281214 | 63127776 | 63128759 | 63318081 | 63318666 |
| SE | 40908 | ENSG00000151883 | PARP8 | chr5 | + | 50741830 | 50741951 | 50668070 | 50668125 | 50744735 | 50744828 |
| SE | 23796 | ENSG00000153071 | DAB2 | chr5 | - | 39381453 | 39381616 | 39376649 | 39377282 | 39388798 | 39388852 |
| SE | 62281 | ENSG00000112031 | MTRF1L | chr6 | - | 152991184 | 152991416 | 152987361 | 152990095 | 152992856 | 152992974 |
| SE | 98928 | ENSG00000102043 | MTMR8 | chrX | - | 64345044 | 64345177 | 64343610 | 64343720 | 64348659 | 64348794 |
| SE | 40504 | ENSG00000285708 | NA | chr3 | - | 71467250 | 71467315 | 71359149 | 71359244 | 71493425 | 71493555 |
| SE | 37281 | ENSG00000174738 | NR1D2 | chr3 | + | 23963223 | 23963330 | 23961976 | 23962605 | 23977222 | 23978497 |
| SE | 52972 | ENSG00000150455 | TIRAP | chr11 | + | 126291424 | 126291569 | 126290802 | 126290961 | 126292476 | 126292687 |
| SE | 78761 | ENSG00000179406 | LINC00174 | chr7 | - | 66467681 | 66467719 | 66450070 | 66450236 | 66476258 | 66476327 |
| SE | 69829 | ENSG00000163803 | PLB1 | chr2 | + | 28605852 | 28605948 | 28602820 | 28602921 | 28606495 | 28606567 |
| SE | 5550 | ENSG00000101052 | IFT52 | chr20 | + | 43618242 | 43618353 | 43613898 | 43613976 | 43618939 | 43619026 |
| SE | 49161 | ENSG00000077044 | DGKD | chr2 | + | 233468422 | 233468553 | 233467085 | 233467203 | 233469370 | 233472096 |
| SE | 54870 | ENSG00000141456 | PELP1 | chr17 | - | 4676752 | 4676812 | 4676356 | 4676507 | 4682501 | 4682573 |
| SE | 76455 | ENSG00000155906 | RMND1 | chr6 | - | 151408642 | 151408900 | 151405719 | 151405836 | 151417278 | 151417399 |
| SE | 109005 | ENSG00000105352 | CEACAM4 | chr19 | - | 41621650 | 41621768 | 41620574 | 41620627 | 41625600 | 41625960 |
| SE | 59974 | ENSG00000076770 | MBNL3 | chrX | - | 132439434 | 132440314 | 132406227 | 132406392 | 132488850 | 132489038 |
| SE | 93870 | ENSG00000116580 | GON4L | chr1 | - | 155832831 | 155832950 | 155831534 | 155831620 | 155853275 | 155853405 |
| SE | 56721 | ENSG00000111737 | RAB35 | chr12 | - | 120108591 | 120108703 | 120108416 | 120108467 | 120116598 | 120117502 |
| SE | 10180 | ENSG00000108846 | ABCC3 | chr17 | + | 50687535 | 50687730 | 50684708 | 50684875 | 50691091 | 50691699 |
| SE | 62420 | ENSG00000113761 | ZNF346 | chr5 | + | 177041777 | 177041870 | 177041125 | 177041229 | 177062057 | 177062151 |
| SE | 35890 | ENSG00000197415 | VEPH1 | chr3 | - | 157489054 | 157489160 | 157470418 | 157470529 | 157495211 | 157495506 |
| SE | 44558 | ENSG00000183020 | AP2A2 | chr11 | + | 1000431 | 1000598 | 994071 | 994245 | 1003721 | 1003804 |
| SE | 96199 | ENSG00000110756 | HPS5 | chr11 | - | 18317750 | 18317907 | 18311516 | 18312024 | 18322132 | 18322178 |
| SE | 44593 | ENSG00000026508 | CD44 | chr11 | + | 35211245 | 35211449 | 35189834 | 35189972 | 35214851 | 35214861 |
| SE | 69902 | ENSG00000105851 | PIK3CG | chr7 | + | 106872536 | 106872602 | 106867549 | 106869556 | 106874699 | 106874803 |
| SE | 10606 | ENSG00000276234 | TADA2A | chr17 | + | 37442563 | 37442652 | 37440504 | 37440662 | 37444695 | 37444768 |
| SE | 34986 | ENSG00000109390 | NDUFC1 | chr4 | - | 139297384 | 139297443 | 139295731 | 139295960 | 139300601 | 139301210 |
| SE | 109149 | ENSG00000185989 | RASA3 | chr13 | - | 114052051 | 114052155 | 114029810 | 114029887 | 114132434 | 114132623 |
| SE | 38052 | ENSG00000215908 | CROCCP2 | chr1 | - | 16642766 | 16642850 | 16633094 | 16633346 | 16643027 | 16643145 |
| SE | 2581 | ENSG00000196684 | HSH2D | chr19 | + | 16155641 | 16155763 | 16154401 | 16154491 | 16157209 | 16157248 |
| SE | 70713 | ENSG00000117362 | APH1A | chr1 | - | 150267715 | 150267789 | 150267355 | 150267478 | 150268697 | 150269023 |
| SE | 70285 | ENSG00000104218 | CSPP1 | chr8 | + | 67162637 | 67162762 | 67161810 | 67161915 | 67163731 | 67163798 |
| SE | 36832 | ENSG00000168038 | ULK4 | chr3 | - | 41705056 | 41705151 | 41681752 | 41681804 | 41705253 | 41705305 |
| SE | 91076 | ENSG00000104164 | BLOC1S6 | chr15 | + | 45603103 | 45603187 | 45592134 | 45592276 | 45605427 | 45605514 |
| SE | 8079 | ENSG00000141252 | VPS53 | chr17 | - | 599944 | 600094 | 586269 | 586364 | 601794 | 601896 |
| SE | 42263 | ENSG00000116001 | TIA1 | chr2 | - | 70228343 | 70228462 | 70227734 | 70227822 | 70229263 | 70229318 |
| SE | 54989 | ENSG00000117859 | OSBPL9 | chr1 | + | 51749501 | 51749588 | 51748368 | 51748398 | 51750144 | 51750195 |
| SE | 13276 | ENSG00000103335 | PIEZO1 | chr16 | - | 88717536 | 88717615 | 88717022 | 88717211 | 88719573 | 88719721 |
| SE | 9379 | ENSG00000121067 | SPOP | chr17 | - | 49636758 | 49636809 | 49622732 | 49622876 | 49668026 | 49668051 |
| SE | 55548 | ENSG00000185379 | RAD51D | chr17 | - | 35106127 | 35106249 | 35103453 | 35103544 | 35106385 | 35106481 |
| SE | 25831 | ENSG00000250334 | LINC00989 | chr4 | + | 79492661 | 79492774 | 79492406 | 79492501 | 79493151 | 79493623 |
| SE | 102513 | ENSG00000166130 | IKBIP | chr12 | - | 98634295 | 98634413 | 98613404 | 98614340 | 98644522 | 98644788 |
| SE | 71008 | ENSG00000115649 | CNPPD1 | chr2 | - | 219174145 | 219174207 | 219173349 | 219173467 | 219174777 | 219174906 |
| SE | 36754 | ENSG00000138231 | DBR1 | chr3 | - | 138163777 | 138163858 | 138163348 | 138163494 | 138167080 | 138167305 |
| SE | 62989 | ENSG00000137497 | NUMA1 | chr11 | - | 72012400 | 72012442 | 72010785 | 72010854 | 72012894 | 72016260 |
| SE | 22093 | ENSG00000111859 | NEDD9 | chr6 | - | 11251563 | 11251726 | 11248932 | 11249214 | 11305991 | 11306155 |
| SE | 69641 | ENSG00000120451 | SNX19 | chr11 | - | 130910017 | 130910137 | 130907952 | 130908083 | 130910269 | 130910370 |
| SE | 96804 | ENSG00000175220 | ARHGAP1 | chr11 | - | 46680204 | 46680282 | 46679647 | 46679776 | 46680486 | 46680563 |
| SE | 5818 | ENSG00000197586 | ENTPD6 | chr20 | + | 25207075 | 25207397 | 25206521 | 25206590 | 25209848 | 25209925 |
| SE | 34931 | ENSG00000076108 | BAZ2A | chr12 | - | 56613014 | 56613233 | 56611772 | 56612246 | 56613952 | 56614138 |
| SE | 83645 | ENSG00000166579 | NDEL1 | chr17 | + | 8460008 | 8460160 | 8454795 | 8454887 | 8466929 | 8468163 |
| SE | 32789 | ENSG00000157601 | MX1 | chr21 | + | 41427205 | 41427304 | 41422755 | 41422911 | 41427754 | 41427866 |
| SE | 102201 | ENSG00000138190 | EXOC6 | chr10 | + | 92935813 | 92935885 | 92934309 | 92934430 | 92940726 | 92940824 |
| SE | 87652 | ENSG00000117676 | RPS6KA1 | chr1 | + | 26561504 | 26561663 | 26561044 | 26561134 | 26571448 | 26571508 |
| SE | 55078 | ENSG00000086200 | IPO11 | chr5 | + | 62476682 | 62476753 | 62474415 | 62474464 | 62483100 | 62483293 |
| SE | 28226 | ENSG00000205937 | RNPS1 | chr16 | - | 2255584 | 2255726 | 2253115 | 2254063 | 2262277 | 2262431 |
| SE | 53731 | ENSG00000151503 | NCAPD3 | chr11 | - | 134210269 | 134210454 | 134209312 | 134209477 | 134220571 | 134220726 |
| SE | 106898 | ENSG00000205885 | C1RL-AS1 | chr12 | + | 7115385 | 7118677 | 7115129 | 7115219 | 7118898 | 7119026 |
| SE | 1407 | ENSG00000100372 | SLC25A17 | chr22 | - | 40776189 | 40776308 | 40773936 | 40774019 | 40777039 | 40777135 |
| SE | 91040 | ENSG00000168522 | FNTA | chr8 | + | 43072180 | 43072307 | 43069554 | 43069659 | 43077215 | 43077226 |
| SE | 479 | ENSG00000142207 | URB1 | chr21 | - | 32366617 | 32366755 | 32363155 | 32363329 | 32368402 | 32368598 |
| SE | 25234 | ENSG00000005100 | DHX33 | chr17 | - | 5455159 | 5455271 | 5453820 | 5453980 | 5455996 | 5456182 |
| SE | 103212 | ENSG00000157625 | TAB3 | chrX | - | 30842965 | 30843049 | 30834050 | 30834152 | 30846550 | 30846644 |
| SE | 81534 | ENSG00000106344 | RBM28 | chr7 | - | 128317956 | 128318106 | 128314763 | 128315020 | 128321265 | 128321424 |
| SE | 41941 | ENSG00000163374 | YY1AP1 | chr1 | - | 155675009 | 155675096 | 155672271 | 155672731 | 155676547 | 155676746 |
| SE | 60991 | ENSG00000109381 | ELF2 | chr4 | - | 139061864 | 139062057 | 139060323 | 139060674 | 139067683 | 139067770 |
| SE | 90434 | ENSG00000165923 | AGBL2 | chr11 | - | 47699445 | 47699553 | 47692102 | 47692256 | 47704542 | 47704728 |
| SE | 56646 | ENSG00000155893 | PXYLP1 | chr3 | + | 141262242 | 141262340 | 141260122 | 141260254 | 141278341 | 141278500 |
| SE | 54184 | ENSG00000175376 | EIF1AD | chr11 | - | 66000302 | 66000505 | 66000052 | 66000161 | 66002039 | 66002160 |
| SE | 80247 | ENSG00000169410 | PTPN9 | chr15 | - | 75527117 | 75527261 | 75524208 | 75524298 | 75578713 | 75578752 |
| SE | 19555 | ENSG00000053770 | AP5M1 | chr14 | + | 57274243 | 57274889 | 57268970 | 57269388 | 57280194 | 57280422 |
| SE | 13134 | ENSG00000118900 | UBN1 | chr16 | + | 4868832 | 4868903 | 4860663 | 4860953 | 4870211 | 4870341 |
| SE | 51724 | ENSG00000037637 | FBXO42 | chr1 | - | 16252287 | 16252404 | 16246839 | 16251785 | 16253095 | 16253152 |
| SE | 82442 | ENSG00000105939 | ZC3HAV1 | chr7 | - | 139055204 | 139055295 | 139053964 | 139054095 | 139064878 | 139064999 |
| SE | 14843 | ENSG00000140479 | PCSK6 | chr15 | - | 101431319 | 101431463 | 101429152 | 101430063 | 101431989 | 101432100 |
| SE | 98452 | ENSG00000177666 | PNPLA2 | chr11 | + | 823997 | 824130 | 823647 | 823855 | 824313 | 824436 |
| SE | 58133 | ENSG00000117523 | PRRC2C | chr1 | + | 171571319 | 171571421 | 171566591 | 171566843 | 171574926 | 171575128 |
| SE | 10 | ENSG00000156299 | TIAM1 | chr21 | - | 31276731 | 31276908 | 31266009 | 31266983 | 31339242 | 31339422 |
| SE | 13979 | ENSG00000103522 | IL21R | chr16 | + | 27403079 | 27403220 | 27402173 | 27402618 | 27430055 | 27430120 |
| SE | 77309 | ENSG00000099968 | BCL2L13 | chr22 | + | 17702242 | 17702386 | 17696140 | 17696210 | 17726676 | 17728699 |
| SE | 13493 | ENSG00000135698 | MPHOSPH6 | chr16 | - | 82151237 | 82151352 | 82149381 | 82149403 | 82151423 | 82151514 |
| SE | 31837 | ENSG00000185278 | ZBTB37 | chr1 | + | 173873466 | 173873566 | 173870196 | 173871148 | 173885635 | 173886636 |
| SE | 5005 | ENSG00000101181 | MTG2 | chr20 | + | 62193415 | 62193624 | 62183027 | 62183057 | 62195801 | 62195949 |
| SE | 50642 | ENSG00000087095 | NLK | chr17 | + | 28185178 | 28185265 | 28172516 | 28172618 | 28191020 | 28191172 |
| SE | 33475 | ENSG00000164327 | RICTOR | chr5 | - | 38978582 | 38978648 | 38975536 | 38975604 | 38981866 | 38982036 |
| SE | 69134 | ENSG00000155252 | PI4K2A | chr10 | + | 97664884 | 97664984 | 97662906 | 97662968 | 97666437 | 97666571 |
| SE | 56432 | ENSG00000050748 | MAPK9 | chr5 | - | 180286974 | 180287067 | 180280439 | 180280608 | 180291847 | 180292053 |
| SE | 81880 | ENSG00000177189 | RPS6KA3 | chrX | - | 20169401 | 20169491 | 20167588 | 20167747 | 20172745 | 20172864 |
| SE | 101272 | ENSG00000134460 | IL2RA | chr10 | - | 6021504 | 6021693 | 6019869 | 6019941 | 6024243 | 6024354 |
| SE | 38095 | ENSG00000160584 | SIK3 | chr11 | - | 116954043 | 116954107 | 116927218 | 116927380 | 116956947 | 116957044 |
| SE | 22953 | ENSG00000170234 | PWWP2A | chr5 | - | 160108544 | 160108614 | 160093757 | 160094065 | 160118804 | 160118833 |
| SE | 90654 | ENSG00000100023 | PPIL2 | chr22 | + | 21675063 | 21675115 | 21672329 | 21672381 | 21681298 | 21681390 |
| SE | 29894 | ENSG00000010165 | EEF1AKNMT | chr1 | + | 171790451 | 171790616 | 171787734 | 171787930 | 171792016 | 171792235 |
| SE | 98610 | ENSG00000049449 | RCN1 | chr11 | + | 32103003 | 32103158 | 32100546 | 32100608 | 32103280 | 32103480 |
| SE | 85099 | ENSG00000173535 | TNFRSF10C | chr8 | + | 23111719 | 23111825 | 23102920 | 23103181 | 23114656 | 23114770 |
| SE | 90226 | ENSG00000172269 | DPAGT1 | chr11 | - | 119097854 | 119098043 | 119096522 | 119097551 | 119098402 | 119098487 |
| SE | 46099 | ENSG00000166912 | MTMR10 | chr15 | - | 30967919 | 30968010 | 30960880 | 30961073 | 30974313 | 30974456 |
| SE | 102536 | ENSG00000197857 | ZNF44 | chr19 | - | 12293231 | 12293375 | 12275955 | 12276082 | 12294691 | 12294794 |
| SE | 18672 | ENSG00000100823 | APEX1 | chr14 | + | 20455913 | 20456101 | 20455225 | 20455703 | 20456667 | 20456683 |
| SE | 75236 | ENSG00000100503 | NIN | chr14 | - | 50821873 | 50822000 | 50806736 | 50806818 | 50830463 | 50830517 |
| SE | 79776 | ENSG00000164808 | SPIDR | chr8 | + | 47729411 | 47729465 | 47713488 | 47713641 | 47735306 | 47735374 |
| SE | 70951 | ENSG00000131781 | FMO5 | chr1 | - | 147201151 | 147201504 | 147190176 | 147190249 | 147208851 | 147209051 |
| SE | 92339 | ENSG00000027697 | IFNGR1 | chr6 | - | 137209849 | 137209972 | 137206962 | 137207077 | 137218671 | 137219244 |
| SE | 105490 | ENSG00000152256 | PDK1 | chr2 | + | 172566855 | 172566933 | 172564977 | 172565073 | 172570725 | 172570820 |
| SE | 8431 | ENSG00000078687 | TNRC6C | chr17 | + | 78079394 | 78079541 | 78075135 | 78075278 | 78086502 | 78086586 |
| SE | 60525 | ENSG00000116984 | MTR | chr1 | + | 236808703 | 236808773 | 236806143 | 236806233 | 236812737 | 236812844 |
| SE | 71076 | ENSG00000134490 | TMEM241 | chr18 | - | 23376826 | 23376941 | 23373711 | 23373763 | 23377645 | 23377671 |
| SE | 105058 | ENSG00000119801 | YPEL5 | chr2 | + | 30148244 | 30148541 | 30146940 | 30147062 | 30158618 | 30158747 |
| SE | 39856 | ENSG00000169239 | CA5B | chrX | + | 15745630 | 15745710 | 15738305 | 15738352 | 15749970 | 15750165 |
| SE | 54916 | ENSG00000010803 | SCMH1 | chr1 | - | 41152598 | 41152741 | 41151613 | 41151684 | 41159724 | 41159933 |
| SE | 3652 | ENSG00000142230 | SAE1 | chr19 | + | 47152897 | 47153040 | 47150201 | 47150375 | 47155113 | 47155177 |
| SE | 51195 | ENSG00000203666 | EFCAB2 | chr1 | + | 245017241 | 245017326 | 245002120 | 245002232 | 245059385 | 245059458 |
| SE | 73924 | ENSG00000155506 | LARP1 | chr5 | + | 154793799 | 154794000 | 154793594 | 154793723 | 154794099 | 154794195 |
| SE | 72657 | ENSG00000153094 | BCL2L11 | chr2 | + | 111153779 | 111153904 | 111150043 | 111150147 | 111164132 | 111164231 |
| SE | 90227 | ENSG00000172269 | DPAGT1 | chr11 | - | 119097854 | 119098043 | 119097141 | 119097297 | 119098402 | 119098487 |
| SE | 56956 | ENSG00000120725 | SIL1 | chr5 | - | 139021170 | 139021292 | 138951787 | 138951884 | 139026800 | 139026992 |
| SE | 37371 | ENSG00000166927 | MS4A7 | chr11 | + | 60385087 | 60385222 | 60383128 | 60383288 | 60394945 | 60395254 |
| SE | 106433 | ENSG00000135297 | MTO1 | chr6 | + | 73482039 | 73482244 | 73480996 | 73481116 | 73482448 | 73482620 |
| SE | 98943 | ENSG00000170291 | ELP5 | chr17 | + | 7256856 | 7257038 | 7254582 | 7254803 | 7258587 | 7258639 |
| SE | 47892 | ENSG00000157500 | APPL1 | chr3 | + | 57253681 | 57253738 | 57252268 | 57252311 | 57256956 | 57257051 |
| SE | 80173 | ENSG00000012124 | CD22 | chr19 | + | 35345601 | 35345720 | 35345050 | 35345126 | 35346150 | 35346235 |
| SE | 39124 | ENSG00000134352 | IL6ST | chr5 | - | 55948101 | 55948982 | 55947492 | 55947589 | 55951463 | 55951604 |
| SE | 93921 | ENSG00000184465 | WDR27 | chr6 | - | 169670568 | 169670693 | 169667981 | 169668185 | 169672254 | 169672396 |
| SE | 27469 | ENSG00000008513 | ST3GAL1 | chr8 | - | 133507198 | 133507321 | 133499134 | 133499189 | 133571692 | 133571887 |
| SE | 102496 | ENSG00000107560 | RAB11FIP2 | chr10 | - | 118033981 | 118034041 | 118015064 | 118015110 | 118038971 | 118039440 |
| SE | 11169 | ENSG00000213918 | DNASE1 | chr16 | + | 3655848 | 3655937 | 3655372 | 3655520 | 3656101 | 3656185 |
| SE | 6604 | ENSG00000134444 | RELCH | chr18 | + | 62284238 | 62284361 | 62282315 | 62282444 | 62285448 | 62285548 |
| SE | 73252 | ENSG00000114956 | DGUOK | chr2 | + | 73946718 | 73946906 | 73926825 | 73927052 | 73958145 | 73958245 |
| SE | 46038 | ENSG00000158517 | NCF1 | chr7 | + | 74782938 | 74783632 | 74780779 | 74780835 | 74785181 | 74785261 |
| SE | 84865 | ENSG00000164074 | ABHD18 | chr4 | + | 128001715 | 128001760 | 127989720 | 127989821 | 128008919 | 128008998 |
| SE | 39855 | ENSG00000169239 | CA5B | chrX | + | 15745630 | 15745710 | 15721496 | 15721592 | 15749970 | 15750165 |
| SE | 67191 | ENSG00000197226 | TBC1D9B | chr5 | - | 179879627 | 179879789 | 179879046 | 179879197 | 179891378 | 179891586 |
| SE | 95458 | ENSG00000180773 | SLC36A4 | chr11 | - | 93184429 | 93184520 | 93182805 | 93182894 | 93185690 | 93185814 |
| SE | 61843 | ENSG00000162929 | KIAA1841 | chr2 | + | 61073457 | 61073551 | 61071605 | 61071792 | 61076919 | 61077158 |
| SE | 7327 | ENSG00000154889 | MPPE1 | chr18 | - | 11905737 | 11905953 | 11896983 | 11897356 | 11906202 | 11906309 |
| SE | 17598 | ENSG00000174373 | RALGAPA1 | chr14 | - | 35627089 | 35627951 | 35625360 | 35625432 | 35634573 | 35634757 |
| SE | 25453 | ENSG00000011243 | AKAP8L | chr19 | - | 15403652 | 15403715 | 15403474 | 15403555 | 15404009 | 15404042 |
| SE | 102736 | ENSG00000040341 | STAU2 | chr8 | - | 73527690 | 73527804 | 73422613 | 73422702 | 73552011 | 73552319 |
| SE | 74553 | ENSG00000114353 | GNAI2 | chr3 | + | 50252396 | 50252538 | 50236203 | 50236453 | 50256191 | 50256320 |
| SE | 82777 | ENSG00000196873 | CBWD3 | chr9 | + | 68298963 | 68299093 | 68297751 | 68297817 | 68299480 | 68299644 |
| SE | 15650 | ENSG00000223509 | NA | chr15 | - | 32524470 | 32524600 | 32522692 | 32523213 | 32528531 | 32528701 |
| SE | 98592 | ENSG00000149269 | PAK1 | chr11 | - | 77397026 | 77397093 | 77392330 | 77392538 | 77411773 | 77411944 |
| SE | 21211 | ENSG00000150403 | TMCO3 | chr13 | + | 113499957 | 113500149 | 113498341 | 113498521 | 113501751 | 113501880 |
| SE | 68349 | ENSG00000125630 | POLR1B | chr2 | + | 112549266 | 112549399 | 112547420 | 112547567 | 112550865 | 112550908 |
| SE | 11273 | ENSG00000125124 | BBS2 | chr16 | - | 56500853 | 56501025 | 56498436 | 56499907 | 56502316 | 56502456 |
| SE | 83293 | ENSG00000003436 | TFPI | chr2 | - | 187529363 | 187529485 | 187513615 | 187513646 | 187554199 | 187554349 |
| SE | 103737 | ENSG00000107937 | GTPBP4 | chr10 | + | 1007017 | 1007128 | 1000947 | 1001013 | 1008957 | 1009035 |
| SE | 14420 | ENSG00000129007 | CALML4 | chr15 | - | 68197439 | 68197628 | 68190704 | 68194112 | 68205120 | 68205151 |
| SE | 14955 | ENSG00000182718 | ANXA2 | chr15 | - | 60390212 | 60390317 | 60386027 | 60386086 | 60396279 | 60396427 |
| SE | 21555 | ENSG00000085382 | HACE1 | chr6 | - | 104811310 | 104811393 | 104796928 | 104797025 | 104833041 | 104833173 |
| SE | 46327 | ENSG00000095564 | BTAF1 | chr10 | + | 91956527 | 91956657 | 91953736 | 91953873 | 91957224 | 91957293 |
| SE | 107381 | ENSG00000168010 | ATG16L2 | chr11 | + | 72823765 | 72823797 | 72822847 | 72822961 | 72824059 | 72824122 |
| SE | 94007 | ENSG00000140988 | RPS2 | chr16 | - | 1964448 | 1964628 | 1964275 | 1964365 | 1964806 | 1964826 |
| SE | 17810 | ENSG00000105669 | COPE | chr19 | - | 18903267 | 18903423 | 18900380 | 18900449 | 18904770 | 18904852 |
| SE | 20197 | ENSG00000137210 | TMEM14B | chr6 | + | 10755141 | 10755232 | 10751132 | 10751234 | 10770076 | 10770204 |
| SE | 62664 | ENSG00000115525 | ST3GAL5 | chr2 | - | 85844395 | 85844521 | 85839205 | 85840392 | 85846376 | 85846563 |
| SE | 99059 | ENSG00000149577 | SIDT2 | chr11 | + | 117186583 | 117186636 | 117185950 | 117186223 | 117187377 | 117187449 |
| SE | 58249 | ENSG00000089127 | OAS1 | chr12 | + | 112911097 | 112911235 | 112906972 | 112907219 | 112916508 | 112916738 |
| SE | 42475 | ENSG00000173083 | HPSE | chr4 | - | 83306202 | 83306317 | 83302149 | 83302268 | 83308844 | 83308951 |
| SE | 102993 | ENSG00000197535 | MYO5A | chr15 | - | 52337809 | 52337884 | 52336462 | 52336556 | 52340195 | 52340394 |
| SE | 75727 | ENSG00000107672 | NSMCE4A | chr10 | - | 121963237 | 121963328 | 121961422 | 121961517 | 121965285 | 121965304 |
| SE | 57313 | ENSG00000181722 | ZBTB20 | chr3 | - | 114380776 | 114380940 | 114380216 | 114380404 | 114389004 | 114389105 |
| SE | 83689 | ENSG00000119321 | FKBP15 | chr9 | - | 113193491 | 113193549 | 113190470 | 113190578 | 113194026 | 113194169 |
| SE | 26962 | ENSG00000172869 | DMXL1 | chr5 | + | 119110150 | 119110283 | 119101934 | 119102006 | 119114474 | 119114541 |
| SE | 56489 | ENSG00000114812 | VIPR1 | chr3 | + | 42534974 | 42535104 | 42532241 | 42532333 | 42536089 | 42536394 |
| SE | 69304 | ENSG00000112367 | FIG4 | chr6 | + | 109695829 | 109695904 | 109691287 | 109691501 | 109715077 | 109715176 |
| SE | 64596 | ENSG00000100644 | HIF1A | chr14 | + | 61720378 | 61720572 | 61695539 | 61695839 | 61721508 | 61721654 |
| SE | 20997 | ENSG00000198399 | ITSN2 | chr2 | - | 24321813 | 24321880 | 24315131 | 24315224 | 24328051 | 24328115 |
| SE | 82135 | ENSG00000132640 | BTBD3 | chr20 | + | 11917777 | 11918011 | 11890816 | 11890954 | 11918333 | 11918463 |
| SE | 11269 | ENSG00000125124 | BBS2 | chr16 | - | 56492523 | 56492582 | 56485589 | 56485738 | 56496966 | 56497079 |
| SE | 105408 | ENSG00000050438 | SLC4A8 | chr12 | + | 51462309 | 51462456 | 51461203 | 51461291 | 51463613 | 51463714 |
| SE | 14078 | ENSG00000130731 | METTL26 | chr16 | - | 634888 | 634956 | 634718 | 634797 | 635280 | 635340 |
| SE | 55796 | ENSG00000144218 | AFF3 | chr2 | - | 99560364 | 99560436 | 99558874 | 99558968 | 99565486 | 99565623 |
| SE | 74401 | ENSG00000168237 | GLYCTK | chr3 | + | 52291744 | 52291922 | 52290959 | 52291111 | 52292259 | 52293475 |
| SE | 44816 | ENSG00000177873 | ZNF619 | chr3 | + | 40483652 | 40483723 | 40482587 | 40482704 | 40486805 | 40488259 |
| SE | 1289 | ENSG00000100239 | PPP6R2 | chr22 | + | 50388985 | 50389111 | 50372071 | 50372150 | 50391866 | 50392015 |
| SE | 106723 | ENSG00000136111 | TBC1D4 | chr13 | - | 75306312 | 75306471 | 75302242 | 75302401 | 75309941 | 75310151 |
| SE | 26109 | ENSG00000164181 | ELOVL7 | chr5 | - | 60802626 | 60802665 | 60799179 | 60799230 | 60843421 | 60843538 |
| SE | 42050 | ENSG00000196352 | CD55 | chr1 | + | 207340390 | 207340508 | 207339396 | 207339417 | 207367369 | 207367487 |
| SE | 14623 | ENSG00000140157 | NIPA2 | chr15 | + | 22839654 | 22839790 | 22838644 | 22838921 | 22845145 | 22845267 |
| SE | 87554 | ENSG00000147050 | KDM6A | chrX | + | 45085864 | 45085979 | 45083459 | 45083608 | 45089742 | 45089930 |
| SE | 89691 | ENSG00000120992 | LYPLA1 | chr8 | - | 54065747 | 54065813 | 54063327 | 54063375 | 54100907 | 54100939 |
| SE | 75170 | ENSG00000072121 | ZFYVE26 | chr14 | - | 67754070 | 67754212 | 67753706 | 67753766 | 67755050 | 67755250 |
| SE | 9949 | ENSG00000109062 | SLC9A3R1 | chr17 | + | 74762011 | 74762173 | 74748627 | 74749287 | 74768146 | 74768236 |
| SE | 81583 | ENSG00000253729 | PRKDC | chr8 | - | 47778458 | 47778660 | 47777685 | 47777874 | 47778727 | 47778799 |
| SE | 74743 | ENSG00000163939 | PBRM1 | chr3 | - | 52641953 | 52642045 | 52634601 | 52634815 | 52644703 | 52644789 |
| SE | 2301 | ENSG00000142002 | DPP9 | chr19 | - | 4722498 | 4722551 | 4719617 | 4719941 | 4723673 | 4723750 |
| SE | 80692 | ENSG00000175471 | MCTP1 | chr5 | - | 94799012 | 94799132 | 94779109 | 94779163 | 94940083 | 94940195 |
| SE | 48290 | ENSG00000229474 | PATL2 | chr15 | - | 44672014 | 44672239 | 44669966 | 44670087 | 44672387 | 44672456 |
| SE | 93337 | ENSG00000095303 | PTGS1 | chr9 | + | 122377898 | 122378015 | 122371612 | 122371834 | 122378432 | 122378573 |
| SE | 9795 | ENSG00000108384 | RAD51C | chr17 | + | 58696692 | 58696859 | 58694930 | 58695189 | 58703195 | 58703329 |
| SE | 67614 | ENSG00000109466 | KLHL2 | chr4 | + | 165305607 | 165305725 | 165299506 | 165299656 | 165310552 | 165310750 |
| SE | 100565 | ENSG00000110011 | DNAJC4 | chr11 | + | 64231031 | 64231147 | 64230651 | 64230940 | 64231870 | 64231964 |
| SE | 10546 | ENSG00000177302 | TOP3A | chr17 | - | 18285141 | 18285307 | 18282697 | 18282841 | 18290556 | 18290686 |
| SE | 49966 | ENSG00000162976 | SLC66A3 | chr2 | + | 11174967 | 11175009 | 11164203 | 11164261 | 11177736 | 11177957 |
| SE | 75930 | ENSG00000173588 | CEP83 | chr12 | - | 94412317 | 94412429 | 94411696 | 94411847 | 94412516 | 94412591 |
| SE | 83701 | ENSG00000177324 | BEND2 | chrX | - | 18195295 | 18195442 | 18191000 | 18191108 | 18201814 | 18201940 |
| SE | 69005 | ENSG00000107521 | HPS1 | chr10 | - | 98435271 | 98435414 | 98433939 | 98434091 | 98435634 | 98435772 |

| | | | | | | | | | |
|---|---|---|---|---|---|---|---|---|---|
| SE | 87804 | ENSG00000111237 | VPS29 | chr12 | - | 110498819 | 110498859 | 110496011 | 110496203 |
| SE | 56343 | ENSG00000117602 | RCAN3 | chr1 | + | 24533082 | 24533254 | 24531217 | 24531391 |
| SE | 50060 | ENSG00000102763 | VWA8 | chr13 | - | 41675214 | 41675296 | 41670945 | 41671147 |
| SE | 109650 | ENSG00000005810 | MYCBP2 | chr13 | - | 77272322 | 77272399 | 77270295 | 77270538 |
| SE | 84408 | ENSG00000188917 | TRMT2B | chrX | - | 101037916 | 101038051 | 101036973 | 101037073 |
| SE | 79167 | ENSG00000134291 | TMEM106C | chr12 | + | 47965281 | 47965345 | 47964208 | 47964423 |
| SE | 68751 | ENSG00000196141 | SPATS2L | chr2 | + | 200383866 | 200384017 | 200329430 | 200329480 |
| SE | 72138 | ENSG00000172936 | MYD88 | chr3 | + | 38140387 | 38140592 | 38139863 | 38139998 |
| SE | 82100 | ENSG00000128891 | CCDC32 | chr15 | - | 40553436 | 40554127 | 40534342 | 40539355 |
| SE | 79512 | ENSG00000146918 | NCAPG2 | chr7 | - | 158690567 | 158690722 | 158689818 | 158689953 |
| SE | 62152 | ENSG00000117625 | RCOR3 | chr1 | + | 211267774 | 211267870 | 211260107 | 211260164 |
| SE | 91053 | ENSG00000104164 | BLOC1S6 | chr15 | + | 45592134 | 45592276 | 45587354 | 45587525 |
| SE | 36970 | ENSG00000153914 | SREK1 | chr5 | + | 66177516 | 66177658 | 66174945 | 66175041 |
| SE | 53088 | ENSG00000103657 | HERC1 | chr15 | - | 63779582 | 63780009 | 63775563 | 63775649 |
| SE | 47497 | ENSG00000143363 | PRUNE1 | chr1 | + | 151025514 | 151025673 | 151017811 | 151017904 |
| SE | 20183 | ENSG00000137210 | TMEM14B | chr6 | + | 10749621 | 10749698 | 10749201 | 10749268 |
| SE | 5344 | ENSG00000101084 | RAB5IF | chr20 | + | 36607889 | 36608000 | 36607714 | 36607818 |
| SE | 43781 | ENSG00000138002 | IFT172 | chr2 | - | 27459708 | 27459829 | 27458778 | 27459522 |
| SE | 29738 | ENSG00000162825 | NBPF20 | chr1 | - | 145394977 | 145395141 | 145393860 | 145393935 |
| SE | 44096 | ENSG00000182923 | CEP63 | chr3 | + | 134559149 | 134559429 | 134551925 | 134552012 |
| SE | 21598 | ENSG00000112234 | FBXL4 | chr6 | - | 98926476 | 98927060 | 98917373 | 98917719 |
| SE | 106041 | ENSG00000204482 | LST1 | chr6 | + | 31587943 | 31587966 | 31587640 | 31587733 |
| SE | 42374 | ENSG00000048740 | CELF2 | chr10 | + | 11321188 | 11321386 | 11314138 | 11314258 |
| SE | 50895 | ENSG00000156515 | HK1 | chr10 | + | 69382486 | 69382791 | 69379861 | 69380095 |
| SE | 48557 | ENSG00000126091 | ST3GAL3 | chr1 | + | 43858134 | 43858308 | 43838218 | 43838311 |
| SE | 19026 | ENSG00000131966 | ACTR10 | chr14 | + | 58219693 | 58219729 | 58215204 | 58215284 |
| SE | 69343 | ENSG00000168958 | MFF | chr2 | + | 227347225 | 227347384 | 227340291 | 227340380 |
| SE | 66475 | ENSG00000131051 | RBM39 | chr20 | - | 35739392 | 35739547 | 35738967 | 35739017 |
| SE | 60942 | ENSG00000182362 | YBEY | chr21 | + | 46291333 | 46291462 | 46286869 | 46287123 |
| SE | 33489 | ENSG00000125037 | EMC3 | chr3 | - | 9981633 | 9981747 | 9977388 | 9977446 |
| SE | 11271 | ENSG00000125124 | BBS2 | chr16 | - | 56500853 | 56500977 | 56499777 | 56499907 |
| SE | 89672 | ENSG00000198851 | CD3E | chr11 | + | 118313706 | 118313874 | 118312152 | 118312170 |
| SE | 20843 | ENSG00000255819 | KLRC4-KLRK1 | chr12 | - | 10389942 | 10390027 | 10388770 | 10388875 |
| SE | 46057 | ENSG00000068097 | HEATR6 | chr17 | - | 60046024 | 60046099 | 60043672 | 60044134 |
| SE | 92698 | ENSG00000180776 | ZDHHC20 | chr13 | - | 21421060 | 21421164 | 21413651 | 21413772 |
| SE | 96790 | ENSG00000259719 | LINC02284 | chr14 | + | 56326179 | 56326225 | 56321485 | 56321611 |
| SE | 105159 | ENSG00000174718 | RESF1 | chr12 | + | 31960776 | 31960871 | 31959369 | 31959491 |
| SE | 36841 | ENSG00000168038 | ULK4 | chr3 | - | 41915976 | 41916052 | 41912806 | 41912899 |
| SE | 72337 | ENSG00000123739 | PLA2G12A | chr4 | - | 109718688 | 109718759 | 109717547 | 109717713 |
| SE | 2293 | ENSG00000142002 | DPP9 | chr19 | - | 4700215 | 4700273 | 4697550 | 4697651 |
| SE | 62857 | ENSG00000121807 | CCR2 | chr3 | + | 46354861 | 46354956 | 46354125 | 46354177 |
| SE | 92490 | ENSG00000130749 | ZC3H4 | chr19 | - | 47110894 | 47110956 | 47094388 | 47094608 |
| SE | 20456 | ENSG00000006744 | ELAC2 | chr17 | - | 12993686 | 12993831 | 12991611 | 12993045 |
| SE | 65325 | ENSG00000285756 | NA | chrX | - | 3822373 | 3822580 | 3818656 | 3818703 |
| SE | 94215 | ENSG00000077463 | SIRT6 | chr19 | - | 4180781 | 4180909 | 4179103 | 4179286 |
| SE | 66353 | ENSG00000197037 | ZSCAN25 | chr7 | + | 99619560 | 99621574 | 99616945 | 99617016 |
| SE | 68085 | ENSG00000135838 | NPL | chr1 | + | 182794355 | 182794439 | 182792231 | 182792286 |
| SE | 97790 | ENSG00000166452 | AKIP1 | chr11 | + | 8917286 | 8917367 | 8914825 | 8914930 |
| SE | 76364 | ENSG00000164674 | SYTL3 | chr6 | + | 158663204 | 158663378 | 158662749 | 158662812 |
| SE | 81699 | ENSG00000171681 | ATF7IP | chr12 | + | 14457206 | 14457295 | 14456560 | 14456634 |
| SE | 41678 | ENSG00000079277 | MKNK1 | chr1 | - | 46582813 | 46582905 | 46580529 | 46580627 |
| SE | 45683 | ENSG00000109323 | MANBA | chr4 | - | 102714437 | 102714561 | 102690595 | 102690771 |
| SE | 62220 | ENSG00000119778 | ATAD2B | chr2 | - | 23863380 | 23863555 | 23857414 | 23857503 |
| SE | 15442 | ENSG00000171763 | SPATA5L1 | chr15 | + | 45407028 | 45407223 | 45405458 | 45405505 |
| SE | 668 | ENSG00000100211 | CBY1 | chr22 | + | 38671069 | 38671188 | 38670766 | 38670989 |
| SE | 24337 | ENSG00000185404 | SP140L | chr2 | + | 230400333 | 230400413 | 230400126 | 230400242 |
| SE | 81748 | ENSG00000176454 | LPCAT4 | chr15 | - | 34362195 | 34362321 | 34360110 | 34360209 |
| SE | 35411 | ENSG00000174891 | RSRC1 | chr3 | + | 158225656 | 158225733 | 158203071 | 158203245 |
| SE | 43325 | ENSG00000135597 | REPS1 | chr6 | - | 138917554 | 138917624 | 138914696 | 138914761 |
| SE | 100189 | ENSG00000078124 | ACER3 | chr11 | + | 77015015 | 77015117 | 76998553 | 76998821 |
| SE | 43483 | ENSG00000018510 | AGPS | chr2 | + | 177442119 | 177442235 | 177440964 | 177441036 |
| SE | 103794 | ENSG00000108175 | ZMIZ1 | chr10 | + | 79162052 | 79162133 | 79139681 | 79139777 |
| SE | 44749 | ENSG00000198265 | HELZ | chr17 | - | 67193966 | 67194042 | 67190156 | 67190355 |
| SE | 93049 | ENSG00000165138 | ANKS6 | chr9 | - | 98780188 | 98780337 | 98778225 | 98778424 |
| SE | 51634 | ENSG00000181523 | SGSH | chr17 | - | 80214614 | 80214783 | 80214289 | 80214328 |
| SE | 6373 | ENSG00000168675 | LDLRAD4 | chr18 | + | 13484036 | 13484119 | 13465014 | 13465111 |
| SE | 106167 | ENSG00000120805 | ARL1 | chr12 | - | 101403141 | 101403255 | 101402864 | 101402946 |
| SE | 79895 | ENSG00000188312 | CENPP | chr9 | + | 92345698 | 92345787 | 92337540 | 92337629 |
| SE | 96914 | ENSG00000100731 | PCNX1 | chr14 | + | 71057508 | 71057724 | 71055503 | 71055562 |
| SE | 85081 | ENSG00000103037 | SETD6 | chr16 | + | 58516807 | 58516928 | 58516477 | 58516672 |
| SE | 43638 | ENSG00000196628 | TCF4 | chr18 | - | 55587044 | 55587136 | 55585279 | 55585352 |
| SE | 49904 | ENSG00000108061 | SHOC2 | chr10 | + | 110985627 | 110985765 | 110964124 | 110965061 |
| SE | 46297 | ENSG00000116830 | TTF2 | chr1 | + | 117084017 | 117084168 | 117081827 | 117081947 |
| SE | 44100 | ENSG00000114982 | KANSL3 | chr2 | - | 96602115 | 96602338 | 96601642 | 96601776 |
| SE | 46329 | ENSG00000095564 | BTAF1 | chr10 | + | 91957224 | 91957293 | 91953736 | 91953873 |
| SE | 76502 | ENSG00000153406 | NMRAL1 | chr16 | - | 4469503 | 4469698 | 4466152 | 4466402 |
| SE | 463 | ENSG00000243927 | MRPS6 | chr21 | + | 34102834 | 34102908 | 34073548 | 34073745 |
| SE | 87322 | ENSG00000215301 | DDX3X | chrX | + | 41343220 | 41343351 | 41342736 | 41342836 |
| SE | 62435 | ENSG00000113761 | ZNF346 | chr5 | + | 177050750 | 177050936 | 177041777 | 177041870 |
| SE | 52952 | ENSG00000144744 | UBA3 | chr3 | - | 69080111 | 69080153 | 69077338 | 69077918 |
| SE | 93588 | ENSG00000165525 | NEMF | chr14 | - | 49838138 | 49838206 | 49834362 | 49834449 |
| SE | 12145 | ENSG00000172775 | PSME3IP1 | chr16 | - | 57175538 | 57175700 | 57173772 | 57173869 |
| SE | 15808 | ENSG00000169375 | SIN3A | chr15 | - | 75400729 | 75400940 | 75400039 | 75400156 |
| SE | 99714 | ENSG00000110075 | PPP6R3 | chr11 | + | 68583042 | 68583129 | 68575957 | 68576043 |
| SE | 102583 | ENSG00000120438 | TCP1 | chr6 | - | 159785899 | 159786020 | 159783989 | 159784067 |
| SE | 105864 | ENSG00000090006 | LTBP4 | chr19 | + | 40617099 | 40617225 | 40616888 | 40617020 |
| SE | 52991 | ENSG00000077585 | GPR137B | chr1 | + | 236183777 | 236183906 | 236168700 | 236168755 |
| SE | 51546 | ENSG00000036549 | ZZZ3 | chr1 | - | 77639448 | 77639602 | 77584516 | 77584655 |
| SE | 45392 | ENSG00000144468 | RHBDD1 | chr2 | + | 226838072 | 226838154 | 226837547 | 226837770 |
| SE | 72303 | ENSG00000059377 | TBXAS1 | chr7 | + | 139911224 | 139911321 | 139875584 | 139875637 |
| SE | 13308 | ENSG00000158805 | ZNF276 | chr16 | + | 89727278 | 89727357 | 89723136 | 89723709 |
| SE | 60408 | ENSG00000196440 | ARMCX4 | chrX | + | 101444051 | 101444155 | 101418909 | 101419000 |
| SE | 97385 | ENSG00000113163 | CERT1 | chr5 | - | 75400204 | 75400291 | 75389591 | 75389687 |
| SE | 102978 | ENSG00000269609 | C10orf95-AS1 | chr10 | + | 102451398 | 102452187 | 102450186 | 102450330 |
| SE | 36627 | ENSG00000169180 | XPO6 | chr16 | - | 28208986 | 28209055 | 28180940 | 28181031 |
| SE | 61969 | ENSG00000163904 | SENP2 | chr3 | + | 185619298 | 185619502 | 185617479 | 185617611 |
| SE | 47541 | ENSG00000066322 | ELOVL1 | chr1 | - | 43366267 | 43366411 | 43365563 | 43365623 |
| SE | 98630 | ENSG00000151498 | ACAD8 | chr11 | + | 134257087 | 134257257 | 134256547 | 134256648 |
| SE | 12989 | ENSG00000122390 | NAA60 | chr16 | + | 3447481 | 3447620 | 3443648 | 3443827 |
| SE | 58739 | ENSG00000172650 | AGAP5 | chr10 | - | 73681007 | 73681341 | 73680070 | 73680106 |
| SE | 96636 | ENSG00000135372 | NAT10 | chr11 | + | 34107052 | 34107319 | 34105616 | 34105792 |
| SE | 39982 | ENSG00000080822 | CLDND1 | chr3 | - | 98521132 | 98521437 | 98518884 | 98518995 |
| SE | 92711 | ENSG00000150776 | NKAPD1 | chr11 | + | 112074886 | 112074916 | 112074315 | 112074358 |
| SE | 109064 | ENSG00000164649 | CDCA7L | chr7 | - | 21940261 | 21940315 | 21916753 | 21916894 |
| SE | 63771 | ENSG00000131748 | STARD3 | chr17 | + | 39657972 | 39658026 | 39657774 | 39657852 |
| SE | 51411 | ENSG00000197302 | ZNF720 | chr16 | + | 31759356 | 31759466 | 31753765 | 31753900 |
| SE | 36512 | ENSG00000168405 | CMAHP | chr6 | - | 25113591 | 25113782 | 25109499 | 25109581 |
| SE | 3304 | ENSG00000076928 | ARHGEF1 | chr19 | + | 41897314 | 41897326 | 41896376 | 41896482 |
| SE | 50582 | ENSG00000139636 | LMBR1L | chr12 | - | 49106653 | 49106746 | 49105923 | 49105957 |
| SE | 20602 | ENSG00000198945 | L3MBTL3 | chr6 | + | 130078557 | 130078634 | 130070846 | 130071127 |
| SE | 71089 | ENSG00000110931 | CAMKK2 | chr12 | - | 121244572 | 121244615 | 121237674 | 121240869 |
| SE | 75169 | ENSG00000072121 | ZFYVE26 | chr14 | - | 67754070 | 67754212 | 67752343 | 67752526 |
| SE | 101878 | ENSG00000173915 | ATP5MK | chr10 | - | 103394198 | 103394394 | 103392370 | 103392466 |
| SE | 16387 | ENSG00000156958 | GALK2 | chr15 | + | 49158844 | 49158968 | 49155770 | 49156016 |
| SE | 7475 | ENSG00000073605 | GSDMB | chr17 | - | 39908175 | 39908214 | 39906960 | 39906987 |
| SE | 104730 | ENSG00000226210 | WASH8P | chr12 | - | 17347 | 17483 | 16968 | 17170 |
| SE | 21668 | ENSG00000130449 | ZSWIM6 | chr5 | + | 61535483 | 61535619 | 61531464 | 61531725 |
| SE | 69518 | ENSG00000168818 | STX18 | chr4 | - | 4425163 | 4425222 | 4423517 | 4423587 |
| SE | 41250 | ENSG00000162819 | BROX | chr1 | + | 222715144 | 222715269 | 222712563 | 222712942 |
| SE | 98058 | ENSG00000186314 | PRELID2 | chr5 | - | 145796441 | 145796547 | 145764930 | 145765000 |
| SE | 73566 | ENSG00000115998 | C2orf42 | chr2 | - | 70182666 | 70182935 | 70181162 | 70181997 |
| SE | 74402 | ENSG00000168237 | GLYCTK | chr3 | + | 52291746 | 52291922 | 52290959 | 52291111 |
| SE | 21205 | ENSG00000120278 | PLEKHG1 | chr6 | + | 150638079 | 150638125 | 150632942 | 150633304 |
| SE | 11039 | ENSG00000132359 | RAP1GAP2 | chr17 | + | 3005340 | 3005440 | 2998220 | 2998376 |
| SE | 67919 | ENSG00000112130 | RNF8 | chr6 | + | 37374619 | 37374709 | 37371511 | 37371574 |
| SE | 49897 | ENSG00000141068 | KSR1 | chr17 | + | 27601892 | 27601958 | 27601359 | 27601401 |
| SE | 27662 | ENSG00000134851 | TMEM165 | chr4 | + | 55417802 | 55417848 | 55416922 | 55417247 |
| SE | 59714 | ENSG00000183530 | PRR14L | chr22 | - | 31738386 | 31738911 | 31725537 | 31725610 |
| SE | 68499 | ENSG00000087087 | SRRT | chr7 | + | 100882457 | 100883090 | 100882052 | 100882241 |
| SE | 44681 | ENSG00000185344 | ATP6V0A2 | chr12 | + | 123726196 | 123726285 | 123723209 | 123724791 |
| SE | 25193 | ENSG00000214253 | FIS1 | chr7 | - | 101244006 | 101244139 | 101240829 | 101240906 |
| SE | 90066 | ENSG00000156787 | TBC1D31 | chr8 | + | 123084161 | 123084340 | 123082701 | 123082817 |
| SE | 53075 | ENSG00000103657 | HERC1 | chr15 | - | 63725291 | 63725513 | 63723181 | 63723355 |
| SE | 1830 | ENSG00000070371 | CLTCL1 | chr22 | - | 19239274 | 19239388 | 19235695 | 19235869 |
| SE | 102832 | ENSG00000148700 | ADD3 | chr10 | + | 110122109 | 110122292 | 110119465 | 110119564 |
| SE | 20137 | ENSG00000112659 | CUL9 | chr6 | + | 43188522 | 43188715 | 43187712 | 43188118 |
| SE | 92067 | ENSG00000108010 | GLRX3 | chr10 | + | 130159745 | 130159834 | 130145210 | 130145319 |
| SE | 12588 | ENSG00000090861 | AARS1 | chr16 | - | 70258032 | 70258217 | 70255719 | 70255836 |
| SE | 103275 | ENSG00000172661 | WASHC2C | chr10 | + | 45787034 | 45787247 | 45786611 | 45786674 |
| SE | 75328 | ENSG00000196262 | PPIA | chr7 | + | 44798747 | 44798814 | 44796680 | 44796793 |
| SE | 64681 | ENSG00000136715 | SAP130 | chr2 | - | 127993186 | 127993308 | 127989563 | 127989866 |
| SE | 89117 | ENSG00000064115 | TM7SF3 | chr12 | - | 26983492 | 26983596 | 26982772 | 26982859 |
| SE | 71253 | ENSG00000236790 | LINC00299 | chr2 | - | 8312725 | 8312849 | 8301808 | 8302425 |
| SE | 62164 | ENSG00000117625 | RCOR3 | chr1 | + | 211304082 | 211304140 | 211295675 | 211295753 |
| SE | 60704 | ENSG00000235750 | KIAA0040 | chr1 | - | 175177610 | 175177684 | 175166561 | 175166737 |
| SE | 80611 | ENSG00000185164 | NOMO2 | chr16 | - | 18527536 | 18527624 | 18524488 | 18524552 |
| SE | 72078 | ENSG00000130414 | NDUFA10 | chr2 | - | 240022171 | 240022340 | 240021116 | 240021412 |
| SE | 54702 | ENSG00000109667 | SLC2A9 | chr4 | - | 9996780 | 9996941 | 9985668 | 9985793 |
| SE | 83502 | ENSG00000106603 | COA1 | chr7 | - | 43711399 | 43711449 | 43665657 | 43665722 |
| SE | 104304 | ENSG00000135655 | USP15 | chr12 | + | 62374389 | 62374535 | 62355330 | 62355475 |
| SE | 103812 | ENSG00000225484 | NUTM2B-AS1 | chr10 | - | 79801949 | 79802003 | 79766289 | 79766324 |
| SE | 3768 | ENSG00000105486 | LIG1 | chr19 | - | 48133980 | 48134066 | 48132981 | 48133097 |
| SE | 68570 | ENSG00000140368 | PSTPIP1 | chr15 | + | 77023569 | 77023667 | 77018456 | 77018531 |
| SE | 97533 | ENSG00000149761 | NUDT22 | chr11 | + | 64227567 | 64227666 | 64226634 | 64227132 |
| SE | 36500 | ENSG00000168405 | CMAHP | chr6 | - | 25084652 | 25084832 | 25084479 | 25084576 |
| SE | 66338 | ENSG00000143207 | COP1 | chr1 | - | 176175909 | 176176007 | 176163814 | 176163891 |
| SE | 77986 | ENSG00000106635 | BCL7B | chr7 | - | 73553547 | 73553669 | 73552166 | 73552242 |
| SE | 4142 | ENSG00000073008 | PVR | chr19 | + | 44658741 | 44658765 | 44657761 | 44657910 |
| SE | 67328 | ENSG00000163728 | TTC14 | chr3 | + | 180604224 | 180604309 | 180603123 | 180603323 |
| SE | 84556 | ENSG00000156170 | NDUFAF6 | chr8 | + | 95036370 | 95036494 | 95035453 | 95035576 |
| SE | 20593 | ENSG00000196367 | TRRAP | chr7 | + | 99032893 | 99033039 | 99031344 | 99031465 |
| SE | 22497 | ENSG00000165097 | KDM1B | chr6 | + | 18213655 | 18213781 | 18211494 | 18212604 |
| SE | 18561 | ENSG00000100697 | DICER1 | chr14 | - | 95129471 | 95129632 | 95126579 | 95126748 |
| SE | 79640 | ENSG00000272333 | KMT2B | chr19 | + | 35728773 | 35728889 | 35728097 | 35728171 |
| SE | 101351 | ENSG00000124795 | DEK | chr6 | - | 18258303 | 18258405 | 18256525 | 18258062 |
| SE | 103859 | ENSG00000138185 | ENTPD1 | chr10 | + | 95823236 | 95823364 | 95756161 | 95756255 |
| SE | 88198 | ENSG00000136279 | DBNL | chr7 | + | 44057972 | 44057992 | 44057781 | 44057859 |
| SE | 12143 | ENSG00000172775 | PSME3IP1 | chr16 | - | 57175538 | 57175700 | 57173772 | 57173869 |
| SE | 28743 | ENSG00000147548 | NSD3 | chr8 | - | 38299443 | 38299590 | 38295795 | 38295952 |
| SE | 95347 | ENSG00000115459 | ELMOD3 | chr2 | + | 85355075 | 85355170 | 85354393 | 85354829 |
| SE | 87423 | ENSG00000021762 | OSBPL5 | chr11 | - | 3120543 | 3120624 | 3119546 | 3119631 |
| SE | 65641 | ENSG00000142856 | ITGB3BP | chr1 | - | 63508527 | 63508570 | 63490082 | 63490218 |
| SE | 30162 | ENSG00000163811 | WDR43 | chr2 | + | 28917892 | 28917995 | 28914068 | 28914208 |
| SE | 5664 | ENSG00000171700 | RGS19 | chr20 | - | 64076856 | 64076954 | 64076524 | 64076646 |
| SE | 69375 | ENSG00000171720 | HDAC3 | chr5 | - | 141634810 | 141634953 | 141629859 | 141630125 |
| SE | 48009 | ENSG00000010810 | FYN | chr6 | - | 111700103 | 111700268 | 111699514 | 111699670 |
| SE | 9713 | ENSG00000141568 | FOXK2 | chr17 | + | 82585251 | 82585271 | 82584012 | 82584188 |
| SE | 43492 | ENSG00000018510 | AGPS | chr2 | + | 177521268 | 177521368 | 177513818 | 177513908 |
| SE | 68503 | ENSG00000087087 | SRRT | chr7 | + | 100888256 | 100888383 | 100888041 | 100888143 |
| SE | 58690 | ENSG00000173409 | ARV1 | chr1 | + | 230999622 | 230999801 | 230997120 | 230997267 |
| SE | 17198 | ENSG00000154719 | MRPL39 | chr21 | - | 25603795 | 25603935 | 25601367 | 25601467 |
| SE | 45822 | ENSG00000169155 | ZBTB43 | chr9 | + | 126808792 | 126808915 | 126805039 | 126805132 |
| SE | 91176 | ENSG00000164941 | INTS8 | chr8 | + | 94856776 | 94856978 | 94853793 | 94853915 |
| SE | 13991 | ENSG00000047578 | KATNIP | chr16 | + | 27654607 | 27654736 | 27648603 | 27648735 |
| SE | 60687 | ENSG00000118961 | LDAH | chr2 | - | 20739970 | 20740205 | 20701569 | 20701652 |
| SE | 71434 | ENSG00000115234 | SNX17 | chr2 | + | 27372622 | 27372740 | 27371268 | 27371343 |
| SE | 39905 | ENSG00000271723 | MROH7-TTC4 | chr1 | + | 54733628 | 54733710 | 54728345 | 54728432 |
| SE | 23005 | ENSG00000171617 | ENC1 | chr5 | - | 74637493 | 74640125 | 74634683 | 74636498 |
| SE | 10757 | ENSG00000154760 | SLFN13 | chr17 | - | 35446839 | 35446932 | 35445447 | 35445693 |
| SE | 81123 | ENSG00000143226 | FCGR2A | chr1 | + | 161506333 | 161506707 | 161505986 | 161506007 |
| SE | 47598 | ENSG00000008869 | HEATR5B | chr2 | - | 37083288 | 37083436 | 37079119 | 37079331 |
| SE | 51373 | ENSG00000066427 | ATXN3 | chr14 | - | 92093745 | 92093831 | 92093251 | 92093318 |
| SE | 7555 | ENSG00000167851 | CD300A | chr17 | + | 74481288 | 74481833 | 74477435 | 74477530 |
| SE | 50272 | ENSG00000134909 | ARHGAP32 | chr11 | - | 129062279 | 129062357 | 128998318 | 128998468 |
| SE | 69395 | ENSG00000135636 | DYSF | chr2 | + | 71668753 | 71668842 | 71667375 | 71667515 |
| SE | 9384 | ENSG00000121067 | SPOP | chr17 | - | 49668026 | 49668078 | 49622732 | 49622876 |
| SE | 45774 | ENSG00000284770 | TBCE | chr1 | + | 235427139 | 235427239 | 235414432 | 235414618 |
| SE | 60233 | ENSG00000065485 | PDIA5 | chr3 | + | 123150233 | 123150364 | 123146098 | 123146259 |
| SE | 93056 | ENSG00000163820 | FYCO1 | chr3 | - | 45919412 | 45921840 | 45917903 | 45919065 |
| SE | 89441 | ENSG00000166946 | CCNDBP1 | chr15 | + | 43185819 | 43185879 | 43185327 | 43185607 |
| SE | 93286 | ENSG00000056586 | RC3H2 | chr9 | - | 122890311 | 122890545 | 122883203 | 122883379 |
| SE | 29949 | ENSG00000136051 | WASHC4 | chr12 | + | 105140293 | 105140401 | 105137885 | 105138011 |
| SE | 6574 | ENSG00000074657 | ZNF532 | chr18 | + | 58934432 | 58934614 | 58918270 | 58920633 |
| SE | 18562 | ENSG00000100697 | DICER1 | chr14 | - | 95131508 | 95131639 | 95130057 | 95130192 |
| SE | 54681 | ENSG00000070010 | UFD1 | chr22 | - | 19475495 | 19475602 | 19475067 | 19475100 |
| SE | 11677 | ENSG00000167207 | NOD2 | chr16 | + | 50719924 | 50720008 | 50716890 | 50716974 |
| SE | 73515 | ENSG00000138380 | CARF | chr2 | + | 202941859 | 202941980 | 202917876 | 202918043 |
| SE | 46808 | ENSG00000059691 | GATB | chr4 | - | 151707987 | 151708101 | 151705184 | 151705269 |
| SE | 80979 | ENSG00000130429 | ARPC1B | chr7 | + | 99376443 | 99376574 | 99374728 | 99374781 |
| SE | 16725 | ENSG00000214013 | GANC | chr15 | + | 42329305 | 42329449 | 42327362 | 42327442 |
| SE | 67466 | ENSG00000162923 | WDR26 | chr1 | - | 224402003 | 224402109 | 224400949 | 224401069 |
| SE | 8408 | ENSG00000173757 | STAT5B | chr17 | - | 42202339 | 42202447 | 42199176 | 42201864 |
| SE | 44853 | ENSG00000143493 | INTS7 | chr1 | - | 211982675 | 211982810 | 211981092 | 211981190 |
| SE | 67984 | ENSG00000197959 | DNM3 | chr1 | + | 172253572 | 172253682 | 172131174 | 172131288 |
| SE | 89717 | ENSG00000215114 | UBXN2B | chr8 | + | 58434857 | 58434948 | 58434394 | 58434504 |
| SE | 29087 | ENSG00000163625 | WDFY3 | chr4 | - | 84896910 | 84897010 | 84860411 | 84860622 |
| SE | 105479 | ENSG00000152256 | PDK1 | chr2 | + | 172562740 | 172562800 | 172562219 | 172562291 |
| SE | 48327 | ENSG00000120798 | NR2C1 | chr12 | - | 95028386 | 95028524 | 95025149 | 95025255 |
| SE | 58380 | ENSG00000169504 | CLIC4 | chr1 | + | 24793176 | 24793213 | 24745381 | 24745625 |
| SE | 77515 | ENSG00000136243 | NUP42 | chr7 | + | 23197172 | 23197247 | 23196679 | 23196766 |
| SE | 109069 | ENSG00000198176 | TFDP1 | chr13 | + | 113631622 | 113631744 | 113610995 | 113611062 |
| SE | 84455 | ENSG00000137522 | RNF121 | chr11 | + | 71978085 | 71978235 | 71960749 | 71960891 |
| SE | 61513 | ENSG00000083520 | DIS3 | chr13 | - | 72780845 | 72781003 | 72778186 | 72778380 |
| SE | 10761 | ENSG00000118046 | STK11 | chr19 | + | 1216267 | 1216398 | 1206815 | 1207203 |
| SE | 91478 | ENSG00000141569 | TRIM65 | chr17 | - | 75892010 | 75892185 | 75889852 | 75891347 |
| SE | 34536 | ENSG00000150779 | TIMM8B | chr11 | - | 112085966 | 112086310 | 112085148 | 112085462 |
| SE | 14583 | ENSG00000140263 | SORD | chr15 | + | 45061066 | 45061226 | 45043256 | 45043421 |
| SE | 47718 | ENSG00000153560 | UBP1 | chr3 | - | 33396744 | 33396878 | 33396161 | 33396280 |
| SE | 76453 | ENSG00000168685 | IL7R | chr5 | + | 35874448 | 35874542 | 35871055 | 35871213 |
| SE | 5336 | ENSG00000171552 | BCL2L1 | chr20 | - | 31666990 | 31667147 | 31664574 | 31666086 |
| SE | 37587 | ENSG00000082074 | FYB1 | chr5 | - | 39153447 | 39153604 | 39141094 | 39141141 |
| SE | 86626 | ENSG00000186312 | CA5BP1 | chrX | + | 15681000 | 15681047 | 15675714 | 15675778 |
| SE | 42976 | ENSG00000121940 | CLCC1 | chr1 | - | 108943835 | 108944057 | 108941404 | 108941498 |

[illegible]

| | | | | | | | | | | | | | | | | | | | | | | | | | |
|---|---|---|---|---|---|---|---|---|---|---|---|---|---|---|---|---|---|---|---|---|---|---|---|---|---|
| SE | 83175 | ENSG00000197965 | MPZL1 | chr1 | + | 167776063 | 167776166 | 167765057 | 167765749 | 167787819 | 167788012 | 2 | 5 | 4 | 1 | 24 | 10 | 6 | 10 | 3 | 10 | 19 | 4 | 1 | 3 |
| SE | 3534 | ENSG00000105483 | CARD8 | chr19 | - | 48223718 | 48223868 | 48221729 | 48221855 | 48231659 | 48232114 | 22 | 28 | 15 | 19 | 27 | 3 | 0 | 0 | 1 | 1 | 10 | 3 | 13 | 7 |
| SE | 1528 | ENSG00000100242 | SUN2 | chr22 | - | 38754641 | 38754690 | 38752506 | 38752665 | 38755762 | 38756000 | 98 | 42 | 93 | 66 | 51 | 53 | 16 | 83 | 75 | 26 | 115 | 61 | 4 | 53 |
| SE | 26188 | ENSG00000059378 | PARP12 | chr7 | - | 140027275 | 140027406 | 140026195 | 140026348 | 140028612 | 140028688 | 174 | 168 | 195 | 224 | 278 | 7 | 1 | 0 | 15 | 11 | 590 | 410 | 130 | 567 |
| SE | 98845 | ENSG00000135365 | PHF21A | chr11 | - | 46118741 | 46118800 | 46092182 | 46092223 | 46120934 | 46121454 | 4 | 0 | 2 | 3 | 2 | 9 | 16 | 15 | 23 | 10 | 18 | 8 | 0 | 5 |
| SE | 75237 | ENSG00000100503 | NIN | chr14 | - | 50821873 | 50822077 | 50805916 | 50806818 | 50830463 | 50830479 | 229 | 153 | 201 | 264 | 140 | 5 | 5 | 7 | 6 | 0 | 57 | 226 | 185 | 127 |
| SE | 86361 | ENSG00000035681 | NSMAF | chr8 | - | 58599969 | 58600021 | 58599749 | 58599870 | 58601284 | 58601348 | 215 | 121 | 175 | 213 | 342 | 2 | 2 | 2 | 3 | 1 | 221 | 190 | 151 | 171 |
| SE | 40397 | ENSG00000163719 | MTMR14 | chr3 | + | 9697710 | 9697866 | 9689963 | 9690143 | 9701789 | 9702216 | 37 | 22 | 42 | 38 | 18 | 16 | 7 | 12 | 19 | 35 | 79 | 42 | 37 | 52 |
| SE | 98109 | ENSG00000171466 | ZNF562 | chr19 | - | 9658008 | 9658135 | 9656546 | 9656653 | 9660719 | 9660874 | 77 | 77 | 49 | 72 | 77 | 3 | 0 | 0 | 0 | 0 | 123 | 79 | 94 | 89 |
| SE | 56415 | ENSG00000104714 | ERICH1 | chr8 | - | 690897 | 691096 | 673766 | 674047 | 692477 | 692612 | 3 | 3 | 8 | 9 | 6 | 64 | 35 | 55 | 87 | 63 | 0 | 2 | 5 | 5 |
| SE | 97315 | ENSG00000070718 | AP3M2 | chr8 | + | 42154477 | 42154536 | 42152945 | 42153105 | 42154615 | 42154677 | 2 | 2 | 2 | 0 | 3 | 12 | 2 | 4 | 7 | 2 | 0 | 0 | 3 | 7 |
| SE | 5527 | ENSG00000125835 | SNRPB | chr20 | - | 2467191 | 2467315 | 2465707 | 2465819 | 2467606 | 2467758 | 15 | 6 | 4 | 15 | 0 | 385 | 218 | 325 | 316 | 312 | 20 | 5 | 0 | 5 |
| SE | 55309 | ENSG00000166938 | DIS3L | chr15 | + | 66306761 | 66306952 | 66294987 | 66295141 | 66308708 | 66308844 | 44 | 25 | 48 | 37 | 28 | 0 | 0 | 0 | 0 | 6 | 4 | 23 | 14 | 29 |
| SE | 45498 | ENSG00000168894 | RNF181 | chr2 | + | 85596856 | 85596931 | 85596518 | 85596649 | 85597444 | 85597604 | 3 | 1 | 0 | 4 | 5 | 2 | 0 | 1 | 3 | 0 | 20 | 3 | 4 | 4 |
| SE | 29719 | ENSG00000138641 | HERC3 | chr4 | + | 88597949 | 88598087 | 88595556 | 88595614 | 88605794 | 88606049 | 61 | 47 | 58 | 89 | 51 | 309 | 237 | 305 | 286 | 219 | 93 | 51 | 57 | 42 |
| SE | 88422 | ENSG00000168172 | HOOK3 | chr8 | + | 42997549 | 42997637 | 42986654 | 42986795 | 43007846 | 43007929 | 164 | 159 | 91 | 181 | 338 | 4 | 1 | 1 | 2 | 5 | 36 | 164 | 219 | 126 |
| SE | 6514 | ENSG00000124155 | PIGT | chr20 | + | 45416516 | 45416694 | 45416096 | 45416343 | 45419294 | 45419395 | 98 | 31 | 58 | 84 | 66 | 2 | 0 | 2 | 0 | 2 | 103 | 98 | 87 | 92 |
| SE | 103250 | ENSG00000122862 | SRGN | chr10 | + | 69097083 | 69097231 | 69088102 | 69088236 | 69103870 | 69104805 | 8736 | 4634 | 5776 | 8440 | 7852 | 32 | 32 | 17 | 24 | 11 | 11832 | 11268 | 10946 | 28467 |
| SE | 47646 | ENSG00000288701 | PRRC2B | chr9 | + | 131479251 | 131479393 | 131477743 | 131477949 | 131481725 | 131481808 | 253 | 128 | 225 | 304 | 164 | 0 | 0 | 0 | 1 | 0 | 276 | 250 | 154 | 201 |
| SE | 22733 | ENSG00000050393 | MCUR1 | chr6 | - | 13791877 | 13791992 | 13786556 | 13790864 | 13798832 | 13798904 | 10 | 11 | 10 | 13 | 19 | 0 | 0 | 0 | 0 | 0 | 1 | 8 | 5 | 4 |
| SE | 27937 | ENSG00000084092 | NOA1 | chr4 | - | 56966619 | 56966736 | 56964405 | 56964526 | 56968383 | 56968515 | 105 | 98 | 162 | 149 | 234 | 3 | 1 | 0 | 2 | 4 | 130 | 109 | 84 | 69 |
| SE | 22096 | ENSG00000111859 | NEDD9 | chr6 | - | 11305065 | 11305347 | 11251563 | 11251726 | 11305991 | 11306155 | 4 | 1 | 0 | 3 | 1 | 2 | 5 | 3 | 8 | 2 | 0 | 1 | 6 | 6 |
| SE | 100488 | ENSG00000213639 | PPP1CB | chr2 | + | 28752037 | 28752176 | 28751738 | 28751820 | 28776850 | 28776982 | 451 | 234 | 337 | 387 | 255 | 0 | 0 | 6 | 2 | 0 | 152 | 227 | 267 | 118 |
| SE | 90732 | ENSG00000168404 | MLKL | chr16 | - | 74678898 | 74678980 | 74675612 | 74675764 | 74685485 | 74685583 | 38 | 27 | 32 | 30 | 39 | 3 | 0 | 0 | 3 | 0 | 40 | 13 | 16 | 43 |
| SE | 100304 | ENSG00000162144 | CYB561A3 | chr11 | - | 61359425 | 61359571 | 61357732 | 61357828 | 61361732 | 61361882 | 0 | 0 | 0 | 8 | 0 | 17 | 10 | 22 | 22 | 20 | 0 | 0 | 0 | 0 |
| SE | 31719 | ENSG00000145012 | LPP | chr3 | + | 188341662 | 188341719 | 188225404 | 188225527 | 188406111 | 188406313 | 3 | 4 | 13 | 21 | 9 | 38 | 15 | 6 | 13 | 9 | 8 | 34 | 27 | 2 |
| SE | 49332 | ENSG00000101452 | DHX35 | chr20 | + | 38972558 | 38972651 | 38969080 | 38969214 | 38983698 | 38983776 | 61 | 52 | 55 | 66 | 54 | 3 | 0 | 1 | 1 | 3 | 60 | 50 | 102 | 84 |
| SE | 92610 | ENSG00000102699 | PARP4 | chr13 | - | 24503644 | 24503777 | 24501632 | 24501834 | 24512705 | 24512778 | 567 | 426 | 471 | 446 | 520 | 7 | 4 | 1 | 3 | 2 | 213 | 401 | 331 | 414 |
| SE | 98264 | ENSG00000160226 | CFAP410 | chr21 | - | 44335757 | 44335804 | 44333032 | 44333262 | 44337648 | 44337667 | 7 | 9 | 10 | 16 | 3 | 3 | 0 | 3 | 0 | 0 | 12 | 10 | 15 | 4 |
| SE | 83440 | ENSG00000122545 | SEPTIN7 | chr7 | + | 35879595 | 35879719 | 35873640 | 35873775 | 35879822 | 35879940 | 15 | 10 | 1 | 16 | 48 | 1412 | 1228 | 1087 | 1542 | 3000 | 0 | 16 | 12 | 2 |
| SE | 36775 | ENSG00000163539 | CLASP2 | chr3 | - | 33644756 | 33644903 | 33632291 | 33632371 | 33684358 | 33684456 | 115 | 56 | 93 | 93 | 144 | 0 | 0 | 0 | 0 | 0 | 29 | 52 | 86 | 49 |
| SE | 62750 | ENSG00000162959 | MEMO1 | chr2 | - | 31883385 | 31883462 | 31869847 | 31869952 | 31891991 | 31892134 | 284 | 199 | 202 | 245 | 352 | 21 | 15 | 17 | 14 | 23 | 391 | 161 | 198 | 301 |
| SE | 41543 | ENSG00000071127 | WDR1 | chr4 | - | 10103895 | 10103986 | 10088663 | 10088741 | 10116112 | 10116234 | 377 | 333 | 347 | 458 | 271 | 10 | 4 | 6 | 17 | 7 | 851 | 676 | 433 | 651 |
| SE | 32447 | ENSG00000234231 | ANAPC1P4 | chr2 | + | 87735673 | 87735765 | 87732386 | 87732621 | 87737717 | 87737852 | 67 | 16 | 85 | 31 | 24 | 1 | 0 | 0 | 0 | 0 | 34 | 153 | 56 | 23 |
| SE | 67052 | ENSG00000170906 | NDUFA3 | chr19 | + | 54106352 | 54106446 | 54105933 | 54106011 | 54106810 | 54106961 | 5 | 7 | 7 | 7 | 10 | 86 | 31 | 57 | 72 | 36 | 2 | 13 | 6 | 9 |
| SE | 54336 | ENSG00000176095 | IP6K1 | chr3 | - | 49747817 | 49748168 | 49738211 | 49738422 | 49786353 | 49786540 | 126 | 50 | 81 | 97 | 40 | 9 | 0 | 2 | 8 | 0 | 80 | 65 | 81 | 52 |
| SE | 849 | ENSG00000242259 | C22orf39 | chr22 | - | 19375630 | 19375792 | 19361726 | 19361931 | 19383619 | 19383705 | 97 | 42 | 66 | 55 | 44 | 0 | 1 | 0 | 0 | 0 | 151 | 65 | 60 | 101 |
| SE | 14061 | ENSG00000140694 | PARN | chr16 | - | 14623923 | 14623953 | 14617589 | 14617650 | 14627105 | 14627187 | 6 | 5 | 8 | 5 | 12 | 302 | 252 | 235 | 292 | 200 | 39 | 1 | 1 | 3 |
| SE | 85358 | ENSG00000102409 | BEX4 | chrX | + | 103215681 | 103215764 | 103215107 | 103215238 | 103216148 | 103217246 | 11 | 11 | 22 | 9 | 4 | 0 | 0 | 2 | 0 | 0 | 36 | 3 | 3 | 23 |
| SE | 41009 | ENSG00000164091 | WDR82 | chr3 | - | 52259196 | 52259266 | 52254433 | 52257519 | 52259716 | 52259872 | 279 | 182 | 283 | 279 | 530 | 0 | 0 | 0 | 0 | 0 | 183 | 201 | 167 | 200 |
| SE | 11439 | ENSG00000167522 | ANKRD11 | chr16 | - | 89316932 | 89317078 | 89305099 | 89305344 | 89418283 | 89418305 | 150 | 89 | 124 | 107 | 125 | 2 | 2 | 2 | 3 | 4 | 364 | 160 | 202 | 212 |
| SE | 87611 | ENSG00000102078 | SLC25A14 | chrX | + | 130349823 | 130349867 | 130349250 | 130349345 | 130350645 | 130350731 | 9 | 6 | 11 | 9 | 9 | 12 | 12 | 9 | 13 | 12 | 11 | 6 | 1 | 4 |
| SE | 50841 | ENSG00000048052 | HDAC9 | chr7 | + | 18502352 | 18502486 | 18496261 | 18496324 | 18502580 | 18502639 | 0 | 2 | 4 | 2 | 5 | 1 | 1 | 0 | 3 | 7 | 9 | 4 | 7 | 0 |
| SE | 78050 | ENSG00000151116 | UEVLD | chr11 | - | 18570213 | 18570377 | 18566346 | 18566482 | 18575346 | 18575412 | 70 | 64 | 40 | 43 | 99 | 4 | 4 | 3 | 2 | 3 | 3 | 40 | 28 | 51 |
| SE | 76914 | ENSG00000167483 | NIBAN3 | chr19 | + | 17530754 | 17530885 | 17527237 | 17527395 | 17532262 | 17532388 | 92 | 62 | 84 | 81 | 22 | 15 | 3 | 17 | 26 | 6 | 35 | 180 | 11 | 48 |
| SE | 58568 | ENSG00000172500 | FIBP | chr11 | - | 65886321 | 65886422 | 65885508 | 65885663 | 65887599 | 65887726 | 140 | 65 | 103 | 110 | 150 | 3 | 1 | 3 | 6 | 10 | 187 | 213 | 81 | 136 |
| SE | 6757 | ENSG00000133313 | CNDP2 | chr18 | + | 74497508 | 74497726 | 74496371 | 74496431 | 74499936 | 74500033 | 3 | 1 | 1 | 2 | 0 | 28 | 22 | 15 | 26 | 7 | 3 | 0 | 0 | 2 |
| SE | 70277 | ENSG00000104218 | CSPP1 | chr8 | + | 67118751 | 67118821 | 67118247 | 67118369 | 67131950 | 67132080 | 31 | 26 | 14 | 31 | 28 | 0 | 0 | 0 | 0 | 0 | 18 | 14 | 16 | 14 |
| SE | 86251 | ENSG00000134516 | DOCK2 | chr5 | + | 169684195 | 169684350 | 169681743 | 169681879 | 169689251 | 169689333 | 1365 | 944 | 1192 | 1125 | 1294 | 12 | 7 | 6 | 5 | 11 | 1801 | 1276 | 1190 | 1586 |
| SE | 39445 | ENSG00000055917 | PUM2 | chr2 | - | 20331716 | 20331937 | 20327309 | 20327378 | 20332966 | 20333063 | 48 | 32 | 43 | 62 | 94 | 21 | 25 | 11 | 5 | 16 | 13 | 42 | 40 | 49 |
| SE | 61494 | ENSG00000124588 | NQO2 | chr6 | + | 3004494 | 3004651 | 2999988 | 3000085 | 3006467 | 3006559 | 18 | 3 | 7 | 7 | 4 | 19 | 6 | 18 | 20 | 4 | 33 | 19 | 13 | 13 |
| SE | 56108 | ENSG00000165689 | ENTR1 | chr9 | - | 136410089 | 136410239 | 136407825 | 136407938 | 136410327 | 136410609 | 1 | 7 | 4 | 9 | 0 | 10 | 1 | 8 | 2 | 1 | 0 | 2 | 1 | 3 |
| SE | 27244 | ENSG00000073331 | ALPK1 | chr4 | + | 112302338 | 112302408 | 112297395 | 112297469 | 112315800 | 112315852 | 1 | 1 | 8 | 0 | 4 | 46 | 46 | 50 | 33 | 38 | 20 | 0 | 0 | 3 |
| SE | 37671 | ENSG00000168615 | ADAM9 | chr8 | + | 39071297 | 39071413 | 39055576 | 39055772 | 39077227 | 39077411 | 63 | 57 | 43 | 67 | 57 | 0 | 4 | 6 | 4 | 1 | 12 | 106 | 85 | 47 |
| SE | 68832 | ENSG00000113456 | RAD1 | chr5 | - | 34913469 | 34913578 | 34911553 | 34911812 | 34913926 | 34914026 | 21 | 18 | 27 | 21 | 56 | 2 | 0 | 0 | 4 | 1 | 43 | 5 | 8 | 10 |
| SE | 43143 | ENSG00000113161 | HMGCR | chr5 | + | 75355055 | 75355214 | 75354502 | 75354697 | 75355364 | 75355522 | 206 | 160 | 163 | 197 | 181 | 42 | 19 | 31 | 54 | 20 | 64 | 38 | 80 | 123 |
| SE | 29877 | ENSG00000154743 | TSEN2 | chr3 | + | 12532661 | 12532773 | 12529761 | 12531659 | 12539160 | 12539591 | 21 | 20 | 48 | 42 | 77 | 0 | 0 | 1 | 6 | 0 | 8 | 10 | 21 | 27 |
| SE | 65511 | ENSG00000140386 | SCAPER | chr15 | - | 76765562 | 76765638 | 76765336 | 76765454 | 76766917 | 76767088 | 64 | 59 | 85 | 104 | 111 | 0 | 8 | 0 | 0 | 1 | 7 | 36 | 33 | 13 |
| SE | 26103 | ENSG00000164181 | ELOVL7 | chr5 | - | 60784093 | 60784212 | 60771902 | 60772093 | 60787333 | 60787431 | 9 | 26 | 7 | 24 | 46 | 55 | 110 | 15 | 143 | 236 | 27 | 25 | 21 | 29 |
| SE | 40501 | ENSG00000114861 | FOXP1 | chr3 | - | 71359149 | 71359244 | 71299694 | 71299880 | 71493425 | 71493555 | 145 | 174 | 197 | 156 | 175 | 7 | 3 | 1 | 4 | 3 | 183 | 175 | 184 | 135 |
| SE | 6555 | ENSG00000154856 | APCDD1 | chr18 | + | 10468468 | 10468652 | 10454634 | 10455039 | 10471529 | 10471740 | 13 | 11 | 12 | 2 | 4 | 0 | 4 | 0 | 0 | 0 | 40 | 4 | 7 | 2 |
| SE | 37351 | ENSG00000075826 | SEC31B | chr10 | - | 100500109 | 100500159 | 100499523 | 100499598 | 100502253 | 100502484 | 12 | 18 | 27 | 36 | 14 | 13 | 8 | 13 | 17 | 6 | 36 | 33 | 35 | 22 |
| SE | 7251 | ENSG00000152242 | C18orf25 | chr18 | + | 46240004 | 46240187 | 46215810 | 46216595 | 46253696 | 46253823 | 17 | 6 | 6 | 6 | 21 | 43 | 48 | 58 | 17 | 42 | 7 | 14 | 17 | 11 |
| SE | 49342 | ENSG00000101452 | DHX35 | chr20 | + | 39025229 | 39025359 | 39023689 | 39023767 | 39030703 | 39030775 | 12 | 7 | 10 | 10 | 10 | 0 | 0 | 0 | 0 | 0 | 6 | 7 | 21 | 7 |

[illegible]

| | | | | | | | | | | | | | | | | | | | | | | | | | | | | | | | | | | | | | | | | | | | | | | |
|---|---|---|---|---|---|---|---|---|---|---|---|---|---|---|---|---|---|---|---|---|---|---|---|---|---|---|---|---|---|---|---|---|---|---|---|---|---|---|---|---|---|---|---|---|---|---|
| SE | 39388 | ENSG00000220804 | LINC01881 | chr2 | + | 242130654 | 242130866 | 242122292 | 242122455 | 242138783 | 242138888 | 23 | 11 | 21 | 26 | 20 | 12 | 14 | 10 | 12 | 72 | 57 | 5 | 4 | 34 | 0 | 11 | 21 | 21 | 2 | 20 | 298 | 149 | 0.434694797 | 1 | 0.489 | 0.282 | 0.512 | 0.52 | 0.122 | 0.722 | 0.106 | 0.087 | 0.895 | 0 | 0.023 |
| SE | 1345 | ENSG00000093000 | NUP50 | chr22 | + | 45183401 | 45183520 | 45181285 | 45181367 | 45184452 | 45184504 | 422 | 337 | 331 | 368 | 714 | 3 | 1 | 1 | 0 | 3 | 174 | 329 | 417 | 353 | 57 | 0 | 0 | 7 | 6 | 2 | 268 | 149 | 0.434896367 | 1 | 0.987 | 0.995 | 0.995 | 1 | 0.992 | 1 | 1 | 0.971 | 0.97 | 0.941 | 0.017 |
| SE | 75043 | ENSG00000130023 | ERMARD | chr6 | + | 169773318 | 169773402 | 169769539 | 169769713 | 169775939 | 169776065 | 11 | 17 | 11 | 21 | 23 | 1 | 3 | 1 | 1 | 0 | 5 | 8 | 6 | 1 | 8 | 0 | 0 | 3 | 2 | 0 | 233 | 149 | 0.434899956 | 1 | 0.876 | 0.784 | 0.876 | 0.931 | 1 | 1 | 1 | 0.561 | 0.242 | 1 | 0.133 |
| SE | 84126 | ENSG00000094841 | UPRT | chrX | + | 75293471 | 75293514 | 75274084 | 75274640 | 75296341 | 75296411 | 64 | 44 | 37 | 50 | 31 | 2 | 1 | 1 | 0 | 2 | 76 | 47 | 39 | 43 | 15 | 0 | 0 | 0 | 0 | 0 | 192 | 149 | 0.434916523 | 1 | 0.961 | 0.972 | 0.966 | 1 | 0.923 | 1 | 1 | 1 | 1 | 1 | -0.036 |
| SE | 9527 | ENSG00000083290 | ULK2 | chr17 | - | 19796094 | 19796282 | 19795621 | 19795725 | 19797395 | 19797682 | 87 | 72 | 65 | 120 | 50 | 3 | 2 | 0 | 1 | 11 | 46 | 77 | 36 | 49 | 62 | 0 | 0 | 14 | 7 | 0 | 298 | 149 | 0.434930445 | 1 | 0.935 | 0.947 | 1 | 0.984 | 0.694 | 1 | 1 | 0.562 | 0.778 | 1 | 0.044 |
| SE | 71296 | ENSG00000064012 | CASP8 | chr2 | + | 201272897 | 201272942 | 201272637 | 201272776 | 201284815 | 201285317 | 154 | 58 | 135 | 111 | 160 | 3 | 3 | 2 | 2 | 2 | 156 | 194 | 258 | 104 | 48 | 0 | 0 | 7 | 0 | 0 | 194 | 149 | 0.434970503 | 1 | 0.975 | 0.937 | 0.981 | 0.977 | 0.984 | 1 | 1 | 0.966 | 1 | 1 | -0.022 |
| SE | 31255 | ENSG00000047365 | ARAP2 | chr4 | - | 36073688 | 36073823 | 36066003 | 36068278 | 36080215 | 36080279 | 364 | 223 | 202 | 296 | 398 | 3 | 7 | 2 | 10 | 4 | 21 | 125 | 99 | 4 | 8 | 0 | 5 | 2 | 0 | 0 | 284 | 149 | 0.435213797 | 1 | 0.985 | 0.944 | 0.981 | 0.94 | 0.981 | 1 | 0.929 | 0.963 | 1 | 1 | -0.012 |
| SE | 43071 | ENSG00000178917 | ZNF852 | chr3 | - | 44502541 | 44502668 | 44499522 | 44500637 | 44510552 | 44510636 | 59 | 66 | 65 | 69 | 97 | 2 | 1 | 1 | 0 | 3 | 55 | 41 | 40 | 33 | 62 | 0 | 0 | 0 | 0 | 0 | 276 | 149 | 0.435415988 | 1 | 0.941 | 0.973 | 0.972 | 1 | 0.946 | 1 | 1 | 1 | 1 | 1 | -0.034 |
| SE | 71024 | ENSG00000163516 | ANKZF1 | chr2 | + | 219231927 | 219232040 | 219229782 | 219229900 | 219232259 | 219232270 | 75 | 49 | 99 | 101 | 117 | 0 | 0 | 0 | 1 | 0 | 87 | 110 | 42 | 104 | 89 | 0 | 0 | 0 | 5 | 4 | 262 | 149 | 0.435446194 | 1 | 1 | 1 | 1 | 0.983 | 1 | 1 | 1 | 1 | 0.922 | 0.927 | 0.027 |
| SE | 82508 | ENSG00000198315 | ZKSCAN8 | chr6 | + | 28148315 | 28148824 | 28144330 | 28144424 | 28149482 | 28149624 | 37 | 28 | 50 | 44 | 34 | 0 | 0 | 0 | 0 | 0 | 21 | 37 | 24 | 17 | 7 | 0 | 3 | 3 | 0 | 0 | 298 | 149 | 0.435461906 | 1 | 1 | 1 | 1 | 1 | 1 | 1 | 0.86 | 0.8 | 1 | 1 | 0.068 |
| SE | 95275 | ENSG00000165219 | GAPVD1 | chr9 | + | 125349389 | 125349519 | 125345889 | 125346941 | 125350294 | 125350404 | 187 | 143 | 192 | 175 | 209 | 0 | 0 | 0 | 0 | 0 | 215 | 168 | 185 | 217 | 126 | 0 | 4 | 0 | 3 | 5 | 279 | 149 | 0.435468233 | 1 | 1 | 1 | 1 | 1 | 1 | 1 | 0.957 | 1 | 0.975 | 0.931 | 0.027 |
| SE | 105614 | ENSG00000139323 | POC1B | chr12 | - | 89472167 | 89472275 | 89466769 | 89466922 | 89491935 | 89492115 | 48 | 28 | 49 | 58 | 54 | 2 | 0 | 1 | 0 | 2 | 52 | 28 | 23 | 30 | 60 | 0 | 6 | 0 | 0 | 0 | 257 | 149 | 0.43549 | 1 | 0.933 | 1 | 0.966 | 1 | 0.94 | 1 | 0.73 | 1 | 1 | 1 | 0.022 |
| SE | 71058 | ENSG00000169914 | OTUD3 | chr1 | + | 19906431 | 19906616 | 19904890 | 19904987 | 19907569 | 19912945 | 28 | 31 | 35 | 26 | 26 | 14 | 11 | 7 | 11 | 10 | 4 | 16 | 12 | 55 | 4 | 10 | 27 | 4 | 12 | 0 | 298 | 149 | 0.435571393 | 1 | 0.5 | 0.585 | 0.714 | 0.542 | 0.565 | 0.167 | 0.229 | 0.6 | 0.696 | 1 | 0.043 |
| SE | 75940 | ENSG00000160185 | UBASH3A | chr21 | + | 42444533 | 42444643 | 42442451 | 42442596 | 42447056 | 42447194 | 19 | 11 | 40 | 55 | 34 | 3 | 1 | 0 | 0 | 0 | 14 | 29 | 3 | 6 | 19 | 0 | 6 | 0 | 0 | 0 | 259 | 149 | 0.435586685 | 1 | 0.785 | 0.864 | 1 | 1 | 1 | 1 | 0.735 | 1 | 1 | 1 | -0.017 |
| SE | 87665 | ENSG00000143569 | UBAP2L | chr1 | + | 154234590 | 154234792 | 154228614 | 154228725 | 154236565 | 154236611 | 670 | 688 | 600 | 808 | 1036 | 2 | 0 | 0 | 0 | 0 | 1686 | 866 | 519 | 890 | 1424 | 0 | 6 | 4 | 1 | 23 | 298 | 149 | 0.435660086 | 1 | 0.994 | 1 | 1 | 1 | 1 | 1 | 0.986 | 0.985 | 0.998 | 0.969 | 0.011 |
| SE | 45856 | ENSG00000062485 | CS | chr12 | - | 56283791 | 56283857 | 56282859 | 56282991 | 56285915 | 56285961 | 352 | 251 | 326 | 390 | 476 | 9 | 1 | 1 | 4 | 2 | 546 | 347 | 262 | 585 | 555 | 7 | 1 | 1 | 5 | 12 | 215 | 149 | 0.435709633 | 1 | 0.964 | 0.994 | 0.996 | 0.985 | 0.994 | 0.982 | 0.996 | 0.995 | 0.988 | 0.97 | 0 |
| SE | 12015 | ENSG00000170100 | ZNF778 | chr16 | + | 89223156 | 89223283 | 89222091 | 89222183 | 89224718 | 89224802 | 21 | 17 | 18 | 23 | 12 | 7 | 2 | 7 | 7 | 0 | 20 | 13 | 17 | 8 | 30 | 0 | 2 | 0 | 2 | 29 | 276 | 149 | 0.435750524 | 1 | 0.618 | 0.821 | 0.581 | 0.639 | 1 | 1 | 0.778 | 1 | 0.683 | 0.358 | -0.032 |
| SE | 67080 | ENSG00000204681 | GABBR1 | chr6 | - | 29630521 | 29630643 | 29629086 | 29629107 | 29631395 | 29631599 | 16 | 17 | 7 | 21 | 7 | 6 | 1 | 0 | 0 | 0 | 36 | 5 | 2 | 11 | 9 | 0 | 0 | 0 | 9 | 0 | 271 | 149 | 0.435797208 | 1 | 0.595 | 0.903 | 1 | 1 | 1 | 1 | 1 | 1 | 0.402 | 1 | 0.019 |
| SE | 75048 | ENSG00000036054 | TBC1D23 | chr3 | + | 100283606 | 100283811 | 100281717 | 100281847 | 100290577 | 100290645 | 153 | 103 | 103 | 98 | 207 | 1 | 0 | 1 | 0 | 0 | 29 | 66 | 50 | 33 | 17 | 0 | 0 | 0 | 6 | 0 | 298 | 149 | 0.435823846 | 1 | 0.987 | 1 | 0.981 | 1 | 1 | 1 | 1 | 1 | 0.733 | 1 | 0.047 |
| SE | 37373 | ENSG00000166927 | MS4A7 | chr11 | + | 60386716 | 60386773 | 60383128 | 60383288 | 60389389 | 60389596 | 175 | 151 | 73 | 134 | 180 | 17 | 3 | 4 | 11 | 9 | 50 | 56 | 27 | 42 | 29 | 15 | 0 | 0 | 2 | 5 | 206 | 149 | 0.435885814 | 1 | 0.882 | 0.973 | 0.93 | 0.898 | 0.935 | 0.707 | 1 | 1 | 0.938 | 0.808 | 0.033 |
| SE | 72691 | ENSG00000104047 | DTWD1 | chr15 | + | 49626743 | 49626893 | 49625112 | 49625431 | 49632158 | 49632302 | 5 | 2 | 3 | 6 | 7 | 50 | 58 | 51 | 70 | 99 | 14 | 1 | 2 | 0 | 0 | 8 | 33 | 16 | 15 | 26 | 298 | 149 | 0.435888689 | 1 | 0.048 | 0.017 | 0.029 | 0.041 | 0.034 | 0.467 | 0.015 | 0.059 | 0 | 0 | -0.074 |
| SE | 14394 | ENSG00000103653 | CSK | chr15 | + | 74801521 | 74801595 | 74800851 | 74801102 | 74801694 | 74801729 | 435 | 237 | 281 | 344 | 226 | 2 | 1 | 0 | 2 | 0 | 403 | 439 | 187 | 242 | 768 | 14 | 0 | 2 | 0 | 6 | 223 | 149 | 0.435902557 | 1 | 0.993 | 0.994 | 1 | 0.991 | 1 | 0.951 | 1 | 0.984 | 1 | 0.988 | 0.011 |
| [illegible] | | | | | | | | | | | | | | | | | | | | | | | | | | | | | | | | | | | | | | | | | | | | | | |

| | | | | | | | | | | | | | | | | | | | | | | | | | | | | | | | | | | | | | | | | | | | | | | |
|---|---|---|---|---|---|---|---|---|---|---|---|---|---|---|---|---|---|---|---|---|---|---|---|---|---|---|---|---|---|---|---|---|---|---|---|---|---|---|---|---|---|---|---|---|---|---|
| SE | 5606 | ENSG00000254806 | SYS1-DBNDD2 | chr20 | + | 45363528 | 45363693 | 45363199 | 45363315 | 45365618 | 45365686 | 34 | 18 | 24 | 33 | 10 | 0 | 5 | 0 | 0 | 0 | 15 | 23 | 21 | 36 | 19 | 0 | 0 | 0 | 0 | 0 | 298 | 149 | 0.453723806 | 1 | 1 | 0.643 | 1 | 1 | 1 | 1 | 1 | 1 | 1 | 1 | -0.071 |
| SE | 61135 | ENSG00000227039 | ITGB2-AS1 | chr21 | + | 44922912 | 44922983 | 44921072 | 44921327 | 44926867 | 44927004 | 14 | 18 | 4 | 11 | 2 | 22 | 40 | 15 | 25 | 5 | 35 | 18 | 13 | 48 | 20 | 57 | 66 | 14 | 86 | 39 | 220 | 149 | 0.453776712 | 1 | 0.301 | 0.234 | 0.153 | 0.23 | 0.213 | 0.294 | 0.156 | 0.386 | 0.274 | 0.258 | -0.047 |
| SE | 98693 | ENSG00000205531 | NAP1L4 | chr11 | - | 2989128 | 2989271 | 2979206 | 2979237 | 2992253 | 2992377 | 13 | 3 | 5 | 11 | 11 | 70 | 30 | 67 | 75 | 41 | 0 | 12 | 3 | 12 | 3 | 56 | 67 | 54 | 29 | 27 | 292 | 149 | 0.453851729 | 1 | 0.087 | 0.049 | 0.037 | 0.07 | 0.12 | 0 | 0.084 | 0.028 | 0.174 | 0.054 | 0.005 |
| SE | 100933 | ENSG00000074319 | TSG101 | chr11 | - | 18480484 | 18480635 | 18469030 | 18469218 | 18481629 | 18481731 | 89 | 26 | 49 | 57 | 37 | 0 | 0 | 0 | 8 | 0 | 51 | 42 | 47 | 53 | 44 | 0 | 0 | 0 | 0 | 0 | 298 | 149 | 0.453912121 | 1 | 1 | 1 | 1 | 0.781 | 1 | 1 | 1 | 1 | 1 | 1 | -0.044 |
| SE | 82431 | ENSG00000157191 | NECAP2 | chr1 | + | 16450124 | 16450241 | 16449092 | 16449201 | 16451837 | 16452015 | 5 | 5 | 8 | 12 | 1 | 147 | 107 | 139 | 130 | 140 | 24 | 3 | 4 | 4 | 0 | 140 | 145 | 57 | 113 | 144 | 266 | 149 | 0.454029708 | 1 | 0.019 | 0.026 | 0.031 | 0.049 | 0.004 | 0.088 | 0.011 | 0.038 | 0.019 | 0 | -0.005 |
| SE | 5971 | ENSG00000170471 | RALGAPB | chr20 | + | 38565358 | 38565478 | 38562531 | 38562697 | 38567095 | 38567232 | 186 | 70 | 141 | 182 | 308 | 1 | 2 | 0 | 0 | 0 | 95 | 130 | 81 | 68 | 39 | 0 | 4 | 3 | 0 | 0 | 269 | 149 | 0.454115522 | 1 | 0.99 | 0.951 | 1 | 1 | 1 | 1 | 0.947 | 0.937 | 1 | 1 | 0.011 |
| SE | 915 | ENSG00000183765 | CHEK2 | chr22 | - | 28696900 | 28696987 | 28695709 | 28695873 | 28699837 | 28699937 | 21 | 9 | 12 | 18 | 51 | 7 | 0 | 0 | 4 | 0 | 6 | 14 | 0 | 18 | 47 | 0 | 0 | 3 | 7 | 5 | 236 | 149 | 0.454378088 | 1 | 0.654 | 1 | 1 | 0.74 | 1 | 1 | 1 | 0 | 0.619 | 0.856 | 0.184 |
| SE | 53933 | ENSG00000138614 | INTS14 | chr15 | - | 65607301 | 65607442 | 65605128 | 65605236 | 65611097 | 65611258 | 24 | 21 | 13 | 22 | 7 | 0 | 0 | 2 | 1 | 3 | 18 | 12 | 6 | 8 | 4 | 0 | 3 | 4 | 0 | 0 | 290 | 149 | 0.45438077 | 1 | 1 | 1 | 0.77 | 0.919 | 0.545 | 1 | 0.673 | 0.435 | 1 | 1 | 0.025 |
| SE | 35901 | ENSG00000146232 | NFKBIE | chr6 | - | 44261625 | 44261848 | 44260450 | 44260539 | 44264981 | 44265593 | 46 | 16 | 31 | 70 | 40 | 0 | 0 | 0 | 5 | 0 | 138 | 62 | 38 | 93 | 52 | 5 | 0 | 0 | 1 | 0 | 298 | 149 | 0.454440009 | 1 | 1 | 1 | 1 | 0.875 | 1 | 0.932 | 1 | 1 | 0.979 | 1 | -0.007 |
| SE | 91067 | ENSG00000104164 | BLOC1S6 | chr15 | + | 45601233 | 45601407 | 45592134 | 45592276 | 45603099 | 45603187 | 2 | 2 | 1 | 7 | 9 | 140 | 123 | 94 | 110 | 156 | 0 | 5 | 4 | 0 | 0 | 62 | 122 | 140 | 66 | 68 | 298 | 149 | 0.454796518 | 1 | 0.007 | 0.008 | 0.005 | 0.031 | 0.028 | 0 | 0.02 | 0.014 | 0 | 0 | 0.009 |
| SE | 86259 | ENSG00000134516 | DOCK2 | chr5 | + | 169712119 | 169712223 | 169711934 | 169712007 | 169714027 | 169714211 | 2233 | 1596 | 1707 | 2061 | 2835 | 19 | 5 | 13 | 10 | 6 | 3314 | 2605 | 2405 | 2757 | 1818 | 30 | 22 | 21 | 8 | 10 | 253 | 149 | 0.454854656 | 1 | 0.986 | 0.995 | 0.987 | 0.992 | 0.996 | 0.985 | 0.986 | 0.985 | 0.995 | 0.991 | 0.003 |
| SE | 6575 | ENSG00000074657 | ZNF532 | chr18 | + | 58934432 | 58934614 | 58918270 | 58920633 | 58953517 | 58953799 | 22 | 22 | 36 | 45 | 35 | 2 | 1 | 2 | 1 | 0 | 38 | 30 | 35 | 56 | 11 | 0 | 0 | 0 | 5 | 0 | 298 | 149 | 0.454878278 | 1 | 0.846 | 0.917 | 0.9 | 0.957 | 1 | 1 | 1 | 1 | 0.848 | 1 | -0.046 |
| SE | 79056 | ENSG00000128536 | CDHR3 | chr7 | + | 105878206 | 105878358 | 105876817 | 105877257 | 105911016 | 105911241 | 2 | 0 | 5 | 0 | 0 | 10 | 4 | 5 | 3 | 2 | 0 | 4 | 4 | 3 | 0 | 6 | 4 | 2 | 0 | 11 | 298 | 149 | 0.455040579 | 1 | 0.091 | 0 | 0.333 | 0 | 0 | 0 | 0.333 | 0.5 | 1 | 0 | -0.282 |
| SE | 38492 | ENSG00000198793 | MTOR | chr1 | - | 11167441 | 11167517 | 11157151 | 11157291 | 11199257 | 11199403 | 72 | 73 | 68 | 88 | 90 | 2 | 3 | 1 | 2 | 0 | 118 | 88 | 98 | 122 | 80 | 0 | 5 | 0 | 0 | 0 | 225 | 149 | 0.455174441 | 1 | 0.96 | 0.942 | 0.978 | 0.967 | 1 | 1 | 0.921 | 1 | 1 | 1 | -0.015 |
| SE | 29084 | ENSG00000163625 | WDFY3 | chr4 | - | 84879405 | 84879494 | 84860563 | 84860622 | 84896910 | 84897010 | 4 | 3 | 3 | 7 | 13 | 25 | 25 | 7 | 22 | 21 | 9 | 12 | 4 | 4 | 20 | 27 | 17 | 29 | 40 | 18 | 238 | 149 | 0.455183598 | 1 | 0.091 | 0.07 | 0.212 | 0.166 | 0.279 | 0.173 | 0.306 | 0.079 | 0.059 | 0.41 | -0.042 |
| SE | 48963 | ENSG00000198821 | CD247 | chr1 | - | 167433023 | 167433059 | 167430639 | 167431746 | 167434019 | 167434066 | 1754 | 621 | 898 | 1027 | 1454 | 13 | 1 | 4 | 6 | 0 | 243 | 479 | 490 | 180 | 410 | 0 | 6 | 7 | 3 | 7 | 185 | 149 | 0.455333184 | 1 | 0.991 | 0.998 | 0.994 | 0.993 | 1 | 1 | 0.985 | 0.983 | 0.98 | 0.979 | 0.01 |
| SE | 45652 | ENSG00000144283 | PKP4 | chr2 | + | 158676738 | 158676867 | 158673882 | 158674000 | 158678580 | 158678654 | 12 | 11 | 15 | 17 | 15 | 42 | 27 | 30 | 29 | 33 | 0 | 27 | 5 | 18 | 35 | 40 | 50 | 28 | 59 | 22 | 278 | 149 | 0.45539035 | 1 | 0.133 | 0.179 | 0.211 | 0.239 | 0.196 | 0 | 0.224 | 0.087 | 0.141 | 0.46 | 0.009 |
| SE | 57951 | ENSG00000242193 | CRYZL2P | chr1 | - | 178022439 | 178022668 | 178006135 | 178008149 | 178026154 | 178026255 | 27 | 14 | 6 | 6 | 4 | 2 | 1 | 0 | 2 | 0 | 10 | 6 | 2 | 37 | 38 | 0 | 0 | 1 | 4 | 1 | 298 | 149 | 0.455740301 | 1 | 0.871 | 0.875 | 1 | 0.6 | 1 | 1 | 1 | 0.5 | 0.822 | 0.95 | 0.015 |
| SE | 35983 | ENSG00000078070 | MCCC1 | chr3 | - | 183092408 | 183092545 | 183086692 | 183086788 | 183094558 | 183094605 | 38 | 44 | 49 | 39 | 37 | 8 | 0 | 3 | 2 | 0 | 77 | 31 | 48 | 26 | 10 | 2 | 0 | 1 | 6 | 0 | 286 | 149 | 0.455815395 | 1 | 0.712 | 1 | 0.895 | 0.91 | 1 | 0.953 | 1 | 0.962 | 0.693 | 1 | -0.018 |
| SE | 108417 | ENSG00000197111 | PCBP2 | chr12 | + | 53460330 | 53460686 | 53455908 | 53456001 | 53461014 | 53461143 | 1 | 9 | 0 | 1 | 0 | 19 | 5 | 11 | 6 | 33 | 0 | 4 | 4 | 4 | 0 | 24 | 12 | 5 | 3 | 25 | 298 | 149 | 0.455886024 | 1 | 0.026 | 0.474 | 0 | 0.077 | 0 | 0 | 0.143 | 0.286 | 0.4 | 0 | -0.05 |
| SE | 6251 | ENSG00000088812 | ATRN | chr20 | + | 3576858 | 3576997 | 3572730 | 3572951 | 3578581 | 3578772 | 68 | 30 | 49 | 49 | 52 | 0 | 0 | 0 | 0 | 0 | 61 | 56 | 39 | 37 | 50 | 0 | 8 | 0 | 0 | 0 | 288 | 149 | 0.455908489 | 1 | 1 | 1 | 1 | 1 | 1 | 1 | 0.784 | 1 | 1 | 1 | 0.043 |
| SE | 16460 | ENSG00000104043 | ATP8B4 | chr15 | - | 50044593 | 50044692 | 50038767 | 50038829 | 50074126 | 50074185 | 31 | 15 | 11 | 28 | 27 | 0 | 0 | 1 | 0 | 0 | 2 | 15 | 20 | 51 | 6 | 0 | 5 | 0 | 0 | 0 | 248 | 149 | 0.455950521 | 1 | 1 | 1 | 0.869 | 1 | 1 | 1 | 0.643 | 1 | 1 | 1 | 0.045 |
| [illegible] | | | | | | | | | | | | | | | | | | | | | | | | | | | | | | | | | | | | | | | | | | | | | | |

| | | | | | | | | | | | | | | | |
|---|---|---|---|---|---|---|---|---|---|---|---|---|---|---|---|
| SE | 35988 | ENSG00000078070 | MCCC1 | chr3 | - | 183094558 | 183094605 | 183092408 | 183092545 | 183099351 | 183099495 | 29 | 30 | 40 | [illegible] |
| SE | 38041 | ENSG00000215908 | CROCCP2 | chr1 | - | 16623869 | 16623997 | 16619844 | 16620024 | 16625774 | 16626005 | 2 | 2 | 0 | [illegible] |
| SE | 10197 | ENSG00000178852 | EFCAB13 | chr17 | + | 47345015 | 47345098 | 47344161 | 47344292 | 47370436 | 47370508 | 4 | 29 | 32 | [illegible] |
| SE | 88922 | ENSG00000176731 | RBIS | chr8 | - | 85219235 | 85219363 | 85217385 | 85217502 | 85220305 | 85220384 | 140 | 4 | 3 | [illegible] |
| SE | 82349 | ENSG00000076555 | ACACB | chr12 | + | 109227370 | 109227489 | 109223814 | 109223904 | 109232668 | 109232806 | 17 | 6 | 9 | [illegible] |
| SE | 23306 | ENSG00000242110 | AMACR | chr5 | - | 34004573 | 34004734 | 33998640 | 33998827 | 34005755 | 34005899 | 33 | 5 | 17 | [illegible] |
| SE | 53607 | ENSG00000213397 | HAUS7 | chrX | - | 153464987 | 153465055 | 153462609 | 153462671 | 153469145 | 153469261 | 33 | 23 | 24 | [illegible] |
| SE | 43331 | ENSG00000135597 | REPS1 | chr6 | - | 138926400 | 138926478 | 138921036 | 138921124 | 138929976 | 138929997 | 52 | 46 | 47 | [illegible] |
| SE | 66392 | ENSG00000128708 | HAT1 | chr2 | + | 171952880 | 171953001 | 171946707 | 171946783 | 171965337 | 171965517 | 242 | 237 | 243 | [illegible] |
| SE | 32888 | ENSG00000068650 | ATP11A | chr13 | + | 112878216 | 112878303 | 112875775 | 112875941 | 112881875 | 112886113 | 22 | 2 | 8 | [illegible] |
| SE | 70952 | ENSG00000131781 | FMO5 | chr1 | - | 147201151 | 147201504 | 147190176 | 147190249 | 147212392 | 147212535 | 25 | 14 | 4 | [illegible] |
| SE | 54858 | ENSG00000133056 | PIK3C2B | chr1 | - | 204459175 | 204459482 | 204457727 | 204457874 | 204459877 | 204459941 | 0 | 0 | 0 | [illegible] |
| SE | 15371 | ENSG00000140521 | POLG | chr15 | - | 89322741 | 89322902 | 89321961 | 89322015 | 89323403 | 89323427 | 201 | 146 | 182 | [illegible] |
| SE | 94123 | ENSG00000139644 | TMBIM6 | chr12 | + | 49741956 | 49742280 | 49741556 | 49741611 | 49752463 | 49752549 | 11 | 5 | 4 | [illegible] |
| SE | 23377 | ENSG00000147044 | CASK | chrX | - | 41739383 | 41739456 | 41671427 | 41671530 | 41745523 | 41745593 | 113 | 88 | 123 | [illegible] |
| SE | 45249 | ENSG00000104419 | NDRG1 | chr8 | - | 133258365 | 133258426 | 133256776 | 133256863 | 133259167 | 133259230 | 204 | 102 | 153 | [illegible] |
| SE | 72935 | ENSG00000048649 | RSF1 | chr11 | - | 77685104 | 77685159 | 77678085 | 77678153 | 77691158 | 77691238 | 35 | 34 | 37 | [illegible] |
| SE | 92118 | ENSG00000233261 | FAM238A | chr10 | + | 26592849 | 26592966 | 26591309 | 26591413 | 26593933 | 26594330 | 2 | 0 | 0 | [illegible] |
| SE | 44164 | ENSG00000008226 | DLEC1 | chr3 | + | 38062580 | 38062801 | 38062168 | 38062368 | 38063840 | 38063919 | 34 | 9 | 13 | [illegible] |
| SE | 58795 | ENSG00000077254 | USP33 | chr1 | - | 77736055 | 77736158 | 77730617 | 77730731 | 77739264 | 77739417 | 142 | 109 | 84 | [illegible] |
| SE | 73958 | ENSG00000166436 | TRIM66 | chr11 | - | 8620049 | 8620124 | 8619382 | 8619535 | 8620445 | 8620572 | 23 | 12 | 31 | [illegible] |
| SE | 83117 | ENSG00000234741 | GAS5 | chr1 | - | 173864483 | 173864704 | 173864256 | 173864304 | 173865470 | 173865547 | 94 | 58 | 125 | [illegible] |
| SE | 73121 | ENSG00000145414 | NAF1 | chr4 | - | 163133153 | 163133256 | 163128668 | 163129348 | 163137198 | 163137250 | 66 | 51 | 47 | [illegible] |
| SE | 30042 | ENSG00000155744 | FAM126B | chr2 | - | 200987364 | 200987532 | 200973717 | 200981871 | 200988261 | 200988421 | 7 | 7 | 19 | [illegible] |
| SE | 22713 | ENSG00000164329 | TENT2 | chr5 | + | 79642843 | 79642910 | 79641104 | 79641196 | 79645122 | 79645192 | 178 | 136 | 139 | [illegible] |
| SE | 47808 | ENSG00000014138 | POLA2 | chr11 | + | 65280991 | 65281147 | 65279537 | 65279626 | 65281669 | 65281732 | 69 | 44 | 82 | [illegible] |
| SE | 12167 | ENSG00000051108 | HERPUD1 | chr16 | + | 56935400 | 56935475 | 56935234 | 56935312 | 56936686 | 56936817 | 294 | 186 | 254 | [illegible] |
| SE | 60570 | ENSG00000155275 | TRMT44 | chr4 | + | 8449668 | 8449888 | 8446475 | 8446590 | 8451959 | 8452028 | 66 | 50 | 55 | [illegible] |
| SE | 47351 | ENSG00000164136 | IL15 | chr4 | + | 141721058 | 141721177 | 141720468 | 141720566 | 141721923 | 141722008 | 0 | 5 | 4 | [illegible] |
| SE | 3571 | ENSG00000186818 | LILRB4 | chr19 | + | 54665129 | 54665180 | 54664798 | 54664849 | 54665814 | 54665931 | 27 | 8 | 13 | [illegible] |
| SE | 37158 | ENSG00000151726 | ACSL1 | chr4 | - | 184776483 | 184776662 | 184773842 | 184773875 | 184776883 | 184776983 | 283 | 164 | 161 | [illegible] |
| SE | 88108 | ENSG00000197265 | GTF2E2 | chr8 | - | 30636929 | 30636960 | 30635031 | 30635123 | 30653432 | 30653602 | 1 | 5 | 6 | [illegible] |
| SE | 86111 | ENSG00000047644 | WWC3 | chrX | + | 10078798 | 10078875 | 10067286 | 10067490 | 10090770 | 10090900 | 18 | 18 | 13 | [illegible] |
| [illegible] | | | | | | | | | | | | | | | |

| | | | | | | | | | | | | | | | |
|---|---|---|---|---|---|---|---|---|---|---|---|---|---|---|---|
| SE | 39072 | ENSG00000172175 | MALT1 | chr18 | + | 58709975 | 58710072 | 58700440 | 58700591 | 58714082 | 58714109 | 103 | 86 | 87 | [illegible] |
| SE | 76423 | ENSG00000168538 | TRAPPC11 | chr4 | + | 183701696 | 183701808 | 183697678 | 183697835 | 183704978 | 183705070 | 123 | 84 | 110 | [illegible] |
| SE | 2498 | ENSG00000160953 | PWWP3A | chr19 | + | 1358587 | 1358693 | 1358393 | 1358464 | 1360135 | 1360606 | 10 | 3 | 1 | [illegible] |
| SE | 11899 | ENSG00000103222 | ABCC1 | chr16 | + | 16079351 | 16079478 | 16076173 | 16076401 | 16086823 | 16086991 | 48 | 23 | 32 | [illegible] |
| SE | 93182 | ENSG00000163558 | PRKCI | chr3 | + | 170267914 | 170268000 | 170263378 | 170263429 | 170270420 | 170270561 | 65 | 78 | 66 | [illegible] |
| SE | 48880 | ENSG00000143514 | TP53BP2 | chr1 | - | 223821219 | 223821367 | 223814175 | 223814353 | 223845653 | 223845794 | 145 | 107 | 117 | [illegible] |
| SE | 10449 | ENSG00000214425 | LRRC37A4P | chr17 | - | 45539162 | 45539243 | 45521306 | 45521425 | 45542763 | 45542835 | 5 | 17 | 27 | [illegible] |
| SE | 90888 | ENSG00000104763 | ASAH1 | chr8 | - | 18083980 | 18084286 | 18075540 | 18075587 | 18084675 | 18084802 | 114 | 47 | 79 | [illegible] |
| SE | 54181 | ENSG00000175376 | EIF1AD | chr11 | - | 66000302 | 66000406 | 66000052 | 66000161 | 66002039 | 66002166 | 86 | 61 | 84 | [illegible] |
| SE | 37618 | ENSG00000082074 | FYB1 | chr5 | - | 39212677 | 39213017 | 39202773 | 39202987 | 39250670 | 39250806 | 25 | 18 | 16 | [illegible] |
| SE | 96655 | ENSG00000277072 | STAG3L2 | chr7 | - | 74887682 | 74887799 | 74885022 | 74885146 | 74888311 | 74888397 | 53 | 27 | 85 | [illegible] |
| SE | 52133 | ENSG00000161835 | TAMALIN | chr12 | + | 52011038 | 52011141 | 52010880 | 52010935 | 52013686 | 52013780 | 18 | 10 | 29 | [illegible] |
| SE | 47457 | ENSG00000083635 | NUFIP1 | chr13 | - | 44959380 | 44959574 | 44949721 | 44949838 | 44965843 | 44965936 | 77 | 67 | 99 | [illegible] |
| SE | 44923 | ENSG00000182095 | TNRC18 | chr7 | - | 5332621 | 5333049 | 5325095 | 5325248 | 5345561 | 5345810 | 77 | 39 | 54 | [illegible] |
| SE | 69312 | ENSG00000112367 | FIG4 | chr6 | + | 109727157 | 109727265 | 109716444 | 109716568 | 109732636 | 109732687 | 91 | 59 | 47 | [illegible] |
| SE | 45366 | ENSG00000143164 | DCAF6 | chr1 | + | 168015780 | 168015951 | 168004532 | 168004793 | 168022987 | 168023047 | 16 | 2 | 17 | [illegible] |
| SE | 79277 | ENSG00000154803 | FLCN | chr17 | - | 17221536 | 17221628 | 17219018 | 17219209 | 17222500 | 17222661 | 64 | 34 | 50 | [illegible] |
| SE | 70753 | ENSG00000040933 | INPP4A | chr2 | + | 98543876 | 98544007 | 98539527 | 98539675 | 98552785 | 98552969 | 196 | 101 | 181 | [illegible] |
| SE | 91676 | ENSG00000092094 | OSGEP | chr14 | - | 20451256 | 20451452 | 20449221 | 20449266 | 20451994 | 20452149 | 9 | 4 | 9 | [illegible] |
| SE | 26267 | ENSG00000137449 | CPEB2 | chr4 | + | 15008337 | 15008427 | 15007304 | 15007595 | 15017187 | 15017278 | 7 | 4 | 0 | [illegible] |
| SE | 42369 | ENSG00000048740 | CELF2 | chr10 | + | 11275056 | 11275120 | 11257737 | 11257872 | 11288417 | 11288552 | 26 | 17 | 12 | [illegible] |
| SE | 61848 | ENSG00000143393 | PI4KB | chr1 | - | 151293399 | 151293462 | 151292601 | 151293033 | 151294017 | 151294138 | 3 | 3 | 2 | [illegible] |
| SE | 12934 | ENSG00000197818 | SLC9A8 | chr20 | + | 49850809 | 49850844 | 49849578 | 49849680 | 49855437 | 49855581 | 16 | 7 | 9 | [illegible] |
| SE | 8991 | ENSG00000161526 | SAP30BP | chr17 | + | 75705610 | 75705693 | 75704755 | 75704814 | 75706007 | 75706092 | 15 | 23 | 11 | [illegible] |
| SE | 58134 | ENSG00000117523 | PRRC2C | chr1 | + | 171571319 | 171571421 | 171568246 | 171568339 | 171574926 | 171575128 | 489 | 395 | 472 | [illegible] |
| SE | 106478 | ENSG00000198270 | TMEM116 | chr12 | - | 111978735 | 111978787 | 111943264 | 111943369 | 111991757 | 111991889 | 6 | 4 | 8 | [illegible] |
| SE | 74759 | ENSG00000163939 | PBRM1 | chr3 | - | 52681703 | 52681820 | 52679581 | 52679726 | 52685748 | 52685794 | 0 | 0 | 2 | [illegible] |
| SE | 70681 | ENSG00000095059 | DHPS | chr19 | - | 12677107 | 12677211 | 12676016 | 12676142 | 12677290 | 12677396 | 209 | 112 | 180 | [illegible] |
| SE | 49605 | ENSG00000076685 | NT5C2 | chr10 | - | 103139405 | 103139479 | 103105705 | 103105801 | 103174857 | 103174982 | 70 | 33 | 56 | [illegible] |
| SE | 28938 | ENSG00000112079 | STK38 | chr6 | - | 36517716 | 36517840 | 36515337 | 36515492 | 36521733 | 36521817 | 2359 | 1702 | 1685 | [illegible] |
| SE | 61599 | ENSG00000061273 | HDAC7 | chr12 | - | 47788834 | 47788951 | 47788095 | 47788164 | 47789260 | 47789348 | 0 | 0 | 0 | [illegible] |
| [illegible] | | | | | | | | | | | | | | | |

| | | | | | | | | | | | | | | | | | | | | | | | | | | | | | | | | | | | | | | | | | | | | | | |
|---|---|---|---|---|---|---|---|---|---|---|---|---|---|---|---|---|---|---|---|---|---|---|---|---|---|---|---|---|---|---|---|---|---|---|---|---|---|---|---|---|---|---|---|---|---|---|
| SE | 26812 | ENSG00000138760 | SCARB2 | chr4 | - | 76176436 | 76176528 | 76175790 | 76175910 | 76180953 | 76181092 | 100 | 68 | 83 | 81 | 242 | 2 | 1 | 0 | 0 | 0 | 117 | 127 | 67 | 143 | 49 | 0 | 5 | 0 | 3 | 0 | 241 | 149 | 0.50838322 | 1 | 0.969 | 0.977 | 1 | 1 | 1 | 1 | 0.94 | 1 | 0.967 | 1 | 0.008 |
| SE | 79732 | ENSG00000168255 | POLR2J3 | chr7 | - | 102545302 | 102545420 | 102544705 | 102544776 | 102566581 | 102566736 | 5 | 2 | 10 | 14 | 10 | 14 | 4 | 3 | 8 | 10 | 19 | 3 | 32 | 15 | 1 | 8 | 4 | 7 | 10 | 7 | 267 | 149 | 0.508389844 | 1 | 0.166 | 0.218 | 0.65 | 0.494 | 0.358 | 0.57 | 0.295 | 0.718 | 0.456 | 0.074 | -0.045 |
| SE | 55311 | ENSG00000166938 | DIS3L | chr15 | + | 66306896 | 66306952 | 66294987 | 66295141 | 66308708 | 66308844 | 30 | 16 | 31 | 24 | 27 | 0 | 0 | 0 | 0 | 6 | 1 | 9 | 9 | 12 | 9 | 0 | 1 | 0 | 0 | 0 | 205 | 149 | 0.508435445 | 1 | 1 | 1 | 1 | 1 | 0.766 | 1 | 0.867 | 1 | 1 | 1 | -0.02 |
| SE | 32746 | ENSG00000163349 | HIPK1 | chr1 | + | 113971823 | 113971954 | 113969955 | 113970197 | 113973023 | 113973571 | 116 | 63 | 108 | 114 | 103 | 1 | 1 | 0 | 0 | 3 | 151 | 90 | 133 | 143 | 41 | 0 | 9 | 0 | 0 | 0 | 280 | 149 | 0.508486582 | 1 | 0.984 | 0.971 | 1 | 1 | 0.948 | 1 | 0.842 | 1 | 1 | 1 | 0.012 |
| SE | 5806 | ENSG00000197586 | ENTPD6 | chr20 | + | 25206521 | 25206590 | 25195700 | 25195867 | 25207075 | 25207397 | 45 | 20 | 29 | 42 | 35 | 13 | 2 | 6 | 20 | 8 | 95 | 44 | 10 | 25 | 34 | 10 | 8 | 0 | 10 | 17 | 218 | 149 | 0.508508887 | 1 | 0.703 | 0.872 | 0.768 | 0.589 | 0.749 | 0.867 | 0.79 | 1 | 0.631 | 0.578 | -0.037 |
| SE | 27438 | ENSG00000136925 | TSTD2 | chr9 | - | 97617756 | 97617877 | 97611573 | 97611699 | 97625680 | 97625775 | 25 | 17 | 40 | 36 | 18 | 3 | 0 | 1 | 1 | 1 | 31 | 25 | 4 | 10 | 14 | 0 | 0 | 3 | 0 | 0 | 270 | 149 | 0.508536395 | 1 | 0.821 | 1 | 0.957 | 0.952 | 0.909 | 1 | 1 | 0.424 | 1 | 1 | 0.043 |
| SE | 5119 | ENSG00000125826 | RBCK1 | chr20 | + | 409880 | 410025 | 408049 | 408779 | 417525 | 417619 | 122 | 77 | 134 | 120 | 156 | 2 | 0 | 3 | 0 | 3 | 208 | 128 | 81 | 172 | 184 | 0 | 0 | 1 | 0 | 0 | 294 | 149 | 0.508595573 | 1 | 0.969 | 1 | 0.958 | 1 | 0.963 | 1 | 1 | 0.976 | 1 | 1 | -0.017 |
| SE | 100513 | ENSG00000054654 | SYNE2 | chr14 | + | 63995043 | 63995202 | 63993834 | 63993969 | 63996946 | 63997158 | 362 | 254 | 209 | 276 | 358 | 8 | 3 | 1 | 4 | 9 | 50 | 142 | 181 | 44 | 40 | 8 | 2 | 2 | 0 | 0 | 298 | 149 | 0.508668907 | 1 | 0.958 | 0.977 | 0.991 | 0.972 | 0.952 | 0.758 | 0.973 | 0.978 | 1 | 1 | 0.028 |
| SE | 80433 | ENSG00000133216 | EPHB2 | chr1 | + | 22892880 | 22893046 | 22882358 | 22882483 | 22895471 | 22895580 | 7 | 2 | 1 | 10 | 1 | 0 | 0 | 0 | 4 | 0 | 69 | 40 | 35 | 51 | 49 | 0 | 0 | 0 | 0 | 0 | 298 | 149 | 0.508703394 | 1 | 1 | 1 | 1 | 0.556 | 1 | 1 | 1 | 1 | 1 | 1 | -0.089 |
| SE | 35660 | ENSG00000010404 | IDS | chrX | - | 149491537 | 149491667 | 149490313 | 149490440 | 149496345 | 149496516 | 51 | 22 | 33 | 45 | 55 | 325 | 182 | 251 | 332 | 312 | 31 | 48 | 55 | 19 | 67 | 209 | 271 | 296 | 271 | 326 | 279 | 149 | 0.508750558 | 1 | 0.077 | 0.061 | 0.066 | 0.068 | 0.086 | 0.073 | 0.086 | 0.09 | 0.036 | 0.099 | -0.005 |
| SE | 43295 | ENSG00000134698 | AGO4 | chr1 | + | 35832436 | 35832570 | 35831811 | 35831931 | 35833989 | 35834174 | 115 | 42 | 68 | 72 | 77 | 0 | 0 | 0 | 0 | 0 | 101 | 65 | 88 | 65 | 44 | 0 | 3 | 1 | 4 | 0 | 283 | 149 | 0.508802687 | 1 | 1 | 1 | 1 | 1 | 1 | 1 | 0.919 | 0.979 | 0.895 | 1 | 0.041 |
| SE | 84502 | ENSG00000128731 | HERC2 | chr15 | - | 28176686 | 28176768 | 28176427 | 28176599 | 28176949 | 28177127 | 68 | 41 | 76 | 68 | 66 | 0 | 1 | 0 | 0 | 0 | 67 | 79 | 47 | 76 | 56 | 0 | 0 | 0 | 8 | 0 | 231 | 149 | 0.508889515 | 1 | 1 | 0.964 | 1 | 1 | 1 | 1 | 1 | 1 | 0.86 | 1 | 0.021 |
| SE | 93467 | ENSG00000106771 | TMEM245 | chr9 | - | 109073247 | 109073438 | 109064476 | 109064567 | 109080838 | 109080943 | 61 | 62 | 102 | 60 | 83 | 1 | 0 | 2 | 3 | 2 | 63 | 65 | 50 | 46 | 19 | 0 | 0 | 0 | 0 | 0 | 298 | 149 | 0.508953703 | 1 | 0.968 | 1 | 0.962 | 0.909 | 0.954 | 1 | 1 | 1 | 1 | 1 | -0.041 |
| SE | 20406 | ENSG00000204623 | ZNRD1ASP | chr6 | - | 30035915 | 30035983 | 30021666 | 30021761 | 30058114 | 30058190 | 13 | 7 | 3 | 9 | 11 | 1 | 2 | 4 | 4 | 3 | 20 | 4 | 1 | 2 | 11 | 0 | 9 | 0 | 0 | 9 | 217 | 149 | 0.508972707 | 1 | 0.899 | 0.706 | 0.34 | 0.607 | 0.716 | 1 | 0.234 | 1 | 1 | 0.456 | -0.084 |
| SE | 82099 | ENSG00000128891 | CCDC32 | chr15 | - | 40553436 | 40554127 | 40531340 | 40532302 | 40557215 | 40557372 | 59 | 52 | 60 | 66 | 93 | 11 | 4 | 8 | 4 | 8 | 95 | 76 | 49 | 118 | 51 | 22 | 9 | 0 | 0 | 8 | 298 | 149 | 0.509004029 | 1 | 0.728 | 0.867 | 0.789 | 0.892 | 0.853 | 0.683 | 0.809 | 1 | 1 | 0.761 | -0.025 |
| SE | 93753 | ENSG00000143409 | MINDY1 | chr1 | - | 151000456 | 151000615 | 150999861 | 150999964 | 151001249 | 151001314 | 184 | 127 | 129 | 201 | 174 | 6 | 1 | 3 | 3 | 13 | 509 | 219 | 124 | 202 | 1132 | 10 | 4 | 8 | 10 | 18 | 298 | 149 | 0.509157192 | 1 | 0.939 | 0.984 | 0.956 | 0.971 | 0.87 | 0.962 | 0.965 | 0.886 | 0.91 | 0.969 | 0.006 |
| SE | 108424 | ENSG00000170653 | ATF7 | chr12 | - | 53517014 | 53517354 | 53507855 | 53508027 | 53523275 | 53523384 | 132 | 102 | 97 | 129 | 147 | 2 | 0 | 2 | 0 | 4 | 284 | 195 | 82 | 221 | 272 | 0 | 0 | 2 | 3 | 0 | 298 | 149 | 0.509563235 | 1 | 0.971 | 1 | 0.96 | 1 | 0.948 | 1 | 1 | 0.953 | 0.974 | 1 | -0.01 |
| SE | 7888 | ENSG00000141580 | WDR45B | chr17 | - | 82630920 | 82631022 | 82627203 | 82627291 | 82643948 | 82644023 | 157 | 101 | 127 | 133 | 305 | 4 | 3 | 1 | 0 | 2 | 248 | 122 | 123 | 191 | 273 | 4 | 2 | 0 | 2 | 0 | 251 | 149 | 0.509681739 | 1 | 0.959 | 0.952 | 0.987 | 1 | 0.989 | 0.974 | 0.973 | 1 | 0.983 | 1 | -0.009 |
| SE | 3640 | ENSG00000239998 | LILRA2 | chr19 | + | 54574300 | 54574582 | 54573840 | 54573912 | 54574730 | 54574910 | 155 | 69 | 63 | 125 | 178 | 1 | 0 | 2 | 1 | 4 | 341 | 221 | 150 | 299 | 336 | 0 | 2 | 0 | 1 | 0 | 298 | 149 | 0.509793686 | 1 | 0.987 | 1 | 0.94 | 0.984 | 0.957 | 1 | 0.982 | 1 | 0.993 | 1 | -0.021 |
| SE | 4451 | ENSG00000181896 | ZNF101 | chr19 | + | 19677863 | 19677986 | 19668850 | 19668966 | 19678725 | 19678786 | 95 | 52 | 153 | 121 | 114 | 4 | 0 | 0 | 4 | 0 | 55 | 88 | 68 | 58 | 54 | 0 | 0 | 0 | 0 | 0 | 272 | 149 | 0.509867748 | 1 | 0.929 | 1 | 1 | 0.943 | 1 | 1 | 1 | 1 | 1 | 1 | -0.026 |
| SE | 93601 | ENSG00000158856 | DMTN | chr8 | + | 22059372 | 22059496 | 22056965 | 22057136 | 22066704 | 22066893 | 12 | 7 | 1 | 20 | 11 | 27 | 16 | 6 | 32 | 7 | 22 | 12 | 8 | 4 | 161 | 33 | 33 | 20 | 38 | 269 | 273 | 149 | 0.510032214 | 1 | 0.195 | 0.193 | 0.083 | 0.254 | 0.462 | 0.267 | 0.166 | 0.179 | 0.054 | 0.246 | 0.055 |
| SE | 19325 | ENSG00000071537 | SEL1L | chr14 | - | 81526732 | 81526964 | 81506073 | 81506241 | 81527700 | 81527738 | 677 | 354 | 363 | 461 | 260 | 2 | 0 | 4 | 1 | 0 | 263 | 423 | 524 | 345 | 164 | 0 | 13 | 2 | 1 | 0 | 298 | 149 | 0.510041588 | 1 | 0.994 | 1 | 0.978 | 0.996 | 1 | 1 | 0.942 | 0.992 | 0.994 | 1 | 0.008 |
| SE | 80804 | ENSG00000151948 | GLT1D1 | chr12 | + | 128875913 | 128876062 | 128853426 | 128853649 | 128888638 | 128888744 | 41 | 27 | 20 | 40 | 32 | 3 | 3 | 3 | 1 | 1 | 148 | 110 | 91 | 122 | 168 | 0 | 2 | 10 | 2 | 9 | 298 | 149 | 0.510094179 | 1 | 0.872 | 0.818 | 0.769 | 0.952 | 0.941 | 1 | 0.965 | 0.82 | 0.968 | 0.903 | -0.061 |
| SE | 75134 | ENSG00000124226 | RNF114 | chr20 | + | 49941582 | 49941711 | 49936376 | 49936552 | 49945381 | 49945488 | 67 | 40 | 67 | 80 | 58 | 1 | 0 | 0 | 3 | 2 | 38 | 67 | 52 | 68 | 78 | 0 | 0 | 2 | 0 | 0 | 278 | 149 | 0.510098858 | 1 | 0.973 | 1 | 1 | 0.935 | 0.94 | 1 | 1 | 0.933 | 1 | 1 | -0.017 |
| SE | 30914 | ENSG00000205726 | ITSN1 | chr21 | + | 33823486 | 33823653 | 33819240 | 33819323 | 33826817 | 33826863 | 11 | 11 | 13 | 20 | 13 | 0 | 0 | 0 | 1 | 0 | 20 | 24 | 6 | 10 | 46 | 0 | 0 | 0 | 4 | 0 | 298 | 149 | 0.510220904 | 1 | 1 | 1 | 1 | 0.909 | 1 | 1 | 1 | 1 | 0.556 | 1 | 0.071 |
| SE | 29665 | ENSG00000155189 | AGPAT5 | chr8 | + | 6747669 | 6747828 | 6732560 | 6732650 | 6755050 | 6755174 | 60 | 50 | 51 | 59 | 86 | 0 | 0 | 0 | 2 | 4 | 41 | 28 | 32 | 14 | 26 | 0 | 4 | 2 | 0 | 0 | 298 | 149 | 0.510308841 | 1 | 1 | 1 | 1 | 0.937 | 0.915 | 1 | 0.778 | 0.889 | 1 | 1 | 0.037 |
| SE | 105862 | ENSG00000090006 | LTBP4 | chr19 | + | 40616888 | 40617042 | 40614179 | 40614446 | 40619346 | 40619493 | 11 | 9 | 12 | 10 | 0 | 6 | 3 | 16 | 4 | 2 | 15 | 5 | 21 | 1 | 2 | 0 | 7 | 10 | 8 | 19 | 298 | 149 | 0.51041032 | 1 | 0.478 | 0.6 | 0.273 | 0.556 | 0 | 1 | 0.263 | 0.512 | 0.059 | 0.05 | 0.005 |
| SE | 107400 | ENSG00000151176 | PLBD2 | chr12 | + | 113384850 | 113384946 | 113384104 | 113384265 | 113385211 | 113385283 | 37 | 22 | 44 | 35 | 27 | 0 | 0 | 5 | 2 | 0 | 45 | 50 | 27 | 23 | 53 | 0 | 0 | 0 | 5 | 0 | 245 | 149 | 0.510453224 | 1 | 1 | 1 | 0.843 | 0.914 | 1 | 1 | 1 | 1 | 0.737 | 1 | 0.004 |
| SE | 58839 | ENSG00000164828 | SUN1 | chr7 | + | 849922 | 850033 | 849519 | 849603 | 851949 | 852043 | 55 | 42 | 43 | 34 | 45 | 0 | 0 | 3 | 0 | 4 | 60 | 29 | 35 | 43 | 29 | 0 | 0 | 1 | 0 | 0 | 260 | 149 | 0.510456824 | 1 | 1 | 1 | 0.891 | 1 | 0.866 | 1 | 1 | 0.953 | 1 | 1 | -0.039 |
| SE | 16198 | ENSG00000140598 | EFL1 | chr15 | - | 82151464 | 82152423 | 82138657 | 82138842 | 82157712 | 82157860 | 140 | 104 | 104 | 133 | 238 | 1 | 1 | 3 | 4 | 0 | 53 | 142 | 82 | 121 | 13 | 0 | 5 | 0 | 0 | 0 | 298 | 149 | 0.510476811 | 1 | 0.986 | 0.981 | 0.945 | 0.943 | 1 | 1 | 0.934 | 1 | 1 | 1 | -0.016 |
| SE | 10687 | ENSG00000278540 | ACACA | chr17 | - | 37210466 | 37210490 | 37207656 | 37207800 | 37221723 | 37221842 | 24 | 4 | 29 | 31 | 32 | 0 | 0 | 3 | 2 | 1 | 16 | 22 | 19 | 9 | 7 | 0 | 1 | 0 | 1 | 0 | 173 | 149 | 0.510536584 | 1 | 1 | 1 | 0.893 | 0.93 | 0.965 | 1 | 0.95 | 1 | 0.886 | 1 | -0.01 |
| SE | 14121 | ENSG00000102897 | LYRM1 | chr16 | + | 20915555 | 20915714 | 20911182 | 20911264 | 20920121 | 20920214 | 147 | 108 | 129 | 145 | 206 | 0 | 0 | 0 | 1 | 0 | 149 | 143 | 134 | 122 | 70 | 0 | 4 | 4 | 0 | 0 | 298 | 149 | 0.510566558 | 1 | 1 | 1 | 1 | 0.986 | 1 | 1 | 0.947 | 0.944 | 1 | 1 | 0.019 |
| SE | 75511 | ENSG00000164366 | CCDC127 | chr5 | - | 216728 | 216859 | 196867 | 205958 | 218092 | 218153 | 60 | 24 | 53 | 29 | 18 | 1 | 0 | 0 | 3 | 0 | 24 | 29 | 36 | 11 | 31 | 0 | 0 | 0 | 0 | 6 | 280 | 149 | 0.510666532 | 1 | 0.97 | 1 | 1 | 0.837 | 1 | 1 | 1 | 1 | 1 | 0.733 | 0.015 |
| SE | 21957 | ENSG00000012660 | ELOVL5 | chr6 | - | 53273219 | 53273344 | 53270592 | 53270727 | 53275089 | 53275261 | 489 | 322 | 460 | 537 | 807 | 13 | 9 | 13 | 20 | 32 | 249 | 418 | 383 | 420 | 188 | 5 | 22 | 13 | 21 | 3 | 274 | 149 | 0.510749506 | 1 | 0.953 | 0.951 | 0.951 | 0.936 | 0.932 | 0.964 | 0.912 | 0.941 | 0.916 | 0.971 | 0.004 |
| SE | 18942 | ENSG00000197324 | LRP10 | chr14 | + | 22873310 | 22873446 | 22872737 | 22872782 | 22875054 | 22875245 | 177 | 66 | 64 | 132 | 86 | 0 | 0 | 3 | 0 | 0 | 48 | 152 | 75 | 78 | 194 | 0 | 0 | 3 | 0 | 5 | 285 | 149 | 0.510775901 | 1 | 1 | 1 | 0.918 | 1 | 1 | 1 | 1 | 0.929 | 1 | 0.953 | 0.007 |
| SE | 90273 | ENSG00000249859 | PVT1 | chr8 | + | 127855154 | 127855319 | 127794532 | 127794734 | 127890588 | 127890998 | 2 | 1 | 1 | 4 | 4 | 22 | 12 | 22 | 19 | 6 | 9 | 0 | 1 | 1 | 5 | 27 | 18 | 24 | 2 | 13 | 298 | 149 | 0.510849342 | 1 | 0.043 | 0.04 | 0.022 | 0.095 | 0.25 | 0.143 | 0 | 0.02 | 0.2 | 0.161 | -0.015 |
| SE | 93961 | ENSG00000114030 | KPNA1 | chr3 | - | 122457725 | 122457848 | 122453869 | 122454001 | 122461223 | 122461318 | 17 | 6 | 10 | 16 | 36 | 70 | 22 | 53 | 46 | 116 | 9 | 18 | 20 | 6 | 0 | 20 | 41 | 28 | 27 | 30 | 272 | 149 | 0.510850525 | 1 | 0.117 | 0.13 | 0.094 | 0.16 | 0.145 | 0.198 | 0.194 | 0.281 | 0.109 | 0 | -0.027 |
| SE | 43970 | ENSG00000108443 | RPS6KB1 | chr17 | + | 59926434 | 59926582 | 59914634 | 59914703 | 59930116 | 59930174 | 433 | 491 | 439 | 491 | 633 | 1 | 0 | 0 | 0 | 0 | 267 | 433 | 684 | 379 | 58 | 3 | 0 | 3 | 6 | 4 | 297 | 149 | 0.510936709 | 1 | 0.995 | 1 | 1 | 1 | 1 | 0.978 | 1 | 0.991 | 0.969 | 0.879 | 0.036 |
| SE | 106494 | ENSG00000185482 | STAC3 | chr12 | - | 57245144 | 57245211 | 57244756 | 57244965 | 57248705 | 57248803 | 7 | 7 | 2 | 2 | 5 | 0 | 0 | 0 | 3 | 0 | 16 | 9 | 11 | 27 | 52 | 0 | 0 | 0 | 0 | 0 | 216 | 149 | 0.511025885 | 1 | 1 | 1 | 1 | 0.315 | 1 | 1 | 1 | 1 | 1 | 1 | -0.137 |
| SE | 46829 | ENSG00000155330 | C16orf87 | chr16 | - | 46809602 | 46809785 | 46796602 | 46803070 | 46831083 | 46831180 | 107 | 90 | 118 | 165 | 139 | 0 | 0 | 0 | 0 | 0 | 22 | 273 | 144 | 90 | 46 | 4 | 4 | 0 | 0 | 0 | 298 | 149 | 0.511034921 | 1 | 1 | 1 | 1 | 1 | 1 | 0.733 | 0.972 | 1 | 1 | 1 | 0.059 |
| SE | 40197 | ENSG00000231607 | DLEU2 | chr13 | - | 50049583 | 50049662 | 50043491 | 50044768 | 50068096 | 50068163 | 52 | 37 | 25 | 62 | 67 | 0 | 2 | 0 | 1 | 0 | 91 | 58 | 55 | 71 | 54 | 0 | 0 | 1 | 0 | 7 | 228 | 149 | 0.511042139 | 1 | 1 | 0.924 | 1 | 0.976 | 1 | 1 | 1 | 0.973 | 1 | 0.834 | 0.019 |
| SE | 60795 | ENSG00000160714 | UBE2Q1 | chr1 | - | 154552092 | 154552183 | 154551770 | 154551819 | 154552403 | 154552464 | 142 | 73 | 149 | 141 | 218 | 0 | 0 | 1 | 0 | 7 | 153 | 149 | 91 | 164 | 263 | 0 | 0 | 4 | 6 | 0 | 240 | 149 | 0.51119963 | 1 | 1 | 1 | 0.989 | 1 | 0.951 | 1 | 1 | 0.934 | 0.944 | 1 | 0.012 |
| SE | 78780 | ENSG00000131023 | LATS1 | chr6 | - | 149683078 | 149684592 | 149679874 | 149680457 | 149695073 | 149695221 | 226 | 216 | 207 | 270 | 413 | 1 | 0 | 0 | 1 | 4 | 156 | 150 | 154 | 85 | 36 | 0 | 6 | 2 | 0 | 0 | 298 | 149 | 0.511236988 | 1 | 0.991 | 1 | 1 | 0.993 | 0.981 | 1 | 0.926 | 0.975 | 1 | 1 | 0.013 |
| SE | 79825 | ENSG00000071243 | ING3 | chr7 | + | 120966625 | 120966697 | 120955558 | 120955624 | 120967528 | 120967648 | 97 | 53 | 57 | 96 | 123 | 0 | 0 | 2 | 0 | 6 | 9 | 39 | 41 | 31 | 26 | 0 | 0 | 0 | 0 | 0 | 221 | 149 | 0.511240685 | 1 | 1 | 1 | 0.951 | 1 | 0.933 | 1 | 1 | 1 | 1 | 1 | -0.023 |
| SE | 66764 | ENSG00000129292 | PHF20L1 | chr8 | + | 132839386 | 132839582 | 132837711 | 132837811 | 132842514 | 132842875 | 202 | 145 | 133 | 256 | 241 | 4 | 1 | 3 | 5 | 3 | 6 | 105 | 57 | 85 | 14 | 0 | 9 | 1 | 0 | 0 | 298 | 149 | 0.51127972 | 1 | 0.962 | 0.986 | 0.957 | 0.962 | 0.976 | 1 | 0.854 | 0.966 | 1 | 1 | 0.005 |
| SE | 11826 | ENSG00000188603 | CLN3 | chr16 | - | 28493494 | 28493664 | 28492019 | 28492215 | 28495125 | 28495545 | 19 | 9 | 11 | 17 | 13 | 0 | 0 | 0 | 0 | 0 | 46 | 19 | 12 | 17 | 51 | 7 | 0 | 0 | 0 | 0 | 298 | 149 | 0.511297062 | 1 | 1 | 1 | 1 | 1 | 1 | 0.767 | 1 | 1 | 1 | 1 | 0.047 |
| SE | 100025 | ENSG00000101972 | STAG2 | chrX | + | 124021366 | 124021431 | 123960518 | 123960866 | 124022530 | 124022671 | 73 | 46 | 46 | 46 | 127 | 75 | 56 | 99 | 118 | 158 | 14 | 23 | 27 | 4 | 0 | 23 | 32 | 20 | 6 | 20 | 214 | 149 | 0.511315069 | 1 | 0.404 | 0.364 | 0.244 | 0.213 | 0.359 | 0.298 | 0.334 | 0.485 | 0.317 | 0 | 0.03 |
| SE | 79656 | ENSG00000167081 | PBX3 | chr9 | + | 125962101 | 125962214 | 125960683 | 125960849 | 125963011 | 125963101 | 23 | 13 | 21 | 26 | 25 | 20 | 16 | 8 | 23 | 7 | 29 | 3 | 17 | 25 | 21 | 26 | 13 | 27 | 2 | 18 | 262 | 149 | 0.511460514 | 1 | 0.395 | 0.316 | 0.599 | 0.391 | 0.67 | 0.388 | 0.116 | 0.264 | 0.877 | 0.399 | 0.065 |
| SE | 101958 | ENSG00000188404 | SELL | chr1 | - | 169701559 | 169701688 | 169696534 | 169696553 | 169703254 | 169703440 | 3486 | 2127 | 3219 | 2554 | 6299 | 15 | 16 | 23 | 14 | 110 | 4066 | 3409 | 2751 | 3619 | 1576 | 25 | 65 | 11 | 12 | 20 | 278 | 149 | 0.511493262 | 1 | 0.992 | 0.986 | 0.987 | 0.99 | 0.968 | 0.989 | 0.966 | 0.993 | 0.994 | 0.977 | 0.001 |
| SE | 73142 | ENSG00000144233 | AMMECR1L | chr2 | - | 127867144 | 127867279 | 127866899 | 127866996 | 127869453 | 127869540 | 11 | 3 | 11 | 9 | 5 | 54 | 48 | 35 | 22 | 60 | 0 | 11 | 8 | 1 | 0 | 32 | 26 | 14 | 21 | 24 | 284 | 149 | 0.511748031 | 1 | 0.097 | 0.032 | 0.142 | 0.177 | 0.042 | 0 | 0.182 | 0.231 | 0.024 | 0 | 0.011 |
| SE | 106760 | ENSG00000078674 | PCM1 | chr8 | + | 17963100 | 17963291 | 17962033 | 17962174 | 17964567 | 17964768 | 535 | 620 | 485 | 579 | 673 | 7 | 17 | 2 | 1 | 14 | 44 | 342 | 314 | 77 | 19 | 0 | 0 | 4 | 2 | 5 | 298 | 149 | 0.511756328 | 1 | 0.974 | 0.948 | 0.992 | 0.997 | 0.96 | 1 | 1 | 0.975 | 0.951 | 0.655 | 0.058 |
| SE | 47542 | ENSG00000144357 | UBR3 | chr2 | + | 169877493 | 169877637 | 169875790 | 169875949 | 169878524 | 169878574 | 205 | 152 | 163 | 178 | 163 | 6 | 5 | 9 | 15 | 7 | 33 | 164 | 110 | 69 | 12 | 0 | 3 | 8 | 17 | 0 | 293 | 149 | 0.51185925 | 1 | 0.946 | 0.939 | 0.902 | 0.858 | 0.922 | 1 | 0.965 | 0.875 | 0.674 | 1 | 0.011 |
| SE | 95299 | ENSG00000172716 | SLFN11 | chr17 | - | 35367602 | 35367700 | 35366946 | 35367063 | 35373473 | 35373695 | 42 | 20 | 35 | 57 | 58 | 5 | 3 | 6 | 5 | 2 | 103 | 68 | 18 | 45 | 29 | 0 | 15 | 0 | 8 | 0 | 247 | 149 | 0.511875783 | 1 | 0.835 | 0.801 | 0.779 | 0.873 | 0.946 | 1 | 0.732 | 1 | 0.772 | 1 | -0.054 |
| SE | 34558 | ENSG00000163755 | HPS3 | chr3 | + | 149160045 | 149160279 | 149158665 | 149158846 | 149162147 | 149162333 | 191 | 106 | 177 | 150 | 257 | 1 | 0 | 1 | 0 | 2 | 27 | 121 | 93 | 61 | 9 | 0 | 6 | 0 | 0 | 0 | 298 | 149 | 0.511934318 | 1 | 0.99 | 1 | 0.989 | 1 | 0.985 | 1 | 0.91 | 1 | 1 | 1 | 0.011 |
| SE | 38154 | ENSG00000138750 | NUP54 | chr4 | - | 76136185 | 76136412 | 76134174 | 76134362 | 76144148 | 76144292 | 179 | 126 | 127 | 171 | 267 | 4 | 6 | 4 | 2 | 2 | 224 | 112 | 126 | 111 | 159 | 0 | 1 | 8 | 4 | 0 | 298 | 149 | 0.511969782 | 1 | 0.957 | 0.913 | 0.941 | 0.977 | 0.985 | 1 | 0.982 | 0.887 | 0.933 | 1 | -0.006 |
| SE | 14222 | ENSG00000167965 | MLST8 | chr16 | + | 2208209 | 2208334 | 2207192 | 2207345 | 2208449 | 2208558 | 25 | 10 | 41 | 23 | 12 | 2 | 8 | 5 | 6 | 1 | 16 | 19 | 10 | 30 | 34 | 0 | 1 | 10 | 5 | 3 | 274 | 149 | 0.511974614 | 1 | 0.872 | 0.405 | 0.817 | 0.676 | 0.867 | 1 | 0.912 | 0.352 | 0.765 | 0.86 | -0.05 |
| SE | 94937 | ENSG00000148180 | GSN | chr9 | + | 121282438 | 121282550 | 121281469 | 121281562 | 121301962 | 121302167 | 0 | 0 | 0 | 0 | 4 | 16 | 7 | 9 | 19 | 5 | 0 | 3 | 0 | 7 | 0 | 30 | 36 | 21 | 79 | 184 | 261 | 149 | 0.511986933 | 1 | 0 | 0 | 0 | 0 | 0.314 | 0 | 0.045 | 0 | 0.048 | 0 | 0.044 |
| SE | 16259 | ENSG00000128881 | TTBK2 | chr15 | - | 42857394 | 42857516 | 42840359 | 42840433 | 42872610 | 42872677 | 3 | 4 | 0 | 0 | 0 | 24 | 33 | 30 | 37 | 18 | 0 | 0 | 0 | 0 | 0 | 14 | 23 | 26 | 7 | 25 | 271 | 149 | 0.512103959 | 1 | 0.064 | 0.062 | 0 | 0 | 0 | 0 | 0 | 0 | 0 | 0 | 0.025 |
| SE | 6168 | ENSG00000132664 | POLR3F | chr20 | + | 18480037 | 18480181 | 18475074 | 18475187 | 18480401 | 18480509 | 44 | 35 | 50 | 29 | 86 | 1 | 4 | 0 | 5 | 15 | 9 | 37 | 32 | 6 | 6 | 0 | 2 | 10 | 7 | 0 | 293 | 149 | 0.512174388 | 1 | 0.957 | 0.817 | 1 | 0.747 | 0.745 | 1 | 0.904 | 0.619 | 0.304 | 1 | 0.088 |
| [illegible] | | | | | | | | | | | | | | | | | | | | | | | | | | | | | | | | | | | | | | | | | | | | | | |

| | | | | | | | | | | | | | | | | | | | | | | | | | | | | | | | | | | | | | | | | | | | | | | |
|---|---|---|---|---|---|---|---|---|---|---|---|---|---|---|---|---|---|---|---|---|---|---|---|---|---|---|---|---|---|---|---|---|---|---|---|---|---|---|---|---|---|---|---|---|---|---|
|SE|92577|ENSG00000137054|POLR1E|chr9|+|37487862|37487939|37486152|37486806|37489314|37489400|88|30|104|78|132|0|0|1|0|0|31|43|41|35|35|0|3|4|0|0|226|149|0.525132298|1|1|1|0.986|1|1|1|0.904|0.871|1|1|0.042|
|SE|62134|ENSG00000182179|UBA7|chr3|-|49810262|49810428|49809977|49810183|49810516|49810672|128|72|76|109|84|2|1|4|0|0|256|165|57|119|342|0|0|0|0|0|298|149|0.52513752|1|0.97|0.973|0.905|1|1|1|1|1|1|1|-0.03|
|SE|87712|ENSG00000242574|HLA-DMB|chr6|-|32935535|32935652|32935341|32935377|32937171|32937456|732|410|609|654|1119|47|26|35|52|93|1402|455|308|776|1987|157|37|15|68|46|266|149|0.525327234|1|0.897|0.898|0.907|0.876|0.871|0.833|0.873|0.92|0.865|0.96|0|
|SE|3610|ENSG00000104974|LILRA1|chr19|+|54594196|54594278|54593631|54593781|54594440|54594476|2|9|3|4|10|48|29|15|32|18|13|5|7|2|41|92|38|13|57|68|231|149|0.525421483|1|0.026|0.167|0.114|0.075|0.264|0.084|0.078|0.258|0.022|0.28|-0.015|
|SE|87849|ENSG00000101752|MIB1|chr18|+|21815615|21815813|21803732|21804014|21819494|21819558|129|65|65|91|186|0|3|0|1|3|31|44|18|14|31|0|0|0|0|0|298|149|0.525433624|1|1|0.915|1|0.978|0.969|1|1|1|1|1|-0.028|
|SE|10931|ENSG00000091592|NLRP1|chr17|-|5515472|5515517|5501813|5501872|5530480|5530704|15|13|13|15|10|0|0|0|1|0|87|11|29|7|55|0|0|0|1|6|194|149|0.525531788|1|1|1|1|0.92|1|1|1|1|0.843|0.876|0.04|
|SE|14917|ENSG00000086666|ZFAND6|chr15|+|80098415|80098578|80059675|80059809|80120327|80120405|108|66|74|92|227|47|31|44|51|17|118|90|39|77|194|7|79|24|23|28|298|149|0.525571627|1|0.535|0.516|0.457|0.474|0.87|0.894|0.363|0.448|0.626|0.776|-0.051|
|SE|21208|ENSG00000120278|PLEKHG1|chr6|+|150821194|150821233|150819678|150819774|150823653|150823676|21|18|23|20|16|0|0|0|2|4|11|18|8|19|15|0|0|0|0|0|188|149|0.525606649|1|1|1|1|0.888|0.76|1|1|1|1|1|-0.07|
|SE|94038|ENSG00000167094|TTC16|chr9|+|127726969|127727112|127726238|127726404|127727269|127727465|19|7|4|20|6|12|4|2|15|5|42|7|5|2|10|17|16|0|8|14|292|149|0.525730232|1|0.447|0.472|0.505|0.405|0.38|0.558|0.183|1|0.113|0.267|0.018|
|SE|96968|ENSG00000183597|TANGO2|chr22|+|20055942|20056013|20053359|20053551|20063337|20063442|30|15|23|32|25|0|0|0|0|0|29|40|32|56|181|0|0|0|0|7|220|149|0.525841264|1|1|1|1|1|1|1|1|1|1|0.946|0.011|
|SE|1511|ENSG00000100307|CBX7|chr22|-|39134400|39134473|39133583|39134048|39138635|39138702|43|16|21|29|22|0|0|1|0|5|10|32|2|13|48|0|0|0|0|0|222|149|0.525859076|1|1|1|0.934|1|0.747|1|1|1|1|1|-0.064|
|SE|53590|ENSG00000115756|HPCAL1|chr2|+|10408561|10408734|10396834|10396920|10419733|10420135|17|7|16|23|14|47|24|28|34|12|6|21|1|24|62|54|44|20|29|46|298|149|0.526119414|1|0.153|0.127|0.222|0.253|0.368|0.053|0.193|0.024|0.293|0.403|0.031|
|SE|77331|ENSG00000214022|REPIN1|chr7|+|150370713|150370884|150369643|150369868|150371227|150371655|3|3|9|0|1|43|15|32|26|19|0|9|3|1|0|37|25|11|22|41|298|149|0.526165742|1|0.034|0.091|0.123|0|0.026|0|0.153|0.12|0.022|0|-0.004|
|SE|62682|ENSG00000115525|ST3GAL5|chr2|-|85852859|85853000|85847860|85848204|85861180|85861231|12|4|11|11|11|100|106|92|111|205|15|7|5|15|17|247|86|58|113|150|290|149|0.526333879|1|0.058|0.019|0.058|0.048|0.027|0.03|0.04|0.042|0.064|0.055|-0.004|
|SE|58701|ENSG00000025039|RRAGD|chr6|-|89380167|89380367|89379223|89379338|89387294|89387590|57|65|50|50|118|0|0|1|0|1|59|31|71|185|63|0|0|0|8|0|298|149|0.526563661|1|1|1|0.962|1|0.983|1|1|1|0.92|1|0.005|
|SE|91158|ENSG00000164941|INTS8|chr8|+|94827262|94827403|94823297|94823561|94827721|94827793|249|150|181|162|414|6|1|0|1|0|37|162|131|114|39|0|2|4|2|0|290|149|0.526595493|1|0.955|0.987|1|0.988|1|1|0.977|0.944|0.967|1|0.008|
|SE|63270|ENSG00000263482|ANTXRLP1|chr10|-|46268776|46268857|46263509|46263608|46269381|46269461|5|11|14|8|7|14|52|44|3|45|0|4|16|2|9|20|0|30|11|2|230|149|0.526606314|1|0.188|0.121|0.171|0.633|0.092|0|1|0.257|0.105|0.745|-0.18|
|SE|53272|ENSG00000178537|SLC25A20|chr3|-|48895754|48895815|48891979|48892072|48898689|48898892|0|0|0|0|0|64|25|38|46|28|0|0|6|0|0|22|59|12|33|25|210|149|0.52662974|1|0|0|0|0|0|0|0|0.262|0|0|-0.052|
|SE|49262|ENSG00000176148|TCP11L1|chr11|+|33043749|33043936|33039571|33039792|33054592|33054725|65|58|48|67|79|0|0|0|2|0|74|76|32|105|63|0|3|0|4|0|298|149|0.52668446|1|1|1|1|0.944|1|1|0.927|1|0.929|1|0.018|
|SE|23013|ENSG00000106123|EPHB6|chr7|+|142861999|142862133|142861051|142861186|142862755|142862788|15|9|32|13|48|5|9|16|18|17|29|20|8|16|46|31|14|2|9|41|283|149|0.526687356|1|0.612|0.345|0.513|0.275|0.598|0.33|0.429|0.678|0.483|0.371|0.01|
|SE|64679|ENSG00000136715|SAP130|chr2|-|127950159|127950408|127949868|127949994|127954985|127955344|59|45|96|94|62|0|0|0|0|0|100|90|90|83|169|0|0|1|0|6|298|149|0.526762701|1|1|1|1|1|1|1|1|0.978|1|0.934|0.018|
|SE|4738|ENSG00000031823|RANBP3|chr19|-|5951392|5951596|5941798|5941835|5957906|5957973|14|8|17|8|24|3|2|1|0|5|14|7|4|7|11|3|0|1|4|2|298|149|0.526881511|1|0.7|0.667|0.895|1|0.706|0.7|1|0.667|0.467|0.733|0.08|
|SE|62408|ENSG00000033867|SLC4A7|chr3|-|27409355|27409530|27404829|27404963|27411641|27411748|71|44|89|64|115|31|26|24|16|59|10|30|33|24|10|5|18|30|0|23|298|149|0.526903395|1|0.534|0.458|0.65|0.667|0.494|0.5|0.455|0.355|1|0.179|0.063|
|SE|11293|ENSG00000140650|PMM2|chr16|+|8802080|8802145|8801798|8801910|8806315|8806407|3|2|0|3|8|4|5|11|9|4|35|0|7|0|0|0|8|13|10|4|214|149|0.527008643|1|0.343|0.218|0|0.188|0.582|1|0|0.273|0|0|0.012|
|SE|40160|ENSG00000138382|METTL5|chr2|-|169821091|169821273|169819560|169819643|169824488|169824640|126|105|113|126|284|0|0|0|0|0|74|117|88|78|35|0|0|0|0|6|298|149|0.527091175|1|1|1|1|1|1|1|1|1|1|0.745|0.051|
|SE|106077|ENSG00000196498|NCOR2|chr12|-|124354477|124354582|124354092|124354196|124354836|124354939|70|53|87|55|17|20|0|0|0|1|73|122|35|73|112|14|10|3|0|0|254|149|0.527128071|1|0.672|1|1|1|0.909|0.754|0.877|0.873|1|1|0.015|
|SE|88371|ENSG00000087088|BAX|chr19|+|48955547|48955599|48954874|48954962|48955686|48955833|65|36|45|47|34|0|1|0|4|0|59|70|41|53|168|9|0|0|0|0|201|149|0.527172297|1|1|0.964|1|0.897|1|0.829|1|1|1|1|0.006|
|SE|35497|ENSG00000051382|PIK3CB|chr3|-|138690999|138691143|138684624|138684803|138694785|138694907|108|99|94|147|80|0|0|0|0|0|67|144|78|170|182|0|0|0|0|8|293|149|0.527178573|1|1|1|1|1|1|1|1|1|1|0.92|0.016|
|SE|40580|ENSG00000163875|MEAF6|chr1|-|37496706|37496736|37496546|37496604|37501803|37501996|1|1|2|2|4|27|8|15|15|31|0|12|0|0|0|16|5|4|16|35|179|149|0.527191993|1|0.03|0.094|0.1|0.1|0.097|0|0.666|0|0|0|-0.049|
|SE|73578|ENSG00000031691|CENPQ|chr6|+|49470158|49470278|49463369|49463453|49470973|49471028|38|29|30|15|77|3|1|1|1|0|21|32|14|13|21|0|0|0|0|0|269|149|0.527237089|1|0.875|0.941|0.943|0.893|1|1|1|1|1|1|-0.07|
|SE|13223|ENSG00000181019|NQO1|chr16|-|69714963|69715077|69713027|69713129|69718122|69718253|9|4|8|1|13|0|0|2|2|0|26|5|11|20|19|0|2|0|2|0|263|149|0.527250085|1|1|1|0.694|0.221|1|1|0.586|1|0.85|1|-0.104|
|SE|10065|ENSG00000108406|DHX40|chr17|+|59570517|59570663|59565612|59565783|59573115|59573235|121|68|101|102|173|0|0|0|0|0|34|108|70|49|18|2|0|0|5|0|295|149|0.527282768|1|1|1|1|1|1|0.896|1|1|0.832|1|0.054|
|SE|6584|ENSG00000074657|ZNF532|chr18|+|58954233|58954334|58953517|58953799|58979054|58979167|9|1|2|6|5|15|18|15|23|15|7|15|4|5|0|41|22|9|8|15|250|149|0.527283543|1|0.263|0.032|0.074|0.135|0.166|0.092|0.289|0.209|0.271|0|-0.038|
|SE|39091|ENSG00000109320|NFKB1|chr4|+|102525511|102525557|102501330|102501788|102529835|102529914|82|63|75|76|137|1|1|3|3|0|85|27|90|108|161|0|1|0|0|0|195|149|0.527297871|1|0.984|0.98|0.95|0.951|1|1|0.954|1|1|1|-0.018|
|SE|81691|ENSG00000171681|ATF7IP|chr12|+|14423908|14425473|14385063|14385400|14434339|14434423|246|181|169|213|275|0|0|0|1|0|45|255|215|97|29|0|0|8|0|0|298|149|0.527343096|1|1|1|1|0.991|1|1|1|0.931|1|1|0.012|
|SE|79521|ENSG00000105778|AVL9|chr7|+|32558928|32559464|32558558|32558628|32570019|32570154|172|118|182|202|258|0|0|0|0|0|69|190|147|176|70|0|0|1|7|0|298|149|0.527359918|1|1|1|1|1|1|1|1|0.987|0.926|1|0.017|
|SE|32958|ENSG00000114127|XRN1|chr3|-|142414134|142414291|142412543|142412663|142417139|142417229|443|434|355|477|753|0|1|0|3|0|42|250|260|33|49|0|7|1|0|0|298|149|0.527363653|1|1|0.995|1|0.988|1|1|0.947|0.992|1|1|0.009|
|SE|59959|ENSG00000117280|RAB29|chr1|-|205774832|205775086|205772495|205772567|205775373|205775449|73|33|40|51|35|2|1|2|0|2|53|79|18|32|39|0|0|0|0|0|298|149|0.527413459|1|0.948|0.943|0.909|1|0.897|1|1|1|1|1|-0.061|
|SE|24006|ENSG00000112983|BRD8|chr5|-|138162053|138162146|138160890|138161068|138163129|138163344|23|29|33|26|30|2|1|3|1|0|10|24|1|7|3|0|4|0|2|0|242|149|0.527414833|1|0.876|0.947|0.871|0.941|1|1|0.787|1|0.683|1|0.033|
|SE|34515|ENSG00000106733|NMRK1|chr9|-|75069894|75070042|75069741|75069813|75077158|75077207|329|294|541|384|405|3|2|2|6|2|200|119|207|77|60|0|0|0|0|7|297|149|0.527425915|1|0.982|0.987|0.993|0.97|0.99|1|1|1|1|0.811|0.022|
|SE|15596|ENSG00000170776|AKAP13|chr15|+|85743491|85743825|85740221|85740272|85744627|85744677|304|163|221|233|266|0|0|0|0|0|234|305|244|199|109|0|0|0|0|10|298|149|0.527511528|1|1|1|1|1|1|1|1|1|1|0.845|0.031|
|SE|62080|ENSG00000263528|IKBKE|chr1|+|206493265|206493378|206491647|206491749|206493919|206493991|36|11|28|36|32|0|1|3|3|0|91|26|7|34|30|0|0|3|5|2|262|149|0.527518947|1|1|0.862|0.841|0.872|1|1|1|0.57|0.795|0.895|0.063|
|SE|46022|ENSG00000173230|GOLGB1|chr3|-|121699811|121699885|121693740|121698929|121702480|121702595|141|125|129|136|128|3|0|0|0|1|15|28|32|8|31|0|6|0|0|0|223|149|0.527534481|1|0.969|1|1|1|0.988|1|0.757|1|1|1|0.04|
|SE|62320|ENSG00000185811|IKZF1|chr7|+|50376532|50376793|50327378|50327757|50387344|50387470|355|211|275|289|162|2|1|2|4|5|152|226|137|182|202|0|6|0|3|0|298|149|0.527538858|1|0.989|0.991|0.986|0.973|0.942|1|0.95|1|0.968|1|-0.007|
|SE|37776|ENSG00000139719|VPS33A|chr12|-|122238586|122238724|122235785|122235923|122239877|122239945|167|129|192|150|416|0|0|1|0|3|230|161|116|172|175|11|0|0|0|0|287|149|0.527566003|1|1|1|0.99|1|0.986|0.916|1|1|1|1|0.012|
|SE|60194|ENSG00000163534|FCRL1|chr1|-|157801914|157802193|157800057|157800085|157807101|157807122|27|15|20|22|12|1|0|7|4|3|9|34|18|6|16|7|0|0|6|0|298|149|0.52761492|1|0.931|1|0.588|0.733|0.667|0.391|1|1|0.333|1|0.039|
|SE|12278|ENSG00000103194|USP10|chr16|+|84744632|84745673|84733434|84733503|84758715|84758807|112|60|113|97|158|0|0|0|0|1|66|102|46|31|84|0|4|0|0|3|298|149|0.527703573|1|1|1|1|1|0.988|1|0.927|1|1|0.933|0.026|
|SE|53516|ENSG00000145390|USP53|chr4|+|119268267|119268420|119267319|119267482|119269690|119269837|92|81|126|98|160|9|5|4|6|13|5|30|16|6|5|0|0|8|1|0|298|149|0.527842576|1|0.836|0.89|0.94|0.891|0.86|1|1|0.5|0.75|1|0.033|
|SE|39|ENSG00000155313|USP25|chr21|+|15849776|15849872|15842397|15842540|15864267|15864446|54|43|49|47|73|281|237|241|244|607|0|54|28|31|0|6|146|206|162|16|245|149|0.527868131|1|0.105|0.099|0.11|0.105|0.068|0|0.184|0.076|0.104|0|0.025|
|SE|75218|ENSG00000100503|NIN|chr14|-|50756491|50758630|50754741|50754867|50759856|50760359|124|113|65|114|137|75|85|126|90|141|14|70|82|33|7|10|106|72|28|0|298|149|0.527931193|1|0.453|0.399|0.205|0.388|0.327|0.412|0.248|0.363|0.371|1|-0.124|
|SE|76818|ENSG00000135317|SNX14|chr6|-|85581195|85581258|85574257|85574378|85593578|85593877|5|2|0|0|0|44|18|25|44|20|0|0|0|0|0|11|25|7|21|10|212|149|0.528018622|1|0.074|0.072|0|0|0|0|0|0|0|0|0.029|
|SE|80127|ENSG00000122515|ZMIZ2|chr7|+|44756974|44757149|44756424|44756539|44757377|44757561|27|13|12|22|22|0|0|0|0|0|58|10|8|15|64|0|3|0|2|0|298|149|0.528057558|1|1|1|1|1|1|1|0.625|1|0.789|1|0.117|
|SE|70499|ENSG00000134086|VHL|chr3|+|10146513|10146636|10141777|10142187|10149786|10150145|183|88|166|148|133|8|2|6|9|4|235|167|147|167|234|0|8|3|18|14|272|149|0.528130784|1|0.926|0.96|0.938|0.9|0.948|1|0.92|0.964|0.836|0.902|0.01|
|SE|2490|ENSG00000141867|BRD4|chr19|-|15268153|15268281|15267415|15267551|15268904|15269042|44|49|52|53|58|759|659|639|815|1639|83|83|45|75|70|1752|1007|923|1067|1936|277|149|0.52813106|1|0.03|0.038|0.042|0.034|0.019|0.025|0.042|0.026|0.036|0.019|0.003|
|SE|30928|ENSG00000105443|CYTH2|chr19|+|48477716|48477719|48474837|48474949|48478068|48478145|9|3|11|13|6|31|10|23|25|15|0|15|13|12|24|12|19|23|24|39|152|149|0.528173877|1|0.222|0.227|0.319|0.338|0.282|0|0.436|0.357|0.329|0.376|-0.022|
|SE|64225|ENSG00000064225|ST3GAL6|chr3|+|98771120|98771190|98770878|98770956|98772812|98772916|18|5|12|8|35|16|20|10|35|13|5|6|23|6|0|12|9|4|9|22|219|149|0.528180819|1|0.434|0.145|0.449|0.135|0.647|0.221|0.312|0.796|0.312|0|0.034|
|SE|76923|ENSG00000167483|NIBAN3|chr19|+|17546685|17546797|17543523|17543631|17549443|17549527|101|104|119|172|116|0|4|0|2|4|272|319|31|154|545|8|13|0|0|1|261|149|0.528337391|1|1|0.937|1|0.98|0.943|0.951|0.933|1|1|0.997|-0.004|
|SE|48168|ENSG00000117543|DPH5|chr1|-|101013709|101013818|100995109|100995149|101021640|101021765|70|44|75|68|157|0|2|1|2|2|78|31|29|29|32|0|0|1|0|0|258|149|0.528337763|1|1|0.927|0.977|0.952|0.978|1|1|0.944|1|1|-0.022|
|SE|36628|ENSG00000169180|XPO6|chr16|-|28208986|28209122|28177219|28177332|28211365|28211550|6|0|5|9|3|4|7|2|13|13|0|6|4|10|5|10|16|18|6|10|285|149|0.528366206|1|0.44|0|0.567|0.266|0.108|0|0.164|0.104|0.466|0.207|0.088|
|SE|9559|ENSG00000100360|IFT27|chr22|-|36767782|36767859|36767062|36767365|36775673|36775699|15|4|10|16|6|0|0|0|0|0|23|8|9|24|17|0|4|0|0|0|226|149|0.528467684|1|1|1|1|1|1|1|0.569|1|1|1|0.086|
|SE|30438|ENSG00000055955|ITIH4|chr3|-|52818051|52818168|52816883|52817058|52818256|52818283|23|8|24|21|17|0|0|0|0|0|5|28|44|70|57|0|0|0|3|2|266|149|0.528512647|1|1|1|1|1|1|1|1|1|0.929|0.941|0.026|
|SE|109319|ENSG00000123104|ITPR2|chr12|-|26475295|26475418|26443542|26443650|26476911|26477007|167|177|158|150|262|0|0|0|0|0|59|139|92|68|13|0|7|0|0|0|272|149|0.52855055|1|1|1|1|1|1|1|0.916|1|1|1|0.017|
|SE|9561|ENSG00000100360|IFT27|chr22|-|36767982|36768099|36767782|36767862|36775673|36776103|0|2|5|0|0|7|8|7|10|3|0|0|6|0|0|20|5|3|6|27|266|149|0.528637397|1|0|0.123|0.286|0|0|0|0|0.528|0|0|-0.024|
|SE|86793|ENSG00000169057|MECP2|chrX|-|154039791|154039903|154032434|154032557|154092183|154092307|7|5|2|2|3|22|19|13|9|4|0|2|0|6|8|32|8|5|5|22|261|149|0.528657585|1|0.154|0.131|0.081|0.113|0.3|0|0.125|0|0.407|0.172|0.015|
|SE|60673|ENSG00000117322|CR2|chr1|+|207471422|207471499|207470996|207471087|207472771|207473179|28|7|29|22|76|0|0|0|0|0|17|17|1|20|36|0|5|0|0|0|226|149|0.528742387|1|1|1|1|1|1|1|0.692|1|1|1|0.062|
|SE|95535|ENSG00000187764|SEMA4D|chr9|-|89388871|89389047|89387385|89387608|89391263|89391415|352|216|444|446|361|0|0|0|0|0|410|386|426|299|746|0|2|0|4|13|298|149|0.52892994|1|1|1|1|1|1|1|0.99|1|0.974|0.966|0.014|
|SE|75581|ENSG00000204568|MRPS18B|chr6|+|30622831|30622898|30619492|30619989|30625501|30626392|39|24|48|43|52|0|0|3|0|3|2|46|20|19|39|0|0|0|0|0|216|149|0.529011461|1|1|1|0.917|1|0.923|1|1|1|1|1|-0.032|
|SE|102071|ENSG00000181192|DHTKD1|chr10|+|12091512|12091684|12088985|12089255|12097683|12097996|37|20|28|31|29|0|0|0|0|6|102|36|24|38|100|0|4|0|0|0|298|149|0.529045939|1|1|1|1|1|0.707|1|0.818|1|1|1|-0.022|
|SE|64217|ENSG00000102524|TNFSF13B|chr13|+|108286802|108286859|108270226|108270424|108303252|108303365|191|161|128|176|213|9|8|1|2|5|176|179|78|119|46|1|10|5|6|0|206|149|0.529255174|1|0.939|0.936|0.989|0.985|0.969|0.992|0.928|0.919|0.935|1|0.009|
|SE|63646|ENSG00000112855|HARS2|chr5|+|140693590|140693665|140691452|140691756|140693934|140693987|66|32|50|56|48|0|0|1|0|0|32|24|30|42|88|0|0|2|0|4|224|149|0.529340472|1|1|1|0.971|1|1|1|1|0.909|1|0.936|0.025|
|SE|61301|ENSG00000067208|EVI5|chr1|-|92625793|92625934|92624175|92624334|92636201|92636336|103|120|83|104|114|3|3|1|0|0|103|110|80|79|72|0|0|1|0|0|290|149|0.529360002|1|0.946|0.954|0.977|1|1|1|1|0.976|1|1|-0.02|
|SE|14079|ENSG00000130731|METTL26|chr16|-|634888|634956|634718|634797|635611|635774|26|10|24|13|16|3|2|1|0|0|10|15|17|37|20|0|0|0|0|0|217|149|0.529374094|1|0.856|0.774|0.943|1|1|1|1|1|1|1|-0.085|
|SE|101681|ENSG00000136935|GOLGA1|chr9|-|124929215|124929290|124928187|124928285|124931315|124931406|128|153|109|152|200|2|1|0|0|0|245|126|165|242|119|9|4|0|0|0|224|149|0.529395907|1|0.977|0.99|1|1|1|0.948|0.954|1|1|1|0.013|
|SE|50601|ENSG00000085871|MGST2|chr4|+|139695196|139695267|139665932|139666077|139703454|139703536|55|28|17|39|32|0|0|0|0|0|49|72|10|73|128|0|0|0|5|2|220|149|0.529461276|1|1|1|1|1|1|1|1|1|0.908|0.977|0.023|
|SE|87177|ENSG00000101849|TBL1X|chrX|+|9653544|9653689|9640272|9640360|9654214|9654322|20|6|13|22|7|18|7|6|31|12|23|35|13|19|39|19|21|22|36|14|294|149|0.529466223|1|0.36|0.303|0.523|0.265|0.228|0.38|0.458|0.23|0.211|0.585|-0.037|
|SE|36915|ENSG00000135622|SEMA4F|chr2|+|74662731|74662825|74657564|74657624|74673456|74673499|12|3|16|18|17|5|3|4|8|7|25|6|4|13|28|25|0|5|3|2|243|149|0.529492811|1|0.595|0.38|0.71|0.58|0.598|0.38|1|0.329|0.727|0.896|-0.094|
|SE|105622|ENSG00000139323|POC1B|chr12|-|89497038|89497342|89492074|89492115|89525119|89525204|16|10|7|20|3|40|30|25|35|23|0|15|7|6|13|15|9|9|16|35|298|149|0.529530397|1|0.167|0.143|0.123|0.222|0.061|0|0.455|0.28|0.158|0.157|-0.067|
|SE|69503|ENSG00000106799|TGFBR1|chr9|+|99132496|99132739|99128854|99129100|99137858|99138089|285|154|157|227|198|2|2|5|2|1|86|105|91|64|21|0|1|0|0|0|298|149|0.529680274|1|0.986|0.975|0.94|0.983|0.99|1|0.981|1|1|1|-0.021|
|SE|87756|ENSG00000063601|MTMR1|chrX|+|150718624|150718700|150699194|150699300|150727214|150727309|82|75|78|80|91|5|0|2|1|0|65|74|52|69|97|0|0|2|0|0|225|149|0.529706321|1|0.916|1|0.963|0.981|1|1|1|0.945|1|1|-0.017|
|SE|1257|ENSG00000100083|GGA1|chr22|+|37625796|37625949|37624968|37625076|37629461|37629526|76|26|79|40|44|4|0|0|0|2|81|41|35|83|132|0|4|3|0|0|298|149|0.52974342|1|0.905|1|1|1|0.917|1|0.837|0.854|1|1|0.026|
|SE|77110|ENSG00000122512|PMS2|chr7|-|6005891|6006031|6003971|6004058|6008996|6009019|67|46|70|77|77|0|2|3|1|0|58|76|92|45|87|0|0|1|0|0|289|149|0.529761455|1|1|0.922|0.923|0.975|1|1|1|0.979|1|1|-0.032|
|SE|14599|ENSG00000137871|ZNF280D|chr15|-|56666394|56666535|56658423|56658486|56666799|56666986|67|67|83|96|138|4|10|3|5|4|6|22|5|11|21|0|5|4|0|0|290|149|0.529863177|1|0.896|0.775|0.934|0.908|0.947|1|0.693|0.391|1|1|0.075|
|SE|6617|ENSG00000154655|L3MBTL4|chr18|-|6171827|6171942|6138193|6138296|6213148|6213259|33|11|9|3|11|0|0|1|0|4|29|14|5|21|37|0|0|0|0|0|264|149|0.529967731|1|1|1|0.836|1|0.608|1|1|1|1|1|-0.111|
|SE|23067|ENSG00000065357|DGKA|chr12|+|55935304|55935404|55932062|55932123|55936422|55936567|5|5|6|9|17|4|4|9|8|0|9|5|11|8|21|0|12|5|16|30|249|149|0.530010267|1|0.428|0.428|0.285|0.402|1|1|0.2|0.568|0.23|0.295|0.05|
|SE|41509|ENSG00000138674|SEC31A|chr4|-|82830936|82830975|82828999|82829058|82842139|82842481|0|3|12|14|3|54|36|53|62|51|0|5|10|13|0|53|30|24|53|11|188|149|0.530060326|1|0|0.062|0.152|0.152|0.045|0|0.117|0.248|0.163|0|-0.023|
|SE|44774|ENSG00000145833|DDX46|chr5|+|134784365|134784541|134782050|134782086|134785464|134785586|126|152|103|134|161|2|0|4|0|2|15|82|83|28|31|0|0|0|0|0|298|149|0.530342155|1|0.969|1|0.928|1|0.976|1|1|1|1|1|-0.025|
|SE|100087|ENSG00000149262|INTS4|chr11|-|77910969|77911152|77907716|77907810|77918820|77918978|18|10|15|10|21|34|20|41|61|30|29|6|11|10|8|31|47|25|41|32|298|149|0.530364867|1|0.209|0.2|0.155|0.076|0.259|0.319|0.06|0.18|0.109|0.111|0.024|
|SE|60777|ENSG00000187741|FANCA|chr16|-|89762637|89762793|89761948|89762022|89764889|89765066|9|3|4|3|5|8|12|6|7|19|0|6|4|8|17|22|6|1|10|19|298|149|0.530368064|1|0.36|0.111|0.25|0.176|0.116|0|0.333|0.667|0.286|0.309|-0.116|
|SE|1562|ENSG00000198089|SFI1|chr22|+|31531057|31531129|31508254|31508376|31546860|31546971|40|30|45|42|38|24|20|29|25|22|25|28|25|47|66|53|17|25|30|16|221|149|0.53037369|1|0.529|0.503|0.511|0.531|0.538|0.241|0.526|0.403|0.514|0.736|0.038|
|SE|63892|ENSG00000073614|KDM5A|chr12|-|297040|297200|295572|295793|306945|307089|465|331|389|397|632|0|0|0|5|0|47|327|284|187|8|7|0|0|0|0|298|149|0.530469408|1|1|1|1|0.975|1|0.77|1|1|1|1|0.041|
|SE|37159|ENSG00000151726|ACSL1|chr4|-|184780331|184780433|184776883|184776983|184783926|184783991|149|128|103|111|142|6|3|2|4|0|273|398|949|522|249|9|5|4|7|9|251|149|0.530482368|1|0.936|0.962|0.968|0.943|1|0.947|0.979|0.993|0.978|0.943|-0.006|
|SE|92852|ENSG00000130958|SLC35D2|chr9|-|96360153|96360221|96336716|96336784|96364463|96364550|23|20|19|17|22|0|0|1|0|0|7|15|24|20|105|0|0|1|0|5|217|149|0.530511474|1|1|1|0.929|1|1|1|1|0.943|1|0.935|0.01|
|SE|6082|ENSG00000105576|TNPO2|chr19|-|12702071|12702147|12701598|12701851|12702822|12702845|23|17|23|27|17|0|0|0|0|0|51|24|15|14|17|0|0|0|0|5|225|149|0.530673444|1|1|1|1|1|1|1|1|1|1|0.692|0.062|
|SE|98733|ENSG00000107290|SETX|chr9|-|132342689|132342799|132336295|132336515|132346260|132346471|387|237|258|394|387|0|0|2|1|0|74|218|225|42|13|0|4|0|5|0|259|149|0.530760767|1|1|1|0.987|0.996|1|1|0.969|1|0.829|1|0.037|
|SE|58154|ENSG00000273136|NBPF26|chr1|+|120822103|120822156|120818122|120818174|120823308|120823360|29|53|14|31|40|6|2|1|0|3|91|30|98|176|24|4|4|9|9|0|202|149|0.530917301|1|0.781|0.951|0.912|1|0.908|0.944|0.847|0.889|0.935|1|-0.013|
|SE|18422|ENSG00000196497|IPO4|chr14|-|24187666|24187796|24187399|24187579|24188215|24188257|18|10|29|15|29|2|4|0|7|8|0|26|22|24|10|1|0|0|6|16|279|149|0.531058423|1|0.828|0.572|1|0.534|0.659|0|1|1|0.681|0.25|0.132|
|SE|49603|ENSG00000076685|NT5C2|chr10|-|103136248|103136372|103106588|103106706|103139405|103139479|7|4|8|9|12|169|152|132|193|294|0|3|27|7|0|357|156|117|295|119|273|149|0.531120333|1|0.022|0.014|0.032|0.025|0.022|0|0.01|0.112|0.013|0|-0.004|
|SE|32369|ENSG00000013441|CLK1|chr2|-|200856890|200856985|200856681|200856811|200857972|200858089|448|294|445|557|1025|0|0|0|0|0|156|235|246|99|72|0|0|0|0|9|244|149|0.531254932|1|1|1|1|1|1|1|1|1|1|0.83|0.034|
|SE|32782|ENSG00000168297|PXK|chr3|+|58420535|58420568|58412900|58412963|58424751|58425175|4|9|6|6|18|86|34|57|73|210|0|7|8|13|6|183|74|67|70|38|182|149|0.531379323|1|0.037|0.178|0.079|0.063|0.066|0|0.072|0.089|0.132|0.114|0.003|
|SE|74383|ENSG00000188846|RPL14|chr3|+|40461607|40461661|40458641|40458736|40461938|40462168|296|182|339|294|512|0|0|1|0|4|118|137|132|113|121|0|0|0|0|11|203|149|0.531421657|1|1|1|0.996|1|0.989|1|1|1|1|0.89|0.019|
|SE|44013|ENSG00000114098|ARMC8|chr3|+|138237481|138237572|138235033|138235114|138239467|138239528|171|141|114|163|204|2|0|0|0|0|65|131|134|130|83|0|6|2|0|0|240|149|0.531457187|1|0.982|1|1|1|1|1|0.931|0.977|1|1|0.015|
|SE|48278|ENSG00000069424|KCNAB2|chr1|+|6097516|6097555|6097268|6097357|6098484|6101178|4|0|0|1|0|210|104|142|224|176|0|0|3|0|17|233|205|93|259|198|188|149|0.531541312|1|0.015|0|0|0.004|0|0|0|0.025|0|0.064|-0.014|
|SE|46003|ENSG00000117620|SLC35A3|chr1|+|100015301|100015420|100011364|100011533|100022385|100022611|84|88|95|105|103|0|1|1|4|6|20|41|55|11|1|9|0|0|0|0|268|149|0.531614087|1|1|0.98|0.981|0.936|0.905|0.553|1|1|1|1|0.05|
|SE|52704|ENSG00000204054|LINC00963|chr9|+|129493415|129493595|129488544|129489299|129503322|129503672|18|13|14|21|0|6|6|2|12|5|17|17|6|18|26|10|11|8|6|0|298|149|0.531732479|1|0.6|0.52|0.778|0.467|0|0.459|0.436|0.273|0.6|1|-0.081|
|SE|106192|ENSG00000135090|TAOK3|chr12|-|118177200|118177329|118172456|118172660|118181370|118181607|164|89|155|111|157|1|0|0|2|2|114|107|175|219|22|0|4|3|1|0|278|149|0.531848659|1|0.989|1|1|0.967|0.977|1|0.935|0.969|0.992|1|0.007|
|SE|85247|ENSG00000147905|ZCCHC7|chr9|+|37305543|37305714|37126311|37126942|37327798|37327834|128|73|102|86|272|2|0|1|0|5|42|85|44|53|32|0|0|0|0|0|298|149|0.531949139|1|0.97|1|0.981|1|0.965|1|1|1|1|1|-0.017|
|SE|5070|ENSG00000101247|NDUFAF5|chr20|+|13816462|13816546|13814076|13814515|13816874|13816957|47|28|22|35|51|2|3|4|6|9|30|42|119|43|8|8|8|6|13|0|233|149|0.532003854|1|0.938|0.856|0.779|0.789|0.784|0.706|0.77|0.927|0.679|1|0.013|
|SE|104443|ENSG00000149100|EIF3M|chr11|+|32594913|32595013|32593768|32593949|32595965|32596010|1687|998|1223|1210|1488|2|0|2|7|5|1234|887|900|869|993|0|17|13|1|3|249|149|0.532057288|1|0.998|1|0.997|0.99|0.994|1|0.969|0.976|0.998|0.995|0.008|
|SE|90103|ENSG00000205038|PKHD1L1|chr8|+|109409864|109409978|109408048|109408206|109412264|109412414|36|31|8|81|69|2|0|0|0|0|120|60|58|29|73|5|0|2|0|0|263|149|0.53207131|1|0.911|1|1|1|1|0.931|1|0.943|1|1|0.007|
|SE|77084|ENSG00000197461|PDGFA|chr7|-|500430|500499|498308|498574|501115|501242|0|8|0|4|2|61|113|23|119|130|16|3|4|0|0|560|150|118|248|836|218|149|0.532072441|1|0|0.046|0|0.022|0.01|0.019|0.013|0.023|0|0|0.005|
|SE|13837|ENSG00000068903|SIRT2|chr19|-|38898378|38898425|38893818|38893867|38899505|38899618|19|3|0|7|2|66|12|30|49|29|5|4|2|15|21|57|89|23|45|172|196|149|0.532083368|1|0.18|0.16|0|0.098|0.05|0.063|0.033|0.062|0.202|0.085|0.009|
|SE|60334|ENSG00000070785|EIF2B3|chr1|-|44881611|44881739|44879817|44880008|44897354|44897444|69|63|71|95|135|1|2|0|1|4|157|109|47|106|80|8|0|1|0|2|277|149|0.53224354|1|0.974|0.944|1|0.981|0.948|0.913|1|0.962|1|0.956|0.003|
|SE|68483|ENSG00000177879|AP3S1|chr5|+|115852978|115853014|115841996|115842106|115866669|115866761|6|2|1|3|7|192|114|107|159|82|0|4|20|5|4|149|220|208|372|77|185|149|0.532291002|1|0.025|0.014|0.007|0.015|0.064|0|0.014|0.072|0.011|0.04|-0.002|
|SE|9342|ENSG00000167280|ENGASE|chr17|+|79083481|79083590|79083019|79083123|79083760|79083931|96|54|104|133|101|1|1|1|4|0|89|60|56|68|77|0|0|0|0|6|258|149|0.532415327|1|0.982|0.969|0.984|0.951|1|1|1|1|1|0.881|0.001|
|SE|5823|ENSG00000197586|ENTPD6|chr20|+|25213262|25213406|25209848|25209925|25214866|25214942|162|92|144|132|172|2|5|1|5|0|214|144|94|86|276|5|0|3|10|0|293|149|0.532438347|1|0.976|0.903|0.987|0.931|1|0.956|1|0.941|0.814|1|0.017|
|SE|84508|ENSG00000128731|HERC2|chr15|-|28196460|28196569|28196214|28196354|28198377|28198503|111|56|186|120|137|0|1|0|0|0|156|159|65|109|98|6|2|0|0|0|258|149|0.532526524|1|1|0.97|1|1|1|0.938|0.979|1|1|1|0.011|
|SE|35140|ENSG00000007923|DNAJC11|chr1|-|6652828|6652951|6651528|6651602|6653910|6654039|77|59|91|74|159|0|0|0|0|2|127|59|43|67|133|0|0|0|0|7|272|149|0.532637806|1|1|1|1|1|0.978|1|1|1|1|0.912|0.013|
|SE|52031|ENSG00000135250|SRPK2|chr7|-|105268805|105268869|105203627|105203785|105388647|105388702|13|1|19|16|27|17|17|9|13|10|3|21|42|19|20|15|15|13|9|17|213|149|0.532638652|1|0.349|0.04|0.596|0.463|0.654|0.123|0.495|0.693|0.596|0.451|-0.051|
|SE|41576|ENSG00000084072|PPIE|chr1|+|39745374|39745498|39743823|39743924|39748902|39749088|172|95|172|187|226|7|1|5|4|5|155|65|59|100|239|13|7|1|0|0|273|149|0.532688491|1|0.931|0.981|0.949|0.962|0.961|0.867|0.835|0.97|1|1|0.022|
|SE|108409|ENSG00000110851|PRDM4|chr12|-|107754828|107754860|107751646|107752209|107756831|107756965|1|3|3|3|6|0|1|0|1|1|0|6|5|6|4|2|1|1|1|0|181|149|0.532709371|1|1|0.712|1|0.712|1|0|0.832|0.805|0.832|1|0.191|
|SE|89280|ENSG00000166169|POLL|chr10|-|101582762|101582891|101580247|101580416|101584601|101584720|10|10|29|33|7|4|0|0|0|0|24|18|11|14|43|0|0|0|0|0|278|149|0.532709972|1|0.573|1|1|1|1|1|1|1|1|1|-0.085|
|SE|96511|ENSG00000150907|FOXO1|chr13|-|40633146|40633248|40560766|40560860|40665582|40665877|10|4|1|5|10|132|90|183|118|95|0|0|11|6|6|67|114|110|45|25|251|149|0.532772165|1|0.043|0.026|0.003|0.025|0.059|0|0|0.056|0.073|0.125|-0.02|
|SE|107841|ENSG00000061936|SFSWAP|chr12|+|131728292|131728428|131711089|131711447|131753122|131753363|84|41|73|90|116|1|1|0|0|0|86|72|104|67|117|0|3|1|3|0|285|149|0.532939457|1|0.978|0.955|1|1|1|1|0.926|0.982|0.921|1|0.021|
|SE|81948|ENSG00000051825|MPHOSPH9|chr12|-|123176687|123176789|123166360|123166789|123179925|123179980|238|175|233|205|289|4|2|1|5|15|38|144|119|61|26|0|1|0|10|0|251|149|0.533114367|1|0.972|0.981|0.993|0.961|0.92|1|0.988|1|0.784|1|0.011|
|SE|55948|ENSG00000187605|TET3|chr2|+|74046277|74048411|74003109|74003166|74073548|74073639|146|95|107|151|78|8|2|9|6|4|319|171|158|206|263|22|6|15|10|8|298|149|0.533180039|1|0.901|0.96|0.856|0.926|0.907|0.879|0.934|0.84|0.912|0.943|0.008|
|SE|96472|ENSG00000108405|P2RX1|chr17|-|3899633|3899761|3898933|3899024|3903201|3903343|74|61|50|123|73|0|0|0|6|0|238|169|121|210|1192|0|0|2|0|18|277|149|0.533366488|1|1|1|1|0.917|1|1|1|0.97|1|0.973|-0.005|
|SE|83060|ENSG00000196305|IARS1|chr9|-|92262135|92262232|92260150|92260234|92262968|92263055|0|0|2|0|1|142|93|173|139|253|10|0|2|2|0|73|115|109|102|16|246|149|0.533372242|1|0|0|0.007|0|0.002|0.077|0|0.011|0.012|0|-0.018|
|SE|33719|ENSG00000155975|VPS37A|chr8|+|17265906|17265981|17246933|17247369|17268257|17268273|35|19|16|16|15|0|0|0|1|0|11|15|20|8|7|0|0|0|4|0|224|149|0.533396009|1|1|1|1|0.914|1|1|1|1|0.571|1|0.069|
|SE|37603|ENSG00000082074|FYB1|chr5|-|39211057|39211091|39202893|39202987|39212677|39213017|2|5|8|2|11|23|18|15|35|28|0|9|4|8|0|13|9|20|25|36|183|149|0.533474068|1|0.066|0.184|0.303|0.044|0.242|0|0.449|0.14|0.207|0|0.009|
|SE|26598|ENSG00000132952|USPL1|chr13|+|30621072|30621239|30617919|30618056|30621763|30621892|25|13|20|27|36|24|16|39|24|31|9|5|16|4|0|0|11|21|6|9|298|149|0.533526554|1|0.342|0.289|0.204|0.36|0.367|1|0.185|0.276|0.25|0|-0.03|
|SE|76072|ENSG00000137692|DCUN1D5|chr11|-|103089226|103089318|103083255|103083326|103091786|103092148|62|39|39|56|65|2|0|1|0|4|28|40|56|27|25|0|0|0|0|0|241|149|0.533641934|1|0.95|1|0.96|1|0.909|1|1|1|1|1|-0.036|
|SE|63292|ENSG00000171132|PRKCE|chr2|+|45840379|45840551|45652400|45652448|45842999|45843063|4|0|6|6|0|33|12|13|23|9|0|8|20|5|5|18|81|45|41|20|298|149|0.533853454|1|0.057|0|0.188|0.115|0|0|0.047|0.182|0.057|0.111|-0.007|
|SE|67582|ENSG00000142188|TMEM50B|chr21|-|33456003|33456097|33455726|33455784|33460412|33460505|23|16|32|20|33|270|181|224|280|139|5|17|24|29|0|205|83|108|130|41|243|149|0.533876772|1|0.05|0.051|0.081|0.042|0.127|0.015|0.112|0.12|0.12|0|-0.003|
|SE|29020|ENSG00000123352|SPATS2|chr12|+|49512861|49512905|49500069|49500205|49514554|49514613|0|0|0|3|3|6|4|13|20|33|0|1|0|0|0|7|8|15|17|7|193|149|0.533915272|1|0|0|0|0.104|0.066|0|0.088|0|0|0|0.016|
|SE|4871|ENSG00000196562|SULF2|chr20|-|47672197|47672393|47666259|47666488|47676493|47676623|237|166|191|104|157|1|0|0|0|0|598|114|70|152|244|11|0|0|2|0|298|149|0.533944008|1|0.992|1|1|1|1|0.965|1|1|0.974|1|0.011|
|SE|106798|ENSG00000173442|EHBP1L1|chr11|+|65583591|65583765|65581210|65581251|65584240|65584398|219|148|177|181|270|4|4|5|7|14|340|224|116|223|442|8|9|2|10|9|298|149|0.534107312|1|0.965|0.949|0.947|0.928|0.906|0.955|0.926|0.967|0.918|0.961|-0.006|
|SE|764|ENSG00000196588|MRTFA|chr22|-|40519463|40519612|40463220|40463286|40533589|40533811|9|7|2|1|14|21|17|20|16|25|0|10|2|1|11|9|8|11|1|15|298|149|0.534117417|1|0.176|0.171|0.048|0.03|0.219|0|0.385|0.083|0.333|0.268|-0.085|
|SE|57782|ENSG00000093010|COMT|chr22|+|19947306|19947490|19941802|19941897|19961198|19961289|3|3|0|0|0|43|11|24|30|31|0|0|0|11|0|14|62|13|22|81|298|149|0.534215202|1|0.034|0.12|0|0|0|0|0|0|0.2|0|-0.009|
|SE|45190|ENSG00000141384|TAF4B|chr18|+|26335174|26335231|26327014|26327140|26357689|26357794|37|36|65|39|98|1|0|0|0|4|58|51|31|30|26|0|0|6|0|0|206|149|0.534242685|1|0.964|1|1|1|0.947|1|1|0.789|1|1|0.024|
|SE|60571|ENSG00000155275|TRMT44|chr4|+|8449697|8449888|8446475|8446590|8451959|8452028|53|39|32|50|88|1|0|0|2|2|96|46|53|72|76|8|0|0|0|0|298|149|0.534426072|1|0.964|1|1|0.926|0.957|0.857|1|1|1|1|-0.002|
|SE|63562|ENSG00000163898|LIPH|chr3|-|185524070|185524160|185519141|185519309|185527483|185527585|5|4|1|6|7|0|1|0|0|0|13|21|6|6|52|0|0|0|0|6|239|149|0.534567035|1|1|0.714|1|1|1|1|1|1|1|0.844|-0.026|
|SE|87294|ENSG00000068394|GPKOW|chrX|-|49113852|49113938|49113406|49113755|49115725|49115795|45|22|31|53|18|0|0|0|0|5|81|67|40|51|33|0|0|0|0|0|235|149|0.534706218|1|1|1|1|1|0.695|1|1|1|1|1|-0.061|
|SE|52569|ENSG00000111445|RFC5|chr12|+|118017994|118018053|118016703|118016892|118019071|118019136|4|1|10|2|5|13|12|15|23|8|8|3|3|10|0|20|25|10|18|31|208|149|0.534791352|1|0.181|0.056|0.323|0.059|0.309|0.223|0.079|0.177|0.285|0|0.033|
|SE|68632|ENSG00000223960|CHROMR|chr2|+|178426365|178426703|178414022|178414113|178430769|178430889|56|32|18|51|30|0|1|0|0|5|19|37|23|35|13|0|0|0|2|0|298|149|0.534804993|1|1|0.941|1|1|0.75|1|1|1|0.897|1|-0.041|
|SE|32555|ENSG00000211456|SACM1L|chr3|+|45735245|45735373|45732052|45732151|45737582|45737652|285|184|206|243|160|20|12|10|14|8|37|95|78|53|8|0|6|9|2|0|277|149|0.534813024|1|0.885|0.892|0.917|0.903|0.915|1|0.895|0.823|0.934|1|-0.028|
|SE|3978|ENSG00000126261|UBA2|chr19|+|34433347|34433412|34430575|34430659|34434867|34434968|345|225|311|356|783|0|0|0|2|0|261|247|206|209|195|8|2|0|0|5|214|149|0.535024139|1|1|1|1|0.992|1|0.958|0.989|1|1|0.964|0.016|
|SE|37608|ENSG00000082074|FYB1|chr5|-|39211057|39211091|39208863|39208964|39212677|39213017|0|0|2|1|4|2|5|1|3|2|0|4|1|1|1|25|4|3|2|10|183|149|0.535070635|1|0|0|0.62|0.213|0.62|0|0.449|0.213|0.289|0.075|0.085|
|SE|66692|ENSG00000137992|DBT|chr1|-|100230732|100230914|100218625|100218747|100235435|100235511|151|116|113|116|184|2|0|3|2|3|13|101|106|59|0|0|0|4|0|2|298|149|0.535254742|1|0.974|1|0.95|0.967|0.968|1|1|0.93|1|0|0.186|
|SE|101790|ENSG00000204161|TMEM273|chr10|-|49165204|49165283|49161598|49161622|49165765|49165796|1|0|3|1|3|64|10|35|67|43|0|0|0|0|0|6|8|11|9|26|228|149|0.535273506|1|0.01|0|0.053|0.01|0.044|0|0|0|0|0|0.023|
|SE|38404|ENSG00000143458|GABPB2|chr1|+|151093191|151093386|151088157|151088297|151097851|151097915|126|76|121|128|224|0|0|0|0|8|90|63|73|88|70|0|0|0|0|0|298|149|0.535336051|1|1|1|1|1|0.933|1|1|1|1|1|-0.013|
|SE|63052|ENSG00000006652|IFRD1|chr7|+|112450507|112450782|112423143|112423432|112455762|112455867|35|32|15|52|50|0|8|1|13|0|135|167|53|99|71|14|6|4|2|9|298|149|0.5353656|1|1|0.667|0.882|0.667|1|0.828|0.933|0.869|0.961|0.798|-0.035|
|SE|8166|ENSG00000086544|ITPKC|chr19|+|40733159|40733364|40729201|40729415|40736985|40737087|27|12|32|32|25|0|0|0|0|0|9|57|23|19|34|0|0|5|0|0|298|149|0.535437715|1|1|1|1|1|1|1|1|0.697|1|1|0.061|
|SE|3909|ENSG00000072958|AP1M1|chr19|+|16210359|16210395|16209029|16209177|16226420|16226547|3|1|4|2|5|153|67|123|165|146|8|2|1|0|0|91|138|63|92|239|185|149|0.53551609|1|0.016|0.012|0.026|0.01|0.027|0.066|0.012|0.013|0|0|0|
|SE|94791|ENSG00000101464|PIGU|chr20|-|34616041|34616139|34588452|34588607|34634614|34634715|47|21|49|39|46|0|0|0|0|0|22|17|19|14|19|0|0|0|0|5|247|149|0.535692043|1|1|1|1|1|1|1|1|1|1|0.696|0.061|
|SE|99328|ENSG00000162231|NXF1|chr11|-|62804081|62804210|62803791|62803978|62805328|62805469|2|0|0|0|2|129|90|82|111|92|8|0|2|3|12|208|169|127|182|217|278|149|0.535724574|1|0.008|0|0|0|0.012|0.02|0|0.008|0.009|0.029|-0.009|
|SE|101622|ENSG00000143702|CEP170|chr1|-|243191017|243191494|243185778|243186422|243199059|243199194|126|100|148|122|188|0|2|0|3|3|14|101|60|17|6|0|0|0|1|0|298|149|0.535802006|1|1|0.962|1|0.953|0.969|1|1|1|0.895|1|-0.002|
|SE|10955|ENSG00000091592|NLRP1|chr17|-|5533303|5533384|5532821|5532984|5536850|5536940|60|26|44|30|60|11|13|12|6|9|117|40|34|33|87|11|19|13|4|10|230|149|0.535816191|1|0.779|0.564|0.704|0.764|0.812|0.873|0.577|0.629|0.842|0.849|-0.029|
|SE|104231|ENSG00000139620|KANSL2|chr12|-|48679035|48679150|48671798|48671962|48679654|48679833|130|76|118|130|137|37|19|33|36|22|165|149|144|146|115|21|61|23|54|55|264|149|0.535882403|1|0.665|0.693|0.669|0.671|0.778|0.816|0.58|0.779|0.604|0.541|0.031|
|SE|45395|ENSG00000144468|RHBDD1|chr2|+|226839408|226839627|226838072|226838154|226864603|226864711|22|22|14|20|44|5|0|2|1|1|20|8|2|25|57|0|0|1|2|5|298|149|0.536243498|1|0.688|1|0.778|0.909|0.957|1|1|0.5|0.862|0.851|0.024|
|SE|66762|ENSG00000129292|PHF20L1|chr8|+|132816887|132817076|132814636|132814889|132817338|132817545|184|184|178|251|242|0|4|1|2|0|81|95|99|60|50|0|5|3|0|0|298|149|0.536296155|1|1|0.958|0.989|0.984|1|1|0.905|0.943|1|1|0.017|
|SE|30202|ENSG00000111266|DUSP16|chr12|-|12500518|12500682|12487027|12487187|12519861|12520000|244|149|295|247|410|1|3|4|5|6|62|39|195|112|36|0|0|8|0|0|298|149|0.536327178|1|0.992|0.961|0.974|0.961|0.972|1|1|0.924|1|1|-0.013|
|SE|25114|ENSG00000153310|CYRIB|chr8|-|129896671|129896854|129879388|129879471|129903311|129903350|17|8|13|4|0|677|462|436|596|542|13|11|8|17|7|402|450|619|620|145|298|149|0.536372741|1|0.012|0.009|0.015|0.003|0|0.016|0.012|0.006|0.014|0.024|-0.007|
|SE|39139|ENSG00000134352|IL6ST|chr5|-|55968275|55968396|55963351|55963506|55969549|55969855|281|349|400|368|665|0|3|0|1|0|76|228|254|119|53|7|3|0|0|0|270|149|0.53650671|1|1|0.985|1|0.995|1|0.857|0.977|1|1|1|0.029|
|SE|168|ENSG00000160194|NDUFV3|chr21|+|42903181|42904276|42896926|42897047|42908863|42909105|111|71|58|76|51|14|10|7|11|13|188|70|38|29|154|6|28|7|10|28|298|149|0.536564912|1|0.799|0.78|0.806|0.776|0.662|0.94|0.556|0.731|0.592|0.733|0.054|
|SE|45992|ENSG00000174175|SELP|chr1|-|169619128|169619219|169617027|169617414|169630071|169630076|60|80|23|150|123|2|2|0|0|3|196|169|74|97|2031|0|0|1|0|0|240|149|0.536567779|1|0.949|0.961|1|1|0.962|1|1|0.979|1|1|-0.021|
|SE|109537|ENSG00000127022|CANX|chr5|+|179704154|179704430|179698980|179699049|179705678|179705852|1|0|6|1|0|17|6|18|3|3|7|0|0|0|12|13|13|14|15|0|298|149|0.536613223|1|0.029|0|0.143|0.143|0|0.212|0|0|0|1|-0.179|
|SE|11839|ENSG00000254788|CKLF-CMTM1|chr16|+|66563121|66563217|66552648|66552793|66566082|66566428|54|19|30|50|31|6|0|1|0|0|14|28|21|28|46|0|4|0|0|0|245|149|0.536621402|1|0.846|1|0.948|1|1|1|0.81|1|1|1|-0.003|
|SE|46173|ENSG00000100280|AP1B1|chr22|-|29329711|29329720|29327679|29328895|29330377|29330532|77|28|48|71|81|73|44|78|50|61|117|77|45|62|408|102|119|43|79|160|158|149|0.53664368|1|0.499|0.375|0.367|0.572|0.556|0.52|0.379|0.497|0.425|0.706|-0.032|
|SE|3299|ENSG00000127445|PIN1|chr19|+|9836822|9836875|9835339|9835402|9838435|9838648|4|0|10|6|1|119|59|91|109|28|8|6|4|2|16|158|122|67|99|200|202|149|0.536915671|1|0.024|0|0.075|0.039|0.026|0.036|0.035|0.042|0.015|0.056|-0.004|
|SE|90823|ENSG00000173064|HECTD4|chr12|-|112247461|112247550|112246900|112247076|112248066|112248171|129|74|88|88|159|4|1|1|2|0|245|121|117|130|203|0|0|1|0|0|238|149|0.536946769|1|0.953|0.979|0.982|0.965|1|1|1|0.987|1|1|-0.022|
|SE|45974|ENSG00000170248|PDCD6IP|chr3|+|33826479|33826580|33825186|33825340|33828837|33828969|140|111|118|156|171|1|2|1|1|0|70|145|112|85|42|0|7|0|0|0|250|149|0.537027335|1|0.988|0.971|0.986|0.989|1|1|0.925|1|1|1|0.002|
|SE|10064|ENSG00000108406|DHX40|chr17|+|59566626|59566794|59565577|59565783|59573115|59573235|68|52|78|79|30|0|0|0|0|0|23|86|68|87|8|2|0|0|5|0|298|149|0.537036346|1|1|1|1|1|1|0.852|1|1|0.897|1|0.05|
|SE|46928|ENSG00000141027|NCOR1|chr17|-|16067893|16068121|16065484|16065691|16070164|16070525|200|109|158|169|290|0|0|0|1|0|106|190|131|118|83|9|0|0|0|0|298|149|0.537160745|1|1|1|1|0.988|1|0.855|1|1|1|1|0.027|
|SE|86332|ENSG00000135048|CEMIP2|chr9|-|71740062|71740232|71734805|71734994|71745017|71745579|440|190|241|288|266|3|0|1|3|0|141|628|880|248|28|0|5|0|1|5|298|149|0.537189571|1|0.987|1|0.992|0.98|1|1|0.984|1|0.992|0.737|0.049|
|SE|89142|ENSG00000111276|CDKN1B|chr12|+|12718824|12718954|12717367|12718314|12721064|12721468|429|316|536|466|658|3|1|3|3|7|238|360|428|231|224|0|10|2|0|3|279|149|0.537195299|1|0.987|0.994|0.99|0.988|0.98|1|0.951|0.991|1|0.976|0.004|
|SE|107867|ENSG00000188906|LRRK2|chr12|+|40225554|40225640|40224976|40225282|40232273|40232383|416|323|205|278|282|4|6|2|1|4|189|207|246|96|30|0|0|1|0|0|235|149|0.537298629|1|0.985|0.972|0.985|0.994|0.978|1|1|0.994|1|1|-0.016|
|SE|60708|ENSG00000241399|CD302|chr2|-|159783358|159783469|159780881|159780998|159798131|159798208|124|111|65|99|190|4|2|1|1|0|100|160|99|80|60|0|0|0|1|0|260|149|0.537313301|1|0.947|0.97|0.974|0.983|1|1|1|1|0.979|1|-0.021|
|SE|48289|ENSG00000229474|PATL2|chr15|-|44672014|44672156|44669966|44670087|44672387|44672456|96|28|52|53|56|3|0|2|2|0|60|232|90|66|170|0|0|0|0|0|291|149|0.537329519|1|0.942|1|0.93|0.931|1|1|1|1|1|1|-0.039|
|SE|49758|ENSG00000077549|CAPZB|chr1|-|19342751|19342864|19338822|19339617|19344357|19344434|4|10|3|13|2|1537|825|985|1110|2342|17|8|4|5|102|2035|1151|908|1137|3549|262|149|0.537496783|1|0.001|0.007|0.002|0.007|0|0.005|0.004|0.002|0.002|0.016|-0.002|
|SE|95845|ENSG00000141551|CSNK1D|chr17|-|82265685|82265796|82255129|82255577|82273305|82273369|192|68|128|172|80|0|0|0|0|0|178|119|251|140|206|0|0|8|4|0|260|149|0.537541887|1|1|1|1|1|1|1|1|0.947|0.953|1|0.02|
|SE|79359|ENSG00000280828|NA|chr7|+|128574719|128574791|128533651|128533730|128579998|128580094|24|13|15|26|28|0|2|3|1|0|104|26|27|90|43|0|0|1|1|0|221|149|0.537550567|1|1|0.814|0.771|0.946|1|1|1|0.948|0.984|1|-0.08|
|SE|40404|ENSG00000008952|SEC62|chr3|+|169988359|169988359|169983160|169983253|169992593|169992721|306|223|324|323|531|0|0|0|4|0|34|143|150|35|8|0|1|6|0|0|269|149|0.537570999|1|1|1|1|0.978|1|1|0.988|0.933|1|1|0.011|
|SE|65316|ENSG00000285756|NA|chrX|-|3818656|3818703|3818441|3818500|3822373|3822580|11|10|22|30|56|28|11|27|33|27|10|14|8|8|10|22|21|2|11|1|196|149|0.537580369|1|0.23|0.409|0.382|0.409|0.612|0.257|0.336|0.753|0.356|0.884|-0.109|
|SE|21295|ENSG00000150867|PIP4K2A|chr10|-|22607926|22608023|22591628|22591781|22609619|22609717|1495|863|1059|1425|1788|1|1|1|1|6|1476|1341|709|974|2494|1|9|10|16|10|246|149|0.537587094|1|0.999|0.998|0.998|0.999|0.994|0.999|0.989|0.977|0.974|0.993|0.011|
|SE|43131|ENSG00000186470|BTN3A2|chr6|+|26368182|26368267|26367974|26368050|26368564|26368912|66|41|71|29|80|100|36|48|34|81|28|59|21|10|127|36|36|15|24|68|234|149|0.537682749|1|0.296|0.42|0.485|0.352|0.386|0.331|0.511|0.471|0.21|0.543|-0.025|
|SE|106414|ENSG00000135297|MTO1|chr6|+|73471459|73471557|73466488|73466606|73473364|73473654|24|17|27|23|39|143|115|163|161|156|24|37|13|19|22|134|169|93|124|71|247|149|0.537817231|1|0.092|0.082|0.091|0.079|0.131|0.098|0.117|0.078|0.085|0.157|-0.012|
|SE|11197|ENSG00000172824|CES4A|chr16|+|67006390|67006519|67005239|67005393|67008973|67009739|1|1|5|2|5|1|3|0|6|3|0|8|1|3|4|4|0|2|3|16|278|149|0.537837765|1|0.349|0.152|1|0.152|0.472|0|1|0.211|0.349|0.118|0.089|
|SE|88675|ENSG00000126216|TUBGCP3|chr13|-|112521662|112521864|112519885|112520021|112522319|112522509|6|5|8|9|11|77|36|38|52|72|0|8|7|3|0|28|30|24|22|28|298|149|0.537926052|1|0.037|0.065|0.095|0.08|0.071|0|0.118|0.127|0.064|0|0.008|
|SE|94610|ENSG00000106829|TLE4|chr9|+|79707122|79707218|79706746|79706899|79708117|79708250|1|2|0|0|6|26|14|23|38|61|10|0|0|0|0|32|26|28|34|119|245|149|0.537937591|1|0.023|0.08|0|0|0.056|0.16|0|0|0|0|0|
|SE|88923|ENSG00000176731|RBIS|chr8|-|85219322|85219363|85217385|85217502|85220305|85220339|12|27|17|13|103|152|129|157|144|286|16|31|18|24|4|214|128|82|93|99|190|149|0.538170271|1|0.058|0.141|0.078|0.066|0.22|0.055|0.16|0.147|0.168|0.031|0|
|SE|100168|ENSG00000078124|ACER3|chr11|+|76985642|76985724|76976288|76976341|76990538|76990574|76|77|31|66|86|2|2|2|1|0|123|83|81|106|63|0|4|0|0|0|231|149|0.538270841|1|0.961|0.961|0.909|0.977|1|1|0.93|1|1|1|-0.024|
|SE|80876|ENSG00000006576|PHTF2|chr7|+|77937709|77937838|77929108|77929327|77940029|77940302|142|152|160|178|154|0|3|2|5|4|42|87|59|44|7|0|5|2|2|0|278|149|0.538272946|1|1|0.964|0.977|0.95|0.954|1|0.903|0.941|0.922|1|0.016|
|SE|45952|ENSG00000117593|DARS2|chr1|+|173833375|173833499|173831534|173831630|173834472|173834519|63|52|56|70|121|3|0|0|0|0|48|28|21|21|10|0|0|0|0|4|273|149|0.538582356|1|0.92|1|1|1|1|1|1|1|1|0.577|0.069|
|SE|22904|ENSG00000140853|NLRC5|chr16|+|57051537|57051621|57047544|57047628|57054750|57054840|164|72|190|157|87|0|0|1|0|0|191|203|143|221|323|0|0|0|0|14|233|149|0.538655482|1|1|1|0.992|1|1|1|1|1|1|0.937|0.011|
|SE|10071|ENSG00000108406|DHX40|chr17|+|59573739|59573967|59573115|59573235|59574187|59574254|169|107|143|154|344|5|0|3|0|5|76|110|131|88|20|0|9|2|0|0|298|149|0.538679168|1|0.944|1|0.96|1|0.972|1|0.859|0.97|1|1|0.009|
|SE|20362|ENSG00000120925|RNF170|chr8|-|42865415|42865489|42861329|42861855|42887727|42887871|40|40|48|27|44|0|1|0|0|6|46|27|17|20|33|0|0|0|0|0|223|149|0.538689722|1|1|0.964|1|1|0.831|1|1|1|1|1|-0.041|
|SE|78655|ENSG00000274272|NA|chr7|-|100577085|100577154|100576693|100577005|100577496|100577558|37|22|39|18|20|0|0|0|0|0|17|68|21|18|45|5|0|0|0|0|218|149|0.538762629|1|1|1|1|1|1|0.699|1|1|1|1|0.06|
|SE|109100|ENSG00000196312|MFSD14C|chr9|-|96942602|96942752|96918642|96918826|96949554|96949585|28|22|19|14|15|14|18|7|8|5|38|20|24|39|20|24|19|12|21|10|298|149|0.538935799|1|0.5|0.379|0.576|0.467|0.6|0.442|0.345|0.5|0.481|0.5|0.051|
|SE|96692|ENSG00000135362|PRR5L|chr11|+|36380519|36380588|36379380|36379467|36381639|36381695|6|5|5|5|5|2|0|1|0|2|2|10|5|4|49|4|0|0|0|0|218|149|0.538938149|1|0.672|1|0.774|1|0.631|0.255|1|1|1|1|-0.036|
|SE|24615|ENSG00000204120|GIGYF2|chr2|+|232787149|232787229|232761395|232761436|232790697|232790915|167|135|174|150|158|0|0|0|0|0|243|137|144|173|73|0|0|10|0|0|298|149|0.539070085|1|1|1|1|1|1|1|1|0.878|1|1|0.024|
|SE|55670|ENSG00000243406|MRPS31P5|chr13|-|52183975|52184114|52180581|52180654|52186682|52186823|35|47|49|29|42|7|7|11|9|11|43|74|89|45|22|10|16|18|10|0|288|149|0.539153466|1|0.721|0.776|0.697|0.625|0.664|0.69|0.705|0.719|0.7|1|-0.066|
|SE|74470|ENSG00000116539|ASH1L|chr1|-|155415743|155415923|155395458|155395553|155438326|155439068|128|116|114|133|171|3|5|0|0|10|48|118|146|109|63|5|2|6|0|0|298|149|0.539161349|1|0.955|0.921|1|1|0.895|0.828|0.967|0.924|1|1|0.01|
|SE|38462|ENSG00000147251|DOCK11|chrX|+|118616201|118616240|118615599|118615711|118618549|118618728|112|86|85|102|207|215|198|161|226|247|13|73|63|12|15|20|123|86|34|20|188|149|0.539367248|1|0.292|0.256|0.295|0.263|0.399|0.34|0.32|0.367|0.219|0.373|-0.023|
|SE|90176|ENSG00000198876|DCAF12|chr9|-|34093285|34093448|34089411|34089590|34096715|34096781|243|169|175|192|258|4|1|1|0|4|243|203|152|277|382|0|9|0|0|0|298|149|0.539415077|1|0.968|0.988|0.989|1|0.97|1|0.919|1|1|1|-0.001|
|SE|82809|ENSG00000136271|DDX56|chr7|-|44572889|44573046|44572573|44572744|44573582|44573664|63|38|51|63|11|0|0|0|0|4|64|34|44|28|79|0|0|0|0|0|298|149|0.539430919|1|1|1|1|1|0.579|1|1|1|1|1|-0.084|
|SE|53796|ENSG00000111653|ING4|chr12|-|6653229|6653396|6652935|6653050|6656726|6656798|214|135|214|225|470|7|8|12|10|12|286|254|103|141|386|25|11|11|3|3|298|149|0.539489831|1|0.939|0.894|0.899|0.918|0.951|0.851|0.92|0.824|0.959|0.985|0.012|
|SE|74702|ENSG00000197548|ATG7|chr3|+|11379971|11380052|11364656|11364734|11426803|11426926|123|65|62|70|84|1|1|0|4|2|130|112|52|121|139|0|0|0|2|0|230|149|0.539551653|1|0.988|0.977|1|0.919|0.965|1|1|1|0.975|1|-0.025|
|SE|76835|ENSG00000146842|TMEM209|chr7|-|130181622|130181719|130178401|130178527|130184183|130184255|142|147|188|158|327|1|8|2|1|1|113|145|111|96|21|0|0|4|3|0|246|149|0.539645206|1|0.989|0.918|0.983|0.99|0.995|1|1|0.944|0.951|1|-0.004|
|SE|29071|ENSG00000163625|WDFY3|chr4|-|84702352|84702506|84696731|84696823|84704337|84704444|57|52|34|57|85|2|1|1|3|0|49|67|110|122|29|0|0|0|0|0|298|149|0.539865086|1|0.934|0.963|0.944|0.905|1|1|1|1|1|1|-0.051|
|SE|62399|ENSG00000122912|SLC25A16|chr10|-|68493448|68493570|68493131|68493198|68503631|68503695|54|64|72|82|75|0|0|2|1|5|24|36|42|56|14|0|0|0|0|0|271|149|0.539881694|1|1|1|0.952|0.978|0.892|1|1|1|1|1|-0.036|
|SE|31074|ENSG00000065882|TBC1D1|chr4|+|38118032|38118192|38102998|38103157|38124961|38125131|137|77|114|107|213|1|0|1|0|6|205|177|164|171|230|0|4|2|0|6|298|149|0.540151641|1|0.986|1|0.983|1|0.947|1|0.957|0.976|1|0.95|0.007|
|SE|62148|ENSG00000162650|ATXN7L2|chr1|+|109489929|109490128|109488846|109489100|109490270|109490392|22|6|20|18|17|0|0|0|0|0|17|63|11|11|15|0|0|0|4|0|298|149|0.54028906|1|1|1|1|1|1|1|1|1|0.579|1|0.084|
|SE|89300|ENSG00000166169|POLL|chr10|-|101585315|101585478|101583507|101583681|101585861|101586156|22|3|25|20|13|0|0|0|0|0|20|27|8|20|11|0|0|0|0|4|298|149|0.540296367|1|1|1|1|1|1|1|1|1|1|0.579|0.084|
|SE|55107|ENSG00000072110|ACTN1|chr14|-|68878988|68879069|68877081|68878523|68879961|68880108|104|132|83|107|180|5|2|1|9|1|254|138|77|199|1224|31|1|5|4|7|230|149|0.540315961|1|0.931|0.977|0.982|0.885|0.991|0.841|0.989|0.909|0.97|0.991|0.013|
|SE|12862|ENSG00000284685|NA|chr16|+|29320927|29321095|29312349|29312589|29326346|29326483|37|10|8|24|5|1|1|0|1|0|[illegible]|11|19|24|19|0|4|0|0|0|298|149|0.540361331|1|0.949|0.833|1|0.923|1|1|0.579|1|1|1|0.025|
|SE|105189|ENSG00000139624|CERS5|chr12|-|50132961|50133064|50129791|50130694|50134399|50134702|0|0|1|0|0|7|3|6|9|20|0|1|0|5|0|26|10|23|31|4|252|149|0.540363304|1|0|1|0.09|0|0|0|0.056|0|0.087|0|-0.011|
|SE|12345|ENSG00000197912|SPG7|chr16|+|89548426|89549713|89548002|89548113|89550493|89550609|1|0|4|3|0|27|19|23|46|20|0|0|3|0|5|55|44|31|79|44|298|149|0.540565508|1|0.018|0|0.08|0.032|0|0|0|0.046|0|0.054|0.006|
|SE|24311|ENSG00000125846|ZNF133|chr20|+|18310168|18310306|18306297|18306393|18315068|18316994|0|1|1|0|4|10|0|3|2|18|0|0|0|0|0|18|3|1|2|3|287|149|0.540703064|1|0|1|0.148|0|0.103|0|0|0|0|0|0.25|
|SE|85025|ENSG00000101901|ALG13|chrX|+|111744667|111744904|111736713|111736879|111752789|111752830|59|55|34|54|58|0|0|0|2|1|31|41|22|20|3|0|0|0|5|0|298|149|0.540946359|1|1|1|1|0.931|0.967|1|1|1|0.667|1|0.046|
|SE|71220|ENSG00000079263|SP140|chr2|+|230250980|230251061|230245862|230245940|230253315|230253417|13|14|25|28|43|0|0|0|0|0|54|26|18|32|26|0|0|5|0|0|230|149|0.540997369|1|1|1|1|1|1|1|1|0.7|1|1|0.06|
|SE|56738|ENSG00000270231|NBPF8|chr1|+|120453908|120454114|120453371|120453444|120455408|120455460|89|116|52|52|208|1|1|0|0|5|99|33|43|28|104|0|0|0|0|5|298|149|0.541049618|1|0.978|0.983|1|1|0.954|1|1|1|1|0.912|0.001|
|SE|85101|ENSG00000147586|MRPS28|chr8|-|80002998|80003120|79918939|79919148|80030053|80030254|38|28|50|22|49|0|0|0|2|0|25|30|14|10|45|0|0|0|4|0|271|149|0.541164701|1|1|1|1|0.858|1|1|1|1|0.579|1|0.056|
|SE|75384|ENSG00000204392|LSM2|chr6|-|31798476|31798507|31797989|31798049|31806074|31806121|56|56|86|55|133|3|1|0|0|10|101|47|32|67|76|0|0|0|0|12|180|149|0.541175596|1|0.939|0.979|1|1|0.917|1|1|1|1|0.84|-0.001|
|SE|95254|ENSG00000165219|GAPVD1|chr9|+|125263550|125263700|125261830|125261959|125298889|125299106|9|1|1|2|0|12|7|18|17|3|12|4|0|0|0|0|21|15|16|30|298|149|0.54131713|1|0.273|0.067|0.027|0.056|0|1|0.087|0|0|0|-0.133|
|SE|55715|ENSG00000116704|SLC35D1|chr1|-|67052011|67052079|67050432|67050504|67052770|67052857|64|34|39|36|65|0|0|0|0|2|36|17|22|29|22|0|5|0|0|0|217|149|0.541339143|1|1|1|1|1|0.957|1|0.7|1|1|1|0.051|
|SE|13560|ENSG00000102908|NFAT5|chr16|+|69568494|69568548|69565093|69566374|69626402|69626528|50|39|32|27|48|22|8|9|19|9|8|33|62|15|13|0|31|17|9|7|203|149|0.541548249|1|0.625|0.782|0.723|0.511|0.797|1|0.439|0.728|0.55|0.577|0.029|
|SE|38169|ENSG00000137074|APTX|chr9|-|32984630|32984857|32924896|32925028|32985970|32986030|63|29|32|32|75|1|0|0|2|3|27|60|39|24|14|0|0|0|0|0|298|149|0.541659683|1|0.969|1|1|0.889|0.926|1|1|1|1|1|-0.043|
|SE|63652|ENSG00000112855|HARS2|chr5|+|140697160|140697406|140696942|140697070|140697568|140697685|122|89|94|101|61|0|0|0|0|0|92|43|40|39|36|0|6|0|0|0|298|149|0.541811934|1|1|1|1|1|1|1|0.782|1|1|1|0.044|
|SE|23955|ENSG00000133835|HSD17B4|chr5|+|119531265|119531404|119529893|119529980|119536422|119536550|295|174|143|220|292|7|4|7|14|15|171|146|154|220|53|6|13|5|0|4|288|149|0.541903174|1|0.956|0.957|0.914|0.89|0.91|0.936|0.853|0.941|1|0.873|0.005|
|SE|47735|ENSG00000145687|SSBP2|chr5|-|81446867|81446922|81440537|81440636|81448789|81448825|11|17|16|17|19|0|0|0|0|0|13|10|2|2|16|0|0|0|0|5|204|149|0.541990364|1|1|1|1|1|1|1|1|1|1|0.7|0.06|
|SE|30662|ENSG00000112893|MAN2A1|chr5|+|109729341|109729513|109716119|109716264|109755328|109755456|495|379|531|474|585|8|1|4|5|2|460|363|342|246|543|0|0|1|0|0|298|149|0.542149242|1|0.969|0.995|0.985|0.979|0.993|1|1|0.994|1|1|-0.015|
|SE|27169|ENSG00000156500|PABIR3|chrX|+|134814770|134814849|134807168|134807708|134821326|134822975|30|16|26|26|60|0|0|0|0|0|8|19|16|25|46|0|0|0|0|7|228|149|0.542383888|1|1|1|1|1|1|1|1|1|1|0.811|0.038|
|SE|97806|ENSG00000011114|BTBD7|chr14|-|93263784|93263993|93261601|93261677|93293857|93294617|167|123|130|135|125|2|3|2|1|0|183|97|133|164|10|0|0|0|0|0|298|149|0.5424116|1|0.977|0.953|0.97|0.985|1|1|1|1|1|1|-0.023|

| | | | | | | | | | | | | | | | | | | | | | | | | | | | | | | | | | | | | | | | | | | | | | | |
|---|---|---|---|---|---|---|---|---|---|---|---|---|---|---|---|---|---|---|---|---|---|---|---|---|---|---|---|---|---|---|---|---|---|---|---|---|---|---|---|---|---|---|---|---|---|---|
| SE | 68026 | ENSG00000172890 | NADSYN1 | chr11 | + | 71481355 | 71481404 | 71480754 | 71480879 | 71481922 | 71482025 | 166 | 110 | 180 | 129 | 183 | 4 | 3 | 1 | 2 | 2 | 227 | 219 | 135 | 150 | 283 | 3 | 3 | 1 | 9 | 2 | 198 | 149 | 0.542421057 | 1 | 0.969 | 0.965 | 0.993 | 0.98 | 0.986 | 0.983 | 0.982 | 0.99 | 0.926 | 0.991 | 0.004 |
| SE | 55429 | ENSG00000116688 | MFN2 | chr1 | + | 11981973 | 11982114 | 11980214 | 11980484 | 11989164 | 11989343 | 185 | 128 | 169 | 128 | 327 | 1 | 6 | 2 | 0 | 1 | 323 | 203 | 161 | 298 | 461 | 1 | 11 | 0 | 2 | 3 | 290 | 149 | 0.542517135 | 1 | 0.99 | 0.916 | 0.977 | 1 | 0.994 | 0.994 | 0.905 | 1 | 0.987 | 0.987 | 0.001 |
| SE | 83356 | ENSG00000087274 | ADD1 | chr4 | + | 2908514 | 2909431 | 2907376 | 2907844 | 2914883 | 2915040 | 646 | 420 | 523 | 579 | 680 | 4 | 5 | 5 | 8 | 1 | 1181 | 926 | 502 | 774 | 1519 | 0 | 7 | 5 | 7 | 8 | 298 | 149 | 0.542639373 | 1 | 0.988 | 0.977 | 0.981 | 0.973 | 0.997 | 1 | 0.985 | 0.98 | 0.982 | 0.99 | -0.004 |
| SE | 16710 | ENSG00000166233 | ARIH1 | chr15 | + | 72570176 | 72570307 | 72567105 | 72567177 | 72572107 | 72572165 | 240 | 178 | 228 | 251 | 412 | 0 | 0 | 1 | 0 | 0 | 117 | 160 | 296 | 236 | 63 | 8 | 0 | 3 | 0 | 0 | 280 | 149 | 0.542644112 | 1 | 1 | 1 | 0.992 | 1 | 1 | 0.886 | 1 | 0.981 | 1 | 1 | 0.025 |
| SE | 88503 | ENSG00000197217 | ENTPD4 | chr8 | - | 23447679 | 23447885 | 23444455 | 23444606 | 23457556 | 23457639 | 138 | 152 | 218 | 177 | 261 | 3 | 6 | 1 | 1 | 0 | 158 | 158 | 184 | 156 | 140 | 0 | 2 | 3 | 2 | 0 | 298 | 149 | 0.542655613 | 1 | 0.958 | 0.927 | 0.991 | 0.989 | 1 | 1 | 0.975 | 0.968 | 0.975 | 1 | -0.011 |
| SE | 43516 | ENSG00000163565 | IFI16 | chr1 | + | 159045296 | 159045464 | 159032523 | 159032691 | 159051678 | 159052098 | 80 | 45 | 50 | 55 | 52 | 105 | 53 | 50 | 86 | 78 | 0 | 94 | 84 | 33 | 0 | 4 | 77 | 16 | 7 | 3 | 298 | 149 | 0.542711648 | 1 | 0.276 | 0.298 | 0.333 | 0.242 | 0.25 | 0 | 0.379 | 0.724 | 0.702 | 0 | -0.081 |
| SE | 61912 | ENSG00000124067 | SLC12A4 | chr16 | - | 67954642 | 67954773 | 67952183 | 67952425 | 67957741 | 67957796 | 27 | 14 | 22 | 25 | 15 | 0 | 1 | 1 | 0 | 0 | 22 | 17 | 7 | 13 | 109 | 0 | 3 | 2 | 0 | 0 | 280 | 149 | 0.542745079 | 1 | 1 | 0.882 | 0.921 | 1 | 1 | 1 | 0.751 | 0.651 | 1 | 1 | 0.08 |
| SE | 34773 | ENSG00000100034 | PPM1F | chr22 | - | 21945842 | 21946108 | 21939531 | 21939680 | 21952791 | 21952831 | 190 | 98 | 86 | 151 | 89 | 0 | 0 | 0 | 0 | 0 | 245 | 141 | 116 | 217 | 317 | 11 | 0 | 1 | 0 | 0 | 298 | 149 | 0.542774509 | 1 | 1 | 1 | 1 | 1 | 1 | 0.918 | 1 | 0.983 | 1 | 1 | 0.02 |
| SE | 102853 | ENSG00000138175 | ARL3 | chr10 | - | 102689892 | 102689943 | 102685815 | 102686001 | 102699372 | 102699489 | 85 | 51 | 92 | 110 | 96 | 0 | 0 | 0 | 0 | 0 | 162 | 97 | 56 | 65 | 76 | 0 | 2 | 1 | 0 | 5 | 200 | 149 | 0.542818296 | 1 | 1 | 1 | 1 | 1 | 1 | 1 | 0.973 | 0.977 | 1 | 0.919 | 0.026 |
| SE | 103725 | ENSG00000136738 | STAM | chr10 | + | 17672974 | 17673078 | 17660463 | 17660548 | 17684674 | 17684750 | 2 | 0 | 0 | 0 | 11 | 45 | 32 | 35 | 71 | 91 | 0 | 0 | 0 | 8 | 0 | 8 | 59 | 44 | 33 | 12 | 253 | 149 | 0.543000409 | 1 | 0.026 | 0 | 0 | 0 | 0.066 | 0 | 0 | 0 | 0.125 | 0 | -0.007 |
| SE | 70698 | ENSG00000114268 | PFKFB4 | chr3 | - | 48548531 | 48548659 | 48547878 | 48548384 | 48549863 | 48549895 | 2 | 13 | 2 | 6 | 16 | 0 | 0 | 0 | 0 | 0 | 10 | 3 | 21 | 9 | 16 | 0 | 3 | 0 | 0 | 0 | 277 | 149 | 0.543014351 | 1 | 1 | 1 | 1 | 1 | 1 | 1 | 0.35 | 1 | 1 | 1 | 0.13 |
| SE | 70680 | ENSG00000095059 | DHPS | chr19 | - | 12677107 | 12677211 | 12676016 | 12676105 | 12677290 | 12677396 | 144 | 65 | 108 | 93 | 170 | 4 | 0 | 0 | 0 | 0 | 84 | 132 | 45 | 78 | 146 | 7 | 0 | 0 | 2 | 0 | 253 | 149 | 0.543038444 | 1 | 0.955 | 1 | 1 | 1 | 1 | 0.876 | 1 | 1 | 0.958 | 1 | 0.024 |
| SE | 101066 | ENSG00000187531 | SIRT7 | chr17 | - | 81916057 | 81916139 | 81915610 | 81915681 | 81917614 | 81917719 | 2 | 0 | 0 | 0 | 2 | 26 | 10 | 13 | 17 | 0 | 0 | 0 | 0 | 5 | 0 | 8 | 46 | 14 | 13 | 37 | 231 | 149 | 0.543169554 | 1 | 0.047 | 0 | 0 | 0 | 1 | 0 | 0 | 0 | 0.199 | 0 | 0.17 |
| SE | 20450 | ENSG00000148814 | LRRC27 | chr10 | + | 132342212 | 132342271 | 132337564 | 132337695 | 132344497 | 132344650 | 4 | 5 | 11 | 20 | 26 | 8 | 2 | 9 | 6 | 12 | 9 | 12 | 0 | 12 | 9 | 9 | 9 | 4 | 14 | 0 | 208 | 149 | 0.543178205 | 1 | 0.264 | 0.642 | 0.467 | 0.705 | 0.608 | 0.417 | 0.489 | 0 | 0.38 | 1 | 0.08 |
| SE | 72959 | ENSG00000204843 | DCTN1 | chr2 | - | 74390582 | 74390728 | 74377999 | 74378245 | 74391793 | 74391843 | 6 | 0 | 1 | 2 | 4 | 62 | 41 | 42 | 58 | 9 | 0 | 10 | 0 | 0 | 5 | 56 | 66 | 51 | 79 | 122 | 295 | 149 | 0.543315971 | 1 | 0.047 | 0 | 0.012 | 0.017 | 0.183 | 0 | 0.071 | 0 | 0 | 0.02 | 0.034 |
| SE | 47498 | ENSG00000143363 | PRUNE1 | chr1 | + | 151025514 | 151025673 | 151018466 | 151018669 | 151027232 | 151027327 | 69 | 74 | 64 | 111 | 195 | 0 | 0 | 2 | 2 | 4 | 167 | 109 | 52 | 104 | 697 | 0 | 0 | 0 | 0 | 4 | 298 | 149 | 0.543383246 | 1 | 1 | 1 | 0.941 | 0.965 | 0.961 | 1 | 1 | 1 | 1 | 0.989 | -0.024 |
| SE | 88704 | ENSG00000183077 | AFMID | chr17 | + | 78205602 | 78205738 | 78205439 | 78205518 | 78205945 | 78206050 | 5 | 2 | 7 | 4 | 11 | 3 | 0 | 0 | 0 | 3 | 16 | 4 | 1 | 4 | 7 | 0 | 0 | 0 | 3 | 0 | 285 | 149 | 0.543400455 | 1 | 0.466 | 1 | 1 | 1 | 0.657 | 1 | 1 | 1 | 0.411 | 1 | -0.058 |
| SE | 7394 | ENSG00000141447 | OSBPL1A | chr18 | - | 24178012 | 24178195 | 24172375 | 24172483 | 24179737 | 24179835 | 33 | 35 | 28 | 33 | 45 | 9 | 0 | 5 | 1 | 0 | 66 | 56 | 52 | 121 | 14 | 12 | 0 | 8 | 9 | 0 | 298 | 149 | 0.543407149 | 1 | 0.647 | 1 | 0.737 | 0.943 | 1 | 0.733 | 1 | 0.765 | 0.871 | 1 | -0.008 |
| SE | 41582 | ENSG00000084072 | PPIE | chr1 | + | 39762516 | 39762651 | 39752909 | 39753052 | 39763688 | 39763914 | 27 | 18 | 15 | 16 | 26 | 15 | 8 | 10 | 18 | 12 | 34 | 44 | 30 | 12 | 60 | 0 | 34 | 21 | 19 | 31 | 284 | 149 | 0.543420022 | 1 | 0.486 | 0.541 | 0.44 | 0.318 | 0.532 | 1 | 0.404 | 0.428 | 0.249 | 0.504 | -0.054 |
| SE | 46702 | ENSG00000171103 | TRMT61B | chr2 | - | 28865016 | 28865119 | 28852407 | 28852499 | 28869578 | 28870295 | 31 | 22 | 34 | 26 | 33 | 0 | 0 | 1 | 0 | 1 | 12 | 17 | 13 | 2 | 26 | 0 | 5 | 0 | 0 | 0 | 252 | 149 | 0.543449062 | 1 | 1 | 1 | 0.953 | 1 | 0.951 | 1 | 0.668 | 1 | 1 | 1 | 0.047 |
| SE | 30103 | ENSG00000113522 | RAD50 | chr5 | + | 132579861 | 132580066 | 132579316 | 132579502 | 132587561 | 132587690 | 249 | 254 | 196 | 263 | 433 | 4 | 0 | 1 | 2 | 7 | 26 | 127 | 82 | 23 | 28 | 0 | 0 | 3 | 0 | 0 | 298 | 149 | 0.543526892 | 1 | 0.969 | 1 | 0.99 | 0.985 | 0.969 | 1 | 1 | 0.932 | 1 | 1 | -0.004 |
| SE | 46317 | ENSG00000093183 | SEC22C | chr3 | - | 42580442 | 42580505 | 42568864 | 42569073 | 42581845 | 42581960 | 0 | 0 | 5 | 7 | 4 | 10 | 6 | 13 | 11 | 1 | 0 | 5 | 1 | 17 | 5 | 12 | 14 | 13 | 6 | 3 | 212 | 149 | 0.543721307 | 1 | 0 | 0 | 0.213 | 0.309 | 0.738 | 0 | 0.201 | 0.051 | 0.666 | 0.539 | -0.039 |
| SE | 1382 | ENSG00000100150 | DEPDC5 | chr22 | + | 31838684 | 31838845 | 31836847 | 31837155 | 31843094 | 31843212 | 39 | 23 | 31 | 40 | 32 | 0 | 0 | 5 | 1 | 0 | 89 | 58 | 19 | 43 | 50 | 0 | 0 | 0 | 0 | 0 | 298 | 149 | 0.543822976 | 1 | 1 | 1 | 0.756 | 0.952 | 1 | 1 | 1 | 1 | 1 | 1 | -0.058 |
| SE | 52767 | ENSG00000198019 | FCGR1B | chr1 | + | 121094673 | 121094958 | 121092995 | 121093250 | 121095871 | 121095934 | 1 | 7 | 3 | 1 | 2 | 31 | 9 | 8 | 13 | 25 | 7 | 0 | 6 | 20 | 0 | 142 | 30 | 12 | 79 | 70 | 298 | 149 | 0.543894653 | 1 | 0.016 | 0.28 | 0.158 | 0.037 | 0.038 | 0.024 | 0 | 0.2 | 0.112 | 0 | 0.039 |
| SE | 24795 | ENSG00000112195 | TREML2 | chr6 | - | 41198108 | 41198429 | 41194424 | 41194833 | 41200953 | 41201149 | 28 | 12 | 30 | 21 | 19 | 0 | 0 | 0 | 3 | 0 | 30 | 30 | 16 | 26 | 29 | 0 | 0 | 1 | 0 | 5 | 298 | 149 | 0.543950698 | 1 | 1 | 1 | 1 | 0.778 | 1 | 1 | 1 | 0.889 | 1 | 0.744 | 0.029 |
| SE | 17097 | ENSG00000215302 | NA | chr15 | + | 30493588 | 30493816 | 30484543 | 30484686 | 30507376 | 30507623 | 161 | 24 | 193 | 172 | 136 | 1 | 0 | 0 | 4 | 3 | 37 | 105 | 133 | 61 | 2 | 0 | 0 | 0 | 0 | 0 | 298 | 149 | 0.543954663 | 1 | 0.988 | 1 | 1 | 0.956 | 0.958 | 1 | 1 | 1 | 1 | 1 | -0.02 |
| SE | 109306 | ENSG00000139597 | N4BP2L1 | chr13 | - | 32404202 | 32404271 | 32400722 | 32403200 | 32407249 | 32407338 | 39 | 27 | 31 | 57 | 42 | 1 | 3 | 1 | 2 | 1 | 59 | 27 | 24 | 78 | 35 | 0 | 3 | 0 | 6 | 0 | 218 | 149 | 0.543957877 | 1 | 0.964 | 0.86 | 0.955 | 0.951 | 0.966 | 1 | 0.86 | 1 | 0.899 | 1 | -0.013 |
| SE | 42231 | ENSG00000100300 | TSPO | chr22 | + | 43159209 | 43159420 | 43151558 | 43151604 | 43161051 | 43161190 | 54 | 16 | 28 | 16 | 17 | 0 | 1 | 4 | 1 | 0 | 88 | 47 | 18 | 41 | 85 | 0 | 3 | 0 | 0 | 0 | 298 | 149 | 0.544001778 | 1 | 1 | 0.889 | 0.778 | 0.889 | 1 | 1 | 0.887 | 1 | 1 | 1 | -0.066 |
| SE | 54102 | ENSG00000115935 | WIPF1 | chr2 | - | 174629665 | 174629868 | 174585522 | 174585611 | 174682773 | 174682916 | 8 | 3 | 4 | 3 | 0 | 2 | 3 | 1 | 11 | 1 | 5 | 11 | 3 | 2 | 0 | 0 | 1 | 1 | 5 | 8 | 298 | 149 | 0.544100416 | 1 | 0.667 | 0.333 | 0.667 | 0.12 | 0 | 1 | 0.846 | 0.6 | 0.167 | 0 | -0.165 |
| SE | 9538 | ENSG00000108932 | SLC16A6 | chr17 | - | 68272638 | 68272767 | 68270838 | 68271654 | 68273926 | 68274070 | 125 | 67 | 82 | 77 | 80 | 2 | 0 | 0 | 1 | 0 | 66 | 147 | 134 | 72 | 22 | 0 | 0 | 0 | 0 | 5 | 278 | 149 | 0.544142925 | 1 | 0.971 | 1 | 1 | 0.976 | 1 | 1 | 1 | | 1 | 0.702 | 0.049 |
| SE | 60355 | ENSG00000172766 | NAA16 | chr13 | + | 41336649 | 41336756 | 41331273 | 41331369 | 41355143 | 41355216 | 47 | 55 | 62 | 63 | 127 | 9 | 3 | 4 | 5 | 8 | 5 | 48 | 14 | 8 | 19 | 3 | 2 | 21 | 0 | 0 | 256 | 149 | 0.544162735 | 1 | 0.752 | 0.914 | 0.9 | 0.88 | 0.902 | 0.492 | 0.933 | 0.28 | 1 | 1 | 0.129 |
| SE | 11237 | ENSG00000102984 | ZNF821 | chr16 | - | 71867911 | 71868037 | 71864137 | 71864242 | 71879906 | 71880023 | 15 | 7 | 7 | 8 | 13 | 1 | 0 | 2 | 2 | 0 | 22 | 22 | 18 | 11 | 45 | 0 | 0 | 0 | 0 | 0 | 275 | 149 | 0.544169411 | 1 | 0.89 | 1 | 0.655 | 0.684 | 1 | 1 | 1 | | 1 | 1 | -0.154 |
| SE | 108969 | ENSG00000111877 | MCM9 | chr6 | - | 118913294 | 118913420 | 118910596 | 118911769 | 118917560 | 118917761 | 123 | 118 | 99 | 129 | 162 | 2 | 0 | 2 | 0 | 4 | 168 | 110 | 133 | 144 | 39 | 0 | 0 | 1 | 0 | 0 | 275 | 149 | 0.54417603 | 1 | 0.971 | 1 | 0.964 | 1 | 0.956 | 1 | 1 | 0.986 | 1 | 1 | -0.019 |
| SE | 44318 | ENSG00000154814 | OXNAD1 | chr3 | + | 16294855 | 16294997 | 16286341 | 16286448 | 16301625 | 16301786 | 408 | 347 | 680 | 548 | 820 | 5 | 2 | 6 | 9 | 15 | 190 | 253 | 390 | 173 | 169 | 0 | 0 | 5 | 11 | 0 | 291 | 149 | 0.54421251 | 1 | 0.977 | 0.989 | 0.983 | 0.969 | 0.966 | 1 | 0.963 | 0.948 | 1 | 1 | -0.005 |
| SE | 97006 | ENSG00000153339 | TRAPPC8 | chr18 | - | 31913368 | 31913522 | 31909666 | 31909760 | 31916271 | 31916446 | 360 | 255 | 312 | 340 | 531 | 9 | 11 | 11 | 6 | 8 | 73 | 230 | 162 | 80 | 81 | 0 | 6 | 0 | 3 | 7 | 298 | 149 | 0.544293197 | 1 | 0.952 | 0.921 | 0.934 | 0.966 | 0.971 | 1 | 0.95 | 1 | 0.93 | 0.853 | 0.002 |
| SE | 47426 | ENSG00000163867 | ZMYM6 | chr1 | - | 35019352 | 35019602 | 35014987 | 35015162 | 35020382 | 35020467 | 144 | 169 | 136 | 158 | 274 | 3 | 0 | 2 | 4 | 1 | 48 | 79 | 95 | 30 | 21 | 0 | 0 | 3 | 0 | 0 | 298 | 149 | 0.544353789 | 1 | 0.96 | 1 | 0.971 | 0.952 | 0.993 | 1 | 1 | 0.941 | 1 | 1 | -0.013 |
| SE | 63040 | ENSG00000285533 | RELA-DT | chr11 | + | 65666130 | 65666217 | 65663426 | 65665543 | 65671422 | 65671704 | 11 | 9 | 13 | 29 | 12 | 0 | 0 | 4 | 1 | 0 | 5 | 13 | 2 | 4 | 12 | 0 | 0 | 0 | 0 | 0 | 236 | 149 | 0.544685289 | 1 | 1 | 1 | 0.672 | 0.948 | 1 | 1 | 1 | 1 | 1 | 1 | -0.076 |
| SE | 18501 | ENSG00000126821 | SGPP1 | chr14 | - | 63698568 | 63698658 | 63684215 | 63686656 | 63727260 | 63728065 | 116 | 63 | 87 | 102 | 94 | 0 | 0 | 2 | 3 | 1 | 19 | 27 | 27 | 7 | 16 | 0 | 0 | 1 | 1 | 0 | 239 | 149 | 0.544801616 | 1 | 0.973 | 1 | 0.964 | 0.955 | 0.983 | 1 | 1 | 0.944 | 0.814 | 1 | 0.023 |
| SE | 67823 | ENSG00000186625 | KATNA1 | chr6 | - | 149632758 | 149632916 | 149623102 | 149623283 | 149638385 | 149638560 | 167 | 108 | 113 | 128 | 170 | 0 | 0 | 1 | 0 | 0 | 213 | 121 | 115 | 100 | 63 | 0 | 0 | 1 | 8 | 0 | 298 | 149 | 0.544807492 | 1 | 1 | 1 | 0.983 | 1 | 1 | 1 | 1 | 0.983 | 0.862 | 1 | 0.028 |
| SE | 60326 | ENSG00000103353 | UBFD1 | chr16 | + | 23566986 | 23567069 | 23562624 | 23562730 | 23570479 | 23570575 | 72 | 40 | 52 | 63 | 62 | 4 | 0 | 0 | 2 | 0 | 27 | 23 | 15 | 34 | 33 | 0 | 0 | 1 | 2 | 0 | 232 | 149 | 0.544860246 | 1 | 0.92 | 1 | 1 | 0.953 | 1 | 1 | 1 | 0.906 | 0.916 | 1 | 0.01 |
| SE | 30050 | ENSG00000155744 | FAM126B | chr2 | - | 201045495 | 201045593 | 201023958 | 201024039 | 201071609 | 201071665 | 65 | 54 | 43 | 77 | 96 | 3 | 0 | 1 | 1 | 1 | 17 | 54 | 66 | 22 | 15 | 0 | 0 | 0 | 0 | 0 | 247 | 149 | 0.544934282 | 1 | 0.929 | 1 | 0.963 | 0.979 | 0.983 | 1 | 1 | 1 | 1 | 1 | -0.029 |
| SE | 64294 | ENSG00000177000 | MTHFR | chr1 | - | 11795097 | 11795348 | 11794728 | 11794863 | 11796205 | 11796399 | 44 | 38 | 34 | 35 | 47 | 0 | 0 | 0 | 0 | 0 | 121 | 66 | 39 | 38 | 156 | 0 | 2 | 0 | 5 | 0 | 298 | 149 | 0.544966016 | 1 | 1 | 1 | 1 | 1 | 1 | 1 | 0.943 | 1 | 0.792 | 1 | 0.053 |
| SE | 91964 | ENSG00000183808 | RBM12B | chr8 | - | 93737306 | 93737355 | 93728154 | 93736438 | 93740628 | 93740696 | 176 | 189 | 153 | 192 | 161 | 2 | 1 | 0 | 1 | 4 | 66 | 99 | 84 | 136 | 19 | 0 | 2 | 6 | 0 | 0 | 198 | 149 | 0.544980557 | 1 | 0.985 | 0.993 | 1 | 0.993 | 0.968 | 1 | 0.974 | 0.913 | 1 | 1 | 0.01 |
| SE | 10670 | ENSG00000214719 | NA | chr17 | + | 30631754 | 30631895 | 30610893 | 30610942 | 30633097 | 30633390 | 45 | 24 | 48 | 32 | 27 | 5 | 3 | 3 | 5 | 5 | 31 | 31 | 27 | 24 | 83 | 0 | 4 | 7 | 1 | 7 | 290 | 149 | 0.545026538 | 1 | 0.822 | 0.804 | 0.892 | 0.767 | 0.735 | 1 | 0.799 | 0.665 | 0.925 | 0.859 | -0.046 |
| SE | 39386 | ENSG00000220804 | LINC01881 | chr2 | + | 242130654 | 242130812 | 242122286 | 242122455 | 242138783 | 242138888 | 26 | 10 | 25 | 26 | 29 | 12 | 14 | 10 | 12 | 72 | 82 | 14 | 2 | 41 | 0 | 11 | 21 | 21 | 2 | 20 | 298 | 149 | 0.545059259 | 1 | 0.52 | 0.263 | 0.556 | 0.52 | 0.168 | 0.788 | 0.25 | 0.045 | 0.911 | 0 | 0.007 |
| SE | 49626 | ENSG00000178927 | CYBC1 | chr17 | - | 82447579 | 82447646 | 82446622 | 82446696 | 82450699 | 82450829 | 11 | 9 | 14 | 7 | 2 | 1 | 0 | 0 | 2 | 0 | 26 | 14 | 8 | 22 | 6 | 0 | 0 | 4 | 0 | 0 | 216 | 149 | 0.545118917 | 1 | 0.884 | 1 | 1 | 0.707 | 1 | 1 | 1 | 0.58 | 1 | 1 | 0.002 |
| SE | 28275 | ENSG00000138443 | ABI2 | chr2 | + | 203394699 | 203394846 | 203391043 | 203391143 | 203395655 | 203395780 | 43 | 30 | 44 | 66 | 86 | 6 | 0 | 2 | 4 | 0 | 79 | 41 | 68 | 80 | 59 | 0 | 0 | 7 | 0 | 11 | 296 | 149 | 0.545147376 | 1 | 0.783 | 1 | 0.917 | 0.893 | 1 | 1 | 1 | 0.83 | 1 | 0.73 | 0.007 |
| SE | 41119 | ENSG00000115523 | GNLY | chr2 | + | 85694915 | 85694996 | 85694357 | 85694470 | 85695319 | 85695423 | 382 | 72 | 73 | 113 | 181 | 2496 | 480 | 702 | 845 | 838 | 110 | 59 | 69 | 34 | 92 | 645 | 335 | 714 | 334 | 509 | 230 | 149 | 0.54520547 | 1 | 0.09 | 0.089 | 0.063 | 0.08 | 0.123 | 0.099 | 0.102 | 0.059 | 0.062 | 0.105 | 0.004 |
| SE | 87673 | ENSG00000143569 | UBAP2L | chr1 | + | 154251480 | 154251653 | 154251040 | 154251318 | 154253899 | 154254089 | 291 | 155 | 268 | 287 | 293 | 0 | 0 | 0 | 1 | 0 | 504 | 364 | 298 | 369 | 584 | 18 | 0 | 0 | 0 | 0 | 298 | 149 | 0.545290215 | 1 | 1 | 1 | 1 | 0.993 | 1 | 0.933 | 1 | 1 | 1 | 1 | 0.012 |
| SE | 63461 | ENSG00000115421 | PAPOLG | chr2 | + | 60791760 | 60791882 | 60786946 | 60787066 | 60792128 | 60792289 | 135 | 72 | 110 | 131 | 306 | 0 | 0 | 0 | 0 | 0 | 49 | 109 | 80 | 33 | 34 | 0 | 6 | 0 | 0 | 0 | 271 | 149 | 0.54530152 | 1 | 1 | 1 | 1 | 1 | 1 | 1 | 0.909 | 1 | 1 | 1 | 0.018 |
| SE | 44738 | ENSG00000198265 | HELZ | chr17 | - | 67137930 | 67138114 | 67128494 | 67128855 | 67145742 | 67145890 | 258 | 188 | 175 | 260 | 269 | 0 | 0 | 0 | 2 | 0 | 76 | 238 | 215 | 109 | 42 | 8 | 0 | 0 | 0 | 0 | 298 | 149 | 0.545351607 | 1 | 1 | 1 | 1 | 0.985 | 1 | 0.826 | 1 | 1 | 1 | 1 | 0.032 |
| SE | 35408 | ENSG00000174891 | RSRC1 | chr3 | + | 158203071 | 158203245 | 158123865 | 158123991 | 158298038 | 158298075 | 148 | 136 | 144 | 186 | 257 | 3 | 3 | 0 | 0 | 3 | 27 | 88 | 95 | 54 | 14 | 0 | 0 | 5 | 0 | 0 | 298 | 149 | 0.545354016 | 1 | 0.961 | 0.958 | 1 | 1 | 0.977 | 1 | 1 | 0.905 | 1 | 1 | -0.002 |
| SE | 106435 | ENSG00000135297 | MTO1 | chr6 | + | 73482448 | 73482620 | 73482028 | 73482244 | 73486597 | 73486761 | 44 | 29 | 41 | 51 | 80 | 2 | 0 | 0 | 0 | 0 | 70 | 29 | 32 | 45 | 9 | 0 | 0 | 2 | 0 | 3 | 298 | 149 | 0.545430164 | 1 | 0.917 | 1 | 1 | 1 | 1 | 1 | 1 | 0.889 | 1 | 0.6 | 0.086 |
| SE | 60271 | ENSG00000082212 | ME2 | chr18 | + | 50917346 | 50917508 | 50912800 | 50912950 | 50918109 | 50918213 | 355 | 287 | 275 | 289 | 937 | 0 | 1 | 0 | 0 | 5 | 263 | 266 | 208 | 345 | 133 | 9 | 0 | 4 | 0 | 0 | 298 | 149 | 0.545480955 | 1 | 1 | 0.993 | 1 | 1 | 0.989 | 0.936 | 1 | 0.963 | 1 | 1 | 0.017 |
| SE | 38474 | ENSG00000147251 | DOCK11 | chrX | + | 118651963 | 118652077 | 118648944 | 118649127 | 118654601 | 118654817 | 683 | 478 | 477 | 550 | 629 | 5 | 0 | 0 | 1 | 0 | 25 | 393 | 297 | 29 | 54 | 0 | 10 | 1 | 0 | 0 | 263 | 149 | 0.545535791 | 1 | 0.987 | 1 | 1 | 0.997 | 1 | 1 | 0.957 | 0.994 | 1 | 1 | 0.007 |
| SE | 96536 | ENSG00000143549 | TPM3 | chr1 | - | 154171412 | 154171488 | 154170648 | 154170711 | 154172907 | 154172978 | 40 | 19 | 26 | 48 | 52 | 8 | 2 | 0 | 0 | 7 | 51 | 58 | 37 | 53 | 167 | 29 | 3 | 1 | 12 | 0 | 225 | 149 | 0.545635486 | 1 | 0.768 | 0.863 | 1 | 1 | 0.831 | 0.538 | 0.928 | 0.961 | 0.745 | 1 | 0.058 |
| SE | 86385 | ENSG00000123562 | MORF4L2 | chrX | - | 103684680 | 103684729 | 103676494 | 103677051 | 103685170 | 103685260 | 49 | 30 | 43 | 51 | 94 | 15 | 5 | 22 | 15 | 19 | 32 | 27 | 31 | 26 | 7 | 6 | 3 | 26 | 6 | 0 | 198 | 149 | 0.545724608 | 1 | 0.711 | 0.819 | 0.595 | 0.719 | 0.788 | 0.801 | 0.871 | 0.473 | 0.765 | 1 | -0.056 |
| SE | 46338 | ENSG00000095564 | BTAF1 | chr10 | + | 92008122 | 92008275 | 91997602 | 91997751 | 92008828 | 92008950 | 753 | 616 | 670 | 674 | 791 | 1 | 1 | 9 | 1 | 8 | 118 | 273 | 420 | 110 | 96 | 0 | 8 | 3 | 1 | 0 | 298 | 149 | 0.545740376 | 1 | 0.997 | 0.997 | 0.974 | 0.997 | 0.98 | 1 | 0.945 | 0.986 | 0.982 | 1 | 0.006 |
| SE | 70634 | ENSG00000188647 | PTAR1 | chr9 | - | 69723325 | 69723630 | 69709521 | 69718684 | 69732138 | 69732352 | 181 | 142 | 133 | 170 | 233 | 6 | 2 | 4 | 6 | 3 | 69 | 74 | 58 | 60 | 5 | 1 | 4 | 7 | 0 | 0 | 298 | 149 | 0.545791292 | 1 | 0.938 | 0.973 | 0.943 | 0.934 | 0.975 | 0.972 | 0.902 | 0.806 | 1 | 1 | 0.017 |
| SE | 24520 | ENSG00000186532 | SMYD4 | chr17 | - | 1799856 | 1801024 | 1787421 | 1787604 | 1804625 | 1804715 | 80 | 57 | 91 | 86 | 88 | 3 | 0 | 1 | 0 | 2 | 171 | 77 | 31 | 109 | 125 | 0 | 0 | 1 | 0 | 0 | 298 | 149 | 0.545844798 | 1 | 0.93 | 1 | 0.978 | 1 | 0.957 | 1 | 1 | 0.939 | 1 | 1 | -0.015 |
| SE | 81610 | ENSG00000253729 | PRKDC | chr8 | - | 47943208 | 47943366 | 47939550 | 47939697 | 47943852 | 47943883 | 237 | 156 | 222 | 259 | 349 | 1 | 0 | 0 | 0 | 0 | 132 | 184 | 154 | 182 | 32 | 0 | 0 | 1 | 7 | 0 | 298 | 149 | 0.545890049 | 1 | 0.992 | 1 | 1 | 1 | 1 | 1 | 1 | 0.987 | 0.929 | 1 | 0.015 |
| SE | 105742 | ENSG00000111142 | METAP2 | chr12 | + | 95494956 | 95495138 | 95494055 | 95494217 | 95496003 | 95496098 | 512 | 390 | 465 | 464 | 780 | 0 | 0 | 1 | 1 | 0 | 120 | 337 | 465 | 251 | 34 | 0 | 8 | 1 | 4 | 0 | 298 | 149 | 0.545939044 | 1 | 1 | 1 | 0.996 | 0.996 | 1 | 1 | 0.955 | 0.996 | 0.969 | 1 | 0.014 |
| SE | 24500 | ENSG00000146067 | FAM193B | chr5 | - | 177531281 | 177531521 | 177524184 | 177525205 | 177532442 | 177532533 | 46 | 35 | 45 | 57 | 41 | 7 | 4 | 6 | 8 | 8 | 40 | 44 | 49 | 69 | 47 | 13 | 28 | 8 | 8 | 0 | 298 | 149 | 0.546041037 | 1 | 0.767 | 0.814 | 0.789 | 0.781 | 0.719 | 0.606 | 0.44 | 0.754 | 0.812 | 1 | 0.052 |
| SE | 77386 | ENSG00000089820 | ARHGAP4 | chrX | - | 153910921 | 153910999 | 153910699 | 153910834 | 153911128 | 153911189 | 323 | 116 | 286 | 240 | 266 | 0 | 0 | 0 | 1 | 1 | 321 | 328 | 240 | 327 | 585 | 0 | 0 | 4 | 15 | 0 | 227 | 149 | 0.546084542 | 1 | 1 | 1 | 1 | 0.994 | 0.994 | 1 | 1 | 0.975 | 0.935 | 1 | 0.016 |
| SE | 79291 | ENSG00000001631 | KRIT1 | chr7 | - | 92234807 | 92234923 | 92226525 | 92226682 | 92235402 | 92235646 | 70 | 58 | 42 | 39 | 95 | 0 | 0 | 0 | 0 | 0 | 8 | 10 | 34 | 21 | 17 | 0 | 4 | 0 | 0 | 0 | 265 | 149 | 0.546177847 | 1 | 1 | 1 | 1 | 1 | 1 | 1 | 0.584 | 1 | 1 | 1 | 0.083 |
| SE | 2786 | ENSG00000011304 | PTBP1 | chr19 | + | 805491 | 805569 | 805012 | 805187 | 806407 | 806556 | 83 | 34 | 80 | 96 | 116 | 33 | 24 | 33 | 61 | 52 | 98 | 71 | 33 | 122 | 141 | 39 | 23 | 50 | 29 | 44 | 227 | 149 | 0.546196201 | 1 | 0.623 | 0.482 | 0.614 | 0.508 | 0.594 | 0.623 | 0.67 | 0.302 | 0.734 | 0.678 | -0.037 |
| SE | 49699 | ENSG00000188732 | FAM221A | chr7 | + | 23688227 | 23688323 | 23684498 | 23684672 | 23689268 | 23689459 | 0 | 1 | 0 | 0 | 0 | 9 | 5 | 10 | 5 | 10 | 0 | 6 | 0 | 0 | 0 | 12 | 13 | 2 | 13 | 9 | 245 | 149 | 0.546209364 | 1 | 0 | 0.108 | 0 | 0 | 0 | 0 | 0.219 | 0 | 0 | 0 | -0.022 |
| SE | 62451 | ENSG00000120265 | PCMT1 | chr6 | + | 149772047 | 149772247 | 149771161 | 149771266 | 149773137 | 149773169 | 17 | 10 | 7 | 3 | 11 | 249 | 158 | 191 | 269 | 273 | 22 | 7 | 23 | 13 | 5 | 505 | 264 | 165 | 331 | 241 | 298 | 149 | 0.546297869 | 1 | 0.033 | 0.031 | 0.018 | 0.006 | 0.02 | 0.021 | 0.013 | 0.065 | 0.019 | 0.01 | -0.004 |
| SE | 66938 | ENSG00000285006 | NA | chr14 | - | 73459033 | 73459305 | 73409975 | 73410068 | 73463474 | 73463642 | 10 | 4 | 5 | 7 | 4 | 8 | 4 | 12 | 0 | 1 | 7 | 3 | 10 | 1 | 16 | 1 | 13 | 0 | 6 | 1 | 298 | 149 | 0.54631687 | 1 | 0.385 | 0.333 | 0.172 | 1 | 0.667 | 0.778 | 0.103 | 1 | 0.077 | 0.889 | -0.058 |
| SE | 21409 | ENSG00000197879 | MYO1C | chr17 | - | 1474609 | 1474690 | 1472122 | 1472228 | 1474811 | 1474858 | 20 | 7 | 20 | 33 | 27 | 1 | 0 | 0 | 0 | 0 | 6 | 44 | 33 | 25 | 175 | 0 | 7 | 0 | 0 | 0 | 230 | 149 | 0.546320949 | 1 | 0.928 | 1 | 1 | 1 | 1 | 1 | 0.803 | 1 | 1 | 1 | 0.025 |
| SE | 72082 | ENSG00000180228 | PRKRA | chr2 | - | 178441609 | 178441704 | 178436144 | 178436319 | 178443266 | 178443384 | 131 | 106 | 124 | 147 | 246 | 0 | 0 | 1 | 5 | 7 | 188 | 115 | 95 | 87 | 71 | 4 | 3 | 2 | 4 | 0 | 244 | 149 | 0.546553424 | 1 | 1 | 1 | 0.987 | 0.947 | 0.955 | 0.966 | 0.959 | 0.967 | 0.93 | 1 | 0.013 |
| SE | 81934 | ENSG00000182165 | TP53TG1 | chr7 | - | 87326262 | 87326449 | 87325136 | 87325444 | 87341522 | 87341684 | 2 | 0 | 5 | 1 | 1 | 31 | 20 | 25 | 28 | 19 | 0 | 0 | 0 | 0 | 0 | 20 | 19 | 10 | 19 | 11 | 298 | 149 | 0.546669303 | 1 | 0.031 | 0 | 0.091 | 0.018 | 0.026 | 0 | 0 | 0 | 0 | 0 | 0.033 |
| SE | 16148 | ENSG00000140400 | MAN2C1 | chr15 | - | 75363209 | 75363290 | 75362641 | 75362748 | 75364487 | 75364665 | 4 | 1 | 10 | 2 | 2 | 3 | 0 | 4 | 3 | 0 | 5 | 9 | 0 | 5 | 9 | 0 | 12 | 9 | 0 | 8 | 230 | 149 | 0.54688113 | 1 | 0.463 | 1 | 0.618 | 0.302 | 1 | 1 | 0.327 | 0 | 1 | 0.422 | 0.127 |
| SE | 96009 | ENSG00000256591 | NA | chr11 | + | 61445940 | 61446231 | 61430149 | 61430182 | 61481781 | 61481918 | 34 | 15 | 25 | 24 | 23 | 0 | 0 | 0 | 0 | 0 | 18 | 35 | 15 | 25 | 28 | 0 | 0 | 0 | 0 | 0 | 298 | 149 | 0.546989452 | 1 | 1 | 1 | 1 | 0.706 | 1 | 1 | 1 | 1 | 1 | 1 | -0.059 |
| SE | 16524 | ENSG00000140367 | UBE2Q2 | chr15 | + | 75877960 | 75878021 | 75876186 | 75876271 | 75883365 | 75883424 | 100 | 62 | 57 | 88 | 80 | 0 | 4 | 1 | 0 | 2 | 47 | 46 | 24 | 25 | 16 | 0 | 0 | 0 | 0 | 0 | 210 | 149 | 0.547007478 | 1 | 1 | 0.917 | 0.976 | 1 | 0.966 | 1 | 1 | 1 | 1 | 1 | -0.028 |
| SE | 53496 | ENSG00000139174 | PRICKLE1 | chr12 | - | 42468625 | 42468829 | 42466193 | 42466380 | 42469449 | 42469587 | 17 | 8 | 12 | 7 | 22 | 0 | 1 | 0 | 1 | 2 | 20 | 38 | 2 | 4 | 75 | 0 | 6 | 0 | 0 | 0 | 298 | 149 | 0.547536237 | 1 | 1 | 0.8 | 1 | 0.778 | 0.846 | 1 | 0.76 | 1 | 1 | 1 | -0.067 |
| SE | 59491 | ENSG00000148660 | CAMK2G | chr10 | - | 73819531 | 73819645 | 73817478 | 73817554 | 73821681 | 73821730 | 14 | 8 | 13 | 6 | 22 | 112 | 92 | 149 | 132 | 158 | 10 | 20 | 16 | 15 | 18 | 223 | 146 | 146 | 171 | 248 | 263 | 149 | 0.547563863 | 1 | 0.066 | 0.047 | 0.047 | 0.025 | 0.073 | 0.025 | 0.072 | 0.058 | 0.047 | 0.039 | 0.003 |
| SE | 75877 | ENSG00000164430 | CGAS | chr6 | - | 73445527 | 73445747 | 73440208 | 73440445 | 73451524 | 73452181 | 207 | 156 | 126 | 182 | 179 | 2 | 0 | 0 | 5 | 2 | 629 | 220 | 172 | 341 | 326 | 0 | 0 | 5 | 5 | 7 | 298 | 149 | 0.547619656 | 1 | 0.981 | 1 | 1 | 0.948 | 0.978 | 1 | 1 | 0.945 | 0.972 | 0.959 | 0.006 |
| SE | 51486 | ENSG00000160087 | UBE2J2 | chr1 | - | 1264782 | 1264830 | 1263345 | 1263386 | 1267861 | 1267992 | 4 | 0 | 1 | 3 | 1 | 65 | 46 | 52 | 56 | 69 | 0 | 4 | 1 | 1 | 0 | 106 | 58 | 20 | 78 | 196 | 197 | 149 | 0.547639353 | 1 | 0.044 | 0 | 0.014 | 0.039 | 0.011 | 0 | 0.05 | 0.036 | 0.01 | 0 | 0.002 |
| SE | 98601 | ENSG00000136527 | TRA2B | chr3 | - | 185923795 | 185923984 | 185922010 | 185922126 | 185925463 | 185925626 | 1638 | 1189 | 1497 | 1790 | 2136 | 2 | 1 | 4 | 6 | 5 | 2085 | 2083 | 1608 | 2455 | 1102 | 18 | 20 | 0 | 13 | 0 | 298 | 149 | 0.547809653 | 1 | 0.998 | 0.998 | 0.995 | 0.993 | 0.995 | 0.983 | 0.981 | 1 | 0.99 | 1 | 0.005 |
| SE | 99540 | ENSG00000188690 | UROS | chr10 | - | 125815033 | 125815130 | 125812213 | 125812288 | 125816436 | 125816525 | 31 | 25 | 36 | 26 | 49 | 0 | 0 | 1 | 1 | 4 | 90 | 32 | 12 | 26 | 62 | 0 | 2 | 0 | 0 | 0 | 246 | 149 | 0.548020384 | 1 | 1 | 1 | 0.956 | 0.94 | 0.881 | 1 | 0.906 | 1 | 1 | 1 | -0.026 |
| SE | 22366 | ENSG00000140455 | USP3 | chr15 | + | 63532646 | 63532707 | 63504510 | 63504830 | 63537024 | 63537056 | 179 | 107 | 116 | 113 | 107 | 2 | 0 | 2 | 0 | 0 | 207 | 173 | 173 | 135 | 164 | 0 | 3 | 0 | 3 | 7 | 210 | 149 | 0.5481112 | 1 | 0.984 | 1 | 0.976 | 1 | 1 | 1 | 0.976 | 1 | 0.97 | 0.943 | 0.014 |
| SE | 55349 | ENSG00000204469 | PRRC2A | chr6 | + | 31636732 | 31636945 | 31636509 | 31636608 | 31637041 | 31637136 | 168 | 44 | 113 | 94 | 47 | 1 | 0 | 2 | 4 | 0 | 71 | 133 | 91 | 101 | 193 | 0 | 0 | 2 | 2 | 0 | 298 | 149 | 0.548187407 | 1 | 0.988 | 1 | 0.966 | 0.922 | 1 | 1 | 1 | 0.958 | 0.962 | 1 | -0.009 |
| SE | 105157 | ENSG00000170946 | DNAJC24 | chr11 | + | 31429482 | 31429606 | 31426286 | 31426355 | 31430270 | 31430703 | 12 | 7 | 16 | 24 | 30 | 42 | 46 | 52 | 42 | 75 | 0 | 5 | 9 | 6 | 0 | 19 | 17 | 4 | 15 | 1 | 273 | 149 | 0.548233761 | 1 | 0.135 | 0.077 | 0.144 | 0.238 | 0.179 | 0 | 0.138 | 0.551 | 0.179 | 0 | -0.019 |
| SE | 80828 | ENSG00000151948 | GLT1D1 | chr12 | + | 128926358 | 128926454 | 128888638 | 128888744 | 128927104 | 128927154 | 65 | 62 | 28 | 53 | 64 | 5 | 0 | 0 | 0 | 0 | 303 | 108 | 82 | 215 | 189 | 0 | 0 | 0 | 0 | 12 | 245 | 149 | 0.548260501 | 1 | 0.888 | 1 | 1 | 1 | 1 | 1 | 1 | 1 | 1 | 0.905 | -0.003 |
| SE | 21215 | ENSG00000150403 | TMCO3 | chr13 | + | 113510237 | 113510424 | 113501751 | 113501880 | 113520615 | 113520661 | 33 | 29 | 39 | 37 | 21 | 1 | 0 | 1 | 3 | 0 | 83 | 43 | 32 | 35 | 100 | 0 | 5 | 2 | 0 | 0 | 298 | 149 | 0.548271196 | 1 | 0.943 | 1 | 0.951 | 0.86 | 1 | 1 | 0.811 | 0.889 | 1 | 1 | 0.011 |
| SE | 5028 | ENSG00000101343 | CRNKL1 | chr20 | - | 20039608 | 20039848 | 20038348 | 20038450 | 20040685 | 20040766 | 236 | 205 | 254 | 265 | 371 | 3 | 3 | 0 | 4 | 8 | 38 | 141 | 234 | 79 | 27 | 0 | 5 | 2 | 5 | 0 | 298 | 149 | 0.54837021 | 1 | 0.975 | 0.972 | 1 | 0.971 | 0.959 | 1 | 0.934 | 0.983 | 0.888 | 1 | 0.014 |
| SE | 61390 | ENSG00000136891 | TEX10 | chr9 | - | 100308499 | 100308681 | 100303631 | 100303842 | 100310298 | 100310379 | 47 | 14 | 42 | 14 | 22 | 2 | 1 | 2 | 4 | 2 | 7 | 24 | 19 | 13 | 0 | 0 | 1 | 6 | 0 | 5 | 298 | 149 | 0.548414324 | 1 | 0.922 | 0.875 | 0.913 | 0.636 | 0.846 | 1 | 0.923 | 0.613 | 1 | 0 | 0.131 |
| SE | 12636 | ENSG00000090857 | PDPR | chr16 | + | 70143509 | 70143658 | 70142470 | 70142686 | 70144420 | 70144533 | 124 | 64 | 124 | 110 | 193 | 4 | 7 | 3 | 0 | 6 | 203 | 148 | 95 | 82 | 388 | 0 | 3 | 0 | 0 | 21 | 298 | 149 | 0.548423503 | 1 | 0.939 | 0.821 | 0.954 | 1 | 0.941 | 1 | 0.961 | 1 | 1 | 0.902 | -0.042 |
| SE | 59127 | ENSG00000047410 | TPR | chr1 | - | 186353687 | 186353850 | 186351975 | 186352110 | 186355409 | 186355558 | 669 | 741 | 555 | 652 | 847 | 3 | 2 | 9 | 0 | 1 | 83 | 134 | 78 | 31 | 23 | 0 | 0 | 6 | 0 | 0 | 298 | 149 | 0.548445059 | 1 | 0.991 | 0.995 | 0.969 | 1 | 0.998 | 1 | 1 | 1 | 0.721 | 1 | 0.046 |
| SE | 90803 | ENSG00000173064 | HECTD4 | chr12 | - | 112169502 | 112169658 | 112167813 | 112167917 | 112170332 | 112170452 | 58 | 29 | 34 | 40 | 45 | 0 | 1 | 0 | 2 | 0 | 39 | 59 | 43 | 35 | 139 | 0 | 1 | 0 | 0 | 5 | 298 | 149 | 0.548846475 | 1 | 1 | 0.935 | 1 | 0.909 | 1 | 1 | 0.967 | 1 | 1 | 0.933 | -0.011 |
| SE | 62623 | ENSG00000033178 | UBA6 | chr4 | - | 67641150 | 67641228 | 67638942 | 67639124 | 67644697 | 67644774 | 183 | 147 | 206 | 227 | 341 | 4 | 0 | 0 | 2 | 0 | 34 | 87 | 62 | 21 | 24 | 0 | 6 | 0 | 0 | 0 | 227 | 149 | 0.548864071 | 1 | 0.968 | 1 | 1 | 0.987 | 1 | 1 | 0.905 | 1 | 1 | 1 | 0.01 |
| SE | 52033 | ENSG00000135250 | SRPK2 | chr7 | - | 105297420 | 105297533 | 105203627 | 105203785 | 105388647 | 105388702 | 0 | 4 | 1 | 1 | 0 | 17 | 17 | 9 | 13 | 10 | 0 | 0 | 0 | 0 | 0 | 15 | 15 | 13 | 9 | 17 | 262 | 149 | 0.548879634 | 1 | 0 | 0.118 | 0.059 | 0.042 | 0 | 0 | 0 | 0 | 0 | 0 | 0.044 |
| SE | 73043 | ENSG00000123607 | TTC21B | chr2 | - | 165945523 | 165945690 | 165943218 | 165943341 | 165949393 | 165949504 | 329 | 217 | 169 | 281 | 332 | 21 | 23 | 6 | 11 | 8 | 73 | 157 | 121 | 51 | 11 | 14 | 0 | 8 | 0 | 0 | 298 | 149 | 0.548939129 | 1 | 0.887 | 0.825 | 0.934 | 0.927 | 0.954 | 0.723 | 1 | 0.883 | 1 | 1 | -0.016 |
| SE | 72723 | ENSG00000172765 | TMCC1 | chr3 | - | 129867009 | 129867098 | 129832757 | 129832826 | 129880308 | 129880559 | 6 | 2 | 3 | 3 | 4 | 40 | 18 | 46 | 30 | 33 | 0 | 0 | 5 | 8 | 0 | 57 | 22 | 24 | 39 | 5 | 238 | 149 | 0.549000381 | 1 | 0.086 | 0.065 | 0.039 | 0.059 | 0.071 | 0 | 0 | 0.115 | 0.114 | 0 | 0.018 |
| SE | 16737 | ENSG00000075131 | TIPIN | chr15 | - | 66349059 | 66349123 | 66341149 | 66341356 | 66349314 | 66349437 | 31 | 32 | 41 | 46 | 53 | 0 | 2 | 3 | 0 | 1 | 27 | 36 | 59 | 27 | 9 | 0 | 0 | 0 | 0 | 0 | 213 | 149 | 0.549010128 | 1 | 1 | 0.918 | 0.905 | 1 | 0.974 | 1 | 1 | 1 | 1 | 1 | -0.041 |
| SE | 38089 | ENSG00000160584 | SIK3 | chr11 | - | 116875373 | 116875451 | 116875157 | 116875267 | 116876252 | 116876363 | 93 | 45 | 68 | 41 | 104 | 0 | 0 | 0 | 0 | 0 | 142 | 103 | 77 | 82 | 118 | 8 | 0 | 0 | 0 | 0 | 227 | 149 | 0.54906592 | 1 | 1 | 1 | 1 | 1 | 1 | 0.921 | 1 | 1 | 1 | 1 | 0.016 |
| SE | 95186 | ENSG00000213930 | GALT | chr9 | + | 34648047 | 34648171 | 34647088 | 34647258 | 34648333 | 34648375 | 25 | 29 | 29 | 18 | 9 | 0 | 0 | 0 | 0 | 0 | 22 | 6 | 8 | 17 | 22 | 0 | 0 | 0 | 0 | 5 | 273 | 149 | 0.549110445 | 1 | 1 | 1 | 1 | 1 | 1 | 1 | 1 | 1 | 1 | 0.706 | 0.059 |
| SE | 19219 | ENSG00000100784 | RPS6KA5 | chr14 | - | 90919279 | 90919466 | 90906148 | 90906299 | 90920205 | 90920309 | 6 | 4 | 5 | 12 | 14 | 78 | 44 | 53 | 63 | 98 | 8 | 5 | 0 | 0 | 0 | 13 | 21 | 40 | 17 | 8 | 298 | 149 | 0.549245341 | 1 | 0.037 | 0.043 | 0.045 | 0.087 | 0.067 | 0.235 | 0.106 | 0 | 0 | 0 | -0.012 |
| SE | 12099 | ENSG00000125149 | C16orf70 | chr16 | + | 67132445 | 67132525 | 67131285 | 67131329 | 67132816 | 67132911 | 102 | 38 | 50 | 64 | 97 | 0 | 0 | 0 | 1 | 0 | 155 | 59 | 69 | 145 | 101 | 5 | 0 | 0 | 0 | 3 | 229 | 149 | 0.549250736 | 1 | 1 | 1 | 1 | 0.977 | 1 | 0.953 | 1 | 1 | 1 | 0.956 | 0.014 |
| SE | 86133 | ENSG00000197713 | RPE | chr2 | + | 210003437 | 210003468 | 210002627 | 210002783 | 210009656 | 210009736 | 11 | 3 | 5 | 10 | 0 | 65 | 39 | 86 | 52 | 17 | 0 | 6 | 3 | 10 | 0 | 48 | 52 | 29 | 50 | 10 | 180 | 149 | 0.549302118 | 1 | 0.123 | 0.06 | 0.046 | 0.137 | 0 | 0 | 0.087 | 0.079 | 0.142 | 0 | 0.012 |
| SE | 19188 | ENSG00000140105 | WARS1 | chr14 | - | 100374135 | 100374244 | 100369086 | 100369258 | 100376259 | 100376334 | 0 | 0 | 30 | 1 | 33 | 75 | 29 | 52 | 42 | 11 | 0 | 9 | 0 | 0 | 68 | 65 | 80 | 75 | 78 | 123 | 258 | 149 | 0.549304753 | 1 | 0 | 0 | 0.25 | 0.014 | 0.634 | 0 | 0.061 | 0 | 0 | 0.242 | 0.119 |
| SE | 69058 | ENSG00000081059 | TCF7 | chr5 | + | 134140795 | 134140888 | 134138950 | 134139038 | 134141020 | 134141350 | 52 | 49 | 83 | 73 | 122 | 2 | 1 | 3 | 2 | 6 | 44 | 7 | 29 | 20 | 51 | 0 | 7 | 0 | 2 | 0 | 242 | 149 | 0.549460676 | 1 | 0.941 | 0.968 | 0.945 | 0.957 | 0.926 | 1 | 0.381 | 1 | 0.86 | 1 | 0.099 |
| SE | 32283 | ENSG00000163964 | PIGX | chr3 | + | 196716857 | 196716921 | 196654526 | 196654647 | 196722414 | 196722556 | 36 | 23 | 17 | 13 | 21 | 0 | 0 | 0 | 0 | 0 | 25 | 4 | 9 | 19 | 8 | 0 | 0 | 0 | 0 | 4 | 213 | 149 | 0.549658111 | 1 | 1 | 1 | 1 | 1 | 1 | 1 | 1 | 1 | 1 | 0.583 | 0.083 |
| SE | 40295 | ENSG00000089682 | RBM41 | chrX | - | 107113396 | 107113468 | 107088734 | 107088839 | 107115351 | 107115556 | 31 | 53 | 38 | 49 | 76 | 11 | 15 | 16 | 27 | 29 | 0 | 15 | 27 | 27 | 0 | 9 | 3 | 3 | 1 | 3 | 221 | 149 | 0.549703168 | 1 | 0.655 | 0.704 | 0.616 | 0.55 | 0.639 | 0 | 0.771 | 0.859 | 0.948 | 0 | 0.117 |
| SE | 5915 | ENSG00000124224 | PPP4R1L | chr20 | - | 58271362 | 58271505 | 58251756 | 58251864 | 58272768 | 58272825 | 35 | 32 | 65 | 28 | 75 | 0 | 1 | 2 | 0 | 2 | 45 | 64 | 67 | 54 | 30 | 0 | 0 | 0 | 0 | 0 | 292 | 149 | 0.549861823 | 1 | 1 | 0.942 | 0.943 | 1 | 0.95 | 1 | 1 | 1 | 1 | 1 | -0.033 |
| SE | 8583 | ENSG00000185359 | HGS | chr17 | + | 81684970 | 81685069 | 81683948 | 81684103 | 81685604 | 81685689 | 2 | 4 | 0 | 0 | 2 | 51 | 26 | 50 | 48 | 21 | 0 | 3 | 2 | 0 | 0 | 49 | 68 | 47 | 59 | 78 | 248 | 149 | 0.54995482 | 1 | 0.023 | 0.085 | 0 | 0 | 0.054 | 0 | 0.026 | 0.025 | 0 | 0 | 0.022 |
| SE | 28894 | ENSG00000164597 | COG5 | chr7 | - | 107281299 | 107281399 | 107258272 | 107258383 | 107283570 | 107283732 | 106 | 115 | 148 | 75 | 82 | 0 | 0 | 1 | 2 | 0 | 57 | 100 | 44 | 89 | 19 | 0 | 3 | 3 | 2 | 0 | 249 | 149 | 0.550026983 | 1 | 1 | 1 | 0.989 | 0.957 | 1 | 1 | 0.952 | 0.898 | 0.964 | 1 | 0.026 |
| SE | 13665 | ENSG00000169189 | NSMCE1 | chr16 | - | 27241798 | 27241889 | 27235215 | 27235299 | 27257434 | 27257581 | 4 | 0 | 7 | 2 | 0 | 104 | 64 | 85 | 104 | 31 | 8 | 2 | 0 | 4 | 0 | 188 | 99 | 46 | 91 | 173 | 240 | 149 | 0.550131674 | 1 | 0.023 | 0 | 0.049 | 0.012 | 0 | 0.026 | 0.012 | 0 | 0.027 | 0 | 0.004 |
| SE | 96938 | ENSG00000089159 | PXN | chr12 | - | 120224642 | 120224727 | 120224150 | 120224377 | 120226352 | 120226795 | 5 | 0 | 4 | 4 | 0 | 7 | 8 | 8 | 10 | 11 | 9 | 3 | 5 | 11 | 12 | 62 | 34 | 29 | 5 | 25 | 234 | 149 | 0.550189097 | 1 | 0.313 | 0 | 0.241 | 0.203 | 0 | 0.085 | 0.053 | 0.099 | 0.583 | 0.234 | -0.059 |
| SE | 21048 | ENSG00000135506 | OS9 | chr12 | + | 57720098 | 57720263 | 57718992 | 57719182 | 57720405 | 57720518 | 751 | 433 | 513 | 625 | 836 | 297 | 252 | 253 | 278 | 438 | 1801 | 913 | 556 | 1061 | 1602 | 798 | 486 | 375 | 558 | 671 | 298 | 149 | 0.550193013 | 1 | 0.558 | 0.462 | 0.503 | 0.529 | 0.488 | 0.53 | 0.484 | 0.426 | 0.487 | 0.544 | 0.014 |
| SE | 11982 | ENSG00000156860 | FBRS | chr16 | + | 30661179 | 30661215 | 30660262 | 30660442 | 30662419 | 30662468 | 22 | 10 | 20 | 26 | 17 | 2 | 0 | 4 | 12 | 19 | 49 | 20 | 26 | 53 | 73 | 15 | 29 | 12 | 28 | 1 | 185 | 149 | 0.550200012 | 1 | 0.899 | 1 | 0.801 | 0.636 | 0.419 | 0.725 | 0.357 | 0.636 | 0.604 | 0.983 | 0.09 |
| SE | 12055 | ENSG00000159720 | ATP6V0D1 | chr16 | - | 67444527 | 67444706 | 67443098 | 67443178 | 67480956 | 67481148 | 321 | 168 | 143 | 233 | 278 | 0 | 0 | 1 | 0 | 2 | 375 | 318 | 224 | 366 | 631 | 0 | 0 | 0 | 0 | 17 | 298 | 149 | 0.5502523 | 1 | 1 | 1 | 0.986 | 1 | 0.986 | 1 | 1 | 1 | 1 | 0.949 | 0.005 |
| SE | 5794 | ENSG00000197586 | ENTPD6 | chr20 | + | 25196153 | 25196254 | 25195791 | 25195867 | 25206521 | 25206590 | 1 | 0 | 1 | 5 | 1 | 38 | 8 | 17 | 30 | 14 | 7 | 0 | 4 | 0 | 0 | 50 | 20 | 10 | 19 | 30 | 250 | 149 | 0.550264233 | 1 | 0.015 | 0 | 0.034 | 0.09 | 0.041 | 0.077 | 0 | 0.193 | 0 | 0 | -0.018 |
| SE | 17231 | ENSG00000197555 | SIPA1L1 | chr14 | + | 71418129 | 71418196 | 71413947 | 71414014 | 71512742 | 71512845 | 24 | 7 | 12 | 13 | 28 | 34 | 26 | 24 | 26 | 28 | 0 | 13 | 13 | 27 | 9 | 32 | 28 | 36 | 15 | 7 | 216 | 149 | 0.550293406 | 1 | 0.327 | 0.157 | 0.256 | 0.256 | 0.408 | 0 | 0.243 | 0.199 | 0.554 | 0.47 | -0.012 |
| SE | 30423 | ENSG00000103502 | CDIPT | chr16 | - | 29862585 | 29862635 | 29861105 | 29861259 | 29862814 | 29863276 | 7 | 3 | 16 | 9 | 14 | 0 | 1 | 1 | 1 | 2 | 20 | 6 | 11 | 18 | 25 | 0 | 5 | 0 | 0 | 0 | 199 | 149 | 0.550416355 | 1 | 1 | 0.692 | 0.923 | 0.871 | 0.84 | 1 | 0.473 | 1 | 1 | 1 | -0.029 |
| SE | 67860 | ENSG00000018280 | SLC11A1 | chr2 | + | 218385146 | 218385266 | 218382272 | 218382375 | 218386634 | 218386741 | 62 | 38 | 20 | 55 | 21 | 2 | 2 | 0 | 3 | 0 | 122 | 150 | 126 | 133 | 154 | 0 | 0 | 0 | 0 | 3 | 269 | 149 | 0.550456664 | 1 | 0.945 | 0.913 | 1 | 0.91 | 1 | 1 | 1 | 1 | 1 | 0.966 | -0.04 |
| SE | 73259 | ENSG00000114956 | DGUOK | chr2 | + | 73950626 | 73950732 | 73946718 | 73946906 | 73958145 | 73958245 | 13 | 4 | 14 | 18 | 29 | 73 | 68 | 73 | 65 | 125 | 13 | 28 | 17 | 12 | 4 | 89 | 99 | 53 | 64 | 136 | 255 | 149 | 0.550491449 | 1 | 0.094 | 0.033 | 0.101 | 0.139 | 0.119 | 0.079 | 0.142 | 0.158 | 0.099 | 0.017 | -0.002 |
| SE | 80513 | ENSG00000179818 | PCBP1-AS1 | chr2 | - | 70032132 | 70032255 | 69996685 | 69996888 | 70051202 | 70051305 | 18 | 17 | 23 | 10 | 14 | 5 | 3 | 9 | 11 | 18 | 0 | 55 | 8 | 24 | 11 | 9 | 13 | 6 | 8 | 7 | 272 | 149 | 0.550492506 | 1 | 0.664 | 0.756 | 0.583 | 0.332 | 0.299 | 0 | 0.699 | 0.422 | 0.622 | 0.463 | 0.086 |
| SE | 54554 | ENSG00000179912 | R3HDM2 | chr12 | - | 57289008 | 57289040 | 57283823 | 57284056 | 57292571 | 57292649 | 105 | 81 | 98 | 110 | 185 | 1 | 3 | 7 | 4 | 18 | 138 | 113 | 66 | 123 | 160 | 8 | 4 | 3 | 6 | 18 | 181 | 149 | 0.550651926 | 1 | 0.989 | 0.957 | 0.92 | 0.958 | 0.894 | 0.934 | 0.959 | 0.948 | 0.944 | 0.88 | 0.011 |
| SE | 86731 | ENSG00000198814 | GK | chrX | + | 30727465 | 30727552 | 30724100 | 30724181 | 30728731 | 30729249 | 23 | 35 | 21 | 36 | 40 | 31 | 17 | 16 | 22 | 48 | 0 | 29 | 17 | 8 | 7 | 11 | 7 | 18 | 21 | 1 | 236 | 149 | 0.550702666 | 1 | 0.319 | 0.565 | 0.453 | 0.508 | 0.345 | 0 | 0.723 | 0.374 | 0.194 | 0.815 | 0.017 |
| SE | 37604 | ENSG00000082074 | FYB1 | chr5 | - | 39211057 | 39211091 | 39202893 | 39202987 | 39219442 | 39219563 | 1 | 2 | 7 | 2 | 14 | 145 | 111 | 72 | 140 | 83 | 4 | 9 | 5 | 10 | 0 | 259 | 109 | 114 | 193 | 56 | 183 | 149 | 0.550791881 | 1 | 0.006 | 0.014 | 0.073 | 0.011 | 0.121 | 0.012 | 0.063 | 0.034 | 0.04 | 0 | 0.015 |
| SE | 12590 | ENSG00000090861 | AARS1 | chr16 | - | 70258052 | 70258217 | 70255727 | 70255836 | 70258979 | 70259186 | 34 | 22 | 37 | 33 | 71 | 9 | 1 | 1 | 12 | 5 | 13 | 25 | 22 | 22 | 57 | 0 | 0 | 0 | 7 | 17 | 298 | 149 | 0.55086404 | 1 | 0.654 | 0.917 | 0.949 | 0.579 | 0.877 | 1 | 1 | 1 | 0.611 | 0.626 | -0.052 |
| SE | 88459 | ENSG00000152601 | MBNL1 | chr3 | + | 152299404 | 152300367 | 152268039 | 152269092 | 152445281 | 152445539 | 677 | 505 | 765 | 789 | 1094 | 5 | 2 | 1 | 6 | 0 | 572 | 634 | 553 | 426 | 664 | 10 | 0 | 5 | 7 | 0 | 298 | 149 | 0.550875377 | 1 | 0.985 | 0.992 | 0.997 | 0.985 | 1 | 0.966 | 1 | 0.982 | 0.968 | 1 | 0.009 |
| SE | 78926 | ENSG00000184903 | IMMP2L | chr7 | - | 111532517 | 111532585 | 111521312 | 111521449 | 111561850 | 111561923 | 3 | 6 | 11 | 5 | 5 | 37 | 11 | 30 | 37 | 17 | 0 | 4 | 0 | 0 | 13 | 8 | 15 | 3 | 3 | 9 | 217 | 149 | 0.55107165 | 1 | 0.053 | 0.272 | 0.201 | 0.085 | 0.168 | 0 | 0.155 | 0 | 0.748 | 0 | -0.025 |
| SE | 6717 | ENSG00000104884 | ERCC2 | chr19 | - | 45355664 | 45355728 | 45354729 | 45354851 | 45357269 | 45357371 | 19 | 14 | 19 | 21 | 6 | 0 | 0 | 0 | 0 | 0 | 30 | 52 | 11 | 11 | 17 | 0 | 2 | 0 | 0 | 4 | 213 | 149 | 0.551089866 | 1 | 1 | 1 | 1 | 1 | 1 | 1 | 0.948 | 1 | 1 | 0.748 | 0.061 |
| SE | 59445 | ENSG00000145246 | ATP10D | chr4 | + | 47554725 | 47554914 | 47546623 | 47546862 | 47557663 | 47558273 | 96 | 66 | 88 | 114 | 142 | 2 | 0 | 2 | 2 | 2 | 57 | 92 | 31 | 91 | 17 | 0 | 0 | 0 | 0 | 0 | 298 | 149 | 0.551122309 | 1 | 0.96 | 1 | 0.957 | 0.966 | 0.973 | 1 | 1 | 1 | 1 | 1 | -0.029 |
| SE | 58593 | ENSG00000186854 | TRABD2A | chr2 | - | 84841860 | 84842007 | 84832054 | 84832145 | 84870224 | 84870785 | 71 | 75 | 177 | 121 | 110 | 2 | 0 | 3 | 1 | 1 | 1 | 28 | 28 | 32 | 26 | 0 | 0 | 0 | 0 | 0 | 296 | 149 | 0.551237111 | 1 | 0.947 | 1 | 0.967 | 0.984 | 0.982 | 1 | 1 | 1 | 1 | 1 | -0.024 |
| SE | 98885 | ENSG00000159352 | PSMD4 | chr1 | + | 151265165 | 151265234 | 151264831 | 151264918 | 151266003 | 151266112 | 142 | 41 | 71 | 100 | 100 | 0 | 1 | 0 | 0 | 0 | 143 | 106 | 81 | 107 | 193 | 8 | 0 | 0 | 0 | 3 | 218 | 149 | 0.551323852 | 1 | 1 | 0.966 | 1 | 1 | 1 | 0.924 | 1 | 1 | 1 | 0.978 | 0.013 |
| SE | 61894 | ENSG00000076242 | MLH1 | chr3 | + | 37025636 | 37026007 | 37012010 | 37012099 | 37028783 | 37028847 | 64 | 33 | 67 | 40 | 73 | 0 | 0 | 3 | 2 | 1 | 71 | 46 | 50 | 53 | 23 | 0 | 0 | 2 | 0 | 0 | 298 | 149 | 0.551397213 | 1 | 1 | 1 | 0.918 | 0.909 | 0.973 | 1 | 1 | 0.926 | 1 | 1 | -0.025 |
| SE | 45771 | ENSG00000285053 | TBCE | chr1 | + | 235401502 | 235401587 | 235380018 | 235380149 | 235414432 | 235414618 | 32 | 23 | 16 | 53 | 36 | 7 | 9 | 5 | 13 | 10 | 34 | 39 | 18 | 29 | 24 | 2 | 19 | 16 | 6 | 0 | 234 | 149 | 0.551404026 | 1 | 0.744 | 0.619 | 0.671 | 0.722 | 0.696 | 0.915 | 0.567 | 0.417 | 0.755 | 1 | -0.04 |
| SE | 86410 | ENSG00000131381 | RBSN | chr3 | - | 15091338 | 15091493 | 15090398 | 15090539 | 15095972 | 15096288 | 2 | 3 | 11 | 11 | 8 | 32 | 20 | 55 | 54 | 39 | 20 | 12 | 4 | 12 | 2 | 73 | 62 | 51 | 97 | 68 | 298 | 149 | 0.551541792 | 1 | 0.03 | 0.07 | 0.091 | 0.092 | 0.093 | 0.12 | 0.088 | 0.038 | 0.058 | 0.014 | 0.012 |
| SE | 33514 | ENSG00000175455 | CCDC14 | chr3 | - | 123931101 | 123931234 | 123913421 | 123915718 | 123933672 | 123933755 | 74 | 63 | 56 | 81 | 88 | 2 | 3 | 0 | 0 | 0 | 27 | 19 | 19 | 2 | 12 | 0 | 4 | 0 | 0 | 0 | 282 | 149 | 0.551588241 | 1 | 0.951 | 0.917 | 1 | 1 | 1 | 1 | 0.715 | 1 | 1 | 1 | 0.031 |
| SE | 20503 | ENSG00000143379 | SETDB1 | chr1 | + | 150962586 | 150962719 | 150960562 | 150961191 | 150962973 | 150963139 | 141 | 84 | 91 | 98 | 104 | 0 | 0 | 0 | 0 | 0 | 183 | 145 | 97 | 74 | 163 | 0 | 2 | 0 | 0 | 6 | 282 | 149 | 0.551651994 | 1 | 1 | 1 | 1 | 1 | 1 | 1 | 0.975 | 1 | 1 | 0.935 | 0.018 |
| SE | 52211 | ENSG00000127463 | EMC1 | chr1 | - | 19238794 | 19238854 | 19238016 | 19238139 | 19239230 | 19239302 | 22 | 12 | 17 | 15 | 15 | 0 | 0 | 0 | 0 | 0 | 10 | 31 | 27 | 17 | 17 | 0 | 0 | 0 | 5 | 0 | 209 | 149 | 0.551759631 | 1 | 1 | 1 | 1 | 1 | 1 | 1 | 1 | 1 | 0.708 | 1 | 0.058 |
| SE | 94087 | ENSG00000136811 | ODF2 | chr9 | + | 128457197 | 128457437 | 128456100 | 128456255 | 128459566 | 128459657 | 6 | 1 | 4 | 6 | 9 | 30 | 15 | 15 | 32 | 3 | 4 | 4 | 6 | 7 | 11 | 19 | 26 | 15 | 32 | 63 | 298 | 149 | 0.551760908 | 1 | 0.091 | 0.032 | 0.118 | 0.086 | 0.6 | 0.095 | 0.071 | 0.167 | 0.099 | 0.08 | 0.083 |
| SE | 29527 | ENSG00000119242 | CCDC92 | chr12 | - | 123956283 | 123956444 | 123944269 | 123944364 | 123972528 | 123972616 | 8 | 2 | 3 | 13 | 11 | 25 | 7 | 24 | 26 | 9 | 9 | 15 | 0 | 0 | 40 | 7 | 23 | 8 | 35 | 107 | 298 | 149 | 0.551779092 | 1 | 0.138 | 0.125 | 0.059 | 0.2 | 0.379 | 0.391 | 0.246 | 0 | 0 | 0.157 | 0.021 |
| SE | 106244 | ENSG00000089154 | GCN1 | chr12 | - | 120184111 | 120184243 | 120178851 | 120178950 | 120184823 | 120184887 | 60 | 43 | 86 | 101 | 109 | 3 | 0 | 0 | 0 | 4 | 102 | 87 | 67 | 60 | 106 | 0 | 0 | 2 | 0 | 0 | 281 | 149 | 0.551796792 | 1 | 0.914 | 1 | 1 | 1 | 0.935 | 1 | 1 | 0.947 | 1 | 1 | -0.02 |
| SE | 75754 | ENSG00000096070 | BRPF3 | chr6 | + | 36211257 | 36211560 | 36210200 | 36210528 | 36217916 | 36218010 | 26 | 12 | 24 | 34 | 40 | 0 | 0 | 0 | 0 | 0 | 59 | 41 | 10 | 3 | 14 | 6 | 0 | 0 | 0 | 0 | 298 | 149 | 0.551800538 | 1 | 1 | | 1 | 1 | 1 | 0.831 | 1 | 1 | 1 | 1 | 0.034 |
| SE | 24041 | ENSG00000122224 | LY9 | chr1 | + | 160816593 | 160816863 | 160814419 | 160814761 | 160818217 | 160818319 | 93 | 39 | 63 | 62 | 113 | 38 | 9 | 35 | 34 | 26 | 5 | 36 | 10 | 17 | 47 | 0 | 11 | 11 | 2 | 17 | 298 | 149 | 0.551824038 | 1 | 0.55 | 0.684 | 0.474 | 0.477 | 0.685 | 1 | 0.621 | 0.312 | 0.81 | 0.58 | -0.091 |
| SE | 56720 | ENSG00000111737 | RAB35 | chr12 | - | 120098810 | 120098935 | 120095098 | 120097373 | 120099029 | 120099106 | 291 | 141 | 192 | 252 | 273 | 5 | 1 | 4 | 4 | 14 | 382 | 240 | 176 | 270 | 307 | 12 | 0 | 0 | 10 | 6 | 274 | 149 | 0.55189938 | 1 | 0.969 | 0.987 | 0.963 | 0.972 | 0.914 | 0.945 | 1 | 1 | 0.936 | 0.965 | -0.008 |
| SE | 41617 | ENSG00000117640 | MTFR1L | chr1 | + | 25826614 | 25826826 | 25826301 | 25826411 | 25829508 | 25829514 | 111 | 49 | 80 | 105 | 82 | 2 | 0 | 1 | 0 | 0 | 142 | 127 | 34 | 56 | 246 | 0 | 0 | 0 | 5 | 4 | 298 | 149 | 0.551907415 | 1 | 0.965 | 1 | 0.976 | 1 | 1 | 1 | 1 | 1 | 0.848 | 0.969 | 0.025 |
| SE | 4565 | ENSG00000123146 | ADGRE5 | chr19 | + | 14396341 | 14396473 | 14390923 | 14391079 | 14397657 | 14397804 | 301 | 155 | 186 | 240 | 209 | 368 | 197 | 263 | 363 | 291 | 395 | 191 | 176 | 245 | 198 | 385 | 369 | 277 | 217 | 524 | 281 | 149 | 0.551909764 | 1 | 0.303 | 0.294 | 0.273 | 0.26 | 0.276 | 0.352 | 0.215 | 0.252 | 0.374 | 0.167 | 0.009 |
| SE | 103611 | ENSG00000078403 | MLLT10 | chr10 | + | 21551815 | 21551921 | 21538832 | 21538912 | 21586293 | 21586348 | 0 | 3 | 3 | 3 | 0 | 36 | 51 | 47 | 59 | 44 | 0 | 6 | 0 | 0 | 0 | 61 | 47 | 50 | 26 | 11 | 255 | 149 | 0.552173093 | 1 | 0 | 0.033 | 0.036 | 0.029 | 0 | 0 | 0.069 | 0 | 0 | 0 | 0.006 |
| SE | 62645 | ENSG00000128815 | WDFY4 | chr10 | + | 48820233 | 48820437 | 48817244 | 48817409 | 48821061 | 48821176 | 59 | 20 | 11 | 53 | 20 | 0 | 0 | 0 | 0 | 0 | 44 | 41 | 30 | 41 | 100 | 0 | 0 | 0 | 0 | 6 | 298 | 149 | 0.552216728 | 1 | 1 | 1 | 1 | 1 | 1 | 1 | 1 | 1 | 1 | 0.893 | 0.021 |
| SE | 82692 | ENSG00000097033 | SH3GLB1 | chr1 | + | 86715723 | 86715865 | 86704569 | 86704971 | 86719506 | 86719635 | 309 | 211 | 228 | 332 | 279 | 3 | 5 | 0 | 2 | 4 | 178 | 199 | 367 | 164 | 120 | 0 | 0 | 3 | 0 | 0 | 291 | 149 | 0.552259815 | 1 | 0.981 | 0.956 | 1 | 0.988 | 0.973 | 1 | 1 | 0.984 | 1 | 1 | -0.017 |
| SE | 95885 | ENSG00000148396 | SEC16A | chr9 | - | 136468414 | 136468512 | 136466956 | 136467083 | 136471974 | 136472111 | 222 | 140 | 189 | 282 | 136 | 0 | 1 | 0 | 0 | 0 | 383 | 410 | 235 | 427 | 495 | 16 | 0 | 3 | 0 | 0 | 247 | 149 | 0.552264519 | 1 | 1 | 0.988 | 1 | 1 | 1 | 0.935 | 1 | 0.979 | 1 | 1 | 0.015 |
| SE | 85259 | ENSG00000176124 | DLEU1 | chr13 | + | 50083141 | 50083194 | 50082168 | 50082557 | 50104703 | 50107218 | 33 | 24 | 28 | 33 | 62 | 27 | 17 | 30 | 29 | 50 | 45 | 38 | 13 | 20 | 18 | 11 | 45 | 37 | 37 | 12 | 202 | 149 | 0.552389944 | 1 | 0.474 | 0.51 | 0.408 | 0.456 | 0.478 | 0.751 | 0.384 | 0.206 | 0.285 | 0.525 | 0.035 |
| SE | 41925 | ENSG00000143110 | C1orf162 | chr1 | + | 111477725 | 111477775 | 111476797 | 111477078 | 111477982 | 111478512 | 139 | 69 | 62 | 88 | 161 | 3 | 2 | 3 | 7 | 13 | 233 | 47 | 75 | 154 | 137 | 0 | 18 | 5 | 16 | 0 | 199 | 149 | 0.552414536 | 1 | 0.972 | 0.963 | 0.939 | 0.904 | 0.903 | 1 | 0.662 | 0.918 | 0.878 | 1 | 0.045 |
| SE | 19339 | ENSG00000100764 | PSMC1 | chr14 | + | 90263800 | 90263847 | 90263317 | 90263442 | 90264040 | 90264169 | 50 | 10 | 27 | 39 | 30 | 0 | 1 | 0 | 6 | 0 | 28 | 27 | 29 | 49 | 5 | 0 | 0 | 0 | 0 | 3 | 196 | 149 | 0.552433575 | 1 | 1 | 0.884 | 1 | 1 | 0.792 | 1 | 1 | 1 | 1 | 0.559 | 0.023 |
| SE | 106219 | ENSG00000070214 | SLC44A1 | chr9 | + | 105299742 | 105299937 | 105299219 | 105299309 | 105309723 | 105309866 | 9 | 1 | 1 | 7 | 9 | 22 | 16 | 23 | 11 | 12 | 0 | 9 | 1 | 18 | 5 | 23 | 10 | 21 | 27 | 42 | 298 | 149 | 0.552483434 | 1 | 0.17 | 0.03 | 0.021 | 0.241 | 0.273 | 0 | 0.31 | 0.023 | 0.25 | 0.056 | 0.019 |
| SE | 5532 | ENSG00000244687 | UBE2V1 | chr20 | - | 50096482 | 50096820 | 50084128 | 50084254 | 50113106 | 50113133 | 73 | 68 | 70 | 69 | 66 | 9 | 3 | 4 | 14 | 4 | 122 | 87 | 88 | 127 | 142 | 5 | 15 | 10 | 2 | 48 | 298 | 149 | 0.552504069 | 1 | 0.802 | 0.919 | 0.897 | 0.711 | 0.892 | 0.924 | 0.744 | 0.815 | 0.969 | 0.597 | 0.034 |
| SE | 76096 | ENSG00000136044 | APPL2 | chr12 | - | 105176875 | 105177016 | 105176034 | 105176082 | 105177225 | 105177262 | 97 | 88 | 93 | 91 | 153 | 1 | 1 | 4 | 4 | 16 | 99 | 66 | 131 | 75 | 13 | 4 | 0 | 1 | 0 | 14 | 290 | 149 | 0.5525571 | 1 | 0.98 | 0.978 | 0.923 | 0.921 | 0.831 | 0.927 | 1 | 0.985 | 1 | 0.323 | 0.08 |
| SE | 1129 | ENSG00000099992 | TBC1D10A | chr22 | - | 30293158 | 30293319 | 30292009 | 30292851 | 30293650 | 30293805 | 6 | 0 | 7 | 8 | 8 | 51 | 21 | 38 | 49 | 46 | 0 | 3 | 2 | 4 | 13 | 6 | 20 | 22 | 27 | 101 | 298 | 149 | 0.552615888 | 1 | 0.056 | 0 | 0.084 | 0.075 | 0.08 | 0 | 0.07 | 0.043 | 0.069 | 0.06 | 0.011 |
| SE | 36361 | ENSG00000173889 | PHC3 | chr3 | - | 170145422 | 170145521 | 170128683 | 170129552 | 170149085 | 170149244 | 114 | 60 | 95 | 126 | 195 | 0 | 0 | 0 | 0 | 0 | 112 | 132 | 39 | 44 | 130 | 0 | 0 | 0 | 7 | 0 | 248 | 149 | 0.552618171 | 1 | 1 | 1 | 1 | 1 | 1 | 1 | 1 | 1 | 0.791 | 1 | 0.042 |
| SE | 63736 | ENSG00000152061 | RABGAP1L | chr1 | + | 174231144 | 174231355 | 174220971 | 174221164 | 174241482 | 174241657 | 768 | 662 | 492 | 742 | 950 | 9 | 9 | 6 | 7 | 3 | 154 | 857 | 566 | 268 | 128 | 5 | 4 | 11 | 4 | 0 | 298 | 149 | 0.552703495 | 1 | 0.977 | 0.974 | 0.976 | 0.981 | 0.994 | 0.939 | 0.991 | 0.963 | 0.971 | 1 | 0.008 |
| SE | 10063 | ENSG00000108406 | DHX40 | chr17 | + | 59566626 | 59566794 | 59565557 | 59565783 | 59570517 | 59570663 | 148 | 112 | 139 | 150 | 157 | 1 | 2 | 4 | 1 | 0 | 52 | 146 | 126 | 117 | 25 | 0 | 0 | 0 | 0 | 0 | 298 | 149 | 0.552722006 | 1 | 0.987 | 0.966 | 0.946 | 0.987 | 1 | 1 | 1 | 1 | 1 | 1 | -0.023 |
| SE | 106799 | ENSG00000173442 | EHBP1L1 | chr11 | + | 65583591 | 65583765 | 65581898 | 65581965 | 65584240 | 65584398 | 223 | 154 | 185 | 185 | 280 | 66 | 18 | 35 | 48 | 51 | 343 | 224 | 117 | 234 | 446 | 69 | 24 | 24 | 76 | 63 | 298 | 149 | 0.552897488 | 1 | 0.628 | 0.811 | 0.725 | 0.658 | 0.733 | 0.713 | 0.824 | 0.709 | 0.606 | 0.78 | -0.015 |
| SE | 107655 | ENSG00000139718 | SETD1B | chr12 | + | 121827518 | 121827650 | 121825199 | 121825366 | 121827734 | 121827854 | 79 | 44 | 125 | 88 | 55 | 0 | 0 | 1 | 0 | 0 | 74 | 141 | 86 | 80 | 162 | 0 | 0 | 0 | 0 | 9 | 281 | 149 | 0.553032083 | 1 | 1 | 1 | 0.985 | 1 | 1 | 1 | 1 | 1 | 1 | 0.905 | 0.016 |
| SE | 1582 | ENSG00000198089 | SFI1 | chr22 | + | 31556941 | 31557059 | 31550253 | 31550348 | 31573057 | 31573214 | 67 | 47 | 74 | 69 | 77 | 0 | 0 | 0 | 0 | 0 | 60 | 73 | 75 | 75 | 143 | 0 | 0 | 0 | 8 | 0 | 267 | 149 | 0.553059294 | 1 | 1 | 1 | 1 | 1 | 1 | 1 | 1 | 1 | 0.84 | 1 | 0.032 |
| SE | 107587 | ENSG00000138777 | PPA2 | chr4 | - | 105453597 | 105453642 | 105449349 | 105449403 | 105473893 | 105473987 | 32 | 26 | 46 | 33 | 53 | 0 | 1 | 1 | 0 | 0 | 24 | 25 | 9 | 26 | 32 | 0 | 6 | 0 | 0 | 0 | 194 | 149 | 0.553070547 | 1 | 1 | 0.952 | 0.972 | 1 | 1 | 1 | 0.762 | 1 | 1 | 1 | 0.032 |
| SE | 13782 | ENSG00000205629 | LCMT1 | chr16 | + | 25174936 | 25175034 | 25170713 | 25170805 | 25178000 | 25178023 | 39 | 40 | 56 | 52 | 99 | 1 | 1 | 0 | 1 | 5 | 99 | 80 | 27 | 40 | 88 | 0 | 0 | 0 | 0 | 0 | 247 | 149 | 0.553111063 | 1 | 0.959 | 0.96 | 1 | 0.969 | 0.923 | 1 | 1 | 1 | 1 | 1 | -0.038 |
| SE | 107507 | ENSG00000123349 | PFDN5 | chr12 | + | 53296453 | 53296551 | 53295541 | 53295639 | 53297849 | 53297924 | 18 | 7 | 25 | 23 | 11 | 208 | 128 | 218 | 168 | 123 | 14 | 12 | 8 | 23 | 26 | 212 | 136 | 153 | 132 | 306 | 247 | 149 | 0.553172426 | 1 | 0.05 | 0.032 | 0.065 | 0.076 | 0.051 | 0.038 | 0.051 | 0.031 | 0.095 | 0.049 | 0.002 |
| SE | 86710 | ENSG00000128654 | MTX2 | chr2 | + | 176329300 | 176329426 | 176328873 | 176328912 | 176337492 | 176337679 | 30 | 28 | 33 | 28 | 51 | 2 | 0 | 0 | 0 | 3 | 42 | 11 | 18 | 15 | 7 | 0 | 0 | 0 | 0 | 5 | 275 | 149 | 0.553185353 | 1 | 0.89 | 1 | 1 | 1 | 0.902 | 1 | 1 | 1 | 1 | 0.431 | 0.072 |
| SE | 1579 | ENSG00000198089 | SFI1 | chr22 | + | 31556941 | 31557059 | 31546860 | 31546971 | 31561289 | 31561333 | 134 | 83 | 109 | 102 | 103 | 0 | 0 | 0 | 0 | 0 | 114 | 88 | 59 | 82 | 173 | 0 | 0 | 0 | 0 | 9 | 267 | 149 | 0.553278735 | 1 | 1 | 1 | 1 | 1 | 1 | 1 | 1 | 1 | 1 | 0.915 | 0.017 |
| SE | 104741 | ENSG00000122694 | GLIPR2 | chr9 | + | 36150871 | 36150949 | 36147785 | 36147894 | 36162361 | 36163902 | 80 | 49 | 57 | 73 | 52 | 0 | 0 | 0 | 0 | 0 | 338 | 128 | 130 | 153 | 338 | 0 | 4 | 3 | 7 | 0 | 227 | 149 | 0.55336459 | 1 | 1 | 1 | 1 | 1 | 1 | 1 | 0.955 | 0.966 | 0.935 | 1 | 0.029 |
| SE | 44753 | ENSG00000175137 | SH3BP5L | chr1 | - | 248812988 | 248813162 | 248810445 | 248812370 | 248814448 | 248814610 | 63 | 82 | 65 | 58 | 33 | 0 | 0 | 0 | 2 | 0 | 67 | 75 | 12 | 69 | 64 | 7 | 0 | 0 | 0 | 0 | 298 | 149 | 0.553423571 | 1 | 1 | 1 | 1 | 0.935 | 1 | 0.827 | 1 | 1 | 1 | 1 | 0.022 |
| SE | 14657 | ENSG00000179361 | ARID3B | chr15 | + | 74591150 | 74591434 | 74589819 | 74590003 | 74591559 | 74591814 | 71 | 32 | 75 | 70 | 104 | 1 | 0 | 0 | 6 | 3 | 72 | 147 | 106 | 89 | 175 | 0 | 7 | 3 | 1 | 6 | 298 | 149 | 0.553496314 | 1 | 0.973 | 1 | 1 | 0.854 | 0.945 | 1 | 0.913 | 0.946 | 0.978 | 0.936 | 0 |
| SE | 79201 | ENSG00000242588 | NA | chr7 | + | 128579998 | 128580094 | 128574750 | 128574791 | 128591849 | 128592017 | 76 | 51 | 63 | 95 | 48 | 17 | 9 | 3 | 13 | 19 | 216 | 102 | 86 | 103 | 299 | 57 | 9 | 5 | 59 | 14 | 245 | 149 | 0.553512909 | 1 | 0.731 | 0.775 | 0.927 | 0.816 | 0.606 | 0.697 | 0.873 | 0.913 | 0.515 | 0.929 | -0.014 |
| SE | 3686 | ENSG00000167613 | LAIR1 | chr19 | - | 54364294 | 54364330 | 54360915 | 54361209 | 54368311 | 54368599 | 36 | 7 | 18 | 15 | 2 | 0 | 0 | 0 | 3 | 0 | 15 | 19 | 8 | 28 | 21 | 0 | 0 | 0 | 0 | 0 | 185 | 149 | 0.553613385 | 1 | 1 | 1 | 1 | 1 | 0.349 | 1 | 1 | 1 | 1 | 1 | -0.13 |
| SE | 51448 | ENSG00000265491 | RNF115 | chr1 | - | 145788907 | 145788966 | 145784538 | 145784596 | 145823771 | 145824095 | 55 | 39 | 49 | 49 | 33 | 0 | 0 | 0 | 1 | 0 | 55 | 38 | 43 | 30 | 49 | 0 | 0 | 0 | 0 | 7 | 208 | 149 | 0.553674581 | 1 | 1 | 1 | 1 | 0.972 | 1 | 1 | 1 | 1 | 1 | 0.834 | 0.028 |
| SE | 101053 | ENSG00000110435 | PDHX | chr11 | + | 34966669 | 34966814 | 34960419 | 34960518 | 34970138 | 34970286 | 202 | 138 | 196 | 232 | 419 | 1 | 0 | 0 | 5 | 0 | 212 | 198 | 94 | 120 | 131 | 0 | 4 | 0 | 0 | 5 | 298 | 149 | 0.553728142 | 1 | 0.99 | 1 | 1 | 0.959 | 1 | 1 | 0.961 | 1 | 1 | 0.929 | 0.012 |
| SE | 95199 | ENSG00000148308 | GTF3C5 | chr9 | + | 133042086 | 133042306 | 133030996 | 133031164 | 133043727 | 133043926 | 36 | 19 | 29 | 27 | 17 | 1 | 2 | 0 | 3 | 0 | 42 | 38 | 24 | 49 | 63 | 0 | 0 | 0 | 0 | 0 | 298 | 149 | 0.553731172 | 1 | 0.947 | 0.826 | 1 | 0.818 | 1 | 1 | 1 | 1 | 1 | 1 | -0.082 |
| SE | 65043 | ENSG00000111275 | ALDH2 | chr12 | + | 111775604 | 111775720 | 111766947 | 111767096 | 111781917 | 111782022 | 5 | 6 | 0 | 5 | 0 | 26 | 15 | 18 | 9 | 9 | 0 | 9 | 5 | 20 | 6 | 39 | 18 | 13 | 37 | 88 | 265 | 149 | 0.553759691 | 1 | 0.098 | 0.184 | 0 | 0.238 | 0 | 0 | 0.219 | 0.178 | 0.233 | 0.037 | -0.029 |
| SE | 60702 | ENSG00000235750 | KIAA0040 | chr1 | - | 175176399 | 175176474 | 175166561 | 175166737 | 175177610 | 175177684 | 3 | 0 | 0 | 3 | 4 | 78 | 36 | 85 | 73 | 66 | 0 | 12 | 0 | 0 | 0 | 59 | 101 | 40 | 49 | 47 | 224 | 149 | 0.553864325 | 1 | 0.025 | 0 | 0 | 0.027 | 0.039 | 0 | 0.073 | 0 | 0 | 0 | 0.004 |
| SE | 37474 | ENSG00000163681 | SLMAP | chr3 | + | 57871635 | 57871698 | 57865241 | 57865292 | 57890040 | 57890100 | 35 | 41 | 43 | 53 | 52 | 7 | 5 | 7 | 12 | 29 | 7 | 64 | 55 | 27 | 6 | 0 | 8 | 9 | 2 | 14 | 212 | 149 | 0.55388794 | 1 | 0.778 | 0.852 | 0.812 | 0.756 | 0.558 | 1 | 0.849 | 0.811 | 0.905 | 0.231 | -0.008 |
| SE | 50378 | ENSG00000058866 | DGKG | chr3 | - | 186207423 | 186207517 | 186188201 | 186188379 | 186211794 | 186211885 | 2 | 1 | 0 | 4 | 0 | 19 | 19 | 11 | 14 | 10 | 0 | 0 | 0 | 0 | 7 | 100 | 21 | 25 | 50 | 43 | 243 | 149 | 0.553997586 | 1 | 0.061 | 0.031 | 0 | 0.149 | 0 | 0 | 0 | 0 | 0.079 | 0 | 0.032 |
| SE | 62442 | ENSG00000113761 | ZNF346 | chr5 | + | 177062057 | 177062151 | 177050702 | 177050936 | 177064511 | 177064601 | 28 | 18 | 38 | 30 | 37 | 0 | 1 | 0 | 0 | 0 | 46 | 52 | 15 | 34 | 108 | 0 | 0 | 0 | 6 | 0 | 243 | 149 | 0.554014 | 1 | 1 | 0.917 | 1 | 1 | 1 | 1 | 1 | 1 | 0.777 | 1 | 0.028 |
| SE | 45090 | ENSG00000100181 | TPTEP1 | chr22 | + | 16647167 | 16647257 | 16638578 | 16638740 | 16647662 | 16647785 | 0 | 3 | 1 | 5 | 0 | 6 | 14 | 0 | 10 | 1 | 0 | 0 | 0 | 0 | 8 | 15 | 13 | 8 | 7 | 39 | 239 | 149 | 0.554147303 | 1 | 0 | 0.118 | 1 | 0.238 | 0 | 0 | 0 | 0 | 0 | 0.113 | 0.249 |
| SE | 70406 | ENSG00000115652 | UXS1 | chr2 | - | 106096717 | 106096821 | 106093307 | 106094157 | 106101057 | 106101118 | 56 | 29 | 54 | 56 | 89 | 1 | 0 | 2 | 2 | 5 | 106 | 43 | 40 | 91 | 148 | 0 | 0 | 0 | 0 | 12 | 253 | 149 | 0.554221946 | 1 | 0.971 | 1 | 0.941 | 0.943 | 0.913 | 1 | 1 | 1 | 1 | 0.879 | -0.022 |
| SE | 20728 | ENSG00000115760 | BIRC6 | chr2 | + | 32490040 | 32490151 | 32488587 | 32488714 | 32491424 | 32491558 | 401 | 317 | 294 | 317 | 499 | 1 | 6 | 2 | 0 | 9 | 24 | 271 | 260 | 109 | 65 | 0 | 1 | 5 | 0 | 0 | 260 | 149 | 0.554427603 | 1 | 0.996 | 0.968 | 0.988 | 1 | 0.969 | 1 | 0.994 | 0.968 | 1 | 1 | -0.008 |
| SE | 88862 | ENSG00000104442 | ARMC1 | chr8 | - | 65605421 | 65605538 | 65605262 | 65605337 | 65613243 | 65613433 | 95 | 64 | 100 | 81 | 177 | 3 | 1 | 4 | 5 | 11 | 38 | 78 | 29 | 48 | 10 | 0 | 2 | 6 | 2 | 0 | 266 | 149 | 0.554479484 | 1 | 0.947 | 0.973 | 0.933 | 0.901 | 0.9 | 1 | 0.956 | 0.73 | 0.931 | 1 | 0.007 |
| SE | 37595 | ENSG00000082074 | FYB1 | chr5 | - | 39208863 | 39208964 | 39202955 | 39202987 | 39250670 | 39250806 | 20 | 4 | 2 | 12 | 9 | 23 | 6 | 8 | 10 | 5 | 11 | 28 | 7 | 2 | 24 | 14 | 13 | 11 | 4 | 20 | 250 | 149 | 0.554666457 | 1 | 0.341 | 0.284 | 0.13 | 0.417 | 0.518 | 0.319 | 0.562 | 0.275 | 0.23 | 0.417 | -0.023 |
| SE | 48607 | ENSG00000131374 | TBC1D5 | chr3 | - | 17518901 | 17518995 | 17508473 | 17508605 | 17586352 | 17586557 | 24 | 24 | 14 | 15 | 18 | 14 | 5 | 10 | 9 | 19 | 35 | 15 | 9 | 16 | 5 | 11 | 9 | 6 | 13 | 0 | 243 | 149 | 0.554841748 | 1 | 0.512 | 0.746 | 0.462 | 0.505 | 0.367 | 0.661 | 0.505 | 0.479 | 0.43 | 1 | -0.097 |
| SE | 79032 | ENSG00000160218 | TRAPPC10 | chr21 | + | 44084121 | 44084263 | 44082787 | 44083302 | 44089832 | 44089933 | 117 | 46 | 104 | 92 | 101 | 0 | 0 | 0 | 1 | 0 | 80 | 91 | 92 | 63 | 68 | 0 | 1 | 2 | 3 | 0 | 291 | 149 | 0.554877123 | 1 | 1 | 1 | 1 | 0.979 | 1 | 1 | 0.979 | 0.959 | 0.915 | 1 | 0.025 |
| SE | 72898 | ENSG00000124831 | LRRFIP1 | chr2 | + | 237708543 | 237708630 | 237692144 | 237692521 | 237748363 | 237748399 | 258 | 162 | 183 | 211 | 96 | 1 | 5 | 1 | 3 | 0 | 119 | 163 | 224 | 259 | 105 | 0 | 4 | 5 | 3 | 0 | 236 | 149 | 0.554896913 | 1 | 0.994 | 0.953 | 0.991 | 0.978 | 1 | 1 | 0.963 | 0.966 | 0.982 | 1 | 0.001 |
| SE | 69835 | ENSG00000163803 | PLB1 | chr2 | + | 28629093 | 28629185 | 28628562 | 28628628 | 28630585 | 28630664 | 34 | 21 | 23 | 42 | 39 | 0 | 0 | 0 | 0 | 0 | 69 | 45 | 43 | 75 | 130 | 0 | 0 | 0 | 8 | 0 | 241 | 149 | 0.554960989 | 1 | 1 | 1 | 1 | 1 | 1 | 1 | 1 | 1 | 0.853 | 1 | 0.029 |
| SE | 54603 | ENSG00000151422 | FER | chr5 | + | 108782662 | 108782755 | 108768092 | 108768238 | 108798123 | 108798389 | 1 | 3 | 0 | 4 | 0 | 18 | 16 | 12 | 19 | 13 | 0 | 0 | 0 | 0 | 8 | 3 | 10 | 3 | 4 | 4 | 242 | 149 | 0.555141156 | 1 | 0.033 | 0.103 | 0 | 0.115 | 0 | 0 | 0 | 0 | 0 | 0.552 | -0.06 |
| SE | 97606 | ENSG00000179832 | MROH1 | chr8 | + | 144242601 | 144242628 | 144242368 | 144242515 | 144243493 | 144243616 | 34 | 37 | 39 | 17 | 35 | 33 | 20 | 35 | 19 | 36 | 10 | 52 | 31 | 53 | 48 | 52 | 11 | 30 | 43 | 84 | 176 | 149 | 0.55515376 | 1 | 0.466 | 0.61 | 0.485 | 0.431 | 0.451 | 0.14 | 0.8 | 0.467 | 0.511 | 0.326 | 0.04 |
| SE | 41444 | ENSG00000284879 | NA | chr2 | + | 87583257 | 87583346 | 87455475 | 87455645 | 87606592 | 87606769 | 42 | 42 | 15 | 25 | 71 | 2 | 4 | 0 | 6 | 0 | 26 | 46 | 38 | 37 | 66 | 8 | 1 | 1 | 2 | 4 | 238 | 149 | 0.555228205 | 1 | 0.929 | 0.868 | 1 | 0.723 | 1 | 0.67 | 0.966 | 0.96 | 0.921 | 0.912 | 0.018 |
| SE | 83731 | ENSG00000118513 | MYB | chr6 | + | 135194355 | 135194460 | 135193837 | 135193918 | 135195747 | 135195917 | 15 | 17 | 20 | 9 | 26 | 0 | 0 | 0 | 0 | 0 | 5 | 2 | 11 | 25 | 4 | 0 | 0 | 0 | 5 | 0 | 254 | 149 | 0.555232568 | 1 | 1 | 1 | 1 | 1 | 1 | 1 | 1 | 1 | 0.746 | 1 | 0.051 |
| SE | 15167 | ENSG00000137842 | TMEM62 | chr15 | + | 43154671 | 43154831 | 43151789 | 43151945 | 43160680 | 43160794 | 125 | 69 | 76 | 78 | 155 | 0 | 0 | 0 | 0 | 0 | 90 | 100 | 112 | 118 | 164 | 0 | 4 | 0 | 4 | 0 | 298 | 149 | 0.555277637 | 1 | 1 | 1 | 1 | 1 | 1 | 1 | 0.926 | 1 | 0.937 | 1 | 0.027 |
| SE | 47312 | ENSG00000133106 | EPSTI1 | chr13 | - | 42889210 | 42889242 | 42886387 | 42888502 | 42895008 | 42895108 | 82 | 48 | 69 | 64 | 70 | 211 | 231 | 203 | 365 | 144 | 589 | 73 | 49 | 256 | 13 | 362 | 500 | 371 | 1058 | 118 | 181 | 149 | 0.55536954 | 1 | 0.242 | 0.146 | 0.219 | 0.126 | 0.286 | 0.573 | 0.107 | 0.098 | 0.166 | 0.083 | -0.002 |
| SE | 20228 | ENSG00000111802 | TDP2 | chr6 | - | 24652982 | 24653153 | 24649978 | 24651069 | 24654411 | 24654530 | 440 | 349 | 408 | 403 | 687 | 6 | 1 | 2 | 1 | 9 | 91 | 263 | 361 | 198 | 33 | 0 | 0 | 0 | 1 | 0 | 298 | 149 | 0.55541668 | 1 | 0.973 | 0.994 | 0.99 | 0.995 | 0.974 | 1 | 1 | 1 | 0.99 | 1 | -0.013 |
| SE | 64927 | ENSG00000204642 | HLA-F | chr6 | + | 29726010 | 29726043 | 29725446 | 29725563 | 29726882 | 29727296 | 191 | 113 | 138 | 160 | 101 | 3 | 6 | 6 | 10 | 6 | 476 | 299 | 208 | 240 | 411 | 7 | 9 | 6 | 3 | 32 | 182 | 149 | 0.555475705 | 1 | 0.981 | 0.939 | 0.95 | 0.929 | 0.932 | 0.982 | 0.965 | 0.966 | 0.985 | 0.913 | -0.016 |
| SE | 64153 | ENSG00000032444 | PNPLA6 | chr19 | + | 7555248 | 7555367 | 7554892 | 7555075 | 7555606 | 7555763 | 36 | 28 | 19 | 30 | 3 | 0 | 0 | 0 | 0 | 0 | 26 | 22 | 5 | 14 | 42 | 0 | 5 | 0 | 0 | 0 | 268 | 149 | 0.555527055 | 1 | 1 | 1 | 1 | 1 | 1 | 1 | 0.71 | 1 | 1 | 1 | 0.058 |
| SE | 46651 | ENSG00000126749 | EMG1 | chr12 | + | 6975228 | 6975378 | 6975089 | 6975148 | 6975695 | 6979936 | 127 | 56 | 130 | 114 | 200 | 7 | 1 | 9 | 2 | 6 | 108 | 47 | 31 | 96 | 53 | 7 | 0 | 3 | 3 | 0 | 298 | 149 | 0.555534553 | 1 | 0.901 | 0.966 | 0.878 | 0.966 | 0.943 | 0.885 | 1 | 0.838 | 0.941 | 1 | -0.002 |
| SE | 80745 | ENSG00000004864 | SLC25A13 | chr7 | - | 96189293 | 96189378 | 96184926 | 96185011 | 96191108 | 96191247 | 7 | 18 | 38 | 4 | 7 | 0 | 0 | 0 | 0 | 0 | 9 | 14 | 3 | 11 | 4 | 4 | 0 | 0 | 0 | 0 | 234 | 149 | 0.555540193 | 1 | 1 | 1 | 1 | 1 | 1 | 0.589 | 1 | 1 | 1 | 1 | 0.082 |
| SE | 65162 | ENSG00000196912 | ANKRD36B | chr2 | - | 97578905 | 97579043 | 97560831 | 97560860 | 97580461 | 97580568 | 139 | 153 | 165 | 126 | 216 | 27 | 13 | 19 | 22 | 23 | 68 | 84 | 70 | 51 | 30 | 4 | 5 | 13 | 6 | 8 | 287 | 149 | 0.555587361 | 1 | 0.728 | 0.859 | 0.818 | 0.748 | 0.83 | 0.898 | 0.897 | 0.737 | 0.815 | 0.661 | -0.005 |
| SE | 42275 | ENSG00000160752 | FDPS | chr1 | + | 155310063 | 155310205 | 155308754 | 155308965 | 155312254 | 155312395 | 186 | 76 | 184 | 163 | 102 | 3 | 1 | 2 | 2 | 0 | 321 | 221 | 81 | 187 | 232 | 0 | 0 | 2 | 2 | 5 | 291 | 149 | 0.55563261 | 1 | 0.969 | 0.975 | 0.979 | 0.977 | 1 | 1 | 1 | 0.954 | 0.98 | 0.96 | 0.001 |
| SE | 89087 | ENSG00000171988 | JMJD1C | chr10 | - | 63264650 | 63264764 | 63217206 | 63217331 | 63268662 | 63268875 | 240 | 246 | 222 | 319 | 587 | 5 | 6 | 0 | 0 | 4 | 86 | 109 | 105 | 75 | 36 | 0 | 9 | 0 | 0 | 0 | 263 | 149 | 0.555638075 | 1 | 0.965 | 0.959 | 1 | 1 | 0.988 | 1 | 0.873 | 1 | 1 | 1 | 0.008 |
| SE | 71399 | ENSG00000122203 | KIAA1191 | chr5 | - | 176359810 | 176359918 | 176352621 | 176352748 | 176361601 | 176361735 | 21 | 7 | 18 | 38 | 9 | 5 | 0 | 0 | 0 | 0 | 4 | 27 | 6 | 14 | 63 | 0 | 1 | 1 | 0 | 0 | 257 | 149 | 0.555639102 | 1 | 0.709 | 1 | 1 | 1 | 1 | 1 | 0.94 | 0.777 | 1 | 1 | -0.002 |
| SE | 55425 | ENSG00000116688 | MFN2 | chr1 | + | 11981809 | 11982114 | 11980298 | 11980484 | 11989164 | 11989343 | 189 | 133 | 173 | 143 | 340 | 1 | 6 | 2 | 0 | 1 | 336 | 204 | 170 | 302 | 512 | 1 | 11 | 0 | 2 | 3 | 298 | 149 | 0.555640868 | 1 | 0.99 | 0.917 | 0.977 | 1 | 0.994 | 0.994 | 0.903 | 1 | 0.987 | 0.988 | 0.001 |
| SE | 100659 | ENSG00000166340 | TPP1 | chr11 | - | 6617353 | 6617428 | 6616974 | 6617153 | 6617625 | 6617776 | 168 | 61 | 103 | 122 | 154 | 3 | 0 | 3 | 0 | 3 | 176 | 167 | 83 | 163 | 475 | 3 | 0 | 1 | 0 | 3 | 224 | 149 | 0.5557402 | 1 | 0.974 | 1 | 0.958 | 1 | 0.972 | 0.975 | 1 | 0.982 | 1 | 1 | -0.011 |
| SE | 18589 | ENSG00000165959 | CLMN | chr14 | - | 95222402 | 95222599 | 95221732 | 95221774 | 95223759 | 95223855 | 0 | 0 | 0 | 0 | 0 | 49 | 26 | 33 | 16 | 53 | 0 | 10 | 0 | 0 | 0 | 45 | 35 | 32 | 26 | 52 | 298 | 149 | 0.555741232 | 1 | 0 | 0 | 0 | 0 | 0 | 0 | 0.125 | 0 | 0 | 0 | -0.025 |
| SE | 103443 | ENSG00000148840 | PPRC1 | chr10 | + | 102138878 | 102138980 | 102137849 | 102138038 | 102139099 | 102142004 | 12 | 6 | 17 | 18 | 11 | 11 | 2 | 2 | 1 | 4 | 49 | 18 | 11 | 26 | 39 | 0 | 11 | 25 | 16 | 7 | 251 | 149 | 0.555795066 | 1 | 0.393 | 0.64 | 0.835 | 0.914 | 0.62 | 1 | 0.493 | 0.207 | 0.491 | 0.768 | 0.089 |
| SE | 11239 | ENSG00000102984 | ZNF821 | chr16 | - | 71881316 | 71881434 | 71879906 | 71880023 | 71883210 | 71883273 | 0 | 1 | 0 | 3 | 2 | 12 | 4 | 8 | 2 | 7 | 0 | 4 | 0 | 2 | 0 | 11 | 14 | 14 | 21 | 9 | 267 | 149 | 0.555805762 | 1 | 0 | 0.122 | 0 | 0.456 | 0.138 | 0 | 0.138 | 0 | 0.05 | 0 | 0.106 |
| SE | 53959 | ENSG00000138101 | DTNB | chr2 | - | 25596085 | 25596240 | 25580720 | 25580826 | 25607235 | 25607293 | 60 | 34 | 73 | 62 | 117 | 0 | 1 | 2 | 3 | 5 | 23 | 47 | 33 | 36 | 57 | 0 | 3 | 0 | 5 | 0 | 298 | 149 | 0.555819865 | 1 | 1 | 0.944 | 0.948 | 0.912 | 0.921 | 1 | 0.887 | 1 | 0.783 | 1 | 0.011 |
| SE | 62078 | ENSG00000263528 | IKBKE | chr1 | + | 206493022 | 206493119 | 206491647 | 206491749 | 206493919 | 206493991 | 52 | 34 | 54 | 51 | 61 | 0 | 1 | 3 | 3 | 0 | 123 | 91 | 28 | 67 | 139 | 0 | 0 | 3 | 5 | 2 | 246 | 149 | 0.555891095 | 1 | 1 | 0.954 | 0.916 | 0.911 | 1 | 1 | 1 | 0.85 | 0.89 | 0.977 | 0.013 |
| SE | 27470 | ENSG00000008513 | ST3GAL1 | chr8 | - | 133539636 | 133539729 | 133499134 | 133499189 | 133545773 | 133545926 | 15 | 11 | 2 | 17 | 37 | 92 | 86 | 90 | 100 | 237 | 0 | 3 | 6 | 16 | 25 | 78 | 89 | 44 | 68 | 116 | 242 | 149 | 0.555997736 | 1 | 0.091 | 0.073 | 0.013 | 0.095 | 0.088 | 0 | 0.02 | 0.077 | 0.127 | 0.117 | 0.004 |
| SE | 82267 | ENSG00000223482 | NUTM2A-AS1 | chr10 | - | 87307956 | 87308060 | 87287958 | 87288495 | 87326629 | 87326672 | 27 | 19 | 4 | 30 | 70 | 75 | 44 | 16 | 76 | 88 | 14 | 26 | 2 | 18 | 80 | 47 | 91 | 20 | 27 | 106 | 253 | 149 | 0.556015677 | 1 | 0.175 | 0.203 | 0.128 | 0.189 | 0.319 | 0.149 | 0.144 | 0.056 | 0.282 | 0.308 | 0.015 |
| SE | 66465 | ENSG00000131051 | RBM39 | chr20 | - | 35729461 | 35729551 | 35729311 | 35729365 | 35731940 | 35732135 | 313 | 286 | 296 | 426 | 352 | 39 | 5 | 8 | 23 | 31 | 67 | 280 | 354 | 276 | 7 | 0 | 10 | 7 | 6 | 15 | 239 | 149 | 0.556245558 | 1 | 0.833 | 0.973 | 0.958 | 0.92 | 0.876 | 1 | 0.946 | 0.969 | 0.966 | 0.225 | 0.091 |
| SE | 45011 | ENSG00000162623 | TYW3 | chr1 | + | 74738689 | 74738788 | 74736188 | 74736622 | 74748750 | 74748822 | 163 | 108 | 111 | 127 | 173 | 3 | 3 | 0 | 2 | 1 | 7 | 83 | 78 | 49 | 8 | 0 | 0 | 0 | 2 | 0 | 248 | 149 | 0.556359898 | 1 | 0.97 | 0.956 | 1 | 0.974 | 0.99 | 1 | 1 | 1 | 0.936 | 1 | -0.009 |
| SE | 83007 | ENSG00000106460 | TMEM106B | chr7 | + | 12224225 | 12224385 | 12218457 | 12218521 | 12229678 | 12229819 | 473 | 334 | 470 | 475 | 972 | 14 | 7 | 8 | 2 | 19 | 62 | 117 | 68 | 22 | 96 | 0 | 2 | 0 | 0 | 9 | 298 | 149 | 0.556658607 | 1 | 0.944 | 0.96 | 0.967 | 0.992 | 0.962 | 1 | 0.967 | 1 | 1 | 0.842 | 0.003 |
| SE | 69981 | ENSG00000135775 | COG2 | chr1 | + | 230688070 | 230688143 | 230686934 | 230687132 | 230688419 | 230688528 | 47 | 22 | 53 | 79 | 80 | 0 | 0 | 3 | 3 | 0 | 64 | 33 | 28 | 36 | 39 | 0 | 0 | 0 | 0 | 0 | 222 | 149 | 0.55666482 | 1 | 1 | 1 | 0.922 | 0.946 | 1 | 1 | 1 | 1 | 1 | 1 | -0.026 |
| SE | 26799 | ENSG00000163281 | GNPDA2 | chr4 | - | 44710952 | 44711137 | 44703245 | 44707926 | 44717112 | 44717295 | 130 | 80 | 104 | 110 | 226 | 2 | 1 | 1 | 5 | 0 | 12 | 43 | 57 | 18 | 23 | 0 | 0 | 1 | 0 | 0 | 298 | 149 | 0.556770139 | 1 | 0.97 | 0.976 | 0.981 | 0.917 | 1 | 1 | 1 | 0.966 | 1 | 1 | -0.024 |
| SE | 48650 | ENSG00000131374 | TBC1D5 | chr3 | - | 17586352 | 17586557 | 17562018 | 17562142 | 17622647 | 17622765 | 29 | 24 | 13 | 21 | 26 | 7 | 3 | 6 | 2 | 9 | 8 | 6 | 11 | 26 | 19 | 8 | 2 | 9 | 5 | 0 | 298 | 149 | 0.556812108 | 1 | 0.674 | 0.8 | 0.52 | 0.84 | 0.591 | 0.333 | 0.6 | 0.379 | 0.722 | 1 | 0.078 |
| SE | 109250 | ENSG00000275066 | SYNRG | chr17 | - | 37570636 | 37570885 | 37566848 | 37568924 | 37571790 | 37571987 | 321 | 197 | 370 | 302 | 482 | 0 | 0 | 4 | 0 | 4 | 114 | 287 | 172 | 158 | 74 | 0 | 5 | 0 | 4 | 0 | 298 | 149 | 0.556815499 | 1 | 1 | 1 | 0.979 | 1 | 0.984 | 1 | 0.966 | 1 | 0.952 | 1 | 0.009 |
| SE | 51078 | ENSG00000169756 | LIMS1 | chr2 | + | 108670780 | 108670847 | 108659604 | 108659764 | 108672324 | 108672445 | 215 | 283 | 128 | 357 | 309 | 9 | 5 | 2 | 9 | 5 | 458 | 319 | 228 | 344 | 463 | 6 | 16 | 9 | 7 | 8 | 216 | 149 | 0.556878554 | 1 | 0.943 | 0.975 | 0.978 | 0.965 | 0.977 | 0.981 | 0.932 | 0.946 | 0.971 | 0.976 | 0.006 |
| SE | 15731 | ENSG00000128394 | APOBEC3F | chr22 | + | 39049424 | 39049581 | 39045183 | 39045542 | 39052073 | 39052287 | 66 | 59 | 57 | 91 | 106 | 6 | 0 | 5 | 4 | 5 | 51 | 98 | 27 | 23 | 57 | 0 | 1 | 0 | 0 | 14 | 298 | 149 | 0.557027706 | 1 | 0.846 | 1 | 0.851 | 0.919 | 0.914 | 1 | 0.98 | 1 | 1 | 0.671 | -0.024 |
| SE | 70424 | ENSG00000132323 | ILKAP | chr2 | - | 238173533 | 238173653 | 238170407 | 238171024 | 238182064 | 238182186 | 67 | 44 | 102 | 75 | 71 | 0 | 2 | 0 | 0 | 0 | 63 | 61 | 82 | 99 | 102 | 1 | 0 | 0 | 0 | 6 | 269 | 149 | 0.55706468 | 1 | 1 | 0.924 | 1 | 1 | 1 | 0.972 | 1 | 1 | 1 | 0.904 | 0.01 |
| SE | 49970 | ENSG00000151690 | MFSD6 | chr2 | + | 190415291 | 190415413 | 190408354 | 190408503 | 190435976 | 190437561 | 14 | 8 | 8 | 10 | 3 | 12 | 2 | 2 | 8 | 1 | 16 | 17 | 12 | 2 | 38 | 4 | 10 | 17 | 2 | 25 | 271 | 149 | 0.557168892 | 1 | 0.391 | 0.687 | 0.687 | 0.407 | 0.623 | 0.687 | 0.483 | 0.28 | 0.355 | 0.455 | 0.107 |
| SE | 94963 | ENSG00000266714 | MYO15B | chr17 | + | 75592427 | 75592541 | 75592237 | 75592301 | 75592678 | 75592840 | 3 | 4 | 4 | 21 | 0 | 2 | 3 | 0 | 0 | 1 | 21 | 7 | 2 | 79 | 32 | 0 | 0 | 5 | 31 | 0 | 263 | 149 | 0.557177965 | 1 | 0.459 | 0.43 | 1 | 1 | 0 | 1 | 1 | 0.185 | 0.591 | 1 | -0.177 |
| SE | 50407 | ENSG00000164306 | PRIMPOL | chr4 | + | 184657081 | 184657320 | 184652022 | 184652100 | 184659339 | 184659437 | 42 | 40 | 43 | 43 | 92 | 0 | 0 | 4 | 2 | 1 | 27 | 25 | 13 | 11 | 39 | 0 | 0 | 2 | 0 | 0 | 298 | 149 | 0.557209059 | 1 | 1 | 1 | 0.843 | 0.915 | 0.979 | 1 | 1 | 0.765 | 1 | 1 | -0.006 |
| SE | 72363 | ENSG00000112877 | CEP72 | chr5 | + | 624470 | 624579 | 620068 | 620261 | 633768 | 633947 | 29 | 12 | 48 | 27 | 29 | 0 | 0 | 1 | 5 | 0 | 36 | 11 | 15 | 11 | 49 | 0 | 0 | 0 | 0 | 0 | 258 | 149 | 0.557287077 | 1 | 1 | 1 | 0.965 | 0.757 | 1 | 1 | 1 | 1 | 1 | 1 | -0.056 |
| SE | 57379 | ENSG00000101040 | ZMYND8 | chr20 | - | 47220257 | 47220324 | 47209213 | 47210897 | 47221313 | 47221474 | 42 | 19 | 29 | 44 | 10 | 1 | 4 | 0 | 8 | 5 | 29 | 45 | 49 | 42 | 1 | 6 | 7 | 8 | 4 | 0 | 216 | 149 | 0.557311007 | 1 | 0.967 | 0.766 | 1 | 0.791 | 0.58 | 0.769 | 0.816 | 0.809 | 0.879 | 1 | -0.034 |
| SE | 91179 | ENSG00000164941 | INTS8 | chr8 | + | 94865505 | 94865690 | 94859510 | 94859632 | 94867139 | 94867196 | 171 | 108 | 112 | 137 | 160 | 1 | 5 | 2 | 2 | 0 | 62 | 97 | 60 | 22 | 7 | 0 | 0 | 0 | 0 | 0 | 298 | 149 | 0.557333351 | 1 | 0.988 | 0.915 | 0.966 | 0.972 | 1 | 1 | 1 | 1 | 1 | 1 | -0.032 |
| SE | 59047 | ENSG00000174021 | GNG5 | chr1 | - | 84501825 | 84501970 | 84498324 | 84498548 | 84506010 | 84506301 | 216 | 124 | 145 | 180 | 114 | 7 | 1 | 1 | 0 | 0 | 338 | 229 | 165 | 248 | 462 | 0 | 0 | 0 | 7 | 0 | 294 | 149 | 0.557451805 | 1 | 0.94 | 0.984 | 0.987 | 1 | 1 | 1 | 1 | 1 | 0.947 | 1 | -0.007 |
| SE | 23779 | ENSG00000113716 | HMGXB3 | chr5 | + | 150026705 | 150026881 | 150024261 | 150024680 | 150027019 | 150027117 | 71 | 32 | 43 | 49 | 34 | 0 | 0 | 0 | 1 | 0 | 48 | 84 | 32 | 39 | 110 | 4 | 0 | 3 | 0 | 0 | 298 | 149 | 0.557507764 | 1 | 1 | 1 | 1 | 0.961 | 1 | 0.857 | 1 | 0.842 | 1 | 1 | 0.052 |
| SE | 87323 | ENSG00000215301 | DDX3X | chrX | + | 41343288 | 41343351 | 41342736 | 41342836 | 41343736 | 41343822 | 196 | 176 | 187 | 248 | 353 | 14 | 16 | 13 | 12 | 22 | 44 | 134 | 157 | 116 | 25 | 6 | 30 | 8 | 1 | 0 | 212 | 149 | 0.55752465 | 1 | 0.908 | 0.885 | 0.91 | 0.936 | 0.919 | 0.838 | 0.758 | 0.932 | 0.988 | 1 | 0.008 |
| SE | 50274 | ENSG00000134909 | ARHGAP32 | chr11 | - | 129123887 | 129123929 | 129123445 | 129123530 | 129124802 | 129124894 | 8 | 16 | 7 | 17 | 28 | 17 | 7 | 2 | 17 | 10 | 33 | 12 | 3 | 20 | 106 | 27 | 13 | 3 | 5 | 30 | 191 | 149 | 0.557554701 | 1 | 0.269 | 0.641 | 0.732 | 0.438 | 0.686 | 0.488 | 0.419 | 0.438 | 0.757 | 0.734 | -0.014 |
| SE | 16862 | ENSG00000138613 | APH1B | chr15 | + | 63287423 | 63287546 | 63286557 | 63286628 | 63302344 | 63302472 | 114 | 113 | 102 | 100 | 94 | 12 | 7 | 11 | 5 | 11 | 108 | 141 | 50 | 96 | 112 | 2 | 23 | 18 | 8 | 5 | 272 | 149 | 0.557576243 | 1 | 0.839 | 0.898 | 0.836 | 0.916 | 0.824 | 0.967 | 0.771 | 0.603 | 0.868 | 0.925 | 0.036 |
| SE | 59026 | ENSG00000171159 | C9orf16 | chr9 | + | 128162003 | 128162115 | 128160459 | 128160486 | 128163442 | 128163737 | 2 | 2 | 0 | 2 | 2 | 66 | 20 | 39 | 45 | 30 | 0 | 0 | 0 | 0 | 0 | 36 | 88 | 29 | 46 | 277 | 261 | 149 | 0.557689282 | 1 | 0.017 | 0.054 | 0 | 0.025 | 0.037 | 0 | 0 | 0 | 0 | 0 | 0.027 |

| | | | | | | | | | | | | | | | | | | | | | | | | | | | | | | | | | | | | | | | | | | | | | | |
|---|---|---|---|---|---|---|---|---|---|---|---|---|---|---|---|---|---|---|---|---|---|---|---|---|---|---|---|---|---|---|---|---|---|---|---|---|---|---|---|---|---|---|---|---|---|---|
| SE | 94998 | ENSG00000096996 | IL12RB1 | chr19 | - | 18072111 | 18072349 | 18070561 | 18071333 | 18073516 | 18073599 | 70 | 34 | 65 | 70 | 35 | 0 | 0 | 0 | 0 | 0 | 50 | 66 | 50 | 38 | 78 | 0 | 0 | 0 | 6 | 0 | 298 | 149 | 0.557814694 | 1 | 1 | 1 | 1 | 1 | 1 | 1 | 1 | 1 | 0.76 | 1 | 0.048 |
| SE | 51921 | ENSG00000074603 | DPP8 | chr15 | - | 65456224 | 65456371 | 65451959 | 65452102 | 65463760 | 65463845 | 182 | 184 | 144 | 175 | 167 | 1 | 3 | 2 | 0 | 5 | 121 | 179 | 275 | 121 | 71 | 0 | 2 | 0 | 0 | 0 | 296 | 149 | 0.557975175 | 1 | 0.989 | 0.969 | 0.973 | 1 | 0.944 | 1 | 0.978 | 1 | 1 | 1 | -0.021 |
| SE | 152 | ENSG00000183527 | PSMG1 | chr21 | - | 39181771 | 39181878 | 39180284 | 39180436 | 39183251 | 39183451 | 28 | 26 | 56 | 40 | 67 | 3 | 0 | 1 | 0 | 0 | 38 | 16 | 27 | 13 | 21 | 0 | 0 | 6 | 0 | 0 | 256 | 149 | 0.558193747 | 1 | 0.845 | 1 | 0.97 | 1 | 1 | 1 | 1 | 0.724 | 1 | 1 | 0.018 |
| SE | 23410 | ENSG00000153113 | CAST | chr5 | + | 96747344 | 96747392 | 96742654 | 96742756 | 96748517 | 96748613 | 201 | 131 | 143 | 148 | 280 | 0 | 1 | 0 | 0 | 0 | 168 | 190 | 113 | 176 | 93 | 0 | 0 | 0 | 0 | 11 | 197 | 149 | 0.55823575 | 1 | 1 | 0.99 | 1 | 1 | 1 | 1 | 1 | 1 | 1 | 0.865 | 0.025 |
| SE | 93238 | ENSG00000124702 | KLHDC3 | chr6 | + | 43017518 | 43017695 | 43017133 | 43017346 | 43017852 | 43017968 | 271 | 111 | 217 | 200 | 154 | 4 | 0 | 1 | 0 | 3 | 198 | 151 | 122 | 127 | 321 | 6 | 2 | 0 | 1 | 0 | 298 | 149 | 0.558415865 | 1 | 0.971 | 1 | 0.991 | 1 | 0.963 | 0.943 | 0.974 | 1 | 0.984 | 1 | 0.005 |
| SE | 4129 | ENSG00000105176 | URI1 | chr19 | + | 29980158 | 29980364 | 29971192 | 29971227 | 29985222 | 29985301 | 2 | 0 | 2 | 0 | 5 | 43 | 30 | 33 | 33 | 48 | 0 | 0 | 0 | 2 | 0 | 7 | 5 | 6 | 8 | 2 | 298 | 149 | 0.55843123 | 1 | 0.023 | 0 | 0.029 | 0 | 0.05 | 0 | 0 | 0 | 0.111 | 0 | -0.002 |
| SE | 37378 | ENSG00000166927 | MS4A7 | chr11 | + | 60389389 | 60389596 | 60383128 | 60383288 | 60394945 | 60395254 | 20 | 7 | 7 | 14 | 13 | 0 | 0 | 0 | 2 | 2 | 15 | 1 | 5 | 4 | 5 | 0 | 0 | 0 | 4 | 0 | 298 | 149 | 0.558501032 | 1 | 1 | 1 | 1 | 0.778 | 0.765 | 1 | 1 | 1 | 0.333 | 1 | 0.042 |
| SE | 31586 | ENSG00000183624 | HMCES | chr3 | + | 129288853 | 129288997 | 129279709 | 129279915 | 129290678 | 129290804 | 139 | 92 | 143 | 129 | 182 | 3 | 1 | 3 | 1 | 0 | 132 | 128 | 169 | 95 | 173 | 0 | 0 | 0 | 0 | 0 | 293 | 149 | 0.55851125 | 1 | 0.959 | 0.979 | 0.96 | 0.985 | 1 | 1 | 1 | 1 | 1 | 1 | -0.023 |
| SE | 28485 | ENSG00000155229 | MMS19 | chr10 | - | 97459361 | 97459526 | 97459222 | 97459282 | 97459658 | 97459741 | 73 | 37 | 68 | 71 | 72 | 0 | 0 | 1 | 3 | 3 | 136 | 43 | 37 | 75 | 106 | 0 | 0 | 0 | 0 | 0 | 298 | 149 | 0.558758905 | 1 | 1 | 1 | 0.971 | 0.922 | 0.923 | 1 | 1 | 1 | 1 | 1 | -0.037 |
| SE | 15307 | ENSG00000114062 | UBE3A | chr15 | - | 25407066 | 25407228 | 25405460 | 25405502 | 25408619 | 25408684 | 26 | 15 | 13 | 15 | 33 | 11 | 8 | 6 | 20 | 8 | 58 | 26 | 3 | 17 | 14 | 3 | 5 | 2 | 13 | 24 | 298 | 149 | 0.558805015 | 1 | 0.542 | 0.484 | 0.52 | 0.273 | 0.673 | 0.906 | 0.722 | 0.429 | 0.395 | 0.226 | -0.037 |
| SE | 72930 | ENSG00000182732 | RGS6 | chr14 | + | 72553289 | 72553356 | 72540040 | 72540094 | 72562416 | 72562589 | 1 | 1 | 0 | 0 | 0 | 4 | 3 | 1 | 5 | 2 | 0 | 0 | 0 | 0 | 7 | 27 | 1 | 1 | 5 | 51 | 216 | 149 | 0.558813535 | 1 | 0.147 | 0.187 | 0 | 0 | 0 | 0 | 0 | 0 | 0 | 0.086 | 0.05 |
| SE | 45417 | ENSG00000142330 | CAPN10 | chr2 | + | 240595933 | 240596152 | 240595023 | 240595304 | 240596318 | 240596521 | 15 | 0 | 14 | 5 | 11 | 20 | 3 | 12 | 7 | 4 | 10 | 1 | 1 | 2 | 22 | 11 | 2 | 0 | 5 | 24 | 298 | 149 | 0.558813871 | 1 | 0.273 | 0 | 0.368 | 0.263 | 0.579 | 0.312 | 0.2 | 1 | 0.167 | 0.314 | -0.102 |
| SE | 89482 | ENSG00000064313 | TAF2 | chr8 | - | 119793365 | 119793451 | 119789591 | 119789746 | 119795531 | 119795631 | 53 | 45 | 45 | 72 | 65 | 0 | 0 | 0 | 0 | 0 | 4 | 52 | 33 | 21 | 34 | 0 | 0 | 6 | 0 | 0 | 235 | 149 | 0.558866413 | 1 | 1 | 1 | 1 | 1 | 1 | 1 | 1 | 0.777 | 1 | 1 | 0.045 |
| SE | 30478 | ENSG00000014824 | SLC30A9 | chr4 | + | 42018388 | 42018505 | 42018110 | 42018170 | 42020415 | 42020515 | 17 | 17 | 14 | 14 | 13 | 125 | 107 | 130 | 78 | 257 | 2 | 10 | 21 | 0 | 0 | 30 | 158 | 75 | 41 | 7 | 266 | 149 | 0.55891835 | 1 | 0.071 | 0.082 | 0.057 | 0.091 | 0.028 | 0.036 | 0.034 | 0.136 | 0 | 0 | 0.025 |
| SE | 83108 | ENSG00000146909 | NOM1 | chr7 | + | 156952473 | 156952598 | 156949711 | 156950724 | 156954102 | 156954298 | 46 | 26 | 50 | 40 | 54 | 2 | 0 | 0 | 1 | 4 | 123 | 29 | 24 | 60 | 11 | 0 | 0 | 0 | 0 | 0 | 274 | 149 | 0.558932492 | 1 | 0.926 | 1 | 1 | 0.956 | 0.88 | 1 | 1 | 1 | 1 | 1 | -0.048 |
| SE | 4780 | ENSG00000064932 | SBNO2 | chr19 | - | 1112188 | 1112301 | 1111995 | 1112067 | 1112401 | 1112537 | 62 | 22 | 31 | 41 | 1 | 0 | 0 | 0 | 0 | 0 | 16 | 68 | 38 | 44 | 58 | 0 | 0 | 2 | 0 | 5 | 262 | 149 | 0.559024012 | 1 | 1 | 1 | 1 | 1 | 1 | 1 | 1 | 0.915 | 1 | 0.868 | 0.043 |
| SE | 72796 | ENSG00000107874 | CUEDC2 | chr10 | - | 102425114 | 102425198 | 102424648 | 102424792 | 102432525 | 102432574 | 92 | 38 | 71 | 77 | 62 | 0 | 0 | 1 | 0 | 0 | 183 | 123 | 63 | 98 | 203 | 0 | 4 | 0 | 1 | 6 | 233 | 149 | 0.559113564 | 1 | 1 | 1 | 0.978 | 1 | 1 | 1 | 0.952 | 1 | 0.984 | 0.956 | 0.017 |
| SE | 63406 | ENSG00000084774 | CAD | chr2 | + | 27239696 | 27239798 | 27239330 | 27239471 | 27240264 | 27240361 | 41 | 15 | 20 | 15 | 47 | 5 | 1 | 1 | 2 | 6 | 20 | 21 | 8 | 17 | 0 | 0 | 0 | 4 | 1 | 15 | 251 | 149 | 0.559164382 | 1 | 0.83 | 0.899 | 0.922 | 0.817 | 0.823 | 1 | 1 | 0.543 | 0.91 | 0 | 0.168 |
| SE | 14080 | ENSG00000130731 | METTL26 | chr16 | - | 634888 | 635063 | 634718 | 634797 | 635280 | 635340 | 43 | 16 | 52 | 28 | 25 | 4 | 8 | 1 | 0 | 0 | 23 | 43 | 29 | 52 | 57 | 0 | 8 | 0 | 0 | 16 | 298 | 149 | 0.559235655 | 1 | 0.843 | 0.5 | 0.963 | 1 | 1 | 1 | 0.729 | 1 | 1 | 0.64 | -0.013 |
| SE | 63350 | ENSG00000151092 | NGLY1 | chr3 | - | 25737333 | 25737455 | 25733871 | 25733982 | 25739576 | 25739799 | 82 | 56 | 60 | 81 | 121 | 1 | 0 | 0 | 2 | 0 | 21 | 31 | 54 | 36 | 3 | 0 | 0 | 0 | 6 | 0 | 271 | 149 | 0.559270445 | 1 | 0.978 | 1 | 1 | 0.957 | 1 | 1 | 1 | 1 | 0.767 | 1 | 0.034 |
| SE | 10544 | ENSG00000177302 | TOP3A | chr17 | - | 18280535 | 18280658 | 18271427 | 18274980 | 18282697 | 18282841 | 82 | 28 | 34 | 33 | 83 | 0 | 0 | 0 | 0 | 2 | 61 | 54 | 47 | 45 | 35 | 0 | 0 | 0 | 0 | 6 | 272 | 149 | 0.559318756 | 1 | 1 | 1 | 1 | 1 | 0.958 | 1 | 1 | 1 | 1 | 0.762 | 0.039 |
| SE | 92461 | ENSG00000100815 | TRIP11 | chr14 | - | 91975171 | 91975286 | 91974626 | 91974743 | 91976107 | 91976189 | 184 | 197 | 195 | 176 | 323 | 3 | 4 | 1 | 2 | 3 | 13 | 24 | 28 | 16 | 13 | 0 | 0 | 0 | 0 | 0 | 264 | 149 | 0.559459373 | 1 | 0.972 | 0.965 | 0.991 | 0.98 | 0.984 | 1 | 1 | 1 | 1 | 1 | -0.022 |
| SE | 60665 | ENSG00000147099 | HDAC8 | chrX | - | 72495155 | 72495268 | 72490928 | 72491006 | 72567888 | 72568030 | 31 | 17 | 22 | 20 | 19 | 3 | 1 | 0 | 2 | 0 | 7 | 28 | 16 | 10 | 15 | 0 | 8 | 0 | 0 | 0 | 262 | 149 | 0.559578805 | 1 | 0.855 | 0.906 | 1 | 0.85 | 1 | 1 | 0.666 | 1 | 1 | 1 | -0.011 |
| SE | 102826 | ENSG00000148700 | ADD3 | chr10 | + | 110105999 | 110106053 | 110100624 | 110100848 | 110112776 | 110112894 | 45 | 35 | 74 | 37 | 55 | 779 | 577 | 833 | 795 | 806 | 6 | 58 | 20 | 19 | 13 | 143 | 683 | 546 | 325 | 180 | 203 | 149 | 0.559604162 | 1 | 0.041 | 0.043 | 0.061 | 0.033 | 0.048 | 0.03 | 0.059 | 0.026 | 0.041 | 0.05 | 0.004 |
| SE | 13085 | ENSG00000131148 | EMC8 | chr16 | - | 85780378 | 85780473 | 85778623 | 85779867 | 85781210 | 85781258 | 55 | 29 | 42 | 49 | 65 | 12 | 2 | 9 | 22 | 27 | 42 | 74 | 48 | 54 | 61 | 5 | 14 | 7 | 9 | 39 | 244 | 149 | 0.559605556 | 1 | 0.737 | 0.899 | 0.74 | 0.576 | 0.595 | 0.837 | 0.763 | 0.807 | 0.786 | 0.489 | -0.027 |
| SE | 75848 | ENSG00000124523 | SIRT5 | chr6 | + | 13600647 | 13600949 | 13599031 | 13599155 | 13611789 | 13615158 | 21 | 7 | 23 | 23 | 5 | 0 | 2 | 0 | 3 | 0 | 18 | 20 | 4 | 23 | 19 | 0 | 0 | 0 | 0 | 0 | 298 | 149 | 0.559621355 | 1 | 1 | 0.636 | 1 | 0.793 | 1 | 1 | 1 | 1 | 1 | 1 | -0.114 |
| SE | 80067 | ENSG00000198198 | SZT2 | chr1 | + | 43416534 | 43416641 | 43404379 | 43404550 | 43419733 | 43419944 | 23 | 19 | 17 | 22 | 17 | 0 | 0 | 0 | 0 | 0 | 61 | 61 | 11 | 10 | 23 | 0 | 0 | 0 | 4 | 0 | 256 | 149 | 0.559685657 | 1 | 1 | 1 | 1 | 1 | 1 | 1 | 1 | 1 | 0.593 | 1 | 0.081 |
| SE | 66927 | ENSG00000152520 | PAN3 | chr13 | + | 28287983 | 28288122 | 28281314 | 28281379 | 28292381 | 28295335 | 152 | 130 | 106 | 131 | 175 | 2 | 0 | 0 | 4 | 2 | 23 | 110 | 115 | 81 | 24 | 0 | 0 | 0 | 2 | 0 | 288 | 149 | 0.559873536 | 1 | 0.975 | 1 | 1 | 0.944 | 0.978 | 1 | 1 | 1 | 0.954 | 1 | -0.011 |
| SE | 1799 | ENSG00000099917 | MED15 | chr22 | + | 20518877 | 20518995 | 20507581 | 20507746 | 20537116 | 20537183 | 6 | 7 | 1 | 3 | 6 | 87 | 37 | 60 | 50 | 30 | 9 | 0 | 8 | 2 | 8 | 76 | 73 | 43 | 83 | 120 | 267 | 149 | 0.559969474 | 1 | 0.037 | 0.095 | 0.009 | 0.032 | 0.1 | 0.062 | 0 | 0.094 | 0.013 | 0.036 | 0.014 |
| SE | 104740 | ENSG00000122694 | GLIPR2 | chr9 | + | 36148546 | 36148650 | 36147785 | 36147894 | 36162361 | 36162910 | 223 | 127 | 132 | 185 | 133 | 0 | 0 | 0 | 0 | 0 | 228 | 245 | 111 | 244 | 374 | 0 | 4 | 3 | 7 | 0 | 253 | 149 | 0.560004238 | 1 | 1 | 1 | 1 | 1 | 1 | 1 | 0.973 | 0.956 | 0.954 | 1 | 0.023 |
| SE | 94654 | ENSG00000011454 | RABGAP1 | chr9 | + | 123101565 | 123101763 | 123099477 | 123099549 | 123103090 | 123103268 | 194 | 188 | 250 | 290 | 374 | 0 | 2 | 0 | 2 | 0 | 191 | 263 | 213 | 172 | 67 | 0 | 11 | 0 | 0 | 0 | 298 | 149 | 0.560013546 | 1 | 1 | 0.979 | 1 | 0.986 | 1 | 1 | 0.923 | 1 | 1 | 1 | 0.008 |
| SE | 48621 | ENSG00000131374 | TBC1D5 | chr3 | - | 17562018 | 17562142 | 17518901 | 17518995 | 17586352 | 17586557 | 22 | 15 | 4 | 24 | 19 | 1 | 2 | 3 | 0 | 6 | 8 | 16 | 10 | 19 | 16 | 8 | 9 | 0 | 2 | 0 | 273 | 149 | 0.560041048 | 1 | 0.923 | 0.804 | 0.421 | 1 | 0.633 | 0.353 | 0.492 | 1 | 0.838 | 1 | 0.02 |
| SE | 63943 | ENSG00000116005 | PCYOX1 | chr2 | + | 70261211 | 70261386 | 70259359 | 70259566 | 70274958 | 70275084 | 93 | 65 | 85 | 86 | 79 | 4 | 0 | 0 | 2 | 2 | 21 | 29 | 35 | 52 | 48 | 0 | 0 | 9 | 0 | 0 | 298 | 149 | 0.560076965 | 1 | 0.921 | 1 | 1 | 0.956 | 0.952 | 1 | 1 | 0.66 | 1 | 1 | 0.034 |
| SE | 12509 | ENSG00000132600 | PRMT7 | chr16 | + | 68347630 | 68347678 | 68346120 | 68346280 | 68348341 | 68348431 | 10 | 10 | 10 | 8 | 3 | 0 | 8 | 0 | 0 | 0 | 3 | 14 | 22 | 0 | 59 | 3 | 0 | 0 | 3 | 0 | 197 | 149 | 0.560087087 | 1 | 1 | 0.486 | 1 | 1 | 1 | 0.431 | 1 | 1 | 0 | 1 | 0.211 |
| SE | 39412 | ENSG00000137393 | RNF144B | chr6 | + | 18427580 | 18427685 | 18399498 | 18399699 | 18439683 | 18439744 | 102 | 128 | 62 | 57 | 105 | 12 | 7 | 4 | 3 | 1 | 101 | 125 | 421 | 86 | 83 | 0 | 14 | 9 | 0 | 13 | 254 | 149 | 0.560113123 | 1 | 0.833 | 0.915 | 0.901 | 0.918 | 0.984 | 1 | 0.84 | 0.965 | 1 | 0.789 | -0.009 |
| SE | 84802 | ENSG00000173681 | BCLAF3 | chrX | - | 19955390 | 19955566 | 19953777 | 19953892 | 19965043 | 19965142 | 148 | 118 | 91 | 158 | 187 | 11 | 5 | 10 | 11 | 7 | 51 | 107 | 152 | 56 | 24 | 11 | 1 | 5 | 5 | 0 | 298 | 149 | 0.560273568 | 1 | 0.871 | 0.922 | 0.82 | 0.878 | 0.93 | 0.699 | 0.982 | 0.938 | 0.848 | 1 | -0.009 |
| SE | 54110 | ENSG00000115935 | WIPF1 | chr2 | - | 174630381 | 174630696 | 174629665 | 174629868 | 174634517 | 174634536 | 24 | 19 | 21 | 25 | 32 | 10 | 2 | 1 | 5 | 0 | 29 | 71 | 91 | 92 | 21 | 7 | 32 | 2 | 21 | 0 | 298 | 149 | 0.560278033 | 1 | 0.545 | 0.826 | 0.913 | 0.714 | 1 | 0.674 | 0.526 | 0.958 | 0.687 | 1 | 0.031 |
| SE | 54880 | ENSG00000158825 | CDA | chr1 | + | 20613841 | 20613899 | 20604927 | 20605039 | 20618451 | 20618903 | 64 | 46 | 60 | 28 | 77 | 10 | 3 | 2 | 12 | 3 | 399 | 212 | 107 | 369 | 189 | 18 | 39 | 13 | 39 | 33 | 207 | 149 | 0.560282659 | 1 | 0.822 | 0.917 | 0.956 | 0.627 | 0.949 | 0.941 | 0.796 | 0.856 | 0.872 | 0.805 | 0 |
| SE | 25127 | ENSG00000153310 | CYRIB | chr8 | - | 129904498 | 129904585 | 129903311 | 129903350 | 129912469 | 129912515 | 179 | 115 | 116 | 153 | 175 | 13 | 4 | 11 | 14 | 15 | 56 | 170 | 209 | 197 | 31 | 13 | 6 | 16 | 9 | 0 | 236 | 149 | 0.560334136 | 1 | 0.897 | 0.948 | 0.869 | 0.873 | 0.88 | 0.731 | 0.947 | 0.892 | 0.933 | 1 | -0.007 |
| [illegible] | | | | | | | | | | | | | | | | | | | | | | | | | | | | | | | | | | | | | | | | | | | | | | |

| | | | | | | | | | | | | | | | | | | | | | | | | | |
|---|---|---|---|---|---|---|---|---|---|---|---|---|---|---|---|---|---|---|---|---|---|---|---|---|---|
| SE | 4827 | ENSG00000078699 | CBFA2T2 | chr20 | + | 33640340 | 33640531 | 33636639 | 33636708 | 33644346 | 33650030 | 158 | 116 | 146 | 137 | 171 | 2 | 0 | 1 | 1 | 4 | 181 | 98 | 84 | 130 |
| SE | 76933 | ENSG00000131018 | SYNE1 | chr6 | - | 152145486 | 152145555 | 152143622 | 152143765 | 152148044 | 152148378 | 98 | 67 | 100 | 82 | 60 | 59 | 29 | 36 | 41 | 45 | 7 | 128 | 101 | 12 |
| SE | 45778 | ENSG00000284770 | TBCE | chr1 | + | 235438768 | 235438922 | 235437321 | 235437474 | 235441813 | 235441823 | 88 | 94 | 75 | 91 | 140 | 0 | 0 | 1 | 5 | 1 | 154 | 88 | 63 | 68 |
| SE | 9269 | ENSG00000179094 | PER1 | chr17 | - | 8145957 | 8146137 | 8144750 | 8144847 | 8146371 | 8146488 | 33 | 12 | 22 | 15 | 13 | 1 | 1 | 0 | 0 | 0 | 7 | 21 | 75 | 8 |
| SE | 14186 | ENSG00000140983 | RHOT2 | chr16 | + | 670674 | 670773 | 670455 | 670557 | 670891 | 670940 | 198 | 75 | 143 | 162 | 116 | 2 | 1 | 3 | 3 | 0 | 179 | 166 | 114 | 113 |
| SE | 106425 | ENSG00000135297 | MTO1 | chr6 | + | 73480996 | 73481116 | 73480674 | 73480805 | 73482039 | 73482244 | 0 | 0 | 2 | 0 | 2 | 70 | 30 | 36 | 50 | 56 | 0 | 0 | 0 | 0 |
| SE | 69429 | ENSG00000123297 | TSFM | chr12 | + | 57792985 | 57793073 | 57787039 | 57787162 | 57796176 | 57796261 | 99 | 49 | 61 | 43 | 63 | 0 | 3 | 5 | 1 | 2 | 60 | 53 | 51 | 36 |
| SE | 8159 | ENSG00000167699 | GLOD4 | chr17 | - | 781862 | 782043 | 778621 | 778744 | 782165 | 782267 | 5 | 0 | 0 | 4 | 0 | 60 | 33 | 35 | 61 | 21 | 2 | 1 | 2 | 0 |
| SE | 100540 | ENSG00000054654 | SYNE2 | chr14 | + | 64128451 | 64128553 | 64126597 | 64126807 | 64129781 | 64129901 | 674 | 455 | 348 | 504 | 353 | 0 | 0 | 0 | 2 | 1 | 82 | 203 | 205 | 59 |
| SE | 9181 | ENSG00000133028 | SCO1 | chr17 | - | 10691871 | 10691964 | 10686726 | 10686842 | 10695740 | 10695831 | 26 | 24 | 32 | 19 | 24 | 0 | 0 | 0 | 0 | 0 | 34 | 21 | 33 | 28 |
| SE | 94357 | ENSG00000021776 | AQR | chr15 | - | 34862866 | 34863041 | 34860041 | 34860155 | 34867523 | 34867609 | 308 | 207 | 245 | 288 | 677 | 2 | 0 | 0 | 0 | 0 | 11 | 231 | 152 | 46 |
| SE | 58794 | ENSG00000077254 | USP33 | chr1 | - | 77734346 | 77734416 | 77730617 | 77730731 | 77739264 | 77739417 | 114 | 79 | 69 | 69 | 97 | 0 | 0 | 0 | 0 | 1 | 21 | 76 | 54 | 64 |
| SE | 95855 | ENSG00000107263 | RAPGEF1 | chr9 | - | 131627912 | 131628096 | 131625921 | 131626422 | 131629101 | 131629254 | 247 | 88 | 148 | 183 | 177 | 1 | 0 | 0 | 8 | 0 | 236 | 242 | 196 | 188 |
| SE | 30693 | ENSG00000132466 | ANKRD17 | chr4 | - | 73139530 | 73140283 | 73135116 | 73135262 | 73141740 | 73141792 | 163 | 127 | 142 | 154 | 94 | 6 | 0 | 3 | 7 | 2 | 119 | 133 | 134 | 90 |
| SE | 27543 | ENSG00000083444 | PLOD1 | chr1 | + | 11957089 | 11957193 | 11956916 | 11957014 | 11957841 | 11957943 | 2 | 0 | 0 | 6 | 0 | 53 | 17 | 32 | 43 | 32 | 0 | 2 | 0 | 4 |
| SE | 5917 | ENSG00000080189 | SLC35C2 | chr20 | - | 46352050 | 46352226 | 46350354 | 46350890 | 46354881 | 46354965 | 184 | 60 | 143 | 124 | 68 | 0 | 0 | 0 | 0 | 0 | 160 | 145 | 56 | 137 |
| SE | 101593 | ENSG00000172671 | ZFAND4 | chr10 | - | 45640240 | 45640421 | 45639705 | 45639963 | 45648293 | 45648534 | 4 | 2 | 3 | 0 | 0 | 30 | 20 | 35 | 36 | 81 | 0 | 0 | 0 | 0 |
| SE | 82274 | ENSG00000136247 | ZDHHC4 | chr7 | + | 6577589 | 6577720 | 6577433 | 6577498 | 6578565 | 6578720 | 15 | 1 | 7 | 4 | 3 | 1 | 0 | 2 | 4 | 0 | 31 | 1 | 1 | 10 |
| SE | 7008 | ENSG00000101782 | RIOK3 | chr18 | + | 23473428 | 23473483 | 23467398 | 23467526 | 23474947 | 23475107 | 76 | 85 | 45 | 91 | 155 | 0 | 0 | 0 | 0 | 0 | 75 | 104 | 116 | 148 |
| SE | 81630 | ENSG00000135093 | USP30 | chr12 | + | 109084952 | 109085073 | 109082694 | 109083062 | 109085666 | 109085922 | 21 | 12 | 31 | 29 | 51 | 0 | 0 | 0 | 0 | 0 | 50 | 20 | 30 | 25 |
| SE | 88036 | ENSG00000156171 | DRAM2 | chr1 | - | 111122615 | 111122643 | 111120515 | 111120693 | 111126226 | 111126294 | 16 | 8 | 10 | 14 | 26 | 7 | 11 | 15 | 11 | 29 | 10 | 2 | 5 | 5 |
| SE | 20558 | ENSG00000108588 | CCDC47 | chr17 | - | 63764015 | 63764190 | 63761229 | 63761351 | 63764739 | 63764793 | 243 | 161 | 181 | 215 | 170 | 1 | 0 | 0 | 0 | 0 | 136 | 164 | 238 | 226 |
| SE | 96894 | ENSG00000100731 | PCNX1 | chr14 | + | 71009633 | 71009724 | 70976941 | 70978648 | 71011491 | 71011549 | 220 | 123 | 189 | 187 | 239 | 0 | 0 | 2 | 1 | 1 | 34 | 250 | 190 | 59 |
| SE | 93606 | ENSG00000158856 | DMTN | chr8 | + | 22080804 | 22080870 | 22080617 | 22080625 | 22081112 | 22081193 | 19 | 30 | 8 | 21 | 17 | 9 | 7 | 4 | 7 | 6 | 50 | 15 | 11 | 7 |
| SE | 55114 | ENSG00000072110 | ACTN1 | chr14 | - | 68909954 | 68910042 | 68909317 | 68909396 | 68912155 | 68912242 | 392 | 361 | 451 | 553 | 746 | 0 | 2 | 3 | 4 | 2 | 1159 | 433 | 444 | 1042 |
| SE | 835 | ENSG00000198911 | SREBF2 | chr22 | + | 41899236 | 41899553 | 41898648 | 41898781 | 41900329 | 41900498 | 0 | 2 | 2 | 0 | 4 | 51 | 25 | 39 | 30 | 34 | 0 | 2 | 1 | 0 |
| SE | 72403 | ENSG00000138835 | RGS3 | chr9 | + | 113582071 | 113582242 | 113580659 | 113581005 | 113583449 | 113583671 | 52 | 34 | 15 | 15 | 6 | 5 | 5 | 4 | 4 | 6 | 15 | 25 | 9 | 11 |
| SE | 37350 | ENSG00000075826 | SEC31B | chr10 | - | 100498704 | 100498804 | 100498028 | 100498207 | 100499159 | 100499258 | 19 | 22 | 36 | 51 | 16 | 7 | 13 | 10 | 5 | 18 | 75 | 42 | 40 | 18 |
| SE | 98349 | ENSG00000213995 | NAXD | chr13 | + | 110625189 | 110625278 | 110624233 | 110624279 | 110627438 | 110627547 | 31 | 22 | 34 | 18 | 32 | 7 | 3 | 12 | 8 | 8 | 20 | 18 | 25 | 57 |
| SE | 55440 | ENSG00000116688 | MFN2 | chr1 | + | 11992554 | 11992690 | 11981809 | 11982114 | 11996155 | 11996318 | 45 | 39 | 45 | 51 | 61 | 0 | 0 | 0 | 0 | 0 | 173 | 41 | 61 | 82 |
| SE | 10749 | ENSG00000108256 | NUFIP2 | chr17 | - | 29285991 | 29287716 | 29267497 | 29267530 | 29293782 | 29294148 | 349 | 332 | 279 | 426 | 497 | 5 | 0 | 4 | 8 | 7 | 135 | 288 | 458 | 192 |
| SE | 101867 | ENSG00000026103 | FAS | chr10 | + | 89010752 | 89010815 | 89010538 | 89010600 | 89011998 | 89012081 | 89 | 78 | 72 | 85 | 81 | 28 | 10 | 29 | 27 | 41 | 0 | 67 | 46 | 14 |
| SE | 52992 | ENSG00000077585 | GPR137B | chr1 | + | 236183777 | 236183906 | 236179878 | 236180028 | 236205125 | 236205198 | 132 | 64 | 58 | 105 | 67 | 0 | 0 | 0 | 0 | 0 | 126 | 101 | 58 | 80 |
| SE | 52567 | ENSG00000111445 | RFC5 | chr12 | + | 118017689 | 118017852 | 118016728 | 118016892 | 118019071 | 118019136 | 2 | 1 | 0 | 4 | 2 | 13 | 12 | 15 | 23 | 8 | 9 | 0 | 3 | 1 |
| SE | 55046 | ENSG00000117713 | ARID1A | chr1 | + | 26762151 | 26762319 | 26761383 | 26761473 | 26762972 | 26763175 | 158 | 76 | 179 | 146 | 119 | 0 | 2 | 0 | 2 | 7 | 310 | 175 | 100 | 154 |
| SE | 77299 | ENSG00000099968 | BCL2L13 | chr22 | + | 17688985 | 17689142 | 17683213 | 17683321 | 17696140 | 17696210 | 277 | 178 | 235 | 229 | 343 | 2 | 3 | 4 | 5 | 7 | 549 | 346 | 198 | 411 |
| SE | 92087 | ENSG00000167325 | RRM1 | chr11 | + | 4119844 | 4119928 | 4111899 | 4112062 | 4121603 | 4121765 | 47 | 22 | 45 | 38 | 48 | 0 | 0 | 1 | 2 | 0 | 5 | 24 | 40 | 41 |
| SE | 28034 | ENSG00000120519 | SLC10A7 | chr4 | - | 146509912 | 146510049 | 146503848 | 146503924 | 146517037 | 146517120 | 163 | 116 | 157 | 125 | 158 | 1 | 0 | 4 | 1 | 2 | 28 | 105 | 87 | 17 |
| SE | 81220 | ENSG00000101307 | SIRPB1 | chr20 | - | 1578337 | 1578694 | 1566152 | 1566267 | 1619868 | 1619996 | 100 | 64 | 38 | 49 | 62 | 0 | 1 | 0 | 0 | 0 | 154 | 146 | 115 | 240 |
| SE | 97535 | ENSG00000149761 | NUDT22 | chr11 | + | 64229246 | 64229344 | 64226634 | 64227132 | 64229477 | 64229539 | 18 | 9 | 11 | 15 | 3 | 0 | 0 | 0 | 0 | 3 | 12 | 31 | 17 | 30 |
| SE | 76363 | ENSG00000164674 | SYTL3 | chr6 | + | 158651752 | 158651842 | 158650013 | 158650078 | 158661268 | 158661385 | 120 | 52 | 56 | 58 | 44 | 3 | 3 | 0 | 0 | 0 | 85 | 176 | 196 | 75 |
| SE | 65729 | ENSG00000138073 | PREB | chr2 | - | 27131671 | 27131831 | 27130755 | 27131508 | 27132009 | 27132082 | 49 | 21 | 35 | 34 | 22 | 1 | 0 | 0 | 0 | 0 | 12 | 31 | 14 | 15 |
| SE | 91111 | ENSG00000173848 | NET1 | chr10 | + | 5452857 | 5452920 | 5452357 | 5452525 | 5453249 | 5453346 | 31 | 23 | 54 | 28 | 91 | 0 | 0 | 0 | 2 | 5 | 82 | 13 | 57 | 46 |
| SE | 91555 | ENSG00000139546 | TARBP2 | chr12 | + | 53503026 | 53503129 | 53502014 | 53502184 | 53503729 | 53503808 | 17 | 2 | 20 | 31 | 17 | 0 | 0 | 2 | 0 | 0 | 16 | 9 | 22 | 16 |
| SE | 41568 | ENSG00000136875 | PRPF4 | chr9 | + | 113283131 | 113283211 | 113282645 | 113282733 | 113283388 | 113283482 | 119 | 70 | 124 | 118 | 127 | 0 | 0 | 0 | 4 | 4 | 183 | 91 | 116 | 74 |
| SE | 58832 | ENSG00000164828 | SUN1 | chr7 | + | 849519 | 849603 | 848417 | 848615 | 851383 | 851482 | 81 | 98 | 84 | 98 | 122 | 2 | 0 | 0 | 0 | 0 | 178 | 92 | 58 | 107 |
| SE | 65174 | ENSG00000268362 | NA | chr19 | + | 24042442 | 24042722 | 24033445 | 24033621 | 24046183 | 24046279 | 1 | 0 | 1 | 3 | 0 | 15 | 7 | 7 | 5 | 9 | 0 | 4 | 0 | 4 |
| SE | 51189 | ENSG00000122986 | HVCN1 | chr12 | - | 110683224 | 110683264 | 110661163 | 110661448 | 110688624 | 110688708 | 201 | 197 | 166 | 199 | 167 | 0 | 0 | 0 | 3 | 0 | 800 | 396 | 105 | 418 |
| SE | 3843 | ENSG00000157540 | DYRK1A | chr21 | + | 37421773 | 37421866 | 37420298 | 37420384 | 37425689 | 37425781 | 45 | 27 | 21 | 47 | 65 | 0 | 1 | 0 | 1 | 0 | 53 | 31 | 10 | 17 |
| SE | 98730 | ENSG00000107290 | SETX | chr9 | - | 132326323 | 132330499 | 132311756 | 132311856 | 132331051 | 132331139 | 776 | 587 | 531 | 733 | 636 | 0 | 0 | 0 | 2 | 0 | 60 | 486 | 456 | 56 |
| SE | 11834 | ENSG00000254788 | CKLF-CMTM1 | chr16 | + | 66558189 | 66558348 | 66552678 | 66552793 | 66566082 | 66566428 | 98 | 43 | 109 | 104 | 60 | 6 | 0 | 1 | 0 | 0 | 82 | 78 | 105 | 93 |
| SE | 83363 | ENSG00000087274 | ADD1 | chr4 | + | 2926638 | 2926675 | 2926013 | 2926112 | 2928170 | 2929307 | 0 | 1 | 2 | 0 | 5 | 390 | 178 | 345 | 344 | 372 | 0 | 3 | 3 | 5 |
| SE | 103252 | ENSG00000172661 | WASHC2C | chr10 | + | 45737982 | 45738045 | 45728861 | 45729026 | 45740072 | 45740246 | 103 | 118 | 91 | 110 | 84 | 0 | 0 | 1 | 0 | 0 | 259 | 90 | 81 | 175 |
| SE | 23254 | ENSG00000108262 | GIT1 | chr17 | - | 29577135 | 29577247 | 29576862 | 29576996 | 29577644 | 29577742 | 62 | 41 | 67 | 104 | 74 | 4 | 0 | 2 | 1 | 0 | 122 | 157 | 62 | 110 |
| SE | 35792 | ENSG00000173226 | IQCB1 | chr3 | - | 121772556 | 121772713 | 121769760 | 121770574 | 121781742 | 121781874 | 110 | 96 | 158 | 163 | 198 | 2 | 1 | 0 | 3 | 2 | 10 | 87 | 80 | 41 |
| SE | 42059 | ENSG00000092871 | RFFL | chr17 | - | 35017522 | 35017606 | 35016369 | 35016580 | 35026373 | 35026568 | 85 | 37 | 79 | 69 | 52 | 2 | 0 | 2 | 2 | 0 | 61 | 76 | 94 | 82 |
| SE | 60589 | ENSG00000070961 | ATP2B1 | chr12 | - | 89601325 | 89601433 | 89599116 | 89599299 | 89603042 | 89603254 | 328 | 206 | 278 | 240 | 488 | 6 | 1 | 3 | 2 | 3 | 23 | 98 | 96 | 8 |
| SE | 40269 | ENSG00000170581 | STAT2 | chr12 | - | 56349514 | 56349636 | 56349344 | 56349509 | 56350411 | 56350432 | 52 | 21 | 27 | 49 | 69 | 0 | 0 | 0 | 0 | 0 | 150 | 27 | 38 | 132 |
| SE | 78805 | ENSG00000135269 | TES | chr7 | + | 116251759 | 116251975 | 116250160 | 116250496 | 116257293 | 116257668 | 285 | 148 | 279 | 222 | 333 | 9 | 2 | 8 | 4 | 7 | 191 | 182 | 107 | 140 |

[illegible]

| | | | | | | | | | | | | | | | |
|---|---|---|---|---|---|---|---|---|---|---|---|---|---|---|---|
| SE | 47831 | ENSG00000187554 | TLR5 | chr1 | - | 223137177 | 223137263 | 223134681 | 223134864 | 223143195 | 223143248 | 12 | 9 | 10 | [illegible] |
| SE | 71218 | ENSG00000079263 | SP140 | chr2 | + | 230248884 | 230248968 | 230245862 | 230245940 | 230253315 | 230253417 | 3 | 7 | 4 | [illegible] |
| SE | 49647 | ENSG00000081760 | AACS | chr12 | + | 125136661 | 125136864 | 125134793 | 125134852 | 125142091 | 125142177 | 18 | 11 | 33 | [illegible] |
| SE | 107378 | ENSG00000168010 | ATG16L2 | chr11 | + | 72822847 | 72822961 | 72822321 | 72822543 | 72824059 | 72824122 | 116 | 116 | 81 | [illegible] |
| SE | 48755 | ENSG00000116473 | RAP1A | chr1 | + | 111706662 | 111706777 | 111704342 | 111704486 | 111709148 | 111709264 | 4 | 10 | 9 | [illegible] |
| SE | 60106 | ENSG00000116288 | PARK7 | chr1 | + | 7977651 | 7977738 | 7970893 | 7970963 | 7984893 | 7985052 | 922 | 657 | 686 | [illegible] |
| SE | 48926 | ENSG00000198408 | OGA | chr10 | - | 101803734 | 101804019 | 101800241 | 101800400 | 101806044 | 101806143 | 1027 | 898 | 927 | [illegible] |
| SE | 61740 | ENSG00000143727 | ACP1 | chr2 | + | 272036 | 272065 | 271865 | 271939 | 272191 | 272305 | 13 | 10 | 29 | [illegible] |
| SE | 2787 | ENSG00000011304 | PTBP1 | chr19 | + | 805512 | 805569 | 805012 | 805187 | 806407 | 806556 | 12 | 8 | 14 | [illegible] |
| SE | 57924 | ENSG00000110200 | ANAPC15 | chr11 | - | 72111411 | 72111496 | 72110543 | 72110603 | 72112649 | 72112754 | 37 | 14 | 10 | [illegible] |
| SE | 32961 | ENSG00000114127 | XRN1 | chr3 | - | 142417139 | 142417229 | 142414134 | 142414291 | 142418503 | 142418609 | 357 | 331 | 253 | [illegible] |
| SE | 48163 | ENSG00000117543 | DPH5 | chr1 | - | 101001466 | 101001587 | 100995109 | 100995149 | 101021640 | 101021765 | 95 | 71 | 87 | [illegible] |
| SE | 95431 | ENSG00000167130 | DOLPP1 | chr9 | + | 129085022 | 129085107 | 129084598 | 129084768 | 129085206 | 129085295 | 14 | 6 | 9 | [illegible] |
| SE | 95261 | ENSG00000165219 | GAPVD1 | chr9 | + | 125295457 | 125295574 | 125261910 | 125261959 | 125298889 | 125299106 | 15 | 11 | 11 | [illegible] |
| SE | 55970 | ENSG00000136161 | RCBTB2 | chr13 | - | 48522307 | 48522403 | 48515585 | 48515741 | 48533027 | 48533080 | 4 | 2 | 2 | [illegible] |
| SE | 47482 | ENSG00000143363 | PRUNE1 | chr1 | + | 151017811 | 151017904 | 151008419 | 151008671 | 151025514 | 151025673 | 19 | 28 | 17 | [illegible] |
| SE | 103290 | ENSG00000196233 | LCOR | chr10 | + | 96858291 | 96858416 | 96833405 | 96833479 | 96907264 | 96907330 | 10 | 6 | 4 | [illegible] |
| SE | 22152 | ENSG00000156639 | ZFAND3 | chr6 | + | 38116571 | 38116739 | 38082391 | 38082457 | 38152234 | 38152844 | 639 | 490 | 451 | [illegible] |
| SE | 79620 | ENSG00000162688 | AGL | chr1 | + | 99870757 | 99870869 | 99870399 | 99870581 | 99874686 | 99874810 | 125 | 109 | 112 | [illegible] |
| SE | 103821 | ENSG00000225484 | NUTM2B-AS1 | chr10 | - | 79812112 | 79812177 | 79808758 | 79808832 | 79814612 | 79814716 | 0 | 0 | 0 | [illegible] |
| SE | 15187 | ENSG00000159459 | UBR1 | chr15 | - | 43022701 | 43022801 | 43021080 | 43021375 | 43024828 | 43024983 | 278 | 180 | 172 | [illegible] |
| SE | 57283 | ENSG00000143119 | CD53 | chr1 | + | 110896652 | 110896733 | 110894959 | 110895055 | 110899123 | 110899922 | 490 | 190 | 325 | [illegible] |
| SE | 87631 | ENSG00000140678 | ITGAX | chr16 | + | 31371333 | 31371497 | 31371083 | 31371214 | 31371629 | 31371784 | 162 | 124 | 136 | [illegible] |
| SE | 56930 | ENSG00000118197 | DDX59 | chr1 | - | 200648438 | 200648567 | 200640810 | 200641185 | 200649073 | 200649149 | 93 | 33 | 58 | [illegible] |
| SE | 73244 | ENSG00000114956 | DGUOK | chr2 | + | 73932597 | 73932683 | 73926896 | 73927052 | 73958145 | 73958245 | 4 | 3 | 6 | [illegible] |
| SE | 71903 | ENSG00000074054 | CLASP1 | chr2 | - | 121530246 | 121530325 | 121503166 | 121503234 | 121605700 | 121606180 | 47 | 42 | 64 | [illegible] |
| SE | 60395 | ENSG00000174796 | THAP6 | chr4 | + | 75521735 | 75521861 | 75516771 | 75516979 | 75542408 | 75542486 | 29 | 52 | 34 | [illegible] |
| SE | 103245 | ENSG00000165732 | DDX21 | chr10 | + | 68977528 | 68977688 | 68974669 | 68974743 | 68978841 | 68978976 | 267 | 274 | 266 | [illegible] |
| SE | 10140 | ENSG00000011260 | UTP18 | chr17 | + | 51273361 | 51273450 | 51268836 | 51268904 | 51275865 | 51275991 | 95 | 81 | 158 | [illegible] |
| SE | 23641 | ENSG00000113263 | ITK | chr5 | + | 157228295 | 157228361 | 157222748 | 157223014 | 157232339 | 157232394 | 643 | 412 | 707 | [illegible] |
| SE | 75281 | ENSG00000024048 | UBR2 | chr6 | + | 42612170 | 42612291 | 42606588 | 42606651 | 42615070 | 42615178 | 262 | 239 | 251 | [illegible] |
| SE | 26450 | ENSG00000083312 | TNPO1 | chr5 | + | 72891809 | 72891896 | 72889785 | 72889957 | 72893138 | 72893246 | 265 | 125 | 181 | [illegible] |
| SE | 24673 | ENSG00000120705 | ETF1 | chr5 | - | 138510564 | 138510629 | 138508668 | 138508816 | 138511044 | 138511200 | 391 | 342 | 361 | [illegible] |

[illegible]

| | | | | | | | | | | | | | | | | | | | | | | | | | | | | | | | | | | | | | | | | | | | | |
|---|---|---|---|---|---|---|---|---|---|---|---|---|---|---|---|---|---|---|---|---|---|---|---|---|---|---|---|---|---|---|---|---|---|---|---|---|---|---|---|---|---|---|---|---|
| SE | 43742 | ENSG00000148019 | CEP78 | chr9 | + | 78246668 | 78246782 | 78243461 | 78243636 | 78248290 | 78248355 | 249 | 97 | 144 | 216 | 226 | 8 | 0 | 2 | 2 | 0 | 61 | 144 | 113 | 63 | 22 | 0 | 0 | 0 | 0 | 0 | 263 | 149 | 0.603045757 | 1 | 0.946 | 1 | 0.976 | 0.984 | 1 | 1 | 1 | 1 | 1 | 1 | -0.019 |
| SE | 50075 | ENSG00000053254 | FOXN3 | chr14 | - | 89190383 | 89190449 | 89180700 | 89180806 | 89280949 | 89281014 | 184 | 112 | 204 | 204 | 330 | 62 | 48 | 87 | 84 | 79 | 238 | 204 | 183 | 229 | 73 | 35 | 54 | 100 | 69 | 40 | 215 | 149 | 0.603050232 | 1 | 0.673 | 0.618 | 0.619 | 0.627 | 0.743 | 0.825 | 0.724 | 0.559 | 0.697 | 0.558 | -0.017 |
| SE | 18630 | ENSG00000256053 | COA8 | chr14 | + | 103581568 | 103581656 | 103571622 | 103571820 | 103587273 | 103587364 | 0 | 5 | 0 | 0 | 0 | 6 | 2 | 7 | 3 | 1 | 0 | 0 | 0 | 7 | 0 | 20 | 2 | 9 | 1 | 3 | 237 | 149 | 0.60309902 | 1 | 0 | 0.611 | 0 | 0 | 0 | 0 | 0 | 0 | 0.815 | 0 | -0.041 |
| SE | 14731 | ENSG00000069966 | GNB5 | chr15 | - | 52147458 | 52147535 | 52141139 | 52141272 | 52149883 | 52149925 | 69 | 31 | 50 | 94 | 70 | 0 | 0 | 0 | 7 | 0 | 62 | 39 | 48 | 28 | 276 | 0 | 0 | 1 | 0 | 0 | 226 | 149 | 0.603251665 | 1 | 1 | 1 | 1 | 0.899 | 1 | 1 | 1 | 0.969 | 1 | 1 | -0.014 |
| SE | 10619 | ENSG00000108599 | AKAP10 | chr17 | - | 19939712 | 19939849 | 19936001 | 19936430 | 19941825 | 19941910 | 190 | 136 | 149 | 172 | 188 | 1 | 0 | 0 | 0 | 0 | 161 | 164 | 150 | 106 | 85 | 3 | 0 | 4 | 2 | 0 | 286 | 149 | 0.603362372 | 1 | 0.99 | 1 | 1 | 1 | 1 | 0.965 | 1 | 0.951 | 0.965 | 1 | 0.022 |
| SE | 60394 | ENSG00000174796 | THAP6 | chr4 | + | 75521735 | 75521861 | 75516771 | 75516979 | 75539858 | 75539959 | 43 | 62 | 46 | 57 | 100 | 2 | 0 | 0 | 3 | 1 | 17 | 25 | 42 | 25 | 16 | 0 | 0 | 0 | 0 | 0 | 275 | 149 | 0.603368769 | 1 | 0.921 | 1 | 1 | 0.911 | 0.982 | 1 | 1 | 1 | 1 | 1 | -0.037 |
| [illegible] | [illegible] | [illegible] | [illegible] | [illegible] | [illegible] | [illegible] | [illegible] | [illegible] | [illegible] | [illegible] | [illegible] | [illegible] | [illegible] | [illegible] | [illegible] | [illegible] | [illegible] | [illegible] | [illegible] | [illegible] | [illegible] | [illegible] | [illegible] | [illegible] | [illegible] | [illegible] | [illegible] | [illegible] | [illegible] | [illegible] | [illegible] | [illegible] | [illegible] | [illegible] | [illegible] | [illegible] | [illegible] | [illegible] | [illegible] | [illegible] | [illegible] | [illegible] | [illegible] | [illegible] |

| | | | | | | | | | | | | | | | |
|---|---|---|---|---|---|---|---|---|---|---|---|---|---|---|---|
| SE | 980 | ENSG00000015475 | BID | chr22 | - | 17750104 | 17750174 | 17739348 | 17739488 | 17774380 | 17774412 | 21 | 7 | 8 | [illegible] |
| SE | 21387 | ENSG00000154309 | DISP1 | chr1 | + | 222830928 | 222831042 | 222827427 | 222827601 | 222835118 | 222835215 | 2 | 2 | 4 | [illegible] |
| SE | 80709 | ENSG00000175471 | MCTP1 | chr5 | - | 95068784 | 95068829 | 95017366 | 95017484 | 95081425 | 95081646 | 3 | 1 | 0 | [illegible] |
| SE | 86691 | ENSG00000147140 | NONO | chrX | + | 71284983 | 71285097 | 71283589 | 71283721 | 71290628 | 71290791 | 8 | 6 | 6 | [illegible] |
| SE | 22712 | ENSG00000164329 | TENT2 | chr5 | + | 79642831 | 79642910 | 79641104 | 79641196 | 79645122 | 79645192 | 229 | 198 | 197 | [illegible] |
| SE | 92844 | ENSG00000130958 | SLC35D2 | chr9 | - | 96345298 | 96345401 | 96343903 | 96343996 | 96364463 | 96364537 | 115 | 72 | 81 | [illegible] |
| SE | 26184 | ENSG00000048140 | TSPAN17 | chr5 | + | 176656894 | 176656956 | 176656699 | 176656816 | 176657517 | 176657618 | 47 | 21 | 55 | [illegible] |
| SE | 109287 | ENSG00000085832 | EPS15 | chr1 | - | 51421785 | 51421858 | 51409534 | 51409696 | 51444888 | 51445045 | 147 | 114 | 102 | [illegible] |
| SE | 1150 | ENSG00000100060 | MFNG | chr22 | - | 37480720 | 37480769 | 37480196 | 37480299 | 37485922 | 37486384 | 238 | 119 | 237 | [illegible] |
| SE | 94092 | ENSG00000136811 | ODF2 | chr9 | + | 128460549 | 128460666 | 128459566 | 128459657 | 128460941 | 128461010 | 47 | 29 | 26 | [illegible] |
| SE | 70210 | ENSG00000085365 | SCAMP1 | chr5 | + | 78418774 | 78418903 | 78416540 | 78416649 | 78459244 | 78459362 | 40 | 28 | 25 | [illegible] |
| SE | 14974 | ENSG00000182718 | ANXA2 | chr15 | - | 60396279 | 60396427 | 60390212 | 60390317 | 60397942 | 60397986 | 2 | 2 | 1 | [illegible] |
| SE | 55005 | ENSG00000138641 | HERC3 | chr4 | + | 88597968 | 88598087 | 88595556 | 88595614 | 88605794 | 88606049 | 36 | 24 | 28 | [illegible] |
| SE | 2472 | ENSG00000130303 | BST2 | chr19 | - | 17404128 | 17404189 | 17403679 | 17403824 | 17405290 | 17405575 | 116 | 57 | 64 | [illegible] |
| SE | 28311 | ENSG00000001036 | FUCA2 | chr6 | - | 143503912 | 143504252 | 143502354 | 143502565 | 143507236 | 143507424 | 259 | 164 | 83 | [illegible] |
| SE | 32923 | ENSG00000075711 | DLG1 | chr3 | - | 197119409 | 197119530 | 197104902 | 197105005 | 197130526 | 197130671 | 118 | 116 | 159 | [illegible] |
| SE | 7647 | ENSG00000184792 | OSBP2 | chr22 | + | 30889489 | 30889636 | 30889176 | 30889234 | 30890727 | 30890973 | 81 | 18 | 14 | [illegible] |
| SE | 96926 | ENSG00000089159 | PXN | chr12 | - | 120215559 | 120215661 | 120215102 | 120215273 | 120221622 | 120221758 | 69 | 27 | 29 | [illegible] |
| SE | 109075 | ENSG00000198176 | TFDP1 | chr13 | + | 113637817 | 113637896 | 113636533 | 113636700 | 113640119 | 113640684 | 37 | 41 | 59 | [illegible] |
| SE | 27228 | ENSG00000163812 | ZDHHC3 | chr3 | - | 44932906 | 44932990 | 44924845 | 44926847 | 44933117 | 44933199 | 12 | 6 | 5 | [illegible] |
| SE | 54404 | ENSG00000141127 | PRPSAP2 | chr17 | + | 18865468 | 18865564 | 18857783 | 18857853 | 18865801 | 18865900 | 141 | 82 | 121 | [illegible] |
| SE | 106400 | ENSG00000089041 | P2RX7 | chr12 | + | 121160901 | 121160974 | 121156078 | 121156147 | 121162423 | 121162520 | 81 | 82 | 67 | [illegible] |
| SE | 15365 | ENSG00000140521 | POLG | chr15 | - | 89320765 | 89321260 | 89319227 | 89319350 | 89321735 | 89321853 | 144 | 92 | 92 | [illegible] |
| SE | 33027 | ENSG00000183323 | CCDC125 | chr5 | - | 69285336 | 69285467 | 69280174 | 69283034 | 69292187 | 69292362 | 91 | 63 | 121 | [illegible] |
| SE | 32250 | ENSG00000156931 | VPS8 | chr3 | + | 184922428 | 184922483 | 184920126 | 184920198 | 184924861 | 184924981 | 29 | 35 | 37 | [illegible] |
| SE | 5347 | ENSG00000101084 | RAB5IF | chr20 | + | 36609600 | 36609758 | 36607714 | 36607818 | 36612009 | 36612557 | 241 | 69 | 143 | [illegible] |
| SE | 74685 | ENSG00000197548 | ATG7 | chr3 | + | 11288645 | 11288740 | 11282192 | 11282438 | 11298685 | 11298855 | 6 | 0 | 0 | [illegible] |
| SE | 13709 | ENSG00000038532 | CLEC16A | chr16 | + | 10977224 | 10977399 | 10972937 | 10973061 | 10979328 | 10979382 | 109 | 79 | 97 | [illegible] |
| SE | 69361 | ENSG00000171720 | HDAC3 | chr5 | - | 141625684 | 141625764 | 141624861 | 141625365 | 141626193 | 141626283 | 151 | 93 | 108 | [illegible] |
| SE | 15460 | ENSG00000140199 | SLC12A6 | chr15 | - | 34238964 | 34239160 | 34238231 | 34238401 | 34240660 | 34240829 | 279 | 210 | 345 | [illegible] |
| SE | 99561 | ENSG00000174483 | BBS1 | chr11 | + | 66523171 | 66523365 | 66521269 | 66521376 | 66523455 | 66523531 | 0 | 3 | 2 | [illegible] |
| SE | 33471 | ENSG00000164327 | RICTOR | chr5 | - | 38952195 | 38952425 | 38949711 | 38950720 | 38952984 | 38953091 | 394 | 333 | 366 | [illegible] |
| SE | 59097 | ENSG00000214941 | ZSWIM7 | chr17 | - | 15977578 | 15977700 | 15976559 | 15977092 | 15977799 | 15977913 | 16 | 10 | 15 | [illegible] |
| SE | 32890 | ENSG00000068650 | ATP11A | chr13 | + | 112880545 | 112880649 | 112875775 | 112875941 | 112881815 | 112887168 | 2 | 3 | 2 | [illegible] |
| SE | 73525 | ENSG00000138380 | CARF | chr2 | + | 202961236 | 202961426 | 202955673 | 202955758 | 202966977 | 202967098 | 60 | 45 | 69 | [illegible] |
| SE | 10362 | ENSG00000197381 | ADARB1 | chr21 | + | 45128401 | 45128573 | 45074579 | 45074793 | 45171609 | 45171684 | 18 | 13 | 14 | [illegible] |
| SE | 35054 | ENSG00000125868 | DSTN | chr20 | + | 17591931 | 17592063 | 17570081 | 17570211 | 17600737 | 17601045 | 3 | 5 | 3 | [illegible] |
| SE | 86219 | ENSG00000010671 | BTK | chrX | - | 101356051 | 101356268 | 101353869 | 101353988 | 101356783 | 101356955 | 86 | 81 | 65 | [illegible] |
| SE | 58155 | ENSG00000273136 | NBPF26 | chr1 | + | 120822103 | 120822267 | 120818122 | 120818174 | 120823308 | 120823360 | 33 | 53 | 20 | [illegible] |
| SE | 54082 | ENSG00000092847 | AGO1 | chr1 | + | 35893673 | 35893810 | 35893096 | 35893278 | 35894036 | 35894171 | 80 | 53 | 53 | [illegible] |
| SE | 81782 | ENSG00000089006 | SNX5 | chr20 | - | 17952586 | 17952710 | 17951499 | 17951595 | 17955364 | 17955475 | 257 | 173 | 223 | [illegible] |
| SE | 10873 | ENSG00000221926 | TRIM16 | chr17 | - | 15632508 | 15632674 | 15631618 | 15631714 | 15636035 | 15636143 | 27 | 8 | 21 | [illegible] |
| SE | 95154 | ENSG00000005238 | FAM214B | chr9 | - | 35108127 | 35108349 | 35107380 | 35107568 | 35108593 | 35108737 | 31 | 8 | 12 | [illegible] |
| SE | 25313 | ENSG00000134644 | PUM1 | chr1 | - | 30992389 | 30992660 | 30981311 | 30981405 | 31028795 | 31028864 | 110 | 62 | 87 | [illegible] |
| SE | 89302 | ENSG00000166169 | POLL | chr10 | - | 101585315 | 101585478 | 101584601 | 101584919 | 101585861 | 101586156 | 30 | 13 | 29 | [illegible] |
| SE | 65805 | ENSG00000116521 | SCAMP3 | chr1 | - | 155261656 | 155261734 | 155260329 | 155260450 | 155262085 | 155262360 | 45 | 13 | 36 | [illegible] |
| SE | 31742 | ENSG00000047849 | MAP4 | chr3 | - | 47921764 | 47921878 | 47915950 | 47917174 | 47928227 | 47928302 | 182 | 108 | 112 | [illegible] |
| SE | 108374 | ENSG00000111554 | MDM1 | chr12 | - | 68316077 | 68316253 | 68314947 | 68315265 | 68316580 | 68316610 | 125 | 120 | 111 | [illegible] |
| SE | 68626 | ENSG00000223960 | CHROMR | chr2 | + | 178426365 | 178426505 | 178413944 | 178414113 | 178430769 | 178430889 | 61 | 36 | 25 | [illegible] |
| SE | 19751 | ENSG00000100614 | PPM1A | chr14 | + | 60250379 | 60250481 | 60249440 | 60249677 | 60282683 | 60283537 | 3 | 2 | 5 | [illegible] |
| SE | 51498 | ENSG00000108021 | TASOR2 | chr10 | + | 5709529 | 5709663 | 5699447 | 5699737 | 5712393 | 5712539 | 100 | 29 | 48 | [illegible] |
| SE | 87209 | ENSG00000086758 | HUWE1 | chrX | - | 53590999 | 53591122 | 53590403 | 53590499 | 53592397 | 53592628 | 601 | 422 | 545 | [illegible] |
| SE | 33525 | ENSG00000082996 | RNF13 | chr3 | + | 149813924 | 149814026 | 149812769 | 149813353 | 149846010 | 149846140 | 3 | 3 | 17 | [illegible] |
| SE | 3179 | ENSG00000076662 | ICAM3 | chr19 | - | 10338681 | 10339245 | 10335670 | 10335976 | 10339538 | 10339619 | 442 | 161 | 289 | [illegible] |
| SE | 16978 | ENSG00000141956 | PRDM15 | chr21 | - | 41834504 | 41834564 | 41828165 | 41828333 | 41835436 | 41835524 | 6 | 1 | 0 | [illegible] |
| SE | 80304 | ENSG00000168137 | SETD5 | chr3 | + | 9468518 | 9468575 | 9464424 | 9464672 | 9470458 | 9470929 | 19 | 21 | 30 | [illegible] |
| SE | 80714 | ENSG00000147535 | PLPP5 | chr8 | - | 38268370 | 38268461 | 38267896 | 38267960 | 38268881 | 38268990 | 42 | 15 | 65 | [illegible] |
| SE | 96421 | ENSG00000159314 | ARHGAP27 | chr17 | - | 45396915 | 45397024 | 45396667 | 45396790 | 45397948 | 45398047 | 350 | 194 | 214 | [illegible] |
| SE | 81970 | ENSG00000051825 | MPHOSPH9 | chr12 | - | 123227462 | 123227616 | 123223037 | 123223127 | 123230260 | 123230446 | 177 | 149 | 213 | [illegible] |
| SE | 85672 | ENSG00000177646 | ACAD9 | chr3 | + | 128884674 | 128884746 | 128879619 | 128879841 | 128893554 | 128893656 | 68 | 48 | 50 | [illegible] |
| SE | 77934 | ENSG00000095015 | MAP3K1 | chr5 | + | 56888225 | 56888357 | 56885931 | 56886063 | 56893530 | 56896152 | 349 | 286 | 295 | [illegible] |
| SE | 21599 | ENSG00000112234 | FBXL4 | chr6 | - | 98927704 | 98927822 | 98926476 | 98927060 | 98934761 | 98934879 | 73 | 50 | 65 | [illegible] |
| SE | 84382 | ENSG00000068323 | TFE3 | chrX | - | 49031396 | 49031544 | 49028725 | 49030601 | 49033464 | 49033540 | 133 | 79 | 114 | [illegible] |
| SE | 71641 | ENSG00000113916 | BCL6 | chr3 | - | 187728359 | 187728544 | 187726730 | 187726898 | 187729049 | 187730021 | 67 | 49 | 54 | [illegible] |
| SE | 27990 | ENSG00000162980 | ARL5A | chr2 | - | 151812356 | 151812440 | 151806820 | 151806972 | 151814168 | 151814316 | 405 | 399 | 278 | [illegible] |
| SE | 1636 | ENSG00000244625 | MIATNB | chr22 | + | 26726643 | 26726794 | 26672766 | 26672987 | 26774155 | 26774217 | 10 | 10 | 14 | [illegible] |
| SE | 10259 | ENSG00000108651 | UTP6 | chr17 | - | 31865365 | 31865438 | 31860903 | 31863516 | 31868045 | 31868112 | 159 | 102 | 131 | [illegible] |
| SE | 34074 | ENSG00000114867 | EIF4G1 | chr3 | + | 184317716 | 184317816 | 184316131 | 184316218 | 184319688 | 184319801 | 107 | 42 | 62 | [illegible] |
| SE | 75258 | ENSG00000175575 | PAAF1 | chr11 | + | 73914412 | 73914504 | 73909398 | 73909593 | 73916544 | 73916616 | 30 | 30 | 22 | [illegible] |
| SE | 103177 | ENSG00000011376 | LARS2 | chr3 | + | 45458742 | 45458886 | 45419668 | 45419729 | 45474242 | 45474350 | 16 | 14 | 26 | [illegible] |
| SE | 52614 | ENSG00000197872 | CYRIA | chr2 | - | 16562004 | 16562141 | 16561455 | 16561533 | 16563988 | 16564094 | 316 | 223 | 175 | [illegible] |
| SE | 27276 | ENSG00000077232 | DNAJC10 | chr2 | + | 182752548 | 182752623 | 182752071 | 182752188 | 182755002 | 182755104 | 0 | 0 | 1 | [illegible] |
| SE | 22552 | ENSG00000167881 | SRP68 | chr17 | - | 76060307 | 76060390 | 76057402 | 76057543 | 76061109 | 76061219 | 379 | 258 | 355 | [illegible] |
| SE | 82097 | ENSG00000128891 | CCDC32 | chr15 | - | 40539238 | 40539355 | 40531340 | 40532302 | 40557215 | 40557372 | 0 | 4 | 6 | [illegible] |
| SE | 10210 | ENSG00000178852 | EFCAB13 | chr17 | + | 47412772 | 47412916 | 47409646 | 47409691 | 47414847 | 47414919 | 25 | 73 | 63 | [illegible] |
| SE | 71766 | ENSG00000021574 | SPAST | chr2 | + | 32126947 | 32127022 | 32116118 | 32116212 | 32128407 | 32128479 | 285 | 286 | 196 | [illegible] |
| SE | 51930 | ENSG00000074603 | DPP8 | chr15 | - | 65480221 | 65480399 | 65478879 | 65479039 | 65485098 | 65485160 | 117 | 99 | 105 | [illegible] |
| SE | 75442 | ENSG00000135049 | AGTPBP1 | chr9 | - | 85660935 | 85660973 | 85657434 | 85657643 | 85669484 | 85669578 | 303 | 220 | 255 | [illegible] |
| SE | 62711 | ENSG00000151779 | NBAS | chr2 | - | 15178987 | 15179116 | 15166913 | 15167323 | 15186741 | 15186880 | 70 | 80 | 82 | [illegible] |
| SE | 28723 | ENSG00000010256 | UQCRC1 | chr3 | - | 48600481 | 48600567 | 48600062 | 48600151 | 48600679 | 48600840 | 274 | 116 | 192 | [illegible] |
| SE | 95096 | ENSG00000148358 | GPR107 | chr9 | + | 130090875 | 130090983 | 130083564 | 130083602 | 130092247 | 130092381 | 108 | 56 | 68 | [illegible] |
| SE | 52457 | ENSG00000072803 | FBXW11 | chr5 | - | 171914342 | 171914405 | 171910571 | 171910797 | 171996889 | 171997051 | 106 | 45 | 55 | [illegible] |
| SE | 53469 | ENSG00000249915 | PDCD6 | chr5 | + | 311292 | 311402 | 304176 | 304221 | 314416 | 314686 | 43 | 27 | 53 | [illegible] |
| SE | 30665 | ENSG00000112893 | MAN2A1 | chr5 | + | 109770354 | 109770541 | 109767534 | 109767708 | 109774787 | 109774965 | 291 | 211 | 282 | [illegible] |
| SE | 14100 | ENSG00000155592 | ZKSCAN2 | chr16 | - | 25243784 | 25244276 | 25236000 | 25240738 | 25246706 | 25247390 | 29 | 14 | 22 | [illegible] |
| SE | 33713 | ENSG00000172046 | USP19 | chr3 | - | 49117058 | 49117361 | 49116726 | 49116943 | 49117436 | 49117570 | 54 | 16 | 29 | [illegible] |
| SE | 64517 | ENSG00000166801 | FAM111A | chr11 | + | 59143496 | 59143695 | 59143082 | 59143304 | 59148835 | 59148953 | 58 | 45 | 40 | [illegible] |
| SE | 65532 | ENSG00000284292 | NA | chr7 | + | 99338180 | 99338285 | 99325878 | 99326004 | 99344292 | 99344515 | 57 | 41 | 59 | [illegible] |
| SE | 50275 | ENSG00000134909 | ARHGAP32 | chr11 | - | 129164318 | 129164427 | 129124802 | 129124894 | 129279145 | 129279324 | 13 | 9 | 6 | [illegible] |
| SE | 82084 | ENSG00000184216 | IRAK1 | chrX | - | 154013042 | 154013343 | 154012528 | 154012678 | 154014041 | 154014278 | 55 | 23 | 37 | [illegible] |
| SE | 56820 | ENSG00000119950 | MXI1 | chr10 | + | 110260998 | 110261145 | 110244827 | 110244857 | 110279179 | 110279294 | 1 | 2 | 0 | [illegible] |
| SE | 17484 | ENSG00000185024 | BRF1 | chr14 | - | 105241264 | 105241414 | 105228819 | 105228913 | 105247070 | 105247193 | 55 | 27 | 30 | [illegible] |
| SE | 73399 | ENSG00000240344 | PPIL3 | chr2 | - | 200887612 | 200887685 | 200885318 | 200885772 | 200888955 | 200889044 | 203 | 99 | 162 | [illegible] |
| SE | 95521 | ENSG00000137145 | DENND4C | chr9 | + | 19357964 | 19358160 | 19356971 | 19357154 | 19360243 | 19360489 | 282 | 197 | 196 | [illegible] |
| SE | 46852 | ENSG00000162894 | FCMR | chr1 | - | 206910209 | 206910340 | 206909724 | 206909868 | 206911729 | 206911952 | 656 | 253 | 582 | [illegible] |
| SE | 62598 | ENSG00000173597 | SULT1B1 | chr4 | - | 69760207 | 69760250 | 69755069 | 69755261 | 69760458 | 69760598 | 13 | 7 | 20 | [illegible] |
| SE | 51237 | ENSG00000146376 | ARHGAP18 | chr6 | - | 129607892 | 129608052 | 129600641 | 129600848 | 129611532 | 129611610 | 160 | 208 | 87 | [illegible] |
| SE | 83818 | ENSG00000167615 | LENG8 | chr19 | + | 54456324 | 54456465 | 54455966 | 54456245 | 54456635 | 54456921 | 153 | 64 | 130 | [illegible] |
| SE | 38046 | ENSG00000215908 | CROCCP2 | chr1 | - | 16633094 | 16633346 | 16632274 | 16632371 | 16635026 | 16635168 | 37 | 16 | 75 | [illegible] |
| SE | 48914 | ENSG00000011295 | TTC19 | chr17 | + | 16024488 | 16024582 | 16006473 | 16006568 | 16025016 | 16025171 | 2 | 2 | 0 | [illegible] |
| SE | 79728 | ENSG00000168255 | POLR2J3 | chr7 | - | 102544705 | 102544776 | 102541500 | 102541662 | 102566996 | 102567070 | 5 | 8 | 15 | [illegible] |
| SE | 101108 | ENSG00000162241 | SLC25A45 | chr11 | - | 65379375 | 65379561 | 65376817 | 65377076 | 65379866 | 65379938 | 53 | 16 | 13 | [illegible] |
| SE | 99043 | ENSG00000087053 | MTMR2 | chr11 | - | 95887611 | 95887759 | 95865600 | 95865676 | 95888155 | 95888261 | 8 | 4 | 4 | [illegible] |
| SE | 18394 | ENSG00000136305 | CIDEB | chr14 | - | 24310406 | 24310548 | 24308478 | 24308830 | 24310648 | 24311395 | 2 | 4 | 4 | [illegible] |
| SE | 33568 | ENSG00000009790 | TRAF3IP3 | chr1 | + | 209772947 | 209773019 | 209763491 | 209763587 | 209775348 | 209775489 | 872 | 649 | 853 | [illegible] |
| SE | 91886 | ENSG00000254995 | STX16-NPEPL1 | chr20 | + | 58691033 | 58691227 | 58673630 | 58673711 | 58691404 | 58691562 | 14 | 14 | 21 | [illegible] |
| SE | 31822 | ENSG00000112701 | SENP6 | chr6 | + | 75647730 | 75647801 | 75634706 | 75634811 | 75659261 | 75659407 | 216 | 144 | 180 | [illegible] |
| SE | 56126 | ENSG00000204406 | MBD5 | chr2 | + | 148054390 | 148054449 | 148021392 | 148021684 | 148178699 | 148178793 | 16 | 46 | 36 | [illegible] |
| SE | 29812 | ENSG00000223547 | ZNF844 | chr19 | + | 12074360 | 12074421 | 12074030 | 12074157 | 12075311 | 12081565 | 20 | 40 | 51 | [illegible] |
| SE | 24080 | ENSG00000146282 | RARS2 | chr6 | - | 87519582 | 87519707 | 87518823 | 87518891 | 87520179 | 87520256 | 156 | 140 | 153 | [illegible] |
| SE | 67555 | ENSG00000115317 | HTRA2 | chr2 | + | 74532618 | 74532714 | 74531855 | 74531925 | 74532819 | 74533152 | 95 | 29 | 68 | [illegible] |
| SE | 73705 | ENSG00000114770 | ABCC5 | chr3 | - | 183949752 | 183949881 | 183947323 | 183947510 | 183949971 | 183950125 | 38 | 19 | 26 | [illegible] |
| SE | 98206 | ENSG00000149273 | RPS3 | chr11 | + | 75402351 | 75402379 | 75401639 | 75401733 | 75404671 | 75404868 | 783 | 386 | 856 | [illegible] |
| SE | 54183 | ENSG00000175376 | EIF1AD | chr11 | - | 66000302 | 66000481 | 65999897 | 66000161 | 66002039 | 66002156 | 117 | 73 | 118 | [illegible] |
| SE | 109070 | ENSG00000198176 | TFDP1 | chr13 | + | 113631622 | 113631744 | 113623179 | 113623286 | 113633119 | 113633285 | 110 | 71 | 99 | [illegible] |
| SE | 32623 | ENSG00000189079 | ARID2 | chr12 | + | 45846855 | 45846937 | 45839328 | 45839496 | 45848835 | 45848970 | 165 | 172 | 136 | [illegible] |
| SE | 82699 | ENSG00000097033 | SH3GLB1 | chr1 | + | 86742207 | 86742436 | 86735078 | 86735179 | 86743127 | 86743268 | 980 | 665 | 672 | [illegible] |
| SE | 49740 | ENSG00000111331 | OAS3 | chr12 | + | 112947945 | 112948099 | 112946644 | 112946981 | 112961070 | 112961246 | 46 | 53 | 48 | [illegible] |
| SE | 95642 | ENSG00000137073 | UBAP2 | chr9 | - | 33948373 | 33948587 | 33944364 | 33944639 | 33953284 | 33953474 | 137 | 90 | 108 | [illegible] |
| SE | 15540 | ENSG00000129003 | VPS13C | chr15 | - | 61954466 | 61954554 | 61951859 | 61951980 | 61958607 | 61958716 | 230 | 239 | 178 | [illegible] |
| SE | 54462 | ENSG00000112303 | VNN2 | chr6 | - | 132752460 | 132752577 | 132751144 | 132751518 | 132755842 | 132756035 | 233 | 90 | 220 | [illegible] |
| SE | 46246 | ENSG00000198301 | SDAD1 | chr4 | - | 75967276 | 75967334 | 75965763 | 75965822 | 75969295 | 75969399 | 270 | 177 | 261 | [illegible] |
| SE | 3157 | ENSG00000213020 | ZNF611 | chr19 | - | 52707468 | 52707542 | 52702812 | 52706864 | 52714014 | 52714141 | 30 | 34 | 31 | [illegible] |
| SE | 100172 | ENSG00000078124 | ACER3 | chr11 | + | 76990538 | 76990574 | 76982677 | 76985724 | 76994155 | 76994226 | 16 | 18 | 6 | [illegible] |
| SE | 1596 | ENSG00000198089 | SFI1 | chr22 | + | 31603743 | 31603819 | 31602606 | 31602785 | 31604308 | 31604404 | 79 | 51 | 94 | [illegible] |
| SE | 65009 | ENSG00000167107 | ACSF2 | chr17 | + | 50464525 | 50464599 | 50464217 | 50464294 | 50471027 | 50471135 | 3 | 4 | 6 | [illegible] |
| SE | 80771 | ENSG00000136280 | CCM2 | chr7 | + | 45068442 | 45068579 | 45064462 | 45064646 | 45069825 | 45069961 | 51 | 20 | 52 | [illegible] |
| SE | 15301 | ENSG00000114062 | UBE3A | chr15 | - | 25405460 | 25405502 | 25373888 | 25375763 | 25438488 | 25439005 | 79 | 69 | 74 | [illegible] |
| SE | 41256 | ENSG00000162819 | BROX | chr1 | + | 222715683 | 222715800 | 222715144 | 222715269 | 222718924 | 222719031 | 96 | 55 | 80 | [illegible] |
| SE | 48305 | ENSG00000229474 | PATL2 | chr15 | - | 44710870 | 44710929 | 44710095 | 44710214 | 44711019 | 44711323 | 34 | 21 | 30 | [illegible] |
| SE | 91350 | ENSG00000167632 | TRAPPC9 | chr8 | - | 140426614 | 140426641 | 140405576 | 140405698 | 140435111 | 140435240 | 84 | 48 | 57 | [illegible] |
| SE | 53802 | ENSG00000111653 | ING4 | chr12 | - | 6657804 | 6657852 | 6656726 | 6656798 | 6663064 | 6663083 | 3 | 0 | 1 | [illegible] |
| SE | 101127 | ENSG00000175224 | ATG13 | chr11 | + | 46621041 | 46621157 | 46617556 | 46617890 | 46630044 | 46630100 | 1 | 1 | 0 | [illegible] |
| SE | 18849 | ENSG00000063761 | ADCK1 | chr14 | + | 77900558 | 77900674 | 77899099 | 77899258 | 77907802 | 77907919 | 4 | 1 | 1 | [illegible] |
| SE | 39108 | ENSG00000116120 | FARSB | chr2 | - | 222624713 | 222624775 | 222624271 | 222624479 | 222628836 | 222628888 | 105 | 86 | 114 | [illegible] |
| SE | 53962 | ENSG00000138101 | DTNB | chr2 | - | 25607235 | 25607321 | 25596085 | 25596240 | 25628170 | 25628384 | 39 | 25 | 67 | [illegible] |
| SE | 38942 | ENSG00000067066 | SP100 | chr2 | + | 230467119 | 230467206 | 230466300 | 230466354 | 230469042 | 230469096 | 247 | 214 | 214 | [illegible] |
| SE | 7289 | ENSG00000101019 | UQCC1 | chr20 | - | 35384037 | 35384133 | 35381917 | 35382025 | 35394091 | 35394196 | 120 | 117 | 82 | [illegible] |
| SE | 81685 | ENSG00000171681 | ATF7IP | chr12 | + | 14420301 | 14420446 | 14365728 | 14365827 | 14423908 | 14424183 | 1 | 1 | 5 | [illegible] |
| SE | 102933 | ENSG00000231131 | LNCAROD | chr10 | - | 52723869 | 52724018 | 52561005 | 52561115 | 52755305 | 52755404 | 0 | 0 | 4 | [illegible] |
| SE | 12933 | ENSG00000197818 | SLC9A8 | chr20 | + | 49850761 | 49850844 | 49849578 | 49849680 | 49855437 | 49855581 | 3 | 2 | 2 | [illegible] |
| SE | 78466 | ENSG00000177156 | TALDO1 | chr11 | + | 752263 | 752473 | 747414 | 747578 | 755878 | 756002 | 2 | 3 | 0 | [illegible] |
| SE | 57264 | ENSG00000143119 | CD53 | chr1 | + | 110894326 | 110894401 | 110891391 | 110891471 | 110897808 | 110897892 | 4 | 0 | 6 | [illegible] |
| SE | 88492 | ENSG00000152601 | MBNL1 | chr3 | + | 152456266 | 152456361 | 152445281 | 152445539 | 152459270 | 152459327 | 196 | 179 | 189 | [illegible] |
| SE | 80516 | ENSG00000179818 | PCBP1-AS1 | chr2 | - | 70051202 | 70051305 | 69996685 | 69996888 | 70053730 | 70053737 | 15 | 4 | 14 | [illegible] |
| SE | 67072 | ENSG00000204681 | GABBR1 | chr6 | - | 29605568 | 29605696 | 29604859 | 29604988 | 29606390 | 29606484 | 108 | 89 | 71 | [illegible] |
| SE | 48471 | ENSG00000259494 | MRPL46 | chr15 | - | 88465586 | 88465773 | 88464702 | 88464876 | 88467149 | 88467388 | 30 | 21 | 43 | [illegible] |
| SE | 96770 | ENSG00000052841 | TTC17 | chr11 | + | 43414589 | 43414776 | 43407352 | 43407577 | 43443324 | 43443584 | 94 | 94 | 96 | [illegible] |
| SE | 80953 | ENSG00000133393 | CEP20 | chr16 | - | 15879803 | 15879888 | 15873484 | 15873627 | 15888557 | 15888603 | 59 | 44 | 55 | [illegible] |
| SE | 5654 | ENSG00000203880 | PCMTD2 | chr20 | + | 64258766 | 64258882 | 64256585 | 64257051 | 64259941 | 64260272 | 3 | 1 | 1 | [illegible] |
| SE | 4766 | ENSG00000104885 | DOT1L | chr19 | + | 2191083 | 2191240 | 2189731 | 2189795 | 2193688 | 2193783 | 29 | 25 | 23 | [illegible] |
| SE | 88977 | ENSG00000165806 | CASP7 | chr10 | + | 113691963 | 113692070 | 113679939 | 113679978 | 113697493 | 113697603 | 2 | 0 | 4 | [illegible] |
| SE | 39900 | ENSG00000271723 | MROH7-TTC4 | chr1 | + | 54731485 | 54731549 | 54728345 | 54728432 | 54737581 | 54737664 | 18 | 17 | 19 | [illegible] |
| SE | 49646 | ENSG00000081760 | AACS | chr12 | + | 125124901 | 125125024 | 125119857 | 125124769 | 125128160 | 125128274 | 11 | 8 | 22 | [illegible] |
| SE | 69536 | ENSG00000177479 | ARIH2 | chr3 | + | 48927461 | 48927813 | 48922747 | 48922811 | 48961611 | 48961679 | 153 | 89 | 116 | [illegible] |
| SE | 29223 | ENSG00000023287 | RB1CC1 | chr8 | - | 52660595 | 52660657 | 52658872 | 52658976 | 52660925 | 52661007 | 191 | 106 | 148 | [illegible] |
| SE | 12238 | ENSG00000205336 | ADGRG1 | chr16 | + | 57647966 | 57648000 | 57646941 | 57647012 | 57650252 | 57650351 | 2 | 3 | 1 | [illegible] |
| SE | 45653 | ENSG00000144283 | PKP4 | chr2 | + | 158678580 | 158678654 | 158673882 | 158674000 | 158680428 | 158680674 | 60 | 41 | 38 | [illegible] |
| SE | 22106 | ENSG00000124784 | RIOK1 | chr6 | + | 7395020 | 7395143 | 7393098 | 7393303 | 7396702 | 7396772 | 84 | 64 | 78 | [illegible] |
| SE | 38598 | ENSG00000122406 | RPL5 | chr1 | + | 92833388 | 92833458 | 92832039 | 92832117 | 92833544 | 92833660 | 1108 | 637 | 1377 | [illegible] |
| SE | 44030 | ENSG00000237943 | PRKCQ-AS1 | chr10 | + | 6584117 | 6584195 | 6583804 | 6583918 | 6585557 | 6585712 | 42 | 40 | 93 | [illegible] |
| SE | 14699 | ENSG00000103591 | AAGAB | chr15 | - | 67220488 | 67220540 | 67209461 | 67209544 | 67231813 | 67231897 | 3 | 2 | 0 | [illegible] |
| SE | 41386 | ENSG00000114796 | KLHL24 | chr3 | + | 183664920 | 183665039 | 183663457 | 183663642 | 183671033 | 183671123 | 154 | 103 | 152 | [illegible] |
| SE | 108116 | ENSG00000101160 | CTSZ | chr20 | - | 58997854 | 58997954 | 58997602 | 58997753 | 58999234 | 58999580 | 13 | 18 | 6 | [illegible] |
| SE | 98485 | ENSG00000151702 | FLI1 | chr11 | + | 128807179 | 128807239 | 128781957 | 128782023 | 128809156 | 128809204 | 312 | 242 | 248 | [illegible] |
| SE | 26579 | ENSG00000006459 | KDM7A | chr7 | - | 140124620 | 140124783 | 140120441 | 140120529 | 140126636 | 140126823 | 226 | 171 | 198 | [illegible] |
| SE | 90072 | ENSG00000156787 | TBC1D31 | chr8 | + | 123134113 | 123134206 | 123130197 | 123130333 | 123140760 | 123140901 | 94 | 66 | 100 | [illegible] |
| SE | 82427 | ENSG00000157191 | NECAP2 | chr1 | + | 16443631 | 16443732 | 16440723 | 16440853 | 16449092 | 16449201 | 236 | 119 | 196 | [illegible] |
| SE | 30321 | ENSG00000249915 | PDCD6 | chr5 | + | 272710 | 272772 | 271620 | 271821 | 304176 | 304221 | 169 | 85 | 99 | [illegible] |
| SE | 15462 | ENSG00000140199 | SLC12A6 | chr15 | - | 34240660 | 34240829 | 34238964 | 34239160 | 34241232 | 34241337 | 238 | 163 | 278 | [illegible] |
| SE | 55467 | ENSG00000100612 | DHRS7 | chr14 | - | 60152938 | 60153178 | 60150064 | 60150187 | 60153958 | 60154005 | 965 | 395 | 553 | [illegible] |
| SE | 50471 | ENSG00000165671 | NSD1 | chr5 | + | 177229764 | 177229886 | 177212110 | 177212195 | 177235820 | 177235945 | 4 | 2 | 4 | [illegible] |
| SE | 15229 | ENSG00000101266 | CSNK2A1 | chr20 | - | 527932 | 528049 | 508450 | 508660 | 543671 | 543712 | 24 | 14 | 15 | [illegible] |
| SE | 63517 | ENSG00000125962 | ARMCX5 | chrX | + | 102601463 | 102601509 | 102599537 | 102599847 | 102601879 | 102604159 | 19 | 12 | 13 | [illegible] |
| SE | 5045 | ENSG00000101126 | ADNP | chr20 | - | 50903888 | 50904001 | 50902016 | 50902109 | 50928650 | 50928732 | 213 | 165 | 193 | [illegible] |
| SE | 31016 | ENSG00000145247 | OCIAD2 | chr4 | - | 48899828 | 48899925 | 48897803 | 48897857 | 48904483 | 48904611 | 158 | 91 | 168 | [illegible] |
| SE | 105451 | ENSG00000179889 | PDXDC1 | chr16 | + | 15001775 | 15001887 | 14998339 | 14998405 | 15004186 | 15004333 | 16 | 15 | 29 | [illegible] |
| SE | 19706 | ENSG00000140090 | SLC24A4 | chr14 | + | 92492826 | 92492924 | 92492174 | 92492240 | 92493475 | 92496252 | 6 | 1 | 1 | [illegible] |
| SE | 38645 | ENSG00000073417 | PDE8A | chr15 | + | 85067013 | 85067204 | 85062496 | 85064426 | 85075861 | 85075918 | 74 | 64 | 49 | [illegible] |
| SE | 81390 | ENSG00000258733 | LINC02328 | chr14 | + | 86041725 | 86041833 | 86035713 | 86035815 | 86070862 | 86070967 | 53 | 17 | 27 | [illegible] |
| SE | 40366 | ENSG00000234663 | LINC01934 | chr2 | + | 181354207 | 181354343 | 181238250 | 181238450 | 181362686 | 181362902 | 46 | 34 | 52 | [illegible] |
| SE | 29399 | ENSG00000127419 | TMEM175 | chr4 | + | 948115 | 948154 | 932479 | 932540 | 950420 | 950518 | 9 | 1 | 3 | [illegible] |
| SE | 93591 | ENSG00000165525 | NEMF | chr14 | - | 49849628 | 49849707 | 49846138 | 49846265 | 49851562 | 49851665 | 3 | 0 | 2 | [illegible] |
| SE | 92351 | ENSG00000165271 | NOL6 | chr9 | - | 33468751 | 33468872 | 33468507 | 33468566 | 33468957 | 33469121 | 25 | 7 | 15 | [illegible] |
| SE | 23044 | ENSG00000152942 | RAD17 | chr5 | + | 69389033 | 69389145 | 69386395 | 69386465 | 69391830 | 69391919 | 179 | 130 | 152 | [illegible] |
| SE | 97377 | ENSG00000113163 | CERT1 | chr5 | - | 75379336 | 75379473 | 75368485 | 75374191 | 75381071 | 75381201 | 149 | 94 | 109 | [illegible] |
| SE | 36646 | ENSG00000274265 | NA | chr1 | + | 149190635 | 149190795 | 149176021 | 149176250 | 149250564 | 149251013 | 3 | 0 | 0 | [illegible] |
| SE | 45209 | ENSG00000121774 | KHDRBS1 | chr1 | + | 32039514 | 32039573 | 32033187 | 32033334 | 32042526 | 32042624 | 176 | 88 | 191 | [illegible] |
| SE | 51864 | ENSG00000180871 | CXCR2 | chr2 | + | 218129313 | 218129365 | 218126027 | 218126353 | 218134776 | 218134846 | 159 | 72 | 83 | [illegible] |
| SE | 6297 | ENSG00000141644 | MBD1 | chr18 | - | 50274185 | 50274353 | 50273563 | 50273863 | 50275128 | 50275245 | 75 | 42 | 49 | [illegible] |
| SE | 72332 | ENSG00000146143 | PRIM2 | chr6 | + | 57507386 | 57507454 | 57382030 | 57382168 | 57532410 | 57532483 | 76 | 38 | 67 | [illegible] |
| SE | 86843 | ENSG00000116754 | SRSF11 | chr1 | + | 70231094 | 70231203 | 70228421 | 70228555 | 70231858 | 70231975 | 11 | 19 | 32 | [illegible] |
| SE | 62149 | ENSG00000162650 | ATXN7L2 | chr1 | + | 109492026 | 109492104 | 109490921 | 109491717 | 109492585 | 109492734 | 1 | 2 | 6 | [illegible] |
| SE | 85688 | ENSG00000177981 | ASB8 | chr12 | - | 48151609 | 48151665 | 48151200 | 48151305 | 48153367 | 48153529 | 1 | 0 | 0 | [illegible] |
| SE | 63112 | ENSG00000115761 | NOL10 | chr2 | - | 10654480 | 10654547 | 10644319 | 10644372 | 10657741 | 10657891 | 95 | 67 | 73 | [illegible] |
| SE | 32089 | ENSG00000242086 | MUC20-OT1 | chr3 | + | 195699029 | 195699207 | 195689338 | 195689438 | 195708133 | 195708268 | 25 | 12 | 7 | [illegible] |
| SE | 44226 | ENSG00000102606 | ARHGEF7 | chr13 | + | 111205288 | 111205373 | 111153713 | 111153991 | 111209871 | 111210002 | 141 | 91 | 130 | [illegible] |
| SE | 33259 | ENSG00000248487 | ABHD14A | chr3 | + | 51980392 | 51980628 | 51978258 | 51978374 | 51980835 | 51981196 | 17 | 12 | 22 | [illegible] |
| SE | 54212 | ENSG00000168395 | ING5 | chr2 | + | 241722938 | 241723074 | 241711977 | 241712071 | 241723209 | 241723271 | 35 | 47 | 41 | [illegible] |
| SE | 49077 | ENSG00000188739 | RBM34 | chr1 | - | 235152705 | 235152765 | 235148403 | 235148447 | 235154880 | 235155112 | 249 | 189 | 191 | [illegible] |
| SE | 35496 | ENSG00000051382 | PIK3CB | chr3 | - | 138688874 | 138688974 | 138684624 | 138684803 | 138694785 | 138694907 | 95 | 86 | 86 | [illegible] |
| SE | 42552 | ENSG00000169241 | SLC50A1 | chr1 | + | 155137560 | 155137722 | 155136298 | 155136376 | 155137978 | 155138098 | 66 | 23 | 51 | [illegible] |
| SE | 62595 | ENSG00000173597 | SULT1B1 | chr4 | - | 69755069 | 69755261 | 69754669 | 69754798 | 69760458 | 69760620 | 414 | 336 | 211 | [illegible] |
| SE | 26511 | ENSG00000114744 | COMMD2 | chr3 | - | 149750677 | 149750851 | 149750256 | 149750382 | 149751402 | 149751485 | 102 | 62 | 86 | [illegible] |
| SE | 106238 | ENSG00000089154 | GCN1 | chr12 | - | 120156460 | 120156604 | 120155591 | 120155719 | 120156911 | 120156992 | 70 | 25 | 97 | [illegible] |
| SE | 26476 | ENSG00000038382 | TRIO | chr5 | + | 14398879 | 14399070 | 14397042 | 14397154 | 14400962 | 14401064 | 112 | 110 | 87 | [illegible] |
| SE | 82819 | ENSG00000155849 | ELMO1 | chr7 | - | 36878009 | 36878117 | 36870392 | 36870475 | 36887559 | 36887672 | 276 | 212 | 297 | [illegible] |
| SE | 11752 | ENSG00000260565 | ERVK13-1 | chr16 | - | 2672706 | 2672859 | 2667087 | 2672570 | 2673282 | 2673387 | 20 | 2 | 6 | [illegible] |
| SE | 64846 | ENSG00000138032 | PPM1B | chr2 | + | 44218479 | 44218537 | 44209209 | 44209327 | 44244227 | 44244383 | 12 | 6 | 5 | [illegible] |
| SE | 3867 | ENSG00000157540 | DYRK1A | chr21 | + | 37456129 | 37456308 | 37421773 | 37421866 | 37472683 | 37472880 | 4 | 2 | 5 | [illegible] |
| SE | 37978 | ENSG00000175348 | TMEM9B | chr11 | - | 8953202 | 8953337 | 8947201 | 8948475 | 8956189 | 8956298 | 211 | 148 | 176 | [illegible] |
| SE | 67501 | ENSG00000068654 | POLR1A | chr2 | - | 86075029 | 86075260 | 86070017 | 86070272 | 86077858 | 86077981 | 114 | 57 | 141 | [illegible] |
| SE | 89745 | ENSG00000156795 | NTAQ1 | chr8 | + | 123436452 | 123436601 | 123429982 | 123430033 | 123437209 | 123437334 | 44 | 43 | 27 | [illegible] |
| SE | 61580 | ENSG00000131876 | SNRPA1 | chr15 | - | 101289813 | 101289922 | 101287655 | 101287702 | 101291961 | 101292040 | 6 | 3 | 7 | [illegible] |
| SE | 27090 | ENSG00000156265 | MAP3K7CL | chr21 | + | 29091500 | 29091576 | 29085812 | 29085917 | 29091667 | 29091761 | 4 | 11 | 8 | [illegible] |
| SE | 93346 | ENSG00000225783 | MIAT | chr22 | + | 26658182 | 26658339 | 26657610 | 26657654 | 26663315 | 26663398 | 0 | 0 | 0 | [illegible] |
| SE | 91557 | ENSG00000139546 | TARBP2 | chr12 | + | 53503026 | 53503212 | 53502014 | 53502184 | 53503729 | 53503808 | 16 | 2 | 29 | [illegible] |
| SE | 66080 | ENSG00000162924 | REL | chr2 | + | 60920573 | 60920642 | 60920040 | 60920109 | 60921762 | 60931610 | 234 | 205 | 140 | [illegible] |
| SE | 82003 | ENSG00000104133 | SPG11 | chr15 | - | 44569397 | 44569505 | 44567423 | 44567592 | 44570524 | 44570652 | 181 | 106 | 141 | [illegible] |
| SE | 42657 | ENSG00000088854 | C20orf194 | chr20 | - | 3287631 | 3287790 | 3281640 | 3281766 | 3294138 | 3294256 | 22 | 31 | 24 | [illegible] |
| SE | 16194 | ENSG00000196547 | MAN2A2 | chr15 | + | 90913272 | 90913374 | 90912876 | 90912991 | 90913613 | 90913755 | 25 | 7 | 15 | [illegible] |
| SE | 31072 | ENSG00000065882 | TBC1D1 | chr4 | + | 38115709 | 38115954 | 38102998 | 38103157 | 38118032 | 38118192 | 260 | 179 | 247 | [illegible] |
| SE | 69739 | ENSG00000149782 | PLCB3 | chr11 | + | 64263497 | 64263597 | 64262646 | 64262808 | 64263690 | 64263795 | 25 | 8 | 23 | [illegible] |
| SE | 12702 | ENSG00000155666 | KDM8 | chr16 | + | 27218960 | 27219110 | 27215944 | 27215989 | 27220392 | 27220485 | 25 | 10 | 26 | [illegible] |
| SE | 84418 | ENSG00000188917 | TRMT2B | chrX | - | 101041316 | 101041371 | 101037916 | 101038051 | 101051250 | 101051366 | 27 | 13 | 19 | [illegible] |
| SE | 10027 | ENSG00000154229 | PRKCA | chr17 | + | 66696426 | 66696487 | 66688950 | 66689047 | 66732687 | 66732825 | 1 | 0 | 0 | [illegible] |
| SE | 10704 | ENSG00000267321 | SNHG30 | chr17 | + | 35573579 | 35573723 | 35568108 | 35568501 | 35573876 | 35574843 | 1 | 0 | 0 | [illegible] |
| SE | 88369 | ENSG00000168300 | PCMTD1 | chr8 | - | 51860860 | 51861246 | 51845660 | 51845763 | 51898929 | 51899113 | 88 | 101 | 85 | [illegible] |
| SE | 68597 | ENSG00000239779 | WBP1 | chr2 | + | 74459477 | 74459562 | 74458449 | 74458671 | 74459642 | 74459745 | 5 | 4 | 16 | [illegible] |
| SE | 2602 | ENSG00000197961 | ZNF121 | chr19 | - | 9568094 | 9568175 | 9560328 | 9567109 | 9569001 | 9569082 | 124 | 118 | 155 | [illegible] |
| SE | 3539 | ENSG00000105483 | CARD8 | chr19 | - | 48238382 | 48238532 | 48232452 | 48234543 | 48240961 | 48241063 | 707 | 443 | 496 | [illegible] |
| SE | 93482 | ENSG00000134077 | THUMPD3 | chr3 | + | 9377818 | 9377888 | 9374515 | 9374646 | 9383198 | 9383309 | 98 | 74 | 99 | [illegible] |
| SE | 61414 | ENSG00000128463 | EMC4 | chr15 | + | 34228436 | 34228589 | 34227694 | 34227846 | 34229752 | 34230156 | 93 | 62 | 91 | [illegible] |

| | | | | | | | | | | | | | | | |
|---|---|---|---|---|---|---|---|---|---|---|---|---|---|---|---|
| SE | 63002 | ENSG00000137497 | NUMA1 | chr11 | - | 72068104 | 72068231 | 72035901 | 72035975 | 72069841 | 72070063 | 13 | 3 | 2 | [illegible] |
| SE | 2354 | ENSG00000099785 | MARCHF2 | chr19 | + | 8426608 | 8426804 | 8421788 | 8422016 | 8430657 | 8430867 | 63 | 46 | 33 | [illegible] |
| SE | 103543 | ENSG00000107862 | GBF1 | chr10 | + | 102368818 | 102368832 | 102368217 | 102368454 | 102369210 | 102369387 | 1 | 1 | 1 | [illegible] |
| SE | 39878 | ENSG00000196821 | ILRUN | chr6 | - | 34646600 | 34646798 | 34606554 | 34606904 | 34654624 | 34654779 | 534 | 271 | 407 | [illegible] |
| SE | 43327 | ENSG00000135597 | REPS1 | chr6 | - | 138917554 | 138917627 | 138914696 | 138914761 | 138920214 | 138920316 | 39 | 41 | 34 | [illegible] |
| SE | 76755 | ENSG00000168036 | CTNNB1 | chr3 | + | 41224953 | 41224991 | 41224525 | 41224753 | 41225102 | 41225207 | 147 | 86 | 85 | [illegible] |
| SE | 43328 | ENSG00000135597 | REPS1 | chr6 | - | 138917554 | 138917627 | 138915857 | 138915976 | 138920214 | 138920316 | 127 | 117 | 127 | [illegible] |
| SE | 16459 | ENSG00000104043 | ATP8B4 | chr15 | - | 50038767 | 50038829 | 50018888 | 50018976 | 50044593 | 50044674 | 22 | 25 | 22 | [illegible] |
| SE | 52341 | ENSG00000142197 | DOP1B | chr21 | + | 36200330 | 36200501 | 36199069 | 36199251 | 36208714 | 36208904 | 42 | 14 | 34 | [illegible] |
| SE | 13668 | ENSG00000090565 | RAB11FIP3 | chr16 | + | 482524 | 482736 | 471294 | 471389 | 488850 | 489000 | 31 | 9 | 14 | [illegible] |
| SE | 50691 | ENSG00000142687 | KIAA0319L | chr1 | - | 35470862 | 35470960 | 35462620 | 35462713 | 35474804 | 35474906 | 58 | 43 | 61 | [illegible] |
| SE | 84462 | ENSG00000137522 | RNF121 | chr11 | + | 71987003 | 71987111 | 71960749 | 71960891 | 71990596 | 71990663 | 37 | 12 | 26 | [illegible] |
| SE | 84862 | ENSG00000164074 | ABHD18 | chr4 | + | 127989720 | 127989821 | 127984338 | 127984423 | 128001715 | 128001760 | 67 | 49 | 79 | [illegible] |
| SE | 66204 | ENSG00000137817 | PARP6 | chr15 | - | 72249244 | 72249314 | 72242619 | 72242699 | 72250019 | 72250092 | 8 | 4 | 12 | [illegible] |
| SE | 108876 | ENSG00000121310 | ECHDC2 | chr1 | - | 52907867 | 52907954 | 52905033 | 52905090 | 52911565 | 52911653 | 57 | 42 | 55 | [illegible] |
| SE | 78528 | ENSG00000164054 | SHISA5 | chr3 | - | 48469727 | 48469843 | 48467875 | 48469573 | 48473009 | 48473103 | 863 | 424 | 644 | [illegible] |
| SE | 91289 | ENSG00000089234 | BRAP | chr12 | - | 111655565 | 111655655 | 111649938 | 111650042 | 111658735 | 111658845 | 128 | 60 | 139 | [illegible] |
| SE | 77115 | ENSG00000164654 | MIOS | chr7 | + | 7568025 | 7568123 | 7566874 | 7567072 | 7572435 | 7572667 | 22 | 12 | 15 | [illegible] |
| SE | 46837 | ENSG00000066827 | ZFAT | chr8 | - | 134608728 | 134608879 | 134601476 | 134602933 | 134637460 | 134637712 | 33 | 13 | 18 | [illegible] |
| SE | 96555 | ENSG00000109321 | AREG | chr4 | + | 74449046 | 74449248 | 74446533 | 74446782 | 74450379 | 74450532 | 57 | 39 | 34 | [illegible] |
| SE | 3619 | ENSG00000104812 | GYS1 | chr19 | - | 48974619 | 48974733 | 48970749 | 48971023 | 48977923 | 48977986 | 95 | 60 | 78 | [illegible] |
| SE | 38984 | ENSG00000004399 | PLXND1 | chr3 | - | 129572841 | 129572941 | 129572608 | 129572748 | 129573593 | 129573745 | 49 | 8 | 25 | [illegible] |
| SE | 84288 | ENSG00000165813 | CCDC186 | chr10 | - | 114135890 | 114135977 | 114134912 | 114135055 | 114136147 | 114136246 | 134 | 151 | 105 | [illegible] |
| SE | 89396 | ENSG00000123124 | WWP1 | chr8 | + | 86435632 | 86435704 | 86435451 | 86435527 | 86438584 | 86438673 | 297 | 225 | 283 | [illegible] |
| SE | 100911 | ENSG00000254732 | NA | chr11 | + | 57789036 | 57789155 | 57742862 | 57742984 | 57791384 | 57791673 | 24 | 21 | 17 | [illegible] |
| SE | 71712 | ENSG00000135976 | ANKRD36 | chr2 | + | 97204190 | 97204263 | 97204067 | 97204096 | 97205939 | 97205968 | 189 | 190 | 150 | [illegible] |
| SE | 66213 | ENSG00000137817 | PARP6 | chr15 | - | 72253713 | 72253762 | 72253436 | 72253504 | 72254454 | 72254520 | 0 | 0 | 0 | [illegible] |
| SE | 48259 | ENSG00000111602 | TIMELESS | chr12 | - | 56421350 | 56421493 | 56420962 | 56421134 | 56421726 | 56421809 | 34 | 33 | 22 | [illegible] |
| SE | 47292 | ENSG00000121957 | GPSM2 | chr1 | + | 108918612 | 108918789 | 108914337 | 108914408 | 108922416 | 108922576 | 22 | 32 | 21 | [illegible] |
| SE | 35198 | ENSG00000101596 | SMCHD1 | chr18 | + | 2775733 | 2775924 | 2772249 | 2772372 | 2777805 | 2777915 | 743 | 681 | 863 | [illegible] |
| SE | 32656 | ENSG00000150995 | ITPR1 | chr3 | + | 4680552 | 4680691 | 4676613 | 4676801 | 4681363 | 4681418 | 182 | 109 | 135 | [illegible] |
| SE | 83411 | ENSG00000146707 | POMZP3 | chr7 | - | 76618496 | 76618615 | 76618182 | 76618300 | 76625521 | 76625683 | 10 | 5 | 6 | [illegible] |
| SE | 77643 | ENSG00000214021 | TTLL3 | chr3 | + | 9816073 | 9816202 | 9813247 | 9813345 | 9818554 | 9818920 | 22 | 15 | 21 | [illegible] |

[illegible]

| | | | | | | | | | | | | | | | | | | | | | | | | | | |
|---|---|---|---|---|---|---|---|---|---|---|---|---|---|---|---|---|---|---|---|---|---|---|---|---|---|---|
| SE | 107500 | ENSG00000123349 | PFDN5 | chr12 | + | 53296243 | 53296275 | 53295838 | 53295941 | 53297849 | 53297924 | 1622 | 887 | 1400 | 1338 | 1648 | 3 | 13 | 11 | 3 | 11 | 1972 | 1168 | 788 | 1165 | [illegible] |
| SE | 73248 | ENSG00000114956 | DGUOK | chr2 | + | 73938909 | 73939022 | 73932597 | 73932683 | 73946718 | 73946906 | 50 | 18 | 38 | 39 | 46 | 13 | 3 | 9 | 2 | 6 | 59 | 53 | 42 | 45 | [illegible] |
| SE | 85689 | ENSG00000177981 | ASB8 | chr12 | - | 48153367 | 48153525 | 48150915 | 48151305 | 48157458 | 48157502 | 88 | 27 | 66 | 73 | 95 | 0 | 0 | 0 | 3 | 0 | 83 | 98 | 82 | 25 | [illegible] |
| SE | 50867 | ENSG00000115568 | ZNF142 | chr2 | - | 218656149 | 218656463 | 218651700 | 218652300 | 218658700 | 218658801 | 14 | 9 | 14 | 13 | 12 | 1 | 0 | 0 | 0 | 0 | 36 | 28 | 21 | 31 | [illegible] |
| SE | 48796 | ENSG00000153922 | CHD1 | chr5 | - | 98901185 | 98901335 | 98900810 | 98901082 | 98902899 | 98902964 | 320 | 297 | 257 | 320 | 415 | 2 | 4 | 2 | 1 | 7 | 72 | 172 | 336 | 64 | [illegible] |
| SE | 68719 | ENSG00000196295 | GARS1-DT | chr7 | - | 30577648 | 30578091 | 30576563 | 30576710 | 30579230 | 30579314 | 108 | 82 | 106 | 142 | 137 | 6 | 0 | 0 | 0 | 0 | 171 | 134 | 94 | 82 | [illegible] |
| SE | 70638 | ENSG00000188647 | PTAR1 | chr9 | - | 69750780 | 69750950 | 69741515 | 69741858 | 69759852 | 69759955 | 152 | 121 | 159 | 146 | 191 | 9 | 3 | 3 | 12 | 5 | 34 | 39 | 37 | 46 | [illegible] |
| SE | 19701 | ENSG00000119599 | DCAF4 | chr14 | + | 72942993 | 72943096 | 72941744 | 72941824 | 72945883 | 72945927 | 44 | 26 | 53 | 48 | 79 | 1 | 0 | 0 | 0 | 0 | 27 | 17 | 43 | 9 | [illegible] |
| SE | 105246 | ENSG00000082898 | XPO1 | chr2 | - | 61488587 | 61488932 | 61488164 | 61488271 | 61490641 | 61490776 | 182 | 125 | 170 | 202 | 402 | 0 | 2 | 2 | 6 | 0 | 34 | 66 | 74 | 52 | [illegible] |
| SE | 23558 | ENSG00000124532 | MRS2 | chr6 | + | 24410726 | 24410799 | 24409460 | 24409573 | 24412221 | 24412395 | 0 | 0 | 0 | 1 | 9 | 34 | 42 | 45 | 38 | 129 | 0 | 0 | 0 | 0 | [illegible] |
| SE | 32635 | ENSG00000177963 | RIC8A | chr11 | + | 212415 | 212511 | 211198 | 211349 | 212614 | 212724 | 172 | 79 | 113 | 155 | 55 | 0 | 0 | 0 | 2 | 0 | 249 | 194 | 115 | 109 | [illegible] |
| SE | 77345 | ENSG00000167380 | ZNF226 | chr19 | + | 44165587 | 44165613 | 44165083 | 44165142 | 44165756 | 44165807 | 1 | 4 | 1 | 2 | 1 | 28 | 19 | 20 | 19 | 10 | 2 | 0 | 1 | 6 | [illegible] |
| SE | 95435 | ENSG00000119383 | PTPA | chr9 | + | 129120512 | 129120610 | 129111313 | 129111631 | 129123051 | 129123138 | 115 | 79 | 129 | 151 | 170 | 2 | 0 | 1 | 0 | 2 | 179 | 200 | 96 | 162 | [illegible] |
| SE | 95545 | ENSG00000187764 | SEMA4D | chr9 | - | 89418362 | 89418437 | 89418055 | 89418118 | 89455887 | 89455953 | 166 | 154 | 184 | 333 | 274 | 123 | 96 | 77 | 157 | 121 | 226 | 122 | 158 | 139 | [illegible] |
| SE | 71857 | ENSG00000105639 | JAK3 | chr19 | - | 17843776 | 17843900 | 17843379 | 17843491 | 17844233 | 17844430 | 120 | 47 | 122 | 116 | 49 | 0 | 0 | 0 | 1 | 0 | 100 | 144 | 50 | 103 | [illegible] |
| SE | 89545 | ENSG00000133317 | LGALS12 | chr11 | + | 63515562 | 63515713 | 63511751 | 63511840 | 63516246 | 63516767 | 8 | 11 | 7 | 17 | 16 | 1 | 0 | 0 | 1 | 3 | 35 | 13 | 8 | 29 | [illegible] |
| SE | 43828 | ENSG00000162946 | DISC1 | chr1 | + | 232008784 | 232008983 | 231958827 | 231958888 | 232026434 | 232026552 | 45 | 62 | 40 | 54 | 56 | 0 | 0 | 0 | 0 | 0 | 93 | 55 | 62 | 47 | [illegible] |
| SE | 67458 | ENSG00000162923 | WDR26 | chr1 | - | 224393827 | 224394013 | 224385145 | 224389860 | 224398514 | 224398593 | 448 | 311 | 304 | 459 | 487 | 4 | 1 | 0 | 3 | 7 | 348 | 642 | 720 | 382 | [illegible] |
| SE | 82554 | ENSG00000106263 | EIF3B | chr7 | + | 2366524 | 2366591 | 2366316 | 2366448 | 2366998 | 2367045 | 161 | 102 | 183 | 189 | 203 | 0 | 0 | 2 | 3 | 2 | 256 | 162 | 126 | 143 | [illegible] |
| SE | 86387 | ENSG00000123562 | MORF4L2 | chrX | - | 103684680 | 103684729 | 103678498 | 103678620 | 103685170 | 103685260 | 50 | 29 | 47 | 50 | 94 | 6 | 8 | 11 | 19 | 12 | 32 | 27 | 31 | 28 | [illegible] |
| SE | 84416 | ENSG00000188917 | TRMT2B | chrX | - | 101041316 | 101041371 | 101036973 | 101037073 | 101051816 | 101052054 | 6 | 8 | 12 | 8 | 15 | 0 | 0 | 0 | 2 | 5 | 13 | 2 | 6 | 1 | [illegible] |
| SE | 79683 | ENSG00000134905 | CARS2 | chr13 | - | 110648317 | 110649297 | 110645966 | 110647239 | 110651033 | 110651100 | 0 | 0 | 0 | 3 | 0 | 127 | 76 | 95 | 95 | 162 | 0 | 1 | 4 | 0 | [illegible] |
| SE | 71267 | ENSG00000121741 | ZMYM2 | chr13 | + | 19970024 | 19970071 | 19965219 | 19965281 | 19993062 | 19993919 | 23 | 20 | 28 | 22 | 51 | 0 | 0 | 3 | 0 | 5 | 7 | 27 | 42 | 22 | [illegible] |
| SE | 21511 | ENSG00000137413 | TAF8 | chr6 | + | 42057388 | 42057513 | 42055951 | 42056014 | 42066311 | 42066459 | 213 | 122 | 175 | 177 | 290 | 0 | 2 | 2 | 0 | 0 | 279 | 207 | 202 | 168 | [illegible] |
| SE | 30072 | ENSG00000115289 | PCGF1 | chr2 | - | 74505951 | 74506057 | 74505736 | 74505770 | 74506731 | 74506884 | 40 | 25 | 17 | 31 | 65 | 5 | 1 | 2 | 1 | 3 | 53 | 12 | 33 | 14 | [illegible] |
| SE | 41110 | ENSG00000122483 | CCDC18 | chr1 | + | 93239309 | 93239473 | 93236247 | 93236390 | 93239682 | 93239896 | 76 | 65 | 73 | 54 | 95 | 1 | 0 | 0 | 0 | 0 | 31 | 34 | 19 | 10 | [illegible] |
| SE | 49588 | ENSG00000076685 | NT5C2 | chr10 | - | 103100751 | 103100943 | 103099925 | 103100019 | 103101044 | 103101102 | 44 | 24 | 35 | 56 | 80 | 156 | 162 | 126 | 182 | 218 | 58 | 79 | 87 | 58 | [illegible] |
| SE | 19201 | ENSG00000140105 | WARS1 | chr14 | - | 100375282 | 100375406 | 100369086 | 100369258 | 100375510 | 100375744 | 102 | 118 | 109 | 97 | 67 | 9 | 10 | 2 | 2 | 6 | 433 | 56 | 43 | 170 | [illegible] |
| SE | 17661 | ENSG00000176681 | LRRC37A | chr17 | + | 46305508 | 46305580 | 46302637 | 46302709 | 46306228 | 46306309 | 4 | 9 | 9 | 7 | 30 | 0 | 0 | 0 | 0 | 5 | 9 | 21 | 1 | 8 | [illegible] |
| SE | 59378 | ENSG00000116127 | ALMS1 | chr2 | + | 73534823 | 73534949 | 73519774 | 73520016 | 73550266 | 73550437 | 105 | 68 | 88 | 77 | 77 | 1 | 2 | 1 | 0 | 4 | 16 | 42 | 54 | 4 | [illegible] |
| SE | 68841 | ENSG00000168769 | TET2 | chr4 | + | 105190359 | 105190486 | 105146292 | 105146979 | 105233896 | 105237351 | 31 | 12 | 13 | 42 | 22 | 26 | 22 | 21 | 21 | 30 | 4 | 12 | 15 | 2 | [illegible] |
| SE | 28024 | ENSG00000138698 | RAP1GDS1 | chr4 | + | 98416744 | 98416888 | 98404476 | 98404602 | 98417366 | 98417498 | 333 | 217 | 173 | 282 | 409 | 1 | 0 | 0 | 2 | 2 | 240 | 253 | 165 | 236 | [illegible] |
| SE | 355 | ENSG00000160294 | MCM3AP | chr21 | - | 46254775 | 46254844 | 46254391 | 46254526 | 46256788 | 46256986 | 78 | 32 | 58 | 73 | 22 | 0 | 0 | 0 | 0 | 0 | 36 | 87 | 56 | 53 | [illegible] |
| SE | 95273 | ENSG00000165219 | GAPVD1 | chr9 | + | 125341176 | 125341264 | 125337220 | 125337591 | 125346818 | 125346941 | 64 | 69 | 88 | 76 | 70 | 0 | 0 | 0 | 0 | 0 | 95 | 56 | 64 | 61 | [illegible] |
| SE | 29105 | ENSG00000061918 | GUCY1B1 | chr4 | + | 155759355 | 155759541 | 155759030 | 155759143 | 155759786 | 155759860 | 3 | 11 | 2 | 24 | 7 | 1 | 3 | 0 | 14 | 10 | 19 | 2 | 14 | 6 | [illegible] |
| SE | 87569 | ENSG00000112159 | MDN1 | chr6 | - | 89690672 | 89690834 | 89688077 | 89688173 | 89692442 | 89693148 | 84 | 31 | 65 | 77 | 48 | 0 | 0 | 0 | 0 | 0 | 89 | 37 | 57 | 55 | [illegible] |
| SE | 90157 | ENSG00000035928 | RFC1 | chr4 | - | 39306591 | 39306701 | 39304813 | 39304928 | 39308635 | 39309032 | 232 | 159 | 235 | 229 | 191 | 0 | 2 | 2 | 4 | 9 | 3 | 116 | 173 | 31 | [illegible] |
| SE | 72805 | ENSG00000188428 | BLOC1S5 | chr6 | - | 8027206 | 8027399 | 8026366 | 8026425 | 8041138 | 8041268 | 21 | 23 | 18 | 43 | 40 | 106 | 65 | 72 | 76 | 142 | 0 | 8 | 33 | 27 | [illegible] |
| SE | 101473 | ENSG00000165309 | ARMC3 | chr10 | + | 22959069 | 22959138 | 22955806 | 22955932 | 22959398 | 22959574 | 14 | 50 | 10 | 67 | 88 | 0 | 2 | 1 | 6 | 9 | 37 | 34 | 16 | 11 | [illegible] |
| SE | 14373 | ENSG00000173517 | PEAK1 | chr15 | - | 77283882 | 77284030 | 77252366 | 77252526 | 77284957 | 77285055 | 33 | 29 | 19 | 17 | 15 | 10 | 9 | 12 | 10 | 3 | 29 | 19 | 26 | 56 | [illegible] |
| SE | 63853 | ENSG00000219626 | FAM228B | chr2 | + | 24080875 | 24080955 | 24076857 | 24076969 | 24095140 | 24095229 | 7 | 11 | 12 | 9 | 10 | 0 | 0 | 0 | 0 | 2 | 7 | 4 | 14 | 24 | [illegible] |
| SE | 96420 | ENSG00000159314 | ARHGAP27 | chr17 | - | 45396486 | 45396585 | 45396206 | 45396284 | 45396667 | 45396790 | 96 | 39 | 64 | 66 | 28 | 1 | 0 | 0 | 0 | 0 | 49 | 61 | 27 | 30 | [illegible] |
| SE | 86071 | ENSG00000164031 | DNAJB14 | chr4 | - | 99930449 | 99930621 | 99922519 | 99923185 | 99946438 | 99946558 | 285 | 197 | 192 | 261 | 325 | 4 | 4 | 2 | 0 | 1 | 48 | 67 | 49 | 17 | [illegible] |
| SE | 100453 | ENSG00000103429 | BFAR | chr16 | + | 14649803 | 14649973 | 14644287 | 14644609 | 14661891 | 14662065 | 16 | 6 | 25 | 16 | 34 | 0 | 1 | 12 | 6 | 3 | 0 | 38 | 12 | 0 | [illegible] |
| SE | 79828 | ENSG00000091106 | NLRC4 | chr2 | - | 32252418 | 32252679 | 32241032 | 32241125 | 32256774 | 32256893 | 122 | 117 | 47 | 52 | 88 | 2 | 0 | 0 | 1 | 0 | 46 | 85 | 33 | 107 | [illegible] |
| SE | 108152 | ENSG00000110911 | SLC11A2 | chr12 | - | 51000562 | 51000633 | 51000312 | 51000419 | 51004787 | 51004907 | 1 | 1 | 0 | 0 | 0 | 28 | 25 | 31 | 20 | 19 | 0 | 0 | 0 | 5 | [illegible] |
| SE | 80834 | ENSG00000151948 | GLT1D1 | chr12 | + | 128927104 | 128927154 | 128888638 | 128888744 | 128945325 | 128945369 | 20 | 16 | 10 | 13 | 13 | 0 | 0 | 0 | 1 | 0 | 39 | 27 | 27 | 22 | [illegible] |
| SE | 29414 | ENSG00000153234 | NR4A2 | chr2 | - | 156326311 | 156326328 | 156326149 | 156326207 | 156326717 | 156326920 | 0 | 0 | 4 | 8 | 2 | 1 | 2 | 8 | 11 | 4 | 0 | 0 | 18 | 30 | [illegible] |
| SE | 2036 | ENSG00000153885 | KCTD15 | chr19 | + | 33799593 | 33799718 | 33798667 | 33798766 | 33800427 | 33800520 | 4 | 0 | 0 | 0 | 0 | 4 | 1 | 2 | 8 | 7 | 0 | 0 | 0 | 0 | [illegible] |
| SE | 32771 | ENSG00000168297 | PXK | chr3 | + | 58391146 | 58391220 | 58390581 | 58390659 | 58394997 | 58395102 | 59 | 30 | 33 | 63 | 125 | 0 | 0 | 1 | 0 | 2 | 118 | 57 | 44 | 96 | [illegible] |
| SE | 46919 | ENSG00000141027 | NCOR1 | chr17 | - | 16034764 | 16034944 | 16029156 | 16032483 | 16039432 | 16039654 | 355 | 175 | 274 | 276 | 528 | 0 | 0 | 0 | 0 | 0 | 159 | 223 | 179 | 265 | [illegible] |
| SE | 95749 | ENSG00000070476 | ZXDC | chr3 | - | 126459652 | 126459737 | 126439631 | 126439727 | 126461534 | 126462220 | 31 | 16 | 12 | 26 | 14 | 0 | 0 | 5 | 0 | 0 | 50 | 21 | 25 | 22 | [illegible] |
| SE | 100577 | ENSG00000250151 | ARPC4-TTLL3 | chr3 | + | 9810620 | 9810709 | 9803842 | 9804013 | 9812942 | 9813071 | 51 | 34 | 49 | 82 | 63 | 0 | 0 | 0 | 0 | 0 | 104 | 84 | 68 | 128 | [illegible] |
| SE | 31352 | ENSG00000124535 | WRNIP1 | chr6 | + | 2768690 | 2768882 | 2765392 | 2766444 | 2770119 | 2770361 | 143 | 74 | 139 | 154 | 194 | 4 | 0 | 0 | 0 | 0 | 128 | 124 | 47 | 122 | [illegible] |
| SE | 63746 | ENSG00000152061 | RABGAP1L | chr1 | + | 174702112 | 174702256 | 174699524 | 174699650 | 174752312 | 174752354 | 213 | 200 | 156 | 220 | 222 | 2 | 4 | 3 | 6 | 0 | 28 | 262 | 278 | 70 | [illegible] |
| SE | 6944 | ENSG00000100983 | GSS | chr20 | - | 34936764 | 34936840 | 34935575 | 34935642 | 34936942 | 34937023 | 24 | 13 | 16 | 18 | 13 | 0 | 0 | 1 | 0 | 4 | 43 | 49 | 26 | 36 | [illegible] |
| SE | 5650 | ENSG00000196700 | ZNF512B | chr20 | - | 63961308 | 63961407 | 63956703 | 63960139 | 63961941 | 63962004 | 17 | 4 | 23 | 14 | 30 | 0 | 0 | 0 | 0 | 0 | 20 | 13 | 13 | 10 | [illegible] |
| SE | 18423 | ENSG00000196497 | IPO4 | chr14 | - | 24187666 | 24187813 | 24187399 | 24187579 | 24188215 | 24188257 | 7 | 4 | 18 | 9 | 18 | 2 | 4 | 0 | 7 | 8 | 0 | 14 | 13 | 10 | [illegible] |
| SE | 86425 | ENSG00000196092 | PAX5 | chr9 | - | 36930887 | 36930944 | 36923354 | 36923484 | 36966548 | 36966724 | 0 | 0 | 0 | 5 | 2 | 26 | 7 | 9 | 31 | 17 | 0 | 0 | 0 | 3 | [illegible] |
| SE | 38146 | ENSG00000198087 | CD2AP | chr6 | + | 47612472 | 47612536 | 47609122 | 47609304 | 47624185 | 47627263 | 155 | 167 | 130 | 170 | 244 | 4 | 1 | 0 | 2 | 0 | 9 | 69 | 92 | 27 | [illegible] |
| SE | 29650 | ENSG00000168310 | IRF2 | chr4 | - | 184418166 | 184418213 | 184398914 | 184399079 | 184418531 | 184418590 | 112 | 73 | 111 | 101 | 268 | 2 | 1 | 1 | 0 | 0 | 179 | 132 | 82 | 109 | [illegible] |
| SE | 37074 | ENSG00000005812 | FBXL3 | chr13 | - | 77021512 | 77021861 | 77018599 | 77018722 | 77026826 | 77027159 | 192 | 92 | 163 | 126 | 144 | 2 | 1 | 1 | 6 | 0 | 30 | 108 | 97 | 28 | [illegible] |
| SE | 86307 | ENSG00000147224 | PRPS1 | chrX | + | 107640903 | 107641000 | 107639294 | 107639478 | 107642365 | 107642490 | 58 | 55 | 79 | 55 | 41 | 1 | 0 | 0 | 0 | 0 | 34 | 68 | 65 | 40 | [illegible] |
| SE | 65298 | ENSG00000149499 | EML3 | chr11 | - | 62606062 | 62606214 | 62605854 | 62605980 | 62606957 | 62607082 | 124 | 38 | 118 | 120 | 127 | 0 | 0 | 0 | 0 | 0 | 111 | 107 | 57 | 86 | [illegible] |
| SE | 1160 | ENSG00000100395 | L3MBTL2 | chr22 | + | 41217122 | 41217202 | 41216138 | 41216262 | 41220733 | 41220868 | 19 | 12 | 25 | 32 | 14 | 0 | 0 | 0 | 0 | 0 | 37 | 37 | 14 | 12 | [illegible] |
| SE | 55260 | ENSG00000135905 | DOCK10 | chr2 | - | 224841803 | 224841896 | 224839953 | 224840072 | 224844752 | 224844839 | 328 | 325 | 370 | 360 | 655 | 2 | 1 | 5 | 1 | 1 | 16 | 227 | 260 | 62 | [illegible] |
| SE | 63644 | ENSG00000112855 | HARS2 | chr5 | + | 140693490 | 140693665 | 140691430 | 140691756 | 140693934 | 140694054 | 48 | 28 | 29 | 38 | 51 | 0 | 0 | 1 | 0 | 0 | 32 | 21 | 34 | 25 | [illegible] |
| SE | 21787 | ENSG00000198625 | MDM4 | chr1 | + | 204537429 | 204537497 | 204532190 | 204532246 | 204538208 | 204538308 | 203 | 166 | 188 | 228 | 276 | 101 | 88 | 78 | 79 | 118 | 238 | 323 | 259 | 338 | [illegible] |
| SE | 109656 | ENSG00000139182 | CLSTN3 | chr12 | + | 7141241 | 7141404 | 7136828 | 7137110 | 7142085 | 7142139 | 78 | 21 | 37 | 58 | 105 | 0 | 0 | 0 | 0 | 0 | 81 | 150 | 53 | 105 | [illegible] |
| SE | 52568 | ENSG00000111445 | RFC5 | chr12 | + | 118017689 | 118018053 | 118016721 | 118016892 | 118019071 | 118019136 | 7 | 2 | 3 | 5 | 7 | 13 | 12 | 15 | 23 | 8 | 17 | 3 | 5 | 16 | [illegible] |
| SE | 77880 | ENSG00000198815 | FOXJ3 | chr1 | - | 42278347 | 42278672 | 42265114 | 42265189 | 42311049 | 42311110 | 322 | 237 | 286 | 309 | 524 | 2 | 0 | 2 | 0 | 0 | 193 | 270 | 233 | 211 | [illegible] |
| SE | 73918 | ENSG00000161692 | DBF4B | chr17 | + | 44732177 | 44732265 | 44730964 | 44731015 | 44734089 | 44734163 | 7 | 2 | 11 | 5 | 13 | 0 | 1 | 1 | 0 | 0 | 22 | 9 | 1 | 9 | [illegible] |
| SE | 55015 | ENSG00000152223 | EPG5 | chr18 | - | 45870566 | 45870742 | 45867562 | 45867748 | 45878375 | 45878448 | 84 | 40 | 41 | 53 | 57 | 0 | 2 | 0 | 0 | 0 | 32 | 25 | 51 | 41 | [illegible] |
| SE | 48217 | ENSG00000141446 | ESCO1 | chr18 | - | 21564202 | 21564317 | 21560858 | 21560990 | 21566145 | 21566206 | 207 | 127 | 157 | 182 | 289 | 1 | 1 | 0 | 1 | 7 | 56 | 33 | 58 | 17 | [illegible] |
| SE | 102076 | ENSG00000181192 | DHTKD1 | chr10 | + | 12100177 | 12100262 | 12097683 | 12097996 | 12101041 | 12101181 | 46 | 34 | 56 | 55 | 83 | 4 | 2 | 1 | 2 | 0 | 81 | 69 | 36 | 48 | [illegible] |
| SE | 54500 | ENSG00000151414 | NEK7 | chr1 | + | 198297126 | 198297240 | 198264124 | 198264235 | 198319411 | 198319522 | 184 | 125 | 140 | 176 | 332 | 7 | 1 | 5 | 4 | 9 | 12 | 72 | 36 | 9 | [illegible] |
| SE | 106844 | ENSG00000123329 | ARHGAP9 | chr12 | - | 57474422 | 57474476 | 57473602 | 57473708 | 57474625 | 57474703 | 119 | 62 | 110 | 128 | 52 | 0 | 0 | 0 | 0 | 0 | 183 | 88 | 98 | 104 | [illegible] |
| SE | 41662 | ENSG00000119185 | ITGB1BP1 | chr2 | - | 9419989 | 9420100 | 9418625 | 9418732 | 9423372 | 9423480 | 32 | 5 | 21 | 11 | 26 | 35 | 19 | 28 | 43 | 3 | 12 | 20 | 28 | 16 | [illegible] |
| SE | 19431 | ENSG00000151445 | VIPAS39 | chr14 | - | 77449292 | 77449357 | 77448493 | 77448550 | 77449713 | 77449752 | 48 | 28 | 53 | 58 | 60 | 0 | 1 | 0 | 0 | 0 | 96 | 60 | 26 | 51 | [illegible] |
| SE | 37980 | ENSG00000175348 | TMEM9B | chr11 | - | 8962091 | 8962183 | 8956189 | 8956298 | 8964889 | 8965011 | 80 | 35 | 62 | 98 | 130 | 0 | 0 | 0 | 0 | 0 | 89 | 83 | 62 | 66 | [illegible] |
| SE | 2649 | ENSG00000129351 | ILF3 | chr19 | + | 10678612 | 10678704 | 10677162 | 10677310 | 10679096 | 10679170 | 374 | 278 | 490 | 438 | 479 | 3 | 2 | 9 | 2 | 10 | 642 | 477 | 382 | 476 | [illegible] |
| SE | 102491 | ENSG00000119979 | DENND10 | chr10 | + | 119129514 | 119129622 | 119120340 | 119120452 | 119132514 | 119132609 | 86 | 79 | 55 | 73 | 96 | 0 | 0 | 0 | 0 | 8 | 48 | 32 | 39 | 58 | [illegible] |
| SE | 70569 | ENSG00000022840 | RNF10 | chr12 | + | 120565071 | 120565189 | 120563747 | 120563943 | 120565427 | 120565527 | 418 | 218 | 349 | 421 | 299 | 2 | 0 | 2 | 1 | 0 | 843 | 660 | 647 | 791 | [illegible] |
| SE | 49624 | ENSG00000178927 | CYBC1 | chr17 | - | 82447579 | 82447621 | 82446622 | 82446696 | 82450699 | 82450737 | 5 | 5 | 9 | 8 | 2 | 1 | 0 | 0 | 2 | 0 | 15 | 20 | 10 | 18 | [illegible] |
| SE | 61521 | ENSG00000138757 | G3BP2 | chr4 | - | 75647028 | 75647157 | 75646337 | 75646456 | 75648638 | 75648741 | 1920 | 1619 | 1630 | 2277 | 2175 | 3 | 1 | 4 | 5 | 4 | 1536 | 1936 | 3285 | 2420 | [illegible] |
| SE | 52634 | ENSG00000197943 | PLCG2 | chr16 | + | 81858262 | 81858356 | 81854443 | 81854587 | 81859115 | 81859163 | 209 | 86 | 122 | 129 | 159 | 6 | 1 | 1 | 2 | 0 | 138 | 232 | 136 | 247 | [illegible] |
| SE | 51043 | ENSG00000112182 | BACH2 | chr6 | - | 90271848 | 90271941 | 90206568 | 90206676 | 90296479 | 90296742 | 12 | 6 | 9 | 12 | 0 | 0 | 1 | 0 | 4 | 2 | 5 | 6 | 3 | 1 | [illegible] |
| SE | 47477 | ENSG00000183023 | SLC8A1 | chr2 | - | 40178386 | 40178493 | 40177751 | 40177855 | 40428472 | 40430271 | 2 | 6 | 2 | 6 | 5 | 70 | 91 | 29 | 43 | 92 | 14 | 3 | 9 | 0 | [illegible] |
| SE | 92929 | ENSG00000131089 | ARHGEF9 | chrX | - | 63722761 | 63722897 | 63706257 | 63706449 | 63724531 | 63724711 | 2 | 3 | 5 | 2 | 7 | 39 | 21 | 38 | 25 | 40 | 0 | 14 | 3 | 2 | [illegible] |
| SE | 21857 | ENSG00000068796 | KIF2A | chr5 | + | 62366413 | 62366481 | 62365242 | 62365353 | 62373686 | 62373837 | 602 | 737 | 494 | 1162 | 1451 | 0 | 7 | 0 | 8 | 0 | 295 | 475 | 418 | 305 | [illegible] |
| SE | 55584 | ENSG00000137275 | RIPK1 | chr6 | + | 3082546 | 3083313 | 3080978 | 3081116 | 3085258 | 3085408 | 59 | 31 | 41 | 77 | 46 | 8 | 8 | 5 | 4 | 3 | 22 | 44 | 36 | 44 | [illegible] |
| SE | 86490 | ENSG00000071889 | FAM3A | chrX | - | 154512313 | 154512431 | 154511867 | 154511871 | 154512822 | 154512936 | 3 | 3 | 15 | 3 | 0 | 48 | 5 | 25 | 18 | 19 | 9 | 7 | 1 | 1 | [illegible] |
| SE | 46423 | ENSG00000162174 | ASRGL1 | chr11 | + | 62356324 | 62356467 | 62337889 | 62338167 | 62356986 | 62357132 | 44 | 31 | 29 | 48 | 48 | 0 | 3 | 0 | 0 | 4 | 139 | 44 | 15 | 54 | [illegible] |
| SE | 77004 | ENSG00000149657 | LSM14B | chr20 | + | 62129784 | 62129952 | 62126225 | 62126439 | 62130529 | 62130605 | 33 | 17 | 30 | 50 | 20 | 0 | 0 | 3 | 3 | 0 | 33 | 24 | 29 | 42 | [illegible] |
| SE | 87380 | ENSG00000134982 | APC | chr5 | + | 112818965 | 112819344 | 112815494 | 112815593 | 112821895 | 112821991 | 144 | 145 | 91 | 164 | 138 | 3 | 0 | 1 | 1 | 3 | 21 | 85 | 39 | 41 | [illegible] |
| SE | 90163 | ENSG00000035928 | RFC1 | chr4 | - | 39312751 | 39312931 | 39311444 | 39311549 | 39316914 | 39317013 | 290 | 256 | 264 | 306 | 444 | 0 | 8 | 0 | 0 | 0 | 0 | 329 | 422 | 114 | [illegible] |
| SE | 94128 | ENSG00000139644 | TMBIM6 | chr12 | + | 49742437 | 49742607 | 49741956 | 49742115 | 49752463 | 49752524 | 2 | 2 | 9 | 4 | 0 | 26 | 7 | 4 | 14 | 7 | 5 | 4 | 5 | 0 | [illegible] |
| SE | 68290 | ENSG00000122085 | MTERF4 | chr2 | - | 241096392 | 241096438 | 241094276 | 241094432 | 241097242 | 241097427 | 20 | 20 | 34 | 31 | 19 | 8 | 8 | 17 | 4 | 19 | 27 | 19 | 22 | 22 | [illegible] |
| SE | 99980 | ENSG00000185187 | SIGIRR | chr11 | - | 409859 | 410027 | 408072 | 408206 | 414822 | 415004 | 108 | 49 | 79 | 82 | 48 | 3 | 1 | 7 | 6 | 7 | 38 | 95 | 71 | 28 | [illegible] |
| SE | 82746 | ENSG00000140396 | NCOA2 | chr8 | - | 70148272 | 70148483 | 70144641 | 70144848 | 70155970 | 70157240 | 185 | 195 | 206 | 223 | 374 | 1 | 1 | 0 | 5 | 3 | 78 | 166 | 221 | 226 | [illegible] |
| SE | 11234 | ENSG00000102984 | ZNF821 | chr16 | - | 71867892 | 71868037 | 71864137 | 71864242 | 71879906 | 71880023 | 16 | 9 | 10 | 15 | 13 | 1 | 0 | 2 | 2 | 0 | 26 | 23 | 17 | 18 | [illegible] |
| SE | 90470 | ENSG00000151461 | UPF2 | chr10 | - | 11955231 | 11955507 | 11952065 | 11952249 | 11956319 | 11956523 | 146 | 124 | 203 | 141 | 235 | 0 | 1 | 0 | 0 | 1 | 30 | 94 | 83 | 9 | [illegible] |
| SE | 13433 | ENSG00000103111 | MON1B | chr16 | + | 77191475 | 77191633 | 77190997 | 77191258 | 77194334 | 77195154 | 103 | 60 | 71 | 87 | 80 | 0 | 1 | 0 | 0 | 1 | 158 | 71 | 55 | 106 | [illegible] |
| SE | 5191 | ENSG00000124214 | STAU1 | chr20 | - | 49153932 | 49154071 | 49151581 | 49151747 | 49174194 | 49174269 | 93 | 90 | 80 | 126 | 144 | 2 | 0 | 0 | 0 | 0 | 110 | 138 | 137 | 108 | [illegible] |
| SE | 4481 | ENSG00000127527 | EPS15L1 | chr19 | - | 16417947 | 16418104 | 16417438 | 16417637 | 16421318 | 16421433 | 160 | 61 | 79 | 150 | 132 | 2 | 1 | 0 | 0 | 0 | 254 | 215 | 117 | 177 | [illegible] |
| SE | 36005 | ENSG00000119725 | ZNF410 | chr14 | + | 73903959 | 73904110 | 73898070 | 73898262 | 73904901 | 73905046 | 202 | 123 | 169 | 147 | 219 | 1 | 2 | 0 | 0 | 4 | 264 | 224 | 148 | 211 | [illegible] |
| SE | 8175 | ENSG00000108799 | EZH1 | chr17 | - | 42713208 | 42713389 | 42712288 | 42712485 | 42717975 | 42718067 | 176 | 82 | 124 | 144 | 251 | 4 | 2 | 1 | 4 | 4 | 172 | 234 | 152 | 145 | [illegible] |
| SE | 91140 | ENSG00000116852 | KIF21B | chr1 | - | 200999348 | 200999466 | 200998383 | 200998575 | 200999882 | 200999964 | 195 | 90 | 166 | 233 | 234 | 0 | 0 | 0 | 2 | 0 | 111 | 291 | 181 | 210 | [illegible] |
| SE | 26954 | ENSG00000144659 | SLC25A38 | chr3 | + | 39394409 | 39394576 | 39391852 | 39392021 | 39396397 | 39396642 | 169 | 89 | 205 | 166 | 135 | 0 | 1 | 0 | 6 | 2 | 77 | 150 | 103 | 73 | [illegible] |
| SE | 3319 | ENSG00000042753 | AP2S1 | chr19 | - | 46839464 | 46839620 | 46838739 | 46838799 | 46845992 | 46846142 | 154 | 81 | 119 | 143 | 83 | 6 | 2 | 1 | 0 | 2 | 229 | 262 | 123 | 103 | [illegible] |
| SE | 65178 | ENSG00000268362 | NA | chr19 | + | 24048936 | 24049014 | 24046183 | 24046279 | 24049404 | 24049548 | 1 | 3 | 4 | 2 | 2 | 20 | 21 | 13 | 22 | 8 | 0 | 0 | 0 | 8 | [illegible] |
| SE | 87544 | ENSG00000147050 | KDM6A | chrX | + | 45062646 | 45062748 | 45061323 | 45061419 | 45063421 | 45063817 | 129 | 94 | 135 | 145 | 185 | 31 | 24 | 28 | 24 | 50 | 56 | 103 | 160 | 162 | [illegible] |
| SE | 84660 | ENSG00000185753 | CXorf38 | chrX | - | 40647006 | 40647141 | 40639008 | 40639128 | 40647304 | 40647521 | 159 | 100 | 122 | 92 | 103 | 6 | 7 | 4 | 2 | 0 | 167 | 102 | 73 | 74 | [illegible] |
| SE | 28592 | ENSG00000105821 | DNAJC2 | chr7 | - | 103327331 | 103327378 | 103326542 | 103326684 | 103327655 | 103327754 | 29 | 31 | 23 | 23 | 24 | 97 | 80 | 72 | 96 | 181 | 9 | 6 | 7 | 9 | [illegible] |
| SE | 4276 | ENSG00000089639 | GMIP | chr19 | - | 19635643 | 19635721 | 19635414 | 19635569 | 19636706 | 19636787 | 62 | 18 | 28 | 39 | 40 | 11 | 2 | 3 | 3 | 3 | 51 | 47 | 49 | 50 | [illegible] |
| SE | 101037 | ENSG00000118276 | B4GALT6 | chr18 | - | 31630958 | 31631146 | 31626988 | 31627121 | 31638643 | 31638760 | 100 | 55 | 45 | 60 | 36 | 0 | 0 | 0 | 0 | 0 | 10 | 40 | 7 | 16 | [illegible] |
| SE | 1780 | ENSG00000100077 | GRK3 | chr22 | + | 25685169 | 25685248 | 25674436 | 25674528 | 25687536 | 25687667 | 75 | 40 | 36 | 59 | 84 | 0 | 0 | 0 | 0 | 0 | 110 | 76 | 39 | 71 | [illegible] |
| SE | 36860 | ENSG00000175792 | RUVBL1 | chr3 | - | 128100594 | 128100744 | 128098881 | 128098945 | 128104772 | 128104924 | 125 | 57 | 111 | 127 | 188 | 3 | 0 | 0 | 4 | 0 | 234 | 128 | 87 | 134 | [illegible] |
| SE | 56287 | ENSG00000134242 | PTPN22 | chr1 | - | 113825141 | 113825172 | 113819298 | 113819654 | 113829591 | 113829707 | 425 | 233 | 282 | 331 | 771 | 9 | 6 | 0 | 4 | 12 | 61 | 182 | 128 | 31 | [illegible] |
| SE | 67050 | ENSG00000170906 | NDUFA3 | chr19 | + | 54105596 | 54105703 | 54103113 | 54103188 | 54105933 | 54106011 | 0 | 0 | 0 | 3 | 0 | 99 | 29 | 62 | 75 | 32 | 0 | 2 | 0 | 8 | [illegible] |
| SE | 102183 | ENSG00000267699 | NA | chr18 | + | 51052634 | 51052725 | 51049294 | 51049324 | 51054780 | 51054993 | 3 | 6 | 3 | 2 | 22 | 208 | 162 | 173 | 148 | 556 | 0 | 1 | 5 | 0 | [illegible] |
| SE | 44537 | ENSG00000076321 | KLHL20 | chr1 | + | 173775633 | 173775842 | 173766145 | 173766289 | 173782123 | 173782230 | 27 | 27 | 22 | 22 | 31 | 1 | 0 | 5 | 2 | 3 | 5 | 39 | 24 | 23 | [illegible] |
| SE | 21562 | ENSG00000085382 | HACE1 | chr6 | - | 104833041 | 104833173 | 104811310 | 104811393 | 104849141 | 104849246 | 28 | 31 | 9 | 17 | 22 | 0 | 0 | 0 | 0 | 1 | 7 | 15 | 12 | 4 | [illegible] |
| SE | 22744 | ENSG00000172262 | ZNF131 | chr5 | + | 43139164 | 43139309 | 43123208 | 43123310 | 43161248 | 43161294 | 188 | 149 | 187 | 237 | 258 | 7 | 13 | 7 | 5 | 10 | 23 | 141 | 150 | 56 | [illegible] |
| SE | 44603 | ENSG00000026025 | VIM | chr10 | + | 17235845 | 17235889 | 17235168 | 17235389 | 17236293 | 17236379 | 6085 | 3794 | 4798 | 5436 | 11256 | 16 | 1 | 4 | 17 | 31 | 14470 | 8589 | 5975 | 9839 | [illegible] |
| SE | 30060 | ENSG00000171298 | GAA | chr17 | + | 80111982 | 80112100 | 80110940 | 80111025 | 80112577 | 80112711 | 62 | 42 | 38 | 56 | 50 | 0 | 0 | 0 | 0 | 0 | 189 | 87 | 66 | 87 | [illegible] |
| SE | 29883 | ENSG00000141562 | NARF | chr17 | + | 82464286 | 82464430 | 82459991 | 82460072 | 82468763 | 82468787 | 156 | 91 | 144 | 201 | 150 | 0 | 0 | 0 | 1 | 4 | 124 | 254 | 127 | 141 | [illegible] |
| SE | 38989 | ENSG00000157216 | SSBP3 | chr1 | - | 54257126 | 54257186 | 54242163 | 54243299 | 54281437 | 54281527 | 104 | 47 | 57 | 81 | 109 | 2 | 4 | 2 | 0 | 0 | 87 | 81 | 39 | 67 | [illegible] |
| SE | 93810 | ENSG00000136869 | TLR4 | chr9 | + | 117708562 | 117708729 | 117708198 | 117708318 | 117712388 | 117715775 | 188 | 70 | 70 | 130 | 95 | 0 | 0 | 0 | 0 | 0 | 78 | 153 | 215 | 108 | [illegible] |
| SE | 104421 | ENSG00000184047 | DIABLO | chr12 | - | 122224644 | 122224748 | 122218265 | 122218397 | 122225964 | 122226039 | 22 | 8 | 21 | 16 | 11 | 1 | 0 | 4 | 0 | 0 | 16 | 15 | 18 | 42 | [illegible] |
| SE | 66077 | ENSG00000162924 | REL | chr2 | + | 60891682 | 60891825 | 60881520 | 60881850 | 60894396 | 60894545 | 123 | 87 | 56 | 99 | 57 | 1 | 0 | 6 | 0 | 0 | 61 | 114 | 124 | 100 | [illegible] |
| SE | 82051 | ENSG00000104133 | SPG11 | chr15 | - | 44620189 | 44620403 | 44615362 | 44615566 | 44621758 | 44621934 | 515 | 420 | 412 | 434 | 755 | 7 | 2 | 2 | 2 | 7 | 129 | 550 | 407 | 362 | [illegible] |
| SE | 2022 | ENSG00000270011 | ZNF559-ZNF177 | chr19 | + | 9339192 | 9339319 | 9338493 | 9338582 | 9341101 | 9341184 | 119 | 70 | 156 | 166 | 97 | 0 | 0 | 3 | 2 | 4 | 37 | 64 | 24 | 21 | [illegible] |
| SE | 63848 | ENSG00000148690 | FRA10AC1 | chr10 | - | 93692008 | 93692093 | 93687403 | 93687449 | 93692645 | 93692729 | 109 | 146 | 112 | 103 | 201 | 4 | 6 | 4 | 5 | 5 | 2 | 47 | 53 | 45 | [illegible] |
| SE | 45687 | ENSG00000109323 | MANBA | chr4 | - | 102722870 | 102723041 | 102714437 | 102714561 | 102723861 | 102723921 | 335 | 159 | 284 | 291 | 477 | 2 | 3 | 2 | 3 | 0 | 298 | 173 | 187 | 267 | [illegible] |
| SE | 104455 | ENSG00000066651 | TRMT11 | chr6 | + | 126006841 | 126006954 | 125999456 | 125999613 | 126008391 | 126008472 | 23 | 22 | 19 | 15 | 24 | 32 | 28 | 39 | 54 | 80 | 0 | 14 | 8 | 6 | [illegible] |
| SE | 1483 | ENSG00000100368 | CSF2RB | chr22 | + | 36930672 | 36930830 | 36930374 | 36930510 | 36932764 | 36932904 | 185 | 85 | 164 | 155 | 106 | 1 | 0 | 2 | 0 | 0 | 231 | 88 | 150 | 230 | [illegible] |
| SE | 59944 | ENSG00000270149 | NA | chr1 | - | 161021009 | 161021112 | 161001284 | 161001353 | 161038879 | 161038944 | 58 | 42 | 35 | 59 | 63 | 2 | 0 | 2 | 3 | 0 | 96 | 45 | 47 | 125 | [illegible] |
| SE | 85103 | ENSG00000147586 | MRPS28 | chr8 | - | 80002998 | 80003180 | 79918716 | 79919148 | 80030053 | 80030232 | 56 | 35 | 70 | 39 | 65 | 0 | 0 | 0 | 2 | 0 | 46 | 63 | 25 | 16 | [illegible] |
| SE | 57471 | ENSG00000115170 | ACVR1 | chr2 | - | 157860032 | 157860179 | 157818384 | 157818559 | 157875795 | 157875820 | 1 | 0 | 0 | 1 | 3 | 3 | 3 | 1 | 4 | 3 | 0 | 0 | 0 | 0 | [illegible] |
| SE | 88097 | ENSG00000139197 | PEX5 | chr12 | + | 7202240 | 7202351 | 7201750 | 7201841 | 7202611 | 7202704 | 33 | 28 | 50 | 27 | 16 | 62 | 23 | 47 | 40 | 86 | 53 | 52 | 8 | 20 | [illegible] |
| SE | 56782 | ENSG00000228956 | SATB1-AS1 | chr3 | + | 18527950 | 18528139 | 18527173 | 18527326 | 18529980 | 18530049 | 9 | 11 | 11 | 8 | 37 | 34 | 37 | 68 | 54 | 171 | 6 | 2 | 4 | 5 | [illegible] |
| SE | 54710 | ENSG00000137965 | IFI44 | chr1 | + | 78660553 | 78660691 | 78659318 | 78659483 | 78662703 | 78662878 | 106 | 175 | 95 | 139 | 138 | 12 | 31 | 15 | 25 | 14 | 85 | 220 | 135 | 126 | [illegible] |
| SE | 15520 | ENSG00000129003 | VPS13C | chr15 | - | 61875731 | 61875845 | 61874875 | 61874951 | 61876972 | 61877054 | 536 | 502 | 402 | 516 | 598 | 0 | 3 | 3 | 6 | 0 | 39 | 113 | 135 | 24 | [illegible] |
| SE | 72750 | ENSG00000240498 | CDKN2B-AS1 | chr9 | + | 22046316 | 22046449 | 22032673 | 22032986 | 22046750 | 22046900 | 14 | 5 | 2 | 9 | 4 | 1 | 0 | 2 | 1 | 0 | 25 | 3 | 4 | 5 | [illegible] |
| SE | 91153 | ENSG00000183049 | CAMK1D | chr10 | + | 12760947 | 12761086 | 12666735 | 12666810 | 12769672 | 12769799 | 156 | 83 | 140 | 120 | 184 | 0 | 0 | 0 | 0 | 0 | 162 | 138 | 86 | 72 | [illegible] |
| SE | 80156 | ENSG00000012124 | CD22 | chr19 | + | 35336035 | 35336341 | 35329186 | 35329230 | 35340880 | 35341138 | 0 | 2 | 2 | 1 | 6 | 5 | 1 | 5 | 8 | 5 | 0 | 9 | 6 | 11 | [illegible] |
| SE | 60509 | ENSG00000115211 | EIF2B4 | chr2 | - | 27369005 | 27369212 | 27368371 | 27368463 | 27369413 | 27369549 | 23 | 19 | 29 | 34 | 37 | 1 | 0 | 0 | 0 | 0 | 53 | 39 | 17 | 34 | [illegible] |
| SE | 82804 | ENSG00000136271 | DDX56 | chr7 | - | 44570757 | 44570877 | 44569983 | 44570128 | 44571491 | 44571736 | 226 | 126 | 193 | 232 | 149 | 0 | 0 | 0 | 0 | 4 | 207 | 133 | 139 | 197 | [illegible] |
| SE | 99310 | ENSG00000101017 | CD40 | chr20 | + | 46126639 | 46126701 | 46123125 | 46123219 | 46128137 | 46128224 | 33 | 18 | 33 | 80 | 95 | 4 | 0 | 3 | 2 | 0 | 92 | 80 | 8 | 40 | [illegible] |
| SE | 36011 | ENSG00000258653 | NA | chr14 | + | 73922492 | 73922697 | 73922065 | 73922206 | 73923394 | 73923522 | 0 | 2 | 6 | 0 | 3 | 52 | 34 | 45 | 45 | 96 | 0 | 0 | 2 | 0 | [illegible] |
| SE | 97452 | ENSG00000142185 | TRPM2 | chr21 | + | 44369176 | 44369343 | 44366753 | 44366934 | 44375832 | 44376013 | 25 | 15 | 17 | 28 | 13 | 0 | 0 | 0 | 0 | 0 | 46 | 16 | 16 | 71 | [illegible] |
| SE | 83509 | ENSG00000196313 | POM121 | chr7 | + | 72913764 | 72913828 | 72890932 | 72891110 | 72926261 | 72926477 | 38 | 25 | 21 | 27 | 64 | 67 | 60 | 77 | 97 | 102 | 41 | 65 | 25 | 29 | [illegible] |
| SE | 62209 | ENSG00000119778 | ATAD2B | chr2 | - | 23799824 | 23799993 | 23798137 | 23798323 | 23810315 | 23810502 | 5 | 2 | 1 | 6 | 0 | 80 | 47 | 72 | 53 | 112 | 0 | 0 | 0 | 0 | [illegible] |
| SE | 89252 | ENSG00000175606 | TMEM70 | chr8 | + | 73979078 | 73979171 | 73978747 | 73978861 | 73981154 | 73981481 | 23 | 20 | 10 | 14 | 13 | 36 | 24 | 41 | 34 | 82 | 34 | 11 | 14 | 8 | [illegible] |
| SE | 31472 | ENSG00000178950 | GAK | chr4 | - | 904636 | 904779 | 898032 | 898158 | 913606 | 913668 | 175 | 102 | 162 | 188 | 190 | 0 | 0 | 1 | 3 | 0 | 232 | 230 | 124 | 210 | [illegible] |
| SE | 89234 | ENSG00000094914 | AAAS | chr12 | - | 53315334 | 53315426 | 53315093 | 53315140 | 53315726 | 53315782 | 99 | 58 | 105 | 93 | 139 | 0 | 1 | 0 | 1 | 0 | 62 | 80 | 59 | 103 | [illegible] |
| SE | 47504 | ENSG00000143363 | PRUNE1 | chr1 | + | 151031371 | 151031464 | 151028785 | 151028944 | 151033805 | 151035681 | 0 | 0 | 4 | 3 | 8 | 55 | 74 | 46 | 77 | 173 | 0 | 0 | 2 | 6 | [illegible] |
| SE | 101129 | ENSG00000175224 | ATG13 | chr11 | + | 46621041 | 46621157 | 46618259 | 46618375 | 46630044 | 46630100 | 1 | 1 | 0 | 5 | 0 | 21 | 13 | 21 | 22 | 15 | 0 | 0 | 0 | 0 | [illegible] |
| SE | 88476 | ENSG00000152601 | MBNL1 | chr3 | + | 152432716 | 152432920 | 152268039 | 152269092 | 152445281 | 152445474 | 942 | 807 | 766 | 1135 | 962 | 5 | 2 | 1 | 6 | 0 | 425 | 1071 | 797 | 452 | [illegible] |
| SE | 102909 | ENSG00000148498 | PARD3 | chr10 | - | 34450316 | 34450448 | 34401825 | 34401917 | 34470084 | 34470263 | 2 | 29 | 6 | 18 | 18 | 0 | 3 | 2 | 5 | 0 | 22 | 34 | 25 | 36 | [illegible] |
| SE | 96960 | ENSG00000183597 | TANGO2 | chr22 | + | 20053359 | 20053551 | 20052464 | 20052584 | 20055942 | 20056013 | 51 | 17 | 32 | 49 | 55 | 42 | 26 | 21 | 21 | 41 | 84 | 64 | 29 | 47 | [illegible] |
| SE | 67737 | ENSG00000004455 | AK2 | chr1 | - | 33012073 | 33012122 | 33010754 | 33010833 | 33013206 | 33013287 | 0 | 0 | 0 | 2 | 0 | 62 | 34 | 60 | 63 | 21 | 0 | 0 | 0 | 2 | [illegible] |
| SE | 97509 | ENSG00000173120 | KDM2A | chr11 | + | 67243008 | 67243092 | 67231565 | 67231960 | 67245188 | 67245458 | 365 | 218 | 397 | 403 | 506 | 7 | 9 | 4 | 5 | 8 | 612 | 432 | 430 | 470 | [illegible] |
| SE | 99976 | ENSG00000185187 | SIGIRR | chr11 | - | 408694 | 408893 | 408072 | 408206 | 409867 | 409977 | 159 | 41 | 115 | 102 | 39 | 18 | 5 | 20 | 6 | 9 | 12 | 78 | 65 | 70 | [illegible] |
| SE | 2539 | ENSG00000104814 | MAP4K1 | chr19 | - | 38609908 | 38610025 | 38608111 | 38608170 | 38611050 | 38611132 | 160 | 92 | 244 | 283 | 568 | 0 | 0 | 0 | 1 | 0 | 329 | 416 | 196 | 178 | [illegible] |
| SE | 102120 | ENSG00000077420 | APBB1IP | chr10 | + | 26541581 | 26541692 | 26533438 | 26533525 | 26560729 | 26560844 | 11 | 5 | 7 | 3 | 14 | 0 | 0 | 0 | 0 | 4 | 27 | 14 | 49 | 51 | [illegible] |
| SE | 12164 | ENSG00000172775 | PSME3IP1 | chr16 | - | 57178805 | 57178852 | 57173727 | 57173869 | 57185820 | 57186031 | 1 | 1 | 1 | 0 | 7 | 88 | 55 | 63 | 72 | 57 | 0 | 4 | 1 | 0 | [illegible] |
| SE | 96004 | ENSG00000256591 | NA | chr11 | + | 61438003 | 61438120 | 61430055 | 61430182 | 61445940 | 61446231 | 3 | 0 | 0 | 3 | 6 | 34 | 15 | 24 | 24 | 23 | 0 | 3 | 3 | 2 | [illegible] |
| SE | 106377 | ENSG00000092531 | SNAP23 | chr15 | + | 42513173 | 42513209 | 42512954 | 42512996 | 42513398 | 42513447 | 3 | 1 | 7 | 3 | 8 | 286 | 319 | 170 | 332 | 581 | 13 | 13 | 8 | 0 | [illegible] |
| SE | 15518 | ENSG00000129003 | VPS13C | chr15 | - | 61864640 | 61864820 | 61863439 | 61863528 | 61868658 | 61868773 | 2 | 7 | 0 | 8 | 15 | 301 | 287 | 223 | 267 | 275 | 0 | 0 | 2 | 0 | [illegible] |
| SE | 68314 | ENSG00000168016 | TRANK1 | chr3 | - | 36842621 | 36842710 | 36838612 | 36838716 | 36846247 | 36846404 | 377 | 230 | 324 | 444 | 394 | 2 | 3 | 2 | 0 | 12 | 313 | 407 | 183 | 296 | [illegible] |
| SE | 86530 | ENSG00000085224 | ATRX | chrX | - | 77717130 | 77717243 | 77698573 | 77698629 | 77785981 | 77786204 | 203 | 192 | 175 | 192 | 223 | 6 | 1 | 0 | 2 | 2 | 53 | 44 | 34 | 25 | [illegible] |
| SE | 100499 | ENSG00000172375 | C2CD2L | chr11 | + | 119112699 | 119112874 | 119112481 | 119112609 | 119113636 | 119113712 | 14 | 2 | 9 | 16 | 11 | 0 | 0 | 0 | 0 | 0 | 13 | 16 | 18 | 22 | [illegible] |
| SE | 9853 | ENSG00000109066 | TMEM104 | chr17 | + | 74785481 | 74785603 | 74777263 | 74777375 | 74788788 | 74788870 | 42 | 25 | 30 | 37 | 50 | 0 | 0 | 1 | 0 | 0 | 84 | 59 | 20 | 45 | [illegible] |
| SE | 71720 | ENSG00000135976 | ANKRD36 | chr2 | + | 97207933 | 97208006 | 97207810 | 97207839 | 97209680 | 97209709 | 77 | 139 | 71 | 116 | 90 | 16 | 21 | 46 | 29 | 22 | 10 | 32 | 30 | 16 | [illegible] |
| SE | 47129 | ENSG00000089012 | SIRPG | chr20 | - | 1636187 | 1636505 | 1630221 | 1630306 | 1649051 | 1649408 | 17 | 7 | 44 | 20 | 47 | 6 | 29 | 22 | 44 | 36 | 6 | 6 | 3 | 11 | [illegible] |
| SE | 19841 | ENSG00000100890 | PRORP | chr14 | + | 35126734 | 35126782 | 35122951 | 35124231 | 35127478 | 35127611 | 110 | 47 | 111 | 86 | 179 | 0 | 1 | 2 | 5 | 16 | 37 | 130 | 87 | 88 | [illegible] |
| SE | 39459 | ENSG00000213024 | NUP62 | chr19 | - | 49918507 | 49918621 | 49909394 | 49909884 | 49929325 | 49929429 | 0 | 3 | 0 | 3 | 2 | 21 | 19 | 38 | 17 | 19 | 0 | 4 | 0 | 0 | [illegible] |
| SE | 47485 | ENSG00000143363 | PRUNE1 | chr1 | + | 151018466 | 151018669 | 151017811 | 151017904 | 151025514 | 151025673 | 21 | 15 | 30 | 35 | 36 | 5 | 0 | 1 | 8 | 1 | 33 | 52 | 6 | 26 | [illegible] |
| SE | 18278 | ENSG00000179364 | PACS2 | chr14 | + | 105348492 | 105348580 | 105314414 | 105315037 | 105352377 | 105352467 | 21 | 14 | 25 | 28 | 24 | 0 | 0 | 0 | 0 | 0 | 34 | 30 | 27 | 19 | [illegible] |
| SE | 108991 | ENSG00000160961 | ZNF333 | chr19 | + | 14701583 | 14701897 | 14699198 | 14699281 | 14705053 | 14705170 | 5 | 0 | 0 | 0 | 0 | 23 | 12 | 22 | 27 | 14 | 0 | 0 | 1 | 4 | [illegible] |
| SE | 86052 | ENSG00000085415 | SEH1L | chr18 | + | 12975732 | 12975882 | 12971152 | 12971251 | 12978751 | 12978892 | 1 | 11 | 1 | 1 | 1 | 52 | 35 | 64 | 82 | 78 | 0 | 0 | 2 | 5 | [illegible] |
| SE | 58191 | ENSG00000172965 | MIR4435-2HG | chr2 | - | 111239801 | 111239996 | 111207813 | 111208661 | 111344060 | 111344237 | 2 | 4 | 1 | 10 | 10 | 16 | 12 | 4 | 23 | 40 | 14 | 0 | 3 | 7 | [illegible] |
| SE | 76927 | ENSG00000167483 | NIBAN3 | chr19 | + | 17549443 | 17549527 | 17546685 | 17546797 | 17551785 | 17552024 | 92 | 91 | 121 | 167 | 95 | 2 | 3 | 1 | 5 | 8 | 266 | 232 | 25 | 139 | [illegible] |
| SE | 74112 | ENSG00000093167 | LRRFIP2 | chr3 | - | 37066223 | 37066325 | 37065809 | 37065942 | 37075023 | 37075116 | 0 | 0 | 0 | 1 | 0 | 46 | 52 | 20 | 40 | 62 | 0 | 2 | 3 | 1 | [illegible] |
| SE | 49876 | ENSG00000109180 | OCIAD1 | chr4 | + | 48832619 | 48832682 | 48831064 | 48831249 | 48842635 | 48842689 | 17 | 9 | 6 | 9 | 10 | 4 | 1 | 2 | 7 | 0 | 38 | 32 | 5 | 14 | [illegible] |
| SE | 12734 | ENSG00000157106 | SMG1 | chr16 | - | 18859021 | 18859181 | 18857785 | 18858290 | 18859555 | 18859703 | 573 | 567 | 512 | 584 | 537 | 0 | 0 | 0 | 4 | 0 | 75 | 521 | 420 | 223 | [illegible] |
| SE | 5270 | ENSG00000185513 | L3MBTL1 | chr20 | + | 43534283 | 43534394 | 43534007 | 43534093 | 43534827 | 43534942 | 15 | 12 | 18 | 14 | 14 | 0 | 0 | 0 | 4 | 0 | 9 | 26 | 3 | 11 | [illegible] |
| SE | 40012 | ENSG00000121289 | CEP89 | chr19 | - | 32926933 | 32926984 | 32926189 | 32926273 | 32931428 | 32931571 | 29 | 25 | 17 | 22 | 24 | 0 | 1 | 0 | 0 | 1 | 18 | 3 | 9 | 12 | [illegible] |
| SE | 12144 | ENSG00000172775 | PSME3IP1 | chr16 | - | 57175538 | 57175700 | 57173772 | 57173869 | 57178709 | 57178852 | 29 | 48 | 22 | 38 | 37 | 0 | 0 | 2 | 4 | 21 | 23 | 67 | 78 | 48 | [illegible] |
| SE | 71301 | ENSG00000064012 | CASP8 | chr2 | + | 201276826 | 201276968 | 201272637 | 201272776 | 201284815 | 201285317 | 328 | 190 | 241 | 239 | 271 | 3 | 3 | 2 | 2 | 2 | 204 | 183 | 203 | 110 | [illegible] |
| SE | 108285 | ENSG00000139132 | FGD4 | chr12 | + | 32601277 | 32601423 | 32598496 | 32598586 | 32607956 | 32608095 | 51 | 34 | 26 | 56 | 75 | 0 | 0 | 0 | 0 | 0 | 87 | 50 | 58 | 50 | [illegible] |
| SE | 106228 | ENSG00000083937 | CHMP2B | chr3 | + | 87240698 | 87240790 | 87227284 | 87227556 | 87245713 | 87245818 | 150 | 112 | 153 | 161 | 149 | 3 | 0 | 0 | 7 | 1 | 54 | 122 | 160 | 121 | [illegible] |
| SE | 42158 | ENSG00000197343 | ZNF655 | chr7 | + | 99562367 | 99562494 | 99560532 | 99560695 | 99563862 | 99564705 | 100 | 51 | 79 | 105 | 169 | 4 | 5 | 8 | 2 | 2 | 126 | 107 | 129 | 117 | [illegible] |
| SE | 30832 | ENSG00000258315 | C17orf49 | chr17 | + | 7016949 | 7017014 | 7016494 | 7016685 | 7017256 | 7017304 | 26 | 9 | 18 | 4 | 25 | 24 | 4 | 11 | 13 | 48 | 29 | 5 | 12 | 2 | [illegible] |
| SE | 16789 | ENSG00000241839 | PLEKHO2 | chr15 | + | 64859893 | 64859998 | 64854920 | 64855037 | 64861476 | 64861575 | 379 | 217 | 231 | 344 | 349 | 0 | 0 | 0 | 0 | 1 | 720 | 348 | 224 | 591 | [illegible] |
| SE | 34054 | ENSG00000114867 | EIF4G1 | chr3 | + | 184316071 | 184316218 | 184315762 | 184315856 | 184317320 | 184317497 | 68 | 51 | 42 | 45 | 38 | 2 | 0 | 1 | 2 | 0 | 33 | 31 | 23 | 18 | [illegible] |
| SE | 33197 | ENSG00000163930 | BAP1 | chr3 | - | 52405764 | 52405912 | 52405109 | 52405294 | 52406252 | 52406322 | 208 | 80 | 165 | 231 | 247 | 0 | 0 | 0 | 0 | 0 | 181 | 213 | 111 | 137 | [illegible] |
| SE | 80422 | ENSG00000126746 | ZNF384 | chr12 | - | 6677166 | 6677259 | 6673215 | 6673440 | 6678126 | 6678460 | 5 | 7 | 13 | 13 | 20 | 66 | 41 | 53 | 67 | 49 | 5 | 13 | 9 | 17 | [illegible] |
| SE | 23797 | ENSG00000153071 | DAB2 | chr5 | - | 39382617 | 39383271 | 39376649 | 39377282 | 39388304 | 39388367 | 120 | 158 | 44 | 153 | 384 | 0 | 2 | 0 | 0 | 0 | 496 | 285 | 169 | 254 | [illegible] |
| SE | 90802 | ENSG00000173064 | HECTD4 | chr12 | - | 112168632 | 112168806 | 112167813 | 112167917 | 112169502 | 112169658 | 1 | 0 | 0 | 2 | 3 | 31 | 14 | 21 | 22 | 18 | 0 | 0 | 0 | 0 | [illegible] |
| SE | 38099 | ENSG00000143811 | PYCR2 | chr1 | - | 225921551 | 225921644 | 225921207 | 225921371 | 225921857 | 225922079 | 113 | 52 | 126 | 115 | 56 | 0 | 0 | 7 | 1 | 1 | 164 | 102 | 26 | 88 | [illegible] |
| SE | 58162 | ENSG00000160688 | FLAD1 | chr1 | + | 154989559 | 154989707 | 154988104 | 154988849 | 154990158 | 154990257 | 73 | 32 | 51 | 43 | 48 | 8 | 3 | 3 | 6 | 3 | 21 | 34 | 10 | 8 | [illegible] |
| SE | 46859 | ENSG00000162894 | FCMR | chr1 | - | 206911729 | 206911952 | 206910209 | 206910340 | 206921817 | 206921867 | 351 | 124 | 327 | 242 | 195 | 4 | 0 | 0 | 2 | 5 | 93 | 260 | 105 | 96 | [illegible] |
| SE | 83146 | ENSG00000188878 | FBF1 | chr17 | - | 75935631 | 75935673 | 75932994 | 75933088 | 75937565 | 75937593 | 13 | 5 | 9 | 6 | 39 | 5 | 3 | 6 | 1 | 6 | 0 | 19 | 1 | 11 | [illegible] |
| SE | 53033 | ENSG00000197183 | NOL4L | chr20 | - | 32474600 | 32474742 | 32456117 | 32456395 | 32511346 | 32511407 | 141 | 98 | 151 | 142 | 84 | 0 | 0 | 0 | 2 | 1 | 9 | 68 | 59 | 83 | [illegible] |
| SE | 50320 | ENSG00000060339 | CCAR1 | chr10 | + | 68722454 | 68722577 | 68721143 | 68721282 | 68736875 | 68736910 | 135 | 130 | 112 | 168 | 235 | 0 | 0 | 0 | 0 | 0 | 144 | 78 | 107 | 84 | [illegible] |
| SE | 45784 | ENSG00000116918 | TSNAX | chr1 | + | 231537212 | 231537327 | 231529254 | 231529359 | 231542480 | 231542611 | 185 | 207 | 278 | 226 | 338 | 0 | 6 | 2 | 2 | 2 | 18 | 106 | 170 | 105 | [illegible] |
| SE | 74720 | ENSG00000112282 | MED23 | chr6 | - | 131607927 | 131608071 | 131606478 | 131606624 | 131610045 | 131610246 | 118 | 97 | 122 | 124 | 221 | 7 | 0 | 1 | 1 | 0 | 92 | 96 | 91 | 47 | [illegible] |
| SE | 35144 | ENSG00000107758 | PPP3CB | chr10 | - | 73444724 | 73444822 | 73436805 | 73438420 | 73446491 | 73446573 | 160 | 165 | 154 | 185 | 273 | 1 | 4 | 5 | 1 | 9 | 234 | 256 | 162 | 206 | [illegible] |
| SE | 4179 | ENSG00000105186 | ANKRD27 | chr19 | - | 32602014 | 32602126 | 32599971 | 32600050 | 32604262 | 32604424 | 100 | 36 | 68 | 65 | 33 | 0 | 1 | 5 | 1 | 0 | 102 | 73 | 35 | 103 | [illegible] |
| SE | 3509 | ENSG00000130755 | GMFG | chr19 | - | 39335434 | 39335554 | 39329543 | 39329626 | 39335973 | 39336028 | 8 | 2 | 13 | 6 | 17 | 0 | 0 | 0 | 0 | 0 | 15 | 6 | 13 | 5 | [illegible] |
| SE | 12512 | ENSG00000132600 | PRMT7 | chr16 | + | 68348341 | 68348431 | 68346144 | 68346280 | 68352247 | 68352409 | 14 | 23 | 9 | 8 | 9 | 3 | 0 | 3 | 10 | 2 | 12 | 4 | 8 | 8 | [illegible] |
| SE | 54234 | ENSG00000143643 | TTC13 | chr1 | - | 230945388 | 230945454 | 230940439 | 230940556 | 230954332 | 230954403 | 117 | 74 | 76 | 97 | 90 | 36 | 12 | 18 | 36 | 20 | 87 | 84 | 99 | 74 | [illegible] |
| SE | 62675 | ENSG00000115525 | ST3GAL5 | chr2 | - | 85848115 | 85848204 | 85847860 | 85847984 | 85861180 | 85861292 | 52 | 58 | 38 | 46 | 81 | 0 | 0 | 0 | 0 | 0 | 89 | 34 | 26 | 58 | [illegible] |
| SE | 6316 | ENSG00000060069 | CTDP1 | chr18 | + | 79728906 | 79729069 | 79717809 | 79718016 | 79736354 | 79736521 | 84 | 52 | 119 | 63 | 79 | 10 | 15 | 22 | 24 | 1 | 103 | 70 | 39 | 58 | [illegible] |
| SE | 83939 | ENSG00000147421 | HMBOX1 | chr8 | + | 28980070 | 28980156 | 28970042 | 28970519 | 29009071 | 29009182 | 208 | 152 | 144 | 201 | 315 | 9 | 13 | 3 | 11 | 15 | 162 | 229 | 187 | 259 | [illegible] |
| SE | 86849 | ENSG00000116754 | SRSF11 | chr1 | + | 70247116 | 70247149 | 70246817 | 70246907 | 70249951 | 70250047 | 19 | 28 | 39 | 25 | 16 | 284 | 282 | 318 | 334 | 410 | 6 | 13 | 19 | 2 | [illegible] |
| SE | 99929 | ENSG00000085733 | CTTN | chr11 | + | 70433624 | 70433718 | 70433100 | 70433278 | 70435025 | 70435152 | 12 | 31 | 4 | 43 | 15 | 0 | 0 | 0 | 1 | 0 | 99 | 57 | 21 | 36 | [illegible] |
| SE | 18156 | ENSG00000100554 | ATP6V1D | chr14 | - | 67345767 | 67345871 | 67343371 | 67343438 | 67347408 | 67347453 | 244 | 190 | 144 | 212 | 358 | 2 | 0 | 1 | 2 | 7 | 77 | 136 | 139 | 130 | [illegible] |
| SE | 92892 | ENSG00000136874 | STX17 | chr9 | + | 99916048 | 99916168 | 99915177 | 99915362 | 99928777 | 99928843 | 3 | 2 | 7 | 2 | 11 | 95 | 128 | 131 | 125 | 180 | 0 | 3 | 0 | 9 | [illegible] |
| SE | 24420 | ENSG00000055163 | CYFIP2 | chr5 | + | 157339056 | 157339256 | 157333326 | 157333446 | 157341069 | 157341094 | 552 | 251 | 526 | 446 | 651 | 2 | 1 | 1 | 4 | 0 | 334 | 557 | 325 | 357 | [illegible] |
| SE | 48156 | ENSG00000117543 | DPH5 | chr1 | - | 101001466 | 101001544 | 100995109 | 100995149 | 101013709 | 101013818 | 50 | 40 | 58 | 73 | 80 | 1 | 2 | 4 | 3 | 3 | 52 | 41 | 53 | 24 | [illegible] |
| SE | 105234 | ENSG00000076067 | RBMS2 | chr12 | + | 56580277 | 56580375 | 56571697 | 56571855 | 56581183 | 56581263 | 5 | 0 | 0 | 0 | 2 | 43 | 36 | 42 | 33 | 27 | 3 | 0 | 0 | 0 | [illegible] |
| SE | 42264 | ENSG00000116001 | TIA1 | chr2 | - | 70229058 | 70229091 | 70227734 | 70227822 | 70229263 | 70229318 | 153 | 107 | 115 | 180 | 240 | 11 | 16 | 5 | 10 | 11 | 61 | 82 | 71 | 85 | [illegible] |
| SE | 54431 | ENSG00000169554 | ZEB2 | chr2 | - | 144424795 | 144424867 | 144403915 | 144404130 | 144429768 | 144430026 | 165 | 100 | 79 | 113 | 100 | 0 | 0 | 0 | 1 | 0 | 136 | 184 | 238 | 99 | [illegible] |
| SE | 82180 | ENSG00000146872 | TLK2 | chr17 | + | 62573214 | 62573367 | 62564759 | 62565137 | 62576708 | 62576775 | 185 | 195 | 198 | 169 | 206 | 0 | 0 | 0 | 4 | 1 | 381 | 192 | 367 | 431 | [illegible] |
| SE | 25201 | ENSG00000160783 | PMF1 | chr1 | + | 156225556 | 156225668 | 156213005 | 156213176 | 156232319 | 156232425 | 12 | 1 | 6 | 10 | 1 | 259 | 109 | 176 | 181 | 201 | 16 | 5 | 11 | 8 | [illegible] |
| SE | 64424 | ENSG00000143919 | CAMKMT | chr2 | + | 44707398 | 44707462 | 44704282 | 44704343 | 44715286 | 44715353 | 14 | 1 | 16 | 9 | 21 | 0 | 0 | 0 | 2 | 0 | 11 | 19 | 10 | 8 | [illegible] |
| SE | 18919 | ENSG00000100461 | RBM23 | chr14 | - | 22905605 | 22905659 | 22905335 | 22905453 | 22906194 | 22906368 | 144 | 43 | 94 | 88 | 127 | 28 | 11 | 29 | 21 | 12 | 107 | 93 | 68 | 95 | [illegible] |
| SE | 65025 | ENSG00000091409 | ITGA6 | chr2 | + | 172467480 | 172467560 | 172465538 | 172465663 | 172474054 | 172474265 | 40 | 20 | 34 | 30 | 18 | 0 | 0 | 2 | 3 | 3 | 36 | 13 | 18 | 17 | [illegible] |
| SE | 62136 | ENSG00000182179 | UBA7 | chr3 | - | 49810516 | 49810672 | 49809977 | 49810183 | 49810751 | 49810832 | 106 | 79 | 72 | 112 | 82 | 0 | 0 | 0 | 0 | 0 | 207 | 119 | 50 | 81 | [illegible] |

| | | | | | | | | | | | | | | |
|---|---|---|---|---|---|---|---|---|---|---|---|---|---|---|
| SE | 36592 | ENSG00000169180 | XPO6 | chr16 | - | 28107521 | 28107677 | 28106382 | 28106497 | 28111816 | 28111890 | 719 | 389 | 777 |
| SE | 90784 | ENSG00000079459 | FDFT1 | chr8 | + | 11808793 | 11808891 | 11803011 | 11803518 | 11809666 | 11809850 | 88 | 46 | 73 |
| SE | 73561 | ENSG00000230006 | ANKRD36BP2 | chr2 | + | 88772972 | 88773044 | 88765806 | 88766196 | 88776500 | 88776573 | 2 | 16 | 27 |
| SE | 75361 | ENSG00000186063 | AIDA | chr1 | - | 222711378 | 222711545 | 222703147 | 222703217 | 222712207 | 222712263 | 2 | 1 | 2 |
| SE | 39463 | ENSG00000213024 | NUP62 | chr19 | - | 49927693 | 49927815 | 49906824 | 49909884 | 49929349 | 49929763 | 86 | 61 | 87 |
| SE | 51624 | ENSG00000121864 | ZNF639 | chr3 | + | 179329933 | 179330055 | 179329633 | 179329728 | 179332988 | 179333123 | 0 | 2 | 8 |
| SE | 85690 | ENSG00000177981 | ASB8 | chr12 | - | 48153367 | 48153529 | 48150917 | 48151305 | 48157458 | 48157477 | 110 | 37 | 86 |
| SE | 100192 | ENSG00000078124 | ACER3 | chr11 | + | 77015015 | 77015659 | 76998762 | 76998821 | 77016674 | 77016779 | 46 | 35 | 22 |
| SE | 80894 | ENSG00000126883 | NUP214 | chr9 | + | 131187288 | 131187364 | 131178310 | 131178410 | 131189052 | 131189131 | 106 | 75 | 120 |
| SE | 10604 | ENSG00000276234 | TADA2A | chr17 | + | 37437737 | 37437829 | 37426949 | 37427009 | 37440504 | 37440579 | 117 | 67 | 96 |
| SE | 60480 | ENSG00000070814 | TCOF1 | chr5 | + | 150383061 | 150383172 | 150379531 | 150379732 | 150387901 | 150388088 | 4 | 2 | 5 |
| SE | 57906 | ENSG00000135899 | SP110 | chr2 | - | 230178156 | 230178255 | 230172843 | 230172959 | 230183571 | 230183640 | 105 | 153 | 110 |
| SE | 5538 | ENSG00000244687 | UBE2V1 | chr20 | - | 50098881 | 50098981 | 50096671 | 50096820 | 50113106 | 50113198 | 2 | 0 | 6 |
| SE | 13072 | ENSG00000123815 | COQ8B | chr19 | - | 40713739 | 40713869 | 40710058 | 40710136 | 40714066 | 40714153 | 0 | 4 | 2 |
| SE | 51063 | ENSG00000112029 | FBXO5 | chr6 | - | 152973045 | 152973136 | 152972271 | 152972454 | 152974906 | 152975621 | 80 | 45 | 82 |
| SE | 40482 | ENSG00000285708 | NA | chr3 | - | 70972554 | 70972676 | 70965889 | 70966056 | 70976940 | 70977042 | 538 | 421 | 717 |
| SE | 1024 | ENSG00000284554 | NA | chr22 | + | 39025556 | 39025671 | 39022821 | 39023014 | 39029362 | 39029519 | 18 | 6 | 3 |
| SE | 18817 | ENSG00000119689 | DLST | chr14 | + | 74901065 | 74901233 | 74900288 | 74900372 | 74902195 | 74902452 | 221 | 128 | 189 |
| SE | 33017 | ENSG00000114316 | USP4 | chr3 | - | 49324893 | 49325039 | 49324696 | 49324763 | 49327685 | 49327816 | 145 | 106 | 127 |
| SE | 51055 | ENSG00000143353 | LYPLAL1 | chr1 | + | 219179399 | 219179426 | 219179146 | 219179246 | 219210531 | 219210647 | 0 | 4 | 4 |
| SE | 65804 | ENSG00000116521 | SCAMP3 | chr1 | - | 155260536 | 155260659 | 155260329 | 155260450 | 155262085 | 155262227 | 59 | 17 | 52 |
| SE | 94858 | ENSG00000167106 | FAM102A | chr9 | - | 127950422 | 127950539 | 127948367 | 127948414 | 127953574 | 127953604 | 337 | 198 | 626 |
| SE | 45645 | ENSG00000144283 | PKP4 | chr2 | + | 158577270 | 158577383 | 158533179 | 158533316 | 158603069 | 158603104 | 17 | 12 | 22 |
| SE | 62243 | ENSG00000197043 | ANXA6 | chr5 | - | 151147883 | 151147926 | 151140152 | 151140243 | 151157679 | 151157747 | 116 | 46 | 72 |
| SE | 84440 | ENSG00000188917 | TRMT2B | chrX | - | 101051643 | 101051733 | 101041316 | 101041371 | 101051816 | 101052054 | 21 | 14 | 15 |
| SE | 44716 | ENSG00000143772 | ITPKB | chr1 | - | 226647166 | 226647380 | 226641920 | 226642125 | 226648671 | 226648771 | 354 | 209 | 406 |
| SE | 33597 | ENSG00000009790 | TRAF3IP3 | chr1 | + | 209780469 | 209780606 | 209779314 | 209779374 | 209782055 | 209782307 | 124 | 84 | 162 |
| SE | 3057 | ENSG00000130520 | LSM4 | chr19 | - | 18310109 | 18310214 | 18309677 | 18309861 | 18312603 | 18312702 | 1 | 0 | 1 |
| SE | 92026 | ENSG00000135315 | CEP162 | chr6 | - | 84217798 | 84217860 | 84215775 | 84215922 | 84219102 | 84219214 | 0 | 0 | 0 |
| SE | 23006 | ENSG00000171617 | ENC1 | chr5 | - | 74637564 | 74637701 | 74634683 | 74636498 | 74640306 | 74640728 | 11 | 3 | 12 |
| SE | 109164 | ENSG00000197442 | MAP3K5 | chr6 | - | 136639542 | 136639638 | 136637324 | 136637406 | 136656306 | 136656460 | 133 | 69 | 83 |
| SE | 19717 | ENSG00000103811 | CTSH | chr15 | - | 78931868 | 78932041 | 78931450 | 78931506 | 78932371 | 78932458 | 7 | 4 | 10 |

| | | | | | | | | | | | | | | | | | | | | | | | | | | | | | | | | | | | | | | | | | | | | | | |
|---|---|---|---|---|---|---|---|---|---|---|---|---|---|---|---|---|---|---|---|---|---|---|---|---|---|---|---|---|---|---|---|---|---|---|---|---|---|---|---|---|---|---|---|---|---|---|
| SE | 62502 | ENSG00000183943 | PRKX | chrX | - | 3641851 | 3641971 | 3626418 | 3626514 | 3655148 | 3655412 | 237 | 112 | 169 | 264 | 179 | 2 | 6 | 16 | 14 | 0 | 327 | 142 | 120 | 220 | 188 | 10 | 1 | 14 | 4 | 6 | 269 | 149 | 0.674668678 | 1 | 0.985 | 0.912 | 0.854 | 0.913 | 1 | 0.948 | 0.987 | 0.826 | 0.968 | 0.946 | -0.002 |
| SE | 22354 | ENSG00000143740 | SNAP47 | chr1 | + | 227747691 | 227748233 | 227728640 | 227728786 | 227758994 | 227759485 | 22 | 8 | 25 | 26 | 18 | 0 | 0 | 0 | 0 | 0 | 11 | 36 | 16 | 20 | 11 | 0 | 5 | 0 | 0 | 0 | 298 | 149 | 0.674701351 | 1 | 1 | | 1 | 1 | | 1 | 0.783 | 1 | 1 | 1 | 0.043 |
| SE | 55695 | ENSG00000135749 | PCNX2 | chr1 | - | 233112844 | 233112966 | 233095754 | 233095863 | 233135012 | 233135190 | 13 | 11 | 29 | 32 | 47 | 15 | 10 | 18 | 17 | 24 | 34 | 5 | 11 | 13 | 12 | 0 | 23 | 15 | 7 | 4 | 271 | 149 | 0.674703886 | 1 | 0.323 | 0.377 | 0.47 | 0.509 | 0.518 | 1 | 0.107 | 0.287 | 0.505 | 0.623 | -0.065 |
| SE | 89712 | ENSG00000215114 | UBXN2B | chr8 | + | 58430518 | 58430669 | 58416849 | 58416953 | 58433159 | 58433243 | 316 | 196 | 251 | 261 | 371 | 0 | 1 | 2 | 1 | 8 | 25 | 218 | 292 | 167 | 24 | 0 | 0 | 1 | 1 | 0 | 298 | 149 | 0.674822931 | 1 | 1 | 0.99 | 0.984 | 0.992 | 0.959 | 1 | 1 | 0.993 | 0.988 | 1 | -0.011 |
| SE | 30641 | ENSG00000101844 | ATG4A | chrX | + | 108128780 | 108128852 | 108126076 | 108126187 | 108131259 | 108131358 | 137 | 142 | 129 | 176 | 326 | 4 | 0 | 5 | 2 | 0 | 154 | 126 | 169 | 193 | 33 | 0 | 0 | 0 | 0 | 0 | 221 | 149 | 0.674831506 | 1 | 0.958 | 1 | 0.946 | 0.983 | 1 | 1 | 1 | 1 | 1 | 1 | -0.023 |
| SE | 72645 | ENSG00000153094 | BCL2L11 | chr2 | + | 111144487 | 111144520 | 111123732 | 111124139 | 111150043 | 111150147 | 0 | 0 | 3 | 0 | 0 | 76 | 22 | 54 | 41 | 70 | 10 | 1 | 3 | 0 | 0 | 107 | 110 | 176 | 87 | 112 | 182 | 149 | 0.674862149 | 1 | 0 | 0 | 0.044 | 0 | 0 | 0.071 | 0.007 | 0.014 | 0 | 0 | -0.01 |
| SE | 17910 | ENSG00000197249 | SERPINA1 | chr14 | - | 94382591 | 94383241 | 94376746 | 94378640 | 94388559 | 94388589 | 415 | 220 | 163 | 241 | 167 | 0 | 0 | 0 | 0 | 0 | 621 | 545 | 302 | 276 | 1000 | 16 | 0 | 2 | 0 | 0 | 298 | 149 | 0.675042223 | 1 | 1 | 1 | 1 | 1 | 1 | 0.951 | 1 | 0.987 | 1 | 1 | 0.012 |
| SE | 30978 | ENSG00000145675 | PIK3R1 | chr5 | + | 68273938 | 68274013 | 68273389 | 68273482 | 68279601 | 68279733 | 316 | 165 | 328 | 346 | 293 | 1 | 0 | 0 | 3 | 0 | 64 | 231 | 442 | 78 | 35 | 0 | 4 | 3 | 3 | 0 | 224 | 149 | 0.675097992 | 1 | 0.995 | 1 | 1 | 0.987 | 1 | 1 | 0.975 | 0.99 | 0.945 | 1 | 0.014 |
| SE | 13016 | ENSG00000140987 | ZSCAN32 | chr16 | - | 3390009 | 3390183 | 3384083 | 3384941 | 3390422 | 3390517 | 17 | 6 | 15 | 13 | 20 | 4 | 0 | 0 | 0 | 0 | 1 | 3 | 4 | 2 | 2 | 0 | 0 | 0 | 0 | 0 | 298 | 149 | 0.675127495 | 1 | 0.68 | 1 | 1 | 1 | 1 | 1 | 1 | 1 | 1 | 1 | -0.064 |
| SE | 58897 | ENSG00000143297 | FCRL5 | chr1 | - | 157524278 | 157524557 | 157521016 | 157521292 | 157527616 | 157527895 | 9 | 17 | 20 | 25 | 10 | 0 | 4 | 0 | 0 | 0 | 48 | 48 | 13 | 19 | 12 | 0 | 0 | 0 | 0 | 0 | 298 | 149 | 0.675205198 | 1 | 1 | 0.68 | 1 | 1 | 1 | 1 | 1 | 1 | 1 | 1 | -0.064 |
| SE | 96586 | ENSG00000198561 | CTNND1 | chr11 | + | 57791384 | 57791673 | 57761796 | 57762046 | 57795576 | 57795729 | 12 | 14 | 5 | 4 | 17 | 0 | 0 | 0 | 0 | 0 | 20 | 0 | 2 | 19 | 32 | 0 | 2 | 0 | 0 | 0 | 298 | 149 | 0.675272287 | 1 | 1 | 1 | 1 | 1 | 1 | 1 | 0 | 1 | 1 | 1 | 0.2 |
| SE | 66210 | ENSG00000137817 | PARP6 | chr15 | - | 72250844 | 72250957 | 72250019 | 72250092 | 72251206 | 72251255 | 47 | 32 | 58 | 72 | 96 | 4 | 14 | 15 | 13 | 49 | 31 | 42 | 59 | 81 | 98 | 18 | 18 | 9 | 27 | 20 | 262 | 149 | 0.675325615 | 1 | 0.87 | 0.565 | 0.687 | 0.759 | 0.527 | 0.495 | 0.57 | 0.789 | 0.63 | 0.736 | 0.038 |
| SE | 8920 | ENSG00000161533 | ACOX1 | chr17 | - | 75973624 | 75973785 | 75960214 | 75960375 | 75978533 | 75978693 | 6 | 12 | 6 | 10 | 1 | 61 | 33 | 37 | 33 | 42 | 6 | 8 | 18 | 16 | 0 | 63 | 63 | 37 | 31 | 65 | 298 | 149 | 0.675345004 | 1 | 0.047 | 0.154 | 0.075 | 0.132 | 0.012 | 0.045 | 0.06 | 0.196 | 0.205 | 0 | -0.017 |
| SE | 106611 | ENSG00000112200 | ZNF451 | chr6 | + | 57128728 | 57128840 | 57124733 | 57124859 | 57133041 | 57133154 | 190 | 145 | 132 | 177 | 272 | 0 | 0 | 0 | 0 | 0 | 53 | 61 | 75 | 42 | 38 | 6 | 0 | 0 | 0 | 0 | 261 | 149 | 0.675425259 | 1 | 1 | 1 | 1 | 1 | 1 | 0.835 | 1 | 1 | 1 | 1 | 0.033 |
| SE | 103260 | ENSG00000172661 | WASHC2C | chr10 | + | 45754485 | 45754545 | 45753179 | 45753237 | 45754935 | 45755113 | 136 | 142 | 158 | 148 | 210 | 0 | 0 | 0 | 0 | 0 | 84 | 157 | 113 | 302 | 110 | 0 | 0 | 0 | 10 | 0 | 209 | 149 | 0.675452259 | 1 | 1 | 1 | 1 | 1 | 1 | 1 | 1 | 1 | 0.956 | 1 | 0.009 |
| SE | 63907 | ENSG00000073614 | KDM5A | chr12 | - | 387247 | 387274 | 385896 | 385974 | 388926 | 389273 | 1 | 0 | 0 | 0 | 0 | 175 | 78 | 170 | 156 | 80 | 0 | 0 | 1 | 10 | 0 | 40 | 112 | 110 | 155 | 35 | 176 | 149 | 0.675512257 | 1 | 0.005 | 0 | 0 | 0 | 0 | 0 | 0 | 0.008 | 0.052 | 0 | -0.011 |
| SE | 9560 | ENSG00000100360 | IFT27 | chr22 | - | 36767782 | 36767862 | 36767305 | 36767365 | 36775673 | 36776119 | 22 | 12 | 17 | 26 | 9 | 0 | 0 | 0 | 0 | 0 | 43 | 13 | 12 | 30 | 44 | 0 | 4 | 0 | 0 | 0 | 229 | 149 | 0.675561744 | 1 | 1 | 1 | 1 | 1 | 1 | 1 | 0.679 | 1 | 1 | 1 | 0.064 |
| SE | 33586 | ENSG00000009790 | TRAF3IP3 | chr1 | + | 209779314 | 209779374 | 209777351 | 209777487 | 209780469 | 209780606 | 34 | 52 | 61 | 44 | 79 | 0 | 0 | 1 | 0 | 3 | 13 | 50 | 18 | 37 | 22 | 0 | 0 | 2 | 0 | 4 | 209 | 149 | 0.675607038 | 1 | 1 | 1 | 0.978 | 1 | 0.949 | 1 | 1 | 0.865 | 1 | 0.797 | 0.053 |
| SE | 78323 | ENSG00000228409 | CCT6P1 | chr7 | + | 65757982 | 65758137 | 65756468 | 65756574 | 65758235 | 65758317 | 65 | 37 | 53 | 45 | 100 | 0 | 0 | 0 | 0 | 7 | 29 | 50 | 126 | 34 | 22 | 0 | 0 | 1 | 0 | 0 | 298 | 149 | 0.675611872 | 1 | 1 | 1 | 1 | 1 | 0.877 | 1 | 1 | 0.984 | 1 | 1 | -0.021 |
| SE | 7546 | ENSG00000167851 | CD300A | chr17 | + | 74474531 | 74474685 | 74466657 | 74466743 | 74481288 | 74481326 | 20 | 3 | 4 | 16 | 0 | 0 | 0 | 0 | 0 | 2 | 14 | 5 | 16 | 13 | 36 | 0 | 0 | 0 | 0 | 0 | 298 | 149 | 0.675612605 | 1 | 1 | 1 | 1 | 1 | 0 | 1 | 1 | 1 | 1 | 1 | -0.2 |
| SE | 101665 | ENSG00000204177 | BMS1P1 | chr10 | + | 46794726 | 46794915 | 46790961 | 46791041 | 46795805 | 46795953 | 15 | 24 | 14 | 19 | 18 | 3 | 3 | 0 | 2 | 0 | 22 | 58 | 18 | 15 | 17 | 10 | 0 | 1 | 0 | 0 | 298 | 149 | 0.675657029 | 1 | 0.714 | 0.8 | 1 | 0.826 | 1 | 0.524 | 1 | 0.9 | 1 | 1 | -0.017 |
| SE | 98284 | ENSG00000068831 | RASGRP2 | chr11 | - | 64730052 | 64730194 | 64728862 | 64729042 | 64735111 | 64735227 | 413 | 278 | 480 | 478 | 560 | 1 | 0 | 1 | 2 | 0 | 304 | 374 | 251 | 294 | 629 | 9 | 0 | 0 | 7 | 0 | 291 | 149 | 0.675675728 | 1 | 0.995 | 1 | 0.996 | 0.992 | 1 | 0.945 | 1 | 1 | 0.956 | 1 | 0.016 |
| SE | 9768 | ENSG00000062716 | VMP1 | chr17 | + | 59811669 | 59811786 | 59808795 | 59808876 | 59817711 | 59817773 | 758 | 486 | 480 | 595 | 837 | 0 | 1 | 0 | 1 | 0 | 748 | 522 | 725 | 561 | 278 | 20 | 0 | 0 | 0 | 0 | 266 | 149 | 0.675700609 | 1 | 1 | 0.996 | 1 | 0.997 | 1 | 0.954 | 1 | 1 | 1 | 1 | 0.008 |
| SE | 96247 | ENSG00000133943 | DGLUCY | chr14 | + | 91173663 | 91173753 | 91173288 | 91173439 | 91175933 | 91176056 | 0 | 0 | 0 | 0 | 0 | 19 | 9 | 31 | 48 | 6 | 0 | 0 | 0 | 0 | 8 | 43 | 18 | 14 | 15 | 46 | 239 | 149 | 0.675728866 | 1 | 0 | 0 | 0 | 0 | 0 | 0 | 0 | 0 | 0 | 0.098 | -0.02 |
| SE | 78413 | ENSG00000161405 | IKZF3 | chr17 | - | 39777884 | 39778010 | 39777650 | 39777767 | 39788257 | 39788374 | 92 | 41 | 58 | 85 | 46 | 262 | 115 | 224 | 234 | 228 | 26 | 102 | 51 | 43 | 35 | 97 | 250 | 154 | 180 | 112 | 275 | 149 | 0.675737305 | 1 | 0.16 | 0.162 | 0.123 | 0.164 | 0.099 | 0.127 | 0.181 | 0.152 | 0.115 | 0.145 | -0.002 |
| SE | 94961 | ENSG00000266714 | MYO15B | chr17 | + | 75590906 | 75591016 | 75590625 | 75590689 | 75591171 | 75591246 | 1 | 2 | 4 | 4 | 2 | 0 | 4 | 3 | 6 | 4 | 0 | 2 | 1 | 11 | 15 | 19 | 9 | 0 | 5 | 0 | 259 | 149 | 0.675838546 | 1 | 1 | 0.223 | 0.434 | 0.277 | 0.223 | 0 | 0.113 | 1 | 0.559 | 1 | -0.103 |
| SE | 18824 | ENSG00000127663 | KDM4B | chr19 | + | 5047475 | 5047669 | 5032865 | 5033031 | 5071009 | 5071059 | 67 | 16 | 51 | 48 | 36 | 0 | 0 | 0 | 0 | 0 | 22 | 77 | 57 | 53 | 124 | 0 | 0 | 6 | 0 | 0 | 298 | 149 | 0.675865168 | 1 | 1 | 1 | 1 | 1 | 1 | 1 | 1 | 0.826 | 1 | 1 | 0.035 |
| SE | 44682 | ENSG00000185344 | ATP6V0A2 | chr12 | + | 123735892 | 123736540 | 123735530 | 123735624 | 123737058 | 123737271 | 5 | 6 | 8 | 7 | 4 | 66 | 31 | 85 | 94 | 81 | 12 | 15 | 19 | 6 | 0 | 144 | 120 | 92 | 135 | 90 | 298 | 149 | 0.675922321 | 1 | 0.036 | 0.088 | 0.045 | 0.036 | 0.024 | 0.04 | 0.059 | 0.094 | 0.022 | 0 | 0.003 |
| SE | 79992 | ENSG00000118058 | KMT2A | chr11 | + | 118520801 | 118520885 | 118519956 | 118520064 | 118521287 | 118521417 | 345 | 179 | 318 | 327 | 211 | 1 | 0 | 0 | 4 | 0 | 110 | 360 | 250 | 191 | 99 | 4 | 5 | 2 | 0 | 0 | 233 | 149 | 0.675926503 | 1 | 0.995 | 1 | 1 | 0.981 | 1 | 0.946 | 0.979 | 0.988 | 1 | 1 | 0.013 |
| SE | 76926 | ENSG00000167483 | NIBAN3 | chr19 | + | 17549443 | 17549527 | 17543523 | 17543631 | 17551785 | 17552024 | 33 | 33 | 36 | 57 | 30 | 3 | 0 | 0 | 2 | 7 | 90 | 77 | 8 | 55 | 157 | 10 | 3 | 0 | 0 | 7 | 233 | 149 | 0.675978836 | 1 | 0.876 | 1 | 1 | 0.948 | 0.733 | 0.852 | 0.943 | 1 | 1 | 0.935 | -0.035 |
| SE | 48526 | ENSG00000111481 | COPZ1 | chr12 | + | 54340546 | 54340615 | 54325082 | 54325181 | 54342205 | 54342287 | 320 | 183 | 269 | 276 | 217 | 2 | 3 | 4 | 7 | 0 | 558 | 424 | 256 | 313 | 633 | 0 | 11 | 0 | 3 | 7 | 218 | 149 | 0.676046602 | 1 | 0.991 | 0.977 | 0.979 | 0.964 | 1 | 1 | 0.963 | 1 | 0.986 | 0.984 | -0.004 |
| SE | 57715 | ENSG00000001497 | LAS1L | chrX | - | 65524563 | 65524614 | 65524055 | 65524262 | 65524964 | 65525050 | 10 | 5 | 14 | 14 | 25 | 36 | 28 | 37 | 26 | 40 | 3 | 18 | 7 | 16 | 11 | 36 | 49 | 10 | 26 | 33 | 200 | 149 | 0.676074314 | 1 | 0.171 | 0.117 | 0.22 | 0.286 | 0.318 | 0.058 | 0.215 | 0.343 | 0.314 | 0.199 | -0.003 |
| SE | 3287 | ENSG00000161671 | EMC10 | chr19 | + | 50481860 | 50481947 | 50480883 | 50480977 | 50482148 | 50483351 | 6 | 4 | 4 | 0 | 0 | 13 | 4 | 19 | 14 | 1 | 0 | 12 | 0 | 4 | 12 | 12 | 26 | 9 | 16 | 30 | 236 | 149 | 0.676119233 | 1 | 0.226 | 0.387 | 0.117 | 0 | 0 | 0 | 0.226 | 0 | 0.136 | 0.202 | 0.033 |
| SE | 47269 | ENSG00000113360 | DROSHA | chr5 | - | 31467141 | 31467166 | 31466181 | 31466281 | 31467938 | 31468006 | 1 | 2 | 4 | 3 | 0 | 79 | 62 | 80 | 53 | 89 | 0 | 4 | 0 | 0 | 0 | 36 | 80 | 26 | 76 | 66 | 174 | 149 | 0.676191385 | 1 | 0.011 | 0.027 | 0.041 | 0.046 | 0 | 0 | 0.041 | 0 | 0 | 0 | 0.017 |
| SE | 49578 | ENSG00000112039 | FANCE | chr6 | + | 35457915 | 35457984 | 35457555 | 35457600 | 35458296 | 35458440 | 12 | 2 | 7 | 6 | 13 | 1 | 0 | 3 | 1 | 0 | 7 | 2 | 7 | 6 | 1 | 0 | 3 | 0 | 0 | 0 | 218 | 149 | 0.676380259 | 1 | 0.891 | 1 | 0.615 | 0.804 | 1 | 1 | 0.313 | 1 | 1 | 1 | -0.001 |
| SE | 4397 | ENSG00000130475 | FCHO1 | chr19 | + | 17766668 | 17766810 | 17764374 | 17764449 | 17770424 | 17770463 | 134 | 60 | 121 | 140 | 107 | 2 | 0 | 0 | 0 | 0 | 95 | 157 | 116 | 126 | 193 | 0 | 0 | 0 | 0 | 9 | 291 | 149 | 0.676385128 | 1 | 0.972 | 1 | 1 | 1 | 1 | 1 | 1 | 1 | 1 | 0.917 | 0.011 |
| SE | 98071 | ENSG00000066926 | FECH | chr18 | - | 57559144 | 57559243 | 57554844 | 57554952 | 57562873 | 57562980 | 51 | 29 | 45 | 46 | 97 | 0 | 0 | 0 | 1 | 0 | 41 | 35 | 23 | 51 | 120 | 1 | 0 | 0 | 6 | 0 | 248 | 149 | 0.676507824 | 1 | 1 | 1 | 1 | 0.965 | 1 | 0.961 | 1 | 1 | 0.836 | 1 | 0.034 |
| SE | 102155 | ENSG00000171130 | ATP6V0E2 | chr7 | + | 149879165 | 149879225 | 149878677 | 149878790 | 149879334 | 149880696 | 2 | 3 | 0 | 0 | 0 | 25 | 4 | 24 | 4 | 11 | 0 | 0 | 0 | 0 | 0 | 16 | 12 | 16 | 3 | 12 | 209 | 149 | 0.676528577 | 1 | 0.054 | 0.348 | 0 | 0 | 0 | 0 | 0 | 0 | 0 | 0 | 0.08 |
| SE | 97362 | ENSG00000110218 | PANX1 | chr11 | + | 94178368 | 94178592 | 94153490 | 94153630 | 94179601 | 94180257 | 71 | 34 | 31 | 69 | 83 | 6 | 0 | 0 | 1 | 0 | 38 | 50 | 57 | 27 | 67 | 0 | 0 | 0 | 0 | 0 | 298 | 149 | 0.67664256 | 1 | 0.855 | 1 | 1 | 0.972 | 1 | 1 | 1 | 1 | 1 | 1 | -0.035 |
| SE | 23432 | ENSG00000172062 | SMN1 | chr5 | + | 70946706 | 70946815 | 70946065 | 70946176 | 70951940 | 70952567 | 1 | 1 | 0 | 2 | 10 | 75 | 46 | 84 | 81 | 129 | 0 | 0 | 1 | 0 | 0 | 18 | 121 | 130 | 70 | 39 | 258 | 149 | 0.676659506 | 1 | 0.008 | 0.012 | 0 | 0.014 | 0.043 | 0 | 0 | 0.004 | 0 | 0 | 0.015 |
| SE | 86976 | ENSG00000116128 | BCL9 | chr1 | + | 147611577 | 147611889 | 147606783 | 147606866 | 147614426 | 147614616 | 27 | 14 | 54 | 21 | 92 | 0 | 0 | 0 | 0 | 0 | 54 | 19 | 15 | 27 | 0 | 0 | 0 | 0 | 0 | 2 | 298 | 149 | 0.676702886 | 1 | 1 | 1 | 1 | 1 | 1 | 1 | 1 | 1 | 1 | 0 | 0.2 |
| SE | 30404 | ENSG00000153064 | BANK1 | chr4 | + | 101829807 | 101830206 | 101790729 | 101790950 | 101855034 | 101855189 | 309 | 290 | 242 | 409 | 666 | 18 | 6 | 4 | 13 | 0 | 105 | 226 | 59 | 46 | 72 | 0 | 5 | 3 | 0 | 5 | 298 | 149 | 0.676747123 | 1 | 0.896 | 0.96 | 0.968 | 0.94 | 1 | 1 | 0.958 | 0.908 | 1 | 0.878 | 0.004 |
| SE | 74792 | ENSG00000178904 | DPY19L3 | chr19 | + | 32432715 | 32432806 | 32411238 | 32411372 | 32436445 | 32436495 | 48 | 30 | 38 | 55 | 60 | 0 | 0 | 1 | 0 | 6 | 8 | 47 | 42 | 22 | 16 | 0 | 1 | 0 | 0 | 0 | 240 | 149 | 0.676876886 | 1 | 1 | 1 | 0.959 | 1 | 0.861 | 1 | 0.967 | 1 | 1 | 1 | -0.029 |
| SE | 62643 | ENSG00000128815 | WDFY4 | chr10 | + | 48810529 | 48810735 | 48807858 | 48807958 | 48811538 | 48811708 | 102 | 62 | 96 | 134 | 129 | 0 | 3 | 0 | 4 | 0 | 179 | 135 | 85 | 87 | 231 | 0 | 1 | 1 | 2 | 0 | 298 | 149 | 0.676982247 | 1 | 1 | 0.912 | 1 | 0.944 | 1 | 1 | 0.985 | 0.977 | 0.956 | 1 | -0.012 |
| SE | 11766 | ENSG00000166548 | TK2 | chr16 | - | 66536963 | 66537017 | 66531379 | 66531469 | 66541878 | 66541953 | 138 | 76 | 78 | 101 | 106 | 3 | 2 | 1 | 1 | 8 | 257 | 154 | 75 | 70 | 281 | 8 | 6 | 4 | 0 | 7 | 203 | 149 | 0.676992898 | 1 | 0.971 | 0.965 | 0.983 | 0.987 | 0.907 | 0.959 | 0.95 | 0.932 | 1 | 0.967 | 0.001 |
| SE | 78580 | ENSG00000088930 | XRN2 | chr20 | + | 21330480 | 21330539 | 21326489 | 21326601 | 21330615 | 21330705 | 89 | 50 | 79 | 66 | 141 | 0 | 0 | 0 | 0 | 0 | 45 | 90 | 127 | 56 | 12 | 0 | 0 | 0 | 7 | 0 | 208 | 149 | 0.677029791 | 1 | 1 | 1 | 1 | 1 | 1 | 1 | 1 | 1 | 0.851 | 1 | 0.03 |
| SE | 64364 | ENSG00000163219 | ARHGAP25 | chr2 | + | 68787839 | 68787956 | 68782232 | 68782320 | 68807275 | 68807480 | 444 | 207 | 334 | 284 | 309 | 2 | 1 | 3 | 0 | 0 | 208 | 290 | 187 | 101 | 190 | 0 | 6 | 3 | 2 | 0 | 266 | 149 | 0.677039564 | 1 | 0.992 | 0.991 | 0.984 | 1 | 1 | 1 | 0.964 | 0.972 | 0.966 | 1 | 0.013 |
| SE | 22587 | ENSG00000127603 | MACF1 | chr1 | + | 39295045 | 39295150 | 39292765 | 39292843 | 39295786 | 39295882 | 106 | 81 | 95 | 94 | 140 | 0 | 0 | 0 | 0 | 0 | 107 | 143 | 74 | 31 | 29 | 0 | 0 | 0 | 5 | 0 | 254 | 149 | 0.677068494 | 1 | 1 | 1 | 1 | 1 | 1 | 1 | 1 | 1 | 0.784 | 1 | 0.043 |
| SE | 21522 | ENSG00000137413 | TAF8 | chr6 | + | 42071346 | 42071468 | 42068464 | 42068607 | 42077099 | 42077239 | 6 | 0 | 2 | 0 | 5 | 70 | 50 | 72 | 87 | 38 | 0 | 8 | 0 | 0 | 8 | 67 | 99 | 51 | 65 | 60 | 271 | 149 | 0.677129736 | 1 | 0.045 | 0 | 0.015 | 0 | 0.067 | 0 | 0.043 | 0 | 0 | 0.068 | 0.003 |
| SE | 23037 | ENSG00000152942 | RAD17 | chr5 | + | 69371453 | 69371557 | 69371034 | 69371171 | 69372033 | 69372217 | 21 | 7 | 22 | 25 | 13 | 86 | 91 | 97 | 104 | 132 | 12 | 10 | 10 | 0 | 0 | 0 | 93 | 56 | 17 | 17 | 253 | 149 | 0.677154599 | 1 | 0.126 | 0.043 | 0.118 | 0.124 | 0.055 | 1 | 0.06 | 0.095 | 0 | 0 | -0.138 |
| SE | 100434 | ENSG00000132275 | RRP8 | chr11 | - | 6600925 | 6601055 | 6600668 | 6600775 | 6601148 | 6601602 | 31 | 11 | 40 | 25 | 42 | 0 | 0 | 0 | 0 | 0 | 9 | 16 | 9 | 15 | 17 | 0 | 4 | 0 | 0 | 0 | 279 | 149 | 0.677198381 | 1 | 1 | 1 | 1 | 1 | 1 | 1 | 0.681 | 1 | 1 | 1 | 0.064 |
| SE | 81460 | ENSG00000106443 | PHF14 | chr7 | + | 11061790 | 11061841 | 11051611 | 11051780 | 11061963 | 11062085 | 51 | 36 | 68 | 42 | 108 | 3 | 0 | 1 | 0 | 0 | 12 | 56 | 11 | 20 | 16 | 0 | 4 | 2 | 0 | 0 | 200 | 149 | 0.677323752 | 1 | 0.927 | 1 | 0.981 | 1 | 1 | 1 | 0.913 | 0.804 | 1 | 1 | 0.038 |
| SE | 48226 | ENSG00000007237 | GAS7 | chr17 | - | 9969676 | 9969762 | 9959201 | 9959255 | 10019776 | 10019897 | 102 | 59 | 60 | 76 | 34 | 0 | 0 | 1 | 0 | 0 | 184 | 134 | 94 | 199 | 276 | 9 | 0 | 2 | 0 | 0 | 235 | 149 | 0.677357999 | 1 | 1 | 1 | 0.974 | 1 | 1 | 0.928 | 1 | 0.968 | 1 | 1 | 0.016 |
| SE | 42442 | ENSG00000163946 | TASOR | chr3 | - | 56663540 | 56663572 | 56660968 | 56661017 | 56666259 | 56666384 | 39 | 30 | 39 | 48 | 109 | 0 | 0 | 0 | 3 | 5 | 8 | 16 | 22 | 29 | 16 | 0 | 0 | 0 | 0 | 0 | 181 | 149 | 0.677388594 | 1 | 1 | 1 | 1 | 0.929 | 0.947 | 1 | 1 | 1 | 1 | 1 | -0.025 |
| SE | 60539 | ENSG00000116984 | MTR | chr1 | + | 236852520 | 236852637 | 236838413 | 236838599 | 236852947 | 236853088 | 111 | 79 | 95 | 97 | 116 | 0 | 0 | 0 | 0 | 0 | 54 | 87 | 83 | 81 | 51 | 0 | 2 | 1 | 5 | 0 | 266 | 149 | 0.677402192 | 1 | 1 | 1 | 1 | 1 | 1 | 1 | 0.961 | 0.979 | 0.901 | 1 | 0.032 |
| SE | 61911 | ENSG00000124067 | SLC12A4 | chr16 | - | 67954103 | 67954219 | 67952241 | 67952425 | 67954642 | 67954773 | 1 | 0 | 3 | 1 | 0 | 19 | 8 | 18 | 14 | 11 | 0 | 0 | 2 | 0 | 0 | 12 | 7 | 6 | 2 | 57 | 265 | 149 | 0.677502071 | 1 | 0.029 | 0 | 0.086 | 0.039 | 0 | 0 | 0 | 0.158 | 0 | 0 | -0.001 |
| SE | 102673 | ENSG00000175029 | CTBP2 | chr10 | - | 125088190 | 125088319 | 125039045 | 125039155 | 125110989 | 125111093 | 4 | 1 | 3 | 7 | 4 | 64 | 28 | 30 | 68 | 38 | 0 | 0 | 0 | 12 | 1 | 2 | 17 | 15 | 14 | 24 | 278 | 149 | 0.67750427 | 1 | 0.032 | 0.019 | 0.051 | 0.052 | 0.053 | 0 | 0 | 0 | 0.315 | 0.022 | -0.026 |
| SE | 35460 | ENSG00000161217 | PCYT1A | chr3 | - | 196239546 | 196239735 | 196234367 | 196238894 | 196241947 | 196242090 | 308 | 191 | 214 | 228 | 367 | 4 | 3 | 4 | 1 | 0 | 406 | 211 | 246 | 280 | 386 | 0 | 0 | 0 | 5 | 0 | 298 | 149 | 0.67762781 | 1 | 0.975 | 0.97 | 0.964 | 0.991 | 1 | 1 | 1 | 1 | 0.966 | 1 | -0.013 |
| SE | 38710 | ENSG00000228794 | LINC01128 | chr1 | + | 841199 | 844498 | 829002 | 829104 | 847653 | 847806 | 11 | 4 | 30 | 13 | 3 | 3 | 2 | 7 | 2 | 7 | 3 | 14 | 0 | 0 | 7 | 0 | 3 | 4 | 4 | 0 | 298 | 149 | 0.677700815 | 1 | 0.647 | 0.5 | 0.682 | 0.765 | 0.176 | 1 | 0.7 | 0 | 0 | 1 | 0.014 |
| SE | 32563 | ENSG00000240682 | ISY1 | chr3 | - | 129158507 | 129158559 | 129156854 | 129156920 | 129159153 | 129159176 | 96 | 68 | 85 | 111 | 86 | 0 | 1 | 0 | 0 | 3 | 241 | 130 | 136 | 167 | 196 | 7 | 0 | 0 | 1 | 3 | 201 | 149 | 0.677775375 | 1 | 1 | 0.981 | 1 | 1 | 0.955 | 0.962 | 1 | 1 | 0.992 | 0.98 | 0 |
| SE | 87560 | ENSG00000112159 | MDN1 | chr6 | - | 89646539 | 89646603 | 89645014 | 89645157 | 89648031 | 89648146 | 79 | 45 | 73 | 54 | 147 | 2 | 2 | 1 | 2 | 2 | 17 | 17 | 31 | 29 | 9 | 0 | 0 | 4 | 0 | 0 | 213 | 149 | 0.677792558 | 1 | 0.965 | 0.94 | 0.981 | 0.95 | 0.981 | 1 | 1 | 0.844 | 1 | 1 | -0.005 |
| SE | 101990 | ENSG00000119541 | VPS4B | chr18 | - | 63397033 | 63397253 | 63393408 | 63393549 | 63400047 | 63400196 | 104 | 70 | 114 | 142 | 205 | 0 | 3 | 2 | 0 | 5 | 48 | 103 | 81 | 25 | 5 | 0 | 3 | 3 | 0 | 0 | 298 | 149 | 0.677841653 | 1 | 1 | 0.921 | 0.966 | 1 | 0.953 | 1 | 0.945 | 0.931 | 1 | 1 | -0.007 |
| SE | 48048 | ENSG00000169299 | PGM2 | chr4 | + | 37844363 | 37844553 | 37840065 | 37840259 | 37845632 | 37845730 | 212 | 135 | 133 | 173 | 229 | 11 | 4 | 8 | 8 | 4 | 47 | 174 | 200 | 193 | 26 | 0 | 21 | 11 | 0 | 0 | 298 | 149 | 0.677922154 | 1 | 0.906 | 0.944 | 0.893 | 0.915 | 0.966 | 1 | 0.806 | 0.901 | 1 | 1 | -0.017 |
| SE | 71948 | ENSG00000197930 | ERO1A | chr14 | - | 52661292 | 52661304 | 52658123 | 52658150 | 52663800 | 52663847 | 648 | 533 | 523 | 683 | 1124 | 0 | 6 | 3 | 9 | 4 | 539 | 761 | 894 | 1126 | 156 | 15 | 0 | 14 | 8 | 0 | 161 | 149 | 0.678013506 | 1 | 1 | 0.988 | 0.994 | 0.986 | 0.996 | 0.971 | 1 | 0.983 | 0.992 | 1 | 0.004 |
| SE | 72096 | ENSG00000122484 | RPAP2 | chr1 | + | 92301475 | 92301590 | 92300193 | 92300239 | 92303976 | 92304075 | 110 | 81 | 78 | 104 | 171 | 2 | 0 | 3 | 2 | 0 | 48 | 24 | 68 | 18 | 20 | 0 | 0 | 1 | 0 | 0 | 264 | 149 | 0.678030549 | 1 | 0.969 | 1 | 0.936 | 0.967 | 1 | 1 | 1 | 0.975 | 1 | 1 | -0.021 |
| SE | 25105 | ENSG00000153310 | CYRIB | chr8 | - | 129879388 | 129879471 | 129871374 | 129871496 | 129903311 | 129903350 | 1284 | 923 | 809 | 1077 | 973 | 19 | 24 | 21 | 42 | 15 | 860 | 909 | 1212 | 1315 | 271 | 14 | 29 | 28 | 27 | 17 | 232 | 149 | 0.678103275 | 1 | 0.977 | 0.961 | 0.961 | 0.943 | 0.977 | 0.975 | 0.953 | 0.965 | 0.969 | 0.911 | 0.009 |
| SE | 103640 | ENSG00000012232 | EXTL3 | chr8 | + | 28713456 | 28713550 | 28701500 | 28701659 | 28715584 | 28715610 | 108 | 32 | 57 | 62 | 52 | 1 | 0 | 0 | 0 | 0 | 51 | 47 | 66 | 34 | 58 | 0 | 0 | 0 | 0 | 6 | 243 | 149 | 0.6781458 | 1 | 0.985 | 1 | 1 | 1 | 1 | 1 | 1 | 1 | 1 | 0.856 | 0.026 |
| SE | 19606 | ENSG00000255185 | PDXDC2P | chr16 | - | 70034170 | 70034360 | 70031899 | 70031968 | 70036431 | 70036578 | 94 | 133 | 103 | 121 | 66 | 0 | 2 | 1 | 2 | 0 | 179 | 204 | 140 | 130 | 143 | 0 | 0 | 5 | 3 | 0 | 298 | 149 | 0.678202383 | 1 | 1 | 0.971 | 0.981 | 0.968 | 1 | 1 | 1 | 0.933 | 0.956 | 1 | 0.006 |
| SE | 30129 | ENSG00000150961 | SEC24D | chr4 | - | 118735726 | 118736569 | 118732732 | 118732912 | 118738260 | 118738379 | 0 | 0 | 1 | 2 | 0 | 107 | 60 | 89 | 82 | 123 | 0 | 2 | 9 | 0 | 0 | 40 | 102 | 81 | 62 | 32 | 298 | 149 | 0.678367715 | 1 | 0 | 0 | 0.006 | 0.012 | 0 | 0 | 0.01 | 0.053 | 0 | 0 | -0.009 |
| SE | 25936 | ENSG00000119522 | DENND1A | chr9 | - | 123387729 | 123387858 | 123383654 | 123383913 | 123403401 | 123403490 | 1 | 2 | 5 | 7 | 2 | 21 | 16 | 19 | 12 | 10 | 11 | 0 | 0 | 17 | 13 | 27 | 20 | 25 | 63 | 88 | 278 | 149 | 0.678372261 | 1 | 0.025 | 0.063 | 0.124 | 0.238 | 0.097 | 0.179 | 0 | 0 | 0.126 | 0.073 | 0.034 |
| SE | 81524 | ENSG00000071462 | BUD23 | chr7 | + | 73693328 | 73693414 | 73692595 | 73692646 | 73697604 | 73697694 | 55 | 14 | 52 | 41 | 34 | 0 | 0 | 0 | 0 | 0 | 26 | 82 | 52 | 42 | 62 | 0 | 0 | 0 | 6 | 0 | 235 | 149 | 0.678387296 | 1 | 1 | 1 | 1 | 1 | 1 | 1 | 1 | 1 | 0.816 | 1 | 0.037 |
| SE | 95782 | ENSG00000148120 | AOPEP | chr9 | + | 94773001 | 94773168 | 94759648 | 94760580 | 94792764 | 94792918 | 27 | 14 | 25 | 13 | 15 | 3 | 1 | 3 | 2 | 3 | 22 | 17 | 14 | 16 | 16 | 0 | 4 | 10 | 1 | 0 | 298 | 149 | 0.67848895 | 1 | 0.818 | 0.875 | 0.806 | 0.765 | 0.714 | 1 | 0.68 | 0.412 | 0.889 | 1 | -0.001 |
| SE | 4309 | ENSG00000182141 | ZNF708 | chr19 | - | 21310500 | 21310627 | 21309245 | 21309341 | 21329209 | 21329410 | 21 | 18 | 19 | 34 | 22 | 11 | 6 | 26 | 14 | 22 | 8 | 10 | 6 | 8 | 0 | 0 | 3 | 9 | 13 | 3 | 276 | 149 | 0.678500642 | 1 | 0.508 | 0.618 | 0.283 | 0.567 | 0.351 | 1 | 0.643 | 0.265 | 0.249 | 0 | 0.034 |
| SE | 64030 | ENSG00000115464 | USP34 | chr2 | - | 61214059 | 61214694 | 61211771 | 61211929 | 61220309 | 61220457 | 272 | 186 | 224 | 303 | 294 | 3 | 0 | 2 | 6 | 0 | 15 | 126 | 137 | 32 | 3 | 0 | 0 | 0 | 2 | 0 | 298 | 149 | 0.678663428 | 1 | 0.978 | 1 | 0.982 | 0.962 | 1 | 1 | 1 | 1 | 0.889 | 1 | 0.007 |
| SE | 106084 | ENSG00000196498 | NCOR2 | chr12 | - | 124372021 | 124372610 | 124363678 | 124363799 | 124378236 | 124378384 | 61 | 53 | 45 | 61 | 72 | 0 | 0 | 0 | 0 | 0 | 133 | 102 | 92 | 107 | 143 | 0 | 0 | 0 | 0 | 8 | 298 | 149 | 0.678673329 | 1 | 1 | 1 | 1 | 1 | 1 | 1 | 1 | 1 | 1 | 0.899 | 0.02 |
| SE | 29130 | ENSG00000248672 | LY75-CD302 | chr2 | - | 159864838 | 159864920 | 159860820 | 159860889 | 159872450 | 159872593 | 192 | 140 | 198 | 132 | 269 | 0 | 1 | 2 | 1 | 1 | 48 | 121 | 104 | 66 | 3 | 6 | 0 | 0 | 0 | 0 | 231 | 149 | 0.678701024 | 1 | 1 | 0.989 | 0.985 | 0.988 | 0.994 | 0.838 | 1 | 1 | 1 | 1 | 0.024 |
| SE | 61523 | ENSG00000138757 | G3BP2 | chr4 | - | 75653982 | 75654081 | 75648638 | 75648741 | 75655065 | 75655246 | 62 | 47 | 54 | 44 | 87 | 450 | 348 | 421 | 563 | 623 | 39 | 50 | 48 | 34 | 50 | 341 | 397 | 777 | 502 | 120 | 248 | 149 | 0.67871136 | 1 | 0.076 | 0.075 | 0.072 | 0.045 | 0.077 | 0.064 | 0.07 | 0.036 | 0.039 | 0.2 | -0.013 |
| SE | 82880 | ENSG00000122643 | NT5C3A | chr7 | - | 33015669 | 33015870 | 33014129 | 33014831 | 33017438 | 33017601 | 582 | 757 | 353 | 973 | 1832 | 0 | 0 | 0 | 4 | 8 | 700 | 893 | 597 | 730 | 476 | 0 | 0 | 0 | 20 | 0 | 298 | 149 | 0.678718668 | 1 | 1 | 1 | 1 | 0.992 | 0.991 | 1 | 1 | 1 | 0.948 | 1 | 0.007 |
| SE | 81210 | ENSG00000110497 | AMBRA1 | chr11 | - | 46508190 | 46508370 | 46494123 | 46494204 | 46541944 | 46542981 | 64 | 20 | 43 | 58 | 45 | 3 | 0 | 0 | 0 | 3 | 29 | 78 | 74 | 26 | 95 | 0 | 0 | 1 | 0 | 0 | 298 | 149 | 0.678752303 | 1 | 0.914 | 1 | 1 | 1 | 0.882 | 1 | 1 | 0.974 | 1 | 1 | -0.036 |
| SE | 52221 | ENSG00000155363 | MOV10 | chr1 | + | 112686965 | 112687106 | 112674736 | 112675049 | 112688327 | 112688483 | 23 | 5 | 9 | 14 | 14 | 0 | 0 | 0 | 0 | 0 | 54 | 21 | 10 | 28 | 62 | 0 | 0 | 2 | 3 | 0 | 290 | 149 | 0.678884035 | 1 | 1 | 1 | 1 | 1 | 1 | 1 | 1 | 0.72 | 0.827 | 1 | 0.091 |
| SE | 10374 | ENSG00000197381 | ADARB1 | chr21 | + | 45184473 | 45184593 | 45183361 | 45183510 | 45184922 | 45185091 | 55 | 45 | 60 | 69 | 89 | 21 | 7 | 33 | 28 | 35 | 20 | 86 | 30 | 51 | 49 | 52 | 23 | 16 | 2 | 32 | 269 | 149 | 0.67906592 | 1 | 0.592 | 0.781 | 0.502 | 0.577 | 0.585 | 0.176 | 0.674 | 0.509 | 0.934 | 0.459 | 0.057 |
| SE | 61085 | ENSG00000198160 | MIER1 | chr1 | + | 66928886 | 66928963 | 66926141 | 66926242 | 66940027 | 66940052 | 28 | 13 | 20 | 26 | 25 | 38 | 45 | 20 | 58 | 34 | 0 | 25 | 31 | 10 | 18 | 5 | 48 | 50 | 5 | 16 | 226 | 149 | 0.679116883 | 1 | 0.327 | 0.16 | 0.397 | 0.228 | 0.326 | 0 | 0.256 | 0.29 | 0.569 | 0.426 | -0.021 |
| SE | 72698 | ENSG00000104047 | DTWD1 | chr15 | + | 49633519 | 49633612 | 49632158 | 49632302 | 49634535 | 49634614 | 2 | 2 | 5 | 2 | 1 | 63 | 62 | 94 | 80 | 150 | 0 | 0 | 0 | 4 | 0 | 23 | 33 | 36 | 40 | 16 | 242 | 149 | 0.679172993 | 1 | 0.019 | 0.019 | 0.032 | 0.015 | 0.004 | 0 | 0 | 0 | 0.058 | 0 | 0.006 |
| SE | 96389 | ENSG00000197563 | PIGN | chr18 | - | 62113133 | 62113316 | 62109833 | 62109973 | 62114560 | 62114626 | 76 | 43 | 80 | 112 | 71 | 0 | 0 | 0 | 0 | 0 | 35 | 83 | 16 | 44 | 22 | 0 | 4 | 2 | 0 | 0 | 298 | 149 | 0.679276032 | 1 | 1 | 1 | 1 | 1 | 1 | 1 | 0.912 | 0.8 | 1 | 1 | 0.058 |
| SE | 31395 | ENSG00000123933 | MXD4 | chr4 | - | 2255154 | 2255351 | 2252407 | 2252522 | 2257981 | 2258011 | 3 | 0 | 5 | 4 | 0 | 159 | 44 | 54 | 75 | 42 | 0 | 2 | 4 | 5 | 0 | 124 | 94 | 66 | 60 | 183 | 298 | 149 | 0.679408039 | 1 | 0.009 | 0 | 0.044 | 0.026 | 0 | 0 | 0.011 | 0.029 | 0.04 | 0 | 0 |
| SE | 57164 | ENSG00000011021 | CLCN6 | chr1 | + | 11826155 | 11826214 | 11824485 | 11824553 | 11827088 | 11827221 | 20 | 18 | 11 | 16 | 25 | 0 | 0 | 0 | 0 | 0 | 76 | 43 | 37 | 37 | 41 | 0 | 6 | 0 | 0 | 0 | 208 | 149 | 0.679533901 | 1 | 1 | 1 | 1 | 1 | 1 | 1 | 0.837 | 1 | 1 | 1 | 0.033 |
| SE | 2710 | ENSG00000011600 | TYROBP | chr19 | - | 35907217 | 35907357 | 35904402 | 35904634 | 35907445 | 35907580 | 1125 | 398 | 407 | 708 | 596 | 3 | 2 | 6 | 4 | 5 | 1487 | 1039 | 581 | 1054 | 1697 | 27 | 3 | 1 | 6 | 1 | 289 | 149 | 0.679565091 | 1 | 0.995 | 0.99 | 0.972 | 0.989 | 0.984 | 0.966 | 0.994 | 0.997 | 0.989 | 0.999 | -0.003 |
| SE | 40222 | ENSG00000084693 | AGBL5 | chr2 | + | 27053895 | 27054059 | 27053401 | 27053573 | 27054629 | 27054807 | 32 | 18 | 25 | 39 | 44 | 0 | 0 | 1 | 0 | 2 | 27 | 50 | 18 | 43 | 122 | 0 | 0 | 3 | 3 | 0 | 298 | 149 | 0.679590749 | 1 | 1 | 1 | 0.926 | 1 | 0.917 | 1 | 1 | 0.75 | 0.878 | 1 | 0.043 |
| SE | 54244 | ENSG00000011405 | PIK3C2A | chr11 | - | 17097056 | 17097264 | 17094260 | 17094385 | 17099859 | 17099969 | 108 | 86 | 76 | 98 | 176 | 0 | 0 | 1 | 4 | 4 | 9 | 39 | 55 | 10 | 9 | 0 | 0 | 3 | 0 | 0 | 298 | 149 | 0.679653775 | 1 | 1 | 1 | 0.974 | 0.925 | 0.957 | 1 | 1 | 0.902 | 1 | 1 | -0.009 |
| SE | 31579 | ENSG00000129071 | MBD4 | chr3 | - | 129432555 | 129432606 | 129431265 | 129431578 | 129433097 | 129433247 | 54 | 56 | 33 | 80 | 149 | 0 | 2 | 0 | 1 | 6 | 35 | 31 | 39 | 34 | 7 | 0 | 4 | 2 | 0 | 0 | 200 | 149 | 0.679656048 | 1 | 1 | 0.954 | 1 | 0.983 | 0.949 | 1 | 0.852 | 0.936 | 1 | 1 | 0.02 |
| SE | 81546 | ENSG00000106348 | IMPDH1 | chr7 | - | 128400831 | 128400891 | 128400332 | 128400539 | 128401014 | 128401116 | 37 | 12 | 24 | 20 | 10 | 0 | 4 | 0 | 0 | 0 | 58 | 32 | 15 | 38 | 78 | 0 | 0 | 0 | 0 | 0 | 209 | 149 | 0.679676249 | 1 | 1 | 0.681 | 1 | 1 | 1 | 1 | 1 | 1 | 1 | 1 | -0.064 |
| SE | 31033 | ENSG00000172795 | DCP2 | chr5 | + | 112981041 | 112981171 | 112976812 | 112976986 | 112985834 | 112985986 | 3 | 0 | 8 | 11 | 5 | 88 | 61 | 65 | 64 | 47 | 0 | 15 | 5 | 3 | 0 | 52 | 70 | 68 | 66 | 41 | 279 | 149 | 0.679768019 | 1 | 0.018 | 0 | 0.062 | 0.084 | 0.054 | 0 | 0.103 | 0.038 | 0.024 | 0 | 0.011 |
| SE | 76174 | ENSG00000132424 | PNISR | chr6 | - | 99403828 | 99403882 | 99402539 | 99402710 | 99404602 | 99404702 | 310 | 406 | 414 | 473 | 438 | 14 | 16 | 9 | 26 | 36 | 117 | 93 | 128 | 67 | 32 | 0 | 3 | 21 | 1 | 0 | 203 | 149 | 0.679828196 | 1 | 0.942 | 0.949 | 0.971 | 0.93 | 0.899 | 1 | 0.958 | 0.817 | 0.98 | 1 | -0.013 |
| SE | 3476 | ENSG00000090924 | PLEKHG2 | chr19 | + | 39422114 | 39422288 | 39421282 | 39421299 | 39422731 | 39423653 | 50 | 31 | 45 | 69 | 48 | 0 | 0 | 0 | 0 | 0 | 158 | 61 | 67 | 59 | 54 | 0 | 0 | 0 | 0 | 6 | 298 | 149 | 0.679863164 | 1 | 1 | 1 | 1 | 1 | 1 | 1 | 1 | 1 | 1 | 0.818 | 0.036 |
| SE | 25445 | ENSG00000011243 | AKAP8L | chr19 | - | 15399301 | 15399410 | 15397713 | 15397855 | 15400793 | 15400864 | 96 | 56 | 59 | 97 | 91 | 0 | 0 | 0 | 0 | 0 | 191 | 73 | 106 | 154 | 236 | 0 | 0 | 2 | 8 | 0 | 258 | 149 | 0.679916471 | 1 | 1 | 1 | 1 | 1 | 1 | 1 | 1 | 0.968 | 0.917 | 1 | 0.023 |
| SE | 58613 | ENSG00000115594 | IL1R1 | chr2 | + | 102157718 | 102157785 | 102153940 | 102154017 | 102164773 | 102164953 | 3 | 7 | 10 | 10 | 3 | 0 | 0 | 0 | 0 | 0 | 5 | 4 | 27 | 35 | 18 | 0 | 0 | 5 | 0 | 0 | 216 | 149 | 0.67998597 | 1 | 1 | 1 | 1 | 1 | 1 | 1 | 1 | 0.788 | 1 | 1 | 0.042 |
| SE | 34927 | ENSG00000076108 | BAZ2A | chr12 | - | 56609735 | 56609937 | 56606632 | 56606733 | 56610113 | 56610215 | 185 | 206 | 204 | 235 | 278 | 0 | 0 | 0 | 0 | 0 | 817 | 439 | 517 | 542 | 728 | 0 | 0 | 3 | 17 | 0 | 298 | 149 | 0.680002164 | 1 | 1 | 1 | 1 | 1 | 1 | 1 | 1 | 0.989 | 0.941 | 1 | 0.014 |
| SE | 67330 | ENSG00000163728 | TTC14 | chr3 | + | 180604477 | 180604607 | 180604224 | 180604309 | 180605765 | 180605837 | 190 | 150 | 208 | 187 | 458 | 3 | 3 | 1 | 0 | 16 | 18 | 63 | 33 | 31 | 19 | 14 | 0 | 0 | 0 | 0 | 279 | 149 | 0.680067775 | 1 | 0.971 | 0.964 | 0.991 | 1 | 0.939 | 0.407 | 1 | 1 | 1 | 1 | 0.092 |
| SE | 82194 | ENSG00000128595 | CALU | chr7 | + | 128754261 | 128754455 | 128748572 | 128748804 | 128754528 | 128754722 | 67 | 47 | 62 | 56 | 49 | 55 | 47 | 76 | 54 | 47 | 73 | 81 | 49 | 87 | 89 | 68 | 106 | 87 | 81 | 60 | 298 | 149 | 0.680113129 | 1 | 0.379 | 0.333 | 0.29 | 0.341 | 0.343 | 0.349 | 0.276 | 0.22 | 0.349 | 0.426 | 0.013 |
| SE | 87576 | ENSG00000112159 | MDN1 | chr6 | - | 89712035 | 89712256 | 89710680 | 89710794 | 89712574 | 89712786 | 106 | 49 | 102 | 87 | 96 | 0 | 0 | 0 | 0 | 0 | 29 | 85 | 77 | 72 | 23 | 4 | 0 | 2 | 0 | 0 | 298 | 149 | 0.680131271 | 1 | 1 | 1 | 1 | 1 | 1 | 0.784 | 1 | 0.951 | 1 | 1 | 0.053 |
| SE | 59730 | ENSG00000149231 | CCDC82 | chr11 | - | 96383961 | 96387613 | 96383268 | 96383473 | 96389843 | 96389929 | 412 | 414 | 313 | 442 | 601 | 2 | 0 | 0 | 0 | 0 | 36 | 118 | 81 | 38 | 42 | 0 | 0 | 7 | 0 | 0 | 298 | 149 | 0.680156818 | 1 | 0.99 | 1 | 1 | 1 | 1 | 1 | 1 | 0.853 | 1 | 1 | 0.027 |
| SE | 59729 | ENSG00000149231 | CCDC82 | chr11 | - | 96383961 | 96386293 | 96380547 | 96383473 | 96389843 | 96389921 | 412 | 414 | 313 | 442 | 601 | 2 | 0 | 0 | 0 | 0 | 36 | 118 | 81 | 38 | 42 | 0 | 0 | 7 | 0 | 0 | 298 | 149 | 0.680157646 | 1 | 0.99 | 1 | 1 | 1 | 1 | 1 | 1 | 0.853 | 1 | 1 | 0.027 |
| SE | 6295 | ENSG00000141644 | MBD1 | chr18 | - | 50274185 | 50274350 | 50273563 | 50273863 | 50275128 | 50275245 | 74 | 42 | 48 | 71 | 9 | 2 | 3 | 1 | 3 | 1 | 47 | 18 | 42 | 41 | 108 | 0 | 5 | 8 | 0 | 0 | 298 | 149 | 0.680253781 | 1 | 0.949 | 0.875 | 0.96 | 0.922 | 0.818 | 1 | 0.643 | 0.724 | 1 | 1 | 0.031 |
| SE | 83898 | ENSG00000196123 | KIAA0895L | chr16 | - | 67181605 | 67181697 | 67180040 | 67180666 | 67183261 | 67184040 | 5 | 6 | 5 | 22 | 26 | 2 | 9 | 9 | 13 | 7 | 107 | 12 | 21 | 48 | 56 | 34 | 14 | 10 | 26 | 53 | 241 | 149 | 0.68027762 | 1 | 0.607 | 0.292 | 0.256 | 0.511 | 0.697 | 0.661 | 0.346 | 0.565 | 0.533 | 0.395 | -0.027 |
| SE | 38887 | ENSG00000111843 | TMEM14C | chr6 | + | 10724555 | 10724633 | 10722914 | 10723241 | 10724960 | 10725037 | 74 | 29 | 55 | 45 | 42 | 0 | 0 | 2 | 0 | 0 | 50 | 35 | 16 | 28 | 39 | 0 | 0 | 0 | 5 | 0 | 227 | 149 | 0.680340171 | 1 | 1 | 1 | 0.948 | 1 | 1 | 1 | 1 | 1 | 0.786 | 1 | 0.032 |
| SE | 108000 | ENSG00000204852 | TCTN1 | chr12 | + | 110644966 | 110645129 | 110640382 | 110640517 | 110647210 | 110647336 | 14 | 12 | 11 | 3 | 6 | 2 | 0 | 1 | 0 | 0 | 6 | 0 | 4 | 11 | 23 | 0 | 2 | 0 | 0 | 0 | 298 | 149 | 0.680455414 | 1 | 0.778 | 1 | 0.846 | 1 | 1 | 1 | 0 | 1 | 1 | 1 | 0.125 |
| SE | 8896 | ENSG00000007168 | PAFAH1B1 | chr17 | + | 2662780 | 2662834 | 2638098 | 2638320 | 2665371 | 2665456 | 3 | 5 | 4 | 0 | 5 | 279 | 214 | 241 | 281 | 323 | 0 | 4 | 4 | 10 | 11 | 352 | 294 | 358 | 349 | 244 | 203 | 149 | 0.680502331 | 1 | 0.008 | 0.017 | 0.012 | 0 | 0.011 | 0 | 0.01 | 0.008 | 0.021 | 0.032 | -0.005 |
| SE | 29834 | ENSG00000129187 | DCTD | chr4 | - | 182916672 | 182916857 | 182916418 | 182916539 | 182917287 | 182917345 | 10 | 4 | 6 | 13 | 5 | 1 | 0 | 0 | 0 | 0 | 8 | 13 | 9 | 0 | 30 | 0 | 0 | 0 | 2 | 0 | 298 | 149 | 0.680612443 | 1 | 0.833 | 1 | 1 | 1 | 1 | 1 | 1 | 1 | 0 | 1 | 0.167 |
| SE | 7120 | ENSG00000168234 | TTC39C | chr18 | + | 24064139 | 24064188 | 24014753 | 24015038 | 24066011 | 24066140 | 114 | 61 | 105 | 141 | 161 | 0 | 0 | 0 | 0 | 0 | 79 | 128 | 97 | 128 | 88 | 0 | 0 | 0 | 8 | 0 | 198 | 149 | 0.68061363 | 1 | 1 | 1 | 1 | 1 | 1 | 1 | 1 | 1 | 0.923 | 1 | 0.015 |
| SE | 73230 | ENSG00000170190 | SLC16A5 | chr17 | + | 75100006 | 75100816 | 75098037 | 75098181 | 75103969 | 75104180 | 8 | 8 | 11 | 6 | 1 | 0 | 0 | 0 | 0 | 0 | 17 | 8 | 6 | 0 | 12 | 0 | 0 | 0 | 2 | 0 | 298 | 149 | 0.680659636 | 1 | 1 | 1 | 1 | 1 | 1 | 1 | 1 | 1 | 0 | 1 | 0.2 |
| SE | 94036 | ENSG00000167094 | TTC16 | chr9 | + | 127726238 | 127726404 | 127724119 | 127724364 | 127727269 | 127727465 | 14 | 8 | 8 | 19 | 10 | 1 | 0 | 0 | 0 | 0 | 17 | 21 | 0 | 8 | 17 | 0 | 0 | 2 | 0 | 0 | 298 | 149 | 0.68066974 | 1 | 0.875 | 1 | 1 | 1 | 1 | 1 | 1 | 0 | 1 | 1 | 0.175 |
| SE | 40812 | ENSG00000055332 | EIF2AK2 | chr2 | - | 37138508 | 37138585 | 37138269 | 37138363 | 37139630 | 37139757 | 248 | 220 | 146 | 297 | 239 | 0 | 0 | 0 | 0 | 0 | 216 | 345 | 277 | 682 | 1 | 0 | 4 | 4 | 9 | 0 | 226 | 149 | 0.680689536 | 1 | 1 | 1 | 1 | 1 | 1 | 1 | 0.983 | 0.979 | 0.98 | 1 | 0.012 |
| SE | 11426 | ENSG00000167522 | ANKRD11 | chr16 | - | 89286038 | 89286186 | 89275092 | 89275191 | 89288527 | 89288670 | 30 | 23 | 20 | 30 | 4 | 0 | 1 | 0 | 2 | 2 | 8 | 53 | 47 | 65 | 29 | 0 | 0 | 5 | 0 | 0 | 297 | 149 | 0.680699836 | 1 | 1 | 0.92 | 1 | 0.883 | 0.501 | 1 | 1 | 0.825 | 1 | 1 | -0.104 |
| SE | 92870 | ENSG00000135018 | UBQLN1 | chr9 | - | 83665029 | 83665145 | 83663874 | 83664043 | 83669184 | 83669327 | 376 | 210 | 371 | 406 | 476 | 2 | 0 | 1 | 14 | 5 | 255 | 276 | 285 | 311 | 152 | 0 | 10 | 4 | 5 | 0 | 265 | 149 | 0.680700699 | 1 | 0.991 | 1 | 0.995 | 0.942 | 0.982 | 1 | 0.939 | 0.976 | 0.972 | 1 | 0.005 |
| SE | 65113 | ENSG00000282572 | NA | chr7 | - | 31059 | 31606 | 20833 | 21029 | 35334 | 35457 | 1 | 2 | 1 | 1 | 0 | 0 | 0 | 0 | 0 | 2 | 14 | 11 | 8 | 29 | 27 | 0 | 0 | 0 | 0 | 1 | 298 | 149 | 0.680738248 | 1 | 1 | 1 | 1 | 1 | 0 | 1 | 1 | 1 | 1 | 0.931 | -0.186 |
| SE | 81307 | ENSG00000141480 | ARRB2 | chr17 | + | 4720392 | 4720472 | 4720215 | 4720299 | 4720585 | 4720640 | 1181 | 625 | 692 | 1018 | 1037 | 0 | 1 | 0 | 0 | 0 | 2376 | 1393 | 1117 | 1528 | 2569 | 23 | 10 | 6 | 15 | 13 | 229 | 149 | 0.680813191 | 1 | 1 | 0.998 | 1 | 1 | 1 | 0.985 | 0.989 | 0.992 | 0.985 | 0.992 | 0.011 |
| SE | 446 | ENSG00000237484 | LINC01684 | chr21 | + | 24488112 | 24488309 | 24485622 | 24485685 | 24526342 | 24526735 | 10 | 4 | 9 | 8 | 13 | 0 | 0 | 0 | 2 | 0 | 12 | 2 | 8 | 0 | 2 | 0 | 0 | 0 | 2 | 0 | 298 | 149 | 0.68084578 | 1 | 1 | 1 | 1 | 0.667 | 1 | 1 | 1 | 1 | 0 | 1 | 0.133 |
| SE | 98201 | ENSG00000149273 | RPS3 | chr11 | + | 75401639 | 75401733 | 75399519 | 75399577 | 75404019 | 75404207 | 12 | 8 | 13 | 13 | 42 | 2 | 0 | 0 | 1 | 0 | 29 | 20 | 14 | 34 | 23 | 0 | 0 | 4 | 0 | 0 | 243 | 149 | 0.680983792 | 1 | 0.786 | 1 | 1 | 0.889 | 1 | 1 | 1 | 0.682 | 1 | 1 | -0.001 |
| SE | 91130 | ENSG00000179950 | PUF60 | chr8 | - | 143820665 | 143820716 | 143818372 | 143818534 | 143821596 | 143821686 | 59 | 23 | 47 | 46 | 44 | 55 | 31 | 32 | 39 | 48 | 88 | 63 | 24 | 34 | 35 | 34 | 60 | 25 | 30 | 52 | 200 | 149 | 0.681110501 | 1 | 0.444 | 0.356 | 0.522 | 0.468 | 0.406 | 0.658 | 0.439 | 0.417 | 0.458 | 0.334 | -0.022 |
| SE | 74583 | ENSG00000128284 | APOL3 | chr22 | - | 36153284 | 36153423 | 36149258 | 36149371 | 36160668 | 36160755 | 2 | 0 | 3 | 0 | 0 | 40 | 28 | 54 | 46 | 16 | 6 | 0 | 0 | 5 | 0 | 87 | 20 | 32 | 38 | 91 | 288 | 149 | 0.681158904 | 1 | 0.025 | 0 | 0.028 | 0 | 0 | 0.034 | 0 | 0 | 0.064 | 0 | -0.009 |
| SE | 63941 | ENSG00000119787 | ATL2 | chr2 | - | 38354066 | 38354177 | 38343267 | 38343512 | 38377142 | 38377262 | 4 | 0 | 0 | 4 | 1 | 66 | 42 | 56 | 58 | 52 | 0 | 0 | 0 | 1 | 0 | 21 | 54 | 56 | 16 | 12 | 260 | 149 | 0.681202512 | 1 | 0.034 | 0 | 0 | 0.038 | 0.011 | 0 | 0 | 0 | 0.035 | 0 | 0.01 |
| SE | 38206 | ENSG00000186501 | TMEM222 | chr1 | + | 27334666 | 27334754 | 27334150 | 27334281 | 27335378 | 27335648 | 1 | 0 | 5 | 2 | 0 | 43 | 17 | 35 | 36 | 23 | 6 | 1 | 0 | 0 | 0 | 24 | 49 | 14 | 26 | 58 | 237 | 149 | 0.68123578 | 1 | 0.014 | 0 | 0.082 | 0.034 | 0 | 0.136 | 0.013 | 0 | 0 | 0 | -0.004 |
| SE | 101554 | ENSG00000088298 | EDEM2 | chr20 | - | 35131641 | 35131783 | 35126250 | 35126375 | 35137879 | 35138005 | 78 | 30 | 73 | 82 | 48 | 5 | 4 | 2 | 8 | 7 | 216 | 125 | 67 | 85 | 236 | 23 | 17 | 3 | 4 | 16 | 291 | 149 | 0.681264373 | 1 | 0.889 | 0.793 | 0.949 | 0.84 | 0.778 | 0.828 | 0.79 | 0.92 | 0.916 | 0.883 | -0.018 |
| SE | 95933 | ENSG00000166199 | ALKBH3 | chr11 | + | 43901515 | 43901725 | 43889761 | 43889828 | 43919037 | 43919042 | 38 | 29 | 38 | 35 | 19 | 3 | 0 | 4 | 0 | 7 | 86 | 54 | 29 | 26 | 29 | 23 | 0 | 1 | 0 | 0 | 298 | 149 | 0.681390667 | 1 | 0.864 | 1 | 0.826 | 1 | 0.576 | 0.652 | 1 | 0.935 | 1 | 1 | -0.064 |
| SE | 59605 | ENSG00000085377 | PREP | chr6 | - | 105376124 | 105376255 | 105373368 | 105373578 | 105377385 | 105377519 | 176 | 106 | 150 | 123 | 225 | 0 | 0 | 0 | 1 | 3 | 112 | 154 | 96 | 156 | 84 | 0 | 0 | 3 | 6 | 0 | 280 | 149 | 0.68153297 | 1 | 1 | 1 | 1 | 0.985 | 0.976 | 1 | 1 | 0.945 | 0.933 | 1 | 0.017 |
| SE | 27303 | ENSG00000136813 | ECPAS | chr9 | - | 111413894 | 111413986 | 111412013 | 111412148 | 111414428 | 111414651 | 253 | 181 | 169 | 241 | 145 | 2 | 0 | 0 | 0 | 0 | 244 | 223 | 298 | 233 | 131 | 0 | 5 | 3 | 4 | 0 | 241 | 149 | 0.681699876 | 1 | 0.987 | 1 | 1 | 1 | 1 | 1 | 0.965 | 0.984 | 0.973 | 1 | 0.013 |
| SE | 307 | ENSG00000157557 | ETS2 | chr21 | + | 38818424 | 38818646 | 38817007 | 38817091 | 38819502 | 38819766 | 182 | 115 | 117 | 117 | 173 | 1 | 0 | 0 | 0 | 0 | 137 | 196 | 564 | 257 | 249 | 0 | 0 | 2 | 10 | 0 | 298 | 149 | 0.681703445 | 1 | 0.989 | 1 | 1 | 1 | 1 | 1 | 1 | 0.993 | 0.928 | 1 | 0.014 |
| SE | 102508 | ENSG00000146826 | TRAPPC14 | chr7 | - | 100156385 | 100156549 | 100155670 | 100155825 | 100156633 | 100156716 | 29 | 16 | 55 | 36 | 45 | 1 | 0 | 0 | 0 | 0 | 67 | 37 | 23 | 23 | 37 | 0 | 5 | 0 | 0 | 0 | 298 | 149 | 0.681807727 | 1 | 1 | 1 | 1 | 1 | 1 | 1 | 0.787 | 1 | 1 | 1 | 0.043 |
| SE | 83010 | ENSG00000106460 | TMEM106B | chr7 | + | 12229678 | 12229819 | 12218457 | 12218521 | 12230388 | 12230438 | 169 | 132 | 193 | 180 | 452 | 0 | 0 | 2 | 2 | 8 | 48 | 47 | 20 | 13 | 11 | 0 | 0 | 3 | 0 | 0 | 290 | 149 | 0.681992107 | 1 | 1 | 1 | 0.98 | 0.979 | 0.967 | 1 | 1 | 0.774 | 1 | 1 | 0.03 |
| SE | 11833 | ENSG00000217555 | CKLF | chr16 | + | 66558189 | 66558348 | 66552568 | 66552793 | 66563121 | 66563217 | 289 | 93 | 254 | 248 | 230 | 60 | 27 | 41 | 64 | 38 | 145 | 188 | 184 | 221 | 173 | 29 | 39 | 21 | 53 | 72 | 298 | 149 | 0.682032494 | 1 | 0.707 | 0.633 | 0.756 | 0.66 | 0.752 | 0.714 | 0.707 | 0.814 | 0.676 | 0.546 | 0.01 |
| SE | 48407 | ENSG00000079335 | CDC14A | chr1 | + | 100455404 | 100455492 | 100442933 | 100442996 | 100462650 | 100462881 | 167 | 149 | 149 | 150 | 243 | 4 | 0 | 2 | 1 | 4 | 22 | 101 | 167 | 52 | 45 | 0 | 0 | 2 | 0 | 0 | 237 | 149 | 0.682147701 | 1 | 0.963 | 1 | 0.979 | 0.99 | 0.974 | 1 | 1 | 0.981 | 1 | 1 | -0.015 |
| SE | 71439 | ENSG00000115234 | SNX17 | chr2 | + | 27375060 | 27375153 | 27374688 | 27374758 | 27375505 | 27375521 | 362 | 162 | 263 | 238 | 405 | 3 | 0 | 2 | 0 | 0 | 258 | 285 | 145 | 228 | 422 | 5 | 0 | 0 | 1 | 9 | 242 | 149 | 0.682343991 | 1 | 0.987 | 1 | 0.988 | 1 | 1 | 0.969 | 1 | 1 | 0.993 | 0.967 | 0.009 |
| SE | 109444 | ENSG00000136146 | MED4 | chr13 | - | 48083370 | 48083428 | 48081644 | 48081731 | 48086281 | 48086452 | 166 | 128 | 170 | 179 | 204 | 3 | 0 | 3 | 1 | 5 | 63 | 144 | 114 | 55 | 35 | 0 | 0 | 2 | 8 | 0 | 207 | 149 | 0.682402436 | 1 | 0.976 | 1 | 0.976 | 0.992 | 0.967 | 1 | 1 | 0.976 | 0.832 | 1 | 0.021 |
| SE | 64215 | ENSG00000117697 | NSL1 | chr1 | - | 212782371 | 212782426 | 212739533 | 212739601 | 212784362 | 212784493 | 159 | 147 | 150 | 140 | 359 | 2 | 0 | 0 | 1 | 9 | 58 | 165 | 80 | 91 | 22 | 0 | 0 | 1 | 3 | 0 | 204 | 149 | 0.682421738 | 1 | 0.983 | 1 | 1 | 0.99 | 0.967 | 1 | 1 | 0.983 | 0.957 | 1 | 0 |
| SE | 67445 | ENSG00000152359 | POC5 | chr5 | - | 75692395 | 75692500 | 75690382 | 75690562 | 75694654 | 75694831 | 36 | 40 | 60 | 44 | 76 | 3 | 1 | 0 | 3 | 5 | 4 | 33 | 13 | 9 | 13 | 0 | 0 | 7 | 1 | 0 | 254 | 149 | 0.682437484 | 1 | 0.876 | 0.959 | 1 | 0.896 | 0.899 | 1 | 1 | 0.521 | 0.841 | 1 | 0.054 |
| SE | 21692 | ENSG00000135341 | MAP3K7 | chr6 | - | 90568557 | 90568623 | 90561621 | 90561667 | 90571696 | 90571807 | 151 | 95 | 119 | 133 | 136 | 3 | 1 | 2 | 0 | 0 | 27 | 73 | 46 | 64 | 28 | 0 | 1 | 4 | 1 | 0 | 215 | 149 | 0.682464497 | 1 | 0.972 | 0.985 | 0.976 | 1 | 1 | 1 | 0.981 | 0.889 | 0.978 | 1 | 0.017 |
| SE | 85108 | ENSG00000147586 | MRPS28 | chr8 | - | 80028694 | 80028796 | 80002998 | 80003180 | 80030035 | 80030289 | 3 | 4 | 5 | 2 | 6 | 25 | 15 | 18 | 20 | 22 | 0 | 8 | 3 | 9 | 1 | 1 | 22 | 31 | 16 | 11 | 251 | 149 | 0.682597885 | 1 | 0.066 | 0.137 | 0.142 | 0.056 | 0.139 | 0 | 0.178 | 0.054 | 0.25 | 0.051 | 0.001 |
| SE | 23279 | ENSG00000145604 | SKP2 | chr5 | + | 36171602 | 36171733 | 36170343 | 36170442 | 36176964 | 36177016 | 178 | 94 | 115 | 131 | 162 | 0 | 0 | 0 | 0 | 0 | 51 | 82 | 76 | 72 | 27 | 6 | 0 | 0 | 0 | 0 | 280 | 149 | 0.682703592 | 1 | 1 | 1 | 1 | 1 | 1 | 0.819 | 1 | 1 | 1 | 1 | 0.036 |
| SE | 19424 | ENSG00000151445 | VIPAS39 | chr14 | - | 77435258 | 77435393 | 77434261 | 77434303 | 77437807 | 77437881 | 41 | 21 | 32 | 40 | 36 | 0 | 0 | 0 | 0 | 5 | 64 | 43 | 42 | 27 | 39 | 0 | 0 | 0 | 0 | 0 | 284 | 149 | 0.682733047 | 1 | 1 | 1 | 1 | 1 | 0.791 | 1 | 1 | 1 | 1 | 1 | -0.042 |
| SE | 30951 | ENSG00000017260 | ATP2C1 | chr3 | + | 131001219 | 131001314 | 130999517 | 130999659 | 131016151 | 131016684 | 125 | 91 | 95 | 82 | 141 | 0 | 0 | 3 | 2 | 4 | 51 | 63 | 108 | 56 | 53 | 5 | 3 | 0 | 0 | 0 | 244 | 149 | 0.68276434 | 1 | 1 | 1 | 0.951 | 0.962 | 0.956 | 0.862 | 0.928 | 1 | 1 | 1 | 0.016 |
| SE | 61031 | ENSG00000066027 | PPP2R5A | chr1 | + | 212329134 | 212329331 | 212285409 | 212286291 | 212333496 | 212333598 | 345 | 134 | 239 | 240 | 409 | 6 | 1 | 3 | 2 | 0 | 81 | 208 | 183 | 240 | 87 | 0 | 0 | 0 | 0 | 0 | 298 | 149 | 0.682798829 | 1 | 0.966 | 0.985 | 0.976 | 0.984 | 1 | 1 | 1 | 1 | 1 | 1 | -0.018 |
| SE | 46815 | ENSG00000197780 | TAF13 | chr1 | - | 109066134 | 109066232 | 109064139 | 109064693 | 109074986 | 109075065 | 221 | 187 | 160 | 287 | 229 | 1 | 1 | 2 | 4 | 3 | 164 | 141 | 293 | 171 | 52 | 0 | 0 | 4 | 1 | 0 | 247 | 149 | 0.682829507 | 1 | 0.993 | 0.991 | 0.98 | 0.977 | 0.979 | 1 | 1 | 0.978 | 0.99 | 1 | -0.01 |
| SE | 43474 | ENSG00000072786 | STK10 | chr5 | - | 172127372 | 172127421 | 172117480 | 172117630 | 172156623 | 172156725 | 563 | 287 | 572 | 627 | 429 | 2 | 0 | 0 | 4 | 0 | 643 | 675 | 420 | 500 | 996 | 0 | 3 | 5 | 16 | 0 | 198 | 149 | 0.682898587 | 1 | 0.995 | 1 | 1 | 0.992 | 1 | 1 | 0.994 | 0.984 | 0.959 | 1 | 0.01 |
| SE | 53974 | ENSG00000188641 | DPYD | chr1 | - | 97515725 | 97515941 | 97450058 | 97450223 | 97549559 | 97549744 | 559 | 511 | 487 | 587 | 670 | 7 | 10 | 2 | 1 | 1 | 289 | 408 | 240 | 337 | 272 | 0 | 3 | 0 | 0 | 0 | 298 | 149 | 0.682905015 | 1 | 0.976 | 0.962 | 0.992 | 0.997 | 0.997 | 1 | 0.986 | 1 | 1 | 1 | -0.012 |
| SE | 10222 | ENSG00000167182 | SP2 | chr17 | + | 47914725 | 47914861 | 47896235 | 47896293 | 47915311 | 47915388 | 1 | 0 | 0 | 1 | 0 | 16 | 4 | 18 | 27 | 7 | 0 | 6 | 0 | 0 | 0 | 2 | 18 | 21 | 21 | 44 | 285 | 149 | 0.683006594 | 1 | 0.032 | 0 | 0 | 0.019 | 0 | 0 | 0.148 | 0 | 0 | 0 | -0.019 |
| SE | 65164 | ENSG00000196912 | ANKRD36B | chr2 | - | 97580461 | 97580568 | 97560831 | 97560860 | 97584943 | 97585117 | 83 | 66 | 97 | 96 | 106 | 51 | 29 | 39 | 70 | 46 | 16 | 49 | 37 | 29 | 17 | 19 | 11 | 7 | 29 | 22 | 256 | 149 | 0.683016622 | 1 | 0.486 | 0.57 | 0.591 | 0.444 | 0.573 | 0.329 | 0.722 | 0.755 | 0.368 | 0.31 | 0.036 |
| SE | 32300 | ENSG00000241553 | ARPC4 | chr3 | + | 9803535 | 9803707 | 9801660 | 9801756 | 9803842 | 9804013 | 25 | 2 | 8 | 19 | 24 | 1027 | 586 | 757 | 915 | 1450 | 23 | 3 | 24 | 33 | 12 | 1604 | 953 | 733 | 1029 | 2549 | 298 | 149 | 0.68305743 | 1 | 0.012 | 0.002 | 0.005 | 0.01 | 0.008 | 0.007 | 0.002 | 0.016 | 0.016 | 0.002 | -0.001 |
| SE | 33284 | ENSG00000214135 | NA | chr3 | - | 197622186 | 197622330 | 197621703 | 197621868 | 197623219 | 197623382 | 45 | 34 | 49 | 64 | 21 | 2 | 5 | 5 | 17 | 3 | 42 | 47 | 47 | 43 | 62 | 7 | 6 | 6 | 16 | 4 | 293 | 149 | 0.683111068 | 1 | 0.92 | 0.776 | 0.833 | 0.657 | 0.781 | 0.753 | 0.799 | 0.799 | 0.577 | 0.887 | 0.03 |
| SE | 38522 | ENSG00000130024 | PHF10 | chr6 | - | 169714732 | 169714842 | 169712385 | 169712539 | 169715707 | 169715857 | 187 | 182 | 274 | 278 | 507 | 2 | 5 | 1 | 2 | 3 | 72 | 102 | 167 | 57 | 101 | 0 | 0 | 1 | 0 | 3 | 259 | 149 | 0.683148863 | 1 | 0.982 | 0.954 | 0.994 | 0.988 | 0.99 | 1 | 1 | 0.99 | 1 | 0.951 | -0.007 |
| SE | 93710 | ENSG00000197329 | PELI1 | chr2 | - | 64104553 | 64104597 | 64100397 | 64100499 | 64104700 | 64104830 | 9 | 9 | 0 | 6 | 9 | 146 | 144 | 105 | 95 | 88 | 0 | 9 | 31 | 15 | 0 | 68 | 150 | 730 | 169 | 19 | 193 | 149 | 0.683183268 | 1 | 0.045 | 0.046 | 0 | 0.046 | 0.073 | 0 | 0.044 | 0.032 | 0.064 | 0 | 0.014 |
| SE | 95888 | ENSG00000080298 | RFX3 | chr9 | - | 3271002 | 3271118 | 3266207 | 3266305 | 3275499 | 3275612 | 31 | 19 | 33 | 45 | 79 | 0 | 0 | 5 | 0 | 0 | 39 | 11 | 41 | 25 | 35 | 0 | 0 | 0 | 0 | 0 | 265 | 149 | 0.683189947 | 1 | 1 | 1 | 0.788 | 1 | 1 | 1 | 1 | 1 | 1 | 1 | -0.042 |
| SE | 80746 | ENSG00000004864 | SLC25A13 | chr7 | - | 96189580 | 96189674 | 96189293 | 96189378 | 96191108 | 96191247 | 9 | 7 | 79 | 17 | 14 | 0 | 8 | 2 | 0 | 2 | 7 | 23 | 17 | 20 | 13 | 0 | 0 | 0 | 8 | 0 | 243 | 149 | 0.683206956 | 1 | 1 | 0.349 | 0.96 | 1 | 0.811 | 1 | 1 | 1 | 0.605 | 1 | -0.097 |
| SE | 82295 | ENSG00000122566 | HNRNPA2B1 | chr7 | - | 26191421 | 26192338 | 26190992 | 26191128 | 26192494 | 26192577 | 1157 | 964 | 801 | 1059 | 898 | 138 | 97 | 161 | 181 | 195 | 306 | 602 | 793 | 944 | 156 | 74 | 125 | 102 | 88 | 38 | 298 | 149 | 0.683319067 | 1 | 0.807 | 0.832 | 0.713 | 0.745 | 0.697 | 0.674 | 0.707 | 0.795 | 0.843 | 0.672 | 0.021 |
| SE | 14517 | ENSG00000185596 | WASH3P | chr15 | + | 101972595 | 101972694 | 101965952 | 101966223 | 101972900 | 101973047 | 123 | 29 | 68 | 51 | 49 | 5 | 1 | 1 | 3 | 0 | 70 | 86 | 36 | 116 | 136 | 9 | 1 | 0 | 0 | 0 | 248 | 149 | 0.683438179 | 1 | 0.937 | 0.946 | 0.976 | 0.911 | 1 | 0.824 | 0.981 | 1 | 1 | 1 | -0.007 |
| SE | 21018 | ENSG00000126214 | KLC1 | chr14 | + | 103687080 | 103687211 | 103679308 | 103679545 | 103700654 | 103700727 | 35 | 14 | 13 | 37 | 24 | 29 | 15 | 24 | 26 | 21 | 21 | 12 | 12 | 46 | 44 | 12 | 23 | 28 | 11 | 31 | 280 | 149 | 0.683526428 | 1 | 0.391 | 0.332 | 0.224 | 0.431 | 0.378 | 0.482 | 0.217 | 0.186 | 0.69 | 0.43 | -0.05 |
| SE | 69088 | ENSG00000136536 | MARCHF7 | chr2 | + | 159743060 | 159743253 | 159729008 | 159729175 | 159745769 | 159745937 | 1031 | 761 | 867 | 973 | 1645 | 3 | 4 | 1 | 4 | 5 | 97 | 496 | 466 | 188 | 38 | 0 | 5 | 7 | 0 | 0 | 298 | 149 | 0.683567996 | 1 | 0.994 | 0.99 | 0.998 | 0.992 | 0.994 | 1 | 0.98 | 0.971 | 1 | 1 | 0.003 |
| SE | 88532 | ENSG00000138378 | STAT4 | chr2 | - | 191068059 | 191068594 | 191066429 | 191066515 | 191069692 | 191069771 | 10 | 6 | 6 | 7 | 1 | 306 | 209 | 242 | 271 | 280 | 0 | 6 | 4 | 3 | 0 | 31 | 134 | 111 | 22 | 39 | 298 | 149 | 0.683708241 | 1 | 0.016 | 0.014 | 0.012 | 0.013 | 0.002 | 0 | 0.022 | 0.018 | 0.064 | 0 | -0.009 |
| SE | 6317 | ENSG00000060069 | CTDP1 | chr18 | + | 79728906 | 79729069 | 79717809 | 79718016 | 79753651 | 79754503 | 49 | 38 | 31 | 29 | 29 | 0 | 2 | 1 | 9 | 3 | 19 | 40 | 29 | 25 | 31 | 0 | 1 | 6 | 2 | 6 | 298 | 149 | 0.683715835 | 1 | 1 | 0.905 | 0.939 | 0.617 | 0.829 | 1 | 0.952 | 0.707 | 0.862 | 0.721 | 0.01 |
| SE | 55647 | ENSG00000114491 | UMPS | chr3 | + | 124742151 | 124742266 | 124738624 | 124738735 | 124743914 | 124744070 | 21 | 17 | 17 | 36 | 40 | 0 | 0 | 0 | 0 | 0 | 0 | 19 | 10 | 4 | 13 | 2 | 0 | 0 | 0 | 0 | 264 | 149 | 0.6838368 | 1 | 1 | 1 | 1 | 1 | 1 | 0 | 1 | 1 | 1 | 1 | 0.2 |
| SE | 71994 | ENSG00000197635 | DPP4 | chr2 | - | 162038948 | 162039021 | 162038301 | 162038422 | 162045531 | 162045612 | 11 | 10 | 18 | 14 | 38 | 0 | 0 | 1 | 4 | 0 | 10 | 5 | 2 | 3 | 1 | 0 | 0 | 0 | 0 | 0 | 222 | 149 | 0.683986321 | 1 | 1 | 1 | 0.924 | 0.701 | 1 | 1 | 1 | 1 | 1 | 1 | -0.075 |
| SE | 88275 | ENSG00000147471 | PLPBP | chr8 | + | 37775917 | 37776016 | 37774935 | 37775481 | 37777972 | 37778080 | 222 | 150 | 201 | 213 | 277 | 4 | 2 | 0 | 0 | 5 | 334 | 120 | 71 | 131 | 298 | 0 | 0 | 0 | 0 | 0 | 248 | 149 | 0.683998646 | 1 | 0.971 | 0.978 | 1 | 1 | 0.971 | 1 | 1 | 1 | 1 | 1 | -0.016 |
| SE | 69397 | ENSG00000135636 | DYSF | chr2 | + | 71669111 | 71669207 | 71667375 | 71667515 | 71669604 | 71669746 | 31 | 41 | 22 | 18 | 44 | 0 | 0 | 0 | 0 | 0 | 119 | 106 | 43 | 89 | 145 | 0 | 0 | 0 | 0 | 9 | 245 | 149 | 0.684005575 | 1 | 1 | 1 | 1 | 1 | 1 | 1 | 1 | 1 | 1 | 0.907 | 0.019 |
| SE | 52752 | ENSG00000136682 | CBWD2 | chr2 | + | 113439019 | 113439117 | 113437790 | 113438022 | 113441350 | 113441437 | 61 | 36 | 63 | 69 | 29 | 106 | 112 | 71 | 111 | 84 | 59 | 67 | 60 | 61 | 37 | 129 | 103 | 111 | 205 | 54 | 247 | 149 | 0.684095598 | 1 | 0.258 | 0.162 | 0.349 | 0.273 | 0.172 | 0.216 | 0.282 | 0.246 | 0.152 | 0.292 | 0.005 |
| SE | 4270 | ENSG00000081665 | ZNF506 | chr19 | - | 19806030 | 19806126 | 19792710 | 19795660 | 19806941 | 19807068 | 71 | 69 | 116 | 124 | 122 | 2 | 2 | 2 | 4 | 3 | 29 | 56 | 50 | 31 | 22 | 0 | 0 | 4 | 0 | 5 | 245 | 149 | 0.684152337 | 1 | 0.956 | 0.955 | 0.972 | 0.95 | 0.961 | 1 | 1 | 0.884 | 1 | 0.728 | 0.036 |
| SE | 22483 | ENSG00000165097 | KDM1B | chr6 | + | 18187791 | 18188002 | 18171362 | 18171479 | 18191196 | 18191381 | 55 | 46 | 31 | 67 | 70 | 0 | 0 | 0 | 0 | 1 | 66 | 53 | 47 | 144 | 34 | 0 | 3 | 0 | 3 | 0 | 298 | 149 | 0.684208334 | 1 | 1 | 1 | 1 | 1 | 0.972 | 1 | 0.898 | 1 | 0.96 | 1 | 0.023 |
| SE | 79789 | ENSG00000106034 | CPED1 | chr7 | + | 121015664 | 121015848 | 120989050 | 120989870 | 121046886 | 121046993 | 42 | 18 | 30 | 30 | 21 | 1 | 4 | 0 | 0 | 0 | 28 | 26 | 10 | 32 | 51 | 0 | 0 | 0 | 0 | 0 | 298 | 149 | 0.684227114 | 1 | 0.955 | 0.692 | 1 | 1 | 1 | 1 | 1 | 1 | 1 | 1 | -0.071 |
| SE | 22083 | ENSG00000111859 | NEDD9 | chr6 | - | 11200270 | 11200421 | 11193590 | 11193692 | 11213280 | 11213727 | 0 | 0 | 0 | 0 | 0 | 72 | 45 | 64 | 69 | 54 | 0 | 0 | 4 | 7 | 0 | 97 | 59 | 81 | 42 | 36 | 298 | 149 | 0.684312038 | 1 | 0 | 0 | 0 | 0 | 0 | 0 | 0 | 0.024 | 0.077 | 0 | -0.02 |
| SE | 75098 | ENSG00000198948 | MFAP3L | chr4 | - | 170006602 | 170006736 | 170004990 | 170006010 | 170026233 | 170026339 | 13 | 20 | 17 | 49 | 42 | 0 | 2 | 0 | 0 | 0 | 41 | 24 | 12 | 35 | 61 | 5 | 0 | 0 | 0 | 0 | 283 | 149 | 0.684348778 | 1 | 1 | 0.84 | 1 | 1 | 1 | 0.812 | 1 | 1 | 1 | 1 | 0.006 |
| SE | 51199 | ENSG00000121486 | TRMT1L | chr1 | - | 185139366 | 185139579 | 185137605 | 185137796 | 185139972 | 185140222 | 371 | 262 | 281 | 241 | 469 | 4 | 3 | 1 | 3 | 2 | 62 | 92 | 111 | 59 | 53 | 0 | 0 | 2 | 0 | 0 | 298 | 149 | 0.684351639 | 1 | 0.979 | 0.978 | 0.993 | 0.976 | 0.992 | 1 | 1 | 0.965 | 1 | 1 | -0.009 |
| SE | 19179 | ENSG00000140105 | WARS1 | chr14 | - | 100374135 | 100374179 | 100361707 | 100361921 | 100375282 | 100375347 | 0 | 0 | 5 | 0 | 0 | 17 | 31 | 10 | 16 | 14 | 0 | 0 | 0 | 0 | 4 | 61 | 11 | 11 | 30 | 53 | 193 | 149 | 0.684423585 | 1 | 0 | 0 | 0.279 | 0 | 0 | 0 | 0 | 0 | 0 | 0.055 | 0.045 |
| SE | 89710 | ENSG00000189233 | NUGGC | chr8 | - | 28067513 | 28067744 | 28064521 | 28064731 | 28068215 | 28068438 | 64 | 23 | 52 | 48 | 56 | 0 | 0 | 8 | 0 | 0 | 85 | 79 | 119 | 56 | 58 | 6 | 0 | 0 | 4 | 0 | 298 | 149 | 0.684536105 | 1 | 1 | 1 | 0.765 | 1 | 1 | 0.876 | 1 | 1 | 0.875 | 1 | 0.003 |
| SE | 31613 | ENSG00000168026 | TTC21A | chr3 | + | 39110028 | 39110139 | 39109084 | 39109214 | 39110850 | 39111017 | 17 | 0 | 14 | 5 | 8 | 0 | 2 | 0 | 0 | 0 | 29 | 11 | 10 | 14 | 10 | 0 | 0 | 0 | 0 | 0 | 260 | 149 | 0.684588602 | 1 | 1 | 0 | 1 | 1 | 1 | 1 | 1 | 1 | 1 | 1 | -0.2 |
| SE | 79943 | ENSG00000158169 | FANCC | chr9 | - | 95186500 | 95186595 | 95172036 | 95172147 | 95240648 | 95240743 | 1 | 1 | 0 | 2 | 2 | 10 | 9 | 4 | 13 | 30 | 0 | 0 | 0 | 0 | 0 | 12 | 2 | 7 | 6 | 3 | 244 | 149 | 0.684593634 | 1 | 0.058 | 0.064 | 0 | 0.086 | 0.039 | 0 | 0 | 0 | 0 | 0 | 0.049 |
| SE | 25116 | ENSG00000153310 | CYRIB | chr8 | - | 129896748 | 129896854 | 129871374 | 129871496 | 129903311 | 129903350 | 12 | 6 | 5 | 2 | 0 | 19 | 24 | 21 | 42 | 15 | 0 | 6 | 5 | 9 | 1 | 14 | 29 | 28 | 27 | 17 | 255 | 149 | 0.68467445 | 1 | 0.27 | 0.127 | 0.122 | 0.027 | 0 | 0 | 0.108 | 0.094 | 0.163 | 0.033 | 0.03 |
| SE | 31323 | ENSG00000143294 | PRCC | chr1 | + | 156791066 | 156791162 | 156786890 | 156787174 | 156791696 | 156791792 | 10 | 2 | 11 | 6 | 17 | 69 | 22 | 47 | 63 | 44 | 8 | 23 | 10 | 21 | 8 | 69 | 47 | 59 | 65 | 118 | 245 | 149 | 0.684696292 | 1 | 0.081 | 0.052 | 0.125 | 0.055 | 0.19 | 0.066 | 0.229 | 0.093 | 0.164 | 0.04 | -0.018 |
| SE | 61389 | ENSG00000163453 | IGFBP7 | chr4 | - | 57032425 | 57032552 | 57030772 | 57031336 | 57033194 | 57033311 | 226 | 65 | 48 | 73 | 102 | 0 | 0 | 0 | 0 | 0 | 350 | 151 | 76 | 248 | 207 | 0 | 0 | 0 | 4 | 7 | 276 | 149 | 0.684710906 | 1 | 1 | 1 | 1 | 1 | 1 | 1 | 1 | 1 | 0.971 | 0.941 | 0.018 |
| SE | 50787 | ENSG00000082512 | TRAF5 | chr1 | + | 211359881 | 211360076 | 211356366 | 211356468 | 211360701 | 211360779 | 171 | 114 | 192 | 175 | 223 | 2 | 0 | 0 | 0 | 2 | 12 | 94 | 94 | 50 | 29 | 0 | 0 | 6 | 0 | 0 | 298 | 149 | 0.684731177 | 1 | 0.977 | 1 | 1 | 1 | 0.982 | 1 | 1 | 0.887 | 1 | 1 | 0.014 |
| SE | 74212 | ENSG00000235257 | ITGA9-AS1 | chr3 | - | 37818809 | 37818930 | 37808494 | 37808880 | 37821013 | 37821100 | 5 | 7 | 2 | 13 | 17 | 3 | 1 | 0 | 11 | 0 | 14 | 9 | 2 | 16 | 27 | 0 | 0 | 3 | 4 | 40 | 270 | 149 | 0.68482336 | 1 | 0.479 | 0.794 | 1 | 0.395 | 1 | 1 | 1 | 0.269 | 0.688 | 0.271 | 0.088 |
| SE | 70542 | ENSG00000073849 | ST6GAL1 | chr3 | + | 187038741 | 187038873 | 186996443 | 186996673 | 187042653 | 187042831 | 271 | 126 | 294 | 288 | 462 | 0 | 0 | 0 | 0 | 0 | 74 | 261 | 91 | 179 | 382 | 0 | 0 | 8 | 0 | 0 | 281 | 149 | 0.684933512 | 1 | 1 | 1 | 1 | 1 | 1 | 1 | 1 | 0.858 | 1 | 1 | 0.028 |
| SE | 20181 | ENSG00000137210 | TMEM14B | chr6 | + | 10749621 | 10749698 | 10749201 | 10749268 | 10755141 | 10755232 | 28 | 13 | 29 | 27 | 19 | 1 | 0 | 3 | 1 | 0 | 38 | 28 | 17 | 15 | 33 | 0 | 1 | 0 | 5 | 1 | 226 | 149 | 0.684972655 | 1 | 0.949 | 1 | 0.864 | 0.947 | 1 | 1 | 0.949 | 1 | 0.664 | 0.956 | 0.038 |
| SE | 2109 | ENSG00000127616 | SMARCA4 | chr19 | + | 11026299 | 11026346 | 11025421 | 11025508 | 11027783 | 11027861 | 66 | 29 | 75 | 89 | 81 | 0 | 0 | 0 | 0 | 0 | 106 | 126 | 77 | 48 | 259 | 0 | 0 | 0 | 7 | 0 | 196 | 149 | 0.684990711 | 1 | 1 | 1 | 1 | 1 | 1 | 1 | 1 | 1 | 0.839 | 1 | 0.032 |
| SE | 72720 | ENSG00000172765 | TMCC1 | chr3 | - | 129860939 | 129861012 | 129848345 | 129848412 | 129880308 | 129880559 | 19 | 29 | 17 | 23 | 37 | 49 | 26 | 49 | 44 | 106 | 36 | 19 | 80 | 33 | 0 | 38 | 51 | 119 | 45 | 13 | 222 | 149 | 0.68511987 | 1 | 0.207 | 0.428 | 0.189 | 0.26 | 0.19 | 0.389 | 0.2 | 0.311 | 0.33 | 0 | 0.009 |
| SE | 98824 | ENSG00000135365 | PHF21A | chr11 | - | 45938156 | 45938312 | 45936493 | 45936569 | 45945839 | 45946003 | 138 | 83 | 91 | 129 | 196 | 3 | 6 | 11 | 8 | 8 | 313 | 179 | 156 | 335 | 224 | 0 | 13 | 15 | 37 | 9 | 298 | 149 | 0.685156353 | 1 | 0.958 | 0.874 | 0.805 | 0.89 | 0.925 | 1 | 0.873 | 0.839 | 0.819 | 0.926 | -0.001 |
| SE | 74695 | ENSG00000197548 | ATG7 | chr3 | + | 11332971 | 11333093 | 11331339 | 11331428 | 11340644 | 11340735 | 131 | 66 | 81 | 95 | 172 | 4 | 3 | 6 | 4 | 2 | 173 | 162 | 57 | 130 | 147 | 15 | 4 | 0 | 3 | 0 | 271 | 149 | 0.685204891 | 1 | 0.947 | 0.924 | 0.881 | 0.929 | 0.979 | 0.864 | 0.957 | 1 | 0.96 | 1 | -0.024 |
| SE | 47645 | ENSG00000288701 | PRRC2B | chr9 | + | 131478473 | 131478619 | 131477743 | 131477949 | 131481725 | 131481808 | 277 | 155 | 210 | 323 | 245 | 0 | 0 | 0 | 1 | 0 | 342 | 329 | 224 | 224 | 340 | 0 | 7 | 1 | 0 | 6 | 295 | 149 | 0.68528156 | 1 | 1 | 1 | 1 | 0.994 | 1 | 1 | 0.96 | 0.991 | 1 | 0.966 | 0.015 |
| SE | 40412 | ENSG00000078304 | PPP2R5C | chr14 | + | 101818991 | 101819085 | 101809904 | 101810036 | 101856685 | 101856885 | 3 | 1 | 5 | 3 | 17 | 339 | 183 | 264 | 290 | 231 | 0 | 6 | 5 | 0 | 0 | 153 | 254 | 207 | 173 | 22 | 243 | 149 | 0.685303298 | 1 | 0.005 | 0.003 | 0.011 | 0.006 | 0.043 | 0 | 0.014 | 0.015 | 0 | 0 | 0.008 |
| SE | 14940 | ENSG00000159216 | RUNX1 | chr21 | - | 34892924 | 34892963 | 34880556 | 34880713 | 35048841 | 35048958 | 21 | 9 | 21 | 20 | 17 | 2 | 0 | 0 | 1 | 3 | 16 | 30 | 26 | 15 | 13 | 0 | 0 | 1 | 0 | 0 | 188 | 149 | 0.685351783 | 1 | 0.893 | 1 | 1 | 0.941 | 0.818 | 1 | 1 | 0.954 | 1 | 1 | -0.06 |
| SE | 6066 | ENSG00000101310 | SEC23B | chr20 | + | 18542295 | 18542402 | 18535652 | 18535742 | 18545955 | 18546033 | 160 | 107 | 122 | 132 | 197 | 2 | 1 | 1 | 1 | 5 | 218 | 176 | 122 | 118 | 203 | 0 | 4 | 0 | 0 | 0 | 256 | 149 | 0.685359438 | 1 | 0.979 | 0.984 | 0.986 | 0.987 | 0.958 | 1 | 0.962 | 1 | 1 | 1 | -0.014 |
| SE | 11394 | ENSG00000074356 | NCBP3 | chr17 | - | 3814321 | 3814483 | 3802157 | 3813279 | 3816115 | 3816270 | 261 | 167 | 284 | 313 | 445 | 0 | 1 | 1 | 0 | 12 | 322 | 233 | 243 | 412 | 103 | 0 | 3 | 1 | 1 | 0 | 298 | 149 | 0.685392152 | 1 | 1 | 0.988 | 0.993 | 1 | 0.949 | 1 | 0.975 | 0.992 | 0.995 | 1 | -0.006 |
| SE | 46220 | ENSG00000114439 | BBX | chr3 | + | 107798522 | 107798720 | 107791239 | 107791299 | 107801094 | 107801281 | 458 | 362 | 401 | 433 | 766 | 0 | 2 | 0 | 0 | 2 | 56 | 447 | 479 | 128 | 12 | 6 | 0 | 0 | 0 | 0 | 298 | 149 | 0.68541186 | 1 | 1 | 0.989 | 1 | 1 | 0.995 | 0.824 | 1 | 1 | 1 | 1 | 0.032 |
| SE | 52734 | ENSG00000143970 | ASXL2 | chr2 | - | 25753533 | 25753639 | 25749695 | 25750413 | 25756017 | 25756114 | 250 | 200 | 230 | 191 | 310 | 11 | 15 | 8 | 19 | 15 | 137 | 234 | 123 | 164 | 48 | 0 | 32 | 8 | 12 | 0 | 255 | 149 | 0.685449265 | 1 | 0.93 | 0.886 | 0.944 | 0.855 | 0.924 | 1 | 0.81 | 0.9 | 0.889 | 1 | -0.012 |
| SE | 31396 | ENSG00000123933 | MXD4 | chr4 | - | 2257981 | 2258011 | 2252407 | 2252522 | 2261724 | 2261799 | 171 | 50 | 56 | 83 | 37 | 0 | 0 | 0 | 0 | 0 | 124 | 106 | 76 | 54 | 209 | 0 | 0 | 0 | 0 | 9 | 179 | 149 | 0.685452174 | 1 | 1 | 1 | 1 | 1 | 1 | 1 | 1 | 1 | 1 | 0.951 | 0.01 |
| SE | 94835 | ENSG00000126882 | FAM78A | chr9 | - | 131263477 | 131263599 | 131258075 | 131261350 | 131275856 | 131276510 | 2 | 3 | 10 | 4 | 4 | 221 | 100 | 143 | 208 | 198 | 0 | 3 | 0 | 0 | 0 | 181 | 145 | 84 | 92 | 162 | 271 | 149 | 0.685471703 | 1 | 0.005 | 0.016 | 0.037 | 0.01 | 0.011 | 0 | 0.011 | 0 | 0 | 0 | 0.014 |
| SE | 37084 | ENSG00000127481 | UBR4 | chr1 | - | 19110095 | 19110223 | 19106836 | 19106966 | 19110379 | 19110464 | 216 | 130 | 203 | 266 | 295 | 0 | 0 | 3 | 0 | 0 | 394 | 376 | 144 | 263 | 361 | 11 | 0 | 0 | 4 | 0 | 277 | 149 | 0.68559639 | 1 | 1 | 1 | 0.973 | 1 | 1 | 0.951 | 1 | 1 | 0.973 | 1 | 0.01 |
| SE | 55989 | ENSG00000121892 | PDS5A | chr4 | - | 39874288 | 39874412 | 39872985 | 39873144 | 39876992 | 39877153 | 498 | 411 | 455 | 501 | 835 | 0 | 0 | 0 | 0 | 4 | 126 | 257 | 284 | 92 | 24 | 0 | 0 | 8 | 2 | 0 | 273 | 149 | 0.685601932 | 1 | 1 | 1 | 1 | 1 | 0.991 | 1 | 1 | 0.951 | 0.962 | 1 | 0.016 |
| SE | 88613 | ENSG00000169139 | UBE2V2 | chr8 | + | 48043032 | 48043181 | 48008430 | 48008470 | 48049852 | 48049978 | 230 | 138 | 160 | 163 | 277 | 2 | 2 | 0 | 1 | 4 | 25 | 114 | 98 | 58 | 4 | 0 | 0 | 0 | 2 | 0 | 298 | 149 | 0.685756604 | 1 | 0.983 | 0.972 | 1 | 0.988 | 0.972 | 1 | 1 | 1 | 0.935 | 1 | -0.004 |
| SE | 18138 | ENSG00000119638 | NEK9 | chr14 | - | 75110320 | 75110371 | 75107342 | 75107487 | 75113338 | 75113403 | 48 | 33 | 71 | 74 | 99 | 0 | 0 | 0 | 0 | 0 | 79 | 32 | 43 | 52 | 44 | 6 | 0 | 0 | 0 | 0 | 200 | 149 | 0.685995432 | 1 | 1 | 1 | 1 | 1 | 1 | 0.907 | 1 | 1 | 1 | 1 | 0.019 |
| SE | 84870 | ENSG00000164074 | ABHD18 | chr4 | + | 128021136 | 128021238 | 128020079 | 128020169 | 128028474 | 128028853 | 109 | 72 | 100 | 124 | 73 | 2 | 3 | 4 | 4 | 3 | 106 | 109 | 115 | 73 | 135 | 0 | 13 | 7 | 0 | 0 | 251 | 149 | 0.686000367 | 1 | 0.97 | 0.934 | 0.937 | 0.948 | 0.935 | 1 | 0.833 | 0.907 | 1 | 1 | -0.003 |
| SE | 4898 | ENSG00000124243 | BCAS4 | chr20 | + | 50875996 | 50876093 | 50841765 | 50841900 | 50876485 | 50876723 | 4 | 1 | 0 | 1 | 0 | 25 | 8 | 22 | 25 | 33 | 2 | 0 | 2 | 0 | 0 | 7 | 34 | 13 | 11 | 21 | 246 | 149 | 0.686014459 | 1 | 0.088 | 0.07 | 0 | 0.024 | 0 | 0.148 | 0 | 0.085 | 0 | 0 | -0.01 |
| SE | 38637 | ENSG00000156875 | MFSD14A | chr1 | + | 100049908 | 100050004 | 100038094 | 100038316 | 100058665 | 100058728 | 121 | 84 | 94 | 106 | 115 | 0 | 1 | 0 | 0 | 0 | 122 | 104 | 105 | 152 | 45 | 0 | 0 | 7 | 1 | 0 | 245 | 149 | 0.686092039 | 1 | 1 | 0.981 | 1 | 1 | 1 | 1 | 1 | 0.901 | 0.989 | 1 | 0.018 |
| SE | 67078 | ENSG00000204681 | GABBR1 | chr6 | - | 29630457 | 29630643 | 29629086 | 29629107 | 29631395 | 29631599 | 14 | 27 | 16 | 28 | 14 | 6 | 1 | 0 | 0 | 0 | 43 | 5 | 4 | 15 | 9 | 0 | 0 | 0 | 9 | 0 | 298 | 149 | 0.68609412 | 1 | 0.538 | 0.931 | 1 | 1 | 1 | 1 | 1 | 1 | 0.455 | 1 | 0.003 |
| SE | 105654 | ENSG00000091039 | OSBPL8 | chr12 | - | 76373343 | 76373433 | 76371447 | 76371584 | 76375272 | 76375370 | 1663 | 1795 | 1047 | 1686 | 2428 | 0 | 0 | 3 | 10 | 0 | 50 | 561 | 535 | 132 | 51 | 5 | 0 | 6 | 0 | 0 | 239 | 149 | 0.686199759 | 1 | 1 | 1 | 0.995 | 0.991 | 1 | 0.862 | 1 | 0.982 | 1 | 1 | 0.028 |
| SE | 8738 | ENSG00000073969 | NSF | chr17 | + | 46755498 | 46755597 | 46755313 | 46755369 | 46755801 | 46755837 | 3 | 0 | 1 | 2 | 0 | 78 | 35 | 41 | 33 | 80 | 0 | 4 | 3 | 0 | 0 | 9 | 70 | 21 | 76 | 6 | 248 | 149 | 0.686204154 | 1 | 0.023 | 0 | 0.014 | 0.035 | 0 | 0 | 0.033 | 0.079 | 0 | 0 | -0.008 |
| SE | 45159 | ENSG00000107099 | DOCK8 | chr9 | + | 336581 | 336718 | 334224 | 334384 | 339005 | 339099 | 1438 | 1065 | 1192 | 1378 | 1780 | 2 | 6 | 3 | 4 | 2 | 2429 | 1853 | 1320 | 1992 | 1374 | 0 | 12 | 9 | 15 | 9 | 286 | 149 | 0.686313461 | 1 | 0.997 | 0.989 | 0.995 | 0.994 | 0.998 | 1 | 0.988 | 0.987 | 0.986 | 0.988 | 0.005 |
| SE | 40086 | ENSG00000125347 | IRF1 | chr5 | - | 132489391 | 132489483 | 132486953 | 132487130 | 132490544 | 132490726 | 12 | 23 | 49 | 50 | 15 | 12 | 1 | 0 | 2 | 0 | 194 | 40 | 56 | 99 | 94 | 0 | 9 | 0 | 8 | 16 | 241 | 149 | 0.686321598 | 1 | 0.382 | 0.934 | 1 | 0.939 | 1 | 1 | 0.733 | 1 | 0.884 | 0.784 | -0.029 |
| SE | 104815 | ENSG00000136404 | TM6SF1 | chr15 | + | 83124671 | 83124776 | 83119577 | 83119681 | 83126754 | 83126818 | 22 | 24 | 6 | 24 | 26 | 0 | 0 | 0 | 0 | 1 | 44 | 15 | 2 | 8 | 2 | 0 | 4 | 0 | 0 | 0 | 254 | 149 | 0.686369606 | 1 | 1 | 1 | 1 | 1 | 0.938 | 1 | 0.687 | 1 | 1 | 1 | 0.05 |
| SE | 86084 | ENSG00000169359 | SLC33A1 | chr3 | - | 155842431 | 155842619 | 155833467 | 155834041 | 155853107 | 155854456 | 92 | 63 | 107 | 67 | 189 | 0 | 0 | 0 | 0 | 0 | 61 | 82 | 37 | 55 | 98 | 0 | 0 | 0 | 6 | 0 | 298 | 149 | 0.686417161 | 1 | 1 | 1 | 1 | 1 | 1 | 1 | 1 | 1 | 0.821 | 1 | 0.036 |
| SE | 29793 | ENSG00000050344 | NFE2L3 | chr7 | + | 26183700 | 26183784 | 26177942 | 26178122 | 26184532 | 26187137 | 125 | 71 | 70 | 72 | 192 | 2 | 0 | 1 | 0 | 0 | 99 | 54 | 28 | 60 | 19 | 3 | 4 | 0 | 0 | 0 | 233 | 149 | 0.686423629 | 1 | 0.976 | 1 | 1 | 0.979 | 1 | 0.955 | 0.896 | 1 | 1 | 1 | 0.021 |
| SE | 8722 | ENSG00000070366 | SMG6 | chr17 | - | 2297242 | 2297353 | 2292870 | 2292977 | 2297862 | 2298055 | 131 | 112 | 111 | 115 | 202 | 1 | 2 | 0 | 0 | 0 | 231 | 160 | 163 | 224 | 324 | 9 | 0 | 0 | 2 | 0 | 260 | 149 | 0.686562112 | 1 | 0.987 | 0.97 | 1 | 1 | 1 | 0.936 | 1 | 1 | 0.985 | 1 | 0.007 |
| SE | 68640 | ENSG00000166295 | ANAPC16 | chr10 | + | 72223887 | 72224056 | 72215999 | 72216072 | 72230365 | 72230440 | 593 | 360 | 680 | 550 | 547 | 10 | 2 | 15 | 12 | 0 | 536 | 515 | 450 | 442 | 670 | 17 | 3 | 12 | 3 | 6 | 298 | 149 | 0.68660918 | 1 | 0.967 | 0.989 | 0.958 | 0.958 | 1 | 0.94 | 0.988 | 0.949 | 0.987 | 0.982 | 0.005 |
| SE | 55615 | ENSG00000149474 | KAT14 | chr20 | + | 18159083 | 18159265 | 18150820 | 18150942 | 18161822 | 18162239 | 98 | 48 | 77 | 67 | 132 | 0 | 0 | 1 | 4 | 2 | 48 | 71 | 31 | 49 | 38 | 0 | 0 | 1 | 6 | 0 | 298 | 149 | 0.686609953 | 1 | 1 | 1 | 0.975 | 0.893 | 0.971 | 1 | 1 | 0.939 | 0.803 | 1 | 0.019 |
| SE | 103151 | ENSG00000186020 | ZNF529 | chr19 | - | 36556103 | 36556197 | 36554669 | 36554796 | 36572332 | 36572392 | 48 | 43 | 55 | 42 | 36 | 0 | 0 | 0 | 0 | 0 | 7 | 31 | 52 | 13 | 15 | 0 | 5 | 0 | 0 | 0 | 243 | 149 | 0.686794589 | 1 | 1 | 1 | 1 | 1 | 1 | 1 | 0.792 | 1 | 1 | 1 | 0.042 |
| SE | 105283 | ENSG00000241370 | RPP21 | chr6 | + | 30345466 | 30345573 | 30345297 | 30345398 | 30346431 | 30346557 | 23 | 9 | 1 | 9 | 14 | 0 | 0 | 0 | 0 | 0 | 0 | 7 | 4 | 5 | 36 | 2 | 0 | 0 | 0 | 0 | 256 | 149 | 0.686805229 | 1 | 1 | 1 | 1 | 1 | 1 | 0 | 1 | 1 | 1 | 1 | 0.2 |
| SE | 20773 | ENSG00000084093 | REST | chr4 | + | 56910629 | 56911536 | 56907875 | 56908213 | 56919786 | 56919870 | 316 | 206 | 241 | 276 | 456 | 2 | 4 | 1 | 2 | 3 | 18 | 215 | 244 | 102 | 47 | 0 | 3 | 0 | 0 | 0 | 298 | 149 | 0.686835126 | 1 | 0.988 | 0.963 | 0.992 | 0.986 | 0.987 | 1 | 0.973 | 1 | 1 | 1 | -0.011 |
| SE | 89575 | ENSG00000095485 | CWF19L1 | chr10 | - | 100260965 | 100261044 | 100256261 | 100256476 | 100261978 | 100262063 | 50 | 54 | 62 | 34 | 40 | 6 | 0 | 0 | 0 | 0 | 39 | 37 | 67 | 59 | 50 | 0 | 0 | 0 | 0 | 4 | 228 | 149 | 0.686947811 | 1 | 0.845 | 1 | 1 | 1 | 1 | 1 | 1 | 1 | 1 | 0.891 | -0.009 |
| SE | 44426 | ENSG00000204435 | CSNK2B | chr6 | + | 31669327 | 31669508 | 31669096 | 31669172 | 31669835 | 31670074 | 266 | 155 | 202 | 241 | 189 | 0 | 0 | 1 | 0 | 0 | 555 | 424 | 285 | 331 | 477 | 0 | 0 | 2 | 6 | 9 | 298 | 149 | 0.686984943 | 1 | 1 | 1 | 0.99 | 1 | 1 | 1 | 1 | 0.986 | 0.965 | 0.964 | 0.015 |
| SE | 21976 | ENSG00000065491 | TBC1D22B | chr6 | + | 37291242 | 37291357 | 37287006 | 37287072 | 37312917 | 37313024 | 61 | 22 | 43 | 58 | 57 | 7 | 3 | 7 | 3 | 17 | 103 | 60 | 33 | 74 | 125 | 14 | 5 | 11 | 7 | 12 | 264 | 149 | 0.686992174 | 1 | 0.831 | 0.805 | 0.776 | 0.916 | 0.654 | 0.806 | 0.871 | 0.629 | 0.856 | 0.855 | -0.007 |
| SE | 3533 | ENSG00000105483 | CARD8 | chr19 | - | 48223718 | 48223868 | 48221729 | 48221855 | 48230437 | 48230498 | 34 | 32 | 24 | 26 | 29 | 97 | 49 | 77 | 105 | 50 | 26 | 15 | 26 | 16 | 8 | 72 | 61 | 52 | 21 | 42 | 298 | 149 | 0.687040791 | 1 | 0.149 | 0.246 | 0.135 | 0.11 | 0.225 | 0.153 | 0.109 | 0.2 | 0.276 | 0.087 | 0.008 |
| SE | 63388 | ENSG00000118263 | KLF7 | chr2 | - | 207123975 | 207124404 | 207088457 | 207088581 | 207165466 | 207165706 | 30 | 25 | 50 | 48 | 71 | 0 | 0 | 0 | 0 | 0 | 38 | 49 | 75 | 58 | 27 | 5 | 0 | 0 | 0 | 0 | 298 | 149 | 0.687106232 | 1 | 1 | 1 | 1 | 1 | 1 | 0.792 | 1 | 1 | 1 | 1 | 0.042 |
| SE | 66428 | ENSG00000102081 | FMR1 | chrX | + | 147938098 | 147938161 | 147937465 | 147937600 | 147940575 | 147940662 | 66 | 60 | 46 | 63 | 84 | 144 | 126 | 149 | 186 | 193 | 8 | 62 | 35 | 31 | 5 | 41 | 53 | 143 | 87 | 21 | 212 | 149 | 0.687219479 | 1 | 0.244 | 0.251 | 0.178 | 0.192 | 0.234 | 0.121 | 0.451 | 0.147 | 0.2 | 0.143 | 0.007 |
| SE | 69946 | ENSG00000108883 | EFTUD2 | chr17 | - | 44853295 | 44853390 | 44852408 | 44852562 | 44853516 | 44853635 | 236 | 123 | 176 | 193 | 161 | 1 | 6 | 0 | 5 | 0 | 259 | 190 | 86 | 164 | 411 | 0 | 0 | 0 | 0 | 6 | 244 | 149 | 0.687246882 | 1 | 0.993 | 0.926 | 1 | 0.959 | 1 | 1 | 1 | 1 | 1 | 0.977 | -0.02 |
| SE | 68022 | ENSG00000172890 | NADSYN1 | chr11 | + | 71473568 | 71473686 | 71473277 | 71473366 | 71474394 | 71474526 | 141 | 45 | 178 | 133 | 84 | 0 | 0 | 1 | 0 | 0 | 124 | 153 | 69 | 197 | 231 | 7 | 2 | 0 | 0 | 0 | 267 | 149 | 0.687284276 | 1 | 1 | 1 | 0.99 | 1 | 1 | 0.908 | 0.977 | 1 | 1 | 1 | 0.021 |
| SE | 69754 | ENSG00000144228 | SPOPL | chr2 | + | 138554438 | 138554535 | 138552553 | 138552681 | 138559021 | 138559199 | 60 | 33 | 33 | 51 | 41 | 124 | 90 | 69 | 125 | 158 | 0 | 28 | 63 | 28 | 0 | 6 | 91 | 105 | 28 | 21 | 246 | 149 | 0.687306521 | 1 | 0.227 | 0.182 | 0.225 | 0.198 | 0.136 | 0 | 0.157 | 0.267 | 0.377 | 0 | 0.033 |
| SE | 96965 | ENSG00000183597 | TANGO2 | chr22 | + | 20055942 | 20056013 | 20052464 | 20052584 | 20063337 | 20063442 | 27 | 22 | 17 | 13 | 23 | 0 | 0 | 0 | 4 | 0 | 33 | 39 | 10 | 41 | 96 | 0 | 0 | 0 | 0 | 0 | 220 | 149 | 0.687315663 | 1 | 1 | 1 | 1 | 0.688 | 1 | 1 | 1 | 1 | 1 | 1 | -0.062 |
| SE | 61761 | ENSG00000158710 | TAGLN2 | chr1 | - | 159919273 | 159919376 | 159918106 | 159918941 | 159919660 | 159919835 | 1740 | 1142 | 1189 | 2479 | 1870 | 1 | 2 | 8 | 0 | 4 | 3471 | 2124 | 1553 | 1793 | 14994 | 28 | 1 | 6 | 20 | 10 | 252 | 149 | 0.687456056 | 1 | 0.999 | 0.997 | 0.989 | 1 | 0.996 | 0.987 | 0.999 | 0.994 | 0.981 | 0.999 | 0.004 |
| SE | 59128 | ENSG00000047410 | TPR | chr1 | - | 186353687 | 186353880 | 186352014 | 186352110 | 186355409 | 186355558 | 342 | 400 | 295 | 331 | 444 | 3 | 2 | 9 | 0 | 1 | 70 | 68 | 48 | 21 | 16 | 0 | 0 | 0 | 6 | 0 | 298 | 149 | 0.687484199 | 1 | 0.983 | 0.99 | 0.942 | 1 | 0.996 | 1 | 1 | 1 | 0.636 | 1 | 0.055 |
| SE | 33102 | ENSG00000178035 | IMPDH2 | chr3 | - | 49025994 | 49026066 | 49025125 | 49025269 | 49026323 | 49026419 | 3 | 0 | 6 | 0 | 0 | 57 | 51 | 84 | 77 | 53 | 0 | 0 | 0 | 0 | 0 | 17 | 38 | 33 | 25 | 59 | 221 | 149 | 0.687516742 | 1 | 0.034 | 0 | 0.046 | 0 | 0 | 0 | 0 | 0 | 0 | 0 | 0.016 |
| SE | 40072 | ENSG00000125347 | IRF1 | chr5 | - | 132485666 | 132485716 | 132484361 | 132484497 | 132486556 | 132486686 | 359 | 228 | 390 | 463 | 392 | 14 | 16 | 17 | 22 | 5 | 1686 | 357 | 661 | 986 | 1030 | 46 | 17 | 11 | 76 | 24 | 199 | 149 | 0.687527428 | 1 | 0.95 | 0.914 | 0.945 | 0.94 | 0.983 | 0.965 | 0.94 | 0.978 | 0.907 | 0.97 | -0.006 |
| SE | 56302 | ENSG00000134242 | PTPN22 | chr1 | - | 113852026 | 113852104 | 113848539 | 113848626 | 113854470 | 113854537 | 906 | 382 | 477 | 632 | 1010 | 35 | 9 | 7 | 12 | 14 | 365 | 410 | 301 | 301 | 264 | 0 | 20 | 4 | 2 | 10 | 227 | 149 | 0.68756165 | 1 | 0.944 | 0.965 | 0.978 | 0.972 | 0.979 | 1 | 0.931 | 0.98 | 0.99 | 0.945 | -0.002 |
| SE | 105605 | ENSG00000139323 | POC1B | chr12 | - | 89470360 | 89470494 | 89466769 | 89466922 | 89472167 | 89472275 | 4 | 5 | 2 | 12 | 15 | 0 | 3 | 0 | 0 | 0 | 9 | 9 | 1 | 12 | 5 | 0 | 0 | 0 | 0 | 0 | 283 | 149 | 0.687752572 | 1 | 1 | 0.467 | 1 | 1 | 1 | 1 | 1 | 1 | 1 | 1 | -0.107 |
| SE | 89977 | ENSG00000153317 | ASAP1 | chr8 | - | 130091972 | 130092143 | 130079901 | 130079971 | 130115627 | 130115735 | 30 | 40 | 49 | 42 | 68 | 0 | 0 | 0 | 0 | 0 | 165 | 91 | 74 | 159 | 187 | 0 | 0 | 7 | 0 | 0 | 298 | 149 | 0.687806107 | 1 | 1 | 1 | 1 | 1 | 1 | 1 | 1 | 0.841 | 1 | 1 | 0.032 |
| SE | 96906 | ENSG00000100731 | PCNX1 | chr14 | + | 71050651 | 71050760 | 71047806 | 71047984 | 71051882 | 71052012 | 336 | 210 | 353 | 331 | 557 | 0 | 0 | 0 | 2 | 5 | 103 | 262 | 381 | 175 | 151 | 8 | 0 | 4 | 0 | 0 | 258 | 149 | 0.68785318 | 1 | 1 | 1 | 1 | 0.99 | 0.985 | 0.881 | 1 | 0.982 | 1 | 1 | 0.022 |
| SE | 11041 | ENSG00000132359 | RAP1GAP2 | chr17 | + | 3005954 | 3006041 | 2998220 | 2998376 | 3008010 | 3008090 | 51 | 41 | 26 | 40 | 6 | 3 | 1 | 1 | 1 | 0 | 66 | 76 | 48 | 61 | 57 | 0 | 2 | 0 | 10 | 0 | 236 | 149 | 0.687878089 | 1 | 0.915 | 0.963 | 0.943 | 0.962 | 1 | 1 | 0.96 | 1 | 0.794 | 1 | 0.006 |

| | | | | | | | | | | | | | | | |
|---|---|---|---|---|---|---|---|---|---|---|---|---|---|---|---|
| SE | 58377 | ENSG00000171105 | INSR | chr19 | - | 7150496 | 7150532 | 7142815 | 7143090 | 7152725 | 7152927 | 0 | 0 | 0 | [illegible] |
| SE | 27390 | ENSG00000181982 | CCDC149 | chr4 | - | 24834947 | 24835032 | 24826939 | 24831650 | 24836435 | 24836508 | 27 | 18 | 26 | [illegible] |
| SE | 66152 | ENSG00000088179 | PTPN4 | chr2 | + | 119885794 | 119885882 | 119882502 | 119882623 | 119900717 | 119900806 | 169 | 115 | 160 | [illegible] |
| SE | 90393 | ENSG00000170633 | RNF34 | chr12 | + | 121402741 | 121402807 | 121400118 | 121400218 | 121416158 | 121416377 | 20 | 8 | 13 | [illegible] |
| SE | 92181 | ENSG00000122692 | SMU1 | chr9 | - | 33060464 | 33060584 | 33057597 | 33057714 | 33062048 | 33062177 | 954 | 655 | 750 | [illegible] |
| SE | 78646 | ENSG00000168014 | C2CD3 | chr11 | - | 74118227 | 74118382 | 74114383 | 74114593 | 74122987 | 74123135 | 76 | 62 | 92 | [illegible] |
| SE | 60499 | ENSG00000115211 | EIF2B4 | chr2 | - | 27367456 | 27367559 | 27367073 | 27367201 | 27369005 | 27369209 | 5 | 7 | 9 | [illegible] |
| SE | 74638 | ENSG00000134440 | NARS1 | chr18 | - | 57620568 | 57620651 | 57615640 | 57615975 | 57621707 | 57621776 | 304 | 223 | 271 | [illegible] |
| SE | 59727 | ENSG00000149231 | CCDC82 | chr11 | - | 96383961 | 96384761 | 96380538 | 96383473 | 96389843 | 96389903 | 414 | 419 | 316 | [illegible] |
| SE | 77284 | ENSG00000113643 | RARS1 | chr5 | + | 168510580 | 168510686 | 168502000 | 168502105 | 168516777 | 168516950 | 62 | 57 | 57 | [illegible] |
| SE | 87134 | ENSG00000104613 | INTS10 | chr8 | + | 19824802 | 19824972 | 19823872 | 19823981 | 19826425 | 19826559 | 183 | 101 | 163 | [illegible] |
| SE | 59936 | ENSG00000159363 | ATP13A2 | chr1 | - | 17002033 | 17002103 | 17000399 | 17000507 | 17002295 | 17002373 | 39 | 16 | 37 | [illegible] |
| SE | 102986 | ENSG00000138111 | MFSD13A | chr10 | + | 102468752 | 102468842 | 102465980 | 102466087 | 102468954 | 102469197 | 9 | 4 | 0 | [illegible] |
| SE | 104696 | ENSG00000211455 | STK38L | chr12 | + | 27317335 | 27317453 | 27315217 | 27315350 | 27317895 | 27318019 | 108 | 105 | 99 | [illegible] |
| SE | 53072 | ENSG00000103657 | HERC1 | chr15 | - | 63723181 | 63723355 | 63718782 | 63718897 | 63725291 | 63725513 | 386 | 278 | 384 | [illegible] |
| SE | 40384 | ENSG00000163719 | MTMR14 | chr3 | + | 9669431 | 9669492 | 9653620 | 9653769 | 9671047 | 9671170 | 167 | 104 | 87 | [illegible] |
| SE | 71685 | ENSG00000135976 | ANKRD36 | chr2 | + | 97124459 | 97124597 | 97122886 | 97122993 | 97127066 | 97127134 | 69 | 59 | 39 | [illegible] |
| SE | 33344 | ENSG00000188313 | PLSCR1 | chr3 | - | 146541133 | 146541180 | 146537995 | 146538044 | 146544466 | 146544521 | 0 | 3 | 0 | [illegible] |
| SE | 88453 | ENSG00000221914 | PPP2R2A | chr8 | + | 26363720 | 26363890 | 26362683 | 26362848 | 26366314 | 26366406 | 196 | 167 | 161 | [illegible] |
| SE | 77601 | ENSG00000127947 | PTPN12 | chr7 | + | 77583554 | 77583650 | 77571152 | 77571186 | 77585542 | 77585571 | 142 | 103 | 63 | [illegible] |
| SE | 51958 | ENSG00000165487 | MICU2 | chr13 | - | 21496051 | 21496160 | 21495160 | 21495318 | 21502925 | 21503097 | 188 | 200 | 172 | [illegible] |
| SE | 78773 | ENSG00000179406 | LINC00174 | chr7 | - | 66489937 | 66490107 | 66487948 | 66488038 | 66491030 | 66491239 | 22 | 20 | 40 | [illegible] |
| [illegible] | | | | | | | | | | | | | | | |

| | | | | | | | | | | | | | | | |
|---|---|---|---|---|---|---|---|---|---|---|---|---|---|---|---|
| SE | 90216 | ENSG00000132294 | EFR3A | chr8 | + | 131996405 | 131996497 | 131987574 | 131987702 | 132000768 | 132000956 | 130 | 96 | 78 | [illegible] |
| SE | 83482 | ENSG00000106603 | COA1 | chr7 | - | 43656032 | 43656153 | 43650611 | 43650712 | 43729428 | 43729505 | 0 | 11 | 12 | [illegible] |
| SE | 2683 | ENSG00000172009 | THOP1 | chr19 | + | 2805015 | 2805176 | 2799688 | 2799791 | 2806916 | 2807052 | 66 | 32 | 39 | [illegible] |
| SE | 29122 | ENSG00000248672 | LY75-CD302 | chr2 | - | 159806972 | 159807140 | 159783358 | 159783469 | 159808448 | 159808571 | 180 | 159 | 144 | [illegible] |
| SE | 100671 | ENSG00000166387 | PPFIBP2 | chr11 | + | 7625776 | 7625891 | 7620934 | 7621027 | 7628284 | 7628346 | 46 | 20 | 66 | [illegible] |
| SE | 89540 | ENSG00000110074 | FOXRED1 | chr11 | + | 126276393 | 126276523 | 126276058 | 126276219 | 126277070 | 126277175 | 26 | 9 | 26 | [illegible] |
| SE | 93555 | ENSG00000095383 | TBC1D2 | chr9 | - | 98203287 | 98203408 | 98201478 | 98201664 | 98210655 | 98210843 | 22 | 7 | 11 | [illegible] |
| SE | 77955 | ENSG00000281103 | TRG-AS1 | chr7 | + | 38346020 | 38346127 | 38341614 | 38341799 | 38350345 | 38350450 | 8 | 17 | 8 | [illegible] |
| SE | 109048 | ENSG00000105507 | CABP5 | chr19 | - | 48041572 | 48041603 | 48040604 | 48040748 | 48043859 | 48044079 | 5 | 16 | 4 | [illegible] |
| SE | 81122 | ENSG00000143226 | FCGR2A | chr1 | + | 161506333 | 161506591 | 161505986 | 161506007 | 161509819 | 161510074 | 537 | 292 | 274 | [illegible] |
| SE | 94312 | ENSG00000132740 | IGHMBP2 | chr11 | + | 68938181 | 68938354 | 68936236 | 68937091 | 68939533 | 68940601 | 27 | 14 | 27 | [illegible] |
| SE | 37996 | ENSG00000095906 | NUBP2 | chr16 | + | 1786756 | 1786955 | 1786536 | 1786655 | 1787676 | 1787777 | 37 | 15 | 39 | [illegible] |
| SE | 68013 | ENSG00000187257 | RSBN1L | chr7 | + | 77749423 | 77750064 | 77736409 | 77736526 | 77765494 | 77765632 | 344 | 220 | 257 | [illegible] |
| SE | 12798 | ENSG00000169217 | CD2BP2 | chr16 | - | 30354183 | 30354322 | 30353900 | 30354058 | 30354603 | 30354707 | 552 | 285 | 503 | [illegible] |
| SE | 104320 | ENSG00000122884 | P4HA1 | chr10 | - | 73044980 | 73045051 | 73030266 | 73030370 | 73046924 | 73047101 | 85 | 83 | 61 | [illegible] |
| SE | 74235 | ENSG00000137478 | FCHSD2 | chr11 | - | 72842620 | 72842841 | 72836744 | 72838874 | 72843150 | 72843328 | 182 | 171 | 212 | [illegible] |
| SE | 83174 | ENSG00000197965 | MPZL1 | chr1 | + | 167773235 | 167773368 | 167765582 | 167765749 | 167787819 | 167787844 | 27 | 25 | 26 | [illegible] |
| SE | 7476 | ENSG00000073605 | GSDMB | chr17 | - | 39908175 | 39908214 | 39906960 | 39906987 | 39909755 | 39909924 | 6 | 3 | 7 | [illegible] |
| SE | 109282 | ENSG00000085832 | EPS15 | chr1 | - | 51405904 | 51406108 | 51403418 | 51403532 | 51408134 | 51408332 | 701 | 419 | 423 | [illegible] |
| SE | 71281 | ENSG00000123143 | PKN1 | chr19 | + | 14469751 | 14469814 | 14469340 | 14469517 | 14469911 | 14469988 | 647 | 318 | 381 | [illegible] |
| SE | 50011 | ENSG00000162298 | SYVN1 | chr11 | - | 65132247 | 65132308 | 65131469 | 65131596 | 65132731 | 65132780 | 14 | 3 | 21 | [illegible] |
| SE | 33711 | ENSG00000172046 | USP19 | chr3 | - | 49116726 | 49116943 | 49116450 | 49116607 | 49117436 | 49117570 | 40 | 24 | 43 | [illegible] |
| SE | 59465 | ENSG00000113312 | TTC1 | chr5 | + | 160051128 | 160051183 | 160043132 | 160043169 | 160064931 | 160065022 | 31 | 25 | 39 | [illegible] |
| SE | 35377 | ENSG00000140262 | TCF12 | chr15 | + | 57001348 | 57001384 | 56921025 | 56921098 | 57063749 | 57063823 | 0 | 4 | 6 | [illegible] |
| SE | 100130 | ENSG00000159063 | ALG8 | chr11 | - | 78107716 | 78107764 | 78106806 | 78106946 | 78109441 | 78109581 | 0 | 0 | 0 | [illegible] |
| SE | 73090 | ENSG00000147036 | LANCL3 | chrX | + | 37668367 | 37668450 | 37667281 | 37667489 | 37675653 | 37684463 | 3 | 3 | 0 | [illegible] |
| SE | 13616 | ENSG00000038358 | EDC4 | chr16 | + | 67882678 | 67882865 | 67882428 | 67882594 | 67883567 | 67883731 | 52 | 36 | 63 | [illegible] |
| SE | 91649 | ENSG00000105339 | DENND3 | chr8 | + | 141163332 | 141163429 | 141155848 | 141155970 | 141165185 | 141165289 | 49 | 44 | 39 | [illegible] |
| SE | 28322 | ENSG00000139505 | MTMR6 | chr13 | - | 25251648 | 25251775 | 25246221 | 25249492 | 25251852 | 25251984 | 263 | 145 | 235 | [illegible] |
| SE | 97802 | ENSG00000011114 | BTBD7 | chr14 | - | 93253646 | 93253790 | 93251462 | 93251652 | 93257194 | 93257355 | 65 | 33 | 41 | [illegible] |
| SE | 10521 | ENSG00000125970 | RALY | chr20 | + | 34031521 | 34031604 | 33993645 | 33994131 | 34072065 | 34072205 | 125 | 91 | 90 | [illegible] |
| SE | 79741 | ENSG00000164808 | SPIDR | chr8 | + | 47293866 | 47294030 | 47291032 | 47291137 | 47396375 | 47396626 | 360 | 281 | 265 | [illegible] |
| SE | 78401 | ENSG00000127980 | PEX1 | chr7 | - | 92501889 | 92502079 | 92501506 | 92501673 | 92503040 | 92503195 | 47 | 38 | 62 | [illegible] |
| SE | 103911 | ENSG00000066084 | DIP2B | chr12 | + | 50731368 | 50731537 | 50728547 | 50728678 | 50732365 | 50732536 | 181 | 105 | 152 | [illegible] |
| SE | 92073 | ENSG00000226752 | CUTALP | chr9 | + | 120844992 | 120845196 | 120844712 | 120844899 | 120847293 | 120847353 | 18 | 49 | 14 | [illegible] |
| SE | 25233 | ENSG00000005100 | DHX33 | chr17 | - | 5453820 | 5453980 | 5453579 | 5453668 | 5455159 | 5455271 | 66 | 23 | 27 | [illegible] |
| SE | 64184 | ENSG00000048991 | R3HDM1 | chr2 | + | 135641535 | 135641604 | 135638895 | 135639122 | 135645378 | 135645527 | 6 | 10 | 7 | [illegible] |
| SE | 38613 | ENSG00000204178 | MACO1 | chr1 | + | 25484126 | 25484274 | 25458390 | 25458892 | 25485612 | 25485795 | 87 | 53 | 64 | [illegible] |
| SE | 79580 | ENSG00000139641 | ESYT1 | chr12 | + | 56138767 | 56138839 | 56138403 | 56138499 | 56142284 | 56142425 | 99 | 75 | 124 | [illegible] |
| SE | 67921 | ENSG00000112130 | RNF8 | chr6 | + | 37376925 | 37377033 | 37371511 | 37371574 | 37390741 | 37391062 | 1 | 6 | 0 | [illegible] |
| SE | 8692 | ENSG00000136485 | DCAF7 | chr17 | + | 63578469 | 63578628 | 63550476 | 63550815 | 63579824 | 63579943 | 382 | 274 | 258 | [illegible] |
| SE | 48391 | ENSG00000162542 | TMCO4 | chr1 | - | 19739823 | 19739960 | 19737371 | 19737456 | 19740776 | 19740941 | 29 | 11 | 14 | [illegible] |
| SE | 109054 | ENSG00000139668 | WDFY2 | chr13 | + | 51727677 | 51727790 | 51703595 | 51703650 | 51739048 | 51739175 | 39 | 29 | 32 | [illegible] |
| SE | 580 | ENSG00000254413 | CHKB-CPT1B | chr22 | - | 50579726 | 50579830 | 50579425 | 50579507 | 50580189 | 50580416 | 188 | 77 | 115 | [illegible] |
| SE | 6250 | ENSG00000088812 | ATRN | chr20 | + | 3575826 | 3575948 | 3572730 | 3572951 | 3578581 | 3578772 | 54 | 54 | 103 | [illegible] |
| SE | 106894 | ENSG00000145050 | MANF | chr3 | + | 51386908 | 51387046 | 51386207 | 51386335 | 51387736 | 51387878 | 5 | 0 | 9 | [illegible] |
| SE | 37190 | ENSG00000033327 | GAB2 | chr11 | - | 78225107 | 78225202 | 78223411 | 78223676 | 78226464 | 78227051 | 155 | 62 | 92 | [illegible] |
| SE | 3921 | ENSG00000171101 | SIGLEC17P | chr19 | + | 51172516 | 51172591 | 51172065 | 51172159 | 51172975 | 51173429 | 36 | 22 | 47 | [illegible] |
| SE | 42996 | ENSG00000065911 | MTHFD2 | chr2 | + | 74201381 | 74201479 | 74198612 | 74198742 | 74205704 | 74205889 | 8 | 3 | 6 | [illegible] |
| SE | 47953 | ENSG00000169398 | PTK2 | chr8 | - | 140803542 | 140803650 | 140800458 | 140800576 | 140818276 | 140818354 | 54 | 102 | 44 | [illegible] |
| SE | 88435 | ENSG00000179715 | PCED1B | chr12 | + | 47223527 | 47223672 | 47216597 | 47216689 | 47235006 | 47235565 | 11 | 9 | 5 | [illegible] |
| SE | 69785 | ENSG00000065675 | PRKCQ | chr10 | - | 6483439 | 6483600 | 6478991 | 6479165 | 6491682 | 6491812 | 144 | 76 | 144 | [illegible] |
| SE | 106146 | ENSG00000089022 | MAPKAPK5 | chr12 | + | 111867571 | 111867669 | 111866155 | 111866231 | 111868752 | 111868861 | 259 | 165 | 286 | [illegible] |
| SE | 107185 | ENSG00000198715 | GLMP | chr1 | - | 156294758 | 156295016 | 156294364 | 156294565 | 156295525 | 156295654 | 46 | 15 | 29 | [illegible] |
| SE | 41804 | ENSG00000136059 | VILL | chr3 | + | 38002395 | 38002575 | 38001701 | 38001860 | 38004254 | 38004399 | 29 | 2 | 40 | [illegible] |
| SE | 60389 | ENSG00000242247 | ARFGAP3 | chr22 | - | 42835361 | 42835493 | 42834241 | 42834325 | 42840943 | 42841016 | 113 | 102 | 96 | [illegible] |
| SE | 42740 | ENSG00000133226 | SRRM1 | chr1 | + | 24648023 | 24648166 | 24647078 | 24647208 | 24648858 | 24648887 | 3 | 6 | 9 | [illegible] |
| SE | 34181 | ENSG00000175193 | PARL | chr3 | - | 183833723 | 183833825 | 183833491 | 183833589 | 183840569 | 183840640 | 267 | 118 | 236 | [illegible] |
| SE | 30161 | ENSG00000163811 | WDR43 | chr2 | + | 28912589 | 28912710 | 28906459 | 28906581 | 28914068 | 28914170 | 188 | 164 | 219 | [illegible] |
| SE | 33551 | ENSG00000164081 | TEX264 | chr3 | + | 51674270 | 51674562 | 51671273 | 51671547 | 51684412 | 51684467 | 34 | 13 | 19 | [illegible] |
| SE | 61418 | ENSG00000053900 | ANAPC4 | chr4 | + | 25413644 | 25413742 | 25409697 | 25409791 | 25414323 | 25414385 | 116 | 116 | 133 | [illegible] |
| SE | 92917 | ENSG00000131089 | ARHGEF9 | chrX | - | 63655493 | 63655737 | 63643979 | 63644048 | 63665885 | 63666017 | 91 | 44 | 96 | [illegible] |
| SE | 88337 | ENSG00000104219 | ZDHHC2 | chr8 | + | 17205654 | 17205775 | 17198380 | 17198413 | 17209931 | 17210058 | 133 | 81 | 104 | [illegible] |
| SE | 75872 | ENSG00000240065 | PSMB9 | chr6 | + | 32857262 | 32857394 | 32856096 | 32856205 | 32858004 | 32858134 | 502 | 209 | 299 | [illegible] |
| SE | 19425 | ENSG00000151445 | VIPAS39 | chr14 | - | 77435843 | 77435919 | 77434261 | 77434303 | 77437807 | 77437881 | 19 | 5 | 19 | [illegible] |
| SE | 89793 | ENSG00000104983 | CCDC61 | chr19 | + | 46003418 | 46003501 | 46003005 | 46003166 | 46006558 | 46006716 | 32 | 17 | 23 | [illegible] |
| SE | 76810 | ENSG00000135317 | SNX14 | chr6 | - | 85567533 | 85567577 | 85557975 | 85558060 | 85572136 | 85572215 | 35 | 22 | 17 | [illegible] |
| SE | 72043 | ENSG00000115053 | NCL | chr2 | - | 231456009 | 231456159 | 231455400 | 231455624 | 231456630 | 231456764 | 763 | 546 | 716 | [illegible] |
| SE | 25723 | ENSG00000247774 | PCED1B-AS1 | chr12 | - | 47210639 | 47210761 | 47209280 | 47209458 | 47216383 | 47216408 | 111 | 51 | 190 | [illegible] |
| SE | 43145 | ENSG00000113161 | HMGCR | chr5 | + | 75358656 | 75358827 | 75356342 | 75356448 | 75359169 | 75359310 | 157 | 98 | 101 | [illegible] |
| SE | 108578 | ENSG00000136169 | SETDB2 | chr13 | + | 49485629 | 49485723 | 49483463 | 49483563 | 49491731 | 49495002 | 116 | 131 | 122 | [illegible] |
| SE | 35906 | ENSG00000187862 | TTC24 | chr1 | + | 156583041 | 156583170 | 156582230 | 156582434 | 156584876 | 156584974 | 11 | 6 | 7 | [illegible] |
| SE | 93602 | ENSG00000158856 | DMTN | chr8 | + | 22059372 | 22059496 | 22057887 | 22057965 | 22066704 | 22066893 | 9 | 7 | 1 | [illegible] |
| SE | 86266 | ENSG00000134516 | DOCK2 | chr5 | + | 169761518 | 169761625 | 169759704 | 169759775 | 169803057 | 169803206 | 1338 | 740 | 794 | [illegible] |
| SE | 60669 | ENSG00000147099 | HDAC8 | chrX | - | 72567888 | 72568030 | 72495155 | 72495268 | 72572056 | 72572095 | 23 | 7 | 18 | [illegible] |
| SE | 48244 | ENSG00000116729 | WLS | chr1 | - | 68193954 | 68194221 | 68159122 | 68159247 | 68232193 | 68232545 | 38 | 29 | 24 | [illegible] |
| SE | 79767 | ENSG00000164808 | SPIDR | chr8 | + | 47727199 | 47727293 | 47713488 | 47713641 | 47735306 | 47735916 | 16 | 18 | 13 | [illegible] |
| SE | 106884 | ENSG00000005379 | TSPOAP1 | chr17 | - | 58326292 | 58326421 | 58324810 | 58325002 | 58326682 | 58326790 | 66 | 15 | 32 | [illegible] |
| SE | 66110 | ENSG00000000938 | FGR | chr1 | - | 27623690 | 27623929 | 27616856 | 27617006 | 27625088 | 27625151 | 333 | 129 | 140 | [illegible] |
| SE | 8057 | ENSG00000141337 | ARSG | chr17 | + | 68368547 | 68368744 | 68356666 | 68356804 | 68385063 | 68385172 | 16 | 10 | 17 | [illegible] |
| SE | 9757 | ENSG00000062716 | VMP1 | chr17 | + | 59764970 | 59765138 | 59735337 | 59735473 | 59773753 | 59773804 | 470 | 327 | 321 | [illegible] |
| SE | 21068 | ENSG00000196182 | STK40 | chr1 | - | 36362558 | 36362656 | 36361220 | 36361340 | 36385722 | 36385924 | 8 | 7 | 5 | [illegible] |
| SE | 13754 | ENSG00000122299 | ZC3H7A | chr16 | - | 11776750 | 11776909 | 11776451 | 11776532 | 11779165 | 11779363 | 310 | 313 | 296 | [illegible] |
| SE | 32569 | ENSG00000113851 | CRBN | chr3 | - | 3153423 | 3153488 | 3150010 | 3151045 | 3156218 | 3156281 | 31 | 19 | 29 | [illegible] |
| SE | 40911 | ENSG00000151883 | PARP8 | chr5 | + | 50750150 | 50750188 | 50667883 | 50668125 | 50759642 | 50759690 | 1728 | 1131 | 1108 | [illegible] |
| SE | 16157 | ENSG00000103196 | CRISPLD2 | chr16 | + | 84845785 | 84845904 | 84838421 | 84838735 | 84850567 | 84850683 | 12 | 4 | 10 | [illegible] |
| SE | 6991 | ENSG00000134779 | TPGS2 | chr18 | - | 36800197 | 36800311 | 36798017 | 36798609 | 36805373 | 36805397 | 217 | 136 | 136 | [illegible] |
| SE | 101389 | ENSG00000148429 | USP6NL | chr10 | - | 11597630 | 11597717 | 11527499 | 11527567 | 11611442 | 11611666 | 13 | 15 | 15 | [illegible] |
| SE | 72668 | ENSG00000284024 | NA | chr10 | + | 14842133 | 14842846 | 14840074 | 14840157 | 14843370 | 14846999 | 51 | 30 | 60 | [illegible] |
| SE | 60514 | ENSG00000116984 | MTR | chr1 | + | 236803427 | 236803642 | 236795259 | 236795737 | 236806143 | 236806233 | 131 | 74 | 140 | [illegible] |
| SE | 24629 | ENSG00000157212 | PAXIP1 | chr7 | - | 154986016 | 154986179 | 154983218 | 154983332 | 154991005 | 154991069 | 8 | 0 | 2 | [illegible] |
| SE | 73441 | ENSG00000198721 | ECI2 | chr6 | - | 4125249 | 4125370 | 4119185 | 4119275 | 4127761 | 4127831 | 47 | 30 | 70 | [illegible] |
| SE | 49165 | ENSG00000066136 | NFYC | chr1 | + | 40723207 | 40723274 | 40691837 | 40691867 | 40738835 | 40738948 | 0 | 0 | 1 | [illegible] |
| SE | 40797 | ENSG00000244682 | FCGR2C | chr1 | + | 161588421 | 161588442 | 161581338 | 161581549 | 161589561 | 161589596 | 130 | 66 | 62 | [illegible] |
| SE | 41724 | ENSG00000116497 | S100PBP | chr1 | + | 32834033 | 32834061 | 32829963 | 32830067 | 32853078 | 32853166 | 3 | 3 | 2 | [illegible] |
| SE | 27028 | ENSG00000198646 | NCOA6 | chr20 | - | 34749402 | 34750519 | 34746806 | 34746928 | 34757219 | 34758104 | 19 | 20 | 29 | [illegible] |
| SE | 106169 | ENSG00000120805 | ARL1 | chr12 | - | 101405843 | 101405981 | 101396142 | 101396577 | 101407641 | 101407746 | 92 | 46 | 79 | [illegible] |
| SE | 101967 | ENSG00000197771 | MCMBP | chr10 | - | 119842471 | 119842595 | 119840842 | 119840960 | 119843247 | 119843426 | 302 | 160 | 183 | [illegible] |
| SE | 65478 | ENSG00000163541 | SUCLG1 | chr2 | - | 84436051 | 84436127 | 84433351 | 84433435 | 84441046 | 84441104 | 3 | 1 | 6 | [illegible] |
| SE | 98662 | ENSG00000107036 | RIC1 | chr9 | + | 5765413 | 5765572 | 5763139 | 5763868 | 5768969 | 5769106 | 154 | 115 | 103 | [illegible] |
| SE | 18969 | ENSG00000015133 | CCDC88C | chr14 | - | 91343598 | 91343657 | 91339883 | 91340024 | 91359641 | 91359711 | 59 | 44 | 50 | [illegible] |
| SE | 11543 | ENSG00000179583 | CIITA | chr16 | + | 10935115 | 10935200 | 10923227 | 10923325 | 10941185 | 10943021 | 19 | 9 | 8 | [illegible] |
| SE | 80913 | ENSG00000067596 | DHX8 | chr17 | + | 43508338 | 43508520 | 43507808 | 43508019 | 43513361 | 43513502 | 226 | 153 | 203 | [illegible] |
| SE | 24557 | ENSG00000172785 | CBWD1 | chr9 | - | 164402 | 164448 | 163977 | 164037 | 172080 | 172172 | 2 | 0 | 0 | [illegible] |
| SE | 57516 | ENSG00000135913 | USP37 | chr2 | - | 218549808 | 218549909 | 218546918 | 218547091 | 218553552 | 218553724 | 102 | 87 | 90 | [illegible] |
| SE | 49461 | ENSG00000115020 | PIKFYVE | chr2 | + | 208277536 | 208277708 | 208276711 | 208276830 | 208298640 | 208298779 | 102 | 63 | 70 | [illegible] |
| SE | 104412 | ENSG00000184047 | DIABLO | chr12 | - | 122218265 | 122218397 | 122216758 | 122216869 | 122224511 | 122224644 | 208 | 92 | 168 | [illegible] |
| SE | 11879 | ENSG00000121281 | ADCY7 | chr16 | + | 50315013 | 50315138 | 50314291 | 50314406 | 50315358 | 50315738 | 551 | 373 | 349 | [illegible] |
| SE | 50873 | ENSG00000115568 | ZNF142 | chr2 | - | 218658979 | 218659130 | 218650358 | 218650526 | 218659368 | 218659476 | 25 | 3 | 14 | [illegible] |
| SE | 33589 | ENSG00000009790 | TRAF3IP3 | chr1 | + | 209779314 | 209779374 | 209778110 | 209778173 | 209780469 | 209780606 | 376 | 334 | 531 | [illegible] |
| SE | 5472 | ENSG00000125885 | MCM8 | chr20 | + | 5963273 | 5963359 | 5958527 | 5958726 | 5967435 | 5967587 | 21 | 23 | 14 | [illegible] |
| SE | 100401 | ENSG00000133812 | SBF2 | chr11 | - | 10042843 | 10042981 | 10031047 | 10031170 | 10193901 | 10193957 | 74 | 82 | 57 | [illegible] |
| SE | 100938 | ENSG00000074319 | TSG101 | chr11 | - | 18514677 | 18514841 | 18509541 | 18509665 | 18519518 | 18519603 | 158 | 91 | 102 | [illegible] |
| SE | 71215 | ENSG00000079263 | SP140 | chr2 | + | 230248884 | 230248968 | 230244987 | 230245071 | 230255451 | 230255532 | 28 | 22 | 23 | [illegible] |
| SE | 107420 | ENSG00000157538 | VPS26C | chr21 | - | 37240495 | 37240639 | 37238204 | 37238609 | 37267237 | 37267318 | 240 | 149 | 202 | [illegible] |
| SE | 35242 | ENSG00000119231 | SENP5 | chr3 | + | 196927795 | 196927879 | 196923413 | 196923551 | 196929632 | 196929683 | 124 | 77 | 105 | [illegible] |
| SE | 39426 | ENSG00000137393 | RNF144B | chr6 | + | 18457154 | 18457359 | 18439683 | 18439744 | 18459606 | 18459751 | 208 | 241 | 119 | [illegible] |
| SE | 52639 | ENSG00000197943 | PLCG2 | chr16 | + | 81905402 | 81905507 | 81900611 | 81900780 | 81907684 | 81907774 | 276 | 171 | 143 | [illegible] |
| SE | 88876 | ENSG00000183255 | PTTG1IP | chr21 | - | 44865394 | 44865447 | 44849585 | 44851627 | 44873501 | 44873690 | 5 | 4 | 4 | [illegible] |
| SE | 27229 | ENSG00000163812 | ZDHHC3 | chr3 | - | 44932906 | 44932990 | 44929305 | 44929436 | 44933117 | 44933199 | 32 | 13 | 16 | [illegible] |
| SE | 4932 | ENSG00000101347 | SAMHD1 | chr20 | - | 36904156 | 36904249 | 36898439 | 36898544 | 36905363 | 36905394 | 3578 | 2733 | 2409 | [illegible] |
| SE | 51259 | ENSG00000144843 | ADPRH | chr3 | + | 119586284 | 119586645 | 119582064 | 119582467 | 119587463 | 119588007 | 34 | 16 | 25 | [illegible] |
| SE | 81194 | ENSG00000176105 | YES1 | chr18 | - | 751704 | 751804 | 745947 | 746051 | 756556 | 756835 | 66 | 49 | 49 | [illegible] |
| SE | 41909 | ENSG00000175203 | DCTN2 | chr12 | - | 57546027 | 57546091 | 57535748 | 57535845 | 57547027 | 57547174 | 54 | 36 | 39 | [illegible] |
| SE | 20263 | ENSG00000118482 | PHF3 | chr6 | + | 63646526 | 63646795 | 63635801 | 63636150 | 63684128 | 63685911 | 16 | 16 | 25 | [illegible] |
| SE | 29110 | ENSG00000061918 | GUCY1B1 | chr4 | + | 155789713 | 155789911 | 155777523 | 155777642 | 155793855 | 155794086 | 54 | 147 | 18 | [illegible] |
| SE | 78647 | ENSG00000168014 | C2CD3 | chr11 | - | 74122987 | 74123135 | 74118227 | 74118382 | 74132843 | 74132972 | 113 | 102 | 118 | [illegible] |
| SE | 87777 | ENSG00000165675 | ENOX2 | chrX | - | 130709242 | 130709331 | 130703119 | 130703254 | 130783546 | 130783690 | 6 | 5 | 12 | [illegible] |
| SE | 18705 | ENSG00000141428 | C18orf21 | chr18 | + | 35972884 | 35972907 | 35972624 | 35972805 | 35974895 | 35975095 | 11 | 13 | 16 | [illegible] |
| SE | 51298 | ENSG00000174780 | SRP72 | chr4 | + | 56483138 | 56483270 | 56478591 | 56478649 | 56484735 | 56484864 | 556 | 336 | 437 | [illegible] |
| SE | 14515 | ENSG00000185596 | WASH3P | chr15 | + | 101972582 | 101972694 | 101966051 | 101966223 | 101972900 | 101973047 | 133 | 23 | 71 | [illegible] |
| SE | 19872 | ENSG00000165501 | LRR1 | chr14 | + | 49607399 | 49608121 | 49602369 | 49602468 | 49614255 | 49614671 | 31 | 9 | 21 | [illegible] |
| SE | 9284 | ENSG00000170037 | CNTROB | chr17 | + | 7934988 | 7935145 | 7934464 | 7934546 | 7936365 | 7936482 | 45 | 44 | 60 | [illegible] |
| SE | 31460 | ENSG00000178950 | GAK | chr4 | - | 851749 | 851974 | 850935 | 851084 | 859361 | 859493 | 122 | 67 | 71 | [illegible] |
| SE | 62535 | ENSG00000004866 | ST7 | chr7 | + | 117170861 | 117170976 | 117138434 | 117138532 | 117189320 | 117189393 | 64 | 48 | 28 | [illegible] |
| SE | 12468 | ENSG00000169896 | ITGAM | chr16 | + | 31265394 | 31265498 | 31261691 | 31261797 | 31265810 | 31265881 | 494 | 130 | 206 | [illegible] |
| SE | 17308 | ENSG00000100445 | SDR39U1 | chr14 | - | 24441677 | 24441795 | 24440782 | 24440926 | 24442177 | 24442260 | 52 | 31 | 42 | [illegible] |
| SE | 22537 | ENSG00000181991 | MRPS11 | chr15 | + | 88476988 | 88477054 | 88475029 | 88475239 | 88477871 | 88477904 | 68 | 26 | 28 | [illegible] |
| SE | 11705 | ENSG00000167978 | SRRM2 | chr16 | + | 2756333 | 2756606 | 2752625 | 2752854 | 2757471 | 2757574 | 344 | 218 | 356 | [illegible] |
| SE | 39022 | ENSG00000003400 | CASP10 | chr2 | + | 201205881 | 201205973 | 201187705 | 201187799 | 201208074 | 201208183 | 82 | 31 | 50 | [illegible] |
| SE | 37605 | ENSG00000082074 | FYB1 | chr5 | - | 39211057 | 39211091 | 39202893 | 39202987 | 39250670 | 39250806 | 1 | 1 | 6 | [illegible] |
| SE | 31081 | ENSG00000108509 | CAMTA2 | chr17 | - | 4979683 | 4980621 | 4978503 | 4978630 | 4981224 | 4981359 | 45 | 21 | 45 | [illegible] |
| SE | 55657 | ENSG00000138119 | MYOF | chr10 | - | 93347616 | 93347782 | 93343855 | 93343932 | 93349807 | 93349969 | 72 | 93 | 42 | [illegible] |
| SE | 45155 | ENSG00000107099 | DOCK8 | chr9 | + | 317042 | 317128 | 311953 | 312166 | 325670 | 325737 | 654 | 418 | 515 | [illegible] |
| SE | 39865 | ENSG00000143457 | GOLPH3L | chr1 | - | 150694107 | 150694173 | 150663631 | 150663763 | 150694655 | 150694850 | 3 | 1 | 9 | [illegible] |
| SE | 13799 | ENSG00000130733 | YIPF2 | chr19 | - | 10924075 | 10924192 | 10923832 | 10923999 | 10925685 | 10925773 | 18 | 6 | 13 | [illegible] |
| SE | 20592 | ENSG00000196367 | TRRAP | chr7 | + | 99030884 | 99031000 | 99030594 | 99030764 | 99031344 | 99031465 | 28 | 21 | 21 | [illegible] |
| SE | 95849 | ENSG00000107263 | RAPGEF1 | chr9 | - | 131596297 | 131596373 | 131588800 | 131588986 | 131598198 | 131598310 | 141 | 52 | 102 | [illegible] |
| SE | 71591 | ENSG00000115504 | EHBP1 | chr2 | + | 62996642 | 62996766 | 62993870 | 62993977 | 63037534 | 63037634 | 84 | 65 | 61 | [illegible] |
| SE | 39673 | ENSG00000181555 | SETD2 | chr3 | - | 47097954 | 47098081 | 47088112 | 47088247 | 47113875 | 47114004 | 49 | 31 | 54 | [illegible] |
| SE | 99031 | ENSG00000087053 | MTMR2 | chr11 | - | 95866402 | 95866486 | 95865600 | 95865676 | 95887611 | 95887684 | 0 | 0 | 0 | [illegible] |
| SE | 87769 | ENSG00000196459 | TRAPPC2 | chrX | - | 13734049 | 13734185 | 13719870 | 13719982 | 13734524 | 13734545 | 23 | 10 | 11 | [illegible] |
| SE | 21932 | ENSG00000112624 | BICRAL | chr6 | + | 42860261 | 42860356 | 42857090 | 42857236 | 42862509 | 42862612 | 297 | 135 | 276 | [illegible] |
| SE | 96601 | ENSG00000108854 | SMURF2 | chr17 | - | 64578491 | 64578576 | 64571797 | 64571956 | 64580788 | 64580991 | 103 | 104 | 124 | [illegible] |
| SE | 100450 | ENSG00000103429 | BFAR | chr16 | + | 14648387 | 14648592 | 14644273 | 14644609 | 14649803 | 14649973 | 147 | 91 | 149 | [illegible] |
| SE | 8490 | ENSG00000177885 | GRB2 | chr17 | - | 75325897 | 75326020 | 75321658 | 75321827 | 75332699 | 75332797 | 629 | 436 | 539 | [illegible] |
| SE | 8877 | ENSG00000153944 | MSI2 | chr17 | + | 57556989 | 57557131 | 57529675 | 57529724 | 57596867 | 57596950 | 5 | 0 | 1 | [illegible] |
| SE | 99699 | ENSG00000110075 | PPP6R3 | chr11 | + | 68564302 | 68564432 | 68558565 | 68558679 | 68567013 | 68567166 | 515 | 329 | 425 | [illegible] |
| SE | 94930 | ENSG00000119403 | PHF19 | chr9 | - | 120861917 | 120862005 | 120861088 | 120861174 | 120862587 | 120862749 | 57 | 48 | 91 | [illegible] |
| SE | 71372 | ENSG00000122203 | KIAA1191 | chr5 | - | 176352621 | 176352748 | 176348249 | 176348356 | 176355570 | 176355749 | 57 | 32 | 27 | [illegible] |
| SE | 69022 | ENSG00000132405 | TBC1D14 | chr4 | + | 7009876 | 7009948 | 7004843 | 7004924 | 7010652 | 7010777 | 53 | 27 | 37 | [illegible] |
| SE | 105484 | ENSG00000152256 | PDK1 | chr2 | + | 172564051 | 172564130 | 172562740 | 172562800 | 172564977 | 172565073 | 1 | 3 | 4 | [illegible] |
| SE | 30214 | ENSG00000132376 | INPP5K | chr17 | - | 1508114 | 1508226 | 1506979 | 1507089 | 1509177 | 1509353 | 215 | 134 | 192 | [illegible] |
| SE | 52675 | ENSG00000198860 | TSEN15 | chr1 | + | 184072050 | 184072194 | 184054727 | 184054863 | 184072826 | 184073568 | 2 | 0 | 4 | [illegible] |
| SE | 28041 | ENSG00000049656 | CLPTM1L | chr5 | - | 1330279 | 1330383 | 1325750 | 1325816 | 1331798 | 1331883 | 232 | 109 | 298 | [illegible] |
| SE | 58633 | ENSG00000134759 | ELP2 | chr18 | + | 36138794 | 36138872 | 36138094 | 36138426 | 36141136 | 36141191 | 156 | 114 | 142 | [illegible] |
| SE | 52453 | ENSG00000072803 | FBXW11 | chr5 | - | 171899003 | 171899094 | 171891466 | 171891604 | 171899913 | 171900100 | 187 | 131 | 155 | [illegible] |
| SE | 97782 | ENSG00000155158 | TTC39B | chr9 | - | 15225916 | 15226012 | 15214138 | 15214249 | 15267913 | 15267948 | 71 | 39 | 103 | [illegible] |
| SE | 66815 | ENSG00000031003 | FAM13B | chr5 | - | 137945901 | 137945997 | 137943132 | 137943216 | 137948954 | 137949184 | 196 | 156 | 135 | [illegible] |
| SE | 46016 | ENSG00000173230 | GOLGB1 | chr3 | - | 121664925 | 121665031 | 121663198 | 121664614 | 121667475 | 121667610 | 101 | 72 | 115 | [illegible] |
| SE | 35910 | ENSG00000187862 | TTC24 | chr1 | + | 156583796 | 156583895 | 156583041 | 156583170 | 156584876 | 156584974 | 5 | 20 | 16 | [illegible] |
| SE | 59924 | ENSG00000170043 | TRAPPC1 | chr17 | - | 7931730 | 7931829 | 7930344 | 7930734 | 7931939 | 7931962 | 7 | 4 | 3 | [illegible] |
| SE | 108949 | ENSG00000136141 | LRCH1 | chr13 | + | 46715559 | 46715664 | 46712524 | 46712597 | 46723220 | 46723330 | 1 | 0 | 0 | [illegible] |
| SE | 95260 | ENSG00000165219 | GAPVD1 | chr9 | + | 125287761 | 125287883 | 125268935 | 125268984 | 125298889 | 125299106 | 7 | 2 | 1 | [illegible] |
| SE | 85351 | ENSG00000137337 | MDC1 | chr6 | - | 30707565 | 30708357 | 30707383 | 30707454 | 30711411 | 30711504 | 27 | 14 | 33 | [illegible] |
| SE | 52679 | ENSG00000198860 | TSEN15 | chr1 | + | 184072156 | 184072298 | 184070615 | 184070732 | 184072826 | 184072986 | 53 | 32 | 55 | [illegible] |
| SE | 95006 | ENSG00000135047 | CTSL | chr9 | + | 87727360 | 87727395 | 87726118 | 87726398 | 87727593 | 87727729 | 0 | 0 | 0 | [illegible] |
| SE | 9239 | ENSG00000178971 | CTC1 | chr17 | - | 8229141 | 8229206 | 8228726 | 8228892 | 8229301 | 8229446 | 38 | 29 | 30 | [illegible] |
| SE | 21232 | ENSG00000136986 | DERL1 | chr8 | - | 123019194 | 123019245 | 123014148 | 123015585 | 123021446 | 123021499 | 109 | 65 | 109 | [illegible] |
| SE | 30617 | ENSG00000131042 | LILRB2 | chr19 | - | 54277888 | 54277939 | 54277549 | 54277597 | 54278259 | 54278544 | 82 | 40 | 29 | [illegible] |
| SE | 48917 | ENSG00000198408 | OGA | chr10 | - | 101790895 | 101791088 | 101787363 | 101787523 | 101792838 | 101792901 | 212 | 231 | 207 | [illegible] |
| SE | 28895 | ENSG00000164597 | COG5 | chr7 | - | 107298141 | 107298346 | 107283570 | 107283732 | 107324439 | 107324521 | 241 | 312 | 279 | [illegible] |
| SE | 11924 | ENSG00000005844 | ITGAL | chr16 | + | 30473964 | 30474083 | 30472657 | 30472898 | 30474195 | 30474298 | 32 | 11 | 29 | [illegible] |
| SE | 82226 | ENSG00000123066 | MED13L | chr12 | - | 116111427 | 116111512 | 116096668 | 116096752 | 116237467 | 116237687 | 435 | 401 | 425 | [illegible] |
| SE | 56409 | ENSG00000151838 | CCDC175 | chr14 | - | 59551354 | 59551436 | 59540674 | 59540746 | 59561118 | 59561228 | 2 | 6 | 1 | [illegible] |
| SE | 62757 | ENSG00000162959 | MEMO1 | chr2 | - | 31932066 | 31932135 | 31920797 | 31920910 | 31943301 | 31943383 | 179 | 87 | 153 | [illegible] |
| SE | 75021 | ENSG00000100814 | CCNB1IP1 | chr14 | - | 20315484 | 20315618 | 20313467 | 20313801 | 20316226 | 20316408 | 2 | 2 | 0 | [illegible] |
| SE | 39142 | ENSG00000134352 | IL6ST | chr5 | - | 55968275 | 55968396 | 55964145 | 55964312 | 55969549 | 55969855 | 590 | 713 | 897 | [illegible] |
| SE | 87427 | ENSG00000125351 | UPF3B | chrX | - | 119843190 | 119843301 | 119841734 | 119841778 | 119845197 | 119845254 | 68 | 50 | 50 | [illegible] |
| SE | 102750 | ENSG00000040341 | STAU2 | chr8 | - | 73688653 | 73688813 | 73672981 | 73673242 | 73738283 | 73738327 | 426 | 383 | 347 | [illegible] |
| SE | 29519 | ENSG00000119242 | CCDC92 | chr12 | - | 123944271 | 123944364 | 123943346 | 123943493 | 123956283 | 123956444 | 52 | 30 | 30 | [illegible] |
| SE | 43621 | ENSG00000196628 | TCF4 | chr18 | - | 55257314 | 55257391 | 55254496 | 55254700 | 55261465 | 55261533 | 16 | 21 | 11 | [illegible] |
| SE | 108338 | ENSG00000111300 | NAA25 | chr12 | - | 112087682 | 112087801 | 112078633 | 112078741 | 112090725 | 112090864 | 30 | 22 | 35 | [illegible] |
| SE | 19200 | ENSG00000140105 | WARS1 | chr14 | - | 100375282 | 100375406 | 100361707 | 100361921 | 100376259 | 100376285 | 17 | 31 | 13 | [illegible] |
| SE | 63202 | ENSG00000118960 | HS1BP3 | chr2 | - | 20638435 | 20638652 | 20624731 | 20624892 | 20645339 | 20645505 | 60 | 24 | 46 | [illegible] |
| SE | 35290 | ENSG00000145016 | RUBCN | chr3 | - | 197691073 | 197691148 | 197684156 | 197684217 | 197693714 | 197693816 | 38 | 31 | 41 | [illegible] |
| SE | 12033 | ENSG00000176715 | ACSF3 | chr16 | + | 89117220 | 89117333 | 89116989 | 89117086 | 89120800 | 89120913 | 0 | 1 | 1 | [illegible] |
| SE | 75244 | ENSG00000065883 | CDK13 | chr7 | + | 40001860 | 40002031 | 39987598 | 39988258 | 40045835 | 40046025 | 434 | 308 | 357 | [illegible] |
| SE | 54754 | ENSG00000083720 | OXCT1 | chr5 | - | 41807330 | 41807438 | 41803068 | 41803163 | 41840450 | 41840511 | 65 | 27 | 79 | [illegible] |
| SE | 82106 | ENSG00000128891 | CCDC32 | chr15 | - | 40557215 | 40557372 | 40553098 | 40554127 | 40562771 | 40563022 | 160 | 99 | 142 | [illegible] |
| SE | 33037 | ENSG00000183323 | CCDC125 | chr5 | - | 69313984 | 69314043 | 69311117 | 69311204 | 69320464 | 69320580 | 31 | 23 | 32 | [illegible] |
| SE | 95054 | ENSG00000198887 | SMC5 | chr9 | + | 70280758 | 70280899 | 70278490 | 70278625 | 70282421 | 70282583 | 219 | 250 | 213 | [illegible] |
| SE | 105520 | ENSG00000084110 | HAL | chr12 | - | 95993771 | 95993838 | 95993450 | 95993488 | 95993925 | 95993998 | 22 | 14 | 10 | [illegible] |
| SE | 84315 | ENSG00000164062 | APEH | chr3 | + | 49676055 | 49676219 | 49675890 | 49675966 | 49676377 | 49676506 | 670 | 359 | 437 | [illegible] |
| SE | 69061 | ENSG00000081059 | TCF7 | chr5 | + | 134144796 | 134144869 | 134143591 | 134143640 | 134146200 | 134148229 | 4 | 0 | 7 | [illegible] |
| SE | 80617 | ENSG00000185164 | NOMO2 | chr16 | - | 18558676 | 18558854 | 18557701 | 18557791 | 18561875 | 18562176 | 0 | 0 | 0 | [illegible] |
| SE | 14160 | ENSG00000005339 | CREBBP | chr16 | - | 3782683 | 3782926 | 3781203 | 3781306 | 3791980 | 3792094 | 111 | 72 | 134 | [illegible] |
| SE | 37888 | ENSG00000196187 | TMEM63A | chr1 | - | 225867111 | 225867163 | 225865896 | 225865967 | 225867887 | 225868030 | 61 | 18 | 51 | [illegible] |
| SE | 70760 | ENSG00000040933 | INPP4A | chr2 | + | 98559462 | 98559495 | 98555552 | 98555808 | 98563464 | 98563637 | 107 | 39 | 70 | [illegible] |
| SE | 85330 | ENSG00000147394 | ZNF185 | chrX | + | 152959698 | 152959896 | 152945266 | 152945464 | 152963838 | 152963949 | 84 | 84 | 33 | [illegible] |
| SE | 28502 | ENSG00000169062 | UPF3A | chr13 | + | 114286301 | 114286400 | 114282836 | 114282943 | 114286518 | 114286629 | 96 | 78 | 111 | [illegible] |
| SE | 32934 | ENSG00000075711 | DLG1 | chr3 | - | 197149742 | 197149796 | 197142717 | 197142768 | 197194424 | 197194589 | 250 | 194 | 248 | [illegible] |
| SE | 97272 | ENSG00000156110 | ADK | chr10 | + | 74600378 | 74600493 | 74589281 | 74589317 | 74670182 | 74670269 | 127 | 195 | 137 | [illegible] |
| SE | 60670 | ENSG00000147099 | HDAC8 | chrX | - | 72568753 | 72568884 | 72495155 | 72495268 | 72572056 | 72572104 | 37 | 28 | 43 | [illegible] |
| SE | 23929 | ENSG00000174989 | FBXW8 | chr12 | + | 116945363 | 116945528 | 116928022 | 116928127 | 116949617 | 116949706 | 52 | 34 | 61 | [illegible] |
| SE | 29518 | ENSG00000109756 | RAPGEF2 | chr4 | + | 159355852 | 159356158 | 159353486 | 159354046 | 159358113 | 159360169 | 41 | 35 | 41 | [illegible] |
| SE | 5693 | ENSG00000188785 | ZNF548 | chr19 | + | 57392800 | 57392948 | 57389964 | 57390114 | 57394187 | 57394223 | 18 | 18 | 34 | [illegible] |
| SE | 69298 | ENSG00000182919 | C11orf54 | chr11 | + | 93757315 | 93757465 | 93754872 | 93755386 | 93759741 | 93759816 | 200 | 104 | 133 | [illegible] |
| SE | 78527 | ENSG00000180233 | ZNRF2 | chr7 | + | 30362376 | 30362456 | 30355727 | 30355833 | 30366034 | 30367689 | 304 | 221 | 201 | [illegible] |
| SE | 19195 | ENSG00000140105 | WARS1 | chr14 | - | 100375282 | 100375350 | 100361707 | 100361921 | 100376259 | 100376307 | 8 | 14 | 9 | [illegible] |
| SE | 40203 | ENSG00000231607 | DLEU2 | chr13 | - | 50075558 | 50075653 | 50068096 | 50068163 | 50125301 | 50125387 | 34 | 22 | 15 | [illegible] |
| SE | 12937 | ENSG00000197818 | SLC9A8 | chr20 | + | 49886751 | 49886898 | 49883845 | 49884066 | 49887828 | 49892241 | 27 | 37 | 39 | [illegible] |
| SE | 56111 | ENSG00000158109 | TPRG1L | chr1 | + | 3625712 | 3625889 | 3625423 | 3625515 | 3627499 | 3627653 | 328 | 128 | 244 | [illegible] |
| SE | 30124 | ENSG00000154144 | TBRG1 | chr11 | + | 124626692 | 124626751 | 124625670 | 124625903 | 124626903 | 124626935 | 8 | 7 | 7 | [illegible] |
| SE | 109536 | ENSG00000127022 | CANX | chr5 | + | 179704154 | 179704430 | 179698976 | 179699102 | 179705678 | 179705852 | 1 | 0 | 7 | [illegible] |
| SE | 56341 | ENSG00000117602 | RCAN3 | chr1 | + | 24531217 | 24531391 | 24514313 | 24514567 | 24535092 | 24535337 | 50 | 99 | 196 | [illegible] |
| SE | 10930 | ENSG00000091592 | NLRP1 | chr17 | - | 5515472 | 5515517 | 5501813 | 5501872 | 5521523 | 5521786 | 15 | 13 | 13 | [illegible] |
| SE | 71572 | ENSG00000166197 | NOLC1 | chr10 | + | 102158048 | 102158214 | 102157430 | 102157555 | 102159192 | 102159308 | 144 | 112 | 153 | [illegible] |
| SE | 2672 | ENSG00000129353 | SLC44A2 | chr19 | + | 10636322 | 10636585 | 10634755 | 10635073 | 10636661 | 10636756 | 94 | 26 | 89 | [illegible] |
| SE | 25117 | ENSG00000153310 | CYRIB | chr8 | - | 129896748 | 129896854 | 129879388 | 129879471 | 129903311 | 129903350 | 10 | 7 | 8 | [illegible] |
| SE | 50848 | ENSG00000048052 | HDAC9 | chr7 | + | 18629349 | 18629481 | 18593907 | 18594029 | 18644670 | 18644793 | 47 | 21 | 18 | [illegible] |
| SE | 77273 | ENSG00000168944 | CEP120 | chr5 | - | 123422466 | 123422575 | 123418358 | 123418515 | 123422949 | 123423324 | 1 | 2 | 2 | [illegible] |
| SE | 39263 | ENSG00000113810 | SMC4 | chr3 | + | 160412529 | 160412849 | 160412325 | 160412453 | 160413472 | 160413583 | 1 | 8 | 1 | [illegible] |

| | | | | | | | | | | | | |
|---|---|---|---|---|---|---|---|---|---|---|---|---|
| SE | 62171 | ENSG00000117625 | RCOR3 | chr1 | + | 211312719 | 211312961 | 211301460 | 211304140 | 211313423 | 211313839 | [illegible] |
| SE | 84654 | ENSG00000185753 | CXorf38 | chrX | - | 40636532 | 40636712 | 40626920 | 40630773 | 40637006 | 40637156 | [illegible] |
| SE | 98979 | ENSG00000255663 | NA | chr11 | + | 114401697 | 114401860 | 114400681 | 114400767 | 114405705 | 114405799 | [illegible] |
| SE | 74419 | ENSG00000163328 | GPR155 | chr2 | - | 174453736 | 174453841 | 174446610 | 174446747 | 174459877 | 174460088 | [illegible] |
| SE | 34868 | ENSG00000117597 | UTP25 | chr1 | + | 209840852 | 209841055 | 209838804 | 209839128 | 209842264 | 209842447 | [illegible] |
| SE | 100944 | ENSG00000074319 | TSG101 | chr11 | - | 18525660 | 18525702 | 18519518 | 18519603 | 18526774 | 18526920 | [illegible] |
| SE | 6763 | ENSG00000133313 | CNDP2 | chr18 | + | 74505848 | 74506011 | 74501089 | 74501472 | 74510812 | 74511013 | [illegible] |
| SE | 106230 | ENSG00000089154 | GCN1 | chr12 | - | 120138322 | 120138415 | 120137900 | 120138044 | 120138694 | 120138856 | [illegible] |
| SE | 42756 | ENSG00000134371 | CDC73 | chr1 | + | 193236255 | 193236356 | 193232992 | 193233154 | 193249729 | 193249871 | [illegible] |
| SE | 8775 | ENSG00000076382 | SPAG5 | chr17 | - | 28584651 | 28584745 | 28584332 | 28584480 | 28585101 | 28585215 | [illegible] |
| SE | 29829 | ENSG00000129187 | DCTD | chr4 | - | 182916418 | 182916539 | 182915460 | 182915575 | 182916672 | 182916857 | [illegible] |
| SE | 76340 | ENSG00000136861 | CDK5RAP2 | chr9 | - | 120453455 | 120453873 | 120447894 | 120448126 | 120458449 | 120458622 | [illegible] |
| SE | 20979 | ENSG00000198399 | ITSN2 | chr2 | - | 24220944 | 24221066 | 24217906 | 24218013 | 24246128 | 24246320 | [illegible] |
| SE | 23434 | ENSG00000172062 | SMN1 | chr5 | + | 70951940 | 70951994 | 70946065 | 70946176 | 70952438 | 70952580 | [illegible] |
| SE | 73463 | ENSG00000003402 | CFLAR | chr2 | + | 201135971 | 201136107 | 201132871 | 201133134 | 201140356 | 201140439 | [illegible] |
| SE | 8945 | ENSG00000132478 | UNK | chr17 | + | 75792312 | 75792478 | 75791759 | 75792019 | 75793442 | 75794124 | [illegible] |
| SE | 97209 | ENSG00000173653 | RCE1 | chr11 | + | 66845499 | 66845562 | 66845165 | 66845237 | 66845859 | 66846528 | [illegible] |
| SE | 88523 | ENSG00000138378 | STAT4 | chr2 | - | 191036163 | 191036299 | 191034547 | 191034597 | 191039198 | 191039297 | [illegible] |
| SE | 30272 | ENSG00000178104 | PDE4DIP | chr1 | + | 148977936 | 148978053 | 148974513 | 148974605 | 148978277 | 148978415 | [illegible] |
| SE | 29086 | ENSG00000163625 | WDFY3 | chr4 | - | 84896910 | 84897010 | 84860411 | 84860622 | 84932269 | 84932363 | [illegible] |
| SE | 53583 | ENSG00000188529 | SRSF10 | chr1 | - | 23978652 | 23978817 | 23977789 | 23978273 | 23980190 | 23980271 | [illegible] |
| SE | 59811 | ENSG00000166377 | ATP9B | chr18 | + | 79253380 | 79253541 | 79213961 | 79214038 | 79303603 | 79303716 | [illegible] |
| SE | 42831 | ENSG00000140575 | IQGAP1 | chr15 | + | 90472839 | 90473010 | 90467449 | 90467592 | 90473714 | 90473798 | [illegible] |
| SE | 51845 | ENSG00000087206 | UIMC1 | chr5 | - | 176969600 | 176969706 | 176958092 | 176958154 | 176970741 | 176970793 | [illegible] |
| SE | 24334 | ENSG00000185404 | SP140L | chr2 | + | 230391714 | 230391807 | 230389918 | 230390023 | 230392086 | 230392229 | [illegible] |
| SE | 35801 | ENSG00000173226 | IQCB1 | chr3 | - | 121797117 | 121797227 | 121795456 | 121795566 | 121807343 | 121807443 | [illegible] |
| SE | 25492 | ENSG00000082397 | EPB41L3 | chr18 | - | 5433902 | 5434121 | 5433468 | 5433556 | 5438034 | 5438110 | [illegible] |
| SE | 68352 | ENSG00000125630 | POLR1B | chr2 | + | 112557909 | 112558081 | 112552644 | 112552816 | 112559292 | 112559395 | [illegible] |
| SE | 40511 | ENSG00000285708 | NA | chr3 | - | 71699613 | 71699708 | 71690009 | 71690165 | 71710411 | 71710484 | [illegible] |
| SE | 2560 | ENSG00000214046 | SMIM7 | chr19 | - | 16654034 | 16654125 | 16646147 | 16647261 | 16659958 | 16660000 | [illegible] |
| SE | 63187 | ENSG00000175662 | TOM1L2 | chr17 | - | 17893660 | 17893810 | 17879626 | 17879743 | 17898595 | 17898674 | [illegible] |
| SE | 30869 | ENSG00000134852 | CLOCK | chr4 | - | 55453676 | 55453824 | 55453053 | 55453129 | 55455896 | 55456003 | [illegible] |
| SE | 91792 | ENSG00000133794 | ARNTL | chr11 | + | 13356738 | 13356777 | 13354300 | 13354448 | 13357054 | 13357096 | [illegible] |
| SE | 90744 | ENSG00000214706 | IFRD2 | chr3 | - | 50289254 | 50289360 | 50288798 | 50288937 | 50289446 | 50289601 | [illegible] |
| SE | 78352 | ENSG00000072609 | CHFR | chr12 | - | 132851617 | 132851734 | 132848640 | 132848724 | 132853430 | 132854104 | [illegible] |
| SE | 42392 | ENSG00000113194 | FAF2 | chr5 | + | 176486354 | 176486489 | 176479187 | 176479256 | 176488950 | 176489027 | [illegible] |
| SE | 68448 | ENSG00000198589 | LRBA | chr4 | - | 150808319 | 150808398 | 150806270 | 150806404 | 150817123 | 150817257 | [illegible] |
| SE | 1325 | ENSG00000100376 | FAM118A | chr22 | + | 45337833 | 45337968 | 45336394 | 45336411 | 45340385 | 45340882 | [illegible] |
| SE | 17529 | ENSG00000087111 | PIGS | chr17 | - | 28556165 | 28556266 | 28554850 | 28555061 | 28556826 | 28556972 | [illegible] |
| SE | 49556 | ENSG00000070269 | TMEM260 | chr14 | + | 56617182 | 56617297 | 56615450 | 56616027 | 56618593 | 56618763 | [illegible] |
| SE | 33071 | ENSG00000088726 | TMEM40 | chr3 | - | 12743899 | 12743989 | 12742453 | 12742507 | 12749759 | 12749840 | [illegible] |
| SE | 105989 | ENSG00000122965 | RBM19 | chr12 | - | 113950082 | 113950154 | 113948832 | 113949036 | 113952511 | 113952590 | [illegible] |
| SE | 72275 | ENSG00000186635 | ARAP1 | chr11 | - | 72704151 | 72704334 | 72702904 | 72703079 | 72705804 | 72705890 | [illegible] |
| SE | 73932 | ENSG00000169967 | MAP3K2 | chr2 | - | 127325727 | 127325807 | 127324173 | 127324241 | 127326686 | 127326817 | [illegible] |
| SE | 44147 | ENSG00000138794 | CASP6 | chr4 | - | 109697621 | 109697768 | 109694580 | 109694700 | 109698299 | 109698342 | [illegible] |
| SE | 26386 | ENSG00000056097 | ZFR | chr5 | - | 32397218 | 32397338 | 32395158 | 32395304 | 32400006 | 32400203 | [illegible] |
| SE | 53005 | ENSG00000158941 | CCAR2 | chr8 | + | 22614389 | 22614503 | 22614091 | 22614314 | 22614837 | 22615001 | [illegible] |
| SE | 98465 | ENSG00000177830 | CHID1 | chr11 | - | 893426 | 893519 | 884067 | 884169 | 899339 | 899401 | [illegible] |
| SE | 5423 | ENSG00000118705 | RPN2 | chr20 | + | 37225687 | 37225802 | 37223877 | 37223969 | 37228549 | 37228744 | [illegible] |
| SE | 88015 | ENSG00000147439 | BIN3 | chr8 | - | 22634412 | 22634525 | 22630441 | 22630578 | 22636524 | 22636586 | [illegible] |
| SE | 25983 | ENSG00000137364 | TPMT | chr6 | - | 18147822 | 18147915 | 18143595 | 18143728 | 18148987 | 18149171 | [illegible] |
| SE | 20230 | ENSG00000170542 | SERPINB9 | chr6 | - | 2893410 | 2893553 | 2891832 | 2891988 | 2895390 | 2895508 | [illegible] |
| SE | 68279 | ENSG00000122085 | MTERF4 | chr2 | - | 241090277 | 241090449 | 241089979 | 241090044 | 241094276 | 241094376 | [illegible] |
| SE | 38814 | ENSG00000229180 | NA | chr7 | - | 66545600 | 66545686 | 66531790 | 66531871 | 66554584 | 66554837 | [illegible] |
| SE | 97702 | ENSG00000170145 | SIK2 | chr11 | + | 111700885 | 111701010 | 111688000 | 111688162 | 111701451 | 111701575 | [illegible] |
| SE | 13224 | ENSG00000174177 | CTU2 | chr16 | + | 88709937 | 88710016 | 88706588 | 88707210 | 88710222 | 88710282 | [illegible] |
| SE | 46350 | ENSG00000028310 | BRD9 | chr5 | - | 869244 | 869394 | 865413 | 865581 | 870472 | 870575 | [illegible] |
| SE | 6478 | ENSG00000171984 | SHLD1 | chr20 | + | 5756711 | 5756848 | 5750452 | 5750479 | 5772861 | 5773043 | [illegible] |
| SE | 40059 | ENSG00000125347 | IRF1 | chr5 | - | 132484361 | 132484497 | 132476631 | 132476661 | 132486556 | 132486686 | [illegible] |
| SE | 107250 | ENSG00000248810 | LINC02432 | chr4 | - | 141324801 | 141324909 | 141323421 | 141323618 | 141332158 | 141332617 | [illegible] |
| SE | 55713 | ENSG00000116704 | SLC35D1 | chr1 | - | 67009084 | 67009167 | 66999349 | 67004448 | 67020368 | 67020447 | [illegible] |
| SE | 57498 | ENSG00000134313 | KIDINS220 | chr2 | - | 8826985 | 8827129 | 8818694 | 8818793 | 8837479 | 8837596 | [illegible] |
| SE | 97988 | ENSG00000197798 | FAM118B | chr11 | + | 126240791 | 126241044 | 126234994 | 126235087 | 126250505 | 126250625 | [illegible] |
| SE | 106819 | ENSG00000139187 | KLRG1 | chr12 | + | 8995118 | 8995288 | 8992069 | 8992310 | 9008974 | 9009075 | [illegible] |
| SE | 79213 | ENSG00000242588 | NA | chr7 | + | 128609486 | 128609579 | 128574750 | 128574791 | 128615859 | 128615970 | [illegible] |
| SE | 1650 | ENSG00000100365 | NCF4 | chr22 | + | 36863146 | 36863225 | 36861005 | 36861203 | 36864044 | 36864129 | [illegible] |
| SE | 70236 | ENSG00000160695 | VPS11 | chr11 | + | 119073197 | 119073399 | 119071595 | 119071843 | 119073799 | 119073951 | [illegible] |
| SE | 46735 | ENSG00000163125 | RPRD2 | chr1 | + | 150444250 | 150444377 | 150443230 | 150443283 | 150446225 | 150446401 | [illegible] |
| SE | 20201 | ENSG00000111845 | PAK1IP1 | chr6 | + | 10697323 | 10697486 | 10694971 | 10695069 | 10702368 | 10702489 | [illegible] |
| SE | 42377 | ENSG00000048740 | CELF2 | chr10 | + | 11321206 | 11321386 | 11314138 | 11314270 | 11325835 | 11325979 | [illegible] |
| SE | 66422 | ENSG00000102081 | FMR1 | chrX | + | 147925539 | 147925633 | 147921932 | 147921985 | 147928658 | 147928807 | [illegible] |
| SE | 1558 | ENSG00000198089 | SFI1 | chr22 | + | 31528689 | 31528863 | 31508254 | 31508376 | 31546860 | 31546971 | [illegible] |
| SE | 45466 | ENSG00000143149 | ALDH9A1 | chr1 | - | 165680486 | 165680683 | 165679441 | 165679582 | 165682106 | 165682241 | [illegible] |
| SE | 25272 | ENSG00000119685 | TTLL5 | chr14 | + | 75757830 | 75757869 | 75752892 | 75752955 | 75764614 | 75764772 | [illegible] |
| SE | 17497 | ENSG00000100577 | GSTZ1 | chr14 | + | 77327911 | 77328037 | 77327471 | 77327552 | 77329122 | 77329201 | [illegible] |
| SE | 67214 | ENSG00000124151 | NCOA3 | chr20 | + | 47652405 | 47652580 | 47650981 | 47651276 | 47652930 | 47653072 | [illegible] |
| SE | 18830 | ENSG00000127663 | KDM4B | chr19 | + | 5110621 | 5110818 | 5082366 | 5082504 | 5131075 | 5131545 | [illegible] |
| SE | 10904 | ENSG00000074370 | ATP2A3 | chr17 | - | 3927980 | 3928038 | 3925345 | 3925438 | 3928662 | 3928780 | [illegible] |
| SE | 40185 | ENSG00000114544 | SLC41A3 | chr3 | - | 126056357 | 126056513 | 126050942 | 126051050 | 126067946 | 126068246 | [illegible] |
| SE | 73676 | ENSG00000151687 | ANKAR | chr2 | + | 189728659 | 189728821 | 189728266 | 189728420 | 189737682 | 189737841 | [illegible] |
| SE | 74582 | ENSG00000128284 | APOL3 | chr22 | - | 36149770 | 36149964 | 36149258 | 36149371 | 36160668 | 36160755 | [illegible] |
| SE | 71134 | ENSG00000197562 | RAB40C | chr16 | + | 618199 | 618260 | 617207 | 617268 | 625431 | 625509 | [illegible] |
| SE | 64935 | ENSG00000170873 | MTSS1 | chr8 | - | 124556231 | 124556354 | 124555741 | 124555904 | 124562781 | 124562992 | [illegible] |
| SE | 51061 | ENSG00000143353 | LYPLAL1 | chr1 | + | 219193129 | 219193251 | 219179399 | 219179426 | 219210531 | 219210647 | [illegible] |
| SE | 73831 | ENSG00000110080 | ST3GAL4 | chr11 | + | 126409267 | 126409411 | 126408306 | 126408496 | 126413504 | 126413648 | [illegible] |
| SE | 57886 | ENSG00000162604 | TM2D1 | chr1 | - | 61694696 | 61694770 | 61683271 | 61683546 | 61700933 | 61701025 | [illegible] |
| SE | 68165 | ENSG00000047230 | CTPS2 | chrX | - | 16691539 | 16691620 | 16689449 | 16689601 | 16693140 | 16693224 | [illegible] |
| SE | 71845 | ENSG00000183486 | MX2 | chr21 | + | 41401969 | 41402128 | 41398984 | 41399019 | 41403266 | 41403343 | [illegible] |
| SE | 11267 | ENSG00000182149 | IST1 | chr16 | + | 71924772 | 71924817 | 71923287 | 71923380 | 71927613 | 71929010 | [illegible] |
| SE | 42667 | ENSG00000088854 | C20orf194 | chr20 | - | 3326196 | 3326284 | 3324891 | 3324968 | 3330645 | 3330682 | [illegible] |
| SE | 91512 | ENSG00000135679 | MDM2 | chr12 | + | 68816811 | 68816945 | 68813553 | 68813628 | 68820324 | 68820374 | [illegible] |
| SE | 58695 | ENSG00000198898 | CAPZA2 | chr7 | + | 116906262 | 116906342 | 116904176 | 116904383 | 116910212 | 116910231 | [illegible] |
| SE | 105485 | ENSG00000152256 | PDK1 | chr2 | + | 172564502 | 172564687 | 172562219 | 172562291 | 172564977 | 172565073 | [illegible] |
| SE | 109122 | ENSG00000006625 | GGCT | chr7 | - | 30500535 | 30500681 | 30496622 | 30497235 | 30504568 | 30504799 | [illegible] |
| SE | 52645 | ENSG00000197943 | PLCG2 | chr16 | + | 81923484 | 81923594 | 81921197 | 81921269 | 81927081 | 81927178 | [illegible] |
| SE | 87633 | ENSG00000140678 | ITGAX | chr16 | + | 31373248 | 31373390 | 31372596 | 31372670 | 31376798 | 31376915 | [illegible] |
| SE | 35602 | ENSG00000114331 | ACAP2 | chr3 | - | 195412845 | 195412916 | 195392089 | 195392147 | 195442794 | 195442905 | [illegible] |
| SE | 36292 | ENSG00000075420 | FNDC3B | chr3 | + | 172307362 | 172307501 | 172298727 | 172298787 | 172310827 | 172310881 | [illegible] |
| SE | 31077 | ENSG00000108509 | CAMTA2 | chr17 | - | 4969499 | 4969520 | 4969149 | 4969337 | 4969629 | 4969701 | [illegible] |
| SE | 5102 | ENSG00000124198 | ARFGEF2 | chr20 | + | 48971119 | 48971354 | 48969146 | 48969277 | 48974765 | 48974874 | [illegible] |
| SE | 62963 | ENSG00000117222 | RBBP5 | chr1 | - | 205098988 | 205099106 | 205097325 | 205097395 | 205099740 | 205099812 | [illegible] |
| SE | 30908 | ENSG00000205726 | ITSN1 | chr21 | + | 33767712 | 33767828 | 33765874 | 33766012 | 33772060 | 33772282 | [illegible] |
| SE | 26620 | ENSG00000164889 | SLC4A2 | chr7 | + | 151074398 | 151074488 | 151074038 | 151074099 | 151074674 | 151074841 | [illegible] |
| SE | 88132 | ENSG00000111364 | DDX55 | chr12 | + | 123617757 | 123617872 | 123616510 | 123616603 | 123618668 | 123618837 | [illegible] |
| SE | 106294 | ENSG00000197057 | DTHD1 | chr4 | + | 36316241 | 36316486 | 36308203 | 36308493 | 36339111 | 36339169 | [illegible] |
| SE | 71650 | ENSG00000163832 | ELP6 | chr3 | - | 47510148 | 47510254 | 47504329 | 47504448 | 47511147 | 47511215 | [illegible] |
| SE | 44353 | ENSG00000142252 | GEMIN7 | chr19 | + | 45079906 | 45080029 | 45079259 | 45079377 | 45090106 | 45090537 | [illegible] |
| SE | 28489 | ENSG00000155229 | MMS19 | chr10 | - | 97468251 | 97468406 | 97467504 | 97467583 | 97469645 | 97469723 | [illegible] |
| SE | 15881 | ENSG00000167491 | GATAD2A | chr19 | + | 19436130 | 19436219 | 19385825 | 19386138 | 19465344 | 19465614 | [illegible] |
| SE | 73139 | ENSG00000144233 | AMMECR1L | chr2 | - | 127865682 | 127865810 | 127861648 | 127865205 | 127866899 | 127866996 | [illegible] |
| SE | 11854 | ENSG00000121274 | TENT4B | chr16 | + | 50223173 | 50223387 | 50217555 | 50217663 | 50224656 | 50224783 | [illegible] |
| SE | 63754 | ENSG00000141429 | GALNT1 | chr18 | + | 35659857 | 35659930 | 35654655 | 35654801 | 35663627 | 35663802 | [illegible] |
| SE | 12498 | ENSG00000132600 | PRMT7 | chr16 | + | 68312041 | 68312176 | 68311018 | 68311099 | 68315896 | 68315957 | [illegible] |
| SE | 89009 | ENSG00000177565 | TBL1XR1 | chr3 | - | 177079826 | 177079945 | 177064919 | 177065022 | 177197120 | 177197184 | [illegible] |
| SE | 68326 | ENSG00000168016 | TRANK1 | chr3 | - | 36892900 | 36892984 | 36892201 | 36892340 | 36899108 | 36899259 | [illegible] |
| SE | 65715 | ENSG00000138434 | ITPRID2 | chr2 | + | 181928160 | 181928278 | 181921947 | 181922412 | 181929560 | 181929784 | [illegible] |
| SE | 95774 | ENSG00000125944 | HNRNPR | chr1 | - | 23333517 | 23333631 | 23323555 | 23323732 | 23337753 | 23337861 | [illegible] |
| SE | 85186 | ENSG00000140332 | TLE3 | chr15 | - | 70054437 | 70054685 | 70053173 | 70053374 | 70055048 | 70055263 | [illegible] |
| SE | 71995 | ENSG00000197635 | DPP4 | chr2 | - | 162039131 | 162039184 | 162038944 | 162039021 | 162045531 | 162045612 | [illegible] |
| SE | 63606 | ENSG00000152127 | MGAT5 | chr2 | + | 134412868 | 134413015 | 134402987 | 134403137 | 134422802 | 134422919 | [illegible] |
| SE | 64947 | ENSG00000163644 | PPM1K | chr4 | - | 88268189 | 88268334 | 88265000 | 88265135 | 88268740 | 88268906 | [illegible] |
| SE | 73533 | ENSG00000138380 | CARF | chr2 | + | 202977268 | 202977332 | 202974333 | 202974496 | 202981554 | 202981685 | [illegible] |
| SE | 67562 | ENSG00000061676 | NCKAP1 | chr2 | - | 182982820 | 182982927 | 182981243 | 182981376 | 182983285 | 182983382 | [illegible] |
| SE | 37856 | ENSG00000114573 | ATP6V1A | chr3 | + | 113780731 | 113780783 | 113778740 | 113778835 | 113781049 | 113781178 | [illegible] |
| SE | 2157 | ENSG00000127511 | SIN3B | chr19 | + | 16862883 | 16862979 | 16862351 | 16862559 | 16863679 | 16863796 | [illegible] |
| SE | 66546 | ENSG00000168438 | CDC40 | chr6 | + | 110217701 | 110217803 | 110215285 | 110215331 | 110219363 | 110219479 | [illegible] |
| SE | 40533 | ENSG00000008128 | CDK11A | chr1 | - | 1712263 | 1712427 | 1709517 | 1709640 | 1716345 | 1716478 | [illegible] |
| SE | 9433 | ENSG00000174238 | PITPNA | chr17 | - | 1541565 | 1541640 | 1538868 | 1538952 | 1548295 | 1548387 | [illegible] |
| SE | 46535 | ENSG00000223745 | CCDC18-AS1 | chr1 | - | 93315148 | 93315269 | 93305387 | 93305470 | 93324634 | 93324691 | [illegible] |
| SE | 99584 | ENSG00000149187 | CELF1 | chr11 | - | 47476845 | 47476959 | 47475335 | 47475521 | 47477296 | 47477425 | [illegible] |
| SE | 12104 | ENSG00000103051 | COG4 | chr16 | - | 70481763 | 70481865 | 70481358 | 70481487 | 70482091 | 70482175 | [illegible] |
| SE | 39241 | ENSG00000197948 | FCHSD1 | chr5 | - | 141647967 | 141648096 | 141647397 | 141647520 | 141648956 | 141649020 | [illegible] |
| SE | 71307 | ENSG00000163754 | GYG1 | chr3 | + | 148992657 | 148992772 | 148991567 | 148991647 | 148994141 | 148994277 | [illegible] |
| SE | 104731 | ENSG00000226210 | WASH8P | chr12 | - | 17722 | 17859 | 16968 | 17170 | 18036 | 18183 | [illegible] |
| SE | 83587 | ENSG00000107897 | ACBD5 | chr10 | - | 27217979 | 27218183 | 27215534 | 27215641 | 27219722 | 27219803 | [illegible] |
| SE | 103044 | ENSG00000099290 | WASHC2A | chr10 | + | 50099977 | 50100064 | 50097674 | 50097802 | 50104041 | 50104143 | [illegible] |
| SE | 107161 | ENSG00000136003 | ISCU | chr12 | + | 108567189 | 108567268 | 108565320 | 108565431 | 108568830 | 108569368 | [illegible] |
| SE | 9503 | ENSG00000154874 | CCDC144B | chr17 | - | 18587665 | 18587879 | 18583340 | 18583523 | 18594720 | 18594862 | [illegible] |
| SE | 28375 | ENSG00000145907 | G3BP1 | chr5 | + | 151800759 | 151800869 | 151799888 | 151800346 | 151803884 | 151803996 | [illegible] |
| SE | 103787 | ENSG00000049323 | LTBP1 | chr2 | + | 33134792 | 33134960 | 33110581 | 33110751 | 33186855 | 33187080 | [illegible] |
| SE | 34578 | ENSG00000009844 | VTA1 | chr6 | + | 142203984 | 142204065 | 142198438 | 142198615 | 142218497 | 142219067 | [illegible] |
| SE | 44400 | ENSG00000134419 | RPS15A | chr16 | - | 18788062 | 18788159 | 18784737 | 18784823 | 18788980 | 18789118 | [illegible] |
| SE | 76449 | ENSG00000168685 | IL7R | chr5 | + | 35867305 | 35867463 | 35860851 | 35860990 | 35871055 | 35871213 | [illegible] |
| SE | 31880 | ENSG00000143669 | LYST | chr1 | - | 235773841 | 235773991 | 235766078 | 235766277 | 235774912 | 235775086 | [illegible] |
| SE | 56272 | ENSG00000248429 | FAM198B-AS1 | chr4 | + | 158176911 | 158177046 | 158172576 | 158172940 | 158179660 | 158179735 | [illegible] |
| SE | 46480 | ENSG00000255154 | HTD2 | chr3 | + | 58315721 | 58315819 | 58310506 | 58310591 | 58316514 | 58316591 | [illegible] |
| SE | 69187 | ENSG00000113712 | CSNK1A1 | chr5 | - | 149515668 | 149515726 | 149513069 | 149513209 | 149517793 | 149517877 | [illegible] |
| SE | 44128 | ENSG00000183826 | BTBD9 | chr6 | - | 38593963 | 38594327 | 38592575 | 38592840 | 38597909 | 38598057 | [illegible] |
| SE | 73504 | ENSG00000138380 | CARF | chr2 | + | 202924296 | 202924415 | 202917876 | 202918043 | 202941859 | 202941980 | [illegible] |
| SE | 54659 | ENSG00000151694 | ADAM17 | chr2 | - | 9536697 | 9536828 | 9527500 | 9527954 | 9543152 | 9543226 | [illegible] |
| SE | 41837 | ENSG00000172456 | FGGY | chr1 | + | 59554123 | 59554227 | 59512310 | 59512439 | 59607802 | 59607910 | [illegible] |
| SE | 93849 | ENSG00000116580 | GON4L | chr1 | - | 155754374 | 155754488 | 155749658 | 155750733 | 155756957 | 155757077 | [illegible] |
| SE | 63802 | ENSG00000181163 | NPM1 | chr5 | + | 171390074 | 171390130 | 171387880 | 171388006 | 171391304 | 171391369 | [illegible] |
| SE | 48705 | ENSG00000143401 | ANP32E | chr1 | - | 150223185 | 150223240 | 150218416 | 150220761 | 150226607 | 150226795 | [illegible] |
| SE | 3837 | ENSG00000157540 | DYRK1A | chr21 | + | 37410569 | 37410670 | 37367556 | 37367628 | 37420298 | 37420384 | [illegible] |
| SE | 67643 | ENSG00000126768 | TIMM17B | chrX | - | 48895351 | 48895454 | 48895037 | 48895101 | 48896758 | 48896858 | [illegible] |
| SE | 57436 | ENSG00000065534 | MYLK | chr3 | - | 123647223 | 123647427 | 123640286 | 123640504 | 123648970 | 123649064 | [illegible] |
| SE | 91734 | ENSG00000182986 | ZNF320 | chr19 | - | 52891228 | 52891346 | 52890240 | 52890328 | 52893778 | 52893856 | [illegible] |
| SE | 52210 | ENSG00000127463 | EMC1 | chr1 | - | 19235129 | 19235252 | 19232935 | 19233135 | 19237141 | 19237238 | [illegible] |
| SE | 42047 | ENSG00000196352 | CD55 | chr1 | + | 207331107 | 207331296 | 207326751 | 207326837 | 207336692 | 207336788 | [illegible] |
| SE | 70378 | ENSG00000132330 | SCLY | chr2 | + | 238068064 | 238068165 | 238064356 | 238064469 | 238073782 | 238073847 | [illegible] |
| SE | 17003 | ENSG00000206149 | HERC2P9 | chr15 | + | 28629265 | 28629441 | 28612058 | 28612173 | 28632636 | 28632807 | [illegible] |
| SE | 83228 | ENSG00000205903 | ZNF316 | chr7 | + | 6642381 | 6642764 | 6641824 | 6641962 | 6642963 | 6643073 | [illegible] |
| SE | 12805 | ENSG00000102882 | MAPK3 | chr16 | - | 30115594 | 30115645 | 30114545 | 30114708 | 30116635 | 30116790 | [illegible] |
| SE | 104642 | ENSG00000079974 | RABL2B | chr22 | - | 50769904 | 50770016 | 50769454 | 50769552 | 50775771 | 50775851 | [illegible] |
| SE | 100410 | ENSG00000167770 | OTUB1 | chr11 | + | 63988626 | 63988752 | 63988336 | 63988398 | 63996529 | 63996648 | [illegible] |
| SE | 15022 | ENSG00000091157 | WDR7 | chr18 | + | 56717963 | 56718159 | 56696241 | 56696462 | 56731382 | 56731597 | [illegible] |
| SE | 102938 | ENSG00000084073 | ZMPSTE24 | chr1 | + | 40262833 | 40262983 | 40260838 | 40260985 | 40267785 | 40267872 | [illegible] |
| SE | 21169 | ENSG00000057663 | ATG5 | chr6 | - | 106217463 | 106217601 | 106201971 | 106202089 | 106248149 | 106248244 | [illegible] |
| SE | 84988 | ENSG00000170832 | USP32 | chr17 | - | 60288522 | 60288682 | 60269449 | 60269557 | 60294682 | 60294801 | [illegible] |
| SE | 25460 | ENSG00000217128 | FNIP1 | chr5 | - | 131698916 | 131699002 | 131679028 | 131679175 | 131704064 | 131704266 | [illegible] |
| SE | 94354 | ENSG00000148090 | AUH | chr9 | - | 91217276 | 91217327 | 91213959 | 91214425 | 91220804 | 91220919 | [illegible] |
| SE | 1727 | ENSG00000100325 | ASCC2 | chr22 | - | 29801993 | 29802208 | 29793359 | 29793676 | 29804637 | 29804824 | [illegible] |
| SE | 64606 | ENSG00000100644 | HIF1A | chr14 | + | 61740542 | 61740627 | 61738086 | 61738373 | 61740754 | 61741188 | [illegible] |
| SE | 92189 | ENSG00000120217 | CD274 | chr9 | + | 5466769 | 5466829 | 5465498 | 5465606 | 5467839 | 5467866 | [illegible] |
| SE | 84517 | ENSG00000128731 | HERC2 | chr15 | - | 28298516 | 28298629 | 28292887 | 28293022 | 28299401 | 28299516 | [illegible] |
| SE | 82992 | ENSG00000265681 | RPL17 | chr18 | - | 49491351 | 49491445 | 49490792 | 49490927 | 49491531 | 49491586 | [illegible] |
| SE | 84947 | ENSG00000139436 | GIT2 | chr12 | - | 109947255 | 109947504 | 109939164 | 109939247 | 109951211 | 109951316 | [illegible] |
| SE | 21181 | ENSG00000143252 | SDHC | chr1 | + | 161339561 | 161339588 | 161328395 | 161328497 | 161340593 | 161340655 | [illegible] |
| SE | 11107 | ENSG00000178188 | SH2B1 | chr16 | + | 28873169 | 28873269 | 28872533 | 28872758 | 28873446 | 28873607 | [illegible] |
| SE | 73893 | ENSG00000123600 | METTL8 | chr2 | - | 171331803 | 171331867 | 171326041 | 171326148 | 171337452 | 171337502 | [illegible] |
| SE | 8953 | ENSG00000108395 | TRIM37 | chr17 | - | 59001597 | 59001741 | 58999380 | 58999459 | 59012327 | 59012446 | [illegible] |
| SE | 78595 | ENSG00000088930 | XRN2 | chr20 | + | 21357735 | 21357792 | 21356585 | 21356665 | 21365420 | 21365489 | [illegible] |
| SE | 185 | ENSG00000160199 | PKNOX1 | chr21 | + | 43010052 | 43010224 | 43007490 | 43007618 | 43013067 | 43013238 | [illegible] |
| SE | 99090 | ENSG00000110274 | CEP164 | chr11 | + | 117338565 | 117338668 | 117335604 | 117335680 | 117344165 | 117344277 | [illegible] |
| SE | 74831 | ENSG00000033100 | CHPF2 | chr7 | + | 151236407 | 151236590 | 151235047 | 151235612 | 151237373 | 151238819 | [illegible] |
| SE | 109538 | ENSG00000127022 | CANX | chr5 | + | 179704215 | 179704430 | 179698934 | 179699102 | 179705678 | 179705852 | [illegible] |
| SE | 108105 | ENSG00000150990 | DHX37 | chr12 | - | 124980489 | 124980838 | 124977341 | 124977490 | 124982510 | 124982623 | [illegible] |
| SE | 59588 | ENSG00000118816 | CCNI | chr4 | - | 77055149 | 77055380 | 77047154 | 77048662 | 77055961 | 77056102 | [illegible] |
| SE | 1163 | ENSG00000100395 | L3MBTL2 | chr22 | + | 41224019 | 41224251 | 41221198 | 41221287 | 41224724 | 41224801 | [illegible] |
| SE | 83974 | ENSG00000068400 | GRIPAP1 | chrX | - | 48998153 | 48998180 | 48997249 | 48997357 | 48999237 | 48999299 | [illegible] |
| SE | 48773 | ENSG00000058804 | NDC1 | chr1 | - | 53800692 | 53800848 | 53796898 | 53797144 | 53803927 | 53804009 | [illegible] |
| SE | 33198 | ENSG00000163930 | BAP1 | chr3 | - | 52407173 | 52407316 | 52406252 | 52406907 | 52407398 | 52407460 | [illegible] |
| SE | 48303 | ENSG00000229474 | PATL2 | chr15 | - | 44710095 | 44710214 | 44676474 | 44676565 | 44711019 | 44711323 | [illegible] |
| SE | 51945 | ENSG00000287856 | NA | chr1 | - | 231370561 | 231370698 | 231367568 | 231367636 | 231373979 | 231374099 | [illegible] |
| SE | 80695 | ENSG00000175471 | MCTP1 | chr5 | - | 94870871 | 94870973 | 94870416 | 94870491 | 94871314 | 94871417 | [illegible] |
| SE | 47728 | ENSG00000145687 | SSBP2 | chr5 | - | 81440557 | 81440636 | 81437429 | 81437458 | 81446867 | 81446922 | [illegible] |
| SE | 39786 | ENSG00000152102 | FAM168B | chr2 | - | 131071854 | 131071938 | 131055552 | 131055695 | 131082576 | 131082657 | [illegible] |
| SE | 68858 | ENSG00000168918 | INPP5D | chr2 | + | 233125747 | 233125919 | 233122106 | 233122257 | 233130507 | 233130648 | [illegible] |
| SE | 10523 | ENSG00000125970 | RALY | chr20 | + | 34065043 | 34065160 | 33993679 | 33994131 | 34072065 | 34072330 | [illegible] |
| SE | 103396 | ENSG00000087338 | GMCL1 | chr2 | + | 69854822 | 69854960 | 69849651 | 69849742 | 69861277 | 69861347 | [illegible] |
| SE | 66985 | ENSG00000135930 | EIF4E2 | chr2 | + | 232557883 | 232558018 | 232556415 | 232556530 | 232566828 | 232566981 | [illegible] |
| SE | 66477 | ENSG00000131051 | RBM39 | chr20 | - | 35740524 | 35740597 | 35738967 | 35739017 | 35740823 | 35740887 | [illegible] |
| SE | 17951 | ENSG00000171469 | ZNF561 | chr19 | - | 9617044 | 9617171 | 9614020 | 9614103 | 9618090 | 9618166 | [illegible] |
| SE | 80882 | ENSG00000126883 | NUP214 | chr9 | + | 131135939 | 131136006 | 131134897 | 131135004 | 131139280 | 131139407 | [illegible] |
| SE | 32651 | ENSG00000150995 | ITPR1 | chr3 | + | 4653841 | 4653886 | 4652122 | 4652218 | 4658123 | 4658278 | [illegible] |
| SE | 25299 | ENSG00000134644 | PUM1 | chr1 | - | 30945345 | 30945483 | 30941150 | 30941272 | 30950132 | 30950261 | [illegible] |
| SE | 7542 | ENSG00000167851 | CD300A | chr17 | + | 74473535 | 74473874 | 74469945 | 74470242 | 74474531 | 74474685 | [illegible] |
| SE | 66819 | ENSG00000031003 | FAM13B | chr5 | - | 137956476 | 137956542 | 137954165 | 137954376 | 137959615 | 137959763 | [illegible] |
| SE | 36252 | ENSG00000168827 | GFM1 | chr3 | + | 158690162 | 158690323 | 158684523 | 158684668 | 158691138 | 158691192 | [illegible] |
| SE | 45064 | ENSG00000007341 | ST7L | chr1 | - | 112591524 | 112591603 | 112583971 | 112584126 | 112597970 | 112598086 | [illegible] |
| SE | 48920 | ENSG00000198408 | OGA | chr10 | - | 101793685 | 101793744 | 101792838 | 101792943 | 101797979 | 101798017 | [illegible] |
| SE | 44939 | ENSG00000138138 | ATAD1 | chr10 | - | 87778520 | 87778731 | 87776320 | 87776427 | 87784469 | 87784670 | [illegible] |
| SE | 78650 | ENSG00000168014 | C2CD3 | chr11 | - | 74161398 | 74161556 | 74139604 | 74139828 | 74168343 | 74168613 | [illegible] |
| SE | 7155 | ENSG00000101639 | CEP192 | chr18 | + | 13037236 | 13037301 | 13030464 | 13030608 | 13042203 | 13042334 | [illegible] |
| SE | 105104 | ENSG00000112297 | CRYBG1 | chr6 | + | 106561341 | 106561500 | 106558485 | 106558625 | 106563763 | 106563926 | [illegible] |
| SE | 40914 | ENSG00000151883 | PARP8 | chr5 | + | 50759642 | 50759732 | 50750150 | 50750188 | 50761820 | 50761898 | [illegible] |
| SE | 52212 | ENSG00000127463 | EMC1 | chr1 | - | 19238794 | 19238857 | 19238016 | 19238139 | 19239230 | 19239302 | [illegible] |
| SE | 31015 | ENSG00000145247 | OCIAD2 | chr4 | - | 48897803 | 48897857 | 48892771 | 48894053 | 48904483 | 48904611 | [illegible] |
| SE | 106151 | ENSG00000089022 | MAPKAPK5 | chr12 | + | 111868752 | 111868861 | 111867571 | 111867669 | 111870270 | 111870360 | [illegible] |
| SE | 1201 | ENSG00000100387 | RBX1 | chr22 | + | 40953554 | 40953633 | 40951346 | 40951476 | 40964046 | 40964117 | [illegible] |
| SE | 41778 | ENSG00000078142 | PIK3C3 | chr18 | + | 41957569 | 41957758 | 41955233 | 41955359 | 41962488 | 41962632 | [illegible] |
| SE | 103564 | ENSG00000150093 | ITGB1 | chr10 | - | 32911885 | 32912124 | 32911447 | 32911670 | 32919884 | 32920084 | [illegible] |
| SE | 21956 | ENSG00000012660 | ELOVL5 | chr6 | - | 53273219 | 53273344 | 53267397 | 53269270 | 53275089 | 53275261 | [illegible] |
| SE | 5056 | ENSG00000177410 | ZFAS1 | chr20 | + | 49280484 | 49280570 | 49279103 | 49279208 | 49289044 | 49289260 | [illegible] |
| SE | 60971 | ENSG00000151693 | ASAP2 | chr2 | + | 9318523 | 9318598 | 9297299 | 9297445 | 9320287 | 9320337 | [illegible] |
| SE | 25719 | ENSG00000247774 | PCED1B-AS1 | chr12 | - | 47210639 | 47210761 | 47205899 | 47206585 | 47216164 | 47216200 | [illegible] |
| SE | 67872 | ENSG00000133627 | ACTR3B | chr7 | + | 152852125 | 152852251 | 152825029 | 152825122 | 152853493 | 152853577 | [illegible] |
| SE | 106457 | ENSG00000198270 | TMEM116 | chr12 | - | 111933885 | 111936830 | 111932584 | 111932659 | 111937159 | 111937243 | [illegible] |
| SE | 88110 | ENSG00000197265 | GTF2E2 | chr8 | - | 30637244 | 30637274 | 30636929 | 30636960 | 30653432 | 30653602 | [illegible] |
| SE | 13140 | ENSG00000118900 | UBN1 | chr16 | + | 4876870 | 4877111 | 4874210 | 4875434 | 4880082 | 4880294 | [illegible] |
| SE | 13891 | ENSG00000069329 | VPS35 | chr16 | - | 46668929 | 46669052 | 46662982 | 46663162 | 46671704 | 46671860 | [illegible] |
| SE | 83614 | ENSG00000130066 | SAT1 | chrX | + | 23783799 | 23783883 | 23783657 | 23783709 | 23785327 | 23785429 | [illegible] |

| | | | | | | | | | | | | | | | | | | | | | | | | | | | | | | | | | | | | | | | | | | | | | | |
|---|---|---|---|---|---|---|---|---|---|---|---|---|---|---|---|---|---|---|---|---|---|---|---|---|---|---|---|---|---|---|---|---|---|---|---|---|---|---|---|---|---|---|---|---|---|---|
| SE | 91205 | ENSG00000288602 | C8orf44-SGK3 | chr8 | + | 66804374 | 66804447 | 66798541 | 66798625 | 66813852 | 66813928 | 119 | 85 | 83 | 104 | 185 | 3 | 0 | 2 | 1 | 2 | 6 | 63 | 76 | 74 | 18 | 0 | 0 | 1 | 6 | 0 | 222 | 149 | 0.727518963 | 1 | 0.964 | 1 | 0.965 | 0.986 | 0.984 | 1 | 1 | 0.981 | 0.892 | 1 | 0.005 |
| SE | 54923 | ENSG00000124731 | TREM1 | chr6 | - | 41280960 | 41281153 | 41273604 | 41276230 | 41282394 | 41282751 | 404 | 242 | 280 | 446 | 687 | 72 | 25 | 17 | 49 | 69 | 966 | 306 | 937 | 525 | 229 | 64 | 64 | 101 | 65 | 28 | 298 | 149 | 0.72753866 | 1 | 0.737 | 0.829 | 0.892 | 0.82 | 0.833 | 0.883 | 0.705 | 0.823 | 0.802 | 0.804 | 0.019 |
| SE | 23176 | ENSG00000113240 | CLK4 | chr5 | - | 178605824 | 178605865 | 178605302 | 178605382 | 178608375 | 178608458 | 17 | 17 | 27 | 21 | 42 | 248 | 191 | 233 | 196 | 446 | 0 | 11 | 13 | 0 | 5 | 30 | 69 | 90 | 20 | 10 | 190 | 149 | 0.727552949 | 1 | 0.051 | 0.065 | 0.083 | 0.078 | 0.069 | 0 | 0.111 | 0.102 | 0 | 0.282 | -0.03 |
| SE | 29824 | ENSG00000129187 | DCTD | chr4 | - | 182914922 | 182915058 | 182894488 | 182894605 | 182915460 | 182915560 | 337 | 181 | 271 | 234 | 393 | 8 | 0 | 4 | 6 | 4 | 309 | 245 | 138 | 105 | 438 | 0 | 12 | 2 | 1 | 3 | 285 | 149 | 0.727601797 | 1 | 0.957 | 1 | 0.973 | 0.953 | 0.981 | 1 | 0.914 | 0.973 | 0.982 | 0.987 | 0.002 |
| SE | 73970 | ENSG00000166436 | TRIM66 | chr11 | - | 8647969 | 8648086 | 8646446 | 8646561 | 8648415 | 8648548 | 51 | 25 | 50 | 62 | 36 | 0 | 0 | 0 | 2 | 0 | 85 | 62 | 42 | 109 | 139 | 0 | 0 | 0 | 8 | 0 | 266 | 149 | 0.727660547 | 1 | 1 | 1 | 1 | 0.946 | 1 | 1 | 1 | 1 | 0.884 | 1 | 0.012 |
| SE | 79017 | ENSG00000160218 | TRAPPC10 | chr21 | + | 44055697 | 44055893 | 44052279 | 44052476 | 44059102 | 44059214 | 429 | 205 | 236 | 302 | 362 | 0 | 2 | 3 | 8 | 0 | 285 | 395 | 310 | 359 | 312 | 0 | 1 | 1 | 4 | 0 | 298 | 149 | 0.727690129 | 1 | 1 | 0.981 | 0.975 | 0.95 | 1 | 1 | 0.995 | 0.994 | 0.978 | 1 | -0.012 |
| SE | 70384 | ENSG00000132330 | SCLY | chr2 | + | 238081708 | 238081836 | 238073782 | 238073847 | 238082044 | 238082202 | 48 | 22 | 56 | 62 | 7 | 0 | 0 | 0 | 0 | 0 | 44 | 21 | 13 | 42 | 51 | 0 | 0 | 0 | 0 | 5 | 277 | 149 | 0.72769706 | 1 | 1 | 1 | 1 | 1 | 1 | 1 | 1 | 1 | 1 | 0.846 | 0.031 |
| SE | 6912 | ENSG00000175354 | PTPN2 | chr18 | - | 12817155 | 12817265 | 12814324 | 12814355 | 12819213 | 12819282 | 252 | 163 | 169 | 256 | 359 | 0 | 2 | 0 | 1 | 0 | 130 | 179 | 197 | 173 | 21 | 0 | 6 | 2 | 0 | 0 | 259 | 149 | 0.727790724 | 1 | 1 | 0.979 | 1 | 0.993 | 1 | 1 | 0.945 | 0.983 | 1 | 1 | 0.009 |
| SE | 27505 | ENSG00000104517 | UBR5 | chr8 | - | 102327741 | 102327896 | 102326545 | 102326663 | 102329089 | 102329200 | 398 | 369 | 360 | 434 | 514 | 5 | 0 | 1 | 2 | 0 | 299 | 452 | 511 | 242 | 138 | 0 | 8 | 7 | 0 | 0 | 298 | 149 | 0.727917935 | 1 | 0.975 | 1 | 0.994 | 0.991 | 1 | 1 | 0.966 | 0.973 | 1 | 1 | 0.004 |
| SE | 25903 | ENSG00000013375 | PGM3 | chr6 | - | 83190808 | 83191014 | 83188613 | 83188798 | 83193178 | 83193231 | 42 | 24 | 43 | 44 | 24 | 4 | 0 | 0 | 0 | 1 | 1 | 11 | 35 | 24 | 30 | 0 | 0 | 4 | 0 | 0 | 298 | 149 | 0.72794935 | 1 | 0.84 | 1 | 1 | 1 | 0.923 | 1 | 1 | 0.814 | 1 | 1 | -0.01 |
| SE | 40087 | ENSG00000125347 | IRF1 | chr5 | - | 132489391 | 132489483 | 132487925 | 132488025 | 132490544 | 132490726 | 297 | 243 | 375 | 419 | 254 | 3 | 3 | 0 | 8 | 2 | 1168 | 211 | 390 | 667 | 677 | 10 | 4 | 0 | 8 | 0 | 241 | 149 | 0.727975671 | 1 | 0.984 | 0.98 | 1 | 0.97 | 0.987 | 0.986 | 0.97 | 1 | 0.981 | 1 | -0.003 |
| SE | 79194 | ENSG00000054277 | OPN3 | chr1 | - | 241600379 | 241600653 | 241597762 | 241597997 | 241604259 | 241604579 | 10 | 3 | 0 | 14 | 2 | 20 | 11 | 8 | 12 | 5 | 0 | 20 | 3 | 4 | 10 | 17 | 15 | 6 | 6 | 19 | 298 | 149 | 0.728005408 | 1 | 0.2 | 0.12 | 0 | 0.368 | 0.167 | 0 | 0.4 | 0.2 | 0.25 | 0.208 | -0.041 |
| SE | 51492 | ENSG00000160087 | UBE2J2 | chr1 | - | 1267861 | 1267992 | 1262852 | 1263386 | 1272852 | 1273471 | 68 | 44 | 50 | 57 | 59 | 0 | 1 | 0 | 0 | 0 | 104 | 53 | 18 | 74 | 190 | 0 | 6 | 0 | 0 | 0 | 280 | 149 | 0.728075906 | 1 | 1 | 0.959 | 1 | 1 | 1 | 1 | 0.825 | 1 | 1 | 1 | 0.027 |
| SE | 3979 | ENSG00000126261 | UBA2 | chr19 | + | 34433347 | 34433412 | 34431860 | 34431931 | 34434867 | 34434968 | 689 | 433 | 631 | 713 | 1477 | 2 | 4 | 1 | 2 | 6 | 437 | 521 | 356 | 391 | 380 | 0 | 0 | 3 | 6 | 10 | 214 | 149 | 0.72815683 | 1 | 0.996 | 0.987 | 0.998 | 0.996 | 0.994 | 1 | 1 | 0.988 | 0.978 | 0.964 | 0.008 |
| SE | 31611 | ENSG00000181789 | COPG1 | chr3 | + | 129268931 | 129269000 | 129268494 | 129268620 | 129271766 | 129271909 | 145 | 125 | 137 | 143 | 102 | 0 | 0 | 1 | 2 | 1 | 230 | 182 | 151 | 157 | 245 | 0 | 0 | 1 | 0 | 12 | 218 | 149 | 0.728224138 | 1 | 1 | 1 | 0.989 | 0.98 | 0.986 | 1 | 1 | 0.99 | 1 | 0.933 | 0.006 |
| SE | 106277 | ENSG00000251301 | LINC02384 | chr12 | - | 68442063 | 68442141 | 68380462 | 68383602 | 68451366 | 68451385 | 95 | 61 | 8 | 12 | 44 | 4 | 0 | 0 | 2 | 0 | 68 | 115 | 28 | 23 | 75 | 0 | 0 | 0 | 0 | 0 | 227 | 149 | 0.72824211 | 1 | 0.94 | 1 | 1 | 0.798 | 1 | 1 | 1 | 1 | 1 | 1 | -0.052 |
| SE | 69504 | ENSG00000106799 | TGFBR1 | chr9 | + | 99132508 | 99132739 | 99128835 | 99129100 | 99137858 | 99138089 | 415 | 215 | 266 | 320 | 310 | 2 | 2 | 5 | 2 | 1 | 167 | 169 | 134 | 106 | 39 | 0 | 1 | 0 | 0 | 0 | 298 | 149 | 0.728247578 | 1 | 0.99 | 0.982 | 0.964 | 0.988 | 0.994 | 1 | 0.988 | 1 | 1 | 1 | -0.014 |
| SE | 39117 | ENSG00000159692 | CTBP1 | chr4 | - | 1241324 | 1241519 | 1238182 | 1238337 | 1248915 | 1248976 | 493 | 211 | 321 | 430 | 377 | 6 | 2 | 6 | 3 | 0 | 262 | 368 | 244 | 196 | 376 | 0 | 2 | 1 | 1 | 9 | 298 | 149 | 0.728504148 | 1 | 0.976 | 0.981 | 0.964 | 0.986 | 1 | 1 | 0.989 | 0.992 | 0.99 | 0.954 | -0.004 |
| SE | 23518 | ENSG00000100284 | TOM1 | chr22 | + | 35330346 | 35330480 | 35327270 | 35327387 | 35331295 | 35331495 | 60 | 25 | 59 | 55 | 43 | 0 | 0 | 0 | 0 | 5 | 80 | 117 | 46 | 90 | 141 | 0 | 0 | 0 | 0 | 0 | 283 | 149 | 0.728526631 | 1 | 1 | 1 | 1 | 1 | 0.819 | 1 | 1 | 1 | 1 | 1 | -0.036 |
| SE | 55314 | ENSG00000166938 | DIS3L | chr15 | + | 66318448 | 66318618 | 66315035 | 66315215 | 66320570 | 66320732 | 78 | 32 | 71 | 85 | 103 | 1 | 0 | 6 | 2 | 0 | 82 | 38 | 35 | 57 | 84 | 5 | 0 | 1 | 2 | 0 | 298 | 149 | 0.728605248 | 1 | 0.975 | 1 | 0.855 | 0.955 | 1 | 0.891 | 1 | 0.946 | 0.934 | 1 | 0.003 |
| SE | 38545 | ENSG00000180098 | TRNAU1AP | chr1 | + | 28561112 | 28561260 | 28560632 | 28560732 | 28561345 | 28561398 | 22 | 6 | 14 | 17 | 16 | 50 | 20 | 53 | 44 | 45 | 5 | 11 | 12 | 20 | 25 | 39 | 36 | 15 | 35 | 63 | 297 | 149 | 0.728639079 | 1 | 0.181 | 0.131 | 0.117 | 0.162 | 0.151 | 0.06 | 0.133 | 0.286 | 0.223 | 0.166 | -0.025 |
| SE | 60421 | ENSG00000159339 | PADI4 | chr1 | + | 17338037 | 17338155 | 17333942 | 17334009 | 17339687 | 17339813 | 121 | 52 | 79 | 145 | 146 | 1 | 0 | 0 | 0 | 0 | 285 | 223 | 199 | 1028 | 618 | 0 | 2 | 0 | 18 | 0 | 267 | 149 | 0.728706704 | 1 | 0.985 | 1 | 1 | 1 | 1 | 1 | 0.984 | 1 | 0.97 | 1 | 0.006 |
| SE | 83365 | ENSG00000166716 | ZNF592 | chr15 | + | 84764706 | 84764815 | 84748591 | 84748664 | 84778182 | 84778312 | 60 | 27 | 42 | 52 | 63 | 0 | 0 | 0 | 0 | 0 | 191 | 78 | 47 | 141 | 137 | 0 | 0 | 1 | 0 | 8 | 258 | 149 | 0.728713055 | 1 | 1 | 1 | 1 | 1 | 1 | 1 | 1 | 0.964 | 1 | 0.908 | 0.026 |
| SE | 5535 | ENSG00000244687 | UBE2V1 | chr20 | - | 50096671 | 50096820 | 50083877 | 50084254 | 50115484 | 50115621 | 221 | 170 | 207 | 205 | 286 | 0 | 1 | 0 | 0 | 0 | 394 | 300 | 218 | 259 | 522 | 0 | 1 | 8 | 1 | 5 | 298 | 149 | 0.728719131 | 1 | 1 | 0.988 | 1 | 1 | 1 | 1 | 0.993 | 0.932 | 0.992 | 0.981 | 0.018 |
| SE | 21911 | ENSG00000109775 | UFSP2 | chr4 | - | 185422484 | 185422553 | 185418586 | 185418770 | 185425865 | 185425978 | 30 | 22 | 32 | 26 | 31 | 0 | 0 | 0 | 0 | 0 | 1 | 21 | 15 | 16 | 3 | 0 | 0 | 4 | 0 | 0 | 218 | 149 | 0.728767813 | 1 | 1 | 1 | 1 | 1 | 1 | 1 | 1 | 0.719 | 1 | 1 | 0.056 |
| SE | 95793 | ENSG00000143614 | GATAD2B | chr1 | - | 153819605 | 153819735 | 153818790 | 153818922 | 153828012 | 153828348 | 289 | 180 | 293 | 309 | 431 | 1 | 0 | 2 | 7 | 2 | 382 | 404 | 221 | 242 | 317 | 10 | 0 | 3 | 0 | 0 | 279 | 149 | 0.728899594 | 1 | 0.994 | 1 | 0.987 | 0.959 | 0.991 | 0.953 | 1 | 0.975 | 1 | 1 | 0.001 |
| SE | 50780 | ENSG00000103351 | CLUAP1 | chr16 | + | 3504719 | 3504831 | 3502043 | 3502084 | 3506330 | 3506415 | 18 | 15 | 18 | 15 | 43 | 0 | 0 | 4 | 0 | 0 | 10 | 1 | 13 | 1 | 4 | 0 | 0 | 0 | 0 | 0 | 261 | 149 | 0.728957736 | 1 | 1 | 1 | 0.72 | 1 | 1 | 1 | 1 | 1 | 1 | 1 | -0.056 |
| SE | 51629 | ENSG00000181523 | SGSH | chr17 | - | 80211868 | 80211923 | 80210396 | 80211011 | 80212070 | 80212218 | 4 | 0 | 4 | 1 | 1 | 11 | 15 | 28 | 16 | 14 | 4 | 0 | 0 | 5 | 0 | 6 | 18 | 11 | 6 | 30 | 204 | 149 | 0.729007117 | 1 | 0.21 | 0 | 0.094 | 0.044 | 0.05 | 0.327 | 0 | 0 | 0.378 | 0 | -0.061 |
| SE | 50024 | ENSG00000165359 | INTS6L | chrX | + | 135556167 | 135556300 | 135551993 | 135552146 | 135569336 | 135569431 | 119 | 115 | 130 | 141 | 267 | 4 | 1 | 2 | 1 | 0 | 63 | 106 | 75 | 78 | 34 | 0 | 0 | 2 | 0 | 0 | 282 | 149 | 0.729007173 | 1 | 0.94 | 0.984 | 0.972 | 0.987 | 1 | 1 | 1 | 0.952 | 1 | 1 | -0.014 |
| SE | 41917 | ENSG00000143110 | C1orf162 | chr1 | + | 111477333 | 111477428 | 111476797 | 111476867 | 111477982 | 111478512 | 112 | 77 | 71 | 110 | 147 | 4 | 0 | 1 | 6 | 14 | 283 | 102 | 142 | 211 | 189 | 13 | 5 | 2 | 13 | 11 | 244 | 149 | 0.729114594 | 1 | 0.945 | 1 | 0.977 | 0.918 | 0.865 | 0.93 | 0.926 | 0.977 | 0.908 | 0.913 | 0.01 |
| SE | 62715 | ENSG00000151779 | NBAS | chr2 | - | 15287072 | 15287183 | 15276850 | 15277101 | 15292536 | 15292766 | 101 | 55 | 73 | 84 | 78 | 0 | 0 | 2 | 2 | 2 | 58 | 48 | 39 | 54 | 30 | 0 | 0 | 0 | 0 | 0 | 260 | 149 | 0.729123838 | 1 | 1 | 1 | 0.954 | 0.96 | 0.957 | 1 | 1 | 1 | 1 | 1 | -0.026 |
| SE | 466 | ENSG00000243927 | MRPS6 | chr21 | + | 34125340 | 34125480 | 34103496 | 34103519 | 34142407 | 34143030 | 112 | 70 | 96 | 110 | 95 | 0 | 0 | 0 | 0 | 0 | 63 | 90 | 30 | 61 | 90 | 6 | 0 | 0 | 0 | 0 | 289 | 149 | 0.729168409 | 1 | 1 | 1 | 1 | 1 | 1 | 0.844 | 1 | 1 | 1 | 1 | 0.031 |
| SE | 37418 | ENSG00000114902 | SPCS1 | chr3 | + | 52706792 | 52706879 | 52706142 | 52706703 | 52707686 | 52708019 | 243 | 131 | 218 | 216 | 89 | 1 | 1 | 4 | 1 | 0 | 266 | 317 | 256 | 233 | 302 | 11 | 0 | 2 | 1 | 0 | 236 | 149 | 0.729244518 | 1 | 0.994 | 0.988 | 0.972 | 0.993 | 1 | 0.939 | 1 | 0.988 | 0.993 | 1 | 0.005 |
| SE | 71275 | ENSG00000121741 | ZMYM2 | chr13 | + | 20062845 | 20062971 | 20061052 | 20061224 | 20064450 | 20064545 | 290 | 237 | 282 | 286 | 453 | 0 | 1 | 4 | 5 | 3 | 142 | 163 | 167 | 57 | 69 | 0 | 0 | 1 | 0 | 0 | 275 | 149 | 0.72925731 | 1 | 1 | 0.992 | 0.974 | 0.969 | 0.988 | 1 | 1 | 0.989 | 1 | 1 | -0.013 |
| SE | 11873 | ENSG00000121281 | ADCY7 | chr16 | + | 50301081 | 50301214 | 50298903 | 50299031 | 50304359 | 50304516 | 91 | 25 | 62 | 92 | 35 | 0 | 0 | 0 | 2 | 0 | 43 | 47 | 42 | 47 | 88 | 5 | 0 | 0 | 0 | 0 | 282 | 149 | 0.729307162 | 1 | 1 | 1 | 1 | 0.96 | 1 | 0.82 | 1 | 1 | 1 | 1 | 0.028 |
| SE | 29964 | ENSG00000143624 | INTS3 | chr1 | + | 153747278 | 153747363 | 153746956 | 153747070 | 153748688 | 153748755 | 79 | 64 | 46 | 81 | 91 | 0 | 0 | 2 | 0 | 1 | 68 | 54 | 25 | 103 | 133 | 5 | 0 | 0 | 8 | 0 | 234 | 149 | 0.729343922 | 1 | 1 | 1 | 0.936 | 1 | 0.983 | 1 | 1 | 1 | 0.891 | 1 | 0.006 |
| SE | 9313 | ENSG00000133030 | MPRIP | chr17 | + | 17176425 | 17176512 | 17173915 | 17174075 | 17177249 | 17177396 | 188 | 101 | 164 | 225 | 157 | 0 | 0 | 0 | 0 | 4 | 110 | 122 | 176 | 102 | 252 | 10 | 0 | 0 | 0 | 0 | 236 | 149 | 0.7293565 | 1 | 1 | 1 | 1 | 1 | 0.961 | 0.874 | 1 | 1 | 1 | 1 | 0.017 |
| SE | 25296 | ENSG00000134644 | PUM1 | chr1 | - | 30941997 | 30942123 | 30941150 | 30941272 | 30950126 | 30950261 | 212 | 137 | 224 | 248 | 167 | 1 | 0 | 0 | 3 | 0 | 208 | 232 | 131 | 181 | 144 | 0 | 0 | 0 | 10 | 0 | 275 | 149 | 0.729401385 | 1 | 0.991 | 1 | 1 | 0.978 | 1 | 1 | 1 | 1 | 0.907 | 1 | 0.012 |
| SE | 51538 | ENSG00000176393 | RNPEP | chr1 | + | 202001658 | 202001767 | 201999901 | 202000015 | 202003236 | 202003318 | 87 | 44 | 62 | 71 | 122 | 0 | 0 | 0 | 0 | 0 | 193 | 91 | 53 | 177 | 328 | 0 | 0 | 2 | 7 | 0 | 258 | 149 | 0.729622945 | 1 | 1 | 1 | 1 | 1 | 1 | 1 | 1 | 0.939 | 0.936 | 1 | 0.025 |
| [illegible] | | | | | | | | | | | | | | | | | | | | | | | | | | | | | | | | | | | | | | | | | | | | | | |

| | | | | | | | | | | | | | | | | | | | | | | | | | | | | | | | | | | | | | | | | | | | | | | |
|---|---|---|---|---|---|---|---|---|---|---|---|---|---|---|---|---|---|---|---|---|---|---|---|---|---|---|---|---|---|---|---|---|---|---|---|---|---|---|---|---|---|---|---|---|---|---|
| SE | 53728 | ENSG00000151503 | NCAPD3 | chr11 | - | 134183012 | 134183153 | 134181076 | 134181184 | 134184636 | 134184752 | 2 | 2 | 2 | 4 | 0 | 22 | 21 | 26 | 21 | 61 | 0 | 0 | 7 | 0 | 0 | 44 | 31 | 10 | 40 | 22 | 290 | 149 | 0.738758575 | 1 | 0.045 | 0.047 | 0.038 | 0.089 | 0 | 0 | 0 | 0.265 | 0 | 0 | -0.009 |
| SE | 60602 | ENSG00000147121 | KRBOX4 | chrX | + | 46462747 | 46462874 | 46450430 | 46450475 | 46463197 | 46463293 | 69 | 38 | 75 | 122 | 87 | 0 | 1 | 0 | 1 | 0 | 82 | 119 | 61 | 81 | 111 | 7 | 0 | 0 | 0 | 0 | 276 | 149 | 0.738765359 | 1 | 1 | 0.954 | 1 | 0.985 | 1 | 0.863 | 1 | 1 | 1 | 1 | 0.015 |
| SE | 65305 | ENSG00000169621 | APLF | chr2 | + | 68513547 | 68513684 | 68502730 | 68502903 | 68526060 | 68526242 | 18 | 18 | 15 | 26 | 45 | 0 | 0 | 0 | 0 | 8 | 39 | 16 | 5 | 16 | 55 | 0 | 9 | 0 | 0 | 3 | 286 | 149 | 0.738808803 | 1 | 1 | 1 | 1 | 1 | 0.746 | 1 | 0.481 | 1 | 1 | 0.905 | 0.072 |
| SE | 83387 | ENSG00000174428 | GTF2IRD2B | chr7 | + | 75123135 | 75123319 | 75120890 | 75121010 | 75125915 | 75125970 | 31 | 20 | 30 | 29 | 33 | 0 | 3 | 1 | 4 | 4 | 72 | 36 | 18 | 32 | 6 | 0 | 1 | 4 | 0 | 7 | 298 | 149 | 0.738897012 | 1 | 1 | 0.769 | 0.938 | 0.784 | 0.805 | 1 | 0.947 | 0.692 | 1 | 0.3 | 0.071 |
| SE | 53891 | ENSG00000166523 | CLEC4E | chr12 | - | 8539854 | 8539947 | 8538953 | 8539306 | 8540760 | 8540873 | 343 | 172 | 100 | 132 | 229 | 0 | 1 | 4 | 0 | 1 | 240 | 315 | 552 | 330 | 160 | 0 | 1 | 8 | 7 | 0 | 242 | 149 | 0.738912292 | 1 | 1 | 0.991 | 0.939 | 1 | 0.993 | 1 | 0.995 | 0.977 | 0.967 | 1 | -0.003 |
| SE | 75459 | ENSG00000197081 | IGF2R | chr6 | + | 160048374 | 160048543 | 160047791 | 160047907 | 160050472 | 160050652 | 974 | 362 | 466 | 601 | 647 | 0 | 0 | 0 | 1 | 0 | 767 | 676 | 743 | 582 | 803 | 13 | 1 | 5 | 3 | 0 | 298 | 149 | 0.738941641 | 1 | 1 | 1 | 1 | 0.997 | 1 | 0.967 | 0.997 | 0.987 | 0.99 | 1 | 0.011 |
| SE | 24215 | ENSG00000113742 | CPEB4 | chr5 | + | 173942883 | 173943049 | 173910522 | 173910604 | 173944966 | 173945140 | 2 | 4 | 3 | 4 | 6 | 17 | 13 | 17 | 18 | 37 | 0 | 10 | 7 | 3 | 0 | 5 | 39 | 34 | 9 | 5 | 298 | 149 | 0.739010378 | 1 | 0.056 | 0.133 | 0.081 | 0.1 | 0.075 | 0 | 0.114 | 0.093 | 0.143 | 0 | 0.019 |
| SE | 97390 | ENSG00000113163 | CERT1 | chr5 | - | 75402971 | 75403058 | 75389591 | 75389687 | 75411010 | 75411103 | 104 | 109 | 70 | 108 | 107 | 5 | 1 | 0 | 1 | 0 | 26 | 71 | 103 | 74 | 45 | 0 | 3 | 0 | 0 | 0 | 236 | 149 | 0.739023069 | 1 | 0.929 | 0.986 | 1 | 0.986 | 1 | 1 | 0.937 | 1 | 1 | 1 | -0.007 |
| SE | 68768 | ENSG00000163872 | YEATS2 | chr3 | + | 183790796 | 183790980 | 183786124 | 183786301 | 183797922 | 183798051 | 93 | 72 | 96 | 102 | 72 | 0 | 0 | 0 | 0 | 0 | 100 | 83 | 64 | 82 | 50 | 0 | 0 | 0 | 6 | 0 | 298 | 149 | 0.739098278 | 1 | 1 | 1 | 1 | 1 | 1 | 1 | 1 | 1 | 0.872 | 1 | 0.026 |
| [illegible] | | | | | | | | | | | | | | | | | | | | | | | | | | | | | | | | | | | | | | | | | | | | | | |

| | | | | | | | | | | | | | | | | | | | | | | | | | | | | | | | | | | | | | | | | | | | | | | |
|---|---|---|---|---|---|---|---|---|---|---|---|---|---|---|---|---|---|---|---|---|---|---|---|---|---|---|---|---|---|---|---|---|---|---|---|---|---|---|---|---|---|---|---|---|---|---|
| SE | 15008 | ENSG00000128915 | ICE2 | chr15 | - | 60466593 | 60466713 | 60456656 | 60456794 | 60468060 | 60468162 | 246 | 165 | 202 | 232 | 342 | 1 | 6 | 4 | 1 | 1 | 6 | 129 | 65 | 23 | 45 | 0 | 0 | 0 | 0 | 0 | 269 | 149 | 0.751488017 | 1 | 0.993 | 0.938 | 0.965 | 0.992 | 0.995 | 1 | 1 | 1 | 1 | 1 | -0.023 |
| SE | 24361 | ENSG00000132780 | NASP | chr1 | + | 45588650 | 45588948 | 45584044 | 45584205 | 45591222 | 45591270 | 0 | 0 | 0 | 0 | 0 | 69 | 56 | 66 | 79 | 28 | 0 | 0 | 0 | 0 | 12 | 67 | 83 | 72 | 133 | 61 | 298 | 149 | 0.751532553 | 1 | 0 | 0 | 0 | 0 | 0 | 0 | 0 | 0 | 0 | 0.09 | -0.018 |
| SE | 98475 | ENSG00000151702 | FLI1 | chr11 | + | 128748236 | 128748290 | 128694091 | 128694276 | 128758114 | 128758326 | 1 | 2 | 0 | 3 | 0 | 435 | 259 | 308 | 311 | 238 | 0 | 0 | 0 | 1 | 17 | 385 | 309 | 222 | 326 | 243 | 203 | 149 | 0.751534097 | 1 | 0.002 | 0.006 | 0 | 0.007 | 0 | 0 | 0 | 0 | 0.002 | 0.049 | -0.007 |
| SE | 71464 | ENSG00000130038 | CRACR2A | chr12 | - | 3638123 | 3638454 | 3633603 | 3633736 | 3641731 | 3641838 | 74 | 50 | 60 | 43 | 55 | 0 | 0 | 0 | 0 | 0 | 59 | 61 | 50 | 37 | 20 | 0 | 0 | 5 | 0 | 0 | 298 | 149 | 0.75158806 | 1 | 1 | 1 | 1 | 1 | 1 | 1 | 1 | 0.833 | 1 | 1 | 0.033 |
| SE | 45808 | ENSG00000108654 | DDX5 | chr17 | - | 64500548 | 64501891 | 64498256 | 64500326 | 64502009 | 64502069 | 5435 | 4131 | 4666 | 6017 | 5552 | 7 | 3 | 9 | 10 | 9 | 1654 | 3322 | 3230 | 2856 | 307 | 22 | 3 | 15 | 11 | 0 | 298 | 149 | 0.751755647 | 1 | 0.997 | 0.999 | 0.996 | 0.997 | 0.997 | 0.974 | 0.998 | 0.991 | 0.992 | 1 | 0.006 |
| SE | 109004 | ENSG00000105352 | CEACAM4 | chr19 | - | 41620574 | 41620627 | 41620210 | 41620242 | 41625600 | 41625960 | 13 | 3 | 7 | 5 | 4 | 0 | 0 | 0 | 0 | 0 | 29 | 17 | 39 | 21 | 14 | 0 | 0 | 5 | 0 | 0 | 202 | 149 | 0.75184574 | 1 | 1 | 1 | 1 | 1 | 1 | 1 | 1 | 0.852 | 1 | 1 | 0.03 |
| SE | 58303 | ENSG00000117262 | GPR89A | chr1 | + | 145646183 | 145646272 | 145645646 | 145645743 | 145647174 | 145647218 | 1 | 3 | 3 | 0 | 1 | 2 | 0 | 0 | 2 | 0 | 13 | 0 | 1 | 7 | 2 | 0 | 7 | 0 | 2 | 0 | 238 | 149 | 0.751871934 | 1 | 0.238 | 1 | 1 | 0 | 1 | 1 | 0 | 1 | 0.687 | 1 | -0.09 |
| SE | 64938 | ENSG00000170873 | MTSS1 | chr8 | - | 124557680 | 124557875 | 124556231 | 124556405 | 124562781 | 124562992 | 100 | 41 | 44 | 88 | 30 | 10 | 0 | 9 | 4 | 0 | 38 | 77 | 51 | 61 | 151 | 0 | 30 | 1 | 2 | 10 | 298 | 149 | 0.751878567 | 1 | 0.833 | 1 | 0.71 | 0.917 | 1 | 1 | 0.562 | 0.962 | 0.938 | 0.883 | 0.023 |
| SE | 64059 | ENSG00000115464 | USP34 | chr2 | - | 61350567 | 61350693 | 61350259 | 61350389 | 61370320 | 61370413 | 450 | 405 | 382 | 450 | 634 | 0 | 2 | 0 | 0 | 0 | 44 | 318 | 260 | 58 | 21 | 0 | 0 | 2 | 5 | 0 | 275 | 149 | 0.751919986 | 1 | 1 | 0.991 | 1 | 1 | 1 | 1 | 1 | 0.986 | 0.863 | 1 | 0.028 |
| SE | 84823 | ENSG00000121390 | PSPC1 | chr13 | - | 19699379 | 19699398 | 19677723 | 19677823 | 19730238 | 19730316 | 1 | 3 | 1 | 7 | 2 | 60 | 28 | 42 | 20 | 60 | 0 | 1 | 1 | 4 | 1 | 6 | 23 | 27 | 14 | 0 | 168 | 149 | 0.751964534 | 1 | 0.015 | 0.087 | 0.021 | 0.237 | 0.029 | 0 | 0.037 | 0.032 | 0.202 | 1 | -0.176 |
| SE | 2385 | ENSG00000170889 | RPS9 | chr19 | + | 54206552 | 54206724 | 54201486 | 54201609 | 54207397 | 54207522 | 92 | 55 | 65 | 47 | 32 | 6 | 14 | 31 | 35 | 18 | 204 | 215 | 104 | 146 | 174 | 56 | 93 | 29 | 69 | 68 | 298 | 149 | 0.752059563 | 1 | 0.885 | 0.663 | 0.512 | 0.402 | 0.471 | 0.646 | 0.536 | 0.642 | 0.514 | 0.561 | 0.007 |
| SE | 90120 | ENSG00000160932 | LY6E | chr8 | + | 143018832 | 143019119 | 143018527 | 143018586 | 143020882 | 143020991 | 1 | 2 | 0 | 4 | 0 | 119 | 76 | 89 | 131 | 28 | 0 | 4 | 1 | 23 | 0 | 435 | 423 | 165 | 376 | 166 | 298 | 149 | 0.752078078 | 1 | 0.004 | 0.013 | 0 | 0.015 | 0 | 0 | 0.005 | 0.003 | 0.03 | 0 | -0.001 |
| SE | 75570 | ENSG00000228716 | DHFR | chr5 | - | 80633876 | 80633992 | 80626225 | 80629165 | 80637882 | 80638009 | 91 | 88 | 164 | 128 | 243 | 0 | 3 | 0 | 0 | 1 | 181 | 153 | 153 | 418 | 127 | 0 | 0 | 0 | 10 | 0 | 265 | 149 | 0.752121722 | 1 | 1 | 0.943 | 1 | 1 | 0.993 | 1 | 1 | 1 | 0.959 | 1 | -0.005 |
| SE | 79275 | ENSG00000128607 | KLHDC10 | chr7 | + | 130129436 | 130129576 | 130127403 | 130127451 | 130130536 | 130135705 | 64 | 46 | 68 | 65 | 59 | 0 | 0 | 0 | 0 | 0 | 44 | 66 | 26 | 49 | 26 | 0 | 0 | 0 | 5 | 0 | 289 | 149 | 0.752181444 | 1 | 1 | 1 | 1 | 1 | 1 | 1 | 1 | 1 | 0.835 | 1 | 0.033 |
| SE | 46806 | ENSG00000059691 | GATB | chr4 | - | 151672761 | 151672896 | 151670503 | 151671302 | 151679812 | 151679891 | 28 | 43 | 27 | 38 | 45 | 1 | 2 | 3 | 0 | 5 | 69 | 32 | 31 | 21 | 67 | 0 | 0 | 1 | 6 | 5 | 284 | 149 | 0.752184205 | 1 | 0.936 | 0.919 | 0.825 | 1 | 0.825 | 1 | 1 | 0.942 | 0.647 | 0.875 | 0.008 |
| SE | 96889 | ENSG00000100731 | PCNX1 | chr14 | + | 70995740 | 70995843 | 70988566 | 70988699 | 71009633 | 71009724 | 15 | 7 | 15 | 12 | 11 | 1 | 2 | 1 | 2 | 0 | 17 | 8 | 11 | 2 | 5 | 0 | 2 | 4 | 0 | 0 | 252 | 149 | 0.75220559 | 1 | 0.899 | 0.674 | 0.899 | 0.78 | 1 | 1 | 0.703 | 0.619 | 1 | 1 | -0.014 |
| SE | 24417 | ENSG00000055163 | CYFIP2 | chr5 | + | 157326170 | 157326267 | 157325481 | 157325638 | 157330741 | 157330850 | 442 | 163 | 331 | 315 | 339 | 7 | 3 | 9 | 4 | 3 | 172 | 358 | 142 | 183 | 267 | 11 | 0 | 7 | 0 | 0 | 246 | 149 | 0.752259498 | 1 | 0.975 | 0.971 | 0.957 | 0.979 | 0.986 | 0.904 | 1 | 0.925 | 1 | 1 | 0.008 |
| SE | 51482 | ENSG00000160087 | UBE2J2 | chr1 | - | 1262169 | 1262390 | 1257207 | 1257310 | 1263345 | 1263386 | 8 | 0 | 0 | 5 | 5 | 120 | 81 | 120 | 143 | 142 | 0 | 0 | 3 | 0 | 11 | 135 | 116 | 67 | 151 | 336 | 298 | 149 | 0.75228016 | 1 | 0.032 | 0 | 0 | 0.017 | 0.017 | 0 | 0 | 0.022 | 0 | 0.016 | 0.006 |
| SE | 76653 | ENSG00000161800 | RACGAP1 | chr12 | - | 50006433 | 50006636 | 50005255 | 50005392 | 50016630 | 50016715 | 70 | 37 | 67 | 98 | 80 | 0 | 2 | 4 | 0 | 0 | 28 | 100 | 94 | 60 | 64 | 0 | 0 | 0 | 1 | 0 | 298 | 149 | 0.752319897 | 1 | 1 | 0.902 | 0.893 | 1 | 1 | 1 | 1 | 1 | 0.968 | 1 | -0.035 |
| SE | 71383 | ENSG00000122203 | KIAA1191 | chr5 | - | 176355570 | 176355749 | 176350612 | 176350737 | 176359480 | 176359567 | 47 | 43 | 48 | 59 | 62 | 4 | 0 | 0 | 1 | 1 | 87 | 82 | 74 | 75 | 218 | 0 | 0 | 0 | 2 | 0 | 298 | 149 | 0.752491476 | 1 | 0.855 | 1 | 1 | 0.967 | 0.969 | 1 | 1 | 1 | 0.949 | 1 | -0.032 |
| SE | 37935 | ENSG00000148843 | PDCD11 | chr10 | + | 103434247 | 103434350 | 103433947 | 103434037 | 103434797 | 103434975 | 104 | 49 | 110 | 99 | 107 | 0 | 0 | 0 | 0 | 0 | 129 | 78 | 73 | 80 | 112 | 0 | 0 | 0 | 0 | 8 | 252 | 149 | 0.75251677 | 1 | 1 | 1 | 1 | 1 | 1 | 1 | 1 | 1 | 1 | 0.892 | 0.022 |
| SE | 44398 | ENSG00000134419 | RPS15A | chr16 | - | 18788062 | 18788105 | 18784737 | 18784823 | 18788980 | 18789118 | 2512 | 1314 | 3112 | 2425 | 5072 | 2 | 0 | 5 | 10 | 0 | 1277 | 1432 | 1389 | 969 | 2159 | 0 | 26 | 6 | 3 | 12 | 192 | 149 | 0.752612577 | 1 | 0.999 | 1 | 0.998 | 0.995 | 1 | 1 | 0.977 | 0.994 | 0.996 | 0.993 | 0.006 |
| SE | 4836 | ENSG00000132670 | PTPRA | chr20 | + | 2955608 | 2955698 | 2947981 | 2948024 | 2964271 | 2964350 | 11 | 10 | 11 | 11 | 36 | 665 | 407 | 387 | 561 | 1143 | 6 | 19 | 11 | 9 | 39 | 753 | 497 | 319 | 505 | 1099 | 239 | 149 | 0.752633314 | 1 | 0.01 | 0.015 | 0.017 | 0.012 | 0.019 | 0.005 | 0.023 | 0.021 | 0.011 | 0.022 | -0.002 |
| SE | 8874 | ENSG00000153944 | MSI2 | chr17 | + | 57370948 | 57370991 | 57262150 | 57262192 | 57401378 | 57401471 | 1 | 2 | 7 | 5 | 3 | 175 | 102 | 153 | 184 | 138 | 0 | 2 | 3 | 1 | 10 | 117 | 148 | 208 | 112 | 48 | 192 | 149 | 0.752657514 | 1 | 0.004 | 0.015 | 0.034 | 0.021 | 0.017 | 0 | 0.01 | 0.011 | 0.007 | 0.139 | -0.015 |
| SE | 68822 | ENSG00000144895 | EIF2A | chr3 | + | 150581617 | 150581746 | 150575648 | 150575762 | 150583199 | 150583265 | 445 | 400 | 501 | 446 | 1251 | 4 | 8 | 5 | 0 | 12 | 52 | 298 | 393 | 147 | 35 | 0 | 7 | 0 | 5 | 0 | 278 | 149 | 0.752827275 | 1 | 0.984 | 0.964 | 0.982 | 1 | 0.982 | 1 | 0.958 | 1 | 0.94 | 1 | 0.003 |
| SE | 65320 | ENSG00000285756 | NA | chrX | - | 3822072 | 3822285 | 3818441 | 3818500 | 3822373 | 3822580 | 30 | 59 | 19 | 26 | 13 | 28 | 11 | 27 | 33 | 27 | 4 | 19 | 7 | 21 | 2 | 22 | 21 | 2 | 11 | 1 | 298 | 149 | 0.752870269 | 1 | 0.349 | 0.728 | 0.26 | 0.283 | 0.194 | 0.083 | 0.311 | 0.636 | 0.488 | 0.5 | -0.041 |
| SE | 84852 | ENSG00000269556 | TMEM185A | chrX | - | 149606730 | 149606809 | 149603986 | 149604070 | 149608626 | 149608829 | 0 | 0 | 0 | 0 | 0 | 28 | 22 | 20 | 36 | 77 | 0 | 5 | 0 | 0 | 0 | 28 | 16 | 16 | 10 | 202 | 228 | 149 | 0.753060554 | 1 | 0 | 0 | 0 | 0 | 0 | 0 | 0.17 | 0 | 0 | 0 | -0.034 |
| SE | 84333 | ENSG00000138593 | SECISBP2L | chr15 | - | 49016847 | 49017015 | 49016559 | 49016701 | 49017547 | 49017628 | 433 | 304 | 315 | 399 | 754 | 1 | 1 | 1 | 0 | 0 | 18 | 327 | 311 | 136 | 16 | 0 | 5 | 3 | 0 | 0 | 298 | 149 | 0.753068218 | 1 | 0.995 | 0.993 | 0.994 | 1 | 1 | 1 | 0.97 | 0.981 | 1 | 1 | 0.006 |
| SE | 18357 | ENSG00000257621 | PSMA3-AS1 | chr14 | - | 58293765 | 58295811 | 58293065 | 58293138 | 58297954 | 58298120 | 8 | 5 | 25 | 7 | 20 | 25 | 12 | 25 | 27 | 8 | 0 | 28 | 1 | 9 | 15 | 19 | 8 | 14 | 7 | 0 | 298 | 149 | 0.75310921 | 1 | 0.138 | 0.172 | 0.333 | 0.115 | 0.556 | 0 | 0.636 | 0.034 | 0.391 | 1 | -0.149 |
| SE | 2213 | ENSG00000006015 | REX1BD | chr19 | + | 18589412 | 18589683 | 18588994 | 18589077 | 18590853 | 18590933 | 86 | 33 | 63 | 56 | 26 | 0 | 0 | 0 | 0 | 0 | 22 | 87 | 27 | 26 | 57 | 0 | 0 | 0 | 0 | 5 | 298 | 149 | 0.753144886 | 1 | 1 | 1 | 1 | 1 | 1 | 1 | 1 | 1 | 1 | 0.851 | 0.03 |
| SE | 32970 | ENSG00000114127 | XRN1 | chr3 | - | 142435950 | 142436056 | 142432660 | 142432893 | 142447869 | 142448028 | 1 | 0 | 4 | 2 | 1 | 390 | 345 | 359 | 355 | 165 | 5 | 10 | 1 | 4 | 0 | 85 | 346 | 367 | 107 | 76 | 255 | 149 | 0.753227696 | 1 | 0.001 | 0 | 0.006 | 0.003 | 0.004 | 0.033 | 0.017 | 0.002 | 0.021 | 0 | -0.012 |
| SE | 65038 | ENSG00000091409 | ITGA6 | chr2 | + | 172489484 | 172489613 | 172488125 | 172488228 | 172491023 | 172491122 | 113 | 94 | 186 | 168 | 255 | 8 | 3 | 23 | 11 | 20 | 16 | 35 | 54 | 70 | 14 | 0 | 0 | 17 | 12 | 0 | 278 | 149 | 0.753254235 | 1 | 0.883 | 0.944 | 0.813 | 0.891 | 0.872 | 1 | 1 | 0.63 | 0.758 | 1 | 0.003 |
| [illegible] | | | | | | | | | | | | | | | | | | | | | | | | | | | | | | | | | | | | | | | | | | | | | | |

| | | | | | | | | | | | | | | | | | | | | | | | | |
|---|---|---|---|---|---|---|---|---|---|---|---|---|---|---|---|---|---|---|---|---|---|---|---|---|
| SE | 96996 | ENSG00000153339 | TRAPPC8 | chr18 | - | 31839311 | 31839457 | 31829196 | 31830989 | 31846715 | 31846817 | 251 | 226 | 215 | 232 | 251 | 1 | 5 | 0 | 0 | 4 | 26 | 131 | 136 | 52 |
| SE | 98252 | ENSG00000173208 | ABCD2 | chr12 | - | 39607598 | 39607714 | 39604761 | 39604930 | 39618676 | 39619803 | 76 | 86 | 100 | 58 | 139 | 2 | 1 | 0 | 0 | 0 | 21 | 35 | 22 | 23 |
| SE | 85713 | ENSG00000072778 | ACADVL | chr17 | + | 7222666 | 7222865 | 7222176 | 7222302 | 7223132 | 7223237 | 330 | 222 | 209 | 301 | 325 | 0 | 0 | 1 | 2 | 0 | 642 | 595 | 346 | 572 |
| SE | 80090 | ENSG00000198198 | SZT2 | chr1 | + | 43443596 | 43443796 | 43443351 | 43443477 | 43445893 | 43445984 | 26 | 5 | 15 | 25 | 19 | 0 | 0 | 0 | 2 | 2 | 30 | 12 | 3 | 9 |
| SE | 1357 | ENSG00000100150 | DEPDC5 | chr22 | + | 31797599 | 31797699 | 31792744 | 31792817 | 31798581 | 31798656 | 37 | 32 | 42 | 29 | 69 | 0 | 2 | 0 | 4 | 0 | 74 | 83 | 26 | 32 |
| SE | 70413 | ENSG00000115652 | UXS1 | chr2 | - | 106125619 | 106125679 | 106122969 | 106123091 | 106129673 | 106129778 | 198 | 125 | 169 | 201 | 271 | 5 | 0 | 0 | 5 | 0 | 277 | 129 | 114 | 222 |
| SE | 73451 | ENSG00000143374 | TARS2 | chr1 | + | 150496481 | 150496628 | 150492184 | 150492489 | 150496809 | 150496908 | 58 | 19 | 48 | 85 | 53 | 10 | 0 | 0 | 1 | 3 | 38 | 50 | 11 | 41 |
| SE | 21043 | ENSG00000135506 | OS9 | chr12 | + | 57695961 | 57696373 | 57695495 | 57695843 | 57715759 | 57715951 | 426 | 309 | 282 | 347 | 419 | 0 | 0 | 2 | 0 | 0 | 711 | 504 | 324 | 475 |
| SE | 45500 | ENSG00000168894 | RNF181 | chr2 | + | 85597103 | 85597178 | 85596518 | 85596649 | 85597444 | 85597604 | 48 | 24 | 51 | 47 | 61 | 2 | 0 | 1 | 3 | 0 | 176 | 50 | 54 | 85 |
| SE | 36429 | ENSG00000121858 | TNFSF10 | chr3 | - | 172511616 | 172511659 | 172505507 | 172506919 | 172514860 | 172514998 | 69 | 42 | 26 | 53 | 56 | 7 | 6 | 3 | 5 | 2 | 49 | 25 | 38 | 54 |
| SE | 61936 | ENSG00000128699 | ORMDL1 | chr2 | - | 189783013 | 189783123 | 189782421 | 189782602 | 189784268 | 189784339 | 73 | 42 | 64 | 81 | 149 | 2 | 4 | 6 | 12 | 3 | 25 | 74 | 46 | 36 |
| SE | 49981 | ENSG00000130772 | MED18 | chr1 | + | 28330596 | 28330735 | 28329039 | 28329180 | 28334416 | 28334585 | 83 | 58 | 52 | 64 | 83 | 0 | 0 | 4 | 0 | 0 | 43 | 54 | 13 | 29 |
| SE | 64871 | ENSG00000115109 | EPB41L5 | chr2 | + | 120074099 | 120074178 | 120073177 | 120073220 | 120075475 | 120075520 | 22 | 21 | 35 | 55 | 75 | 0 | 3 | 3 | 3 | 0 | 4 | 29 | 1 | 5 |
| SE | 60090 | ENSG00000075651 | PLD1 | chr3 | - | 171733443 | 171733509 | 171726003 | 171726076 | 171734864 | 171734970 | 64 | 44 | 32 | 40 | 65 | 6 | 0 | 0 | 0 | 0 | 42 | 72 | 58 | 52 |
| SE | 98484 | ENSG00000151702 | FLI1 | chr11 | + | 128805365 | 128805431 | 128781957 | 128782023 | 128809156 | 128809204 | 171 | 84 | 138 | 119 | 117 | 6 | 6 | 11 | 29 | 12 | 144 | 136 | 67 | 117 |
| SE | 8114 | ENSG00000141252 | VPS53 | chr17 | - | 696824 | 697004 | 661808 | 661895 | 697417 | 697484 | 2 | 4 | 1 | 0 | 4 | 53 | 38 | 30 | 47 | 77 | 0 | 0 | 5 | 0 |
| SE | 50807 | ENSG00000117481 | NSUN4 | chr1 | + | 46341322 | 46341497 | 46340819 | 46340919 | 46344800 | 46345144 | 1 | 1 | 4 | 4 | 0 | 27 | 4 | 20 | 18 | 17 | 0 | 0 | 6 | 2 |
| SE | 90244 | ENSG00000066855 | MTFR1 | chr8 | + | 65689565 | 65689599 | 65682352 | 65682451 | 65693643 | 65693759 | 4 | 1 | 2 | 0 | 0 | 24 | 26 | 14 | 28 | 31 | 0 | 0 | 2 | 0 |
| SE | 88921 | ENSG00000176731 | RBIS | chr8 | - | 85219235 | 85219360 | 85217385 | 85217502 | 85220305 | 85220353 | 89 | 3 | 1 | 60 | 0 | 152 | 129 | 157 | 144 | 286 | 68 | 0 | 0 | 0 |
| SE | 728 | ENSG00000099991 | CABIN1 | chr22 | + | 24041138 | 24041273 | 24038347 | 24038461 | 24042903 | 24043084 | 45 | 27 | 51 | 63 | 81 | 28 | 1 | 6 | 11 | 13 | 78 | 57 | 43 | 102 |
| SE | 8679 | ENSG00000198909 | MAP3K3 | chr17 | + | 63672844 | 63672914 | 63666985 | 63667060 | 63681765 | 63681899 | 2 | 0 | 0 | 0 | 0 | 299 | 161 | 190 | 229 | 439 | 0 | 0 | 0 | 9 |
| SE | 30883 | ENSG00000003056 | M6PR | chr12 | - | 8946715 | 8946824 | 8946228 | 8946405 | 8949487 | 8949645 | 3 | 0 | 0 | 0 | 4 | 222 | 230 | 290 | 351 | 121 | 0 | 0 | 6 | 1 |
| SE | 84450 | ENSG00000189376 | C8orf76 | chr8 | - | 123231299 | 123231757 | 123219966 | 123220297 | 123237797 | 123237941 | 93 | 50 | 45 | 53 | 98 | 1 | 3 | 1 | 6 | 1 | 72 | 108 | 56 | 90 |
| SE | 68857 | ENSG00000168918 | INPP5D | chr2 | + | 233125744 | 233125919 | 233122106 | 233122257 | 233130507 | 233130648 | 583 | 291 | 425 | 555 | 511 | 0 | 0 | 0 | 1 | 0 | 717 | 716 | 383 | 547 |
| SE | 51154 | ENSG00000165458 | INPPL1 | chr11 | + | 72238041 | 72238175 | 72237123 | 72237796 | 72238262 | 72239105 | 67 | 39 | 40 | 49 | 10 | 1 | 0 | 0 | 2 | 0 | 29 | 69 | 32 | 49 |
| SE | 47871 | ENSG00000146247 | PHIP | chr6 | - | 78965944 | 78966056 | 78965702 | 78965764 | 78969834 | 78969917 | 209 | 164 | 208 | 240 | 247 | 2 | 5 | 3 | 1 | 1 | 32 | 37 | 46 | 12 |
| SE | 7748 | ENSG00000167863 | ATP5PD | chr17 | - | 75040091 | 75040163 | 75039208 | 75039271 | 75042180 | 75042277 | 849 | 544 | 849 | 828 | 977 | 5 | 4 | 4 | 2 | 7 | 1448 | 1080 | 701 | 971 |
| SE | 34869 | ENSG00000117597 | UTP25 | chr1 | + | 209840852 | 209841055 | 209838804 | 209839128 | 209851203 | 209857565 | 55 | 35 | 48 | 63 | 127 | 0 | 0 | 0 | 4 | 2 | 51 | 27 | 33 | 58 |
| SE | 90844 | ENSG00000129696 | TTI2 | chr8 | - | 33511966 | 33512712 | 33509745 | 33509932 | 33513081 | 33513135 | 36 | 17 | 24 | 20 | 51 | 0 | 0 | 0 | 2 | 3 | 67 | 10 | 18 | 11 |
| SE | 40302 | ENSG00000234663 | LINC01934 | chr2 | + | 181195346 | 181195472 | 181123931 | 181123980 | 181226470 | 181226539 | 7 | 4 | 9 | 10 | 1 | 39 | 28 | 24 | 40 | 15 | 0 | 12 | 4 | 4 |
| SE | 57233 | ENSG00000178691 | SUZ12 | chr17 | + | 31982998 | 31983104 | 31976520 | 31976614 | 31988319 | 31988497 | 419 | 376 | 384 | 388 | 652 | 8 | 10 | 16 | 17 | 29 | 115 | 182 | 199 | 75 |
| SE | 45091 | ENSG00000100181 | TPTEP1 | chr22 | + | 16647167 | 16647257 | 16638578 | 16638740 | 16648526 | 16648830 | 0 | 2 | 1 | 3 | 0 | 7 | 21 | 0 | 10 | 3 | 0 | 0 | 0 | 0 |
| SE | 70635 | ENSG00000188647 | PTAR1 | chr9 | - | 69723325 | 69723630 | 69709524 | 69718568 | 69732138 | 69732352 | 92 | 68 | 78 | 90 | 148 | 0 | 0 | 1 | 6 | 0 | 12 | 56 | 34 | 41 |
| SE | 95640 | ENSG00000137073 | UBAP2 | chr9 | - | 33944364 | 33944639 | 33943419 | 33943589 | 33953284 | 33953474 | 29 | 8 | 24 | 32 | 23 | 0 | 0 | 4 | 0 | 0 | 10 | 14 | 14 | 40 |
| SE | 41068 | ENSG00000160741 | CRTC2 | chr1 | - | 153955064 | 153955166 | 153954872 | 153954989 | 153958344 | 153958612 | 69 | 42 | 57 | 67 | 39 | 0 | 0 | 0 | 0 | 0 | 79 | 114 | 65 | 26 |
| SE | 59323 | ENSG00000196591 | HDAC2 | chr6 | - | 113953276 | 113953418 | 113946007 | 113949260 | 113956012 | 113956151 | 461 | 329 | 427 | 431 | 414 | 1 | 0 | 1 | 3 | 0 | 127 | 203 | 164 | 263 |
| SE | 56515 | ENSG00000105676 | ARMC6 | chr19 | + | 19037551 | 19037707 | 19033584 | 19033930 | 19042710 | 19042877 | 1 | 6 | 0 | 1 | 7 | 20 | 3 | 6 | 17 | 5 | 0 | 0 | 0 | 9 |
| SE | 55109 | ENSG00000072110 | ACTN1 | chr14 | - | 68882872 | 68883055 | 68882457 | 68882592 | 68884167 | 68884308 | 334 | 261 | 280 | 343 | 490 | 0 | 0 | 0 | 0 | 0 | 765 | 379 | 413 | 977 |
| SE | 26904 | ENSG00000160613 | PCSK7 | chr11 | - | 117207653 | 117207735 | 117207066 | 117207160 | 117207969 | 117208064 | 11 | 3 | 4 | 22 | 23 | 48 | 32 | 42 | 40 | 38 | 10 | 21 | 4 | 24 |
| SE | 38194 | ENSG00000137074 | APTX | chr9 | - | 32997242 | 32997348 | 32989758 | 32989895 | 33001566 | 33001583 | 5 | 7 | 6 | 1 | 7 | 20 | 8 | 11 | 20 | 4 | 0 | 6 | 6 | 29 |
| SE | 104540 | ENSG00000075303 | SLC25A40 | chr7 | - | 87843753 | 87843863 | 87841632 | 87841714 | 87846948 | 87847122 | 268 | 237 | 235 | 234 | 319 | 4 | 0 | 0 | 0 | 3 | 223 | 231 | 183 | 252 |
| SE | 75949 | ENSG00000147383 | NSDHL | chrX | + | 152867570 | 152867673 | 152865818 | 152865961 | 152868783 | 152869363 | 70 | 16 | 33 | 42 | 114 | 0 | 0 | 0 | 4 | 2 | 52 | 59 | 27 | 21 |
| SE | 34930 | ENSG00000076108 | BAZ2A | chr12 | - | 56610408 | 56610539 | 56610116 | 56610215 | 56611572 | 56611633 | 407 | 294 | 401 | 460 | 345 | 0 | 0 | 0 | 0 | 0 | 1376 | 787 | 846 | 895 |
| SE | 33561 | ENSG00000009790 | TRAF3IP3 | chr1 | + | 209759981 | 209760384 | 209759033 | 209759134 | 209762514 | 209762662 | 642 | 444 | 532 | 432 | 357 | 0 | 1 | 0 | 0 | 0 | 612 | 766 | 426 | 294 |
| SE | 32391 | ENSG00000232112 | TMA7 | chr3 | + | 48440540 | 48440628 | 48440283 | 48440458 | 48443847 | 48444203 | 380 | 215 | 348 | 377 | 452 | 0 | 0 | 0 | 3 | 0 | 288 | 272 | 150 | 176 |
| SE | 20479 | ENSG00000164080 | RAD54L2 | chr3 | + | 51637081 | 51637503 | 51635592 | 51635789 | 51638143 | 51638321 | 41 | 13 | 28 | 34 | 31 | 0 | 1 | 0 | 0 | 0 | 53 | 65 | 30 | 16 |
| SE | 52926 | ENSG00000159023 | EPB41 | chr1 | + | 29112367 | 29112448 | 29109335 | 29109437 | 29115698 | 29115702 | 712 | 480 | 837 | 808 | 1251 | 4 | 3 | 8 | 12 | 1 | 624 | 560 | 623 | 666 |
| SE | 48590 | ENSG00000150681 | RGS18 | chr1 | + | 192161452 | 192161487 | 192160377 | 192160439 | 192181291 | 192181454 | 34 | 76 | 23 | 114 | 94 | 575 | 1216 | 358 | 1269 | 2456 | 39 | 25 | 39 | 7 |
| SE | 96381 | ENSG00000197563 | PIGN | chr18 | - | 62070389 | 62070506 | 62044256 | 62045979 | 62072672 | 62072725 | 0 | 0 | 0 | 7 | 0 | 7 | 9 | 8 | 19 | 17 | 0 | 4 | 0 | 0 |
| SE | 41584 | ENSG00000118308 | IRAG2 | chr12 | + | 25061591 | 25061653 | 25052511 | 25052956 | 25062819 | 25062900 | 183 | 136 | 133 | 177 | 255 | 210 | 170 | 161 | 206 | 318 | 87 | 135 | 107 | 92 |
| SE | 60088 | ENSG00000075651 | PLD1 | chr3 | - | 171688675 | 171688876 | 171687370 | 171687584 | 171692331 | 171692442 | 42 | 18 | 20 | 27 | 35 | 0 | 0 | 0 | 0 | 0 | 6 | 24 | 25 | 78 |
| SE | 59028 | ENSG00000171159 | C9orf16 | chr9 | + | 128162003 | 128162952 | 128160313 | 128160486 | 128163442 | 128163890 | 3 | 4 | 5 | 1 | 2 | 66 | 20 | 39 | 45 | 30 | 0 | 9 | 3 | 0 |
| SE | 82112 | ENSG00000136048 | DRAM1 | chr12 | + | 101914173 | 101914232 | 101901290 | 101901433 | 101920108 | 101920201 | 6 | 8 | 9 | 6 | 24 | 2 | 12 | 1 | 3 | 3 | 33 | 22 | 11 | 48 |
| SE | 99420 | ENSG00000198730 | CTR9 | chr11 | + | 10774011 | 10774169 | 10773126 | 10773273 | 10775206 | 10775303 | 58 | 77 | 50 | 62 | 68 | 0 | 0 | 0 | 0 | 0 | 18 | 100 | 65 | 47 |
| SE | 24211 | ENSG00000113742 | CPEB4 | chr5 | + | 173932449 | 173932500 | 173910522 | 173910604 | 173943025 | 173943049 | 15 | 9 | 11 | 24 | 11 | 3 | 0 | 0 | 0 | 4 | 12 | 16 | 13 | 6 |
| SE | 96597 | ENSG00000198561 | CTNND1 | chr11 | + | 57815914 | 57816001 | 57815393 | 57815500 | 57816296 | 57819178 | 3 | 0 | 0 | 5 | 1 | 97 | 84 | 67 | 83 | 158 | 0 | 0 | 0 | 0 |
| SE | 108568 | ENSG00000185189 | NRBP2 | chr8 | - | 143839508 | 143839825 | 143839313 | 143839408 | 143839928 | 143840030 | 31 | 20 | 29 | 36 | 24 | 0 | 0 | 0 | 0 | 0 | 9 | 27 | 24 | 32 |
| SE | 72003 | ENSG00000139629 | GALNT6 | chr12 | - | 51383460 | 51383582 | 51379545 | 51379884 | 51390849 | 51390937 | 20 | 5 | 9 | 7 | 9 | 0 | 1 | 1 | 1 | 0 | 4 | 6 | 10 | 6 |
| SE | 108864 | ENSG00000125304 | TM9SF2 | chr13 | + | 99539445 | 99539557 | 99537738 | 99537863 | 99540713 | 99540793 | 1037 | 734 | 730 | 926 | 1348 | 0 | 1 | 0 | 1 | 4 | 955 | 699 | 677 | 1077 |
| SE | 102944 | ENSG00000233695 | GAS6-AS1 | chr13 | + | 113834577 | 113834764 | 113833491 | 113833579 | 113838076 | 113838247 | 1 | 2 | 6 | 7 | 3 | 1 | 0 | 0 | 4 | 1 | 50 | 18 | 0 | 38 |
| SE | 75783 | ENSG00000204231 | RXRB | chr6 | - | 33199168 | 33199416 | 33198307 | 33198464 | 33200241 | 33200614 | 37 | 15 | 24 | 29 | 17 | 4 | 1 | 4 | 1 | 0 | 11 | 23 | 24 | 17 |
| SE | 17646 | ENSG00000100796 | PPP4R3A | chr14 | - | 91485631 | 91485730 | 91481575 | 91482193 | 91490746 | 91490802 | 592 | 479 | 492 | 590 | 673 | 0 | 0 | 0 | 0 | 0 | 537 | 573 | 626 | 551 |
| SE | 42658 | ENSG00000088854 | C20orf194 | chr20 | - | 3290128 | 3290217 | 3287631 | 3287790 | 3294138 | 3294256 | 19 | 29 | 25 | 18 | 13 | 0 | 0 | 1 | 0 | 2 | 80 | 18 | 20 | 19 |
| SE | 9548 | ENSG00000266967 | AARSD1 | chr17 | - | 42953723 | 42953778 | 42951799 | 42951894 | 42954875 | 42954967 | 36 | 15 | 30 | 44 | 19 | 13 | 28 | 17 | 14 | 5 | 50 | 43 | 28 | 51 |
| [illegible] | | | | | | | | | | | | | | | | | | | | | | | | |

| | | | | | | | | | | | | | | | | | | | | | | | | | | | | | | | | | | | | | | | | | | | | | | |
|---|---|---|---|---|---|---|---|---|---|---|---|---|---|---|---|---|---|---|---|---|---|---|---|---|---|---|---|---|---|---|---|---|---|---|---|---|---|---|---|---|---|---|---|---|---|---|
| SE | 50765 | ENSG00000112531 | QKI | chr6 | + | 163457380 | 163457551 | 163455278 | 163455421 | 163478779 | 163478896 | 21 | 24 | 30 | 22 | 46 | 212 | 175 | 121 | 156 | 191 | 37 | 31 | 32 | 46 | 58 | 346 | 223 | 288 | 307 | 149 | 298 | 149 | 0.77399872 | 1 | 0.047 | 0.064 | 0.11 | 0.066 | 0.107 | 0.051 | 0.065 | 0.053 | 0.07 | 0.163 | -0.002 |
| SE | 4985 | ENSG00000000419 | DPM1 | chr20 | - | 50940864 | 50940933 | 50934904 | 50936262 | 50942030 | 50942126 | 417 | 338 | 276 | 329 | 292 | 1 | 3 | 2 | 3 | 9 | 130 | 207 | 405 | 268 | 21 | 0 | 0 | 6 | 0 | 0 | 218 | 149 | 0.774008658 | 1 | 0.997 | 0.987 | 0.99 | 0.987 | 0.957 | 1 | 1 | 0.979 | 1 | 1 | -0.012 |
| SE | 47021 | ENSG00000175309 | PHYKPL | chr5 | - | 178222354 | 178222492 | 178215275 | 178215430 | 178222851 | 178222934 | 46 | 26 | 36 | 29 | 39 | 0 | 0 | 0 | 1 | 0 | 22 | 31 | 26 | 22 | 53 | 0 | 0 | 0 | 0 | 5 | 287 | 149 | 0.774062891 | 1 | 1 | 1 | 1 | 0.938 | 1 | 1 | 1 | 1 | 1 | 0.846 | 0.018 |
| SE | 63285 | ENSG00000177084 | POLE | chr12 | - | 132660968 | 132661164 | 132659294 | 132659509 | 132661526 | 132661684 | 84 | 47 | 74 | 99 | 70 | 0 | 0 | 0 | 0 | 0 | 132 | 84 | 75 | 231 | 259 | 8 | 0 | 0 | 0 | 0 | 298 | 149 | 0.774111323 | 1 | 1 | 1 | 1 | 1 | 1 | 0.892 | 1 | 1 | 1 | 1 | 0.022 |
| SE | 10608 | ENSG00000276234 | TADA2A | chr17 | + | 37444695 | 37444768 | 37440504 | 37440662 | 37458523 | 37458587 | 15 | 14 | 13 | 19 | 11 | 1 | 3 | 0 | 0 | 0 | 5 | 6 | 2 | 18 | 17 | 3 | 0 | 0 | 0 | 0 | 222 | 149 | 0.774158786 | 1 | 0.91 | 0.758 | 1 | 1 | 1 | 0.528 | 1 | 1 | 1 | 1 | 0.028 |
| SE | 108642 | ENSG00000136104 | RNASEH2B | chr13 | + | 50960144 | 50960171 | 50949462 | 50949505 | 50969931 | 50970363 | 3 | 0 | 2 | 3 | 6 | 84 | 87 | 71 | 84 | 134 | 0 | 11 | 0 | 0 | 0 | 73 | 69 | 29 | 27 | 27 | 176 | 149 | 0.774160153 | 1 | 0.029 | 0 | 0.023 | 0.029 | 0.037 | 0 | 0.119 | 0 | 0 | 0 | 0 |
| SE | 72886 | ENSG00000272752 | STAG3L5P-PVRIG2P-PILRB | chr7 | + | 100353372 | 100353483 | 100353209 | 100353270 | 100355142 | 100355240 | 22 | 12 | 6 | 25 | 3 | 2 | 2 | 10 | 5 | 4 | 15 | 12 | 9 | 16 | 22 | 22 | 0 | 7 | 8 | 5 | 260 | 149 | 0.774200653 | 1 | 0.863 | 0.775 | 0.256 | 0.741 | 0.301 | 0.281 | 1 | 0.424 | 0.534 | 0.716 | -0.004 |
| SE | 50317 | ENSG00000148335 | NTMT1 | chr9 | + | 129632649 | 129632865 | 129626162 | 129626295 | 129634053 | 129634221 | 18 | 10 | 12 | 22 | 14 | 6 | 0 | 4 | 0 | 0 | 51 | 27 | 30 | 36 | 56 | 12 | 2 | 0 | 0 | 4 | 298 | 149 | 0.774339361 | 1 | 0.6 | 1 | 0.6 | 1 | 1 | 0.68 | 0.871 | 1 | 1 | 0.875 | -0.045 |
| SE | 89811 | ENSG00000122882 | ECD | chr10 | - | 73163732 | 73163950 | 73160433 | 73160551 | 73167865 | 73168008 | 145 | 92 | 108 | 142 | 250 | 1 | 9 | 2 | 7 | 1 | 108 | 116 | 144 | 148 | 24 | 0 | 0 | 13 | 2 | 0 | 298 | 149 | 0.774366117 | 1 | 0.986 | 0.836 | 0.964 | 0.91 | 0.992 | 1 | 1 | 0.847 | 0.974 | 1 | -0.027 |
| SE | 44918 | ENSG00000162739 | SLAMF6 | chr1 | - | 160496060 | 160496393 | 160491124 | 160491388 | 160523143 | 160523255 | 461 | 135 | 287 | 279 | 416 | 10 | 1 | 6 | 1 | 7 | 127 | 309 | 119 | 156 | 98 | 0 | 5 | 9 | 0 | 0 | 298 | 149 | 0.774445291 | 1 | 0.958 | 0.985 | 0.96 | 0.993 | 0.967 | 1 | 0.969 | 0.869 | 1 | 1 | 0.005 |
| SE | 72280 | ENSG00000186635 | ARAP1 | chr11 | - | 72732514 | 72732597 | 72726619 | 72727172 | 72752327 | 72752408 | 48 | 25 | 26 | 49 | 20 | 7 | 3 | 1 | 3 | 1 | 59 | 59 | 22 | 36 | 42 | 0 | 0 | 14 | 0 | 12 | 232 | 149 | 0.774688591 | 1 | 0.815 | 0.843 | 0.943 | 0.913 | 0.928 | 1 | 1 | 0.502 | 1 | 0.692 | 0.05 |
| SE | 84215 | ENSG00000214783 | POLR2J4 | chr7 | - | 43988085 | 43988247 | 43987677 | 43987888 | 44014605 | 44014783 | 14 | 3 | 10 | 11 | 7 | 0 | 0 | 0 | 0 | 0 | 11 | 15 | 8 | 25 | 27 | 0 | 0 | 0 | 4 | 0 | 298 | 149 | 0.774715665 | 1 | 1 | 1 | 1 | 1 | 1 | 1 | 1 | 1 | 0.758 | 1 | 0.048 |
| SE | 95658 | ENSG00000137073 | UBAP2 | chr9 | - | 33971650 | 33971737 | 33963725 | 33963791 | 33973182 | 33973237 | 60 | 61 | 63 | 87 | 168 | 4 | 3 | 2 | 1 | 0 | 100 | 84 | 43 | 98 | 117 | 0 | 4 | 4 | 3 | 0 | 236 | 149 | 0.774739906 | 1 | 0.904 | 0.928 | 0.952 | 0.982 | 1 | 1 | 0.93 | 0.872 | 0.954 | 1 | 0.002 |
| SE | 14828 | ENSG00000069943 | PIGB | chr15 | + | 55324779 | 55324834 | 55321272 | 55321390 | 55327530 | 55327635 | 5 | 5 | 3 | 4 | 10 | 18 | 16 | 10 | 23 | 55 | 0 | 0 | 1 | 5 | 7 | 6 | 10 | 4 | 4 | 5 | 204 | 149 | 0.774775045 | 1 | 0.169 | 0.186 | 0.18 | 0.113 | 0.117 | 0 | 0 | 0.154 | 0.477 | 0.506 | -0.074 |
| SE | 64926 | ENSG00000204642 | HLA-F | chr6 | + | 29725992 | 29726043 | 29725533 | 29725563 | 29726882 | 29727296 | 8 | 3 | 2 | 4 | 9 | 3 | 6 | 6 | 10 | 6 | 3 | 7 | 1 | 10 | 17 | 7 | 9 | 6 | 3 | 32 | 200 | 149 | 0.774830869 | 1 | 0.665 | 0.271 | 0.199 | 0.23 | 0.528 | 0.242 | 0.367 | 0.11 | 0.713 | 0.284 | 0.035 |
| SE | 86931 | ENSG00000122122 | SASH3 | chrX | + | 129792988 | 129793139 | 129792626 | 129792836 | 129793641 | 129795201 | 544 | 249 | 340 | 415 | 290 | 0 | 0 | 0 | 0 | 0 | 306 | 456 | 195 | 357 | 530 | 13 | 0 | 0 | 0 | 0 | 298 | 149 | 0.77486709 | 1 | 1 | 1 | 1 | 1 | 1 | 0.922 | 1 | 1 | 1 | 1 | 0.016 |
| SE | 62389 | ENSG00000051596 | THOC3 | chr5 | - | 175964950 | 175965155 | 175961272 | 175961434 | 175967110 | 175967267 | 67 | 43 | 50 | 69 | 38 | 0 | 1 | 0 | 0 | 0 | 66 | 33 | 44 | 42 | 66 | 5 | 0 | 0 | 0 | 0 | 298 | 149 | 0.774875509 | 1 | 1 | 0.956 | 1 | 1 | 1 | 0.868 | 1 | 1 | 1 | 1 | 0.018 |
| SE | 76900 | ENSG00000204590 | GNL1 | chr6 | - | 30552466 | 30552661 | 30547351 | 30547530 | 30553083 | 30553179 | 125 | 66 | 133 | 132 | 78 | 0 | 0 | 0 | 0 | 0 | 97 | 139 | 58 | 87 | 192 | 0 | 0 | 4 | 3 | 0 | 298 | 149 | 0.775281584 | 1 | 1 | 1 | 1 | 1 | 1 | 1 | 1 | 0.879 | 0.935 | 1 | 0.037 |
| SE | 22367 | ENSG00000140455 | USP3 | chr15 | + | 63532646 | 63532707 | 63504510 | 63504830 | 63553714 | 63553798 | 25 | 14 | 21 | 11 | 8 | 2 | 3 | 0 | 0 | 0 | 31 | 33 | 22 | 20 | 29 | 0 | 0 | 0 | 0 | 0 | 210 | 149 | 0.775394613 | 1 | 0.899 | 0.768 | 1 | 1 | 1 | 1 | 1 | 1 | 1 | 1 | -0.067 |
| SE | 16205 | ENSG00000140598 | EFL1 | chr15 | - | 82252690 | 82252775 | 82241269 | 82241403 | 82259087 | 82259155 | 175 | 125 | 111 | 109 | 151 | 8 | 6 | 6 | 8 | 10 | 63 | 141 | 78 | 102 | 18 | 0 | 12 | 8 | 7 | 0 | 234 | 149 | 0.775414677 | 1 | 0.933 | 0.93 | 0.922 | 0.897 | 0.906 | 1 | 0.882 | 0.861 | 0.903 | 1 | -0.012 |
| SE | 16844 | ENSG00000151575 | TEX9 | chr15 | + | 56388471 | 56388520 | 56383951 | 56384031 | 56389317 | 56389400 | 13 | 19 | 15 | 12 | 36 | 0 | 3 | 1 | 0 | 2 | 2 | 6 | 9 | 2 | 9 | 0 | 0 | 1 | 0 | 0 | 198 | 149 | 0.775424326 | 1 | 1 | 0.827 | 0.919 | 1 | 0.931 | 1 | 1 | 0.871 | 1 | 1 | -0.039 |
| SE | 57281 | ENSG00000143119 | CD53 | chr1 | + | 110896652 | 110896733 | 110894326 | 110894401 | 110899123 | 110899922 | 12 | 3 | 10 | 8 | 21 | 14 | 14 | 11 | 30 | 31 | 2 | 35 | 26 | 21 | 19 | 24 | 22 | 26 | 35 | 59 | 230 | 149 | 0.775470897 | 1 | 0.357 | 0.122 | 0.371 | 0.147 | 0.305 | 0.051 | 0.508 | 0.393 | 0.28 | 0.173 | -0.021 |
| SE | 69659 | ENSG00000143390 | RFX5 | chr1 | - | 151346488 | 151346606 | 151346204 | 151346333 | 151347210 | 151347252 | 304 | 166 | 238 | 209 | 337 | 0 | 0 | 0 | 0 | 0 | 299 | 267 | 155 | 278 | 363 | 4 | 0 | 0 | 0 | 8 | 267 | 149 | 0.775505929 | 1 | 1 | 1 | 1 | 1 | 1 | 0.977 | 1 | 1 | 1 | 0.962 | 0.012 |
| SE | 65538 | ENSG00000284292 | NA | chr7 | + | 99348851 | 99348959 | 99338180 | 99338285 | 99353908 | 99354121 | 40 | 39 | 36 | 39 | 22 | 0 | 0 | 2 | 0 | 0 | 78 | 67 | 47 | 30 | 72 | 0 | 0 | 5 | 0 | 0 | 257 | 149 | 0.775528077 | 1 | 1 | 1 | 0.913 | 1 | 1 | 1 | 1 | 0.845 | 1 | 1 | 0.014 |
| SE | 49300 | ENSG00000120029 | ARMH3 | chr10 | - | 102001970 | 102002072 | 101995296 | 101995355 | 102006539 | 102006633 | 121 | 70 | 84 | 123 | 122 | 4 | 3 | 3 | 3 | 1 | 42 | 143 | 84 | 66 | 145 | 0 | 3 | 8 | 5 | 0 | 251 | 149 | 0.775552023 | 1 | 0.947 | 0.933 | 0.943 | 0.961 | 0.986 | 1 | 0.966 | 0.862 | 0.887 | 1 | 0.011 |
| SE | 59678 | ENSG00000152683 | SLC30A6 | chr2 | + | 32174062 | 32174147 | 32171286 | 32171373 | 32184272 | 32184338 | 24 | 22 | 23 | 23 | 20 | 0 | 0 | 0 | 0 | 4 | 9 | 1 | 16 | 5 | 3 | 0 | 0 | 0 | 0 | 0 | 234 | 149 | 0.775633243 | 1 | 1 | 1 | 1 | 1 | 0.761 | 1 | 1 | 1 | 1 | 1 | -0.048 |
| SE | 30619 | ENSG00000131042 | LILRB2 | chr19 | - | 54280263 | 54280299 | 54279790 | 54280075 | 54280960 | 54281097 | 94 | 60 | 35 | 99 | 19 | 0 | 0 | 2 | 0 | 0 | 205 | 140 | 76 | 128 | 269 | 0 | 0 | 7 | 5 | 0 | 185 | 149 | 0.775653058 | 1 | 1 | 1 | 0.934 | 1 | 1 | 1 | 1 | 0.897 | 0.954 | 1 | 0.017 |
| SE | 40736 | ENSG00000073792 | IGF2BP2 | chr3 | - | 185672540 | 185672669 | 185658340 | 185658409 | 185675295 | 185675431 | 29 | 36 | 4 | 50 | 88 | 7 | 1 | 6 | 5 | 16 | 64 | 41 | 34 | 129 | 200 | 20 | 15 | 3 | 19 | 9 | 278 | 149 | 0.775675501 | 1 | 0.689 | 0.951 | 0.263 | 0.843 | 0.747 | 0.632 | 0.594 | 0.859 | 0.784 | 0.923 | -0.06 |
| SE | 107499 | ENSG00000123349 | PFDN5 | chr12 | + | 53296243 | 53296275 | 53295508 | 53295639 | 53297849 | 53297924 | 607 | 355 | 528 | 516 | 640 | 208 | 128 | 218 | 168 | 123 | 841 | 542 | 339 | 477 | 863 | 212 | 136 | 153 | 132 | 306 | 181 | 149 | 0.77568576 | 1 | 0.706 | 0.695 | 0.666 | 0.717 | 0.811 | 0.766 | 0.766 | 0.646 | 0.748 | 0.699 | -0.006 |
| SE | 31030 | ENSG00000038219 | BOD1L1 | chr4 | - | 13615311 | 13615502 | 13614195 | 13614810 | 13619942 | 13620067 | 204 | 207 | 164 | 204 | 225 | 0 | 3 | 2 | 0 | 4 | 34 | 149 | 102 | 44 | 42 | 0 | 0 | 1 | 0 | 0 | 298 | 149 | 0.775703017 | 1 | 1 | 0.972 | 0.976 | 1 | 0.966 | 1 | 1 | 0.981 | 1 | 1 | -0.013 |
| SE | 56398 | ENSG00000151838 | CCDC175 | chr14 | - | 59540674 | 59540746 | 59538704 | 59538840 | 59543343 | 59543454 | 37 | 56 | 14 | 63 | 100 | 0 | 0 | 0 | 0 | 0 | 415 | 36 | 31 | 64 | 192 | 6 | 0 | 3 | 0 | 0 | 221 | 149 | 0.775757105 | 1 | 1 | 1 | 1 | 1 | 1 | 0.979 | 1 | 0.874 | 1 | 1 | 0.029 |
| SE | 96964 | ENSG00000183597 | TANGO2 | chr22 | + | 20055942 | 20056013 | 20052464 | 20052584 | 20061529 | 20061576 | 54 | 30 | 35 | 30 | 57 | 3 | 4 | 1 | 4 | 6 | 76 | 59 | 23 | 69 | 198 | 0 | 5 | 2 | 7 | 15 | 220 | 149 | 0.775807668 | 1 | 0.924 | 0.836 | 0.96 | 0.836 | 0.865 | 1 | 0.889 | 0.886 | 0.87 | 0.899 | -0.025 |
| SE | 26489 | ENSG00000112685 | EXOC2 | chr6 | - | 556483 | 556564 | 555953 | 556013 | 562783 | 562845 | 203 | 140 | 205 | 197 | 239 | 0 | 2 | 2 | 0 | 0 | 145 | 127 | 109 | 190 | 54 | 0 | 5 | 2 | 3 | 0 | 230 | 149 | 0.775808267 | 1 | 1 | 0.978 | 0.985 | 1 | 1 | 1 | 0.943 | 0.972 | 0.976 | 1 | 0.014 |
| SE | 80126 | ENSG00000122515 | ZMIZ2 | chr7 | + | 44756946 | 44757149 | 44756424 | 44756539 | 44757377 | 44757561 | 69 | 29 | 29 | 50 | 34 | 0 | 0 | 0 | 0 | 0 | 73 | 21 | 21 | 31 | 168 | 0 | 3 | 0 | 2 | 0 | 298 | 149 | 0.775841007 | 1 | 1 | 1 | 1 | 1 | 1 | 1 | 0.778 | 1 | 0.886 | 1 | 0.067 |
| SE | 36695 | ENSG00000158062 | UBXN11 | chr1 | - | 26300925 | 26301024 | 26297961 | 26298062 | 26301693 | 26301722 | 106 | 61 | 59 | 79 | 52 | 51 | 54 | 43 | 66 | 22 | 227 | 121 | 85 | 113 | 257 | 131 | 40 | 67 | 101 | 166 | 248 | 149 | 0.775841127 | 1 | 0.555 | 0.404 | 0.452 | 0.418 | 0.587 | 0.51 | 0.645 | 0.433 | 0.402 | 0.482 | -0.011 |
| SE | 23980 | ENSG00000277149 | TYW1B | chr7 | - | 72728821 | 72728931 | 72713620 | 72713798 | 72744483 | 72744601 | 38 | 30 | 13 | 14 | 23 | 0 | 0 | 0 | 0 | 0 | 28 | 3 | 22 | 15 | 24 | 0 | 0 | 4 | 0 | 0 | 259 | 149 | 0.775871252 | 1 | 1 | 1 | 1 | 1 | 1 | 1 | 1 | 0.76 | 1 | 1 | 0.048 |
| SE | 75578 | ENSG00000135678 | CPM | chr12 | - | 68885791 | 68885889 | 68871783 | 68871956 | 68932677 | 68932840 | 32 | 4 | 9 | 27 | 19 | 4 | 3 | 4 | 1 | 2 | 4 | 5 | 27 | 18 | 6 | 0 | 0 | 2 | 9 | 6 | 247 | 149 | 0.775876392 | 1 | 0.828 | 0.446 | 0.576 | 0.942 | 0.851 | 1 | 1 | 0.891 | 0.547 | 0.376 | -0.034 |
| SE | 29449 | ENSG00000130338 | TULP4 | chr6 | + | 158282262 | 158282378 | 158232235 | 158232303 | 158312050 | 158312358 | 59 | 64 | 56 | 47 | 69 | 5 | 7 | 10 | 15 | 0 | 63 | 48 | 78 | 64 | 47 | 4 | 18 | 3 | 13 | 0 | 265 | 149 | 0.775993921 | 1 | 0.869 | 0.837 | 0.759 | 0.638 | 1 | 0.899 | 0.6 | 0.936 | 0.735 | 1 | -0.013 |
| SE | 64240 | ENSG00000175104 | TRAF6 | chr11 | - | 36497107 | 36497266 | 36494975 | 36495047 | 36498489 | 36498640 | 173 | 96 | 90 | 161 | 239 | 0 | 2 | 1 | 0 | 0 | 151 | 178 | 215 | 181 | 93 | 0 | 0 | 5 | 3 | 0 | 298 | 149 | 0.776040709 | 1 | 1 | 0.96 | 0.978 | 1 | 1 | 1 | 1 | 0.956 | 0.968 | 1 | 0.003 |
| SE | 27321 | ENSG00000166913 | YWHAB | chr20 | + | 44887647 | 44887742 | 44885710 | 44885886 | 44901530 | 44901756 | 92 | 46 | 40 | 59 | 83 | 631 | 413 | 422 | 568 | 335 | 190 | 63 | 26 | 107 | 83 | 873 | 464 | 541 | 839 | 516 | 244 | 149 | 0.776178001 | 1 | 0.082 | 0.064 | 0.055 | 0.06 | 0.131 | 0.117 | 0.077 | 0.029 | 0.072 | 0.089 | 0.002 |
| SE | 22278 | ENSG00000159082 | SYNJ1 | chr21 | - | 32657715 | 32657872 | 32657002 | 32657120 | 32664912 | 32665071 | 52 | 48 | 70 | 66 | 43 | 5 | 3 | 2 | 5 | 3 | 31 | 37 | 33 | 26 | 25 | 0 | 1 | 4 | 6 | 0 | 298 | 149 | 0.776227862 | 1 | 0.839 | 0.889 | 0.946 | 0.868 | 0.878 | 1 | 0.949 | 0.805 | 0.684 | 1 | -0.004 |
| SE | 30422 | ENSG00000103502 | CDIPT | chr16 | - | 29859441 | 29859523 | 29858356 | 29859334 | 29860580 | 29860662 | 114 | 53 | 82 | 93 | 99 | 4 | 0 | 0 | 4 | 6 | 153 | 120 | 58 | 81 | 160 | 0 | 6 | 3 | 2 | 6 | 231 | 149 | 0.776269041 | 1 | 0.948 | 1 | 1 | 0.937 | 0.914 | 1 | 0.928 | 0.926 | 0.963 | 0.945 | 0.007 |
| SE | 99367 | ENSG00000156599 | ZDHHC5 | chr11 | + | 57690030 | 57690437 | 57688507 | 57688665 | 57692610 | 57692702 | 243 | 93 | 228 | 200 | 245 | 2 | 0 | 2 | 0 | 0 | 298 | 252 | 160 | 188 | 250 | 0 | 0 | 6 | 6 | 0 | 298 | 149 | 0.776291701 | 1 | 0.984 | 1 | 0.983 | 1 | 1 | 1 | 1 | 0.93 | 0.94 | 1 | 0.019 |
| SE | 54752 | ENSG00000083720 | OXCT1 | chr5 | - | 41805566 | 41805681 | 41803068 | 41803163 | 41840450 | 41840511 | 54 | 36 | 52 | 50 | 56 | 1 | 0 | 0 | 0 | 0 | 11 | 40 | 29 | 22 | 44 | 0 | 0 | 0 | 4 | 0 | 264 | 149 | 0.776291882 | 1 | 0.968 | 1 | 1 | 1 | 1 | 1 | 1 | 1 | 0.756 | 1 | 0.042 |
| SE | 53090 | ENSG00000103657 | HERC1 | chr15 | - | 63780461 | 63780552 | 63775187 | 63775649 | 63833821 | 63833915 | 5 | 5 | 2 | 4 | 4 | 18 | 15 | 18 | 11 | 12 | 0 | 2 | 6 | 2 | 13 | 20 | 18 | 15 | 6 | 9 | 240 | 149 | 0.776365722 | 1 | 0.147 | 0.171 | 0.065 | 0.184 | 0.171 | 0 | 0.065 | 0.199 | 0.171 | 0.473 | -0.034 |
| SE | 1033 | ENSG00000183741 | CBX6 | chr22 | - | 38871901 | 38871945 | 38871691 | 38871757 | 38872121 | 38872216 | 128 | 74 | 90 | 104 | 49 | 0 | 0 | 1 | 0 | 0 | 96 | 74 | 64 | 87 | 160 | 0 | 0 | 2 | 1 | 7 | 193 | 149 | 0.776387555 | 1 | 1 | 1 | 0.986 | 1 | 1 | 1 | 1 | 0.961 | 0.985 | 0.946 | 0.019 |
| SE | 83474 | ENSG00000106603 | COA1 | chr7 | - | 43650611 | 43650712 | 43647227 | 43648652 | 43665657 | 43665722 | 15 | 13 | 16 | 13 | 10 | 9 | 4 | 4 | 11 | 20 | 14 | 0 | 12 | 13 | 8 | 0 | 2 | 16 | 6 | 4 | 250 | 149 | 0.776398005 | 1 | 0.498 | 0.66 | 0.704 | 0.413 | 0.23 | 1 | 0 | 0.309 | 0.564 | 0.544 | 0.018 |
| SE | 38797 | ENSG00000165113 | GKAP1 | chr9 | - | 83792950 | 83793017 | 83788600 | 83788678 | 83799184 | 83799328 | 7 | 6 | 3 | 8 | 23 | 6 | 8 | 9 | 7 | 13 | 23 | 3 | 1 | 2 | 0 | 0 | 0 | 7 | 4 | 3 | 216 | 149 | 0.776442352 | 1 | 0.446 | 0.341 | 0.187 | 0.441 | 0.55 | 1 | 1 | 0.09 | 0.256 | 0 | -0.076 |
| SE | 87761 | ENSG00000063601 | MTMR1 | chrX | + | 150751029 | 150751240 | 150750729 | 150750843 | 150755688 | 150755865 | 14 | 4 | 7 | 8 | 2 | 64 | 41 | 70 | 56 | 59 | 0 | 13 | 6 | 11 | 0 | 84 | 47 | 29 | 28 | 46 | 298 | 149 | 0.776538949 | 1 | 0.099 | 0.047 | 0.048 | 0.067 | 0.017 | 0 | 0.121 | 0.094 | 0.164 | 0 | -0.02 |
| SE | 73580 | ENSG00000129472 | RAB2B | chr14 | - | 21468356 | 21468449 | 21463655 | 21463767 | 21474866 | 21474934 | 66 | 38 | 43 | 59 | 39 | 0 | 0 | 0 | 2 | 0 | 86 | 48 | 51 | 82 | 89 | 0 | 0 | 0 | 6 | 0 | 242 | 149 | 0.776723266 | 1 | 1 | 1 | 1 | 0.948 | 1 | 1 | 1 | 1 | 0.894 | 1 | 0.011 |
| SE | 54962 | ENSG00000169991 | IFFO2 | chr1 | - | 18919677 | 18919773 | 18916902 | 18917042 | 18921060 | 18921121 | 21 | 11 | 39 | 23 | 12 | 0 | 2 | 3 | 0 | 0 | 10 | 7 | 16 | 18 | 18 | 0 | 0 | 0 | 0 | 0 | 245 | 149 | 0.776772136 | 1 | 1 | 0.77 | 0.888 | 1 | 1 | 1 | 1 | 1 | 1 | 1 | -0.068 |
| SE | 22717 | ENSG00000164329 | TENT2 | chr5 | + | 79656957 | 79657001 | 79648616 | 79648693 | 79668891 | 79669028 | 64 | 62 | 60 | 74 | 97 | 0 | 0 | 0 | 0 | 0 | 12 | 61 | 56 | 29 | 16 | 0 | 0 | 0 | 0 | 4 | 193 | 149 | 0.77678163 | 1 | 1 | 1 | 1 | 1 | 1 | 1 | 1 | 1 | 1 | 0.755 | 0.049 |
| SE | 47128 | ENSG00000089012 | SIRPG | chr20 | - | 1635266 | 1635599 | 1630221 | 1630306 | 1649051 | 1649408 | 58 | 12 | 71 | 43 | 88 | 6 | 29 | 22 | 44 | 36 | 8 | 38 | 13 | 15 | 27 | 7 | 17 | 8 | 0 | 21 | 298 | 149 | 0.777055236 | 1 | 0.829 | 0.171 | 0.617 | 0.328 | 0.55 | 0.364 | 0.528 | 0.448 | 1 | 0.391 | -0.047 |
| SE | 57406 | ENSG00000101040 | ZMYND8 | chr20 | - | 47310055 | 47310204 | 47298728 | 47298872 | 47347855 | 47347926 | 93 | 122 | 81 | 97 | 121 | 0 | 0 | 6 | 0 | 0 | 195 | 112 | 94 | 142 | 126 | 0 | 0 | 0 | 0 | 0 | 298 | 149 | 0.777062692 | 1 | 1 | 1 | 0.871 | 1 | 1 | 1 | 1 | 1 | 1 | 1 | -0.026 |
| SE | 34099 | ENSG00000135083 | CCNJL | chr5 | - | 160276268 | 160276412 | 160259468 | 160259771 | 160280524 | 160280738 | 2 | 0 | 1 | 0 | 0 | 6 | 2 | 4 | 7 | 3 | 0 | 0 | 1 | 0 | 4 | 8 | 4 | 9 | 1 | 6 | 293 | 149 | 0.777125368 | 1 | 0.145 | 0 | 0.113 | 0 | 0 | 0 | 0 | 0.053 | 0 | 0.253 | -0.01 |
| SE | 30434 | ENSG00000164038 | SLC9B2 | chr4 | - | 103043295 | 103043445 | 103031699 | 103031808 | 103044889 | 103044991 | 23 | 8 | 35 | 9 | 25 | 0 | 0 | 0 | 0 | 4 | 18 | 24 | 5 | 8 | 23 | 0 | 0 | 2 | 0 | 0 | 298 | 149 | 0.777172028 | 1 | 1 | 1 | 1 | 1 | 0.758 | 1 | 1 | 0.556 | 1 | 1 | 0.04 |
| SE | 65569 | ENSG00000147526 | TACC1 | chr8 | + | 38788703 | 38788819 | 38787235 | 38787743 | 38825307 | 38825368 | 78 | 19 | 64 | 86 | 30 | 2 | 2 | 0 | 2 | 0 | 25 | 50 | 42 | 61 | 15 | 0 | 0 | 0 | 0 | 3 | 265 | 149 | 0.777234306 | 1 | 0.956 | 0.842 | 1 | 0.96 | 1 | 1 | 1 | 1 | 1 | 0.738 | 0.004 |
| SE | 10215 | ENSG00000167182 | SP2 | chr17 | + | 47897814 | 47897927 | 47896181 | 47896293 | 47915311 | 47915388 | 6 | 6 | 4 | 15 | 26 | 16 | 4 | 18 | 27 | 7 | 42 | 9 | 0 | 10 | 24 | 2 | 18 | 21 | 21 | 44 | 262 | 149 | 0.777271336 | 1 | 0.176 | 0.46 | 0.112 | 0.24 | 0.679 | 0.923 | 0.221 | 0 | 0.213 | 0.237 | 0.015 |
| SE | 61863 | ENSG00000143393 | PI4KB | chr1 | - | 151324744 | 151324917 | 151315572 | 151316509 | 151327270 | 151327377 | 52 | 18 | 24 | 45 | 45 | 29 | 11 | 27 | 8 | 13 | 77 | 43 | 19 | 29 | 90 | 14 | 28 | 12 | 22 | 55 | 298 | 149 | 0.77730579 | 1 | 0.473 | 0.45 | 0.308 | 0.738 | 0.634 | 0.733 | 0.434 | 0.442 | 0.397 | 0.45 | 0.029 |
| SE | 14231 | ENSG00000142192 | APP | chr21 | - | 26000014 | 26000182 | 25982343 | 25982477 | 26021839 | 26022042 | 463 | 351 | 375 | 633 | 419 | 8 | 12 | 8 | 9 | 6 | 1375 | 638 | 389 | 1167 | 2227 | 28 | 2 | 17 | 27 | 21 | 298 | 149 | 0.777311596 | 1 | 0.967 | 0.936 | 0.959 | 0.972 | 0.972 | 0.961 | 0.994 | 0.92 | 0.956 | 0.981 | -0.001 |
| SE | 99305 | ENSG00000101017 | CD40 | chr20 | + | 46121723 | 46121898 | 46118372 | 46118394 | 46122232 | 46122358 | 14 | 5 | 23 | 23 | 21 | 1 | 0 | 0 | 2 | 0 | 11 | 25 | 5 | 10 | 55 | 0 | 4 | 0 | 0 | 0 | 298 | 149 | 0.777312592 | 1 | 0.875 | 1 | 1 | 0.852 | 1 | 1 | 0.758 | 1 | 1 | 1 | -0.006 |
| SE | 81932 | ENSG00000105819 | PMPCB | chr7 | + | 103307595 | 103307708 | 103304410 | 103304490 | 103308951 | 103309095 | 201 | 146 | 245 | 193 | 385 | 61 | 52 | 55 | 51 | 96 | 54 | 78 | 130 | 84 | 9 | 7 | 20 | 73 | 79 | 0 | 262 | 149 | 0.777328071 | 1 | 0.652 | 0.615 | 0.717 | 0.683 | 0.695 | 0.814 | 0.689 | 0.503 | 0.377 | 1 | -0.004 |
| SE | 72089 | ENSG00000180228 | PRKRA | chr2 | - | 178445425 | 178445535 | 178444421 | 178444500 | 178447504 | 178447586 | 9 | 3 | 9 | 9 | 13 | 75 | 37 | 71 | 65 | 111 | 4 | 13 | 3 | 3 | 0 | 83 | 14 | 26 | 47 | 14 | 259 | 149 | 0.777333167 | 1 | 0.065 | 0.045 | 0.068 | 0.074 | 0.063 | 0.027 | 0.348 | 0.062 | 0.035 | 0 | -0.031 |
| SE | 20831 | ENSG00000107929 | LARP4B | chr10 | - | 844976 | 845055 | 842931 | 843068 | 863742 | 863883 | 244 | 161 | 231 | 287 | 346 | 2 | 0 | 4 | 5 | 0 | 310 | 285 | 120 | 407 | 216 | 0 | 0 | 0 | 0 | 13 | 228 | 149 | 0.777398594 | 1 | 0.988 | 1 | 0.974 | 0.974 | 1 | 1 | 1 | 1 | 1 | 0.916 | 0.004 |
| SE | 65951 | ENSG00000123240 | OPTN | chr10 | + | 13108961 | 13109015 | 13108137 | 13108289 | 13109111 | 13109288 | 49 | 19 | 42 | 37 | 13 | 13 | 6 | 41 | 13 | 14 | 41 | 36 | 36 | 12 | 2 | 0 | 11 | 19 | 6 | 21 | 203 | 149 | 0.777444146 | 1 | 0.735 | 0.699 | 0.429 | 0.676 | 0.405 | 1 | 0.706 | 0.582 | 0.595 | 0.065 | -0.001 |
| SE | 43107 | ENSG00000119048 | UBE2B | chr5 | + | 134376668 | 134376694 | 134374382 | 134374463 | 134380718 | 134380808 | 442 | 276 | 345 | 348 | 610 | 3 | 3 | 2 | 0 | 2 | 141 | 242 | 391 | 336 | 165 | 1 | 0 | 9 | 6 | 0 | 175 | 149 | 0.777478248 | 1 | 0.992 | 0.987 | 0.993 | 1 | 0.996 | 0.992 | 1 | 0.974 | 0.979 | 1 | 0.005 |
| SE | 39448 | ENSG00000055917 | PUM2 | chr2 | - | 20332966 | 20333063 | 20327309 | 20327378 | 20350596 | 20350702 | 24 | 27 | 15 | 8 | 18 | 132 | 107 | 80 | 128 | 100 | 6 | 3 | 13 | 17 | 7 | 72 | 66 | 50 | 75 | 16 | 246 | 149 | 0.77758863 | 1 | 0.099 | 0.133 | 0.102 | 0.036 | 0.098 | 0.048 | 0.027 | 0.136 | 0.121 | 0.209 | -0.015 |
| SE | 76324 | ENSG00000073050 | XRCC1 | chr19 | - | 43560909 | 43561020 | 43554645 | 43554804 | 43574909 | 43575002 | 40 | 23 | 35 | 39 | 27 | 2 | 0 | 0 | 0 | 0 | 102 | 51 | 34 | 71 | 69 | 7 | 0 | 0 | 0 | 0 | 260 | 149 | 0.777604464 | 1 | 0.92 | 1 | 1 | 1 | 1 | 0.893 | 1 | 1 | 1 | 1 | 0.005 |
| SE | 106262 | ENSG00000116198 | CEP104 | chr1 | - | 3826707 | 3826744 | 3826369 | 3826436 | 3829265 | 3829373 | 41 | 34 | 41 | 38 | 60 | 1 | 5 | 0 | 0 | 1 | 36 | 24 | 23 | 56 | 43 | 0 | 0 | 0 | 0 | 0 | 186 | 149 | 0.777607351 | 1 | 0.97 | 0.845 | 1 | 1 | 0.98 | 1 | 1 | 1 | 1 | 1 | -0.041 |
| SE | 100648 | ENSG00000166340 | TPP1 | chr11 | - | 6616004 | 6616074 | 6615441 | 6615562 | 6616314 | 6616479 | 335 | 162 | 184 | 255 | 187 | 0 | 3 | 0 | 0 | 0 | 500 | 359 | 202 | 358 | 1056 | 6 | 6 | 0 | 6 | 2 | 219 | 149 | 0.777608205 | 1 | 1 | 0.974 | 1 | 1 | 1 | 0.983 | 0.976 | 1 | 0.976 | 0.997 | 0.008 |
| SE | 41632 | ENSG00000183258 | DDX41 | chr5 | - | 177513010 | 177513082 | 177512495 | 177512876 | 177513352 | 177513484 | 148 | 81 | 100 | 134 | 113 | 0 | 0 | 1 | 0 | 0 | 275 | 192 | 124 | 197 | 332 | 0 | 4 | 2 | 7 | 0 | 221 | 149 | 0.77770755 | 1 | 1 | 1 | 0.985 | 1 | 1 | 1 | 0.97 | 0.977 | 0.95 | 1 | 0.018 |
| SE | 4282 | ENSG00000053501 | USE1 | chr19 | + | 17215801 | 17215851 | 17215356 | 17215507 | 17215991 | 17216070 | 37 | 20 | 55 | 32 | 29 | 5 | 0 | 0 | 0 | 0 | 49 | 33 | 13 | 12 | 30 | 0 | 0 | 0 | 0 | 0 | 199 | 149 | 0.777754426 | 1 | 0.847 | 1 | 1 | 1 | 1 | 1 | 1 | 1 | 1 | 1 | -0.031 |
| SE | 26146 | ENSG00000213445 | SIPA1 | chr11 | + | 65642249 | 65642377 | 65640800 | 65641600 | 65642462 | 65642639 | 248 | 75 | 191 | 234 | 192 | 1 | 2 | 6 | 3 | 0 | 76 | 300 | 93 | 179 | 236 | 0 | 2 | 2 | 9 | 0 | 277 | 149 | 0.777827824 | 1 | 0.993 | 0.953 | 0.945 | 0.977 | 1 | 1 | 0.988 | 0.962 | 0.915 | 1 | 0.001 |

[illegible]

| | | | | | | | | | | | | | | |
|---|---|---|---|---|---|---|---|---|---|---|---|---|---|---|
| SE | 14339 | ENSG00000007545 | CRAMP1 | chr16 | + | 1662760 | 1662835 | 1662466 | 1662671 | 1665056 | 1665138 | 61 | 18 | 44 |
| SE | 91259 | ENSG00000154114 | TBCEL | chr11 | + | 121036504 | 121036612 | 121035240 | 121036314 | 121045673 | 121045823 | 6 | 10 | 9 |
| SE | 37673 | ENSG00000168615 | ADAM9 | chr8 | + | 39076146 | 39076275 | 39071297 | 39071403 | 39077227 | 39077411 | 5 | 5 | 3 |
| SE | 82257 | ENSG00000223482 | NUTM2A-AS1 | chr10 | - | 87241181 | 87245855 | 87208185 | 87208512 | 87246114 | 87246212 | 31 | 16 | 11 |
| SE | 105673 | ENSG00000156502 | SUPV3L1 | chr10 | + | 69199103 | 69199197 | 69198371 | 69198552 | 69200279 | 69200353 | 59 | 53 | 55 |
| SE | 96592 | ENSG00000198561 | CTNND1 | chr11 | + | 57794009 | 57794081 | 57791384 | 57791673 | 57795576 | 57795729 | 67 | 82 | 41 |
| SE | 29778 | ENSG00000101445 | PPP1R16B | chr20 | + | 38905968 | 38906094 | 38902667 | 38902792 | 38906978 | 38907054 | 238 | 92 | 158 |
| SE | 64131 | ENSG00000168906 | MAT2A | chr2 | + | 85541254 | 85541377 | 85541082 | 85541160 | 85541632 | 85541745 | 746 | 574 | 622 |
| SE | 92817 | ENSG00000156471 | PTDSS1 | chr8 | + | 96287021 | 96287146 | 96284108 | 96284153 | 96295097 | 96295256 | 373 | 186 | 431 |
| SE | 108070 | ENSG00000004700 | RECQL | chr12 | - | 21486478 | 21486585 | 21483375 | 21483574 | 21490198 | 21490378 | 358 | 320 | 270 |
| SE | 51687 | ENSG00000198837 | DENND4B | chr1 | - | 153934759 | 153934964 | 153934134 | 153934302 | 153936059 | 153936188 | 36 | 32 | 26 |
| SE | 67569 | ENSG00000076003 | MCM6 | chr2 | - | 135865012 | 135865163 | 135862606 | 135862748 | 135866131 | 135866277 | 257 | 152 | 239 |
| SE | 57936 | ENSG00000168439 | STIP1 | chr11 | + | 64197853 | 64197974 | 64197492 | 64197595 | 64199939 | 64200008 | 234 | 212 | 174 |
| SE | 62647 | ENSG00000128815 | WDFY4 | chr10 | + | 48830699 | 48830885 | 48828777 | 48828896 | 48832572 | 48832709 | 247 | 189 | 169 |
| SE | 5185 | ENSG00000101189 | MRGBP | chr20 | + | 62798974 | 62799049 | 62798586 | 62798668 | 62799455 | 62801729 | 48 | 11 | 27 |
| SE | 27267 | ENSG00000077232 | DNAJC10 | chr2 | + | 182736248 | 182736386 | 182731029 | 182731107 | 182739544 | 182739709 | 46 | 47 | 48 |
| SE | 37083 | ENSG00000127481 | UBR4 | chr1 | - | 19101519 | 19101641 | 19100375 | 19100573 | 19104083 | 19104257 | 280 | 186 | 279 |
| SE | 100139 | ENSG00000159063 | ALG8 | chr11 | - | 78112649 | 78112770 | 78109441 | 78109581 | 78114265 | 78114392 | 43 | 22 | 35 |
| SE | 54921 | ENSG00000256235 | SMIM3 | chr5 | + | 150782473 | 150782590 | 150778804 | 150778972 | 150795429 | 150796734 | 0 | 0 | 4 |
| SE | 62139 | ENSG00000182179 | UBA7 | chr3 | - | 49812647 | 49812738 | 49812407 | 49812543 | 49813061 | 49813168 | 83 | 38 | 90 |
| SE | 106979 | ENSG00000111271 | ACAD10 | chr12 | + | 111733922 | 111734068 | 111729805 | 111729956 | 111736830 | 111737004 | 27 | 12 | 27 |
| SE | 27146 | ENSG00000178562 | CD28 | chr2 | + | 203726632 | 203726989 | 203706523 | 203706617 | 203729647 | 203729772 | 136 | 119 | 234 |
| SE | 24477 | ENSG00000197321 | SVIL | chr10 | - | 29530606 | 29530668 | 29529704 | 29529844 | 29550596 | 29551263 | 39 | 27 | 39 |
| SE | 52189 | ENSG00000248710 | NA | chr3 | - | 160319759 | 160319939 | 160307662 | 160307781 | 160356012 | 160356150 | 58 | 97 | 81 |
| SE | 15809 | ENSG00000169375 | SIN3A | chr15 | - | 75401851 | 75401970 | 75400039 | 75400156 | 75407054 | 75407144 | 106 | 70 | 104 |
| SE | 36360 | ENSG00000173889 | PHC3 | chr3 | - | 170136418 | 170136665 | 170128683 | 170129552 | 170149085 | 170149244 | 211 | 174 | 192 |
| SE | 17220 | ENSG00000126804 | ZBTB1 | chr14 | + | 64516599 | 64516751 | 64504725 | 64504946 | 64521486 | 64521606 | 74 | 22 | 23 |
| SE | 82245 | ENSG00000128604 | IRF5 | chr7 | + | 128947022 | 128947056 | 128946500 | 128946562 | 128947229 | 128947535 | 60 | 18 | 29 |
| SE | 39074 | ENSG00000172175 | MALT1 | chr18 | + | 58710920 | 58710953 | 58709321 | 58710072 | 58714082 | 58714109 | 20 | 31 | 28 |
| SE | 86828 | ENSG00000005889 | ZFX | chrX | + | 24175388 | 24175443 | 24173606 | 24173723 | 24179182 | 24179550 | 5 | 0 | 2 |
| SE | 80729 | ENSG00000163636 | PSMD6 | chr3 | - | 64022738 | 64022801 | 64022317 | 64022523 | 64023274 | 64023456 | 1 | 2 | 0 |
| SE | 10732 | ENSG00000006125 | AP2B1 | chr17 | + | 35670856 | 35670898 | 35657598 | 35657791 | 35671753 | 35671900 | 81 | 45 | 92 |
| SE | 39524 | ENSG00000186517 | ARHGAP30 | chr1 | - | 161052625 | 161052797 | 161052439 | 161052543 | 161053257 | 161053385 | 1034 | 586 | 780 |
| SE | 39310 | ENSG00000116514 | RNF19B | chr1 | - | 32938396 | 32938528 | 32936453 | 32937309 | 32942251 | 32942459 | 113 | 76 | 97 |
| SE | 35896 | ENSG00000197415 | VEPH1 | chr3 | - | 157499394 | 157499675 | 157495211 | 157495506 | 157533512 | 157533616 | 0 | 0 | 0 |
| SE | 42156 | ENSG00000197343 | ZNF655 | chr7 | + | 99561887 | 99562494 | 99560532 | 99560695 | 99563862 | 99564705 | 214 | 111 | 170 |
| SE | 75095 | ENSG00000188554 | NBR1 | chr17 | + | 43203680 | 43203786 | 43202654 | 43202712 | 43209900 | 43211688 | 278 | 122 | 217 |
| SE | 26537 | ENSG00000138303 | ASCC1 | chr10 | - | 72210731 | 72210831 | 72203426 | 72203524 | 72213186 | 72213277 | 140 | 111 | 157 |
| SE | 635 | ENSG00000100129 | EIF3L | chr22 | + | 37870175 | 37870347 | 37863271 | 37863345 | 37875840 | 37876011 | 738 | 465 | 672 |
| SE | 43941 | ENSG00000077782 | FGFR1 | chr8 | - | 38426121 | 38426245 | 38423312 | 38424699 | 38427920 | 38428093 | 43 | 34 | 24 |
| SE | 54553 | ENSG00000179912 | R3HDM2 | chr12 | - | 57288856 | 57288910 | 57283823 | 57284056 | 57289008 | 57289040 | 2 | 2 | 1 |
| SE | 24834 | ENSG00000170085 | SIMC1 | chr5 | + | 176322272 | 176322425 | 176313690 | 176313845 | 176324628 | 176324757 | 53 | 23 | 51 |
| SE | 50896 | ENSG00000156515 | HK1 | chr10 | + | 69384332 | 69384481 | 69382486 | 69382791 | 69384795 | 69384915 | 562 | 466 | 439 |
| SE | 38490 | ENSG00000198793 | MTOR | chr1 | - | 11150125 | 11150226 | 11146675 | 11146791 | 11157151 | 11157291 | 87 | 51 | 71 |
| SE | 80118 | ENSG00000197024 | ZNF398 | chr7 | + | 149166157 | 149166284 | 149153944 | 149154340 | 149166816 | 149166930 | 132 | 85 | 129 |
| SE | 97267 | ENSG00000156110 | ADK | chr10 | + | 74527759 | 74528093 | 74525255 | 74525426 | 74589281 | 74589317 | 7 | 5 | 3 |
| SE | 57070 | ENSG00000003756 | RBM5 | chr3 | + | 50115427 | 50115607 | 50112613 | 50113544 | 50115905 | 50115943 | 398 | 315 | 334 |
| SE | 56608 | ENSG00000075151 | EIF4G3 | chr1 | - | 20892618 | 20892729 | 20886200 | 20886371 | 20893516 | 20893636 | 52 | 7 | 20 |
| SE | 67156 | ENSG00000152117 | SMPD4BP | chr2 | + | 131516856 | 131516957 | 131516313 | 131516404 | 131517726 | 131517890 | 6 | 6 | 22 |
| SE | 40860 | ENSG00000117020 | AKT3 | chr1 | - | 243739522 | 243739613 | 243695590 | 243695716 | 243843124 | 243843282 | 2 | 1 | 3 |
| SE | 72530 | ENSG00000135426 | TESPA1 | chr12 | - | 54961167 | 54961267 | 54948014 | 54950390 | 54962430 | 54963238 | 139 | 91 | 284 |
| SE | 32554 | ENSG00000211456 | SACM1L | chr3 | + | 45735234 | 45735373 | 45732052 | 45732151 | 45737582 | 45737652 | 535 | 355 | 412 |
| SE | 8169 | ENSG00000108799 | EZH1 | chr17 | - | 42704601 | 42704683 | 42703192 | 42703820 | 42705087 | 42705183 | 86 | 71 | 102 |
| SE | 71394 | ENSG00000122203 | KIAA1191 | chr5 | - | 176359480 | 176359567 | 176352621 | 176352748 | 176359810 | 176359918 | 26 | 18 | 30 |
| SE | 6741 | ENSG00000166401 | SERPINB8 | chr18 | + | 63983578 | 63983721 | 63981720 | 63981838 | 63985092 | 63985245 | 277 | 315 | 228 |
| SE | 23719 | ENSG00000124782 | RREB1 | chr6 | + | 7228996 | 7231907 | 7226466 | 7226656 | 7246423 | 7247221 | 89 | 63 | 104 |
| SE | 101174 | ENSG00000133816 | MICAL2 | chr11 | + | 12255642 | 12255750 | 12249183 | 12249246 | 12256784 | 12256909 | 2 | 3 | 8 |
| SE | 18109 | ENSG00000075413 | MARK3 | chr14 | + | 103487378 | 103487707 | 103480386 | 103480490 | 103491776 | 103492034 | 7 | 4 | 1 |
| SE | 106947 | ENSG00000127334 | DYRK2 | chr12 | + | 67649796 | 67649945 | 67648744 | 67649182 | 67657105 | 67665406 | 88 | 28 | 96 |
| SE | 2794 | ENSG00000011304 | PTBP1 | chr19 | + | 807868 | 807902 | 799412 | 799443 | 808359 | 808452 | 95 | 42 | 78 |
| SE | 19718 | ENSG00000103811 | CTSH | chr15 | - | 78931900 | 78932038 | 78931036 | 78931505 | 78932371 | 78932458 | 1 | 3 | 1 |
| SE | 71445 | ENSG00000123983 | ACSL3 | chr2 | + | 222892263 | 222892301 | 222887829 | 222887888 | 222900673 | 222900780 | 0 | 0 | 0 |
| SE | 96961 | ENSG00000183597 | TANGO2 | chr22 | + | 20053436 | 20053551 | 20052464 | 20052584 | 20055942 | 20056013 | 96 | 39 | 69 |
| SE | 100354 | ENSG00000174842 | GLMN | chr1 | - | 92266699 | 92266741 | 92266418 | 92266492 | 92267912 | 92268002 | 32 | 16 | 26 |
| SE | 17326 | ENSG00000258461 | NA | chr15 | + | 42385678 | 42385956 | 42384482 | 42384552 | 42386166 | 42386285 | 0 | 0 | 0 |
| SE | 51645 | ENSG00000081154 | PCNP | chr3 | + | 101579789 | 101580004 | 101574179 | 101574279 | 101585436 | 101585511 | 428 | 382 | 423 |
| SE | 84540 | ENSG00000288640 | NA | chr7 | + | 112475429 | 112475676 | 112472765 | 112472861 | 112484811 | 112484943 | 88 | 92 | 54 |
| SE | 9295 | ENSG00000187838 | PLSCR3 | chr17 | - | 7393143 | 7393364 | 7392790 | 7392952 | 7393466 | 7393482 | 76 | 32 | 90 |
| SE | 66488 | ENSG00000155561 | NUP205 | chr7 | + | 135643191 | 135643358 | 135638556 | 135638683 | 135644894 | 135644978 | 143 | 92 | 141 |
| SE | 40192 | ENSG00000231607 | DLEU2 | chr13 | - | 50029277 | 50029492 | 50027089 | 50027406 | 50044650 | 50044768 | 68 | 73 | 38 |
| SE | 2566 | ENSG00000214046 | SMIM7 | chr19 | - | 16659958 | 16660000 | 16659394 | 16659447 | 16660084 | 16660114 | 147 | 67 | 141 |
| SE | 3748 | ENSG00000187116 | LILRA5 | chr19 | - | 54312536 | 54312621 | 54311863 | 54312148 | 54313016 | 54313074 | 87 | 28 | 38 |
| SE | 72279 | ENSG00000186635 | ARAP1 | chr11 | - | 72714151 | 72714321 | 72712437 | 72712568 | 72721849 | 72722028 | 300 | 189 | 163 |
| SE | 28111 | ENSG00000125386 | FAM193A | chr4 | + | 2715321 | 2715493 | 2699679 | 2700544 | 2716022 | 2716104 | 1 | 2 | 0 |
| SE | 77810 | ENSG00000134250 | NOTCH2 | chr1 | - | 119968076 | 119968232 | 119967432 | 119967621 | 119969510 | 119969744 | 304 | 183 | 217 |
| SE | 15696 | ENSG00000174485 | DENND4A | chr15 | - | 65752378 | 65752628 | 65741714 | 65741784 | 65756139 | 65756472 | 280 | 258 | 258 |
| SE | 58048 | ENSG00000082701 | GSK3B | chr3 | - | 119882636 | 119882834 | 119876412 | 119876508 | 119905754 | 119905852 | 0 | 0 | 2 |
| SE | 6660 | ENSG00000141627 | DYM | chr18 | - | 49286433 | 49286616 | 49272177 | 49272303 | 49331863 | 49332006 | 110 | 67 | 101 |
| SE | 24010 | ENSG00000112983 | BRD8 | chr5 | - | 138164086 | 138164133 | 138163129 | 138163344 | 138164319 | 138164413 | 166 | 120 | 171 |
| SE | 105266 | ENSG00000082898 | XPO1 | chr2 | - | 61529065 | 61529159 | 61526419 | 61526521 | 61533771 | 61533903 | 12 | 2 | 12 |
| SE | 75129 | ENSG00000248124 | RRN3P1 | chr16 | - | 21810774 | 21810838 | 21809645 | 21809715 | 21811777 | 21811837 | 38 | 19 | 24 |
| SE | 14632 | ENSG00000140157 | NIPA2 | chr15 | + | 22853211 | 22853268 | 22851041 | 22851870 | 22858539 | 22858630 | 62 | 36 | 37 |
| SE | 43598 | ENSG00000058673 | ZC3H11A | chr1 | + | 203802956 | 203803016 | 203801574 | 203801628 | 203816925 | 203817124 | 230 | 153 | 211 |
| SE | 76809 | ENSG00000135317 | SNX14 | chr6 | - | 85567533 | 85567577 | 85549722 | 85549879 | 85572136 | 85572215 | 33 | 22 | 17 |
| SE | 63005 | ENSG00000137497 | NUMA1 | chr11 | - | 72068495 | 72068617 | 72068104 | 72068231 | 72069841 | 72069911 | 0 | 0 | 2 |
| SE | 105677 | ENSG00000196922 | ZNF252P | chr8 | - | 144996506 | 144996607 | 144994478 | 144994592 | 144999602 | 144999769 | 20 | 18 | 15 |
| SE | 66084 | ENSG00000155366 | RHOC | chr1 | - | 112705099 | 112705168 | 112703492 | 112703806 | 112707077 | 112707114 | 69 | 33 | 25 |
| SE | 100619 | ENSG00000132254 | ARFIP2 | chr11 | - | 6478578 | 6478652 | 6478083 | 6478198 | 6478737 | 6478959 | 3 | 1 | 3 |
| SE | 55675 | ENSG00000272886 | DCP1A | chr3 | - | 53338090 | 53338145 | 53319406 | 53319473 | 53342143 | 53342271 | 2 | 0 | 7 |
| SE | 12614 | ENSG00000168872 | DDX19A | chr16 | + | 70356845 | 70356906 | 70356111 | 70356247 | 70361417 | 70361496 | 15 | 8 | 12 |
| SE | 85322 | ENSG00000147394 | ZNF185 | chrX | + | 152928574 | 152928661 | 152922719 | 152922809 | 152931671 | 152931776 | 26 | 47 | 14 |
| SE | 1029 | ENSG00000244509 | APOBEC3C | chr22 | + | 39015594 | 39015751 | 39014256 | 39014379 | 39017765 | 39018045 | 937 | 384 | 528 |
| SE | 1634 | ENSG00000244625 | MIATNB | chr22 | + | 26726643 | 26726720 | 26717252 | 26717459 | 26774155 | 26774217 | 0 | 3 | 2 |
| SE | 107267 | ENSG00000186815 | TPCN1 | chr12 | + | 113294105 | 113294228 | 113293268 | 113293349 | 113295959 | 113296012 | 4 | 5 | 10 |
| SE | 63053 | ENSG00000006652 | IFRD1 | chr7 | + | 112450577 | 112450782 | 112422886 | 112423432 | 112455762 | 112455855 | 18 | 10 | 14 |
| SE | 95805 | ENSG00000167721 | TSR1 | chr17 | - | 2334471 | 2334896 | 2333389 | 2333716 | 2335259 | 2335394 | 97 | 67 | 104 |
| SE | 46404 | ENSG00000168724 | DNAJC21 | chr5 | + | 34933814 | 34933908 | 34929558 | 34929916 | 34935709 | 34935833 | 57 | 41 | 53 |
| SE | 71926 | ENSG00000157637 | SLC38A10 | chr17 | - | 81270917 | 81271024 | 81260237 | 81260394 | 81275968 | 81276151 | 292 | 155 | 190 |
| SE | 65312 | ENSG00000285756 | NA | chrX | - | 3818441 | 3818500 | 3817527 | 3817778 | 3828505 | 3829387 | 15 | 18 | 27 |
| SE | 69984 | ENSG00000164938 | TP53INP1 | chr8 | - | 94932069 | 94932107 | 94925971 | 94930728 | 94939859 | 94940220 | 123 | 96 | 172 |
| SE | 3138 | ENSG00000213762 | ZNF134 | chr19 | + | 57618791 | 57618877 | 57614255 | 57614503 | 57619411 | 57619508 | 2 | 0 | 0 |
| SE | 38086 | ENSG00000160584 | SIK3 | chr11 | - | 116867945 | 116868089 | 116863667 | 116863818 | 116870140 | 116870251 | 58 | 31 | 64 |
| SE | 36185 | ENSG00000116731 | PRDM2 | chr1 | + | 13742004 | 13742157 | 13731017 | 13731117 | 13749360 | 13749463 | 84 | 63 | 109 |
| SE | 74052 | ENSG00000111011 | RSRC2 | chr12 | - | 122514711 | 122514728 | 122511108 | 122511188 | 122515104 | 122515227 | 7 | 7 | 20 |
| SE | 82752 | ENSG00000140396 | NCOA2 | chr8 | - | 70216659 | 70216764 | 70174755 | 70174859 | 70296743 | 70296800 | 20 | 10 | 22 |
| SE | 8087 | ENSG00000141252 | VPS53 | chr17 | - | 627173 | 627316 | 623532 | 623674 | 653290 | 653410 | 89 | 60 | 82 |
| SE | 99645 | ENSG00000110075 | PPP6R3 | chr11 | + | 68505144 | 68505249 | 68460730 | 68460827 | 68519500 | 68519651 | 34 | 12 | 28 |
| SE | 41691 | ENSG00000247828 | TMEM161B-AS1 | chr5 | + | 88282714 | 88282866 | 88269467 | 88282106 | 88285743 | 88285851 | 34 | 27 | 26 |
| SE | 17328 | ENSG00000258461 | NA | chr15 | + | 42388927 | 42389096 | 42387752 | 42387886 | 42392638 | 42392722 | 33 | 56 | 23 |
| SE | 93090 | ENSG00000113924 | HGD | chr3 | - | 120644318 | 120644443 | 120641588 | 120641693 | 120646972 | 120647052 | 2 | 1 | 0 |
| SE | 86868 | ENSG00000090316 | MAEA | chr4 | + | 1327626 | 1327703 | 1322380 | 1322503 | 1332756 | 1332854 | 138 | 36 | 77 |
| SE | 19880 | ENSG00000152240 | HAUS1 | chr18 | + | 46119925 | 46120060 | 46105193 | 46105368 | 46122466 | 46122590 | 92 | 40 | 75 |
| SE | 74247 | ENSG00000137478 | FCHSD2 | chr11 | - | 72902542 | 72902638 | 72889828 | 72889945 | 72921827 | 72921950 | 429 | 336 | 307 |
| SE | 46943 | ENSG00000141027 | NCOR1 | chr17 | - | 16121051 | 16121269 | 16119422 | 16119485 | 16126081 | 16126206 | 210 | 161 | 173 |
| SE | 64214 | ENSG00000117697 | NSL1 | chr1 | - | 212766188 | 212766273 | 212739533 | 212739601 | 212782371 | 212782426 | 26 | 32 | 20 |
| SE | 26733 | ENSG00000125089 | SH3TC1 | chr4 | + | 8209664 | 8209822 | 8205166 | 8205366 | 8212700 | 8212828 | 32 | 8 | 17 |
| SE | 73412 | ENSG00000196290 | NIF3L1 | chr2 | + | 200899384 | 200899468 | 200895263 | 200895390 | 200903493 | 200903896 | 28 | 5 | 19 |
| SE | 34512 | ENSG00000116350 | SRSF4 | chr1 | - | 29168502 | 29168638 | 29160374 | 29160517 | 29181645 | 29181890 | 1 | 0 | 0 |
| SE | 8590 | ENSG00000185359 | HGS | chr17 | + | 81692384 | 81692553 | 81691446 | 81691571 | 81693502 | 81693581 | 1 | 0 | 2 |
| SE | 48188 | ENSG00000006831 | ADIPOR2 | chr12 | + | 1772841 | 1772961 | 1754257 | 1754514 | 1777853 | 1777915 | 200 | 151 | 160 |
| SE | 4838 | ENSG00000132670 | PTPRA | chr20 | + | 2964271 | 2964350 | 2947981 | 2948024 | 2975214 | 2975241 | 308 | 170 | 219 |
| SE | 9287 | ENSG00000170037 | CNTROB | chr17 | + | 7948486 | 7949137 | 7948156 | 7948327 | 7949384 | 7949918 | 26 | 21 | 29 |
| SE | 78598 | ENSG00000083168 | KAT6A | chr8 | - | 41942792 | 41943000 | 41940841 | 41941444 | 41943747 | 41943979 | 358 | 237 | 335 |
| SE | 82181 | ENSG00000146872 | TLK2 | chr17 | + | 62574344 | 62574410 | 62573214 | 62573367 | 62576708 | 62576775 | 5 | 0 | 4 |
| SE | 7397 | ENSG00000141447 | OSBPL1A | chr18 | - | 24181144 | 24181279 | 24172375 | 24172483 | 24196124 | 24196200 | 16 | 20 | 9 |
| SE | 39494 | ENSG00000010322 | NISCH | chr3 | + | 52477573 | 52477642 | 52476297 | 52476599 | 52478096 | 52478282 | 126 | 58 | 153 |
| SE | 26390 | ENSG00000056097 | ZFR | chr5 | - | 32403905 | 32404097 | 32403105 | 32403397 | 32406773 | 32407021 | 345 | 234 | 287 |
| SE | 15996 | ENSG00000068305 | MEF2A | chr15 | + | 99598429 | 99598511 | 99566482 | 99566599 | 99632977 | 99633173 | 90 | 131 | 72 |
| SE | 79865 | ENSG00000106617 | PRKAG2 | chr7 | - | 151786469 | 151786541 | 151781151 | 151781431 | 151876506 | 151876848 | 31 | 41 | 34 |
| SE | 95139 | ENSG00000165282 | PIGO | chr9 | - | 35093420 | 35093580 | 35093209 | 35093290 | 35093900 | 35094024 | 46 | 37 | 43 |
| SE | 40284 | ENSG00000271383 | NBPF19 | chr1 | + | 149477944 | 149478047 | 149475759 | 149476107 | 149480141 | 149480214 | 67 | 118 | 90 |
| SE | 43249 | ENSG00000181666 | ZNF875 | chr19 | + | 37359425 | 37359546 | 37347776 | 37347872 | 37362108 | 37362146 | 0 | 0 | 0 |
| SE | 12938 | ENSG00000167397 | VKORC1 | chr16 | - | 31092782 | 31092864 | 31090841 | 31091342 | 31093311 | 31093421 | 3 | 0 | 1 |
| SE | 28280 | ENSG00000138443 | ABI2 | chr2 | + | 203395658 | 203395780 | 203394699 | 203394846 | 203402575 | 203402734 | 26 | 12 | 25 |
| SE | 60778 | ENSG00000187741 | FANCA | chr16 | - | 89778340 | 89778462 | 89775741 | 89775815 | 89778800 | 89778850 | 0 | 0 | 0 |
| SE | 82465 | ENSG00000119471 | HSDL2 | chr9 | + | 112416840 | 112416944 | 112405623 | 112405722 | 112418859 | 112418958 | 41 | 18 | 20 |
| SE | 69582 | ENSG00000134262 | AP4B1 | chr1 | - | 113902637 | 113902734 | 113901754 | 113901885 | 113904604 | 113904844 | 22 | 8 | 17 |
| SE | 72923 | ENSG00000182732 | RGS6 | chr14 | + | 72495151 | 72495262 | 72478267 | 72478329 | 72510153 | 72510279 | 18 | 41 | 9 |
| SE | 57196 | ENSG00000084112 | SSH1 | chr12 | - | 108792285 | 108792829 | 108778190 | 108789244 | 108798999 | 108799200 | 104 | 58 | 64 |
| SE | 42955 | ENSG00000121940 | CLCC1 | chr1 | - | 108938165 | 108938308 | 108937076 | 108937418 | 108939635 | 108939782 | 6 | 8 | 12 |
| SE | 22550 | ENSG00000167881 | SRP68 | chr17 | - | 76043828 | 76043958 | 76040902 | 76040978 | 76045291 | 76045386 | 239 | 185 | 255 |
| SE | 21238 | ENSG00000055208 | TAB2 | chr6 | + | 149342684 | 149342833 | 149317711 | 149318015 | 149369908 | 149370075 | 12 | 20 | 39 |
| SE | 51813 | ENSG00000181135 | ZNF707 | chr8 | + | 143692244 | 143692330 | 143691599 | 143691713 | 143693670 | 143693703 | 5 | 0 | 0 |
| SE | 109176 | ENSG00000197442 | MAP3K5 | chr6 | - | 136720449 | 136720589 | 136705109 | 136705133 | 136791709 | 136792477 | 252 | 141 | 222 |
| SE | 92843 | ENSG00000130958 | SLC35D2 | chr9 | - | 96345298 | 96345401 | 96343903 | 96343996 | 96360153 | 96360192 | 115 | 74 | 85 |
| SE | 36380 | ENSG00000155621 | C9orf85 | chr9 | + | 71971504 | 71971618 | 71947005 | 71947112 | 71982656 | 71983250 | 23 | 17 | 18 |
| SE | 99884 | ENSG00000167323 | STIM1 | chr11 | + | 4084672 | 4084765 | 4083262 | 4083498 | 4086476 | 4086543 | 0 | 1 | 5 |
| SE | 38589 | ENSG00000132879 | FBXO44 | chr1 | + | 11658532 | 11658628 | 11658266 | 11658393 | 11658735 | 11658871 | 32 | 13 | 45 |
| SE | 16718 | ENSG00000214013 | GANC | chr15 | + | 42278481 | 42278590 | 42276271 | 42276410 | 42287690 | 42287722 | 120 | 70 | 45 |
| SE | 61042 | ENSG00000066027 | PPP2R5A | chr1 | + | 212348388 | 212348497 | 212347346 | 212347406 | 212356625 | 212356676 | 187 | 98 | 123 |
| SE | 19983 | ENSG00000112584 | FAM120B | chr6 | + | 170358225 | 170358318 | 170348150 | 170348323 | 170388286 | 170388493 | 73 | 40 | 72 |
| SE | 5592 | ENSG00000126001 | CEP250 | chr20 | + | 35497718 | 35498067 | 35494523 | 35494657 | 35498594 | 35498716 | 15 | 3 | 23 |
| SE | 44554 | ENSG00000183020 | AP2A2 | chr11 | + | 972061 | 972255 | 959436 | 959505 | 977094 | 977120 | 37 | 21 | 39 |
| SE | 50609 | ENSG00000133275 | CSNK1G2 | chr19 | + | 1978304 | 1978345 | 1969507 | 1969959 | 1978441 | 1978511 | 221 | 117 | 203 |
| SE | 67809 | ENSG00000055044 | NOP58 | chr2 | + | 202287659 | 202287724 | 202284344 | 202284481 | 202290322 | 202290457 | 227 | 140 | 275 |
| SE | 17082 | ENSG00000206190 | ATP10A | chr15 | - | 25691714 | 25691791 | 25687702 | 25687828 | 25701915 | 25702100 | 30 | 19 | 32 |
| SE | 90774 | ENSG00000274536 | MIR223HG | chrX | + | 66017064 | 66017189 | 66015460 | 66015551 | 66018823 | 66020422 | 33 | 16 | 3 |
| SE | 46251 | ENSG00000135148 | TRAFD1 | chr12 | + | 112130510 | 112130569 | 112125537 | 112125618 | 112134737 | 112134873 | 286 | 157 | 221 |
| SE | 38959 | ENSG00000101457 | DNTTIP1 | chr20 | + | 45795344 | 45795443 | 45793920 | 45794017 | 45801073 | 45801142 | 121 | 90 | 112 |
| SE | 83310 | ENSG00000182487 | NCF1B | chr7 | + | 73231810 | 73231890 | 73229566 | 73230260 | 73234613 | 73234718 | 97 | 52 | 78 |
| SE | 39518 | ENSG00000118260 | CREB1 | chr2 | + | 207577109 | 207577224 | 207576630 | 207576693 | 207577504 | 207577655 | 9 | 25 | 7 |
| SE | 82327 | ENSG00000198742 | SMURF1 | chr7 | - | 99047683 | 99047882 | 99045697 | 99045801 | 99049562 | 99049709 | 69 | 55 | 87 |
| SE | 54446 | ENSG00000164305 | CASP3 | chr4 | - | 184631036 | 184631157 | 184627695 | 184629501 | 184631764 | 184631940 | 185 | 130 | 141 |
| SE | 102161 | ENSG00000163249 | CCNYL1 | chr2 | + | 207747046 | 207747213 | 207742222 | 207742342 | 207750956 | 207751114 | 96 | 71 | 50 |
| SE | 62165 | ENSG00000117625 | RCOR3 | chr1 | + | 211304082 | 211304140 | 211295675 | 211295753 | 211313423 | 211314591 | 108 | 92 | 106 |
| SE | 92643 | ENSG00000156990 | RPUSD3 | chr3 | - | 9838984 | 9839171 | 9837848 | 9838207 | 9840183 | 9840307 | 60 | 31 | 64 |
| SE | 1349 | ENSG00000100150 | DEPDC5 | chr22 | + | 31764974 | 31765060 | 31760643 | 31760702 | 31766584 | 31766668 | 122 | 53 | 96 |
| SE | 67956 | ENSG00000123374 | CDK2 | chr12 | + | 55969474 | 55969576 | 55968777 | 55968948 | 55971043 | 55971087 | 20 | 14 | 21 |
| SE | 3465 | ENSG00000105197 | TIMM50 | chr19 | + | 39488538 | 39488645 | 39488005 | 39488217 | 39489718 | 39489768 | 77 | 25 | 69 |
| SE | 11948 | ENSG00000005844 | ITGAL | chr16 | + | 30489253 | 30492069 | 30489081 | 30489155 | 30494211 | 30494363 | 1167 | 643 | 670 |
| SE | 82981 | ENSG00000186088 | GSAP | chr7 | - | 77353571 | 77353641 | 77352943 | 77353026 | 77355212 | 77355430 | 310 | 198 | 203 |
| SE | 106900 | ENSG00000205885 | C1RL-AS1 | chr12 | + | 7118898 | 7119026 | 7115129 | 7115219 | 7119499 | 7119553 | 16 | 5 | 6 |
| SE | 19341 | ENSG00000167705 | RILP | chr17 | - | 1648349 | 1648495 | 1647834 | 1647957 | 1648798 | 1649032 | 17 | 16 | 16 |
| SE | 10609 | ENSG00000276234 | TADA2A | chr17 | + | 37444695 | 37444768 | 37442563 | 37442652 | 37458523 | 37458587 | 74 | 44 | 69 |
| SE | 80128 | ENSG00000122515 | ZMIZ2 | chr7 | + | 44757042 | 44757149 | 44756424 | 44756539 | 44757377 | 44757561 | 26 | 13 | 8 |
| SE | 7672 | ENSG00000258315 | C17orf49 | chr17 | + | 7015773 | 7015875 | 7015039 | 7015156 | 7016494 | 7016685 | 237 | 112 | 180 |
| SE | 99664 | ENSG00000110075 | PPP6R3 | chr11 | + | 68544837 | 68545024 | 68519500 | 68519651 | 68551120 | 68551127 | 24 | 12 | 20 |
| SE | 95112 | ENSG00000096872 | IFT74 | chr9 | + | 27048147 | 27048274 | 27047273 | 27047371 | 27055608 | 27055772 | 75 | 65 | 59 |
| SE | 92787 | ENSG00000134333 | LDHA | chr11 | + | 18399430 | 18399548 | 18396553 | 18396968 | 18400836 | 18400872 | 1837 | 1257 | 1635 |
| SE | 6151 | ENSG00000282826 | FRG1CP | chr20 | - | 28591393 | 28591519 | 28590556 | 28590614 | 28600391 | 28600461 | 105 | 189 | 162 |
| SE | 42081 | ENSG00000026751 | SLAMF7 | chr1 | + | 160748193 | 160748514 | 160739056 | 160739356 | 160752185 | 160752248 | 183 | 71 | 137 |
| SE | 56534 | ENSG00000103489 | XYLT1 | chr16 | - | 17198211 | 17198414 | 17158828 | 17158909 | 17200481 | 17200654 | 53 | 44 | 49 |
| SE | 47943 | ENSG00000169398 | PTK2 | chr8 | - | 140735250 | 140735455 | 140717597 | 140717709 | 140739017 | 140739107 | 63 | 100 | 30 |
| SE | 21662 | ENSG00000130449 | ZSWIM6 | chr5 | + | 61494259 | 61494410 | 61472680 | 61473037 | 61521262 | 61521442 | 74 | 77 | 71 |
| SE | 30118 | ENSG00000154144 | TBRG1 | chr11 | + | 124624930 | 124625001 | 124622835 | 124623233 | 124626903 | 124627050 | 15 | 11 | 27 |
| SE | 102824 | ENSG00000148700 | ADD3 | chr10 | + | 110079239 | 110079297 | 110077004 | 110077118 | 110100624 | 110100704 | 120 | 96 | 118 |
| SE | 93228 | ENSG00000110455 | ACCS | chr11 | + | 44073446 | 44073517 | 44067627 | 44067915 | 44074611 | 44074681 | 7 | 3 | 6 |
| SE | 71571 | ENSG00000166197 | NOLC1 | chr10 | + | 102158048 | 102158214 | 102157018 | 102157074 | 102159192 | 102159308 | 59 | 47 | 65 |
| SE | 89476 | ENSG00000064313 | TAF2 | chr8 | - | 119761607 | 119761772 | 119760454 | 119760738 | 119762414 | 119762534 | 1 | 3 | 6 |
| SE | 44002 | ENSG00000114098 | ARMC8 | chr3 | + | 138209816 | 138209893 | 138187247 | 138187599 | 138221925 | 138221997 | 76 | 37 | 62 |
| SE | 86284 | ENSG00000134516 | DOCK2 | chr5 | + | 170081841 | 170081984 | 170078974 | 170079146 | 170082795 | 170082934 | 196 | 89 | 159 |
| SE | 95510 | ENSG00000137145 | DENND4C | chr9 | + | 19305351 | 19305527 | 19300186 | 19300331 | 19316416 | 19316517 | 478 | 311 | 329 |
| SE | 59634 | ENSG00000163466 | ARPC2 | chr2 | + | 218217371 | 218217544 | 218217140 | 218217254 | 218225919 | 218225954 | 485 | 277 | 346 |
| SE | 23360 | ENSG00000147044 | CASK | chrX | - | 41542690 | 41542806 | 41534892 | 41534973 | 41553718 | 41553915 | 70 | 54 | 94 |
| SE | 84586 | ENSG00000078589 | P2RY10 | chrX | + | 78947814 | 78947863 | 78945390 | 78945495 | 78960507 | 78963727 | 37 | 35 | 30 |
| SE | 79154 | ENSG00000145191 | EIF2B5 | chr3 | + | 184137619 | 184138075 | 184136611 | 184136736 | 184138165 | 184138246 | 168 | 68 | 177 |
| SE | 28067 | ENSG00000121350 | PYROXD1 | chr12 | + | 21456864 | 21457086 | 21455994 | 21456095 | 21461024 | 21461061 | 13 | 17 | 36 |
| SE | 10948 | ENSG00000091592 | NLRP1 | chr17 | - | 5521523 | 5521786 | 5515472 | 5515517 | 5530480 | 5530704 | 93 | 41 | 105 |
| SE | 6453 | ENSG00000134758 | RNF138 | chr18 | + | 32123517 | 32123574 | 32113744 | 32113860 | 32124733 | 32124845 | 257 | 213 | 263 |
| SE | 86146 | ENSG00000017483 | SLC38A5 | chrX | - | 48462232 | 48462291 | 48462006 | 48462144 | 48462897 | 48462980 | 34 | 11 | 27 |
| SE | 17469 | ENSG00000159110 | IFNAR2 | chr21 | + | 33260596 | 33260727 | 33252661 | 33252830 | 33262560 | 33262756 | 281 | 186 | 216 |
| SE | 1428 | ENSG00000100201 | DDX17 | chr22 | - | 38489227 | 38489458 | 38488025 | 38488115 | 38492055 | 38492115 | 23 | 30 | 37 |
| SE | 72198 | ENSG00000145425 | RPS3A | chr4 | + | 151100974 | 151101085 | 151100484 | 151100588 | 151102870 | 151103079 | 5807 | 3435 | 6067 |
| SE | 55827 | ENSG00000143641 | GALNT2 | chr1 | + | 230265240 | 230265367 | 230262921 | 230263005 | 230274444 | 230274564 | 395 | 209 | 224 |
| SE | 27075 | ENSG00000002549 | LAP3 | chr4 | + | 17595409 | 17595534 | 17588818 | 17588977 | 17598455 | 17598558 | 107 | 153 | 116 |
| SE | 78408 | ENSG00000127980 | PEX1 | chr7 | - | 92518994 | 92519078 | 92518140 | 92518255 | 92522101 | 92522245 | 90 | 61 | 114 |
| SE | 20323 | ENSG00000111913 | RIPOR2 | chr6 | - | 24870865 | 24870889 | 24869093 | 24869147 | 24872880 | 24872959 | 1534 | 1171 | 1679 |
| SE | 58592 | ENSG00000186854 | TRABD2A | chr2 | - | 84839148 | 84839323 | 84835918 | 84838241 | 84870224 | 84870785 | 26 | 33 | 94 |
| SE | 9216 | ENSG00000184381 | PLA2G6 | chr22 | - | 38114433 | 38114637 | 38113486 | 38113654 | 38115526 | 38115681 | 3 | 0 | 0 |
| SE | 96222 | ENSG00000110768 | GTF2H1 | chr11 | + | 18360614 | 18360707 | 18358524 | 18358640 | 18365782 | 18365898 | 161 | 140 | 184 |
| SE | 38393 | ENSG00000143458 | GABPB2 | chr1 | + | 151073038 | 151073157 | 151070577 | 151070934 | 151088189 | 151088297 | 19 | 7 | 8 |
| SE | 68017 | ENSG00000172890 | NADSYN1 | chr11 | + | 71458427 | 71458556 | 71455109 | 71455170 | 71463431 | 71463485 | 74 | 46 | 62 |
| SE | 79807 | ENSG00000136286 | MYO1G | chr7 | - | 44966072 | 44966280 | 44965636 | 44965860 | 44966671 | 44966838 | 662 | 280 | 391 |
| SE | 93690 | ENSG00000198780 | FAM169A | chr5 | - | 74838964 | 74839050 | 74834425 | 74834597 | 74840073 | 74840173 | 164 | 39 | 40 |
| SE | 44192 | ENSG00000113583 | C5orf15 | chr5 | - | 133967395 | 133967459 | 133959777 | 133960020 | 133968445 | 133968625 | 0 | 0 | 2 |
| SE | 31430 | ENSG00000155508 | CNOT8 | chr5 | + | 154865191 | 154865385 | 154858538 | 154858768 | 154870660 | 154870704 | 283 | 224 | 264 |
| SE | 64373 | ENSG00000264538 | SUZ12P1 | chr17 | + | 30735032 | 30735097 | 30734062 | 30734943 | 30743300 | 30743369 | 48 | 52 | 69 |
| SE | 24101 | ENSG00000143889 | HNRNPLL | chr2 | - | 38583845 | 38583926 | 38581912 | 38581985 | 38585643 | 38585881 | 93 | 44 | 59 |
| SE | 90655 | ENSG00000100023 | PPIL2 | chr22 | + | 21685637 | 21685726 | 21684752 | 21684913 | 21686482 | 21686558 | 18 | 7 | 14 |
| SE | 102122 | ENSG00000077420 | APBB1IP | chr10 | + | 26560104 | 26560203 | 26541581 | 26541692 | 26560729 | 26560844 | 533 | 335 | 425 |
| SE | 8330 | ENSG00000159259 | CHAF1B | chr21 | + | 36411462 | 36411604 | 36409373 | 36409465 | 36412883 | 36413315 | 7 | 13 | 20 |
| SE | 62688 | ENSG00000115525 | ST3GAL5 | chr2 | - | 85852859 | 85853099 | 85851263 | 85851703 | 85861180 | 85861292 | 11 | 4 | 10 |
| SE | 7533 | ENSG00000169718 | DUS1L | chr17 | - | 82058340 | 82058408 | 82057881 | 82058254 | 82058780 | 82058818 | 84 | 61 | 112 |
| SE | 100067 | ENSG00000171204 | TMEM126B | chr11 | + | 85631686 | 85631808 | 85628592 | 85628688 | 85634085 | 85634176 | 117 | 43 | 81 |
| SE | 82228 | ENSG00000123066 | MED13L | chr12 | - | 116230432 | 116230532 | 116111427 | 116111512 | 116237467 | 116237569 | 5 | 1 | 1 |
| SE | 79985 | ENSG00000118058 | KMT2A | chr11 | + | 118507507 | 118507609 | 118506263 | 118506646 | 118509135 | 118509200 | 40 | 14 | 34 |
| SE | 28477 | ENSG00000104972 | LILRB1 | chr19 | + | 54633637 | 54633688 | 54633015 | 54633318 | 54633973 | 54634021 | 76 | 27 | 23 |
| SE | 25235 | ENSG00000005100 | DHX33 | chr17 | - | 5463528 | 5463689 | 5462318 | 5462546 | 5468570 | 5468982 | 46 | 28 | 29 |
| SE | 57239 | ENSG00000143815 | LBR | chr1 | - | 225411336 | 225411440 | 225410290 | 225410416 | 225412453 | 225412645 | 750 | 372 | 720 |
| SE | 19242 | ENSG00000140943 | MBTPS1 | chr16 | - | 84074596 | 84074741 | 84069865 | 84070038 | 84081746 | 84081908 | 83 | 35 | 66 |
| SE | 57916 | ENSG00000135899 | SP110 | chr2 | - | 230216780 | 230216928 | 230214949 | 230215118 | 230221704 | 230221784 | 330 | 312 | 343 |
| SE | 86371 | ENSG00000035681 | NSMAF | chr8 | - | 58637233 | 58637389 | 58635467 | 58635546 | 58642983 | 58643073 | 16 | 5 | 5 |
| SE | 85072 | ENSG00000147459 | DOCK5 | chr8 | + | 25388808 | 25388941 | 25382673 | 25382778 | 25389090 | 25389232 | 4 | 0 | 2 |

| | | | | | | | | | | | | | | | | | | | | | | | | | | | | | | | | | | | | | | | | | | | | | | |
|---|---|---|---|---|---|---|---|---|---|---|---|---|---|---|---|---|---|---|---|---|---|---|---|---|---|---|---|---|---|---|---|---|---|---|---|---|---|---|---|---|---|---|---|---|---|---|
| SE | 22597 | ENSG00000127603 | MACF1 | chr1 | + | 39331202 | 39336653 | 39328600 | 39328706 | 39337181 | 39337331 | 288 | 164 | 225 | 260 | 208 | 3 | 0 | 2 | 9 | 10 | 28 | 334 | 245 | 36 | 33 | 0 | 1 | 0 | 3 | 12 | 298 | 149 | 0.797302873 | 1 | 0.98 | 1 | 0.983 | 0.935 | 0.912 | 1 | 0.994 | 1 | 0.857 | 0.579 | 0.076 |
| SE | 36213 | ENSG00000118855 | MFSD1 | chr3 | + | 158813964 | 158814067 | 158809001 | 158809287 | 158819648 | 158819706 | 691 | 574 | 311 | 764 | 1173 | 5 | 2 | 2 | 0 | 2 | 728 | 410 | 472 | 612 | 1617 | 13 | 0 | 2 | 0 | 11 | 252 | 149 | 0.797339317 | 1 | 0.988 | 0.994 | 0.989 | 1 | 0.997 | 0.971 | 1 | 0.993 | 1 | 0.989 | 0.003 |
| SE | 96232 | ENSG00000133943 | DGLUCY | chr14 | + | 91147780 | 91147858 | 91114062 | 91114283 | 91157638 | 91157690 | 20 | 29 | 19 | 21 | 28 | 41 | 25 | 38 | 33 | 51 | 31 | 29 | 11 | 36 | 63 | 52 | 42 | 31 | 101 | 57 | 227 | 149 | 0.797353829 | 1 | 0.243 | 0.432 | 0.247 | 0.295 | 0.265 | 0.281 | 0.312 | 0.189 | 0.19 | 0.42 | 0.018 |
| SE | 10548 | ENSG00000177302 | TOP3A | chr17 | - | 18292644 | 18292852 | 18285406 | 18285520 | 18294702 | 18294785 | 89 | 74 | 70 | 84 | 206 | 0 | 1 | 1 | 0 | 0 | 273 | 107 | 92 | 119 | 239 | 0 | 1 | 0 | 0 | 9 | 298 | 149 | 0.797446518 | 1 | 1 | 0.974 | 0.972 | 1 | 1 | 1 | 0.982 | 1 | 1 | 0.93 | 0.007 |
| SE | 57187 | ENSG00000197323 | TRIM33 | chr1 | - | 114430797 | 114430912 | 114427747 | 114427894 | 114433616 | 114433733 | 311 | 326 | 247 | 342 | 501 | 6 | 14 | 3 | 12 | 24 | 47 | 239 | 260 | 89 | 78 | 0 | 3 | 18 | 5 | 3 | 264 | 149 | 0.797497 | 1 | 0.967 | 0.929 | 0.979 | 0.941 | 0.922 | 1 | 0.978 | 0.891 | 0.909 | 0.936 | 0.005 |
| SE | 20043 | ENSG00000079482 | OPHN1 | chrX | - | 68119247 | 68119332 | 68113180 | 68113239 | 68192918 | 68192993 | 17 | 17 | 3 | 26 | 26 | 1 | 1 | 0 | 0 | 3 | 138 | 25 | 26 | 32 | 100 | 0 | 0 | 2 | 3 | 0 | 234 | 149 | 0.797586172 | 1 | 0.915 | 0.915 | 1 | 1 | 0.847 | 1 | 1 | 0.892 | 0.872 | 1 | -0.017 |
| SE | 30637 | ENSG00000189308 | LIN54 | chr4 | - | 82978882 | 82979006 | 82970326 | 82970469 | 82984823 | 82984876 | 60 | 65 | 66 | 60 | 98 | 0 | 0 | 0 | 0 | 0 | 101 | 99 | 165 | 86 | 75 | 0 | 0 | 0 | 6 | 0 | 273 | 149 | 0.797668616 | 1 | 1 | 1 | 1 | 1 | 1 | 1 | 1 | 1 | 0.887 | 1 | 0.023 |
| SE | 8550 | ENSG00000067191 | CACNB1 | chr17 | - | 39184783 | 39184864 | 39184324 | 39184383 | 39185130 | 39185150 | 25 | 5 | 5 | 11 | 13 | 0 | 0 | 0 | 0 | 0 | 13 | 21 | 22 | 8 | 17 | 0 | 4 | 0 | 0 | 0 | 230 | 149 | 0.797681463 | 1 | 1 | 1 | 1 | 1 | 1 | 1 | 0.773 | 1 | 1 | 1 | 0.045 |
| SE | 35663 | ENSG00000010404 | IDS | chrX | - | 149499228 | 149499354 | 149498106 | 149498307 | 149500948 | 149501037 | 5 | 1 | 2 | 13 | 14 | 594 | 371 | 403 | 500 | 816 | 8 | 19 | 16 | 6 | 7 | 642 | 553 | 539 | 558 | 712 | 275 | 149 | 0.797695818 | 1 | 0.005 | 0.001 | 0.003 | 0.014 | 0.009 | 0.007 | 0.018 | 0.016 | 0.006 | 0.005 | -0.004 |
| SE | 51249 | ENSG00000164022 | AIMP1 | chr4 | + | 106320797 | 106320889 | 106316551 | 106316594 | 106324984 | 106325118 | 1 | 2 | 0 | 1 | 4 | 34 | 22 | 29 | 28 | 22 | 0 | 0 | 0 | 0 | 0 | 18 | 40 | 32 | 14 | 13 | 241 | 149 | 0.797706039 | 1 | 0.018 | 0.053 | 0 | 0.022 | 0.101 | 0 | 0 | 0 | 0 | 0 | 0.039 |
| SE | 51375 | ENSG00000066427 | ATXN3 | chr14 | - | 92093745 | 92093850 | 92093251 | 92093318 | 92096092 | 92096137 | 99 | 47 | 55 | 76 | 72 | 33 | 27 | 26 | 40 | 59 | 22 | 59 | 39 | 39 | 12 | 11 | 37 | 53 | 44 | 0 | 254 | 149 | 0.797763909 | 1 | 0.638 | 0.505 | 0.554 | 0.527 | 0.417 | 0.54 | 0.483 | 0.302 | 0.342 | 1 | -0.005 |
| SE | 109304 | ENSG00000139597 | N4BP2L1 | chr13 | - | 32403655 | 32403761 | 32400752 | 32403200 | 32404202 | 32404271 | 0 | 1 | 6 | 2 | 0 | 39 | 32 | 33 | 48 | 33 | 0 | 0 | 0 | 0 | 0 | 34 | 26 | 25 | 57 | 24 | 255 | 149 | 0.797774306 | 1 | 0 | 0.018 | 0.096 | 0.024 | 0 | 0 | 0 | 0 | 0 | 0 | 0.028 |
| [illegible] | | | | | | | | | | | | | | | | | | | | | | | | | | | | | | | | | | | | | | | | | | | | | | |

| | | | | | | | | | | | | | | | | | | | | | | | | | | | | | | | | | | | | | | | | | | | | | | |
|---|---|---|---|---|---|---|---|---|---|---|---|---|---|---|---|---|---|---|---|---|---|---|---|---|---|---|---|---|---|---|---|---|---|---|---|---|---|---|---|---|---|---|---|---|---|---|
| SE | 15166 | ENSG00000137842 | TMEM62 | chr15 | + | 43151789 | 43151945 | 43149028 | 43149151 | 43154671 | 43154831 | 87 | 41 | 40 | 70 | 122 | 0 | 0 | 0 | 0 | 0 | 65 | 75 | 64 | 60 | 87 | 0 | 0 | 0 | 0 | 7 | 298 | 149 | 0.809728888 | 1 | 1 | 1 | 1 | 1 | 1 | 1 | 1 | 1 | 1 | 0.861 | 0.028 |
| SE | 103008 | ENSG00000213047 | DENND1B | chr1 | - | 197583151 | 197583253 | 197552254 | 197553112 | 197607072 | 197607174 | 69 | 43 | 34 | 66 | 58 | 0 | 0 | 0 | 0 | 0 | 36 | 52 | 42 | 9 | 19 | 0 | 6 | 0 | 0 | 0 | 251 | 149 | 0.809730264 | 1 | 1 | 1 | 1 | 1 | 1 | 1 | 0.837 | 1 | 1 | 1 | 0.033 |
| SE | 96205 | ENSG00000117751 | PPP1R8 | chr1 | + | 27838698 | 27838852 | 27832755 | 27832816 | 27843185 | 27843330 | 117 | 62 | 78 | 79 | 62 | 2 | 0 | 0 | 0 | 4 | 101 | 46 | 60 | 71 | 40 | 0 | 0 | 0 | 0 | 0 | 298 | 149 | 0.809748824 | 1 | 0.967 | 1 | 1 | 1 | 0.886 | 1 | 1 | 1 | 1 | 1 | -0.029 |
| SE | 19129 | ENSG00000183576 | SETD3 | chr14 | - | 99461191 | 99461340 | 99459112 | 99459185 | 99463485 | 99463578 | 1228 | 671 | 850 | 1038 | 1660 | 15 | 6 | 7 | 4 | 2 | 1363 | 743 | 514 | 780 | 1114 | 11 | 10 | 9 | 1 | 0 | 298 | 149 | 0.80988361 | 1 | 0.976 | 0.982 | 0.984 | 0.992 | 0.998 | 0.984 | 0.974 | 0.966 | 0.997 | 1 | 0.002 |
| SE | 65494 | ENSG00000121989 | ACVR2A | chr2 | + | 147898240 | 147898330 | 147896300 | 147896508 | 147899457 | 147899567 | 2 | 2 | 4 | 6 | 6 | 228 | 205 | 144 | 325 | 543 | 4 | 0 | 8 | 4 | 20 | 632 | 191 | 100 | 263 | 981 | 239 | 149 | 0.809945323 | 1 | 0.005 | 0.006 | 0.017 | 0.011 | 0.007 | 0.004 | 0 | 0.048 | 0.009 | 0.013 | -0.006 |
| SE | 38897 | ENSG00000151466 | SCLT1 | chr4 | - | 128936654 | 128936851 | 128891058 | 128891137 | 128946006 | 128946152 | 46 | 70 | 29 | 69 | 107 | 0 | 0 | 4 | 0 | 0 | 2 | 24 | 27 | 13 | 53 | 0 | 0 | 0 | 0 | 0 | 298 | 149 | 0.809976464 | 1 | 1 | 1 | 0.784 | 1 | 1 | 1 | 1 | 1 | 1 | 1 | -0.043 |
| SE | 49226 | ENSG00000142864 | SERBP1 | chr1 | - | 67420008 | 67420186 | 67415165 | 67415339 | 67424199 | 67424277 | 1688 | 1319 | 1481 | 1683 | 3179 | 9 | 1 | 4 | 5 | 5 | 1554 | 1451 | 1762 | 1543 | 556 | 0 | 11 | 9 | 12 | 3 | 298 | 149 | 0.809995493 | 1 | 0.989 | 0.998 | 0.995 | 0.994 | 0.997 | 1 | 0.985 | 0.99 | 0.985 | 0.989 | 0.005 |
| SE | 68737 | ENSG00000196295 | GARS1-DT | chr7 | - | 30584710 | 30584808 | 30579230 | 30579314 | 30585329 | 30585459 | 50 | 27 | 43 | 41 | 60 | 33 | 18 | 25 | 45 | 32 | 72 | 54 | 38 | 43 | 82 | 32 | 35 | 19 | 48 | 89 | 247 | 149 | 0.810021417 | 1 | 0.478 | 0.475 | 0.509 | 0.355 | 0.531 | 0.576 | 0.482 | 0.547 | 0.351 | 0.357 | 0.007 |
| SE | 7078 | ENSG00000175387 | SMAD2 | chr18 | - | 47930360 | 47930446 | 47896520 | 47896809 | 47930799 | 47930837 | 75 | 47 | 73 | 75 | 39 | 18 | 2 | 4 | 9 | 1 | 18 | 25 | 38 | 25 | 50 | 0 | 23 | 8 | 0 | 0 | 235 | 149 | 0.810045419 | 1 | 0.725 | 0.937 | 0.92 | 0.841 | 0.961 | 1 | 0.408 | 0.751 | 1 | 1 | 0.045 |
| SE | 74652 | ENSG00000114383 | TUSC2 | chr3 | - | 50327027 | 50327219 | 50326356 | 50326477 | 50327953 | 50328251 | 0 | 0 | 1 | 1 | 0 | 13 | 7 | 11 | 11 | 3 | 2 | 0 | 0 | 0 | 0 | 0 | 8 | 9 | 4 | 28 | 298 | 149 | 0.810086847 | 1 | 0 | 0 | 0.043 | 0.043 | 0 | 1 | 0 | 0 | 0 | 0 | -0.183 |
| SE | 91889 | ENSG00000254995 | STX16-NPEPL1 | chr20 | + | 58691404 | 58691562 | 58691033 | 58691227 | 58691723 | 58691780 | 18 | 31 | 21 | 41 | 35 | 0 | 0 | 0 | 0 | 0 | 29 | 65 | 58 | 36 | 10 | 4 | 0 | 0 | 0 | 0 | 298 | 149 | 0.810126629 | 1 | 1 | 1 | 1 | 1 | 1 | 0.784 | 1 | 1 | 1 | 1 | 0.043 |
| SE | 41315 | ENSG00000137411 | VARS2 | chr6 | + | 30923352 | 30923505 | 30923103 | 30923231 | 30924353 | 30924560 | 38 | 26 | 60 | 29 | 51 | 0 | 0 | 0 | 4 | 0 | 26 | 49 | 14 | 16 | 29 | 0 | 0 | 0 | 0 | 0 | 298 | 149 | 0.810159265 | 1 | 1 | 1 | 1 | 0.784 | 1 | 1 | 1 | 1 | 1 | 1 | -0.043 |
| SE | 54040 | ENSG00000180353 | HCLS1 | chr3 | - | 121647318 | 121647371 | 121644817 | 121644928 | 121657278 | 121657352 | 493 | 462 | 459 | 542 | 799 | 0 | 1 | 1 | 3 | 0 | 1301 | 644 | 594 | 816 | 1248 | 0 | 9 | 3 | 12 | 3 | 202 | 149 | 0.810185253 | 1 | 1 | 0.997 | 0.997 | 0.993 | 1 | 1 | 0.981 | 0.993 | 0.98 | 0.997 | 0.007 |
| SE | 109346 | ENSG00000102710 | SUPT20H | chr13 | - | 37033448 | 37033588 | 37031738 | 37031895 | 37040404 | 37040458 | 289 | 200 | 267 | 302 | 444 | 2 | 2 | 4 | 8 | 4 | 108 | 164 | 149 | 111 | 53 | 0 | 5 | 5 | 0 | 0 | 289 | 149 | 0.810242137 | 1 | 0.987 | 0.981 | 0.972 | 0.951 | 0.983 | 1 | 0.944 | 0.939 | 1 | 1 | -0.002 |
| SE | 53098 | ENSG00000006756 | ARSD | chrX | - | 2917803 | 2918227 | 2910658 | 2910793 | 2920600 | 2920723 | 34 | 17 | 23 | 13 | 29 | 0 | 0 | 0 | 0 | 4 | 23 | 27 | 6 | 50 | 28 | 0 | 0 | 0 | 2 | 0 | 298 | 149 | 0.810246805 | 1 | 1 | 1 | 1 | 1 | 0.784 | 1 | 1 | 1 | 0.926 | 1 | -0.028 |
| SE | 6957 | ENSG00000100983 | GSS | chr20 | - | 34954268 | 34954547 | 34951761 | 34951860 | 34955726 | 34955798 | 5 | 0 | 0 | 0 | 0 | 20 | 2 | 15 | 16 | 11 | 0 | 0 | 0 | 0 | 0 | 29 | 9 | 18 | 11 | 10 | 298 | 149 | 0.810250071 | 1 | 0.111 | 0 | 0 | 0 | 0 | 0 | 0 | 0 | 0 | 0 | 0.022 |
| SE | 105991 | ENSG00000122965 | RBM19 | chr12 | - | 113960058 | 113960178 | 113959864 | 113959903 | 113962231 | 113962414 | 196 | 120 | 202 | 225 | 192 | 0 | 0 | 0 | 2 | 0 | 317 | 275 | 203 | 232 | 366 | 8 | 0 | 4 | 0 | 0 | 269 | 149 | 0.810283686 | 1 | 1 | 1 | 1 | 0.984 | 1 | 0.956 | 1 | 0.966 | 1 | 1 | 0.012 |
| SE | 17650 | ENSG00000100949 | RABGGTA | chr14 | - | 24270836 | 24270998 | 24270333 | 24270458 | 24271112 | 24271169 | 35 | 5 | 39 | 23 | 11 | 0 | 0 | 0 | 0 | 0 | 14 | 83 | 23 | 19 | 44 | 0 | 5 | 0 | 0 | 0 | 298 | 149 | 0.81032645 | 1 | 1 | 1 | 1 | 1 | 1 | 1 | 0.892 | 1 | 1 | 1 | 0.022 |
| SE | 52104 | ENSG00000158006 | PAFAH2 | chr1 | - | 25976681 | 25976773 | 25974479 | 25974650 | 25982363 | 25982477 | 128 | 77 | 74 | 118 | 85 | 1 | 1 | 1 | 1 | 3 | 30 | 96 | 76 | 53 | 38 | 0 | 0 | 8 | 0 | 0 | 241 | 149 | 0.810374635 | 1 | 0.988 | 0.979 | 0.979 | 0.986 | 0.946 | 1 | 1 | 0.855 | 1 | 1 | 0.005 |
| SE | 70497 | ENSG00000170266 | GLB1 | chr3 | - | 33077093 | 33077237 | 33072543 | 33072713 | 33096718 | 33096792 | 4 | 1 | 3 | 11 | 2 | 6 | 10 | 5 | 7 | 9 | 11 | 7 | 2 | 6 | 0 | 10 | 12 | 2 | 10 | 27 | 293 | 149 | 0.81039522 | 1 | 0.253 | 0.048 | 0.234 | 0.444 | 0.102 | 0.359 | 0.229 | 0.337 | 0.234 | 0 | -0.016 |
| SE | 50880 | ENSG00000170017 | ALCAM | chr3 | + | 105533602 | 105533690 | 105532001 | 105532066 | 105534662 | 105534845 | 117 | 103 | 109 | 102 | 264 | 0 | 0 | 0 | 0 | 0 | 48 | 88 | 95 | 23 | 36 | 0 | 0 | 0 | 4 | 0 | 237 | 149 | 0.810412856 | 1 | 1 | 1 | 1 | 1 | 1 | 1 | 1 | 1 | 0.783 | 1 | 0.043 |
| SE | 23950 | ENSG00000133835 | HSD17B4 | chr5 | + | 119495715 | 119495813 | 119492099 | 119493946 | 119496542 | 119496646 | 3 | 0 | 0 | 0 | 4 | 231 | 161 | 141 | 194 | 344 | 0 | 0 | 4 | 11 | 0 | 99 | 166 | 100 | 161 | 55 | 247 | 149 | 0.810430859 | 1 | 0.008 | 0 | 0 | 0 | 0.007 | 0 | 0 | 0.024 | 0.04 | 0 | -0.01 |
| SE | 65444 | ENSG00000166164 | BRD7 | chr16 | - | 50334710 | 50334895 | 50333573 | 50333697 | 50339975 | 50340086 | 218 | 243 | 178 | 284 | 264 | 0 | 0 | 0 | 0 | 0 | 421 | 256 | 445 | 495 | 163 | 14 | 0 | 0 | 0 | 0 | 298 | 149 | 0.810472697 | 1 | 1 | 1 | 1 | 1 | 1 | 0.938 | 1 | 1 | 1 | 1 | 0.012 |
| SE | 91579 | ENSG00000070501 | POLB | chr8 | + | 42369848 | 42369988 | 42362611 | 42362698 | 42371562 | 42371765 | 82 | 68 | 64 | 82 | 75 | 0 | 2 | 1 | 3 | 0 | 58 | 35 | 71 | 40 | 7 | 2 | 0 | 0 | 0 | 0 | 289 | 149 | 0.81055626 | 1 | 1 | 0.946 | 0.971 | 0.934 | 1 | 0.937 | 1 | 1 | 1 | 1 | -0.017 |
| SE | 14736 | ENSG00000185658 | BRWD1 | chr21 | - | 39210785 | 39210929 | 39209994 | 39210147 | 39212665 | 39212707 | 217 | 278 | 275 | 267 | 443 | 12 | 13 | 13 | 15 | 15 | 100 | 82 | 58 | 98 | 115 | 0 | 22 | 14 | 0 | 0 | 293 | 149 | 0.810563783 | 1 | 0.902 | 0.916 | 0.915 | 0.901 | 0.938 | 1 | 0.655 | 0.678 | 1 | 1 | 0.048 |
| SE | 88032 | ENSG00000156171 | DRAM2 | chr1 | - | 111119614 | 111119681 | 111118864 | 111118897 | 111119876 | 111119959 | 3 | 4 | 2 | 5 | 3 | 223 | 108 | 126 | 154 | 265 | 0 | 1 | 4 | 0 | 0 | 92 | 105 | 52 | 74 | 24 | 216 | 149 | 0.810584144 | 1 | 0.009 | 0.025 | 0.011 | 0.022 | 0.008 | 0 | 0.007 | 0.05 | 0 | 0 | 0.004 |
| SE | 53822 | ENSG00000173575 | CHD2 | chr15 | + | 92978147 | 92978383 | 92974878 | 92974950 | 92979134 | 92979213 | 458 | 361 | 493 | 548 | 489 | 0 | 0 | 0 | 0 | 3 | 246 | 637 | 954 | 417 | 126 | 0 | 0 | 5 | 11 | 0 | 298 | 149 | 0.810586988 | 1 | 1 | 1 | 1 | 1 | 0.988 | 1 | 1 | 0.99 | 0.95 | 1 | 0.01 |
| SE | 80369 | ENSG00000204371 | EHMT2 | chr6 | - | 31888968 | 31889070 | 31888598 | 31888747 | 31889227 | 31889342 | 111 | 62 | 83 | 90 | 92 | 39 | 15 | 65 | 41 | 50 | 71 | 135 | 20 | 73 | 83 | 38 | 51 | 10 | 44 | 38 | 251 | 149 | 0.810611599 | 1 | 0.628 | 0.71 | 0.431 | 0.566 | 0.522 | 0.526 | 0.611 | 0.543 | 0.496 | 0.565 | 0.023 |
| SE | 384 | ENSG00000237945 | LINC00649 | chr21 | + | 33945809 | 33945895 | 33931159 | 33931742 | 33969236 | 33969358 | 23 | 7 | 32 | 39 | 54 | 4 | 0 | 0 | 0 | 0 | 10 | 74 | 7 | 9 | 9 | 0 | 0 | 0 | 0 | 0 | 235 | 149 | 0.810634083 | 1 | 0.785 | 1 | 1 | 1 | 1 | 1 | 1 | 1 | 1 | 1 | -0.043 |
| SE | 36690 | ENSG00000158062 | UBXN11 | chr1 | - | 26296918 | 26296995 | 26285822 | 26286037 | 26297426 | 26297481 | 165 | 129 | 88 | 138 | 57 | 0 | 0 | 0 | 0 | 0 | 471 | 161 | 150 | 260 | 438 | 13 | 1 | 0 | 0 | 0 | 226 | 149 | 0.810669531 | 1 | 1 | 1 | 1 | 1 | 1 | 0.96 | 0.991 | 1 | 1 | 1 | 0.01 |
| SE | 2656 | ENSG00000129351 | ILF3 | chr19 | + | 10682536 | 10682704 | 10681218 | 10682192 | 10683136 | 10683262 | 330 | 184 | 363 | 310 | 206 | 0 | 0 | 0 | 0 | 0 | 485 | 299 | 283 | 304 | 468 | 0 | 3 | 2 | 8 | 0 | 298 | 149 | 0.810725076 | 1 | 1 | 1 | 1 | 1 | 1 | 1 | 0.98 | 0.986 | 0.95 | 1 | 0.017 |
| SE | 58142 | ENSG00000117523 | PRRC2C | chr1 | + | 171591151 | 171591200 | 171588378 | 171588505 | 171591586 | 171592223 | 49 | 19 | 28 | 52 | 66 | 138 | 93 | 103 | 138 | 188 | 0 | 28 | 47 | 22 | 0 | 20 | 69 | 37 | 26 | 11 | 198 | 149 | 0.810726143 | 1 | 0.211 | 0.133 | 0.17 | 0.221 | 0.209 | 0 | 0.234 | 0.489 | 0.389 | 0 | -0.034 |
| SE | 96583 | ENSG00000198561 | CTNND1 | chr11 | + | 57789036 | 57789155 | 57761796 | 57762119 | 57791384 | 57791673 | 24 | 23 | 16 | 20 | 22 | 6 | 3 | 0 | 1 | 4 | 18 | 12 | 5 | 19 | 42 | 27 | 0 | 0 | 15 | 0 | 268 | 149 | 0.810732941 | 1 | 0.69 | 0.81 | 1 | 0.917 | 0.754 | 0.27 | 1 | 1 | 0.413 | 1 | 0.098 |
| SE | 91104 | ENSG00000107864 | CPEB3 | chr10 | - | 92143028 | 92143118 | 92111075 | 92111194 | 92180962 | 92181019 | 28 | 24 | 16 | 32 | 18 | 5 | 0 | 1 | 2 | 0 | 4 | 43 | 15 | 18 | 20 | 0 | 4 | 0 | 3 | 0 | 239 | 149 | 0.810910296 | 1 | 0.777 | 1 | 0.909 | 0.909 | 1 | 1 | 0.87 | 1 | 0.789 | 1 | -0.013 |
| SE | 24212 | ENSG00000113742 | CPEB4 | chr5 | + | 173932449 | 173932500 | 173910522 | 173910604 | 173944966 | 173945092 | 42 | 23 | 24 | 49 | 39 | 17 | 13 | 17 | 18 | 37 | 17 | 40 | 36 | 32 | 6 | 5 | 39 | 34 | 9 | 5 | 200 | 149 | 0.810991918 | 1 | 0.648 | 0.569 | 0.513 | 0.67 | 0.44 | 0.717 | 0.433 | 0.441 | 0.726 | 0.472 | 0.01 |
| SE | 53893 | ENSG00000112033 | PPARD | chr6 | + | 35397497 | 35397593 | 35347066 | 35347150 | 35410986 | 35411217 | 3 | 3 | 6 | 1 | 0 | 35 | 33 | 60 | 47 | 37 | 0 | 4 | 2 | 1 | 8 | 47 | 26 | 33 | 78 | 118 | 245 | 149 | 0.811020591 | 1 | 0.05 | 0.052 | 0.057 | 0.013 | 0 | 0 | 0.086 | 0.036 | 0.008 | 0.04 | 0 |
| SE | 16297 | ENSG00000255571 | MIR9-3HG | chr15 | + | 89389344 | 89389482 | 89387430 | 89387881 | 89395027 | 89395040 | 25 | 7 | 14 | 12 | 17 | 1 | 0 | 0 | 0 | 0 | 6 | 28 | 44 | 5 | 2 | 0 | 4 | 0 | 0 | 0 | 287 | 149 | 0.811094482 | 1 | 0.928 | 1 | 1 | 1 | 1 | 1 | 0.784 | 1 | 1 | 1 | 0.029 |
| SE | 46241 | ENSG00000139645 | ANKRD52 | chr12 | - | 56254864 | 56254952 | 56254577 | 56254720 | 56255783 | 56255984 | 26 | 7 | 17 | 22 | 3 | 0 | 0 | 0 | 0 | 0 | 11 | 20 | 6 | 11 | 26 | 0 | 0 | 3 | 0 | 0 | 237 | 149 | 0.81110892 | 1 | 1 | 1 | 1 | 1 | 1 | 1 | 1 | 0.557 | 1 | 1 | 0.089 |
| SE | 36427 | ENSG00000121858 | TNFSF10 | chr3 | - | 172509216 | 172509321 | 172505507 | 172506919 | 172514860 | 172514998 | 101 | 124 | 88 | 100 | 233 | 7 | 6 | 3 | 5 | 2 | 126 | 48 | 66 | 135 | 6 | 1 | 1 | 4 | 10 | 2 | 254 | 149 | 0.811117935 | 1 | 0.894 | 0.924 | 0.945 | 0.921 | 0.986 | 0.987 | 0.966 | 0.906 | 0.888 | 0.638 | 0.057 |
| SE | 102073 | ENSG00000181192 | DHTKD1 | chr10 | + | 12094072 | 12094271 | 12088985 | 12089255 | 12097683 | 12097958 | 41 | 19 | 34 | 40 | 53 | 0 | 0 | 0 | 0 | 6 | 117 | 33 | 21 | 22 | 65 | 0 | 4 | 0 | 0 | 0 | 298 | 149 | 0.811147197 | 1 | 1 | 1 | 1 | 1 | 0.815 | 1 | 0.805 | 1 | 1 | 1 | 0.002 |
| SE | 98724 | ENSG00000107290 | SETX | chr9 | - | 132283263 | 132283413 | 132281466 | 132281574 | 132286422 | 132286494 | 420 | 265 | 324 | 378 | 652 | 0 | 11 | 0 | 0 | 3 | 55 | 235 | 207 | 76 | 15 | 0 | 0 | 0 | 0 | 0 | 298 | 149 | 0.811213987 | 1 | 1 | 0.923 | 1 | 1 | 0.991 | 1 | 1 | 1 | 1 | 1 | -0.017 |
| SE | 100107 | ENSG00000166333 | ILK | chr11 | + | 6608045 | 6608211 | 6603815 | 6604360 | 6608393 | 6608489 | 651 | 447 | 414 | 724 | 503 | 10 | 7 | 10 | 6 | 4 | 1312 | 724 | 499 | 734 | 5819 | 3 | 11 | 4 | 14 | 83 | 298 | 149 | 0.811217264 | 1 | 0.97 | 0.97 | 0.954 | 0.984 | 0.984 | 0.995 | 0.971 | 0.984 | 0.963 | 0.972 | -0.005 |
| SE | 3305 | ENSG00000076928 | ARHGEF1 | chr19 | + | 41897314 | 41897326 | 41896376 | 41896482 | 41898441 | 41898587 | 21 | 23 | 37 | 29 | 16 | 39 | 22 | 42 | 38 | 19 | 47 | 51 | 59 | 34 | 22 | 72 | 34 | 36 | 75 | 76 | 161 | 149 | 0.811238559 | 1 | 0.333 | 0.492 | 0.449 | 0.414 | 0.438 | 0.377 | 0.581 | 0.603 | 0.296 | 0.211 | 0.012 |
| SE | 1159 | ENSG00000100395 | L3MBTL2 | chr22 | + | 41213892 | 41214026 | 41209695 | 41209933 | 41219418 | 41219536 | 69 | 33 | 68 | 81 | 46 | 0 | 0 | 0 | 0 | 0 | 14 | 26 | 28 | 39 | 88 | 0 | 0 | 4 | 0 | 0 | 283 | 149 | 0.811276056 | 1 | 1 | 1 | 1 | 1 | 1 | 1 | 1 | 0.787 | 1 | 1 | 0.043 |
| SE | 69956 | ENSG00000108883 | EFTUD2 | chr17 | - | 44879555 | 44879638 | 44875933 | 44876100 | 44880553 | 44880644 | 269 | 161 | 189 | 237 | 244 | 1 | 2 | 0 | 0 | 1 | 359 | 302 | 132 | 184 | 441 | 0 | 0 | 8 | 3 | 3 | 232 | 149 | 0.811361998 | 1 | 0.994 | 0.981 | 1 | 1 | 0.994 | 1 | 1 | 0.914 | 0.975 | 0.99 | 0.018 |
| SE | 106485 | ENSG00000176915 | ANKLE2 | chr12 | - | 132733995 | 132734071 | 132730232 | 132730270 | 132734384 | 132734575 | 6 | 10 | 8 | 17 | 10 | 77 | 68 | 81 | 91 | 79 | 15 | 36 | 7 | 4 | 5 | 146 | 77 | 77 | 90 | 73 | 225 | 149 | 0.811367988 | 1 | 0.049 | 0.089 | 0.061 | 0.11 | 0.077 | 0.064 | 0.236 | 0.057 | 0.029 | 0.043 | -0.009 |
| SE | 52737 | ENSG00000143970 | ASXL2 | chr2 | - | 25786894 | 25786960 | 25771439 | 25771540 | 25799384 | 25799535 | 1 | 0 | 2 | 1 | 0 | 39 | 29 | 25 | 30 | 36 | 0 | 5 | 0 | 1 | 0 | 28 | 28 | 30 | 22 | 30 | 215 | 149 | 0.811457262 | 1 | 0.017 | 0 | 0.053 | 0.023 | 0 | 0 | 0.11 | 0 | 0.031 | 0 | -0.01 |
| SE | 3154 | ENSG00000213020 | ZNF611 | chr19 | - | 52707420 | 52707542 | 52706653 | 52706864 | 52714014 | 52714141 | 19 | 25 | 27 | 29 | 15 | 19 | 23 | 19 | 39 | 16 | 18 | 17 | 4 | 4 | 0 | 0 | 19 | 13 | 6 | 2 | 271 | 149 | 0.811507012 | 1 | 0.355 | 0.374 | 0.439 | 0.29 | 0.34 | 1 | 0.33 | 0.145 | 0.268 | 0 | 0.011 |
| SE | 95479 | ENSG00000095319 | NUP188 | chr9 | + | 128999171 | 128999317 | 128998537 | 128998623 | 128999623 | 128999805 | 152 | 90 | 144 | 122 | 131 | 0 | 0 | 0 | 6 | 3 | 192 | 131 | 61 | 132 | 181 | 0 | 0 | 4 | 6 | 0 | 295 | 149 | 0.811513055 | 1 | 1 | 1 | 1 | 0.911 | 0.957 | 1 | 1 | 0.885 | 0.917 | 1 | 0.013 |
| SE | 20702 | ENSG00000115760 | BIRC6 | chr2 | + | 32436052 | 32436184 | 32433643 | 32433804 | 32439507 | 32439686 | 532 | 502 | 379 | 559 | 616 | 1 | 2 | 0 | 5 | 0 | 71 | 602 | 613 | 304 | 63 | 0 | 4 | 3 | 6 | 0 | 281 | 149 | 0.811562231 | 1 | 0.996 | 0.993 | 1 | 0.983 | 1 | 1 | 0.988 | 0.991 | 0.964 | 1 | 0.006 |
| SE | 64585 | ENSG00000135624 | CCT7 | chr2 | + | 73244543 | 73244715 | 73242972 | 73243129 | 73247761 | 73247789 | 419 | 213 | 400 | 402 | 478 | 0 | 0 | 0 | 0 | 0 | 432 | 364 | 195 | 330 | 356 | 0 | 0 | 4 | 8 | 0 | 298 | 149 | 0.811611919 | 1 | 1 | 1 | 1 | 1 | 1 | 1 | 1 | 0.961 | 0.954 | 1 | 0.017 |
| SE | 60618 | ENSG00000111196 | MAGOHB | chr12 | - | 10609097 | 10609383 | 10607853 | 10607936 | 10609830 | 10609941 | 34 | 60 | 35 | 66 | 87 | 54 | 57 | 72 | 67 | 88 | 61 | 52 | 17 | 6 | 7 | 5 | 50 | 31 | 41 | 10 | 298 | 149 | 0.811739704 | 1 | 0.239 | 0.345 | 0.196 | 0.33 | 0.331 | 0.859 | 0.342 | 0.215 | 0.068 | 0.259 | -0.06 |
| SE | 35305 | ENSG00000165288 | BRWD3 | chrX | - | 80707426 | 80707503 | 80704677 | 80704846 | 80709427 | 80709577 | 141 | 130 | 135 | 137 | 349 | 2 | 0 | 0 | 1 | 0 | 34 | 67 | 84 | 30 | 6 | 0 | 6 | 0 | 0 | 0 | 226 | 149 | 0.811766714 | 1 | 0.979 | 1 | 1 | 0.989 | 1 | 1 | 0.88 | 1 | 1 | 1 | 0.018 |
| SE | 26734 | ENSG00000125089 | SH3TC1 | chr4 | + | 8209747 | 8209822 | 8205166 | 8205366 | 8212700 | 8212828 | 27 | 6 | 11 | 16 | 5 | 0 | 0 | 0 | 0 | 0 | 5 | 8 | 9 | 6 | 32 | 0 | 1 | 1 | 0 | 3 | 224 | 149 | 0.811814009 | 1 | 1 | 1 | 1 | 1 | 1 | 1 | 0.842 | 0.857 | 1 | 0.876 | 0.085 |
| SE | 76238 | ENSG00000166278 | C2 | chr6 | + | 31944979 | 31945029 | 31944726 | 31944853 | 31945177 | 31945391 | 19 | 4 | 5 | 5 | 6 | 0 | 0 | 0 | 0 | 0 | 45 | 27 | 13 | 50 | 20 | 0 | 0 | 0 | 5 | 0 | 199 | 149 | 0.811891911 | 1 | 1 | 1 | 1 | 1 | 1 | 1 | 1 | 1 | 0.882 | 1 | 0.024 |
| SE | 29177 | ENSG00000227355 | NA | chr9 | + | 121451974 | 121452173 | 121375230 | 121375325 | 121462656 | 121462770 | 2 | 7 | 1 | 5 | 6 | 15 | 25 | 5 | 21 | 18 | 10 | 6 | 1 | 9 | 10 | 47 | 39 | 0 | 50 | 68 | 298 | 149 | 0.811932967 | 1 | 0.062 | 0.123 | 0.091 | 0.106 | 0.143 | 0.096 | 0.071 | 1 | 0.083 | 0.068 | -0.159 |
| SE | 43805 | ENSG00000184588 | PDE4B | chr1 | + | 66368786 | 66368969 | 66367942 | 66368065 | 66372312 | 66372738 | 253 | 246 | 362 | 327 | 642 | 4 | 5 | 4 | 2 | 10 | 382 | 496 | 697 | 426 | 149 | 3 | 3 | 10 | 11 | 0 | 298 | 149 | 0.811934998 | 1 | 0.969 | 0.961 | 0.978 | 0.988 | 0.97 | 0.985 | 0.988 | 0.972 | 0.951 | 1 | -0.006 |
| SE | 6299 | ENSG00000141644 | MBD1 | chr18 | - | 50274976 | 50275045 | 50274185 | 50274350 | 50275128 | 50275155 | 39 | 21 | 34 | 46 | 36 | 0 | 0 | 0 | 0 | 0 | 19 | 42 | 31 | 23 | 46 | 0 | 0 | 0 | 0 | 6 | 218 | 149 | 0.811958926 | 1 | 1 | 1 | 1 | 1 | 1 | 1 | 1 | 1 | 1 | 0.84 | 0.032 |
| SE | 92075 | ENSG00000226752 | CUTALP | chr9 | + | 120846681 | 120846803 | 120844992 | 120845196 | 120847293 | 120847353 | 0 | 8 | 6 | 0 | 3 | 6 | 12 | 5 | 7 | 43 | 0 | 3 | 4 | 6 | 0 | 7 | 22 | 11 | 3 | 10 | 271 | 149 | 0.812004404 | 1 | 0 | 0.268 | 0.398 | 0 | 0.037 | 0 | 0.07 | 0.167 | 0.524 | 0 | -0.012 |
| SE | 48018 | ENSG00000010810 | FYN | chr6 | - | 111740980 | 111741093 | 111719804 | 111720062 | 111846588 | 111846629 | 61 | 15 | 28 | 34 | 49 | 1 | 4 | 0 | 0 | 0 | 19 | 24 | 50 | 16 | 22 | 0 | 0 | 3 | 0 | 0 | 262 | 149 | 0.812047491 | 1 | 0.972 | 0.681 | 1 | 1 | 1 | 1 | 1 | 0.905 | 1 | 1 | -0.05 |
| SE | 106797 | ENSG00000173442 | EHBP1L1 | chr11 | + | 65581538 | 65583765 | 65581210 | 65581373 | 65584240 | 65584398 | 509 | 358 | 396 | 443 | 456 | 6 | 0 | 1 | 1 | 9 | 953 | 539 | 311 | 609 | 1203 | 1 | 1 | 6 | 4 | 3 | 298 | 149 | 0.812156632 | 1 | 0.977 | 1 | 0.995 | 0.996 | 0.962 | 0.998 | 0.996 | 0.963 | 0.987 | 0.995 | -0.002 |
| SE | 104837 | ENSG00000205356 | TECPR1 | chr7 | - | 98245921 | 98246165 | 98244884 | 98245067 | 98251418 | 98251548 | 74 | 42 | 74 | 60 | 52 | 0 | 2 | 1 | 4 | 0 | 144 | 58 | 47 | 56 | 130 | 0 | 0 | 0 | 0 | 0 | 298 | 149 | 0.812227967 | 1 | 1 | 0.913 | 0.974 | 0.882 | 1 | 1 | 1 | 1 | 1 | 1 | -0.046 |
| SE | 34919 | ENSG00000172673 | THEMIS | chr6 | - | 127900841 | 127901081 | 127829475 | 127829934 | 127918427 | 127918540 | 103 | 82 | 94 | 66 | 84 | 0 | 0 | 2 | 4 | 0 | 34 | 67 | 25 | 11 | 3 | 0 | 0 | 0 | 0 | 0 | 298 | 149 | 0.812242775 | 1 | 1 | 1 | 0.959 | 0.892 | 1 | 1 | 1 | 1 | 1 | 1 | -0.03 |
| SE | 52482 | ENSG00000173209 | AHSA2P | chr2 | + | 61183546 | 61183662 | 61181319 | 61181405 | 61184711 | 61184800 | 14 | 19 | 20 | 14 | 28 | 376 | 354 | 368 | 478 | 332 | 11 | 9 | 20 | 22 | 5 | 245 | 400 | 312 | 272 | 152 | 265 | 149 | 0.812289864 | 1 | 0.021 | 0.029 | 0.03 | 0.016 | 0.045 | 0.025 | 0.012 | 0.035 | 0.043 | 0.018 | 0.002 |
| SE | 105548 | ENSG00000012822 | CALCOCO1 | chr12 | - | 53715792 | 53716047 | 53715199 | 53715325 | 53716259 | 53716415 | 430 | 262 | 364 | 541 | 460 | 0 | 0 | 0 | 1 | 0 | 1021 | 734 | 528 | 578 | 1570 | 8 | 5 | 3 | 6 | 7 | 298 | 149 | 0.81231403 | 1 | 1 | 1 | 1 | 0.996 | 1 | 0.985 | 0.987 | 0.989 | 0.98 | 0.991 | 0.013 |
| SE | 14067 | ENSG00000140694 | PARN | chr16 | - | 14627268 | 14627336 | 14627105 | 14627187 | 14628171 | 14628251 | 413 | 361 | 319 | 391 | 412 | 5 | 4 | 0 | 5 | 6 | 508 | 558 | 316 | 399 | 235 | 0 | 0 | 4 | 8 | 5 | 217 | 149 | 0.812408058 | 1 | 0.983 | 0.984 | 1 | 0.982 | 0.979 | 1 | 1 | 0.982 | 0.972 | 0.97 | 0.001 |
| SE | 16222 | ENSG00000077238 | IL4R | chr16 | + | 27360939 | 27361020 | 27360765 | 27360815 | 27362251 | 27362373 | 3 | 2 | 5 | 1 | 0 | 181 | 90 | 150 | 156 | 175 | 5 | 0 | 0 | 6 | 11 | 242 | 277 | 162 | 210 | 268 | 230 | 149 | 0.812410267 | 1 | 0.011 | 0.014 | 0.021 | 0.004 | 0 | 0.013 | 0 | 0 | 0.018 | 0.026 | -0.001 |
| SE | 34797 | ENSG00000136152 | COG3 | chr13 | + | 45496161 | 45496312 | 45493346 | 45493486 | 45503243 | 45503349 | 244 | 154 | 190 | 245 | 314 | 7 | 3 | 9 | 4 | 15 | 33 | 211 | 164 | 117 | 25 | 9 | 4 | 8 | 0 | 0 | 298 | 149 | 0.812539816 | 1 | 0.946 | 0.963 | 0.913 | 0.968 | 0.913 | 0.647 | 0.963 | 0.911 | 1 | 1 | 0.036 |
| SE | 4913 | ENSG00000258366 | RTEL1 | chr20 | + | 63660675 | 63660796 | 63659232 | 63659504 | 63661297 | 63661332 | 0 | 0 | 0 | 0 | 0 | 23 | 9 | 17 | 30 | 34 | 0 | 4 | 0 | 0 | 2 | 10 | 31 | 26 | 33 | 10 | 270 | 149 | 0.812582265 | 1 | 0 | 0 | 0 | 0 | 0 | 0 | 0.066 | 0 | 0 | 0.099 | -0.033 |
| SE | 74182 | ENSG00000116698 | SMG7 | chr1 | + | 183547102 | 183547252 | 183545965 | 183546337 | 183549207 | 183549288 | 12 | 9 | 14 | 24 | 22 | 84 | 55 | 91 | 96 | 138 | 33 | 25 | 29 | 31 | 4 | 218 | 123 | 101 | 115 | 188 | 298 | 149 | 0.812602106 | 1 | 0.067 | 0.076 | 0.071 | 0.111 | 0.074 | 0.07 | 0.092 | 0.126 | 0.119 | 0.011 | -0.004 |
| SE | 68740 | ENSG00000196295 | GARS1-DT | chr7 | - | 30585329 | 30585479 | 30579230 | 30579314 | 30594710 | 30594740 | 93 | 71 | 76 | 132 | 89 | 0 | 0 | 0 | 0 | 0 | 107 | 139 | 80 | 134 | 202 | 0 | 7 | 0 | 0 | 0 | 298 | 149 | 0.812719051 | 1 | 1 | 1 | 1 | 1 | 1 | 1 | 0.908 | 1 | 1 | 1 | 0.018 |
| SE | 8554 | ENSG00000067191 | CACNB1 | chr17 | - | 39185949 | 39186104 | 39184783 | 39184864 | 39186495 | 39186572 | 4 | 0 | 0 | 0 | 1 | 13 | 7 | 6 | 12 | 5 | 0 | 0 | 0 | 0 | 0 | 12 | 23 | 13 | 6 | 9 | 298 | 149 | 0.812777541 | 1 | 0.133 | 0 | 0 | 0 | 0.091 | 0 | 0 | 0 | 0 | 0 | 0.045 |
| SE | 59635 | ENSG00000163466 | ARPC2 | chr2 | + | 218217462 | 218217544 | 218217146 | 218217254 | 218225919 | 218225954 | 615 | 330 | 418 | 494 | 285 | 1 | 0 | 2 | 4 | 0 | 878 | 681 | 548 | 826 | 709 | 0 | 16 | 0 | 0 | 0 | 231 | 149 | 0.812793281 | 1 | 0.997 | 1 | 0.993 | 0.988 | 1 | 1 | 0.965 | 1 | 1 | 1 | 0.003 |
| SE | 69834 | ENSG00000163803 | PLB1 | chr2 | + | 28625056 | 28625108 | 28620671 | 28620978 | 28626427 | 28626508 | 23 | 10 | 13 | 18 | 24 | 0 | 0 | 0 | 0 | 0 | 13 | 36 | 11 | 55 | 66 | 0 | 0 | 0 | 6 | 0 | 201 | 149 | 0.812924418 | 1 | 1 | 1 | 1 | 1 | 1 | 1 | 1 | 1 | 0.872 | 1 | 0.026 |
| SE | 841 | ENSG00000242259 | C22orf39 | chr22 | + | 19356889 | 19357051 | 19356229 | 19356288 | 19359335 | 19359484 | 65 | 37 | 61 | 55 | 14 | 1 | 0 | 4 | 1 | 0 | 85 | 87 | 26 | 35 | 62 | 4 | 0 | 0 | 0 | 0 | 298 | 149 | 0.812966561 | 1 | 0.97 | 1 | 0.884 | 0.965 | 1 | 0.914 | 1 | 1 | 1 | 1 | -0.019 |
| SE | 1676 | ENSG00000100281 | HMGXB4 | chr22 | + | 35263981 | 35264096 | 35263795 | 35263874 | 35264647 | 35265567 | 38 | 18 | 27 | 18 | 59 | 50 | 21 | 37 | 28 | 65 | 44 | 41 | 44 | 27 | 6 | 43 | 45 | 51 | 57 | 3 | 264 | 149 | 0.813022137 | 1 | 0.3 | 0.326 | 0.292 | 0.266 | 0.339 | 0.366 | 0.34 | 0.327 | 0.211 | 0.53 | -0.05 |
| SE | 4839 | ENSG00000132670 | PTPRA | chr20 | + | 2964271 | 2964350 | 2947981 | 2948024 | 2986764 | 2986849 | 306 | 170 | 220 | 288 | 557 | 1 | 3 | 0 | 0 | 0 | 297 | 215 | 129 | 219 | 395 | 0 | 0 | 0 | 14 | 0 | 228 | 149 | 0.813081565 | 1 | 0.995 | 0.974 | 1 | 1 | 1 | 1 | 1 | 1 | 0.911 | 1 | 0.012 |
| SE | 46934 | ENSG00000141027 | NCOR1 | chr17 | - | 16075533 | 16075702 | 16073428 | 16073569 | 16079963 | 16080064 | 467 | 323 | 367 | 457 | 365 | 0 | 0 | 0 | 1 | 0 | 245 | 538 | 461 | 403 | 196 | 10 | 3 | 0 | 0 | 0 | 298 | 149 | 0.813152033 | 1 | 1 | 1 | 1 | 0.996 | 1 | 0.925 | 0.989 | 1 | 1 | 1 | 0.016 |
| SE | 85793 | ENSG00000258529 | NA | chr11 | - | 111844600 | 111844723 | 111840654 | 111840809 | 111853379 | 111853485 | 66 | 29 | 77 | 45 | 106 | 0 | 0 | 0 | 1 | 0 | 106 | 68 | 84 | 27 | 18 | 0 | 0 | 0 | 4 | 0 | 272 | 149 | 0.813158808 | 1 | 1 | 1 | 1 | 0.961 | 1 | 1 | 1 | 1 | 0.787 | 1 | 0.035 |
| SE | 99901 | ENSG00000073910 | FRY | chr13 | + | 32147838 | 32147947 | 32147281 | 32147385 | 32149747 | 32149834 | 95 | 74 | 64 | 86 | 123 | 0 | 0 | 0 | 2 | 0 | 85 | 106 | 95 | 111 | 36 | 7 | 1 | 0 | 0 | 0 | 258 | 149 | 0.813179997 | 1 | 1 | 1 | 1 | 0.961 | 1 | 0.875 | 0.984 | 1 | 1 | 1 | 0.02 |
| SE | 101801 | ENSG00000090621 | PABPC4 | chr1 | - | 39565105 | 39565366 | 39564685 | 39564773 | 39567750 | 39567846 | 193 | 138 | 233 | 180 | 289 | 0 | 0 | 2 | 0 | 0 | 174 | 145 | 154 | 82 | 200 | 0 | 0 | 3 | 5 | 0 | 298 | 149 | 0.813231849 | 1 | 1 | 1 | 0.983 | 1 | 1 | 1 | 1 | 0.963 | 0.891 | 1 | 0.026 |
| SE | 20267 | ENSG00000118482 | PHF3 | chr6 | + | 63646618 | 63646795 | 63635835 | 63636150 | 63684128 | 63685911 | 6 | 0 | 5 | 6 | 2 | 8 | 8 | 10 | 7 | 3 | 12 | 4 | 5 | 0 | 0 | 10 | 11 | 3 | 3 | 1 | 298 | 149 | 0.813254473 | 1 | 0.273 | 0 | 0.2 | 0.3 | 0.25 | 0.375 | 0.154 | 0.455 | 0 | 0 | 0.008 |
| SE | 48863 | ENSG00000184677 | ZBTB40 | chr1 | + | 22502298 | 22502441 | 22491399 | 22491533 | 22506048 | 22506241 | 54 | 30 | 71 | 63 | 49 | 3 | 3 | 0 | 2 | 5 | 68 | 51 | 44 | 27 | 47 | 0 | 5 | 5 | 0 | 3 | 292 | 149 | 0.813275157 | 1 | 0.902 | 0.836 | 1 | 0.941 | 0.833 | 1 | 0.839 | 0.818 | 1 | 0.889 | -0.007 |
| SE | 43869 | ENSG00000153898 | MCOLN2 | chr1 | - | 84958528 | 84958702 | 84952445 | 84952530 | 84965548 | 84965708 | 30 | 40 | 26 | 35 | 28 | 4 | 0 | 0 | 0 | 0 | 25 | 70 | 62 | 57 | 12 | 0 | 0 | 0 | 0 | 0 | 298 | 149 | 0.813374298 | 1 | 0.789 | 1 | 1 | 1 | 1 | 1 | 1 | 1 | 1 | 1 | -0.042 |
| SE | 82850 | ENSG00000146776 | ATXN7L1 | chr7 | - | 105610528 | 105610603 | 105604771 | 105607890 | 105613861 | 105614816 | 23 | 30 | 18 | 39 | 16 | 0 | 0 | 0 | 0 | 0 | 22 | 19 | 7 | 23 | 12 | 4 | 0 | 0 | 0 | 0 | 224 | 149 | 0.813386537 | 1 | 1 | 1 | 1 | 1 | 1 | 0.785 | 1 | 1 | 1 | 1 | 0.043 |
| SE | 108964 | ENSG00000102543 | CDADC1 | chr13 | + | 49265878 | 49266003 | 49259345 | 49259523 | 49267489 | 49267525 | 16 | 9 | 9 | 18 | 12 | 31 | 29 | 21 | 25 | 19 | 0 | 10 | 43 | 12 | 7 | 32 | 29 | 44 | 37 | 0 | 274 | 149 | 0.813410428 | 1 | 0.219 | 0.144 | 0.189 | 0.281 | 0.256 | 0 | 0.158 | 0.347 | 0.15 | 1 | -0.113 |
| SE | 66274 | ENSG00000267281 | ATF7-NPFF | chr12 | - | 53534501 | 53534659 | 53532509 | 53532623 | 53537414 | 53537552 | 21 | 27 | 21 | 24 | 48 | 2 | 0 | 1 | 2 | 0 | 93 | 45 | 22 | 55 | 78 | 0 | 0 | 0 | 2 | 0 | 298 | 149 | 0.813468777 | 1 | 0.84 | 1 | 0.913 | 0.857 | 1 | 1 | 1 | 1 | 0.932 | 1 | -0.064 |
| SE | 91763 | ENSG00000143748 | NVL | chr1 | - | 224313033 | 224313151 | 224311799 | 224311857 | 224317693 | 224317793 | 5 | 2 | 4 | 0 | 7 | 50 | 40 | 47 | 47 | 116 | 0 | 0 | 3 | 10 | 0 | 6 | 44 | 52 | 26 | 20 | 267 | 149 | 0.813502163 | 1 | 0.053 | 0.027 | 0.045 | 0 | 0.033 | 0 | 0 | 0.031 | 0.177 | 0 | -0.01 |
| SE | 75276 | ENSG00000024048 | UBR2 | chr6 | + | 42592150 | 42592229 | 42573733 | 42573993 | 42603587 | 42603718 | 68 | 49 | 50 | 59 | 40 | 9 | 5 | 9 | 6 | 3 | 42 | 51 | 53 | 54 | 41 | 0 | 2 | 26 | 2 | 7 | 228 | 149 | 0.813508348 | 1 | 0.832 | 0.865 | 0.784 | 0.865 | 0.897 | 1 | 0.943 | 0.571 | 0.946 | 0.793 | -0.002 |
| SE | 70233 | ENSG00000160695 | VPS11 | chr11 | + | 119071595 | 119071843 | 119070233 | 119070397 | 119073197 | 119073399 | 112 | 65 | 115 | 99 | 138 | 3 | 13 | 3 | 3 | 0 | 130 | 102 | 79 | 108 | 115 | 0 | 1 | 9 | 10 | 4 | 298 | 149 | 0.813519263 | 1 | 0.949 | 0.714 | 0.95 | 0.943 | 1 | 1 | 0.981 | 0.814 | 0.844 | 0.935 | -0.004 |
| SE | 103183 | ENSG00000011376 | LARS2 | chr3 | + | 45513134 | 45513235 | 45500441 | 45500579 | 45516093 | 45516276 | 64 | 53 | 59 | 62 | 91 | 0 | 0 | 0 | 1 | 0 | 91 | 67 | 44 | 66 | 40 | 0 | 0 | 0 | 6 | 0 | 250 | 149 | 0.813561148 | 1 | 1 | 1 | 1 | 0.974 | 1 | 1 | 1 | 1 | 0.868 | 1 | 0.021 |
| SE | 89088 | ENSG00000171988 | JMJD1C | chr10 | - | 63264650 | 63264764 | 63217206 | 63217331 | 63380317 | 63380482 | 130 | 140 | 126 | 160 | 338 | 3 | 1 | 1 | 4 | 4 | 35 | 43 | 50 | 10 | 22 | 0 | 6 | 1 | 0 | 0 | 263 | 149 | 0.813563539 | 1 | 0.961 | 0.988 | 0.986 | 0.958 | 0.98 | 1 | 0.802 | 0.966 | 1 | 1 | 0.021 |
| SE | 23428 | ENSG00000148110 | MFSD14B | chr9 | + | 94456239 | 94456337 | 94453957 | 94454070 | 94458321 | 94458464 | 274 | 119 | 213 | 224 | 384 | 27 | 6 | 19 | 15 | 49 | 536 | 354 | 173 | 276 | 629 | 45 | 33 | 15 | 41 | 52 | 247 | 149 | 0.813612907 | 1 | 0.86 | 0.923 | 0.871 | 0.9 | 0.825 | 0.878 | 0.866 | 0.874 | 0.802 | 0.879 | 0.016 |
| SE | 46212 | ENSG00000114439 | BBX | chr3 | + | 107661827 | 107661944 | 107645835 | 107645909 | 107710451 | 107710622 | 8 | 5 | 0 | 4 | 0 | 104 | 91 | 89 | 82 | 101 | 0 | 8 | 4 | 0 | 0 | 66 | 88 | 93 | 39 | 23 | 266 | 149 | 0.813872472 | 1 | 0.041 | 0.03 | 0 | 0.027 | 0 | 0 | 0.048 | 0.024 | 0 | 0 | 0.005 |
| SE | 80280 | ENSG00000168137 | SETD5 | chr3 | + | 9429826 | 9430051 | 9428822 | 9429009 | 9433844 | 9433950 | 68 | 33 | 59 | 66 | 80 | 46 | 25 | 22 | 23 | 14 | 63 | 68 | 55 | 57 | 27 | 40 | 44 | 10 | 13 | 24 | 298 | 149 | 0.81387491 | 1 | 0.425 | 0.398 | 0.573 | 0.589 | 0.741 | 0.441 | 0.436 | 0.733 | 0.687 | 0.36 | 0.014 |
| SE | 60 | ENSG00000160285 | LSS | chr21 | - | 46196201 | 46196267 | 46195675 | 46195756 | 46205835 | 46205941 | 41 | 11 | 24 | 32 | 9 | 0 | 0 | 0 | 2 | 0 | 20 | 62 | 15 | 25 | 92 | 0 | 0 | 1 | 4 | 0 | 215 | 149 | 0.813881149 | 1 | 1 | 1 | 1 | 0.917 | 1 | 1 | 1 | 0.912 | 0.812 | 1 | 0.039 |
| SE | 107685 | ENSG00000128585 | MKLN1 | chr7 | + | 131429032 | 131429145 | 131414644 | 131414710 | 131437784 | 131437997 | 442 | 463 | 278 | 459 | 453 | 1 | 1 | 0 | 3 | 0 | 149 | 413 | 462 | 159 | 22 | 0 | 0 | 5 | 6 | 0 | 262 | 149 | 0.813904815 | 1 | 0.996 | 0.996 | 1 | 0.989 | 1 | 1 | 1 | 0.981 | 0.938 | 1 | 0.012 |
| SE | 94645 | ENSG00000011454 | RABGAP1 | chr9 | + | 123070349 | 123070424 | 123065347 | 123065601 | 123073551 | 123073677 | 47 | 34 | 41 | 56 | 50 | 0 | 0 | 0 | 0 | 0 | 22 | 22 | 23 | 30 | 54 | 0 | 4 | 0 | 0 | 0 | 224 | 149 | 0.813943763 | 1 | 1 | 1 | 1 | 1 | 1 | 1 | 0.785 | 1 | 1 | 1 | 0.043 |
| SE | 45357 | ENSG00000115267 | IFIH1 | chr2 | - | 162278204 | 162278328 | 162277414 | 162277693 | 162279995 | 162280112 | 147 | 245 | 126 | 213 | 183 | 0 | 0 | 0 | 4 | 0 | 75 | 131 | 73 | 128 | 8 | 0 | 4 | 2 | 0 | 0 | 273 | 149 | 0.81403529 | 1 | 1 | 1 | 1 | 0.967 | 1 | 1 | 0.947 | 0.952 | 1 | 1 | 0.014 |
| SE | 14536 | ENSG00000185043 | CIB1 | chr15 | - | 90232218 | 90232327 | 90231356 | 90231507 | 90233668 | 90233703 | 418 | 144 | 300 | 264 | 153 | 0 | 0 | 0 | 0 | 0 | 306 | 445 | 240 | 246 | 409 | 0 | 0 | 2 | 0 | 11 | 258 | 149 | 0.814107856 | 1 | 1 | 1 | 1 | 1 | 1 | 1 | 1 | 0.986 | 1 | 0.956 | 0.012 |
| SE | 8854 | ENSG00000136436 | CALCOCO2 | chr17 | + | 48860313 | 48860449 | 48856091 | 48856187 | 48862275 | 48862304 | 986 | 926 | 931 | 1029 | 1918 | 3 | 3 | 9 | 12 | 8 | 1118 | 980 | 914 | 992 | 393 | 0 | 6 | 3 | 7 | 0 | 285 | 149 | 0.814124015 | 1 | 0.994 | 0.994 | 0.982 | 0.978 | 0.992 | 1 | 0.988 | 0.994 | 0.987 | 1 | -0.006 |
| SE | 106465 | ENSG00000198270 | TMEM116 | chr12 | - | 111937159 | 111937243 | 111932584 | 111932659 | 111938160 | 111938210 | 29 | 23 | 27 | 19 | 64 | 0 | 4 | 0 | 0 | 0 | 8 | 19 | 18 | 26 | 26 | 0 | 0 | 0 | 0 | 0 | 233 | 149 | 0.814217782 | 1 | 1 | 0.786 | 1 | 1 | 1 | 1 | 1 | 1 | 1 | 1 | -0.043 |
| SE | 62413 | ENSG00000033867 | SLC4A7 | chr3 | - | 27431297 | 27431669 | 27424036 | 27424152 | 27433915 | 27434104 | 0 | 0 | 0 | 2 | 4 | 43 | 23 | 43 | 38 | 52 | 0 | 5 | 0 | 0 | 0 | 25 | 20 | 18 | 3 | 5 | 298 | 149 | 0.814444274 | 1 | 0 | 0 | 0 | 0.026 | 0.037 | 0 | 0.111 | 0 | 0 | 0 | -0.01 |
| SE | 10012 | ENSG00000141540 | TTYH2 | chr17 | + | 74249343 | 74249399 | 74247957 | 74249080 | 74249935 | 74250014 | 23 | 10 | 25 | 18 | 6 | 1 | 1 | 0 | 3 | 0 | 41 | 14 | 14 | 5 | 80 | 0 | 0 | 1 | 2 | 2 | 205 | 149 | 0.814509911 | 1 | 0.944 | 0.879 | 1 | 0.813 | 1 | 1 | 1 | 0.911 | 0.645 | 0.967 | 0.023 |
| SE | 98716 | ENSG00000107290 | SETX | chr9 | - | 132266126 | 132266249 | 132261355 | 132264985 | 132269614 | 132269702 | 7 | 5 | 2 | 2 | 8 | 189 | 152 | 174 | 203 | 320 | 0 | 0 | 2 | 4 | 0 | 27 | 151 | 58 | 28 | 19 | 272 | 149 | 0.81452164 | 1 | 0.02 | 0.018 | 0.006 | 0.005 | 0.014 | 0 | 0 | 0.019 | 0.073 | 0 | -0.006 |
| SE | 98983 | ENSG00000076043 | REXO2 | chr11 | + | 114440655 | 114440739 | 114439434 | 114439675 | 114445978 | 114446087 | 22 | 9 | 21 | 19 | 4 | 0 | 0 | 0 | 0 | 0 | 23 | 18 | 18 | 8 | 15 | 4 | 0 | 0 | 0 | 0 | 233 | 149 | 0.814616123 | 1 | 1 | 1 | 1 | 1 | 1 | 0.786 | 1 | 1 | 1 | 1 | 0.043 |
| SE | 78719 | ENSG00000166886 | NAB2 | chr12 | + | 57093406 | 57093598 | 57093062 | 57093195 | 57094611 | 57095476 | 15 | 9 | 13 | 23 | 9 | 4 | 0 | 0 | 2 | 0 | 23 | 3 | 1 | 52 | 93 | 0 | 0 | 4 | 0 | 0 | 298 | 149 | 0.814688251 | 1 | 0.652 | 1 | 1 | 0.852 | 1 | 1 | 1 | 0.111 | 1 | 1 | 0.079 |
| SE | 51429 | ENSG00000107745 | MICU1 | chr10 | - | 72477492 | 72477603 | 72477173 | 72477256 | 72508154 | 72508269 | 0 | 4 | 0 | 0 | 3 | 129 | 108 | 73 | 116 | 313 | 9 | 0 | 4 | 5 | 0 | 240 | 144 | 79 | 172 | 500 | 260 | 149 | 0.814720118 | 1 | 0 | 0.021 | 0 | 0 | 0.005 | 0.021 | 0 | 0.028 | 0.016 | 0 | -0.008 |
| SE | 58221 | ENSG00000172965 | MIR4435-2HG | chr2 | - | 111469602 | 111469987 | 111429442 | 111429620 | 111494884 | 111495025 | 3 | 0 | 4 | 4 | 0 | 65 | 73 | 44 | 70 | 91 | 8 | 0 | 1 | 1 | 4 | 243 | 122 | 43 | 115 | 344 | 298 | 149 | 0.814747091 | 1 | 0.023 | 0 | 0.043 | 0.028 | 0 | 0.016 | 0 | 0.011 | 0.004 | 0.006 | 0.011 |
| SE | 2233 | ENSG00000161091 | MFSD12 | chr19 | - | 3546254 | 3546425 | 3546073 | 3546168 | 3547271 | 3547331 | 25 | 17 | 32 | 31 | 19 | 1 | 0 | 0 | 3 | 1 | 25 | 23 | 28 | 23 | 14 | 0 | 0 | 0 | 1 | 0 | 298 | 149 | 0.814802152 | 1 | 0.926 | 1 | 1 | 0.838 | 0.905 | 1 | 1 | 1 | 0.92 | 1 | -0.05 |
| SE | 16195 | ENSG00000196547 | MAN2A2 | chr15 | + | 90913272 | 90913406 | 90912876 | 90912991 | 90913613 | 90913755 | 60 | 25 | 37 | 53 | 5 | 3 | 2 | 0 | 0 | 0 | 100 | 104 | 69 | 120 | 298 | 0 | 0 | 3 | 0 | 0 | 283 | 149 | 0.814865273 | 1 | 0.913 | 0.868 | 1 | 1 | 1 | 1 | 1 | 0.924 | 1 | 1 | -0.029 |
| SE | 23106 | ENSG00000111912 | NCOA7 | chr6 | + | 125928173 | 125928247 | 125927662 | 125927758 | 125928635 | 125929027 | 134 | 151 | 106 | 151 | 83 | 0 | 0 | 0 | 1 | 0 | 27 | 119 | 158 | 248 | 12 | 0 | 2 | 3 | 5 | 0 | 223 | 149 | 0.814892084 | 1 | 1 | 1 | 1 | 0.99 | 1 | 1 | 0.975 | 0.972 | 0.971 | 1 | 0.014 |
| SE | 48578 | ENSG00000126091 | ST3GAL3 | chr1 | + | 43921442 | 43921638 | 43920781 | 43920928 | 43930131 | 43931057 | 4 | 0 | 0 | 0 | 0 | 7 | 5 | 5 | 11 | 4 | 0 | 0 | 0 | 0 | 0 | 4 | 13 | 4 | 2 | 81 | 298 | 149 | 0.814894224 | 1 | 0.222 | 0 | 0 | 0 | 0 | 0 | 0 | 0 | 0 | 0 | 0.044 |
| SE | 103430 | ENSG00000107669 | ATE1 | chr10 | - | 121922348 | 121922411 | 121913789 | 121913893 | 121924265 | 121924329 | 142 | 144 | 128 | 187 | 395 | 2 | 1 | 0 | 2 | 6 | 143 | 99 | 143 | 185 | 164 | 0 | 0 | 2 | 0 | 4 | 212 | 149 | 0.814905515 | 1 | 0.98 | 0.99 | 1 | 0.985 | 0.979 | 1 | 1 | 0.98 | 1 | 0.966 | -0.002 |
| SE | 11848 | ENSG00000121274 | TENT4B | chr16 | + | 50153746 | 50154259 | 50152910 | 50153032 | 50211322 | 50211446 | 50 | 31 | 32 | 68 | 6 | 0 | 0 | 0 | 0 | 0 | 52 | 32 | 42 | 46 | 24 | 0 | 4 | 1 | 0 | 0 | 298 | 149 | 0.814988912 | 1 | 1 | 1 | 1 | 1 | 1 | 1 | 0.8 | 0.955 | 1 | 1 | 0.049 |
| SE | 49882 | ENSG00000109180 | OCIAD1 | chr4 | + | 48849946 | 48850082 | 48848398 | 48848446 | 48857212 | 48857252 | 553 | 430 | 455 | 516 | 975 | 0 | 0 | 1 | 4 | 1 | 634 | 378 | 303 | 415 | 526 | 14 | 0 | 4 | 0 | 0 | 285 | 149 | 0.815041366 | 1 | 1 | 1 | 0.996 | 0.985 | 0.998 | 0.959 | 1 | 0.975 | 1 | 1 | 0.009 |
| SE | 45697 | ENSG00000049541 | RFC2 | chr7 | - | 74246661 | 74246763 | 74243145 | 74243246 | 74249738 | 74249780 | 26 | 14 | 26 | 31 | 57 | 0 | 0 | 0 | 0 | 0 | 51 | 16 | 21 | 25 | 30 | 0 | 0 | 0 | 4 | 0 | 251 | 149 | 0.815107314 | 1 | 1 | 1 | 1 | 1 | 1 | 1 | 1 | 1 | 0.788 | 1 | 0.042 |
| SE | 41578 | ENSG00000084072 | PPIE | chr1 | + | 39745413 | 39745498 | 39743823 | 39743924 | 39748902 | 39749088 | 60 | 33 | 67 | 58 | 91 | 7 | 1 | 5 | 4 | 5 | 74 | 26 | 28 | 30 | 76 | 13 | 7 | 1 | 0 | 0 | 234 | 149 | 0.815142452 | 1 | 0.845 | 0.955 | 0.895 | 0.902 | 0.921 | 0.784 | 0.703 | 0.947 | 1 | 1 | 0.017 |
| SE | 63234 | ENSG00000068784 | SRBD1 | chr2 | - | 45562652 | 45562756 | 45553622 | 45553730 | 45573206 | 45573342 | 85 | 57 | 60 | 56 | 164 | 0 | 0 | 4 | 2 | 0 | 80 | 62 | 17 | 14 | 20 | 0 | 0 | 0 | 0 | 0 | 253 | 149 | 0.815199082 | 1 | 1 | 1 | 0.898 | 0.943 | 1 | 1 | 1 | 1 | 1 | 1 | -0.032 |
| SE | 8901 | ENSG00000007168 | PAFAH1B1 | chr17 | + | 2676504 | 2676606 | 2672303 | 2674288 | 2680163 | 2680320 | 428 | 267 | 393 | 436 | 644 | 2 | 0 | 3 | 1 | 0 | 304 | 438 | 379 | 524 | 320 | 7 | 7 | 0 | 1 | 0 | 251 | 149 | 0.815252644 | 1 | 0.992 | 1 | 0.987 | 0.996 | 1 | 0.963 | 0.974 | 1 | 0.997 | 1 | 0.008 |
| SE | 90730 | ENSG00000168404 | MLKL | chr16 | - | 74676300 | 74676425 | 74675617 | 74675764 | 74685485 | 74685583 | 36 | 24 | 24 | 37 | 20 | 3 | 0 | 0 | 3 | 0 | 56 | 45 | 86 | 86 | 16 | 0 | 3 | 0 | 2 | 0 | 274 | 149 | 0.815404863 | 1 | 0.867 | 1 | 1 | 0.87 | 1 | 1 | 0.891 | 1 | 0.959 | 1 | -0.023 |
| SE | 71162 | ENSG00000117758 | STX12 | chr1 | + | 27811991 | 27812060 | 27810245 | 27810289 | 27812162 | 27812268 | 0 | 0 | 0 | 6 | 0 | 222 | 136 | 108 | 179 | 216 | 7 | 6 | 2 | 2 | 0 | 165 | 238 | 152 | 219 | 76 | 218 | 149 | 0.81541033 | 1 | 0 | 0 | 0 | 0.022 | 0 | 0.028 | 0.017 | 0.009 | 0.006 | 0 | -0.008 |
| SE | 23361 | ENSG00000147044 | CASK | chrX | - | 41542690 | 41542806 | 41534892 | 41534973 | 41559778 | 41559847 | 39 | 26 | 47 | 37 | 58 | 0 | 1 | 4 | 0 | 0 | 11 | 34 | 26 | 26 | 6 | 0 | 0 | 0 | 0 | 0 | 265 | 149 | 0.815415954 | 1 | 1 | 0.936 | 0.869 | 1 | 1 | 1 | 1 | 1 | 1 | 1 | -0.039 |
| SE | 64019 | ENSG00000115464 | USP34 | chr2 | - | 61203139 | 61203263 | 61192900 | 61192980 | 61204255 | 61204380 | 125 | 92 | 194 | 212 | 482 | 5 | 1 | 4 | 0 | 2 | 12 | 79 | 53 | 38 | 9 | 0 | 0 | 0 | 0 | 0 | 273 | 149 | 0.81545796 | 1 | 0.932 | 0.98 | 0.964 | 1 | 0.992 | 1 | 1 | 1 | 1 | 1 | -0.026 |
| SE | 70719 | ENSG00000183283 | DAZAP2 | chr12 | + | 51240870 | 51241039 | 51240342 | 51240461 | 51242329 | 51242816 | 161 | 79 | 114 | 141 | 119 | 2 | 1 | 1 | 4 | 11 | 250 | 187 | 179 | 290 | 364 | 0 | 11 | 11 | 7 | 2 | 298 | 149 | 0.815464374 | 1 | 0.976 | 0.975 | 0.983 | 0.946 | 0.844 | 1 | 0.895 | 0.891 | 0.954 | 0.989 | -0.001 |
| SE | 73334 | ENSG00000115970 | THADA | chr2 | - | 43538428 | 43538566 | 43527878 | 43527988 | 43541158 | 43541303 | 10 | 2 | 1 | 0 | 0 | 79 | 56 | 71 | 71 | 94 | 0 | 6 | 4 | 0 | 0 | 33 | 43 | 42 | 71 | 7 | 287 | 149 | 0.81551791 | 1 | 0.062 | 0.018 | 0.007 | 0 | 0 | 0 | 0.068 | 0.047 | 0 | 0 | -0.006 |
| SE | 11936 | ENSG00000005844 | ITGAL | chr16 | + | 30479090 | 30479208 | 30475512 | 30475580 | 30481438 | 30481584 | 246 | 94 | 149 | 183 | 118 | 53 | 19 | 21 | 21 | 27 | 270 | 308 | 172 | 201 | 468 | 53 | 59 | 40 | 22 | 85 | 267 | 149 | 0.81557663 | 1 | 0.721 | 0.734 | 0.798 | 0.829 | 0.709 | 0.74 | 0.744 | 0.706 | 0.836 | 0.754 | 0.002 |
| SE | 89265 | ENSG00000104325 | DECR1 | chr8 | + | 90015540 | 90015666 | 90001487 | 90001561 | 90017123 | 90017326 | 1 | 7 | 5 | 9 | 0 | 90 | 52 | 76 | 74 | 68 | 0 | 5 | 6 | 23 | 0 | 168 | 95 | 114 | 111 | 51 | 275 | 149 | 0.815585014 | 1 | 0.006 | 0.068 | 0.034 | 0.062 | 0 | 0 | 0.028 | 0.028 | 0.101 | 0 | 0.003 |
| SE | 67017 | ENSG00000112658 | SRF | chr6 | + | 43176547 | 43176667 | 43173846 | 43174113 | 43178293 | 43178485 | 26 | 14 | 19 | 27 | 16 | 0 | 0 | 0 | 4 | 0 | 32 | 15 | 16 | 27 | 53 | 0 | 0 | 0 | 0 | 0 | 269 | 149 | 0.815617868 | 1 | 1 | 1 | 1 | 0.789 | 1 | 1 | 1 | 1 | 1 | 1 | -0.042 |
| SE | 10910 | ENSG00000074370 | ATP2A3 | chr17 | - | 3935191 | 3935277 | 3930300 | 3930434 | 3936266 | 3936469 | 218 | 83 | 245 | 279 | 89 | 0 | 0 | 0 | 3 | 0 | 275 | 417 | 204 | 256 | 1654 | 9 | 0 | 0 | 4 | 0 | 235 | 149 | 0.815681384 | 1 | 1 | 1 | 1 | 0.983 | 1 | 0.951 | 1 | 1 | 0.976 | 1 | 0.011 |
| SE | 102118 | ENSG00000077420 | APBB1IP | chr10 | + | 26513538 | 26513660 | 26511746 | 26511906 | 26533438 | 26533525 | 1439 | 1228 | 1078 | 1377 | 1754 | 4 | 8 | 12 | 7 | 9 | 2240 | 1533 | 1123 | 1777 | 2361 | 14 | 10 | 10 | 12 | 0 | 271 | 149 | 0.815743797 | 1 | 0.995 | 0.988 | 0.98 | 0.991 | 0.991 | 0.989 | 0.988 | 0.984 | 0.988 | 1 | -0.001 |
| SE | 18524 | ENSG00000203485 | INF2 | chr14 | + | 104715283 | 104715340 | 104714202 | 104714856 | 104718794 | 104719603 | 83 | 60 | 71 | 80 | 117 | 6 | 2 | 2 | 3 | 4 | 148 | 23 | 28 | 31 | 282 | 0 | 10 | 1 | 1 | 11 | 206 | 149 | 0.815749318 | 1 | 0.909 | 0.956 | 0.963 | 0.951 | 0.955 | 1 | 0.625 | 0.953 | 0.957 | 0.949 | 0.05 |
| SE | 31962 | ENSG00000137502 | RAB30 | chr11 | - | 83060076 | 83060178 | 82997223 | 82997324 | 83071690 | 83071829 | 1 | 0 | 1 | 3 | 0 | 34 | 27 | 35 | 33 | 26 | 0 | 0 | 8 | 0 | 0 | 65 | 58 | 66 | 36 | 1 | 251 | 149 | 0.815770373 | 1 | 0.017 | 0 | 0.017 | 0.051 | 0 | 0 | 0 | 0.067 | 0 | 0 | 0.004 |
| SE | 33311 | ENSG00000166128 | RAB8B | chr15 | + | 63223847 | 63223979 | 63189602 | 63189748 | 63244755 | 63244816 | 45 | 20 | 35 | 31 | 29 | 637 | 535 | 398 | 562 | 403 | 0 | 24 | 52 | 13 | 0 | 73 | 513 | 568 | 157 | 73 | 281 | 149 | 0.815796141 | 1 | 0.036 | 0.019 | 0.045 | 0.028 | 0.037 | 0 | 0.024 | 0.046 | 0.042 | 0 | 0.011 |
| SE | 61153 | ENSG00000111711 | GOLT1B | chr12 | + | 21508382 | 21508561 | 21501792 | 21501948 | 21512294 | 21512376 | 103 | 59 | 85 | 88 | 226 | 1 | 0 | 0 | 0 | 0 | 8 | 56 | 14 | 30 | 31 | 0 | 4 | 0 | 1 | 0 | 298 | 149 | 0.815867787 | 1 | 0.981 | 1 | 1 | 1 | 1 | 1 | 0.875 | 1 | 0.938 | 1 | 0.034 |
| SE | 49621 | ENSG00000178927 | CYBC1 | chr17 | - | 82447579 | 82447621 | 82445863 | 82445960 | 82449169 | 82449292 | 19 | 9 | 6 | 9 | 9 | 4 | 0 | 0 | 0 | 0 | 19 | 13 | 4 | 6 | 22 | 0 | 0 | 0 | 0 | 0 | 191 | 149 | 0.815907047 | 1 | 0.787 | 1 | 1 | 1 | 1 | 1 | 1 | 1 | 1 | 1 | -0.043 |
| SE | 21688 | ENSG00000135341 | MAP3K7 | chr6 | - | 90552048 | 90552179 | 90550467 | 90550549 | 90553457 | 90553586 | 224 | 142 | 169 | 169 | 322 | 0 | 0 | 0 | 0 | 0 | 41 | 114 | 89 | 62 | 28 | 0 | 0 | 0 | 0 | 4 | 280 | 149 | 0.815938312 | 1 | 1 | 1 | 1 | 1 | 1 | 1 | 1 | 1 | 1 | 0.788 | 0.042 |
| SE | 72455 | ENSG00000163482 | STK36 | chr2 | + | 218679167 | 218679261 | 218676028 | 218676278 | 218679559 | 218679617 | 2 | 1 | 1 | 14 | 0 | 1 | 0 | 0 | 0 | 7 | 48 | 5 | 0 | 7 | 6 | 0 | 0 | 2 | 0 | 3 | 243 | 149 | 0.815975826 | 1 | 0.551 | 1 | 1 | 1 | 0 | 1 | 1 | 0 | 1 | 0.551 | 0 |
| SE | 107056 | ENSG00000111652 | COPS7A | chr12 | + | 6727925 | 6728001 | 6724286 | 6724818 | 6728222 | 6728311 | 36 | 21 | 43 | 39 | 24 | 6 | 1 | 0 | 7 | 11 | 41 | 26 | 14 | 21 | 52 | 0 | 2 | 10 | 3 | 6 | 225 | 149 | 0.81598186 | 1 | 0.799 | 0.933 | 1 | 0.787 | 0.591 | 1 | 0.896 | 0.481 | 0.823 | 0.852 | 0.012 |
| SE | 58326 | ENSG00000116667 | C1orf21 | chr1 | + | 184507597 | 184507682 | 184477385 | 184477603 | 184590738 | 184590815 | 96 | 25 | 26 | 84 | 91 | 2 | 3 | 0 | 3 | 0 | 46 | 56 | 51 | 17 | 53 | 0 | 0 | 0 | 0 | 8 | 234 | 149 | 0.816018852 | 1 | 0.968 | 0.841 | 1 | 0.947 | 1 | 1 | 1 | 1 | 1 | 0.808 | -0.01 |
| SE | 21058 | ENSG00000196182 | STK40 | chr1 | - | 36343363 | 36343448 | 36339623 | 36341973 | 36343859 | 36343979 | 89 | 40 | 56 | 106 | 34 | 0 | 0 | 0 | 1 | 0 | 135 | 89 | 58 | 75 | 376 | 0 | 0 | 2 | 0 | 9 | 234 | 149 | 0.816053854 | 1 | 1 | 1 | 1 | 0.985 | 1 | 1 | 1 | 0.949 | 1 | 0.964 | 0.014 |
| SE | 64621 | ENSG00000148737 | TCF7L2 | chr10 | + | 113089399 | 113089540 | 113040024 | 113040126 | 113141183 | 113141316 | 4 | 1 | 0 | 1 | 0 | 22 | 36 | 13 | 18 | 33 | 0 | 0 | 0 | 9 | 3 | 152 | 25 | 18 | 99 | 53 | 290 | 149 | 0.816118483 | 1 | 0.085 | 0.014 | 0 | 0.028 | 0 | 0 | 0 | 0 | 0.045 | 0.028 | 0.011 |
| SE | 12637 | ENSG00000090857 | PDPR | chr16 | + | 70146133 | 70146228 | 70144420 | 70144533 | 70148463 | 70148553 | 77 | 41 | 55 | 77 | 89 | 1 | 0 | 2 | 0 | 0 | 90 | 66 | 65 | 64 | 127 | 0 | 5 | 0 | 0 | 1 | 244 | 149 | 0.816216256 | 1 | 0.979 | 1 | 0.944 | 1 | 1 | 1 | 0.89 | 1 | 1 | 0.987 | 0.009 |
| SE | 14859 | ENSG00000185418 | TARS3 | chr15 | - | 101705682 | 101705747 | 101703858 | 101703937 | 101711879 | 101712001 | 22 | 20 | 35 | 36 | 24 | 0 | 0 | 1 | 4 | 0 | 19 | 35 | 13 | 21 | 51 | 0 | 0 | 3 | 0 | 0 | 214 | 149 | 0.816252594 | 1 | 1 | 1 | 0.961 | 0.862 | 1 | 1 | 1 | 0.751 | 1 | 1 | 0.014 |
| SE | 11885 | ENSG00000103222 | ABCC1 | chr16 | + | 16014490 | 16014628 | 16009775 | 16009901 | 16016495 | 16016621 | 258 | 103 | 262 | 274 | 382 | 2 | 1 | 0 | 2 | 3 | 317 | 339 | 301 | 290 | 594 | 0 | 0 | 1 | 12 | 0 | 287 | 149 | 0.8162801 | 1 | 0.985 | 0.982 | 1 | 0.986 | 0.985 | 1 | 1 | 0.994 | 0.926 | 1 | 0.004 |
| SE | 107103 | ENSG00000187109 | NAP1L1 | chr12 | - | 76068908 | 76068994 | 76067370 | 76067473 | 76084566 | 76084666 | 1171 | 1304 | 1194 | 1681 | 2717 | 8 | 4 | 0 | 2 | 0 | 927 | 948 | 995 | 469 | 2188 | 0 | 9 | 3 | 10 | 4 | 235 | 149 | 0.816381959 | 1 | 0.989 | 0.995 | 1 | 0.998 | 1 | 1 | 0.985 | 0.995 | 0.967 | 0.997 | 0.008 |
| SE | 16345 | ENSG00000140497 | SCAMP2 | chr15 | - | 74848599 | 74848645 | 74845472 | 74845593 | 74851352 | 74851481 | 187 | 78 | 133 | 152 | 172 | 0 | 0 | 0 | 0 | 0 | 262 | 207 | 127 | 170 | 397 | 0 | 0 | 0 | 0 | 13 | 195 | 149 | 0.816434561 | 1 | 1 | 1 | 1 | 1 | 1 | 1 | 1 | 1 | 1 | 0.959 | 0.008 |
| SE | 82193 | ENSG00000128595 | CALU | chr7 | + | 128748572 | 128748804 | 128739291 | 128739432 | 128758870 | 128759037 | 37 | 35 | 46 | 50 | 14 | 0 | 0 | 0 | 0 | 0 | 46 | 66 | 80 | 79 | 39 | 0 | 0 | 5 | 0 | 0 | 298 | 149 | 0.816464698 | 1 | 1 | 1 | 1 | 1 | 1 | 1 | 1 | 0.889 | 1 | 1 | 0.022 |
| SE | 28151 | ENSG00000267364 | NA | chr17 | + | 35321734 | 35321865 | 35313495 | 35313644 | 35323147 | 35323260 | 11 | 0 | 21 | 5 | 3 | 10 | 2 | 1 | 5 | 0 | 3 | 9 | 4 | 0 | 6 | 0 | 2 | 6 | 4 | 0 | 280 | 149 | 0.816582163 | 1 | 0.369 | 0 | 0.918 | 0.347 | 1 | 1 | 0.705 | 0.262 | 0 | 1 | -0.067 |
| SE | 90587 | ENSG00000138413 | IDH1 | chr2 | - | 208251429 | 208251567 | 208247381 | 208248660 | 208253885 | 208253959 | 269 | 201 | 179 | 230 | 332 | 0 | 0 | 0 | 1 | 1 | 767 | 517 | 230 | 470 | 302 | 0 | 0 | 0 | 13 | 0 | 287 | 149 | 0.816586051 | 1 | 1 | 1 | 1 | 0.992 | 0.994 | 1 | 1 | 1 | 0.949 | 1 | 0.007 |
| SE | 87451 | ENSG00000100483 | VCPKMT | chr14 | - | 50112614 | 50112719 | 50108632 | 50109713 | 50114284 | 50114404 | 24 | 28 | 28 | 26 | 65 | 7 | 2 | 6 | 2 | 4 | 32 | 39 | 76 | 29 | 5 | 0 | 8 | 5 | 0 | 10 | 254 | 149 | 0.816671251 | 1 | 0.668 | 0.891 | 0.732 | 0.884 | 0.905 | 1 | 0.741 | 0.899 | 1 | 0.227 | 0.043 |
| SE | 8547 | ENSG00000189159 | JPT1 | chr17 | - | 75149090 | 75149185 | 75148528 | 75148671 | 75154341 | 75154675 | 0 | 0 | 0 | 0 | 0 | 150 | 56 | 62 | 112 | 22 | 0 | 0 | 0 | 0 | 14 | 168 | 139 | 123 | 191 | 201 | 244 | 149 | 0.816780978 | 1 | 0 | 0 | 0 | 0 | 0 | 0 | 0 | 0 | 0 | 0.041 | -0.008 |
| SE | 35000 | ENSG00000105993 | DNAJB6 | chr7 | + | 157382245 | 157382377 | 157367372 | 157367483 | 157409794 | 157410001 | 461 | 323 | 316 | 546 | 398 | 5 | 1 | 2 | 3 | 3 | 597 | 511 | 545 | 478 | 1040 | 0 | 1 | 2 | 7 | 12 | 281 | 149 | 0.816797577 | 1 | 0.98 | 0.994 | 0.988 | 0.99 | 0.986 | 1 | 0.996 | 0.993 | 0.973 | 0.979 | -0.001 |
| SE | 65374 | ENSG00000087008 | ACOX3 | chr4 | - | 8374977 | 8375152 | 8366281 | 8367080 | 8381491 | 8381607 | 10 | 12 | 8 | 9 | 14 | 0 | 1 | 3 | 0 | 0 | 9 | 3 | 0 | 8 | 44 | 0 | 0 | 1 | 0 | 0 | 298 | 149 | 0.816984157 | 1 | 1 | 0.857 | 0.571 | 1 | 1 | 1 | 1 | 0 | 1 | 1 | 0.086 |
| SE | 54406 | ENSG00000141127 | PRPSAP2 | chr17 | + | 18865468 | 18865564 | 18859339 | 18859475 | 18865801 | 18865900 | 139 | 83 | 123 | 99 | 180 | 0 | 0 | 0 | 0 | 0 | 166 | 145 | 85 | 99 | 148 | 0 | 0 | 0 | 7 | 0 | 245 | 149 | 0.816987468 | 1 | 1 | 1 | 1 | 1 | 1 | 1 | 1 | 1 | 0.896 | 1 | 0.021 |
| SE | 16526 | ENSG00000140367 | UBE2Q2 | chr15 | + | 75879097 | 75879188 | 75873427 | 75873568 | 75883365 | 75883424 | 247 | 156 | 170 | 191 | 247 | 4 | 3 | 3 | 7 | 13 | 65 | 88 | 73 | 65 | 21 | 0 | 2 | 8 | 4 | 0 | 240 | 149 | 0.816998424 | 1 | 0.975 | 0.97 | 0.972 | 0.944 | 0.922 | 1 | 0.965 | 0.85 | 0.91 | 1 | 0.012 |
| SE | 31977 | ENSG00000082293 | COL19A1 | chr6 | + | 70023626 | 70023680 | 69962825 | 69962870 | 70034244 | 70034298 | 22 | 23 | 35 | 21 | 29 | 0 | 0 | 1 | 0 | 0 | 16 | 54 | 6 | 5 | 11 | 0 | 0 | 0 | 3 | 0 | 203 | 149 | 0.817003454 | 1 | 1 | 1 | 0.963 | 1 | 1 | 1 | 1 | 1 | 0.55 | 1 | 0.083 |
| SE | 3547 | ENSG00000105483 | CARD8 | chr19 | - | 48246074 | 48246381 | 48240961 | 48241063 | 48249522 | 48249633 | 33 | 0 | 18 | 23 | 27 | 202 | 154 | 164 | 221 | 177 | 26 | 3 | 9 | 8 | 20 | 126 | 121 | 87 | 80 | 146 | 298 | 149 | 0.817079323 | 1 | 0.076 | 0 | 0.052 | 0.049 | 0.071 | 0.094 | 0.012 | 0.049 | 0.048 | 0.064 | -0.004 |
| SE | 28173 | ENSG00000138796 | HADH | chr4 | + | 108027938 | 108028069 | 108026063 | 108027760 | 108033175 | 108033292 | 6 | 7 | 8 | 14 | 4 | 46 | 40 | 64 | 54 | 39 | 0 | 15 | 8 | 4 | 0 | 59 | 25 | 15 | 27 | 56 | 280 | 149 | 0.817132299 | 1 | 0.065 | 0.085 | 0.062 | 0.121 | 0.052 | 0 | 0.242 | 0.221 | 0.073 | 0 | -0.03 |
| SE | 13055 | ENSG00000131652 | THOC6 | chr16 | + | 3026062 | 3026166 | 3025923 | 3025988 | 3026250 | 3026288 | 61 | 39 | 50 | 55 | 84 | 0 | 0 | 0 | 0 | 0 | 72 | 36 | 22 | 41 | 81 | 5 | 0 | 0 | 0 | 0 | 253 | 149 | 0.81713935 | 1 | 1 | 1 | 1 | 1 | 1 | 0.895 | 1 | 1 | 1 | 1 | 0.021 |
| SE | 79736 | ENSG00000168255 | POLR2J3 | chr7 | - | 102566996 | 102568184 | 102541500 | 102541662 | 102569834 | 102569924 | 11 | 13 | 16 | 25 | 40 | 0 | 1 | 0 | 0 | 4 | 57 | 28 | 14 | 43 | 45 | 0 | 0 | 0 | 0 | 0 | 298 | 149 | 0.817287887 | 1 | 1 | 0.867 | 1 | 1 | 0.833 | 1 | 1 | 1 | 1 | 1 | -0.06 |
| SE | 90575 | ENSG00000198105 | ZNF248 | chr10 | - | 37856434 | 37856532 | 37856295 | 37856337 | 37857184 | 37857197 | 41 | 16 | 30 | 13 | 27 | 1 | 4 | 3 | 4 | 8 | 6 | 10 | 2 | 2 | 0 | 0 | 1 | 2 | 0 | 6 | 247 | 149 | 0.817313567 | 1 | 0.961 | 0.707 | 0.858 | 0.662 | 0.671 | 1 | 0.858 | 0.376 | 1 | 0 | 0.125 |
| SE | 12249 | ENSG00000059122 | FLYWCH1 | chr16 | + | 2915407 | 2915476 | 2914260 | 2914289 | 2929612 | 2930010 | 0 | 2 | 1 | 0 | 2 | 14 | 17 | 6 | 22 | 6 | 0 | 0 | 0 | 0 | 0 | 4 | 32 | 14 | 24 | 41 | 218 | 149 | 0.817368356 | 1 | 0 | 0.074 | 0.102 | 0 | 0.186 | 0 | 0 | 0 | 0 | 0 | 0.072 |
| SE | 87986 | ENSG00000198373 | WWP2 | chr16 | + | 69871803 | 69871931 | 69840125 | 69840263 | 69888038 | 69888158 | 73 | 43 | 55 | 56 | 60 | 0 | 0 | 0 | 0 | 0 | 110 | 87 | 47 | 92 | 142 | 0 | 0 | 1 | 7 | 0 | 277 | 149 | 0.817400279 | 1 | 1 | 1 | 1 | 1 | 1 | 1 | 1 | 0.962 | 0.876 | 1 | 0.032 |
| SE | 58848 | ENSG00000164828 | SUN1 | chr7 | + | 890927 | 891066 | 887369 | 887519 | 895333 | 896435 | 32 | 14 | 19 | 45 | 23 | 0 | 0 | 0 | 0 | 0 | 29 | 10 | 23 | 22 | 28 | 4 | 0 | 0 | 0 | 0 | 288 | 149 | 0.817407362 | 1 | 1 | 1 | 1 | 1 | 1 | 0.79 | 1 | 1 | 1 | 1 | 0.042 |
| SE | 56925 | ENSG00000269713 | NBPF9 | chr1 | - | 149092754 | 149092785 | 149090752 | 149090894 | 149098437 | 149098706 | 5 | 3 | 10 | 1 | 5 | 5 | 7 | 16 | 9 | 4 | 8 | 4 | 0 | 1 | 4 | 16 | 4 | 3 | 8 | 0 | 180 | 149 | 0.817457506 | 1 | 0.453 | 0.262 | 0.341 | 0.084 | 0.509 | 0.293 | 0.453 | 0 | 0.094 | 1 | -0.038 |
| SE | 23921 | ENSG00000124177 | CHD6 | chr20 | - | 41514804 | 41514952 | 41512845 | 41512995 | 41533049 | 41533285 | 57 | 57 | 84 | 69 | 59 | 0 | 4 | 0 | 0 | 0 | 44 | 89 | 85 | 79 | 18 | 0 | 0 | 5 | 0 | 0 | 297 | 149 | 0.817529615 | 1 | 1 | 1 | 0.913 | 1 | 1 | 1 | 1 | 0.895 | 1 | 1 | 0.004 |
| SE | 6655 | ENSG00000141627 | DYM | chr18 | - | 49118743 | 49118926 | 49097401 | 49097515 | 49163684 | 49163787 | 242 | 196 | 261 | 234 | 540 | 1 | 0 | 1 | 1 | 1 | 191 | 123 | 199 | 213 | 160 | 0 | 7 | 0 | 0 | 0 | 298 | 149 | 0.817575129 | 1 | 0.992 | 1 | 0.992 | 0.992 | 0.996 | 1 | 0.898 | 1 | 1 | 1 | 0.015 |
| SE | 89148 | ENSG00000235531 | MSC-AS1 | chr8 | + | 71965408 | 71965534 | 71962922 | 71963048 | 72052538 | 72052616 | 8 | 19 | 17 | 14 | 12 | 0 | 2 | 0 | 0 | 0 | 36 | 6 | 18 | 3 | 93 | 0 | 0 | 0 | 0 | 6 | 275 | 149 | 0.817612928 | 1 | 1 | 0.837 | 1 | 1 | 1 | 1 | 1 | 1 | 1 | 0.894 | -0.011 |
| SE | 62170 | ENSG00000117625 | RCOR3 | chr1 | + | 211312719 | 211312961 | 211295675 | 211295753 | 211313423 | 211313839 | 39 | 35 | 30 | 51 | 75 | 2 | 5 | 1 | 4 | 3 | 44 | 68 | 23 | 28 | 63 | 8 | 0 | 0 | 0 | 7 | 298 | 149 | 0.817692531 | 1 | 0.907 | 0.778 | 0.938 | 0.864 | 0.926 | 0.733 | 1 | 1 | 1 | 0.818 | -0.028 |
| SE | 17588 | ENSG00000012983 | MAP4K5 | chr14 | - | 50438091 | 50438094 | 50437893 | 50438008 | 50440012 | 50440073 | 48 | 52 | 32 | 62 | 118 | 4 | 13 | 8 | 5 | 0 | 14 | 24 | 29 | 17 | 2 | 1 | 6 | 0 | 7 | 0 | 152 | 149 | 0.817724398 | 1 | 0.922 | 0.797 | 0.797 | 0.924 | 1 | 0.932 | 0.797 | 1 | 0.704 | 1 | 0.001 |
| SE | 44417 | ENSG00000056050 | HPF1 | chr4 | - | 169748743 | 169748842 | 169741956 | 169742107 | 169753675 | 169753835 | 22 | 15 | 53 | 37 | 60 | 2 | 0 | 0 | 0 | 3 | 1 | 24 | 7 | 11 | 4 | 0 | 0 | 0 | 0 | 0 | 248 | 149 | 0.817736438 | 1 | 0.869 | 1 | 1 | 1 | 0.923 | 1 | 1 | 1 | 1 | 1 | -0.042 |
| SE | 95322 | ENSG00000165699 | TSC1 | chr9 | - | 132910570 | 132910689 | 132907300 | 132907370 | 132911001 | 132911113 | 25 | 26 | 28 | 27 | 26 | 0 | 0 | 0 | 4 | 0 | 17 | 19 | 38 | 27 | 33 | 0 | 0 | 0 | 0 | 0 | 268 | 149 | 0.817773572 | 1 | 1 | 1 | 1 | 0.79 | 1 | 1 | 1 | 1 | 1 | 1 | -0.042 |
| SE | 74888 | ENSG00000187037 | GPR141 | chr7 | + | 37685459 | 37685583 | 37683765 | 37683903 | 37709717 | 37709799 | 62 | 28 | 29 | 29 | 73 | 1 | 4 | 0 | 0 | 0 | 17 | 26 | 40 | 40 | 16 | 0 | 0 | 0 | 0 | 0 | 273 | 149 | 0.817825734 | 1 | 0.971 | 0.793 | 1 | 1 | 1 | 1 | 1 | 1 | 1 | 1 | -0.047 |
| SE | 20184 | ENSG00000137210 | TMEM14B | chr6 | + | 10751132 | 10751234 | 10749201 | 10749268 | 10755141 | 10755232 | 19 | 10 | 30 | 27 | 9 | 1 | 0 | 3 | 1 | 0 | 36 | 25 | 19 | 31 | 40 | 0 | 1 | 0 | 5 | 1 | 251 | 149 | 0.817881038 | 1 | 0.919 | 1 | 0.856 | 0.941 | 1 | 1 | 0.937 | 1 | 0.786 | 0.96 | 0.007 |
| SE | 47459 | ENSG00000083635 | NUFIP1 | chr13 | - | 44979189 | 44979266 | 44965843 | 44965936 | 44979889 | 44979952 | 85 | 107 | 99 | 68 | 144 | 0 | 2 | 0 | 0 | 0 | 112 | 85 | 106 | 142 | 57 | 0 | 6 | 0 | 3 | 0 | 226 | 149 | 0.818023844 | 1 | 1 | 0.972 | 1 | 1 | 1 | 1 | 0.903 | 1 | 0.969 | 1 | 0.02 |
| SE | 33873 | ENSG00000132153 | DHX30 | chr3 | + | 47818021 | 47818117 | 47810656 | 47810711 | 47840876 | 47841178 | 26 | 16 | 32 | 32 | 55 | 0 | 0 | 0 | 0 | 0 | 30 | 11 | 25 | 17 | 45 | 0 | 0 | 4 | 0 | 0 | 245 | 149 | 0.818041093 | 1 | 1 | 1 | 1 | 1 | 1 | 1 | 1 | 0.792 | 1 | 1 | 0.042 |
| SE | 34745 | ENSG00000145431 | PDGFC | chr4 | - | 156767772 | 156767876 | 156762802 | 156763206 | 156767961 | 156767990 | 27 | 23 | 17 | 42 | 29 | 2 | 1 | 0 | 2 | 0 | 98 | 15 | 50 | 35 | 72 | 0 | 0 | 0 | 0 | 0 | 253 | 149 | 0.818059814 | 1 | 0.888 | 0.931 | 1 | 0.925 | 1 | 1 | 1 | 1 | 1 | 1 | -0.051 |
| SE | 85635 | ENSG00000108679 | LGALS3BP | chr17 | - | 78979445 | 78979586 | 78977139 | 78977214 | 78979823 | 78979923 | 0 | 0 | 0 | 0 | 0 | 50 | 20 | 37 | 56 | 13 | 13 | 0 | 0 | 0 | 0 | 123 | 146 | 47 | 249 | 105 | 290 | 149 | 0.818083471 | 1 | 0 | 0 | 0 | 0 | 0 | 0.052 | 0 | 0 | 0 | 0 | -0.01 |
| SE | 26508 | ENSG00000114744 | COMMD2 | chr3 | - | 149750256 | 149750382 | 149741389 | 149741718 | 149750677 | 149750851 | 14 | 5 | 19 | 12 | 23 | 155 | 134 | 104 | 142 | 304 | 8 | 9 | 15 | 11 | 0 | 109 | 79 | 114 | 89 | 75 | 275 | 149 | 0.81809183 | 1 | 0.047 | 0.02 | 0.09 | 0.044 | 0.039 | 0.038 | 0.058 | 0.067 | 0.063 | 0 | 0.003 |
| SE | 96397 | ENSG00000197563 | PIGN | chr18 | - | 62157128 | 62157227 | 62148213 | 62148253 | 62157686 | 62157808 | 41 | 50 | 33 | 49 | 51 | 0 | 0 | 0 | 0 | 0 | 4 | 15 | 25 | 31 | 22 | 0 | 0 | 4 | 0 | 0 | 248 | 149 | 0.818126603 | 1 | 1 | 1 | 1 | 1 | 1 | 1 | 1 | 0.79 | 1 | 1 | 0.042 |
| SE | 1134 | ENSG00000128294 | TPST2 | chr22 | - | 26543107 | 26543277 | 26540788 | 26541718 | 26544603 | 26544675 | 0 | 0 | 0 | 0 | 0 | 30 | 14 | 17 | 31 | 2 | 0 | 0 | 1 | 2 | 4 | 22 | 40 | 3 | 18 | 70 | 298 | 149 | 0.818165502 | 1 | 0 | 0 | 0 | 0 | 0 | 0 | 0 | 0.143 | 0.053 | 0.028 | -0.045 |
| SE | 8491 | ENSG00000177885 | GRB2 | chr17 | - | 75393550 | 75393765 | 75332699 | 75332797 | 75405488 | 75405678 | 529 | 350 | 395 | 526 | 307 | 0 | 0 | 1 | 0 | 0 | 1004 | 690 | 444 | 853 | 1524 | 13 | 0 | 4 | 0 | 0 | 298 | 149 | 0.818270292 | 1 | 1 | 1 | 0.995 | 1 | 1 | 0.975 | 1 | 0.982 | 1 | 1 | 0.008 |
| SE | 77966 | ENSG00000173200 | PARP15 | chr3 | + | 122615285 | 122615320 | 122613040 | 122613268 | 122615778 | 122615857 | 13 | 8 | 10 | 10 | 10 | 8 | 5 | 2 | 25 | 7 | 0 | 20 | 8 | 0 | 3 | 4 | 14 | 0 | 7 | 0 | 184 | 149 | 0.81830155 | 1 | 0.568 | 0.564 | 0.802 | 0.245 | 0.536 | 0 | 0.536 | 1 | 0 | 1 | 0.036 |
| SE | 96228 | ENSG00000133943 | DGLUCY | chr14 | + | 91145220 | 91145313 | 91114369 | 91114528 | 91157638 | 91157690 | 27 | 9 | 17 | 23 | 25 | 0 | 8 | 6 | 16 | 11 | 39 | 8 | 6 | 26 | 60 | 10 | 8 | 1 | 15 | 18 | 242 | 149 | 0.818383973 | 1 | 1 | 0.409 | 0.636 | 0.47 | 0.583 | 0.706 | 0.381 | 0.787 | 0.516 | 0.672 | 0.007 |
| SE | 39952 | ENSG00000164253 | WDR41 | chr5 | - | 77449759 | 77449870 | 77440812 | 77440997 | 77451290 | 77451353 | 248 | 108 | 131 | 183 | 318 | 0 | 0 | 0 | 0 | 0 | 64 | 105 | 65 | 159 | 31 | 4 | 0 | 2 | 0 | 0 | 260 | 149 | 0.818432582 | 1 | 1 | 1 | 1 | 1 | 1 | 0.902 | 1 | 0.949 | 1 | 1 | 0.03 |
| SE | 22970 | ENSG00000198612 | COPS8 | chr2 | + | 237089861 | 237089994 | 237088604 | 237088653 | 237094089 | 237094186 | 167 | 68 | 129 | 127 | 180 | 4 | 2 | 2 | 14 | 4 | 115 | 52 | 60 | 76 | 76 | 3 | 4 | 6 | 2 | 0 | 282 | 149 | 0.818432898 | 1 | 0.957 | 0.947 | 0.971 | 0.827 | 0.96 | 0.953 | 0.873 | 0.841 | 0.953 | 1 | 0.008 |
| SE | 48921 | ENSG00000198408 | OGA | chr10 | - | 101793912 | 101793998 | 101792838 | 101792901 | 101797979 | 101798154 | 229 | 157 | 217 | 202 | 342 | 1 | 0 | 0 | 0 | 0 | 38 | 128 | 205 | 115 | 18 | 0 | 3 | 4 | 0 | 0 | 235 | 149 | 0.818437464 | 1 | 0.993 | 1 | 1 | 1 | 1 | 1 | 0.964 | 0.97 | 1 | 1 | 0.012 |
| SE | 32937 | ENSG00000075711 | DLG1 | chr3 | - | 197161639 | 197161738 | 197142717 | 197142768 | 197194424 | 197194589 | 43 | 18 | 39 | 38 | 144 | 30 | 19 | 24 | 28 | 45 | 0 | 32 | 36 | 19 | 7 | 6 | 12 | 39 | 10 | 0 | 248 | 149 | 0.818625697 | 1 | 0.463 | 0.363 | 0.494 | 0.449 | 0.658 | 0 | 0.616 | 0.357 | 0.533 | 1 | -0.016 |
| SE | 61470 | ENSG00000096080 | MRPS18A | chr6 | - | 43678517 | 43678625 | 43675201 | 43675617 | 43681088 | 43681120 | 106 | 62 | 109 | 85 | 159 | 1 | 0 | 3 | 2 | 3 | 101 | 182 | 73 | 136 | 152 | 0 | 2 | 0 | 0 | 0 | 257 | 149 | 0.818642757 | 1 | 0.984 | 1 | 0.955 | 0.961 | 0.968 | 1 | 0.981 | 1 | 1 | 1 | -0.023 |
| SE | 74881 | ENSG00000162585 | FAAP20 | chr1 | - | 2192646 | 2192975 | 2189579 | 2189781 | 2193638 | 2193910 | 3 | 0 | 8 | 4 | 0 | 34 | 3 | 15 | 16 | 3 | 11 | 2 | 2 | 0 | 0 | 0 | 16 | 1 | 25 | 16 | 298 | 149 | 0.818646971 | 1 | 0.042 | 0 | 0.211 | 0.111 | 0 | 1 | 0.059 | 0.5 | 0 | 0 | -0.239 |
| SE | 104066 | ENSG00000072694 | FCGR2B | chr1 | + | 161675256 | 161675313 | 161673959 | 161674073 | 161677327 | 161677365 | 15 | 4 | 8 | 25 | 19 | 7 | 3 | 12 | 2 | 15 | 12 | 18 | 1 | 0 | 9 | 0 | 11 | 2 | 10 | 0 | 206 | 149 | 0.818802788 | 1 | 0.608 | 0.491 | 0.325 | 0.9 | 0.478 | 1 | 0.542 | 0.266 | 0 | 1 | -0.001 |
| SE | 69913 | ENSG00000064995 | TAF11 | chr6 | - | 34880288 | 34880376 | 34879966 | 34880063 | 34882931 | 34883080 | 180 | 123 | 145 | 188 | 169 | 0 | 0 | 0 | 1 | 8 | 217 | 194 | 108 | 140 | 227 | 0 | 0 | 0 | 0 | 0 | 237 | 149 | 0.818810943 | 1 | 1 | 1 | 1 | 0.992 | 0.93 | 1 | 1 | 1 | 1 | 1 | -0.016 |
| SE | 36623 | ENSG00000169180 | XPO6 | chr16 | - | 28177219 | 28177332 | 28175897 | 28176095 | 28211365 | 28211476 | 442 | 275 | 456 | 405 | 1137 | 2 | 0 | 4 | 2 | 13 | 620 | 426 | 710 | 501 | 493 | 0 | 0 | 3 | 0 | 0 | 262 | 149 | 0.818825903 | 1 | 0.992 | 1 | 0.985 | 0.991 | 0.98 | 1 | 1 | 0.993 | 1 | 1 | -0.009 |
| SE | 40174 | ENSG00000115306 | SPTBN1 | chr2 | + | 54637712 | 54637803 | 54632565 | 54632768 | 54642982 | 54643129 | 637 | 501 | 1032 | 741 | 1265 | 0 | 0 | 1 | 0 | 0 | 1092 | 675 | 534 | 812 | 725 | 18 | 0 | 0 | 2 | 0 | 240 | 149 | 0.818906828 | 1 | 1 | 1 | 0.998 | 1 | 1 | 0.974 | 1 | 1 | 0.996 | 1 | 0.006 |
| SE | 73351 | ENSG00000115604 | IL18R1 | chr2 | + | 102373936 | 102373993 | 102371952 | 102372118 | 102375906 | 102376063 | 54 | 30 | 43 | 63 | 66 | 78 | 62 | 69 | 143 | 105 | 0 | 20 | 174 | 6 | 8 | 28 | 27 | 193 | 27 | 0 | 206 | 149 | 0.819111387 | 1 | 0.334 | 0.259 | 0.311 | 0.242 | 0.313 | 0 | 0.349 | 0.395 | 0.138 | 1 | -0.085 |
| SE | 76263 | ENSG00000154839 | SKA1 | chr18 | + | 50380125 | 50380250 | 50375819 | 50375918 | 50382128 | 50382226 | 6 | 1 | 12 | 18 | 26 | 0 | 1 | 0 | 1 | 0 | 71 | 8 | 40 | 42 | 25 | 0 | 0 | 2 | 3 | 0 | 274 | 149 | 0.819189089 | 1 | 1 | 0.352 | 1 | 0.907 | 1 | 1 | 1 | 0.916 | 0.884 | 1 | -0.108 |
| SE | 74168 | ENSG00000116698 | SMG7 | chr1 | + | 183473720 | 183473898 | 183472620 | 183472649 | 183501130 | 183501267 | 5 | 5 | 0 | 9 | 3 | 5 | 0 | 3 | 3 | 0 | 4 | 7 | 5 | 2 | 11 | 3 | 1 | 2 | 13 | 0 | 298 | 149 | 0.819265112 | 1 | 0.333 | 1 | 0 | 0.6 | 1 | 0.4 | 0.778 | 0.556 | 0.071 | 1 | 0.026 |
| SE | 87382 | ENSG00000134982 | APC | chr5 | + | 112823306 | 112823360 | 112821948 | 112821991 | 112827107 | 112827247 | 12 | 19 | 13 | 9 | 17 | 48 | 58 | 28 | 69 | 76 | 9 | 0 | 7 | 3 | 5 | 1 | 38 | 17 | 19 | 10 | 203 | 149 | 0.819290105 | 1 | 0.155 | 0.194 | 0.254 | 0.087 | 0.141 | 0.869 | 0 | 0.232 | 0.104 | 0.268 | -0.128 |
| SE | 41142 | ENSG00000009780 | FAM76A | chr1 | + | 27730231 | 27730333 | 27727471 | 27727536 | 27732602 | 27732657 | 0 | 0 | 3 | 0 | 2 | 37 | 18 | 34 | 15 | 34 | 0 | 0 | 0 | 7 | 0 | 39 | 24 | 26 | 25 | 11 | 251 | 149 | 0.819318753 | 1 | 0 | 0 | 0.05 | 0 | 0.034 | 0 | 0 | 0 | 0.143 | 0 | -0.012 |
| SE | 64465 | ENSG00000233369 | GTF2IP4 | chr7 | + | 73184660 | 73184719 | 73182076 | 73182148 | 73190065 | 73190137 | 85 | 44 | 61 | 51 | 86 | 0 | 1 | 0 | 0 | 0 | 87 | 48 | 57 | 21 | 76 | 0 | 0 | 0 | 4 | 0 | 208 | 149 | 0.819371089 | 1 | 1 | 0.969 | 1 | 1 | 1 | 1 | 1 | 1 | 0.79 | 1 | 0.036 |
| SE | 55838 | ENSG00000188186 | LAMTOR4 | chr7 | + | 100152715 | 100152757 | 100149356 | 100149579 | 100153399 | 100153517 | 5 | 0 | 7 | 2 | 6 | 66 | 39 | 49 | 53 | 8 | 0 | 8 | 19 | 2 | 11 | 69 | 72 | 30 | 92 | 162 | 191 | 149 | 0.819621705 | 1 | 0.056 | 0 | 0.1 | 0.029 | 0.369 | 0 | 0.08 | 0.331 | 0.017 | 0.05 | 0.015 |
| SE | 105220 | ENSG00000102967 | DHODH | chr16 | + | 72015683 | 72015858 | 72014450 | 72014672 | 72017023 | 72017079 | 1 | 0 | 0 | 2 | 0 | 20 | 20 | 23 | 34 | 32 | 0 | 4 | 0 | 0 | 5 | 106 | 37 | 12 | 21 | 22 | 298 | 149 | 0.819725152 | 1 | 0.024 | 0 | 0 | 0.029 | 0 | 0 | 0.051 | 0 | 0 | 0.102 | -0.02 |
| SE | 109169 | ENSG00000197442 | MAP3K5 | chr6 | - | 136650983 | 136651091 | 136637324 | 136637406 | 136656306 | 136656460 | 79 | 76 | 64 | 71 | 192 | 0 | 1 | 0 | 6 | 0 | 69 | 117 | 85 | 95 | 62 | 0 | 4 | 0 | 0 | 0 | 257 | 149 | 0.819748478 | 1 | 1 | 0.978 | 1 | 0.873 | 1 | 1 | 0.944 | 1 | 1 | 1 | -0.019 |
| SE | 71491 | ENSG00000211460 | TSN | chr2 | + | 121757239 | 121757434 | 121755662 | 121755845 | 121758709 | 121758806 | 49 | 11 | 22 | 24 | 22 | 1 | 0 | 0 | 3 | 1 | 39 | 22 | 43 | 42 | 4 | 0 | 0 | 0 | 0 | 0 | 298 | 149 | 0.819787327 | 1 | 0.961 | 1 | 1 | 0.8 | 0.917 | 1 | 1 | 1 | 1 | 1 | -0.064 |
| SE | 95934 | ENSG00000166199 | ALKBH3 | chr11 | + | 43901515 | 43901725 | 43892040 | 43892129 | 43919037 | 43919042 | 119 | 72 | 110 | 75 | 99 | 1 | 1 | 1 | 0 | 9 | 271 | 135 | 72 | 98 | 136 | 0 | 1 | 7 | 7 | 0 | 298 | 149 | 0.819797177 | 1 | 0.983 | 0.973 | 0.982 | 1 | 0.846 | 1 | 0.985 | 0.837 | 0.875 | 1 | 0.017 |
| SE | 66934 | ENSG00000167658 | EEF2 | chr19 | - | 3979807 | 3980066 | 3979328 | 3979436 | 3980513 | 3980709 | 5635 | 2802 | 6553 | 5319 | 7911 | 1 | 2 | 0 | 0 | 0 | 3876 | 4412 | 3062 | 2229 | 6911 | 10 | 6 | 0 | 23 | 20 | 298 | 149 | 0.819847592 | 1 | 1 | 0.999 | 1 | 1 | 1 | 0.995 | 0.997 | 1 | 0.98 | 0.994 | 0.007 |
| SE | 75943 | ENSG00000196730 | DAPK1 | chr9 | + | 87643375 | 87643468 | 87641968 | 87642058 | 87645894 | 87646014 | 184 | 247 | 115 | 153 | 385 | 3 | 2 | 0 | 0 | 4 | 507 | 149 | 141 | 168 | 387 | 0 | 7 | 0 | 7 | 0 | 242 | 149 | 0.819955935 | 1 | 0.974 | 0.987 | 1 | 1 | 0.983 | 1 | 0.929 | 1 | 0.937 | 1 | 0.016 |
| SE | 48595 | ENSG00000131374 | TBC1D5 | chr3 | - | 17340654 | 17340695 | 17307991 | 17308134 | 17372074 | 17372134 | 1 | 0 | 0 | 2 | 5 | 167 | 90 | 119 | 162 | 185 | 0 | 2 | 4 | 8 | 0 | 157 | 117 | 125 | 173 | 51 | 190 | 149 | 0.820013889 | 1 | 0.005 | 0 | 0 | 0.01 | 0.021 | 0 | 0.013 | 0.024 | 0.035 | 0 | -0.007 |
| SE | 99891 | ENSG00000122741 | DCAF10 | chr9 | + | 37860047 | 37860193 | 37854818 | 37854982 | 37861139 | 37861441 | 61 | 47 | 73 | 69 | 88 | 0 | 0 | 0 | 0 | 0 | 14 | 46 | 66 | 46 | 30 | 0 | 0 | 0 | 0 | 4 | 295 | 149 | 0.820113942 | 1 | 1 | 1 | 1 | 1 | 1 | 1 | 1 | 1 | 1 | 0.791 | 0.042 |
| SE | 43312 | ENSG00000009413 | REV3L | chr6 | - | 111367114 | 111368028 | 111363852 | 111363978 | 111372595 | 111376757 | 63 | 51 | 60 | 88 | 61 | 0 | 0 | 0 | 6 | 0 | 11 | 37 | 8 | 10 | 1 | 0 | 0 | 0 | 0 | 0 | 298 | 149 | 0.820178529 | 1 | 1 | 1 | 1 | 0.88 | 1 | 1 | 1 | 1 | 1 | 1 | -0.024 |
| SE | 62778 | ENSG00000163785 | RYK | chr3 | - | 134175608 | 134175768 | 134159236 | 134159373 | 134175929 | 134176039 | 281 | 202 | 258 | 281 | 405 | 0 | 0 | 0 | 2 | 1 | 147 | 197 | 135 | 150 | 68 | 0 | 0 | 7 | 0 | 0 | 298 | 149 | 0.820299104 | 1 | 1 | 1 | 1 | 0.986 | 0.995 | 1 | 1 | 0.906 | 1 | 1 | 0.015 |
| SE | 5208 | ENSG00000124207 | CSE1L | chr20 | + | 49092045 | 49092127 | 49090936 | 49091022 | 49094139 | 49094286 | 185 | 150 | 134 | 137 | 288 | 0 | 1 | 1 | 3 | 2 | 36 | 65 | 32 | 19 | 21 | 0 | 3 | 0 | 2 | 0 | 231 | 149 | 0.820352948 | 1 | 1 | 0.99 | 0.989 | 0.967 | 0.989 | 1 | 0.933 | 1 | 0.86 | 1 | 0.028 |
| SE | 71918 | ENSG00000157637 | SLC38A10 | chr17 | - | 81250069 | 81250093 | 81246884 | 81247061 | 81251492 | 81251612 | 6 | 6 | 3 | 2 | 0 | 105 | 41 | 40 | 57 | 24 | 6 | 0 | 0 | 4 | 15 | 72 | 81 | 62 | 93 | 123 | 173 | 149 | 0.82058 | 1 | 0.047 | 0.112 | 0.061 | 0.029 | 0 | 0.067 | 0 | 0 | 0.036 | 0.095 | 0.01 |
| SE | 56119 | ENSG00000134070 | IRAK2 | chr3 | + | 10200368 | 10200515 | 10177837 | 10178020 | 10209588 | 10209692 | 37 | 9 | 23 | 38 | 9 | 1 | 0 | 3 | 1 | 0 | 37 | 21 | 19 | 72 | 4 | 0 | 0 | 0 | 0 | 0 | 296 | 149 | 0.820615977 | 1 | 0.949 | 1 | 0.794 | 0.95 | 1 | 1 | 1 | 1 | 1 | 1 | -0.061 |
| SE | 13356 | ENSG00000140950 | MEAK7 | chr16 | - | 84497418 | 84497647 | 84495682 | 84495913 | 84497933 | 84498111 | 0 | 3 | 0 | 0 | 2 | 12 | 3 | 8 | 19 | 35 | 0 | 0 | 0 | 0 | 1 | 1 | 37 | 19 | 28 | 16 | 298 | 149 | 0.820625182 | 1 | 0 | 0.333 | 0 | 0 | 0.028 | 0 | 0 | 0 | 0 | 0.03 | 0.066 |
| SE | 83780 | ENSG00000111412 | SPRING1 | chr12 | + | 116720295 | 116720447 | 116719762 | 116719876 | 116723066 | 116723223 | 190 | 95 | 138 | 191 | 163 | 2 | 0 | 0 | 2 | 9 | 66 | 197 | 69 | 68 | 131 | 2 | 1 | 2 | 6 | 0 | 298 | 149 | 0.82062819 | 1 | 0.979 | 1 | 1 | 0.979 | 0.901 | 0.943 | 0.99 | 0.945 | 0.85 | 1 | 0.026 |
| SE | 10108 | ENSG00000108433 | GOSR2 | chr17 | + | 46931104 | 46931207 | 46929519 | 46929584 | 46932066 | 46932199 | 119 | 90 | 108 | 140 | 130 | 0 | 0 | 0 | 2 | 0 | 104 | 84 | 69 | 69 | 126 | 0 | 0 | 0 | 6 | 0 | 252 | 149 | 0.820636331 | 1 | 1 | 1 | 1 | 0.976 | 1 | 1 | 1 | 1 | 0.872 | 1 | 0.021 |
| SE | 45572 | ENSG00000117335 | CD46 | chr1 | + | 207770320 | 207770362 | 207767204 | 207767823 | 207783291 | 207783330 | 620 | 429 | 564 | 636 | 866 | 6 | 2 | 7 | 8 | 13 | 132 | 364 | 368 | 290 | 97 | 0 | 13 | 1 | 2 | 0 | 191 | 149 | 0.82072517 | 1 | 0.988 | 0.994 | 0.984 | 0.984 | 0.981 | 1 | 0.956 | 0.997 | 0.991 | 1 | -0.003 |
| SE | 63693 | ENSG00000131061 | ZNF341 | chr20 | + | 33781290 | 33781387 | 33770083 | 33770292 | 33783731 | 33783864 | 23 | 7 | 21 | 24 | 14 | 0 | 1 | 1 | 0 | 0 | 61 | 16 | 4 | 27 | 31 | 2 | 3 | 0 | 0 | 0 | 246 | 149 | 0.820755589 | 1 | 1 | 0.809 | 0.927 | 1 | 1 | 0.949 | 0.764 | 1 | 1 | 1 | 0.005 |
| SE | 75455 | ENSG00000197081 | IGF2R | chr6 | + | 160029549 | 160029655 | 160027184 | 160027314 | 160032550 | 160032713 | 504 | 148 | 235 | 342 | 271 | 0 | 0 | 0 | 0 | 0 | 243 | 346 | 245 | 233 | 410 | 0 | 0 | 6 | 8 | 0 | 255 | 149 | 0.820839737 | 1 | 1 | 1 | 1 | 1 | 1 | 1 | 1 | 0.96 | 0.945 | 1 | 0.019 |
| SE | 52703 | ENSG00000204054 | LINC00963 | chr9 | + | 129490843 | 129491128 | 129488806 | 129489299 | 129503322 | 129504274 | 3 | 0 | 4 | 2 | 0 | 6 | 6 | 2 | 12 | 5 | 0 | 0 | 6 | 1 | 12 | 10 | 11 | 8 | 6 | 0 | 298 | 149 | 0.820857814 | 1 | 0.2 | 0 | 0.5 | 0.077 | 0 | 0 | 0 | 0.273 | 0.077 | 1 | -0.115 |
| SE | 30386 | ENSG00000120137 | PANK3 | chr5 | - | 168559031 | 168559157 | 168548494 | 168557621 | 168561392 | 168561516 | 137 | 104 | 142 | 125 | 330 | 0 | 0 | 0 | 0 | 0 | 11 | 44 | 45 | 45 | 22 | 0 | 4 | 1 | 0 | 0 | 275 | 149 | 0.820893543 | 1 | 1 | 1 | 1 | 1 | 1 | 1 | 0.856 | 0.961 | 1 | 1 | 0.037 |
| SE | 4448 | ENSG00000105058 | FAM32A | chr19 | + | 16190519 | 16190573 | 16189387 | 16189511 | 16190886 | 16191378 | 50 | 31 | 68 | 96 | 143 | 0 | 0 | 0 | 0 | 0 | 104 | 88 | 69 | 89 | 98 | 0 | 0 | 0 | 0 | 7 | 203 | 149 | 0.82104395 | 1 | 1 | 1 | 1 | 1 | 1 | 1 | 1 | 1 | 1 | 0.911 | 0.018 |
| SE | 24999 | ENSG00000099940 | SNAP29 | chr22 | + | 20883470 | 20883569 | 20881048 | 20881134 | 20887678 | 20887819 | 224 | 208 | 122 | 270 | 239 | 0 | 0 | 0 | 0 | 0 | 724 | 211 | 187 | 374 | 724 | 9 | 0 | 0 | 7 | 0 | 248 | 149 | 0.82113731 | 1 | 1 | 1 | 1 | 1 | 1 | 0.98 | 1 | 1 | 0.97 | 1 | 0.01 |
| SE | 90944 | ENSG00000185803 | SLC52A2 | chr8 | + | 144360589 | 144360713 | 144359622 | 144360493 | 144360802 | 144360879 | 7 | 3 | 1 | 4 | 2 | 3 | 0 | 0 | 0 | 0 | 22 | 13 | 2 | 3 | 15 | 0 | 0 | 0 | 0 | 0 | 273 | 149 | 0.821245429 | 1 | 0.56 | 1 | 1 | 1 | 1 | 1 | 1 | 1 | 1 | 1 | -0.088 |
| SE | 4950 | ENSG00000101347 | SAMHD1 | chr20 | - | 36935028 | 36935189 | 36930759 | 36930875 | 36941038 | 36941111 | 3629 | 2726 | 2437 | 3217 | 3840 | 5 | 1 | 6 | 6 | 2 | 6273 | 2143 | 1048 | 3386 | 2324 | 5 | 5 | 15 | 0 | 12 | 298 | 149 | 0.821283208 | 1 | 0.997 | 0.999 | 0.995 | 0.996 | 0.999 | 0.998 | 0.995 | 0.972 | 1 | 0.99 | 0.006 |
| SE | 68707 | ENSG00000196295 | GARS1-DT | chr7 | - | 30552099 | 30552179 | 30551317 | 30551479 | 30561465 | 30561487 | 21 | 21 | 11 | 16 | 21 | 0 | 0 | 0 | 0 | 0 | 20 | 8 | 6 | 9 | 32 | 0 | 0 | 3 | 0 | 0 | 229 | 149 | 0.821298226 | 1 | 1 | 1 | 1 | 1 | 1 | 1 | 1 | 0.565 | 1 | 1 | 0.087 |
| SE | 59796 | ENSG00000166377 | ATP9B | chr18 | + | 79077981 | 79078157 | 79069284 | 79069529 | 79096475 | 79096481 | 5 | 2 | 6 | 1 | 0 | 21 | 12 | 13 | 21 | 10 | 0 | 0 | 3 | 0 | 14 | 68 | 19 | 18 | 15 | 10 | 298 | 149 | 0.821317518 | 1 | 0.106 | 0.077 | 0.188 | 0.023 | 0 | 0 | 0 | 0.077 | 0 | 0.412 | -0.019 |
| SE | 107714 | ENSG00000177575 | CD163 | chr12 | - | 7481160 | 7481256 | 7479859 | 7479996 | 7482642 | 7482762 | 157 | 86 | 58 | 164 | 131 | 5 | 3 | 2 | 9 | 1 | 611 | 982 | 631 | 394 | 206 | 21 | 25 | 0 | 31 | 0 | 245 | 149 | 0.82134666 | 1 | 0.95 | 0.946 | 0.946 | 0.917 | 0.988 | 0.947 | 0.96 | 1 | 0.885 | 1 | -0.009 |
| SE | 19972 | ENSG00000136758 | YME1L1 | chr10 | - | 27153226 | 27153281 | 27148905 | 27149040 | 27154177 | 27154371 | 3 | 6 | 1 | 2 | 2 | 137 | 99 | 132 | 152 | 191 | 0 | 0 | 0 | 0 | 0 | 28 | 178 | 129 | 103 | 48 | 204 | 149 | 0.821382974 | 1 | 0.016 | 0.042 | 0.006 | 0.01 | 0.008 | 0 | 0 | 0 | 0 | 0 | 0.016 |
| SE | 51537 | ENSG00000176393 | RNPEP | chr1 | + | 202001375 | 202001488 | 201999901 | 202000015 | 202001658 | 202001722 | 239 | 139 | 171 | 212 | 353 | 0 | 0 | 0 | 0 | 0 | 431 | 306 | 238 | 386 | 666 | 13 | 0 | 0 | 0 | 0 | 262 | 149 | 0.821414653 | 1 | 1 | 1 | 1 | 1 | 1 | 0.95 | 1 | 1 | 1 | 1 | 0.01 |
| SE | 99224 | ENSG00000086848 | ALG9 | chr11 | - | 111805093 | 111805315 | 111786283 | 111786520 | 111809642 | 111809709 | 0 | 7 | 6 | 2 | 6 | 28 | 22 | 43 | 30 | 53 | 0 | 0 | 10 | 17 | 1 | 30 | 62 | 51 | 68 | 40 | 298 | 149 | 0.821442058 | 1 | 0 | 0.137 | 0.065 | 0.032 | 0.054 | 0 | 0 | 0.089 | 0.111 | 0.012 | 0.015 |

| | | | | | | | | | | | | | | | | | | | | | | | | | | | | | | | | | | | | | | | | | | | | | | |
|---|---|---|---|---|---|---|---|---|---|---|---|---|---|---|---|---|---|---|---|---|---|---|---|---|---|---|---|---|---|---|---|---|---|---|---|---|---|---|---|---|---|---|---|---|---|---|
| SE | 82057 | ENSG00000104133 | SPG11 | chr15 | - | 44629232 | 44629388 | 44627635 | 44628844 | 44633504 | 44633637 | 588 | 351 | 426 | 523 | 913 | 0 | 0 | 0 | 0 | 0 | 146 | 474 | 412 | 336 | 25 | 0 | 0 | 0 | 10 | 0 | 298 | 149 | 0.821481421 | 1 | 1 | 1 | 1 | 1 | 1 | 1 | 1 | 1 | 0.944 | 1 | 0.011 |
| SE | 79311 | ENSG00000001631 | KRIT1 | chr7 | - | 92245426 | 92245595 | 92242033 | 92242137 | 92245905 | 92246100 | 2 | 1 | 13 | 13 | 1 | 0 | 0 | 3 | 0 | 2 | 7 | 3 | 7 | 3 | 2 | 6 | 0 | 0 | 0 | 0 | 298 | 149 | 0.821519759 | 1 | 1 | 1 | 0.684 | 1 | 0.2 | 0.368 | 1 | 1 | 1 | 1 | -0.097 |
| SE | 84603 | ENSG00000245958 | SEPTIN7P14 | chr4 | + | 119488644 | 119488717 | 119462895 | 119462979 | 119493332 | 119493437 | 25 | 21 | 13 | 25 | 39 | 60 | 51 | 52 | 62 | 87 | 18 | 15 | 19 | 13 | 29 | 49 | 55 | 72 | 21 | 53 | 222 | 149 | 0.821566073 | 1 | 0.219 | 0.217 | 0.144 | 0.213 | 0.231 | 0.198 | 0.155 | 0.15 | 0.294 | 0.269 | -0.008 |
| SE | 1049 | ENSG00000100099 | HPS4 | chr22 | - | 26472831 | 26472939 | 26472301 | 26472418 | 26476992 | 26477011 | 88 | 30 | 91 | 51 | 77 | 0 | 0 | 0 | 0 | 0 | 41 | 72 | 36 | 47 | 29 | 0 | 0 | 1 | 4 | 0 | 257 | 149 | 0.821583379 | 1 | 1 | 1 | 1 | 1 | 1 | 1 | 1 | 0.954 | 0.872 | 1 | 0.035 |
| SE | 106695 | ENSG00000141376 | BCAS3 | chr17 | + | 60688931 | 60689046 | 60683981 | 60684036 | 60689685 | 60689761 | 5 | 10 | 2 | 7 | 19 | 107 | 57 | 70 | 61 | 87 | 17 | 0 | 8 | 4 | 10 | 87 | 89 | 55 | 31 | 65 | 264 | 149 | 0.821704735 | 1 | 0.026 | 0.09 | 0.016 | 0.061 | 0.11 | 0.099 | 0 | 0.076 | 0.068 | 0.08 | -0.004 |
| SE | 46637 | ENSG00000141556 | TBCD | chr17 | + | 82937270 | 82937360 | 82932657 | 82932735 | 82938048 | 82938136 | 52 | 17 | 34 | 25 | 39 | 0 | 0 | 2 | 0 | 3 | 30 | 28 | 23 | 46 | 77 | 0 | 0 | 0 | 0 | 0 | 239 | 149 | 0.821768692 | 1 | 1 | 1 | 0.914 | 1 | 0.89 | 1 | 1 | 1 | 1 | 1 | -0.039 |
| SE | 44023 | ENSG00000143179 | UCK2 | chr1 | + | 165896189 | 165896332 | 165891225 | 165891322 | 165903181 | 165903279 | 33 | 33 | 44 | 47 | 60 | 1 | 0 | 2 | 2 | 0 | 69 | 47 | 29 | 66 | 45 | 0 | 0 | 0 | 0 | 0 | 292 | 149 | 0.82177281 | 1 | 0.944 | 1 | 0.918 | 0.923 | 1 | 1 | 1 | 1 | 1 | 1 | -0.043 |
| SE | 26515 | ENSG00000204859 | ZBTB48 | chr1 | + | 6585918 | 6586030 | 6582057 | 6582299 | 6587204 | 6587291 | 7 | 12 | 4 | 8 | 24 | 0 | 0 | 0 | 3 | 0 | 9 | 19 | 4 | 5 | 32 | 0 | 0 | 3 | 0 | 0 | 261 | 149 | 0.821996993 | 1 | 1 | 1 | 1 | 0.604 | 1 | 1 | 1 | 0.432 | 1 | 1 | 0.034 |
| SE | 72079 | ENSG00000180228 | PRKRA | chr2 | - | 178436144 | 178436319 | 178431291 | 178432254 | 178441609 | 178441704 | 207 | 184 | 221 | 192 | 434 | 6 | 0 | 0 | 6 | 0 | 217 | 165 | 199 | 140 | 93 | 0 | 0 | 0 | 0 | 0 | 298 | 149 | 0.822057524 | 1 | 0.945 | 1 | 1 | 0.941 | 1 | 1 | 1 | 1 | 1 | 1 | -0.023 |
| SE | 20844 | ENSG00000255819 | KLRC4-KLRK1 | chr12 | - | 10389942 | 10390027 | 10388770 | 10388875 | 10407635 | 10407766 | 545 | 252 | 289 | 448 | 422 | 55 | 26 | 40 | 84 | 83 | 58 | 231 | 166 | 136 | 22 | 29 | 25 | 24 | 9 | 1 | 234 | 149 | 0.822103824 | 1 | 0.863 | 0.861 | 0.821 | 0.773 | 0.764 | 0.56 | 0.855 | 0.815 | 0.906 | 0.933 | 0.003 |
| SE | 25953 | ENSG00000119522 | DENND1A | chr9 | - | 123769513 | 123769563 | 123757702 | 123757822 | 123878950 | 123879021 | 18 | 18 | 17 | 25 | 40 | 0 | 0 | 0 | 0 | 0 | 49 | 40 | 23 | 46 | 61 | 0 | 0 | 0 | 0 | 7 | 199 | 149 | 0.822104974 | 1 | 1 | 1 | 1 | 1 | 1 | 1 | 1 | 1 | 1 | 0.867 | 0.027 |
| SE | 64597 | ENSG00000100644 | HIF1A | chr14 | + | 61720381 | 61720572 | 61695512 | 61695839 | 61721508 | 61721654 | 406 | 329 | 309 | 439 | 583 | 0 | 0 | 0 | 0 | 0 | 287 | 522 | 531 | 223 | 189 | 0 | 0 | 10 | 5 | 0 | 298 | 149 | 0.822113395 | 1 | 1 | 1 | 1 | 1 | 1 | 1 | 1 | 0.964 | 0.957 | 1 | 0.016 |
| SE | 53032 | ENSG00000197183 | NOL4L | chr20 | - | 32456117 | 32456395 | 32453575 | 32453761 | 32474600 | 32474735 | 89 | 44 | 84 | 88 | 51 | 0 | 0 | 0 | 0 | 0 | 21 | 33 | 61 | 84 | 92 | 0 | 0 | 0 | 5 | 0 | 298 | 149 | 0.822138238 | 1 | 1 | 1 | 1 | 1 | 1 | 1 | 1 | 1 | 0.894 | 1 | 0.021 |
| SE | 69349 | ENSG00000123472 | ATPAF1 | chr1 | - | 46643193 | 46643301 | 46632736 | 46635970 | 46645160 | 46645256 | 159 | 144 | 170 | 150 | 250 | 0 | 1 | 0 | 0 | 8 | 194 | 134 | 157 | 203 | 213 | 0 | 0 | 3 | 0 | 0 | 257 | 149 | 0.822147624 | 1 | 1 | 0.988 | 1 | 1 | 0.948 | 1 | 1 | 0.968 | 1 | 1 | -0.006 |
| SE | 46911 | ENSG00000055609 | KMT2C | chr7 | - | 152265037 | 152265209 | 152262701 | 152263130 | 152273704 | 152273867 | 400 | 344 | 345 | 495 | 341 | 1 | 0 | 0 | 0 | 2 | 262 | 402 | 362 | 281 | 299 | 0 | 12 | 0 | 0 | 0 | 298 | 149 | 0.822203973 | 1 | 0.995 | 1 | 1 | 1 | 0.988 | 1 | 0.944 | 1 | 1 | 1 | 0.008 |
| SE | 9971 | ENSG00000170921 | TANC2 | chr17 | + | 63151269 | 63151380 | 63099174 | 63099357 | 63193990 | 63194082 | 3 | 1 | 0 | 1 | 3 | 45 | 10 | 19 | 24 | 31 | 0 | 0 | 2 | 0 | 6 | 8 | 9 | 21 | 27 | 0 | 260 | 149 | 0.822216606 | 1 | 0.037 | 0.054 | 0 | 0.023 | 0.053 | 0 | 0 | 0.052 | 0 | 1 | -0.177 |
| SE | 79739 | ENSG00000164808 | SPIDR | chr8 | + | 47284027 | 47284094 | 47279861 | 47280017 | 47291032 | 47291137 | 164 | 84 | 110 | 136 | 87 | 1 | 1 | 0 | 0 | 0 | 173 | 207 | 120 | 127 | 92 | 0 | 0 | 1 | 0 | 7 | 216 | 149 | 0.822244546 | 1 | 0.991 | 0.983 | 1 | 1 | 1 | 1 | 1 | 0.988 | 1 | 0.901 | 0.017 |
| SE | 18805 | ENSG00000119689 | DLST | chr14 | + | 74885585 | 74885634 | 74882590 | 74882624 | 74889094 | 74889147 | 159 | 43 | 107 | 131 | 28 | 0 | 0 | 0 | 0 | 0 | 73 | 138 | 77 | 96 | 245 | 0 | 7 | 0 | 0 | 4 | 198 | 149 | 0.822298843 | 1 | 1 | 1 | 1 | 1 | 1 | 1 | 0.937 | 1 | 1 | 0.979 | 0.017 |
| SE | 56224 | ENSG00000106462 | EZH2 | chr7 | - | 148836821 | 148837000 | 148832633 | 148832750 | 148846496 | 148846598 | 4 | 7 | 7 | 6 | 4 | 6 | 2 | 1 | 3 | 6 | 17 | 5 | 5 | 9 | 10 | 0 | 6 | 33 | 25 | 0 | 298 | 149 | 0.822312799 | 1 | 0.25 | 0.636 | 0.778 | 0.5 | 0.25 | 1 | 0.294 | 0.07 | 0.153 | 1 | -0.021 |
| SE | 50778 | ENSG00000257698 | GIHCG | chr12 | - | 57935469 | 57935592 | 57931294 | 57931737 | 57935838 | 57936166 | 21 | 8 | 15 | 7 | 6 | 0 | 0 | 0 | 3 | 0 | 3 | 4 | 3 | 13 | 3 | 0 | 0 | 1 | 0 | 0 | 272 | 149 | 0.822354268 | 1 | 1 | 1 | 1 | 0.561 | 1 | 1 | 1 | 0.622 | 1 | 1 | -0.012 |
| SE | 73917 | ENSG00000161692 | DBF4B | chr17 | + | 44729904 | 44730096 | 44722879 | 44723022 | 44730964 | 44731015 | 20 | 11 | 37 | 17 | 18 | 0 | 0 | 0 | 0 | 0 | 49 | 26 | 2 | 33 | 27 | 0 | 0 | 1 | 3 | 0 | 298 | 149 | 0.822406712 | 1 | 1 | 1 | 1 | 1 | 1 | 1 | 1 | 0.5 | 0.846 | 1 | 0.131 |
| SE | 45975 | ENSG00000170248 | PDCD6IP | chr3 | + | 33826479 | 33826580 | 33825186 | 33825340 | 33828852 | 33828969 | 367 | 288 | 321 | 381 | 568 | 0 | 4 | 4 | 6 | 2 | 203 | 301 | 263 | 232 | 83 | 0 | 4 | 2 | 0 | 0 | 250 | 149 | 0.822474239 | 1 | 1 | 0.977 | 0.98 | 0.974 | 0.994 | 1 | 0.978 | 0.987 | 1 | 1 | -0.008 |
| SE | 39398 | ENSG00000160712 | IL6R | chr1 | + | 154435968 | 154436110 | 154434989 | 154435156 | 154445046 | 154445094 | 208 | 162 | 217 | 179 | 165 | 4 | 0 | 1 | 6 | 0 | 221 | 124 | 71 | 158 | 185 | 0 | 0 | 0 | 0 | 0 | 291 | 149 | 0.822487962 | 1 | 0.964 | 1 | 0.991 | 0.939 | 1 | 1 | 1 | 1 | 1 | 1 | -0.021 |
| SE | 101861 | ENSG00000144815 | NXPE3 | chr3 | + | 101807052 | 101807126 | 101801234 | 101801989 | 101816794 | 101816904 | 121 | 115 | 118 | 161 | 118 | 6 | 0 | 3 | 3 | 2 | 104 | 95 | 146 | 142 | 68 | 9 | 0 | 3 | 0 | 0 | 223 | 149 | 0.822489415 | 1 | 0.931 | 1 | 0.963 | 0.973 | 0.975 | 0.885 | 1 | 0.97 | 1 | 1 | -0.003 |
| SE | 44565 | ENSG00000026508 | CD44 | chr11 | + | 35198120 | 35198246 | 35189834 | 35190065 | 35214851 | 35214914 | 21 | 11 | 5 | 7 | 27 | 560 | 408 | 542 | 461 | 519 | 44 | 23 | 20 | 14 | 0 | 1081 | 515 | 558 | 618 | 653 | 275 | 149 | 0.822578133 | 1 | 0.02 | 0.014 | 0.005 | 0.008 | 0.027 | 0.022 | 0.024 | 0.019 | 0.012 | 0 | -0.001 |
| SE | 68770 | ENSG00000100523 | DDHD1 | chr14 | - | 53051843 | 53051927 | 53036744 | 53046949 | 53054437 | 53054629 | 44 | 16 | 37 | 29 | 33 | 15 | 16 | 14 | 8 | 5 | 20 | 13 | 15 | 9 | 2 | 7 | 6 | 15 | 2 | 0 | 233 | 149 | 0.822662156 | 1 | 0.652 | 0.39 | 0.628 | 0.699 | 0.808 | 0.646 | 0.581 | 0.39 | 0.742 | 1 | -0.036 |
| SE | 31546 | ENSG00000171608 | PIK3CD | chr1 | + | 9716958 | 9717108 | 9716275 | 9716619 | 9718693 | 9718915 | 117 | 31 | 74 | 99 | 84 | 0 | 0 | 0 | 0 | 0 | 86 | 77 | 45 | 83 | 83 | 7 | 0 | 0 | 0 | 0 | 298 | 149 | 0.822707634 | 1 | 1 | 1 | 1 | 1 | 1 | 0.86 | 1 | 1 | 1 | 1 | 0.028 |
| SE | 31583 | ENSG00000129071 | MBD4 | chr3 | - | 129436460 | 129437308 | 129434061 | 129434136 | 129437719 | 129437950 | 434 | 333 | 378 | 464 | 776 | 2 | 5 | 5 | 4 | 1 | 183 | 389 | 355 | 377 | 120 | 0 | 0 | 3 | 2 | 0 | 298 | 149 | 0.822756203 | 1 | 0.991 | 0.971 | 0.974 | 0.983 | 0.997 | 1 | 1 | 0.983 | 0.99 | 1 | -0.011 |
| SE | 84189 | ENSG00000137642 | SORL1 | chr11 | + | 121605401 | 121605571 | 121605112 | 121605239 | 121606844 | 121606957 | 1171 | 598 | 918 | 1001 | 1565 | 0 | 1 | 2 | 0 | 0 | 426 | 428 | 882 | 568 | 393 | 0 | 0 | 4 | 3 | 10 | 298 | 149 | 0.822839086 | 1 | 1 | 0.997 | 0.996 | 1 | 1 | 1 | 1 | 0.991 | 0.99 | 0.952 | 0.012 |
| SE | 46200 | ENSG00000117533 | VAMP4 | chr1 | - | 171719170 | 171719218 | 171710713 | 171710814 | 171728523 | 171728570 | 22 | 26 | 30 | 32 | 45 | 0 | 0 | 0 | 0 | 0 | 5 | 14 | 9 | 12 | 16 | 3 | 0 | 0 | 0 | 0 | 197 | 149 | 0.822857698 | 1 | 1 | 1 | 1 | 1 | 1 | 0.558 | 1 | 1 | 1 | 1 | 0.088 |
| SE | 23441 | ENSG00000166888 | STAT6 | chr12 | - | 57099293 | 57099440 | 57099014 | 57099078 | 57099766 | 57099794 | 930 | 573 | 802 | 983 | 1385 | 5 | 1 | 2 | 2 | 4 | 1599 | 1108 | 564 | 948 | 2098 | 0 | 11 | 7 | 2 | 7 | 296 | 149 | 0.822884363 | 1 | 0.989 | 0.997 | 0.995 | 0.996 | 0.994 | 1 | 0.981 | 0.976 | 0.996 | 0.993 | 0.005 |
| SE | 86003 | ENSG00000244754 | N4BP2L2 | chr13 | - | 32522181 | 32522270 | 32517874 | 32518003 | 32527407 | 32527532 | 263 | 161 | 184 | 219 | 543 | 4 | 2 | 0 | 3 | 0 | 79 | 118 | 110 | 41 | 146 | 0 | 0 | 0 | 6 | 0 | 238 | 149 | 0.823009668 | 1 | 0.976 | 0.981 | 1 | 0.979 | 1 | 1 | 1 | 1 | 0.811 | 1 | 0.025 |
| [illegible] | | | | | | | | | | | | | | | | | | | | | | | | | | | | | | | | | | | | | | | | | | | | | | |

| | | | | | | | | | | | |
|---|---|---|---|---|---|---|---|---|---|---|---|
| SE | 14059 | ENSG00000140694 | PARN | chr16 | - | 14617589 | 14617650 | 14610643 | 14610809 | 14627105 | 14627136 |
| SE | 45739 | ENSG00000058600 | POLR3E | chr16 | + | 22305378 | 22305494 | 22305155 | 22305206 | 22308147 | 22308225 |
| SE | 29073 | ENSG00000163625 | WDFY3 | chr4 | - | 84726860 | 84726911 | 84724425 | 84724594 | 84733381 | 84733609 |
| SE | 29016 | ENSG00000123352 | SPATS2 | chr12 | + | 49486249 | 49486348 | 49484589 | 49484669 | 49489464 | 49489573 |
| SE | 58169 | ENSG00000172965 | MIR4435-2HG | chr2 | - | 111207719 | 111208661 | 111196998 | 111197194 | 111344060 | 111344237 |
| SE | 50182 | ENSG00000185808 | PIGP | chr21 | - | 37067261 | 37067380 | 37062845 | 37065712 | 37069551 | 37069624 |
| SE | 7931 | ENSG00000108479 | GALK1 | chr17 | - | 75762703 | 75763439 | 75758590 | 75758599 | 75763896 | 75764086 |
| SE | 15871 | ENSG00000186654 | PRR5 | chr22 | + | 44732250 | 44732391 | 44730883 | 44731821 | 44736771 | 44737662 |
| SE | 22151 | ENSG00000156639 | ZFAND3 | chr6 | + | 38116571 | 38116739 | 38061592 | 38061775 | 38152234 | 38152844 |
| SE | 6052 | ENSG00000149639 | SOGA1 | chr20 | - | 36817406 | 36817469 | 36815125 | 36816307 | 36829053 | 36829212 |
| SE | 103308 | ENSG00000134452 | FBH1 | chr10 | + | 5891018 | 5891116 | 5890279 | 5890346 | 5903019 | 5903175 |
| SE | 35887 | ENSG00000197415 | VEPH1 | chr3 | - | 157413880 | 157414090 | 157381155 | 157381376 | 157428321 | 157428488 |
| SE | 24418 | ENSG00000055163 | CYFIP2 | chr5 | + | 157327972 | 157328049 | 157325481 | 157325638 | 157330741 | 157330850 |
| SE | 35055 | ENSG00000125868 | DSTN | chr20 | + | 17604554 | 17604631 | 17600737 | 17601045 | 17607036 | 17608242 |
| SE | 60103 | ENSG00000116288 | PARK7 | chr1 | + | 7965323 | 7965425 | 7962762 | 7962875 | 7969344 | 7969404 |
| SE | 89253 | ENSG00000104325 | DECR1 | chr8 | + | 90005291 | 90005369 | 90001475 | 90001561 | 90005447 | 90005845 |
| SE | 100002 | ENSG00000116898 | MRPS15 | chr1 | - | 36460691 | 36460776 | 36457922 | 36457981 | 36461263 | 36461312 |
| SE | 79781 | ENSG00000198846 | TOX | chr8 | - | 58959942 | 58960008 | 58939301 | 58939544 | 59118885 | 59119147 |
| SE | 99583 | ENSG00000149187 | CELF1 | chr11 | - | 47476845 | 47476956 | 47475335 | 47475521 | 47477296 | 47477425 |
| SE | 52492 | ENSG00000283199 | C13orf46 | chr13 | - | 113968466 | 113968514 | 113964926 | 113964994 | 113968594 | 113968760 |
| SE | 98003 | ENSG00000205758 | CRYZL1 | chr21 | - | 33631485 | 33631557 | 33624682 | 33624760 | 33641680 | 33641720 |
| SE | 92608 | ENSG00000102699 | PARP4 | chr13 | - | 24500315 | 24500382 | 24499300 | 24499376 | 24501632 | 24501834 |
| SE | 42710 | ENSG00000066557 | LRRC40 | chr1 | - | 70181085 | 70181209 | 70178850 | 70178993 | 70184784 | 70184914 |
| SE | 43914 | ENSG00000165029 | ABCA1 | chr9 | - | 104786872 | 104786976 | 104786297 | 104786390 | 104787919 | 104788054 |
| SE | 79150 | ENSG00000104299 | INTS9 | chr8 | - | 28859435 | 28859563 | 28850008 | 28850273 | 28889873 | 28889909 |
| SE | 20822 | ENSG00000119899 | SLC17A5 | chr6 | - | 73638411 | 73638499 | 73636620 | 73636707 | 73641690 | 73641924 |
| SE | 26095 | ENSG00000285000 | NA | chr5 | + | 77039530 | 77039719 | 77036555 | 77036720 | 77046346 | 77046677 |
| SE | 6600 | ENSG00000134444 | RELCH | chr18 | + | 62280359 | 62280442 | 62275373 | 62275473 | 62280645 | 62280709 |
| SE | 64192 | ENSG00000171824 | EXOSC10 | chr1 | - | 11077329 | 11077443 | 11076841 | 11076948 | 11077600 | 11077651 |
| SE | 84073 | ENSG00000197157 | SND1 | chr7 | + | 128089488 | 128089692 | 128086470 | 128087051 | 128091836 | 128091881 |
| SE | 38053 | ENSG00000215908 | CROCCP2 | chr1 | - | 16642766 | 16642850 | 16635026 | 16635168 | 16643027 | 16643145 |
| SE | 38155 | ENSG00000138750 | NUP54 | chr4 | - | 76144148 | 76144292 | 76136185 | 76136412 | 76144389 | 76144473 |
| SE | 17925 | ENSG00000197249 | SERPINA1 | chr14 | - | 94388559 | 94388660 | 94382591 | 94383241 | 94388800 | 94388847 |
| SE | 5510 | ENSG00000101294 | HM13 | chr20 | + | 31554745 | 31554829 | 31550063 | 31550121 | 31559610 | 31559647 |
| SE | 29768 | ENSG00000133065 | SLC41A1 | chr1 | - | 205794869 | 205795018 | 205789093 | 205791718 | 205795343 | 205795478 |
| SE | 31009 | ENSG00000163694 | RBM47 | chr4 | - | 40437770 | 40438924 | 40432650 | 40432862 | 40466576 | 40466699 |
| SE | 92361 | ENSG00000143437 | ARNT | chr1 | - | 150842423 | 150842468 | 150836279 | 150836493 | 150846262 | 150846307 |
| SE | 66857 | ENSG00000082781 | ITGB5 | chr3 | - | 124817620 | 124817710 | 124809021 | 124809156 | 124821312 | 124821458 |
| SE | 33046 | ENSG00000239697 | TNFSF12 | chr17 | + | 7553586 | 7553825 | 7550942 | 7550978 | 7556777 | 7556902 |
| SE | 40060 | ENSG00000125347 | IRF1 | chr5 | - | 132484361 | 132484497 | 132481608 | 132484075 | 132485666 | 132485716 |
| SE | 76269 | ENSG00000065427 | KARS1 | chr16 | - | 75640183 | 75640349 | 75636453 | 75636547 | 75641563 | 75641723 |
| SE | 55863 | ENSG00000177200 | CHD9 | chr16 | + | 53262986 | 53263097 | 53255599 | 53255779 | 53267293 | 53267490 |
| SE | 78101 | ENSG00000141258 | SGSM2 | chr17 | + | 2367270 | 2367405 | 2365214 | 2365341 | 2371261 | 2371415 |
| SE | 29373 | ENSG00000101161 | PRPF6 | chr20 | + | 64011284 | 64011503 | 64010199 | 64010318 | 64016722 | 64016845 |
| SE | 52691 | ENSG00000196235 | SUPT5H | chr19 | + | 39458293 | 39458305 | 39457674 | 39457740 | 39458817 | 39458887 |
| SE | 44590 | ENSG00000026508 | CD44 | chr11 | + | 35209964 | 35210054 | 35189834 | 35190065 | 35214851 | 35214914 |
| SE | 106579 | ENSG00000166012 | TAF1D | chr11 | - | 93731149 | 93731216 | 93730676 | 93730712 | 93734624 | 93734660 |
| SE | 36747 | ENSG00000173193 | PARP14 | chr3 | + | 122703741 | 122703978 | 122699389 | 122701635 | 122708189 | 122708268 |
| SE | 63298 | ENSG00000138463 | SLC49A4 | chr3 | + | 122826799 | 122827065 | 122806856 | 122806950 | 122833316 | 122833446 |
| SE | 61067 | ENSG00000235194 | PPP1R3E | chr14 | - | 23300106 | 23300325 | 23299400 | 23299541 | 23300757 | 23300864 |
| SE | 18636 | ENSG00000256500 | NA | chr14 | + | 103662115 | 103662194 | 103657545 | 103657776 | 103662701 | 103662927 |
| SE | 42020 | ENSG00000163931 | TKT | chr3 | - | 53237796 | 53237921 | 53234982 | 53235174 | 53240250 | 53240348 |
| SE | 95373 | ENSG00000115459 | ELMOD3 | chr2 | + | 85371439 | 85371562 | 85371085 | 85371209 | 85377343 | 85377474 |
| SE | 106052 | ENSG00000130177 | CDC16 | chr13 | + | 114236644 | 114236699 | 114234886 | 114235132 | 114236798 | 114236896 |
| SE | 35062 | ENSG00000156030 | MIDEAS | chr14 | - | 73727457 | 73727524 | 73726829 | 73726972 | 73729639 | 73729985 |
| SE | 56204 | ENSG00000124942 | AHNAK | chr11 | - | 62536468 | 62536567 | 62535944 | 62536098 | 62546659 | 62546796 |
| SE | 72076 | ENSG00000130414 | NDUFA10 | chr2 | - | 240018554 | 240018639 | 240011616 | 240014860 | 240021196 | 240021412 |
| SE | 68023 | ENSG00000172890 | NADSYN1 | chr11 | + | 71476032 | 71477438 | 71474394 | 71474526 | 71478394 | 71478469 |
| SE | 101214 | ENSG00000150433 | TMEM218 | chr11 | - | 125102131 | 125102351 | 125101200 | 125101303 | 125102733 | 125102809 |
| SE | 53209 | ENSG00000083544 | TDRD3 | chr13 | + | 60484692 | 60484757 | 60483774 | 60483846 | 60485798 | 60485948 |
| SE | 97344 | ENSG00000229124 | VIM-AS1 | chr10 | - | 17216166 | 17216274 | 17214238 | 17215364 | 17229344 | 17229985 |
| SE | 105093 | ENSG00000112297 | CRYBG1 | chr6 | + | 106530175 | 106530315 | 106527304 | 106527470 | 106539402 | 106539529 |
| SE | 19906 | ENSG00000100890 | PRORP | chr14 | + | 35123504 | 35124231 | 35122758 | 35122910 | 35126734 | 35126782 |
| SE | 55500 | ENSG00000124181 | PLCG1 | chr20 | + | 41137520 | 41137858 | 41136959 | 41137035 | 41159605 | 41159758 |
| SE | 38766 | ENSG00000148399 | DPH7 | chr9 | - | 137577469 | 137577603 | 137576308 | 137576402 | 137578624 | 137578889 |
| SE | 82233 | ENSG00000123066 | MED13L | chr12 | - | 116232058 | 116232156 | 116230532 | 116230622 | 116236820 | 116236890 |
| SE | 27732 | ENSG00000152818 | UTRN | chr6 | + | 144438744 | 144438895 | 144437564 | 144437746 | 144440351 | 144440471 |
| SE | 89085 | ENSG00000171988 | JMJD1C | chr10 | - | 63219877 | 63219983 | 63217206 | 63217331 | 63380317 | 63380482 |
| SE | 80943 | ENSG00000100997 | ABHD12 | chr20 | - | 25320198 | 25320318 | 25309445 | 25309575 | 25323324 | 25323430 |
| SE | 42364 | ENSG00000048740 | CELF2 | chr10 | + | 11249152 | 11249201 | 11217424 | 11217507 | 11257737 | 11257872 |
| SE | 54661 | ENSG00000151694 | ADAM17 | chr2 | - | 9543152 | 9543285 | 9527500 | 9527954 | 9555508 | 9555774 |
| SE | 84013 | ENSG00000111581 | NUP107 | chr12 | + | 68691967 | 68692112 | 68690467 | 68690746 | 68696818 | 68696910 |
| SE | 104551 | ENSG00000150045 | KLRF1 | chr12 | + | 9841811 | 9841951 | 9833302 | 9833452 | 9842320 | 9842433 |
| SE | 72835 | ENSG00000129158 | SERGEF | chr11 | - | 17988596 | 17988755 | 17878207 | 17878244 | 17992930 | 17992993 |
| SE | 21433 | ENSG00000164180 | TMEM161B | chr5 | - | 88240812 | 88240916 | 88228440 | 88228528 | 88268720 | 88268799 |
| SE | 94141 | ENSG00000139644 | TMBIM6 | chr12 | + | 49758351 | 49758480 | 49758226 | 49758275 | 49758682 | 49758707 |
| SE | 51766 | ENSG00000114841 | DNAH1 | chr3 | + | 52398031 | 52398162 | 52397706 | 52397877 | 52398849 | 52399201 |
| SE | 7733 | ENSG00000108389 | MTMR4 | chr17 | - | 58503743 | 58503898 | 58494931 | 58496330 | 58504049 | 58504220 |
| SE | 13564 | ENSG00000102908 | NFAT5 | chr16 | + | 69647027 | 69647586 | 69626402 | 69626528 | 69653235 | 69653428 |
| SE | 48646 | ENSG00000131374 | TBC1D5 | chr3 | - | 17586352 | 17586557 | 17518901 | 17518995 | 17622647 | 17622765 |
| SE | 109305 | ENSG00000139597 | N4BP2L1 | chr13 | - | 32403655 | 32403761 | 32400752 | 32403200 | 32404320 | 32404397 |
| SE | 86529 | ENSG00000085224 | ATRX | chrX | - | 77699509 | 77699547 | 77698573 | 77698629 | 77717130 | 77717243 |
| SE | 96994 | ENSG00000153339 | TRAPPC8 | chr18 | - | 31832083 | 31832173 | 31829196 | 31830989 | 31839311 | 31839457 |
| SE | 81968 | ENSG00000051825 | MPHOSPH9 | chr12 | - | 123227462 | 123227572 | 123223037 | 123223127 | 123230260 | 123230522 |
| SE | 20271 | ENSG00000118482 | PHF3 | chr6 | + | 63678827 | 63679020 | 63646526 | 63646795 | 63679999 | 63680161 |
| SE | 86045 | ENSG00000156313 | RPGR | chrX | - | 38317810 | 38317893 | 38317315 | 38317465 | 38318828 | 38318987 |
| SE | 75818 | ENSG00000153147 | SMARCA5 | chr4 | + | 143540362 | 143540495 | 143538785 | 143538938 | 143543508 | 143543657 |
| SE | 92968 | ENSG00000108669 | CYTH1 | chr17 | - | 78698268 | 78698380 | 78680970 | 78681042 | 78698819 | 78698968 |
| SE | 100319 | ENSG00000081237 | PTPRC | chr1 | + | 198694017 | 198694140 | 198692346 | 198692373 | 198696711 | 198696909 |
| SE | 71415 | ENSG00000171033 | PKIA | chr8 | + | 78572810 | 78572939 | 78516339 | 78516468 | 78598357 | 78598535 |
| SE | 15190 | ENSG00000159459 | UBR1 | chr15 | - | 43025380 | 43025429 | 43024828 | 43024983 | 43026560 | 43026663 |
| SE | 50164 | ENSG00000025708 | TYMP | chr22 | - | 50527164 | 50527283 | 50526575 | 50526738 | 50527587 | 50527717 |
| SE | 47339 | ENSG00000166669 | ATF7IP2 | chr16 | + | 10472109 | 10472183 | 10440363 | 10440462 | 10473478 | 10473534 |
| SE | 35482 | ENSG00000019169 | MARCO | chr2 | + | 118991776 | 118991875 | 118990588 | 118990633 | 118992431 | 118992476 |
| SE | 98497 | ENSG00000215252 | GOLGA8B | chr15 | - | 34552894 | 34553091 | 34551137 | 34551276 | 34553744 | 34553956 |
| SE | 101817 | ENSG00000138286 | FAM149B1 | chr10 | + | 73228059 | 73228184 | 73210250 | 73210438 | 73230421 | 73230525 |
| SE | 55358 | ENSG00000131127 | ZNF141 | chr4 | + | 344334 | 344430 | 343781 | 343908 | 372663 | 373007 |
| SE | 61787 | ENSG00000104343 | UBE2W | chr8 | - | 73825146 | 73825249 | 73810473 | 73810629 | 73830380 | 73830472 |
| SE | 13854 | ENSG00000102893 | PHKB | chr16 | + | 47580294 | 47580358 | 47547432 | 47547548 | 47587667 | 47587758 |
| SE | 80912 | ENSG00000067596 | DHX8 | chr17 | + | 43499955 | 43500103 | 43498861 | 43498959 | 43504643 | 43504825 |
| SE | 11297 | ENSG00000140650 | PMM2 | chr16 | + | 8804766 | 8804843 | 8801798 | 8801910 | 8806315 | 8806407 |
| SE | 11886 | ENSG00000103222 | ABCC1 | chr16 | + | 16014490 | 16014628 | 16009775 | 16009901 | 16033108 | 16033170 |
| SE | 56403 | ENSG00000151838 | CCDC175 | chr14 | - | 59543343 | 59543454 | 59540674 | 59540746 | 59545162 | 59545299 |
| SE | 36269 | ENSG00000169715 | MT1E | chr16 | + | 56626465 | 56626531 | 56625474 | 56625879 | 56626879 | 56627112 |
| SE | 11458 | ENSG00000187555 | USP7 | chr16 | - | 8900529 | 8900630 | 8899120 | 8899188 | 8900989 | 8901057 |
| SE | 67708 | ENSG00000145332 | KLHL8 | chr4 | - | 87183202 | 87183389 | 87178476 | 87178620 | 87220417 | 87220525 |
| SE | 51507 | ENSG00000108021 | TASOR2 | chr10 | + | 5714134 | 5714207 | 5712822 | 5712918 | 5717658 | 5717750 |
| SE | 46034 | ENSG00000158517 | NCF1 | chr7 | + | 74779256 | 74779422 | 74777268 | 74777347 | 74780779 | 74780802 |
| SE | 9064 | ENSG00000215041 | NEURL4 | chr17 | - | 7327164 | 7327230 | 7326710 | 7327009 | 7327439 | 7327500 |
| SE | 11916 | ENSG00000070770 | CSNK2A2 | chr16 | - | 58167206 | 58167308 | 58166583 | 58166684 | 58167684 | 58167795 |
| SE | 31001 | ENSG00000113504 | SLC12A7 | chr5 | - | 1088305 | 1088360 | 1086902 | 1087033 | 1093532 | 1093575 |
| SE | 93258 | ENSG00000148908 | RGS10 | chr10 | - | 119515508 | 119515652 | 119499816 | 119500259 | 119526031 | 119526118 |
| SE | 50608 | ENSG00000134574 | DDB2 | chr11 | + | 47237618 | 47237712 | 47235269 | 47235412 | 47237836 | 47237900 |
| SE | 40257 | ENSG00000170581 | STAT2 | chr12 | - | 56348751 | 56348851 | 56348528 | 56348623 | 56349425 | 56349509 |
| SE | 69854 | ENSG00000107581 | EIF3A | chr10 | - | 119065398 | 119065570 | 119061223 | 119061328 | 119069445 | 119069654 |
| SE | 12754 | ENSG00000089280 | FUS | chr16 | + | 31190274 | 31190398 | 31190039 | 31190141 | 31190741 | 31190842 |
| SE | 89288 | ENSG00000166169 | POLL | chr10 | - | 101583507 | 101583681 | 101582762 | 101582891 | 101587245 | 101587406 |
| SE | 69906 | ENSG00000105851 | PIK3CG | chr7 | + | 106884154 | 106884266 | 106883032 | 106883163 | 106886134 | 106886292 |
| SE | 31213 | ENSG00000025293 | PHF20 | chr20 | + | 35847349 | 35847434 | 35842572 | 35842744 | 35863012 | 35863400 |
| SE | 39299 | ENSG00000141293 | SKAP1 | chr17 | - | 48184722 | 48184847 | 48182393 | 48182457 | 48345904 | 48345965 |
| SE | 98934 | ENSG00000175115 | PACS1 | chr11 | + | 66221153 | 66221247 | 66220630 | 66220791 | 66227503 | 66227584 |
| SE | 17434 | ENSG00000140563 | MCTP2 | chr15 | + | 94345128 | 94345164 | 94340812 | 94340924 | 94356136 | 94356301 |
| SE | 74269 | ENSG00000137959 | IFI44L | chr1 | + | 78635341 | 78635489 | 78628950 | 78628999 | 78637031 | 78637203 |
| SE | 75871 | ENSG00000240065 | PSMB9 | chr6 | + | 32857262 | 32857394 | 32854191 | 32854289 | 32858004 | 32858134 |
| SE | 3859 | ENSG00000157540 | DYRK1A | chr21 | + | 37428866 | 37428908 | 37425689 | 37425781 | 37472683 | 37472880 |
| SE | 22829 | ENSG00000107968 | MAP3K8 | chr10 | + | 30437175 | 30437406 | 30434183 | 30434378 | 30438915 | 30439274 |
| SE | 102580 | ENSG00000120438 | TCP1 | chr6 | - | 159783940 | 159784067 | 159780934 | 159781110 | 159785899 | 159785997 |
| SE | 35559 | ENSG00000182606 | TRAK1 | chr3 | + | 42199176 | 42199253 | 42194803 | 42194941 | 42200817 | 42201054 |
| SE | 96334 | ENSG00000166181 | API5 | chr11 | + | 43326506 | 43326611 | 43323429 | 43323636 | 43327788 | 43327878 |
| SE | 100178 | ENSG00000078124 | ACER3 | chr11 | + | 76997205 | 76997273 | 76989512 | 76990574 | 76998762 | 76998821 |
| SE | 50245 | ENSG00000126012 | KDM5C | chrX | - | 53196685 | 53197044 | 53195915 | 53196054 | 53197770 | 53197876 |
| SE | 70953 | ENSG00000131781 | FMO5 | chr1 | - | 147208851 | 147209051 | 147190176 | 147190249 | 147212392 | 147212535 |
| SE | 57929 | ENSG00000168439 | STIP1 | chr11 | + | 64194188 | 64194330 | 64193077 | 64193287 | 64194478 | 64194620 |
| SE | 86237 | ENSG00000184634 | MED12 | chrX | + | 71132844 | 71132956 | 71132376 | 71132538 | 71133122 | 71133212 |
| SE | 13976 | ENSG00000170540 | ARL6IP1 | chr16 | - | 18797924 | 18798044 | 18795175 | 18795581 | 18798700 | 18798834 |
| SE | 60117 | ENSG00000122223 | CD244 | chr1 | - | 160836194 | 160836254 | 160834050 | 160834116 | 160838450 | 160838518 |
| SE | 56076 | ENSG00000160679 | CHTOP | chr1 | + | 153642245 | 153642429 | 153638083 | 153638448 | 153645063 | 153646302 |
| SE | 98770 | ENSG00000157600 | TMEM164 | chrX | + | 110067346 | 110067396 | 110003500 | 110004164 | 110109079 | 110109146 |
| SE | 72413 | ENSG00000114978 | MOB1A | chr2 | - | 74159090 | 74159254 | 74152527 | 74156645 | 74165217 | 74165351 |
| SE | 90459 | ENSG00000070081 | NUCB2 | chr11 | + | 17301743 | 17301870 | 17296103 | 17296211 | 17309571 | 17309675 |
| SE | 74351 | ENSG00000187118 | CMC1 | chr3 | + | 28316752 | 28316859 | 28316352 | 28316423 | 28319508 | 28319776 |
| SE | 103298 | ENSG00000196233 | LCOR | chr10 | + | 96949007 | 96949295 | 96907667 | 96907747 | 96952102 | 96952196 |
| SE | 78981 | ENSG00000183735 | TBK1 | chr12 | + | 64457750 | 64458018 | 64455839 | 64455957 | 64460188 | 64460329 |
| SE | 18032 | ENSG00000139990 | DCAF5 | chr14 | - | 69118138 | 69118275 | 69116365 | 69116495 | 69119193 | 69119230 |
| SE | 4532 | ENSG00000131355 | ADGRE3 | chr19 | - | 14668801 | 14668852 | 14663417 | 14663540 | 14674745 | 14674844 |
| SE | 46783 | ENSG00000096060 | FKBP5 | chr6 | - | 35580035 | 35580221 | 35573584 | 35575942 | 35587033 | 35587117 |
| SE | 62554 | ENSG00000092330 | TINF2 | chr14 | - | 24241019 | 24241116 | 24239642 | 24240875 | 24241674 | 24241776 |
| SE | 23720 | ENSG00000124782 | RREB1 | chr6 | + | 7228996 | 7231907 | 7226466 | 7226656 | 7248510 | 7249274 |
| SE | 19329 | ENSG00000182979 | MTA1 | chr14 | + | 105450260 | 105450324 | 105450057 | 105450184 | 105454192 | 105454298 |
| SE | 84653 | ENSG00000131196 | NFATC1 | chr18 | + | 79486247 | 79486937 | 79467449 | 79467582 | 79527527 | 79527741 |
| SE | 16214 | ENSG00000077238 | IL4R | chr16 | + | 27334315 | 27334417 | 27330065 | 27330198 | 27340185 | 27340234 |
| SE | 27142 | ENSG00000137806 | NDUFAF1 | chr15 | - | 41388447 | 41388522 | 41387217 | 41387593 | 41394858 | 41395044 |
| SE | 17932 | ENSG00000197249 | SERPINA1 | chr14 | - | 94388559 | 94388663 | 94382591 | 94383241 | 94388800 | 94389010 |
| SE | 72943 | ENSG00000048649 | RSF1 | chr11 | - | 77725544 | 77725699 | 77700720 | 77702495 | 77740730 | 77740936 |
| SE | 1293 | ENSG00000100239 | PPP6R2 | chr22 | + | 50406688 | 50406875 | 50393892 | 50394135 | 50414551 | 50414689 |
| SE | 55960 | ENSG00000136161 | RCBTB2 | chr13 | - | 48510628 | 48510771 | 48502723 | 48502914 | 48511769 | 48511877 |
| SE | 96754 | ENSG00000183431 | SF3A3 | chr1 | - | 37978719 | 37978827 | 37976883 | 37976953 | 37978987 | 37979055 |
| SE | 63793 | ENSG00000119866 | BCL11A | chr2 | - | 60508605 | 60508824 | 60468731 | 60468833 | 60545970 | 60546300 |
| SE | 20108 | ENSG00000078319 | PMS2P1 | chr7 | - | 100325690 | 100325779 | 100320438 | 100321084 | 100328835 | 100329019 |
| SE | 41910 | ENSG00000175203 | DCTN2 | chr12 | - | 57546027 | 57546096 | 57535494 | 57535845 | 57547027 | 57547079 |
| SE | 69622 | ENSG00000156711 | MAPK13 | chr6 | + | 36136018 | 36136048 | 36135752 | 36135861 | 36136483 | 36136515 |
| SE | 81084 | ENSG00000140750 | ARHGAP17 | chr16 | - | 24927683 | 24927732 | 24919394 | 24920260 | 24930783 | 24931160 |
| SE | 15872 | ENSG00000186654 | PRR5 | chr22 | + | 44735026 | 44735162 | 44732250 | 44732391 | 44736771 | 44736786 |
| SE | 4399 | ENSG00000130475 | FCHO1 | chr19 | + | 17772644 | 17772741 | 17772456 | 17772555 | 17774238 | 17774241 |
| SE | 97796 | ENSG00000196914 | ARHGEF12 | chr11 | + | 120409393 | 120409450 | 120407737 | 120407823 | 120420752 | 120420851 |
| SE | 95860 | ENSG00000107263 | RAPGEF1 | chr9 | - | 131638634 | 131638791 | 131629101 | 131629254 | 131643238 | 131643391 |
| SE | 14914 | ENSG00000086666 | ZFAND6 | chr15 | + | 80075163 | 80075242 | 80059567 | 80059809 | 80120327 | 80120498 |
| SE | 97070 | ENSG00000175216 | CKAP5 | chr11 | - | 46816197 | 46816404 | 46811006 | 46811178 | 46818309 | 46818503 |
| SE | 62323 | ENSG00000185811 | IKZF1 | chr7 | + | 50382539 | 50382707 | 50327637 | 50327757 | 50387344 | 50387470 |
| SE | 59502 | ENSG00000148660 | CAMK2G | chr10 | - | 73839538 | 73839601 | 73837467 | 73837511 | 73842168 | 73842211 |
| SE | 70303 | ENSG00000133059 | DSTYK | chr1 | - | 205148204 | 205148339 | 205142504 | 205147745 | 205150679 | 205150794 |
| SE | 16347 | ENSG00000140497 | SCAMP2 | chr15 | - | 74848599 | 74848701 | 74843731 | 74845593 | 74851352 | 74851481 |
| SE | 99811 | ENSG00000115368 | WDR75 | chr2 | + | 189470075 | 189470245 | 189469343 | 189469439 | 189470818 | 189470878 |
| SE | 10993 | ENSG00000083454 | P2RX5 | chr17 | - | 3690955 | 3691027 | 3690604 | 3690680 | 3691643 | 3691773 |
| SE | 35715 | ENSG00000124802 | EEF1E1 | chr6 | - | 8095377 | 8095495 | 8089984 | 8090281 | 8097266 | 8097311 |
| SE | 43065 | ENSG00000101474 | APMAP | chr20 | - | 24983902 | 24984019 | 24973644 | 24973737 | 24992593 | 24992642 |
| SE | 89571 | ENSG00000095485 | CWF19L1 | chr10 | - | 100260217 | 100260319 | 100253420 | 100253539 | 100260965 | 100261028 |
| SE | 43612 | ENSG00000168765 | GSTM4 | chr1 | + | 109657589 | 109657671 | 109657214 | 109657279 | 109657771 | 109657794 |
| SE | 34969 | ENSG00000117133 | RPF1 | chr1 | + | 84495881 | 84496063 | 84495372 | 84495455 | 84496243 | 84496370 |
| SE | 25432 | ENSG00000288674 | NA | chr1 | + | 226881887 | 226882048 | 226875364 | 226875550 | 226883704 | 226883919 |
| SE | 107003 | ENSG00000133858 | ZFC3H1 | chr12 | - | 71642435 | 71642559 | 71638417 | 71638515 | 71644094 | 71644318 |
| SE | 41877 | ENSG00000089327 | FXYD5 | chr19 | + | 35156843 | 35156974 | 35155550 | 35155611 | 35157420 | 35157501 |
| SE | 74452 | ENSG00000116209 | TMEM59 | chr1 | - | 54047266 | 54047372 | 54043372 | 54043525 | 54052999 | 54053192 |
| SE | 7576 | ENSG00000175155 | YPEL2 | chr17 | + | 59353214 | 59353526 | 59331654 | 59331824 | 59388326 | 59388370 |
| SE | 109189 | ENSG00000124496 | TRERF1 | chr6 | - | 42268153 | 42269848 | 42265750 | 42265797 | 42300637 | 42300749 |
| SE | 76781 | ENSG00000135317 | SNX14 | chr6 | - | 85533598 | 85533800 | 85530191 | 85530275 | 85536791 | 85536924 |
| SE | 57900 | ENSG00000135899 | SP110 | chr2 | - | 230171695 | 230171767 | 230170620 | 230170761 | 230172065 | 230172174 |
| SE | 71830 | ENSG00000183486 | MX2 | chr21 | + | 41363268 | 41363389 | 41362006 | 41362055 | 41376835 | 41377155 |
| SE | 8831 | ENSG00000109083 | IFT20 | chr17 | - | 28332145 | 28332382 | 28331401 | 28331987 | 28335339 | 28335435 |
| SE | 43558 | ENSG00000110172 | CHORDC1 | chr11 | - | 90214017 | 90214175 | 90211214 | 90211318 | 90215173 | 90215230 |
| SE | 7015 | ENSG00000101782 | RIOK3 | chr18 | + | 23479325 | 23479424 | 23477178 | 23477268 | 23481171 | 23481500 |
| SE | 64402 | ENSG00000101577 | LPIN2 | chr18 | - | 2928590 | 2928660 | 2927721 | 2927811 | 2929064 | 2929158 |
| SE | 100397 | ENSG00000133812 | SBF2 | chr11 | - | 9992414 | 9992543 | 9989496 | 9989595 | 9992989 | 9993103 |
| SE | 106359 | ENSG00000076513 | ANKRD13A | chr12 | + | 110033796 | 110033957 | 110030644 | 110030758 | 110036260 | 110036328 |
| SE | 8999 | ENSG00000108469 | RECQL5 | chr17 | - | 75650516 | 75650718 | 75631491 | 75631668 | 75651185 | 75651265 |
| SE | 94106 | ENSG00000113734 | BNIP1 | chr5 | + | 173151565 | 173151694 | 173146865 | 173146958 | 173154321 | 173154413 |
| SE | 57825 | ENSG00000090104 | RGS1 | chr1 | + | 192576284 | 192576365 | 192575772 | 192575929 | 192576773 | 192576835 |
| SE | 84978 | ENSG00000170832 | USP32 | chr17 | - | 60211012 | 60211118 | 60209369 | 60209543 | 60211375 | 60211514 |
| SE | 88698 | ENSG00000183077 | AFMID | chr17 | + | 78204827 | 78204900 | 78202702 | 78202751 | 78205945 | 78206050 |
| SE | 109520 | ENSG00000115561 | CHMP3 | chr2 | - | 86529217 | 86529397 | 86510357 | 86510479 | 86563303 | 86563479 |
| SE | 30620 | ENSG00000131042 | LILRB2 | chr19 | - | 54280462 | 54280544 | 54279790 | 54280075 | 54280960 | 54281097 |
| SE | 71222 | ENSG00000079263 | SP140 | chr2 | + | 230250980 | 230251061 | 230248884 | 230248968 | 230253315 | 230253417 |
| SE | 68446 | ENSG00000198589 | LRBA | chr4 | - | 150597082 | 150597115 | 150590712 | 150590859 | 150599006 | 150599131 |
| SE | 95480 | ENSG00000095319 | NUP188 | chr9 | + | 128999623 | 128999805 | 128999171 | 128999317 | 129001528 | 129001729 |
| SE | 86489 | ENSG00000071889 | FAM3A | chrX | - | 154512313 | 154512411 | 154511847 | 154511871 | 154512822 | 154512936 |
| SE | 24192 | ENSG00000104731 | KLHDC4 | chr16 | - | 87748672 | 87748809 | 87730551 | 87730644 | 87755193 | 87755254 |
| SE | 103284 | ENSG00000196233 | LCOR | chr10 | + | 96833405 | 96833479 | 96832259 | 96832399 | 96907264 | 96907330 |
| SE | 100679 | ENSG00000166387 | PPFIBP2 | chr11 | + | 7639731 | 7639870 | 7635551 | 7635593 | 7641478 | 7641620 |
| SE | 101398 | ENSG00000134461 | ANKRD16 | chr10 | - | 5883968 | 5884077 | 5882702 | 5883167 | 5885722 | 5885765 |
| SE | 42597 | ENSG00000078140 | UBE2K | chr4 | + | 39777681 | 39777810 | 39774833 | 39774933 | 39778359 | 39778384 |
| SE | 36259 | ENSG00000069849 | ATP1B3 | chr3 | + | 141916375 | 141916440 | 141915969 | 141916020 | 141921976 | 141922063 |
| SE | 31491 | ENSG00000161013 | MGAT4B | chr5 | - | 179800483 | 179800597 | 179800183 | 179800259 | 179800906 | 179800939 |
| SE | 98691 | ENSG00000205531 | NAP1L4 | chr11 | - | 2989128 | 2989271 | 2979206 | 2979237 | 2991793 | 2992102 |
| SE | 9339 | ENSG00000068079 | IFI35 | chr17 | + | 43013775 | 43013882 | 43013475 | 43013662 | 43014107 | 43014456 |
| SE | 90773 | ENSG00000174744 | BRMS1 | chr11 | - | 66340966 | 66341046 | 66340773 | 66340870 | 66341205 | 66341333 |
| SE | 103263 | ENSG00000172661 | WASHC2C | chr10 | + | 45769454 | 45769618 | 45765678 | 45765810 | 45773255 | 45773358 |
| SE | 19592 | ENSG00000029364 | SLC39A9 | chr14 | + | 69424093 | 69424202 | 69398689 | 69399465 | 69442068 | 69442266 |
| SE | 32772 | ENSG00000168297 | PXK | chr3 | + | 58391772 | 58391847 | 58390581 | 58390659 | 58394997 | 58395102 |
| SE | 83641 | ENSG00000166579 | NDEL1 | chr17 | + | 8445710 | 8445864 | 8444259 | 8444357 | 8446753 | 8446780 |
| SE | 17397 | ENSG00000100473 | COCH | chr14 | + | 30884909 | 30885041 | 30884552 | 30884656 | 30885393 | 30885620 |
| SE | 5400 | ENSG00000078747 | ITCH | chr20 | + | 34480598 | 34480732 | 34479629 | 34479789 | 34481065 | 34481176 |
| SE | 77495 | ENSG00000160799 | CCDC12 | chr3 | - | 46923606 | 46923668 | 46923142 | 46923363 | 46925455 | 46925535 |
| SE | 70686 | ENSG00000095059 | DHPS | chr19 | - | 12679800 | 12679922 | 12679622 | 12679719 | 12680195 | 12680281 |
| SE | 4760 | ENSG00000141905 | NFIC | chr19 | + | 3456549 | 3456635 | 3452481 | 3452666 | 3462751 | 3462900 |
| SE | 103259 | ENSG00000172661 | WASHC2C | chr10 | + | 45753179 | 45753237 | 45752587 | 45752706 | 45754485 | 45754545 |
| SE | 70184 | ENSG00000115282 | TTC31 | chr2 | + | 74490024 | 74490127 | 74483321 | 74483410 | 74490243 | 74490473 |
| SE | 80914 | ENSG00000067596 | DHX8 | chr17 | + | 43513361 | 43513502 | 43508338 | 43508520 | 43517166 | 43517322 |
| SE | 72660 | ENSG00000153094 | BCL2L11 | chr2 | + | 111161418 | 111161550 | 111153779 | 111153904 | 111164132 | 111164231 |
| SE | 38961 | ENSG00000101457 | DNTTIP1 | chr20 | + | 45801998 | 45802057 | 45801401 | 45801458 | 45803332 | 45803378 |
| SE | 50556 | ENSG00000204131 | NHSL2 | chrX | + | 72137093 | 72137225 | 72134508 | 72134704 | 72138440 | 72140771 |
| SE | 17478 | ENSG00000270580 | PKD1P6-NPIPP1 | chr16 | - | 15105548 | 15105818 | 15104722 | 15104758 | 15106114 | 15106150 |
| SE | 967 | ENSG00000188677 | PARVB | chr22 | + | 44119037 | 44119140 | 44100052 | 44100123 | 44131486 | 44131627 |
| SE | 2674 | ENSG00000129353 | SLC44A2 | chr19 | + | 10638226 | 10638315 | 10638022 | 10638093 | 10643278 | 10643528 |

| | | | | | | | | | | | | | | |
|---|---|---|---|---|---|---|---|---|---|---|---|---|---|---|
| SE | 106845 | ENSG00000123329 | ARHGAP9 | chr12 | - | 57474422 | 57474476 | 57474041 | 57474176 | 57474625 | 57474703 | 408 | 195 | 327 |
| SE | 25931 | ENSG00000113319 | RASGRF2 | chr5 | + | 81108995 | 81109078 | 81094855 | 81094992 | 81113537 | 81113920 | 10 | 21 | 41 |
| SE | 68082 | ENSG00000135838 | NPL | chr1 | + | 182791247 | 182791330 | 182789772 | 182789805 | 182792231 | 182792286 | 0 | 4 | 0 |
| SE | 17435 | ENSG00000140563 | MCTP2 | chr15 | + | 94358481 | 94358612 | 94356136 | 94356301 | 94367604 | 94367791 | 238 | 125 | 118 |
| SE | 76042 | ENSG00000137312 | FLOT1 | chr6 | - | 30741791 | 30741867 | 30741189 | 30741333 | 30742146 | 30742203 | 37 | 7 | 13 |
| SE | 95098 | ENSG00000148358 | GPR107 | chr9 | + | 130099456 | 130099532 | 130092247 | 130092381 | 130100628 | 130100702 | 170 | 79 | 81 |
| SE | 19715 | ENSG00000103811 | CTSH | chr15 | - | 78929411 | 78929493 | 78923780 | 78927781 | 78931450 | 78931506 | 295 | 317 | 373 |
| SE | 23631 | ENSG00000135074 | ADAM19 | chr5 | - | 157518822 | 157518888 | 157513423 | 157513505 | 157519838 | 157520031 | 116 | 68 | 122 |
| SE | 86233 | ENSG00000184634 | MED12 | chrX | + | 71125350 | 71125495 | 71124975 | 71125146 | 71125662 | 71125713 | 76 | 49 | 66 |
| SE | 4649 | ENSG00000142347 | MYO1F | chr19 | - | 8553349 | 8553437 | 8551665 | 8552164 | 8554476 | 8554564 | 401 | 189 | 262 |
| SE | 51877 | ENSG00000147576 | ADHFE1 | chr8 | + | 66445217 | 66445414 | 66444593 | 66444748 | 66447263 | 66447341 | 21 | 24 | 45 |
| SE | 78789 | ENSG00000106245 | BUD31 | chr7 | + | 99410033 | 99410169 | 99408968 | 99409245 | 99411063 | 99411186 | 195 | 123 | 148 |
| SE | 58202 | ENSG00000172965 | MIR4435-2HG | chr2 | - | 111248147 | 111248241 | 111208755 | 111208936 | 111344060 | 111344237 | 23 | 15 | 6 |
| SE | 20110 | ENSG00000078319 | PMS2P1 | chr7 | - | 100325690 | 100325868 | 100320438 | 100321084 | 100328835 | 100329019 | 8 | 9 | 31 |
| SE | 47537 | ENSG00000066322 | ELOVL1 | chr1 | - | 43364323 | 43364460 | 43363397 | 43364137 | 43365563 | 43365623 | 334 | 146 | 185 |
| SE | 36034 | ENSG00000145819 | ARHGAP26 | chr5 | + | 142915541 | 142915684 | 142913198 | 142913293 | 142932046 | 142932125 | 5 | 4 | 3 |
| SE | 103457 | ENSG00000107959 | PITRM1 | chr10 | - | 3156929 | 3157064 | 3155590 | 3155729 | 3157434 | 3157531 | 453 | 327 | 531 |
| SE | 5546 | ENSG00000101052 | IFT52 | chr20 | + | 43603759 | 43603889 | 43596434 | 43596522 | 43604182 | 43604258 | 103 | 107 | 98 |
| SE | 48002 | ENSG00000010810 | FYN | chr6 | - | 111696276 | 111696456 | 111694627 | 111694704 | 111702884 | 111703034 | 933 | 506 | 673 |
| SE | 96470 | ENSG00000110514 | MADD | chr11 | + | 47326543 | 47326552 | 47324470 | 47324577 | 47326737 | 47326807 | 2 | 1 | 2 |
| SE | 24150 | ENSG00000164574 | GALNT10 | chr5 | + | 154218028 | 154218149 | 154190747 | 154191025 | 154294815 | 154294918 | 3 | 0 | 0 |
| SE | 49778 | ENSG00000116337 | AMPD2 | chr1 | + | 109620913 | 109621266 | 109619836 | 109620278 | 109625302 | 109625433 | 8 | 3 | 9 |

[illegible]

| | | | | | | | | | | | | | | | | | | | | | | | | | | | | | | | | | | | | | | | | | | | | | | |
|---|---|---|---|---|---|---|---|---|---|---|---|---|---|---|---|---|---|---|---|---|---|---|---|---|---|---|---|---|---|---|---|---|---|---|---|---|---|---|---|---|---|---|---|---|---|---|
| SE | 103036 | ENSG00000099290 | WASHC2A | chr10 | + | 50087274 | 50087322 | 50085496 | 50085558 | 50090775 | 50090886 | 211 | 147 | 142 | 212 | 226 | 0 | 0 | 0 | 0 | 0 | 607 | 273 | 239 | 372 | 236 | 0 | 10 | 0 | 0 | 0 | 197 | 149 | 0.858284638 | 1 | 1 | 1 | 1 | 1 | 1 | 1 | 0.954 | 1 | 1 | 1 | 0.009 |
| SE | 22474 | ENSG00000259040 | BLOC1S5-TXNDC5 | chr6 | - | 7891620 | 7891736 | 7889494 | 7889581 | 7895105 | 7895202 | 349 | 131 | 942 | 416 | 247 | 0 | 0 | 0 | 0 | 0 | 202 | 956 | 1909 | 192 | 353 | 0 | 4 | 4 | 0 | 10 | 265 | 149 | 0.858331511 | 1 | 1 | 1 | 1 | 1 | 1 | 1 | 0.993 | 0.996 | 1 | 0.952 | 0.012 |
| SE | 60226 | ENSG00000286129 | NA | chr22 | + | 22308696 | 22308819 | 22307532 | 22307591 | 22309735 | 22309883 | 15 | 8 | 62 | 22 | 36 | 0 | 0 | 2 | 0 | 0 | 229 | 7 | 25 | 33 | 50 | 2 | 0 | 1 | 3 | 0 | 272 | 149 | 0.858338047 | 1 | 1 | 1 | 0.944 | 1 | 1 | 0.984 | 1 | 0.932 | 0.858 | 1 | 0.034 |
| SE | 77996 | ENSG00000146729 | NIPSNAP2 | chr7 | + | 55982209 | 55982280 | 55978906 | 55981567 | 55983727 | 55983812 | 259 | 190 | 249 | 264 | 472 | 4 | 4 | 2 | 3 | 0 | 172 | 268 | 197 | 238 | 50 | 0 | 0 | 0 | 1 | 0 | 220 | 149 | 0.858339952 | 1 | 0.978 | 0.97 | 0.988 | 0.983 | 1 | | 1 | 1 | 0.994 | 1 | -0.015 |
| SE | 39221 | ENSG00000150753 | CCT5 | chr5 | + | 10262480 | 10262618 | 10258385 | 10258535 | 10263133 | 10263314 | 411 | 293 | 330 | 430 | 383 | 0 | 1 | 1 | 0 | 5 | 467 | 267 | 177 | 441 | 207 | 0 | 5 | 3 | 0 | 4 | 287 | 149 | 0.85836181 | 1 | 1 | 0.993 | 0.994 | 1 | 0.975 | 1 | 0.965 | 0.968 | 1 | 0.964 | 0.013 |
| SE | 17820 | ENSG00000155465 | SLC7A7 | chr14 | - | 22775832 | 22775936 | 22775443 | 22775540 | 22776194 | 22776232 | 571 | 245 | 328 | 439 | 437 | 0 | 1 | 1 | 0 | 3 | 711 | 498 | 237 | 391 | 930 | 4 | 11 | 1 | 2 | 4 | 253 | 149 | 0.858418742 | 1 | 1 | 0.993 | 0.995 | 1 | 0.988 | 0.991 | 0.964 | 0.993 | 0.991 | 0.993 | 0.009 |
| SE | 36364 | ENSG00000173889 | PHC3 | chr3 | - | 170145422 | 170145521 | 170136365 | 170136665 | 170149085 | 170149244 | 347 | 214 | 332 | 429 | 633 | 17 | 6 | 8 | 4 | 10 | 327 | 352 | 219 | 219 | 506 | 0 | 22 | 12 | 3 | 1 | 248 | 149 | 0.858459499 | 1 | 0.925 | 0.955 | 0.961 | 0.985 | 0.974 | 1 | 0.906 | 0.916 | 0.978 | 0.997 | 0.001 |
| SE | 63812 | ENSG00000181163 | NPM1 | chr5 | + | 171400838 | 171400925 | 171400152 | 171400210 | 171405301 | 171405403 | 1621 | 1167 | 2092 | 1674 | 3786 | 30 | 9 | 23 | 19 | 50 | 223 | 781 | 1000 | 661 | 57 | 1 | 14 | 7 | 11 | 4 | 236 | 149 | 0.858551983 | 1 | 0.972 | 0.988 | 0.983 | 0.982 | 0.98 | 0.993 | 0.972 | 0.989 | 0.974 | 0.9 | 0.015 |
| SE | 65299 | ENSG00000149499 | EML3 | chr11 | - | 62607665 | 62607821 | 62606957 | 62607099 | 62608200 | 62608296 | 244 | 156 | 232 | 204 | 318 | 1 | 3 | 0 | 0 | 8 | 401 | 207 | 118 | 174 | 502 | 0 | 3 | 0 | 3 | 0 | 298 | 149 | 0.858679102 | 1 | 0.992 | 0.963 | 1 | 1 | 0.952 | 1 | 0.972 | 1 | 0.967 | 1 | -0.006 |
| SE | 38158 | ENSG00000138750 | NUP54 | chr4 | - | 76145554 | 76145691 | 76144389 | 76144473 | 76148307 | 76148377 | 5 | 1 | 1 | 6 | 10 | 27 | 17 | 35 | 35 | 33 | 24 | 2 | 0 | 3 | 0 | 5 | 20 | 13 | 11 | 15 | 286 | 149 | 0.858719899 | 1 | 0.088 | 0.03 | 0.015 | 0.082 | 0.136 | 0.714 | 0.05 | 0 | 0.124 | 0 | -0.107 |
| SE | 63503 | ENSG00000166478 | ZNF143 | chr11 | + | 9472676 | 9472769 | 9471301 | 9471420 | 9473940 | 9473954 | 133 | 79 | 96 | 113 | 126 | 2 | 0 | 1 | 0 | 0 | 74 | 102 | 118 | 87 | 26 | 0 | 8 | 0 | 0 | 0 | 242 | 149 | 0.858743794 | 1 | 0.976 | 1 | 0.983 | 1 | 1 | 1 | 0.887 | 1 | 1 | 1 | 0.014 |
| SE | 82372 | ENSG00000257923 | CUX1 | chr7 | + | 102097363 | 102097501 | 102070338 | 102070417 | 102104335 | 102104459 | 400 | 308 | 304 | 430 | 540 | 0 | 2 | 1 | 0 | 0 | 1430 | 647 | 497 | 832 | 2223 | 11 | 9 | 0 | 1 | 7 | 287 | 149 | 0.858751272 | 1 | 1 | 0.988 | 0.994 | 1 | 1 | 0.985 | 0.974 | 1 | 0.998 | 0.994 | 0.006 |
| SE | 107464 | ENSG00000100599 | RIN3 | chr14 | + | 92641237 | 92641329 | 92615406 | 92615479 | 92651581 | 92651943 | 238 | 131 | 118 | 216 | 345 | 2 | 0 | 0 | 0 | 2 | 332 | 326 | 160 | 306 | 554 | 0 | 0 | 0 | 0 | 12 | 241 | 149 | 0.858813577 | 1 | 0.987 | 1 | 1 | 1 | 0.991 | 1 | 1 | 1 | 1 | 0.966 | 0.002 |
| SE | 1537 | ENSG00000100055 | CYTH4 | chr22 | + | 37301934 | 37302031 | 37300881 | 37301019 | 37303253 | 37303295 | 0 | 1 | 0 | 0 | 0 | 160 | 70 | 119 | 134 | 190 | 12 | 0 | 0 | 0 | 0 | 207 | 211 | 141 | 127 | 289 | 246 | 149 | 0.858905636 | 1 | 0 | 0.009 | 0 | 0 | 0 | 0.034 | 0 | 0 | 0 | 0 | -0.005 |
| SE | 34172 | ENSG00000163660 | CCNL1 | chr3 | - | 157157010 | 157157083 | 157153035 | 157153156 | 157158865 | 157158975 | 102 | 94 | 114 | 136 | 181 | 74 | 40 | 61 | 69 | 80 | 45 | 59 | 112 | 44 | 18 | 29 | 29 | 69 | 23 | 5 | 222 | 149 | 0.858952108 | 1 | 0.481 | 0.612 | 0.556 | 0.57 | 0.603 | 0.51 | 0.577 | 0.521 | 0.562 | 0.707 | -0.011 |
| SE | 14891 | ENSG00000166855 | CLPX | chr15 | - | 65156843 | 65156932 | 65155691 | 65155856 | 65157745 | 65157910 | 196 | 143 | 146 | 161 | 191 | 0 | 2 | 0 | 7 | 0 | 48 | 168 | 171 | 127 | 29 | 0 | 0 | 0 | 0 | 0 | 238 | 149 | 0.858953137 | 1 | 1 | 0.978 | 1 | 0.935 | 1 | 1 | 1 | 1 | 1 | 1 | -0.017 |
| SE | 87988 | ENSG00000198373 | WWP2 | chr16 | + | 69930129 | 69930258 | 69929447 | 69929529 | 69931151 | 69931227 | 249 | 110 | 205 | 261 | 271 | 0 | 0 | 0 | 0 | 0 | 228 | 226 | 139 | 229 | 301 | 0 | 0 | 0 | 3 | 7 | 278 | 149 | 0.858977621 | 1 | 1 | 1 | 1 | 1 | 1 | 1 | 1 | 1 | 0.976 | 0.958 | 0.013 |
| SE | 52746 | ENSG00000163166 | IWS1 | chr2 | - | 127498139 | 127498237 | 127495997 | 127496148 | 127502814 | 127502872 | 89 | 136 | 70 | 99 | 118 | 15 | 11 | 20 | 12 | 12 | 55 | 82 | 85 | 46 | 4 | 0 | 15 | 18 | 17 | 0 | 247 | 149 | 0.859008781 | 1 | 0.782 | 0.882 | 0.679 | 0.833 | 0.856 | 1 | 0.767 | 0.74 | 0.62 | 1 | -0.019 |
| SE | 80313 | ENSG00000241973 | PI4KA | chr22 | - | 20727773 | 20727864 | 20727229 | 20727397 | 20729312 | 20729506 | 188 | 105 | 208 | 188 | 228 | 1 | 0 | 0 | 0 | 0 | 312 | 203 | 138 | 133 | 337 | 0 | 6 | 1 | 3 | 0 | 240 | 149 | 0.859025594 | 1 | 0.992 | 1 | 1 | 1 | 1 | 1 | 0.955 | 0.988 | 0.965 | 1 | 0.017 |
| SE | 71309 | ENSG00000163754 | GYG1 | chr3 | + | 148996301 | 148996476 | 148994141 | 148994277 | 148996741 | 148996892 | 258 | 171 | 214 | 215 | 194 | 0 | 0 | 0 | 1 | 0 | 272 | 405 | 231 | 565 | 330 | 0 | 10 | 0 | 0 | 0 | 298 | 149 | 0.859053008 | 1 | 1 | 1 | 1 | 0.991 | 1 | 1 | 0.953 | 1 | 1 | 1 | 0.008 |
| SE | 93356 | ENSG00000127084 | FGD3 | chr9 | + | 93013851 | 93013998 | 93011213 | 93011272 | 93015736 | 93015829 | 625 | 334 | 540 | 572 | 842 | 3 | 1 | 0 | 0 | 0 | 609 | 432 | 375 | 459 | 643 | 12 | 1 | 0 | 0 | 0 | 296 | 149 | 0.859090572 | 1 | 0.991 | 0.994 | 1 | 1 | 1 | 0.962 | 0.995 | 1 | 1 | 1 | 0.006 |
| SE | 14146 | ENSG00000005339 | CREBBP | chr16 | - | 3740398 | 3740549 | 3736507 | 3736815 | 3744893 | 3744957 | 377 | 189 | 290 | 241 | 231 | 0 | 0 | 0 | 0 | 0 | 398 | 285 | 385 | 559 | 360 | 8 | 4 | 0 | 0 | 0 | 298 | 149 | 0.859204603 | 1 | 1 | 1 | 1 | 1 | 1 | 0.961 | 0.973 | 1 | 1 | 1 | 0.013 |
| SE | 60789 | ENSG00000160714 | UBE2Q1 | chr1 | - | 154550937 | 154551004 | 154548576 | 154550469 | 154551396 | 154551492 | 341 | 236 | 331 | 342 | 834 | 0 | 1 | 0 | 1 | 0 | 456 | 371 | 327 | 376 | 710 | 6 | 0 | 0 | 0 | 10 | 216 | 149 | 0.859219937 | 1 | 1 | 0.994 | 1 | 0.996 | 1 | 0.981 | 1 | 1 | 1 | 0.98 | 0.006 |
| SE | 90973 | ENSG00000132669 | RIN2 | chr20 | + | 19815222 | 19815298 | 19799621 | 19799747 | 19889565 | 19890334 | 4 | 0 | 0 | 1 | 2 | 17 | 11 | 11 | 4 | 5 | 0 | 9 | 0 | 0 | 0 | 28 | 25 | 17 | 18 | 34 | 225 | 149 | 0.859232953 | 1 | 0.135 | 0 | 0 | 0.142 | 0.209 | 0 | 0.193 | 0 | 0 | 0 | 0.059 |
| SE | 75476 | ENSG00000197081 | IGF2R | chr6 | + | 160079579 | 160079787 | 160078200 | 160078362 | 160080128 | 160080275 | 760 | 226 | 373 | 466 | 355 | 0 | 0 | 0 | 0 | 0 | 409 | 438 | 448 | 447 | 572 | 0 | 1 | 0 | 11 | 0 | 298 | 149 | 0.859238046 | 1 | 1 | 1 | 1 | 1 | 1 | 1 | 0.995 | 1 | 0.953 | 1 | 0.01 |
| SE | 25203 | ENSG00000160783 | PMF1 | chr1 | + | 156232319 | 156232425 | 156212992 | 156213176 | 156233627 | 156233652 | 349 | 177 | 272 | 272 | 281 | 6 | 0 | 4 | 2 | 3 | 631 | 502 | 384 | 429 | 879 | 0 | 0 | 4 | 0 | 20 | 255 | 149 | 0.85925165 | 1 | 0.971 | 1 | 0.975 | 0.988 | 0.982 | 1 | 1 | 0.982 | 1 | 0.963 | -0.006 |
| SE | 92878 | ENSG00000135018 | UBQLN1 | chr9 | - | 83678440 | 83678599 | 83666349 | 83669327 | 83679774 | 83680037 | 483 | 322 | 448 | 521 | 816 | 2 | 0 | 0 | 0 | 0 | 485 | 470 | 382 | 457 | 307 | 0 | 0 | 0 | 0 | 10 | 298 | 149 | 0.859263188 | 1 | 0.992 | 1 | 1 | 1 | 1 | 1 | 1 | 1 | 1 | 0.939 | 0.011 |
| SE | 42008 | ENSG00000137996 | RTCA | chr1 | + | 100267165 | 100267245 | 100266523 | 100266624 | 100268151 | 100268295 | 2 | 1 | 1 | 5 | 2 | 52 | 25 | 36 | 30 | 20 | 0 | 0 | 12 | 2 | 0 | 37 | 28 | 34 | 41 | 13 | 229 | 149 | 0.859264641 | 1 | 0.024 | 0.025 | 0.018 | 0.098 | 0.061 | 0 | 0 | 0.187 | 0.031 | 0 | 0.002 |
| SE | 107924 | ENSG00000111335 | OAS2 | chr12 | + | 112979718 | 112979789 | 112978423 | 112978785 | 112987037 | 112988803 | 7 | 0 | 2 | 7 | 0 | 181 | 127 | 138 | 221 | 171 | 2 | 4 | 1 | 10 | 0 | 526 | 362 | 108 | 505 | 115 | 220 | 149 | 0.859305039 | 1 | 0.026 | 0 | 0.01 | 0.021 | 0 | 0.003 | 0.007 | 0.006 | 0.013 | 0 | 0.006 |
| SE | 20167 | ENSG00000090615 | GOLGA3 | chr12 | - | 132827606 | 132827713 | 132822271 | 132822311 | 132828802 | 132828858 | 14 | 11 | 17 | 35 | 8 | 16 | 5 | 12 | 2 | 6 | 29 | 29 | 10 | 0 | 14 | 0 | 3 | 6 | 8 | 35 | 256 | 149 | 0.859331074 | 1 | 0.337 | 0.561 | 0.452 | 0.911 | 0.437 | 1 | 0.849 | 0.492 | 0 | 0.189 | 0.034 |
| SE | 28388 | ENSG00000204842 | ATXN2 | chr12 | - | 111479069 | 111479102 | 111470557 | 111470742 | 111482786 | 111482831 | 0 | 0 | 0 | 1 | 0 | 2 | 1 | 3 | 3 | 1 | 0 | 0 | 1 | 1 | 3 | 3 | 3 | 4 | 10 | 25 | 182 | 149 | 0.859348076 | 1 | 0 | 0 | 0 | 0.214 | 0 | 0 | 0 | 0.17 | 0.076 | 0.089 | -0.024 |
| SE | 41235 | ENSG00000119782 | FKBP1B | chr2 | + | 24054103 | 24054192 | 24053901 | 24053949 | 24060813 | 24060926 | 0 | 0 | 2 | 3 | 4 | 2 | 1 | 5 | 6 | 6 | 0 | 2 | 0 | 0 | 14 | 12 | 8 | 7 | 4 | 33 | 238 | 149 | 0.859390603 | 1 | 0 | 0 | 0.2 | 0.238 | 0.294 | 0 | 0.135 | 0 | 0 | 0.21 | 0.077 |
| SE | 15886 | ENSG00000167491 | GATAD2A | chr19 | + | 19492305 | 19492438 | 19465339 | 19465614 | 19492580 | 19492596 | 635 | 346 | 502 | 624 | 553 | 5 | 0 | 10 | 13 | 4 | 1020 | 780 | 609 | 545 | 1325 | 7 | 6 | 1 | 7 | 26 | 282 | 149 | 0.859395825 | 1 | 0.985 | 1 | 0.964 | 0.962 | 0.986 | 0.987 | 0.986 | 0.997 | 0.976 | 0.964 | -0.003 |
| SE | 100102 | ENSG00000149262 | INTS4 | chr11 | - | 77987603 | 77987698 | 77981458 | 77981576 | 77991107 | 77991299 | 3 | 3 | 1 | 0 | 13 | 89 | 51 | 84 | 91 | 92 | 5 | 3 | 7 | 3 | 0 | 64 | 96 | 70 | 106 | 69 | 244 | 149 | 0.859443659 | 1 | 0.02 | 0.035 | 0.007 | 0 | 0.079 | 0.046 | 0.019 | 0.058 | 0.017 | 0 | 0 |
| SE | 67464 | ENSG00000162923 | WDR26 | chr1 | - | 224398514 | 224398593 | 224397726 | 224398226 | 224398888 | 224399034 | 857 | 567 | 646 | 872 | 793 | 0 | 0 | 0 | 0 | 1 | 602 | 970 | 1215 | 814 | 475 | 0 | 1 | 1 | 14 | 0 | 228 | 149 | 0.859451233 | 1 | 1 | 1 | 1 | 1 | 0.998 | 1 | 0.998 | 0.999 | 0.974 | 1 | 0.005 |
| SE | 40171 | ENSG00000115306 | SPTBN1 | chr2 | + | 54621399 | 54621512 | 54617607 | 54617688 | 54622299 | 54622487 | 227 | 182 | 342 | 279 | 481 | 0 | 0 | 0 | 0 | 0 | 421 | 196 | 163 | 352 | 340 | 0 | 5 | 0 | 6 | 0 | 262 | 149 | 0.859506113 | 1 | 1 | 1 | 1 | 1 | 1 | 1 | 0.957 | 1 | 0.971 | 1 | 0.014 |
| SE | 39902 | ENSG00000271723 | MROH7-TTC4 | chr1 | + | 54731485 | 54731700 | 54728345 | 54728432 | 54737581 | 54737664 | 43 | 46 | 50 | 53 | 93 | 0 | 3 | 1 | 2 | 1 | 40 | 40 | 36 | 25 | 51 | 0 | 0 | 1 | 0 | 5 | 298 | 149 | 0.859626802 | 1 | 1 | 0.885 | 0.962 | 0.93 | 0.979 | 1 | 1 | 0.947 | 1 | 0.836 | -0.005 |
| SE | 8189 | ENSG00000108799 | EZH1 | chr17 | - | 42724304 | 42724424 | 42722794 | 42722915 | 42727634 | 42727763 | 271 | 205 | 305 | 314 | 514 | 19 | 9 | 8 | 14 | 12 | 411 | 339 | 297 | 231 | 305 | 13 | 16 | 7 | 2 | 23 | 269 | 149 | 0.859704866 | 1 | 0.888 | 0.927 | 0.955 | 0.926 | 0.96 | 0.946 | 0.921 | 0.959 | 0.985 | 0.88 | -0.007 |
| SE | 38512 | ENSG00000117298 | ECE1 | chr1 | - | 21233557 | 21233661 | 21227930 | 21228041 | 21235849 | 21235927 | 339 | 253 | 327 | 318 | 518 | 0 | 0 | 1 | 2 | 0 | 764 | 566 | 496 | 568 | 1225 | 8 | 0 | 1 | 9 | 3 | 253 | 149 | 0.859746987 | 1 | 1 | 1 | 0.995 | 0.989 | 1 | 0.983 | 1 | 0.997 | 0.974 | 0.996 | 0.007 |
| SE | 95318 | ENSG00000165699 | TSC1 | chr9 | - | 132903650 | 132903817 | 132902604 | 132902787 | 132904410 | 132904454 | 61 | 34 | 110 | 102 | 87 | 1 | 2 | 2 | 2 | 0 | 91 | 111 | 67 | 76 | 71 | 0 | 1 | 0 | 2 | 4 | 298 | 149 | 0.859773967 | 1 | 0.968 | 0.895 | 0.965 | 0.962 | 1 | 1 | 0.982 | 1 | 0.95 | 0.899 | -0.008 |
| SE | 66967 | ENSG00000286239 | NA | chr2 | - | 223894233 | 223894339 | 223884647 | 223884749 | 223895503 | 223895630 | 228 | 187 | 180 | 226 | 176 | 5 | 0 | 1 | 4 | 1 | 177 | 201 | 242 | 270 | 72 | 0 | 5 | 5 | 0 | 0 | 255 | 149 | 0.859882812 | 1 | 0.964 | 1 | 0.991 | 0.971 | 0.99 | 1 | 0.959 | 0.966 | 1 | 1 | -0.002 |
| SE | 47784 | ENSG00000154358 | OBSCN | chr1 | + | 228292522 | 228292786 | 228291897 | 228292161 | 228293352 | 228293616 | 24 | 30 | 79 | 68 | 42 | 3 | 4 | 12 | 6 | 2 | 51 | 5 | 28 | 51 | 36 | 19 | 4 | 4 | 0 | 0 | 298 | 149 | 0.859936287 | 1 | 0.8 | 0.789 | 0.767 | 0.85 | 0.913 | 0.573 | 0.385 | 0.778 | 1 | 1 | 0.077 |
| SE | 14089 | ENSG00000197006 | METTL9 | chr16 | + | 21612644 | 21612835 | 21599557 | 21599898 | 21617864 | 21618009 | 854 | 617 | 578 | 586 | 1334 | 19 | 5 | 12 | 5 | 8 | 1501 | 756 | 558 | 1085 | 954 | 0 | 23 | 23 | 11 | 3 | 298 | 149 | 0.85998261 | 1 | 0.957 | 0.984 | 0.96 | 0.983 | 0.988 | 1 | 0.943 | 0.924 | 0.98 | 0.994 | 0.006 |
| SE | 100244 | ENSG00000084234 | APLP2 | chr11 | + | 130122304 | 130122513 | 130110537 | 130110661 | 130126699 | 130126830 | 542 | 430 | 379 | 373 | 418 | 0 | 0 | 0 | 0 | 0 | 1979 | 1234 | 390 | 1026 | 1071 | 0 | 2 | 0 | 0 | 14 | 298 | 149 | 0.860006832 | 1 | 1 | 1 | 1 | 1 | 1 | 1 | 0.997 | 1 | 1 | 0.975 | 0.006 |
| SE | 5632 | ENSG00000124209 | RAB22A | chr20 | + | 58353272 | 58353344 | 58343717 | 58343799 | 58353431 | 58353538 | 72 | 73 | 105 | 69 | 163 | 0 | 2 | 0 | 3 | 2 | 67 | 88 | 64 | 83 | 27 | 0 | 0 | 0 | 0 | 0 | 221 | 149 | 0.860021063 | 1 | 1 | 0.961 | 1 | 0.939 | 0.982 | 1 | 1 | 1 | 1 | 1 | -0.024 |
| SE | 79362 | ENSG00000280828 | NA | chr7 | + | 128578889 | 128578933 | 128533651 | 128533730 | 128579998 | 128580094 | 36 | 17 | 21 | 46 | 26 | 0 | 2 | 3 | 1 | 0 | 175 | 60 | 79 | 89 | 148 | 0 | 0 | 1 | 1 | 0 | 193 | 149 | 0.86004403 | 1 | 1 | 0.868 | 0.844 | 0.973 | 1 | 1 | 1 | 0.984 | 0.986 | 1 | -0.057 |
| SE | 59966 | ENSG00000103671 | TRIP4 | chr15 | + | 64414084 | 64414211 | 64409612 | 64409828 | 64418540 | 64418728 | 222 | 156 | 145 | 171 | 305 | 6 | 1 | 7 | 9 | 15 | 175 | 184 | 151 | 151 | 50 | 6 | 4 | 2 | 6 | 9 | 276 | 149 | 0.860074019 | 1 | 0.952 | 0.988 | 0.918 | 0.911 | 0.917 | 0.94 | 0.961 | 0.976 | 0.931 | 0.75 | 0.026 |
| SE | 3636 | ENSG00000239998 | LILRA2 | chr19 | + | 54573697 | 54573912 | 54573027 | 54573435 | 54574075 | 54574111 | 96 | 40 | 23 | 77 | 18 | 1 | 3 | 3 | 1 | 0 | 131 | 134 | 40 | 114 | 178 | 0 | 2 | 1 | 0 | 11 | 298 | 149 | 0.860083183 | 1 | 0.98 | 0.87 | 0.793 | 0.975 | 1 | 1 | 0.971 | 0.952 | 1 | 0.89 | -0.039 |
| SE | 66457 | ENSG00000131051 | RBM39 | chr20 | - | 35724569 | 35724722 | 35721739 | 35721877 | 35725037 | 35725063 | 2375 | 1877 | 2279 | 2918 | 3622 | 0 | 0 | 0 | 3 | 0 | 877 | 1881 | 2307 | 1341 | 794 | 14 | 0 | 0 | 0 | 0 | 298 | 149 | 0.860120736 | 1 | 1 | 1 | 1 | 0.998 | 1 | 0.969 | 1 | 1 | 1 | 1 | 0.006 |
| SE | 18912 | ENSG00000100804 | PSMB5 | chr14 | - | 23027744 | 23027829 | 23025857 | 23026375 | 23033367 | 23033674 | 9 | 0 | 3 | 4 | 0 | 32 | 17 | 25 | 25 | 23 | 5 | 2 | 1 | 0 | 8 | 0 | 24 | 3 | 6 | 64 | 234 | 149 | 0.860140227 | 1 | 0.152 | 0 | 0.071 | 0.092 | 0 | 1 | 0.05 | 0.175 | 0 | 0.074 | -0.197 |
| SE | 75321 | ENSG00000204256 | BRD2 | chr6 | + | 32977756 | 32978005 | 32976561 | 32977570 | 32980341 | 32980464 | 1294 | 840 | 1119 | 1294 | 1073 | 0 | 0 | 0 | 1 | 0 | 2293 | 1421 | 1231 | 1608 | 1916 | 13 | 5 | 0 | 13 | 11 | 298 | 149 | 0.86018575 | 1 | 1 | 1 | 1 | 0.998 | 1 | 0.989 | 0.993 | 1 | 0.984 | 0.989 | 0.009 |
| SE | 93475 | ENSG00000136827 | TOR1A | chr9 | - | 129818177 | 129818297 | 129812964 | 129814222 | 129818519 | 129818647 | 0 | 1 | 0 | 1 | 3 | 165 | 136 | 171 | 187 | 289 | 0 | 14 | 0 | 0 | 0 | 384 | 297 | 119 | 334 | 344 | 269 | 149 | 0.860285655 | 1 | 0 | 0.004 | 0 | 0.003 | 0.006 | 0 | 0.025 | 0 | 0 | 0 | -0.002 |
| SE | 32019 | ENSG00000100003 | SEC14L2 | chr22 | + | 30415758 | 30415865 | 30410595 | 30410679 | 30415947 | 30416007 | 3 | 4 | 30 | 20 | 23 | 0 | 0 | 0 | 0 | 0 | 2 | 38 | 57 | 17 | 341 | 0 | 0 | 0 | 0 | 8 | 256 | 149 | 0.860308924 | 1 | 1 | 1 | 1 | 1 | 1 | 1 | 1 | 1 | 1 | 0.961 | 0.008 |
| SE | 76614 | ENSG00000112695 | COX7A2 | chr6 | - | 75240300 | 75240385 | 75237674 | 75237988 | 75243716 | 75243788 | 438 | 234 | 394 | 385 | 328 | 0 | 0 | 0 | 0 | 0 | 479 | 407 | 314 | 461 | 587 | 0 | 0 | 0 | 0 | 12 | 234 | 149 | 0.860583629 | 1 | 1 | 1 | 1 | 1 | 1 | 1 | 1 | 1 | 1 | 0.969 | 0.006 |
| SE | 15207 | ENSG00000101266 | CSNK2A1 | chr20 | - | 492253 | 492364 | 489502 | 489881 | 495718 | 495802 | 389 | 301 | 417 | 418 | 535 | 0 | 0 | 0 | 0 | 0 | 325 | 442 | 368 | 380 | 324 | 10 | 0 | 0 | 0 | 0 | 260 | 149 | 0.860600015 | 1 | 1 | 1 | 1 | 1 | 1 | 0.949 | 1 | 1 | 1 | 1 | 0.01 |
| SE | 693 | ENSG00000186575 | NF2 | chr22 | + | 29642201 | 29642285 | 29639089 | 29639212 | 29654656 | 29654725 | 196 | 177 | 174 | 276 | 200 | 0 | 0 | 0 | 0 | 0 | 470 | 281 | 188 | 264 | 466 | 0 | 0 | 3 | 4 | 8 | 233 | 149 | 0.860624262 | 1 | 1 | 1 | 1 | 1 | 1 | 1 | 1 | 0.976 | 0.977 | 0.974 | 0.015 |
| SE | 51041 | ENSG00000112182 | BACH2 | chr6 | - | 90252512 | 90252590 | 90206568 | 90206681 | 90271848 | 90271941 | 22 | 27 | 31 | 23 | 52 | 12 | 16 | 35 | 29 | 28 | 0 | 66 | 35 | 11 | 3 | 8 | 14 | 2 | 9 | 6 | 227 | 149 | 0.860714859 | 1 | 0.546 | 0.526 | 0.368 | 0.342 | 0.549 | 0 | 0.756 | 0.92 | 0.445 | 0.247 | -0.007 |
| SE | 55111 | ENSG00000072110 | ACTN1 | chr14 | - | 68893654 | 68893747 | 68892052 | 68892283 | 68902476 | 68902562 | 157 | 144 | 156 | 249 | 238 | 0 | 0 | 1 | 0 | 0 | 531 | 214 | 200 | 476 | 1985 | 0 | 0 | 4 | 0 | 18 | 242 | 149 | 0.860728341 | 1 | 1 | 1 | 0.99 | 1 | 1 | 1 | 1 | 0.969 | 1 | 0.985 | 0.007 |
| SE | 108306 | ENSG00000115310 | RTN4 | chr2 | - | 54974694 | 54974764 | 54973820 | 54973867 | 54982514 | 54982653 | 1241 | 881 | 800 | 1165 | 1239 | 4 | 1 | 2 | 0 | 3 | 2717 | 1179 | 836 | 1909 | 2758 | 0 | 6 | 15 | 0 | 12 | 219 | 149 | 0.860841183 | 1 | 0.995 | 0.998 | 0.996 | 1 | 0.996 | 1 | 0.993 | 0.974 | 1 | 0.994 | 0.005 |
| SE | 90346 | ENSG00000249859 | PVT1 | chr8 | + | 128070159 | 128070272 | 128049366 | 128049748 | 128096517 | 128096654 | 8 | 8 | 12 | 4 | 2 | 0 | 0 | 2 | 1 | 3 | 12 | 4 | 6 | 1 | 0 | 0 | 3 | 0 | 0 | 4 | 262 | 149 | 0.860926123 | 1 | 1 | 1 | 0.773 | 0.695 | 0.275 | 1 | 0.431 | 1 | 1 | 0 | 0.062 |
| SE | 81605 | ENSG00000253729 | PRKDC | chr8 | - | 47918276 | 47918383 | 47915327 | 47915418 | 47927193 | 47927353 | 457 | 355 | 467 | 380 | 528 | 0 | 1 | 0 | 2 | 0 | 237 | 557 | 352 | 481 | 30 | 0 | 0 | 0 | 11 | 0 | 256 | 149 | 0.860997516 | 1 | 1 | 0.995 | 1 | 0.991 | 1 | 1 | 1 | 1 | 0.962 | 1 | 0.005 |
| SE | 51983 | ENSG00000071054 | MAP4K4 | chr2 | + | 101863820 | 101864051 | 101859642 | 101859864 | 101864929 | 101865003 | 0 | 2 | 0 | 0 | 0 | 42 | 38 | 32 | 50 | 24 | 0 | 6 | 6 | 0 | 0 | 40 | 67 | 62 | 82 | 33 | 298 | 149 | 0.86100964 | 1 | 0 | 0.026 | 0 | 0 | 0 | 0 | 0.043 | 0.046 | 0 | 0 | -0.013 |
| SE | 14922 | ENSG00000086666 | ZFAND6 | chr15 | + | 80098415 | 80098578 | 80075163 | 80075315 | 80120327 | 80120405 | 98 | 63 | 69 | 83 | 228 | 0 | 1 | 0 | 1 | 4 | 147 | 77 | 37 | 78 | 180 | 7 | 0 | 0 | 0 | 0 | 298 | 149 | 0.861020584 | 1 | 1 | 0.969 | 1 | 0.976 | 0.966 | 0.913 | 1 | 1 | 1 | 1 | 0 |
| SE | 52800 | ENSG00000005175 | RPAP3 | chr12 | - | 47679492 | 47679594 | 47670106 | 47670345 | 47679703 | 47679774 | 156 | 127 | 140 | 156 | 276 | 2 | 5 | 2 | 12 | 12 | 25 | 64 | 31 | 6 | 0 | 0 | 2 | 4 | 0 | 3 | 251 | 149 | 0.861058801 | 1 | 0.979 | 0.938 | 0.977 | 0.885 | 0.932 | 1 | 0.95 | 0.821 | 1 | 0 | 0.188 |
| SE | 44055 | ENSG00000054118 | THRAP3 | chr1 | + | 36225411 | 36225463 | 36224432 | 36224505 | 36259381 | 36259484 | 12 | 5 | 12 | 6 | 4 | 183 | 99 | 175 | 174 | 126 | 0 | 16 | 23 | 15 | 0 | 255 | 256 | 278 | 162 | 181 | 201 | 149 | 0.861100747 | 1 | 0.046 | 0.036 | 0.048 | 0.025 | 0.023 | 0 | 0.044 | 0.058 | 0.064 | 0 | 0.002 |
| SE | 661 | ENSG00000100216 | TOMM22 | chr22 | + | 38682322 | 38682441 | 38681956 | 38682095 | 38682878 | 38682996 | 474 | 205 | 427 | 461 | 271 | 0 | 0 | 0 | 0 | 0 | 530 | 481 | 273 | 288 | 502 | 11 | 0 | 0 | 0 | 0 | 268 | 149 | 0.861127482 | 1 | 1 | 1 | 1 | 1 | 1 | 0.964 | 1 | 1 | 1 | 1 | 0.007 |
| SE | 2404 | ENSG00000126247 | CAPNS1 | chr19 | + | 36146246 | 36146312 | 36145975 | 36146054 | 36149577 | 36149636 | 1083 | 514 | 570 | 697 | 1227 | 0 | 0 | 0 | 0 | 0 | 2124 | 997 | 689 | 958 | 3563 | 24 | 0 | 0 | 0 | 0 | 215 | 149 | 0.861328796 | 1 | 1 | 1 | 1 | 1 | 1 | 0.984 | 1 | 1 | 1 | 1 | 0.003 |
| SE | 10738 | ENSG00000286065 | SLFN12L | chr17 | - | 35479116 | 35480195 | 35478074 | 35478185 | 35522278 | 35522969 | 101 | 65 | 83 | 74 | 67 | 13 | 11 | 10 | 1 | 8 | 49 | 64 | 80 | 28 | 35 | 11 | 11 | 7 | 1 | 0 | 298 | 149 | 0.861361987 | 1 | 0.795 | 0.747 | 0.806 | 0.974 | 0.807 | 0.69 | 0.744 | 0.851 | 0.933 | 1 | -0.018 |
| SE | 65489 | ENSG00000125875 | TBC1D20 | chr20 | - | 444253 | 444406 | 441856 | 442043 | 445049 | 445130 | 0 | 0 | 0 | 0 | 0 | 92 | 56 | 70 | 89 | 94 | 11 | 0 | 0 | 0 | 0 | 88 | 69 | 55 | 91 | 203 | 298 | 149 | 0.861370385 | 1 | 0 | 0 | 0 | 0 | 0 | 0.059 | 0 | 0 | 0 | 0 | -0.012 |
| SE | 10422 | ENSG00000126581 | BECN1 | chr17 | - | 42811654 | 42811797 | 42810133 | 42810928 | 42823747 | 42823879 | 134 | 81 | 118 | 121 | 119 | 0 | 0 | 0 | 0 | 0 | 306 | 132 | 107 | 123 | 300 | 0 | 2 | 1 | 0 | 5 | 292 | 149 | 0.861394097 | 1 | 1 | 1 | 1 | 1 | 1 | 1 | 0.971 | 0.982 | 1 | 0.968 | 0.016 |
| SE | 23905 | ENSG00000196911 | KPNA5 | chr6 | + | 116698703 | 116698798 | 116692292 | 116692392 | 116702018 | 116702150 | 318 | 238 | 327 | 306 | 468 | 8 | 1 | 5 | 0 | 0 | 93 | 161 | 117 | 72 | 186 | 0 | 3 | 2 | 0 | 0 | 244 | 149 | 0.861408558 | 1 | 0.96 | 0.993 | 0.976 | 1 | 1 | 1 | 0.97 | 0.973 | 1 | 1 | -0.003 |
| SE | 11963 | ENSG00000005844 | ITGAL | chr16 | + | 30496096 | 30496294 | 30494712 | 30494850 | 30499073 | 30499234 | 1111 | 518 | 537 | 737 | 506 | 0 | 0 | 0 | 0 | 0 | 1052 | 1161 | 573 | 697 | 1355 | 1 | 0 | 1 | 4 | 15 | 298 | 149 | 0.861479118 | 1 | 1 | 1 | 1 | 1 | 1 | 0.998 | 1 | 0.997 | 0.989 | 0.978 | 0.008 |
| SE | 73163 | ENSG00000285437 | POLR2J3 | chr7 | - | 102568009 | 102568184 | 102565975 | 102567083 | 102569834 | 102569924 | 21 | 10 | 18 | 10 | 28 | 63 | 89 | 98 | 179 | 183 | 20 | 23 | 12 | 38 | 37 | 326 | 150 | 179 | 184 | 132 | 298 | 149 | 0.86157511 | 1 | 0.143 | 0.053 | 0.084 | 0.027 | 0.071 | 0.03 | 0.071 | 0.032 | 0.094 | 0.123 | 0.006 |
| SE | 83897 | ENSG00000196123 | KIAA0895L | chr16 | - | 67181557 | 67181697 | 67180040 | 67180666 | 67183261 | 67183952 | 7 | 9 | 4 | 24 | 29 | 2 | 9 | 9 | 13 | 7 | 119 | 20 | 24 | 48 | 40 | 34 | 14 | 10 | 26 | 53 | 289 | 149 | 0.861592578 | 1 | 0.643 | 0.34 | 0.186 | 0.488 | 0.681 | 0.643 | 0.424 | 0.553 | 0.488 | 0.28 | -0.01 |
| SE | 45157 | ENSG00000107099 | DOCK8 | chr9 | + | 328021 | 328171 | 317042 | 317128 | 332397 | 332478 | 390 | 242 | 368 | 366 | 395 | 10 | 0 | 0 | 0 | 0 | 625 | 512 | 382 | 682 | 469 | 0 | 0 | 0 | 0 | 0 | 298 | 149 | 0.861604973 | 1 | 0.951 | 1 | 1 | 1 | 1 | 1 | 1 | 1 | 1 | 1 | -0.01 |
| SE | 97595 | ENSG00000139722 | VPS37B | chr12 | - | 122868479 | 122868562 | 122865329 | 122867607 | 122895951 | 122896124 | 180 | 107 | 100 | 120 | 224 | 1 | 0 | 0 | 1 | 0 | 267 | 238 | 274 | 212 | 354 | 0 | 0 | 2 | 9 | 0 | 232 | 149 | 0.861629094 | 1 | 0.991 | 1 | 1 | 0.987 | 1 | 1 | 1 | 0.989 | 0.938 | 1 | 0.01 |
| SE | 97743 | ENSG00000130309 | COLGALT1 | chr19 | + | 17560347 | 17560465 | 17559310 | 17559421 | 17567405 | 17567540 | 167 | 81 | 112 | 162 | 94 | 0 | 0 | 0 | 0 | 0 | 203 | 262 | 119 | 193 | 536 | 0 | 0 | 0 | 9 | 0 | 267 | 149 | 0.861715613 | 1 | 1 | 1 | 1 | 1 | 1 | 1 | 1 | 1 | 0.923 | 1 | 0.015 |
| SE | 62508 | ENSG00000049167 | ERCC8 | chr5 | - | 60892036 | 60893643 | 60890623 | 60891086 | 60898275 | 60898400 | 0 | 0 | 1 | 0 | 1 | 46 | 27 | 30 | 24 | 36 | 0 | 8 | 0 | 0 | 0 | 41 | 45 | 17 | 14 | 4 | 298 | 149 | 0.861732628 | 1 | 0 | 0 | 0.016 | 0 | 0.014 | 0 | 0.082 | 0 | 0 | 0 | -0.01 |
| SE | 80908 | ENSG00000067596 | DHX8 | chr17 | + | 43492680 | 43493040 | 43492182 | 43492292 | 43493444 | 43493589 | 239 | 162 | 173 | 248 | 262 | 0 | 0 | 0 | 0 | 0 | 545 | 319 | 297 | 321 | 283 | 0 | 0 | 0 | 10 | 0 | 298 | 149 | 0.861745329 | 1 | 1 | 1 | 1 | 1 | 1 | 1 | 1 | 1 | 0.941 | 1 | 0.012 |
| SE | 98605 | ENSG00000136527 | TRA2B | chr3 | - | 185926600 | 185926734 | 185925084 | 185925626 | 185937824 | 185937987 | 553 | 325 | 499 | 573 | 769 | 3 | 6 | 16 | 7 | 4 | 366 | 668 | 441 | 620 | 369 | 0 | 0 | 12 | 5 | 14 | 283 | 149 | 0.861781782 | 1 | 0.99 | 0.966 | 0.943 | 0.977 | 0.99 | 1 | 1 | 0.951 | 0.985 | 0.933 | -0.001 |
| SE | 76417 | ENSG00000168538 | TRAPPC11 | chr4 | + | 183682731 | 183682825 | 183680119 | 183680267 | 183683974 | 183684006 | 177 | 158 | 120 | 146 | 216 | 3 | 0 | 0 | 2 | 5 | 18 | 103 | 122 | 60 | 4 | 0 | 0 | 5 | 2 | 0 | 243 | 149 | 0.861788427 | 1 | 0.973 | 1 | 1 | 0.978 | 0.964 | 1 | 1 | 0.937 | 0.948 | 1 | 0.006 |
| SE | 88497 | ENSG00000152601 | MBNL1 | chr3 | + | 152458129 | 152458193 | 152456266 | 152456361 | 152459270 | 152459327 | 8 | 4 | 4 | 10 | 8 | 338 | 300 | 327 | 438 | 526 | 0 | 1 | 1 | 6 | 0 | 97 | 208 | 216 | 138 | 126 | 213 | 149 | 0.861934628 | 1 | 0.016 | 0.009 | 0.008 | 0.016 | 0.011 | 0 | 0.003 | 0.003 | 0.03 | 0 | 0.005 |
| SE | 15228 | ENSG00000101266 | CSNK2A1 | chr20 | - | 527932 | 528049 | 505117 | 505229 | 543671 | 543712 | 16 | 4 | 7 | 2 | 5 | 4 | 2 | 1 | 3 | 4 | 15 | 1 | 0 | 3 | 3 | 0 | 2 | 2 | 5 | 0 | 266 | 149 | 0.861979059 | 1 | 0.691 | 0.528 | 0.797 | 0.272 | 0.412 | 1 | 0.219 | 0 | 0.252 | 1 | 0.046 |
| SE | 96821 | ENSG00000147874 | HAUS6 | chr9 | - | 19089411 | 19089559 | 19087090 | 19087156 | 19093170 | 19093213 | 108 | 119 | 110 | 143 | 194 | 1 | 0 | 0 | 2 | 5 | 35 | 112 | 108 | 55 | 36 | 0 | 6 | 1 | 0 | 0 | 297 | 149 | 0.862071678 | 1 | 0.982 | 1 | 1 | 0.973 | 0.951 | 1 | 0.904 | 0.982 | 1 | 1 | 0.004 |
| SE | 42056 | ENSG00000092871 | RFFL | chr17 | - | 35014739 | 35014763 | 35005989 | 35012149 | 35016369 | 35016580 | 114 | 72 | 96 | 84 | 70 | 4 | 0 | 0 | 3 | 0 | 73 | 73 | 113 | 124 | 106 | 0 | 0 | 0 | 3 | 5 | 173 | 149 | 0.86209276 | 1 | 0.961 | 1 | 1 | 0.96 | 1 | 1 | 1 | 1 | 0.973 | 0.948 | 0 |
| SE | 64866 | ENSG00000118640 | VAMP8 | chr2 | + | 85579741 | 85579874 | 85579008 | 85579167 | 85581575 | 85582031 | 8 | 3 | 2 | 1 | 30 | 959 | 436 | 598 | 657 | 1030 | 0 | 3 | 0 | 2 | 5 | 1051 | 778 | 285 | 626 | 1173 | 282 | 149 | 0.862104028 | 1 | 0.004 | 0.004 | 0.002 | 0.001 | 0.015 | 0 | 0.002 | 0 | 0.002 | 0.002 | 0.004 |
| SE | 107842 | ENSG00000061936 | SFSWAP | chr12 | + | 131764455 | 131764686 | 131756472 | 131756644 | 131766117 | 131766308 | 190 | 136 | 212 | 230 | 298 | 11 | 10 | 11 | 18 | 15 | 249 | 219 | 238 | 253 | 490 | 22 | 22 | 13 | 24 | 2 | 298 | 149 | 0.862176174 | 1 | 0.896 | 0.872 | 0.906 | 0.865 | 0.909 | 0.85 | 0.833 | 0.902 | 0.841 | 0.992 | 0.006 |
| SE | 57411 | ENSG00000101040 | ZMYND8 | chr20 | - | 47349860 | 47349988 | 47347855 | 47347926 | 47356656 | 47356889 | 0 | 0 | 0 | 0 | 6 | 34 | 46 | 26 | 43 | 40 | 0 | 10 | 0 | 1 | 0 | 94 | 57 | 36 | 64 | 22 | 277 | 149 | 0.862220445 | 1 | 0 | 0 | 0 | 0 | 0.075 | 0 | 0.086 | 0 | 0.008 | 0 | -0.004 |
| SE | 76924 | ENSG00000167483 | NIBAN3 | chr19 | + | 17546685 | 17546797 | 17543523 | 17543631 | 17551785 | 17551960 | 35 | 38 | 31 | 60 | 51 | 3 | 0 | 0 | 2 | 7 | 69 | 139 | 12 | 64 | 237 | 10 | 3 | 0 | 0 | 7 | 261 | 149 | 0.86234477 | 1 | 0.869 | 1 | 1 | 0.945 | 0.806 | 0.798 | 0.964 | 1 | 1 | 0.951 | -0.019 |
| SE | 12168 | ENSG00000051108 | HERPUD1 | chr16 | + | 56935403 | 56935475 | 56935234 | 56935312 | 56936686 | 56936817 | 121 | 86 | 127 | 122 | 181 | 13 | 9 | 3 | 12 | 10 | 132 | 190 | 165 | 126 | 163 | 9 | 3 | 2 | 15 | 32 | 221 | 149 | 0.862358865 | 1 | 0.863 | 0.866 | 0.966 | 0.873 | 0.924 | 0.908 | 0.977 | 0.982 | 0.85 | 0.774 | 0 |
| SE | 108383 | ENSG00000111554 | MDM1 | chr12 | - | 68331106 | 68331221 | 68325004 | 68327510 | 68332227 | 68332257 | 41 | 35 | 22 | 54 | 34 | 0 | 1 | 0 | 6 | 0 | 8 | 32 | 19 | 45 | 2 | 13 | 0 | 0 | 0 | 0 | 264 | 149 | 0.86237253 | 1 | 1 | 0.952 | 1 | 0.836 | 1 | 0.258 | 1 | 1 | 1 | 1 | 0.106 |
| SE | 13989 | ENSG00000047578 | KATNIP | chr16 | + | 27648603 | 27648735 | 27631064 | 27631162 | 27677728 | 27677996 | 32 | 15 | 23 | 28 | 54 | 6 | 3 | 2 | 1 | 6 | 83 | 42 | 51 | 41 | 82 | 5 | 4 | 3 | 13 | 9 | 281 | 149 | 0.862473692 | 1 | 0.739 | 0.726 | 0.859 | 0.937 | 0.827 | 0.898 | 0.848 | 0.9 | 0.626 | 0.829 | -0.003 |
| SE | 45906 | ENSG00000162434 | JAK1 | chr1 | - | 64956933 | 64957017 | 64886258 | 64886341 | 64966332 | 64966448 | 0 | 0 | 1 | 0 | 0 | 248 | 173 | 187 | 217 | 87 | 0 | 12 | 2 | 0 | 0 | 252 | 206 | 244 | 214 | 131 | 233 | 149 | 0.862526574 | 1 | 0 | 0 | 0.003 | 0 | 0 | 0 | 0.036 | 0.005 | 0 | 0 | -0.008 |
| SE | 93234 | ENSG00000110455 | ACCS | chr11 | + | 44082064 | 44082192 | 44081178 | 44081320 | 44083168 | 44083208 | 0 | 3 | 4 | 11 | 9 | 32 | 6 | 6 | 47 | 18 | 6 | 0 | 8 | 3 | 0 | 0 | 1 | 10 | 30 | 12 | 277 | 149 | 0.862563108 | 1 | 0 | 0.212 | 0.264 | 0.112 | 0.212 | 1 | 0 | 0.301 | 0.051 | 0 | -0.11 |
| SE | 10661 | ENSG00000161395 | PGAP3 | chr17 | - | 39685921 | 39686019 | 39684596 | 39684749 | 39687833 | 39688001 | 60 | 37 | 71 | 39 | 39 | 0 | 0 | 0 | 0 | 0 | 24 | 41 | 14 | 10 | 80 | 0 | 0 | 0 | 0 | 5 | 247 | 149 | 0.862578032 | 1 | 1 | 1 | 1 | 1 | 1 | 1 | 1 | 1 | 1 | 0.906 | 0.019 |
| SE | 27311 | ENSG00000136813 | ECPAS | chr9 | - | 111472896 | 111473000 | 111451424 | 111451555 | 111483984 | 111484057 | 26 | 26 | 20 | 25 | 52 | 2 | 0 | 1 | 2 | 0 | 33 | 26 | 32 | 38 | 18 | 0 | 0 | 0 | 0 | 0 | 253 | 149 | 0.862636439 | 1 | 0.884 | 1 | 0.922 | 0.88 | 1 | 1 | 1 | 1 | 1 | 1 | -0.063 |
| SE | 45648 | ENSG00000144283 | PKP4 | chr2 | + | 158624877 | 158625427 | 158621230 | 158621421 | 158631752 | 158631941 | 38 | 42 | 58 | 65 | 116 | 3 | 0 | 1 | 0 | 0 | 34 | 65 | 82 | 135 | 149 | 0 | 1 | 2 | 0 | 4 | 298 | 149 | 0.862674527 | 1 | 0.864 | 1 | 0.967 | 1 | 1 | 1 | 0.97 | 0.953 | 1 | 0.949 | -0.008 |
| SE | 47281 | ENSG00000113360 | DROSHA | chr5 | - | 31531449 | 31531525 | 31530797 | 31530924 | 31531989 | 31532074 | 33 | 28 | 35 | 36 | 36 | 4 | 6 | 1 | 2 | 7 | 7 | 66 | 26 | 33 | 34 | 8 | 3 | 3 | 6 | 0 | 225 | 149 | 0.862722887 | 1 | 0.845 | 0.756 | 0.959 | 0.923 | 0.773 | 0.367 | 0.936 | 0.852 | 0.785 | 1 | 0.063 |
| SE | 11708 | ENSG00000167978 | SRRM2 | chr16 | + | 2758984 | 2759047 | 2758469 | 2758547 | 2759139 | 2759172 | 742 | 550 | 667 | 704 | 1126 | 1 | 1 | 1 | 1 | 0 | 1588 | 782 | 863 | 1215 | 1319 | 6 | 1 | 2 | 5 | 18 | 212 | 149 | 0.862731309 | 1 | 0.998 | 0.997 | 0.998 | 0.998 | 1 | 0.995 | 0.998 | 0.997 | 0.994 | 0.981 | 0.005 |
| SE | 53485 | ENSG00000115350 | POLE4 | chr2 | + | 74959705 | 74959756 | 74959340 | 74959425 | 74960104 | 74960146 | 1 | 6 | 1 | 0 | 0 | 68 | 33 | 45 | 56 | 80 | 0 | 0 | 0 | 0 | 6 | 89 | 77 | 40 | 56 | 126 | 200 | 149 | 0.862814658 | 1 | 0.011 | 0.119 | 0.016 | 0 | 0 | 0 | 0 | 0 | 0 | 0.034 | 0.022 |
| SE | 28051 | ENSG00000114738 | MAPKAPK3 | chr3 | + | 50642252 | 50642332 | 50641706 | 50641771 | 50646739 | 50646825 | 161 | 101 | 119 | 163 | 97 | 0 | 0 | 0 | 0 | 0 | 375 | 161 | 110 | 307 | 456 | 0 | 0 | 0 | 0 | 12 | 229 | 149 | 0.86295316 | 1 | 1 | 1 | 1 | 1 | 1 | 1 | 1 | 1 | 1 | 0.961 | 0.008 |
| SE | 104887 | ENSG00000142166 | IFNAR1 | chr21 | + | 33340998 | 33341174 | 33335523 | 33335647 | 33343267 | 33343422 | 425 | 307 | 415 | 443 | 733 | 3 | 4 | 2 | 4 | 0 | 257 | 303 | 386 | 366 | 38 | 0 | 1 | 1 | 0 | 0 | 298 | 149 | 0.86297791 | 1 | 0.986 | 0.975 | 0.99 | 0.982 | 1 | 1 | 0.993 | 0.995 | 1 | 1 | -0.011 |
| SE | 31055 | ENSG00000065882 | TBC1D1 | chr4 | + | 38020590 | 38020695 | 38018353 | 38018443 | 38021585 | 38021718 | 231 | 147 | 203 | 190 | 199 | 0 | 0 | 0 | 0 | 0 | 509 | 247 | 247 | 315 | 436 | 0 | 7 | 0 | 4 | 0 | 254 | 149 | 0.862984951 | 1 | 1 | 1 | 1 | 1 | 1 | 1 | 0.954 | 1 | 0.979 | 1 | 0.013 |
| SE | 70666 | ENSG00000168214 | RBPJ | chr4 | + | 26430418 | 26430522 | 26429897 | 26430053 | 26430691 | 26430703 | 250 | 181 | 185 | 235 | 403 | 3 | 0 | 1 | 4 | 7 | 24 | 141 | 149 | 111 | 9 | 0 | 0 | 0 | 8 | 0 | 253 | 149 | 0.863013917 | 1 | 0.98 | 1 | 0.991 | 0.972 | 0.971 | 1 | 1 | 1 | 0.891 | 1 | 0.005 |
| SE | 107354 | ENSG00000110921 | MVK | chr12 | + | 109579164 | 109579282 | 109575997 | 109576145 | 109579801 | 109579946 | 0 | 0 | 5 | 2 | 0 | 23 | 10 | 20 | 21 | 11 | 0 | 0 | 0 | 0 | 0 | 16 | 17 | 3 | 2 | 6 | 267 | 149 | 0.863118041 | 1 | 0 | 0 | 0.122 | 0.05 | 0 | 0 | 0 | 0 | 0 | 0 | 0.034 |
| SE | 59237 | ENSG00000117153 | KLHL12 | chr1 | - | 202896853 | 202896960 | 202895521 | 202895717 | 202909009 | 202909124 | 131 | 65 | 101 | 124 | 112 | 0 | 0 | 1 | 0 | 0 | 113 | 92 | 79 | 51 | 79 | 0 | 2 | 4 | 0 | 0 | 256 | 149 | 0.863132871 | 1 | 1 | 1 | 0.983 | 1 | 1 | 1 | 0.964 | 0.92 | 1 | 1 | 0.02 |
| SE | 109134 | ENSG00000185989 | RASA3 | chr13 | - | 114018103 | 114018252 | 114017236 | 114017351 | 114021403 | 114021508 | 345 | 139 | 247 | 240 | 213 | 0 | 0 | 0 | 0 | 0 | 247 | 209 | 197 | 137 | 441 | 0 | 0 | 0 | 0 | 10 | 298 | 149 | 0.863154423 | 1 | 1 | 1 | 1 | 1 | 1 | 1 | 1 | 1 | 1 | 0.957 | 0.009 |
| SE | 15665 | ENSG00000092439 | TRPM7 | chr15 | - | 50614122 | 50614263 | 50613706 | 50613841 | 50619744 | 50619798 | 252 | 262 | 269 | 279 | 319 | 1 | 0 | 0 | 3 | 0 | 12 | 211 | 175 | 74 | 27 | 0 | 2 | 1 | 5 | 0 | 290 | 149 | 0.863164081 | 1 | 0.992 | 1 | 1 | 0.98 | 1 | 1 | 0.982 | 0.989 | 0.884 | 1 | 0.023 |
| SE | 37495 | ENSG00000163681 | SLMAP | chr3 | + | 57925844 | 57925934 | 57922888 | 57923023 | 57927295 | 57927393 | 179 | 178 | 105 | 206 | 317 | 62 | 36 | 69 | 55 | 95 | 8 | 57 | 35 | 56 | 17 | 0 | 97 | 79 | 26 | 0 | 239 | 149 | 0.863204567 | 1 | 0.643 | 0.755 | 0.487 | 0.7 | 0.675 | 1 | 0.268 | 0.216 | 0.573 | 1 | 0.041 |
| SE | 54089 | ENSG00000115935 | WIPF1 | chr2 | - | 174575203 | 174575380 | 174571168 | 174572446 | 174585522 | 174585611 | 402 | 221 | 344 | 362 | 423 | 0 | 0 | 0 | 0 | 0 | 291 | 333 | 280 | 336 | 674 | 9 | 0 | 0 | 2 | 0 | 298 | 149 | 0.863253985 | 1 | 1 | 1 | 1 | 1 | 1 | 0.942 | 1 | 1 | 0.988 | 1 | 0.014 |
| SE | 55815 | ENSG00000106615 | RHEB | chr7 | - | 151484736 | 151484804 | 151477332 | 151477415 | 151490942 | 151491014 | 296 | 206 | 200 | 289 | 280 | 0 | 0 | 0 | 1 | 0 | 606 | 390 | 273 | 584 | 795 | 0 | 0 | 0 | 3 | 15 | 217 | 149 | 0.863362044 | 1 | 1 | 1 | 1 | 0.995 | 1 | 1 | 1 | 1 | 0.993 | 0.973 | 0.006 |
| SE | 37199 | ENSG00000149480 | MTA2 | chr11 | - | 62596636 | 62596825 | 62596457 | 62596532 | 62597315 | 62597415 | 564 | 267 | 469 | 480 | 432 | 0 | 0 | 0 | 0 | 0 | 668 | 664 | 312 | 453 | 561 | 0 | 5 | 0 | 0 | 9 | 298 | 149 | 0.863425835 | 1 | 1 | 1 | 1 | 1 | 1 | 1 | 0.985 | 1 | 1 | 0.969 | 0.009 |
| SE | 103557 | ENSG00000150093 | ITGB1 | chr10 | - | 32901511 | 32901635 | 32887331 | 32889002 | 32908367 | 32908534 | 691 | 486 | 351 | 835 | 606 | 0 | 0 | 0 | 0 | 12 | 70 | 400 | 595 | 216 | 132 | 0 | 0 | 0 | 0 | 0 | 273 | 149 | 0.863455849 | 1 | 1 | 1 | 1 | 1 | 0.965 | 1 | 1 | 1 | 1 | 1 | -0.007 |
| SE | 52540 | ENSG00000029534 | ANK1 | chr8 | - | 41672515 | 41672912 | 41672353 | 41672426 | 41684543 | 41684690 | 5 | 4 | 1 | 1 | 7 | 0 | 6 | 1 | 1 | 3 | 0 | 7 | 6 | 4 | 91 | 9 | 2 | 4 | 0 | 46 | 298 | 149 | 0.863483495 | 1 | 1 | 0.25 | 0.333 | 0.333 | 0.538 | 0 | 0.636 | 0.429 | 1 | 0.497 | -0.022 |
| SE | 63612 | ENSG00000138029 | HADHB | chr2 | + | 26269952 | 26269997 | 26263379 | 26263479 | 26273650 | 26273750 | 202 | 143 | 175 | 179 | 231 | 0 | 2 | 1 | 0 | 0 | 277 | 250 | 146 | 236 | 375 | 6 | 0 | 0 | 0 | 8 | 194 | 149 | 0.863545441 | 1 | 1 | 0.982 | 0.993 | 1 | 1 | 0.973 | 1 | 1 | 1 | 0.973 | 0.006 |
| SE | 26406 | ENSG00000165494 | PCF11 | chr11 | + | 83165599 | 83166714 | 83164206 | 83164401 | 83167124 | 83167308 | 535 | 395 | 458 | 533 | 675 | 0 | 0 | 0 | 0 | 0 | 11 | 285 | 254 | 110 | 12 | 0 | 4 | 0 | 3 | 0 | 298 | 149 | 0.863630648 | 1 | 1 | 1 | 1 | 1 | 1 | 1 | 0.973 | 1 | 0.948 | 1 | 0.016 |
| SE | 57931 | ENSG00000168439 | STIP1 | chr11 | + | 64194478 | 64194620 | 64193077 | 64193287 | 64195644 | 64195811 | 153 | 116 | 132 | 208 | 249 | 0 | 0 | 0 | 0 | 0 | 275 | 107 | 104 | 131 | 180 | 10 | 0 | 0 | 0 | 0 | 291 | 149 | 0.863713454 | 1 | 1 | 1 | 1 | 1 | 1 | 0.934 | 1 | 1 | 1 | 1 | 0.013 |
| SE | 85519 | ENSG00000139083 | ETV6 | chr12 | + | 11716586 | 11716743 | 11650010 | 11650160 | 11752449 | 11752875 | 1 | 0 | 0 | 0 | 0 | 84 | 58 | 76 | 84 | 145 | 12 | 0 | 0 | 0 | 0 | 166 | 120 | 121 | 93 | 127 | 298 | 149 | 0.863782798 | 1 | 0.006 | 0 | 0 | 0 | 0 | 0.035 | 0 | 0 | 0 | 0 | -0.006 |
| SE | 62282 | ENSG00000112031 | MTRF1L | chr6 | - | 152994512 | 152994676 | 152992856 | 152992974 | 152995135 | 152995294 | 156 | 160 | 128 | 179 | 229 | 2 | 0 | 0 | 2 | 0 | 187 | 266 | 181 | 265 | 126 | 0 | 7 | 0 | 2 | 0 | 298 | 149 | 0.86380965 | 1 | 0.975 | 1 | 1 | 0.978 | 1 | 1 | 0.95 | 1 | 0.985 | 1 | 0.004 |
| SE | 29210 | ENSG00000101109 | STK4 | chr20 | + | 44997168 | 44997306 | 44995089 | 44995257 | 45000391 | 45000520 | 1809 | 1275 | 1685 | 1982 | 1987 | 2 | 2 | 0 | 4 | 3 | 1479 | 1342 | 1735 | 1332 | 561 | 5 | 5 | 19 | 3 | 0 | 287 | 149 | 0.863893462 | 1 | 0.998 | 0.997 | 1 | 0.996 | 0.997 | 0.994 | 0.993 | 0.979 | 0.996 | 1 | 0.005 |
| SE | 108522 | ENSG00000065615 | CYB5R4 | chr6 | + | 83921081 | 83921175 | 83919396 | 83919454 | 83922437 | 83922470 | 348 | 285 | 283 | 296 | 470 | 0 | 0 | 0 | 0 | 0 | 80 | 241 | 308 | 105 | 29 | 0 | 7 | 0 | 1 | 0 | 243 | 149 | 0.863923143 | 1 | 1 | 1 | 1 | 1 | 1 | 1 | 0.955 | 1 | 0.985 | 1 | 0.012 |
| SE | 87206 | ENSG00000086758 | HUWE1 | chrX | - | 53564573 | 53564722 | 53562830 | 53562929 | 53565066 | 53565239 | 491 | 427 | 455 | 584 | 321 | 0 | 1 | 1 | 1 | 0 | 920 | 562 | 467 | 683 | 623 | 0 | 6 | 1 | 8 | 0 | 298 | 149 | 0.863933086 | 1 | 1 | 0.995 | 0.996 | 0.997 | 1 | 1 | 0.979 | 0.996 | 0.977 | 1 | 0.007 |
| SE | 31560 | ENSG00000163913 | IFT122 | chr3 | + | 129461227 | 129461304 | 129451913 | 129451998 | 129463559 | 129463626 | 14 | 6 | 7 | 18 | 22 | 10 | 5 | 6 | 1 | 11 | 65 | 15 | 12 | 23 | 55 | 22 | 9 | 10 | 16 | 11 | 226 | 149 | 0.864006127 | 1 | 0.48 | 0.442 | 0.435 | 0.922 | 0.569 | 0.661 | 0.524 | 0.442 | 0.487 | 0.767 | -0.007 |
| SE | 109325 | ENSG00000123104 | ITPR2 | chr12 | - | 26495151 | 26495260 | 26494152 | 26494340 | 26550246 | 26550355 | 349 | 294 | 171 | 290 | 411 | 0 | 0 | 0 | 0 | 0 | 52 | 196 | 263 | 63 | 8 | 0 | 0 | 7 | 0 | 0 | 258 | 149 | 0.864013564 | 1 | 1 | 1 | 1 | 1 | 1 | 1 | 1 | 0.956 | 1 | 1 | 0.009 |
| SE | 9139 | ENSG00000006695 | COX10 | chr17 | + | 14191988 | 14192221 | 14159876 | 14159947 | 14206809 | 14207268 | 61 | 26 | 60 | 60 | 72 | 1 | 0 | 1 | 5 | 0 | 65 | 37 | 42 | 22 | 95 | 0 | 5 | 1 | 0 | 0 | 298 | 149 | 0.864042793 | 1 | 0.968 | 1 | 0.968 | 0.857 | 1 | 1 | 0.787 | 0.955 | 1 | 1 | 0.01 |
| SE | 80054 | ENSG00000140287 | HDC | chr15 | - | 50248244 | 50248343 | 50243142 | 50243244 | 50252429 | 50252520 | 25 | 15 | 29 | 34 | 26 | 3 | 0 | 2 | 4 | 0 | 11 | 31 | 2 | 10 | 4 | 0 | 0 | 2 | 5 | 0 | 248 | 149 | 0.864112293 | 1 | 0.834 | 1 | 0.897 | 0.836 | 1 | 1 | 1 | 0.375 | 0.546 | 1 | 0.129 |
| SE | 46734 | ENSG00000163125 | RPRD2 | chr1 | + | 150441880 | 150441958 | 150440922 | 150441023 | 150443230 | 150443283 | 308 | 222 | 206 | 236 | 406 | 122 | 94 | 121 | 187 | 184 | 244 | 310 | 156 | 202 | 152 | 111 | 170 | 77 | 121 | 81 | 227 | 149 | 0.864115967 | 1 | 0.624 | 0.608 | 0.528 | 0.453 | 0.592 | 0.591 | 0.545 | 0.571 | 0.523 | 0.552 | 0.005 |
| SE | 11889 | ENSG00000103222 | ABCC1 | chr16 | + | 16016495 | 16016621 | 16014490 | 16014628 | 16033108 | 16033170 | 180 | 74 | 171 | 177 | 247 | 4 | 2 | 4 | 8 | 2 | 163 | 208 | 175 | 223 | 370 | 0 | 9 | 11 | 2 | 6 | 275 | 149 | 0.864150538 | 1 | 0.961 | 0.952 | 0.959 | 0.923 | 0.985 | 1 | 0.926 | 0.896 | 0.984 | 0.971 | 0.001 |
| SE | 106987 | ENSG00000111271 | ACAD10 | chr12 | + | 111749010 | 111749103 | 111748316 | 111748475 | 111749172 | 111749769 | 0 | 0 | 0 | 1 | 0 | 10 | 5 | 24 | 29 | 12 | 0 | 0 | 0 | 0 | 7 | 47 | 7 | 9 | 14 | 29 | 242 | 149 | 0.864183623 | 1 | 0 | 0 | 0 | 0.021 | 0 | 0 | 0 | 0 | 0 | 0.129 | -0.022 |
| SE | 43500 | ENSG00000153563 | CD8A | chr2 | - | 86789639 | 86789750 | 86788529 | 86788560 | 86790327 | 86790681 | 135 | 56 | 107 | 128 | 57 | 0 | 0 | 0 | 0 | 0 | 72 | 165 | 69 | 92 | 172 | 3 | 2 | 2 | 0 | 0 | 260 | 149 | 0.864189376 | 1 | 1 | 1 | 1 | 1 | 1 | 0.932 | 0.979 | 0.952 | 1 | 1 | 0.027 |
| SE | 38477 | ENSG00000147251 | DOCK11 | chrX | + | 118675935 | 118676049 | 118671022 | 118671145 | 118676590 | 118676737 | 908 | 655 | 641 | 892 | 1209 | 2 | 2 | 8 | 8 | 7 | 80 | 493 | 466 | 62 | 64 | 0 | 1 | 2 | 0 | 0 | 263 | 149 | 0.864276192 | 1 | 0.996 | 0.995 | 0.978 | 0.984 | 0.99 | 1 | 0.996 | 0.992 | 1 | 1 | -0.009 |
| SE | 17352 | ENSG00000133961 | NUMB | chr14 | - | 73297210 | 73297285 | 73292733 | 73292874 | 73323129 | 73323204 | 445 | 358 | 283 | 319 | 458 | 3 | 1 | 4 | 1 | 0 | 768 | 329 | 456 | 484 | 437 | 0 | 6 | 1 | 3 | 7 | 224 | 149 | 0.86443702 | 1 | 0.99 | 0.996 | 0.979 | 0.995 | 1 | 1 | 0.973 | 0.997 | 0.991 | 0.976 | 0.005 |
| SE | 45411 | ENSG00000070190 | DAPP1 | chr4 | + | 99863769 | 99863855 | 99863009 | 99863072 | 99866033 | 99866121 | 296 | 520 | 202 | 608 | 598 | 0 | 0 | 0 | 0 | 0 | 634 | 618 | 278 | 531 | 283 | 0 | 0 | 4 | 9 | 0 | 235 | 149 | 0.864485683 | 1 | 1 | 1 | 1 | 1 | 1 | 1 | 1 | 0.978 | 0.974 | 1 | 0.01 |
| SE | 19412 | ENSG00000092203 | TOX4 | chr14 | + | 21482551 | 21482611 | 21477495 | 21477564 | 21487450 | 21487693 | 2 | 3 | 11 | 4 | 2 | 237 | 148 | 218 | 304 | 233 | 0 | 0 | 2 | 0 | 0 | 408 | 298 | 273 | 221 | 384 | 209 | 149 | 0.864495211 | 1 | 0.006 | 0.014 | 0.035 | 0.009 | 0.006 | 0 | 0 | 0.005 | 0 | 0 | 0.013 |
| SE | 32991 | ENSG00000114316 | USP4 | chr3 | - | 49278813 | 49278902 | 49277143 | 49278451 | 49280743 | 49280847 | 302 | 208 | 249 | 291 | 402 | 9 | 9 | 5 | 13 | 12 | 502 | 293 | 309 | 341 | 423 | 14 | 7 | 6 | 12 | 23 | 238 | 149 | 0.864531268 | 1 | 0.955 | 0.935 | 0.969 | 0.933 | 0.954 | 0.957 | 0.963 | 0.97 | 0.947 | 0.92 | -0.002 |
| SE | 40075 | ENSG00000125347 | IRF1 | chr5 | - | 132486250 | 132486373 | 132483761 | 132484075 | 132486556 | 132486639 | 616 | 378 | 594 | 791 | 682 | 10 | 0 | 6 | 5 | 2 | 2000 | 414 | 846 | 991 | 1263 | 0 | 2 | 2 | 4 | 0 | 272 | 149 | 0.86456921 | 1 | 0.971 | 1 | 0.982 | 0.989 | 0.995 | 1 | 0.991 | 0.996 | 0.993 | 1 | -0.009 |
| SE | 106489 | ENSG00000176915 | ANKLE2 | chr12 | - | 132743153 | 132743276 | 132741418 | 132741485 | 132747831 | 132748020 | 165 | 107 | 156 | 179 | 190 | 1 | 0 | 0 | 2 | 0 | 229 | 198 | 218 | 172 | 250 | 0 | 10 | 0 | 0 | 0 | 272 | 149 | 0.864623291 | 1 | 0.989 | 1 | 1 | 0.98 | 1 | 1 | 0.916 | 1 | 1 | 1 | 0.011 |
| SE | 21572 | ENSG00000198833 | UBE2J1 | chr6 | - | 89342323 | 89342455 | 89338458 | 89338543 | 89352538 | 89352722 | 515 | 496 | 620 | 578 | 747 | 0 | 0 | 0 | 1 | 0 | 1583 | 1200 | 1569 | 1305 | 930 | 0 | 16 | 0 | 0 | 0 | 281 | 149 | 0.864630184 | 1 | 1 | 1 | 1 | 0.997 | 1 | 1 | 0.975 | 1 | 1 | 1 | 0.004 |
| SE | 97703 | ENSG00000170145 | SIK2 | chr11 | + | 111701451 | 111701575 | 111700885 | 111701010 | 111703202 | 111703423 | 92 | 51 | 70 | 100 | 73 | 0 | 0 | 0 | 0 | 0 | 155 | 94 | 49 | 74 | 81 | 0 | 6 | 0 | 0 | 0 | 273 | 149 | 0.864683679 | 1 | 1 | 1 | 1 | 1 | 1 | 1 | 0.895 | 1 | 1 | 1 | 0.021 |
| SE | 87944 | ENSG00000104361 | NIPAL2 | chr8 | - | 98217038 | 98217286 | 98212404 | 98212501 | 98222478 | 98222600 | 14 | 1 | 4 | 19 | 7 | 42 | 13 | 33 | 28 | 42 | 0 | 11 | 5 | 7 | 0 | 4 | 33 | 11 | 22 | 2 | 298 | 149 | 0.86479698 | 1 | 0.143 | 0.037 | 0.057 | 0.253 | 0.077 | 0 | 0.143 | 0.185 | 0.137 | 0 | 0.02 |
| SE | 28901 | ENSG00000145495 | MARCHF6 | chr5 | + | 10381799 | 10381943 | 10378758 | 10378832 | 10390331 | 10390491 | 243 | 197 | 213 | 274 | 249 | 2 | 0 | 8 | 0 | 0 | 332 | 263 | 283 | 219 | 455 | 0 | 12 | 1 | 0 | 0 | 293 | 149 | 0.864805202 | 1 | 0.984 | 1 | 0.931 | 1 | 1 | 1 | 0.918 | 0.993 | 1 | 1 | 0.001 |
| SE | 46920 | ENSG00000141027 | NCOR1 | chr17 | - | 16048844 | 16048985 | 16046950 | 16047093 | 16057513 | 16057730 | 134 | 113 | 152 | 156 | 127 | 0 | 0 | 1 | 0 | 0 | 88 | 114 | 96 | 115 | 59 | 0 | 0 | 2 | 4 | 0 | 290 | 149 | 0.864925542 | 1 | 1 | 1 | 0.987 | 1 | 1 | 1 | 1 | 0.961 | 0.937 | 1 | 0.018 |
| SE | 41198 | ENSG00000146416 | AIG1 | chr6 | + | 143284109 | 143284225 | 143165081 | 143165183 | 143333281 | 143333293 | 42 | 24 | 25 | 36 | 60 | 0 | 1 | 8 | 0 | 1 | 35 | 16 | 19 | 50 | 94 | 0 | 0 | 3 | 7 | 0 | 265 | 149 | 0.865068072 | 1 | 1 | 0.931 | 0.637 | 1 | 0.971 | 1 | 1 | 0.781 | 0.801 | 1 | -0.009 |
| SE | 37633 | ENSG00000082074 | FYB1 | chr5 | - | 39250670 | 39250806 | 39208863 | 39208964 | 39270568 | 39270644 | 34 | 2 | 2 | 5 | 3 | 5 | 2 | 0 | 1 | 1 | 7 | 22 | 10 | 4 | 12 | 0 | 3 | 3 | 0 | 3 | 285 | 149 | 0.865084608 | 1 | 0.78 | 0.343 | 1 | 0.723 | 0.611 | 1 | 0.793 | 0.635 | 1 | 0.677 | -0.13 |
| SE | 99094 | ENSG00000110274 | CEP164 | chr11 | + | 117373750 | 117373831 | 117371079 | 117371466 | 117375707 | 117375791 | 79 | 46 | 53 | 79 | 91 | 2 | 0 | 3 | 10 | 0 | 99 | 89 | 82 | 63 | 123 | 0 | 2 | 1 | 6 | 8 | 230 | 149 | 0.865089388 | 1 | 0.962 | 1 | 0.92 | 0.837 | 1 | 1 | 0.966 | 0.982 | 0.872 | 0.909 | -0.002 |
| SE | 12642 | ENSG00000090857 | PDPR | chr16 | + | 70153390 | 70153573 | 70148463 | 70148553 | 70156474 | 70156614 | 104 | 63 | 93 | 85 | 89 | 7 | 1 | 9 | 7 | 6 | 88 | 157 | 55 | 83 | 337 | 0 | 16 | 0 | 2 | 43 | 298 | 149 | 0.865114914 | 1 | 0.881 | 0.969 | 0.838 | 0.859 | 0.881 | 1 | 0.831 | 1 | 0.954 | 0.797 | -0.031 |
| SE | 84943 | ENSG00000139436 | GIT2 | chr12 | - | 109945259 | 109945349 | 109939164 | 109939247 | 109951166 | 109951316 | 25 | 21 | 24 | 24 | 25 | 1 | 0 | 4 | 5 | 1 | 60 | 26 | 31 | 32 | 27 | 8 | 0 | 2 | 4 | 0 | 239 | 149 | 0.865197169 | 1 | 0.94 | 1 | 0.789 | 0.75 | 0.94 | 0.824 | 1 | 0.906 | 0.833 | 1 | -0.029 |
| SE | 85084 | ENSG00000164162 | ANAPC10 | chr4 | - | 145064571 | 145064692 | 144994574 | 144995603 | 145081659 | 145081750 | 84 | 44 | 67 | 83 | 225 | 3 | 5 | 1 | 4 | 9 | 48 | 50 | 78 | 43 | 15 | 17 | 0 | 2 | 0 | 0 | 270 | 149 | 0.865261132 | 1 | 0.939 | 0.829 | 0.974 | 0.92 | 0.932 | 0.609 | 1 | 0.956 | 1 | 1 | 0.006 |
| SE | 35778 | ENSG00000188687 | SLC4A5 | chr2 | - | 74328119 | 74328186 | 74314944 | 74315025 | 74334026 | 74334139 | 10 | 3 | 4 | 9 | 4 | 0 | 2 | 2 | 0 | 0 | 3 | 12 | 13 | 6 | 5 | 0 | 4 | 2 | 0 | 0 | 216 | 149 | 0.865300037 | 1 | 1 | 0.509 | 0.58 | 1 | 1 | 1 | 0.674 | 0.818 | 1 | 1 | -0.081 |
| SE | 20590 | ENSG00000196367 | TRRAP | chr7 | + | 99030594 | 99030757 | 99012070 | 99014955 | 99030884 | 99031000 | 10 | 8 | 9 | 8 | 14 | 0 | 1 | 0 | 4 | 0 | 16 | 2 | 5 | 7 | 2 | 0 | 0 | 2 | 2 | 0 | 298 | 149 | 0.865306585 | 1 | 1 | 0.8 | 1 | 0.5 | 1 | 1 | 1 | 0.556 | 0.636 | 1 | 0.022 |
| SE | 64028 | ENSG00000115464 | USP34 | chr2 | - | 61214059 | 61214218 | 61211771 | 61211929 | 61220309 | 61220457 | 211 | 151 | 164 | 232 | 219 | 3 | 0 | 2 | 6 | 0 | 15 | 78 | 97 | 33 | 3 | 0 | 0 | 0 | 2 | 0 | 298 | 149 | 0.865383828 | 1 | 0.972 | 1 | 0.976 | 0.951 | 1 | 1 | 1 | 1 | 0.892 | 1 | 0.001 |
| SE | 48643 | ENSG00000131374 | TBC1D5 | chr3 | - | 17586352 | 17586557 | 17508548 | 17508605 | 17622647 | 17622765 | 18 | 15 | 20 | 12 | 27 | 23 | 18 | 9 | 22 | 43 | 11 | 9 | 8 | 22 | 3 | 24 | 20 | 17 | 18 | 0 | 298 | 149 | 0.865462781 | 1 | 0.281 | 0.294 | 0.526 | 0.214 | 0.239 | 0.186 | 0.184 | 0.19 | 0.379 | 1 | -0.077 |
| SE | 73507 | ENSG00000138380 | CARF | chr2 | + | 202924356 | 202924415 | 202917876 | 202918043 | 202941859 | 202941980 | 15 | 9 | 17 | 8 | 22 | 51 | 46 | 59 | 44 | 34 | 5 | 12 | 4 | 11 | 1 | 35 | 25 | 23 | 13 | 9 | 208 | 149 | 0.865525515 | 1 | 0.174 | 0.123 | 0.171 | 0.115 | 0.317 | 0.093 | 0.256 | 0.111 | 0.377 | 0.074 | -0.002 |
| SE | 4765 | ENSG00000104885 | DOT1L | chr19 | + | 2191011 | 2191240 | 2189731 | 2189795 | 2193688 | 2193783 | 76 | 47 | 63 | 100 | 54 | 0 | 0 | 0 | 0 | 1 | 213 | 112 | 59 | 144 | 158 | 0 | 0 | 1 | 0 | 6 | 298 | 149 | 0.865530389 | 1 | 1 | 1 | 1 | 1 | 0.964 | 1 | 1 | 0.967 | 1 | 0.929 | 0.014 |
| SE | 44484 | ENSG00000090266 | NDUFB2 | chr7 | + | 140700873 | 140700959 | 140696708 | 140696842 | 140702865 | 140703010 | 0 | 0 | 6 | 1 | 0 | 89 | 20 | 31 | 50 | 30 | 0 | 0 | 0 | 0 | 0 | 1 | 15 | 31 | 29 | 78 | 235 | 149 | 0.865638003 | 1 | 0 | 0 | 0.109 | 0.013 | 0 | 0 | 0 | 0 | 0 | 0 | 0.024 |
| SE | 18217 | ENSG00000089916 | GPATCH2L | chr14 | + | 76180763 | 76180849 | 76177987 | 76178042 | 76195877 | 76195972 | 194 | 152 | 183 | 209 | 360 | 0 | 0 | 1 | 0 | 2 | 138 | 184 | 161 | 142 | 192 | 8 | 0 | 0 | 1 | 0 | 235 | 149 | 0.865647978 | 1 | 1 | 1 | 0.991 | 1 | 0.991 | 0.916 | 1 | 1 | 0.989 | 1 | 0.015 |
| SE | 102295 | ENSG00000100225 | FBXO7 | chr22 | + | 32495492 | 32495530 | 32487744 | 32493281 | 32498143 | 32498608 | 352 | 209 | 353 | 370 | 699 | 5 | 3 | 3 | 7 | 2 | 553 | 404 | 424 | 474 | 850 | 10 | 6 | 3 | 0 | 6 | 187 | 149 | 0.865652441 | 1 | 0.982 | 0.982 | 0.989 | 0.977 | 0.996 | 0.978 | 0.982 | 0.991 | 1 | 0.991 | -0.003 |
| SE | 44980 | ENSG00000138798 | EGF | chr4 | + | 109993246 | 109993369 | 109988583 | 109988709 | 109994732 | 109994880 | 31 | 23 | 2 | 57 | 45 | 0 | 0 | 0 | 2 | 0 | 86 | 85 | 23 | 57 | 214 | 0 | 6 | 0 | 0 | 0 | 272 | 149 | 0.865652468 | 1 | 1 | 1 | 1 | 0.94 | 1 | 1 | 0.886 | 1 | 1 | 1 | 0.011 |
| SE | 86245 | ENSG00000134516 | DOCK2 | chr5 | + | 169670541 | 169670597 | 169669287 | 169669328 | 169671077 | 169671174 | 480 | 310 | 293 | 328 | 382 | 1 | 0 | 1 | 1 | 1 | 634 | 464 | 524 | 523 | 335 | 2 | 8 | 3 | 7 | 0 | 205 | 149 | 0.865681687 | 1 | 0.997 | 1 | 0.995 | 0.996 | 0.996 | 0.996 | 0.977 | 0.992 | 0.982 | 1 | 0.007 |
| SE | 27914 | ENSG00000118564 | FBXL5 | chr4 | - | 15644492 | 15644708 | 15640787 | 15640883 | 15655203 | 15655341 | 691 | 532 | 387 | 595 | 586 | 1 | 2 | 1 | 1 | 0 | 539 | 497 | 622 | 771 | 97 | 0 | 2 | 7 | 6 | 0 | 298 | 149 | 0.865717547 | 1 | 0.997 | 0.993 | 0.995 | 0.997 | 1 | 1 | 0.992 | 0.978 | 0.985 | 1 | 0.005 |
| SE | 51149 | ENSG00000165458 | INPPL1 | chr11 | + | 72230992 | 72231189 | 72230795 | 72230898 | 72231497 | 72231615 | 215 | 83 | 147 | 178 | 151 | 0 | 0 | 0 | 0 | 0 | 241 | 263 | 138 | 230 | 430 | 5 | 0 | 4 | 1 | 0 | 298 | 149 | 0.865767451 | 1 | 1 | 1 | 1 | 1 | 1 | 0.96 | 1 | 0.945 | 0.991 | 1 | 0.021 |
| SE | 96695 | ENSG00000135362 | PRR5L | chr11 | + | 36393845 | 36394011 | 36376380 | 36376704 | 36400996 | 36401264 | 262 | 34 | 43 | 159 | 93 | 20 | 15 | 8 | 7 | 0 | 13 | 142 | 30 | 14 | 81 | 0 | 14 | 0 | 1 | 14 | 298 | 149 | 0.865796485 | 1 | 0.868 | 0.531 | 0.729 | 0.919 | 1 | 1 | 0.835 | 1 | 0.875 | 0.743 | -0.081 |
| SE | 98413 | ENSG00000175334 | BANF1 | chr11 | + | 66002843 | 66003070 | 66002255 | 66002513 | 66003234 | 66003335 | 1 | 0 | 9 | 3 | 1 | 55 | 25 | 36 | 99 | 36 | 0 | 0 | 4 | 3 | 8 | 54 | 57 | 28 | 79 | 118 | 298 | 149 | 0.865851868 | 1 | 0.009 | 0 | 0.111 | 0.015 | 0.014 | 0 | 0 | 0.067 | 0.019 | 0.033 | 0.006 |
| SE | 30898 | ENSG00000145220 | LYAR | chr4 | - | 4286518 | 4286572 | 4283620 | 4283795 | 4290035 | 4290154 | 157 | 57 | 97 | 103 | 63 | 1 | 1 | 0 | 5 | 0 | 23 | 88 | 63 | 36 | 6 | 0 | 0 | 0 | 0 | 0 | 203 | 149 | 0.865861312 | 1 | 0.991 | 0.977 | 1 | 0.938 | 1 | 1 | 1 | 1 | 1 | 1 | -0.019 |
| SE | 103507 | ENSG00000107862 | GBF1 | chr10 | + | 102352457 | 102352518 | 102351842 | 102351951 | 102353599 | 102353654 | 93 | 63 | 97 | 98 | 186 | 2 | 2 | 5 | 1 | 7 | 233 | 103 | 84 | 146 | 197 | 0 | 4 | 4 | 13 | 0 | 210 | 149 | 0.865979459 | 1 | 0.971 | 0.957 | 0.932 | 0.986 | 0.95 | 1 | 0.948 | 0.937 | 0.888 | 1 | 0.005 |
| SE | 84258 | ENSG00000172943 | PHF8 | chrX | - | 53987765 | 53987944 | 53987077 | 53987163 | 53992735 | 53992839 | 373 | 275 | 277 | 315 | 377 | 1 | 0 | 0 | 0 | 0 | 664 | 340 | 306 | 507 | 561 | 0 | 5 | 6 | 0 | 0 | 298 | 149 | 0.865987965 | 1 | 0.995 | 1 | 1 | 1 | 1 | 1 | 0.971 | 0.962 | 1 | 1 | 0.012 |
| SE | 5162 | ENSG00000088827 | SIGLEC1 | chr20 | - | 3694662 | 3694923 | 3694220 | 3694532 | 3696585 | 3696888 | 12 | 22 | 13 | 63 | 7 | 0 | 0 | 0 | 0 | 0 | 758 | 315 | 91 | 714 | 60 | 0 | 4 | 0 | 7 | 0 | 298 | 149 | 0.866118735 | 1 | 1 | 1 | 1 | 1 | 1 | 1 | 0.975 | 1 | 0.981 | 1 | 0.009 |
| SE | 351 | ENSG00000160294 | MCM3AP | chr21 | - | 46240810 | 46241017 | 46236828 | 46236979 | 46243464 | 46243722 | 177 | 72 | 129 | 131 | 94 | 2 | 0 | 0 | 0 | 0 | 216 | 127 | 134 | 229 | 272 | 8 | 0 | 0 | 0 | 0 | 298 | 149 | 0.866141619 | 1 | 0.978 | 1 | 1 | 1 | 1 | 0.931 | 1 | 1 | 1 | 1 | 0.009 |
| SE | 18306 | ENSG00000100938 | GMPR2 | chr14 | + | 24233478 | 24233593 | 24232917 | 24233340 | 24235736 | 24235816 | 43 | 37 | 55 | 53 | 73 | 4 | 0 | 1 | 1 | 0 | 10 | 83 | 28 | 48 | 48 | 0 | 0 | 0 | 0 | 0 | 264 | 149 | 0.866151607 | 1 | 0.859 | 1 | 0.969 | 0.968 | 1 | 1 | 1 | 1 | 1 | 1 | -0.041 |
| SE | 41935 | ENSG00000163374 | YY1AP1 | chr1 | - | 155672520 | 155672731 | 155670319 | 155670464 | 155675009 | 155675051 | 167 | 106 | 149 | 171 | 257 | 0 | 1 | 0 | 2 | 3 | 335 | 286 | 175 | 252 | 205 | 0 | 0 | 6 | 0 | 4 | 298 | 149 | 0.866167038 | 1 | 1 | 0.981 | 1 | 0.977 | 0.977 | 1 | 1 | 0.936 | 1 | 0.962 | 0.007 |
| SE | 67205 | ENSG00000124151 | NCOA3 | chr20 | + | 47633495 | 47633636 | 47627921 | 47628023 | 47634047 | 47634195 | 332 | 220 | 348 | 296 | 769 | 0 | 0 | 0 | 0 | 0 | 164 | 333 | 311 | 252 | 64 | 8 | 0 | 1 | 0 | 0 | 290 | 149 | 0.866182976 | 1 | 1 | 1 | 1 | 1 | 1 | 0.913 | 1 | 0.994 | 1 | 1 | 0.019 |
| SE | 53814 | ENSG00000173575 | CHD2 | chr15 | + | 92946037 | 92946082 | 92945820 | 92945865 | 92946247 | 92946340 | 173 | 163 | 167 | 185 | 368 | 0 | 3 | 2 | 0 | 7 | 65 | 194 | 284 | 168 | 29 | 0 | 2 | 0 | 0 | 0 | 194 | 149 | 0.866242017 | 1 | 1 | 0.977 | 0.985 | 1 | 0.976 | 1 | 0.987 | 1 | 1 | 1 | -0.01 |
| SE | 106823 | ENSG00000100982 | PCIF1 | chr20 | + | 45939214 | 45939339 | 45938980 | 45939123 | 45940474 | 45940612 | 217 | 88 | 136 | 151 | 148 | 0 | 0 | 0 | 2 | 0 | 181 | 239 | 142 | 161 | 435 | 0 | 0 | 8 | 0 | 0 | 274 | 149 | 0.86624713 | 1 | 1 | 1 | 1 | 0.976 | 1 | 1 | 1 | 0.906 | 1 | 1 | 0.014 |
| SE | 18291 | ENSG00000140153 | WDR20 | chr14 | + | 102197754 | 102197847 | 102194937 | 102195120 | 102208602 | 102208902 | 3 | 17 | 6 | 7 | 10 | 24 | 5 | 10 | 19 | 30 | 20 | 0 | 12 | 19 | 0 | 10 | 13 | 21 | 0 | 10 | 242 | 149 | 0.866438091 | 1 | 0.071 | 0.677 | 0.27 | 0.185 | 0.17 | 0.552 | 0 | 0.26 | 1 | 0 | -0.088 |
| SE | 75117 | ENSG00000114423 | CBLB | chr3 | - | 105839422 | 105839470 | 105776395 | 105776542 | 105853413 | 105853664 | 1 | 1 | 0 | 2 | 3 | 175 | 117 | 135 | 118 | 186 | 0 | 2 | 1 | 5 | 0 | 26 | 58 | 99 | 24 | 20 | 197 | 149 | 0.866474153 | 1 | 0.004 | 0.006 | 0 | 0.013 | 0.012 | 0 | 0.025 | 0.008 | 0.136 | 0 | -0.027 |
| SE | 74022 | ENSG00000051620 | HEBP2 | chr6 | + | 138412082 | 138412256 | 138405970 | 138406033 | 138412879 | 138413154 | 0 | 0 | 0 | 0 | 6 | 1 | 11 | 5 | 6 | 16 | 0 | 0 | 0 | 0 | 0 | 5 | 9 | 15 | 1 | 7 | 298 | 149 | 0.866592864 | 1 | 0 | 0 | 0 | 0 | 0.158 | 0 | 0 | 0 | 0 | 0 | 0.032 |
| SE | 98481 | ENSG00000151702 | FLI1 | chr11 | + | 128772781 | 128772985 | 128768117 | 128768272 | 128781957 | 128782023 | 1271 | 702 | 953 | 1240 | 1580 | 0 | 0 | 2 | 2 | 6 | 1125 | 1154 | 602 | 1020 | 881 | 0 | 14 | 2 | 0 | 0 | 298 | 149 | 0.866655065 | 1 | 1 | 1 | 0.996 | 0.997 | 0.992 | 1 | 0.976 | 0.993 | 1 | 1 | 0.003 |
| SE | 56265 | ENSG00000143995 | MEIS1 | chr2 | + | 66512148 | 66512294 | 66464108 | 66464220 | 66547942 | 66548019 | 44 | 93 | 26 | 265 | 39 | 4 | 1 | 0 | 9 | 14 | 279 | 124 | 125 | 214 | 301 | 7 | 14 | 11 | 16 | 8 | 295 | 149 | 0.866657099 | 1 | 0.847 | 0.979 | 1 | 0.937 | 0.585 | 0.953 | 0.817 | 0.852 | 0.871 | 0.95 | -0.019 |
| SE | 91337 | ENSG00000120868 | APAF1 | chr12 | + | 98725414 | 98725540 | 98723638 | 98723764 | 98727172 | 98727191 | 250 | 245 | 190 | 222 | 474 | 0 | 1 | 0 | 2 | 0 | 139 | 210 | 81 | 296 | 23 | 0 | 0 | 3 | 4 | 0 | 275 | 149 | 0.86675501 | 1 | 1 | 0.993 | 1 | 0.984 | 1 | 1 | 1 | 0.936 | 0.976 | 1 | 0.013 |
| SE | 100348 | ENSG00000081237 | PTPRC | chr1 | + | 198742231 | 198742367 | 198735126 | 198735252 | 198744053 | 198744203 | 1328 | 915 | 905 | 1283 | 2097 | 0 | 5 | 1 | 4 | 0 | 102 | 679 | 489 | 83 | 101 | 0 | 0 | 0 | 1 | 6 | 285 | 149 | 0.866765197 | 1 | 1 | 0.99 | 0.998 | 0.994 | 1 | 1 | 1 | 1 | 0.977 | 0.898 | 0.021 |
| SE | 69614 | ENSG00000163412 | EIF4E3 | chr3 | - | 71696459 | 71696520 | 71693874 | 71693941 | 71725191 | 71725402 | 117 | 125 | 143 | 176 | 316 | 0 | 0 | 0 | 0 | 0 | 283 | 190 | 247 | 357 | 113 | 10 | 0 | 0 | 0 | 0 | 210 | 149 | 0.866845822 | 1 | 1 | 1 | 1 | 1 | 1 | 0.953 | 1 | 1 | 1 | 1 | 0.009 |
| SE | 105268 | ENSG00000082898 | XPO1 | chr2 | - | 61534422 | 61534518 | 61533771 | 61533903 | 61537561 | 61538356 | 0 | 0 | 1 | 2 | 5 | 334 | 259 | 307 | 321 | 240 | 0 | 2 | 0 | 2 | 7 | 55 | 197 | 184 | 153 | 33 | 245 | 149 | 0.866850798 | 1 | 0 | 0 | 0.002 | 0.004 | 0.013 | 0 | 0.006 | 0 | 0.008 | 0.114 | -0.022 |
| SE | 79034 | ENSG00000160218 | TRAPPC10 | chr21 | + | 44086801 | 44086960 | 44084121 | 44084263 | 44087698 | 44087928 | 499 | 219 | 367 | 489 | 356 | 3 | 0 | 0 | 0 | 0 | 499 | 442 | 381 | 394 | 402 | 0 | 0 | 0 | 0 | 11 | 298 | 149 | 0.867039549 | 1 | 0.988 | 1 | 1 | 1 | 1 | 1 | 1 | 1 | 1 | 0.948 | 0.008 |
| SE | 53406 | ENSG00000151304 | SRFBP1 | chr5 | + | 122019259 | 122019341 | 121994598 | 121994670 | 122020087 | 122020802 | 87 | 71 | 75 | 77 | 258 | 1 | 1 | 0 | 0 | 0 | 4 | 44 | 42 | 43 | 20 | 0 | 3 | 2 | 0 | 0 | 231 | 149 | 0.867179236 | 1 | 0.982 | 0.979 | 1 | 1 | 1 | 1 | 0.904 | 0.931 | 1 | 1 | 0.025 |
| SE | 54236 | ENSG00000143643 | TTC13 | chr1 | - | 230956420 | 230956560 | 230954332 | 230954403 | 230958223 | 230958299 | 18 | 4 | 3 | 6 | 16 | 189 | 111 | 98 | 169 | 156 | 7 | 13 | 0 | 0 | 24 | 138 | 155 | 123 | 106 | 169 | 289 | 149 | 0.867187841 | 1 | 0.047 | 0.018 | 0.016 | 0.018 | 0.05 | 0.025 | 0.041 | 0 | 0 | 0.068 | 0.003 |
| SE | 30952 | ENSG00000017260 | ATP2C1 | chr3 | + | 131001219 | 131001344 | 130999517 | 130999659 | 131016151 | 131016669 | 150 | 113 | 123 | 110 | 169 | 0 | 0 | 3 | 2 | 4 | 80 | 71 | 123 | 75 | 66 | 5 | 3 | 0 | 0 | 0 | 274 | 149 | 0.867266379 | 1 | 1 | 1 | 0.957 | 0.968 | 0.958 | 0.897 | 0.928 | 1 | 1 | 1 | 0.012 |
| SE | 73740 | ENSG00000119772 | DNMT3A | chr2 | - | 25239314 | 25239513 | 25239129 | 25239215 | 25240301 | 25240450 | 4 | 4 | 0 | 5 | 9 | 37 | 31 | 56 | 48 | 54 | 6 | 1 | 0 | 2 | 3 | 27 | 9 | 31 | 22 | 4 | 298 | 149 | 0.867463918 | 1 | 0.051 | 0.061 | 0 | 0.05 | 0.077 | 0.1 | 0.053 | 0 | 0.043 | 0.273 | -0.046 |
| SE | 67618 | ENSG00000175215 | CTDSP2 | chr12 | - | 57826345 | 57826402 | 57824226 | 57824319 | 57826995 | 57827097 | 487 | 252 | 446 | 513 | 379 | 0 | 0 | 0 | 0 | 0 | 461 | 478 | 283 | 288 | 924 | 0 | 0 | 0 | 11 | 0 | 206 | 149 | 0.867477549 | 1 | 1 | 1 | 1 | 1 | 1 | 1 | 1 | 1 | 0.95 | 1 | 0.01 |
| SE | 24056 | ENSG00000161960 | EIF4A1 | chr17 | + | 7578119 | 7578244 | 7577826 | 7577916 | 7578341 | 7578630 | 132 | 65 | 132 | 150 | 202 | 0 | 1 | 0 | 1 | 2 | 365 | 169 | 154 | 223 | 308 | 0 | 0 | 7 | 3 | 0 | 274 | 149 | 0.867579963 | 1 | 1 | 0.972 | 1 | 0.988 | 0.982 | 1 | 1 | 0.923 | 0.976 | 1 | 0.009 |
| SE | 95417 | ENSG00000148356 | LRSAM1 | chr9 | + | 127487536 | 127487763 | 127485735 | 127485835 | 127489443 | 127489518 | 28 | 19 | 22 | 13 | 7 | 0 | 0 | 0 | 1 | 0 | 72 | 36 | 24 | 37 | 82 | 6 | 0 | 0 | 0 | 0 | 298 | 149 | 0.867583963 | 1 | 1 | 1 | 1 | 0.867 | 1 | 0.857 | 1 | 1 | 1 | 1 | 0.002 |
| SE | 9399 | ENSG00000132383 | RPA1 | chr17 | + | 1872433 | 1872526 | 1853100 | 1853189 | 1875660 | 1875705 | 241 | 154 | 222 | 205 | 325 | 2 | 3 | 1 | 4 | 6 | 167 | 142 | 122 | 130 | 118 | 9 | 3 | 0 | 0 | 0 | 242 | 149 | 0.867592786 | 1 | 0.987 | 0.969 | 0.993 | 0.969 | 0.971 | 0.92 | 0.967 | 1 | 1 | 1 | 0 |
| SE | 103790 | ENSG00000049323 | LTBP1 | chr2 | + | 33280038 | 33280158 | 33275800 | 33275923 | 33293159 | 33293282 | 119 | 173 | 24 | 311 | 311 | 0 | 0 | 0 | 0 | 0 | 577 | 150 | 137 | 184 | 2240 | 0 | 5 | 4 | 0 | 4 | 269 | 149 | 0.867618037 | 1 | 1 | 1 | 1 | 1 | 1 | 1 | 0.943 | 0.95 | 1 | 0.997 | 0.022 |
| SE | 16152 | ENSG00000140400 | MAN2C1 | chr15 | - | 75365600 | 75365698 | 75364487 | 75364665 | 75366521 | 75366592 | 2 | 0 | 0 | 0 | 0 | 139 | 45 | 154 | 120 | 58 | 0 | 8 | 0 | 1 | 5 | 239 | 147 | 169 | 90 | 177 | 247 | 149 | 0.867707398 | 1 | 0.009 | 0 | 0 | 0 | 0 | 0 | 0.032 | 0 | 0.007 | 0.017 | -0.009 |
| SE | 79144 | ENSG00000104299 | INTS9 | chr8 | - | 28812326 | 28812461 | 28796543 | 28796655 | 28813491 | 28813612 | 79 | 54 | 76 | 63 | 181 | 0 | 1 | 0 | 1 | 2 | 35 | 34 | 50 | 45 | 69 | 0 | 3 | 1 | 2 | 0 | 284 | 149 | 0.867823369 | 1 | 1 | 0.966 | 1 | 0.971 | 0.979 | 1 | 0.856 | 0.963 | 0.922 | 1 | 0.035 |
| SE | 44066 | ENSG00000054118 | THRAP3 | chr1 | + | 36296582 | 36296770 | 36293850 | 36293935 | 36300885 | 36301084 | 1226 | 834 | 1015 | 969 | 2180 | 5 | 1 | 2 | 1 | 4 | 1869 | 1043 | 1200 | 1296 | 995 | 13 | 10 | 3 | 1 | 0 | 298 | 149 | 0.867957227 | 1 | 0.992 | 0.998 | 0.996 | 0.998 | 0.996 | 0.986 | 0.981 | 0.995 | 0.998 | 1 | 0.004 |
| SE | 32995 | ENSG00000114316 | USP4 | chr3 | - | 49280743 | 49280847 | 49278813 | 49278902 | 49283986 | 49284136 | 304 | 177 | 236 | 258 | 390 | 0 | 0 | 0 | 2 | 0 | 452 | 290 | 250 | 269 | 318 | 7 | 4 | 2 | 0 | 0 | 253 | 149 | 0.868023892 | 1 | 1 | 1 | 1 | 0.987 | 1 | 0.974 | 0.977 | 0.987 | 1 | 1 | 0.01 |
| SE | 69064 | ENSG00000081059 | TCF7 | chr5 | + | 134145259 | 134145340 | 134143591 | 134143640 | 134146223 | 134146303 | 13 | 3 | 11 | 5 | 23 | 171 | 132 | 335 | 223 | 344 | 0 | 9 | 6 | 3 | 0 | 59 | 69 | 60 | 63 | 46 | 230 | 149 | 0.86803058 | 1 | 0.047 | 0.015 | 0.021 | 0.014 | 0.042 | 0 | 0.078 | 0.061 | 0.03 | 0 | -0.006 |
| SE | 20575 | ENSG00000196367 | TRRAP | chr7 | + | 98890334 | 98890445 | 98881974 | 98882024 | 98892423 | 98892528 | 331 | 214 | 285 | 352 | 590 | 1 | 1 | 1 | 5 | 9 | 523 | 277 | 314 | 389 | 513 | 0 | 0 | 0 | 11 | 0 | 260 | 149 | 0.868119561 | 1 | 0.995 | 0.992 | 0.994 | 0.976 | 0.974 | 1 | 1 | 1 | 0.953 | 1 | -0.004 |
| SE | 86163 | ENSG00000113845 | TIMMDC1 | chr3 | + | 119503953 | 119504021 | 119498580 | 119498927 | 119513640 | 119513719 | 24 | 16 | 28 | 35 | 40 | 0 | 0 | 1 | 0 | 5 | 38 | 14 | 8 | 63 | 45 | 0 | 0 | 0 | 0 | 7 | 217 | 149 | 0.868139219 | 1 | 1 | 0.917 | 1 | 0.828 | 1 | 1 | 1 | 1 | 0.861 | 1 | -0.023 |
| SE | 91762 | ENSG00000143748 | NVL | chr1 | - | 224311799 | 224311857 | 224307990 | 224308263 | 224317693 | 224317793 | 80 | 69 | 79 | 78 | 186 | 3 | 9 | 10 | 7 | 5 | 55 | 86 | 89 | 58 | 21 | 0 | 5 | 12 | 7 | 0 | 207 | 149 | 0.868155133 | 1 | 0.95 | 0.847 | 0.85 | 0.889 | 0.964 | 1 | 0.925 | 0.842 | 0.856 | 1 | -0.025 |
| SE | 79550 | ENSG00000117868 | ESYT2 | chr7 | - | 158773340 | 158773396 | 158767653 | 158767774 | 158788003 | 158788093 | 500 | 302 | 533 | 471 | 943 | 1 | 2 | 0 | 0 | 0 | 349 | 277 | 248 | 359 | 255 | 0 | 10 | 1 | 0 | 0 | 205 | 149 | 0.868198827 | 1 | 0.997 | 0.991 | 1 | 1 | 1 | 1 | 0.953 | 0.994 | 1 | 1 | 0.008 |
| SE | 51406 | ENSG00000197302 | ZNF720 | chr16 | + | 31722625 | 31722752 | 31713233 | 31713435 | 31723257 | 31723353 | 47 | 28 | 27 | 53 | 31 | 1 | 0 | 1 | 0 | 3 | 19 | 21 | 12 | 8 | 12 | 0 | 0 | 2 | 0 | 0 | 276 | 149 | 0.868313573 | 1 | 0.962 | 1 | 0.936 | 1 | 0.848 | 1 | 1 | 0.764 | 1 | 1 | -0.004 |
| SE | 40283 | ENSG00000271383 | NBPF19 | chr1 | + | 149477944 | 149478047 | 149475759 | 149476107 | 149478879 | 149479094 | 125 | 168 | 138 | 152 | 164 | 0 | 0 | 0 | 0 | 0 | 253 | 131 | 185 | 170 | 91 | 9 | 0 | 2 | 0 | 0 | 252 | 149 | 0.868358741 | 1 | 1 | 1 | 1 | 1 | 1 | 0.943 | 1 | 0.982 | 1 | 1 | 0.015 |
| SE | 85307 | ENSG00000054983 | GALC | chr14 | - | 87986968 | 87987056 | 87986488 | 87986602 | 87988143 | 87988207 | 5 | 8 | 0 | 2 | 0 | 63 | 38 | 34 | 43 | 39 | 0 | 5 | 7 | 0 | 0 | 16 | 55 | 33 | 27 | 28 | 237 | 149 | 0.868362163 | 1 | 0.048 | 0.117 | 0 | 0.028 | 0 | 0 | 0.054 | 0.118 | 0 | 0 | 0.004 |
| SE | 32134 | ENSG00000084463 | WBP11 | chr12 | - | 14801877 | 14802032 | 14801319 | 14801428 | 14803351 | 14803486 | 2 | 1 | 0 | 4 | 0 | 110 | 50 | 92 | 90 | 118 | 6 | 12 | 2 | 3 | 0 | 295 | 167 | 163 | 190 | 120 | 298 | 149 | 0.868622088 | 1 | 0.009 | 0.01 | 0 | 0.022 | 0 | 0.01 | 0.035 | 0.006 | 0.008 | 0 | -0.004 |
| SE | 8404 | ENSG00000175106 | TVP23C | chr17 | - | 15540432 | 15540561 | 15502263 | 15503232 | 15545784 | 15545916 | 16 | 9 | 0 | 37 | 0 | 2 | 0 | 7 | 0 | 1 | 5 | 6 | 17 | 38 | 0 | 0 | 2 | 3 | 0 | 4 | 278 | 149 | 0.868683975 | 1 | 0.811 | 1 | 0 | 1 | 0 | 1 | 0.617 | 0.752 | 1 | 0 | -0.112 |
| SE | 92598 | ENSG00000102699 | PARP4 | chr13 | - | 24459971 | 24460136 | 24459263 | 24459310 | 24469023 | 24469110 | 684 | 467 | 534 | 559 | 763 | 1 | 0 | 4 | 0 | 0 | 253 | 486 | 402 | 469 | 52 | 0 | 5 | 2 | 5 | 0 | 298 | 149 | 0.868791768 | 1 | 0.997 | 1 | 0.985 | 1 | 1 | 1 | 0.98 | 0.99 | 0.979 | 1 | 0.007 |
| SE | 18123 | ENSG00000075413 | MARK3 | chr14 | + | 103498501 | 103498528 | 103491538 | 103492034 | 103500155 | 103500200 | 1 | 1 | 0 | 3 | 0 | 15 | 16 | 12 | 16 | 2 | 0 | 0 | 1 | 0 | 0 | 79 | 7 | 12 | 25 | 18 | 176 | 149 | 0.868954426 | 1 | 0.053 | 0.05 | 0 | 0.137 | 0 | 0 | 0 | 0.066 | 0 | 0 | 0.035 |
| SE | 36527 | ENSG00000169230 | PRELID1 | chr5 | + | 177305914 | 177305984 | 177304624 | 177304850 | 177306097 | 177306176 | 377 | 397 | 319 | 446 | 298 | 0 | 0 | 0 | 0 | 0 | 1719 | 585 | 565 | 885 | 1339 | 0 | 0 | 0 | 14 | 9 | 219 | 149 | 0.868961338 | 1 | 1 | 1 | 1 | 1 | 1 | 1 | 1 | 1 | 0.977 | 0.99 | 0.007 |
| SE | 31370 | ENSG00000115073 | ACTR1B | chr2 | - | 97658426 | 97658643 | 97658223 | 97658316 | 97658878 | 97659003 | 82 | 29 | 88 | 86 | 49 | 0 | 0 | 0 | 4 | 2 | 59 | 46 | 31 | 41 | 70 | 5 | 0 | 2 | 0 | 0 | 298 | 149 | 0.868975392 | 1 | 1 | 1 | 1 | 0.915 | 0.925 | 0.855 | 1 | 0.886 | 1 | 1 | 0.02 |
| SE | 48852 | ENSG00000163563 | MNDA | chr1 | + | 158843954 | 158844122 | 158843278 | 158843415 | 158845586 | 158846003 | 2123 | 1830 | 1398 | 1958 | 4676 | 0 | 10 | 0 | 0 | 2 | 605 | 1879 | 1556 | 1381 | 319 | 0 | 0 | 0 | 0 | 10 | 298 | 149 | 0.868987899 | 1 | 1 | 0.989 | 1 | 1 | 0.999 | 1 | 1 | 1 | 1 | 0.941 | 0.009 |
| SE | 18739 | ENSG00000080824 | HSP90AA1 | chr14 | - | 102085757 | 102086124 | 102085297 | 102085431 | 102086216 | 102086378 | 2699 | 1836 | 2358 | 3054 | 3869 | 0 | 0 | 0 | 0 | 0 | 250 | 1762 | 2106 | 725 | 117 | 10 | 0 | 0 | 0 | 0 | 298 | 149 | 0.869002915 | 1 | 1 | 1 | 1 | 1 | 1 | 0.926 | 1 | 1 | 1 | 1 | 0.015 |
| SE | 73661 | ENSG00000081019 | RSBN1 | chr1 | - | 113797362 | 113798036 | 113777670 | 113777808 | 113811709 | 113812040 | 150 | 76 | 142 | 133 | 261 | 2 | 0 | 0 | 5 | 0 | 35 | 135 | 149 | 73 | 54 | 0 | 0 | 0 | 0 | 0 | 298 | 149 | 0.869014243 | 1 | 0.974 | 1 | 1 | 0.93 | 1 | 1 | 1 | 1 | 1 | 1 | -0.019 |
| SE | 89744 | ENSG00000156795 | NTAQ1 | chr8 | + | 123435441 | 123435542 | 123429982 | 123430033 | 123436452 | 123436601 | 4 | 1 | 11 | 11 | 9 | 19 | 27 | 20 | 11 | 26 | 0 | 0 | 10 | 4 | 7 | 11 | 9 | 16 | 8 | 0 | 250 | 149 | 0.869140848 | 1 | 0.111 | 0.022 | 0.247 | 0.373 | 0.171 | 0 | 0 | 0.271 | 0.23 | 1 | -0.115 |
| SE | 106762 | ENSG00000078674 | PCM1 | chr8 | + | 17969576 | 17969748 | 17966979 | 17967170 | 17972328 | 17972687 | 506 | 415 | 479 | 422 | 745 | 0 | 4 | 2 | 1 | 1 | 8 | 123 | 125 | 14 | 8 | 0 | 0 | 5 | 0 | 0 | 298 | 149 | 0.869254751 | 1 | 1 | 0.981 | 0.992 | 0.995 | 0.997 | 1 | 1 | 0.926 | 1 | 1 | 0.008 |
| SE | 8696 | ENSG00000136485 | DCAF7 | chr17 | + | 63583501 | 63583711 | 63579824 | 63579943 | 63585210 | 63585251 | 242 | 120 | 243 | 216 | 204 | 0 | 0 | 6 | 2 | 0 | 193 | 225 | 102 | 166 | 310 | 0 | 0 | 0 | 0 | 0 | 298 | 149 | 0.869355499 | 1 | 1 | 1 | 0.953 | 0.982 | 1 | 1 | 1 | 1 | 1 | 1 | -0.013 |
| SE | 101879 | ENSG00000173915 | ATP5MK | chr10 | - | 103394198 | 103394394 | 103392370 | 103392466 | 103396408 | 103396466 | 57 | 81 | 49 | 81 | 1 | 551 | 239 | 378 | 378 | 284 | 56 | 42 | 27 | 0 | 85 | 302 | 317 | 229 | 343 | 244 | 298 | 149 | 0.869426548 | 1 | 0.049 | 0.145 | 0.061 | 0.097 | 0.002 | 0.085 | 0.062 | 0.056 | 0 | 0.148 | 0.001 |
| SE | 65510 | ENSG00000140386 | SCAPER | chr15 | - | 76753807 | 76753948 | 76733228 | 76733384 | 76764960 | 76765072 | 124 | 222 | 195 | 211 | 173 | 0 | 0 | 1 | 0 | 0 | 49 | 28 | 106 | 46 | 14 | 0 | 0 | 5 | 0 | 0 | 290 | 149 | 0.86943872 | 1 | 1 | 1 | 1 | 0.991 | 1 | 1 | 1 | 0.916 | 1 | 1 | 0.015 |
| SE | 51454 | ENSG00000172432 | GTPBP2 | chr6 | - | 43625362 | 43625560 | 43624887 | 43625062 | 43625755 | 43625864 | 283 | 328 | 221 | 353 | 361 | 0 | 0 | 0 | 0 | 0 | 784 | 419 | 360 | 559 | 1590 | 12 | 0 | 0 | 0 | 0 | 298 | 149 | 0.869534377 | 1 | 1 | 1 | 1 | 1 | 1 | 0.97 | 1 | 1 | 1 | 1 | 0.006 |
| SE | 66432 | ENSG00000102081 | FMR1 | chrX | + | 147943130 | 147943326 | 147940575 | 147940662 | 147944943 | 147945051 | 284 | 252 | 230 | 283 | 337 | 1 | 0 | 4 | 2 | 1 | 100 | 160 | 223 | 184 | 38 | 0 | 0 | 2 | 5 | 0 | 298 | 149 | 0.86953472 | 1 | 0.993 | 1 | 0.966 | 0.986 | 0.994 | 1 | 1 | 0.982 | 0.948 | 1 | 0.002 |
| SE | 2910 | ENSG00000090554 | FLT3LG | chr19 | + | 49478908 | 49479047 | 49476422 | 49476566 | 49480297 | 49480476 | 96 | 47 | 142 | 85 | 69 | 9 | 5 | 6 | 12 | 6 | 65 | 29 | 52 | 19 | 32 | 0 | 6 | 3 | 10 | 0 | 288 | 149 | 0.869569146 | 1 | 0.847 | 0.829 | 0.924 | 0.786 | 0.856 | 1 | 0.714 | 0.9 | 0.496 | 1 | 0.026 |
| SE | 99523 | ENSG00000188690 | UROS | chr10 | - | 125789282 | 125789363 | 125788585 | 125789005 | 125794879 | 125794978 | 9 | 5 | 4 | 0 | 18 | 67 | 35 | 43 | 44 | 79 | 8 | 1 | 6 | 10 | 7 | 41 | 80 | 28 | 19 | 82 | 230 | 149 | 0.869609239 | 1 | 0.08 | 0.085 | 0.057 | 0 | 0.129 | 0.112 | 0.008 | 0.122 | 0.254 | 0.052 | -0.039 |
| SE | 91599 | ENSG00000104687 | GSR | chr8 | - | 30692968 | 30693055 | 30689160 | 30689319 | 30696379 | 30696479 | 249 | 146 | 213 | 250 | 215 | 1 | 3 | 0 | 0 | 0 | 438 | 454 | 198 | 292 | 450 | 4 | 0 | 0 | 0 | 10 | 236 | 149 | 0.869654834 | 1 | 0.994 | 0.968 | 1 | 1 | 1 | 0.986 | 1 | 1 | 1 | 0.966 | 0.002 |
| SE | 316 | ENSG00000182670 | TTC3 | chr21 | + | 37129002 | 37129063 | 37126079 | 37126143 | 37132681 | 37132766 | 330 | 297 | 345 | 429 | 773 | 1 | 0 | 5 | 0 | 3 | 166 | 238 | 164 | 87 | 66 | 0 | 4 | 5 | 0 | 0 | 210 | 149 | 0.869849519 | 1 | 0.996 | 1 | 0.98 | 1 | 0.995 | 1 | 0.977 | 0.959 | 1 | 1 | 0.007 |
| SE | 8977 | ENSG00000161526 | SAP30BP | chr17 | + | 75668048 | 75668169 | 75667250 | 75667478 | 75668515 | 75668625 | 2 | 2 | 0 | 5 | 9 | 221 | 144 | 189 | 219 | 388 | 14 | 1 | 6 | 2 | 0 | 385 | 315 | 184 | 402 | 448 | 270 | 149 | 0.869849888 | 1 | 0.005 | 0.008 | 0 | 0.012 | 0.013 | 0.02 | 0.002 | 0.018 | 0.003 | 0 | -0.001 |
| SE | 33449 | ENSG00000156976 | EIF4A2 | chr3 | + | 186787494 | 186787584 | 186787126 | 186787264 | 186787802 | 186787882 | 2936 | 1775 | 2673 | 2832 | 6477 | 7 | 6 | 14 | 8 | 15 | 542 | 1528 | 1837 | 1063 | 183 | 0 | 0 | 2 | 11 | 5 | 239 | 149 | 0.869856075 | 1 | 0.996 | 0.995 | 0.992 | 0.995 | 0.996 | 1 | 1 | 0.998 | 0.984 | 0.958 | 0.007 |
| SE | 5077 | ENSG00000022277 | RTF2 | chr20 | + | 56472325 | 56472415 | 56468584 | 56468766 | 56473300 | 56473360 | 3 | 0 | 6 | 0 | 4 | 231 | 124 | 167 | 159 | 114 | 0 | 1 | 6 | 15 | 2 | 268 | 213 | 207 | 278 | 331 | 239 | 149 | 0.869899462 | 1 | 0.008 | 0 | 0.022 | 0 | 0.021 | 0 | 0.003 | 0.018 | 0.033 | 0.004 | -0.001 |
| SE | 12500 | ENSG00000132600 | PRMT7 | chr16 | + | 68312041 | 68312176 | 68311062 | 68311111 | 68315896 | 68315957 | 3 | 2 | 11 | 6 | 2 | 0 | 3 | 1 | 5 | 2 | 7 | 3 | 8 | 7 | 2 | 0 | 0 | 2 | 11 | 24 | 284 | 149 | 0.870130334 | 1 | 1 | 0.259 | 0.852 | 0.386 | 0.344 | 1 | 1 | 0.677 | 0.25 | 0.042 | -0.026 |
| SE | 75202 | ENSG00000204410 | MSH5 | chr6 | + | 31744189 | 31744299 | 31743903 | 31744025 | 31744545 | 31744581 | 25 | 22 | 24 | 13 | 43 | 0 | 0 | 0 | 1 | 5 | 69 | 47 | 16 | 30 | 50 | 5 | 0 | 0 | 0 | 2 | 259 | 149 | 0.870234735 | 1 | 1 | 1 | 1 | 0.882 | 0.832 | 0.888 | 1 | 1 | 1 | 0.935 | -0.022 |
| SE | 26492 | ENSG00000112685 | EXOC2 | chr6 | - | 619429 | 619543 | 617710 | 617835 | 629834 | 629961 | 197 | 93 | 202 | 186 | 242 | 0 | 1 | 0 | 0 | 0 | 28 | 98 | 92 | 123 | 38 | 0 | 0 | 6 | 0 | 0 | 263 | 149 | 0.870282657 | 1 | 1 | 0.981 | 1 | 1 | 1 | 1 | 1 | 0.897 | 1 | 1 | 0.017 |
| SE | 17110 | ENSG00000276550 | HERC2P2 | chr15 | + | 22547880 | 22548072 | 22546937 | 22547089 | 22554384 | 22554572 | 47 | 26 | 50 | 42 | 25 | 1 | 0 | 1 | 0 | 2 | 19 | 61 | 13 | 4 | 9 | 0 | 4 | 0 | 1 | 0 | 298 | 149 | 0.87031179 | 1 | 0.959 | 1 | 0.962 | 1 | 0.862 | 1 | 0.884 | 1 | 0.667 | 1 | 0.046 |
| SE | 106438 | ENSG00000135297 | MTO1 | chr6 | + | 73486597 | 73486761 | 73482028 | 73482244 | 73492233 | 73492352 | 7 | 2 | 4 | 9 | 15 | 3 | 0 | 1 | 0 | 0 | 17 | 12 | 6 | 9 | 3 | 0 | 0 | 0 | 0 | 3 | 298 | 149 | 0.870413368 | 1 | 0.538 | 1 | 0.667 | 1 | 1 | 1 | 1 | 1 | 1 | 0.333 | -0.026 |
| SE | 87256 | ENSG00000046651 | OFD1 | chrX | + | 13763744 | 13763855 | 13762343 | 13762444 | 13767126 | 13767284 | 149 | 147 | 184 | 155 | 197 | 31 | 41 | 20 | 31 | 38 | 88 | 73 | 76 | 65 | 3 | 0 | 46 | 34 | 26 | 0 | 260 | 149 | 0.870429507 | 1 | 0.734 | 0.673 | 0.841 | 0.741 | 0.748 | 1 | 0.476 | 0.562 | 0.589 | 1 | 0.022 |

| | | | | | | | | | | | | | | | | | | | | | | | | | | | | | | | | | | | | | | | | | | | | | | |
|---|---|---|---|---|---|---|---|---|---|---|---|---|---|---|---|---|---|---|---|---|---|---|---|---|---|---|---|---|---|---|---|---|---|---|---|---|---|---|---|---|---|---|---|---|---|---|
| SE | 100639 | ENSG00000179532 | DNHD1 | chr11 | + | 6548644 | 6548933 | 6548209 | 6548402 | 6556682 | 6558297 | 7 | 5 | 7 | 1 | 5 | 3 | 0 | 3 | 0 | 2 | 24 | 30 | 8 | 2 | 9 | 0 | 12 | 2 | 8 | 0 | 298 | 149 | 0.870461637 | 1 | 0.538 | 1 | 0.538 | 1 | 0.556 | 1 | 0.556 | 0.667 | 0.111 | 1 | 0.06 |
| SE | 28185 | ENSG00000164190 | NIPBL | chr5 | + | 36962122 | 36962274 | 36961483 | 36961583 | 36970875 | 36971036 | 445 | 371 | 373 | 411 | 585 | 0 | 0 | 0 | 1 | 0 | 76 | 213 | 167 | 113 | 96 | 0 | 0 | 4 | 3 | 0 | 298 | 149 | 0.870572508 | 1 | 1 | 1 | 1 | 0.995 | 1 | 1 | 1 | 0.954 | 0.95 | 1 | 0.018 |
| SE | 59344 | ENSG00000168216 | LMBRD1 | chr6 | - | 69708849 | 69709231 | 69701888 | 69701953 | 69713644 | 69713797 | 4 | 8 | 3 | 3 | 1 | 138 | 133 | 130 | 135 | 193 | 0 | 1 | 1 | 1 | 0 | 72 | 59 | 65 | 44 | 31 | 298 | 149 | 0.870575383 | 1 | 0.014 | 0.029 | 0.011 | 0.011 | 0.003 | 0 | 0.008 | 0.008 | 0.011 | 0 | 0.008 |
| SE | 88359 | ENSG00000166167 | BTRC | chr10 | + | 101531236 | 101531333 | 101526012 | 101526199 | 101532294 | 101532432 | 166 | 136 | 148 | 156 | 181 | 1 | 2 | 2 | 5 | 0 | 51 | 179 | 124 | 91 | 49 | 0 | 0 | 6 | 2 | 0 | 246 | 149 | 0.870696778 | 1 | 0.99 | 0.976 | 0.978 | 0.95 | 1 | 1 | 1 | 0.926 | 0.965 | 1 | 0.001 |
| SE | 82882 | ENSG00000122643 | NT5C3A | chr7 | - | 33019634 | 33019724 | 33017203 | 33017601 | 33021271 | 33021357 | 434 | 448 | 277 | 582 | 977 | 0 | 0 | 0 | 2 | 2 | 475 | 635 | 332 | 438 | 287 | 0 | 4 | 0 | 15 | 0 | 239 | 149 | 0.870747501 | 1 | 1 | 1 | 1 | 0.995 | 0.997 | 1 | 0.99 | 1 | 0.948 | 1 | 0.011 |
| SE | 52927 | ENSG00000159023 | EPB41 | chr1 | + | 29115698 | 29115803 | 29112367 | 29112448 | 29116818 | 29117048 | 596 | 408 | 774 | 701 | 1270 | 0 | 0 | 1 | 2 | 9 | 493 | 462 | 570 | 516 | 610 | 0 | 4 | 0 | 0 | 11 | 254 | 149 | 0.870779145 | 1 | 1 | 1 | 0.998 | 0.995 | 0.988 | 1 | 0.985 | 1 | 1 | 0.97 | 0.005 |
| SE | 96351 | ENSG00000149428 | HYOU1 | chr11 | - | 119051034 | 119051173 | 119049776 | 119049837 | 119051437 | 119051625 | 434 | 160 | 394 | 387 | 409 | 0 | 0 | 0 | 0 | 0 | 711 | 406 | 407 | 404 | 595 | 0 | 2 | 0 | 10 | 2 | 288 | 149 | 0.870792067 | 1 | 1 | 1 | 1 | 1 | 1 | 1 | 0.991 | 1 | 0.954 | 0.994 | 0.012 |
| SE | 77035 | ENSG00000014123 | UFL1 | chr6 | + | 96525296 | 96525394 | 96524381 | 96524410 | 96526320 | 96526435 | 360 | 354 | 372 | 323 | 573 | 1 | 0 | 10 | 2 | 0 | 37 | 190 | 110 | 67 | 42 | 0 | 0 | 0 | 0 | 0 | 247 | 149 | 0.87091174 | 1 | 0.995 | 1 | 0.957 | 0.99 | 1 | 1 | 1 | 1 | 1 | 1 | -0.012 |
| SE | 52837 | ENSG00000111348 | ARHGDIB | chr12 | - | 14956175 | 14956333 | 14950531 | 14950724 | 14961536 | 14961605 | 47 | 35 | 58 | 39 | 83 | 3587 | 2361 | 2650 | 2923 | 6559 | 139 | 58 | 58 | 69 | 97 | 5538 | 3895 | 3086 | 4525 | 7830 | 298 | 149 | 0.871012128 | 1 | 0.007 | 0.007 | 0.011 | 0.007 | 0.006 | 0.012 | 0.007 | 0.009 | 0.008 | 0.006 | -0.001 |
| SE | 23898 | ENSG00000196911 | KPNA5 | chr6 | + | 116681446 | 116681552 | 116681186 | 116681338 | 116689319 | 116689453 | 3 | 1 | 0 | 3 | 0 | 13 | 7 | 20 | 9 | 9 | 0 | 4 | 2 | 0 | 0 | 6 | 9 | 6 | 6 | 9 | 255 | 149 | 0.87113725 | 1 | 0.119 | 0.077 | 0 | 0.163 | 0 | 0 | 0.206 | 0.163 | 0 | 0 | -0.002 |
| SE | 57377 | ENSG00000101040 | ZMYND8 | chr20 | - | 47212641 | 47212725 | 47209213 | 47210897 | 47220257 | 47220324 | 32 | 30 | 24 | 51 | 41 | 25 | 8 | 17 | 21 | 2 | 64 | 59 | 53 | 63 | 13 | 21 | 26 | 35 | 19 | 0 | 233 | 149 | 0.871170318 | 1 | 0.45 | 0.706 | 0.474 | 0.608 | 0.929 | 0.661 | 0.592 | 0.492 | 0.68 | 1 | -0.052 |
| SE | 70672 | ENSG00000115762 | PLEKHB2 | chr2 | + | 131132901 | 131132991 | 131130720 | 131130760 | 131140166 | 131140275 | 570 | 417 | 447 | 583 | 1021 | 4 | 6 | 13 | 14 | 4 | 1041 | 434 | 372 | 591 | 696 | 21 | 3 | 2 | 15 | 0 | 239 | 149 | 0.871249598 | 1 | 0.989 | 0.977 | 0.955 | 0.963 | 0.994 | 0.969 | 0.989 | 0.991 | 0.961 | 1 | -0.006 |
| SE | 106356 | ENSG00000076513 | ANKRD13A | chr12 | + | 110030644 | 110030758 | 110028946 | 110029635 | 110033796 | 110033931 | 519 | 347 | 456 | 492 | 835 | 2 | 0 | 3 | 3 | 9 | 893 | 564 | 510 | 647 | 932 | 0 | 2 | 10 | 0 | 10 | 263 | 149 | 0.871285368 | 1 | 0.993 | 1 | 0.989 | 0.989 | 0.981 | 1 | 0.994 | 0.967 | 1 | 0.981 | 0.002 |
| SE | 12179 | ENSG00000051108 | HERPUD1 | chr16 | + | 56942131 | 56942237 | 56939236 | 56939359 | 56943125 | 56943297 | 87 | 47 | 76 | 78 | 54 | 1 | 0 | 0 | 5 | 0 | 176 | 158 | 174 | 100 | 105 | 0 | 9 | 0 | 0 | 0 | 255 | 149 | 0.871410799 | 1 | 0.981 | 1 | 1 | 0.901 | 1 | 1 | 0.911 | 1 | 1 | 1 | -0.006 |
| SE | 108323 | ENSG00000089248 | ERP29 | chr12 | + | 112019755 | 112019894 | 112013425 | 112013609 | 112022149 | 112023168 | 324 | 171 | 244 | 252 | 177 | 66 | 31 | 45 | 50 | 15 | 400 | 400 | 343 | 354 | 575 | 80 | 82 | 56 | 83 | 59 | 288 | 149 | 0.871429263 | 1 | 0.717 | 0.741 | 0.737 | 0.723 | 0.859 | 0.721 | 0.716 | 0.76 | 0.688 | 0.834 | 0.012 |
| SE | 18444 | ENSG00000100911 | PSME2 | chr14 | - | 24145057 | 24145184 | 24144399 | 24144468 | 24145226 | 24145273 | 452 | 368 | 464 | 473 | 530 | 0 | 0 | 0 | 0 | 1 | 1447 | 546 | 365 | 749 | 927 | 0 | 0 | 0 | 0 | 14 | 276 | 149 | 0.871442411 | 1 | 1 | 1 | 1 | 1 | 0.997 | 1 | 1 | 1 | 1 | 0.973 | 0.005 |
| SE | 39701 | ENSG00000198734 | F5 | chr1 | - | 169523800 | 169523904 | 169523196 | 169523352 | 169524836 | 169524908 | 97 | 68 | 36 | 64 | 123 | 0 | 0 | 0 | 1 | 2 | 22 | 181 | 148 | 69 | 54 | 0 | 0 | 4 | 2 | 0 | 253 | 149 | 0.871476013 | 1 | 1 | 1 | 1 | 0.974 | 0.973 | 1 | 1 | 0.956 | 0.953 | 1 | 0.008 |
| SE | 34007 | ENSG00000060971 | ACAA1 | chr3 | - | 38127785 | 38127866 | 38125825 | 38126700 | 38129289 | 38129388 | 381 | 264 | 331 | 294 | 314 | 0 | 0 | 0 | 0 | 4 | 728 | 438 | 300 | 472 | 975 | 8 | 3 | 6 | 0 | 0 | 230 | 149 | 0.871492957 | 1 | 1 | 1 | 1 | 1 | 0.981 | 0.983 | 0.99 | 0.97 | 1 | 1 | 0.008 |
| SE | 20034 | ENSG00000139974 | SLC38A6 | chr14 | + | 61048150 | 61048297 | 61046106 | 61046167 | 61050511 | 61050636 | 1 | 8 | 2 | 1 | 19 | 20 | 11 | 20 | 16 | 35 | 0 | 7 | 12 | 1 | 2 | 14 | 8 | 4 | 12 | 17 | 296 | 149 | 0.871694026 | 1 | 0.025 | 0.268 | 0.048 | 0.031 | 0.215 | 0 | 0.306 | 0.602 | 0.04 | 0.056 | -0.083 |
| SE | 67810 | ENSG00000055044 | NOP58 | chr2 | + | 202292776 | 202292903 | 202291124 | 202291270 | 202297378 | 202297513 | 248 | 182 | 297 | 265 | 379 | 9 | 0 | 0 | 0 | 0 | 53 | 237 | 262 | 131 | 18 | 0 | 0 | 1 | 0 | 0 | 276 | 149 | 0.871716746 | 1 | 0.937 | 1 | 1 | 1 | 1 | 1 | 1 | 0.993 | 1 | 1 | -0.011 |
| SE | 31537 | ENSG00000155903 | RASA2 | chr3 | + | 141579436 | 141579489 | 141576999 | 141577106 | 141580367 | 141580445 | 0 | 5 | 2 | 2 | 0 | 208 | 122 | 200 | 161 | 302 | 0 | 3 | 0 | 4 | 0 | 19 | 63 | 101 | 29 | 7 | 202 | 149 | 0.871734861 | 1 | 0 | 0.029 | 0.007 | 0.009 | 0 | 0 | 0.034 | 0 | 0.092 | 0 | -0.016 |
| SE | 32130 | ENSG00000084463 | WBP11 | chr12 | - | 14796806 | 14797003 | 14793730 | 14793922 | 14799634 | 14799728 | 42 | 42 | 37 | 66 | 101 | 0 | 0 | 0 | 0 | 0 | 91 | 76 | 54 | 56 | 28 | 0 | 5 | 2 | 0 | 0 | 298 | 149 | 0.871775299 | 1 | 1 | 1 | 1 | 1 | 1 | 1 | 0.884 | 0.931 | 1 | 1 | 0.037 |
| SE | 37102 | ENSG00000127481 | UBR4 | chr1 | - | 19168026 | 19168184 | 19167021 | 19167231 | 19169434 | 19169532 | 300 | 117 | 255 | 245 | 231 | 0 | 0 | 0 | 2 | 0 | 176 | 284 | 248 | 317 | 399 | 7 | 0 | 0 | 4 | 0 | 298 | 149 | 0.871805321 | 1 | 1 | 1 | 1 | 0.984 | 1 | 0.926 | 1 | 1 | 0.975 | 1 | 0.017 |
| SE | 20995 | ENSG00000198399 | ITSN2 | chr2 | - | 24313459 | 24313523 | 24312211 | 24312375 | 24315131 | 24315224 | 346 | 253 | 231 | 289 | 688 | 2 | 6 | 3 | 3 | 5 | 39 | 220 | 162 | 140 | 88 | 0 | 5 | 0 | 0 | 0 | 213 | 149 | 0.871962484 | 1 | 0.992 | 0.967 | 0.982 | 0.985 | 0.99 | 1 | 0.969 | 1 | 1 | 1 | -0.011 |
| SE | 38366 | ENSG00000186074 | CD300LF | chr17 | - | 74705203 | 74705377 | 74704477 | 74704816 | 74712823 | 74712978 | 3 | 3 | 8 | 8 | 6 | 157 | 71 | 82 | 148 | 85 | 27 | 3 | 2 | 6 | 12 | 309 | 60 | 68 | 174 | 330 | 298 | 149 | 0.872048133 | 1 | 0.009 | 0.021 | 0.047 | 0.026 | 0.034 | 0.042 | 0.024 | 0.014 | 0.017 | 0.018 | 0.004 |
| SE | 23448 | ENSG00000166888 | STAT6 | chr12 | - | 57106527 | 57106580 | 57105467 | 57106339 | 57107230 | 57107314 | 419 | 263 | 277 | 345 | 325 | 0 | 0 | 0 | 1 | 0 | 785 | 471 | 220 | 463 | 872 | 10 | 0 | 5 | 3 | 0 | 202 | 149 | 0.872119848 | 1 | 1 | 1 | 1 | 0.996 | 1 | 0.983 | 1 | 0.97 | 0.991 | 1 | 0.01 |
| SE | 43977 | ENSG00000108443 | RPS6KB1 | chr17 | + | 59936214 | 59936277 | 59935192 | 59935300 | 59936463 | 59936541 | 122 | 112 | 106 | 127 | 176 | 0 | 0 | 0 | 0 | 8 | 49 | 96 | 102 | 19 | 65 | 0 | 0 | 2 | 0 | 0 | 212 | 149 | 0.872120515 | 1 | 1 | 1 | 1 | 1 | 0.939 | 1 | 1 | 0.973 | 1 | 1 | -0.007 |
| SE | 26319 | ENSG00000113300 | CNOT6 | chr5 | + | 180567848 | 180568003 | 180567087 | 180567242 | 180569109 | 180569340 | 220 | 148 | 192 | 169 | 318 | 0 | 4 | 2 | 3 | 0 | 52 | 166 | 109 | 153 | 37 | 4 | 0 | 1 | 0 | 0 | 298 | 149 | 0.872154645 | 1 | 1 | 0.949 | 0.98 | 0.966 | 1 | 0.867 | 1 | 0.982 | 1 | 1 | 0.009 |
| SE | 42644 | ENSG00000008130 | NADK | chr1 | - | 1756778 | 1756882 | 1756502 | 1756608 | 1757180 | 1757268 | 1 | 0 | 3 | 0 | 0 | 104 | 39 | 80 | 104 | 105 | 0 | 0 | 1 | 10 | 0 | 95 | 94 | 41 | 107 | 190 | 253 | 149 | 0.872156607 | 1 | 0.006 | 0 | 0.022 | 0 | 0 | 0 | 0 | 0.014 | 0.052 | 0 | -0.008 |
| SE | 23074 | ENSG00000065357 | DGKA | chr12 | + | 55937977 | 55938052 | 55935335 | 55937090 | 55938510 | 55938558 | 42 | 43 | 110 | 74 | 99 | 0 | 1 | 1 | 5 | 0 | 59 | 55 | 51 | 48 | 47 | 0 | 4 | 1 | 0 | 1 | 224 | 149 | 0.872301575 | 1 | 1 | 0.966 | 0.987 | 0.908 | 1 | 1 | 0.901 | 0.971 | 1 | 0.969 | 0.004 |
| SE | 59965 | ENSG00000103671 | TRIP4 | chr15 | + | 64406329 | 64406459 | 64400742 | 64400821 | 64409612 | 64409684 | 180 | 137 | 151 | 152 | 169 | 3 | 3 | 7 | 1 | 1 | 162 | 170 | 101 | 104 | 16 | 0 | 0 | 5 | 7 | 0 | 279 | 149 | 0.872310468 | 1 | 0.97 | 0.961 | 0.92 | 0.988 | 0.989 | 1 | 1 | 0.915 | 0.888 | 1 | 0.005 |
| [illegible] | | | | | | | | | | | | | | | | | | | | | | | | | | | | | | | | | | | | | | | | | | | | | | |

| | | | | | | | | | | | | | | | | | | | | | | | | | | | | | | | | | |
|---|---|---|---|---|---|---|---|---|---|---|---|---|---|---|---|---|---|---|---|---|---|---|---|---|---|---|---|---|---|---|---|---|---|
| SE | 39166 | ENSG00000165240 | ATP7A | chrX | + | 78003072 | 78003236 | 77998477 | 77998684 | 78009101 | 78009263 | 108 | 108 | 71 | 109 | 157 | 0 | 0 | 0 | 1 | 0 | 2 | 37 | 82 | 58 | 4 | 0 | 3 | 0 | 3 | 0 | 298 | 149 |
| SE | 81429 | ENSG00000011275 | RNF216 | chr7 | - | 5721032 | 5721172 | 5716715 | 5716766 | 5725323 | 5725438 | 327 | 225 | 344 | 366 | 321 | 3 | 1 | 0 | 1 | 1 | 422 | 344 | 364 | 327 | 503 | 0 | 0 | 3 | 0 | 13 | 289 | 149 |
| SE | 15884 | ENSG00000167491 | GATAD2A | chr19 | + | 19436130 | 19436219 | 19427695 | 19427992 | 19465339 | 19465577 | 9 | 4 | 17 | 9 | 2 | 2 | 5 | 9 | 10 | 4 | 20 | 19 | 6 | 25 | 45 | 29 | 20 | 13 | 8 | 10 | 238 | 149 |
| SE | 89562 | ENSG00000095485 | CWF19L1 | chr10 | - | 100236849 | 100236969 | 100235666 | 100235764 | 100238021 | 100238231 | 212 | 183 | 188 | 198 | 190 | 7 | 11 | 9 | 7 | 20 | 499 | 364 | 371 | 442 | 316 | 44 | 13 | 15 | 17 | 18 | 269 | 149 |
| SE | 31713 | ENSG00000145012 | LPP | chr3 | + | 188225404 | 188225527 | 188208197 | 188208318 | 188406111 | 188406260 | 48 | 20 | 11 | 21 | 26 | 2 | 1 | 6 | 0 | 0 | 63 | 19 | 26 | 21 | 42 | 10 | 0 | 2 | 0 | 0 | 272 | 149 |
| SE | 7276 | ENSG00000186481 | ANKRD20A5P | chr18 | + | 14242139 | 14242419 | 14234686 | 14234815 | 14243011 | 14243223 | 6 | 1 | 11 | 3 | 6 | 9 | 4 | 4 | 12 | 3 | 0 | 17 | 7 | 0 | 5 | 5 | 6 | 3 | 13 | 0 | 298 | 149 |
| SE | 84193 | ENSG00000137642 | SORL1 | chr11 | + | 121611075 | 121611158 | 121608103 | 121608176 | 121612735 | 121612832 | 1595 | 813 | 1211 | 1239 | 1536 | 3 | 0 | 1 | 0 | 0 | 716 | 536 | 1036 | 721 | 355 | 0 | 1 | 14 | 5 | 0 | 232 | 149 |
| SE | 67372 | ENSG00000187688 | TRPV2 | chr17 | + | 16420114 | 16420248 | 16417561 | 16417868 | 16422598 | 16422889 | 277 | 150 | 192 | 295 | 203 | 0 | 0 | 0 | 0 | 0 | 485 | 276 | 121 | 206 | 428 | 0 | 2 | 0 | 1 | 9 | 283 | 149 |
| SE | 83695 | ENSG00000119321 | FKBP15 | chr9 | - | 113207211 | 113207296 | 113206010 | 113206578 | 113211476 | 113211592 | 166 | 118 | 116 | 168 | 128 | 2 | 5 | 1 | 0 | 0 | 212 | 153 | 56 | 136 | 139 | 0 | 0 | 0 | 0 | 0 | 234 | 149 |
| SE | 46079 | ENSG00000068097 | HEATR6 | chr17 | - | 60073961 | 60074159 | 60073845 | 60073886 | 60076129 | 60076237 | 8 | 5 | 2 | 8 | 4 | 18 | 3 | 3 | 5 | 2 | 9 | 3 | 3 | 4 | 0 | 0 | 13 | 2 | 2 | 6 | 298 | 149 |
| SE | 78064 | ENSG00000127952 | STYXL1 | chr7 | - | 76028641 | 76028703 | 76021850 | 76021992 | 76030420 | 76030524 | 80 | 40 | 64 | 34 | 139 | 0 | 0 | 0 | 0 | 2 | 111 | 73 | 30 | 73 | 56 | 8 | 0 | 0 | 0 | 0 | 211 | 149 |
| SE | 62003 | ENSG00000044115 | CTNNA1 | chr5 | + | 138932577 | 138932712 | 138930829 | 138930935 | 138933801 | 138934213 | 299 | 127 | 129 | 217 | 238 | 0 | 0 | 0 | 0 | 0 | 607 | 215 | 131 | 303 | 479 | 0 | 7 | 0 | 3 | 0 | 284 | 149 |
| SE | 27118 | ENSG00000070756 | PABPC1 | chr8 | - | 100706650 | 100706805 | 100705588 | 100705673 | 100709132 | 100709223 | 2677 | 1464 | 2576 | 2705 | 1954 | 4 | 2 | 19 | 7 | 14 | 2692 | 1726 | 1977 | 1618 | 3192 | 14 | 22 | 7 | 8 | 7 | 298 | 149 |
| SE | 43390 | ENSG00000198791 | CNOT7 | chr8 | - | 17243441 | 17243545 | 17242991 | 17243185 | 17245035 | 17245247 | 7 | 0 | 12 | 6 | 9 | 729 | 603 | 584 | 769 | 1195 | 0 | 19 | 3 | 2 | 0 | 528 | 537 | 582 | 537 | 158 | 253 | 149 |
| SE | 57998 | ENSG00000070882 | OSBPL3 | chr7 | - | 24806775 | 24806902 | 24804314 | 24804437 | 24809806 | 24809951 | 140 | 96 | 100 | 127 | 73 | 6 | 0 | 0 | 3 | 0 | 27 | 115 | 64 | 52 | 19 | 0 | 5 | 0 | 0 | 3 | 276 | 149 |
| SE | 20766 | ENSG00000111897 | SERINC1 | chr6 | - | 122451887 | 122452057 | 122447120 | 122447265 | 122453769 | 122453907 | 358 | 264 | 326 | 387 | 843 | 24 | 11 | 5 | 23 | 28 | 115 | 198 | 291 | 127 | 13 | 0 | 11 | 30 | 0 | 0 | 298 | 149 |
| SE | 6674 | ENSG00000141627 | DYM | chr18 | - | 49393644 | 49393742 | 49391592 | 49391645 | 49430254 | 49430447 | 1 | 2 | 4 | 4 | 2 | 87 | 45 | 39 | 83 | 99 | 0 | 5 | 1 | 0 | 5 | 66 | 71 | 54 | 65 | 33 | 247 | 149 |
| SE | 89502 | ENSG00000156162 | DPY19L4 | chr8 | + | 94756035 | 94756159 | 94739644 | 94739790 | 94761699 | 94761834 | 93 | 59 | 57 | 59 | 44 | 4 | 4 | 8 | 7 | 1 | 35 | 25 | 21 | 21 | 5 | 0 | 4 | 1 | 0 | 18 | 273 | 149 |
| SE | 34928 | ENSG00000076108 | BAZ2A | chr12 | - | 56609735 | 56609946 | 56606632 | 56606733 | 56610113 | 56610215 | 636 | 548 | 682 | 767 | 714 | 0 | 0 | 0 | 0 | 0 | 2370 | 1297 | 1391 | 1513 | 1985 | 0 | 0 | 3 | 17 | 0 | 298 | 149 |
| SE | 50054 | ENSG00000137628 | DDX60 | chr4 | - | 168287047 | 168287203 | 168285392 | 168285498 | 168288173 | 168288315 | 130 | 136 | 133 | 183 | 367 | 1 | 4 | 1 | 2 | 0 | 39 | 104 | 39 | 71 | 15 | 0 | 1 | 0 | 0 | 0 | 298 | 149 |
| SE | 67457 | ENSG00000162923 | WDR26 | chr1 | - | 224393827 | 224394013 | 224385145 | 224389860 | 224398096 | 224398226 | 1182 | 803 | 786 | 1138 | 1297 | 10 | 13 | 19 | 18 | 35 | 923 | 1579 | 1771 | 1001 | 560 | 13 | 10 | 61 | 22 | 2 | 298 | 149 |
| SE | 84097 | ENSG00000197892 | KIF13B | chr8 | - | 29196186 | 29196199 | 29190996 | 29191057 | 29245345 | 29245439 | 116 | 98 | 62 | 115 | 105 | 1 | 0 | 0 | 1 | 3 | 117 | 133 | 92 | 117 | 40 | 0 | 0 | 4 | 4 | 0 | 162 | 149 |
| SE | 94740 | ENSG00000106948 | AKNA | chr9 | - | 114381059 | 114381446 | 114376465 | 114377532 | 114394356 | 114394405 | 611 | 266 | 507 | 551 | 311 | 3 | 1 | 4 | 4 | 2 | 522 | 738 | 493 | 372 | 788 | 4 | 0 | 3 | 5 | 8 | 298 | 149 |
| SE | 1687 | ENSG00000185721 | DRG1 | chr22 | + | 31421307 | 31421399 | 31420255 | 31420425 | 31423279 | 31423319 | 16 | 3 | 3 | 3 | 2 | 129 | 72 | 122 | 118 | 203 | 0 | 7 | 2 | 0 | 17 | 140 | 180 | 81 | 102 | 121 | 241 | 149 |
| SE | 68344 | ENSG00000125630 | POLR1B | chr2 | + | 112545830 | 112545922 | 112542188 | 112542671 | 112547011 | 112547179 | 3 | 0 | 2 | 0 | 0 | 20 | 9 | 24 | 26 | 3 | 0 | 0 | 0 | 0 | 0 | 9 | 10 | 7 | 13 | 2 | 241 | 149 |
| SE | 8260 | ENSG00000108312 | UBTF | chr17 | - | 44213217 | 44213282 | 44212818 | 44212939 | 44215653 | 44215809 | 523 | 310 | 556 | 522 | 1075 | 4 | 3 | 1 | 0 | 9 | 494 | 592 | 316 | 379 | 722 | 0 | 4 | 7 | 0 | 8 | 214 | 149 |
| SE | 49094 | ENSG00000072736 | NFATC3 | chr16 | + | 68183239 | 68183366 | 68181474 | 68181530 | 68190767 | 68191775 | 595 | 334 | 463 | 511 | 1054 | 2 | 9 | 2 | 5 | 5 | 276 | 535 | 300 | 376 | 264 | 0 | 6 | 6 | 0 | 3 | 276 | 149 |
| SE | 27412 | ENSG00000164125 | GASK1B | chr4 | - | 158155610 | 158155825 | 158130785 | 158131012 | 158170465 | 158171599 | 290 | 178 | 175 | 166 | 215 | 0 | 0 | 1 | 0 | 8 | 88 | 458 | 244 | 210 | 44 | 0 | 0 | 2 | 0 | 0 | 298 | 149 |
| SE | 69664 | ENSG00000184708 | EIF4ENIF1 | chr22 | - | 31450288 | 31450360 | 31449347 | 31449531 | 31454143 | 31454376 | 89 | 54 | 81 | 67 | 128 | 0 | 4 | 1 | 5 | 3 | 46 | 74 | 28 | 58 | 110 | 0 | 3 | 0 | 0 | 7 | 221 | 149 |
| SE | 12348 | ENSG00000197912 | SPG7 | chr16 | + | 89549499 | 89549713 | 89548002 | 89548113 | 89550493 | 89550609 | 0 | 0 | 2 | 2 | 0 | 27 | 19 | 23 | 46 | 20 | 0 | 0 | 3 | 0 | 5 | 55 | 44 | 31 | 79 | 44 | 298 | 149 |
| SE | 2489 | ENSG00000141867 | BRD4 | chr19 | - | 15268153 | 15268240 | 15267513 | 15267551 | 15268904 | 15269042 | 35 | 31 | 30 | 39 | 47 | 759 | 659 | 639 | 815 | 1639 | 76 | 38 | 37 | 46 | 70 | 1752 | 1007 | 923 | 1067 | 1936 | 236 | 149 |
| SE | 73109 | ENSG00000057252 | SOAT1 | chr1 | + | 179323436 | 179323495 | 179302676 | 179302802 | 179335505 | 179335657 | 351 | 210 | 220 | 270 | 330 | 7 | 4 | 2 | 0 | 2 | 124 | 203 | 139 | 162 | 69 | 0 | 6 | 3 | 0 | 0 | 208 | 149 |
| SE | 14279 | ENSG00000103249 | CLCN7 | chr16 | - | 1452754 | 1452893 | 1451622 | 1451716 | 1453833 | 1453894 | 310 | 217 | 267 | 264 | 341 | 0 | 0 | 0 | 0 | 0 | 415 | 445 | 177 | 319 | 617 | 0 | 0 | 1 | 9 | 0 | 288 | 149 |

[illegible]

| | | | | | | | | | | | | | | | |
|---|---|---|---|---|---|---|---|---|---|---|---|---|---|---|---|
| SE | 37779 | ENSG00000139719 | VPS33A | chr12 | - | 122242381 | 122242508 | 122239877 | 122239945 | 122249870 | 122250045 | 85 | 45 | 99 | [illegible] |
| SE | 54673 | ENSG00000070010 | UFD1 | chr22 | - | 19471686 | 19471808 | 19467402 | 19468003 | 19479082 | 19479144 | 191 | 103 | 207 | [illegible] |
| SE | 18182 | ENSG00000072042 | RDH11 | chr14 | - | 67692437 | 67692593 | 67691139 | 67691244 | 67692933 | 67693052 | 264 | 219 | 146 | [illegible] |
| SE | 86352 | ENSG00000176871 | WSB2 | chr12 | - | 118038287 | 118038388 | 118036337 | 118036510 | 118042840 | 118042878 | 200 | 129 | 136 | [illegible] |
| SE | 38273 | ENSG00000134684 | YARS1 | chr1 | - | 32797591 | 32797638 | 32791161 | 32791254 | 32797762 | 32797843 | 11 | 2 | 7 | [illegible] |
| SE | 70524 | ENSG00000167850 | CD300C | chr17 | - | 74544608 | 74544947 | 74542860 | 74542987 | 74545721 | 74546115 | 121 | 74 | 60 | [illegible] |
| SE | 7737 | ENSG00000108389 | MTMR4 | chr17 | - | 58514991 | 58515083 | 58512851 | 58512941 | 58516579 | 58516618 | 7 | 0 | 3 | [illegible] |
| SE | 17424 | ENSG00000126822 | PLEKHG3 | chr14 | + | 64737861 | 64738224 | 64737355 | 64737375 | 64738741 | 64738855 | 5 | 4 | 3 | [illegible] |
| SE | 76035 | ENSG00000137312 | FLOT1 | chr6 | - | 30741189 | 30741333 | 30740678 | 30740798 | 30742146 | 30742307 | 339 | 149 | 239 | [illegible] |
| SE | 92693 | ENSG00000180776 | ZDHHC20 | chr13 | - | 21402796 | 21402866 | 21391721 | 21391854 | 21413651 | 21413772 | 164 | 94 | 148 | [illegible] |
| SE | 100101 | ENSG00000149262 | INTS4 | chr11 | - | 77987528 | 77987698 | 77981458 | 77981576 | 77991107 | 77991299 | 5 | 7 | 13 | [illegible] |
| SE | 63841 | ENSG00000075292 | ZNF638 | chr2 | + | 71428546 | 71428651 | 71426459 | 71427414 | 71431326 | 71431428 | 203 | 194 | 196 | [illegible] |
| SE | 66684 | ENSG00000005483 | KMT2E | chr7 | + | 105108543 | 105108658 | 105107899 | 105107925 | 105108941 | 105109228 | 5 | 3 | 4 | [illegible] |
| SE | 46766 | ENSG00000117115 | PADI2 | chr1 | - | 17084598 | 17084702 | 17083725 | 17083837 | 17086520 | 17086699 | 48 | 14 | 42 | [illegible] |
| SE | 25095 | ENSG00000153310 | CYRIB | chr8 | - | 129849242 | 129849369 | 129846803 | 129846874 | 129850834 | 129850914 | 1059 | 638 | 655 | [illegible] |
| SE | 63370 | ENSG00000151092 | NGLY1 | chr3 | - | 25777700 | 25777840 | 25764128 | 25764311 | 25778573 | 25778688 | 1 | 6 | 0 | [illegible] |
| SE | 21960 | ENSG00000012660 | ELOVL5 | chr6 | - | 53277645 | 53277743 | 53276178 | 53276256 | 53291775 | 53291963 | 0 | 1 | 1 | [illegible] |
| SE | 31417 | ENSG00000155508 | CNOT8 | chr5 | + | 154863206 | 154863395 | 154858536 | 154858801 | 154865191 | 154865240 | 216 | 157 | 220 | [illegible] |
| SE | 17566 | ENSG00000197102 | DYNC1H1 | chr14 | + | 102024475 | 102024580 | 102022750 | 102022880 | 102026573 | 102026707 | 1 | 0 | 4 | [illegible] |
| SE | 55101 | ENSG00000072110 | ACTN1 | chr14 | - | 68878457 | 68878520 | 68877081 | 68877240 | 68878988 | 68879069 | 1 | 4 | 0 | [illegible] |
| SE | 106089 | ENSG00000196498 | NCOR2 | chr12 | - | 124385744 | 124385887 | 124372021 | 124372610 | 124398118 | 124398181 | 103 | 72 | 72 | [illegible] |
| SE | 27306 | ENSG00000136813 | ECPAS | chr9 | - | 111414428 | 111414651 | 111413894 | 111413986 | 111416271 | 111416352 | 246 | 182 | 185 | [illegible] |

[illegible]

| | | | | | | | | | | | |
|---|---|---|---|---|---|---|---|---|---|---|---|
| SE | 53467 | ENSG00000249915 | PDCD6 | chr5 | + | 306601 | 306760 | 304176 | 304221 | 314416 | 314974 |
| SE | 25066 | ENSG00000171206 | TRIM8 | chr10 | + | 102656746 | 102656920 | 102656269 | 102656385 | 102657033 | 102657101 |
| SE | 51587 | ENSG00000135821 | GLUL | chr1 | - | 182386361 | 182386402 | 182385698 | 182385887 | 182387130 | 182387292 |
| SE | 52013 | ENSG00000123636 | BAZ2B | chr2 | - | 159453612 | 159453801 | 159446781 | 159446975 | 159478574 | 159478721 |
| SE | 83972 | ENSG00000068400 | GRIPAP1 | chrX | - | 48997249 | 48997357 | 48993427 | 48993578 | 48999237 | 48999299 |
| SE | 43471 | ENSG00000072786 | STK10 | chr5 | - | 172096425 | 172096560 | 172093411 | 172093960 | 172105655 | 172105737 |
| SE | 5390 | ENSG00000078747 | ITCH | chr20 | + | 34424479 | 34424525 | 34413741 | 34413879 | 34438473 | 34438501 |
| SE | 94783 | ENSG00000119314 | PTBP3 | chr9 | - | 112275843 | 112276013 | 112268048 | 112268195 | 112297840 | 112297916 |
| SE | 44243 | ENSG00000102606 | ARHGEF7 | chr13 | + | 111274730 | 111274790 | 111273813 | 111273952 | 111275531 | 111275678 |
| SE | 32510 | ENSG00000213585 | VDAC1 | chr5 | - | 133975870 | 133976021 | 133973790 | 133973848 | 133980728 | 133980956 |
| SE | 2855 | ENSG00000099860 | GADD45B | chr19 | + | 2476528 | 2476630 | 2476126 | 2476402 | 2477028 | 2477126 |
| SE | 103581 | ENSG00000150093 | ITGB1 | chr10 | - | 32936349 | 32936423 | 32935491 | 32935558 | 32958144 | 32958251 |
| SE | 8207 | ENSG00000033627 | ATP6V0A1 | chr17 | + | 42495033 | 42495188 | 42494333 | 42494473 | 42495625 | 42495716 |
| SE | 54902 | ENSG00000156983 | BRPF1 | chr3 | + | 9745811 | 9745930 | 9745572 | 9745709 | 9746299 | 9746454 |
| SE | 75215 | ENSG00000100503 | NIN | chr14 | - | 50743415 | 50743529 | 50741581 | 50741728 | 50747991 | 50748105 |
| SE | 14050 | ENSG00000140694 | PARN | chr16 | - | 14591912 | 14592033 | 14586317 | 14586361 | 14593300 | 14593378 |
| SE | 101734 | ENSG00000196693 | ZNF33B | chr10 | - | 42632294 | 42632439 | 42631807 | 42632024 | 42638473 | 42638551 |
| SE | 57475 | ENSG00000154845 | PPP4R1 | chr18 | - | 9549196 | 9549338 | 9546790 | 9547952 | 9550051 | 9550186 |
| SE | 28666 | ENSG00000002586 | CD99 | chrX | + | 2733356 | 2733374 | 2726259 | 2726373 | 2738202 | 2738256 |
| SE | 83584 | ENSG00000107897 | ACBD5 | chr10 | - | 27210813 | 27211081 | 27208245 | 27208445 | 27215534 | 27215641 |
| SE | 93190 | ENSG00000139112 | GABARAPL1 | chr12 | + | 10214668 | 10214779 | 10212896 | 10213219 | 10218062 | 10218141 |
| SE | 51750 | ENSG00000114841 | DNAH1 | chr3 | + | 52351961 | 52352103 | 52350507 | 52350590 | 52353102 | 52353301 |
| SE | 34002 | ENSG00000060971 | ACAA1 | chr3 | - | 38126161 | 38126700 | 38125825 | 38125881 | 38127785 | 38127866 |
| SE | 14855 | ENSG00000185418 | TARS3 | chr15 | - | 101699330 | 101699435 | 101685895 | 101686062 | 101701085 | 101701184 |
| SE | 33859 | ENSG00000136522 | MRPL47 | chr3 | - | 179602651 | 179602737 | 179601729 | 179601790 | 179604526 | 179604640 |
| SE | 18311 | ENSG00000100938 | GMPR2 | chr14 | + | 24235736 | 24235820 | 24233478 | 24233598 | 24235966 | 24236005 |
| SE | 70571 | ENSG00000022840 | RNF10 | chr12 | + | 120566824 | 120566980 | 120565427 | 120565529 | 120571190 | 120571291 |
| SE | 89084 | ENSG00000171988 | JMJD1C | chr10 | - | 63219877 | 63219983 | 63217206 | 63217331 | 63268662 | 63268875 |
| SE | 28699 | ENSG00000112249 | ASCC3 | chr6 | - | 100725538 | 100725703 | 100718074 | 100718251 | 100766564 | 100766705 |
| SE | 99235 | ENSG00000137720 | C11orf1 | chr11 | + | 111883135 | 111883202 | 111882367 | 111882598 | 111883770 | 111883805 |
| SE | 60001 | ENSG00000137496 | IL18BP | chr11 | + | 71999227 | 71999662 | 71998908 | 71999019 | 71999926 | 71999938 |
| SE | 15098 | ENSG00000128739 | SNRPN | chr15 | + | 24974310 | 24974456 | 24967931 | 24968082 | 24975357 | 24975509 |
| SE | 18569 | ENSG00000168411 | RFWD3 | chr16 | - | 74636345 | 74636577 | 74632522 | 74632673 | 74644361 | 74644453 |
| SE | 93613 | ENSG00000187239 | FNBP1 | chr9 | - | 129923843 | 129923996 | 129908889 | 129908999 | 129924959 | [illegible] |
| SE | 3789 | ENSG00000090372 | STRN4 | chr19 | - | 46733038 | 46733236 | 46730731 | 46730873 | 46736822 | [illegible] |
| SE | 99427 | ENSG00000110321 | EIF4G2 | chr11 | - | 10801660 | 10801774 | 10800961 | 10801087 | 10802048 | [illegible] |
| SE | 27896 | ENSG00000150977 | RILPL2 | chr12 | - | 123423043 | 123423157 | 123415038 | 123415921 | 123436081 | [illegible] |
| SE | 52558 | ENSG00000153283 | CD96 | chr3 | + | 111606699 | 111606792 | 111598119 | 111598210 | 111623753 | [illegible] |
| SE | 18033 | ENSG00000139990 | DCAF5 | chr14 | - | 69118138 | 69118278 | 69116365 | 69116495 | 69119193 | [illegible] |
| SE | 36183 | ENSG00000116731 | PRDM2 | chr1 | + | 13732778 | 13732882 | 13730999 | 13731117 | 13749360 | [illegible] |
| SE | 101249 | ENSG00000166224 | SGPL1 | chr10 | + | 70869791 | 70869897 | 70868344 | 70868433 | 70875401 | [illegible] |
| SE | 84042 | ENSG00000182220 | ATP6AP2 | chrX | + | 40597252 | 40597344 | 40591233 | 40591365 | 40597526 | [illegible] |
| SE | 33089 | ENSG00000144741 | SLC25A26 | chr3 | + | 66356030 | 66356084 | 66346363 | 66346408 | 66362859 | [illegible] |
| SE | 57378 | ENSG00000101040 | ZMYND8 | chr20 | - | 47212641 | 47212725 | 47209213 | 47210897 | 47221313 | [illegible] |
| SE | 105513 | ENSG00000153179 | RASSF3 | chr12 | + | 64688215 | 64688453 | 64684786 | 64684894 | 64691469 | [illegible] |
| SE | 99841 | ENSG00000013725 | CD6 | chr11 | + | 61013918 | 61014014 | 61013422 | 61013563 | 61017478 | [illegible] |
| SE | 59711 | ENSG00000183530 | PRR14L | chr22 | - | 31688155 | 31688227 | 31681346 | 31685803 | 31701655 | [illegible] |
| SE | 57658 | ENSG00000144535 | DIS3L2 | chr2 | + | 232198526 | 232198776 | 232163458 | 232163632 | 232210325 | [illegible] |
| SE | 19849 | ENSG00000100519 | PSMC6 | chr14 | + | 52718228 | 52718352 | 52718080 | 52718142 | 52718976 | [illegible] |
| SE | 98033 | ENSG00000085117 | CD82 | chr11 | + | 44587709 | 44587886 | 44587474 | 44587556 | 44594642 | [illegible] |
| SE | 75268 | ENSG00000100504 | PYGL | chr14 | - | 50913028 | 50913130 | 50912155 | 50912303 | 50914700 | [illegible] |
| SE | 12876 | ENSG00000176953 | NFATC2IP | chr16 | + | 28954564 | 28954682 | 28952131 | 28952204 | 28963704 | [illegible] |
| SE | 30782 | ENSG00000102796 | DHRS12 | chr13 | - | 51771820 | 51771911 | 51767994 | 51769293 | 51773929 | [illegible] |
| SE | 8640 | ENSG00000261371 | PECAM1 | chr17 | - | 64350379 | 64350433 | 64348259 | 64348322 | 64352389 | [illegible] |
| SE | 43410 | ENSG00000132849 | PATJ | chr1 | + | 61766278 | 61766473 | 61763012 | 61763179 | 61769282 | [illegible] |
| SE | 80687 | ENSG00000175471 | MCTP1 | chr5 | - | 94714776 | 94714886 | 94710817 | 94710927 | 94779109 | [illegible] |
| SE | 79958 | ENSG00000042980 | ADAM28 | chr8 | + | 24320235 | 24320307 | 24313387 | 24313580 | 24321217 | [illegible] |
| SE | 56503 | ENSG00000176853 | FAM91A1 | chr8 | + | 123789612 | 123789745 | 123787663 | 123787750 | 123798089 | [illegible] |
| SE | 37585 | ENSG00000082074 | FYB1 | chr5 | - | 39139232 | 39139252 | 39138656 | 39138691 | 39141094 | [illegible] |
| [illegible] | | | | | | | | | | | |

| | | | | | | | | | | | | | | | |
|---|---|---|---|---|---|---|---|---|---|---|---|---|---|---|---|
| SE | 19327 | ENSG00000182979 | MTA1 | chr14 | + | 105445666 | 105445852 | 105445417 | 105445511 | 105449358 | 105449409 | 1 | 0 | 2 | [illegible] |
| SE | 67479 | ENSG00000162923 | WDR26 | chr1 | - | 224431681 | 224431781 | 224429296 | 224431581 | 224433683 | 224433850 | 444 | 306 | 326 | [illegible] |
| SE | 81579 | ENSG00000165055 | METTL2B | chr7 | + | 128498035 | 128498142 | 128493803 | 128493943 | 128500902 | 128500968 | 75 | 35 | 69 | [illegible] |
| SE | 79486 | ENSG00000145725 | PPIP5K2 | chr5 | + | 103183233 | 103183407 | 103180020 | 103180188 | 103184671 | 103184744 | 9 | 14 | 2 | [illegible] |
| SE | 5049 | ENSG00000101126 | ADNP | chr20 | - | 50904765 | 50904849 | 50903888 | 50904001 | 50928650 | 50928825 | 33 | 16 | 14 | [illegible] |
| SE | 75726 | ENSG00000107672 | NSMCE4A | chr10 | - | 121961840 | 121962107 | 121961422 | 121961517 | 121963237 | 121963328 | 2 | 2 | 5 | [illegible] |
| SE | 93084 | ENSG00000113924 | HGD | chr3 | - | 120638454 | 120638581 | 120633146 | 120633328 | 120641588 | 120641693 | 18 | 16 | 9 | [illegible] |
| SE | 51651 | ENSG00000081154 | PCNP | chr3 | + | 101585436 | 101585511 | 101579588 | 101580004 | 101592626 | 101592771 | 181 | 163 | 174 | [illegible] |
| SE | 7909 | ENSG00000159842 | ABR | chr17 | - | 1083519 | 1083627 | 1079329 | 1079390 | 1091664 | 1091850 | 333 | 199 | 182 | [illegible] |
| SE | 29160 | ENSG00000144043 | TEX261 | chr2 | - | 70991829 | 70991983 | 70989576 | 70989816 | 70993695 | 70993775 | 69 | 49 | 80 | [illegible] |
| SE | 16354 | ENSG00000140497 | SCAMP2 | chr15 | - | 74854580 | 74854649 | 74852068 | 74854119 | 74873198 | 74873302 | 268 | 109 | 203 | [illegible] |
| SE | 10260 | ENSG00000108651 | UTP6 | chr17 | - | 31866654 | 31866715 | 31865365 | 31865438 | 31868045 | 31868112 | 11 | 21 | 18 | [illegible] |
| SE | 64362 | ENSG00000163219 | ARHGAP25 | chr2 | + | 68787839 | 68787956 | 68774800 | 68775420 | 68807272 | 68807480 | 282 | 89 | 139 | [illegible] |
| SE | 72306 | ENSG00000059377 | TBXAS1 | chr7 | + | 139953367 | 139953456 | 139936190 | 139936307 | 139957633 | 139957764 | 185 | 102 | 110 | [illegible] |
| SE | 67601 | ENSG00000008083 | JARID2 | chr6 | + | 15505164 | 15505295 | 15504499 | 15504592 | 15507135 | 15507254 | 0 | 0 | 0 | [illegible] |
| SE | 99616 | ENSG00000109919 | MTCH2 | chr11 | - | 47628952 | 47629046 | 47627079 | 47627127 | 47630554 | 47630614 | 124 | 65 | 109 | [illegible] |
| SE | 76796 | ENSG00000135317 | SNX14 | chr6 | - | 85557975 | 85558060 | 85549722 | 85549879 | 85565331 | 85565419 | 257 | 154 | 162 | [illegible] |
| SE | 14744 | ENSG00000185658 | BRWD1 | chr21 | - | 39224407 | 39224469 | 39215236 | 39215362 | 39225085 | 39225197 | 97 | 84 | 82 | [illegible] |
| SE | 71225 | ENSG00000079263 | SP140 | chr2 | + | 230253315 | 230253417 | 230248884 | 230248968 | 230255451 | 230255532 | 31 | 54 | 48 | [illegible] |
| SE | 91275 | ENSG00000154114 | TBCEL | chr11 | + | 121057507 | 121057614 | 121055051 | 121055308 | 121058344 | 121058471 | 2 | 3 | 2 | [illegible] |
| SE | 765 | ENSG00000196588 | MRTFA | chr22 | - | 40594673 | 40594735 | 40552105 | 40552367 | 40636477 | 40636685 | 44 | 11 | 38 | [illegible] |
| SE | 32370 | ENSG00000013441 | CLK1 | chr2 | - | 200856890 | 200856985 | 200856681 | 200856811 | 200859679 | 200859746 | 444 | 289 | 441 | [illegible] |
| SE | 39014 | ENSG00000003400 | CASP10 | chr2 | + | 201195841 | 201195948 | 201192983 | 201193119 | 201209069 | 201209562 | 77 | 74 | 75 | [illegible] |
| SE | 5172 | ENSG00000088888 | MAVS | chr20 | + | 3861331 | 3861504 | 3857634 | 3857809 | 3862253 | 3862413 | 125 | 59 | 97 | [illegible] |
| SE | 77600 | ENSG00000127947 | PTPN12 | chr7 | + | 77581426 | 77581503 | 77571077 | 77571186 | 77585542 | 77585581 | 214 | 156 | 114 | [illegible] |
| SE | 9238 | ENSG00000178971 | CTC1 | chr17 | - | 8229141 | 8229206 | 8228502 | 8228629 | 8229301 | 8229446 | 22 | 15 | 23 | [illegible] |
| SE | 56567 | ENSG00000253710 | ALG11 | chr13 | + | 52018912 | 52019143 | 52012390 | 52012462 | 52024005 | 52024937 | 61 | 25 | 46 | [illegible] |
| SE | 74346 | ENSG00000187118 | CMC1 | chr3 | + | 28294433 | 28294526 | 28291800 | 28291954 | 28316332 | 28316407 | 3 | 5 | 1 | [illegible] |
| SE | 81132 | ENSG00000241468 | ATP5MF | chr7 | - | 99459146 | 99459263 | 99458189 | 99458355 | 99460085 | 99460193 | 802 | 388 | 591 | [illegible] |
| SE | 79680 | ENSG00000134905 | CARS2 | chr13 | - | 110646504 | 110646631 | 110645966 | 110646090 | 110647100 | 110647239 | 1 | 3 | 4 | [illegible] |
| SE | 69253 | ENSG00000168385 | SEPTIN2 | chr2 | + | 241317498 | 241317599 | 241315288 | 241315610 | 241324215 | 241324241 | 38 | 23 | 37 | [illegible] |
| SE | 71333 | ENSG00000135631 | RAB11FIP5 | chr2 | - | 73079650 | 73081663 | 73075992 | 73076182 | 73088049 | 73088749 | 3 | 0 | 0 | [illegible] |
| SE | 92861 | ENSG00000147872 | PLIN2 | chr9 | - | 19126113 | 19126309 | 19123312 | 19123647 | 19126396 | 19126448 | 180 | 86 | 121 | [illegible] |
| [illegible] | | | | | | | | | | | | | | | |

| | | | | | | | | | | | | | | | | | | | | | | | | | | | | | | | | | | | | | | | | | | | | | | |
|---|---|---|---|---|---|---|---|---|---|---|---|---|---|---|---|---|---|---|---|---|---|---|---|---|---|---|---|---|---|---|---|---|---|---|---|---|---|---|---|---|---|---|---|---|---|---|
| SE | 84292 | ENSG00000165813 | CCDC186 | chr10 | - | 114173140 | 114173265 | 114161607 | 114163329 | 114174014 | 114174202 | 49 | 33 | 24 | 32 | 27 | 28 | 21 | 23 | 37 | 34 | 5 | 31 | 17 | 5 | 0 | 0 | 19 | 11 | 3 | 11 | 274 | 149 | 0.9779937 | 1 | 0.488 | 0.461 | 0.362 | 0.32 | 0.302 | 1 | 0.47 | 0.457 | 0.475 | 0 | -0.094 |
| SE | 26399 | ENSG00000138744 | NAAA | chr4 | - | 75918760 | 75918789 | 75917466 | 75917773 | 75919908 | 75919975 | 31 | 7 | 11 | 21 | 12 | 26 | 17 | 26 | 27 | 9 | 29 | 1 | 5 | 0 | 59 | 25 | 5 | 0 | 14 | 29 | 178 | 149 | 0.978644191 | 1 | 0.5 | 0.256 | 0.262 | 0.394 | 0.527 | 0.493 | 0.143 | 1 | 0 | 0.63 | -0.065 |
| SE | 30123 | ENSG00000154144 | TBRG1 | chr11 | + | 124626472 | 124626609 | 124625670 | 124625903 | 124626903 | 124627050 | 17 | 15 | 14 | 34 | 15 | 20 | 17 | 21 | 19 | 67 | 18 | 15 | 7 | 5 | 32 | 18 | 12 | 25 | 19 | 25 | 286 | 149 | 0.979068906 | 1 | 0.307 | 0.315 | 0.258 | 0.482 | 0.104 | 0.343 | 0.394 | 0.127 | 0.121 | 0.4 | 0.016 |
| SE | 32821 | ENSG00000088387 | DOCK9 | chr13 | - | 98825871 | 98825975 | 98824397 | 98824504 | 98826829 | 98826887 | 0 | 0 | 2 | 4 | 9 | 43 | 40 | 72 | 70 | 33 | 0 | 1 | 7 | 0 | 0 | 12 | 12 | 29 | 28 | 2 | 253 | 149 | 0.979192287 | 1 | 0 | 0 | 0.016 | 0.033 | 0.138 | 0 | 0.047 | 0.124 | 0 | 0 | 0.003 |
| SE | 96233 | ENSG00000133943 | DGLUCY | chr14 | + | 91147780 | 91147858 | 91114369 | 91114528 | 91157638 | 91157690 | 20 | 30 | 19 | 21 | 26 | 0 | 8 | 6 | 16 | 11 | 31 | 29 | 13 | 33 | 63 | 10 | 8 | 1 | 15 | 18 | 227 | 149 | 0.979272415 | 1 | 1 | 0.711 | 0.675 | 0.463 | 0.608 | 0.67 | 0.704 | 0.895 | 0.591 | 0.697 | -0.02 |
| SE | 73059 | ENSG00000115993 | TRAK2 | chr2 | - | 201401017 | 201401094 | 201399376 | 201399493 | 201407402 | 201407597 | 196 | 128 | 154 | 188 | 170 | 24 | 6 | 13 | 10 | 24 | 70 | 258 | 131 | 133 | 38 | 0 | 18 | 15 | 12 | 11 | 226 | 149 | 0.979573356 | 1 | 0.843 | 0.934 | 0.886 | 0.925 | 0.824 | 1 | 0.904 | 0.852 | 0.88 | 0.695 | 0.016 |
| SE | 12363 | ENSG00000072864 | NDE1 | chr16 | + | 15672689 | 15672771 | 15667285 | 15667439 | 15677800 | 15677949 | 1 | 8 | 0 | 11 | 0 | 223 | 151 | 207 | 213 | 232 | 5 | 6 | 4 | 2 | 9 | 307 | 192 | 221 | 187 | 462 | 231 | 149 | 0.979682467 | 1 | 0.003 | 0.033 | 0 | 0.032 | 0 | 0.01 | 0.02 | 0.012 | 0.007 | 0.012 | 0.001 |
| SE | 49483 | ENSG00000115020 | PIKFYVE | chr2 | + | 208330522 | 208330694 | 208329841 | 208329913 | 208335305 | 208335419 | 72 | 27 | 58 | 49 | 147 | 0 | 0 | 0 | 1 | 0 | 4 | 24 | 45 | 11 | 11 | 0 | 1 | 4 | 0 | 0 | 298 | 149 | 0.97970522 | 1 | 1 | 1 | 1 | 0.961 | 1 | 1 | 0.923 | 0.849 | 1 | 1 | 0.038 |
| SE | 25402 | ENSG00000186704 | DTX2P1 | chr7 | + | 76995454 | 76995595 | 76990268 | 76990369 | 77000308 | 77000388 | 10 | 6 | 3 | 6 | 10 | 6 | 6 | 6 | 12 | 0 | 9 | 10 | 4 | 2 | 5 | 15 | 2 | 2 | 6 | 1 | 290 | 149 | 0.979850427 | 1 | 0.461 | 0.339 | 0.204 | 0.204 | 1 | 0.236 | 0.72 | 0.507 | 0.146 | 0.72 | -0.024 |
| SE | 28330 | ENSG00000141385 | AFG3L2 | chr18 | - | 12356693 | 12356831 | 12353004 | 12353158 | 12358669 | 12358943 | 103 | 94 | 130 | 141 | 176 | 2 | 1 | 0 | 2 | 3 | 239 | 156 | 184 | 146 | 153 | 0 | 3 | 2 | 4 | 0 | 287 | 149 | 0.979928067 | 1 | 0.964 | 0.98 | 1 | 0.973 | 0.968 | 1 | 0.964 | 0.979 | 0.95 | 1 | -0.002 |
| SE | 37852 | ENSG00000114126 | TFDP2 | chr3 | - | 142054114 | 142054149 | 142005440 | 142005544 | 142093060 | 142093127 | 66 | 25 | 45 | 34 | 54 | 116 | 48 | 61 | 82 | 107 | 35 | 28 | 34 | 20 | 36 | 51 | 76 | 64 | 69 | 30 | 184 | 149 | 0.980860368 | 1 | 0.315 | 0.297 | 0.374 | 0.251 | 0.29 | 0.357 | 0.23 | 0.301 | 0.19 | 0.493 | -0.009 |
| SE | 20982 | ENSG00000198399 | ITSN2 | chr2 | - | 24271765 | 24271941 | 24270670 | 24270768 | 24286211 | 24286351 | 95 | 80 | 81 | 108 | 90 | 2 | 0 | 0 | 0 | 4 | 7 | 48 | 33 | 23 | 13 | 0 | 0 | 0 | 0 | 0 | 298 | 149 | 0.98104031 | 1 | 0.96 | 1 | 1 | 1 | 0.918 | 1 | 1 | 1 | 1 | 1 | -0.024 |
| SE | 9242 | ENSG00000178971 | CTC1 | chr17 | - | 8231275 | 8231469 | 8230562 | 8230651 | 8231725 | 8231815 | 159 | 102 | 110 | 200 | 148 | 1 | 2 | 2 | 3 | 2 | 155 | 135 | 108 | 127 | 202 | 0 | 2 | 5 | 3 | 0 | 298 | 149 | 0.981362989 | 1 | 0.988 | 0.962 | 0.965 | 0.971 | 0.974 | 1 | 0.971 | 0.915 | 0.955 | 1 | 0.004 |
| SE | 62861 | ENSG00000112335 | SNX3 | chr6 | - | 108222233 | 108222320 | 108214497 | 108214622 | 108222949 | 108223045 | 3 | 2 | 2 | 3 | 17 | 607 | 437 | 392 | 621 | 708 | 9 | 25 | 8 | 20 | 0 | 1588 | 876 | 641 | 1282 | 2398 | 236 | 149 | 0.981679771 | 1 | 0.003 | 0.003 | 0.003 | 0.003 | 0.015 | 0.004 | 0.018 | 0.008 | 0.01 | 0 | -0.003 |
| SE | 65675 | ENSG00000151502 | VPS26B | chr11 | + | 134234896 | 134235053 | 134224670 | 134225345 | 134239990 | 134240155 | 316 | 192 | 287 | 367 | 546 | 1 | 1 | 0 | 0 | 1 | 399 | 393 | 179 | 256 | 664 | 14 | 0 | 1 | 0 | 0 | 298 | 149 | 0.981893713 | 1 | 0.994 | 0.99 | 1 | 1 | 0.996 | 0.934 | 1 | 0.989 | 1 | 1 | 0.011 |
| SE | 17465 | ENSG00000159110 | IFNAR2 | chr21 | + | 33246256 | 33246482 | 33244950 | 33245074 | 33246717 | 33246890 | 7 | 15 | 21 | 18 | 14 | 501 | 407 | 455 | 426 | 509 | 13 | 12 | 20 | 11 | 10 | 467 | 340 | 483 | 386 | 343 | 298 | 149 | 0.983122362 | 1 | 0.007 | 0.018 | 0.023 | 0.021 | 0.014 | 0.014 | 0.017 | 0.02 | 0.014 | 0.014 | 0.001 |
| SE | 53248 | ENSG00000138688 | KIAA1109 | chr4 | + | 122209158 | 122209337 | 122207540 | 122207638 | 122209817 | 122209924 | 369 | 366 | 318 | 382 | 864 | 0 | 1 | 0 | 2 | 2 | 46 | 135 | 119 | 77 | 34 | 0 | 0 | 6 | 0 | 0 | 298 | 149 | 0.983457431 | 1 | 1 | 0.995 | 1 | 0.99 | 0.995 | 1 | 1 | 0.908 | 1 | 1 | 0.014 |
| SE | 4341 | ENSG00000064607 | SUGP2 | chr19 | - | 19008316 | 19008428 | 19004167 | 19004646 | 19009854 | 19010342 | 233 | 163 | 210 | 266 | 106 | 3 | 0 | 1 | 3 | 0 | 579 | 339 | 395 | 474 | 328 | 10 | 0 | 4 | 0 | 0 | 261 | 149 | 0.983479052 | 1 | 0.978 | 1 | 0.992 | 0.981 | 1 | 0.971 | 1 | 0.983 | 1 | 1 | -0.001 |
| SE | 36426 | ENSG00000121858 | TNFSF10 | chr3 | - | 172509216 | 172509321 | 172505507 | 172506919 | 172511616 | 172511659 | 420 | 467 | 290 | 341 | 683 | 0 | 0 | 1 | 1 | 0 | 326 | 217 | 194 | 379 | 67 | 0 | 0 | 10 | 3 | 0 | 254 | 149 | 0.984572686 | 1 | 1 | 1 | 0.994 | 0.995 | 1 | 1 | 1 | 0.919 | 0.987 | 1 | 0.017 |
| SE | 89905 | ENSG00000104375 | STK3 | chr8 | - | 98638459 | 98638560 | 98595329 | 98596169 | 98706466 | 98706634 | 8 | 0 | 1 | 1 | 4 | 63 | 58 | 28 | 66 | 71 | 0 | 0 | 0 | 9 | 0 | 53 | 34 | 20 | 14 | 30 | 250 | 149 | 0.985062601 | 1 | 0.07 | 0 | 0.021 | 0.009 | 0.032 | 0 | 0 | 0 | 0.277 | 0 | -0.029 |
| SE | 41944 | ENSG00000163374 | YY1AP1 | chr1 | - | 155679927 | 155680008 | 155679408 | 155679512 | 155680415 | 155680456 | 0 | 1 | 8 | 2 | 8 | 131 | 81 | 98 | 122 | 197 | 0 | 16 | 0 | 3 | 6 | 264 | 143 | 100 | 167 | 260 | 230 | 149 | 0.985377942 | 1 | 0 | 0.008 | 0.05 | 0.011 | 0.026 | 0 | 0.068 | 0 | 0.012 | 0.015 | 0 |
| SE | 55971 | ENSG00000136161 | RCBTB2 | chr13 | - | 48522307 | 48522403 | 48521897 | 48521962 | 48524658 | 48524757 | 33 | 31 | 25 | 45 | 64 | 15 | 7 | 1 | 4 | 2 | 8 | 21 | 22 | 17 | 25 | 0 | 4 | 6 | 5 | 0 | 245 | 149 | 0.985875539 | 1 | 0.572 | 0.729 | 0.938 | 0.872 | 0.951 | 1 | 0.761 | 0.69 | 0.674 | 1 | -0.013 |
| SE | 61800 | ENSG00000196975 | ANXA4 | chr2 | + | 69803392 | 69803515 | 69788053 | 69788141 | 69804532 | 69804627 | 9 | 3 | 12 | 7 | 14 | 36 | 25 | 20 | 30 | 25 | 12 | 10 | 16 | 38 | 8 | 83 | 51 | 21 | 43 | 69 | 272 | 149 | 0.985908961 | 1 | 0.12 | 0.062 | 0.247 | 0.113 | 0.235 | 0.073 | 0.097 | 0.294 | 0.326 | 0.06 | -0.015 |
| SE | 26633 | ENSG00000124279 | FASTKD3 | chr5 | - | 7861582 | 7861773 | 7861148 | 7861263 | 7862822 | 7862997 | 41 | 36 | 49 | 40 | 92 | 4 | 0 | 1 | 1 | 0 | 8 | 23 | 45 | 6 | 4 | 0 | 0 | 0 | 0 | 0 | 298 | 149 | 0.986310996 | 1 | 0.837 | 1 | 0.961 | 0.952 | 1 | 1 | 1 | 1 | 1 | 1 | -0.05 |
| SE | 80691 | ENSG00000175471 | MCTP1 | chr5 | - | 94799012 | 94799132 | 94779109 | 94779163 | 94868332 | 94868452 | 120 | 110 | 50 | 139 | 138 | 15 | 37 | 4 | 35 | 36 | 53 | 65 | 53 | 62 | 42 | 2 | 15 | 5 | 24 | 26 | 269 | 149 | 0.986848932 | 1 | 0.816 | 0.622 | 0.874 | 0.687 | 0.68 | 0.936 | 0.706 | 0.854 | 0.589 | 0.472 | 0.024 |
| SE | 14480 | ENSG00000213614 | HEXA | chr15 | - | 72370430 | 72370694 | 72356524 | 72356617 | 72375719 | 72375972 | 3 | 0 | 2 | 1 | 5 | 163 | 70 | 91 | 119 | 81 | 2 | 10 | 0 | 1 | 5 | 200 | 140 | 107 | 155 | 223 | 298 | 149 | 0.986873874 | 1 | 0.009 | 0 | 0.011 | 0.004 | 0.03 | 0.005 | 0.034 | 0 | 0.003 | 0.011 | 0 |
| SE | 10869 | ENSG00000108474 | PIGL | chr17 | + | 16342572 | 16342720 | 16326484 | 16326905 | 16347544 | 16347674 | 37 | 38 | 38 | 38 | 45 | 4 | 4 | 8 | 11 | 3 | 10 | 30 | 41 | 25 | 10 | 6 | 3 | 3 | 0 | 6 | 297 | 149 | 0.987096293 | 1 | 0.823 | 0.827 | 0.704 | 0.634 | 0.883 | 0.455 | 0.834 | 0.873 | 1 | 0.455 | 0.051 |
| SE | 10908 | ENSG00000074370 | ATP2A3 | chr17 | - | 3928239 | 3928327 | 3925290 | 3925441 | 3928662 | 3928780 | 13 | 3 | 11 | 10 | 1 | 133 | 44 | 103 | 166 | 43 | 18 | 33 | 7 | 2 | 39 | 121 | 263 | 88 | 162 | 780 | 237 | 149 | 0.987181618 | 1 | 0.058 | 0.041 | 0.063 | 0.036 | 0.014 | 0.086 | 0.073 | 0.048 | 0.008 | 0.03 | -0.007 |
| SE | 99093 | ENSG00000110274 | CEP164 | chr11 | + | 117363428 | 117363506 | 117362403 | 117362538 | 117371079 | 117371466 | 26 | 8 | 36 | 39 | 25 | 20 | 17 | 18 | 30 | 24 | 52 | 47 | 29 | 27 | 89 | 54 | 31 | 16 | 27 | 91 | 227 | 149 | 0.987260773 | 1 | 0.46 | 0.236 | 0.568 | 0.46 | 0.406 | 0.387 | 0.499 | 0.543 | 0.396 | 0.391 | -0.017 |
| SE | 58006 | ENSG00000070882 | OSBPL3 | chr7 | - | 24863199 | 24863292 | 24861612 | 24861769 | 24863495 | 24863599 | 32 | 20 | 37 | 39 | 19 | 24 | 6 | 12 | 33 | 30 | 18 | 8 | 16 | 12 | 13 | 0 | 10 | 16 | 13 | 13 | 242 | 149 | 0.987504151 | 1 | 0.451 | 0.672 | 0.655 | 0.421 | 0.281 | 1 | 0.33 | 0.381 | 0.362 | 0.381 | 0.005 |
| SE | 93231 | ENSG00000110455 | ACCS | chr11 | + | 44073446 | 44073535 | 44071255 | 44071315 | 44074611 | 44074681 | 13 | 3 | 6 | 15 | 2 | 13 | 1 | 7 | 13 | 3 | 6 | 6 | 6 | 14 | 7 | 11 | 6 | 5 | 21 | 1 | 238 | 149 | 0.987772391 | 1 | 0.385 | 0.653 | 0.349 | 0.419 | 0.294 | 0.255 | 0.385 | 0.429 | 0.294 | 0.814 | -0.015 |
| SE | 60512 | ENSG00000143382 | ADAMTSL4 | chr1 | + | 150556620 | 150556793 | 150556161 | 150556366 | 150556938 | 150556976 | 29 | 28 | 14 | 36 | 18 | 1 | 0 | 1 | 1 | 4 | 74 | 18 | 22 | 62 | 68 | 0 | 3 | 1 | 3 | 2 | 298 | 149 | 0.987838642 | 1 | 0.935 | 1 | 0.875 | 0.947 | 0.692 | 1 | 0.75 | 0.917 | 0.912 | 0.944 | -0.015 |

| | | | | | | | | | | | | | | | | | | | | | | | | | | | | | |
|---|---|---|---|---|---|---|---|---|---|---|---|---|---|---|---|---|---|---|---|---|---|---|---|---|---|---|---|---|---|
| SE | 395 | ENSG00000159086 | PAXBP1 | chr21 | - | 32755229 | 32755353 | 32751118 | 32751218 | 32759079 | 32759269 | 94 | 52 | 77 | 131 | 132 | 0 | 1 | 0 | 0 | 0 | 18 | 57 | 25 | 7 | 3 | 0 | 0 | 0 | 0 | 0 | 273 | 149 |
| SE | 396 | ENSG00000159086 | PAXBP1 | chr21 | - | 32759079 | 32759269 | 32755212 | 32755353 | 32759776 | 32759994 | 84 | 62 | 90 | 141 | 130 | 0 | 0 | 0 | 0 | 6 | 0 | 39 | 24 | 5 | 1 | 0 | 0 | 0 | 0 | 0 | 298 | 149 |
| SE | 397 | ENSG00000159086 | PAXBP1 | chr21 | - | 32761058 | 32761162 | 32759401 | 32759994 | 32762095 | 32762317 | 180 | 158 | 209 | 234 | 185 | 0 | 0 | 0 | 0 | 1 | 32 | 133 | 78 | 58 | 35 | 0 | 0 | 0 | 0 | 0 | 253 | 149 |
| SE | 401 | ENSG00000154642 | C21orf91 | chr21 | - | 17811355 | 17811471 | 17796722 | 17797118 | 17818191 | 17818325 | 6 | 0 | 0 | 0 | 0 | 149 | 136 | 148 | 147 | 403 | 0 | 0 | 0 | 0 | 0 | 21 | 68 | 58 | 32 | 21 | 265 | 149 |
| SE | 406 | ENSG00000275895 | U2AF1L5 | chr21 | - | 6493043 | 6493110 | 6492130 | 6492197 | 6495992 | 6496080 | 0 | 0 | 1 | 2 | 1 | 0 | 0 | 0 | 1 | 1 | 0 | 0 | 0 | 1 | 0 | 0 | 0 | 0 | 0 | 0 | 216 | 149 |
| SE | 407 | ENSG00000278932 | NA | chr21 | + | 9325604 | 9325823 | 9325038 | 9325223 | 9327239 | 9327298 | 3 | 2 | 4 | 6 | 5 | 4 | 0 | 2 | 0 | 0 | 4 | 1 | 0 | 0 | 0 | 0 | 0 | 0 | 0 | 0 | 298 | 149 |
| SE | 410 | ENSG00000205930 | C21orf62-AS1 | chr21 | + | 32784796 | 32784919 | 32772099 | 32772308 | 32788611 | 32788701 | 6 | 23 | 21 | 17 | 34 | 0 | 0 | 0 | 0 | 0 | 0 | 16 | 3 | 15 | 39 | 0 | 0 | 1 | 0 | 0 | 272 | 149 |
| SE | 411 | ENSG00000205930 | C21orf62-AS1 | chr21 | + | 32788611 | 32788701 | 32784796 | 32784919 | 32796938 | 32797042 | 5 | 11 | 18 | 10 | 11 | 0 | 0 | 2 | 0 | 0 | 0 | 8 | 2 | 10 | 17 | 0 | 0 | 0 | 0 | 0 | 239 | 149 |
| SE | 412 | ENSG00000205930 | C21orf62-AS1 | chr21 | + | 32788611 | 32788701 | 32784796 | 32784919 | 32851748 | 32852490 | 2 | 3 | 5 | 3 | 9 | 0 | 0 | 2 | 3 | 0 | 0 | 0 | 2 | 6 | 4 | 0 | 0 | 0 | 0 | 0 | 239 | 149 |
| SE | 416 | ENSG00000156261 | CCT8 | chr21 | - | 29064967 | 29065105 | 29063351 | 29063530 | 29066715 | 29066777 | 664 | 714 | 596 | 659 | 1332 | 0 | 1 | 0 | 0 | 0 | 234 | 486 | 826 | 526 | 32 | 0 | 0 | 0 | 0 | 0 | 287 | 149 |
| SE | 417 | ENSG00000156261 | CCT8 | chr21 | - | 29066890 | 29067071 | 29066715 | 29066777 | 29067555 | 29067705 | 505 | 437 | 433 | 402 | 868 | 0 | 0 | 0 | 0 | 0 | 83 | 261 | 448 | 241 | 18 | 0 | 0 | 0 | 2 | 0 | 298 | 149 |
| SE | 418 | ENSG00000156261 | CCT8 | chr21 | - | 29069422 | 29069502 | 29067555 | 29067705 | 29070246 | 29070337 | 274 | 218 | 248 | 199 | 351 | 0 | 0 | 0 | 1 | 0 | 32 | 187 | 154 | 128 | 11 | 0 | 0 | 0 | 0 | 0 | 229 | 149 |
| SE | 419 | ENSG00000156261 | CCT8 | chr21 | - | 29070246 | 29070337 | 29067555 | 29067705 | 29073530 | 29073565 | 138 | 87 | 107 | 86 | 48 | 0 | 0 | 0 | 1 | 0 | 11 | 82 | 105 | 63 | 4 | 0 | 0 | 0 | 0 | 0 | 240 | 149 |
| SE | 420 | ENSG00000156261 | CCT8 | chr21 | - | 29070246 | 29070337 | 29069422 | 29069502 | 29071951 | 29071988 | 166 | 107 | 132 | 102 | 154 | 0 | 0 | 0 | 0 | 0 | 20 | 116 | 110 | 83 | 11 | 0 | 0 | 2 | 0 | 0 | 240 | 149 |
| SE | 421 | ENSG00000156261 | CCT8 | chr21 | - | 29071951 | 29071988 | 29070246 | 29070337 | 29073259 | 29073351 | 0 | 1 | 0 | 0 | 1 | 2 | 1 | 1 | 1 | 0 | 0 | 0 | 2 | 0 | 0 | 0 | 2 | 1 | 0 | 0 | 186 | 149 |
| SE | 422 | ENSG00000156261 | CCT8 | chr21 | - | 29071951 | 29071988 | 29070246 | 29070337 | 29073530 | 29073797 | 1 | 1 | 0 | 0 | 1 | 167 | 106 | 126 | 103 | 69 | 0 | 0 | 2 | 0 | 0 | 11 | 100 | 113 | 69 | 4 | 186 | 149 |
| SE | 423 | ENSG00000156304 | SCAF4 | chr21 | - | 31690796 | 31690953 | 31688306 | 31688464 | 31691816 | 31691930 | 196 | 109 | 156 | 200 | 262 | 0 | 0 | 1 | 0 | 0 | 149 | 153 | 156 | 152 | 132 | 0 | 0 | 0 | 0 | 0 | 298 | 149 |
| SE | 425 | ENSG00000156304 | SCAF4 | chr21 | - | 31705422 | 31705467 | 31703764 | 31703926 | 31706273 | 31706357 | 323 | 230 | 224 | 282 | 386 | 0 | 3 | 0 | 0 | 0 | 290 | 320 | 248 | 172 | 213 | 0 | 0 | 0 | 0 | 0 | 194 | 149 |
| SE | 431 | ENSG00000237484 | LINC01684 | chr21 | + | 24441168 | 24441353 | 24438252 | 24438433 | 24441794 | 24441913 | 10 | 2 | 0 | 5 | 11 | 6 | 1 | 0 | 0 | 0 | 10 | 6 | 3 | 0 | 8 | 9 | 2 | 0 | 0 | 0 | 298 | 149 |
| SE | 433 | ENSG00000237484 | LINC01684 | chr21 | + | 24441794 | 24441913 | 24428728 | 24428904 | 24443963 | 24444128 | 7 | 7 | 4 | 10 | 10 | 0 | 9 | 0 | 2 | 0 | 0 | 1 | 0 | 0 | 5 | 0 | 0 | 0 | 0 | 0 | 268 | 149 |
| SE | 434 | ENSG00000237484 | LINC01684 | chr21 | + | 24441794 | 24441913 | 24428728 | 24428904 | 24485622 | 24485685 | 5 | 6 | 9 | 6 | 13 | 0 | 0 | 4 | 4 | 0 | 0 | 1 | 0 | 0 | 0 | 0 | 0 | 0 | 0 | 0 | 268 | 149 |
| SE | 435 | ENSG00000237484 | LINC01684 | chr21 | + | 24443557 | 24443743 | 24441794 | 24441913 | 24443963 | 24444128 | 0 | 1 | 0 | 0 | 0 | 7 | 6 | 0 | 9 | 2 | 0 | 0 | 0 | 3 | 10 | 0 | 0 | 0 | 0 | 5 | 298 | 149 |
| SE | 437 | ENSG00000237484 | LINC01684 | chr21 | + | 24443963 | 24444128 | 24441794 | 24441913 | 24480382 | 24480703 | 7 | 6 | 0 | 9 | 2 | 0 | 2 | 0 | 0 | 0 | 0 | 0 | 0 | 0 | 5 | 0 | 0 | 0 | 0 | 0 | 298 | 149 |
| SE | 438 | ENSG00000237484 | LINC01684 | chr21 | + | 24443963 | 24444128 | 24441794 | 24441913 | 24485622 | 24485685 | 7 | 6 | 0 | 9 | 2 | 4 | 4 | 5 | 2 | 5 | 0 | 0 | 0 | 0 | 5 | 0 | 0 | 0 | 0 | 0 | 298 | 149 |
| SE | 439 | ENSG00000237484 | LINC01684 | chr21 | + | 24443963 | 24444128 | 24441794 | 24441913 | 24526342 | 24526892 | 7 | 6 | 0 | 11 | 2 | 0 | 3 | 0 | 1 | 1 | 0 | 0 | 0 | 0 | 17 | 0 | 1 | 0 | 0 | 0 | 298 | 149 |
| SE | 441 | ENSG00000237484 | LINC01684 | chr21 | + | 24485622 | 24485685 | 24441779 | 24441913 | 24488112 | 24488309 | 4 | 3 | 1 | 2 | 5 | 0 | 1 | 0 | 0 | 0 | 12 | 0 | 0 | 0 | 2 | 0 | 0 | 0 | 0 | 0 | 212 | 149 |
| SE | 442 | ENSG00000237484 | LINC01684 | chr21 | + | 24485622 | 24485685 | 24441779 | 24441913 | 24526342 | 24526892 | 0 | 0 | 0 | 2 | 1 | 0 | 3 | 0 | 1 | 1 | 0 | 0 | 0 | 2 | 0 | 0 | 1 | 0 | 0 | 0 | 212 | 149 |
| SE | 443 | ENSG00000237484 | LINC01684 | chr21 | + | 24485622 | 24485685 | 24443963 | 24444128 | 24526342 | 24526892 | 0 | 0 | 0 | 2 | 0 | 0 | 0 | 0 | 2 | 0 | 0 | 0 | 0 | 2 | 0 | 0 | 0 | 0 | 0 | 12 | 212 | 149 |
| SE | 445 | ENSG00000237484 | LINC01684 | chr21 | + | 24488112 | 24488309 | 24441794 | 24441913 | 24526342 | 24526735 | 6 | 2 | 6 | 1 | 9 | 0 | 3 | 0 | 1 | 1 | 0 | 2 | 8 | 0 | 0 | 0 | 1 | 0 | 0 | 0 | 298 | 149 |
| SE | 449 | ENSG00000237484 | LINC01684 | chr21 | + | 24522888 | 24522999 | 24441794 | 24441913 | 24526342 | 24526892 | 0 | 0 | 0 | 0 | 2 | 0 | 3 | 0 | 1 | 1 | 0 | 0 | 0 | 0 | 0 | 0 | 1 | 0 | 0 | 0 | 260 | 149 |
| SE | 450 | ENSG00000237484 | LINC01684 | chr21 | + | 24522888 | 24522999 | 24488112 | 24488309 | 24526342 | 24526892 | 0 | 0 | 0 | 0 | 2 | 6 | 1 | 6 | 1 | 9 | 0 | 0 | 0 | 0 | 0 | 0 | 2 | 8 | 0 | 0 | 260 | 149 |
| SE | 451 | ENSG00000159131 | GART | chr21 | - | 33505560 | 33505702 | 33504411 | 33504527 | 33509782 | 33509920 | 52 | 29 | 48 | 37 | 91 | 0 | 0 | 0 | 0 | 2 | 29 | 23 | 17 | 22 | 17 | 0 | 0 | 0 | 0 | 0 | 291 | 149 |
| SE | 454 | ENSG00000159131 | GART | chr21 | - | 33520905 | 33521015 | 33520363 | 33520562 | 33522187 | 33522282 | 45 | 23 | 36 | 34 | 33 | 1 | 0 | 0 | 0 | 0 | 74 | 28 | 12 | 13 | 40 | 0 | 1 | 0 | 0 | 0 | 259 | 149 |
| SE | 455 | ENSG00000159131 | GART | chr21 | - | 33528166 | 33528335 | 33524008 | 33525000 | 33528518 | 33528604 | 251 | 160 | 201 | 207 | 287 | 0 | 0 | 2 | 0 | 0 | 231 | 119 | 81 | 154 | 136 | 0 | 0 | 0 | 0 | 0 | 298 | 149 |
| SE | 456 | ENSG00000159131 | GART | chr21 | - | 33528849 | 33528937 | 33528514 | 33528604 | 33530758 | 33530884 | 166 | 95 | 134 | 117 | 236 | 0 | 0 | 0 | 0 | 0 | 213 | 140 | 75 | 98 | 95 | 0 | 0 | 0 | 0 | 1 | 237 | 149 |
| SE | 457 | ENSG00000159131 | GART | chr21 | - | 33529566 | 33529768 | 33528849 | 33528937 | 33530758 | 33530884 | 0 | 0 | 0 | 0 | 0 | 122 | 70 | 86 | 80 | 116 | 0 | 0 | 2 | 0 | 0 | 165 | 97 | 42 | 65 | 71 | 298 | 149 |
| SE | 458 | ENSG00000159131 | GART | chr21 | - | 33530758 | 33530884 | 33528849 | 33528937 | 33531488 | 33531557 | 284 | 239 | 221 | 284 | 359 | 0 | 1 | 0 | 1 | 3 | 549 | 301 | 216 | 335 | 346 | 0 | 7 | 2 | 0 | 0 | 275 | 149 |
| SE | 459 | ENSG00000159131 | GART | chr21 | - | 33531488 | 33531557 | 33530709 | 33530884 | 33532344 | 33532456 | 214 | 200 | 168 | 245 | 291 | 0 | 0 | 0 | 0 | 0 | 434 | 244 | 212 | 329 | 308 | 0 | 3 | 0 | 0 | 0 | 218 | 149 |
| SE | 460 | ENSG00000159131 | GART | chr21 | - | 33534578 | 33534753 | 33532344 | 33532456 | 33535224 | 33535320 | 262 | 242 | 249 | 280 | 361 | 0 | 0 | 0 | 0 | 0 | 457 | 324 | 246 | 396 | 341 | 0 | 4 | 0 | 0 | 0 | 298 | 149 |
| SE | 462 | ENSG00000243927 | MRPS6 | chr21 | + | 34102834 | 34102908 | 34073223 | 34073268 | 34125340 | 34125480 | 0 | 0 | 2 | 0 | 0 | 1 | 1 | 3 | 2 | 2 | 0 | 0 | 0 | 0 | 7 | 0 | 0 | 0 | 0 | 11 | 223 | 149 |
| SE | 464 | ENSG00000243927 | MRPS6 | chr21 | + | 34103496 | 34103519 | 34102834 | 34102908 | 34125340 | 34125480 | 14 | 7 | 7 | 11 | 13 | 1 | 0 | 2 | 3 | 0 | 0 | 12 | 2 | 4 | 3 | 0 | 0 | 0 | 2 | 7 | 172 | 149 |
| SE | 467 | ENSG00000275464 | NA | chr21 | - | 5134360 | 5134479 | 5133901 | 5134120 | 5135088 | 5135154 | 1 | 0 | 5 | 2 | 0 | 8 | 0 | 8 | 10 | 2 | 0 | 0 | 0 | 0 | 0 | 0 | 10 | 3 | 0 | 0 | 268 | 149 |
| SE | 468 | ENSG00000275464 | NA | chr21 | - | 5143221 | 5143292 | 5142531 | 5142668 | 5144090 | 5144234 | 45 | 8 | 45 | 37 | 0 | 1 | 0 | 0 | 0 | 0 | 16 | 38 | 14 | 24 | 54 | 0 | 0 | 0 | 0 | 0 | 220 | 149 |
| SE | 471 | ENSG00000142207 | URB1 | chr21 | - | 32319216 | 32319414 | 32317675 | 32317917 | 32320530 | 32320640 | 52 | 26 | 45 | 53 | 26 | 0 | 0 | 0 | 1 | 0 | 37 | 27 | 14 | 21 | 24 | 0 | 0 | 0 | 2 | 0 | 298 | 149 |
| SE | 473 | ENSG00000142207 | URB1 | chr21 | - | 32345373 | 32345575 | 32344569 | 32344756 | 32346955 | 32347811 | 19 | 23 | 38 | 38 | 16 | 1 | 0 | 1 | 0 | 0 | 33 | 31 | 12 | 11 | 12 | 0 | 0 | 0 | 0 | 0 | 298 | 149 |
| SE | 477 | ENSG00000142207 | URB1 | chr21 | - | 32353932 | 32354103 | 32352709 | 32352906 | 32354858 | 32354997 | 36 | 15 | 40 | 43 | 31 | 0 | 3 | 0 | 0 | 0 | 32 | 20 | 25 | 17 | 29 | 0 | 0 | 0 | 2 | 0 | 298 | 149 |
| SE | 481 | ENSG00000142207 | URB1 | chr21 | - | 32378444 | 32378541 | 32375397 | 32375483 | 32383421 | 32383554 | 44 | 35 | 94 | 84 | 67 | 0 | 0 | 0 | 2 | 0 | 82 | 39 | 13 | 17 | 25 | 0 | 0 | 1 | 0 | 0 | 246 | 149 |
| SE | 489 | ENSG00000280071 | GATD3B | chr21 | - | 5127897 | 5127950 | 5125878 | 5125992 | 5128213 | 5128273 | 17 | 11 | 20 | 15 | 0 | 0 | 0 | 1 | 0 | 0 | 52 | 15 | 6 | 13 | 5 | 0 | 0 | 0 | 0 | 0 | 202 | 149 |
| SE | 490 | ENSG00000281903 | LINC02246 | chr21 | - | 14853273 | 14853431 | 14825522 | 14825624 | 14857541 | 14857708 | 6 | 4 | 1 | 8 | 6 | 2 | 2 | 1 | 6 | 10 | 2 | 10 | 1 | 3 | 0 | 0 | 0 | 1 | 1 | 0 | 298 | 149 |
| SE | 491 | ENSG00000281903 | LINC02246 | chr21 | - | 14902898 | 14903015 | 14857541 | 14857708 | 14913462 | 14913522 | 0 | 0 | 0 | 0 | 0 | 0 | 1 | 0 | 0 | 0 | 0 | 0 | 0 | 0 | 7 | 0 | 0 | 0 | 0 | 0 | 266 | 149 |
| SE | 552 | ENSG00000100221 | JOSD1 | chr22 | - | 38688934 | 38689129 | 38685542 | 38688001 | 38689295 | 38689424 | 158 | 57 | 101 | 162 | 142 | 0 | 1 | 0 | 0 | 0 | 77 | 153 | 167 | 137 | 228 | 0 | 0 | 0 | 0 | 0 | 298 | 149 |
| SE | 559 | ENSG00000187860 | CCDC157 | chr22 | + | 30370325 | 30370950 | 30369431 | 30369603 | 30371649 | 30371727 | 4 | 3 | 2 | 1 | 0 | 0 | 0 | 1 | 0 | 0 | 0 | 0 | 0 | 0 | 2 | 0 | 0 | 0 | 0 | 0 | 298 | 149 |
| SE | 560 | ENSG00000187860 | CCDC157 | chr22 | + | 30372074 | 30372286 | 30370325 | 30371727 | 30373596 | 30373764 | 0 | 6 | 3 | 0 | 0 | 3 | 0 | 1 | 1 | 0 | 0 | 3 | 0 | 0 | 2 | 0 | 0 | 0 | 0 | 5 | 298 | 149 |
| SE | 561 | ENSG00000187860 | CCDC157 | chr22 | + | 30375478 | 30375663 | 30373922 | 30374091 | 30376258 | 30376347 | 4 | 0 | 0 | 2 | 7 | 0 | 0 | 0 | 0 | 2 | 0 | 12 | 0 | 2 | 18 | 0 | 0 | 0 | 4 | 0 | 298 | 149 |
| SE | 562 | ENSG00000187860 | CCDC157 | chr22 | + | 30375485 | 30375663 | 30373922 | 30374091 | 30376258 | 30376347 | 4 | 0 | 0 | 2 | 7 | 0 | 0 | 0 | 0 | 2 | 0 | 19 | 0 | 2 | 10 | 0 | 0 | 0 | 4 | 0 | 298 | 149 |
| SE | 568 | ENSG00000100228 | RAB36 | chr22 | + | 23159162 | 23159253 | 23158897 | 23158979 | 23160878 | 23160998 | 1 | 0 | 0 | 3 | 2 | 0 | 0 | 0 | 0 | 0 | 12 | 0 | 0 | 1 | 0 | 0 | 3 | 0 | 0 | 0 | 240 | 149 |
| SE | 575 | ENSG00000100299 | ARSA | chr22 | - | 50627555 | 50627792 | 50627165 | 50627406 | 50627990 | 50628159 | 24 | 10 | 23 | 10 | 16 | 0 | 0 | 0 | 1 | 0 | 22 | 35 | 20 | 7 | 20 | 0 | 0 | 0 | 0 | 0 | 298 | 149 |
| SE | 576 | ENSG00000254413 | CHKB-CPT1B | chr22 | - | 50570292 | 50570406 | 50569575 | 50569668 | 50570890 | 50571043 | 37 | 11 | 11 | 45 | 61 | 0 | 1 | 0 | 1 | 0 | 54 | 60 | 82 | 60 | 46 | 0 | 0 | 1 | 4 | 0 | 263 | 149 |
| SE | 577 | ENSG00000254413 | CHKB-CPT1B | chr22 | - | 50576856 | 50577034 | 50576535 | 50576637 | 50577323 | 50577463 | 18 | 20 | 4 | 17 | 19 | 0 | 0 | 0 | 0 | 0 | 39 | 45 | 38 | 35 | 47 | 0 | 1 | 0 | 0 | 0 | 298 | 149 |

[illegible]

| | | | | | | | | | | | | | | | | | | | | | | | | | | | | | | | | | | | | | | | | | | | | | | |
|---|---|---|---|---|---|---|---|---|---|---|---|---|---|---|---|---|---|---|---|---|---|---|---|---|---|---|---|---|---|---|---|---|---|---|---|---|---|---|---|---|---|---|---|---|---|---|
| SE | 947 | ENSG00000100418 | DESI1 | chr22 | - | 41604043 | 41604153 | 41603258 | 41603381 | 41620751 | 41621043 | 81 | 53 | 69 | 52 | 76 | 0 | 0 | 0 | 0 | 0 | 132 | 83 | 61 | 138 | 161 | 0 | 0 | 2 | 0 | 0 | 259 | 149 | 1 | 1 | 1 | 1 | 1 | 1 | 1 | 1 | 1 | 0.946 | 1 | 1 | 0.011 |
| SE | 948 | ENSG00000100418 | DESI1 | chr22 | - | 41607261 | 41607331 | 41603258 | 41603381 | 41607839 | 41607861 | 19 | 4 | 19 | 11 | 17 | 0 | 0 | 0 | 0 | 0 | 9 | 12 | 8 | 8 | 21 | 0 | 0 | 2 | 0 | 0 | 219 | 149 | 1 | 1 | 1 | 1 | 1 | 1 | 1 | 1 | 1 | 0.731 | 1 | 1 | 0.054 |
| SE | 949 | ENSG00000100418 | DESI1 | chr22 | - | 41607261 | 41607331 | 41603258 | 41603381 | 41620751 | 41621043 | 1 | 0 | 0 | 0 | 0 | 0 | 0 | 0 | 0 | 0 | 0 | 2 | 0 | 0 | 0 | 0 | 0 | 2 | 0 | 0 | 219 | 149 | 1 | 1 | 1 | NA | NA | NA | NA | NA | 1 | 0 | NA | NA | 0.5 |
| SE | 950 | ENSG00000100418 | DESI1 | chr22 | - | 41607261 | 41607331 | 41604043 | 41604153 | 41607839 | 41607861 | 62 | 15 | 57 | 48 | 47 | 0 | 0 | 0 | 0 | 0 | 18 | 55 | 35 | 61 | 92 | 0 | 0 | 0 | 3 | 0 | 219 | 149 | 1 | 1 | 1 | 1 | 1 | 1 | 1 | 1 | 1 | 1 | 0.933 | 1 | 0.013 |
| SE | 952 | ENSG00000100418 | DESI1 | chr22 | - | 41607839 | 41607861 | 41603258 | 41603381 | 41620751 | 41621043 | 4 | 1 | 4 | 1 | 0 | 0 | 0 | 0 | 0 | 0 | 11 | 4 | 2 | 5 | 0 | 0 | 0 | 2 | 0 | 0 | 171 | 149 | 1 | 1 | 1 | 1 | 1 | 1 | NA | 1 | 1 | 0.466 | 1 | NA | 0.133 |
| SE | 953 | ENSG00000100418 | DESI1 | chr22 | - | 41607839 | 41607861 | 41604043 | 41604153 | 41620751 | 41621043 | 4 | 1 | 4 | 1 | 0 | 0 | 1 | 0 | 0 | 2 | 11 | 4 | 0 | 8 | 0 | 0 | 0 | 0 | 12 | 0 | 171 | 149 | 1 | 1 | 1 | 0.466 | 1 | 1 | 0 | 1 | 1 | NA | 0.367 | NA | -0.096 |
| SE | 954 | ENSG00000100418 | DESI1 | chr22 | - | 41607839 | 41607861 | 41607261 | 41607331 | 41620751 | 41621043 | 35 | 9 | 43 | 35 | 29 | 2 | 0 | 0 | 0 | 0 | 28 | 47 | 21 | 44 | 60 | 0 | 0 | 0 | 2 | 0 | 171 | 149 | 1 | 1 | 0.938 | 1 | 1 | 1 | 1 | 1 | 1 | 1 | 0.95 | 1 | -0.002 |
| SE | 955 | ENSG00000100417 | PMM1 | chr22 | - | 41577807 | 41577923 | 41576899 | 41577440 | 41578805 | 41578881 | 37 | 8 | 33 | 48 | 23 | 0 | 0 | 0 | 0 | 0 | 29 | 54 | 15 | 34 | 58 | 0 | 1 | 1 | 0 | 0 | 265 | 149 | 1 | 1 | 1 | 1 | 1 | 1 | 1 | 1 | 0.968 | 0.894 | 1 | 1 | 0.028 |
| SE | 958 | ENSG00000100417 | PMM1 | chr22 | - | 41586296 | 41586430 | 41586075 | 41586193 | 41589718 | 41589796 | 0 | 0 | 0 | 0 | 0 | 4 | 0 | 6 | 3 | 3 | 0 | 0 | 0 | 0 | 11 | 0 | 3 | 1 | 5 | 6 | 283 | 149 | 1 | 1 | 0 | NA | 0 | 0 | 0 | NA | 0 | 0 | 0 | 0.491 | -0.123 |
| SE | 959 | ENSG00000100417 | PMM1 | chr22 | - | 41589051 | 41589163 | 41586075 | 41586193 | 41589718 | 41589847 | 0 | 0 | 0 | 0 | 3 | 4 | 0 | 6 | 3 | 3 | 0 | 0 | 14 | 5 | 13 | 0 | 3 | 1 | 5 | 6 | 261 | 149 | 1 | 1 | 0 | NA | 0 | 0 | 0.363 | NA | 0 | 0.889 | 0.363 | 0.553 | -0.361 |
| SE | 960 | ENSG00000100413 | POLR3H | chr22 | - | 41532603 | 41532745 | 41532093 | 41532157 | 41540698 | 41540795 | 35 | 7 | 29 | 34 | 75 | 0 | 1 | 0 | 0 | 0 | 52 | 7 | 15 | 31 | 61 | 0 | 0 | 0 | 1 | 0 | 291 | 149 | 1 | 1 | 1 | 0.782 | 1 | 1 | 1 | 1 | 1 | 1 | 0.941 | 1 | -0.032 |
| SE | 961 | ENSG00000100413 | POLR3H | chr22 | - | 41532658 | 41532745 | 41532093 | 41532157 | 41540698 | 41540795 | 72 | 46 | 67 | 91 | 123 | 0 | 1 | 0 | 0 | 0 | 102 | 36 | 56 | 78 | 112 | 0 | 0 | 0 | 1 | 0 | 236 | 149 | 1 | 1 | 1 | 0.967 | 1 | 1 | 1 | 1 | 1 | 1 | 0.98 | 1 | -0.003 |
| SE | 963 | ENSG00000100413 | POLR3H | chr22 | - | 41540698 | 41540795 | 41532603 | 41532745 | 41543990 | 41544121 | 93 | 37 | 82 | 84 | 129 | 0 | 0 | 0 | 3 | 0 | 112 | 29 | 41 | 60 | 96 | 0 | 0 | 0 | 0 | 0 | 246 | 149 | 1 | 1 | 1 | 1 | 1 | 0.944 | 1 | 1 | 1 | 1 | 1 | 1 | -0.011 |
| SE | 964 | ENSG00000183506 | PI4KAP2 | chr22 | - | 21476643 | 21476764 | 21475991 | 21476160 | 21477789 | 21477915 | 22 | 21 | 34 | 46 | 30 | 0 | 0 | 1 | 0 | 0 | 39 | 72 | 31 | 14 | 98 | 0 | 0 | 0 | 0 | 0 | 270 | 149 | 1 | 1 | 1 | 1 | 0.949 | 1 | 1 | 1 | 1 | 1 | 1 | 1 | -0.01 |
| SE | 966 | ENSG00000188677 | PARVB | chr22 | + | 44100052 | 44100123 | 44093927 | 44094017 | 44119037 | 44119140 | 53 | 49 | 41 | 83 | 59 | 0 | 0 | 0 | 0 | 0 | 183 | 74 | 58 | 41 | 820 | 0 | 0 | 0 | 0 | 5 | 220 | 149 | 1 | 1 | 1 | 1 | 1 | 1 | 1 | 1 | 1 | 1 | 1 | 0.991 | 0.002 |
| SE | 968 | ENSG00000188677 | PARVB | chr22 | + | 44131486 | 44131627 | 44119037 | 44119140 | 44132893 | 44132965 | 86 | 119 | 64 | 188 | 200 | 1 | 0 | 0 | 0 | 0 | 429 | 190 | 107 | 128 | 2475 | 0 | 0 | 0 | 0 | 0 | 290 | 149 | 1 | 1 | 0.978 | 1 | 1 | 1 | 1 | 1 | 1 | 1 | 1 | 1 | -0.004 |
| SE | 969 | ENSG00000188677 | PARVB | chr22 | + | 44151482 | 44151551 | 44147860 | 44147922 | 44157981 | 44158083 | 58 | 56 | 39 | 115 | 106 | 0 | 2 | 0 | 0 | 2 | 325 | 116 | 80 | 77 | 1415 | 0 | 0 | 0 | 0 | 0 | 218 | 149 | 1 | 1 | 1 | 0.95 | 1 | 1 | 0.973 | 1 | 1 | 1 | 1 | 1 | -0.015 |
| SE | 970 | ENSG00000188677 | PARVB | chr22 | + | 44163857 | 44163930 | 44157981 | 44158083 | 44168601 | 44168923 | 82 | 72 | 54 | 176 | 112 | 0 | 0 | 0 | 2 | 0 | 505 | 115 | 106 | 101 | 2274 | 9 | 0 | 0 | 0 | 0 | 222 | 149 | 1 | 1 | 1 | 1 | 1 | 0.983 | 1 | 0.974 | 1 | 1 | 1 | 1 | 0.002 |
| SE | 971 | ENSG00000184164 | CRELD2 | chr22 | + | 49919729 | 49919840 | 49919229 | 49919312 | 49920155 | 49920247 | 44 | 24 | 43 | 34 | 34 | 0 | 1 | 0 | 3 | 0 | 84 | 25 | 38 | 33 | 51 | 0 | 0 | 4 | 0 | 0 | 260 | 149 | 1 | 1 | 1 | 0.932 | 1 | 0.867 | 1 | 1 | 1 | 0.845 | 1 | 1 | -0.009 |
| SE | 973 | ENSG00000184164 | CRELD2 | chr22 | + | 49923233 | 49923317 | 49922611 | 49922707 | 49924359 | 49924455 | 56 | 22 | 47 | 45 | 55 | 0 | 0 | 0 | 0 | 0 | 65 | 50 | 49 | 39 | 93 | 0 | 0 | 1 | 0 | 0 | 233 | 149 | 1 | 1 | 1 | 1 | 1 | 1 | 1 | 1 | 1 | 0.969 | 1 | 1 | 0.006 |
| SE | 974 | ENSG00000184164 | CRELD2 | chr22 | + | 49923233 | 49923317 | 49922611 | 49922707 | 49925416 | 49925557 | 18 | 11 | 4 | 6 | 8 | 0 | 0 | 0 | 2 | 0 | 20 | 12 | 13 | 4 | 11 | 0 | 0 | 0 | 0 | 0 | 233 | 149 | 1 | 1 | 1 | 1 | 1 | 0.657 | 1 | 1 | 1 | 1 | 1 | 1 | -0.069 |
| SE | 975 | ENSG00000184164 | CRELD2 | chr22 | + | 49924359 | 49924455 | 49922611 | 49922707 | 49925416 | 49925557 | 41 | 31 | 55 | 47 | 54 | 0 | 0 | 0 | 2 | 0 | 66 | 52 | 43 | 33 | 77 | 0 | 0 | 0 | 0 | 0 | 245 | 149 | 1 | 1 | 1 | 1 | 1 | 0.935 | 1 | 1 | 1 | 1 | 1 | 1 | -0.013 |
| SE | 979 | ENSG00000015475 | BID | chr22 | - | 17743802 | 17744013 | 17739348 | 17739488 | 17774380 | 17774495 | 76 | 45 | 41 | 65 | 49 | 0 | 0 | 0 | 0 | 0 | 92 | 87 | 48 | 84 | 165 | 0 | 3 | 2 | 0 | 0 | 298 | 149 | 1 | 1 | 1 | 1 | 1 | 1 | 1 | 1 | 0.935 | 0.923 | 1 | 1 | 0.028 |
| SE | 982 | ENSG00000100401 | RANGAP1 | chr22 | - | 41252871 | 41252991 | 41251006 | 41251109 | 41254307 | 41254494 | 92 | 53 | 101 | 99 | 92 | 0 | 0 | 0 | 1 | 0 | 79 | 114 | 69 | 32 | 117 | 4 | 0 | 0 | 0 | 0 | 269 | 149 | 1 | 1 | 1 | 1 | 1 | 0.982 | 1 | 0.916 | 1 | 1 | 1 | 1 | 0.013 |
| SE | 983 | ENSG00000100401 | RANGAP1 | chr22 | - | 41256190 | 41256290 | 41256020 | 41256105 | 41256710 | 41256824 | 185 | 143 | 176 | 269 | 141 | 0 | 0 | 0 | 2 | 0 | 337 | 255 | 217 | 233 | 274 | 0 | 0 | 0 | 0 | 0 | 249 | 149 | 1 | 1 | 1 | 1 | 1 | 0.988 | 1 | 1 | 1 | 1 | 1 | 1 | -0.002 |
| SE | 985 | ENSG00000100401 | RANGAP1 | chr22 | - | 41264663 | 41264843 | 41261445 | 41261580 | 41268096 | 41268156 | 77 | 34 | 67 | 96 | 45 | 0 | 0 | 1 | 0 | 0 | 80 | 77 | 43 | 58 | 161 | 0 | 0 | 0 | 0 | 0 | 298 | 149 | 1 | 1 | 1 | 1 | 0.971 | 1 | 1 | 1 | 1 | 1 | 1 | 1 | -0.006 |
| SE | 986 | ENSG00000100401 | RANGAP1 | chr22 | - | 41284900 | 41285153 | 41280932 | 41281082 | 41285543 | 41285965 | 3 | 1 | 0 | 1 | 1 | 2 | 0 | 2 | 7 | 1 | 0 | 0 | 0 | 0 | 0 | 0 | 0 | 3 | 1 | 1 | 298 | 149 | 1 | 1 | 0.429 | 1 | 0 | 0.067 | 0.333 | NA | NA | 0 | 0 | 0 | 0.366 |
| SE | 987 | ENSG00000128185 | DGCR6L | chr22 | - | 20316118 | 20316219 | 20315335 | 20315476 | 20319638 | 20319799 | 63 | 18 | 78 | 55 | 47 | 0 | 0 | 2 | 1 | 0 | 23 | 42 | 51 | 69 | 92 | 0 | 0 | 0 | 2 | 0 | 250 | 149 | 1 | 1 | 1 | 1 | 0.959 | 0.97 | 1 | 1 | 1 | 1 | 0.954 | 1 | -0.005 |
| SE | 994 | ENSG00000243156 | MICAL3 | chr22 | - | 17803810 | 17803861 | 17791201 | 17791301 | 17808843 | 17808937 | 4 | 0 | 1 | 0 | 0 | 7 | 4 | 10 | 12 | 5 | 0 | 2 | 0 | 0 | 0 | 0 | 31 | 3 | 13 | 4 | 200 | 149 | 1 | 1 | 0.299 | 0 | 0.069 | 0 | 0 | NA | 0.046 | 0 | 0 | 0 | 0.062 |
| SE | 1001 | ENSG00000243156 | MICAL3 | chr22 | - | 17831853 | 17832107 | 17822029 | 17822170 | 17841821 | 17842017 | 14 | 9 | 8 | 9 | 2 | 0 | 0 | 1 | 0 | 0 | 26 | 71 | 5 | 17 | 27 | 0 | 0 | 0 | 1 | 0 | 298 | 149 | 1 | 1 | 1 | 1 | 0.8 | 1 | 1 | 1 | 1 | 1 | 0.895 | 1 | -0.019 |
| SE | 1003 | ENSG00000243156 | MICAL3 | chr22 | - | 17884274 | 17884358 | 17871836 | 17872023 | 17885877 | 17886051 | 1 | 0 | 0 | 0 | 0 | 19 | 15 | 44 | 24 | 37 | 0 | 3 | 1 | 0 | 0 | 19 | 70 | 32 | 24 | 123 | 233 | 149 | 1 | 1 | 0.033 | 0 | 0 | 0 | 0 | 0 | 0.027 | 0.02 | 0 | 0 | -0.003 |
| SE | 1004 | ENSG00000243156 | MICAL3 | chr22 | - | 17893807 | 17893904 | 17891484 | 17891632 | 17895283 | 17895288 | 57 | 33 | 46 | 32 | 40 | 0 | 0 | 1 | 0 | 0 | 59 | 92 | 29 | 57 | 108 | 0 | 0 | 0 | 0 | 0 | 246 | 149 | 1 | 1 | 1 | 1 | 0.965 | 1 | 1 | 1 | 1 | 1 | 1 | 1 | -0.007 |
| SE | 1005 | ENSG00000243156 | MICAL3 | chr22 | - | 17896723 | 17896981 | 17896245 | 17896361 | 17899447 | 17899548 | 51 | 54 | 62 | 78 | 36 | 0 | 0 | 0 | 0 | 2 | 121 | 168 | 58 | 102 | 136 | 0 | 0 | 0 | 0 | 0 | 298 | 149 | 1 | 1 | 1 | 1 | 1 | 1 | 0.9 | 1 | 1 | 1 | 1 | 1 | -0.02 |
| SE | 1008 | ENSG00000243156 | MICAL3 | chr22 | - | 17902630 | 17902747 | 17900841 | 17900997 | 17904631 | 17904839 | 20 | 9 | 23 | 20 | 18 | 0 | 0 | 0 | 0 | 0 | 39 | 65 | 39 | 21 | 120 | 0 | 0 | 1 | 0 | 0 | 266 | 149 | 1 | 1 | 1 | 1 | 1 | 1 | 1 | 1 | 1 | 0.956 | 1 | 1 | 0.009 |
| SE | 1009 | ENSG00000131100 | ATP6V1E1 | chr22 | - | 17598193 | 17598288 | 17594528 | 17594616 | 17600026 | 17600095 | 561 | 396 | 420 | 511 | 551 | 0 | 0 | 1 | 1 | 0 | 624 | 477 | 469 | 544 | 965 | 0 | 0 | 2 | 0 | 0 | 244 | 149 | 1 | 1 | 1 | 1 | 0.996 | 0.997 | 1 | 1 | 1 | 0.993 | 1 | 1 | 0 |
| SE | 1011 | ENSG00000131100 | ATP6V1E1 | chr22 | - | 17619033 | 17619127 | 17613210 | 17613320 | 17619460 | 17619526 | 0 | 0 | 0 | 0 | 0 | 406 | 308 | 288 | 389 | 529 | 0 | 0 | 0 | 0 | 12 | 648 | 535 | 472 | 484 | 941 | 243 | 149 | 1 | 1 | 0 | 0 | 0 | 0 | 0 | 0 | 0 | 0 | 0 | 0.008 | -0.002 |
| SE | 1012 | ENSG00000131100 | ATP6V1E1 | chr22 | - | 17619460 | 17619526 | 17612975 | 17613320 | 17628602 | 17628733 | 573 | 414 | 423 | 584 | 713 | 5 | 4 | 4 | 4 | 2 | 989 | 707 | 634 | 635 | 1284 | 0 | 0 | 5 | 0 | 10 | 215 | 149 | 1 | 1 | 0.988 | 0.986 | 0.987 | 0.99 | 0.996 | 1 | 1 | 0.989 | 1 | 0.989 | -0.006 |
| SE | 1014 | ENSG00000187051 | RPS19BP1 | chr22 | - | 39530463 | 39530554 | 39529819 | 39529917 | 39532394 | 39532523 | 2 | 0 | 0 | 0 | 0 | 57 | 18 | 28 | 30 | 18 | 0 | 0 | 0 | 0 | 0 | 36 | 31 | 17 | 23 | 14 | 240 | 149 | 1 | 1 | 0.021 | 0 | 0 | 0 | 0 | 0 | 0 | 0 | 0 | 0 | 0.004 |
| SE | 1016 | ENSG00000100335 | MIEF1 | chr22 | + | 39503695 | 39503877 | 39502251 | 39502437 | 39504202 | 39504302 | 0 | 0 | 0 | 0 | 0 | 21 | 13 | 21 | 31 | 10 | 1 | 0 | 0 | 0 | 0 | 34 | 22 | 38 | 18 | 45 | 298 | 149 | 1 | 1 | 0 | 0 | 0 | 0 | 0 | 0.014 | 0 | 0 | 0 | 0 | -0.003 |
| SE | 1021 | ENSG00000100324 | TAB1 | chr22 | + | 39417710 | 39417849 | 39416790 | 39416877 | 39418731 | 39418845 | 63 | 59 | 65 | 99 | 65 | 1 | 0 | 0 | 0 | 0 | 141 | 141 | 44 | 91 | 218 | 0 | 0 | 0 | 2 | 0 | 288 | 149 | 1 | 1 | 0.97 | 1 | 1 | 1 | 1 | 1 | 1 | 1 | 0.959 | 1 | 0.002 |
| SE | 1025 | ENSG00000284554 | NA | chr22 | + | 39025556 | 39025671 | 39022821 | 39023014 | 39031693 | 39031973 | 2 | 1 | 1 | 1 | 1 | 16 | 10 | 8 | 10 | 8 | 0 | 0 | 0 | 3 | 0 | 15 | 7 | 6 | 10 | 0 | 264 | 149 | 1 | 1 | 0.066 | 0.053 | 0.066 | 0.053 | 0.066 | 0 | 0 | 0 | 0.145 | NA | 0.025 |
| SE | 1028 | ENSG00000284554 | NA | chr22 | + | 39029362 | 39029519 | 39025556 | 39025671 | 39031693 | 39031973 | 112 | 44 | 72 | 60 | 83 | 1 | 0 | 1 | 0 | 0 | 116 | 120 | 41 | 59 | 74 | 0 | 0 | 0 | 0 | 0 | 298 | 149 | 1 | 1 | 0.982 | 1 | 0.973 | 1 | 1 | 1 | 1 | 1 | 1 | 1 | -0.009 |
| SE | 1031 | ENSG00000128383 | APOBEC3A | chr22 | + | 38962097 | 38962213 | 38961386 | 38961681 | 38962494 | 38963183 | 21 | 41 | 29 | 44 | 26 | 0 | 0 | 0 | 0 | 0 | 307 | 114 | 105 | 369 | 46 | 1 | 0 | 0 | 0 | 0 | 265 | 149 | 1 | 1 | 1 | 1 | 1 | 1 | 1 | 0.994 | 1 | 1 | 1 | 1 | 0.001 |
| SE | 1037 | ENSG00000182057 | OGFRP1 | chr22 | + | 42274655 | 42274821 | 42269784 | 42270025 | 42274987 | 42275033 | 0 | 0 | 0 | 4 | 0 | 0 | 0 | 1 | 0 | 2 | 0 | 3 | 0 | 0 | 0 | 0 | 0 | 3 | 7 | 0 | 298 | 149 | 1 | 1 | NA | NA | 0 | 1 | 0 | NA | 1 | 0 | 0 | NA | 0 |
| SE | 1043 | ENSG00000100099 | HPS4 | chr22 | - | 26465454 | 26465551 | 26463916 | 26464826 | 26466050 | 26466104 | 68 | 27 | 25 | 40 | 60 | 0 | 0 | 0 | 0 | 0 | 23 | 69 | 41 | 12 | 46 | 0 | 0 | 1 | 0 | 0 | 246 | 149 | 1 | 1 | 1 | 1 | 1 | 1 | 1 | 1 | 1 | 0.961 | 1 | 1 | 0.008 |
| SE | 1045 | ENSG00000100099 | HPS4 | chr22 | - | 26468550 | 26468623 | 26465454 | 26466262 | 26470718 | 26470813 | 50 | 22 | 44 | 40 | 62 | 0 | 1 | 0 | 0 | 0 | 34 | 46 | 21 | 30 | 10 | 0 | 0 | 0 | 0 | 0 | 222 | 149 | 1 | 1 | 1 | 0.937 | 1 | 1 | 1 | 1 | 1 | 1 | 1 | 1 | -0.013 |
| SE | 1046 | ENSG00000100099 | HPS4 | chr22 | - | 26471212 | 26471288 | 26470718 | 26470813 | 26472301 | 26472418 | 1 | 1 | 1 | 1 | 2 | 39 | 17 | 51 | 28 | 47 | 0 | 1 | 0 | 0 | 2 | 42 | 40 | 33 | 52 | 6 | 225 | 149 | 1 | 1 | 0.017 | 0.037 | 0.013 | 0.023 | 0.027 | 0 | 0.016 | 0 | 0 | 0.181 | -0.016 |
| SE | 1048 | ENSG00000100099 | HPS4 | chr22 | - | 26471212 | 26471388 | 26470718 | 26470922 | 26472301 | 26472418 | 5 | 4 | 0 | 2 | 8 | 4 | 1 | 2 | 2 | 0 | 0 | 3 | 1 | 0 | 6 | 0 | 0 | 0 | 0 | 0 | 298 | 149 | 1 | 1 | 0.385 | 0.667 | 0 | 0.333 | 1 | NA | 1 | 1 | NA | 1 | -0.523 |
| SE | 1050 | ENSG00000100099 | HPS4 | chr22 | - | 26472831 | 26472939 | 26472301 | 26472418 | 26481721 | 26482240 | 27 | 6 | 44 | 28 | 31 | 0 | 0 | 0 | 0 | 0 | 0 | 37 | 13 | 32 | 14 | 0 | 0 | 2 | 0 | 0 | 257 | 149 | 1 | 1 | 1 | 1 | 1 | 1 | 1 | NA | 1 | 0.79 | 1 | 1 | 0.052 |
| SE | 1053 | ENSG00000100099 | HPS4 | chr22 | - | 26482745 | 26482873 | 26481834 | 26482240 | 26483764 | 26483828 | 6 | 0 | 3 | 4 | 1 | 6 | 4 | 6 | 5 | 1 | 0 | 0 | 0 | 0 | 0 | 0 | 19 | 6 | 25 | 24 | 277 | 149 | 1 | 1 | 0.35 | 0 | 0.212 | 0.301 | 0.35 | NA | 0 | 0 | 0 | 0 | 0.243 |
| SE | 1056 | ENSG00000253352 | TUG1 | chr22 | + | 30972047 | 30972172 | 30971196 | 30971610 | 30972854 | 30973587 | 3 | 7 | 11 | 15 | 18 | 0 | 0 | 1 | 0 | 8 | 0 | 6 | 4 | 1 | 8 | 0 | 0 | 0 | 0 | 0 | 274 | 149 | 1 | 1 | 1 | 1 | 0.857 | 1 | 0.55 | NA | 1 | 1 | 1 | 1 | -0.119 |
| SE | 1062 | ENSG00000284431 | NA | chr22 | + | 40349831 | 40350035 | 40346528 | 40346711 | 40358863 | 40359035 | 35 | 25 | 61 | 36 | 48 | 0 | 0 | 0 | 1 | 0 | 85 | 45 | 37 | 74 | 70 | 0 | 0 | 0 | 0 | 0 | 298 | 149 | 1 | 1 | 1 | 1 | 1 | 0.947 | 1 | 1 | 1 | 1 | 1 | 1 | -0.011 |
| SE | 1065 | ENSG00000284431 | NA | chr22 | + | 40402138 | 40402205 | 40400695 | 40400813 | 40404246 | 40404455 | 67 | 73 | 67 | 44 | 75 | 0 | 2 | 0 | 0 | 0 | 67 | 48 | 26 | 30 | 116 | 0 | 0 | 0 | 0 | 0 | 216 | 149 | 1 | 1 | 1 | 0.962 | 1 | 1 | 1 | 1 | 1 | 1 | 1 | 1 | -0.008 |
| SE | 1068 | ENSG00000250479 | CHCHD10 | chr22 | - | 23767373 | 23767540 | 23766127 | 23766275 | 23767833 | 23767920 | 4 | 0 | 1 | 2 | 0 | 0 | 0 | 0 | 0 | 0 | 0 | 0 | 1 | 0 | 11 | 0 | 1 | 0 | 0 | 0 | 298 | 149 | 1 | 1 | 1 | NA | 1 | 1 | NA | NA | 0 | 1 | NA | 1 | 0.333 |
| SE | 1069 | ENSG00000250479 | CHCHD10 | chr22 | - | 23767373 | 23767593 | 23766127 | 23766275 | 23767833 | 23767942 | 14 | 4 | 5 | 7 | 0 | 0 | 0 | 0 | 0 | 0 | 8 | 0 | 1 | 7 | 24 | 0 | 1 | 0 | 0 | 0 | 298 | 149 | 1 | 1 | 1 | 1 | 1 | 1 | NA | 1 | 0 | 1 | 1 | 1 | 0.2 |
| SE | 1073 | ENSG00000100330 | MTMR3 | chr22 | + | 30002879 | 30002993 | 29998760 | 29998857 | 30007113 | 30007319 | 171 | 79 | 95 | 104 | 181 | 0 | 0 | 0 | 1 | 0 | 169 | 223 | 194 | 235 | 224 | 0 | 0 | 0 | 0 | 0 | 263 | 149 | 1 | 1 | 1 | 1 | 1 | 0.983 | 1 | 1 | 1 | 1 | 1 | 1 | -0.003 |
| SE | 1077 | ENSG00000100330 | MTMR3 | chr22 | + | 30007900 | 30008032 | 30007113 | 30007319 | 30009017 | 30009129 | 251 | 138 | 167 | 216 | 197 | 0 | 0 | 0 | 0 | 0 | 541 | 374 | 339 | 492 | 426 | 0 | 0 | 0 | 1 | 0 | 281 | 149 | 1 | 1 | 1 | 1 | 1 | 1 | 1 | 1 | 1 | 1 | 0.996 | 1 | 0.001 |
| SE | 1078 | ENSG00000100330 | MTMR3 | chr22 | + | 30009017 | 30009129 | 30002879 | 30002993 | 30013355 | 30013541 | 0 | 0 | 0 | 3 | 4 | 0 | 0 | 0 | 0 | 0 | 27 | 9 | 2 | 14 | 7 | 0 | 0 | 0 | 5 | 0 | 261 | 149 | 1 | 1 | NA | NA | NA | 1 | 1 | 1 | 1 | 1 | 0.615 | 1 | 0.077 |
| SE | 1083 | ENSG00000100330 | MTMR3 | chr22 | + | 30016527 | 30016698 | 30009017 | 30009129 | 30017926 | 30018072 | 269 | 106 | 132 | 175 | 280 | 0 | 0 | 0 | 0 | 0 | 256 | 319 | 237 | 280 | 375 | 0 | 0 | 0 | 4 | 0 | 298 | 149 | 1 | 1 | 1 | 1 | 1 | 1 | 1 | 1 | 1 | 1 | 0.972 | 1 | 0.006 |
| SE | 1084 | ENSG00000100330 | MTMR3 | chr22 | + | 30017926 | 30018072 | 30009017 | 30009129 | 30019479 | 30020884 | 98 | 44 | 54 | 83 | 54 | 0 | 0 | 0 | 0 | 0 | 112 | 164 | 158 | 166 | 193 | 0 | 5 | 0 | 0 | 0 | 295 | 149 | 1 | 1 | 1 | 1 | 1 | 1 | 1 | 1 | 0.943 | 1 | 1 | 1 | 0.011 |
| SE | 1086 | ENSG00000100330 | MTMR3 | chr22 | + | 30022608 | 30022697 | 30019479 | 30020884 | 30025629 | 30026495 | 72 | 40 | 48 | 61 | 69 | 0 | 0 | 0 | 0 | 0 | 78 | 107 | 70 | 95 | 88 | 0 | 1 | 2 | 0 | 0 | 238 | 149 | 1 | 1 | 1 | 1 | 1 | 1 | 1 | 1 | 0.985 | 0.956 | 1 | 1 | 0.012 |
| SE | 1088 | ENSG00000185651 | UBE2L3 | chr22 | + | 21592860 | 21592956 | 21567713 | 21567771 | 21610856 | 21611043 | 294 | 205 | 203 | 175 | 297 | 0 | 0 | 1 | 0 | 0 | 737 | 337 | 284 | 446 | 473 | 0 | 0 | 0 | 0 | 0 | 245 | 149 | 1 | 1 | 1 | 1 | 0.992 | 1 | 1 | 1 | 1 | 1 | 1 | 1 | -0.002 |
| SE | 1089 | ENSG00000100030 | MAPK1 | chr22 | - | 21772872 | 21772982 | 21769039 | 21769320 | 21788256 | 21788388 | 649 | 284 | 458 | 537 | 549 | 0 | 0 | 0 | 0 | 0 | 599 | 492 | 325 | 546 | 671 | 0 | 0 | 0 | 4 | 0 | 259 | 149 | 1 | 1 | 1 | 1 | 1 | 1 | 1 | 1 | 1 | 1 | 0.987 | 1 | 0.003 |
| SE | 1091 | ENSG00000100030 | MAPK1 | chr22 | - | 21788256 | 21788388 | 21772872 | 21772982 | 21799011 | 21799128 | 423 | 230 | 336 | 374 | 395 | 4 | 2 | 1 | 0 | 2 | 402 | 392 | 276 | 403 | 504 | 0 | 0 | 0 | 0 | 4 | 281 | 149 | 1 | 1 | 0.982 | 0.984 | 0.994 | 1 | 0.991 | 1 | 1 | 1 | 1 | 0.985 | -0.007 |
| SE | 1092 | ENSG00000100030 | MAPK1 | chr22 | - | 21788693 | 21788808 | 21772872 | 21772982 | 21799011 | 21799128 | 426 | 176 | 300 | 388 | 472 | 4 | 2 | 1 | 0 | 2 | 341 | 346 | 265 | 294 | 432 | 0 | 0 | 0 | 0 | 4 | 264 | 149 | 1 | 1 | 0.984 | 0.98 | 0.994 | 1 | 0.993 | 1 | 1 | 1 | 1 | 0.984 | -0.007 |
| SE | 1094 | ENSG00000100030 | MAPK1 | chr22 | - | 21799011 | 21799128 | 21788693 | 21788808 | 21805849 | 21806039 | 806 | 366 | 567 | 655 | 775 | 1 | 0 | 0 | 0 | 0 | 668 | 591 | 488 | 613 | 811 | 0 | 0 | 0 | 0 | 0 | 266 | 149 | 1 | 1 | 0.998 | 1 | 1 | 1 | 1 | 1 | 1 | 1 | 1 | 1 | 0 |
| SE | 1098 | ENSG00000100031 | GGT1 | chr22 | + | 24608989 | 24609231 | 24607953 | 24608023 | 24609964 | 24610033 | 0 | 0 | 0 | 2 | 0 | 1 | 2 | 5 | 3 | 1 | 0 | 0 | 0 | 5 | 10 | 0 | 2 | 0 | 0 | 20 | 298 | 149 | 1 | 1 | 0 | 0 | 0 | 0.25 | 0 | NA | 0 | NA | 1 | 0.2 | -0.35 |
| SE | 1099 | ENSG00000100031 | GGT1 | chr22 | + | 24609961 | 24610033 | 24607953 | 24608023 | 24610357 | 24610531 | 1 | 3 | 3 | 0 | 1 | 0 | 0 | 0 | 0 | 3 | 0 | 12 | 0 | 1 | 9 | 0 | 0 | 0 | 5 | 0 | 221 | 149 | 1 | 1 | 1 | 1 | 1 | NA | 0.183 | NA | 1 | NA | 0.119 | 1 | 0.089 |
| SE | 1100 | ENSG00000100031 | GGT1 | chr22 | + | 24609964 | 24610033 | 24607953 | 24608023 | 24610357 | 24610531 | 1 | 3 | 6 | 2 | 1 | 0 | 0 | 0 | 0 | 3 | 0 | 11 | 0 | 0 | 29 | 0 | 0 | 0 | 5 | 0 | 218 | 149 | 1 | 1 | 1 | 1 | 1 | 1 | 0.186 | NA | 1 | NA | 0 | 1 | 0.171 |
| SE | 1102 | ENSG00000100031 | GGT1 | chr22 | + | 24620912 | 24621070 | 24620327 | 24620520 | 24623106 | 24623256 | 3 | 6 | 9 | 17 | 8 | 0 | 0 | 1 | 0 | 0 | 14 | 37 | 23 | 10 | 30 | 0 | 0 | 1 | 0 | 0 | 298 | 149 | 1 | 1 | 1 | 1 | 0.818 | 1 | 1 | 1 | 1 | 0.92 | 1 | 1 | -0.02 |
| SE | 1113 | ENSG00000100014 | SPECC1L | chr22 | + | 24330255 | 24330431 | 24328845 | 24328919 | 24334409 | 24334573 | 144 | 146 | 139 | 245 | 320 | 0 | 0 | 0 | 1 | 1 | 225 | 140 | 165 | 161 | 122 | 0 | 1 | 0 | 0 | 0 | 298 | 149 | 1 | 1 | 1 | 1 | 1 | 0.992 | 0.994 | 1 | 0.986 | 1 | 1 | 1 | 0 |
| SE | 1114 | ENSG00000100014 | SPECC1L | chr22 | + | 24330255 | 24330431 | 24328845 | 24328919 | 24347085 | 24347176 | 51 | 56 | 36 | 109 | 104 | 0 | 0 | 0 | 0 | 0 | 100 | 60 | 59 | 56 | 50 | 0 | 0 | 1 | 0 | 0 | 298 | 149 | 1 | 1 | 1 | 1 | 1 | 1 | 1 | 1 | 1 | 0.967 | 1 | 1 | 0.007 |
| SE | 1115 | ENSG00000100014 | SPECC1L | chr22 | + | 24334409 | 24334573 | 24328845 | 24328919 | 24347085 | 24347176 | 0 | 0 | 0 | 3 | 4 | 0 | 0 | 0 | 0 | 0 | 0 | 1 | 2 | 10 | 0 | 0 | 0 | 1 | 0 | 0 | 298 | 149 | 1 | 1 | NA | NA | NA | 1 | 1 | NA | 1 | 0.5 | 1 | NA | 0.167 |
| SE | 1116 | ENSG00000100014 | SPECC1L | chr22 | + | 24334409 | 24334573 | 24328845 | 24330255 | 24347085 | 24347176 | 93 | 90 | 103 | 139 | 219 | 0 | 0 | 0 | 1 | 0 | 125 | 80 | 108 | 115 | 72 | 0 | 0 | 0 | 0 | 0 | 298 | 149 | 1 | 1 | 1 | 1 | 1 | 0.986 | 1 | 1 | 1 | 1 | 1 | 1 | -0.003 |
| SE | 1122 | ENSG00000128340 | RAC2 | chr22 | - | 37226670 | 37226803 | 37225269 | 37226039 | 37231230 | 37231390 | 1227 | 538 | 946 | 1141 | 931 | 1 | 0 | 0 | 5 | 0 | 862 | 1118 | 671 | 901 | 1492 | 0 | 0 | 0 | 0 | 0 | 282 | 149 | 1 | 1 | 0.998 | 1 | 1 | 0.992 | 1 | 1 | 1 | 1 | 1 | 1 | -0.002 |
| SE | 1124 | ENSG00000128340 | RAC2 | chr22 | - | 37231931 | 37231994 | 37231230 | 37231390 | 37232800 | 37232918 | 2689 | 1106 | 1726 | 2079 | 1462 | 2 | 2 | 5 | 4 | 0 | 2449 | 2431 | 1321 | 2347 | 3685 | 0 | 2 | 2 | 8 | 19 | 212 | 149 | 1 | 1 | 0.999 | 0.997 | 0.996 | 0.997 | 1 | 1 | 0.999 | 0.998 | 0.995 | 0.993 | 0.001 |
| SE | 1125 | ENSG00000128340 | RAC2 | chr22 | - | 37231931 | 37231994 | 37231230 | 37231390 | 37241586 | 37241658 | 644 | 221 | 404 | 490 | 411 | 0 | 0 | 0 | 1 | 0 | 358 | 541 | 304 | 408 | 687 | 0 | 0 | 0 | 0 | 0 | 212 | 149 | 1 | 1 | 1 | 1 | 1 | 0.997 | 1 | 1 | 1 | 1 | 1 | 1 | -0.001 |
| SE | 1126 | ENSG00000128340 | RAC2 | chr22 | - | 37232800 | 37232918 | 37231230 | 37231390 | 37241586 | 37241658 | 611 | 227 | 354 | 476 | 197 | 0 | 0 | 0 | 1 | 0 | 595 | 576 | 313 | 536 | 836 | 0 | 0 | 0 | 0 | 0 | 267 | 149 | 1 | 1 | 1 | 1 | 1 | 0.996 | 1 | 1 | 1 | 1 | 1 | 1 | -0.001 |
| SE | 1127 | ENSG00000128340 | RAC2 | chr22 | - | 37232800 | 37232918 | 37231931 | 37231994 | 37241586 | 37241658 | 2654 | 1111 | 1672 | 2062 | 1246 | 0 | 1 | 1 | 1 | 0 | 2686 | 2464 | 1328 | 2467 | 3815 | 0 | 0 | 0 | 0 | 0 | 267 | 149 | 1 | 1 | 1 | 0.998 | 0.999 | 0.999 | 1 | 1 | 1 | 1 | 1 | 1 | -0.001 |
| SE | 1128 | ENSG00000128340 | RAC2 | chr22 | - | 37244113 | 37244202 | 37241586 | 37241658 | 37244289 | 37244441 | 495 | 202 | 310 | 407 | 173 | 0 | 1 | 0 | 0 | 0 | 460 | 485 | 297 | 423 | 712 | 0 | 0 | 0 | 0 | 0 | 238 | 149 | 1 | 1 | 1 | 0.992 | 1 | 1 | 1 | 1 | 1 | 1 | 1 | 1 | -0.002 |
| SE | 1133 | ENSG00000128294 | TPST2 | chr22 | - | 26532694 | 26532745 | 26528213 | 26528262 | 26536287 | 26536486 | 160 | 87 | 135 | 215 | 209 | 2 | 0 | 0 | 0 | 0 | 156 | 161 | 78 | 100 | 597 | 0 | 0 | 0 | 0 | 0 | 200 | 149 | 1 | 1 | 0.983 | 1 | 1 | 1 | 1 | 1 | 1 | 1 | 1 | 1 | -0.003 |
| SE | 1136 | ENSG00000128294 | TPST2 | chr22 | - | 26544603 | 26544675 | 26540788 | 26541718 | 26590052 | 26590125 | 30 | 14 | 15 | 30 | 2 | 3 | 0 | 0 | 0 | 0 | 6 | 40 | 3 | 18 | 67 | 0 | 0 | 0 | 0 | 0 | 221 | 149 | 1 | 1 | 0.871 | 1 | 1 | 1 | 1 | 1 | 1 | 1 | 1 | 1 | -0.026 |
| SE | 1138 | ENSG00000100104 | SRRD | chr22 | + | 26488389 | 26488488 | 26487981 | 26488288 | 26490043 | 26490198 | 91 | 61 | 96 | 83 | 100 | 0 | 3 | 0 | 0 | 0 | 115 | 121 | 113 | 96 | 151 | 0 | 0 | 0 | 0 | 0 | 248 | 149 | 1 | 1 | 1 | 0.924 | 1 | 1 | 1 | 1 | 1 | 1 | 1 | 1 | -0.015 |
| SE | 1140 | ENSG00000128203 | ASPHD2 | chr22 | + | 26442458 | 26442572 | 26433391 | 26434501 | 26443096 | 26445015 | 60 | 46 | 50 | 43 | 12 | 0 | 0 | 0 | 2 | 0 | 8 | 41 | 27 | 32 | 119 | 0 | 2 | 0 | 0 | 1 | 263 | 149 | 1 | 1 | 1 | 1 | 1 | 1 | 1 | 1 | 0.921 | 1 | 1 | 0.985 | 0.019 |
| SE | 1144 | ENSG00000205853 | RFPL3S | chr22 | - | 32362971 | 32363058 | 32359885 | 32360839 | 32367689 | 32367714 | 0 | 1 | 0 | 5 | 2 | 0 | 0 | 0 | 2 | 0 | 0 | 0 | 1 | 0 | 0 | 0 | 0 | 0 | 0 | 0 | 236 | 149 | 1 | 1 | NA | 1 | NA | 0.612 | 1 | NA | NA | 1 | NA | NA | -0.129 |
| SE | 1148 | ENSG00000100060 | MFNG | chr22 | - | 37480196 | 37480299 | 37479344 | 37479498 | 37485922 | 37486379 | 82 | 45 | 91 | 75 | 64 | 0 | 0 | 1 | 1 | 0 | 81 | 63 | 34 | 59 | 191 | 0 | 0 | 1 | 0 | 0 | 252 | 149 | 1 | 1 | 1 | 1 | 0.982 | 0.978 | 1 | 1 | 1 | 0.953 | 1 | 1 | 0.001 |
| SE | 1157 | ENSG00000100395 | L3MBTL2 | chr22 | + | 41213850 | 41214026 | 41209695 | 41209933 | 41219418 | 41219536 | 0 | 0 | 2 | 0 | 0 | 0 | 0 | 0 | 0 | 0 | 0 | 0 | 4 | 1 | 0 | 0 | 0 | 4 | 0 | 0 | 298 | 149 | 1 | 1 | NA | NA | 1 | NA | NA | NA | NA | 0.333 | 1 | NA | 0.334 |
| SE | 1164 | ENSG00000100395 | L3MBTL2 | chr22 | + | 41227574 | 41227679 | 41227088 | 41227323 | 41227803 | 41227869 | 0 | 1 | 0 | 0 | 0 | 22 | 10 | 37 | 18 | 24 | 0 | 0 | 0 | 0 | 0 | 16 | 30 | 10 | 7 | 10 | 254 | 149 | 1 | 1 | 0 | 0.055 | 0 | 0 | 0 | 0 | 0 | 0 | 0 | 0 | 0.011 |
| SE | 1166 | ENSG00000182841 | RRP7BP | chr22 | - | 42576980 | 42577098 | 42576511 | 42576609 | 42577843 | 42577962 | 111 | 62 | 98 | 105 | 48 | 0 | 0 | 0 | 1 | 0 | 97 | 52 | 43 | 102 | 131 | 0 | 0 | 0 | 4 | 0 | 267 | 149 | 1 | 1 | 1 | 1 | 1 | 0.983 | 1 | 1 | 1 | 1 | 0.934 | 1 | 0.01 |
| SE | 1169 | ENSG00000169635 | HIC2 | chr22 | + | 21442758 | 21442857 | 21417370 | 21417560 | 21444921 | 21449434 | 0 | 0 | 3 | 0 | 0 | 0 | 0 | 0 | 0 | 0 | 0 | 0 | 0 | 0 | 0 | 5 | 0 | 0 | 1 | 1 | 248 | 149 | 1 | 1 | NA | NA | 1 | NA | NA | 0 | NA | NA | 0 | 0 | 1 |
| SE | 1171 | ENSG00000100241 | SBF1 | chr22 | - | 50466149 | 50466258 | 50465960 | 50466074 | 50466349 | 50466482 | 121 | 40 | 79 | 85 | 62 | 0 | 0 | 0 | 0 | 4 | 51 | 103 | 83 | 22 | 159 | 0 | 0 | 0 | 0 | 0 | 258 | 149 | 1 | 1 | 1 | 1 | 1 | 1 | 0.9 | 1 | 1 | 1 | 1 | 1 | -0.02 |
| SE | 1172 | ENSG00000196236 | XPNPEP3 | chr22 | + | 40860677 | 40860793 | 40857096 | 40857245 | 40861617 | 40862770 | 8 | 5 | 1 | 0 | 2 | 3 | 3 | 3 | 1 | 0 | 0 | 8 | 0 | 0 | 0 | 0 | 0 | 8 | 11 | 0 | 265 | 149 | 1 | 1 | 0.6 | 0.484 | 0.158 | 0 | 1 | NA | 1 | 0 | 0 | NA | 0.115 |
| SE | 1174 | ENSG00000196236 | XPNPEP3 | chr22 | + | 40881769 | 40882177 | 40868998 | 40869115 | 40886312 | 40886515 | 55 | 25 | 49 | 51 | 76 | 2 | 0 | 0 | 0 | 0 | 100 | 37 | 52 | 33 | 95 | 0 | 0 | 0 | 0 | 0 | 298 | 149 | 1 | 1 | 0.932 | 1 | 1 | 1 | 1 | 1 | 1 | 1 | 1 | 1 | -0.014 |
| SE | 1175 | ENSG00000196236 | XPNPEP3 | chr22 | + | 40914238 | 40914324 | 40909121 | 40909235 | 40922332 | 40922513 | 48 | 39 | 44 | 57 | 43 | 0 | 0 | 0 | 0 | 0 | 81 | 61 | 35 | 27 | 54 | 0 | 0 | 2 | 0 | 0 | 235 | 149 | 1 | 1 | 1 | 1 | 1 | 1 | 1 | 1 | 1 | 0.917 | 1 | 1 | 0.017 |
| SE | 1176 | ENSG00000196236 | XPNPEP3 | chr22 | + | 40922332 | 40922513 | 40914238 | 40914324 | 40924361 | 40924482 | 58 | 41 | 60 | 64 | 50 | 1 | 0 | 0 | 0 | 0 | 84 | 80 | 26 | 32 | 52 | 0 | 0 | 0 | 0 | 0 | 298 | 149 | 1 | 1 | 0.967 | 1 | 1 | 1 | 1 | 1 | 1 | 1 | 1 | 1 | -0.007 |
| SE | 1179 | ENSG00000100351 | GRAP2 | chr22 | + | 39947092 | 39947184 | 39901083 | 39901330 | 39955818 | 39955910 | 610 | 528 | 451 | 923 | 759 | 7 | 1 | 3 | 7 | 4 | 1375 | 861 | 439 | 759 | 3802 | 5 | 0 | 0 | 0 | 7 | 241 | 149 | 1 | 1 | 0.982 | 0.997 | 0.989 | 0.988 | 0.992 | 0.994 | 1 | 1 | 1 | 0.997 | -0.009 |
| SE | 1181 | ENSG00000100351 | GRAP2 | chr22 | + | 39947092 | 39947184 | 39901083 | 39901330 | 39965989 | 39966158 | 124 | 74 | 119 | 115 | 115 | 0 | 2 | 0 | 1 | 0 | 72 | 66 | 46 | 47 | 56 | 0 | 0 | 0 | 0 | 0 | 241 | 149 | 1 | 1 | 1 | 0.958 | 1 | 0.986 | 1 | 1 | 1 | 1 | 1 | 1 | -0.011 |
| SE | 1182 | ENSG00000100351 | GRAP2 | chr22 | + | 39947092 | 39947184 | 39926590 | 39926853 | 39955818 | 39955910 | 376 | 415 | 242 | 751 | 580 | 1 | 0 | 0 | 2 | 0 | 1267 | 729 | 344 | 630 | 3653 | 0 | 0 | 0 | 0 | 0 | 241 | 149 | 1 | 1 | 0.996 | 1 | 1 | 0.996 | 1 | 1 | 1 | 1 | 1 | 1 | -0.002 |
| SE | 1183 | ENSG00000100351 | GRAP2 | chr22 | + | 39955818 | 39955910 | 39901083 | 39901330 | 39960054 | 39960174 | 265 | 278 | 191 | 532 | 518 | 1 | 0 | 4 | 0 | 3 | 972 | 662 | 248 | 429 | 2675 | 8 | 0 | 0 | 2 | 4 | 241 | 149 | 1 | 1 | 0.994 | 1 | 0.967 | 1 | 0.991 | 0.987 | 1 | 1 | 0.993 | 0.998 | -0.005 |
| SE | 1184 | ENSG00000100351 | GRAP2 | chr22 | + | 39955818 | 39955910 | 39901083 | 39901330 | 39965989 | 39966109 | 2 | 0 | 4 | 6 | 3 | 0 | 2 | 0 | 1 | 0 | 5 | 0 | 0 | 0 | 4 | 0 | 0 | 0 | 0 | 0 | 241 | 149 | 1 | 1 | 1 | 0 | 1 | 0.788 | 1 | 1 | NA | NA | NA | 1 | -0.242 |
| SE | 1187 | ENSG00000100351 | GRAP2 | chr22 | + | 39960054 | 39960174 | 39901083 | 39901330 | 39965989 | 39966109 | 362 | 389 | 252 | 827 | 577 | 0 | 2 | 0 | 1 | 0 | 1101 | 968 | 340 | 731 | 3936 | 0 | 0 | 0 | 0 | 0 | 269 | 149 | 1 | 1 | 1 | 0.991 | 1 | 0.998 | 1 | 1 | 1 | 1 | 1 | 1 | -0.002 |
| SE | 1189 | ENSG00000100351 | GRAP2 | chr22 | + | 39960054 | 39960174 | 39946816 | 39947184 | 39969410 | 39969533 | 34 | 32 | 21 | 41 | 45 | 2 | 0 | 0 | 0 | 0 | 65 | 62 | 27 | 32 | 233 | 0 | 0 | 0 | 0 | 0 | 269 | 149 | 1 | 1 | 0.904 | 1 | 1 | 1 | 1 | 1 | 1 | 1 | 1 | 1 | -0.019 |
| SE | 1190 | ENSG00000100351 | GRAP2 | chr22 | + | 39960054 | 39960174 | 39955818 | 39955910 | 39965989 | 39966109 | 885 | 940 | 583 | 1843 | 1515 | 5 | 1 | 1 | 2 | 3 | 2958 | 2179 | 835 | 1578 | 9275 | 0 | 0 | 0 | 0 | 0 | 269 | 149 | 1 | 1 | 0.99 | 0.998 | 0.997 | 0.998 | 0.996 | 1 | 1 | 1 | 1 | 1 | -0.004 |
| SE | 1192 | ENSG00000100351 | GRAP2 | chr22 | + | 39968041 | 39968272 | 39960054 | 39960174 | 39969410 | 39969533 | 97 | 62 | 62 | 196 | 80 | 0 | 0 | 0 | 0 | 0 | 346 | 231 | 115 | 170 | 1216 | 0 | 1 | 0 | 0 | 0 | 298 | 149 | 1 | 1 | 1 | 1 | 1 | 1 | 1 | 1 | 0.991 | 1 | 1 | 1 | 0.002 |
| SE | 1194 | ENSG00000100227 | POLDIP3 | chr22 | - | 42591954 | 42592084 | 42583720 | 42585968 | 42595536 | 42595603 | 94 | 54 | 126 | 97 | 95 | 1 | 0 | 0 | 0 | 0 | 110 | 142 | 79 | 114 | 117 | 0 | 0 | 0 | 2 | 0 | 279 | 149 | 1 | 1 | 0.98 | 1 | 1 | 1 | 1 | 1 | 1 | 1 | 0.968 | 1 | 0.002 |
| SE | 1195 | ENSG00000100227 | POLDIP3 | chr22 | - | 42595536 | 42595603 | 42591954 | 42592084 | 42596185 | 42596365 | 40 | 9 | 59 | 40 | 59 | 1 | 0 | 0 | 0 | 0 | 65 | 35 | 19 | 39 | 50 | 0 | 0 | 0 | 0 | 0 | 216 | 149 | 1 | 1 | 0.965 | 1 | 1 | 1 | 1 | 1 | 1 | 1 | 1 | 1 | -0.007 |
| SE | 1196 | ENSG00000100227 | POLDIP3 | chr22 | - | 42595536 | 42595614 | 42591954 | 42592084 | 42596185 | 42596365 | 199 | 94 | 204 | 161 | 181 | 1 | 0 | 0 | 0 | 0 | 219 | 229 | 142 | 175 | 207 | 0 | 0 | 0 | 0 | 0 | 227 | 149 | 1 | 1 | 0.992 | 1 | 1 | 1 | 1 | 1 | 1 | 1 | 1 | 1 | -0.002 |
| SE | 1202 | ENSG00000100387 | RBX1 | chr22 | + | 40953554 | 40953633 | 40951346 | 40951476 | 40967798 | 40967884 | 185 | 128 | 149 | 160 | 71 | 0 | 0 | 1 | 1 | 0 | 330 | 277 | 110 | 291 | 334 | 0 | 0 | 1 | 0 | 0 | 228 | 149 | 1 | 1 | 1 | 1 | 0.99 | 0.991 | 1 | 1 | 1 | 0.986 | 1 | 1 | -0.001 |
| SE | 1203 | ENSG00000100387 | RBX1 | chr22 | + | 40964046 | 40964117 | 40951377 | 40951476 | 40967798 | 40967884 | 225 | 182 | 186 | 214 | 120 | 0 | 0 | 1 | 1 | 0 | 446 | 278 | 146 | 339 | 494 | 0 | 0 | 1 | 0 | 0 | 220 | 149 | 1 | 1 | 1 | 1 | 0.992 | 0.993 | 1 | 1 | 1 | 0.99 | 1 | 1 | -0.001 |
| SE | 1204 | ENSG00000100387 | RBX1 | chr22 | + | 40964046 | 40964117 | 40953554 | 40953633 | 40967798 | 40967884 | 660 | 546 | 527 | 696 | 456 | 3 | 2 | 0 | 1 | 1 | 1365 | 902 | 410 | 1063 | 1378 | 0 | 3 | 0 | 2 | 4 | 220 | 149 | 1 | 1 | 0.993 | 0.995 | 1 | 0.998 | 0.997 | 1 | 0.995 | 1 | 0.997 | 0.996 | -0.001 |
| SE | 1205 | ENSG00000100387 | RBX1 | chr22 | + | 40964046 | 40964117 | 40953554 | 40953633 | 40972475 | 40973309 | 224 | 200 | 194 | 283 | 219 | 0 | 0 | 0 | 0 | 2 | 517 | 350 | 137 | 366 | 523 | 0 | 0 | 0 | 0 | 0 | 220 | 149 | 1 | 1 | 1 | 1 | 1 | 1 | 0.987 | 1 | 1 | 1 | 1 | 1 | -0.003 |
| SE | 1206 | ENSG00000100387 | RBX1 | chr22 | + | 40967798 | 40967884 | 40953554 | 40953633 | 40972475 | 40973309 | 136 | 82 | 91 | 139 | 119 | 0 | 0 | 0 | 0 | 2 | 128 | 170 | 71 | 187 | 208 | 0 | 0 | 0 | 0 | 0 | 235 | 149 | 1 | 1 | 1 | 1 | 1 | 1 | 0.974 | 1 | 1 | 1 | 1 | 1 | -0.005 |
| SE | 1208 | ENSG00000235513 | L3MBTL2-AS1 | chr22 | - | 41213629 | 41214703 | 41207591 | 41209951 | 41217031 | 41217910 | 8 | 5 | 0 | 3 | 10 | 0 | 0 | 0 | 1 | 0 | 7 | 0 | 0 | 2 | 0 | 2 | 1 | 2 | 0 | 0 | 298 | 149 | 1 | 1 | 1 | 1 | NA | 0.6 | 1 | 0.636 | 0 | 0 | 1 | NA | 0.491 |
| SE | 1209 | ENSG00000235513 | L3MBTL2-AS1 | chr22 | - | 41213795 | 41214703 | 41208551 | 41209951 | 41217031 | 41217310 | 0 | 2 | 0 | 3 | 4 | 0 | 0 | 0 | 1 | 0 | 0 | 0 | 0 | 0 | 0 | 2 | 1 | 2 | 0 | 0 | 298 | 149 | 1 | 1 | NA | 1 | NA | 0.6 | 1 | 0 | 0 | 0 | NA | NA | 0.867 |
| SE | 1210 | ENSG00000100393 | EP300 | chr22 | + | 41117186 | 41117821 | 41092591 | 41093098 | 41125863 | 41126040 | 344 | 190 | 269 | 300 | 259 | 0 | 0 | 0 | 1 | 0 | 327 | 328 | 282 | 409 | 260 | 1 | 0 | 0 | 0 | 0 | 298 | 149 | 1 | 1 | 1 | 1 | 1 | 0.993 | 1 | 0.994 | 1 | 1 | 1 | 1 | 0 |
| SE | 1211 | ENSG00000100393 | EP300 | chr22 | + | 41131387 | 41131633 | 41129889 | 41130003 | 41135812 | 41135906 | 353 | 214 | 266 | 376 | 343 | 0 | 1 | 0 | 0 | 0 | 503 | 353 | 316 | 498 | 314 | 0 | 0 | 3 | 3 | 0 | 298 | 149 | 1 | 1 | 1 | 0.991 | 1 | 1 | 1 | 1 | 1 | 0.981 | 0.988 | 1 | 0.004 |
| SE | 1212 | ENSG00000100393 | EP300 | chr22 | + | 41137652 | 41137790 | 41135812 | 41135906 | 41140139 | 41140257 | 506 | 435 | 458 | 533 | 548 | 0 | 0 | 0 | 0 | 0 | 1060 | 666 | 658 | 938 | 521 | 5 | 0 | 0 | 6 | 6 | 287 | 149 | 1 | 1 | 1 | 1 | 1 | 1 | 1 | 0.991 | 1 | 1 | 0.988 | 0.978 | 0.009 |
| SE | 1213 | ENSG00000100393 | EP300 | chr22 | + | 41141047 | 41141222 | 41140139 | 41140257 | 41146738 | 41146809 | 635 | 503 | 550 | 636 | 961 | 3 | 0 | 0 | 1 | 0 | 1393 | 770 | 873 | 1093 | 754 | 0 | 0 | 0 | 0 | 0 | 298 | 149 | 1 | 1 | 0.991 | 1 | 1 | 0.997 | 1 | 1 | 1 | 1 | 1 | 1 | -0.002 |
| SE | 1215 | ENSG00000100393 | EP300 | chr22 | + | 41149037 | 41149175 | 41147836 | 41147946 | 41151832 | 41152012 | 134 | 83 | 130 | 120 | 136 | 0 | 0 | 3 | 0 | 0 | 192 | 183 | 125 | 261 | 127 | 0 | 0 | 0 | 0 | 0 | 287 | 149 | 1 | 1 | 1 | 1 | 0.957 | 1 | 1 | 1 | 1 | 1 | 1 | 1 | -0.009 |
| SE | 1217 | ENSG00000100393 | EP300 | chr22 | + | 41152205 | 41152350 | 41151832 | 41152012 | 41154994 | 41155113 | 465 | 342 | 346 | 483 | 490 | 0 | 0 | 0 | 0 | 0 | 854 | 501 | 419 | 901 | 370 | 0 | 0 | 0 | 4 | 0 | 294 | 149 | 1 | 1 | 1 | 1 | 1 | 1 | 1 | 1 | 1 | 1 | 0.991 | 1 | 0.002 |
| SE | 1218 | ENSG00000100393 | EP300 | chr22 | + | 41166598 | 41166666 | 41164052 | 41164130 | 41168448 | 41168599 | 391 | 277 | 354 | 409 | 564 | 0 | 0 | 0 | 1 | 0 | 393 | 386 | 394 | 546 | 317 | 0 | 0 | 0 | 0 | 0 | 217 | 149 | 1 | 1 | 1 | 1 | 1 | 0.996 | 1 | 1 | 1 | 1 | 1 | 1 | -0.001 |
| SE | 1219 | ENSG00000100393 | EP300 | chr22 | + | 41168720 | 41168867 | 41168448 | 41168599 | 41169502 | 41169616 | 490 | 333 | 332 | 428 | 540 | 0 | 1 | 0 | 0 | 0 | 593 | 376 | 454 | 548 | 283 | 0 | 0 | 0 | 0 | 0 | 296 | 149 | 1 | 1 | 1 | 0.994 | 1 | 1 | 1 | 1 | 1 | 1 | 1 | 1 | -0.001 |
| SE | 1220 | ENSG00000100393 | EP300 | chr22 | + | 41169502 | 41169616 | 41168720 | 41168867 | 41170405 | 41170571 | 335 | 325 | 295 | 435 | 581 | 1 | 1 | 0 | 1 | 2 | 897 | 411 | 463 | 642 | 307 | 6 | 0 | 0 | 3 | 0 | 263 | 149 | 1 | 1 | 0.995 | 0.995 | 1 | 0.996 | 0.994 | 0.988 | 1 | 1 | 0.992 | 1 | 0 |
| SE | 1221 | ENSG00000100393 | EP300 | chr22 | + | 41170405 | 41170571 | 41169502 | 41169616 | 41172498 | 41172663 | 678 | 665 | 518 | 723 | 1002 | 0 | 2 | 1 | 5 | 4 | 1825 | 743 | 1037 | 1305 | 690 | 0 | 0 | 3 | 1 | 0 | 298 | 149 | 1 | 1 | 1 | 0.994 | 0.996 | 0.986 | 0.992 | 1 | 1 | 0.994 | 0.998 | 1 | -0.005 |
| SE | 1233 | ENSG00000100124 | ANKRD54 | chr22 | - | 37833683 | 37833750 | 37832992 | 37833206 | 37833916 | 37834728 | 1 | 1 | 2 | 7 | 11 | 0 | 0 | 0 | 0 | 0 | 0 | 8 | 0 | 0 | 0 | 0 | 1 | 0 | 0 | 0 | 216 | 149 | 1 | 1 | 1 | 1 | 1 | 1 | 1 | NA | 0.847 | NA | NA | NA | 0.153 |
| SE | 1234 | ENSG00000100124 | ANKRD54 | chr22 | - | 37833683 | 37833750 | 37832992 | 37833206 | 37838499 | 37838593 | 1 | 1 | 2 | 5 | 7 | 0 | 0 | 0 | 4 | 0 | 0 | 8 | 0 | 0 | 0 | 0 | 0 | 0 | 0 | 0 | 216 | 149 | 1 | 1 | 1 | 1 | 1 | 0.463 | 1 | NA | 1 | NA | NA | NA | -0.107 |
| SE | 1235 | ENSG00000100124 | ANKRD54 | chr22 | - | 37833683 | 37833755 | 37832957 | 37833206 | 37833916 | 37834728 | 2 | 1 | 2 | 9 | 7 | 0 | 0 | 0 | 0 | 0 | 5 | 8 | 2 | 0 | 2 | 0 | 1 | 0 | 0 | 0 | 221 | 149 | 1 | 1 | 1 | 1 | 1 | 1 | 1 | 1 | 0.844 | 1 | NA | 1 | 0.039 |
| SE | 1236 | ENSG00000100124 | ANKRD54 | chr22 | - | 37833683 | 37833755 | 37832957 | 37833206 | 37838499 | 37838593 | 9 | 7 | 7 | 7 | 23 | 0 | 0 | 0 | 4 | 0 | 0 | 8 | 1 | 0 | 2 | 0 | 0 | 0 | 0 | 0 | 221 | 149 | 1 | 1 | 1 | 1 | 1 | 0.541 | 1 | NA | 1 | 1 | NA | 1 | -0.092 |
| SE | 1240 | ENSG00000100154 | TTC28 | chr22 | - | 27990788 | 27990812 | 27989877 | 27990007 | 27992586 | 27992663 | 0 | 0 | 2 | 0 | 1 | 0 | 0 | 0 | 1 | 0 | 0 | 0 | 0 | 0 | 0 | 0 | 0 | 0 | 0 | 5 | 173 | 149 | 1 | 1 | NA | NA | 1 | 0 | 1 | NA | NA | NA | NA | 0 | 0.667 |
| SE | 1241 | ENSG00000100097 | LGALS1 | chr22 | + | 37678375 | 37678392 | 37676985 | 37677065 | 37678482 | 37678654 | 0 | 0 | 0 | 0 | 1 | 967 | 313 | 579 | 708 | 356 | 0 | 0 | 0 | 0 | 0 | 1128 | 927 | 396 | 754 | 1443 | 166 | 149 | 1 | 1 | 0 | 0 | 0 | 0 | 0.003 | 0 | 0 | 0 | 0 | 0 | 0.001 |
| SE | 1242 | ENSG00000100097 | LGALS1 | chr22 | + | 37678375 | 37678654 | 37676998 | 37677065 | 37679602 | 37679802 | 762 | 216 | 523 | 539 | 280 | 8 | 2 | 2 | 0 | 0 | 845 | 763 | 342 | 508 | 1102 | 0 | 8 | 0 | 4 | 5 | 298 | 149 | 1 | 1 | 0.979 | 0.982 | 0.992 | 1 | 1 | 1 | 0.979 | 1 | 0.984 | 0.991 | 0 |
| SE | 1243 | ENSG00000100097 | LGALS1 | chr22 | + | 37678482 | 37678654 | 37676855 | 37677065 | 37679602 | 37679640 | 1729 | 529 | 1101 | 1247 | 635 | 8 | 2 | 2 | 0 | 0 | 1952 | 1689 | 737 | 1255 | 2545 | 0 | 8 | 0 | 4 | 5 | 298 | 149 | 1 | 1 | 0.991 | 0.992 | 0.996 | 1 | 1 | 1 | 0.991 | 1 | 0.994 | 0.996 | 0 |
| SE | 1245 | ENSG00000100083 | GGA1 | chr22 | + | 37613023 | 37613191 | 37608844 | 37608903 | 37614193 | 37614274 | 0 | 0 | 2 | 0 | 0 | 0 | 0 | 0 | 0 | 0 | 0 | 0 | 0 | 0 | 0 | 0 | 1 | 1 | 0 | 0 | 298 | 149 | 1 | 1 | NA | NA | 1 | NA | NA | NA | 0 | 0 | NA | NA | 1 |
| SE | 1246 | ENSG00000100083 | GGA1 | chr22 | + | 37613023 | 37613191 | 37608959 | 37609154 | 37614189 | 37614274 | 7 | 1 | 10 | 1 | 3 | 0 | 0 | 1 | 2 | 0 | 8 | 8 | 5 | 11 | 27 | 0 | 0 | 0 | 0 | 0 | 298 | 149 | 1 | 1 | 1 | 1 | 0.833 | 0.2 | 1 | 1 | 1 | 1 | 1 | 1 | -0.193 |
| SE | 1249 | ENSG00000100083 | GGA1 | chr22 | + | 37613032 | 37613191 | 37609059 | 37609154 | 37614189 | 37614274 | 7 | 1 | 10 | 1 | 3 | 0 | 0 | 1 | 2 | 0 | 8 | 6 | 5 | 7 | 27 | 0 | 0 | 0 | 0 | 0 | 298 | 149 | 1 | 1 | 1 | 1 | 0.833 | 0.2 | 1 | 1 | 1 | 1 | 1 | 1 | -0.193 |
| SE | 1250 | ENSG00000100083 | GGA1 | chr22 | + | 37613118 | 37613191 | 37608976 | 37609154 | 37614189 | 37614195 | 1 | 0 | 8 | 0 | 0 | 0 | 0 | 1 | 2 | 0 | 4 | 5 | 5 | 7 | 22 | 0 | 0 | 0 | 0 | 0 | 222 | 149 | 1 | 1 | 1 | NA | 0.843 | 0 | NA | 1 | 1 | 1 | 1 | 1 | -0.386 |
| SE | 1252 | ENSG00000100083 | GGA1 | chr22 | + | 37614189 | 37614274 | 37608724 | 37608903 | 37616921 | 37616997 | 44 | 20 | 49 | 33 | 8 | 0 | 0 | 1 | 0 | 0 | 63 | 46 | 25 | 72 | 86 | 0 | 0 | 0 | 0 | 0 | 234 | 149 | 1 | 1 | 1 | 1 | 0.969 | 1 | 1 | 1 | 1 | 1 | 1 | 1 | -0.006 |
| SE | 1253 | ENSG00000100083 | GGA1 | chr22 | + | 37614193 | 37614274 | 37608834 | 37608903 | 37616921 | 37616997 | 31 | 17 | 25 | 21 | 5 | 0 | 0 | 1 | 0 | 0 | 38 | 38 | 22 | 51 | 74 | 0 | 0 | 0 | 0 | 0 | 230 | 149 | 1 | 1 | 1 | 1 | 0.942 | 1 | 1 | 1 | 1 | 1 | 1 | 1 | -0.012 |
| SE | 1254 | ENSG00000100083 | GGA1 | chr22 | + | 37616921 | 37616997 | 37614132 | 37614274 | 37618447 | 37618515 | 57 | 23 | 66 | 34 | 15 | 0 | 0 | 0 | 1 | 0 | 67 | 71 | 40 | 67 | 122 | 0 | 0 | 0 | 0 | 0 | 225 | 149 | 1 | 1 | 1 | 1 | 1 | 0.957 | 1 | 1 | 1 | 1 | 1 | 1 | -0.009 |
| SE | 1255 | ENSG00000100083 | GGA1 | chr22 | + | 37616921 | 37617048 | 37614189 | 37614274 | 37618447 | 37618546 | 23 | 12 | 39 | 16 | 6 | 0 | 0 | 0 | 1 | 0 | 40 | 30 | 24 | 53 | 49 | 0 | 0 | 0 | 0 | 0 | 276 | 149 | 1 | 1 | 1 | 1 | 1 | 0.896 | 1 | 1 | 1 | 1 | 1 | 1 | -0.021 |
| SE | 1258 | ENSG00000100083 | GGA1 | chr22 | + | 37629997 | 37630170 | 37629461 | 37629526 | 37630902 | 37631099 | 51 | 31 | 59 | 42 | 18 | 0 | 0 | 0 | 0 | 0 | 146 | 78 | 49 | 41 | 163 | 0 | 0 | 0 | 2 | 0 | 298 | 149 | 1 | 1 | 1 | 1 | 1 | 1 | 1 | 1 | 1 | 1 | 0.911 | 1 | 0.018 |
| SE | 1259 | ENSG00000100083 | GGA1 | chr22 | + | 37632404 | 37632515 | 37631995 | 37632165 | 37632600 | 37632677 | 29 | 20 | 18 | 20 | 21 | 0 | 0 | 0 | 2 | 0 | 40 | 42 | 26 | 37 | 36 | 0 | 0 | 0 | 0 | 0 | 260 | 149 | 1 | 1 | 1 | 1 | 1 | 0.851 | 1 | 1 | 1 | 1 | 1 | 1 | -0.03 |
| SE | 1265 | ENSG00000100095 | SEZ6L | chr22 | + | 26298983 | 26299169 | 26296887 | 26297080 | 26305978 | 26306144 | 0 | 0 | 0 | 10 | 0 | 0 | 0 | 1 | 0 | 0 | 0 | 0 | 0 | 0 | 5 | 0 | 0 | 0 | 0 | 0 | 298 | 149 | 1 | 1 | NA | NA | 0 | 1 | NA | NA | NA | NA | NA | 1 | -0.5 |
| SE | 1266 | ENSG00000100095 | SEZ6L | chr22 | + | 26351051 | 26351243 | 26347718 | 26347913 | 26365371 | 26365566 | 3 | 2 | 3 | 3 | 8 | 0 | 0 | 1 | 0 | 0 | 6 | 0 | 7 | 0 | 0 | 0 | 0 | 0 | 0 | 0 | 298 | 149 | 1 | 1 | 1 | 1 | 0.6 | 1 | 1 | 1 | NA | 1 | NA | NA | -0.08 |
| SE | 1267 | ENSG00000100095 | SEZ6L | chr22 | + | 26365371 | 26365566 | 26351051 | 26351243 | 26375574 | 26375689 | 4 | 2 | 0 | 2 | 11 | 0 | 0 | 1 | 0 | 2 | 0 | 0 | 0 | 3 | 0 | 0 | 0 | 0 | 0 | 0 | 298 | 149 | 1 | 1 | 1 | 1 | 0 | 1 | 0.733 | NA | NA | NA | 1 | NA | -0.253 |
| SE | 1269 | ENSG00000100095 | SEZ6L | chr22 | + | 26373450 | 26373483 | 26365371 | 26365566 | 26375574 | 26375689 | 12 | 3 | 0 | 2 | 7 | 1 | 1 | 0 | 1 | 9 | 4 | 0 | 4 | 8 | 9 | 0 | 0 | 0 | 3 | 0 | 182 | 149 | 1 | 1 | 0.908 | 0.711 | NA | 0.621 | 0.389 | NA | NA | 1 | 0.686 | 1 | -0.238 |
| SE | 1270 | ENSG00000100095 | SEZ6L | chr22 | + | 26373450 | 26373483 | 26365371 | 26365566 | 26375577 | 26375689 | 0 | 1 | 0 | 2 | 1 | 1 | 0 | 0 | 2 | 0 | 0 | 0 | 2 | 0 | 0 | 0 | 0 | 0 | 0 | 0 | 182 | 149 | 1 | 1 | 0 | 1 | NA | 0.45 | 1 | NA | NA | 1 | NA | NA | -0.387 |
| SE | 1272 | ENSG00000186951 | PPARA | chr22 | + | 46176752 | 46176836 | 46151883 | 46151970 | 46188722 | 46188822 | 11 | 2 | 5 | 4 | 0 | 2 | 0 | 0 | 0 | 0 | 4 | 4 | 8 | 6 | 0 | 0 | 0 | 0 | 0 | 0 | 233 | 149 | 1 | 1 | 0.779 | 1 | 1 | 1 | NA | 1 | 1 | 1 | 1 | NA | -0.055 |
| SE | 1273 | ENSG00000186951 | PPARA | chr22 | + | 46176752 | 46176836 | 46151883 | 46151970 | 46198341 | 46198410 | 8 | 2 | 5 | 3 | 0 | 0 | 0 | 0 | 1 | 0 | 0 | 7 | 16 | 1 | 0 | 0 | 0 | 2 | 1 | 0 | 233 | 149 | 1 | 1 | 1 | 1 | 1 | 0.657 | NA | NA | 1 | 0.836 | 0.39 | NA | 0.172 |
| SE | 1274 | ENSG00000186951 | PPARA | chr22 | + | 46188722 | 46188822 | 46151887 | 46151970 | 46198341 | 46198591 | 2 | 3 | 3 | 0 | 0 | 0 | 0 | 0 | 1 | 0 | 1 | 0 | 0 | 0 | 0 | 0 | 0 | 2 | 1 | 0 | 249 | 149 | 1 | 1 | 1 | 1 | 1 | 0 | NA | 1 | NA | 0 | 0 | NA | 0.417 |
| SE | 1275 | ENSG00000186951 | PPARA | chr22 | + | 46188722 | 46188822 | 46151887 | 46151970 | 46198344 | 46198410 | 6 | 1 | 2 | 2 | 1 | 1 | 0 | 0 | 0 | 0 | 23 | 17 | 3 | 4 | 0 | 0 | 0 | 4 | 0 | 0 | 249 | 149 | 1 | 1 | 0.782 | 1 | 1 | 1 | 1 | 1 | 1 | 0.31 | 1 | NA | 0.129 |
| SE | 1281 | ENSG00000186951 | PPARA | chr22 | + | 46231791 | 46232239 | 46219811 | 46220014 | 46235132 | 46235403 | 41 | 26 | 36 | 47 | 37 | 1 | 0 | 0 | 0 | 0 | 33 | 32 | 15 | 37 | 0 | 0 | 0 | 0 | 0 | 0 | 298 | 149 | 1 | 1 | 0.953 | 1 | 1 | 1 | 1 | 1 | 1 | 1 | 1 | NA | -0.009 |
| SE | 1282 | ENSG00000188511 | C22orf34 | chr22 | - | 49416601 | 49416735 | 49414523 | 49416035 | 49416814 | 49416928 | 8 | 5 | 1 | 4 | 4 | 0 | 6 | 3 | 1 | 2 | 16 | 33 | 10 | 0 | 0 | 4 | 7 | 2 | 0 | 0 | 283 | 149 | 1 | 1 | 1 | 0.305 | 0.149 | 0.678 | 0.513 | 0.678 | 0.713 | 0.725 | NA | NA | -0.176 |
| SE | 1283 | ENSG00000188511 | C22orf34 | chr22 | - | 49440875 | 49441212 | 49416814 | 49416928 | 49657404 | 49657457 | 9 | 0 | 1 | 4 | 1 | 4 | 5 | 4 | 0 | 2 | 0 | 1 | 4 | 0 | 0 | 0 | 18 | 1 | 0 | 0 | 298 | 149 | 1 | 1 | 0.529 | 0 | 0.111 | 1 | 0.2 | NA | 0.027 | 0.667 | NA | NA | 0.021 |
| SE | 1284 | ENSG00000188511 | C22orf34 | chr22 | - | 49624446 | 49624681 | 49624166 | 49624253 | 49624857 | 49624936 | 1 | 6 | 0 | 1 | 3 | 2 | 0 | 0 | 0 | 0 | 0 | 0 | 0 | 0 | 2 | 0 | 0 | 0 | 0 | 0 | 298 | 149 | 1 | 1 | 0.2 | 1 | NA | 1 | 1 | NA | NA | NA | NA | 1 | -0.2 |
| SE | 1285 | ENSG00000188511 | C22orf34 | chr22 | - | 49624857 | 49624936 | 49624369 | 49624681 | 49657404 | 49657542 | 1 | 3 | 0 | 0 | 0 | 0 | 0 | 0 | 0 | 0 | 0 | 0 | 0 | 0 | 0 | 0 | 0 | 0 | 0 | 2 | 228 | 149 | 1 | 1 | 1 | 1 | NA | NA | NA | NA | NA | NA | NA | 0 | 1 |
| SE | 1286 | ENSG00000100362 | PVALB | chr22 | - | 36811474 | 36811524 | 36800700 | 36800918 | 36813645 | 36813755 | 2 | 1 | 0 | 1 | 1 | 18 | 7 | 9 | 12 | 69 | 5 | 0 | 0 | 0 | 10 | 143 | 30 | 5 | 4 | 362 | 199 | 149 | 1 | 1 | 0.077 | 0.097 | 0 | 0.059 | 0.011 | 0.026 | 0 | 0 | 0 | 0.02 | 0.04 |
| SE | 1287 | ENSG00000197182 | MIRLET7BHG | chr22 | + | 46097925 | 46098010 | 46053868 | 46053920 | 46098493 | 46098558 | 2 | 5 | 4 | 5 | 2 | 0 | 2 | 0 | 0 | 0 | 0 | 2 | 0 | 6 | 0 | 0 | 8 | 0 | 0 | 0 | 234 | 149 | 1 | 1 | 1 | 0.614 | 1 | 1 | 1 | NA | 0.137 | NA | 1 | NA | 0.354 |
| SE | 1294 | ENSG00000100239 | PPP6R2 | chr22 | + | 50418866 | 50418979 | 50416091 | 50416157 | 50419345 | 50419462 | 49 | 37 | 47 | 40 | 38 | 0 | 0 | 0 | 1 | 0 | 55 | 24 | 43 | 47 | 133 | 0 | 0 | 0 | 0 | 0 | 262 | 149 | 1 | 1 | 1 | 1 | 1 | 0.958 | 1 | 1 | 1 | 1 | 1 | 1 | -0.008 |
| SE | 1297 | ENSG00000100239 | PPP6R2 | chr22 | + | 50438579 | 50438762 | 50438173 | 50438298 | 50439700 | 50439857 | 158 | 86 | 171 | 167 | 88 | 0 | 0 | 0 | 0 | 0 | 160 | 250 | 128 | 127 | 283 | 0 | 0 | 0 | 2 | 0 | 298 | 149 | 1 | 1 | 1 | 1 | 1 | 1 | 1 | 1 | 1 | 1 | 0.969 | 1 | 0.006 |
| SE | 1298 | ENSG00000100239 | PPP6R2 | chr22 | + | 50440800 | 50441026 | 50439960 | 50440049 | 50443865 | 50444117 | 34 | 17 | 33 | 32 | 15 | 0 | 0 | 0 | 0 | 0 | 16 | 18 | 6 | 20 | 10 | 0 | 0 | 0 | 1 | 0 | 298 | 149 | 1 | 1 | 1 | 1 | 1 | 1 | 1 | 1 | 1 | 1 | 0.909 | 1 | 0.018 |
| SE | 1299 | ENSG00000100239 | PPP6R2 | chr22 | + | 50440821 | 50441026 | 50439960 | 50440049 | 50443865 | 50444117 | 86 | 43 | 88 | 100 | 41 | 0 | 0 | 0 | 0 | 0 | 67 | 105 | 36 | 74 | 79 | 0 | 0 | 0 | 1 | 0 | 298 | 149 | 1 | 1 | 1 | 1 | 1 | 1 | 1 | 1 | 1 | 1 | 0.974 | 1 | 0.005 |
| SE | 1302 | ENSG00000138942 | RNF185 | chr22 | + | 31192683 | 31192702 | 31160182 | 31160303 | 31195468 | 31195581 | 4 | 10 | 1 | 3 | 2 | 2 | 0 | 0 | 0 | 0 | 19 | 12 | 2 | 6 | 0 | 0 | 0 | 0 | 0 | 0 | 168 | 149 | 1 | 1 | 0.639 | 1 | 1 | 1 | 1 | 1 | 1 | 1 | 1 | NA | -0.072 |
| SE | 1305 | ENSG00000138942 | RNF185 | chr22 | + | 31196935 | 31196990 | 31195468 | 31195581 | 31201497 | 31201615 | 34 | 26 | 28 | 24 | 11 | 2 | 0 | 0 | 0 | 0 | 24 | 44 | 26 | 44 | 24 | 3 | 0 | 0 | 0 | 0 | 204 | 149 | 1 | 1 | 0.925 | 1 | 1 | 1 | 1 | 0.854 | 1 | 1 | 1 | 1 | 0.014 |
| SE | 1306 | ENSG00000138942 | RNF185 | chr22 | + | 31196935 | 31197157 | 31192683 | 31192702 | 31201497 | 31201615 | 0 | 0 | 1 | 2 | 0 | 1 | 0 | 0 | 2 | 0 | 0 | 0 | 0 | 3 | 0 | 0 | 0 | 0 | 0 | 15 | 298 | 149 | 1 | 1 | 0 | NA | 1 | 0.333 | NA | NA | NA | NA | 1 | 0 | -0.056 |
| SE | 1310 | ENSG00000100376 | FAM118A | chr22 | + | 45321601 | 45321793 | 45318479 | 45318640 | 45322041 | 45322215 | 0 | 0 | 2 | 0 | 0 | 0 | 1 | 0 | 1 | 0 | 0 | 0 | 0 | 10 | 0 | 0 | 0 | 0 | 0 | 0 | 298 | 149 | 1 | 1 | NA | 0 | 1 | 0 | NA | NA | NA | NA | 1 | NA | -0.667 |
| SE | 1311 | ENSG00000100376 | FAM118A | chr22 | + | 45321601 | 45321793 | 45318479 | 45318640 | 45322370 | 45322383 | 0 | 0 | 2 | 3 | 4 | 4 | 0 | 7 | 5 | 2 | 0 | 0 | 0 | 5 | 0 | 0 | 5 | 3 | 3 | 0 | 298 | 149 | 1 | 1 | 0 | NA | 0.125 | 0.231 | 0.5 | NA | 0 | 0 | 0.455 | NA | 0.062 |
| SE | 1312 | ENSG00000100376 | FAM118A | chr22 | + | 45321601 | 45321793 | 45318790 | 45318960 | 45322370 | 45322383 | 0 | 0 | 0 | 3 | 4 | 4 | 1 | 1 | 3 | 0 | 0 | 0 | 0 | 5 | 0 | 0 | 0 | 0 | 0 | 0 | 298 | 149 | 1 | 1 | 0 | 0 | 0 | 0.333 | 1 | NA | NA | NA | 1 | NA | -0.733 |
| SE | 1313 | ENSG00000100376 | FAM118A | chr22 | + | 45322041 | 45322215 | 45318479 | 45318640 | 45322370 | 45322383 | 0 | 1 | 2 | 5 | 2 | 4 | 0 | 7 | 5 | 2 | 0 | 0 | 1 | 5 | 0 | 0 | 5 | 3 | 3 | 0 | 298 | 149 | 1 | 1 | 0 | 1 | 0.125 | 0.333 | 0.333 | NA | 0 | 0.143 | 0.455 | NA | 0.159 |
| SE | 1314 | ENSG00000100376 | FAM118A | chr22 | + | 45322041 | 45322215 | 45321601 | 45321793 | 45322370 | 45322383 | 0 | 0 | 2 | 4 | 2 | 0 | 0 | 0 | 3 | 4 | 0 | 0 | 1 | 15 | 0 | 0 | 0 | 0 | 5 | 0 | 298 | 149 | 1 | 1 | NA | NA | 1 | 0.4 | 0.2 | NA | NA | | 0.6 | NA | -0.267 |
| SE | 1317 | ENSG00000100376 | FAM118A | chr22 | + | 45330602 | 45330731 | 45323174 | 45323427 | 45335349 | 45335382 | 0 | 0 | 0 | 0 | 5 | 0 | 0 | 1 | 0 | 0 | 0 | 11 | 3 | 0 | 0 | 12 | 0 | 0 | 0 | 3 | 278 | 149 | 1 | 1 | NA | NA | 0 | NA | 1 | 0 | 1 | 1 | NA | 0 | 0 |
| SE | 1318 | ENSG00000100376 | FAM118A | chr22 | + | 45330602 | 45330731 | 45323174 | 45323427 | 45340385 | 45341955 | 0 | 0 | 1 | 0 | 5 | 0 | 0 | 0 | 0 | 0 | 0 | 11 | 0 | 0 | 0 | 0 | 1 | 0 | 0 | 0 | 278 | 149 | 1 | 1 | NA | NA | 1 | NA | 1 | NA | 0.855 | NA | NA | NA | 0.145 |
| SE | 1320 | ENSG00000100376 | FAM118A | chr22 | + | 45332424 | 45332710 | 45330456 | 45330731 | 45335349 | 45335382 | 184 | 21 | 52 | 350 | 101 | 0 | 0 | 0 | 0 | 0 | 90 | 59 | 9 | 342 | 39 | 0 | 0 | 3 | 0 | 0 | 298 | 149 | 1 | 1 | 1 | 1 | 1 | 1 | 1 | 1 | 1 | 0.6 | 1 | 1 | 0.08 |
| SE | 1322 | ENSG00000100376 | FAM118A | chr22 | + | 45335349 | 45335382 | 45330456 | 45330731 | 45340385 | 45340761 | 0 | 0 | 0 | 0 | 0 | 0 | 0 | 1 | 0 | 0 | 0 | 0 | 3 | 0 | 0 | 0 | 0 | 0 | 0 | 0 | 182 | 149 | 1 | 1 | NA | NA | 0 | NA | NA | NA | NA | 1 | NA | NA | -1 |
| SE | 1323 | ENSG00000100376 | FAM118A | chr22 | + | 45335349 | 45335382 | 45332424 | 45332710 | 45336327 | 45336410 | 119 | 16 | 30 | 206 | 93 | 1 | 0 | 0 | 0 | 0 | 54 | 53 | 10 | 271 | 34 | 0 | 0 | 0 | 0 | 0 | 182 | 149 | 1 | 1 | 0.99 | 1 | 1 | 1 | 1 | 1 | 1 | 1 | 1 | 1 | -0.002 |
| SE | 1324 | ENSG00000100376 | FAM118A | chr22 | + | 45336327 | 45336411 | 45335215 | 45335382 | 45340385 | 45340761 | 229 | 38 | 86 | 484 | 155 | 0 | 0 | 1 | 0 | 0 | 121 | 119 | 29 | 571 | 94 | 0 | 0 | 0 | 0 | 1 | 233 | 149 | 1 | 1 | 1 | 1 | 0.982 | 1 | 1 | 1 | 1 | 1 | 1 | 0.984 | 0 |
| SE | 1326 | ENSG00000205593 | DENND6B | chr22 | - | 50314399 | 50314494 | 50313813 | 50314272 | 50314604 | 50314700 | 39 | 19 | 33 | 33 | 28 | 0 | 0 | 0 | 0 | 0 | 58 | 49 | 52 | 13 | 110 | 0 | 0 | 1 | 0 | 0 | 244 | 149 | 1 | 1 | 1 | 1 | 1 | 1 | 1 | 1 | 1 | 0.969 | 1 | 1 | 0.006 |
| SE | 1334 | ENSG00000093000 | NUP50 | chr22 | + | 45171808 | 45172037 | 45171599 | 45171683 | 45175893 | 45176080 | 0 | 0 | 2 | 2 | 3 | 252 | 179 | 218 | 216 | 450 | 0 | 4 | 6 | 3 | 0 | 151 | 196 | 123 | 190 | 80 | 298 | 149 | 1 | 1 | 0 | 0 | 0.005 | 0.005 | 0.003 | 0 | 0.01 | 0.024 | 0.008 | 0 | -0.006 |
| SE | 1338 | ENSG00000093000 | NUP50 | chr22 | + | 45178237 | 45178900 | 45171599 | 45171683 | 45183401 | 45183520 | 20 | 22 | 18 | 12 | 26 | 0 | 0 | 0 | 0 | 0 | 24 | 29 | 45 | 33 | 14 | 0 | 3 | 0 | 0 | 0 | 298 | 149 | 1 | 1 | 1 | 1 | 1 | 1 | 1 | 1 | 0.829 | 1 | 1 | 1 | 0.034 |
| SE | 1341 | ENSG00000093000 | NUP50 | chr22 | + | 45181285 | 45181367 | 45171599 | 45171683 | 45183401 | 45183515 | 142 | 144 | 135 | 136 | 320 | 0 | 0 | 0 | 0 | 0 | 64 | 128 | 189 | 127 | 19 | 0 | 3 | 0 | 0 | 0 | 231 | 149 | 1 | 1 | 1 | 1 | 1 | 1 | 1 | 1 | 0.965 | 1 | 1 | 1 | 0.007 |
| SE | 1342 | ENSG00000093000 | NUP50 | chr22 | + | 45181285 | 45181367 | 45171599 | 45171683 | 45184452 | 45184504 | 3 | 4 | 2 | 1 | 0 | 0 | 0 | 0 | 0 | 0 | 0 | 0 | 21 | 4 | 2 | 0 | 0 | 4 | 0 | 0 | 231 | 149 | 1 | 1 | 1 | 1 | 1 | 1 | NA | NA | NA | 0.772 | 1 | 1 | 0.076 |
| SE | 1343 | ENSG00000093000 | NUP50 | chr22 | + | 45181285 | 45181367 | 45177698 | 45178900 | 45183401 | 45183515 | 374 | 295 | 313 | 327 | 616 | 0 | 0 | 0 | 0 | 0 | 125 | 265 | 314 | 264 | 47 | 0 | 0 | 2 | 0 | 0 | 231 | 149 | 1 | 1 | 1 | 1 | 1 | 1 | 1 | 1 | 1 | 0.99 | 1 | 1 | 0.002 |
| SE | 1344 | ENSG00000093000 | NUP50 | chr22 | + | 45183401 | 45183520 | 45171599 | 45171683 | 45184452 | 45184504 | 144 | 109 | 85 | 121 | 219 | 0 | 0 | 0 | 0 | 0 | 53 | 131 | 132 | 129 | 24 | 0 | 0 | 4 | 0 | 0 | 268 | 149 | 1 | 1 | 1 | 1 | 1 | 1 | 1 | 1 | 1 | 0.948 | 1 | 1 | 0.01 |
| SE | 1348 | ENSG00000100150 | DEPDC5 | chr22 | + | 31760655 | 31760702 | 31758526 | 31758633 | 31764974 | 31765060 | 93 | 40 | 77 | 103 | 132 | 5 | 0 | 0 | 0 | 0 | 133 | 126 | 61 | 109 | 140 | 0 | 5 | 0 | 0 | 0 | 196 | 149 | 1 | 1 | 0.934 | 1 | 1 | 1 | 1 | 1 | 0.95 | 1 | 1 | 1 | -0.003 |
| SE | 1353 | ENSG00000100150 | DEPDC5 | chr22 | + | 31783906 | 31783985 | 31778098 | 31778168 | 31784813 | 31784875 | 100 | 76 | 95 | 105 | 103 | 0 | 0 | 0 | 0 | 1 | 212 | 145 | 92 | 173 | 214 | 0 | 0 | 0 | 0 | 0 | 228 | 149 | 1 | 1 | 1 | 1 | 1 | 1 | 0.985 | 1 | 1 | 1 | 1 | 1 | -0.003 |
| SE | 1354 | ENSG00000100150 | DEPDC5 | chr22 | + | 31784813 | 31784875 | 31783906 | 31783985 | 31792032 | 31792102 | 135 | 79 | 124 | 122 | 100 | 2 | 0 | 1 | 0 | 0 | 257 | 154 | 97 | 207 | 311 | 0 | 0 | 0 | 0 | 0 | 211 | 149 | 1 | 1 | 0.979 | 1 | 0.989 | 1 | 1 | 1 | 1 | 1 | 1 | 1 | -0.006 |
| SE | 1355 | ENSG00000100150 | DEPDC5 | chr22 | + | 31784813 | 31785107 | 31783906 | 31783985 | 31792032 | 31792102 | 30 | 37 | 47 | 43 | 63 | 2 | 0 | 1 | 0 | 0 | 88 | 52 | 31 | 57 | 93 | 0 | 0 | 0 | 0 | 0 | 298 | 149 | 1 | 1 | 0.882 | 1 | 0.959 | 1 | 1 | 1 | 1 | 1 | 1 | 1 | -0.032 |
| SE | 1356 | ENSG00000100150 | DEPDC5 | chr22 | + | 31792032 | 31792102 | 31784813 | 31784875 | 31792744 | 31792768 | 125 | 64 | 96 | 103 | 96 | 0 | 2 | 0 | 0 | 0 | 223 | 154 | 84 | 179 | 261 | 0 | 1 | 1 | 0 | 3 | 219 | 149 | 1 | 1 | 1 | 0.956 | 1 | 1 | 1 | 1 | 0.991 | 0.983 | 1 | 0.983 | 0 |
| SE | 1358 | ENSG00000100150 | DEPDC5 | chr22 | + | 31797599 | 31797703 | 31792686 | 31792817 | 31798581 | 31798656 | 113 | 83 | 96 | 92 | 127 | 0 | 2 | 0 | 4 | 0 | 171 | 151 | 65 | 93 | 144 | 0 | 0 | 0 | 0 | 3 | 253 | 149 | 1 | 1 | 1 | 0.961 | 1 | 0.931 | 1 | 1 | 1 | 1 | 1 | 0.966 | -0.015 |
| SE | 1359 | ENSG00000100150 | DEPDC5 | chr22 | + | 31798581 | 31798656 | 31797599 | 31797703 | 31802703 | 31802738 | 108 | 66 | 84 | 85 | 104 | 0 | 0 | 3 | 0 | 0 | 152 | 145 | 71 | 98 | 133 | 0 | 0 | 0 | 0 | 0 | 224 | 149 | 1 | 1 | 1 | 1 | 0.949 | 1 | 1 | 1 | 1 | 1 | 1 | 1 | -0.01 |
| SE | 1360 | ENSG00000100150 | DEPDC5 | chr22 | + | 31804161 | 31804223 | 31802703 | 31802838 | 31804841 | 31804915 | 59 | 44 | 40 | 60 | 94 | 0 | 0 | 0 | 0 | 0 | 23 | 96 | 47 | 64 | 64 | 0 | 3 | 0 | 0 | 0 | 211 | 149 | 1 | 1 | 1 | 1 | 1 | 1 | 1 | 1 | 0.958 | 1 | 1 | 1 | 0.008 |
| SE | 1361 | ENSG00000100150 | DEPDC5 | chr22 | + | 31804841 | 31804915 | 31804161 | 31804223 | 31806121 | 31806191 | 66 | 59 | 57 | 69 | 107 | 0 | 0 | 0 | 1 | 0 | 68 | 95 | 68 | 107 | 89 | 0 | 0 | 0 | 0 | 0 | 223 | 149 | 1 | 1 | 1 | 1 | 1 | 0.979 | 1 | 1 | 1 | 1 | 1 | 1 | -0.004 |
| SE | 1362 | ENSG00000100150 | DEPDC5 | chr22 | + | 31806121 | 31806191 | 31804841 | 31804915 | 31809610 | 31809647 | 63 | 58 | 66 | 69 | 134 | 0 | 0 | 1 | 0 | 0 | 86 | 109 | 73 | 129 | 98 | 0 | 0 | 0 | 0 | 0 | 219 | 149 | 1 | 1 | 1 | 1 | 0.978 | 1 | 1 | 1 | 1 | 1 | 1 | 1 | -0.004 |
| SE | 1363 | ENSG00000100150 | DEPDC5 | chr22 | + | 31809610 | 31809647 | 31806121 | 31806191 | 31810520 | 31810641 | 55 | 42 | 67 | 54 | 140 | 2 | 1 | 1 | 0 | 0 | 90 | 79 | 55 | 117 | 84 | 0 | 0 | 0 | 2 | 0 | 186 | 149 | 1 | 1 | 0.957 | 0.971 | 0.982 | 1 | 1 | 1 | 1 | 1 | 0.979 | 1 | -0.014 |
| SE | 1364 | ENSG00000100150 | DEPDC5 | chr22 | + | 31814991 | 31815212 | 31810520 | 31810641 | 31819021 | 31819040 | 73 | 56 | 77 | 73 | 162 | 0 | 0 | 0 | 2 | 0 | 164 | 92 | 69 | 118 | 159 | 0 | 0 | 0 | 0 | 0 | 298 | 149 | 1 | 1 | 1 | 1 | 1 | 0.948 | 1 | 1 | 1 | 1 | 1 | 1 | -0.01 |
| SE | 1366 | ENSG00000100150 | DEPDC5 | chr22 | + | 31822692 | 31822790 | 31821501 | 31821637 | 31833914 | 31833980 | 38 | 37 | 42 | 36 | 28 | 0 | 0 | 0 | 0 | 0 | 220 | 112 | 57 | 83 | 144 | 0 | 0 | 1 | 3 | 0 | 247 | 149 | 1 | 1 | 1 | 1 | 1 | 1 | 1 | 1 | 1 | 0.972 | 0.943 | 1 | 0.017 |
| SE | 1367 | ENSG00000100150 | DEPDC5 | chr22 | + | 31831063 | 31831225 | 31819021 | 31819225 | 31833914 | 31833980 | 2 | 0 | 0 | 1 | 1 | 13 | 4 | 8 | 6 | 4 | 0 | 1 | 0 | 8 | 0 | 16 | 1 | 4 | 12 | 10 | 298 | 149 | 1 | 1 | 0.071 | 0 | 0 | 0.077 | 0.111 | 0 | 0.333 | 0 | 0.25 | 0 | -0.065 |
| SE | 1368 | ENSG00000100150 | DEPDC5 | chr22 | + | 31833914 | 31833980 | 31819021 | 31819225 | 31836971 | 31837155 | 13 | 5 | 14 | 11 | 1 | 0 | 0 | 0 | 0 | 1 | 0 | 5 | 5 | 15 | 2 | 0 | 0 | 0 | 0 | 0 | 215 | 149 | 1 | 1 | 1 | 1 | 1 | 1 | 0.409 | NA | 1 | 1 | 1 | 1 | -0.118 |
| SE | 1369 | ENSG00000100150 | DEPDC5 | chr22 | + | 31833914 | 31833980 | 31822692 | 31822790 | 31836971 | 31837155 | 17 | 6 | 16 | 15 | 11 | 0 | 0 | 0 | 1 | 0 | 53 | 40 | 26 | 28 | 6 | 0 | 0 | 0 | 0 | 0 | 215 | 149 | 1 | 1 | 1 | 1 | 1 | 0.912 | 1 | 1 | 1 | 1 | 1 | 1 | -0.018 |
| SE | 1370 | ENSG00000100150 | DEPDC5 | chr22 | + | 31833914 | 31833980 | 31822692 | 31822790 | 31836998 | 31837155 | 23 | 17 | 16 | 27 | 23 | 1 | 0 | 0 | 1 | 0 | 98 | 54 | 37 | 17 | 50 | 0 | 1 | 0 | 0 | 0 | 215 | 149 | 1 | 1 | 0.941 | 1 | 1 | 1 | 1 | 1 | 0.974 | 1 | 1 | 1 | -0.007 |
| SE | 1371 | ENSG00000100150 | DEPDC5 | chr22 | + | 31833914 | 31833980 | 31822692 | 31822790 | 31838684 | 31838845 | 12 | 2 | 5 | 6 | 8 | 0 | 0 | 0 | 0 | 0 | 53 | 35 | 19 | 10 | 6 | 0 | 3 | 0 | 0 | 0 | 215 | 149 | 1 | 1 | 1 | 1 | 1 | 1 | 1 | 1 | 0.89 | 1 | 1 | 1 | 0.022 |
| SE | 1372 | ENSG00000100150 | DEPDC5 | chr22 | + | 31836971 | 31837155 | 31822692 | 31822790 | 31838684 | 31838845 | 29 | 16 | 18 | 24 | 23 | 0 | 0 | 0 | 0 | 0 | 45 | 49 | 9 | 24 | 26 | 0 | 3 | 0 | 0 | 0 | 298 | 149 | 1 | 1 | 1 | 1 | 1 | 1 | 1 | 1 | 0.891 | 1 | 1 | 1 | 0.022 |
| SE | 1373 | ENSG00000100150 | DEPDC5 | chr22 | + | 31836971 | 31837155 | 31833914 | 31833980 | 31838684 | 31838845 | 37 | 20 | 33 | 34 | 27 | 1 | 0 | 0 | 1 | 0 | 45 | 54 | 17 | 47 | 26 | 0 | 0 | 0 | 0 | 0 | 298 | 149 | 1 | 1 | 0.949 | 1 | 1 | 0.944 | 1 | 1 | 1 | 1 | 1 | 1 | -0.021 |
| SE | 1374 | ENSG00000100150 | DEPDC5 | chr22 | + | 31836971 | 31837155 | 31833914 | 31833980 | 31843094 | 31843212 | 8 | 4 | 20 | 12 | 4 | 0 | 0 | 2 | 0 | 0 | 0 | 5 | 8 | 23 | 0 | 0 | 0 | 0 | 0 | 0 | 298 | 149 | 1 | 1 | 1 | 1 | 0.833 | 1 | 1 | NA | 1 | 1 | 1 | NA | -0.033 |
| SE | 1375 | ENSG00000100150 | DEPDC5 | chr22 | + | 31836998 | 31837155 | 31833865 | 31833980 | 31838684 | 31838845 | 46 | 32 | 31 | 47 | 42 | 1 | 0 | 0 | 1 | 0 | 111 | 70 | 29 | 37 | 78 | 0 | 0 | 0 | 0 | 0 | 298 | 149 | 1 | 1 | 0.958 | 1 | 1 | 0.959 | 1 | 1 | 1 | 1 | 1 | 1 | -0.017 |
| SE | 1376 | ENSG00000100150 | DEPDC5 | chr22 | + | 31836998 | 31837155 | 31833865 | 31833980 | 31843094 | 31843212 | 17 | 16 | 18 | 25 | 19 | 0 | 0 | 2 | 0 | 0 | 66 | 21 | 20 | 13 | 52 | 0 | 0 | 0 | 0 | 0 | 298 | 149 | 1 | 1 | 1 | 1 | 0.818 | 1 | 1 | 1 | 1 | 1 | 1 | 1 | -0.036 |
| SE | 1377 | ENSG00000100150 | DEPDC5 | chr22 | + | 31837093 | 31837155 | 31833848 | 31833980 | 31838684 | 31838721 | 21 | 7 | 11 | 13 | 19 | 1 | 0 | 0 | 1 | 0 | 40 | 20 | 4 | 11 | 6 | 0 | 0 | 0 | 0 | 0 | 211 | 149 | 1 | 1 | 0.937 | 1 | 1 | 0.902 | 1 | 1 | 1 | 1 | 1 | 1 | -0.032 |
| SE | 1378 | ENSG00000100150 | DEPDC5 | chr22 | + | 31837093 | 31837155 | 31833848 | 31833980 | 31843094 | 31843212 | 0 | 0 | 5 | 4 | 0 | 0 | 0 | 2 | 0 | 0 | 0 | 0 | 0 | 2 | 0 | 0 | 0 | 0 | 0 | 0 | 211 | 149 | 1 | 1 | NA | NA | 0.638 | 1 | NA | NA | NA | NA | 1 | NA | -0.181 |
| SE | 1381 | ENSG00000100150 | DEPDC5 | chr22 | + | 31838684 | 31838845 | 31833865 | 31833980 | 31843094 | 31843212 | 7 | 6 | 11 | 16 | 9 | 0 | 0 | 2 | 0 | 0 | 35 | 11 | 9 | 18 | 24 | 0 | 0 | 0 | 0 | 0 | 298 | 149 | 1 | 1 | 1 | 1 | 0.733 | 1 | 1 | 1 | 1 | 1 | 1 | 1 | -0.053 |
| SE | 1384 | ENSG00000100150 | DEPDC5 | chr22 | + | 31857444 | 31857553 | 31846833 | 31846967 | 31870589 | 31870744 | 40 | 22 | 41 | 31 | 22 | 2 | 0 | 0 | 1 | 0 | 25 | 32 | 32 | 28 | 81 | 0 | 0 | 0 | 0 | 0 | 258 | 149 | 1 | 1 | 0.92 | 1 | 1 | 0.947 | 1 | 1 | 1 | 1 | 1 | 1 | -0.027 |
| SE | 1386 | ENSG00000100150 | DEPDC5 | chr22 | + | 31876086 | 31876265 | 31874272 | 31874405 | 31879524 | 31879752 | 1 | 8 | 9 | 7 | 5 | 0 | 0 | 0 | 1 | 0 | 14 | 11 | 3 | 10 | 41 | 0 | 0 | 1 | 0 | 0 | 298 | 149 | 1 | 1 | 1 | 1 | 1 | 0.778 | 1 | 1 | 1 | 0.6 | 1 | 1 | 0.036 |
| SE | 1387 | ENSG00000100150 | DEPDC5 | chr22 | + | 31876156 | 31876265 | 31873826 | 31874405 | 31879524 | 31879752 | 15 | 15 | 28 | 14 | 8 | 0 | 0 | 0 | 1 | 0 | 19 | 23 | 16 | 33 | 71 | 0 | 0 | 1 | 0 | 0 | 258 | 149 | 1 | 1 | 1 | 1 | 1 | 0.89 | 1 | 1 | 1 | 0.902 | 1 | 1 | -0.002 |
| SE | 1389 | ENSG00000100150 | DEPDC5 | chr22 | + | 31895358 | 31895954 | 31893581 | 31893751 | 31901741 | 31901802 | 2 | 0 | 3 | 1 | 0 | 2 | 0 | 0 | 1 | 0 | 0 | 14 | 14 | 6 | 27 | 0 | 3 | 0 | 0 | 0 | 298 | 149 | 1 | 1 | 0.333 | NA | 1 | 0.333 | NA | NA | 0.7 | 1 | 1 | 1 | -0.37 |
| SE | 1395 | ENSG00000183172 | SMDT1 | chr22 | + | 42081924 | 42082065 | 42079690 | 42079905 | 42083118 | 42083353 | 277 | 113 | 252 | 201 | 474 | 0 | 0 | 2 | 0 | 0 | 141 | 209 | 141 | 84 | 250 | 0 | 5 | 0 | 0 | 0 | 290 | 149 | 1 | 1 | 1 | 1 | 0.985 | 1 | 1 | 1 | 0.956 | 1 | 1 | 1 | 0.006 |
| SE | 1396 | ENSG00000183172 | SMDT1 | chr22 | + | 42081924 | 42082065 | 42079699 | 42079954 | 42083118 | 42083353 | 340 | 145 | 345 | 307 | 521 | 0 | 0 | 2 | 0 | 0 | 169 | 291 | 158 | 103 | 386 | 0 | 7 | 0 | 0 | 0 | 290 | 149 | 1 | 1 | 1 | 1 | 1 | 1 | 1 | 1 | 0.955 | 1 | 1 | 1 | 0.009 |
| SE | 1399 | ENSG00000100106 | TRIOBP | chr22 | + | 37713209 | 37713411 | 37710426 | 37710566 | 37715762 | 37715934 | 5 | 1 | 2 | 0 | 0 | 0 | 0 | 2 | 0 | 0 | 11 | 5 | 0 | 7 | 0 | 0 | 5 | 0 | 0 | 0 | 298 | 149 | 1 | 1 | 1 | 1 | 0.333 | NA | NA | 1 | 0.333 | NA | 1 | NA | 0 |
| SE | 1400 | ENSG00000100106 | TRIOBP | chr22 | + | 37715762 | 37715934 | 37713209 | 37713411 | 37723184 | 37726503 | 4 | 3 | 6 | 1 | 0 | 0 | 0 | 2 | 0 | 0 | 11 | 0 | 0 | 3 | 0 | 0 | 0 | 0 | 0 | 9 | 298 | 149 | 1 | 1 | 1 | 1 | 1 | 1 | NA | 1 | NA | NA | 1 | 0 | 0.333 |
| SE | 1402 | ENSG00000100106 | TRIOBP | chr22 | + | 37750641 | 37750730 | 37746321 | 37746420 | 37751771 | 37751828 | 0 | 0 | 0 | 0 | 0 | 26 | 8 | 13 | 18 | 4 | 0 | 0 | 0 | 1 | 0 | 53 | 24 | 7 | 40 | 32 | 238 | 149 | 1 | 1 | 0 | 0 | 0 | 0 | 0 | 0 | 0 | 0 | 0.015 | 0 | -0.003 |
| SE | 1404 | ENSG00000100106 | TRIOBP | chr22 | + | 37769027 | 37769187 | 37768073 | 37768176 | 37769261 | 37769375 | 153 | 58 | 87 | 143 | 115 | 0 | 0 | 0 | 0 | 4 | 154 | 191 | 87 | 288 | 302 | 0 | 0 | 0 | 0 | 0 | 298 | 149 | 1 | 1 | 1 | 1 | 1 | 1 | 0.935 | 1 | 1 | 1 | 1 | 1 | -0.013 |
| SE | 1405 | ENSG00000100106 | TRIOBP | chr22 | + | 37769261 | 37769375 | 37768073 | 37768176 | 37771649 | 37771736 | 21 | 15 | 17 | 25 | 24 | 0 | 0 | 0 | 0 | 0 | 30 | 57 | 22 | 70 | 94 | 0 | 0 | 1 | 0 | 0 | 263 | 149 | 1 | 1 | 1 | 1 | 1 | 1 | 1 | 1 | 1 | 1 | 0.975 | 1 | 0.005 |
| SE | 1406 | ENSG00000100372 | SLC25A17 | chr22 | - | 40773936 | 40774019 | 40769629 | 40770981 | 40777039 | 40777135 | 49 | 19 | 38 | 36 | 100 | 0 | 2 | 0 | 0 | 0 | 70 | 40 | 16 | 15 | 75 | 0 | 0 | 0 | 0 | 0 | 232 | 149 | 1 | 1 | 1 | 0.859 | 1 | 1 | 1 | 1 | 1 | 1 | 1 | 1 | -0.028 |
| SE | 1408 | ENSG00000100372 | SLC25A17 | chr22 | - | 40779757 | 40779894 | 40779008 | 40779125 | 40792524 | 40792676 | 1 | 5 | 1 | 5 | 0 | 14 | 6 | 20 | 21 | 37 | 0 | 12 | 0 | 0 | 14 | 4 | 0 | 4 | 0 | 0 | 286 | 149 | 1 | 1 | 0.036 | 0.303 | 0.025 | 0.11 | 0 | NA | 0.61 | NA | 0 | 1 | -0.442 |
| SE | 1409 | ENSG00000100372 | SLC25A17 | chr22 | - | 40779757 | 40779894 | 40779008 | 40779125 | 40794513 | 40794580 | 2 | 2 | 0 | 3 | 0 | 0 | 1 | 0 | 0 | 0 | 0 | 4 | 0 | 0 | 13 | 4 | 0 | 0 | 0 | 0 | 286 | 149 | 1 | 1 | 1 | 0.51 | NA | 1 | NA | 0 | 1 | NA | NA | 1 | 0.17 |
| SE | 1411 | ENSG00000100372 | SLC25A17 | chr22 | - | 40792524 | 40792676 | 40779008 | 40779125 | 40794513 | 40794580 | 28 | 20 | 34 | 43 | 73 | 0 | 1 | 0 | 0 | 0 | 10 | 12 | 0 | 10 | 11 | 4 | 0 | 0 | 0 | 0 | 298 | 149 | 1 | 1 | 1 | 0.909 | 1 | 1 | 1 | 0.556 | 1 | NA | 1 | 1 | 0.093 |
| SE | 1413 | ENSG00000100372 | SLC25A17 | chr22 | - | 40792524 | 40792676 | 40779008 | 40779125 | 40819194 | 40819350 | 14 | 6 | 20 | 21 | 37 | 0 | 1 | 1 | 0 | 0 | 0 | 4 | 0 | 4 | 0 | 0 | 0 | 0 | 0 | 4 | 298 | 149 | 1 | 1 | 1 | 0.75 | 0.909 | 1 | 1 | NA | 1 | NA | 1 | 0 | 0.265 |
| SE | 1414 | ENSG00000100372 | SLC25A17 | chr22 | - | 40792524 | 40792676 | 40779757 | 40779894 | 40794513 | 40794580 | 14 | 17 | 15 | 24 | 36 | 1 | 0 | 0 | 0 | 0 | 10 | 16 | 0 | 6 | 12 | 0 | 0 | 0 | 0 | 0 | 298 | 149 | 1 | 1 | 0.875 | 1 | 1 | 1 | 1 | 1 | 1 | NA | 1 | 1 | -0.025 |
| SE | 1415 | ENSG00000100372 | SLC25A17 | chr22 | - | 40794513 | 40794580 | 40779008 | 40779125 | 40797283 | 40797336 | 1 | 1 | 1 | 0 | 0 | 1 | 0 | 0 | 0 | 0 | 4 | 0 | 0 | 0 | 0 | 0 | 0 | 0 | 0 | 0 | 216 | 149 | 1 | 1 | 0.408 | 1 | 1 | NA | NA | 1 | NA | NA | NA | NA | -0.197 |
| SE | 1417 | ENSG00000100372 | SLC25A17 | chr22 | - | 40794513 | 40794580 | 40792524 | 40792676 | 40797253 | 40797336 | 7 | 9 | 7 | 10 | 20 | 0 | 0 | 0 | 3 | 0 | 4 | 8 | 0 | 0 | 11 | 0 | 0 | 0 | 0 | 0 | 216 | 149 | 1 | 1 | 1 | 1 | 1 | 0.697 | 1 | 1 | 1 | NA | NA | 1 | -0.061 |
| SE | 1420 | ENSG00000100372 | SLC25A17 | chr22 | - | 40797253 | 40797336 | 40794513 | 40794580 | 40799022 | 40799083 | 1 | 4 | 0 | 3 | 2 | 13 | 8 | 8 | 14 | 14 | 0 | 0 | 0 | 0 | 2 | 10 | 0 | 1 | 6 | 0 | 232 | 149 | 1 | 1 | 0.047 | 0.243 | 0 | 0.121 | 0.084 | 0 | NA | 0 | 0 | 1 | -0.151 |
| SE | 1422 | ENSG00000100372 | SLC25A17 | chr22 | - | 40797283 | 40797336 | 40794513 | 40794580 | 40799022 | 40799083 | 3 | 4 | 3 | 4 | 4 | 13 | 8 | 8 | 14 | 14 | 0 | 0 | 0 | 0 | 2 | 10 | 0 | 1 | 6 | 0 | 202 | 149 | 1 | 1 | 0.145 | 0.269 | 0.217 | 0.174 | 0.174 | 0 | NA | 0 | 0 | 1 | -0.054 |
| SE | 1424 | ENSG00000100372 | SLC25A17 | chr22 | - | 40799022 | 40799083 | 40779008 | 40779125 | 40819194 | 40819326 | 10 | 4 | 7 | 9 | 3 | 0 | 1 | 1 | 0 | 0 | 0 | 2 | 0 | 0 | 22 | 0 | 0 | 0 | 0 | 4 | 210 | 149 | 1 | 1 | 1 | 0.739 | 0.832 | 1 | 1 | NA | 1 | NA | NA | 0.796 | 0.016 |
| SE | 1427 | ENSG00000100372 | SLC25A17 | chr22 | - | 40817202 | 40817377 | 40799022 | 40799083 | 40819194 | 40819338 | 0 | 0 | 2 | 0 | 0 | 19 | 7 | 9 | 15 | 3 | 0 | 0 | 0 | 0 | 0 | 4 | 2 | 0 | 0 | 24 | 298 | 149 | 1 | 1 | 0 | 0 | 0.1 | 0 | 0 | 0 | 0 | NA | NA | 0 | 0.02 |
| SE | 1429 | ENSG00000100201 | DDX17 | chr22 | - | 38494629 | 38494802 | 38494020 | 38494131 | 38495795 | 38495937 | 1031 | 868 | 1192 | 1295 | 1635 | 0 | 0 | 0 | 0 | 2 | 1198 | 1124 | 808 | 1094 | 867 | 0 | 0 | 0 | 0 | 0 | 298 | 149 | 1 | 1 | 1 | 1 | 1 | 1 | 0.998 | 1 | 1 | 1 | 1 | 1 | 0 |

| | | | | | | | | | | | | | | | | | | | | | | | | | | | | | | | | | | | | | | | | | | | | | | | | |
|---|---|---|---|---|---|---|---|---|---|---|---|---|---|---|---|---|---|---|---|---|---|---|---|---|---|---|---|---|---|---|---|---|---|---|---|---|---|---|---|---|---|---|---|---|---|---|---|---|
| SE | 1430 | ENSG00000100201 | DDX17 | chr22 | - | 38494885 | 38495046 | 38494309 | 38494802 | 38495795 | 38495937 | 3180 | 2192 | 3238 | 3387 | 4023 | 0 | 0 | 0 | 0 | 0 | 2650 | 2488 | 1978 | 2239 | 2534 | 0 | 0 | 0 | 1 | 0 | 298 | 149 | 1 | 1 | 1 | 1 | 1 | 1 | 1 | 1 | 1 | 1 | 1 | 1 | 0.999 | 1 | 0 |
| SE | 1432 | ENSG00000100201 | DDX17 | chr22 | - | 38498084 | 38498150 | 38495795 | 38495937 | 38498439 | 38498573 | 2675 | 2010 | 2969 | 2828 | 3467 | 2 | 0 | 0 | 0 | 0 | 2128 | 2139 | 1788 | 2088 | 2308 | 0 | 0 | 0 | 0 | 0 | 215 | 149 | 1 | 1 | 0.999 | 1 | 1 | 1 | 1 | 1 | 1 | 1 | 1 | 1 | 1 | 1 | 0 |
| SE | 1433 | ENSG00000100201 | DDX17 | chr22 | - | 38498439 | 38498573 | 38498084 | 38498150 | 38499399 | 38499499 | 3275 | 2545 | 3477 | 3425 | 4952 | 0 | 0 | 0 | 0 | 0 | 2857 | 2750 | 2274 | 2823 | 2943 | 0 | 0 | 0 | 4 | 0 | 283 | 149 | 1 | 1 | 1 | 1 | 1 | 1 | 1 | 1 | 1 | 1 | 1 | 1 | 0.997 | 1 | 0.001 |
| SE | 1434 | ENSG00000100201 | DDX17 | chr22 | - | 38501129 | 38501280 | 38499399 | 38499499 | 38505950 | 38506294 | 1728 | 1287 | 1872 | 1942 | 2225 | 0 | 0 | 0 | 2 | 1 | 1311 | 1524 | 1230 | 1380 | 1599 | 0 | 0 | 0 | 0 | 2 | 298 | 149 | 1 | 1 | 1 | 1 | 1 | 0.998 | 0.999 | 1 | 0.999 | 1 | 1 | 1 | 1 | 0.998 | 0 |
| SE | 1436 | ENSG00000075234 | TTC38 | chr22 | + | 46281598 | 46281718 | 46278585 | 46278661 | 46283972 | 46284032 | 232 | 70 | 90 | 117 | 130 | 0 | 0 | 0 | 0 | 0 | 199 | 155 | 98 | 126 | 167 | 0 | 5 | 0 | 0 | 0 | 269 | 149 | 1 | 1 | 1 | 1 | 1 | 1 | 1 | 1 | 1 | 1 | 0.945 | 1 | 1 | 1 | 0.011 |
| SE | 1438 | ENSG00000075234 | TTC38 | chr22 | + | 46285240 | 46285279 | 46283972 | 46284032 | 46287072 | 46287154 | 277 | 29 | 64 | 116 | 118 | 0 | 0 | 1 | 0 | 0 | 95 | 124 | 69 | 55 | 128 | 0 | 0 | 0 | 0 | 0 | 188 | 149 | 1 | 1 | 1 | 1 | 0.981 | 1 | 1 | 1 | 1 | 1 | 1 | 1 | 1 | 1 | -0.004 |
| SE | 1440 | ENSG00000100347 | SAMM50 | chr22 | + | 43968730 | 43968818 | 43964451 | 43964553 | 43972235 | 43972342 | 252 | 158 | 228 | 231 | 343 | 0 | 6 | 0 | 1 | 0 | 180 | 284 | 99 | 143 | 216 | 0 | 0 | 1 | 0 | 0 | 237 | 149 | 1 | 1 | 1 | 0.943 | 1 | 0.993 | 1 | 1 | 1 | 1 | 1 | 0.984 | 1 | 1 | -0.01 |
| SE | 1441 | ENSG00000100347 | SAMM50 | chr22 | + | 43973235 | 43973323 | 43972870 | 43973001 | 43976054 | 43976183 | 120 | 68 | 143 | 137 | 172 | 0 | 0 | 0 | 0 | 0 | 209 | 114 | 66 | 78 | 159 | 0 | 0 | 0 | 2 | 0 | 237 | 149 | 1 | 1 | 1 | 1 | 1 | 1 | 1 | 1 | 1 | 1 | 1 | 1 | 0.961 | 1 | 0.008 |
| SE | 1442 | ENSG00000100347 | SAMM50 | chr22 | + | 43976054 | 43976183 | 43973235 | 43973323 | 43976749 | 43976821 | 161 | 75 | 179 | 173 | 180 | 1 | 0 | 0 | 0 | 2 | 234 | 131 | 112 | 115 | 169 | 0 | 0 | 0 | 0 | 0 | 278 | 149 | 1 | 1 | 0.989 | 1 | 1 | 1 | 0.98 | 1 | 0.98 | 1 | 1 | 1 | 1 | 1 | -0.006 |
| SE | 1444 | ENSG00000186654 | PRR5 | chr22 | + | 44725243 | 44725292 | 44714590 | 44714671 | 44726576 | 44726634 | 40 | 9 | 21 | 31 | 16 | 0 | 0 | 2 | 0 | 0 | 0 | 12 | 18 | 3 | 14 | 0 | 1 | 3 | 0 | 0 | 198 | 149 | 1 | 1 | 1 | 1 | 0.888 | 1 | 1 | NA | 1 | NA | 0.9 | 0.819 | 1 | 1 | 0.048 |
| SE | 1445 | ENSG00000248405 | PRR5-ARHGAP8 | chr22 | + | 44731729 | 44731821 | 44726576 | 44726634 | 44732250 | 44732391 | 52 | 8 | 39 | 45 | 32 | 0 | 0 | 1 | 0 | 0 | 3 | 19 | 4 | 4 | 34 | 0 | 0 | 0 | 0 | 0 | 241 | 149 | 1 | 1 | 1 | 1 | 0.96 | 1 | 1 | 1 | 1 | 1 | 1 | 1 | 1 | 1 | -0.008 |
| SE | 1446 | ENSG00000248405 | PRR5-ARHGAP8 | chr22 | + | 44731729 | 44731821 | 44726576 | 44726634 | 44786456 | 44786606 | 6 | 2 | 7 | 8 | 7 | 0 | 0 | 0 | 0 | 0 | 0 | 3 | 1 | 1 | 0 | 0 | 2 | 14 | 0 | 6 | 241 | 149 | 1 | 1 | 1 | 1 | 1 | 1 | 1 | NA | 1 | NA | 0.481 | 0.042 | 1 | 0 | 0.619 |
| SE | 1447 | ENSG00000241484 | ARHGAP8 | chr22 | + | 44802076 | 44802164 | 44786456 | 44786606 | 44808306 | 44808438 | 4 | 6 | 2 | 0 | 0 | 0 | 0 | 0 | 0 | 0 | 0 | 4 | 34 | 10 | 0 | 0 | 3 | 10 | 0 | 0 | 237 | 149 | 1 | 1 | 1 | 1 | 1 | NA | NA | NA | | NA | 0.456 | 0.681 | 1 | NA | 0.288 |
| SE | 1448 | ENSG00000248405 | PRR5-ARHGAP8 | chr22 | + | 44802076 | 44802164 | 44786456 | 44786606 | 44814671 | 44814758 | 1 | 2 | 0 | 0 | 0 | 0 | 0 | 0 | 1 | 0 | 0 | 1 | 13 | 3 | 0 | 0 | 0 | 1 | 0 | 6 | 237 | 149 | 1 | 1 | 1 | 1 | NA | 0 | NA | NA | | NA | 1 | 0.891 | 1 | 0 | -0.056 |
| SE | 1449 | ENSG00000248405 | PRR5-ARHGAP8 | chr22 | + | 44808306 | 44808438 | 44786456 | 44786606 | 44814671 | 44814758 | 6 | 0 | 1 | 2 | 0 | 0 | 0 | 0 | 1 | 0 | 0 | 35 | 44 | 4 | 0 | 0 | 0 | 1 | 0 | 6 | 281 | 149 | 1 | 1 | 1 | NA | 1 | 0.515 | NA | NA | | NA | 1 | 0.959 | 1 | 0 | 0.099 |
| SE | 1476 | ENSG00000226328 | NUP50-DT | chr22 | - | 45138823 | 45138916 | 45133019 | 45134797 | 45152404 | 45152623 | 18 | 10 | 14 | 17 | 19 | 0 | 0 | 1 | 0 | 0 | 40 | 19 | 2 | 5 | 12 | 3 | 0 | 0 | 0 | 0 | 242 | 149 | 1 | 1 | 1 | 1 | 0.896 | 1 | 1 | 0.891 | 1 | 0.891 | 1 | 1 | 1 | 1 | 0.001 |
| SE | 1477 | ENSG00000226328 | NUP50-DT | chr22 | - | 45138823 | 45138916 | 45133019 | 45134797 | 45163576 | 45163781 | 10 | 10 | 13 | 17 | 17 | 1 | 0 | 0 | 0 | 0 | 40 | 19 | 2 | 4 | 3 | 0 | 0 | 0 | 0 | 0 | 242 | 149 | 1 | 1 | 0.86 | 1 | 1 | 1 | 1 | 1 | 1 | 1 | 1 | 1 | 1 | 1 | -0.028 |
| SE | 1478 | ENSG00000100368 | CSF2RB | chr22 | + | 36922035 | 36922283 | 36913627 | 36913677 | 36923243 | 36923367 | 313 | 205 | 316 | 282 | 374 | 0 | 1 | 0 | 0 | 0 | 549 | 179 | 265 | 358 | 366 | 0 | 3 | 0 | 0 | 0 | 298 | 149 | 1 | 1 | 1 | 0.99 | 1 | 1 | 1 | 1 | 1 | 1 | 0.968 | 1 | 1 | 1 | 0.004 |
| SE | 1479 | ENSG00000100368 | CSF2RB | chr22 | + | 36925986 | 36926177 | 36923243 | 36923367 | 36929401 | 36929559 | 148 | 70 | 133 | 125 | 50 | 0 | 0 | 0 | 0 | 0 | 232 | 99 | 97 | 173 | 189 | 0 | 1 | 0 | 3 | 0 | 298 | 149 | 1 | 1 | 1 | 1 | 1 | 1 | 1 | 1 | 1 | 1 | 0.98 | 1 | 0.966 | 1 | 0.011 |
| SE | 1480 | ENSG00000100368 | CSF2RB | chr22 | + | 36925986 | 36926177 | 36923243 | 36923367 | 36932764 | 36932904 | 84 | 27 | 79 | 58 | 24 | 3 | 0 | 0 | 0 | 0 | 135 | 44 | 40 | 81 | 81 | 0 | 0 | 0 | 0 | 0 | 298 | 149 | 1 | 1 | 0.933 | 1 | 1 | 1 | 1 | 1 | 1 | 1 | 1 | 1 | 1 | 1 | -0.013 |
| SE | 1481 | ENSG00000100368 | CSF2RB | chr22 | + | 36930374 | 36930510 | 36929638 | 36929807 | 36932764 | 36932904 | 65 | 22 | 50 | 34 | 5 | 0 | 0 | 1 | 0 | 0 | 133 | 34 | 35 | 43 | 53 | 0 | 0 | 0 | 0 | 0 | 285 | 149 | 1 | 1 | 1 | 1 | 0.963 | 1 | 1 | 1 | 1 | 1 | 1 | 1 | 1 | 1 | -0.007 |
| SE | 1484 | ENSG00000100368 | CSF2RB | chr22 | + | 36935350 | 36935441 | 36933831 | 36933994 | 36935629 | 36935687 | 126 | 43 | 82 | 109 | 15 | 0 | 0 | 0 | 0 | 0 | 73 | 30 | 83 | 68 | 100 | 0 | 0 | 0 | 1 | 0 | 240 | 149 | 1 | 1 | 1 | 1 | 1 | 1 | 1 | 1 | 1 | 1 | 1 | 1 | 0.977 | 1 | 0.005 |
| [illegible] | | | | | | | | | | | | | | | | | | | | | | | | | | | | | | | | | | | | | | | | | | | | | | | | |

| | | | | | | | | | | | | | | |
|---|---|---|---|---|---|---|---|---|---|---|---|---|---|---|
| SE | 2007 | ENSG00000270011 | ZNF559-ZNF177 | chr19 | + | 9330114 | 9330189 | 9324711 | 9324780 | 9337795 | 9338017 | 0 | 0 | 1 |
| SE | 2008 | ENSG00000270011 | ZNF559-ZNF177 | chr19 | + | 9330114 | 9330189 | 9324711 | 9324780 | 9338493 | 9338582 | 0 | 0 | 2 |
| SE | 2011 | ENSG00000270011 | ZNF559-ZNF177 | chr19 | + | 9333014 | 9333113 | 9324694 | 9324780 | 9337795 | 9338017 | 1 | 1 | 5 |
| SE | 2012 | ENSG00000270011 | ZNF559-ZNF177 | chr19 | + | 9333014 | 9333113 | 9324694 | 9324780 | 9338493 | 9338582 | 0 | 1 | 3 |
| SE | 2016 | ENSG00000270011 | ZNF559-ZNF177 | chr19 | + | 9337795 | 9338017 | 9330114 | 9330189 | 9338493 | 9338582 | 18 | 5 | 16 |
| SE | 2017 | ENSG00000270011 | ZNF559-ZNF177 | chr19 | + | 9337795 | 9338017 | 9333014 | 9333113 | 9338493 | 9338582 | 18 | 5 | 16 |
| SE | 2018 | ENSG00000270011 | ZNF559-ZNF177 | chr19 | + | 9338493 | 9338582 | 9324694 | 9324780 | 9341101 | 9341184 | 13 | 9 | 11 |
| SE | 2021 | ENSG00000270011 | ZNF559-ZNF177 | chr19 | + | 9338493 | 9338582 | 9337795 | 9337858 | 9339192 | 9339319 | 82 | 52 | 105 |
| SE | 2029 | ENSG00000270011 | ZNF559-ZNF177 | chr19 | + | 9371609 | 9371752 | 9364861 | 9364948 | 9376315 | 9376423 | 4 | 1 | 2 |
| SE | 2030 | ENSG00000270011 | ZNF559-ZNF177 | chr19 | + | 9378961 | 9379088 | 9378258 | 9378344 | 9379526 | 9379569 | 5 | 11 | 7 |
| SE | 2032 | ENSG00000189042 | ZNF567 | chr19 | + | 36689400 | 36689497 | 36687627 | 36687686 | 36694801 | 36694876 | 13 | 11 | 12 |
| SE | 2033 | ENSG00000189042 | ZNF567 | chr19 | + | 36694801 | 36694876 | 36689396 | 36689497 | 36712385 | 36712512 | 23 | 20 | 25 |
| SE | 2034 | ENSG00000189042 | ZNF567 | chr19 | + | 36718947 | 36720561 | 36712780 | 36712867 | 36723188 | 36723346 | 51 | 39 | 60 |
| SE | 2039 | ENSG00000268500 | SIGLEC5 | chr19 | - | 51627461 | 51627746 | 51627148 | 51627248 | 51627833 | 51628091 | 32 | 18 | 6 |
| SE | 2040 | ENSG00000099331 | MYO9B | chr19 | + | 17145396 | 17145491 | 17101655 | 17102557 | 17152643 | 17152706 | 372 | 207 | 324 |
| SE | 2041 | ENSG00000099331 | MYO9B | chr19 | + | 17152643 | 17152706 | 17145396 | 17145491 | 17153966 | 17154066 | 229 | 130 | 200 |
| SE | 2042 | ENSG00000099331 | MYO9B | chr19 | + | 17156908 | 17157038 | 17154314 | 17154415 | 17159394 | 17159484 | 555 | 256 | 405 |
| SE | 2044 | ENSG00000099331 | MYO9B | chr19 | + | 17162349 | 17162466 | 17159394 | 17159484 | 17162987 | 17163122 | 474 | 211 | 381 |
| SE | 2046 | ENSG00000099331 | MYO9B | chr19 | + | 17167942 | 17168064 | 17162987 | 17163122 | 17172335 | 17172477 | 458 | 203 | 366 |
| SE | 2047 | ENSG00000099331 | MYO9B | chr19 | + | 17175662 | 17175741 | 17172758 | 17172963 | 17180926 | 17181040 | 371 | 177 | 278 |
| SE | 2048 | ENSG00000099331 | MYO9B | chr19 | + | 17175662 | 17175741 | 17172758 | 17172963 | 17183828 | 17183868 | 87 | 35 | 63 |
| SE | 2049 | ENSG00000099331 | MYO9B | chr19 | + | 17177796 | 17177886 | 17175662 | 17175741 | 17180926 | 17181040 | 2 | 0 | 0 |
| SE | 2051 | ENSG00000099331 | MYO9B | chr19 | + | 17180926 | 17181040 | 17172758 | 17172963 | 17183828 | 17183868 | 125 | 81 | 102 |
| SE | 2053 | ENSG00000099331 | MYO9B | chr19 | + | 17180926 | 17181040 | 17177796 | 17177886 | 17183828 | 17183868 | 126 | 81 | 101 |
| SE | 2054 | ENSG00000099331 | MYO9B | chr19 | + | 17183828 | 17183868 | 17175662 | 17175741 | 17184864 | 17184898 | 65 | 22 | 53 |
| SE | 2055 | ENSG00000099331 | MYO9B | chr19 | + | 17184864 | 17184987 | 17175662 | 17175741 | 17185920 | 17186001 | 126 | 61 | 96 |
| SE | 2056 | ENSG00000099331 | MYO9B | chr19 | + | 17185920 | 17186001 | 17184864 | 17184987 | 17187934 | 17187939 | 343 | 189 | 250 |
| SE | 2057 | ENSG00000099331 | MYO9B | chr19 | + | 17187934 | 17188045 | 17185920 | 17186001 | 17191096 | 17191219 | 445 | 260 | 358 |
| SE | 2058 | ENSG00000099331 | MYO9B | chr19 | + | 17197791 | 17197858 | 17194555 | 17195473 | 17198183 | 17198308 | 158 | 81 | 87 |
| SE | 2059 | ENSG00000099331 | MYO9B | chr19 | + | 17198183 | 17198308 | 17194555 | 17195473 | 17200292 | 17200426 | 139 | 82 | 107 |
| SE | 2060 | ENSG00000099331 | MYO9B | chr19 | + | 17198183 | 17198308 | 17197791 | 17197858 | 17200638 | 17200829 | 124 | 64 | 64 |
| SE | 2061 | ENSG00000099331 | MYO9B | chr19 | + | 17200292 | 17200426 | 17198117 | 17198308 | 17200638 | 17200829 | 420 | 239 | 320 |
| SE | 2104 | ENSG00000127616 | SMARCA4 | chr19 | + | 10980776 | 10980852 | 10960991 | 10961174 | 10984123 | 10984373 | 1 | 0 | 0 |
| SE | 2106 | ENSG00000127616 | SMARCA4 | chr19 | + | 10989316 | 10989443 | 10987665 | 10987924 | 10991149 | 10991323 | 100 | 71 | 117 |
| SE | 2108 | ENSG00000127616 | SMARCA4 | chr19 | + | 11010380 | 11010531 | 11007901 | 11008023 | 11012948 | 11013112 | 217 | 115 | 179 |
| SE | 2110 | ENSG00000127616 | SMARCA4 | chr19 | + | 11026299 | 11026346 | 11025421 | 11025619 | 11027783 | 11027861 | 8 | 0 | 8 |
| SE | 2118 | ENSG00000127616 | SMARCA4 | chr19 | + | 11056216 | 11056382 | 11041306 | 11041560 | 11058254 | 11058363 | 0 | 0 | 0 |
| SE | 2119 | ENSG00000127616 | SMARCA4 | chr19 | + | 11056216 | 11056382 | 11041306 | 11041560 | 11058257 | 11058363 | 0 | 0 | 2 |
| SE | 2120 | ENSG00000127616 | SMARCA4 | chr19 | + | 11058787 | 11058889 | 11058254 | 11058363 | 11059752 | 11059845 | 217 | 145 | 213 |
| SE | 2123 | ENSG00000213889 | PPM1N | chr19 | + | 45499948 | 45500066 | 45499609 | 45499725 | 45500455 | 45500561 | 9 | 0 | 2 |
| SE | 2124 | ENSG00000213889 | PPM1N | chr19 | + | 45500455 | 45500561 | 45498606 | 45500066 | 45500649 | 45500710 | 21 | 13 | 6 |
| SE | 2126 | ENSG00000213889 | PPM1N | chr19 | + | 45500680 | 45500710 | 45500455 | 45500561 | 45502016 | 45502310 | 4 | 10 | 2 |
| SE | 2127 | ENSG00000213889 | PPM1N | chr19 | + | 45500689 | 45500710 | 45500455 | 45500561 | 45502016 | 45502310 | 4 | 9 | 2 |
| SE | 2139 | ENSG00000167470 | MIDN | chr19 | + | 1251561 | 1251649 | 1249887 | 1250529 | 1251838 | 1251901 | 68 | 25 | 55 |
| SE | 2143 | ENSG00000131351 | HAUS8 | chr19 | - | 17052824 | 17052966 | 17049728 | 17050176 | 17055860 | 17056002 | 65 | 25 | 34 |
| SE | 2144 | ENSG00000131351 | HAUS8 | chr19 | - | 17062697 | 17062779 | 17059996 | 17060089 | 17069030 | 17069086 | 26 | 4 | 17 |
| SE | 2146 | ENSG00000131351 | HAUS8 | chr19 | - | 17069030 | 17069086 | 17059996 | 17060092 | 17073273 | 17073335 | 10 | 4 | 2 |
| SE | 2150 | ENSG00000160111 | CPAMD8 | chr19 | - | 16928008 | 16928234 | 16925195 | 16925372 | 16928941 | 16929240 | 26 | 16 | 16 |
| SE | 2152 | ENSG00000160111 | CPAMD8 | chr19 | - | 16947073 | 16947227 | 16945548 | 16945679 | 16951968 | 16952200 | 6 | 16 | 19 |
| SE | 2154 | ENSG00000160111 | CPAMD8 | chr19 | - | 16976001 | 16976151 | 16975096 | 16975258 | 16977367 | 16977540 | 5 | 1 | 6 |
| SE | 2155 | ENSG00000160111 | CPAMD8 | chr19 | - | 17002265 | 17002350 | 17000413 | 17000522 | 17004272 | 17004386 | 2 | 5 | 13 |
| SE | 2156 | ENSG00000127511 | SIN3B | chr19 | + | 16862351 | 16862559 | 16854142 | 16854261 | 16863679 | 16863796 | 102 | 44 | 94 |
| SE | 2158 | ENSG00000104936 | DMPK | chr19 | - | 45770970 | 45771060 | 45769708 | 45770640 | 45771567 | 45771665 | 2 | 3 | 1 |
| SE | 2159 | ENSG00000104936 | DMPK | chr19 | - | 45770970 | 45771396 | 45769792 | 45770640 | 45771567 | 45771665 | 7 | 8 | 4 |
| SE | 2163 | ENSG00000104936 | DMPK | chr19 | - | 45780322 | 45780381 | 45779777 | 45779869 | 45782192 | 45782552 | 15 | 2 | 7 |
| SE | 2164 | ENSG00000079432 | CIC | chr19 | + | 42287809 | 42287975 | 42287544 | 42287727 | 42289180 | 42289406 | 176 | 109 | 131 |
| SE | 2165 | ENSG00000079432 | CIC | chr19 | + | 42288887 | 42289090 | 42287809 | 42287975 | 42289180 | 42289406 | 83 | 42 | 129 |
| SE | 2166 | ENSG00000079432 | CIC | chr19 | + | 42289847 | 42289951 | 42289180 | 42289406 | 42290232 | 42291466 | 77 | 60 | 91 |
| SE | 2167 | ENSG00000079432 | CIC | chr19 | + | 42291557 | 42291745 | 42290232 | 42291466 | 42292085 | 42292207 | 53 | 15 | 28 |
| SE | 2168 | ENSG00000079432 | CIC | chr19 | + | 42292299 | 42292466 | 42292085 | 42292207 | 42292565 | 42292859 | 102 | 51 | 46 |
| SE | 2170 | ENSG00000079805 | DNM2 | chr19 | + | 10759737 | 10759811 | 10718054 | 10718403 | 10772478 | 10772628 | 373 | 166 | 331 |
| SE | 2171 | ENSG00000079805 | DNM2 | chr19 | + | 10772478 | 10772628 | 10759737 | 10759811 | 10775702 | 10775815 | 581 | 268 | 549 |
| SE | 2172 | ENSG00000079805 | DNM2 | chr19 | + | 10782959 | 10783120 | 10777117 | 10777216 | 10786563 | 10786706 | 736 | 297 | 601 |
| SE | 2173 | ENSG00000079805 | DNM2 | chr19 | + | 10786563 | 10786706 | 10782959 | 10783120 | 10793719 | 10793751 | 638 | 297 | 570 |
| SE | 2176 | ENSG00000079805 | DNM2 | chr19 | + | 10796060 | 10796199 | 10795371 | 10795439 | 10798485 | 10798572 | 583 | 342 | 459 |
| SE | 2177 | ENSG00000079805 | DNM2 | chr19 | + | 10797379 | 10797518 | 10795371 | 10795439 | 10798485 | 10798572 | 391 | 247 | 448 |
| SE | 2180 | ENSG00000079805 | DNM2 | chr19 | + | 10802287 | 10802358 | 10798485 | 10798572 | 10805915 | 10805967 | 806 | 450 | 716 |
| SE | 2181 | ENSG00000079805 | DNM2 | chr19 | + | 10802287 | 10802358 | 10798485 | 10798572 | 10812263 | 10812300 | 353 | 213 | 347 |
| SE | 2183 | ENSG00000079805 | DNM2 | chr19 | + | 10805915 | 10805967 | 10798485 | 10798572 | 10812263 | 10812300 | 11 | 12 | 11 |
| SE | 2184 | ENSG00000079805 | DNM2 | chr19 | + | 10805915 | 10805967 | 10802287 | 10802358 | 10812263 | 10812300 | 360 | 182 | 296 |
| SE | 2187 | ENSG00000176024 | ZNF613 | chr19 | + | 51929731 | 51929824 | 51927146 | 51927540 | 51936027 | 51936125 | 8 | 1 | 0 |
| SE | 2188 | ENSG00000176024 | ZNF613 | chr19 | + | 51929731 | 51929896 | 51927447 | 51927540 | 51936027 | 51936235 | 20 | 13 | 5 |
| SE | 2189 | ENSG00000176024 | ZNF613 | chr19 | + | 51936027 | 51936235 | 51929731 | 51929896 | 51940208 | 51940335 | 15 | 12 | 2 |
| SE | 2190 | ENSG00000161551 | ZNF577 | chr19 | - | 51878388 | 51878515 | 51877281 | 51877377 | 51880322 | 51880401 | 29 | 26 | 36 |
| SE | 2201 | ENSG00000214189 | ZNF788P | chr19 | + | 12110633 | 12110694 | 12092467 | 12092470 | 12135774 | 12135927 | 0 | 0 | 0 |
| SE | 2202 | ENSG00000214189 | ZNF788P | chr19 | + | 12110633 | 12110694 | 12110304 | 12110431 | 12111245 | 12113048 | 0 | 1 | 0 |
| SE | 2203 | ENSG00000219665 | ZNF433-AS1 | chr19 | + | 11990168 | 11990381 | 11987766 | 11987823 | 12001773 | 12001822 | 0 | 0 | 1 |
| SE | 2205 | ENSG00000219665 | ZNF433-AS1 | chr19 | + | 12001773 | 12001822 | 11987616 | 11987823 | 12035553 | 12035772 | 8 | 2 | 3 |
| SE | 2207 | ENSG00000197054 | ZNF763 | chr19 | + | 11977370 | 11977431 | 11976821 | 11977164 | 11978115 | 11978275 | 25 | 25 | 36 |
| SE | 2211 | ENSG00000267179 | NA | chr19 | + | 11961660 | 11961758 | 11956504 | 11956642 | 11977037 | 11977164 | 1 | 2 | 0 |
| SE | 2217 | ENSG00000221983 | UBA52 | chr19 | + | 18573292 | 18573403 | 18571856 | 18571909 | 18574869 | 18574972 | 2 | 3 | 1 |
| SE | 2218 | ENSG00000221983 | UBA52 | chr19 | + | 18573661 | 18573748 | 18573264 | 18573403 | 18574869 | 18574972 | 3858 | 1639 | 4183 |
| SE | 2219 | ENSG00000105701 | FKBP8 | chr19 | - | 18533269 | 18533347 | 18532663 | 18532795 | 18537600 | 18537704 | 340 | 118 | 292 |
| SE | 2220 | ENSG00000105701 | FKBP8 | chr19 | - | 18537600 | 18537773 | 18533269 | 18533347 | 18539370 | 18539459 | 195 | 76 | 192 |
| SE | 2221 | ENSG00000105701 | FKBP8 | chr19 | - | 18538215 | 18538433 | 18537600 | 18537773 | 18539370 | 18539459 | 292 | 134 | 258 |
| SE | 2222 | ENSG00000105701 | FKBP8 | chr19 | - | 18538215 | 18538436 | 18537600 | 18537773 | 18539370 | 18539459 | 410 | 173 | 334 |
| SE | 2224 | ENSG00000105701 | FKBP8 | chr19 | - | 18539370 | 18539459 | 18537600 | 18537773 | 18541678 | 18541995 | 0 | 0 | 0 |
| SE | 2225 | ENSG00000105701 | FKBP8 | chr19 | - | 18539370 | 18539459 | 18538215 | 18538436 | 18541678 | 18541995 | 62 | 29 | 47 |
| SE | 2226 | ENSG00000105701 | FKBP8 | chr19 | - | 18539550 | 18539720 | 18539370 | 18539459 | 18541678 | 18541995 | 840 | 403 | 693 |
| SE | 2237 | ENSG00000105643 | ARRDC2 | chr19 | + | 18007589 | 18007842 | 18001131 | 18001573 | 18008710 | 18008777 | 0 | 0 | 0 |
| SE | 2240 | ENSG00000189068 | VSTM1 | chr19 | - | 54042168 | 54042196 | 54041915 | 54041953 | 54051409 | 54051448 | 7 | 10 | 8 |
| SE | 2241 | ENSG00000189068 | VSTM1 | chr19 | - | 54042226 | 54042369 | 54041915 | 54041953 | 54051409 | 54051448 | 35 | 26 | 11 |
| SE | 2243 | ENSG00000189068 | VSTM1 | chr19 | - | 54051409 | 54051448 | 54042226 | 54042369 | 54058305 | 54058423 | 80 | 71 | 24 |
| SE | 2247 | ENSG00000189068 | VSTM1 | chr19 | - | 54058305 | 54058590 | 54051409 | 54051448 | 54063743 | 54063897 | 62 | 53 | 15 |
| SE | 2248 | ENSG00000189068 | VSTM1 | chr19 | - | 54058696 | 54058732 | 54058305 | 54058590 | 54063743 | 54063897 | 51 | 17 | 19 |
| SE | 2251 | ENSG00000161860 | SYCE2 | chr19 | - | 12900003 | 12900120 | 12898785 | 12899385 | 12900459 | 12900648 | 0 | 0 | 2 |
| SE | 2252 | ENSG00000105607 | GCDH | chr19 | + | 12891486 | 12891522 | 12891159 | 12891395 | 12891830 | 12891974 | 46 | 20 | 50 |
| SE | 2253 | ENSG00000105607 | GCDH | chr19 | + | 12891486 | 12891660 | 12891270 | 12891395 | 12891830 | 12891974 | 36 | 18 | 42 |
| SE | 2256 | ENSG00000105607 | GCDH | chr19 | + | 12892386 | 12892434 | 12892115 | 12892178 | 12893482 | 12893493 | 0 | 0 | 1 |
| SE | 2257 | ENSG00000196466 | ZNF799 | chr19 | - | 12398133 | 12398251 | 12393296 | 12393423 | 12401067 | 12401268 | 0 | 0 | 0 |
| SE | 2260 | ENSG00000234773 | NA | chr19 | + | 12207331 | 12207708 | 12195014 | 12195143 | 12234712 | 12234824 | 0 | 2 | 3 |
| SE | 2261 | ENSG00000234773 | NA | chr19 | + | 12234712 | 12234824 | 12195016 | 12195143 | 12237089 | 12237767 | 5 | 7 | 39 |
| SE | 2270 | ENSG00000105619 | TFPT | chr19 | - | 54110050 | 54110121 | 54108325 | 54108395 | 54115246 | 54115288 | 10 | 3 | 10 |
| SE | 2271 | ENSG00000105619 | TFPT | chr19 | - | 54114441 | 54114700 | 54108325 | 54108395 | 54115246 | 54115285 | 4 | 17 | 4 |
| SE | 2274 | ENSG00000104824 | HNRNPL | chr19 | - | 38838893 | 38839015 | 38838396 | 38838598 | 38840487 | 38840559 | 366 | 187 | 347 |
| SE | 2275 | ENSG00000104824 | HNRNPL | chr19 | - | 38840095 | 38840376 | 38838396 | 38838598 | 38840487 | 38840559 | 270 | 162 | 235 |
| SE | 2276 | ENSG00000104824 | HNRNPL | chr19 | - | 38840095 | 38840376 | 38838893 | 38839015 | 38840487 | 38840559 | 471 | 261 | 458 |
| SE | 2278 | ENSG00000104824 | HNRNPL | chr19 | - | 38845852 | 38846090 | 38843370 | 38843914 | 38847315 | 38847434 | 382 | 188 | 312 |
| SE | 2282 | ENSG00000180448 | ARHGAP45 | chr19 | + | 1073947 | 1074014 | 1073582 | 1073746 | 1074103 | 1074241 | 916 | 487 | 934 |
| SE | 2283 | ENSG00000180448 | ARHGAP45 | chr19 | + | 1077856 | 1078045 | 1074798 | 1074879 | 1079702 | 1079840 | 753 | 338 | 619 |
| SE | 2284 | ENSG00000180448 | ARHGAP45 | chr19 | + | 1080463 | 1080547 | 1080254 | 1080379 | 1080681 | 1080786 | 393 | 153 | 362 |
| SE | 2285 | ENSG00000180448 | ARHGAP45 | chr19 | + | 1084237 | 1084346 | 1082840 | 1083353 | 1085659 | 1085996 | 376 | 150 | 286 |
| SE | 2287 | ENSG00000188321 | ZNF559 | chr19 | + | 9337714 | 9338017 | 9333014 | 9333113 | 9338493 | 9338582 | 18 | 5 | 16 |
| SE | 2288 | ENSG00000188321 | ZNF559 | chr19 | + | 9337795 | 9337865 | 9324694 | 9324780 | 9338493 | 9338582 | 11 | 4 | 9 |
| SE | 2292 | ENSG00000142002 | DPP9 | chr19 | - | 4697550 | 4697651 | 4695377 | 4695555 | 4700215 | 4700273 | 136 | 65 | 98 |
| SE | 2294 | ENSG00000142002 | DPP9 | chr19 | - | 4700215 | 4700277 | 4697550 | 4697651 | 4702026 | 4702155 | 99 | 49 | 52 |
| SE | 2295 | ENSG00000142002 | DPP9 | chr19 | - | 4703885 | 4704054 | 4702602 | 4702716 | 4704130 | 4704151 | 103 | 44 | 80 |
| SE | 2296 | ENSG00000142002 | DPP9 | chr19 | - | 4714080 | 4714337 | 4705857 | 4705970 | 4719850 | 4719941 | 116 | 44 | 86 |
| SE | 2302 | ENSG00000126453 | BCL2L12 | chr19 | + | 49667018 | 49667161 | 49666684 | 49666799 | 49668850 | 49668937 | 31 | 4 | 13 |
| SE | 2304 | ENSG00000126453 | BCL2L12 | chr19 | + | 49667021 | 49667161 | 49666684 | 49666799 | 49668850 | 49668937 | 21 | 0 | 6 |
| SE | 2310 | ENSG00000105699 | LSR | chr19 | + | 35258944 | 35259064 | 35250314 | 35250659 | 35266358 | 35266532 | 15 | 10 | 17 |
| SE | 2311 | ENSG00000105699 | LSR | chr19 | + | 35261924 | 35261981 | 35258944 | 35259064 | 35262545 | 35262692 | 0 | 0 | 3 |
| SE | 2313 | ENSG00000105699 | LSR | chr19 | + | 35262545 | 35262692 | 35258944 | 35259064 | 35266358 | 35266532 | 29 | 12 | 32 |
| SE | 2317 | ENSG00000099875 | MKNK2 | chr19 | - | 2046185 | 2046283 | 2043502 | 2043582 | 2046366 | 2046468 | 531 | 258 | 427 |
| SE | 2318 | ENSG00000257103 | LSM14A | chr19 | + | 34195218 | 34195320 | 34194477 | 34194641 | 34196633 | 34196666 | 0 | 0 | 2 |
| SE | 2319 | ENSG00000257103 | LSM14A | chr19 | + | 34196633 | 34196763 | 34194477 | 34194641 | 34208928 | 34209051 | 770 | 471 | 727 |
| SE | 2320 | ENSG00000257103 | LSM14A | chr19 | + | 34196633 | 34196763 | 34194477 | 34194641 | 34215595 | 34215661 | 354 | 214 | 327 |
| SE | 2321 | ENSG00000257103 | LSM14A | chr19 | + | 34208928 | 34209051 | 34194477 | 34194641 | 34215595 | 34215661 | 33 | 50 | 38 |
| SE | 2322 | ENSG00000257103 | LSM14A | chr19 | + | 34208928 | 34209051 | 34196633 | 34196763 | 34215123 | 34215300 | 738 | 470 | 680 |
| SE | 2323 | ENSG00000257103 | LSM14A | chr19 | + | 34208928 | 34209051 | 34196633 | 34196763 | 34215595 | 34215661 | 316 | 236 | 326 |
| SE | 2324 | ENSG00000257103 | LSM14A | chr19 | + | 34215123 | 34215300 | 34196633 | 34196763 | 34215595 | 34215661 | 226 | 172 | 188 |
| SE | 2326 | ENSG00000257103 | LSM14A | chr19 | + | 34215595 | 34215661 | 34208928 | 34209051 | 34219390 | 34219573 | 59 | 58 | 60 |
| SE | 2330 | ENSG00000189144 | ZNF573 | chr19 | - | 37740640 | 37740710 | 37739905 | 37740194 | 37770004 | 37770097 | 1 | 1 | 0 |
| SE | 2331 | ENSG00000189144 | ZNF573 | chr19 | - | 37758475 | 37758596 | 37738286 | 37740194 | 37759046 | 37759139 | 0 | 3 | 0 |
| SE | 2332 | ENSG00000189144 | ZNF573 | chr19 | - | 37758475 | 37758596 | 37738286 | 37740194 | 37770004 | 37770097 | 0 | 2 | 1 |
| SE | 2334 | ENSG00000189144 | ZNF573 | chr19 | - | 37759046 | 37759139 | 37738286 | 37740194 | 37770004 | 37770097 | 2 | 5 | 17 |
| SE | 2335 | ENSG00000189144 | ZNF573 | chr19 | - | 37759046 | 37759139 | 37738286 | 37740194 | 37771563 | 37771696 | 0 | 0 | 5 |
| SE | 2336 | ENSG00000189144 | ZNF573 | chr19 | - | 37759046 | 37759139 | 37758475 | 37758596 | 37770004 | 37770097 | 2 | 6 | 12 |
| SE | 2337 | ENSG00000189144 | ZNF573 | chr19 | - | 37770004 | 37770097 | 37738301 | 37740194 | 37771563 | 37771690 | 38 | 35 | 47 |
| SE | 2338 | ENSG00000189144 | ZNF573 | chr19 | - | 37770004 | 37770097 | 37759046 | 37759139 | 37771563 | 37771690 | 25 | 32 | 50 |
| SE | 2339 | ENSG00000189144 | ZNF573 | chr19 | - | 37770004 | 37770097 | 37759046 | 37759139 | 37773660 | 37773751 | 0 | 1 | 5 |
| SE | 2341 | ENSG00000189144 | ZNF573 | chr19 | - | 37771563 | 37771690 | 37770004 | 37770097 | 37773660 | 37773751 | 28 | 24 | 39 |
| SE | 2342 | ENSG00000189144 | ZNF573 | chr19 | - | 37771563 | 37771696 | 37759046 | 37759139 | 37773660 | 37773745 | 44 | 30 | 50 |
| SE | 2344 | ENSG00000189144 | ZNF573 | chr19 | - | 37772937 | 37772982 | 37771563 | 37771690 | 37773660 | 37773751 | 4 | 1 | 4 |
| SE | 2346 | ENSG00000189144 | ZNF573 | chr19 | - | 37773660 | 37773751 | 37771563 | 37771690 | 37779543 | 37779553 | 1 | 0 | 1 |
| SE | 2355 | ENSG00000099785 | MARCHF2 | chr19 | + | 8426608 | 8426804 | 8421788 | 8422016 | 8438387 | 8439004 | 39 | 20 | 10 |
| SE | 2356 | ENSG00000099785 | MARCHF2 | chr19 | + | 8430657 | 8430867 | 8421788 | 8422016 | 8438387 | 8439017 | 37 | 21 | 31 |
| SE | 2357 | ENSG00000099785 | MARCHF2 | chr19 | + | 8430657 | 8430867 | 8426608 | 8426804 | 8438387 | 8439017 | 62 | 47 | 54 |
| SE | 2360 | ENSG00000104880 | ARHGEF18 | chr19 | + | 7362780 | 7362905 | 7348936 | 7349241 | 7372811 | 7373071 | 419 | 368 | 572 |
| SE | 2361 | ENSG00000104880 | ARHGEF18 | chr19 | + | 7372811 | 7373071 | 7362780 | 7362905 | 7375719 | 7375870 | 372 | 273 | 529 |
| SE | 2363 | ENSG00000104880 | ARHGEF18 | chr19 | + | 7378393 | 7378451 | 7376642 | 7376757 | 7379121 | 7379166 | 262 | 174 | 309 |
| SE | 2364 | ENSG00000104880 | ARHGEF18 | chr19 | + | 7379121 | 7379166 | 7378393 | 7378451 | 7380916 | 7380994 | 275 | 188 | 303 |
| SE | 2365 | ENSG00000104880 | ARHGEF18 | chr19 | + | 7380916 | 7380994 | 7379121 | 7379166 | 7382791 | 7382894 | 308 | 240 | 360 |
| SE | 2366 | ENSG00000104880 | ARHGEF18 | chr19 | + | 7440188 | 7440482 | 7383061 | 7383203 | 7441652 | 7441765 | 274 | 166 | 284 |
| SE | 2367 | ENSG00000104880 | ARHGEF18 | chr19 | + | 7440343 | 7440482 | 7383061 | 7383203 | 7441652 | 7441765 | 587 | 393 | 653 |
| SE | 2368 | ENSG00000104880 | ARHGEF18 | chr19 | + | 7444203 | 7444454 | 7441911 | 7442052 | 7447042 | 7447168 | 714 | 515 | 809 |
| SE | 2369 | ENSG00000104880 | ARHGEF18 | chr19 | + | 7447042 | 7447168 | 7444203 | 7444454 | 7451148 | 7451266 | 497 | 305 | 555 |
| SE | 2370 | ENSG00000104880 | ARHGEF18 | chr19 | + | 7456326 | 7456403 | 7453466 | 7453715 | 7458511 | 7458690 | 552 | 353 | 615 |
| SE | 2371 | ENSG00000104880 | ARHGEF18 | chr19 | + | 7458511 | 7458690 | 7456326 | 7456403 | 7459902 | 7459994 | 649 | 475 | 801 |
| SE | 2372 | ENSG00000104880 | ARHGEF18 | chr19 | + | 7458511 | 7458690 | 7456326 | 7456403 | 7462151 | 7462334 | 402 | 265 | 430 |
| SE | 2373 | ENSG00000104880 | ARHGEF18 | chr19 | + | 7459902 | 7459994 | 7456326 | 7456403 | 7462151 | 7462334 | 171 | 130 | 153 |
| SE | 2374 | ENSG00000104880 | ARHGEF18 | chr19 | + | 7459902 | 7459994 | 7458511 | 7458690 | 7462151 | 7462334 | 419 | 338 | 524 |
| SE | 2375 | ENSG00000104880 | ARHGEF18 | chr19 | + | 7462151 | 7462334 | 7459902 | 7459994 | 7463817 | 7463955 | 462 | 341 | 512 |
| SE | 2376 | ENSG00000104880 | ARHGEF18 | chr19 | + | 7463817 | 7463955 | 7462151 | 7462334 | 7464559 | 7464687 | 271 | 170 | 249 |
| SE | 2377 | ENSG00000104880 | ARHGEF18 | chr19 | + | 7468824 | 7469131 | 7467213 | 7467684 | 7469903 | 7470029 | 119 | 56 | 137 |
| SE | 2379 | ENSG00000170889 | RPS9 | chr19 | + | 54201486 | 54201609 | 54201132 | 54201281 | 54207397 | 54207647 | 1633 | 994 | 2020 |
| SE | 2380 | ENSG00000170889 | RPS9 | chr19 | + | 54206275 | 54206462 | 54201132 | 54201281 | 54207397 | 54207647 | 2277 | 1230 | 2901 |
| SE | 2381 | ENSG00000170889 | RPS9 | chr19 | + | 54206275 | 54206462 | 54201486 | 54201609 | 54206552 | 54206565 | 2567 | 1410 | 3395 |
| SE | 2383 | ENSG00000170889 | RPS9 | chr19 | + | 54206275 | 54206724 | 54201159 | 54201281 | 54207397 | 54207644 | 104 | 59 | 71 |
| SE | 2387 | ENSG00000167608 | TMC4 | chr19 | - | 54160466 | 54160545 | 54160107 | 54160374 | 54160877 | 54161033 | 14 | 0 | 8 |
| SE | 2388 | ENSG00000167608 | TMC4 | chr19 | - | 54160466 | 54160610 | 54160107 | 54160374 | 54160877 | 54160972 | 11 | 0 | 2 |
| SE | 2391 | ENSG00000105618 | PRPF31 | chr19 | + | 54118240 | 54118455 | 54115778 | 54115797 | 54118572 | 54118633 | 196 | 153 | 223 |
| SE | 2392 | ENSG00000105618 | PRPF31 | chr19 | + | 54118247 | 54118455 | 54115753 | 54115797 | 54118572 | 54118633 | 192 | 148 | 222 |
| SE | 2393 | ENSG00000105618 | PRPF31 | chr19 | + | 54118270 | 54118455 | 54115409 | 54115797 | 54118572 | 54118633 | 253 | 192 | 303 |
| SE | 2394 | ENSG00000105618 | PRPF31 | chr19 | + | 54128304 | 54128377 | 54128072 | 54128200 | 54129056 | 54129185 | 124 | 60 | 127 |
| SE | 2395 | ENSG00000204604 | ZNF468 | chr19 | - | 52845178 | 52845296 | 52838535 | 52842151 | 52849086 | 52849213 | 44 | 29 | 15 |
| SE | 2396 | ENSG00000204604 | ZNF468 | chr19 | - | 52846557 | 52846611 | 52841935 | 52842151 | 52849086 | 52849213 | 0 | 0 | 2 |
| SE | 2397 | ENSG00000204604 | ZNF468 | chr19 | - | 52846557 | 52846611 | 52845236 | 52845296 | 52849086 | 52849213 | 0 | 0 | 0 |
| SE | 2398 | ENSG00000204604 | ZNF468 | chr19 | - | 52853871 | 52854107 | 52849086 | 52849213 | 52854257 | 52854345 | 14 | 5 | 10 |
| SE | 2399 | ENSG00000204604 | ZNF468 | chr19 | - | 52853871 | 52854126 | 52849086 | 52849213 | 52854257 | 52854345 | 23 | 13 | 6 |
| SE | 2400 | ENSG00000269825 | NA | chr19 | - | 52660761 | 52660843 | 52655560 | 52655687 | 52690442 | 52690533 | 32 | 32 | 39 |
| SE | 2402 | ENSG00000126247 | CAPNS1 | chr19 | + | 36142651 | 36142741 | 36142299 | 36142333 | 36142908 | 36142966 | 2229 | 930 | 1367 |
| SE | 2403 | ENSG00000126247 | CAPNS1 | chr19 | + | 36146195 | 36146312 | 36145975 | 36146054 | 36149577 | 36149580 | 2674 | 1207 | 1476 |
| SE | 2406 | ENSG00000161277 | THAP8 | chr19 | - | 36039943 | 36040136 | 36039322 | 36039718 | 36054134 | 36054285 | 11 | 14 | 13 |
| SE | 2412 | ENSG00000239382 | ALKBH6 | chr19 | - | 36011403 | 36011464 | 36010289 | 36011045 | 36013020 | 36013089 | 44 | 20 | 43 |
| SE | 2413 | ENSG00000239382 | ALKBH6 | chr19 | - | 36013000 | 36013089 | 36011403 | 36011464 | 36013343 | 36013422 | 11 | 8 | 13 |
| SE | 2414 | ENSG00000239382 | ALKBH6 | chr19 | - | 36013020 | 36013089 | 36011179 | 36011464 | 36013343 | 36013422 | 45 | 33 | 36 |
| SE | 2422 | ENSG00000167642 | SPINT2 | chr19 | + | 38283626 | 38283797 | 38264572 | 38264998 | 38287875 | 38287905 | 53 | 46 | 54 |
| SE | 2429 | ENSG00000132024 | CC2D1A | chr19 | + | 13918069 | 13918194 | 13913403 | 13913638 | 13918503 | 13918576 | 69 | 6 | 27 |
| SE | 2431 | ENSG00000132024 | CC2D1A | chr19 | + | 13926516 | 13926590 | 13923694 | 13923811 | 13926666 | 13926725 | 83 | 35 | 71 |
| SE | 2432 | ENSG00000132024 | CC2D1A | chr19 | + | 13927174 | 13927265 | 13926977 | 13927077 | 13927892 | 13928030 | 38 | 28 | 52 |
| SE | 2434 | ENSG00000104979 | C19orf53 | chr19 | + | 13775470 | 13775613 | 13774651 | 13774707 | 13778051 | 13778461 | 1 | 0 | 0 |
| SE | 2435 | ENSG00000104979 | C19orf53 | chr19 | + | 13775476 | 13775613 | 13774651 | 13774707 | 13778051 | 13778207 | 1 | 0 | 0 |
| SE | 2437 | ENSG00000104957 | CCDC130 | chr19 | + | 13757417 | 13757473 | 13756196 | 13756279 | 13757785 | 13757846 | 143 | 89 | 177 |
| SE | 2444 | ENSG00000088247 | KHSRP | chr19 | - | 6420072 | 6420144 | 6419202 | 6419260 | 6420421 | 6420471 | 311 | 143 | 298 |
| SE | 2448 | ENSG00000105127 | AKAP8 | chr19 | - | 15368234 | 15368322 | 15362109 | 15362251 | 15370145 | 15370179 | 85 | 78 | 107 |
| SE | 2449 | ENSG00000105127 | AKAP8 | chr19 | - | 15368234 | 15368339 | 15362109 | 15362251 | 15370145 | 15370179 | 30 | 29 | 36 |
| SE | 2454 | ENSG00000105127 | AKAP8 | chr19 | - | 15374602 | 15374635 | 15373716 | 15374065 | 15379712 | 15379785 | 9 | 14 | 18 |
| SE | 2467 | ENSG00000141971 | MVB12A | chr19 | + | 17413096 | 17413104 | 17406108 | 17406296 | 17413220 | 17413305 | 6 | 9 | 4 |
| SE | 2468 | ENSG00000141971 | MVB12A | chr19 | + | 17413220 | 17413305 | 17406144 | 17406296 | 17420312 | 17420411 | 0 | 0 | 0 |
| SE | 2471 | ENSG00000141971 | MVB12A | chr19 | + | 17424620 | 17424677 | 17424005 | 17424067 | 17424930 | 17424989 | 38 | 12 | 17 |
| SE | 2483 | ENSG00000160877 | NACC1 | chr19 | + | 13117993 | 13118131 | 13116861 | 13116935 | 13135199 | 13135704 | 27 | 10 | 12 |
| SE | 2484 | ENSG00000141867 | BRD4 | chr19 | - | 15244709 | 15244762 | 15244230 | 15244600 | 15254151 | 15254262 | 359 | 228 | 287 |
| SE | 2485 | ENSG00000141867 | BRD4 | chr19 | - | 15256063 | 15256263 | 15255296 | 15255592 | 15256963 | 15257173 | 161 | 107 | 189 |
| SE | 2486 | ENSG00000141867 | BRD4 | chr19 | - | 15263419 | 15263548 | 15256963 | 15257173 | 15264403 | 15264620 | 229 | 108 | 212 |
| SE | 2488 | ENSG00000141867 | BRD4 | chr19 | - | 15267415 | 15267551 | 15265353 | 15265643 | 15272814 | 15273133 | 591 | 434 | 533 |
| SE | 2491 | ENSG00000141867 | BRD4 | chr19 | - | 15268904 | 15269042 | 15265353 | 15265643 | 15272814 | 15272981 | 469 | 328 | 445 |
| SE | 2492 | ENSG00000141867 | BRD4 | chr19 | - | 15268904 | 15269042 | 15267415 | 15267551 | 15272814 | 15272981 | 1227 | 985 | 1082 |
| SE | 2493 | ENSG00000141867 | BRD4 | chr19 | - | 15272814 | 15273133 | 15268904 | 15269042 | 15332289 | 15332443 | 515 | 365 | 495 |
| SE | 2494 | ENSG00000141867 | BRD4 | chr19 | - | 15275656 | 15275839 | 15272814 | 15273133 | 15331835 | 15331907 | 0 | 0 | 0 |
| SE | 2495 | ENSG00000141867 | BRD4 | chr19 | - | 15275656 | 15275839 | 15272814 | 15273133 | 15332289 | 15332545 | 0 | 0 | 1 |
| SE | 2500 | ENSG00000160953 | PWWP3A | chr19 | + | 1370641 | 1371078 | 1369595 | 1369646 | 1373071 | 1373160 | 87 | 31 | 40 |
| SE | 2502 | ENSG00000160953 | PWWP3A | chr19 | + | 1373071 | 1373160 | 1370641 | 1371078 | 1376518 | 1376865 | 34 | 17 | 34 |
| SE | 2507 | ENSG00000011451 | WIZ | chr19 | - | 15431011 | 15431182 | 15429585 | 15430089 | 15436805 | 15437129 | 8 | 0 | 5 |
| SE | 2508 | ENSG00000011451 | WIZ | chr19 | - | 15438577 | 15440715 | 15436805 | 15437129 | 15442675 | 15442748 | 2 | 4 | 10 |
| SE | 2513 | ENSG00000182472 | CAPN12 | chr19 | - | 38733702 | 38733781 | 38731106 | 38731223 | 38734812 | 38734870 | 5 | 5 | 1 |
| SE | 2514 | ENSG00000182472 | CAPN12 | chr19 | - | 38733702 | 38733781 | 38731106 | 38731223 | 38735369 | 38735405 | 4 | 5 | 1 |
| SE | 2515 | ENSG00000182472 | CAPN12 | chr19 | - | 38733702 | 38733781 | 38731106 | 38731223 | 38735501 | 38735544 | 3 | 5 | 1 |
| SE | 2516 | ENSG00000182472 | CAPN12 | chr19 | - | 38733702 | 38733940 | 38731106 | 38731223 | 38734812 | 38734870 | 4 | 5 | 4 |
| SE | 2517 | ENSG00000182472 | CAPN12 | chr19 | - | 38734141 | 38734204 | 38733702 | 38733781 | 38734318 | 38734389 | 1 | 2 | 0 |
| SE | 2518 | ENSG00000182472 | CAPN12 | chr19 | - | 38734318 | 38734389 | 38733702 | 38733781 | 38734812 | 38734870 | 0 | 0 | 2 |
| SE | 2520 | ENSG00000182472 | CAPN12 | chr19 | - | 38734812 | 38734870 | 38731106 | 38731223 | 38735369 | 38735410 | 0 | 0 | 1 |
| SE | 2522 | ENSG00000182472 | CAPN12 | chr19 | - | 38734812 | 38734870 | 38733702 | 38733781 | 38735369 | 38735410 | 2 | 0 | 0 |
| SE | 2525 | ENSG00000182472 | CAPN12 | chr19 | - | 38735369 | 38735429 | 38731106 | 38731223 | 38735501 | 38735544 | 0 | 0 | 0 |
| SE | 2526 | ENSG00000182472 | CAPN12 | chr19 | - | 38735369 | 38735429 | 38734565 | 38734870 | 38735501 | 38735544 | 0 | 0 | 3 |
| SE | 2527 | ENSG00000178982 | EIF3K | chr19 | + | 38620336 | 38620435 | 38619185 | 38619327 | 38624076 | 38624197 | 2516 | 1056 | 2210 |
| SE | 2532 | ENSG00000104814 | MAP4K1 | chr19 | - | 38589177 | 38589275 | 38587640 | 38587817 | 38593281 | 38593337 | 1 | 2 | 0 |
| SE | 2533 | ENSG00000104814 | MAP4K1 | chr19 | - | 38592449 | 38592497 | 38587640 | 38587817 | 38593281 | 38593337 | 0 | 0 | 2 |
| SE | 2534 | ENSG00000104814 | MAP4K1 | chr19 | - | 38593281 | 38593337 | 38587640 | 38587817 | 38595484 | 38595543 | 41 | 14 | 33 |
| SE | 2535 | ENSG00000104814 | MAP4K1 | chr19 | - | 38597485 | 38597594 | 38597325 | 38597384 | 38599924 | 38599985 | 203 | 85 | 192 |
| SE | 2537 | ENSG00000104814 | MAP4K1 | chr19 | - | 38605567 | 38605730 | 38605408 | 38605491 | 38606172 | 38606215 | 73 | 19 | 65 |
| SE | 2538 | ENSG00000104814 | MAP4K1 | chr19 | - | 38609595 | 38609674 | 38608111 | 38608170 | 38609908 | 38610025 | 275 | 195 | 342 |
| SE | 2540 | ENSG00000104814 | MAP4K1 | chr19 | - | 38614244 | 38614292 | 38614042 | 38614085 | 38614389 | 38614445 | 418 | 243 | 355 |

| | | | | | | | | | | | |
|---|---|---|---|---|---|---|---|---|---|---|---|
| SE | 2543 | ENSG00000104814 | MAP4K1 | chr19 | - | 38614389 | 38614445 | 38614066 | 38614292 | 38616194 | [illegible] |
| SE | 2544 | ENSG00000104814 | MAP4K1 | chr19 | - | 38617353 | 38617444 | 38616177 | 38616259 | 38617567 | [illegible] |
| SE | 2558 | ENSG00000214046 | SMIM7 | chr19 | - | 16652604 | 16652961 | 16645660 | 16647261 | 16659958 | [illegible] |
| SE | 2561 | ENSG00000214046 | SMIM7 | chr19 | - | 16654034 | 16654125 | 16652355 | 16652961 | 16659958 | [illegible] |
| SE | 2571 | ENSG00000126457 | PRMT1 | chr19 | + | 49683926 | 49684069 | 49682195 | 49682259 | 49684753 | [illegible] |
| SE | 2572 | ENSG00000126457 | PRMT1 | chr19 | + | 49684753 | 49684841 | 49683926 | 49684069 | 49684921 | [illegible] |
| SE | 2573 | ENSG00000126457 | PRMT1 | chr19 | + | 49684921 | 49685037 | 49683926 | 49684069 | 49686092 | [illegible] |
| SE | 2574 | ENSG00000196684 | HSH2D | chr19 | + | 16148723 | 16148800 | 16143735 | 16143774 | 16151446 | [illegible] |
| SE | 2576 | ENSG00000196684 | HSH2D | chr19 | + | 16151446 | 16151578 | 16148723 | 16148800 | 16152551 | [illegible] |
| SE | 2579 | ENSG00000196684 | HSH2D | chr19 | + | 16154398 | 16154491 | 16152551 | 16153208 | 16157209 | [illegible] |
| SE | 2580 | ENSG00000196684 | HSH2D | chr19 | + | 16154401 | 16154491 | 16153031 | 16153208 | 16157209 | [illegible] |
| SE | 2582 | ENSG00000186272 | ZNF17 | chr19 | + | 57413595 | 57413636 | 57411160 | 57411406 | 57417911 | [illegible] |
| SE | 2583 | ENSG00000186272 | ZNF17 | chr19 | + | 57413595 | 57413636 | 57411160 | 57411406 | 57419634 | [illegible] |
| SE | 2584 | ENSG00000186272 | ZNF17 | chr19 | + | 57417911 | 57418038 | 57411178 | 57411406 | 57419634 | [illegible] |
| SE | 2588 | ENSG00000197128 | ZNF772 | chr19 | - | 57475023 | 57475146 | 57469585 | 57474421 | 57475659 | [illegible] |
| SE | 2589 | ENSG00000197128 | ZNF772 | chr19 | - | 57475032 | 57475146 | 57474360 | 57474421 | 57475659 | [illegible] |
| SE | 2590 | ENSG00000197128 | ZNF772 | chr19 | - | 57475047 | 57475146 | 57474360 | 57474421 | 57475659 | [illegible] |
| SE | 2591 | ENSG00000197128 | ZNF772 | chr19 | - | 57475659 | 57475786 | 57469585 | 57474421 | 57476633 | [illegible] |
| SE | 2600 | ENSG00000185869 | ZNF829 | chr19 | - | 36915129 | 36915251 | 36914964 | 36915022 | 36916010 | [illegible] |
| SE | 2603 | ENSG00000197961 | ZNF121 | chr19 | - | 9569001 | 9569082 | 9568094 | 9568175 | 9584460 | [illegible] |
| SE | 2604 | ENSG00000197961 | ZNF121 | chr19 | - | 9584185 | 9584348 | 9569001 | 9569082 | 9584460 | [illegible] |
| SE | 2608 | ENSG00000160570 | DEDD2 | chr19 | - | 42201988 | 42202072 | 42199792 | 42199829 | 42209699 | [illegible] |

[illegible]

| | | | | | | | | | | | | | | | |
|---|---|---|---|---|---|---|---|---|---|---|---|---|---|---|---|
| SE | 3023 | ENSG00000213096 | ZNF254 | chr19 | + | 24105939 | 24106066 | 24087183 | 24087337 | 24106547 | 24106643 | 26 | 23 | 22 | [illegible] |
| SE | 3024 | ENSG00000213096 | ZNF254 | chr19 | + | 24106547 | 24106643 | 24087159 | 24087337 | 24126253 | 24126320 | 30 | 17 | 6 | [illegible] |
| SE | 3025 | ENSG00000131845 | ZNF304 | chr19 | + | 57352654 | 57352735 | 57351325 | 57351697 | 57353724 | 57353851 | 4 | 0 | 0 | [illegible] |
| SE | 3026 | ENSG00000131845 | ZNF304 | chr19 | + | 57353724 | 57353851 | 57351270 | 57351697 | 57356029 | 57358780 | 21 | 11 | 16 | [illegible] |
| SE | 3028 | ENSG00000174917 | MICOS13 | chr19 | - | 5678774 | 5678902 | 5678420 | 5678648 | 5679344 | 5679396 | 0 | 0 | 1 | [illegible] |
| SE | 3029 | ENSG00000007255 | TRAPPC6A | chr19 | - | 45164163 | 45164247 | 45163915 | 45164009 | 45164852 | 45164970 | 92 | 20 | 93 | [illegible] |
| SE | 3032 | ENSG00000115268 | RPS15 | chr19 | + | 1438806 | 1438892 | 1438384 | 1438428 | 1440018 | 1440241 | 5638 | 2389 | 5837 | [illegible] |
| SE | 3033 | ENSG00000115268 | RPS15 | chr19 | + | 1438810 | 1438892 | 1438357 | 1438428 | 1440018 | 1440253 | 3403 | 1405 | 3319 | [illegible] |
| SE | 3034 | ENSG00000115268 | RPS15 | chr19 | + | 1439605 | 1439826 | 1438806 | 1438892 | 1440018 | 1440253 | 0 | 0 | 2 | [illegible] |
| SE | 3036 | ENSG00000115268 | RPS15 | chr19 | + | 1439715 | 1439826 | 1438806 | 1438892 | 1440018 | 1440233 | 0 | 0 | 1 | [illegible] |
| SE | 3043 | ENSG00000178078 | STAP2 | chr19 | - | 4326941 | 4327007 | 4325395 | 4325545 | 4327123 | 4327226 | 0 | 0 | 8 | [illegible] |
| SE | 3044 | ENSG00000178078 | STAP2 | chr19 | - | 4328674 | 4328809 | 4327315 | 4327385 | 4329960 | 4330061 | 0 | 1 | 2 | [illegible] |
| SE | 3046 | ENSG00000104973 | MED25 | chr19 | + | 49819171 | 49819296 | 49818570 | 49818616 | 49828448 | 49828547 | 27 | 7 | 33 | [illegible] |
| SE | 3047 | ENSG00000104973 | MED25 | chr19 | + | 49836225 | 49836406 | 49835726 | 49835945 | 49838579 | 49840383 | 11 | 5 | 4 | [illegible] |
| SE | 3050 | ENSG00000104960 | PTOV1 | chr19 | + | 49860267 | 49860336 | 49859985 | 49860183 | 49860613 | 49860725 | 119 | 90 | 127 | [illegible] |
| SE | 3054 | ENSG00000105655 | ISYNA1 | chr19 | - | 18436972 | 18437105 | 18436329 | 18436479 | 18437598 | 18437760 | 4 | 0 | 1 | [illegible] |
| SE | 3055 | ENSG00000105655 | ISYNA1 | chr19 | - | 18436972 | 18437105 | 18436683 | 18436877 | 18437859 | 18437988 | 2 | 0 | 3 | [illegible] |
| SE | 3068 | ENSG00000171049 | FPR2 | chr19 | + | 51762416 | 51762535 | 51761197 | 51761230 | 51768644 | 51769068 | 1 | 0 | 1 | [illegible] |
| SE | 3071 | ENSG00000105492 | SIGLEC6 | chr19 | - | 51527746 | 51527828 | 51517818 | 51520255 | 51529723 | 51529981 | 5 | 0 | 7 | [illegible] |
| SE | 3072 | ENSG00000105492 | SIGLEC6 | chr19 | - | 51528159 | 51528253 | 51519729 | 51520255 | 51529723 | 51529981 | 1 | 5 | 8 | [illegible] |
| SE | 3073 | ENSG00000105492 | SIGLEC6 | chr19 | - | 51528159 | 51528253 | 51527746 | 51527828 | 51528368 | 51528447 | 1 | 1 | 1 | [illegible] |
| SE | 3074 | ENSG00000105492 | SIGLEC6 | chr19 | - | 51528159 | 51528253 | 51527746 | 51527828 | 51529723 | 51529981 | 2 | 9 | 10 | [illegible] |
| SE | 3075 | ENSG00000105492 | SIGLEC6 | chr19 | - | 51529723 | 51529981 | 51519729 | 51520255 | 51530436 | 51530484 | 5 | 1 | 1 | [illegible] |
| SE | 3076 | ENSG00000105492 | SIGLEC6 | chr19 | - | 51529723 | 51529981 | 51519729 | 51520255 | 51530680 | 51530959 | 4 | 0 | 7 | [illegible] |
| SE | 3078 | ENSG00000105492 | SIGLEC6 | chr19 | - | 51530436 | 51530484 | 51529389 | 51529981 | 51530680 | 51530926 | 7 | 0 | 0 | [illegible] |
| SE | 3080 | ENSG00000105492 | SIGLEC6 | chr19 | - | 51530436 | 51530517 | 51529723 | 51529981 | 51530680 | 51530959 | 3 | 3 | 0 | [illegible] |
| SE | 3082 | ENSG00000248098 | BCKDHA | chr19 | + | 41414048 | 41414157 | 41410636 | 41411009 | 41419134 | 41419296 | 94 | 36 | 82 | [illegible] |
| SE | 3084 | ENSG00000248098 | BCKDHA | chr19 | + | 41419134 | 41419296 | 41414048 | 41414157 | 41422163 | 41422207 | 89 | 32 | 124 | [illegible] |
| SE | 3085 | ENSG00000248098 | BCKDHA | chr19 | + | 41422922 | 41423169 | 41422628 | 41422770 | 41424437 | 41425001 | 64 | 33 | 58 | [illegible] |
| SE | 3086 | ENSG00000248098 | BCKDHA | chr19 | + | 41422997 | 41423169 | 41422628 | 41422770 | 41424437 | 41425002 | 173 | 82 | 169 | [illegible] |
| SE | 3090 | ENSG00000187266 | EPOR | chr19 | - | 11382846 | 11382949 | 11381929 | 11382105 | 11383096 | 11383232 | 0 | 2 | 0 | [illegible] |
| SE | 3091 | ENSG00000187266 | EPOR | chr19 | - | 11382846 | 11382982 | 11381929 | 11382105 | 11383096 | 11383232 | 0 | 3 | 0 | [illegible] |
| SE | 3093 | ENSG00000167600 | CYP2S1 | chr19 | + | 41194543 | 41194709 | 41193218 | 41193441 | 41197778 | 41197928 | 6 | 10 | 3 | [illegible] |
| SE | 3094 | ENSG00000167600 | CYP2S1 | chr19 | + | 41194543 | 41194709 | 41193218 | 41193441 | 41201230 | 41201372 | 1 | 1 | 2 | [illegible] |
| SE | 3099 | ENSG00000167600 | CYP2S1 | chr19 | + | 41203449 | 41203637 | 41201230 | 41201372 | 41205957 | 41205975 | 11 | 19 | 6 | [illegible] |
| SE | 3100 | ENSG00000198182 | ZNF607 | chr19 | - | 37707913 | 37708009 | 37696372 | 37699895 | 37709695 | 37709822 | 12 | 12 | 18 | [illegible] |
| SE | 3102 | ENSG00000198182 | ZNF607 | chr19 | - | 37707913 | 37708012 | 37696370 | 37699895 | 37709695 | 37709822 | 13 | 4 | 9 | [illegible] |
| SE | 3107 | ENSG00000180479 | ZNF571 | chr19 | - | 37583970 | 37584097 | 37563823 | 37566291 | 37586667 | 37586745 | 22 | 16 | 32 | [illegible] |
| SE | 3108 | ENSG00000180479 | ZNF571 | chr19 | - | 37594219 | 37594319 | 37586667 | 37586745 | 37594740 | 37594755 | 0 | 2 | 4 | [illegible] |
| SE | 3112 | ENSG00000171817 | ZNF540 | chr19 | + | 37601006 | 37601105 | 37599625 | 37599752 | 37611512 | 37611577 | 5 | 1 | 2 | [illegible] |
| SE | 3113 | ENSG00000171817 | ZNF540 | chr19 | + | 37601009 | 37601105 | 37598375 | 37598456 | 37611512 | 37611690 | 4 | 0 | 2 | [illegible] |
| SE | 3114 | ENSG00000171817 | ZNF540 | chr19 | + | 37601009 | 37601105 | 37599625 | 37599752 | 37611512 | 37611690 | 6 | 3 | 6 | [illegible] |
| SE | 3123 | ENSG00000188227 | ZNF793 | chr19 | + | 37520183 | 37520312 | 37508263 | 37508403 | 37522531 | 37522647 | 7 | 6 | 2 | [illegible] |
| SE | 3124 | ENSG00000188227 | ZNF793 | chr19 | + | 37522531 | 37522647 | 37508264 | 37508403 | 37523409 | 37523454 | 6 | 0 | 9 | [illegible] |
| SE | 3125 | ENSG00000188227 | ZNF793 | chr19 | + | 37522531 | 37522647 | 37520183 | 37520312 | 37532355 | 37532482 | 4 | 1 | 1 | [illegible] |
| SE | 3126 | ENSG00000188227 | ZNF793 | chr19 | + | 37523409 | 37523454 | 37522531 | 37522647 | 37532355 | 37532482 | 14 | 3 | 12 | [illegible] |
| SE | 3128 | ENSG00000266916 | ZNF793-AS1 | chr19 | - | 37505845 | 37506040 | 37497158 | 37497592 | 37506838 | 37507087 | 0 | 2 | 0 | [illegible] |
| SE | 3129 | ENSG00000121417 | ZNF211 | chr19 | + | 57633712 | 57633905 | 57633215 | 57633436 | 57634022 | 57634061 | 15 | 14 | 7 | [illegible] |
| SE | 3130 | ENSG00000121417 | ZNF211 | chr19 | + | 57633712 | 57633905 | 57633215 | 57633436 | 57634628 | 57634755 | 0 | 5 | 3 | [illegible] |
| SE | 3131 | ENSG00000121417 | ZNF211 | chr19 | + | 57633785 | 57633905 | 57633180 | 57633436 | 57634022 | 57634061 | 20 | 10 | 8 | [illegible] |
| SE | 3132 | ENSG00000121417 | ZNF211 | chr19 | + | 57633785 | 57633905 | 57633180 | 57633436 | 57634628 | 57634755 | 2 | 1 | 2 | [illegible] |
| SE | 3133 | ENSG00000121417 | ZNF211 | chr19 | + | 57634022 | 57634061 | 57633167 | 57633436 | 57634628 | 57634755 | 5 | 9 | 5 | [illegible] |
| SE | 3135 | ENSG00000121417 | ZNF211 | chr19 | + | 57634022 | 57634061 | 57633712 | 57633905 | 57634628 | 57634755 | 19 | 12 | 12 | [illegible] |
| SE | 3136 | ENSG00000121417 | ZNF211 | chr19 | + | 57634628 | 57634755 | 57634022 | 57634061 | 57640703 | 57642226 | 62 | 29 | 48 | [illegible] |
| SE | 3140 | ENSG00000251369 | ZNF550 | chr19 | - | 57543650 | 57544583 | 57541839 | 57543243 | 57546456 | 57547993 | 22 | 10 | 44 | [illegible] |
| SE | 3141 | ENSG00000251369 | ZNF550 | chr19 | - | 57544236 | 57544583 | 57541839 | 57543243 | 57546456 | 57547993 | 19 | 6 | 38 | [illegible] |
| SE | 3142 | ENSG00000251369 | ZNF550 | chr19 | - | 57552626 | 57552722 | 57546456 | 57547993 | 57556230 | 57556357 | 13 | 21 | 30 | [illegible] |
| SE | 3143 | ENSG00000198538 | ZNF28 | chr19 | - | 52805144 | 52805262 | 52801309 | 52801702 | 52808006 | 52808133 | 18 | 14 | 19 | [illegible] |
| SE | 3144 | ENSG00000198538 | ZNF28 | chr19 | - | 52805144 | 52805262 | 52801309 | 52801702 | 52817943 | 52818031 | 8 | 6 | 13 | [illegible] |
| SE | 3145 | ENSG00000198538 | ZNF28 | chr19 | - | 52805144 | 52805262 | 52801456 | 52801660 | 52808006 | 52808133 | 10 | 7 | 8 | [illegible] |
| SE | 3146 | ENSG00000198538 | ZNF28 | chr19 | - | 52808006 | 52808133 | 52797407 | 52801702 | 52817943 | 52818031 | 33 | 26 | 15 | [illegible] |
| SE | 3149 | ENSG00000189190 | ZNF600 | chr19 | - | 52774574 | 52774701 | 52764549 | 52767772 | 52778825 | 52778907 | 304 | 188 | 79 | [illegible] |
| SE | 3150 | ENSG00000189190 | ZNF600 | chr19 | - | 52781316 | 52781464 | 52778825 | 52778907 | 52786594 | 52786724 | 52 | 15 | 0 | [illegible] |
| SE | 3151 | ENSG00000189190 | ZNF600 | chr19 | - | 52781316 | 52781464 | 52780827 | 52781055 | 52786594 | 52786724 | 95 | 38 | 3 | [illegible] |
| SE | 3152 | ENSG00000213020 | ZNF611 | chr19 | - | 52707391 | 52707538 | 52704731 | 52706864 | 52714014 | 52714141 | 4 | 1 | 0 | [illegible] |
| SE | 3153 | ENSG00000213020 | ZNF611 | chr19 | - | 52707420 | 52707542 | 52706653 | 52706864 | 52711206 | 52711304 | 8 | 10 | 7 | [illegible] |
| SE | 3155 | ENSG00000213020 | ZNF611 | chr19 | - | 52707420 | 52707542 | 52706653 | 52706864 | 52715831 | 52715913 | 5 | 8 | 7 | [illegible] |
| SE | 3156 | ENSG00000213020 | ZNF611 | chr19 | - | 52707468 | 52707542 | 52702812 | 52706864 | 52711206 | 52711304 | 12 | 17 | 9 | [illegible] |
| SE | 3158 | ENSG00000213020 | ZNF611 | chr19 | - | 52707468 | 52707542 | 52702812 | 52706864 | 52715831 | 52715913 | 7 | 13 | 8 | [illegible] |
| SE | 3159 | ENSG00000213020 | ZNF611 | chr19 | - | 52708176 | 52708326 | 52706621 | 52706864 | 52714014 | 52714141 | 0 | 0 | 2 | [illegible] |
| SE | 3160 | ENSG00000213020 | ZNF611 | chr19 | - | 52711206 | 52711304 | 52706655 | 52706864 | 52714014 | 52714141 | 5 | 4 | 1 | [illegible] |
| SE | 3161 | ENSG00000213020 | ZNF611 | chr19 | - | 52711206 | 52711304 | 52707420 | 52707542 | 52714014 | 52714141 | 8 | 7 | 2 | [illegible] |
| SE | 3163 | ENSG00000213020 | ZNF611 | chr19 | - | 52714014 | 52714141 | 52707420 | 52707542 | 52715831 | 52715913 | 67 | 57 | 78 | [illegible] |
| SE | 3166 | ENSG00000213020 | ZNF611 | chr19 | - | 52716110 | 52716233 | 52715831 | 52715913 | 52735000 | 52735038 | 6 | 2 | 0 | [illegible] |
| SE | 3167 | ENSG00000213020 | ZNF611 | chr19 | - | 52723870 | 52723989 | 52721152 | 52721267 | 52728729 | 52728831 | 0 | 5 | 0 | [illegible] |
| SE | 3168 | ENSG00000213020 | ZNF611 | chr19 | - | 52728729 | 52728831 | 52715831 | 52715913 | 52729905 | 52730005 | 62 | 32 | 60 | [illegible] |
| SE | 3170 | ENSG00000213020 | ZNF611 | chr19 | - | 52728729 | 52728831 | 52716110 | 52716233 | 52735000 | 52735034 | 2 | 1 | 2 | [illegible] |
| SE | 3171 | ENSG00000213020 | ZNF611 | chr19 | - | 52728923 | 52728944 | 52728729 | 52728831 | 52729905 | 52730005 | 0 | 3 | 0 | [illegible] |
| SE | 3172 | ENSG00000213020 | ZNF611 | chr19 | - | 52729905 | 52730005 | 52715831 | 52715913 | 52735000 | 52735011 | 0 | 1 | 0 | [illegible] |
| SE | 3173 | ENSG00000213020 | ZNF611 | chr19 | - | 52729905 | 52730005 | 52728729 | 52728831 | 52735000 | 52735011 | 33 | 17 | 35 | [illegible] |
| SE | 3174 | ENSG00000076662 | ICAM3 | chr19 | - | 10334527 | 10334782 | 10334159 | 10334408 | 10335670 | 10335773 | 894 | 368 | 798 | [illegible] |
| SE | 3180 | ENSG00000076662 | ICAM3 | chr19 | - | 10339017 | 10339245 | 10338681 | 10338863 | 10339538 | 10339624 | 3 | 1 | 3 | [illegible] |
| SE | 3182 | ENSG00000161847 | RAVER1 | chr19 | - | 10321047 | 10321259 | 10320654 | 10320951 | 10321530 | 10321618 | 33 | 6 | 19 | [illegible] |
| SE | 3186 | ENSG00000220201 | ZGLP1 | chr19 | - | 10305389 | 10305483 | 10304802 | 10305208 | 10305845 | 10305952 | 13 | 3 | 1 | [illegible] |
| SE | 3187 | ENSG00000220201 | ZGLP1 | chr19 | - | 10309348 | 10309448 | 10308184 | 10308550 | 10309556 | 10309783 | 0 | 0 | 3 | [illegible] |
| SE | 3191 | ENSG00000188033 | ZNF490 | chr19 | - | 12582849 | 12582910 | 12576099 | 12581724 | 12583429 | 12583556 | 66 | 43 | 42 | [illegible] |
| SE | 3192 | ENSG00000188033 | ZNF490 | chr19 | - | 12583429 | 12583556 | 12582849 | 12582910 | 12609157 | 12609202 | 45 | 36 | 43 | [illegible] |
| SE | 3194 | ENSG00000128000 | ZNF780B | chr19 | - | 40047374 | 40047470 | 40028259 | 40036626 | 40048669 | 40048796 | 61 | 63 | 70 | [illegible] |
| SE | 3195 | ENSG00000128000 | ZNF780B | chr19 | - | 40047405 | 40047470 | 40033578 | 40036626 | 40048669 | 40048796 | 29 | 32 | 41 | [illegible] |
| SE | 3196 | ENSG00000128000 | ZNF780B | chr19 | - | 40047405 | 40047473 | 40036530 | 40036626 | 40048669 | 40048796 | 12 | 12 | 15 | [illegible] |
| SE | 3197 | ENSG00000128000 | ZNF780B | chr19 | - | 40048917 | 40049056 | 40048669 | 40048796 | 40050323 | 40050377 | 0 | 4 | 1 | [illegible] |
| SE | 3198 | ENSG00000128000 | ZNF780B | chr19 | - | 40050323 | 40050377 | 40048669 | 40048796 | 40055415 | 40055576 | 53 | 48 | 40 | [illegible] |
| SE | 3199 | ENSG00000128000 | ZNF780B | chr19 | - | 40050323 | 40050377 | 40048669 | 40048796 | 40056188 | 40056231 | 44 | 43 | 47 | [illegible] |
| SE | 3200 | ENSG00000128000 | ZNF780B | chr19 | - | 40055415 | 40055576 | 40048669 | 40048796 | 40056188 | 40056231 | 3 | 1 | 4 | [illegible] |
| SE | 3203 | ENSG00000013275 | PSMC4 | chr19 | + | 39972145 | 39972244 | 39971164 | 39971238 | 39972368 | 39972555 | 116 | 37 | 81 | [illegible] |
| SE | 3204 | ENSG00000013275 | PSMC4 | chr19 | + | 39972145 | 39972244 | 39971164 | 39971238 | 39972461 | 39972555 | 19 | 12 | 23 | [illegible] |
| SE | 3205 | ENSG00000013275 | PSMC4 | chr19 | + | 39980069 | 39980146 | 39979816 | 39979984 | 39980285 | 39980454 | 182 | 92 | 147 | [illegible] |
| SE | 3206 | ENSG00000160007 | ARHGAP35 | chr19 | + | 46937263 | 46937408 | 46918487 | 46922356 | 46987988 | 46988066 | 147 | 118 | 182 | [illegible] |
| SE | 3208 | ENSG00000071655 | MBD3 | chr19 | - | 1581091 | 1581269 | 1577745 | 1578534 | 1582621 | 1582712 | 131 | 41 | 99 | [illegible] |
| SE | 3209 | ENSG00000071655 | MBD3 | chr19 | - | 1582621 | 1582712 | 1581091 | 1581269 | 1584539 | 1584677 | 203 | 67 | 149 | [illegible] |
| SE | 3210 | ENSG00000071655 | MBD3 | chr19 | - | 1583252 | 1583380 | 1582621 | 1582712 | 1584539 | 1584573 | 0 | 0 | 1 | [illegible] |
| SE | 3212 | ENSG00000071626 | DAZAP1 | chr19 | + | 1418665 | 1418731 | 1418203 | 1418370 | 1421150 | 1421258 | 216 | 154 | 183 | [illegible] |
| SE | 3213 | ENSG00000071626 | DAZAP1 | chr19 | + | 1422347 | 1422396 | 1421147 | 1421258 | 1425877 | 1425960 | 296 | 225 | 270 | [illegible] |
| SE | 3216 | ENSG00000071626 | DAZAP1 | chr19 | + | 1428841 | 1428995 | 1422347 | 1422396 | 1430221 | 1430362 | 53 | 26 | 64 | [illegible] |
| SE | 3218 | ENSG00000071626 | DAZAP1 | chr19 | + | 1429966 | 1429996 | 1426348 | 1428995 | 1430221 | 1430362 | 270 | 130 | 210 | [illegible] |
| SE | 3219 | ENSG00000071626 | DAZAP1 | chr19 | + | 1430221 | 1430362 | 1422347 | 1422396 | 1432513 | 1432690 | 70 | 24 | 71 | [illegible] |
| SE | 3220 | ENSG00000071626 | DAZAP1 | chr19 | + | 1433738 | 1433827 | 1432513 | 1432690 | 1434736 | 1435684 | 2 | 1 | 1 | [illegible] |
| SE | 3221 | ENSG00000071626 | DAZAP1 | chr19 | + | 1433738 | 1433827 | 1432513 | 1432690 | 1434739 | 1435687 | 1 | 4 | 1 | [illegible] |
| SE | 3222 | ENSG00000115286 | NDUFS7 | chr19 | + | 1386517 | 1386639 | 1383917 | 1383942 | 1387810 | 1387847 | 0 | 0 | 0 | [illegible] |
| SE | 3223 | ENSG00000115286 | NDUFS7 | chr19 | + | 1388524 | 1388593 | 1387738 | 1387847 | 1388832 | 1388938 | 47 | 12 | 23 | [illegible] |
| SE | 3224 | ENSG00000063245 | EPN1 | chr19 | + | 55677549 | 55677734 | 55676624 | 55677218 | 55678526 | 55678855 | 2 | 1 | 3 | [illegible] |
| SE | 3226 | ENSG00000063245 | EPN1 | chr19 | + | 55694725 | 55694983 | 55692950 | 55693037 | 55695147 | 55695441 | 64 | 22 | 55 | [illegible] |
| SE | 3228 | ENSG00000179954 | SSC5D | chr19 | + | 55497879 | 55498197 | 55494609 | 55494783 | 55499815 | 55500412 | 0 | 0 | 0 | [illegible] |
| SE | 3229 | ENSG00000197483 | ZNF628 | chr19 | + | 55479833 | 55479917 | 55476616 | 55476807 | 55481200 | 55484487 | 2 | 0 | 0 | [illegible] |
| SE | 3230 | ENSG00000063241 | ISOC2 | chr19 | - | 55455635 | 55455845 | 55455259 | 55455330 | 55456348 | 55456489 | 14 | 6 | 18 | [illegible] |
| SE | 3231 | ENSG00000063241 | ISOC2 | chr19 | - | 55455635 | 55455845 | 55455259 | 55455378 | 55456348 | 55456489 | 10 | 5 | 8 | [illegible] |
| SE | 3233 | ENSG00000108106 | UBE2S | chr19 | - | 55404060 | 55404147 | 55401578 | 55401762 | 55404287 | 55404478 | 1 | 0 | 0 | [illegible] |
| SE | 3235 | ENSG00000133247 | KMT5C | chr19 | + | 55342214 | 55342380 | 55341615 | 55342046 | 55342741 | 55342851 | 3 | 5 | 6 | [illegible] |
| SE | 3236 | ENSG00000133247 | KMT5C | chr19 | + | 55343977 | 55343997 | 55343679 | 55343843 | 55346212 | 55346349 | 3 | 2 | 7 | [illegible] |
| SE | 3240 | ENSG00000180061 | TMEM150B | chr19 | - | 55322647 | 55322743 | 55320968 | 55321093 | 55325271 | 55325305 | 1 | 10 | 0 | [illegible] |
| SE | 3244 | ENSG00000221946 | FXYD7 | chr19 | + | 35151439 | 35151482 | 35151253 | 35151328 | 35151632 | 35151673 | 2 | 4 | 13 | [illegible] |
| SE | 3247 | ENSG00000271109 | NA | chr19 | - | 34850668 | 34850794 | 34849045 | 34849549 | 34860533 | 34860715 | 4 | 2 | 5 | [illegible] |
| SE | 3248 | ENSG00000271109 | NA | chr19 | - | 34860173 | 34860302 | 34849045 | 34849549 | 34860533 | 34860715 | 12 | 5 | 39 | [illegible] |
| SE | 3249 | ENSG00000271109 | NA | chr19 | - | 34860173 | 34860302 | 34850668 | 34850794 | 34860533 | 34860715 | 18 | 12 | 18 | [illegible] |
| SE | 3250 | ENSG00000245680 | ZNF585B | chr19 | - | 37189660 | 37189753 | 37181578 | 37187244 | 37190023 | 37190150 | 42 | 17 | 29 | [illegible] |
| SE | 3251 | ENSG00000245680 | ZNF585B | chr19 | - | 37190023 | 37190150 | 37181578 | 37187244 | 37198938 | 37199008 | 14 | 3 | 8 | [illegible] |
| SE | 3252 | ENSG00000245680 | ZNF585B | chr19 | - | 37190023 | 37190150 | 37181578 | 37187244 | 37207039 | 37207254 | 14 | 9 | 19 | [illegible] |
| SE | 3253 | ENSG00000245680 | ZNF585B | chr19 | - | 37190023 | 37190150 | 37181578 | 37187244 | 37210458 | 37210505 | 0 | 0 | 3 | [illegible] |
| SE | 3254 | ENSG00000245680 | ZNF585B | chr19 | - | 37195880 | 37196019 | 37190023 | 37190150 | 37207039 | 37207254 | 0 | 0 | 0 | [illegible] |
| SE | 3255 | ENSG00000245680 | ZNF585B | chr19 | - | 37198938 | 37199054 | 37187079 | 37187244 | 37207039 | 37207254 | 5 | 7 | 1 | [illegible] |
| SE | 3257 | ENSG00000245680 | ZNF585B | chr19 | - | 37198938 | 37199054 | 37190023 | 37190150 | 37199399 | 37199497 | 17 | 3 | 8 | [illegible] |
| SE | 3258 | ENSG00000245680 | ZNF585B | chr19 | - | 37198938 | 37199054 | 37190023 | 37190150 | 37207039 | 37207254 | 19 | 9 | 8 | [illegible] |
| SE | 3259 | ENSG00000245680 | ZNF585B | chr19 | - | 37198938 | 37199054 | 37190023 | 37190150 | 37210458 | 37210505 | 12 | 3 | 7 | [illegible] |
| SE | 3260 | ENSG00000245680 | ZNF585B | chr19 | - | 37199399 | 37199497 | 37190072 | 37190150 | 37207039 | 37207254 | 4 | 0 | 2 | [illegible] |
| SE | 3261 | ENSG00000245680 | ZNF585B | chr19 | - | 37199399 | 37199497 | 37198938 | 37199054 | 37207039 | 37207254 | 8 | 0 | 3 | [illegible] |
| SE | 3262 | ENSG00000245680 | ZNF585B | chr19 | - | 37207039 | 37207254 | 37181578 | 37187244 | 37210458 | 37210531 | 17 | 6 | 7 | [illegible] |
| SE | 3263 | ENSG00000245680 | ZNF585B | chr19 | - | 37207039 | 37207254 | 37190023 | 37190150 | 37210440 | 37210484 | 27 | 10 | 21 | [illegible] |
| SE | 3264 | ENSG00000245680 | ZNF585B | chr19 | - | 37207039 | 37207254 | 37190023 | 37190150 | 37210458 | 37210531 | 31 | 15 | 24 | [illegible] |
| SE | 3265 | ENSG00000245680 | ZNF585B | chr19 | - | 37207039 | 37207254 | 37198938 | 37199054 | 37210458 | 37210531 | 24 | 12 | 7 | [illegible] |
| SE | 3267 | ENSG00000196967 | ZNF585A | chr19 | - | 37150834 | 37150879 | 37115035 | 37115165 | 37155864 | 37155957 | 0 | 0 | 0 | [illegible] |
| SE | 3268 | ENSG00000196967 | ZNF585A | chr19 | - | 37165594 | 37165696 | 37156228 | 37156355 | 37169838 | 37170054 | 35 | 9 | 17 | [illegible] |
| SE | 3279 | ENSG00000087076 | HSD17B14 | chr19 | - | 48813432 | 48813552 | 48813017 | 48813348 | 48813662 | 48813730 | 2 | 0 | 3 | [illegible] |
| SE | 3280 | ENSG00000087076 | HSD17B14 | chr19 | - | 48813455 | 48813552 | 48813017 | 48813348 | 48813662 | 48813730 | 3 | 0 | 4 | [illegible] |
| SE | 3281 | ENSG00000087076 | HSD17B14 | chr19 | - | 48831667 | 48831759 | 48815036 | 48815141 | 48832665 | 48832732 | 0 | 0 | 2 | [illegible] |
| SE | 3282 | ENSG00000105552 | BCAT2 | chr19 | - | 48796146 | 48796329 | 48795099 | 48795464 | 48796427 | 48796502 | 0 | 0 | 2 | [illegible] |
| SE | 3284 | ENSG00000105552 | BCAT2 | chr19 | - | 48806999 | 48807074 | 48800186 | 48800297 | 48810983 | 48811027 | 0 | 0 | 0 | [illegible] |
| SE | 3286 | ENSG00000105552 | BCAT2 | chr19 | - | 48807755 | 48808236 | 48806999 | 48807074 | 48810983 | 48811027 | 1 | 0 | 1 | [illegible] |
| SE | 3300 | ENSG00000127445 | PIN1 | chr19 | + | 9838011 | 9838075 | 9835345 | 9835402 | 9838435 | 9838648 | 0 | 2 | 0 | [illegible] |
| SE | 3303 | ENSG00000076928 | ARHGEF1 | chr19 | + | 41896376 | 41896482 | 41895348 | 41895486 | 41897314 | 41897326 | 531 | 288 | 523 | [illegible] |
| SE | 3308 | ENSG00000076928 | ARHGEF1 | chr19 | + | 41900835 | 41900933 | 41898423 | 41898587 | 41901886 | 41902033 | 0 | 0 | 1 | [illegible] |
| SE | 3309 | ENSG00000076928 | ARHGEF1 | chr19 | + | 41902783 | 41902898 | 41902532 | 41902658 | 41903306 | 41903407 | 1025 | 415 | 785 | [illegible] |
| SE | 3310 | ENSG00000076928 | ARHGEF1 | chr19 | + | 41903655 | 41903784 | 41903306 | 41903407 | 41904034 | 41904110 | 91 | 49 | 77 | [illegible] |
| SE | 3311 | ENSG00000076928 | ARHGEF1 | chr19 | + | 41903706 | 41903784 | 41903306 | 41903407 | 41904034 | 41904110 | 519 | 298 | 537 | [illegible] |
| SE | 3313 | ENSG00000076928 | ARHGEF1 | chr19 | + | 41905174 | 41905261 | 41904948 | 41905036 | 41905759 | 41905827 | 656 | 279 | 534 | [illegible] |
| SE | 3315 | ENSG00000076928 | ARHGEF1 | chr19 | + | 41906702 | 41906803 | 41905938 | 41906025 | 41907104 | 41907448 | 225 | 101 | 265 | [illegible] |
| SE | 3321 | ENSG00000198521 | ZNF43 | chr19 | - | 21819094 | 21819221 | 21817887 | 21817986 | 21836035 | 21836182 | 60 | 28 | 64 | [illegible] |
| SE | 3322 | ENSG00000198521 | ZNF43 | chr19 | - | 21819094 | 21819221 | 21817887 | 21817986 | 21843144 | 21843326 | 31 | 28 | 53 | [illegible] |
| SE | 3323 | ENSG00000198521 | ZNF43 | chr19 | - | 21819094 | 21819221 | 21817887 | 21817986 | 21845391 | 21845506 | 29 | 24 | 50 | [illegible] |
| SE | 3325 | ENSG00000198521 | ZNF43 | chr19 | - | 21843144 | 21843326 | 21817887 | 21817986 | 21845391 | 21845506 | 5 | 9 | 1 | [illegible] |
| SE | 3327 | ENSG00000198521 | ZNF43 | chr19 | - | 21843144 | 21843326 | 21819094 | 21819221 | 21845391 | 21845506 | 7 | 13 | 4 | [illegible] |
| SE | 3328 | ENSG00000198521 | ZNF43 | chr19 | - | 21843144 | 21843326 | 21819094 | 21819221 | 21845429 | 21845506 | 2 | 5 | 5 | [illegible] |
| SE | 3330 | ENSG00000198521 | ZNF43 | chr19 | - | 21845391 | 21845506 | 21843144 | 21843326 | 21850937 | 21851027 | 9 | 10 | 5 | [illegible] |
| SE | 3331 | ENSG00000198521 | ZNF43 | chr19 | - | 21845429 | 21845506 | 21819094 | 21819221 | 21851904 | 21851968 | 0 | 2 | 1 | [illegible] |
| SE | 3332 | ENSG00000198521 | ZNF43 | chr19 | - | 21845429 | 21845506 | 21843144 | 21843326 | 21851904 | 21851968 | 0 | 2 | 1 | [illegible] |
| SE | 3334 | ENSG00000198521 | ZNF43 | chr19 | - | 21850937 | 21851027 | 21843144 | 21843326 | 21851904 | 21852017 | 0 | 0 | 1 | [illegible] |
| SE | 3335 | ENSG00000198521 | ZNF43 | chr19 | - | 21850937 | 21851027 | 21845391 | 21845506 | 21851904 | 21852017 | 3 | 2 | 5 | [illegible] |
| SE | 3337 | ENSG00000198521 | ZNF43 | chr19 | - | 21850937 | 21851162 | 21845391 | 21845506 | 21851904 | 21852028 | 3 | 3 | 6 | [illegible] |
| SE | 3338 | ENSG00000198521 | ZNF43 | chr19 | - | 21851162 | 21851360 | 21845391 | 21845506 | 21851904 | 21852029 | 1 | 0 | 1 | [illegible] |
| SE | 3341 | ENSG00000012061 | ERCC1 | chr19 | - | 45413962 | 45414034 | 45413433 | 45413745 | 45416820 | 45416897 | 67 | 35 | 65 | [illegible] |
| SE | 3342 | ENSG00000012061 | ERCC1 | chr19 | - | 45414860 | 45414960 | 45413435 | 45413745 | 45416820 | 45416859 | 68 | 49 | 60 | [illegible] |
| SE | 3343 | ENSG00000012061 | ERCC1 | chr19 | - | 45414860 | 45414960 | 45413962 | 45414034 | 45416820 | 45416859 | 110 | 79 | 105 | [illegible] |
| SE | 3344 | ENSG00000012061 | ERCC1 | chr19 | - | 45416820 | 45416897 | 45409618 | 45409725 | 45419097 | 45419186 | 23 | 15 | 20 | [illegible] |
| SE | 3345 | ENSG00000012061 | ERCC1 | chr19 | - | 45419097 | 45419197 | 45416490 | 45416897 | 45420323 | 45420427 | 128 | 76 | 110 | [illegible] |
| SE | 3346 | ENSG00000012061 | ERCC1 | chr19 | - | 45419097 | 45419197 | 45416490 | 45416897 | 45421177 | 45421393 | 64 | 44 | 54 | [illegible] |
| SE | 3347 | ENSG00000012061 | ERCC1 | chr19 | - | 45419410 | 45419489 | 45419097 | 45419197 | 45420323 | 45420382 | 1 | 0 | 0 | [illegible] |
| SE | 3348 | ENSG00000012061 | ERCC1 | chr19 | - | 45420323 | 45420427 | 45416490 | 45416897 | 45421177 | 45421393 | 39 | 27 | 46 | [illegible] |
| SE | 3349 | ENSG00000012061 | ERCC1 | chr19 | - | 45420323 | 45420427 | 45419097 | 45419197 | 45421177 | 45421393 | 101 | 55 | 101 | [illegible] |
| SE | 3351 | ENSG00000012061 | ERCC1 | chr19 | - | 45423269 | 45423381 | 45421177 | 45421393 | 45449172 | 45449244 | 47 | 32 | 44 | [illegible] |
| SE | 3352 | ENSG00000161677 | JOSD2 | chr19 | - | 50507573 | 50507699 | 50506377 | 50506572 | 50510285 | 50510445 | 24 | 3 | 19 | [illegible] |
| SE | 3353 | ENSG00000187912 | CLEC17A | chr19 | + | 14595273 | 14595315 | 14594758 | 14594800 | 14596875 | 14597013 | 23 | 3 | 20 | [illegible] |
| SE | 3354 | ENSG00000187912 | CLEC17A | chr19 | + | 14600030 | 14600182 | 14599653 | 14599812 | 14606992 | 14607102 | 36 | 15 | 33 | [illegible] |
| SE | 3357 | ENSG00000130164 | LDLR | chr19 | + | 11100222 | 11100345 | 11089461 | 11089615 | 11102663 | 11102786 | 71 | 28 | 37 | [illegible] |
| SE | 3363 | ENSG00000130164 | LDLR | chr19 | + | 11116858 | 11116998 | 11116093 | 11116212 | 11123173 | 11123344 | 43 | 5 | 29 | [illegible] |
| SE | 3364 | ENSG00000130164 | LDLR | chr19 | + | 11120091 | 11120233 | 11111513 | 11111639 | 11123173 | 11123344 | 0 | 0 | 0 | [illegible] |
| SE | 3365 | ENSG00000130164 | LDLR | chr19 | + | 11120091 | 11120233 | 11116093 | 11116212 | 11123173 | 11123344 | 0 | 0 | 2 | [illegible] |
| SE | 3368 | ENSG00000130164 | LDLR | chr19 | + | 11123173 | 11123344 | 11120369 | 11120522 | 11128007 | 11128085 | 94 | 36 | 86 | [illegible] |
| SE | 3370 | ENSG00000232677 | LINC00665 | chr19 | - | 36316177 | 36316257 | 36316038 | 36316070 | 36321299 | 36321357 | 2 | 0 | 0 | [illegible] |
| SE | 3371 | ENSG00000232677 | LINC00665 | chr19 | - | 36316177 | 36316348 | 36315173 | 36316070 | 36321299 | 36321357 | 2 | 0 | 0 | [illegible] |
| SE | 3372 | ENSG00000232677 | LINC00665 | chr19 | - | 36321992 | 36322030 | 36321029 | 36321357 | 36330252 | 36330492 | 3 | 0 | 0 | [illegible] |
| SE | 3373 | ENSG00000232677 | LINC00665 | chr19 | - | 36322129 | 36322222 | 36321992 | 36322030 | 36330252 | 36330492 | 7 | 1 | 2 | [illegible] |
| SE | 3374 | ENSG00000232677 | LINC00665 | chr19 | - | 36322129 | 36322225 | 36321992 | 36322030 | 36330252 | 36330492 | 6 | 1 | 2 | [illegible] |
| SE | 3375 | ENSG00000232677 | LINC00665 | chr19 | - | 36322129 | 36322384 | 36321992 | 36322030 | 36330252 | 36330492 | 6 | 1 | 2 | [illegible] |
| SE | 3376 | ENSG00000232677 | LINC00665 | chr19 | - | 36326052 | 36327225 | 36324925 | 36325764 | 36330252 | 36330492 | 4 | 0 | 0 | [illegible] |
| SE | 3377 | ENSG00000232677 | LINC00665 | chr19 | - | 36326289 | 36327225 | 36324923 | 36325764 | 36330252 | 36330492 | 3 | 0 | 0 | [illegible] |
| SE | 3378 | ENSG00000232677 | LINC00665 | chr19 | - | 36326481 | 36327225 | 36324925 | 36325764 | 36330252 | 36330492 | 3 | 0 | 0 | [illegible] |
| SE | 3381 | ENSG00000254004 | ZNF260 | chr19 | - | 36525154 | 36525373 | 36510686 | 36515699 | 36528218 | 36528268 | 32 | 17 | 19 | [illegible] |
| SE | 3382 | ENSG00000105518 | TMEM205 | chr19 | - | 11345251 | 11345415 | 11342775 | 11343120 | 11345519 | 11345628 | 77 | 17 | 34 | [illegible] |
| SE | 3387 | ENSG00000181007 | ZFP82 | chr19 | - | 36409780 | 36409867 | 36407886 | 36408013 | 36418491 | 36418644 | 20 | 7 | 8 | [illegible] |
| SE | 3388 | ENSG00000142065 | ZFP14 | chr19 | - | 36362111 | 36362238 | 36360434 | 36360533 | 36367883 | 36367971 | 85 | 60 | 57 | [illegible] |
| SE | 3393 | ENSG00000177025 | C19orf18 | chr19 | - | 57961390 | 57961551 | 57958436 | 57958717 | 57972462 | 57972504 | 0 | 3 | 0 | [illegible] |
| SE | 3394 | ENSG00000177025 | C19orf18 | chr19 | - | 57966529 | 57966632 | 57961390 | 57961551 | 57972462 | 57972504 | 1 | 4 | 6 | [illegible] |
| SE | 3395 | ENSG00000177025 | C19orf18 | chr19 | - | 57972462 | 57972504 | 57966529 | 57966632 | 57974098 | 57974203 | 0 | 3 | 2 | [illegible] |
| SE | 3397 | ENSG00000173480 | ZNF417 | chr19 | - | 57912059 | 57912186 | 57908479 | 57910114 | 57916378 | 57916591 | 17 | 10 | 25 | [illegible] |
| SE | 3398 | ENSG00000173480 | ZNF417 | chr19 | - | 57912059 | 57912189 | 57905773 | 57910114 | 57916378 | 57916592 | 29 | 8 | 20 | [illegible] |
| SE | 3399 | ENSG00000173480 | ZNF417 | chr19 | - | 57915337 | 57915531 | 57912059 | 57912186 | 57916378 | 57916577 | 0 | 0 | 1 | [illegible] |
| SE | 3400 | ENSG00000173480 | ZNF417 | chr19 | - | 57915337 | 57915531 | 57912059 | 57912189 | 57916378 | 57916577 | 0 | 0 | 0 | [illegible] |
| SE | 3401 | ENSG00000173480 | ZNF417 | chr19 | - | 57915564 | 57915665 | 57912059 | 57912186 | 57916378 | 57916592 | 7 | 0 | 0 | [illegible] |
| SE | 3402 | ENSG00000173480 | ZNF417 | chr19 | - | 57915564 | 57915665 | 57912059 | 57912189 | 57916378 | 57916592 | 3 | 0 | 0 | [illegible] |
| SE | 3405 | ENSG00000204514 | ZNF814 | chr19 | - | 57875155 | 57875226 | 57868457 | 57868579 | 57876915 | 57877042 | 2 | 2 | 6 | [illegible] |
| SE | 3406 | ENSG00000204514 | ZNF814 | chr19 | - | 57876915 | 57877042 | 57850314 | 57850561 | 57888766 | 57888780 | 7 | 19 | 14 | [illegible] |
| SE | 3410 | ENSG00000267106 | ZNF561-AS1 | chr19 | + | 9626783 | 9626860 | 9621486 | 9621645 | 9632227 | 9632417 | 5 | 0 | 4 | [illegible] |
| SE | 3411 | ENSG00000267106 | ZNF561-AS1 | chr19 | + | 9626783 | 9627806 | 9621266 | 9621350 | 9632227 | 9632417 | 1 | 3 | 1 | [illegible] |
| SE | 3412 | ENSG00000267106 | ZNF561-AS1 | chr19 | + | 9626783 | 9627806 | 9621511 | 9621645 | 9632227 | 9632417 | 3 | 0 | 2 | [illegible] |
| SE | 3413 | ENSG00000267106 | ZNF561-AS1 | chr19 | + | 9627715 | 9627806 | 9621248 | 9621350 | 9632227 | 9632417 | 2 | 0 | 1 | [illegible] |
| SE | 3414 | ENSG00000267106 | ZNF561-AS1 | chr19 | + | 9627715 | 9627806 | 9621502 | 9621645 | 9632227 | 9632417 | 1 | 0 | 1 | [illegible] |
| SE | 3415 | ENSG00000267106 | ZNF561-AS1 | chr19 | + | 9627715 | 9627806 | 9626783 | 9626860 | 9632227 | 9632417 | 1 | 0 | 2 | [illegible] |
| SE | 3416 | ENSG00000127452 | FBXL12 | chr19 | - | 9813230 | 9813274 | 9810266 | 9811717 | 9814137 | 9814342 | 0 | 4 | 3 | [illegible] |
| SE | 3418 | ENSG00000127452 | FBXL12 | chr19 | - | 9813230 | 9813274 | 9810266 | 9811717 | 9818727 | 9819079 | 1 | 2 | 3 | [illegible] |
| SE | 3419 | ENSG00000127452 | FBXL12 | chr19 | - | 9814137 | 9814339 | 9811155 | 9811717 | 9818544 | 9818813 | 1 | 0 | 0 | [illegible] |
| SE | 3420 | ENSG00000127452 | FBXL12 | chr19 | - | 9814137 | 9814342 | 9811307 | 9811717 | 9818544 | 9818617 | 1 | 0 | 0 | [illegible] |
| SE | 3423 | ENSG00000127452 | FBXL12 | chr19 | - | 9818544 | 9818617 | 9810269 | 9811717 | 9818727 | 9818816 | 18 | 5 | 9 | [illegible] |
| SE | 3424 | ENSG00000127452 | FBXL12 | chr19 | - | 9818544 | 9818617 | 9810269 | 9811717 | 9818753 | 9818785 | 12 | 2 | 13 | [illegible] |
| SE | 3425 | ENSG00000127452 | FBXL12 | chr19 | - | 9818544 | 9818617 | 9813230 | 9813274 | 9818727 | 9818816 | 10 | 5 | 10 | [illegible] |
| SE | 3426 | ENSG00000130818 | ZNF426 | chr19 | - | 9530984 | 9531067 | 9523222 | 9529636 | 9532844 | 9532925 | 37 | 48 | 43 | [illegible] |
| SE | 3427 | ENSG00000130818 | ZNF426 | chr19 | - | 9533839 | 9533966 | 9532844 | 9532925 | 9535187 | 9535279 | 45 | 40 | 72 | [illegible] |
| SE | 3428 | ENSG00000130818 | ZNF426 | chr19 | - | 9535187 | 9535279 | 9533839 | 9533966 | 9536207 | 9536356 | 47 | 44 | 83 | [illegible] |
| SE | 3429 | ENSG00000177045 | SIX5 | chr19 | - | 45766349 | 45767155 | 45764784 | 45766111 | 45768041 | 45769252 | 0 | 1 | 1 | [illegible] |
| SE | 3430 | ENSG00000125755 | SYMPK | chr19 | - | 45827836 | 45827918 | 45827509 | 45827623 | 45828969 | 45829205 | 94 | 30 | 85 | [illegible] |
| SE | 3431 | ENSG00000125755 | SYMPK | chr19 | - | 45838460 | 45838615 | 45835077 | 45835228 | 45842249 | 45842489 | 139 | 108 | 193 | [illegible] |
| SE | 3432 | ENSG00000125755 | SYMPK | chr19 | - | 45842249 | 45842489 | 45838460 | 45838615 | 45844029 | 45844200 | 131 | 73 | 138 | [illegible] |
| SE | 3433 | ENSG00000125755 | SYMPK | chr19 | - | 45854174 | 45854507 | 45852481 | 45852535 | 45863057 | 45863142 | 72 | 36 | 31 | [illegible] |
| SE | 3435 | ENSG00000125731 | SH2D3A | chr19 | - | 6759593 | 6759670 | 6754830 | 6755315 | 6760637 | 6760987 | 13 | 25 | 43 | [illegible] |
| SE | 3436 | ENSG00000125731 | SH2D3A | chr19 | - | 6759593 | 6759670 | 6754830 | 6755315 | 6763679 | 6763816 | 4 | 12 | 11 | [illegible] |

| | | | | | | | | | | | | | | | | | | | | | | | | | |
|---|---|---|---|---|---|---|---|---|---|---|---|---|---|---|---|---|---|---|---|---|---|---|---|---|---|
| SE | 3437 | ENSG00000125731 | SH2D3A | chr19 | - | 6760637 | 6760987 | 6754830 | 6755315 | 6763679 | 6763816 | 8 | 2 | 28 | 9 | 12 | 2 | 0 | 0 | 0 | 0 | 8 | 9 | 1 | 0 |
| SE | 3438 | ENSG00000125731 | SH2D3A | chr19 | - | 6760637 | 6760987 | 6757329 | 6759670 | 6763679 | 6763816 | 20 | 15 | 62 | 38 | 19 | 1 | 0 | 1 | 1 | 0 | 28 | 26 | 17 | 11 |
| SE | 3439 | ENSG00000125731 | SH2D3A | chr19 | - | 6763679 | 6763816 | 6760637 | 6760987 | 6767386 | 6767425 | 16 | 4 | 34 | 15 | 14 | 0 | 0 | 0 | 2 | 0 | 8 | 16 | 3 | 0 |
| SE | 3444 | ENSG00000125734 | GPR108 | chr19 | - | 6735621 | 6735704 | 6734182 | 6734307 | 6736591 | 6736711 | 64 | 20 | 44 | 35 | 51 | 0 | 0 | 1 | 0 | 0 | 34 | 74 | 42 | 27 |
| SE | 3445 | ENSG00000125734 | GPR108 | chr19 | - | 6735907 | 6735958 | 6734182 | 6734307 | 6736591 | 6736711 | 25 | 14 | 10 | 6 | 3 | 0 | 0 | 1 | 0 | 0 | 4 | 15 | 13 | 0 |
| SE | 3447 | ENSG00000205744 | DENND1C | chr19 | - | 6471264 | 6471305 | 6470294 | 6470366 | 6471405 | 6471496 | 81 | 31 | 50 | 97 | 44 | 0 | 0 | 2 | 1 | 0 | 139 | 122 | 67 | 102 |
| SE | 3450 | ENSG00000205744 | DENND1C | chr19 | - | 6479986 | 6479990 | 6479858 | 6479902 | 6481678 | 6481774 | 1 | 0 | 0 | 1 | 3 | 0 | 0 | 1 | 0 | 0 | 0 | 0 | 0 | 1 |
| SE | 3451 | ENSG00000205744 | DENND1C | chr19 | - | 6479986 | 6480051 | 6479858 | 6479902 | 6481678 | 6481787 | 40 | 30 | 36 | 49 | 41 | 0 | 0 | 1 | 0 | 0 | 41 | 115 | 44 | 61 |
| SE | 3453 | ENSG00000230510 | PPP5D1 | chr19 | - | 46576598 | 46576717 | 46526924 | 46527072 | 46600882 | 46601200 | 1 | 0 | 0 | 1 | 0 | 0 | 0 | 0 | 0 | 0 | 0 | 0 | 0 | 0 |
| SE | 3454 | ENSG00000230510 | PPP5D1 | chr19 | - | 46589145 | 46589235 | 46526924 | 46527072 | 46600882 | 46601200 | 1 | 0 | 0 | 0 | 0 | 0 | 0 | 0 | 0 | 0 | 0 | 0 | 0 | 0 |
| SE | 3460 | ENSG00000105197 | TIMM50 | chr19 | + | 39482884 | 39482963 | 39481882 | 39482033 | 39483134 | 39483156 | 5 | 0 | 3 | 8 | 2 | 0 | 0 | 0 | 0 | 0 | 9 | 0 | 1 | 0 |
| SE | 3464 | ENSG00000105197 | TIMM50 | chr19 | + | 39485543 | 39485602 | 39483134 | 39483156 | 39485687 | 39485807 | 39 | 10 | 55 | 39 | 57 | 1 | 0 | 0 | 0 | 0 | 74 | 40 | 41 | 28 |
| SE | 3466 | ENSG00000105193 | RPS16 | chr19 | - | 39433493 | 39433569 | 39433206 | 39433418 | 39433664 | 39433761 | 3290 | 1118 | 3999 | 3068 | 2611 | 1 | 0 | 0 | 1 | 2 | 1183 | 2108 | 1416 | 960 |
| SE | 3467 | ENSG00000105193 | RPS16 | chr19 | - | 39433517 | 39433569 | 39433259 | 39433418 | 39433664 | 39433761 | 3290 | 1112 | 4000 | 3066 | 2604 | 1 | 0 | 0 | 1 | 2 | 1183 | 2109 | 1411 | 960 |
| SE | 3468 | ENSG00000105193 | RPS16 | chr19 | - | 39433521 | 39433569 | 39433136 | 39433418 | 39433664 | 39433761 | 7114 | 2524 | 8391 | 6725 | 5749 | 1 | 0 | 0 | 1 | 2 | 2615 | 4322 | 3137 | 2200 |
| SE | 3469 | ENSG00000090924 | PLEKHG2 | chr19 | + | 39420750 | 39420852 | 39420625 | 39420659 | 39421074 | 39421113 | 30 | 7 | 17 | 13 | 16 | 0 | 0 | 0 | 2 | 0 | 76 | 17 | 25 | 31 |
| SE | 3477 | ENSG00000063322 | MED29 | chr19 | + | 39392463 | 39392522 | 39391322 | 39391638 | 39393552 | 39393637 | 128 | 69 | 99 | 102 | 189 | 0 | 0 | 2 | 0 | 1 | 85 | 104 | 83 | 118 |
| SE | 3480 | ENSG00000006712 | PAF1 | chr19 | - | 39388338 | 39388467 | 39386693 | 39386799 | 39388559 | 39388676 | 269 | 170 | 267 | 294 | 369 | 0 | 0 | 0 | 0 | 0 | 554 | 335 | 326 | 343 |
| SE | 3481 | ENSG00000006712 | PAF1 | chr19 | - | 39388559 | 39388676 | 39386693 | 39386799 | 39388761 | 39388865 | 108 | 84 | 91 | 112 | 74 | 0 | 0 | 0 | 0 | 0 | 191 | 172 | 78 | 124 |
| SE | 3484 | ENSG00000179134 | SAMD4B | chr19 | + | 39348287 | 39348389 | 39342467 | 39342576 | 39354005 | 39354066 | 1 | 1 | 0 | 3 | 0 | 5 | 0 | 3 | 1 | 2 | 0 | 0 | 0 | 0 |
| SE | 3485 | ENSG00000179134 | SAMD4B | chr19 | + | 39351816 | 39352374 | 39348287 | 39348389 | 39354005 | 39354066 | 1 | 6 | 1 | 1 | 3 | 0 | 1 | 0 | 2 | 0 | 0 | 0 | 0 | 9 |
| SE | 3486 | ENSG00000179134 | SAMD4B | chr19 | + | 39354005 | 39354066 | 39342395 | 39342576 | 39356688 | 39357089 | 7 | 1 | 7 | 8 | 5 | 3 | 0 | 0 | 0 | 0 | 0 | 2 | 5 | 7 |
| SE | 3488 | ENSG00000179134 | SAMD4B | chr19 | + | 39379965 | 39380084 | 39378503 | 39378589 | 39380586 | 39380755 | 72 | 26 | 45 | 47 | 35 | 1 | 0 | 0 | 0 | 0 | 23 | 47 | 44 | 55 |
| SE | 3492 | ENSG00000130755 | GMFG | chr19 | - | 39329543 | 39329626 | 39328358 | 39329073 | 39335260 | 39335310 | 334 | 183 | 280 | 349 | 494 | 2 | 0 | 0 | 0 | 0 | 371 | 386 | 249 | 460 |
| SE | 3493 | ENSG00000130755 | GMFG | chr19 | - | 39329543 | 39329626 | 39328358 | 39329073 | 39335973 | 39336037 | 332 | 183 | 270 | 342 | 493 | 0 | 1 | 2 | 0 | 0 | 371 | 382 | 241 | 442 |
| SE | 3496 | ENSG00000130755 | GMFG | chr19 | - | 39333076 | 39333126 | 39328999 | 39329073 | 39335260 | 39335310 | 450 | 229 | 427 | 416 | 449 | 2 | 0 | 0 | 0 | 0 | 436 | 418 | 391 | 486 |
| SE | 3497 | ENSG00000130755 | GMFG | chr19 | - | 39333076 | 39333126 | 39328999 | 39329073 | 39335973 | 39336005 | 32 | 3 | 39 | 6 | 16 | 0 | 1 | 2 | 0 | 0 | 12 | 3 | 3 | 15 |
| SE | 3500 | ENSG00000130755 | GMFG | chr19 | - | 39333076 | 39333126 | 39329543 | 39329626 | 39335973 | 39336005 | 482 | 355 | 529 | 532 | 518 | 0 | 0 | 0 | 0 | 0 | 795 | 606 | 482 | 748 |
| SE | 3503 | ENSG00000130755 | GMFG | chr19 | - | 39335260 | 39335310 | 39329543 | 39329626 | 39335973 | 39336017 | 1 | 0 | 5 | 2 | 0 | 0 | 0 | 0 | 0 | 0 | 0 | 1 | 0 | 14 |
| SE | 3506 | ENSG00000130755 | GMFG | chr19 | - | 39335434 | 39335531 | 39329543 | 39329626 | 39335973 | 39336005 | 187 | 104 | 184 | 206 | 142 | 0 | 0 | 0 | 0 | 0 | 288 | 227 | 172 | 179 |
| SE | 3513 | ENSG00000130669 | PAK4 | chr19 | + | 39168229 | 39168312 | 39125797 | 39125919 | 39169531 | 39169757 | 4 | 2 | 12 | 13 | 4 | 3 | 4 | 1 | 2 | 0 | 13 | 6 | 5 | 0 |
| SE | 3514 | ENSG00000130669 | PAK4 | chr19 | + | 39168624 | 39168840 | 39168229 | 39168302 | 39169531 | 39169757 | 0 | 0 | 0 | 2 | 0 | 4 | 0 | 11 | 3 | 1 | 0 | 0 | 0 | 0 |
| SE | 3515 | ENSG00000130669 | PAK4 | chr19 | + | 39168624 | 39168840 | 39168229 | 39168312 | 39169531 | 39169757 | 0 | 0 | 0 | 1 | 0 | 0 | 1 | 3 | 4 | 0 | 0 | 0 | 0 | 0 |
| SE | 3516 | ENSG00000130669 | PAK4 | chr19 | + | 39172917 | 39173106 | 39169531 | 39169757 | 39173575 | 39174010 | 2 | 3 | 0 | 4 | 0 | 0 | 0 | 0 | 3 | 0 | 0 | 6 | 2 | 14 |
| SE | 3517 | ENSG00000130669 | PAK4 | chr19 | + | 39172917 | 39173376 | 39169531 | 39169757 | 39173575 | 39174010 | 6 | 3 | 2 | 8 | 0 | 0 | 0 | 0 | 3 | 0 | 0 | 7 | 2 | 14 |
| SE | 3518 | ENSG00000130669 | PAK4 | chr19 | + | 39174930 | 39175064 | 39173575 | 39174010 | 39175311 | 39175438 | 21 | 7 | 21 | 24 | 22 | 0 | 0 | 0 | 0 | 0 | 10 | 12 | 20 | 17 |
| SE | 3523 | ENSG00000105483 | CARD8 | chr19 | - | 48186549 | 48186765 | 48180769 | 48183092 | 48190495 | 48190600 | 0 | 0 | 2 | 0 | 0 | 1 | 0 | 0 | 0 | 0 | 0 | 0 | 3 | 0 |
| SE | 3527 | ENSG00000105483 | CARD8 | chr19 | - | 48218870 | 48219012 | 48215339 | 48215384 | 48230437 | 48230498 | 197 | 95 | 136 | 190 | 120 | 1 | 0 | 1 | 0 | 1 | 68 | 65 | 63 | 50 |
| SE | 3528 | ENSG00000105483 | CARD8 | chr19 | - | 48218870 | 48219012 | 48215339 | 48215384 | 48231659 | 48231810 | 196 | 95 | 136 | 189 | 120 | 0 | 0 | 0 | 1 | 0 | 68 | 65 | 63 | 50 |
| SE | 3529 | ENSG00000105483 | CARD8 | chr19 | - | 48221729 | 48221855 | 48215137 | 48215384 | 48230437 | 48230647 | 119 | 65 | 92 | 123 | 66 | 1 | 0 | 1 | 0 | 1 | 93 | 67 | 68 | 24 |
| SE | 3530 | ENSG00000105483 | CARD8 | chr19 | - | 48221729 | 48221855 | 48215137 | 48215384 | 48231659 | 48231810 | 34 | 19 | 17 | 22 | 19 | 0 | 0 | 0 | 1 | 0 | 21 | 14 | 23 | 3 |
| SE | 3531 | ENSG00000105483 | CARD8 | chr19 | - | 48221729 | 48221855 | 48218870 | 48219012 | 48230437 | 48230647 | 325 | 224 | 269 | 376 | 236 | 1 | 0 | 1 | 1 | 0 | 186 | 217 | 175 | 115 |
| SE | 3532 | ENSG00000105483 | CARD8 | chr19 | - | 48221729 | 48221855 | 48218870 | 48219012 | 48231659 | 48231810 | 235 | 176 | 197 | 273 | 187 | 0 | 0 | 1 | 0 | 0 | 114 | 161 | 123 | 94 |
| SE | 3540 | ENSG00000105483 | CARD8 | chr19 | - | 48240961 | 48241063 | 48234277 | 48234543 | 48249522 | 48249614 | 264 | 196 | 223 | 296 | 237 | 0 | 0 | 0 | 1 | 0 | 219 | 147 | 128 | 137 |
| SE | 3541 | ENSG00000105483 | CARD8 | chr19 | - | 48240961 | 48241063 | 48234277 | 48234543 | 48249750 | 48249816 | 206 | 120 | 199 | 213 | 219 | 0 | 0 | 0 | 0 | 0 | 195 | 108 | 134 | 138 |
| SE | 3542 | ENSG00000105483 | CARD8 | chr19 | - | 48240961 | 48241063 | 48234277 | 48234543 | 48255791 | 48255871 | 108 | 71 | 98 | 119 | 112 | 1 | 0 | 0 | 0 | 0 | 127 | 58 | 72 | 66 |
| SE | 3543 | ENSG00000105483 | CARD8 | chr19 | - | 48240961 | 48241063 | 48238261 | 48238532 | 48242539 | 48242644 | 305 | 176 | 225 | 296 | 313 | 2 | 0 | 0 | 0 | 0 | 246 | 186 | 175 | 103 |
| SE | 3550 | ENSG00000105483 | CARD8 | chr19 | - | 48246074 | 48246381 | 48242539 | 48242644 | 48249750 | 48249847 | 4 | 2 | 0 | 12 | 12 | 12 | 2 | 2 | 5 | 9 | 18 | 0 | 0 | 4 |
| SE | 3551 | ENSG00000105483 | CARD8 | chr19 | - | 48249522 | 48249633 | 48232452 | 48234543 | 48249750 | 48249810 | 111 | 52 | 98 | 125 | 60 | 0 | 0 | 0 | 0 | 0 | 52 | 74 | 60 | 42 |
| SE | 3552 | ENSG00000105483 | CARD8 | chr19 | - | 48249522 | 48249633 | 48232452 | 48234543 | 48255791 | 48255871 | 0 | 0 | 0 | 3 | 0 | 1 | 0 | 0 | 0 | 0 | 6 | 0 | 0 | 0 |
| SE | 3556 | ENSG00000105483 | CARD8 | chr19 | - | 48249522 | 48249633 | 48246074 | 48246381 | 48249750 | 48249810 | 133 | 52 | 105 | 141 | 76 | 0 | 0 | 0 | 3 | 0 | 69 | 75 | 66 | 50 |
| SE | 3557 | ENSG00000105483 | CARD8 | chr19 | - | 48249750 | 48249847 | 48234402 | 48234543 | 48255791 | 48255871 | 40 | 18 | 35 | 31 | 85 | 1 | 0 | 0 | 0 | 0 | 36 | 32 | 23 | 27 |
| SE | 3560 | ENSG00000105483 | CARD8 | chr19 | - | 48249750 | 48249878 | 48234402 | 48234543 | 48255791 | 48255911 | 0 | 0 | 0 | 1 | 3 | 1 | 0 | 0 | 0 | 0 | 0 | 0 | 1 | 0 |
| SE | 3563 | ENSG00000077312 | SNRPA | chr19 | + | 40757331 | 40757504 | 40751202 | 40751481 | 40759430 | 40759610 | 194 | 152 | 744 | 807 | 986 | 1 | 0 | 2 | 3 | 1 | 1363 | 1234 | 814 | 1037 |
| SE | 3564 | ENSG00000077312 | SNRPA | chr19 | + | 40757331 | 40757504 | 40751202 | 40751481 | 40759439 | 40759579 | 172 | 139 | 396 | 444 | 634 | 0 | 0 | 0 | 0 | 0 | 717 | 615 | 422 | 527 |
| SE | 3566 | ENSG00000077312 | SNRPA | chr19 | + | 40758783 | 40758869 | 40757331 | 40757504 | 40759430 | 40759579 | 1 | 0 | 0 | 0 | 0 | 22 | 14 | 348 | 368 | 352 | 0 | 0 | 0 | 0 |

[illegible]

| | | | | | | | | | | | | | | | | | | | | | | | | | | | | | | | | | | | | | | | | | | | | | | |
|---|---|---|---|---|---|---|---|---|---|---|---|---|---|---|---|---|---|---|---|---|---|---|---|---|---|---|---|---|---|---|---|---|---|---|---|---|---|---|---|---|---|---|---|---|---|---|
| SE | 3986 | ENSG00000126261 | UBA2 | chr19 | + | 34454443 | 34454556 | 34454259 | 34454353 | 34464025 | 34464119 | 231 | 146 | 171 | 215 | 357 | 0 | 0 | 0 | 0 | 0 | 66 | 124 | 152 | 96 | 28 | 4 | 0 | 0 | 0 | 0 | 262 | 149 | 1 | 1 | 1 | 1 | 1 | 1 | 1 | 0.904 | 1 | 1 | 1 | 1 | 0.019 |
| SE | 3987 | ENSG00000126261 | UBA2 | chr19 | + | 34458768 | 34458924 | 34454259 | 34454353 | 34464025 | 34464131 | 6 | 2 | 4 | 3 | 4 | 0 | 0 | 0 | 0 | 0 | 44 | 2 | 3 | 0 | 6 | 4 | 0 | 0 | 0 | 0 | 298 | 149 | 1 | 1 | 1 | 1 | 1 | 1 | 1 | 0.846 | 1 | 1 | NA | 1 | 0.038 |
| SE | 3989 | ENSG00000126261 | UBA2 | chr19 | + | 34458768 | 34458924 | 34454443 | 34454556 | 34464025 | 34464131 | 174 | 123 | 185 | 184 | 359 | 1 | 1 | 0 | 1 | 2 | 119 | 88 | 90 | 61 | 57 | 0 | 0 | 0 | 0 | 0 | 298 | 149 | 1 | 1 | 0.989 | 0.984 | 1 | 0.989 | 0.989 | 1 | 1 | 1 | 1 | 1 | -0.01 |
| SE | 3990 | ENSG00000126261 | UBA2 | chr19 | + | 34460469 | 34460566 | 34454443 | 34454556 | 34464025 | 34464119 | 203 | 163 | 196 | 199 | 324 | 1 | 1 | 0 | 1 | 2 | 67 | 105 | 148 | 102 | 41 | 0 | 0 | 0 | 0 | 0 | 246 | 149 | 1 | 1 | 0.992 | 0.99 | 1 | 0.992 | 0.99 | 1 | 1 | 1 | 1 | 1 | -0.007 |
| SE | 3992 | ENSG00000126261 | UBA2 | chr19 | + | 34464025 | 34464131 | 34460469 | 34460566 | 34466877 | 34467014 | 556 | 450 | 583 | 589 | 1020 | 1 | 0 | 0 | 1 | 0 | 210 | 410 | 448 | 402 | 92 | 0 | 0 | 0 | 0 | 0 | 255 | 149 | 1 | 1 | 0.997 | 1 | 1 | 0.997 | 1 | 1 | 1 | 1 | 1 | 1 | -0.001 |
| SE | 3994 | ENSG00000104859 | CLASRP | chr19 | + | 45052070 | 45052168 | 45039520 | 45040311 | 45052790 | 45052853 | 85 | 43 | 49 | 86 | 58 | 0 | 0 | 0 | 0 | 2 | 189 | 103 | 83 | 123 | 240 | 0 | 0 | 0 | 0 | 0 | 247 | 149 | 1 | 1 | 1 | 1 | 1 | 1 | 0.946 | 1 | 1 | 1 | 1 | 1 | -0.011 |
| SE | 3995 | ENSG00000104859 | CLASRP | chr19 | + | 45068014 | 45068058 | 45067336 | 45067594 | 45069065 | 45069124 | 2 | 0 | 0 | 3 | 0 | 0 | 0 | 0 | 2 | 0 | 3 | 3 | 3 | 2 | 10 | 0 | 0 | 0 | 0 | 0 | 193 | 149 | 1 | 1 | 1 | NA | NA | 0.537 | NA | 1 | 1 | 1 | 1 | 1 | -0.232 |
| SE | 3997 | ENSG00000104859 | CLASRP | chr19 | + | 45070536 | 45070561 | 45069838 | 45070104 | 45070802 | 45070855 | 47 | 27 | 58 | 52 | 33 | 0 | 0 | 0 | 0 | 0 | 147 | 57 | 72 | 111 | 137 | 0 | 2 | 0 | 0 | 0 | 174 | 149 | 1 | 1 | 1 | 1 | 1 | 1 | 1 | 1 | 0.961 | 1 | 1 | 1 | 0.008 |
| SE | 4000 | ENSG00000105379 | ETFB | chr19 | - | 51350328 | 51350391 | 51346899 | 51347058 | 51353131 | 51353290 | 126 | 81 | 133 | 144 | 73 | 0 | 0 | 0 | 0 | 0 | 236 | 166 | 136 | 175 | 225 | 0 | 0 | 0 | 2 | 0 | 212 | 149 | 1 | 1 | 1 | 1 | 1 | 1 | 1 | 1 | 1 | 1 | 0.984 | 1 | 0.003 |
| SE | 4001 | ENSG00000105379 | ETFB | chr19 | - | 51353131 | 51353290 | 51350186 | 51350391 | 51354149 | 51354308 | 261 | 177 | 248 | 304 | 200 | 0 | 1 | 0 | 0 | 0 | 449 | 325 | 297 | 371 | 466 | 0 | 4 | 2 | 0 | 0 | 298 | 149 | 1 | 1 | 1 | 0.989 | 1 | 1 | 1 | 1 | 0.976 | 0.987 | 1 | 1 | 0.005 |
| SE | 4009 | ENSG00000189013 | KIR2DL4 | chr19 | + | 54813124 | 54813228 | 54808832 | 54808883 | 54813689 | 54813742 | 0 | 0 | 0 | 1 | 0 | 1 | 0 | 0 | 0 | 0 | 0 | 2 | 0 | 0 | 0 | 0 | 0 | 0 | 0 | 0 | 253 | 149 | 1 | 1 | 0 | NA | NA | 1 | NA | NA | 1 | NA | NA | NA | -0.5 |
| SE | 4010 | ENSG00000189013 | KIR2DL4 | chr19 | + | 54813124 | 54813295 | 54808832 | 54808883 | 54813689 | 54813742 | 0 | 0 | 0 | 1 | 0 | 1 | 0 | 0 | 0 | 0 | 0 | 2 | 0 | 0 | 0 | 0 | 0 | 0 | 0 | 0 | 298 | 149 | 1 | 1 | 0 | NA | NA | 1 | NA | NA | 1 | NA | NA | NA | -0.5 |
| SE | 4011 | ENSG00000105369 | CD79A | chr19 | + | 41879534 | 41879653 | 41878989 | 41879289 | 41880669 | 41880738 | 420 | 140 | 371 | 389 | 407 | 0 | 0 | 1 | 0 | 0 | 348 | 890 | 210 | 213 | 715 | 0 | 0 | 0 | 0 | 0 | 268 | 149 | 1 | 1 | 1 | 1 | 0.995 | 1 | 1 | 1 | 1 | 1 | 1 | 1 | -0.001 |
| SE | 4013 | ENSG00000213753 | CENPBD1P1 | chr19 | + | 58581244 | 58583172 | 58575437 | 58575588 | 58598966 | 58605223 | 28 | 27 | 36 | 44 | 10 | 1 | 0 | 0 | 0 | 0 | 13 | 29 | 15 | 10 | 56 | 0 | 0 | 3 | 0 | 0 | 298 | 149 | 1 | 1 | 0.933 | 1 | 1 | 1 | 1 | 1 | 1 | 0.714 | 1 | 1 | 0.044 |
| SE | 4017 | ENSG00000104856 | RELB | chr19 | + | 45025605 | 45025737 | 45025328 | 45025420 | 45028887 | 45028992 | 109 | 81 | 98 | 63 | 125 | 0 | 0 | 0 | 0 | 0 | 484 | 86 | 122 | 285 | 370 | 0 | 0 | 0 | 3 | 0 | 281 | 149 | 1 | 1 | 1 | 1 | 1 | 1 | 1 | 1 | 1 | 1 | 0.981 | 1 | 0.004 |
| SE | 4018 | ENSG00000104856 | RELB | chr19 | + | 45032533 | 45032749 | 45028887 | 45028992 | 45034243 | 45034312 | 80 | 56 | 79 | 50 | 90 | 1 | 0 | 1 | 2 | 0 | 292 | 92 | 59 | 129 | 142 | 17 | 3 | 6 | 4 | 4 | 298 | 149 | 1 | 1 | 0.976 | 1 | 0.975 | 0.926 | 1 | 0.896 | 0.939 | 0.831 | 0.942 | 0.947 | 0.064 |
| SE | 4019 | ENSG00000104856 | RELB | chr19 | + | 45034243 | 45034528 | 45032423 | 45032749 | 45037404 | 45037568 | 30 | 19 | 37 | 25 | 38 | 0 | 0 | 0 | 0 | 0 | 90 | 41 | 49 | 61 | 49 | 0 | 4 | 0 | 0 | 0 | 298 | 149 | 1 | 1 | 1 | 1 | 1 | 1 | 1 | 1 | 0.837 | 1 | 1 | 1 | 0.033 |
| SE | 4020 | ENSG00000142556 | ZNF614 | chr19 | - | 52017116 | 52017359 | 52012764 | 52013252 | 52018007 | 52018103 | 10 | 16 | 19 | 26 | 22 | 0 | 0 | 0 | 0 | 0 | 0 | 5 | 3 | 4 | 0 | 0 | 0 | 0 | 6 | 0 | 298 | 149 | 1 | 1 | 1 | 1 | 1 | 1 | 1 | NA | 1 | 1 | 0.25 | NA | 0.25 |
| SE | 4021 | ENSG00000142556 | ZNF614 | chr19 | - | 52017575 | 52017697 | 52017000 | 52017359 | 52018007 | 52018103 | 4 | 7 | 11 | 10 | 11 | 10 | 16 | 19 | 26 | 21 | 0 | 3 | 1 | 0 | 0 | 0 | 5 | 3 | 4 | 0 | 271 | 149 | 1 | 1 | 0.18 | 0.194 | 0.241 | 0.175 | 0.224 | NA | 0.248 | 0.155 | 0 | NA | 0.068 |
| SE | 4022 | ENSG00000198453 | ZNF568 | chr19 | + | 36922585 | 36922846 | 36917578 | 36917648 | 36925199 | 36925258 | 35 | 39 | 22 | 52 | 40 | 7 | 8 | 8 | 8 | 12 | 0 | 16 | 14 | 9 | 3 | 0 | 13 | 6 | 7 | 1 | 298 | 149 | 1 | 1 | 0.714 | 0.709 | 0.579 | 0.765 | 0.625 | NA | 0.381 | 0.538 | 0.391 | 0.6 | 0.201 |
| SE | 4023 | ENSG00000198453 | ZNF568 | chr19 | + | 36925199 | 36925258 | 36917578 | 36917648 | 36936745 | 36936872 | 12 | 11 | 13 | 14 | 22 | 1 | 0 | 0 | 1 | 0 | 0 | 11 | 12 | 3 | 1 | 0 | 0 | 0 | 0 | 0 | 208 | 149 | 1 | 1 | 0.896 | 1 | 1 | 0.909 | 1 | NA | 1 | 1 | 1 | 1 | -0.039 |
| SE | 4024 | ENSG00000198453 | ZNF568 | chr19 | + | 36931552 | 36931645 | 36925199 | 36925258 | 36936745 | 36936824 | 0 | 0 | 2 | 0 | 5 | 22 | 27 | 19 | 39 | 32 | 0 | 9 | 1 | 2 | 0 | 0 | 10 | 19 | 3 | 4 | 242 | 149 | 1 | 1 | 0 | 0 | 0.061 | 0 | 0.088 | NA | 0.357 | 0.031 | 0.291 | 0 | -0.14 |
| SE | 4026 | ENSG00000198453 | ZNF568 | chr19 | + | 36976326 | 36976367 | 36974419 | 36974466 | 36991750 | 36991846 | 0 | 0 | 0 | 2 | 0 | 0 | 0 | 0 | 0 | 0 | 0 | 0 | 0 | 0 | 0 | 0 | 0 | 0 | 5 | 0 | 190 | 149 | 1 | 1 | NA | NA | NA | 1 | NA | NA | NA | NA | 0 | NA | 1 |
| SE | 4027 | ENSG00000198453 | ZNF568 | chr19 | + | 36991175 | 36991299 | 36974419 | 36974466 | 36991750 | 36991846 | 15 | 0 | 1 | 1 | 7 | 0 | 0 | 0 | 0 | 0 | 0 | 0 | 0 | 0 | 0 | 0 | 0 | 0 | 5 | 0 | 273 | 149 | 1 | 1 | 1 | NA | 1 | 1 | 1 | NA | NA | NA | 0 | NA | 1 |
| SE | 4031 | ENSG00000268163 | NA | chr19 | - | 57475659 | 57475786 | 57456125 | 57456182 | 57476633 | 57476672 | 2 | 1 | 1 | 1 | 9 | 8 | 0 | 2 | 3 | 0 | 0 | 0 | 4 | 0 | 0 | 0 | 0 | 3 | 0 | 0 | 276 | 149 | 1 | 1 | 0.119 | 1 | 0.213 | 0.153 | 1 | NA | NA | 0.419 | NA | NA | 0.078 |
| SE | 4032 | ENSG00000197619 | ZNF615 | chr19 | - | 52000258 | 52000378 | 51991337 | 51994837 | 52001812 | 52001908 | 2 | 3 | 4 | 23 | 6 | 1 | 2 | 1 | 2 | 0 | 0 | 1 | 2 | 0 | 0 | 0 | 0 | 0 | 0 | 0 | 269 | 149 | 1 | 1 | 0.526 | 0.454 | 0.689 | 0.864 | 1 | NA | 1 | 1 | NA | NA | -0.293 |
| SE | 4033 | ENSG00000197619 | ZNF615 | chr19 | - | 52000345 | 52000378 | 51991331 | 51994837 | 52001812 | 52001908 | 12 | 14 | 7 | 19 | 17 | 1 | 2 | 1 | 2 | 0 | 0 | 6 | 9 | 3 | 0 | 0 | 0 | 0 | 0 | 0 | 182 | 149 | 1 | 1 | 0.908 | 0.851 | 0.851 | 0.886 | 1 | NA | 1 | 1 | 1 | NA | -0.101 |
| SE | 4035 | ENSG00000197619 | ZNF615 | chr19 | - | 52003696 | 52003900 | 52002154 | 52002281 | 52007292 | 52007330 | 10 | 16 | 17 | 11 | 18 | 0 | 0 | 0 | 0 | 5 | 22 | 10 | 13 | 6 | 0 | 0 | 0 | 0 | 0 | 0 | 298 | 149 | 1 | 1 | 1 | 1 | 1 | 1 | 0.643 | 1 | 1 | 1 | 1 | NA | -0.071 |
| SE | 4036 | ENSG00000197619 | ZNF615 | chr19 | - | 52003696 | 52003900 | 52002154 | 52002281 | 52008140 | 52008163 | 10 | 19 | 17 | 10 | 20 | 1 | 1 | 0 | 0 | 0 | 11 | 13 | 9 | 6 | 0 | 0 | 0 | 0 | 0 | 0 | 298 | 149 | 1 | 1 | 0.833 | 0.905 | 1 | 1 | 1 | 1 | 1 | 1 | 1 | NA | -0.052 |
| SE | 4037 | ENSG00000197619 | ZNF615 | chr19 | - | 52004416 | 52004723 | 52003696 | 52003900 | 52007292 | 52007330 | 2 | 3 | 3 | 2 | 0 | 1 | 2 | 4 | 3 | 2 | 0 | 0 | 0 | 9 | 0 | 11 | 2 | 4 | 0 | 0 | 298 | 149 | 1 | 1 | 0.5 | 0.429 | 0.273 | 0.25 | 0 | 0 | 0 | 0 | 1 | NA | 0.04 |
| SE | 4038 | ENSG00000197619 | ZNF615 | chr19 | - | 52004416 | 52004723 | 52003696 | 52003900 | 52008140 | 52008201 | 2 | 2 | 0 | 1 | 0 | 1 | 5 | 4 | 2 | 4 | 0 | 0 | 0 | 3 | 0 | 0 | 5 | 0 | 0 | 0 | 298 | 149 | 1 | 1 | 0.5 | 0.167 | 0 | 0.2 | 0 | NA | 0 | NA | 1 | NA | -0.327 |
| SE | 4039 | ENSG00000197619 | ZNF615 | chr19 | - | 52004416 | 52004849 | 52003848 | 52003900 | 52007292 | 52007330 | 2 | 3 | 0 | 1 | 0 | 1 | 2 | 4 | 3 | 2 | 0 | 0 | 0 | 3 | 0 | 11 | 2 | 4 | 0 | 0 | 298 | 149 | 1 | 1 | 0.5 | 0.429 | 0 | 0.143 | 0 | 0 | 0 | 0 | 1 | NA | -0.036 |
| SE | 4041 | ENSG00000197619 | ZNF615 | chr19 | - | 52007292 | 52007330 | 52003696 | 52003900 | 52008140 | 52008200 | 1 | 2 | 4 | 3 | 2 | 1 | 5 | 4 | 2 | 4 | 11 | 2 | 4 | 0 | 0 | 0 | 5 | 0 | 0 | 0 | 187 | 149 | 1 | 1 | 0.443 | 0.242 | 0.443 | 0.544 | 0.285 | 1 | 0.242 | 1 | NA | NA | -0.356 |
| SE | 4043 | ENSG00000126267 | COX6B1 | chr19 | + | 35651232 | 35651349 | 35648322 | 35648403 | 35654570 | 35654671 | 515 | 206 | 436 | 439 | 433 | 3 | 1 | 2 | 2 | 3 | 474 | 519 | 298 | 414 | 773 | 0 | 3 | 0 | 4 | 0 | 266 | 149 | 1 | 1 | 0.99 | 0.991 | 0.992 | 0.992 | 0.988 | 1 | 0.99 | 1 | 0.983 | 1 | -0.004 |
| SE | 4044 | ENSG00000126267 | COX6B1 | chr19 | + | 35654570 | 35654671 | 35651232 | 35651349 | 35658593 | 35658726 | 752 | 285 | 584 | 621 | 698 | 2 | 1 | 7 | 1 | 0 | 644 | 654 | 412 | 633 | 940 | 2 | 6 | 5 | 0 | 4 | 250 | 149 | 1 | 1 | 0.996 | 0.994 | 0.98 | 0.997 | 1 | 0.995 | 0.985 | 0.98 | 1 | 0.993 | 0.003 |
| SE | 4045 | ENSG00000186230 | ZNF749 | chr19 | + | 57441884 | 57442011 | 57435324 | 57435593 | 57443290 | 57447101 | 6 | 3 | 11 | 8 | 6 | 0 | 4 | 3 | 7 | 0 | 0 | 1 | 0 | 0 | 12 | 0 | 0 | 1 | 3 | 0 | 276 | 149 | 1 | 1 | 1 | 0.288 | 0.664 | 0.382 | 1 | NA | 1 | 0 | 0 | 1 | 0.167 |
| SE | 4046 | ENSG00000186230 | ZNF749 | chr19 | + | 57441884 | 57442011 | 57435346 | 57435667 | 57443290 | 57447101 | 4 | 2 | 10 | 7 | 6 | 0 | 0 | 1 | 1 | 0 | 0 | 1 | 0 | 0 | 12 | 0 | 0 | 0 | 0 | 0 | 276 | 149 | 1 | 1 | 1 | 1 | 0.844 | 0.791 | 1 | NA | 1 | NA | NA | 1 | -0.073 |
| SE | 4052 | ENSG00000249115 | HAUS5 | chr19 | + | 35617123 | 35617193 | 35615226 | 35615386 | 35617286 | 35617369 | 20 | 6 | 18 | 16 | 5 | 3 | 0 | 0 | 0 | 0 | 0 | 27 | 14 | 4 | 52 | 0 | 0 | 0 | 0 | 0 | 219 | 149 | 1 | 1 | 0.819 | 1 | 1 | 1 | 1 | NA | 1 | 1 | 1 | 1 | -0.036 |
| SE | 4053 | ENSG00000249115 | HAUS5 | chr19 | + | 35617286 | 35617369 | 35617123 | 35617193 | 35617854 | 35617912 | 42 | 7 | 27 | 25 | 16 | 2 | 2 | 0 | 0 | 0 | 0 | 46 | 14 | 15 | 99 | 0 | 0 | 0 | 0 | 0 | 232 | 149 | 1 | 1 | 0.931 | 0.692 | 1 | 1 | 1 | NA | 1 | 1 | 1 | 1 | -0.075 |
| SE | 4054 | ENSG00000249115 | HAUS5 | chr19 | + | 35618401 | 35618465 | 35618070 | 35618195 | 35618568 | 35618699 | 6 | 4 | 3 | 6 | 1 | 0 | 0 | 1 | 0 | 0 | 0 | 12 | 5 | 8 | 16 | 0 | 0 | 0 | 0 | 0 | 213 | 149 | 1 | 1 | 1 | 1 | 0.677 | 1 | 1 | NA | 1 | 1 | 1 | 1 | -0.065 |
| SE | 4056 | ENSG00000248385 | TARM1 | chr19 | - | 54075882 | 54075918 | 54074823 | 54075114 | 54081306 | 54081365 | 1 | 0 | 0 | 0 | 0 | 0 | 0 | 0 | 0 | 0 | 0 | 7 | 3 | 20 | 0 | 0 | 4 | 0 | 0 | 7 | 185 | 149 | 1 | 1 | 1 | NA | NA | NA | NA | NA | 0.585 | 1 | 1 | 0 | 0.354 |
| SE | 4057 | ENSG00000269533 | NA | chr19 | + | 57397047 | 57397174 | 57394187 | 57394223 | 57410879 | 57411167 | 50 | 35 | 62 | 42 | 27 | 0 | 1 | 0 | 0 | 0 | 24 | 66 | 45 | 8 | 6 | 0 | 0 | 0 | 1 | 0 | 276 | 149 | 1 | 1 | 1 | 0.95 | 1 | 1 | 1 | 1 | 1 | 1 | 0.812 | 1 | 0.028 |
| SE | 4058 | ENSG00000267254 | ZNF790-AS1 | chr19 | + | 36801202 | 36801240 | 36797501 | 36797811 | 36801875 | 36801933 | 7 | 3 | 1 | 7 | 0 | 1 | 0 | 0 | 0 | 0 | 0 | 11 | 6 | 4 | 6 | 0 | 6 | 1 | 5 | 0 | 187 | 149 | 1 | 1 | 0.848 | 1 | 1 | 1 | NA | NA | 0.594 | 0.827 | 0.389 | 1 | 0.26 |
| SE | 4059 | ENSG00000267254 | ZNF790-AS1 | chr19 | + | 36801875 | 36801933 | 36797515 | 36797811 | 36823730 | 36823821 | 1 | 0 | 1 | 0 | 0 | 1 | 0 | 0 | 0 | 0 | 0 | 6 | 0 | 5 | 2 | 0 | 0 | 0 | 0 | 0 | 207 | 149 | 1 | 1 | 0.419 | NA | 1 | NA | NA | NA | 1 | NA | 1 | 1 | -0.29 |
| SE | 4060 | ENSG00000267254 | ZNF790-AS1 | chr19 | + | 36812509 | 36812606 | 36801875 | 36801933 | 36823730 | 36823821 | 0 | 0 | 1 | 1 | 0 | 1 | 1 | 1 | 0 | 0 | 0 | 0 | 7 | 0 | 0 | 0 | 0 | 0 | 0 | 2 | 246 | 149 | 1 | 1 | 0 | 0 | 0.377 | 1 | NA | NA | NA | 1 | NA | 0 | -0.156 |
| SE | 4061 | ENSG00000267254 | ZNF790-AS1 | chr19 | + | 36812509 | 36812606 | 36801875 | 36801933 | 36827649 | 36828115 | 0 | 0 | 0 | 1 | 0 | 0 | 0 | 0 | 0 | 0 | 0 | 0 | 7 | 0 | 0 | 0 | 0 | 0 | 4 | 12 | 246 | 149 | 1 | 1 | NA | NA | NA | 1 | NA | NA | NA | 1 | 0 | 0 | 0.667 |
| SE | 4063 | ENSG00000267254 | ZNF790-AS1 | chr19 | + | 36823730 | 36823821 | 36801875 | 36801933 | 36827649 | 36828101 | 2 | 1 | 1 | 0 | 0 | 0 | 0 | 0 | 0 | 0 | 0 | 0 | 2 | 2 | 2 | 0 | 0 | 0 | 4 | 12 | 240 | 149 | 1 | 1 | 1 | 1 | 1 | NA | NA | NA | NA | 1 | 0.237 | 0.094 | 0.556 |
| SE | 4065 | ENSG00000099326 | MZF1 | chr19 | - | 58570993 | 58571429 | 58570343 | 58570527 | 58573054 | 58573298 | 10 | 5 | 23 | 7 | 7 | 1 | 0 | 0 | 0 | 0 | 8 | 26 | 1 | 4 | 24 | 0 | 0 | 0 | 0 | 0 | 298 | 149 | 1 | 1 | 0.833 | 1 | 1 | 1 | 1 | 1 | 1 | 1 | 1 | 1 | -0.033 |
| SE | 4069 | ENSG00000105771 | SMG9 | chr19 | - | 43739672 | 43740218 | 43738121 | 43738217 | 43747441 | 43747539 | 4 | 0 | 0 | 5 | 0 | 0 | 0 | 0 | 0 | 0 | 0 | 0 | 0 | 7 | 1 | 0 | 0 | 0 | 0 | 1 | 298 | 149 | 1 | 1 | 1 | NA | NA | 1 | NA | NA | NA | NA | 1 | 0.333 | 0.334 |
| SE | 4070 | ENSG00000105771 | SMG9 | chr19 | - | 43740106 | 43740218 | 43737849 | 43738217 | 43747441 | 43747539 | 26 | 22 | 29 | 52 | 26 | 0 | 0 | 0 | 0 | 0 | 42 | 28 | 10 | 31 | 52 | 0 | 0 | 0 | 0 | 1 | 261 | 149 | 1 | 1 | 1 | 1 | 1 | 1 | 1 | 1 | 1 | 1 | 1 | 0.967 | 0.007 |
| SE | 4077 | ENSG00000267767 | LINC01801 | chr19 | - | 34829163 | 34829253 | 34818586 | 34818766 | 34830405 | 34830526 | 7 | 0 | 4 | 4 | 0 | 0 | 0 | 0 | 1 | 0 | 0 | 0 | 1 | 0 | 0 | 0 | 1 | 0 | 0 | 0 | 239 | 149 | 1 | 1 | 1 | NA | 1 | 0.714 | NA | NA | 0 | 1 | NA | NA | 0.405 |
| SE | 4080 | ENSG00000011422 | PLAUR | chr19 | - | 43652224 | 43652371 | 43648578 | 43649143 | 43665315 | 43665459 | 118 | 73 | 100 | 143 | 181 | 0 | 0 | 0 | 0 | 0 | 229 | 144 | 218 | 229 | 241 | 0 | 0 | 0 | 3 | 0 | 296 | 149 | 1 | 1 | 1 | 1 | 1 | 1 | 1 | 1 | 1 | 1 | 0.975 | 1 | 0.005 |
| SE | 4082 | ENSG00000011422 | PLAUR | chr19 | - | 43655438 | 43655573 | 43648578 | 43649143 | 43665315 | 43665459 | 7 | 2 | 4 | 9 | 9 | 0 | 0 | 0 | 0 | 0 | 39 | 15 | 17 | 33 | 14 | 0 | 0 | 0 | 3 | 0 | 284 | 149 | 1 | 1 | 1 | 1 | 1 | 1 | 1 | 1 | 1 | 1 | 0.852 | 1 | 0.03 |
| SE | 4084 | ENSG00000011422 | PLAUR | chr19 | - | 43655438 | 43655573 | 43652224 | 43652371 | 43665315 | 43665459 | 70 | 29 | 44 | 80 | 53 | 0 | 0 | 4 | 0 | 0 | 133 | 100 | 96 | 140 | 147 | 0 | 0 | 1 | 0 | 0 | 284 | 149 | 1 | 1 | 1 | 1 | 0.852 | 1 | 1 | 1 | 1 | 0.981 | 1 | 1 | -0.026 |
| SE | 4085 | ENSG00000011422 | PLAUR | chr19 | - | 43656478 | 43656640 | 43648578 | 43649143 | 43665315 | 43665459 | 202 | 129 | 120 | 190 | 297 | 0 | 0 | 0 | 0 | 0 | 412 | 288 | 291 | 365 | 498 | 0 | 0 | 0 | 3 | 0 | 298 | 149 | 1 | 1 | 1 | 1 | 1 | 1 | 1 | 1 | 1 | 1 | 0.984 | 1 | 0.003 |
| SE | 4086 | ENSG00000011422 | PLAUR | chr19 | - | 43656478 | 43656640 | 43652224 | 43652371 | 43665315 | 43665459 | 213 | 130 | 129 | 196 | 310 | 0 | 0 | 4 | 0 | 0 | 440 | 296 | 295 | 380 | 484 | 0 | 0 | 1 | 0 | 0 | 298 | 149 | 1 | 1 | 1 | 1 | 0.942 | 1 | 1 | 1 | 1 | 0.993 | 1 | 1 | -0.01 |
| SE | 4089 | ENSG00000168813 | ZNF507 | chr19 | + | 32347244 | 32347338 | 32345609 | 32345783 | 32352828 | 32354957 | 37 | 39 | 45 | 44 | 69 | 7 | 1 | 3 | 0 | 3 | 21 | 21 | 50 | 47 | 0 | 0 | 0 | 1 | 0 | 0 | 243 | 149 | 1 | 1 | 0.764 | 0.96 | 0.902 | 1 | 0.934 | 1 | 1 | 0.968 | 1 | NA | -0.08 |
| SE | 4090 | ENSG00000168813 | ZNF507 | chr19 | + | 32356615 | 32356733 | 32352828 | 32354957 | 32360503 | 32360618 | 119 | 66 | 106 | 126 | 168 | 0 | 0 | 2 | 0 | 0 | 18 | 78 | 102 | 24 | 0 | 0 | 0 | 1 | 0 | 0 | 267 | 149 | 1 | 1 | 1 | 1 | 0.967 | 1 | 1 | 1 | 1 | 1 | 1 | NA | -0.007 |
| SE | 4091 | ENSG00000168813 | ZNF507 | chr19 | + | 32360503 | 32360618 | 32356615 | 32356733 | 32382466 | 32382601 | 120 | 58 | 111 | 121 | 227 | 0 | 1 | 0 | 1 | 0 | 11 | 74 | 79 | 17 | 6 | 0 | 0 | 0 | 0 | 0 | 264 | 149 | 1 | 1 | 1 | 0.97 | 1 | 0.986 | 1 | 1 | 1 | 1 | 1 | 1 | -0.009 |
| SE | 4095 | ENSG00000105048 | TNNT1 | chr19 | - | 55146433 | 55146466 | 55145399 | 55145565 | 55146680 | 55146705 | 2 | 2 | 0 | 0 | 0 | 1 | 0 | 0 | 0 | 0 | 6 | 0 | 4 | 1 | 12 | 0 | 0 | 0 | 0 | 0 | 182 | 149 | 1 | 1 | 0.621 | 1 | NA | NA | NA | 1 | NA | 1 | 1 | 1 | -0.19 |
| SE | 4099 | ENSG00000105738 | SIPA1L3 | chr19 | + | 38088720 | 38088851 | 38081255 | 38083099 | 38099961 | 38100150 | 76 | 40 | 55 | 75 | 49 | 0 | 0 | 0 | 0 | 0 | 97 | 142 | 56 | 62 | 162 | 5 | 0 | 0 | 0 | 0 | 280 | 149 | 1 | 1 | 1 | 1 | 1 | 1 | 1 | 0.912 | 1 | 1 | 1 | 1 | 0.018 |
| SE | 4100 | ENSG00000105738 | SIPA1L3 | chr19 | + | 38119305 | 38119882 | 38110226 | 38110384 | 38130497 | 38130772 | 158 | 72 | 151 | 159 | 145 | 0 | 0 | 0 | 0 | 0 | 269 | 178 | 92 | 152 | 310 | 0 | 3 | 0 | 0 | 0 | 298 | 149 | 1 | 1 | 1 | 1 | 1 | 1 | 1 | 1 | 0.967 | 1 | 1 | 1 | 0.007 |
| SE | 4103 | ENSG00000105738 | SIPA1L3 | chr19 | + | 38162252 | 38162371 | 38152839 | 38152967 | 38164478 | 38164906 | 38 | 25 | 20 | 33 | 28 | 1 | 0 | 0 | 0 | 0 | 109 | 51 | 43 | 26 | 126 | 0 | 3 | 0 | 0 | 0 | 268 | 149 | 1 | 1 | 0.955 | 1 | 1 | 1 | 1 | 1 | 0.904 | 1 | 1 | 1 | 0.01 |
| SE | 4106 | ENSG00000269858 | EGLN2 | chr19 | + | 40806554 | 40806674 | 40799544 | 40801415 | 40807483 | 40807551 | 63 | 22 | 61 | 40 | 14 | 0 | 0 | 0 | 0 | 0 | 8 | 19 | 11 | 40 | 59 | 0 | 0 | 0 | 0 | 4 | 269 | 149 | 1 | 1 | 1 | 1 | 1 | 1 | 1 | 1 | 1 | 1 | 1 | 0.891 | 0.022 |
| SE | 4108 | ENSG00000269858 | EGLN2 | chr19 | + | 40807137 | 40807274 | 40801289 | 40801415 | 40807483 | 40807551 | 129 | 69 | 134 | 95 | 45 | 0 | 0 | 0 | 0 | 0 | 117 | 45 | 46 | 63 | 133 | 0 | 0 | 0 | 0 | 4 | 286 | 149 | 1 | 1 | 1 | 1 | 1 | 1 | 1 | 1 | 1 | 1 | 1 | 0.945 | 0.011 |
| SE | 4109 | ENSG00000269858 | EGLN2 | chr19 | + | 40807137 | 40807274 | 40801289 | 40801415 | 40807808 | 40807936 | 7 | 1 | 5 | 3 | 0 | 1 | 0 | 3 | 3 | 0 | 0 | 0 | 6 | 8 | 3 | 4 | 0 | 0 | 21 | 6 | 286 | 149 | 1 | 1 | 0.785 | 1 | 0.465 | 0.343 | NA | 0 | NA | 1 | 0.166 | 0.207 | 0.305 |
| SE | 4110 | ENSG00000269858 | EGLN2 | chr19 | + | 40807137 | 40807274 | 40804469 | 40806674 | 40807483 | 40807551 | 211 | 103 | 219 | 180 | 102 | 0 | 0 | 0 | 2 | 0 | 134 | 92 | 72 | 82 | 178 | 0 | 0 | 0 | 0 | 0 | 286 | 149 | 1 | 1 | 1 | 1 | 1 | 0.979 | 1 | 1 | 1 | 1 | 1 | 1 | -0.004 |
| SE | 4114 | ENSG00000269858 | EGLN2 | chr19 | + | 40807483 | 40807551 | 40804072 | 40807274 | 40807808 | 40807913 | 233 | 147 | 229 | 205 | 96 | 0 | 0 | 0 | 0 | 0 | 193 | 97 | 82 | 92 | 243 | 0 | 0 | 1 | 0 | 0 | 217 | 149 | 1 | 1 | 1 | 1 | 1 | 1 | 1 | 1 | 1 | 0.983 | 1 | 1 | 0.003 |
| SE | 4122 | ENSG00000105204 | DYRK1B | chr19 | - | 39826951 | 39826987 | 39826671 | 39826867 | 39827284 | 39827425 | 2 | 0 | 3 | 1 | 0 | 7 | 3 | 10 | 11 | 2 | 0 | 3 | 0 | 1 | 0 | 0 | 2 | 0 | 3 | 1 | 185 | 149 | 1 | 1 | 0.187 | 0 | 0.195 | 0.068 | 0 | NA | 0.547 | NA | 0.212 | 0 | -0.163 |
| SE | 4123 | ENSG00000105204 | DYRK1B | chr19 | - | 39830374 | 39830563 | 39829879 | 39830027 | 39830663 | 39830783 | 96 | 65 | 79 | 92 | 60 | 0 | 0 | 0 | 0 | 2 | 97 | 81 | 35 | 71 | 101 | 0 | 0 | 0 | 0 | 0 | 298 | 149 | 1 | 1 | 1 | 1 | 1 | 1 | 0.938 | 1 | 1 | 1 | 1 | 1 | -0.012 |
| SE | 4124 | ENSG00000105204 | DYRK1B | chr19 | - | 39831804 | 39831968 | 39830663 | 39830783 | 39834022 | 39834152 | 30 | 11 | 28 | 20 | 0 | 0 | 0 | 1 | 0 | 0 | 24 | 17 | 21 | 18 | 16 | 0 | 0 | 0 | 0 | 0 | 298 | 149 | 1 | 1 | 1 | 1 | 0.933 | 1 | NA | 1 | 1 | 1 | 1 | 1 | -0.017 |
| SE | 4127 | ENSG00000105176 | URI1 | chr19 | + | 29971192 | 29971227 | 29942238 | 29942664 | 30005360 | 30005436 | 2 | 0 | 0 | 0 | 2 | 2 | 0 | 0 | 0 | 0 | 0 | 0 | 0 | 0 | 0 | 0 | 2 | 0 | 1 | 0 | 184 | 149 | 1 | 1 | 0.447 | NA | NA | NA | 1 | NA | 0 | NA | 0 | NA | 0.724 |
| SE | 4131 | ENSG00000105176 | URI1 | chr19 | + | 30011093 | 30011236 | 30009004 | 30009353 | 30012284 | 30012531 | 954 | 582 | 816 | 847 | 1127 | 2 | 1 | 0 | 1 | 0 | 334 | 537 | 494 | 440 | 210 | 0 | 4 | 0 | 4 | 0 | 292 | 149 | 1 | 1 | 0.996 | 0.997 | 1 | 0.998 | 1 | 1 | 0.986 | 1 | 0.982 | 1 | 0.005 |
| SE | 4132 | ENSG00000105176 | URI1 | chr19 | + | 30014180 | 30014222 | 30012422 | 30012531 | 30014886 | 30015026 | 1 | 0 | 1 | 0 | 0 | 52 | 24 | 42 | 43 | 86 | 0 | 0 | 0 | 0 | 0 | 12 | 25 | 18 | 35 | 0 | 191 | 149 | 1 | 1 | 0.015 | 0 | 0.018 | 0 | 0 | 0 | 0 | 0 | 0 | NA | 0.007 |
| SE | 4134 | ENSG00000171570 | RAB4B-EGLN2 | chr19 | + | 40783920 | 40784075 | 40780070 | 40780499 | 40786664 | 40786760 | 179 | 87 | 137 | 132 | 55 | 0 | 0 | 0 | 1 | 0 | 161 | 149 | 72 | 91 | 193 | 0 | 0 | 0 | 0 | 0 | 298 | 149 | 1 | 1 | 1 | 1 | 1 | 0.985 | 1 | 1 | 1 | 1 | 1 | 1 | -0.003 |
| SE | 4135 | ENSG00000269547 | NA | chr19 | - | 38915840 | 38915906 | 38915403 | 38915749 | 38916036 | 38916129 | 20 | 4 | 18 | 6 | 9 | 0 | 1 | 0 | 0 | 0 | 31 | 6 | 6 | 10 | 9 | 0 | 0 | 0 | 0 | 0 | 215 | 149 | 1 | 1 | 1 | 0.735 | 1 | 1 | 1 | 1 | 1 | 1 | 1 | 1 | -0.053 |
| SE | 4137 | ENSG00000269547 | NA | chr19 | - | 38918093 | 38918139 | 38917920 | 38918008 | 38918421 | 38918530 | 13 | 6 | 16 | 6 | 20 | 1 | 0 | 0 | 0 | 0 | 24 | 4 | 10 | 6 | 19 | 0 | 0 | 0 | 0 | 0 | 195 | 149 | 1 | 1 | 0.909 | 1 | 1 | 1 | 1 | 1 | 1 | 1 | 1 | 1 | -0.018 |
| SE | 4141 | ENSG00000073008 | PVR | chr19 | + | 44657761 | 44657910 | 44653899 | 44654017 | 44661291 | 44661323 | 14 | 18 | 13 | 22 | 8 | 0 | 0 | 0 | 0 | 0 | 36 | 7 | 17 | 47 | 32 | 0 | 0 | 1 | 0 | 0 | 298 | 149 | 1 | 1 | 1 | 1 | 1 | 1 | 1 | 1 | 1 | 0.895 | 1 | 1 | 0.021 |
| SE | 4144 | ENSG00000007129 | CEACAM21 | chr19 | + | 41577199 | 41577559 | 41576165 | 41576338 | 41579352 | 41579628 | 15 | 7 | 12 | 7 | 13 | 0 | 0 | 0 | 0 | 0 | 10 | 34 | 8 | 35 | 22 | 0 | 0 | 0 | 1 | 0 | 298 | 149 | 1 | 1 | 1 | 1 | 1 | 1 | 1 | 1 | 1 | 1 | 0.946 | 1 | 0.011 |
| SE | 4145 | ENSG00000007129 | CEACAM21 | chr19 | + | 41579269 | 41579628 | 41577199 | 41577559 | 41584349 | 41584443 | 50 | 36 | 69 | 66 | 101 | 0 | 0 | 0 | 0 | 0 | 31 | 27 | 13 | 45 | 21 | 0 | 0 | 0 | 3 | 0 | 298 | 149 | 1 | 1 | 1 | 1 | 1 | 1 | 1 | 1 | 1 | 1 | 0.882 | 1 | 0.024 |
| SE | 4146 | ENSG00000007129 | CEACAM21 | chr19 | + | 41579352 | 41579628 | 41577199 | 41577559 | 41584349 | 41584443 | 24 | 14 | 35 | 26 | 28 | 0 | 0 | 0 | 0 | 0 | 15 | 26 | 5 | 57 | 18 | 0 | 0 | 0 | 3 | 0 | 298 | 149 | 1 | 1 | 1 | 1 | 1 | 1 | 1 | 1 | 1 | 1 | 0.905 | 1 | 0.019 |
| SE | 4147 | ENSG00000007129 | CEACAM21 | chr19 | + | 41586153 | 41586195 | 41585839 | 41585871 | 41586463 | 41586844 | 11 | 1 | 2 | 5 | 8 | 7 | 0 | 0 | 0 | 0 | 0 | 9 | 2 | 14 | 4 | 0 | 5 | 3 | 10 | 6 | 191 | 149 | 1 | 1 | 0.551 | 1 | 1 | 1 | 1 | NA | 0.584 | 0.342 | 0.522 | 0.342 | 0.463 |
| SE | 4148 | ENSG00000167637 | ZNF283 | chr19 | + | 43827775 | 43827940 | 43827373 | 43827444 | 43828201 | 43828281 | 28 | 8 | 9 | 26 | 9 | 5 | 1 | 8 | 5 | 2 | 8 | 0 | 9 | 2 | 14 | 0 | 0 | 5 | 0 | 0 | 298 | 149 | 1 | 1 | 0.737 | 0.8 | 0.36 | 0.722 | 0.692 | 1 | NA | 0.474 | 1 | 1 | -0.206 |
| SE | 4149 | ENSG00000167637 | ZNF283 | chr19 | + | 43831317 | 43831381 | 43828195 | 43828281 | 43835504 | 43835592 | 25 | 11 | 25 | 21 | 33 | 0 | 0 | 2 | 1 | 1 | 5 | 5 | 11 | 5 | 7 | 0 | 0 | 0 | 0 | 0 | 213 | 149 | 1 | 1 | 1 | 1 | 0.897 | 0.936 | 0.958 | 1 | 1 | 1 | 1 | 1 | -0.042 |
| SE | 4150 | ENSG00000167637 | ZNF283 | chr19 | + | 43831317 | 43831381 | 43828195 | 43828281 | 43846938 | 43848809 | 8 | 3 | 10 | 4 | 10 | 0 | 0 | 0 | 1 | 0 | 0 | 0 | 4 | 0 | 0 | 0 | 0 | 0 | 0 | 0 | 213 | 149 | 1 | 1 | 1 | 1 | 1 | 0.737 | 1 | NA | NA | 1 | NA | NA | -0.053 |
| SE | 4152 | ENSG00000167637 | ZNF283 | chr19 | + | 43835504 | 43835592 | 43828198 | 43828281 | 43846938 | 43848898 | 2 | 0 | 4 | 1 | 1 | 0 | 0 | 0 | 1 | 0 | 0 | 1 | 0 | 0 | 0 | 0 | 0 | 0 | 0 | 0 | 237 | 149 | 1 | 1 | 1 | NA | 1 | 0.386 | 1 | NA | 1 | NA | NA | NA | -0.153 |
| SE | 4153 | ENSG00000167637 | ZNF283 | chr19 | + | 43835504 | 43835592 | 43831317 | 43831381 | 43837052 | 43837132 | 33 | 28 | 28 | 32 | 61 | 0 | 1 | 0 | 0 | 0 | 5 | 27 | 13 | 5 | 14 | 0 | 0 | 0 | 0 | 0 | 237 | 149 | 1 | 1 | 1 | 0.946 | 1 | 1 | 1 | 1 | 1 | 1 | 1 | 1 | -0.011 |
| SE | 4154 | ENSG00000167637 | ZNF283 | chr19 | + | 43835504 | 43835592 | 43831317 | 43831381 | 43846938 | 43848898 | 15 | 8 | 13 | 15 | 25 | 0 | 0 | 0 | 0 | 2 | 0 | 7 | 4 | 0 | 7 | 0 | 0 | 0 | 0 | 0 | 237 | 149 | 1 | 1 | 1 | 1 | 1 | 1 | 0.887 | NA | 1 | 1 | NA | 1 | -0.023 |
| SE | 4156 | ENSG00000167637 | ZNF283 | chr19 | + | 43837052 | 43837132 | 43835504 | 43835592 | 43846938 | 43852017 | 9 | 7 | 5 | 12 | 13 | 2 | 0 | 2 | 2 | 1 | 5 | 6 | 5 | 5 | 0 | 0 | 1 | 0 | 0 | 0 | 229 | 149 | 1 | 1 | 0.745 | 1 | 0.619 | 0.796 | 0.894 | 1 | 0.796 | 1 | 1 | NA | -0.138 |
| SE | 4157 | ENSG00000167637 | ZNF283 | chr19 | + | 43837052 | 43837179 | 43831317 | 43831381 | 43846938 | 43848809 | 6 | 4 | 0 | 3 | 2 | 0 | 0 | 0 | 0 | 2 | 0 | 5 | 0 | 5 | 0 | 0 | 0 | 0 | 0 | 0 | 276 | 149 | 1 | 1 | 1 | 1 | NA | 1 | 0.351 | NA | 1 | NA | 1 | NA | -0.162 |
| SE | 4159 | ENSG00000104825 | NFKBIB | chr19 | + | 38907220 | 38907309 | 38905201 | 38905535 | 38907398 | 38907649 | 105 | 54 | 112 | 138 | 81 | 1 | 2 | 0 | 0 | 0 | 194 | 213 | 129 | 205 | 288 | 0 | 0 | 0 | 2 | 0 | 238 | 149 | 1 | 1 | 0.985 | 0.944 | 1 | 1 | 1 | 1 | 1 | 1 | 0.985 | 1 | -0.011 |
| SE | 4171 | ENSG00000130726 | TRIM28 | chr19 | + | 58549330 | 58549650 | 58548987 | 58549240 | 58549736 | 58549860 | 270 | 90 | 304 | 246 | 184 | 0 | 0 | 0 | 0 | 0 | 187 | 189 | 142 | 159 | 298 | 0 | 0 | 0 | 1 | 0 | 298 | 149 | 1 | 1 | 1 | 1 | 1 | 1 | 1 | 1 | 1 | 1 | 0.988 | 1 | 0.002 |
| SE | 4172 | ENSG00000267041 | ZNF850 | chr19 | - | 36761642 | 36761738 | 36743479 | 36750804 | 36762304 | 36762431 | 13 | 14 | 16 | 18 | 9 | 0 | 0 | 2 | 3 | 1 | 8 | 10 | 15 | 20 | 0 | 0 | 0 | 1 | 0 | 0 | 245 | 149 | 1 | 1 | 1 | 1 | 0.83 | 0.785 | 0.846 | 1 | 1 | 0.901 | 1 | NA | -0.083 |
| SE | 4180 | ENSG00000105186 | ANKRD27 | chr19 | - | 32605834 | 32605954 | 32604262 | 32604424 | 32607634 | 32607832 | 49 | 25 | 46 | 39 | 23 | 0 | 0 | 2 | 0 | 0 | 45 | 46 | 29 | 44 | 34 | 0 | 0 | 0 | 0 | 0 | 269 | 149 | 1 | 1 | 1 | 1 | 0.927 | 1 | 1 | 1 | 1 | 1 | 1 | 1 | -0.015 |
| SE | 4181 | ENSG00000105186 | ANKRD27 | chr19 | - | 32607634 | 32607832 | 32605834 | 32605954 | 32615657 | 32615780 | 60 | 40 | 98 | 63 | 60 | 0 | 1 | 0 | 0 | 0 | 104 | 72 | 32 | 76 | 140 | 0 | 0 | 0 | 0 | 0 | 298 | 149 | 1 | 1 | 1 | 0.952 | 1 | 1 | 1 | 1 | 1 | 1 | 1 | 1 | -0.01 |
| SE | 4182 | ENSG00000105186 | ANKRD27 | chr19 | - | 32615657 | 32615780 | 32607634 | 32607832 | 32617588 | 32617633 | 50 | 37 | 88 | 54 | 47 | 0 | 1 | 0 | 1 | 1 | 99 | 50 | 32 | 61 | 137 | 0 | 0 | 1 | 4 | 0 | 272 | 149 | 1 | 1 | 1 | 0.953 | 1 | 0.967 | 0.963 | 1 | 1 | 0.946 | 0.893 | 1 | 0.009 |
| SE | 4183 | ENSG00000105186 | ANKRD27 | chr19 | - | 32642023 | 32642145 | 32640306 | 32640385 | 32643122 | 32643199 | 127 | 83 | 112 | 113 | 112 | 1 | 0 | 1 | 0 | 0 | 258 | 104 | 117 | 194 | 221 | 0 | 1 | 0 | 1 | 0 | 271 | 149 | 1 | 1 | 0.986 | 1 | 0.984 | 1 | 1 | 1 | 0.983 | 1 | 0.991 | 1 | -0.001 |
| SE | 4184 | ENSG00000105186 | ANKRD27 | chr19 | - | 32643122 | 32643199 | 32642023 | 32642145 | 32643286 | 32643300 | 121 | 84 | 95 | 102 | 105 | 0 | 0 | 0 | 0 | 0 | 160 | 70 | 89 | 142 | 247 | 0 | 0 | 0 | 2 | 0 | 226 | 149 | 1 | 1 | 1 | 1 | 1 | 1 | 1 | 1 | 1 | 1 | 0.979 | 1 | 0.004 |
| SE | 4187 | ENSG00000105186 | ANKRD27 | chr19 | - | 32643571 | 32643631 | 32643430 | 32643484 | 32649681 | 32649792 | 47 | 15 | 20 | 45 | 27 | 1 | 1 | 0 | 0 | 0 | 36 | 26 | 29 | 35 | 96 | 0 | 0 | 0 | 0 | 0 | 209 | 149 | 1 | 1 | 0.971 | 0.914 | 1 | 1 | 1 | 1 | 1 | 1 | 1 | 1 | -0.023 |
| SE | 4188 | ENSG00000105186 | ANKRD27 | chr19 | - | 32644324 | 32644479 | 32643430 | 32643484 | 32649681 | 32649792 | 10 | 15 | 6 | 7 | 3 | 1 | 1 | 0 | 0 | 0 | 13 | 7 | 10 | 4 | 21 | 0 | 0 | 0 | 0 | 0 | 298 | 149 | 1 | 1 | 0.833 | 0.882 | 1 | 1 | 1 | 1 | 1 | 1 | 1 | 1 | -0.057 |
| SE | 4189 | ENSG00000105186 | ANKRD27 | chr19 | - | 32644324 | 32644479 | 32643571 | 32643631 | 32646458 | 32646615 | 147 | 95 | 107 | 113 | 202 | 0 | 0 | 0 | 0 | 0 | 99 | 93 | 97 | 118 | 155 | 0 | 0 | 0 | 1 | 0 | 298 | 149 | 1 | 1 | 1 | 1 | 1 | 1 | 1 | 1 | 1 | 1 | 0.983 | 1 | 0.003 |
| SE | 4190 | ENSG00000105186 | ANKRD27 | chr19 | - | 32644324 | 32644479 | 32643571 | 32643631 | 32649681 | 32649792 | 94 | 63 | 64 | 77 | 116 | 0 | 0 | 0 | 0 | 0 | 60 | 53 | 67 | 76 | 85 | 0 | 0 | 0 | 0 | 2 | 298 | 149 | 1 | 1 | 1 | 1 | 1 | 1 | 1 | 1 | 1 | 1 | 1 | 0.955 | 0.009 |
| SE | 4191 | ENSG00000105186 | ANKRD27 | chr19 | - | 32646458 | 32646615 | 32643571 | 32643631 | 32649681 | 32649792 | 81 | 38 | 69 | 55 | 93 | 0 | 0 | 0 | 0 | 0 | 60 | 34 | 31 | 38 | 29 | 0 | 0 | 0 | 0 | 2 | 298 | 149 | 1 | 1 | 1 | 1 | 1 | 1 | 1 | 1 | 1 | 1 | 1 | 0.879 | 0.024 |
| SE | 4193 | ENSG00000105186 | ANKRD27 | chr19 | - | 32646458 | 32646615 | 32644396 | 32644476 | 32649681 | 32649792 | 81 | 38 | 69 | 57 | 93 | 0 | 0 | 1 | 0 | 0 | 60 | 34 | 31 | 37 | 29 | 0 | 0 | 0 | 0 | 0 | 298 | 149 | 1 | 1 | 1 | 1 | 0.972 | 1 | 1 | 1 | 1 | 1 | 1 | 1 | -0.006 |
| SE | 4194 | ENSG00000105186 | ANKRD27 | chr19 | - | 32649681 | 32649792 | 32646458 | 32646615 | 32658913 | 32659045 | 111 | 52 | 99 | 77 | 128 | 6 | 0 | 0 | 0 | 0 | 80 | 57 | 38 | 51 | 64 | 0 | 0 | 0 | 0 | 0 | 260 | 149 | 1 | 1 | 0.914 | 1 | 1 | 1 | 1 | 1 | 1 | 1 | 1 | 1 | -0.017 |
| SE | 4200 | ENSG00000083807 | SLC27A5 | chr19 | - | 58483531 | 58483646 | 58481063 | 58481187 | 58499122 | 58499263 | 2 | 2 | 0 | 0 | 1 | 1 | 0 | 0 | 0 | 0 | 0 | 0 | 1 | 0 | 0 | 0 | 0 | 0 | 0 | 0 | 264 | 149 | 1 | 1 | 0.53 | 1 | NA | NA | 1 | NA | NA | 1 | NA | NA | -0.157 |
| SE | 4202 | ENSG00000086967 | MYBPC2 | chr19 | + | 50436616 | 50436734 | 50436011 | 50436160 | 50437472 | 50437521 | 2 | 0 | 2 | 0 | 0 | 0 | 0 | 0 | 0 | 0 | 0 | 3 | 0 | 0 | 0 | 14 | 0 | 0 | 0 | 12 | 267 | 149 | 1 | 1 | 1 | NA | 1 | NA | NA | 0 | 1 | NA | NA | 0 | 0.667 |
| SE | 4205 | ENSG00000187994 | RINL | chr19 | - | 38871077 | 38871227 | 38870569 | 38870992 | 38871646 | 38871711 | 106 | 26 | 64 | 51 | 48 | 2 | 0 | 1 | 0 | 0 | 48 | 112 | 29 | 54 | 141 | 0 | 0 | 0 | 0 | 0 | 298 | 149 | 1 | 1 | 0.964 | 1 | 0.97 | 1 | 1 | 1 | 1 | 1 | 1 | 1 | -0.013 |
| SE | 4207 | ENSG00000187994 | RINL | chr19 | - | 38873885 | 38873988 | 38871465 | 38871870 | 38876330 | 38876474 | 108 | 22 | 32 | 65 | 43 | 3 | 1 | 0 | 1 | 0 | 73 | 112 | 11 | 36 | 54 | 0 | 0 | 0 | 0 | 0 | 252 | 149 | 1 | 1 | 0.955 | 0.929 | 1 | 0.975 | 1 | 1 | 1 | 1 | 1 | 1 | -0.028 |
| SE | 4208 | ENSG00000187994 | RINL | chr19 | - | 38876330 | 38876474 | 38873750 | 38873988 | 38876692 | 38876781 | 75 | 17 | 30 | 42 | 23 | 0 | 0 | 0 | 1 | 0 | 55 | 79 | 8 | 22 | 53 | 0 | 0 | 0 | 0 | 0 | 293 | 149 | 1 | 1 | 1 | 1 | 1 | 0.955 | 1 | 1 | 1 | 1 | 1 | 1 | -0.009 |
| SE | 4209 | ENSG00000187994 | RINL | chr19 | - | 38876330 | 38876490 | 38873885 | 38873988 | 38876692 | 38876781 | 92 | 28 | 30 | 65 | 28 | 0 | 0 | 0 | 1 | 0 | 77 | 110 | 8 | 40 | 88 | 0 | 0 | 0 | 0 | 0 | 298 | 149 | 1 | 1 | 1 | 1 | 1 | 0.97 | 1 | 1 | 1 | 1 | 1 | 1 | -0.006 |
| SE | 4212 | ENSG00000064547 | LPAR2 | chr19 | - | 19627753 | 19627969 | 19627396 | 19627504 | 19628091 | 19628167 | 1 | 3 | 5 | 2 | 0 | 0 | 1 | 6 | 2 | 0 | 0 | 1 | 1 | 0 | 7 | 0 | 2 | 0 | 0 | 0 | 298 | 149 | 1 | 1 | 1 | 0.6 | 0.294 | 0.333 | NA | NA | 0.2 | 1 | NA | 1 | -0.177 |
| SE | 4224 | ENSG00000161249 | DMKN | chr19 | - | 35506125 | 35506185 | 35505938 | 35505986 | 35508204 | 35508240 | 0 | 0 | 2 | 0 | 0 | 2 | 0 | 0 | 2 | 0 | 0 | 0 | 3 | 0 | 0 | 0 | 0 | 0 | 0 | 0 | 209 | 149 | 1 | 1 | 0 | NA | 1 | 0 | NA | NA | NA | 1 | NA | NA | -0.667 |
| SE | 4225 | ENSG00000161249 | DMKN | chr19 | - | 35506125 | 35506185 | 35505938 | 35505986 | 35509910 | 35509961 | 0 | 0 | 2 | 1 | 1 | 0 | 1 | 0 | 3 | 0 | 0 | 0 | 3 | 0 | 0 | 0 | 2 | 2 | 0 | 0 | 209 | 149 | 1 | 1 | NA | 0 | 1 | 0.192 | 1 | NA | 0 | 0.517 | NA | NA | 0.29 |
| SE | 4226 | ENSG00000161249 | DMKN | chr19 | - | 35506478 | 35506665 | 35506125 | 35506185 | 35509910 | 35509961 | 0 | 0 | 0 | 0 | 0 | 0 | 0 | 0 | 1 | 1 | 0 | 0 | 0 | 0 | 5 | 0 | 0 | 0 | 0 | 0 | 298 | 149 | 1 | 1 | NA | NA | NA | 0 | 0 | NA | NA | NA | NA | 1 | -1 |
| SE | 4228 | ENSG00000161249 | DMKN | chr19 | - | 35507457 | 35507511 | 35505938 | 35505986 | 35508204 | 35508240 | 0 | 0 | 0 | 0 | 0 | 2 | 0 | 0 | 2 | 0 | 0 | 2 | 1 | 0 | 0 | 0 | 0 | 0 | 0 | 0 | 203 | 149 | 1 | 1 | 0 | NA | NA | 0 | NA | NA | 1 | 1 | NA | NA | -1 |
| SE | 4229 | ENSG00000161249 | DMKN | chr19 | - | 35507457 | 35507511 | 35505938 | 35505986 | 35509910 | 35509961 | 0 | 0 | 0 | 0 | 0 | 0 | 1 | 0 | 3 | 0 | 0 | 0 | 4 | 0 | 0 | 0 | 2 | 2 | 0 | 0 | 203 | 149 | 1 | 1 | NA | 0 | NA | 0 | NA | NA | 0 | 0.595 | NA | NA | -0.297 |
| SE | 4231 | ENSG00000161249 | DMKN | chr19 | - | 35507457 | 35507511 | 35506125 | 35506185 | 35509910 | 35509961 | 0 | 0 | 0 | 0 | 0 | 0 | 0 | 0 | 1 | 1 | 0 | 0 | 5 | 0 | 0 | 0 | 0 | 0 | 0 | 0 | 203 | 149 | 1 | 1 | NA | NA | NA | 0 | 0 | NA | NA | 1 | NA | NA | -1 |
| SE | 4232 | ENSG00000161249 | DMKN | chr19 | - | 35508204 | 35508240 | 35505938 | 35505986 | 35509910 | 35509961 | 2 | 0 | 0 | 2 | 0 | 0 | 1 | 0 | 3 | 0 | 0 | 0 | 0 | 0 | 0 | 0 | 2 | 2 | 0 | 0 | 185 | 149 | 1 | 1 | 1 | 0 | NA | 0.349 | NA | NA | 0 | 0 | NA | NA | 0.45 |
| SE | 4235 | ENSG00000161249 | DMKN | chr19 | - | 35508204 | 35508240 | 35507457 | 35507511 | 35509910 | 35509961 | 0 | 0 | 1 | 0 | 0 | 0 | 0 | 0 | 0 | 0 | 0 | 2 | 1 | 0 | 0 | 0 | 0 | 4 | 0 | 0 | 185 | 149 | 1 | 1 | NA | NA | 1 | NA | NA | | 1 | 0.168 | NA | NA | 0.416 |
| SE | 4248 | ENSG00000089335 | ZNF302 | chr19 | + | 34679827 | 34679959 | 34678736 | 34678813 | 34682776 | 34682897 | 1 | 0 | 0 | 0 | 2 | 46 | 45 | 90 | 54 | 130 | 0 | 0 | 0 | 0 | 0 | 0 | 28 | 20 | 4 | 5 | 281 | 149 | 1 | 1 | 0.011 | 0 | 0 | 0 | 0.008 | NA | 0 | 0 | 0 | 0 | 0.004 |
| SE | 4252 | ENSG00000089335 | ZNF302 | chr19 | + | 34683151 | 34683238 | 34682776 | 34682897 | 34684251 | 34686394 | 34 | 79 | 66 | 59 | 79 | 2 | 3 | 2 | 1 | 4 | 0 | 25 | 4 | 0 | 5 | 0 | 0 | 3 | 1 | 0 | 236 | 149 | 1 | 1 | 0.915 | 0.943 | 0.954 | 0.974 | 0.926 | NA | 1 | 0.457 | 0 | 1 | 0.328 |
| SE | 4253 | ENSG00000089335 | ZNF302 | chr19 | + | 34683154 | 34683238 | 34681282 | 34682897 | 34684251 | 34684464 | 76 | 111 | 109 | 91 | 159 | 2 | 3 | 2 | 1 | 4 | 7 | 64 | 19 | 0 | 0 | 0 | 0 | 3 | 1 | 0 | 233 | 149 | 1 | 1 | 0.96 | 0.959 | 0.972 | 0.983 | 0.962 | 1 | 1 | 0.802 | 0 | NA | 0.267 |
| SE | 4257 | ENSG00000126249 | PDCD2L | chr19 | + | 34421518 | 34421667 | 34413736 | 34413847 | 34425989 | 34426150 | 63 | 32 | 44 | 66 | 69 | 1 | 0 | 0 | 0 | 0 | 26 | 60 | 19 | 8 | 14 | 0 | 0 | 0 | 0 | 0 | 298 | 149 | 1 | 1 | 0.969 | 1 | 1 | 1 | 1 | 1 | 1 | 1 | 1 | 1 | -0.006 |
| SE | 4258 | ENSG00000283201 | NA | chr19 | - | 23232166 | 23232293 | 23231265 | 23231361 | 23260863 | 23260934 | 1 | 2 | 5 | 6 | 0 | 0 | 0 | 0 | 0 | 0 | 0 | 0 | 1 | 2 | 0 | 0 | 0 | 7 | 0 | 0 | 276 | 149 | 1 | 1 | 1 | 1 | 1 | 1 | NA | NA | NA | 0.072 | 1 | NA | 0.464 |
| SE | 4259 | ENSG00000284428 | NA | chr19 | - | 23270217 | 23270292 | 23261619 | 23261744 | 23274075 | 23274196 | 2 | 3 | 2 | 6 | 4 | 10 | 3 | 4 | 0 | 0 | 0 | 5 | 3 | 0 | 10 | 0 | 0 | 1 | 3 | 3 | 224 | 149 | 1 | 1 | 0.117 | 0.399 | 0.25 | 1 | 1 | NA | 1 | 0.666 | 0 | 0.689 | -0.036 |
| SE | 4261 | ENSG00000131944 | FAAP24 | chr19 | + | 32973425 | 32973562 | 32972220 | 32972346 | 32974059 | 32974212 | 1 | 1 | 3 | 2 | 2 | 0 | 0 | 1 | 0 | 2 | 0 | 9 | 8 | 4 | 31 | 0 | 0 | 0 | 0 | 0 | 286 | 149 | 1 | 1 | 1 | 1 | 0.61 | 1 | 0.343 | NA | 1 | 1 | 1 | 1 | -0.209 |
| SE | 4267 | ENSG00000081665 | ZNF506 | chr19 | - | 19791561 | 19791731 | 19786561 | 19786854 | 19792410 | 19792484 | 9 | 3 | 7 | 5 | 14 | 1 | 0 | 0 | 1 | 0 | 0 | 12 | 7 | 13 | 0 | 0 | 1 | 0 | 0 | 0 | 298 | 149 | 1 | 1 | 0.818 | 1 | 1 | 0.714 | 1 | NA | 0.857 | 1 | 1 | NA | -0.046 |
| SE | 4271 | ENSG00000081665 | ZNF506 | chr19 | - | 19806030 | 19806126 | 19792710 | 19795660 | 19821600 | 19821709 | 20 | 28 | 38 | 41 | 40 | 1 | 0 | 0 | 1 | 2 | 0 | 10 | 16 | 5 | 0 | 2 | 0 | 1 | 0 | 0 | 245 | 149 | 1 | 1 | 0.924 | 1 | 1 | 0.961 | 0.924 | 0 | 1 | 0.907 | 1 | NA | 0.235 |
| SE | 4272 | ENSG00000081665 | ZNF506 | chr19 | - | 19806941 | 19807068 | 19794551 | 19795660 | 19821600 | 19821709 | 24 | 15 | 45 | 39 | 11 | 1 | 0 | 0 | 1 | 2 | 10 | 40 | 10 | 22 | 22 | 2 | 0 | 1 | 0 | 0 | 276 | 149 | 1 | 1 | 0.928 | 1 | 1 | 0.955 | 0.748 | 0.73 | 1 | 0.844 | 1 | 1 | 0.011 |
| SE | 4273 | ENSG00000081665 | ZNF506 | chr19 | - | 19806941 | 19807068 | 19806030 | 19806126 | 19821600 | 19821709 | 100 | 60 | 132 | 141 | 153 | 0 | 0 | 0 | 2 | 0 | 58 | 107 | 55 | 65 | 65 | 0 | 0 | 0 | 0 | 0 | 276 | 149 | 1 | 1 | 1 | 1 | 1 | 0.974 | 1 | 1 | 1 | 1 | 1 | 1 | -0.005 |
| SE | 4274 | ENSG00000089639 | GMIP | chr19 | - | 19634506 | 19634703 | 19633802 | 19634190 | 19634791 | 19634929 | 294 | 148 | 197 | 238 | 174 | 2 | 0 | 2 | 0 | 0 | 362 | 258 | 214 | 315 | 482 | 0 | 0 | 4 | 0 | 0 | 298 | 149 | 1 | 1 | 0.987 | 1 | 0.98 | 1 | 1 | 1 | 1 | 0.964 | 1 | 1 | 0.001 |
| SE | 4278 | ENSG00000089639 | GMIP | chr19 | - | 19641809 | 19641867 | 19640445 | 19640571 | 19641975 | 19642051 | 244 | 123 | 230 | 198 | 208 | 1 | 2 | 0 | 0 | 0 | 490 | 454 | 293 | 352 | 591 | 0 | 3 | 1 | 1 | 9 | 207 | 149 | 1 | 1 | 0.994 | 0.978 | 1 | 1 | 1 | 1 | 0.991 | 0.995 | 0.996 | 0.979 | 0.002 |
| SE | 4283 | ENSG00000053501 | USE1 | chr19 | + | 17217452 | 17217462 | 17216168 | 17216321 | 17218363 | 17218391 | 48 | 23 | 43 | 66 | 19 | 2 | 0 | 0 | 0 | 0 | 65 | 35 | 21 | 19 | 27 | 0 | 0 | 0 | 0 | 0 | 159 | 149 | 1 | 1 | 0.957 | 1 | 1 | 1 | 1 | 1 | 1 | 1 | 1 | 1 | -0.009 |
| SE | 4284 | ENSG00000053501 | USE1 | chr19 | + | 17219212 | 17219387 | 17218363 | 17218391 | 17219630 | 17219640 | 70 | 35 | 63 | 87 | 44 | 0 | 0 | 0 | 0 | 0 | 99 | 46 | 23 | 58 | 60 | 0 | 0 | 0 | 3 | 0 | 298 | 149 | 1 | 1 | 1 | 1 | 1 | 1 | 1 | 1 | 1 | 1 | 0.906 | 1 | 0.019 |
| SE | 4289 | ENSG00000261824 | LINC00662 | chr19 | - | 27727816 | 27727979 | 27721701 | 27721815 | 27732795 | 27732967 | 3 | 0 | 0 | 1 | 4 | 0 | 0 | 0 | 1 | 0 | 0 | 0 | 0 | 0 | 0 | 0 | 1 | 0 | 0 | 0 | 298 | 149 | 1 | 1 | 1 | NA | NA | 0.333 | 1 | NA | 0 | NA | NA | NA | 0.778 |
| SE | 4295 | ENSG00000261824 | LINC00662 | chr19 | - | 27746020 | 27746340 | 27732795 | 27732967 | 27792999 | 27793232 | 10 | 5 | 0 | 5 | 9 | 0 | 0 | 0 | 0 | 0 | 0 | 3 | 0 | 2 | 0 | 0 | 0 | 0 | 2 | 0 | 298 | 149 | 1 | 1 | 1 | 1 | NA | 1 | 1 | NA | 1 | NA | 0.333 | NA | 0.334 |
| SE | 4296 | ENSG00000261824 | LINC00662 | chr19 | - | 27746080 | 27746340 | 27732795 | 27732967 | 27792999 | 27793232 | 9 | 7 | 1 | 9 | 20 | 0 | 0 | 0 | 0 | 0 | 0 | 4 | 0 | 2 | 0 | 0 | 0 | 0 | 2 | 0 | 298 | 149 | 1 | 1 | 1 | 1 | 1 | 1 | 1 | NA | 1 | NA | 0.333 | NA | 0.334 |
| SE | 4298 | ENSG00000261824 | LINC00662 | chr19 | - | 27746097 | 27746340 | 27731331 | 27732967 | 27792999 | 27793138 | 15 | 6 | 4 | 9 | 16 | 0 | 0 | 0 | 0 | 0 | 0 | 8 | 1 | 12 | 22 | 0 | 0 | 0 | 2 | 0 | 298 | 149 | 1 | 1 | 1 | 1 | 1 | 1 | 1 | NA | 1 | 1 | 0.75 | 1 | 0.062 |
| SE | 4300 | ENSG00000261824 | LINC00662 | chr19 | - | 27771663 | 27771989 | 27746097 | 27746340 | 27792999 | 27793232 | 0 | 6 | 0 | 0 | 0 | 9 | 5 | 0 | 5 | 9 | 0 | 0 | 0 | 0 | 0 | 0 | 3 | 0 | 2 | 0 | 298 | 149 | 1 | 1 | 0 | 0.375 | NA | 0 | 0 | NA | 0 | NA | 0 | NA | 0.094 |
| SE | 4303 | ENSG00000261824 | LINC00662 | chr19 | - | 27778710 | 27779043 | 27774166 | 27774252 | 27792999 | 27793232 | 2 | 8 | 2 | 2 | 2 | 2 | 0 | 0 | 1 | 0 | 8 | 0 | 3 | 5 | 8 | 0 | 0 | 1 | 0 | 0 | 298 | 149 | 1 | 1 | 0.333 | 1 | 1 | 0.5 | 1 | 1 | NA | 0.6 | 1 | 1 | -0.133 |
| SE | 4304 | ENSG00000261824 | LINC00662 | chr19 | - | 27778710 | 27779043 | 27774743 | 27774792 | 27792999 | 27793232 | 2 | 8 | 2 | 2 | 2 | 1 | 0 | 2 | 0 | 0 | 8 | 0 | 3 | 5 | 8 | 0 | 0 | 0 | 0 | 0 | 298 | 149 | 1 | 1 | 0.5 | 1 | 0.333 | 1 | 1 | 1 | NA | 1 | 1 | 1 | -0.233 |
| SE | 4305 | ENSG00000182141 | ZNF708 | chr19 | - | 21306817 | 21306941 | 21294542 | 21294739 | 21309245 | 21309341 | 2 | 3 | 0 | 0 | 0 | 31 | 28 | 41 | 63 | 61 | 0 | 0 | 0 | 0 | 0 | 0 | 16 | 21 | 0 | 16 | 273 | 149 | 1 | 1 | 0.034 | 0.055 | 0 | 0 | 0 | NA | 0 | 0 | NA | 0 | 0.018 |
| SE | 4306 | ENSG00000182141 | ZNF708 | chr19 | - | 21309245 | 21309341 | 21291159 | 21294739 | 21327959 | 21328141 | 24 | 21 | 29 | 51 | 44 | 0 | 2 | 0 | 0 | 1 | 0 | 12 | 18 | 0 | 15 | 0 | 2 | 0 | 0 | 0 | 245 | 149 | 1 | 1 | 1 | 0.865 | 1 | 1 | 0.964 | NA | 0.785 | 1 | NA | 1 | 0.037 |
| SE | 4307 | ENSG00000182141 | ZNF708 | chr19 | - | 21309245 | 21309341 | 21291159 | 21294739 | 21329209 | 21329336 | 32 | 24 | 48 | 59 | 56 | 2 | 0 | 0 | 3 | 1 | 0 | 14 | 19 | 13 | 12 | 0 | 2 | 0 | 0 | 0 | 245 | 149 | 1 | 1 | 0.907 | 1 | 1 | 0.923 | 0.971 | NA | 0.81 | 1 | 1 | 1 | 0.008 |
| SE | 4310 | ENSG00000182141 | ZNF708 | chr19 | - | 21327959 | 21328141 | 21292603 | 21294739 | 21329209 | 21329425 | 9 | 8 | 3 | 5 | 7 | 2 | 0 | 0 | 3 | 1 | 7 | 12 | 7 | 0 | 0 | 0 | 2 | 0 | 0 | 0 | 298 | 149 | 1 | 1 | 0.692 | 1 | 1 | 0.455 | 0.778 | 1 | 0.75 | 1 | NA | NA | -0.132 |
| SE | 4312 | ENSG00000182141 | ZNF708 | chr19 | - | 21327959 | 21328141 | 21310500 | 21310627 | 21329209 | 21329425 | 12 | 6 | 5 | 10 | 12 | 6 | 3 | 4 | 10 | 0 | 7 | 13 | 11 | 7 | 0 | 4 | 4 | 2 | 0 | 0 | 298 | 149 | 1 | 1 | 0.5 | 0.5 | 0.385 | 0.333 | 1 | 0.467 | 0.619 | 0.733 | 1 | NA | -0.161 |
| SE | 4313 | ENSG00000213999 | MEF2B | chr19 | - | 19146272 | 19146384 | 19145566 | 19146022 | 19146554 | 19146648 | 2 | 0 | 0 | 2 | 7 | 0 | 1 | 0 | 0 | 0 | 0 | 0 | 0 | 0 | 7 | 0 | 0 | 0 | 0 | 0 | 261 | 149 | 1 | 1 | 1 | 0 | NA | 1 | 1 | NA | NA | NA | NA | 1 | -0.25 |
| SE | 4316 | ENSG00000213999 | MEF2B | chr19 | - | 19157024 | 19157142 | 19150681 | 19150764 | 19170204 | 19170289 | 0 | 0 | 1 | 1 | 0 | 0 | 0 | 0 | 0 | 0 | 0 | 0 | 4 | 3 | 13 | 0 | 4 | 2 | 0 | 0 | 267 | 149 | 1 | 1 | NA | NA | 1 | 1 | NA | NA | 0 | 0.527 | 1 | 1 | 0.368 |
| SE | 4319 | ENSG00000269110 | NA | chr19 | - | 20646278 | 20646405 | 20645683 | 20645779 | 20661443 | 20661583 | 42 | 43 | 45 | 51 | 42 | 1 | 0 | 0 | 0 | 0 | 3 | 18 | 18 | 32 | 2 | 0 | 0 | 0 | 0 | 0 | 276 | 149 | 1 | 1 | 0.958 | 1 | 1 | 1 | 1 | 1 | 1 | 1 | 1 | 1 | -0.008 |
| SE | 4326 | ENSG00000187187 | ZNF546 | chr19 | + | 39998247 | 39998332 | 39997860 | 39997915 | 40006095 | 40006182 | 8 | 3 | 3 | 2 | 0 | 3 | 2 | 0 | 2 | 5 | 0 | 0 | 0 | 0 | 0 | 0 | 7 | 2 | 3 | 0 | 234 | 149 | 1 | 1 | 0.629 | 0.489 | 1 | 0.389 | 0 | NA | 0 | 0 | 0 | NA | 0.501 |
| SE | 4327 | ENSG00000187187 | ZNF546 | chr19 | + | 39998247 | 39998410 | 39997860 | 39997915 | 40006095 | 40006182 | 13 | 4 | 7 | 2 | 3 | 3 | 2 | 0 | 2 | 5 | 0 | 5 | 0 | 0 | 0 | 0 | 7 | 2 | 3 | 0 | 298 | 149 | 1 | 1 | 0.684 | 0.5 | 1 | 0.333 | 0.231 | NA | 0.263 | 0 | 0 | NA | 0.462 |
| SE | 4329 | ENSG00000187187 | ZNF546 | chr19 | + | 40006095 | 40006182 | 39997860 | 39997915 | 40007273 | 40007400 | 12 | 10 | 5 | 15 | 13 | 0 | 0 | 1 | 0 | 0 | 0 | 11 | 11 | 6 | 0 | 0 | 0 | 0 | 0 | 0 | 236 | 149 | 1 | 1 | 1 | 1 | 0.759 | 1 | 1 | NA | 1 | 1 | 1 | NA | -0.048 |
| SE | 4334 | ENSG00000064607 | SUGP2 | chr19 | - | 18993647 | 18993740 | 18990889 | 18991149 | 18994365 | 18995280 | 19 | 11 | 25 | 16 | 28 | 0 | 0 | 0 | 0 | 0 | 0 | 21 | 19 | 28 | 6 | 0 | 0 | 1 | 5 | 0 | 242 | 149 | 1 | 1 | 1 | 1 | 1 | 1 | 1 | NA | 1 | 0.921 | 0.775 | 1 | 0.076 |
| SE | 4337 | ENSG00000064607 | SUGP2 | chr19 | - | 19001612 | 19001674 | 18994365 | 18995280 | 19004167 | 19004273 | 84 | 76 | 74 | 78 | 55 | 0 | 0 | 0 | 3 | 0 | 82 | 81 | 73 | 128 | 45 | 1 | 0 | 5 | 0 | 0 | 211 | 149 | 1 | 1 | 1 | 1 | 1 | 0.948 | 1 | 0.983 | 1 | 0.912 | 1 | 1 | 0.011 |
| SE | 4339 | ENSG00000064607 | SUGP2 | chr19 | - | 19004167 | 19004646 | 19001612 | 19001674 | 19008316 | 19008428 | 198 | 158 | 181 | 217 | 108 | 2 | 0 | 0 | 0 | 0 | 395 | 231 | 314 | 354 | 279 | 0 | 0 | 0 | 0 | 0 | 298 | 149 | 1 | 1 | 0.98 | 1 | 1 | 1 | 1 | 1 | 1 | 1 | 1 | 1 | -0.004 |
| SE | 4342 | ENSG00000064607 | SUGP2 | chr19 | - | 19009854 | 19010342 | 19008316 | 19008428 | 19019108 | 19019229 | 278 | 219 | 303 | 328 | 214 | 0 | 0 | 0 | 0 | 0 | 599 | 400 | 381 | 497 | 357 | 0 | 7 | 1 | 0 | 0 | 298 | 149 | 1 | 1 | 1 | 1 | 1 | 1 | 1 | 1 | 0.966 | 0.995 | 1 | 1 | 0.008 |
| SE | 4344 | ENSG00000064607 | SUGP2 | chr19 | - | 19030950 | 19031082 | 19024618 | 19026226 | 19033436 | 19033490 | 227 | 155 | 203 | 215 | 135 | 0 | 0 | 0 | 0 | 0 | 248 | 263 | 255 | 363 | 247 | 0 | 1 | 0 | 5 | 0 | 281 | 149 | 1 | 1 | 1 | 1 | 1 | 1 | 1 | 1 | 0.993 | 1 | 0.975 | 1 | 0.006 |
| SE | 4345 | ENSG00000064607 | SUGP2 | chr19 | - | 19030950 | 19031082 | 19024618 | 19026226 | 19033761 | 19033816 | 200 | 146 | 186 | 194 | 134 | 0 | 0 | 0 | 0 | 0 | 186 | 252 | 233 | 330 | 236 | 0 | 0 | 0 | 1 | 0 | 281 | 149 | 1 | 1 | 1 | 1 | 1 | 1 | 1 | 1 | 1 | 1 | 0.994 | 1 | 0.001 |
| SE | 4346 | ENSG00000064607 | SUGP2 | chr19 | - | 19030950 | 19031107 | 19024618 | 19026226 | 19033761 | 19034023 | 188 | 142 | 177 | 185 | 125 | 0 | 0 | 0 | 0 | 0 | 190 | 250 | 233 | 320 | 221 | 0 | 0 | 0 | 1 | 0 | 298 | 149 | 1 | 1 | 1 | 1 | 1 | 1 | 1 | 1 | 1 | 1 | 0.994 | 1 | 0.001 |
| SE | 4347 | ENSG00000130311 | DDA1 | chr19 | + | 17315933 | 17315995 | 17314337 | 17314389 | 17319545 | 17319725 | 144 | 88 | 109 | 132 | 154 | 0 | 0 | 0 | 0 | 4 | 274 | 189 | 120 | 184 | 233 | 0 | 0 | 0 | 2 | 0 | 211 | 149 | 1 | 1 | 1 | 1 | 1 | 1 | 0.965 | 1 | 1 | 1 | 0.985 | 1 | -0.004 |
| SE | 4348 | ENSG00000197372 | ZNF675 | chr19 | - | 23656721 | 23656822 | 23654545 | 23654706 | 23662113 | 23662209 | 2 | 6 | 4 | 7 | 2 | 30 | 32 | 26 | 16 | 20 | 0 | 9 | 0 | 7 | 3 | 0 | 1 | 0 | 0 | 0 | 250 | 149 | 1 | 1 | 0.038 | 0.101 | 0.084 | 0.207 | 0.056 | NA | 0.843 | NA | 1 | 1 | -0.85 |
| SE | 4350 | ENSG00000130511 | SSBP4 | chr19 | + | 18425908 | 18426083 | 18419623 | 18419707 | 18427350 | 18427423 | 0 | 0 | 0 | 0 | 0 | 71 | 14 | 26 | 33 | 8 | 0 | 3 | 0 | 0 | 0 | 52 | 48 | 18 | 10 | 31 | 298 | 149 | 1 | 1 | 0 | 0 | 0 | 0 | 0 | 0 | 0.03 | 0 | 0 | 0 | -0.006 |
| SE | 4354 | ENSG00000197013 | ZNF429 | chr19 | + | 21505984 | 21506080 | 21505603 | 21505774 | 21518574 | 21518988 | 0 | 4 | 0 | 2 | 2 | 2 | 0 | 1 | 0 | 0 | 0 | 0 | 0 | 0 | 5 | 0 | 0 | 1 | 1 | 0 | 245 | 149 | 1 | 1 | 0 | 1 | 0 | 1 | 1 | NA | NA | 0 | 0 | 1 | 0.267 |
| SE | 4355 | ENSG00000197013 | ZNF429 | chr19 | + | 21505984 | 21506080 | 21505603 | 21505774 | 21529657 | 21529784 | 0 | 4 | 0 | 1 | 2 | 4 | 2 | 14 | 8 | 2 | 0 | 1 | 0 | 0 | 5 | 0 | 4 | 0 | 0 | 0 | 245 | 149 | 1 | 1 | 0 | 0.549 | 0 | 0.071 | 0.378 | NA | 0.132 | NA | NA | 1 | -0.366 |
| SE | 4356 | ENSG00000197013 | ZNF429 | chr19 | + | 21529083 | 21529279 | 21505660 | 21505774 | 21529657 | 21529784 | 0 | 4 | 0 | 3 | 9 | 4 | 2 | 14 | 8 | 2 | 0 | 0 | 0 | 6 | 0 | 0 | 4 | 0 | 0 | 0 | 298 | 149 | 1 | 1 | 0 | 0.5 | 0 | 0.158 | 0.692 | NA | 0 | NA | 1 | NA | -0.23 |
| SE | 4357 | ENSG00000197013 | ZNF429 | chr19 | + | 21529657 | 21529784 | 21505601 | 21505774 | 21530588 | 21530684 | 44 | 27 | 24 | 32 | 48 | 2 | 0 | 0 | 0 | 0 | 13 | 15 | 4 | 10 | 0 | 0 | 0 | 0 | 0 | 0 | 276 | 149 | 1 | 1 | 0.922 | 1 | 1 | 1 | 1 | 1 | 1 | 1 | 1 | NA | -0.016 |
| SE | 4358 | ENSG00000197013 | ZNF429 | chr19 | + | 21530588 | 21530684 | 21529657 | 21529784 | 21536279 | 21536499 | 69 | 58 | 38 | 58 | 100 | 0 | 2 | 0 | 0 | 0 | 13 | 18 | 8 | 16 | 7 | 0 | 0 | 1 | 0 | 0 | 245 | 149 | 1 | 1 | 1 | 0.946 | 1 | 1 | 1 | 1 | 1 | 0.83 | 1 | 1 | 0.023 |
| SE | 4365 | ENSG00000167232 | ZNF91 | chr19 | - | 23342169 | 23342265 | 23339230 | 23339260 | 23373741 | 23373837 | 4 | 1 | 3 | 6 | 15 | 0 | 0 | 0 | 0 | 5 | 0 | 0 | 0 | 0 | 0 | 0 | 4 | 0 | 0 | 0 | 245 | 149 | 1 | 1 | 1 | 1 | 1 | 1 | 0.646 | NA | 0 | NA | NA | NA | 0.929 |
| SE | 4366 | ENSG00000167232 | ZNF91 | chr19 | - | 23373741 | 23373837 | 23338748 | 23339054 | 23395324 | 23395457 | 1 | 1 | 2 | 1 | 0 | 0 | 0 | 2 | 0 | 0 | 0 | 0 | 2 | 0 | 0 | 0 | 0 | 0 | 0 | 0 | 245 | 149 | 1 | 1 | 1 | 1 | 0.378 | 1 | NA | NA | NA | 1 | NA | NA | -0.155 |
| SE | 4367 | ENSG00000167232 | ZNF91 | chr19 | - | 23373741 | 23373837 | 23357695 | 23362725 | 23374637 | 23374764 | 88 | 61 | 66 | 79 | 80 | 2 | 0 | 1 | 2 | 0 | 9 | 46 | 34 | 24 | 20 | 0 | 3 | 2 | 0 | 0 | 245 | 149 | 1 | 1 | 0.964 | 1 | 0.976 | 0.96 | 1 | 1 | 0.903 | 0.912 | 1 | 1 | 0.017 |
| SE | 4368 | ENSG00000167232 | ZNF91 | chr19 | - | 23373741 | 23373837 | 23357695 | 23362725 | 23395324 | 23395457 | 33 | 22 | 13 | 28 | 32 | 3 | 1 | 0 | 0 | 0 | 1 | 5 | 8 | 0 | 11 | 0 | 0 | 0 | 0 | 0 | 245 | 149 | 1 | 1 | 0.87 | 0.93 | 1 | 1 | 1 | 1 | 1 | 1 | NA | 1 | -0.04 |
| SE | 4369 | ENSG00000167232 | ZNF91 | chr19 | - | 23374637 | 23374764 | 23359402 | 23362725 | 23395324 | 23395446 | 54 | 41 | 57 | 67 | 19 | 3 | 1 | 0 | 0 | 0 | 14 | 57 | 52 | 16 | 29 | 0 | 0 | 0 | 0 | 0 | 276 | 149 | 1 | 1 | 0.907 | 0.957 | 1 | 1 | 1 | 1 | 1 | 1 | 1 | 1 | -0.027 |
| SE | 4370 | ENSG00000167232 | ZNF91 | chr19 | - | 23374637 | 23374764 | 23373741 | 23373837 | 23395324 | 23395446 | 129 | 98 | 137 | 141 | 80 | 1 | 1 | 3 | 1 | 0 | 32 | 127 | 124 | 51 | 53 | 0 | 0 | 2 | 0 | 0 | 276 | 149 | 1 | 1 | 0.986 | 0.981 | 0.961 | 0.987 | 1 | 1 | 1 | 0.971 | 1 | 1 | -0.011 |
| SE | 4372 | ENSG00000118620 | ZNF430 | chr19 | + | 21022788 | 21022881 | 21020633 | 21020803 | 21056630 | 21060050 | 23 | 11 | 26 | 20 | 11 | 0 | 1 | 0 | 0 | 0 | 8 | 15 | 29 | 11 | 6 | 0 | 1 | 0 | 0 | 0 | 242 | 149 | 1 | 1 | 1 | 0.871 | 1 | 1 | 1 | 1 | 0.902 | 1 | 1 | 1 | -0.006 |
| SE | 4373 | ENSG00000118620 | ZNF430 | chr19 | + | 21033455 | 21033582 | 21022788 | 21022881 | 21034088 | 21036802 | 91 | 91 | 93 | 92 | 98 | 0 | 0 | 0 | 0 | 2 | 50 | 82 | 98 | 53 | 27 | 0 | 0 | 0 | 0 | 0 | 276 | 149 | 1 | 1 | 1 | 1 | 1 | 1 | 0.964 | 1 | 1 | 1 | 1 | 1 | -0.007 |
| SE | 4380 | ENSG00000064545 | TMEM161A | chr19 | - | 19134783 | 19134887 | 19132879 | 19133210 | 19138425 | 19138447 | 3 | 0 | 1 | 5 | 0 | 0 | 0 | 0 | 0 | 0 | 0 | 6 | 0 | 6 | 4 | 0 | 0 | 1 | 0 | 0 | 253 | 149 | 1 | 1 | 1 | NA | 1 | 1 | NA | NA | 1 | 0 | 1 | 1 | 0.25 |
| SE | 4381 | ENSG00000064545 | TMEM161A | chr19 | - | 19134783 | 19134916 | 19133129 | 19133210 | 19138425 | 19138467 | 2 | 0 | 1 | 8 | 0 | 0 | 0 | 0 | 0 | 0 | 0 | 2 | 0 | 3 | 4 | 0 | 0 | 1 | 0 | 0 | 282 | 149 | 1 | 1 | 1 | NA | 1 | 1 | NA | NA | 1 | 0 | 1 | 1 | 0.25 |
| SE | 4388 | ENSG00000229676 | ZNF492 | chr19 | + | 22653919 | 22654015 | 22653306 | 22653433 | 22663799 | 22667671 | 0 | 4 | 1 | 1 | 2 | 0 | 0 | 0 | 0 | 0 | 0 | 2 | 0 | 0 | 0 | 0 | 0 | 2 | 0 | 0 | 245 | 149 | 1 | 1 | NA | 1 | 1 | 1 | 1 | NA | 1 | 0 | NA | NA | 0.5 |
| SE | 4392 | ENSG00000130475 | FCHO1 | chr19 | + | 17754591 | 17754621 | 17754316 | 17754361 | 17755117 | 17755191 | 5 | 2 | 13 | 2 | 0 | 1 | 1 | 8 | 5 | 0 | 0 | 9 | 1 | 6 | 10 | 0 | 3 | 7 | 10 | 23 | 179 | 149 | 1 | 1 | 0.806 | 0.625 | 0.575 | 0.25 | NA | NA | 0.714 | 0.106 | 0.333 | 0.266 | 0.209 |
| SE | 4394 | ENSG00000130475 | FCHO1 | chr19 | + | 17755075 | 17755191 | 17747717 | 17747814 | 17762761 | 17762853 | 26 | 16 | 39 | 35 | 11 | 3 | 0 | 0 | 0 | 0 | 56 | 39 | 21 | 10 | 55 | 0 | 0 | 0 | 0 | 0 | 265 | 149 | 1 | 1 | 0.83 | 1 | 1 | 1 | 1 | 1 | 1 | 1 | 1 | 1 | -0.034 |
| SE | 4395 | ENSG00000130475 | FCHO1 | chr19 | + | 17755117 | 17755191 | 17754316 | 17754361 | 17762761 | 17762805 | 25 | 17 | 45 | 35 | 11 | 0 | 0 | 0 | 1 | 0 | 56 | 42 | 28 | 15 | 78 | 0 | 0 | 0 | 0 | 0 | 223 | 149 | 1 | 1 | 1 | 1 | 1 | 0.959 | 1 | 1 | 1 | 1 | 1 | 1 | -0.008 |
| SE | 4401 | ENSG00000130475 | FCHO1 | chr19 | + | 17776246 | 17776271 | 17775982 | 17776161 | 17776634 | 17776686 | 60 | 30 | 43 | 44 | 39 | 0 | 0 | 0 | 1 | 0 | 39 | 75 | 19 | 44 | 21 | 0 | 0 | 0 | 0 | 0 | 174 | 149 | 1 | 1 | 1 | 1 | 1 | 0.974 | 1 | 1 | 1 | 1 | 1 | 1 | -0.005 |
| SE | 4402 | ENSG00000130475 | FCHO1 | chr19 | + | 17776634 | 17776686 | 17776156 | 17776271 | 17778136 | 17778228 | 67 | 23 | 43 | 53 | 55 | 0 | 0 | 0 | 1 | 0 | 36 | 73 | 30 | 46 | 48 | 0 | 3 | 0 | 0 | 0 | 201 | 149 | 1 | 1 | 1 | 1 | 1 | 0.975 | 1 | 1 | 0.947 | 1 | 1 | 1 | 0.006 |
| SE | 4404 | ENSG00000130475 | FCHO1 | chr19 | + | 17778608 | 17778884 | 17778136 | 17778228 | 17781230 | 17781343 | 40 | 8 | 27 | 43 | 42 | 0 | 0 | 0 | 0 | 0 | 22 | 32 | 32 | 8 | 60 | 0 | 0 | 0 | 1 | 0 | 298 | 149 | 1 | 1 | 1 | 1 | 1 | 1 | 1 | 1 | 1 | 1 | 0.8 | 1 | 0.04 |
| SE | 4406 | ENSG00000130475 | FCHO1 | chr19 | + | 17784724 | 17784924 | 17784102 | 17784235 | 17786573 | 17786629 | 91 | 46 | 68 | 64 | 35 | 1 | 1 | 0 | 0 | 0 | 40 | 98 | 36 | 61 | 122 | 3 | 0 | 0 | 0 | 0 | 298 | 149 | 1 | 1 | 0.978 | 0.958 | 1 | 1 | 1 | 0.87 | 1 | 1 | 1 | 1 | 0.013 |
| SE | 4408 | ENSG00000184635 | ZNF93 | chr19 | + | 19914733 | 19914822 | 19900982 | 19901091 | 19915279 | 19915406 | 0 | 0 | 0 | 1 | 0 | 3 | 0 | 4 | 6 | 1 | 0 | 0 | 0 | 0 | 0 | 0 | 0 | 2 | 0 | 2 | 238 | 149 | 1 | 1 | 0 | NA | 0 | 0.094 | 0 | NA | NA | 0 | NA | 0 | 0.024 |
| SE | 4410 | ENSG00000184635 | ZNF93 | chr19 | + | 19926100 | 19926221 | 19916559 | 19916655 | 19933181 | 19933274 | 0 | 0 | 3 | 0 | 0 | 6 | 1 | 17 | 30 | 5 | 0 | 0 | 0 | 0 | 0 | 10 | 15 | 6 | 4 | 6 | 270 | 149 | 1 | 1 | 0 | 0 | 0.089 | 0 | 0 | 0 | 0 | 0 | 0 | 0 | 0.018 |
| SE | 4412 | ENSG00000105662 | CRTC1 | chr19 | + | 18744107 | 18744155 | 18742909 | 18743026 | 18745822 | 18745960 | 0 | 1 | 1 | 0 | 1 | 21 | 9 | 11 | 22 | 18 | 0 | 17 | 0 | 0 | 0 | 0 | 10 | 9 | 8 | 26 | 197 | 149 | 1 | 1 | 0 | 0.078 | 0.064 | 0 | 0.04 | NA | 0.563 | 0 | 0 | 0 | -0.104 |
| SE | 4414 | ENSG00000105662 | CRTC1 | chr19 | + | 18768484 | 18768793 | 18765403 | 18765528 | 18775640 | 18775821 | 3 | 11 | 4 | 5 | 7 | 0 | 0 | 0 | 0 | 0 | 0 | 2 | 1 | 2 | 11 | 0 | 0 | 1 | 0 | 0 | 298 | 149 | 1 | 1 | 1 | 1 | 1 | 1 | 1 | NA | 1 | 0.333 | 1 | 1 | 0.167 |
| SE | 4415 | ENSG00000105662 | CRTC1 | chr19 | + | 18771441 | 18771546 | 18768484 | 18768793 | 18775640 | 18775821 | 0 | 2 | 1 | 0 | 0 | 0 | 0 | 0 | 1 | 0 | 8 | 7 | 0 | 0 | 0 | 0 | 0 | 0 | 0 | 0 | 254 | 149 | 1 | 1 | NA | 1 | 1 | 0 | NA | 1 | 1 | NA | NA | NA | -0.333 |
| SE | 4416 | ENSG00000105662 | CRTC1 | chr19 | + | 18774899 | 18774986 | 18765403 | 18765528 | 18775640 | 18775821 | 9 | 2 | 3 | 3 | 1 | 0 | 0 | 0 | 0 | 0 | 0 | 10 | 2 | 1 | 4 | 0 | 0 | 1 | 0 | 0 | 236 | 149 | 1 | 1 | 1 | 1 | 1 | 1 | 1 | NA | 1 | 0.558 | 1 | 1 | 0.111 |
| SE | 4417 | ENSG00000105662 | CRTC1 | chr19 | + | 18774899 | 18774986 | 18771441 | 18771546 | 18775640 | 18775821 | 19 | 4 | 4 | 7 | 15 | 0 | 1 | 0 | 0 | 0 | 4 | 15 | 2 | 3 | 30 | 0 | 0 | 0 | 0 | 0 | 236 | 149 | 1 | 1 | 1 | 0.716 | 1 | 1 | 1 | 1 | 1 | 1 | 1 | 1 | -0.057 |
| SE | 4418 | ENSG00000105726 | ATP13A1 | chr19 | - | 19654542 | 19654700 | 19653968 | 19654144 | 19655118 | 19655239 | 120 | 56 | 124 | 120 | 97 | 0 | 0 | 0 | 0 | 0 | 138 | 147 | 81 | 127 | 234 | 0 | 0 | 3 | 0 | 4 | 298 | 149 | 1 | 1 | 1 | 1 | 1 | 1 | 1 | 1 | 1 | 0.931 | 1 | 0.967 | 0.02 |
| SE | 4423 | ENSG00000105072 | C19orf44 | chr19 | + | 16503064 | 16503380 | 16500791 | 16501551 | 16506700 | 16506774 | 7 | 1 | 5 | 6 | 13 | 1 | 0 | 0 | 0 | 0 | 24 | 5 | 5 | 6 | 21 | 0 | 0 | 0 | 0 | 0 | 298 | 149 | 1 | 1 | 0.778 | 1 | 1 | 1 | 1 | 1 | 1 | 1 | 1 | 1 | -0.044 |
| SE | 4426 | ENSG00000105072 | C19orf44 | chr19 | + | 16514496 | 16514663 | 16509498 | 16509988 | 16517229 | 16517341 | 1 | 0 | 1 | 0 | 0 | 0 | 2 | 0 | 0 | 3 | 11 | 0 | 6 | 0 | 14 | 0 | 0 | 0 | 0 | 0 | 298 | 149 | 1 | 1 | 1 | 0 | 1 | NA | 0 | 1 | NA | 1 | NA | 1 | -0.5 |
| SE | 4430 | ENSG00000105705 | SUGP1 | chr19 | - | 19296988 | 19297344 | 19280184 | 19280291 | 19302264 | 19302388 | 146 | 86 | 127 | 101 | 213 | 0 | 0 | 0 | 0 | 4 | 148 | 135 | 45 | 122 | 151 | 0 | 0 | 0 | 0 | 0 | 298 | 149 | 1 | 1 | 1 | 1 | 1 | 1 | 0.964 | 1 | 1 | 1 | 1 | 1 | -0.007 |
| SE | 4431 | ENSG00000105705 | SUGP1 | chr19 | - | 19302264 | 19302388 | 19296988 | 19297344 | 19303347 | 19303406 | 173 | 114 | 154 | 119 | 351 | 1 | 0 | 0 | 0 | 0 | 177 | 166 | 70 | 131 | 238 | 0 | 0 | 0 | 0 | 0 | 273 | 149 | 1 | 1 | 0.99 | 1 | 1 | 1 | 1 | 1 | 1 | 1 | 1 | 1 | -0.002 |
| SE | 4432 | ENSG00000105705 | SUGP1 | chr19 | - | 19302264 | 19302443 | 19296988 | 19297344 | 19303347 | 19303448 | 97 | 53 | 71 | 57 | 168 | 1 | 0 | 0 | 0 | 0 | 73 | 64 | 32 | 70 | 89 | 0 | 0 | 0 | 0 | 0 | 298 | 149 | 1 | 1 | 0.98 | 1 | 1 | 1 | 1 | 1 | 1 | 1 | 1 | 1 | -0.004 |
| SE | 4434 | ENSG00000105705 | SUGP1 | chr19 | - | 19303723 | 19304043 | 19303347 | 19303406 | 19305848 | 19306076 | 0 | 0 | 0 | 0 | 7 | 0 | 0 | 0 | 0 | 2 | 3 | 0 | 4 | 0 | 2 | 0 | 3 | 0 | 5 | 0 | 298 | 149 | 1 | 1 | NA | NA | NA | NA | 0.636 | 1 | 0 | 1 | 0 | 1 | 0.036 |
| SE | 4436 | ENSG00000105705 | SUGP1 | chr19 | - | 19310096 | 19310200 | 19305645 | 19306076 | 19316421 | 19316522 | 119 | 73 | 99 | 106 | 408 | 2 | 0 | 1 | 0 | 4 | 221 | 178 | 106 | 247 | 232 | 1 | 0 | 0 | 0 | 1 | 253 | 149 | 1 | 1 | 0.972 | 1 | 0.983 | 1 | 0.984 | 0.992 | 1 | 1 | 1 | 0.993 | -0.009 |
| SE | 4440 | ENSG00000186010 | NDUFA13 | chr19 | + | 19527280 | 19527352 | 19526024 | 19526260 | 19527700 | 19527770 | 560 | 290 | 464 | 479 | 274 | 1 | 0 | 0 | 0 | 0 | 801 | 532 | 362 | 508 | 788 | 0 | 1 | 0 | 0 | 9 | 221 | 149 | 1 | 1 | 0.997 | 1 | 1 | 1 | 1 | 1 | 0.997 | 1 | 1 | 0.983 | 0.003 |
| SE | 4441 | ENSG00000186010 | NDUFA13 | chr19 | + | 19527280 | 19527372 | 19526181 | 19526260 | 19527700 | 19527770 | 231 | 123 | 196 | 180 | 61 | 1 | 0 | 0 | 0 | 0 | 433 | 210 | 149 | 253 | 302 | 0 | 1 | 0 | 0 | 9 | 241 | 149 | 1 | 1 | 0.993 | 1 | 1 | 1 | 1 | 1 | 0.992 | 1 | 1 | 0.954 | 0.009 |
| SE | 4442 | ENSG00000267419 | ZNF56 | chr19 | + | 19788949 | 19789072 | 19786608 | 19786691 | 19789679 | 19789821 | 8 | 4 | 2 | 0 | 14 | 1 | 0 | 0 | 0 | 0 | 0 | 0 | 3 | 0 | 0 | 0 | 0 | 2 | 0 | 0 | 272 | 149 | 1 | 1 | 0.814 | 1 | 1 | NA | 1 | NA | NA | 0.451 | NA | NA | 0.502 |
| SE | 4443 | ENSG00000267419 | ZNF56 | chr19 | + | 19789679 | 19789821 | 19786608 | 19786691 | 19806280 | 19806376 | 1 | 0 | 0 | 0 | 0 | 0 | 0 | 0 | 1 | 0 | 0 | 0 | 2 | 0 | 0 | 0 | 0 | 0 | 0 | 0 | 291 | 149 | 1 | 1 | 1 | NA | NA | 0 | NA | NA | NA | 1 | NA | NA | -0.5 |
| SE | 4444 | ENSG00000105058 | FAM32A | chr19 | + | 16185623 | 16185765 | 16185409 | 16185516 | 16190886 | 16191448 | 68 | 59 | 36 | 59 | 64 | 0 | 0 | 0 | 1 | 0 | 173 | 98 | 54 | 138 | 120 | 0 | 0 | 0 | 0 | 0 | 291 | 149 | 1 | 1 | 1 | 1 | 1 | 0.968 | 1 | 1 | 1 | 1 | 1 | 1 | -0.006 |
| SE | 4446 | ENSG00000105058 | FAM32A | chr19 | + | 16189387 | 16189511 | 16185623 | 16185765 | 16190886 | 16191378 | 0 | 1 | 2 | 0 | 6 | 7 | 3 | 2 | 0 | 5 | 0 | 0 | 0 | 0 | 7 | 12 | 12 | 2 | 3 | 3 | 273 | 149 | 1 | 1 | 0 | 0.154 | 0.353 | NA | 0.396 | 0 | 0 | 0 | 0 | 0.56 | 0.114 |
| SE | 4449 | ENSG00000074855 | ANO8 | chr19 | - | 17327437 | 17327569 | 17327234 | 17327345 | 17327688 | 17327880 | 30 | 13 | 15 | 23 | 4 | 0 | 0 | 0 | 4 | 0 | 0 | 11 | 23 | 11 | 28 | 0 | 0 | 0 | 0 | 0 | 281 | 149 | 1 | 1 | 1 | 1 | 1 | 0.753 | 1 | NA | 1 | 1 | 1 | 1 | -0.049 |
| SE | 4450 | ENSG00000181896 | ZNF101 | chr19 | + | 19672209 | 19672358 | 19668900 | 19668966 | 19677863 | 19677990 | 0 | 0 | 0 | 0 | 2 | 104 | 47 | 149 | 126 | 96 | 0 | 0 | 0 | 0 | 0 | 44 | 94 | 74 | 58 | 58 | 298 | 149 | 1 | 1 | 0 | 0 | 0 | 0 | 0.01 | 0 | 0 | 0 | 0 | 0 | 0.002 |
| SE | 4452 | ENSG00000181896 | ZNF101 | chr19 | + | 19677863 | 19677986 | 19668850 | 19668966 | 19679180 | 19680032 | 87 | 40 | 134 | 114 | 96 | 2 | 2 | 1 | 0 | 0 | 43 | 72 | 67 | 50 | 50 | 0 | 0 | 0 | 3 | 0 | 272 | 149 | 1 | 1 | 0.96 | 0.916 | 0.987 | 1 | 1 | 1 | 1 | 1 | 0.901 | 1 | -0.008 |
| SE | 4453 | ENSG00000181896 | ZNF101 | chr19 | + | 19677863 | 19677990 | 19668795 | 19668966 | 19678725 | 19678786 | 162 | 88 | 279 | 228 | 233 | 4 | 0 | 0 | 4 | 0 | 78 | 165 | 106 | 106 | 80 | 0 | 0 | 0 | 0 | 0 | 276 | 149 | 1 | 1 | 0.956 | 1 | 1 | 0.969 | 1 | 1 | 1 | 1 | 1 | 1 | -0.015 |
| SE | 4454 | ENSG00000181896 | ZNF101 | chr19 | + | 19677863 | 19677990 | 19668795 | 19668966 | 19679180 | 19680329 | 97 | 45 | 136 | 122 | 101 | 2 | 2 | 1 | 0 | 0 | 70 | 109 | 74 | 51 | 56 | 0 | 0 | 0 | 3 | 0 | 276 | 149 | 1 | 1 | 0.963 | 0.924 | 0.987 | 1 | 1 | 1 | 1 | 1 | 0.902 | 1 | -0.006 |
| SE | 4455 | ENSG00000181896 | ZNF101 | chr19 | + | 19678725 | 19678786 | 19668795 | 19668966 | 19679180 | 19679523 | 64 | 45 | 95 | 66 | 159 | 2 | 2 | 1 | 0 | 0 | 39 | 53 | 54 | 28 | 28 | 0 | 0 | 0 | 3 | 0 | 210 | 149 | 1 | 1 | 0.958 | 0.941 | 0.985 | 1 | 1 | 1 | 1 | 1 | 0.869 | 1 | 0.003 |
| SE | 4456 | ENSG00000181896 | ZNF101 | chr19 | + | 19678725 | 19678786 | 19676140 | 19677986 | 19679180 | 19679523 | 68 | 58 | 114 | 71 | 177 | 0 | 0 | 0 | 2 | 0 | 51 | 73 | 59 | 36 | 32 | 0 | 0 | 4 | 0 | 0 | 210 | 149 | 1 | 1 | 1 | 1 | 1 | 0.962 | 1 | 1 | 1 | 0.913 | 1 | 1 | 0.01 |
| SE | 4467 | ENSG00000160117 | ANKLE1 | chr19 | + | 17284088 | 17284266 | 17283224 | 17283962 | 17285430 | 17285590 | 8 | 0 | 0 | 0 | 0 | 0 | 0 | 0 | 0 | 0 | 8 | 5 | 0 | 19 | 0 | 11 | 0 | 0 | 2 | 0 | 298 | 149 | 1 | 1 | 1 | NA | NA | NA | NA | 0.267 | 1 | NA | 0.826 | NA | 0.302 |
| SE | 4469 | ENSG00000266913 | LINC01841 | chr19 | - | 14320239 | 14320351 | 14305457 | 14305675 | 14325307 | 14325431 | 6 | 4 | 3 | 1 | 5 | 0 | 0 | 1 | 0 | 0 | 0 | 7 | 1 | 6 | 4 | 0 | 0 | 0 | 0 | 0 | 261 | 149 | 1 | 1 | 1 | 1 | 0.631 | 1 | 1 | NA | 1 | 1 | 1 | 1 | -0.074 |
| SE | 4471 | ENSG00000266913 | LINC01841 | chr19 | - | 14347314 | 14347409 | 14325351 | 14325431 | 14369942 | 14370196 | 5 | 1 | 4 | 6 | 8 | 0 | 0 | 0 | 0 | 0 | 0 | 11 | 2 | 4 | 1 | 0 | 0 | 0 | 3 | 0 | 244 | 149 | 1 | 1 | 1 | 1 | 1 | 1 | 1 | NA | 1 | 1 | 0.449 | 1 | 0.138 |
| SE | 4472 | ENSG00000266913 | LINC01841 | chr19 | - | 14366866 | 14366960 | 14347314 | 14347409 | 14369942 | 14370196 | 4 | 1 | 0 | 1 | 0 | 3 | 0 | 2 | 2 | 4 | 0 | 3 | 0 | 0 | 0 | 0 | 3 | 0 | 0 | 0 | 243 | 149 | 1 | 1 | 0.45 | 1 | 0 | 0.235 | 0 | NA | 0.38 | NA | NA | NA | -0.043 |
| SE | 4478 | ENSG00000127527 | EPS15L1 | chr19 | - | 16395343 | 16395467 | 16393950 | 16394001 | 16402320 | 16402485 | 110 | 70 | 75 | 114 | 94 | 2 | 0 | 0 | 1 | 0 | 107 | 79 | 75 | 109 | 221 | 0 | 0 | 0 | 0 | 0 | 273 | 149 | 1 | 1 | 0.968 | 1 | 1 | 0.984 | 1 | 1 | 1 | 1 | 1 | 1 | -0.01 |
| SE | 4480 | ENSG00000127527 | EPS15L1 | chr19 | - | 16417551 | 16417637 | 16413772 | 16413845 | 16421318 | 16421476 | 93 | 41 | 48 | 80 | 86 | 0 | 1 | 0 | 0 | 0 | 146 | 113 | 61 | 86 | 297 | 0 | 0 | 0 | 0 | 0 | 235 | 149 | 1 | 1 | 1 | 0.963 | 1 | 1 | 1 | 1 | 1 | 1 | 1 | 1 | -0.007 |
| SE | 4482 | ENSG00000127527 | EPS15L1 | chr19 | - | 16425082 | 16425316 | 16421318 | 16421476 | 16428701 | 16428761 | 171 | 79 | 119 | 134 | 218 | 2 | 0 | 0 | 0 | 0 | 158 | 214 | 142 | 141 | 423 | 0 | 0 | 0 | 0 | 0 | 298 | 149 | 1 | 1 | 0.977 | 1 | 1 | 1 | 1 | 1 | 1 | 1 | 1 | 1 | -0.005 |
| SE | 4483 | ENSG00000127527 | EPS15L1 | chr19 | - | 16436936 | 16436999 | 16434364 | 16434490 | 16440861 | 16440909 | 45 | 34 | 47 | 55 | 68 | 0 | 0 | 1 | 2 | 2 | 49 | 54 | 50 | 51 | 173 | 0 | 0 | 0 | 0 | 0 | 212 | 149 | 1 | 1 | 1 | 1 | 0.971 | 0.951 | 0.96 | 1 | 1 | 1 | 1 | 1 | -0.024 |
| SE | 4484 | ENSG00000127527 | EPS15L1 | chr19 | - | 16437769 | 16437865 | 16436595 | 16436999 | 16440861 | 16440909 | 109 | 46 | 95 | 148 | 140 | 0 | 0 | 0 | 0 | 0 | 182 | 178 | 106 | 142 | 333 | 0 | 0 | 0 | 3 | 0 | 245 | 149 | 1 | 1 | 1 | 1 | 1 | 1 | 1 | 1 | 1 | 1 | 0.966 | 1 | 0.007 |
| SE | 4485 | ENSG00000127527 | EPS15L1 | chr19 | - | 16440861 | 16440909 | 16437769 | 16437865 | 16441891 | 16441981 | 76 | 34 | 73 | 113 | 103 | 0 | 0 | 0 | 0 | 0 | 95 | 92 | 87 | 138 | 280 | 0 | 0 | 0 | 2 | 0 | 197 | 149 | 1 | 1 | 1 | 1 | 1 | 1 | 1 | 1 | 1 | 1 | 0.981 | 1 | 0.004 |
| SE | 4486 | ENSG00000127527 | EPS15L1 | chr19 | - | 16440884 | 16440909 | 16437769 | 16437865 | 16441891 | 16441981 | 10 | 5 | 11 | 12 | 9 | 0 | 0 | 0 | 0 | 0 | 5 | 3 | 16 | 23 | 56 | 0 | 0 | 0 | 2 | 0 | 174 | 149 | 1 | 1 | 1 | 1 | 1 | 1 | 1 | 1 | 1 | 1 | 0.908 | 1 | 0.018 |
| SE | 4489 | ENSG00000198551 | ZNF627 | chr19 | + | 11564060 | 11564184 | 11559373 | 11559616 | 11564444 | 11564492 | 0 | 1 | 2 | 2 | 0 | 3 | 3 | 2 | 0 | 0 | 0 | 2 | 0 | 6 | 0 | 0 | 0 | 0 | 0 | 0 | 273 | 149 | 1 | 1 | 0 | 0.154 | 0.353 | 1 | NA | NA | 1 | NA | 1 | NA | -0.623 |
| SE | 4490 | ENSG00000198551 | ZNF627 | chr19 | + | 11564060 | 11564184 | 11559373 | 11559616 | 11575249 | 11575353 | 0 | 2 | 0 | 1 | 0 | 0 | 0 | 0 | 0 | 0 | 0 | 0 | 0 | 0 | 0 | 0 | 0 | 2 | 0 | 0 | 273 | 149 | 1 | 1 | NA | 1 | NA | 1 | NA | NA | NA | 0 | NA | NA | 1 |
| SE | 4491 | ENSG00000198551 | ZNF627 | chr19 | + | 11564444 | 11564492 | 11559373 | 11559616 | 11575249 | 11575353 | 3 | 3 | 3 | 0 | 0 | 0 | 0 | 0 | 0 | 0 | 0 | 0 | 0 | 0 | 0 | 0 | 0 | 2 | 0 | 0 | 197 | 149 | 1 | 1 | 1 | 1 | 1 | NA | NA | NA | NA | 0 | NA | NA | 1 |
| SE | 4492 | ENSG00000198551 | ZNF627 | chr19 | + | 11564444 | 11564492 | 11564060 | 11564184 | 11575249 | 11575353 | 0 | 0 | 3 | 1 | 0 | 0 | 1 | 0 | 0 | 0 | 0 | 2 | 0 | 6 | 0 | 0 | 0 | 0 | 0 | 0 | 197 | 149 | 1 | 1 | NA | 0 | 1 | 1 | NA | NA | 1 | NA | 1 | NA | -0.333 |
| SE | 4493 | ENSG00000198551 | ZNF627 | chr19 | + | 11604209 | 11604364 | 11597461 | 11597630 | 11614526 | 11614653 | 3 | 0 | 0 | 0 | 0 | 6 | 1 | 7 | 2 | 4 | 0 | 0 | 0 | 0 | 0 | 0 | 3 | 4 | 2 | 0 | 298 | 149 | 1 | 1 | 0.2 | 0 | 0 | 0 | 0 | NA | 0 | 0 | 0 | NA | 0.04 |
| SE | 4494 | ENSG00000198551 | ZNF627 | chr19 | + | 11614826 | 11614887 | 11614526 | 11614653 | 11616694 | 11616773 | 33 | 12 | 32 | 14 | 42 | 0 | 0 | 1 | 0 | 0 | 6 | 23 | 29 | 5 | 2 | 0 | 0 | 0 | 0 | 0 | 210 | 149 | 1 | 1 | 1 | 1 | 0.958 | 1 | 1 | 1 | 1 | 1 | 1 | 1 | -0.008 |
| SE | 4499 | ENSG00000105135 | ILVBL | chr19 | - | 15116000 | 15116071 | 15114983 | 15115921 | 15116151 | 15116321 | 7 | 1 | 17 | 5 | 1 | 1 | 0 | 0 | 0 | 0 | 8 | 11 | 2 | 5 | 6 | 0 | 0 | 0 | 0 | 0 | 220 | 149 | 1 | 1 | 0.826 | 1 | 1 | 1 | 1 | 1 | 1 | 1 | 1 | 1 | -0.035 |
| SE | 4500 | ENSG00000105135 | ILVBL | chr19 | - | 15116407 | 15116522 | 15115884 | 15116321 | 15117880 | 15117931 | 62 | 21 | 38 | 22 | 12 | 0 | 0 | 0 | 1 | 0 | 13 | 36 | 5 | 28 | 21 | 0 | 0 | 0 | 0 | 0 | 264 | 149 | 1 | 1 | 1 | 1 | 1 | 0.925 | 1 | 1 | 1 | 1 | 1 | 1 | -0.015 |

| | | | | | | | | | | | | | | |
|---|---|---|---|---|---|---|---|---|---|---|---|---|---|---|
| SE | 4509 | ENSG00000167460 | TPM4 | chr19 | + | 16081440 | 16081538 | 16080764 | 16081343 | 16081912 | 16082046 | 0 | 0 | 0 |
| SE | 4512 | ENSG00000167460 | TPM4 | chr19 | + | 16086422 | 16086540 | 16081448 | 16082046 | 16088026 | 16088097 | 1069 | 1253 | 796 |
| SE | 4513 | ENSG00000167460 | TPM4 | chr19 | + | 16088026 | 16088097 | 16086293 | 16086540 | 16089044 | 16089120 | 1537 | 1828 | 1112 |
| SE | 4514 | ENSG00000167460 | TPM4 | chr19 | + | 16093039 | 16093132 | 16089044 | 16089120 | 16093535 | 16093598 | 1 | 2 | 5 |
| SE | 4515 | ENSG00000167460 | TPM4 | chr19 | + | 16093535 | 16093598 | 16088026 | 16089120 | 16093683 | 16093746 | 1561 | 1832 | 1146 |
| SE | 4516 | ENSG00000167460 | TPM4 | chr19 | + | 16093683 | 16093753 | 16088026 | 16089120 | 16101263 | 16101409 | 253 | 237 | 172 |
| SE | 4517 | ENSG00000167460 | TPM4 | chr19 | + | 16095263 | 16095357 | 16093683 | 16093753 | 16101263 | 16102875 | 0 | 0 | 1 |
| SE | 4518 | ENSG00000167460 | TPM4 | chr19 | + | 16095263 | 16095893 | 16093683 | 16093753 | 16101263 | 16102948 | 0 | 0 | 1 |
| SE | 4519 | ENSG00000167461 | RAB8A | chr19 | + | 16121749 | 16121810 | 16118225 | 16118286 | 16125469 | 16125547 | 827 | 645 | 571 |
| SE | 4520 | ENSG00000167461 | RAB8A | chr19 | + | 16127436 | 16127526 | 16125469 | 16125547 | 16128025 | 16128091 | 1526 | 1174 | 1223 |
| SE | 4525 | ENSG00000131355 | ADGRE3 | chr19 | - | 14654981 | 14655165 | 14651084 | 14651204 | 14661962 | 14662118 | 69 | 55 | 29 |
| SE | 4526 | ENSG00000131355 | ADGRE3 | chr19 | - | 14658479 | 14658550 | 14651084 | 14651204 | 14661962 | 14662118 | 38 | 38 | 27 |
| SE | 4527 | ENSG00000131355 | ADGRE3 | chr19 | - | 14658512 | 14658550 | 14651084 | 14651204 | 14661962 | 14662118 | 39 | 39 | 29 |
| SE | 4529 | ENSG00000131355 | ADGRE3 | chr19 | - | 14658512 | 14658550 | 14654981 | 14655165 | 14661962 | 14662118 | 72 | 67 | 42 |
| SE | 4533 | ENSG00000127507 | ADGRE2 | chr19 | - | 14746231 | 14746323 | 14732391 | 14736244 | 14746895 | 14746962 | 57 | 23 | 18 |
| SE | 4534 | ENSG00000127507 | ADGRE2 | chr19 | - | 14746231 | 14746323 | 14743615 | 14743784 | 14746895 | 14746962 | 156 | 90 | 46 |
| SE | 4535 | ENSG00000127507 | ADGRE2 | chr19 | - | 14751435 | 14751671 | 14746895 | 14746962 | 14764432 | 14764610 | 79 | 47 | 31 |
| SE | 4536 | ENSG00000127507 | ADGRE2 | chr19 | - | 14752328 | 14752526 | 14751435 | 14751671 | 14754953 | 14755127 | 79 | 72 | 40 |
| SE | 4537 | ENSG00000127507 | ADGRE2 | chr19 | - | 14752328 | 14752526 | 14751435 | 14751671 | 14756237 | 14756345 | 29 | 23 | 26 |
| SE | 4538 | ENSG00000127507 | ADGRE2 | chr19 | - | 14752328 | 14752526 | 14751435 | 14751671 | 14764432 | 14764610 | 27 | 21 | 26 |
| SE | 4539 | ENSG00000127507 | ADGRE2 | chr19 | - | 14754953 | 14755127 | 14751435 | 14751671 | 14756237 | 14756345 | 6 | 5 | 0 |
| SE | 4545 | ENSG00000127507 | ADGRE2 | chr19 | - | 14756237 | 14756345 | 14751435 | 14751671 | 14764432 | 14764610 | 72 | 57 | 43 |
| SE | 4547 | ENSG00000127507 | ADGRE2 | chr19 | - | 14764432 | 14764610 | 14751435 | 14751671 | 14765319 | 14765397 | 76 | 55 | 41 |
| SE | 4555 | ENSG00000127507 | ADGRE2 | chr19 | - | 14774255 | 14774306 | 14773937 | 14774054 | 14776725 | 14776756 | 39 | 13 | 12 |
| SE | 4559 | ENSG00000123146 | ADGRE5 | chr19 | + | 14388449 | 14388500 | 14381155 | 14381545 | 14390923 | 14391079 | 71 | 13 | 55 |
| SE | 4560 | ENSG00000123146 | ADGRE5 | chr19 | + | 14388701 | 14388818 | 14381415 | 14381545 | 14390923 | 14391079 | 724 | 289 | 368 |
| SE | 4561 | ENSG00000123146 | ADGRE5 | chr19 | + | 14388701 | 14388818 | 14388449 | 14388500 | 14390923 | 14391079 | 1128 | 426 | 580 |
| SE | 4562 | ENSG00000123146 | ADGRE5 | chr19 | + | 14390923 | 14391079 | 14388701 | 14388818 | 14397657 | 14397804 | 1176 | 532 | 676 |
| SE | 4563 | ENSG00000123146 | ADGRE5 | chr19 | + | 14391390 | 14391492 | 14390923 | 14391079 | 14397657 | 14397804 | 1 | 0 | 0 |
| SE | 4564 | ENSG00000123146 | ADGRE5 | chr19 | + | 14396341 | 14396473 | 14390923 | 14391079 | 14397076 | 14397223 | 356 | 187 | 175 |
| SE | 4568 | ENSG00000123146 | ADGRE5 | chr19 | + | 14397657 | 14397804 | 14390923 | 14391079 | 14397888 | 14397911 | 1355 | 701 | 897 |
| SE | 4569 | ENSG00000123146 | ADGRE5 | chr19 | + | 14397888 | 14397935 | 14390923 | 14391079 | 14398061 | 14398139 | 348 | 234 | 249 |
| SE | 4570 | ENSG00000123146 | ADGRE5 | chr19 | + | 14397888 | 14397935 | [illegible] | [illegible] | [illegible] | [illegible] | [illegible] | [illegible] | [illegible] |

| | | | | | | | | | | | | | | | |
|---|---|---|---|---|---|---|---|---|---|---|---|---|---|---|---|
| SE | 4991 | ENSG00000000419 | DPM1 | chr20 | - | 50941128 | 50941209 | 50940864 | 50940933 | 50945736 | 50945762 | 3 | 10 | 7 | [illegible] |
| SE | 4992 | ENSG00000000419 | DPM1 | chr20 | - | 50941128 | 50941209 | 50940864 | 50940955 | 50942030 | 50942126 | 2 | 4 | 3 | [illegible] |
| SE | 4993 | ENSG00000000419 | DPM1 | chr20 | - | 50942030 | 50942126 | 50936147 | 50936262 | 50945736 | 50945762 | 175 | 120 | 115 | [illegible] |
| SE | 4994 | ENSG00000000419 | DPM1 | chr20 | - | 50942030 | 50942126 | 50940864 | 50940933 | 50945736 | 50945762 | 390 | 280 | 259 | [illegible] |
| SE | 4997 | ENSG00000000419 | DPM1 | chr20 | - | 50948628 | 50948662 | 50945736 | 50945923 | 50955185 | 50955285 | 287 | 205 | 206 | [illegible] |
| SE | 4998 | ENSG00000089048 | ESF1 | chr20 | - | 13718907 | 13718984 | 13717367 | 13717514 | 13728377 | 13728465 | 67 | 29 | 34 | [illegible] |
| SE | 4999 | ENSG00000089048 | ESF1 | chr20 | - | 13769906 | 13770021 | 13766776 | 13766924 | 13771330 | 13771483 | 94 | 123 | 53 | [illegible] |
| SE | 5001 | ENSG00000101224 | CDC25B | chr20 | + | 3801721 | 3801802 | 3801253 | 3801388 | 3801923 | 3802100 | 481 | 246 | 554 | [illegible] |
| SE | 5003 | ENSG00000101224 | CDC25B | chr20 | + | 3804568 | 3804680 | 3803403 | 3803537 | 3804820 | 3805218 | 518 | 216 | 573 | [illegible] |
| SE | 5004 | ENSG00000101181 | MTG2 | chr20 | + | 62188789 | 62188805 | 62183050 | 62183057 | 62193415 | 62193624 | 2 | 0 | 0 | [illegible] |
| SE | 5006 | ENSG00000101181 | MTG2 | chr20 | + | 62195801 | 62195949 | 62183027 | 62183057 | 62198633 | 62198852 | 2 | 0 | 4 | [illegible] |
| SE | 5009 | ENSG00000088833 | NSFL1C | chr20 | - | 1445665 | 1445830 | 1442161 | 1443911 | 1452492 | 1452630 | 298 | 156 | 235 | [illegible] |
| SE | 5010 | ENSG00000088833 | NSFL1C | chr20 | - | 1445665 | 1446149 | 1443734 | 1443911 | 1452492 | 1452630 | 207 | 99 | 158 | [illegible] |
| SE | 5011 | ENSG00000088833 | NSFL1C | chr20 | - | 1452492 | 1452630 | 1445665 | 1445830 | 1453030 | 1453140 | 206 | 169 | 176 | [illegible] |
| SE | 5012 | ENSG00000088833 | NSFL1C | chr20 | - | 1453030 | 1453140 | 1445665 | 1445830 | 1454966 | 1455132 | 0 | 2 | 2 | [illegible] |
| SE | 5013 | ENSG00000088833 | NSFL1C | chr20 | - | 1454212 | 1454305 | 1453030 | 1453140 | 1454966 | 1455037 | 245 | 212 | 180 | [illegible] |
| SE | 5014 | ENSG00000088833 | NSFL1C | chr20 | - | 1455713 | 1455719 | 1454966 | 1455132 | 1458199 | 1458274 | 0 | 6 | 0 | [illegible] |
| SE | 5019 | ENSG00000088833 | NSFL1C | chr20 | - | 1464328 | 1464426 | 1458029 | 1458274 | 1473791 | 1473842 | 104 | 71 | 79 | [illegible] |
| SE | 5020 | ENSG00000088833 | NSFL1C | chr20 | - | 1472714 | 1473012 | 1458199 | 1458274 | 1473791 | 1473842 | 10 | 12 | 0 | [illegible] |
| SE | 5021 | ENSG00000088833 | NSFL1C | chr20 | - | 1472714 | 1473012 | 1464328 | 1464426 | 1473791 | 1473842 | 21 | 17 | 2 | [illegible] |
| SE | 5022 | ENSG00000088833 | NSFL1C | chr20 | - | 1473117 | 1473260 | 1472714 | 1473012 | 1473791 | 1473842 | 18 | 9 | 1 | [illegible] |
| SE | 5023 | ENSG00000101236 | RNF24 | chr20 | - | 3948236 | 3948279 | 3945176 | 3945218 | 3963874 | 3964024 | 52 | 45 | 41 | [illegible] |
| SE | 5024 | ENSG00000101236 | RNF24 | chr20 | - | 3963874 | 3964024 | 3948236 | 3948279 | 3974331 | 3974387 | 47 | 40 | 44 | [illegible] |
| SE | 5026 | ENSG00000101343 | CRNKL1 | chr20 | - | 20037322 | 20037534 | 20036005 | 20036362 | 20038348 | 20038452 | 134 | 77 | 96 | [illegible] |
| SE | 5027 | ENSG00000101343 | CRNKL1 | chr20 | - | 20037322 | 20037571 | 20034367 | 20036362 | 20038348 | 20038450 | 207 | 132 | 184 | [illegible] |
| SE | 5030 | ENSG00000101343 | CRNKL1 | chr20 | - | 20040685 | 20040766 | 20038348 | 20038450 | 20041565 | 20041625 | 67 | 64 | 70 | [illegible] |
| SE | 5032 | ENSG00000101343 | CRNKL1 | chr20 | - | 20043491 | 20043662 | 20042324 | 20042516 | 20045307 | 20045486 | 235 | 126 | 322 | [illegible] |
| SE | 5033 | ENSG00000173418 | NAA20 | chr20 | + | 20018361 | 20018517 | 20017309 | 20017449 | 20022455 | 20022480 | 1 | 0 | 1 | [illegible] |
| SE | 5035 | ENSG00000173418 | NAA20 | chr20 | + | 20025668 | 20025767 | 20017358 | 20017449 | 20026783 | 20026919 | 116 | 82 | 103 | [illegible] |
| SE | 5036 | ENSG00000173418 | NAA20 | chr20 | + | 20025668 | 20025767 | 20017358 | 20017449 | 20032507 | 20032653 | 0 | 0 | 0 | [illegible] |
| SE | 5037 | ENSG00000173418 | NAA20 | chr20 | + | 20026783 | 20026919 | 20017309 | 20017449 | 20032507 | 20032653 | 88 | 60 | 70 | [illegible] |
| SE | 5038 | ENSG00000173418 | NAA20 | chr20 | + | 20026783 | 20026919 | 20025668 | 20025767 | 20032507 | 20032653 | 274 | 220 | 252 | [illegible] |
| SE | 5042 | ENSG00000101126 | ADNP | chr20 | - | 50902016 | 50902109 | 50888915 | 50894512 | 50930825 | 50931364 | 124 | 118 | 156 | [illegible] |
| SE | 5044 | ENSG00000101126 | ADNP | chr20 | - | 50903888 | 50904001 | 50888917 | 50894512 | 50930825 | 50931421 | 4 | 4 | 9 | [illegible] |
| SE | 5046 | ENSG00000101126 | ADNP | chr20 | - | 50903888 | 50904001 | 50902016 | 50902109 | 50930825 | 50931421 | 194 | 152 | 176 | [illegible] |
| SE | 5047 | ENSG00000101126 | ADNP | chr20 | - | 50904765 | 50904849 | 50903888 | 50904001 | 50909499 | 50909791 | 30 | 10 | 12 | [illegible] |
| SE | 5051 | ENSG00000101126 | ADNP | chr20 | - | 50918738 | 50918781 | 50903888 | 50904001 | 50928650 | 50928825 | 0 | 2 | 0 | [illegible] |
| SE | 5053 | ENSG00000101126 | ADNP | chr20 | - | 50928650 | 50928825 | 50888917 | 50894512 | 50930825 | 50931375 | 9 | 0 | 4 | [illegible] |
| SE | 5055 | ENSG00000101126 | ADNP | chr20 | - | 50928650 | 50928825 | 50904765 | 50904849 | 50930825 | 50931375 | 12 | 6 | 6 | [illegible] |
| SE | 5061 | ENSG00000101247 | NDUFAF5 | chr20 | + | 13793159 | 13793227 | 13788588 | 13788652 | 13798460 | 13798500 | 5 | 5 | 2 | [illegible] |
| SE | 5071 | ENSG00000124103 | FAM209A | chr20 | + | 56518086 | 56518297 | 56517186 | 56517201 | 56525803 | 56526075 | 328 | 179 | 260 | [illegible] |
| SE | 5072 | ENSG00000124103 | FAM209A | chr20 | + | 56524889 | 56525057 | 56518086 | 56518297 | 56525803 | 56526075 | 1 | 2 | 4 | [illegible] |
| SE | 5073 | ENSG00000022277 | RTF2 | chr20 | + | 56470599 | 56470751 | 56468685 | 56468766 | 56471613 | 56471669 | 3 | 0 | 1 | [illegible] |
| SE | 5074 | ENSG00000022277 | RTF2 | chr20 | + | 56470599 | 56470941 | 56468657 | 56468766 | 56471613 | 56471669 | 3 | 0 | 1 | [illegible] |
| SE | 5075 | ENSG00000022277 | RTF2 | chr20 | + | 56471613 | 56471669 | 56468678 | 56468766 | 56472325 | 56472415 | 0 | 0 | 2 | [illegible] |
| SE | 5076 | ENSG00000022277 | RTF2 | chr20 | + | 56471613 | 56471669 | 56468678 | 56468766 | 56473300 | 56473360 | 1 | 2 | 0 | [illegible] |
| SE | 5078 | ENSG00000022277 | RTF2 | chr20 | + | 56472325 | 56472415 | 56471613 | 56471669 | 56473300 | 56473360 | 4 | 0 | 6 | [illegible] |
| SE | 5079 | ENSG00000022277 | RTF2 | chr20 | + | 56474677 | 56474771 | 56473300 | 56473395 | 56476984 | 56477032 | 408 | 274 | 373 | [illegible] |
| SE | 5080 | ENSG00000022277 | RTF2 | chr20 | + | 56513314 | 56513428 | 56484110 | 56484189 | 56516934 | 56516989 | 459 | 396 | 427 | [illegible] |
| SE | 5082 | ENSG00000101138 | CSTF1 | chr20 | + | 56397206 | 56397484 | 56395520 | 56395721 | 56398966 | 56399161 | 63 | 45 | 57 | [illegible] |
| SE | 5083 | ENSG00000101138 | CSTF1 | chr20 | + | 56397643 | 56397841 | 56397206 | 56397484 | 56398966 | 56399161 | 226 | 133 | 219 | [illegible] |
| SE | 5084 | ENSG00000087586 | AURKA | chr20 | - | 56373407 | 56373556 | 56369664 | 56370659 | 56381432 | 56381571 | 25 | 14 | 28 | [illegible] |
| SE | 5086 | ENSG00000087586 | AURKA | chr20 | - | 56388686 | 56388784 | 56388155 | 56388202 | 56390555 | 56390665 | 0 | 0 | 0 | [illegible] |
| SE | 5087 | ENSG00000087586 | AURKA | chr20 | - | 56388686 | 56388784 | 56388155 | 56388202 | 56392153 | 56392215 | 0 | 0 | 0 | [illegible] |
| SE | 5090 | ENSG00000087586 | AURKA | chr20 | - | 56390555 | 56390665 | 56388155 | 56388202 | 56392153 | 56392215 | 0 | 0 | 1 | [illegible] |
| SE | 5091 | ENSG00000087586 | AURKA | chr20 | - | 56390555 | 56390665 | 56388155 | 56388202 | 56392167 | 56392202 | 0 | 0 | 1 | [illegible] |
| SE | 5094 | ENSG00000124198 | ARFGEF2 | chr20 | + | 48941198 | 48941229 | 48921710 | 48922010 | 48951322 | 48951469 | 2 | 0 | 2 | [illegible] |
| SE | 5095 | ENSG00000124198 | ARFGEF2 | chr20 | + | 48941863 | 48941987 | 48941198 | 48941229 | 48951322 | 48951469 | 104 | 57 | 103 | [illegible] |
| SE | 5096 | ENSG00000124198 | ARFGEF2 | chr20 | + | 48953555 | 48953790 | 48952704 | 48952884 | 48963829 | 48963898 | 199 | 104 | 195 | [illegible] |
| SE | 5097 | ENSG00000124198 | ARFGEF2 | chr20 | + | 48963829 | 48963898 | 48953555 | 48953790 | 48965871 | 48966023 | 95 | 48 | 108 | [illegible] |
| SE | 5101 | ENSG00000124198 | ARFGEF2 | chr20 | + | 48971119 | 48971354 | 48969146 | 48969277 | 48972325 | 48972425 | 123 | 109 | 103 | [illegible] |
| SE | 5103 | ENSG00000124198 | ARFGEF2 | chr20 | + | 48972325 | 48972425 | 48969146 | 48969277 | 48974765 | 48974874 | 2 | 0 | 0 | [illegible] |
| SE | 5108 | ENSG00000124198 | ARFGEF2 | chr20 | + | 48976015 | 48976199 | 48973144 | 48973284 | 48984728 | 48984840 | 95 | 67 | 78 | [illegible] |
| SE | 5109 | ENSG00000124198 | ARFGEF2 | chr20 | + | 48988303 | 48988388 | 48985407 | 48985613 | 48988490 | 48988662 | 253 | 220 | 231 | [illegible] |
| SE | 5110 | ENSG00000124198 | ARFGEF2 | chr20 | + | 48998192 | 48998233 | 48995782 | 48995882 | 48998335 | 48998505 | 38 | 29 | 42 | [illegible] |
| SE | 5111 | ENSG00000124198 | ARFGEF2 | chr20 | + | 49018883 | 49018998 | 49017495 | 49017550 | 49023050 | 49023181 | 143 | 105 | 171 | [illegible] |
| SE | 5113 | ENSG00000124198 | ARFGEF2 | chr20 | + | 49028529 | 49028668 | 49025312 | 49025481 | 49032048 | 49032166 | 104 | 58 | 134 | [illegible] |
| SE | 5114 | ENSG00000124198 | ARFGEF2 | chr20 | + | 49032048 | 49032166 | 49028529 | 49028668 | 49033022 | 49033199 | 118 | 72 | 142 | [illegible] |
| SE | 5127 | ENSG00000125841 | NRSN2 | chr20 | + | 355325 | 355349 | 349637 | 349832 | 359258 | 359405 | 0 | 0 | 0 | [illegible] |
| SE | 5132 | ENSG00000196476 | C20orf96 | chr20 | - | 275967 | 276086 | 270862 | 271267 | 276792 | 276879 | 18 | 10 | 12 | [illegible] |
| SE | 5134 | ENSG00000131043 | AAR2 | chr20 | + | 36236881 | 36237037 | 36236130 | 36236782 | 36239820 | 36240625 | 2 | 0 | 2 | [illegible] |
| SE | 5135 | ENSG00000131043 | AAR2 | chr20 | + | 36236881 | 36237037 | 36236477 | 36236503 | 36239820 | 36240625 | 2 | 0 | 3 | [illegible] |
| SE | 5136 | ENSG00000131043 | AAR2 | chr20 | + | 36236889 | 36237037 | 36236467 | 36236503 | 36239820 | 36240625 | 2 | 0 | 2 | [illegible] |
| SE | 5138 | ENSG00000131043 | AAR2 | chr20 | + | 36246200 | 36246278 | 36244696 | 36244926 | 36255577 | 36256914 | 0 | 0 | 0 | [illegible] |
| SE | 5160 | ENSG00000088827 | SIGLEC1 | chr20 | - | 3689154 | 3689227 | 3686969 | 3688619 | 3689599 | 3689702 | 10 | 9 | 14 | [illegible] |
| SE | 5161 | ENSG00000088827 | SIGLEC1 | chr20 | - | 3693446 | 3693698 | 3692861 | 3693131 | 3694220 | 3694532 | 5 | 4 | 5 | [illegible] |
| SE | 5164 | ENSG00000088827 | SIGLEC1 | chr20 | - | 3703196 | 3703451 | 3701341 | 3701641 | 3703824 | 3704091 | 17 | 21 | 9 | [illegible] |
| SE | 5165 | ENSG00000088827 | SIGLEC1 | chr20 | - | 3707079 | 3707237 | 3706346 | 3706706 | 3712469 | 3712600 | 4 | 5 | 3 | [illegible] |
| SE | 5166 | ENSG00000088888 | MAVS | chr20 | + | 3854557 | 3854741 | 3846798 | 3846903 | 3857634 | 3857809 | 109 | 67 | 132 | [illegible] |
| SE | 5167 | ENSG00000088888 | MAVS | chr20 | + | 3854557 | 3854741 | 3846798 | 3846903 | 3861331 | 3861504 | 32 | 7 | 28 | [illegible] |
| SE | 5168 | ENSG00000088888 | MAVS | chr20 | + | 3857634 | 3857809 | 3846833 | 3846903 | 3861331 | 3861504 | 60 | 36 | 44 | [illegible] |
| SE | 5169 | ENSG00000088888 | MAVS | chr20 | + | 3857634 | 3857809 | 3854557 | 3854741 | 3861331 | 3861504 | 137 | 96 | 149 | [illegible] |
| SE | 5170 | ENSG00000088888 | MAVS | chr20 | + | 3857634 | 3857809 | 3854557 | 3854741 | 3862253 | 3862413 | 85 | 61 | 113 | [illegible] |
| SE | 5171 | ENSG00000088888 | MAVS | chr20 | + | 3861331 | 3861504 | 3854557 | 3854741 | 3862253 | 3862413 | 65 | 23 | 54 | [illegible] |
| SE | 5173 | ENSG00000275491 | LINC01730 | chr20 | + | 3809896 | 3810014 | 3808356 | 3808602 | 3811529 | 3811741 | 10 | 7 | 10 | [illegible] |
| SE | 5174 | ENSG00000101220 | C20orf27 | chr20 | - | 3754396 | 3754519 | 3753507 | 3754158 | 3755468 | 3755607 | 80 | 32 | 63 | [illegible] |
| SE | 5177 | ENSG00000101191 | DIDO1 | chr20 | - | 62895048 | 62895165 | 62894809 | 62894914 | 62896232 | 62896392 | 198 | 155 | 198 | [illegible] |
| SE | 5178 | ENSG00000101191 | DIDO1 | chr20 | - | 62905886 | 62906100 | 62896530 | 62896996 | 62907146 | 62907359 | 395 | 393 | 416 | [illegible] |
| SE | 5179 | ENSG00000101191 | DIDO1 | chr20 | - | 62909698 | 62910020 | 62907146 | 62907359 | 62926431 | 62926469 | 461 | 339 | 430 | [illegible] |
| SE | 5180 | ENSG00000101191 | DIDO1 | chr20 | - | 62910773 | 62911614 | 62909698 | 62910020 | 62926431 | 62926469 | 439 | 357 | 432 | [illegible] |
| SE | 5181 | ENSG00000101191 | DIDO1 | chr20 | - | 62914209 | 62914406 | 62910773 | 62911614 | 62926431 | 62926469 | 534 | 345 | 349 | [illegible] |
| SE | 5182 | ENSG00000101191 | DIDO1 | chr20 | - | 62914209 | 62914406 | 62910773 | 62911614 | 62926438 | 62926471 | 330 | 201 | 323 | [illegible] |
| SE | 5183 | ENSG00000060491 | OGFR | chr20 | + | 62809584 | 62809663 | 62805885 | 62808325 | 62811461 | 62811610 | 119 | 64 | 114 | [illegible] |
| SE | 5184 | ENSG00000060491 | OGFR | chr20 | + | 62810498 | 62810565 | 62809584 | 62809663 | 62811461 | 62811610 | 397 | 203 | 277 | [illegible] |
| SE | 5186 | ENSG00000124214 | STAU1 | chr20 | - | 49117776 | 49118096 | 49117125 | 49117248 | 49119981 | 49120128 | 59 | 36 | 54 | [illegible] |
| SE | 5187 | ENSG00000124214 | STAU1 | chr20 | - | 49118332 | 49118408 | 49117776 | 49118096 | 49119981 | 49120128 | 221 | 115 | 182 | [illegible] |
| SE | 5188 | ENSG00000124214 | STAU1 | chr20 | - | 49123091 | 49123229 | 49119981 | 49120128 | 49124374 | 49124587 | 101 | 69 | 119 | [illegible] |
| SE | 5189 | ENSG00000124214 | STAU1 | chr20 | - | 49123091 | 49123235 | 49119981 | 49120128 | 49124374 | 49124587 | 170 | 146 | 204 | [illegible] |
| SE | 5190 | ENSG00000124214 | STAU1 | chr20 | - | 49151581 | 49151747 | 49135832 | 49135931 | 49153932 | 49154071 | 181 | 164 | 213 | [illegible] |
| SE | 5199 | ENSG00000124207 | CSE1L | chr20 | + | 49067189 | 49067280 | 49066364 | 49066510 | 49068714 | 49068822 | 155 | 96 | 175 | [illegible] |
| SE | 5200 | ENSG00000124207 | CSE1L | chr20 | + | 49072285 | 49072453 | 49070204 | 49070297 | 49072567 | 49072697 | 277 | 211 | 262 | [illegible] |
| SE | 5201 | ENSG00000124207 | CSE1L | chr20 | + | 49072567 | 49072697 | 49070204 | 49070297 | 49074784 | 49074850 | 139 | 144 | 164 | [illegible] |
| SE | 5202 | ENSG00000124207 | CSE1L | chr20 | + | 49076979 | 49077064 | 49075317 | 49075520 | 49078560 | 49078622 | 259 | 167 | 201 | [illegible] |
| SE | 5203 | ENSG00000124207 | CSE1L | chr20 | + | 49078560 | 49078622 | 49076979 | 49077064 | 49084025 | 49084162 | 251 | 162 | 223 | [illegible] |
| SE | 5205 | ENSG00000124207 | CSE1L | chr20 | + | 49085282 | 49085386 | 49084025 | 49084162 | 49089246 | 49089397 | 69 | 96 | 75 | [illegible] |
| SE | 5207 | ENSG00000124207 | CSE1L | chr20 | + | 49088008 | 49088106 | 49085282 | 49085386 | 49089246 | 49089397 | 162 | 128 | 135 | [illegible] |
| SE | 5210 | ENSG00000175063 | UBE2C | chr20 | + | 45813436 | 45813464 | 45812575 | 45812796 | 45814383 | 45814470 | 3 | 0 | 6 | [illegible] |
| SE | 5211 | ENSG00000175063 | UBE2C | chr20 | + | 45813436 | 45813464 | 45813005 | 45813290 | 45814383 | 45814470 | 0 | 8 | 3 | [illegible] |
| SE | 5213 | ENSG00000175063 | UBE2C | chr20 | + | 45814383 | 45814470 | 45813436 | 45813464 | 45815540 | 45815745 | 4 | 8 | 12 | [illegible] |
| SE | 5235 | ENSG00000101265 | RASSF2 | chr20 | - | 4788216 | 4788268 | 4787632 | 4787701 | 4789595 | 4789697 | 86 | 55 | 65 | [illegible] |
| SE | 5236 | ENSG00000101265 | RASSF2 | chr20 | - | 4788216 | 4788268 | 4787632 | 4787754 | 4789595 | 4789697 | 515 | 283 | 339 | [illegible] |
| SE | 5237 | ENSG00000101265 | RASSF2 | chr20 | - | 4789595 | 4789697 | 4787632 | 4787754 | 4790450 | 4790611 | 214 | 95 | 120 | [illegible] |
| SE | 5238 | ENSG00000101265 | RASSF2 | chr20 | - | 4792538 | 4792627 | 4790450 | 4790611 | 4795814 | 4795966 | 579 | 305 | 388 | [illegible] |
| SE | 5240 | ENSG00000226648 | PLCG1-AS1 | chr20 | - | 41101171 | 41101289 | 41100657 | 41100792 | 41133486 | 41133645 | 0 | 3 | 2 | [illegible] |
| SE | 5241 | ENSG00000125971 | DYNLRB1 | chr20 | + | 34526267 | 34526343 | 34516413 | 34516461 | 34534627 | 34534795 | 140 | 86 | 70 | [illegible] |
| SE | 5242 | ENSG00000125971 | DYNLRB1 | chr20 | + | 34529851 | 34530234 | 34526267 | 34526343 | 34534627 | 34534795 | 2 | 1 | 0 | [illegible] |
| SE | 5259 | ENSG00000132646 | PCNA | chr20 | - | 5118609 | 5118677 | 5117469 | 5117664 | 5118768 | 5118866 | 514 | 316 | 500 | [illegible] |
| SE | 5260 | ENSG00000132646 | PCNA | chr20 | - | 5118768 | 5118866 | 5117469 | 5117664 | 5119577 | 5119914 | 57 | 24 | 45 | [illegible] |
| SE | 5263 | ENSG00000185513 | L3MBTL1 | chr20 | + | 43515008 | 43515159 | 43514407 | 43514776 | 43515918 | 43516177 | 6 | 5 | 2 | [illegible] |
| SE | 5266 | ENSG00000185513 | L3MBTL1 | chr20 | + | 43518606 | 43518703 | 43516092 | 43516177 | 43528656 | 43528745 | 0 | 0 | 0 | [illegible] |
| SE | 5267 | ENSG00000185513 | L3MBTL1 | chr20 | + | 43530797 | 43530889 | 43530283 | 43530419 | 43532772 | 43532924 | 0 | 0 | 1 | [illegible] |
| SE | 5268 | ENSG00000185513 | L3MBTL1 | chr20 | + | 43532772 | 43532924 | 43530283 | 43530419 | 43533341 | 43533418 | 6 | 3 | 4 | [illegible] |
| SE | 5269 | ENSG00000185513 | L3MBTL1 | chr20 | + | 43534264 | 43534394 | 43534007 | 43534093 | 43534827 | 43534942 | 5 | 8 | 6 | [illegible] |
| SE | 5271 | ENSG00000185513 | L3MBTL1 | chr20 | + | 43535836 | 43535936 | 43534283 | 43534942 | 43536096 | 43536294 | 5 | 11 | 7 | [illegible] |
| SE | 5273 | ENSG00000185513 | L3MBTL1 | chr20 | + | 43540752 | 43540815 | 43538974 | 43540308 | 43548146 | 43550941 | 4 | 2 | 7 | [illegible] |
| SE | 5274 | ENSG00000101421 | CHMP4B | chr20 | + | 33850951 | 33851066 | 33848466 | 33848644 | 33852076 | 33852203 | 1015 | 680 | 958 | [illegible] |
| SE | 5275 | ENSG00000101421 | CHMP4B | chr20 | + | 33852076 | 33852203 | 33850951 | 33851066 | 33853495 | 33854366 | 676 | 471 | 629 | [illegible] |
| SE | 5284 | ENSG00000198026 | ZNF335 | chr20 | - | 45957585 | 45957680 | 45953688 | 45953948 | 45957834 | 45957928 | 29 | 12 | 13 | [illegible] |
| SE | 5285 | ENSG00000198026 | ZNF335 | chr20 | - | 45960448 | 45960525 | 45960207 | 45960368 | 45960615 | 45960732 | 74 | 59 | 82 | [illegible] |
| SE | 5290 | ENSG00000100979 | PLTP | chr20 | - | 45909515 | 45909671 | 45907840 | 45907904 | 45909941 | 45910070 | 9 | 0 | 0 | [illegible] |
| SE | 5294 | ENSG00000101473 | ACOT8 | chr20 | - | 45848449 | 45848675 | 45842022 | 45844420 | 45856782 | 45856924 | 53 | 20 | 43 | [illegible] |
| SE | 5296 | ENSG00000101473 | ACOT8 | chr20 | - | 45853922 | 45853979 | 45848532 | 45848675 | 45855158 | 45855292 | 0 | 0 | 1 | [illegible] |
| SE | 5298 | ENSG00000101473 | ACOT8 | chr20 | - | 45855158 | 45855292 | 45848449 | 45848675 | 45856782 | 45856924 | 33 | 19 | 24 | [illegible] |
| SE | 5302 | ENSG00000124104 | SNX21 | chr20 | + | 45840321 | 45840458 | 45835032 | 45835116 | 45840638 | 45841090 | 0 | 1 | 0 | [illegible] |
| SE | 5303 | ENSG00000124104 | SNX21 | chr20 | + | 45840447 | 45840458 | 45834576 | 45835116 | 45840638 | 45840907 | 1 | 2 | 0 | [illegible] |
| SE | 5306 | ENSG00000125900 | SIRPD | chr20 | - | 1551690 | 1552038 | 1537154 | 1537313 | 1557580 | 1557705 | 2 | 1 | 4 | [illegible] |
| SE | 5309 | ENSG00000125818 | PSMF1 | chr20 | + | 1125497 | 1125650 | 1118597 | 1118902 | 1127425 | 1127508 | 466 | 279 | 363 | [illegible] |
| SE | 5312 | ENSG00000125818 | PSMF1 | chr20 | + | 1164317 | 1164476 | 1135177 | 1135306 | 1186339 | 1186495 | 0 | 0 | 0 | [illegible] |
| SE | 5313 | ENSG00000125818 | PSMF1 | chr20 | + | 1164317 | 1164476 | 1163129 | 1163183 | 1165028 | 1165059 | 321 | 198 | 308 | [illegible] |
| SE | 5319 | ENSG00000088970 | KIZ | chr20 | + | 21132096 | 21132159 | 21126006 | 21126204 | 21136389 | 21136449 | 98 | 90 | 77 | [illegible] |
| SE | 5322 | ENSG00000088970 | KIZ | chr20 | + | 21136389 | 21136552 | 21132096 | 21132159 | 21145564 | 21145654 | 173 | 197 | 171 | [illegible] |
| SE | 5326 | ENSG00000088970 | KIZ | chr20 | + | 21145564 | 21145654 | 21136389 | 21136552 | 21161870 | 21162507 | 120 | 122 | 117 | [illegible] |
| SE | 5327 | ENSG00000088970 | KIZ | chr20 | + | 21161870 | 21162507 | 21145564 | 21145654 | 21162849 | 21163159 | 132 | 112 | 152 | [illegible] |
| SE | 5328 | ENSG00000088970 | KIZ | chr20 | + | 21205490 | 21205584 | 21162849 | 21163159 | 21229010 | 21229115 | 36 | 25 | 35 | [illegible] |
| SE | 5329 | ENSG00000088970 | KIZ | chr20 | + | 21214534 | 21214700 | 21205490 | 21205584 | 21229010 | 21229115 | 25 | 17 | 24 | [illegible] |
| SE | 5333 | ENSG00000088970 | KIZ | chr20 | + | 21232733 | 21232830 | 21229010 | 21229115 | 21244244 | 21244288 | 96 | 40 | 92 | [illegible] |
| SE | 5334 | ENSG00000088970 | KIZ | chr20 | + | 21244244 | 21244288 | 21232733 | 21232830 | 21246478 | 21246618 | 49 | 19 | 48 | [illegible] |
| SE | 5335 | ENSG00000171552 | BCL2L1 | chr20 | - | 31666990 | 31667147 | 31664574 | 31666086 | 31719344 | 31719417 | 0 | 0 | 0 | [illegible] |
| SE | 5342 | ENSG00000101084 | RAB5IF | chr20 | + | 36607714 | 36607818 | 36605778 | 36606065 | 36609600 | 36609721 | 364 | 128 | 237 | [illegible] |
| SE | 5348 | ENSG00000232907 | DLGAP4-AS1 | chr20 | - | 36508811 | 36508825 | 36507701 | 36508274 | 36510879 | 36511032 | 0 | 0 | 0 | [illegible] |
| SE | 5349 | ENSG00000232907 | DLGAP4-AS1 | chr20 | - | 36510879 | 36511032 | 36507701 | 36508274 | 36571440 | 36571543 | 0 | 1 | 0 | [illegible] |
| SE | 5351 | ENSG00000232907 | DLGAP4-AS1 | chr20 | - | 36527723 | 36527924 | 36525500 | 36525566 | 36571440 | 36571543 | 1 | 0 | 1 | [illegible] |
| SE | 5352 | ENSG00000182541 | LIMK2 | chr22 | + | 31225719 | 31225819 | 31212238 | 31212424 | 31258290 | 31258426 | 26 | 18 | 23 | [illegible] |
| SE | 5354 | ENSG00000182541 | LIMK2 | chr22 | + | 31259888 | 31260077 | 31258290 | 31258426 | 31262133 | 31262239 | 55 | 21 | 45 | [illegible] |
| SE | 5356 | ENSG00000182541 | LIMK2 | chr22 | + | 31268143 | 31268200 | 31267775 | 31267907 | 31271135 | 31271201 | 63 | 31 | 35 | [illegible] |
| SE | 5359 | ENSG00000125991 | ERGIC3 | chr20 | + | 35542322 | 35542393 | 35542037 | 35542185 | 35542512 | 35542600 | 222 | 76 | 331 | [illegible] |
| SE | 5360 | ENSG00000125991 | ERGIC3 | chr20 | + | 35542512 | 35542600 | 35542191 | 35542393 | 35542821 | 35542941 | 354 | 172 | 446 | [illegible] |
| SE | 5362 | ENSG00000125991 | ERGIC3 | chr20 | + | 35549162 | 35549207 | 35548807 | 35548865 | 35555043 | 35555075 | 0 | 0 | 0 | [illegible] |
| SE | 5363 | ENSG00000125991 | ERGIC3 | chr20 | + | 35554371 | 35554386 | 35548807 | 35548865 | 35555043 | 35555075 | 0 | 1 | 0 | [illegible] |
| SE | 5366 | ENSG00000101150 | TPD52L2 | chr20 | + | 63875815 | 63875875 | 63873667 | 63873816 | 63889189 | 63889238 | 30 | 12 | 12 | [illegible] |
| SE | 5367 | ENSG00000101150 | TPD52L2 | chr20 | + | 63882718 | 63882820 | 63873667 | 63873816 | 63889189 | 63889238 | 156 | 89 | 111 | [illegible] |
| SE | 5368 | ENSG00000101150 | TPD52L2 | chr20 | + | 63882718 | 63882820 | 63875815 | 63875875 | 63889189 | 63889238 | 267 | 154 | 185 | [illegible] |
| SE | 5370 | ENSG00000101150 | TPD52L2 | chr20 | + | 63887563 | 63887605 | 63882718 | 63882820 | 63889189 | 63889238 | 0 | 1 | 0 | [illegible] |
| SE | 5371 | ENSG00000101150 | TPD52L2 | chr20 | + | 63889189 | 63889238 | 63882718 | 63882820 | 63889849 | 63890424 | 331 | 178 | 224 | [illegible] |
| SE | 5375 | ENSG00000089177 | KIF16B | chr20 | - | 16356329 | 16356452 | 16335925 | 16336015 | 16370585 | 16370636 | 24 | 11 | 30 | [illegible] |
| SE | 5378 | ENSG00000089177 | KIF16B | chr20 | - | 16428952 | 16429004 | 16427103 | 16427241 | 16429862 | 16429982 | 38 | 40 | 47 | [illegible] |
| SE | 5379 | ENSG00000089177 | KIF16B | chr20 | - | 16515547 | 16515664 | 16512825 | 16512923 | 16526091 | 16526205 | 71 | 41 | 61 | [illegible] |
| SE | 5380 | ENSG00000078747 | ITCH | chr20 | + | 34365459 | 34365576 | 34363305 | 34363349 | 34369393 | 34369470 | 4 | 2 | 1 | [illegible] |
| SE | 5381 | ENSG00000078747 | ITCH | chr20 | + | 34369393 | 34369470 | 34363240 | 34363349 | 34393790 | 34393881 | 201 | 94 | 173 | [illegible] |
| SE | 5382 | ENSG00000078747 | ITCH | chr20 | + | 34392736 | 34392914 | 34369393 | 34369470 | 34393769 | 34393881 | 2 | 1 | 2 | [illegible] |
| SE | 5385 | ENSG00000078747 | ITCH | chr20 | + | 34393790 | 34393881 | 34369393 | 34369470 | 34408650 | 34408792 | 452 | 223 | 377 | [illegible] |
| SE | 5389 | ENSG00000078747 | ITCH | chr20 | + | 34417116 | 34417239 | 34413741 | 34413879 | 34424479 | 34424525 | 7 | 5 | 2 | [illegible] |
| SE | 5391 | ENSG00000078747 | ITCH | chr20 | + | 34438473 | 34438631 | 34413741 | 34413879 | 34440154 | 34440344 | 193 | 133 | 207 | [illegible] |
| SE | 5392 | ENSG00000078747 | ITCH | chr20 | + | 34442207 | 34442303 | 34440154 | 34440344 | 34445286 | 34445461 | 615 | 409 | 528 | [illegible] |
| SE | 5393 | ENSG00000078747 | ITCH | chr20 | + | 34457389 | 34457474 | 34449410 | 34449480 | 34462092 | 34462221 | 303 | 196 | 293 | [illegible] |
| SE | 5394 | ENSG00000078747 | ITCH | chr20 | + | 34470047 | 34470120 | 34462092 | 34462221 | 34471443 | 34471515 | 359 | 186 | 282 | [illegible] |
| SE | 5397 | ENSG00000078747 | ITCH | chr20 | + | 34479629 | 34479789 | 34471443 | 34471515 | 34480598 | 34480732 | 405 | 252 | 300 | [illegible] |
| SE | 5398 | ENSG00000078747 | ITCH | chr20 | + | 34479629 | 34479789 | 34471443 | 34471515 | 34481065 | 34481176 | 1 | 2 | 4 | [illegible] |
| SE | 5399 | ENSG00000078747 | ITCH | chr20 | + | 34480598 | 34480732 | 34471443 | 34471515 | 34481065 | 34481176 | 374 | 257 | 289 | [illegible] |
| SE | 5401 | ENSG00000078747 | ITCH | chr20 | + | 34489265 | 34489386 | 34481065 | 34481206 | 34489821 | 34489926 | 530 | 350 | 404 | [illegible] |
| SE | 5402 | ENSG00000078747 | ITCH | chr20 | + | 34489821 | 34489926 | 34489265 | 34489386 | 34492500 | 34492597 | 604 | 432 | 473 | [illegible] |
| SE | 5403 | ENSG00000130584 | ZBTB46 | chr20 | - | 63769249 | 63769383 | 63768072 | 63768118 | 63775677 | 63775962 | 0 | 1 | 0 | [illegible] |
| SE | 5408 | ENSG00000149636 | DSN1 | chr20 | - | 36771424 | 36771473 | 36770872 | 36771193 | 36773661 | 36773720 | 12 | 1 | 4 | [illegible] |
| SE | 5410 | ENSG00000198053 | SIRPA | chr20 | + | 1922312 | 1922645 | 1921394 | 1921712 | 1924763 | 1924877 | 397 | 257 | 228 | [illegible] |
| SE | 5412 | ENSG00000198053 | SIRPA | chr20 | + | 1927874 | 1927899 | 1924763 | 1924877 | 1934714 | 1934754 | 229 | 198 | 119 | [illegible] |
| SE | 5413 | ENSG00000118705 | RPN2 | chr20 | + | 37198396 | 37198492 | 37184179 | 37184373 | 37199049 | 37199225 | 506 | 163 | 408 | [illegible] |
| SE | 5415 | ENSG00000118705 | RPN2 | chr20 | + | 37198396 | 37198492 | 37184179 | 37184373 | 37204766 | 37204901 | 175 | 73 | 187 | [illegible] |
| SE | 5419 | ENSG00000118705 | RPN2 | chr20 | + | 37203884 | 37203960 | 37184179 | 37184373 | 37204766 | 37204901 | 77 | 54 | 96 | [illegible] |
| SE | 5420 | ENSG00000118705 | RPN2 | chr20 | + | 37203884 | 37203960 | 37198396 | 37198492 | 37204766 | 37204901 | 78 | 52 | 95 | [illegible] |
| SE | 5421 | ENSG00000118705 | RPN2 | chr20 | + | 37204766 | 37204901 | 37184179 | 37184373 | 37207272 | 37207443 | 204 | 75 | 182 | [illegible] |
| SE | 5422 | ENSG00000118705 | RPN2 | chr20 | + | 37213759 | 37213865 | 37210046 | 37210165 | 37223877 | 37223969 | 854 | 464 | 777 | [illegible] |
| SE | 5424 | ENSG00000118705 | RPN2 | chr20 | + | 37229972 | 37230059 | 37228549 | 37228744 | 37232295 | 37232391 | 606 | 297 | 518 | [illegible] |
| SE | 5425 | ENSG00000118705 | RPN2 | chr20 | + | 37229972 | 37230059 | 37228549 | 37228744 | 37234019 | 37234095 | 293 | 145 | 235 | [illegible] |
| SE | 5426 | ENSG00000118705 | RPN2 | chr20 | + | 37232295 | 37232391 | 37228549 | 37228744 | 37234019 | 37234095 | 244 | 97 | 226 | [illegible] |
| SE | 5428 | ENSG00000118705 | RPN2 | chr20 | + | 37232295 | 37232391 | 37229972 | 37230059 | 37236579 | 37236709 | 216 | 105 | 191 | [illegible] |
| SE | 5429 | ENSG00000118705 | RPN2 | chr20 | + | 37234019 | 37234095 | 37229869 | 37230059 | 37236579 | 37236709 | 171 | 105 | 149 | [illegible] |
| SE | 5430 | ENSG00000118705 | RPN2 | chr20 | + | 37234019 | 37234095 | 37232295 | 37232391 | 37236579 | 37236709 | 532 | 253 | 491 | [illegible] |
| SE | 5431 | ENSG00000118705 | RPN2 | chr20 | + | 37236579 | 37236709 | 37234019 | 37234095 | 37241302 | 37241417 | 558 | 271 | 534 | [illegible] |
| SE | 5434 | ENSG00000126005 | MMP24OS | chr20 | - | 35236080 | 35236191 | 35224743 | 35224952 | 35272004 | 35272145 | 2 | 8 | 1 | [illegible] |
| SE | 5435 | ENSG00000126005 | MMP24OS | chr20 | - | 35236080 | 35236191 | 35225599 | 35225997 | 35272004 | 35272145 | 2 | 10 | 1 | [illegible] |
| SE | 5436 | ENSG00000126005 | MMP24OS | chr20 | - | 35276155 | 35276314 | 35272004 | 35272145 | 35277389 | 35277932 | 0 | 1 | 4 | [illegible] |
| SE | 5437 | ENSG00000126005 | MMP24OS | chr20 | - | 35276155 | 35276314 | 35272004 | 35272145 | 35278008 | 35278103 | 0 | 1 | 3 | [illegible] |
| SE | 5438 | ENSG00000126005 | MMP24OS | chr20 | - | 35277389 | 35277932 | 35272004 | 35272145 | 35278008 | 35278027 | 13 | 3 | 6 | [illegible] |
| SE | 5440 | ENSG00000101367 | MAPRE1 | chr20 | + | 32825924 | 32826048 | 32819953 | 32820028 | 32833716 | 32833862 | 748 | 474 | 654 | [illegible] |
| SE | 5441 | ENSG00000101367 | MAPRE1 | chr20 | + | 32833716 | 32833862 | 32825924 | 32826048 | 32836633 | 32836841 | 1163 | 791 | 1012 | [illegible] |
| SE | 5442 | ENSG00000101367 | MAPRE1 | chr20 | + | 32833716 | 32833862 | 32825924 | 32826048 | 32839734 | 32839856 | 583 | 355 | 507 | [illegible] |
| SE | 5443 | ENSG00000101367 | MAPRE1 | chr20 | + | 32836633 | 32836841 | 32825924 | 32826048 | 32839734 | 32839856 | 582 | 484 | 516 | [illegible] |
| SE | 5451 | ENSG00000285382 | NA | chr20 | - | 32656575 | 32657968 | 32651732 | 32651981 | 32704022 | 32704071 | 2 | 0 | 0 | [illegible] |
| SE | 5452 | ENSG00000285382 | NA | chr20 | - | 32704439 | 32704489 | 32704022 | 32704071 | 32706582 | 32706620 | 43 | 35 | 33 | [illegible] |
| SE | 5453 | ENSG00000285382 | NA | chr20 | - | 32704813 | 32704904 | 32704439 | 32704489 | 32706582 | 32706620 | 185 | 121 | 159 | [illegible] |
| SE | 5454 | ENSG00000285382 | NA | chr20 | - | 32728088 | 32728139 | 32727892 | 32727995 | 32743307 | 32743567 | 12 | 7 | 18 | [illegible] |
| SE | 5455 | ENSG00000285382 | NA | chr20 | - | 32728088 | 32728142 | 32727892 | 32727995 | 32743307 | 32743523 | 23 | 14 | 36 | [illegible] |
| SE | 5458 | ENSG00000101346 | POFUT1 | chr20 | + | 32216608 | 32216721 | 32215268 | 32215451 | 32228262 | 32228455 | 97 | 39 | 93 | [illegible] |
| SE | 5459 | ENSG00000101337 | TM9SF4 | chr20 | + | 32133012 | 32133126 | 32109713 | 32109755 | 32136073 | 32136173 | 130 | 91 | 89 | [illegible] |
| SE | 5462 | ENSG00000101337 | TM9SF4 | chr20 | + | 32136073 | 32136173 | 32133012 | 32133126 | 32141496 | 32141665 | 215 | 138 | 203 | [illegible] |
| SE | 5465 | ENSG00000101337 | TM9SF4 | chr20 | + | 32141765 | 32141895 | 32141496 | 32141665 | 32142981 | 32143105 | 405 | 306 | 326 | [illegible] |
| SE | 5466 | ENSG00000101337 | TM9SF4 | chr20 | + | 32145090 | 32145209 | 32142981 | 32143105 | 32145311 | 32145423 | 219 | 126 | 162 | [illegible] |
| SE | 5473 | ENSG00000125885 | MCM8 | chr20 | + | 5972006 | 5972037 | 5967829 | 5968025 | 5972575 | 5972726 | 4 | 0 | 4 | [illegible] |
| SE | 5474 | ENSG00000125885 | MCM8 | chr20 | + | 5972006 | 5972037 | 5967829 | 5968025 | 5973055 | 5973196 | 34 | 17 | 25 | [illegible] |
| SE | 5475 | ENSG00000125885 | MCM8 | chr20 | + | 5972006 | 5972037 | 5967829 | 5968025 | 5977875 | 5978017 | 4 | 2 | 4 | [illegible] |
| SE | 5476 | ENSG00000125885 | MCM8 | chr20 | + | 5972575 | 5972726 | 5967829 | 5968025 | 5973055 | 5973196 | 0 | 0 | 2 | [illegible] |
| SE | 5477 | ENSG00000125885 | MCM8 | chr20 | + | 5972575 | 5972726 | 5972006 | 5972037 | 5973055 | 5973196 | 0 | 0 | 2 | [illegible] |
| SE | 5478 | ENSG00000125885 | MCM8 | chr20 | + | 5973055 | 5973196 | 5967829 | 5968025 | 5977875 | 5978017 | 21 | 23 | 24 | [illegible] |
| SE | 5479 | ENSG00000125885 | MCM8 | chr20 | + | 5973055 | 5973196 | 5972006 | 5972037 | 5977875 | 5978017 | 51 | 39 | 46 | [illegible] |
| SE | 5480 | ENSG00000125885 | MCM8 | chr20 | + | 5977875 | 5978017 | 5973055 | 5973196 | 5982969 | 5983165 | 56 | 60 | 71 | [illegible] |
| SE | 5481 | ENSG00000125885 | MCM8 | chr20 | + | 5984780 | 5985000 | 5982969 | 5983165 | 5985921 | 5986131 | 33 | 37 | 39 | [illegible] |
| SE | 5482 | ENSG00000130589 | HELZ2 | chr20 | - | 63561600 | 63561745 | 63561349 | 63561466 | 63561822 | 63561984 | 124 | 43 | 81 | [illegible] |
| SE | 5501 | ENSG00000101294 | HM13 | chr20 | + | 31544946 | 31545035 | 31538178 | 31538261 | 31549028 | 31549114 | 163 | 64 | 102 | [illegible] |
| SE | 5503 | ENSG00000101294 | HM13 | chr20 | + | 31544946 | 31545035 | 31538178 | 31538359 | 31549028 | 31549114 | 41 | 27 | 29 | [illegible] |
| SE | 5504 | ENSG00000101294 | HM13 | chr20 | + | 31544946 | 31545035 | 31538178 | 31538359 | 31550063 | 31550121 | 1 | 4 | 0 | [illegible] |
| SE | 5505 | ENSG00000101294 | HM13 | chr20 | + | 31544946 | 31545035 | 31538178 | 31538554 | 31550063 | 31550121 | 4 | 2 | 5 | [illegible] |
| SE | 5506 | ENSG00000101294 | HM13 | chr20 | + | 31549028 | 31549114 | 31538178 | 31538261 | 31550063 | 31550121 | 1 | 1 | 1 | [illegible] |
| SE | 5507 | ENSG00000101294 | HM13 | chr20 | + | 31549028 | 31549114 | 31538178 | 31538359 | 31550063 | 31550121 | 1 | 1 | 1 | [illegible] |

| | | | | | | | | | | | | | | |
|---|---|---|---|---|---|---|---|---|---|---|---|---|---|---|
| SE | 5508 | ENSG00000101294 | HM13 | chr20 | + | 31549028 | 31549114 | 31544946 | 31545035 | 31550063 | 31550121 | 62 | 25 | [illegible] |
| SE | 5512 | ENSG00000101294 | HM13 | chr20 | + | 31564118 | 31564218 | 31561633 | 31561736 | 31566209 | 31566295 | 0 | 1 | [illegible] |
| SE | 5514 | ENSG00000101294 | HM13 | chr20 | + | 31566209 | 31566295 | 31561633 | 31561736 | 31567639 | 31567740 | 70 | 15 | [illegible] |
| SE | 5515 | ENSG00000101294 | HM13 | chr20 | + | 31566209 | 31566295 | 31561633 | 31561736 | 31568077 | 31568211 | 73 | 16 | [illegible] |
| SE | 5518 | ENSG00000101294 | HM13 | chr20 | + | 31567639 | 31567740 | 31561633 | 31561736 | 31569119 | 31569215 | 0 | 0 | [illegible] |
| SE | 5519 | ENSG00000101294 | HM13 | chr20 | + | 31567639 | 31567740 | 31566209 | 31566295 | 31569119 | 31569215 | 3 | 0 | [illegible] |
| SE | 5528 | ENSG00000244687 | UBE2V1 | chr20 | - | 50084128 | 50084254 | 50081123 | 50082914 | 50096671 | 50096820 | 524 | 353 | [illegible] |
| SE | 5529 | ENSG00000244687 | UBE2V1 | chr20 | - | 50084128 | 50084254 | 50081123 | 50082914 | 50113106 | 50113172 | 329 | 196 | [illegible] |
| SE | 5530 | ENSG00000244687 | UBE2V1 | chr20 | - | 50096482 | 50096534 | 50084128 | 50084254 | 50096671 | 50096820 | 4 | 1 | [illegible] |
| SE | 5531 | ENSG00000244687 | UBE2V1 | chr20 | - | 50096482 | 50096534 | 50094914 | 50095382 | 50096671 | 50096820 | 2 | 0 | [illegible] |
| SE | 5533 | ENSG00000244687 | UBE2V1 | chr20 | - | 50096671 | 50096820 | 50081123 | 50082914 | 50113106 | 50113125 | 71 | 67 | [illegible] |
| SE | 5536 | ENSG00000244687 | UBE2V1 | chr20 | - | 50096671 | 50096820 | 50083877 | 50084254 | 50115698 | 50115954 | 208 | 166 | [illegible] |
| SE | 5537 | ENSG00000244687 | UBE2V1 | chr20 | - | 50098881 | 50098964 | 50096671 | 50096820 | 50113106 | 50113167 | 1 | 0 | [illegible] |
| SE | 5540 | ENSG00000244687 | UBE2V1 | chr20 | - | 50111288 | 50111562 | 50096671 | 50096820 | 50113106 | 50113145 | 0 | 0 | [illegible] |
| SE | 5541 | ENSG00000244687 | UBE2V1 | chr20 | - | 50115484 | 50115621 | 50084128 | 50084254 | 50115698 | 50115954 | 1 | 1 | [illegible] |
| SE | 5542 | ENSG00000244687 | UBE2V1 | chr20 | - | 50115484 | 50115621 | 50096671 | 50096820 | 50115698 | 50115954 | 14 | 4 | [illegible] |
| SE | 5544 | ENSG00000101057 | MYBL2 | chr20 | + | 43682793 | 43682886 | 43673805 | 43673899 | 43686851 | 43687072 | 9 | 1 | [illegible] |
| SE | 5551 | ENSG00000101052 | IFT52 | chr20 | + | 43620856 | 43620925 | 43618939 | 43619026 | 43623890 | 43624045 | 88 | 66 | [illegible] |
| SE | 5552 | ENSG00000101052 | IFT52 | chr20 | + | 43642478 | 43642624 | 43637144 | 43637253 | 43646935 | 43647222 | 66 | 60 | [illegible] |
| SE | 5553 | ENSG00000223891 | OSER1-DT | chr20 | + | 44214887 | 44215003 | 44210942 | 44211077 | 44224820 | 44225969 | 6 | 0 | [illegible] |
| SE | 5554 | ENSG00000223891 | OSER1-DT | chr20 | + | 44214887 | 44215003 | 44210994 | 44211356 | 44224820 | 44225969 | 7 | 0 | [illegible] |
| SE | 5556 | ENSG00000223891 | OSER1-DT | chr20 | + | 44217366 | 44217682 | 44214887 | 44215003 | 44224820 | 44225972 | 12 | 3 | [illegible] |
| SE | 5557 | ENSG00000223891 | OSER1-DT | chr20 | + | 44223718 | 44223872 | 44214887 | 44215003 | 44224820 | 44225071 | 0 | 0 | [illegible] |
| SE | 5558 | ENSG00000223891 | OSER1-DT | chr20 | + | 44223718 | 44223872 | 44217366 | 44217682 | 44224820 | 44225071 | 0 | 0 | [illegible] |
| SE | 5560 | ENSG00000160410 | SHKBP1 | chr19 | + | 40578122 | 40578212 | 40577556 | 40577630 | 40578461 | 40578542 | 42 | 31 | [illegible] |
| SE | 5561 | ENSG00000160410 | SHKBP1 | chr19 | + | 40578153 | 40578212 | 40577556 | 40577630 | 40578461 | 40578542 | 139 | 72 | [illegible] |
| SE | 5562 | ENSG00000160410 | SHKBP1 | chr19 | + | 40580565 | 40580656 | 40580166 | 40580485 | 40580745 | 40580838 | 221 | 78 | [illegible] |
| SE | 5564 | ENSG00000160410 | SHKBP1 | chr19 | + | 40586773 | 40586944 | 40580745 | 40580936 | 40588623 | 40588779 | 85 | 27 | [illegible] |
| SE | 5565 | ENSG00000160410 | SHKBP1 | chr19 | + | 40586773 | 40586944 | 40583600 | 40583717 | 40588623 | 40588779 | 160 | 48 | [illegible] |
| SE | 5573 | ENSG00000088340 | FER1L4 | chr20 | - | 35560122 | 35560364 | 35559155 | 35559627 | 35560539 | 35560638 | 0 | 0 | [illegible] |
| SE | 5574 | ENSG00000088340 | FER1L4 | chr20 | - | 35560162 | 35560364 | 35559340 | 35559627 | 35560539 | 35560638 | 0 | 0 | [illegible] |
| SE | 5577 | ENSG00000088340 | FER1L4 | chr20 | - | 35578835 | 35578967 | 35576785 | 35576828 | 35579055 | 35579114 | 0 | 0 | [illegible] |
| SE | 5582 | ENSG00000126001 | CEP250 | chr20 | + | 35458303 | 35458374 | 35455163 | 35455242 | 35459982 | 35460105 | 2 | 2 | [illegible] |
| SE | 5583 | ENSG00000126001 | CEP250 | chr20 | + | 35458303 | 35458374 | 35455174 | 35455237 | 35459982 | 35460105 | 1 | 1 | [illegible] |
| SE | 5585 | ENSG00000126001 | CEP250 | chr20 | + | 35469889 | 35469986 | 35467303 | 35467555 | 35472049 | 35472151 | 10 | 28 | [illegible] |
| SE | 5586 | ENSG00000126001 | CEP250 | chr20 | + | 35474793 | 35474852 | 35474000 | 35474052 | 35475501 | 35475646 | 2 | 2 | [illegible] |
| SE | 5588 | ENSG00000126001 | CEP250 | chr20 | + | 35477870 | 35478101 | 35476448 | 35476595 | 35479230 | 35479424 | 13 | 9 | [illegible] |
| SE | 5590 | ENSG00000126001 | CEP250 | chr20 | + | 35494523 | 35494657 | 35493428 | 35493572 | 35496576 | 35496715 | 90 | 85 | [illegible] |
| SE | 5594 | ENSG00000126001 | CEP250 | chr20 | + | 35500048 | 35500169 | 35498594 | 35498716 | 35507737 | 35507851 | 12 | 5 | [illegible] |
| SE | 5595 | ENSG00000126001 | CEP250 | chr20 | + | 35507737 | 35507851 | 35500048 | 35500169 | 35508034 | 35508190 | 47 | 23 | [illegible] |
| SE | 5597 | ENSG00000126001 | CEP250 | chr20 | + | 35508942 | 35509044 | 35508034 | 35508190 | 35509997 | 35510054 | 89 | 38 | [illegible] |
| SE | 5602 | ENSG00000167633 | KIR3DL1 | chr19 | + | 54821564 | 54821858 | 54819712 | 54820012 | 54825027 | 54825078 | 30 | 20 | [illegible] |
| SE | 5603 | ENSG00000167633 | KIR3DL1 | chr19 | + | 54825027 | 54825078 | 54821564 | 54821858 | 54829360 | 54829465 | 23 | 12 | [illegible] |
| SE | 5605 | ENSG00000215440 | NPEPL1 | chr20 | + | 58713916 | 58714093 | 58713419 | 58713543 | 58714559 | 58714670 | 54 | 36 | [illegible] |
| SE | 5609 | ENSG00000183647 | ZNF530 | chr19 | + | 57600727 | 57600779 | 57599884 | 57600134 | 57604276 | 57604406 | 3 | 0 | [illegible] |
| SE | 5612 | ENSG00000131067 | GGT7 | chr20 | - | 34851230 | 34851368 | 34849960 | 34850060 | 34852154 | 34852272 | 13 | 10 | [illegible] |
| SE | 5613 | ENSG00000131067 | GGT7 | chr20 | - | 34862813 | 34862965 | 34861444 | 34861562 | 34863312 | 34863548 | 8 | 8 | [illegible] |
| SE | 5614 | ENSG00000131067 | GGT7 | chr20 | - | 34863679 | 34863730 | 34863312 | 34863548 | 34872646 | 34872822 | 0 | 0 | [illegible] |
| SE | 5619 | ENSG00000213742 | ZNF337-AS1 | chr20 | + | 25674896 | 25675078 | 25673851 | 25674007 | 25676935 | 25677052 | 6 | 1 | [illegible] |
| SE | 5622 | ENSG00000101413 | RPRD1B | chr20 | + | 38040434 | 38040564 | 38033745 | 38034098 | 38057531 | 38057644 | 25 | 7 | [illegible] |
| SE | 5623 | ENSG00000101413 | RPRD1B | chr20 | + | 38048347 | 38048481 | 38033724 | 38034098 | 38057531 | 38057644 | 66 | 34 | [illegible] |
| SE | 5625 | ENSG00000101413 | RPRD1B | chr20 | + | 38066080 | 38066256 | 38059393 | 38059520 | 38089725 | 38089839 | 122 | 119 | [illegible] |
| SE | 5626 | ENSG00000124205 | EDN3 | chr20 | + | 59322371 | 59322417 | 59321016 | 59321193 | 59324330 | 59324474 | 0 | 1 | [illegible] |
| SE | 5627 | ENSG00000124205 | EDN3 | chr20 | + | 59322371 | 59322452 | 59321016 | 59321193 | 59324330 | 59325992 | 0 | 0 | [illegible] |
| SE | 5628 | ENSG00000124205 | EDN3 | chr20 | + | 59322371 | 59322482 | 59321016 | 59321193 | 59324330 | 59324572 | 0 | 0 | [illegible] |
| SE | 5631 | ENSG00000124209 | RAB22A | chr20 | + | 58343717 | 58343799 | 58311042 | 58311122 | 58353272 | 58353344 | 83 | 68 | [illegible] |
| SE | 5633 | ENSG00000124209 | RAB22A | chr20 | + | 58354155 | 58354265 | 58353431 | 58353538 | 58359605 | 58367507 | 159 | 147 | [illegible] |
| SE | 5634 | ENSG00000080839 | RBL1 | chr20 | - | 37003701 | 37003866 | 36996348 | 36998929 | 37007410 | 37007559 | 49 | 29 | [illegible] |
| SE | 5635 | ENSG00000080839 | RBL1 | chr20 | - | 37007410 | 37007559 | 37003573 | 37003866 | 37018278 | 37018369 | 81 | 37 | [illegible] |
| SE | 5636 | ENSG00000080839 | RBL1 | chr20 | - | 37020658 | 37020730 | 37018278 | 37018369 | 37022649 | 37022826 | 69 | 70 | [illegible] |
| SE | 5638 | ENSG00000080839 | RBL1 | chr20 | - | 37035241 | 37035508 | 37022649 | 37022826 | 37040152 | 37040285 | 63 | 40 | [illegible] |
| SE | 5639 | ENSG00000080839 | RBL1 | chr20 | - | 37040152 | 37040285 | 37035241 | 37035508 | 37044085 | 37044250 | 126 | 57 | [illegible] |
| SE | 5640 | ENSG00000080839 | RBL1 | chr20 | - | 37055552 | 37055656 | 37047052 | 37047190 | 37056145 | 37056258 | 98 | 72 | [illegible] |
| SE | 5641 | ENSG00000080839 | RBL1 | chr20 | - | 37065423 | 37065473 | 37062083 | 37062270 | 37066723 | 37066884 | 99 | 76 | [illegible] |
| SE | 5642 | ENSG00000080839 | RBL1 | chr20 | - | 37066992 | 37067021 | 37066723 | 37066884 | 37067232 | 37067297 | 18 | 7 | [illegible] |
| SE | 5643 | ENSG00000080839 | RBL1 | chr20 | - | 37066992 | 37067021 | 37066723 | 37066884 | 37067985 | 37068055 | 20 | 8 | [illegible] |
| SE | 5644 | ENSG00000080839 | RBL1 | chr20 | - | 37066992 | 37067121 | 37066723 | 37066884 | 37067232 | 37067297 | 193 | 140 | [illegible] |
| SE | 5645 | ENSG00000080839 | RBL1 | chr20 | - | 37066992 | 37067121 | 37066723 | 37066884 | 37067985 | 37068186 | 126 | 107 | [illegible] |
| SE | 5646 | ENSG00000080839 | RBL1 | chr20 | - | 37067232 | 37067297 | 37066723 | 37066884 | 37067985 | 37068186 | 23 | 18 | [illegible] |
| SE | 5647 | ENSG00000080839 | RBL1 | chr20 | - | 37067232 | 37067297 | 37066992 | 37067021 | 37067985 | 37068186 | 23 | 18 | [illegible] |
| SE | 5649 | ENSG00000080839 | RBL1 | chr20 | - | 37081887 | 37082029 | 37068086 | 37068186 | 37088988 | 37089122 | 3 | 2 | [illegible] |
| SE | 5651 | ENSG00000196700 | ZNF512B | chr20 | - | 63967380 | 63967523 | 63966875 | 63967004 | 63967829 | 63967952 | 21 | 14 | [illegible] |
| SE | 5653 | ENSG00000203880 | PCMTD2 | chr20 | + | 64258766 | 64258882 | 64255843 | 64255870 | 64259941 | 64260272 | 5 | 0 | [illegible] |
| SE | 5656 | ENSG00000203880 | PCMTD2 | chr20 | + | 64259941 | 64260272 | 64255694 | 64255870 | 64273220 | 64275653 | 17 | 9 | [illegible] |
| SE | 5657 | ENSG00000203880 | PCMTD2 | chr20 | + | 64264428 | 64264531 | 64259941 | 64260272 | 64267886 | 64268010 | 62 | 35 | [illegible] |
| SE | 5658 | ENSG00000203880 | PCMTD2 | chr20 | + | 64265257 | 64265429 | 64264428 | 64264531 | 64267886 | 64268010 | 344 | 230 | [illegible] |
| SE | 5659 | ENSG00000203880 | PCMTD2 | chr20 | + | 64267886 | 64268010 | 64259941 | 64260272 | 64273220 | 64273545 | 161 | 123 | [illegible] |
| SE | 5660 | ENSG00000203880 | PCMTD2 | chr20 | + | 64267886 | 64268010 | 64265257 | 64265429 | 64273220 | 64273545 | 294 | 206 | [illegible] |
| SE | 5661 | ENSG00000203880 | PCMTD2 | chr20 | + | 64267967 | 64268010 | 64265257 | 64265429 | 64273220 | 64274139 | 85 | 46 | [illegible] |
| SE | 5662 | ENSG00000171700 | RGS19 | chr20 | - | 64076524 | 64076646 | 64074466 | 64074541 | 64079872 | 64079988 | 81 | 34 | [illegible] |
| SE | 5665 | ENSG00000125510 | OPRL1 | chr20 | + | 64086504 | 64086599 | 64080195 | 64080352 | 64091965 | 64092116 | 0 | 0 | [illegible] |
| SE | 5666 | ENSG00000125510 | OPRL1 | chr20 | + | 64086504 | 64086599 | 64080195 | 64080352 | 64092687 | 64092953 | 0 | 0 | [illegible] |
| SE | 5667 | ENSG00000125510 | OPRL1 | chr20 | + | 64091965 | 64092116 | 64080081 | 64080352 | 64092687 | 64092953 | 2 | 0 | [illegible] |
| SE | 5668 | ENSG00000125510 | OPRL1 | chr20 | + | 64091965 | 64092116 | 64086504 | 64086599 | 64092687 | 64092953 | 2 | 0 | [illegible] |
| SE | 5669 | ENSG00000125510 | OPRL1 | chr20 | + | 64092687 | 64092953 | 64080195 | 64080352 | 64097801 | 64098157 | 4 | 4 | [illegible] |
| SE | 5676 | ENSG00000064787 | BCAS1 | chr20 | - | 53957431 | 53957497 | 53953431 | 53953695 | 53985286 | 53985499 | 1 | 0 | [illegible] |
| SE | 5678 | ENSG00000064787 | BCAS1 | chr20 | - | 53966905 | 53967073 | 53953431 | 53953695 | 53985286 | 53985499 | 3 | 0 | [illegible] |
| SE | 5683 | ENSG00000064787 | BCAS1 | chr20 | - | 53975388 | 53975430 | 53966905 | 53967073 | 53985286 | 53985499 | 0 | 0 | [illegible] |
| SE | 5684 | ENSG00000064787 | BCAS1 | chr20 | - | 53992511 | 53992646 | 53985286 | 53985499 | 53995011 | 53995056 | 2 | 0 | [illegible] |
| SE | 5686 | ENSG00000184402 | SS18L1 | chr20 | + | 62154280 | 62154465 | 62143794 | 62143889 | 62158671 | 62158942 | 5 | 2 | [illegible] |
| SE | 5690 | ENSG00000188785 | ZNF548 | chr19 | + | 57390721 | 57390845 | 57390004 | 57390114 | 57392800 | 57392948 | 13 | 12 | [illegible] |
| SE | 5692 | ENSG00000188785 | ZNF548 | chr19 | + | 57392787 | 57392948 | 57390745 | 57390845 | 57394187 | 57394223 | 14 | 17 | [illegible] |
| SE | 5697 | ENSG00000188785 | ZNF548 | chr19 | + | 57397047 | 57397174 | 57394042 | 57394223 | 57398429 | 57398436 | 66 | 37 | [illegible] |
| SE | 5698 | ENSG00000101079 | NDRG3 | chr20 | - | 36654800 | 36654839 | 36651765 | 36653701 | 36656359 | 36656411 | 0 | 0 | [illegible] |
| SE | 5699 | ENSG00000101079 | NDRG3 | chr20 | - | 36660336 | 36660384 | 36656496 | 36656532 | 36665045 | 36665097 | 96 | 32 | [illegible] |
| SE | 5700 | ENSG00000101079 | NDRG3 | chr20 | - | 36680815 | 36680902 | 36671340 | 36671397 | 36682517 | 36682578 | 138 | 64 | [illegible] |
| SE | 5701 | ENSG00000101079 | NDRG3 | chr20 | - | 36687491 | 36687612 | 36684412 | 36684475 | 36688678 | 36688784 | 187 | 87 | [illegible] |
| SE | 5702 | ENSG00000101079 | NDRG3 | chr20 | - | 36688678 | 36688784 | 36684412 | 36684475 | 36706971 | 36707007 | 83 | 39 | [illegible] |
| SE | 5718 | ENSG00000198276 | UCKL1 | chr20 | - | 63942469 | 63942492 | 63941109 | 63941208 | 63944396 | 63944458 | 3 | 0 | [illegible] |
| SE | 5720 | ENSG00000198276 | UCKL1 | chr20 | - | 63943652 | 63943669 | 63942469 | 63942492 | 63944396 | 63944458 | 0 | 3 | [illegible] |
| SE | 5721 | ENSG00000198276 | UCKL1 | chr20 | - | 63948224 | 63948475 | 63946452 | 63946643 | 63956259 | 63956341 | 0 | 1 | [illegible] |
| SE | 5722 | ENSG00000020256 | ZFP64 | chr20 | - | 52165863 | 52166025 | 52164694 | 52164757 | 52186831 | 52187065 | 60 | 37 | [illegible] |
| SE | 5730 | ENSG00000154930 | ACSS1 | chr20 | - | 25013535 | 25013662 | 25012600 | 25012664 | 25013960 | 25014073 | 66 | 27 | [illegible] |
| SE | 5731 | ENSG00000154930 | ACSS1 | chr20 | - | 25015137 | 25015230 | 25013960 | 25014073 | 25020009 | 25020147 | 94 | 27 | [illegible] |
| SE | 5734 | ENSG00000154930 | ACSS1 | chr20 | - | 25021388 | 25021536 | 25015137 | 25015230 | 25030758 | 25030958 | 0 | 2 | [illegible] |
| SE | 5735 | ENSG00000154930 | ACSS1 | chr20 | - | 25022939 | 25023092 | 25021388 | 25021536 | 25023465 | 25023641 | 193 | 74 | [illegible] |
| SE | 5739 | ENSG00000154930 | ACSS1 | chr20 | - | 25030758 | 25030958 | 25013960 | 25014073 | 25048084 | 25048181 | 42 | 27 | [illegible] |
| SE | 5740 | ENSG00000154930 | ACSS1 | chr20 | - | 25030758 | 25030958 | 25013960 | 25014073 | 25057768 | 25058139 | 0 | 1 | [illegible] |
| SE | 5742 | ENSG00000154930 | ACSS1 | chr20 | - | 25048084 | 25048181 | 25013960 | 25014073 | 25057768 | 25058139 | 25 | 8 | [illegible] |
| SE | 5743 | ENSG00000154930 | ACSS1 | chr20 | - | 25048084 | 25048181 | 25030758 | 25030958 | 25057768 | 25058139 | 66 | 35 | [illegible] |
| SE | 5756 | ENSG00000131044 | TTLL9 | chr20 | + | 31938181 | 31938263 | 31937395 | 31937509 | 31939141 | 31939266 | 0 | 0 | [illegible] |
| SE | 5769 | ENSG00000101365 | IDH3B | chr20 | - | 2659698 | 2659793 | 2659524 | 2659585 | 2660262 | 2660308 | 69 | 40 | [illegible] |
| SE | 5770 | ENSG00000101365 | IDH3B | chr20 | - | 2659698 | 2660176 | 2659524 | 2659585 | 2660262 | 2660359 | 309 | 157 | [illegible] |
| SE | 5771 | ENSG00000101365 | IDH3B | chr20 | - | 2660029 | 2660176 | 2659698 | 2659793 | 2660262 | 2660308 | 316 | 154 | [illegible] |
| SE | 5772 | ENSG00000101365 | IDH3B | chr20 | - | 2660067 | 2660176 | 2659698 | 2659793 | 2660262 | 2660309 | 182 | 96 | [illegible] |
| SE | 5773 | ENSG00000101365 | IDH3B | chr20 | - | 2660456 | 2660590 | 2660262 | 2660365 | 2660696 | 2660829 | 210 | 127 | [illegible] |
| SE | 5774 | ENSG00000101365 | IDH3B | chr20 | - | 2663659 | 2663758 | 2663445 | 2663566 | 2663924 | 2664005 | 299 | 116 | [illegible] |
| SE | 5777 | ENSG00000101193 | GID8 | chr20 | + | 62941490 | 62941620 | 62938146 | 62938253 | 62942986 | 62943183 | 115 | 89 | [illegible] |
| SE | 5778 | ENSG00000101193 | GID8 | chr20 | + | 62943494 | 62943692 | 62942986 | 62943183 | 62944738 | 62948475 | 432 | 285 | [illegible] |
| SE | 5788 | ENSG00000149646 | CNBD2 | chr20 | + | 35987394 | 35987533 | 35984626 | 35984778 | 35995037 | 35995152 | 0 | 0 | [illegible] |
| SE | 5793 | ENSG00000197586 | ENTPD6 | chr20 | + | 25196153 | 25196254 | 25195791 | 25195867 | 25197104 | 25197226 | 0 | 0 | [illegible] |
| SE | 5795 | ENSG00000197586 | ENTPD6 | chr20 | + | 25196153 | 25196254 | 25195791 | 25195867 | 25207075 | 25207397 | 0 | 0 | [illegible] |
| SE | 5796 | ENSG00000197586 | ENTPD6 | chr20 | + | 25196153 | 25196254 | 25195791 | 25195867 | 25207078 | 25207397 | 0 | 0 | [illegible] |
| SE | 5797 | ENSG00000197586 | ENTPD6 | chr20 | + | 25196153 | 25196254 | 25195791 | 25195867 | 25209848 | 25209925 | 1 | 0 | [illegible] |
| SE | 5799 | ENSG00000197586 | ENTPD6 | chr20 | + | 25197104 | 25197226 | 25195791 | 25195867 | 25207075 | 25207397 | 3 | 0 | [illegible] |
| SE | 5800 | ENSG00000197586 | ENTPD6 | chr20 | + | 25197104 | 25197226 | 25195791 | 25195867 | 25207078 | 25207397 | 3 | 0 | [illegible] |
| SE | 5801 | ENSG00000197586 | ENTPD6 | chr20 | + | 25197104 | 25197226 | 25195791 | 25195867 | 25209848 | 25209925 | 3 | 0 | [illegible] |
| SE | 5802 | ENSG00000197586 | ENTPD6 | chr20 | + | 25197104 | 25197226 | 25196153 | 25196254 | 25206521 | 25206590 | 3 | 2 | [illegible] |
| SE | 5803 | ENSG00000197586 | ENTPD6 | chr20 | + | 25197104 | 25197226 | 25196153 | 25196254 | 25207075 | 25207397 | 0 | 0 | [illegible] |
| SE | 5804 | ENSG00000197586 | ENTPD6 | chr20 | + | 25197104 | 25197226 | 25196153 | 25196254 | 25207078 | 25207397 | 0 | 0 | [illegible] |
| SE | 5805 | ENSG00000197586 | ENTPD6 | chr20 | + | 25197104 | 25197226 | 25196153 | 25196254 | 25209848 | 25209925 | 0 | 0 | [illegible] |
| SE | 5807 | ENSG00000197586 | ENTPD6 | chr20 | + | 25206521 | 25206590 | 25195700 | 25195867 | 25207078 | 25207397 | 43 | 13 | [illegible] |
| SE | 5809 | ENSG00000197586 | ENTPD6 | chr20 | + | 25206521 | 25206590 | 25196153 | 25196254 | 25207075 | 25207397 | 22 | 16 | [illegible] |
| SE | 5810 | ENSG00000197586 | ENTPD6 | chr20 | + | 25206521 | 25206590 | 25196153 | 25196254 | 25207078 | 25207397 | 15 | 7 | [illegible] |
| SE | 5811 | ENSG00000197586 | ENTPD6 | chr20 | + | 25206521 | 25206590 | 25196153 | 25196254 | 25209848 | 25209925 | 2 | 0 | [illegible] |
| SE | 5812 | ENSG00000197586 | ENTPD6 | chr20 | + | 25206521 | 25206590 | 25197104 | 25197226 | 25207075 | 25207397 | 23 | 17 | [illegible] |
| SE | 5814 | ENSG00000197586 | ENTPD6 | chr20 | + | 25206521 | 25206590 | 25197104 | 25197226 | 25209848 | 25209925 | 1 | 0 | [illegible] |
| SE | 5816 | ENSG00000197586 | ENTPD6 | chr20 | + | 25207075 | 25207397 | 25196153 | 25196254 | 25209848 | 25209925 | 23 | 14 | [illegible] |
| SE | 5817 | ENSG00000197586 | ENTPD6 | chr20 | + | 25207075 | 25207397 | 25197104 | 25197226 | 25209848 | 25209925 | 23 | 14 | [illegible] |
| SE | 5820 | ENSG00000197586 | ENTPD6 | chr20 | + | 25207078 | 25207397 | 25196153 | 25196254 | 25209848 | 25209925 | 23 | 14 | [illegible] |
| SE | 5821 | ENSG00000197586 | ENTPD6 | chr20 | + | 25207078 | 25207397 | 25197104 | 25197226 | 25209848 | 25209925 | 23 | 14 | [illegible] |
| SE | 5824 | ENSG00000197586 | ENTPD6 | chr20 | + | 25218549 | 25218614 | 25217501 | 25217581 | 25221231 | 25221333 | 67 | 31 | [illegible] |
| SE | 5825 | ENSG00000197586 | ENTPD6 | chr20 | + | 25218549 | 25218614 | 25217501 | 25217581 | 25222837 | 25222978 | 36 | 22 | [illegible] |
| SE | 5826 | ENSG00000197586 | ENTPD6 | chr20 | + | 25221231 | 25221333 | 25217501 | 25217581 | 25222837 | 25222970 | 22 | 18 | [illegible] |
| SE | 5832 | ENSG00000100994 | PYGB | chr20 | + | 25276645 | 25276757 | 25274591 | 25274723 | 25277243 | 25277326 | 146 | 79 | [illegible] |
| SE | 5833 | ENSG00000100994 | PYGB | chr20 | + | 25292405 | 25292613 | 25290480 | 25290622 | 25294157 | 25294292 | 168 | 91 | [illegible] |
| SE | 5834 | ENSG00000218018 | RBM38-AS1 | chr20 | - | 57377551 | 57377727 | 57377139 | 57377177 | 57392278 | 57392626 | 0 | 0 | [illegible] |
| SE | 5835 | ENSG00000101194 | SLC17A9 | chr20 | + | 62953210 | 62953272 | 62952706 | 62952889 | 62956764 | 62956962 | 0 | 1 | [illegible] |
| SE | 5842 | ENSG00000101391 | CDK5RAP1 | chr20 | - | 33360350 | 33360491 | 33358838 | 33359123 | 33366858 | 33367005 | 222 | 157 | [illegible] |
| SE | 5843 | ENSG00000101391 | CDK5RAP1 | chr20 | - | 33372641 | 33372697 | 33370498 | 33370629 | 33374114 | 33374212 | 17 | 18 | [illegible] |
| SE | 5845 | ENSG00000101391 | CDK5RAP1 | chr20 | - | 33374114 | 33374212 | 33370498 | 33370629 | 33392141 | 33392242 | 2 | 0 | [illegible] |
| SE | 5847 | ENSG00000101391 | CDK5RAP1 | chr20 | - | 33379460 | 33379691 | 33374114 | 33374212 | 33392141 | 33392242 | 48 | 20 | [illegible] |
| SE | 5848 | ENSG00000101391 | CDK5RAP1 | chr20 | - | 33383479 | 33383521 | 33379460 | 33379691 | 33385649 | 33385770 | 1 | 1 | [illegible] |
| SE | 5849 | ENSG00000101391 | CDK5RAP1 | chr20 | - | 33385649 | 33385770 | 33370498 | 33370629 | 33392141 | 33392242 | 2 | 6 | [illegible] |
| SE | 5850 | ENSG00000101391 | CDK5RAP1 | chr20 | - | 33385649 | 33385770 | 33374114 | 33374212 | 33392141 | 33392242 | 2 | 7 | [illegible] |
| SE | 5851 | ENSG00000101391 | CDK5RAP1 | chr20 | - | 33385649 | 33385770 | 33379460 | 33379691 | 33392141 | 33392242 | 34 | 26 | [illegible] |
| SE | 5854 | ENSG00000101391 | CDK5RAP1 | chr20 | - | 33392141 | 33392242 | 33379460 | 33379691 | 33401427 | 33401452 | 0 | 0 | [illegible] |
| SE | 5855 | ENSG00000101391 | CDK5RAP1 | chr20 | - | 33392141 | 33392242 | 33387322 | 33387533 | 33394031 | 33394066 | 73 | 45 | [illegible] |
| SE | 5856 | ENSG00000101391 | CDK5RAP1 | chr20 | - | 33394031 | 33394066 | 33387322 | 33387533 | 33395012 | 33395116 | 7 | 9 | [illegible] |
| SE | 5857 | ENSG00000101391 | CDK5RAP1 | chr20 | - | 33394031 | 33394066 | 33392141 | 33392242 | 33396760 | 33396936 | 17 | 15 | [illegible] |
| SE | 5860 | ENSG00000101391 | CDK5RAP1 | chr20 | - | 33395012 | 33395116 | 33394031 | 33394066 | 33401427 | 33401505 | 27 | 11 | [illegible] |
| SE | 5862 | ENSG00000101391 | CDK5RAP1 | chr20 | - | 33396760 | 33396936 | 33394031 | 33394066 | 33401427 | 33401517 | 0 | 0 | [illegible] |
| SE | 5863 | ENSG00000101391 | CDK5RAP1 | chr20 | - | 33396760 | 33396936 | 33395012 | 33395116 | 33401427 | 33401517 | 29 | 17 | [illegible] |
| SE | 5865 | ENSG00000101391 | CDK5RAP1 | chr20 | - | 33396760 | 33397084 | 33394031 | 33394066 | 33401427 | 33401519 | 24 | 13 | [illegible] |
| SE | 5866 | ENSG00000101391 | CDK5RAP1 | chr20 | - | 33396760 | 33397084 | 33395012 | 33395116 | 33401427 | 33401519 | 53 | 30 | [illegible] |
| SE | 5882 | ENSG00000132819 | RBM38 | chr20 | + | 57392396 | 57392492 | 57391452 | 57391795 | 57392653 | 57392777 | 1 | 0 | [illegible] |
| SE | 5885 | ENSG00000132819 | RBM38 | chr20 | + | 57399927 | 57399968 | 57393278 | 57393333 | 57407542 | 57409333 | 0 | 0 | [illegible] |
| SE | 5888 | ENSG00000101425 | BPI | chr20 | + | 38307566 | 38307681 | 38304149 | 38304353 | 38308929 | 38309058 | 32 | 23 | [illegible] |
| SE | 5889 | ENSG00000101425 | BPI | chr20 | + | 38308929 | 38309058 | 38307566 | 38307681 | 38310490 | 38310652 | 31 | 17 | [illegible] |
| SE | 5890 | ENSG00000101425 | BPI | chr20 | + | 38311873 | 38311937 | 38310490 | 38310652 | 38318412 | 38318476 | 11 | 9 | [illegible] |
| SE | 5891 | ENSG00000101425 | BPI | chr20 | + | 38323869 | 38324046 | 38320182 | 38320274 | 38324773 | 38324833 | 40 | 23 | [illegible] |
| SE | 5892 | ENSG00000101425 | BPI | chr20 | + | 38323921 | 38324046 | 38320182 | 38320274 | 38324773 | 38324833 | 12 | 7 | [illegible] |
| SE | 5893 | ENSG00000101425 | BPI | chr20 | + | 38327587 | 38327655 | 38326264 | 38326432 | 38334429 | 38334493 | 0 | 0 | [illegible] |
| SE | 5894 | ENSG00000101425 | BPI | chr20 | + | 38331047 | 38331090 | 38327587 | 38327655 | 38334429 | 38334493 | 14 | 1 | [illegible] |
| SE | 5895 | ENSG00000124228 | DDX27 | chr20 | + | 49222956 | 49223016 | 49221451 | 49221598 | 49223267 | 49223433 | 78 | 79 | [illegible] |
| SE | 5896 | ENSG00000124228 | DDX27 | chr20 | + | 49224774 | 49224991 | 49223267 | 49223433 | 49226429 | 49226466 | 0 | 0 | [illegible] |
| SE | 5897 | ENSG00000124228 | DDX27 | chr20 | + | 49224944 | 49224991 | 49223267 | 49223433 | 49226429 | 49226535 | 5 | 10 | [illegible] |
| SE | 5898 | ENSG00000124228 | DDX27 | chr20 | + | 49225112 | 49225199 | 49224774 | 49224991 | 49226429 | 49226466 | 112 | 65 | [illegible] |
| SE | 5899 | ENSG00000124228 | DDX27 | chr20 | + | 49226429 | 49226535 | 49225112 | 49225199 | 49228714 | 49228888 | 165 | 87 | [illegible] |
| SE | 5900 | ENSG00000124228 | DDX27 | chr20 | + | 49228714 | 49228888 | 49226429 | 49226535 | 49230198 | 49230349 | 105 | 45 | [illegible] |
| SE | 5901 | ENSG00000124228 | DDX27 | chr20 | + | 49230198 | 49230349 | 49228714 | 49228888 | 49233305 | 49233405 | 44 | 16 | [illegible] |
| SE | 5903 | ENSG00000124228 | DDX27 | chr20 | + | 49234934 | 49235088 | 49233567 | 49233709 | 49236149 | 49236231 | 118 | 94 | [illegible] |
| SE | 5904 | ENSG00000124228 | DDX27 | chr20 | + | 49238948 | 49239055 | 49233837 | 49236510 | 49239235 | 49239338 | 198 | 120 | [illegible] |
| SE | 5905 | ENSG00000124228 | DDX27 | chr20 | + | 49242082 | 49242206 | 49241892 | 49241987 | 49242593 | 49242681 | 139 | 68 | [illegible] |
| SE | 5906 | ENSG00000101004 | NINL | chr20 | - | 25476042 | 25477089 | 25469990 | 25470095 | 25478922 | 25479206 | 13 | 1 | [illegible] |
| SE | 5908 | ENSG00000101004 | NINL | chr20 | - | 25498209 | 25498346 | 25496662 | 25496803 | 25500839 | 25501010 | 9 | 4 | [illegible] |
| SE | 5914 | ENSG00000124224 | PPP4R1L | chr20 | - | 58271362 | 58271505 | 58248114 | 58248273 | 58272768 | 58272825 | 27 | 19 | [illegible] |
| SE | 5916 | ENSG00000124224 | PPP4R1L | chr20 | - | 58286313 | 58286446 | 58272768 | 58272875 | 58309225 | 58309270 | 35 | 30 | [illegible] |
| SE | 5919 | ENSG00000080189 | SLC35C2 | chr20 | - | 46355801 | 46355874 | 46355072 | 46355241 | 46356573 | 46356637 | 35 | 16 | [illegible] |
| SE | 5941 | ENSG00000170471 | RALGAPB | chr20 | + | 38497352 | 38497516 | 38492929 | 38493132 | 38509076 | 38509208 | 193 | 143 | [illegible] |
| SE | 5942 | ENSG00000170471 | RALGAPB | chr20 | + | 38497352 | 38497516 | 38492929 | 38493132 | 38516191 | 38516370 | 193 | 143 | [illegible] |
| SE | 5944 | ENSG00000170471 | RALGAPB | chr20 | + | 38497479 | 38497516 | 38492929 | 38493132 | 38509076 | 38509208 | 1 | 0 | [illegible] |
| SE | 5945 | ENSG00000170471 | RALGAPB | chr20 | + | 38497479 | 38497516 | 38492929 | 38493132 | 38516191 | 38516370 | 1 | 0 | [illegible] |
| SE | 5946 | ENSG00000170471 | RALGAPB | chr20 | + | 38499446 | 38499633 | 38492929 | 38493132 | 38509076 | 38509208 | 61 | 38 | [illegible] |
| SE | 5947 | ENSG00000170471 | RALGAPB | chr20 | + | 38499446 | 38499633 | 38492929 | 38493132 | 38516191 | 38516370 | 3 | 0 | [illegible] |
| SE | 5949 | ENSG00000170471 | RALGAPB | chr20 | + | 38499446 | 38499633 | 38497352 | 38497516 | 38516191 | 38516370 | 55 | 34 | [illegible] |
| SE | 5950 | ENSG00000170471 | RALGAPB | chr20 | + | 38509076 | 38509208 | 38492929 | 38493132 | 38516191 | 38516370 | 139 | 93 | [illegible] |
| SE | 5951 | ENSG00000170471 | RALGAPB | chr20 | + | 38509076 | 38509208 | 38497352 | 38497516 | 38516191 | 38516370 | 138 | 93 | [illegible] |
| SE | 5952 | ENSG00000170471 | RALGAPB | chr20 | + | 38509076 | 38509208 | 38499446 | 38499633 | 38516191 | 38516370 | 196 | 131 | [illegible] |
| SE | 5953 | ENSG00000170471 | RALGAPB | chr20 | + | 38517505 | 38517654 | 38516191 | 38516370 | 38517783 | 38518000 | 268 | 166 | [illegible] |
| SE | 5954 | ENSG00000170471 | RALGAPB | chr20 | + | 38517783 | 38518000 | 38517505 | 38517654 | 38521496 | 38521698 | 263 | 193 | [illegible] |
| SE | 5955 | ENSG00000170471 | RALGAPB | chr20 | + | 38521496 | 38521698 | 38517783 | 38518000 | 38524777 | 38524945 | 217 | 165 | [illegible] |
| SE | 5957 | ENSG00000170471 | RALGAPB | chr20 | + | 38531166 | 38531231 | 38525894 | 38526042 | 38532729 | 38532859 | 154 | 105 | [illegible] |
| SE | 5961 | ENSG00000170471 | RALGAPB | chr20 | + | 38541040 | 38541192 | 38539775 | 38539958 | 38548688 | 38548795 | 72 | 39 | [illegible] |
| SE | 5962 | ENSG00000170471 | RALGAPB | chr20 | + | 38546242 | 38546430 | 38541040 | 38541192 | 38548688 | 38548795 | 165 | 146 | [illegible] |
| SE | 5964 | ENSG00000170471 | RALGAPB | chr20 | + | 38548688 | 38548795 | 38541040 | 38541192 | 38551070 | 38551223 | 93 | 95 | [illegible] |
| SE | 5966 | ENSG00000170471 | RALGAPB | chr20 | + | 38553866 | 38554076 | 38551070 | 38551223 | 38558294 | 38558453 | 400 | 340 | [illegible] |
| SE | 5967 | ENSG00000170471 | RALGAPB | chr20 | + | 38562531 | 38562697 | 38558294 | 38558453 | 38574149 | 38574298 | 85 | 58 | [illegible] |
| SE | 5969 | ENSG00000170471 | RALGAPB | chr20 | + | 38565355 | 38565478 | 38562531 | 38562697 | 38570768 | 38570847 | 50 | 27 | [illegible] |
| SE | 5970 | ENSG00000170471 | RALGAPB | chr20 | + | 38565355 | 38565478 | 38562531 | 38562697 | 38574149 | 38574298 | 50 | 27 | [illegible] |
| SE | 5972 | ENSG00000170471 | RALGAPB | chr20 | + | 38565358 | 38565478 | 38562531 | 38562697 | 38570768 | 38570847 | 50 | 20 | [illegible] |
| SE | 5973 | ENSG00000170471 | RALGAPB | chr20 | + | 38565358 | 38565478 | 38562531 | 38562697 | 38574149 | 38574298 | 50 | 20 | [illegible] |
| SE | 5974 | ENSG00000170471 | RALGAPB | chr20 | + | 38567095 | 38567232 | 38562531 | 38562697 | 38570768 | 38570847 | 1 | 3 | [illegible] |
| SE | 5975 | ENSG00000170471 | RALGAPB | chr20 | + | 38567095 | 38567232 | 38562531 | 38562697 | 38574149 | 38574298 | 2 | 2 | [illegible] |
| SE | 5976 | ENSG00000170471 | RALGAPB | chr20 | + | 38567095 | 38567232 | 38565355 | 38565478 | 38570768 | 38570847 | 142 | 58 | [illegible] |
| SE | 5977 | ENSG00000170471 | RALGAPB | chr20 | + | 38567095 | 38567232 | 38565355 | 38565478 | 38574149 | 38574298 | 143 | 57 | [illegible] |
| SE | 5979 | ENSG00000170471 | RALGAPB | chr20 | + | 38569887 | 38569996 | 38567095 | 38567232 | 38570768 | 38570847 | 168 | 127 | [illegible] |
| SE | 5981 | ENSG00000170471 | RALGAPB | chr20 | + | 38570768 | 38570847 | 38562531 | 38562697 | 38574149 | 38574298 | 132 | 88 | [illegible] |
| SE | 5982 | ENSG00000170471 | RALGAPB | chr20 | + | 38570768 | 38570847 | 38565355 | 38565478 | 38574149 | 38574298 | 132 | 88 | [illegible] |
| SE | 5983 | ENSG00000170471 | RALGAPB | chr20 | + | 38570768 | 38570847 | 38567095 | 38567232 | 38574149 | 38574298 | 132 | 89 | [illegible] |
| SE | 5984 | ENSG00000170471 | RALGAPB | chr20 | + | 38570768 | 38570847 | 38568466 | 38569996 | 38574149 | 38574298 | 268 | 190 | [illegible] |
| SE | 5985 | ENSG00000170471 | RALGAPB | chr20 | + | 38574149 | 38574298 | 38570768 | 38570847 | 38574773 | 38575070 | 314 | 253 | [illegible] |
| SE | 5996 | ENSG00000062598 | ELMO2 | chr20 | - | 46394404 | 46394503 | 46394048 | 46394089 | 46406547 | 46406632 | 87 | 32 | [illegible] |
| SE | 5999 | ENSG00000062598 | ELMO2 | chr20 | - | 46398696 | 46398771 | 46393980 | 46394089 | 46406547 | 46406559 | 1 | 0 | [illegible] |
| SE | 6003 | ENSG00000174365 | SNHG11 | chr20 | + | 38447929 | 38448093 | 38446886 | 38447623 | 38448661 | 38448730 | 2 | 2 | [illegible] |
| SE | 6004 | ENSG00000174365 | SNHG11 | chr20 | + | 38447929 | 38448093 | 38446886 | 38447623 | 38448665 | 38448730 | 1 | 1 | [illegible] |
| SE | 6009 | ENSG00000229976 | NA | chr20 | + | 39763954 | 39764004 | 39757201 | 39757480 | 39765587 | 39765658 | 3 | 1 | [illegible] |
| SE | 6013 | ENSG00000124191 | TOX2 | chr20 | + | 44064776 | 44064857 | 44054298 | 44054526 | 44065711 | 44066107 | 0 | 0 | [illegible] |
| SE | 6014 | ENSG00000124191 | TOX2 | chr20 | + | 44065711 | 44066107 | 44054298 | 44054526 | 44066729 | 44066857 | 0 | 0 | [illegible] |
| SE | 6017 | ENSG00000130684 | ZNF337 | chr20 | - | 25685995 | 25686122 | 25685566 | 25685662 | 25686390 | 25686466 | 54 | 39 | [illegible] |
| SE | 6018 | ENSG00000130684 | ZNF337 | chr20 | - | 25685995 | 25686249 | 25685566 | 25685662 | 25686390 | 25686466 | 33 | 22 | [illegible] |
| SE | 6020 | ENSG00000196756 | SNHG17 | chr20 | - | 38422086 | 38422241 | 38421005 | 38421098 | 38425934 | 38426048 | 3 | 0 | [illegible] |
| SE | 6023 | ENSG00000196756 | SNHG17 | chr20 | - | 38422091 | 38422241 | 38421005 | 38421098 | 38425934 | 38426048 | 2 | 2 | [illegible] |

| | | | | | | | | | | | | | | | | | | | | | | | | | |
|---|---|---|---|---|---|---|---|---|---|---|---|---|---|---|---|---|---|---|---|---|---|---|---|---|---|
| SE | 6025 | ENSG00000196756 | SNHG17 | chr20 | - | 38422095 | 38422241 | 38421005 | 38421098 | 38425934 | 38426048 | 1 | 0 | 7 | [illegible] | [illegible] | [illegible] | [illegible] | [illegible] | [illegible] | [illegible] | [illegible] | [illegible] | [illegible] | [illegible] |
| SE | 6028 | ENSG00000196756 | SNHG17 | chr20 | - | 38422290 | 38422368 | 38421005 | 38421098 | 38425934 | 38426503 | 1 | 0 | 0 | [illegible] | [illegible] | [illegible] | [illegible] | [illegible] | [illegible] | [illegible] | [illegible] | [illegible] | [illegible] | [illegible] |
| SE | 6031 | ENSG00000196756 | SNHG17 | chr20 | - | 38425934 | 38426048 | 38421005 | 38422241 | 38426418 | 38426503 | 3 | 1 | 5 | [illegible] | [illegible] | [illegible] | [illegible] | [illegible] | [illegible] | [illegible] | [illegible] | [illegible] | [illegible] | [illegible] |
| SE | 6032 | ENSG00000196756 | SNHG17 | chr20 | - | 38425934 | 38426052 | 38421005 | 38421098 | 38426418 | 38426503 | 0 | 0 | 0 | 3 | 0 | 6 | 3 | 8 | 14 | 2 | 0 | 0 | 1 | 0 |
| SE | 6033 | ENSG00000196756 | SNHG17 | chr20 | - | 38425934 | 38426052 | 38421005 | 38422241 | 38426418 | 38426503 | 1 | 0 | 4 | 1 | 0 | 1 | 3 | 5 | 1 | 0 | 5 | 0 | 1 | 0 |
| SE | 6035 | ENSG00000196756 | SNHG17 | chr20 | - | 38427679 | 38427766 | 38425934 | 38426503 | 38428550 | 38429805 | 1 | 0 | 0 | 3 | 0 | 0 | 0 | 0 | 0 | 0 | 0 | 0 | 0 | 4 |
| SE | 6037 | ENSG00000196756 | SNHG17 | chr20 | - | 38427679 | 38427766 | 38425934 | 38426503 | 38431040 | 38431044 | 1 | 0 | 0 | 2 | 0 | 7 | 8 | 12 | 15 | 8 | 0 | 0 | 0 | 4 |
| SE | 6038 | ENSG00000196756 | SNHG17 | chr20 | - | 38427679 | 38427847 | 38426418 | 38426503 | 38431040 | 38431112 | 1 | 0 | 0 | 2 | 0 | 7 | 8 | 12 | 15 | 8 | 0 | 0 | 0 | 4 |
| SE | 6041 | ENSG00000196756 | SNHG17 | chr20 | - | 38434503 | 38434626 | 38430779 | 38431112 | 38435196 | 38435273 | 3 | 4 | 0 | 3 | 10 | 0 | 0 | 0 | 0 | 0 | 44 | 24 | 4 | 9 |
| SE | 6042 | ENSG00000196756 | SNHG17 | chr20 | - | 38434519 | 38434626 | 38431040 | 38431112 | 38435196 | 38435352 | 1 | 0 | 0 | 2 | 6 | 0 | 0 | 0 | 0 | 0 | 31 | 8 | 1 | 1 |
| SE | 6043 | ENSG00000101442 | ACTR5 | chr20 | + | 38750009 | 38750239 | 38748459 | 38748853 | 38752130 | 38752300 | 39 | 17 | 27 | 37 | 27 | 4 | 0 | 0 | 0 | 0 | 35 | 31 | 20 | 29 |
| SE | 6046 | ENSG00000196839 | ADA | chr20 | - | 44623006 | 44623078 | 44622828 | 44622930 | 44629046 | 44629169 | 7 | 12 | 3 | 12 | 15 | 0 | 0 | 0 | 0 | 0 | 3 | 9 | 3 | 3 |
| SE | 6047 | ENSG00000196839 | ADA | chr20 | - | 44624201 | 44624329 | 44622828 | 44622930 | 44626455 | 44626491 | 5 | 0 | 7 | 2 | 6 | 0 | 0 | 0 | 0 | 0 | 0 | 4 | 4 | 1 |
| SE | 6049 | ENSG00000196839 | ADA | chr20 | - | 44626455 | 44626599 | 44622828 | 44622930 | 44629046 | 44629169 | 97 | 34 | 47 | 65 | 43 | 0 | 0 | 0 | 0 | 0 | 47 | 65 | 41 | 61 |
| SE | 6051 | ENSG00000149639 | SOGA1 | chr20 | - | 36812703 | 36812844 | 36809981 | 36810107 | 36815125 | 36816281 | 16 | 4 | 9 | 20 | 2 | 0 | 0 | 0 | 2 | 0 | 48 | 11 | 10 | 6 |
| SE | 6054 | ENSG00000132792 | CTNNBL1 | chr20 | + | 37727301 | 37727402 | 37693954 | 37694152 | 37732878 | 37733067 | 1 | 0 | 0 | 0 | 0 | 26 | 47 | 46 | 72 | 39 | 0 | 0 | 0 | 2 |
| SE | 6055 | ENSG00000132792 | CTNNBL1 | chr20 | + | 37765196 | 37765290 | 37757558 | 37757656 | 37767952 | 37768044 | 189 | 166 | 185 | 206 | 235 | 0 | 0 | 0 | 2 | 1 | 391 | 218 | 153 | 253 |
| SE | 6057 | ENSG00000132792 | CTNNBL1 | chr20 | + | 37832247 | 37832345 | 37802866 | 37803048 | 37840101 | 37840199 | 2 | 0 | 0 | 0 | 0 | 107 | 92 | 81 | 115 | 98 | 0 | 0 | 0 | 0 |
| SE | 6058 | ENSG00000101310 | SEC23B | chr20 | + | 18508090 | 18508297 | 18507543 | 18507669 | 18510821 | 18510991 | 1 | 0 | 0 | 1 | 2 | 2 | 0 | 4 | 5 | 4 | 6 | 10 | 1 | 0 |
| SE | 6059 | ENSG00000101310 | SEC23B | chr20 | + | 18509615 | 18509943 | 18507939 | 18507990 | 18510821 | 18511014 | 0 | 0 | 0 | 0 | 0 | 44 | 22 | 25 | 36 | 14 | 0 | 0 | 1 | 0 |
| SE | 6060 | ENSG00000101310 | SEC23B | chr20 | + | 18524934 | 18525020 | 18524432 | 18524669 | 18525787 | 18525932 | 225 | 164 | 200 | 222 | 152 | 0 | 1 | 0 | 0 | 0 | 428 | 219 | 232 | 308 |
| SE | 6061 | ENSG00000101310 | SEC23B | chr20 | + | 18525787 | 18525932 | 18524934 | 18525020 | 18526372 | 18526531 | 306 | 225 | 257 | 292 | 211 | 0 | 1 | 1 | 0 | 0 | 529 | 269 | 339 | 384 |
| SE | 6062 | ENSG00000101310 | SEC23B | chr20 | + | 18527495 | 18527611 | 18526372 | 18526531 | 18530679 | 18530803 | 222 | 122 | 175 | 166 | 126 | 1 | 0 | 0 | 0 | 0 | 241 | 157 | 156 | 208 |
| SE | 6063 | ENSG00000101310 | SEC23B | chr20 | + | 18530679 | 18530803 | 18527495 | 18527611 | 18532663 | 18532744 | 306 | 177 | 233 | 249 | 279 | 0 | 0 | 0 | 0 | 0 | 474 | 249 | 246 | 240 |
| SE | 6064 | ENSG00000101310 | SEC23B | chr20 | + | 18532663 | 18532744 | 18530679 | 18530803 | 18535652 | 18535742 | 328 | 190 | 274 | 266 | 390 | 1 | 0 | 0 | 0 | 0 | 522 | 304 | 243 | 230 |
| SE | 6065 | ENSG00000101310 | SEC23B | chr20 | + | 18542295 | 18542402 | 18535652 | 18535742 | 18543018 | 18543172 | 262 | 168 | 221 | 235 | 265 | 1 | 1 | 0 | 0 | 1 | 460 | 340 | 245 | 237 |
| SE | 6068 | ENSG00000101310 | SEC23B | chr20 | + | 18543018 | 18543172 | 18542295 | 18542402 | 18545955 | 18546033 | 192 | 105 | 165 | 168 | 147 | 0 | 1 | 0 | 2 | 0 | 321 | 257 | 193 | 188 |
| SE | 6069 | ENSG00000101310 | SEC23B | chr20 | + | 18545955 | 18546033 | 18535652 | 18535742 | 18548608 | 18548770 | 109 | 50 | 74 | 72 | 74 | 0 | 0 | 2 | 0 | 0 | 79 | 89 | 57 | 60 |
| SE | 6070 | ENSG00000101310 | SEC23B | chr20 | + | 18551088 | 18551175 | 18548608 | 18548770 | 18554234 | 18554390 | 157 | 62 | 109 | 126 | 131 | 0 | 0 | 2 | 1 | 1 | 150 | 173 | 115 | 149 |
| SE | 6071 | ENSG00000124102 | PI3 | chr20 | + | 45175860 | 45176136 | 45174901 | 45175001 | 45176369 | 45176544 | 6 | 0 | 4 | 24 | 19 | 0 | 0 | 0 | 2 | 0 | 0 | 0 | 22 | 20 |
| SE | 6076 | ENSG00000089057 | SLC23A2 | chr20 | - | 4869905 | 4870053 | 4867769 | 4867875 | 4873935 | 4874092 | 31 | 16 | 13 | 27 | 38 | 1 | 0 | 0 | 0 | 0 | 23 | 8 | 18 | 33 |
| SE | 6078 | ENSG00000089057 | SLC23A2 | chr20 | - | 4884752 | 4884823 | 4874575 | 4874696 | 4899554 | 4899712 | 1 | 3 | 1 | 7 | 0 | 0 | 2 | 0 | 3 | 0 | 12 | 6 | 5 | 14 |
| SE | 6080 | ENSG00000089057 | SLC23A2 | chr20 | - | 4899554 | 4899712 | 4885820 | 4885909 | 4902441 | 4902558 | 76 | 44 | 46 | 70 | 58 | 0 | 0 | 0 | 1 | 0 | 80 | 73 | 51 | 57 |
| SE | 6081 | ENSG00000089057 | SLC23A2 | chr20 | - | 4932454 | 4932716 | 4912879 | 4912978 | 4970792 | 4970915 | 103 | 31 | 62 | 72 | 82 | 0 | 1 | 0 | 0 | 0 | 183 | 89 | 130 | 108 |
| SE | 6083 | ENSG00000105576 | TNPO2 | chr19 | - | 12702071 | 12702177 | 12701751 | 12701851 | 12702822 | 12702918 | 17 | 12 | 15 | 14 | 0 | 0 | 0 | 0 | 0 | 0 | 25 | 19 | 10 | 3 |
| SE | 6084 | ENSG00000105576 | TNPO2 | chr19 | - | 12703713 | 12703801 | 12703427 | 12703526 | 12705239 | 12705398 | 46 | 33 | 43 | 41 | 20 | 0 | 0 | 0 | 0 | 0 | 50 | 49 | 33 | 34 |
| SE | 6085 | ENSG00000105576 | TNPO2 | chr19 | - | 12705491 | 12705599 | 12705239 | 12705398 | 12705681 | 12705768 | 53 | 46 | 52 | 59 | 57 | 1 | 0 | 0 | 0 | 0 | 78 | 31 | 31 | 42 |
| SE | 6086 | ENSG00000105576 | TNPO2 | chr19 | - | 12705681 | 12705768 | 12705491 | 12705599 | 12706195 | 12706367 | 29 | 32 | 33 | 62 | 37 | 0 | 0 | 0 | 0 | 0 | 60 | 22 | 16 | 27 |
| SE | 6087 | ENSG00000105576 | TNPO2 | chr19 | - | 12705681 | 12705830 | 12705491 | 12705599 | 12706195 | 12706367 | 23 | 24 | 20 | 31 | 27 | 0 | 0 | 0 | 0 | 0 | 32 | 17 | 9 | 15 |
| SE | 6089 | ENSG00000105576 | TNPO2 | chr19 | - | 12711295 | 12711461 | 12710620 | 12710773 | 12711552 | 12711613 | 97 | 59 | 92 | 99 | 116 | 0 | 1 | 1 | 0 | 0 | 58 | 129 | 60 | 61 |
| SE | 6090 | ENSG00000105576 | TNPO2 | chr19 | - | 12711552 | 12711613 | 12710620 | 12710773 | 12714820 | 12714939 | 13 | 17 | 23 | 24 | 5 | 0 | 0 | 0 | 0 | 0 | 13 | 11 | 4 | 15 |
| SE | 6091 | ENSG00000171456 | ASXL1 | chr20 | + | 32360709 | 32360806 | 32360016 | 32360586 | 32366383 | 32366466 | 0 | 1 | 1 | 2 | 5 | 1 | 0 | 0 | 0 | 0 | 0 | 0 | 4 | 10 |
| SE | 6092 | ENSG00000171456 | ASXL1 | chr20 | + | 32360793 | 32360806 | 32358965 | 32359851 | 32366383 | 32366466 | 0 | 0 | 0 | 0 | 0 | 0 | 0 | 0 | 1 | 0 | 0 | 0 | 1 | 0 |
| SE | 6093 | ENSG00000171456 | ASXL1 | chr20 | + | 32360793 | 32360806 | 32360347 | 32360586 | 32366383 | 32366466 | 0 | 0 | 1 | 0 | 0 | 1 | 0 | 0 | 0 | 0 | 0 | 0 | 1 | 0 |
| SE | 6098 | ENSG00000171456 | ASXL1 | chr20 | + | 32431582 | 32431679 | 32428763 | 32431484 | 32432879 | 32432895 | 129 | 103 | 188 | 173 | 122 | 2 | 0 | 0 | 0 | 0 | 238 | 280 | 183 | 212 |
| SE | 6134 | ENSG00000131069 | ACSS2 | chr20 | + | 34914095 | 34914117 | 34913752 | 34913825 | 34914322 | 34914437 | 16 | 14 | 13 | 7 | 19 | 0 | 1 | 0 | 0 | 0 | 48 | 11 | 14 | 30 |
| SE | 6135 | ENSG00000131069 | ACSS2 | chr20 | + | 34914095 | 34914171 | 34913752 | 34913825 | 34914322 | 34914437 | 47 | 23 | 39 | 38 | 33 | 0 | 1 | 0 | 0 | 0 | 152 | 47 | 42 | 81 |
| SE | 6137 | ENSG00000131069 | ACSS2 | chr20 | + | 34915217 | 34915256 | 34914322 | 34914437 | 34919434 | 34919572 | 1 | 0 | 0 | 0 | 0 | 23 | 11 | 19 | 34 | 13 | 0 | 0 | 0 | 0 |
| SE | 6138 | ENSG00000131069 | ACSS2 | chr20 | + | 34921785 | 34921866 | 34921543 | 34921600 | 34923322 | 34923431 | 47 | 22 | 24 | 26 | 28 | 1 | 0 | 0 | 0 | 4 | 110 | 41 | 31 | 57 |
| SE | 6140 | ENSG00000131069 | ACSS2 | chr20 | + | 34926876 | 34926951 | 34926104 | 34926281 | 34927086 | 34927205 | 59 | 39 | 56 | 67 | 112 | 0 | 0 | 0 | 0 | 0 | 200 | 98 | 94 | 121 |
| SE | 6142 | ENSG00000282826 | FRG1CP | chr20 | - | 28583018 | 28583110 | 28581918 | 28582029 | 28586298 | 28586403 | 81 | 125 | 149 | 144 | 208 | 0 | 0 | 0 | 1 | 0 | 45 | 30 | 89 | 72 |
| SE | 6143 | ENSG00000282826 | FRG1CP | chr20 | - | 28583869 | 28583946 | 28581918 | 28582029 | 28586298 | 28586403 | 9 | 14 | 15 | 18 | 25 | 0 | 0 | 0 | 1 | 0 | 1 | 2 | 6 | 0 |
| SE | 6145 | ENSG00000282826 | FRG1CP | chr20 | - | 28588642 | 28588757 | 28586298 | 28586403 | 28590556 | 28590614 | 162 | 282 | 253 | 285 | 438 | 0 | 0 | 1 | 0 | 0 | 90 | 103 | 195 | 147 |
| SE | 6146 | ENSG00000282826 | FRG1CP | chr20 | - | 28588642 | 28588757 | 28586298 | 28586403 | 28600391 | 28600461 | 87 | 140 | 131 | 146 | 173 | 2 | 1 | 0 | 0 | 0 | 35 | 69 | 114 | 70 |
| SE | 6160 | ENSG00000286235 | NA | chr20 | + | 6015360 | 6015490 | 6009774 | 6009912 | 6031284 | 6031370 | 123 | 77 | 102 | 112 | 340 | 0 | 0 | 1 | 1 | 1 | 93 | 119 | 64 | 104 |
| SE | 6161 | ENSG00000286235 | NA | chr20 | + | 6015360 | 6015490 | 6009774 | 6009912 | 6032011 | 6032080 | 73 | 35 | 42 | 45 | 113 | 0 | 0 | 0 | 1 | 1 | 31 | 58 | 35 | 63 |
| SE | 6162 | ENSG00000286235 | NA | chr20 | + | 6031284 | 6031370 | 6009774 | 6009912 | 6032011 | 6032080 | 34 | 26 | 38 | 51 | 80 | 0 | 0 | 0 | 0 | 1 | 48 | 46 | 16 | 44 |
| SE | 6163 | ENSG00000088766 | CRLS1 | chr20 | + | 6031284 | 6031370 | 6015360 | 6015490 | 6032011 | 6032080 | 88 | 66 | 99 | 121 | 306 | 1 | 0 | 2 | 1 | 0 | 110 | 107 | 47 | 80 |
| SE | 6167 | ENSG00000132664 | POLR3F | chr20 | + | 18475074 | 18475187 | 18473390 | 18473458 | 18480401 | 18480509 | 25 | 18 | 19 | 13 | 33 | 0 | 0 | 0 | 0 | 0 | 5 | 13 | 41 | 18 |
| SE | 6169 | ENSG00000132664 | POLR3F | chr20 | + | 18481618 | 18481810 | 18480401 | 18480509 | 18483480 | 18484643 | 83 | 85 | 83 | 88 | 141 | 1 | 2 | 4 | 3 | 3 | 17 | 78 | 132 | 42 |
| SE | 6173 | ENSG00000088766 | CRLS1 | chr20 | + | 6023379 | 6023422 | 6015360 | 6015490 | 6026170 | 6026238 | 4 | 0 | 1 | 3 | 4 | 0 | 0 | 1 | 0 | 12 | 0 | 0 | 3 | 4 |
| SE | 6175 | ENSG00000088766 | CRLS1 | chr20 | + | 6026170 | 6026238 | 6015360 | 6015490 | 6028671 | 6028725 | 0 | 0 | 1 | 0 | 12 | 3 | 0 | 0 | 0 | 4 | 0 | 0 | 6 | 0 |
| SE | 6177 | ENSG00000088766 | CRLS1 | chr20 | + | 6026170 | 6026238 | 6023379 | 6023422 | 6031284 | 6031370 | 7 | 0 | 2 | 2 | 10 | 4 | 3 | 0 | 1 | 2 | 0 | 2 | 5 | 2 |
| SE | 6179 | ENSG00000088766 | CRLS1 | chr20 | + | 6028671 | 6028725 | 6015360 | 6015490 | 6032011 | 6032080 | 3 | 0 | 0 | 0 | 4 | 1 | 0 | 2 | 1 | 0 | 6 | 1 | 1 | 1 |
| SE | 6180 | ENSG00000088766 | CRLS1 | chr20 | + | 6028671 | 6028725 | 6026170 | 6026238 | 6031284 | 6031370 | 0 | 0 | 2 | 0 | 2 | 5 | 0 | 2 | 3 | 0 | 6 | 0 | 3 | 0 |
| SE | 6186 | ENSG00000125772 | GPCPD1 | chr20 | - | 5565016 | 5565078 | 5561464 | 5561530 | 5566732 | 5566772 | 264 | 283 | 280 | 288 | 477 | 0 | 1 | 0 | 0 | 0 | 29 | 235 | 340 | 168 |
| SE | 6188 | ENSG00000125772 | GPCPD1 | chr20 | - | 5567482 | 5567560 | 5566732 | 5566772 | 5570146 | 5570239 | 223 | 196 | 195 | 210 | 384 | 0 | 1 | 0 | 2 | 5 | 19 | 160 | 279 | 116 |
| SE | 6189 | ENSG00000125772 | GPCPD1 | chr20 | - | 5567796 | 5567842 | 5567482 | 5567560 | 5570146 | 5570239 | 0 | 0 | 3 | 0 | 4 | 149 | 125 | 135 | 121 | 245 | 0 | 3 | 5 | 0 |
| SE | 6190 | ENSG00000125772 | GPCPD1 | chr20 | - | 5570146 | 5570239 | 5567482 | 5567560 | 5573914 | 5573954 | 278 | 222 | 228 | 223 | 453 | 0 | 0 | 0 | 4 | 0 | 15 | 207 | 236 | 115 |
| SE | 6191 | ENSG00000125772 | GPCPD1 | chr20 | - | 5580007 | 5580131 | 5578379 | 5578611 | 5584280 | 5584322 | 588 | 603 | 493 | 500 | 889 | 2 | 1 | 0 | 0 | 1 | 261 | 567 | 792 | 347 |
| SE | 6193 | ENSG00000125772 | GPCPD1 | chr20 | - | 5593326 | 5593411 | 5584280 | 5584322 | 5598724 | 5598821 | 175 | 126 | 143 | 125 | 294 | 1 | 0 | 1 | 0 | 2 | 55 | 139 | 163 | 100 |
| SE | 6194 | ENSG00000125772 | GPCPD1 | chr20 | - | 5604363 | 5604440 | 5598724 | 5598821 | 5610841 | 5610924 | 121 | 97 | 130 | 103 | 216 | 0 | 1 | 0 | 1 | 0 | 62 | 158 | 128 | 128 |
| SE | 6196 | ENSG00000116017 | ARID3A | chr19 | + | 961761 | 961940 | 960091 | 960164 | 964247 | 964366 | 2 | 0 | 0 | 0 | 0 | 36 | 21 | 49 | 48 | 34 | 0 | 0 | 4 | 0 |
| SE | 6197 | ENSG00000116017 | ARID3A | chr19 | + | 964832 | 965080 | 964247 | 964431 | 966571 | 966868 | 57 | 48 | 76 | 101 | 46 | 1 | 0 | 0 | 0 | 0 | 170 | 106 | 70 | 180 |
| SE | 6198 | ENSG00000116017 | ARID3A | chr19 | + | 964832 | 965080 | 964247 | 964431 | 968404 | 968503 | 41 | 31 | 52 | 69 | 31 | 0 | 0 | 0 | 0 | 0 | 122 | 82 | 57 | 133 |
| SE | 6200 | ENSG00000116017 | ARID3A | chr19 | + | 966571 | 966868 | 964832 | 965080 | 968404 | 968503 | 37 | 30 | 44 | 78 | 40 | 0 | 0 | 0 | 2 | 0 | 98 | 73 | 35 | 87 |
| SE | 6204 | ENSG00000125844 | RRBP1 | chr20 | - | 17615925 | 17616009 | 17615430 | 17615529 | 17618595 | 17618679 | 42 | 33 | 23 | 27 | 23 | 0 | 0 | 0 | 0 | 0 | 51 | 34 | 60 | 85 |
| SE | 6206 | ENSG00000125844 | RRBP1 | chr20 | - | 17627347 | 17627382 | 17625511 | 17625602 | 17627503 | 17627682 | 43 | 18 | 35 | 21 | 19 | 1 | 0 | 0 | 0 | 0 | 27 | 75 | 51 | 40 |
| SE | 6207 | ENSG00000125844 | RRBP1 | chr20 | - | 17627503 | 17627682 | 17627347 | 17627382 | 17629822 | 17629961 | 60 | 23 | 52 | 46 | 19 | 0 | 0 | 1 | 0 | 0 | 35 | 96 | 54 | 66 |
| SE | 6208 | ENSG00000125844 | RRBP1 | chr20 | - | 17636576 | 17636729 | 17635545 | 17635664 | 17641796 | 17641919 | 201 | 69 | 99 | 157 | 75 | 3 | 0 | 0 | 0 | 0 | 237 | 217 | 191 | 168 |
| SE | 6209 | ENSG00000125844 | RRBP1 | chr20 | - | 17658595 | 17660528 | 17642978 | 17643127 | 17679998 | 17680075 | 103 | 55 | 58 | 77 | 72 | 0 | 0 | 0 | 0 | 0 | 244 | 129 | 155 | 188 |
| SE | 6211 | ENSG00000088876 | ZNF343 | chr20 | - | 2486640 | 2486763 | 2481822 | 2484656 | 2492698 | 2492825 | 3 | 6 | 3 | 8 | 19 | 16 | 19 | 12 | 16 | 12 | 24 | 3 | 0 | 0 |
| SE | 6214 | ENSG00000088876 | ZNF343 | chr20 | - | 2495414 | 2495524 | 2493777 | 2494044 | 2500655 | 2500742 | 0 | 2 | 0 | 0 | 0 | 30 | 20 | 22 | 19 | 10 | 0 | 0 | 0 | 0 |

| | | | | | | | | | | | | | | | | | | | | | | | | | | | | | | | | | | | | | | | | | | | | | | |
|---|---|---|---|---|---|---|---|---|---|---|---|---|---|---|---|---|---|---|---|---|---|---|---|---|---|---|---|---|---|---|---|---|---|---|---|---|---|---|---|---|---|---|---|---|---|---|
| SE | 6497 | ENSG00000176014 | TUBB6 | chr18 | + | 12308686 | 12308795 | 12308231 | 12308349 | 12310942 | 12311053 | 81 | 13 | 41 | 36 | 31 | 0 | 0 | 0 | 0 | 0 | 197 | 65 | 27 | 32 | 99 | 0 | 3 | 0 | 0 | 0 | 258 | 149 | 1 | 1 | 1 | 1 | 1 | 1 | 1 | 1 | 0.926 | 1 | 1 | 1 | 0.015 |
| SE | 6500 | ENSG00000176014 | TUBB6 | chr18 | + | 12320628 | 12320744 | 12310942 | 12311053 | 12325066 | 12325197 | 0 | 0 | 0 | 1 | 0 | 28 | 1 | 11 | 24 | 6 | 0 | 0 | 0 | 0 | 0 | 70 | 22 | 13 | 24 | 12 | 265 | 149 | 1 | 1 | 0 | 0 | 0 | 0.023 | 0 | 0 | 0 | 0 | 0 | 0 | 0.005 |
| SE | 6502 | ENSG00000105429 | MEGF8 | chr19 | + | 42357403 | 42357584 | 42356773 | 42356981 | 42358143 | 42358307 | 8 | 0 | 2 | 3 | 0 | 0 | 0 | 0 | 2 | 0 | 0 | 18 | 0 | 1 | 7 | 0 | 0 | 0 | 0 | 0 | 298 | 149 | 1 | 1 | 1 | NA | 1 | 0.429 | NA | NA | 1 | NA | 1 | 1 | -0.19 |
| SE | 6503 | ENSG00000105429 | MEGF8 | chr19 | + | 42358786 | 42358954 | 42358143 | 42358307 | 42359097 | 42359242 | 4 | 0 | 9 | 0 | 0 | 0 | 0 | 0 | 0 | 0 | 0 | 2 | 2 | 2 | 0 | 11 | 0 | 0 | 0 | 11 | 298 | 149 | 1 | 1 | 1 | NA | 1 | NA | NA | 0 | 1 | | 1 | 0 | 0.4 |
| SE | 6505 | ENSG00000167088 | SNRPD1 | chr18 | + | 21622724 | 21622801 | 21612313 | 21612443 | 21623747 | 21623939 | 488 | 401 | 494 | 489 | 650 | 3 | 0 | 3 | 0 | 0 | 1014 | 435 | 612 | 700 | 414 | 10 | 0 | 1 | 1 | 0 | 226 | 149 | 1 | 1 | 0.991 | 1 | 0.991 | 1 | 1 | 0.985 | 1 | 0.998 | 0.998 | 1 | 0 |
| SE | 6506 | ENSG00000167088 | SNRPD1 | chr18 | + | 21623747 | 21623939 | 21622724 | 21622801 | 21629061 | 21629479 | 794 | 696 | 747 | 812 | 1155 | 0 | 0 | 0 | 0 | 0 | 1833 | 646 | 1051 | 1236 | 702 | 0 | 0 | 1 | 0 | 0 | 298 | 149 | 1 | 1 | 1 | 1 | 1 | 1 | 1 | 1 | 1 | 0.998 | 1 | 1 | 0 |
| SE | 6507 | ENSG00000167088 | SNRPD1 | chr18 | + | 21623874 | 21623939 | 21622724 | 21622801 | 21629061 | 21629109 | 0 | 5 | 11 | 1 | 12 | 0 | 0 | 0 | 0 | 0 | 0 | 0 | 4 | 6 | 0 | 0 | 0 | 1 | 0 | 0 | 214 | 149 | 1 | 1 | NA | 1 | 1 | 1 | 1 | NA | NA | 0.736 | 1 | NA | 0.132 |
| SE | 6515 | ENSG00000124155 | PIGT | chr20 | + | 45416516 | 45416694 | 45416096 | 45416343 | 45419503 | 45419590 | 91 | 29 | 48 | 72 | 62 | 0 | 0 | 0 | 0 | 0 | 88 | 92 | 73 | 74 | 144 | 0 | 0 | 3 | 0 | 0 | 298 | 149 | 1 | 1 | 1 | 1 | 1 | 1 | 1 | 1 | 1 | 0.924 | 1 | 1 | 0.015 |
| SE | 6530 | ENSG00000124155 | PIGT | chr20 | + | 45419294 | 45419395 | 45416083 | 45416343 | 45419503 | 45419590 | 92 | 34 | 78 | 60 | 71 | 0 | 0 | 0 | 0 | 0 | 68 | 147 | 91 | 86 | 145 | 0 | 0 | 3 | 0 | 0 | 250 | 149 | 1 | 1 | 1 | 1 | 1 | 1 | 1 | 1 | 1 | 0.948 | 1 | 1 | 0.01 |
| SE | 6538 | ENSG00000124155 | PIGT | chr20 | + | 45420135 | 45420223 | 45419472 | 45419590 | 45420331 | 45420424 | 363 | 139 | 228 | 216 | 236 | 0 | 0 | 0 | 0 | 0 | 421 | 438 | 251 | 246 | 495 | 0 | 0 | 3 | 0 | 0 | 237 | 149 | 1 | 1 | 1 | 1 | 1 | 1 | 1 | 1 | 1 | 0.981 | 1 | 1 | 0.004 |
| SE | 6539 | ENSG00000124155 | PIGT | chr20 | + | 45420152 | 45420223 | 45419503 | 45419590 | 45420331 | 45420429 | 172 | 62 | 107 | 101 | 132 | 0 | 0 | 0 | 0 | 0 | 228 | 197 | 112 | 114 | 242 | 0 | 0 | 3 | 0 | 0 | 220 | 149 | 1 | 1 | 1 | 1 | 1 | 1 | 1 | 1 | 1 | 0.962 | 1 | 1 | 0.008 |
| SE | 6540 | ENSG00000124155 | PIGT | chr20 | + | 45421382 | 45421533 | 45420527 | 45420693 | 45424215 | 45424381 | 35 | 6 | 28 | 26 | 23 | 0 | 0 | 0 | 0 | 0 | 47 | 48 | 22 | 39 | 85 | 0 | 2 | 0 | 0 | 0 | 298 | 149 | 1 | 1 | 1 | 1 | 1 | 1 | 1 | 1 | 0.923 | 1 | 1 | 1 | 0.015 |
| SE | 6541 | ENSG00000124155 | PIGT | chr20 | + | 45421382 | 45421583 | 45420507 | 45420693 | 45424215 | 45424262 | 74 | 25 | 63 | 60 | 46 | 0 | 0 | 0 | 0 | 0 | 76 | 92 | 69 | 70 | 164 | 0 | 2 | 0 | 0 | 0 | 298 | 149 | 1 | 1 | 1 | 1 | 1 | 1 | 1 | 1 | 0.958 | 1 | 1 | 1 | 0.008 |
| SE | 6542 | ENSG00000124155 | PIGT | chr20 | + | 45421382 | 45421664 | 45420527 | 45420693 | 45424215 | 45424381 | 35 | 6 | 28 | 25 | 23 | 0 | 0 | 0 | 0 | 0 | 47 | 48 | 22 | 39 | 85 | 0 | 2 | 0 | 0 | 0 | 298 | 149 | 1 | 1 | 1 | 1 | 1 | 1 | 1 | 1 | 0.923 | 1 | 1 | 1 | 0.015 |
| SE | 6545 | ENSG00000124155 | PIGT | chr20 | + | 45424495 | 45424579 | 45423103 | 45424381 | 45427421 | 45427545 | 43 | 5 | 28 | 41 | 19 | 1 | 0 | 0 | 0 | 0 | 17 | 51 | 66 | 41 | 90 | 0 | 0 | 0 | 0 | 0 | 233 | 149 | 1 | 1 | 0.965 | 1 | 1 | 1 | 1 | 1 | 1 | 1 | 1 | 1 | -0.007 |
| SE | 6554 | ENSG00000134265 | NAPG | chr18 | + | 10534465 | 10534496 | 10533535 | 10533553 | 10539761 | 10539871 | 138 | 61 | 70 | 82 | 46 | 1 | 0 | 0 | 0 | 0 | 37 | 54 | 57 | 62 | 20 | 0 | 0 | 0 | 0 | 0 | 180 | 149 | 1 | 1 | 0.991 | 1 | 1 | 1 | 1 | 1 | 1 | 1 | 1 | 1 | -0.002 |
| SE | 6558 | ENSG00000074657 | ZNF532 | chr18 | + | 58865303 | 58865385 | 58862599 | 58863166 | 58865469 | 58865579 | 7 | 0 | 4 | 11 | 8 | 0 | 0 | 0 | 0 | 0 | 0 | 9 | 5 | 1 | 0 | 0 | 2 | 1 | 0 | 0 | 231 | 149 | 1 | 1 | 1 | NA | 1 | 1 | 1 | NA | 0.744 | 0.763 | 1 | NA | 0.164 |
| SE | 6559 | ENSG00000074657 | ZNF532 | chr18 | + | 58865303 | 58865385 | 58862599 | 58863166 | 58918270 | 58918661 | 0 | 0 | 0 | 2 | 0 | 0 | 0 | 0 | 0 | 1 | 0 | 0 | 0 | 0 | 0 | 0 | 0 | 0 | 3 | 0 | 231 | 149 | 1 | 1 | NA | NA | NA | 1 | 0 | NA | NA | NA | 0 | NA | 0.5 |
| SE | 6560 | ENSG00000074657 | ZNF532 | chr18 | + | 58865303 | 58865385 | 58863561 | 58863686 | 58865469 | 58865579 | 7 | 0 | 4 | 10 | 8 | 0 | 0 | 0 | 0 | 0 | 0 | 11 | 5 | 1 | 0 | 0 | 0 | 0 | 1 | 0 | 231 | 149 | 1 | 1 | 1 | NA | 1 | 1 | 1 | NA | 1 | 1 | 0.392 | NA | 0.203 |
| SE | 6561 | ENSG00000074657 | ZNF532 | chr18 | + | 58865303 | 58865385 | 58863603 | 58863750 | 58865469 | 58865579 | 8 | 0 | 4 | 10 | 9 | 0 | 1 | 0 | 1 | 0 | 0 | 14 | 5 | 1 | 0 | 4 | 0 | 0 | 2 | 0 | 231 | 149 | 1 | 1 | 1 | 0 | 1 | 0.866 | 1 | 0 | 1 | 1 | 0.244 | NA | 0.212 |
| SE | 6564 | ENSG00000074657 | ZNF532 | chr18 | + | 58865303 | 58865385 | 58864876 | 58864986 | 58865469 | 58865579 | 7 | 0 | 4 | 10 | 8 | 0 | 1 | 0 | 1 | 0 | 0 | 9 | 5 | 1 | 0 | 0 | 0 | 0 | 4 | 0 | 231 | 149 | 1 | 1 | 1 | 0 | 1 | 0.866 | 1 | NA | 1 | 1 | 0.139 | NA | 0.06 |
| SE | 6565 | ENSG00000074657 | ZNF532 | chr18 | + | 58865303 | 58865385 | 58864876 | 58864986 | 58918270 | 58918661 | 0 | 0 | 0 | 1 | 0 | 0 | 0 | 0 | 0 | 0 | 0 | 0 | 0 | 0 | 0 | 0 | 0 | 1 | 0 | 0 | 231 | 149 | 1 | 1 | NA | NA | NA | 1 | NA | NA | NA | 0 | NA | NA | 1 |
| SE | 6566 | ENSG00000074657 | ZNF532 | chr18 | + | 58865469 | 58865579 | 58862947 | 58863166 | 58918270 | 58918492 | 21 | 5 | 21 | 28 | 30 | 0 | 0 | 0 | 0 | 1 | 6 | 25 | 20 | 31 | 14 | 0 | 0 | 0 | 3 | 0 | 259 | 149 | 1 | 1 | 1 | 1 | 1 | 1 | 0.945 | 1 | 1 | 1 | 0.856 | 1 | 0.018 |
| SE | 6567 | ENSG00000074657 | ZNF532 | chr18 | + | 58865469 | 58865579 | 58864092 | 58864235 | 58918270 | 58918492 | 21 | 5 | 21 | 28 | 30 | 0 | 2 | 0 | 0 | 0 | 6 | 23 | 19 | 31 | 14 | 0 | 0 | 0 | 0 | 0 | 259 | 149 | 1 | 1 | 1 | 0.59 | 1 | 1 | 1 | 1 | 1 | 1 | 1 | 1 | -0.082 |
| SE | 6568 | ENSG00000074657 | ZNF532 | chr18 | + | 58865469 | 58865579 | 58864820 | 58864986 | 58918270 | 58918492 | 21 | 6 | 21 | 29 | 30 | 0 | 0 | 0 | 0 | 0 | 6 | 23 | 19 | 35 | 14 | 0 | 0 | 1 | 0 | 0 | 259 | 149 | 1 | 1 | 1 | 1 | 1 | 1 | 1 | 1 | 1 | 0.916 | 1 | 1 | 0.017 |
| SE | 6569 | ENSG00000074657 | ZNF532 | chr18 | + | 58865469 | 58865579 | 58864876 | 58865385 | 58918270 | 58918492 | 28 | 5 | 26 | 40 | 52 | 0 | 0 | 0 | 1 | 0 | 6 | 35 | 26 | 35 | 14 | 0 | 0 | 0 | 0 | 0 | 259 | 149 | 1 | 1 | 1 | 1 | 1 | 0.958 | 1 | 1 | 1 | 1 | 1 | 1 | -0.008 |
| SE | 6570 | ENSG00000074657 | ZNF532 | chr18 | + | 58896424 | 58896622 | 58865553 | 58865579 | 58918270 | 58918608 | 0 | 0 | 1 | 3 | 1 | 22 | 5 | 22 | 34 | 37 | 0 | 0 | 4 | 0 | 0 | 8 | 30 | 20 | 31 | 21 | 298 | 149 | 1 | 1 | 0 | 0 | 0.022 | 0.042 | 0.013 | 0 | 0 | 0.091 | 0 | 0 | -0.003 |
| SE | 6571 | ENSG00000074657 | ZNF532 | chr18 | + | 58918270 | 58920633 | 58865469 | 58865579 | 58939444 | 58939621 | 23 | 5 | 30 | 34 | 38 | 0 | 0 | 1 | 0 | 0 | 14 | 30 | 26 | 33 | 21 | 0 | 0 | 0 | 0 | 0 | 298 | 149 | 1 | 1 | 1 | 1 | 0.938 | 1 | 1 | 1 | 1 | 1 | 1 | 1 | -0.012 |
| SE | 6577 | ENSG00000074657 | ZNF532 | chr18 | + | 58939444 | 58939621 | 58918270 | 58920633 | 58953517 | 58953799 | 4 | 3 | 12 | 2 | 5 | 2 | 1 | 2 | 1 | 0 | 18 | 8 | 11 | 12 | 0 | 0 | 0 | 0 | 5 | 0 | 298 | 149 | 1 | 1 | 0.5 | 0.6 | 0.75 | 0.5 | 1 | 1 | 1 | 1 | 0.545 | NA | -0.216 |
| SE | 6578 | ENSG00000074657 | ZNF532 | chr18 | + | 58939444 | 58939621 | 58934231 | 58934614 | 58948066 | 58948081 | 42 | 29 | 32 | 53 | 63 | 4 | 0 | 0 | 1 | 0 | 69 | 34 | 27 | 99 | 15 | 0 | 0 | 0 | 0 | 0 | 298 | 149 | 1 | 1 | 0.84 | 1 | 1 | 0.964 | 1 | 1 | 1 | 1 | 1 | 1 | -0.039 |
| SE | 6579 | ENSG00000074657 | ZNF532 | chr18 | + | 58939444 | 58939621 | 58934231 | 58934614 | 58953517 | 58953799 | 20 | 10 | 19 | 24 | 10 | 0 | 0 | 0 | 2 | 0 | 18 | 17 | 13 | 37 | 10 | 0 | 2 | 0 | 0 | 0 | 298 | 149 | 1 | 1 | 1 | 1 | 1 | 0.857 | 1 | 1 | 0.81 | 1 | 1 | 1 | 0.009 |
| SE | 6581 | ENSG00000074657 | ZNF532 | chr18 | + | 58948066 | 58948229 | 58934432 | 58934614 | 58953517 | 58953799 | 64 | 39 | 33 | 57 | 50 | 0 | 0 | 0 | 2 | 0 | 108 | 51 | 54 | 87 | 15 | 0 | 2 | 0 | 0 | 0 | 298 | 149 | 1 | 1 | 1 | 1 | 1 | 0.934 | 1 | 1 | 0.927 | 1 | 1 | 1 | 0.001 |
| SE | 6585 | ENSG00000074657 | ZNF532 | chr18 | + | 58981469 | 58981617 | 58979054 | 58979167 | 58983971 | 58984094 | 32 | 33 | 22 | 40 | 20 | 0 | 0 | 0 | 0 | 0 | 23 | 39 | 19 | 20 | 13 | 0 | 3 | 0 | 0 | 0 | 297 | 149 | 1 | 1 | 1 | 1 | 1 | 1 | 1 | 1 | 0.867 | 1 | 1 | 1 | 0.027 |
| [illegible] | [illegible] | [illegible] | [illegible] | [illegible] | [illegible] | [illegible] | [illegible] | [illegible] | [illegible] | [illegible] | [illegible] | [illegible] | [illegible] | [illegible] | [illegible] | [illegible] | [illegible] | [illegible] | [illegible] | [illegible] | [illegible] | [illegible] | [illegible] | [illegible] | [illegible] | [illegible] | [illegible] | [illegible] | [illegible] | [illegible] | [illegible] | [illegible] | [illegible] | [illegible] | [illegible] | [illegible] | [illegible] | [illegible] | [illegible] | [illegible] | [illegible] | [illegible] | [illegible] | [illegible] | [illegible] | [illegible] |

| | | | | | | | | | | | | | | |
|---|---|---|---|---|---|---|---|---|---|---|---|---|---|---|
| SE | 7094 | ENSG00000134046 | MBD2 | chr18 | - | 54166075 | 54166166 | 54164522 | 54164700 | 54188873 | 54189011 | 991 | 885 | 771 |
| SE | 7097 | ENSG00000134046 | MBD2 | chr18 | - | 54188873 | 54189011 | 54166075 | 54166166 | 54204997 | 54205021 | 695 | 585 | 579 |
| SE | 7098 | ENSG00000134046 | MBD2 | chr18 | - | 54188873 | 54189011 | 54166075 | 54166166 | 54224017 | 54224669 | 262 | 223 | 195 |
| SE | 7099 | ENSG00000134046 | MBD2 | chr18 | - | 54204997 | 54205157 | 54166075 | 54166166 | 54224017 | 54224617 | 489 | 378 | 434 |
| SE | 7100 | ENSG00000134046 | MBD2 | chr18 | - | 54204997 | 54205157 | 54188873 | 54189011 | 54224017 | 54224617 | 923 | 741 | 820 |
| SE | 7107 | ENSG00000105426 | PTPRS | chr19 | - | 5216719 | 5216767 | 5215497 | 5215595 | 5218419 | 5218532 | 7 | 7 | 1 |
| SE | 7110 | ENSG00000105426 | PTPRS | chr19 | - | 5229490 | 5229684 | 5225726 | 5225844 | 5231309 | 5231615 | 4 | 0 | 1 |
| SE | 7112 | ENSG00000105426 | PTPRS | chr19 | - | 5273441 | 5273583 | 5265007 | 5265196 | 5274198 | 5274344 | 39 | 44 | 56 |
| SE | 7121 | ENSG00000168234 | TTC39C | chr18 | + | 24066011 | 24066140 | 24014753 | 24015038 | 24080584 | 24080779 | 16 | 12 | 7 |
| SE | 7124 | ENSG00000168234 | TTC39C | chr18 | + | 24114553 | 24114647 | 24082912 | 24083081 | 24123833 | 24123937 | 49 | 46 | 66 |
| SE | 7126 | ENSG00000168234 | TTC39C | chr18 | + | 24123833 | 24123943 | 24118124 | 24118232 | 24128885 | 24128983 | 71 | 46 | 90 |
| SE | 7127 | ENSG00000168234 | TTC39C | chr18 | + | 24125426 | 24125550 | 24123595 | 24123943 | 24128885 | 24128983 | 199 | 95 | 203 |
| SE | 7128 | ENSG00000168234 | TTC39C | chr18 | + | 24128885 | 24128983 | 24125426 | 24125550 | 24130312 | 24130417 | 257 | 137 | 243 |
| SE | 7129 | ENSG00000168234 | TTC39C | chr18 | + | 24131881 | 24131920 | 24130312 | 24130417 | 24132484 | 24132509 | 251 | 189 | 262 |
| SE | 7133 | ENSG00000101773 | RBBP8 | chr18 | + | 22968805 | 22968918 | 22949617 | 22949713 | 22975152 | 22975219 | 71 | 72 | 33 |
| SE | 7134 | ENSG00000101773 | RBBP8 | chr18 | + | 22975152 | 22975219 | 22949617 | 22949713 | 22982217 | 22982393 | 26 | 29 | 15 |
| SE | 7135 | ENSG00000101773 | RBBP8 | chr18 | + | 22977789 | 22977887 | 22975180 | 22975219 | 22982217 | 22982393 | 6 | 2 | 1 |
| SE | 7136 | ENSG00000101773 | RBBP8 | chr18 | + | 22993720 | 22993847 | 22992747 | 22993639 | 22996373 | 22996462 | 150 | 130 | 123 |
| SE | 7137 | ENSG00000101773 | RBBP8 | chr18 | + | 23006362 | 23006432 | 23001570 | 23001729 | 23016827 | 23016924 | 26 | 38 | 26 |
| SE | 7138 | ENSG00000101773 | RBBP8 | chr18 | + | 23016827 | 23016924 | 23006362 | 23006432 | 23022128 | 23022270 | 29 | 33 | 28 |
| SE | 7139 | ENSG00000101773 | RBBP8 | chr18 | + | 23022128 | 23022270 | 23016795 | 23016924 | 23026142 | 23026370 | 37 | 30 | 29 |
| SE | 7140 | ENSG00000141425 | RPRD1A | chr18 | - | 35988731 | 35988835 | 35985337 | 35985386 | 35990525 | 35990595 | 0 | 0 | 0 |
| SE | 7142 | ENSG00000141425 | RPRD1A | chr18 | - | 36040806 | 36040865 | 36033707 | 36033837 | 36062977 | 36063075 | 15 | 8 | 14 |
| SE | 7144 | ENSG00000141425 | RPRD1A | chr18 | - | 36062977 | 36063075 | 36033751 | 36033837 | 36067253 | 36067576 | 2 | 0 | 1 |
| SE | 7146 | ENSG00000101639 | CEP192 | chr18 | + | 12999420 | 12999588 | 12991361 | 12991437 | 13001456 | 13001582 | 217 | 109 | 171 |
| SE | 7147 | ENSG00000101639 | CEP192 | chr18 | + | 13000311 | 13000486 | 12999420 | 12999588 | 13001456 | 13001582 | 3 | 0 | 0 |
| SE | 7148 | ENSG00000101639 | CEP192 | chr18 | + | 13001456 | 13001582 | 12999420 | 12999588 | 13008455 | 13008631 | 363 | 170 | 244 |
| SE | 7149 | ENSG00000101639 | CEP192 | chr18 | + | 13012972 | 13013025 | 13008455 | 13008631 | 13015327 | 13015448 | 259 | 126 | 232 |
| SE | 7150 | ENSG00000101639 | CEP192 | chr18 | + | 13015327 | 13015448 | 13008455 | 13008631 | 13017187 | 13017336 | 269 | 166 | 166 |
| SE | 7153 | ENSG00000101639 | CEP192 | chr18 | + | 13037236 | 13037301 | 13030464 | 13030608 | 13038514 | 13038579 | 57 | 24 | 46 |
| SE | 7157 | ENSG00000101639 | CEP192 | chr18 | + | 13038369 | 13038579 | 13030464 | 13030608 | 13042203 | 13042334 | 2 | 2 | 2 |

[illegible]

| | | | | | | | | | | | | | | |
|---|---|---|---|---|---|---|---|---|---|---|---|---|---|---|
| SE | 7643 | ENSG00000270647 | TAF15 | chr17 | + | 35844476 | 35845038 | 35844279 | 35844368 | 35846905 | 35847234 | 405 | 334 | 405 |
| SE | 7649 | ENSG00000184792 | OSBP2 | chr22 | + | 30893816 | 30894001 | 30893637 | 30893733 | 30905836 | 30906069 | 68 | 12 | 6 |
| SE | 7650 | ENSG00000170296 | GABARAP | chr17 | - | 7241341 | 7241460 | 7240007 | 7240919 | 7242240 | 7242358 | 1058 | 772 | 825 |
| SE | 7653 | ENSG00000004975 | DVL2 | chr17 | - | 7230727 | 7230797 | 7230284 | 7230430 | 7234068 | 7234223 | 55 | 30 | 49 |
| SE | 7658 | ENSG00000161944 | ASGR2 | chr17 | - | 7114614 | 7114811 | 7114116 | 7114310 | 7114946 | 7115015 | 25 | 6 | 6 |
| SE | 7659 | ENSG00000161944 | ASGR2 | chr17 | - | 7114614 | 7114811 | 7114116 | 7114310 | 7115053 | 7115522 | 17 | 4 | 2 |
| SE | 7661 | ENSG00000132514 | CLEC10A | chr17 | - | 7075975 | 7076071 | 7075730 | 7075885 | 7076732 | 7076804 | 11 | 33 | 11 |
| SE | 7662 | ENSG00000132514 | CLEC10A | chr17 | - | 7075984 | 7076071 | 7074794 | 7075885 | 7076732 | 7076804 | 34 | 46 | 30 |
| SE | 7663 | ENSG00000132514 | CLEC10A | chr17 | - | 7075984 | 7076152 | 7075730 | 7075885 | 7076732 | 7076804 | 28 | 52 | 24 |
| SE | 7664 | ENSG00000070413 | DGCR2 | chr22 | - | 19041057 | 19041294 | 19036281 | 19039121 | 19041806 | 19041959 | 74 | 27 | 57 |
| SE | 7665 | ENSG00000070413 | DGCR2 | chr22 | - | 19041057 | 19041959 | 19037836 | 19039121 | 19048439 | 19048643 | 115 | 46 | 123 |
| SE | 7666 | ENSG00000070413 | DGCR2 | chr22 | - | 19041806 | 19041959 | 19036281 | 19039121 | 19048439 | 19048643 | 90 | 39 | 108 |
| SE | 7667 | ENSG00000070413 | DGCR2 | chr22 | - | 19048439 | 19048643 | 19041057 | 19041959 | 19056985 | 19057053 | 169 | 70 | 166 |
| SE | 7668 | ENSG00000070413 | DGCR2 | chr22 | - | 19056985 | 19057162 | 19048378 | 19048643 | 19063201 | 19063277 | 194 | 82 | 130 |
| SE | 7669 | ENSG00000070413 | DGCR2 | chr22 | - | 19060565 | 19060955 | 19056985 | 19057162 | 19063201 | 19063278 | 0 | 0 | 6 |
| SE | 7670 | ENSG00000070413 | DGCR2 | chr22 | - | 19063201 | 19063278 | 19056985 | 19057162 | 19064847 | 19065058 | 168 | 70 | 108 |
| SE | 7679 | ENSG00000108839 | ALOX12 | chr17 | + | 7006485 | 7006607 | 7005857 | 7006027 | 7009746 | 7009847 | 29 | 57 | 9 |
| SE | 7681 | ENSG00000168517 | HEXIM2 | chr17 | + | 45162487 | 45162635 | 45161874 | 45162031 | 45162749 | 45162859 | 12 | 0 | 9 |
| SE | 7684 | ENSG00000170004 | CHD3 | chr17 | + | 7893804 | 7893935 | 7893285 | 7893569 | 7894126 | 7894265 | 183 | 137 | 278 |
| SE | 7687 | ENSG00000170004 | CHD3 | chr17 | + | 7897970 | 7898102 | 7897082 | 7897294 | 7898495 | 7898595 | 415 | 358 | 656 |
| SE | 7688 | ENSG00000170004 | CHD3 | chr17 | + | 7898495 | 7898595 | 7897970 | 7898102 | 7899010 | 7899202 | 308 | 236 | 509 |
| SE | 7689 | ENSG00000170004 | CHD3 | chr17 | + | 7899895 | 7900033 | 7899342 | 7899543 | 7900289 | 7900411 | 544 | 449 | 767 |
| SE | 7690 | ENSG00000170004 | CHD3 | chr17 | + | 7899901 | 7900033 | 7899342 | 7899543 | 7900289 | 7900411 | 249 | 203 | 311 |
| SE | 7691 | ENSG00000170004 | CHD3 | chr17 | + | 7900289 | 7900411 | 7899342 | 7899543 | 7900557 | 7900731 | 202 | 127 | 255 |
| SE | 7692 | ENSG00000170004 | CHD3 | chr17 | + | 7901243 | 7901375 | 7900851 | 7900993 | 7902609 | 7902727 | 314 | 224 | 509 |
| SE | 7695 | ENSG00000170004 | CHD3 | chr17 | + | 7906868 | 7907031 | 7906552 | 7906697 | 7907125 | 7907247 | 547 | 400 | 663 |
| SE | 7696 | ENSG00000170004 | CHD3 | chr17 | + | 7907352 | 7907488 | 7907125 | 7907247 | 7907893 | 7908019 | 186 | 173 | 251 |
| SE | 7699 | ENSG00000170004 | CHD3 | chr17 | + | 7907890 | 7908019 | 7907352 | 7907488 | 7909142 | 7909338 | 0 | 1 | 3 |
| SE | 7700 | ENSG00000170004 | CHD3 | chr17 | + | 7907893 | 7908019 | 7907352 | 7907488 | 7909142 | 7909338 | 50 | 49 | 80 |
| SE | 7701 | ENSG00000170004 | CHD3 | chr17 | + | 7910427 | 7910591 | 7909142 | 7909338 | 7910846 | 7910973 | 130 | 95 | 135 |
| SE | 7702 | ENSG00000170004 | CHD3 | chr17 | + | 7910438 | 7910591 | 7909142 | 7909338 | 7910846 | 7910895 | 52 | 38 | 59 |
| SE | 7703 | ENSG00000170004 | CHD3 | chr17 | + | 7910846 | 7910973 | 7909500 | 7910591 | 7911463 | 7911750 | 139 | 103 | 148 |
| SE | 7710 | ENSG00000136492 | BRIP1 | chr17 | - | 61685835 | 61686165 | 61679138 | 61684140 | 61693429 | 61693512 | 4 | 6 | 7 |
| SE | 7711 | ENSG00000136492 | BRIP1 | chr17 | - | 61687112 | 61687237 | 61685835 | 61686165 | 61693429 | 61693512 | 0 | 1 | 0 |
| SE | 7714 | ENSG00000136492 | BRIP1 | chr17 | - | 61778119 | 61778179 | 61776400 | 61776502 | 61778672 | 61778778 | 0 | 0 | 0 |
| SE | 7716 | ENSG00000136492 | BRIP1 | chr17 | - | 61778119 | 61778179 | 61776400 | 61776562 | 61778672 | 61778778 | 1 | 0 | 1 |
| SE | 7718 | ENSG00000136492 | BRIP1 | chr17 | - | 61778672 | 61778778 | 61776400 | 61776502 | 61780260 | 61780401 | 2 | 1 | 1 |
| SE | 7720 | ENSG00000136492 | BRIP1 | chr17 | - | 61778672 | 61778778 | 61778119 | 61778179 | 61780260 | 61780401 | 2 | 0 | 1 |
| SE | 7721 | ENSG00000136492 | BRIP1 | chr17 | - | 61780260 | 61780401 | 61776400 | 61776562 | 61780839 | 61781005 | 12 | 15 | 10 |
| SE | 7722 | ENSG00000136492 | BRIP1 | chr17 | - | 61783818 | 61783907 | 61780839 | 61781005 | 61784269 | 61784424 | 0 | 1 | 0 |
| SE | 7723 | ENSG00000136492 | BRIP1 | chr17 | - | 61795731 | 61796540 | 61793596 | 61793729 | 61799099 | 61799299 | 0 | 0 | 0 |
| SE | 7728 | ENSG00000136492 | BRIP1 | chr17 | - | 61861446 | 61861565 | 61859795 | 61859907 | 61863283 | 61863500 | 9 | 14 | 16 |
| SE | 7729 | ENSG00000136492 | BRIP1 | chr17 | - | 61861446 | 61861569 | 61859795 | 61859907 | 61861909 | 61862075 | 9 | 14 | 16 |
| SE | 7730 | ENSG00000136492 | BRIP1 | chr17 | - | 61861446 | 61861569 | 61859795 | 61859907 | 61863283 | 61863326 | 11 | 16 | 25 |
| SE | 7731 | ENSG00000136492 | BRIP1 | chr17 | - | 61861909 | 61862075 | 61859795 | 61859907 | 61863283 | 61863409 | 0 | 0 | 0 |
| SE | 7735 | ENSG00000108389 | MTMR4 | chr17 | - | 58504778 | 58504974 | 58504302 | 58504488 | 58505471 | 58505583 | 90 | 52 | 98 |
| SE | 7738 | ENSG00000108506 | INTS2 | chr17 | - | 61869272 | 61869380 | 61869033 | 61869139 | 61869736 | 61869988 | 35 | 23 | 28 |
| SE | 7739 | ENSG00000108506 | INTS2 | chr17 | - | 61881006 | 61881171 | 61877886 | 61878088 | 61884900 | 61885005 | 48 | 20 | 39 |
| SE | 7740 | ENSG00000108506 | INTS2 | chr17 | - | 61889785 | 61889894 | 61884900 | 61885005 | 61891512 | 61891689 | 32 | 38 | 55 |
| SE | 7742 | ENSG00000108506 | INTS2 | chr17 | - | 61904459 | 61904585 | 61897667 | 61897739 | 61907407 | 61907634 | 53 | 17 | 31 |
| SE | 7743 | ENSG00000108506 | INTS2 | chr17 | - | 61919399 | 61919513 | 61911939 | 61912070 | 61921724 | 61921827 | 58 | 55 | 56 |
| SE | 7744 | ENSG00000108506 | INTS2 | chr17 | - | 61921221 | 61921342 | 61919399 | 61919513 | 61921724 | 61921827 | 1 | 0 | 1 |
| SE | 7745 | ENSG00000108506 | INTS2 | chr17 | - | 61921724 | 61921827 | 61919399 | 61919513 | 61926351 | 61926662 | 34 | 33 | 28 |
| SE | 7746 | ENSG00000108506 | INTS2 | chr17 | - | 61924960 | 61925099 | 61921724 | 61921827 | 61926351 | 61926662 | 70 | 54 | 63 |
| SE | 7749 | ENSG00000167863 | ATP5PD | chr17 | - | 75042180 | 75042277 | 75040091 | 75040163 | 75046924 | 75046950 | 407 | 260 | 392 |
| SE | 7760 | ENSG00000161513 | FDXR | chr17 | - | 74869949 | 74870102 | 74866783 | 74866879 | 74872035 | 74872133 | 0 | 0 | 0 |
| SE | 7762 | ENSG00000109118 | PHF12 | chr17 | - | 28907589 | 28907672 | 28906855 | 28906994 | 28908782 | 28908881 | 220 | 133 | 183 |
| SE | 7765 | ENSG00000109118 | PHF12 | chr17 | - | 28908782 | 28908881 | 28906855 | 28906994 | 28911111 | 28911237 | 0 | 0 | 0 |
| SE | 7767 | ENSG00000109118 | PHF12 | chr17 | - | 28910225 | 28910369 | 28906855 | 28906994 | 28911111 | 28911174 | 67 | 76 | 82 |
| SE | 7768 | ENSG00000109118 | PHF12 | chr17 | - | 28910225 | 28910369 | 28908782 | 28908881 | 28911111 | 28911174 | 199 | 180 | 195 |
| SE | 7774 | ENSG00000109118 | PHF12 | chr17 | - | 28921687 | 28921808 | 28919142 | 28919275 | 28923908 | 28923947 | 256 | 138 | 252 |
| SE | 7775 | ENSG00000109118 | PHF12 | chr17 | - | 28923908 | 28924302 | 28921687 | 28921808 | 28926990 | 28927063 | 332 | 164 | 280 |
| SE | 7776 | ENSG00000109118 | PHF12 | chr17 | - | 28923908 | 28924302 | 28921687 | 28921808 | 28950064 | 28950246 | 83 | 51 | 83 |
| SE | 7777 | ENSG00000109118 | PHF12 | chr17 | - | 28923908 | 28927063 | 28921687 | 28921808 | 28950064 | 28950246 | 164 | 85 | 138 |
| SE | 7780 | ENSG00000109118 | PHF12 | chr17 | - | 28926990 | 28927063 | 28923908 | 28924302 | 28950064 | 28950246 | 317 | 147 | 250 |
| SE | 7781 | ENSG00000109118 | PHF12 | chr17 | - | 28944502 | 28944569 | 28926990 | 28927063 | 28950064 | 28950662 | 5 | 0 | 0 |
| SE | 7782 | ENSG00000109118 | PHF12 | chr17 | - | 28950064 | 28950246 | 28923908 | 28927063 | 28950894 | 28950904 | 416 | 202 | 349 |
| SE | 7784 | ENSG00000278053 | DDX52 | chr17 | - | 37618291 | 37618384 | 37609738 | 37614353 | 37619767 | 37619839 | 92 | 126 | 112 |
| SE | 7788 | ENSG00000278053 | DDX52 | chr17 | - | 37630029 | 37630173 | 37628560 | 37628672 | 37632112 | 37632298 | 129 | 114 | 98 |
| SE | 7789 | ENSG00000278053 | DDX52 | chr17 | - | 37632112 | 37632298 | 37630029 | 37630173 | 37633287 | 37633418 | 169 | 156 | 149 |
| SE | 7791 | ENSG00000278053 | DDX52 | chr17 | - | 37635550 | 37635604 | 37633287 | 37633418 | 37642109 | 37642308 | 3 | 0 | 4 |
| SE | 7793 | ENSG00000005961 | ITGA2B | chr17 | - | 44374353 | 44374470 | 44372180 | 44372423 | 44374658 | 44374760 | 73 | 38 | 18 |
| SE | 7794 | ENSG00000005961 | ITGA2B | chr17 | - | 44374658 | 44374760 | 44374353 | 44374470 | 44374997 | 44375111 | 65 | 46 | 10 |
| SE | 7795 | ENSG00000005961 | ITGA2B | chr17 | - | 44379688 | 44379814 | 44378642 | 44378710 | 44380001 | 44380153 | 118 | 69 | 33 |
| SE | 7797 | ENSG00000005961 | ITGA2B | chr17 | - | 44385163 | 44385202 | 44384947 | 44385076 | 44385281 | 44385335 | 13 | 5 | 0 |
| SE | 7798 | ENSG00000005961 | ITGA2B | chr17 | - | 44385281 | 44385335 | 44384947 | 44385076 | 44385550 | 44385809 | 61 | 39 | 9 |
| SE | 7802 | ENSG00000030582 | GRN | chr17 | + | 44345565 | 44345605 | 44345264 | 44345334 | 44349157 | 44349302 | 3 | 0 | 0 |
| SE | 7803 | ENSG00000030582 | GRN | chr17 | + | 44345565 | 44345605 | 44345264 | 44345334 | 44349161 | 44349302 | 0 | 1 | 0 |
| SE | 7806 | ENSG00000030582 | GRN | chr17 | + | 44345609 | 44345741 | 44345261 | 44345334 | 44349161 | 44349302 | 0 | 2 | 0 |
| SE | 7807 | ENSG00000030582 | GRN | chr17 | + | 44346088 | 44346228 | 44345301 | 44345334 | 44349157 | 44349275 | 4 | 1 | 1 |
| SE | 7808 | ENSG00000030582 | GRN | chr17 | + | 44346088 | 44346228 | 44345301 | 44345605 | 44349157 | 44349275 | 5 | 0 | 1 |
| SE | 7810 | ENSG00000030582 | GRN | chr17 | + | 44349624 | 44349751 | 44349425 | 44349551 | 44350227 | 44350340 | 162 | 68 | 94 |
| SE | 7811 | ENSG00000030582 | GRN | chr17 | + | 44349666 | 44349751 | 44349425 | 44349551 | 44350227 | 44350283 | 717 | 289 | 382 |
| SE | 7813 | ENSG00000030582 | GRN | chr17 | + | 44351549 | 44351795 | 44351362 | 44351460 | 44352014 | 44352248 | 2550 | 1126 | 1395 |
| SE | 7814 | ENSG00000030582 | GRN | chr17 | + | 44352014 | 44352248 | 44351549 | 44351795 | 44352340 | 44352571 | 1568 | 620 | 787 |
| SE | 7819 | ENSG00000087995 | METTL2A | chr17 | + | 62444836 | 62444943 | 62440616 | 62440756 | 62447700 | 62447766 | 65 | 27 | 31 |
| SE | 7820 | ENSG00000087995 | METTL2A | chr17 | + | 62447700 | 62447766 | 62444836 | 62444943 | 62448574 | 62448914 | 38 | 19 | 33 |
| SE | 7830 | ENSG00000108829 | LRRC59 | chr17 | - | 50385117 | 50385291 | 50381237 | 50383235 | 50388059 | 50388132 | 248 | 137 | 167 |
| SE | 7831 | ENSG00000108829 | LRRC59 | chr17 | - | 50388059 | 50388132 | 50385117 | 50385291 | 50392397 | 50392502 | 252 | 155 | 188 |
| SE | 7834 | ENSG00000154920 | EME1 | chr17 | + | 50378594 | 50378681 | 50376065 | 50376193 | 50378773 | 50378895 | 1 | 5 | 0 |
| SE | 7835 | ENSG00000154920 | EME1 | chr17 | + | 50378599 | 50378681 | 50376065 | 50376193 | 50378773 | 50378895 | 2 | 0 | 0 |
| SE | 7843 | ENSG00000108819 | PPP1R9B | chr17 | - | 50139262 | 50139316 | 50135967 | 50136197 | 50143597 | 50143718 | 87 | 43 | 73 |
| SE | 7846 | ENSG00000108819 | PPP1R9B | chr17 | - | 50139428 | 50139581 | 50139262 | 50139316 | 50143597 | 50143718 | 1023 | 783 | 827 |
| SE | 7850 | ENSG00000108819 | PPP1R9B | chr17 | - | 50140092 | 50140228 | 50139428 | 50139581 | 50143597 | 50143718 | 560 | 340 | 484 |
| SE | 7853 | ENSG00000108819 | PPP1R9B | chr17 | - | 50141268 | 50141373 | 50139428 | 50139581 | 50143597 | 50143718 | 292 | 176 | 185 |
| SE | 7854 | ENSG00000108819 | PPP1R9B | chr17 | - | 50141268 | 50141373 | 50140092 | 50140228 | 50143597 | 50143718 | 863 | 561 | 572 |
| SE | 7856 | ENSG00000167100 | SAMD14 | chr17 | - | 50125090 | 50125360 | 50124919 | 50124971 | 50129516 | 50129754 | 2 | 4 | 0 |
| SE | 7857 | ENSG00000108825 | PTGES3L-AARSD1 | chr17 | - | 42971606 | 42971696 | 42970288 | 42970342 | 42979169 | 42979268 | 3 | 1 | 1 |
| SE | 7859 | ENSG00000267060 | PTGES3L | chr17 | - | 42979549 | 42979663 | 42979377 | 42979444 | 42980045 | 42980080 | 2 | 7 | 3 |
| SE | 7860 | ENSG00000176845 | METRNL | chr17 | + | 83084937 | 83085323 | 83079608 | 83079985 | 83093166 | 83093226 | 213 | 62 | 80 |
| SE | 7862 | ENSG00000100425 | BRD1 | chr22 | - | 49787389 | 49787522 | 49777677 | 49777813 | 49794033 | 49794294 | 38 | 17 | 35 |
| SE | 7866 | ENSG00000100425 | BRD1 | chr22 | - | 49822950 | 49824331 | 49804203 | 49804360 | 49827496 | 49827512 | 66 | 50 | 69 |
| SE | 7867 | ENSG00000175711 | B3GNTL1 | chr17 | - | 82960358 | 82960486 | 82957358 | 82957494 | 82961056 | 82961332 | 1 | 0 | 2 |
| SE | 7868 | ENSG00000175711 | B3GNTL1 | chr17 | - | 82965650 | 82965746 | 82961056 | 82961181 | 83005114 | 83005210 | 26 | 13 | 18 |
| SE | 7871 | ENSG00000175711 | B3GNTL1 | chr17 | - | 83005574 | 83005671 | 83005114 | 83005210 | 83014453 | 83014519 | 0 | 0 | 0 |
| SE | 7872 | ENSG00000175711 | B3GNTL1 | chr17 | - | 83014453 | 83014519 | 83005114 | 83005210 | 83035034 | 83035099 | 40 | 17 | 32 |
| SE | 7875 | ENSG00000141560 | FN3KRP | chr17 | + | 82720249 | 82720363 | 82718905 | 82719057 | 82722803 | 82722886 | 11 | 9 | 9 |
| SE | 7876 | ENSG00000141560 | FN3KRP | chr17 | + | 82720271 | 82720363 | 82718905 | 82719057 | 82722803 | 82722886 | 38 | 34 | 41 |
| SE | 7878 | ENSG00000141560 | FN3KRP | chr17 | + | 82720760 | 82720965 | 82720271 | 82720363 | 82722803 | 82722886 | 3 | 2 | 0 |
| SE | 7882 | ENSG00000141580 | WDR45B | chr17 | - | 82619042 | 82619094 | 82617295 | 82617397 | 82621608 | 82621799 | 43 | 33 | 43 |
| SE | 7883 | ENSG00000141580 | WDR45B | chr17 | - | 82619042 | 82619128 | 82617295 | 82617397 | 82621608 | 82621799 | 188 | 138 | 220 |
| SE | 7884 | ENSG00000141580 | WDR45B | chr17 | - | 82625388 | 82625483 | 82621608 | 82621799 | 82627203 | 82627291 | 214 | 158 | 196 |
| SE | 7885 | ENSG00000141580 | WDR45B | chr17 | - | 82627203 | 82627291 | 82621608 | 82621799 | 82643948 | 82644023 | 0 | 0 | 0 |
| SE | 7886 | ENSG00000141580 | WDR45B | chr17 | - | 82627203 | 82627291 | 82625388 | 82625483 | 82630920 | 82631022 | 196 | 132 | 173 |
| SE | 7889 | ENSG00000153933 | DGKE | chr17 | + | 56845689 | 56845809 | 56844018 | 56844178 | 56847921 | 56848065 | 84 | 68 | 89 |
| SE | 7890 | ENSG00000153933 | DGKE | chr17 | + | 56847359 | 56847429 | 56845689 | 56845809 | 56847515 | 56847658 | 0 | 0 | 2 |
| SE | 7891 | ENSG00000153933 | DGKE | chr17 | + | 56847921 | 56848065 | 56845689 | 56845809 | 56848695 | 56848853 | 62 | 41 | 65 |
| SE | 7892 | ENSG00000153933 | DGKE | chr17 | + | 56848695 | 56848853 | 56847921 | 56848065 | 56849180 | 56849232 | 79 | 77 | 87 |
| SE | 7893 | ENSG00000153933 | DGKE | chr17 | + | 56856511 | 56856625 | 56849180 | 56849232 | 56858593 | 56858665 | 98 | 76 | 82 |
| SE | 7895 | ENSG00000214226 | C17orf67 | chr17 | - | 56824738 | 56824853 | 56815755 | 56816010 | 56832897 | 56833342 | 0 | 0 | 3 |
| SE | 7896 | ENSG00000214226 | C17orf67 | chr17 | - | 56825080 | 56825321 | 56824738 | 56824853 | 56832897 | 56833342 | 7 | 6 | 11 |
| SE | 7899 | ENSG00000159842 | ABR | chr17 | - | 1056034 | 1056109 | 1050536 | 1050634 | 1056997 | 1057102 | 373 | 242 | 232 |
| SE | 7900 | ENSG00000159842 | ABR | chr17 | - | 1056034 | 1056109 | 1050536 | 1050634 | 1057969 | 1058045 | 129 | 100 | 96 |
| SE | 7901 | ENSG00000159842 | ABR | chr17 | - | 1056997 | 1057102 | 1050553 | 1050634 | 1057969 | 1058045 | 313 | 177 | 174 |
| SE | 7902 | ENSG00000159842 | ABR | chr17 | - | 1056997 | 1057102 | 1055282 | 1056109 | 1057969 | 1058045 | 558 | 312 | 305 |
| SE | 7903 | ENSG00000159842 | ABR | chr17 | - | 1058744 | 1058867 | 1057969 | 1058045 | 1067076 | 1067242 | 465 | 252 | 276 |
| SE | 7904 | ENSG00000159842 | ABR | chr17 | - | 1058744 | 1058867 | 1057969 | 1058045 | 1069968 | 1069983 | 233 | 135 | 128 |
| SE | 7905 | ENSG00000159842 | ABR | chr17 | - | 1067076 | 1067242 | 1057969 | 1058045 | 1069968 | 1069983 | 174 | 68 | 113 |
| SE | 7907 | ENSG00000159842 | ABR | chr17 | - | 1073624 | 1073677 | 1072613 | 1072754 | 1079329 | 1079390 | 333 | 166 | 181 |
| SE | 7910 | ENSG00000159842 | ABR | chr17 | - | 1100636 | 1100735 | 1091664 | 1091850 | 1125182 | 1125367 | 124 | 54 | 73 |
| SE | 7911 | ENSG00000159842 | ABR | chr17 | - | 1100636 | 1100735 | 1091664 | 1091850 | 1228792 | 1229738 | 101 | 47 | 56 |
| SE | 7912 | ENSG00000159842 | ABR | chr17 | - | 1125182 | 1125367 | 1091664 | 1091850 | 1228792 | 1229004 | 18 | 15 | 6 |
| SE | 7913 | ENSG00000159842 | ABR | chr17 | - | 1125182 | 1125367 | 1100636 | 1100735 | 1228792 | 1229004 | 41 | 22 | 22 |
| SE | 7916 | ENSG00000108953 | YWHAE | chr17 | - | 1354210 | 1354347 | 1344274 | 1345499 | 1361091 | 1361294 | 1168 | 925 | 892 |
| SE | 7917 | ENSG00000108953 | YWHAE | chr17 | - | 1354210 | 1354347 | 1344274 | 1345499 | 1364858 | 1365058 | 640 | 457 | 458 |
| SE | 7919 | ENSG00000108953 | YWHAE | chr17 | - | 1361091 | 1361298 | 1354210 | 1354347 | 1361901 | 1362008 | 1610 | 1481 | 1224 |
| SE | 7923 | ENSG00000108953 | YWHAE | chr17 | - | 1361901 | 1362008 | 1361091 | 1361298 | 1364858 | 1365058 | 1397 | 1345 | 1034 |
| SE | 7927 | ENSG00000108953 | YWHAE | chr17 | - | 1364858 | 1365058 | 1361670 | 1362008 | 1400046 | 1400206 | 1125 | 1032 | 783 |
| SE | 7929 | ENSG00000177370 | TIMM22 | chr17 | + | 998778 | 998975 | 997128 | 997380 | 999511 | 999584 | 136 | 80 | 111 |
| SE | 7932 | ENSG00000108479 | GALK1 | chr17 | - | 75763580 | 75763667 | 75762757 | 75763439 | 75763896 | 75764086 | 1 | 0 | 0 |
| SE | 7933 | ENSG00000198862 | LTN1 | chr21 | - | 28932469 | 28932664 | 28931154 | 28931322 | 28935108 | 28935329 | 106 | 95 | 67 |
| SE | 7934 | ENSG00000198862 | LTN1 | chr21 | - | 28935108 | 28935329 | 28932469 | 28932664 | 28936525 | 28936697 | 161 | 101 | 99 |
| SE | 7935 | ENSG00000198862 | LTN1 | chr21 | - | 28943666 | 28943904 | 28943261 | 28943336 | 28944382 | 28944596 | 265 | 160 | 187 |
| SE | 7936 | ENSG00000198862 | LTN1 | chr21 | - | 28944382 | 28944596 | 28943666 | 28943904 | 28945806 | 28945951 | 143 | 93 | 114 |
| SE | 7938 | ENSG00000198862 | LTN1 | chr21 | - | 28958385 | 28958539 | 28956761 | 28956948 | 28959457 | 28959697 | 45 | 31 | 28 |
| SE | 7939 | ENSG00000198862 | LTN1 | chr21 | - | 28964576 | 28964766 | 28960516 | 28960706 | 28965864 | 28965906 | 4 | 5 | 7 |
| SE | 7940 | ENSG00000198862 | LTN1 | chr21 | - | 28964576 | 28964766 | 28961316 | 28961533 | 28965864 | 28965906 | 3 | 5 | 7 |
| SE | 7941 | ENSG00000198862 | LTN1 | chr21 | - | 28965864 | 28965906 | 28960516 | 28960706 | 28966369 | 28967179 | 92 | 57 | 58 |
| SE | 7943 | ENSG00000198862 | LTN1 | chr21 | - | 28971270 | 28971444 | 28970551 | 28970742 | 28981118 | 28981299 | 142 | 105 | 142 |
| SE | 7944 | ENSG00000198862 | LTN1 | chr21 | - | 28971270 | 28971444 | 28970551 | 28970742 | 28982315 | 28982368 | 72 | 51 | 77 |
| SE | 7945 | ENSG00000198862 | LTN1 | chr21 | - | 28981118 | 28981299 | 28970551 | 28970742 | 28982315 | 28982368 | 65 | 66 | 63 |
| SE | 7946 | ENSG00000198862 | LTN1 | chr21 | - | 28981118 | 28981299 | 28971270 | 28971444 | 28982315 | 28982368 | 133 | 119 | 125 |
| SE | 7947 | ENSG00000198862 | LTN1 | chr21 | - | 28982315 | 28982368 | 28970551 | 28970742 | 28984691 | 28984922 | 38 | 29 | 55 |
| SE | 7948 | ENSG00000108448 | TRIM16L | chr17 | + | 18699359 | 18699457 | 18699153 | 18699275 | 18701356 | 18701445 | 14 | 3 | 0 |
| SE | 7950 | ENSG00000108448 | TRIM16L | chr17 | + | 18731069 | 18731235 | 18727524 | 18727758 | 18732038 | 18732134 | 29 | 7 | 22 |
| SE | 7955 | ENSG00000184640 | SEPTIN9 | chr17 | + | 77322788 | 77322948 | 77320312 | 77320348 | 77402058 | 77402415 | 0 | 0 | 0 |
| SE | 7956 | ENSG00000184640 | SEPTIN9 | chr17 | + | 77402058 | 77402703 | 77319517 | 77320348 | 77482143 | 77482335 | 104 | 50 | 87 |
| SE | 7958 | ENSG00000184640 | SEPTIN9 | chr17 | + | 77496213 | 77496389 | 77492979 | 77493076 | 77497314 | 77497366 | 4 | 1 | 5 |
| SE | 7959 | ENSG00000276293 | PIP4K2B | chr17 | - | 38778333 | 38778372 | 38777686 | 38777800 | 38779382 | 38779529 | 114 | 110 | 145 |
| SE | 7962 | ENSG00000276293 | PIP4K2B | chr17 | - | 38792725 | 38792882 | 38786822 | 38786920 | 38799983 | 38800126 | 2 | 3 | 1 |
| SE | 7963 | ENSG00000167447 | SMG8 | chr17 | + | 59209899 | 59211810 | 59209399 | 59209511 | 59212340 | 59212486 | 55 | 21 | 38 |
| SE | 7967 | ENSG00000126561 | STAT5A | chr17 | + | 42289369 | 42289539 | 42288500 | 42288598 | 42289865 | 42290022 | 34 | 17 | 53 |
| SE | 7970 | ENSG00000126561 | STAT5A | chr17 | + | 42289401 | 42289539 | 42288055 | 42288598 | 42289865 | 42290022 | 57 | 27 | 84 |
| SE | 7971 | ENSG00000126561 | STAT5A | chr17 | + | 42289401 | 42289539 | 42288055 | 42288598 | 42291971 | 42292061 | 22 | 11 | 34 |
| SE | 7972 | ENSG00000126561 | STAT5A | chr17 | + | 42289401 | 42289539 | 42288055 | 42288598 | 42295618 | 42295793 | 22 | 11 | 34 |
| SE | 7973 | ENSG00000126561 | STAT5A | chr17 | + | 42289429 | 42289539 | 42288495 | 42288598 | 42289865 | 42290022 | 31 | 15 | 38 |
| SE | 7976 | ENSG00000126561 | STAT5A | chr17 | + | 42289865 | 42290022 | 42288468 | 42288598 | 42291971 | 42292061 | 100 | 55 | 113 |
| SE | 7977 | ENSG00000126561 | STAT5A | chr17 | + | 42289865 | 42290022 | 42288468 | 42288598 | 42295618 | 42295793 | 11 | 2 | 7 |
| SE | 7978 | ENSG00000126561 | STAT5A | chr17 | + | 42289865 | 42290022 | 42289212 | 42289539 | 42291971 | 42292061 | 136 | 73 | 166 |
| SE | 7980 | ENSG00000126561 | STAT5A | chr17 | + | 42291971 | 42292061 | 42288055 | 42288598 | 42295618 | 42295793 | 70 | 51 | 67 |
| SE | 7981 | ENSG00000126561 | STAT5A | chr17 | + | 42291971 | 42292061 | 42289212 | 42289539 | 42295618 | 42295793 | 70 | 51 | 67 |
| SE | 7984 | ENSG00000126561 | STAT5A | chr17 | + | 42307401 | 42307496 | 42306113 | 42306447 | 42307592 | 42307723 | 226 | 173 | 263 |
| SE | 7985 | ENSG00000126561 | STAT5A | chr17 | + | 42309046 | 42309098 | 42308177 | 42308333 | 42309376 | 42309484 | 200 | 141 | 236 |
| SE | 7987 | ENSG00000126858 | RHOT1 | chr17 | + | 32171042 | 32171101 | 32142453 | 32142729 | 32173830 | 32173912 | 68 | 42 | 48 |
| SE | 7988 | ENSG00000126858 | RHOT1 | chr17 | + | 32171046 | 32171101 | 32142559 | 32142729 | 32173830 | 32173912 | 18 | 9 | 12 |
| SE | 7990 | ENSG00000126858 | RHOT1 | chr17 | + | 32173830 | 32173912 | 32171042 | 32171101 | 32175961 | 32176015 | 33 | 21 | 31 |
| SE | 7991 | ENSG00000126858 | RHOT1 | chr17 | + | 32175318 | 32175362 | 32173830 | 32173912 | 32175961 | 32176015 | 322 | 204 | 199 |
| SE | 7993 | ENSG00000126858 | RHOT1 | chr17 | + | 32176160 | 32176213 | 32175961 | 32176015 | 32182756 | 32182865 | 299 | 186 | 187 |
| SE | 7994 | ENSG00000126858 | RHOT1 | chr17 | + | 32182756 | 32182865 | 32176160 | 32176213 | 32183170 | 32183272 | 324 | 209 | 200 |
| SE | 7996 | ENSG00000126858 | RHOT1 | chr17 | + | 32202769 | 32202900 | 32200955 | 32201056 | 32203889 | 32203973 | 239 | 191 | 161 |
| SE | 7997 | ENSG00000126858 | RHOT1 | chr17 | + | 32203889 | 32203973 | 32202769 | 32202900 | 32206909 | 32207029 | 281 | 207 | 176 |
| SE | 7998 | ENSG00000126858 | RHOT1 | chr17 | + | 32206909 | 32207029 | 32203889 | 32203973 | 32208106 | 32208309 | 357 | 291 | 246 |
| SE | 8003 | ENSG00000126858 | RHOT1 | chr17 | + | 32211115 | 32211238 | 32208106 | 32208309 | 32253084 | 32253374 | 8 | 5 | 8 |
| SE | 8005 | ENSG00000126858 | RHOT1 | chr17 | + | 32221048 | 32221186 | 32211115 | 32211238 | 32224615 | 32225503 | 1 | 0 | 1 |
| SE | 8009 | ENSG00000129197 | RPAIN | chr17 | + | 5425970 | 5426082 | 5422768 | 5422829 | 5426235 | 5426299 | 69 | 36 | 76 |
| SE | 8023 | ENSG00000073350 | LLGL2 | chr17 | + | 75533641 | 75533705 | 75525778 | 75525825 | 75543396 | 75543501 | 0 | 0 | 0 |
| SE | 8024 | ENSG00000073350 | LLGL2 | chr17 | + | 75538530 | 75538636 | 75525788 | 75525825 | 75543396 | 75543501 | 7 | 0 | 3 |
| SE | 8025 | ENSG00000073350 | LLGL2 | chr17 | + | 75538530 | 75538636 | 75533641 | 75533705 | 75543396 | 75543501 | 6 | 3 | 4 |
| SE | 8026 | ENSG00000073350 | LLGL2 | chr17 | + | 75564352 | 75564507 | 75563751 | 75563806 | 75568475 | 75568693 | 40 | 15 | 20 |
| SE | 8030 | ENSG00000073350 | LLGL2 | chr17 | + | 75574452 | 75574668 | 75574212 | 75574260 | 75574870 | 75575121 | 10 | 3 | 6 |
| SE | 8031 | ENSG00000073350 | LLGL2 | chr17 | + | 75574609 | 75574668 | 75574110 | 75574260 | 75574870 | 75574921 | 3 | 1 | 1 |
| SE | 8032 | ENSG00000073350 | LLGL2 | chr17 | + | 75574609 | 75574668 | 75574110 | 75574260 | 75574873 | 75574906 | 3 | 0 | 0 |
| SE | 8035 | ENSG00000182173 | TSEN54 | chr17 | + | 75518994 | 75519047 | 75517556 | 75517655 | 75521408 | 75521510 | 62 | 23 | 60 |
| SE | 8036 | ENSG00000182173 | TSEN54 | chr17 | + | 75519002 | 75519047 | 75517556 | 75517655 | 75521408 | 75521510 | 12 | 4 | 14 |
| SE | 8038 | ENSG00000182173 | TSEN54 | chr17 | + | 75521675 | 75523335 | 75521408 | 75521510 | 75524261 | 75524723 | 0 | 0 | 0 |
| SE | 8039 | ENSG00000182173 | TSEN54 | chr17 | + | 75521704 | 75522333 | 75521408 | 75521510 | 75523274 | 75523335 | 87 | 41 | 84 |
| SE | 8040 | ENSG00000182173 | TSEN54 | chr17 | + | 75521704 | 75522521 | 75521408 | 75521510 | 75523274 | 75523335 | 91 | 38 | 67 |
| SE | 8041 | ENSG00000182173 | TSEN54 | chr17 | + | 75521704 | 75522535 | 75521408 | 75521510 | 75523274 | 75523335 | 76 | 37 | 66 |
| SE | 8042 | ENSG00000182173 | TSEN54 | chr17 | + | 75521704 | 75523335 | 75521408 | 75521510 | 75524261 | 75524420 | 76 | 37 | 66 |
| SE | 8043 | ENSG00000182173 | TSEN54 | chr17 | + | 75522006 | 75522333 | 75521408 | 75521510 | 75523274 | 75523287 | 11 | 4 | 20 |
| SE | 8044 | ENSG00000182173 | TSEN54 | chr17 | + | 75523249 | 75523335 | 75521704 | 75522521 | 75523662 | 75523779 | 14 | 6 | 25 |
| SE | 8045 | ENSG00000182173 | TSEN54 | chr17 | + | 75523274 | 75523335 | 75521408 | 75521510 | 75524261 | 75524693 | 0 | 0 | 0 |
| SE | 8046 | ENSG00000182173 | TSEN54 | chr17 | + | 75523274 | 75523335 | 75521704 | 75522521 | 75523662 | 75523779 | 20 | 2 | 18 |
| SE | 8047 | ENSG00000182173 | TSEN54 | chr17 | + | 75523662 | 75523779 | 75521408 | 75523335 | 75524261 | 75524420 | 66 | 21 | 72 |
| SE | 8049 | ENSG00000196704 | AMZ2 | chr17 | + | 68249010 | 68249096 | 68205480 | 68206238 | 68250187 | 68250470 | 3 | 1 | 6 |
| SE | 8053 | ENSG00000196704 | AMZ2 | chr17 | + | 68250793 | 68250967 | 68249199 | 68250470 | 68251049 | 68251065 | 336 | 188 | 275 |
| SE | 8056 | ENSG00000141337 | ARSG | chr17 | + | 68343603 | 68343791 | 68306942 | 68307711 | 68347124 | 68347172 | 71 | 46 | 55 |
| SE | 8058 | ENSG00000141337 | ARSG | chr17 | + | 68370443 | 68370524 | 68368547 | 68368744 | 68385063 | 68385172 | 96 | 37 | 53 |
| SE | 8062 | ENSG00000185262 | UBALD2 | chr17 | + | 76265906 | 76265969 | 76265347 | 76265625 | 76270193 | 76270383 | 97 | 32 | 105 |
| SE | 8063 | ENSG00000100348 | TXN2 | chr22 | - | 36468601 | 36468717 | 36467883 | 36467917 | 36476732 | 36476856 | 1 | 1 | 4 |
| SE | 8064 | ENSG00000100348 | TXN2 | chr22 | - | 36476732 | 36476856 | 36467045 | 36467917 | 36480574 | 36480748 | 605 | 360 | 541 |
| SE | 8067 | ENSG00000214087 | ARL16 | chr17 | - | 81683012 | 81683126 | 81682033 | 81682149 | 81683535 | 81683594 | 37 | 11 | 14 |
| SE | 8072 | ENSG00000141252 | VPS53 | chr17 | - | 519566 | 519703 | 519122 | 519298 | 519825 | 519930 | 2 | 3 | 0 |
| SE | 8073 | ENSG00000141252 | VPS53 | chr17 | - | 525871 | 525993 | 521600 | 521738 | 532841 | 532911 | 0 | 0 | 0 |
| SE | 8074 | ENSG00000141252 | VPS53 | chr17 | - | 532841 | 532911 | 521600 | 521738 | 537027 | 537176 | 85 | 48 | 86 |
| SE | 8075 | ENSG00000141252 | VPS53 | chr17 | - | 551871 | 551950 | 536365 | 537176 | 553379 | 553462 | 101 | 46 | 92 |
| SE | 8077 | ENSG00000141252 | VPS53 | chr17 | - | 560425 | 560573 | 547671 | 553462 | 562502 | 562745 | 112 | 45 | 82 |
| SE | 8080 | ENSG00000141252 | VPS53 | chr17 | - | 601794 | 601896 | 584774 | 586364 | 623532 | 623669 | 148 | 160 | 140 |
| SE | 8081 | ENSG00000141252 | VPS53 | chr17 | - | 622888 | 622976 | 601794 | 601896 | 623532 | 623674 | 3 | 0 | 0 |
| SE | 8082 | ENSG00000141252 | VPS53 | chr17 | - | 623532 | 623669 | 601794 | 601896 | 627173 | 627316 | 93 | 103 | 87 |
| SE | 8084 | ENSG00000141252 | VPS53 | chr17 | - | 623532 | 623674 | 600870 | 601896 | 627173 | 627316 | 183 | 165 | 168 |
| SE | 8085 | ENSG00000141252 | VPS53 | chr17 | - | 627173 | 627316 | 623532 | 623674 | 628087 | 628170 | 229 | 192 | 218 |
| SE | 8086 | ENSG00000141252 | VPS53 | chr17 | - | 627173 | 627316 | 623532 | 623674 | 631549 | 631628 | 93 | 64 | 85 |
| SE | 8088 | ENSG00000141252 | VPS53 | chr17 | - | 627225 | 627316 | 623532 | 623674 | 628087 | 628231 | 109 | 92 | 102 |
| SE | 8089 | ENSG00000141252 | VPS53 | chr17 | - | 628087 | 628231 | 623532 | 623674 | 631549 | 631628 | 65 | 50 | 81 |
| SE | 8090 | ENSG00000141252 | VPS53 | chr17 | - | 628087 | 628231 | 623532 | 623674 | 653290 | 653410 | 3 | 1 | 2 |
| SE | 8092 | ENSG00000141252 | VPS53 | chr17 | - | 628087 | 628231 | 627173 | 627316 | 653290 | 653410 | 146 | 134 | 137 |
| SE | 8094 | ENSG00000141252 | VPS53 | chr17 | - | 631549 | 631628 | 627173 | 627316 | 653290 | 653410 | 33 | 17 | 24 |
| SE | 8095 | ENSG00000141252 | VPS53 | chr17 | - | 631549 | 631628 | 628087 | 628231 | 653290 | 653410 | 91 | 68 | 109 |
| SE | 8097 | ENSG00000141252 | VPS53 | chr17 | - | 655837 | 655953 | 651059 | 653410 | 661808 | 661895 | 128 | 79 | 85 |
| SE | 8098 | ENSG00000141252 | VPS53 | chr17 | - | 655837 | 655953 | 651059 | 653410 | 697417 | 697484 | 48 | 31 | 30 |
| SE | 8101 | ENSG00000141252 | VPS53 | chr17 | - | 661808 | 661895 | 653290 | 653410 | 697417 | 697484 | 22 | 23 | 15 |
| SE | 8108 | ENSG00000141252 | VPS53 | chr17 | - | 685898 | 686005 | 661808 | 661895 | 697417 | 697484 | 0 | 0 | 0 |

| | | | | | | | | | | |
|---|---|---|---|---|---|---|---|---|---|---|
| SE | 8110 | ENSG00000141252 | VPS53 | chr17 | - | 695077 | 695225 | 661808 | 661895 | [illegible] |
| SE | 8112 | ENSG00000141252 | VPS53 | chr17 | - | 695077 | 695225 | 670545 | 670807 | [illegible] |
| SE | 8113 | ENSG00000141252 | VPS53 | chr17 | - | 695077 | 695225 | 670545 | 670807 | [illegible] |
| SE | 8115 | ENSG00000141252 | VPS53 | chr17 | - | 696824 | 697004 | 670545 | 670807 | [illegible] |
| SE | 8116 | ENSG00000141252 | VPS53 | chr17 | - | 696824 | 697004 | 695077 | 695225 | [illegible] |
| SE | 8117 | ENSG00000141252 | VPS53 | chr17 | - | 697203 | 697294 | 696863 | 697004 | [illegible] |
| SE | 8118 | ENSG00000141252 | VPS53 | chr17 | - | 697417 | 697484 | 695077 | 695225 | [illegible] |
| SE | 8119 | ENSG00000141252 | VPS53 | chr17 | - | 710532 | 710613 | 698556 | 699380 | [illegible] |
| SE | 8121 | ENSG00000120071 | KANSL1 | chr17 | - | 46032046 | 46032299 | 46029915 | 46031703 | [illegible] |
| SE | 8122 | ENSG00000120071 | KANSL1 | chr17 | - | 46033079 | 46033192 | 46032046 | 46032299 | [illegible] |
| SE | 8123 | ENSG00000120071 | KANSL1 | chr17 | - | 46033402 | 46033460 | 46033044 | 46033192 | [illegible] |
| SE | 8124 | ENSG00000120071 | KANSL1 | chr17 | - | 46033402 | 46033469 | 46033079 | 46033192 | [illegible] |
| SE | 8125 | ENSG00000120071 | KANSL1 | chr17 | - | 46034160 | 46034279 | 46033402 | 46033460 | [illegible] |
| SE | 8126 | ENSG00000120071 | KANSL1 | chr17 | - | 46034160 | 46034282 | 46033402 | 46033460 | [illegible] |
| SE | 8127 | ENSG00000120071 | KANSL1 | chr17 | - | 46034160 | 46034285 | 46033402 | 46033460 | [illegible] |
| SE | 8134 | ENSG00000120071 | KANSL1 | chr17 | - | 46156773 | 46156895 | 46094559 | 46094701 | [illegible] |
| SE | 8138 | ENSG00000120071 | KANSL1 | chr17 | - | 46170854 | 46172232 | 46092869 | 46094701 | [illegible] |
| SE | 8140 | ENSG00000073536 | NLE1 | chr17 | - | 35134942 | 35135059 | 35133338 | 35133498 | [illegible] |
| SE | 8141 | ENSG00000073536 | NLE1 | chr17 | - | 35137542 | 35137640 | 35137000 | 35137193 | [illegible] |
| SE | 8142 | ENSG00000073536 | NLE1 | chr17 | - | 35139234 | 35139314 | 35137813 | 35137890 | [illegible] |
| SE | 8143 | ENSG00000073536 | NLE1 | chr17 | - | 35139819 | 35140066 | 35139234 | 35139314 | [illegible] |
| SE | 8144 | ENSG00000073536 | NLE1 | chr17 | - | 35139848 | 35140066 | 35139234 | 35139314 | [illegible] |
| [illegible] | | | | | | | | | | |

[illegible]

| | | | | | | | | | | | | | | | |
|---|---|---|---|---|---|---|---|---|---|---|---|---|---|---|---|
| SE | 9103 | ENSG00000129657 | SEC14L1 | chr17 | + | 77142649 | 77142750 | 77141069 | 77141107 | 77143566 | 77143659 | 118 | 99 | 111 | [illegible] |
| SE | 9104 | ENSG00000129657 | SEC14L1 | chr17 | + | 77193420 | 77193549 | 77191180 | 77191312 | 77194676 | 77194733 | 574 | 422 | 413 | [illegible] |
| SE | 9109 | ENSG00000129657 | SEC14L1 | chr17 | + | 77213917 | 77214019 | 77213313 | 77213492 | 77216548 | 77216802 | 156 | 62 | 72 | [illegible] |
| SE | 9111 | ENSG00000234912 | SNHG20 | chr17 | + | 77093243 | 77093347 | 77088748 | 77089271 | 77099017 | 77099174 | 11 | 5 | 16 | [illegible] |
| SE | 9112 | ENSG00000129667 | RHBDF2 | chr17 | - | 76477656 | 76477785 | 76477179 | 76477298 | 76478805 | 76479009 | 196 | 130 | 140 | [illegible] |
| SE | 9115 | ENSG00000129667 | RHBDF2 | chr17 | - | 76487973 | 76488132 | 76487711 | 76487851 | 76493061 | 76493154 | 26 | 10 | 11 | [illegible] |
| SE | 9116 | ENSG00000129667 | RHBDF2 | chr17 | - | 76487973 | 76488132 | 76487711 | 76487851 | 76501352 | 76501412 | 32 | 9 | 10 | [illegible] |
| SE | 9117 | ENSG00000129667 | RHBDF2 | chr17 | - | 76488014 | 76488132 | 76487711 | 76487909 | 76493061 | 76493154 | 3 | 0 | 1 | [illegible] |
| SE | 9121 | ENSG00000070526 | ST6GALNAC1 | chr17 | - | 76637599 | 76637736 | 76629011 | 76629711 | 76643507 | 76643786 | 0 | 3 | 2 | [illegible] |
| SE | 9125 | ENSG00000163597 | SNHG16 | chr17 | + | 76557969 | 76558279 | 76557765 | 76557857 | 76558373 | 76558600 | 0 | 1 | 0 | [illegible] |
| SE | 9131 | ENSG00000163597 | SNHG16 | chr17 | + | 76558944 | 76559043 | 76558373 | 76558600 | 76561287 | 76561398 | 81 | 19 | 80 | [illegible] |
| SE | 9137 | ENSG00000006695 | COX10 | chr17 | + | 14076734 | 14077056 | 14074322 | 14074456 | 14102117 | 14102242 | 151 | 102 | 132 | [illegible] |
| SE | 9140 | ENSG00000006695 | COX10 | chr17 | + | 14191988 | 14192221 | 14159876 | 14159947 | 14209508 | 14209965 | 48 | 20 | 52 | [illegible] |
| SE | 9141 | ENSG00000006695 | COX10 | chr17 | + | 14209508 | 14209965 | 14191988 | 14192221 | 14211287 | 14211508 | 4 | 1 | 7 | [illegible] |
| SE | 9143 | ENSG00000006695 | COX10 | chr17 | + | 14215809 | 14215937 | 14211287 | 14211508 | 14216684 | 14217665 | 2 | 0 | 0 | [illegible] |
| SE | 9144 | ENSG00000236088 | COX10-AS1 | chr17 | - | 13880301 | 13880371 | 13776879 | 13777349 | 13892724 | 13892839 | 8 | 3 | 0 | [illegible] |
| SE | 9145 | ENSG00000236088 | COX10-AS1 | chr17 | - | 13898608 | 13898732 | 13892724 | 13892842 | 14034007 | 14034416 | 0 | 0 | 0 | [illegible] |
| SE | 9146 | ENSG00000236088 | COX10-AS1 | chr17 | - | 14033551 | 14033661 | 14028765 | 14030571 | 14034007 | 14034416 | 1 | 3 | 3 | [illegible] |
| SE | 9147 | ENSG00000236088 | COX10-AS1 | chr17 | - | 14033551 | 14033661 | 14028896 | 14031093 | 14034007 | 14034416 | 1 | 3 | 3 | [illegible] |
| SE | 9155 | ENSG00000065559 | MAP2K4 | chr17 | + | 12032265 | 12032298 | 12020833 | 12021001 | 12054888 | 12054991 | 0 | 2 | 0 | [illegible] |
| SE | 9157 | ENSG00000065559 | MAP2K4 | chr17 | + | 12054888 | 12054991 | 12020869 | 12021001 | 12095574 | 12095694 | 5 | 0 | 5 | [illegible] |
| SE | 9158 | ENSG00000065559 | MAP2K4 | chr17 | + | 12081355 | 12081530 | 12020869 | 12021001 | 12095574 | 12095694 | 73 | 58 | 77 | [illegible] |
| SE | 9161 | ENSG00000065559 | MAP2K4 | chr17 | + | 12107789 | 12107909 | 12095574 | 12095694 | 12110374 | 12110426 | 179 | 166 | 157 | [illegible] |
| SE | 9163 | ENSG00000065559 | MAP2K4 | chr17 | + | 12139838 | 12139884 | 12129138 | 12129287 | 12141146 | 12141260 | 101 | 80 | 105 | [illegible] |
| SE | 9166 | ENSG00000154957 | ZNF18 | chr17 | - | 11993501 | 11993654 | 11992539 | 11992911 | 11997430 | 11997475 | 0 | 0 | 3 | [illegible] |
| SE | 9167 | ENSG00000154957 | ZNF18 | chr17 | - | 11995557 | 11995679 | 11992442 | 11992911 | 11996809 | 11997110 | 0 | 0 | 2 | [illegible] |
| SE | 9175 | ENSG00000263400 | TMEM220-AS1 | chr17 | + | 10754228 | 10754362 | 10736601 | 10736638 | 10802398 | 10802508 | 1 | 0 | 0 | [illegible] |
| SE | 9177 | ENSG00000187824 | TMEM220 | chr17 | - | 10723269 | 10723329 | 10713294 | 10715588 | 10725010 | 10725134 | 9 | 8 | 18 | [illegible] |
| SE | 9180 | ENSG00000133028 | SCO1 | chr17 | - | 10691589 | 10691649 | 10686726 | 10686842 | 10691871 | 10691999 | 0 | 0 | 1 | [illegible] |
| SE | 9182 | ENSG00000133028 | SCO1 | chr17 | - | 10692763 | 10692961 | 10691871 | 10691964 | 10695740 | 10695831 | 142 | 84 | 99 | [illegible] |
| SE | 9183 | ENSG00000133028 | SCO1 | chr17 | - | 10692856 | 10692961 | 10691871 | 10691964 | 10695740 | 10695831 | 63 | 17 | 34 | [illegible] |
| SE | 9184 | ENSG00000109063 | MYH3 | chr17 | - | 10634016 | 10634182 | 10633590 | 10633715 | 10634839 | 10635023 | 19 | 7 | 17 | [illegible] |
| SE | 9185 | ENSG00000109063 | MYH3 | chr17 | - | 10647181 | 10647280 | 10645928 | 10646032 | 10648556 | 10648649 | 1 | 3 | 8 | [illegible] |
| SE | 9186 | ENSG00000109063 | MYH3 | chr17 | - | 10647362 | 10647426 | 10647181 | 10647280 | 10648556 | 10648649 | 19 | 2 | 5 | [illegible] |
| SE | 9193 | ENSG00000124215 | CDH26 | chr20 | + | 60001342 | 60001411 | 59999585 | 59999663 | 60002812 | 60002866 | 2 | 0 | 1 | [illegible] |
| SE | 9194 | ENSG00000124215 | CDH26 | chr20 | + | 60002812 | 60002866 | 59999585 | 59999663 | 60006712 | 60006787 | 2 | 0 | 1 | [illegible] |
| SE | 9205 | ENSG00000141576 | RNF157 | chr17 | - | 76156209 | 76156321 | 76155561 | 76155734 | 76158392 | 76158501 | 19 | 8 | 52 | [illegible] |
| SE | 9206 | ENSG00000141576 | RNF157 | chr17 | - | 76156209 | 76156370 | 76155561 | 76155734 | 76158392 | 76158523 | 8 | 6 | 30 | [illegible] |
| SE | 9207 | ENSG00000141576 | RNF157 | chr17 | - | 76166460 | 76166527 | 76165501 | 76165545 | 76167008 | 76167082 | 75 | 53 | 196 | [illegible] |
| SE | 9208 | ENSG00000276231 | PIK3R6 | chr17 | - | [illegible] | [illegible] | [illegible] | [illegible] | [illegible] | [illegible] | [illegible] | [illegible] | [illegible] | [illegible] |
| SE | 9209 | ENSG00000276231 | PIK3R6 | chr17 | - | [illegible] | [illegible] | [illegible] | [illegible] | [illegible] | [illegible] | [illegible] | [illegible] | [illegible] | [illegible] |
| SE | 9210 | ENSG00000276231 | PIK3R6 | chr17 | - | [illegible] | [illegible] | [illegible] | [illegible] | [illegible] | [illegible] | [illegible] | [illegible] | [illegible] | [illegible] |
| SE | 9212 | ENSG00000276231 | PIK3R6 | chr17 | - | [illegible] | [illegible] | [illegible] | [illegible] | [illegible] | [illegible] | [illegible] | [illegible] | [illegible] | [illegible] |
| SE | 9214 | ENSG00000276231 | PIK3R6 | chr17 | - | [illegible] | [illegible] | [illegible] | [illegible] | [illegible] | [illegible] | [illegible] | [illegible] | [illegible] | [illegible] |
| SE | 9215 | ENSG00000184381 | PLA2G6 | chr22 | - | [illegible] | [illegible] | [illegible] | [illegible] | [illegible] | [illegible] | [illegible] | [illegible] | [illegible] | [illegible] |
| SE | 9218 | ENSG00000184381 | PLA2G6 | chr22 | - | [illegible] | [illegible] | [illegible] | [illegible] | [illegible] | [illegible] | [illegible] | [illegible] | [illegible] | [illegible] |
| SE | 9219 | ENSG00000184381 | PLA2G6 | chr22 | - | [illegible] | [illegible] | [illegible] | [illegible] | [illegible] | [illegible] | [illegible] | [illegible] | [illegible] | [illegible] |
| SE | 9221 | ENSG00000184381 | PLA2G6 | chr22 | - | [illegible] | [illegible] | [illegible] | [illegible] | [illegible] | [illegible] | [illegible] | [illegible] | [illegible] | [illegible] |
| SE | 9222 | ENSG00000184381 | PLA2G6 | chr22 | - | [illegible] | [illegible] | [illegible] | [illegible] | [illegible] | [illegible] | [illegible] | [illegible] | [illegible] | [illegible] |
| SE | 9223 | ENSG00000184381 | PLA2G6 | chr22 | - | [illegible] | [illegible] | [illegible] | [illegible] | [illegible] | [illegible] | [illegible] | [illegible] | [illegible] | [illegible] |
| SE | 9224 | ENSG00000184381 | PLA2G6 | chr22 | - | [illegible] | [illegible] | [illegible] | [illegible] | [illegible] | [illegible] | [illegible] | [illegible] | [illegible] | [illegible] |
| SE | 9225 | ENSG00000184381 | PLA2G6 | chr22 | - | [illegible] | [illegible] | [illegible] | [illegible] | [illegible] | [illegible] | [illegible] | [illegible] | [illegible] | [illegible] |
| SE | 9227 | ENSG00000184381 | PLA2G6 | chr22 | - | [illegible] | [illegible] | [illegible] | [illegible] | [illegible] | [illegible] | [illegible] | [illegible] | [illegible] | [illegible] |
| SE | 9229 | ENSG00000184381 | PLA2G6 | chr22 | - | [illegible] | [illegible] | [illegible] | [illegible] | [illegible] | [illegible] | [illegible] | [illegible] | [illegible] | [illegible] |
| SE | 9232 | ENSG00000184381 | PLA2G6 | chr22 | - | [illegible] | [illegible] | [illegible] | [illegible] | [illegible] | [illegible] | [illegible] | [illegible] | [illegible] | [illegible] |
| SE | 9235 | ENSG00000178921 | PFAS | chr17 | + | [illegible] | [illegible] | [illegible] | [illegible] | [illegible] | [illegible] | [illegible] | [illegible] | [illegible] | [illegible] |
| SE | 9248 | ENSG00000178971 | CTC1 | chr17 | - | [illegible] | [illegible] | [illegible] | [illegible] | [illegible] | [illegible] | [illegible] | [illegible] | [illegible] | [illegible] |
| SE | 9251 | ENSG00000283403 | NA | chr19 | - | [illegible] | [illegible] | [illegible] | [illegible] | [illegible] | [illegible] | [illegible] | [illegible] | [illegible] | [illegible] |
| SE | 9254 | ENSG00000178999 | AURKB | chr17 | - | [illegible] | [illegible] | [illegible] | [illegible] | [illegible] | [illegible] | [illegible] | [illegible] | [illegible] | [illegible] |
| SE | 9255 | ENSG00000178999 | AURKB | chr17 | - | [illegible] | [illegible] | [illegible] | [illegible] | [illegible] | [illegible] | [illegible] | [illegible] | [illegible] | [illegible] |
| SE | 9257 | ENSG00000178999 | AURKB | chr17 | - | [illegible] | [illegible] | [illegible] | [illegible] | [illegible] | [illegible] | [illegible] | [illegible] | [illegible] | [illegible] |
| SE | 9258 | ENSG00000178999 | AURKB | chr17 | - | [illegible] | [illegible] | [illegible] | [illegible] | [illegible] | [illegible] | [illegible] | [illegible] | [illegible] | [illegible] |
| SE | 9259 | ENSG00000178999 | AURKB | chr17 | - | [illegible] | [illegible] | [illegible] | [illegible] | [illegible] | [illegible] | [illegible] | [illegible] | [illegible] | [illegible] |
| SE | 9260 | ENSG00000179029 | TMEM107 | chr17 | - | [illegible] | [illegible] | [illegible] | [illegible] | [illegible] | [illegible] | [illegible] | [illegible] | [illegible] | [illegible] |
| SE | 9263 | ENSG00000179094 | PER1 | chr17 | - | [illegible] | [illegible] | [illegible] | [illegible] | [illegible] | [illegible] | [illegible] | [illegible] | [illegible] | [illegible] |
| SE | 9265 | ENSG00000179094 | PER1 | chr17 | - | [illegible] | [illegible] | [illegible] | [illegible] | [illegible] | [illegible] | [illegible] | [illegible] | [illegible] | [illegible] |
| SE | 9266 | ENSG00000179094 | PER1 | chr17 | - | [illegible] | [illegible] | [illegible] | [illegible] | [illegible] | [illegible] | [illegible] | [illegible] | [illegible] | [illegible] |
| SE | 9271 | ENSG00000179094 | PER1 | chr17 | - | [illegible] | [illegible] | [illegible] | [illegible] | [illegible] | [illegible] | [illegible] | [illegible] | [illegible] | [illegible] |
| SE | 9272 | ENSG00000179094 | PER1 | chr17 | - | [illegible] | [illegible] | [illegible] | [illegible] | [illegible] | [illegible] | [illegible] | [illegible] | [illegible] | [illegible] |
| SE | 9273 | ENSG00000179094 | PER1 | chr17 | - | [illegible] | [illegible] | [illegible] | [illegible] | [illegible] | [illegible] | [illegible] | [illegible] | [illegible] | [illegible] |
| SE | 9274 | ENSG00000179094 | PER1 | chr17 | - | [illegible] | [illegible] | [illegible] | [illegible] | [illegible] | [illegible] | [illegible] | [illegible] | [illegible] | [illegible] |
| SE | 9275 | ENSG00000179094 | PER1 | chr17 | - | [illegible] | [illegible] | [illegible] | [illegible] | [illegible] | [illegible] | [illegible] | [illegible] | [illegible] | [illegible] |
| SE | 9276 | ENSG00000179094 | PER1 | chr17 | - | [illegible] | [illegible] | [illegible] | [illegible] | [illegible] | [illegible] | [illegible] | [illegible] | [illegible] | [illegible] |
| SE | 9281 | ENSG00000063244 | U2AF2 | chr19 | + | [illegible] | [illegible] | [illegible] | [illegible] | [illegible] | [illegible] | [illegible] | [illegible] | [illegible] | [illegible] |
| SE | 9294 | ENSG00000187838 | PLSCR3 | chr17 | - | [illegible] | [illegible] | [illegible] | [illegible] | [illegible] | [illegible] | [illegible] | [illegible] | [illegible] | [illegible] |
| SE | 9296 | ENSG00000187838 | PLSCR3 | chr17 | - | [illegible] | [illegible] | [illegible] | [illegible] | [illegible] | [illegible] | [illegible] | [illegible] | [illegible] | [illegible] |
| SE | 9297 | ENSG00000133030 | MPRIP | chr17 | + | [illegible] | [illegible] | [illegible] | [illegible] | [illegible] | [illegible] | [illegible] | [illegible] | [illegible] | [illegible] |
| SE | 9298 | ENSG00000133030 | MPRIP | chr17 | + | [illegible] | [illegible] | [illegible] | [illegible] | [illegible] | [illegible] | [illegible] | [illegible] | [illegible] | [illegible] |
| SE | 9299 | ENSG00000133030 | MPRIP | chr17 | + | [illegible] | [illegible] | [illegible] | [illegible] | [illegible] | [illegible] | [illegible] | [illegible] | [illegible] | [illegible] |
| SE | 9304 | ENSG00000133030 | MPRIP | chr17 | + | [illegible] | [illegible] | [illegible] | [illegible] | [illegible] | [illegible] | [illegible] | [illegible] | [illegible] | [illegible] |
| SE | 9305 | ENSG00000133030 | MPRIP | chr17 | + | [illegible] | [illegible] | [illegible] | [illegible] | [illegible] | [illegible] | [illegible] | [illegible] | [illegible] | [illegible] |
| SE | 9306 | ENSG00000133030 | MPRIP | chr17 | + | [illegible] | [illegible] | [illegible] | [illegible] | [illegible] | [illegible] | [illegible] | [illegible] | [illegible] | [illegible] |
| SE | 9309 | ENSG00000133030 | MPRIP | chr17 | + | [illegible] | [illegible] | [illegible] | [illegible] | [illegible] | [illegible] | [illegible] | [illegible] | [illegible] | [illegible] |
| SE | 9312 | ENSG00000133030 | MPRIP | chr17 | + | [illegible] | [illegible] | [illegible] | [illegible] | [illegible] | [illegible] | [illegible] | [illegible] | [illegible] | [illegible] |
| SE | 9314 | ENSG00000133030 | MPRIP | chr17 | + | [illegible] | [illegible] | [illegible] | [illegible] | [illegible] | [illegible] | [illegible] | [illegible] | [illegible] | [illegible] |
| SE | 9316 | ENSG00000133030 | MPRIP | chr17 | + | [illegible] | [illegible] | [illegible] | [illegible] | [illegible] | [illegible] | [illegible] | [illegible] | [illegible] | [illegible] |
| SE | 9317 | ENSG00000126351 | THRA | chr17 | + | [illegible] | [illegible] | [illegible] | [illegible] | [illegible] | [illegible] | [illegible] | [illegible] | [illegible] | [illegible] |
| SE | 9322 | ENSG00000167740 | CYB5D2 | chr17 | + | [illegible] | [illegible] | [illegible] | [illegible] | [illegible] | [illegible] | [illegible] | [illegible] | [illegible] | [illegible] |
| SE | 9323 | ENSG00000108439 | PNPO | chr17 | + | [illegible] | [illegible] | [illegible] | [illegible] | [illegible] | [illegible] | [illegible] | [illegible] | [illegible] | [illegible] |
| SE | 9325 | ENSG00000108439 | PNPO | chr17 | + | [illegible] | [illegible] | [illegible] | [illegible] | [illegible] | [illegible] | [illegible] | [illegible] | [illegible] | [illegible] |
| SE | 9326 | ENSG00000108439 | PNPO | chr17 | + | [illegible] | [illegible] | [illegible] | [illegible] | [illegible] | [illegible] | [illegible] | [illegible] | [illegible] | [illegible] |
| SE | 9327 | ENSG00000108439 | PNPO | chr17 | + | [illegible] | [illegible] | [illegible] | [illegible] | [illegible] | [illegible] | [illegible] | [illegible] | [illegible] | [illegible] |
| SE | 9329 | ENSG00000108439 | PNPO | chr17 | + | [illegible] | [illegible] | [illegible] | [illegible] | [illegible] | [illegible] | [illegible] | [illegible] | [illegible] | [illegible] |
| SE | 9330 | ENSG00000108439 | PNPO | chr17 | + | [illegible] | [illegible] | [illegible] | [illegible] | [illegible] | [illegible] | [illegible] | [illegible] | [illegible] | [illegible] |
| SE | 9333 | ENSG00000108774 | RAB5C | chr17 | - | [illegible] | [illegible] | [illegible] | [illegible] | [illegible] | [illegible] | [illegible] | [illegible] | [illegible] | [illegible] |
| SE | 9334 | ENSG00000108774 | RAB5C | chr17 | - | [illegible] | [illegible] | [illegible] | [illegible] | [illegible] | [illegible] | [illegible] | [illegible] | [illegible] | [illegible] |
| SE | 9335 | ENSG00000108774 | RAB5C | chr17 | - | [illegible] | [illegible] | [illegible] | [illegible] | [illegible] | [illegible] | [illegible] | [illegible] | [illegible] | [illegible] |
| SE | 9337 | ENSG00000108774 | RAB5C | chr17 | - | [illegible] | [illegible] | [illegible] | [illegible] | [illegible] | [illegible] | [illegible] | [illegible] | [illegible] | [illegible] |
| SE | 9338 | ENSG00000108774 | RAB5C | chr17 | - | [illegible] | [illegible] | [illegible] | [illegible] | [illegible] | [illegible] | [illegible] | [illegible] | [illegible] | [illegible] |
| SE | 9341 | ENSG00000167280 | ENGASE | chr17 | + | [illegible] | [illegible] | [illegible] | [illegible] | [illegible] | [illegible] | [illegible] | [illegible] | [illegible] | [illegible] |
| SE | 9343 | ENSG00000131469 | RPL27 | chr17 | + | [illegible] | [illegible] | [illegible] | [illegible] | [illegible] | [illegible] | [illegible] | [illegible] | [illegible] | [illegible] |
| SE | 9345 | ENSG00000131469 | RPL27 | chr17 | + | [illegible] | [illegible] | [illegible] | [illegible] | [illegible] | [illegible] | [illegible] | [illegible] | [illegible] | [illegible] |
| SE | 9346 | ENSG00000259207 | ITGB3 | chr17 | + | [illegible] | [illegible] | [illegible] | [illegible] | [illegible] | [illegible] | [illegible] | [illegible] | [illegible] | [illegible] |
| SE | 9347 | ENSG00000259207 | ITGB3 | chr17 | + | [illegible] | [illegible] | [illegible] | [illegible] | [illegible] | [illegible] | [illegible] | [illegible] | [illegible] | [illegible] |
| SE | 9348 | ENSG00000259207 | ITGB3 | chr17 | + | [illegible] | [illegible] | [illegible] | [illegible] | [illegible] | [illegible] | [illegible] | [illegible] | [illegible] | [illegible] |
| SE | 9349 | ENSG00000141026 | MED9 | chr17 | + | [illegible] | [illegible] | [illegible] | [illegible] | [illegible] | [illegible] | [illegible] | [illegible] | [illegible] | [illegible] |
| SE | 9351 | ENSG00000141030 | COPS3 | chr17 | - | [illegible] | [illegible] | [illegible] | [illegible] | [illegible] | [illegible] | [illegible] | [illegible] | [illegible] | [illegible] |
| SE | 9352 | ENSG00000141030 | COPS3 | chr17 | - | [illegible] | [illegible] | [illegible] | [illegible] | [illegible] | [illegible] | [illegible] | [illegible] | [illegible] | [illegible] |
| SE | 9356 | ENSG00000141030 | COPS3 | chr17 | - | [illegible] | [illegible] | [illegible] | [illegible] | [illegible] | [illegible] | [illegible] | [illegible] | [illegible] | [illegible] |
| SE | 9357 | ENSG00000141030 | COPS3 | chr17 | - | [illegible] | [illegible] | [illegible] | [illegible] | [illegible] | [illegible] | [illegible] | [illegible] | [illegible] | [illegible] |
| SE | 9358 | ENSG00000141030 | COPS3 | chr17 | - | [illegible] | [illegible] | [illegible] | [illegible] | [illegible] | [illegible] | [illegible] | [illegible] | [illegible] | [illegible] |
| SE | 9359 | ENSG00000141030 | COPS3 | chr17 | - | [illegible] | [illegible] | [illegible] | [illegible] | [illegible] | [illegible] | [illegible] | [illegible] | [illegible] | [illegible] |
| SE | 9363 | ENSG00000264187 | NA | chr17 | - | [illegible] | [illegible] | [illegible] | [illegible] | [illegible] | [illegible] | [illegible] | [illegible] | [illegible] | [illegible] |
| SE | 9365 | ENSG00000128438 | TBC1D27P | chr17 | - | [illegible] | [illegible] | [illegible] | [illegible] | [illegible] | [illegible] | [illegible] | [illegible] | [illegible] | [illegible] |
| SE | 9370 | ENSG00000121067 | SPOP | chr17 | - | [illegible] | [illegible] | [illegible] | [illegible] | [illegible] | [illegible] | [illegible] | [illegible] | [illegible] | [illegible] |
| SE | 9371 | ENSG00000121067 | SPOP | chr17 | - | [illegible] | [illegible] | [illegible] | [illegible] | [illegible] | [illegible] | [illegible] | [illegible] | [illegible] | [illegible] |
| SE | 9372 | ENSG00000121067 | SPOP | chr17 | - | [illegible] | [illegible] | [illegible] | [illegible] | [illegible] | [illegible] | [illegible] | [illegible] | [illegible] | [illegible] |
| SE | 9375 | ENSG00000121067 | SPOP | chr17 | - | [illegible] | [illegible] | [illegible] | [illegible] | [illegible] | [illegible] | [illegible] | [illegible] | [illegible] | [illegible] |
| SE | 9376 | ENSG00000121067 | SPOP | chr17 | - | [illegible] | [illegible] | [illegible] | [illegible] | [illegible] | [illegible] | [illegible] | [illegible] | [illegible] | [illegible] |
| SE | 9378 | ENSG00000121067 | SPOP | chr17 | - | [illegible] | [illegible] | [illegible] | [illegible] | [illegible] | [illegible] | [illegible] | [illegible] | [illegible] | [illegible] |
| SE | 9382 | ENSG00000121067 | SPOP | chr17 | - | [illegible] | [illegible] | [illegible] | [illegible] | [illegible] | [illegible] | [illegible] | [illegible] | [illegible] | [illegible] |
| SE | 9383 | ENSG00000121067 | SPOP | chr17 | - | [illegible] | [illegible] | [illegible] | [illegible] | [illegible] | [illegible] | [illegible] | [illegible] | [illegible] | [illegible] |
| SE | 9386 | ENSG00000121067 | SPOP | chr17 | - | [illegible] | [illegible] | [illegible] | [illegible] | [illegible] | [illegible] | [illegible] | [illegible] | [illegible] | [illegible] |
| SE | 9392 | ENSG00000080986 | NDC80 | chr18 | + | [illegible] | [illegible] | [illegible] | [illegible] | [illegible] | [illegible] | [illegible] | [illegible] | [illegible] | [illegible] |
| SE | 9397 | ENSG00000132383 | RPA1 | chr17 | + | [illegible] | [illegible] | [illegible] | [illegible] | [illegible] | [illegible] | [illegible] | [illegible] | [illegible] | [illegible] |
| SE | 9398 | ENSG00000132383 | RPA1 | chr17 | + | [illegible] | [illegible] | [illegible] | [illegible] | [illegible] | [illegible] | [illegible] | [illegible] | [illegible] | [illegible] |
| SE | 9400 | ENSG00000132383 | RPA1 | chr17 | + | [illegible] | [illegible] | [illegible] | [illegible] | [illegible] | [illegible] | [illegible] | [illegible] | [illegible] | [illegible] |
| SE | 9401 | ENSG00000132383 | RPA1 | chr17 | + | [illegible] | [illegible] | [illegible] | [illegible] | [illegible] | [illegible] | [illegible] | [illegible] | [illegible] | [illegible] |
| SE | 9402 | ENSG00000132383 | RPA1 | chr17 | + | [illegible] | [illegible] | [illegible] | [illegible] | [illegible] | [illegible] | [illegible] | [illegible] | [illegible] | [illegible] |
| SE | 9403 | ENSG00000132383 | RPA1 | chr17 | + | [illegible] | [illegible] | [illegible] | [illegible] | [illegible] | [illegible] | [illegible] | [illegible] | [illegible] | [illegible] |
| SE | 9405 | ENSG00000105053 | VRK3 | chr19 | - | [illegible] | [illegible] | [illegible] | [illegible] | [illegible] | [illegible] | [illegible] | [illegible] | [illegible] | [illegible] |
| SE | 9407 | ENSG00000105053 | VRK3 | chr19 | - | [illegible] | [illegible] | [illegible] | [illegible] | [illegible] | [illegible] | [illegible] | [illegible] | [illegible] | [illegible] |
| SE | 9408 | ENSG00000105053 | VRK3 | chr19 | - | [illegible] | [illegible] | [illegible] | [illegible] | [illegible] | [illegible] | [illegible] | [illegible] | [illegible] | [illegible] |
| SE | 9413 | ENSG00000105053 | VRK3 | chr19 | - | [illegible] | [illegible] | [illegible] | [illegible] | [illegible] | [illegible] | [illegible] | [illegible] | [illegible] | [illegible] |
| SE | 9414 | ENSG00000105053 | VRK3 | chr19 | - | [illegible] | [illegible] | [illegible] | [illegible] | [illegible] | [illegible] | [illegible] | [illegible] | [illegible] | [illegible] |
| SE | 9415 | ENSG00000105053 | VRK3 | chr19 | - | [illegible] | [illegible] | [illegible] | [illegible] | [illegible] | [illegible] | [illegible] | [illegible] | [illegible] | [illegible] |
| SE | 9416 | ENSG00000105053 | VRK3 | chr19 | - | [illegible] | [illegible] | [illegible] | [illegible] | [illegible] | [illegible] | [illegible] | [illegible] | [illegible] | [illegible] |
| SE | 9417 | ENSG00000105053 | VRK3 | chr19 | - | [illegible] | [illegible] | [illegible] | [illegible] | [illegible] | [illegible] | [illegible] | [illegible] | [illegible] | [illegible] |
| SE | 9421 | ENSG00000174231 | PRPF8 | chr17 | - | [illegible] | [illegible] | [illegible] | [illegible] | [illegible] | [illegible] | [illegible] | [illegible] | [illegible] | [illegible] |
| SE | 9422 | ENSG00000174231 | PRPF8 | chr17 | - | [illegible] | [illegible] | [illegible] | [illegible] | [illegible] | [illegible] | [illegible] | [illegible] | [illegible] | [illegible] |
| SE | 9424 | ENSG00000174231 | PRPF8 | chr17 | - | [illegible] | [illegible] | [illegible] | [illegible] | [illegible] | [illegible] | [illegible] | [illegible] | [illegible] | [illegible] |
| SE | 9425 | ENSG00000174231 | PRPF8 | chr17 | - | [illegible] | [illegible] | [illegible] | [illegible] | [illegible] | [illegible] | [illegible] | [illegible] | [illegible] | [illegible] |
| SE | 9427 | ENSG00000130402 | ACTN4 | chr19 | + | [illegible] | [illegible] | [illegible] | [illegible] | [illegible] | [illegible] | [illegible] | [illegible] | [illegible] | [illegible] |
| SE | 9428 | ENSG00000130402 | ACTN4 | chr19 | + | [illegible] | [illegible] | [illegible] | [illegible] | [illegible] | [illegible] | [illegible] | [illegible] | [illegible] | [illegible] |
| SE | 9429 | ENSG00000130402 | ACTN4 | chr19 | + | [illegible] | [illegible] | [illegible] | [illegible] | [illegible] | [illegible] | [illegible] | [illegible] | [illegible] | [illegible] |
| SE | 9430 | ENSG00000130402 | ACTN4 | chr19 | + | [illegible] | [illegible] | [illegible] | [illegible] | [illegible] | [illegible] | [illegible] | [illegible] | [illegible] | [illegible] |
| SE | 9435 | ENSG00000174238 | PITPNA | chr17 | - | [illegible] | [illegible] | [illegible] | [illegible] | [illegible] | [illegible] | [illegible] | [illegible] | [illegible] | [illegible] |
| SE | 9437 | ENSG00000174238 | PITPNA | chr17 | - | [illegible] | [illegible] | [illegible] | [illegible] | [illegible] | [illegible] | [illegible] | [illegible] | [illegible] | [illegible] |
| SE | 9438 | ENSG00000174238 | PITPNA | chr17 | - | [illegible] | [illegible] | [illegible] | [illegible] | [illegible] | [illegible] | [illegible] | [illegible] | [illegible] | [illegible] |
| SE | 9442 | ENSG00000244045 | TMEM199 | chr17 | + | [illegible] | [illegible] | [illegible] | [illegible] | [illegible] | [illegible] | [illegible] | [illegible] | [illegible] | [illegible] |
| SE | 9443 | ENSG00000004142 | POLDIP2 | chr17 | - | [illegible] | [illegible] | [illegible] | [illegible] | [illegible] | [illegible] | [illegible] | [illegible] | [illegible] | [illegible] |
| SE | 9444 | ENSG00000004142 | POLDIP2 | chr17 | - | [illegible] | [illegible] | [illegible] | [illegible] | [illegible] | [illegible] | [illegible] | [illegible] | [illegible] | [illegible] |
| SE | 9445 | ENSG00000004142 | POLDIP2 | chr17 | - | [illegible] | [illegible] | [illegible] | [illegible] | [illegible] | [illegible] | [illegible] | [illegible] | [illegible] | [illegible] |
| SE | 9453 | ENSG00000129245 | FXR2 | chr17 | - | [illegible] | [illegible] | [illegible] | [illegible] | [illegible] | [illegible] | [illegible] | [illegible] | [illegible] | [illegible] |
| SE | 9455 | ENSG00000129245 | FXR2 | chr17 | - | [illegible] | [illegible] | [illegible] | [illegible] | [illegible] | [illegible] | [illegible] | [illegible] | [illegible] | [illegible] |
| SE | 9456 | ENSG00000129245 | FXR2 | chr17 | - | [illegible] | [illegible] | [illegible] | [illegible] | [illegible] | [illegible] | [illegible] | [illegible] | [illegible] | [illegible] |
| SE | 9458 | ENSG00000161956 | SENP3 | chr17 | + | [illegible] | [illegible] | [illegible] | [illegible] | [illegible] | [illegible] | [illegible] | [illegible] | [illegible] | [illegible] |
| SE | 9459 | ENSG00000277957 | SENP3-EIF4A1 | chr17 | + | [illegible] | [illegible] | [illegible] | [illegible] | [illegible] | [illegible] | [illegible] | [illegible] | [illegible] | [illegible] |
| SE | 9460 | ENSG00000277957 | SENP3-EIF4A1 | chr17 | + | [illegible] | [illegible] | [illegible] | [illegible] | [illegible] | [illegible] | [illegible] | [illegible] | [illegible] | [illegible] |
| SE | 9461 | ENSG00000161960 | EIF4A1 | chr17 | + | [illegible] | [illegible] | [illegible] | [illegible] | [illegible] | [illegible] | [illegible] | [illegible] | [illegible] | [illegible] |
| SE | 9464 | ENSG00000277957 | SENP3-EIF4A1 | chr17 | + | [illegible] | [illegible] | [illegible] | [illegible] | [illegible] | [illegible] | [illegible] | [illegible] | [illegible] | [illegible] |
| SE | 9465 | ENSG00000277957 | SENP3-EIF4A1 | chr17 | + | [illegible] | [illegible] | [illegible] | [illegible] | [illegible] | [illegible] | [illegible] | [illegible] | [illegible] | [illegible] |
| SE | 9466 | ENSG00000161960 | EIF4A1 | chr17 | + | [illegible] | [illegible] | [illegible] | [illegible] | [illegible] | [illegible] | [illegible] | [illegible] | [illegible] | [illegible] |
| SE | 9467 | ENSG00000277957 | SENP3-EIF4A1 | chr17 | + | [illegible] | [illegible] | [illegible] | [illegible] | [illegible] | [illegible] | [illegible] | [illegible] | [illegible] | [illegible] |
| SE | 9468 | ENSG00000277957 | SENP3-EIF4A1 | chr17 | + | [illegible] | [illegible] | [illegible] | [illegible] | [illegible] | [illegible] | [illegible] | [illegible] | [illegible] | [illegible] |
| SE | 9469 | ENSG00000161960 | EIF4A1 | chr17 | + | [illegible] | [illegible] | [illegible] | [illegible] | [illegible] | [illegible] | [illegible] | [illegible] | [illegible] | [illegible] |
| SE | 9470 | ENSG00000277957 | SENP3-EIF4A1 | chr17 | + | [illegible] | [illegible] | [illegible] | [illegible] | [illegible] | [illegible] | [illegible] | [illegible] | [illegible] | [illegible] |
| SE | 9471 | ENSG00000161960 | EIF4A1 | chr17 | + | [illegible] | [illegible] | [illegible] | [illegible] | [illegible] | [illegible] | [illegible] | [illegible] | [illegible] | [illegible] |
| SE | 9475 | ENSG00000248871 | TNFSF12-TNFSF13 | chr17 | + | [illegible] | [illegible] | [illegible] | [illegible] | [illegible] | [illegible] | [illegible] | [illegible] | [illegible] | [illegible] |
| SE | 9477 | ENSG00000248871 | TNFSF12-TNFSF13 | chr17 | + | [illegible] | [illegible] | [illegible] | [illegible] | [illegible] | [illegible] | [illegible] | [illegible] | [illegible] | [illegible] |
| SE | 9479 | ENSG00000248871 | TNFSF12-TNFSF13 | chr17 | + | [illegible] | [illegible] | [illegible] | [illegible] | [illegible] | [illegible] | [illegible] | [illegible] | [illegible] | [illegible] |
| SE | 9481 | ENSG00000273018 | FAM106A | chr17 | - | [illegible] | [illegible] | [illegible] | [illegible] | [illegible] | [illegible] | [illegible] | [illegible] | [illegible] | [illegible] |
| SE | 9484 | ENSG00000273018 | FAM106A | chr17 | - | [illegible] | [illegible] | [illegible] | [illegible] | [illegible] | [illegible] | [illegible] | [illegible] | [illegible] | [illegible] |
| SE | 9485 | ENSG00000273018 | FAM106A | chr17 | - | [illegible] | [illegible] | [illegible] | [illegible] | [illegible] | [illegible] | [illegible] | [illegible] | [illegible] | [illegible] |
| SE | 9489 | ENSG00000273018 | FAM106A | chr17 | - | [illegible] | [illegible] | [illegible] | [illegible] | [illegible] | [illegible] | [illegible] | [illegible] | [illegible] | [illegible] |
| SE | 9490 | ENSG00000273018 | FAM106A | chr17 | - | [illegible] | [illegible] | [illegible] | [illegible] | [illegible] | [illegible] | [illegible] | [illegible] | [illegible] | [illegible] |
| SE | 9491 | ENSG00000273018 | FAM106A | chr17 | - | [illegible] | [illegible] | [illegible] | [illegible] | [illegible] | [illegible] | [illegible] | [illegible] | [illegible] | [illegible] |
| SE | 9494 | ENSG00000273018 | FAM106A | chr17 | - | [illegible] | [illegible] | [illegible] | [illegible] | [illegible] | [illegible] | [illegible] | [illegible] | [illegible] | [illegible] |
| SE | 9497 | ENSG00000171928 | TVP23B | chr17 | + | [illegible] | [illegible] | [illegible] | [illegible] | [illegible] | [illegible] | [illegible] | [illegible] | [illegible] | [illegible] |
| SE | 9501 | ENSG00000154874 | CCDC144B | chr17 | - | [illegible] | [illegible] | [illegible] | [illegible] | [illegible] | [illegible] | [illegible] | [illegible] | [illegible] | [illegible] |
| SE | 9502 | ENSG00000154874 | CCDC144B | chr17 | - | [illegible] | [illegible] | [illegible] | [illegible] | [illegible] | [illegible] | [illegible] | [illegible] | [illegible] | [illegible] |
| SE | 9504 | ENSG00000154874 | CCDC144B | chr17 | - | [illegible] | [illegible] | [illegible] | [illegible] | [illegible] | [illegible] | [illegible] | [illegible] | [illegible] | [illegible] |
| SE | 9505 | ENSG00000154874 | CCDC144B | chr17 | - | [illegible] | [illegible] | [illegible] | [illegible] | [illegible] | [illegible] | [illegible] | [illegible] | [illegible] | [illegible] |
| SE | 9506 | ENSG00000154874 | CCDC144B | chr17 | - | [illegible] | [illegible] | [illegible] | [illegible] | [illegible] | [illegible] | [illegible] | [illegible] | [illegible] | [illegible] |
| SE | 9510 | ENSG00000171916 | LGALS9C | chr17 | + | [illegible] | [illegible] | [illegible] | [illegible] | [illegible] | [illegible] | [illegible] | [illegible] | [illegible] | [illegible] |
| SE | 9512 | ENSG00000171916 | LGALS9C | chr17 | + | [illegible] | [illegible] | [illegible] | [illegible] | [illegible] | [illegible] | [illegible] | [illegible] | [illegible] | [illegible] |
| SE | 9513 | ENSG00000171916 | LGALS9C | chr17 | + | [illegible] | [illegible] | [illegible] | [illegible] | [illegible] | [illegible] | [illegible] | [illegible] | [illegible] | [illegible] |
| SE | 9516 | ENSG00000181610 | MRPS23 | chr17 | - | [illegible] | [illegible] | [illegible] | [illegible] | [illegible] | [illegible] | [illegible] | [illegible] | [illegible] | [illegible] |
| SE | 9521 | ENSG00000083290 | ULK2 | chr17 | - | [illegible] | [illegible] | [illegible] | [illegible] | [illegible] | [illegible] | [illegible] | [illegible] | [illegible] | [illegible] |
| SE | 9522 | ENSG00000083290 | ULK2 | chr17 | - | [illegible] | [illegible] | [illegible] | [illegible] | [illegible] | [illegible] | [illegible] | [illegible] | [illegible] | [illegible] |
| SE | 9523 | ENSG00000083290 | ULK2 | chr17 | - | [illegible] | [illegible] | [illegible] | [illegible] | [illegible] | [illegible] | [illegible] | [illegible] | [illegible] | [illegible] |
| SE | 9525 | ENSG00000083290 | ULK2 | chr17 | - | [illegible] | [illegible] | [illegible] | [illegible] | [illegible] | [illegible] | [illegible] | [illegible] | [illegible] | [illegible] |
| SE | 9526 | ENSG00000083290 | ULK2 | chr17 | - | [illegible] | [illegible] | [illegible] | [illegible] | [illegible] | [illegible] | [illegible] | [illegible] | [illegible] | [illegible] |
| SE | 9528 | ENSG00000083290 | ULK2 | chr17 | - | [illegible] | [illegible] | [illegible] | [illegible] | [illegible] | [illegible] | [illegible] | [illegible] | [illegible] | [illegible] |
| SE | 9530 | ENSG00000083290 | ULK2 | chr17 | - | [illegible] | [illegible] | [illegible] | [illegible] | [illegible] | [illegible] | [illegible] | [illegible] | [illegible] | [illegible] |
| SE | 9531 | ENSG00000083290 | ULK2 | chr17 | - | [illegible] | [illegible] | [illegible] | [illegible] | [illegible] | [illegible] | [illegible] | [illegible] | [illegible] | [illegible] |
| SE | 9532 | ENSG00000083290 | ULK2 | chr17 | - | [illegible] | [illegible] | [illegible] | [illegible] | [illegible] | [illegible] | [illegible] | [illegible] | [illegible] | [illegible] |
| SE | 9533 | ENSG00000083290 | ULK2 | chr17 | - | [illegible] | [illegible] | [illegible] | [illegible] | [illegible] | [illegible] | [illegible] | [illegible] | [illegible] | [illegible] |
| SE | 9537 | ENSG00000108932 | SLC16A6 | chr17 | - | [illegible] | [illegible] | [illegible] | [illegible] | [illegible] | [illegible] | [illegible] | [illegible] | [illegible] | [illegible] |
| SE | 9540 | ENSG00000108932 | SLC16A6 | chr17 | - | [illegible] | [illegible] | [illegible] | [illegible] | [illegible] | [illegible] | [illegible] | [illegible] | [illegible] | [illegible] |
| SE | 9542 | ENSG00000108932 | SLC16A6 | chr17 | - | [illegible] | [illegible] | [illegible] | [illegible] | [illegible] | [illegible] | [illegible] | [illegible] | [illegible] | [illegible] |
| SE | 9550 | ENSG00000266967 | AARSD1 | chr17 | - | [illegible] | [illegible] | [illegible] | [illegible] | [illegible] | [illegible] | [illegible] | [illegible] | [illegible] | [illegible] |
| SE | 9551 | ENSG00000266967 | AARSD1 | chr17 | - | [illegible] | [illegible] | [illegible] | [illegible] | [illegible] | [illegible] | [illegible] | [illegible] | [illegible] | [illegible] |
| SE | 9556 | ENSG00000100360 | IFT27 | chr22 | - | [illegible] | [illegible] | [illegible] | [illegible] | [illegible] | [illegible] | [illegible] | [illegible] | [illegible] | [illegible] |
| SE | 9563 | ENSG00000130844 | ZNF331 | chr19 | + | [illegible] | [illegible] | [illegible] | [illegible] | [illegible] | [illegible] | [illegible] | [illegible] | [illegible] | [illegible] |
| SE | 9568 | ENSG00000130844 | ZNF331 | chr19 | + | [illegible] | [illegible] | [illegible] | [illegible] | [illegible] | [illegible] | [illegible] | [illegible] | [illegible] | [illegible] |
| SE | 9569 | ENSG00000213246 | SUPT4H1 | chr17 | - | [illegible] | [illegible] | [illegible] | [illegible] | [illegible] | [illegible] | [illegible] | [illegible] | [illegible] | [illegible] |
| SE | 9570 | ENSG00000213246 | SUPT4H1 | chr17 | - | [illegible] | [illegible] | [illegible] | [illegible] | [illegible] | [illegible] | [illegible] | [illegible] | [illegible] | [illegible] |
| SE | 9571 | ENSG00000213246 | SUPT4H1 | chr17 | - | [illegible] | [illegible] | [illegible] | [illegible] | [illegible] | [illegible] | [illegible] | [illegible] | [illegible] | [illegible] |
| SE | 9573 | ENSG00000108604 | SMARCD2 | chr17 | - | [illegible] | [illegible] | [illegible] | [illegible] | [illegible] | [illegible] | [illegible] | [illegible] | [illegible] | [illegible] |
| SE | 9575 | ENSG00000108604 | SMARCD2 | chr17 | - | [illegible] | [illegible] | [illegible] | [illegible] | [illegible] | [illegible] | [illegible] | [illegible] | [illegible] | [illegible] |
| SE | 9576 | ENSG00000108604 | SMARCD2 | chr17 | - | [illegible] | [illegible] | [illegible] | [illegible] | [illegible] | [illegible] | [illegible] | [illegible] | [illegible] | [illegible] |
| SE | 9580 | ENSG00000141298 | SSH2 | chr17 | - | [illegible] | [illegible] | [illegible] | [illegible] | [illegible] | [illegible] | [illegible] | [illegible] | [illegible] | [illegible] |
| SE | 9581 | ENSG00000141298 | SSH2 | chr17 | - | [illegible] | [illegible] | [illegible] | [illegible] | [illegible] | [illegible] | [illegible] | [illegible] | [illegible] | [illegible] |
| SE | 9582 | ENSG00000141298 | SSH2 | chr17 | - | [illegible] | [illegible] | [illegible] | [illegible] | [illegible] | [illegible] | [illegible] | [illegible] | [illegible] | [illegible] |
| SE | 9583 | ENSG00000141298 | SSH2 | chr17 | - | [illegible] | [illegible] | [illegible] | [illegible] | [illegible] | [illegible] | [illegible] | [illegible] | [illegible] | [illegible] |
| SE | 9584 | ENSG00000141298 | SSH2 | chr17 | - | [illegible] | [illegible] | [illegible] | [illegible] | [illegible] | [illegible] | [illegible] | [illegible] | [illegible] | [illegible] |
| SE | 9586 | ENSG00000141298 | SSH2 | chr17 | - | [illegible] | [illegible] | [illegible] | [illegible] | [illegible] | [illegible] | [illegible] | [illegible] | [illegible] | [illegible] |
| SE | 9587 | ENSG00000141298 | SSH2 | chr17 | - | [illegible] | [illegible] | [illegible] | [illegible] | [illegible] | [illegible] | [illegible] | [illegible] | [illegible] | [illegible] |
| SE | 9588 | ENSG00000141298 | SSH2 | chr17 | - | [illegible] | [illegible] | [illegible] | [illegible] | [illegible] | [illegible] | [illegible] | [illegible] | [illegible] | [illegible] |
| SE | 9589 | ENSG00000141298 | SSH2 | chr17 | - | [illegible] | [illegible] | [illegible] | [illegible] | [illegible] | [illegible] | [illegible] | [illegible] | [illegible] | [illegible] |
| SE | 9590 | ENSG00000141298 | SSH2 | chr17 | - | [illegible] | [illegible] | [illegible] | [illegible] | [illegible] | [illegible] | [illegible] | [illegible] | [illegible] | [illegible] |
| SE | 9591 | ENSG00000141298 | SSH2 | chr17 | - | [illegible] | [illegible] | [illegible] | [illegible] | [illegible] | [illegible] | [illegible] | [illegible] | [illegible] | [illegible] |
| SE | 9592 | ENSG00000141298 | SSH2 | chr17 | - | [illegible] | [illegible] | [illegible] | [illegible] | [illegible] | [illegible] | [illegible] | [illegible] | [illegible] | [illegible] |
| SE | 9593 | ENSG00000141298 | SSH2 | chr17 | - | [illegible] | [illegible] | [illegible] | [illegible] | [illegible] | [illegible] | [illegible] | [illegible] | [illegible] | [illegible] |
| SE | 9594 | ENSG00000141298 | SSH2 | chr17 | - | [illegible] | [illegible] | [illegible] | [illegible] | [illegible] | [illegible] | [illegible] | [illegible] | [illegible] | [illegible] |
| SE | 9595 | ENSG00000141298 | SSH2 | chr17 | - | [illegible] | [illegible] | [illegible] | [illegible] | [illegible] | [illegible] | [illegible] | [illegible] | [illegible] | [illegible] |
| SE | 9596 | ENSG00000141298 | SSH2 | chr17 | - | [illegible] | [illegible] | [illegible] | [illegible] | [illegible] | [illegible] | [illegible] | [illegible] | [illegible] | [illegible] |
| SE | 9597 | ENSG00000141298 | SSH2 | chr17 | - | [illegible] | [illegible] | [illegible] | [illegible] | [illegible] | [illegible] | [illegible] | [illegible] | [illegible] | [illegible] |
| SE | 9598 | ENSG00000141298 | SSH2 | chr17 | - | [illegible] | [illegible] | [illegible] | [illegible] | [illegible] | [illegible] | [illegible] | [illegible] | [illegible] | [illegible] |
| SE | 9601 | ENSG00000141298 | SSH2 | chr17 | - | [illegible] | [illegible] | [illegible] | [illegible] | [illegible] | [illegible] | [illegible] | [illegible] | [illegible] | [illegible] |
| SE | 9617 | ENSG00000173821 | RNF213 | chr17 | + | [illegible] | [illegible] | [illegible] | [illegible] | [illegible] | [illegible] | [illegible] | [illegible] | [illegible] | [illegible] |
| SE | 9619 | ENSG00000173821 | RNF213 | chr17 | + | [illegible] | [illegible] | [illegible] | [illegible] | [illegible] | [illegible] | [illegible] | [illegible] | [illegible] | [illegible] |

| | | | | | | | | | | | | | | | |
|---|---|---|---|---|---|---|---|---|---|---|---|---|---|---|---|
| SE | 9620 | ENSG00000173821 | RNF213 | chr17 | + | 80291627 | 80291827 | 80290569 | 80290728 | 80294719 | 80295003 | 1823 | 1307 | 1489 | [illegible] |
| SE | 9622 | ENSG00000173821 | RNF213 | chr17 | + | 80309017 | 80309171 | 80307127 | 80307201 | 80313011 | 80313167 | 1275 | 798 | 996 | [illegible] |
| SE | 9623 | ENSG00000173821 | RNF213 | chr17 | + | 80325029 | 80325198 | 80319189 | 80319312 | 80327815 | 80327989 | 1182 | 795 | 1106 | [illegible] |
| SE | 9624 | ENSG00000173821 | RNF213 | chr17 | + | 80339200 | 80340356 | 80337832 | 80337997 | 80343131 | 80343325 | 835 | 438 | 770 | [illegible] |
| SE | 9625 | ENSG00000173821 | RNF213 | chr17 | + | 80343856 | 80344015 | 80343131 | 80343325 | 80344677 | 80348286 | 945 | 629 | 806 | [illegible] |
| SE | 9627 | ENSG00000173821 | RNF213 | chr17 | + | 80349769 | 80349906 | 80344677 | 80348286 | 80351684 | 80351803 | 356 | 137 | 308 | [illegible] |
| SE | 9628 | ENSG00000173821 | RNF213 | chr17 | + | 80350300 | 80350396 | 80344677 | 80348286 | 80351684 | 80351803 | 511 | 362 | 351 | [illegible] |
| SE | 9629 | ENSG00000173821 | RNF213 | chr17 | + | 80350300 | 80350396 | 80349769 | 80349906 | 80351684 | 80351803 | 948 | 618 | 712 | [illegible] |
| SE | 9630 | ENSG00000173821 | RNF213 | chr17 | + | 80351684 | 80351803 | 80344677 | 80348286 | 80358287 | 80358479 | 4 | 0 | 0 | [illegible] |
| SE | 9631 | ENSG00000173821 | RNF213 | chr17 | + | 80352939 | 80353059 | 80351684 | 80351803 | 80353511 | 80353666 | 1354 | 915 | 738 | [illegible] |
| SE | 9632 | ENSG00000173821 | RNF213 | chr17 | + | 80358287 | 80358479 | 80354440 | 80354576 | 80360060 | 80360206 | 969 | 498 | 787 | [illegible] |
| SE | 9633 | ENSG00000173821 | RNF213 | chr17 | + | 80361733 | 80361888 | 80360060 | 80360206 | 80363101 | 80363314 | 580 | 297 | 418 | [illegible] |
| SE | 9634 | ENSG00000173821 | RNF213 | chr17 | + | 80363101 | 80363314 | 80361733 | 80361888 | 80363608 | 80363790 | 661 | 295 | 479 | [illegible] |
| SE | 9635 | ENSG00000173821 | RNF213 | chr17 | + | 80363608 | 80363790 | 80363101 | 80363314 | 80364432 | 80364553 | 721 | 343 | 535 | [illegible] |
| SE | 9636 | ENSG00000173821 | RNF213 | chr17 | + | 80369767 | 80369867 | 80369501 | 80369671 | 80371873 | 80371985 | 1006 | 739 | 757 | [illegible] |
| SE | 9637 | ENSG00000173821 | RNF213 | chr17 | + | 80376881 | 80376963 | 80376300 | 80376543 | 80377761 | 80377796 | 876 | 543 | 767 | [illegible] |
| SE | 9638 | ENSG00000173821 | RNF213 | chr17 | + | 80379619 | 80379714 | 80377761 | 80377796 | 80380830 | 80380987 | 720 | 469 | 718 | [illegible] |
| SE | 9639 | ENSG00000173821 | RNF213 | chr17 | + | 80381546 | 80383070 | 80380830 | 80380987 | 80385537 | 80385621 | 288 | 144 | 185 | [illegible] |
| SE | 9640 | ENSG00000173821 | RNF213 | chr17 | + | 80382978 | 80383070 | 80381546 | 80381727 | 80383676 | 80383928 | 724 | 462 | 541 | [illegible] |
| SE | 9641 | ENSG00000173821 | RNF213 | chr17 | + | 80383676 | 80383928 | 80381546 | 80383070 | 80385038 | 80385171 | 1205 | 811 | 909 | [illegible] |
| SE | 9642 | ENSG00000173821 | RNF213 | chr17 | + | 80383676 | 80383928 | 80381546 | 80383070 | 80385537 | 80385621 | 523 | 347 | 414 | [illegible] |
| SE | 9643 | ENSG00000173821 | RNF213 | chr17 | + | 80385038 | 80385171 | 80381546 | 80383070 | 80385537 | 80385621 | 773 | 535 | 585 | [illegible] |
| SE | 9645 | ENSG00000173821 | RNF213 | chr17 | + | 80386689 | 80386891 | 80386249 | 80386430 | 80388611 | 80388689 | 1062 | 596 | 793 | [illegible] |
| SE | 9646 | ENSG00000173821 | RNF213 | chr17 | + | 80389172 | 80389367 | 80388611 | 80388689 | 80389827 | 80389917 | 856 | 482 | 702 | [illegible] |
| SE | 9647 | ENSG00000173821 | RNF213 | chr17 | + | 80390011 | 80390196 | 80389827 | 80389917 | 80393344 | 80395253 | 865 | 548 | 749 | [illegible] |
| SE | 9653 | ENSG00000087157 | PGS1 | chr17 | + | 78398251 | 78398351 | 78396307 | 78396385 | 78399347 | 78399537 | 138 | 64 | 135 | [illegible] |
| SE | 9663 | ENSG00000181045 | SLC26A11 | chr17 | + | 80222654 | 80222847 | 80221345 | 80221794 | 80223179 | 80223255 | 7 | 2 | 14 | [illegible] |
| SE | 9664 | ENSG00000181045 | SLC26A11 | chr17 | + | 80222654 | 80222847 | 80221345 | 80221794 | 80223251 | 80223272 | 22 | 5 | 24 | [illegible] |
| SE | 9665 | ENSG00000181045 | SLC26A11 | chr17 | + | 80227817 | 80227960 | 80225581 | 80225916 | 80236927 | 80237010 | 57 | 18 | 57 | [illegible] |
| SE | 9666 | ENSG00000181045 | SLC26A11 | chr17 | + | 80237307 | 80237439 | 80236927 | 80237103 | 80237521 | 80237594 | 0 | 1 | 0 | [illegible] |
| SE | 9668 | ENSG00000055483 | USP36 | chr17 | - | 78796316 | 78796494 | 78787465 | 78787658 | 78797785 | 78797902 | 0 | 0 | 4 | [illegible] |
| SE | 9669 | ENSG00000055483 | USP36 | chr17 | - | 78796316 | 78796494 | 78787465 | 78787658 | 78798399 | 78798455 | 0 | 0 | 5 | [illegible] |
| [illegible] | [illegible] | [illegible] | [illegible] | [illegible] | [illegible] | [illegible] | [illegible] | [illegible] | [illegible] | [illegible] | [illegible] | [illegible] | [illegible] | [illegible] | [illegible] |

| | | | | | | | | | | | | | | | | | | | | | | | | |
|---|---|---|---|---|---|---|---|---|---|---|---|---|---|---|---|---|---|---|---|---|---|---|---|---|
| SE | 10070 | ENSG00000108406 | DHX40 | chr17 | + | 59573115 | 59573235 | 59570517 | 59570663 | 59573739 | 59573756 | 208 | 125 | 176 | 191 | 322 | 0 | 1 | 0 | 0 | 0 | 75 | 160 | 138 | 104 | [illegible] |
| SE | 10072 | ENSG00000108406 | DHX40 | chr17 | + | 59575339 | 59575471 | 59574187 | 59574254 | 59577265 | 59577365 | 194 | 114 | 183 | 152 | 398 | 0 | 0 | 0 | 0 | 2 | 38 | 175 | 140 | 153 | [illegible] |
| SE | 10077 | ENSG00000108423 | TUBD1 | chr17 | - | 59863663 | 59863847 | 59859478 | 59860424 | 59866608 | 59866749 | 57 | 25 | 43 | 33 | 77 | 4 | 0 | 5 | 0 | 5 | 0 | 16 | 4 | 0 | [illegible] |
| SE | 10078 | ENSG00000108423 | TUBD1 | chr17 | - | 59863663 | 59863847 | 59859478 | 59860424 | 59874538 | 59874703 | 30 | 12 | 23 | 19 | 53 | 0 | 0 | 1 | 1 | 0 | 0 | 12 | 3 | 0 | [illegible] |
| SE | 10079 | ENSG00000108423 | TUBD1 | chr17 | - | 59863663 | 59863847 | 59859478 | 59860424 | 59878102 | 59878334 | 30 | 11 | 23 | 23 | 51 | 0 | 0 | 1 | 2 | 1 | 0 | 9 | 3 | 0 | [illegible] |
| SE | 10080 | ENSG00000108423 | TUBD1 | chr17 | - | 59863663 | 59863847 | 59859478 | 59860424 | 59886082 | 59886230 | 30 | 11 | 24 | 18 | 50 | 1 | 0 | 0 | 0 | 0 | 0 | 9 | 3 | 0 | [illegible] |
| SE | 10081 | ENSG00000108423 | TUBD1 | chr17 | - | 59866595 | 59866749 | 59860033 | 59860424 | 59874538 | 59874703 | 18 | 2 | 6 | 17 | 13 | 0 | 0 | 1 | 1 | 0 | 7 | 0 | 5 | 3 | [illegible] |
| SE | 10083 | ENSG00000108423 | TUBD1 | chr17 | - | 59866608 | 59866749 | 59859478 | 59860424 | 59874538 | 59874703 | 22 | 2 | 11 | 16 | 18 | 0 | 0 | 1 | 1 | 0 | 7 | 0 | 5 | 5 | [illegible] |
| SE | 10084 | ENSG00000108423 | TUBD1 | chr17 | - | 59866608 | 59866749 | 59859478 | 59860424 | 59878102 | 59878334 | 10 | 3 | 10 | 6 | 10 | 0 | 0 | 1 | 2 | 1 | 0 | 11 | 3 | 4 | [illegible] |
| SE | 10085 | ENSG00000108423 | TUBD1 | chr17 | - | 59866608 | 59866749 | 59859478 | 59860424 | 59886082 | 59886230 | 4 | 0 | 7 | 0 | 8 | 1 | 0 | 0 | 0 | 0 | 0 | 0 | 0 | 2 | [illegible] |
| SE | 10087 | ENSG00000108423 | TUBD1 | chr17 | - | 59866608 | 59866749 | 59863663 | 59863847 | 59878102 | 59878334 | 33 | 17 | 25 | 20 | 32 | 0 | 0 | 0 | 5 | 2 | 0 | 16 | 4 | 2 | [illegible] |
| SE | 10088 | ENSG00000108423 | TUBD1 | chr17 | - | 59866608 | 59866749 | 59863663 | 59863847 | 59886082 | 59886230 | 27 | 14 | 22 | 14 | 30 | 0 | 0 | 1 | 0 | 1 | 0 | 5 | 1 | 0 | [illegible] |
| SE | 10089 | ENSG00000108423 | TUBD1 | chr17 | - | 59874538 | 59874703 | 59859478 | 59860424 | 59878102 | 59878334 | 39 | 23 | 20 | 24 | 29 | 0 | 0 | 1 | 2 | 1 | 0 | 7 | 13 | 6 | [illegible] |
| SE | 10090 | ENSG00000108423 | TUBD1 | chr17 | - | 59874538 | 59874703 | 59863663 | 59863847 | 59878102 | 59878334 | 39 | 24 | 19 | 24 | 33 | 0 | 0 | 0 | 5 | 2 | 0 | 10 | 13 | 6 | [illegible] |
| SE | 10092 | ENSG00000108423 | TUBD1 | chr17 | - | 59880893 | 59881110 | 59878102 | 59878334 | 59886082 | 59886230 | 50 | 50 | 62 | 58 | 94 | 1 | 0 | 0 | 6 | 4 | 0 | 29 | 17 | 4 | [illegible] |
| SE | 10093 | ENSG00000108423 | TUBD1 | chr17 | - | 59884660 | 59884756 | 59866608 | 59866749 | 59886082 | 59886230 | 2 | 0 | 0 | 0 | 4 | 0 | 0 | 2 | 0 | 3 | 0 | 0 | 0 | 3 | [illegible] |
| SE | 10094 | ENSG00000108423 | TUBD1 | chr17 | - | 59884660 | 59884756 | 59880893 | 59881110 | 59886082 | 59886230 | 4 | 2 | 0 | 0 | 4 | 29 | 30 | 51 | 39 | 44 | 0 | 0 | 0 | 3 | [illegible] |
| SE | 10096 | ENSG00000108423 | TUBD1 | chr17 | - | 59886082 | 59886230 | 59878300 | 59878334 | 59892696 | 59892893 | 12 | 6 | 22 | 17 | 10 | 0 | 0 | 2 | 1 | 0 | 0 | 8 | 5 | 0 | [illegible] |
| SE | 10102 | ENSG00000108433 | GOSR2 | chr17 | + | 46929519 | 46929584 | 46923074 | 46923221 | 46931098 | 46931207 | 150 | 78 | 127 | 147 | 93 | 0 | 0 | 0 | 0 | 2 | 86 | 139 | 80 | 104 | [illegible] |
| SE | 10103 | ENSG00000108433 | GOSR2 | chr17 | + | 46929523 | 46929584 | 46923137 | 46923221 | 46931098 | 46931207 | 36 | 22 | 28 | 48 | 35 | 0 | 0 | 0 | 0 | 2 | 34 | 46 | 42 | 36 | [illegible] |
| SE | 10104 | ENSG00000108433 | GOSR2 | chr17 | + | 46931098 | 46931207 | 46929519 | 46929584 | 46932066 | 46932199 | 265 | 165 | 220 | 286 | 209 | 0 | 0 | 0 | 2 | 0 | 198 | 226 | 163 | 162 | [illegible] |
| SE | 10110 | ENSG00000108433 | GOSR2 | chr17 | + | 46932066 | 46932199 | 46929523 | 46929584 | 46935028 | 46935154 | 54 | 58 | 55 | 49 | 49 | 1 | 0 | 2 | 0 | 1 | 54 | 48 | 16 | 26 | [illegible] |
| SE | 10111 | ENSG00000108433 | GOSR2 | chr17 | + | 46932066 | 46932199 | 46929523 | 46929584 | 46938598 | 46938704 | 3 | 1 | 3 | 5 | 3 | 0 | 0 | 0 | 1 | 0 | 6 | 11 | 10 | 18 | [illegible] |
| SE | 10112 | ENSG00000108433 | GOSR2 | chr17 | + | 46932066 | 46932199 | 46931098 | 46931207 | 46935028 | 46935154 | 193 | 160 | 170 | 212 | 193 | 0 | 0 | 1 | 0 | 0 | 170 | 143 | 95 | 101 | [illegible] |
| SE | 10113 | ENSG00000108433 | GOSR2 | chr17 | + | 46935028 | 46935169 | 46929519 | 46929584 | 46938598 | 46938602 | 75 | 62 | 106 | 120 | 132 | 0 | 0 | 0 | 1 | 0 | 169 | 122 | 48 | 77 | [illegible] |
| SE | 10117 | ENSG00000108433 | GOSR2 | chr17 | + | 46948565 | 46948657 | 46938598 | 46938704 | 46948942 | 46949347 | 3 | 0 | 1 | 4 | 8 | 1 | 0 | 1 | 1 | 2 | 25 | 9 | 2 | 7 | [illegible] |
| SE | 10119 | ENSG00000108433 | GOSR2 | chr17 | + | 46948942 | 46949347 | 46938598 | 46938704 | 46955500 | 46955712 | 1 | 0 | 1 | 1 | 2 | 0 | 1 | 1 | 1 | 0 | 0 | 2 | 0 | 10 | [illegible] |
| SE | 10120 | ENSG00000108433 | GOSR2 | chr17 | + | 46948942 | 46949347 | 46938598 | 46938704 | 46963696 | 46965039 | 1 | 0 | 1 | 1 | 2 | 0 | 1 | 1 | 2 | 0 | 0 | 2 | 0 | 3 | [illegible] |
| SE | 10122 | ENSG00000108433 | GOSR2 | chr17 | + | 46955500 | 46955712 | 46938598 | 46938704 | 46963696 | 46965039 | 0 | 3 | 2 | 1 | 0 | 0 | 1 | 1 | 2 | 0 | 0 | 0 | 3 | 4 | [illegible] |
| SE | 10124 | ENSG00000108433 | GOSR2 | chr17 | + | 46955500 | 46955712 | 46948942 | 46949347 | 46963696 | 46965039 | 0 | 2 | 1 | 0 | 0 | 0 | 0 | 0 | 0 | 0 | 0 | 0 | 0 | 8 | [illegible] |
| SE | 10127 | ENSG00000108433 | GOSR2 | chr17 | + | 46963696 | 46965039 | 46948942 | 46949347 | 46966533 | 46966851 | 2 | 2 | 5 | 3 | 10 | 0 | 0 | 0 | 0 | 0 | 10 | 4 | 0 | 1 | [illegible] |
| SE | 10131 | ENSG00000198231 | DDX42 | chr17 | + | 63774188 | 63774376 | 63773883 | 63774091 | 63787033 | 63787270 | 74 | 22 | 69 | 49 | 4 | 0 | 0 | 0 | 1 | 2 | 61 | 90 | 53 | 60 | [illegible] |
| SE | 10133 | ENSG00000198231 | DDX42 | chr17 | + | 63787033 | 63787270 | 63774169 | 63774376 | 63792411 | 63792504 | 203 | 119 | 208 | 163 | 129 | 1 | 0 | 0 | 0 | 0 | 104 | 213 | 176 | 192 | [illegible] |
| SE | 10134 | ENSG00000198231 | DDX42 | chr17 | + | 63799479 | 63799523 | 63798037 | 63798099 | 63800467 | 63800617 | 2 | 3 | 2 | 8 | 7 | 2 | 5 | 2 | 2 | 12 | 0 | 5 | 8 | 3 | [illegible] |
| SE | 10136 | ENSG00000198231 | DDX42 | chr17 | + | 63816867 | 63816966 | 63815562 | 63815673 | 63817693 | 63818029 | 328 | 282 | 360 | 347 | 368 | 0 | 0 | 0 | 1 | 0 | 208 | 381 | 280 | 270 | [illegible] |
| SE | 10138 | ENSG00000011260 | UTP18 | chr17 | + | 51266181 | 51266280 | 51263273 | 51263386 | 51268836 | 51268904 | 140 | 128 | 158 | 223 | 248 | 0 | 0 | 0 | 0 | 0 | 175 | 128 | 58 | 175 | [illegible] |
| SE | 10139 | ENSG00000011260 | UTP18 | chr17 | + | 51268836 | 51268904 | 51266181 | 51266280 | 51273361 | 51273450 | 104 | 95 | 152 | 188 | 175 | 0 | 0 | 0 | 0 | 0 | 118 | 94 | 39 | 156 | [illegible] |
| SE | 10143 | ENSG00000011260 | UTP18 | chr17 | + | 51280388 | 51280479 | 51275865 | 51275991 | 51285244 | 51285368 | 43 | 24 | 29 | 35 | 95 | 0 | 0 | 0 | 0 | 0 | 19 | 38 | 31 | 63 | [illegible] |
| SE | 10146 | ENSG00000011260 | UTP18 | chr17 | + | 51288547 | 51288591 | 51288028 | 51288203 | 51296964 | 51297003 | 1 | 0 | 2 | 5 | 1 | 3 | 0 | 0 | 1 | 0 | 0 | 2 | 0 | 2 | [illegible] |
| SE | 10147 | ENSG00000011260 | UTP18 | chr17 | + | 51293902 | 51294045 | 51288028 | 51288203 | 51296964 | 51297003 | 108 | 122 | 126 | 125 | 203 | 3 | 0 | 0 | 1 | 0 | 58 | 76 | 74 | 83 | [illegible] |
| SE | 10148 | ENSG00000011260 | UTP18 | chr17 | + | 51293902 | 51294045 | 51288547 | 51288591 | 51296964 | 51297003 | 38 | 41 | 42 | 38 | 53 | 1 | 0 | 0 | 0 | 0 | 10 | 24 | 20 | 36 | [illegible] |
| SE | 10151 | ENSG00000073584 | SMARCE1 | chr17 | - | 40630207 | 40630286 | 40627780 | 40629568 | 40630713 | 40630924 | 1 | 1 | 3 | 6 | 4 | 1 | 0 | 0 | 0 | 0 | 16 | 5 | 10 | 1 | [illegible] |
| SE | 10152 | ENSG00000073584 | SMARCE1 | chr17 | - | 40630713 | 40630924 | 40624961 | 40628993 | 40631591 | 40631693 | 446 | 480 | 483 | 507 | 620 | 0 | 0 | 0 | 0 | 0 | 321 | 500 | 630 | 619 | [illegible] |
| SE | 10153 | ENSG00000073584 | SMARCE1 | chr17 | - | 40630713 | 40630924 | 40627760 | 40629568 | 40631591 | 40631693 | 272 | 321 | 323 | 341 | 452 | 0 | 0 | 0 | 0 | 0 | 257 | 365 | 462 | 409 | [illegible] |
| SE | 10154 | ENSG00000073584 | SMARCE1 | chr17 | - | 40631591 | 40631693 | 40629334 | 40630924 | 40632194 | 40632367 | 475 | 485 | 525 | 520 | 817 | 0 | 0 | 0 | 0 | 0 | 431 | 598 | 729 | 604 | [illegible] |
| SE | 10155 | ENSG00000073584 | SMARCE1 | chr17 | - | 40631591 | 40631711 | 40630713 | 40630924 | 40632194 | 40632367 | 219 | 262 | 266 | 280 | 387 | 0 | 0 | 0 | 0 | 0 | 243 | 301 | 389 | 339 | [illegible] |
| SE | 10156 | ENSG00000073584 | SMARCE1 | chr17 | - | 40632194 | 40632367 | 40630713 | 40631693 | 40635930 | 40636098 | 471 | 364 | 482 | 494 | 754 | 5 | 0 | 0 | 1 | 0 | 347 | 558 | 589 | 440 | [illegible] |
| SE | 10158 | ENSG00000073584 | SMARCE1 | chr17 | - | 40642117 | 40642559 | 40637491 | 40637572 | 40645575 | 40645619 | 71 | 51 | 75 | 101 | 99 | 0 | 0 | 2 | 1 | 0 | 25 | 82 | 100 | 56 | [illegible] |
| SE | 10159 | ENSG00000073584 | SMARCE1 | chr17 | - | 40642454 | 40642559 | 40637095 | 40637572 | 40645575 | 40645619 | 204 | 151 | 194 | 238 | 248 | 0 | 0 | 2 | 1 | 0 | 87 | 189 | 212 | 164 | [illegible] |
| SE | 10160 | ENSG00000073584 | SMARCE1 | chr17 | - | 40642454 | 40642559 | 40637095 | 40637572 | 40645795 | 40645847 | 93 | 77 | 95 | 109 | 118 | 0 | 1 | 0 | 0 | 0 | 40 | 74 | 91 | 88 | [illegible] |
| SE | 10163 | ENSG00000073584 | SMARCE1 | chr17 | - | 40645575 | 40645619 | 40637491 | 40637572 | 40645795 | 40645836 | 5 | 5 | 11 | 7 | 6 | 0 | 1 | 0 | 0 | 0 | 9 | 0 | 6 | 3 | [illegible] |
| SE | 10165 | ENSG00000073584 | SMARCE1 | chr17 | - | 40645575 | 40645619 | 40640573 | 40642559 | 40645795 | 40645836 | 116 | 80 | 110 | 135 | 141 | 0 | 0 | 2 | 0 | 0 | 56 | 115 | 129 | 79 | [illegible] |
| SE | 10166 | ENSG00000073584 | SMARCE1 | chr17 | - | 40645575 | 40645619 | 40640573 | 40642559 | 40647772 | 40647833 | 40 | 25 | 26 | 28 | 41 | 2 | 0 | 0 | 0 | 0 | 22 | 33 | 33 | 20 | [illegible] |
| SE | 10169 | ENSG00000073584 | SMARCE1 | chr17 | - | 40645795 | 40645847 | 40642895 | 40645619 | 40647772 | 40647814 | 83 | 55 | 93 | 115 | 108 | 0 | 0 | 0 | 0 | 0 | 25 | 89 | 97 | 57 | [illegible] |
| SE | 10170 | ENSG00000005884 | ITGA3 | chr17 | + | 50068055 | 50068305 | 50064527 | 50064607 | 50070843 | 50070930 | 0 | 9 | 12 | 22 | 1 | 0 | 0 | 1 | 0 | 0 | 0 | 9 | 14 | 0 | [illegible] |
| SE | 10171 | ENSG00000005884 | ITGA3 | chr17 | + | 50081096 | 50081193 | 50080316 | 50080375 | 50081309 | 50081380 | 0 | 0 | 2 | 0 | 0 | 6 | 1 | 6 | 2 | 0 | 0 | 0 | 0 | 3 | [illegible] |
| SE | 10175 | ENSG00000108846 | ABCC3 | chr17 | + | 50660922 | 50661114 | 50659181 | 50659368 | 50667553 | 50667757 | 15 | 12 | 4 | 12 | 5 | 0 | 0 | 0 | 0 | 2 | 0 | 8 | 6 | 6 | [illegible] |
| SE | 10177 | ENSG00000108846 | ABCC3 | chr17 | + | 50665152 | 50665245 | 50659181 | 50659368 | 50667553 | 50667594 | 9 | 8 | 4 | 22 | 38 | 0 | 0 | 0 | 0 | 2 | 70 | 16 | 25 | 5 | [illegible] |
| SE | 10178 | ENSG00000108846 | ABCC3 | chr17 | + | 50667862 | 50668009 | 50667553 | 50667757 | 50668429 | 50668517 | 28 | 51 | 9 | 123 | 99 | 0 | 0 | 0 | 0 | 0 | 280 | 64 | 72 | 75 | [illegible] |
| SE | 10181 | ENSG00000005243 | COPZ2 | chr17 | - | 48027675 | 48027789 | 48026166 | 48026475 | 48028471 | 48028510 | 1 | 0 | 0 | 6 | 0 | 4 | 0 | 1 | 0 | 6 | 4 | 2 | 3 | 0 | [illegible] |
| SE | 10182 | ENSG00000005243 | COPZ2 | chr17 | - | 48028471 | 48028510 | 48026166 | 48026475 | 48029124 | 48029176 | 7 | 0 | 1 | 0 | 6 | 0 | 0 | 0 | 1 | 0 | 0 | 9 | 2 | 1 | [illegible] |
| SE | 10184 | ENSG00000005243 | COPZ2 | chr17 | - | 48032155 | 48032233 | 48029124 | 48029176 | 48033210 | 48033252 | 2 | 0 | 0 | 3 | 3 | 0 | 0 | 0 | 0 | 0 | 0 | 2 | 0 | 0 | [illegible] |
| SE | 10185 | ENSG00000005243 | COPZ2 | chr17 | - | 48032685 | 48032741 | 48029124 | 48029176 | 48033210 | 48033282 | 5 | 2 | 0 | 1 | 1 | 0 | 0 | 0 | 0 | 0 | 5 | 12 | 8 | 6 | [illegible] |
| SE | 10186 | ENSG00000005243 | COPZ2 | chr17 | - | 48032685 | 48032741 | 48029124 | 48029176 | 48033862 | 48033944 | 0 | 0 | 0 | 0 | 6 | 0 | 0 | 0 | 1 | 0 | 0 | 5 | 0 | 0 | [illegible] |
| SE | 10188 | ENSG00000005243 | COPZ2 | chr17 | - | 48033210 | 48033302 | 48029124 | 48029176 | 48033862 | 48033872 | 5 | 0 | 1 | 4 | 0 | 0 | 0 | 0 | 1 | 0 | 0 | 1 | 0 | 5 | [illegible] |
| SE | 10192 | ENSG00000178852 | EFCAB13 | chr17 | + | 47324453 | 47324523 | 47323954 | 47324074 | 47326225 | 47326387 | 1 | 3 | 4 | 13 | 8 | 0 | 0 | 2 | 2 | 0 | 0 | 1 | 5 | 2 | [illegible] |
| SE | 10193 | ENSG00000178852 | EFCAB13 | chr17 | + | 47326225 | 47326387 | 47324453 | 47324523 | 47328268 | 47328383 | 13 | 51 | 53 | 78 | 118 | 0 | 0 | 1 | 0 | 0 | 26 | 32 | 27 | 7 | [illegible] |
| SE | 10194 | ENSG00000178852 | EFCAB13 | chr17 | + | 47328268 | 47328383 | 47326225 | 47326387 | 47335195 | 47335276 | 30 | 118 | 104 | 122 | 273 | 0 | 0 | 2 | 0 | 0 | 74 | 74 | 33 | 25 | [illegible] |
| SE | 10196 | ENSG00000178852 | EFCAB13 | chr17 | + | 47335195 | 47335356 | 47328047 | 47328383 | 47341920 | 47342032 | 43 | 131 | 131 | 148 | 326 | 0 | 0 | 0 | 3 | 0 | 72 | 100 | 31 | 50 | [illegible] |
| SE | 10198 | ENSG00000178852 | EFCAB13 | chr17 | + | 47345015 | 47345098 | 47344161 | 47344292 | 47374471 | 47374639 | 4 | 15 | 21 | 19 | 56 | 0 | 0 | 0 | 0 | 2 | 40 | 19 | 3 | 3 | [illegible] |
| SE | 10199 | ENSG00000178852 | EFCAB13 | chr17 | + | 47347807 | 47347951 | 47345015 | 47345098 | 47361377 | 47361521 | 0 | 11 | 0 | 14 | 4 | 12 | 6 | 0 | 21 | 3 | 0 | 0 | 0 | 3 | [illegible] |
| SE | 10200 | ENSG00000178852 | EFCAB13 | chr17 | + | 47347807 | 47347951 | 47345015 | 47345098 | 47370436 | 47370508 | 0 | 9 | 0 | 2 | 4 | 0 | 15 | 16 | 24 | 75 | 0 | 0 | 0 | 1 | [illegible] |
| SE | 10203 | ENSG00000178852 | EFCAB13 | chr17 | + | 47361377 | 47361521 | 47347807 | 47347951 | 47370436 | 47370508 | 0 | 52 | 0 | 86 | 0 | 0 | 2 | 0 | 0 | 0 | 0 | 46 | 0 | 45 | [illegible] |
| SE | 10204 | ENSG00000178852 | EFCAB13 | chr17 | + | 47370436 | 47370508 | 47344161 | 47344292 | 47374471 | 47374638 | 0 | 54 | 28 | 79 | 49 | 0 | 0 | 0 | 0 | 2 | 0 | 22 | 7 | 35 | [illegible] |
| SE | 10205 | ENSG00000178852 | EFCAB13 | chr17 | + | 47370436 | 47370508 | 47345015 | 47345098 | 47374471 | 47374638 | 0 | 69 | 44 | 103 | 118 | 0 | 1 | 5 | 2 | 16 | 0 | 25 | 10 | 35 | [illegible] |
| SE | 10208 | ENSG00000178852 | EFCAB13 | chr17 | + | 47391436 | 47391580 | 47377653 | 47377903 | 47394024 | 47394099 | 7 | 14 | 24 | 37 | 51 | 0 | 1 | 0 | 0 | 1 | 0 | 4 | 7 | 0 | [illegible] |
| SE | 10209 | ENSG00000178852 | EFCAB13 | chr17 | + | 47409646 | 47409691 | 47404561 | 47404633 | 47414847 | 47414919 | 0 | 48 | 18 | 20 | 63 | 0 | 0 | 0 | 0 | 0 | 0 | 53 | 14 | 0 | [illegible] |
| SE | 10211 | ENSG00000178852 | EFCAB13 | chr17 | + | 47414847 | 47414919 | 47409646 | 47409691 | 47429817 | 47429961 | 9 | 16 | 5 | 38 | 27 | 0 | 0 | 0 | 2 | 0 | 18 | 20 | 5 | 16 | [illegible] |
| SE | 10212 | ENSG00000178852 | EFCAB13 | chr17 | + | 47429817 | 47429961 | 47414847 | 47414919 | 47440430 | 47441312 | 0 | 7 | 7 | 13 | 31 | 0 | 0 | 1 | 2 | 3 | 12 | 19 | 7 | 4 | [illegible] |
| SE | 10213 | ENSG00000167182 | SP2 | chr17 | + | 47897814 | 47897927 | 47896181 | 47896293 | 47909663 | 47909730 | 0 | 0 | 0 | 2 | 0 | 0 | 0 | 0 | 0 | 0 | 0 | 0 | 0 | 0 | [illegible] |
| SE | 10214 | ENSG00000167182 | SP2 | chr17 | + | 47897814 | 47897927 | 47896181 | 47896293 | 47914725 | 47914861 | 0 | 2 | 2 | 0 | 0 | 1 | 0 | 0 | 1 | 1 | 0 | 0 | 0 | 0 | [illegible] |
| SE | 10217 | ENSG00000167182 | SP2 | chr17 | + | 47909663 | 47909730 | 47896261 | 47896293 | 47914725 | 47914861 | 0 | 0 | 0 | 0 | 0 | 1 | 0 | 0 | 1 | 1 | 0 | 0 | 0 | 0 | [illegible] |
| SE | 10219 | ENSG00000167182 | SP2 | chr17 | + | 47909663 | 47909730 | 47897814 | 47897927 | 47914725 | 47914861 | 0 | 0 | 0 | 0 | 0 | 0 | 2 | 2 | 0 | 0 | 0 | 0 | 0 | 0 | [illegible] |
| SE | 10224 | ENSG00000167182 | SP2 | chr17 | + | 47914725 | 47914861 | 47909663 | 47909730 | 47915311 | 47915388 | 0 | 0 | 0 | 0 | 0 | 0 | 0 | 0 | 2 | 1 | 0 | 5 | 0 | 0 | [illegible] |
| SE | 10225 | ENSG00000167182 | SP2 | chr17 | + | 47916155 | 47917130 | 47915311 | 47915388 | 47922961 | 47923274 | 59 | 31 | 50 | 93 | 59 | 0 | 0 | 0 | 0 | 0 | 86 | 82 | 49 | 96 | [illegible] |
| SE | 10226 | ENSG00000108960 | MMD | chr17 | - | 55401468 | 55401538 | 55392621 | 55394534 | 55403766 | 55403868 | 225 | 302 | 106 | 425 | 274 | 0 | 0 | 0 | 2 | 0 | 377 | 290 | 200 | 412 | [illegible] |
| SE | 10227 | ENSG00000108960 | MMD | chr17 | - | 55401468 | 55401538 | 55392621 | 55394534 | 55407745 | 55407820 | 27 | 34 | 17 | 81 | 59 | 0 | 1 | 0 | 1 | 0 | 65 | 74 | 20 | 54 | [illegible] |
| SE | 10229 | ENSG00000108960 | MMD | chr17 | - | 55403766 | 55403868 | 55392621 | 55394534 | 55407745 | 55407820 | 224 | 309 | 100 | 406 | 234 | 0 | 1 | 0 | 1 | 0 | 384 | 244 | 182 | 385 | [illegible] |
| SE | 10230 | ENSG00000108960 | MMD | chr17 | - | 55403766 | 55403868 | 55392621 | 55394534 | 55411256 | 55411417 | 0 | 5 | 2 | 7 | 0 | 0 | 0 | 0 | 1 | 0 | 0 | 2 | 11 | 17 | [illegible] |
| SE | 10231 | ENSG00000108960 | MMD | chr17 | - | 55403766 | 55403868 | 55401468 | 55401538 | 55407745 | 55407820 | 421 | 574 | 185 | 739 | 447 | 0 | 0 | 0 | 0 | 5 | 696 | 451 | 358 | 734 | [illegible] |
| SE | 10232 | ENSG00000108960 | MMD | chr17 | - | 55403766 | 55403868 | 55401468 | 55401538 | 55411256 | 55411417 | 178 | 250 | 87 | 310 | 189 | 1 | 2 | 0 | 2 | 0 | 286 | 186 | 162 | 341 | [illegible] |
| SE | 10235 | ENSG00000108960 | MMD | chr17 | - | 55407745 | 55407820 | 55401468 | 55401538 | 55411256 | 55411417 | 50 | 63 | 12 | 74 | 47 | 1 | 2 | 0 | 2 | 0 | 125 | 35 | 36 | 69 | [illegible] |
| SE | 10237 | ENSG00000108960 | MMD | chr17 | - | 55411256 | 55411417 | 55403766 | 55403868 | 55414150 | 55414232 | 134 | 220 | 86 | 292 | 137 | 0 | 0 | 0 | 2 | 0 | 322 | 191 | 129 | 249 | [illegible] |
| SE | 10238 | ENSG00000123159 | GIPC1 | chr19 | - | 14478683 | 14478765 | 14477759 | 14478567 | 14479411 | 14479524 | 48 | 21 | 44 | 49 | 47 | 0 | 0 | 0 | 0 | 0 | 28 | 66 | 36 | 43 | [illegible] |
| SE | 10241 | ENSG00000123159 | GIPC1 | chr19 | - | 14482688 | 14483006 | 14480592 | 14480778 | 14492856 | 14492912 | 35 | 15 | 23 | 27 | 33 | 0 | 0 | 1 | 0 | 0 | 16 | 23 | 10 | 8 | [illegible] |
| SE | 10244 | ENSG00000123159 | GIPC1 | chr19 | - | 14491655 | 14491743 | 14480592 | 14480778 | 14492856 | 14492912 | 29 | 12 | 18 | 13 | 24 | 0 | 0 | 1 | 0 | 0 | 20 | 13 | 6 | 14 | [illegible] |
| SE | 10246 | ENSG00000123159 | GIPC1 | chr19 | - | 14491655 | 14491743 | 14482688 | 14483006 | 14492856 | 14492912 | 39 | 14 | 25 | 27 | 27 | 2 | 0 | 1 | 0 | 0 | 8 | 14 | 13 | 4 | [illegible] |
| SE | 10250 | ENSG00000123159 | GIPC1 | chr19 | - | 14492856 | 14492912 | 14491655 | 14491743 | 14496036 | 14496052 | 21 | 11 | 20 | 15 | 32 | 2 | 1 | 1 | 0 | 0 | 10 | 18 | 14 | 6 | [illegible] |
| SE | 10253 | ENSG00000267248 | NA | chr17 | - | 60128955 | 60129090 | 60126534 | 60126865 | 60132435 | 60132512 | 2 | 0 | 0 | 1 | 0 | 0 | 0 | 0 | 0 | 0 | 0 | 0 | 0 | 1 | [illegible] |
| SE | 10254 | ENSG00000267248 | NA | chr17 | - | 60128955 | 60129090 | 60126534 | 60126865 | 60134755 | 60135525 | 0 | 0 | 0 | 2 | 0 | 0 | 0 | 0 | 0 | 0 | 0 | 0 | 0 | 0 | [illegible] |
| SE | 10255 | ENSG00000267248 | NA | chr17 | - | 60132435 | 60132512 | 60126530 | 60126865 | 60134755 | 60134841 | 3 | 0 | 0 | 0 | 0 | 0 | 0 | 0 | 0 | 0 | 0 | 0 | 1 | 6 | [illegible] |
| SE | 10256 | ENSG00000267248 | NA | chr17 | - | 60132435 | 60132512 | 60128955 | 60129090 | 60134755 | 60134841 | 5 | 0 | 0 | 0 | 0 | 0 | 0 | 0 | 1 | 0 | 0 | 0 | 1 | 1 | [illegible] |
| SE | 10257 | ENSG00000263715 | LINC02210-CRHR1 | chr17 | + | 45672826 | 45672902 | 45629903 | 45630158 | 45807009 | 45807097 | 0 | 0 | 1 | 0 | 0 | 0 | 0 | 0 | 0 | 0 | 0 | 0 | 0 | 0 | [illegible] |
| SE | 10261 | ENSG00000108651 | UTP6 | chr17 | - | 31873377 | 31873487 | 31868045 | 31868112 | 31873672 | 31873753 | 261 | 155 | 196 | 230 | 186 | 0 | 0 | 2 | 0 | 0 | 138 | 211 | 84 | 168 | [illegible] |
| SE | 10262 | ENSG00000108651 | UTP6 | chr17 | - | 31875233 | 31875413 | 31873672 | 31873753 | 31878249 | 31878327 | 165 | 120 | 158 | 206 | 141 | 1 | 1 | 0 | 1 | 2 | 91 | 134 | 64 | 141 | [illegible] |
| SE | 10265 | ENSG00000108651 | UTP6 | chr17 | - | 31885979 | 31886061 | 31884423 | 31884505 | 31887235 | 31887313 | 169 | 158 | 120 | 190 | 201 | 0 | 2 | 0 | 0 | 0 | 199 | 79 | 103 | 173 | [illegible] |
| SE | 10266 | ENSG00000108651 | UTP6 | chr17 | - | 31889284 | 31889403 | 31887235 | 31887313 | 31892259 | 31892323 | 240 | 170 | 230 | 351 | 318 | 0 | 2 | 2 | 3 | 0 | 697 | 175 | 172 | 238 | [illegible] |
| SE | 10267 | ENSG00000108651 | UTP6 | chr17 | - | 31892259 | 31892323 | 31889284 | 31889403 | 31892746 | 31892794 | 164 | 104 | 141 | 220 | 186 | 1 | 0 | 0 | 0 | 0 | 378 | 93 | 81 | 134 | [illegible] |
| SE | 10276 | ENSG00000125648 | SLC25A23 | chr19 | - | 6455951 | 6456092 | 6454558 | 6454717 | 6456419 | 6456531 | 3 | 0 | 0 | 0 | 2 | 8 | 3 | 8 | 12 | 1 | 0 | 2 | 0 | 0 | [illegible] |
| SE | 10277 | ENSG00000176208 | ATAD5 | chr17 | + | 30860432 | 30860612 | 30858160 | 30858323 | 30865703 | 30865800 | 40 | 36 | 39 | 62 | 58 | 0 | 0 | 0 | 1 | 0 | 8 | 25 | 18 | 33 | [illegible] |
| SE | 10278 | ENSG00000176208 | ATAD5 | chr17 | + | 30865703 | 30865800 | 30860432 | 30860612 | 30868332 | 30868412 | 32 | 37 | 26 | 43 | 39 | 2 | 0 | 0 | 0 | 0 | 10 | 8 | 7 | 12 | [illegible] |
| SE | 10279 | ENSG00000176208 | ATAD5 | chr17 | + | 30876373 | 30876550 | 30869495 | 30869646 | 30877415 | 30877549 | 39 | 22 | 26 | 31 | 55 | 0 | 0 | 1 | 0 | 0 | 32 | 24 | 18 | 17 | [illegible] |
| SE | 10280 | ENSG00000176208 | ATAD5 | chr17 | + | 30879422 | 30879487 | 30878002 | 30878096 | 30887191 | 30887372 | 21 | 14 | 12 | 7 | 25 | 0 | 0 | 1 | 0 | 0 | 15 | 35 | 5 | 0 | [illegible] |
| SE | 10281 | ENSG00000176208 | ATAD5 | chr17 | + | 30893293 | 30894150 | 30892606 | 30892788 | 30894563 | 30894722 | 26 | 17 | 17 | 23 | 30 | 0 | 0 | 0 | 0 | 4 | 0 | 13 | 15 | 2 | [illegible] |
| SE | 10284 | ENSG00000108771 | DHX58 | chr17 | - | 42111724 | 42111893 | 42111295 | 42111497 | 42112112 | 42112205 | 65 | 60 | 84 | 74 | 52 | 0 | 0 | 0 | 2 | 0 | 241 | 150 | 42 | 220 | [illegible] |
| SE | 10285 | ENSG00000108771 | DHX58 | chr17 | - | 42111745 | 42111893 | 42111295 | 42111497 | 42112112 | 42112205 | 19 | 23 | 41 | 41 | 19 | 0 | 0 | 0 | 2 | 0 | 140 | 82 | 19 | 150 | [illegible] |
| SE | 10293 | ENSG00000168259 | DNAJC7 | chr17 | - | 41987818 | 41987910 | 41983562 | 41983636 | 41988731 | 41988896 | 465 | 412 | 429 | 417 | 882 | 0 | 0 | 0 | 0 | 0 | 1007 | 349 | 662 | 752 | [illegible] |
| SE | 10294 | ENSG00000168259 | DNAJC7 | chr17 | - | 41988731 | 41988896 | 41986565 | 41987910 | 41989403 | 41989557 | 557 | 456 | 556 | 551 | 1050 | 0 | 2 | 0 | 0 | 0 | 1266 | 505 | 865 | 977 | [illegible] |
| SE | 10296 | ENSG00000168259 | DNAJC7 | chr17 | - | 41992798 | 41992867 | 41990263 | 41990382 | 41994869 | 41994944 | 0 | 0 | 4 | 0 | 0 | 167 | 127 | 139 | 158 | 346 | 0 | 0 | 0 | 0 | [illegible] |
| SE | 10299 | ENSG00000168259 | DNAJC7 | chr17 | - | 42000481 | 42000570 | 41997114 | 41997239 | 42017339 | 42017345 | 214 | 134 | 155 | 136 | 129 | 0 | 0 | 0 | 0 | 0 | 262 | 164 | 176 | 256 | [illegible] |
| SE | 10310 | ENSG00000108848 | LUC7L3 | chr17 | + | 50723818 | 50723965 | 50719628 | 50719831 | 50736959 | 50737026 | 0 | 0 | 1 | 0 | 0 | 337 | 243 | 366 | 342 | 211 | 0 | 0 | 0 | 0 | [illegible] |
| SE | 10313 | ENSG00000108848 | LUC7L3 | chr17 | + | 50737286 | 50737479 | 50736959 | 50737026 | 50738140 | 50738199 | 94 | 59 | 75 | 81 | 63 | 0 | 0 | 0 | 2 | 0 | 31 | 25 | 23 | 17 | [illegible] |
| SE | 10319 | ENSG00000108848 | LUC7L3 | chr17 | + | 50741101 | 50741246 | 50740266 | 50740345 | 50741656 | 50741673 | 1665 | 1516 | 1843 | 1801 | 1295 | 0 | 0 | 0 | 0 | 0 | 515 | 692 | 537 | 488 | [illegible] |
| SE | 10320 | ENSG00000108848 | LUC7L3 | chr17 | + | 50741656 | 50741731 | 50740266 | 50740345 | 50743705 | 50743792 | 127 | 140 | 165 | 130 | 260 | 0 | 0 | 0 | 0 | 0 | 41 | 43 | 54 | 32 | [illegible] |
| SE | 10321 | ENSG00000108848 | LUC7L3 | chr17 | + | 50744651 | 50744813 | 50743705 | 50743810 | 50745719 | 50746003 | 416 | 575 | 537 | 498 | 1032 | 1 | 1 | 0 | 0 | 0 | 200 | 235 | 256 | 216 | [illegible] |
| SE | 10323 | ENSG00000108848 | LUC7L3 | chr17 | + | 50749218 | 50749344 | 50748388 | 50748416 | 50750500 | 50750550 | 35 | 27 | 39 | 52 | 34 | 2 | 2 | 0 | 1 | 2 | 0 | 18 | 6 | 0 | [illegible] |
| SE | 10324 | ENSG00000108848 | LUC7L3 | chr17 | + | 50749223 | 50749344 | 50746541 | 50746702 | 50750746 | 50751013 | 0 | 5 | 1 | 6 | 2 | 2 | 0 | 2 | 3 | 2 | 0 | 0 | 1 | 0 | [illegible] |
| SE | 10326 | ENSG00000108848 | LUC7L3 | chr17 | + | 50750500 | 50750641 | 50746541 | 50746702 | 50750746 | 50750852 | 373 | 417 | 488 | 474 | 706 | 2 | 0 | 2 | 3 | 2 | 32 | 221 | 181 | 91 | [illegible] |
| SE | 10327 | ENSG00000108848 | LUC7L3 | chr17 | + | 50751296 | 50751362 | 50750746 | 50750852 | 50752199 | 50752585 | 8 | 4 | 4 | 9 | 12 | 7 | 4 | 7 | 5 | 9 | 0 | 9 | 8 | 0 | [illegible] |
| SE | 10334 | ENSG00000188825 | LINC00910 | chr17 | - | 43356391 | 43356530 | 43341177 | 43341459 | 43377670 | 43377813 | 0 | 0 | 0 | 1 | 0 | 0 | 0 | 0 | 1 | 0 | 0 | 12 | 1 | 0 | [illegible] |
| SE | 10335 | ENSG00000188825 | LINC00910 | chr17 | - | 43377670 | 43377813 | 43356391 | 43356530 | 43379122 | 43379262 | 0 | 1 | 1 | 1 | 3 | 0 | 0 | 1 | 0 | 0 | 0 | 4 | 2 | 0 | [illegible] |
| SE | 10336 | ENSG00000188825 | LINC00910 | chr17 | - | 43379122 | 43379262 | 43375927 | 43377813 | 43381272 | 43381660 | 6 | 1 | 1 | 1 | 12 | 3 | 0 | 0 | 0 | 1 | 0 | 5 | 2 | 0 | [illegible] |
| SE | 10338 | ENSG00000188825 | LINC00910 | chr17 | - | 43387935 | 43388057 | 43381272 | 43381660 | 43388288 | 43388331 | 1 | 0 | 3 | 0 | 0 | 8 | 3 | 4 | 5 | 15 | 0 | 0 | 0 | 0 | [illegible] |
| SE | 10340 | ENSG00000198496 | NBR2 | chr17 | + | 43132598 | 43132723 | 43131207 | 43131270 | 43132955 | 43133137 | 5 | 0 | 2 | 1 | 1 | 4 | 1 | 1 | 1 | 2 | 0 | 0 | 0 | 0 | [illegible] |
| SE | 10341 | ENSG00000198496 | NBR2 | chr17 | + | 43132955 | 43133137 | 43131207 | 43131270 | 43138155 | 43138360 | 4 | 1 | 1 | 1 | 2 | 0 | 2 | 0 | 0 | 0 | 0 | 4 | 0 | 2 | [illegible] |
| SE | 10343 | ENSG00000198496 | NBR2 | chr17 | + | 43138155 | 43138360 | 43131207 | 43131270 | 43138656 | 43138922 | 0 | 2 | 0 | 0 | 0 | 11 | 12 | 14 | 12 | 15 | 0 | 0 | 0 | 0 | [illegible] |
| SE | 10345 | ENSG00000228782 | MRPL45P2 | chr17 | - | 47486762 | 47486884 | 47482991 | 47483090 | 47490165 | 47490295 | 0 | 0 | 1 | 0 | 2 | 5 | 12 | 15 | 5 | 5 | 0 | 0 | 0 | 0 | [illegible] |
| SE | 10347 | ENSG00000131467 | PSME3 | chr17 | + | 42833775 | 42833939 | 42833417 | 42833673 | 42834093 | 42834256 | 0 | 0 | 0 | 0 | 0 | 0 | 1 | 0 | 0 | 0 | 0 | 0 | 0 | 0 | [illegible] |
| SE | 10348 | ENSG00000131467 | PSME3 | chr17 | + | 42833775 | 42833939 | 42833417 | 42833673 | 42834343 | 42834376 | 3 | 0 | 3 | 0 | 0 | 97 | 47 | 70 | 99 | 69 | 0 | 2 | 0 | 0 | [illegible] |
| SE | 10350 | ENSG00000131467 | PSME3 | chr17 | + | 42834093 | 42834256 | 42833775 | 42833939 | 42834343 | 42834376 | 2 | 0 | 1 | 1 | 2 | 3 | 0 | 3 | 0 | 0 | 0 | 2 | 1 | 7 | [illegible] |
| SE | 10354 | ENSG00000131467 | PSME3 | chr17 | + | 42838092 | 42838205 | 42837648 | 42837697 | 42838944 | 42839012 | 156 | 92 | 111 | 133 | 210 | 0 | 0 | 0 | 0 | 0 | 256 | 163 | 105 | 186 | [illegible] |
| SE | 10356 | ENSG00000131467 | PSME3 | chr17 | + | 42838730 | 42838799 | 42837648 | 42837697 | 42838944 | 42838976 | 182 | 113 | 161 | 146 | 156 | 0 | 0 | 0 | 0 | 0 | 232 | 205 | 133 | 206 | [illegible] |
| SE | 10357 | ENSG00000131467 | PSME3 | chr17 | + | 42838730 | 42838799 | 42838092 | 42838205 | 42838944 | 42838976 | 403 | 246 | 352 | 369 | 480 | 4 | 0 | 0 | 0 | 1 | 565 | 435 | 295 | 454 | [illegible] |
| SE | 10358 | ENSG00000131467 | PSME3 | chr17 | + | 42839111 | 42839166 | 42838944 | 42839012 | 42839293 | 42839303 | 374 | 212 | 304 | 311 | 401 | 0 | 0 | 1 | 0 | 0 | 482 | 347 | 268 | 320 | [illegible] |
| SE | 10363 | ENSG00000197381 | ADARB1 | chr21 | + | 45132214 | 45132306 | 45128401 | 45128573 | 45133558 | 45133682 | 0 | 0 | 0 | 2 | 0 | 1 | 0 | 0 | 0 | 3 | 0 | 0 | 2 | 0 | [illegible] |
| SE | 10365 | ENSG00000197381 | ADARB1 | chr21 | + | 45133558 | 45133682 | 45128401 | 45128573 | 45134714 | 45134832 | 1 | 0 | 0 | 0 | 0 | 14 | 4 | 13 | 10 | 25 | 0 | 0 | 0 | 1 | [illegible] |
| SE | 10370 | ENSG00000197381 | ADARB1 | chr21 | + | 45144630 | 45144962 | 45134714 | 45134832 | 45171609 | 45171759 | 2 | 1 | 0 | 0 | 1 | 12 | 4 | 16 | 20 | 16 | 10 | 0 | 3 | 0 | [illegible] |
| SE | 10371 | ENSG00000197381 | ADARB1 | chr21 | + | 45171609 | 45171684 | 45128401 | 45128573 | 45175729 | 45176126 | 20 | 22 | 34 | 37 | 39 | 1 | 0 | 0 | 0 | 0 | 55 | 37 | 23 | 59 | [illegible] |
| SE | 10372 | ENSG00000197381 | ADARB1 | chr21 | + | 45175729 | 45176664 | 45171609 | 45171684 | 45180329 | 45180444 | 43 | 38 | 69 | 68 | 75 | 0 | 2 | 0 | 0 | 2 | 92 | 74 | 35 | 60 | [illegible] |
| SE | 10373 | ENSG00000197381 | ADARB1 | chr21 | + | 45181479 | 45181516 | 45180435 | 45180444 | 45182584 | 45182753 | 0 | 0 | 0 | 2 | 1 | 32 | 20 | 40 | 47 | 45 | 0 | 0 | 0 | 0 | [illegible] |
| SE | 10375 | ENSG00000197381 | ADARB1 | chr21 | + | 45204554 | 45204736 | 45184922 | 45185091 | 45220835 | 45221014 | 41 | 29 | 50 | 56 | 44 | 0 | 0 | 0 | 1 | 2 | 50 | 31 | 17 | 20 | [illegible] |
| SE | 10376 | ENSG00000126246 | IGFLR1 | chr19 | - | 35739709 | 35739769 | 35739214 | 35739626 | 35740379 | 35740549 | 22 | 9 | 14 | 18 | 15 | 0 | 2 | 0 | 0 | 0 | 24 | 11 | 10 | 15 | [illegible] |
| SE | 10378 | ENSG00000126246 | IGFLR1 | chr19 | - | 35739709 | 35740086 | 35739256 | 35739626 | 35740379 | 35740564 | 69 | 14 | 32 | 44 | 18 | 0 | 2 | 0 | 0 | 0 | 50 | 28 | 19 | 32 | [illegible] |
| SE | 10380 | ENSG00000126246 | IGFLR1 | chr19 | - | 35740379 | 35740564 | 35739709 | 35740086 | 35741023 | 35741223 | 26 | 5 | 16 | 14 | 26 | 0 | 0 | 1 | 1 | 0 | 18 | 20 | 12 | 3 | [illegible] |
| SE | 10386 | ENSG00000108773 | KAT2A | chr17 | - | 42117906 | 42118017 | 42117677 | 42117814 | 42118296 | 42118403 | 39 | 16 | 43 | 23 | 21 | 0 | 0 | 1 | 0 | 3 | 21 | 51 | 13 | 19 | [illegible] |
| SE | 10387 | ENSG00000108773 | KAT2A | chr17 | - | 42120705 | 42120829 | 42120224 | 42120370 | 42120965 | 42121058 | 55 | 29 | 38 | 73 | 39 | 0 | 0 | 0 | 1 | 0 | 63 | 31 | 10 | 51 | [illegible] |
| SE | 10391 | ENSG00000239704 | CDRT4 | chr17 | - | 15440207 | 15440285 | 15436014 | 15438200 | 15503097 | 15503608 | 10 | 9 | 5 | 6 | 14 | 1 | 0 | 1 | 2 | 0 | 0 | 1 | 10 | 0 | [illegible] |
| SE | 10394 | ENSG00000159210 | SNF8 | chr17 | - | 48931642 | 48931717 | 48929315 | 48930612 | 48936169 | 48936242 | 32 | 18 | 26 | 41 | 31 | 2 | 0 | 0 | 0 | 0 | 99 | 29 | 56 | 76 | [illegible] |
| SE | 10395 | ENSG00000159210 | SNF8 | chr17 | - | 48933204 | 48933346 | 48929315 | 48930612 | 48936169 | 48936242 | 76 | 51 | 67 | 83 | 84 | 2 | 0 | 0 | 0 | 0 | 151 | 76 | 61 | 119 | [illegible] |
| SE | 10396 | ENSG00000159210 | SNF8 | chr17 | - | 48933204 | 48933346 | 48929315 | 48930612 | 48937019 | 48937124 | 0 | 1 | 1 | 0 | 0 | 0 | 1 | 0 | 0 | 0 | 15 | 0 | 0 | 9 | [illegible] |
| SE | 10397 | ENSG00000159210 | SNF8 | chr17 | - | 48933204 | 48933346 | 48931642 | 48931717 | 48936169 | 48936242 | 187 | 117 | 129 | 160 | 210 | 0 | 0 | 0 | 4 | 0 | 384 | 152 | 145 | 223 | [illegible] |
| SE | 10398 | ENSG00000159210 | SNF8 | chr17 | - | 48934478 | 48934576 | 48933204 | 48933346 | 48936169 | 48936242 | 0 | 0 | 0 | 0 | 0 | 83 | 59 | 72 | 91 | 84 | 0 | 0 | 1 | 0 | [illegible] |
| SE | 10399 | ENSG00000159210 | SNF8 | chr17 | - | 48936130 | 48936242 | 48933204 | 48933346 | 48937019 | 48937124 | 95 | 90 | 109 | 134 | 140 | 0 | 1 | 0 | 0 | 0 | 368 | 203 | 108 | 233 | [illegible] |
| SE | 10400 | ENSG00000159210 | SNF8 | chr17 | - | 48936169 | 48936242 | 48929315 | 48930612 | 48937019 | 48937124 | 97 | 90 | 109 | 133 | 134 | 0 | 1 | 0 | 0 | 0 | 368 | 200 | 108 | 232 | [illegible] |
| SE | 10401 | ENSG00000159210 | SNF8 | chr17 | - | 48936169 | 48936242 | 48933204 | 48933346 | 48937019 | 48937124 | 176 | 149 | 180 | 223 | 212 | 0 | 1 | 0 | 0 | 0 | 532 | 283 | 181 | 346 | [illegible] |
| SE | 10405 | ENSG00000159210 | SNF8 | chr17 | - | 48940923 | 48941062 | 48936655 | 48937124 | 48943924 | 48943975 | 284 | 227 | 274 | 335 | 313 | 0 | 0 | 0 | 2 | 0 | 784 | 439 | 243 | 503 | [illegible] |
| SE | 10409 | ENSG00000282851 | BISPR | chr19 | + | 17413220 | 17413305 | 17406076 | 17406296 | 17415199 | 17415734 | 1 | 0 | 0 | 2 | 0 | 1 | 0 | 4 | 2 | 1 | 0 | 6 | 3 | 0 | [illegible] |
| SE | 10410 | ENSG00000187607 | ZNF286A | chr17 | + | 15700047 | 15700366 | 15699576 | 15699777 | 15701151 | 15701240 | 14 | 13 | 11 | 24 | 10 | 3 | 0 | 1 | 0 | 0 | 15 | 25 | 15 | 0 | [illegible] |
| SE | 10411 | ENSG00000187607 | ZNF286A | chr17 | + | 15700134 | 15700366 | 15699576 | 15699777 | 15701151 | 15701240 | 17 | 10 | 9 | 21 | 11 | 3 | 0 | 1 | 0 | 0 | 30 | 37 | 5 | 0 | [illegible] |
| SE | 10412 | ENSG00000187607 | ZNF286A | chr17 | + | 15700207 | 15700366 | 15699743 | 15699777 | 15701151 | 15701240 | 14 | 8 | 8 | 20 | 12 | 3 | 0 | 1 | 0 | 0 | 15 | 25 | 5 | 0 | [illegible] |
| SE | 10413 | ENSG00000187607 | ZNF286A | chr17 | + | 15706386 | 15706501 | 15701151 | 15701240 | 15708154 | 15708247 | 66 | 44 | 74 | 59 | 71 | 0 | 1 | 2 | 0 | 0 | 0 | 39 | 34 | 9 | [illegible] |
| SE | 10416 | ENSG00000173786 | CNP | chr17 | + | 41968067 | 41968740 | 41966794 | 41966887 | 41971891 | 41972031 | 133 | 73 | 91 | 138 | 60 | 0 | 2 | 0 | 2 | 1 | 287 | 156 | 68 | 160 | [illegible] |
| SE | 10417 | ENSG00000173786 | CNP | chr17 | + | 41968072 | 41968740 | 41966814 | 41966887 | 41971891 | 41972031 | 121 | 62 | 80 | 126 | 56 | 0 | 2 | 0 | 2 | 1 | 287 | 127 | 63 | 148 | [illegible] |
| SE | 10418 | ENSG00000173786 | CNP | chr17 | + | 41971891 | 41972031 | 41968067 | 41968740 | 41973474 | 41973817 | 273 | 205 | 223 | 349 | 311 | 0 | 2 | 0 | 0 | 0 | 854 | 354 | 216 | 609 | [illegible] |
| SE | 10425 | ENSG00000126581 | BECN1 | chr17 | - | 42812230 | 42812356 | 42811654 | 42811797 | 42813947 | 42813989 | 3 | 0 | 1 | 0 | 6 | 220 | 126 | 169 | 209 | 221 | 0 | 3 | 4 | 1 | [illegible] |
| SE | 10426 | ENSG00000126581 | BECN1 | chr17 | - | 42812230 | 42812356 | 42811654 | 42811797 | 42814523 | 42815014 | 0 | 0 | 1 | 0 | 0 | 1 | 0 | 2 | 0 | 0 | 0 | 0 | 2 | 0 | [illegible] |
| SE | 10428 | ENSG00000126581 | BECN1 | chr17 | - | 42813947 | 42814008 | 42811654 | 42811797 | 42814523 | 42814673 | 263 | 155 | 191 | 244 | 266 | 1 | 0 | 2 | 0 | 0 | 334 | 207 | 130 | 200 | [illegible] |
| SE | 10429 | ENSG00000126581 | BECN1 | chr17 | - | 42814523 | 42814673 | 42810309 | 42810928 | 42815907 | 42816054 | 139 | 101 | 103 | 139 | 171 | 0 | 0 | 0 | 0 | 0 | 316 | 163 | 246 | 231 | [illegible] |
| SE | 10435 | ENSG00000126581 | BECN1 | chr17 | - | 42815907 | 42816054 | 42813547 | 42814673 | 42823747 | 42823879 | 120 | 87 | 88 | 116 | 157 | 0 | 0 | 0 | 0 | 0 | 213 | 131 | 135 | 144 | [illegible] |
| SE | 10436 | ENSG00000126581 | BECN1 | chr17 | - | 42818220 | 42818415 | 42810669 | 42810928 | 42823747 | 42823879 | 0 | 0 | 0 | 1 | 0 | 0 | 0 | 0 | 0 | 0 | 5 | 4 | 1 | 2 | [illegible] |
| SE | 10437 | ENSG00000126581 | BECN1 | chr17 | - | 42818220 | 42818415 | 42814598 | 42814673 | 42823747 | 42823879 | 2 | 0 | 1 | 2 | 0 | 0 | 0 | 0 | 0 | 0 | 4 | 4 | 5 | 14 | [illegible] |
| SE | 10438 | ENSG00000126581 | BECN1 | chr17 | - | 42818220 | 42818415 | 42815669 | 42816054 | 42823747 | 42823879 | 130 | 87 | 88 | 153 | 223 | 0 | 0 | 0 | 0 | 0 | 275 | 185 | 155 | 124 | [illegible] |
| SE | 10440 | ENSG00000126581 | BECN1 | chr17 | - | 42818543 | 42819609 | 42818220 | 42818415 | 42823747 | 42823879 | 189 | 126 | 145 | 177 | 230 | 0 | 0 | 0 | 0 | 0 | 401 | 258 | 209 | 305 | [illegible] |
| SE | 10441 | ENSG00000126581 | BECN1 | chr17 | - | 42818786 | 42818877 | 42818543 | 42818680 | 42823747 | 42819609 | 249 | 153 | 212 | 240 | 199 | 2 | 0 | 0 | 0 | 2 | 507 | 400 | 239 | 319 | [illegible] |
| SE | 10442 | ENSG00000126581 | BECN1 | chr17 | - | 42819547 | 42819609 | 42818220 | 42818415 | 42823747 | 42823879 | 0 | 0 | 1 | 0 | 1 | 0 | 0 | 0 | 0 | 0 | 0 | 4 | 0 | 2 | [illegible] |
| SE | 10443 | ENSG00000126581 | BECN1 | chr17 | - | 42820773 | 42820841 | 42818543 | 42819609 | 42823747 | 42823879 | 287 | 165 | 299 | 309 | 244 | 0 | 1 | 1 | 0 | 1 | 359 | 420 | 286 | 301 | [illegible] |
| SE | 10448 | ENSG00000214425 | LRRC37A4P | chr17 | - | 45521306 | 45521425 | 45510243 | 45510293 | 45539162 | 45539243 | 14 | 27 | 39 | 44 | 38 | 2 | 2 | 0 | 0 | 0 | 18 | 42 | 49 | 45 | [illegible] |
| SE | 10450 | ENSG00000214425 | LRRC37A4P | chr17 | - | 45539891 | 45539963 | 45539162 | 45539243 | 45542763 | 45542835 | 22 | 24 | 26 | 35 | 45 | 0 | 1 | 0 | 0 | 0 | 39 | 29 | 37 | 19 | [illegible] |
| SE | 10455 | ENSG00000141698 | NT5C3B | chr17 | - | 41827425 | 41827626 | 41825056 | 41825657 | 41830800 | 41830890 | 20 | 15 | 36 | 26 | 23 | 0 | 1 | 0 | 0 | 0 | 25 | 58 | 21 | 50 | [illegible] |
| SE | 10456 | ENSG00000141698 | NT5C3B | chr17 | - | 41828789 | 41828952 | 41825199 | 41825657 | 41830800 | 41830890 | 34 | 30 | 54 | 56 | 49 | 0 | 1 | 0 | 0 | 0 | 11 | 72 | 16 | 88 | [illegible] |
| SE | 10458 | ENSG00000141698 | NT5C3B | chr17 | - | 41830800 | 41830890 | 41828789 | 41828952 | 41832391 | 41832477 | 43 | 37 | 83 | 83 | 77 | 2 | 0 | 0 | 2 | 0 | 20 | 116 | 29 | 121 | [illegible] |
| SE | 10459 | ENSG00000141698 | NT5C3B | chr17 | - | 41832607 | 41832703 | 41832391 | 41832477 | 41835069 | 41835116 | 0 | 3 | 2 | 0 | 0 | 26 | 14 | 37 | 30 | 50 | 0 | 0 | 0 | 0 | [illegible] |
| SE | 10460 | ENSG00000141698 | NT5C3B | chr17 | - | 41835587 | 41835642 | 41835202 | 41835272 | 41835858 | 41836227 | 0 | 0 | 0 | 1 | 2 | 3 | 3 | 3 | 6 | 15 | 0 | 5 | 0 | 9 | [illegible] |
| SE | 10461 | ENSG00000141698 | NT5C3B | chr17 | - | 41835587 | 41835682 | 41835202 | 41835272 | 41835858 | 41835957 | 0 | 0 | 0 | 2 | 2 | 3 | 3 | 3 | 6 | 15 | 0 | 6 | 0 | 8 | [illegible] |
| SE | 10465 | ENSG00000266088 | NA | chr17 | + | 40523464 | 40523655 | 40520714 | 40520824 | 40526443 | 40527000 | 22 | 27 | 18 | 38 | 25 | 0 | 0 | 0 | 0 | 1 | 39 | 24 | 8 | 27 | [illegible] |
| SE | 10468 | ENSG00000186566 | GPATCH8 | chr17 | - | 44405920 | 44406051 | 44395280 | 44401453 | 44424348 | 44424492 | 384 | 280 | 344 | 449 | 627 | 0 | 0 | 0 | 1 | 0 | 391 | 465 | 442 | 407 | [illegible] |
| SE | 10469 | ENSG00000186566 | GPATCH8 | chr17 | - | 44407216 | 44407266 | 44405920 | 44406051 | 44424348 | 44424492 | 4 | 0 | 0 | 2 | 0 | 243 | 170 | 205 | 252 | 449 | 0 | 0 | 0 | 0 | [illegible] |
| SE | 10470 | ENSG00000186566 | GPATCH8 | chr17 | - | 44424348 | 44424492 | 44405920 | 44406051 | 44435064 | 44435151 | 445 | 305 | 407 | 464 | 823 | 1 | 0 | 0 | 1 | 0 | 374 | 459 | 386 | 461 | [illegible] |
| SE | 10471 | ENSG00000186566 | GPATCH8 | chr17 | - | 44435064 | 44435151 | 44424348 | 44424492 | 44436477 | 44436545 | 361 | 242 | 359 | 374 | 654 | 0 | 0 | 1 | 0 | 0 | 304 | 459 | 380 | 415 | [illegible] |
| SE | 10472 | ENSG00000186566 | GPATCH8 | chr17 | - | 44435064 | 44435151 | 44424348 | 44424492 | 44464471 | 44464544 | 115 | 68 | 102 | 100 | 215 | 0 | 0 | 0 | 0 | 0 | 83 | 110 | 105 | 105 | [illegible] |
| SE | 10473 | ENSG00000186566 | GPATCH8 | chr17 | - | 44436477 | 44436545 | 44424348 | 44424492 | 44464471 | 44464544 | 71 | 27 | 69 | 90 | 132 | 0 | 0 | 0 | 0 | 0 | 76 | 77 | 38 | 72 | [illegible] |
| SE | 10475 | ENSG00000186566 | GPATCH8 | chr17 | - | 44436477 | 44436545 | 44435064 | 44435151 | 44464471 | 44464544 | 283 | 163 | 284 | 341 | 502 | 2 | 0 | 0 | 0 | 0 | 273 | 332 | 223 | 325 | [illegible] |
| SE | 10481 | ENSG00000186566 | GPATCH8 | chr17 | - | 44464471 | 44464544 | 44438670 | 44438715 | 44474828 | 44474903 | 49 | 39 | 44 | 70 | 97 | 0 | 0 | 0 | 1 | 0 | 46 | 62 | 35 | 33 | [illegible] |
| SE | 10486 | ENSG00000186566 | GPATCH8 | chr17 | - | 44474828 | 44474903 | 44464471 | 44464544 | 44503325 | 44503389 | 151 | 65 | 154 | 184 | 255 | 0 | 0 | 0 | 1 | 0 | 149 | 176 | 111 | 143 | [illegible] |
| SE | 10490 | ENSG00000101407 | TTI1 | chr20 | - | 38019004 | 38019144 | 38011514 | 38013857 | 38033403 | 38033461 | 1 | 0 | 4 | 0 | 0 | 27 | 5 | 24 | 18 | 35 | 7 | 8 | 2 | 0 | [illegible] |
| SE | 10496 | ENSG00000141696 | P3H4 | chr17 | - | 41803286 | 41803431 | 41801946 | 41802979 | 41806795 | 41806848 | 20 | 1 | 8 | 12 | 6 | 0 | 0 | 0 | 2 | 0 | 6 | 9 | 6 | 7 | [illegible] |
| SE | 10497 | ENSG00000141696 | P3H4 | chr17 | - | 41807858 | 41808004 | 41806795 | 41806879 | 41809705 | 41809834 | 7 | 0 | 8 | 11 | 8 | 0 | 0 | 1 | 0 | 0 | 0 | 3 | 0 | 2 | [illegible] |
| SE | 10498 | ENSG00000141696 | P3H4 | chr17 | - | 41807858 | 41808004 | 41806795 | 41806879 | 41810862 | 41811034 | 9 | 0 | 3 | 8 | 4 | 0 | 1 | 0 | 0 | 0 | 0 | 3 | 0 | 2 | [illegible] |
| SE | 10499 | ENSG00000141696 | P3H4 | chr17 | - | 41809705 | 41809834 | 41806795 | 41806879 | 41810862 | 41811034 | 0 | 0 | 5 | 4 | 0 | 0 | 1 | 0 | 0 | 0 | 0 | 4 | 7 | 0 | [illegible] |
| SE | 10500 | ENSG00000141696 | P3H4 | chr17 | - | 41809705 | 41809834 | 41807858 | 41808004 | 41810862 | 41811034 | 1 | 0 | 9 | 7 | 4 | 3 | 0 | 0 | 0 | 0 | 0 | 4 | 7 | 0 | [illegible] |
| SE | 10501 | ENSG00000172057 | ORMDL3 | chr17 | - | 39924754 | 39924805 | 39924029 | 39924225 | 39925168 | 39925371 | 0 | 0 | 0 | 0 | 0 | 0 | 1 | 0 | 0 | 0 | 0 | 7 | 0 | 0 | [illegible] |
| SE | 10502 | ENSG00000172057 | ORMDL3 | chr17 | - | 39925168 | 39925371 | 39924029 | 39924220 | 39925622 | 39926841 | 0 | 0 | 1 | 2 | 0 | 1 | 2 | 0 | 0 | 0 | 0 | 2 | 3 | 0 | [illegible] |
| SE | 10503 | ENSG00000172057 | ORMDL3 | chr17 | - | 39925168 | 39925371 | 39924029 | 39924225 | 39925622 | 39926841 | 0 | 1 | 1 | 2 | 0 | 0 | 0 | 0 | 1 | 0 | 0 | 0 | 3 | 0 | [illegible] |
| SE | 10504 | ENSG00000167914 | GSDMA | chr17 | + | 39973809 | 39973830 | 39972586 | 39972613 | 39974899 | 39975014 | 0 | 0 | 0 | 0 | 0 | 4 | 1 | 0 | 0 | 0 | 0 | 0 | 0 | 0 | [illegible] |
| SE | 10505 | ENSG00000167914 | GSDMA | chr17 | + | 39974272 | 39974427 | 39973809 | 39973830 | 39974899 | 39975014 | 1 | 2 | 4 | 2 | 9 | 0 | 0 | 0 | 0 | 0 | 0 | 0 | 0 | 3 | [illegible] |
| SE | 10515 | ENSG00000091536 | MYO15A | chr17 | + | 18166360 | 18166521 | 18163741 | 18163838 | 18167589 | 18167723 | 0 | 0 | 1 | 1 | 0 | 0 | 0 | 0 | 0 | 0 | 0 | 0 | 0 | 4 | [illegible] |
| SE | 10520 | ENSG00000125970 | RALY | chr20 | + | 34031521 | 34031604 | 33993645 | 33994131 | 34065043 | 34065160 | 25 | 7 | 8 | 14 | 3 | 0 | 0 | 1 | 0 | 0 | 24 | 25 | 13 | 29 | [illegible] |
| SE | 10527 | ENSG00000125970 | RALY | chr20 | + | 34075873 | 34076040 | 34073818 | 34073866 | 34076701 | 34076815 | 276 | 79 | 154 | 231 | 124 | 0 | 0 | 0 | 0 | 6 | 187 | 133 | 126 | 205 | [illegible] |
| SE | 10528 | ENSG00000131242 | RAB11FIP4 | chr17 | + | 31509076 | 31509481 | 31488117 | 31488272 | 31517650 | 31517708 | 0 | 0 | 0 | 0 | 0 | 9 | 6 | 5 | 11 | 1 | 0 | 0 | 0 | 1 | [illegible] |
| SE | 10529 | ENSG00000131242 | RAB11FIP4 | chr17 | + | 31517650 | 31517877 | 31487999 | 31488272 | 31521165 | 31521184 | 79 | 41 | 55 | 60 | 49 | 0 | 0 | 0 | 0 | 0 | 82 | 73 | 39 | 61 | [illegible] |
| SE | 10532 | ENSG00000131242 | RAB11FIP4 | chr17 | + | 31521165 | 31521360 | 31517650 | 31517877 | 31521914 | 31522049 | 175 | 92 | 133 | 119 | 136 | 0 | 0 | 0 | 3 | 0 | 156 | 179 | 104 | 168 | [illegible] |
| SE | 10534 | ENSG00000131242 | RAB11FIP4 | chr17 | + | 31525089 | 31525230 | 31523511 | 31523611 | 31527841 | 31527893 | 49 | 21 | 35 | 38 | 32 | 0 | 0 | 0 | 0 | 0 | 15 | 19 | 39 | 32 | [illegible] |
| SE | 10537 | ENSG00000213297 | ZNF625-ZNF20 | chr19 | - | 12145257 | 12146224 | 12142640 | 12142736 | 12147394 | 12147455 | 17 | 2 | 4 | 7 | 12 | 0 | 0 | 0 | 0 | 1 | 4 | 5 | 2 | 6 | [illegible] |
| SE | 10542 | ENSG00000177302 | TOP3A | chr17 | - | 18277674 | 18278357 | 18271427 | 18274980 | 18280535 | 18280658 | 110 | 56 | 71 | 82 | 86 | 0 | 0 | 0 | 0 | 0 | 118 | 100 | 67 | 56 | [illegible] |
| SE | 10547 | ENSG00000177302 | TOP3A | chr17 | - | 18285406 | 18285520 | 18285141 | 18285307 | 18290556 | 18290686 | 82 | 52 | 73 | 82 | 32 | 0 | 0 | 0 | 0 | 0 | 114 | 131 | 53 | 47 | [illegible] |
| SE | 10550 | ENSG00000177302 | TOP3A | chr17 | - | 18292644 | 18292852 | 18290841 | 18291027 | 18299558 | 18299633 | 46 | 15 | 31 | 41 | 29 | 1 | 0 | 0 | 0 | 1 | 43 | 66 | 29 | 35 | [illegible] |
| SE | 10551 | ENSG00000177302 | TOP3A | chr17 | - | 18294702 | 18294785 | 18290841 | 18291027 | 18299558 | 18299633 | 28 | 32 | 21 | 22 | 84 | 1 | 0 | 0 | 0 | 1 | 120 | 29 | 25 | 62 | [illegible] |
| SE | 10552 | ENSG00000177302 | TOP3A | chr17 | - | 18294702 | 18294785 | 18292644 | 18292852 | 18299558 | 18299633 | 116 | 102 | 87 | 103 | 282 | 0 | 1 | 0 | 0 | 0 | 371 | 130 | 117 | 171 | [illegible] |
| SE | 10553 | ENSG00000177302 | TOP3A | chr17 | - | 18299558 | 18299633 | 18294702 | 18294785 | 18301884 | 18301985 | 144 | 111 | 103 | 131 | 366 | 1 | 0 | 2 | 0 | 2 | 371 | 148 | 121 | 186 | [illegible] |
| SE | 10554 | ENSG00000177302 | TOP3A | chr17 | - | 18301884 | 18301985 | 18299558 | 18299633 | 18302263 | 18302361 | 104 | 65 | 82 | 90 | 240 | 0 | 0 | 0 | 0 | 0 | 258 | 138 | 83 | 126 | [illegible] |
| SE | 10557 | ENSG00000177302 | TOP3A | chr17 | - | 18307472 | 18307570 | 18306890 | 18306966 | 18308412 | 18308424 | 1 | 0 | 0 | 1 | 3 | 0 | 0 | 0 | 0 | 0 | 0 | 0 | 0 | 0 | [illegible] |
| SE | 10559 | ENSG00000177302 | TOP3A | chr17 | - | 18308350 | 18308424 | 18306549 | 18306966 | 18308881 | 18308941 | 126 | 79 | 62 | 100 | 177 | 1 | 1 | 0 | 0 | 3 | 251 | 144 | 110 | 106 | [illegible] |
| SE | 10560 | ENSG00000177302 | TOP3A | chr17 | - | 18308412 | 18308424 | 18306890 | 18306966 | 18308881 | 18308941 | 2 | 2 | 2 | 2 | 0 | 1 | 1 | 0 | 0 | 3 | 9 | 3 | 0 | 2 | [illegible] |

| | | | | | | | | | | | | | | | | | | | | | | | | | | | | | | | | | | | | | | | | | | | | | | |
|---|---|---|---|---|---|---|---|---|---|---|---|---|---|---|---|---|---|---|---|---|---|---|---|---|---|---|---|---|---|---|---|---|---|---|---|---|---|---|---|---|---|---|---|---|---|---|
| SE | 10566 | ENSG00000181481 | RNF135 | chr17 | + | 30987943 | 30988106 | 30984616 | 30984760 | 30993839 | 30993904 | 55 | 25 | 24 | 37 | 44 | 0 | 0 | 1 | 0 | 0 | 69 | 68 | 25 | 23 | 76 | 0 | 1 | 0 | 0 | 0 | 298 | 149 | 1 | 1 | 1 | 1 | 0.923 | 1 | 1 | 1 | 0.971 | 1 | 1 | 1 | -0.01 |
| SE | 10570 | ENSG00000188895 | MSL1 | chr17 | + | 40126182 | 40126406 | 40121970 | 40123380 | 40129244 | 40129260 | 727 | 625 | 891 | 926 | 1765 | 0 | 1 | 0 | 2 | 0 | 1159 | 873 | 755 | 1222 | 1161 | 0 | 0 | 1 | 0 | 2 | 298 | 149 | 1 | 1 | 1 | 0.997 | 1 | 0.996 | 1 | 1 | 1 | 0.997 | 1 | 0.997 | 0 |
| SE | 10573 | ENSG00000188895 | MSL1 | chr17 | + | 40133533 | 40133658 | 40133041 | 40133109 | 40133826 | 40133886 | 340 | 313 | 347 | 407 | 768 | 0 | 0 | 2 | 2 | 0 | 509 | 405 | 423 | 596 | 424 | 0 | 0 | 2 | 2 | 0 | 274 | 149 | 1 | 1 | 1 | 1 | 0.99 | 0.991 | 1 | 1 | 1 | 0.991 | 0.994 | 1 | -0.001 |
| SE | 10574 | ENSG00000188895 | MSL1 | chr17 | + | 40133533 | 40133658 | 40133041 | 40133109 | 40134278 | 40134450 | 188 | 130 | 167 | 204 | 334 | 0 | 0 | 0 | 0 | 0 | 293 | 165 | 192 | 300 | 166 | 0 | 0 | 0 | 1 | 0 | 274 | 149 | 1 | 1 | 1 | 1 | 1 | 1 | 1 | 1 | 1 | 1 | 0.994 | 1 | 0.001 |
| SE | 10575 | ENSG00000188895 | MSL1 | chr17 | + | 40133826 | 40133899 | 40133041 | 40133109 | 40134278 | 40134450 | 42 | 49 | 48 | 63 | 59 | 0 | 0 | 0 | 0 | 0 | 63 | 38 | 70 | 56 | 47 | 0 | 0 | 0 | 1 | 0 | 222 | 149 | 1 | 1 | 1 | 1 | 1 | 1 | 1 | 1 | 1 | 1 | 0.974 | 1 | 0.005 |
| SE | 10577 | ENSG00000108306 | FBXL20 | chr17 | - | 39261451 | 39261567 | 39259130 | 39259296 | 39264174 | 39264387 | 71 | 56 | 56 | 82 | 170 | 1 | 0 | 0 | 0 | 0 | 135 | 101 | 87 | 76 | 56 | 0 | 0 | 0 | 0 | 0 | 265 | 149 | 1 | 1 | 0.976 | 1 | 1 | 1 | 1 | 1 | 1 | 1 | 1 | 1 | -0.005 |
| SE | 10579 | ENSG00000108306 | FBXL20 | chr17 | - | 39274969 | 39275100 | 39270795 | 39270856 | 39281388 | 39281463 | 90 | 69 | 80 | 135 | 116 | 0 | 0 | 0 | 0 | 0 | 107 | 80 | 88 | 111 | 50 | 0 | 0 | 3 | 2 | 0 | 280 | 149 | 1 | 1 | 1 | 1 | 1 | 1 | 1 | 1 | 1 | 0.94 | 0.967 | 1 | 0.019 |
| SE | 10581 | ENSG00000108306 | FBXL20 | chr17 | - | 39282728 | 39282855 | 39281388 | 39281463 | 39285477 | 39285573 | 85 | 58 | 92 | 91 | 127 | 2 | 0 | 0 | 0 | 0 | 112 | 89 | 74 | 129 | 73 | 0 | 0 | 0 | 0 | 0 | 276 | 149 | 1 | 1 | 0.958 | 1 | 1 | 1 | 1 | 1 | 1 | 1 | 1 | 1 | -0.008 |
| SE | 10582 | ENSG00000108306 | FBXL20 | chr17 | - | 39282728 | 39282855 | 39281388 | 39281463 | 39297126 | 39297195 | 47 | 33 | 49 | 47 | 56 | 1 | 0 | 0 | 0 | 0 | 75 | 34 | 38 | 50 | 26 | 0 | 0 | 0 | 0 | 0 | 276 | 149 | 1 | 1 | 0.962 | 1 | 1 | 1 | 1 | 1 | 1 | 1 | 1 | 1 | -0.008 |
| SE | 10583 | ENSG00000108306 | FBXL20 | chr17 | - | 39285477 | 39285573 | 39281388 | 39281463 | 39297126 | 39297195 | 33 | 21 | 23 | 26 | 44 | 1 | 0 | 0 | 0 | 0 | 40 | 51 | 19 | 51 | 19 | 0 | 0 | 0 | 0 | 0 | 245 | 149 | 1 | 1 | 0.953 | 1 | 1 | 1 | 1 | 1 | 1 | 1 | 1 | 1 | -0.009 |
| SE | 10585 | ENSG00000108306 | FBXL20 | chr17 | - | 39301000 | 39301075 | 39298989 | 39299084 | 39303584 | 39303639 | 147 | 105 | 116 | 146 | 117 | 0 | 0 | 2 | 1 | 3 | 220 | 148 | 169 | 251 | 145 | 0 | 0 | 3 | 2 | 0 | 224 | 149 | 1 | 1 | 1 | 1 | 0.975 | 0.99 | 0.963 | 1 | 1 | 0.974 | 0.988 | 1 | -0.007 |
| SE | 10588 | ENSG00000274211 | SOCS7 | chr17 | + | 38365307 | 38365409 | 38364751 | 38364856 | 38366286 | 38366417 | 0 | 0 | 0 | 1 | 0 | 11 | 5 | 8 | 9 | 1 | 0 | 2 | 1 | 0 | 0 | 0 | 9 | 5 | 6 | 5 | 251 | 149 | 1 | 1 | 0 | 0 | 0 | 0.062 | 0 | NA | 0.117 | 0.106 | 0 | 0 | -0.043 |
| SE | 10590 | ENSG00000274211 | SOCS7 | chr17 | + | 38375839 | 38375962 | 38367881 | 38368050 | 38377713 | 38377842 | 0 | 1 | 0 | 0 | 0 | 55 | 37 | 29 | 53 | 33 | 0 | 3 | 0 | 0 | 0 | 46 | 36 | 42 | 43 | 77 | 272 | 149 | 1 | 1 | 0 | 0.015 | 0 | 0 | 0 | 0 | 0.044 | 0 | 0 | 0 | -0.006 |
| SE | 10592 | ENSG00000274211 | SOCS7 | chr17 | + | 38395847 | 38395998 | 38395308 | 38395444 | 38399512 | 38399648 | 80 | 57 | 52 | 90 | 71 | 0 | 0 | 0 | 0 | 2 | 128 | 92 | 42 | 83 | 153 | 0 | 0 | 0 | 0 | 0 | 298 | 149 | 1 | 1 | 1 | 1 | 1 | 1 | 0.947 | 1 | 1 | 1 | 1 | 1 | -0.011 |
| SE | 10594 | ENSG00000283103 | NA | chr19 | - | 58322361 | 58322494 | 58314787 | 58315036 | 58325274 | 58325361 | 0 | 0 | 0 | 2 | 0 | 26 | 3 | 16 | 10 | 4 | 0 | 0 | 0 | 0 | 0 | 12 | 12 | 2 | 0 | 7 | 282 | 149 | 1 | 1 | 0 | 0 | 0 | 0.096 | 0 | 0 | 0 | 0 | NA | 0 | 0.019 |
| SE | 10595 | ENSG00000283103 | NA | chr19 | - | 58325274 | 58325361 | 58314670 | 58315036 | 58326822 | 58326859 | 29 | 3 | 17 | 10 | 4 | 10 | 1 | 3 | 10 | 2 | 12 | 12 | 2 | 0 | 7 | 0 | 0 | 0 | 0 | 9 | 236 | 149 | 1 | 1 | 0.647 | 0.654 | 0.782 | 0.387 | 0.558 | 1 | 1 | 1 | NA | 0.329 | -0.227 |
| SE | 10598 | ENSG00000125691 | RPL23 | chr17 | - | 38850361 | 38850475 | 38847859 | 38850214 | 38852603 | 38852732 | 18644 | 11612 | 20251 | 17552 | 22310 | 4 | 2 | 13 | 4 | 6 | 15481 | 10415 | 11110 | 9605 | 15208 | 11 | 3 | 5 | 9 | 6 | 263 | 149 | 1 | 1 | 1 | 1 | 0.999 | 1 | 1 | 0.999 | 0.999 | 0.999 | 0.998 | 0.999 | 0.001 |
| SE | 10599 | ENSG00000125691 | RPL23 | chr17 | - | 38853021 | 38853105 | 38850080 | 38852732 | 38853697 | 38853721 | 14347 | 8772 | 15741 | 13850 | 14897 | 0 | 0 | 0 | 1 | 0 | 12179 | 8330 | 8668 | 7008 | 11146 | 0 | 0 | 0 | 0 | 0 | 233 | 149 | 1 | 1 | 1 | 1 | 1 | 1 | 1 | 1 | 1 | 1 | 1 | 1 | 0 |
| SE | 10601 | ENSG00000276234 | TADA2A | chr17 | + | 37423508 | 37423615 | 37411268 | 37411390 | 37437737 | 37437955 | 35 | 27 | 27 | 30 | 38 | 0 | 1 | 0 | 2 | 0 | 13 | 14 | 15 | 20 | 12 | 0 | 0 | 0 | 0 | 0 | 256 | 149 | 1 | 1 | 1 | 0.94 | 1 | 0.897 | 1 | 1 | 1 | 1 | 1 | 1 | -0.033 |
| SE | 10602 | ENSG00000276234 | TADA2A | chr17 | + | 37426949 | 37427009 | 37411268 | 37411390 | 37437737 | 37437829 | 67 | 30 | 46 | 44 | 80 | 0 | 1 | 0 | 2 | 0 | 44 | 32 | 23 | 20 | 18 | 0 | 0 | 0 | 0 | 0 | 209 | 149 | 1 | 1 | 1 | 0.955 | 1 | 0.94 | 1 | 1 | 1 | 1 | 1 | 1 | -0.021 |
| SE | 10603 | ENSG00000276234 | TADA2A | chr17 | + | 37426949 | 37427009 | 37423508 | 37423615 | 37437737 | 37437829 | 101 | 47 | 64 | 61 | 109 | 0 | 0 | 1 | 2 | 0 | 35 | 46 | 31 | 26 | 38 | 0 | 0 | 3 | 0 | 0 | 209 | 149 | 1 | 1 | 1 | 1 | 0.979 | 0.956 | 1 | 1 | 1 | 0.88 | 1 | 1 | 0.011 |
| SE | 10607 | ENSG00000276234 | TADA2A | chr17 | + | 37442563 | 37442652 | 37440504 | 37440662 | 37458523 | 37458587 | 63 | 28 | 52 | 45 | 74 | 1 | 3 | 0 | 0 | 0 | 13 | 29 | 16 | 26 | 35 | 3 | 0 | 0 | 0 | 0 | 238 | 149 | 1 | 1 | 0.975 | 0.854 | 1 | 1 | 1 | 0.731 | 1 | 1 | 1 | 1 | 0.02 |
| SE | 10610 | ENSG00000276234 | TADA2A | chr17 | + | 37458523 | 37458587 | 37444695 | 37444768 | 37462077 | 37462121 | 34 | 24 | 39 | 27 | 20 | 0 | 4 | 0 | 1 | 0 | 60 | 7 | 3 | 24 | 29 | 0 | 0 | 0 | 0 | 0 | 213 | 149 | 1 | 1 | 1 | 0.808 | 1 | 0.95 | 1 | 1 | 1 | 1 | 1 | 1 | -0.048 |
| SE | 10611 | ENSG00000276234 | TADA2A | chr17 | + | 37462077 | 37462121 | 37458523 | 37458587 | 37465430 | 37465471 | 28 | 22 | 42 | 28 | 30 | 1 | 0 | 0 | 0 | 0 | 31 | 20 | 9 | 25 | 28 | 0 | 0 | 0 | 0 | 0 | 193 | 149 | 1 | 1 | 0.956 | 1 | 1 | 1 | 1 | 1 | 1 | 1 | 1 | 1 | -0.009 |
| SE | 10612 | ENSG00000276234 | TADA2A | chr17 | + | 37471093 | 37471137 | 37470399 | 37470532 | 37474555 | 37474629 | 48 | 32 | 34 | 43 | 55 | 1 | 2 | 0 | 1 | 0 | 58 | 23 | 12 | 34 | 3 | 0 | 5 | 0 | 0 | 0 | 193 | 149 | 1 | 1 | 0.974 | 0.925 | 1 | 0.971 | 1 | 1 | 0.78 | 1 | 1 | 1 | 0.018 |
| SE | 10613 | ENSG00000276234 | TADA2A | chr17 | + | 37474555 | 37474629 | 37471093 | 37471137 | 37476796 | 37476850 | 97 | 49 | 49 | 81 | 85 | 4 | 0 | 0 | 0 | 0 | 87 | 35 | 24 | 49 | 35 | 0 | 0 | 0 | 0 | 0 | 223 | 149 | 1 | 1 | 0.942 | 1 | 1 | 1 | 1 | 1 | 1 | 1 | 1 | 1 | -0.012 |
| SE | 10615 | ENSG00000108599 | AKAP10 | chr17 | - | 19924407 | 19924517 | 19920035 | 19920118 | 19931804 | 19931978 | 230 | 173 | 158 | 200 | 287 | 1 | 1 | 1 | 3 | 1 | 79 | 227 | 205 | 164 | 33 | 0 | 4 | 2 | 0 | 0 | 259 | 149 | 1 | 1 | 0.992 | 0.99 | 0.989 | 0.975 | 0.994 | 1 | 0.97 | 0.983 | 1 | 1 | -0.003 |
| SE | 10618 | ENSG00000108599 | AKAP10 | chr17 | - | 19936285 | 19936430 | 19924407 | 19924517 | 19939712 | 19939849 | 147 | 106 | 123 | 142 | 150 | 0 | 0 | 0 | 0 | 0 | 125 | 124 | 131 | 92 | 58 | 0 | 0 | 0 | 5 | 0 | 294 | 149 | 1 | [illegible] | [illegible] | [illegible] | [illegible] | [illegible] | [illegible] | [illegible] | [illegible] | [illegible] | [illegible] | [illegible] | [illegible] |
| SE | 10620 | ENSG00000108599 | AKAP10 | chr17 | - | 19939712 | 19939849 | 19936001 | 19936430 | 19947406 | 19947475 | 142 | 103 | 117 | 134 | 138 | 0 | 1 | 0 | 1 | 0 | 102 | 106 | 115 | 75 | 58 | [illegible] | [illegible] | [illegible] | [illegible] | [illegible] | [illegible] | [illegible] | [illegible] | [illegible] | [illegible] | [illegible] | [illegible] | [illegible] | [illegible] | [illegible] | [illegible] | [illegible] | [illegible] | [illegible] | [illegible] |
| SE | 10622 | ENSG00000108599 | AKAP10 | chr17 | - | 19940886 | 19941010 | 19939712 | 19939849 | 19947406 | 19947505 | 71 | 41 | 70 | 51 | 48 | 0 | 0 | 0 | 3 | 0 | 63 | 49 | 37 | 46 | 18 | [illegible] | [illegible] | [illegible] | [illegible] | [illegible] | [illegible] | [illegible] | [illegible] | [illegible] | [illegible] | [illegible] | [illegible] | [illegible] | [illegible] | [illegible] | [illegible] | [illegible] | [illegible] | [illegible] | [illegible] |
| SE | 10623 | ENSG00000108599 | AKAP10 | chr17 | - | 19941825 | 19941910 | 19936001 | 19936430 | 19947406 | 19947475 | 113 | 78 | 76 | 82 | 128 | 0 | 1 | 0 | 1 | 0 | 57 | 91 | 84 | 60 | 35 | 0 | 0 | 0 | 0 | 0 | 234 | 149 | 1 | [illegible] | [illegible] | [illegible] | [illegible] | [illegible] | [illegible] | [illegible] | [illegible] | [illegible] | [illegible] | [illegible] | [illegible] |
| SE | 10624 | ENSG00000108599 | AKAP10 | chr17 | - | 19941825 | 19941910 | 19939712 | 19939849 | 19947406 | 19947475 | 160 | 110 | 108 | 122 | 178 | 0 | 0 | 0 | 3 | 0 | 113 | 149 | 114 | 89 | 62 | 0 | 0 | 0 | 0 | 0 | 234 | 149 | 1 | [illegible] | [illegible] | [illegible] | [illegible] | [illegible] | [illegible] | [illegible] | [illegible] | [illegible] | [illegible] | [illegible] | [illegible] |
| SE | 10625 | ENSG00000108599 | AKAP10 | chr17 | - | 19941825 | 19941910 | 19940886 | 19941010 | 19947406 | 19947475 | 208 | 132 | 136 | 135 | 209 | 0 | 0 | 0 | 0 | 0 | 96 | 168 | 127 | 101 | 78 | 0 | 0 | 1 | 0 | 0 | 234 | 149 | 1 | [illegible] | [illegible] | [illegible] | [illegible] | [illegible] | [illegible] | [illegible] | [illegible] | [illegible] | [illegible] | [illegible] | [illegible] |
| SE | 10626 | ENSG00000108599 | AKAP10 | chr17 | - | 19947406 | 19947505 | 19941825 | 19941910 | 19958013 | 19958567 | 244 | 194 | 186 | 196 | 315 | 1 | 0 | 0 | 0 | 0 | 131 | 205 | 173 | 108 | 94 | 0 | 0 | 0 | 0 | 0 | 248 | 149 | 1 | [illegible] | [illegible] | [illegible] | [illegible] | [illegible] | [illegible] | [illegible] | [illegible] | [illegible] | [illegible] | [illegible] | [illegible] |
| SE | 10627 | ENSG00000108599 | AKAP10 | chr17 | - | 19962839 | 19962966 | 19958424 | 19958571 | 19968413 | 19968461 | 32 | 17 | 15 | 30 | 35 | 0 | 0 | 0 | 2 | 0 | 18 | 38 | 18 | 7 | 0 | 0 | 0 | 0 | 0 | 0 | 276 | 149 | 1 | [illegible] | [illegible] | [illegible] | [illegible] | [illegible] | [illegible] | [illegible] | [illegible] | [illegible] | [illegible] | [illegible] | [illegible] |
| SE | 10628 | ENSG00000108599 | AKAP10 | chr17 | - | 19962839 | 19962991 | 19958443 | 19958571 | 19968413 | 19968461 | 36 | 20 | 18 | 31 | 40 | 0 | 0 | 0 | 2 | 0 | 18 | 49 | 21 | 7 | 0 | 0 | 0 | 0 | 0 | 0 | 298 | 149 | 1 | [illegible] | [illegible] | [illegible] | [illegible] | [illegible] | [illegible] | [illegible] | [illegible] | [illegible] | [illegible] | [illegible] | [illegible] |
| SE | 10629 | ENSG00000108599 | AKAP10 | chr17 | - | 19962839 | 19963022 | 19958013 | 19958571 | 19968413 | 19968461 | 54 | 35 | 34 | 48 | 70 | 0 | 0 | 0 | 2 | 0 | 23 | 55 | 28 | 18 | 0 | 0 | 0 | 0 | 0 | 0 | 298 | 149 | 1 | [illegible] | [illegible] | [illegible] | [illegible] | [illegible] | [illegible] | [illegible] | [illegible] | [illegible] | [illegible] | [illegible] | [illegible] |
| SE | 10630 | ENSG00000108599 | AKAP10 | chr17 | - | 19967918 | 19968034 | 19962859 | 19963022 | 19968413 | 19968461 | 0 | 0 | 1 | 0 | 0 | 21 | 16 | 17 | 18 | 30 | 0 | 0 | 0 | 4 | 0 | 5 | 12 | 7 | 11 | 0 | 265 | 149 | 1 | [illegible] | [illegible] | [illegible] | [illegible] | [illegible] | [illegible] | [illegible] | [illegible] | [illegible] | [illegible] | [illegible] | [illegible] |
| SE | 10631 | ENSG00000108599 | AKAP10 | chr17 | - | 19968413 | 19968461 | 19962839 | 19963022 | 19977591 | 19977679 | 15 | 16 | 17 | 21 | 32 | 0 | 2 | 0 | 0 | 0 | 5 | 22 | 5 | 16 | 0 | 0 | 0 | 0 | 0 | 0 | 197 | 149 | 1 | [illegible] | [illegible] | [illegible] | [illegible] | [illegible] | [illegible] | [illegible] | [illegible] | [illegible] | [illegible] | [illegible] | [illegible] |
| SE | 10633 | ENSG00000277791 | PSMB3 | chr17 | + | 38760430 | 38760608 | 38755882 | 38755990 | 38764118 | 38764225 | 282 | 126 | 281 | 304 | 397 | 0 | 0 | 3 | 0 | 0 | 411 | 419 | 163 | 297 | 515 | 0 | 2 | 0 | 0 | 0 | 298 | 149 | 1 | [illegible] | [illegible] | [illegible] | [illegible] | [illegible] | [illegible] | [illegible] | [illegible] | [illegible] | [illegible] | [illegible] | [illegible] |
| SE | 10636 | ENSG00000277791 | PSMB3 | chr17 | + | 38762410 | 38762505 | 38755882 | 38755990 | 38764118 | 38764225 | 250 | 89 | 179 | 158 | 227 | 0 | 0 | 3 | 0 | 0 | 322 | 300 | 158 | 270 | 373 | 0 | 2 | 0 | 0 | 0 | 244 | 149 | 1 | [illegible] | [illegible] | [illegible] | [illegible] | [illegible] | [illegible] | [illegible] | [illegible] | [illegible] | [illegible] | [illegible] | [illegible] |
| SE | 10637 | ENSG00000277791 | PSMB3 | chr17 | + | 38762410 | 38762505 | 38760384 | 38760608 | 38764118 | 38764225 | 634 | 220 | 452 | 422 | 585 | 1 | 0 | 1 | 0 | 0 | 654 | 663 | 308 | 598 | 922 | 0 | 1 | 0 | 0 | 0 | 244 | 149 | 1 | [illegible] | [illegible] | [illegible] | [illegible] | [illegible] | [illegible] | [illegible] | [illegible] | [illegible] | [illegible] | [illegible] | [illegible] |
| SE | 10639 | ENSG00000171475 | WIPF2 | chr17 | + | 40256390 | 40256522 | 40219303 | 40219492 | 40260534 | 40260627 | 80 | 42 | 63 | 66 | 51 | 0 | 0 | 0 | 0 | 0 | 158 | 65 | 79 | 94 | 121 | 0 | 3 | 0 | 0 | 0 | 281 | 149 | 1 | [illegible] | [illegible] | [illegible] | [illegible] | [illegible] | [illegible] | [illegible] | [illegible] | [illegible] | [illegible] | [illegible] | [illegible] |
| SE | 10645 | ENSG00000171475 | WIPF2 | chr17 | + | 40264489 | 40265146 | 40260534 | 40260667 | 40277082 | 40277184 | 1 | 1 | 6 | 2 | 0 | 0 | 0 | 0 | 0 | 0 | 47 | 6 | 11 | 14 | 0 | 0 | 0 | 0 | 0 | 0 | 298 | 149 | 1 | [illegible] | [illegible] | [illegible] | [illegible] | [illegible] | [illegible] | [illegible] | [illegible] | [illegible] | [illegible] | [illegible] | [illegible] |
| SE | 10646 | ENSG00000171475 | WIPF2 | chr17 | + | 40273789 | 40273999 | 40260534 | 40260667 | 40277082 | 40277132 | 88 | 56 | 63 | 83 | 91 | 0 | 0 | 0 | 0 | 0 | 70 | 69 | 76 | 105 | 99 | 0 | 0 | 0 | 3 | 0 | 298 | 149 | 1 | [illegible] | [illegible] | [illegible] | [illegible] | [illegible] | [illegible] | [illegible] | [illegible] | [illegible] | [illegible] | [illegible] | [illegible] |
| SE | 10647 | ENSG00000171475 | WIPF2 | chr17 | + | 40273789 | 40273999 | 40264489 | 40265146 | 40277082 | 40277132 | 121 | 69 | 110 | 112 | 138 | 0 | 0 | 0 | 0 | 0 | 110 | 115 | 101 | 146 | 145 | 0 | 0 | 0 | 3 | 0 | 298 | 149 | 1 | [illegible] | [illegible] | [illegible] | [illegible] | [illegible] | [illegible] | [illegible] | [illegible] | [illegible] | [illegible] | [illegible] | [illegible] |
| SE | 10648 | ENSG00000273559 | CWC25 | chr17 | - | 38806296 | 38806395 | 38802699 | 38802861 | 38806764 | 38806976 | 123 | 107 | 123 | 110 | 77 | 1 | 2 | 0 | 1 | 0 | 107 | 102 | 177 | 153 | 95 | 0 | 0 | 2 | 4 | 0 | 248 | 149 | 1 | [illegible] | [illegible] | [illegible] | [illegible] | [illegible] | [illegible] | [illegible] | [illegible] | [illegible] | [illegible] | [illegible] | [illegible] |
| SE | 10653 | ENSG00000161395 | PGAP3 | chr17 | - | 39673050 | 39673255 | 39671121 | 39672866 | 39673513 | 39673650 | 34 | 2 | 13 | 27 | 3 | 0 | 0 | 4 | 0 | 2 | 0 | 7 | 14 | 3 | 10 | 0 | 0 | 0 | 0 | 0 | 298 | 149 | 1 | [illegible] | [illegible] | [illegible] | [illegible] | [illegible] | [illegible] | [illegible] | [illegible] | [illegible] | [illegible] | [illegible] | [illegible] |
| SE | 10654 | ENSG00000161395 | PGAP3 | chr17 | - | 39673050 | 39673255 | 39671121 | 39672866 | 39673992 | 39674054 | 19 | 2 | 12 | 19 | 0 | 0 | 0 | 0 | 0 | 0 | 0 | 3 | 8 | 3 | 2 | 0 | 0 | 2 | 0 | 0 | 298 | 149 | 1 | [illegible] | [illegible] | [illegible] | [illegible] | [illegible] | [illegible] | [illegible] | [illegible] | [illegible] | [illegible] | [illegible] | [illegible] |
| SE | 10655 | ENSG00000161395 | PGAP3 | chr17 | - | 39673513 | 39673650 | 39671121 | 39672866 | 39673992 | 39674054 | 13 | 3 | 11 | 9 | 10 | 0 | 0 | 0 | 0 | 0 | 0 | 11 | 0 | 0 | 6 | 0 | 0 | 2 | 0 | 0 | 286 | 149 | 1 | [illegible] | [illegible] | [illegible] | [illegible] | [illegible] | [illegible] | [illegible] | [illegible] | [illegible] | [illegible] | [illegible] | [illegible] |
| SE | 10657 | ENSG00000161395 | PGAP3 | chr17 | - | 39673992 | 39674054 | 39671121 | 39672866 | 39684596 | 39684749 | 0 | 0 | 0 | 0 | 0 | 2 | 0 | 0 | 0 | 0 | 0 | 0 | 0 | 2 | 0 | 0 | 0 | 0 | 0 | 0 | 211 | 149 | 1 | [illegible] | [illegible] | [illegible] | [illegible] | [illegible] | [illegible] | [illegible] | [illegible] | [illegible] | [illegible] | [illegible] | [illegible] |
| SE | 10660 | ENSG00000161395 | PGAP3 | chr17 | - | 39684596 | 39684749 | 39674616 | 39674679 | 39685921 | 39686019 | 69 | 39 | 72 | 44 | 53 | 0 | 1 | 0 | 0 | 0 | 12 | 43 | 13 | 15 | 66 | 0 | 0 | 0 | 0 | 0 | 298 | 149 | 1 | [illegible] | [illegible] | [illegible] | [illegible] | [illegible] | [illegible] | [illegible] | [illegible] | [illegible] | [illegible] | [illegible] | [illegible] |
| SE | 10663 | ENSG00000108344 | PSMD3 | chr17 | + | 39984293 | 39984484 | 39980806 | 39981190 | 39986574 | 39986712 | 198 | 68 | 141 | 129 | 66 | 0 | 0 | 0 | 0 | 0 | 125 | 213 | 162 | 141 | 177 | 0 | 2 | 0 | 0 | 0 | 298 | 149 | 1 | [illegible] | [illegible] | [illegible] | [illegible] | [illegible] | [illegible] | [illegible] | [illegible] | [illegible] | [illegible] | [illegible] | [illegible] |
| SE | 10664 | ENSG00000108344 | PSMD3 | chr17 | + | 39988682 | 39988819 | 39986574 | 39986712 | 39989738 | 39989793 | 139 | 62 | 119 | 129 | 162 | 0 | 0 | 0 | 0 | 0 | 123 | 163 | 69 | 115 | 219 | 0 | 0 | 0 | 0 | 0 | 286 | 149 | 1 | [illegible] | [illegible] | [illegible] | [illegible] | [illegible] | [illegible] | [illegible] | [illegible] | [illegible] | [illegible] | [illegible] | [illegible] |
| SE | 10665 | ENSG00000108344 | PSMD3 | chr17 | + | 39996182 | 39996338 | 39995423 | 39995527 | 39997329 | 39997380 | 277 | 127 | 184 | 225 | 199 | 0 | 0 | 0 | 0 | 0 | 312 | 307 | 232 | 224 | 425 | 0 | 0 | 0 | 7 | 0 | 298 | 149 | 1 | [illegible] | [illegible] | [illegible] | [illegible] | [illegible] | [illegible] | [illegible] | [illegible] | [illegible] | [illegible] | [illegible] | [illegible] |
| SE | 10668 | ENSG00000214719 | NA | chr17 | + | 30607551 | 30607665 | 30600891 | 30601012 | 30608429 | 30608562 | 59 | 34 | 57 | 37 | 44 | 2 | 0 | 2 | 0 | 0 | 38 | 46 | 41 | 35 | 85 | 0 | 3 | 0 | 0 | 0 | 263 | 149 | 1 | [illegible] | [illegible] | [illegible] | [illegible] | [illegible] | [illegible] | [illegible] | [illegible] | [illegible] | [illegible] | [illegible] | [illegible] |
| SE | 10675 | ENSG00000278540 | ACACA | chr17 | - | 37111530 | 37111643 | 37097829 | 37097984 | 37121354 | 37121490 | 19 | 21 | 31 | 17 | 20 | 0 | 0 | 1 | 0 | 0 | 23 | 15 | 15 | 19 | 8 | 0 | 0 | 0 | 0 | 0 | 262 | 149 | 1 | [illegible] | [illegible] | [illegible] | [illegible] | [illegible] | [illegible] | [illegible] | [illegible] | [illegible] | [illegible] | [illegible] | [illegible] |
| SE | 10676 | ENSG00000278540 | ACACA | chr17 | - | 37113087 | 37113265 | 37111530 | 37111643 | 37121354 | 37121490 | 33 | 29 | 41 | 25 | 21 | 1 | 0 | 0 | 0 | 0 | 25 | 63 | 42 | 63 | 11 | 0 | 0 | 0 | 0 | 0 | 298 | 149 | 1 | [illegible] | [illegible] | [illegible] | [illegible] | [illegible] | [illegible] | [illegible] | [illegible] | [illegible] | [illegible] | [illegible] | [illegible] |
| SE | 10677 | ENSG00000278540 | ACACA | chr17 | - | 37122530 | 37122627 | 37121354 | 37121490 | 37125697 | 37125794 | 19 | 10 | 21 | 22 | 24 | 0 | 0 | 0 | 0 | 0 | 7 | 12 | 11 | 10 | 45 | 0 | 0 | 3 | 0 | 0 | 246 | 149 | 1 | [illegible] | [illegible] | [illegible] | [illegible] | [illegible] | [illegible] | [illegible] | [illegible] | [illegible] | [illegible] | [illegible] | [illegible] |
| SE | 10678 | ENSG00000278540 | ACACA | chr17 | - | 37122530 | 37122627 | 37121354 | 37121490 | 37129364 | 37129485 | 9 | 6 | 13 | 13 | 18 | 0 | 0 | 0 | 0 | 0 | 3 | 4 | 8 | 7 | 18 | 0 | 1 | 0 | 0 | 0 | 246 | 149 | 1 | [illegible] | [illegible] | [illegible] | [illegible] | [illegible] | [illegible] | [illegible] | [illegible] | [illegible] | [illegible] | [illegible] | [illegible] |
| SE | 10679 | ENSG00000278540 | ACACA | chr17 | - | 37125697 | 37125794 | 37121354 | 37121490 | 37129364 | 37129485 | 6 | 4 | 5 | 2 | 3 | 0 | 0 | 0 | 0 | 0 | 3 | 1 | 1 | 3 | 11 | 0 | 1 | 0 | 0 | 0 | 246 | 149 | 1 | [illegible] | [illegible] | [illegible] | [illegible] | [illegible] | [illegible] | [illegible] | [illegible] | [illegible] | [illegible] | [illegible] | [illegible] |
| SE | 10680 | ENSG00000278540 | ACACA | chr17 | - | 37125697 | 37125794 | 37122530 | 37122627 | 37129364 | 37129485 | 15 | 8 | 13 | 12 | 10 | 0 | 0 | 0 | 0 | 1 | 7 | 17 | 4 | 4 | 33 | 0 | 0 | 0 | 0 | 0 | 246 | 149 | 1 | [illegible] | [illegible] | [illegible] | [illegible] | [illegible] | [illegible] | [illegible] | [illegible] | [illegible] | [illegible] | [illegible] | [illegible] |
| SE | 10682 | ENSG00000278540 | ACACA | chr17 | - | 37179259 | 37179406 | 37161599 | 37162050 | 37181200 | 37181356 | 47 | 17 | 39 | 19 | 70 | 0 | 0 | 0 | 0 | 0 | 12 | 51 | 35 | 25 | 16 | 0 | 0 | 0 | 0 | 0 | 296 | 149 | 1 | [illegible] | [illegible] | [illegible] | [illegible] | [illegible] | [illegible] | [illegible] | [illegible] | [illegible] | [illegible] | [illegible] | [illegible] |
| SE | 10683 | ENSG00000278540 | ACACA | chr17 | - | 37192089 | 37192305 | 37191119 | 37191275 | 37193373 | 37193415 | 77 | 38 | 62 | 84 | 54 | 1 | 0 | 0 | 0 | 0 | 133 | 64 | 92 | 82 | 35 | 0 | 0 | 0 | 0 | 0 | 298 | 149 | 1 | [illegible] | [illegible] | [illegible] | [illegible] | [illegible] | [illegible] | [illegible] | [illegible] | [illegible] | [illegible] | [illegible] | [illegible] |
| SE | 10684 | ENSG00000278540 | ACACA | chr17 | - | 37200138 | 37200183 | 37193373 | 37193415 | 37200426 | 37200483 | 33 | 24 | 31 | 30 | 17 | 0 | 0 | 0 | 0 | 0 | 36 | 21 | 34 | 38 | 8 | 0 | 0 | 1 | 0 | 0 | 194 | 149 | 1 | [illegible] | [illegible] | [illegible] | [illegible] | [illegible] | [illegible] | [illegible] | [illegible] | [illegible] | [illegible] | [illegible] | [illegible] |
| SE | 10685 | ENSG00000278540 | ACACA | chr17 | - | 37200426 | 37200483 | 37193373 | 37193415 | 37205764 | 37205812 | 7 | 4 | 14 | 2 | 10 | 0 | 0 | 1 | 0 | 0 | 0 | 3 | 8 | 3 | 0 | 0 | 0 | 0 | 0 | 0 | 206 | 149 | 1 | [illegible] | [illegible] | [illegible] | [illegible] | [illegible] | [illegible] | [illegible] | [illegible] | [illegible] | [illegible] | [illegible] | [illegible] |
| SE | 10686 | ENSG00000278540 | ACACA | chr17 | - | 37206782 | 37206879 | 37205764 | 37205872 | 37207656 | 37207800 | 19 | 19 | 39 | 29 | 41 | 0 | 0 | 2 | 0 | 0 | 25 | 45 | 41 | 22 | 24 | 0 | 0 | 0 | 0 | 0 | 246 | 149 | 1 | [illegible] | [illegible] | [illegible] | [illegible] | [illegible] | [illegible] | [illegible] | [illegible] | [illegible] | [illegible] | [illegible] | [illegible] |
| SE | 10688 | ENSG00000278540 | ACACA | chr17 | - | 37224991 | 37225105 | 37223511 | 37223601 | 37226338 | 37226452 | 66 | 37 | 52 | 66 | 74 | 0 | 0 | 3 | 1 | 0 | 48 | 63 | 40 | 40 | 45 | 0 | 0 | 2 | 2 | 0 | 263 | 149 | 1 | [illegible] | [illegible] | [illegible] | [illegible] | [illegible] | [illegible] | [illegible] | [illegible] | [illegible] | [illegible] | [illegible] | [illegible] |
| SE | 10689 | ENSG00000278540 | ACACA | chr17 | - | 37234974 | 37235099 | 37226338 | 37226452 | 37240475 | 37240564 | 76 | 36 | 50 | 55 | 104 | 0 | 0 | 0 | 0 | 0 | 58 | 52 | 38 | 57 | 46 | 0 | 0 | 2 | 0 | 0 | 274 | 149 | 1 | [illegible] | [illegible] | [illegible] | [illegible] | [illegible] | [illegible] | [illegible] | [illegible] | [illegible] | [illegible] | [illegible] | [illegible] |
| SE | 10690 | ENSG00000278540 | ACACA | chr17 | - | 37241952 | 37242053 | 37240475 | 37240564 | 37243370 | 37243559 | 56 | 31 | 27 | 45 | 57 | 0 | 0 | 0 | 1 | 0 | 98 | 58 | 25 | 53 | 50 | 0 | 0 | 0 | 0 | 0 | 250 | 149 | 1 | [illegible] | [illegible] | [illegible] | [illegible] | [illegible] | [illegible] | [illegible] | [illegible] | [illegible] | [illegible] | [illegible] | [illegible] |
| SE | 10691 | ENSG00000278540 | ACACA | chr17 | - | 37246825 | 37246976 | 37245079 | 37245214 | 37248010 | 37248156 | 24 | 17 | 31 | 47 | 47 | 0 | 0 | 0 | 0 | 0 | 3 | 35 | 33 | 23 | 27 | 0 | 0 | 1 | 0 | 0 | 298 | 149 | 1 | [illegible] | [illegible] | [illegible] | [illegible] | [illegible] | [illegible] | [illegible] | [illegible] | [illegible] | [illegible] | [illegible] | [illegible] |
| SE | 10693 | ENSG00000278540 | ACACA | chr17 | - | 37252885 | 37253036 | 37252004 | 37252108 | 37257702 | 37257866 | 48 | 58 | 68 | 54 | 34 | 0 | 0 | 0 | 0 | 3 | 44 | 92 | 55 | 84 | 23 | 0 | 0 | 0 | 0 | 0 | 298 | 149 | 1 | [illegible] | [illegible] | [illegible] | [illegible] | [illegible] | [illegible] | [illegible] | [illegible] | [illegible] | [illegible] | [illegible] | [illegible] |
| SE | 10697 | ENSG00000278540 | ACACA | chr17 | - | 37345252 | 37345382 | 37339803 | 37339850 | 37365442 | 37365750 | 0 | 1 | 0 | 0 | 0 | 0 | 0 | 0 | 0 | 0 | 0 | 0 | 0 | 0 | 0 | 0 | 0 | 0 | 1 | 0 | 279 | 149 | 1 | [illegible] | [illegible] | [illegible] | [illegible] | [illegible] | [illegible] | [illegible] | [illegible] | [illegible] | [illegible] | [illegible] | [illegible] |
| SE | 10699 | ENSG00000223704 | LINC01422 | chr22 | + | 26877328 | 26877413 | 26861981 | 26862105 | 26884847 | 26885691 | 9 | 9 | 9 | 2 | 3 | 1 | 1 | 0 | 1 | 0 | 0 | 5 | 8 | 8 | 2 | 0 | 0 | 0 | 0 | 0 | 234 | 149 | 1 | [illegible] | [illegible] | [illegible] | [illegible] | [illegible] | [illegible] | [illegible] | [illegible] | [illegible] | [illegible] | [illegible] | [illegible] |
| SE | 10705 | ENSG00000275832 | ARHGAP23 | chr17 | + | 38477578 | 38477705 | 38471862 | 38472006 | 38477827 | 38477896 | 0 | 0 | 4 | 1 | 0 | 0 | 0 | 0 | 0 | 0 | 0 | 0 | 1 | 0 | 0 | 0 | 6 | 0 | 0 | 0 | 276 | 149 | 1 | [illegible] | [illegible] | [illegible] | [illegible] | [illegible] | [illegible] | [illegible] | [illegible] | [illegible] | [illegible] | [illegible] | [illegible] |
| SE | 10707 | ENSG00000275832 | ARHGAP23 | chr17 | + | 38498413 | 38498510 | 38497784 | 38497826 | 38500596 | 38500628 | 2 | 1 | 0 | 0 | 0 | 1 | 0 | 0 | 0 | 0 | 6 | 0 | 0 | 10 | 0 | 0 | 0 | 0 | 0 | 0 | 246 | 149 | 1 | [illegible] | [illegible] | [illegible] | [illegible] | [illegible] | [illegible] | [illegible] | [illegible] | [illegible] | [illegible] | [illegible] | [illegible] |
| SE | 10709 | ENSG00000274487 | NPEPPSP1 | chr17 | - | 38208994 | 38209116 | 38205634 | 38205742 | 38219045 | 38219123 | 3 | 0 | 3 | 0 | 3 | 0 | 0 | 0 | 0 | 2 | 12 | 0 | 0 | 0 | 8 | 0 | 0 | 0 | 0 | 0 | 271 | 149 | 1 | [illegible] | [illegible] | [illegible] | [illegible] | [illegible] | [illegible] | [illegible] | [illegible] | [illegible] | [illegible] | [illegible] | [illegible] |
| SE | 10717 | ENSG00000278845 | MRPL45 | chr17 | + | 38298448 | 38298626 | 38297022 | 38297249 | 38299350 | 38299468 | 85 | 59 | 92 | 85 | 176 | 0 | 0 | 0 | 0 | 0 | 83 | 72 | 52 | 50 | 118 | 0 | 1 | 0 | 0 | 0 | 298 | 149 | 1 | [illegible] | [illegible] | [illegible] | [illegible] | [illegible] | [illegible] | [illegible] | [illegible] | [illegible] | [illegible] | [illegible] | [illegible] |
| SE | 10718 | ENSG00000278845 | MRPL45 | chr17 | + | 38306532 | 38306631 | 38299350 | 38299468 | 38318686 | 38318735 | 105 | 75 | 108 | 106 | 294 | 2 | 0 | 0 | 0 | 0 | 127 | 79 | 48 | 35 | 57 | 0 | 1 | 0 | 0 | 0 | 248 | 149 | 1 | [illegible] | [illegible] | [illegible] | [illegible] | [illegible] | [illegible] | [illegible] | [illegible] | [illegible] | [illegible] | [illegible] | [illegible] |
| SE | 10719 | ENSG00000278845 | MRPL45 | chr17 | + | 38320617 | 38320767 | 38318686 | 38318735 | 38322125 | 38322299 | 164 | 108 | 163 | 147 | 597 | 4 | 1 | 1 | 0 | 0 | 174 | 79 | 67 | 47 | 109 | 0 | 0 | 0 | 0 | 0 | 298 | 149 | 1 | [illegible] | [illegible] | [illegible] | [illegible] | [illegible] | [illegible] | [illegible] | [illegible] | [illegible] | [illegible] | [illegible] | [illegible] |
| SE | 10720 | ENSG00000277972 | CISD3 | chr17 | + | 38731319 | 38731439 | 38730358 | 38730795 | 38733275 | 38735044 | 60 | 29 | 44 | 76 | 22 | 0 | 0 | 0 | 0 | 0 | 44 | 36 | 31 | 39 | 70 | 0 | 0 | 2 | 0 | 0 | 269 | 149 | 1 | [illegible] | [illegible] | [illegible] | [illegible] | [illegible] | [illegible] | [illegible] | [illegible] | [illegible] | [illegible] | [illegible] | [illegible] |
| SE | 10725 | ENSG00000006125 | AP2B1 | chr17 | + | 35588624 | 35588733 | 35587415 | 35587597 | 35594007 | 35594067 | 0 | 0 | 0 | 1 | 0 | 21 | 14 | 14 | 12 | 18 | 0 | 0 | 0 | 0 | 0 | 39 | 32 | 44 | 14 | 12 | 258 | 149 | 1 | [illegible] | [illegible] | [illegible] | [illegible] | [illegible] | [illegible] | [illegible] | [illegible] | [illegible] | [illegible] | [illegible] | [illegible] |
| SE | 10727 | ENSG00000006125 | AP2B1 | chr17 | + | 35600985 | 35601071 | 35598229 | 35598335 | 35605704 | 35605840 | 4 | 4 | 4 | 6 | 9 | 312 | 210 | 272 | 263 | 434 | 0 | 0 | 6 | 0 | 0 | 293 | 232 | 207 | 225 | 330 | 235 | 149 | 1 | [illegible] | [illegible] | [illegible] | [illegible] | [illegible] | [illegible] | [illegible] | [illegible] | [illegible] | [illegible] | [illegible] | [illegible] |
| SE | 10729 | ENSG00000006125 | AP2B1 | chr17 | + | 35641876 | 35641975 | 35639594 | 35639760 | 35650529 | 35650789 | 735 | 457 | 537 | 664 | 765 | 0 | 0 | 0 | 3 | 1 | 700 | 402 | 388 | 430 | 1048 | 4 | 0 | 0 | 2 | 0 | 248 | 149 | 1 | [illegible] | [illegible] | [illegible] | [illegible] | [illegible] | [illegible] | [illegible] | [illegible] | [illegible] | [illegible] | [illegible] | [illegible] |
| SE | 10730 | ENSG00000006125 | AP2B1 | chr17 | + | 35650529 | 35650789 | 35641876 | 35641975 | 35657598 | 35657791 | 672 | 419 | 507 | 553 | 683 | 0 | 0 | 0 | 0 | 0 | 640 | 350 | 351 | 429 | 953 | 0 | 0 | 0 | 2 | 0 | 298 | 149 | 1 | [illegible] | [illegible] | [illegible] | [illegible] | [illegible] | [illegible] | [illegible] | [illegible] | [illegible] | [illegible] | [illegible] | [illegible] |
| SE | 10733 | ENSG00000006125 | AP2B1 | chr17 | + | 35682694 | 35682824 | 35674175 | 35674321 | 35709223 | 35709308 | 370 | 220 | 320 | 355 | 555 | 0 | 0 | 0 | 0 | 4 | 388 | 280 | 186 | 266 | 584 | 0 | 0 | 0 | 0 | 0 | 279 | 149 | 1 | [illegible] | [illegible] | [illegible] | [illegible] | [illegible] | [illegible] | [illegible] | [illegible] | [illegible] | [illegible] | [illegible] | [illegible] |
| SE | 10734 | ENSG00000006125 | AP2B1 | chr17 | + | 35710233 | 35710320 | 35709223 | 35709308 | 35717194 | 35717349 | 496 | 305 | 362 | 451 | 882 | 0 | 0 | 0 | 2 | 0 | 619 | 381 | 235 | 282 | 587 | 0 | 0 | 0 | 0 | 0 | 236 | 149 | 1 | [illegible] | [illegible] | [illegible] | [illegible] | [illegible] | [illegible] | [illegible] | [illegible] | [illegible] | [illegible] | [illegible] | [illegible] |
| SE | 10735 | ENSG00000006125 | AP2B1 | chr17 | + | 35717194 | 35717349 | 35710233 | 35710320 | 35723624 | 35724086 | 513 | 321 | 403 | 482 | 1059 | 0 | 2 | 0 | 0 | 0 | 643 | 492 | 299 | 338 | 798 | 0 | 0 | 0 | 0 | 0 | 298 | 149 | 1 | [illegible] | [illegible] | [illegible] | [illegible] | [illegible] | [illegible] | [illegible] | [illegible] | [illegible] | [illegible] | [illegible] | [illegible] |
| SE | 10736 | ENSG00000166582 | CENPV | chr17 | - | 16349139 | 16349333 | 16348615 | 16348685 | 16349930 | 16349956 | 0 | 1 | 3 | 6 | 2 | 8 | 3 | 3 | 5 | 3 | 0 | 2 | 0 | 9 | 8 | 0 | 5 | 2 | 0 | 8 | 298 | 149 | 1 | [illegible] | [illegible] | [illegible] | [illegible] | [illegible] | [illegible] | [illegible] | [illegible] | [illegible] | [illegible] | [illegible] | [illegible] |
| SE | 10737 | ENSG00000286065 | SLFN12L | chr17 | - | 35478074 | 35478185 | 35474903 | 35475485 | 35522278 | 35522969 | 137 | 94 | 91 | 64 | 80 | 0 | 1 | 1 | 0 | 4 | 50 | 71 | 87 | 59 | 17 | 0 | 0 | 0 | 0 | 0 | 260 | 149 | 1 | [illegible] | [illegible] | [illegible] | [illegible] | [illegible] | [illegible] | [illegible] | [illegible] | [illegible] | [illegible] | [illegible] | [illegible] |
| SE | 10740 | ENSG00000215067 | ALOX12-AS1 | chr17 | - | 7002483 | 7002620 | 6999146 | 6999586 | 7009795 | 7009889 | 21 | 15 | 17 | 16 | 47 | 0 | 0 | 0 | 1 | 0 | 21 | 23 | 11 | 6 | 9 | 0 | 0 | 0 | 0 | 0 | 286 | 149 | 1 | [illegible] | [illegible] | [illegible] | [illegible] | [illegible] | [illegible] | [illegible] | [illegible] | [illegible] | [illegible] | [illegible] | [illegible] |
| SE | 10742 | ENSG00000215067 | ALOX12-AS1 | chr17 | - | 7009795 | 7009889 | 7001131 | 7002620 | 7010058 | 7010137 | 15 | 13 | 8 | 11 | 34 | 0 | 0 | 0 | 0 | 1 | 29 | 12 | 10 | 4 | 9 | 0 | 0 | 0 | 0 | 0 | 243 | 149 | 1 | [illegible] | [illegible] | [illegible] | [illegible] | [illegible] | [illegible] | [illegible] | [illegible] | [illegible] | [illegible] | [illegible] | [illegible] |
| SE | 10744 | ENSG00000215067 | ALOX12-AS1 | chr17 | - | 7010058 | 7010137 | 7002483 | 7002620 | 7012195 | 7012331 | 1 | 0 | 0 | 0 | 3 | 11 | 12 | 21 | 20 | 45 | 0 | 0 | 0 | 0 | 0 | 8 | 33 | 12 | 25 | 17 | 228 | 149 | 1 | [illegible] | [illegible] | [illegible] | [illegible] | [illegible] | [illegible] | [illegible] | [illegible] | [illegible] | [illegible] | [illegible] | [illegible] |
| SE | 10746 | ENSG00000131899 | LLGL1 | chr17 | + | 18241450 | 18241715 | 18240577 | 18240873 | 18241884 | 18241999 | 41 | 10 | 24 | 32 | 4 | 0 | 0 | 0 | 0 | 0 | 39 | 28 | 17 | 22 | 61 | 0 | 0 | 0 | 0 | 0 | 298 | 149 | 1 | [illegible] | [illegible] | [illegible] | [illegible] | [illegible] | [illegible] | [illegible] | [illegible] | [illegible] | [illegible] | [illegible] | [illegible] |
| SE | 10748 | ENSG00000108256 | NUFIP2 | chr17 | - | 29267497 | 29267530 | 29255838 | 29264591 | 29285991 | 29287716 | 257 | 278 | 191 | 305 | 467 | 0 | 1 | 0 | 2 | 0 | 79 | 190 | 328 | 122 | 15 | 0 | 0 | 0 | 0 | 0 | 182 | 149 | 1 | [illegible] | [illegible] | [illegible] | [illegible] | [illegible] | [illegible] | [illegible] | [illegible] | [illegible] | [illegible] | [illegible] | [illegible] |
| SE | 10758 | ENSG00000154760 | SLFN13 | chr17 | - | 35447267 | 35447414 | 35444614 | 35445693 | 35448721 | 35448764 | 81 | 14 | 31 | 37 | 16 | 0 | 0 | 0 | 0 | 0 | 14 | 28 | 40 | 80 | 2 | 0 | 0 | 1 | 3 | 0 | 296 | 149 | 1 | [illegible] | [illegible] | [illegible] | [illegible] | [illegible] | [illegible] | [illegible] | [illegible] | [illegible] | [illegible] | [illegible] | [illegible] |
| SE | 10759 | ENSG00000236320 | SLFN14 | chr17 | - | 35554575 | 35554704 | 35552729 | 35553444 | 35557002 | 35558098 | 10 | 52 | 3 | 46 | 17 | 0 | 0 | 0 | 0 | 0 | 44 | 56 | 29 | 61 | 41 | 0 | 0 | 1 | 1 | 0 | 278 | 149 | 1 | [illegible] | [illegible] | [illegible] | [illegible] | [illegible] | [illegible] | [illegible] | [illegible] | [illegible] | [illegible] | [illegible] | [illegible] |
| SE | 10762 | ENSG00000118046 | STK11 | chr19 | + | 1220372 | 1220505 | 1219323 | 1219413 | 1220580 | 1220617 | 173 | 105 | 181 | 197 | 177 | 0 | 0 | 1 | 0 | 4 | 109 | 162 | 53 | 68 | 215 | 0 | 0 | 0 | 3 | 0 | 282 | 149 | 1 | [illegible] | [illegible] | [illegible] | [illegible] | [illegible] | [illegible] | [illegible] | [illegible] | [illegible] | [illegible] | [illegible] | [illegible] |
| SE | 10763 | ENSG00000118046 | STK11 | chr19 | + | 1223637 | 1226663 | 1223147 | 1223172 | 1227592 | 1228095 | 54 | 25 | 35 | 51 | 30 | 0 | 1 | 0 | 2 | 2 | 21 | 34 | 9 | 36 | 55 | 0 | 0 | 0 | 0 | 0 | 298 | 149 | 1 | [illegible] | [illegible] | [illegible] | [illegible] | [illegible] | [illegible] | [illegible] | [illegible] | [illegible] | [illegible] | [illegible] | [illegible] |
| SE | 10764 | ENSG00000118046 | STK11 | chr19 | + | 1226453 | 1226663 | 1222984 | 1223172 | 1227592 | 1228431 | 167 | 63 | 99 | 100 | 88 | 0 | 1 | 0 | 2 | 2 | 46 | 102 | 30 | 52 | 162 | 0 | 0 | 0 | 0 | 0 | 298 | 149 | 1 | [illegible] | [illegible] | [illegible] | [illegible] | [illegible] | [illegible] | [illegible] | [illegible] | [illegible] | [illegible] | [illegible] | [illegible] |
| SE | 10765 | ENSG00000172171 | TEFM | chr17 | - | 30900412 | 30900521 | 30898985 | 30899606 | 30904065 | 30904529 | 15 | 12 | 10 | 9 | 10 | 0 | 6 | 2 | 2 | 3 | 0 | 23 | 0 | 2 | 0 | 0 | 0 | 0 | 1 | 0 | 258 | 149 | 1 | [illegible] | [illegible] | [illegible] | [illegible] | [illegible] | [illegible] | [illegible] | [illegible] | [illegible] | [illegible] | [illegible] | [illegible] |
| SE | 10766 | ENSG00000172171 | TEFM | chr17 | - | 30900412 | 30900562 | 30898985 | 30899606 | 30904065 | 30904529 | 21 | 28 | 29 | 27 | 49 | 0 | 6 | 2 | 2 | 3 | 4 | 23 | 5 | 9 | 0 | 0 | 0 | 1 | 0 | 0 | 298 | 149 | 1 | [illegible] | [illegible] | [illegible] | [illegible] | [illegible] | [illegible] | [illegible] | [illegible] | [illegible] | [illegible] | [illegible] | [illegible] |
| SE | 10767 | ENSG00000132205 | EMILIN2 | chr18 | + | 2847808 | 2847931 | 2847005 | 2847322 | 2884963 | 2885139 | 223 | 125 | 89 | 169 | 159 | 0 | 0 | 0 | 0 | 0 | 912 | 276 | 161 | 416 | 614 | 6 | 0 | 0 | 0 | 0 | 272 | 149 | 1 | [illegible] | [illegible] | [illegible] | [illegible] | [illegible] | [illegible] | [illegible] | [illegible] | [illegible] | [illegible] | [illegible] | [illegible] |
| SE | 10768 | ENSG00000132205 | EMILIN2 | chr18 | + | 2890560 | 2892486 | 2884963 | 2885139 | 2906782 | 2907085 | 313 | 213 | 153 | 261 | 251 | 2 | 0 | 0 | 0 | 0 | 1197 | 348 | 182 | 466 | 626 | 0 | 2 | 0 | 7 | 0 | 298 | 149 | 1 | [illegible] | [illegible] | [illegible] | [illegible] | [illegible] | [illegible] | [illegible] | [illegible] | [illegible] | [illegible] | [illegible] | [illegible] |
| SE | 10772 | ENSG00000108578 | BLMH | chr17 | - | 30271270 | 30271388 | 30266878 | 30266954 | 30272560 | 30272628 | 147 | 126 | 147 | 172 | 246 | 0 | 0 | 0 | 0 | 0 | 278 | 147 | 191 | 133 | 122 | 0 | 0 | 1 | 0 | 0 | 267 | 149 | 1 | [illegible] | [illegible] | [illegible] | [illegible] | [illegible] | [illegible] | [illegible] | [illegible] | [illegible] | [illegible] | [illegible] | [illegible] |
| SE | 10773 | ENSG00000108578 | BLMH | chr17 | - | 30285387 | 30285480 | 30274037 | 30274197 | 30286813 | 30286851 | 307 | 196 | 285 | 206 | 393 | 1 | 0 | 0 | 0 | 0 | 272 | 241 | 303 | 244 | 166 | 0 | 0 | 0 | 0 | 0 | 242 | 149 | 1 | [illegible] | [illegible] | [illegible] | [illegible] | [illegible] | [illegible] | [illegible] | [illegible] | [illegible] | [illegible] | [illegible] | [illegible] |
| SE | 10775 | ENSG00000105556 | MIER2 | chr19 | - | 311844 | 311939 | 308800 | 308925 | 312190 | 312272 | 37 | 23 | 22 | 33 | 29 | 0 | 0 | 0 | 0 | 0 | 40 | 37 | 25 | 18 | 62 | 0 | 0 | 0 | 1 | 0 | 244 | 149 | 1 | [illegible] | [illegible] | [illegible] | [illegible] | [illegible] | [illegible] | [illegible] | [illegible] | [illegible] | [illegible] | [illegible] | [illegible] |
| SE | 10778 | ENSG00000176658 | MYO1D | chr17 | - | 32654476 | 32654621 | 32653842 | 32653947 | 32659114 | 32659338 | 32 | 4 | 63 | 53 | 44 | 0 | 0 | 0 | 1 | 0 | 4 | 17 | 32 | 8 | 21 | 0 | 0 | 0 | 0 | 0 | 294 | 149 | 1 | [illegible] | [illegible] | [illegible] | [illegible] | [illegible] | [illegible] | [illegible] | [illegible] | [illegible] | [illegible] | [illegible] | [illegible] |
| SE | 10779 | ENSG00000176658 | MYO1D | chr17 | - | 32745521 | 32745689 | 32745210 | 32745285 | 32748935 | 32749006 | 0 | 0 | 0 | 0 | 0 | 14 | 7 | 25 | 19 | 21 | 0 | 0 | 1 | 0 | 0 | 0 | 5 | 16 | 1 | 6 | 298 | 149 | 1 | [illegible] | [illegible] | [illegible] | [illegible] | [illegible] | [illegible] | [illegible] | [illegible] | [illegible] | [illegible] | [illegible] | [illegible] |
| SE | 10780 | ENSG00000108582 | CPD | chr17 | + | 30384988 | 30385236 | 30378926 | 30379726 | 30420840 | 30420983 | 222 | 119 | 148 | 178 | 137 | 0 | 0 | 0 | 1 | 0 | 82 | 161 | 300 | 176 | 38 | 0 | 0 | 0 | 0 | 0 | 298 | 149 | 1 | [illegible] | [illegible] | [illegible] | [illegible] | [illegible] | [illegible] | [illegible] | [illegible] | [illegible] | [illegible] | [illegible] | [illegible] |
| SE | 10781 | ENSG00000108582 | CPD | chr17 | + | 30421663 | 30421833 | 30420840 | 30420983 | 30422673 | 30423023 | 321 | 140 | 210 | 256 | 500 | 0 | 0 | 0 | 1 | 2 | 49 | 372 | 286 | 111 | 41 | 0 | 0 | 3 | 0 | 0 | 298 | 149 | 1 | [illegible] | [illegible] | [illegible] | [illegible] | [illegible] | [illegible] | [illegible] | [illegible] | [illegible] | [illegible] | [illegible] | [illegible] |
| SE | 10785 | ENSG00000108582 | CPD | chr17 | + | 30451756 | 30451846 | 30445690 | 30446020 | 30455338 | 30455470 | 211 | 93 | 132 | 187 | 252 | 0 | 1 | 2 | 0 | 0 | 17 | 186 | 246 | 69 | 1 | 0 | 0 | 0 | 0 | 0 | 239 | 149 | 1 | [illegible] | [illegible] | [illegible] | [illegible] | [illegible] | [illegible] | [illegible] | [illegible] | [illegible] | [illegible] | [illegible] | [illegible] |
| SE | 10786 | ENSG00000108582 | CPD | chr17 | + | 30461179 | 30461311 | 30456461 | 30456526 | 30461876 | 30462062 | 226 | 104 | 143 | 181 | 203 | 0 | 0 | 0 | 0 | 0 | 19 | 220 | 246 | 50 | 0 | 0 | 0 | 3 | 0 | 0 | 281 | 149 | 1 | [illegible] | [illegible] | [illegible] | [illegible] | [illegible] | [illegible] | [illegible] | [illegible] | [illegible] | [illegible] | [illegible] | [illegible] |
| SE | 10788 | ENSG00000274180 | NATD1 | chr17 | - | 21244105 | 21244224 | 21238869 | 21243429 | 21253158 | 21253410 | 11 | 3 | 5 | 10 | 0 | 0 | 0 | 0 | 0 | 0 | 0 | 10 | 3 | 10 | 13 | 0 | 0 | 0 | 0 | 0 | 268 | 149 | 1 | [illegible] | [illegible] | [illegible] | [illegible] | [illegible] | [illegible] | [illegible] | [illegible] | [illegible] | [illegible] | [illegible] | [illegible] |
| SE | 10795 | ENSG00000154898 | CCDC144CP | chr17 | + | 20392363 | 20392584 | 20391474 | 20391768 | 20400520 | 20400563 | 6 | 10 | 1 | 8 | 18 | 1 | 0 | 0 | 1 | 0 | 0 | 4 | 0 | 0 | 0 | 0 | 0 | 0 | 0 | 0 | 298 | 149 | 1 | [illegible] | [illegible] | [illegible] | [illegible] | [illegible] | [illegible] | [illegible] | [illegible] | [illegible] | [illegible] | [illegible] | [illegible] |
| SE | 10796 | ENSG00000154898 | CCDC144CP | chr17 | + | 20394020 | 20394230 | 20392363 | 20392584 | 20400520 | 20400563 | 0 | 0 | 0 | 0 | 2 | 0 | 1 | 0 | 4 | 6 | 0 | 1 | 0 | 0 | 0 | 0 | 3 | 0 | 0 | 0 | 298 | 149 | 1 | [illegible] | [illegible] | [illegible] | [illegible] | [illegible] | [illegible] | [illegible] | [illegible] | [illegible] | [illegible] | [illegible] | [illegible] |
| SE | 10798 | ENSG00000091542 | ALKBH5 | chr17 | + | 18194954 | 18195035 | 18183827 | 18185013 | 18206814 | 18206970 | 274 | 100 | 216 | 247 | 106 | 1 | 0 | 0 | 0 | 2 | 229 | 267 | 201 | 202 | 437 | 0 | 3 | 0 | 0 | 0 | 230 | 149 | 1 | [illegible] | [illegible] | [illegible] | [illegible] | [illegible] | [illegible] | [illegible] | [illegible] | [illegible] | [illegible] | [illegible] | [illegible] |
| SE | 10803 | ENSG00000177731 | FLII | chr17 | - | 18245353 | 18245419 | 18244814 | 18245272 | 18245554 | 18245660 | 286 | 163 | 216 | 241 | 253 | 0 | 0 | 0 | 1 | 0 | 314 | 269 | 165 | 205 | 507 | 0 | 0 | 0 | 0 | 0 | 215 | 149 | 1 | [illegible] | [illegible] | [illegible] | [illegible] | [illegible] | [illegible] | [illegible] | [illegible] | [illegible] | [illegible] | [illegible] | [illegible] |
| SE | 10804 | ENSG00000177731 | FLII | chr17 | - | 18245743 | 18245850 | 18245353 | 18245660 | 18245933 | 18245989 | 369 | 177 | 233 | 318 | 322 | 0 | 0 | 0 | 1 | 2 | 445 | 370 | 188 | 276 | 713 | 0 | 0 | 0 | 0 | 0 | 256 | 149 | 1 | [illegible] | [illegible] | [illegible] | [illegible] | [illegible] | [illegible] | [illegible] | [illegible] | [illegible] | [illegible] | [illegible] | [illegible] |
| SE | 10805 | ENSG00000177731 | FLII | chr17 | - | 18247656 | 18247848 | 18247168 | 18247357 | 18247928 | 18248033 | 328 | 147 | 282 | 270 | 152 | 0 | 0 | 7 | 0 | 0 | 301 | 380 | 165 | 219 | 922 | 0 | 0 | 6 | 0 | 4 | 298 | 149 | 1 | [illegible] | [illegible] | [illegible] | [illegible] | [illegible] | [illegible] | [illegible] | [illegible] | [illegible] | [illegible] | [illegible] | [illegible] |
| SE | 10806 | ENSG00000177731 | FLII | chr17 | - | 18248799 | 18248883 | 18248549 | 18248721 | 18249126 | 18249201 | 575 | 394 | 487 | 562 | 603 | 2 | 1 | 0 | 0 | 0 | 1230 | 868 | 537 | 787 | 2099 | 0 | 0 | 7 | 0 | 12 | 233 | 149 | 1 | [illegible] | [illegible] | [illegible] | [illegible] | [illegible] | [illegible] | [illegible] | [illegible] | [illegible] | [illegible] | [illegible] | [illegible] |
| SE | 10807 | ENSG00000177731 | FLII | chr17 | - | 18249126 | 18249201 | 18248549 | 18248721 | 18249325 | 18249408 | 201 | 111 | 190 | 198 | 263 | 0 | 0 | 0 | 0 | 0 | 428 | 286 | 233 | 180 | 838 | 0 | 0 | 0 | 1 | 0 | 224 | 149 | 1 | [illegible] | [illegible] | [illegible] | [illegible] | [illegible] | [illegible] | [illegible] | [illegible] | [illegible] | [illegible] | [illegible] | [illegible] |
| SE | 10808 | ENSG00000177731 | FLII | chr17 | - | 18249325 | 18249408 | 18249126 | 18249201 | 18250837 | 18251017 | 515 | 326 | 472 | 528 | 513 | 0 | 1 | 4 | 1 | 2 | 1060 | 755 | 521 | 612 | 2020 | 0 | 2 | 0 | 0 | 0 | 232 | 149 | 1 | [illegible] | [illegible] | [illegible] | [illegible] | [illegible] | [illegible] | [illegible] | [illegible] | [illegible] | [illegible] | [illegible] | [illegible] |
| SE | 10809 | ENSG00000177731 | FLII | chr17 | - | 18249325 | 18249408 | 18249126 | 18249201 | 18251264 | 18251477 | 194 | 140 | 162 | 200 | 200 | 0 | 0 | 0 | 0 | 0 | 458 | 339 | 200 | 295 | 785 | 0 | 0 | 0 | 1 | 0 | 232 | 149 | 1 | [illegible] | [illegible] | [illegible] | [illegible] | [illegible] | [illegible] | [illegible] | [illegible] | [illegible] | [illegible] | [illegible] | [illegible] |
| SE | 10810 | ENSG00000177731 | FLII | chr17 | - | 18250837 | 18251017 | 18249126 | 18249201 | 18251264 | 18251477 | 225 | 110 | 167 | 179 | 195 | 0 | 0 | 0 | 0 | 0 | 269 | 239 | 145 | 180 | 581 | 0 | 0 | 0 | 1 | 0 | 298 | 149 | 1 | [illegible] | [illegible] | [illegible] | [illegible] | [illegible] | [illegible] | [illegible] | [illegible] | [illegible] | [illegible] | [illegible] | [illegible] |
| SE | 10811 | ENSG00000177731 | FLII | chr17 | - | 18250837 | 18251017 | 18249325 | 18249408 | 18251264 | 18251477 | 547 | 295 | 473 | 508 | 507 | 0 | 0 | 0 | 0 | 0 | 879 | 657 | 466 | 497 | 1821 | 8 | 4 | 0 | 0 | 5 | 298 | 149 | 1 | [illegible] | [illegible] | [illegible] | [illegible] | [illegible] | [illegible] | [illegible] | [illegible] | [illegible] | [illegible] | [illegible] | [illegible] |
| SE | 10812 | ENSG00000177731 | FLII | chr17 | - | 18253543 | 18253719 | 18253300 | 18253458 | 18254078 | 18254178 | 224 | 104 | 177 | 147 | 137 | 3 | 0 | 0 | 0 | 0 | 204 | 166 | 102 | 91 | 486 | 0 | 0 | 1 | 4 | 0 | 298 | 149 | 1 | [illegible] | [illegible] | [illegible] | [illegible] | [illegible] | [illegible] | [illegible] | [illegible] | [illegible] | [illegible] | [illegible] | [illegible] |
| SE | 10813 | ENSG00000177731 | FLII | chr17 | - | 18253543 | 18254182 | 18253300 | 18253458 | 18254520 | 18254638 | 233 | 111 | 184 | 206 | 145 | 0 | 0 | 0 | 0 | 0 | 192 | 243 | 121 | 119 | 510 | 0 | 0 | 0 | 3 | 0 | 298 | 149 | 1 | [illegible] | [illegible] | [illegible] | [illegible] | [illegible] | [illegible] | [illegible] | [illegible] | [illegible] | [illegible] | [illegible] | [illegible] |
| SE | 10814 | ENSG00000177731 | FLII | chr17 | - | 18253598 | 18253719 | 18253300 | 18253458 | 18254078 | 18254182 | 84 | 43 | 63 | 50 | 46 | 3 | 0 | 0 | 0 | 0 | 71 | 52 | 31 | 43 | 183 | 0 | 0 | 1 | 4 | 0 | 270 | 149 | 1 | [illegible] | [illegible] | [illegible] | [illegible] | [illegible] | [illegible] | [illegible] | [illegible] | [illegible] | [illegible] | [illegible] | [illegible] |
| SE | 10815 | ENSG00000177731 | FLII | chr17 | - | 18254078 | 18254182 | 18253356 | 18253458 | 18254520 | 18254619 | 106 | 55 | 79 | 116 | 58 | 0 | 0 | 0 | 0 | 0 | 70 | 140 | 55 | 79 | 226 | 0 | 0 | 0 | 1 | 0 | 253 | 149 | 1 | [illegible] | [illegible] | [illegible] | [illegible] | [illegible] | [illegible] | [illegible] | [illegible] | [illegible] | [illegible] | [illegible] | [illegible] |
| SE | 10818 | ENSG00000177731 | FLII | chr17 | - | 18255182 | 18255263 | 18254768 | 18254854 | 18256525 | 18256597 | 168 | 94 | 137 | 163 | 152 | 1 | 0 | 0 | 0 | 0 | 225 | 234 | 141 | 228 | 521 | 0 | 3 | 0 | 1 | 0 | 230 | 149 | 1 | [illegible] | [illegible] | [illegible] | [illegible] | [illegible] | [illegible] | [illegible] | [illegible] | [illegible] | [illegible] | [illegible] | [illegible] |
| SE | 10826 | ENSG00000109046 | WSB1 | chr17 | + | 27297186 | 27297308 | 27294079 | 27294435 | 27301787 | 27301956 | 2 | 0 | 0 | 2 | 0 | 149 | 97 | 109 | 122 | 98 | 0 | 0 | 0 | 0 | 0 | 93 | 78 | 60 | 78 | 37 | 271 | 149 | 1 | [illegible] | [illegible] | [illegible] | [illegible] | [illegible] | [illegible] | [illegible] | [illegible] | [illegible] | [illegible] | [illegible] | [illegible] |
| SE | 10834 | ENSG00000109046 | WSB1 | chr17 | + | 27304779 | 27304911 | 27302434 | 27303635 | 27306781 | 27306858 | 1093 | 960 | 786 | 919 | 1269 | 1 | 0 | 2 | 0 | 1 | 1087 | 797 | 770 | 965 | 551 | 0 | 1 | 0 | 0 | 0 | 281 | 149 | 1 | [illegible] | [illegible] | [illegible] | [illegible] | [illegible] | [illegible] | [illegible] | [illegible] | [illegible] | [illegible] | [illegible] | [illegible] |
| SE | 10838 | ENSG00000142494 | SLC47A1 | chr17 | + | 19551423 | 19551468 | 19549634 | 19549677 | 19566789 | 19566859 | 0 | 1 | 0 | 0 | 0 | 0 | 0 | 0 | 0 | 0 | 0 | 0 | 0 | 0 | 0 | 5 | 0 | 0 | 0 | 0 | 194 | 149 | 1 | [illegible] | [illegible] | [illegible] | [illegible] | [illegible] | [illegible] | [illegible] | [illegible] | [illegible] | [illegible] | [illegible] | [illegible] |
| SE | 10840 | ENSG00000142494 | SLC47A1 | chr17 | + | 19567095 | 19567228 | 19566789 | 19566859 | 19571477 | 19571572 | 0 | 4 | 0 | 3 | 5 | 0 | 2 | 0 | 0 | 0 | 0 | 0 | 0 | 11 | 14 | 0 | 0 | 0 | 0 | 0 | 282 | 149 | 1 | [illegible] | [illegible] | [illegible] | [illegible] | [illegible] | [illegible] | [illegible] | [illegible] | [illegible] | [illegible] | [illegible] | [illegible] |
| SE | 10841 | ENSG00000142494 | SLC47A1 | chr17 | + | 19571477 | 19571572 | 19567095 | 19567228 | 19572779 | 19572861 | 4 | 5 | 4 | 5 | 5 | 0 | 1 | 2 | 0 | 0 | 0 | 0 | 0 | 3 | 7 | 0 | 0 | 0 | 0 | 0 | 244 | 149 | 1 | [illegible] | [illegible] | [illegible] | [illegible] | [illegible] | [illegible] | [illegible] | [illegible] | [illegible] | [illegible] | [illegible] | [illegible] |
| SE | 10842 | ENSG00000142494 | SLC47A1 | chr17 | + | 19572779 | 19572861 | 19567095 | 19567228 | 19577326 | 19577514 | 0 | 4 | 7 | 1 | 1 | 0 | 0 | 0 | 0 | 1 | 5 | 0 | 0 | 8 | 0 | 0 | 0 | 0 | 0 | 0 | 231 | 149 | 1 | [illegible] | [illegible] | [illegible] | [illegible] | [illegible] | [illegible] | [illegible] | [illegible] | [illegible] | [illegible] | [illegible] | [illegible] |
| SE | 10845 | ENSG00000029725 | RABEP1 | chr17 | + | 5308693 | 5308822 | 5282283 | 5282520 | 5331948 | 5332152 | 146 | 134 | 130 | 143 | 144 | 1 | 1 | 0 | 0 | 0 | 87 | 190 | 174 | 110 | 56 | 0 | 6 | 1 | 0 | 0 | 278 | 149 | 1 | [illegible] | [illegible] | [illegible] | [illegible] | [illegible] | [illegible] | [illegible] | [illegible] | [illegible] | [illegible] | [illegible] | [illegible] |
| SE | 10847 | ENSG00000029725 | RABEP1 | chr17 | + | 5335183 | 5335344 | 5331948 | 5332152 | 5346789 | 5346925 | 285 | 313 | 242 | 330 | 303 | 0 | 0 | 0 | 1 | 0 | 163 | 453 | 452 | 314 | 175 | 0 | 0 | 3 | 1 | 0 | 298 | 149 | 1 | [illegible] | [illegible] | [illegible] | [illegible] | [illegible] | [illegible] | [illegible] | [illegible] | [illegible] | [illegible] | [illegible] | [illegible] |
| SE | 10848 | ENSG00000029725 | RABEP1 | chr17 | + | 5338018 | 5338138 | 5331948 | 5332152 | 5346789 | 5346925 | 191 | 165 | 149 | 203 | 386 | 0 | 0 | 2 | 1 | 0 | 41 | 198 | 128 | 79 | 6 | 0 | 0 | 3 | 1 | 0 | 269 | 149 | 1 | [illegible] | [illegible] | [illegible] | [illegible] | [illegible] | [illegible] | [illegible] | [illegible] | [illegible] | [illegible] | [illegible] | [illegible] |
| SE | 10849 | ENSG00000029725 | RABEP1 | chr17 | + | 5338018 | 5338138 | 5335183 | 5335344 | 5346789 | 5346925 | 363 | 289 | 274 | 363 | 608 | 1 | 0 | 0 | 0 | 0 | 61 | 348 | 207 | 147 | 9 | 0 | 0 | 0 | 0 | 0 | 269 | 149 | 1 | [illegible] | [illegible] | [illegible] | [illegible] | [illegible] | [illegible] | [illegible] | [illegible] | [illegible] | [illegible] | [illegible] | [illegible] |
| SE | 10852 | ENSG00000029725 | RABEP1 | chr17 | + | 5361207 | 5361675 | 5354358 | 5354490 | 5362911 | 5363016 | 533 | 354 | 366 | 509 | 340 | 1 | 0 | 0 | 0 | 0 | 155 | 407 | 272 | 211 | 169 | 0 | 0 | 0 | 0 | 0 | 298 | 149 | 1 | [illegible] | [illegible] | [illegible] | [illegible] | [illegible] | [illegible] | [illegible] | [illegible] | [illegible] | [illegible] | [illegible] | [illegible] |
| SE | 10854 | ENSG00000029725 | RABEP1 | chr17 | + | 5365121 | 5365238 | 5362911 | 5363016 | 5368369 | 5368468 | 154 | 117 | 153 | 171 | 115 | 1 | 0 | 0 | 0 | 0 | 77 | 122 | 117 | 98 | 56 | 0 | 0 | 0 | 0 | 0 | 266 | 149 | 1 | [illegible] | [illegible] | [illegible] | [illegible] | [illegible] | [illegible] | [illegible] | [illegible] | [illegible] | [illegible] | [illegible] | [illegible] |
| SE | 10855 | ENSG00000029725 | RABEP1 | chr17 | + | 5377115 | 5377305 | 5373313 | 5373454 | 5378176 | 5378232 | 158 | 131 | 105 | 154 | 147 | 2 | 0 | 0 | 0 | 0 | 12 | 128 | 85 | 53 | 11 | 0 | 0 | 0 | 0 | 0 | 298 | 149 | 1 | [illegible] | [illegible] | [illegible] | [illegible] | [illegible] | [illegible] | [illegible] | [illegible] | [illegible] | [illegible] | [illegible] | [illegible] |
| SE | 10856 | ENSG00000029725 | RABEP1 | chr17 | + | 5380363 | 5380462 | 5378176 | 5378232 | 5381388 | 5381505 | 166 | 123 | 154 | 145 | 173 | 5 | 9 | 2 | 2 | 2 | 13 | 159 | 97 | 35 | 0 | 0 | 8 | 3 | 0 | 0 | 248 | 149 | 1 | [illegible] | [illegible] | [illegible] | [illegible] | [illegible] | [illegible] | [illegible] | [illegible] | [illegible] | [illegible] | [illegible] | [illegible] |
| SE | 10857 | ENSG00000029725 | RABEP1 | chr17 | + | 5380363 | 5380462 | 5378176 | 5378232 | 5383121 | 5386340 | 32 | 18 | 17 | 33 | 46 | 4 | 1 | 1 | 2 | 5 | 0 | 50 | 39 | 2 | 0 | 0 | 10 | 2 | 2 | 0 | 248 | 149 | 1 | [illegible] | [illegible] | [illegible] | [illegible] | [illegible] | [illegible] | [illegible] | [illegible] | [illegible] | [illegible] | [illegible] | [illegible] |
| SE | 10858 | ENSG00000029725 | RABEP1 | chr17 | + | 5381388 | 5381505 | 5378176 | 5378232 | 5383121 | 5383223 | 181 | 147 | 173 | 173 | 206 | 4 | 1 | 1 | 2 | 5 | 0 | 141 | 92 | 35 | 0 | 0 | 10 | 2 | 2 | 0 | 266 | 149 | 1 | [illegible] | [illegible] | [illegible] | [illegible] | [illegible] | [illegible] | [illegible] | [illegible] | [illegible] | [illegible] | [illegible] | [illegible] |
| SE | 10859 | ENSG00000029725 | RABEP1 | chr17 | + | 5381388 | 5381505 | 5380363 | 5380462 | 5383121 | 5383223 | 307 | 242 | 311 | 289 | 354 | 0 | 2 | 4 | 5 | 23 | 6 | 239 | 156 | 68 | 0 | 0 | 5 | 11 | 0 | 0 | 266 | 149 | 1 | [illegible] | [illegible] | [illegible] | [illegible] | [illegible] | [illegible] | [illegible] | [illegible] | [illegible] | [illegible] | [illegible] | [illegible] |
| SE | 10861 | ENSG00000167840 | ZNF232 | chr17 | - | 5108925 | 5109052 | 5105540 | 5106506 | 5109393 | 5109868 | 8 | 0 | 2 | 2 | 5 | 1 | 0 | 0 | 1 | 4 | 0 | 0 | 2 | 3 | 0 | 5 | 0 | 0 | 0 | 0 | 276 | 149 | 1 | [illegible] | [illegible] | [illegible] | [illegible] | [illegible] | [illegible] | [illegible] | [illegible] | [illegible] | [illegible] | [illegible] | [illegible] |
| SE | 10862 | ENSG00000167840 | ZNF232 | chr17 | - | 5108925 | 5109052 | 5105634 | 5106533 | 5109393 | 5109868 | 10 | 5 | 3 | 4 | 6 | 5 | 7 | 0 | 2 | 9 | 0 | 1 | 2 | 0 | 0 | 0 | 0 | 0 | 0 | 0 | 276 | 149 | 1 | [illegible] | [illegible] | [illegible] | [illegible] | [illegible] | [illegible] | [illegible] | [illegible] | [illegible] | [illegible] | [illegible] | [illegible] |
| SE | 10863 | ENSG00000167840 | ZNF232 | chr17 | - | 5111075 | 5111219 | 5108925 | 5109868 | 5111799 | 5111815 | 12 | 5 | 9 | 8 | 5 | 1 | 0 | 4 | 0 | 0 | 0 | 26 | 3 | 1 | 7 | 0 | 0 | 0 | 0 | 0 | 293 | 149 | 1 | [illegible] | [illegible] | [illegible] | [illegible] | [illegible] | [illegible] | [illegible] | [illegible] | [illegible] | [illegible] | [illegible] | [illegible] |
| SE | 10864 | ENSG00000167840 | ZNF232 | chr17 | - | 5111075 | 5111219 | 5109393 | 5109768 | 5111799 | 5111815 | 7 | 0 | 4 | 3 | 0 | 0 | 0 | 0 | 0 | 0 | 0 | 21 | 4 | 13 | 5 | 0 | 0 | 3 | 2 | 0 | 293 | 149 | 1 | [illegible] | [illegible] | [illegible] | [illegible] | [illegible] | [illegible] | [illegible] | [illegible] | [illegible] | [illegible] | [illegible] | [illegible] |
| SE | 10865 | ENSG00000189152 | GRAPL | chr17 | + | 19144102 | 19144170 | 19138548 | 19138588 | 19158318 | 19158783 | 0 | 0 | 0 | 3 | 0 | 0 | 5 | 0 | 0 | 0 | 0 | 0 | 0 | 0 | 0 | 2 | 0 | 0 | 0 | 0 | 217 | 149 | 1 | [illegible] | [illegible] | [illegible] | [illegible] | [illegible] | [illegible] | [illegible] | [illegible] | [illegible] | [illegible] | [illegible] | [illegible] |
| SE | 10866 | ENSG00000108474 | PIGL | chr17 | + | 16233970 | 16234070 | 16217190 | 16217461 | 16299887 | 16299978 | 39 | 28 | 56 | 39 | 39 | 0 | 0 | 0 | 0 | 0 | 23 | 33 | 30 | 22 | 62 | 0 | 0 | 0 | 4 | 0 | 249 | 149 | 1 | [illegible] | [illegible] | [illegible] | [illegible] | [illegible] | [illegible] | [illegible] | [illegible] | [illegible] | [illegible] | [illegible] | [illegible] |
| SE | 10867 | ENSG00000108474 | PIGL | chr17 | + | 16279656 | 16279812 | 16233970 | 16234070 | 16284020 | 16284068 | 0 | 0 | 0 | 0 | 5 | 2 | 5 | 3 | 0 | 3 | 15 | 0 | 0 | 0 | 0 | 0 | 3 | 0 | 2 | 0 | 298 | 149 | 1 | [illegible] | [illegible] | [illegible] | [illegible] | [illegible] | [illegible] | [illegible] | [illegible] | [illegible] | [illegible] | [illegible] | [illegible] |
| SE | 10868 | ENSG00000108474 | PIGL | chr17 | + | 16316680 | 16316712 | 16313546 | 16313614 | 16317774 | 16317908 | 3 | 3 | 4 | 3 | 1 | 1 | 0 | 0 | 0 | 0 | 0 | 5 | 3 | 1 | 6 | 0 | 0 | 0 | 0 | 0 | 181 | 149 | 1 | [illegible] | [illegible] | [illegible] | [illegible] | [illegible] | [illegible] | [illegible] | [illegible] | [illegible] | [illegible] | [illegible] | [illegible] |
| SE | 10870 | ENSG00000181350 | LRRC75A | chr17 | - | 16446879 | 16447031 | 16441576 | 16444131 | 16447844 | 16448294 | 1 | 0 | 1 | 0 | 0 | 3 | 3 | 2 | 1 | 0 | 0 | 0 | 0 | 0 | 0 | 0 | 1 | 0 | 0 | 0 | 298 | 149 | 1 | [illegible] | [illegible] | [illegible] | [illegible] | [illegible] | [illegible] | [illegible] | [illegible] | [illegible] | [illegible] | [illegible] | [illegible] |
| SE | 10872 | ENSG00000251537 | NA | chr17 | - | 15607573 | 15607683 | 15605222 | 15605423 | 15612700 | 15612823 | 0 | 0 | 0 | 0 | 0 | 1 | 0 | 0 | 0 | 0 | 0 | 2 | 0 | 0 | 0 | 0 | 0 | 0 | 0 | 0 | 259 | 149 | 1 | [illegible] | [illegible] | [illegible] | [illegible] | [illegible] | [illegible] | [illegible] | [illegible] | [illegible] | [illegible] | [illegible] | [illegible] |
| SE | 10875 | ENSG00000198920 | KIAA0753 | chr17 | - | 6590509 | 6590578 | 6589806 | 6590003 | 6594971 | 6595053 | 1 | 12 | 14 | 10 | 13 | 0 | 2 | 0 | 0 | 0 | 0 | 10 | 6 | 14 | 0 | 0 | 0 | 0 | 0 | 0 | 218 | 149 | 1 | [illegible] | [illegible] | [illegible] | [illegible] | [illegible] | [illegible] | [illegible] | [illegible] | [illegible] | [illegible] | [illegible] | [illegible] |
| SE | 10876 | ENSG00000198920 | KIAA0753 | chr17 | - | 6590509 | 6590630 | 6589778 | 6590003 | 6594971 | 6594999 | 25 | 17 | 23 | 35 | 31 | 0 | 2 | 0 | 0 | 0 | 4 | 30 | 2 | 35 | 13 | 0 | 0 | 0 | 0 | 0 | 270 | 149 | 1 | [illegible] | [illegible] | [illegible] | [illegible] | [illegible] | [illegible] | [illegible] | [illegible] | [illegible] | [illegible] | [illegible] | [illegible] |
| SE | 10877 | ENSG00000198920 | KIAA0753 | chr17 | - | 6599236 | 6599320 | 6596157 | 6596343 | 6606872 | 6606928 | 7 | 10 | 7 | 11 | 3 | 3 | 0 | 0 | 0 | 0 | 0 | 7 | 3 | 13 | 9 | 0 | 0 | 0 | 0 | 0 | 233 | 149 | 1 | [illegible] | [illegible] | [illegible] | [illegible] | [illegible] | [illegible] | [illegible] | [illegible] | [illegible] | [illegible] | [illegible] | [illegible] |
| SE | 10880 | ENSG00000198920 | KIAA0753 | chr17 | - | 6620787 | 6620998 | 6611918 | 6612148 | 6622881 | 6623097 | 29 | 30 | 40 | 35 | 45 | 0 | 0 | 0 | 0 | 0 | 27 | 20 | 24 | 46 | 0 | 0 | 0 | 1 | 0 | 0 | 298 | 149 | 1 | [illegible] | [illegible] | [illegible] | [illegible] | [illegible] | [illegible] | [illegible] | [illegible] | [illegible] | [illegible] | [illegible] | [illegible] |
| SE | 10881 | ENSG00000198920 | KIAA0753 | chr17 | - | 6624754 | 6624861 | 6623508 | 6623571 | 6628116 | 6628487 | 7 | 2 | 7 | 5 | 11 | 1 | 1 | 0 | 0 | 0 | 0 | 4 | 4 | 2 | 0 | 0 | 0 | 3 | 1 | 0 | 256 | 149 | 1 | [illegible] | [illegible] | [illegible] | [illegible] | [illegible] | [illegible] | [illegible] | [illegible] | [illegible] | [illegible] | [illegible] | [illegible] |
| SE | 10882 | ENSG00000198920 | KIAA0753 | chr17 | - | 6624754 | 6624861 | 6623936 | 6624006 | 6628116 | 6628487 | 6 | 4 | 2 | 5 | 14 | 0 | 0 | 0 | 3 | 0 | 0 | 2 | 4 | 2 | 0 | 0 | 0 | 0 | 1 | 2 | 256 | 149 | 1 | [illegible] | [illegible] | [illegible] | [illegible] | [illegible] | [illegible] | [illegible] | [illegible] | [illegible] | [illegible] | [illegible] | [illegible] |
| SE | 10883 | ENSG00000198920 | KIAA0753 | chr17 | - | 6635010 | 6635171 | 6628116 | 6628741 | 6640636 | 6640711 | 22 | 14 | 22 | 17 | 13 | 0 | 0 | 0 | 1 | 0 | 29 | 31 | 18 | 9 | 11 | 0 | 0 | 0 | 0 | 0 | 298 | 149 | 1 | [illegible] | [illegible] | [illegible] | [illegible] | [illegible] | [illegible] | [illegible] | [illegible] | [illegible] | [illegible] | [illegible] | [illegible] |
| SE | 10884 | ENSG00000198920 | KIAA0753 | chr17 | - | 6635400 | 6635634 | 6635010 | 6635171 | 6640636 | 6640662 | 1 | 0 | 2 | 4 | 2 | 1 | 4 | 6 | 4 | 2 | 0 | 0 | 4 | 1 | 0 | 9 | 5 | 4 | 1 | 0 | 298 | 149 | 1 | [illegible] | [illegible] | [illegible] | [illegible] | [illegible] | [illegible] | [illegible] | [illegible] | [illegible] | [illegible] | [illegible] | [illegible] |
| SE | 10885 | ENSG00000262304 | NA | chr17 | - | 3609308 | 3610972 | 3592066 | 3592383 | 3621236 | 3621412 | 0 | 0 | 0 | 0 | 0 | 1 | 0 | 2 | 1 | 0 | 0 | 0 | 0 | 6 | 0 | 0 | 0 | 0 | 2 | 0 | 298 | 149 | 1 | [illegible] | [illegible] | [illegible] | [illegible] | [illegible] | [illegible] | [illegible] | [illegible] | [illegible] | [illegible] | [illegible] | [illegible] |
| SE | 10891 | ENSG00000129250 | KIF1C | chr17 | + | 4999012 | 4999046 | 4997953 | 4998156 | 4999849 | 4999970 | 0 | 0 | 1 | 0 | 0 | 6 | 2 | 7 | 13 | 0 | 0 | 0 | 0 | 0 | 0 | 20 | 20 | 6 | 2 | 20 | 183 | 149 | 1 | [illegible] | [illegible] | [illegible] | [illegible] | [illegible] | [illegible] | [illegible] | [illegible] | [illegible] | [illegible] | [illegible] | [illegible] |
| SE | 10893 | ENSG00000129250 | KIF1C | chr17 | + | 5002058 | 5002124 | 5001221 | 5001401 | 5002463 | 5002642 | 17 | 24 | 16 | 15 | 21 | 0 | 1 | 0 | 0 | 0 | 10 | 40 | 7 | 29 | 70 | 0 | 0 | 0 | 0 | 0 | 215 | 149 | 1 | [illegible] | [illegible] | [illegible] | [illegible] | [illegible] | [illegible] | [illegible] | [illegible] | [illegible] | [illegible] | [illegible] | [illegible] |
| SE | 10895 | ENSG00000176974 | SHMT1 | chr17 | - | 18333165 | 18333288 | 18330554 | 18330671 | 18340042 | 18340255 | 15 | 2 | 14 | 8 | 11 | 2 | 0 | 1 | 0 | 0 | 25 | 41 | 8 | 24 | 31 | 0 | 0 | 3 | 1 | 0 | 272 | 149 | 1 | [illegible] | [illegible] | [illegible] | [illegible] | [illegible] | [illegible] | [illegible] | [illegible] | [illegible] | [illegible] | [illegible] | [illegible] |
| SE | 10897 | ENSG00000176974 | SHMT1 | chr17 | - | 18348324 | 18348440 | 18347495 | 18347656 | 18353671 | 18353817 | 43 | 11 | 63 | 25 | 47 | 2 | 0 | 0 | 0 | 0 | 45 | 82 | 39 | 39 | 38 | 0 | 0 | 0 | 0 | 0 | 265 | 149 | 1 | [illegible] | [illegible] | [illegible] | [illegible] | [illegible] | [illegible] | [illegible] | [illegible] | [illegible] | [illegible] | [illegible] | [illegible] |
| SE | 10898 | ENSG00000176974 | SHMT1 | chr17 | - | 18360413 | 18360552 | 18355978 | 18356000 | 18363371 | 18363542 | 1 | 0 | 0 | 0 | 0 | 17 | 7 | 11 | 13 | 0 | 0 | 4 | 0 | 0 | 0 | 6 | 39 | 5 | 30 | 4 | 288 | 149 | 1 | [illegible] | [illegible] | [illegible] | [illegible] | [illegible] | [illegible] | [illegible] | [illegible] | [illegible] | [illegible] | [illegible] | [illegible] |
| SE | 10899 | ENSG00000074370 | ATP2A3 | chr17 | - | 3927980 | 3928038 | 3925290 | 3925441 | 3928226 | 3928327 | 5 | 0 | 0 | 1 | 0 | 3 | 0 | 1 | 2 | 0 | 0 | 1 | 7 | 0 | 0 | 0 | 0 | 1 | 9 | 13 | 207 | 149 | 1 | [illegible] | [illegible] | [illegible] | [illegible] | [illegible] | [illegible] | [illegible] | [illegible] | [illegible] | [illegible] | [illegible] | [illegible] |
| SE | 10901 | ENSG00000074370 | ATP2A3 | chr17 | - | 3927980 | 3928038 | 3925290 | 3925441 | 3928662 | 3928780 | 3 | 0 | 0 | 1 | 0 | 133 | 44 | 103 | 166 | 43 | 0 | 1 | 7 | 0 | 0 | 121 | 263 | 88 | 162 | 780 | 207 | 149 | 1 | [illegible] | [illegible] | [illegible] | [illegible] | [illegible] | [illegible] | [illegible] | [illegible] | [illegible] | [illegible] | [illegible] | [illegible] |
| SE | 10902 | ENSG00000074370 | ATP2A3 | chr17 | - | 3927980 | 3928038 | 3925345 | 3925438 | 3928226 | 3928327 | 0 | 0 | 0 | 0 | 0 | 0 | 0 | 1 | 0 | 0 | 0 | 0 | 0 | 0 | 8 | 0 | 0 | 0 | 0 | 0 | 207 | 149 | 1 | [illegible] | [illegible] | [illegible] | [illegible] | [illegible] | [illegible] | [illegible] | [illegible] | [illegible] | [illegible] | [illegible] | [illegible] |
| SE | 10903 | ENSG00000074370 | ATP2A3 | chr17 | - | 3927980 | 3928038 | 3925345 | 3925438 | 3928239 | 3928327 | 0 | 0 | 0 | 0 | 0 | 0 | 0 | 0 | 1 | 0 | 0 | 0 | 0 | 1 | 8 | 0 | 7 | 1 | 4 | 0 | 207 | 149 | 1 | [illegible] | [illegible] | [illegible] | [illegible] | [illegible] | [illegible] | [illegible] | [illegible] | [illegible] | [illegible] | [illegible] | [illegible] |
| SE | 10911 | ENSG00000074370 | ATP2A3 | chr17 | - | 3942605 | 3942731 | 3941435 | 3941654 | 3943390 | 3943522 | 490 | 204 | 476 | 581 | 457 | 0 | 0 | 0 | 0 | 0 | 631 | 593 | 325 | 444 | 2388 | 0 | 0 | 3 | 0 | 0 | 275 | 149 | 1 | [illegible] | [illegible] | [illegible] | [illegible] | [illegible] | [illegible] | [illegible] | [illegible] | [illegible] | [illegible] | [illegible] | [illegible] |
| SE | 10912 | ENSG00000074370 | ATP2A3 | chr17 | - | 3945059 | 3945148 | 3944703 | 3944806 | 3947390 | 3947855 | 71 | 15 | 53 | 93 | 50 | 0 | 0 | 0 | 0 | 0 | 26 | 146 | 55 | 62 | 597 | 0 | 0 | 0 | 0 | 1 | 238 | 149 | 1 | [illegible] | [illegible] | [illegible] | [illegible] | [illegible] | [illegible] | [illegible] | [illegible] | [illegible] | [illegible] | [illegible] | [illegible] |
| SE | 10914 | ENSG00000074370 | ATP2A3 | chr17 | - | 3951580 | 3951685 | 3951250 | 3951389 | 3953692 | 3953710 | 285 | 130 | 241 | 260 | 93 | 0 | 0 | 0 | 0 | 0 | 269 | 447 | 189 | 280 | 1540 | 0 | 0 | 0 | 0 | 10 | 254 | 149 | 1 | [illegible] | [illegible] | [illegible] | [illegible] | [illegible] | [illegible] | [illegible] | [illegible] | [illegible] | [illegible] | [illegible] | [illegible] |
| SE | 10918 | ENSG00000074370 | ATP2A3 | chr17 | - | 3953692 | 3953710 | 3951580 | 3951685 | 3964173 | 3964236 | 6 | 1 | 1 | 5 | 0 | 0 | 0 | 0 | 0 | 0 | 1 | 1 | 0 | 0 | 16 | 12 | 0 | 0 | 0 | 0 | 167 | 149 | 1 | [illegible] | [illegible] | [illegible] | [illegible] | [illegible] | [illegible] | [illegible] | [illegible] | [illegible] | [illegible] | [illegible] | [illegible] |
| SE | 10919 | ENSG00000074370 | ATP2A3 | chr17 | - | 3953692 | 3953710 | 3953280 | 3953429 | 3964173 | 3964236 | 87 | 55 | 55 | 61 | 16 | 0 | 0 | 1 | 1 | 0 | 85 | 86 | 45 | 36 | 326 | 0 | 3 | 0 | 0 | 0 | 167 | 149 | 1 | [illegible] | [illegible] | [illegible] | [illegible] | [illegible] | [illegible] | [illegible] | [illegible] | [illegible] | [illegible] | [illegible] | [illegible] |
| SE | 10921 | ENSG00000196267 | ZNF836 | chr19 | - | 52169647 | 52169781 | 52168057 | 52168152 | 52171335 | 52171621 | 77 | 67 | 115 | 84 | 48 | 2 | 0 | 0 | 0 | 0 | 23 | 62 | 41 | 30 | 43 | 0 | 4 | 0 | 0 | 0 | 283 | 149 | 1 | [illegible] | [illegible] | [illegible] | [illegible] | [illegible] | [illegible] | [illegible] | [illegible] | [illegible] | [illegible] | [illegible] | [illegible] |
| SE | 10923 | ENSG00000091592 | NLRP1 | chr17 | - | 5501736 | 5501872 | 5499426 | 5499950 | 5521523 | 5521786 | 10 | 11 | 10 | 22 | 3 | 1 | 0 | 0 | 0 | 0 | 20 | 5 | 8 | 34 | 27 | 3 | 0 | 0 | 0 | 0 | 285 | 149 | 1 | [illegible] | [illegible] | [illegible] | [illegible] | [illegible] | [illegible] | [illegible] | [illegible] | [illegible] | [illegible] | [illegible] | [illegible] |
| SE | 10924 | ENSG00000091592 | NLRP1 | chr17 | - | 5501736 | 5501872 | 5499426 | 5499950 | 5530480 | 5530704 | 10 | 11 | 10 | 22 | 3 | 1 | 0 | 1 | 0 | 0 | 20 | 5 | 7 | 35 | 27 | 0 | 0 | 0 | 0 | 0 | 285 | 149 | 1 | [illegible] | [illegible] | [illegible] | [illegible] | [illegible] | [illegible] | [illegible] | [illegible] | [illegible] | [illegible] | [illegible] | [illegible] |
| SE | 10926 | ENSG00000091592 | NLRP1 | chr17 | - | 5515472 | 5515517 | 5499427 | 5499950 | 5521523 | 5521786 | 2 | 0 | 0 | 1 | 0 | 0 | 0 | 0 | 0 | 0 | 2 | 0 | 0 | 3 | 6 | 3 | 0 | 0 | 0 | 0 | 194 | 149 | 1 | [illegible] | [illegible] | [illegible] | [illegible] | [illegible] | [illegible] | [illegible] | [illegible] | [illegible] | [illegible] | [illegible] | [illegible] |
| SE | 10927 | ENSG00000091592 | NLRP1 | chr17 | - | 5515472 | 5515517 | 5499427 | 5499950 | 5530480 | 5530704 | 2 | 0 | 0 | 0 | 0 | 1 | 0 | 0 | 0 | 0 | 2 | 0 | 0 | 6 | 0 | 0 | 0 | 0 | 0 | 0 | 194 | 149 | 1 | [illegible] | [illegible] | [illegible] | [illegible] | [illegible] | [illegible] | [illegible] | [illegible] | [illegible] | [illegible] | [illegible] | [illegible] |
| SE | 10929 | ENSG00000091592 | NLRP1 | chr17 | - | 5515472 | 5515517 | 5501813 | 5501872 | 5520880 | 5521012 | 15 | 13 | 13 | 15 | 10 | 1 | 0 | 0 | 0 | 0 | 87 | 11 | 23 | 7 | 55 | 0 | 0 | 0 | 0 | 0 | 194 | 149 | 1 | [illegible] | [illegible] | [illegible] | [illegible] | [illegible] | [illegible] | [illegible] | [illegible] | [illegible] | [illegible] | [illegible] | [illegible] |
| SE | 10934 | ENSG00000091592 | NLRP1 | chr17 | - | 5515472 | 5515517 | 5514117 | 5515073 | 5517745 | 5517887 | 138 | 118 | 191 | 168 | 126 | 0 | 0 | 0 | 0 | 0 | 388 | 175 | 144 | 206 | 330 | 0 | 0 | 0 | 0 | 0 | 194 | 149 | 1 | [illegible] | [illegible] | [illegible] | [illegible] | [illegible] | [illegible] | [illegible] | [illegible] | [illegible] | [illegible] | [illegible] | [illegible] |
| SE | 10935 | ENSG00000091592 | NLRP1 | chr17 | - | 5517745 | 5517887 | 5499426 | 5499950 | 5521523 | 5521786 | 51 | 68 | 76 | 51 | 52 | 0 | 0 | 0 | 0 | 0 | 183 | 87 | 94 | 66 | 201 | 3 | 0 | 0 | 0 | 0 | 291 | 149 | 1 | [illegible] | [illegible] | [illegible] | [illegible] | [illegible] | [illegible] | [illegible] | [illegible] | [illegible] | [illegible] | [illegible] | [illegible] |
| SE | 10936 | ENSG00000091592 | NLRP1 | chr17 | - | 5517745 | 5517887 | 5499426 | 5499950 | 5530480 | 5530704 | 1 | 8 | 6 | 4 | 6 | 1 | 0 | 1 | 0 | 0 | 36 | 8 | 4 | 14 | 14 | 0 | 0 | 0 | 0 | 0 | 291 | 149 | 1 | [illegible] | [illegible] | [illegible] | [illegible] | [illegible] | [illegible] | [illegible] | [illegible] | [illegible] | [illegible] | [illegible] | [illegible] |
| SE | 10937 | ENSG00000091592 | NLRP1 | chr17 | - | 5517745 | 5517887 | 5501399 | 5501872 | 5520880 | 5521012 | 375 | 346 | 416 | 412 | 334 | 1 | 0 | 0 | 0 | 0 | 1050 | 451 | 390 | 630 | 1042 | 0 | 0 | 0 | 0 | 0 | 291 | 149 | 1 | [illegible] | [illegible] | [illegible] | [illegible] | [illegible] | [illegible] | [illegible] | [illegible] | [illegible] | [illegible] | [illegible] | [illegible] |
| SE | 10938 | ENSG00000091592 | NLRP1 | chr17 | - | 5517745 | 5517887 | 5501399 | 5501872 | 5521523 | 5521786 | 215 | 246 | 273 | 244 | 187 | 0 | 0 | 0 | 1 | 0 | 753 | 284 | 292 | 385 | 716 | 0 | 0 | 0 | 0 | 6 | 291 | 149 | 1 | [illegible] | [illegible] | [illegible] | [illegible] | [illegible] | [illegible] | [illegible] | [illegible] | [illegible] | [illegible] | [illegible] | [illegible] |
| SE | 10939 | ENSG00000091592 | NLRP1 | chr17 | - | 5517745 | 5517887 | 5501399 | 5501872 | 5530480 | 5530704 | 165 | 186 | 203 | 197 | 141 | 0 | 0 | 0 | 1 | 0 | 606 | 205 | 202 | 333 | 524 | 0 | 0 | 0 | 1 | 6 | 291 | 149 | 1 | [illegible] | [illegible] | [illegible] | [illegible] | [illegible] | [illegible] | [illegible] | [illegible] | [illegible] | [illegible] | [illegible] | [illegible] |
| SE | 10940 | ENSG00000091592 | NLRP1 | chr17 | - | 5517745 | 5517887 | 5515472 | 5515517 | 5520880 | 5521012 | 356 | 277 | 392 | 368 | 334 | 0 | 0 | 0 | 0 | 0 | 786 | 390 | 308 | 468 | 839 | 8 | 1 | 0 | 0 | 0 | 291 | 149 | 1 | [illegible] | [illegible] | [illegible] | [illegible] | [illegible] | [illegible] | [illegible] | [illegible] | [illegible] | [illegible] | [illegible] | [illegible] |
| SE | 10942 | ENSG00000091592 | NLRP1 | chr17 | - | 5517745 | 5517887 | 5515472 | 5515517 | 5530480 | 5530704 | 146 | 116 | 179 | 153 | 141 | 0 | 0 | 0 | 0 | 0 | 347 | 144 | 121 | 173 | 314 | 0 | 0 | 6 | 0 | 0 | 291 | 149 | 1 | [illegible] | [illegible] | [illegible] | [illegible] | [illegible] | [illegible] | [illegible] | [illegible] | [illegible] | [illegible] | [illegible] | [illegible] |
| SE | 10943 | ENSG00000091592 | NLRP1 | chr17 | - | 5520880 | 5521012 | 5501399 | 5501872 | 5521523 | 5521786 | 128 | 78 | 123 | 118 | 106 | 0 | 0 | 0 | 1 | 0 | 235 | 130 | 117 | 86 | 221 | 0 | 0 | 0 | 0 | 6 | 281 | 149 | 1 | [illegible] | [illegible] | [illegible] | [illegible] | [illegible] | [illegible] | [illegible] | [illegible] | [illegible] | [illegible] | [illegible] | [illegible] |
| SE | 10946 | ENSG00000091592 | NLRP1 | chr17 | - | 5521523 | 5521786 | 5499426 | 5499950 | 5530480 | 5530704 | 93 | 41 | 105 | 106 | 103 | 1 | 0 | 1 | 0 | 0 | 219 | 101 | 93 | 80 | 276 | 0 | 0 | 0 | 0 | 0 | 298 | 149 | 1 | [illegible] | [illegible] | [illegible] | [illegible] | [illegible] | [illegible] | [illegible] | [illegible] | [illegible] | [illegible] | [illegible] | [illegible] |
| SE | 10947 | ENSG00000091592 | NLRP1 | chr17 | - | 5521523 | 5521786 | 5501399 | 5501872 | 5530480 | 5530704 | 93 | 41 | 105 | 107 | 103 | 0 | 0 | 0 | 1 | 0 | 216 | 101 | 94 | 80 | 282 | 0 | 0 | 0 | 1 | 6 | 298 | 149 | 1 | [illegible] | [illegible] | [illegible] | [illegible] | [illegible] | [illegible] | [illegible] | [illegible] | [illegible] | [illegible] | [illegible] | [illegible] |
| SE | 10950 | ENSG00000091592 | NLRP1 | chr17 | - | 5530480 | 5530704 | 5521523 | 5521786 | 5532821 | 5532984 | 241 | 155 | 252 | 323 | 325 | 0 | 0 | 4 | 0 | 0 | 652 | 253 | 256 | 234 | 614 | 0 | 5 | 2 | 0 | 0 | 298 | 149 | 1 | [illegible] | [illegible] | [illegible] | [illegible] | [illegible] | [illegible] | [illegible] | [illegible] | [illegible] | [illegible] | [illegible] | [illegible] |
| SE | 10951 | ENSG00000091592 | NLRP1 | chr17 | - | 5532821 | 5532984 | 5530480 | 5530704 | 5536850 | 5536940 | 159 | 127 | 159 | 223 | 231 | 0 | 0 | 0 | 0 | 0 | 447 | 171 | 176 | 158 | 348 | 0 | 0 | 1 | 0 | 0 | 298 | 149 | 1 | [illegible] | [illegible] | [illegible] | [illegible] | [illegible] | [illegible] | [illegible] | [illegible] | [illegible] | [illegible] | [illegible] | [illegible] |
| SE | 10952 | ENSG00000091592 | NLRP1 | chr17 | - | 5533291 | 5533384 | 5532821 | 5532984 | 5533896 | 5533988 | 65 | 42 | 61 | 64 | 22 | 2 | 1 | 0 | 0 | 0 | 119 | 38 | 50 | 52 | 142 | 0 | 0 | 1 | 0 | 0 | 242 | 149 | 1 | [illegible] | [illegible] | [illegible] | [illegible] | [illegible] | [illegible] | [illegible] | [illegible] | [illegible] | [illegible] | [illegible] | [illegible] |
| SE | 10954 | ENSG00000091592 | NLRP1 | chr17 | - | 5533303 | 5533384 | 5532821 | 5532984 | 5533896 | 5533988 | 157 | 85 | 143 | 155 | 106 | 2 | 1 | 1 | 0 | 0 | 310 | 96 | 103 | 142 | 327 | 0 | 0 | 1 | 0 | 0 | 230 | 149 | 1 | [illegible] | [illegible] | [illegible] | [illegible] | [illegible] | [illegible] | [illegible] | [illegible] | [illegible] | [illegible] | [illegible] | [illegible] |
| SE | 10957 | ENSG00000091592 | NLRP1 | chr17 | - | 5533896 | 5533988 | 5532821 | 5532984 | 5539414 | 5539585 | 5 | 3 | 4 | 4 | 0 | 0 | 0 | 0 | 0 | 0 | 10 | 4 | 4 | 7 | 0 | 21 | 0 | 2 | 0 | 0 | 241 | 149 | 1 | [illegible] | [illegible] | [illegible] | [illegible] | [illegible] | [illegible] | [illegible] | [illegible] | [illegible] | [illegible] | [illegible] | [illegible] |
| SE | 10958 | ENSG00000091592 | NLRP1 | chr17 | - | 5533896 | 5533988 | 5532821 | 5532984 | 5541856 | 5542027 | 2 | 1 | 2 | 0 | 0 | 0 | 1 | 0 | 0 | 0 | 0 | 0 | 1 | 0 | 0 | 0 | 0 | 0 | 0 | 0 | 241 | 149 | 1 | [illegible] | [illegible] | [illegible] | [illegible] | [illegible] | [illegible] | [illegible] | [illegible] | [illegible] | [illegible] | [illegible] | [illegible] |
| SE | 10959 | ENSG00000091592 | NLRP1 | chr17 | - | 5533896 | 5533988 | 5533291 | 5533384 | 5536850 | 5536940 | 169 | 109 | 205 | 206 | 91 | 2 | 0 | 2 | 0 | 0 | 430 | 124 | 149 | 246 | 491 | 0 | 0 | 0 | 2 | 0 | 241 | 149 | 1 | [illegible] | [illegible] | [illegible] | [illegible] | [illegible] | [illegible] | [illegible] | [illegible] | [illegible] | [illegible] | [illegible] | [illegible] |
| SE | 10962 | ENSG00000091592 | NLRP1 | chr17 | - | 5539414 | 5539585 | 5532821 | 5532984 | 5541856 | 5542027 | 62 | 35 | 79 | 65 | 26 | 0 | 1 | 0 | 0 | 0 | 170 | 78 | 56 | 83 | 160 | 0 | 0 | 0 | 0 | 0 | 298 | 149 | 1 | [illegible] | [illegible] | [illegible] | [illegible] | [illegible] | [illegible] | [illegible] | [illegible] | [illegible] | [illegible] | [illegible] | [illegible] |
| SE | 10963 | ENSG00000091592 | NLRP1 | chr17 | - | 5539414 | 5539585 | 5533896 | 5533988 | 5541856 | 5542027 | 68 | 37 | 83 | 72 | 26 | 0 | 0 | 1 | 0 | 0 | 164 | 85 | 61 | 90 | 167 | 0 | 0 | 0 | 0 | 0 | 298 | 149 | 1 | [illegible] | [illegible] | [illegible] | [illegible] | [illegible] | [illegible] | [illegible] | [illegible] | [illegible] | [illegible] | [illegible] | [illegible] |
| SE | 10964 | ENSG00000091592 | NLRP1 | chr17 | - | 5553385 | 5553556 | 5541856 | 5542027 | 5558338 | 5560043 | 190 | 154 | 271 | 267 | 188 | 0 | 1 | 0 | 0 | 0 | 384 | 185 | 244 | 319 | 581 | 0 | 0 | 0 | 0 | 0 | 298 | 149 | 1 | [illegible] | [illegible] | [illegible] | [illegible] | [illegible] | [illegible] | [illegible] | [illegible] | [illegible] | [illegible] | [illegible] | [illegible] |
| SE | 10965 | ENSG00000091592 | NLRP1 | chr17 | - | 5558338 | 5560043 | 5553385 | 5553556 | 5581858 | 5582062 | 268 | 229 | 317 | 341 | 265 | 1 | 0 | 0 | 0 | 3 | 1114 | 318 | 396 | 555 | 1113 | 0 | 0 | 3 | 0 | 0 | 298 | 149 | 1 | [illegible] | [illegible] | [illegible] | [illegible] | [illegible] | [illegible] | [illegible] | [illegible] | [illegible] | [illegible] | [illegible] | [illegible] |
| SE | 10966 | ENSG00000091592 | NLRP1 | chr17 | - | 5582669 | 5582846 | 5581858 | 5582062 | 5583686 | 5583957 | 385 | 214 | 350 | 405 | 310 | 0 | 0 | 0 | 2 | 0 | 958 | 484 | 415 | 494 | 1095 | 0 | 0 | 0 | 0 | 0 | 298 | 149 | 1 | [illegible] | [illegible] | [illegible] | [illegible] | [illegible] | [illegible] | [illegible] | [illegible] | [illegible] | [illegible] | [illegible] | [illegible] |
| SE | 10967 | ENSG00000091592 | NLRP1 | chr17 | - | 5582669 | 5582846 | 5581858 | 5582062 | 5584292 | 5584448 | 147 | 67 | 124 | 162 | 123 | 0 | 0 | 0 | 0 | 0 | 464 | 195 | 195 | 253 | 547 | 0 | 0 | 1 | 0 | 0 | 298 | 149 | 1 | [illegible] | [illegible] | [illegible] | [illegible] | [illegible] | [illegible] | [illegible] | [illegible] | [illegible] | [illegible] | [illegible] | [illegible] |
| SE | 10968 | ENSG00000091592 | NLRP1 | chr17 | - | 5583686 | 5583959 | 5581858 | 5582062 | 5584292 | 5584448 | 24 | 3 | 11 | 13 | 33 | 0 | 0 | 0 | 0 | 0 | 0 | 8 | 4 | 11 | 19 | 0 | 0 | 1 | 0 | 0 | 298 | 149 | 1 | [illegible] | [illegible] | [illegible] | [illegible] | [illegible] | [illegible] | [illegible] | [illegible] | [illegible] | [illegible] | [illegible] | [illegible] |
| SE | 10969 | ENSG00000091592 | NLRP1 | chr17 | - | 5583686 | 5583959 | 5582669 | 5582846 | 5584292 | 5584448 | 262 | 150 | 237 | 254 | 225 | 0 | 0 | 0 | 0 | 5 | 494 | 297 | 224 | 252 | 567 | 0 | 0 | 0 | 0 | 0 | 298 | 149 | 1 | [illegible] | [illegible] | [illegible] | [illegible] | [illegible] | [illegible] | [illegible] | [illegible] | [illegible] | [illegible] | [illegible] | [illegible] |
| SE | 10970 | ENSG00000091592 | NLRP1 | chr17 | - | 5583686 | 5584103 | 5581858 | 5582062 | 5584292 | 5584512 | 37 | 9 | 31 | 24 | 31 | 0 | 0 | 0 | 0 | 5 | 44 | 24 | 29 | 35 | 40 | 0 | 0 | 1 | 0 | 0 | 298 | 149 | 1 | [illegible] | [illegible] | [illegible] | [illegible] | [illegible] | [illegible] | [illegible] | [illegible] | [illegible] | [illegible] | [illegible] | [illegible] |
| SE | 10971 | ENSG00000091592 | NLRP1 | chr17 | - | 5583686 | 5584103 | 5582669 | 5582846 | 5584292 | 5584512 | 275 | 156 | 257 | 265 | 223 | 0 | 0 | 0 | 0 | 5 | 538 | 313 | 249 | 276 | 588 | 0 | 0 | 0 | 0 | 0 | 298 | 149 | 1 | [illegible] | [illegible] | [illegible] | [illegible] | [illegible] | [illegible] | [illegible] | [illegible] | [illegible] | [illegible] | [illegible] | [illegible] |
| SE | 10973 | ENSG00000274114 | ALOX15P1 | chr17 | + | 6657861 | 6657984 | 6657644 | 6657783 | 6659924 | 6660057 | 2 | 2 | 2 | 5 | 0 | 0 | 0 | 0 | 1 | 4 | 0 | 2 | 2 | 0 | 0 | 0 | 0 | 0 | 0 | 0 | 272 | 149 | 1 | [illegible] | [illegible] | [illegible] | [illegible] | [illegible] | [illegible] | [illegible] | [illegible] | [illegible] | [illegible] | [illegible] | [illegible] |
| SE | 10977 | ENSG00000274114 | ALOX15P1 | chr17 | + | 6658753 | 6658914 | 6657861 | 6657984 | 6659924 | 6660057 | 0 | 0 | 1 | 2 | 7 | 0 | 0 | 0 | 0 | 0 | 0 | 0 | 0 | 0 | 0 | 0 | 2 | 0 | 0 | 0 | 298 | 149 | 1 | [illegible] | [illegible] | [illegible] | [illegible] | [illegible] | [illegible] | [illegible] | [illegible] | [illegible] | [illegible] | [illegible] | [illegible] |
| SE | 10979 | ENSG00000083454 | P2RX5 | chr17 | - | 3681895 | 3681978 | 3679589 | 3679784 | 3688011 | 3688105 | 226 | 116 | 232 | 249 | 287 | 0 | 0 | 0 | 0 | 0 | 224 | 564 | 140 | 99 | 454 | 0 | 0 | 1 | 0 | 0 | 232 | 149 | 1 | [illegible] | [illegible] | [illegible] | [illegible] | [illegible] | [illegible] | [illegible] | [illegible] | [illegible] | [illegible] | [illegible] | [illegible] |
| SE | 10980 | ENSG00000083454 | P2RX5 | chr17 | - | 3688011 | 3688105 | 3681895 | 3681978 | 3690069 | 3690150 | 83 | 49 | 76 | 93 | 59 | 0 | 0 | 0 | 0 | 0 | 76 | 214 | 68 | 27 | 164 | 0 | 2 | 0 | 0 | 0 | 243 | 149 | 1 | [illegible] | [illegible] | [illegible] | [illegible] | [illegible] | [illegible] | [illegible] | [illegible] | [illegible] | [illegible] | [illegible] | [illegible] |
| SE | 10982 | ENSG00000083454 | P2RX5 | chr17 | - | 3688625 | 3688676 | 3688011 | 3688105 | 3690069 | 3690150 | 35 | 15 | 56 | 49 | 81 | 0 | 1 | 1 | 0 | 2 | 68 | 81 | 29 | 44 | 120 | 0 | 4 | 1 | 2 | 5 | 200 | 149 | 1 | [illegible] | [illegible] | [illegible] | [illegible] | [illegible] | [illegible] | [illegible] | [illegible] | [illegible] | [illegible] | [illegible] | [illegible] |
| SE | 10987 | ENSG00000083454 | P2RX5 | chr17 | - | 3689491 | 3689630 | 3688011 | 3688105 | 3690069 | 3690150 | 95 | 31 | 81 | 100 | 97 | 0 | 1 | 1 | 0 | 2 | 49 | 165 | 46 | 51 | 153 | 0 | 4 | 1 | 2 | 5 | 288 | 149 | 1 | [illegible] | [illegible] | [illegible] | [illegible] | [illegible] | [illegible] | [illegible] | [illegible] | [illegible] | [illegible] | [illegible] | [illegible] |
| SE | 10989 | ENSG00000083454 | P2RX5 | chr17 | - | 3689491 | 3689630 | 3688625 | 3688676 | 3690426 | 3690523 | 18 | 10 | 19 | 22 | 14 | 0 | 0 | 0 | 0 | 0 | 0 | 62 | 13 | 4 | 110 | 0 | 3 | 0 | 0 | 0 | 288 | 149 | 1 | [illegible] | [illegible] | [illegible] | [illegible] | [illegible] | [illegible] | [illegible] | [illegible] | [illegible] | [illegible] | [illegible] | [illegible] |
| SE | 10991 | ENSG00000083454 | P2RX5 | chr17 | - | 3690069 | 3690150 | 3688625 | 3688676 | 3690426 | 3690523 | 94 | 40 | 92 | 111 | 185 | 0 | 0 | 0 | 0 | 0 | 77 | 265 | 44 | 60 | 222 | 0 | 3 | 0 | 0 | 0 | 230 | 149 | 1 | [illegible] | [illegible] | [illegible] | [illegible] | [illegible] | [illegible] | [illegible] | [illegible] | [illegible] | [illegible] | [illegible] | [illegible] |
| SE | 10992 | ENSG00000083454 | P2RX5 | chr17 | - | 3690069 | 3690150 | 3689491 | 3689630 | 3690426 | 3690523 | 185 | 67 | 164 | 208 | 270 | 0 | 0 | 0 | 0 | 0 | 106 | 414 | 80 | 95 | 330 | 0 | 0 | 2 | 0 | 0 | 230 | 149 | 1 | [illegible] | [illegible] | [illegible] | [illegible] | [illegible] | [illegible] | [illegible] | [illegible] | [illegible] | [illegible] | [illegible] | [illegible] |
| SE | 10997 | ENSG00000004660 | CAMKK1 | chr17 | - | 3881626 | 3881648 | 3880345 | 3880434 | 3882256 | 3882370 | 0 | 5 | 0 | 3 | 0 | 0 | 0 | 0 | 0 | 1 | 0 | 6 | 1 | 0 | 9 | 0 | 0 | 0 | 0 | 0 | 171 | 149 | 1 | [illegible] | [illegible] | [illegible] | [illegible] | [illegible] | [illegible] | [illegible] | [illegible] | [illegible] | [illegible] | [illegible] | [illegible] |
| SE | 10998 | ENSG00000004660 | CAMKK1 | chr17 | - | 3881626 | 3881648 | 3880345 | 3880434 | 3882527 | 3882564 | 3 | 4 | 13 | 5 | 0 | 2 | 1 | 0 | 2 | 0 | 0 | 4 | 9 | 28 | 27 | 0 | 0 | 0 | 0 | 0 | 171 | 149 | 1 | [illegible] | [illegible] | [illegible] | [illegible] | [illegible] | [illegible] | [illegible] | [illegible] | [illegible] | [illegible] | [illegible] | [illegible] |
| SE | 10999 | ENSG00000004660 | CAMKK1 | chr17 | - | 3882256 | 3882370 | 3880345 | 3880434 | 3882527 | 3882564 | 3 | 0 | 0 | 3 | 2 | 2 | 1 | 0 | 2 | 0 | 0 | 6 | 0 | 0 | 4 | 0 | 0 | 0 | 0 | 0 | 263 | 149 | 1 | [illegible] | [illegible] | [illegible] | [illegible] | [illegible] | [illegible] | [illegible] | [illegible] | [illegible] | [illegible] | [illegible] | [illegible] |
| SE | 11000 | ENSG00000004660 | CAMKK1 | chr17 | - | 3882256 | 3882370 | 3881626 | 3881648 | 3882527 | 3882564 | 3 | 5 | 0 | 5 | 1 | 3 | 5 | 13 | 7 | 0 | 0 | 10 | 0 | 0 | 7 | 0 | 4 | 9 | 30 | 29 | 263 | 149 | 1 | [illegible] | [illegible] | [illegible] | [illegible] | [illegible] | [illegible] | [illegible] | [illegible] | [illegible] | [illegible] | [illegible] | [illegible] |
| SE | 11001 | ENSG00000004660 | CAMKK1 | chr17 | - | 3882527 | 3882564 | 3881626 | 3881648 | 3883428 | 3883480 | 0 | 0 | 6 | 3 | 0 | 0 | 0 | 0 | 0 | 0 | 0 | 0 | 2 | 23 | 17 | 0 | 4 | 0 | 2 | 0 | 186 | 149 | 1 | [illegible] | [illegible] | [illegible] | [illegible] | [illegible] | [illegible] | [illegible] | [illegible] | [illegible] | [illegible] | [illegible] | [illegible] |
| SE | 11003 | ENSG00000108559 | NUP88 | chr17 | - | 5386707 | 5386826 | 5384832 | 5386269 | 5386983 | 5387069 | 235 | 170 | 188 | 203 | 710 | 0 | 2 | 0 | 0 | 0 | 161 | 193 | 243 | 205 | 56 | 0 | 0 | 0 | 0 | 0 | 268 | 149 | 1 | [illegible] | [illegible] | [illegible] | [illegible] | [illegible] | [illegible] | [illegible] | [illegible] | [illegible] | [illegible] | [illegible] | [illegible] |
| SE | 11004 | ENSG00000108559 | NUP88 | chr17 | - | 5387385 | 5387466 | 5386983 | 5387110 | 5387604 | 5387670 | 238 | 155 | 171 | 184 | 565 | 0 | 0 | 0 | 0 | 0 | 117 | 135 | 169 | 189 | 70 | 0 | 3 | 0 | 0 | 0 | 230 | 149 | 1 | [illegible] | [illegible] | [illegible] | [illegible] | [illegible] | [illegible] | [illegible] | [illegible] | [illegible] | [illegible] | [illegible] | [illegible] |
| SE | 11005 | ENSG00000108559 | NUP88 | chr17 | - | 5387778 | 5387904 | 5387385 | 5387670 | 5388801 | 5388850 | 180 | 123 | 178 | 172 | 392 | 1 | 1 | 0 | 0 | 0 | 107 | 137 | 139 | 143 | 48 | 0 | 0 | 0 | 3 | 0 | 275 | 149 | 1 | [illegible] | [illegible] | [illegible] | [illegible] | [illegible] | [illegible] | [illegible] | [illegible] | [illegible] | [illegible] | [illegible] | [illegible] |
| SE | 11006 | ENSG00000108559 | NUP88 | chr17 | - | 5387778 | 5387952 | 5387604 | 5387670 | 5388801 | 5388960 | 72 | 59 | 76 | 65 | 172 | 1 | 1 | 0 | 0 | 0 | 35 | 75 | 51 | 57 | 6 | 0 | 0 | 0 | 3 | 0 | 298 | 149 | 1 | [illegible] | [illegible] | [illegible] | [illegible] | [illegible] | [illegible] | [illegible] | [illegible] | [illegible] | [illegible] | [illegible] | [illegible] |

| | | | | | | | | | | | | | | | | | | | | | | | | | | | | | | | | | | | | | | | | | | | | | | |
|---|---|---|---|---|---|---|---|---|---|---|---|---|---|---|---|---|---|---|---|---|---|---|---|---|---|---|---|---|---|---|---|---|---|---|---|---|---|---|---|---|---|---|---|---|---|---|
| SE | 11007 | ENSG00000108559 | NUP88 | chr17 | - | 5388801 | 5388960 | 5387778 | 5387904 | 5391560 | 5391662 | 230 | 128 | 207 | 232 | 516 | 1 | 4 | 1 | 1 | 0 | 190 | 161 | 235 | 199 | 83 | 0 | 0 | 0 | 1 | 0 | 298 | 149 | 1 | 1 | 0.991 | 0.941 | 0.99 | 0.991 | 1 | 1 | 1 | 1 | 0.99 | 1 | -0.015 |
| SE | 11008 | ENSG00000108559 | NUP88 | chr17 | - | 5394890 | 5394981 | 5391474 | 5391662 | 5399551 | 5399633 | 204 | 110 | 204 | 173 | 465 | 3 | 0 | 0 | 0 | 0 | 101 | 179 | 195 | 160 | 91 | 0 | 3 | 2 | 0 | 0 | 240 | 149 | 1 | 1 | 0.977 | 1 | 1 | 1 | 1 | 1 | 0.974 | 0.984 | 1 | 1 | 0.004 |
| SE | 11009 | ENSG00000108559 | NUP88 | chr17 | - | 5399551 | 5399650 | 5394890 | 5394981 | 5404098 | 5404246 | 172 | 65 | 142 | 117 | 322 | 1 | 0 | 0 | 0 | 0 | 94 | 134 | 119 | 70 | 60 | 0 | 0 | 2 | 0 | 0 | 248 | 149 | 1 | 1 | 0.99 | 1 | 1 | 1 | 1 | 1 | 1 | 0.973 | 1 | 1 | 0.003 |
| SE | 11010 | ENSG00000108559 | NUP88 | chr17 | - | 5399551 | 5399650 | 5394890 | 5394981 | 5405056 | 5405243 | 102 | 52 | 95 | 79 | 198 | 0 | 0 | 0 | 0 | 3 | 37 | 89 | 100 | 49 | 55 | 0 | 0 | 2 | 0 | 0 | 248 | 149 | 1 | 1 | 1 | 1 | 1 | 1 | 0.975 | 1 | 1 | 0.968 | 1 | 1 | 0.001 |
| SE | 11011 | ENSG00000108559 | NUP88 | chr17 | - | 5404098 | 5404246 | 5394890 | 5394981 | 5405056 | 5405243 | 63 | 21 | 63 | 46 | 98 | 0 | 0 | 0 | 0 | 3 | 29 | 23 | 26 | 34 | 12 | 0 | 0 | 2 | 0 | 0 | 297 | 149 | 1 | 1 | 1 | 1 | 1 | 1 | 0.942 | 1 | 1 | 0.867 | 1 | 1 | 0.015 |
| SE | 11013 | ENSG00000108559 | NUP88 | chr17 | - | 5410702 | 5410789 | 5408732 | 5408909 | 5414008 | 5414134 | 173 | 103 | 180 | 173 | 385 | 2 | 0 | 0 | 0 | 0 | 179 | 174 | 155 | 207 | 129 | 0 | 2 | 0 | 0 | 0 | 236 | 149 | 1 | 1 | 0.982 | 1 | 1 | 1 | 1 | 1 | 0.982 | 1 | 1 | 1 | 0 |
| SE | 11016 | ENSG00000261879 | ZNF594-DT | chr17 | + | 5209749 | 5209895 | 5192026 | 5192163 | 5214049 | 5214209 | 5 | 0 | 7 | 1 | 7 | 0 | 0 | 0 | 2 | 0 | 0 | 0 | 0 | 1 | 8 | 0 | 0 | 0 | 0 | 0 | 295 | 149 | 1 | 1 | 1 | NA | 1 | 0.202 | 1 | NA | NA | NA | 1 | 1 | -0.2 |
| SE | 11017 | ENSG00000261879 | ZNF594-DT | chr17 | + | 5214049 | 5214209 | 5192026 | 5192163 | 5215146 | 5215293 | 13 | 7 | 19 | 14 | 32 | 0 | 0 | 0 | 2 | 0 | 1 | 6 | 22 | 11 | 18 | 0 | 0 | 0 | 0 | 0 | 298 | 149 | 1 | 1 | 1 | 1 | 1 | 0.778 | 1 | 1 | 1 | 1 | 1 | 1 | -0.044 |
| SE | 11018 | ENSG00000261879 | ZNF594-DT | chr17 | + | 5223316 | 5223484 | 5215146 | 5215293 | 5233823 | 5233942 | 14 | 9 | 23 | 13 | 18 | 0 | 1 | 0 | 0 | 0 | 11 | 11 | 11 | 9 | 31 | 0 | 0 | 0 | 0 | 0 | 298 | 149 | 1 | 1 | 1 | 0.818 | 1 | 1 | 1 | 1 | 1 | 1 | 1 | 1 | -0.036 |
| SE | 11019 | ENSG00000261879 | ZNF594-DT | chr17 | + | 5223316 | 5223484 | 5215146 | 5215293 | 5239906 | 5240109 | 14 | 5 | 16 | 13 | 18 | 0 | 0 | 0 | 1 | 0 | 8 | 10 | 12 | 9 | 31 | 0 | 0 | 0 | 0 | 0 | 298 | 149 | 1 | 1 | 1 | 1 | 1 | 0.867 | 1 | 1 | 1 | 1 | 1 | 1 | -0.027 |
| SE | 11026 | ENSG00000241525 | NA | chr17 | - | 413128 | 413413 | 404534 | 404592 | 413978 | 414023 | 2 | 3 | 4 | 0 | 0 | 0 | 3 | 3 | 1 | 0 | 0 | 0 | 0 | 1 | 18 | 0 | 8 | 0 | 0 | 0 | 298 | 149 | 1 | 1 | 1 | 0.333 | 0.4 | 0 | NA | NA | 0 | NA | 1 | 1 | -0.233 |
| SE | 11030 | ENSG00000072849 | DERL2 | chr17 | - | 5481295 | 5481389 | 5481100 | 5481142 | 5482808 | 5482879 | 78 | 53 | 78 | 54 | 120 | 0 | 2 | 0 | 0 | 0 | 44 | 57 | 48 | 47 | 44 | 0 | 0 | 0 | 0 | 0 | 243 | 149 | 1 | 1 | 1 | 0.942 | 1 | 1 | 1 | 1 | 1 | 1 | 1 | 1 | -0.012 |
| SE | 11031 | ENSG00000072849 | DERL2 | chr17 | - | 5485150 | 5485216 | 5480053 | 5480144 | 5486068 | 5486148 | 16 | 5 | 17 | 10 | 11 | 0 | 0 | 1 | 0 | 0 | 8 | 18 | 12 | 8 | 9 | 0 | 0 | 0 | 0 | 0 | 215 | 149 | 1 | 1 | 1 | 1 | 0.922 | 1 | 1 | 1 | 1 | 1 | 1 | 1 | -0.016 |
| SE | 11032 | ENSG00000072849 | DERL2 | chr17 | - | 5485150 | 5485216 | 5482269 | 5482882 | 5486068 | 5486148 | 96 | 65 | 98 | 79 | 126 | 0 | 1 | 0 | 0 | 0 | 55 | 72 | 74 | 53 | 78 | 0 | 3 | 0 | 0 | 0 | 215 | 149 | 1 | 1 | 1 | 0.978 | 1 | 1 | 1 | 1 | 0.943 | 1 | 1 | 1 | 0.007 |
| SE | 11033 | ENSG00000132359 | RAP1GAP2 | chr17 | + | 2905283 | 2905368 | 2800514 | 2800550 | 2957758 | 2957794 | 110 | 50 | 36 | 81 | 15 | 1 | 0 | 0 | 0 | 0 | 124 | 64 | 55 | 54 | 119 | 0 | 0 | 0 | 0 | 0 | 234 | 149 | 1 | 1 | 0.986 | 1 | 1 | 1 | 1 | 1 | 1 | 1 | 1 | 1 | -0.003 |
| SE | 11034 | ENSG00000132359 | RAP1GAP2 | chr17 | + | 2957758 | 2957794 | 2905283 | 2905368 | 2963429 | 2963462 | 75 | 32 | 24 | 64 | 17 | 1 | 0 | 0 | 0 | 0 | 68 | 39 | 42 | 27 | 92 | 0 | 0 | 0 | 0 | 0 | 185 | 149 | 1 | 1 | 0.984 | 1 | 1 | 1 | 1 | 1 | 1 | 1 | 1 | 1 | -0.003 |
| SE | 11036 | ENSG00000132359 | RAP1GAP2 | chr17 | + | 2965539 | 2965643 | 2963855 | 2964068 | 2980286 | 2980365 | 363 | 192 | 131 | 290 | 107 | 0 | 0 | 0 | 0 | 0 | 397 | 333 | 218 | 338 | 355 | 0 | 9 | 0 | 0 | 0 | 253 | 149 | 1 | 1 | 1 | 1 | 1 | 1 | 1 | 1 | 0.956 | 1 | 1 | 1 | 0.009 |
| SE | 11037 | ENSG00000132359 | RAP1GAP2 | chr17 | + | 2980286 | 2980365 | 2965539 | 2965643 | 2981194 | 2981248 | 365 | 220 | 124 | 318 | 134 | 0 | 0 | 0 | 0 | 0 | 516 | 396 | 215 | 350 | 340 | 0 | 0 | 0 | 6 | 0 | 228 | 149 | 1 | 1 | 1 | 1 | 1 | 1 | 1 | 1 | 1 | 1 | 0.974 | 1 | 0.005 |
| SE | 11038 | ENSG00000132359 | RAP1GAP2 | chr17 | + | 2991296 | 2991397 | 2984982 | 2985066 | 2995336 | 2995466 | 544 | 258 | 214 | 395 | 262 | 0 | 0 | 2 | 0 | 0 | 471 | 395 | 317 | 379 | 502 | 0 | 0 | 2 | 0 | 0 | 250 | 149 | 1 | 1 | 1 | 1 | 1 | 0.992 | 1 | 1 | 1 | 0.99 | 1 | 1 | 0 |
| SE | 11043 | ENSG00000132359 | RAP1GAP2 | chr17 | + | 3020476 | 3020595 | 3008010 | 3008145 | 3026007 | 3026121 | 103 | 24 | 34 | 55 | 59 | 0 | 0 | 0 | 0 | 0 | 208 | 67 | 67 | 98 | 91 | 0 | 5 | 0 | 0 | 0 | 268 | 149 | 1 | 1 | 1 | 1 | 1 | 1 | 1 | 1 | 0.882 | 1 | 1 | 1 | 0.024 |
| SE | 11044 | ENSG00000091640 | SPAG7 | chr17 | - | 4960233 | 4960318 | 4960021 | 4960111 | 4960458 | 4960547 | 397 | 217 | 286 | 365 | 345 | 0 | 1 | 2 | 1 | 6 | 880 | 435 | 315 | 352 | 753 | 0 | 0 | 5 | 5 | 0 | 234 | 149 | 1 | 1 | 1 | 0.993 | 0.989 | 0.996 | 0.973 | 1 | 1 | 0.976 | 0.978 | 1 | -0.001 |
| SE | 11045 | ENSG00000185722 | ANKFY1 | chr17 | - | 4165397 | 4167911 | 4163956 | 4164173 | 4169197 | 4169288 | 14 | 7 | 32 | 19 | 17 | 0 | 1 | 0 | 0 | 0 | 37 | 8 | 6 | 22 | 11 | 0 | 2 | 1 | 0 | 0 | 298 | 149 | 1 | 1 | 1 | 0.778 | 1 | 1 | 1 | 1 | 0.667 | 0.75 | 1 | 1 | 0.072 |
| SE | 11047 | ENSG00000185722 | ANKFY1 | chr17 | - | 4179720 | 4179877 | 4178856 | 4179057 | 4181253 | 4181372 | 160 | 102 | 116 | 157 | 81 | 0 | 0 | 0 | 0 | 0 | 190 | 157 | 81 | 227 | 141 | 0 | 0 | 0 | 2 | 0 | 298 | 149 | 1 | 1 | 1 | 1 | 1 | 1 | 1 | 1 | 1 | 1 | 0.983 | 1 | 0.003 |
| SE | 11055 | ENSG00000267508 | ZNF285 | chr19 | - | 44394519 | 44394540 | 44392205 | 44392466 | 44397198 | 44397256 | 2 | 0 | 0 | 2 | 2 | 2 | 2 | 0 | 0 | 0 | 0 | 14 | 1 | 0 | 0 | 0 | 0 | 0 | 3 | 0 | 170 | 149 | 1 | 1 | 0.467 | 0 | NA | 1 | 1 | NA | 1 | 1 | 0 | NA | -0.05 |
| SE | 11056 | ENSG00000257950 | P2RX5-TAX1BP3 | chr17 | - | 3669057 | 3672493 | 3664678 | 3664798 | 3679589 | 3679784 | 3 | 0 | 0 | 2 | 5 | 0 | 0 | 0 | 0 | 0 | 0 | 11 | 0 | 0 | 21 | 0 | 3 | 1 | 0 | 0 | 298 | 149 | 1 | 1 | 1 | NA | NA | 1 | 1 | NA | 0.647 | 0 | NA | 1 | 0.451 |
| SE | 11058 | ENSG00000172530 | BANP | chr16 | + | 87952180 | 87952273 | 87951444 | 87951515 | 87981035 | 87981127 | 0 | 0 | 1 | 1 | 0 | 3 | 0 | 0 | 0 | 2 | 0 | 0 | 0 | 1 | 0 | 0 | 0 | 1 | 0 | 0 | 242 | 149 | 1 | 1 | 0 | NA | 1 | 1 | 0 | NA | NA | 0 | 1 | NA | 0 |
| SE | 11060 | ENSG00000172530 | BANP | chr16 | + | 87975047 | 87975185 | 87951431 | 87951515 | 87981035 | 87981049 | 120 | 73 | 67 | 100 | 96 | 3 | 0 | 0 | 0 | 2 | 135 | 91 | 63 | 66 | 89 | 0 | 0 | 1 | 0 | 0 | 287 | 149 | 1 | 1 | 0.954 | 1 | 1 | 1 | 0.961 | 1 | 1 | 0.97 | 1 | 1 | -0.011 |
| SE | 11061 | ENSG00000172530 | BANP | chr16 | + | 87975047 | 87975185 | 87952180 | 87952273 | 87981035 | 87981049 | 94 | 49 | 51 | 56 | 79 | 0 | 0 | 0 | 0 | 0 | 133 | 57 | 45 | 41 | 68 | 0 | 0 | 0 | 1 | 0 | 287 | 149 | 1 | 1 | 1 | 1 | 1 | 1 | 1 | 1 | 1 | 1 | 0.955 | 1 | 0.009 |
| SE | 11062 | ENSG00000172530 | BANP | chr16 | + | 87975047 | 87975203 | 87951431 | 87951515 | 87981035 | 87981127 | 41 | 21 | 19 | 43 | 27 | 3 | 0 | 0 | 0 | 2 | 2 | 36 | 19 | 27 | 34 | 0 | 0 | 1 | 0 | 0 | 298 | 149 | 1 | 1 | 0.872 | 1 | 1 | 1 | 0.871 | 1 | 1 | 0.905 | 1 | 1 | -0.032 |
| SE | 11065 | ENSG00000172530 | BANP | chr16 | + | 87978439 | 87978692 | 87975047 | 87975203 | 87981035 | 87981102 | 0 | 0 | 0 | 0 | 0 | 6 | 3 | 3 | 3 | 9 | 0 | 7 | 0 | 0 | 7 | 0 | 4 | 0 | 11 | 6 | 298 | 149 | 1 | 1 | 0 | 0 | 0 | 0 | 0 | NA | 0.467 | NA | 0 | 0.368 | -0.278 |
| SE | 11066 | ENSG00000172530 | BANP | chr16 | + | 87984059 | 87984259 | 87981035 | 87981127 | 88004294 | 88004301 | 186 | 127 | 177 | 125 | 286 | 0 | 1 | 0 | 1 | 0 | 236 | 152 | 137 | 133 | 250 | 0 | 0 | 0 | 0 | 0 | 298 | 149 | 1 | 1 | 1 | 0.984 | 1 | 0.984 | 1 | 1 | 1 | 1 | 1 | 1 | -0.006 |
| SE | 11067 | ENSG00000172530 | BANP | chr16 | + | 87984083 | 87984259 | 87981035 | 87981127 | 88004294 | 88004411 | 30 | 25 | 24 | 15 | 19 | 0 | 1 | 0 | 1 | 0 | 36 | 35 | 28 | 24 | 62 | 0 | 0 | 0 | 0 | 0 | 298 | 149 | 1 | 1 | 1 | 0.926 | 1 | 0.882 | 1 | 1 | 1 | 1 | 1 | 1 | -0.038 |
| SE | 11068 | ENSG00000172530 | BANP | chr16 | + | 88003481 | 88003563 | 87984059 | 87984259 | 88003810 | 88003931 | 0 | 2 | 2 | 1 | 2 | 0 | 0 | 0 | 0 | 0 | 0 | 0 | 0 | 0 | 0 | 0 | 0 | 0 | 0 | 3 | 231 | 149 | 1 | 1 | NA | 1 | 1 | 1 | 1 | NA | NA | NA | NA | 0 | 1 |
| SE | 11069 | ENSG00000172530 | BANP | chr16 | + | 88003481 | 88003563 | 87984059 | 87984259 | 88004294 | 88004391 | 0 | 2 | 0 | 1 | 0 | 30 | 25 | 24 | 15 | 19 | 0 | 0 | 0 | 0 | 0 | 36 | 35 | 28 | 24 | 62 | 231 | 149 | 1 | 1 | 0 | 0.049 | 0 | 0.041 | 0 | 0 | 0 | 0 | 0 | 0 | 0.018 |
| SE | 11071 | ENSG00000172530 | BANP | chr16 | + | 88006089 | 88006265 | 87984059 | 87984259 | 88018427 | 88018455 | 50 | 16 | 35 | 40 | 21 | 0 | 1 | 0 | 0 | 0 | 33 | 25 | 16 | 17 | 68 | 0 | 0 | 0 | 0 | 0 | 298 | 149 | 1 | 1 | 1 | 0.889 | 1 | 1 | 1 | 1 | 1 | 1 | 1 | 1 | -0.022 |
| SE | 11072 | ENSG00000172530 | BANP | chr16 | + | 88018427 | 88018667 | 87984059 | 87984259 | 88033108 | 88033219 | 0 | 1 | 1 | 1 | 0 | 0 | 0 | 0 | 1 | 0 | 0 | 2 | 0 | 0 | 0 | 0 | 0 | 0 | 0 | 0 | 298 | 149 | 1 | 1 | NA | 1 | 1 | 0.333 | NA | NA | 1 | NA | NA | NA | -0.222 |
| SE | 11073 | ENSG00000172530 | BANP | chr16 | + | 88027482 | 88027650 | 88018427 | 88018667 | 88033108 | 88033219 | 149 | 31 | 81 | 92 | 60 | 0 | 0 | 1 | 1 | 0 | 111 | 68 | 66 | 51 | 166 | 0 | 2 | 0 | 0 | 0 | 298 | 149 | 1 | 1 | 1 | 1 | 0.976 | 0.979 | 1 | 1 | 0.944 | 1 | 1 | 1 | 0.002 |
| SE | 11074 | ENSG00000172530 | BANP | chr16 | + | 88065266 | 88065332 | 88037972 | 88038011 | 88072068 | 88072212 | 6 | 0 | 1 | 5 | 2 | 8 | 7 | 3 | 6 | 2 | 16 | 7 | 2 | 0 | 0 | 0 | 1 | 3 | 4 | 0 | 215 | 149 | 1 | 1 | 0.342 | 0 | 0.188 | 0.366 | 0.409 | 1 | 0.829 | 0.316 | 0 | NA | -0.275 |
| SE | 11075 | ENSG00000172530 | BANP | chr16 | + | 88072068 | 88072212 | 88018427 | 88018667 | 88076589 | 88076661 | 3 | 8 | 14 | 10 | 0 | 0 | 0 | 0 | 1 | 0 | 0 | 5 | 7 | 0 | 5 | 0 | 0 | 0 | 0 | 0 | 293 | 149 | 1 | 1 | 1 | 1 | 1 | 0.836 | NA | NA | 1 | 1 | NA | 1 | -0.041 |
| SE | 11076 | ENSG00000172530 | BANP | chr16 | + | 88072068 | 88072212 | 88037972 | 88038011 | 88076589 | 88076661 | 11 | 15 | 17 | 12 | 2 | 0 | 0 | 0 | 0 | 0 | 0 | 6 | 10 | 4 | 0 | 3 | 0 | 0 | 0 | 0 | 293 | 149 | 1 | 1 | 1 | 1 | 1 | 1 | 1 | 0 | 1 | 1 | 1 | NA | 0.25 |
| SE | 11077 | ENSG00000167377 | ZNF23 | chr16 | - | 71450589 | 71450715 | 71447596 | 71449882 | 71453242 | 71453350 | 0 | 0 | 1 | 0 | 0 | 1 | 0 | 3 | 0 | 0 | 0 | 0 | 0 | 0 | 0 | 0 | 0 | 1 | 0 | 0 | 275 | 149 | 1 | 1 | 0 | NA | 0.153 | NA | NA | NA | NA | 0 | NA | NA | 0.076 |
| SE | 11078 | ENSG00000167377 | ZNF23 | chr16 | - | 71453242 | 71453350 | 71447596 | 71449882 | 71456763 | 71456828 | 1 | 0 | 4 | 0 | 0 | 0 | 0 | 0 | 1 | 0 | 0 | 0 | 1 | 0 | 0 | 0 | 0 | 0 | 0 | 0 | 257 | 149 | 1 | 1 | 1 | NA | 1 | 0 | NA | NA | NA | 1 | NA | NA | -0.333 |
| SE | 11081 | ENSG00000167377 | ZNF23 | chr16 | - | 71454041 | 71454168 | 71453242 | 71453350 | 71456763 | 71456828 | 13 | 21 | 24 | 16 | 22 | 0 | 0 | 2 | 0 | 0 | 6 | 7 | 3 | 0 | 4 | 0 | 0 | 0 | 0 | 0 | 276 | 149 | 1 | 1 | 1 | 1 | 0.866 | 1 | 1 | 1 | 1 | 1 | NA | 1 | -0.027 |
| SE | 11082 | ENSG00000167377 | ZNF23 | chr16 | - | 71454041 | 71454218 | 71453242 | 71453350 | 71456763 | 71456828 | 14 | 20 | 20 | 17 | 22 | 0 | 0 | 2 | 0 | 0 | 6 | 8 | 3 | 0 | 4 | 0 | 0 | 0 | 0 | 0 | 298 | 149 | 1 | 1 | 1 | 1 | 0.833 | 1 | 1 | 1 | | 1 | NA | 1 | -0.033 |
| SE | 11083 | ENSG00000167377 | ZNF23 | chr16 | - | 71456071 | 71456121 | 71454041 | 71454168 | 71456763 | 71456828 | 2 | 1 | 5 | 3 | 1 | 1 | 2 | 5 | 0 | 0 | 0 | 3 | 0 | 1 | 0 | 0 | 0 | 0 | 0 | 0 | 199 | 149 | 1 | 1 | 0.6 | 0.272 | 0.428 | 1 | 1 | NA | 1 | NA | 1 | NA | -0.34 |
| SE | 11084 | ENSG00000167377 | ZNF23 | chr16 | - | 71456071 | 71456121 | 71454041 | 71454218 | 71456763 | 71456828 | 1 | 0 | 6 | 3 | 1 | 2 | 0 | 0 | 1 | 0 | 0 | 3 | 0 | 0 | 0 | 0 | 0 | 0 | 0 | 0 | 199 | 149 | 1 | 1 | 0.272 | NA | 1 | 0.692 | 1 | NA | 1 | NA | NA | NA | -0.259 |
| SE | 11085 | ENSG00000167377 | ZNF23 | chr16 | - | 71461658 | 71462004 | 71456781 | 71456828 | 71462209 | 71462392 | 4 | 1 | 4 | 0 | 0 | 4 | 2 | 2 | 1 | 2 | 0 | 0 | 3 | 0 | 0 | 0 | 3 | 0 | 0 | 0 | 298 | 149 | 1 | 1 | 0.333 | 0.2 | 0.5 | 0 | 0 | NA | 0 | 1 | NA | NA | -0.293 |
| SE | 11091 | ENSG00000135709 | KIAA0513 | chr16 | + | 85072924 | 85072998 | 85071782 | 85071882 | 85075843 | 85075914 | 138 | 141 | 67 | 117 | 205 | 0 | 0 | 0 | 0 | 0 | 295 | 148 | 83 | 209 | 588 | 0 | 0 | 0 | 3 | 5 | 223 | 149 | 1 | 1 | 1 | 1 | 1 | 1 | 1 | 1 | 1 | 1 | 0.979 | 0.987 | 0.007 |
| SE | 11092 | ENSG00000135709 | KIAA0513 | chr16 | + | 85078414 | 85078455 | 85077424 | 85077632 | 85078924 | 85079003 | 119 | 130 | 72 | 94 | 105 | 0 | 0 | 1 | 0 | 0 | 260 | 81 | 57 | 206 | 447 | 0 | 0 | 0 | 3 | 0 | 190 | 149 | 1 | 1 | 1 | 1 | 0.983 | 1 | 1 | 1 | 1 | 1 | 0.982 | 1 | 0 |
| SE | 11095 | ENSG00000178188 | SH2B1 | chr16 | + | 28861617 | 28861747 | 28846642 | 28846827 | 28863667 | 28863754 | 0 | 0 | 0 | 2 | 0 | 0 | 1 | 1 | 0 | 1 | 0 | 0 | 0 | 0 | 0 | 0 | 0 | 1 | 0 | 0 | 279 | 149 | 1 | 1 | NA | 0 | 0 | 1 | 0 | NA | NA | 0 | NA | NA | 0.25 |
| SE | 11096 | ENSG00000178188 | SH2B1 | chr16 | + | 28863289 | 28863345 | 28861617 | 28861747 | 28863667 | 28863754 | 0 | 0 | 0 | 0 | 0 | 0 | 0 | 0 | 2 | 0 | 0 | 0 | 0 | 3 | 0 | 0 | 0 | 0 | 0 | 0 | 205 | 149 | 1 | 1 | NA | NA | NA | 0 | NA | NA | NA | NA | 1 | NA | -1 |
| SE | 11099 | ENSG00000178188 | SH2B1 | chr16 | + | 28866011 | 28867033 | 28863756 | 28863945 | 28867373 | 28867432 | 3 | 0 | 0 | 6 | 15 | 3 | 1 | 2 | 1 | 2 | 5 | 0 | 0 | 8 | 0 | 0 | 0 | 2 | 0 | 0 | 298 | 149 | 1 | 1 | 0.333 | 0 | 0 | 0.75 | 0.789 | 1 | NA | 0 | 1 | NA | -0.292 |
| SE | 11102 | ENSG00000178188 | SH2B1 | chr16 | + | 28869005 | 28869097 | 28867153 | 28867432 | 28869207 | 28869383 | 177 | 131 | 222 | 223 | 345 | 0 | 2 | 4 | 0 | 0 | 131 | 214 | 84 | 122 | 274 | 0 | 0 | 0 | 0 | 0 | 241 | 149 | 1 | 1 | 1 | 0.976 | 0.972 | 1 | 1 | 1 | 1 | 1 | 1 | 1 | -0.01 |
| SE | 11104 | ENSG00000178188 | SH2B1 | chr16 | + | 28872533 | 28872705 | 28872189 | 28872401 | 28873169 | 28873269 | 92 | 49 | 126 | 97 | 51 | 0 | 0 | 0 | 0 | 0 | 102 | 125 | 45 | 60 | 160 | 0 | 0 | 3 | 0 | 0 | 298 | 149 | 1 | 1 | 1 | 1 | 1 | 1 | 1 | 1 | 1 | 0.882 | 1 | 1 | 0.024 |
| SE | 11105 | ENSG00000178188 | SH2B1 | chr16 | + | 28872533 | 28872758 | 28872189 | 28872401 | 28873169 | 28873269 | 45 | 23 | 78 | 60 | 35 | 0 | 0 | 0 | 0 | 0 | 33 | 59 | 24 | 14 | 64 | 0 | 0 | 3 | 0 | 0 | 298 | 149 | 1 | 1 | 1 | 1 | 1 | 1 | 1 | 1 | 1 | 0.8 | 1 | 1 | 0.04 |
| SE | 11106 | ENSG00000178188 | SH2B1 | chr16 | + | 28873169 | 28873269 | 28872533 | 28872705 | 28873446 | 28873607 | 71 | 35 | 84 | 60 | 28 | 0 | 1 | 0 | 0 | 0 | 100 | 78 | 34 | 73 | 138 | 0 | 0 | 0 | 0 | 0 | 249 | 149 | 1 | 1 | 1 | 0.954 | 1 | 1 | 1 | 1 | 1 | 1 | 1 | 1 | -0.009 |
| SE | 11108 | ENSG00000205456 | TP53TG3D | chr16 | + | 32254393 | 32254683 | 32253820 | 32253987 | 32254890 | 32254935 | 4 | 0 | 0 | 3 | 1 | 1 | 0 | 0 | 0 | 0 | 0 | 0 | 0 | 2 | 2 | 0 | 0 | 0 | 0 | 0 | 298 | 149 | 1 | 1 | 0.667 | NA | NA | 1 | 1 | NA | NA | NA | 1 | 1 | -0.111 |
| SE | 11113 | ENSG00000142512 | SIGLEC10 | chr19 | - | 51417081 | 51417358 | 51416713 | 51416950 | 51417544 | 51417591 | 39 | 52 | 21 | 68 | 42 | 0 | 0 | 0 | 4 | 0 | 44 | 55 | 48 | 58 | 116 | 0 | 2 | 0 | 0 | 0 | 298 | 149 | 1 | 1 | 1 | 1 | 1 | 0.895 | 1 | 1 | 0.932 | 1 | 1 | 1 | -0.007 |
| SE | 11115 | ENSG00000142512 | SIGLEC10 | chr19 | - | 51417081 | 51417465 | 51416665 | 51416950 | 51417544 | 51417581 | 63 | 52 | 36 | 80 | 49 | 0 | 0 | 0 | 4 | 0 | 93 | 76 | 62 | 76 | 149 | 0 | 2 | 0 | 0 | 0 | 298 | 149 | 1 | 1 | 1 | 1 | 1 | 0.909 | 1 | 1 | 0.95 | 1 | 1 | 1 | -0.008 |
| SE | 11117 | ENSG00000185947 | ZNF267 | chr16 | + | 31875202 | 31875302 | 31873830 | 31873969 | 31884497 | 31884624 | 2 | 1 | 0 | 2 | 1 | 75 | 63 | 41 | 76 | 73 | 0 | 0 | 3 | 0 | 0 | 22 | 81 | 80 | 53 | 17 | 249 | 149 | 1 | 1 | 0.016 | 0.009 | 0 | 0.016 | 0.008 | 0 | 0 | 0.022 | 0 | 0 | 0.005 |
| SE | 11118 | ENSG00000185947 | ZNF267 | chr16 | + | 31890206 | 31890250 | 31885160 | 31885256 | 31914475 | 31914558 | 0 | 0 | 5 | 0 | 5 | 233 | 209 | 194 | 308 | 320 | 0 | 0 | 4 | 0 | 0 | 151 | 160 | 210 | 94 | 126 | 193 | 149 | 1 | 1 | 0 | 0 | 0.02 | 0 | 0.012 | 0 | 0 | 0.014 | 0 | 0 | 0.004 |
| SE | 11121 | ENSG00000008517 | IL32 | chr16 | + | 3065780 | 3065826 | 3065311 | 3065494 | 3067376 | 3067415 | 509 | 100 | 349 | 345 | 300 | 1 | 0 | 0 | 0 | 0 | 185 | 270 | 169 | 88 | 325 | 0 | 0 | 0 | 0 | 0 | 195 | 149 | 1 | 1 | 0.997 | 1 | 1 | 1 | 1 | 1 | 1 | 1 | 1 | 1 | -0.001 |
| SE | 11122 | ENSG00000008517 | IL32 | chr16 | + | 3065780 | 3065826 | 3065662 | 3065710 | 3067376 | 3067415 | 521 | 99 | 354 | 368 | 309 | 1 | 0 | 3 | 1 | 0 | 185 | 282 | 176 | 90 | 331 | 0 | 4 | 0 | 0 | 0 | 195 | 149 | 1 | 1 | 0.997 | 1 | 0.989 | 0.996 | 1 | 1 | 0.982 | 1 | 1 | 1 | 0 |
| SE | 11123 | ENSG00000008517 | IL32 | chr16 | + | 3065783 | 3065826 | 3065296 | 3065494 | 3067376 | 3067415 | 515 | 100 | 362 | 349 | 298 | 1 | 0 | 0 | 0 | 0 | 185 | 276 | 170 | 90 | 344 | 0 | 0 | 0 | 0 | 0 | 192 | 149 | 1 | 1 | 0.998 | 1 | 1 | 1 | 1 | 1 | 1 | 1 | 1 | 1 | 0 |
| SE | 11124 | ENSG00000008517 | IL32 | chr16 | + | 3065783 | 3065826 | 3065609 | 3065710 | 3067376 | 3067415 | 613 | 118 | 427 | 412 | 377 | 1 | 0 | 3 | 1 | 0 | 268 | 328 | 198 | 97 | 398 | 0 | 4 | 0 | 0 | 0 | 192 | 149 | 1 | 1 | 0.998 | 1 | 0.991 | 0.997 | 1 | 1 | 0.985 | 1 | 1 | 1 | 0 |
| SE | 11126 | ENSG00000008517 | IL32 | chr16 | + | 3067553 | 3067613 | 3067376 | 3067415 | 3068179 | 3068239 | 505 | 113 | 375 | 351 | 333 | 0 | 0 | 0 | 0 | 0 | 213 | 231 | 146 | 85 | 306 | 0 | 0 | 1 | 0 | 0 | 209 | 149 | 1 | 1 | 1 | 1 | 1 | 1 | 1 | 1 | 1 | 0.99 | 1 | 1 | 0.002 |
| SE | 11127 | ENSG00000008517 | IL32 | chr16 | + | 3067983 | 3068010 | 3067376 | 3067415 | 3068179 | 3068239 | 66 | 15 | 69 | 65 | 43 | 0 | 0 | 0 | 0 | 0 | 9 | 30 | 13 | 16 | 72 | 0 | 0 | 1 | 0 | 0 | 176 | 149 | 1 | 1 | 1 | 1 | 1 | 1 | 1 | 1 | 1 | 0.917 | 1 | 1 | 0.017 |
| SE | 11128 | ENSG00000008517 | IL32 | chr16 | + | 3067983 | 3068010 | 3067376 | 3067613 | 3068179 | 3068239 | 640 | 139 | 567 | 497 | 403 | 0 | 3 | 1 | 0 | 0 | 299 | 421 | 216 | 144 | 398 | 0 | 1 | 3 | 0 | 0 | 176 | 149 | 1 | 1 | 1 | 0.975 | 0.998 | 1 | 1 | 1 | 0.997 | 0.984 | 1 | 1 | -0.002 |
| SE | 11129 | ENSG00000008517 | IL32 | chr16 | + | 3068179 | 3068239 | 3067803 | 3068010 | 3068989 | 3069109 | 609 | 147 | 632 | 521 | 555 | 1 | 0 | 1 | 1 | 0 | 244 | 445 | 220 | 133 | 374 | 0 | 0 | 0 | 2 | 0 | 209 | 149 | 1 | 1 | 0.998 | 1 | 0.998 | 0.997 | 1 | 1 | 1 | 1 | 0.979 | 1 | 0.003 |
| SE | 11141 | ENSG00000277481 | PKD1L3 | chr16 | - | 71947491 | 71947591 | 71944029 | 71944170 | 71949782 | 71950017 | 0 | 0 | 1 | 2 | 5 | 0 | 0 | 0 | 0 | 0 | 0 | 0 | 0 | 3 | 0 | 0 | 0 | 0 | 0 | 12 | 249 | 149 | 1 | 1 | NA | NA | 1 | 1 | 1 | NA | NA | NA | 1 | 0 | 0.5 |
| SE | 11142 | ENSG00000277481 | PKD1L3 | chr16 | - | 71951563 | 71951744 | 71950117 | 71950310 | 71952893 | 71953093 | 0 | 0 | 0 | 0 | 0 | 0 | 0 | 2 | 1 | 0 | 4 | 4 | 2 | 0 | 17 | 0 | 0 | 1 | 6 | 0 | 298 | 149 | 1 | 1 | NA | NA | 0 | 0 | NA | 1 | 1 | 0.5 | 0 | 1 | -0.7 |
| SE | 11143 | ENSG00000124120 | TTPAL | chr20 | + | 44478623 | 44478647 | 44475887 | 44475991 | 44479984 | 44480444 | 5 | 4 | 3 | 2 | 0 | 5 | 2 | 6 | 4 | 2 | 7 | 5 | 0 | 7 | 0 | 9 | 0 | 0 | 6 | 0 | 173 | 149 | 1 | 1 | 0.463 | 0.633 | 0.301 | 0.301 | 0 | 0.401 | 1 | NA | 0.501 | NA | -0.294 |
| SE | 11144 | ENSG00000124120 | TTPAL | chr20 | + | 44484336 | 44484428 | 44479984 | 44480444 | 44486595 | 44486706 | 5 | 5 | 19 | 12 | 10 | 0 | 0 | 0 | 0 | 5 | 0 | 21 | 8 | 17 | 26 | 0 | 0 | 1 | 0 | 0 | 241 | 149 | 1 | 1 | 1 | 1 | 1 | 1 | 0.553 | NA | 1 | 0.832 | 1 | 1 | -0.047 |
| SE | 11145 | ENSG00000124120 | TTPAL | chr20 | + | 44484336 | 44484530 | 44479984 | 44480444 | 44486595 | 44486706 | 34 | 32 | 58 | 49 | 88 | 0 | 0 | 0 | 0 | 5 | 53 | 42 | 30 | 45 | 38 | 0 | 0 | 1 | 0 | 0 | 298 | 149 | 1 | 1 | 1 | 1 | 1 | 1 | 0.898 | 1 | 1 | 0.938 | 1 | 1 | -0.008 |
| SE | 11146 | ENSG00000124120 | TTPAL | chr20 | + | 44484336 | 44484530 | 44479984 | 44480444 | 44489262 | 44494603 | 6 | 8 | 18 | 15 | 14 | 0 | 0 | 1 | 0 | 0 | 0 | 25 | 9 | 29 | 34 | 0 | 0 | 0 | 0 | 0 | 298 | 149 | 1 | 1 | 1 | 1 | 0.9 | 1 | 1 | NA | 1 | 1 | 1 | 1 | -0.02 |
| SE | 11147 | ENSG00000124120 | TTPAL | chr20 | + | 44486595 | 44486706 | 44479984 | 44480444 | 44489262 | 44489453 | 40 | 25 | 55 | 39 | 98 | 0 | 0 | 1 | 0 | 0 | 38 | 29 | 28 | 43 | 7 | 0 | 0 | 0 | 0 | 0 | 260 | 149 | 1 | 1 | 1 | 1 | 0.969 | 1 | 1 | 1 | 1 | 1 | 1 | 1 | -0.006 |
| SE | 11149 | ENSG00000205832 | C16orf96 | chr16 | + | 4574971 | 4575058 | 4574708 | 4574789 | 4575173 | 4576635 | 0 | 0 | 0 | 0 | 2 | 0 | 0 | 0 | 0 | 1 | 0 | 6 | 0 | 0 | 0 | 0 | 0 | 0 | 0 | 0 | 236 | 149 | 1 | 1 | NA | NA | NA | NA | 0.558 | NA | 1 | NA | NA | NA | -0.442 |
| SE | 11151 | ENSG00000103404 | USP31 | chr16 | - | 23082437 | 23082557 | 23078978 | 23080171 | 23084859 | 23084989 | 18 | 8 | 12 | 21 | 30 | 1 | 0 | 0 | 0 | 0 | 58 | 20 | 10 | 23 | 52 | 0 | 0 | 0 | 2 | 0 | 269 | 149 | 1 | 1 | 0.909 | 1 | 1 | 1 | 1 | 1 | 1 | 1 | 0.864 | 1 | 0.009 |
| SE | 11152 | ENSG00000103404 | USP31 | chr16 | - | 23102318 | 23102463 | 23090623 | 23090804 | 23105440 | 23105576 | 71 | 40 | 46 | 62 | 54 | 0 | 0 | 0 | 0 | 1 | 101 | 52 | 22 | 50 | 171 | 0 | 0 | 0 | 0 | 0 | 294 | 149 | 1 | 1 | 1 | 1 | 1 | 1 | 0.965 | 1 | 1 | 1 | 1 | 1 | -0.007 |
| SE | 11154 | ENSG00000168488 | ATXN2L | chr16 | + | 28826239 | 28826390 | 28825769 | 28825841 | 28829400 | 28829492 | 499 | 360 | 474 | 538 | 523 | 0 | 0 | 0 | 0 | 0 | 933 | 575 | 411 | 584 | 937 | 0 | 0 | 2 | 0 | 0 | 298 | 149 | 1 | 1 | 1 | 1 | 1 | 1 | 1 | 1 | 1 | 0.99 | 1 | 1 | 0.002 |
| SE | 11157 | ENSG00000168488 | ATXN2L | chr16 | + | 28832186 | 28832399 | 28830961 | 28831072 | 28833058 | 28833354 | 9 | 2 | 8 | 11 | 8 | 0 | 0 | 0 | 0 | 0 | 0 | 13 | 6 | 18 | 27 | 0 | 0 | 0 | 2 | 0 | 298 | 149 | 1 | 1 | 1 | 1 | 1 | 1 | 1 | NA | 1 | 1 | 0.818 | 1 | 0.045 |
| SE | 11158 | ENSG00000168488 | ATXN2L | chr16 | + | 28832204 | 28832399 | 28830961 | 28831072 | 28833058 | 28833354 | 94 | 23 | 100 | 98 | 83 | 0 | 0 | 0 | 0 | 0 | 171 | 84 | 71 | 64 | 206 | 0 | 0 | 0 | 2 | 0 | 298 | 149 | 1 | 1 | 1 | 1 | 1 | 1 | 1 | 1 | 1 | 1 | 0.941 | 1 | 0.012 |
| SE | 11161 | ENSG00000168488 | ATXN2L | chr16 | + | 28832816 | 28832887 | 28832186 | 28832399 | 28833058 | 28833354 | 92 | 59 | 100 | 107 | 172 | 0 | 0 | 0 | 0 | 0 | 222 | 137 | 84 | 146 | 146 | 0 | 3 | 1 | 4 | 0 | 220 | 149 | 1 | 1 | 1 | 1 | 1 | 1 | 1 | 1 | 0.969 | 0.983 | 0.961 | 1 | 0.017 |
| SE | 11164 | ENSG00000168488 | ATXN2L | chr16 | + | 28836325 | 28836490 | 28835932 | 28836122 | 28836750 | 28837237 | 0 | 0 | 1 | 2 | 3 | 2 | 0 | 1 | 0 | 0 | 1 | 0 | 1 | 0 | 8 | 3 | 8 | 6 | 0 | 20 | 298 | 149 | 1 | 1 | 0 | NA | 0.333 | 1 | 1 | 0.143 | 0 | 0.077 | NA | 0.167 | 0.487 |
| SE | 11165 | ENSG00000168488 | ATXN2L | chr16 | + | 28836325 | 28836490 | 28835932 | 28836176 | 28836727 | 28837237 | 0 | 0 | 3 | 0 | 0 | 0 | 0 | 0 | 0 | 0 | 0 | 1 | 1 | 0 | 0 | 0 | 0 | 3 | 0 | 0 | 298 | 149 | 1 | 1 | NA | NA | 1 | NA | NA | NA | 1 | 0.143 | NA | NA | 0.428 |
| SE | 11166 | ENSG00000168488 | ATXN2L | chr16 | + | 28836325 | 28836490 | 28835932 | 28836176 | 28836750 | 28837237 | 0 | 0 | 0 | 0 | 0 | 4 | 0 | 0 | 0 | 2 | 0 | 1 | 2 | 0 | 0 | 8 | 0 | 3 | 0 | 0 | 298 | 149 | 1 | 1 | 0 | NA | NA | NA | 0 | 0 | 1 | 0.25 | NA | NA | -0.417 |
| SE | 11167 | ENSG00000168488 | ATXN2L | chr16 | + | 28836448 | 28836485 | 28835932 | 28836122 | 28836750 | 28837237 | 0 | 0 | 0 | 0 | 0 | 2 | 0 | 1 | 0 | 0 | 0 | 0 | 1 | 1 | 0 | 3 | 8 | 6 | 0 | 20 | 186 | 149 | 1 | 1 | 0 | NA | 0 | NA | NA | 0 | 0 | 0.118 | 1 | 0 | -0.224 |
| SE | 11168 | ENSG00000213918 | DNASE1 | chr16 | + | 3651316 | 3651921 | 3642891 | 3643036 | 3654960 | 3655044 | 6 | 0 | 1 | 5 | 1 | 8 | 2 | 4 | 8 | 5 | 0 | 0 | 2 | 3 | 3 | 0 | 6 | 0 | 9 | 12 | 298 | 149 | 1 | 1 | 0.273 | 0 | 0.111 | 0.238 | 0.091 | NA | 0 | 1 | 0.143 | 0.111 | -0.171 |
| SE | 11171 | ENSG00000213918 | DNASE1 | chr16 | + | 3656963 | 3657111 | 3656637 | 3656753 | 3657186 | 3657341 | 3 | 5 | 7 | 1 | 0 | 0 | 0 | 0 | 0 | 0 | 0 | 2 | 13 | 5 | 4 | 0 | 0 | 0 | 7 | 0 | 297 | 149 | 1 | 1 | 1 | 1 | 1 | 1 | NA | NA | 1 | 1 | 0.264 | 1 | 0.184 |
| SE | 11172 | ENSG00000213918 | DNASE1 | chr16 | + | 3656998 | 3657111 | 3656637 | 3656753 | 3657186 | 3657341 | 11 | 5 | 8 | 15 | 0 | 0 | 0 | 0 | 0 | 0 | 3 | 12 | 2 | 12 | 10 | 0 | 0 | 0 | 7 | 0 | 262 | 149 | 1 | 1 | 1 | 1 | 1 | 1 | NA | 1 | 1 | 1 | 0.494 | 1 | 0.101 |
| SE | 11174 | ENSG00000102878 | HSF4 | chr16 | + | 67167302 | 67167402 | 67167119 | 67167222 | 67167474 | 67167599 | 3 | 4 | 0 | 8 | 0 | 3 | 0 | 0 | 0 | 0 | 0 | 4 | 1 | 1 | 0 | 0 | 0 | 0 | 0 | 0 | 249 | 149 | 1 | 1 | 0.374 | 1 | NA | 1 | NA | NA | 1 | 1 | 1 | NA | -0.209 |
| SE | 11181 | ENSG00000102878 | HSF4 | chr16 | + | 67169278 | 67169348 | 67169035 | 67169101 | 67169630 | 67169730 | 9 | 1 | 5 | 6 | 2 | 0 | 0 | 0 | 0 | 1 | 0 | 0 | 1 | 5 | 16 | 0 | 0 | 0 | 0 | 0 | 219 | 149 | 1 | 1 | 1 | 1 | 1 | 1 | 0.576 | NA | NA | 1 | 1 | 1 | -0.085 |
| SE | 11183 | ENSG00000067955 | CBFB | chr16 | + | 67036638 | 67036755 | 67029726 | 67029813 | 67082212 | 67082308 | 138 | 101 | 155 | 187 | 127 | 1 | 0 | 0 | 0 | 0 | 95 | 152 | 170 | 194 | 65 | 0 | 0 | 0 | 0 | 0 | 266 | 149 | 1 | 1 | 0.987 | 1 | 1 | 1 | 1 | 1 | 1 | 1 | 1 | 1 | -0.003 |
| SE | 11184 | ENSG00000067955 | CBFB | chr16 | + | 67036638 | 67036755 | 67029726 | 67030177 | 67066681 | 67066798 | 247 | 187 | 198 | 268 | 211 | 1 | 1 | 0 | 1 | 0 | 153 | 220 | 194 | 253 | 110 | 0 | 0 | 3 | 0 | 0 | 266 | 149 | 1 | 1 | 0.993 | 0.991 | 1 | 0.993 | 1 | 1 | 0.976 | 1 | 1 | 1 | 0 |
| SE | 11186 | ENSG00000067955 | CBFB | chr16 | + | 67066681 | 67066798 | 67029726 | 67029813 | 67082212 | 67082234 | 342 | 229 | 238 | 320 | 278 | 1 | 0 | 0 | 0 | 0 | 271 | 284 | 370 | 326 | 102 | 0 | 0 | 0 | 0 | 0 | 266 | 149 | 1 | 1 | 0.995 | 1 | 1 | 1 | 1 | 1 | 1 | 1 | 1 | 1 | -0.001 |
| SE | 11188 | ENSG00000067955 | CBFB | chr16 | + | 67066681 | 67066798 | 67036638 | 67036755 | 67082212 | 67082234 | 573 | 412 | 429 | 581 | 479 | 1 | 2 | 3 | 3 | 4 | 418 | 481 | 532 | 571 | 212 | 0 | 0 | 1 | 0 | 0 | 266 | 149 | 1 | 1 | 0.997 | 0.991 | 0.988 | 0.991 | 0.985 | 1 | 1 | 0.997 | 1 | 1 | -0.009 |
| SE | 11190 | ENSG00000067955 | CBFB | chr16 | + | 67082212 | 67082308 | 67066681 | 67066798 | 67098709 | 67101040 | 383 | 285 | 301 | 383 | 346 | 0 | 1 | 2 | 0 | 0 | 266 | 343 | 371 | 380 | 153 | 0 | 0 | 3 | 0 | 0 | 245 | 149 | 1 | 1 | 1 | 0.994 | 0.989 | 1 | 1 | 1 | 1 | 0.987 | 1 | 1 | -0.001 |
| SE | 11192 | ENSG00000067955 | CBFB | chr16 | + | 67082212 | 67082339 | 67066681 | 67066798 | 67098709 | 67098811 | 401 | 306 | 333 | 412 | 344 | 0 | 1 | 2 | 0 | 0 | 358 | 325 | 357 | 422 | 169 | 0 | 0 | 3 | 0 | 0 | 276 | 149 | 1 | 1 | 1 | 0.994 | 0.989 | 1 | 1 | 1 | 1 | 0.985 | 1 | 1 | 0 |
| SE | 11193 | ENSG00000067955 | CBFB | chr16 | + | 67093413 | 67093629 | 67082212 | 67082339 | 67098709 | 67100093 | 0 | 0 | 1 | 0 | 0 | 79 | 62 | 99 | 91 | 96 | 0 | 0 | 0 | 0 | 0 | 106 | 39 | 38 | 62 | 20 | 298 | 149 | 1 | 1 | 0 | 0 | 0.005 | 0 | 0 | 0 | 0 | 0 | 0 | 0 | 0.001 |
| SE | 11195 | ENSG00000172824 | CES4A | chr16 | + | 67004792 | 67004873 | 67004083 | 67004224 | 67005239 | 67005393 | 5 | 1 | 4 | 6 | 11 | 1 | 0 | 1 | 0 | 1 | 34 | 10 | 13 | 4 | 25 | 0 | 0 | 0 | 0 | 2 | 230 | 149 | 1 | 1 | 0.764 | 1 | 0.722 | 1 | 0.877 | 1 | 1 | 1 | 1 | 0.89 | -0.105 |
| SE | 11196 | ENSG00000172824 | CES4A | chr16 | + | 67006390 | 67006519 | 67005239 | 67005393 | 67006744 | 67006817 | 0 | 1 | 5 | 2 | 5 | 1 | 0 | 0 | 0 | 0 | 0 | 14 | 6 | 6 | 4 | 0 | 0 | 5 | 4 | 0 | 278 | 149 | 1 | 1 | 0 | 1 | 1 | 1 | 1 | NA | 1 | 0.391 | 0.446 | 1 | 0.091 |
| SE | 11198 | ENSG00000172824 | CES4A | chr16 | + | 67006744 | 67006817 | 67004792 | 67006519 | 67008973 | 67009735 | 1 | 1 | 3 | 2 | 0 | 1 | 0 | 0 | 0 | 0 | 14 | 6 | 7 | 5 | 5 | 0 | 0 | 0 | 0 | 0 | 222 | 149 | 1 | 1 | 0.402 | 1 | 1 | 1 | NA | 1 | 1 | 1 | 1 | 1 | -0.149 |
| SE | 11199 | ENSG00000172824 | CES4A | chr16 | + | 67006744 | 67006817 | 67005239 | 67005393 | 67008973 | 67009735 | 2 | 1 | 3 | 2 | 0 | 1 | 3 | 0 | 6 | 3 | 14 | 0 | 7 | 6 | 5 | 4 | 0 | 2 | 3 | 16 | 222 | 149 | 1 | 1 | 0.573 | 0.183 | 1 | 0.183 | 0 | 0.701 | NA | 0.701 | 0.573 | 0.173 | -0.149 |
| SE | 11201 | ENSG00000172831 | CES2 | chr16 | + | 66938036 | 66938241 | 66934443 | 66935711 | 66939216 | 66939358 | 38 | 7 | 31 | 19 | 12 | 0 | 0 | 0 | 0 | 0 | 33 | 66 | 7 | 24 | 55 | 0 | 4 | 0 | 0 | 0 | 298 | 149 | 1 | 1 | 1 | 1 | 1 | 1 | 1 | 1 | 0.892 | 1 | 1 | 1 | 0.022 |
| SE | 11202 | ENSG00000172831 | CES2 | chr16 | + | 66938036 | 66938241 | 66934470 | 66935976 | 66939216 | 66939358 | 19 | 6 | 20 | 17 | 9 | 1 | 0 | 0 | 0 | 0 | 19 | 44 | 6 | 19 | 55 | 0 | 0 | 0 | 0 | 0 | 298 | 149 | 1 | 1 | 0.905 | 1 | 1 | 1 | 1 | 1 | 1 | 1 | 1 | 1 | -0.019 |
| SE | 11203 | ENSG00000172831 | CES2 | chr16 | + | 66938036 | 66938241 | 66934470 | 66935976 | 66940221 | 66940355 | 0 | 1 | 2 | 3 | 0 | 0 | 1 | 0 | 0 | 0 | 0 | 0 | 1 | 6 | 10 | 0 | 0 | 0 | 3 | 0 | 298 | 149 | 1 | 1 | NA | 0.333 | 1 | 1 | NA | NA | NA | 1 | 0.5 | 1 | -0.056 |
| SE | 11204 | ENSG00000172831 | CES2 | chr16 | + | 66939216 | 66939358 | 66934470 | 66935976 | 66940221 | 66940355 | 16 | 2 | 11 | 13 | 13 | 0 | 1 | 0 | 0 | 0 | 13 | 14 | 12 | 15 | 4 | 0 | 0 | 0 | 3 | 0 | 291 | 149 | 1 | 1 | 1 | 0.506 | 1 | 1 | 1 | 1 | 1 | 1 | 0.719 | 1 | -0.043 |
| SE | 11206 | ENSG00000172831 | CES2 | chr16 | + | 66941123 | 66941222 | 66940436 | 66940695 | 66941505 | 66941646 | 41 | 38 | 51 | 54 | 6 | 0 | 0 | 0 | 2 | 0 | 52 | 68 | 49 | 29 | 66 | 0 | 0 | 0 | 0 | 0 | 248 | 149 | 1 | 1 | 1 | 1 | 1 | 0.942 | 1 | 1 | 1 | 1 | 1 | 1 | -0.012 |
| SE | 11207 | ENSG00000172831 | CES2 | chr16 | + | 66941767 | 66941848 | 66941505 | 66941646 | 66942104 | 66942249 | 83 | 54 | 104 | 70 | 77 | 0 | 0 | 1 | 0 | 1 | 99 | 76 | 50 | 58 | 101 | 0 | 3 | 0 | 0 | 0 | 230 | 149 | 1 | 1 | 1 | 1 | 0.985 | 1 | 0.98 | 1 | 0.943 | 1 | 1 | 1 | 0.004 |
| SE | 11208 | ENSG00000172831 | CES2 | chr16 | + | 66941767 | 66941848 | 66941505 | 66941646 | 66942647 | 66942737 | 27 | 14 | 41 | 28 | 23 | 1 | 0 | 0 | 0 | 0 | 37 | 20 | 6 | 7 | 26 | 0 | 0 | 0 | 0 | 0 | 230 | 149 | 1 | 1 | 0.946 | 1 | 1 | 1 | 1 | 1 | 1 | 1 | 1 | 1 | -0.011 |
| SE | 11209 | ENSG00000172831 | CES2 | chr16 | + | 66942104 | 66942249 | 66941505 | 66941646 | 66942647 | 66942737 | 62 | 51 | 99 | 57 | 60 | 1 | 0 | 0 | 0 | 0 | 117 | 90 | 63 | 57 | 96 | 0 | 0 | 0 | 0 | 0 | 294 | 149 | 1 | 1 | 0.969 | 1 | 1 | 1 | 1 | 1 | 1 | 1 | 1 | 1 | -0.006 |
| SE | 11211 | ENSG00000172831 | CES2 | chr16 | + | 66942647 | 66942737 | 66942104 | 66942249 | 66943298 | 66943371 | 55 | 42 | 63 | 50 | 40 | 1 | 0 | 0 | 0 | 0 | 93 | 76 | 32 | 73 | 65 | 0 | 0 | 0 | 2 | 0 | 239 | 149 | 1 | 1 | 1 | 1 | 1 | 1 | 1 | 1 | 1 | 1 | 0.958 | 1 | 0.008 |
| SE | 11212 | ENSG00000172831 | CES2 | chr16 | + | 66942647 | 66942785 | 66942104 | 66942249 | 66943298 | 66943371 | 118 | 91 | 164 | 119 | 92 | 0 | 0 | 0 | 0 | 0 | 205 | 110 | 106 | 151 | 193 | 0 | 0 | 0 | 2 | 0 | 287 | 149 | 1 | 1 | 1 | 1 | 1 | 1 | 1 | 1 | 1 | 1 | 0.975 | 1 | 0.005 |
| SE | 11214 | ENSG00000140829 | DHX38 | chr16 | + | 72094396 | 72094454 | 72093923 | 72094051 | 72096138 | 72096480 | 0 | 1 | 0 | 0 | 0 | 208 | 140 | 183 | 227 | 152 | 0 | 0 | 0 | 0 | 0 | 578 | 282 | 228 | 400 | 582 | 207 | 149 | 1 | 1 | 0 | 0.005 | 0 | 0 | 0 | 0 | 0 | 0 | 0 | 0 | 0.001 |
| SE | 11215 | ENSG00000140829 | DHX38 | chr16 | + | 72096821 | 72097009 | 72096138 | 72096480 | 72098644 | 72098792 | 156 | 153 | 191 | 203 | 217 | 0 | 0 | 0 | 0 | 0 | 425 | 245 | 229 | 320 | 660 | 0 | 0 | 0 | 1 | 0 | 298 | 149 | 1 | 1 | 1 | 1 | 1 | 1 | 1 | 1 | 1 | 1 | 0.994 | 1 | 0.001 |
| SE | 11217 | ENSG00000140829 | DHX38 | chr16 | + | 72098644 | 72098792 | 72096138 | 72096480 | 72099926 | 72099045 | 87 | 38 | 76 | 93 | 91 | 0 | 0 | 0 | 0 | 0 | 172 | 132 | 88 | 163 | 234 | 0 | 0 | 3 | 0 | 0 | 297 | 149 | 1 | 1 | 1 | 1 | 1 | 1 | 1 | 1 | 1 | 0.936 | 1 | 1 | 0.013 |
| SE | 11218 | ENSG00000140829 | DHX38 | chr16 | + | 72098644 | 72098792 | 72096821 | 72097009 | 72099203 | 72099280 | 7 | 2 | 3 | 6 | 2 | 0 | 0 | 0 | 0 | 0 | 11 | 16 | 2 | 27 | 32 | 0 | 0 | 1 | 0 | 0 | 297 | 149 | 1 | 1 | 1 | 1 | 1 | 1 | 1 | 1 | 1 | 0.501 | 1 | 1 | 0.1 |
| SE | 11222 | ENSG00000140829 | DHX38 | chr16 | + | 72104485 | 72104626 | 72103945 | 72104131 | 72105026 | 72105137 | 90 | 50 | 94 | 80 | 69 | 0 | 0 | 1 | 0 | 0 | 99 | 71 | 45 | 86 | 100 | 0 | 0 | 0 | 0 | 0 | 290 | 149 | 1 | 1 | 1 | 1 | 0.98 | 1 | 1 | 1 | 1 | 1 | 1 | 1 | -0.004 |
| SE | 11224 | ENSG00000140829 | DHX38 | chr16 | + | 72108799 | 72108925 | 72108472 | 72108607 | 72109414 | 72109510 | 193 | 100 | 212 | 209 | 138 | 0 | 0 | 0 | 0 | 0 | 396 | 240 | 182 | 215 | 326 | 0 | 0 | 1 | 0 | 0 | 275 | 149 | 1 | 1 | 1 | 1 | 1 | 1 | 1 | 1 | 1 | 0.99 | 1 | 1 | 0.002 |
| SE | 11227 | ENSG00000102984 | ZNF821 | chr16 | - | 71864137 | 71864242 | 71859679 | 71860672 | 71867892 | 71868037 | 2 | 2 | 9 | 7 | 0 | 0 | 0 | 0 | 0 | 0 | 11 | 6 | 10 | 23 | 0 | 0 | 2 | 0 | 0 | 0 | 254 | 149 | 1 | 1 | 1 | 1 | 1 | 1 | NA | 1 | 0.638 | 1 | 1 | NA | 0.091 |
| SE | 11228 | ENSG00000102984 | ZNF821 | chr16 | - | 71864137 | 71864242 | 71859679 | 71860672 | 71867911 | 71868037 | 1 | 2 | 8 | 4 | 0 | 0 | 0 | 0 | 1 | 0 | 11 | 9 | 11 | 23 | 2 | 0 | 0 | 0 | 0 | 6 | 254 | 149 | 1 | 1 | 1 | 1 | 1 | 0.701 | NA | 1 | 1 | 1 | 1 | 0.164 | 0.092 |
| SE | 11229 | ENSG00000102984 | ZNF821 | chr16 | - | 71864137 | 71864242 | 71859679 | 71860672 | 71879906 | 71880023 | 2 | 2 | 10 | 5 | 0 | 0 | 0 | 0 | 0 | 2 | 11 | 6 | 9 | 23 | 0 | 2 | 0 | 2 | 0 | 0 | 254 | 149 | 1 | 1 | 1 | 1 | 1 | 1 | 0 | 0.763 | 1 | 0.725 | 1 | NA | -0.072 |
| SE | 11232 | ENSG00000102984 | ZNF821 | chr16 | - | 71864902 | 71865048 | 71864137 | 71864242 | 71879906 | 71880023 | 12 | 6 | 11 | 15 | 12 | 1 | 0 | 2 | 2 | 0 | 10 | 25 | 13 | 14 | 22 | 0 | 0 | 0 | 0 | 0 | 295 | 149 | 1 | 1 | 0.858 | 1 | 0.735 | 0.791 | 1 | 1 | 1 | 1 | 1 | 1 | -0.123 |
| SE | 11233 | ENSG00000102984 | ZNF821 | chr16 | - | 71867892 | 71868037 | 71859684 | 71860672 | 71879906 | 71880023 | 15 | 9 | 9 | 11 | 13 | 0 | 0 | 0 | 0 | 2 | 26 | 25 | 16 | 18 | 43 | 2 | 0 | 2 | 0 | 0 | 294 | 149 | 1 | 1 | 1 | 1 | 1 | 1 | 0.767 | 0.868 | 1 | 0.802 | 1 | 1 | 0.019 |
| SE | 11236 | ENSG00000102984 | ZNF821 | chr16 | - | 71867911 | 71868037 | 71859679 | 71860672 | 71879906 | 71880023 | 15 | 7 | 7 | 8 | 13 | 0 | 0 | 0 | 0 | 2 | 22 | 19 | 16 | 11 | 49 | 2 | 0 | 2 | 0 | 0 | 275 | 149 | 1 | 1 | 1 | 1 | 1 | 1 | 0.779 | 0.856 | 1 | 0.813 | 1 | 1 | 0.022 |
| SE | 11241 | ENSG00000182149 | IST1 | chr16 | + | 71903164 | 71903258 | 71894720 | 71894858 | 71915625 | 71915728 | 6 | 0 | 2 | 0 | 0 | 0 | 2 | 0 | 1 | 3 | 0 | 0 | 0 | 1 | 0 | 0 | 0 | 4 | 1 | 7 | 243 | 149 | 1 | 1 | 1 | 0 | 1 | 0 | 0 | NA | NA | 0 | 0.38 | 0 | 0.273 |
| SE | 11242 | ENSG00000182149 | IST1 | chr16 | + | 71903164 | 71903258 | 71895550 | 71895589 | 71903589 | 71903676 | 0 | 0 | 1 | 0 | 3 | 0 | 0 | 0 | 0 | 0 | 0 | 0 | 0 | 0 | 0 | 6 | 0 | 0 | 0 | 0 | 243 | 149 | 1 | 1 | NA | NA | 1 | NA | | 0 | NA | NA | NA | NA | 1 |
| SE | 11244 | ENSG00000182149 | IST1 | chr16 | + | 71903589 | 71903676 | 71895552 | 71895589 | 71915625 | 71915728 | 0 | 0 | 0 | 0 | 0 | 107 | 60 | 120 | 90 | 64 | 6 | 0 | 0 | 0 | 0 | 182 | 111 | 132 | 135 | 103 | 236 | 149 | 1 | 1 | 0 | 0 | 0 | 0 | 0 | 0.02 | 0 | 0 | 0 | 0 | -0.004 |
| SE | 11246 | ENSG00000182149 | IST1 | chr16 | + | 71915601 | 71915728 | 71895493 | 71895589 | 71916461 | 71916642 | 130 | 118 | 123 | 131 | 198 | 2 | 1 | 0 | 3 | 0 | 212 | 118 | 147 | 103 | 153 | 0 | 0 | 0 | 0 | 0 | 276 | 149 | 1 | 1 | 0.972 | 0.985 | 1 | 0.959 | 1 | 1 | 1 | 1 | 1 | 1 | -0.017 |
| SE | 11247 | ENSG00000182149 | IST1 | chr16 | + | 71915625 | 71915728 | 71895507 | 71895589 | 71916461 | 71916493 | 232 | 173 | 240 | 216 | 262 | 2 | 1 | 0 | 3 | 0 | 367 | 212 | 265 | 210 | 256 | 0 | 0 | 0 | 0 | 0 | 252 | 149 | 1 | 1 | 0.986 | 0.99 | 1 | 0.977 | 1 | 1 | 1 | 1 | 1 | 1 | -0.009 |
| SE | 11248 | ENSG00000182149 | IST1 | chr16 | + | 71916461 | 71916642 | 71895493 | 71895589 | 71920738 | 71920822 | 4 | 2 | 0 | 3 | 0 | 0 | 0 | 2 | 0 | 0 | 0 | 0 | 0 | 2 | 0 | 0 | 0 | 0 | 0 | 0 | 298 | 149 | 1 | 1 | 1 | 1 | 0 | 1 | NA | NA | NA | NA | 1 | NA | -0.25 |
| SE | 11250 | ENSG00000182149 | IST1 | chr16 | + | 71917046 | 71917134 | 71916461 | 71916642 | 71920738 | 71920822 | 552 | 358 | 480 | 605 | 592 | 2 | 1 | 0 | 0 | 0 | 1051 | 624 | 657 | 723 | 846 | 0 | 0 | 0 | 2 | 0 | 237 | 149 | 1 | 1 | 0.994 | 0.996 | 1 | 1 | 1 | 1 | 1 | 1 | 0.996 | 1 | -0.001 |
| SE | 11251 | ENSG00000182149 | IST1 | chr16 | + | 71917046 | 71917134 | 71916461 | 71916642 | 71922473 | 71922540 | 139 | 100 | 123 | 157 | 134 | 0 | 0 | 0 | 0 | 0 | 287 | 172 | 187 | 173 | 190 | 0 | 4 | 0 | 3 | 0 | 237 | 149 | 1 | 1 | 1 | 1 | 1 | 1 | 1 | 1 | 0.964 | 1 | 0.973 | 1 | 0.013 |
| SE | 11253 | ENSG00000182149 | IST1 | chr16 | + | 71920738 | 71920822 | 71917046 | 71917134 | 71921342 | 71921400 | 534 | 321 | 456 | 546 | 599 | 0 | 0 | 0 | 0 | 4 | 806 | 514 | 572 | 666 | 765 | 0 | 0 | 0 | 0 | 0 | 233 | 149 | 1 | 1 | 1 | 1 | 1 | 1 | 0.99 | 1 | 1 | 1 | 1 | 1 | -0.002 |
| SE | 11254 | ENSG00000182149 | IST1 | chr16 | + | 71921342 | 71921453 | 71917046 | 71917134 | 71922473 | 71922540 | 309 | 230 | 311 | 299 | 714 | 0 | 1 | 0 | 0 | 0 | 360 | 343 | 248 | 252 | 441 | 0 | 0 | 0 | 0 | 0 | 260 | 149 | 1 | 1 | 1 | 0.992 | 1 | 1 | 1 | 1 | 1 | 1 | 1 | 1 | -0.002 |
| SE | 11255 | ENSG00000182149 | IST1 | chr16 | + | 71923287 | 71923380 | 71922473 | 71922680 | 71924090 | 71924281 | 208 | 132 | 291 | 233 | 501 | 0 | 0 | 0 | 0 | 0 | 237 | 276 | 123 | 204 | 423 | 0 | 0 | 0 | 2 | 0 | 242 | 149 | 1 | 1 | 1 | 1 | 1 | 1 | 1 | 1 | 1 | 1 | 0.984 | 1 | 0.003 |
| SE | 11257 | ENSG00000182149 | IST1 | chr16 | + | 71923287 | 71923380 | 71922473 | 71922680 | 71927613 | 71927890 | 213 | 129 | 289 | 233 | 503 | 0 | 1 | 1 | 0 | 0 | 237 | 270 | 122 | 201 | 406 | 0 | 0 | 0 | 0 | 0 | 242 | 149 | 1 | 1 | 1 | 0.988 | 0.994 | 1 | 1 | 1 | 1 | 1 | 1 | 1 | -0.004 |
| SE | 11258 | ENSG00000182149 | IST1 | chr16 | + | 71923287 | 71923384 | 71922473 | 71922680 | 71927613 | 71928974 | 208 | 129 | 292 | 231 | 501 | 0 | 1 | 1 | 0 | 0 | 237 | 270 | 121 | 201 | 406 | 0 | 0 | 0 | 0 | 0 | 246 | 149 | 1 | 1 | 1 | 0.987 | 0.994 | 1 | 1 | 1 | 1 | 1 | 1 | 1 | -0.004 |
| SE | 11260 | ENSG00000182149 | IST1 | chr16 | + | 71924090 | 71924281 | 71922626 | 71922680 | 71927613 | 71928049 | 0 | 1 | 0 | 0 | 0 | 0 | 1 | 1 | 0 | 0 | 0 | 0 | 0 | 2 | 0 | 0 | 0 | 0 | 0 | 0 | 298 | 149 | 1 | 1 | NA | 0.333 | 0 | NA | NA | NA | NA | NA | 1 | NA | -0.834 |
| SE | 11262 | ENSG00000182149 | IST1 | chr16 | + | 71924090 | 71924281 | 71923287 | 71923380 | 71924772 | 71924817 | 1 | 3 | 2 | 2 | 1 | 4 | 0 | 4 | 4 | 3 | 0 | 6 | 4 | 3 | 17 | 0 | 0 | 7 | 2 | 0 | 298 | 149 | 1 | 1 | 0.111 | 1 | 0.2 | 0.2 | 0.143 | NA | 1 | 0.222 | 0.429 | 1 | -0.332 |
| SE | 11263 | ENSG00000182149 | IST1 | chr16 | + | 71924090 | 71924281 | 71923287 | 71923380 | 71927613 | 71928049 | 0 | 4 | 2 | 2 | 1 | 5 | 0 | 0 | 2 | 3 | 0 | 6 | 2 | 3 | 17 | 0 | 0 | 1 | 0 | 0 | 298 | 149 | 1 | 1 | 0 | 1 | 1 | 0.333 | 0.143 | NA | 1 | 0.5 | 1 | 1 | -0.38 |
| SE | 11264 | ENSG00000182149 | IST1 | chr16 | + | 71924768 | 71924817 | 71922473 | 71922680 | 71927613 | 71927890 | 127 | 74 | 110 | 119 | 226 | 0 | 1 | 1 | 0 | 0 | 77 | 102 | 129 | 97 | 113 | 0 | 0 | 0 | 0 | 0 | 198 | 149 | 1 | 1 | 1 | 0.982 | 0.988 | 1 | 1 | 1 | 1 | 1 | 1 | 1 | -0.006 |
| SE | 11265 | ENSG00000182149 | IST1 | chr16 | + | 71924768 | 71924817 | 71923287 | 71923380 | 71927613 | 71927890 | 333 | 154 | 241 | 257 | 436 | 5 | 0 | 0 | 2 | 3 | 196 | 200 | 245 | 179 | 225 | 0 | 0 | 1 | 0 | 0 | 198 | 149 | 1 | 1 | 0.98 | 1 | 1 | 0.99 | 0.991 | 1 | 1 | 0.995 | 1 | 1 | -0.007 |
| SE | 11266 | ENSG00000182149 | IST1 | chr16 | + | 71924768 | 71924817 | 71924090 | 71924281 | 71927613 | 71927890 | 125 | 76 | 106 | 121 | 228 | 0 | 1 | 0 | 0 | 0 | 96 | 121 | 131 | 104 | 111 | 0 | 0 | 0 | 0 | 0 | 198 | 149 | 1 | 1 | 1 | 0.983 | 1 | 1 | 1 | 1 | 1 | 1 | 1 | 1 | -0.003 |
| SE | 11268 | ENSG00000182149 | IST1 | chr16 | + | 71924772 | 71924817 | 71924090 | 71924281 | 71927613 | 71929010 | 111 | 56 | 87 | 102 | 170 | 0 | 1 | 0 | 0 | 0 | 66 | 78 | 109 | 67 | 105 | 0 | 0 | 0 | 0 | 0 | 194 | 149 | 1 | 1 | 1 | 0.977 | 1 | 1 | 1 | 1 | 1 | 1 | 1 | 1 | -0.005 |
| SE | 11270 | ENSG00000125124 | BBS2 | chr16 | - | 56497742 | 56497880 | 56496416 | 56497079 | 56498436 | 56498568 | 211 | 175 | 238 | 232 | 263 | 2 | 0 | 0 | 0 | 0 | 131 | 142 | 62 | 81 | 34 | 0 | 0 | 0 | 0 | 0 | 287 | 149 | 1 | 1 | 0.982 | 1 | 1 | 1 | 1 | 1 | 1 | 1 | 1 | 1 | -0.004 |
| SE | 11276 | ENSG00000125124 | BBS2 | chr16 | - | 56501352 | 56501497 | 56500638 | 56501025 | 56501944 | 56502079 | 50 | 18 | 41 | 38 | 47 | 0 | 0 | 0 | 0 | 1 | 13 | 22 | 21 | 1 | 16 | 0 | 0 | 0 | 0 | 0 | 294 | 149 | 1 | 1 | 1 | 1 | 1 | 1 | 0.96 | 1 | 1 | 1 | 1 | 1 | -0.008 |
| SE | 11277 | ENSG00000125124 | BBS2 | chr16 | - | 56501352 | 56501497 | 56500638 | 56501025 | 56502316 | 56502456 | 102 | 52 | 91 | 74 | 142 | 0 | 1 | 2 | 1 | 0 | 19 | 43 | 34 | 24 | 31 | 0 | 0 | 2 | 0 | 0 | 294 | 149 | 1 | 1 | 1 | 0.963 | 0.958 | 0.974 | 1 | 1 | 1 | 0.896 | 1 | 1 | 0 |
| SE | 11278 | ENSG00000125124 | BBS2 | chr16 | - | 56501352 | 56502076 | 56500853 | 56501025 | 56502316 | 56502456 | 51 | 22 | 43 | 37 | 50 | 0 | 1 | 2 | 1 | 0 | 13 | 21 | 21 | 1 | 16 | 0 | 0 | 2 | 0 | 0 | 298 | 149 | 1 | 1 | 1 | 0.917 | 0.915 | 0.949 | 1 | 1 | 1 | 0.84 | 1 | 1 | -0.012 |
| SE | 11279 | ENSG00000125124 | BBS2 | chr16 | - | 56501944 | 56502079 | 56500853 | 56501025 | 56502316 | 56502456 | 3 | 1 | 7 | 2 | 1 | 0 | 1 | 2 | 1 | 0 | 0 | 1 | 4 | 0 | 0 | 0 | 0 | 2 | 0 | 0 | 284 | 149 | 1 | 1 | 1 | 0.344 | 0.647 | 0.512 | 1 | NA | 1 | 0.512 | NA | NA | -0.055 |
| SE | 11285 | ENSG00000125124 | BBS2 | chr16 | - | 56509356 | 56509494 | 56506119 | 56506224 | 56509956 | 56510034 | 0 | 0 | 2 | 6 | 0 | 131 | 100 | 131 | 115 | 182 | 0 | 0 | 0 | 0 | 0 | 56 | 29 | 58 | 38 | 33 | 287 | 149 | 1 | 1 | 0 | 0 | 0.008 | 0.026 | 0 | 0 | 0 | 0 | 0 | 0 | 0.007 |
| SE | 11290 | ENSG00000140650 | PMM2 | chr16 | + | 8802080 | 8802145 | 8801798 | 8801910 | 8802260 | 8802389 | 3 | 2 | 0 | 5 | 8 | 1 | 0 | 0 | 0 | 0 | 30 | 0 | 16 | 0 | 0 | 0 | 0 | 0 | 0 | 0 | 214 | 149 | 1 | 1 | 0.676 | 1 | NA | 1 | 1 | 1 | NA | 1 | NA | NA | -0.081 |
| SE | 11291 | ENSG00000140650 | PMM2 | chr16 | + | 8802080 | 8802145 | 8801798 | 8801910 | 8802289 | 8802389 | 3 | 2 | 0 | 3 | 8 | 0 | 0 | 2 | 0 | 0 | 30 | 0 | 7 | 0 | 0 | 0 | 0 | 1 | 0 | 0 | 214 | 149 | 1 | 1 | 1 | 1 | 0 | 1 | 1 | 1 | NA | 0.83 | NA | NA | -0.115 |
| SE | 11295 | ENSG00000140650 | PMM2 | chr16 | + | 8802260 | 8802389 | 8802080 | 8802145 | 8804766 | 8804843 | 3 | 1 | 0 | 2 | 0 | 1 | 3 | 0 | 2 | 0 | 16 | 7 | 16 | 4 | 12 | 11 | 0 | 4 | 0 | 0 | 278 | 149 | 1 | 1 | 0.617 | 0.152 | NA | 0.349 | NA | 0.438 | 1 | 0.682 | | 1 | -0.451 |
| SE | 11296 | ENSG00000140650 | PMM2 | chr16 | + | 8802289 | 8802389 | 8801798 | 8801910 | 8811078 | 8811178 | 0 | 0 | 1 | 3 | 0 | 0 | 0 | 0 | 1 | 3 | 0 | 0 | 0 | 0 | 0 | 11 | 0 | 0 | 4 | 0 | 249 | 149 | 1 | 1 | NA | NA | 1 | 0.642 | 0 | 0 | NA | NA | 0 | NA | 0.547 |
| SE | 11298 | ENSG00000140650 | PMM2 | chr16 | + | 8804766 | 8804843 | 8801798 | 8801910 | 8811078 | 8811111 | 43 | 27 | 20 | 41 | 45 | 0 | 0 | 0 | 1 | 3 | 40 | 41 | 20 | 33 | 42 | 11 | 0 | 0 | 4 | 0 | 226 | 149 | 1 | 1 | 1 | 1 | 1 | 0.964 | 0.908 | 0.706 | 1 | 1 | 0.845 | 1 | 0.064 |
| SE | 11299 | ENSG00000140650 | PMM2 | chr16 | + | 8804766 | 8804843 | 8802080 | 8802145 | 8806315 | 8806407 | 37 | 38 | 24 | 34 | 45 | 0 | 0 | 0 | 0 | 0 | 56 | 37 | 32 | 43 | 30 | 5 | 0 | 0 | 4 | 0 | 226 | 149 | 1 | 1 | 1 | 1 | 1 | 1 | 1 | 0.881 | 1 | 1 | 1 | 1 | 0.024 |
| SE | 11300 | ENSG00000140650 | PMM2 | chr16 | + | 8804766 | 8804843 | 8802260 | 8802389 | 8811078 | 8811111 | 3 | 0 | 1 | 0 | 0 | 0 | 0 | 1 | 3 | 0 | 16 | 0 | 2 | 0 | 0 | 0 | 0 | 0 | 0 | 0 | 226 | 149 | 1 | 1 | 1 | NA | 0.397 | 0 | NA | 1 | NA | 1 | NA | NA | -0.534 |
| SE | 11303 | ENSG00000140650 | PMM2 | chr16 | + | 8806315 | 8806407 | 8804766 | 8804843 | 8811078 | 8811111 | 113 | 87 | 119 | 110 | 155 | 0 | 0 | 0 | 0 | 0 | 237 | 132 | 98 | 121 | 229 | 0 | 5 | 0 | 0 | 0 | 241 | 149 | 1 | 1 | 1 | 1 | 1 | 1 | 1 | 1 | 0.942 | 1 | 1 | 1 | 0.012 |
| SE | 11308 | ENSG00000140650 | PMM2 | chr16 | + | 8850932 | 8851050 | 8849848 | 8849993 | 8855033 | 8855217 | 0 | 0 | 0 | 0 | 0 | 4 | 3 | 0 | 0 | 0 | 0 | 0 | 0 | 0 | 3 | 0 | 0 | 0 | 1 | 5 | 267 | 149 | 1 | 1 | 0 | 0 | NA | NA | NA | NA | NA | NA | 0 | 0.251 | -0.126 |
| SE | 11309 | ENSG00000140650 | PMM2 | chr16 | + | 8853525 | 8853636 | 8849848 | 8849993 | 8855033 | 8856245 | 0 | 0 | 4 | 3 | 8 | 4 | 3 | 0 | 0 | 0 | 0 | 5 | 5 | 13 | 10 | 0 | 0 | 0 | 1 | 5 | 260 | 149 | 1 | 1 | 0 | 0 | 1 | 1 | 1 | NA | 1 | 1 | 0.882 | 0.534 | -0.254 |
| SE | 11310 | ENSG00000122257 | RBBP6 | chr16 | + | 24558420 | 24558534 | 24556307 | 24556447 | 24559504 | 24559677 | 0 | 0 | 0 | 1 | 1 | 85 | 78 | 94 | 115 | 204 | 0 | 0 | 0 | 0 | 0 | 43 | 68 | 41 | 22 | 13 | 263 | 149 | 1 | 1 | 0 | 0 | 0 | 0.005 | 0.003 | 0 | 0 | 0 | 0 | 0 | 0.002 |
| SE | 11313 | ENSG00000122257 | RBBP6 | chr16 | + | 24567791 | 24567893 | 24567142 | 24567505 | 24570875 | 24572231 | 15 | 16 | 12 | 10 | 9 | 0 | 0 | 0 | 0 | 0 | 6 | 4 | 1 | 7 | 0 | 0 | 1 | 1 | 0 | 0 | 251 | 149 | 1 | 1 | 1 | 1 | 1 | 1 | 1 | 1 | 0.704 | 0.372 | 1 | NA | 0.231 |
| SE | 11314 | ENSG00000122257 | RBBP6 | chr16 | + | 24568744 | 24570499 | 24567142 | 24567505 | 24570875 | 24572231 | 154 | 134 | 140 | 162 | 222 | 0 | 0 | 0 | 0 | 0 | 42 | 76 | 104 | 24 | 23 | 0 | 1 | 1 | 0 | 0 | 298 | 149 | 1 | 1 | 1 | 1 | 1 | 1 | 1 | 1 | 0.974 | 0.981 | 1 | 1 | 0.009 |
| SE | 11315 | ENSG00000122257 | RBBP6 | chr16 | + | 24568744 | 24570499 | 24567791 | 24567893 | 24570875 | 24572231 | 186 | 148 | 180 | 200 | 272 | 0 | 0 | 0 | 0 | 0 | 16 | 98 | 83 | 29 | 12 | 0 | 0 | 1 | 0 | 0 | 298 | 149 | 1 | 1 | 1 | 1 | 1 | 1 | 1 | 1 | 1 | 0.976 | 1 | 1 | 0.005 |
| SE | 11321 | ENSG00000162004 | CCDC78 | chr16 | - | 723856 | 723936 | 722921 | 723161 | 724105 | 724205 | 2 | 0 | 3 | 5 | 0 | 0 | 0 | 0 | 0 | 0 | 0 | 0 | 3 | 0 | 0 | 0 | 0 | 2 | 0 | 0 | 229 | 149 | 1 | 1 | 1 | NA | 1 | 1 | NA | NA | NA | 0.494 | NA | NA | 0.506 |
| SE | 11322 | ENSG00000162004 | CCDC78 | chr16 | - | 725412 | 725580 | 722581 | 725293 | 725793 | 725880 | 1 | 5 | 3 | 6 | 10 | 1 | 0 | 0 | 0 | 0 | 0 | 0 | 0 | 8 | 18 | 0 | 0 | 0 | 4 | 0 | 298 | 149 | 1 | 1 | 0.333 | 1 | 1 | 1 | 1 | NA | NA | NA | 0.5 | 1 | 0.117 |
| SE | 11323 | ENSG00000162004 | CCDC78 | chr16 | - | 725412 | 725591 | 724983 | 725293 | 725793 | 725855 | 0 | 0 | 0 | 3 | 5 | 1 | 0 | 0 | 0 | 0 | 0 | 0 | 0 | 8 | 9 | 0 | 0 | 0 | 4 | 0 | 298 | 149 | 1 | 1 | 0 | NA | NA | 1 | 1 | NA | NA | NA | 0.5 | 1 | -0.083 |
| SE | 11324 | ENSG00000162004 | CCDC78 | chr16 | - | 725445 | 725580 | 724906 | 725293 | 725793 | 725880 | 1 | 5 | 3 | 3 | 0 | 1 | 0 | 0 | 0 | 0 | 0 | 0 | 0 | 0 | 9 | 0 | 0 | 0 | 4 | 0 | 284 | 149 | 1 | 1 | 0.344 | 1 | 1 | 1 | NA | NA | NA | NA | 0 | 1 | 0.336 |
| SE | 11325 | ENSG00000162004 | CCDC78 | chr16 | - | 725965 | 726085 | 724105 | 725880 | 726614 | 726880 | 1 | 5 | 4 | 4 | 0 | 1 | 0 | 0 | 0 | 0 | 9 | 3 | 0 | 4 | 0 | 0 | 0 | 0 | 0 | 0 | 269 | 149 | 1 | 1 | 0.356 | 1 | 1 | 1 | NA | 1 | 1 | NA | 1 | NA | -0.161 |
| SE | 11329 | ENSG00000033011 | ALG1 | chr16 | + | 5079063 | 5079106 | 5078756 | 5078878 | 5079747 | 5079807 | 3 | 1 | 2 | 9 | 4 | 2 | 1 | 1 | 1 | 1 | 0 | 1 | 4 | 0 | 0 | 15 | 4 | 3 | 0 | 0 | 192 | 149 | 1 | 1 | 0.538 | 0.437 | 0.608 | 0.875 | 0.756 | 0 | 0 | 0.206 | 1 | NA | 0.341 |
| SE | 11330 | ENSG00000033011 | ALG1 | chr16 | + | 5079065 | 5079102 | 5078756 | 5078878 | 5079747 | 5079807 | 4 | 0 | 2 | 1 | 0 | 2 | 1 | 1 | 1 | 1 | 17 | 3 | 0 | 0 | 16 | 15 | 4 | 3 | 0 | 0 | 186 | 149 | 1 | 1 | 0.616 | 0 | 0.616 | 0.445 | 0 | 0.476 | 0.375 | 0 | NA | 1 | -0.127 |
| SE | 11332 | ENSG00000091262 | ABCC6 | chr16 | - | 16155791 | 16155854 | 16154872 | 16155031 | 16157662 | 16157809 | 0 | 1 | 0 | 1 | 0 | 0 | 2 | 0 | 0 | 0 | 6 | 1 | 0 | 5 | 0 | 0 | 0 | 0 | 0 | 0 | 212 | 149 | 1 | 1 | NA | 0.26 | NA | 1 | NA | 1 | 1 | NA | 1 | NA | -0.37 |
| SE | 11333 | ENSG00000091262 | ABCC6 | chr16 | - | 16162992 | 16163192 | 16161437 | 16161564 | 16165622 | 16165933 | 0 | 1 | 3 | 0 | 0 | 0 | 2 | 0 | 0 | 0 | 14 | 0 | 2 | 5 | 0 | 14 | 0 | 0 | 0 | 0 | 298 | 149 | 1 | 1 | NA | 0.2 | 1 | NA | NA | 0.333 | NA | 1 | 1 | NA | -0.178 |
| SE | 11334 | ENSG00000091262 | ABCC6 | chr16 | - | 16177451 | 16177626 | 16175910 | 16175986 | 16178797 | 16178965 | 2 | 6 | 2 | 5 | 1 | 0 | 0 | 0 | 0 | 0 | 0 | 8 | 3 | 1 | 19 | 0 | 0 | 1 | 0 | 0 | 298 | 149 | 1 | 1 | 1 | 1 | 1 | 1 | 1 | NA | 1 | 0.6 | 1 | 1 | 0.1 |
| SE | 11335 | ENSG00000091262 | ABCC6 | chr16 | - | 16192829 | 16192922 | 16190163 | 16190367 | 16198020 | 16198182 | 3 | 0 | 0 | 4 | 4 | 0 | 0 | 0 | 0 | 0 | 6 | 2 | 1 | 0 | 10 | 6 | 0 | 0 | 2 | 0 | 242 | 149 | 1 | 1 | 1 | NA | NA | 1 | 1 | 0.381 | 1 | 1 | 0 | 1 | 0.324 |
| SE | 11336 | ENSG00000091262 | ABCC6 | chr16 | - | 16220272 | 16220388 | 16219821 | 16219947 | 16221648 | 16221831 | 1 | 3 | 0 | 0 | 2 | 0 | 1 | 0 | 0 | 0 | 8 | 0 | 1 | 0 | 7 | 0 | 3 | 0 | 0 | 0 | 265 | 149 | 1 | 1 | 1 | 0.628 | NA | NA | 1 | 1 | 0 | 1 | NA | 1 | 0.126 |
| SE | 11338 | ENSG00000175595 | ERCC4 | chr16 | + | 13924529 | 13924637 | 13922030 | 13922211 | 13926560 | 13926756 | 0 | 2 | 0 | 0 | 0 | 26 | 9 | 6 | 17 | 35 | 0 | 0 | 0 | 0 | 0 | 0 | 11 | 15 | 4 | 0 | 257 | 149 | 1 | 1 | 0 | 0.114 | 0 | 0 | 0 | NA | 0 | 0 | 0 | NA | 0.023 |
| SE | 11339 | ENSG00000175595 | ERCC4 | chr16 | + | 13926153 | 13926291 | 13922030 | 13922211 | 13926560 | 13926756 | 2 | 0 | 0 | 0 | 0 | 26 | 9 | 6 | 17 | 35 | 0 | 0 | 2 | 0 | 0 | 0 | 11 | 15 | 4 | 0 | 287 | 149 | 1 | 1 | 0.038 | 0 | 0 | 0 | 0 | NA | 0 | 0.065 | 0 | NA | -0.014 |
| SE | 11340 | ENSG00000175595 | ERCC4 | chr16 | + | 13928027 | 13928235 | 13924529 | 13926756 | 13930709 | 13930890 | 54 | 35 | 43 | 48 | 52 | 0 | 1 | 2 | 0 | 1 | 28 | 56 | 31 | 37 | 5 | 0 | 0 | 0 | 0 | 0 | 298 | 149 | 1 | 1 | 1 | 0.946 | 0.915 | 1 | 0.963 | 1 | 1 | 1 | 1 | 1 | -0.035 |
| SE | 11341 | ENSG00000175595 | ERCC4 | chr16 | + | 13928121 | 13928235 | 13926551 | 13926756 | 13930709 | 13930890 | 26 | 22 | 27 | 28 | 25 | 0 | 1 | 2 | 0 | 1 | 28 | 48 | 27 | 25 | 5 | 0 | 0 | 0 | 0 | 0 | 263 | 149 | 1 | 1 | 1 | 0.926 | 0.884 | 1 | 0.934 | 1 | 1 | 1 | 1 | 1 | -0.051 |
| SE | 11347 | ENSG00000186153 | WWOX | chr16 | + | 78117213 | 78117414 | 78114975 | 78115154 | 78118954 | 78120464 | 0 | 0 | 1 | 1 | 3 | 4 | 4 | 15 | 0 | 7 | 0 | 0 | 1 | 0 | 0 | 0 | 0 | 6 | 0 | 0 | 298 | 149 | 1 | 1 | 0 | 0 | 0.032 | 1 | 0.176 | NA | NA | 0.077 | NA | NA | 0.165 |
| SE | 11350 | ENSG00000186153 | WWOX | chr16 | + | 78179707 | 78179786 | 78164182 | 78164289 | 78386859 | 78388566 | 1 | 0 | 0 | 0 | 0 | 13 | 4 | 7 | 7 | 17 | 0 | 0 | 0 | 0 | 0 | 26 | 41 | 11 | 21 | 15 | 228 | 149 | 1 | 1 | 0.048 | 0 | 0 | 0 | 0 | 0 | 0 | 0 | 0 | 0 | 0.01 |
| SE | 11358 | ENSG00000186153 | WWOX | chr16 | + | 78756939 | 78757015 | 78535746 | 78535842 | 79211607 | 79212667 | 2 | 0 | 0 | 0 | 0 | 0 | 0 | 2 | 0 | 0 | 0 | 8 | 1 | 0 | 0 | 0 | 0 | 0 | 0 | 0 | 225 | 149 | 1 | 1 | 1 | NA | 0 | NA | NA | NA | 1 | 1 | NA | NA | -0.5 |
| SE | 11359 | ENSG00000186153 | WWOX | chr16 | + | 79040103 | 79040245 | 78756939 | 78757015 | 79211607 | 79212637 | 0 | 0 | 1 | 0 | 0 | 2 | 0 | 0 | 0 | 0 | 0 | 0 | 0 | 0 | 0 | 0 | 12 | 0 | 0 | 0 | 291 | 149 | 1 | 1 | 0 | NA | 1 | NA | NA | NA | 0 | NA | NA | NA | 0.5 |
| SE | 11364 | ENSG00000284428 | NA | chr19 | - | 23272618 | 23272719 | 23269762 | 23270292 | 23274075 | 23274196 | 0 | 0 | 2 | 0 | 2 | 1 | 1 | 0 | 5 | 10 | 0 | 0 | 4 | 0 | 0 | 0 | 10 | 0 | 0 | 10 | 250 | 149 | 1 | 1 | 0 | 0 | 1 | 0 | 0.107 | NA | 0 | 1 | NA | 0 | -0.112 |
| SE | 11365 | ENSG00000284428 | NA | chr19 | - | 23272618 | 23272723 | 23261619 | 23261744 | 23274075 | 23274228 | 0 | 0 | 0 | 1 | 0 | 10 | 3 | 4 | 0 | 0 | 0 | 0 | 0 | 0 | 0 | 0 | 0 | 1 | 3 | 3 | 254 | 149 | 1 | 1 | 0 | 0 | 0 | 1 | NA | NA | NA | 0 | 0 | 0 | 0.25 |
| SE | 11372 | ENSG00000166747 | AP1G1 | chr16 | - | 71734608 | 71734707 | 71728999 | 71733159 | 71738941 | 71739102 | 266 | 185 | 232 | 261 | 308 | 0 | 4 | 0 | 2 | 0 | 94 | 148 | 293 | 142 | 45 | 0 | 0 | 1 | 1 | 0 | 248 | 149 | 1 | 1 | 1 | 0.965 | 1 | 0.987 | 1 | 1 | 1 | 0.994 | 0.988 | 1 | -0.006 |
| SE | 11373 | ENSG00000166747 | AP1G1 | chr16 | - | 71739233 | 71739341 | 71738941 | 71739102 | 71745143 | 71745270 | 212 | 163 | 133 | 184 | 186 | 0 | 0 | 0 | 0 | 0 | 110 | 185 | 320 | 175 | 58 | 0 | 0 | 1 | 0 | 0 | 257 | 149 | 1 | 1 | 1 | 1 | 1 | 1 | 1 | 1 | 1 | 0.995 | 1 | 1 | 0.001 |
| SE | 11375 | ENSG00000166747 | AP1G1 | chr16 | - | 71749893 | 71749983 | 71748250 | 71748378 | 71750209 | 71750332 | 239 | 147 | 172 | 199 | 286 | 0 | 0 | 0 | 1 | 0 | 88 | 112 | 187 | 114 | 56 | 0 | 0 | 0 | 0 | 0 | 239 | 149 | 1 | 1 | 1 | 1 | 1 | 0.992 | 1 | 1 | 1 | 1 | 1 | 1 | -0.002 |
| SE | 11378 | ENSG00000166747 | AP1G1 | chr16 | - | 71765488 | 71765584 | 71764645 | 71764726 | 71767876 | 71767885 | 112 | 78 | 70 | 94 | 113 | 0 | 1 | 0 | 0 | 0 | 18 | 74 | 113 | 80 | 6 | 0 | 0 | 0 | 0 | 0 | 245 | 149 | 1 | 1 | 1 | 0.979 | 1 | 1 | 1 | 1 | 1 | 1 | 1 | 1 | -0.004 |
| SE | 11379 | ENSG00000166747 | AP1G1 | chr16 | - | 71765488 | 71765584 | 71764645 | 71764726 | 71769622 | 71769699 | 243 | 142 | 139 | 189 | 224 | 0 | 0 | 0 | 1 | 0 | 56 | 173 | 241 | 152 | 30 | 0 | 0 | 0 | 0 | 0 | 245 | 149 | 1 | 1 | 1 | 1 | 1 | 0.991 | 1 | 1 | 1 | 1 | 1 | 1 | -0.002 |
| SE | 11382 | ENSG00000166747 | AP1G1 | chr16 | - | 71769622 | 71769699 | 71767876 | 71767885 | 71771155 | 71771252 | 36 | 21 | 10 | 25 | 31 | 0 | 0 | 0 | 0 | 0 | 0 | 38 | 23 | 25 | 3 | 0 | 0 | 2 | 0 | 0 | 226 | 149 | 1 | 1 | 1 | 1 | 1 | 1 | 1 | NA | 1 | 0.883 | 1 | 1 | 0.029 |
| SE | 11383 | ENSG00000166747 | AP1G1 | chr16 | - | 71771155 | 71771252 | 71769618 | 71769699 | 71773220 | 71773362 | 236 | 121 | 167 | 181 | 266 | 1 | 0 | 0 | 0 | 0 | 69 | 151 | 213 | 127 | 45 | 0 | 0 | 0 | 0 | 0 | 246 | 149 | 1 | 1 | 0.993 | 1 | 1 | 1 | 1 | 1 | 1 | 1 | 1 | 1 | -0.001 |
| SE | 11385 | ENSG00000166747 | AP1G1 | chr16 | - | 71773220 | 71773362 | 71771139 | 71771252 | 71774467 | 71774592 | 269 | 165 | 206 | 236 | 306 | 0 | 0 | 0 | 0 | 2 | 49 | 161 | 246 | 168 | 54 | 0 | 4 | 0 | 0 | 0 | 291 | 149 | 1 | 1 | 1 | 1 | 1 | 1 | 0.987 | 1 | 0.954 | 1 | 1 | 1 | 0.007 |
| SE | 11386 | ENSG00000166747 | AP1G1 | chr16 | - | 71806685 | 71806728 | 71789278 | 71789482 | 71807800 | 71807838 | 9 | 6 | 10 | 5 | 7 | 6 | 4 | 2 | 3 | 9 | 0 | 8 | 5 | 1 | 0 | 0 | 6 | 2 | 2 | 0 | 192 | 149 | 1 | 1 | 0.538 | 0.538 | 0.795 | 0.564 | 0.376 | NA | 0.509 | 0.66 | 0.28 | NA | 0.079 |
| SE | 11387 | ENSG00000166747 | AP1G1 | chr16 | - | 71806685 | 71806728 | 71789278 | 71789482 | 71808014 | 71808150 | 1 | 2 | 2 | 0 | 3 | 1 | 0 | 1 | 1 | 3 | 0 | 4 | 3 | 6 | 0 | 0 | 0 | 0 | 3 | 0 | 192 | 149 | 1 | 1 | 0.437 | 1 | 0.608 | 0 | 0.437 | NA | 1 | 1 | 0.608 | NA | -0.373 |
| SE | 11388 | ENSG00000166747 | AP1G1 | chr16 | - | 71807800 | 71807838 | 71789319 | 71789482 | 71808014 | 71808150 | 2 | 2 | 1 | 2 | 7 | 1 | 0 | 1 | 1 | 3 | 0 | 6 | 0 | 0 | 0 | 0 | 0 | 0 | 3 | 0 | 187 | 149 | 1 | 1 | 0.614 | 1 | 0.443 | 0.614 | 0.65 | NA | 1 | NA | 0 | NA | 0.164 |
| SE | 11389 | ENSG00000166747 | AP1G1 | chr16 | - | 71807800 | 71807838 | 71806685 | 71806728 | 71808014 | 71808150 | 0 | 1 | 4 | 4 | 4 | 0 | 0 | 0 | 0 | 0 | 0 | 4 | 0 | 0 | 5 | 0 | 0 | 0 | 6 | 0 | 187 | 149 | 1 | 1 | NA | 1 | 1 | 1 | 1 | NA | 1 | NA | 0 | 1 | 0.333 |
| SE | 11390 | ENSG00000166747 | AP1G1 | chr16 | - | 71808014 | 71808150 | 71789278 | 71789482 | 71808504 | 71808677 | 6 | 0 | 1 | 2 | 7 | 2 | 0 | 0 | 1 | 0 | 0 | 1 | 0 | 5 | 0 | 0 | 1 | 3 | 0 | 2 | 285 | 149 | 1 | 1 | 0.611 | NA | 1 | 0.511 | 1 | NA | 0.343 | 0 | 1 | 0 | 0.445 |
| SE | 11395 | ENSG00000074356 | NCBP3 | chr17 | - | 3816115 | 3816270 | 3802157 | 3813279 | 3818262 | 3818572 | 199 | 110 | 166 | 175 | 251 | 0 | 0 | 0 | 0 | 0 | 180 | 175 | 118 | 250 | 82 | 0 | 0 | 3 | 0 | 0 | 298 | 149 | 1 | 1 | 1 | 1 | 1 | 1 | 1 | 1 | 1 | 0.952 | 1 | 1 | 0.01 |
| SE | 11396 | ENSG00000074356 | NCBP3 | chr17 | - | 3821248 | 3821352 | 3818262 | 3818572 | 3821952 | 3822052 | 145 | 176 | 156 | 220 | 201 | 0 | 0 | 0 | 0 | 0 | 285 | 214 | 252 | 373 | 155 | 1 | 0 | 3 | 0 | 0 | 253 | 149 | 1 | 1 | 1 | 1 | 1 | 1 | 1 | 0.994 | 1 | 0.98 | 1 | 1 | 0.005 |
| SE | 11398 | ENSG00000074356 | NCBP3 | chr17 | - | 3829242 | 3829368 | 3826086 | 3826215 | 3840099 | 3840205 | 251 | 195 | 120 | 210 | 231 | 3 | 0 | 1 | 0 | 0 | 280 | 258 | 205 | 436 | 141 | 0 | 0 | 0 | 2 | 0 | 275 | 149 | 1 | 1 | 0.978 | 1 | 0.985 | 1 | 1 | 1 | 1 | 1 | 0.992 | 1 | -0.006 |
| SE | 11399 | ENSG00000074356 | NCBP3 | chr17 | - | 3829242 | 3829368 | 3826086 | 3826215 | 3843085 | 3843151 | 132 | 112 | 71 | 108 | 128 | 2 | 0 | 0 | 0 | 0 | 168 | 177 | 124 | 251 | 85 | 0 | 0 | 0 | 0 | 0 | 275 | 149 | 1 | 1 | 0.973 | 1 | 1 | 1 | 1 | 1 | 1 | 1 | 1 | 1 | -0.005 |
| SE | 11400 | ENSG00000074356 | NCBP3 | chr17 | - | 3840099 | 3840205 | 3826086 | 3826215 | 3843085 | 3843151 | 42 | 18 | 28 | 32 | 35 | 2 | 0 | 0 | 0 | 0 | 70 | 20 | 26 | 49 | 0 | 0 | 0 | 0 | 0 | 0 | 255 | 149 | 1 | 1 | 0.925 | 1 | 1 | 1 | 1 | 1 | 1 | 1 | 1 | NA | -0.015 |
| SE | 11403 | ENSG00000040199 | PHLPP2 | chr16 | - | 71656570 | 71656681 | 71655239 | 71655434 | 71658232 | 71658363 | 40 | 28 | 15 | 21 | 43 | 0 | 0 | 2 | 2 | 0 | 0 | 6 | 25 | 28 | 3 | 0 | 0 | 0 | 0 | 0 | 260 | 149 | 1 | 1 | 1 | 1 | 0.811 | 0.857 | 1 | NA | 1 | 1 | 1 | 1 | -0.066 |
| SE | 11405 | ENSG00000040199 | PHLPP2 | chr16 | - | 71684475 | 71684601 | 71681750 | 71681905 | 71690518 | 71690709 | 38 | 17 | 25 | 39 | 36 | 1 | 2 | 7 | 9 | 1 | 0 | 12 | 21 | 1 | 0 | 0 | 8 | 1 | 9 | 0 | 275 | 149 | 1 | 1 | 0.954 | 0.822 | 0.659 | 0.701 | 0.951 | NA | 0.448 | 0.919 | 0.057 | NA | 0.343 |
| SE | 11407 | ENSG00000040199 | PHLPP2 | chr16 | - | 71702597 | 71702731 | 71690518 | 71690709 | 71723768 | 71723812 | 10 | 10 | 17 | 14 | 17 | 0 | 0 | 0 | 0 | 0 | 8 | 6 | 3 | 1 | 7 | 0 | 2 | 0 | 0 | 0 | 283 | 149 | 1 | 1 | 1 | 1 | 1 | 1 | 1 | 1 | 0.612 | 1 | 1 | 1 | 0.078 |
| SE | 11408 | ENSG00000040199 | PHLPP2 | chr16 | - | 71714511 | 71714801 | 71690518 | 71690709 | 71723768 | 71723812 | 7 | 2 | 1 | 3 | 3 | 0 | 0 | 0 | 0 | 0 | 0 | 5 | 1 | 3 | 0 | 0 | 2 | 0 | 0 | 0 | 298 | 149 | 1 | 1 | 1 | 1 | 1 | 1 | 1 | NA | 0.556 | 1 | 1 | NA | 0.148 |
| SE | 11409 | ENSG00000040199 | PHLPP2 | chr16 | - | 71714511 | 71714801 | 71702597 | 71702731 | 71723768 | 71723812 | 11 | 4 | 11 | 9 | 15 | 2 | 0 | 1 | 1 | 0 | 0 | 17 | 7 | 3 | 7 | 0 | 0 | 0 | 0 | 0 | 298 | 149 | 1 | 1 | 0.733 | 1 | 0.846 | 0.818 | 1 | NA | 1 | 1 | 1 | 1 | -0.121 |
| SE | 11410 | ENSG00000261611 | NA | chr16 | - | 71482081 | 71482143 | 71478878 | 71479005 | 71484588 | 71484748 | 1 | 7 | 12 | 6 | 2 | 1 | 3 | 1 | 1 | 0 | 0 | 5 | 0 | 2 | 4 | 0 | 0 | 0 | 0 | 0 | 211 | 149 | 1 | 1 | 0.414 | 0.622 | 0.894 | 0.809 | 1 | NA | 1 | NA | 1 | 1 | -0.252 |
| SE | 11413 | ENSG00000242612 | DECR2 | chr16 | + | 405512 | 405665 | 404955 | 405024 | 406345 | 406397 | 0 | 0 | 1 | 0 | 0 | 4 | 0 | 3 | 0 | 0 | 0 | 0 | 0 | 0 | 0 | 18 | 0 | 0 | 9 | 0 | 298 | 149 | 1 | 1 | 0 | NA | 0.143 | NA | NA | 0 | NA | NA | 0 | NA | 0.071 |
| SE | 11419 | ENSG00000103202 | NME4 | chr16 | + | 398207 | 398394 | 397191 | 397313 | 398989 | 399123 | 0 | 1 | 0 | 2 | 0 | 7 | 0 | 0 | 0 | 0 | 0 | 0 | 0 | 0 | 0 | 0 | 0 | 0 | 0 | 4 | 298 | 149 | 1 | 1 | 0 | 1 | NA | 1 | NA | NA | NA | NA | NA | 0 | 0.667 |
| SE | 11420 | ENSG00000103202 | NME4 | chr16 | + | 399378 | 399480 | 398207 | 399123 | 399626 | 399739 | 162 | 25 | 56 | 57 | 57 | 0 | 0 | 1 | 3 | 0 | 81 | 27 | 39 | 48 | 259 | 0 | 0 | 0 | 0 | 0 | 251 | 149 | 1 | 1 | 1 | 1 | 0.971 | 0.919 | 1 | 1 | 1 | 1 | 1 | 1 | -0.022 |
| SE | 11423 | ENSG00000129925 | PGAP6 | chr16 | - | 377670 | 377848 | 377377 | 377585 | 384255 | 384396 | 28 | 8 | 14 | 13 | 0 | 0 | 1 | 0 | 0 | 0 | 23 | 31 | 16 | 18 | 28 | 0 | 0 | 0 | 0 | 0 | 298 | 149 | 1 | 1 | 1 | 0.8 | 1 | 1 | NA | 1 | 1 | 1 | 1 | 1 | -0.05 |
| SE | 11424 | ENSG00000086504 | MRPL28 | chr16 | - | 368500 | 368635 | 368327 | 368414 | 369067 | 369220 | 155 | 67 | 118 | 137 | 125 | 0 | 0 | 0 | 0 | 0 | 273 | 237 | 181 | 282 | 303 | 0 | 0 | 1 | 0 | 0 | 284 | 149 | 1 | 1 | 1 | 1 | 1 | 1 | 1 | 1 | 1 | 0.99 | 1 | 1 | 0.002 |
| SE | 11427 | ENSG00000167522 | ANKRD11 | chr16 | - | 89288527 | 89288670 | 89286038 | 89286186 | 89305205 | 89305344 | 30 | 23 | 19 | 27 | 4 | 0 | 0 | 0 | 0 | 0 | 8 | 52 | 40 | 65 | 29 | 0 | 1 | 0 | 0 | 0 | 292 | 149 | 1 | 1 | 1 | 1 | 1 | 1 | 1 | 1 | 0.964 | 1 | 1 | 1 | 0.007 |
| SE | 11428 | ENSG00000167522 | ANKRD11 | chr16 | - | 89290624 | 89291183 | 89288527 | 89288670 | 89305205 | 89305239 | 189 | 105 | 161 | 173 | 129 | 0 | 1 | 0 | 1 | 0 | 270 | 168 | 172 | 182 | 297 | 0 | 1 | 0 | 0 | 0 | 298 | 149 | 1 | 1 | 1 | 0.981 | 1 | 0.989 | 1 | 1 | 0.988 | 1 | 1 | 1 | -0.004 |
| SE | 11430 | ENSG00000167522 | ANKRD11 | chr16 | - | 89291680 | 89291777 | 89291012 | 89291183 | 89316932 | 89317075 | 2 | 0 | 7 | 3 | 7 | 1 | 0 | 0 | 0 | 0 | 8 | 3 | 13 | 3 | 12 | 0 | 0 | 0 | 4 | 0 | 246 | 149 | 1 | 1 | 0.548 | NA | 1 | 1 | 1 | 1 | 1 | 1 | 0.312 | 1 | 0.025 |
| SE | 11432 | ENSG00000167522 | ANKRD11 | chr16 | - | 89305205 | 89305344 | 89270816 | 89270909 | 89418283 | 89418368 | 2 | 2 | 3 | 3 | 4 | 0 | 0 | 0 | 0 | 1 | 18 | 3 | 16 | 2 | 0 | 0 | 0 | 0 | 0 | 0 | 288 | 149 | 1 | 1 | 1 | 1 | 1 | 1 | 0.674 | 1 | 1 | 1 | 1 | NA | -0.065 |
| SE | 11433 | ENSG00000167522 | ANKRD11 | chr16 | - | 89305205 | 89305344 | 89290971 | 89291183 | 89316932 | 89317019 | 134 | 85 | 134 | 125 | 122 | 1 | 0 | 0 | 0 | 0 | 333 | 146 | 206 | 177 | 337 | 0 | 0 | 0 | 4 | 0 | 288 | 149 | 1 | 1 | 0.986 | 1 | 1 | 1 | 1 | 1 | 1 | 1 | 0.958 | 1 | 0.006 |
| SE | 11434 | ENSG00000167522 | ANKRD11 | chr16 | - | 89305205 | 89305344 | 89291680 | 89291777 | 89316932 | 89317019 | 107 | 60 | 99 | 91 | 99 | 0 | 0 | 0 | 0 | 3 | 253 | 124 | 146 | 145 | 260 | 0 | 0 | 0 | 0 | 0 | 288 | 149 | 1 | 1 | 1 | 1 | 1 | 1 | 0.945 | 1 | 1 | 1 | 1 | 1 | -0.011 |
| SE | 11436 | ENSG00000167522 | ANKRD11 | chr16 | - | 89305205 | 89305344 | 89298867 | 89300962 | 89316932 | 89317019 | 130 | 65 | 90 | 112 | 92 | 0 | 1 | 0 | 0 | 0 | 286 | 143 | 161 | 156 | 354 | 0 | 0 | 0 | 1 | 0 | 288 | 149 | 1 | 1 | 1 | 0.971 | 1 | 1 | 1 | 1 | 1 | 1 | 0.988 | 1 | -0.003 |
| SE | 11440 | ENSG00000167522 | ANKRD11 | chr16 | - | 89395696 | 89396061 | 89361494 | 89361677 | 89418283 | 89418499 | 1 | 0 | 1 | 0 | 0 | 0 | 0 | 1 | 0 | 2 | 13 | 0 | 0 | 0 | 0 | 0 | 0 | 5 | 0 | 0 | 298 | 149 | 1 | 1 | 1 | NA | 0.333 | NA | 0 | 1 | NA | 0 | NA | NA | -0.056 |
| SE | 11442 | ENSG00000167522 | ANKRD11 | chr16 | - | 89412449 | 89412563 | 89392375 | 89392679 | 89418283 | 89418499 | 0 | 0 | 0 | 0 | 0 | 0 | 0 | 0 | 1 | 0 | 0 | 0 | 0 | 2 | 0 | 0 | 0 | 1 | 0 | 0 | 263 | 149 | 1 | 1 | NA | NA | NA | 0 | NA | NA | NA | 0 | 1 | NA | -0.5 |
| SE | 11443 | ENSG00000167522 | ANKRD11 | chr16 | - | 89412449 | 89412563 | 89395696 | 89396061 | 89418283 | 89418499 | 0 | 0 | 0 | 0 | 0 | 1 | 0 | 1 | 0 | 0 | 0 | 0 | 0 | 2 | 0 | 13 | 0 | 0 | 0 | 0 | 263 | 149 | 1 | 1 | 0 | NA | 0 | NA | NA | 0 | NA | NA | 1 | NA | -0.5 |
| SE | 11444 | ENSG00000167522 | ANKRD11 | chr16 | - | 89418283 | 89418368 | 89316932 | 89317075 | 89431255 | 89431326 | 31 | 22 | 28 | 33 | 28 | 0 | 0 | 1 | 0 | 0 | 74 | 14 | 22 | 37 | 57 | 0 | 3 | 0 | 0 | 0 | 234 | 149 | 1 | 1 | 1 | 1 | 0.947 | 1 | 1 | 1 | 0.748 | 1 | 1 | 1 | 0.04 |
| SE | 11447 | ENSG00000187555 | USP7 | chr16 | - | 8895641 | 8895741 | 8895030 | 8895150 | 8896998 | 8897099 | 495 | 367 | 441 | 417 | 449 | 0 | 1 | 0 | 0 | 0 | 371 | 348 | 498 | 507 | 204 | 0 | 0 | 0 | 0 | 0 | 249 | 149 | 1 | 1 | 1 | 0.995 | 1 | 1 | 1 | 1 | 1 | 1 | 1 | 1 | -0.001 |
| SE | 11449 | ENSG00000187555 | USP7 | chr16 | - | 8898530 | 8898639 | 8898359 | 8898437 | 8900529 | 8900630 | 192 | 147 | 163 | 209 | 241 | 0 | 1 | 0 | 0 | 0 | 190 | 161 | 235 | 235 | 42 | 0 | 0 | 0 | 0 | 0 | 258 | 149 | 1 | 1 | 1 | 0.988 | 1 | 1 | 1 | 1 | 1 | 1 | 1 | 1 | -0.002 |
| SE | 11451 | ENSG00000187555 | USP7 | chr16 | - | 8899120 | 8899188 | 8898530 | 8898639 | 8900529 | 8900630 | 180 | 123 | 131 | 187 | 118 | 1 | 0 | 3 | 0 | 0 | 53 | 168 | 162 | 210 | 64 | 0 | 0 | 0 | 5 | 0 | 217 | 149 | 1 | 1 | 0.992 | 1 | 0.968 | 1 | 1 | 1 | 1 | 1 | 0.966 | 1 | -0.001 |
| SE | 11453 | ENSG00000187555 | USP7 | chr16 | - | 8899603 | 8899757 | 8898530 | 8898639 | 8900529 | 8900630 | 358 | 244 | 349 | 348 | 511 | 1 | 0 | 3 | 0 | 0 | 288 | 325 | 348 | 269 | 73 | 0 | 0 | 0 | 5 | 0 | 298 | 149 | 1 | 1 | 0.994 | 1 | 0.983 | 1 | 1 | 1 | 1 | 1 | 0.964 | 1 | 0.003 |
| SE | 11454 | ENSG00000187555 | USP7 | chr16 | - | 8899603 | 8899757 | 8898530 | 8898639 | 8900989 | 8901057 | 7 | 5 | 6 | 7 | 4 | 2 | 3 | 1 | 0 | 1 | 10 | 35 | 16 | 30 | 0 | 6 | 3 | 2 | 3 | 0 | 298 | 149 | 1 | 1 | 0.636 | 0.455 | 0.75 | 1 | 0.667 | 0.455 | 0.854 | 0.8 | 0.833 | NA | -0.034 |
| SE | 11455 | ENSG00000187555 | USP7 | chr16 | - | 8899603 | 8899757 | 8899120 | 8899188 | 8900529 | 8900630 | 698 | 502 | 633 | 725 | 845 | 2 | 1 | 4 | 4 | 0 | 474 | 579 | 694 | 541 | 143 | 0 | 0 | 1 | 6 | 0 | 298 | 149 | 1 | 1 | 0.994 | 0.996 | 0.988 | 0.989 | 1 | 1 | 1 | 0.997 | 0.978 | 1 | -0.002 |

| | | | | | | | | | | | | | | | |
|---|---|---|---|---|---|---|---|---|---|---|---|---|---|---|---|
| SE | 11456 | ENSG00000187555 | USP7 | chr16 | - | 8899603 | 8899757 | 8899120 | 8899188 | 8900989 | 8901057 | 347 | 263 | 290 | [illegible] |
| SE | 11462 | ENSG00000187555 | USP7 | chr16 | - | 8900989 | 8901057 | 8900529 | 8900630 | 8901141 | 8901219 | 494 | 356 | 446 | [illegible] |
| SE | 11464 | ENSG00000187555 | USP7 | chr16 | - | 8903267 | 8903402 | 8902138 | 8902482 | 8904434 | 8904522 | 488 | 352 | 477 | [illegible] |
| SE | 11465 | ENSG00000187555 | USP7 | chr16 | - | 8904434 | 8904524 | 8903267 | 8903402 | 8905186 | 8905331 | 96 | 55 | 85 | [illegible] |
| SE | 11466 | ENSG00000187555 | USP7 | chr16 | - | 8904434 | 8904565 | 8903267 | 8903402 | 8905186 | 8905331 | 540 | 344 | 492 | [illegible] |
| SE | 11467 | ENSG00000187555 | USP7 | chr16 | - | 8905186 | 8905331 | 8904434 | 8904565 | 8906425 | 8906582 | 1213 | 952 | 1219 | [illegible] |
| SE | 11468 | ENSG00000187555 | USP7 | chr16 | - | 8915444 | 8915525 | 8915253 | 8915344 | 8916501 | 8916556 | 574 | 477 | 504 | [illegible] |
| SE | 11472 | ENSG00000187555 | USP7 | chr16 | - | 8917025 | 8917156 | 8916501 | 8916556 | 8919030 | 8919139 | 623 | 471 | 580 | [illegible] |
| SE | 11473 | ENSG00000187555 | USP7 | chr16 | - | 8919030 | 8919139 | 8917025 | 8917156 | 8920358 | 8920447 | 741 | 580 | 658 | [illegible] |
| SE | 11475 | ENSG00000187555 | USP7 | chr16 | - | 8921156 | 8921295 | 8920358 | 8920447 | 8923353 | 8923413 | 438 | 378 | 396 | [illegible] |
| SE | 11476 | ENSG00000187555 | USP7 | chr16 | - | 8923214 | 8923413 | 8921156 | 8921295 | 8930292 | 8930397 | 787 | 563 | 762 | [illegible] |
| SE | 11477 | ENSG00000187555 | USP7 | chr16 | - | 8923353 | 8923413 | 8921156 | 8921295 | 8930292 | 8930397 | 95 | 57 | 96 | [illegible] |
| SE | 11479 | ENSG00000187555 | USP7 | chr16 | - | 8930292 | 8930397 | 8923214 | 8923413 | 8963206 | 8963422 | 410 | 276 | 409 | [illegible] |
| SE | 11485 | ENSG00000153048 | CARHSP1 | chr16 | - | 8860104 | 8860218 | 8858912 | 8859335 | 8868965 | 8869006 | 0 | 0 | 3 | [illegible] |
| SE | 11487 | ENSG00000153048 | CARHSP1 | chr16 | - | 8868048 | 8868160 | 8859170 | 8859335 | 8868531 | 8868719 | 1 | 1 | 0 | [illegible] |
| SE | 11488 | ENSG00000166743 | ACSM1 | chr16 | - | 20625422 | 20625522 | 20624095 | 20624215 | 20627188 | 20627316 | 0 | 0 | 1 | [illegible] |
| SE | 11491 | ENSG00000166743 | ACSM1 | chr16 | - | 20636738 | 20636840 | 20627188 | 20627316 | 20640460 | 20640584 | 0 | 0 | 0 | [illegible] |
| SE | 11492 | ENSG00000166743 | ACSM1 | chr16 | - | 20636738 | 20637451 | 20627255 | 20627316 | 20640460 | 20640584 | 1 | 0 | 0 | [illegible] |
| SE | 11493 | ENSG00000166743 | ACSM1 | chr16 | - | 20637370 | 20637451 | 20627188 | 20627316 | 20640460 | 20640584 | 1 | 0 | 0 | [illegible] |
| SE | 11495 | ENSG00000166743 | ACSM1 | chr16 | - | 20637370 | 20637451 | 20636738 | 20636840 | 20640460 | 20640584 | 1 | 2 | 0 | [illegible] |
| SE | 11496 | ENSG00000166743 | ACSM1 | chr16 | - | 20640460 | 20640584 | 20627188 | 20627316 | 20661793 | 20661873 | 2 | 0 | 0 | [illegible] |
| SE | 11498 | ENSG00000166743 | ACSM1 | chr16 | - | 20669826 | 20669986 | 20661793 | 20661873 | 20671530 | 20671671 | 5 | 0 | 2 | [illegible] |
| SE | 11499 | ENSG00000166743 | ACSM1 | chr16 | - | 20669826 | 20669990 | 20661793 | 20661873 | 20671530 | 20671671 | 1 | 0 | 0 | [illegible] |
| SE | 11512 | ENSG00000090581 | GNPTG | chr16 | + | 1357699 | 1357832 | 1352238 | 1352306 | 1361742 | 1361797 | 0 | 0 | 0 | [illegible] |
| SE | 11513 | ENSG00000090581 | GNPTG | chr16 | + | 1361694 | 1361797 | 1352238 | 1352306 | 1361871 | 1361955 | 23 | 1 | 22 | [illegible] |
| SE | 11514 | ENSG00000090581 | GNPTG | chr16 | + | 1361742 | 1361797 | 1352070 | 1352306 | 1361871 | 1361955 | 48 | 12 | 40 | [illegible] |
| SE | 11528 | ENSG00000179583 | CIITA | chr16 | + | 10895668 | 10895764 | 10895281 | 10895428 | 10898669 | 10898732 | 354 | 253 | 212 | [illegible] |
| SE | 11529 | ENSG00000179583 | CIITA | chr16 | + | 10901513 | 10901558 | 10898921 | 10899002 | 10902657 | 10902801 | 84 | 90 | 49 | [illegible] |
| SE | 11532 | ENSG00000179583 | CIITA | chr16 | + | 10906498 | 10908149 | 10904743 | 10904812 | 10908982 | 10909187 | 54 | 22 | 31 | [illegible] |
| SE | 11534 | ENSG00000179583 | CIITA | chr16 | + | 10906498 | 10909187 | 10904743 | 10904812 | 10910187 | 10910259 | 149 | 64 | 84 | [illegible] |
| SE | 11535 | ENSG00000179583 | CIITA | chr16 | + | 10908982 | 10909187 | 10904743 | 10904812 | 10910187 | 10910259 | 98 | 44 | 57 | [illegible] |
| SE | 11536 | ENSG00000179583 | CIITA | chr16 | + | 10909028 | 10909187 | 10904743 | 10904812 | 10910187 | 10910259 | 98 | 45 | 57 | [illegible] |
| SE | 11540 | ENSG00000179583 | CIITA | chr16 | + | 10923227 | 10923325 | 10922406 | 10922490 | 10935115 | 10935200 | 71 | 46 | 59 | [illegible] |
| SE | 11541 | ENSG00000179583 | CIITA | chr16 | + | 10923227 | 10923325 | 10922406 | 10922490 | 10941185 | 10943021 | 59 | 32 | 54 | [illegible] |
| SE | 11542 | ENSG00000179583 | CIITA | chr16 | + | 10935115 | 10935200 | 10922406 | 10922490 | 10941185 | 10943021 | 0 | 0 | 0 | [illegible] |
| SE | 11547 | ENSG00000103494 | RPGRIP1L | chr16 | - | 53622218 | 53622356 | 53619024 | 53619208 | 53636438 | 53636512 | 0 | 0 | 0 | [illegible] |
| SE | 11549 | ENSG00000103494 | RPGRIP1L | chr16 | - | 53638309 | 53638411 | 53637694 | 53637854 | 53641032 | 53641116 | 1 | 9 | 16 | [illegible] |
| SE | 11550 | ENSG00000103494 | RPGRIP1L | chr16 | - | 53645624 | 53646003 | 53641284 | 53641475 | 53648963 | 53649115 | 21 | 10 | 16 | [illegible] |
| SE | 11551 | ENSG00000103494 | RPGRIP1L | chr16 | - | 53658771 | 53658878 | 53658413 | 53658464 | 53664869 | 53665009 | 19 | 35 | 21 | [illegible] |
| SE | 11552 | ENSG00000103494 | RPGRIP1L | chr16 | - | 53671509 | 53671583 | 53664869 | 53665009 | 53672869 | 53673016 | 7 | 27 | 24 | [illegible] |
| SE | 11555 | ENSG00000125814 | NAPB | chr20 | - | 23395138 | 23395185 | 23394921 | 23394999 | 23397071 | 23397188 | 14 | 3 | 9 | [illegible] |
| SE | 11556 | ENSG00000125814 | NAPB | chr20 | - | 23395138 | 23395185 | 23394921 | 23394999 | 23402992 | 23403072 | 6 | 4 | 2 | [illegible] |
| SE | 11558 | ENSG00000125814 | NAPB | chr20 | - | 23397071 | 23397188 | 23395138 | 23395185 | 23402992 | 23403072 | 21 | 4 | 14 | [illegible] |
| SE | 11559 | ENSG00000125814 | NAPB | chr20 | - | 23397071 | 23397200 | 23395138 | 23395185 | 23402992 | 23403072 | 6 | 0 | 6 | [illegible] |
| SE | 11563 | ENSG00000125166 | GOT2 | chr16 | - | 58716662 | 58716813 | 58716013 | 58716179 | 58718195 | 58718300 | 188 | 74 | 258 | [illegible] |
| SE | 11564 | ENSG00000125166 | GOT2 | chr16 | - | 58719195 | 58719255 | 58718216 | 58718688 | 58722149 | 58722278 | 114 | 82 | 136 | [illegible] |
| SE | 11568 | ENSG00000103042 | SLC38A7 | chr16 | - | 58671044 | 58671244 | 58670112 | 58670167 | 58672095 | 58672243 | 3 | 3 | 1 | [illegible] |
| SE | 11569 | ENSG00000103042 | SLC38A7 | chr16 | - | 58672095 | 58672243 | 58670112 | 58670167 | 58675939 | 58676022 | 4 | 4 | 5 | [illegible] |
| SE | 11570 | ENSG00000103042 | SLC38A7 | chr16 | - | 58675212 | 58675342 | 58672095 | 58672243 | 58675939 | 58676054 | 0 | 0 | 1 | [illegible] |
| SE | 11571 | ENSG00000103042 | SLC38A7 | chr16 | - | 58678332 | 58678474 | 58677325 | 58677424 | 58678695 | 58678763 | 4 | 8 | 13 | [illegible] |
| SE | 11572 | ENSG00000103042 | SLC38A7 | chr16 | - | 58678695 | 58678894 | 58678332 | 58678474 | 58679856 | 58679873 | 14 | 4 | 7 | [illegible] |
| SE | 11612 | ENSG00000089505 | CMTM1 | chr16 | + | 66569935 | 66570094 | 66566438 | 66566945 | 66577103 | 66577202 | 0 | 0 | 1 | [illegible] |
| SE | 11613 | ENSG00000089505 | CMTM1 | chr16 | + | 66571136 | 66571270 | 66569935 | 66570094 | 66577103 | 66577202 | 0 | 0 | 10 | [illegible] |
| SE | 11626 | ENSG00000153066 | TXNDC11 | chr16 | - | 11684164 | 11684245 | 11679082 | 11679837 | 11687856 | 11687966 | 155 | 96 | 184 | [illegible] |
| SE | 11628 | ENSG00000153066 | TXNDC11 | chr16 | - | 11688302 | 11688445 | 11687856 | 11687966 | 11698124 | 11698325 | 191 | 84 | 195 | [illegible] |
| SE | 11629 | ENSG00000153066 | TXNDC11 | chr16 | - | 11691289 | 11692082 | 11687856 | 11687966 | 11698124 | 11698325 | 101 | 49 | 111 | [illegible] |
| SE | 11631 | ENSG00000153066 | TXNDC11 | chr16 | - | 11698124 | 11698325 | 11691289 | 11692082 | 11700451 | 11700564 | 332 | 162 | 400 | [illegible] |
| SE | 11632 | ENSG00000153066 | TXNDC11 | chr16 | - | 11698124 | 11698325 | 11691289 | 11692082 | 11733981 | 11733999 | 100 | 46 | 107 | [illegible] |
| SE | 11633 | ENSG00000153066 | TXNDC11 | chr16 | - | 11700451 | 11700564 | 11691289 | 11692082 | 11721576 | 11721670 | 103 | 52 | 101 | [illegible] |
| SE | 11634 | ENSG00000153066 | TXNDC11 | chr16 | - | 11700451 | 11700564 | 11691289 | 11692082 | 11733981 | 11733999 | 0 | 0 | 4 | [illegible] |
| SE | 11635 | ENSG00000153066 | TXNDC11 | chr16 | - | 11700451 | 11700564 | 11698124 | 11698325 | 11721313 | 11721435 | 147 | 63 | 192 | [illegible] |
| SE | 11637 | ENSG00000153066 | TXNDC11 | chr16 | - | 11700451 | 11700564 | 11698124 | 11698325 | 11733981 | 11733999 | 147 | 62 | 194 | [illegible] |
| SE | 11638 | ENSG00000153066 | TXNDC11 | chr16 | - | 11703669 | 11703750 | 11700451 | 11700564 | 11721576 | 11721670 | 0 | 1 | 1 | [illegible] |
| SE | 11639 | ENSG00000153066 | TXNDC11 | chr16 | - | 11721313 | 11721435 | 11698124 | 11698325 | 11721576 | 11721670 | 1 | 0 | 0 | [illegible] |
| SE | 11640 | ENSG00000153066 | TXNDC11 | chr16 | - | 11721313 | 11721435 | 11700451 | 11700564 | 11721576 | 11721670 | 1 | 1 | 2 | [illegible] |
| SE | 11641 | ENSG00000153066 | TXNDC11 | chr16 | - | 11721576 | 11721670 | 11691289 | 11692082 | 11733981 | 11733999 | 30 | 11 | 28 | [illegible] |
| SE | 11642 | ENSG00000153066 | TXNDC11 | chr16 | - | 11721576 | 11721670 | 11698133 | 11698325 | 11733981 | 11733999 | 47 | 18 | 49 | [illegible] |
| SE | 11643 | ENSG00000153066 | TXNDC11 | chr16 | - | 11721576 | 11721670 | 11698133 | 11698325 | 11736016 | 11736130 | 13 | 4 | 22 | [illegible] |
| SE | 11644 | ENSG00000153066 | TXNDC11 | chr16 | - | 11721576 | 11721670 | 11700451 | 11700564 | 11730644 | 11730774 | 287 | 145 | 332 | [illegible] |
| SE | 11645 | ENSG00000153066 | TXNDC11 | chr16 | - | 11721576 | 11721670 | 11700451 | 11700564 | 11733981 | 11733999 | 179 | 84 | 175 | [illegible] |
| SE | 11646 | ENSG00000153066 | TXNDC11 | chr16 | - | 11730644 | 11730774 | 11700451 | 11700564 | 11733981 | 11734079 | 69 | 43 | 98 | [illegible] |
| SE | 11649 | ENSG00000153066 | TXNDC11 | chr16 | - | 11733981 | 11734079 | 11698124 | 11698325 | 11736016 | 11736233 | 40 | 19 | 49 | [illegible] |
| SE | 11653 | ENSG00000103549 | RNF40 | chr16 | + | 30764937 | 30765059 | 30764178 | 30764385 | 30765180 | 30765327 | 140 | 63 | 125 | [illegible] |
| SE | 11654 | ENSG00000103549 | RNF40 | chr16 | + | 30766162 | 30766282 | 30765424 | 30765499 | 30766378 | 30766558 | 314 | 153 | 264 | [illegible] |
| SE | 11656 | ENSG00000103549 | RNF40 | chr16 | + | 30769185 | 30769398 | 30768837 | 30768987 | 30769474 | 30769600 | 343 | 179 | 231 | [illegible] |
| SE | 11657 | ENSG00000103549 | RNF40 | chr16 | + | 30769185 | 30769600 | 30768837 | 30768987 | 30771832 | 30771973 | 315 | 178 | 211 | [illegible] |
| SE | 11658 | ENSG00000103549 | RNF40 | chr16 | + | 30769474 | 30769600 | 30768837 | 30768987 | 30771832 | 30771973 | 96 | 55 | 54 | [illegible] |
| SE | 11659 | ENSG00000099385 | BCL7C | chr16 | - | 30892839 | 30892948 | 30892585 | 30892747 | 30893211 | 30893290 | 97 | 44 | 77 | [illegible] |
| SE | 11663 | ENSG00000121270 | ABCC11 | chr16 | - | 48196231 | 48196321 | 48193878 | 48193982 | 48197970 | 48198067 | 0 | 1 | 0 | [illegible] |
| SE | 11669 | ENSG00000167207 | NOD2 | chr16 | + | 50716586 | 50716670 | 50710557 | 50712373 | 50719924 | 50720008 | 2 | 0 | 4 | [illegible] |
| SE | 11670 | ENSG00000167207 | NOD2 | chr16 | + | 50716586 | 50716670 | 50710557 | 50712373 | 50722621 | 50722705 | 1 | 0 | 4 | [illegible] |
| SE | 11672 | ENSG00000167207 | NOD2 | chr16 | + | 50716890 | 50716974 | 50716586 | 50716670 | 50719924 | 50720008 | 17 | 24 | 41 | [illegible] |
| SE | 11674 | ENSG00000167207 | NOD2 | chr16 | + | 50719924 | 50720008 | 50710557 | 50712373 | 50722621 | 50722705 | 9 | 4 | 4 | [illegible] |
| SE | 11678 | ENSG00000162066 | AMDHD2 | chr16 | + | 2520768 | 2520905 | 2520364 | 2520541 | 2520983 | 2521123 | 25 | 8 | 22 | [illegible] |
| SE | 11680 | ENSG00000162066 | AMDHD2 | chr16 | + | 2527572 | 2527615 | 2520983 | 2521123 | 2527772 | 2527836 | 4 | 0 | 4 | [illegible] |
| SE | 11682 | ENSG00000162066 | AMDHD2 | chr16 | + | 2528649 | 2528718 | 2528351 | 2528559 | 2528993 | 2529060 | 8 | 0 | 12 | [illegible] |
| SE | 11683 | ENSG00000162066 | AMDHD2 | chr16 | + | 2528993 | 2529095 | 2528451 | 2528718 | 2529474 | 2529517 | 13 | 1 | 20 | [illegible] |
| SE | 11694 | ENSG00000140995 | DEF8 | chr16 | + | 89950066 | 89950306 | 89949416 | 89949513 | 89954242 | 89954376 | 0 | 0 | 0 | [illegible] |
| SE | 11695 | ENSG00000140995 | DEF8 | chr16 | + | 89950066 | 89950328 | 89949416 | 89949513 | 89954242 | 89954376 | 0 | 0 | 0 | [illegible] |
| SE | 11696 | ENSG00000140995 | DEF8 | chr16 | + | 89952557 | 89952682 | 89949416 | 89949513 | 89954242 | 89954376 | 0 | 0 | 0 | [illegible] |
| SE | 11697 | ENSG00000140995 | DEF8 | chr16 | + | 89952561 | 89952682 | 89949416 | 89949513 | 89954242 | 89954376 | 0 | 0 | 0 | [illegible] |
| SE | 11698 | ENSG00000140995 | DEF8 | chr16 | + | 89954242 | 89954376 | 89948696 | 89948814 | 89955168 | 89955266 | 33 | 12 | 24 | [illegible] |
| SE | 11700 | ENSG00000140995 | DEF8 | chr16 | + | 89954288 | 89954376 | 89948754 | 89948814 | 89955168 | 89955266 | 27 | 6 | 9 | [illegible] |
| SE | 11703 | ENSG00000140995 | DEF8 | chr16 | + | 89959013 | 89959155 | 89957252 | 89957660 | 89960930 | 89960959 | 169 | 89 | 160 | [illegible] |
| SE | 11706 | ENSG00000167978 | SRRM2 | chr16 | + | 2756333 | 2756606 | 2752637 | 2752846 | 2757471 | 2757574 | 345 | 229 | 358 | [illegible] |
| SE | 11707 | ENSG00000167978 | SRRM2 | chr16 | + | 2757471 | 2757579 | 2752637 | 2752846 | 2757780 | 2757945 | 282 | 199 | 258 | [illegible] |
| SE | 11709 | ENSG00000167978 | SRRM2 | chr16 | + | 2759568 | 2759661 | 2758984 | 2759047 | 2760300 | 2760499 | 116 | 97 | 133 | [illegible] |
| SE | 11713 | ENSG00000100266 | PACSIN2 | chr22 | - | 42882183 | 42882304 | 42879047 | 42879169 | 42884385 | 42884561 | 262 | 273 | 201 | [illegible] |
| SE | 11714 | ENSG00000100266 | PACSIN2 | chr22 | - | 42888642 | 42888798 | 42884385 | 42884561 | 42893456 | 42893613 | 60 | 91 | 56 | [illegible] |
| SE | 11715 | ENSG00000100266 | PACSIN2 | chr22 | - | 42890946 | 42891182 | 42888642 | 42888798 | 42893456 | 42893613 | 231 | 224 | 170 | [illegible] |
| SE | 11716 | ENSG00000100266 | PACSIN2 | chr22 | - | 42909426 | 42909565 | 42893456 | 42893613 | 42912020 | 42912157 | 0 | 0 | 0 | [illegible] |
| SE | 11718 | ENSG00000100266 | PACSIN2 | chr22 | - | 42912020 | 42912157 | 42893456 | 42893613 | 43015020 | 43015098 | 134 | 91 | 113 | [illegible] |
| SE | 11721 | ENSG00000010361 | FUZ | chr19 | - | 49808114 | 49808159 | 49806884 | 49807374 | 49808413 | 49808491 | 1 | 0 | 3 | [illegible] |
| SE | 11722 | ENSG00000010361 | FUZ | chr19 | - | 49809377 | 49809575 | 49809162 | 49809258 | 49811362 | 49811467 | 12 | 1 | 22 | [illegible] |
| SE | 11723 | ENSG00000010361 | FUZ | chr19 | - | 49812250 | 49812335 | 49811630 | 49811699 | 49812614 | 49812736 | 10 | 9 | 8 | [illegible] |
| SE | 11724 | ENSG00000010361 | FUZ | chr19 | - | 49812250 | 49812335 | 49811630 | 49811699 | 49812637 | 49812736 | 4 | 6 | 4 | [illegible] |
| SE | 11727 | ENSG00000010361 | FUZ | chr19 | - | 49812446 | 49812736 | 49812250 | 49812335 | 49813132 | 49813235 | 5 | 5 | 5 | [illegible] |
| SE | 11729 | ENSG00000010361 | FUZ | chr19 | - | 49812614 | 49812736 | 49812250 | 49812335 | 49813132 | 49813249 | 12 | 6 | 8 | [illegible] |
| SE | 11735 | ENSG00000260565 | ERVK13-1 | chr16 | - | 2669410 | 2669561 | 2660355 | 2663407 | 2671344 | 2671405 | 5 | 12 | 10 | [illegible] |
| SE | 11737 | ENSG00000260565 | ERVK13-1 | chr16 | - | 2669410 | 2669561 | 2660698 | 2663430 | 2671344 | 2671405 | 3 | 4 | 4 | [illegible] |
| SE | 11741 | ENSG00000260565 | ERVK13-1 | chr16 | - | 2670863 | 2670974 | 2669410 | 2669561 | 2671344 | 2672273 | 0 | 0 | 0 | [illegible] |
| SE | 11742 | ENSG00000260565 | ERVK13-1 | chr16 | - | 2670863 | 2670974 | 2669410 | 2669561 | 2671594 | 2672273 | 0 | 0 | 0 | [illegible] |
| SE | 11743 | ENSG00000260565 | ERVK13-1 | chr16 | - | 2671023 | 2671231 | 2669410 | 2669561 | 2671344 | 2672273 | 0 | 0 | 1 | [illegible] |
| SE | 11753 | ENSG00000260565 | ERVK13-1 | chr16 | - | 2672706 | 2672859 | 2669410 | 2672562 | 2673282 | 2673387 | 10 | 2 | 1 | [illegible] |
| SE | 11759 | ENSG00000166548 | TK2 | chr16 | - | 66517788 | 66517877 | 66513730 | 66513811 | 66528993 | 66529067 | 47 | 38 | 28 | [illegible] |
| SE | 11760 | ENSG00000166548 | TK2 | chr16 | - | 66528993 | 66529067 | 66517788 | 66517877 | 66531379 | 66531469 | 205 | 115 | 124 | [illegible] |
| SE | 11761 | ENSG00000166548 | TK2 | chr16 | - | 66531379 | 66531469 | 66517788 | 66517877 | 66541878 | 66541953 | 1 | 0 | 1 | [illegible] |
| SE | 11762 | ENSG00000166548 | TK2 | chr16 | - | 66531379 | 66531469 | 66517788 | 66517877 | 66548977 | 66549009 | 0 | 0 | 0 | [illegible] |
| SE | 11763 | ENSG00000166548 | TK2 | chr16 | - | 66531379 | 66531469 | 66528993 | 66529067 | 66536963 | 66537017 | 187 | 105 | 104 | [illegible] |
| SE | 11768 | ENSG00000166548 | TK2 | chr16 | - | 66541878 | 66541953 | 66517788 | 66517877 | 66548977 | 66549009 | 16 | 16 | 14 | [illegible] |
| SE | 11774 | ENSG00000166548 | TK2 | chr16 | - | 66546146 | 66546226 | 66541878 | 66541953 | 66548977 | 66549009 | 1 | 0 | 0 | [illegible] |
| SE | 11775 | ENSG00000166548 | TK2 | chr16 | - | 66547970 | 66548065 | 66536963 | 66537017 | 66548977 | 66549009 | 2 | 0 | 0 | [illegible] |
| SE | 11780 | ENSG00000166548 | TK2 | chr16 | - | 66548977 | 66549009 | 66541878 | 66541953 | 66549937 | 66550005 | 31 | 29 | 18 | [illegible] |
| SE | 11781 | ENSG00000166548 | TK2 | chr16 | - | 66550252 | 66550327 | 66548977 | 66549009 | 66551411 | 66551718 | 0 | 2 | 0 | [illegible] |
| SE | 11793 | ENSG00000261656 | BEAN1-AS1 | chr16 | - | 66476677 | 66476722 | 66475601 | 66475678 | 66477943 | 66478117 | 1 | 0 | 0 | [illegible] |
| SE | 11806 | ENSG00000188603 | CLN3 | chr16 | - | 28484005 | 28484118 | 28482104 | 28482672 | 28487455 | 28487541 | 18 | 3 | 12 | [illegible] |
| SE | 11807 | ENSG00000188603 | CLN3 | chr16 | - | 28486346 | 28486490 | 28484005 | 28484118 | 28486577 | 28486650 | 47 | 11 | 6 | [illegible] |
| SE | 11809 | ENSG00000188603 | CLN3 | chr16 | - | 28486346 | 28486524 | 28484005 | 28484118 | 28486577 | 28487541 | 35 | 4 | 3 | [illegible] |
| SE | 11812 | ENSG00000188603 | CLN3 | chr16 | - | 28487455 | 28487541 | 28484005 | 28484118 | 28488590 | 28488662 | 0 | 2 | 3 | [illegible] |
| SE | 11830 | ENSG00000140931 | CMTM3 | chr16 | + | 66609434 | 66609530 | 66608308 | 66608464 | 66609882 | 66610003 | 101 | 33 | 98 | [illegible] |
| SE | 11835 | ENSG00000254788 | CKLF-CMTM1 | chr16 | + | 66558189 | 66558348 | 66552678 | 66552793 | 66569935 | 66570086 | 96 | 38 | 104 | [illegible] |
| SE | 11836 | ENSG00000254788 | CKLF-CMTM1 | chr16 | + | 66558189 | 66558348 | 66552678 | 66552793 | 66577103 | 66577202 | 98 | 38 | 104 | [illegible] |
| SE | 11837 | ENSG00000254788 | CKLF-CMTM1 | chr16 | + | 66563071 | 66563217 | 66552586 | 66552793 | 66569935 | 66570086 | 5 | 1 | 6 | [illegible] |
| SE | 11838 | ENSG00000254788 | CKLF-CMTM1 | chr16 | + | 66563071 | 66563217 | 66558189 | 66558348 | 66569935 | 66570086 | 1 | 0 | 1 | [illegible] |
| SE | 11840 | ENSG00000254788 | CKLF-CMTM1 | chr16 | + | 66563121 | 66563217 | 66552648 | 66552793 | 66577103 | 66577202 | 39 | 19 | 29 | [illegible] |
| SE | 11844 | ENSG00000217555 | CKLF | chr16 | + | 66563071 | 66563217 | 66552562 | 66552793 | 66566082 | 66566251 | 20 | 1 | 7 | [illegible] |
| SE | 11845 | ENSG00000217555 | CKLF | chr16 | + | 66563071 | 66563217 | 66558189 | 66558348 | 66566082 | 66566251 | 16 | 0 | 2 | [illegible] |
| SE | 11849 | ENSG00000121274 | TENT4B | chr16 | + | 50211322 | 50211446 | 50153305 | 50154259 | 50214220 | 50214267 | 109 | 63 | 82 | [illegible] |
| SE | 11850 | ENSG00000121274 | TENT4B | chr16 | + | 50214220 | 50214267 | 50211322 | 50211446 | 50216074 | 50216195 | 109 | 45 | 98 | [illegible] |
| SE | 11851 | ENSG00000121274 | TENT4B | chr16 | + | 50217555 | 50217663 | 50216074 | 50216195 | 50222305 | 50222434 | 224 | 170 | 200 | [illegible] |
| SE | 11855 | ENSG00000121274 | TENT4B | chr16 | + | 50223173 | 50223387 | 50222305 | 50222434 | 50224656 | 50224783 | 329 | 180 | 246 | [illegible] |
| SE | 11857 | ENSG00000121274 | TENT4B | chr16 | + | 50223173 | 50223400 | 50222305 | 50222434 | 50224656 | 50224783 | 168 | 108 | 134 | [illegible] |
| SE | 11858 | ENSG00000121274 | TENT4B | chr16 | + | 50225097 | 50225285 | 50224890 | 50224994 | 50227838 | 50228003 | 190 | 129 | 146 | [illegible] |
| SE | 11859 | ENSG00000121274 | TENT4B | chr16 | + | 50225238 | 50225285 | 50224890 | 50224994 | 50227838 | 50228003 | 37 | 22 | 23 | [illegible] |
| SE | 11861 | ENSG00000121281 | ADCY7 | chr16 | + | 50287911 | 50288350 | 50246136 | 50246203 | 50292675 | 50292825 | 0 | 2 | 1 | [illegible] |
| SE | 11862 | ENSG00000121281 | ADCY7 | chr16 | + | 50287911 | 50288350 | 50266550 | 50266587 | 50291735 | 50291897 | 16 | 6 | 10 | [illegible] |
| SE | 11863 | ENSG00000121281 | ADCY7 | chr16 | + | 50287911 | 50288350 | 50266550 | 50266587 | 50292675 | 50292825 | 16 | 6 | 9 | [illegible] |
| SE | 11864 | ENSG00000121281 | ADCY7 | chr16 | + | 50287911 | 50288350 | 50266550 | 50266680 | 50292675 | 50292825 | 19 | 10 | 8 | [illegible] |
| SE | 11866 | ENSG00000121281 | ADCY7 | chr16 | + | 50289220 | 50289313 | 50287881 | 50288350 | 50290456 | 50290518 | 0 | 0 | 0 | [illegible] |
| SE | 11867 | ENSG00000121281 | ADCY7 | chr16 | + | 50290456 | 50290660 | 50287910 | 50288350 | 50291735 | 50291897 | 79 | 23 | 39 | [illegible] |
| SE | 11868 | ENSG00000121281 | ADCY7 | chr16 | + | 50290456 | 50290660 | 50287910 | 50288350 | 50292675 | 50292825 | 19 | 6 | 12 | [illegible] |
| SE | 11869 | ENSG00000121281 | ADCY7 | chr16 | + | 50291735 | 50291897 | 50266550 | 50266587 | 50292675 | 50292825 | 44 | 14 | 36 | [illegible] |
| SE | 11870 | ENSG00000121281 | ADCY7 | chr16 | + | 50291735 | 50291897 | 50287910 | 50288350 | 50292675 | 50292825 | 43 | 14 | 37 | [illegible] |
| SE | 11871 | ENSG00000121281 | ADCY7 | chr16 | + | 50291735 | 50291897 | 50290456 | 50290660 | 50292675 | 50292825 | 103 | 32 | 63 | [illegible] |
| SE | 11872 | ENSG00000121281 | ADCY7 | chr16 | + | 50300714 | 50300873 | 50298903 | 50299031 | 50301081 | 50301124 | 441 | 202 | 301 | [illegible] |
| SE | 11875 | ENSG00000121281 | ADCY7 | chr16 | + | 50308326 | 50308411 | 50307049 | 50307147 | 50308666 | 50308792 | 212 | 95 | 110 | [illegible] |
| SE | 11876 | ENSG00000121281 | ADCY7 | chr16 | + | 50309547 | 50309646 | 50308666 | 50308792 | 50310686 | 50310880 | 215 | 84 | 131 | [illegible] |
| SE | 11878 | ENSG00000121281 | ADCY7 | chr16 | + | 50312889 | 50313036 | 50312035 | 50312191 | 50313957 | 50314062 | 478 | 365 | 332 | [illegible] |
| SE | 11881 | ENSG00000103222 | ABCC1 | chr16 | + | 16009775 | 16009901 | 16007815 | 16007992 | 16016495 | 16016621 | 70 | 24 | 47 | [illegible] |
| SE | 11882 | ENSG00000103222 | ABCC1 | chr16 | + | 16009775 | 16009901 | 16007815 | 16007992 | 16033108 | 16033170 | 70 | 23 | 47 | [illegible] |
| SE | 11883 | ENSG00000103222 | ABCC1 | chr16 | + | 16014490 | 16014628 | 16007815 | 16007992 | 16016495 | 16016621 | 123 | 51 | 132 | [illegible] |
| SE | 11884 | ENSG00000103222 | ABCC1 | chr16 | + | 16014490 | 16014628 | 16007815 | 16007992 | 16033108 | 16033170 | 4 | 3 | 5 | [illegible] |
| SE | 11887 | ENSG00000103222 | ABCC1 | chr16 | + | 16016495 | 16016621 | 16007815 | 16007992 | 16033108 | 16033170 | 54 | 22 | 39 | [illegible] |
| SE | 11890 | ENSG00000103222 | ABCC1 | chr16 | + | 16033108 | 16033170 | 16014490 | 16014628 | 16036471 | 16036603 | 41 | 19 | 30 | [illegible] |
| SE | 11891 | ENSG00000103222 | ABCC1 | chr16 | + | 16044449 | 16044680 | 16036471 | 16036603 | 16045835 | 16046013 | 152 | 112 | 158 | [illegible] |
| SE | 11892 | ENSG00000103222 | ABCC1 | chr16 | + | 16056091 | 16056295 | 16052723 | 16052816 | 16068155 | 16068302 | 158 | 78 | 152 | [illegible] |
| SE | 11894 | ENSG00000103222 | ABCC1 | chr16 | + | 16071641 | 16071729 | 16056091 | 16056295 | 16090404 | 16090588 | 0 | 0 | 0 | [illegible] |
| SE | 11896 | ENSG00000103222 | ABCC1 | chr16 | + | 16076325 | 16076401 | 16071641 | 16071729 | 16090404 | 16090588 | 17 | 10 | 27 | [illegible] |
| SE | 11897 | ENSG00000103222 | ABCC1 | chr16 | + | 16079351 | 16079478 | 16071641 | 16071729 | 16090404 | 16090588 | 2 | 1 | 0 | [illegible] |
| SE | 11898 | ENSG00000103222 | ABCC1 | chr16 | + | 16079351 | 16079478 | 16076173 | 16076401 | 16083365 | 16083542 | 128 | 55 | 83 | [illegible] |
| SE | 11902 | ENSG00000103222 | ABCC1 | chr16 | + | 16083365 | 16083542 | 16076173 | 16076401 | 16090404 | 16090588 | 0 | 1 | 0 | [illegible] |
| SE | 11904 | ENSG00000103222 | ABCC1 | chr16 | + | 16083365 | 16083542 | 16079351 | 16079478 | 16090404 | 16090588 | 84 | 33 | 55 | [illegible] |
| SE | 11906 | ENSG00000103222 | ABCC1 | chr16 | + | 16086823 | 16086991 | 16079351 | 16079478 | 16090404 | 16090588 | 82 | 35 | 75 | [illegible] |
| SE | 11907 | ENSG00000103222 | ABCC1 | chr16 | + | 16086823 | 16086991 | 16083365 | 16083542 | 16090404 | 16090588 | 115 | 70 | 101 | [illegible] |
| SE | 11908 | ENSG00000103222 | ABCC1 | chr16 | + | 16098872 | 16098902 | 16090404 | 16090588 | 16102626 | 16102717 | 2 | 0 | 0 | [illegible] |
| SE | 11909 | ENSG00000103222 | ABCC1 | chr16 | + | 16105399 | 16105460 | 16102626 | 16102717 | 16106737 | 16106873 | 0 | 2 | 0 | [illegible] |
| SE | 11912 | ENSG00000103222 | ABCC1 | chr16 | + | 16131788 | 16131935 | 16121974 | 16122174 | 16134349 | 16134508 | 47 | 20 | 33 | [illegible] |
| SE | 11917 | ENSG00000070770 | CSNK2A2 | chr16 | - | 58168609 | 58168693 | 58167684 | 58167795 | 58174450 | 58174510 | 292 | 219 | 306 | [illegible] |
| SE | 11919 | ENSG00000070770 | CSNK2A2 | chr16 | - | 58168609 | 58168693 | 58167684 | 58167795 | 58196732 | 58196844 | 107 | 78 | 113 | [illegible] |
| SE | 11921 | ENSG00000070770 | CSNK2A2 | chr16 | - | 58174450 | 58174510 | 58167684 | 58167795 | 58196732 | 58196844 | 1 | 0 | 0 | [illegible] |
| SE | 11922 | ENSG00000070770 | CSNK2A2 | chr16 | - | 58174450 | 58174510 | 58168609 | 58168693 | 58184259 | 58184310 | 236 | 194 | 265 | [illegible] |
| SE | 11923 | ENSG00000070770 | CSNK2A2 | chr16 | - | 58174450 | 58174510 | 58168609 | 58168693 | 58196732 | 58196844 | 60 | 58 | 57 | [illegible] |
| SE | 11925 | ENSG00000005844 | ITGAL | chr16 | + | 30474195 | 30474287 | 30472744 | 30472898 | 30475288 | 30475400 | 281 | 113 | 150 | [illegible] |
| SE | 11927 | ENSG00000005844 | ITGAL | chr16 | + | 30474195 | 30474298 | 30472710 | 30472898 | 30475288 | 30475400 | 308 | 129 | 159 | [illegible] |
| SE | 11928 | ENSG00000005844 | ITGAL | chr16 | + | 30474195 | 30474298 | 30472710 | 30472898 | 30475305 | 30475400 | 677 | 289 | 354 | [illegible] |
| SE | 11929 | ENSG00000005844 | ITGAL | chr16 | + | 30475288 | 30475400 | 30474195 | 30474298 | 30479090 | 30479190 | 1 | 6 | 4 | [illegible] |
| SE | 11930 | ENSG00000005844 | ITGAL | chr16 | + | 30475305 | 30475400 | 30474000 | 30474298 | 30479090 | 30479174 | 103 | 53 | 48 | [illegible] |
| SE | 11931 | ENSG00000005844 | ITGAL | chr16 | + | 30475305 | 30475400 | 30474000 | 30474298 | 30481438 | 30481584 | 103 | 53 | 47 | [illegible] |
| SE | 11932 | ENSG00000005844 | ITGAL | chr16 | + | 30475512 | 30475580 | 30475288 | 30475400 | 30479090 | 30479174 | 832 | 342 | 501 | [illegible] |
| SE | 11933 | ENSG00000005844 | ITGAL | chr16 | + | 30475512 | 30475580 | 30475288 | 30475400 | 30481438 | 30481584 | 436 | 183 | 244 | [illegible] |
| SE | 11934 | ENSG00000005844 | ITGAL | chr16 | + | 30479090 | 30479208 | 30474195 | 30474298 | 30481438 | 30481584 | 5 | 1 | 5 | [illegible] |
| SE | 11937 | ENSG00000005844 | ITGAL | chr16 | + | 30479090 | 30479208 | 30475512 | 30475580 | 30483826 | 30483959 | 236 | 93 | 145 | [illegible] |
| SE | 11939 | ENSG00000005844 | ITGAL | chr16 | + | 30479330 | 30479461 | 30479089 | 30479208 | 30483826 | 30483959 | 635 | 244 | 353 | [illegible] |
| SE | 11940 | ENSG00000005844 | ITGAL | chr16 | + | 30481438 | 30481584 | 30475512 | 30475580 | 30483826 | 30483959 | 1471 | 669 | 845 | [illegible] |
| SE | 11941 | ENSG00000005844 | ITGAL | chr16 | + | 30481438 | 30481584 | 30475512 | 30475580 | 30489081 | 30489155 | 53 | 19 | 23 | [illegible] |
| SE | 11942 | ENSG00000005844 | ITGAL | chr16 | + | 30481438 | 30481584 | 30479090 | 30479208 | 30483826 | 30483959 | 1428 | 651 | 829 | [illegible] |
| SE | 11943 | ENSG00000005844 | ITGAL | chr16 | + | 30481438 | 30481584 | 30479330 | 30479461 | 30483826 | 30483959 | 2566 | 1178 | 1476 | [illegible] |
| SE | 11944 | ENSG00000005844 | ITGAL | chr16 | + | 30483826 | 30483959 | 30481438 | 30481584 | 30489081 | 30489155 | 1293 | 581 | 779 | [illegible] |
| SE | 11945 | ENSG00000005844 | ITGAL | chr16 | + | 30484112 | 30484263 | 30483826 | 30483959 | 30489081 | 30489155 | 2229 | 1102 | 1186 | [illegible] |
| SE | 11946 | ENSG00000005844 | ITGAL | chr16 | + | 30489253 | 30489386 | 30488999 | 30489155 | 30494211 | 30494363 | 2505 | 1407 | 1447 | [illegible] |
| SE | 11947 | ENSG00000005844 | ITGAL | chr16 | + | 30489253 | 30489386 | 30488999 | 30489155 | 30496435 | 30496566 | 1163 | 639 | 664 | [illegible] |
| SE | 11951 | ENSG00000005844 | ITGAL | chr16 | + | 30494211 | 30494363 | 30489253 | 30489386 | 30496435 | 30496566 | 1344 | 772 | 784 | [illegible] |
| SE | 11952 | ENSG00000005844 | ITGAL | chr16 | + | 30494211 | 30494363 | 30489253 | 30489386 | 30499073 | 30499234 | 1344 | 772 | 783 | [illegible] |
| SE | 11953 | ENSG00000005844 | ITGAL | chr16 | + | 30494712 | 30494850 | 30494211 | 30494363 | 30496096 | 30496201 | 2360 | 1168 | 1214 | [illegible] |
| SE | 11954 | ENSG00000005844 | ITGAL | chr16 | + | 30494712 | 30494850 | 30494211 | 30494363 | 30496435 | 30496566 | 1259 | 655 | 681 | [illegible] |
| SE | 11955 | ENSG00000005844 | ITGAL | chr16 | + | 30494712 | 30494850 | 30494211 | 30494363 | 30499073 | 30499234 | 1255 | 652 | 681 | [illegible] |
| SE | 11956 | ENSG00000005844 | ITGAL | chr16 | + | 30496096 | 30496201 | 30494211 | 30494363 | 30496435 | 30496566 | 0 | 0 | 1 | [illegible] |
| SE | 11958 | ENSG00000005844 | ITGAL | chr16 | + | 30496096 | 30496294 | 30489253 | 30489386 | 30496435 | 30496566 | 595 | 264 | 306 | [illegible] |
| SE | 11959 | ENSG00000005844 | ITGAL | chr16 | + | 30496096 | 30496294 | 30489253 | 30489386 | 30499073 | 30499234 | 1 | 2 | 1 | [illegible] |
| SE | 11961 | ENSG00000005844 | ITGAL | chr16 | + | 30496096 | 30496294 | 30494211 | 30494363 | 30499073 | 30499234 | 0 | 0 | 1 | [illegible] |
| SE | 11964 | ENSG00000005844 | ITGAL | chr16 | + | 30496096 | 30496566 | 30494211 | 30494363 | 30499073 | 30499234 | 687 | 318 | 456 | [illegible] |
| SE | 11965 | ENSG00000005844 | ITGAL | chr16 | + | 30496096 | 30496566 | 30494712 | 30494850 | 30499073 | 30499234 | 1798 | 836 | 992 | [illegible] |
| SE | 11966 | ENSG00000005844 | ITGAL | chr16 | + | 30496435 | 30496566 | 30489253 | 30489386 | 30499073 | 30499234 | 665 | 316 | 446 | [illegible] |
| SE | 11967 | ENSG00000005844 | ITGAL | chr16 | + | 30496435 | 30496566 | 30494211 | 30494363 | 30499073 | 30499234 | 664 | 316 | 447 | [illegible] |
| SE | 11969 | ENSG00000005844 | ITGAL | chr16 | + | 30496435 | 30496566 | 30496096 | 30496294 | 30499073 | 30499234 | 1258 | 578 | 751 | [illegible] |
| SE | 11970 | ENSG00000005844 | ITGAL | chr16 | + | 30499120 | 30499489 | 30496431 | 30496566 | 30504174 | 30504219 | 741 | 366 | 468 | [illegible] |
| SE | 11974 | ENSG00000005844 | ITGAL | chr16 | + | 30505246 | 30505300 | 30504174 | 30504264 | 30505388 | 30505462 | 819 | 379 | 430 | [illegible] |
| SE | 11975 | ENSG00000005844 | ITGAL | chr16 | + | 30505388 | 30505462 | 30504174 | 30504264 | 30510360 | 30510395 | 9 | 2 | 1 | [illegible] |
| SE | 11976 | ENSG00000005844 | ITGAL | chr16 | + | 30506714 | 30506856 | 30505386 | 30505462 | 30510360 | 30510395 | 1915 | 857 | 1054 | [illegible] |
| SE | 11977 | ENSG00000005844 | ITGAL | chr16 | + | 30510880 | 30510960 | 30510360 | 30510471 | 30511049 | 30511136 | 1545 | 669 | 807 | [illegible] |
| SE | 11978 | ENSG00000005844 | ITGAL | chr16 | + | 30510892 | 30510960 | 30510360 | 30510471 | 30511049 | 30511136 | 415 | 198 | 240 | [illegible] |
| SE | 11979 | ENSG00000005844 | ITGAL | chr16 | + | 30513770 | 30513846 | 30511049 | 30511136 | 30516972 | 30517070 | 742 | 370 | 395 | [illegible] |
| SE | 11980 | ENSG00000005844 | ITGAL | chr16 | + | 30516972 | 30517086 | 30513770 | 30513846 | 30517648 | 30517705 | 557 | 264 | 261 | [illegible] |
| SE | 11981 | ENSG00000005844 | ITGAL | chr16 | + | 30517796 | 30517895 | 30517648 | 30517705 | 30518623 | 30518719 | 395 | 136 | 169 | [illegible] |
| SE | 11985 | ENSG00000156860 | FBRS | chr16 | + | 30665305 | 30665401 | 30665034 | 30665079 | 30665637 | 30665706 | 143 | 67 | 93 | [illegible] |
| SE | 11988 | ENSG00000197162 | ZNF785 | chr16 | - | 30579794 | 30579847 | 30573739 | 30573891 | 30582510 | 30582804 | 3 | 2 | 2 | [illegible] |
| SE | 11990 | ENSG00000169951 | ZNF764 | chr16 | - | 30557732 | 30557846 | 30553763 | 30556110 | 30557986 | 30558263 | 9 | 5 | 23 | [illegible] |
| SE | 11994 | ENSG00000007516 | BAIAP3 | chr16 | + | 1346846 | 1346955 | 1346604 | 1346684 | 1347297 | 1347369 | 11 | 1 | 5 | [illegible] |
| SE | 11995 | ENSG00000007516 | BAIAP3 | chr16 | + | 1346933 | 1346955 | 1346604 | 1346684 | 1347297 | 1347369 | 3 | 0 | 0 | [illegible] |
| SE | 11996 | ENSG00000103275 | UBE2I | chr16 | + | 1312346 | 1312558 | 1309677 | 1309773 | 1314019 | 1314096 | 2 | 1 | 6 | [illegible] |
| SE | 11998 | ENSG00000103275 | UBE2I | chr16 | + | 1312346 | 1312667 | 1309655 | 1309773 | 1314019 | 1314096 | 2 | 1 | 0 | [illegible] |
| SE | 11999 | ENSG00000103275 | UBE2I | chr16 | + | 1312346 | 1312667 | 1309802 | 1310253 | 1314019 | 1314096 | 0 | 0 | 0 | [illegible] |
| SE | 12001 | ENSG00000103275 | UBE2I | chr16 | + | 1314019 | 1314096 | 1309635 | 1309773 | 1315653 | 1315730 | 40 | 27 | 34 | [illegible] |

| | | | | | | | | | | | | | | | | | | | | | | | | | | | | | | | | | | | | | | | | | | | | | | |
|---|---|---|---|---|---|---|---|---|---|---|---|---|---|---|---|---|---|---|---|---|---|---|---|---|---|---|---|---|---|---|---|---|---|---|---|---|---|---|---|---|---|---|---|---|---|---|
| SE | 12002 | ENSG00000103275 | UBE2I | chr16 | + | 1314292 | 1314376 | 1309647 | 1309773 | 1315653 | 1315726 | 364 | 225 | 288 | 340 | 639 | 0 | 0 | 1 | 0 | 0 | 740 | 511 | 389 | 456 | 723 | 0 | 0 | 0 | 0 | 0 | 233 | 149 | 1 | 1 | 1 | 1 | 0.995 | 1 | 1 | 1 | 1 | 1 | 1 | 1 | -0.001 |
| SE | 12004 | ENSG00000103275 | UBE2I | chr16 | + | 1319701 | 1319862 | 1315653 | 1315726 | 1320173 | 1320199 | 0 | 0 | 6 | 4 | 3 | 659 | 334 | 504 | 582 | 1004 | 7 | 1 | 3 | 7 | 12 | 1234 | 821 | 654 | 813 | 1237 | 298 | 149 | 1 | 1 | 0 | 0 | 0.006 | 0.003 | 0.001 | 0.003 | 0.001 | 0.002 | 0.004 | 0.005 | -0.001 |
| SE | 12005 | ENSG00000103275 | UBE2I | chr16 | + | 1320437 | 1320517 | 1317166 | 1320283 | 1324729 | 1324753 | 768 | 547 | 625 | 789 | 975 | 0 | 0 | 0 | 1 | 0 | 1802 | 1065 | 831 | 1223 | 1833 | 0 | 0 | 0 | 0 | 0 | 229 | 149 | 1 | 1 | 1 | 1 | 1 | 0.998 | 1 | 1 | 1 | 1 | 1 | 1 | 0 |
| SE | 12018 | ENSG00000101624 | CEP76 | chr18 | - | 12686261 | 12686450 | 12680661 | 12680828 | 12691358 | 12691487 | 26 | 18 | 28 | 39 | 18 | 0 | 1 | 0 | 0 | 0 | 14 | 10 | 5 | 26 | 0 | 0 | 0 | 0 | 0 | 0 | 298 | 149 | 1 | 1 | 1 | 0.9 | 1 | 1 | 1 | 1 | 1 | 1 | 1 | NA | -0.02 |
| SE | 12021 | ENSG00000101624 | CEP76 | chr18 | - | 12695253 | 12695351 | 12691232 | 12691487 | 12697222 | 12697408 | 32 | 13 | 30 | 24 | 63 | 0 | 0 | 1 | 0 | 0 | 0 | 23 | 7 | 29 | 12 | 0 | 0 | 0 | 0 | 0 | 247 | 149 | 1 | 1 | 1 | 1 | 0.948 | 1 | 1 | NA | 1 | 1 | 1 | 1 | -0.01 |
| SE | 12025 | ENSG00000176715 | ACSF3 | chr16 | + | 89098590 | 89098763 | 89093851 | 89093992 | 89100661 | 89100767 | 3 | 3 | 10 | 8 | 7 | 0 | 0 | 0 | 0 | 0 | 0 | 3 | 0 | 18 | 27 | 0 | 12 | 0 | 0 | 0 | 298 | 149 | 1 | 1 | 1 | 1 | 1 | 1 | 1 | NA | 0.111 | NA | 1 | 1 | 0.296 |
| SE | 12027 | ENSG00000176715 | ACSF3 | chr16 | + | 89100661 | 89101347 | 89093854 | 89093992 | 89102603 | 89102648 | 35 | 15 | 36 | 41 | 23 | 0 | 0 | 0 | 2 | 0 | 33 | 44 | 23 | 28 | 37 | 0 | 0 | 0 | 0 | 0 | 298 | 149 | 1 | 1 | 1 | 1 | 1 | 0.911 | 1 | 1 | 1 | 1 | 1 | 1 | -0.018 |
| SE | 12029 | ENSG00000176715 | ACSF3 | chr16 | + | 89112091 | 89112246 | 89102603 | 89102759 | 89120800 | 89120913 | 83 | 23 | 70 | 80 | 64 | 0 | 0 | 0 | 0 | 0 | 110 | 85 | 75 | 57 | 165 | 0 | 1 | 0 | 0 | 0 | 298 | 149 | 1 | 1 | 1 | 1 | 1 | 1 | 1 | 1 | 0.977 | 1 | 1 | 1 | 0.005 |
| SE | 12035 | ENSG00000176715 | ACSF3 | chr16 | + | 89120800 | 89120913 | 89112091 | 89112246 | 89133135 | 89133262 | 67 | 42 | 66 | 81 | 55 | 0 | 0 | 0 | 0 | 0 | 199 | 123 | 94 | 100 | 243 | 0 | 0 | 0 | 4 | 0 | 262 | 149 | 1 | 1 | 1 | 1 | 1 | 1 | 1 | 1 | 1 | 1 | 0.934 | 1 | 0.013 |
| SE | 12036 | ENSG00000176715 | ACSF3 | chr16 | + | 89141066 | 89141276 | 89133135 | 89133262 | 89145266 | 89145401 | 1 | 2 | 0 | 0 | 0 | 89 | 63 | 61 | 54 | 80 | 0 | 0 | 0 | 0 | 5 | 148 | 89 | 52 | 105 | 149 | 298 | 149 | 1 | 1 | 0.006 | 0.016 | 0 | 0 | 0 | 0 | 0 | 0 | 0 | 0.017 | 0.001 |
| SE | 12038 | ENSG00000103018 | CYB5B | chr16 | + | 69459092 | 69459121 | 69448114 | 69448144 | 69462429 | 69462902 | 220 | 113 | 132 | 194 | 132 | 1 | 0 | 1 | 0 | 0 | 116 | 183 | 80 | 73 | 119 | 0 | 0 | 0 | 0 | 0 | 178 | 149 | 1 | 1 | 0.995 | 1 | 0.991 | 1 | 1 | 1 | 1 | 1 | 1 | 1 | -0.003 |
| SE | 12040 | ENSG00000213380 | COG8 | chr16 | - | 69330812 | 69331095 | 69326427 | 69329179 | 69332713 | 69332882 | 77 | 28 | 49 | 58 | 47 | 0 | 0 | 0 | 0 | 0 | 39 | 70 | 25 | 31 | 86 | 0 | 0 | 0 | 1 | 3 | 298 | 149 | 1 | 1 | 1 | 1 | 1 | 1 | 1 | 1 | 1 | 1 | 0.939 | 0.935 | 0.025 |
| SE | 12044 | ENSG00000132612 | VPS4A | chr16 | + | 69318822 | 69318942 | 69318649 | 69318711 | 69319386 | 69319543 | 225 | 78 | 172 | 196 | 267 | 0 | 0 | 0 | 0 | 0 | 304 | 287 | 175 | 192 | 484 | 0 | 2 | 0 | 0 | 0 | 269 | 149 | 1 | 1 | 1 | 1 | 1 | 1 | 1 | 1 | 0.988 | 1 | 1 | 1 | 0.002 |
| SE | 12045 | ENSG00000183889 | NA | chr16 | + | 16318550 | 16318734 | 16317895 | 16318079 | 16318922 | 16319038 | 3 | 4 | 2 | 4 | 0 | 0 | 3 | 0 | 0 | 0 | 15 | 8 | 0 | 0 | 8 | 0 | 0 | 0 | 6 | 2 | 298 | 149 | 1 | 1 | 1 | 0.4 | 1 | 1 | NA | 1 | 1 | NA | 0 | 0.667 | 0.183 |
| SE | 12047 | ENSG00000183889 | NA | chr16 | + | 16331853 | 16331980 | 16331552 | 16331763 | 16333646 | 16333884 | 6 | 2 | 5 | 6 | 6 | 0 | 0 | 0 | 2 | 0 | 8 | 2 | 0 | 0 | 10 | 0 | 0 | 0 | 0 | 0 | 276 | 149 | 1 | 1 | 1 | 1 | 1 | 0.618 | 1 | 1 | 1 | NA | NA | 1 | -0.076 |
| SE | 12048 | ENSG00000183889 | NA | chr16 | + | 16335860 | 16335992 | 16333646 | 16333884 | 16340305 | 16340434 | 0 | 0 | 0 | 3 | 0 | 4 | 2 | 1 | 0 | 0 | 0 | 0 | 0 | 0 | 0 | 0 | 0 | 7 | 0 | 0 | 281 | 149 | 1 | 1 | 0 | 0 | 0 | 1 | NA | NA | NA | 0 | NA | NA | 0.25 |
| SE | 12049 | ENSG00000183889 | NA | chr16 | + | 16335860 | 16336137 | 16333736 | 16333884 | 16340305 | 16340501 | 0 | 1 | 1 | 3 | 1 | 4 | 2 | 1 | 0 | 0 | 0 | 0 | 0 | 0 | 0 | 0 | 0 | 7 | 0 | 0 | 298 | 149 | 1 | 1 | 0 | 0.2 | 0.333 | 1 | 1 | NA | NA | 0 | NA | NA | 0.507 |
| SE | 12050 | ENSG00000183889 | NA | chr16 | + | 16336051 | 16336137 | 16333646 | 16333884 | 16340305 | 16340434 | 8 | 11 | 14 | 19 | 7 | 4 | 2 | 1 | 0 | 0 | 37 | 0 | 1 | 22 | 20 | 0 | 0 | 7 | 0 | 0 | 235 | 149 | 1 | 1 | 0.559 | 0.777 | 0.899 | 1 | 1 | 1 | NA | 0.083 | 1 | 1 | 0.076 |
| SE | 12051 | ENSG00000183889 | NA | chr16 | + | 16347402 | 16347510 | 16344307 | 16344452 | 16348526 | 16348587 | 6 | 1 | 9 | 11 | 1 | 0 | 0 | 1 | 0 | 0 | 26 | 15 | 5 | 0 | 0 | 0 | 0 | 0 | 0 | 0 | 257 | 149 | 1 | 1 | 1 | 1 | 0.839 | 1 | 1 | 1 | 1 | 1 | NA | NA | -0.032 |
| SE | 12052 | ENSG00000183889 | NA | chr16 | + | 16348526 | 16348587 | 16344307 | 16344452 | 16348696 | 16348732 | 5 | 0 | 2 | 5 | 0 | 0 | 0 | 0 | 0 | 2 | 0 | 0 | 3 | 0 | 0 | 0 | 0 | 0 | 0 | 0 | 210 | 149 | 1 | 1 | 1 | NA | 1 | 1 | 0 | NA | NA | 1 | NA | NA | -0.25 |
| SE | 12053 | ENSG00000183889 | NA | chr16 | + | 16349228 | 16349298 | 16348696 | 16348732 | 16350091 | 16350203 | 4 | 1 | 1 | 5 | 0 | 6 | 0 | 4 | 11 | 2 | 5 | 1 | 8 | 0 | 0 | 16 | 15 | 3 | 0 | 0 | 219 | 149 | 1 | 1 | 0.312 | 1 | 0.145 | 0.236 | 0 | 0.175 | 0.043 | 0.645 | NA | NA | 0.051 |
| SE | 12056 | ENSG00000159720 | ATP6V0D1 | chr16 | - | 67445811 | 67445965 | 67444527 | 67444706 | 67453543 | 67453715 | 1 | 0 | 0 | 0 | 2 | 426 | 178 | 236 | 312 | 405 | 0 | 0 | 0 | 0 | 0 | 480 | 478 | 254 | 493 | 1008 | 298 | 149 | 1 | 1 | 0.001 | 0 | 0 | 0 | 0.002 | 0 | 0 | 0 | 0 | 0 | 0.001 |
| [illegible] | | | | | | | | | | | | | | | | | | | | | | | | | | | | | | | | | | | | | | | | | | | | | | |

| | | | | | | | | | | | | | | | | |
|---|---|---|---|---|---|---|---|---|---|---|---|---|---|---|---|---|
| SE | 12545 | ENSG00000103148 | NPRL3 | chr16 | - | 130521 | 130591 | 112621 | 112775 | 138149 | 138334 | 6 | 1 | 3 | 5 | 0 |
| SE | 12554 | ENSG00000214078 | CPNE1 | chr20 | - | 35626734 | 35626826 | 35626043 | 35626381 | 35627279 | 35627399 | 4 | 0 | 6 | 0 | 4 |
| SE | 12555 | ENSG00000214078 | CPNE1 | chr20 | - | 35627279 | 35627413 | 35626562 | 35626803 | 35630438 | 35630456 | 291 | 118 | 330 | 177 | 283 |
| SE | 12556 | ENSG00000214078 | CPNE1 | chr20 | - | 35630438 | 35630490 | 35627279 | 35627413 | 35630740 | 35630795 | 480 | 212 | 507 | 277 | 657 |
| SE | 12557 | ENSG00000214078 | CPNE1 | chr20 | - | 35630438 | 35630490 | 35627279 | 35627413 | 35630900 | 35631034 | 99 | 38 | 110 | 31 | 55 |
| SE | 12558 | ENSG00000214078 | CPNE1 | chr20 | - | 35630740 | 35630795 | 35627279 | 35627413 | 35630900 | 35631034 | 50 | 34 | 45 | 39 | 40 |
| SE | 12560 | ENSG00000214078 | CPNE1 | chr20 | - | 35630900 | 35631034 | 35630438 | 35630490 | 35631113 | 35631173 | 148 | 110 | 203 | 121 | 416 |
| SE | 12567 | ENSG00000214078 | CPNE1 | chr20 | - | 35658929 | 35658982 | 35632794 | 35632923 | 35664878 | 35664932 | 5 | 2 | 7 | 10 | 9 |
| SE | 12573 | ENSG00000214078 | CPNE1 | chr20 | - | 35658929 | 35659014 | 35632794 | 35632923 | 35664878 | 35664932 | 9 | 3 | 9 | 12 | 14 |
| SE | 12574 | ENSG00000244462 | RBM12 | chr20 | - | 35658929 | 35659014 | 35648924 | 35655344 | 35664759 | 35664900 | 731 | 427 | 742 | 746 | 579 |
| SE | 12578 | ENSG00000087263 | OGFOD1 | chr16 | + | 56454773 | 56454921 | 56453262 | 56453408 | 56458547 | 56458901 | 0 | 0 | 0 | 0 | 1 |
| SE | 12579 | ENSG00000087263 | OGFOD1 | chr16 | + | 56462533 | 56462634 | 56458128 | 56458594 | 56466151 | 56466179 | 118 | 84 | 131 | 106 | 229 |
| SE | 12580 | ENSG00000087263 | OGFOD1 | chr16 | + | 56466875 | 56466967 | 56466151 | 56466268 | 56467164 | 56467198 | 145 | 69 | 131 | 152 | 260 |
| SE | 12583 | ENSG00000034713 | GABARAPL2 | chr16 | + | 75568036 | 75568209 | 75566851 | 75566907 | 75577278 | 75577530 | 738 | 719 | 662 | 996 | 683 |
| SE | 12596 | ENSG00000090861 | AARS1 | chr16 | - | 70265537 | 70265662 | 70261043 | 70261157 | 70267658 | 70267796 | 78 | 49 | 70 | 70 | 60 |
| SE | 12599 | ENSG00000157353 | FCSK | chr16 | + | 70457912 | 70457975 | 70454594 | 70454630 | 70463168 | 70463272 | 0 | 0 | 0 | 0 | 2 |
| SE | 12603 | ENSG00000157353 | FCSK | chr16 | + | 70467885 | 70467966 | 70466881 | 70467471 | 70468848 | 70468968 | 8 | 0 | 3 | 5 | 0 |
| SE | 12604 | ENSG00000157353 | FCSK | chr16 | + | 70469151 | 70469323 | 70468848 | 70468968 | 70470313 | 70470426 | 6 | 6 | 2 | 4 | 0 |
| SE | 12605 | ENSG00000157353 | FCSK | chr16 | + | 70476186 | 70476334 | 70475647 | 70475767 | 70478271 | 70478459 | 0 | 0 | 3 | 0 | 0 |
| SE | 12607 | ENSG00000157350 | ST3GAL2 | chr16 | - | 70388366 | 70388546 | 70383189 | 70383235 | 70394981 | 70395175 | 186 | 104 | 169 | 187 | 112 |
| SE | 12610 | ENSG00000168872 | DDX19A | chr16 | + | 70350556 | 70350605 | 70346860 | 70347048 | 70355484 | 70355535 | 86 | 55 | 94 | 106 | 138 |
| SE | 12611 | ENSG00000168872 | DDX19A | chr16 | + | 70356111 | 70356247 | 70355484 | 70355535 | 70361417 | 70361498 | 214 | 126 | 206 | 183 | 398 |
| SE | 12612 | ENSG00000168872 | DDX19A | chr16 | + | 70356128 | 70356247 | 70355484 | 70355535 | 70361417 | 70361510 | 88 | 57 | 91 | 92 | 150 |
| SE | 12615 | ENSG00000168872 | DDX19A | chr16 | + | 70360776 | 70360894 | 70356111 | 70356247 | 70361417 | 70361510 | 0 | 0 | 0 | 0 | 10 |
| SE | 12620 | ENSG00000168872 | DDX19A | chr16 | + | 70364542 | 70364645 | 70361417 | 70361510 | 70365016 | 70365131 | 97 | 64 | 97 | 90 | 194 |
| SE | 12621 | ENSG00000168872 | DDX19A | chr16 | + | 70365016 | 70365131 | 70361417 | 70361510 | 70366084 | 70366184 | 32 | 23 | 37 | 30 | 64 |
| SE | 12623 | ENSG00000261777 | DDX19A-DT | chr16 | - | 70341939 | 70342099 | 70333782 | 70333960 | 70346198 | 70346747 | 5 | 9 | 9 | 13 | 18 |
| SE | 12627 | ENSG00000090857 | PDPR | chr16 | + | 70120460 | 70120719 | 70114343 | 70114440 | 70127259 | 70127393 | 99 | 54 | 82 | 110 | 95 |
| SE | 12628 | ENSG00000090857 | PDPR | chr16 | + | 70130422 | 70130544 | 70128783 | 70129122 | 70131301 | 70131419 | 66 | 24 | 97 | 56 | 93 |
| SE | 12635 | ENSG00000090857 | PDPR | chr16 | + | 70138898 | 70139023 | 70136193 | 70136386 | 70142233 | 70142389 | 158 | 83 | 148 | 158 | 110 |
| SE | 12638 | ENSG00000090857 | PDPR | chr16 | + | 70146133 | 70146228 | 70144420 | 70144533 | 70156474 | 70156614 | 25 | 14 | 24 | 23 | 47 |
| SE | 12639 | ENSG00000090857 | PDPR | chr16 | + | 70147540 | 70147673 | 70146133 | 70146228 | 70148463 | 70148553 | 0 | 0 | 0 | 0 | 3 |
| SE | 12640 | ENSG00000090857 | PDPR | chr16 | + | 70148463 | 70148553 | 70144420 | 70144533 | 70156474 | 70156859 | 6 | 0 | 6 | 6 | 6 |
| SE | 12641 | ENSG00000090857 | PDPR | chr16 | + | 70148463 | 70148553 | 70145803 | 70146228 | 70156474 | 70156859 | 46 | 22 | 28 | 44 | 41 |
| SE | 12645 | ENSG00000196712 | NF1 | chr17 | + | 31181709 | 31181785 | 31181421 | 31181489 | 31182507 | 31182665 | 94 | 60 | 43 | 63 | 72 |
| SE | 12646 | ENSG00000196712 | NF1 | chr17 | + | 31221849 | 31221929 | 31219004 | 31219118 | 31223443 | 31223567 | 102 | 65 | 55 | 80 | 92 |
| SE | 12647 | ENSG00000196712 | NF1 | chr17 | + | 31223443 | 31223567 | 31221849 | 31221929 | 31225094 | 31225250 | 155 | 92 | 74 | 121 | 147 |
| SE | 12648 | ENSG00000196712 | NF1 | chr17 | + | 31252937 | 31253000 | 31248983 | 31249119 | 31257686 | 31257726 | 10 | 4 | 1 | 0 | 5 |
| SE | 12649 | ENSG00000196712 | NF1 | chr17 | + | 31252937 | 31253000 | 31248983 | 31249119 | 31258343 | 31258502 | 37 | 29 | 22 | 18 | 23 |
| SE | 12650 | ENSG00000196712 | NF1 | chr17 | + | 31257686 | 31257726 | 31248983 | 31249119 | 31258343 | 31258502 | 0 | 1 | 0 | 0 | 0 |
| SE | 12653 | ENSG00000196712 | NF1 | chr17 | + | 31327498 | 31327839 | 31325819 | 31326252 | 31330295 | 31330498 | 158 | 126 | 155 | 131 | 270 |
| SE | 12655 | ENSG00000196712 | NF1 | chr17 | + | 31334663 | 31334730 | 31330328 | 31330498 | 31334837 | 31335031 | 2 | 0 | 0 | 0 | 0 |
| SE | 12656 | ENSG00000196712 | NF1 | chr17 | + | 31336634 | 31336914 | 31336332 | 31336473 | 31337367 | 31337582 | 104 | 64 | 90 | 109 | 145 |
| SE | 12657 | ENSG00000196712 | NF1 | chr17 | + | 31338001 | 31338139 | 31337816 | 31337880 | 31338703 | 31338805 | 39 | 25 | 23 | 44 | 64 |
| SE | 12659 | ENSG00000196712 | NF1 | chr17 | + | 31338024 | 31338139 | 31337762 | 31337880 | 31338703 | 31338805 | 113 | 95 | 100 | 155 | 146 |
| SE | 12662 | ENSG00000196712 | NF1 | chr17 | + | 31338703 | 31338805 | 31338001 | 31338139 | 31340504 | 31340645 | 103 | 65 | 90 | 101 | 111 |
| SE | 12665 | ENSG00000196712 | NF1 | chr17 | + | 31356459 | 31356582 | 31352256 | 31352414 | 31356959 | 31357090 | 103 | 97 | 98 | 85 | 129 |
| SE | 12667 | ENSG00000196712 | NF1 | chr17 | + | 31358479 | 31358622 | 31356959 | 31357090 | 31358968 | 31359015 | 41 | 32 | 29 | 51 | 120 |
| SE | 12668 | ENSG00000196712 | NF1 | chr17 | + | 31367224 | 31367278 | 31360486 | 31360703 | 31374012 | 31374433 | 8 | 0 | 0 | 0 | 0 |
| SE | 12669 | ENSG00000196712 | NF1 | chr17 | + | 31379105 | 31379357 | 31378890 | 31379023 | 31381872 | 31382106 | 0 | 0 | 1 | 0 | 0 |
| SE | 12674 | ENSG00000140905 | GCSH | chr16 | - | 81084462 | 81084594 | 81081944 | 81082963 | 81090600 | 81090680 | 26 | 14 | 31 | 37 | 27 |
| SE | 12679 | ENSG00000140905 | GCSH | chr16 | - | 81090996 | 81091107 | 81090600 | 81090680 | 81096130 | 81096318 | 0 | 0 | 3 | 1 | 0 |
| SE | 12680 | ENSG00000260643 | NA | chr16 | - | 81057904 | 81058064 | 81053664 | 81054094 | 81084462 | 81084594 | 0 | 0 | 0 | 0 | 2 |
| SE | 12690 | ENSG00000177398 | UMODL1 | chr21 | + | 42128003 | 42128145 | 42127677 | 42127831 | 42129712 | 42129797 | 3 | 0 | 0 | 0 | 0 |
| SE | 12691 | ENSG00000213599 | SLX1A-SULT1A3 | chr16 | + | 30201195 | 30201293 | 30200980 | 30201106 | 30202689 | 30202816 | 0 | 1 | 1 | 1 | 0 |
| SE | 12697 | ENSG00000155666 | KDM8 | chr16 | + | 27214875 | 27215008 | 27210092 | 27210621 | 27215944 | 27215989 | 20 | 4 | 25 | 4 | 13 |
| SE | 12699 | ENSG00000155666 | KDM8 | chr16 | + | 27215944 | 27215989 | 27210092 | 27210621 | 27218960 | 27219046 | 2 | 1 | 1 | 0 | 0 |
| SE | 12700 | ENSG00000155666 | KDM8 | chr16 | + | 27218960 | 27219046 | 27215944 | 27215989 | 27220392 | 27220485 | 10 | 3 | 13 | 2 | 7 |
| SE | 12704 | ENSG00000155666 | KDM8 | chr16 | + | 27220392 | 27220485 | 27218960 | 27219110 | 27220565 | 27220859 | 20 | 10 | 11 | 5 | 6 |
| SE | 12708 | ENSG00000067836 | ROGDI | chr16 | - | 4801049 | 4801321 | 4800497 | 4800578 | 4801502 | 4801585 | 0 | 4 | 0 | 3 | 0 |
| SE | 12710 | ENSG00000067836 | ROGDI | chr16 | - | 4802381 | 4802453 | 4801028 | 4801585 | 4802526 | 4802615 | 28 | 13 | 14 | 34 | 16 |
| SE | 12712 | ENSG00000103227 | LMF1 | chr16 | - | 910930 | 911079 | 893006 | 893072 | 934243 | 934254 | 7 | 12 | 18 | 21 | 23 |
| SE | 12714 | ENSG00000103227 | LMF1 | chr16 | - | 931641 | 931793 | 910930 | 911079 | 934243 | 934254 | 1 | 2 | 0 | 1 | 4 |
| SE | 12715 | ENSG00000103227 | LMF1 | chr16 | - | 933990 | 934145 | 911015 | 911079 | 934243 | 934254 | 6 | 0 | 15 | 8 | 0 |
| SE | 12718 | ENSG00000181938 | GINS3 | chr16 | + | 58395101 | 58395218 | 58392393 | 58392787 | 58403097 | 58403331 | 0 | 0 | 0 | 0 | 0 |
| SE | 12722 | ENSG00000157106 | SMG1 | chr16 | - | 18815174 | 18815281 | 18811947 | 18812127 | 18815439 | 18815651 | 317 | 247 | 307 | 330 | 599 |
| SE | 12723 | ENSG00000157106 | SMG1 | chr16 | - | 18819501 | 18819654 | 18817290 | 18817470 | 18828030 | 18828168 | 190 | 141 | 147 | 155 | 203 |
| SE | 12724 | ENSG00000157106 | SMG1 | chr16 | - | 18829925 | 18830115 | 18829285 | 18829755 | 18830218 | 18830369 | 341 | 317 | 334 | 308 | 438 |
| SE | 12729 | ENSG00000157106 | SMG1 | chr16 | - | 18838356 | 18838466 | 18837252 | 18837443 | 18838550 | 18838689 | 248 | 197 | 207 | 206 | 184 |
| SE | 12730 | ENSG00000157106 | SMG1 | chr16 | - | 18839697 | 18839946 | 18838460 | 18838689 | 18841564 | 18841794 | 248 | 181 | 272 | 204 | 225 |
| SE | 12731 | ENSG00000157106 | SMG1 | chr16 | - | 18845428 | 18845651 | 18842207 | 18842454 | 18847452 | 18847607 | 317 | 294 | 298 | 353 | 364 |
| SE | 12732 | ENSG00000157106 | SMG1 | chr16 | - | 18847452 | 18847607 | 18845428 | 18845651 | 18847815 | 18848033 | 228 | 231 | 245 | 270 | 365 |
| SE | 12733 | ENSG00000157106 | SMG1 | chr16 | - | 18854655 | 18854904 | 18853582 | 18853867 | 18858169 | 18858290 | 264 | 258 | 271 | 284 | 326 |
| SE | 12735 | ENSG00000157106 | SMG1 | chr16 | - | 18868522 | 18868719 | 18868189 | 18868354 | 18869103 | 18869303 | 272 | 226 | 272 | 287 | 386 |
| SE | 12736 | ENSG00000157106 | SMG1 | chr16 | - | 18868632 | 18868719 | 18868189 | 18868354 | 18869103 | 18869303 | 90 | 96 | 76 | 88 | 141 |
| SE | 12737 | ENSG00000157106 | SMG1 | chr16 | - | 18871363 | 18871482 | 18870800 | 18870888 | 18872183 | 18872345 | 371 | 335 | 328 | 397 | 782 |
| SE | 12738 | ENSG00000157106 | SMG1 | chr16 | - | 18877130 | 18877232 | 18875101 | 18876393 | 18879494 | 18879719 | 316 | 308 | 315 | 235 | 415 |
| SE | 12739 | ENSG00000157106 | SMG1 | chr16 | - | 18879494 | 18879719 | 18877130 | 18877232 | 18882164 | 18882338 | 198 | 198 | 197 | 181 | 249 |
| SE | 12740 | ENSG00000157106 | SMG1 | chr16 | - | 18885540 | 18885666 | 18885089 | 18885162 | 18889371 | 18889585 | 241 | 158 | 166 | 232 | 310 |
| SE | 12743 | ENSG00000157106 | SMG1 | chr16 | - | 18899986 | 18900044 | 18896792 | 18896956 | 18911362 | 18911713 | 45 | 35 | 23 | 38 | 47 |
| SE | 12744 | ENSG00000157106 | SMG1 | chr16 | - | 18899986 | 18900044 | 18896792 | 18896956 | 18925678 | 18925750 | 43 | 33 | 25 | 40 | 42 |
| SE | 12747 | ENSG00000089280 | FUS | chr16 | + | 31182512 | 31182664 | 31182397 | 31182422 | 31183857 | 31184002 | 326 | 174 | 313 | 371 | 93 |
| SE | 12748 | ENSG00000089280 | FUS | chr16 | + | 31182512 | 31182664 | 31182397 | 31182422 | 31183860 | 31184002 | 255 | 137 | 240 | 261 | 79 |
| SE | 12751 | ENSG00000089280 | FUS | chr16 | + | 31188324 | 31188357 | 31184938 | 31185179 | 31189122 | 31189226 | 440 | 373 | 402 | 336 | 387 |
| SE | 12752 | ENSG00000089280 | FUS | chr16 | + | 31189119 | 31189226 | 31188324 | 31188357 | 31189664 | 31189794 | 771 | 628 | 831 | 604 | 766 |
| SE | 12753 | ENSG00000089280 | FUS | chr16 | + | 31189122 | 31189226 | 31184938 | 31188357 | 31189664 | 31189794 | 2020 | 1562 | 1988 | 1559 | 1857 |
| SE | 12760 | ENSG00000007541 | PIGQ | chr16 | + | 574527 | 574763 | 574065 | 574438 | 575838 | 575970 | 3 | 3 | 3 | 4 | 0 |
| SE | 12764 | ENSG00000007541 | PIGQ | chr16 | + | 580857 | 580972 | 580182 | 580263 | 582882 | 584109 | 8 | 0 | 5 | 10 | 9 |
| SE | 12766 | ENSG00000007541 | PIGQ | chr16 | + | 582247 | 582309 | 580182 | 580263 | 582882 | 582893 | 1 | 0 | 0 | 0 | 0 |
| SE | 12770 | ENSG00000186260 | MRTFB | chr16 | + | 14148666 | 14148729 | 14140543 | 14140760 | 14149285 | 14149536 | 0 | 0 | 0 | 0 | 0 |
| SE | 12772 | ENSG00000186260 | MRTFB | chr16 | + | 14252364 | 14252502 | 14251861 | 14252023 | 14258100 | 14258161 | 10 | 11 | 14 | 12 | 7 |
| SE | 12775 | ENSG00000168802 | CHTF8 | chr16 | - | 69121052 | 69121170 | 69118009 | 69120649 | 69132483 | 69132573 | 104 | 59 | 56 | 75 | 97 |
| SE | 12777 | ENSG00000168802 | CHTF8 | chr16 | - | 69121435 | 69121493 | 69118009 | 69120649 | 69132483 | 69132567 | 20 | 34 | 21 | 32 | 33 |
| SE | 12780 | ENSG00000132207 | SLX1A | chr16 | + | 30194885 | 30195227 | 30193886 | 30194791 | 30196973 | 30197101 | 25 | 1 | 6 | 9 | 0 |
| SE | 12782 | ENSG00000213658 | LAT | chr16 | + | 28985853 | 28985888 | 28985712 | 28985740 | 28986134 | 28986216 | 122 | 66 | 122 | 114 | 71 |
| SE | 12783 | ENSG00000213658 | LAT | chr16 | + | 28986134 | 28986216 | 28985841 | 28985888 | 28986381 | 28986399 | 84 | 72 | 108 | 132 | 52 |
| SE | 12785 | ENSG00000213658 | LAT | chr16 | + | 28986539 | 28986569 | 28986098 | 28986446 | 28986656 | 28986697 | 279 | 187 | 295 | 403 | 194 |
| SE | 12787 | ENSG00000213658 | LAT | chr16 | + | 28989773 | 28989839 | 28987565 | 28989589 | 28989932 | 28990012 | 620 | 370 | 585 | 628 | 436 |
| SE | 12788 | ENSG00000213658 | LAT | chr16 | + | 28989773 | 28989839 | 28987565 | 28989589 | 28990188 | 28990513 | 51 | 25 | 36 | 49 | 29 |
| SE | 12789 | ENSG00000213658 | LAT | chr16 | + | 28989932 | 28990019 | 28987565 | 28989589 | 28990188 | 28990296 | 130 | 79 | 145 | 182 | 80 |
| SE | 12790 | ENSG00000213658 | LAT | chr16 | + | 28989932 | 28990019 | 28989576 | 28989839 | 28990188 | 28990296 | 703 | 430 | 695 | 772 | 491 |
| SE | 12793 | ENSG00000102879 | CORO1A | chr16 | + | 30185208 | 30185407 | 30183601 | 30183725 | 30186597 | 30186720 | 1367 | 467 | 1087 | 1053 | 471 |
| SE | 12794 | ENSG00000102879 | CORO1A | chr16 | + | 30186597 | 30186720 | 30185208 | 30185407 | 30186815 | 30186938 | 2071 | 714 | 1600 | 1541 | 698 |
| SE | 12795 | ENSG00000102879 | CORO1A | chr16 | + | 30186815 | 30186945 | 30185208 | 30186720 | 30187038 | 30187083 | 2518 | 894 | 1917 | 1917 | 778 |
| SE | 12796 | ENSG00000102879 | CORO1A | chr16 | + | 30187941 | 30188087 | 30187724 | 30187829 | 30188191 | 30188249 | 2759 | 1114 | 2253 | 2205 | 1611 |
| SE | 12797 | ENSG00000102879 | CORO1A | chr16 | + | 30188360 | 30188576 | 30188191 | 30188249 | 30188859 | 30188964 | 2717 | 1058 | 2005 | 2025 | 1506 |
| SE | 12799 | ENSG00000160014 | CALM3 | chr19 | + | 46605826 | 46605857 | 46601275 | 46601437 | 46608196 | 46608327 | 579 | 416 | 427 | 676 | 280 |
| SE | 12800 | ENSG00000160014 | CALM3 | chr19 | + | 46605993 | 46606185 | 46605826 | 46605857 | 46608196 | 46608340 | 2 | 5 | 2 | 4 | 3 |
| SE | 12804 | ENSG00000261416 | CORO1A-AS1 | chr16 | - | 30184148 | 30184331 | 30183504 | 30183771 | 30184503 | 30184523 | 6 | 1 | 5 | 3 | 0 |
| SE | 12809 | ENSG00000102882 | MAPK3 | chr16 | - | 30118098 | 30118163 | 30117669 | 30117784 | 30118348 | 30118538 | 5 | 0 | 18 | 3 | 4 |
| SE | 12812 | ENSG00000102886 | GDPD3 | chr16 | - | 30108372 | 30108432 | 30104843 | 30105009 | 30111387 | 30111396 | 3 | 0 | 1 | 0 | 0 |
| SE | 12813 | ENSG00000149923 | PPP4C | chr16 | + | 30076314 | 30076475 | 30075977 | 30076094 | 30081258 | 30081310 | 230 | 94 | 147 | 133 | 157 |
| SE | 12821 | ENSG00000169592 | INO80E | chr16 | + | 29996807 | 29996860 | 29996243 | 29996617 | 30000757 | 30000812 | 240 | 125 | 209 | 181 | 280 |
| SE | 12822 | ENSG00000169592 | INO80E | chr16 | + | 30000928 | 30001040 | 30000757 | 30000836 | 30001211 | 30001229 | 93 | 81 | 112 | 100 | 144 |
| SE | 12823 | ENSG00000169592 | INO80E | chr16 | + | 30000928 | 30001040 | 30000757 | 30000836 | 30005220 | 30005717 | 86 | 62 | 86 | 79 | 142 |
| SE | 12824 | ENSG00000169592 | INO80E | chr16 | + | 30001211 | 30001530 | 30000757 | 30000836 | 30005220 | 30005323 | 25 | 15 | 22 | 13 | 33 |
| SE | 12826 | ENSG00000169592 | INO80E | chr16 | + | 30001211 | 30004657 | 30000757 | 30000836 | 30005220 | 30005774 | 5 | 1 | 8 | 3 | 0 |
| SE | 12830 | ENSG00000169592 | INO80E | chr16 | + | 30002435 | 30003013 | 30001413 | 30001530 | 30003435 | 30003526 | 11 | 17 | 7 | 6 | 0 |
| SE | 12833 | ENSG00000169592 | INO80E | chr16 | + | 30004567 | 30004657 | 30001211 | 30001530 | 30005367 | 30005435 | 5 | 4 | 1 | 0 | 0 |
| SE | 12838 | ENSG00000288632 | NA | chr16 | - | 29554866 | 29554992 | 29553085 | 29553183 | 29558664 | 29558878 | 31 | 8 | 33 | 23 | 60 |
| SE | 12840 | ENSG00000288632 | NA | chr16 | - | 29569270 | 29569328 | 29566085 | 29566249 | 29613102 | 29613717 | 13 | 15 | 15 | 36 | 4 |
| SE | 12841 | ENSG00000169203 | NPIPB12 | chr16 | - | 29488022 | 29488091 | 29487338 | 29487399 | 29491466 | 29491662 | 6 | 1 | 2 | 3 | 6 |
| SE | 12845 | ENSG00000169203 | NPIPB12 | chr16 | - | 29500142 | 29500274 | 29495478 | 29495607 | 29505472 | 29505630 | 0 | 0 | 0 | 0 | 0 |
| SE | 12846 | ENSG00000261740 | BOLA2-SMG1P6 | chr16 | - | 29446801 | 29446922 | 29444397 | 29444499 | 29454015 | 29454113 | 27 | 21 | 36 | 48 | 17 |
| SE | 12847 | ENSG00000261740 | BOLA2-SMG1P6 | chr16 | - | 29446801 | 29447026 | 29444397 | 29444499 | 29454015 | 29454113 | 37 | 24 | 34 | 56 | 23 |
| SE | 12848 | ENSG00000261740 | BOLA2-SMG1P6 | chr16 | - | 29450111 | 29450276 | 29446801 | 29447026 | 29452108 | 29454113 | 0 | 0 | 0 | 0 | 0 |
| SE | 12849 | ENSG00000261740 | BOLA2-SMG1P6 | chr16 | - | 29450111 | 29450276 | 29446801 | 29447026 | 29454015 | 29454113 | 1 | 0 | 0 | 0 | 0 |
| SE | 12850 | ENSG00000261740 | BOLA2-SMG1P6 | chr16 | - | 29452108 | 29454113 | 29446801 | 29447026 | 29454190 | 29454356 | 9 | 5 | 7 | 5 | 4 |
| SE | 12851 | ENSG00000261740 | BOLA2-SMG1P6 | chr16 | - | 29454015 | 29454113 | 29446801 | 29446922 | 29454190 | 29454347 | 16 | 11 | 19 | 15 | 8 |
| SE | 12852 | ENSG00000261740 | BOLA2-SMG1P6 | chr16 | - | 29454015 | 29454113 | 29446801 | 29447026 | 29454190 | 29454347 | 22 | 13 | 15 | 21 | 14 |
| SE | 12853 | ENSG00000181031 | RPH3AL | chr17 | - | 247110 | 247285 | 219622 | 219736 | 281767 | 281854 | 2 | 0 | 6 | 5 | 3 |
| SE | 12858 | ENSG00000121413 | ZSCAN18 | chr19 | - | 58086905 | 58087008 | 58086173 | 58086266 | 58087315 | 58087404 | 19 | 17 | 23 | 23 | 8 |
| SE | 12861 | ENSG00000121413 | ZSCAN18 | chr19 | - | 58093227 | 58093398 | 58089864 | 58090386 | 58098114 | 58098214 | 3 | 0 | 1 | 0 | 0 |
| SE | 12863 | ENSG00000103472 | RRN3P2 | chr16 | + | 29076655 | 29076761 | 29074975 | 29075121 | 29077900 | 29077957 | 62 | 66 | 50 | 85 | 124 |
| SE | 12864 | ENSG00000103472 | RRN3P2 | chr16 | + | 29077900 | 29077957 | 29074975 | 29075121 | 29084638 | 29084768 | 4 | 2 | 1 | 3 | 1 |
| SE | 12866 | ENSG00000103472 | RRN3P2 | chr16 | + | 29086213 | 29086275 | 29085216 | 29085276 | 29090739 | 29090838 | 8 | 19 | 16 | 24 | 20 |
| SE | 12867 | ENSG00000103472 | RRN3P2 | chr16 | + | 29094237 | 29094330 | 29090739 | 29090838 | 29096130 | 29096261 | 100 | 80 | 67 | 61 | 80 |
| SE | 12868 | ENSG00000103472 | RRN3P2 | chr16 | + | 29094237 | 29094408 | 29090739 | 29090838 | 29096130 | 29096261 | 45 | 46 | 40 | 45 | 59 |
| SE | 12870 | ENSG00000103472 | RRN3P2 | chr16 | + | 29097131 | 29097316 | 29096130 | 29096261 | 29099104 | 29099213 | 88 | 92 | 70 | 80 | 91 |
| SE | 12871 | ENSG00000103472 | RRN3P2 | chr16 | + | 29097192 | 29097316 | 29096130 | 29096261 | 29099104 | 29099213 | 72 | 54 | 56 | 66 | 87 |
| SE | 12872 | ENSG00000103472 | RRN3P2 | chr16 | + | 29101986 | 29102139 | 29099104 | 29099213 | 29104074 | 29104162 | 131 | 127 | 118 | 139 | 114 |
| SE | 12874 | ENSG00000176953 | NFATC2IP | chr16 | + | 28954564 | 28954682 | 28952131 | 28952204 | 28955977 | 28956058 | 138 | 116 | 122 | 190 | 158 |
| SE | 12877 | ENSG00000176953 | NFATC2IP | chr16 | + | 28955977 | 28956058 | 28954564 | 28954682 | 28956150 | 28956337 | 96 | 88 | 84 | 125 | 130 |
| SE | 12879 | ENSG00000176953 | NFATC2IP | chr16 | + | 28956150 | 28956337 | 28954564 | 28954682 | 28958990 | 28959100 | 2 | 0 | 0 | 2 | 0 |
| SE | 12880 | ENSG00000176953 | NFATC2IP | chr16 | + | 28956150 | 28956337 | 28954564 | 28954682 | 28963704 | 28967097 | 3 | 0 | 0 | 0 | 0 |
| SE | 12881 | ENSG00000176953 | NFATC2IP | chr16 | + | 28958716 | 28958861 | 28954564 | 28954682 | 28958990 | 28959018 | 72 | 69 | 91 | 64 | 70 |
| SE | 12882 | ENSG00000176953 | NFATC2IP | chr16 | + | 28958716 | 28958861 | 28956150 | 28956337 | 28958990 | 28959018 | 123 | 104 | 168 | 104 | 104 |
| SE | 12883 | ENSG00000176953 | NFATC2IP | chr16 | + | 28958990 | 28959100 | 28954564 | 28954682 | 28963704 | 28964279 | 51 | 35 | 47 | 37 | 26 |
| SE | 12884 | ENSG00000176953 | NFATC2IP | chr16 | + | 28958990 | 28959100 | 28956150 | 28956337 | 28963704 | 28964279 | 51 | 35 | 46 | 39 | 26 |
| SE | 12885 | ENSG00000177455 | CD19 | chr16 | + | 28932345 | 28932612 | 28931967 | 28932088 | 28932910 | 28933114 | 17 | 10 | 22 | 12 | 15 |
| SE | 12886 | ENSG00000177455 | CD19 | chr16 | + | 28935443 | 28935554 | 28933233 | 28933509 | 28936511 | 28936600 | 23 | 15 | 31 | 16 | 41 |
| SE | 12889 | ENSG00000177548 | RABEP2 | chr16 | - | 28905866 | 28905878 | 28905703 | 28905759 | 28906018 | 28906196 | 20 | 6 | 14 | 7 | 9 |
| SE | 12890 | ENSG00000177548 | RABEP2 | chr16 | - | 28906018 | 28906196 | 28905866 | 28905878 | 28908608 | 28908764 | 53 | 31 | 54 | 33 | 24 |
| SE | 12891 | ENSG00000177548 | RABEP2 | chr16 | - | 28911083 | 28911179 | 28910286 | 28910986 | 28914235 | 28914586 | 30 | 10 | 29 | 22 | 10 |
| SE | 12893 | ENSG00000177548 | RABEP2 | chr16 | - | 28924402 | 28924615 | 28919687 | 28919943 | 28925102 | 28925177 | 33 | 23 | 27 | 36 | 25 |
| SE | 12894 | ENSG00000178952 | TUFM | chr16 | - | 28844229 | 28844334 | 28843949 | 28844101 | 28844418 | 28844551 | 543 | 313 | 418 | 641 | 350 |
| SE | 12898 | ENSG00000176476 | SGF29 | chr16 | + | 28590630 | 28590666 | 28590116 | 28590442 | 28590772 | 28590935 | 104 | 43 | 105 | 98 | 146 |
| SE | 12902 | ENSG00000261832 | NA | chr16 | - | 28476250 | 28476375 | 28466719 | 28466902 | 28477463 | 28477635 | 1 | 0 | 0 | 0 | 0 |
| SE | 12905 | ENSG00000140740 | UQCRC2 | chr16 | + | 21957234 | 21957318 | 21953287 | 21953456 | 21957416 | 21957566 | 571 | 270 | 486 | 460 | 436 |
| SE | 12906 | ENSG00000140740 | UQCRC2 | chr16 | + | 21959338 | 21959555 | 21958534 | 21958599 | 21961327 | 21961465 | 16 | 26 | 11 | 60 | 70 |
| SE | 12909 | ENSG00000140740 | UQCRC2 | chr16 | + | 21962459 | 21962516 | 21958534 | 21958599 | 21962760 | 21962819 | 459 | 249 | 376 | 366 | 562 |
| SE | 12910 | ENSG00000140740 | UQCRC2 | chr16 | + | 21968627 | 21968685 | 21965407 | 21965505 | 21971524 | 21971620 | 443 | 251 | 406 | 417 | 516 |
| SE | 12911 | ENSG00000140740 | UQCRC2 | chr16 | + | 21971524 | 21971620 | 21968461 | 21968685 | 21971922 | 21972122 | 609 | 372 | 566 | 561 | 560 |
| SE | 12913 | ENSG00000185710 | SMG1P4 | chr16 | - | 21890763 | 21890882 | 21890202 | 21890290 | 21891591 | 21891747 | 159 | 54 | 0 | 88 | 2 |
| SE | 12917 | ENSG00000187193 | MT1X | chr16 | + | 56683164 | 56683230 | 56682469 | 56682568 | 56683957 | 56684196 | 107 | 28 | 141 | 58 | 51 |
| SE | 12918 | ENSG00000187193 | MT1X | chr16 | + | 56683164 | 56683386 | 56682480 | 56682568 | 56683957 | 56684196 | 43 | 12 | 24 | 18 | 9 |
| SE | 12919 | ENSG00000197863 | ZNF790 | chr19 | - | 36823284 | 36823380 | 36817427 | 36820114 | 36823666 | 36823790 | 25 | 10 | 16 | 32 | 18 |
| SE | 12920 | ENSG00000198417 | MT1F | chr16 | + | 56658674 | 56658740 | 56657958 | 56658086 | 56659072 | 56659303 | 29 | 18 | 20 | 22 | 37 |
| SE | 12921 | ENSG00000198417 | MT1F | chr16 | + | 56658674 | 56658782 | 56658011 | 56658086 | 56659072 | 56659305 | 12 | 1 | 9 | 2 | 0 |
| SE | 12922 | ENSG00000108798 | ABI3 | chr17 | + | 49216530 | 49216680 | 49210410 | 49210841 | 49217738 | 49217915 | 196 | 66 | 76 | 135 | 77 |
| SE | 12923 | ENSG00000108798 | ABI3 | chr17 | + | 49216530 | 49216698 | 49210706 | 49210841 | 49217738 | 49217915 | 406 | 153 | 165 | 273 | 149 |
| SE | 12924 | ENSG00000108798 | ABI3 | chr17 | + | 49220168 | 49220326 | 49219857 | 49219953 | 49222090 | 49222225 | 148 | 64 | 47 | 88 | 40 |
| SE | 12926 | ENSG00000197818 | SLC9A8 | chr20 | + | 49815007 | 49815189 | 49812712 | 49812948 | 49823060 | 49823141 | 84 | 58 | 80 | 51 | 37 |
| SE | 12927 | ENSG00000197818 | SLC9A8 | chr20 | + | 49823060 | 49823141 | 49815007 | 49815189 | 49839540 | 49839599 | 72 | 55 | 65 | 55 | 47 |
| SE | 12929 | ENSG00000197818 | SLC9A8 | chr20 | + | 49845035 | 49845119 | 49839540 | 49839599 | 49849578 | 49849680 | 49 | 43 | 53 | 54 | 59 |
| SE | 12930 | ENSG00000197818 | SLC9A8 | chr20 | + | 49845035 | 49845119 | 49839540 | 49839599 | 49855437 | 49855581 | 16 | 12 | 16 | 21 | 22 |
| SE | 12931 | ENSG00000197818 | SLC9A8 | chr20 | + | 49849578 | 49849680 | 49839540 | 49839599 | 49855437 | 49855581 | 6 | 7 | 9 | 14 | 4 |
| SE | 12936 | ENSG00000197818 | SLC9A8 | chr20 | + | 49877980 | 49878063 | 49874704 | 49874821 | 49880923 | 49881035 | 44 | 40 | 54 | 38 | 39 |
| SE | 12942 | ENSG00000255439 | NA | chr16 | - | 31084118 | 31084335 | 31083438 | 31083809 | 31084557 | 31084703 | 1 | 0 | 2 | 0 | 0 |
| SE | 12946 | ENSG00000260083 | MIR762HG | chr16 | + | 30888713 | 30888958 | 30875778 | 30876118 | 30894547 | 30895136 | 1 | 0 | 0 | 0 | 0 |
| SE | 12948 | ENSG00000196118 | CCDC189 | chr16 | - | 30759011 | 30759101 | 30757422 | 30757654 | 30759177 | 30759247 | 0 | 6 | 0 | 0 | 0 |
| SE | 12949 | ENSG00000196118 | CCDC189 | chr16 | - | 30759011 | 30759247 | 30757435 | 30757654 | 30759341 | 30759458 | 2 | 0 | 1 | 0 | 1 |
| SE | 12950 | ENSG00000196118 | CCDC189 | chr16 | - | 30759177 | 30759247 | 30757422 | 30757654 | 30759341 | 30759444 | 0 | 1 | 1 | 0 | 9 |
| SE | 12951 | ENSG00000196118 | CCDC189 | chr16 | - | 30759341 | 30759444 | 30757422 | 30757654 | 30760283 | 30760668 | 1 | 1 | 1 | 0 | 0 |
| SE | 12952 | ENSG00000196118 | CCDC189 | chr16 | - | 30759341 | 30759458 | 30757422 | 30757654 | 30760283 | 30760419 | 0 | 0 | 1 | 0 | 0 |
| SE | 12953 | ENSG00000196118 | CCDC189 | chr16 | - | 30759341 | 30759724 | 30757466 | 30757654 | 30760283 | 30760453 | 1 | 0 | 0 | 2 | 0 |
| SE | 12954 | ENSG00000196118 | CCDC189 | chr16 | - | 30759653 | 30759724 | 30759341 | 30759458 | 30760283 | 30760453 | 1 | 0 | 0 | 2 | 0 |
| SE | 12955 | ENSG00000196118 | CCDC189 | chr16 | - | 30759653 | 30759809 | 30759341 | 30759458 | 30760283 | 30760668 | 0 | 0 | 0 | 0 | 0 |
| SE | 12956 | ENSG00000196118 | CCDC189 | chr16 | - | 30760605 | 30760668 | 30760036 | 30760453 | 30761197 | 30761229 | 2 | 5 | 3 | 4 | 0 |
| SE | 12959 | ENSG00000103415 | HMOX2 | chr16 | + | 4488643 | 4488742 | 4476339 | 4476487 | 4505483 | 4505610 | 0 | 0 | 0 | 0 | 0 |
| SE | 12962 | ENSG00000103415 | HMOX2 | chr16 | + | 4505483 | 4505610 | 4476397 | 4476487 | 4506894 | 4507012 | 268 | 125 | 191 | 211 | 222 |
| SE | 12963 | ENSG00000103423 | DNAJA3 | chr16 | + | 4431598 | 4431672 | 4425864 | 4426092 | 4434383 | 4434517 | 1 | 0 | 0 | 0 | 0 |
| SE | 12964 | ENSG00000103423 | DNAJA3 | chr16 | + | 4434383 | 4434517 | 4425859 | 4426092 | 4437401 | 4437485 | 90 | 62 | 98 | 77 | 101 |
| SE | 12972 | ENSG00000262468 | LINC01569 | chr16 | - | 4246972 | 4247061 | 4245900 | 4246860 | 4251321 | 4252034 | 6 | 1 | 4 | 9 | 2 |
| SE | 12979 | ENSG00000126602 | TRAP1 | chr16 | - | 3685995 | 3686136 | 3679718 | 3679790 | 3689054 | 3689137 | 207 | 118 | 199 | 219 | 203 |
| SE | 12983 | ENSG00000126602 | TRAP1 | chr16 | - | 3710384 | 3710453 | 3690826 | 3690985 | 3717420 | 3717512 | 0 | 0 | 1 | 0 | 0 |
| SE | 12984 | ENSG00000167984 | NLRC3 | chr16 | - | 3544245 | 3544329 | 3543424 | 3543508 | 3548134 | 3548218 | 138 | 62 | 148 | 134 | 37 |
| SE | 12986 | ENSG00000167984 | NLRC3 | chr16 | - | 3554241 | 3554325 | 3552195 | 3552279 | 3556910 | 3556994 | 61 | 27 | 60 | 57 | 48 |
| SE | 12987 | ENSG00000167984 | NLRC3 | chr16 | - | 3555780 | 3555916 | 3554241 | 3554325 | 3556910 | 3556994 | 4 | 1 | 4 | 8 | 2 |
| SE | 12990 | ENSG00000122390 | NAA60 | chr16 | + | 3447481 | 3447620 | 3443687 | 3443893 | 3448470 | 3448540 | 5 | 4 | 1 | 0 | 2 |
| SE | 12992 | ENSG00000122390 | NAA60 | chr16 | + | 3448470 | 3448540 | 3443677 | 3443827 | 3476221 | 3476337 | 13 | 22 | 29 | 28 | 23 |
| SE | 12997 | ENSG00000122390 | NAA60 | chr16 | + | 3467676 | 3468245 | 3458075 | 3458181 | 3476221 | 3476299 | 0 | 1 | 0 | 1 | 0 |
| SE | 13002 | ENSG00000122390 | NAA60 | chr16 | + | 3476221 | 3476337 | 3457984 | 3458181 | 3479470 | 3479591 | 183 | 96 | 140 | 176 | 85 |
| SE | 13003 | ENSG00000122390 | NAA60 | chr16 | + | 3476221 | 3476337 | 3457984 | 3458181 | 3482501 | 3482598 | 31 | 20 | 20 | 31 | 3 |
| SE | 13004 | ENSG00000122390 | NAA60 | chr16 | + | 3479470 | 3479600 | 3448470 | 3448540 | 3482501 | 3482540 | 103 | 44 | 72 | 100 | 131 |
| SE | 13005 | ENSG00000122390 | NAA60 | chr16 | + | 3479470 | 3479600 | 3457992 | 3458181 | 3482501 | 3482540 | 104 | 44 | 70 | 102 | 131 |
| SE | 13006 | ENSG00000122390 | NAA60 | chr16 | + | 3479470 | 3479600 | 3476221 | 3476337 | 3482501 | 3482540 | 258 | 128 | 191 | 248 | 216 |
| SE | 13007 | ENSG00000122390 | NAA60 | chr16 | + | 3482501 | 3482598 | 3479470 | 3479600 | 3483362 | 3483385 | 183 | 90 | 122 | 144 | 278 |
| SE | 13008 | ENSG00000122390 | NAA60 | chr16 | + | 3482501 | 3482598 | 3479470 | 3479600 | 3484690 | 3485083 | 51 | 28 | 40 | 57 | 96 |
| SE | 13009 | ENSG00000122390 | NAA60 | chr16 | + | 3484690 | 3485083 | 3482501 | 3482598 | 3485466 | 3485578 | 3 | 0 | 3 | 6 | 0 |
| SE | 13010 | ENSG00000122390 | NAA60 | chr16 | + | 3484761 | 3485061 | 3482501 | 3482598 | 3485466 | 3485604 | 12 | 4 | 7 | 5 | 5 |
| SE | 13011 | ENSG00000122390 | NAA60 | chr16 | + | 3484761 | 3485083 | 3482501 | 3482598 | 3485466 | 3485786 | 3 | 0 | 4 | 4 | 0 |
| SE | 13012 | ENSG00000140987 | ZSCAN32 | chr16 | - | 3384458 | 3384941 | 3382080 | 3383711 | 3390009 | 3390133 | 28 | 27 | 33 | 39 | 51 |
| SE | 13019 | ENSG00000140987 | ZSCAN32 | chr16 | - | 3390422 | 3390517 | 3383700 | 3384941 | 3393648 | 3393814 | 8 | 4 | 9 | 6 | 9 |
| SE | 13020 | ENSG00000140987 | ZSCAN32 | chr16 | - | 3390422 | 3390517 | 3383700 | 3384941 | 3397191 | 3397744 | 4 | 0 | 0 | 0 | 3 |
| SE | 13021 | ENSG00000140987 | ZSCAN32 | chr16 | - | 3390422 | 3390517 | 3390009 | 3390133 | 3393648 | 3393814 | 7 | 5 | 18 | 13 | 14 |
| SE | 13022 | ENSG00000140987 | ZSCAN32 | chr16 | - | 3390422 | 3390517 | 3390009 | 3390133 | 3397191 | 3397744 | 2 | 1 | 6 | 2 | 3 |
| SE | 13024 | ENSG00000140987 | ZSCAN32 | chr16 | - | 3390422 | 3390517 | 3390009 | 3390183 | 3397191 | 3397744 | 1 | 0 | 0 | 0 | 3 |
| SE | 13027 | ENSG00000140987 | ZSCAN32 | chr16 | - | 3393648 | 3393814 | 3390009 | 3390133 | 3397191 | 3397744 | 15 | 4 | 9 | 7 | 17 |
| SE | 13030 | ENSG00000140987 | ZSCAN32 | chr16 | - | 3397191 | 3397744 | 3384458 | 3384941 | 3400944 | 3400976 | 29 | 11 | 36 | 14 | 7 |
| SE | 13032 | ENSG00000140993 | TIGD7 | chr16 | - | 3304763 | 3304944 | 3301678 | 3302009 | 3305211 | 3305430 | 1 | 0 | 0 | 0 | 0 |
| SE | 13035 | ENSG00000006194 | ZNF263 | chr16 | + | 3285680 | 3285754 | 3283505 | 3284205 | 3286022 | 3286149 | 19 | 13 | 10 | 12 | 3 |
| SE | 13037 | ENSG00000006194 | ZNF263 | chr16 | + | 3288453 | 3288570 | 3286022 | 3286149 | 3289392 | 3289875 | 48 | 22 | 24 | 21 | 27 |
| SE | 13040 | ENSG00000085644 | ZNF213 | chr16 | + | 3138974 | 3139098 | 3138417 | 3138541 | 3140688 | 3141532 | 9 | 4 | 11 | 8 | 4 |
| SE | 13041 | ENSG00000263072 | ZNF213-AS1 | chr16 | - | 3124486 | 3124898 | 3114967 | 3116519 | 3129247 | 3129653 | 3 | 5 | 1 | 5 | 0 |
| SE | 13042 | ENSG00000263072 | ZNF213-AS1 | chr16 | - | 3129247 | 3129310 | 3114967 | 3116519 | 3129460 | 3129653 | 2 | 0 | 0 | 0 | 0 |
| SE | 13046 | ENSG00000263072 | ZNF213-AS1 | chr16 | - | 3132849 | 3132945 | 3131746 | 3132103 | 3134629 | 3134873 | 8 | 5 | 8 | 10 | 6 |
| SE | 13047 | ENSG00000263072 | ZNF213-AS1 | chr16 | - | 3132849 | 3133194 | 3131746 | 3132103 | 3134629 | 3134863 | 9 | 6 | 11 | 12 | 6 |
| SE | 13056 | ENSG00000131652 | THOC6 | chr16 | + | 3027169 | 3027280 | 3027010 | 3027073 | 3027365 | 3027500 | 83 | 26 | 64 | 43 | 47 |
| SE | 13063 | ENSG00000123815 | COQ8B | chr19 | - | 40703714 | 40703855 | 40703368 | 40703622 | 40705095 | 40705174 | 34 | 5 | 29 | 43 | 4 |
| SE | 13064 | ENSG00000123815 | COQ8B | chr19 | - | 40703714 | 40703855 | 40703368 | 40703622 | 40710058 | 40710136 | 13 | 1 | 12 | 13 | 0 |
| SE | 13066 | ENSG00000123815 | COQ8B | chr19 | - | 40705095 | 40705181 | 40703714 | 40703855 | 40710058 | 40710136 | 13 | 3 | 15 | 22 | 4 |
| SE | 13067 | ENSG00000123815 | COQ8B | chr19 | - | 40705095 | 40705447 | 40703540 | 40703622 | 40710058 | 40710136 | 15 | 7 | 11 | 12 | 2 |

| | | | | | | | | | | | | | | | | | | | | | | | | | | | | | | | | | | | | | | | | | | | | | | |
|---|---|---|---|---|---|---|---|---|---|---|---|---|---|---|---|---|---|---|---|---|---|---|---|---|---|---|---|---|---|---|---|---|---|---|---|---|---|---|---|---|---|---|---|---|---|---|
| SE | 13068 | ENSG00000123815 | COQ8B | chr19 | - | 40705095 | 40705447 | 40703540 | 40703855 | 40710058 | 40710136 | 30 | 10 | 26 | 37 | 6 | 0 | 0 | 0 | 0 | 0 | 14 | 38 | 17 | 30 | 37 | 0 | 0 | 0 | 3 | 0 | 298 | 149 | 1 | 1 | 1 | 1 | 1 | 1 | 1 | 1 | 1 | 1 | 0.833 | 1 | 0.033 |
| SE | 13074 | ENSG00000123815 | COQ8B | chr19 | - | 40714066 | 40714133 | 40707631 | 40710136 | 40714277 | 40714397 | 37 | 16 | 35 | 36 | 15 | 2 | 0 | 0 | 0 | 0 | 25 | 50 | 17 | 18 | 57 | 0 | 0 | 0 | 0 | 0 | 216 | 149 | 1 | 1 | 0.927 | 1 | 1 | 1 | 1 | 1 | 1 | 1 | 1 | 1 | -0.015 |
| SE | 13075 | ENSG00000123815 | COQ8B | chr19 | - | 40714066 | 40714153 | 40710058 | 40710136 | 40714277 | 40714397 | 6 | 3 | 8 | 3 | 0 | 2 | 0 | 0 | 0 | 0 | 9 | 29 | 4 | 5 | 22 | 0 | 0 | 0 | 0 | 0 | 236 | 149 | 1 | 1 | 0.654 | 1 | 1 | 1 | NA | 1 | 1 | 1 | 1 | 1 | -0.087 |
| SE | 13078 | ENSG00000123815 | COQ8B | chr19 | - | 40714530 | 40714635 | 40714277 | 40714397 | 40715056 | 40715078 | 30 | 8 | 18 | 20 | 24 | 0 | 0 | 1 | 0 | 0 | 23 | 41 | 14 | 26 | 56 | 0 | 0 | 0 | 0 | 0 | 254 | 149 | 1 | 1 | 1 | 1 | 0.913 | 1 | 1 | 1 | 1 | 1 | 1 | 1 | -0.017 |
| SE | 13081 | ENSG00000123815 | COQ8B | chr19 | - | 40716586 | 40716703 | 40714530 | 40714635 | 40716861 | 40716882 | 6 | 3 | 2 | 2 | 0 | 1 | 0 | 2 | 0 | 0 | 0 | 3 | 0 | 6 | 0 | 0 | 5 | 0 | 0 | 0 | 266 | 149 | 1 | 1 | 0.771 | 1 | 0.359 | 1 | NA | NA | 0.252 | NA | 1 | NA | 0.156 |
| SE | 13084 | ENSG00000183793 | NPIPA5 | chr16 | - | 15366652 | 15366760 | 15365573 | 15365634 | 15369711 | 15369856 | 3 | 10 | 6 | 1 | 51 | 2 | 0 | 0 | 0 | 0 | 9 | 0 | 0 | 13 | 2 | 0 | 0 | 0 | 0 | 0 | 257 | 149 | 1 | 1 | 0.465 | 1 | 1 | 1 | 1 | 1 | NA | NA | 1 | 1 | -0.107 |
| SE | 13086 | ENSG00000131148 | EMC8 | chr16 | - | 85781210 | 85781280 | 85778623 | 85779867 | 85788973 | 85789050 | 22 | 8 | 19 | 31 | 30 | 2 | 0 | 0 | 0 | 0 | 5 | 26 | 11 | 15 | 45 | 0 | 0 | 0 | 0 | 0 | 219 | 149 | 1 | 1 | 0.882 | 1 | 1 | 1 | 1 | 1 | 1 | 1 | 1 | 1 | -0.024 |
| SE | 13092 | ENSG00000130304 | SLC27A1 | chr19 | + | 17483881 | 17484121 | 17470488 | 17470707 | 17486562 | 17486957 | 0 | 0 | 0 | 0 | 0 | 6 | 0 | 0 | 2 | 0 | 0 | 0 | 0 | 0 | 7 | 0 | 13 | 0 | 2 | 5 | 298 | 149 | 1 | 1 | 0 | NA | NA | 0 | NA | NA | 0 | NA | 0 | 0.412 | -0.137 |
| SE | 13094 | ENSG00000130304 | SLC27A1 | chr19 | + | 17488847 | 17488939 | 17487459 | 17487529 | 17489007 | 17489065 | 64 | 21 | 34 | 29 | 10 | 0 | 0 | 1 | 1 | 0 | 101 | 29 | 19 | 63 | 92 | 0 | 0 | 0 | 0 | 0 | 241 | 149 | 1 | 1 | 1 | 1 | 0.955 | 0.947 | 1 | 1 | 1 | 1 | 1 | 1 | -0.02 |
| SE | 13095 | ENSG00000130304 | SLC27A1 | chr19 | + | 17500711 | 17500876 | 17500494 | 17500632 | 17501272 | 17501419 | 41 | 15 | 26 | 13 | 10 | 0 | 0 | 0 | 0 | 0 | 12 | 35 | 0 | 37 | 33 | 0 | 0 | 1 | 0 | 0 | 298 | 149 | 1 | 1 | 1 | 1 | 1 | 1 | 1 | 1 | 1 | 0 | 1 | 1 | 0.2 |
| SE | 13096 | ENSG00000130304 | SLC27A1 | chr19 | + | 17500711 | 17500876 | 17500494 | 17500632 | 17504454 | 17504900 | 24 | 11 | 17 | 11 | 6 | 0 | 0 | 0 | 0 | 0 | 4 | 23 | 0 | 29 | 21 | 0 | 0 | 0 | 2 | 0 | 298 | 149 | 1 | 1 | 1 | 1 | 1 | 1 | 1 | 1 | 1 | NA | 0.879 | 1 | 0.03 |
| SE | 13097 | ENSG00000130304 | SLC27A1 | chr19 | + | 17501272 | 17501419 | 17500494 | 17500632 | 17504454 | 17504986 | 5 | 5 | 10 | 2 | 0 | 0 | 0 | 0 | 0 | 0 | 8 | 13 | 8 | 0 | 12 | 0 | 0 | 0 | 2 | 0 | 296 | 149 | 1 | 1 | 1 | 1 | 1 | 1 | NA | 1 | 1 | 1 | 0 | 1 | 0.2 |
| SE | 13099 | ENSG00000129993 | CBFA2T3 | chr16 | - | 88880719 | 88880788 | 88879269 | 88879460 | 88881290 | 88881489 | 12 | 6 | 6 | 18 | 2 | 0 | 0 | 0 | 0 | 0 | 0 | 12 | 1 | 26 | 18 | 0 | 0 | 0 | 1 | 0 | 218 | 149 | 1 | 1 | 1 | 1 | 1 | 1 | 1 | NA | 1 | 1 | 0.947 | 1 | 0.013 |
| SE | 13100 | ENSG00000129993 | CBFA2T3 | chr16 | - | 88880987 | 88881148 | 88880719 | 88880788 | 88881290 | 88881313 | 2 | 0 | 0 | 5 | 0 | 7 | 2 | 2 | 11 | 3 | 0 | 3 | 1 | 2 | 6 | 0 | 7 | 0 | 9 | 17 | 298 | 149 | 1 | 1 | 0.125 | 0 | 0 | 0.185 | 0 | NA | 0.176 | 1 | 0.1 | 0.15 | -0.294 |
| SE | 13115 | ENSG00000167264 | DUS2 | chr16 | + | 68054573 | 68054617 | 68053563 | 68053655 | 68056363 | 68056376 | 51 | 61 | 50 | 101 | 51 | 0 | 0 | 0 | 0 | 0 | 133 | 102 | 94 | 42 | 148 | 0 | 0 | 0 | 4 | 0 | 193 | 149 | 1 | 1 | 1 | 1 | 1 | 1 | 1 | 1 | 1 | 1 | 0.89 | 1 | 0.022 |
| SE | 13116 | ENSG00000167264 | DUS2 | chr16 | + | 68056363 | 68056424 | 68053563 | 68053655 | 68061065 | 68061113 | 22 | 16 | 17 | 25 | 34 | 0 | 0 | 0 | 0 | 0 | 38 | 18 | 21 | 48 | 43 | 0 | 2 | 0 | 0 | 0 | 210 | 149 | 1 | 1 | 1 | 1 | 1 | 1 | 1 | 1 | 0.865 | 1 | 1 | 1 | 0.027 |
| SE | 13118 | ENSG00000167264 | DUS2 | chr16 | + | 68066565 | 68066636 | 68066316 | 68066382 | 68070939 | 68071108 | 30 | 17 | 22 | 35 | 31 | 0 | 0 | 0 | 0 | 2 | 92 | 38 | 8 | 42 | 66 | 0 | 0 | 0 | 0 | 0 | 220 | 149 | 1 | 1 | 1 | 1 | 1 | 1 | 0.913 | 1 | 1 | 1 | 1 | 1 | -0.017 |
| SE | 13119 | ENSG00000167264 | DUS2 | chr16 | + | 68070133 | 68070220 | 68066565 | 68066636 | 68070939 | 68071108 | 48 | 30 | 59 | 71 | 81 | 2 | 0 | 0 | 0 | 0 | 63 | 73 | 67 | 62 | 126 | 0 | 0 | 0 | 0 | 0 | 236 | 149 | 1 | 1 | 0.938 | 1 | 1 | 1 | 1 | 1 | 1 | 1 | 1 | 1 | -0.012 |
| SE | 13121 | ENSG00000167264 | DUS2 | chr16 | + | 68075354 | 68075504 | 68074033 | 68074155 | 68078444 | 68078518 | 34 | 18 | 24 | 16 | 18 | 0 | 0 | 0 | 0 | 0 | 35 | 48 | 32 | 6 | 64 | 0 | 0 | 0 | 0 | 3 | 298 | 149 | 1 | 1 | 1 | 1 | 1 | 1 | 1 | 1 | 1 | 1 | 1 | 0.914 | 0.017 |
| SE | 13123 | ENSG00000118900 | UBN1 | chr16 | + | 4852878 | 4853166 | 4846664 | 4848393 | 4857989 | 4858076 | 106 | 73 | 79 | 131 | 86 | 0 | 0 | 0 | 0 | 0 | 220 | 184 | 161 | 262 | 349 | 0 | 0 | 0 | 6 | 0 | 298 | 149 | 1 | 1 | 1 | 1 | 1 | 1 | 1 | 1 | 1 | 1 | 0.956 | 1 | 0.009 |
| SE | 13124 | ENSG00000118900 | UBN1 | chr16 | + | 4852878 | 4853166 | 4847480 | 4848210 | 4857989 | 4858076 | 114 | 76 | 85 | 144 | 81 | 0 | 0 | 2 | 0 | 0 | 256 | 179 | 159 | 273 | 360 | 0 | 0 | 0 | 0 | 4 | 298 | 149 | 1 | 1 | 1 | 1 | 0.955 | 1 | 1 | 1 | 1 | 1 | 1 | 0.978 | -0.005 |
| SE | 13127 | ENSG00000118900 | UBN1 | chr16 | + | 4858877 | 4859159 | 4858573 | 4858663 | 4860663 | 4860953 | 0 | 0 | 0 | 0 | 0 | 0 | 1 | 0 | 0 | 5 | 0 | 3 | 0 | 0 | 7 | 19 | 4 | 14 | 15 | 10 | 298 | 149 | 1 | 1 | NA | 0 | NA | NA | 0 | 0 | 0.273 | 0 | 0 | 0.259 | -0.106 |
| SE | 13129 | ENSG00000118900 | UBN1 | chr16 | + | 4859864 | 4859968 | 4858877 | 4859159 | 4860663 | 4860953 | 400 | 388 | 347 | 425 | 561 | 0 | 0 | 0 | 0 | 0 | 1369 | 665 | 798 | 997 | 1363 | 0 | 3 | 0 | 0 | 7 | 253 | 149 | 1 | 1 | 1 | 1 | 1 | 1 | 1 | 1 | 0.992 | | 1 | 0.991 | 0.003 |
| SE | 13130 | ENSG00000118900 | UBN1 | chr16 | + | 4860663 | 4860953 | 4858573 | 4858663 | 4868832 | 4868903 | 1 | 1 | 0 | 0 | 5 | 0 | 0 | 0 | 0 | 0 | 19 | 4 | 17 | 20 | 10 | 0 | 6 | 2 | 0 | 0 | 298 | 149 | 1 | 1 | 1 | 1 | NA | NA | 1 | 1 | 0.25 | 0.81 | 1 | 1 | 0.188 |
| SE | 13131 | ENSG00000118900 | UBN1 | chr16 | + | 4860663 | 4860953 | 4859864 | 4859968 | 4868832 | 4868903 | 211 | 204 | 183 | 205 | 319 | 0 | 0 | 0 | 0 | 0 | 748 | 385 | 461 | 489 | 770 | 0 | 0 | 1 | 0 | 0 | 298 | 149 | 1 | 1 | 1 | 1 | 1 | 1 | 1 | 1 | 1 | 0.996 | 1 | 1 | 0.001 |
| SE | 13133 | ENSG00000118900 | UBN1 | chr16 | + | 4860663 | 4861102 | 4859864 | 4859968 | 4868832 | 4868903 | 301 | 239 | 230 | 252 | 355 | 0 | 0 | 0 | 0 | 0 | 812 | 455 | 535 | 593 | 973 | 0 | 0 | 1 | 0 | 0 | 298 | 149 | 1 | 1 | 1 | 1 | 1 | 1 | 1 | 1 | 1 | 0.996 | 1 | 1 | 0.001 |
| SE | 13136 | ENSG00000118900 | UBN1 | chr16 | + | 4870211 | 4870341 | 4860744 | 4861102 | 4870515 | 4870572 | 153 | 98 | 109 | 180 | 105 | 0 | 0 | 0 | 0 | 0 | 602 | 289 | 249 | 303 | 416 | 0 | 0 | 0 | 4 | 0 | 279 | 149 | 1 | 1 | 1 | 1 | 1 | 1 | 1 | 1 | 1 | 1 | 0.976 | 1 | 0.005 |
| SE | 13138 | ENSG00000118900 | UBN1 | chr16 | + | 4873030 | 4873073 | 4872883 | 4872934 | 4874210 | 4874968 | 122 | 48 | 106 | 107 | 140 | 0 | 0 | 0 | 0 | 0 | 61 | 112 | 96 | 165 | 183 | 0 | 0 | 0 | 6 | 0 | 192 | 149 | 1 | 1 | 1 | 1 | 1 | 1 | 1 | 1 | 1 | 1 | 0.955 | 1 | 0.009 |
| SE | 13141 | ENSG00000118900 | UBN1 | chr16 | + | 4877022 | 4877111 | 4875271 | 4875434 | 4877384 | 4877474 | 15 | 1 | 7 | 8 | 6 | 0 | 2 | 2 | 0 | 0 | 0 | 13 | 6 | 6 | 5 | 0 | 5 | 2 | 0 | 0 | 238 | 149 | 1 | 1 | 1 | 0.238 | 0.687 | 1 | 1 | NA | 0.619 | 0.653 | 1 | 1 | -0.033 |
| SE | 13142 | ENSG00000118900 | UBN1 | chr16 | + | 4877384 | 4877474 | 4874210 | 4875434 | 4880082 | 4880445 | 6 | 6 | 6 | 10 | 0 | 0 | 0 | 0 | 1 | 0 | 0 | 10 | 6 | 4 | 8 | 0 | 0 | 0 | 5 | 0 | 239 | 149 | 1 | 1 | 1 | 1 | 1 | 0.862 | NA | NA | 1 | 1 | 0.333 | 1 | 0.132 |
| SE | 13143 | ENSG00000118900 | UBN1 | chr16 | + | 4877384 | 4877474 | 4876870 | 4877111 | 4880082 | 4880445 | 29 | 5 | 16 | 28 | 9 | 5 | 1 | 1 | 1 | 6 | 0 | 25 | 9 | 7 | 13 | 0 | 0 | 2 | 1 | 0 | 239 | 149 | 1 | 1 | 0.783 | 0.757 | 0.909 | 0.946 | 0.483 | NA | 1 | 0.737 | 0.814 | 1 | -0.112 |
| SE | 13153 | ENSG00000103121 | CMC2 | chr16 | - | 80981805 | 80981877 | 80980769 | 80980879 | 81006733 | 81006873 | 13 | 8 | 12 | 14 | 17 | 0 | 0 | 0 | 0 | 0 | 10 | 43 | 32 | 24 | 95 | 0 | 0 | 0 | 2 | 0 | 221 | 149 | 1 | 1 | 1 | 1 | 1 | 1 | 1 | 1 | 1 | 1 | 0.89 | 1 | 0.022 |
| SE | 13154 | ENSG00000103121 | CMC2 | chr16 | - | 80988539 | 80988596 | 80981805 | 80981877 | 80991753 | 80991957 | 3 | 0 | 1 | 0 | 0 | 2 | 0 | 0 | 1 | 0 | 0 | 5 | 0 | 1 | 3 | 4 | 1 | 1 | 0 | 0 | 206 | 149 | 1 | 1 | 0.52 | NA | 1 | 0 | NA | 0 | 0.783 | 0 | 1 | 1 | -0.05 |
| SE | 13155 | ENSG00000103121 | CMC2 | chr16 | - | 80988539 | 80988596 | 80981805 | 80981877 | 80997313 | 80997429 | 6 | 0 | 1 | 0 | 0 | 87 | 60 | 59 | 104 | 65 | 0 | 5 | 0 | 0 | 6 | 122 | 142 | 76 | 179 | 239 | 206 | 149 | 1 | 1 | 0.048 | 0 | 0.012 | 0 | 0 | 0 | 0.025 | 0 | 0 | 0.018 | 0.003 |
| SE | 13156 | ENSG00000103121 | CMC2 | chr16 | - | 80991753 | 80991957 | 80981805 | 80981877 | 80996922 | 80997020 | 2 | 0 | 0 | 1 | 0 | 0 | 0 | 0 | 1 | 0 | 8 | 1 | 1 | 1 | 0 | 2 | 0 | 0 | 6 | 2 | 298 | 149 | 1 | 1 | 1 | NA | NA | 0.333 | NA | 0.667 | 1 | 1 | 0.077 | 0 | 0.118 |
| SE | 13158 | ENSG00000103121 | CMC2 | chr16 | - | 80991753 | 80991957 | 80988539 | 80988596 | 80997313 | 80997429 | 0 | 0 | 0 | 1 | 0 | 3 | 0 | 0 | 1 | 0 | 16 | 3 | 0 | 1 | 0 | 0 | 0 | 0 | 1 | 3 | 298 | 149 | 1 | 1 | 0 | NA | NA | 0.333 | NA | 1 | 1 | NA | 0.333 | 0 | -0.417 |
| SE | 13160 | ENSG00000103121 | CMC2 | chr16 | - | 80996922 | 80997020 | 80991753 | 80991957 | 80997313 | 80997429 | 1 | 0 | 0 | 1 | 4 | 0 | 0 | 0 | 1 | 0 | 4 | 0 | 6 | 2 | 0 | 16 | 3 | 0 | 0 | 0 | 247 | 149 | 1 | 1 | 1 | NA | NA | 0.376 | 1 | 0.131 | 0 | 1 | 1 | NA | 0.259 |
| SE | 13161 | ENSG00000103121 | CMC2 | chr16 | - | 80997313 | 80997429 | 80976049 | 80976179 | 81006733 | 81006854 | 38 | 12 | 20 | 41 | 1 | 0 | 0 | 0 | 0 | 0 | 40 | 64 | 28 | 100 | 54 | 4 | 0 | 0 | 0 | 0 | 265 | 149 | 1 | 1 | 1 | 1 | 1 | 1 | 1 | 0.849 | 1 | 1 | 1 | 1 | 0.03 |
| SE | 13162 | ENSG00000103121 | CMC2 | chr16 | - | 80997313 | 80997429 | 80980769 | 80980879 | 81006733 | 81006854 | 34 | 13 | 23 | 36 | 1 | 0 | 0 | 0 | 0 | 0 | 38 | 71 | 26 | 82 | 51 | 0 | 0 | 0 | 2 | 0 | 265 | 149 | 1 | 1 | 1 | 1 | 1 | 1 | 1 | 1 | 1 | 1 | 0.958 | 1 | 0.008 |
| SE | 13164 | ENSG00000103121 | CMC2 | chr16 | - | 80998284 | 80998351 | 80997313 | 80997429 | 81001247 | 81001334 | 23 | 12 | 0 | 12 | 12 | 3 | 1 | 1 | 1 | 9 | 0 | 16 | 8 | 14 | 38 | 0 | 8 | 1 | 3 | 3 | 216 | 149 | 1 | 1 | 0.841 | 0.892 | 0 | 0.892 | 0.479 | NA | 0.58 | 0.847 | 0.763 | 0.897 | -0.151 |
| SE | 13166 | ENSG00000103121 | CMC2 | chr16 | - | 80998284 | 80998354 | 80997313 | 80997429 | 81001247 | 81001334 | 21 | 12 | 0 | 11 | 12 | 3 | 1 | 1 | 1 | 9 | 0 | 16 | 8 | 11 | 39 | 0 | 8 | 1 | 3 | 3 | 219 | 149 | 1 | 1 | 0.826 | 0.891 | 0 | 0.882 | 0.476 | NA | 0.576 | 0.845 | 0.714 | 0.898 | -0.143 |
| SE | 13170 | ENSG00000103121 | CMC2 | chr16 | - | 81001247 | 81001334 | 80998284 | 80998351 | 81006733 | 81006858 | 2 | 0 | 2 | 3 | 0 | 2 | 0 | 1 | 0 | 0 | 0 | 1 | 0 | 5 | 0 | 0 | 4 | 0 | 2 | 0 | 236 | 149 | 1 | 1 | 0.387 | NA | 0.558 | 1 | NA | NA | 0.136 | NA | 0.612 | NA | 0.274 |
| SE | 13171 | ENSG00000103121 | CMC2 | chr16 | - | 81001247 | 81001334 | 80998284 | 80998354 | 81006733 | 81006858 | 0 | 0 | 2 | 2 | 0 | 0 | 0 | 4 | 0 | 0 | 0 | 1 | 0 | 0 | 1 | 0 | 0 | 0 | 0 | 0 | 236 | 149 | 1 | 1 | NA | NA | 0.24 | 1 | NA | NA | 1 | NA | NA | 1 | -0.38 |
| SE | 13175 | ENSG00000140718 | FTO | chr16 | + | 53810139 | 53810217 | 53703962 | 53704229 | 53825863 | 53826491 | 21 | 16 | 29 | 33 | 47 | 0 | 0 | 0 | 0 | 0 | 14 | 28 | 36 | 34 | 18 | 0 | 2 | 0 | 0 | 0 | 227 | 149 | 1 | 1 | 1 | 1 | 1 | 1 | 1 | 1 | 0.902 | 1 | 1 | 1 | 0.02 |
| SE | 13177 | ENSG00000140718 | FTO | chr16 | + | 53825863 | 53826491 | 53810139 | 53810217 | 53844154 | 53844298 | 90 | 65 | 106 | 115 | 131 | 0 | 0 | 0 | 0 | 0 | 97 | 114 | 79 | 60 | 51 | 0 | 0 | 0 | 0 | 1 | 298 | 149 | 1 | 1 | 1 | 1 | 1 | 1 | 1 | 1 | 1 | 1 | 1 | 0.962 | 0.008 |
| SE | 13178 | ENSG00000140718 | FTO | chr16 | + | 53844154 | 53844298 | 53825863 | 53826491 | 53873785 | 53873865 | 158 | 102 | 176 | 163 | 205 | 0 | 0 | 0 | 0 | 1 | 257 | 204 | 136 | 95 | 97 | 0 | 0 | 1 | 0 | 0 | 293 | 149 | 1 | 1 | 1 | 1 | 1 | 1 | 0.99 | 1 | 1 | 0.986 | 1 | 1 | 0.001 |
| SE | 13179 | ENSG00000140718 | FTO | chr16 | + | 53873785 | 53873865 | 53844154 | 53844298 | 53879843 | 53879987 | 219 | 151 | 210 | 215 | 254 | 0 | 0 | 0 | 1 | 0 | 329 | 257 | 200 | 208 | 244 | 0 | 0 | 0 | 0 | 0 | 229 | 149 | 1 | 1 | 1 | 1 | 1 | 0.993 | 1 | 1 | 1 | 1 | 1 | 1 | -0.001 |
| SE | 13180 | ENSG00000140718 | FTO | chr16 | + | 53879843 | 53879987 | 53873785 | 53873865 | 53888831 | 53888951 | 347 | 259 | 257 | 288 | 363 | 0 | 0 | 4 | 0 | 2 | 431 | 325 | 269 | 254 | 386 | 0 | 0 | 0 | 0 | 0 | 293 | 149 | 1 | 1 | 1 | 1 | 0.97 | 1 | 0.989 | 1 | 1 | 1 | 1 | 1 | -0.008 |
| SE | 13181 | ENSG00000140718 | FTO | chr16 | + | 53888831 | 53888951 | 53879843 | 53879987 | 53933984 | 53934109 | 274 | 217 | 198 | 235 | 309 | 0 | 0 | 0 | 0 | 0 | 288 | 280 | 231 | 142 | 329 | 0 | 0 | 4 | 0 | 0 | 269 | 149 | 1 | 1 | 1 | 1 | 1 | 1 | 1 | 1 | 1 | 0.97 | 1 | 1 | 0.006 |
| SE | 13182 | ENSG00000140718 | FTO | chr16 | + | 53956682 | 53956789 | 53933984 | 53934109 | 53965720 | 53965858 | 0 | 0 | 0 | 0 | 9 | 4 | 2 | 0 | 0 | 0 | 0 | 0 | 0 | 1 | 0 | 0 | 3 | 1 | 0 | 0 | 256 | 149 | 1 | 1 | 0 | 0 | NA | NA | 1 | NA | 0 | 0 | 1 | NA | 0 |
| SE | 13183 | ENSG00000140718 | FTO | chr16 | + | 53984937 | 53984954 | 53934029 | 53934109 | 54111761 | 54114463 | 2 | 0 | 0 | 0 | 0 | 61 | 40 | 72 | 69 | 102 | 0 | 0 | 0 | 0 | 0 | 33 | 75 | 37 | 20 | 42 | 166 | 149 | 1 | 1 | 0.029 | 0 | 0 | 0 | 0 | 0 | 0 | 0 | 0 | 0 | 0.006 |
| SE | 13191 | ENSG00000103035 | PSMD7 | chr16 | + | 74304302 | 74304394 | 74302211 | 74302292 | 74305288 | 74305888 | 537 | 401 | 569 | 584 | 1211 | 1 | 0 | 0 | 1 | 0 | 1221 | 717 | 759 | 916 | 577 | 0 | 1 | 0 | 12 | 0 | 241 | 149 | 1 | 1 | 0.997 | 1 | 1 | 0.997 | 1 | 1 | 0.998 | 1 | 0.979 | 1 | 0.003 |
| SE | 13193 | ENSG00000103005 | USB1 | chr16 | + | 58014272 | 58014294 | 58009928 | 58010112 | 58017333 | 58017439 | 3 | 0 | 4 | 2 | 6 | 2 | 0 | 4 | 2 | 1 | 0 | 0 | 0 | 6 | 0 | 0 | 3 | 5 | 3 | 6 | 171 | 149 | 1 | 1 | 0.567 | NA | 0.466 | 0.466 | 0.839 | NA | 0 | 0 | 0.635 | 0 | 0.426 |
| SE | 13195 | ENSG00000103005 | USB1 | chr16 | + | 58017333 | 58017439 | 58013628 | 58014326 | 58018971 | 58019036 | 78 | 48 | 52 | 57 | 74 | 0 | 0 | 0 | 0 | 2 | 125 | 61 | 47 | 93 | 35 | 0 | 1 | 0 | 0 | 0 | 255 | 149 | 1 | 1 | 1 | 1 | 1 | 1 | 0.956 | 1 | 0.973 | 1 | 1 | 1 | -0.003 |
| SE | 13196 | ENSG00000103005 | USB1 | chr16 | + | 58018971 | 58019055 | 58015112 | 58017439 | 58020140 | 58020542 | 61 | 33 | 47 | 48 | 54 | 0 | 0 | 0 | 0 | 0 | 64 | 69 | 59 | 102 | 61 | 0 | 0 | 0 | 1 | 0 | 233 | 149 | 1 | 1 | 1 | 1 | 1 | 1 | 1 | 1 | 1 | 1 | 0.985 | 1 | 0.003 |
| SE | 13198 | ENSG00000140836 | ZFHX3 | chr16 | - | 72957426 | 72960194 | 72950468 | 72950965 | 73058529 | 73059012 | 24 | 18 | 24 | 33 | 5 | 0 | 0 | 0 | 0 | 0 | 50 | 11 | 10 | 9 | 27 | 0 | 0 | 1 | 0 | 0 | 298 | 149 | 1 | 1 | 1 | 1 | 1 | 1 | 1 | 1 | 1 | 0.833 | 1 | 1 | 0.033 |
| SE | 13201 | ENSG00000184517 | ZFP1 | chr16 | + | 75166769 | 75166896 | 75152908 | 75152966 | 75169252 | 75169402 | 20 | 15 | 12 | 21 | 24 | 0 | 0 | 1 | 2 | 0 | 8 | 23 | 9 | 16 | 0 | 0 | 0 | 5 | 0 | 0 | 276 | 149 | 1 | 1 | 1 | 1 | 0.866 | 0.85 | 1 | 1 | 1 | 0.493 | 1 | NA | 0.07 |
| SE | 13202 | ENSG00000184517 | ZFP1 | chr16 | + | 75166769 | 75166913 | 75152908 | 75152966 | 75169252 | 75169326 | 8 | 14 | 7 | 7 | 11 | 0 | 0 | 1 | 2 | 0 | 4 | 10 | 5 | 7 | 0 | 0 | 0 | 5 | 0 | 0 | 293 | 149 | 1 | 1 | 1 | 1 | 0.781 | 0.64 | 1 | 1 | 1 | 0.337 | 1 | NA | 0.05 |
| SE | 13203 | ENSG00000259768 | NA | chr16 | + | 72713332 | 72713416 | 72665122 | 72665333 | 72745404 | 72745466 | 1 | 2 | 3 | 0 | 0 | 0 | 1 | 0 | 0 | 0 | 6 | 0 | 6 | 2 | 0 | 0 | 0 | 0 | 0 | 0 | 233 | 149 | 1 | 1 | 1 | 0.561 | 1 | NA | NA | 1 | NA | 1 | 1 | NA | -0.146 |
| SE | 13204 | ENSG00000259768 | NA | chr16 | + | 72713946 | 72713989 | 72713332 | 72713416 | 72745404 | 72745466 | 0 | 0 | 1 | 1 | 2 | 0 | 1 | 2 | 0 | 0 | 6 | 3 | 9 | 7 | 8 | 0 | 0 | 0 | 2 | 0 | 192 | 149 | 1 | 1 | NA | 0 | 0.28 | 1 | 1 | 1 | 1 | 1 | 0.731 | 1 | -0.376 |
| SE | 13205 | ENSG00000259768 | NA | chr16 | + | 72745404 | 72745466 | 72713946 | 72713989 | 72781727 | 72781819 | 0 | 0 | 1 | 1 | 0 | 0 | 0 | 0 | 0 | 0 | 0 | 3 | 6 | 9 | 9 | 0 | 0 | 0 | 0 | 2 | 211 | 149 | 1 | 1 | NA | NA | 1 | 1 | NA | NA | 1 | 1 | 1 | 0.761 | 0.06 |
| SE | 13206 | ENSG00000259768 | NA | chr16 | + | 72747120 | 72747182 | 72745404 | 72745466 | 72781727 | 72781819 | 0 | 1 | 0 | 1 | 0 | 0 | 0 | 2 | 0 | 0 | 0 | 0 | 0 | 0 | 0 | 0 | 0 | 6 | 9 | 9 | 211 | 149 | 1 | 1 | NA | 1 | 0 | 1 | NA | NA | NA | 0 | 0 | 0 | 0.667 |
| SE | 13207 | ENSG00000259768 | NA | chr16 | + | 72781727 | 72781819 | 72745404 | 72745466 | 72787162 | 72791624 | 0 | 0 | 3 | 1 | 0 | 0 | 0 | 0 | 0 | 0 | 0 | 0 | 6 | 7 | 0 | 0 | 0 | 3 | 0 | 0 | 241 | 149 | 1 | 1 | NA | NA | 1 | 1 | NA | NA | NA | 0.553 | 1 | NA | 0.224 |
| SE | 13213 | ENSG00000103226 | NOMO3 | chr16 | + | 16251962 | 16252100 | 16250927 | 16251080 | 16252432 | 16252522 | 7 | 5 | 7 | 16 | 12 | 0 | 1 | 0 | 0 | 1 | 33 | 19 | 8 | 6 | 46 | 0 | 1 | 0 | 0 | 0 | 287 | 149 | 1 | 1 | 1 | 0.722 | 1 | 1 | 0.862 | 1 | 0.908 | 1 | 1 | 1 | -0.065 |
| SE | 13215 | ENSG00000103226 | NOMO3 | chr16 | + | 16261501 | 16261676 | 16256007 | 16256158 | 16263073 | 16263215 | 158 | 92 | 116 | 249 | 148 | 0 | 0 | 0 | 0 | 0 | 229 | 173 | 209 | 144 | 279 | 0 | 0 | 0 | 3 | 0 | 298 | 149 | 1 | 1 | 1 | 1 | 1 | 1 | 1 | 1 | 1 | 1 | 0.96 | 1 | 0.008 |
| SE | 13220 | ENSG00000181019 | NQO1 | chr16 | - | 69713027 | 69713129 | 69709400 | 69711281 | 69714963 | 69715077 | 11 | 2 | 8 | 4 | 5 | 2 | 0 | 0 | 0 | 0 | 26 | 4 | 4 | 21 | 0 | 0 | 0 | 0 | 0 | 0 | 251 | 149 | 1 | 1 | 0.766 | 1 | 1 | 1 | 1 | 1 | 1 | 1 | 1 | NA | -0.047 |
| SE | 13226 | ENSG00000140876 | NUDT7 | chr16 | + | 77725430 | 77725584 | 77722491 | 77722617 | 77741581 | 77742258 | 22 | 9 | 17 | 27 | 18 | 0 | 0 | 2 | 0 | 3 | 0 | 7 | 17 | 0 | 2 | 0 | 0 | 0 | 0 | 0 | 298 | 149 | 1 | 1 | 1 | 1 | 0.81 | 1 | 0.75 | NA | 1 | 1 | NA | 1 | -0.088 |
| SE | 13228 | ENSG00000140876 | NUDT7 | chr16 | + | 77735827 | 77735986 | 77725430 | 77725584 | 77736649 | 77736743 | 8 | 2 | 7 | 7 | 18 | 1 | 0 | 0 | 0 | 5 | 0 | 0 | 4 | 5 | 0 | 0 | 0 | 0 | 4 | 0 | 298 | 149 | 1 | 1 | 0.8 | 1 | 1 | 1 | 0.643 | NA | NA | 1 | 0.385 | NA | 0.196 |
| SE | 13229 | ENSG00000140876 | NUDT7 | chr16 | + | 77735827 | 77735986 | 77725430 | 77725584 | 77741581 | 77742260 | 10 | 1 | 6 | 9 | 20 | 9 | 3 | 10 | 9 | 10 | 0 | 0 | 12 | 5 | 0 | 0 | 2 | 5 | 0 | 0 | 298 | 149 | 1 | 1 | 0.357 | 0.143 | 0.231 | 0.333 | 0.5 | NA | 0 | 0.545 | 1 | NA | -0.202 |
| SE | 13230 | ENSG00000140876 | NUDT7 | chr16 | + | 77736649 | 77736743 | 77725430 | 77725584 | 77741581 | 77741969 | 4 | 0 | 0 | 0 | 6 | 9 | 3 | 10 | 9 | 10 | 0 | 0 | 0 | 4 | 0 | 0 | 2 | 5 | 0 | 0 | 243 | 149 | 1 | 1 | 0.214 | 0 | 0 | 0 | 0.269 | NA | 0 | 0 | 1 | NA | -0.237 |
| SE | 13231 | ENSG00000140876 | NUDT7 | chr16 | + | 77736649 | 77736743 | 77735827 | 77735986 | 77741581 | 77741969 | 5 | 1 | 3 | 1 | 4 | 4 | 0 | 2 | 3 | 5 | 0 | 0 | 0 | 0 | 0 | 0 | 0 | 8 | 0 | 0 | 243 | 149 | 1 | 1 | 0.434 | 1 | 0.479 | 0.17 | 0.329 | NA | NA | 0 | NA | NA | 0.482 |
| SE | 13233 | ENSG00000153774 | CFDP1 | chr16 | - | 75301756 | 75301905 | 75293907 | 75294042 | 75305023 | 75305438 | 0 | 0 | 4 | 0 | 0 | 21 | 18 | 21 | 21 | 29 | 0 | 0 | 0 | 0 | 0 | 16 | 7 | 9 | 27 | 14 | 298 | 149 | 1 | 1 | 0 | 0 | 0.087 | 0 | 0 | 0 | 0 | 0 | 0 | 0 | 0.017 |
| SE | 13234 | ENSG00000153774 | CFDP1 | chr16 | - | 75362874 | 75363035 | 75305023 | 75305182 | 75391379 | 75391508 | 1 | 0 | 1 | 1 | 0 | 2 | 0 | 4 | 3 | 0 | 0 | 0 | 0 | 0 | 0 | 0 | 2 | 3 | 2 | 0 | 298 | 149 | 1 | 1 | 0.2 | NA | 0.111 | 0.143 | NA | NA | 0 | 0 | 0 | NA | 0.151 |
| SE | 13235 | ENSG00000153774 | CFDP1 | chr16 | - | 75362874 | 75363035 | 75305023 | 75305182 | 75395089 | 75395211 | 0 | 0 | 1 | 1 | 0 | 100 | 99 | 85 | 99 | 126 | 0 | 1 | 0 | 0 | 0 | 36 | 23 | 67 | 77 | 57 | 298 | 149 | 1 | 1 | 0 | 0 | 0.006 | 0.005 | 0 | 0 | 0.021 | 0 | 0 | 0 | -0.002 |
| SE | 13237 | ENSG00000153774 | CFDP1 | chr16 | - | 75391379 | 75391508 | 75362874 | 75363035 | 75395089 | 75395136 | 8 | 1 | 3 | 8 | 8 | 0 | 0 | 0 | 0 | 0 | 31 | 12 | 7 | 2 | 0 | 0 | 1 | 0 | 0 | 0 | 278 | 149 | 1 | 1 | 1 | 1 | 1 | 1 | 1 | 1 | 0.865 | 1 | 1 | NA | 0.034 |
| SE | 13239 | ENSG00000153774 | CFDP1 | chr16 | - | 75411824 | 75411952 | 75394681 | 75395209 | 75412534 | 75412754 | 163 | 164 | 143 | 155 | 222 | 0 | 0 | 1 | 0 | 2 | 130 | 96 | 123 | 197 | 52 | 0 | 2 | 0 | 0 | 0 | 277 | 149 | 1 | 1 | 1 | 1 | 0.987 | 1 | 0.984 | 1 | 0.963 | 1 | 1 | 1 | 0.002 |
| SE | 13275 | ENSG00000103335 | PIEZO1 | chr16 | - | 88716805 | 88716898 | 88716558 | 88716731 | 88717022 | 88717211 | 210 | 94 | 133 | 165 | 223 | 0 | 1 | 0 | 0 | 0 | 232 | 202 | 150 | 181 | 433 | 0 | 0 | 0 | 0 | 0 | 242 | 149 | 1 | 1 | 1 | 0.983 | 1 | 1 | 1 | 1 | 1 | 1 | 1 | 1 | -0.003 |
| SE | 13279 | ENSG00000141012 | GALNS | chr16 | - | 88818006 | 88818124 | 88813733 | 88814525 | 88822588 | 88822710 | 34 | 3 | 28 | 37 | 12 | 0 | 0 | 0 | 0 | 0 | 61 | 70 | 13 | 39 | 70 | 0 | 1 | 0 | 2 | 0 | 267 | 149 | 1 | 1 | 1 | 1 | 1 | 1 | 1 | 1 | 0.975 | 1 | 0.916 | 1 | 0.022 |
| SE | 13282 | ENSG00000141012 | GALNS | chr16 | - | 88843029 | 88843199 | 88842705 | 88842829 | 88856757 | 88856826 | 2 | 1 | 1 | 0 | 2 | 2 | 0 | 0 | 4 | 1 | 0 | 4 | 2 | 4 | 0 | 0 | 2 | 4 | 4 | 15 | 298 | 149 | 1 | 1 | 0.333 | 1 | 1 | 0 | 0.5 | NA | 0.5 | 0.2 | 0.333 | 0 | 0.308 |
| SE | 13283 | ENSG00000051523 | CYBA | chr16 | - | 88646519 | 88646603 | 88646131 | 88646197 | 88646754 | 88646838 | 4 | 0 | 2 | 5 | 1 | 966 | 318 | 481 | 600 | 485 | 8 | 6 | 1 | 6 | 0 | 743 | 721 | 391 | 688 | 1017 | 233 | 149 | 1 | 1 | 0.003 | 0 | 0.003 | 0.005 | 0.001 | 0.007 | 0.005 | 0.002 | 0.006 | 0 | -0.002 |
| SE | 13284 | ENSG00000051523 | CYBA | chr16 | - | 88646754 | 88646838 | 88645086 | 88646197 | 88647100 | 88647175 | 1380 | 446 | 688 | 849 | 588 | 4 | 0 | 0 | 0 | 0 | 958 | 1078 | 565 | 941 | 1472 | 0 | 0 | 0 | 1 | 0 | 233 | 149 | 1 | 1 | 0.995 | 1 | 1 | 1 | 1 | 1 | 1 | 1 | 0.998 | 1 | -0.001 |
| SE | 13285 | ENSG00000237440 | ZNF737 | chr19 | - | 20553708 | 20553835 | 20552474 | 20552570 | 20565637 | 20565781 | 53 | 27 | 52 | 30 | 16 | 0 | 0 | 5 | 0 | 0 | 0 | 19 | 20 | 11 | 1 | 0 | 0 | 1 | 0 | 0 | 276 | 149 | 1 | 1 | 1 | 1 | 0.849 | 1 | 1 | NA | 1 | 0.915 | 1 | 1 | -0.009 |
| SE | 13292 | ENSG00000102974 | CTCF | chr16 | + | 67611950 | 67612121 | 67610589 | 67611613 | 67616744 | 67616878 | 674 | 459 | 628 | 579 | 1287 | 0 | 0 | 0 | 0 | 2 | 387 | 456 | 316 | 395 | 233 | 0 | 0 | 0 | 2 | 0 | 298 | 149 | 1 | 1 | 1 | 1 | 1 | 1 | 0.997 | 1 | 1 | 1 | 0.99 | 1 | 0.001 |
| SE | 13293 | ENSG00000102974 | CTCF | chr16 | + | 67616744 | 67616878 | 67611950 | 67612121 | 67620696 | 67620817 | 613 | 347 | 529 | 545 | 844 | 0 | 0 | 0 | 0 | 0 | 312 | 496 | 311 | 307 | 153 | 0 | 0 | 0 | 2 | 0 | 283 | 149 | 1 | 1 | 1 | 1 | 1 | 1 | 1 | 1 | 1 | 1 | 0.988 | 1 | 0.002 |
| SE | 13294 | ENSG00000102974 | CTCF | chr16 | + | 67620696 | 67620817 | 67614962 | 67616878 | 67621441 | 67621591 | 474 | 228 | 419 | 435 | 512 | 0 | 0 | 0 | 2 | 0 | 205 | 398 | 245 | 263 | 94 | 0 | 2 | 0 | 2 | 0 | 270 | 149 | 1 | 1 | 1 | 1 | 1 | 0.992 | 1 | 1 | 0.991 | 1 | 0.986 | 1 | 0.003 |
| SE | 13298 | ENSG00000102974 | CTCF | chr16 | + | 67621441 | 67621591 | 67620696 | 67620817 | 67626554 | 67626715 | 385 | 205 | 367 | 360 | 504 | 0 | 0 | 0 | 1 | 0 | 166 | 266 | 214 | 276 | 92 | 0 | 0 | 0 | 1 | 0 | 298 | 149 | 1 | 1 | 1 | 1 | 1 | 0.994 | 1 | 1 | 1 | 1 | 0.993 | 1 | 0 |
| SE | 13300 | ENSG00000102974 | CTCF | chr16 | + | 67628369 | 67628552 | 67626508 | 67626715 | 67629397 | 67629533 | 298 | 168 | 254 | 294 | 398 | 0 | 0 | 1 | 0 | 0 | 210 | 260 | 273 | 322 | 176 | 0 | 0 | 2 | 0 | 0 | 298 | 149 | 1 | 1 | 1 | 1 | 0.992 | 1 | 1 | 1 | 1 | 0.986 | 1 | 1 | 0.001 |
| SE | 13301 | ENSG00000102974 | CTCF | chr16 | + | 67635514 | 67635632 | 67629397 | 67629533 | 67636689 | 67636851 | 0 | 0 | 0 | 0 | 3 | 219 | 161 | 208 | 192 | 417 | 0 | 0 | 2 | 0 | 0 | 237 | 181 | 212 | 155 | 159 | 267 | 149 | 1 | 1 | 0 | 0 | 0 | 0 | 0.004 | 0 | 0 | 0.005 | 0 | 0 | 0 |
| SE | 13302 | ENSG00000103264 | FBXO31 | chr16 | - | 87358410 | 87358497 | 87347173 | 87347250 | 87360294 | 87360366 | 2 | 0 | 2 | 2 | 0 | 33 | 3 | 23 | 21 | 9 | 0 | 5 | 5 | 4 | 5 | 0 | 17 | 21 | 10 | 24 | 236 | 149 | 1 | 1 | 0.037 | 0 | 0.052 | 0.057 | 0 | NA | 0.157 | 0.131 | 0.202 | 0.116 | -0.122 |
| SE | 13304 | ENSG00000158805 | ZNF276 | chr16 | + | 89723136 | 89723183 | 89722530 | 89722834 | 89723259 | 89723709 | 45 | 21 | 64 | 51 | 9 | 2 | 0 | 1 | 0 | 0 | 0 | 22 | 10 | 13 | 8 | 0 | 0 | 6 | 0 | 0 | 196 | 149 | 1 | 1 | 0.945 | 1 | 0.98 | 1 | 1 | NA | 1 | 0.559 | 1 | 1 | 0.095 |
| SE | 13305 | ENSG00000158805 | ZNF276 | chr16 | + | 89723136 | 89723709 | 89722530 | 89722834 | 89727278 | 89727357 | 67 | 41 | 74 | 72 | 32 | 0 | 0 | 2 | 0 | 0 | 35 | 100 | 27 | 55 | 90 | 0 | 0 | 0 | 0 | 0 | 298 | 149 | 1 | 1 | 1 | 1 | 0.949 | 1 | 1 | 1 | 1 | 1 | 1 | 1 | -0.01 |
| SE | 13306 | ENSG00000158805 | ZNF276 | chr16 | + | 89723259 | 89723709 | 89722530 | 89722834 | 89727278 | 89727325 | 58 | 35 | 66 | 63 | 27 | 0 | 0 | 2 | 0 | 0 | 35 | 100 | 32 | 54 | 81 | 0 | 0 | 0 | 0 | 0 | 298 | 149 | 1 | 1 | 1 | 1 | 0.943 | 1 | 1 | 1 | 1 | 1 | 1 | 1 | -0.011 |
| SE | 13313 | ENSG00000103257 | SLC7A5 | chr16 | - | 87851723 | 87851849 | 87841049 | 87841155 | 87868884 | 87869507 | 7 | 7 | 12 | 21 | 9 | 0 | 0 | 0 | 0 | 0 | 8 | 28 | 77 | 18 | 16 | 0 | 0 | 3 | 0 | 0 | 275 | 149 | 1 | 1 | 1 | 1 | 1 | 1 | 1 | 1 | 1 | 0.933 | 1 | 1 | 0.013 |
| SE | 13316 | ENSG00000039523 | RIPOR1 | chr16 | + | 67529636 | 67529859 | 67528880 | 67528914 | 67538423 | 67538550 | 0 | 0 | 0 | 0 | 0 | 6 | 1 | 3 | 0 | 0 | 0 | 0 | 2 | 0 | 0 | 0 | 1 | 4 | 1 | 10 | 298 | 149 | 1 | 1 | 0 | 0 | 0 | NA | NA | NA | 0 | 0.2 | 0 | 0 | -0.05 |
| SE | 13317 | ENSG00000039523 | RIPOR1 | chr16 | + | 67538423 | 67538550 | 67528816 | 67528914 | 67538989 | 67539068 | 6 | 1 | 3 | 0 | 0 | 1 | 0 | 0 | 0 | 0 | 0 | 1 | 4 | 1 | 10 | 0 | 0 | 0 | 0 | 0 | 276 | 149 | 1 | 1 | 0.764 | 1 | 1 | NA | NA | NA | 1 | 1 | 1 | 1 | -0.079 |
| SE | 13318 | ENSG00000039523 | RIPOR1 | chr16 | + | 67538423 | 67538550 | 67529790 | 67530087 | 67538989 | 67539068 | 2 | 0 | 0 | 0 | 0 | 0 | 0 | 0 | 0 | 0 | 0 | 0 | 0 | 0 | 0 | 0 | 0 | 4 | 0 | 0 | 276 | 149 | 1 | 1 | 1 | NA | NA | NA | NA | NA | NA | 0 | NA | NA | 1 |
| SE | 13319 | ENSG00000039523 | RIPOR1 | chr16 | + | 67538659 | 67538824 | 67538423 | 67538550 | 67538989 | 67539068 | 43 | 23 | 23 | 30 | 8 | 2 | 0 | 0 | 0 | 0 | 50 | 24 | 37 | 14 | 46 | 0 | 0 | 0 | 0 | 0 | 298 | 149 | 1 | 1 | 0.915 | 1 | 1 | 1 | 1 | 1 | 1 | 1 | 1 | 1 | -0.017 |
| SE | 13320 | ENSG00000039523 | RIPOR1 | chr16 | + | 67538671 | 67538824 | 67538077 | 67538550 | 67538989 | 67538996 | 51 | 30 | 29 | 47 | 8 | 2 | 0 | 0 | 0 | 0 | 67 | 28 | 48 | 19 | 90 | 0 | 0 | 0 | 0 | 0 | 298 | 149 | 1 | 1 | 0.927 | 1 | 1 | 1 | 1 | 1 | 1 | 1 | 1 | 1 | -0.015 |
| SE | 13322 | ENSG00000177663 | IL17RA | chr22 | + | 17097061 | 17097086 | 17084958 | 17085229 | 17097796 | 17097943 | 191 | 138 | 98 | 105 | 212 | 1 | 0 | 0 | 0 | 0 | 222 | 157 | 96 | 232 | 295 | 0 | 0 | 0 | 0 | 5 | 174 | 149 | 1 | 1 | 0.994 | 1 | 1 | 1 | 1 | 1 | 1 | 1 | 1 | 0.981 | 0.003 |
| SE | 13325 | ENSG00000177663 | IL17RA | chr22 | + | 17105590 | 17105602 | 17104725 | 17104810 | 17105852 | 17105954 | 360 | 215 | 222 | 277 | 339 | 3 | 0 | 2 | 0 | 0 | 920 | 472 | 249 | 537 | 948 | 11 | 5 | 0 | 0 | 0 | 161 | 149 | 1 | 1 | 0.991 | 1 | 0.99 | 1 | 1 | 0.987 | 0.989 | 1 | 1 | 1 | 0.001 |
| SE | 13327 | ENSG00000103248 | MTHFSD | chr16 | - | 86546558 | 86546649 | 86542100 | 86542213 | 86548034 | 86548111 | 32 | 17 | 23 | 18 | 34 | 0 | 1 | 0 | 0 | 0 | 18 | 29 | 6 | 6 | 42 | 0 | 1 | 0 | 0 | 0 | 240 | 149 | 1 | 1 | 1 | 0.913 | 1 | 1 | 1 | 1 | 0.947 | 1 | 1 | 1 | -0.007 |
| SE | 13328 | ENSG00000103248 | MTHFSD | chr16 | - | 86546558 | 86546649 | 86542100 | 86542213 | 86548463 | 86548577 | 57 | 34 | 65 | 50 | 82 | 0 | 0 | 0 | 2 | 0 | 73 | 64 | 13 | 15 | 66 | 0 | 0 | 1 | 0 | 0 | 240 | 149 | 1 | 1 | 1 | 1 | 1 | 0.939 | 1 | 1 | 1 | 0.89 | 1 | 1 | 0.01 |
| SE | 13329 | ENSG00000103248 | MTHFSD | chr16 | - | 86548034 | 86548111 | 86542100 | 86542213 | 86548463 | 86548577 | 4 | 5 | 2 | 0 | 3 | 0 | 0 | 0 | 2 | 0 | 0 | 2 | 1 | 0 | 0 | 0 | 0 | 1 | 0 | 0 | 226 | 149 | 1 | 1 | 1 | 1 | 1 | 0 | 1 | NA | 1 | 0.397 | NA | NA | 0.102 |
| SE | 13331 | ENSG00000103248 | MTHFSD | chr16 | - | 86548034 | 86548111 | 86547145 | 86547568 | 86548463 | 86548577 | 4 | 4 | 3 | 0 | 10 | 0 | 0 | 0 | 0 | 3 | 0 | 1 | 1 | 0 | 0 | 0 | 0 | 0 | 0 | 0 | 226 | 149 | 1 | 1 | 1 | 1 | 1 | NA | 0.687 | NA | 1 | 1 | NA | NA | -0.078 |
| SE | 13336 | ENSG00000103248 | MTHFSD | chr16 | - | 86554644 | 86554748 | 86552032 | 86552146 | 86555168 | 86555203 | 19 | 8 | 21 | 10 | 18 | 0 | 0 | 0 | 3 | 0 | 20 | 29 | 5 | 7 | 22 | 0 | 1 | 0 | 0 | 0 | 253 | 149 | 1 | 1 | 1 | 1 | 1 | 0.663 | 1 | 1 | 0.945 | 1 | 1 | 1 | -0.056 |
| SE | 13337 | ENSG00000103248 | MTHFSD | chr16 | - | 86554644 | 86554751 | 86551972 | 86552146 | 86555168 | 86555178 | 19 | 11 | 24 | 15 | 24 | 0 | 0 | 0 | 3 | 0 | 25 | 24 | 6 | 11 | 31 | 0 | 1 | 0 | 0 | 0 | 256 | 149 | 1 | 1 | 1 | 1 | 1 | 0.744 | 1 | 1 | 0.933 | 1 | 1 | 1 | -0.038 |
| SE | 13341 | ENSG00000103091 | WDR59 | chr16 | - | 74889702 | 74889815 | 74888168 | 74888319 | 74892483 | 74892565 | 53 | 40 | 79 | 57 | 71 | 0 | 1 | 2 | 2 | 0 | 69 | 44 | 32 | 57 | 75 | 0 | 0 | 0 | 0 | 0 | 262 | 149 | 1 | 1 | 1 | 0.958 | 0.957 | 0.942 | 1 | 1 | 1 | 1 | 1 | 1 | -0.029 |
| SE | 13346 | ENSG00000103091 | WDR59 | chr16 | - | 74938149 | 74938266 | 74923925 | 74924003 | 74942737 | 74942826 | 78 | 40 | 74 | 76 | 69 | 0 | 1 | 1 | 0 | 0 | 94 | 44 | 41 | 48 | 105 | 0 | 0 | 0 | 0 | 0 | 266 | 149 | 1 | 1 | 1 | 0.957 | 0.976 | 1 | 1 | 1 | 1 | 1 | 1 | 1 | -0.013 |
| SE | 13347 | ENSG00000103091 | WDR59 | chr16 | - | 74938149 | 74938266 | 74923925 | 74924003 | 74948518 | 74948556 | 39 | 20 | 42 | 43 | 28 | 0 | 0 | 0 | 0 | 0 | 39 | 14 | 27 | 14 | 47 | 0 | 0 | 0 | 0 | 2 | 266 | 149 | 1 | 1 | 1 | 1 | 1 | 1 | 1 | 1 | 1 | 1 | 1 | 0.929 | 0.014 |
| SE | 13348 | ENSG00000103091 | WDR59 | chr16 | - | 74942737 | 74942826 | 74923925 | 74924003 | 74948518 | 74948556 | 29 | 20 | 37 | 44 | 45 | 0 | 0 | 0 | 0 | 0 | 21 | 30 | 23 | 22 | 61 | 0 | 0 | 0 | 0 | 2 | 238 | 149 | 1 | 1 | 1 | 1 | 1 | 1 | 1 | 1 | 1 | 1 | 1 | 0.95 | 0.01 |
| SE | 13349 | ENSG00000103091 | WDR59 | chr16 | - | 74942737 | 74942826 | 74938149 | 74938266 | 74948518 | 74948556 | 65 | 39 | 67 | 75 | 86 | 0 | 0 | 1 | 0 | 0 | 76 | 60 | 36 | 53 | 119 | 0 | 0 | 0 | 0 | 0 | 238 | 149 | 1 | 1 | 1 | 1 | 0.977 | 1 | 1 | 1 | 1 | 1 | 1 | 1 | -0.005 |
| SE | 13350 | ENSG00000103091 | WDR59 | chr16 | - | 74942737 | 74942826 | 74938149 | 74938266 | 74951457 | 74951543 | 6 | 7 | 9 | 5 | 15 | 0 | 0 | 0 | 1 | 0 | 29 | 14 | 2 | 12 | 7 | 0 | 0 | 0 | 0 | 0 | 238 | 149 | 1 | 1 | 1 | 1 | 1 | 0.758 | 1 | 1 | 1 | 1 | 1 | 1 | -0.048 |
| SE | 13351 | ENSG00000103091 | WDR59 | chr16 | - | 74948518 | 74948556 | 74942737 | 74942826 | 74949717 | 74949798 | 83 | 45 | 76 | 98 | 124 | 0 | 0 | 1 | 0 | 0 | 47 | 67 | 48 | 57 | 137 | 0 | 0 | 0 | 0 | 0 | 187 | 149 | 1 | 1 | 1 | 1 | 0.984 | 1 | 1 | 1 | 1 | 1 | 1 | 1 | -0.003 |
| SE | 13352 | ENSG00000103091 | WDR59 | chr16 | - | 74949717 | 74949798 | 74942737 | 74942826 | 74951457 | 74951543 | 36 | 29 | 32 | 39 | 38 | 0 | 0 | 0 | 0 | 0 | 32 | 28 | 18 | 28 | 44 | 0 | 0 | 0 | 3 | 0 | 230 | 149 | 1 | 1 | 1 | 1 | 1 | 1 | 1 | 1 | 1 | 1 | 0.858 | 1 | 0.028 |
| SE | 13354 | ENSG00000103091 | WDR59 | chr16 | - | 74959349 | 74959592 | 74956483 | 74956610 | 74965772 | 74965822 | 0 | 0 | 0 | 0 | 0 | 20 | 14 | 27 | 35 | 8 | 0 | 0 | 0 | 5 | 0 | 13 | 31 | 17 | 31 | 39 | 298 | 149 | 1 | 1 | 0 | 0 | 0 | 0 | 0 | 0 | 0 | 0 | 0.075 | 0 | -0.015 |
| SE | 13358 | ENSG00000140950 | MEAK7 | chr16 | - | 84502720 | 84502862 | 84497933 | 84498111 | 84503957 | 84504147 | 4 | 2 | 1 | 1 | 0 | 0 | 0 | 2 | 0 | 0 | 0 | 4 | 0 | 0 | 0 | 0 | 0 | 0 | 0 | 0 | 291 | 149 | 1 | 1 | 1 | 1 | 0.204 | 1 | NA | NA | 1 | NA | NA | NA | -0.199 |
| SE | 13359 | ENSG00000140950 | MEAK7 | chr16 | - | 84502720 | 84502862 | 84497933 | 84498111 | 84504600 | 84504621 | 4 | 2 | 1 | 1 | 0 | 0 | 1 | 0 | 2 | 12 | 0 | 4 | 0 | 0 | 0 | 0 | 0 | 4 | 2 | 6 | 291 | 149 | 1 | 1 | 1 | 0.506 | 1 | 0.204 | 0 | NA | 1 | 0 | 0 | 0 | 0.292 |
| SE | 13360 | ENSG00000140950 | MEAK7 | chr16 | - | 84503957 | 84504147 | 84497933 | 84498111 | 84504600 | 84504647 | 0 | 0 | 4 | 0 | 0 | 0 | 1 | 0 | 2 | 12 | 0 | 0 | 0 | 0 | 0 | 0 | 0 | 4 | 2 | 6 | 298 | 149 | 1 | 1 | NA | 0 | 1 | 0 | 0 | NA | NA | 0 | 0 | 0 | 0.25 |
| SE | 13363 | ENSG00000005194 | CIAPIN1 | chr16 | - | 57431150 | 57431266 | 57430257 | 57430339 | 57432486 | 57432560 | 64 | 54 | 63 | 58 | 160 | 0 | 0 | 1 | 3 | 0 | 71 | 48 | 40 | 60 | 81 | 0 | 0 | 0 | 0 | 0 | 265 | 149 | 1 | 1 | 1 | 1 | 0.973 | 0.916 | 1 | 1 | 1 | 1 | 1 | 1 | -0.022 |
| SE | 13364 | ENSG00000005194 | CIAPIN1 | chr16 | - | 57432486 | 57432560 | 57431150 | 57431266 | 57434043 | 57434212 | 78 | 66 | 97 | 79 | 205 | 2 | 0 | 0 | 0 | 0 | 86 | 75 | 52 | 51 | 94 | 0 | 0 | 0 | 0 | 0 | 223 | 149 | 1 | 1 | 0.963 | 1 | 1 | 1 | 1 | 1 | 1 | 1 | 1 | 1 | -0.007 |
| SE | 13365 | ENSG00000005194 | CIAPIN1 | chr16 | - | 57434043 | 57434212 | 57432486 | 57432560 | 57436655 | 57436732 | 145 | 87 | 146 | 123 | 261 | 0 | 0 | 0 | 0 | 0 | 150 | 125 | 70 | 75 | 153 | 0 | 0 | 1 | 0 | 0 | 298 | 149 | 1 | 1 | 1 | 1 | 1 | 1 | 1 | 1 | 1 | 0.972 | 1 | 1 | 0.006 |
| SE | 13366 | ENSG00000005194 | CIAPIN1 | chr16 | - | 57436655 | 57436732 | 57433909 | 57434212 | 57439181 | 57439295 | 104 | 57 | 90 | 82 | 129 | 0 | 0 | 0 | 0 | 3 | 98 | 125 | 51 | 74 | 113 | 0 | 0 | 2 | 0 | 0 | 226 | 149 | 1 | 1 | 1 | 1 | 1 | 1 | 0.966 | 1 | 1 | 0.944 | 1 | 1 | 0.004 |
| SE | 13371 | ENSG00000074755 | ZZEF1 | chr17 | - | 4016322 | 4016466 | 4014346 | 4014515 | 4017835 | 4017971 | 108 | 58 | 77 | 110 | 56 | 0 | 0 | 0 | 0 | 0 | 183 | 82 | 90 | 95 | 122 | 0 | 0 | 0 | 3 | 0 | 293 | 149 | 1 | 1 | 1 | 1 | 1 | 1 | 1 | 1 | 1 | 1 | 0.942 | 1 | 0.012 |
| SE | 13372 | ENSG00000074755 | ZZEF1 | chr17 | - | 4016632 | 4016774 | 4016322 | 4016466 | 4017370 | 4017504 | 0 | 0 | 0 | 1 | 0 | 105 | 52 | 72 | 81 | 81 | 0 | 0 | 0 | 0 | 0 | 145 | 77 | 76 | 88 | 92 | 291 | 149 | 1 | 1 | 0 | 0 | 0 | 0.006 | 0 | 0 | 0 | 0 | 0 | 0 | 0.001 |
| SE | 13373 | ENSG00000074755 | ZZEF1 | chr17 | - | 4017370 | 4017730 | 4016322 | 4016466 | 4017835 | 4017971 | 212 | 91 | 140 | 160 | 173 | 0 | 0 | 2 | 0 | 0 | 208 | 181 | 111 | 209 | 187 | 0 | 0 | 0 | 0 | 0 | 298 | 149 | 1 | 1 | 1 | 1 | 0.972 | 1 | 1 | 1 | 1 | 1 | 1 | 1 | -0.006 |
| SE | 13374 | ENSG00000074755 | ZZEF1 | chr17 | - | 4021128 | 4021320 | 4019668 | 4019769 | 4022708 | 4022828 | 231 | 103 | 231 | 223 | 201 | 0 | 0 | 0 | 0 | 0 | 242 | 321 | 193 | 204 | 296 | 7 | 0 | 0 | 0 | 0 | 298 | 149 | 1 | 1 | 1 | 1 | 1 | 1 | 1 | 0.945 | 1 | 1 | 1 | 1 | 0.011 |
| SE | 13375 | ENSG00000074755 | ZZEF1 | chr17 | - | 4022708 | 4022828 | 4019668 | 4019769 | 4024918 | 4025118 | 59 | 37 | 65 | 53 | 25 | 0 | 0 | 0 | 0 | 3 | 116 | 71 | 41 | 106 | 45 | 0 | 0 | 0 | 0 | 0 | 269 | 149 | 1 | 1 | 1 | 1 | 1 | 1 | 0.822 | 1 | 1 | 1 | 1 | 1 | -0.036 |
| SE | 13376 | ENSG00000074755 | ZZEF1 | chr17 | - | 4022708 | 4022828 | 4019668 | 4019769 | 4032125 | 4032258 | 2 | 0 | 6 | 5 | 2 | 0 | 0 | 0 | 0 | 0 | 0 | 0 | 0 | 1 | 0 | 0 | 2 | 0 | 0 | 0 | 269 | 149 | 1 | 1 | 1 | NA | 1 | 1 | 1 | NA | 0 | NA | 1 | NA | 0.5 |
| SE | 13377 | ENSG00000074755 | ZZEF1 | chr17 | - | 4024918 | 4025118 | 4019668 | 4019769 | 4032125 | 4032258 | 76 | 49 | 58 | 74 | 111 | 0 | 0 | 0 | 0 | 0 | 141 | 101 | 79 | 108 | 169 | 0 | 2 | 0 | 0 | 0 | 298 | 149 | 1 | 1 | 1 | 1 | 1 | 1 | 1 | 1 | 0.962 | 1 | 1 | 1 | 0.008 |
| SE | 13380 | ENSG00000074755 | ZZEF1 | chr17 | - | 4050780 | 4051043 | 4049707 | 4049859 | 4051970 | 4052136 | 187 | 79 | 145 | 159 | 127 | 0 | 0 | 0 | 0 | 0 | 254 | 229 | 122 | 131 | 224 | 6 | 0 | 0 | 0 | 0 | 298 | 149 | 1 | 1 | 1 | 1 | 1 | 1 | 1 | 0.955 | 1 | 1 | 1 | 1 | 0.009 |
| SE | 13381 | ENSG00000074755 | ZZEF1 | chr17 | - | 4051970 | 4052136 | 4050780 | 4051043 | 4054056 | 4054195 | 182 | 73 | 172 | 155 | 183 | 0 | 0 | 0 | 0 | 4 | 268 | 266 | 183 | 187 | 297 | 0 | 0 | 0 | 0 | 0 | 298 | 149 | 1 | 1 | 1 | 1 | 1 | 1 | 0.958 | 1 | 1 | 1 | 1 | 1 | -0.008 |
| SE | 13382 | ENSG00000074755 | ZZEF1 | chr17 | - | 4054056 | 4054195 | 4051970 | 4052136 | 4057993 | 4058155 | 82 | 32 | 92 | 90 | 92 | 0 | 0 | 0 | 0 | 0 | 166 | 155 | 147 | 134 | 165 | 0 | 0 | 0 | 3 | 0 | 288 | 149 | 1 | 1 | 1 | 1 | 1 | 1 | 1 | 1 | 1 | 1 | 0.959 | 1 | 0.008 |
| SE | 13386 | ENSG00000074755 | ZZEF1 | chr17 | - | 4057993 | 4058155 | 4056215 | 4056345 | 4059170 | 4059290 | 321 | 223 | 299 | 314 | 529 | 0 | 0 | 0 | 0 | 0 | 609 | 489 | 348 | 428 | 559 | 0 | 1 | 0 | 8 | 0 | 298 | 149 | 1 | 1 | 1 | 1 | 1 | 1 | 1 | 1 | 0.996 | 1 | 0.964 | 1 | 0.008 |
| SE | 13387 | ENSG00000074755 | ZZEF1 | chr17 | - | 4062752 | 4062917 | 4059170 | 4059290 | 4064360 | 4064534 | 176 | 144 | 193 | 232 | 191 | 0 | 3 | 0 | 0 | 0 | 266 | 233 | 167 | 308 | 305 | 0 | 0 | 0 | 0 | 0 | 298 | 149 | 1 | 1 | 1 | 0.96 | 1 | 1 | 1 | 1 | 1 | 1 | 1 | 1 | -0.008 |
| SE | 13388 | ENSG00000074755 | ZZEF1 | chr17 | - | 4070683 | 4070924 | 4067162 | 4067242 | 4074149 | 4074351 | 122 | 98 | 98 | 116 | 126 | 1 | 0 | 0 | 0 | 0 | 152 | 137 | 80 | 163 | 269 | 0 | 0 | 5 | 0 | 0 | 298 | 149 | 1 | 1 | 0.984 | 1 | 1 | 1 | 1 | 1 | 1 | 0.889 | 1 | 1 | 0.019 |
| SE | 13393 | ENSG00000074755 | ZZEF1 | chr17 | - | 4074149 | 4074351 | 4072607 | 4072756 | 4075096 | 4075178 | 311 | 191 | 241 | 243 | 389 | 0 | 0 | 0 | 0 | 1 | 488 | 410 | 267 | 302 | 431 | 0 | 0 | 0 | 0 | 0 | 298 | 149 | 1 | 1 | 1 | 1 | 1 | 1 | 0.995 | 1 | 1 | 1 | 1 | 1 | -0.001 |
| SE | 13394 | ENSG00000074755 | ZZEF1 | chr17 | - | 4077882 | 4078042 | 4076867 | 4076989 | 4081375 | 4081490 | 158 | 75 | 169 | 137 | 102 | 0 | 0 | 0 | 0 | 3 | 223 | 194 | 97 | 121 | 260 | 0 | 0 | 0 | 0 | 3 | 298 | 149 | 1 | 1 | 1 | 1 | 1 | 1 | 0.944 | 1 | 1 | 1 | 1 | 0.977 | -0.007 |
| SE | 13395 | ENSG00000074755 | ZZEF1 | chr17 | - | 4086485 | 4086655 | 4085669 | 4085803 | 4088677 | 4088893 | 97 | 80 | 101 | 88 | 112 | 0 | 1 | 0 | 0 | 0 | 170 | 173 | 113 | 190 | 204 | 0 | 0 | 0 | 0 | 0 | 298 | 149 | 1 | 1 | 1 | 0.976 | 1 | 1 | 1 | 1 | 1 | 1 | 1 | 1 | -0.005 |
| SE | 13396 | ENSG00000074755 | ZZEF1 | chr17 | - | 4087433 | 4087534 | 4086485 | 4086655 | 4088677 | 4088893 | 117 | 80 | 122 | 119 | 122 | 0 | 0 | 0 | 0 | 0 | 223 | 134 | 123 | 171 | 152 | 0 | 0 | 1 | 0 | 0 | 250 | 149 | 1 | 1 | 1 | 1 | 1 | 1 | 1 | 1 | 1 | 0.987 | 1 | 1 | 0.003 |
| SE | 13397 | ENSG00000074755 | ZZEF1 | chr17 | - | 4102316 | 4102415 | 4096608 | 4096700 | 4104632 | 4104811 | 163 | 122 | 145 | 140 | 178 | 1 | 1 | 0 | 2 | 0 | 397 | 255 | 202 | 263 | 339 | 0 | 4 | 0 | 2 | 0 | 248 | 149 | 1 | 1 | 0.99 | 0.987 | 1 | 0.977 | 1 | 1 | 0.975 | 1 | 1 | 1 | -0.004 |
| SE | 13399 | ENSG00000140968 | IRF8 | chr16 | + | 85900927 | 85901246 | 85899162 | 85899223 | 85903014 | 85903156 | 1 | 0 | 2 | 1 | 0 | 84 | 54 | 58 | 55 | 11 | 0 | 0 | 0 | 0 | 0 | 145 | 91 | 35 | 87 | 193 | 298 | 149 | 1 | 1 | 0.006 | 0 | 0.017 | 0.009 | 0 | 0 | 0 | 0 | 0 | 0 | 0.006 |
| SE | 13402 | ENSG00000135686 | KLHL36 | chr16 | + | 84659759 | 84659917 | 84656870 | 84657944 | 84661577 | 84662309 | 94 | 33 | 90 | 91 | 40 | 1 | 0 | 0 | 1 | 0 | 71 | 64 | 46 | 61 | 88 | 4 | 0 | 0 | 0 | 0 | 298 | 149 | 1 | 1 | 0.979 | 1 | 1 | 0.978 | 1 | 0.899 | 1 | 1 | 1 | 1 | 0.012 |
| SE | 13404 | ENSG00000260630 | SNAI3-AS1 | chr16 | + | 88671565 | 88671693 | 88671276 | 88671465 | 88671812 | 88672122 | 1 | 0 | 0 | 0 | 0 | 6 | 4 | 3 | 3 | 0 | 0 | 0 | 0 | 0 | 0 | 0 | 2 | 0 | 0 | 0 | 277 | 149 | 1 | 1 | 0.082 | 0 | 0 | 0 | NA | NA | 0 | NA | NA | NA | 0.021 |
| SE | 13405 | ENSG00000172840 | PDP2 | chr16 | + | 66883014 | 66883102 | 66880518 | 66880640 | 66884230 | 66884594 | 1 | 1 | 4 | 2 | 4 | 11 | 4 | 16 | 13 | 10 | 0 | 0 | 0 | 0 | 0 | 0 | 2 | 8 | 4 | 0 | 237 | 149 | 1 | 1 | 0.054 | 0.136 | 0.136 | 0.088 | 0.201 | NA | 0 | 0 | 0 | NA | 0.123 |
| SE | 13406 | ENSG00000172840 | PDP2 | chr16 | + | 66883014 | 66883102 | 66880548 | 66880664 | 66884230 | 66884594 | 1 | 1 | 5 | 0 | 0 | 2 | 2 | 0 | 1 | 0 | 0 | 0 | 0 | 0 | 0 | 0 | 0 | 2 | 0 | 5 | 237 | 149 | 1 | 1 | 0.239 | 0.239 | 1 | 0 | NA | NA | NA | 0 | NA | 0 | 0.369 |
| SE | 13423 | ENSG00000140948 | ZCCHC14 | chr16 | - | 87420606 | 87420716 | 87419782 | 87419877 | 87423809 | 87423881 | 84 | 46 | 102 | 101 | 300 | 0 | 0 | 0 | 1 | 0 | 31 | 64 | 57 | 71 | 81 | 0 | 1 | 0 | 0 | 0 | 259 | 149 | 1 | 1 | 1 | 1 | 1 | 0.983 | 1 | 1 | 0.974 | 1 | 1 | 1 | 0.002 |
| SE | 13424 | ENSG00000140948 | ZCCHC14 | chr16 | - | 87423809 | 87423881 | 87420606 | 87420716 | 87460007 | 87460131 | 11 | 5 | 10 | 6 | 30 | 0 | 0 | 1 | 0 | 0 | 6 | 10 | 6 | 21 | 10 | 0 | 0 | 0 | 0 | 0 | 221 | 149 | 1 | 1 | 1 | 1 | 0.871 | 1 | 1 | 1 | 1 | 1 | 1 | 1 | -0.026 |
| SE | 13425 | ENSG00000140948 | ZCCHC14 | chr16 | - | 87433127 | 87433201 | 87423809 | 87423881 | 87460007 | 87460131 | 60 | 25 | 55 | 72 | 186 | 1 | 0 | 0 | 0 | 5 | 18 | 37 | 41 | 40 | 54 | 0 | 0 | 1 | 0 | 0 | 223 | 149 | 1 | 1 | 0.976 | 1 | 1 | 1 | 0.961 | 1 | 1 | 0.965 | 1 | 1 | -0.006 |
| SE | 13428 | ENSG00000140941 | MAP1LC3B | chr16 | + | 87398814 | 87398870 | 87391597 | 87392467 | 87402174 | 87402281 | 243 | 168 | 191 | 321 | 283 | 1 | 0 | 2 | 0 | 0 | 248 | 325 | 285 | 231 | 625 | 0 | 0 | 2 | 6 | 0 | 205 | 149 | 1 | 1 | 0.994 | 1 | 0.986 | 1 | 1 | 1 | 1 | 0.99 | 0.965 | 1 | 0.005 |
| SE | 13430 | ENSG00000140941 | MAP1LC3B | chr16 | + | 87402174 | 87402281 | 87398814 | 87398870 | 87402922 | 87403778 | 772 | 507 | 521 | 928 | 1147 | 0 | 0 | 0 | 0 | 0 | 663 | 905 | 708 | 721 | 1687 | 0 | 4 | 0 | 3 | 0 | 256 | 149 | 1 | 1 | 1 | 1 | 1 | 1 | 1 | 1 | 0.992 | 1 | 0.993 | 1 | 0.003 |
| SE | 13431 | ENSG00000103175 | WFDC1 | chr16 | + | 84318271 | 84318355 | 84312960 | 84313153 | 84319430 | 84319571 | 1 | 0 | 0 | 0 | 0 | 0 | 0 | 0 | 0 | 0 | 0 | 0 | 2 | 4 | 20 | 0 | 0 | 0 | 0 | 1 | 233 | 149 | 1 | 1 | 1 | NA | NA | NA | NA | NA | NA | 1 | 1 | 0.927 | 0.024 |
| SE | 13432 | ENSG00000103111 | MON1B | chr16 | + | 77191475 | 77191633 | 77190997 | 77191258 | 77193450 | 77193606 | 197 | 118 | 134 | 153 | 116 | 0 | 0 | 0 | 0 | 2 | 188 | 128 | 71 | 147 | 191 | 0 | 0 | 0 | 0 | 0 | 298 | 149 | 1 | 1 | 1 | 1 | 1 | 1 | 0.967 | 1 | 1 | 1 | 1 | 1 | -0.007 |
| SE | 13437 | ENSG00000141098 | GFOD2 | chr16 | - | 67677765 | 67677857 | 67675899 | 67676053 | 67685456 | 67685612 | 0 | 0 | 1 | 0 | 0 | 4 | 2 | 3 | 3 | 9 | 0 | 0 | 0 | 0 | 0 | 0 | 0 | 2 | 4 | 4 | 241 | 149 | 1 | 1 | 0 | 0 | 0.171 | 0 | 0 | NA | NA | 0 | 0 | 0 | 0.034 |
| SE | 13441 | ENSG00000167508 | MVD | chr16 | - | 88660747 | 88660841 | 88660519 | 88660637 | 88663010 | 88663148 | 0 | 0 | 0 | 3 | 6 | 0 | 0 | 2 | 0 | 0 | 0 | 0 | 0 | 0 | 0 | 0 | 0 | 0 | 1 | 0 | 243 | 149 | 1 | 1 | NA | NA | 0 | 1 | 1 | NA | NA | NA | 0 | NA | 0.667 |
| SE | 13445 | ENSG00000131149 | GSE1 | chr16 | + | 85633913 | 85634132 | 85613321 | 85613398 | 85648551 | 85648751 | 8 | 5 | 7 | 3 | 4 | 6 | 0 | 1 | 0 | 0 | 14 | 0 | 4 | 6 | 24 | 10 | 0 | 0 | 5 | 0 | 298 | 149 | 1 | 1 | 0.4 | 1 | 0.778 | 1 | 1 | 0.412 | NA | 1 | 0.375 | 1 | 0.139 |
| SE | 13447 | ENSG00000131149 | GSE1 | chr16 | + | 85661145 | 85661765 | 85657276 | 85657604 | 85662980 | 85663093 | 64 | 25 | 35 | 50 | 25 | 0 | 0 | 0 | 0 | 0 | 84 | 70 | 31 | 36 | 90 | 0 | 0 | 0 | 0 | 2 | 298 | 149 | 1 | 1 | 1 | 1 | 1 | 1 | 1 | 1 | 1 | 1 | 1 | 0.957 | 0.009 |
| SE | 13448 | ENSG00000131149 | GSE1 | chr16 | + | 85665014 | 85665128 | 85663343 | 85663614 | 85665975 | 85666347 | 113 | 72 | 108 | 108 | 60 | 0 | 0 | 1 | 0 | 1 | 102 | 136 | 81 | 153 | 220 | 6 | 0 | 0 | 0 | 0 | 263 | 149 | 1 | 1 | 1 | 1 | 0.984 | 1 | 0.971 | 0.906 | 1 | 1 | 1 | 1 | 0.01 |
| SE | 13449 | ENSG00000131149 | GSE1 | chr16 | + | 85665975 | 85666347 | 85662290 | 85663614 | 85668139 | 85668424 | 76 | 47 | 56 | 46 | 26 | 0 | 0 | 0 | 0 | 0 | 144 | 93 | 88 | 95 | 183 | 0 | 0 | 0 | 0 | 1 | 298 | 149 | 1 | 1 | 1 | 1 | 1 | 1 | 1 | 1 | 1 | 1 | 1 | 0.989 | 0.002 |
| SE | 13450 | ENSG00000131149 | GSE1 | chr16 | + | 85672181 | 85672309 | 85670994 | 85671098 | 85672404 | 85672893 | 0 | 0 | 4 | 0 | 0 | 125 | 90 | 68 | 106 | 54 | 0 | 0 | 0 | 1 | 4 | 102 | 81 | 77 | 130 | 120 | 277 | 149 | 1 | 1 | 0 | 0 | 0.031 | 0 | 0 | 0 | 0 | 0 | 0.004 | 0.018 | 0.002 |
| SE | 13451 | ENSG00000205078 | SYCE1L | chr16 | + | 77211212 | 77211276 | 77208209 | 77208269 | 77212129 | 77212199 | 0 | 0 | 0 | 0 | 0 | 1 | 0 | 0 | 0 | 0 | 0 | 0 | 0 | 5 | 0 | 0 | 0 | 0 | 0 | 0 | 213 | 149 | 1 | 1 | 0 | NA | NA | NA | NA | NA | NA | NA | 1 | NA | -1 |
| SE | 13462 | ENSG00000259972 | NA | chr16 | - | 74306254 | 74306421 | 74305255 | 74305337 | 74332566 | 74332613 | 0 | 1 | 0 | 0 | 0 | 0 | 0 | 1 | 3 | 0 | 0 | 0 | 0 | 0 | 0 | 0 | 3 | 3 | 0 | 0 | 298 | 149 | 1 | 1 | NA | 1 | 0 | 0 | NA | NA | 0 | 0 | NA | NA | 0.333 |
| SE | 13489 | ENSG00000135736 | CCDC102A | chr16 | - | 57518067 | 57518277 | 57516292 | 57516463 | 57518624 | 57518741 | 45 | 18 | 32 | 31 | 26 | 0 | 0 | 0 | 1 | 0 | 22 | 58 | 16 | 12 | 47 | 0 | 0 | 0 | 0 | 0 | 298 | 149 | 1 | 1 | 1 | 1 | 1 | 0.939 | 1 | 1 | 1 | 1 | 1 | 1 | -0.012 |
| SE | 13490 | ENSG00000135736 | CCDC102A | chr16 | - | 57518624 | 57518741 | 57516292 | 57516463 | 57521067 | 57521176 | 30 | 19 | 31 | 24 | 18 | 0 | 0 | 0 | 0 | 0 | 22 | 45 | 13 | 10 | 31 | 0 | 0 | 1 | 0 | 0 | 266 | 149 | 1 | 1 | 1 | 1 | 1 | 1 | 1 | 1 | 1 | 0.879 | 1 | 1 | 0.024 |
| SE | 13491 | ENSG00000135736 | CCDC102A | chr16 | - | 57525900 | 57526127 | 57521067 | 57521176 | 57528592 | 57529324 | 31 | 1 | 26 | 35 | 1 | 0 | 0 | 0 | 0 | 0 | 0 | 5 | 6 | 2 | 24 | 0 | 0 | 0 | 0 | 8 | 298 | 149 | 1 | 1 | 1 | 1 | 1 | 1 | 1 | NA | 1 | 1 | 1 | 0.6 | 0.1 |
| SE | 13494 | ENSG00000135698 | MPHOSPH6 | chr16 | - | 82151423 | 82151514 | 82149308 | 82149403 | 82164081 | 82164194 | 147 | 120 | 114 | 166 | 220 | 0 | 0 | 1 | 0 | 3 | 31 | 81 | 72 | 30 | 12 | 0 | 0 | 1 | 0 | 0 | 240 | 149 | 1 | 1 | 1 | 1 | 0.986 | 1 | 0.979 | 1 | 1 | 0.978 | 1 | 1 | -0.003 |
| SE | 13495 | ENSG00000135698 | MPHOSPH6 | chr16 | - | 82162211 | 82162313 | 82151166 | 82151514 | 82164081 | 82164194 | 1 | 0 | 0 | 1 | 0 | 88 | 69 | 73 | 104 | 144 | 0 | 1 | 0 | 0 | 0 | 6 | 39 | 27 | 23 | 6 | 251 | 149 | 1 | 1 | 0.007 | 0 | 0 | 0.006 | 0 | 0 | 0.015 | 0 | 0 | 0 | 0 |
| SE | 13496 | ENSG00000135698 | MPHOSPH6 | chr16 | - | 82164081 | 82164194 | 82151083 | 82151514 | 82167539 | 82167597 | 75 | 45 | 69 | 86 | 108 | 0 | 1 | 0 | 0 | 0 | 11 | 33 | 15 | 27 | 6 | 0 | 0 | 0 | 0 | 0 | 262 | 149 | 1 | 1 | 1 | 0.962 | 1 | 1 | 1 | 1 | 1 | 1 | 1 | 1 | -0.008 |
| SE | 13497 | ENSG00000135698 | MPHOSPH6 | chr16 | - | 82164651 | 82164741 | 82164081 | 82164194 | 82167539 | 82167597 | 3 | 0 | 0 | 2 | 0 | 0 | 1 | 4 | 6 | 5 | 0 | 0 | 6 | 0 | 0 | 5 | 15 | 9 | 5 | 0 | 239 | 149 | 1 | 1 | 1 | 1 | 0 | 0.172 | 0 | 0 | 0 | 0.294 | 0 | NA | 0.161 |
| SE | 13498 | ENSG00000135698 | MPHOSPH6 | chr16 | - | 82164651 | 82164741 | 82164081 | 82164194 | 82170124 | 82170193 | 1 | 0 | 0 | 2 | 0 | 32 | 46 | 23 | 43 | 40 | 0 | 0 | 6 | 0 | 0 | 5 | 28 | 19 | 1 | 0 | 239 | 149 | 1 | 1 | 0.019 | 0 | 0 | 0.028 | 0 | 0 | 0 | 0.164 | 0 | NA | -0.032 |
| SE | 13499 | ENSG00000135698 | MPHOSPH6 | chr16 | - | 82164651 | 82164940 | 82164074 | 82164194 | 82170124 | 82170218 | 1 | 0 | 1 | 2 | 0 | 32 | 46 | 23 | 43 | 40 | 0 | 0 | 6 | 0 | 0 | 5 | 28 | 19 | 1 | 0 | 298 | 149 | 1 | 1 | 0.015 | 0 | 0.021 | 0.023 | 0 | 0 | 0 | 0.136 | 0 | NA | -0.022 |
| SE | 13500 | ENSG00000135698 | MPHOSPH6 | chr16 | - | 82167539 | 82167597 | 82164081 | 82164194 | 82170124 | 82170213 | 0 | 1 | 4 | 5 | 5 | 32 | 46 | 23 | 43 | 40 | 5 | 15 | 9 | 5 | 0 | 5 | 28 | 19 | 1 | 0 | 207 | 149 | 1 | 1 | 0 | 0.015 | 0.111 | 0.077 | 0.083 | 0.419 | 0.278 | 0.254 | 0.783 | NA | -0.376 |
| SE | 13502 | ENSG00000131153 | GINS2 | chr16 | - | 85678539 | 85678666 | 85676197 | 85678337 | 85681581 | 85681681 | 7 | 7 | 26 | 17 | 28 | 0 | 0 | 0 | 0 | 0 | 28 | 14 | 41 | 95 | 39 | 0 | 0 | 1 | 0 | 0 | 276 | 149 | 1 | 1 | 1 | 1 | 1 | 1 | 1 | 1 | 1 | 0.957 | 1 | 1 | 0.009 |
| SE | 13511 | ENSG00000196696 | PDXDC2P-NPIPB14P | chr16 | - | 69978221 | 69978282 | 69978076 | 69978112 | 69982349 | 69982494 | 22 | 45 | 29 | 39 | 49 | 0 | 1 | 3 | 0 | 0 | 25 | 66 | 30 | 51 | 23 | 0 | 0 | 0 | 0 | 0 | 210 | 149 | 1 | 1 | 1 | 0.97 | 0.873 | 1 | 1 | 1 | 1 | 1 | 1 | 1 | -0.031 |
| SE | 13513 | ENSG00000159761 | C16orf86 | chr16 | + | 67667226 | 67667525 | 67666836 | 67667072 | 67667877 | 67668098 | 2 | 2 | 2 | 1 | 0 | 0 | 0 | 3 | 0 | 0 | 19 | 5 | 0 | 0 | 18 | 0 | 3 | 0 | 0 | 0 | 298 | 149 | 1 | 1 | 1 | 1 | 0.25 | 1 | NA | 1 | 0.455 | NA | NA | 1 | -0.006 |
| SE | 13514 | ENSG00000159761 | C16orf86 | chr16 | + | 67667295 | 67667525 | 67666813 | 67667072 | 67667877 | 67668098 | 2 | 2 | 2 | 2 | 0 | 0 | 0 | 3 | 0 | 0 | 19 | 9 | 0 | 0 | 18 | 0 | 3 | 0 | 0 | 0 | 298 | 149 | 1 | 1 | 1 | 1 | 0.25 | 1 | NA | 1 | 0.6 | NA | NA | 1 | -0.054 |
| SE | 13517 | ENSG00000159593 | NAE1 | chr16 | - | 66805911 | 66806026 | 66802877 | 66803118 | 66808520 | 66808548 | 85 | 38 | 55 | 57 | 156 | 1 | 0 | 0 | 1 | 0 | 0 | 17 | 28 | 2 | 0 | 0 | 0 | 0 | 0 | 0 | 264 | 149 | 1 | 1 | 0.98 | 1 | 1 | 0.97 | 1 | NA | 1 | 1 | 1 | NA | -0.01 |
| SE | 13518 | ENSG00000159593 | NAE1 | chr16 | - | 66808520 | 66808613 | 66805911 | 66806026 | 66808988 | 66809075 | 237 | 164 | 198 | 218 | 377 | 0 | 0 | 0 | 0 | 0 | 14 | 94 | 81 | 12 | 14 | 0 | 3 | 0 | 0 | 0 | 242 | 149 | 1 | 1 | 1 | 1 | 1 | 1 | 1 | 1 | 0.951 | 1 | 1 | 1 | 0.01 |
| SE | 13519 | ENSG00000159593 | NAE1 | chr16 | - | 66816580 | 66816672 | 66813786 | 66813846 | 66816964 | 66817028 | 209 | 171 | 266 | 206 | 295 | 0 | 0 | 0 | 0 | 1 | 62 | 124 | 133 | 73 | 13 | 0 | 0 | 0 | 0 | 0 | 241 | 149 | 1 | 1 | 1 | 1 | 1 | 1 | 0.995 | 1 | 1 | 1 | 1 | 1 | -0.001 |
| SE | 13520 | ENSG00000159593 | NAE1 | chr16 | - | 66816964 | 66817028 | 66816580 | 66816672 | 66817424 | 66817487 | 241 | 181 | 269 | 217 | 349 | 1 | 1 | 0 | 0 | 0 | 68 | 132 | 179 | 93 | 24 | 0 | 0 | 0 | 0 | 0 | 213 | 149 | 1 | 1 | 0.994 | 0.992 | 1 | 1 | 1 | 1 | 1 | 1 | 1 | 1 | -0.003 |
| SE | 13521 | ENSG00000159593 | NAE1 | chr16 | - | 66821449 | 66821559 | 66818527 | 66818637 | 66823226 | 66823262 | 237 | 168 | 261 | 273 | 347 | 1 | 0 | 0 | 0 | 0 | 28 | 156 | 168 | 76 | 37 | 0 | 0 | 0 | 0 | 0 | 259 | 149 | 1 | 1 | 0.993 | 1 | 1 | 1 | 1 | 1 | 1 | 1 | 1 | 1 | -0.001 |
| SE | 13522 | ENSG00000159593 | NAE1 | chr16 | - | 66823226 | 66823262 | 66821449 | 66821559 | 66823528 | 66823600 | 57 | 31 | 51 | 65 | 59 | 0 | 0 | 0 | 0 | 0 | 4 | 40 | 21 | 15 | 0 | 0 | 0 | 2 | 0 | 0 | 185 | 149 | 1 | 1 | 1 | 1 | 1 | 1 | 1 | 1 | 1 | 0.894 | 1 | NA | 0.026 |
| SE | 13523 | ENSG00000159593 | NAE1 | chr16 | - | 66823226 | 66823306 | 66821445 | 66821559 | 66823528 | 66823600 | 151 | 100 | 166 | 177 | 214 | 0 | 0 | 0 | 0 | 0 | 16 | 90 | 85 | 39 | 4 | 0 | 0 | 2 | 0 | 0 | 229 | 149 | 1 | 1 | 1 | 1 | 1 | 1 | 1 | 1 | 1 | 0.965 | 1 | 1 | 0.007 |
| SE | 13524 | ENSG00000159593 | NAE1 | chr16 | - | 66823226 | 66823306 | 66821445 | 66821559 | 66824854 | 66824885 | 74 | 44 | 74 | 87 | 106 | 0 | 0 | 1 | 0 | 0 | 11 | 59 | 38 | 21 | 0 | 0 | 0 | 0 | 0 | 0 | 229 | 149 | 1 | 1 | 1 | 1 | 0.98 | 1 | 1 | 1 | 1 | 1 | 1 | NA | -0.004 |
| SE | 13526 | ENSG00000159593 | NAE1 | chr16 | - | 66823528 | 66823600 | 66821445 | 66821559 | 66824854 | 66824885 | 12 | 22 | 15 | 10 | 20 | 0 | 0 | 1 | 0 | 0 | 0 | 6 | 8 | 5 | 0 | 0 | 0 | 0 | 0 | 0 | 221 | 149 | 1 | 1 | 1 | 1 | 0.91 | 1 | 1 | NA | 1 | 1 | 1 | NA | -0.018 |
| SE | 13527 | ENSG00000159593 | NAE1 | chr16 | - | 66823528 | 66823600 | 66823226 | 66823306 | 66824854 | 66824885 | 89 | 76 | 103 | 92 | 128 | 0 | 1 | 0 | 0 | 0 | 2 | 34 | 45 | 23 | 0 | 0 | 0 | 0 | 0 | 0 | 221 | 149 | 1 | 1 | 1 | 0.981 | 1 | 1 | 1 | 1 | 1 | 1 | 1 | NA | -0.004 |
| SE | 13528 | ENSG00000159593 | NAE1 | chr16 | - | 66824854 | 66824885 | 66823341 | 66823600 | 66826522 | 66826583 | 62 | 61 | 70 | 61 | 73 | 0 | 4 | 0 | 7 | 1 | 0 | 28 | 28 | 16 | 0 | 3 | 2 | 0 | 1 | 0 | 180 | 149 | 1 | 1 | 1 | 0.927 | 1 | 0.878 | 0.984 | 0 | 0.921 | 1 | 0.93 | NA | 0.245 |
| SE | 13532 | ENSG00000159593 | NAE1 | chr16 | - | 66827934 | 66828077 | 66826676 | 66826780 | 66830846 | 66830976 | 5 | 1 | 0 | 0 | 0 | 4 | 6 | 9 | 12 | 6 | 0 | 0 | 2 | 0 | 0 | 11 | 0 | 8 | 2 | 0 | 292 | 149 | 1 | 1 | 0.389 | 0.078 | 0 | 0 | 0 | 0 | NA | 0.113 | 0 | NA | 0.056 |
| SE | 13533 | ENSG00000159593 | NAE1 | chr16 | - | 66827979 | 66828077 | 66826676 | 66826780 | 66830846 | 66830976 | 9 | 1 | 1 | 0 | 0 | 4 | 6 | 9 | 12 | 6 | 0 | 0 | 4 | 0 | 0 | 11 | 0 | 8 | 2 | 0 | 247 | 149 | 1 | 1 | 0.576 | 0.091 | 0.063 | 0 | 0 | 0 | NA | 0.232 | 0 | NA | 0.069 |
| SE | 13534 | ENSG00000159593 | NAE1 | chr16 | - | 66827979 | 66828142 | 66826676 | 66826780 | 66830846 | 66830917 | 4 | 0 | 1 | 0 | 0 | 4 | 6 | 9 | 12 | 6 | 0 | 0 | 2 | 0 | 0 | 11 | 0 | 8 | 2 | 0 | 298 | 149 | 1 | 1 | 0.333 | 0 | 0.053 | 0 | 0 | 0 | NA | 0.111 | 0 | NA | 0.04 |
| SE | 13540 | ENSG00000103047 | TANGO6 | chr16 | + | 68900433 | 68900546 | 68880547 | 68880630 | 68907442 | 68907575 | 93 | 61 | 101 | 66 | 51 | 0 | 0 | 0 | 0 | 3 | 161 | 88 | 43 | 55 | 97 | 0 | 0 | 0 | 0 | 0 | 262 | 149 | 1 | 1 | 1 | 1 | 1 | 1 | 0.906 | 1 | 1 | 1 | 1 | 1 | -0.019 |
| SE | 13541 | ENSG00000103047 | TANGO6 | chr16 | + | 68902327 | 68902504 | 68880547 | 68880630 | 68907442 | 68907533 | 25 | 16 | 36 | 26 | 41 | 0 | 0 | 0 | 0 | 3 | 17 | 47 | 0 | 18 | 64 | 0 | 0 | 0 | 0 | 0 | 298 | 149 | 1 | 1 | 1 | 1 | 1 | 1 | 0.872 | 1 | 1 | NA | 1 | 1 | -0.026 |
| SE | 13543 | ENSG00000103047 | TANGO6 | chr16 | + | 68907442 | 68907575 | 68902327 | 68902504 | 68909210 | 68909402 | 52 | 40 | 58 | 68 | 76 | 0 | 0 | 0 | 0 | 0 | 21 | 74 | 9 | 37 | 110 | 0 | 0 | 0 | 3 | 0 | 282 | 149 | 1 | 1 | 1 | 1 | 1 | 1 | 1 | 1 | 1 | 1 | 0.867 | 1 | 0.027 |
| SE | 13544 | ENSG00000103047 | TANGO6 | chr16 | + | 68930237 | 68930295 | 68927567 | 68928083 | 68974027 | 68974168 | 189 | 131 | 143 | 145 | 206 | 0 | 0 | 2 | 0 | 0 | 237 | 179 | 149 | 228 | 322 | 0 | 0 | 0 | 0 | 4 | 207 | 149 | 1 | 1 | 1 | 1 | 0.981 | 1 | 1 | 1 | 1 | 1 | 1 | 0.983 | 0 |
| SE | 13546 | ENSG00000102977 | ACD | chr16 | - | 67657985 | 67658362 | 67657761 | 67657853 | 67658719 | 67658816 | 23 | 3 | 21 | 23 | 11 | 0 | 0 | 0 | 0 | 0 | 33 | 5 | 5 | 0 | 16 | 0 | 0 | 1 | 0 | 0 | 298 | 149 | 1 | 1 | 1 | 1 | 1 | 1 | 1 | 1 | 1 | 0.714 | NA | 1 | 0.072 |
| SE | 13550 | ENSG00000118557 | PMFBP1 | chr16 | - | 72122913 | 72122988 | 72112156 | 72112845 | 72123545 | 72123649 | 1 | 0 | 1 | 0 | 0 | 0 | 0 | 0 | 0 | 0 | 0 | 4 | 2 | 0 | 0 | 0 | 0 | 0 | 3 | 0 | 224 | 149 | 1 | 1 | 1 | NA | 1 | NA | NA | NA | 1 | 1 | 0 | NA | 0.333 |
| SE | 13551 | ENSG00000118557 | PMFBP1 | chr16 | - | 72124766 | 72124934 | 72123545 | 72123649 | 72125237 | 72125405 | 3 | 0 | 4 | 2 | 0 | 0 | 0 | 2 | 0 | 0 | 0 | 4 | 3 | 16 | 0 | 0 | 0 | 0 | 0 | 0 | 298 | 149 | 1 | 1 | 1 | NA | 0.5 | 1 | NA | NA | 1 | 1 | 1 | NA | -0.167 |
| SE | 13552 | ENSG00000118557 | PMFBP1 | chr16 | - | 72128376 | 72128569 | 72125967 | 72126132 | 72128656 | 72128794 | 0 | 0 | 0 | 0 | 0 | 6 | 2 | 4 | 1 | 1 | 0 | 1 | 0 | 0 | 0 | 0 | 0 | 2 | 7 | 0 | 298 | 149 | 1 | 1 | 0 | 0 | 0 | 0 | 0 | NA | 1 | 0 | 0 | NA | -0.333 |
| SE | 13553 | ENSG00000118557 | PMFBP1 | chr16 | - | 72150607 | 72150829 | 72140411 | 72140582 | 72154210 | 72154459 | 2 | 1 | 1 | 0 | 0 | 1 | 0 | 0 | 0 | 0 | 0 | 4 | 1 | 0 | 0 | 0 | 0 | 0 | 1 | 0 | 298 | 149 | 1 | 1 | 0.5 | 1 | 1 | NA | NA | NA | 1 | 1 | 0 | NA | 0.167 |
| SE | 13554 | ENSG00000118557 | PMFBP1 | chr16 | - | 72154210 | 72154459 | 72140411 | 72140582 | 72164763 | 72164916 | 1 | 0 | 0 | 0 | 0 | 0 | 0 | 0 | 0 | 0 | 0 | 0 | 0 | 1 | 0 | 0 | 0 | 1 | 0 | 0 | 298 | 149 | 1 | 1 | 1 | NA | NA | NA | NA | NA | NA | 0 | 1 | NA | 0.5 |
| SE | 13556 | ENSG00000141101 | NOB1 | chr16 | - | 69749558 | 69749630 | 69749212 | 69749338 | 69752240 | 69752253 | 64 | 43 | 85 | 83 | 76 | 0 | 0 | 0 | 0 | 0 | 184 | 113 | 41 | 59 | 157 | 0 | 1 | 0 | 1 | 0 | 221 | 149 | 1 | 1 | 1 | 1 | 1 | 1 | 1 | 1 | 0.987 | 1 | 0.975 | 1 | 0.008 |
| SE | 13566 | ENSG00000102908 | NFAT5 | chr16 | + | 69695135 | 69695379 | 69691748 | 69694239 | 69696359 | 69697390 | 155 | 170 | 102 | 173 | 215 | 0 | 0 | 0 | 2 | 0 | 54 | 154 | 190 | 42 | 37 | 0 | 1 | 5 | 0 | 0 | 298 | 149 | 1 | 1 | 1 | 1 | 1 | 0.977 | 1 | 1 | 0.987 | 0.95 | 1 | 1 | 0.008 |
| SE | 13573 | ENSG00000167261 | DPEP2 | chr16 | - | 67989322 | 67989398 | 67987393 | 67987760 | 67990820 | 67990939 | 9 | 6 | 7 | 8 | 25 | 0 | 0 | 0 | 0 | 3 | 6 | 4 | 13 | 18 | 24 | 0 | 0 | 2 | 0 | 0 | 225 | 149 | 1 | 1 | 1 | 1 | 1 | 1 | 0.847 | 1 | 1 | 0.811 | 1 | 1 | 0.007 |
| SE | 13575 | ENSG00000167261 | DPEP2 | chr16 | - | 67990820 | 67990939 | 67989322 | 67989398 | 67999374 | 67999512 | 1 | 0 | 1 | 2 | 5 | 1 | 0 | 0 | 0 | 0 | 0 | 8 | 2 | 2 | 0 | 0 | 0 | 2 | 0 | 0 | 268 | 149 | 1 | 1 | 0.357 | NA | 1 | 1 | 1 | NA | 1 | 0.357 | 1 | NA | 0.054 |
| SE | 13576 | ENSG00000167261 | DPEP2 | chr16 | - | 67990820 | 67990939 | 67989322 | 67989398 | 67999395 | 67999518 | 1 | 0 | 1 | 1 | 0 | 0 | 0 | 5 | 0 | 0 | 0 | 4 | 2 | 5 | 0 | 0 | 3 | 1 | 0 | 2 | 268 | 149 | 1 | 1 | 1 | NA | 0.1 | 1 | NA | NA | 0.426 | 0.527 | 1 | 0 | 0.212 |
| SE | 13577 | ENSG00000167261 | DPEP2 | chr16 | - | 67990820 | 67990939 | 67989322 | 67989398 | 68000447 | 68000586 | 8 | 0 | 0 | 1 | 0 | 0 | 0 | 0 | 0 | 1 | 0 | 4 | 2 | 2 | 0 | 0 | 0 | 2 | 0 | 0 | 268 | 149 | 1 | 1 | 1 | NA | NA | 1 | 0 | NA | 1 | 0.357 | 1 | NA | -0.119 |
| SE | 13578 | ENSG00000167261 | DPEP2 | chr16 | - | 67990820 | 67990997 | 67989322 | 67989398 | 67999374 | 67999484 | 2 | 0 | 1 | 1 | 0 | 1 | 0 | 0 | 0 | 0 | 0 | 7 | 2 | 6 | 0 | 0 | 0 | 2 | 0 | 0 | 298 | 149 | 1 | 1 | 0.5 | NA | 1 | 1 | NA | NA | 1 | 0.333 | 1 | NA | 0.056 |
| SE | 13579 | ENSG00000167261 | DPEP2 | chr16 | - | 67990820 | 67990997 | 67989322 | 67989398 | 68000447 | 68000564 | 1 | 0 | 3 | 1 | 0 | 0 | 0 | 0 | 0 | 1 | 0 | 4 | 2 | 2 | 0 | 0 | 0 | 2 | 0 | 0 | 298 | 149 | 1 | 1 | 1 | NA | 1 | 1 | 0 | NA | 1 | 0.333 | 1 | NA | -0.028 |
| SE | 13581 | ENSG00000167261 | DPEP2 | chr16 | - | 67991114 | 67991184 | 67990820 | 67990939 | 67992063 | 67992193 | 1 | 1 | 0 | 2 | 11 | 0 | 0 | 0 | 2 | 2 | 24 | 0 | 0 | 1 | 27 | 0 | 0 | 3 | 0 | 0 | 219 | 149 | 1 | 1 | 1 | 1 | NA | 0.405 | 0.789 | 1 | NA | 0 | 1 | 1 | 0.049 |
| SE | 13582 | ENSG00000167261 | DPEP2 | chr16 | - | 67991114 | 67991184 | 67990820 | 67990939 | 67999374 | 67999484 | 1 | 1 | 1 | 2 | 11 | 0 | 0 | 1 | 1 | 5 | 14 | 0 | 0 | 1 | 36 | 0 | 4 | 0 | 0 | 0 | 219 | 149 | 1 | 1 | 1 | 1 | 0.405 | 0.576 | 0.599 | 1 | 0 | NA | 1 | 1 | -0.034 |
| SE | 13583 | ENSG00000167261 | DPEP2 | chr16 | - | 67991114 | 67991184 | 67990820 | 67990939 | 67999395 | 67999518 | 1 | 1 | 0 | 2 | 11 | 0 | 0 | 1 | 0 | 0 | 14 | 0 | 0 | 1 | 27 | 0 | 0 | 0 | 3 | 0 | 219 | 149 | 1 | 1 | 1 | 1 | 0 | 1 | 1 | 1 | NA | NA | 0.185 | 1 | 0.072 |
| SE | 13584 | ENSG00000167261 | DPEP2 | chr16 | - | 67991114 | 67991184 | 67990820 | 67990939 | 68000447 | 68000564 | 3 | 1 | 0 | 2 | 11 | 9 | 0 | 0 | 0 | 0 | 28 | 0 | 0 | 1 | 27 | 0 | 0 | 0 | 0 | 0 | 219 | 149 | 1 | 1 | 0.185 | 1 | NA | 1 | 1 | 1 | NA | NA | 1 | 1 | -0.204 |
| SE | 13585 | ENSG00000167261 | DPEP2 | chr16 | - | 67991114 | 67991184 | 67990820 | 67990997 | 67991837 | 67991979 | 57 | 24 | 31 | 36 | 39 | 1 | 0 | 0 | 0 | 0 | 39 | 23 | 34 | 37 | 67 | 0 | 0 | 0 | 0 | 0 | 219 | 149 | 1 | 1 | 0.975 | 1 | 1 | 1 | 1 | 1 | 1 | 1 | 1 | 1 | -0.005 |
| SE | 13587 | ENSG00000167261 | DPEP2 | chr16 | - | 67991114 | 67991184 | 67990820 | 67990997 | 68000447 | 68000564 | 17 | 10 | 13 | 11 | 5 | 0 | 0 | 3 | 0 | 0 | 17 | 10 | 18 | 7 | 22 | 0 | 0 | 0 | 0 | 0 | 219 | 149 | 1 | 1 | 1 | 1 | 0.747 | 1 | 1 | 1 | 1 | 1 | 1 | 1 | -0.051 |
| SE | 13588 | ENSG00000167261 | DPEP2 | chr16 | - | 67991837 | 67991979 | 67990820 | 67990939 | 67992063 | 67992193 | 40 | 16 | 42 | 45 | 43 | 0 | 0 | 0 | 2 | 2 | 64 | 21 | 36 | 39 | 50 | 0 | 0 | 3 | 0 | 0 | 291 | 149 | 1 | 1 | 1 | 1 | 1 | 0.92 | 0.917 | 1 | 1 | 0.86 | 1 | 1 | -0.005 |
| SE | 13591 | ENSG00000167261 | DPEP2 | chr16 | - | 67992063 | 67992193 | 67990820 | 67990939 | 68000447 | 68000586 | 2 | 4 | 4 | 11 | 6 | 9 | 0 | 0 | 0 | 0 | 0 | 0 | 3 | 0 | 0 | 0 | 0 | 0 | 0 | 0 | 279 | 149 | 1 | 1 | 0.106 | 1 | 1 | 1 | 1 | NA | NA | 1 | NA | NA | -0.179 |
| SE | 13592 | ENSG00000167261 | DPEP2 | chr16 | - | 67992063 | 67992193 | 67991114 | 67991184 | 67999374 | 67999512 | 13 | 12 | 13 | 25 | 29 | 0 | 2 | 2 | 0 | 0 | 19 | 0 | 17 | 8 | 55 | 0 | 0 | 2 | 0 | 9 | 279 | 149 | 1 | 1 | 1 | 0.762 | 0.776 | 1 | 1 | NA | | 0.819 | 1 | 0.765 | 0.012 |
| SE | 13593 | ENSG00000167261 | DPEP2 | chr16 | - | 67992063 | 67992193 | 67991114 | 67991184 | 68000447 | 68000586 | 2 | 4 | 4 | 9 | 4 | 2 | 0 | 0 | 1 | 3 | 10 | 0 | 3 | 0 | 0 | 14 | 0 | 0 | 0 | 0 | 279 | 149 | 1 | 1 | 0.348 | 1 | 1 | 0.828 | 0.416 | 0.276 | NA | 1 | NA | NA | 0.08 |
| SE | 13594 | ENSG00000167261 | DPEP2 | chr16 | - | 67992509 | 67992636 | 67992063 | 67992193 | 67992949 | 67993257 | 13 | 8 | 15 | 22 | 15 | 1 | 0 | 2 | 0 | 0 | 0 | 5 | 7 | 0 | 19 | 0 | 0 | 1 | 0 | 0 | 276 | 149 | 1 | 1 | 0.875 | 1 | 0.802 | 1 | 1 | NA | 1 | 0.791 | NA | 1 | 0.005 |
| SE | 13596 | ENSG00000167261 | DPEP2 | chr16 | - | 67992509 | 67992636 | 67992063 | 67992193 | 67999395 | 67999518 | 12 | 5 | 10 | 19 | 9 | 4 | 0 | 2 | 1 | 15 | 0 | 2 | 11 | 0 | 19 | 0 | 2 | 3 | 2 | 5 | 276 | 149 | 1 | 1 | 0.618 | 1 | 0.73 | 0.911 | 0.245 | NA | 0.351 | 0.664 | 0 | 0.672 | 0.279 |
| SE | 13597 | ENSG00000167261 | DPEP2 | chr16 | - | 67992949 | 67993257 | 67991642 | 67992193 | 67999266 | 67999748 | 1 | 0 | 2 | 0 | 0 | 1 | 0 | 0 | 0 | 0 | 0 | 0 | 1 | 0 | 0 | 7 | 0 | 0 | 2 | 0 | 298 | 149 | 1 | 1 | 0.333 | NA | 1 | NA | NA | 0 | NA | 1 | 0 | NA | 0.333 |
| SE | 13599 | ENSG00000167261 | DPEP2 | chr16 | - | 67992949 | 67993257 | 67991642 | 67992193 | 67999395 | 67999518 | 8 | 3 | 8 | 0 | 18 | 4 | 0 | 2 | 1 | 15 | 0 | 4 | 4 | 5 | 19 | 0 | 2 | 3 | 2 | 5 | 298 | 149 | 1 | 1 | 0.5 | 1 | 0.667 | 0 | 0.375 | NA | 0.5 | 0.4 | 0.556 | 0.655 | -0.019 |
| SE | 13600 | ENSG00000167261 | DPEP2 | chr16 | - | 67992949 | 67993257 | 67991642 | 67992193 | 68000447 | 68000581 | 4 | 0 | 3 | 9 | 5 | 2 | 4 | 4 | 9 | 4 | 11 | 0 | 1 | 4 | 0 | 0 | 0 | 0 | 0 | 0 | 298 | 149 | 1 | 1 | 0.5 | 0 | 0.273 | 0.333 | 0.385 | 1 | NA | 1 | 1 | NA | -0.702 |
| SE | 13601 | ENSG00000167261 | DPEP2 | chr16 | - | 67992949 | 67993257 | 67992509 | 67992636 | 67999374 | 67999484 | 13 | 5 | 7 | 4 | 11 | 0 | 0 | 0 | 0 | 1 | 0 | 7 | 8 | 3 | 0 | 0 | 0 | 4 | 0 | 0 | 298 | 149 | 1 | 1 | 1 | 1 | 1 | 1 | 0.846 | NA | 1 | 0.5 | 1 | NA | 0.136 |
| SE | 13602 | ENSG00000167261 | DPEP2 | chr16 | - | 67992949 | 67993257 | 67992509 | 67992636 | 67999395 | 67999518 | 8 | 6 | 11 | 3 | 24 | 0 | 0 | 0 | 0 | 0 | 0 | 7 | 4 | 5 | 19 | 0 | 0 | 5 | 0 | 0 | 298 | 149 | 1 | 1 | 1 | 1 | 1 | 1 | 1 | NA | 1 | 0.286 | 1 | 1 | 0.178 |
| SE | 13603 | ENSG00000167261 | DPEP2 | chr16 | - | 67999266 | 67999748 | 67992063 | 67992193 | 68000447 | 68000586 | 1 | 2 | 1 | 0 | 0 | 2 | 4 | 4 | 9 | 4 | 7 | 0 | 0 | 2 | 0 | 0 | 0 | 0 | 0 | 0 | 298 | 149 | 1 | 1 | 0.2 | 0.2 | 0.111 | 0 | 0 | 1 | NA | NA | 1 | NA | -0.898 |
| SE | 13604 | ENSG00000167261 | DPEP2 | chr16 | - | 67999266 | 67999748 | 67992949 | 67993257 | 68000447 | 68000586 | 0 | 2 | 1 | 0 | 0 | 3 | 0 | 1 | 9 | 5 | 0 | 0 | 0 | 0 | 0 | 11 | 0 | 0 | 4 | 0 | 298 | 149 | 1 | 1 | 0 | 1 | 0.333 | 0 | 0 | 0 | NA | NA | 0 | NA | 0.267 |
| SE | 13608 | ENSG00000087253 | LPCAT2 | chr16 | + | 55532823 | 55532882 | 55531913 | 55531974 | 55537577 | 55537632 | 33 | 27 | 24 | 30 | 87 | 0 | 0 | 1 | 0 | 0 | 45 | 20 | 28 | 42 | 0 | 0 | 0 | 0 | 0 | 0 | 208 | 149 | 1 | 1 | 1 | 1 | 0.945 | 1 | 1 | 1 | 1 | 1 | 1 | NA | -0.011 |
| SE | 13610 | ENSG00000087253 | LPCAT2 | chr16 | + | 55549276 | 55549402 | 55545484 | 55545817 | 55550948 | 55551102 | 100 | 111 | 68 | 90 | 129 | 0 | 0 | 0 | 0 | 0 | 115 | 103 | 96 | 174 | 54 | 0 | 0 | 1 | 2 | 0 | 275 | 149 | 1 | 1 | 1 | 1 | 1 | 1 | 1 | 1 | 1 | 0.981 | 0.979 | 1 | 0.008 |
| SE | 13611 | ENSG00000087253 | LPCAT2 | chr16 | + | 55574630 | 55574729 | 55550948 | 55551102 | 55579108 | 55579244 | 121 | 121 | 100 | 119 | 167 | 0 | 3 | 0 | 0 | 0 | 114 | 132 | 157 | 282 | 61 | 0 | 0 | 0 | 0 | 0 | 248 | 149 | 1 | 1 | 1 | 0.96 | 1 | 1 | 1 | 1 | 1 | 1 | 1 | 1 | -0.008 |
| SE | 13614 | ENSG00000038358 | EDC4 | chr16 | + | 67881496 | 67881533 | 67881264 | 67881417 | 67881667 | 67881845 | 109 | 40 | 79 | 87 | 75 | 0 | 0 | 0 | 0 | 0 | 111 | 72 | 78 | 96 | 172 | 0 | 0 | 1 | 0 | 0 | 186 | 149 | 1 | 1 | 1 | 1 | 1 | 1 | 1 | 1 | 1 | 0.984 | 1 | 1 | 0.003 |
| SE | 13615 | ENSG00000038358 | EDC4 | chr16 | + | 67881667 | 67882109 | 67881496 | 67881533 | 67882211 | 67882327 | 125 | 53 | 103 | 102 | 83 | 0 | 0 | 0 | 0 | 0 | 124 | 102 | 111 | 130 | 174 | 0 | 3 | 0 | 0 | 0 | 298 | 149 | 1 | 1 | 1 | 1 | 1 | 1 | 1 | 1 | 0.944 | 1 | 1 | 1 | 0.011 |
| SE | 13619 | ENSG00000038358 | EDC4 | chr16 | + | 67883114 | 67883177 | 67882957 | 67883006 | 67883567 | 67883731 | 12 | 3 | 7 | 5 | 0 | 0 | 0 | 1 | 0 | 0 | 0 | 5 | 8 | 11 | 0 | 0 | 0 | 4 | 0 | 0 | 212 | 149 | 1 | 1 | 1 | 1 | 0.831 | 1 | NA | NA | 1 | 1 | 0.659 | NA | 0.071 |
| SE | 13620 | ENSG00000213398 | LCAT | chr16 | - | 67940736 | 67940831 | 67939881 | 67940478 | 67942362 | 67942587 | 0 | 0 | 0 | 2 | 0 | 5 | 4 | 3 | 10 | 0 | 0 | 0 | 8 | 2 | 0 | 0 | 5 | 0 | 0 | 8 | 244 | 149 | 1 | 1 | 0 | 0 | 0 | 0.109 | NA | NA | 0 | 1 | 1 | 0 | -0.473 |

| | | | | | | | | | | | | | | | | | | | | | | | | | |
|---|---|---|---|---|---|---|---|---|---|---|---|---|---|---|---|---|---|---|---|---|---|---|---|---|---|
| SE | 13642 | ENSG00000270049 | ATP6V0D1-DT | chr16 | + | 67492024 | 67492121 | 67484109 | 67484328 | 67504419 | 67505914 | 1 | 3 | 6 | 6 | 10 | 0 | 0 | 0 | 2 | 0 | 10 | 7 | 1 | 0 |
| SE | 13644 | ENSG00000102898 | NUTF2 | chr16 | + | 67847277 | 67847456 | 67846934 | 67846985 | 67868339 | 67868411 | 0 | 1 | 1 | 0 | 0 | 0 | 0 | 0 | 2 | 0 | 0 | 0 | 0 | 0 |
| SE | 13645 | ENSG00000102898 | NUTF2 | chr16 | + | 67865101 | 67865229 | 67846922 | 67846985 | 67868339 | 67868411 | 201 | 72 | 170 | 172 | 253 | 0 | 0 | 0 | 2 | 0 | 299 | 321 | 112 | 153 |
| SE | 13646 | ENSG00000102898 | NUTF2 | chr16 | + | 67865101 | 67865229 | 67847160 | 67847456 | 67868339 | 67868411 | 181 | 70 | 150 | 144 | 243 | 0 | 0 | 1 | 0 | 0 | 286 | 298 | 99 | 132 |
| SE | 13647 | ENSG00000197170 | PSMD12 | chr17 | - | 67344605 | 67344780 | 67342185 | 67342263 | 67345744 | 67345857 | 388 | 290 | 336 | 391 | 520 | 0 | 0 | 0 | 1 | 0 | 122 | 299 | 427 | 191 |
| SE | 13648 | ENSG00000197170 | PSMD12 | chr17 | - | 67345744 | 67345857 | 67344605 | 67344780 | 67347115 | 67347250 | 290 | 196 | 257 | 283 | 406 | 0 | 0 | 0 | 4 | 0 | 70 | 250 | 296 | 132 |
| SE | 13649 | ENSG00000197170 | PSMD12 | chr17 | - | 67350228 | 67350336 | 67348549 | 67348654 | 67357302 | 67357431 | 283 | 204 | 239 | 283 | 340 | 0 | 0 | 1 | 1 | 0 | 131 | 232 | 359 | 274 |
| SE | 13650 | ENSG00000197170 | PSMD12 | chr17 | - | 67355340 | 67355422 | 67350257 | 67350336 | 67357302 | 67357578 | 5 | 0 | 4 | 0 | 0 | 180 | 112 | 141 | 163 | 163 | 0 | 1 | 0 | 0 |
| SE | 13651 | ENSG00000197170 | PSMD12 | chr17 | - | 67357302 | 67357431 | 67350228 | 67350336 | 67366411 | 67366605 | 163 | 108 | 133 | 152 | 142 | 0 | 0 | 0 | 0 | 0 | 60 | 114 | 190 | 105 |
| SE | 13654 | ENSG00000197170 | PSMD12 | chr17 | - | 67362775 | 67362892 | 67361094 | 67361261 | 67366411 | 67366547 | 0 | 0 | 2 | 0 | 0 | 0 | 0 | 1 | 0 | 0 | 0 | 1 | 0 | 0 |
| SE | 13659 | ENSG00000140848 | CPNE2 | chr16 | + | 57134774 | 57134826 | 57127848 | 57127903 | 57146084 | 57146321 | 29 | 16 | 17 | 34 | 40 | 1 | 0 | 0 | 0 | 0 | 108 | 44 | 11 | 159 |
| SE | 13660 | ENSG00000140848 | CPNE2 | chr16 | + | 57134774 | 57134826 | 57129051 | 57129150 | 57146084 | 57146321 | 0 | 0 | 2 | 3 | 0 | 0 | 0 | 0 | 0 | 0 | 0 | 0 | 4 | 6 |
| SE | 13661 | ENSG00000140848 | CPNE2 | chr16 | + | 57137148 | 57137282 | 57134774 | 57134826 | 57146084 | 57146321 | 68 | 38 | 33 | 59 | 35 | 0 | 0 | 1 | 3 | 0 | 125 | 49 | 15 | 198 |
| SE | 13662 | ENSG00000169189 | NSMCE1 | chr16 | - | 27225725 | 27225846 | 27224993 | 27225236 | 27226719 | 27226836 | 153 | 62 | 117 | 119 | 219 | 0 | 0 | 1 | 0 | 0 | 131 | 118 | 30 | 99 |
| SE | 13663 | ENSG00000169189 | NSMCE1 | chr16 | - | 27241352 | 27241484 | 27235177 | 27235299 | 27241798 | 27241889 | 0 | 0 | 2 | 2 | 0 | 4 | 0 | 6 | 2 | 4 | 0 | 0 | 0 | 0 |
| SE | 13664 | ENSG00000169189 | NSMCE1 | chr16 | - | 27241352 | 27241889 | 27235177 | 27235299 | 27257434 | 27257581 | 0 | 0 | 5 | 2 | 0 | 104 | 64 | 85 | 104 | 31 | 0 | 7 | 0 | 0 |
| SE | 13669 | ENSG00000090565 | RAB11FIP3 | chr16 | + | 491133 | 491268 | 488850 | 489000 | 496823 | 496859 | 16 | 2 | 7 | 6 | 7 | 6 | 6 | 2 | 6 | 1 | 0 | 3 | 4 | 9 |
| SE | 13670 | ENSG00000090565 | RAB11FIP3 | chr16 | + | 491133 | 491268 | 488850 | 489000 | 503003 | 503097 | 13 | 0 | 3 | 1 | 2 | 1 | 0 | 0 | 2 | 0 | 0 | 2 | 1 | 5 |
| SE | 13671 | ENSG00000090565 | RAB11FIP3 | chr16 | + | 496823 | 496859 | 488850 | 489000 | 503003 | 503063 | 18 | 10 | 2 | 18 | 3 | 1 | 0 | 0 | 2 | 0 | 0 | 15 | 15 | 15 |
| SE | 13672 | ENSG00000090565 | RAB11FIP3 | chr16 | + | 496823 | 496859 | 491133 | 491268 | 503003 | 503063 | 13 | 7 | 3 | 17 | 7 | 0 | 0 | 0 | 1 | 0 | 0 | 8 | 11 | 13 |
| SE | 13673 | ENSG00000090565 | RAB11FIP3 | chr16 | + | 503003 | 503097 | 496823 | 496859 | 510659 | 510800 | 9 | 6 | 1 | 5 | 6 | 0 | 0 | 0 | 0 | 0 | 0 | 3 | 5 | 1 |
| SE | 13675 | ENSG00000006007 | GDE1 | chr16 | - | 19504880 | 19505092 | 19501692 | 19503617 | 19507686 | 19507779 | 302 | 193 | 165 | 230 | 434 | 0 | 0 | 0 | 0 | 0 | 681 | 368 | 355 | 400 |
| SE | 13676 | ENSG00000006007 | GDE1 | chr16 | - | 19504880 | 19505092 | 19501692 | 19503617 | 19510838 | 19510944 | 88 | 46 | 58 | 80 | 110 | 0 | 1 | 0 | 0 | 0 | 110 | 75 | 69 | 87 |
| SE | 13678 | ENSG00000006007 | GDE1 | chr16 | - | 19507686 | 19507779 | 19501692 | 19503617 | 19510838 | 19510944 | 73 | 52 | 42 | 64 | 99 | 0 | 1 | 0 | 0 | 0 | 237 | 84 | 110 | 124 |
| SE | 13679 | ENSG00000006007 | GDE1 | chr16 | - | 19507686 | 19507779 | 19504880 | 19505092 | 19510838 | 19510944 | 287 | 200 | 148 | 215 | 425 | 4 | 1 | 1 | 3 | 2 | 802 | 376 | 394 | 438 |
| SE | 13680 | ENSG00000006007 | GDE1 | chr16 | - | 19517013 | 19517185 | 19510838 | 19510944 | 19521703 | 19522123 | 159 | 95 | 89 | 105 | 272 | 0 | 0 | 0 | 0 | 0 | 413 | 228 | 215 | 212 |
| SE | 13681 | ENSG00000006007 | GDE1 | chr16 | - | 19517013 | 19517189 | 19510838 | 19510944 | 19521703 | 19522098 | 173 | 101 | 98 | 122 | 277 | 0 | 0 | 0 | 0 | 0 | 424 | 257 | 233 | 218 |
| SE | 13682 | ENSG00000077235 | GTF3C1 | chr16 | - | 27462293 | 27462486 | 27460612 | 27461562 | 27463540 | 27463592 | 89 | 36 | 47 | 62 | 31 | 1 | 0 | 0 | 0 | 0 | 13 | 51 | 24 | 27 |
| SE | 13683 | ENSG00000077235 | GTF3C1 | chr16 | - | 27462293 | 27462486 | 27460612 | 27461562 | 27464319 | 27464409 | 52 | 12 | 23 | 24 | 15 | 0 | 0 | 1 | 0 | 0 | 13 | 25 | 15 | 14 |
| SE | 13685 | ENSG00000077235 | GTF3C1 | chr16 | - | 27463540 | 27463592 | 27462293 | 27462486 | 27464319 | 27464409 | 40 | 24 | 23 | 39 | 17 | 0 | 0 | 0 | 0 | 0 | 0 | 29 | 7 | 12 |
| SE | 13690 | ENSG00000077235 | GTF3C1 | chr16 | - | 27488553 | 27488635 | 27488218 | 27488415 | 27489042 | 27489123 | 192 | 114 | 133 | 191 | 99 | 0 | 0 | 0 | 0 | 0 | 217 | 266 | 155 | 254 |
| SE | 13693 | ENSG00000077235 | GTF3C1 | chr16 | - | 27498629 | 27498733 | 27495210 | 27495492 | 27501190 | 27501344 | 114 | 54 | 84 | 81 | 143 | 0 | 0 | 0 | 0 | 0 | 155 | 87 | 57 | 73 |
| SE | 13694 | ENSG00000077235 | GTF3C1 | chr16 | - | 27506846 | 27507156 | 27505898 | 27506116 | 27511748 | 27511901 | 83 | 53 | 70 | 99 | 93 | 0 | 0 | 0 | 0 | 0 | 112 | 101 | 69 | 70 |
| SE | 13696 | ENSG00000077235 | GTF3C1 | chr16 | - | 27528597 | 27528721 | 27511748 | 27511901 | 27533290 | 27533387 | 227 | 143 | 182 | 272 | 207 | 0 | 0 | 0 | 0 | 0 | 455 | 300 | 160 | 220 |
| SE | 13697 | ENSG00000077235 | GTF3C1 | chr16 | - | 27533290 | 27533387 | 27528597 | 27528721 | 27537783 | 27537927 | 184 | 116 | 164 | 233 | 175 | 0 | 0 | 0 | 0 | 3 | 402 | 258 | 148 | 215 |
| SE | 13698 | ENSG00000077235 | GTF3C1 | chr16 | - | 27545313 | 27545523 | 27538179 | 27538356 | 27549669 | 27549913 | 195 | 118 | 199 | 176 | 272 | 0 | 0 | 0 | 0 | 0 | 274 | 319 | 167 | 200 |
| SE | 13699 | ENSG00000140961 | OSGIN1 | chr16 | + | 83951127 | 83951287 | 83949066 | 83949506 | 83957639 | 83957738 | 2 | 0 | 4 | 1 | 0 | 0 | 3 | 0 | 0 | 0 | 0 | 1 | 0 | 0 |
| SE | 13705 | ENSG00000141086 | CTRL | chr16 | - | 67931097 | 67931201 | 67930919 | 67930999 | 67931800 | 67931862 | 0 | 4 | 5 | 0 | 0 | 0 | 0 | 0 | 0 | 0 | 10 | 0 | 0 | 0 |
| SE | 13707 | ENSG00000038532 | CLEC16A | chr16 | + | 10962454 | 10962588 | 10957781 | 10957910 | 10969160 | 10969309 | 136 | 93 | 146 | 182 | 298 | 0 | 0 | 0 | 0 | 0 | 340 | 256 | 199 | 320 |
| SE | 13711 | ENSG00000038532 | CLEC16A | chr16 | + | 10977224 | 10977399 | 10972937 | 10973061 | 11003073 | 11003257 | 69 | 48 | 50 | 90 | 142 | 0 | 1 | 0 | 0 | 0 | 96 | 60 | 60 | 106 |
| SE | 13713 | ENSG00000038532 | CLEC16A | chr16 | + | 10979328 | 10979382 | 10972937 | 10973061 | 11003073 | 11003257 | 0 | 0 | 0 | 2 | 2 | 0 | 1 | 0 | 0 | 0 | 0 | 2 | 2 | 0 |
| SE | 13714 | ENSG00000038532 | CLEC16A | chr16 | + | 10979328 | 10979382 | 10977224 | 10977399 | 10982877 | 10982991 | 66 | 56 | 76 | 103 | 136 | 2 | 0 | 0 | 2 | 0 | 219 | 195 | 123 | 170 |
| SE | 13715 | ENSG00000038532 | CLEC16A | chr16 | + | 10979328 | 10979382 | 10977224 | 10977399 | 11003073 | 11003257 | 13 | 9 | 16 | 27 | 35 | 0 | 0 | 0 | 0 | 0 | 17 | 39 | 20 | 26 |
| SE | 13716 | ENSG00000038532 | CLEC16A | chr16 | + | 10982877 | 10982991 | 10972937 | 10973061 | 11003073 | 11003257 | 68 | 62 | 103 | 126 | 128 | 0 | 1 | 0 | 0 | 0 | 313 | 221 | 143 | 213 |
| SE | 13717 | ENSG00000038532 | CLEC16A | chr16 | + | 10982877 | 10982991 | 10977224 | 10977399 | 11003073 | 11003257 | 70 | 62 | 103 | 127 | 123 | 0 | 0 | 0 | 0 | 0 | 313 | 221 | 141 | 213 |
| SE | 13718 | ENSG00000038532 | CLEC16A | chr16 | + | 10982877 | 10982991 | 10977224 | 10977399 | 11020192 | 11020325 | 6 | 3 | 1 | 2 | 2 | 1 | 0 | 0 | 0 | 0 | 0 | 0 | 3 | 1 |
| SE | 13719 | ENSG00000038532 | CLEC16A | chr16 | + | 10982877 | 10982991 | 10979328 | 10979382 | 11003073 | 11003257 | 123 | 111 | 165 | 205 | 233 | 0 | 0 | 0 | 0 | 2 | 525 | 372 | 246 | 367 |
| SE | 13720 | ENSG00000038532 | CLEC16A | chr16 | + | 11003073 | 11003257 | 10977224 | 10977399 | 11020192 | 11020325 | 22 | 24 | 43 | 42 | 53 | 1 | 0 | 0 | 0 | 0 | 53 | 58 | 50 | 91 |
| SE | 13722 | ENSG00000038532 | CLEC16A | chr16 | + | 11003073 | 11003305 | 10977224 | 10977399 | 11020192 | 11020325 | 23 | 13 | 17 | 9 | 38 | 1 | 0 | 0 | 0 | 0 | 16 | 28 | 14 | 15 |
| SE | 13724 | ENSG00000038532 | CLEC16A | chr16 | + | 11020192 | 11020325 | 10982877 | 10982991 | 11024820 | 11024919 | 19 | 15 | 16 | 26 | 22 | 0 | 0 | 0 | 0 | 0 | 13 | 28 | 25 | 45 |
| SE | 13726 | ENSG00000038532 | CLEC16A | chr16 | + | 11039753 | 11039876 | 11024817 | 11024921 | 11042253 | 11042363 | 129 | 71 | 114 | 120 | 107 | 3 | 0 | 0 | 0 | 0 | 110 | 115 | 130 | 181 |
| SE | 13727 | ENSG00000038532 | CLEC16A | chr16 | + | 11051512 | 11051641 | 11047291 | 11047342 | 11060901 | 11061022 | 133 | 72 | 73 | 108 | 80 | 0 | 0 | 0 | 0 | 0 | 135 | 78 | 114 | 136 |
| SE | 13728 | ENSG00000038532 | CLEC16A | chr16 | + | 11060901 | 11061022 | 11051512 | 11051641 | 11120614 | 11120766 | 83 | 44 | 62 | 81 | 35 | 0 | 0 | 0 | 0 | 0 | 52 | 39 | 89 | 107 |
| SE | 13741 | ENSG00000122299 | ZC3H7A | chr16 | - | 11761467 | 11761511 | 11758430 | 11758539 | 11761909 | 11762043 | 39 | 50 | 50 | 55 | 48 | 11 | 0 | 3 | 8 | 3 | 59 | 49 | 58 | 32 |
| SE | 13743 | ENSG00000122299 | ZC3H7A | chr16 | - | 11763477 | 11763659 | 11762670 | 11762747 | 11765052 | 11765153 | 320 | 230 | 296 | 303 | 458 | 0 | 2 | 0 | 0 | 0 | 278 | 269 | 353 | 227 |
| SE | 13744 | ENSG00000122299 | ZC3H7A | chr16 | - | 11765052 | 11765153 | 11763477 | 11763659 | 11765488 | 11765561 | 220 | 167 | 201 | 232 | 327 | 0 | 2 | 0 | 0 | 0 | 228 | 226 | 267 | 168 |
| SE | 13746 | ENSG00000122299 | ZC3H7A | chr16 | - | 11769030 | 11769095 | 11768314 | 11768501 | 11774235 | 11774519 | 86 | 73 | 88 | 77 | 130 | 1 | 0 | 0 | 0 | 0 | 94 | 65 | 166 | 84 |
| SE | 13747 | ENSG00000122299 | ZC3H7A | chr16 | - | 11770782 | 11770987 | 11768314 | 11768501 | 11774235 | 11774292 | 153 | 105 | 165 | 159 | 318 | 1 | 0 | 0 | 0 | 0 | 143 | 226 | 242 | 154 |
| SE | 13748 | ENSG00000122299 | ZC3H7A | chr16 | - | 11770782 | 11770987 | 11769030 | 11769095 | 11774235 | 11774292 | 262 | 222 | 292 | 289 | 513 | 0 | 0 | 0 | 0 | 0 | 243 | 324 | 433 | 234 |
| SE | 13749 | ENSG00000122299 | ZC3H7A | chr16 | - | 11776319 | 11776358 | 11774979 | 11775013 | 11776451 | 11776532 | 217 | 162 | 200 | 231 | 281 | 0 | 4 | 2 | 0 | 2 | 229 | 258 | 301 | 312 |
| SE | 13750 | ENSG00000122299 | ZC3H7A | chr16 | - | 11776319 | 11776358 | 11774979 | 11775013 | 11779165 | 11779363 | 34 | 6 | 24 | 22 | 39 | 0 | 0 | 0 | 0 | 0 | 5 | 55 | 25 | 10 |
| SE | 13751 | ENSG00000122299 | ZC3H7A | chr16 | - | 11776451 | 11776532 | 11776319 | 11776358 | 11776750 | 11776909 | 279 | 268 | 265 | 322 | 438 | 0 | 0 | 0 | 0 | 0 | 281 | 287 | 444 | 421 |
| SE | 13755 | ENSG00000122299 | ZC3H7A | chr16 | - | 11779165 | 11779363 | 11776750 | 11776909 | 11781424 | 11781464 | 331 | 253 | 356 | 304 | 741 | 0 | 0 | 0 | 1 | 0 | 270 | 285 | 419 | 289 |
| SE | 13757 | ENSG00000234719 | NPIPB2 | chr16 | - | 11929384 | 11929445 | 11928623 | 11929275 | 11933516 | 11933661 | 3 | 2 | 3 | 11 | 8 | 0 | 0 | 0 | 1 | 0 | 0 | 0 | 3 | 5 |
| SE | 13758 | ENSG00000234719 | NPIPB2 | chr16 | - | 11930449 | 11930551 | 11929384 | 11929445 | 11933516 | 11933661 | 0 | 0 | 1 | 1 | 0 | 0 | 2 | 2 | 6 | 7 | 0 | 0 | 0 | 0 |
| SE | 13760 | ENSG00000234719 | NPIPB2 | chr16 | - | 11956143 | 11956237 | 11937539 | 11937668 | 11976567 | 11976638 | 1 | 0 | 3 | 2 | 0 | 1 | 2 | 1 | 0 | 0 | 11 | 5 | 0 | 0 |
| SE | 13762 | ENSG00000264772 | NA | chr17 | + | 7580465 | 7580585 | 7577568 | 7580327 | 7580710 | 7580783 | 1155 | 574 | 648 | 796 | 738 | 0 | 0 | 0 | 3 | 1 | 2607 | 1115 | 687 | 1344 |
| SE | 13763 | ENSG00000264772 | NA | chr17 | + | 7580710 | 7580783 | 7580465 | 7580585 | 7580895 | 7581066 | 1265 | 608 | 695 | 822 | 754 | 0 | 0 | 0 | 0 | 0 | 2689 | 1158 | 694 | 1280 |
| SE | 13764 | ENSG00000264772 | NA | chr17 | + | 7580895 | 7581066 | 7580710 | 7580783 | 7581377 | 7582024 | 1514 | 688 | 803 | 946 | 797 | 0 | 0 | 0 | 0 | 0 | 2795 | 1464 | 884 | 1570 |
| SE | 13768 | ENSG00000171241 | SHCBP1 | chr16 | - | 46595551 | 46595670 | 46584002 | 46584089 | 46599830 | 46599962 | 8 | 8 | 26 | 15 | 10 | 0 | 0 | 0 | 0 | 0 | 7 | 19 | 44 | 26 |
| SE | 13769 | ENSG00000171241 | SHCBP1 | chr16 | - | 46604227 | 46604461 | 46603593 | 46604143 | 46608296 | 46608389 | 8 | 10 | 23 | 11 | 20 | 0 | 0 | 0 | 0 | 0 | 6 | 29 | 14 | 55 |
| SE | 13771 | ENSG00000171241 | SHCBP1 | chr16 | - | 46618204 | 46618372 | 46617633 | 46617749 | 46621256 | 46621368 | 1 | 4 | 7 | 0 | 5 | 0 | 0 | 0 | 0 | 0 | 0 | 4 | 23 | 11 |
| SE | 13777 | ENSG00000205629 | LCMT1 | chr16 | + | 25161101 | 25161204 | 25151530 | 25151615 | 25164597 | 25164718 | 132 | 77 | 111 | 128 | 136 | 0 | 2 | 0 | 0 | 0 | 161 | 143 | 76 | 116 |
| SE | 13780 | ENSG00000205629 | LCMT1 | chr16 | + | 25170713 | 25170805 | 25169111 | 25169213 | 25178000 | 25178226 | 41 | 8 | 20 | 26 | 37 | 0 | 0 | 0 | 0 | 0 | 21 | 35 | 12 | 24 |
| SE | 13781 | ENSG00000205629 | LCMT1 | chr16 | + | 25174936 | 25175034 | 25169111 | 25169213 | 25178000 | 25178023 | 13 | 13 | 14 | 15 | 44 | 0 | 0 | 0 | 0 | 0 | 38 | 27 | 12 | 17 |
| SE | 13784 | ENSG00000129636 | ITFG1 | chr16 | - | 47237964 | 47238008 | 47218867 | 47218946 | 47258631 | 47258722 | 116 | 100 | 69 | 119 | 230 | 0 | 1 | 0 | 0 | 1 | 104 | 127 | 108 | 93 |
| SE | 13785 | ENSG00000129636 | ITFG1 | chr16 | - | 47313728 | 47313823 | 47311239 | 47311412 | 47365787 | 47365869 | 426 | 372 | 294 | 350 | 412 | 2 | 0 | 0 | 0 | 1 | 341 | 352 | 342 | 381 |
| SE | 13786 | ENSG00000129636 | ITFG1 | chr16 | - | 47454012 | 47454158 | 47452732 | 47452790 | 47459102 | 47459130 | 732 | 573 | 496 | 689 | 951 | 0 | 1 | 0 | 0 | 2 | 838 | 679 | 593 | 553 |
| SE | 13791 | ENSG00000161999 | JMJD8 | chr16 | - | 684065 | 684155 | 683860 | 683909 | 684233 | 684334 | 5 | 0 | 0 | 5 | 0 | 2 | 0 | 0 | 0 | 0 | 0 | 7 | 1 | 2 |
| SE | 13792 | ENSG00000166123 | GPT2 | chr16 | + | 46885466 | 46885577 | 46884693 | 46884958 | 46897647 | 46897737 | 0 | 0 | 0 | 1 | 0 | 1 | 0 | 3 | 1 | 0 | 0 | 0 | 0 | 0 |
| SE | 13793 | ENSG00000166123 | GPT2 | chr16 | + | 46900681 | 46900790 | 46897647 | 46897737 | 46906841 | 46906975 | 2 | 0 | 10 | 3 | 0 | 0 | 0 | 0 | 0 | 0 | 0 | 0 | 9 | 0 |
| SE | 13794 | ENSG00000166123 | GPT2 | chr16 | + | 46918620 | 46918757 | 46915297 | 46916707 | 46922241 | 46922416 | 5 | 3 | 9 | 5 | 0 | 0 | 0 | 0 | 0 | 0 | 36 | 3 | 0 | 2 |
| SE | 13800 | ENSG00000130733 | YIPF2 | chr19 | - | 10925685 | 10925773 | 10923841 | 10924192 | 10927629 | 10927716 | 31 | 9 | 13 | 17 | 8 | 0 | 0 | 2 | 0 | 0 | 7 | 18 | 15 | 9 |
| SE | 13809 | ENSG00000102910 | LONP2 | chr16 | + | 48262777 | 48262872 | 48261423 | 48261587 | 48270015 | 48270274 | 44 | 54 | 275 | 250 | 317 | 0 | 0 | 0 | 0 | 1 | 256 | 200 | 279 | 150 |
| SE | 13810 | ENSG00000102910 | LONP2 | chr16 | + | 48296014 | 48296165 | 48277337 | 48277479 | 48299661 | 48299788 | 383 | 267 | 359 | 318 | 654 | 1 | 0 | 0 | 0 | 0 | 405 | 364 | 341 | 341 |
| SE | 13811 | ENSG00000102910 | LONP2 | chr16 | + | 48296084 | 48296165 | 48277337 | 48277479 | 48299661 | 48299788 | 137 | 108 | 128 | 104 | 211 | 1 | 0 | 0 | 0 | 0 | 125 | 110 | 126 | 124 |
| SE | 13812 | ENSG00000102910 | LONP2 | chr16 | + | 48299661 | 48299788 | 48296014 | 48296165 | 48303171 | 48303283 | 346 | 248 | 323 | 277 | 502 | 4 | 0 | 1 | 0 | 0 | 364 | 339 | 310 | 280 |
| SE | 13813 | ENSG00000102910 | LONP2 | chr16 | + | 48303171 | 48303305 | 48299661 | 48299788 | 48334215 | 48334358 | 285 | 157 | 212 | 254 | 320 | 0 | 0 | 1 | 0 | 0 | 265 | 273 | 188 | 177 |
| SE | 13814 | ENSG00000102910 | LONP2 | chr16 | + | 48336392 | 48336498 | 48334215 | 48334358 | 48347506 | 48347616 | 0 | 0 | 2 | 0 | 0 | 60 | 64 | 57 | 79 | 102 | 0 | 0 | 0 | 0 |
| SE | 13816 | ENSG00000102910 | LONP2 | chr16 | + | 48347506 | 48347714 | 48334215 | 48334358 | 48351580 | 48351685 | 61 | 64 | 57 | 79 | 102 | 3 | 0 | 0 | 1 | 0 | 67 | 69 | 37 | 72 |
| SE | 13817 | ENSG00000102910 | LONP2 | chr16 | + | 48348099 | 48348290 | 48334215 | 48334358 | 48351580 | 48351685 | 162 | 106 | 150 | 156 | 108 | 3 | 0 | 0 | 1 | 0 | 187 | 150 | 149 | 142 |
| SE | 13818 | ENSG00000102910 | LONP2 | chr16 | + | 48348099 | 48348290 | 48347506 | 48347714 | 48351580 | 48351685 | 329 | 188 | 266 | 289 | 212 | 1 | 0 | 0 | 0 | 0 | 324 | 248 | 245 | 255 |
| SE | 13819 | ENSG00000102910 | LONP2 | chr16 | + | 48356664 | 48356710 | 48351580 | 48351798 | 48361476 | 48361836 | 3 | 3 | 6 | 9 | 4 | 3 | 0 | 0 | 0 | 0 | 0 | 0 | 3 | 2 |
| SE | 13828 | ENSG00000068903 | SIRT2 | chr19 | - | 38880820 | 38880897 | 38880684 | 38880736 | 38881099 | 38881155 | 71 | 20 | 56 | 64 | 55 | 0 | 0 | 0 | 0 | 0 | 111 | 90 | 37 | 53 |
| SE | 13829 | ENSG00000068903 | SIRT2 | chr19 | - | 38889253 | 38889323 | 38889086 | 38889155 | 38889688 | 38889745 | 0 | 0 | 0 | 0 | 0 | 54 | 16 | 28 | 45 | 33 | 0 | 0 | 1 | 0 |
| SE | 13832 | ENSG00000068903 | SIRT2 | chr19 | - | 38893413 | 38893527 | 38890102 | 38890144 | 38893818 | 38893867 | 170 | 52 | 93 | 138 | 86 | 0 | 0 | 0 | 0 | 0 | 173 | 212 | 104 | 104 |
| SE | 13833 | ENSG00000068903 | SIRT2 | chr19 | - | 38893818 | 38893867 | 38890102 | 38890144 | 38899505 | 38899518 | 5 | 3 | 3 | 9 | 3 | 0 | 2 | 0 | 0 | 0 | 5 | 0 | 3 | 7 |
| SE | 13840 | ENSG00000068903 | SIRT2 | chr19 | - | 38898793 | 38898918 | 38898378 | 38898425 | 38899505 | 38899714 | 5 | 2 | 0 | 0 | 1 | 12 | 4 | 0 | 2 | 1 | 0 | 1 | 0 | 0 |
| SE | 13845 | ENSG00000102893 | PHKB | chr16 | + | 47462340 | 47462452 | 47461349 | 47461426 | 47463881 | 47463993 | 0 | 1 | 2 | 0 | 0 | 12 | 11 | 10 | 8 | 9 | 0 | 0 | 0 | 0 |
| SE | 13846 | ENSG00000102893 | PHKB | chr16 | + | 47462340 | 47462452 | 47461349 | 47461426 | 47497398 | 47497488 | 0 | 1 | 2 | 1 | 0 | 23 | 46 | 23 | 41 | 11 | 0 | 0 | 0 | 0 |
| SE | 13850 | ENSG00000102893 | PHKB | chr16 | + | 47499755 | 47499894 | 47497398 | 47497488 | 47502990 | 47503090 | 400 | 390 | 285 | 446 | 535 | 1 | 0 | 0 | 0 | 0 | 622 | 628 | 364 | 379 |
| SE | 13851 | ENSG00000102893 | PHKB | chr16 | + | 47502990 | 47503090 | 47499755 | 47499894 | 47511664 | 47511731 | 248 | 206 | 175 | 278 | 386 | 3 | 0 | 0 | 0 | 3 | 461 | 420 | 233 | 239 |
| SE | 13862 | ENSG00000102893 | PHKB | chr16 | + | 47698447 | 47698588 | 47696380 | 47696488 | 47699228 | 47699745 | 240 | 280 | 245 | 340 | 370 | 0 | 0 | 1 | 2 | 0 | 130 | 173 | 221 | 244 |
| SE | 13866 | ENSG00000174628 | IQCK | chr16 | + | 19730429 | 19730494 | 19717694 | 19718487 | 19733697 | 19733827 | 1 | 3 | 3 | 4 | 2 | 0 | 0 | 0 | 0 | 0 | 0 | 0 | 0 | 0 |
| SE | 13869 | ENSG00000174628 | IQCK | chr16 | + | 19763847 | 19764112 | 19735352 | 19735450 | 19788837 | 19788922 | 1 | 0 | 6 | 0 | 5 | 0 | 0 | 0 | 0 | 0 | 0 | 1 | 0 | 0 |
| SE | 13871 | ENSG00000103319 | EEF2K | chr16 | + | 22250653 | 22250691 | 22248754 | 22248815 | 22251150 | 22251322 | 26 | 19 | 30 | 26 | 24 | 0 | 1 | 0 | 0 | 0 | 44 | 28 | 15 | 43 |
| SE | 13872 | ENSG00000103319 | EEF2K | chr16 | + | 22257642 | 22257770 | 22257252 | 22257385 | 22258493 | 22258695 | 43 | 22 | 30 | 47 | 34 | 0 | 0 | 0 | 0 | 0 | 46 | 40 | 29 | 34 |
| SE | 13873 | ENSG00000103319 | EEF2K | chr16 | + | 22266389 | 22266524 | 22264817 | 22264880 | 22266687 | 22266876 | 39 | 32 | 64 | 32 | 38 | 1 | 0 | 1 | 1 | 0 | 52 | 42 | 22 | 13 |
| SE | 13875 | ENSG00000180096 | SEPTIN1 | chr16 | - | 30379434 | 30379534 | 30379017 | 30379183 | 30379765 | 30380033 | 132 | 75 | 145 | 112 | 123 | 0 | 1 | 0 | 0 | 0 | 40 | 92 | 80 | 71 |
| SE | 13876 | ENSG00000180096 | SEPTIN1 | chr16 | - | 30379434 | 30379534 | 30379017 | 30379183 | 30379931 | 30380033 | 510 | 275 | 471 | 484 | 432 | 0 | 0 | 0 | 3 | 0 | 266 | 323 | 205 | 141 |
| SE | 13877 | ENSG00000180096 | SEPTIN1 | chr16 | - | 30381126 | 30381244 | 30379931 | 30380033 | 30381759 | 30381883 | 193 | 100 | 177 | 164 | 196 | 0 | 0 | 1 | 3 | 0 | 80 | 170 | 65 | 56 |
| SE | 13878 | ENSG00000180096 | SEPTIN1 | chr16 | - | 30381126 | 30381473 | 30379765 | 30380033 | 30381759 | 30381883 | 546 | 278 | 506 | 501 | 416 | 0 | 0 | 1 | 3 | 0 | 297 | 405 | 170 | 125 |
| SE | 13879 | ENSG00000180096 | SEPTIN1 | chr16 | - | 30381338 | 30381473 | 30381126 | 30381244 | 30381759 | 30381883 | 667 | 340 | 603 | 625 | 516 | 0 | 0 | 0 | 1 | 0 | 330 | 482 | 196 | 138 |
| SE | 13881 | ENSG00000180096 | SEPTIN1 | chr16 | - | 30382092 | 30382365 | 30381842 | 30381883 | 30382524 | 30382598 | 153 | 52 | 169 | 124 | 103 | 0 | 0 | 0 | 0 | 0 | 69 | 152 | 52 | 47 |
| SE | 13882 | ENSG00000180096 | SEPTIN1 | chr16 | - | 30382524 | 30382598 | 30382092 | 30382365 | 30382734 | 30382813 | 59 | 27 | 68 | 59 | 38 | 0 | 1 | 1 | 0 | 0 | 8 | 52 | 21 | 12 |
| SE | 13886 | ENSG00000103512 | NOMO1 | chr16 | + | 14889095 | 14889215 | 14886760 | 14886862 | 14894997 | 14895090 | 133 | 76 | 125 | 142 | 104 | 0 | 0 | 0 | 0 | 0 | 172 | 164 | 138 | 106 |
| SE | 13890 | ENSG00000069329 | VPS35 | chr16 | - | 46662982 | 46663162 | 46662242 | 46662482 | 46668929 | 46669052 | 381 | 286 | 281 | 367 | 538 | 1 | 1 | 0 | 1 | 0 | 103 | 248 | 176 | 288 |
| SE | 13893 | ENSG00000069329 | VPS35 | chr16 | - | 46674563 | 46674660 | 46674313 | 46674462 | 46676582 | 46676692 | 619 | 416 | 510 | 620 | 825 | 0 | 0 | 0 | 1 | 0 | 232 | 458 | 202 | 346 |
| SE | 13894 | ENSG00000069329 | VPS35 | chr16 | - | 46677314 | 46677398 | 46676504 | 46676692 | 46678942 | 46679043 | 479 | 322 | 397 | 455 | 651 | 0 | 0 | 0 | 0 | 0 | 111 | 345 | 124 | 203 |
| SE | 13895 | ENSG00000069329 | VPS35 | chr16 | - | 46678942 | 46679156 | 46677314 | 46677398 | 46680670 | 46680853 | 870 | 778 | 703 | 844 | 1567 | 2 | 1 | 0 | 0 | 0 | 438 | 699 | 518 | 680 |
| SE | 13896 | ENSG00000069329 | VPS35 | chr16 | - | 46682477 | 46682666 | 46682078 | 46682175 | 46683507 | 46683606 | 0 | 0 | 0 | 0 | 0 | 182 | 104 | 113 | 167 | 80 | 0 | 0 | 0 | 1 |
| SE | 13907 | ENSG00000067365 | METTL22 | chr16 | + | 8623852 | 8624118 | 8621709 | 8621775 | 8625495 | 8625713 | 5 | 2 | 2 | 0 | 0 | 4 | 5 | 4 | 16 | 0 | 14 | 4 | 12 | 3 |
| SE | 13909 | ENSG00000067365 | METTL22 | chr16 | + | 8635038 | 8635079 | 8631246 | 8632138 | 8635167 | 8635282 | 7 | 1 | 3 | 3 | 3 | 0 | 1 | 0 | 0 | 0 | 8 | 0 | 0 | 1 |
| SE | 13910 | ENSG00000067365 | METTL22 | chr16 | + | 8635167 | 8635312 | 8628729 | 8629110 | 8639090 | 8639162 | 22 | 27 | 25 | 21 | 31 | 0 | 0 | 1 | 0 | 0 | 77 | 16 | 17 | 29 |
| SE | 13915 | ENSG00000069345 | DNAJA2 | chr16 | - | 46964610 | 46964807 | 46959274 | 46959419 | 46967512 | 46967646 | 542 | 435 | 540 | 640 | 1050 | 0 | 0 | 3 | 0 | 1 | 875 | 564 | 524 | 689 |
| SE | 13917 | ENSG00000069345 | DNAJA2 | chr16 | - | 46967512 | 46967646 | 46959274 | 46959419 | 46968083 | 46968164 | 435 | 292 | 376 | 492 | 724 | 0 | 0 | 0 | 0 | 0 | 649 | 449 | 422 | 591 |
| SE | 13919 | ENSG00000069345 | DNAJA2 | chr16 | - | 46968083 | 46968164 | 46967512 | 46967646 | 46971348 | 46971572 | 642 | 426 | 585 | 748 | 1097 | 0 | 0 | 0 | 0 | 0 | 1065 | 691 | 634 | 844 |
| SE | 13920 | ENSG00000069345 | DNAJA2 | chr16 | - | 46971348 | 46971572 | 46968083 | 46968164 | 46971895 | 46971955 | 545 | 405 | 461 | 680 | 859 | 0 | 0 | 0 | 1 | 0 | 907 | 558 | 606 | 651 |
| SE | 13921 | ENSG00000069345 | DNAJA2 | chr16 | - | 46971348 | 46971659 | 46968083 | 46968164 | 46971895 | 46971955 | 337 | 248 | 305 | 399 | 617 | 0 | 0 | 0 | 1 | 0 | 550 | 363 | 326 | 408 |
| SE | 13922 | ENSG00000069345 | DNAJA2 | chr16 | - | 46971895 | 46971955 | 46971348 | 46971572 | 46973494 | 46973664 | 214 | 144 | 162 | 290 | 218 | 0 | 0 | 0 | 0 | 0 | 348 | 187 | 277 | 254 |
| SE | 13930 | ENSG00000131797 | CLUHP3 | chr16 | + | 31701662 | 31701858 | 31700589 | 31700738 | 31703155 | 31703367 | 12 | 7 | 6 | 10 | 0 | 0 | 5 | 5 | 4 | 0 | 8 | 7 | 6 | 5 |
| SE | 13933 | ENSG00000131797 | CLUHP3 | chr16 | + | 31705348 | 31705472 | 31704219 | 31704390 | 31705671 | 31705792 | 33 | 25 | 62 | 62 | 82 | 1 | 0 | 0 | 0 | 0 | 22 | 38 | 45 | 74 |
| SE | 13934 | ENSG00000131797 | CLUHP3 | chr16 | + | 31705671 | 31705792 | 31705348 | 31705472 | 31705889 | 31706034 | 37 | 26 | 81 | 84 | 102 | 0 | 0 | 0 | 0 | 0 | 26 | 40 | 67 | 74 |
| SE | 13936 | ENSG00000131797 | CLUHP3 | chr16 | + | 31705889 | 31705994 | 31705669 | 31705792 | 31706890 | 31707351 | 7 | 8 | 22 | 36 | 19 | 0 | 0 | 0 | 0 | 3 | 14 | 8 | 14 | 10 |
| SE | 13938 | ENSG00000131797 | CLUHP3 | chr16 | + | 31705889 | 31706034 | 31705671 | 31705792 | 31706890 | 31707422 | 23 | 19 | 45 | 70 | 44 | 0 | 0 | 0 | 0 | 3 | 18 | 15 | 42 | 28 |
| SE | 13939 | ENSG00000131797 | CLUHP3 | chr16 | + | 31706420 | 31706454 | 31705889 | 31705994 | 31706890 | 31707351 | 0 | 0 | 0 | 0 | 2 | 0 | 4 | 1 | 2 | 2 | 0 | 0 | 0 | 0 |
| SE | 13942 | ENSG00000262246 | CORO7 | chr16 | - | 4357967 | 4358103 | 4356431 | 4357259 | 4358366 | 4358450 | 306 | 155 | 309 | 241 | 188 | 3 | 0 | 0 | 0 | 0 | 430 | 564 | 332 | 436 |
| SE | 13945 | ENSG00000103426 | CORO7-PAM16 | chr16 | - | 4388544 | 4388631 | 4387985 | 4388068 | 4395288 | 4395339 | 77 | 30 | 64 | 73 | 16 | 0 | 0 | 0 | 0 | 0 | 74 | 97 | 49 | 69 |
| SE | 13946 | ENSG00000103426 | CORO7-PAM16 | chr16 | - | 4412355 | 4412430 | 4408180 | 4408251 | 4413307 | 4413404 | 238 | 90 | 155 | 155 | 56 | 0 | 0 | 0 | 0 | 0 | 256 | 224 | 148 | 197 |
| SE | 13947 | ENSG00000262246 | CORO7 | chr16 | - | 4413307 | 4413404 | 4412355 | 4412430 | 4416458 | 4416595 | 148 | 57 | 79 | 100 | 35 | 0 | 1 | 0 | 1 | 0 | 147 | 142 | 103 | 113 |
| SE | 13950 | ENSG00000243716 | NPIPB5 | chr16 | + | 22516764 | 22517102 | 22513763 | 22513921 | 22519137 | 22519235 | 2 | 1 | 0 | 2 | 3 | 2 | 1 | 2 | 0 | 0 | 0 | 1 | 0 | 0 |
| SE | 13951 | ENSG00000243716 | NPIPB5 | chr16 | + | 22519137 | 22519269 | 22513763 | 22513921 | 22523819 | 22523908 | 2 | 1 | 2 | 0 | 0 | 1 | 0 | 0 | 0 | 2 | 11 | 0 | 1 | 0 |
| SE | 13952 | ENSG00000243716 | NPIPB5 | chr16 | + | 22519137 | 22519414 | 22513759 | 22513921 | 22523819 | 22523948 | 2 | 1 | 2 | 0 | 0 | 1 | 0 | 0 | 0 | 2 | 11 | 0 | 1 | 0 |
| SE | 13954 | ENSG00000243716 | NPIPB5 | chr16 | + | 22530942 | 22531044 | 22527742 | 22527973 | 22532059 | 22532120 | 0 | 1 | 0 | 0 | 0 | 1 | 0 | 0 | 0 | 0 | 0 | 2 | 0 | 1 |
| SE | 13956 | ENSG00000141646 | SMAD4 | chr18 | + | 51049294 | 51049324 | 51048685 | 51048860 | 51054780 | 51054993 | 258 | 187 | 199 | 170 | 648 | 0 | 0 | 0 | 0 | 2 | 119 | 155 | 165 | 102 |
| SE | 13958 | ENSG00000141646 | SMAD4 | chr18 | + | 51058339 | 51058456 | 51058124 | 51058244 | 51059865 | 51059916 | 244 | 104 | 234 | 217 | 325 | 0 | 0 | 0 | 0 | 0 | 102 | 139 | 216 | 91 |
| SE | 13961 | ENSG00000141646 | SMAD4 | chr18 | + | 51067018 | 51067187 | 51065422 | 51065606 | 51076637 | 51076776 | 170 | 88 | 166 | 135 | 198 | 0 | 0 | 0 | 1 | 0 | 61 | 55 | 137 | 41 |
| SE | 13964 | ENSG00000080603 | SRCAP | chr16 | + | 30709850 | 30710128 | 30709512 | 30709735 | 30710753 | 30710847 | 344 | 209 | 326 | 351 | 440 | 0 | 0 | 0 | 0 | 0 | 526 | 507 | 272 | 478 |
| SE | 13965 | ENSG00000080603 | SRCAP | chr16 | + | 30711834 | 30712157 | 30711570 | 30711744 | 30712261 | 30712439 | 961 | 777 | 688 | 1066 | 1325 | 0 | 0 | 0 | 2 | 0 | 2583 | 1312 | 850 | 1371 |
| SE | 13967 | ENSG00000080603 | SRCAP | chr16 | + | 30716065 | 30716202 | 30713518 | 30713711 | 30716292 | 30716479 | 221 | 125 | 186 | 295 | 268 | 1 | 0 | 0 | 0 | 0 | 260 | 270 | 159 | 215 |
| SE | 13968 | ENSG00000080603 | SRCAP | chr16 | + | 30716292 | 30716479 | 30713518 | 30713711 | 30720161 | 30720331 | 108 | 36 | 75 | 78 | 75 | 0 | 0 | 0 | 0 | 0 | 123 | 63 | 28 | 46 |
| SE | 13972 | ENSG00000080603 | SRCAP | chr16 | + | 30722962 | 30723229 | 30722121 | 30722286 | 30723583 | 30725082 | 87 | 29 | 63 | 62 | 47 | 0 | 0 | 0 | 0 | 0 | 133 | 78 | 50 | 87 |
| SE | 13973 | ENSG00000080603 | SRCAP | chr16 | + | 30729369 | 30729572 | 30728965 | 30729231 | 30733279 | 30733449 | 209 | 95 | 201 | 201 | 108 | 0 | 0 | 0 | 1 | 0 | 111 | 256 | 118 | 152 |
| SE | 13974 | ENSG00000080603 | SRCAP | chr16 | + | 30736540 | 30736624 | 30736199 | 30736394 | 30737048 | 30739733 | 135 | 70 | 134 | 184 | 109 | 0 | 0 | 1 | 0 | 0 | 244 | 188 | 115 | 123 |
| SE | 13975 | ENSG00000170540 | ARL6IP1 | chr16 | - | 18795463 | 18795581 | 18794598 | 18794683 | 18797924 | 18797952 | 818 | 418 | 725 | 731 | 916 | 0 | 0 | 0 | 1 | 0 | 328 | 730 | 602 | 475 |
| SE | 13977 | ENSG00000170540 | ARL6IP1 | chr16 | - | 18798700 | 18798834 | 18795175 | 18795581 | 18801430 | 18801544 | 105 | 64 | 110 | 71 | 74 | 0 | 0 | 0 | 0 | 0 | 29 | 73 | 45 | 42 |
| SE | 13982 | ENSG00000103522 | IL21R | chr16 | + | 27446006 | 27446088 | 27445176 | 27445276 | 27448533 | 27448674 | 151 | 68 | 79 | 104 | 59 | 0 | 1 | 0 | 0 | 0 | 71 | 100 | 70 | 57 |
| SE | 13984 | ENSG00000153443 | UBALD1 | chr16 | - | 4610492 | 4610555 | 4609367 | 4609908 | 4614677 | 4614872 | 1 | 0 | 0 | 2 | 0 | 0 | 0 | 0 | 0 | 0 | 0 | 4 | 4 | 0 |
| SE | 13987 | ENSG00000047578 | KATNIP | chr16 | + | 27631064 | 27631162 | 27628587 | 27628830 | 27648603 | 27648735 | 75 | 45 | 53 | 88 | 87 | 0 | 0 | 0 | 0 | 0 | 120 | 115 | 96 | 120 |
| SE | 13992 | ENSG00000047578 | KATNIP | chr16 | + | 27654607 | 27654736 | 27648603 | 27648735 | 27681398 | 27681530 | 16 | 9 | 25 | 24 | 48 | 0 | 0 | 0 | 0 | 3 | 62 | 55 | 34 | 47 |
| SE | 13993 | ENSG00000047578 | KATNIP | chr16 | + | 27677728 | 27677996 | 27648603 | 27648735 | 27681398 | 27681530 | 25 | 18 | 48 | 36 | 50 | 0 | 0 | 0 | 0 | 3 | 94 | 36 | 43 | 58 |
| SE | 13994 | ENSG00000047578 | KATNIP | chr16 | + | 27677728 | 27677996 | 27654607 | 27654736 | 27681398 | 27681530 | 34 | 25 | 52 | 37 | 69 | 0 | 0 | 0 | 0 | 0 | 63 | 65 | 41 | 51 |
| SE | 14001 | ENSG00000047578 | KATNIP | chr16 | + | 27740040 | 27740920 | 27721557 | 27721695 | 27751718 | 27751924 | 39 | 21 | 23 | 41 | 49 | 0 | 0 | 0 | 0 | 0 | 56 | 41 | 33 | 22 |
| SE | 14002 | ENSG00000047578 | KATNIP | chr16 | + | 27740040 | 27740920 | 27721557 | 27721695 | 27754172 | 27754251 | 39 | 21 | 23 | 41 | 49 | 0 | 0 | 0 | 0 | 0 | 56 | 43 | 33 | 22 |
| SE | 14003 | ENSG00000047578 | KATNIP | chr16 | + | 27749583 | 27750306 | 27721557 | 27721695 | 27751718 | 27751924 | 52 | 51 | 32 | 44 | 78 | 0 | 0 | 0 | 0 | 0 | 123 | 63 | 63 | 74 |
| SE | 14004 | ENSG00000047578 | KATNIP | chr16 | + | 27749583 | 27750306 | 27740040 | 27740920 | 27751718 | 27751924 | 90 | 74 | 66 | 73 | 95 | 0 | 0 | 0 | 0 | 0 | 179 | 131 | 91 | 142 |
| SE | 14005 | ENSG00000047578 | KATNIP | chr16 | + | 27751718 | 27751924 | 27721557 | 27721695 | 27754172 | 27754251 | 16 | 15 | 32 | 14 | 20 | 0 | 0 | 0 | 0 | 0 | 20 | 26 | 24 | 17 |
| SE | 14006 | ENSG00000047578 | KATNIP | chr16 | + | 27751718 | 27751924 | 27740040 | 27740920 | 27754172 | 27754251 | 16 | 15 | 32 | 14 | 20 | 0 | 0 | 0 | 0 | 0 | 20 | 23 | 24 | 17 |
| SE | 14007 | ENSG00000047578 | KATNIP | chr16 | + | 27754172 | 27754251 | 27751718 | 27751924 | 27761412 | 27761590 | 28 | 21 | 35 | 20 | 26 | 0 | 0 | 1 | 0 | 0 | 29 | 36 | 26 | 27 |
| SE | 14008 | ENSG00000047578 | KATNIP | chr16 | + | 27769860 | 27770018 | 27766308 | 27766474 | 27771587 | 27771652 | 40 | 16 | 20 | 25 | 4 | 0 | 1 | 0 | 0 | 0 | 45 | 39 | 23 | 36 |
| SE | 14009 | ENSG00000185716 | MOSMO | chr16 | + | 22064337 | 22064438 | 22008208 | 22008407 | 22075486 | 22075699 | 0 | 0 | 0 | 1 | 0 | 17 | 12 | 33 | 28 | 8 | 0 | 0 | 0 | 0 |
| SE | 14010 | ENSG00000185716 | MOSMO | chr16 | + | 22083675 | 22083868 | 22075486 | 22075699 | 22085207 | 22085275 | 1 | 0 | 0 | 3 | 4 | 1 | 0 | 2 | 3 | 3 | 0 | 4 | 0 | 0 |
| SE | 14011 | ENSG00000185716 | MOSMO | chr16 | + | 22085207 | 22085275 | 22075486 | 22075699 | 22085643 | 22085750 | 2 | 1 | 2 | 8 | 6 | 6 | 7 | 5 | 10 | 25 | 0 | 7 | 1 | 1 |
| SE | 14014 | ENSG00000090905 | TNRC6A | chr16 | + | 24730252 | 24730300 | 24729650 | 24729846 | 24776932 | 24777027 | 10 | 13 | 8 | 6 | 6 | 0 | 0 | 0 | 2 | 0 | 0 | 5 | 22 | 10 |
| SE | 14015 | ENSG00000090905 | TNRC6A | chr16 | + | 24750725 | 24750813 | 24729650 | 24729846 | 24776932 | 24777358 | 0 | 0 | 0 | 2 | 0 | 0 | 0 | 0 | 2 | 0 | 2 | 7 | 0 | 2 |
| SE | 14016 | ENSG00000090905 | TNRC6A | chr16 | + | 24750725 | 24750813 | 24730252 | 24730300 | 24758338 | 24758360 | 33 | 37 | 37 | 51 | 25 | 1 | 1 | 0 | 1 | 0 | 16 | 23 | 49 | 24 |
| SE | 14017 | ENSG00000090905 | TNRC6A | chr16 | + | 24750725 | 24750813 | 24730252 | 24730300 | 24776932 | 24777358 | 9 | 5 | 8 | 9 | 7 | 0 | 0 | 0 | 2 | 0 | 0 | 4 | 14 | 8 |
| SE | 14018 | ENSG00000090905 | TNRC6A | chr16 | + | 24758338 | 24758360 | 24730252 | 24730300 | 24776932 | 24777358 | 4 | 3 | 5 | 2 | 1 | 0 | 0 | 0 | 2 | 0 | 7 | 5 | 5 | 5 |
| SE | 14019 | ENSG00000090905 | TNRC6A | chr16 | + | 24758338 | 24758360 | 24750725 | 24750813 | 24776932 | 24777358 | 25 | 26 | 30 | 32 | 15 | 0 | 0 | 0 | 4 | 0 | 18 | 21 | 26 | 22 |
| SE | 14024 | ENSG00000090905 | TNRC6A | chr16 | + | 24804704 | 24804851 | 24804176 | 24804319 | 24805013 | 24805151 | 1 | 2 | 0 | 0 | 0 | 160 | 140 | 185 | 186 | 200 | 0 | 0 | 0 | 0 |
| SE | 14025 | ENSG00000090905 | TNRC6A | chr16 | + | 24805013 | 24805151 | 24804176 | 24804319 | 24805604 | 24805733 | 192 | 177 | 238 | 212 | 235 | 0 | 0 | 0 | 0 | 1 | 32 | 143 | 170 | 114 |
| SE | 14026 | ENSG00000090905 | TNRC6A | chr16 | + | 24805604 | 24805733 | 24804333 | 24805151 | 24806205 | 24806283 | 92 | 71 | 88 | 80 | 91 | 0 | 0 | 2 | 0 | 2 | 8 | 43 | 64 | 37 |
| SE | 14027 | ENSG00000090905 | TNRC6A | chr16 | + | 24805604 | 24805733 | 24804333 | 24805151 | 24806573 | 24806784 | 32 | 34 | 55 | 22 | 41 | 0 | 0 | 0 | 0 | 0 | 0 | 26 | 26 | 19 |
| SE | 14029 | ENSG00000090905 | TNRC6A | chr16 | + | 24805673 | 24805733 | 24805013 | 24805151 | 24806573 | 24806754 | 5 | 1 | 1 | 6 | 4 | 0 | 0 | 0 | 0 | 0 | 0 | 4 | 4 | 0 |
| SE | 14030 | ENSG00000090905 | TNRC6A | chr16 | + | 24806205 | 24806283 | 24804333 | 24805151 | 24806573 | 24806754 | 37 | 27 | 22 | 25 | 10 | 0 | 0 | 0 | 0 | 0 | 18 | 25 | 30 | 26 |
| SE | 14031 | ENSG00000090905 | TNRC6A | chr16 | + | 24806205 | 24806283 | 24805013 | 24805733 | 24806573 | 24806754 | 100 | 64 | 58 | 85 | 59 | 0 | 0 | 1 | 0 | 0 | 32 | 44 | 67 | 44 |
| SE | 14032 | ENSG00000090905 | TNRC6A | chr16 | + | 24815146 | 24815305 | 24809349 | 24809481 | 24816815 | 24816956 | 127 | 121 | 167 | 161 | 279 | 0 | 0 | 0 | 2 | 0 | 47 | 110 | 122 | 41 |
| SE | 14033 | ENSG00000090905 | TNRC6A | chr16 | + | 24815354 | 24815475 | 24809349 | 24809481 | 24816815 | 24816956 | 12 | 3 | 4 | 5 | 4 | 0 | 0 | 0 | 2 | 0 | 7 | 9 | 5 | 8 |
| SE | 14035 | ENSG00000090905 | TNRC6A | chr16 | + | 24818592 | 24818700 | 24816584 | 24816956 | 24820138 | 24820180 | 186 | 162 | 245 | 227 | 319 | 4 | 2 | 0 | 2 | 0 | 29 | 192 | 209 | 117 |
| SE | 14036 | ENSG00000140694 | PARN | chr16 | - | 14482269 | 14482388 | 14446956 | 14447081 | 14482637 | 14482827 | 0 | 1 | 0 | 3 | 3 | 74 | 83 | 83 | 68 | 91 | 0 | 0 | 0 | 2 |
| SE | 14038 | ENSG00000140694 | PARN | chr16 | - | 14531053 | 14531174 | 14482637 | 14482827 | 14552020 | 14552095 | 0 | 0 | 2 | 3 | 4 | 108 | 75 | 118 | 104 | 114 | 0 | 0 | 0 | 0 |
| SE | 14041 | ENSG00000140694 | PARN | chr16 | - | 14558372 | 14558489 | 14555653 | 14555709 | 14580873 | 14580943 | 0 | 0 | 2 | 8 | 0 | 145 | 80 | 144 | 119 | 143 | 0 | 0 | 0 | 0 |
| SE | 14042 | ENSG00000140694 | PARN | chr16 | - | 14578530 | 14578628 | 14555653 | 14555709 | 14580873 | 14580943 | 0 | 1 | 3 | 0 | 0 | 145 | 80 | 144 | 119 | 143 | 0 | 0 | 0 | 0 |
| SE | 14044 | ENSG00000140694 | PARN | chr16 | - | 14584748 | 14584791 | 14584346 | 14584422 | 14593300 | 14593378 | 87 | 68 | 91 | 68 | 133 | 0 | 1 | 4 | 0 | 0 | 104 | 99 | 101 | 59 |
| SE | 14047 | ENSG00000140694 | PARN | chr16 | - | 14586317 | 14586361 | 14584748 | 14584791 | 14599903 | 14599960 | 28 | 17 | 22 | 30 | 37 | 0 | 0 | 0 | 0 | 0 | 5 | 10 | 23 | 25 |
| SE | 14048 | ENSG00000140694 | PARN | chr16 | - | 14588097 | 14588172 | 14586317 | 14586361 | 14593300 | 14593378 | 0 | 0 | 0 | 0 | 0 | 166 | 140 | 180 | 179 | 409 | 0 | 0 | 0 | 5 |
| SE | 14049 | ENSG00000140694 | PARN | chr16 | - | 14591912 | 14592033 | 14584748 | 14584791 | 14593300 | 14593378 | 2 | 0 | 0 | 0 | 1 | 2 | 12 | 1 | 0 | 6 | 0 | 1 | 1 | 0 |
| SE | 14051 | ENSG00000140694 | PARN | chr16 | - | 14591912 | 14593378 | 14584748 | 14584791 | 14599903 | 14599960 | 72 | 57 | 78 | 86 | 178 | 0 | 0 | 0 | 0 | 0 | 97 | 47 | 58 | 62 |
| SE | 14053 | ENSG00000140694 | PARN | chr16 | - | 14593300 | 14593378 | 14584748 | 14584791 | 14599903 | 14599960 | 68 | 67 | 78 | 85 | 174 | 0 | 0 | 0 | 0 | 0 | 121 | 54 | 58 | 62 |
| SE | 14054 | ENSG00000140694 | PARN | chr16 | - | 14593300 | 14593378 | 14586317 | 14586361 | 14599903 | 14599960 | 213 | 179 | 240 | 236 | 569 | 1 | 0 | 2 | 2 | 2 | 254 | 202 | 178 | 182 |
| SE | 14055 | ENSG00000140694 | PARN | chr16 | - | 14604145 | 14604223 | 14599903 | 14599960 | 14606483 | 14606526 | 76 | 63 | 88 | 82 | 173 | 3 | 0 | 0 | 0 | 0 | 101 | 62 | 64 | 74 |
| SE | 14056 | ENSG00000140694 | PARN | chr16 | - | 14604145 | 14604226 | 14599903 | 14599960 | 14606483 | 14606526 | 270 | 219 | 269 | 249 | 482 | 3 | 0 | 0 | 0 | 0 | 412 | 308 | 318 | 312 |
| SE | 14057 | ENSG00000140694 | PARN | chr16 | - | 14606483 | 14606526 | 14604145 | 14604226 | 14608280 | 14608319 | 218 | 183 | 215 | 188 | 351 | 2 | 2 | 1 | 1 | 0 | 320 | 267 | 256 | 273 |
| SE | 14058 | ENSG00000140694 | PARN | chr16 | - | 14617589 | 14617650 | 14610643 | 14610809 | 14623923 | 14623953 | 58 | 31 | 34 | 30 | 90 | 0 | 0 | 1 | 0 | 0 | 51 | 24 | 45 | 27 |
| SE | 14060 | ENSG00000140694 | PARN | chr16 | - | 14623923 | 14623953 | 14610643 | 14610809 | 14627105 | 14627187 | 0 | 0 | 1 | 1 | 0 | 2 | 7 | 10 | 2 | 3 | 0 | 0 | 1 | 0 |
| SE | 14062 | ENSG00000140694 | PARN | chr16 | - | 14627105 | 14627136 | 14617589 | 14617650 | 14627268 | 14627336 | 44 | 31 | 39 | 43 | 27 | 0 | 0 | 0 | 1 | 0 | 19 | 40 | 32 | 19 |
| SE | 14063 | ENSG00000140694 | PARN | chr16 | - | 14627105 | 14627174 | 14617589 | 14617650 | 14627268 | 14627326 | 111 | 89 | 89 | 117 | 82 | 0 | 0 | 0 | 1 | 0 | 106 | 125 | 77 | 92 |
| SE | 14064 | ENSG00000140694 | PARN | chr16 | - | 14627105 | 14627187 | 14617589 | 14617650 | 14627268 | 14627336 | 427 | 355 | 308 | 428 | 384 | 0 | 0 | 0 | 1 | 0 | 480 | 524 | 333 | 382 |
| SE | 14065 | ENSG00000140694 | PARN | chr16 | - | 14627105 | 14627187 | 14617589 | 14617650 | 14628171 | 14628251 | 131 | 97 | 94 | 125 | 101 | 0 | 0 | 0 | 0 | 0 | 107 | 129 | 83 | 110 |
| SE | 14066 | ENSG00000140694 | PARN | chr16 | - | 14627268 | 14627336 | 14617589 | 14617650 | 14628171 | 14628251 | 105 | 92 | 93 | 81 | 116 | 0 | 0 | 0 | 0 | 0 | 112 | 161 | 63 | 116 |
| SE | 14082 | ENSG00000130731 | METTL26 | chr16 | - | 635280 | 635340 | 634663 | 634797 | 635611 | 635774 | 6 | 4 | 13 | 5 | 3 | 3 | 2 | 1 | 0 | 0 | 0 | 11 | 3 | 3 |
| SE | 14083 | ENSG00000130731 | METTL26 | chr16 | - | 635280 | 635340 | 634663 | 634797 | 636093 | 636278 | 1 | 4 | 2 | 0 | 0 | 2 | 1 | 0 | 3 | 0 | 0 | 4 | 0 | 0 |
| SE | 14085 | ENSG00000130731 | METTL26 | chr16 | - | 635517 | 635774 | 634736 | 634797 | 636093 | 636260 | 5 | 3 | 6 | 4 | 0 | 2 | 1 | 0 | 3 | 0 | 0 | 4 | 2 | 2 |
| SE | 14086 | ENSG00000130731 | METTL26 | chr16 | - | 635517 | 635774 | 635280 | 635340 | 636093 | 636260 | 6 | 8 | 6 | 6 | 0 | 2 | 0 | 2 | 4 | 0 | 0 | 4 | 2 | 2 |
| SE | 14087 | ENSG00000130731 | METTL26 | chr16 | - | 635611 | 635774 | 634718 | 634797 | 636093 | 636235 | 8 | 5 | 7 | 4 | 0 | 2 | 1 | 0 | 3 | 0 | 0 | 4 | 2 | 2 |
| SE | 14090 | ENSG00000197006 | METTL9 | chr16 | + | 21612644 | 21612835 | 21599717 | 21599828 | 21617864 | 21618009 | 663 | 500 | 442 | 446 | 1168 | 0 | 0 | 1 | 0 | 0 | 1275 | 632 | 432 | 908 |
| SE | 14091 | ENSG00000197006 | METTL9 | chr16 | + | 21617864 | 21618074 | 21599591 | 21599898 | 21624930 | 21625114 | 795 | 364 | 587 | 558 | 951 | 0 | 1 | 1 | 0 | 0 | 1035 | 917 | 625 | 1013 |
| SE | 14093 | ENSG00000197006 | METTL9 | chr16 | + | 21624930 | 21625115 | 21612699 | 21612835 | 21655229 | 21656119 | 176 | 91 | 159 | 127 | 196 | 0 | 0 | 1 | 0 | 0 | 332 | 273 | 197 | 303 |
| SE | 14101 | ENSG00000155592 | ZKSCAN2 | chr16 | - | 25246706 | 25247390 | 25236000 | 25240738 | 25251908 | 25252035 | 12 | 8 | 14 | 10 | 15 | 0 | 0 | 0 | 1 | 0 | 13 | 13 | 22 | 20 |
| SE | 14102 | ENSG00000155592 | ZKSCAN2 | chr16 | - | 25251908 | 25252035 | 25246270 | 25247390 | 25252945 | 25253037 | 19 | 11 | 21 | 12 | 18 | 2 | 0 | 0 | 0 | 0 | 13 | 18 | 28 | 24 |
| SE | 14103 | ENSG00000155592 | ZKSCAN2 | chr16 | - | 25251908 | 25252046 | 25246706 | 25247390 | 25252945 | 25253037 | 7 | 6 | 10 | 4 | 15 | 2 | 0 | 0 | 0 | 0 | 13 | 13 | 14 | 13 |
| SE | 14104 | ENSG00000261448 | NA | chr16 | + | 17137515 | 17137614 | 17134503 | 17134550 | 17138241 | 17138736 | 2 | 1 | 0 | 0 | 2 | 1 | 1 | 0 | 1 | 0 | 0 | 0 | 0 | 0 |
| SE | 14106 | ENSG00000102897 | LYRM1 | chr16 | + | 20902485 | 20902717 | 20900449 | 20900535 | 20911182 | 20911264 | 0 | 1 | 2 | 2 | 0 | 0 | 0 | 0 | 0 | 1 | 0 | 0 | 2 | 0 |
| SE | 14108 | ENSG00000102897 | LYRM1 | chr16 | + | 20902485 | 20902717 | 20900449 | 20900535 | 20920121 | 20920214 | 0 | 3 | 0 | 7 | 0 | 1 | 3 | 2 | 0 | 7 | 0 | 2 | 0 | 0 |
| SE | 14109 | ENSG00000102897 | LYRM1 | chr16 | + | 20902485 | 20902717 | 20900665 | 20900889 | 20911182 | 20911264 | 1 | 1 | 2 | 5 | 0 | 2 | 0 | 1 | 0 | 0 | 0 | 0 | 2 | 0 |
| SE | 14111 | ENSG00000102897 | LYRM1 | chr16 | + | 20902485 | 20902717 | 20900665 | 20900889 | 20920121 | 20920214 | 1 | 3 | 0 | 10 | 0 | 0 | 1 | 2 | 0 | 0 | 0 | 2 | 0 | 0 |
| SE | 14112 | ENSG00000102897 | LYRM1 | chr16 | + | 20911182 | 20911264 | 20900449 | 20900535 | 20915555 | 20915714 | 2 | 8 | 2 | 7 | 2 | 15 | 14 | 17 | 0 | 11 | 0 | 0 | 7 | 4 |
| SE | 14113 | ENSG00000102897 | LYRM1 | chr16 | + | 20911182 | 20911264 | 20900449 | 20900535 | 20920121 | 20920214 | 0 | 0 | 0 | 0 | 0 | 1 | 3 | 2 | 0 | 7 | 0 | 0 | 4 | 0 |
| SE | 14114 | ENSG00000102897 | LYRM1 | chr16 | + | 20911182 | 20911264 | 20900770 | 20900889 | 20915555 | 20915714 | 4 | 8 | 3 | 7 | 1 | 1 | 1 | 3 | 9 | 0 | 0 | 0 | 7 | 4 |
| SE | 14115 | ENSG00000102897 | LYRM1 | chr16 | + | 20911182 | 20911264 | 20900770 | 20900889 | 20920121 | 20920214 | 0 | 0 | 1 | 0 | 0 | 0 | 1 | 2 | 0 | 0 | 0 | 0 | 4 | 0 |
| SE | 14117 | ENSG00000102897 | LYRM1 | chr16 | + | 20911182 | 20911264 | 20902485 | 20902717 | 20920121 | 20920214 | 0 | 1 | 2 | 2 | 0 | 0 | 3 | 0 | 7 | 0 | 0 | 0 | 4 | 0 |
| SE | 14122 | ENSG00000102897 | LYRM1 | chr16 | + | 20920121 | 20920214 | 20915555 | 20915714 | 20923999 | 20924116 | 210 | 159 | 221 | 201 | 304 | 1 | 0 | 0 | 0 | 1 | 196 | 219 | 175 | 191 |
| SE | 14124 | ENSG00000102858 | MGRN1 | chr16 | + | 4668264 | 4668312 | 4665098 | 4665151 | 4671390 | 4671459 | 191 | 101 | 136 | 176 | 228 | 0 | 0 | 0 | 0 | 0 | 156 | 257 | 141 | 261 |

| | | | | | | | | | | | | | | | | | | | | | | | | | | | | | | | | | | | | | | | | | | | | | | |
|---|---|---|---|---|---|---|---|---|---|---|---|---|---|---|---|---|---|---|---|---|---|---|---|---|---|---|---|---|---|---|---|---|---|---|---|---|---|---|---|---|---|---|---|---|---|---|
| SE | 14126 | ENSG00000102858 | MGRN1 | chr16 | + | 4677462 | 4677572 | 4673497 | 4673657 | 4681549 | 4681776 | 95 | 36 | 60 | 71 | 47 | 2 | 0 | 0 | 0 | 0 | 59 | 100 | 64 | 97 | 119 | 0 | 5 | 0 | 1 | 0 | 259 | 149 | 1 | 1 | 0.965 | 1 | 1 | 1 | 1 | 1 | 0.92 | 1 | 0.982 | 1 | 0.013 |
| SE | 14128 | ENSG00000102858 | MGRN1 | chr16 | + | 4682822 | 4682946 | 4681347 | 4681776 | 4683223 | 4683269 | 137 | 65 | 106 | 121 | 148 | 0 | 0 | 0 | 0 | 0 | 121 | 96 | 73 | 142 | 118 | 0 | 0 | 0 | 0 | 1 | 273 | 149 | 1 | 1 | 1 | 1 | 1 | 1 | 1 | 1 | 1 | 1 | 1 | 0.985 | 0.003 |
| SE | 14130 | ENSG00000205746 | PKD1P4 | chr16 | - | 18344820 | 18344915 | 18344139 | 18344283 | 18345377 | 18345511 | 0 | 0 | 0 | 0 | 0 | 0 | 0 | 0 | 4 | 0 | 0 | 0 | 0 | 2 | 0 | 0 | 6 | 0 | 0 | 0 | 244 | 149 | 1 | 1 | NA | NA | NA | 0 | NA | NA | 0 | NA | 1 | NA | -0.5 |
| SE | 14136 | ENSG00000007376 | RPUSD1 | chr16 | - | 786826 | 786928 | 784973 | 786377 | 787076 | 787179 | 44 | 26 | 29 | 27 | 57 | 2 | 0 | 0 | 0 | 0 | 26 | 41 | 18 | 24 | 28 | 0 | 0 | 0 | 0 | 0 | 251 | 149 | 1 | 1 | 0.929 | 1 | 1 | 1 | 1 | 1 | 1 | 1 | 1 | 1 | -0.014 |
| SE | 14137 | ENSG00000007376 | RPUSD1 | chr16 | - | 787067 | 787179 | 786826 | 786928 | 787353 | 787477 | 14 | 4 | 9 | 15 | 3 | 2 | 0 | 0 | 0 | 0 | 15 | 1 | 3 | 10 | 18 | 0 | 1 | 0 | 0 | 0 | 261 | 149 | 1 | 1 | 0.8 | 1 | 1 | 1 | 1 | 1 | 0.363 | 1 | 1 | 1 | 0.087 |
| SE | 14138 | ENSG00000007376 | RPUSD1 | chr16 | - | 787076 | 787179 | 786193 | 786928 | 787353 | 787469 | 23 | 16 | 23 | 28 | 24 | 2 | 0 | 0 | 0 | 0 | 23 | 19 | 14 | 11 | 34 | 0 | 1 | 0 | 0 | 0 | 252 | 149 | 1 | 1 | 0.872 | 1 | 1 | 1 | 1 | 1 | 0.918 | 1 | 1 | 1 | -0.009 |
| SE | 14139 | ENSG00000007376 | RPUSD1 | chr16 | - | 787555 | 787645 | 787353 | 787477 | 788255 | 788334 | 2 | 2 | 7 | 2 | 1 | 1 | 0 | 0 | 2 | 0 | 0 | 1 | 1 | 0 | 0 | 0 | 0 | 2 | 0 | 0 | 239 | 149 | 1 | 1 | 0.555 | 1 | 1 | 0.384 | 1 | NA | 1 | 0.238 | NA | NA | 0.169 |
| SE | 14140 | ENSG00000007376 | RPUSD1 | chr16 | - | 787555 | 787744 | 787343 | 787477 | 788255 | 788329 | 7 | 5 | 9 | 9 | 0 | 1 | 0 | 0 | 2 | 0 | 7 | 5 | 2 | 0 | 11 | 0 | 0 | 2 | 0 | 0 | 298 | 149 | 1 | 1 | 0.778 | 1 | 1 | 0.692 | NA | 1 | 1 | 0.333 | NA | 1 | 0.034 |
| SE | 14147 | ENSG00000005339 | CREBBP | chr16 | - | 3744893 | 3744961 | 3740398 | 3740549 | 3749626 | 3749683 | 184 | 87 | 153 | 119 | 96 | 0 | 0 | 0 | 0 | 0 | 152 | 99 | 136 | 194 | 106 | 0 | 0 | 2 | 0 | 0 | 217 | 149 | 1 | 1 | 1 | 1 | 1 | 1 | 1 | 1 | 1 | 0.979 | 1 | 1 | 0.004 |
| SE | 14148 | ENSG00000005339 | CREBBP | chr16 | - | 3745276 | 3745354 | 3744893 | 3744961 | 3749626 | 3749683 | 260 | 132 | 199 | 149 | 204 | 0 | 0 | 0 | 0 | 0 | 163 | 180 | 150 | 245 | 213 | 0 | 2 | 0 | 0 | 0 | 227 | 149 | 1 | 1 | 1 | 1 | 1 | 1 | 1 | 1 | 0.983 | 1 | 1 | 1 | 0.003 |
| SE | 14149 | ENSG00000005339 | CREBBP | chr16 | - | 3749626 | 3749683 | 3745276 | 3745354 | 3751725 | 3751806 | 143 | 68 | 130 | 99 | 150 | 0 | 0 | 0 | 0 | 0 | 99 | 98 | 85 | 125 | 112 | 0 | 3 | 0 | 0 | 0 | 206 | 149 | 1 | 1 | 1 | 1 | 1 | 1 | 1 | 1 | 0.959 | 1 | 1 | 1 | 0.008 |
| SE | 14150 | ENSG00000005339 | CREBBP | chr16 | - | 3751725 | 3751806 | 3749626 | 3749683 | 3757287 | 3757291 | 236 | 143 | 200 | 189 | 191 | 0 | 0 | 0 | 0 | 0 | 323 | 223 | 253 | 253 | 283 | 0 | 0 | 0 | 0 | 2 | 230 | 149 | 1 | 1 | 1 | 1 | 1 | 1 | 1 | 1 | 1 | 1 | 1 | 0.989 | 0.002 |
| SE | 14151 | ENSG00000005339 | CREBBP | chr16 | - | 3751725 | 3751836 | 3749626 | 3749683 | 3757287 | 3757376 | 22 | 14 | 27 | 14 | 31 | 0 | 0 | 0 | 0 | 0 | 74 | 13 | 33 | 37 | 30 | 0 | 0 | 0 | 0 | 2 | 260 | 149 | 1 | 1 | 1 | 1 | 1 | 1 | 1 | 1 | 1 | 1 | 1 | 0.896 | 0.021 |
| SE | 14152 | ENSG00000005339 | CREBBP | chr16 | - | 3757287 | 3757376 | 3751725 | 3751806 | 3757808 | 3758048 | 397 | 222 | 308 | 346 | 335 | 0 | 0 | 0 | 0 | 0 | 461 | 329 | 367 | 360 | 423 | 0 | 7 | 0 | 3 | 0 | 238 | 149 | 1 | 1 | 1 | 1 | 1 | 1 | 1 | 1 | 0.967 | 1 | 0.987 | 1 | 0.009 |
| SE | 14154 | ENSG00000005339 | CREBBP | chr16 | - | 3773750 | 3773930 | 3770569 | 3770986 | 3774568 | 3774693 | 332 | 146 | 233 | 265 | 445 | 0 | 0 | 0 | 0 | 1 | 620 | 382 | 387 | 345 | 522 | 0 | 0 | 2 | 0 | 0 | 298 | 149 | 1 | 1 | 1 | 1 | 1 | 1 | 0.996 | 1 | 1 | 0.99 | 1 | 1 | 0.001 |
| SE | 14156 | ENSG00000005339 | CREBBP | chr16 | - | 3778699 | 3778817 | 3778010 | 3778182 | 3780731 | 3780867 | 895 | 504 | 660 | 859 | 921 | 0 | 0 | 0 | 0 | 1 | 1783 | 1118 | 1039 | 1161 | 1414 | 0 | 3 | 0 | 0 | 0 | 267 | 149 | 1 | 1 | 1 | 1 | 1 | 1 | 0.998 | 1 | 0.995 | 1 | 1 | 1 | 0.001 |
| SE | 14157 | ENSG00000005339 | CREBBP | chr16 | - | 3778699 | 3778817 | 3778010 | 3778182 | 3781203 | 3781306 | 352 | 226 | 296 | 371 | 437 | 0 | 0 | 0 | 0 | 2 | 672 | 393 | 381 | 409 | 507 | 0 | 0 | 0 | 0 | 0 | 267 | 149 | 1 | 1 | 1 | 1 | 1 | 1 | 0.992 | 1 | 1 | 1 | 1 | 1 | -0.002 |
| SE | 14158 | ENSG00000005339 | CREBBP | chr16 | - | 3780731 | 3780878 | 3778010 | 3778182 | 3781203 | 3781306 | 151 | 94 | 132 | 160 | 121 | 0 | 0 | 0 | 0 | 2 | 378 | 194 | 212 | 332 | 342 | 0 | 0 | 0 | 0 | 0 | 296 | 149 | 1 | 1 | 1 | 1 | 1 | 1 | 0.968 | 1 | 1 | 1 | 1 | 1 | -0.006 |
| SE | 14162 | ENSG00000005339 | CREBBP | chr16 | - | 3791980 | 3792094 | 3781203 | 3781306 | 3793385 | 3793626 | 147 | 74 | 166 | 167 | 246 | 1 | 0 | 0 | 0 | 1 | 223 | 139 | 92 | 139 | 151 | 0 | 0 | 0 | 7 | 0 | 263 | 149 | 1 | 1 | 0.988 | 1 | 1 | 1 | 0.993 | 1 | 1 | 1 | 0.918 | 1 | 0.013 |
| SE | 14164 | ENSG00000005339 | CREBBP | chr16 | - | 3810602 | 3810779 | 3793385 | 3793626 | 3850296 | 3851009 | 488 | 284 | 458 | 535 | 331 | 0 | 0 | 1 | 1 | 0 | 723 | 527 | 517 | 639 | 644 | 11 | 0 | 1 | 1 | 0 | 298 | 149 | 1 | 1 | 1 | 1 | 0.996 | 0.996 | 1 | 0.97 | 1 | 0.996 | 0.997 | 1 | 0.006 |
| SE | 14165 | ENSG00000005339 | CREBBP | chr16 | - | 3850296 | 3851009 | 3810602 | 3810779 | 3879831 | 3880120 | 442 | 233 | 347 | 419 | 197 | 2 | 0 | 1 | 0 | 0 | 607 | 437 | 459 | 485 | 550 | 0 | 1 | 1 | 0 | 0 | 298 | 149 | 1 | 1 | 0.991 | 1 | 0.994 | 1 | 1 | 1 | 0.995 | 0.996 | 1 | 1 | -0.001 |
| SE | 14171 | ENSG00000180747 | SMG1P3 | chr16 | - | 21447927 | 21448087 | 21446682 | 21447197 | 21448461 | 21448609 | 6 | 4 | 2 | 6 | 6 | 0 | 0 | 0 | 0 | 0 | 13 | 10 | 6 | 12 | 0 | 1 | 0 | 0 | 0 | 0 | 298 | 149 | 1 | 1 | 1 | 1 | 1 | 1 | 1 | 0.867 | 1 | 1 | 1 | NA | 0.033 |
| SE | 14172 | ENSG00000180747 | SMG1P3 | chr16 | - | 21467091 | 21467316 | 21464715 | 21464817 | 21469746 | 21469919 | 25 | 17 | 21 | 18 | 29 | 0 | 0 | 2 | 0 | 0 | 33 | 34 | 21 | 26 | 13 | 0 | 0 | 0 | 0 | 0 | 298 | 149 | 1 | 1 | 1 | 1 | 0.84 | 1 | 1 | 1 | 1 | 1 | 1 | 1 | -0.032 |
| SE | 14173 | ENSG00000180747 | SMG1P3 | chr16 | - | 21473123 | 21473249 | 21472672 | 21472745 | 21519827 | 21520444 | 4 | 2 | 3 | 2 | 0 | 1 | 1 | 0 | 2 | 0 | 0 | 1 | 4 | 5 | 0 | 0 | 5 | 0 | 0 | 0 | 275 | 149 | 1 | 1 | 0.684 | 0.52 | 1 | 0.351 | NA | NA | 0.098 | 1 | 1 | NA | -0.061 |
| SE | 14175 | ENSG00000180747 | SMG1P3 | chr16 | - | 21476598 | 21476812 | 21473123 | 21473249 | 21519827 | 21520444 | 9 | 8 | 5 | 2 | 15 | 2 | 0 | 0 | 1 | 0 | 0 | 23 | 1 | 14 | 17 | 0 | 0 | 0 | 0 | 0 | 298 | 149 | 1 | 1 | 0.692 | 1 | 1 | 0.5 | 1 | NA | 1 | 1 | 1 | 1 | -0.162 |
| SE | 14177 | ENSG00000180747 | SMG1P3 | chr16 | - | 21484371 | 21484535 | 21479441 | 21479578 | 21487577 | 21487616 | 25 | 305 | 392 | 44 | 369 | 0 | 0 | 0 | 0 | 0 | 992 | 685 | 461 | 1366 | 271 | 0 | 1 | 0 | 0 | 0 | 298 | 149 | 1 | 1 | 1 | 1 | 1 | 1 | 1 | 1 | 0.997 | 1 | 1 | 1 | 0.001 |
| SE | 14178 | ENSG00000180747 | SMG1P3 | chr16 | - | 21484371 | 21484535 | 21479441 | 21479578 | 21501776 | 21501886 | 8 | 224 | 247 | 12 | 275 | 0 | 0 | 0 | 0 | 0 | 719 | 457 | 289 | 945 | 189 | 0 | 1 | 0 | 0 | 0 | 298 | 149 | 1 | 1 | 1 | 1 | 1 | 1 | 1 | 1 | 0.996 | 1 | 1 | 1 | 0.001 |
| SE | 14179 | ENSG00000180747 | SMG1P3 | chr16 | - | 21487577 | 21487616 | 21479544 | 21479578 | 21501776 | 21501886 | 0 | 2 | 0 | 0 | 0 | 0 | 0 | 0 | 0 | 0 | 0 | 0 | 0 | 0 | 0 | 0 | 1 | 0 | 0 | 0 | 188 | 149 | 1 | 1 | NA | 1 | NA | NA | NA | NA | 0 | NA | NA | NA | 1 |
| SE | 14180 | ENSG00000180747 | SMG1P3 | chr16 | - | 21487577 | 21487616 | 21484371 | 21484535 | 21501776 | 21501886 | 0 | 64 | 95 | 4 | 64 | 0 | 11 | 3 | 0 | 0 | 260 | 150 | 112 | 313 | 43 | 47 | 9 | 4 | 24 | 0 | 188 | 149 | 1 | 1 | NA | 0.822 | 0.962 | 1 | 1 | 0.814 | 0.93 | 0.957 | 0.912 | 1 | 0.023 |
| SE | 14182 | ENSG00000140983 | RHOT2 | chr16 | + | 668487 | 668569 | 668131 | 668411 | 668655 | 668699 | 117 | 50 | 89 | 104 | 33 | 1 | 0 | 0 | 0 | 0 | 70 | 115 | 108 | 69 | 215 | 0 | 4 | 0 | 0 | 0 | 231 | 149 | 1 | 1 | 0.987 | 1 | 1 | 1 | 1 | 1 | 0.949 | 1 | 1 | 1 | 0.008 |
| SE | 14185 | ENSG00000140983 | RHOT2 | chr16 | + | 670674 | 670773 | 670455 | 670557 | 670888 | 671000 | 109 | 49 | 79 | 85 | 73 | 0 | 0 | 1 | 0 | 0 | 92 | 99 | 70 | 66 | 85 | 0 | 0 | 0 | 0 | 0 | 248 | 149 | 1 | 1 | 1 | 1 | 0.979 | 1 | 1 | 1 | 1 | 1 | 1 | 1 | -0.004 |
| SE | 14195 | ENSG00000064270 | ATP2C2 | chr16 | + | 84410703 | 84410765 | 84410567 | 84410603 | 84415482 | 84415526 | 0 | 0 | 0 | 1 | 0 | 0 | 0 | 0 | 0 | 0 | 0 | 0 | 4 | 50 | 10 | 0 | 0 | 2 | 0 | 0 | 211 | 149 | 1 | 1 | NA | NA | NA | 1 | NA | NA | NA | 0.585 | 1 | 1 | 0.138 |
| SE | 14196 | ENSG00000064270 | ATP2C2 | chr16 | + | 84440856 | 84440958 | 84439426 | 84439524 | 84442509 | 84442599 | 0 | 0 | 0 | 2 | 0 | 0 | 0 | 0 | 0 | 0 | 0 | 3 | 4 | 54 | 1 | 0 | 0 | 0 | 7 | 0 | 251 | 149 | 1 | 1 | NA | NA | NA | 1 | NA | NA | 1 | 1 | 0.821 | 1 | 0.045 |
| SE | 14198 | ENSG00000064270 | ATP2C2 | chr16 | + | 84459505 | 84459592 | 84459269 | 84459386 | 84460653 | 84460801 | 0 | 0 | 0 | 0 | 0 | 0 | 0 | 0 | 1 | 0 | 0 | 0 | 3 | 0 | 0 | 0 | 2 | 0 | 10 | 0 | 236 | 149 | 1 | 1 | NA | NA | NA | 0 | NA | NA | 0 | 1 | 0 | NA | -0.333 |
| SE | 14199 | ENSG00000196678 | ERI2 | chr16 | - | 20790849 | 20790932 | 20789478 | 20789557 | 20799262 | 20799351 | 0 | 0 | 0 | 1 | 4 | 0 | 0 | 0 | 0 | 0 | 10 | 0 | 1 | 0 | 0 | 0 | 6 | 0 | 0 | 0 | 232 | 149 | 1 | 1 | NA | NA | NA | 1 | 1 | 1 | 0 | 1 | NA | NA | 0.333 |
| SE | 14201 | ENSG00000196678 | ERI2 | chr16 | - | 20805714 | 20805945 | 20803602 | 20803670 | 20806407 | 20806446 | 14 | 2 | 4 | 9 | 12 | 7 | 2 | 1 | 3 | 0 | 0 | 6 | 1 | 0 | 0 | 0 | 0 | 0 | 3 | 0 | 298 | 149 | 1 | 1 | 0.5 | 0.333 | 0.667 | 0.6 | 1 | NA | 1 | 1 | 0 | NA | -0.047 |
| SE | 14203 | ENSG00000103199 | ZNF500 | chr16 | - | 4762270 | 4762335 | 4760491 | 4760588 | 4762572 | 4762713 | 13 | 4 | 13 | 32 | 2 | 0 | 0 | 0 | 1 | 0 | 11 | 20 | 5 | 13 | 15 | 0 | 0 | 0 | 0 | 0 | 214 | 149 | 1 | 1 | 1 | 1 | 1 | 0.957 | 1 | 1 | 1 | 1 | 1 | 1 | -0.009 |
| SE | 14205 | ENSG00000103381 | CPPED1 | chr16 | - | 12781184 | 12781403 | 12704623 | 12705049 | 12803706 | 12803851 | 244 | 182 | 123 | 195 | 192 | 1 | 0 | 0 | 0 | 0 | 435 | 240 | 178 | 354 | 266 | 0 | 0 | 0 | 0 | 0 | 298 | 149 | 1 | 1 | 0.992 | 1 | 1 | 1 | 1 | 1 | 1 | 1 | 1 | 1 | -0.002 |
| SE | 14206 | ENSG00000162104 | ADCY9 | chr16 | - | 3992145 | 3992270 | 3989088 | 3989096 | 3993405 | 3993510 | 64 | 8 | 19 | 43 | 74 | 0 | 0 | 0 | 2 | 0 | 27 | 26 | 25 | 22 | 94 | 0 | 0 | 0 | 0 | 1 | 274 | 149 | 1 | 1 | 1 | 1 | 1 | 0.921 | 1 | 1 | 1 | 1 | 1 | 0.981 | -0.012 |
| SE | 14207 | ENSG00000162104 | ADCY9 | chr16 | - | 3992145 | 3992363 | 3988993 | 3989096 | 3993405 | 3993510 | 106 | 17 | 29 | 77 | 107 | 0 | 0 | 0 | 2 | 0 | 44 | 42 | 27 | 34 | 135 | 0 | 0 | 0 | 0 | 1 | 298 | 149 | 1 | 1 | 1 | 1 | 1 | 0.951 | 1 | 1 | 1 | 1 | 1 | 0.985 | -0.007 |
| SE | 14212 | ENSG00000140990 | NDUFB10 | chr16 | + | 1961152 | 1961258 | 1959551 | 1959754 | 1961496 | 1961636 | 74 | 53 | 65 | 44 | 33 | 0 | 0 | 0 | 1 | 1 | 230 | 68 | 68 | 113 | 191 | 3 | 0 | 0 | 0 | 0 | 255 | 149 | 1 | 1 | 1 | 1 | 1 | 0.963 | 0.951 | 0.978 | 1 | 1 | 1 | 1 | -0.013 |
| SE | 14213 | ENSG00000140990 | NDUFB10 | chr16 | + | 1961152 | 1961291 | 1959537 | 1959754 | 1961496 | 1961611 | 260 | 172 | 195 | 193 | 227 | 0 | 0 | 0 | 1 | 1 | 486 | 292 | 235 | 356 | 494 | 3 | 0 | 0 | 0 | 0 | 288 | 149 | 1 | 1 | 1 | 1 | 1 | 0.99 | 0.992 | 0.988 | 1 | 1 | 1 | 1 | -0.001 |
| SE | 14215 | ENSG00000156858 | PRR14 | chr16 | + | 30651595 | 30651668 | 30650716 | 30650806 | 30651795 | 30651964 | 15 | 2 | 11 | 10 | 4 | 0 | 0 | 0 | 3 | 0 | 4 | 4 | 4 | 17 | 14 | 0 | 0 | 0 | 0 | 0 | 222 | 149 | 1 | 1 | 1 | 1 | 1 | 0.691 | 1 | 1 | 1 | 1 | 1 | 1 | -0.062 |
| SE | 14220 | ENSG00000167965 | MLST8 | chr16 | + | 2207856 | 2208024 | 2207196 | 2207345 | 2208209 | 2208328 | 0 | 0 | 2 | 0 | 0 | 23 | 8 | 31 | 19 | 7 | 0 | 0 | 0 | 0 | 0 | 16 | 21 | 5 | 29 | 30 | 298 | 149 | 1 | 1 | 0 | 0 | 0.031 | 0 | 0 | 0 | 0 | 0 | 0 | 0 | 0.006 |
| SE | 14221 | ENSG00000167965 | MLST8 | chr16 | + | 2207856 | 2208024 | 2207196 | 2207345 | 2208449 | 2209412 | 0 | 2 | 0 | 0 | 0 | 2 | 8 | 5 | 6 | 1 | 0 | 0 | 0 | 0 | 0 | 0 | 1 | 10 | 5 | 3 | 298 | 149 | 1 | 1 | 0 | 0.111 | 0 | 0 | 0 | NA | 0 | 0 | 0 | 0 | 0.022 |
| SE | 14224 | ENSG00000142192 | APP | chr21 | - | 25897572 | 25897673 | 25891721 | 25891868 | 25954589 | 25954689 | 422 | 331 | 258 | 455 | 777 | 0 | 0 | 0 | 0 | 0 | 1024 | 416 | 276 | 827 | 1717 | 7 | 0 | 0 | 0 | 0 | 250 | 149 | 1 | 1 | 1 | 1 | 1 | 1 | 1 | 0.989 | 1 | 1 | 1 | 1 | 0.002 |
| SE | 14226 | ENSG00000142192 | APP | chr21 | - | 25911740 | 25911962 | 25897572 | 25897673 | 25954589 | 25954689 | 493 | 383 | 308 | 499 | 511 | 1 | 0 | 0 | 0 | 0 | 1209 | 531 | 433 | 1125 | 2357 | 0 | 0 | 1 | 0 | 0 | 298 | 149 | 1 | 1 | 0.996 | 1 | 1 | 1 | 1 | 1 | 1 | 0.995 | 1 | 1 | 0 |
| SE | 14227 | ENSG00000142192 | APP | chr21 | - | 25975069 | 25975228 | 25955626 | 25955755 | 25982343 | 25982477 | 203 | 141 | 123 | 256 | 338 | 0 | 0 | 0 | 0 | 0 | 394 | 197 | 136 | 352 | 917 | 0 | 7 | 0 | 0 | 0 | 298 | 149 | 1 | 1 | 1 | 1 | 1 | 1 | 1 | 1 | 0.934 | 1 | 1 | 1 | 0.013 |
| SE | 14228 | ENSG00000142192 | APP | chr21 | - | 25975953 | 25976028 | 25975069 | 25975228 | 25982343 | 25982477 | 221 | 163 | 177 | 290 | 317 | 0 | 0 | 0 | 1 | 0 | 554 | 254 | 182 | 447 | 1045 | 0 | 4 | 0 | 1 | 0 | 224 | 149 | 1 | 1 | 1 | 1 | 1 | 0.995 | 1 | 1 | 0.977 | 1 | 0.997 | 1 | 0.004 |
| SE | 14232 | ENSG00000142192 | APP | chr21 | - | 26000014 | 26000182 | 25997187 | 25997416 | 26021839 | 26022042 | 352 | 217 | 257 | 493 | 332 | 2 | 0 | 0 | 0 | 0 | 1008 | 401 | 328 | 898 | 1791 | 0 | 0 | 0 | 0 | 0 | 298 | 149 | 1 | 1 | 0.989 | 1 | 1 | 1 | 1 | 1 | 1 | 1 | 1 | 1 | -0.002 |
| SE | 14233 | ENSG00000142192 | APP | chr21 | - | 26089942 | 26090072 | 26053235 | 26053348 | 26111978 | 26112146 | 288 | 180 | 220 | 325 | 372 | 0 | 0 | 0 | 0 | 0 | 808 | 403 | 285 | 605 | 1467 | 0 | 0 | 1 | 0 | 0 | 279 | 149 | 1 | 1 | 1 | 1 | 1 | 1 | 1 | 1 | 1 | 0.993 | 1 | 1 | 0.001 |
| SE | 14235 | ENSG00000103326 | CAPN15 | chr16 | + | 536028 | 536142 | 533945 | 533998 | 546816 | 548287 | 17 | 0 | 10 | 7 | 3 | 2 | 0 | 0 | 0 | 0 | 0 | 3 | 4 | 1 | 0 | 0 | 0 | 1 | 0 | 8 | 263 | 149 | 1 | 1 | 0.828 | NA | 1 | 1 | 1 | NA | 1 | 0.694 | 1 | 0 | 0.283 |
| SE | 14236 | ENSG00000125877 | ITPA | chr20 | + | 3213168 | 3213226 | 3208867 | 3209033 | 3213984 | 3214058 | 5 | 5 | 3 | 13 | 0 | 0 | 0 | 5 | 0 | 0 | 0 | 3 | 0 | 2 | 0 | 0 | 7 | 0 | 0 | 0 | 207 | 149 | 1 | 1 | 1 | 1 | 0.302 | 1 | NA | NA | 0 | NA | 1 | NA | 0.326 |
| SE | 14238 | ENSG00000125877 | ITPA | chr20 | + | 3213168 | 3213226 | 3209512 | 3209566 | 3213984 | 3214058 | 3 | 2 | 2 | 8 | 0 | 0 | 0 | 1 | 0 | 0 | 0 | 8 | 0 | 0 | 0 | 0 | 0 | 0 | 0 | 0 | 207 | 149 | 1 | 1 | 1 | 1 | 0.59 | 1 | NA | NA | 1 | NA | NA | NA | -0.103 |
| SE | 14244 | ENSG00000260807 | CEROX1 | chr16 | - | 979600 | 979758 | 979024 | 979110 | 981144 | 981298 | 3 | 3 | 3 | 2 | 6 | 0 | 1 | 0 | 0 | 0 | 0 | 0 | 3 | 0 | 0 | 0 | 0 | 0 | 0 | 0 | 298 | 149 | 1 | 1 | 1 | 0.6 | 1 | 1 | 1 | NA | NA | 1 | NA | NA | -0.08 |
| SE | 14245 | ENSG00000260807 | CEROX1 | chr16 | - | 979600 | 979763 | 979024 | 979110 | 981144 | 981293 | 3 | 3 | 3 | 2 | 6 | 0 | 1 | 0 | 0 | 0 | 0 | 0 | 3 | 0 | 0 | 0 | 0 | 0 | 0 | 0 | 298 | 149 | 1 | 1 | 1 | 0.6 | 1 | 1 | 1 | NA | NA | 1 | NA | NA | -0.08 |
| SE | 14247 | ENSG00000233024 | NPIPA9 | chr16 | - | 18372577 | 18372854 | 18368257 | 18368386 | 18374831 | 18374979 | 0 | 1 | 0 | 0 | 0 | 0 | 0 | 0 | 0 | 0 | 0 | 0 | 0 | 0 | 0 | 0 | 0 | 0 | 0 | 5 | 298 | 149 | 1 | 1 | NA | 1 | NA | NA | NA | NA | NA | NA | NA | 0 | 1 |
| SE | 14248 | ENSG00000233024 | NPIPA9 | chr16 | - | 18372722 | 18372854 | 18368257 | 18368386 | 18374831 | 18375069 | 0 | 1 | 0 | 0 | 0 | 0 | 0 | 0 | 0 | 0 | 0 | 0 | 0 | 0 | 0 | 0 | 0 | 0 | 0 | 5 | 281 | 149 | 1 | 1 | NA | 1 | NA | NA | NA | NA | NA | NA | NA | 0 | 1 |
| SE | 14250 | ENSG00000059145 | UNKL | chr16 | - | 1385207 | 1385385 | 1371518 | 1371611 | 1390631 | 1390694 | 43 | 20 | 32 | 30 | 25 | 0 | 0 | 2 | 0 | 0 | 37 | 29 | 8 | 6 | 22 | 2 | 2 | 1 | 0 | 0 | 298 | 149 | 1 | 1 | 1 | 1 | 0.889 | 1 | 1 | 0.902 | 0.879 | 0.8 | 1 | 1 | 0.062 |
| [illegible] | | | | | | | | | | | | | | | | | | | | | | | | | | | | | | | | | | | | | | | | | | | | | | |

| | | | | | | | | | | | | | | | |
|---|---|---|---|---|---|---|---|---|---|---|---|---|---|---|---|
| SE | 14617 | ENSG00000243708 | PLA2G4B | chr15 | + | 41845637 | 41845775 | 41845202 | 41845320 | 41845942 | 41846047 | 41 | 22 | 27 | [illegible] |
| SE | 14618 | ENSG00000168970 | JMJD7-PLA2G4B | chr15 | + | 41847336 | 41847523 | 41846668 | 41846835 | 41847648 | 41848143 | 41 | 29 | 47 | [illegible] |
| SE | 14625 | ENSG00000140157 | NIPA2 | chr15 | + | 22839654 | 22839790 | 22838675 | 22839174 | 22845145 | 22845267 | 27 | 10 | 17 | [illegible] |
| SE | 14631 | ENSG00000140157 | NIPA2 | chr15 | + | 22851638 | 22851870 | 22838688 | 22839174 | 22858539 | 22858630 | 9 | 2 | 1 | [illegible] |
| SE | 14637 | ENSG00000260007 | NA | chr15 | - | 68199540 | 68199681 | 68197528 | 68197628 | 68218535 | 68218573 | 11 | 2 | 3 | [illegible] |
| SE | 14658 | ENSG00000179361 | ARID3B | chr15 | + | 74593134 | 74593236 | 74591559 | 74591814 | 74595610 | 74598129 | 27 | 5 | 26 | [illegible] |
| SE | 14659 | ENSG00000179361 | ARID3B | chr15 | + | 74593137 | 74593236 | 74591559 | 74591814 | 74595610 | 74598131 | 47 | 21 | 45 | [illegible] |
| SE | 14660 | ENSG00000140488 | CELF6 | chr15 | - | 72288537 | 72288618 | 72288307 | 72288451 | 72288867 | 72288930 | 0 | 3 | 1 | [illegible] |
| SE | 14661 | ENSG00000140488 | CELF6 | chr15 | - | 72288867 | 72288930 | 72288537 | 72288618 | 72289137 | 72289287 | 0 | 2 | 0 | [illegible] |
| SE | 14662 | ENSG00000140488 | CELF6 | chr15 | - | 72289938 | 72290018 | 72289626 | 72289770 | 72290126 | 72290255 | 6 | 1 | 5 | [illegible] |
| SE | 14663 | ENSG00000140488 | CELF6 | chr15 | - | 72292197 | 72292265 | 72290126 | 72290255 | 72304745 | 72304794 | 0 | 0 | 0 | [illegible] |
| SE | 14665 | ENSG00000273025 | NA | chr15 | - | 72267474 | 72267671 | 72266745 | 72266822 | 72271022 | 72271104 | 146 | 72 | 95 | [illegible] |
| SE | 14666 | ENSG00000156232 | WHAMM | chr15 | + | 82816691 | 82816842 | 82813102 | 82813276 | 82817919 | 82818089 | 141 | 76 | 172 | [illegible] |
| SE | 14667 | ENSG00000156232 | WHAMM | chr15 | + | 82823099 | 82823287 | 82819322 | 82819488 | 82826409 | 82826496 | 176 | 172 | 190 | [illegible] |
| SE | 14668 | ENSG00000156232 | WHAMM | chr15 | + | 82826750 | 82826846 | 82826409 | 82826496 | 82830598 | 82831079 | 240 | 173 | 249 | [illegible] |
| SE | 14671 | ENSG00000250988 | SNHG21 | chr15 | + | 82756729 | 82756785 | 82754628 | 82754784 | 82756903 | 82756939 | 1 | 5 | 5 | [illegible] |
| SE | 14683 | ENSG00000166794 | PPIB | chr15 | - | 64159266 | 64159311 | 64156724 | 64156909 | 64160103 | 64160197 | 1 | 2 | 0 | [illegible] |
| SE | 14684 | ENSG00000166794 | PPIB | chr15 | - | 64160103 | 64160193 | 64156724 | 64156909 | 64162040 | 64162154 | 449 | 238 | 531 | [illegible] |
| SE | 14685 | ENSG00000166794 | PPIB | chr15 | - | 64160103 | 64160197 | 64156724 | 64156909 | 64162040 | 64162154 | 1455 | 723 | 1530 | [illegible] |
| SE | 14686 | ENSG00000166794 | PPIB | chr15 | - | 64162040 | 64162154 | 64156724 | 64160197 | 64162851 | 64163019 | 1623 | 770 | 1650 | [illegible] |
| SE | 14688 | ENSG00000182511 | FES | chr15 | + | 90889305 | 90889443 | 90887186 | 90887370 | 90889516 | 90889636 | 152 | 73 | 94 | [illegible] |
| SE | 14690 | ENSG00000182511 | FES | chr15 | + | 90890091 | 90890278 | 90889839 | 90889962 | 90890365 | 90890484 | 47 | 24 | 36 | [illegible] |
| SE | 14693 | ENSG00000182511 | FES | chr15 | + | 90893099 | 90893194 | 90892706 | 90892825 | 90893290 | 90893414 | 88 | 24 | 52 | [illegible] |
| SE | 14698 | ENSG00000103591 | AAGAB | chr15 | - | 67203547 | 67203597 | 67200666 | 67202898 | 67204043 | 67204148 | 143 | 89 | 116 | [illegible] |
| SE | 14700 | ENSG00000103591 | AAGAB | chr15 | - | 67231813 | 67231897 | 67209461 | 67209544 | 67235978 | 67236068 | 167 | 113 | 124 | [illegible] |
| SE | 14706 | ENSG00000259363 | NA | chr15 | + | 99812972 | 99813086 | 99807435 | 99808302 | 99876952 | 99877148 | 12 | 0 | 3 | [illegible] |
| SE | 14707 | ENSG00000259363 | NA | chr15 | + | 99839082 | 99839134 | 99812972 | 99813086 | 99876952 | 99877148 | 0 | 0 | 0 | [illegible] |
| SE | 14708 | ENSG00000259363 | NA | chr15 | + | 99880676 | 99880827 | 99876952 | 99877148 | 99882967 | 99883926 | 18 | 10 | 6 | [illegible] |
| SE | 14715 | ENSG00000140545 | MFGE8 | chr15 | - | 88899655 | 88899811 | 88898682 | 88899532 | 88901550 | 88901735 | 39 | 15 | 42 | [illegible] |
| SE | 14716 | ENSG00000140545 | MFGE8 | chr15 | - | 88903936 | 88904162 | 88902182 | 88902231 | 88905756 | 88905901 | 3 | 2 | 3 | [illegible] |
| SE | 14718 | ENSG00000140545 | MFGE8 | chr15 | - | 88912123 | 88912176 | 88909791 | 88909923 | 88913246 | 88913379 | 2 | 0 | 5 | [illegible] |
| SE | 14720 | ENSG00000104064 | GABPB1 | chr15 | - | 50282278 | 50282391 | 50278541 | 50278784 | 50286067 | 50286183 | 2 | 0 | 0 | [illegible] |
| SE | 14721 | ENSG00000104064 | GABPB1 | chr15 | - | 50286067 | 50286183 | 50275388 | 50278784 | 50289482 | 50289668 | 108 | 80 | 89 | [illegible] |
| SE | 14723 | ENSG00000104064 | GABPB1 | chr15 | - | 50309690 | 50309798 | 50303965 | 50304133 | 50354984 | 50355153 | 78 | 52 | 87 | [illegible] |
| SE | 14724 | ENSG00000104064 | GABPB1 | chr15 | - | 50314415 | 50314670 | 50309690 | 50309798 | 50354984 | 50355191 | 1 | 2 | 1 | [illegible] |
| SE | 14725 | ENSG00000104064 | GABPB1 | chr15 | - | 50346371 | 50346395 | 50309690 | 50309798 | 50354984 | 50355202 | 5 | 1 | 10 | [illegible] |
| SE | 14728 | ENSG00000069966 | GNB5 | chr15 | - | 52133377 | 52133469 | 52128195 | 52128244 | 52135612 | 52135756 | 36 | 23 | 38 | [illegible] |
| SE | 14729 | ENSG00000069966 | GNB5 | chr15 | - | 52135612 | 52135756 | 52133377 | 52133469 | 52141139 | 52141211 | 49 | 35 | 45 | [illegible] |
| SE | 14732 | ENSG00000069966 | GNB5 | chr15 | - | 52149883 | 52149925 | 52147166 | 52147535 | 52153939 | 52154076 | 50 | 25 | 44 | [illegible] |
| SE | 14733 | ENSG00000185658 | BRWD1 | chr21 | - | 39202324 | 39202545 | 39200218 | 39200386 | 39206107 | 39206274 | 106 | 95 | 139 | [illegible] |
| SE | 14734 | ENSG00000185658 | BRWD1 | chr21 | - | 39206107 | 39206274 | 39202324 | 39202545 | 39213480 | 39213553 | 64 | 47 | 83 | [illegible] |
| SE | 14735 | ENSG00000185658 | BRWD1 | chr21 | - | 39209994 | 39210147 | 39206107 | 39206274 | 39212665 | 39212707 | 248 | 180 | 201 | [illegible] |
| SE | 14738 | ENSG00000185658 | BRWD1 | chr21 | - | 39212665 | 39212707 | 39206107 | 39206274 | 39213480 | 39213553 | 57 | 54 | 59 | [illegible] |
| SE | 14739 | ENSG00000185658 | BRWD1 | chr21 | - | 39218151 | 39218272 | 39215236 | 39215362 | 39218504 | 39218660 | 347 | 304 | 386 | [illegible] |
| SE | 14745 | ENSG00000185658 | BRWD1 | chr21 | - | 39224407 | 39224469 | 39215236 | 39215362 | 39228499 | 39228582 | 0 | 0 | 0 | [illegible] |
| SE | 14746 | ENSG00000185658 | BRWD1 | chr21 | - | 39224407 | 39224469 | 39218504 | 39218660 | 39228499 | 39228582 | 102 | 103 | 113 | [illegible] |
| SE | 14747 | ENSG00000185658 | BRWD1 | chr21 | - | 39225085 | 39225197 | 39215236 | 39215362 | 39228499 | 39228582 | 95 | 56 | 77 | [illegible] |
| SE | 14748 | ENSG00000185658 | BRWD1 | chr21 | - | 39225085 | 39225197 | 39224407 | 39224469 | 39228499 | 39228582 | 226 | 162 | 207 | [illegible] |
| SE | 14749 | ENSG00000185658 | BRWD1 | chr21 | - | 39228499 | 39228582 | 39225085 | 39225197 | 39232176 | 39232284 | 103 | 86 | 78 | [illegible] |
| SE | 14750 | ENSG00000185658 | BRWD1 | chr21 | - | 39229311 | 39229436 | 39228499 | 39228582 | 39232176 | 39232284 | 226 | 174 | 222 | [illegible] |
| SE | 14751 | ENSG00000185658 | BRWD1 | chr21 | - | 39238478 | 39238573 | 39236594 | 39236784 | 39247700 | 39247832 | 207 | 165 | 195 | [illegible] |
| SE | 14752 | ENSG00000185658 | BRWD1 | chr21 | - | 39247700 | 39247832 | 39238478 | 39238573 | 39250795 | 39250889 | 249 | 195 | 229 | [illegible] |
| SE | 14753 | ENSG00000185658 | BRWD1 | chr21 | - | 39250795 | 39250889 | 39238478 | 39238573 | 39255644 | 39255828 | 56 | 43 | 59 | [illegible] |
| SE | 14754 | ENSG00000185658 | BRWD1 | chr21 | - | 39255644 | 39255828 | 39250795 | 39250889 | 39258486 | 39258672 | 182 | 132 | 182 | [illegible] |
| SE | 14755 | ENSG00000185658 | BRWD1 | chr21 | - | 39255664 | 39255828 | 39250795 | 39250889 | 39258486 | 39258672 | 71 | 75 | 83 | [illegible] |
| SE | 14756 | ENSG00000185658 | BRWD1 | chr21 | - | 39278742 | 39278813 | 39274373 | 39274472 | 39280147 | 39280213 | 95 | 79 | 87 | [illegible] |
| SE | 14757 | ENSG00000185658 | BRWD1 | chr21 | - | 39278742 | 39278813 | 39277250 | 39277351 | 39280147 | 39280213 | 352 | 288 | 283 | [illegible] |
| SE | 14758 | ENSG00000185658 | BRWD1 | chr21 | - | 39296264 | 39296363 | 39295742 | 39295903 | 39298431 | 39298582 | 155 | 93 | 138 | [illegible] |
| SE | 14760 | ENSG00000103994 | ZNF106 | chr15 | - | 42424833 | 42425025 | 42423997 | 42424060 | 42428017 | 42428104 | 208 | 105 | 157 | [illegible] |
| SE | 14761 | ENSG00000103994 | ZNF106 | chr15 | - | 42428017 | 42428134 | 42424833 | 42425025 | 42435383 | 42435518 | 237 | 152 | 165 | [illegible] |
| SE | 14762 | ENSG00000103994 | ZNF106 | chr15 | - | 42435383 | 42435518 | 42428017 | 42428134 | 42437231 | 42437377 | 396 | 273 | 244 | [illegible] |
| SE | 14763 | ENSG00000103994 | ZNF106 | chr15 | - | 42438611 | 42438667 | 42437231 | 42437377 | 42439032 | 42439552 | 387 | 297 | 275 | [illegible] |
| SE | 14764 | ENSG00000103994 | ZNF106 | chr15 | - | 42442072 | 42442414 | 42439032 | 42439813 | 42444201 | 42444262 | 340 | 239 | 308 | [illegible] |
| SE | 14765 | ENSG00000103994 | ZNF106 | chr15 | - | 42444201 | 42444262 | 42441740 | 42442414 | 42444826 | 42444981 | 302 | 172 | 213 | [illegible] |
| SE | 14766 | ENSG00000103994 | ZNF106 | chr15 | - | 42444826 | 42444981 | 42441740 | 42442414 | 42446588 | 42446598 | 135 | 100 | 109 | [illegible] |
| SE | 14767 | ENSG00000103994 | ZNF106 | chr15 | - | 42448071 | 42448705 | 42446588 | 42446658 | 42449770 | 42451954 | 301 | 206 | 210 | [illegible] |
| SE | 14769 | ENSG00000103994 | ZNF106 | chr15 | - | 42449770 | 42451954 | 42446588 | 42446658 | 42466052 | 42466114 | 0 | 1 | 5 | [illegible] |
| SE | 14770 | ENSG00000103994 | ZNF106 | chr15 | - | 42449770 | 42451954 | 42448071 | 42448705 | 42456957 | 42457158 | 164 | 84 | 122 | [illegible] |
| SE | 14774 | ENSG00000103994 | ZNF106 | chr15 | - | 42456957 | 42457158 | 42449770 | 42451954 | 42466052 | 42466114 | 269 | 218 | 221 | [illegible] |
| SE | 14775 | ENSG00000103994 | ZNF106 | chr15 | - | 42466052 | 42466114 | 42456957 | 42457158 | 42472235 | 42472321 | 385 | 307 | 302 | [illegible] |
| SE | 14776 | ENSG00000140264 | SERF2 | chr15 | + | 43791975 | 43792383 | 43785409 | 43785534 | 43792974 | 43793083 | 810 | 516 | 590 | [illegible] |
| SE | 14777 | ENSG00000140264 | SERF2 | chr15 | + | 43793709 | 43793936 | 43792974 | 43793083 | 43794373 | 43796089 | 1984 | 1182 | 1304 | [illegible] |
| SE | 14781 | ENSG00000140264 | SERF2 | chr15 | + | 43801517 | 43801569 | 43801131 | 43801187 | 43801710 | 43804427 | 53 | 35 | 53 | [illegible] |
| SE | 14787 | ENSG00000169918 | OTUD7A | chr15 | - | 31649678 | 31649829 | 31570017 | 31570197 | 31655095 | 31655250 | 0 | 1 | 0 | [illegible] |
| SE | 14792 | ENSG00000092445 | TYRO3 | chr15 | + | 41567359 | 41567537 | 41564783 | 41565141 | 41568216 | 41568265 | 2 | 0 | 0 | [illegible] |
| SE | 14795 | ENSG00000062524 | LTK | chr15 | - | 41505914 | 41506005 | 41505712 | 41505777 | 41507094 | 41507174 | 33 | 6 | 22 | [illegible] |
| SE | 14796 | ENSG00000062524 | LTK | chr15 | - | 41507513 | 41507657 | 41507094 | 41507290 | 41508068 | 41508221 | 32 | 16 | 20 | [illegible] |
| SE | 14798 | ENSG00000062524 | LTK | chr15 | - | 41507561 | 41507657 | 41507094 | 41507290 | 41508068 | 41508221 | 28 | 18 | 21 | [illegible] |
| SE | 14799 | ENSG00000062524 | LTK | chr15 | - | 41511163 | 41511346 | 41509030 | 41509129 | 41511421 | 41511578 | 1 | 0 | 2 | [illegible] |
| SE | 14801 | ENSG00000137815 | RTF1 | chr15 | + | 41438320 | 41438431 | 41417103 | 41417313 | 41452900 | 41452942 | 65 | 62 | 62 | [illegible] |
| SE | 14802 | ENSG00000137815 | RTF1 | chr15 | + | 41440235 | 41440333 | 41438320 | 41438431 | 41452900 | 41452994 | 0 | 0 | 0 | [illegible] |
| SE | 14804 | ENSG00000137815 | RTF1 | chr15 | + | 41466140 | 41466252 | 41464770 | 41464885 | 41470256 | 41470392 | 156 | 110 | 168 | [illegible] |
| SE | 14805 | ENSG00000137815 | RTF1 | chr15 | + | 41471171 | 41471349 | 41470256 | 41470392 | 41474619 | 41474702 | 203 | 133 | 234 | [illegible] |
| SE | 14806 | ENSG00000137815 | RTF1 | chr15 | + | 41479102 | 41479198 | 41478484 | 41478625 | 41480213 | 41480290 | 268 | 145 | 212 | [illegible] |
| SE | 14807 | ENSG00000137804 | NUSAP1 | chr15 | + | 41342385 | 41342454 | 41332155 | 41333050 | 41349097 | 41349241 | 29 | 15 | 56 | [illegible] |
| SE | 14809 | ENSG00000137804 | NUSAP1 | chr15 | + | 41365401 | 41365589 | 41358148 | 41358258 | 41371526 | 41371684 | 17 | 15 | 48 | [illegible] |
| SE | 14810 | ENSG00000137804 | NUSAP1 | chr15 | + | 41365404 | 41365589 | 41358148 | 41358258 | 41371526 | 41371684 | 25 | 17 | 58 | [illegible] |
| SE | 14811 | ENSG00000137804 | NUSAP1 | chr15 | + | 41365815 | 41365863 | 41365404 | 41365589 | 41371526 | 41371684 | 0 | 1 | 0 | [illegible] |
| SE | 14815 | ENSG00000104147 | OIP5 | chr15 | - | 41313272 | 41313354 | 41309272 | 41309849 | 41319657 | 41319780 | 0 | 0 | 2 | [illegible] |
| SE | 14816 | ENSG00000104147 | OIP5 | chr15 | - | 41319657 | 41319780 | 41313272 | 41313354 | 41331914 | 41331981 | 1 | 0 | 2 | [illegible] |
| SE | 14818 | ENSG00000247556 | OIP5-AS1 | chr15 | + | 41290304 | 41290416 | 41286010 | 41286100 | 41298040 | 41298069 | 2 | 0 | 0 | [illegible] |
| SE | 14819 | ENSG00000137876 | RSL24D1 | chr15 | - | 55192402 | 55192833 | 55190974 | 55191047 | 55196809 | 55196914 | 301 | 174 | 274 | [illegible] |
| SE | 14820 | ENSG00000137876 | RSL24D1 | chr15 | - | 55192710 | 55192833 | 55190974 | 55191047 | 55196809 | 55196937 | 300 | 173 | 272 | [illegible] |
| SE | 14821 | ENSG00000137876 | RSL24D1 | chr15 | - | 55192719 | 55192833 | 55190974 | 55191047 | 55196809 | 55196902 | 732 | 396 | 663 | [illegible] |
| SE | 14823 | ENSG00000137876 | RSL24D1 | chr15 | - | 55194152 | 55194183 | 55192719 | 55192833 | 55194295 | 55194359 | 0 | 2 | 0 | [illegible] |
| SE | 14825 | ENSG00000137876 | RSL24D1 | chr15 | - | 55194295 | 55194359 | 55192719 | 55192833 | 55196809 | 55196958 | 1 | 0 | 0 | [illegible] |
| SE | 14826 | ENSG00000137876 | RSL24D1 | chr15 | - | 55195303 | 55195591 | 55194295 | 55194359 | 55196809 | 55196958 | 0 | 0 | 0 | [illegible] |
| SE | 14827 | ENSG00000069943 | PIGB | chr15 | + | 55324779 | 55324834 | 55319272 | 55319413 | 55327530 | 55327635 | 2 | 3 | 1 | [illegible] |
| SE | 14830 | ENSG00000069943 | PIGB | chr15 | + | 55327530 | 55327635 | 55321269 | 55321390 | 55329723 | 55329854 | 30 | 23 | 30 | [illegible] |
| SE | 14831 | ENSG00000069943 | PIGB | chr15 | + | 55329723 | 55329854 | 55321269 | 55321390 | 55333866 | 55334007 | 35 | 26 | 38 | [illegible] |
| SE | 14832 | ENSG00000069943 | PIGB | chr15 | + | 55333866 | 55334007 | 55329723 | 55329854 | 55339266 | 55339289 | 56 | 42 | 55 | [illegible] |
| SE | 14835 | ENSG00000140479 | PCSK6 | chr15 | - | 101305105 | 101305355 | 101300749 | 101300869 | 101307212 | 101307325 | 7 | 6 | 0 | [illegible] |
| SE | 14839 | ENSG00000140479 | PCSK6 | chr15 | - | 101364998 | 101365044 | 101346372 | 101347777 | 101366195 | 101366332 | 0 | 0 | 0 | [illegible] |
| SE | 14840 | ENSG00000140479 | PCSK6 | chr15 | - | 101364998 | 101365044 | 101355239 | 101356039 | 101366195 | 101366332 | 0 | 0 | 0 | [illegible] |
| SE | 14844 | ENSG00000140479 | PCSK6 | chr15 | - | 101431989 | 101432100 | 101429986 | 101430063 | 101443555 | 101443660 | 13 | 13 | 3 | [illegible] |
| SE | 14845 | ENSG00000140479 | PCSK6 | chr15 | - | 101443555 | 101443660 | 101431989 | 101432100 | 101489373 | 101489697 | 5 | 14 | 2 | [illegible] |
| SE | 14846 | ENSG00000185418 | TARS3 | chr15 | - | 101655866 | 101656020 | 101654393 | 101654730 | 101656921 | 101657036 | 0 | 0 | 0 | [illegible] |
| SE | 14847 | ENSG00000185418 | TARS3 | chr15 | - | 101657784 | 101657857 | 101656921 | 101657036 | 101661711 | 101661816 | 35 | 34 | 38 | [illegible] |
| SE | 14848 | ENSG00000185418 | TARS3 | chr15 | - | 101671485 | 101671586 | 101661711 | 101661816 | 101671670 | 101671748 | 60 | 39 | 33 | [illegible] |
| SE | 14849 | ENSG00000185418 | TARS3 | chr15 | - | 101684074 | 101684237 | 101675599 | 101675737 | 101685895 | 101686062 | 90 | 67 | 73 | [illegible] |
| SE | 14850 | ENSG00000185418 | TARS3 | chr15 | - | 101684074 | 101684237 | 101675599 | 101675737 | 101699330 | 101699435 | 48 | 40 | 34 | [illegible] |
| SE | 14851 | ENSG00000185418 | TARS3 | chr15 | - | 101685895 | 101686062 | 101675599 | 101675737 | 101699330 | 101699435 | 21 | 10 | 14 | [illegible] |
| SE | 14852 | ENSG00000185418 | TARS3 | chr15 | - | 101685895 | 101686062 | 101675599 | 101675737 | 101701085 | 101701184 | 59 | 49 | 63 | [illegible] |
| SE | 14853 | ENSG00000185418 | TARS3 | chr15 | - | 101685895 | 101686062 | 101684074 | 101684237 | 101699330 | 101699435 | 62 | 37 | 53 | [illegible] |
| SE | 14856 | ENSG00000185418 | TARS3 | chr15 | - | 101703858 | 101703937 | 101702238 | 101702385 | 101708792 | 101708828 | 68 | 65 | 64 | [illegible] |
| SE | 14857 | ENSG00000185418 | TARS3 | chr15 | - | 101703858 | 101703937 | 101702238 | 101702385 | 101711879 | 101712001 | 68 | 65 | 65 | [illegible] |
| SE | 14858 | ENSG00000185418 | TARS3 | chr15 | - | 101705682 | 101705747 | 101703858 | 101703937 | 101708792 | 101708828 | 107 | 76 | 122 | [illegible] |
| SE | 14860 | ENSG00000185418 | TARS3 | chr15 | - | 101708792 | 101708910 | 101702238 | 101702385 | 101711879 | 101712001 | 22 | 41 | 39 | [illegible] |
| SE | 14861 | ENSG00000185418 | TARS3 | chr15 | - | 101708792 | 101708910 | 101703858 | 101703937 | 101711879 | 101712001 | 22 | 41 | 39 | [illegible] |
| SE | 14862 | ENSG00000185418 | TARS3 | chr15 | - | 101708792 | 101708910 | 101705682 | 101705747 | 101711879 | 101712001 | 105 | 99 | 127 | [illegible] |
| SE | 14864 | ENSG00000185418 | TARS3 | chr15 | - | 101714839 | 101714963 | 101711879 | 101712001 | 101721125 | 101721322 | 45 | 66 | 62 | [illegible] |
| SE | 14865 | ENSG00000184277 | TM2D3 | chr15 | - | 101626447 | 101626563 | 101621643 | 101621755 | 101632942 | 101633026 | 2 | 7 | 0 | [illegible] |
| SE | 14866 | ENSG00000184277 | TM2D3 | chr15 | - | 101651207 | 101651391 | 101650069 | 101650161 | 101651695 | 101651998 | 0 | 1 | 0 | [illegible] |
| SE | 14867 | ENSG00000184277 | TM2D3 | chr15 | - | 101651695 | 101651773 | 101650003 | 101650161 | 101652270 | 101652381 | 12 | 4 | 4 | [illegible] |
| SE | 14869 | ENSG00000128989 | ARPP19 | chr15 | - | 52564189 | 52564246 | 52557099 | 52557222 | 52568288 | 52568832 | 5 | 5 | 8 | [illegible] |
| SE | 14871 | ENSG00000128989 | ARPP19 | chr15 | - | 52564189 | 52564246 | 52557099 | 52557222 | 52569115 | 52569167 | 6 | 5 | 8 | [illegible] |
| SE | 14873 | ENSG00000128989 | ARPP19 | chr15 | - | 52568847 | 52568902 | 52557099 | 52557222 | 52569111 | 52569167 | 101 | 52 | 79 | [illegible] |
| SE | 14875 | ENSG00000128989 | ARPP19 | chr15 | - | 52568847 | 52568902 | 52564189 | 52564246 | 52569115 | 52569154 | 7 | 5 | 7 | [illegible] |
| SE | 14876 | ENSG00000128833 | MYO5C | chr15 | - | 52208553 | 52208643 | 52205815 | 52205966 | 52211729 | 52211884 | 3 | 5 | 1 | [illegible] |
| SE | 14878 | ENSG00000128833 | MYO5C | chr15 | - | 52232621 | 52232685 | 52229132 | 52229313 | 52235669 | 52235763 | 1 | 1 | 4 | [illegible] |
| SE | 14889 | ENSG00000166855 | CLPX | chr15 | - | 65153546 | 65153639 | 65152429 | 65152536 | 65154781 | 65155081 | 231 | 270 | 306 | [illegible] |
| SE | 14892 | ENSG00000166855 | CLPX | chr15 | - | 65162603 | 65162645 | 65158574 | 65158751 | 65164028 | 65164188 | 182 | 177 | 200 | [illegible] |
| SE | 14895 | ENSG00000090487 | SPG21 | chr15 | - | 64969254 | 64969362 | 64965319 | 64965460 | 64970113 | 64970222 | 803 | 436 | 543 | [illegible] |
| SE | 14896 | ENSG00000090487 | SPG21 | chr15 | - | 64969254 | 64969362 | 64965319 | 64965460 | 64974601 | 64974747 | 307 | 158 | 219 | [illegible] |
| SE | 14897 | ENSG00000090487 | SPG21 | chr15 | - | 64970113 | 64970222 | 64965319 | 64965460 | 64974601 | 64974747 | 229 | 135 | 157 | [illegible] |
| SE | 14898 | ENSG00000090487 | SPG21 | chr15 | - | 64970113 | 64970222 | 64969254 | 64969362 | 64974601 | 64974747 | 724 | 413 | 488 | [illegible] |
| SE | 14901 | ENSG00000090487 | SPG21 | chr15 | - | 64980863 | 64981025 | 64976472 | 64976555 | 64983506 | 64983593 | 303 | 105 | 214 | [illegible] |
| SE | 14902 | ENSG00000090487 | SPG21 | chr15 | - | 64980863 | 64981025 | 64976472 | 64976555 | 64989664 | 64989946 | 195 | 68 | 139 | [illegible] |
| SE | 14903 | ENSG00000090487 | SPG21 | chr15 | - | 64983506 | 64983593 | 64976472 | 64976555 | 64989664 | 64989815 | 27 | 9 | 13 | [illegible] |
| SE | 14905 | ENSG00000090487 | SPG21 | chr15 | - | 64987171 | 64987261 | 64983506 | 64983593 | 64988618 | 64988816 | 0 | 1 | 0 | [illegible] |
| SE | 14906 | ENSG00000090487 | SPG21 | chr15 | - | 64988618 | 64988816 | 64983506 | 64983593 | 64989405 | 64989496 | 7 | 0 | 4 | [illegible] |
| SE | 14908 | ENSG00000086666 | ZFAND6 | chr15 | + | 80073265 | 80073330 | 80059713 | 80059809 | 80075163 | 80075315 | 0 | 1 | 0 | [illegible] |
| SE | 14912 | ENSG00000086666 | ZFAND6 | chr15 | + | 80073265 | 80073330 | 80072587 | 80072647 | 80098415 | 80098578 | 0 | 1 | 2 | [illegible] |
| SE | 14913 | ENSG00000086666 | ZFAND6 | chr15 | + | 80073265 | 80073330 | 80072587 | 80072647 | 80120327 | 80120498 | 2 | 0 | 0 | [illegible] |
| SE | 14915 | ENSG00000086666 | ZFAND6 | chr15 | + | 80075163 | 80075315 | 80072589 | 80072647 | 80098415 | 80098578 | 2 | 1 | 0 | [illegible] |
| SE | 14916 | ENSG00000086666 | ZFAND6 | chr15 | + | 80075163 | 80075315 | 80073265 | 80073330 | 80098415 | 80098578 | 2 | 3 | 0 | [illegible] |
| SE | 14918 | ENSG00000086666 | ZFAND6 | chr15 | + | 80098415 | 80098578 | 80060423 | 80060740 | 80120327 | 80120405 | 98 | 62 | 71 | [illegible] |
| SE | 14923 | ENSG00000086666 | ZFAND6 | chr15 | + | 80111427 | 80111559 | 80098415 | 80098578 | 80120327 | 80120440 | 0 | 0 | 1 | [illegible] |
| SE | 14924 | ENSG00000086666 | ZFAND6 | chr15 | + | 80112775 | 80112854 | 80111427 | 80111559 | 80120327 | 80120404 | 0 | 0 | 0 | [illegible] |
| SE | 14925 | ENSG00000086666 | ZFAND6 | chr15 | + | 80115026 | 80115070 | 80059634 | 80059809 | 80120327 | 80120498 | 1 | 0 | 0 | [illegible] |
| SE | 14926 | ENSG00000086666 | ZFAND6 | chr15 | + | 80122699 | 80122800 | 80121711 | 80121820 | 80131179 | 80131233 | 492 | 258 | 352 | [illegible] |
| SE | 14927 | ENSG00000086666 | ZFAND6 | chr15 | + | 80131179 | 80131293 | 80122699 | 80122800 | 80137479 | 80137678 | 358 | 207 | 270 | [illegible] |
| SE | 14929 | ENSG00000159216 | RUNX1 | chr21 | - | 34799300 | 34799462 | 34787800 | 34792610 | 34834409 | 34834601 | 102 | 65 | 116 | [illegible] |
| SE | 14932 | ENSG00000159216 | RUNX1 | chr21 | - | 34859473 | 34859578 | 34799300 | 34799462 | 34880556 | 34880713 | 132 | 71 | 116 | [illegible] |
| SE | 14933 | ENSG00000159216 | RUNX1 | chr21 | - | 34859473 | 34859578 | 34799300 | 34799462 | 34892924 | 34892963 | 20 | 5 | 27 | [illegible] |
| SE | 14934 | ENSG00000159216 | RUNX1 | chr21 | - | 34880556 | 34880713 | 34799300 | 34799462 | 34892924 | 34892963 | 2 | 3 | 1 | [illegible] |
| SE | 14936 | ENSG00000159216 | RUNX1 | chr21 | - | 34880556 | 34880713 | 34859473 | 34859578 | 35048841 | 35048958 | 134 | 82 | 119 | [illegible] |
| SE | 14939 | ENSG00000159216 | RUNX1 | chr21 | - | 34892924 | 34892963 | 34859473 | 34859578 | 35048841 | 35048958 | 21 | 7 | 20 | [illegible] |
| SE | 14942 | ENSG00000180953 | ST20 | chr15 | - | 79907628 | 79907684 | 79898846 | 79899125 | 79914587 | 79915256 | 10 | 3 | 19 | [illegible] |
| SE | 14943 | ENSG00000180953 | ST20 | chr15 | - | 79907628 | 79907684 | 79898846 | 79899125 | 79923451 | 79923702 | 8 | 5 | 8 | [illegible] |
| SE | 14944 | ENSG00000259332 | ST20-MTHFS | chr15 | - | 79889092 | 79889354 | 79845149 | 79845442 | 79907628 | 79907684 | 68 | 35 | 51 | [illegible] |
| SE | 14945 | ENSG00000259332 | ST20-MTHFS | chr15 | - | 79907628 | 79907684 | 79889092 | 79889354 | 79923451 | 79923744 | 0 | 0 | 0 | [illegible] |
| SE | 14947 | ENSG00000182718 | ANXA2 | chr15 | - | 60351723 | 60351819 | 60351191 | 60351251 | 60352382 | 60352476 | 1881 | 1312 | 1186 | [illegible] |
| SE | 14948 | ENSG00000182718 | ANXA2 | chr15 | - | 60364428 | 60364523 | 60358458 | 60361054 | 60386027 | 60386086 | 993 | 728 | 712 | [illegible] |
| SE | 14949 | ENSG00000182718 | ANXA2 | chr15 | - | 60374618 | 60374682 | 60364428 | 60364523 | 60382341 | 60382441 | 1 | 0 | 2 | [illegible] |
| SE | 14950 | ENSG00000182718 | ANXA2 | chr15 | - | 60382341 | 60382441 | 60364428 | 60364523 | 60386027 | 60386086 | 1398 | 845 | 796 | [illegible] |
| SE | 14959 | ENSG00000182718 | ANXA2 | chr15 | - | 60393012 | 60393440 | 60386027 | 60386086 | 60397257 | 60397338 | 0 | 0 | 1 | [illegible] |
| SE | 14960 | ENSG00000182718 | ANXA2 | chr15 | - | 60393012 | 60393440 | 60386027 | 60386086 | 60397942 | 60397986 | 0 | 0 | 1 | [illegible] |
| SE | 14962 | ENSG00000182718 | ANXA2 | chr15 | - | 60393037 | 60393440 | 60386027 | 60386086 | 60397942 | 60397986 | 0 | 0 | 0 | [illegible] |
| SE | 14966 | ENSG00000182718 | ANXA2 | chr15 | - | 60396150 | 60396427 | 60386027 | 60386086 | 60397942 | 60397986 | 2 | 8 | 2 | [illegible] |
| SE | 14967 | ENSG00000182718 | ANXA2 | chr15 | - | 60396150 | 60396427 | 60390212 | 60390317 | 60397257 | 60397338 | 0 | 2 | 2 | [illegible] |
| SE | 14972 | ENSG00000182718 | ANXA2 | chr15 | - | 60396279 | 60396427 | 60386027 | 60386086 | 60397942 | 60397986 | 2 | 1 | 2 | [illegible] |
| SE | 14973 | ENSG00000182718 | ANXA2 | chr15 | - | 60396279 | 60396427 | 60390212 | 60390317 | 60397257 | 60397338 | 0 | 3 | 2 | [illegible] |
| SE | 14975 | ENSG00000182718 | ANXA2 | chr15 | - | 60396372 | 60396427 | 60386027 | 60386086 | 60397942 | 60397986 | 0 | 0 | 0 | [illegible] |
| SE | 14976 | ENSG00000182718 | ANXA2 | chr15 | - | 60396372 | 60396427 | 60390212 | 60390317 | 60397942 | 60397986 | 0 | 0 | 0 | [illegible] |
| SE | 14980 | ENSG00000182718 | ANXA2 | chr15 | - | 60397257 | 60397338 | 60390212 | 60390317 | 60397942 | 60397964 | 0 | 1 | 2 | [illegible] |
| SE | 14982 | ENSG00000182718 | ANXA2 | chr15 | - | 60397257 | 60397338 | 60395644 | 60396427 | 60397942 | 60397964 | 0 | 3 | 2 | [illegible] |
| SE | 14983 | ENSG00000267827 | NA | chr19 | - | 52169647 | 52169785 | 52168057 | 52168152 | 52171335 | 52171469 | 43 | 31 | 61 | [illegible] |
| SE | 14984 | ENSG00000250021 | ARPIN-AP3S2 | chr15 | - | 89837614 | 89837722 | 89834307 | 89835643 | 89871474 | 89871546 | 165 | 112 | 181 | [illegible] |
| SE | 14985 | ENSG00000250021 | ARPIN-AP3S2 | chr15 | - | 89871474 | 89871546 | 89837614 | 89837722 | 89888520 | 89888632 | 152 | 93 | 162 | [illegible] |
| SE | 14986 | ENSG00000250021 | ARPIN-AP3S2 | chr15 | - | 89871474 | 89871546 | 89837614 | 89837722 | 89889048 | 89889140 | 78 | 50 | 81 | [illegible] |
| SE | 14987 | ENSG00000250021 | ARPIN-AP3S2 | chr15 | - | 89888520 | 89888632 | 89837614 | 89837722 | 89889048 | 89889140 | 55 | 37 | 49 | [illegible] |
| SE | 14988 | ENSG00000250021 | ARPIN-AP3S2 | chr15 | - | 89888520 | 89888632 | 89871474 | 89871546 | 89889048 | 89889140 | 130 | 79 | 129 | [illegible] |
| SE | 14991 | ENSG00000037757 | MRI1 | chr19 | + | 13765953 | 13766065 | 13764870 | 13765109 | 13768358 | 13768680 | 11 | 6 | 7 | [illegible] |
| SE | 14996 | ENSG00000157823 | AP3S2 | chr15 | - | 89878264 | 89878300 | 89871474 | 89871546 | 89889048 | 89889140 | 3 | 0 | 0 | [illegible] |
| SE | 15000 | ENSG00000128915 | ICE2 | chr15 | - | 60428428 | 60428687 | 60419608 | 60423762 | 60431933 | 60431984 | 144 | 108 | 138 | [illegible] |
| SE | 15001 | ENSG00000128915 | ICE2 | chr15 | - | 60431933 | 60431984 | 60428428 | 60428687 | 60436142 | 60436227 | 153 | 118 | 137 | [illegible] |
| SE | 15002 | ENSG00000128915 | ICE2 | chr15 | - | 60436142 | 60436227 | 60431933 | 60431984 | 60442415 | 60442545 | 234 | 168 | 195 | [illegible] |
| SE | 15003 | ENSG00000128915 | ICE2 | chr15 | - | 60442415 | 60442545 | 60436142 | 60436227 | 60447969 | 60448145 | 224 | 141 | 160 | [illegible] |
| SE | 15004 | ENSG00000128915 | ICE2 | chr15 | - | 60447969 | 60448145 | 60436142 | 60436227 | 60448847 | 60449841 | 99 | 96 | 145 | [illegible] |
| SE | 15007 | ENSG00000128915 | ICE2 | chr15 | - | 60456656 | 60456794 | 60455325 | 60455442 | 60466593 | 60466713 | 236 | 162 | 174 | [illegible] |
| SE | 15009 | ENSG00000128915 | ICE2 | chr15 | - | 60466593 | 60466713 | 60456656 | 60456794 | 60476062 | 60476167 | 125 | 68 | 84 | [illegible] |
| SE | 15010 | ENSG00000128915 | ICE2 | chr15 | - | 60468060 | 60468162 | 60456656 | 60456794 | 60476062 | 60476167 | 35 | 33 | 51 | [illegible] |
| SE | 15011 | ENSG00000128915 | ICE2 | chr15 | - | 60468060 | 60468162 | 60466593 | 60466713 | 60476062 | 60476167 | 152 | 115 | 154 | [illegible] |
| SE | 15012 | ENSG00000128915 | ICE2 | chr15 | - | 60468060 | 60468322 | 60456656 | 60456794 | 60476062 | 60476167 | 69 | 46 | 43 | [illegible] |
| SE | 15013 | ENSG00000128915 | ICE2 | chr15 | - | 60468060 | 60468322 | 60466593 | 60466713 | 60476062 | 60476167 | 188 | 135 | 154 | [illegible] |
| SE | 15014 | ENSG00000128915 | ICE2 | chr15 | - | 60477936 | 60478069 | 60476062 | 60476167 | 60478567 | 60478637 | 83 | 49 | 72 | [illegible] |
| SE | 15015 | ENSG00000128915 | ICE2 | chr15 | - | 60477936 | 60478069 | 60476062 | 60476167 | 60479002 | 60479073 | 121 | 69 | 92 | [illegible] |
| SE | 15016 | ENSG00000128915 | ICE2 | chr15 | - | 60478567 | 60478637 | 60477936 | 60478069 | 60479002 | 60479112 | 2 | 1 | 3 | [illegible] |
| SE | 15017 | ENSG00000128915 | ICE2 | chr15 | - | 60478567 | 60478650 | 60476062 | 60476167 | 60479002 | 60479107 | 1 | 1 | 0 | [illegible] |
| SE | 15018 | ENSG00000128915 | ICE2 | chr15 | - | 60478567 | 60478650 | 60477936 | 60478069 | 60479002 | 60479107 | 2 | 1 | 3 | [illegible] |
| SE | 15019 | ENSG00000128915 | ICE2 | chr15 | - | 60478567 | 60478660 | 60477936 | 60478069 | 60479002 | 60479103 | 2 | 1 | 3 | [illegible] |
| SE | 15020 | ENSG00000128915 | ICE2 | chr15 | - | 60478567 | 60478660 | 60477936 | 60478069 | 60479071 | 60479132 | 2 | 1 | 3 | [illegible] |
| SE | 15021 | ENSG00000091157 | WDR7 | chr18 | + | 56694949 | 56695198 | 56694618 | 56694760 | 56696241 | 56696462 | 270 | 225 | 202 | [illegible] |
| SE | 15024 | ENSG00000091157 | WDR7 | chr18 | + | 56816030 | 56816144 | 56781532 | 56781656 | 56879943 | 56880165 | 155 | 90 | 122 | [illegible] |
| SE | 15028 | ENSG00000091157 | WDR7 | chr18 | + | 57020744 | 57020849 | 56962429 | 56962529 | 57027003 | 57029792 | 112 | 80 | 101 | [illegible] |
| SE | 15032 | ENSG00000121057 | AKAP1 | chr17 | + | 57107945 | 57108048 | 57105440 | 57107178 | 57110024 | 57110158 | 1 | 0 | 0 | [illegible] |
| SE | 15039 | ENSG00000185880 | TRIM69 | chr15 | + | 44758612 | 44758803 | 44756367 | 44756463 | 44759639 | 44759662 | 0 | 0 | 0 | [illegible] |
| SE | 15040 | ENSG00000185880 | TRIM69 | chr15 | + | 44758612 | 44758854 | 44756367 | 44756463 | 44759639 | 44759662 | 3 | 6 | 7 | [illegible] |
| SE | 15041 | ENSG00000185880 | TRIM69 | chr15 | + | 44758620 | 44758854 | 44756367 | 44756463 | 44759639 | 44759662 | 3 | 6 | 7 | [illegible] |
| SE | 15042 | ENSG00000166783 | MARF1 | chr16 | - | 15598853 | 15599024 | 15594386 | 15596937 | 15600427 | 15600553 | 173 | 118 | 192 | [illegible] |
| SE | 15043 | ENSG00000166783 | MARF1 | chr16 | - | 15598853 | 15599117 | 15596634 | 15596937 | 15600427 | 15600460 | 56 | 59 | 85 | [illegible] |
| SE | 15045 | ENSG00000166783 | MARF1 | chr16 | - | 15604167 | 15604398 | 15601990 | 15602203 | 15608290 | 15608518 | 308 | 248 | 288 | [illegible] |
| SE | 15046 | ENSG00000166783 | MARF1 | chr16 | - | 15610974 | 15611108 | 15609522 | 15609725 | 15611591 | 15611734 | 367 | 297 | 375 | [illegible] |
| SE | 15047 | ENSG00000166783 | MARF1 | chr16 | - | 15611591 | 15611734 | 15610974 | 15611108 | 15612556 | 15612777 | 324 | 267 | 388 | [illegible] |
| SE | 15049 | ENSG00000166783 | MARF1 | chr16 | - | 15617051 | 15617171 | 15615829 | 15616005 | 15617298 | 15617535 | 363 | 311 | 322 | [illegible] |
| SE | 15050 | ENSG00000166783 | MARF1 | chr16 | - | 15620450 | 15620531 | 15617298 | 15617535 | 15624768 | 15624927 | 92 | 59 | 67 | [illegible] |
| SE | 15051 | ENSG00000166783 | MARF1 | chr16 | - | 15621732 | 15621911 | 15620450 | 15620531 | 15624768 | 15624927 | 148 | 102 | 111 | [illegible] |
| SE | 15053 | ENSG00000166783 | MARF1 | chr16 | - | 15622933 | 15623123 | 15621732 | 15621911 | 15624768 | 15624927 | 269 | 223 | 277 | [illegible] |
| SE | 15054 | ENSG00000166783 | MARF1 | chr16 | - | 15622933 | 15623123 | 15621732 | 15621911 | 15625015 | 15625173 | 176 | 141 | 192 | [illegible] |
| SE | 15055 | ENSG00000166783 | MARF1 | chr16 | - | 15622933 | 15623126 | 15621732 | 15621911 | 15624768 | 15624927 | 205 | 153 | 211 | [illegible] |
| SE | 15056 | ENSG00000166783 | MARF1 | chr16 | - | 15624768 | 15624927 | 15621732 | 15621911 | 15625015 | 15625173 | 124 | 124 | 126 | [illegible] |
| SE | 15058 | ENSG00000166783 | MARF1 | chr16 | - | 15625015 | 15625173 | 15622933 | 15623123 | 15631380 | 15631498 | 3 | 3 | 12 | [illegible] |
| SE | 15060 | ENSG00000166783 | MARF1 | chr16 | - | 15625371 | 15625797 | 15625015 | 15625173 | 15631380 | 15631498 | 236 | 161 | 189 | [illegible] |
| SE | 15062 | ENSG00000166783 | MARF1 | chr16 | - | 15625371 | 15625800 | 15625015 | 15625173 | 15631380 | 15631498 | 236 | 163 | 190 | [illegible] |
| SE | 15064 | ENSG00000166783 | MARF1 | chr16 | - | 15630331 | 15630504 | 15625371 | 15625797 | 15631380 | 15631498 | 120 | 54 | 121 | [illegible] |
| SE | 15065 | ENSG00000166783 | MARF1 | chr16 | - | 15630331 | 15630504 | 15625371 | 15625800 | 15631380 | 15631498 | 158 | 74 | 149 | [illegible] |
| SE | 15066 | ENSG00000166783 | MARF1 | chr16 | - | 15631380 | 15631498 | 15625015 | 15625173 | 15633616 | 15633834 | 257 | 160 | 231 | [illegible] |
| SE | 15069 | ENSG00000137877 | SPTBN5 | chr15 | - | 41853581 | 41853787 | 41853257 | 41853447 | 41854049 | 41854205 | 19 | 0 | 4 | [illegible] |
| SE | 15070 | ENSG00000137877 | SPTBN5 | chr15 | - | 41868397 | 41868601 | 41868068 | 41868218 | 41869840 | 41870020 | 2 | 0 | 0 | [illegible] |
| SE | 15071 | ENSG00000137877 | SPTBN5 | chr15 | - | 41879733 | 41879864 | 41879259 | 41879499 | 41881033 | 41881234 | 3 | 0 | 2 | [illegible] |
| SE | 15072 | ENSG00000137877 | SPTBN5 | chr15 | - | 41880159 | 41880312 | 41879733 | 41879864 | 41881033 | 41881234 | 4 | 6 | 4 | [illegible] |
| SE | 15073 | ENSG00000137877 | SPTBN5 | chr15 | - | 41887927 | 41888085 | 41887212 | 41887441 | 41890088 | 41890205 | 1 | 4 | 9 | [illegible] |

| | | | | | | | | | | | | | | | | | | | | | | | | | |
|---|---|---|---|---|---|---|---|---|---|---|---|---|---|---|---|---|---|---|---|---|---|---|---|---|---|
| SE | 15074 | ENSG00000168970 | JMJD7-PLA2G4B | chr15 | + | 41834739 | 41834893 | 41828094 | 41828188 | 41834969 | 41835223 | 16 | 13 | 28 | [illegible] | [illegible] | [illegible] | [illegible] | [illegible] | [illegible] | [illegible] | [illegible] | [illegible] | [illegible] | [illegible] |
| SE | 15075 | ENSG00000168970 | JMJD7-PLA2G4B | chr15 | + | 41836780 | 41836940 | 41836474 | 41836551 | 41837067 | 41840230 | 29 | 7 | 38 | 43 | 25 | 0 | 0 | 0 | 0 | 0 | 55 | 27 | 27 | 11 |
| SE | 15077 | ENSG00000168970 | JMJD7-PLA2G4B | chr15 | + | 41843125 | 41843223 | 41842192 | 41842591 | 41843675 | 41843811 | 6 | 5 | 14 | 6 | 1 | 5 | 1 | 4 | 14 | 7 | 0 | 6 | 0 | 4 |
| SE | 15078 | ENSG00000243789 | JMJD7 | chr15 | + | 41828393 | 41828501 | 41828106 | 41828188 | 41834739 | 41834893 | 0 | 0 | 1 | 0 | 0 | 7 | 7 | 10 | 8 | 1 | 0 | 0 | 0 | 0 |
| SE | 15079 | ENSG00000243789 | JMJD7 | chr15 | + | 41829103 | 41829292 | 41828112 | 41828188 | 41834739 | 41834804 | 0 | 0 | 1 | 2 | 8 | 7 | 7 | 10 | 8 | 1 | 0 | 2 | 9 | 0 |
| SE | 15091 | ENSG00000128739 | SNRPN | chr15 | + | 24967931 | 24968082 | 24962113 | 24962209 | 24974397 | 24974456 | 188 | 137 | 253 | 233 | 170 | 0 | 0 | 0 | 1 | 0 | 133 | 210 | 90 | 55 |
| SE | 15092 | ENSG00000128739 | SNRPN | chr15 | + | 24967931 | 24968082 | 24962113 | 24962209 | 24974641 | 24974957 | 182 | 132 | 243 | 224 | 175 | 0 | 0 | 0 | 4 | 0 | 125 | 187 | 86 | 49 |
| SE | 15093 | ENSG00000128739 | SNRPN | chr15 | + | 24967931 | 24968082 | 24962113 | 24962209 | 24976876 | 24977029 | 181 | 132 | 242 | 225 | 166 | 0 | 0 | 0 | 0 | 0 | 129 | 187 | 86 | 49 |
| SE | 15095 | ENSG00000128739 | SNRPN | chr15 | + | 24974287 | 24974456 | 24967931 | 24968082 | 24976304 | 24976416 | 6 | 2 | 4 | 2 | 11 | 0 | 0 | 0 | 1 | 0 | 0 | 4 | 5 | 1 |
| SE | 15096 | ENSG00000128739 | SNRPN | chr15 | + | 24974310 | 24974456 | 24962113 | 24962209 | 24974641 | 24974957 | 4 | 1 | 8 | 9 | 5 | 0 | 0 | 0 | 4 | 0 | 0 | 0 | 10 | 0 |
| SE | 15099 | ENSG00000128739 | SNRPN | chr15 | + | 24974310 | 24974456 | 24967931 | 24968082 | 24976304 | 24976327 | 278 | 175 | 293 | 312 | 230 | 0 | 0 | 0 | 1 | 0 | 192 | 259 | 105 | 106 |
| SE | 15101 | ENSG00000128739 | SNRPN | chr15 | + | 24974386 | 24974456 | 24967931 | 24968082 | 24976304 | 24976416 | 3 | 2 | 3 | 8 | 5 | 0 | 0 | 0 | 1 | 0 | 0 | 4 | 2 | 0 |
| SE | 15103 | ENSG00000128739 | SNRPN | chr15 | + | 24974397 | 24974456 | 24967931 | 24968082 | 24976304 | 24976416 | 0 | 3 | 1 | 4 | 2 | 0 | 0 | 0 | 1 | 0 | 0 | 4 | 1 | 4 |
| SE | 15105 | ENSG00000128739 | SNRPN | chr15 | + | 24974641 | 24974957 | 24967931 | 24968082 | 24976304 | 24976416 | 1 | 0 | 1 | 0 | 9 | 0 | 0 | 0 | 1 | 0 | 0 | 0 | 0 | 2 |
| SE | 15107 | ENSG00000128739 | SNRPN | chr15 | + | 24974641 | 24974957 | 24974310 | 24974456 | 24976304 | 24976416 | 2 | 0 | 6 | 6 | 5 | 4 | 2 | 3 | 1 | 11 | 0 | 0 | 2 | 2 |
| SE | 15108 | ENSG00000128739 | SNRPN | chr15 | + | 24975357 | 24975509 | 24967931 | 24968082 | 24976304 | 24976327 | 733 | 478 | 552 | 671 | 772 | 0 | 0 | 0 | 1 | 0 | 417 | 723 | 416 | 372 |
| SE | 15110 | ENSG00000128739 | SNRPN | chr15 | + | 24975357 | 24975509 | 24974287 | 24974456 | 24976304 | 24976327 | 1210 | 791 | 979 | 1166 | 1350 | 4 | 2 | 3 | 1 | 11 | 770 | 1200 | 695 | 636 |
| SE | 15111 | ENSG00000128739 | SNRPN | chr15 | + | 24975357 | 24975509 | 24974641 | 24974957 | 24976304 | 24976327 | 741 | 491 | 559 | 673 | 779 | 0 | 0 | 0 | 0 | 0 | 421 | 746 | 429 | 380 |
| SE | 15112 | ENSG00000128739 | SNRPN | chr15 | + | 24976304 | 24976416 | 24967931 | 24968082 | 24976876 | 24977029 | 75 | 68 | 98 | 96 | 128 | 0 | 0 | 0 | 1 | 0 | 51 | 99 | 40 | 22 |
| SE | 15113 | ENSG00000128739 | SNRPN | chr15 | + | 24976304 | 24976416 | 24975357 | 24975509 | 24976876 | 24977029 | 807 | 545 | 647 | 758 | 890 | 0 | 0 | 1 | 0 | 0 | 463 | 821 | 456 | 394 |
| SE | 15114 | ENSG00000168803 | ADAL | chr15 | + | 43331899 | 43332000 | 43330671 | 43330859 | 43333310 | 43333434 | 9 | 7 | 12 | 5 | 7 | 0 | 0 | 2 | 0 | 2 | 0 | 1 | 0 | 0 |
| SE | 15116 | ENSG00000168803 | ADAL | chr15 | + | 43334944 | 43335170 | 43333310 | 43333434 | 43335695 | 43335826 | 36 | 33 | 29 | 24 | 33 | 1 | 0 | 0 | 0 | 0 | 13 | 10 | 4 | 25 |
| SE | 15117 | ENSG00000168803 | ADAL | chr15 | + | 43335695 | 43335826 | 43331899 | 43332000 | 43336625 | 43336686 | 16 | 21 | 11 | 14 | 10 | 0 | 1 | 0 | 0 | 0 | 0 | 0 | 17 | 0 |
| SE | 15118 | ENSG00000168803 | ADAL | chr15 | + | 43335695 | 43335826 | 43334937 | 43335170 | 43336625 | 43336686 | 49 | 47 | 35 | 32 | 35 | 0 | 0 | 0 | 1 | 0 | 13 | 5 | 21 | 20 |
| SE | 15119 | ENSG00000168803 | ADAL | chr15 | + | 43336625 | 43336686 | 43335695 | 43335826 | 43340264 | 43340352 | 24 | 22 | 14 | 15 | 15 | 0 | 0 | 4 | 0 | 0 | 0 | 0 | 26 | 5 |
| SE | 15120 | ENSG00000168803 | ADAL | chr15 | + | 43347015 | 43347096 | 43345853 | 43345980 | 43348906 | 43349045 | 21 | 10 | 8 | 6 | 13 | 3 | 3 | 1 | 1 | 2 | 19 | 7 | 9 | 8 |
| SE | 15121 | ENSG00000168803 | ADAL | chr15 | + | 43350940 | 43351041 | 43348906 | 43349045 | 43351750 | 43353898 | 15 | 6 | 8 | 15 | 7 | 1 | 0 | 0 | 0 | 1 | 0 | 6 | 8 | 16 |
| SE | 15123 | ENSG00000198838 | RYR3 | chr15 | + | 33728856 | 33729026 | 33726385 | 33726506 | 33731473 | 33731694 | 0 | 0 | 0 | 1 | 0 | 0 | 0 | 0 | 0 | 0 | 0 | 0 | 0 | 0 |
| SE | 15132 | ENSG00000104093 | DMXL2 | chr15 | - | 51451644 | 51451697 | 51450128 | 51450346 | 51456065 | 51456193 | 7 | 15 | 8 | 10 | 13 | 0 | 4 | 0 | 0 | 2 | 0 | 3 | 5 | 4 |
| SE | 15134 | ENSG00000104093 | DMXL2 | chr15 | - | 51455150 | 51455228 | 51450128 | 51450346 | 51456065 | 51456193 | 60 | 72 | 50 | 82 | 45 | 0 | 4 | 0 | 0 | 2 | 7 | 39 | 19 | 0 |
| SE | 15138 | ENSG00000104093 | DMXL2 | chr15 | - | 51464674 | 51464876 | 51463075 | 51463496 | 51466183 | 51466311 | 121 | 140 | 95 | 94 | 57 | 0 | 1 | 5 | 1 | 0 | 12 | 73 | 85 | 3 |
| SE | 15139 | ENSG00000104093 | DMXL2 | chr15 | - | 51465565 | 51465651 | 51463075 | 51463496 | 51466183 | 51466311 | 66 | 72 | 50 | 65 | 107 | 0 | 1 | 5 | 1 | 0 | 4 | 51 | 15 | 16 |
| SE | 15141 | ENSG00000104093 | DMXL2 | chr15 | - | 51471222 | 51471401 | 51466183 | 51466311 | 51474343 | 51474592 | 359 | 334 | 209 | 287 | 340 | 1 | 0 | 0 | 3 | 6 | 27 | 230 | 174 | 37 |
| SE | 15142 | ENSG00000104093 | DMXL2 | chr15 | - | 51478270 | 51478347 | 51476588 | 51476716 | 51479947 | 51480139 | 124 | 153 | 74 | 106 | 155 | 0 | 0 | 2 | 0 | 2 | 15 | 77 | 29 | 19 |
| SE | 15143 | ENSG00000104093 | DMXL2 | chr15 | - | 51478270 | 51478347 | 51476588 | 51476719 | 51479947 | 51480139 | 202 | 203 | 105 | 149 | 191 | 2 | 3 | 2 | 0 | 0 | 24 | 95 | 56 | 22 |
| SE | 15144 | ENSG00000104093 | DMXL2 | chr15 | - | 51479947 | 51480139 | 51476588 | 51476716 | 51480541 | 51481623 | 107 | 132 | 73 | 89 | 138 | 0 | 0 | 0 | 0 | 0 | 8 | 48 | 26 | 2 |
| SE | 15145 | ENSG00000104093 | DMXL2 | chr15 | - | 51486072 | 51486337 | 51480541 | 51481623 | 51487953 | 51488119 | 246 | 294 | 148 | 220 | 306 | 0 | 0 | 0 | 0 | 3 | 16 | 138 | 98 | 11 |
| SE | 15147 | ENSG00000104093 | DMXL2 | chr15 | - | 51495023 | 51495134 | 51491577 | 51491747 | 51507133 | 51507253 | 106 | 83 | 43 | 82 | 76 | 0 | 0 | 0 | 0 | 0 | 4 | 82 | 38 | 5 |
| SE | 15148 | ENSG00000104093 | DMXL2 | chr15 | - | 51498551 | 51500231 | 51491577 | 51491747 | 51507133 | 51507253 | 8 | 8 | 1 | 8 | 7 | 0 | 0 | 0 | 0 | 0 | 0 | 8 | 18 | 2 |
| SE | 15149 | ENSG00000104093 | DMXL2 | chr15 | - | 51498551 | 51500231 | 51495023 | 51495134 | 51507133 | 51507253 | 110 | 83 | 39 | 63 | 82 | 0 | 2 | 0 | 1 | 1 | 0 | 36 | 35 | 2 |
| SE | 15151 | ENSG00000104093 | DMXL2 | chr15 | - | 51535662 | 51535784 | 51517077 | 51517167 | 51536165 | 51536862 | 290 | 285 | 156 | 226 | 345 | 0 | 0 | 0 | 0 | 3 | 82 | 135 | 87 | 64 |
| SE | 15152 | ENSG00000104093 | DMXL2 | chr15 | - | 51538212 | 51538452 | 51537487 | 51537759 | 51542332 | 51542507 | 196 | 254 | 124 | 208 | 193 | 0 | 0 | 0 | 1 | 0 | 42 | 98 | 103 | 70 |
| SE | 15153 | ENSG00000104093 | DMXL2 | chr15 | - | 51542332 | 51542507 | 51538212 | 51538452 | 51545582 | 51545766 | 195 | 252 | 129 | 211 | 114 | 1 | 2 | 0 | 0 | 0 | 45 | 135 | 118 | 59 |
| SE | 15154 | ENSG00000104093 | DMXL2 | chr15 | - | 51564124 | 51564260 | 51563380 | 51563447 | 51565087 | 51565166 | 167 | 174 | 102 | 151 | 213 | 1 | 0 | 0 | 0 | 0 | 45 | 101 | 72 | 38 |
| SE | 15155 | ENSG00000104093 | DMXL2 | chr15 | - | 51565087 | 51565166 | 51564124 | 51564260 | 51568486 | 51568558 | 163 | 150 | 88 | 109 | 152 | 0 | 1 | 0 | 0 | 0 | 43 | 121 | 61 | 55 |
| SE | 15156 | ENSG00000104093 | DMXL2 | chr15 | - | 51568486 | 51568558 | 51564124 | 51564260 | 51576055 | 51576181 | 57 | 53 | 35 | 34 | 15 | 0 | 4 | 0 | 0 | 0 | 12 | 27 | 10 | 17 |
| SE | 15157 | ENSG00000104093 | DMXL2 | chr15 | - | 51576055 | 51576181 | 51547229 | 51547408 | 51622458 | 51622635 | 25 | 13 | 27 | 20 | 19 | 0 | 1 | 0 | 0 | 0 | 14 | 65 | 11 | 13 |
| SE | 15158 | ENSG00000137842 | TMEM62 | chr15 | + | 43134256 | 43134368 | 43133161 | 43133402 | 43135511 | 43135649 | 15 | 3 | 10 | 10 | 13 | 0 | 0 | 0 | 0 | 0 | 4 | 23 | 8 | 23 |
| SE | 15161 | ENSG00000137842 | TMEM62 | chr15 | + | 43138573 | 43138619 | 43135511 | 43135649 | 43148754 | 43148879 | 7 | 4 | 5 | 0 | 3 | 18 | 4 | 7 | 7 | 8 | 0 | 5 | 9 | 10 |
| SE | 15168 | ENSG00000137842 | TMEM62 | chr15 | + | 43169592 | 43169677 | 43160393 | 43160794 | 43181180 | 43181299 | 38 | 21 | 28 | 35 | 54 | 0 | 0 | 0 | 0 | 0 | 12 | 47 | 64 | 40 |
| SE | 15170 | ENSG00000137842 | TMEM62 | chr15 | + | 43181180 | 43181299 | 43169592 | 43169677 | 43184259 | 43184385 | 16 | 8 | 24 | 21 | 58 | 0 | 0 | 0 | 0 | 0 | 9 | 32 | 41 | 35 |
| SE | 15175 | ENSG00000171953 | ATPAF2 | chr17 | - | 18021744 | 18021857 | 18021122 | 18021238 | 18024623 | 18024704 | 62 | 47 | 73 | 41 | 61 | 1 | 0 | 0 | 0 | 0 | 82 | 121 | 58 | 98 |
| SE | 15176 | ENSG00000171953 | ATPAF2 | chr17 | - | 18024623 | 18024704 | 18021744 | 18021857 | 18026318 | 18026416 | 43 | 41 | 59 | 36 | 51 | 0 | 0 | 0 | 0 | 0 | 139 | 104 | 50 | 88 |
| SE | 15178 | ENSG00000171953 | ATPAF2 | chr17 | - | 18028614 | 18028659 | 18026318 | 18026416 | 18038880 | 18039050 | 28 | 16 | 9 | 18 | 10 | 0 | 1 | 0 | 0 | 0 | 47 | 22 | 7 | 25 |
| SE | 15179 | ENSG00000159459 | UBR1 | chr15 | - | 42977879 | 42977947 | 42976716 | 42976867 | 42983896 | 42983993 | 94 | 56 | 83 | 90 | 134 | 0 | 0 | 0 | 0 | 0 | 9 | 60 | 27 | 16 |
| SE | 15181 | ENSG00000159459 | UBR1 | chr15 | - | 42983896 | 42983993 | 42977879 | 42977947 | 42988818 | 42988967 | 28 | 17 | 25 | 28 | 29 | 0 | 1 | 0 | 0 | 0 | 0 | 30 | 19 | 4 |
| SE | 15182 | ENSG00000159459 | UBR1 | chr15 | - | 42984886 | 42984942 | 42977879 | 42977947 | 42988818 | 42988967 | 13 | 7 | 11 | 15 | 18 | 0 | 1 | 0 | 0 | 0 | 0 | 12 | 7 | 8 |
| SE | 15183 | ENSG00000159459 | UBR1 | chr15 | - | 42984886 | 42984942 | 42983896 | 42983993 | 42988818 | 42988967 | 51 | 21 | 42 | 58 | 77 | 3 | 2 | 0 | 2 | 0 | 0 | 23 | 17 | 20 |
| SE | 15184 | ENSG00000159459 | UBR1 | chr15 | - | 42998167 | 42998265 | 42990029 | 42990120 | 43002554 | 43002704 | 70 | 55 | 77 | 87 | 103 | 2 | 0 | 0 | 1 | 0 | 0 | 54 | 47 | 15 |
| SE | 15185 | ENSG00000159459 | UBR1 | chr15 | - | 43002554 | 43002704 | 42998167 | 42998265 | 43003836 | 43003930 | 105 | 71 | 79 | 90 | 137 | 3 | 0 | 0 | 0 | 0 | 20 | 87 | 49 | 20 |
| SE | 15186 | ENSG00000159459 | UBR1 | chr15 | - | 43017094 | 43017181 | 43015655 | 43015869 | 43021274 | 43021375 | 201 | 128 | 138 | 176 | 204 | 1 | 0 | 0 | 0 | 0 | 43 | 154 | 126 | 99 |
| SE | 15188 | ENSG00000159459 | UBR1 | chr15 | - | 43024828 | 43024983 | 43022701 | 43022801 | 43026560 | 43026663 | 138 | 73 | 102 | 96 | 74 | 0 | 0 | 2 | 0 | 0 | 58 | 83 | 80 | 44 |
| SE | 15194 | ENSG00000159459 | UBR1 | chr15 | - | 43047160 | 43047289 | 43043213 | 43043395 | 43048391 | 43048491 | 152 | 85 | 94 | 120 | 127 | 1 | 0 | 0 | 0 | 0 | 0 | 88 | 101 | 40 |
| SE | 15195 | ENSG00000159459 | UBR1 | chr15 | - | 43048391 | 43048491 | 43047160 | 43047289 | 43054741 | 43054899 | 152 | 80 | 81 | 103 | 114 | 10 | 5 | 1 | 8 | 11 | 0 | 71 | 81 | 22 |
| SE | 15196 | ENSG00000159459 | UBR1 | chr15 | - | 43048391 | 43048491 | 43047160 | 43047289 | 43056343 | 43056442 | 57 | 38 | 35 | 41 | 54 | 0 | 0 | 1 | 0 | 0 | 0 | 20 | 40 | 20 |
| SE | 15197 | ENSG00000159459 | UBR1 | chr15 | - | 43054741 | 43054899 | 43047160 | 43047289 | 43056343 | 43056442 | 177 | 121 | 109 | 136 | 184 | 0 | 0 | 1 | 0 | 0 | 24 | 158 | 136 | 62 |
| SE | 15198 | ENSG00000159459 | UBR1 | chr15 | - | 43054741 | 43054899 | 43048391 | 43048491 | 43056343 | 43056442 | 264 | 160 | 157 | 190 | 237 | 2 | 1 | 1 | 0 | 4 | 24 | 193 | 162 | 65 |
| SE | 15199 | ENSG00000159459 | UBR1 | chr15 | - | 43058340 | 43058429 | 43056343 | 43056442 | 43059084 | 43059192 | 116 | 98 | 95 | 111 | 166 | 0 | 0 | 0 | 0 | 0 | 37 | 70 | 110 | 56 |
| SE | 15200 | ENSG00000159459 | UBR1 | chr15 | - | 43074978 | 43075089 | 43070794 | 43070925 | 43082637 | 43082716 | 273 | 212 | 198 | 243 | 321 | 0 | 0 | 0 | 0 | 4 | 83 | 250 | 256 | 185 |

| | | | | | | | | | | | | | | | | | | | | | | | | | | | | | | | | | | | | | | | | | | | | | | |
|---|---|---|---|---|---|---|---|---|---|---|---|---|---|---|---|---|---|---|---|---|---|---|---|---|---|---|---|---|---|---|---|---|---|---|---|---|---|---|---|---|---|---|---|---|---|---|
| SE | 15511 | ENSG00000166664 | CHRFAM7A | chr15 | - | 30371097 | 30371184 | 30367417 | 30367527 | 30372146 | 30372341 | 2 | 1 | 0 | 0 | 0 | 0 | 0 | 0 | 0 | 0 | 0 | 0 | 1 | 0 | 0 | 0 | 0 | 0 | 2 | 0 | 236 | 149 | 1 | 1 | 1 | 1 | NA | NA | NA | NA | NA | 1 | 0 | NA | 0.5 |
| SE | 15513 | ENSG00000166664 | CHRFAM7A | chr15 | - | 30383318 | 30383382 | 30380352 | 30380399 | 30386862 | 30386987 | 6 | 4 | 0 | 4 | 0 | 0 | 0 | 0 | 1 | 0 | 0 | 0 | 6 | 0 | 0 | 0 | 0 | 0 | 2 | 0 | 213 | 149 | 1 | 1 | 1 | 1 | NA | 0.737 | NA | NA | NA | 1 | 0 | NA | 0.412 |
| SE | 15514 | ENSG00000129003 | VPS13C | chr15 | - | 61856285 | 61856409 | 61854870 | 61854954 | 61863439 | 61863528 | 619 | 652 | 476 | 594 | 868 | 2 | 0 | 3 | 1 | 2 | 35 | 109 | 179 | 15 | 9 | 0 | 0 | 0 | 0 | 0 | 273 | 149 | 1 | 1 | 0.994 | 1 | 0.989 | 0.997 | 0.996 | 1 | 1 | 1 | 1 | 1 | -0.005 |
| SE | 15515 | ENSG00000129003 | VPS13C | chr15 | - | 61856285 | 61856409 | 61854870 | 61854954 | 61868658 | 61868773 | 290 | 323 | 226 | 280 | 483 | 0 | 0 | 0 | 0 | 2 | 33 | 57 | 84 | 3 | 5 | 0 | 0 | 1 | 0 | 0 | 273 | 149 | 1 | 1 | 1 | 1 | 1 | 1 | 0.992 | 1 | 1 | 0.979 | 1 | 1 | 0.003 |
| SE | 15516 | ENSG00000129003 | VPS13C | chr15 | - | 61863439 | 61863528 | 61854870 | 61854954 | 61868658 | 61868773 | 141 | 137 | 102 | 123 | 139 | 0 | 0 | 0 | 0 | 2 | 9 | 36 | 40 | 4 | 6 | 0 | 0 | 1 | 0 | 0 | 238 | 149 | 1 | 1 | 1 | 1 | 1 | 1 | 0.978 | 1 | 1 | 0.962 | 1 | 1 | 0.003 |
| SE | 15517 | ENSG00000129003 | VPS13C | chr15 | - | 61863439 | 61863528 | 61856285 | 61856409 | 61868658 | 61868773 | 439 | 428 | 335 | 406 | 515 | 0 | 0 | 0 | 0 | 1 | 11 | 88 | 130 | 16 | 10 | 0 | 0 | 0 | 0 | 0 | 238 | 149 | 1 | 1 | 1 | 1 | 1 | 1 | 0.997 | 1 | 1 | 1 | 1 | 1 | -0.001 |
| SE | 15519 | ENSG00000129003 | VPS13C | chr15 | - | 61873245 | 61873409 | 61871988 | 61872034 | 61874875 | 61874951 | 487 | 490 | 417 | 482 | 653 | 1 | 0 | 0 | 0 | 0 | 31 | 130 | 128 | 27 | 34 | 0 | 0 | 0 | 0 | 0 | 298 | 149 | 1 | 1 | 0.996 | 1 | 1 | 1 | 1 | 1 | 1 | 1 | 1 | 1 | -0.001 |
| SE | 15521 | ENSG00000129003 | VPS13C | chr15 | - | 61878606 | 61878746 | 61876972 | 61877054 | 61880608 | 61880722 | 452 | 447 | 315 | 432 | 404 | 0 | 0 | 0 | 0 | 6 | 36 | 131 | 161 | 27 | 16 | 0 | 0 | 0 | 0 | 0 | 289 | 149 | 1 | 1 | 1 | 1 | 1 | 1 | 0.972 | 1 | 1 | 1 | 1 | 1 | -0.006 |
| SE | 15523 | ENSG00000129003 | VPS13C | chr15 | - | 61890164 | 61890400 | 61884127 | 61884269 | 61907263 | 61907390 | 579 | 587 | 390 | 578 | 646 | 0 | 0 | 0 | 0 | 2 | 61 | 228 | 170 | 40 | 6 | 0 | 0 | 0 | 0 | 0 | 298 | 149 | 1 | 1 | 1 | 1 | 1 | 1 | 0.994 | 1 | 1 | 1 | 1 | 1 | -0.001 |
| SE | 15524 | ENSG00000129003 | VPS13C | chr15 | - | 61907263 | 61907390 | 61890164 | 61890400 | 61908991 | 61909125 | 774 | 554 | 536 | 587 | 773 | 1 | 1 | 0 | 0 | 0 | 65 | 245 | 172 | 42 | 22 | 0 | 0 | 0 | 0 | 0 | 276 | 149 | 1 | 1 | 0.998 | 0.997 | 1 | 1 | 1 | 1 | 1 | 1 | 1 | 1 | -0.001 |
| SE | 15525 | ENSG00000129003 | VPS13C | chr15 | - | 61908991 | 61909125 | 61906642 | 61907390 | 61910176 | 61910305 | 1141 | 977 | 867 | 885 | 1134 | 0 | 0 | 0 | 0 | 4 | 161 | 360 | 349 | 58 | 81 | 0 | 0 | 3 | 0 | 0 | 283 | 149 | 1 | 1 | 1 | 1 | 1 | 1 | 0.993 | 1 | 1 | 0.984 | 1 | 1 | 0.002 |
| SE | 15526 | ENSG00000129003 | VPS13C | chr15 | - | 61920497 | 61920647 | 61920066 | 61920331 | 61921946 | 61922033 | 884 | 747 | 641 | 740 | 622 | 1 | 2 | 0 | 1 | 0 | 89 | 283 | 272 | 38 | 52 | 0 | 0 | 0 | 0 | 0 | 298 | 149 | 1 | 1 | 0.998 | 0.995 | 1 | 0.997 | 1 | 1 | 1 | 1 | 1 | 1 | -0.002 |
| SE | 15527 | ENSG00000129003 | VPS13C | chr15 | - | 61925455 | 61925548 | 61922396 | 61922762 | 61927090 | 61927320 | 252 | 165 | 151 | 159 | 292 | 0 | 0 | 2 | 0 | 2 | 6 | 49 | 54 | 0 | 8 | 0 | 0 | 4 | 0 | 0 | 242 | 149 | 1 | 1 | 1 | 1 | 0.979 | 1 | 0.989 | 1 | 1 | 0.893 | NA | 1 | 0.02 |
| SE | 15528 | ENSG00000129003 | VPS13C | chr15 | - | 61929500 | 61929748 | 61927090 | 61927320 | 61931089 | 61931259 | 487 | 343 | 382 | 417 | 713 | 2 | 4 | 0 | 3 | 2 | 39 | 154 | 69 | 35 | 57 | 0 | 1 | 0 | 0 | 0 | 298 | 149 | 1 | 1 | 0.992 | 0.977 | 1 | 0.986 | 0.994 | 1 | 0.987 | 1 | 1 | 1 | -0.008 |
| SE | 15529 | ENSG00000129003 | VPS13C | chr15 | - | 61931089 | 61931259 | 61927090 | 61927320 | 61936596 | 61936750 | 3 | 5 | 0 | 4 | 2 | 0 | 0 | 2 | 0 | 0 | 0 | 1 | 0 | 0 | 0 | 0 | 0 | 0 | 0 | 0 | 298 | 149 | 1 | 1 | 1 | 1 | 0 | 1 | 1 | NA | 1 | NA | NA | NA | -0.2 |
| SE | 15530 | ENSG00000129003 | VPS13C | chr15 | - | 61934218 | 61934331 | 61931089 | 61931259 | 61936596 | 61936750 | 625 | 473 | 438 | 495 | 744 | 1 | 1 | 0 | 1 | 0 | 21 | 170 | 150 | 26 | 48 | 0 | 0 | 0 | 0 | 0 | 262 | 149 | 1 | 1 | 0.997 | 0.996 | 1 | 0.996 | 1 | 1 | 1 | 1 | 1 | 1 | -0.002 |
| SE | 15532 | ENSG00000129003 | VPS13C | chr15 | - | 61945714 | 61945882 | 61941762 | 61942067 | 61946306 | 61946410 | 774 | 701 | 615 | 679 | 1057 | 0 | 0 | 0 | 1 | 0 | 37 | 219 | 191 | 16 | 33 | 0 | 0 | 0 | 0 | 0 | 298 | 149 | 1 | 1 | 1 | 1 | 1 | 0.997 | 1 | 1 | 1 | 1 | 1 | 1 | -0.001 |
| SE | 15533 | ENSG00000129003 | VPS13C | chr15 | - | 61946306 | 61946410 | 61945714 | 61945882 | 61947192 | 61947309 | 698 | 600 | 505 | 587 | 521 | 4 | 2 | 0 | 4 | 0 | 50 | 206 | 171 | 18 | 44 | 0 | 1 | 0 | 0 | 0 | 253 | 149 | 1 | 1 | 0.99 | 0.994 | 1 | 0.989 | 1 | 1 | 0.992 | 1 | 1 | 1 | -0.004 |
| SE | 15534 | ENSG00000129003 | VPS13C | chr15 | - | 61950357 | 61950417 | 61949442 | 61949605 | 61950944 | 61951024 | 438 | 361 | 339 | 382 | 465 | 10 | 0 | 1 | 2 | 3 | 27 | 190 | 109 | 12 | 26 | 0 | 3 | 0 | 0 | 0 | 209 | 149 | 1 | 1 | 0.969 | 1 | 0.996 | 0.993 | 0.991 | 1 | 0.978 | 1 | 1 | 1 | -0.006 |
| SE | 15535 | ENSG00000129003 | VPS13C | chr15 | - | 61951823 | 61951980 | 61950944 | 61951024 | 61954420 | 61954554 | 769 | 627 | 542 | 677 | 959 | 1 | 1 | 0 | 3 | 0 | 72 | 299 | 199 | 16 | 80 | 0 | 0 | 1 | 0 | 0 | 298 | 149 | 1 | 1 | 0.997 | 0.997 | 1 | 0.991 | 1 | 1 | 1 | 0.99 | 1 | 1 | -0.001 |
| SE | 15536 | ENSG00000129003 | VPS13C | chr15 | - | 61951823 | 61951980 | 61950944 | 61951024 | 61958607 | 61958653 | 421 | 353 | 300 | 356 | 490 | 0 | 0 | 0 | 2 | 0 | 41 | 168 | 91 | 14 | 35 | 0 | 0 | 0 | 0 | 0 | 298 | 149 | 1 | 1 | 1 | 1 | 1 | 0.989 | 1 | 1 | 1 | 1 | 1 | 1 | -0.002 |
| SE | 15537 | ENSG00000129003 | VPS13C | chr15 | - | 61954420 | 61954554 | 61950944 | 61951024 | 61958607 | 61958653 | 268 | 290 | 204 | 289 | 338 | 0 | 0 | 0 | 2 | 0 | 38 | 113 | 110 | 5 | 25 | 0 | 0 | 0 | 0 | 0 | 283 | 149 | 1 | 1 | 1 | 1 | 1 | 0.987 | 1 | 1 | 1 | 1 | 1 | 1 | -0.003 |
| SE | 15538 | ENSG00000129003 | VPS13C | chr15 | - | 61954420 | 61954554 | 61951823 | 61951980 | 61958607 | 61958653 | 619 | 564 | 445 | 607 | 811 | 4 | 1 | 0 | 0 | 8 | 69 | 246 | 217 | 8 | 70 | 0 | 3 | 0 | 1 | 0 | 283 | 149 | 1 | 1 | 0.988 | 0.997 | 1 | 1 | 0.982 | 1 | 0.977 | 1 | 0.808 | 1 | 0.036 |
| SE | 15539 | ENSG00000129003 | VPS13C | chr15 | - | 61954466 | 61954554 | 61950944 | 61951024 | 61958607 | 61958716 | 225 | 236 | 175 | 233 | 271 | 0 | 0 | 0 | 2 | 0 | 43 | 108 | 99 | 3 | 16 | 0 | 0 | 0 | 0 | 0 | 237 | 149 | 1 | 1 | 1 | 1 | 1 | 0.987 | 1 | 1 | 1 | 1 | 1 | 1 | -0.003 |
| SE | 15541 | ENSG00000129003 | VPS13C | chr15 | - | 61962748 | 61962852 | 61962370 | 61962538 | 61963834 | 61963951 | 1402 | 1300 | 944 | 1203 | 915 | 3 | 7 | 0 | 5 | 0 | 188 | 673 | 470 | 74 | 121 | 0 | 0 | 0 | 0 | 0 | 253 | 149 | 1 | 1 | 0.996 | 0.991 | 1 | 0.993 | 1 | 1 | 1 | 1 | 1 | 1 | -0.004 |
| SE | 15542 | ENSG00000129003 | VPS13C | chr15 | - | 61963834 | 61963951 | 61962748 | 61962852 | 61964698 | 61964861 | 1122 | 936 | 730 | 939 | 905 | 0 | 0 | 0 | 0 | 0 | 100 | 510 | 315 | 41 | 86 | 0 | 0 | 1 | 0 | 0 | 266 | 149 | 1 | 1 | 1 | 1 | 1 | 1 | 1 | 1 | 1 | 0.994 | 1 | 1 | 0.001 |
| SE | 15543 | ENSG00000129003 | VPS13C | chr15 | - | 61977081 | 61977199 | 61974287 | 61974417 | 61978625 | 61978749 | 634 | 508 | 463 | 470 | 868 | 5 | 5 | 0 | 0 | 4 | 27 | 237 | 125 | 37 | 34 | 0 | 0 | 0 | 0 | 0 | 267 | 149 | 1 | 1 | 0.986 | 0.983 | 1 | 1 | 0.992 | 1 | 1 | 1 | 1 | 1 | -0.008 |
| SE | 15544 | ENSG00000129003 | VPS13C | chr15 | - | 61982458 | 61982573 | 61981341 | 61981478 | 61983819 | 61984012 | 652 | 611 | 478 | 624 | 880 | 1 | 0 | 0 | 0 | 0 | 46 | 272 | 176 | 40 | 32 | 0 | 0 | 0 | 0 | 0 | 264 | 149 | 1 | 1 | 0.997 | 1 | 1 | 1 | 1 | 1 | 1 | 1 | 1 | 1 | -0.001 |
| SE | 15547 | ENSG00000129003 | VPS13C | chr15 | - | 62000563 | 62000626 | 61991672 | 61991802 | 62007307 | 62007479 | 903 | 958 | 654 | 883 | 836 | 3 | 0 | 1 | 2 | 0 | 72 | 468 | 318 | 69 | 38 | 0 | 0 | 0 | 0 | 0 | 212 | 149 | 1 | 1 | 0.995 | 1 | 0.998 | 0.997 | 1 | 1 | 1 | 1 | 1 | 1 | -0.002 |
| SE | 15548 | ENSG00000129003 | VPS13C | chr15 | - | 62007307 | 62007479 | 62000563 | 62000626 | 62008654 | 62008761 | 1843 | 1893 | 1308 | 1629 | 1651 | 1 | 3 | 0 | 4 | 0 | 417 | 901 | 572 | 207 | 400 | 0 | 0 | 0 | 0 | 0 | 298 | 149 | 1 | 1 | 0.999 | 0.997 | 1 | 0.995 | 1 | 1 | 1 | 1 | 1 | 1 | -0.002 |
| SE | 15549 | ENSG00000129003 | VPS13C | chr15 | - | 62020478 | 62020538 | 62013932 | 62013992 | 62023410 | 62023520 | 1024 | 977 | 721 | 852 | 1185 | 0 | 0 | 0 | 1 | 0 | 133 | 449 | 287 | 77 | 79 | 0 | 0 | 0 | 0 | 0 | 209 | 149 | 1 | 1 | 1 | 1 | 1 | 0.998 | 1 | 1 | 1 | 1 | 1 | 1 | 0 |
| SE | 15551 | ENSG00000129003 | VPS13C | chr15 | - | 62023779 | 62023845 | 62023410 | 62023520 | 62033440 | 62033542 | 0 | 0 | 0 | 3 | 2 | 348 | 273 | 215 | 288 | 378 | 0 | 0 | 2 | 0 | 0 | 52 | 116 | 112 | 22 | 64 | 215 | 149 | 1 | 1 | 0 | 0 | 0 | 0.007 | 0.004 | 0 | 0 | 0.012 | 0 | 0 | 0 |
| SE | 15552 | ENSG00000129003 | VPS13C | chr15 | - | 62028357 | 62028420 | 62023410 | 62023520 | 62033440 | 62033542 | 0 | 1 | 2 | 0 | 6 | 348 | 273 | 215 | 288 | 378 | 0 | 0 | 0 | 0 | 0 | 52 | 116 | 112 | 22 | 64 | 212 | 149 | 1 | 1 | 0 | 0.003 | 0.006 | 0 | 0.011 | 0 | 0 | 0 | 0 | 0 | 0.004 |
| SE | 15553 | ENSG00000129003 | VPS13C | chr15 | - | 62028357 | 62028420 | 62023779 | 62023845 | 62033440 | 62033542 | 0 | 1 | 0 | 0 | 2 | 0 | 0 | 0 | 3 | 0 | 0 | 0 | 0 | 0 | 0 | 0 | 0 | 2 | 0 | 0 | 212 | 149 | 1 | 1 | NA | 1 | NA | 0 | 1 | NA | NA | 0 | NA | NA | 0.667 |
| SE | 15554 | ENSG00000129003 | VPS13C | chr15 | - | 62041323 | 62041366 | 62034956 | 62035052 | 62044211 | 62044255 | 352 | 297 | 203 | 324 | 323 | 0 | 1 | 0 | 0 | 0 | 107 | 173 | 94 | 27 | 68 | 0 | 0 | 0 | 0 | 0 | 192 | 149 | 1 | 1 | 1 | 0.996 | 1 | 1 | 1 | 1 | 1 | 1 | 1 | 1 | -0.001 |
| SE | 15555 | ENSG00000184056 | VPS33B | chr15 | - | 91003084 | 91003131 | 91002049 | 91002182 | 91004876 | 91004931 | 45 | 29 | 50 | 48 | 47 | 2 | 0 | 0 | 0 | 0 | 25 | 71 | 56 | 61 | 74 | 0 | 0 | 0 | 0 | 0 | 196 | 149 | 1 | 1 | 0.945 | 1 | 1 | 1 | 1 | 1 | 1 | 1 | 1 | 1 | -0.011 |
| SE | 15556 | ENSG00000184056 | VPS33B | chr15 | - | 91006371 | 91006445 | 91005972 | 91006059 | 91006651 | 91006729 | 69 | 48 | 56 | 40 | 63 | 1 | 2 | 0 | 0 | 2 | 54 | 49 | 46 | 34 | 57 | 0 | 0 | 0 | 0 | 0 | 223 | 149 | 1 | 1 | 0.979 | 0.941 | 1 | 1 | 0.955 | 1 | 1 | 1 | 1 | 1 | -0.025 |
| SE | 15557 | ENSG00000184056 | VPS33B | chr15 | - | 91006371 | 91006445 | 91005972 | 91006059 | 91006949 | 91007046 | 27 | 17 | 26 | 23 | 32 | 0 | 1 | 2 | 0 | 0 | 50 | 40 | 43 | 28 | 28 | 0 | 0 | 2 | 1 | 0 | 223 | 149 | 1 | 1 | 1 | 0.919 | 0.897 | 1 | 1 | 1 | 1 | 0.935 | 0.949 | 1 | -0.014 |
| SE | 15560 | ENSG00000184056 | VPS33B | chr15 | - | 91006651 | 91006729 | 91005972 | 91006059 | 91006949 | 91007046 | 13 | 12 | 5 | 8 | 8 | 0 | 1 | 2 | 0 | 0 | 5 | 0 | 3 | 0 | 0 | 0 | 0 | 2 | 1 | 0 | 227 | 149 | 1 | 1 | 1 | 0.887 | 0.621 | 1 | 1 | 1 | NA | 0.496 | 0 | NA | 0.403 |
| SE | 15563 | ENSG00000184056 | VPS33B | chr15 | - | 91006949 | 91007046 | 91006371 | 91006445 | 91007468 | 91007573 | 37 | 35 | 36 | 29 | 36 | 0 | 0 | 0 | 0 | 0 | 76 | 47 | 57 | 28 | 65 | 0 | 0 | 2 | 0 | 0 | 246 | 149 | 1 | 1 | 1 | 1 | 1 | 1 | 1 | 1 | 1 | 0.945 | 1 | 1 | 0.011 |
| SE | 15564 | ENSG00000184056 | VPS33B | chr15 | - | 91013803 | 91013871 | 91009800 | 91009846 | 91014383 | 91014433 | 28 | 11 | 35 | 31 | 22 | 0 | 0 | 0 | 0 | 0 | 49 | 33 | 39 | 19 | 34 | 0 | 0 | 1 | 0 | 0 | 217 | 149 | 1 | 1 | 1 | 1 | 1 | 1 | 1 | 1 | 1 | 0.964 | 1 | 1 | 0.007 |
| SE | 15566 | ENSG00000184056 | VPS33B | chr15 | - | 91016962 | 91017024 | 91014383 | 91014433 | 91022153 | 91022596 | 6 | 2 | 10 | 8 | 12 | 0 | 0 | 0 | 0 | 0 | 4 | 3 | 6 | 11 | 8 | 0 | 0 | 1 | 0 | 0 | 211 | 149 | 1 | 1 | 1 | 1 | 1 | 1 | 1 | 1 | 1 | 0.809 | 1 | 1 | 0.038 |
| SE | 15568 | ENSG00000184056 | VPS33B | chr15 | - | 91017804 | 91017885 | 91014383 | 91014433 | 91022153 | 91022513 | 8 | 6 | 12 | 7 | 2 | 0 | 0 | 0 | 0 | 0 | 16 | 17 | 11 | 2 | 4 | 0 | 0 | 1 | 0 | 0 | 230 | 149 | 1 | 1 | 1 | 1 | 1 | 1 | 1 | 1 | 1 | 0.877 | 1 | 1 | 0.025 |
| SE | 15569 | ENSG00000184056 | VPS33B | chr15 | - | 91017804 | 91017885 | 91016962 | 91017024 | 91022153 | 91022513 | 21 | 14 | 39 | 19 | 5 | 0 | 0 | 0 | 0 | 0 | 51 | 50 | 40 | 8 | 62 | 4 | 0 | 0 | 0 | 0 | 230 | 149 | 1 | 1 | 1 | 1 | 1 | 1 | 1 | 0.892 | 1 | 1 | 1 | 1 | 0.022 |
| SE | 15573 | ENSG00000170776 | AKAP13 | chr15 | + | 85533583 | 85533880 | 85521427 | 85521575 | 85543771 | 85543955 | 150 | 106 | 158 | 171 | 200 | 0 | 0 | 2 | 0 | 0 | 93 | 212 | 137 | 127 | 133 | 0 | 0 | 0 | 0 | 0 | 298 | 149 | 1 | 1 | 1 | 1 | 0.975 | 1 | 1 | 1 | 1 | 1 | 1 | 1 | -0.005 |
| SE | 15578 | ENSG00000170776 | AKAP13 | chr15 | + | 85669721 | 85669830 | 85664562 | 85664755 | 85682157 | 85682212 | 833 | 652 | 524 | 685 | 1062 | 1 | 0 | 0 | 0 | 0 | 775 | 1072 | 1151 | 1204 | 250 | 0 | 0 | 0 | 0 | 0 | 258 | 149 | 1 | 1 | 0.998 | 1 | 1 | 1 | 1 | 1 | 1 | 1 | 1 | 1 | 0 |
| SE | 15579 | ENSG00000170776 | AKAP13 | chr15 | + | 85684740 | 85684873 | 85682157 | 85682212 | 85693276 | 85693451 | 534 | 456 | 359 | 439 | 674 | 0 | 0 | 0 | 0 | 0 | 744 | 829 | 846 | 876 | 187 | 0 | 0 | 0 | 3 | 0 | 282 | 149 | 1 | 1 | 1 | 1 | 1 | 1 | 1 | 1 | 1 | 1 | 0.994 | 1 | 0.001 |
| SE | 15580 | ENSG00000170776 | AKAP13 | chr15 | + | 85693276 | 85693451 | 85684740 | 85684873 | 85708018 | 85708086 | 910 | 566 | 699 | 736 | 1053 | 0 | 0 | 0 | 0 | 0 | 766 | 1025 | 861 | 740 | 159 | 0 | 1 | 0 | 0 | 0 | 298 | 149 | 1 | 1 | 1 | 1 | 1 | 1 | 1 | 1 | 0.998 | 1 | 1 | 1 | 0 |
| SE | 15581 | ENSG00000170776 | AKAP13 | chr15 | + | 85708018 | 85708086 | 85693098 | 85693451 | 85710578 | 85710645 | 1148 | 598 | 850 | 912 | 1359 | 0 | 0 | 0 | 1 | 0 | 492 | 1129 | 794 | 675 | 127 | 0 | 0 | 0 | 3 | 0 | 217 | 149 | 1 | 1 | 1 | 1 | 1 | 0.998 | 1 | 1 | 1 | 1 | 0.994 | 1 | 0.001 |
| SE | 15582 | ENSG00000170776 | AKAP13 | chr15 | + | 85710578 | 85710645 | 85708018 | 85708086 | 85715787 | 85715901 | 921 | 493 | 655 | 753 | 970 | 2 | 0 | 0 | 0 | 0 | 385 | 891 | 600 | 534 | 115 | 0 | 0 | 0 | 0 | 0 | 216 | 149 | 1 | 1 | 0.997 | 1 | 1 | 1 | 1 | 1 | 1 | 1 | 1 | 1 | -0.001 |
| SE | 15583 | ENSG00000170776 | AKAP13 | chr15 | + | 85710581 | 85710645 | 85708018 | 85708086 | 85715787 | 85715923 | 274 | 136 | 211 | 207 | 234 | 2 | 0 | 0 | 0 | 0 | 112 | 250 | 174 | 205 | 49 | 0 | 0 | 0 | 0 | 0 | 213 | 149 | 1 | 1 | 0.99 | 1 | 1 | 1 | 1 | 1 | 1 | 1 | 1 | 1 | -0.002 |
| SE | 15585 | ENSG00000170776 | AKAP13 | chr15 | + | 85718006 | 85718159 | 85717289 | 85717402 | 85719075 | 85719326 | 1312 | 940 | 916 | 1093 | 1421 | 0 | 0 | 0 | 0 | 2 | 997 | 1313 | 1279 | 1179 | 209 | 0 | 0 | 0 | 0 | 0 | 298 | 149 | 1 | 1 | 1 | 1 | 1 | 1 | 0.997 | 1 | 1 | 1 | 1 | 1 | -0.001 |
| SE | 15586 | ENSG00000170776 | AKAP13 | chr15 | + | 85718012 | 85718159 | 85717151 | 85717402 | 85719075 | 85719326 | 1394 | 963 | 1014 | 1145 | 1522 | 0 | 0 | 0 | 0 | 2 | 1024 | 1400 | 1470 | 1428 | 253 | 0 | 0 | 0 | 0 | 0 | 296 | 149 | 1 | 1 | 1 | 1 | 1 | 1 | 0.997 | 1 | 1 | 1 | 1 | 1 | -0.001 |
| SE | 15587 | ENSG00000170776 | AKAP13 | chr15 | + | 85727380 | 85727463 | 85727065 | 85727247 | 85730512 | 85730707 | 870 | 597 | 553 | 668 | 1095 | 0 | 0 | 0 | 2 | 0 | 857 | 933 | 972 | 887 | 168 | 0 | 0 | 0 | 0 | 0 | 232 | 149 | 1 | 1 | 1 | 1 | 1 | 0.995 | 1 | 1 | 1 | 1 | 1 | 1 | -0.001 |
| SE | 15589 | ENSG00000170776 | AKAP13 | chr15 | + | 85735559 | 85735630 | 85734991 | 85735150 | 85736089 | 85736134 | 984 | 664 | 572 | 812 | 928 | 0 | 3 | 3 | 5 | 0 | 1402 | 1127 | 1263 | 1114 | 330 | 0 | 4 | 11 | 1 | 0 | 220 | 149 | 1 | 1 | 1 | 0.993 | 0.992 | 0.991 | 1 | 1 | 0.995 | 0.987 | 0.999 | 1 | -0.001 |
| SE | 15590 | ENSG00000170776 | AKAP13 | chr15 | + | 85736089 | 85736134 | 85735559 | 85735630 | 85740221 | 85740272 | 433 | 284 | 281 | 374 | 353 | 0 | 0 | 0 | 0 | 1 | 368 | 445 | 448 | 325 | 93 | 0 | 0 | 0 | 0 | 0 | 194 | 149 | 1 | 1 | 1 | 1 | 1 | 1 | 0.996 | 1 | 1 | 1 | 1 | 1 | -0.001 |
| SE | 15591 | ENSG00000170776 | AKAP13 | chr15 | + | 85740221 | 85740272 | 85736089 | 85736134 | 85741045 | 85741495 | 404 | 235 | 294 | 339 | 337 | 4 | 2 | 5 | 1 | 3 | 266 | 411 | 348 | 227 | 84 | 0 | 3 | 1 | 1 | 0 | 200 | 149 | 1 | 1 | 0.987 | 0.989 | 0.978 | 0.996 | 0.988 | 1 | 0.99 | 0.996 | 0.994 | 1 | -0.008 |
| SE | 15597 | ENSG00000170776 | AKAP13 | chr15 | + | 85743491 | 85743825 | 85741045 | 85741495 | 85744627 | 85744677 | 474 | 219 | 311 | 351 | 324 | 1 | 0 | 0 | 0 | 0 | 296 | 424 | 337 | 254 | 170 | 0 | 0 | 1 | 4 | 0 | 298 | 149 | 1 | 1 | 0.996 | 1 | 1 | 1 | 1 | 1 | 1 | 0.994 | 0.969 | 1 | 0.007 |
| SE | 15599 | ENSG00000169032 | MAP2K1 | chr15 | + | 66481754 | 66481879 | 66444655 | 66444707 | 66484889 | 66485191 | 187 | 99 | 154 | 197 | 150 | 0 | 0 | 0 | 0 | 0 | 231 | 216 | 191 | 240 | 358 | 5 | 0 | 0 | 0 | 0 | 274 | 149 | 1 | 1 | 1 | 1 | 1 | 1 | 1 | 0.962 | 1 | 1 | 1 | 1 | 0.008 |

| | | | | | | | | | | | | | | | | | | | | | | | | | | | | | | | | | | | | | | | | | | | | | | |
|---|---|---|---|---|---|---|---|---|---|---|---|---|---|---|---|---|---|---|---|---|---|---|---|---|---|---|---|---|---|---|---|---|---|---|---|---|---|---|---|---|---|---|---|---|---|---|
| SE | 15966 | ENSG00000137845 | ADAM10 | chr15 | - | 58679123 | 58679282 | 58665096 | 58665197 | 58682195 | 58682314 | 1218 | 1274 | 848 | 1126 | 1986 | 1 | 0 | 1 | 1 | 0 | 384 | 741 | 684 | 227 | 160 | 0 | 0 | 0 | 0 | 0 | 298 | 149 | 1 | 1 | 0.998 | 1 | 0.998 | 0.998 | 1 | 1 | 1 | 1 | 1 | 1 | -0.001 |
| SE | 15967 | ENSG00000137845 | ADAM10 | chr15 | - | 58682195 | 58682314 | 58665145 | 58665197 | 58717576 | 58717727 | 603 | 562 | 413 | 600 | 602 | 1 | 1 | 0 | 0 | 0 | 123 | 301 | 311 | 119 | 78 | 0 | 0 | 6 | 0 | 0 | 268 | 149 | 1 | 1 | 0.997 | 0.997 | 1 | 1 | 1 | 1 | 1 | 0.966 | 1 | 1 | 0.006 |
| SE | 15968 | ENSG00000137845 | ADAM10 | chr15 | - | 58682195 | 58682314 | 58679123 | 58679282 | 58717576 | 58717727 | 1231 | 1240 | 839 | 1141 | 1609 | 0 | 0 | 10 | 0 | 2 | 330 | 660 | 621 | 216 | 158 | 0 | 0 | 0 | 0 | 0 | 268 | 149 | 1 | 1 | 1 | 1 | 0.979 | 1 | 0.998 | 1 | 1 | 1 | 1 | 1 | -0.005 |
| SE | 15969 | ENSG00000137845 | ADAM10 | chr15 | - | 58698279 | 58698404 | 58665181 | 58665197 | 58717576 | 58717727 | 13 | 3 | 8 | 7 | 7 | 1 | 1 | 0 | 0 | 0 | 0 | 19 | 13 | 0 | 4 | 0 | 0 | 6 | 0 | 0 | 274 | 149 | 1 | 1 | 0.876 | 0.62 | 1 | 1 | 1 | NA | 1 | 0.541 | NA | 1 | 0.052 |
| SE | 15972 | ENSG00000137845 | ADAM10 | chr15 | - | 58717576 | 58717727 | 58682195 | 58682314 | 58749479 | 58749707 | 969 | 888 | 611 | 869 | 927 | 0 | 1 | 0 | 0 | 0 | 182 | 442 | 436 | 181 | 102 | 0 | 0 | 0 | 0 | 0 | 298 | 149 | 1 | 1 | 1 | 0.998 | 1 | 1 | 1 | 1 | 1 | 1 | 1 | 1 | 0 |
| SE | 15977 | ENSG00000185787 | MORF4L1 | chr15 | + | 78887268 | 78887349 | 78886140 | 78886227 | 78890988 | 78891014 | 1197 | 920 | 1041 | 1266 | 1416 | 3 | 2 | 2 | 5 | 1 | 2347 | 1419 | 1675 | 1659 | 1076 | 0 | 5 | 2 | 4 | 4 | 230 | 149 | 1 | 1 | 0.996 | 0.997 | 0.997 | 0.994 | 0.999 | 1 | 0.995 | 0.998 | 0.996 | 0.994 | 0 |
| SE | 15978 | ENSG00000185787 | MORF4L1 | chr15 | + | 78892211 | 78892313 | 78890988 | 78891572 | 78893538 | 78893590 | 2833 | 1977 | 2578 | 3042 | 5392 | 1 | 0 | 0 | 0 | 2 | 3196 | 2779 | 2556 | 3041 | 1505 | 0 | 3 | 0 | 0 | 0 | 251 | 149 | 1 | 1 | 0.999 | 1 | 1 | 1 | 0.999 | 1 | 0.998 | 1 | 1 | 1 | 0 |
| SE | 15979 | ENSG00000185787 | MORF4L1 | chr15 | + | 78893538 | 78893627 | 78892211 | 78892313 | 78894057 | 78894134 | 4418 | 3230 | 3943 | 4748 | 9265 | 0 | 1 | 2 | 0 | 0 | 5132 | 4191 | 3984 | 4719 | 2474 | 0 | 4 | 0 | 0 | 0 | 238 | 149 | 1 | 1 | 1 | 1 | 0.999 | 1 | 1 | 1 | 0.998 | 1 | 1 | 1 | 0 |
| SE | 15980 | ENSG00000185787 | MORF4L1 | chr15 | + | 78893538 | 78893627 | 78892211 | 78892313 | 78894819 | 78894904 | 1725 | 1340 | 1570 | 1879 | 4126 | 0 | 0 | 0 | 0 | 0 | 1957 | 1728 | 1500 | 1924 | 1011 | 0 | 0 | 0 | 5 | 0 | 238 | 149 | 1 | 1 | 1 | 1 | 1 | 1 | 1 | 1 | 1 | 1 | 0.996 | 1 | 0.001 |
| SE | 15981 | ENSG00000185787 | MORF4L1 | chr15 | + | 78894057 | 78894230 | 78892211 | 78892313 | 78894819 | 78894904 | 366 | 219 | 312 | 423 | 400 | 0 | 0 | 0 | 0 | 0 | 417 | 282 | 453 | 377 | 167 | 0 | 0 | 0 | 5 | 0 | 298 | 149 | 1 | 1 | 1 | 1 | 1 | 1 | 1 | 1 | 1 | 1 | 0.974 | 1 | 0.005 |
| SE | 15984 | ENSG00000101230 | ISM1 | chr20 | + | 13279633 | 13279898 | 13270503 | 13270743 | 13288539 | 13288683 | 5 | 2 | 8 | 6 | 6 | 0 | 0 | 4 | 0 | 0 | 0 | 0 | 3 | 0 | 0 | 0 | 0 | 0 | 0 | 0 | 298 | 149 | 1 | 1 | 1 | 1 | 0.5 | 1 | 1 | NA | NA | 1 | NA | NA | -0.1 |
| SE | 15986 | ENSG00000235568 | NFAM1 | chr22 | - | 42386988 | 42387078 | 42380406 | 42385220 | 42397857 | 42397956 | 129 | 73 | 56 | 93 | 112 | 0 | 0 | 1 | 0 | 0 | 223 | 92 | 68 | 144 | 260 | 0 | 0 | 0 | 2 | 2 | 239 | 149 | 1 | 1 | 1 | 1 | 0.972 | 1 | 1 | 1 | 1 | 1 | 0.978 | 0.988 | 0.001 |
| SE | 15987 | ENSG00000235568 | NFAM1 | chr22 | - | 42397857 | 42397956 | 42386988 | 42387078 | 42409434 | 42409547 | 253 | 138 | 113 | 179 | 130 | 0 | 0 | 0 | 1 | 0 | 623 | 223 | 146 | 296 | 584 | 0 | 0 | 0 | 0 | 0 | 248 | 149 | 1 | 1 | 1 | 1 | 1 | 0.991 | 1 | 1 | 1 | 1 | 1 | 1 | -0.002 |
| SE | 16000 | ENSG00000068305 | MEF2A | chr15 | + | 99645560 | 99645764 | 99632977 | 99633173 | 99671322 | 99671454 | 228 | 269 | 210 | 238 | 455 | 0 | 0 | 0 | 2 | 0 | 134 | 275 | 237 | 214 | 92 | 0 | 1 | 0 | 0 | 0 | 298 | 149 | 1 | 1 | 1 | 1 | 1 | 0.983 | 1 | 1 | 0.993 | 1 | 1 | 1 | -0.002 |
| SE | 16001 | ENSG00000068305 | MEF2A | chr15 | + | 99645560 | 99645764 | 99632977 | 99633173 | 99671519 | 99671657 | 204 | 232 | 162 | 232 | 455 | 1 | 0 | 2 | 0 | 0 | 119 | 205 | 159 | 170 | 53 | 4 | 0 | 0 | 0 | 0 | 298 | 149 | 1 | 1 | 0.99 | 1 | 0.976 | 1 | 1 | 0.937 | 1 | 1 | 1 | 1 | 0.006 |
| SE | 16002 | ENSG00000068305 | MEF2A | chr15 | + | 99671322 | 99671454 | 99632977 | 99633173 | 99671519 | 99671657 | 0 | 0 | 0 | 2 | 2 | 1 | 0 | 2 | 0 | 0 | 0 | 1 | 0 | 0 | 0 | 4 | 0 | 0 | 0 | 0 | 281 | 149 | 1 | 1 | 0 | NA | 0 | 1 | 1 | 0 | 1 | NA | NA | NA | 0 |
| SE | 16003 | ENSG00000068305 | MEF2A | chr15 | + | 99671322 | 99671454 | 99632977 | 99633173 | 99674392 | 99674612 | 73 | 69 | 84 | 73 | 233 | 1 | 0 | 0 | 0 | 0 | 72 | 117 | 127 | 104 | 47 | 0 | 0 | 0 | 0 | 0 | 281 | 149 | 1 | 1 | 0.975 | 1 | 1 | 1 | 1 | 1 | 1 | 1 | 1 | 1 | -0.005 |
| SE | 16005 | ENSG00000068305 | MEF2A | chr15 | + | 99671519 | 99671657 | 99632977 | 99633173 | 99674392 | 99674612 | 41 | 61 | 44 | 63 | 107 | 1 | 0 | 0 | 0 | 0 | 15 | 35 | 23 | 26 | 17 | 0 | 0 | 0 | 0 | 0 | 287 | 149 | 1 | 1 | 0.955 | 1 | 1 | 1 | 1 | 1 | 1 | 1 | 1 | 1 | -0.009 |
| SE | 16008 | ENSG00000068305 | MEF2A | chr15 | + | 99690240 | 99690428 | 99675398 | 99675458 | 99710633 | 99710760 | 112 | 118 | 116 | 112 | 149 | 3 | 0 | 0 | 0 | 0 | 74 | 149 | 146 | 219 | 43 | 0 | 0 | 1 | 0 | 0 | 298 | 149 | 1 | 1 | 0.949 | 1 | 1 | 1 | 1 | 1 | 1 | 0.986 | 1 | 1 | -0.007 |
| SE | 16010 | ENSG00000068305 | MEF2A | chr15 | + | 99706728 | 99706855 | 99675398 | 99675458 | 99710633 | 99710760 | 141 | 119 | 92 | 117 | 190 | 3 | 0 | 0 | 0 | 0 | 79 | 152 | 130 | 135 | 24 | 0 | 0 | 1 | 0 | 0 | 276 | 149 | 1 | 1 | 0.962 | 1 | 1 | 1 | 1 | 1 | 1 | 0.986 | 1 | 1 | -0.005 |
| SE | 16011 | ENSG00000068305 | MEF2A | chr15 | + | 99706728 | 99706855 | 99675398 | 99675458 | 99712389 | 99713324 | 5 | 6 | 2 | 0 | 5 | 0 | 0 | 0 | 0 | 6 | 14 | 8 | 4 | 5 | 3 | 0 | 0 | 0 | 0 | 0 | 276 | 149 | 1 | 1 | 1 | 1 | 1 | NA | 0.31 | 1 | 1 | 1 | 1 | 1 | -0.172 |
| SE | 16012 | ENSG00000068305 | MEF2A | chr15 | + | 99706728 | 99706855 | 99690240 | 99690428 | 99710633 | 99710760 | 227 | 206 | 174 | 250 | 339 | 0 | 0 | 2 | 4 | 0 | 135 | 301 | 233 | 255 | 79 | 0 | 0 | 1 | 0 | 0 | 276 | 149 | 1 | 1 | 1 | 1 | 0.979 | 0.971 | 1 | 1 | 1 | 0.992 | 1 | 1 | -0.008 |
| SE | 16013 | ENSG00000068305 | MEF2A | chr15 | + | 99710633 | 99710760 | 99675398 | 99675458 | 99712389 | 99712835 | 161 | 131 | 112 | 169 | 245 | 0 | 0 | 0 | 0 | 6 | 140 | 212 | 171 | 168 | 44 | 0 | 0 | 0 | 0 | 0 | 276 | 149 | 1 | 1 | 1 | 1 | 1 | 1 | 0.957 | 1 | 1 | 1 | 1 | 1 | -0.009 |
| SE | 16014 | ENSG00000068305 | MEF2A | chr15 | + | 99710633 | 99710760 | 99706594 | 99706855 | 99712389 | 99712835 | 298 | 254 | 218 | 298 | 441 | 0 | 3 | 1 | 0 | 0 | 211 | 383 | 308 | 311 | 76 | 0 | 5 | 2 | 0 | 0 | 276 | 149 | 1 | 1 | 1 | 0.979 | 0.992 | 1 | 1 | 1 | 0.976 | 0.988 | 1 | 1 | 0.001 |
| SE | 16033 | ENSG00000258476 | LINC02207 | chr15 | + | 93865342 | 93865648 | 93863033 | 93863216 | 93867399 | 93867722 | 3 | 0 | 0 | 0 | 0 | 0 | 0 | 0 | 1 | 0 | 0 | 2 | 8 | 2 | 0 | 0 | 0 | 0 | 0 | 0 | 298 | 149 | 1 | 1 | 1 | NA | NA | 0 | NA | NA | 1 | 1 | 1 | NA | -0.5 |
| SE | 16036 | ENSG00000258476 | LINC02207 | chr15 | + | 93865373 | 93865648 | 93863007 | 93863216 | 93867399 | 93867722 | 14 | 0 | 2 | 5 | 7 | 0 | 0 | 0 | 1 | 0 | 11 | 26 | 16 | 14 | 0 | 0 | 0 | 0 | 0 | 0 | 298 | 149 | 1 | 1 | 1 | NA | 1 | 0.714 | 1 | 1 | 1 | 1 | 1 | NA | -0.072 |
| SE | 16038 | ENSG00000258476 | LINC02207 | chr15 | + | 93869690 | 93869833 | 93865342 | 93865648 | 93878033 | 93878352 | 0 | 0 | 2 | 0 | 0 | 3 | 0 | 0 | 0 | 4 | 0 | 0 | 6 | 0 | 0 | 0 | 6 | 4 | 0 | 0 | 292 | 149 | 1 | 1 | 0 | NA | 1 | NA | 0 | NA | 0 | 0.434 | NA | NA | 0.116 |
| SE | 16042 | ENSG00000048471 | SNX29 | chr16 | + | 11999296 | 11999358 | 11976733 | 11976813 | 12002990 | 12003043 | 23 | 25 | 28 | 15 | 26 | 0 | 0 | 0 | 0 | 0 | 61 | 31 | 46 | 65 | 59 | 0 | 0 | 0 | 2 | 0 | 211 | 149 | 1 | 1 | 1 | 1 | 1 | 1 | 1 | 1 | 1 | 1 | 0.958 | 1 | 0.008 |
| SE | 16043 | ENSG00000048471 | SNX29 | chr16 | + | 12002990 | 12003043 | 11999296 | 11999358 | 12027319 | 12027444 | 24 | 25 | 40 | 26 | 47 | 0 | 0 | 0 | 0 | 1 | 104 | 40 | 64 | 97 | 64 | 0 | 0 | 0 | 0 | 5 | 202 | 149 | 1 | 1 | 1 | 1 | 1 | 1 | 0.972 | 1 | 1 | 1 | 1 | 0.904 | 0.014 |
| SE | 16044 | ENSG00000048471 | SNX29 | chr16 | + | 12027319 | 12027444 | 12002990 | 12003043 | 12042896 | 12043077 | 49 | 40 | 56 | 54 | 93 | 0 | 0 | 2 | 0 | 0 | 191 | 73 | 105 | 210 | 140 | 0 | 4 | 0 | 0 | 0 | 274 | 149 | 1 | 1 | 1 | 1 | 0.938 | 1 | 1 | 1 | 0.908 | 1 | 1 | 1 | 0.006 |
| SE | 16045 | ENSG00000048471 | SNX29 | chr16 | + | 12029612 | 12029734 | 12027319 | 12027444 | 12042896 | 12042997 | 0 | 0 | 0 | 0 | 0 | 34 | 21 | 27 | 33 | 47 | 0 | 0 | 1 | 0 | 0 | 97 | 45 | 62 | 128 | 86 | 271 | 149 | 1 | 1 | 0 | 0 | 0 | 0 | 0 | 0 | 0 | 0.009 | 0 | 0 | -0.002 |
| SE | 16046 | ENSG00000048471 | SNX29 | chr16 | + | 12042896 | 12043077 | 12027319 | 12027444 | 12046383 | 12046454 | 41 | 30 | 52 | 49 | 58 | 0 | 0 | 3 | 0 | 0 | 134 | 70 | 85 | 165 | 146 | 0 | 0 | 0 | 0 | 0 | 298 | 149 | 1 | 1 | 1 | 1 | 0.897 | 1 | 1 | 1 | 1 | 1 | 1 | 1 | -0.021 |
| SE | 16047 | ENSG00000048471 | SNX29 | chr16 | + | 12046383 | 12046454 | 12027319 | 12027444 | 12048371 | 12048620 | 17 | 16 | 27 | 22 | 49 | 0 | 0 | 1 | 0 | 0 | 131 | 50 | 47 | 57 | 118 | 0 | 4 | 0 | 0 | 0 | 220 | 149 | 1 | 1 | 1 | 1 | 0.948 | 1 | 1 | 1 | 0.894 | 1 | 1 | 1 | 0.011 |
| SE | 16049 | ENSG00000048471 | SNX29 | chr16 | + | 12078832 | 12078915 | 12069056 | 12069132 | 12126632 | 12126696 | 85 | 51 | 79 | 87 | 37 | 0 | 0 | 0 | 0 | 0 | 202 | 224 | 139 | 210 | 165 | 0 | 4 | 0 | 0 | 0 | 232 | 149 | 1 | 1 | 1 | 1 | 1 | 1 | 1 | 1 | 0.973 | 1 | 1 | 1 | 0.005 |
| SE | 16051 | ENSG00000048471 | SNX29 | chr16 | + | 12356162 | 12356279 | 12277932 | 12278036 | 12398445 | 12398501 | 77 | 40 | 79 | 94 | 125 | 0 | 0 | 0 | 0 | 0 | 116 | 92 | 73 | 131 | 97 | 0 | 1 | 0 | 0 | 0 | 266 | 149 | 1 | 1 | 1 | 1 | 1 | 1 | 1 | 1 | 0.981 | 1 | 1 | 1 | 0.004 |
| SE | 16052 | ENSG00000048471 | SNX29 | chr16 | + | 12398445 | 12398501 | 12356162 | 12356279 | 12403447 | 12403529 | 54 | 29 | 58 | 72 | 97 | 0 | 0 | 0 | 0 | 0 | 125 | 71 | 64 | 100 | 96 | 0 | 0 | 0 | 2 | 0 | 205 | 149 | 1 | 1 | 1 | 1 | 1 | 1 | 1 | 1 | 1 | 1 | 0.973 | 1 | 0.005 |
| SE | 16053 | ENSG00000048471 | SNX29 | chr16 | + | 12477718 | 12477859 | 12403447 | 12403529 | 12524701 | 12524841 | 79 | 58 | 103 | 105 | 269 | 3 | 0 | 0 | 0 | 0 | 341 | 124 | 107 | 135 | 216 | 0 | 3 | 0 | 0 | 0 | 290 | 149 | 1 | 1 | 0.931 | 1 | 1 | 1 | 1 | 1 | 0.955 | 1 | 1 | 1 | -0.005 |
| SE | 16057 | ENSG00000169612 | RAMAC | chr15 | + | 82989009 | 82989188 | 82987335 | 82987384 | 82989880 | 82991057 | 601 | 354 | 455 | 410 | 449 | 0 | 0 | 1 | 0 | 1 | 597 | 614 | 464 | 539 | 342 | 0 | 0 | 0 | 0 | 0 | 298 | 149 | 1 | 1 | 1 | 1 | 0.996 | 1 | 0.996 | 1 | 1 | 1 | 1 | 1 | -0.002 |
| SE | 16064 | ENSG00000128908 | INO80 | chr15 | - | 40983761 | 40983921 | 40982861 | 40983077 | 40984196 | 40984352 | 229 | 102 | 132 | 214 | 230 | 0 | 2 | 0 | 0 | 0 | 138 | 225 | 166 | 138 | 176 | 0 | 0 | 0 | 0 | 0 | 298 | 149 | 1 | 1 | 1 | 0.962 | 1 | 1 | 1 | 1 | 1 | 1 | 1 | 1 | -0.008 |
| SE | 16065 | ENSG00000128908 | INO80 | chr15 | - | 41005592 | 41005687 | 40997528 | 40997601 | 41016087 | 41016215 | 87 | 60 | 102 | 100 | 76 | 0 | 0 | 0 | 1 | 0 | 108 | 112 | 101 | 107 | 95 | 0 | 5 | 0 | 0 | 0 | 244 | 149 | 1 | 1 | 1 | 1 | 1 | 0.984 | 1 | 1 | 0.932 | 1 | 1 | 1 | 0.01 |
| SE | 16067 | ENSG00000128908 | INO80 | chr15 | - | 41020899 | 41021125 | 41016087 | 41016215 | 41027595 | 41027736 | 122 | 103 | 134 | 115 | 86 | 0 | 0 | 0 | 0 | 0 | 155 | 169 | 139 | 133 | 96 | 0 | 6 | 0 | 0 | 0 | 298 | 149 | 1 | 1 | 1 | 1 | 1 | 1 | 1 | 1 | 0.934 | 1 | 1 | 1 | 0.013 |
| SE | 16069 | ENSG00000128908 | INO80 | chr15 | - | 41023225 | 41023346 | 41020899 | 41021125 | 41027595 | 41027599 | 1 | 0 | 0 | 0 | 0 | 98 | 93 | 109 | 97 | 74 | 0 | 0 | 0 | 0 | 0 | 151 | 140 | 124 | 94 | 76 | 270 | 149 | 1 | 1 | 0.006 | 0 | 0 | 0 | 0 | 0 | 0 | 0 | 0 | 0 | 0.001 |
| SE | 16072 | ENSG00000128908 | INO80 | chr15 | - | 41044903 | 41045075 | 41027595 | 41027736 | 41047407 | 41047501 | 135 | 86 | 115 | 118 | 104 | 2 | 0 | 0 | 0 | 0 | 185 | 176 | 137 | 155 | 95 | 0 | 0 | 0 | 0 | 0 | 298 | 149 | 1 | 1 | 0.971 | 1 | 1 | 1 | 1 | 1 | 1 | 1 | 1 | 1 | -0.006 |
| SE | 16073 | ENSG00000128908 | INO80 | chr15 | - | 41048211 | 41048276 | 41047407 | 41047501 | 41049286 | 41049420 | 96 | 37 | 74 | 77 | 85 | 0 | 0 | 0 | 0 | 0 | 60 | 70 | 49 | 72 | 87 | 0 | 4 | 0 | 0 | 0 | 214 | 149 | 1 | 1 | 1 | 1 | 1 | 1 | 1 | 1 | 0.924 | 1 | 1 | 1 | 0.015 |
| SE | 16074 | ENSG00000128908 | INO80 | chr15 | - | 41053928 | 41054014 | 41049934 | 41050102 | 41055246 | 41055364 | 316 | 260 | 233 | 249 | 649 | 2 | 0 | 0 | 1 | 1 | 474 | 390 | 317 | 338 | 186 | 0 | 0 | 0 | 0 | 0 | 235 | 149 | 1 | 1 | 0.99 | 1 | 1 | 0.994 | 0.998 | 1 | 1 | 1 | 1 | 1 | -0.004 |
| SE | 16075 | ENSG00000128908 | INO80 | chr15 | - | 41055246 | 41055364 | 41049934 | 41050102 | 41056621 | 41056706 | 65 | 38 | 72 | 47 | 72 | 0 | 1 | 0 | 0 | 0 | 69 | 98 | 58 | 69 | 35 | 2 | 0 | 2 | 0 | 0 | 267 | 149 | 1 | 1 | 1 | 0.955 | 1 | 1 | 1 | 0.951 | 1 | 0.942 | 1 | 1 | 0.012 |
| SE | 16076 | ENSG00000128908 | INO80 | chr15 | - | 41055246 | 41055364 | 41053928 | 41054014 | 41058638 | 41058781 | 131 | 105 | 112 | 112 | 219 | 0 | 0 | 0 | 1 | 0 | 171 | 182 | 151 | 158 | 71 | 0 | 0 | 0 | 0 | 0 | 267 | 149 | 1 | 1 | 1 | 1 | 1 | 0.984 | 1 | 1 | 1 | 1 | 1 | 1 | -0.003 |
| SE | 16077 | ENSG00000128908 | INO80 | chr15 | - | 41056621 | 41056706 | 41055246 | 41055364 | 41058638 | 41058781 | 191 | 141 | 183 | 161 | 245 | 3 | 0 | 0 | 0 | 2 | 301 | 252 | 171 | 178 | 98 | 0 | 0 | 1 | 0 | 0 | 234 | 149 | 1 | 1 | 0.976 | 1 | 1 | 1 | 0.987 | 1 | 1 | 0.991 | 1 | 1 | -0.006 |
| SE | 16079 | ENSG00000128908 | INO80 | chr15 | - | 41058638 | 41058781 | 41056621 | 41056706 | 41069569 | 41069665 | 112 | 88 | 107 | 101 | 146 | 4 | 0 | 0 | 0 | 0 | 191 | 142 | 111 | 107 | 45 | 0 | 0 | 0 | 0 | 0 | 292 | 149 | 1 | 1 | 0.935 | 1 | 1 | 1 | 1 | 1 | 1 | 1 | 1 | 1 | -0.013 |
| SE | 16080 | ENSG00000128908 | INO80 | chr15 | - | 41059866 | 41059926 | 41056621 | 41056706 | 41069569 | 41069659 | 43 | 45 | 39 | 41 | 81 | 4 | 0 | 0 | 0 | 0 | 127 | 56 | 63 | 77 | 52 | 0 | 0 | 0 | 0 | 0 | 209 | 149 | 1 | 1 | 0.885 | 1 | 1 | 1 | 1 | 1 | 1 | 1 | 1 | 1 | -0.023 |
| SE | 16083 | ENSG00000128908 | INO80 | chr15 | - | 41069569 | 41069665 | 41059866 | 41059926 | 41071848 | 41072058 | 78 | 54 | 75 | 81 | 130 | 0 | 0 | 0 | 0 | 2 | 68 | 122 | 79 | 81 | 18 | 0 | 0 | 0 | 0 | 0 | 245 | 149 | 1 | 1 | 1 | 1 | 1 | 1 | 0.975 | 1 | 1 | 1 | 1 | 1 | -0.005 |
| SE | 16084 | ENSG00000128908 | INO80 | chr15 | - | 41070466 | 41070547 | 41059866 | 41059926 | 41071848 | 41072015 | 52 | 40 | 50 | 63 | 74 | 0 | 0 | 0 | 0 | 2 | 124 | 81 | 70 | 73 | 11 | 0 | 0 | 0 | 0 | 0 | 230 | 149 | 1 | 1 | 1 | 1 | 1 | 1 | 0.96 | 1 | 1 | 1 | 1 | 1 | -0.008 |
| SE | 16086 | ENSG00000128908 | INO80 | chr15 | - | 41071848 | 41072058 | 41070466 | 41070547 | 41073427 | 41073495 | 221 | 146 | 204 | 226 | 352 | 0 | 1 | 0 | 0 | 0 | 391 | 289 | 261 | 291 | 117 | 0 | 0 | 2 | 0 | 0 | 298 | 149 | 1 | 1 | 1 | 0.986 | 1 | 1 | 1 | 1 | 1 | 0.985 | 1 | 1 | 0 |
| SE | 16087 | ENSG00000128908 | INO80 | chr15 | - | 41074369 | 41074565 | 41073427 | 41073495 | 41079700 | 41079904 | 235 | 177 | 195 | 266 | 299 | 1 | 0 | 0 | 0 | 0 | 391 | 308 | 287 | 320 | 164 | 0 | 1 | 0 | 0 | 0 | 298 | 149 | 1 | 1 | 0.992 | 1 | 1 | 1 | 1 | 1 | 0.994 | 1 | 1 | 1 | 0 |
| SE | 16088 | ENSG00000128908 | INO80 | chr15 | - | 41079700 | 41079904 | 41074369 | 41074565 | 41081019 | 41081073 | 269 | 175 | 230 | 269 | 336 | 0 | 0 | 1 | 0 | 0 | 343 | 332 | 283 | 372 | 236 | 0 | 0 | 0 | 3 | 0 | 298 | 149 | 1 | 1 | 1 | 1 | 0.991 | 1 | 1 | 1 | 1 | 1 | 0.984 | 1 | 0.001 |
| SE | 16089 | ENSG00000128908 | INO80 | chr15 | - | 41081019 | 41081073 | 41079700 | 41079904 | 41085368 | 41085583 | 146 | 77 | 123 | 124 | 213 | 1 | 0 | 0 | 2 | 1 | 149 | 159 | 152 | 209 | 120 | 0 | 0 | 2 | 3 | 0 | 203 | 149 | 1 | 1 | 0.991 | 1 | 1 | 0.978 | 0.994 | 1 | 1 | 0.982 | 0.981 | 1 | 0 |
| SE | 16090 | ENSG00000128908 | INO80 | chr15 | - | 41087561 | 41087682 | 41085368 | 41085583 | 41092026 | 41092182 | 232 | 144 | 209 | 207 | 400 | 0 | 1 | 0 | 0 | 3 | 332 | 178 | 170 | 280 | 167 | 0 | 0 | 1 | 4 | 0 | 270 | 149 | 1 | 1 | 1 | 0.988 | 1 | 1 | 0.987 | 1 | 1 | 0.989 | 0.975 | 1 | 0.002 |
| SE | 16091 | ENSG00000128908 | INO80 | chr15 | - | 41095600 | 41095668 | 41092026 | 41092182 | 41095758 | 41095928 | 197 | 117 | 188 | 166 | 316 | 0 | 0 | 0 | 0 | 0 | 336 | 173 | 181 | 241 | 121 | 0 | 0 | 0 | 4 | 0 | 217 | 149 | 1 | 1 | 1 | 1 | 1 | 1 | 1 | 1 | 1 | 1 | 0.976 | 1 | 0.005 |
| SE | 16095 | ENSG00000128928 | IVD | chr15 | + | 40407635 | 40407725 | 40405484 | 40405971 | 40407938 | 40407990 | 54 | 12 | 43 | 20 | 20 | 0 | 0 | 0 | 0 | 0 | 35 | 65 | 26 | 2 | 95 | 0 | 0 | 0 | 1 | 0 | 239 | 149 | 1 | 1 | 1 | 1 | 1 | 1 | 1 | 1 | 1 | 1 | 0.555 | 1 | 0.089 |
| [illegible] | | | | | | | | | | | | | | | | | | | | | | | | | | | | | | | | | | | | | | | | | | | | | | |

| | | | | | | | | | | | | | | | | | | | | | | | | | | | | | | | | | | | | | | | | | | | | | | |
|---|---|---|---|---|---|---|---|---|---|---|---|---|---|---|---|---|---|---|---|---|---|---|---|---|---|---|---|---|---|---|---|---|---|---|---|---|---|---|---|---|---|---|---|---|---|---|
| SE | 16452 | ENSG00000104043 | ATP8B4 | chr15 | - | 49996676 | 49996759 | 49981205 | 49981294 | 50002152 | 50002223 | 15 | 25 | 11 | 18 | 14 | 0 | 1 | 1 | 0 | 0 | 7 | 7 | 11 | 27 | 10 | 0 | 0 | 0 | 0 | 0 | 232 | 149 | 1 | 1 | 1 | 0.941 | 0.876 | 1 | 1 | 1 | 1 | 1 | 1 | 1 | -0.037 |
| SE | 16454 | ENSG00000104043 | ATP8B4 | chr15 | - | 50002152 | 50002223 | 49987390 | 49987549 | 50010844 | 50010917 | 2 | 6 | 1 | 3 | 4 | 0 | 2 | 0 | 0 | 0 | 16 | 12 | 2 | 20 | 0 | 0 | 0 | 2 | 2 | 0 | 220 | 149 | 1 | 1 | 1 | 0.67 | 1 | | 1 | 1 | 1 | 0.404 | 0.871 | NA | 0.115 |
| SE | 16455 | ENSG00000104043 | ATP8B4 | chr15 | - | 50010844 | 50010917 | 50002152 | 50002223 | 50038767 | 50038829 | 41 | 41 | 15 | 32 | 34 | 0 | 0 | 0 | 1 | 0 | 23 | 34 | 25 | 69 | 12 | 0 | 0 | 0 | 0 | 0 | 222 | 149 | 1 | 1 | 1 | 1 | 1 | 0.956 | 1 | 1 | 1 | 1 | 1 | 1 | -0.009 |
| SE | 16457 | ENSG00000104043 | ATP8B4 | chr15 | - | 50018888 | 50018976 | 50010844 | 50010917 | 50044593 | 50044692 | 0 | 0 | 0 | 2 | 0 | 0 | 4 | 0 | 1 | 0 | 0 | 0 | 0 | 4 | 0 | 0 | 0 | 0 | 0 | 0 | 237 | 149 | 1 | 1 | NA | 0 | NA | 0.557 | NA | NA | NA | NA | 1 | NA | -0.722 |
| SE | 16458 | ENSG00000104043 | ATP8B4 | chr15 | - | 50038767 | 50038829 | 50010844 | 50010917 | 50044593 | 50044674 | 52 | 32 | 26 | 56 | 40 | 0 | 4 | 0 | 1 | 0 | 32 | 38 | 31 | 90 | 34 | 0 | 0 | 0 | 0 | 0 | 211 | 149 | 1 | 1 | 1 | 0.85 | 1 | 0.975 | 1 | 1 | 1 | 1 | 1 | 1 | -0.035 |
| SE | 16461 | ENSG00000104043 | ATP8B4 | chr15 | - | 50047350 | 50047464 | 50044593 | 50044692 | 50074126 | 50074185 | 97 | 90 | 90 | 136 | 100 | 1 | 0 | 0 | 0 | 0 | 93 | 91 | 46 | 200 | 105 | 0 | 0 | 0 | 1 | 0 | 263 | 149 | 1 | 1 | 0.982 | 1 | 1 | 1 | 1 | 1 | 1 | 1 | 0.991 | 1 | -0.002 |
| SE | 16462 | ENSG00000104043 | ATP8B4 | chr15 | - | 50106938 | 50107008 | 50074126 | 50074185 | 50119122 | 50119218 | 44 | 37 | 29 | 51 | 24 | 0 | 1 | 1 | 0 | 0 | 38 | 28 | 13 | 48 | 9 | 0 | 0 | 0 | 0 | 0 | 219 | 149 | 1 | 1 | 1 | 0.962 | 0.952 | 1 | 1 | 1 | 1 | 1 | 1 | 1 | -0.017 |
| SE | 16463 | ENSG00000285331 | NA | chr15 | - | 56847323 | 56847481 | 56827523 | 56827651 | 56890013 | 56890167 | 0 | 0 | 3 | 3 | 2 | 0 | 0 | 1 | 0 | 0 | 0 | 0 | 2 | 3 | 0 | 0 | 0 | 0 | 0 | 0 | 298 | 149 | 1 | 1 | NA | NA | 0.6 | 1 | 1 | NA | NA | 1 | 1 | NA | -0.133 |
| SE | 16464 | ENSG00000285331 | NA | chr15 | - | 56849955 | 56850186 | 56847323 | 56847481 | 56890013 | 56890167 | 2 | 0 | 0 | 4 | 0 | 0 | 0 | 1 | 0 | 0 | 0 | 0 | 0 | 0 | 0 | 0 | 0 | 2 | 3 | 0 | 298 | 149 | 1 | 1 | 1 | NA | 0 | 1 | NA | NA | NA | 0 | 0 | NA | 0.667 |
| SE | 16465 | ENSG00000285331 | NA | chr15 | - | 56905747 | 56905865 | 56890013 | 56890167 | 56917064 | 56917212 | 0 | 1 | 0 | 1 | 0 | 2 | 1 | 1 | 3 | 0 | 0 | 0 | 0 | 0 | 0 | 0 | 4 | 2 | 0 | 0 | 267 | 149 | 1 | 1 | 0 | 0.358 | 0 | 0.157 | NA | NA | 0 | 0 | NA | NA | 0.129 |
| SE | 16466 | ENSG00000285331 | NA | chr15 | - | 56917064 | 56917212 | 56890013 | 56890167 | 56917539 | 56917648 | 33 | 5 | 18 | 16 | 19 | 0 | 0 | 2 | 1 | 0 | 12 | 45 | 6 | 13 | 2 | 0 | 0 | 0 | 0 | 0 | 297 | 149 | 1 | 1 | 1 | 1 | 0.819 | 0.889 | 1 | 1 | 1 | 1 | 1 | 1 | -0.058 |
| SE | 16475 | ENSG00000166035 | LIPC | chr15 | + | 58541784 | 58541967 | 58538332 | 58538517 | 58542533 | 58542651 | 20 | 8 | 2 | 6 | 6 | 0 | 0 | 0 | 0 | 0 | 33 | 0 | 4 | 8 | 25 | 0 | 0 | 0 | 3 | 0 | 298 | 149 | 1 | 1 | 1 | 1 | 1 | 1 | 1 | 1 | NA | 1 | 0.571 | 1 | 0.107 |
| SE | 16476 | ENSG00000247240 | UBL7-AS1 | chr15 | + | 74466035 | 74466130 | 74461278 | 74462027 | 74479059 | 74479222 | 3 | 3 | 6 | 5 | 20 | 2 | 0 | 0 | 0 | 0 | 27 | 3 | 4 | 6 | 2 | 0 | 0 | 0 | 0 | 0 | 244 | 149 | 1 | 1 | 0.478 | 1 | 1 | 1 | 1 | 1 | 1 | 1 | 1 | 1 | -0.104 |
| SE | 16477 | ENSG00000247240 | UBL7-AS1 | chr15 | + | 74466035 | 74466130 | 74461297 | 74462098 | 74479059 | 74479222 | 6 | 6 | 11 | 4 | 18 | 1 | 0 | 0 | 1 | 0 | 15 | 6 | 4 | 7 | 2 | 0 | 0 | 0 | 0 | 0 | 244 | 149 | 1 | 1 | 0.786 | 1 | 1 | 0.71 | 1 | 1 | 1 | 1 | 1 | 1 | -0.101 |
| SE | 16479 | ENSG00000167173 | C15orf39 | chr15 | + | 75201960 | 75202059 | 75198865 | 75199161 | 75205998 | 75206112 | 33 | 11 | 27 | 33 | 3 | 0 | 2 | 0 | 0 | 0 | 44 | 9 | 16 | 43 | 67 | 0 | 0 | 0 | 0 | 0 | 248 | 149 | 1 | 1 | 1 | 0.768 | 1 | 1 | 1 | 1 | 1 | 1 | 1 | 1 | -0.046 |
| SE | 16483 | ENSG00000072071 | ADGRL1 | chr19 | - | 14164620 | 14164705 | 14162947 | 14163406 | 14167015 | 14167030 | 1 | 0 | 1 | 0 | 0 | 1 | 0 | 3 | 0 | 3 | 0 | 9 | 0 | 0 | 0 | 0 | 0 | 0 | 0 | 0 | 234 | 149 | 1 | 1 | 0.389 | NA | 0.175 | NA | 0 | NA | 1 | NA | NA | NA | -0.812 |
| SE | 16484 | ENSG00000072071 | ADGRL1 | chr19 | - | 14164620 | 14164705 | 14162947 | 14163406 | 14170681 | 14170734 | 1 | 0 | 0 | 0 | 0 | 3 | 1 | 5 | 2 | 2 | 0 | 9 | 0 | 0 | 0 | 0 | 26 | 0 | 6 | 0 | 234 | 149 | 1 | 1 | 0.175 | 0 | 0 | 0 | 0 | NA | 0.181 | NA | 0 | NA | -0.056 |
| SE | 16485 | ENSG00000072071 | ADGRL1 | chr19 | - | 14167015 | 14167030 | 14162605 | 14163406 | 14170681 | 14170734 | 1 | 0 | 2 | 0 | 2 | 3 | 1 | 5 | 2 | 2 | 0 | 0 | 0 | 0 | 0 | 0 | 26 | 0 | 6 | 0 | 164 | 149 | 1 | 1 | 0.232 | 0 | 0.267 | 0 | 0.476 | NA | 0 | NA | 0 | NA | 0.195 |
| SE | 16487 | ENSG00000140474 | ULK3 | chr15 | - | 74837750 | 74837798 | 74837368 | 74837435 | 74838151 | 74838173 | 30 | 12 | 18 | 41 | 30 | 0 | 0 | 2 | 0 | 0 | 15 | 41 | 17 | 18 | 66 | 0 | 0 | 0 | 0 | 0 | 197 | 149 | 1 | 1 | 1 | 1 | 0.872 | 1 | 1 | 1 | 1 | 1 | 1 | 1 | -0.026 |
| SE | 16488 | ENSG00000140474 | ULK3 | chr15 | - | 74837756 | 74837798 | 74837368 | 74837435 | 74838151 | 74838192 | 4 | 14 | 49 | 9 | 2 | 0 | 0 | 2 | 0 | 0 | 21 | 1 | 13 | 8 | 18 | 0 | 0 | 0 | 0 | 0 | 191 | 149 | 1 | 1 | 1 | 1 | 0.95 | 1 | 1 | 1 | 1 | 1 | 1 | 1 | -0.01 |
| SE | 16490 | ENSG00000140474 | ULK3 | chr15 | - | 74838151 | 74838192 | 74837750 | 74837798 | 74838265 | 74838344 | 29 | 19 | 38 | 28 | 19 | 1 | 0 | 2 | 0 | 1 | 15 | 32 | 15 | 8 | 46 | 0 | 0 | 7 | 0 | 0 | 190 | 149 | 1 | 1 | 0.958 | 1 | 0.937 | 1 | 0.937 | 1 | 1 | 0.627 | 1 | 1 | 0.041 |
| SE | 16491 | ENSG00000140474 | ULK3 | chr15 | - | 74838151 | 74838192 | 74837750 | 74837798 | 74838439 | 74838504 | 5 | 3 | 13 | 5 | 15 | 0 | 0 | 0 | 0 | 0 | 6 | 19 | 9 | 4 | 11 | 0 | 3 | 0 | 0 | 0 | 190 | 149 | 1 | 1 | 1 | 1 | 1 | 1 | 1 | 1 | 0.832 | 1 | 1 | 1 | 0.034 |
| SE | 16492 | ENSG00000140474 | ULK3 | chr15 | - | 74838265 | 74838344 | 74837750 | 74837798 | 74838439 | 74838504 | 19 | 5 | 20 | 24 | 11 | 0 | 0 | 0 | 0 | 0 | 21 | 16 | 13 | 7 | 34 | 0 | 3 | 0 | 0 | 0 | 228 | 149 | 1 | 1 | 1 | 1 | 1 | 1 | 1 | 1 | 0.777 | 1 | 1 | 1 | 0.045 |
| SE | 16503 | ENSG00000140367 | UBE2Q2 | chr15 | + | 75854385 | 75854487 | 75843306 | 75843846 | 75859877 | 75859885 | 188 | 132 | 154 | 169 | 191 | 0 | 0 | 0 | 0 | 0 | 225 | 131 | 140 | 41 | 59 | 0 | 0 | 0 | 3 | 0 | 251 | 149 | 1 | 1 | 1 | 1 | 1 | 1 | 1 | 1 | 1 | 1 | 0.89 | 1 | 0.022 |
| SE | 16505 | ENSG00000140367 | UBE2Q2 | chr15 | + | 75868950 | 75869010 | 75854385 | 75854487 | 75873427 | 75873568 | 102 | 88 | 89 | 97 | 128 | 0 | 0 | 0 | 1 | 0 | 104 | 45 | 85 | 44 | 47 | 0 | 0 | 0 | 0 | 0 | 209 | 149 | 1 | 1 | 1 | 1 | 1 | 0.986 | 1 | 1 | 1 | 1 | 1 | 1 | -0.003 |
| SE | 16507 | ENSG00000140367 | UBE2Q2 | chr15 | + | 75868950 | 75869010 | 75859877 | 75859982 | 75876186 | 75876271 | 94 | 78 | 82 | 113 | 133 | 0 | 0 | 0 | 0 | 0 | 190 | 65 | 68 | 41 | 79 | 0 | 0 | 2 | 0 | 0 | 209 | 149 | 1 | 1 | 1 | 1 | 1 | 1 | 1 | 1 | 1 | 0.96 | 1 | 1 | 0.008 |
| SE | 16510 | ENSG00000140367 | UBE2Q2 | chr15 | + | 75873427 | 75873568 | 75868950 | 75869010 | 75877960 | 75878021 | 224 | 185 | 195 | 240 | 354 | 0 | 0 | 0 | 0 | 1 | 224 | 183 | 196 | 107 | 140 | 0 | 0 | 2 | 0 | 0 | 290 | 149 | 1 | 1 | 1 | 1 | 1 | 1 | 0.995 | 1 | 1 | 0.981 | 1 | 1 | 0.003 |
| SE | 16513 | ENSG00000140367 | UBE2Q2 | chr15 | + | 75876186 | 75876271 | 75868950 | 75869010 | 75877960 | 75878021 | 166 | 152 | 123 | 140 | 199 | 0 | 0 | 0 | 0 | 1 | 60 | 70 | 63 | 48 | 41 | 0 | 0 | 2 | 0 | 0 | 234 | 149 | 1 | 1 | 1 | 1 | 1 | 1 | 0.992 | 1 | 1 | 0.953 | 1 | 1 | 0.008 |
| SE | 16514 | ENSG00000140367 | UBE2Q2 | chr15 | + | 75876186 | 75876271 | 75868950 | 75869010 | 75879097 | 75879188 | 6 | 7 | 4 | 8 | 5 | 0 | 0 | 0 | 0 | 0 | 0 | 2 | 6 | 0 | 0 | 0 | 2 | 0 | 0 | 10 | 234 | 149 | 1 | 1 | 1 | 1 | 1 | 1 | 1 | NA | 0.389 | 1 | NA | 0 | 0.537 |
| SE | 16515 | ENSG00000140367 | UBE2Q2 | chr15 | + | 75876186 | 75876271 | 75868950 | 75869010 | 75883365 | 75883424 | 3 | 6 | 0 | 7 | 2 | 2 | 5 | 1 | 1 | 3 | 0 | 0 | 3 | 0 | 0 | 8 | 0 | 0 | 0 | 0 | 234 | 149 | 1 | 1 | 0.489 | 0.433 | 0 | 0.817 | 0.298 | 0 | NA | 1 | NA | NA | -0.093 |
| SE | 16520 | ENSG00000140367 | UBE2Q2 | chr15 | + | 75877960 | 75878021 | 75868950 | 75869010 | 75883365 | 75883424 | 1 | 0 | 0 | 0 | 0 | 2 | 5 | 1 | 1 | 3 | 0 | 0 | 0 | 0 | 0 | 8 | 0 | 0 | 0 | 0 | 210 | 149 | 1 | 1 | 0.262 | 0 | 0 | 0 | 0 | 0 | NA | NA | NA | NA | 0.052 |
| SE | 16522 | ENSG00000140367 | UBE2Q2 | chr15 | + | 75877960 | 75878021 | 75873427 | 75873568 | 75883365 | 75883424 | 1 | 2 | 5 | 2 | 5 | 4 | 3 | 3 | 7 | 13 | 0 | 1 | 4 | 0 | 0 | 0 | 2 | 8 | 4 | 0 | 210 | 149 | 1 | 1 | 0.151 | 0.321 | 0.542 | 0.169 | 0.214 | NA | 0.262 | 0.262 | 0 | NA | 0.105 |
| SE | 16523 | ENSG00000140367 | UBE2Q2 | chr15 | + | 75877960 | 75878021 | 75876186 | 75876271 | 75879097 | 75879188 | 375 | 315 | 263 | 319 | 382 | 4 | 1 | 4 | 1 | 3 | 106 | 179 | 138 | 101 | 93 | 0 | 2 | 3 | 0 | 2 | 210 | 149 | 1 | 1 | 0.985 | 0.996 | 0.979 | 0.996 | 0.989 | 1 | 0.984 | 0.97 | 1 | 0.971 | 0.004 |
| SE | 16527 | ENSG00000140367 | UBE2Q2 | chr15 | + | 75879097 | 75879188 | 75876186 | 75876271 | 75883365 | 75883424 | 243 | 147 | 169 | 180 | 239 | 0 | 4 | 1 | 0 | 2 | 51 | 87 | 64 | 65 | 16 | 0 | 0 | 0 | 0 | 0 | 240 | 149 | 1 | 1 | 1 | 0.958 | 0.991 | 1 | 0.987 | | 1 | 1 | 1 | 1 | -0.013 |
| SE | 16529 | ENSG00000140367 | UBE2Q2 | chr15 | + | 75883365 | 75883424 | 75873427 | 75873568 | 75896994 | 75897061 | 3 | 2 | 3 | 4 | 7 | 1 | 0 | 0 | 0 | 0 | 0 | 2 | 3 | 0 | 0 | 0 | 0 | 0 | 0 | 0 | 208 | 149 | 1 | 1 | 0.682 | 1 | 1 | 1 | 1 | NA | 1 | 1 | NA | NA | -0.064 |
| SE | 16530 | ENSG00000140367 | UBE2Q2 | chr15 | + | 75883365 | 75883424 | 75877960 | 75878021 | 75890434 | 75890483 | 99 | 63 | 56 | 74 | 34 | 0 | 1 | 0 | 0 | 0 | 6 | 37 | 35 | 18 | 9 | 0 | 0 | 0 | 0 | 0 | 208 | 149 | 1 | 1 | 1 | 0.978 | 1 | 1 | 1 | 1 | 1 | 1 | 1 | 1 | -0.004 |
| SE | 16531 | ENSG00000140367 | UBE2Q2 | chr15 | + | 75890434 | 75890483 | 75883365 | 75883424 | 75890918 | 75891014 | 431 | 258 | 284 | 301 | 336 | 0 | 0 | 1 | 0 | 2 | 56 | 136 | 127 | 95 | 23 | 0 | 0 | 0 | 0 | 0 | 198 | 149 | 1 | 1 | 1 | 1 | 0.995 | 1 | 0.992 | 1 | 1 | 1 | 1 | 1 | -0.003 |
| SE | 16532 | ENSG00000140367 | UBE2Q2 | chr15 | + | 75890434 | 75890483 | 75883365 | 75883424 | 75896994 | 75897061 | 129 | 91 | 91 | 95 | 97 | 0 | 2 | 1 | 0 | 9 | 4 | 50 | 32 | 26 | 0 | 0 | 0 | 1 | 0 | 0 | 198 | 149 | 1 | 1 | 1 | 0.972 | 0.986 | 1 | 0.89 | 1 | 1 | 0.96 | 1 | NA | -0.02 |
| SE | 16533 | ENSG00000140367 | UBE2Q2 | chr15 | + | 75890918 | 75891014 | 75883365 | 75883424 | 75896994 | 75897061 | 53 | 24 | 45 | 43 | 31 | 0 | 2 | 1 | 0 | 9 | 0 | 25 | 5 | 2 | 6 | 0 | 0 | 1 | 0 | 0 | 245 | 149 | 1 | 1 | 1 | 0.879 | 0.965 | 1 | 0.677 | NA | 1 | 0.753 | 1 | 1 | -0.034 |
| SE | 16534 | ENSG00000140367 | UBE2Q2 | chr15 | + | 75890918 | 75891014 | 75883365 | 75883424 | 75899426 | 75899650 | 0 | 0 | 0 | 0 | 2 | 3 | 3 | 0 | 8 | 4 | 0 | 0 | 0 | 0 | 0 | 5 | 0 | 0 | 6 | 0 | 245 | 149 | 1 | 1 | 0 | 0 | NA | 0 | 0.233 | 0 | NA | NA | 0 | NA | 0.058 |
| SE | 16536 | ENSG00000140367 | UBE2Q2 | chr15 | + | 75896994 | 75897061 | 75883365 | 75883424 | 75899426 | 75899494 | 47 | 37 | 48 | 36 | 76 | 3 | 3 | 0 | 8 | 4 | 12 | 10 | 18 | 1 | 0 | 5 | 0 | 0 | 6 | 0 | 216 | 149 | 1 | 1 | 0.915 | 0.895 | 1 | 0.756 | 0.929 | 0.623 | 1 | 1 | 0.103 | NA | 0.217 |
| SE | 16537 | ENSG00000140367 | UBE2Q2 | chr15 | + | 75896994 | 75897061 | 75890918 | 75891014 | 75899426 | 75899494 | 235 | 139 | 184 | 159 | 248 | 0 | 0 | 0 | 2 | 0 | 15 | 71 | 62 | 28 | 12 | 0 | 0 | 0 | 0 | 0 | 216 | 149 | 1 | 1 | 1 | 1 | 1 | 0.982 | 1 | 1 | 1 | 1 | 1 | 1 | -0.004 |
| SE | 16538 | ENSG00000278202 | NA | chr15 | + | 82432575 | 82432631 | 82431829 | 82431949 | 82432816 | 82432897 | 1 | 8 | 2 | 5 | 0 | 1 | 6 | 1 | 7 | 0 | 0 | 0 | 2 | 0 | 0 | 0 | 0 | 2 | 0 | 0 | 205 | 149 | 1 | 1 | 0.421 | 0.492 | 0.592 | 0.342 | NA | NA | NA | 0.421 | NA | NA | 0.041 |
| SE | 16539 | ENSG00000138623 | SEMA7A | chr15 | - | 74418267 | 74418309 | 74417876 | 74417969 | 74418800 | 74418952 | 7 | 3 | 1 | 2 | 0 | 0 | 0 | 1 | 0 | 0 | 0 | 3 | 0 | 0 | 0 | 0 | 0 | 0 | 0 | 0 | 191 | 149 | 1 | 1 | 1 | 1 | 0.438 | 1 | NA | NA | 1 | NA | NA | NA | -0.14 |
| SE | 16544 | ENSG00000166839 | ANKDD1A | chr15 | + | 64926900 | 64926999 | 64926065 | 64926170 | 64930821 | 64930920 | 0 | 0 | 9 | 1 | 0 | 0 | 0 | 0 | 0 | 0 | 0 | 7 | 4 | 3 | 2 | 0 | 4 | 0 | 0 | 0 | 248 | 149 | 1 | 1 | NA | NA | 1 | 1 | NA | NA | 0.513 | 1 | 1 | 1 | 0.122 |
| SE | 16546 | ENSG00000166839 | ANKDD1A | chr15 | + | 64947403 | 64947593 | 64943483 | 64943582 | 64949840 | 64949972 | 2 | 5 | 2 | 4 | 8 | 0 | 0 | 0 | 0 | 0 | 13 | 1 | 13 | 6 | 0 | 0 | 0 | 4 | 0 | 0 | 298 | 149 | 1 | 1 | 1 | 1 | 1 | 1 | 1 | 1 | 1 | 0.619 | 1 | NA | 0.095 |
| SE | 16548 | ENSG00000137764 | MAP2K5 | chr15 | + | 67563282 | 67563350 | 67550033 | 67550082 | 67580753 | 67580823 | 130 | 107 | 157 | 135 | 180 | 0 | 0 | 0 | 1 | 0 | 155 | 179 | 93 | 92 | 256 | 0 | 0 | 0 | 0 | 0 | 217 | 149 | 1 | 1 | 1 | 1 | 1 | 0.989 | 1 | 1 | 1 | 1 | 1 | 1 | -0.002 |
| SE | 16550 | ENSG00000137764 | MAP2K5 | chr15 | + | 67585889 | 67585930 | 67563282 | 67563350 | 67586845 | 67586913 | 30 | 37 | 37 | 44 | 52 | 0 | 0 | 3 | 1 | 1 | 73 | 59 | 30 | 53 | 57 | 0 | 0 | 0 | 5 | 0 | 190 | 149 | 1 | 1 | 1 | 1 | 0.906 | 0.972 | 0.976 | 1 | 1 | 1 | 0.893 | 1 | -0.008 |
| SE | 16551 | ENSG00000137764 | MAP2K5 | chr15 | + | 67585889 | 67585930 | 67580753 | 67580823 | 67592925 | 67592974 | 33 | 36 | 45 | 43 | 36 | 0 | 0 | 0 | 0 | 1 | 27 | 68 | 19 | 15 | 61 | 0 | 0 | 0 | 0 | 0 | 190 | 149 | 1 | 1 | 1 | 1 | 1 | 1 | 0.966 | 1 | 1 | 1 | 1 | 1 | -0.007 |
| SE | 16552 | ENSG00000137764 | MAP2K5 | chr15 | + | 67586845 | 67586913 | 67585741 | 67585930 | 67592925 | 67592974 | 157 | 150 | 176 | 173 | 210 | 0 | 0 | 1 | 0 | 0 | 281 | 193 | 161 | 209 | 304 | 0 | 0 | 0 | 0 | 0 | 217 | 149 | 1 | 1 | 1 | 1 | 0.992 | 1 | 1 | 1 | 1 | 1 | 1 | 1 | -0.002 |
| SE | 16553 | ENSG00000137764 | MAP2K5 | chr15 | + | 67600684 | 67600749 | 67592925 | 67592974 | 67630887 | 67630927 | 181 | 174 | 211 | 210 | 254 | 0 | 0 | 0 | 0 | 0 | 282 | 190 | 150 | 241 | 335 | 0 | 4 | 0 | 0 | 0 | 214 | 149 | 1 | 1 | 1 | 1 | 1 | 1 | 1 | 1 | 0.971 | 1 | 1 | 1 | 0.006 |
| SE | 16555 | ENSG00000137764 | MAP2K5 | chr15 | + | 67642416 | 67642497 | 67641444 | 67641658 | 67646230 | 67646299 | 4 | 4 | 2 | 3 | 0 | 2 | 0 | 1 | 2 | 0 | 1 | 1 | 0 | 0 | 0 | 0 | 0 | 1 | 6 | 0 | 230 | 149 | 1 | 1 | 0.564 | 1 | 0.564 | 0.493 | NA | 1 | 1 | 0 | 0 | NA | 0.155 |
| SE | 16557 | ENSG00000137764 | MAP2K5 | chr15 | + | 67670419 | 67670447 | 67664596 | 67664645 | 67679695 | 67679956 | 1 | 0 | 0 | 0 | 0 | 1 | 0 | 2 | 0 | 0 | 0 | 0 | 0 | 0 | 0 | 0 | 0 | 0 | 0 | 3 | 177 | 149 | 1 | 1 | 0.457 | NA | 0 | NA | NA | NA | NA | NA | NA | 0 | 0.229 |
| SE | 16558 | ENSG00000137764 | MAP2K5 | chr15 | + | 67670419 | 67670447 | 67664596 | 67664645 | 67692478 | 67692552 | 3 | 0 | 1 | 0 | 0 | 53 | 32 | 56 | 65 | 93 | 0 | 0 | 2 | 0 | 0 | 55 | 35 | 21 | 39 | 97 | 177 | 149 | 1 | 1 | 0.045 | 0 | 0.015 | 0 | 0 | 0 | 0 | 0.074 | 0 | 0 | -0.003 |
| SE | 16560 | ENSG00000137764 | MAP2K5 | chr15 | + | 67693517 | 67693568 | 67664596 | 67664645 | 67703336 | 67703408 | 28 | 7 | 27 | 36 | 33 | 0 | 0 | 0 | 1 | 0 | 47 | 29 | 14 | 13 | 57 | 4 | 0 | 0 | 0 | 0 | 200 | 149 | 1 | 1 | 1 | 1 | 1 | 0.964 | 1 | 0.897 | 1 | 1 | 1 | 1 | 0.013 |
| SE | 16561 | ENSG00000137764 | MAP2K5 | chr15 | + | 67703336 | 67703408 | 67693517 | 67693568 | 67727915 | 67727945 | 68 | 33 | 100 | 87 | 111 | 1 | 1 | 0 | 0 | 0 | 106 | 65 | 35 | 43 | 148 | 0 | 0 | 0 | 0 | 0 | 221 | 149 | 1 | 1 | 0.979 | 0.957 | 1 | 1 | 1 | 1 | 1 | 1 | 1 | 1 | -0.013 |
| SE | 16566 | ENSG00000137764 | MAP2K5 | chr15 | + | 67748568 | 67748601 | 67727915 | 67727945 | 67769601 | 67769663 | 9 | 4 | 12 | 5 | 19 | 0 | 0 | 0 | 1 | 0 | 0 | 12 | 4 | 6 | 42 | 0 | 0 | 0 | 0 | 0 | 182 | 149 | 1 | 1 | 1 | 1 | 1 | 0.804 | 1 | NA | 1 | 1 | 1 | 1 | -0.039 |
| SE | 16567 | ENSG00000137764 | MAP2K5 | chr15 | + | 67769601 | 67769663 | 67748568 | 67748601 | 67772706 | 67772752 | 108 | 49 | 81 | 84 | 108 | 0 | 0 | 0 | 0 | 5 | 51 | 86 | 35 | 52 | 174 | 0 | 0 | 1 | 0 | 0 | 211 | 149 | 1 | 1 | 1 | 1 | 1 | 1 | 0.938 | 1 | 1 | 0.961 | 1 | 1 | -0.005 |
| SE | 16568 | ENSG00000137764 | MAP2K5 | chr15 | + | 67772706 | 67772752 | 67748568 | 67748601 | 67806645 | 67806732 | 47 | 23 | 37 | 32 | 60 | 1 | 3 | 0 | 0 | 0 | 33 | 46 | 19 | 31 | 26 | 0 | 0 | 0 | 0 | 0 | 195 | 149 | 1 | 1 | 0.973 | 0.854 | 1 | 1 | 1 | 1 | 1 | 1 | 1 | 1 | -0.035 |
| SE | 16569 | ENSG00000179151 | EDC3 | chr15 | - | 74635408 | 74635626 | 74630557 | 74632946 | 74640465 | 74640619 | 93 | 45 | 86 | 106 | 111 | 0 | 0 | 1 | 0 | 0 | 42 | 89 | 23 | 65 | 81 | 0 | 0 | 0 | 0 | 0 | 298 | 149 | 1 | 1 | 1 | 1 | 0.977 | 1 | 1 | 1 | 1 | 1 | 1 | 1 | -0.005 |
| SE | 16570 | ENSG00000179151 | EDC3 | chr15 | - | 74640465 | 74640619 | 74635180 | 74635626 | 74655732 | 74655893 | 123 | 39 | 100 | 138 | 154 | 0 | 0 | 1 | 2 | 0 | 100 | 157 | 27 | 63 | 125 | 0 | 0 | 0 | 0 | 0 | 298 | 149 | 1 | 1 | 1 | 1 | 0.98 | 0.972 | 1 | 1 | 1 | 1 | 1 | 1 | -0.01 |
| SE | 16571 | ENSG00000179151 | EDC3 | chr15 | - | 74655732 | 74656068 | 74640465 | 74640619 | 74671454 | 74671774 | 114 | 47 | 152 | 173 | 150 | 0 | 0 | 1 | 0 | 0 | 181 | 199 | 59 | 110 | 201 | 0 | 0 | 0 | 0 | 0 | 298 | 149 | 1 | 1 | 1 | 1 | 0.987 | 1 | 1 | 1 | 1 | 1 | 1 | 1 | -0.003 |
| SE | 16572 | ENSG00000179151 | EDC3 | chr15 | - | 74671454 | 74671774 | 74655732 | 74656068 | 74674960 | 74675142 | 99 | 46 | 141 | 151 | 110 | 0 | 0 | 0 | 1 | 0 | 196 | 150 | 56 | 98 | 194 | 0 | 0 | 0 | 0 | 0 | 298 | 149 | 1 | 1 | 1 | 1 | 1 | 0.987 | 1 | 1 | 1 | 1 | 1 | 1 | -0.003 |
| SE | 16575 | ENSG00000179151 | EDC3 | chr15 | - | 74676940 | 74677022 | 74674960 | 74675142 | 74695879 | 74695987 | 0 | 3 | 2 | 2 | 2 | 10 | 5 | 13 | 12 | 6 | 0 | 0 | 0 | 8 | 0 | 0 | 24 | 9 | 4 | 0 | 231 | 149 | 1 | 1 | 0 | 0.279 | 0.09 | 0.097 | 0.177 | NA | 0 | 0 | 0.563 | NA | -0.059 |
| SE | 16577 | ENSG00000179151 | EDC3 | chr15 | - | 74687090 | 74687179 | 74674960 | 74675142 | 74692885 | 74693018 | 11 | 1 | 2 | 1 | 3 | 0 | 0 | 0 | 0 | 4 | 6 | 7 | 2 | 0 | 5 | 0 | 0 | 0 | 0 | 0 | 238 | 149 | 1 | 1 | 1 | 1 | 1 | 1 | 0.32 | 1 | 1 | 1 | NA | 1 | -0.136 |
| SE | 16581 | ENSG00000179151 | EDC3 | chr15 | - | 74692885 | 74693018 | 74674960 | 74675142 | 74695879 | 74696020 | 0 | 0 | 0 | 0 | 4 | 10 | 5 | 13 | 12 | 6 | 0 | 0 | 0 | 0 | 0 | 0 | 24 | 9 | 4 | 0 | 282 | 149 | 1 | 1 | 0 | 0 | 0 | 0 | 0.26 | NA | 0 | 0 | 0 | NA | 0.052 |
| SE | 16586 | ENSG00000138621 | PPCDC | chr15 | + | 75028246 | 75028453 | 75023585 | 75023626 | 75043440 | 75043536 | 12 | 5 | 13 | 12 | 6 | 3 | 0 | 0 | 0 | 0 | 41 | 13 | 3 | 6 | 33 | 0 | 0 | 0 | 0 | 0 | 298 | 149 | 1 | 1 | 0.667 | 1 | 1 | 1 | 1 | 1 | 1 | 1 | 1 | 1 | -0.067 |
| SE | 16587 | ENSG00000138621 | PPCDC | chr15 | + | 75043440 | 75043536 | 75028246 | 75028453 | 75044385 | 75044514 | 42 | 39 | 38 | 18 | 21 | 1 | 0 | 1 | 0 | 0 | 53 | 23 | 41 | 17 | 102 | 0 | 0 | 0 | 1 | 0 | 245 | 149 | 1 | 1 | 0.962 | 1 | 0.959 | 1 | 1 | 1 | 1 | 1 | 0.912 | 1 | 0.002 |
| SE | 16589 | ENSG00000138621 | PPCDC | chr15 | + | 75044385 | 75044514 | 75028248 | 75028453 | 75048552 | 75048721 | 36 | 30 | 25 | 22 | 10 | 0 | 1 | 0 | 3 | 0 | 48 | 26 | 19 | 33 | 89 | 0 | 0 | 1 | 0 | 0 | 278 | 149 | 1 | 1 | 1 | 0.941 | 1 | 0.797 | 1 | 1 | 1 | 0.911 | 1 | 1 | -0.035 |
| SE | 16591 | ENSG00000136381 | IREB2 | chr15 | + | 78462921 | 78463087 | 78439794 | 78439881 | 78465250 | 78465264 | 468 | 358 | 390 | 354 | 473 | 0 | 0 | 2 | 0 | 0 | 275 | 445 | 502 | 308 | 185 | 0 | 0 | 2 | 1 | 0 | 298 | 149 | 1 | 1 | 1 | 1 | 0.99 | 1 | 1 | 1 | 1 | 0.992 | 0.994 | 1 | 0.001 |
| SE | 16592 | ENSG00000136381 | IREB2 | chr15 | + | 78465250 | 78465388 | 78462921 | 78463087 | 78466270 | 78466436 | 397 | 304 | 314 | 300 | 272 | 0 | 0 | 0 | 0 | 4 | 161 | 325 | 412 | 216 | 171 | 0 | 0 | 0 | 4 | 0 | 287 | 149 | 1 | 1 | 1 | 1 | 1 | 1 | 0.972 | 1 | 1 | 1 | 0.966 | 1 | 0.001 |
| SE | 16593 | ENSG00000136381 | IREB2 | chr15 | + | 78465250 | 78465388 | 78462921 | 78463087 | 78470531 | 78470601 | 221 | 181 | 175 | 170 | 188 | 1 | 0 | 0 | 0 | 0 | 120 | 206 | 261 | 138 | 93 | 0 | 0 | 0 | 0 | 0 | 287 | 149 | 1 | 1 | 0.991 | 1 | 1 | 1 | 1 | 1 | 1 | 1 | 1 | 1 | -0.002 |
| SE | 16594 | ENSG00000136381 | IREB2 | chr15 | + | 78466270 | 78466489 | 78462921 | 78463087 | 78470531 | 78470601 | 138 | 101 | 134 | 145 | 121 | 1 | 0 | 0 | 0 | 0 | 44 | 86 | 120 | 46 | 19 | 0 | 0 | 0 | 0 | 0 | 298 | 149 | 1 | 1 | 0.986 | 1 | 1 | 1 | 1 | 1 | 1 | 1 | 1 | 1 | -0.003 |
| SE | 16596 | ENSG00000136381 | IREB2 | chr15 | + | 78470531 | 78470601 | 78466270 | 78466489 | 78471740 | 78471801 | 213 | 155 | 187 | 198 | 180 | 5 | 5 | 4 | 2 | 3 | 68 | 146 | 175 | 58 | 45 | 13 | 0 | 1 | 0 | 0 | 219 | 149 | 1 | 1 | 0.967 | 0.955 | 0.97 | 0.985 | 0.976 | 0.781 | 1 | 0.992 | 1 | 1 | 0.016 |
| SE | 16598 | ENSG00000136381 | IREB2 | chr15 | + | 78483317 | 78483434 | 78478296 | 78478397 | 78484760 | 78484920 | 129 | 98 | 104 | 132 | 347 | 0 | 0 | 0 | 0 | 3 | 23 | 130 | 72 | 40 | 8 | 0 | 2 | 0 | 0 | 0 | 266 | 149 | 1 | 1 | 1 | 1 | 1 | 1 | 0.985 | 1 | 0.973 | 1 | 1 | 1 | 0.002 |
| SE | 16599 | ENSG00000136381 | IREB2 | chr15 | + | 78488179 | 78488336 | 78487732 | 78487817 | 78488646 | 78488771 | 241 | 231 | 243 | 202 | 301 | 0 | 0 | 0 | 0 | 0 | 56 | 113 | 223 | 99 | 5 | 0 | 0 | 1 | 0 | 0 | 298 | 149 | 1 | 1 | 1 | 1 | 1 | 1 | 1 | 1 | 1 | 1 | 0.98 | 1 | 0.004 |
| SE | 16600 | ENSG00000136381 | IREB2 | chr15 | + | 78488646 | 78488771 | 78488179 | 78488336 | 78490421 | 78490526 | 255 | 231 | 271 | 231 | 272 | 0 | 0 | 0 | 0 | 0 | 20 | 143 | 214 | 109 | 7 | 0 | 2 | 0 | 0 | 0 | 274 | 149 | 1 | 1 | 1 | 1 | 1 | 1 | 1 | 1 | 0.975 | 1 | 1 | 1 | 0.005 |
| SE | 16602 | ENSG00000140284 | SLC27A2 | chr15 | + | 50198234 | 50198482 | 50197499 | 50197709 | 50202486 | 50202645 | 0 | 0 | 0 | 0 | 0 | 2 | 1 | 2 | 1 | 0 | 0 | 1 | 0 | 0 | 0 | 0 | 4 | 1 | 9 | 0 | 298 | 149 | 1 | 1 | 0 | 0 | 0 | 0 | NA | NA | 0.111 | 0 | 0 | NA | -0.037 |
| SE | 16604 | ENSG00000138629 | UBL7 | chr15 | - | 74451435 | 74451520 | 74445976 | 74446227 | 74456551 | 74456671 | 0 | 0 | 0 | 0 | 0 | 1 | 0 | 3 | 0 | 0 | 4 | 0 | 0 | 0 | 0 | 0 | 4 | 0 | 0 | 0 | 234 | 149 | 1 | 1 | 0 | NA | 0 | NA | NA | 1 | NA | NA | NA | NA | -1 |
| SE | 16606 | ENSG00000138629 | UBL7 | chr15 | - | 74451435 | 74451520 | 74450801 | 74450859 | 74456551 | 74456671 | 35 | 13 | 29 | 47 | 67 | 1 | 1 | 0 | 1 | 2 | 24 | 31 | 9 | 24 | 67 | 2 | 6 | 0 | 0 | 0 | 234 | 149 | 1 | 1 | 0.957 | 0.892 | 1 | 0.968 | 0.955 | 0.884 | 0.767 | 1 | 1 | 1 | 0.024 |
| SE | 16607 | ENSG00000138629 | UBL7 | chr15 | - | 74452295 | 74452378 | 74450801 | 74450859 | 74456551 | 74456671 | 53 | 30 | 43 | 52 | 51 | 1 | 1 | 0 | 1 | 2 | 27 | 33 | 33 | 24 | 66 | 2 | 6 | 0 | 0 | 0 | 232 | 149 | 1 | 1 | 0.971 | 0.951 | 1 | 0.971 | 0.942 | 0.897 | 0.779 | 1 | 1 | 1 | 0.032 |
| SE | 16608 | ENSG00000138629 | UBL7 | chr15 | - | 74452295 | 74452378 | 74451435 | 74451520 | 74456551 | 74456671 | 119 | 50 | 80 | 105 | 109 | 2 | 0 | 0 | 0 | 0 | 98 | 78 | 51 | 42 | 172 | 0 | 0 | 0 | 0 | 0 | 232 | 149 | 1 | 1 | 0.974 | 1 | 1 | 1 | 1 | 1 | 1 | 1 | 1 | 1 | -0.005 |
| SE | 16609 | ENSG00000138629 | UBL7 | chr15 | - | 74456551 | 74456671 | 74452295 | 74452378 | 74458683 | 74458896 | 193 | 70 | 126 | 151 | 149 | 2 | 0 | 0 | 0 | 0 | 133 | 92 | 93 | 71 | 258 | 0 | 0 | 0 | 0 | 0 | 269 | 149 | 1 | 1 | 0.982 | 1 | 1 | 1 | 1 | 1 | 1 | 1 | 1 | 1 | -0.004 |
| SE | 16615 | ENSG00000140382 | HMG20A | chr15 | + | 77471782 | 77471814 | 77470909 | 77471042 | 77477554 | 77477630 | 78 | 74 | 89 | 80 | 135 | 1 | 0 | 0 | 1 | 0 | 96 | 52 | 65 | 63 | 35 | 0 | 0 | 0 | 0 | 0 | 181 | 149 | 1 | 1 | 0.985 | 1 | 1 | 0.985 | 1 | 1 | 1 | 1 | 1 | 1 | -0.006 |
| SE | 16616 | ENSG00000140382 | HMG20A | chr15 | + | 77477554 | 77477630 | 77471782 | 77471814 | 77478294 | 77478510 | 95 | 89 | 102 | 97 | 181 | 0 | 0 | 0 | 1 | 0 | 84 | 58 | 87 | 56 | 38 | 0 | 0 | 0 | 0 | 0 | 225 | 149 | 1 | 1 | 1 | 1 | 1 | 0.985 | 1 | 1 | 1 | 1 | 1 | 1 | -0.003 |
| SE | 16617 | ENSG00000067141 | NEO1 | chr15 | + | 73273810 | 73274005 | 73272454 | 73272562 | 73278130 | 73278199 | 36 | 23 | 53 | 24 | 69 | 0 | 0 | 0 | 1 | 0 | 0 | 5 | 5 | 2 | 18 | 0 | 0 | 0 | 0 | 0 | 298 | 149 | 1 | 1 | 1 | 1 | 1 | 0.923 | 1 | NA | 1 | 1 | 1 | 1 | -0.015 |
| SE | 16618 | ENSG00000067141 | NEO1 | chr15 | + | 73274691 | 73274724 | 73273810 | 73274005 | 73278130 | 73278199 | 3 | 0 | 2 | 5 | 0 | 12 | 10 | 22 | 12 | 29 | 0 | 0 | 3 | 0 | 0 | 0 | 5 | 3 | 0 | 0 | 182 | 149 | 1 | 1 | 0.17 | 0 | 0.069 | 0.254 | 0 | NA | 0 | 0.45 | NA | NA | -0.126 |
| SE | 16619 | ENSG00000067141 | NEO1 | chr15 | + | 73293389 | 73293548 | 73289073 | 73289238 | 73298347 | 73298611 | 7 | 14 | 14 | 28 | 29 | 2 | 4 | 7 | 2 | 5 | 7 | 0 | 18 | 0 | 14 | 0 | 5 | 0 | 0 | 8 | 298 | 149 | 1 | 1 | 0.636 | 0.636 | 0.5 | 0.875 | 0.744 | 1 | 0 | 1 | NA | 0.467 | 0.061 |
| SE | 16627 | ENSG00000248508 | SRP14-AS1 | chr15 | + | 40046861 | 40047221 | 40045960 | 40046069 | 40056688 | 40057003 | 1 | 0 | 5 | 1 | 7 | 4 | 5 | 6 | 10 | 2 | 0 | 16 | 0 | 0 | 0 | 19 | 0 | 3 | 6 | 0 | 298 | 149 | 1 | 1 | 0.111 | 0 | 0.294 | 0.048 | 0.636 | 0 | 1 | 0 | 0 | NA | -0.032 |
| SE | 16628 | ENSG00000248508 | SRP14-AS1 | chr15 | + | 40056452 | 40056641 | 40045960 | 40046069 | 40056729 | 40057000 | 5 | 0 | 6 | 2 | 6 | 0 | 0 | 0 | 1 | 0 | 16 | 7 | 0 | 11 | 0 | 0 | 0 | 0 | 0 | 0 | 298 | 149 | 1 | 1 | 1 | NA | 1 | 0.5 | 1 | 1 | 1 | NA | 1 | NA | -0.125 |
| SE | 16629 | ENSG00000248508 | SRP14-AS1 | chr15 | + | 40056452 | 40056641 | 40045960 | 40046069 | 40059132 | 40059289 | 5 | 0 | 3 | 2 | 4 | 2 | 0 | 1 | 0 | 0 | 10 | 3 | 0 | 15 | 0 | 0 | 0 | 0 | 0 | 0 | 298 | 149 | 1 | 1 | 0.556 | NA | 0.6 | 1 | 1 | 1 | 1 | NA | 1 | NA | -0.211 |
| SE | 16630 | ENSG00000248508 | SRP14-AS1 | chr15 | + | 40056452 | 40056641 | 40045960 | 40046069 | 40060096 | 40060263 | 5 | 0 | 3 | 2 | 10 | 0 | 0 | 0 | 0 | 0 | 10 | 4 | 0 | 10 | 0 | 0 | 0 | 0 | 0 | 7 | 298 | 149 | 1 | 1 | 1 | NA | 1 | 1 | 1 | 1 | 1 | NA | 1 | 0 | 0.25 |
| SE | 16631 | ENSG00000248508 | SRP14-AS1 | chr15 | + | 40059132 | 40059289 | 40045960 | 40046069 | 40060096 | 40060263 | 2 | 0 | 1 | 0 | 0 | 0 | 0 | 0 | 0 | 0 | 0 | 0 | 0 | 3 | 0 | 0 | 0 | 0 | 0 | 7 | 298 | 149 | 1 | 1 | 1 | NA | 1 | NA | NA | NA | NA | NA | NA | 0 | 1 |
| SE | 16632 | ENSG00000248508 | SRP14-AS1 | chr15 | + | 40059132 | 40059289 | 40056452 | 40056641 | 40060096 | 40060263 | 0 | 0 | 0 | 0 | 0 | 0 | 0 | 0 | 0 | 6 | 0 | 0 | 0 | 5 | 0 | 0 | 1 | 0 | 0 | 0 | 298 | 149 | 1 | 1 | NA | NA | NA | NA | 0 | NA | 0 | NA | 1 | NA | -0.5 |
| SE | 16634 | ENSG00000248508 | SRP14-AS1 | chr15 | + | 40060096 | 40060263 | 40045960 | 40046069 | 40064460 | 40064601 | 0 | 0 | 0 | 0 | 5 | 0 | 3 | 0 | 1 | 0 | 0 | 0 | 0 | 0 | 7 | 0 | 1 | 0 | 0 | 0 | 298 | 149 | 1 | 1 | NA | 0 | NA | 0 | 1 | NA | NA | NA | NA | 1 | -0.667 |
| SE | 16635 | ENSG00000248508 | SRP14-AS1 | chr15 | + | 40064295 | 40064601 | 40045960 | 40046069 | 40065220 | 40065705 | 3 | 5 | 1 | 9 | 15 | 7 | 4 | 2 | 2 | 16 | 0 | 6 | 0 | 0 | 3 | 0 | 9 | 0 | 0 | 0 | 298 | 149 | 1 | 1 | 0.176 | 0.385 | 0.2 | 0.692 | 0.319 | NA | 0.25 | NA | NA | 1 | -0.271 |
| SE | 16636 | ENSG00000248508 | SRP14-AS1 | chr15 | + | 40064460 | 40064601 | 40045960 | 40046069 | 40065220 | 40065705 | 3 | 7 | 1 | 9 | 15 | 7 | 4 | 2 | 2 | 16 | 0 | 4 | 0 | 0 | 0 | 0 | 9 | 0 | 0 | 0 | 290 | 149 | 1 | 1 | 0.18 | 0.473 | 0.204 | 0.698 | 0.325 | NA | 0.186 | NA | NA | NA | 0.19 |
| SE | 16659 | ENSG00000174444 | RPL4 | chr15 | - | 66500292 | 66500376 | 66500055 | 66500176 | 66500948 | 66501105 | 12187 | 7612 | 17237 | 14121 | 32289 | 0 | 0 | 0 | 0 | 0 | 12142 | 9214 | 9609 | 7255 | 14817 | 0 | 3 | 0 | 0 | 0 | 233 | 149 | 1 | 1 | 1 | 1 | 1 | 1 | 1 | 1 | 0.999 | 1 | 1 | 1 | 0 |
| SE | 16660 | ENSG00000174444 | RPL4 | chr15 | - | 66501787 | 66501912 | 66500948 | 66501504 | 66502611 | 66502750 | 15659 | 8907 | 21174 | 16159 | 32095 | 0 | 0 | 1 | 4 | 12 | 11990 | 9317 | 9453 | 6573 | 15748 | 0 | 0 | 0 | 0 | 4 | 274 | 149 | 1 | 1 | 1 | 1 | 1 | 1 | 0.999 | 1 | 1 | 1 | 1 | 1 | 0 |
| SE | 16661 | ENSG00000174444 | RPL4 | chr15 | - | 66502611 | 66502871 | 66501787 | 66501912 | 66503357 | 66503529 | 6093 | 3376 | 7861 | 5838 | 10332 | 0 | 0 | 0 | 1 | 0 | 3882 | 3394 | 3271 | 2267 | 5585 | 0 | 0 | 0 | 0 | 0 | 298 | 149 | 1 | 1 | 1 | 1 | 1 | 1 | 1 | 1 | 1 | 1 | 1 | 1 | 0 |
| SE | 16662 | ENSG00000174444 | RPL4 | chr15 | - | 66502611 | 66503164 | 66501787 | 66501912 | 66503357 | 66503529 | 7749 | 4123 | 9667 | 7237 | 12371 | 0 | 0 | 0 | 1 | 0 | 4642 | 4302 | 4027 | 2909 | 7193 | 0 | 0 | 0 | 0 | 0 | 298 | 149 | 1 | 1 | 1 | 1 | 1 | 1 | 1 | 1 | 1 | 1 | 1 | 1 | 0 |
| SE | 16663 | ENSG00000180304 | OAZ2 | chr15 | - | 64689083 | 64689169 | 64687572 | 64688834 | 64690373 | 64690549 | 186 | 134 | 123 | 161 | 171 | 4 | 0 | 1 | 1 | 2 | 239 | 173 | 104 | 152 | 223 | 0 | 0 | 0 | 0 | 0 | 235 | 149 | 1 | 1 | 0.967 | 1 | 0.987 | 0.99 | 0.982 | 1 | 1 | 1 | 1 | 1 | -0.015 |
| SE | 16666 | ENSG00000180304 | OAZ2 | chr15 | - | 64693074 | 64693181 | 64691433 | 64691591 | 64695963 | 64696050 | 1 | 0 | 2 | 0 | 0 | 0 | 8 | 0 | 0 | 0 | 0 | 0 | 0 | 10 | 0 | 0 | 0 | 4 | 0 | 0 | 256 | 149 | 1 | 1 | 1 | 0 | 1 | NA | NA | NA | NA | 0 | 1 | NA | 0.167 |
| SE | 16669 | ENSG00000180304 | OAZ2 | chr15 | - | 64695963 | 64696050 | 64693074 | 64693181 | 64703028 | 64703218 | 0 | 0 | 0 | 0 | 0 | 1 | 0 | 0 | 0 | 9 | 0 | 0 | 0 | 7 | 0 | 0 | 1 | 2 | 0 | 9 | 236 | 149 | 1 | 1 | 0 | NA | NA | NA | 0 | NA | 0 | 0 | 1 | 0 | -0.25 |
| SE | 16671 | ENSG00000103599 | IQCH | chr15 | + | 67344062 | 67344191 | 67261271 | 67261394 | 67357344 | 67357421 | 0 | 0 | 0 | 0 | 0 | 0 | 2 | 0 | 0 | 1 | 0 | 1 | 0 | 0 | 0 | 0 | 0 | 0 | 0 | 0 | 278 | 149 | 1 | 1 | NA | 0 | NA | NA | 0 | NA | 1 | NA | NA | NA | -1 |
| SE | 16672 | ENSG00000103599 | IQCH | chr15 | + | 67359846 | 67359885 | 67357344 | 67357421 | 67372110 | 67372444 | 1 | 1 | 1 | 0 | 0 | 2 | 0 | 0 | 0 | 0 | 0 | 0 | 0 | 0 | 1 | 0 | 0 | 0 | 0 | 0 | 188 | 149 | 1 | 1 | 0.284 | 1 | 1 | NA | NA | NA | NA | NA | NA | 1 | -0.239 |
| SE | 16678 | ENSG00000140464 | PML | chr15 | + | 73998003 | 73998476 | 73994672 | 73994941 | 74022827 | 74023408 | 29 | 5 | 21 | 32 | 3 | 0 | 0 | 0 | 0 | 0 | 103 | 56 | 22 | 57 | 67 | 0 | 0 | 0 | 1 | 0 | 298 | 149 | 1 | 1 | 1 | 1 | 1 | 1 | 1 | 1 | 1 | 1 | 0.966 | 1 | 0.007 |
| SE | 16682 | ENSG00000140464 | PML | chr15 | + | 74033155 | 74033414 | 74032571 | 74032715 | 74034477 | 74034530 | 52 | 35 | 64 | 52 | 40 | 0 | 1 | 0 | 0 | 0 | 75 | 55 | 32 | 132 | 77 | 0 | 0 | 0 | 0 | 0 | 298 | 149 | 1 | 1 | 1 | 0.946 | 1 | 1 | 1 | 1 | 1 | 1 | 1 | 1 | -0.011 |
| SE | 16684 | ENSG00000140464 | PML | chr15 | + | 74033155 | 74033422 | 74032571 | 74032715 | 74034477 | 74034530 | 25 | 14 | 20 | 18 | 21 | 0 | 1 | 0 | 0 | 0 | 43 | 22 | 12 | 43 | 16 | 0 | 0 | 0 | 0 | 0 | 298 | 149 | 1 | 1 | 1 | 0.875 | 1 | 1 | 1 | 1 | 1 | 1 | 1 | 1 | -0.025 |
| SE | 16693 | ENSG00000137841 | PLCB2 | chr15 | - | 40292338 | 40292443 | 40291873 | 40292158 | 40292925 | 40293025 | 338 | 184 | 212 | 256 | 274 | 0 | 0 | 0 | 0 | 0 | 339 | 353 | 187 | 272 | 524 | 0 | 1 | 4 | 0 | 0 | 254 | 149 | 1 | 1 | 1 | 1 | 1 | 1 | 1 | 1 | 0.995 | 0.965 | 1 | 1 | 0.008 |
| SE | 16694 | ENSG00000137841 | PLCB2 | chr15 | - | 40296521 | 40296634 | 40296295 | 40296392 | 40296745 | 40296908 | 350 | 230 | 272 | 295 | 414 | 8 | 0 | 0 | 0 | 1 | 515 | 309 | 232 | 386 | 606 | 0 | 0 | 1 | 3 | 3 | 262 | 149 | 1 | 1 | 0.961 | 1 | 1 | 1 | 0.996 | 1 | 1 | 0.992 | 0.987 | 0.991 | -0.003 |
| SE | 16696 | ENSG00000137841 | PLCB2 | chr15 | - | 40296745 | 40296908 | 40296295 | 40296392 | 40297520 | 40297605 | 201 | 110 | 107 | 187 | 104 | 0 | 0 | 0 | 0 | 0 | 361 | 189 | 146 | 307 | 425 | 0 | 1 | 0 | 8 | 0 | 298 | 149 | 1 | 1 | 1 | 1 | 1 | 1 | 1 | 1 | 0.99 | 1 | 0.95 | 1 | 0.012 |
| SE | 16697 | ENSG00000137841 | PLCB2 | chr15 | - | 40296757 | 40296908 | 40296295 | 40296392 | 40297520 | 40297605 | 194 | 110 | 107 | 187 | 103 | 0 | 0 | 0 | 0 | 0 | 361 | 189 | 146 | 304 | 437 | 0 | 1 | 0 | 8 | 0 | 298 | 149 | 1 | 1 | 1 | 1 | 1 | 1 | 1 | 1 | 0.99 | 1 | 0.95 | 1 | 0.012 |
| SE | 16698 | ENSG00000137841 | PLCB2 | chr15 | - | 40298797 | 40298964 | 40298561 | 40298708 | 40299127 | 40299228 | 918 | 604 | 713 | 1122 | 1307 | 0 | 0 | 2 | 0 | 1 | 1737 | 1140 | 817 | 1526 | 2195 | 0 | 0 | 2 | 0 | 0 | 298 | 149 | 1 | 1 | 1 | 1 | 0.994 | 1 | 0.998 | 1 | 1 | 0.995 | 1 | 1 | -0.001 |
| SE | 16699 | ENSG00000137841 | PLCB2 | chr15 | - | 40299127 | 40299228 | 40298797 | 40298964 | 40301956 | 40302032 | 544 | 326 | 427 | 654 | 763 | 0 | 0 | 0 | 0 | 0 | 872 | 656 | 434 | 787 | 1340 | 13 | 0 | 0 | 0 | 0 | 250 | 149 | 1 | 1 | 1 | 1 | 1 | 1 | 1 | 0.976 | 1 | 1 | 1 | 1 | 0.005 |
| SE | 16700 | ENSG00000137841 | PLCB2 | chr15 | - | 40302135 | 40302189 | 40301956 | 40302032 | 40302269 | 40302349 | 347 | 150 | 199 | 297 | 283 | 0 | 0 | 0 | 2 | 0 | 413 | 317 | 166 | 271 | 660 | 0 | 2 | 0 | 8 | 0 | 203 | 149 | 1 | 1 | 1 | 1 | 1 | 0.991 | 1 | 1 | 0.991 | 1 | 0.961 | 1 | 0.008 |
| SE | 16701 | ENSG00000137841 | PLCB2 | chr15 | - | 40302468 | 40302609 | 40302269 | 40302349 | 40303287 | 40303356 | 326 | 145 | 219 | 241 | 263 | 2 | 0 | 0 | 2 | 0 | 448 | 309 | 225 | 295 | 561 | 0 | 3 | 0 | 4 | 0 | 290 | 149 | 1 | 1 | 0.988 | 1 | 1 | 0.984 | 1 | 1 | 0.981 | 1 | 0.974 | 1 | 0.003 |
| SE | 16704 | ENSG00000166233 | ARIH1 | chr15 | + | 72518066 | 72518134 | 72496790 | 72496858 | 72555270 | 72555363 | 3 | 2 | 7 | 3 | 3 | 0 | 0 | 0 | 1 | 0 | 0 | 0 | 17 | 2 | 0 | 0 | 0 | 0 | 0 | 0 | 217 | 149 | 1 | 1 | 1 | 1 | 1 | 0.673 | 1 | NA | NA | 1 | 1 | NA | -0.065 |
| SE | 16706 | ENSG00000166233 | ARIH1 | chr15 | + | 72555270 | 72555363 | 72518066 | 72518134 | 72561482 | 72561547 | 0 | 4 | 0 | 2 | 2 | 0 | 0 | 0 | 0 | 0 | 1 | 3 | 1 | 4 | 0 | 0 | 0 | 1 | 0 | 0 | 242 | 149 | 1 | 1 | NA | 1 | NA | 1 | 1 | 1 | 1 | 0.381 | 1 | NA | 0.155 |
| SE | 16707 | ENSG00000166233 | ARIH1 | chr15 | + | 72555851 | 72555907 | 72555270 | 72555363 | 72561482 | 72561547 | 217 | 101 | 182 | 218 | 288 | 1 | 1 | 0 | 1 | 0 | 80 | 151 | 178 | 114 | 42 | 1 | 0 | 0 | 0 | 0 | 205 | 149 | 1 | 1 | 0.994 | 0.987 | 1 | 0.994 | 1 | 0.983 | 1 | 1 | 1 | 1 | -0.002 |
| SE | 16711 | ENSG00000166233 | ARIH1 | chr15 | + | 72572262 | 72572380 | 72572125 | 72572165 | 72580730 | 72580991 | 0 | 1 | 1 | 0 | 0 | 141 | 81 | 91 | 122 | 225 | 0 | 3 | 0 | 0 | 0 | 58 | 89 | 105 | 73 | 38 | 267 | 149 | 1 | 1 | 0 | 0.007 | 0.006 | 0 | 0 | 0 | 0.018 | 0 | 0 | 0 | -0.001 |
| SE | 16716 | ENSG00000214013 | GANC | chr15 | + | 42278481 | 42278590 | 42276271 | 42276410 | 42280916 | 42281112 | 66 | 52 | 39 | 53 | 106 | 0 | 0 | 0 | 1 | 0 | 100 | 49 | 25 | 89 | 58 | 0 | 0 | 0 | 0 | 0 | 258 | 149 | 1 | 1 | 1 | 1 | 1 | 0.968 | 1 | 1 | 1 | 1 | 1 | 1 | -0.006 |
| SE | 16717 | ENSG00000214013 | GANC | chr15 | + | 42278481 | 42278590 | 42276271 | 42276410 | 42283610 | 42283902 | 63 | 36 | 24 | 45 | 76 | 0 | 0 | 0 | 2 | 0 | 85 | 36 | 21 | 86 | 50 | 0 | 0 | 0 | 0 | 0 | 258 | 149 | 1 | 1 | 1 | 1 | 1 | 0.929 | 1 | 1 | 1 | 1 | 1 | 1 | -0.014 |
| SE | 16719 | ENSG00000214013 | GANC | chr15 | + | 42280916 | 42281133 | 42276347 | 42276410 | 42283610 | 42284389 | 37 | 21 | 26 | 24 | 57 | 0 | 0 | 0 | 2 | 0 | 0 | 26 | 15 | 11 | 36 | 0 | 0 | 0 | 0 | 0 | 298 | 149 | 1 | 1 | 1 | 1 | 1 | 0.857 | 1 | NA | 1 | 1 | 1 | 1 | -0.029 |
| SE | 16726 | ENSG00000214013 | GANC | chr15 | + | 42329305 | 42329449 | 42327362 | 42327442 | 42338388 | 42338490 | 31 | 32 | 34 | 48 | 58 | 0 | 0 | 0 | 0 | 0 | 65 | 34 | 22 | 47 | 28 | 0 | 0 | 2 | 0 | 0 | 293 | 149 | 1 | 1 | 1 | 1 | 1 | 1 | 1 | 1 | 1 | 0.848 | 1 | 1 | 0.03 |
| SE | 16727 | ENSG00000214013 | GANC | chr15 | + | 42330575 | 42330672 | 42327362 | 42327442 | 42338388 | 42338490 | 25 | 12 | 36 | 22 | 44 | 0 | 0 | 0 | 0 | 0 | 13 | 52 | 36 | 40 | 11 | 0 | 0 | 2 | 0 | 0 | 246 | 149 | 1 | 1 | 1 | 1 | 1 | 1 | 1 | 1 | 1 | 0.916 | 1 | 1 | 0.017 |
| SE | 16729 | ENSG00000214013 | GANC | chr15 | + | 42340689 | 42340754 | 42339668 | 42339912 | 42343077 | 42343154 | 26 | 34 | 28 | 40 | 15 | 0 | 0 | 0 | 1 | 0 | 41 | 38 | 33 | 61 | 19 | 0 | 0 | 0 | 0 | 0 | 214 | 149 | 1 | 1 | 1 | 1 | 1 | 0.965 | 1 | 1 | 1 | 1 | 1 | 1 | -0.007 |
| SE | 16732 | ENSG00000261801 | LOXL1-AS1 | chr15 | - | 73916342 | 73916447 | 73912184 | 73912303 | 73919837 | 73919870 | 2 | 2 | 4 | 5 | 2 | 0 | 0 | 3 | 0 | 0 | 0 | 0 | 1 | 0 | 0 | 0 | 0 | 4 | 0 | 0 | 254 | 149 | 1 | 1 | 1 | 1 | 0.439 | 1 | 1 | NA | NA | 0.128 | NA | NA | 0.76 |
| SE | 16733 | ENSG00000261801 | LOXL1-AS1 | chr15 | - | 73919837 | 73920038 | 73916303 | 73916447 | 73927776 | 73927940 | 2 | 2 | 3 | 3 | 2 | 1 | 0 | 0 | 0 | 0 | 0 | 0 | 1 | 0 | 0 | 0 | 0 | 0 | 0 | 0 | 298 | 149 | 1 | 1 | 0.5 | 1 | 1 | 1 | 1 | NA | NA | 1 | NA | NA | -0.1 |
| SE | 16738 | ENSG00000075131 | TIPIN | chr15 | - | 66349059 | 66349123 | 66341149 | 66341356 | 66351524 | 66351600 | 15 | 22 | 17 | 21 | 22 | 0 | 0 | 2 | 0 | 0 | 7 | 15 | 29 | 7 | 0 | 0 | 0 | 0 | 0 | 0 | 213 | 149 | 1 | 1 | 1 | 1 | 0.856 | 1 | 1 | 1 | 1 | 1 | 1 | NA | -0.029 |
| SE | 16739 | ENSG00000075131 | TIPIN | chr15 | - | 66349314 | 66349437 | 66341149 | 66341356 | 66351524 | 66351600 | 26 | 33 | 44 | 31 | 41 | 0 | 0 | 2 | 0 | 0 | 63 | 16 | 26 | 42 | 20 | 0 | 0 | 0 | 0 | 0 | 272 | 149 | 1 | 1 | 1 | 1 | 0.923 | 1 | 1 | 1 | 1 | 1 | 1 | 1 | -0.015 |
| SE | 16740 | ENSG00000075131 | TIPIN | chr15 | - | 66349314 | 66349437 | 66349059 | 66349123 | 66351524 | 66351600 | 41 | 41 | 65 | 54 | 71 | 0 | 0 | 0 | 0 | 0 | 83 | 35 | 56 | 63 | 29 | 0 | 0 | 0 | 1 | 0 | 272 | 149 | 1 | 1 | 1 | 1 | 1 | 1 | 1 | 1 | 1 | 1 | 0.972 | 1 | 0.006 |
| SE | 16742 | ENSG00000075131 | TIPIN | chr15 | - | 66352128 | 66352207 | 66349314 | 66349437 | 66352814 | 66352955 | 16 | 14 | 15 | 14 | 9 | 0 | 0 | 0 | 1 | 0 | 10 | 9 | 6 | 22 | 4 | 0 | 0 | 0 | 0 | 0 | 228 | 149 | 1 | 1 | 1 | 1 | 1 | 0.901 | 1 | 1 | 1 | 1 | 1 | 1 | -0.02 |
| SE | 16743 | ENSG00000125107 | CNOT1 | chr16 | - | 58525988 | 58526138 | 58525178 | 58525359 | 58528474 | 58528648 | 675 | 342 | 545 | 545 | 1010 | 0 | 0 | 0 | 1 | 0 | 348 | 426 | 403 | 438 | 110 | 0 | 0 | 0 | 0 | 0 | 298 | 149 | 1 | 1 | 1 | 1 | 1 | 0.996 | 1 | 1 | 1 | 1 | 1 | 1 | -0.001 |
| SE | 16744 | ENSG00000125107 | CNOT1 | chr16 | - | 58527922 | 58528105 | 58525988 | 58526138 | 58528474 | 58528648 | 1 | 0 | 0 | 2 | 0 | 279 | 127 | 222 | 241 | 475 | 0 | 0 | 0 | 0 | 0 | 160 | 206 | 160 | 184 | 33 | 298 | 149 | 1 | 1 | 0.002 | 0 | 0 | 0.004 | 0 | 0 | 0 | 0 | 0 | 0 | 0.001 |
| SE | 16745 | ENSG00000125107 | CNOT1 | chr16 | - | 58531957 | 58532075 | 58530245 | 58530347 | 58532231 | 58532395 | 292 | 197 | 259 | 251 | 330 | 0 | 0 | 0 | 0 | 0 | 97 | 172 | 184 | 225 | 48 | 0 | 4 | 0 | 0 | 0 | 267 | 149 | 1 | 1 | 1 | 1 | 1 | 1 | 1 | 1 | 0.96 | 1 | 1 | 1 | 0.008 |
| SE | 16746 | ENSG00000125107 | CNOT1 | chr16 | - | 58542230 | 58542335 | 58541500 | 58541620 | 58542427 | 58542568 | 229 | 190 | 265 | 278 | 255 | 0 | 0 | 1 | 0 | 0 | 82 | 280 | 121 | 144 | 28 | 0 | 0 | 0 | 0 | 0 | 254 | 149 | 1 | 1 | 1 | 1 | 1 | 0.994 | 1 | 1 | 1 | 1 | 1 | 1 | -0.001 |
| SE | 16749 | ENSG00000125107 | CNOT1 | chr16 | - | 58547185 | 58547296 | 58546671 | 58546749 | 58547565 | 58547682 | 768 | 541 | 640 | 648 | 1602 | 0 | 0 | 0 | 2 | 0 | 434 | 637 | 454 | 591 | 56 | 0 | 0 | 0 | 0 | 0 | 260 | 149 | 1 | 1 | 1 | 1 | 1 | 1 | 0.998 | 1 | 1 | 1 | 1 | 1 | 0 |
| SE | 16750 | ENSG00000125107 | CNOT1 | chr16 | - | 58547185 | 58547296 | 58546671 | 58546749 | 58549718 | 58549898 | 573 | 381 | 495 | 487 | 1329 | 0 | 1 | 0 | 1 | 0 | 354 | 473 | 354 | 444 | 32 | 0 | 3 | 0 | 2 | 0 | 260 | 149 | 1 | 1 | 1 | 0.995 | 1 | 0.996 | 1 | 1 | 0.989 | 1 | 0.992 | 1 | 0.002 |
| SE | 16751 | ENSG00000125107 | CNOT1 | chr16 | - | 58547565 | 58547682 | 58546671 | 58546749 | 58549718 | 58549898 | 239 | 152 | 176 | 223 | 806 | 0 | 1 | 0 | 1 | 0 | 98 | 199 | 100 | 240 | 44 | 0 | 3 | 0 | 2 | 0 | 266 | 149 | 1 | 1 | 1 | 0.988 | 1 | 0.992 | 1 | 1 | 0.974 | 1 | 0.985 | 1 | 0.004 |
| SE | 16752 | ENSG00000125107 | CNOT1 | chr16 | - | 58547565 | 58547682 | 58547185 | 58547296 | 58549718 | 58549898 | 443 | 316 | 325 | 389 | 1098 | 3 | 3 | 0 | 0 | 3 | 178 | 369 | 201 | 393 | 68 | 0 | 0 | 1 | 6 | 0 | 266 | 149 | 1 | 1 | 0.988 | 0.983 | 1 | 1 | 0.995 | 1 | 1 | 0.991 | 0.973 | 1 | 0 |
| SE | 16754 | ENSG00000125107 | CNOT1 | chr16 | - | 58551588 | 58551819 | 58549718 | 58549898 | 58553781 | 58553860 | 392 | 319 | 356 | 403 | 816 | 0 | 0 | 0 | 0 | 0 | 235 | 428 | 256 | 293 | 72 | 0 | 0 | 2 | 0 | 0 | 298 | 149 | 1 | 1 | 1 | 1 | 1 | 1 | 1 | 1 | 1 | 0.985 | 1 | 1 | 0.003 |
| SE | 16756 | ENSG00000125107 | CNOT1 | chr16 | - | 58555250 | 58555537 | 58551602 | 58551819 | 58555783 | 58555908 | 459 | 366 | 400 | 495 | 416 | 0 | 0 | 0 | 0 | 0 | 151 | 486 | 375 | 480 | 48 | 0 | 2 | 0 | 0 | 0 | 298 | 149 | 1 | 1 | 1 | 1 | 1 | 1 | 1 | 1 | 0.992 | 1 | 1 | 1 | 0.002 |
| SE | 16757 | ENSG00000125107 | CNOT1 | chr16 | - | 58556846 | 58556990 | 58555783 | 58555908 | 58558472 | 58558674 | 210 | 214 | 179 | 206 | 203 | 0 | 0 | 0 | 0 | 1 | 118 | 219 | 134 | 208 | 34 | 0 | 0 | 0 | 0 | 0 | 293 | 149 | 1 | 1 | 1 | 1 | 1 | 1 | 0.99 | 1 | 1 | 1 | 1 | 1 | -0.002 |
| SE | 16758 | ENSG00000125107 | CNOT1 | chr16 | - | 58556846 | 58556990 | 58555783 | 58555908 | 58560211 | 58560362 | 196 | 193 | 160 | 186 | 197 | 0 | 4 | 0 | 0 | 0 | 115 | 197 | 123 | 193 | 21 | 0 | 0 | 3 | 0 | 0 | 293 | 149 | 1 | 1 | 1 | 0.961 | 1 | 1 | 1 | 1 | 1 | 0.954 | 1 | 1 | 0.001 |
| SE | 16759 | ENSG00000125107 | CNOT1 | chr16 | - | 58556846 | 58556993 | 58555783 | 58555908 | 58558472 | 58558490 | 413 | 356 | 342 | 445 | 309 | 0 | 0 | 0 | 0 | 1 | 223 | 453 | 310 | 413 | 42 | 0 | 0 | 0 | 0 | 0 | 296 | 149 | 1 | 1 | 1 | 1 | 1 | 1 | 0.994 | 1 | 1 | 1 | 1 | 1 | -0.001 |
| SE | 16760 | ENSG00000125107 | CNOT1 | chr16 | - | 58556846 | 58556993 | 58555783 | 58555908 | 58560211 | 58560362 | 204 | 198 | 161 | 198 | 200 | 0 | 4 | 0 | 0 | 0 | 115 | 211 | 162 | 186 | 25 | 0 | 0 | 3 | 0 | 0 | 296 | 149 | 1 | 1 | 1 | 0.961 | 1 | 1 | 1 | 1 | 1 | 0.965 | 1 | 1 | -0.001 |
| SE | 16761 | ENSG00000125107 | CNOT1 | chr16 | - | 58556861 | 58556993 | 58555783 | 58555908 | 58558472 | 58558617 | 285 | 227 | 241 | 330 | 161 | 0 | 0 | 0 | 0 | 1 | 134 | 348 | 236 | 352 | 29 | 0 | 0 | 0 | 0 | 0 | 281 | 149 | 1 | 1 | 1 | 1 | 1 | 1 | 0.988 | 1 | 1 | 1 | 1 | 1 | -0.002 |
| SE | 16762 | ENSG00000125107 | CNOT1 | chr16 | - | 58556861 | 58556993 | 58555783 | 58555908 | 58560211 | 58560362 | 78 | 70 | 59 | 84 | 53 | 0 | 4 | 0 | 0 | 0 | 26 | 109 | 81 | 130 | 12 | 0 | 0 | 3 | 0 | 0 | 281 | 149 | 1 | 1 | 1 | 0.903 | 1 | 1 | 1 | 1 | 1 | 0.935 | 1 | 1 | -0.006 |
| SE | 16763 | ENSG00000125107 | CNOT1 | chr16 | - | 58558472 | 58558674 | 58555783 | 58555908 | 58560211 | 58560362 | 231 | 110 | 180 | 175 | 301 | 0 | 4 | 0 | 0 | 0 | 64 | 120 | 99 | 91 | 43 | 0 | 0 | 3 | 0 | 0 | 298 | 149 | 1 | 1 | 1 | 0.932 | 1 | 1 | 1 | 1 | 1 | 0.943 | 1 | 1 | -0.002 |
| SE | 16766 | ENSG00000125107 | CNOT1 | chr16 | - | 58574608 | 58574760 | 58560211 | 58560362 | 58575006 | 58575063 | 379 | 202 | 318 | 331 | 396 | 0 | 1 | 0 | 0 | 0 | 86 | 230 | 239 | 231 | 50 | 0 | 0 | 0 | 0 | 0 | 298 | 149 | 1 | 1 | 1 | 0.99 | 1 | 1 | 1 | 1 | 1 | 1 | 1 | 1 | -0.002 |
| SE | 16767 | ENSG00000125107 | CNOT1 | chr16 | - | 58576462 | 58576582 | 58575006 | 58575129 | 58578698 | 58578939 | 877 | 577 | 684 | 797 | 652 | 4 | 2 | 0 | 3 | 3 | 542 | 801 | 561 | 696 | 224 | 0 | 4 | 2 | 6 | 0 | 269 | 149 | 1 | 1 | 0.992 | 0.994 | 1 | 0.993 | 0.992 | 1 | 0.991 | 0.994 | 0.985 | 1 | 0 |
| SE | 16768 | ENSG00000125107 | CNOT1 | chr16 | - | 58578698 | 58578939 | 58576462 | 58576582 | 58580632 | 58580760 | 709 | 443 | 552 | 642 | 777 | 2 | 0 | 0 | 1 | 0 | 385 | 584 | 435 | 538 | 172 | 0 | 0 | 0 | 2 | 0 | 298 | 149 | 1 | 1 | 0.994 | 1 | 1 | 0.997 | 1 | 1 | 1 | 1 | 0.993 | 1 | 0 |
| SE | 16769 | ENSG00000125107 | CNOT1 | chr16 | - | 58582792 | 58582903 | 58581344 | 58581515 | 58585337 | 58585506 | 467 | 400 | 443 | 495 | 713 | 0 | 0 | 0 | 1 | 0 | 346 | 463 | 307 | 370 | 96 | 0 | 0 | 0 | 0 | 0 | 260 | 149 | 1 | 1 | 1 | 1 | 1 | 0.996 | 1 | 1 | 1 | 1 | 1 | 1 | -0.001 |
| SE | 16770 | ENSG00000125107 | CNOT1 | chr16 | - | 58583055 | 58583182 | 58582792 | 58582903 | 58585337 | 58585506 | 711 | 530 | 613 | 692 | 703 | 0 | 0 | 0 | 3 | 0 | 332 | 581 | 403 | 523 | 220 | 0 | 0 | 0 | 0 | 0 | 276 | 149 | 1 | 1 | 1 | 1 | 1 | 0.992 | 1 | 1 | 1 | 1 | 1 | 1 | -0.002 |
| SE | 16771 | ENSG00000125107 | CNOT1 | chr16 | - | 58585337 | 58585506 | 58583055 | 58583182 | 58586544 | 58586748 | 605 | 438 | 512 | 600 | 521 | 2 | 0 | 0 | 0 | 0 | 308 | 523 | 366 | 515 | 199 | 8 | 2 | 1 | 0 | 0 | 298 | 149 | 1 | 1 | 0.993 | 1 | 1 | 1 | 1 | 0.951 | 0.992 | 0.995 | 1 | 1 | 0.011 |
| SE | 16772 | ENSG00000125107 | CNOT1 | chr16 | - | 58586544 | 58586748 | 58585337 | 58585506 | 58587344 | 58587413 | 258 | 185 | 219 | 265 | 269 | 0 | 0 | 1 | 0 | 0 | 175 | 254 | 191 | 275 | 107 | 0 | 0 | 0 | 0 | 0 | 298 | 149 | 1 | 1 | 1 | 1 | 0.991 | 1 | 1 | 1 | 1 | 1 | 1 | 1 | -0.002 |
| SE | 16773 | ENSG00000125107 | CNOT1 | chr16 | - | 58587200 | 58587255 | 58586544 | 58586748 | 58587344 | 58587413 | 310 | 146 | 251 | 280 | 499 | 0 | 0 | 1 | 1 | 0 | 173 | 265 | 154 | 271 | 73 | 0 | 0 | 0 | 0 | 0 | 204 | 149 | 1 | 1 | 1 | 1 | 0.995 | 0.995 | 1 | 1 | 1 | 1 | 1 | 1 | -0.002 |
| SE | 16774 | ENSG00000125107 | CNOT1 | chr16 | - | 58587779 | 58587878 | 58587344 | 58587413 | 58588798 | 58588906 | 518 | 247 | 453 | 484 | 533 | 0 | 0 | 1 | 0 | 0 | 127 | 477 | 235 | 284 | 164 | 0 | 1 | 0 | 0 | 0 | 248 | 149 | 1 | 1 | 1 | 1 | 0.996 | 1 | 1 | 1 | 0.997 | 1 | 1 | 1 | 0 |
| SE | 16775 | ENSG00000125107 | CNOT1 | chr16 | - | 58588798 | 58588906 | 58587779 | 58587878 | 58599235 | 58599511 | 613 | 313 | 526 | 570 | 489 | 0 | 0 | 0 | 0 | 0 | 184 | 586 | 291 | 334 | 226 | 0 | 0 | 2 | 0 | 0 | 257 | 149 | 1 | 1 | 1 | 1 | 1 | 1 | 1 | 1 | 1 | 0.988 | 1 | 1 | 0.002 |
| SE | 16776 | ENSG00000125107 | CNOT1 | chr16 | - | 58599235 | 58599511 | 58588798 | 58588906 | 58629727 | 58629826 | 354 | 188 | 313 | 333 | 267 | 0 | 0 | 1 | 0 | 0 | 116 | 305 | 191 | 227 | 147 | 0 | 0 | 1 | 0 | 0 | 298 | 149 | 1 | 1 | 1 | 1 | 0.994 | 1 | 1 | 1 | 1 | 0.99 | 1 | 1 | 0.001 |
| SE | 16777 | ENSG00000125107 | CNOT1 | chr16 | - | 58623281 | 58623409 | 58588820 | 58588906 | 58629727 | 58629885 | 2 | 4 | 12 | 14 | 2 | 0 | 0 | 1 | 0 | 0 | 22 | 5 | 4 | 1 | 7 | 0 | 0 | 1 | 0 | 0 | 277 | 149 | 1 | 1 | 1 | 1 | 0.866 | 1 | 1 | 1 | 1 | 0.683 | 1 | 1 | 0.037 |
| SE | 16778 | ENSG00000125107 | CNOT1 | chr16 | - | 58623281 | 58623409 | 58599262 | 58599511 | 58624652 | 58624794 | 7 | 1 | 14 | 7 | 2 | 0 | 3 | 7 | 3 | 3 | 2 | 31 | 9 | 7 | 0 | 0 | 1 | 3 | 2 | 0 | 277 | 149 | 1 | 1 | 1 | 0.152 | 0.518 | 0.557 | 0.264 | 1 | 0.943 | 0.617 | 0.653 | NA | -0.305 |
| SE | 16779 | ENSG00000125107 | CNOT1 | chr16 | - | 58623281 | 58623409 | 58599262 | 58599511 | 58624672 | 58624794 | 2 | 1 | 4 | 4 | 2 | 4 | 2 | 2 | 5 | 2 | 2 | 23 | 8 | 6 | 0 | 0 | 0 | 0 | 0 | 0 | 277 | 149 | 1 | 1 | 0.212 | 0.212 | 0.518 | 0.301 | 0.35 | 1 | 1 | 1 | 1 | NA | -0.681 |
| SE | 16788 | ENSG00000241839 | PLEKHO2 | chr15 | + | 64854920 | 64855037 | 64848592 | 64848742 | 64859893 | 64859998 | 417 | 226 | 264 | 361 | 246 | 0 | 0 | 0 | 0 | 0 | 854 | 457 | 240 | 834 | 767 | 0 | 0 | 0 | 1 | 0 | 266 | 149 | 1 | 1 | 1 | 1 | 1 | 1 | 1 | 1 | 1 | 1 | 0.998 | 1 | 0 |
| SE | 16794 | ENSG00000268895 | A1BG-AS1 | chr19 | + | 58353320 | 58353474 | 58347750 | 58347844 | 58353713 | 58353857 | 7 | 2 | 5 | 8 | 6 | 1 | 1 | 0 | 0 | 0 | 0 | 4 | 5 | 10 | 9 | 0 | 0 | 0 | 0 | 2 | 298 | 149 | 1 | 1 | 0.778 | 0.5 | 1 | 1 | 1 | NA | 1 | 1 | 1 | 0.692 | -0.067 |
| SE | 16795 | ENSG00000268895 | A1BG-AS1 | chr19 | + | 58353378 | 58353474 | 58347752 | 58347844 | 58353713 | 58353857 | 7 | 2 | 5 | 8 | 6 | 1 | 1 | 0 | 0 | 0 | 0 | 4 | 5 | 11 | 9 | 0 | 0 | 0 | 0 | 2 | 245 | 149 | 1 | 1 | 0.81 | 0.549 | 1 | 1 | 1 | NA | 1 | 1 | 1 | 0.732 | -0.061 |
| SE | 16800 | ENSG00000276141 | WHAMMP3 | chr15 | + | 22674420 | 22674586 | 22672689 | 22672859 | 22678196 | 22678383 | 67 | 18 | 75 | 55 | 43 | 0 | 1 | 0 | 0 | 0 | 7 | 26 | 26 | 8 | 3 | 0 | 0 | 3 | 0 | 0 | 298 | 149 | 1 | 1 | 1 | 0.9 | 1 | 1 | 1 | 1 | 1 | 0.812 | 1 | 1 | 0.018 |
| SE | 16801 | ENSG00000276141 | WHAMMP3 | chr15 | + | 22676804 | 22676934 | 22672689 | 22672859 | 22678196 | 22678383 | 11 | 6 | 14 | 10 | 19 | 0 | 1 | 0 | 0 | 0 | 0 | 8 | 11 | 6 | 9 | 0 | 0 | 3 | 0 | 0 | 279 | 149 | 1 | 1 | 1 | 0.762 | 1 | 1 | 1 | NA | 1 | 0.662 | 1 | 1 | 0.037 |
| SE | 16802 | ENSG00000276141 | WHAMMP3 | chr15 | + | 22676804 | 22676934 | 22674420 | 22674586 | 22678196 | 22678383 | 24 | 15 | 14 | 21 | 36 | 27 | 14 | 23 | 4 | 11 | 0 | 14 | 13 | 13 | 12 | 0 | 6 | 22 | 5 | 0 | 279 | 149 | 1 | 1 | 0.322 | 0.364 | 0.245 | 0.737 | 0.636 | NA | 0.555 | 0.24 | 0.581 | 1 | -0.133 |
| SE | 16804 | ENSG00000137814 | HAUS2 | chr15 | + | 42558197 | 42558290 | 42548827 | 42548965 | 42561269 | 42561402 | 8 | 11 | 13 | 16 | 21 | 0 | 0 | 0 | 0 | 0 | 19 | 22 | 15 | 31 | 0 | 0 | 0 | 0 | 3 | 0 | 242 | 149 | 1 | 1 | 1 | 1 | 1 | 1 | 1 | 1 | 1 | 1 | 0.864 | NA | 0.034 |
| SE | 16805 | ENSG00000137814 | HAUS2 | chr15 | + | 42559338 | 42559408 | 42548827 | 42548965 | 42561269 | 42561402 | 35 | 25 | 24 | 31 | 89 | 0 | 0 | 0 | 0 | 0 | 1 | 34 | 22 | 44 | 17 | 0 | 0 | 0 | 3 | 0 | 219 | 149 | 1 | 1 | 1 | 1 | 1 | 1 | 1 | 1 | 1 | 1 | 0.909 | 1 | 0.018 |
| SE | 16806 | ENSG00000137814 | HAUS2 | chr15 | + | 42559338 | 42559408 | 42558197 | 42558290 | 42560781 | 42560870 | 36 | 20 | 33 | 27 | 56 | 0 | 0 | 0 | 0 | 1 | 49 | 40 | 28 | 42 | 16 | 0 | 0 | 0 | 0 | 3 | 219 | 149 | 1 | 1 | 1 | 1 | 1 | 1 | 0.974 | 1 | 1 | 1 | 1 | 0.784 | 0.038 |
| SE | 16812 | ENSG00000137814 | HAUS2 | chr15 | + | 42561269 | 42561402 | 42559338 | 42559408 | 42563748 | 42563857 | 157 | 135 | 148 | 102 | 393 | 0 | 0 | 0 | 0 | 4 | 78 | 133 | 145 | 157 | 44 | 0 | 0 | 0 | 0 | 0 | 282 | 149 | 1 | 1 | 1 | 1 | 1 | 1 | 0.981 | 1 | 1 | 1 | 1 | 1 | -0.004 |
| SE | 16817 | ENSG00000166200 | COPS2 | chr15 | - | 49139527 | 49139653 | 49137347 | 49137437 | 49144226 | 49144304 | 504 | 479 | 461 | 507 | 1213 | 0 | 0 | 0 | 0 | 9 | 25 | 344 | 166 | 57 | 26 | 0 | 0 | 0 | 0 | 0 | 275 | 149 | 1 | 1 | 1 | 1 | 1 | 1 | 0.986 | 1 | 1 | 1 | 1 | 1 | -0.003 |
| SE | 16818 | ENSG00000166200 | COPS2 | chr15 | - | 49139527 | 49139653 | 49137347 | 49137458 | 49144226 | 49144304 | 398 | 370 | 363 | 371 | 988 | 0 | 0 | 1 | 4 | 0 | 18 | 227 | 120 | 51 | 20 | 0 | 0 | 0 | 0 | 0 | 275 | 149 | 1 | 1 | 1 | 1 | 0.995 | 0.98 | 1 | 1 | 1 | 1 | 1 | 1 | -0.005 |
| SE | 16822 | ENSG00000137878 | GCOM1 | chr15 | + | 57621607 | 57621700 | 57618032 | 57618188 | 57625778 | 57625892 | 21 | 16 | 12 | 52 | 20 | 0 | 0 | 0 | 0 | 1 | 47 | 184 | 12 | 72 | 513 | 0 | 0 | 0 | 0 | 0 | 242 | 149 | 1 | 1 | 1 | 1 | 1 | 1 | 0.925 | 1 | 1 | 1 | 1 | 1 | -0.015 |
| SE | 16825 | ENSG00000137878 | GCOM1 | chr15 | + | 57625778 | 57625892 | 57621607 | 57621700 | 57629701 | 57629854 | 23 | 20 | 5 | 62 | 32 | 0 | 2 | 0 | 0 | 0 | 56 | 252 | 25 | 102 | 701 | 0 | 1 | 0 | 0 | 0 | 263 | 149 | 1 | 1 | 1 | 0.85 | 1 | 1 | 1 | 1 | 0.993 | 1 | 1 | 1 | -0.029 |
| SE | 16829 | ENSG00000137878 | GCOM1 | chr15 | + | 57661449 | 57661533 | 57646136 | 57646669 | 57684401 | 57684674 | 0 | 1 | 0 | 0 | 0 | 0 | 0 | 0 | 0 | 0 | 0 | 6 | 0 | 1 | 0 | 0 | 2 | 0 | 0 | 0 | 233 | 149 | 1 | 1 | NA | 1 | NA | NA | NA | NA | 0.657 | NA | 1 | NA | 0.171 |
| SE | 16831 | ENSG00000137878 | GCOM1 | chr15 | + | 57674967 | 57675068 | 57661449 | 57661533 | 57684401 | 57684674 | 33 | 22 | 10 | 42 | 46 | 0 | 0 | 0 | 0 | 0 | 73 | 117 | 22 | 31 | 408 | 0 | 2 | 0 | 1 | 0 | 250 | 149 | 1 | 1 | 1 | 1 | 1 | 1 | 1 | 1 | 0.972 | 1 | 0.949 | 1 | 0.016 |
| SE | 16840 | ENSG00000137878 | GCOM1 | chr15 | + | 57708713 | 57709358 | 57706491 | 57706955 | 57711983 | 57712188 | 214 | 108 | 168 | 170 | 306 | 0 | 0 | 0 | 1 | 0 | 111 | 116 | 95 | 143 | 52 | 0 | 1 | 0 | 2 | 0 | 298 | 149 | 1 | 1 | 1 | 1 | 1 | 0.988 | 1 | 1 | 0.983 | 1 | 0.973 | 1 | 0.006 |
| SE | 16841 | ENSG00000137878 | GCOM1 | chr15 | + | 57709184 | 57709358 | 57706491 | 57706955 | 57711983 | 57712188 | 156 | 86 | 111 | 121 | 211 | 0 | 0 | 0 | 1 | 0 | 45 | 75 | 65 | 114 | 50 | 0 | 1 | 0 | 2 | 0 | 298 | 149 | 1 | 1 | 1 | 1 | 1 | 0.984 | 1 | 1 | 0.974 | 1 | 0.966 | 1 | 0.009 |
| SE | 16845 | ENSG00000151575 | TEX9 | chr15 | + | 56389317 | 56389400 | 56383951 | 56384031 | 56391242 | 56391385 | 5 | 14 | 3 | 8 | 12 | 1 | 0 | 0 | 0 | 2 | 10 | 2 | 8 | 12 | 13 | 0 | 0 | 1 | 0 | 0 | 232 | 149 | 1 | 1 | 0.763 | 1 | 1 | 1 | 0.794 | 1 | 1 | 0.837 | 1 | 1 | -0.056 |
| SE | 16846 | ENSG00000151575 | TEX9 | chr15 | + | 56389317 | 56389400 | 56383951 | 56384031 | 56412301 | 56412436 | 0 | 3 | 1 | 0 | 2 | 0 | 1 | 0 | 0 | 0 | 0 | 0 | 1 | 0 | 0 | 0 | 0 | 0 | 0 | 4 | 232 | 149 | 1 | 1 | NA | 0.658 | 1 | NA | 1 | NA | NA | 1 | NA | 0 | 0.386 |
| SE | 16847 | ENSG00000151575 | TEX9 | chr15 | + | 56391242 | 56391418 | 56383951 | 56384031 | 56412301 | 56412436 | 3 | 0 | 4 | 0 | 8 | 0 | 1 | 0 | 0 | 0 | 0 | 0 | 1 | 0 | 0 | 0 | 0 | 0 | 0 | 4 | 298 | 149 | 1 | 1 | 1 | 0 | 1 | NA | 1 | NA | NA | 1 | NA | 0 | 0.25 |
| SE | 16848 | ENSG00000151575 | TEX9 | chr15 | + | 56391242 | 56391418 | 56389317 | 56389400 | 56394164 | 56394247 | 41 | 49 | 30 | 46 | 68 | 0 | 0 | 1 | 0 | 0 | 60 | 45 | 22 | 33 | 55 | 0 | 0 | 0 | 0 | 0 | 298 | 149 | 1 | 1 | 1 | 1 | 0.938 | 1 | 1 | 1 | 1 | 1 | 1 | 1 | -0.012 |
| SE | 16850 | ENSG00000151575 | TEX9 | chr15 | + | 56427604 | 56427739 | 56412301 | 56412436 | 56428366 | 56428473 | 4 | 15 | 19 | 12 | 26 | 2 | 0 | 0 | 0 | 0 | 0 | 16 | 2 | 0 | 17 | 0 | 1 | 0 | 0 | 0 | 284 | 149 | 1 | 1 | 0.512 | 1 | 1 | 1 | 1 | NA | 0.894 | 1 | NA | 1 | -0.062 |
| SE | 16854 | ENSG00000174442 | ZWILCH | chr15 | + | 66518878 | 66519078 | 66515525 | 66515644 | 66520589 | 66520660 | 14 | 9 | 14 | 19 | 61 | 0 | 0 | 0 | 0 | 2 | 0 | 30 | 17 | 5 | 19 | 0 | 0 | 0 | 0 | 0 | 298 | 149 | 1 | 1 | 1 | 1 | 1 | 1 | 0.938 | NA | 1 | 1 | 1 | 1 | -0.012 |
| SE | 16856 | ENSG00000174442 | ZWILCH | chr15 | + | 66527289 | 66527383 | 66523510 | 66523748 | 66527856 | 66527912 | 14 | 3 | 6 | 5 | 18 | 0 | 0 | 1 | 0 | 0 | 0 | 0 | 4 | 0 | 0 | 0 | 0 | 0 | 0 | 0 | 243 | 149 | 1 | 1 | 1 | 1 | 0.786 | 1 | 1 | NA | NA | 1 | NA | NA | -0.043 |
| SE | 16857 | ENSG00000174442 | ZWILCH | chr15 | + | 66529493 | 66529573 | 66528851 | 66528957 | 66532246 | 66532403 | 28 | 14 | 21 | 35 | 47 | 0 | 0 | 1 | 0 | 0 | 0 | 12 | 26 | 2 | 8 | 0 | 0 | 0 | 0 | 0 | 229 | 149 | 1 | 1 | 1 | 1 | 0.932 | 1 | 1 | NA | 1 | 1 | 1 | 1 | -0.014 |
| SE | 16858 | ENSG00000174442 | ZWILCH | chr15 | + | 66540097 | 66540210 | 66537167 | 66537263 | 66546590 | 66546705 | 65 | 29 | 34 | 47 | 53 | 3 | 0 | 0 | 0 | 0 | 17 | 17 | 39 | 15 | 19 | 0 | 0 | 0 | 0 | 0 | 262 | 149 | 1 | 1 | 0.925 | 1 | 1 | 1 | 1 | 1 | 1 | 1 | 1 | 1 | -0.015 |
| SE | 16860 | ENSG00000138613 | APH1B | chr15 | + | 63286557 | 63286628 | 63279160 | 63279331 | 63302344 | 63302472 | 34 | 27 | 22 | 22 | 37 | 0 | 1 | 0 | 0 | 0 | 25 | 62 | 41 | 44 | 34 | 0 | 0 | 2 | 0 | 0 | 220 | 149 | 1 | 1 | 1 | 0.948 | 1 | 1 | 1 | 1 | 1 | 0.933 | 1 | 1 | 0.003 |
| SE | 16861 | ENSG00000138613 | APH1B | chr15 | + | 63286557 | 63286628 | 63279160 | 63279331 | 63305613 | 63305735 | 23 | 20 | 11 | 19 | 27 | 1 | 0 | 0 | 0 | 0 | 23 | 50 | 26 | 36 | 29 | 0 | 0 | 0 | 0 | 0 | 220 | 149 | 1 | 1 | 0.94 | 1 | 1 | 1 | 1 | 1 | 1 | 1 | 1 | 1 | -0.012 |
| SE | 16864 | ENSG00000138613 | APH1B | chr15 | + | 63302344 | 63302472 | 63279160 | 63279331 | 63305613 | 63305735 | 15 | 10 | 12 | 9 | 13 | 1 | 0 | 0 | 0 | 0 | 23 | 35 | 9 | 20 | 19 | 0 | 0 | 0 | 0 | 0 | 277 | 149 | 1 | 1 | 0.89 | 1 | 1 | 1 | 1 | 1 | 1 | 1 | 1 | 1 | -0.022 |
| SE | 16867 | ENSG00000166734 | GOLM2 | chr15 | + | 44319406 | 44319553 | 44289158 | 44289356 | 44322964 | 44323019 | 0 | 1 | 1 | 1 | 4 | 180 | 115 | 120 | 120 | 90 | 0 | 7 | 1 | 0 | 0 | 111 | 150 | 149 | 135 | 105 | 296 | 149 | 1 | 1 | 0 | 0.004 | 0.004 | 0.004 | 0.022 | 0 | 0.023 | 0.003 | 0 | 0 | 0.002 |
| SE | 16869 | ENSG00000166734 | GOLM2 | chr15 | + | 44328684 | 44328787 | 44322964 | 44323019 | 44331987 | 44332059 | 220 | 183 | 187 | 168 | 200 | 4 | 0 | 0 | 0 | 0 | 247 | 168 | 189 | 199 | 159 | 0 | 1 | 0 | 2 | 0 | 252 | 149 | 1 | 1 | 0.97 | 1 | 1 | 1 | 1 | 1 | 0.99 | 1 | 0.983 | 1 | -0.001 |
| SE | 16871 | ENSG00000166734 | GOLM2 | chr15 | + | 44338236 | 44338317 | 44337762 | 44337907 | 44379689 | 44379788 | 714 | 534 | 579 | 725 | 2006 | 2 | 0 | 1 | 3 | 8 | 504 | 544 | 585 | 584 | 198 | 0 | 8 | 6 | 3 | 0 | 230 | 149 | 1 | 1 | 0.996 | 1 | 0.997 | 0.994 | 0.994 | 1 | 0.978 | 0.984 | 0.992 | 1 | 0.005 |
| SE | 16875 | ENSG00000051180 | RAD51 | chr15 | + | 40706176 | 40706294 | 40701063 | 40701201 | 40709024 | 40709116 | 9 | 4 | 12 | 25 | 16 | 0 | 0 | 0 | 0 | 0 | 69 | 11 | 30 | 50 | 0 | 0 | 0 | 2 | 3 | 0 | 267 | 149 | 1 | 1 | 1 | 1 | 1 | 1 | 1 | 1 | 1 | 0.893 | 0.903 | NA | 0.051 |
| SE | 16877 | ENSG00000051180 | RAD51 | chr15 | + | 40709024 | 40709116 | 40701063 | 40701201 | 40718804 | 40718827 | 0 | 0 | 7 | 0 | 3 | 0 | 0 | 0 | 0 | 0 | 0 | 20 | 3 | 9 | 0 | 0 | 0 | 0 | 2 | 0 | 241 | 149 | 1 | 1 | NA | NA | 1 | NA | 1 | NA | 1 | 1 | 0.736 | NA | 0.088 |
| SE | 16878 | ENSG00000051180 | RAD51 | chr15 | + | 40718804 | 40718899 | 40709024 | 40709116 | 40728710 | 40728824 | 3 | 1 | 18 | 8 | 16 | 0 | 1 | 1 | 7 | 0 | 26 | 37 | 13 | 26 | 0 | 22 | 0 | 6 | 3 | 0 | 244 | 149 | 1 | 1 | 1 | 0.379 | 0.917 | 0.411 | 1 | 0.419 | 1 | 0.57 | 0.841 | NA | 0.034 |
| SE | 16879 | ENSG00000051180 | RAD51 | chr15 | + | 40729504 | 40729634 | 40728710 | 40728824 | 40729852 | 40729974 | 4 | 0 | 12 | 4 | 6 | 0 | 0 | 0 | 0 | 0 | 54 | 32 | 6 | 35 | 4 | 0 | 0 | 0 | 0 | 7 | 279 | 149 | 1 | 1 | 1 | NA | 1 | 1 | 1 | 1 | 1 | 1 | 1 | 0.234 | 0.153 |
| SE | 16883 | ENSG00000131473 | ACLY | chr17 | - | 41868708 | 41868785 | 41866916 | 41867904 | 41869042 | 41869125 | 952 | 542 | 600 | 883 | 1170 | 0 | 0 | 1 | 2 | 2 | 1230 | 765 | 613 | 873 | 1075 | 2 | 0 | 1 | 1 | 0 | 226 | 149 | 1 | 1 | 1 | 1 | 0.997 | 0.997 | 0.997 | 0.998 | 1 | 0.998 | 0.998 | 1 | -0.001 |
| SE | 16884 | ENSG00000131473 | ACLY | chr17 | - | 41869042 | 41869125 | 41866916 | 41867904 | 41871688 | 41871832 | 3 | 1 | 2 | 7 | 2 | 0 | 0 | 2 | 0 | 2 | 10 | 0 | 1 | 1 | 0 | 0 | 3 | 1 | 4 | 0 | 232 | 149 | 1 | 1 | 1 | 1 | 0.391 | 1 | 0.391 | 1 | 0 | 0.391 | 0.138 | NA | 0.374 |
| SE | 16886 | ENSG00000131473 | ACLY | chr17 | - | 41887598 | 41887703 | 41886111 | 41886308 | 41892278 | 41892447 | 346 | 145 | 233 | 259 | 195 | 1 | 0 | 0 | 0 | 0 | 262 | 379 | 205 | 262 | 485 | 0 | 0 | 0 | 0 | 0 | 254 | 149 | 1 | 1 | 0.995 | 1 | 1 | 1 | 1 | 1 | 1 | 1 | 1 | 1 | -0.001 |
| SE | 16888 | ENSG00000131473 | ACLY | chr17 | - | 41896619 | 41896649 | 41893032 | 41893174 | 41898630 | 41898785 | 38 | 14 | 40 | 31 | 32 | 0 | 0 | 0 | 0 | 0 | 81 | 37 | 9 | 29 | 48 | 0 | 4 | 0 | 0 | 0 | 179 | 149 | 1 | 1 | 1 | 1 | 1 | 1 | 1 | 1 | 0.885 | 1 | 1 | 1 | 0.023 |
| SE | 16889 | ENSG00000131473 | ACLY | chr17 | - | 41897748 | 41897839 | 41893032 | 41893174 | 41898630 | 41898785 | 131 | 61 | 95 | 103 | 60 | 0 | 0 | 0 | 0 | 0 | 163 | 193 | 85 | 101 | 207 | 0 | 4 | 0 | 0 | 0 | 240 | 149 | 1 | 1 | 1 | 1 | 1 | 1 | 1 | 1 | 0.968 | 1 | 1 | 1 | 0.006 |
| SE | 16890 | ENSG00000131473 | ACLY | chr17 | - | 41897748 | 41897839 | 41896619 | 41896649 | 41898630 | 41898785 | 329 | 116 | 229 | 240 | 152 | 1 | 0 | 0 | 0 | 4 | 396 | 371 | 199 | 239 | 534 | 0 | 0 | 3 | 0 | 0 | 240 | 149 | 1 | 1 | 0.995 | 1 | 1 | 1 | 0.959 | 1 | 1 | 0.976 | 1 | 1 | -0.004 |
| SE | 16891 | ENSG00000131473 | ACLY | chr17 | - | 41898630 | 41898785 | 41897748 | 41897839 | 41904728 | 41904790 | 172 | 71 | 127 | 135 | 97 | 0 | 0 | 0 | 2 | 0 | 218 | 214 | 92 | 146 | 266 | 0 | 0 | 0 | 0 | 0 | 298 | 149 | 1 | 1 | 1 | 1 | 1 | 0.971 | 1 | 1 | 1 | 1 | 1 | 1 | -0.006 |
| SE | 16892 | ENSG00000131473 | ACLY | chr17 | - | 41901695 | 41901813 | 41898630 | 41898785 | 41904728 | 41904790 | 365 | 221 | 298 | 388 | 194 | 0 | 1 | 2 | 1 | 0 | 414 | 468 | 183 | 276 | 543 | 0 | 5 | 1 | 0 | 5 | 267 | 149 | 1 | 1 | 1 | 0.992 | 0.988 | 0.995 | 1 | 1 | 0.981 | 0.99 | 1 | 0.984 | 0.004 |
| SE | 16893 | ENSG00000131473 | ACLY | chr17 | - | 41905521 | 41905658 | 41904728 | 41904790 | 41918879 | 41918951 | 146 | 101 | 120 | 167 | 78 | 0 | 0 | 0 | 0 | 0 | 171 | 168 | 120 | 139 | 216 | 0 | 0 | 1 | 0 | 0 | 286 | 149 | 1 | 1 | 1 | 1 | 1 | 1 | 1 | 1 | 1 | 0.984 | 1 | 1 | 0.003 |
| SE | 16894 | ENSG00000170113 | NIPA1 | chr15 | + | 22812162 | 22812253 | 22810748 | 22810796 | 22814082 | 22814153 | 11 | 1 | 8 | 4 | 2 | 0 | 0 | 0 | 0 | 2 | 0 | 0 | 0 | 9 | 9 | 0 | 0 | 0 | 0 | 0 | 240 | 149 | 1 | 1 | 1 | 1 | 1 | 1 | 0.383 | NA | NA | NA | 1 | 1 | -0.123 |
| SE | 16895 | ENSG00000170113 | NIPA1 | chr15 | + | 22812162 | 22812253 | 22810748 | 22810796 | 22820312 | 22820473 | 20 | 4 | 16 | 7 | 7 | 0 | 0 | 0 | 1 | 0 | 0 | 5 | 3 | 3 | 19 | 0 | 0 | 1 | 3 | 0 | 240 | 149 | 1 | 1 | 1 | 1 | 1 | 0.813 | 1 | NA | 1 | 0.651 | 0.383 | 1 | 0.204 |
| SE | 16896 | ENSG00000170113 | NIPA1 | chr15 | + | 22813632 | 22813714 | 22812162 | 22812253 | 22814082 | 22814153 | 5 | 0 | 0 | 0 | 0 | 3 | 0 | 5 | 1 | 0 | 0 | 2 | 0 | 0 | 0 | 0 | 0 | 0 | 6 | 7 | 231 | 149 | 1 | 1 | 0.518 | NA | 0 | 0 | NA | NA | 1 | NA | 0 | 0 | -0.161 |
| SE | 16897 | ENSG00000170113 | NIPA1 | chr15 | + | 22813632 | 22813714 | 22812162 | 22812253 | 22820312 | 22820473 | 1 | 0 | 0 | 0 | 0 | 12 | 3 | 13 | 4 | 5 | 0 | 0 | 0 | 0 | 0 | 0 | 5 | 3 | 0 | 20 | 231 | 149 | 1 | 1 | 0.051 | 0 | 0 | 0 | 0 | NA | 1 | 0 | NA | 0 | 0.01 |
| SE | 16898 | ENSG00000170113 | NIPA1 | chr15 | + | 22814082 | 22814153 | 22810748 | 22810796 | 22820312 | 22820473 | 4 | 2 | 3 | 0 | 2 | 0 | 0 | 0 | 1 | 0 | 0 | 3 | 1 | 0 | 0 | 0 | 0 | 1 | 3 | 0 | 220 | 149 | 1 | 1 | 1 | 1 | 1 | 0 | 1 | NA | 1 | 0.404 | 0 | NA | 0.332 |
| SE | 16899 | ENSG00000170113 | NIPA1 | chr15 | + | 22814082 | 22814153 | 22812162 | 22812253 | 22820312 | 22820473 | 7 | 2 | 8 | 1 | 0 | 12 | 3 | 13 | 4 | 5 | 0 | 3 | 1 | 6 | 0 | 0 | 5 | 3 | 0 | 20 | 220 | 149 | 1 | 1 | 0.283 | 0.311 | 0.294 | 0.145 | 0 | NA | 0.289 | 0.184 | 1 | 0 | -0.162 |
| SE | 16901 | ENSG00000170113 | NIPA1 | chr15 | + | 22819287 | 22819416 | 22814082 | 22814153 | 22820312 | 22820473 | 1 | 0 | 0 | 0 | 1 | 7 | 2 | 7 | 1 | 2 | 0 | 0 | 0 | 0 | 0 | 0 | 6 | 1 | 0 | 0 | 278 | 149 | 1 | 1 | 0.071 | 0 | 0 | 0 | 0.211 | NA | 0 | 0 | NA | NA | 0.056 |
| SE | 16902 | ENSG00000141564 | RPTOR | chr17 | + | 80625690 | 80625793 | 80544818 | 80545791 | 80643727 | 80643810 | 158 | 109 | 169 | 157 | 202 | 0 | 0 | 0 | 0 | 1 | 212 | 191 | 114 | 150 | 294 | 0 | 0 | 0 | 0 | 0 | 252 | 149 | 1 | 1 | 1 | 1 | 1 | 1 | 0.992 | 1 | 1 | 1 | 1 | 1 | -0.002 |
| SE | 16906 | ENSG00000141564 | RPTOR | chr17 | + | 80707840 | 80707999 | 80643727 | 80643810 | 80730559 | 80730706 | 86 | 75 | 90 | 91 | 82 | 0 | 1 | 1 | 0 | 0 | 97 | 122 | 73 | 118 | 210 | 0 | 0 | 0 | 0 | 0 | 298 | 149 | 1 | 1 | 1 | 0.974 | 0.978 | 1 | 1 | 1 | 1 | 1 | 1 | 1 | -0.01 |
| SE | 16907 | ENSG00000141564 | RPTOR | chr17 | + | 80730559 | 80730706 | 80707840 | 80707999 | 80754009 | 80754104 | 73 | 49 | 70 | 74 | 45 | 1 | 0 | 0 | 0 | 0 | 63 | 112 | 62 | 48 | 153 | 0 | 0 | 3 | 0 | 0 | 296 | 149 | 1 | 1 | 0.974 | 1 | 1 | 1 | 1 | 1 | 1 | 0.912 | 1 | 1 | 0.012 |
| SE | 16908 | ENSG00000141564 | RPTOR | chr17 | + | 80730559 | 80730706 | 80708932 | 80709114 | 80754009 | 80754104 | 61 | 31 | 52 | 56 | 49 | 0 | 0 | 1 | 0 | 0 | 32 | 85 | 61 | 35 | 64 | 0 | 0 | 0 | 0 | 0 | 296 | 149 | 1 | 1 | 1 | 1 | 0.963 | 1 | 1 | 1 | 1 | 1 | 1 | 1 | -0.007 |
| SE | 16910 | ENSG00000141564 | RPTOR | chr17 | + | 80823078 | 80823223 | 80822200 | 80822301 | 80837921 | 80837997 | 96 | 49 | 70 | 100 | 88 | 2 | 1 | 0 | 0 | 0 | 59 | 65 | 55 | 37 | 65 | 0 | 0 | 0 | 0 | 0 | 294 | 149 | 1 | 1 | 0.961 | 0.961 | 1 | 1 | 1 | 1 | 1 | 1 | 1 | 1 | -0.016 |
| SE | 16911 | ENSG00000141564 | RPTOR | chr17 | + | 80857789 | 80857900 | 80855463 | 80855547 | 80880414 | 80880489 | 41 | 11 | 43 | 41 | 34 | 0 | 0 | 0 | 0 | 0 | 6 | 51 | 28 | 30 | 94 | 0 | 0 | 1 | 0 | 0 | 260 | 149 | 1 | 1 | 1 | 1 | 1 | 1 | 1 | 1 | 1 | 0.941 | 1 | 1 | 0.012 |
| SE | 16912 | ENSG00000141564 | RPTOR | chr17 | + | 80880414 | 80880489 | 80857789 | 80857900 | 80891719 | 80891837 | 9 | 2 | 7 | 9 | 5 | 0 | 0 | 0 | 0 | 0 | 0 | 11 | 8 | 4 | 35 | 0 | 0 | 2 | 0 | 0 | 224 | 149 | 1 | 1 | 1 | 1 | 1 | 1 | 1 | NA | 1 | 0.727 | 1 | 1 | 0.068 |
| SE | 16914 | ENSG00000141564 | RPTOR | chr17 | + | 80940495 | 80940601 | 80925369 | 80925480 | 80945666 | 80945781 | 113 | 42 | 79 | 80 | 61 | 1 | 0 | 0 | 0 | 0 | 86 | 75 | 35 | 57 | 97 | 0 | 0 | 4 | 0 | 0 | 255 | 149 | 1 | 1 | 0.985 | 1 | 1 | 1 | 1 | 1 | 1 | 0.836 | 1 | 1 | 0.03 |
| SE | 16915 | ENSG00000141564 | RPTOR | chr17 | + | 80957623 | 80957730 | 80949442 | 80949547 | 80960077 | 80960205 | 53 | 15 | 42 | 42 | 39 | 0 | 0 | 0 | 0 | 0 | 50 | 62 | 26 | 25 | 45 | 0 | 0 | 0 | 1 | 0 | 256 | 149 | 1 | 1 | 1 | 1 | 1 | 1 | 1 | 1 | 1 | 1 | 0.936 | 1 | 0.013 |
| SE | 16918 | ENSG00000170298 | LGALS9B | chr17 | - | 20455302 | 20455398 | 20453571 | 20453607 | 20456560 | 20456671 | 34 | 1 | 5 | 6 | 0 | 7 | 0 | 0 | 0 | 0 | 0 | 0 | 2 | 0 | 0 | 0 | 0 | 0 | 0 | 0 | 245 | 149 | 1 | 1 | 0.747 | 1 | 1 | 1 | NA | NA | NA | 1 | NA | NA | -0.063 |
| SE | 16919 | ENSG00000170298 | LGALS9B | chr17 | - | 20457377 | 20457563 | 20456560 | 20456671 | 20458182 | 20458260 | 9 | 1 | 2 | 3 | 0 | 36 | 3 | 5 | 7 | 6 | 5 | 0 | 0 | 0 | 0 | 0 | 0 | 2 | 0 | 0 | 298 | 149 | 1 | 1 | 0.111 | 0.143 | 0.167 | 0.176 | 0 | 1 | NA | 0 | NA | NA | -0.381 |
| SE | 16920 | ENSG00000170298 | LGALS9B | chr17 | - | 20460351 | 20460443 | 20458182 | 20458384 | 20467431 | 20467535 | 114 | 5 | 2 | 17 | 5 | 6 | 0 | 0 | 0 | 0 | 0 | 0 | 2 | 0 | 0 | 5 | 0 | 0 | 0 | 0 | 241 | 149 | 1 | 1 | 0.922 | 1 | 1 | 1 | 1 | 0 | NA | 1 | NA | NA | 0.484 |
| SE | 16924 | ENSG00000171606 | ZNF274 | chr19 | + | 58185711 | 58185838 | 58183493 | 58183998 | 58186946 | 58187023 | 93 | 45 | 104 | 105 | 98 | 1 | 4 | 1 | 2 | 2 | 62 | 82 | 86 | 67 | 81 | 0 | 3 | 1 | 1 | 0 | 276 | 149 | 1 | 1 | 0.98 | 0.859 | 0.983 | 0.966 | 0.964 | 1 | 0.937 | 0.979 | 0.973 | 1 | -0.027 |
| SE | 16929 | ENSG00000171606 | ZNF274 | chr19 | + | 58206719 | 58207202 | 58186946 | 58187042 | 58212160 | 58213552 | 48 | 30 | 62 | 64 | 72 | 0 | 0 | 1 | 0 | 0 | 49 | 55 | 56 | 30 | 54 | 0 | 0 | 0 | 0 | 0 | 298 | 149 | 1 | 1 | 1 | 1 | 0.969 | 1 | 1 | 1 | 1 | 1 | 1 | 1 | -0.006 |
| SE | 16931 | ENSG00000171606 | ZNF274 | chr19 | + | 58211559 | 58211686 | 58206719 | 58207202 | 58212160 | 58213552 | 27 | 24 | 46 | 39 | 23 | 0 | 0 | 3 | 0 | 0 | 28 | 35 | 21 | 23 | 15 | 0 | 0 | 0 | 0 | 0 | 276 | 149 | 1 | 1 | 1 | 1 | 0.892 | 1 | 1 | 1 | 1 | 1 | 1 | 1 | -0.022 |
| SE | 16932 | ENSG00000154237 | LRRK1 | chr15 | + | 100920030 | 100920255 | 100919326 | 100919451 | 100924510 | 100924729 | 0 | 0 | 0 | 0 | 0 | 5 | 1 | 0 | 0 | 0 | 4 | 0 | 4 | 0 | 0 | 0 | 0 | 6 | 2 | 4 | 298 | 149 | 1 | 1 | 0 | 0 | NA | NA | NA | 1 | NA | 0.25 | 0 | 0 | -0.312 |

| | | | | | | | | | | | | | | | |
|---|---|---|---|---|---|---|---|---|---|---|---|---|---|---|---|
| SE | 16933 | ENSG00000154237 | LRRK1 | chr15 | + | 101014315 | 101014428 | 101012007 | 101012145 | 101015325 | 101015402 | 36 | 75 | 38 | [illegible] |
| SE | 16936 | ENSG00000154237 | LRRK1 | chr15 | + | 101022382 | 101022597 | 101021052 | 101021182 | 101024799 | 101024949 | 0 | 4 | 2 | [illegible] |
| SE | 16937 | ENSG00000154237 | LRRK1 | chr15 | + | 101045980 | 101046152 | 101028955 | 101029232 | 101048493 | 101048657 | 39 | 41 | 29 | [illegible] |
| SE | 16939 | ENSG00000154237 | LRRK1 | chr15 | + | 101048493 | 101048657 | 101028955 | 101029232 | 101049643 | 101049783 | 46 | 23 | 27 | [illegible] |
| SE | 16941 | ENSG00000154237 | LRRK1 | chr15 | + | 101054945 | 101055223 | 101051710 | 101051960 | 101056855 | 101057050 | 9 | 8 | 20 | [illegible] |
| SE | 16942 | ENSG00000154237 | LRRK1 | chr15 | + | 101062380 | 101062690 | 101061170 | 101061288 | 101065351 | 101066205 | 6 | 4 | 0 | [illegible] |
| SE | 16948 | ENSG00000172575 | RASGRP1 | chr15 | - | 38494381 | 38494767 | 38488102 | 38490688 | 38502311 | 38502421 | 173 | 127 | 157 | [illegible] |
| SE | 16950 | ENSG00000172575 | RASGRP1 | chr15 | - | 38498793 | 38498946 | 38490553 | 38490688 | 38502311 | 38502421 | 6 | 7 | 4 | [illegible] |
| SE | 16952 | ENSG00000172575 | RASGRP1 | chr15 | - | 38498793 | 38498946 | 38494381 | 38494767 | 38502311 | 38502421 | 104 | 52 | 116 | [illegible] |
| SE | 16954 | ENSG00000172575 | RASGRP1 | chr15 | - | 38500102 | 38500139 | 38490553 | 38490688 | 38502311 | 38502421 | 3 | 2 | 5 | [illegible] |
| SE | 16955 | ENSG00000172575 | RASGRP1 | chr15 | - | 38500102 | 38500139 | 38494381 | 38494767 | 38502311 | 38502421 | 4 | 4 | 15 | [illegible] |
| SE | 16956 | ENSG00000172575 | RASGRP1 | chr15 | - | 38500102 | 38500139 | 38498793 | 38498946 | 38502311 | 38502421 | 10 | 18 | 35 | [illegible] |
| SE | 16959 | ENSG00000172575 | RASGRP1 | chr15 | - | 38502311 | 38502421 | 38500102 | 38500139 | 38507725 | 38508001 | 8 | 18 | 15 | [illegible] |
| SE | 16960 | ENSG00000172575 | RASGRP1 | chr15 | - | 38502311 | 38502421 | 38500102 | 38500139 | 38511603 | 38511705 | 7 | 10 | 11 | [illegible] |
| SE | 16963 | ENSG00000172575 | RASGRP1 | chr15 | - | 38507725 | 38508001 | 38500102 | 38500139 | 38511603 | 38511705 | 261 | 152 | 298 | [illegible] |
| SE | 16964 | ENSG00000172575 | RASGRP1 | chr15 | - | 38507725 | 38508001 | 38502311 | 38502421 | 38511603 | 38511705 | 262 | 161 | 303 | [illegible] |
| SE | 16966 | ENSG00000172575 | RASGRP1 | chr15 | - | 38518291 | 38518423 | 38516196 | 38516350 | 38519308 | 38519364 | 432 | 296 | 478 | [illegible] |
| SE | 16967 | ENSG00000172575 | RASGRP1 | chr15 | - | 38518291 | 38518423 | 38516196 | 38516350 | 38559820 | 38560005 | 157 | 76 | 135 | [illegible] |
| SE | 16972 | ENSG00000141956 | PRDM15 | chr21 | - | 41821066 | 41821230 | 41820094 | 41820174 | 41821902 | 41822037 | 61 | 22 | 55 | [illegible] |
| SE | 16974 | ENSG00000141956 | PRDM15 | chr21 | - | 41823317 | 41823449 | 41821902 | 41822037 | 41828165 | 41828281 | 17 | 15 | 13 | [illegible] |
| SE | 16975 | ENSG00000141956 | PRDM15 | chr21 | - | 41825959 | 41826054 | 41821902 | 41822037 | 41828165 | 41828281 | 7 | 14 | 8 | [illegible] |
| SE | 16976 | ENSG00000141956 | PRDM15 | chr21 | - | 41825959 | 41826054 | 41823317 | 41823449 | 41828165 | 41828281 | 36 | 29 | 23 | [illegible] |
| SE | 16979 | ENSG00000141956 | PRDM15 | chr21 | - | 41839622 | 41839853 | 41837933 | 41838063 | 41847089 | 41847191 | 24 | 22 | 26 | [illegible] |
| SE | 16982 | ENSG00000248905 | FMN1 | chr15 | - | 32888171 | 32888292 | 32857014 | 32857107 | 32898833 | 32898893 | 27 | 94 | 82 | [illegible] |
| SE | 16983 | ENSG00000248905 | FMN1 | chr15 | - | 32898833 | 32898893 | 32888171 | 32888292 | 32899978 | 32900125 | 18 | 62 | 62 | [illegible] |
| SE | 16984 | ENSG00000248905 | FMN1 | chr15 | - | 32901910 | 32902040 | 32899794 | 32900125 | 32908489 | 32908578 | 18 | 117 | 59 | [illegible] |
| SE | 16985 | ENSG00000248905 | FMN1 | chr15 | - | 32908489 | 32908578 | 32901910 | 32902040 | 32926173 | 32926260 | 6 | 23 | 24 | [illegible] |
| [illegible] | | | | | | | | | | | | | | | |

| | | | | | | | | | | | | | | |
|---|---|---|---|---|---|---|---|---|---|---|---|---|---|---|
| SE | 17451 | ENSG00000140563 | MCTP2 | chr15 | + | 94458136 | 94458246 | 94442918 | 94442960 | 94470332 | 94470442 | 231 | 97 | 130 |
| SE | 17452 | ENSG00000140563 | MCTP2 | chr15 | + | 94470332 | 94470442 | 94458136 | 94458246 | 94476695 | 94476793 | 163 | 69 | 89 |
| SE | 17453 | ENSG00000198133 | TMEM229B | chr14 | - | 67503486 | 67503551 | 67486999 | 67487172 | 67515085 | 67515182 | 0 | 0 | 1 |
| SE | 17456 | ENSG00000159110 | IFNAR2 | chr21 | + | 33241839 | 33241977 | 33229937 | 33230216 | 33243672 | 33243714 | 317 | 232 | 279 |
| SE | 17457 | ENSG00000159110 | IFNAR2 | chr21 | + | 33241839 | 33241977 | 33229937 | 33230216 | 33244950 | 33245074 | 16 | 7 | 15 |
| SE | 17458 | ENSG00000159110 | IFNAR2 | chr21 | + | 33241839 | 33241977 | 33230408 | 33230557 | 33244950 | 33245074 | 1 | 1 | 1 |
| SE | 17459 | ENSG00000159110 | IFNAR2 | chr21 | + | 33241885 | 33241977 | 33229900 | 33230216 | 33243672 | 33243714 | 305 | 221 | 265 |
| SE | 17460 | ENSG00000159110 | IFNAR2 | chr21 | + | 33241885 | 33241977 | 33229900 | 33230216 | 33244950 | 33245074 | 6 | 0 | 2 |
| SE | 17461 | ENSG00000159110 | IFNAR2 | chr21 | + | 33241885 | 33241977 | 33230408 | 33230557 | 33244950 | 33245074 | 0 | 0 | 1 |
| SE | 17462 | ENSG00000159110 | IFNAR2 | chr21 | + | 33243672 | 33243714 | 33229900 | 33230216 | 33244950 | 33245074 | 56 | 45 | 54 |
| SE | 17463 | ENSG00000159110 | IFNAR2 | chr21 | + | 33243672 | 33243714 | 33241839 | 33241977 | 33244950 | 33245074 | 380 | 267 | 324 |
| SE | 17466 | ENSG00000159110 | IFNAR2 | chr21 | + | 33248708 | 33248854 | 33246717 | 33246890 | 33252661 | 33252830 | 772 | 643 | 709 |
| SE | 17468 | ENSG00000159110 | IFNAR2 | chr21 | + | 33252661 | 33252830 | 33248708 | 33248854 | 33262792 | 33262855 | 446 | 330 | 388 |
| SE | 17479 | ENSG00000270580 | PKD1P6-NPIPP1 | chr16 | - | 15107329 | 15107437 | 15106259 | 15106320 | 15110413 | 15110558 | 94 | 69 | 76 |
| SE | 17480 | ENSG00000270580 | PKD1P6-NPIPP1 | chr16 | - | 15127469 | 15127596 | 15125690 | 15125807 | 15127686 | 15127897 | 2 | 4 | 9 |
| SE | 17481 | ENSG00000270580 | PKD1P6-NPIPP1 | chr16 | - | 15130634 | 15130791 | 15130201 | 15130454 | 15131086 | 15131601 | 18 | 3 | 15 |
| SE | 17482 | ENSG00000185024 | BRF1 | chr14 | - | 105226250 | 105226290 | 105226068 | 105226161 | 105226633 | 105226760 | 58 | 21 | 62 |
| SE | 17483 | ENSG00000185024 | BRF1 | chr14 | - | 105228819 | 105228913 | 105226633 | 105226760 | 105247070 | 105247193 | 12 | 6 | 18 |
| SE | 17488 | ENSG00000100580 | TMED8 | chr14 | - | 77343696 | 77343823 | 77343177 | 77343483 | 77346348 | 77346478 | 66 | 74 | 98 |
| SE | 17490 | ENSG00000100580 | TMED8 | chr14 | - | 77346348 | 77346478 | 77343177 | 77343483 | 77351672 | 77351751 | 12 | 6 | 7 |
| SE | 17492 | ENSG00000100577 | GSTZ1 | chr14 | + | 77322546 | 77322732 | 77320995 | 77321183 | 77324869 | 77324921 | 0 | 0 | 1 |
| SE | 17494 | ENSG00000100577 | GSTZ1 | chr14 | + | 77324869 | 77324921 | 77320998 | 77321183 | 77326837 | 77326905 | 14 | 2 | 15 |
| SE | 17513 | ENSG00000172590 | MRPL52 | chr14 | + | 22833417 | 22833482 | 22830186 | 22830254 | 22834171 | 22834345 | 30 | 17 | 33 |
| SE | 17515 | ENSG00000172590 | MRPL52 | chr14 | + | 22833417 | 22833489 | 22830186 | 22830254 | 22834171 | 22834346 | 4 | 5 | 6 |
| SE | 17517 | ENSG00000125952 | MAX | chr14 | - | 65065536 | 65065602 | 65060917 | 65062002 | 65077912 | 65078036 | 0 | 0 | 1 |
| SE | 17518 | ENSG00000125952 | MAX | chr14 | - | 65069693 | 65070369 | 65065536 | 65065602 | 65077912 | 65078036 | 0 | 0 | 1 |
| SE | 17521 | ENSG00000125952 | MAX | chr14 | - | 65077562 | 65077643 | 65077384 | 65077428 | 65077912 | 65078036 | 0 | 1 | 4 |
| SE | 17523 | ENSG00000125952 | MAX | chr14 | - | 65077912 | 65078036 | 65069693 | 65070369 | 65093707 | 65093815 | 462 | 388 | 367 |
| SE | 17537 | ENSG00000100813 | ACIN1 | chr14 | - | 23062167 | 23062275 | 23061297 | 23061622 | 23062415 | 23062523 | 384 | 252 | 229 |
| SE | 17538 | ENSG00000100813 | ACIN1 | chr14 | - | 23062415 | 23062523 | 23061297 | 23061622 | 23063435 | 23063577 | 0 | 0 | 0 |
| SE | 17539 | ENSG00000100813 | ACIN1 | chr14 | - | 23062928 | 23063074 | 23062415 | 23062523 | 23063435 | 23063577 | 401 | 263 | 373 |
| SE | 17540 | ENSG00000100813 | ACIN1 | chr14 | - | 23065965 | 23066008 | 23064354 | 23064488 | 23067313 | 23068671 | 97 | 72 | 105 |
| SE | 17541 | ENSG00000100813 | ACIN1 | chr14 | - | 23065965 | 23066008 | 23064354 | 23064488 | 23069475 | 23069617 | 242 | 152 | 189 |
| SE | 17542 | ENSG00000100813 | ACIN1 | chr14 | - | 23067313 | 23068671 | 23064354 | 23064488 | 23069475 | 23069525 | 3 | 5 | 13 |
| SE | 17546 | ENSG00000100813 | ACIN1 | chr14 | - | 23076469 | 23076548 | 23069475 | 23069617 | 23078150 | 23078266 | 6 | 7 | 4 |
| SE | 17547 | ENSG00000100813 | ACIN1 | chr14 | - | 23078819 | 23079038 | 23077796 | 23078266 | 23079546 | 23079593 | 265 | 149 | 260 |
| SE | 17548 | ENSG00000100813 | ACIN1 | chr14 | - | 23081747 | 23081836 | 23079546 | 23080809 | 23090521 | 23090633 | 301 | 277 | 311 |
| SE | 17550 | ENSG00000100813 | ACIN1 | chr14 | - | 23090521 | 23090633 | 23081747 | 23081836 | 23093478 | 23093544 | 209 | 179 | 176 |
| SE | 17551 | ENSG00000100813 | ACIN1 | chr14 | - | 23093478 | 23093544 | 23090521 | 23090633 | 23094974 | 23095136 | 306 | 235 | 277 |
| SE | 17552 | ENSG00000131943 | C19orf12 | chr19 | - | 29705273 | 29705394 | 29702841 | 29702977 | 29708253 | 29708423 | 3 | 0 | 0 |
| SE | 17553 | ENSG00000131943 | C19orf12 | chr19 | - | 29708253 | 29708423 | 29698936 | 29702977 | 29714906 | 29715060 | 12 | 2 | 2 |
| SE | 17554 | ENSG00000131943 | C19orf12 | chr19 | - | 29708253 | 29708423 | 29698936 | 29702977 | 29715124 | 29715224 | 21 | 5 | 3 |
| SE | 17555 | ENSG00000197102 | DYNC1H1 | chr14 | + | 101975711 | 101975799 | 101964572 | 101964947 | 101979318 | 101979492 | 189 | 102 | 183 |
| SE | 17556 | ENSG00000197102 | DYNC1H1 | chr14 | + | 101975713 | 101975799 | 101964687 | 101964947 | 101979318 | 101979492 | 96 | 52 | 94 |
| SE | 17557 | ENSG00000197102 | DYNC1H1 | chr14 | + | 101994985 | 101995096 | 101994672 | 101994849 | 101995180 | 101995300 | 214 | 109 | 236 |
| SE | 17558 | ENSG00000197102 | DYNC1H1 | chr14 | + | 101995180 | 101995300 | 101994985 | 101995096 | 101997034 | 101997274 | 369 | 228 | 347 |
| SE | 17559 | ENSG00000197102 | DYNC1H1 | chr14 | + | 101996807 | 101996900 | 101995180 | 101995300 | 101997034 | 101997274 | 2 | 5 | 5 |
| SE | 17560 | ENSG00000197102 | DYNC1H1 | chr14 | + | 102000594 | 102000717 | 102000285 | 102000399 | 102000953 | 102001064 | 0 | 0 | 0 |
| SE | 17561 | ENSG00000197102 | DYNC1H1 | chr14 | + | 102008177 | 102008337 | 102007007 | 102007108 | 102009842 | 102010086 | 384 | 214 | 305 |
| SE | 17562 | ENSG00000197102 | DYNC1H1 | chr14 | + | 102014007 | 102014247 | 102012313 | 102012470 | 102015104 | 102015332 | 4 | 0 | 1 |
| SE | 17563 | ENSG00000197102 | DYNC1H1 | chr14 | + | 102022750 | 102022880 | 102019892 | 102020056 | 102026573 | 102026707 | 365 | 197 | 258 |
| SE | 17565 | ENSG00000197102 | DYNC1H1 | chr14 | + | 102024284 | 102024349 | 102022750 | 102022880 | 102026573 | 102026707 | 0 | 0 | 0 |
| SE | 17567 | ENSG00000197102 | DYNC1H1 | chr14 | + | 102024475 | 102024580 | 102024284 | 102024349 | 102026573 | 102026707 | 1 | 0 | 0 |
| SE | 17568 | ENSG00000197102 | DYNC1H1 | chr14 | + | 102027173 | 102027288 | 102026573 | 102026707 | 102027382 | 102027544 | 667 | 421 | 526 |
| SE | 17571 | ENSG00000197102 | DYNC1H1 | chr14 | + | 102036488 | 102036642 | 102034324 | 102034452 | 102038459 | 102038606 | 559 | 308 | 456 |
| SE | 17572 | ENSG00000197102 | DYNC1H1 | chr14 | + | 102038459 | 102038606 | 102036488 | 102036642 | 102038697 | 102038848 | 374 | 243 | 320 |
| SE | 17573 | ENSG00000197102 | DYNC1H1 | chr14 | + | 102040235 | 102040410 | 102039490 | 102039732 | 102040597 | 102040673 | 228 | 99 | 194 |
| SE | 17574 | ENSG00000197102 | DYNC1H1 | chr14 | + | 102040235 | 102040410 | 102039490 | 102039732 | 102041573 | 102041734 | 104 | 42 | 88 |
| SE | 17575 | ENSG00000197102 | DYNC1H1 | chr14 | + | 102040235 | 102040439 | 102039637 | 102039732 | 102040597 | 102040673 | 106 | 42 | 88 |
| SE | 17576 | ENSG00000197102 | DYNC1H1 | chr14 | + | 102040597 | 102040673 | 102039637 | 102039732 | 102041573 | 102041734 | 72 | 45 | 61 |
| SE | 17578 | ENSG00000197102 | DYNC1H1 | chr14 | + | 102041573 | 102041734 | 102040597 | 102040673 | 102042012 | 102042124 | 183 | 107 | 178 |
| SE | 17579 | ENSG00000197102 | DYNC1H1 | chr14 | + | 102044594 | 102044698 | 102044273 | 102044491 | 102047816 | 102048028 | 298 | 199 | 206 |
| SE | 17581 | ENSG00000012983 | MAP4K5 | chr14 | - | 50423120 | 50423176 | 50418500 | 50420106 | 50425906 | 50425977 | 108 | 86 | 82 |
| SE | 17582 | ENSG00000012983 | MAP4K5 | chr14 | - | 50425906 | 50425977 | 50423120 | 50423176 | 50428661 | 50428754 | 124 | 114 | 115 |
| SE | 17583 | ENSG00000012983 | MAP4K5 | chr14 | - | 50428661 | 50428754 | 50425906 | 50425977 | 50429191 | 50429256 | 113 | 110 | 118 |
| SE | 17584 | ENSG00000012983 | MAP4K5 | chr14 | - | 50429191 | 50429256 | 50428661 | 50428754 | 50434393 | 50434571 | 49 | 50 | 61 |
| SE | 17585 | ENSG00000012983 | MAP4K5 | chr14 | - | 50429191 | 50429260 | 50428661 | 50428754 | 50434393 | 50434571 | 94 | 89 | 114 |
| SE | 17586 | ENSG00000012983 | MAP4K5 | chr14 | - | 50434393 | 50434571 | 50429191 | 50429260 | 50434961 | 50435065 | 112 | 79 | 108 |
| SE | 17587 | ENSG00000012983 | MAP4K5 | chr14 | - | 50434961 | 50435065 | 50434393 | 50434571 | 50437475 | 50437534 | 100 | 56 | 80 |
| SE | 17589 | ENSG00000012983 | MAP4K5 | chr14 | - | 50445040 | 50445194 | 50443938 | 50444036 | 50446078 | 50446121 | 171 | 133 | 165 |
| SE | 17590 | ENSG00000012983 | MAP4K5 | chr14 | - | 50456515 | 50456594 | 50448773 | 50448832 | 50462664 | 50462781 | 164 | 122 | 121 |
| SE | 17591 | ENSG00000012983 | MAP4K5 | chr14 | - | 50482360 | 50482416 | 50475076 | 50475149 | 50485577 | 50485642 | 58 | 39 | 30 |
| SE | 17595 | ENSG00000174373 | RALGAPA1 | chr14 | - | 35542026 | 35542029 | 35538356 | 35539690 | 35549109 | 35549234 | 1 | 0 | 0 |
| SE | 17597 | ENSG00000174373 | RALGAPA1 | chr14 | - | 35595633 | 35595789 | 35572559 | 35572718 | 35605585 | 35605709 | 140 | 119 | 145 |
| SE | 17599 | ENSG00000174373 | RALGAPA1 | chr14 | - | 35634573 | 35634757 | 35627089 | 35627951 | 35635463 | 35635598 | 140 | 86 | 155 |
| SE | 17600 | ENSG00000174373 | RALGAPA1 | chr14 | - | 35635463 | 35635598 | 35634573 | 35634757 | 35651804 | 35651873 | 111 | 60 | 111 |
| SE | 17601 | ENSG00000174373 | RALGAPA1 | chr14 | - | 35659137 | 35659196 | 35655806 | 35655915 | 35664641 | 35664767 | 99 | 78 | 102 |
| SE | 17602 | ENSG00000174373 | RALGAPA1 | chr14 | - | 35659137 | 35659196 | 35655806 | 35655915 | 35671388 | 35671517 | 46 | 34 | 55 |
| SE | 17603 | ENSG00000174373 | RALGAPA1 | chr14 | - | 35664641 | 35664767 | 35655806 | 35655915 | 35671388 | 35671517 | 38 | 34 | 36 |
| SE | 17604 | ENSG00000174373 | RALGAPA1 | chr14 | - | 35664641 | 35664767 | 35659137 | 35659196 | 35671388 | 35671517 | 91 | 78 | 84 |
| SE | 17605 | ENSG00000174373 | RALGAPA1 | chr14 | - | 35677949 | 35678102 | 35674515 | 35674709 | 35683808 | 35683985 | 123 | 67 | 122 |
| SE | 17606 | ENSG00000174373 | RALGAPA1 | chr14 | - | 35688458 | 35688540 | 35686541 | 35686623 | 35689835 | 35690003 | 2 | 1 | 4 |
| SE | 17607 | ENSG00000174373 | RALGAPA1 | chr14 | - | 35700161 | 35700302 | 35688458 | 35690003 | 35721687 | 35721849 | 0 | 0 | 1 |
| SE | 17608 | ENSG00000174373 | RALGAPA1 | chr14 | - | 35728361 | 35728510 | 35725023 | 35725153 | 35738512 | 35738650 | 291 | 256 | 280 |
| SE | 17610 | ENSG00000174373 | RALGAPA1 | chr14 | - | 35756792 | 35756908 | 35752023 | 35752162 | 35760828 | 35760857 | 187 | 120 | 150 |
| SE | 17611 | ENSG00000174373 | RALGAPA1 | chr14 | - | 35760828 | 35761006 | 35756792 | 35756908 | 35762709 | 35762753 | 151 | 119 | 161 |
| SE | 17612 | ENSG00000174373 | RALGAPA1 | chr14 | - | 35762709 | 35762753 | 35760828 | 35761006 | 35770941 | 35770999 | 120 | 96 | 141 |
| SE | 17613 | ENSG00000174373 | RALGAPA1 | chr14 | - | 35770941 | 35770999 | 35762709 | 35762753 | 35775005 | 35775055 | 105 | 70 | 131 |
| SE | 17618 | ENSG00000101384 | JAG1 | chr20 | - | 10643777 | 10643863 | 10642487 | 10642601 | 10644356 | 10644384 | 3 | 1 | 5 |
| SE | 17619 | ENSG00000101384 | JAG1 | chr20 | - | 10646938 | 10647103 | 10645721 | 10646084 | 10647959 | 10648028 | 8 | 10 | 7 |
| SE | 17620 | ENSG00000092036 | HAUS4 | chr14 | - | 22947867 | 22948013 | 22947600 | 22947731 | 22950313 | 22950410 | 168 | 94 | 151 |
| SE | 17621 | ENSG00000092036 | HAUS4 | chr14 | - | 22950313 | 22950410 | 22947600 | 22947731 | 22951554 | 22951689 | 54 | 29 | 49 |
| SE | 17632 | ENSG00000100796 | PPP4R3A | chr14 | - | 91462734 | 91462877 | 91462048 | 91462239 | 91465249 | 91465419 | 1007 | 978 | 807 |
| SE | 17633 | ENSG00000100796 | PPP4R3A | chr14 | - | 91465249 | 91465419 | 91462734 | 91462877 | 91470836 | 91470995 | 816 | 679 | 637 |
| SE | 17635 | ENSG00000100796 | PPP4R3A | chr14 | - | 91470836 | 91470995 | 91465249 | 91465419 | 91473032 | 91473135 | 531 | 368 | 426 |
| SE | 17638 | ENSG00000100796 | PPP4R3A | chr14 | - | 91476407 | 91476524 | 91474640 | 91475966 | 91481575 | 91481998 | 169 | 155 | 150 |
| SE | 17639 | ENSG00000100796 | PPP4R3A | chr14 | - | 91476908 | 91476986 | 91474640 | 91475966 | 91481575 | 91481873 | 246 | 219 | 183 |
| SE | 17640 | ENSG00000100796 | PPP4R3A | chr14 | - | 91476908 | 91476986 | 91474640 | 91475966 | 91485631 | 91485730 | 1 | 2 | 0 |
| SE | 17641 | ENSG00000100796 | PPP4R3A | chr14 | - | 91476908 | 91476986 | 91476407 | 91476524 | 91481575 | 91481873 | 1000 | 865 | 806 |
| SE | 17642 | ENSG00000100796 | PPP4R3A | chr14 | - | 91481575 | 91482193 | 91475810 | 91475966 | 91485631 | 91485730 | 410 | 360 | 327 |
| SE | 17643 | ENSG00000100796 | PPP4R3A | chr14 | - | 91481575 | 91482193 | 91476908 | 91476986 | 91485631 | 91485730 | 813 | 716 | 659 |
| SE | 17648 | ENSG00000100949 | RABGGTA | chr14 | - | 24269079 | 24269163 | 24268826 | 24268993 | 24269490 | 24269694 | 44 | 17 | 25 |
| SE | 17649 | ENSG00000100949 | RABGGTA | chr14 | - | 24270836 | 24270947 | 24270333 | 24270458 | 24271112 | 24271169 | 54 | 8 | 46 |
| SE | 17656 | ENSG00000100916 | BRMS1L | chr14 | + | 35829794 | 35829907 | 35826440 | 35826658 | 35831409 | 35831500 | 7 | 19 | 4 |
| SE | 17657 | ENSG00000100916 | BRMS1L | chr14 | + | 35831409 | 35831500 | 35829794 | 35829907 | 35832977 | 35833105 | 20 | 28 | 25 |
| SE | 17658 | ENSG00000100916 | BRMS1L | chr14 | + | 35832977 | 35833105 | 35831409 | 35831500 | 35862589 | 35862686 | 20 | 35 | 29 |
| SE | 17659 | ENSG00000100916 | BRMS1L | chr14 | + | 35834843 | 35834923 | 35832977 | 35833105 | 35858576 | 35858681 | 18 | 14 | 8 |
| SE | 17660 | ENSG00000100916 | BRMS1L | chr14 | + | 35834843 | 35834923 | 35832977 | 35833105 | 35862589 | 35862686 | 49 | 50 | 44 |
| SE | 17663 | ENSG00000176681 | LRRC37A | chr17 | + | 46330449 | 46331981 | 46328391 | 46328510 | 46335544 | 46335594 | 0 | 0 | 2 |
| SE | 17665 | ENSG00000176681 | LRRC37A | chr17 | + | 46337295 | 46337350 | 46335544 | 46335594 | 46337428 | 46337568 | 36 | 22 | 85 |
| SE | 17667 | ENSG00000100906 | NFKBIA | chr14 | - | 35402393 | 35402534 | 35401539 | 35402060 | 35403149 | 35403360 | 1244 | 1110 | 1603 |
| SE | 17668 | ENSG00000100906 | NFKBIA | chr14 | - | 35402393 | 35402859 | 35401925 | 35402060 | 35403149 | 35403238 | 1358 | 1151 | 1791 |
| SE | 17670 | ENSG00000100906 | NFKBIA | chr14 | - | 35402770 | 35402859 | 35402393 | 35402663 | 35403149 | 35403360 | 812 | 460 | 1136 |
| SE | 17675 | ENSG00000126787 | DLGAP5 | chr14 | - | 55148593 | 55148684 | 55148111 | 55148473 | 55150798 | 55150848 | 0 | 0 | 0 |
| SE | 17676 | ENSG00000126787 | DLGAP5 | chr14 | - | 55191068 | 55191176 | 55188941 | 55189180 | 55191462 | 55191608 | 0 | 0 | 2 |
| SE | 17679 | ENSG00000119682 | AREL1 | chr14 | - | 74667464 | 74667594 | 74667318 | 74667377 | 74669648 | 74669774 | 156 | 90 | 142 |
| SE | 17681 | ENSG00000119682 | AREL1 | chr14 | - | 74669946 | 74670126 | 74669648 | 74669774 | 74671407 | 74671483 | 108 | 62 | 85 |
| SE | 17683 | ENSG00000119682 | AREL1 | chr14 | - | 74669946 | 74670871 | 74669648 | 74669774 | 74671407 | 74671483 | 201 | 116 | 152 |
| SE | 17684 | ENSG00000119682 | AREL1 | chr14 | - | 74670761 | 74670871 | 74669648 | 74669774 | 74671407 | 74671483 | 90 | 54 | 68 |
| SE | 17687 | ENSG00000119682 | AREL1 | chr14 | - | 74674033 | 74674111 | 74673076 | 74673218 | 74675698 | 74675708 | 132 | 84 | 132 |
| SE | 17688 | ENSG00000119682 | AREL1 | chr14 | - | 74675698 | 74675946 | 74673076 | 74673218 | 74676140 | 74676321 | 136 | 78 | 96 |
| SE | 17689 | ENSG00000119682 | AREL1 | chr14 | - | 74676582 | 74676752 | 74676140 | 74676321 | 74683295 | 74683533 | 252 | 144 | 186 |
| SE | 17690 | ENSG00000119682 | AREL1 | chr14 | - | 74678178 | 74678329 | 74676582 | 74676752 | 74683295 | 74683533 | 0 | 0 | 2 |
| SE | 17691 | ENSG00000119682 | AREL1 | chr14 | - | 74683295 | 74683533 | 74676582 | 74676752 | 74684453 | 74684680 | 182 | 90 | 138 |
| SE | 17692 | ENSG00000119682 | AREL1 | chr14 | - | 74683295 | 74683533 | 74676582 | 74676752 | 74685599 | 74685660 | 114 | 46 | 90 |
| SE | 17693 | ENSG00000119682 | AREL1 | chr14 | - | 74684453 | 74684680 | 74676582 | 74676752 | 74685599 | 74685646 | 93 | 58 | 84 |
| SE | 17698 | ENSG00000119682 | AREL1 | chr14 | - | 74698763 | 74698856 | 74692040 | 74692328 | 74701668 | 74701774 | 2 | 2 | 8 |
| SE | 17700 | ENSG00000119682 | AREL1 | chr14 | - | 74701668 | 74701774 | 74692040 | 74692328 | 74712932 | 74713087 | 0 | 0 | 0 |
| SE | 17701 | ENSG00000119682 | AREL1 | chr14 | - | 74701668 | 74701774 | 74698763 | 74698856 | 74712932 | 74713087 | 1 | 2 | 1 |
| SE | 17703 | ENSG00000100749 | VRK1 | chr14 | + | 96826836 | 96826950 | 96797402 | 96797447 | 96833466 | 96833631 | 0 | 0 | 0 |
| SE | 17707 | ENSG00000100749 | VRK1 | chr14 | + | 96846094 | 96846164 | 96837761 | 96837817 | 96847256 | 96847344 | 432 | 277 | 316 |
| SE | 17709 | ENSG00000100749 | VRK1 | chr14 | + | 96846102 | 96846164 | 96837761 | 96837817 | 96847256 | 96847344 | 110 | 75 | 80 |
| SE | 17715 | ENSG00000100749 | VRK1 | chr14 | + | 96856129 | 96856250 | 96855223 | 96855350 | 96856527 | 96856586 | 156 | 89 | 114 |
| SE | 17716 | ENSG00000100749 | VRK1 | chr14 | + | 96856129 | 96856250 | 96855223 | 96855352 | 96856527 | 96856586 | 157 | 92 | 109 |
| SE | 17717 | ENSG00000100749 | VRK1 | chr14 | + | 96856527 | 96856586 | 96855223 | 96855350 | 96860646 | 96860735 | 0 | 0 | 0 |
| SE | 17718 | ENSG00000100749 | VRK1 | chr14 | + | 96858892 | 96859027 | 96856527 | 96856586 | 96860556 | 96860735 | 1 | 0 | 0 |
| SE | 17724 | ENSG00000100749 | VRK1 | chr14 | + | 96876029 | 96876120 | 96860556 | 96860735 | 96931330 | 96931375 | 43 | 34 | 40 |
| SE | 17725 | ENSG00000100749 | VRK1 | chr14 | + | 96876029 | 96876120 | 96860556 | 96860735 | 96943516 | 96943680 | 43 | 34 | 40 |
| SE | 17727 | ENSG00000100749 | VRK1 | chr14 | + | 96876032 | 96876120 | 96860556 | 96860735 | 96931330 | 96931556 | 0 | 1 | 1 |
| SE | 17729 | ENSG00000100749 | VRK1 | chr14 | + | 96877501 | 96877573 | 96876029 | 96876120 | 96881176 | 96881391 | 3 | 0 | 11 |
| SE | 17730 | ENSG00000100749 | VRK1 | chr14 | + | 96877505 | 96877573 | 96876029 | 96876120 | 96881176 | 96881606 | 3 | 0 | 10 |
| SE | 17732 | ENSG00000100749 | VRK1 | chr14 | + | 96931330 | 96931450 | 96860556 | 96860735 | 96943516 | 96943680 | 7 | 0 | 1 |
| SE | 17734 | ENSG00000100749 | VRK1 | chr14 | + | 96931330 | 96931556 | 96860556 | 96860735 | 96943516 | 96943680 | 7 | 0 | 1 |
| SE | 17738 | ENSG00000100934 | SEC23A | chr14 | - | 39045162 | 39045324 | 39042785 | 39042872 | 39048651 | 39048729 | 297 | 180 | 284 |
| SE | 17740 | ENSG00000100934 | SEC23A | chr14 | - | 39061764 | 39061871 | 39048651 | 39048729 | 39063323 | 39063413 | 97 | 68 | 98 |
| SE | 17741 | ENSG00000100934 | SEC23A | chr14 | - | 39064912 | 39064993 | 39063323 | 39063413 | 39067172 | 39067296 | 174 | 135 | 171 |
| SE | 17743 | ENSG00000100934 | SEC23A | chr14 | - | 39086928 | 39087008 | 39085761 | 39085906 | 39093186 | 39093244 | 19 | 15 | 13 |
| SE | 17744 | ENSG00000100934 | SEC23A | chr14 | - | 39091476 | 39091713 | 39086928 | 39087008 | 39093186 | 39093244 | 152 | 113 | 121 |
| SE | 17745 | ENSG00000100934 | SEC23A | chr14 | - | 39092540 | 39092627 | 39091476 | 39091713 | 39093186 | 39093244 | 192 | 119 | 163 |
| SE | 17748 | ENSG00000006432 | MAP3K9 | chr14 | - | 70735960 | 70736029 | 70734385 | 70734498 | 70738244 | 70738398 | 1 | 0 | 0 |
| SE | 17750 | ENSG00000006432 | MAP3K9 | chr14 | - | 70749932 | 70750081 | 70742350 | 70742591 | 70761001 | 70761182 | 10 | 3 | 6 |
| SE | 17751 | ENSG00000006432 | MAP3K9 | chr14 | - | 70761001 | 70761182 | 70749932 | 70750081 | 70800666 | 70801080 | 17 | 4 | 13 |
| SE | 17763 | ENSG00000258689 | LINC01269 | chr14 | + | 70703694 | 70703787 | 70699994 | 70700200 | 70712009 | 70712153 | 0 | 0 | 1 |
| SE | 17764 | ENSG00000133997 | MED6 | chr14 | - | 70591265 | 70591381 | 70585755 | 70585783 | 70592879 | 70592988 | 222 | 157 | 196 |
| SE | 17767 | ENSG00000133997 | MED6 | chr14 | - | 70597617 | 70597777 | 70596211 | 70596702 | 70600615 | 70600643 | 96 | 96 | 140 |
| SE | 17772 | ENSG00000092200 | RPGRIP1 | chr14 | + | 21324874 | 21325070 | 21324568 | 21324725 | 21325231 | 21325383 | 0 | 2 | 1 |
| SE | 17773 | ENSG00000092200 | RPGRIP1 | chr14 | + | 21334601 | 21334705 | 21330248 | 21330387 | 21343035 | 21343165 | 3 | 4 | 3 |
| SE | 17774 | ENSG00000092200 | RPGRIP1 | chr14 | + | 21334604 | 21334705 | 21330248 | 21330387 | 21343035 | 21343228 | 8 | 10 | 12 |
| SE | 17775 | ENSG00000092200 | RPGRIP1 | chr14 | + | 21334604 | 21334705 | 21330248 | 21330387 | 21345112 | 21345197 | 5 | 6 | 5 |
| SE | 17776 | ENSG00000092200 | RPGRIP1 | chr14 | + | 21343035 | 21343228 | 21330248 | 21330387 | 21345112 | 21345197 | 7 | 4 | 4 |
| SE | 17777 | ENSG00000092200 | RPGRIP1 | chr14 | + | 21343035 | 21343228 | 21334604 | 21334705 | 21345112 | 21345197 | 8 | 8 | 12 |
| SE | 17779 | ENSG00000100490 | CDKL1 | chr14 | - | 50334564 | 50334621 | 50331925 | 50332432 | 50338946 | 50339029 | 20 | 19 | 18 |
| SE | 17780 | ENSG00000100490 | CDKL1 | chr14 | - | 50344985 | 50345058 | 50342131 | 50342222 | 50359027 | 50359149 | 18 | 11 | 6 |
| SE | 17781 | ENSG00000100490 | CDKL1 | chr14 | - | 50359027 | 50359149 | 50342131 | 50342222 | 50395700 | 50396076 | 0 | 1 | 3 |
| SE | 17782 | ENSG00000100490 | CDKL1 | chr14 | - | 50363220 | 50363314 | 50359027 | 50359149 | 50363844 | 50363976 | 2 | 0 | 0 |
| SE | 17783 | ENSG00000100485 | SOS2 | chr14 | - | 50120274 | 50120384 | 50117129 | 50118853 | 50129960 | 50130002 | 122 | 101 | 123 |
| SE | 17784 | ENSG00000100485 | SOS2 | chr14 | - | 50134122 | 50134239 | 50130500 | 50130762 | 50138611 | 50138784 | 368 | 285 | 295 |
| SE | 17787 | ENSG00000100485 | SOS2 | chr14 | - | 50145476 | 50145596 | 50139941 | 50140059 | 50150007 | 50150230 | 102 | 69 | 70 |
| SE | 17788 | ENSG00000100485 | SOS2 | chr14 | - | 50153069 | 50153173 | 50150007 | 50150230 | 50156998 | 50157121 | 410 | 354 | 293 |
| SE | 17789 | ENSG00000100485 | SOS2 | chr14 | - | 50156998 | 50157121 | 50153069 | 50153173 | 50159430 | 50160086 | 225 | 185 | 161 |
| SE | 17791 | ENSG00000100485 | SOS2 | chr14 | - | 50174453 | 50174552 | 50161481 | 50161609 | 50180571 | 50180654 | 183 | 164 | 145 |
| SE | 17792 | ENSG00000100485 | SOS2 | chr14 | - | 50180571 | 50180682 | 50174453 | 50174552 | 50182462 | 50182606 | 349 | 279 | 233 |
| SE | 17794 | ENSG00000100485 | SOS2 | chr14 | - | 50188496 | 50188700 | 50182462 | 50182606 | 50199690 | 50199855 | 464 | 351 | 293 |
| SE | 17795 | ENSG00000100485 | SOS2 | chr14 | - | 50199690 | 50199855 | 50188496 | 50188700 | 50200952 | 50201084 | 560 | 360 | 347 |
| SE | 17796 | ENSG00000100485 | SOS2 | chr14 | - | 50204283 | 50204409 | 50200952 | 50201084 | 50231196 | 50231254 | 115 | 69 | 73 |
| SE | 17800 | ENSG00000140092 | FBLN5 | chr14 | - | 91939849 | 91939950 | 91936946 | 91937201 | 91940564 | 91940616 | 0 | 0 | 0 |
| SE | 17804 | ENSG00000165929 | TC2N | chr14 | - | 91792366 | 91792558 | 91787512 | 91787627 | 91798298 | 91798399 | 484 | 456 | 649 |
| SE | 17805 | ENSG00000165929 | TC2N | chr14 | - | 91797784 | 91797901 | 91787512 | 91787627 | 91798298 | 91798399 | 571 | 528 | 821 |
| SE | 17807 | ENSG00000165929 | TC2N | chr14 | - | 91797784 | 91797901 | 91792366 | 91792558 | 91798988 | 91799064 | 602 | 571 | 861 |
| SE | 17809 | ENSG00000165929 | TC2N | chr14 | - | 91798298 | 91798399 | 91797784 | 91797901 | 91798988 | 91798998 | 850 | 753 | 1197 |
| SE | 17812 | ENSG00000105669 | COPE | chr19 | - | 18906959 | 18907112 | 18905575 | 18905629 | 18910970 | 18911071 | 328 | 127 | 255 |
| SE | 17813 | ENSG00000105669 | COPE | chr19 | - | 18912983 | 18913046 | 18910566 | 18911071 | 18919222 | 18919354 | 255 | 156 | 200 |
| SE | 17815 | ENSG00000140006 | WDR89 | chr14 | - | 63641298 | 63641407 | 63624927 | 63625033 | 63641803 | 63641853 | 0 | 4 | 3 |
| SE | 17816 | ENSG00000157227 | MMP14 | chr14 | + | 22843021 | 22843172 | 22842409 | 22842717 | 22843256 | 22843418 | 0 | 0 | 0 |
| SE | 17817 | ENSG00000155465 | SLC7A7 | chr14 | - | 22773932 | 22774116 | 22773221 | 22773716 | 22812899 | 22813440 | 236 | 117 | 130 |
| SE | 17819 | ENSG00000155465 | SLC7A7 | chr14 | - | 22775443 | 22775540 | 22773932 | 22774116 | 22775832 | 22775936 | 478 | 193 | 268 |
| SE | 17821 | ENSG00000155465 | SLC7A7 | chr14 | - | 22778792 | 22778937 | 22776194 | 22776318 | 22779925 | 22780051 | 317 | 122 | 153 |
| SE | 17822 | ENSG00000155465 | SLC7A7 | chr14 | - | 22779925 | 22780051 | 22778792 | 22778937 | 22812899 | 22813440 | 267 | 102 | 133 |
| SE | 17823 | ENSG00000155465 | SLC7A7 | chr14 | - | 22779925 | 22780051 | 22778792 | 22778937 | 22815319 | 22815435 | 157 | 60 | 69 |
| SE | 17824 | ENSG00000155465 | SLC7A7 | chr14 | - | 22812899 | 22813440 | 22778792 | 22778937 | 22815319 | 22815435 | 141 | 59 | 96 |
| SE | 17825 | ENSG00000155465 | SLC7A7 | chr14 | - | 22812899 | 22813440 | 22779925 | 22780051 | 22815319 | 22815435 | 254 | 102 | 163 |
| SE | 17827 | ENSG00000155465 | SLC7A7 | chr14 | - | 22814694 | 22814883 | 22813307 | 22813440 | 22815313 | 22815467 | 0 | 2 | 0 |
| SE | 17830 | ENSG00000155465 | SLC7A7 | chr14 | - | 22815319 | 22815451 | 22812899 | 22813440 | 22819637 | 22819756 | 143 | 59 | 95 |
| SE | 17831 | ENSG00000155465 | SLC7A7 | chr14 | - | 22815319 | 22815451 | 22812899 | 22813440 | 22821121 | 22821232 | 146 | 66 | 96 |
| SE | 17832 | ENSG00000155465 | SLC7A7 | chr14 | - | 22819637 | 22819756 | 22815319 | 22815451 | 22821121 | 22821440 | 0 | 1 | 2 |
| SE | 17833 | ENSG00000155465 | SLC7A7 | chr14 | - | 22819637 | 22819756 | 22815319 | 22815451 | 22823023 | 22823072 | 2 | 1 | 2 |
| SE | 17834 | ENSG00000155465 | SLC7A7 | chr14 | - | 22821121 | 22821232 | 22815319 | 22815451 | 22822918 | 22822963 | 4 | 8 | 1 |
| SE | 17835 | ENSG00000155465 | SLC7A7 | chr14 | - | 22821121 | 22821232 | 22815319 | 22815451 | 22823023 | 22823072 | 3 | 8 | 3 |
| SE | 17836 | ENSG00000155465 | SLC7A7 | chr14 | - | 22821121 | 22821440 | 22815319 | 22815451 | 22823023 | 22823072 | 3 | 8 | 3 |
| SE | 17837 | ENSG00000155465 | SLC7A7 | chr14 | - | 22821121 | 22821440 | 22819637 | 22819756 | 22823023 | 22823072 | 0 | 0 | 0 |
| SE | 17840 | ENSG00000151327 | FAM177A1 | chr14 | + | 35050783 | 35050868 | 35046603 | 35046628 | 35053277 | 35053451 | 2 | 1 | 0 |
| SE | 17841 | ENSG00000151327 | FAM177A1 | chr14 | + | 35053277 | 35053451 | 35046271 | 35046628 | 35077149 | 35077216 | 185 | 148 | 131 |
| SE | 17842 | ENSG00000151327 | FAM177A1 | chr14 | + | 35053277 | 35053451 | 35046271 | 35046628 | 35078926 | 35079012 | 12 | 18 | 8 |
| SE | 17843 | ENSG00000151327 | FAM177A1 | chr14 | + | 35077149 | 35077216 | 35046271 | 35046628 | 35078926 | 35079012 | 156 | 170 | 119 |
| SE | 17844 | ENSG00000151327 | FAM177A1 | chr14 | + | 35077149 | 35077216 | 35053199 | 35053451 | 35078926 | 35079012 | 329 | 307 | 242 |
| SE | 17845 | ENSG00000151327 | FAM177A1 | chr14 | + | 35078926 | 35079024 | 35077149 | 35077216 | 35081021 | 35081103 | 288 | 255 | 208 |
| SE | 17847 | ENSG00000151327 | FAM177A1 | chr14 | + | 35106366 | 35106484 | 35099675 | 35102729 | 35109894 | 35109960 | 1 | 0 | 0 |
| SE | 17848 | ENSG00000151327 | FAM177A1 | chr14 | + | 35109894 | 35109960 | 35099675 | 35102729 | 35110569 | 35110726 | 2 | 5 | 0 |
| SE | 17849 | ENSG00000151327 | FAM177A1 | chr14 | + | 35110569 | 35110642 | 35102680 | 35102729 | 35112863 | 35113130 | 2 | 0 | 0 |
| SE | 17851 | ENSG00000089775 | ZBTB25 | chr14 | - | 64499440 | 64499563 | 64490360 | 64490540 | 64502900 | 64503127 | 11 | 9 | 14 |
| SE | 17852 | ENSG00000089775 | ZBTB25 | chr14 | - | 64499440 | 64499563 | 64490360 | 64490540 | 64503660 | 64503838 | 11 | 9 | 13 |
| SE | 17853 | ENSG00000089775 | ZBTB25 | chr14 | - | 64502568 | 64502815 | 64499440 | 64499563 | 64502900 | 64503127 | 7 | 7 | 14 |
| SE | 17854 | ENSG00000089775 | ZBTB25 | chr14 | - | 64502900 | 64503127 | 64490360 | 64490540 | 64503660 | 64503838 | 11 | 11 | 19 |
| SE | 17855 | ENSG00000089775 | ZBTB25 | chr14 | - | 64502900 | 64503127 | 64490360 | 64490540 | 64504486 | 64504584 | 11 | 10 | 18 |
| SE | 17856 | ENSG00000089775 | ZBTB25 | chr14 | - | 64502900 | 64503127 | 64499440 | 64499563 | 64503660 | 64503838 | 0 | 1 | 4 |
| SE | 17858 | ENSG00000092199 | HNRNPC | chr14 | - | 21211405 | 21211566 | 21209146 | 21211306 | 21211809 | 21211903 | 1801 | 1130 | 1451 |
| SE | 17859 | ENSG00000092199 | HNRNPC | chr14 | - | 21233952 | 21234229 | 21212959 | 21213117 | 21269297 | 21269400 | 424 | 217 | 433 |
| SE | 17860 | ENSG00000092199 | HNRNPC | chr14 | - | 21233952 | 21234229 | 21230897 | 21231072 | 21269297 | 21269400 | 2396 | 1490 | 2359 |
| SE | 17861 | ENSG00000092199 | HNRNPC | chr14 | - | 21234768 | 21234831 | 21233952 | 21234229 | 21263310 | 21263336 | 0 | 0 | 0 |
| SE | 17862 | ENSG00000092199 | HNRNPC | chr14 | - | 21236372 | 21236460 | 21233952 | 21234229 | 21263310 | 21263336 | 4 | 2 | 6 |
| SE | 17863 | ENSG00000092199 | HNRNPC | chr14 | - | 21254546 | 21254593 | 21233994 | 21234229 | 21263310 | 21263336 | 1 | 0 | 0 |
| SE | 17865 | ENSG00000092199 | HNRNPC | chr14 | - | 21263310 | 21263336 | 21233952 | 21234229 | 21268622 | 21268651 | 446 | 214 | 415 |
| SE | 17869 | ENSG00000092199 | HNRNPC | chr14 | - | 21263666 | 21263829 | 21263310 | 21263582 | 21269297 | 21269406 | 189 | 151 | 176 |
| SE | 17874 | ENSG00000187097 | ENTPD5 | chr14 | - | 73971851 | 73971908 | 73970009 | 73970125 | 73973876 | 73973978 | 12 | 14 | 19 |
| SE | 17876 | ENSG00000187097 | ENTPD5 | chr14 | - | 73973876 | 73973978 | 73972883 | 73973024 | 73974923 | 73974985 | 43 | 55 | 37 |
| SE | 17878 | ENSG00000187097 | ENTPD5 | chr14 | - | 73975935 | 73976015 | 73974923 | 73974985 | 73977023 | 73977059 | 9 | 17 | 4 |
| SE | 17879 | ENSG00000187097 | ENTPD5 | chr14 | - | 73976323 | 73976412 | 73974923 | 73974985 | 73977023 | 73977059 | 11 | 12 | 7 |
| SE | 17881 | ENSG00000187097 | ENTPD5 | chr14 | - | 74007695 | 74007812 | 73987885 | 73988172 | 74011090 | 74011150 | 0 | 0 | 0 |
| SE | 17882 | ENSG00000187097 | ENTPD5 | chr14 | - | 74007695 | 74007812 | 73996516 | 73996776 | 74011090 | 74011150 | 0 | 0 | 0 |
| SE | 17883 | ENSG00000129467 | ADCY4 | chr14 | - | 24318653 | 24318778 | 24318348 | 24318568 | 24319097 | 24319212 | 3 | 2 | 4 |
| SE | 17886 | ENSG00000129467 | ADCY4 | chr14 | - | 24321715 | 24321976 | 24319741 | 24319888 | 24322065 | 24322708 | 0 | 0 | 3 |
| SE | 17894 | ENSG00000129484 | PARP2 | chr14 | + | 20351046 | 20351122 | 20350525 | 20350622 | 20352244 | 20352347 | 26 | 29 | 41 |
| SE | 17895 | ENSG00000129484 | PARP2 | chr14 | + | 20354084 | 20354247 | 20352244 | 20352347 | 20354834 | 20354947 | 29 | 28 | 29 |

| | | | | | | | | | | | | | | | |
|---|---|---|---|---|---|---|---|---|---|---|---|---|---|---|---|
| SE | 17896 | ENSG00000129484 | PARP2 | chr14 | + | 20354808 | 20354947 | 20354084 | 20354247 | 20355751 | 20355815 | 53 | 29 | 38 | [illegible] |
| SE | 17897 | ENSG00000129484 | PARP2 | chr14 | + | 20356589 | 20356689 | 20356306 | 20356434 | 20357050 | 20357149 | 42 | 36 | 41 | [illegible] |
| SE | 17898 | ENSG00000129484 | PARP2 | chr14 | + | 20357395 | 20357520 | 20357050 | 20357149 | 20357637 | 20357904 | 18 | 16 | 13 | [illegible] |
| SE | 17899 | ENSG00000197249 | SERPINA1 | chr14 | - | 94379463 | 94379611 | 94376746 | 94378640 | 94380870 | 94381141 | 2498 | 1496 | 1152 | [illegible] |
| SE | 17901 | ENSG00000197249 | SERPINA1 | chr14 | - | 94379463 | 94379611 | 94376746 | 94378640 | 94388559 | 94388602 | 1274 | 743 | 568 | [illegible] |
| SE | 17902 | ENSG00000197249 | SERPINA1 | chr14 | - | 94379463 | 94379611 | 94376746 | 94378640 | 94390456 | 94390637 | 1273 | 745 | 570 | [illegible] |
| SE | 17903 | ENSG00000197249 | SERPINA1 | chr14 | - | 94380650 | 94381141 | 94379463 | 94379611 | 94382591 | 94383246 | 2251 | 1513 | 1059 | [illegible] |
| SE | 17904 | ENSG00000197249 | SERPINA1 | chr14 | - | 94380870 | 94381141 | 94376746 | 94378640 | 94382591 | 94383241 | 2249 | 1512 | 1058 | [illegible] |
| SE | 17905 | ENSG00000197249 | SERPINA1 | chr14 | - | 94380870 | 94381141 | 94376746 | 94378640 | 94388559 | 94388589 | 0 | 1 | 0 | [illegible] |
| SE | 17906 | ENSG00000197249 | SERPINA1 | chr14 | - | 94380870 | 94381141 | 94376746 | 94378640 | 94390456 | 94390637 | 0 | 1 | 0 | [illegible] |
| SE | 17907 | ENSG00000197249 | SERPINA1 | chr14 | - | 94380870 | 94381141 | 94379112 | 94379611 | 94382591 | 94383241 | 3474 | 2264 | 1642 | [illegible] |
| SE | 17908 | ENSG00000197249 | SERPINA1 | chr14 | - | 94380870 | 94381141 | 94379112 | 94379611 | 94388559 | 94388589 | 1225 | 753 | 584 | [illegible] |
| SE | 17909 | ENSG00000197249 | SERPINA1 | chr14 | - | 94380870 | 94381141 | 94379112 | 94379611 | 94390456 | 94390637 | 1225 | 753 | 584 | [illegible] |
| SE | 17911 | ENSG00000197249 | SERPINA1 | chr14 | - | 94382591 | 94383241 | 94376746 | 94378640 | 94390456 | 94390637 | 1097 | 529 | 550 | [illegible] |
| SE | 17912 | ENSG00000197249 | SERPINA1 | chr14 | - | 94382591 | 94383241 | 94379112 | 94379611 | 94388559 | 94388589 | 414 | 216 | 163 | [illegible] |
| SE | 17913 | ENSG00000197249 | SERPINA1 | chr14 | - | 94382591 | 94383241 | 94379112 | 94379611 | 94390456 | 94390637 | 1096 | 525 | 550 | [illegible] |
| SE | 17914 | ENSG00000197249 | SERPINA1 | chr14 | - | 94382591 | 94383241 | 94380870 | 94381141 | 94388559 | 94388589 | 2663 | 1727 | 1223 | [illegible] |
| SE | 17915 | ENSG00000197249 | SERPINA1 | chr14 | - | 94382591 | 94383241 | 94380870 | 94381141 | 94390456 | 94390637 | 3345 | 2036 | 1610 | [illegible] |
| SE | 17916 | ENSG00000197249 | SERPINA1 | chr14 | - | 94384033 | 94384130 | 94382768 | 94383379 | 94388559 | 94388663 | 0 | 0 | 0 | [illegible] |
| SE | 17917 | ENSG00000197249 | SERPINA1 | chr14 | - | 94384033 | 94384130 | 94382768 | 94383379 | 94390456 | 94390635 | 0 | 0 | 1 | [illegible] |
| SE | 17918 | ENSG00000197249 | SERPINA1 | chr14 | - | 94384033 | 94384130 | 94383170 | 94383241 | 94388559 | 94388663 | 0 | 0 | 0 | [illegible] |
| SE | 17919 | ENSG00000197249 | SERPINA1 | chr14 | - | 94384033 | 94384130 | 94383170 | 94383241 | 94390456 | 94390635 | 0 | 0 | 0 | [illegible] |
| SE | 17920 | ENSG00000197249 | SERPINA1 | chr14 | - | 94384033 | 94384130 | 94383628 | 94383647 | 94388559 | 94388663 | 0 | 0 | 0 | [illegible] |
| SE | 17921 | ENSG00000197249 | SERPINA1 | chr14 | - | 94384033 | 94384130 | 94383628 | 94383647 | 94390456 | 94390635 | 0 | 0 | 0 | [illegible] |
| SE | 17922 | ENSG00000197249 | SERPINA1 | chr14 | - | 94388559 | 94388660 | 94378372 | 94378640 | 94390456 | 94390662 | 61 | 33 | 21 | [illegible] |
| SE | 17926 | ENSG00000197249 | SERPINA1 | chr14 | - | 94388559 | 94388660 | 94382591 | 94383241 | 94388818 | 94389010 | 353 | 204 | 141 | [illegible] |
| SE | 17929 | ENSG00000197249 | SERPINA1 | chr14 | - | 94388559 | 94388663 | 94376746 | 94378640 | 94390456 | 94390650 | 72 | 32 | 25 | [illegible] |
| SE | 17933 | ENSG00000197249 | SERPINA1 | chr14 | - | 94388559 | 94388663 | 94382591 | 94383241 | 94388818 | 94389010 | 366 | 210 | 147 | [illegible] |
| SE | 17936 | ENSG00000197249 | SERPINA1 | chr14 | - | 94388559 | 94388663 | 94383628 | 94383647 | 94390456 | 94390650 | 72 | 36 | 27 | [illegible] |
| SE | 17937 | ENSG00000197249 | SERPINA1 | chr14 | - | 94388559 | 94388663 | 94384033 | 94384130 | 94390456 | 94390650 | 72 | 32 | 25 | [illegible] |
| SE | 17944 | ENSG00000197249 | SERPINA1 | chr14 | - | 94388818 | 94389010 | 94388559 | 94388663 | 94390419 | 94390635 | 46 | 23 | 14 | [illegible] |
| SE | 17947 | ENSG00000171469 | ZNF561 | chr19 | - | 9614020 | 9614103 | 9607325 | 9611336 | 9617048 | 9617171 | 49 | 41 | 37 | [illegible] |
| [illegible] | [illegible] | [illegible] | [illegible] | [illegible] | [illegible] | [illegible] | [illegible] | [illegible] | [illegible] | [illegible] | [illegible] | [illegible] | [illegible] | [illegible] | [illegible] |

| | | | | | | | | | | | | | | | |
|---|---|---|---|---|---|---|---|---|---|---|---|---|---|---|---|
| SE | 18297 | ENSG00000100711 | ZFYVE21 | chr14 | + | 103727558 | 103727914 | 103726791 | 103726842 | 103728907 | 103728983 | 21 | 12 | 11 | [illegible] |
| SE | 18298 | ENSG00000100711 | ZFYVE21 | chr14 | + | 103727745 | 103727914 | 103726791 | 103726842 | 103728907 | 103728983 | 35 | 18 | 14 | [illegible] |
| SE | 18300 | ENSG00000100711 | ZFYVE21 | chr14 | + | 103729792 | 103729846 | 103729090 | 103729182 | 103732619 | 103732762 | 2 | 0 | 0 | [illegible] |
| SE | 18301 | ENSG00000100938 | GMPR2 | chr14 | + | 24233218 | 24233340 | 24232421 | 24232788 | 24233478 | 24233586 | 91 | 49 | 92 | [illegible] |
| SE | 18308 | ENSG00000100938 | GMPR2 | chr14 | + | 24233478 | 24233598 | 24232900 | 24233340 | 24235736 | 24235810 | 119 | 73 | 132 | [illegible] |
| SE | 18312 | ENSG00000100938 | GMPR2 | chr14 | + | 24237041 | 24237152 | 24235966 | 24236140 | 24237244 | 24237351 | 91 | 60 | 85 | [illegible] |
| SE | 18313 | ENSG00000100938 | GMPR2 | chr14 | + | 24237070 | 24237152 | 24235966 | 24236140 | 24237244 | 24237287 | 172 | 101 | 169 | [illegible] |
| SE | 18317 | ENSG00000100575 | TIMM9 | chr14 | - | 58411906 | 58411971 | 58410842 | 58410938 | 58427053 | 58427187 | 99 | 59 | 100 | [illegible] |
| SE | 18319 | ENSG00000100575 | TIMM9 | chr14 | - | 58424007 | 58424095 | 58410842 | 58410938 | 58427053 | 58427114 | 16 | 16 | 13 | [illegible] |
| SE | 18321 | ENSG00000100575 | TIMM9 | chr14 | - | 58424007 | 58424095 | 58411906 | 58411971 | 58427391 | 58427527 | 43 | 33 | 55 | [illegible] |
| SE | 18322 | ENSG00000100575 | TIMM9 | chr14 | - | 58427053 | 58427242 | 58424007 | 58424095 | 58427391 | 58427527 | 78 | 50 | 80 | [illegible] |
| SE | 18324 | ENSG00000032219 | ARID4A | chr14 | + | 58301579 | 58301690 | 58299536 | 58299860 | 58304943 | 58304990 | 232 | 194 | 190 | [illegible] |
| SE | 18325 | ENSG00000032219 | ARID4A | chr14 | + | 58303537 | 58303761 | 58301579 | 58301690 | 58304943 | 58305009 | 0 | 0 | 2 | [illegible] |
| SE | 18327 | ENSG00000032219 | ARID4A | chr14 | + | 58304943 | 58305009 | 58301579 | 58301690 | 58306021 | 58306112 | 210 | 190 | 181 | [illegible] |
| SE | 18330 | ENSG00000032219 | ARID4A | chr14 | + | 58330002 | 58330169 | 58329527 | 58329604 | 58344694 | 58344767 | 306 | 210 | 249 | [illegible] |
| SE | 18331 | ENSG00000032219 | ARID4A | chr14 | + | 58344694 | 58344767 | 58330002 | 58330169 | 58346410 | 58346505 | 158 | 127 | 149 | [illegible] |
| SE | 18332 | ENSG00000032219 | ARID4A | chr14 | + | 58346410 | 58346505 | 58344694 | 58344767 | 58347019 | 58347117 | 158 | 139 | 153 | [illegible] |
| SE | 18334 | ENSG00000032219 | ARID4A | chr14 | + | 58351072 | 58351323 | 58347646 | 58347878 | 58353657 | 58353855 | 105 | 132 | 93 | [illegible] |
| SE | 18335 | ENSG00000032219 | ARID4A | chr14 | + | 58360900 | 58361042 | 58359131 | 58359216 | 58364169 | 58365300 | 94 | 121 | 102 | [illegible] |
| SE | 18336 | ENSG00000032219 | ARID4A | chr14 | + | 58366023 | 58366068 | 58365517 | 58365622 | 58366882 | 58367029 | 16 | 8 | 12 | [illegible] |
| SE | 18337 | ENSG00000032219 | ARID4A | chr14 | + | 58366023 | 58366230 | 58365517 | 58365622 | 58366882 | 58367029 | 198 | 166 | 202 | [illegible] |
| SE | 18338 | ENSG00000257621 | PSMA3-AS1 | chr14 | - | 58273978 | 58274010 | 58265364 | 58267393 | 58285590 | 58285756 | 11 | 8 | 23 | [illegible] |

[illegible]

| | | | | | | | | | | | | | | | | | | | | | | | | | | | | | | | | | | | | | | | | | | | | | | |
|---|---|---|---|---|---|---|---|---|---|---|---|---|---|---|---|---|---|---|---|---|---|---|---|---|---|---|---|---|---|---|---|---|---|---|---|---|---|---|---|---|---|---|---|---|---|---|
| SE | 18778 | ENSG00000160190 | SLC37A1 | chr21 | + | 42542403 | 42542480 | 42539511 | 42539647 | 42543435 | 42543602 | 42 | 27 | 44 | 46 | 26 | 0 | 0 | 0 | 0 | 0 | 61 | 42 | 26 | 31 | 184 | 0 | 0 | 0 | 4 | 0 | 226 | 149 | 1 | 1 | 1 | 1 | 1 | 1 | 1 | 1 | 1 | 1 | 0.836 | 1 | 0.033 |
| SE | 18779 | ENSG00000160190 | SLC37A1 | chr21 | + | 42558957 | 42559089 | 42554061 | 42554142 | 42562077 | 42562168 | 57 | 26 | 44 | 74 | 39 | 0 | 0 | 1 | 0 | 0 | 61 | 52 | 29 | 55 | 155 | 0 | 0 | 0 | 0 | 0 | 281 | 149 | 1 | 1 | 1 | 1 | 0.959 | 1 | 1 | 1 | 1 | 1 | 1 | 1 | -0.008 |
| SE | 18780 | ENSG00000160190 | SLC37A1 | chr21 | + | 42564707 | 42564793 | 42563814 | 42563877 | 42565826 | 42565875 | 28 | 32 | 34 | 50 | 31 | 1 | 1 | 0 | 0 | 2 | 69 | 41 | 43 | 53 | 144 | 0 | 0 | 0 | 5 | 0 | 235 | 149 | 1 | 1 | 0.947 | 0.953 | 1 | 1 | 0.908 | 1 | 1 | 1 | 0.87 | 1 | -0.012 |
| SE | 18787 | ENSG00000119684 | MLH3 | chr14 | - | 75033418 | 75033490 | 75032067 | 75032179 | 75038339 | 75038412 | 17 | 13 | 23 | 17 | 16 | 4 | 4 | 4 | 5 | 5 | 0 | 1 | 25 | 7 | 0 | 0 | 4 | 4 | 0 | 0 | 221 | 149 | 1 | 1 | 0.741 | 0.687 | 0.795 | 0.696 | 0.683 | NA | 0.144 | 0.808 | 1 | NA | 0.07 |
| SE | 18791 | ENSG00000119684 | MLH3 | chr14 | - | 75038339 | 75038412 | 75032067 | 75032179 | 75046375 | 75046641 | 4 | 0 | 3 | 3 | 1 | 0 | 0 | 1 | 0 | 0 | 0 | 0 | 5 | 0 | 0 | 0 | 0 | 0 | 0 | 0 | 222 | 149 | 1 | 1 | 1 | NA | 0.668 | 1 | 1 | NA | NA | 1 | NA | NA | -0.083 |
| SE | 18792 | ENSG00000119684 | MLH3 | chr14 | - | 75038339 | 75038412 | 75033418 | 75033490 | 75039910 | 75040015 | 13 | 17 | 17 | 26 | 17 | 0 | 1 | 0 | 0 | 0 | 9 | 8 | 18 | 17 | 0 | 0 | 0 | 0 | 0 | 0 | 222 | 149 | 1 | 1 | 1 | 0.919 | 1 | 1 | 1 | 1 | 1 | 1 | 1 | NA | -0.016 |
| SE | 18793 | ENSG00000119684 | MLH3 | chr14 | - | 75038339 | 75038412 | 75033418 | 75033490 | 75041614 | 75041700 | 8 | 8 | 13 | 10 | 15 | 0 | 0 | 0 | 0 | 3 | 0 | 2 | 13 | 11 | 0 | 0 | 0 | 0 | 0 | 0 | 222 | 149 | 1 | 1 | 1 | 1 | 1 | 1 | 0.77 | NA | 1 | 1 | 1 | NA | -0.046 |
| SE | 18795 | ENSG00000119684 | MLH3 | chr14 | - | 75039910 | 75040015 | 75033418 | 75033490 | 75041614 | 75041700 | 4 | 10 | 6 | 9 | 19 | 0 | 0 | 0 | 0 | 3 | 0 | 0 | 3 | 0 | 0 | 0 | 0 | 0 | 0 | 0 | 254 | 149 | 1 | 1 | 1 | 1 | 1 | 1 | 0.788 | NA | NA | 1 | NA | NA | -0.042 |
| SE | 18796 | ENSG00000119684 | MLH3 | chr14 | - | 75039910 | 75040015 | 75038339 | 75038412 | 75041614 | 75041700 | 12 | 21 | 14 | 27 | 32 | 3 | 3 | 7 | 3 | 5 | 9 | 12 | 13 | 11 | 0 | 0 | 2 | 4 | 0 | 0 | 254 | 149 | 1 | 1 | 0.701 | 0.804 | 0.54 | 0.841 | 0.79 | 1 | 0.779 | 0.656 | 1 | NA | -0.124 |
| SE | 18797 | ENSG00000119684 | MLH3 | chr14 | - | 75039910 | 75040015 | 75038339 | 75038412 | 75046375 | 75047201 | 8 | 11 | 6 | 17 | 8 | 1 | 0 | 2 | 1 | 0 | 0 | 12 | 7 | 5 | 0 | 0 | 0 | 1 | 0 | 0 | 254 | 149 | 1 | 1 | 0.824 | 1 | 0.638 | 0.909 | 1 | NA | 1 | 0.804 | 1 | NA | -0.06 |
| SE | 18798 | ENSG00000119684 | MLH3 | chr14 | - | 75041614 | 75041700 | 75038339 | 75038412 | 75046375 | 75046444 | 1 | 2 | 2 | 3 | 0 | 1 | 0 | 2 | 1 | 0 | 0 | 2 | 4 | 0 | 0 | 0 | 0 | 1 | 0 | 0 | 235 | 149 | 1 | 1 | 0.388 | 1 | 0.388 | 0.655 | NA | NA | 1 | 0.717 | NA | NA | -0.251 |
| SE | 18800 | ENSG00000119684 | MLH3 | chr14 | - | 75042378 | 75042477 | 75041614 | 75041700 | 75046375 | 75046388 | 41 | 35 | 46 | 26 | 82 | 0 | 0 | 0 | 0 | 0 | 0 | 19 | 12 | 0 | 0 | 0 | 0 | 2 | 0 | 0 | 248 | 149 | 1 | 1 | 1 | 1 | 1 | 1 | 1 | NA | 1 | 0.783 | NA | NA | 0.109 |
| SE | 18802 | ENSG00000119689 | DLST | chr14 | + | 74882590 | 74882624 | 74881890 | 74882016 | 74885585 | 74885634 | 116 | 26 | 78 | 90 | 14 | 4 | 1 | 0 | 1 | 0 | 39 | 92 | 49 | 68 | 182 | 0 | 1 | 0 | 5 | 0 | 183 | 149 | 1 | 1 | 0.959 | 0.955 | 1 | 0.987 | 1 | 1 | 0.987 | 1 | 0.917 | 1 | -0.001 |
| SE | 18803 | ENSG00000119689 | DLST | chr14 | + | 74882590 | 74882624 | 74881890 | 74882016 | 74889094 | 74889147 | 0 | 0 | 0 | 0 | 0 | 0 | 1 | 0 | 0 | 0 | 0 | 7 | 0 | 4 | 2 | 0 | 3 | 13 | 6 | 0 | 183 | 149 | 1 | 1 | NA | 0 | NA | NA | NA | NA | 0.655 | 0 | 0.352 | 1 | -0.502 |
| SE | 18806 | ENSG00000119689 | DLST | chr14 | + | 74889094 | 74889147 | 74885585 | 74885634 | 74889274 | 74889349 | 183 | 60 | 108 | 151 | 65 | 1 | 0 | 0 | 0 | 0 | 120 | 172 | 79 | 102 | 311 | 0 | 0 | 0 | 0 | 0 | 202 | 149 | 1 | 1 | 0.993 | 1 | 1 | 1 | 1 | 1 | 1 | 1 | 1 | 1 | -0.001 |
| SE | 18807 | ENSG00000119689 | DLST | chr14 | + | 74889094 | 74889147 | 74885585 | 74885634 | 74889896 | 74889952 | 47 | 14 | 40 | 38 | 8 | 0 | 0 | 0 | 1 | 0 | 32 | 50 | 26 | 42 | 102 | 0 | 0 | 0 | 0 | 0 | 202 | 149 | 1 | 1 | 1 | 1 | 1 | 0.966 | 1 | 1 | 1 | 1 | 1 | 1 | -0.007 |
| SE | 18808 | ENSG00000119689 | DLST | chr14 | + | 74889274 | 74889349 | 74885585 | 74885634 | 74889896 | 74889952 | 94 | 44 | 62 | 75 | 91 | 0 | 0 | 0 | 1 | 0 | 37 | 76 | 41 | 43 | 134 | 0 | 0 | 0 | 0 | 0 | 224 | 149 | 1 | 1 | 1 | 1 | 1 | 0.98 | 1 | 1 | 1 | 1 | 1 | 1 | -0.004 |
| SE | 18809 | ENSG00000119689 | DLST | chr14 | + | 74889274 | 74889349 | 74889094 | 74889147 | 74889896 | 74889952 | 240 | 89 | 148 | 206 | 151 | 0 | 0 | 0 | 0 | 0 | 177 | 231 | 120 | 184 | 345 | 0 | 0 | 7 | 0 | 0 | 224 | 149 | 1 | 1 | 1 | 1 | 1 | 1 | 1 | 1 | 1 | 0.919 | 1 | 1 | 0.016 |
| SE | 18810 | ENSG00000119689 | DLST | chr14 | + | 74889877 | 74889952 | 74889274 | 74889349 | 74891055 | 74891193 | 31 | 22 | 41 | 12 | 36 | 1 | 1 | 0 | 0 | 0 | 40 | 15 | 32 | 47 | 34 | 0 | 0 | 0 | 0 | 1 | 224 | 149 | 1 | 1 | 0.954 | 0.936 | 1 | 1 | 1 | 1 | 1 | 1 | 1 | 0.958 | -0.014 |
| SE | 18811 | ENSG00000119689 | DLST | chr14 | + | 74889896 | 74889952 | 74889274 | 74889349 | 74891055 | 74891167 | 175 | 85 | 130 | 130 | 172 | 1 | 1 | 0 | 0 | 0 | 117 | 152 | 90 | 117 | 298 | 0 | 0 | 0 | 0 | 1 | 205 | 149 | 1 | 1 | 0.992 | 0.984 | 1 | 1 | 1 | 1 | 1 | 1 | 1 | 0.995 | -0.004 |
| SE | 18813 | ENSG00000119689 | DLST | chr14 | + | 74898368 | 74898499 | 74893347 | 74893424 | 74899922 | 74899996 | 177 | 100 | 174 | 156 | 188 | 0 | 0 | 0 | 3 | 0 | 175 | 275 | 168 | 218 | 399 | 0 | 0 | 0 | 6 | 0 | 280 | 149 | 1 | 1 | 1 | 1 | 1 | 0.965 | 1 | 1 | 1 | 1 | 0.951 | 1 | 0.003 |
| SE | 18814 | ENSG00000119689 | DLST | chr14 | + | 74898368 | 74898499 | 74893347 | 74893424 | 74900288 | 74900372 | 2 | 2 | 12 | 8 | 2 | 0 | 0 | 0 | 1 | 0 | 11 | 18 | 7 | 43 | 23 | 0 | 0 | 0 | 1 | 0 | 280 | 149 | 1 | 1 | 1 | 1 | 1 | 0.81 | 1 | 1 | 1 | 1 | 0.958 | 1 | -0.03 |
| SE | 18815 | ENSG00000119689 | DLST | chr14 | + | 74899922 | 74899996 | 74893347 | 74893424 | 74900288 | 74900372 | 39 | 33 | 47 | 53 | 26 | 0 | 0 | 0 | 1 | 0 | 60 | 62 | 35 | 71 | 112 | 0 | 0 | 0 | 1 | 0 | 223 | 149 | 1 | 1 | 1 | 1 | 1 | 0.973 | 1 | 1 | 1 | 1 | 0.979 | 1 | -0.001 |
| SE | 18818 | ENSG00000125520 | SLC2A4RG | chr20 | + | 63741868 | 63742056 | 63741369 | 63741479 | 63742129 | 63742166 | 98 | 55 | 108 | 93 | 73 | 0 | 0 | 1 | 0 | 0 | 22 | 113 | 58 | 80 | 98 | 0 | 0 | 0 | 0 | 0 | 298 | 149 | 1 | 1 | 1 | 1 | 0.982 | 1 | 1 | 1 | 1 | 1 | 1 | 1 | -0.004 |
| SE | 18825 | ENSG00000127663 | KDM4B | chr19 | + | 5047475 | 5047669 | 5041136 | 5041251 | 5071009 | 5071059 | 178 | 76 | 171 | 147 | 118 | 0 | 0 | 1 | 0 | 0 | 259 | 234 | 164 | 144 | 344 | 0 | 0 | 1 | 0 | 0 | 298 | 149 | 1 | 1 | 1 | 1 | 0.988 | 1 | 1 | 1 | 1 | 0.988 | 1 | 1 | 0 |
| SE | 18826 | ENSG00000127663 | KDM4B | chr19 | + | 5047475 | 5047669 | 5041136 | 5041251 | 5077366 | 5077470 | 114 | 62 | 123 | 102 | 83 | 0 | 0 | 0 | 0 | 0 | 241 | 163 | 114 | 102 | 226 | 0 | 0 | 0 | 0 | 6 | 298 | 149 | 1 | 1 | 1 | 1 | 1 | 1 | 1 | 1 | 1 | 1 | 1 | 0.95 | 0.01 |
| SE | 18827 | ENSG00000127663 | KDM4B | chr19 | + | 5071009 | 5071059 | 5041136 | 5041251 | 5077366 | 5077470 | 4 | 0 | 5 | 3 | 1 | 0 | 0 | 0 | 0 | 0 | 3 | 4 | 2 | 7 | 26 | 0 | 0 | 0 | 0 | 6 | 199 | 149 | 1 | 1 | 1 | NA | 1 | 1 | 1 | 1 | 1 | 1 | 1 | 0.764 | 0.047 |
| SE | 18829 | ENSG00000127663 | KDM4B | chr19 | + | 5077366 | 5077470 | 5047475 | 5047669 | 5082366 | 5082437 | 17 | 6 | 21 | 16 | 9 | 0 | 0 | 0 | 5 | 0 | 0 | 17 | 17 | 22 | 47 | 0 | 0 | 4 | 0 | 0 | 253 | 149 | 1 | 1 | 1 | 1 | 1 | 0.653 | 1 | NA | 1 | 0.715 | 1 | 1 | 0.002 |
| SE | 18846 | ENSG00000063761 | ADCK1 | chr14 | + | 77827853 | 77827971 | 77822434 | 77822518 | 77859075 | 77859279 | 0 | 0 | 0 | 2 | 2 | 4 | 1 | 6 | 8 | 2 | 3 | 2 | 0 | 0 | 0 | 0 | 18 | 7 | 4 | 0 | 267 | 149 | 1 | 1 | 0 | 0 | 0 | 0.122 | 0.358 | 1 | 0.058 | 0 | 0 | NA | -0.169 |
| SE | 18848 | ENSG00000063761 | ADCK1 | chr14 | + | 77887090 | 77887249 | 77859075 | 77859279 | 77899099 | 77899258 | 17 | 10 | 12 | 14 | 5 | 0 | 0 | 0 | 2 | 0 | 34 | 24 | 10 | 19 | 17 | 0 | 0 | 0 | 0 | 0 | 298 | 149 | 1 | 1 | 1 | 1 | 1 | 0.778 | 1 | 1 | 1 | 1 | 1 | 1 | -0.044 |
| SE | 18852 | ENSG00000119705 | SLIRP | chr14 | + | 77710554 | 77710665 | 77708113 | 77708208 | 77710837 | 77710896 | 2 | 2 | 2 | 0 | 2 | 76 | 56 | 85 | 83 | 81 | 0 | 0 | 0 | 0 | 0 | 100 | 73 | 73 | 59 | 20 | 260 | 149 | 1 | 1 | 0.015 | 0.02 | 0.013 | 0 | 0.014 | 0 | 0 | 0 | 0 | 0 | 0.012 |

[illegible]

| | | | | | | | | | | | | | | |
|---|---|---|---|---|---|---|---|---|---|---|---|---|---|---|
| SE | 19271 | ENSG00000187621 | TCL6 | chr14 | + | 95670441 | 95670577 | 95668125 | 95669617 | 95671330 | 95671487 | 28 | 17 | 12 |
| SE | 19272 | ENSG00000259084 | NA | chr14 | + | 95688178 | 95688274 | 95671330 | 95671487 | 95690735 | 95690906 | 7 | 0 | 0 |
| SE | 19274 | ENSG00000127152 | BCL11B | chr14 | - | 99231344 | 99231557 | 99169286 | 99176195 | 99271160 | 99272197 | 47 | 39 | 55 |
| SE | 19275 | ENSG00000127152 | BCL11B | chr14 | - | 99257470 | 99257839 | 99169286 | 99176195 | 99271160 | 99271485 | 171 | 81 | 189 |
| SE | 19276 | ENSG00000127152 | BCL11B | chr14 | - | 99257470 | 99257839 | 99231344 | 99231557 | 99271160 | 99271485 | 166 | 109 | 274 |
| SE | 19280 | ENSG00000099899 | TRMT2A | chr22 | - | 20115265 | 20115447 | 20114964 | 20115079 | 20115671 | 20115688 | 73 | 40 | 42 |
| SE | 19281 | ENSG00000099899 | TRMT2A | chr22 | - | 20115265 | 20115447 | 20114964 | 20115079 | 20116037 | 20116612 | 25 | 16 | 19 |
| SE | 19282 | ENSG00000099899 | TRMT2A | chr22 | - | 20115671 | 20115780 | 20114964 | 20115079 | 20116037 | 20116100 | 67 | 14 | 48 |
| SE | 19286 | ENSG00000284930 | NA | chr14 | - | 73462528 | 73462689 | 73458849 | 73459305 | 73463474 | 73463618 | 99 | 75 | 46 |
| SE | 19293 | ENSG00000162086 | ZNF75A | chr16 | + | 3312676 | 3312768 | 3310610 | 3310946 | 3316911 | 3317022 | 3 | 3 | 4 |
| SE | 19294 | ENSG00000162086 | ZNF75A | chr16 | + | 3312676 | 3312768 | 3310610 | 3310946 | 3317189 | 3317252 | 3 | 1 | 1 |
| SE | 19297 | ENSG00000162086 | ZNF75A | chr16 | + | 3316911 | 3317022 | 3310610 | 3310946 | 3317189 | 3317252 | 17 | 27 | 18 |
| SE | 19298 | ENSG00000162086 | ZNF75A | chr16 | + | 3316911 | 3317022 | 3312676 | 3312768 | 3317189 | 3317252 | 18 | 32 | 25 |
| SE | 19299 | ENSG00000162086 | ZNF75A | chr16 | + | 3316911 | 3317022 | 3313048 | 3313175 | 3317189 | 3317252 | 39 | 42 | 39 |
| SE | 19301 | ENSG00000119698 | PPP4R4 | chr14 | + | 94240672 | 94240795 | 94237564 | 94237686 | 94241787 | 94241957 | 2 | 0 | 0 |
| SE | 19304 | ENSG00000198752 | CDC42BPB | chr14 | - | 102959630 | 102959710 | 102954601 | 102954688 | 102963060 | 102963155 | 30 | 38 | 32 |
| SE | 19305 | ENSG00000156414 | TDRD9 | chr14 | + | 103963078 | 103963176 | 103955663 | 103955770 | 103965332 | 103965481 | 20 | 8 | 19 |
| SE | 19308 | ENSG00000156414 | TDRD9 | chr14 | + | 103994518 | 103994603 | 103994331 | 103994386 | 103995749 | 103995807 | 15 | 12 | 5 |
| SE | 19309 | ENSG00000156414 | TDRD9 | chr14 | + | 103998623 | 103998728 | 103995749 | 103995807 | 104004237 | 104004335 | 19 | 5 | 7 |
| SE | 19310 | ENSG00000156414 | TDRD9 | chr14 | + | 104007159 | 104007204 | 104006781 | 104006845 | 104008412 | 104008466 | 11 | 12 | 6 |
| SE | 19312 | ENSG00000156414 | TDRD9 | chr14 | + | 104018091 | 104018192 | 104015980 | 104016088 | 104024568 | 104024680 | 4 | 7 | 9 |
| SE | 19315 | ENSG00000156414 | TDRD9 | chr14 | + | 104026046 | 104026136 | 104025563 | 104025776 | 104026678 | 104026939 | 1 | 2 | 5 |
| SE | 19316 | ENSG00000156414 | TDRD9 | chr14 | + | 104031107 | 104031263 | 104026678 | 104026939 | 104032016 | 104032052 | 15 | 6 | 6 |
| SE | 19317 | ENSG00000156414 | TDRD9 | chr14 | + | 104034959 | 104035056 | 104033959 | 104034069 | 104040195 | 104040334 | 17 | 10 | 12 |
| SE | 19318 | ENSG00000156414 | TDRD9 | chr14 | + | 104049864 | 104049937 | 104049424 | 104049680 | 104051980 | 104052281 | 0 | 1 | 0 |
| SE | 19319 | ENSG00000071537 | SEL1L | chr14 | - | 81484224 | 81484397 | 81479611 | 81479740 | 81485671 | 81485746 | 484 | 235 | 270 |
| SE | 19320 | ENSG00000071537 | SEL1L | chr14 | - | 81487389 | 81487538 | 81486288 | 81486454 | 81489251 | 81489314 | 197 | 100 | 129 |
| SE | 19321 | ENSG00000071537 | SEL1L | chr14 | - | 81487854 | 81487942 | 81487389 | 81487538 | 81489251 | 81489314 | 312 | 170 | 182 |
| SE | 19323 | ENSG00000071537 | SEL1L | chr14 | - | 81499458 | 81499518 | 81498357 | 81498494 | 81499608 | 81499662 | 184 | 111 | 127 |
| SE | 19324 | ENSG00000071537 | SEL1L | chr14 | - | 81504200 | 81504306 | 81502720 | 81502883 | 81506073 | 81506155 | 327 | 207 | 218 |
| SE | 19331 | ENSG00000182979 | MTA1 | chr14 | + | 105458269 | 105458372 | 105454192 | 105454310 | 105460357 | 105460457 | 147 | 82 | 154 |
| SE | 19333 | ENSG00000182979 | MTA1 | chr14 | + | 105465093 | 105465183 | 105464462 | 105464863 | 105466425 | 105466578 | 72 | 35 | 43 |
| SE | 19334 | ENSG00000182979 | MTA1 | chr14 | + | 105466706 | 105466742 | 105465333 | 105466578 | 105469466 | 105469498 | 48 | 23 | 63 |
| SE | 19336 | ENSG00000125384 | PTGER2 | chr14 | + | 52315234 | 52315391 | 52314304 | 52314501 | 52327220 | 52327509 | 132 | 77 | 89 |
| SE | 19338 | ENSG00000100764 | PSMC1 | chr14 | + | 90263661 | 90263847 | 90263317 | 90263442 | 90264040 | 90264169 | 230 | 115 | 171 |
| SE | 19340 | ENSG00000167705 | RILP | chr17 | - | 1646905 | 1646989 | 1646149 | 1646619 | 1647834 | 1647957 | 39 | 22 | 22 |
| SE | 19349 | ENSG00000089737 | DDX24 | chr14 | - | 94054995 | 94055184 | 94052997 | 94053127 | 94057821 | 94057897 | 685 | 438 | 713 |
| SE | 19350 | ENSG00000089737 | DDX24 | chr14 | - | 94054995 | 94055184 | 94052997 | 94053127 | 94060097 | 94060613 | 524 | 329 | 526 |
| SE | 19351 | ENSG00000089737 | DDX24 | chr14 | - | 94054995 | 94057897 | 94052997 | 94053127 | 94060097 | 94060613 | 668 | 414 | 726 |
| SE | 19352 | ENSG00000089737 | DDX24 | chr14 | - | 94057821 | 94057897 | 94052997 | 94053127 | 94060097 | 94060613 | 94 | 57 | 122 |
| SE | 19353 | ENSG00000089737 | DDX24 | chr14 | - | 94057821 | 94057897 | 94054995 | 94055184 | 94060097 | 94060613 | 251 | 165 | 309 |
| SE | 19354 | ENSG00000089737 | DDX24 | chr14 | - | 94060097 | 94060613 | 94054995 | 94057897 | 94060912 | 94061066 | 576 | 361 | 729 |
| SE | 19357 | ENSG00000119720 | NRDE2 | chr14 | - | 90286353 | 90286492 | 90278557 | 90279133 | 90288216 | 90288885 | 61 | 20 | 22 |
| SE | 19358 | ENSG00000119720 | NRDE2 | chr14 | - | 90290220 | 90290607 | 90288216 | 90289145 | 90292696 | 90292872 | 32 | 25 | 43 |
| SE | 19359 | ENSG00000119720 | NRDE2 | chr14 | - | 90292696 | 90292872 | 90290220 | 90290607 | 90298259 | 90298380 | 25 | 27 | 33 |
| SE | 19361 | ENSG00000119720 | NRDE2 | chr14 | - | 90301773 | 90301803 | 90301238 | 90301372 | 90303934 | 90304382 | 0 | 0 | 0 |
| SE | 19372 | ENSG00000100811 | YY1 | chr14 | + | 100274697 | 100274758 | 100262303 | 100262466 | 100277417 | 100278142 | 136 | 87 | 132 |
| SE | 19375 | ENSG00000070367 | EXOC5 | chr14 | - | 57233987 | 57234032 | 57233742 | 57233883 | 57235710 | 57235820 | 120 | 104 | 73 |
| SE | 19376 | ENSG00000070367 | EXOC5 | chr14 | - | 57235710 | 57235820 | 57233987 | 57234032 | 57237337 | 57237358 | 209 | 163 | 138 |
| SE | 19377 | ENSG00000070367 | EXOC5 | chr14 | - | 57239594 | 57239659 | 57237337 | 57237366 | 57244164 | 57244359 | 179 | 157 | 130 |
| SE | 19378 | ENSG00000070367 | EXOC5 | chr14 | - | 57239594 | 57239659 | 57237337 | 57237366 | 57246710 | 57246858 | 53 | 38 | 39 |
| SE | 19379 | ENSG00000070367 | EXOC5 | chr14 | - | 57244164 | 57244359 | 57237337 | 57237366 | 57246710 | 57246827 | 123 | 114 | 93 |
| SE | 19381 | ENSG00000070367 | EXOC5 | chr14 | - | 57244164 | 57244359 | 57243247 | 57243636 | 57246710 | 57246827 | 125 | 113 | 95 |
| SE | 19382 | ENSG00000258875 | NA | chr14 | + | 91243912 | 91243981 | 91242934 | 91243013 | 91244354 | 91244499 | 1 | 0 | 5 |
| SE | 19384 | ENSG00000129460 | NGDN | chr14 | + | 23477199 | 23477356 | 23476165 | 23476407 | 23477502 | 23477538 | 147 | 107 | 160 |
| SE | 19388 | ENSG00000100629 | CEP128 | chr14 | - | 80527168 | 80527261 | 80526868 | 80526982 | 80530808 | 80530886 | 0 | 2 | 0 |
| SE | 19389 | ENSG00000100629 | CEP128 | chr14 | - | 80784894 | 80785545 | 80777881 | 80778046 | 80792759 | 80793110 | 25 | 21 | 48 |
| SE | 19390 | ENSG00000100629 | CEP128 | chr14 | - | 80831142 | 80831294 | 80792759 | 80793110 | 80836204 | 80836336 | 46 | 65 | 58 |
| SE | 19391 | ENSG00000100629 | CEP128 | chr14 | - | 80836204 | 80836337 | 80792759 | 80793110 | 80838203 | 80838278 | 21 | 34 | 32 |
| SE | 19392 | ENSG00000100629 | CEP128 | chr14 | - | 80862756 | 80862873 | 80840681 | 80840768 | 80895717 | 80895790 | 35 | 19 | 36 |
| SE | 19393 | ENSG00000100629 | CEP128 | chr14 | - | 80939384 | 80939540 | 80916400 | 80916562 | 80958177 | 80958285 | 17 | 12 | 16 |
| SE | 19394 | ENSG00000100629 | CEP128 | chr14 | - | 80942283 | 80942380 | 80939384 | 80939540 | 80958177 | 80958285 | 0 | 2 | 0 |
| SE | 19395 | ENSG00000100629 | CEP128 | chr14 | - | 80958177 | 80958285 | 80916400 | 80916562 | 80959431 | 80959484 | 4 | 0 | 4 |
| SE | 19396 | ENSG00000100629 | CEP128 | chr14 | - | 80958177 | 80958285 | 80942283 | 80942380 | 80959431 | 80959484 | 3 | 0 | 2 |
| SE | 19400 | ENSG00000197119 | SLC25A29 | chr14 | - | 100295625 | 100295697 | 100293293 | 100293377 | 100298841 | 100298885 | 1 | 0 | 0 |
| SE | 19401 | ENSG00000197119 | SLC25A29 | chr14 | - | 100295625 | 100295993 | 100293293 | 100293377 | 100298841 | 100298885 | 8 | 0 | 0 |
| SE | 19406 | ENSG00000151320 | AKAP6 | chr14 | + | 32628036 | 32628098 | 32600628 | 32600792 | 32678310 | 32678459 | 0 | 0 | 1 |
| SE | 19410 | ENSG00000092203 | TOX4 | chr14 | + | 21477495 | 21477564 | 21477121 | 21477284 | 21487450 | 21487602 | 231 | 148 | 219 |
| SE | 19411 | ENSG00000092203 | TOX4 | chr14 | + | 21477495 | 21477592 | 21477237 | 21477284 | 21487450 | 21487693 | 2 | 1 | 4 |
| SE | 19413 | ENSG00000092203 | TOX4 | chr14 | + | 21492295 | 21492376 | 21489172 | 21489403 | 21492507 | 21492606 | 632 | 444 | 556 |
| SE | 19414 | ENSG00000092203 | TOX4 | chr14 | + | 21495228 | 21495392 | 21492507 | 21493257 | 21496545 | 21496603 | 287 | 153 | 163 |
| SE | 19417 | ENSG00000198894 | CIPC | chr14 | + | 77099883 | 77099961 | 77098311 | 77098361 | 77105656 | 77105844 | 0 | 2 | 0 |
| SE | 19419 | ENSG00000198894 | CIPC | chr14 | + | 77109811 | 77109981 | 77105656 | 77105844 | 77113424 | 77113522 | 58 | 30 | 44 |
| SE | 19421 | ENSG00000151445 | VIPAS39 | chr14 | - | 77429680 | 77429767 | 77429005 | 77429095 | 77433841 | 77433931 | 92 | 48 | 79 |
| SE | 19422 | ENSG00000151445 | VIPAS39 | chr14 | - | 77434261 | 77434303 | 77433841 | 77433931 | 77435258 | 77435393 | 53 | 28 | 42 |
| SE | 19426 | ENSG00000151445 | VIPAS39 | chr14 | - | 77435843 | 77435919 | 77435170 | 77435393 | 77437807 | 77437881 | 43 | 18 | 31 |
| SE | 19427 | ENSG00000151445 | VIPAS39 | chr14 | - | 77437807 | 77437881 | 77435843 | 77435919 | 77441065 | 77441093 | 91 | 40 | 80 |
| SE | 19432 | ENSG00000151445 | VIPAS39 | chr14 | - | 77449713 | 77449752 | 77449292 | 77449357 | 77453298 | 77453401 | 2 | 3 | 5 |
| SE | 19437 | ENSG00000119636 | BBOF1 | chr14 | + | 74022915 | 74023144 | 74019348 | 74019534 | 74029183 | 74029249 | 13 | 4 | 0 |
| SE | 19438 | ENSG00000119636 | BBOF1 | chr14 | + | 74034027 | 74034171 | 74029183 | 74029249 | 74040564 | 74040645 | 11 | 23 | 24 |
| SE | 19439 | ENSG00000119636 | BBOF1 | chr14 | + | 74046059 | 74046130 | 74040564 | 74040645 | 74047929 | 74048074 | 2 | 9 | 8 |
| SE | 19441 | ENSG00000119636 | BBOF1 | chr14 | + | 74047929 | 74048074 | 74040564 | 74040645 | 74055583 | 74055685 | 0 | 4 | 5 |
| SE | 19442 | ENSG00000119636 | BBOF1 | chr14 | + | 74048345 | 74048453 | 74047929 | 74048074 | 74049701 | 74050195 | 0 | 0 | 1 |
| SE | 19443 | ENSG00000119636 | BBOF1 | chr14 | + | 74049701 | 74050195 | 74047929 | 74048074 | 74055583 | 74055685 | 7 | 1 | 9 |
| SE | 19444 | ENSG00000119636 | BBOF1 | chr14 | + | 74055583 | 74055685 | 74049701 | 74050195 | 74056905 | 74056980 | 6 | 2 | 12 |
| SE | 19450 | ENSG00000198668 | CALM1 | chr14 | + | 90403861 | 90403968 | 90401258 | 90401402 | 90404378 | 90404496 | 4906 | 3225 | 4168 |
| SE | 19451 | ENSG00000198668 | CALM1 | chr14 | + | 90404378 | 90404514 | 90403631 | 90403968 | 90404688 | 90404746 | 3946 | 2434 | 3365 |
| SE | 19452 | ENSG00000198668 | CALM1 | chr14 | + | 90404398 | 90404514 | 90403861 | 90403968 | 90404688 | 90404833 | 744 | 379 | 599 |
| SE | 19461 | ENSG00000125746 | EML2 | chr19 | - | 45624718 | 45624818 | 45621206 | 45621332 | 45629950 | 45630046 | 1 | 1 | 5 |
| SE | 19462 | ENSG00000125746 | EML2 | chr19 | - | 45624718 | 45624818 | 45621482 | 45621637 | 45626704 | 45626752 | 90 | 43 | 75 |
| SE | 19464 | ENSG00000125746 | EML2 | chr19 | - | 45628697 | 45628795 | 45626704 | 45626839 | 45629950 | 45630046 | 2 | 0 | 2 |
| SE | 19465 | ENSG00000125746 | EML2 | chr19 | - | 45629950 | 45630046 | 45626704 | 45626839 | 45632860 | 45632971 | 98 | 42 | 58 |
| SE | 19466 | ENSG00000125746 | EML2 | chr19 | - | 45632860 | 45633139 | 45629963 | 45630046 | 45634321 | 45634496 | 48 | 20 | 36 |
| SE | 19467 | ENSG00000139977 | NAA30 | chr14 | + | 57390956 | 57391728 | 57390585 | 57390705 | 57396751 | 57396875 | 73 | 20 | 32 |
| SE | 19468 | ENSG00000139977 | NAA30 | chr14 | + | 57396751 | 57396875 | 57390956 | 57391728 | 57409378 | 57409931 | 44 | 16 | 28 |
| SE | 19470 | ENSG00000139946 | PELI2 | chr14 | + | 56178334 | 56178464 | 56119108 | 56119210 | 56279675 | 56279777 | 131 | 83 | 89 |
| SE | 19471 | ENSG00000139946 | PELI2 | chr14 | + | 56279675 | 56279777 | 56178334 | 56178464 | 56288436 | 56288634 | 217 | 131 | 160 |
| SE | 19472 | ENSG00000033170 | FUT8 | chr14 | + | 65430215 | 65430309 | 65411862 | 65411993 | 65455620 | 65455718 | 0 | 0 | 2 |
| SE | 19473 | ENSG00000033170 | FUT8 | chr14 | + | 65430215 | 65430309 | 65413068 | 65413214 | 65455620 | 65455718 | 0 | 0 | 4 |
| SE | 19475 | ENSG00000033170 | FUT8 | chr14 | + | 65455620 | 65455718 | 65412729 | 65413214 | 65561336 | 65561652 | 31 | 18 | 33 |
| SE | 19476 | ENSG00000033170 | FUT8 | chr14 | + | 65455620 | 65455718 | 65412729 | 65413214 | 65615977 | 65616093 | 3 | 2 | 5 |
| SE | 19477 | ENSG00000033170 | FUT8 | chr14 | + | 65479706 | 65479784 | 65455620 | 65455718 | 65561336 | 65561685 | 0 | 2 | 0 |
| SE | 19478 | ENSG00000033170 | FUT8 | chr14 | + | 65516242 | 65516388 | 65455620 | 65455718 | 65561336 | 65561610 | 0 | 0 | 1 |
| SE | 19479 | ENSG00000033170 | FUT8 | chr14 | + | 65561336 | 65561766 | 65455620 | 65455718 | 65615977 | 65616093 | 140 | 91 | 144 |
| SE | 19480 | ENSG00000033170 | FUT8 | chr14 | + | 65615977 | 65616093 | 65561336 | 65561766 | 65616210 | 65616373 | 186 | 155 | 200 |
| SE | 19483 | ENSG00000033170 | FUT8 | chr14 | + | 65645959 | 65646117 | 65629491 | 65629606 | 65669242 | 65669480 | 0 | 0 | 0 |
| SE | 19485 | ENSG00000033170 | FUT8 | chr14 | + | 65646519 | 65646598 | 65629491 | 65629606 | 65669242 | 65669480 | 0 | 1 | 0 |
| SE | 19486 | ENSG00000033170 | FUT8 | chr14 | + | 65646519 | 65646598 | 65645959 | 65646117 | 65669242 | 65669480 | 0 | 1 | 0 |
| SE | 19487 | ENSG00000033170 | FUT8 | chr14 | + | 65721774 | 65722021 | 65669242 | 65669480 | 65724146 | 65724323 | 116 | 80 | 166 |
| SE | 19488 | ENSG00000033170 | FUT8 | chr14 | + | 65721774 | 65722021 | 65669242 | 65669480 | 65733230 | 65733381 | 48 | 34 | 78 |
| SE | 19489 | ENSG00000033170 | FUT8 | chr14 | + | 65724146 | 65724323 | 65669242 | 65669480 | 65733230 | 65733381 | 40 | 28 | 41 |
| SE | 19490 | ENSG00000033170 | FUT8 | chr14 | + | 65724146 | 65724323 | 65669242 | 65669480 | 65742092 | 65742936 | 4 | 0 | 0 |
| SE | 19491 | ENSG00000033170 | FUT8 | chr14 | + | 65724146 | 65724323 | 65721774 | 65722021 | 65733230 | 65733381 | 108 | 74 | 129 |
| SE | 19492 | ENSG00000033170 | FUT8 | chr14 | + | 65733230 | 65733381 | 65669242 | 65669480 | 65742092 | 65742936 | 51 | 75 | 87 |
| SE | 19500 | ENSG00000161558 | TMEM143 | chr19 | - | 48360071 | 48360183 | 48345159 | 48345354 | 48363290 | 48363674 | 5 | 2 | 5 |
| SE | 19507 | ENSG00000100479 | POLE2 | chr14 | - | 49655670 | 49655843 | 49655004 | 49655094 | 49663314 | 49663387 | 10 | 12 | 10 |
| SE | 19508 | ENSG00000100479 | POLE2 | chr14 | - | 49663314 | 49663387 | 49655670 | 49655843 | 49664625 | 49664663 | 13 | 18 | 19 |
| SE | 19509 | ENSG00000100479 | POLE2 | chr14 | - | 49666329 | 49666413 | 49665093 | 49665163 | 49669523 | 49669598 | 23 | 35 | 25 |
| SE | 19511 | ENSG00000100479 | POLE2 | chr14 | - | 49674122 | 49674216 | 49669523 | 49669598 | 49674349 | 49674427 | 25 | 29 | 27 |
| SE | 19514 | ENSG00000134815 | DHX34 | chr19 | + | 47376914 | 47376989 | 47376442 | 47376560 | 47377099 | 47377206 | 5 | 0 | 4 |
| SE | 19516 | ENSG00000080815 | PSEN1 | chr14 | + | 73137124 | 73137223 | 73136434 | 73136583 | 73147794 | 73147876 | 0 | 0 | 2 |
| SE | 19517 | ENSG00000080815 | PSEN1 | chr14 | + | 73138106 | 73138152 | 73136446 | 73136579 | 73147794 | 73147876 | 1 | 2 | 3 |
| SE | 19518 | ENSG00000080815 | PSEN1 | chr14 | + | 73138111 | 73138152 | 73136476 | 73136579 | 73147794 | 73147876 | 1 | 2 | 3 |
| SE | 19520 | ENSG00000080815 | PSEN1 | chr14 | + | 73138111 | 73138152 | 73137124 | 73137223 | 73147794 | 73147876 | 2 | 2 | 3 |
| SE | 19522 | ENSG00000080815 | PSEN1 | chr14 | + | 73146077 | 73146126 | 73138111 | 73138152 | 73147794 | 73147876 | 2 | 0 | 0 |
| SE | 19523 | ENSG00000080815 | PSEN1 | chr14 | + | 73147942 | 73148106 | 73147794 | 73147876 | 73170796 | 73171047 | 127 | 77 | 70 |
| SE | 19524 | ENSG00000080815 | PSEN1 | chr14 | + | 73147966 | 73148094 | 73147491 | 73147876 | 73170796 | 73170895 | 334 | 214 | 272 |
| SE | 19525 | ENSG00000080815 | PSEN1 | chr14 | + | 73147966 | 73148106 | 73147794 | 73147876 | 73170796 | 73171007 | 288 | 184 | 206 |
| SE | 19526 | ENSG00000080815 | PSEN1 | chr14 | + | 73147977 | 73148094 | 73147794 | 73147876 | 73170796 | 73170998 | 142 | 86 | 107 |
| SE | 19528 | ENSG00000080815 | PSEN1 | chr14 | + | 73186852 | 73186920 | 73173565 | 73173707 | 73192643 | 73192864 | 185 | 128 | 152 |
| SE | 19529 | ENSG00000080815 | PSEN1 | chr14 | + | 73192643 | 73192864 | 73186852 | 73186920 | 73198030 | 73198129 | 297 | 220 | 184 |
| SE | 19530 | ENSG00000080815 | PSEN1 | chr14 | + | 73192643 | 73192864 | 73186852 | 73186920 | 73206385 | 73206472 | 81 | 55 | 78 |
| SE | 19531 | ENSG00000080815 | PSEN1 | chr14 | + | 73198030 | 73198129 | 73186852 | 73186920 | 73206385 | 73206472 | 353 | 283 | 198 |
| SE | 19533 | ENSG00000080815 | PSEN1 | chr14 | + | 73206385 | 73206472 | 73192643 | 73192864 | 73211768 | 73211853 | 115 | 109 | 83 |
| SE | 19534 | ENSG00000080815 | PSEN1 | chr14 | + | 73208823 | 73208915 | 73206385 | 73206472 | 73211768 | 73211942 | 1 | 2 | 0 |
| SE | 19535 | ENSG00000080815 | PSEN1 | chr14 | + | 73211768 | 73211942 | 73206152 | 73206472 | 73217125 | 73217158 | 575 | 519 | 406 |
| SE | 19536 | ENSG00000187790 | FANCM | chr14 | + | 45137068 | 45137241 | 45135929 | 45136539 | 45140631 | 45140709 | 9 | 3 | 4 |
| SE | 19537 | ENSG00000187790 | FANCM | chr14 | + | 45140631 | 45140709 | 45137068 | 45137241 | 45148836 | 45148995 | 11 | 3 | 6 |
| SE | 19538 | ENSG00000187790 | FANCM | chr14 | + | 45151396 | 45151528 | 45148836 | 45148995 | 45154696 | 45154822 | 16 | 15 | 15 |
| SE | 19539 | ENSG00000187790 | FANCM | chr14 | + | 45153919 | 45154052 | 45151396 | 45151528 | 45154696 | 45154822 | 24 | 27 | 32 |
| SE | 19540 | ENSG00000187790 | FANCM | chr14 | + | 45153919 | 45154052 | 45151396 | 45151528 | 45164358 | 45164565 | 17 | 15 | 24 |
| SE | 19541 | ENSG00000187790 | FANCM | chr14 | + | 45154696 | 45154822 | 45151396 | 45151528 | 45164358 | 45164565 | 1 | 0 | 1 |
| SE | 19542 | ENSG00000187790 | FANCM | chr14 | + | 45154696 | 45154822 | 45153919 | 45154052 | 45164358 | 45164565 | 5 | 10 | 9 |
| SE | 19543 | ENSG00000187790 | FANCM | chr14 | + | 45155372 | 45155459 | 45154696 | 45154822 | 45164358 | 45164565 | 32 | 25 | 19 |
| SE | 19544 | ENSG00000187790 | FANCM | chr14 | + | 45159095 | 45159280 | 45155372 | 45155459 | 45164358 | 45164565 | 41 | 36 | 37 |
| SE | 19545 | ENSG00000187790 | FANCM | chr14 | + | 45159095 | 45159280 | 45155372 | 45155459 | 45166949 | 45167163 | 19 | 15 | 21 |
| SE | 19546 | ENSG00000187790 | FANCM | chr14 | + | 45164358 | 45164565 | 45155372 | 45155459 | 45166949 | 45167163 | 20 | 13 | 23 |
| SE | 19547 | ENSG00000187790 | FANCM | chr14 | + | 45164358 | 45164565 | 45159095 | 45159280 | 45166949 | 45167163 | 42 | 34 | 38 |
| SE | 19548 | ENSG00000187790 | FANCM | chr14 | + | 45170588 | 45170746 | 45166949 | 45167163 | 45173054 | 45173210 | 49 | 39 | 48 |
| SE | 19549 | ENSG00000187790 | FANCM | chr14 | + | 45195480 | 45195585 | 45188801 | 45189362 | 45196171 | 45196547 | 0 | 0 | 2 |
| SE | 19550 | ENSG00000187790 | FANCM | chr14 | + | 45196171 | 45196547 | 45188801 | 45189362 | 45198643 | 45198935 | 30 | 15 | 30 |
| SE | 19552 | ENSG00000178386 | ZNF223 | chr19 | + | 44060454 | 44060581 | 44055108 | 44055191 | 44060748 | 44060841 | 7 | 7 | 16 |
| SE | 19554 | ENSG00000053770 | AP5M1 | chr14 | + | 57273708 | 57273750 | 57269055 | 57269388 | 57274243 | 57274889 | 0 | 0 | 1 |
| SE | 19556 | ENSG00000053770 | AP5M1 | chr14 | + | 57280194 | 57280422 | 57274243 | 57274889 | 57282088 | 57282228 | 294 | 169 | 175 |
| SE | 19557 | ENSG00000053770 | AP5M1 | chr14 | + | 57282088 | 57282228 | 57280194 | 57280422 | 57282933 | 57283019 | 277 | 171 | 170 |
| SE | 19559 | ENSG00000198718 | TOGARAM1 | chr14 | + | 44999362 | 44999497 | 44962189 | 44964467 | 45004060 | 45004288 | 38 | 14 | 25 |
| SE | 19560 | ENSG00000198718 | TOGARAM1 | chr14 | + | 45004060 | 45004366 | 44999362 | 44999497 | 45006007 | 45006267 | 79 | 47 | 66 |
| SE | 19561 | ENSG00000198718 | TOGARAM1 | chr14 | + | 45006007 | 45006267 | 45004060 | 45004366 | 45008912 | 45009145 | 84 | 52 | 66 |
| SE | 19562 | ENSG00000198718 | TOGARAM1 | chr14 | + | 45006007 | 45006267 | 45004060 | 45004366 | 45011974 | 45012075 | 47 | 38 | 48 |
| SE | 19563 | ENSG00000198718 | TOGARAM1 | chr14 | + | 45008912 | 45009145 | 45004060 | 45004366 | 45011974 | 45012075 | 20 | 7 | 4 |
| SE | 19566 | ENSG00000198718 | TOGARAM1 | chr14 | + | 45025782 | 45025872 | 45011886 | 45012075 | 45027298 | 45027474 | 58 | 55 | 42 |
| SE | 19568 | ENSG00000198718 | TOGARAM1 | chr14 | + | 45032222 | 45032376 | 45028175 | 45028329 | 45043685 | 45043791 | 45 | 48 | 55 |
| SE | 19569 | ENSG00000198718 | TOGARAM1 | chr14 | + | 45043685 | 45043791 | 45032222 | 45032376 | 45046544 | 45046703 | 23 | 21 | 27 |
| SE | 19571 | ENSG00000198718 | TOGARAM1 | chr14 | + | 45046544 | 45046703 | 45044634 | 45044870 | 45052435 | 45052562 | 18 | 20 | 21 |
| SE | 19577 | ENSG00000140044 | JDP2 | chr14 | + | 75437897 | 75438121 | 75428021 | 75428252 | 75461425 | 75461530 | 37 | 31 | 19 |
| SE | 19578 | ENSG00000165898 | ISCA2 | chr14 | + | 74494049 | 74494152 | 74493755 | 74493845 | 74494825 | 74495130 | 36 | 25 | 25 |
| SE | 19582 | ENSG00000119681 | LTBP2 | chr14 | - | 74507178 | 74507310 | 74506697 | 74506823 | 74508603 | 74508729 | 3 | 5 | 0 |
| SE | 19583 | ENSG00000119681 | LTBP2 | chr14 | - | 74507972 | 74508095 | 74506697 | 74506823 | 74508603 | 74508729 | 1 | 0 | 6 |
| SE | 19585 | ENSG00000175985 | PLEKHD1 | chr14 | + | 69500870 | 69500947 | 69500576 | 69500666 | 69501733 | 69501825 | 0 | 0 | 0 |
| SE | 19586 | ENSG00000119723 | COQ6 | chr14 | + | 73955450 | 73955509 | 73953434 | 73953569 | 73955804 | 73955928 | 33 | 18 | 38 |
| SE | 19587 | ENSG00000119723 | COQ6 | chr14 | + | 73958146 | 73958277 | 73955329 | 73955928 | 73958970 | 73959078 | 36 | 25 | 24 |
| SE | 19590 | ENSG00000119723 | COQ6 | chr14 | + | 73961736 | 73961903 | 73959414 | 73959522 | 73962969 | 73963108 | 9 | 8 | 21 |
| SE | 19591 | ENSG00000119723 | COQ6 | chr14 | + | 73961736 | 73961903 | 73961454 | 73961570 | 73962969 | 73963108 | 47 | 18 | 62 |
| SE | 19595 | ENSG00000029364 | SLC39A9 | chr14 | + | 69442068 | 69442266 | 69424093 | 69424202 | 69454811 | 69454897 | 122 | 75 | 113 |
| SE | 19596 | ENSG00000029364 | SLC39A9 | chr14 | + | 69453240 | 69453309 | 69424093 | 69424202 | 69454811 | 69454897 | 25 | 8 | 13 |
| SE | 19598 | ENSG00000029364 | SLC39A9 | chr14 | + | 69458362 | 69458729 | 69455731 | 69455866 | 69461696 | 69462381 | 45 | 26 | 37 |
| SE | 19604 | ENSG00000255185 | PDXDC2P | chr16 | - | 70018046 | 70018097 | 70014103 | 70014167 | 70021823 | 70021949 | 61 | 74 | 38 |
| SE | 19605 | ENSG00000255185 | PDXDC2P | chr16 | - | 70031899 | 70031968 | 70030987 | 70031066 | 70034170 | 70034360 | 88 | 102 | 71 |
| SE | 19607 | ENSG00000255185 | PDXDC2P | chr16 | - | 70038904 | 70038985 | 70036431 | 70036578 | 70042350 | 70042416 | 55 | 52 | 33 |
| SE | 19608 | ENSG00000255185 | PDXDC2P | chr16 | - | 70042350 | 70042416 | 70038904 | 70038985 | 70042929 | 70043003 | 28 | 22 | 14 |
| SE | 19609 | ENSG00000255185 | PDXDC2P | chr16 | - | 70064670 | 70064776 | 70042929 | 70043003 | 70065755 | 70065785 | 0 | 0 | 0 |
| SE | 19610 | ENSG00000203546 | NA | chr14 | - | 31343895 | 31344065 | 31337475 | 31337614 | 31347747 | 31347907 | 10 | 4 | 17 |
| SE | 19611 | ENSG00000203546 | NA | chr14 | - | 31345086 | 31345276 | 31343895 | 31344065 | 31347747 | 31347907 | 26 | 28 | 26 |
| SE | 19613 | ENSG00000203546 | NA | chr14 | - | 31364188 | 31364298 | 31358893 | 31359057 | 31371809 | 31371909 | 73 | 35 | 47 |
| SE | 19616 | ENSG00000203546 | NA | chr14 | - | 31398672 | 31398781 | 31395198 | 31395348 | 31400300 | 31400512 | 64 | 48 | 39 |
| SE | 19617 | ENSG00000203546 | NA | chr14 | - | 31400300 | 31400512 | 31398672 | 31398781 | 31402849 | 31403049 | 112 | 94 | 78 |
| SE | 19619 | ENSG00000203546 | NA | chr14 | - | 31428143 | 31428188 | 31402849 | 31403049 | 31453274 | 31453344 | 0 | 0 | 2 |
| SE | 19620 | ENSG00000129521 | EGLN3 | chr14 | - | 33926959 | 33927033 | 33924226 | 33925922 | 33929075 | 33929212 | 5 | 21 | 11 |
| SE | 19626 | ENSG00000237356 | NA | chr14 | + | 53188031 | 53188153 | 53170433 | 53170501 | 53189868 | 53189936 | 2 | 0 | 0 |
| SE | 19628 | ENSG00000237356 | NA | chr14 | + | 53313715 | 53313790 | 53201429 | 53201483 | 53318030 | 53318138 | 0 | 0 | 1 |
| SE | 19629 | ENSG00000237356 | NA | chr14 | + | 53313715 | 53313790 | 53217954 | 53218055 | 53318030 | 53318138 | 0 | 0 | 0 |
| SE | 19631 | ENSG00000237356 | NA | chr14 | + | 53510724 | 53510879 | 53369451 | 53369501 | 53687198 | 53688970 | 1 | 0 | 2 |
| SE | 19636 | ENSG00000119711 | ALDH6A1 | chr14 | - | 74065180 | 74065360 | 74064821 | 74064920 | 74066704 | 74066886 | 74 | 37 | 98 |
| SE | 19638 | ENSG00000119661 | DNAL1 | chr14 | + | 73651166 | 73651327 | 73645016 | 73645042 | 73653169 | 73653215 | 1 | 0 | 0 |
| SE | 19639 | ENSG00000119661 | DNAL1 | chr14 | + | 73653169 | 73653215 | 73644990 | 73645042 | 73654846 | 73654885 | 0 | 2 | 2 |
| SE | 19642 | ENSG00000119661 | DNAL1 | chr14 | + | 73654846 | 73654885 | 73653169 | 73653215 | 73661986 | 73662042 | 2 | 0 | 0 |
| SE | 19643 | ENSG00000119661 | DNAL1 | chr14 | + | 73658846 | 73658956 | 73654846 | 73654885 | 73661986 | 73662042 | 25 | 17 | 24 |
| SE | 19646 | ENSG00000119661 | DNAL1 | chr14 | + | 73672944 | 73673091 | 73671541 | 73671597 | 73687258 | 73687385 | 0 | 0 | 2 |
| SE | 19647 | ENSG00000119661 | DNAL1 | chr14 | + | 73689374 | 73689515 | 73687258 | 73687385 | 73695901 | 73695942 | 50 | 25 | 33 |
| SE | 19649 | ENSG00000081177 | EXD2 | chr14 | + | 69203916 | 69204000 | 69191528 | 69191591 | 69209423 | 69209803 | 9 | 5 | 4 |
| SE | 19651 | ENSG00000081177 | EXD2 | chr14 | + | 69203916 | 69204000 | 69191765 | 69192039 | 69209423 | 69209803 | 4 | 6 | 2 |
| SE | 19653 | ENSG00000081177 | EXD2 | chr14 | + | 69209423 | 69209803 | 69191510 | 69191591 | 69228815 | 69229072 | 16 | 10 | 14 |
| SE | 19654 | ENSG00000081177 | EXD2 | chr14 | + | 69209423 | 69209803 | 69203916 | 69204000 | 69228815 | 69229072 | 21 | 13 | 17 |
| SE | 19655 | ENSG00000081177 | EXD2 | chr14 | + | 69210111 | 69210201 | 69209423 | 69209803 | 69228815 | 69229072 | 1 | 0 | 1 |
| SE | 19656 | ENSG00000081177 | EXD2 | chr14 | + | 69234699 | 69235031 | 69230471 | 69230598 | 69236045 | 69236152 | 24 | 14 | 26 |
| SE | 19657 | ENSG00000081177 | EXD2 | chr14 | + | 69234773 | 69235031 | 69230471 | 69230598 | 69236045 | 69236152 | 9 | 8 | 16 |
| SE | 19658 | ENSG00000205659 | LIN52 | chr14 | + | 74091231 | 74091306 | 74084795 | 74084993 | 74095947 | 74095985 | 31 | 18 | 47 |
| SE | 19659 | ENSG00000205659 | LIN52 | chr14 | + | 74091237 | 74091306 | 74084962 | 74084993 | 74095947 | 74095985 | 17 | 8 | 26 |
| SE | 19660 | ENSG00000205659 | LIN52 | chr14 | + | 74101154 | 74101238 | 74097793 | 74097860 | 74198921 | 74200414 | 36 | 21 | 49 |
| SE | 19663 | ENSG00000119688 | ABCD4 | chr14 | - | 74292550 | 74292642 | 74292286 | 74292376 | 74292747 | 74292826 | 28 | 20 | 22 |
| SE | 19665 | ENSG00000119688 | ABCD4 | chr14 | - | 74293153 | 74293279 | 74292747 | 74292869 | 74295147 | 74295198 | 12 | 1 | 8 |
| SE | 19666 | ENSG00000119688 | ABCD4 | chr14 | - | 74293153 | 74293310 | 74292747 | 74292869 | 74295147 | 74295198 | 12 | 1 | 8 |
| SE | 19669 | ENSG00000119688 | ABCD4 | chr14 | - | 74297638 | 74298069 | 74296332 | 74296449 | 74300149 | 74300268 | 10 | 0 | 4 |
| SE | 19671 | ENSG00000119688 | ABCD4 | chr14 | - | 74297929 | 74298069 | 74296332 | 74296449 | 74300149 | 74300268 | 27 | 13 | 23 |
| SE | 19672 | ENSG00000119688 | ABCD4 | chr14 | - | 74299547 | 74299675 | 74296332 | 74296449 | 74300149 | 74300268 | 6 | 11 | 17 |
| SE | 19674 | ENSG00000119688 | ABCD4 | chr14 | - | 74299547 | 74299675 | 74297835 | 74298069 | 74302874 | 74302910 | 34 | 24 | 39 |
| SE | 19675 | ENSG00000119688 | ABCD4 | chr14 | - | 74300149 | 74300268 | 74297638 | 74298069 | 74302874 | 74302918 | 39 | 8 | 14 |
| SE | 19680 | ENSG00000258989 | NA | chr14 | + | 61547742 | 61547886 | 61530406 | 61530595 | 61657604 | 61657964 | 580 | 266 | 422 |
| SE | 19681 | ENSG00000100647 | SUSD6 | chr14 | + | 69703394 | 69703592 | 69658512 | 69658713 | 69704603 | 69704742 | 334 | 108 | 216 |
| SE | 19682 | ENSG00000100647 | SUSD6 | chr14 | + | 69704603 | 69704742 | 69658512 | 69658713 | 69708676 | 69709104 | 117 | 64 | 97 |
| SE | 19689 | ENSG00000050130 | JKAMP | chr14 | + | 59484989 | 59485116 | 59484442 | 59484593 | 59486712 | 59486804 | 6 | 4 | 6 |
| SE | 19690 | ENSG00000050130 | JKAMP | chr14 | + | 59484989 | 59485116 | 59484442 | 59484593 | 59486730 | 59486804 | 3 | 1 | 0 |
| SE | 19691 | ENSG00000050130 | JKAMP | chr14 | + | 59486680 | 59486804 | 59484568 | 59484593 | 59487673 | 59487828 | 58 | 31 | 51 |
| SE | 19692 | ENSG00000050130 | JKAMP | chr14 | + | 59486712 | 59486804 | 59484516 | 59484593 | 59487673 | 59487828 | 79 | 39 | 67 |
| SE | 19693 | ENSG00000050130 | JKAMP | chr14 | + | 59486730 | 59486804 | 59484502 | 59484593 | 59487673 | 59487828 | 44 | 19 | 37 |
| SE | 19694 | ENSG00000050130 | JKAMP | chr14 | + | 59498726 | 59498908 | 59495017 | 59495087 | 59501190 | 59501247 | 68 | 37 | 45 |
| SE | 19702 | ENSG00000258760 | NA | chr14 | - | 65269444 | 65269512 | 65247637 | 65247723 | 65318490 | 65318790 | 0 | 1 | 0 |
| SE | 19705 | ENSG00000140090 | SLC24A4 | chr14 | + | 92447370 | 92447424 | 92445316 | 92445342 | 92449073 | 92449159 | 14 | 19 | 9 |
| SE | 19709 | ENSG00000103811 | CTSH | chr15 | - | 78926174 | 78926254 | 78925333 | 78925440 | 78927712 | 78927781 | 0 | 0 | 0 |
| SE | 19710 | ENSG00000103811 | CTSH | chr15 | - | 78926562 | 78926684 | 78925333 | 78925440 | 78926781 | 78926868 | 1 | 0 | 2 |
| SE | 19711 | ENSG00000103811 | CTSH | chr15 | - | 78926781 | 78926830 | 78925333 | 78925440 | 78927712 | 78927781 | 1 | 0 | 0 |
| SE | 19712 | ENSG00000103811 | CTSH | chr15 | - | 78926781 | 78926868 | 78925333 | 78925440 | 78927712 | 78927781 | 0 | 0 | 0 |
| SE | 19713 | ENSG00000103811 | CTSH | chr15 | - | 78926781 | 78926939 | 78925333 | 78925440 | 78927712 | 78927781 | 0 | 0 | 0 |

| | | | |
|---|---|---|---|
| [illegible] | [illegible] | [illegible] | [illegible] |

| | | | | | | | | | | | | | | |
|---|---|---|---|---|---|---|---|---|---|---|---|---|---|---|
| SE | 20234 | ENSG00000004809 | SLC22A16 | chr6 | - | 110438719 | 110438847 | 110435851 | 110435961 | 110442243 | 110442775 | 8 | 0 | 2 |
| SE | 20235 | ENSG00000004809 | SLC22A16 | chr6 | - | 110446872 | 110446990 | 110442243 | 110442775 | 110456899 | 110457017 | 1 | 2 | 0 |
| SE | 20236 | ENSG00000004809 | SLC22A16 | chr6 | - | 110456537 | 110456803 | 110446872 | 110446990 | 110456899 | 110457017 | 6 | 1 | 0 |
| SE | 20238 | ENSG00000106305 | AIMP2 | chr7 | + | 6014901 | 6015052 | 6009254 | 6009378 | 6015145 | 6015352 | 0 | 1 | 1 |
| SE | 20239 | ENSG00000106305 | AIMP2 | chr7 | + | 6015145 | 6015352 | 6009271 | 6009498 | 6017813 | 6018045 | 96 | 42 | 52 |
| SE | 20241 | ENSG00000124789 | NUP153 | chr6 | - | 17624560 | 17624833 | 17616526 | 17616695 | 17625807 | 17626164 | 244 | 172 | 220 |
| SE | 20242 | ENSG00000124789 | NUP153 | chr6 | - | 17624560 | 17624833 | 17616526 | 17616695 | 17628654 | 17629539 | 124 | 65 | 86 |
| SE | 20243 | ENSG00000124789 | NUP153 | chr6 | - | 17625807 | 17626164 | 17616526 | 17616695 | 17628654 | 17629539 | 196 | 166 | 182 |
| SE | 20246 | ENSG00000124789 | NUP153 | chr6 | - | 17637152 | 17637770 | 17632649 | 17632844 | 17646066 | 17646154 | 91 | 121 | 93 |
| SE | 20247 | ENSG00000124789 | NUP153 | chr6 | - | 17639938 | 17640064 | 17637152 | 17637770 | 17646066 | 17646154 | 414 | 308 | 397 |
| SE | 20249 | ENSG00000124789 | NUP153 | chr6 | - | 17647806 | 17647905 | 17646066 | 17646154 | 17649162 | 17649300 | 427 | 286 | 362 |
| SE | 20250 | ENSG00000124789 | NUP153 | chr6 | - | 17649162 | 17649300 | 17646066 | 17646154 | 17661652 | 17661779 | 139 | 99 | 125 |
| SE | 20251 | ENSG00000124789 | NUP153 | chr6 | - | 17651833 | 17651926 | 17649162 | 17649300 | 17661652 | 17661779 | 1 | 1 | 1 |
| SE | 20253 | ENSG00000124789 | NUP153 | chr6 | - | 17662017 | 17662070 | 17649162 | 17649300 | 17665238 | 17665385 | 128 | 103 | 136 |
| SE | 20254 | ENSG00000124789 | NUP153 | chr6 | - | 17665238 | 17665385 | 17662017 | 17662070 | 17668974 | 17669028 | 392 | 366 | 421 |
| SE | 20257 | ENSG00000124789 | NUP153 | chr6 | - | 17669292 | 17669337 | 17668974 | 17669028 | 17674904 | 17675033 | 31 | 36 | 49 |
| SE | 20258 | ENSG00000124789 | NUP153 | chr6 | - | 17669429 | 17669546 | 17669292 | 17669337 | 17674904 | 17675033 | 432 | 371 | 434 |
| SE | 20261 | ENSG00000118482 | PHF3 | chr6 | + | 63646526 | 63646795 | 63635801 | 63636150 | 63678827 | 63679020 | 16 | 18 | 30 |
| SE | 20264 | ENSG00000118482 | PHF3 | chr6 | + | 63646526 | 63646795 | 63635801 | 63636150 | 63691736 | 63692043 | 14 | 18 | 22 |
| SE | 20265 | ENSG00000118482 | PHF3 | chr6 | + | 63646526 | 63646795 | 63635801 | 63636150 | 63694580 | 63694764 | 14 | 16 | 22 |
| SE | 20268 | ENSG00000118482 | PHF3 | chr6 | + | 63646618 | 63646795 | 63635835 | 63636150 | 63691736 | 63692043 | 4 | 2 | 2 |
| SE | 20269 | ENSG00000118482 | PHF3 | chr6 | + | 63646618 | 63646795 | 63635835 | 63636150 | 63694580 | 63694690 | 4 | 0 | 2 |
| SE | 20273 | ENSG00000118482 | PHF3 | chr6 | + | 63679999 | 63680161 | 63635822 | 63636150 | 63691736 | 63692043 | 55 | 33 | 44 |
| SE | 20274 | ENSG00000118482 | PHF3 | chr6 | + | 63679999 | 63680161 | 63635822 | 63636150 | 63694580 | 63694690 | 46 | 27 | 40 |
| SE | 20276 | ENSG00000118482 | PHF3 | chr6 | + | 63679999 | 63680161 | 63646525 | 63646795 | 63691736 | 63692043 | 101 | 67 | 94 |
| SE | 20277 | ENSG00000118482 | PHF3 | chr6 | + | 63679999 | 63680161 | 63646525 | 63646795 | 63694580 | 63694690 | 92 | 61 | 90 |
| SE | 20278 | ENSG00000118482 | PHF3 | chr6 | + | 63684128 | 63685911 | 63635819 | 63636150 | 63691736 | 63692043 | 269 | 294 | 254 |
| SE | 20280 | ENSG00000118482 | PHF3 | chr6 | + | 63684128 | 63685911 | 63646525 | 63646795 | 63691736 | 63692043 | 263 | 286 | 247 |
| SE | 20281 | ENSG00000118482 | PHF3 | chr6 | + | 63684128 | 63685911 | 63646525 | 63646795 | 63694580 | 63694690 | 11 | 4 | 10 |
| SE | 20284 | ENSG00000118482 | PHF3 | chr6 | + | 63691736 | 63692043 | 63635801 | 63636150 | 63694580 | 63694690 | 260 | 236 | 231 |
| SE | 20285 | ENSG00000118482 | PHF3 | chr6 | + | 63691736 | 63692043 | 63646525 | 63646795 | 63694580 | 63694690 | 259 | 237 | 231 |
| SE | 20286 | ENSG00000118482 | PHF3 | chr6 | + | 63691736 | 63692043 | 63679999 | 63680161 | 63694580 | 63694690 | 271 | 242 | 235 |

[illegible]

| | | | | | | | | | | | | | | | | | | | | | | | | | | | | | | | | | | | | | | | | | | | | | | |
|---|---|---|---|---|---|---|---|---|---|---|---|---|---|---|---|---|---|---|---|---|---|---|---|---|---|---|---|---|---|---|---|---|---|---|---|---|---|---|---|---|---|---|---|---|---|---|
| SE | 20686 | ENSG00000166024 | R3HCC1L | chr10 | + | 98211306 | 98211348 | 98208100 | 98209899 | 98231511 | 98231687 | 6 | 6 | 5 | 3 | 6 | 67 | 45 | 46 | 55 | 50 | 0 | 1 | 0 | 5 | 0 | 0 | 19 | 66 | 3 | 0 | 191 | 149 | 1 | 1 | 0.065 | 0.094 | 0.078 | 0.041 | 0.086 | NA | 0.039 | 0 | 0.565 | NA | -0.129 |
| SE | 20687 | ENSG00000166024 | R3HCC1L | chr10 | + | 98211306 | 98211348 | 98208100 | 98209899 | 98234445 | 98234516 | 3 | 3 | 4 | 4 | 5 | 5 | 4 | 12 | 7 | 3 | 0 | 3 | 0 | 0 | 0 | 0 | 2 | 12 | 2 | 0 | 191 | 149 | 1 | 1 | 0.319 | 0.369 | 0.206 | 0.308 | 0.565 | NA | 0.539 | 0 | 0 | NA | 0.174 |
| SE | 20688 | ENSG00000166024 | R3HCC1L | chr10 | + | 98231511 | 98231687 | 98163286 | 98163397 | 98234445 | 98234516 | 59 | 72 | 53 | 65 | 83 | 0 | 0 | 0 | 0 | 0 | 0 | 67 | 87 | 15 | 4 | 0 | 0 | 0 | 0 | 12 | 298 | 149 | 1 | 1 | 1 | 1 | 1 | 1 | 1 | NA | 1 | 1 | 1 | 0.143 | 0.214 |
| SE | 20689 | ENSG00000166024 | R3HCC1L | chr10 | + | 98231511 | 98231687 | 98208100 | 98209899 | 98234445 | 98234516 | 126 | 117 | 99 | 120 | 133 | 5 | 4 | 12 | 7 | 3 | 0 | 86 | 153 | 18 | 4 | 0 | 2 | 12 | 2 | 0 | 298 | 149 | 1 | 1 | 0.926 | 0.936 | 0.805 | 0.896 | 0.957 | NA | 0.956 | 0.864 | 0.818 | 1 | -0.005 |
| SE | 20690 | ENSG00000166024 | R3HCC1L | chr10 | + | 98231511 | 98231687 | 98211306 | 98211348 | 98234445 | 98234516 | 62 | 78 | 54 | 67 | 84 | 0 | 3 | 0 | 3 | 0 | 0 | 68 | 87 | 20 | 4 | 0 | 3 | 0 | 0 | 0 | 298 | 149 | 1 | 1 | 1 | 0.929 | 1 | 0.918 | 1 | NA | 0.919 | 1 | 1 | 1 | -0.01 |
| SE | 20691 | ENSG00000166024 | R3HCC1L | chr10 | + | 98234445 | 98234516 | 98208100 | 98209899 | 98235424 | 98235520 | 40 | 42 | 41 | 28 | 20 | 0 | 0 | 1 | 1 | 0 | 8 | 16 | 53 | 15 | 0 | 0 | 0 | 1 | 0 | 0 | 220 | 149 | 1 | 1 | 1 | 1 | 0.965 | 0.95 | 1 | 1 | 1 | 0.973 | 1 | NA | -0.01 |
| SE | 20692 | ENSG00000165025 | SYK | chr9 | + | 90864588 | 90864667 | 90862205 | 90862344 | 90865047 | 90865097 | 418 | 298 | 339 | 400 | 536 | 0 | 0 | 0 | 0 | 0 | 1232 | 727 | 418 | 852 | 1183 | 5 | 0 | 0 | 0 | 0 | 228 | 149 | 1 | 1 | 1 | 1 | 1 | 1 | 1 | 0.994 | 1 | 1 | 1 | 1 | 0.001 |
| SE | 20693 | ENSG00000165025 | SYK | chr9 | + | 90865047 | 90865097 | 90862205 | 90862344 | 90867130 | 90867199 | 69 | 58 | 51 | 81 | 101 | 0 | 0 | 0 | 0 | 0 | 275 | 153 | 59 | 151 | 215 | 6 | 0 | 0 | 0 | 0 | 199 | 149 | 1 | 1 | 1 | 1 | 1 | 1 | 1 | 0.972 | 1 | 1 | 1 | 1 | 0.006 |
| SE | 20695 | ENSG00000165025 | SYK | chr9 | + | 90887748 | 90887889 | 90878763 | 90878953 | 90888514 | 90888627 | 676 | 531 | 514 | 575 | 804 | 0 | 0 | 0 | 0 | 0 | 2897 | 1272 | 619 | 1559 | 1926 | 0 | 5 | 0 | 0 | 0 | 290 | 149 | 1 | 1 | 1 | 1 | 1 | 1 | 1 | 1 | 0.992 | 1 | 1 | 1 | 0.002 |
| SE | 20696 | ENSG00000165025 | SYK | chr9 | + | 90888514 | 90888627 | 90887748 | 90887889 | 90895527 | 90898549 | 657 | 472 | 484 | 590 | 859 | 3 | 1 | 1 | 3 | 1 | 2300 | 1069 | 574 | 1388 | 1877 | 0 | 8 | 0 | 9 | 0 | 262 | 149 | 1 | 1 | 0.992 | 0.996 | 0.996 | 0.991 | 0.998 | 1 | 0.987 | 1 | 0.989 | 1 | -0.001 |
| SE | 20697 | ENSG00000111325 | OGFOD2 | chr12 | + | 122975810 | 122975867 | 122975219 | 122975383 | 122976653 | 122976767 | 6 | 1 | 7 | 2 | 2 | 1 | 0 | 2 | 0 | 0 | 0 | 3 | 3 | 2 | 29 | 0 | 0 | 0 | 1 | 0 | 206 | 149 | 1 | 1 | 0.813 | 1 | 0.717 | 1 | 1 | NA | 1 | 1 | 0.591 | 1 | 0.008 |
| SE | 20698 | ENSG00000115760 | BIRC6 | chr2 | + | 32401462 | 32401623 | 32401162 | 32401385 | 32406498 | 32406557 | 494 | 320 | 400 | 473 | 571 | 5 | 3 | 1 | 1 | 4 | 165 | 560 | 424 | 204 | 167 | 0 | 4 | 1 | 0 | 0 | 298 | 149 | 1 | 1 | 0.98 | 0.982 | 0.995 | 0.996 | 0.986 | 1 | 0.986 | 0.995 | 1 | 1 | -0.008 |
| SE | 20699 | ENSG00000115760 | BIRC6 | chr2 | + | 32406498 | 32406557 | 32401462 | 32401623 | 32414768 | 32416163 | 286 | 211 | 274 | 297 | 320 | 0 | 0 | 0 | 0 | 0 | 46 | 342 | 200 | 148 | 38 | 0 | 0 | 1 | 0 | 0 | 208 | 149 | 1 | 1 | 1 | 1 | 1 | 1 | 1 | 1 | 1 | 0.993 | 1 | 1 | 0.001 |
| SE | 20700 | ENSG00000115760 | BIRC6 | chr2 | + | 32414768 | 32416163 | 32406498 | 32406557 | 32429145 | 32429295 | 320 | 262 | 337 | 329 | 424 | 0 | 0 | 0 | 0 | 0 | 69 | 381 | 223 | 135 | 20 | 0 | 0 | 1 | 0 | 0 | 298 | 149 | 1 | 1 | 1 | 1 | 1 | 1 | 1 | 1 | 1 | 0.991 | 1 | 1 | 0.002 |
| SE | 20703 | ENSG00000115760 | BIRC6 | chr2 | + | 32439507 | 32439686 | 32436052 | 32436184 | 32441328 | 32441435 | 816 | 753 | 624 | 828 | 892 | 0 | 3 | 0 | 0 | 0 | 136 | 875 | 890 | 385 | 87 | 0 | 0 | 0 | 0 | 0 | 298 | 149 | 1 | 1 | 1 | 0.992 | 1 | 1 | 1 | 1 | 1 | 1 | 1 | 1 | -0.002 |
| SE | 20705 | ENSG00000115760 | BIRC6 | chr2 | + | 32448794 | 32448928 | 32445520 | 32445668 | 32453807 | 32453942 | 212 | 122 | 177 | 189 | 362 | 0 | 2 | 0 | 1 | 2 | 94 | 191 | 116 | 64 | 32 | 0 | 0 | 3 | 0 | 0 | 283 | 149 | 1 | 1 | 1 | 0.97 | 1 | 0.99 | 0.99 | 1 | 1 | 0.953 | 1 | 1 | -0.001 |
| SE | 20707 | ENSG00000115760 | BIRC6 | chr2 | + | 32453807 | 32453942 | 32445520 | 32445668 | 32463193 | 32463381 | 62 | 38 | 57 | 77 | 161 | 0 | 0 | 8 | 4 | 0 | 0 | 50 | 33 | 11 | 0 | 0 | 0 | 1 | 0 | 0 | 284 | 149 | 1 | 1 | 1 | 1 | 0.789 | 0.91 | 1 | NA | 1 | 0.945 | 1 | NA | -0.042 |
| SE | 20708 | ENSG00000115760 | BIRC6 | chr2 | + | 32453807 | 32453942 | 32448794 | 32448928 | 32463193 | 32463381 | 190 | 91 | 138 | 188 | 360 | 0 | 0 | 0 | 1 | 0 | 58 | 150 | 106 | 53 | 22 | 0 | 0 | 2 | 0 | 0 | 284 | 149 | 1 | 1 | 1 | 1 | 1 | 0.99 | 1 | 1 | 1 | 0.965 | 1 | 1 | 0.005 |
| SE | 20710 | ENSG00000115760 | BIRC6 | chr2 | + | 32453807 | 32453954 | 32448794 | 32448928 | 32463193 | 32463381 | 161 | 73 | 114 | 137 | 262 | 0 | 0 | 0 | 1 | 0 | 74 | 126 | 96 | 51 | 55 | 0 | 0 | 2 | 0 | 0 | 296 | 149 | 1 | 1 | 1 | 1 | 1 | 0.986 | 1 | 1 | 1 | 0.96 | 1 | 1 | 0.005 |
| SE | 20711 | ENSG00000115760 | BIRC6 | chr2 | + | 32463193 | 32463381 | 32453807 | 32453942 | 32464508 | 32464823 | 135 | 74 | 129 | 142 | 259 | 0 | 0 | 0 | 0 | 0 | 7 | 104 | 91 | 25 | 2 | 0 | 0 | 1 | 0 | 0 | 298 | 149 | 1 | 1 | 1 | 1 | 1 | 1 | 1 | 1 | 1 | 0.978 | 1 | 1 | 0.004 |
| SE | 20713 | ENSG00000115760 | BIRC6 | chr2 | + | 32464508 | 32464823 | 32453807 | 32453942 | 32467524 | 32467739 | 1 | 2 | 4 | 4 | 0 | 1 | 0 | 0 | 2 | 3 | 10 | 26 | 21 | 5 | 0 | 0 | 0 | 0 | 6 | 0 | 298 | 149 | 1 | 1 | 0.333 | 1 | 1 | 0.5 | 0 | 1 | 1 | 1 | 0.294 | NA | -0.257 |
| SE | 20714 | ENSG00000115760 | BIRC6 | chr2 | + | 32464508 | 32464823 | 32453807 | 32453954 | 32467524 | 32467739 | 1 | 3 | 4 | 4 | 0 | 1 | 0 | 0 | 0 | 0 | 10 | 33 | 25 | 5 | 0 | 0 | 0 | 0 | 0 | 0 | 298 | 149 | 1 | 1 | 0.333 | 1 | 1 | 1 | NA | 1 | 1 | 1 | 1 | NA | -0.167 |
| SE | 20715 | ENSG00000115760 | BIRC6 | chr2 | + | 32464508 | 32464823 | 32463193 | 32463381 | 32465064 | 32465164 | 288 | 207 | 267 | 269 | 311 | 1 | 1 | 4 | 1 | 3 | 62 | 231 | 227 | 62 | 56 | 0 | 4 | 0 | 0 | 0 | 298 | 149 | 1 | 1 | 0.993 | 0.99 | 0.971 | 0.993 | 0.981 | 1 | 0.967 | 1 | 1 | 1 | -0.008 |
| SE | 20718 | ENSG00000115760 | BIRC6 | chr2 | + | 32467524 | 32467739 | 32464508 | 32464823 | 32467902 | 32468111 | 278 | 253 | 293 | 275 | 400 | 0 | 0 | 0 | 0 | 0 | 111 | 315 | 311 | 94 | 45 | 0 | 6 | 0 | 0 | 0 | 298 | 149 | 1 | 1 | 1 | 1 | 1 | 1 | 1 | 1 | 0.963 | 1 | 1 | 1 | 0.007 |
| SE | 20719 | ENSG00000115760 | BIRC6 | chr2 | + | 32467524 | 32467739 | 32465064 | 32465164 | 32467902 | 32468111 | 498 | 384 | 449 | 446 | 594 | 0 | 0 | 1 | 0 | 0 | 145 | 403 | 471 | 141 | 100 | 0 | 0 | 2 | 0 | 0 | 298 | 149 | 1 | 1 | 1 | 1 | 0.996 | 1 | 1 | 1 | 1 | 0.992 | 1 | 1 | 0.001 |
| SE | 20720 | ENSG00000115760 | BIRC6 | chr2 | + | 32468436 | 32468783 | 32467902 | 32468111 | 32469394 | 32469614 | 291 | 263 | 287 | 312 | 436 | 0 | 1 | 0 | 1 | 0 | 70 | 276 | 297 | 108 | 31 | 0 | 0 | 0 | 0 | 0 | 298 | 149 | 1 | 1 | 1 | 0.992 | 1 | 0.994 | 1 | 1 | 1 | 1 | 1 | 1 | -0.003 |
| SE | 20721 | ENSG00000115760 | BIRC6 | chr2 | + | 32469394 | 32469614 | 32468436 | 32468783 | 32470167 | 32470301 | 308 | 283 | 267 | 285 | 291 | 0 | 0 | 0 | 0 | 0 | 59 | 296 | 255 | 100 | 23 | 0 | 0 | 5 | 0 | 0 | 298 | 149 | 1 | 1 | 1 | 1 | 1 | 1 | 1 | 1 | 1 | 0.962 | 1 | 1 | 0.008 |
| SE | 20723 | ENSG00000115760 | BIRC6 | chr2 | + | 32473111 | 32473239 | 32471013 | 32471151 | 32476212 | 32476344 | 369 | 326 | 320 | 298 | 518 | 1 | 0 | 0 | 0 | 2 | 31 | 342 | 356 | 68 | 10 | 0 | 0 | 0 | 0 | 0 | 277 | 149 | 1 | 1 | 0.995 | 1 | 1 | 1 | 0.993 | 1 | 1 | 1 | 1 | 1 | -0.002 |
| SE | 20726 | ENSG00000115760 | BIRC6 | chr2 | + | 32481319 | 32481453 | 32479461 | 32479617 | 32482428 | 32482582 | 642 | 446 | 376 | 496 | 635 | 0 | 3 | 0 | 0 | 3 | 95 | 571 | 574 | 258 | 155 | 0 | 2 | 0 | 2 | 0 | 283 | 149 | 1 | 1 | 1 | 0.987 | 1 | 1 | 0.991 | 1 | 0.993 | 1 | 0.985 | 1 | 0 |
| SE | 20729 | ENSG00000115760 | BIRC6 | chr2 | + | 32491424 | 32491558 | 32488587 | 32488714 | 32493539 | 32493667 | 211 | 143 | 143 | 156 | 108 | 0 | 1 | 0 | 0 | 0 | 61 | 90 | 145 | 65 | 43 | 0 | 0 | 0 | 0 | 0 | 283 | 149 | 1 | 1 | 1 | 0.987 | 1 | 1 | 1 | 1 | 1 | 1 | 1 | 1 | -0.003 |
| SE | 20731 | ENSG00000115760 | BIRC6 | chr2 | + | 32502794 | 32502891 | 32501712 | 32501888 | 32503041 | 32503236 | 282 | 210 | 219 | 293 | 319 | 0 | 0 | 0 | 0 | 0 | 93 | 264 | 197 | 22 | 10 | 0 | 2 | 0 | 0 | 0 | 246 | 149 | 1 | 1 | 1 | 1 | 1 | 1 | 1 | 1 | 0.988 | 1 | 1 | 1 | 0.002 |
| SE | 20732 | ENSG00000115760 | BIRC6 | chr2 | + | 32509737 | 32509994 | 32507979 | 32508259 | 32510525 | 32510634 | 198 | 116 | 156 | 165 | 244 | 2 | 0 | 0 | 0 | 0 | 12 | 140 | 157 | 21 | 0 | 0 | 0 | 0 | 0 | 0 | 298 | 149 | 1 | 1 | 0.98 | 1 | 1 | 1 | 1 | 1 | 1 | 1 | 1 | NA | -0.004 |

[illegible]

| | | | | | | | | | | | | | | |
|---|---|---|---|---|---|---|---|---|---|---|---|---|---|---|
| SE | 21225 | ENSG00000272325 | NUDT3 | chr6 | - | 34341861 | 34341972 | 34295640 | 34295685 | 34392263 | 34392669 | 453 | 382 | 387 |
| SE | 21233 | ENSG00000136986 | DERL1 | chr8 | - | 123019194 | 123019305 | 123013169 | 123015585 | 123021446 | 123021499 | 526 | 347 | 339 |
| SE | 21234 | ENSG00000136986 | DERL1 | chr8 | - | 123021446 | 123021499 | 123019194 | 123019245 | 123022683 | 123022779 | 130 | 72 | 108 |
| SE | 21235 | ENSG00000136986 | DERL1 | chr8 | - | 123024985 | 123025050 | 123021446 | 123021499 | 123041969 | 123042122 | 0 | 0 | 1 |
| SE | 21239 | ENSG00000055208 | TAB2 | chr6 | + | 149342684 | 149342833 | 149317711 | 149318015 | 149378017 | 149379518 | 3 | 4 | 6 |
| SE | 21240 | ENSG00000055208 | TAB2 | chr6 | + | 149342684 | 149342833 | 149318581 | 149318782 | 149369908 | 149370075 | 9 | 17 | 35 |
| SE | 21241 | ENSG00000055208 | TAB2 | chr6 | + | 149342684 | 149342833 | 149318581 | 149318782 | 149378017 | 149379518 | 0 | 1 | 2 |
| SE | 21243 | ENSG00000055208 | TAB2 | chr6 | + | 149369908 | 149370099 | 149317711 | 149318015 | 149378017 | 149379518 | 361 | 331 | 357 |
| SE | 21244 | ENSG00000055208 | TAB2 | chr6 | + | 149369908 | 149370099 | 149318581 | 149318782 | 149378017 | 149379518 | 345 | 316 | 327 |
| SE | 21245 | ENSG00000055208 | TAB2 | chr6 | + | 149369908 | 149370099 | 149342656 | 149342833 | 149378017 | 149379518 | 354 | 332 | 362 |
| SE | 21246 | ENSG00000055208 | TAB2 | chr6 | + | 149378017 | 149379518 | 149369908 | 149370099 | 149397603 | 149397764 | 811 | 663 | 814 |
| SE | 21248 | ENSG00000055208 | TAB2 | chr6 | + | 149397603 | 149397764 | 149378017 | 149379518 | 149397968 | 149398062 | 773 | 609 | 850 |
| SE | 21249 | ENSG00000055208 | TAB2 | chr6 | + | 149397603 | 149397764 | 149378017 | 149379518 | 149409576 | 149411395 | 466 | 348 | 487 |
| SE | 21250 | ENSG00000055208 | TAB2 | chr6 | + | 149397968 | 149398062 | 149378017 | 149379518 | 149409576 | 149411395 | 0 | 0 | 0 |
| SE | 21251 | ENSG00000055208 | TAB2 | chr6 | + | 149397968 | 149398062 | 149397603 | 149397764 | 149399103 | 149399184 | 581 | 524 | 679 |
| SE | 21252 | ENSG00000055208 | TAB2 | chr6 | + | 149397968 | 149398062 | 149397603 | 149397764 | 149409576 | 149411395 | 206 | 208 | 254 |
| SE | 21253 | ENSG00000055208 | TAB2 | chr6 | + | 149399103 | 149399184 | 149397603 | 149397764 | 149409576 | 149409617 | 80 | 80 | 110 |
| SE | 21254 | ENSG00000055208 | TAB2 | chr6 | + | 149399103 | 149399184 | 149397691 | 149398062 | 149409576 | 149409617 | 456 | 405 | 542 |
| SE | 21257 | ENSG00000231426 | FILNC1 | chr6 | + | 139856763 | 139856819 | 139839686 | 139839758 | 139859284 | 139859808 | 1 | 0 | 1 |
| SE | 21260 | ENSG00000155111 | CDK19 | chr6 | - | 110626775 | 110626845 | 110623289 | 110623362 | 110627001 | 110627145 | 93 | 80 | 81 |
| SE | 21265 | ENSG00000155111 | CDK19 | chr6 | - | 110670430 | 110670541 | 110667433 | 110667574 | 110746125 | 110746201 | 91 | 62 | 58 |
| SE | 21266 | ENSG00000155111 | CDK19 | chr6 | - | 110746125 | 110746201 | 110670430 | 110670541 | 110773875 | 110773937 | 46 | 34 | 30 |
| SE | 21271 | ENSG00000150867 | PIP4K2A | chr10 | - | 22550658 | 22550772 | 22541803 | 22542047 | 22567850 | 22567889 | 2901 | 2075 | 1846 |
| SE | 21272 | ENSG00000150867 | PIP4K2A | chr10 | - | 22550658 | 22550772 | 22541803 | 22542047 | 22573310 | 22573457 | 1014 | 826 | 607 |
| SE | 21273 | ENSG00000150867 | PIP4K2A | chr10 | - | 22567850 | 22567889 | 22541803 | 22542047 | 22573310 | 22573457 | 329 | 233 | 238 |
| SE | 21274 | ENSG00000150867 | PIP4K2A | chr10 | - | 22567850 | 22567889 | 22550658 | 22550772 | 22573310 | 22573457 | 2170 | 1453 | 1437 |
| SE | 21279 | ENSG00000150867 | PIP4K2A | chr10 | - | 22573310 | 22573457 | 22550658 | 22550772 | 22591628 | 22591675 | 1546 | 998 | 1065 |
| SE | 21281 | ENSG00000150867 | PIP4K2A | chr10 | - | 22573310 | 22573457 | 22550658 | 22550772 | 22609619 | 22609717 | 18 | 7 | 4 |
| SE | 21282 | ENSG00000150867 | PIP4K2A | chr10 | - | 22573310 | 22573457 | 22550658 | 22550772 | 22714182 | 22714578 | 8 | 5 | 3 |
| SE | 21283 | ENSG00000150867 | PIP4K2A | chr10 | - | 22573310 | 22573457 | 22567850 | 22567889 | 22591628 | 22591675 | 3640 | 2376 | 2439 |
| SE | 21284 | ENSG00000150867 | PIP4K2A | chr10 | - | 22573310 | 22573457 | 22567850 | 22567889 | 22607926 | 22608023 | 2194 | 1424 | 1439 |
| SE | 21285 | ENSG00000150867 | PIP4K2A | chr10 | - | 22573310 | 22573457 | 22569012 | 22569054 | 22591628 | 22591675 | 1552 | 1005 | 1088 |
| SE | 21287 | ENSG00000150867 | PIP4K2A | chr10 | - | 22591628 | 22591781 | 22550658 | 22550772 | 22609619 | 22609717 | 2 | 5 | 1 |
| SE | 21288 | ENSG00000150867 | PIP4K2A | chr10 | - | 22591628 | 22591781 | 22550658 | 22550772 | 22714182 | 22714578 | 1 | 5 | 0 |
| SE | 21289 | ENSG00000150867 | PIP4K2A | chr10 | - | 22591628 | 22591781 | 22567850 | 22567889 | 22607926 | 22608023 | 1069 | 634 | 790 |
| SE | 21292 | ENSG00000150867 | PIP4K2A | chr10 | - | 22591628 | 22591781 | 22573310 | 22573457 | 22714182 | 22714578 | 1539 | 994 | 1064 |
| SE | 21293 | ENSG00000150867 | PIP4K2A | chr10 | - | 22607926 | 22608023 | 22550658 | 22550772 | 22609619 | 22609717 | 436 | 236 | 271 |
| SE | 21296 | ENSG00000150867 | PIP4K2A | chr10 | - | 22609619 | 22609717 | 22550658 | 22550772 | 22714182 | 22714578 | 286 | 181 | 171 |
| SE | 21297 | ENSG00000150867 | PIP4K2A | chr10 | - | 22609619 | 22609717 | 22573310 | 22573457 | 22714182 | 22714578 | 295 | 183 | 174 |
| SE | 21298 | ENSG00000150867 | PIP4K2A | chr10 | - | 22609619 | 22609717 | 22591628 | 22591781 | 22714182 | 22714578 | 285 | 182 | 172 |
| SE | 21299 | ENSG00000150867 | PIP4K2A | chr10 | - | 22646218 | 22646272 | 22609619 | 22609717 | 22659609 | 22659730 | 1 | 0 | 1 |
| SE | 21301 | ENSG00000129534 | MIS18BP1 | chr14 | - | 45217018 | 45217179 | 45210110 | 45210528 | 45218281 | 45218432 | 432 | 360 | 242 |
| SE | 21303 | ENSG00000129534 | MIS18BP1 | chr14 | - | 45227662 | 45227814 | 45226742 | 45226836 | 45231140 | 45231298 | 853 | 793 | 434 |
| SE | 21304 | ENSG00000129534 | MIS18BP1 | chr14 | - | 45231140 | 45231298 | 45227662 | 45227814 | 45232508 | 45232627 | 617 | 616 | 314 |
| SE | 21305 | ENSG00000129534 | MIS18BP1 | chr14 | - | 45231140 | 45231298 | 45227662 | 45227814 | 45232732 | 45232820 | 840 | 822 | 420 |
| SE | 21306 | ENSG00000129534 | MIS18BP1 | chr14 | - | 45232508 | 45232627 | 45227662 | 45227814 | 45232732 | 45232820 | 6 | 1 | 10 |
| SE | 21308 | ENSG00000129534 | MIS18BP1 | chr14 | - | 45232732 | 45232820 | 45231140 | 45231298 | 45235813 | 45235944 | 305 | 298 | 176 |
| SE | 21309 | ENSG00000129534 | MIS18BP1 | chr14 | - | 45235813 | 45235944 | 45232732 | 45232820 | 45237647 | 45237721 | 532 | 537 | 289 |
| SE | 21311 | ENSG00000129534 | MIS18BP1 | chr14 | - | 45242760 | 45242874 | 45237647 | 45237721 | 45253034 | 45253198 | 13 | 16 | 11 |
| SE | 21312 | ENSG00000129534 | MIS18BP1 | chr14 | - | 45242760 | 45242874 | 45242033 | 45242518 | 45246742 | 45247377 | 544 | 485 | 322 |
| SE | 21313 | ENSG00000129534 | MIS18BP1 | chr14 | - | 45246742 | 45247377 | 45242760 | 45242874 | 45253034 | 45253198 | 321 | 299 | 208 |
| SE | 21315 | ENSG00000126860 | EVI2A | chr17 | - | 31320372 | 31320513 | 31317559 | 31319023 | 31320956 | 31321032 | 18 | 8 | 15 |
| SE | 21317 | ENSG00000137055 | PLAA | chr9 | - | 26907833 | 26907998 | 26903371 | 26906076 | 26910337 | 26910439 | 162 | 137 | 131 |
| SE | 21318 | ENSG00000137055 | PLAA | chr9 | - | 26910337 | 26910439 | 26907609 | 26907998 | 26913878 | 26913947 | 139 | 104 | 135 |
| SE | 21320 | ENSG00000137055 | PLAA | chr9 | - | 26917096 | 26917165 | 26913878 | 26913947 | 26919309 | 26919461 | 196 | 159 | 182 |
| SE | 21322 | ENSG00000137055 | PLAA | chr9 | - | 26935012 | 26935197 | 26928307 | 26928408 | 26946896 | 26946983 | 104 | 72 | 71 |
| SE | 21323 | ENSG00000137055 | PLAA | chr9 | - | 26935012 | 26935206 | 26928307 | 26928408 | 26946896 | 26947233 | 121 | 90 | 88 |
| SE | 21341 | ENSG00000157017 | GHRL | chr3 | - | 10289761 | 10289875 | 10286703 | 10286812 | 10292841 | 10292917 | 1 | 2 | 1 |
| SE | 21343 | ENSG00000157017 | GHRL | chr3 | - | 10289761 | 10289878 | 10286703 | 10286812 | 10292841 | 10292917 | 1 | 2 | 1 |
| SE | 21351 | ENSG00000056972 | TRAF3IP2 | chr6 | - | 111595702 | 111595820 | 111591544 | 111592094 | 111601195 | 111601318 | 0 | 0 | 0 |
| SE | 21355 | ENSG00000056972 | TRAF3IP2 | chr6 | - | 111601195 | 111601318 | 111595702 | 111595820 | 111605775 | 111605844 | 0 | 0 | 2 |
| SE | 21358 | ENSG00000206560 | ANKRD28 | chr3 | - | 15677479 | 15677562 | 15676973 | 15677056 | 15678208 | 15678354 | 87 | 77 | 67 |
| SE | 21359 | ENSG00000206560 | ANKRD28 | chr3 | - | 15713526 | 15713641 | 15712139 | 15712222 | 15720914 | 15721122 | 59 | 71 | 48 |
| SE | 21361 | ENSG00000206560 | ANKRD28 | chr3 | - | 15724381 | 15724524 | 15720914 | 15721127 | 15735409 | 15735497 | 126 | 102 | 94 |
| SE | 21362 | ENSG00000206560 | ANKRD28 | chr3 | - | 15735409 | 15735497 | 15724381 | 15724524 | 15751749 | 15751820 | 23 | 21 | 19 |
| SE | 21364 | ENSG00000206560 | ANKRD28 | chr3 | - | 15737032 | 15737233 | 15735409 | 15735497 | 15766233 | 15766312 | 98 | 67 | 59 |
| SE | 21365 | ENSG00000206560 | ANKRD28 | chr3 | - | 15751749 | 15751820 | 15735409 | 15735497 | 15766233 | 15766312 | 41 | 55 | 37 |
| SE | 21367 | ENSG00000206560 | ANKRD28 | chr3 | - | 15766233 | 15766312 | 15751749 | 15751820 | 15795222 | 15795306 | 116 | 145 | 105 |
| SE | 21368 | ENSG00000206560 | ANKRD28 | chr3 | - | 15795222 | 15795306 | 15766233 | 15766312 | 15814239 | 15814309 | 49 | 61 | 43 |
| SE | 21370 | ENSG00000206560 | ANKRD28 | chr3 | - | 15814239 | 15814309 | 15766233 | 15766312 | 15859376 | 15859469 | 0 | 0 | 0 |
| SE | 21371 | ENSG00000206560 | ANKRD28 | chr3 | - | 15814239 | 15814309 | 15795222 | 15795306 | 15859376 | 15859469 | 9 | 2 | 4 |
| SE | 21375 | ENSG00000154309 | DISP1 | chr1 | + | 222823855 | 222823945 | 222815058 | 222815078 | 222827427 | 222827601 | 5 | 6 | 8 |
| SE | 21376 | ENSG00000154309 | DISP1 | chr1 | + | 222823855 | 222823945 | 222815058 | 222815078 | 222835118 | 222835215 | 6 | 6 | 6 |
| SE | 21377 | ENSG00000154309 | DISP1 | chr1 | + | 222823855 | 222823945 | 222815077 | 222815171 | 222827427 | 222827601 | 5 | 6 | 8 |
| SE | 21378 | ENSG00000154309 | DISP1 | chr1 | + | 222823855 | 222823945 | 222815077 | 222815171 | 222835118 | 222835215 | 6 | 6 | 6 |
| SE | 21393 | ENSG00000154309 | DISP1 | chr1 | + | 222835118 | 222835215 | 222823855 | 222823945 | 222872270 | 222872432 | 0 | 2 | 2 |
| SE | 21394 | ENSG00000154309 | DISP1 | chr1 | + | 222835118 | 222835215 | 222823855 | 222823945 | 222899780 | 222899865 | 0 | 2 | 3 |
| SE | 21395 | ENSG00000154309 | DISP1 | chr1 | + | 222836995 | 222837066 | 222835118 | 222835215 | 222837218 | 222837380 | 12 | 17 | 25 |
| SE | 21397 | ENSG00000154309 | DISP1 | chr1 | + | 222872270 | 222872432 | 222823883 | 222823945 | 222899780 | 222899865 | 0 | 0 | 0 |
| SE | 21398 | ENSG00000154309 | DISP1 | chr1 | + | 222872270 | 222872432 | 222835118 | 222835215 | 222899780 | 222899865 | 0 | 0 | 1 |
| SE | 21401 | ENSG00000154309 | DISP1 | chr1 | + | 222899780 | 222899865 | 222815063 | 222815078 | 222942806 | 222943332 | 3 | 0 | 1 |
| SE | 21402 | ENSG00000154309 | DISP1 | chr1 | + | 222899780 | 222899865 | 222835118 | 222835215 | 222942806 | 222943332 | 3 | 0 | 3 |
| SE | 21403 | ENSG00000154309 | DISP1 | chr1 | + | 222899780 | 222899865 | 222872270 | 222872432 | 222942806 | 222943332 | 3 | 0 | 1 |
| SE | 21408 | ENSG00000154309 | DISP1 | chr1 | + | 222994884 | 222994982 | 222991854 | 222992110 | 223002384 | 223005882 | 6 | 3 | 9 |
| SE | 21410 | ENSG00000197879 | MYO1C | chr17 | - | 1486951 | 1487098 | 1484147 | 1484303 | 1491599 | 1491610 | 0 | 0 | 3 |
| SE | 21412 | ENSG00000280739 | EIF1B-AS1 | chr3 | - | 40248183 | 40248471 | 40244451 | 40244638 | 40301236 | 40301396 | 2 | 1 | 5 |
| SE | 21416 | ENSG00000112299 | VNN1 | chr6 | - | 132693023 | 132693315 | 132692222 | 132692584 | 132693989 | 132694182 | 172 | 69 | 18 |
| SE | 21422 | ENSG00000164180 | TMEM161B | chr5 | - | 88197668 | 88197765 | 88195046 | 88196488 | 88198975 | 88199069 | 60 | 25 | 47 |
| SE | 21426 | ENSG00000164180 | TMEM161B | chr5 | - | 88221300 | 88221471 | 88220614 | 88220719 | 88225768 | 88225866 | 0 | 0 | 2 |
| SE | 21427 | ENSG00000164180 | TMEM161B | chr5 | - | 88221300 | 88221471 | 88220614 | 88220719 | 88228444 | 88228528 | 1 | 0 | 1 |
| SE | 21429 | ENSG00000164180 | TMEM161B | chr5 | - | 88221673 | 88221760 | 88220562 | 88220719 | 88228444 | 88228528 | 11 | 3 | 1 |
| SE | 21431 | ENSG00000164180 | TMEM161B | chr5 | - | 88225768 | 88225866 | 88221300 | 88221471 | 88228444 | 88228528 | 16 | 15 | 23 |
| SE | 21432 | ENSG00000164180 | TMEM161B | chr5 | - | 88225768 | 88225866 | 88221673 | 88221760 | 88228444 | 88228528 | 34 | 21 | 42 |
| SE | 21434 | ENSG00000164180 | TMEM161B | chr5 | - | 88240812 | 88240916 | 88228440 | 88228528 | 88268787 | 88269070 | 50 | 31 | 58 |
| SE | 21454 | ENSG00000126777 | KTN1 | chr14 | + | 55601756 | 55601880 | 55580253 | 55580354 | 55612018 | 55612571 | 14 | 5 | 16 |
| SE | 21456 | ENSG00000126777 | KTN1 | chr14 | + | 55629956 | 55630097 | 55627911 | 55628028 | 55633234 | 55633341 | 307 | 318 | 284 |
| SE | 21457 | ENSG00000126777 | KTN1 | chr14 | + | 55637197 | 55637364 | 55636448 | 55636536 | 55637778 | 55637847 | 305 | 235 | 227 |
| SE | 21458 | ENSG00000126777 | KTN1 | chr14 | + | 55646972 | 55647007 | 55641691 | 55641760 | 55648024 | 55648088 | 165 | 156 | 150 |
| SE | 21459 | ENSG00000126777 | KTN1 | chr14 | + | 55650568 | 55650637 | 55650327 | 55650418 | 55651286 | 55651375 | 107 | 82 | 88 |
| SE | 21461 | ENSG00000126777 | KTN1 | chr14 | + | 55651286 | 55651375 | 55650327 | 55650418 | 55651889 | 55651927 | 3 | 3 | 4 |
| SE | 21463 | ENSG00000126777 | KTN1 | chr14 | + | 55663954 | 55664041 | 55661273 | 55661612 | 55667240 | 55667330 | 148 | 152 | 98 |
| SE | 21464 | ENSG00000126777 | KTN1 | chr14 | + | 55663954 | 55664041 | 55661273 | 55661612 | 55670728 | 55670809 | 77 | 94 | 47 |
| SE | 21465 | ENSG00000126777 | KTN1 | chr14 | + | 55667240 | 55667330 | 55661521 | 55661612 | 55670728 | 55670809 | 322 | 251 | 238 |
| SE | 21466 | ENSG00000126777 | KTN1 | chr14 | + | 55667240 | 55667330 | 55663342 | 55664041 | 55670728 | 55670809 | 254 | 195 | 199 |
| SE | 21467 | ENSG00000126777 | KTN1 | chr14 | + | 55671784 | 55671877 | 55671565 | 55671655 | 55672629 | 55672697 | 407 | 353 | 332 |
| SE | 21468 | ENSG00000126777 | KTN1 | chr14 | + | 55672629 | 55672701 | 55671565 | 55671877 | 55672928 | 55673012 | 295 | 291 | 243 |
| SE | 21470 | ENSG00000126777 | KTN1 | chr14 | + | 55680664 | 55680741 | 55679662 | 55679685 | 55684098 | 55684253 | 1 | 0 | 1 |
| SE | 21471 | ENSG00000126777 | KTN1 | chr14 | + | 55680685 | 55680741 | 55679564 | 55679685 | 55684098 | 55684579 | 5 | 5 | 6 |
| SE | 21475 | ENSG00000018189 | RUFY3 | chr4 | + | 70741626 | 70741680 | 70722010 | 70722751 | 70762518 | 70762692 | 1 | 0 | 1 |
| SE | 21476 | ENSG00000018189 | RUFY3 | chr4 | + | 70741626 | 70741680 | 70734433 | 70734593 | 70762518 | 70762692 | 1 | 0 | 2 |
| SE | 21477 | ENSG00000018189 | RUFY3 | chr4 | + | 70762518 | 70762692 | 70721963 | 70722751 | 70763551 | 70763669 | 62 | 48 | 57 |
| SE | 21478 | ENSG00000018189 | RUFY3 | chr4 | + | 70778368 | 70778438 | 70775167 | 70775233 | 70783090 | 70783183 | 125 | 140 | 106 |
| SE | 21479 | ENSG00000018189 | RUFY3 | chr4 | + | 70793764 | 70793904 | 70789494 | 70789592 | 70794794 | 70794894 | 49 | 50 | 44 |
| SE | 21480 | ENSG00000018189 | RUFY3 | chr4 | + | 70793784 | 70793904 | 70789494 | 70789592 | 70794794 | 70794894 | 64 | 63 | 70 |
| SE | 21482 | ENSG00000018189 | RUFY3 | chr4 | + | 70794794 | 70794894 | 70789494 | 70789592 | 70802955 | 70802983 | 3 | 2 | 0 |
| SE | 21483 | ENSG00000018189 | RUFY3 | chr4 | + | 70800140 | 70800205 | 70789494 | 70789592 | 70802955 | 70802983 | 13 | 21 | 5 |
| SE | 21490 | ENSG00000018189 | RUFY3 | chr4 | + | 70804347 | 70804416 | 70802982 | 70803059 | 70806515 | 70806553 | 29 | 21 | 22 |
| SE | 21497 | ENSG00000187630 | DHRS4L2 | chr14 | + | 23995031 | 23995133 | 23990181 | 23990359 | 24001383 | 24001517 | 3 | 0 | 2 |
| SE | 21498 | ENSG00000187630 | DHRS4L2 | chr14 | + | 24000862 | 24001084 | 23990290 | 23990359 | 24001383 | 24001517 | 5 | 3 | 5 |
| SE | 21501 | ENSG00000213204 | NA | chr6 | + | 87488057 | 87488102 | 87477361 | 87477539 | 87500507 | 87500667 | 11 | 6 | 3 |
| SE | 21502 | ENSG00000213204 | NA | chr6 | + | 87488057 | 87488102 | 87477361 | 87477539 | 87501157 | 87501310 | 3 | 3 | 0 |
| SE | 21504 | ENSG00000213204 | NA | chr6 | + | 87500507 | 87500667 | 87488057 | 87488102 | 87501157 | 87501310 | 199 | 172 | 130 |
| SE | 21505 | ENSG00000164414 | SLC35A1 | chr6 | + | 87506381 | 87506448 | 87501157 | 87501310 | 87508419 | 87508596 | 80 | 39 | 61 |
| SE | 21506 | ENSG00000213204 | NA | chr6 | + | 87506880 | 87507061 | 87506381 | 87506448 | 87508419 | 87508596 | 3 | 2 | 1 |
| SE | 21507 | ENSG00000164414 | SLC35A1 | chr6 | + | 87508419 | 87508596 | 87506381 | 87506448 | 87509040 | 87509175 | 122 | 58 | 89 |
| SE | 21509 | ENSG00000111911 | HINT3 | chr6 | + | 125972258 | 125972328 | 125966886 | 125967004 | 125974846 | 125974973 | 12 | 11 | 14 |
| SE | 21510 | ENSG00000111911 | HINT3 | chr6 | + | 125974846 | 125974973 | 125972258 | 125972328 | 125977643 | 125980244 | 41 | 45 | 34 |
| SE | 21512 | ENSG00000137413 | TAF8 | chr6 | + | 42057388 | 42057513 | 42055951 | 42056014 | 42068464 | 42068607 | 143 | 98 | 127 |
| SE | 21513 | ENSG00000137413 | TAF8 | chr6 | + | 42057388 | 42057513 | 42055951 | 42056014 | 42077099 | 42077239 | 142 | 96 | 124 |
| SE | 21514 | ENSG00000137413 | TAF8 | chr6 | + | 42066311 | 42066459 | 42055951 | 42056014 | 42068464 | 42068607 | 92 | 44 | 82 |
| SE | 21515 | ENSG00000137413 | TAF8 | chr6 | + | 42066311 | 42066459 | 42055951 | 42056014 | 42077099 | 42077239 | 1 | 2 | 2 |
| SE | 21517 | ENSG00000137413 | TAF8 | chr6 | + | 42066311 | 42066459 | 42057388 | 42057513 | 42077099 | 42077239 | 72 | 26 | 51 |
| SE | 21518 | ENSG00000137413 | TAF8 | chr6 | + | 42068464 | 42068607 | 42055951 | 42056014 | 42077099 | 42077239 | 69 | 44 | 71 |
| SE | 21519 | ENSG00000137413 | TAF8 | chr6 | + | 42068464 | 42068607 | 42057388 | 42057513 | 42077099 | 42077239 | 70 | 46 | 73 |
| SE | 21520 | ENSG00000137413 | TAF8 | chr6 | + | 42068464 | 42068607 | 42066311 | 42066459 | 42077099 | 42077239 | 161 | 86 | 149 |
| SE | 21521 | ENSG00000137413 | TAF8 | chr6 | + | 42068503 | 42068607 | 42066311 | 42066459 | 42077099 | 42077239 | 52 | 48 | 65 |
| SE | 21523 | ENSG00000108468 | CBX1 | chr17 | - | 48075988 | 48076178 | 48075005 | 48075100 | 48076864 | 48077041 | 155 | 138 | 120 |
| SE | 21524 | ENSG00000108468 | CBX1 | chr17 | - | 48076000 | 48076178 | 48074925 | 48075100 | 48076864 | 48077041 | 200 | 169 | 154 |
| SE | 21525 | ENSG00000108468 | CBX1 | chr17 | - | 48076864 | 48077041 | 48075988 | 48076178 | 48101267 | 48101287 | 199 | 157 | 149 |
| SE | 21533 | ENSG00000093134 | VNN3 | chr6 | - | 132724605 | 132724979 | 132722925 | 132723907 | 132734252 | 132734383 | 27 | 19 | 17 |
| SE | 21534 | ENSG00000093134 | VNN3 | chr6 | - | 132726721 | 132726838 | 132724605 | 132724979 | 132728740 | 132728979 | 4 | 2 | 3 |
| SE | 21535 | ENSG00000093134 | VNN3 | chr6 | - | 132726721 | 132726838 | 132724605 | 132724979 | 132734252 | 132734386 | 4 | 2 | 0 |
| SE | 21537 | ENSG00000093134 | VNN3 | chr6 | - | 132726721 | 132727012 | 132724605 | 132724979 | 132734252 | 132734383 | 5 | 2 | 2 |
| SE | 21539 | ENSG00000093134 | VNN3 | chr6 | - | 132728740 | 132728933 | 132726721 | 132727012 | 132734252 | 132734383 | 18 | 6 | 16 |
| SE | 21540 | ENSG00000093134 | VNN3 | chr6 | - | 132728740 | 132728979 | 132724605 | 132724979 | 132734252 | 132734383 | 5 | 2 | 11 |
| SE | 21541 | ENSG00000093134 | VNN3 | chr6 | - | 132728740 | 132728979 | 132726721 | 132726838 | 132734252 | 132734383 | 5 | 1 | 8 |
| SE | 21542 | ENSG00000093134 | VNN3 | chr6 | - | 132728740 | 132728979 | 132726721 | 132727012 | 132734252 | 132734383 | 19 | 4 | 15 |
| SE | 21543 | ENSG00000093134 | VNN3 | chr6 | - | 132731374 | 132731527 | 132728740 | 132728979 | 132734252 | 132734333 | 0 | 3 | 0 |
| SE | 21550 | ENSG00000198746 | GPATCH3 | chr1 | - | 26892410 | 26892538 | 26890487 | 26891226 | 26892669 | 26892791 | 69 | 23 | 35 |
| SE | 21551 | ENSG00000198746 | GPATCH3 | chr1 | - | 26894235 | 26894410 | 26893388 | 26893448 | 26897300 | 26897725 | 51 | 29 | 57 |
| SE | 21553 | ENSG00000085382 | HACE1 | chr6 | - | 104750340 | 104750472 | 104744511 | 104744610 | 104771192 | 104771389 | 64 | 42 | 49 |
| SE | 21554 | ENSG00000085382 | HACE1 | chr6 | - | 104780352 | 104780457 | 104777205 | 104777317 | 104784085 | 104784173 | 3 | 0 | 3 |
| SE | 21556 | ENSG00000085382 | HACE1 | chr6 | - | 104811310 | 104811393 | 104796928 | 104797025 | 104843222 | 104843298 | 10 | 6 | 10 |
| SE | 21557 | ENSG00000085382 | HACE1 | chr6 | - | 104811310 | 104811393 | 104796928 | 104797025 | 104849141 | 104849246 | 10 | 5 | 8 |
| SE | 21560 | ENSG00000085382 | HACE1 | chr6 | - | 104833041 | 104833173 | 104796928 | 104797025 | 104849141 | 104849246 | 2 | 5 | 4 |
| SE | 21563 | ENSG00000085382 | HACE1 | chr6 | - | 104843222 | 104843298 | 104796928 | 104797025 | 104849141 | 104849246 | 13 | 9 | 5 |
| SE | 21564 | ENSG00000085382 | HACE1 | chr6 | - | 104843222 | 104843298 | 104811310 | 104811393 | 104849141 | 104849246 | 11 | 8 | 7 |
| SE | 21566 | ENSG00000085382 | HACE1 | chr6 | - | 104849141 | 104849246 | 104833041 | 104833173 | 104850906 | 104850996 | 32 | 15 | 5 |
| SE | 21567 | ENSG00000085382 | HACE1 | chr6 | - | 104850906 | 104850996 | 104849141 | 104849246 | 104852316 | 104852371 | 45 | 16 | 13 |
| SE | 21570 | ENSG00000198833 | UBE2J1 | chr6 | - | 89338204 | 89338310 | 89335301 | 89335431 | 89338458 | 89338543 | 655 | 705 | 937 |
| SE | 21571 | ENSG00000198833 | UBE2J1 | chr6 | - | 89342323 | 89342455 | 89338458 | 89338543 | 89343682 | 89343756 | 839 | 802 | 1036 |
| SE | 21574 | ENSG00000198833 | UBE2J1 | chr6 | - | 89343682 | 89343756 | 89342323 | 89342455 | 89352538 | 89352722 | 369 | 322 | 459 |
| SE | 21575 | ENSG00000075568 | TMEM131 | chr2 | - | 97766113 | 97766263 | 97762034 | 97762200 | 97766477 | 97766602 | 146 | 118 | 140 |
| SE | 21576 | ENSG00000075568 | TMEM131 | chr2 | - | 97766477 | 97766602 | 97762034 | 97762200 | 97775842 | 97776018 | 3 | 1 | 1 |
| SE | 21579 | ENSG00000075568 | TMEM131 | chr2 | - | 97775842 | 97776018 | 97772296 | 97772424 | 97793394 | 97793553 | 60 | 56 | 43 |
| SE | 21580 | ENSG00000075568 | TMEM131 | chr2 | - | 97792385 | 97792984 | 97775842 | 97776018 | 97793394 | 97793553 | 200 | 121 | 164 |
| SE | 21583 | ENSG00000075568 | TMEM131 | chr2 | - | 97805087 | 97805205 | 97802651 | 97802790 | 97805375 | 97805451 | 172 | 115 | 173 |
| SE | 21584 | ENSG00000075568 | TMEM131 | chr2 | - | 97805087 | 97805205 | 97802651 | 97802790 | 97805550 | 97805703 | 111 | 70 | 103 |
| SE | 21585 | ENSG00000075568 | TMEM131 | chr2 | - | 97805375 | 97805451 | 97802651 | 97802790 | 97805550 | 97805703 | 33 | 28 | 28 |
| SE | 21586 | ENSG00000075568 | TMEM131 | chr2 | - | 97805375 | 97805451 | 97805087 | 97805205 | 97805550 | 97805703 | 94 | 73 | 98 |
| SE | 21587 | ENSG00000075568 | TMEM131 | chr2 | - | 97809687 | 97809774 | 97805550 | 97805703 | 97811127 | 97811232 | 308 | 253 | 276 |
| SE | 21588 | ENSG00000075568 | TMEM131 | chr2 | - | 97818612 | 97818721 | 97815198 | 97815307 | 97833364 | 97833426 | 314 | 220 | 213 |
| SE | 21589 | ENSG00000075568 | TMEM131 | chr2 | - | 97833364 | 97833426 | 97818612 | 97818721 | 97837076 | 97837157 | 61 | 29 | 33 |
| SE | 21590 | ENSG00000075568 | TMEM131 | chr2 | - | 97834620 | 97834677 | 97833364 | 97833426 | 97837076 | 97837157 | 51 | 46 | 36 |
| SE | 21591 | ENSG00000075568 | TMEM131 | chr2 | - | 97834774 | 97834925 | 97834620 | 97834677 | 97837076 | 97837157 | 294 | 236 | 223 |
| SE | 21594 | ENSG00000112234 | FBXL4 | chr6 | - | 98899267 | 98899481 | 98880552 | 98880624 | 98905425 | 98905670 | 138 | 96 | 152 |
| SE | 21596 | ENSG00000112234 | FBXL4 | chr6 | - | 98917373 | 98917719 | 98905425 | 98905670 | 98927704 | 98927822 | 41 | 31 | 35 |
| SE | 21597 | ENSG00000112234 | FBXL4 | chr6 | - | 98926476 | 98927060 | 98905425 | 98905670 | 98927704 | 98927822 | 56 | 29 | 60 |
| SE | 21600 | ENSG00000112234 | FBXL4 | chr6 | - | 98934761 | 98934879 | 98927704 | 98927822 | 98947805 | 98947946 | 38 | 29 | 37 |
| SE | 21604 | ENSG00000283352 | NA | chr6 | - | 57926425 | 57926510 | 57924416 | 57924591 | 57930103 | 57930291 | 9 | 10 | 12 |
| SE | 21606 | ENSG00000283352 | NA | chr6 | - | 57961151 | 57961224 | 57959027 | 57959111 | 57961320 | 57961382 | 25 | 15 | 22 |
| SE | 21607 | ENSG00000143507 | DUSP10 | chr1 | - | 221706094 | 221706466 | 221701423 | 221702677 | 221738933 | 221739787 | 40 | 31 | 51 |
| SE | 21608 | ENSG00000143507 | DUSP10 | chr1 | - | 221706094 | 221706466 | 221701423 | 221702677 | 221741980 | 221742089 | 4 | 8 | 23 |
| SE | 21609 | ENSG00000143507 | DUSP10 | chr1 | - | 221738933 | 221739787 | 221701423 | 221702677 | 221741980 | 221742089 | 75 | 46 | 42 |
| SE | 21611 | ENSG00000129515 | SNX6 | chr14 | - | 34575755 | 34575842 | 34561092 | 34563175 | 34581560 | 34581600 | 378 | 247 | 257 |
| SE | 21612 | ENSG00000129515 | SNX6 | chr14 | - | 34575755 | 34575842 | 34567853 | 34568013 | 34581560 | 34581600 | 1106 | 769 | 757 |
| SE | 21614 | ENSG00000129515 | SNX6 | chr14 | - | 34586229 | 34586305 | 34581560 | 34581600 | 34593044 | 34593150 | 595 | 438 | 449 |
| SE | 21615 | ENSG00000129515 | SNX6 | chr14 | - | 34603347 | 34603471 | 34597549 | 34597645 | 34604134 | 34604174 | 287 | 212 | 187 |
| SE | 21616 | ENSG00000129515 | SNX6 | chr14 | - | 34603347 | 34603471 | 34597549 | 34597645 | 34605595 | 34605717 | 922 | 655 | 743 |
| SE | 21619 | ENSG00000129515 | SNX6 | chr14 | - | 34604134 | 34604268 | 34597549 | 34597645 | 34605595 | 34605717 | 11 | 8 | 7 |
| SE | 21623 | ENSG00000079950 | STX7 | chr6 | - | 132469950 | 132470047 | 132468402 | 132468475 | 132470573 | 132470626 | 612 | 523 | 404 |
| SE | 21624 | ENSG00000079950 | STX7 | chr6 | - | 132471462 | 132471600 | 132470573 | 132470626 | 132475592 | 132475662 | 281 | 178 | 182 |
| SE | 21625 | ENSG00000079950 | STX7 | chr6 | - | 132472281 | 132472375 | 132471462 | 132471600 | 132475592 | 132475662 | 747 | 676 | 575 |
| SE | 21627 | ENSG00000281706 | LINC01012 | chr6 | + | 27707338 | 27707416 | 27702313 | 27702417 | 27710123 | 27710222 | 0 | 0 | 1 |
| SE | 21629 | ENSG00000281706 | LINC01012 | chr6 | + | 27710680 | 27710889 | 27702313 | 27702417 | 27713003 | 27713352 | 0 | 0 | 4 |
| SE | 21630 | ENSG00000258864 | NA | chr5 | + | 112827928 | 112828006 | 112827212 | 112827247 | 112828855 | 112828972 | 128 | 103 | 92 |
| SE | 21631 | ENSG00000258864 | NA | chr5 | + | 112864620 | 112864732 | 112864456 | 112864528 | 112867403 | 112867582 | 494 | 350 | 351 |
| SE | 21633 | ENSG00000230148 | HOXB-AS1 | chr17 | + | 48545120 | 48545357 | 48544350 | 48544412 | 48545494 | 48545564 | 0 | 15 | 6 |
| SE | 21634 | ENSG00000230148 | HOXB-AS1 | chr17 | + | 48548374 | 48548463 | 48545494 | 48545564 | 48549888 | 48549946 | 2 | 10 | 9 |
| SE | 21636 | ENSG00000118181 | RPS25 | chr11 | - | 119017957 | 119018053 | 119017361 | 119017545 | 119018281 | 119018343 | 7188 | 3952 | 8406 |
| SE | 21638 | ENSG00000130349 | MTRES1 | chr6 | + | 107030024 | 107030107 | 107028568 | 107028797 | 107039748 | 107040230 | 8 | 5 | 1 |
| SE | 21640 | ENSG00000163092 | XIRP2 | chr2 | + | 167239854 | 167239965 | 167218165 | 167218300 | 167240663 | 167240736 | 2 | 5 | 2 |
| SE | 21641 | ENSG00000163092 | XIRP2 | chr2 | + | 167242568 | 167251947 | 167241776 | 167241910 | 167254031 | 167254165 | 0 | 0 | 0 |
| SE | 21644 | ENSG00000221988 | PPT2 | chr6 | + | 32154586 | 32154777 | 32154094 | 32154394 | 32155767 | 32155783 | 1 | 1 | 1 |
| SE | 21649 | ENSG00000231889 | TRAF3IP2-AS1 | chr6 | + | 111484743 | 111484861 | 111483561 | 111483665 | 111490967 | 111493003 | 1 | 1 | 1 |
| SE | 21651 | ENSG00000231889 | TRAF3IP2-AS1 | chr6 | + | 111484743 | 111484861 | 111483561 | 111483665 | 111574620 | 111574802 | 3 | 2 | 6 |
| SE | 21652 | ENSG00000231889 | TRAF3IP2-AS1 | chr6 | + | 111574620 | 111574658 | 111483611 | 111483665 | 111576361 | 111576753 | 1 | 0 | 1 |
| SE | 21653 | ENSG00000231889 | TRAF3IP2-AS1 | chr6 | + | 111574620 | 111574802 | 111483540 | 111483665 | 111576361 | 111576470 | 11 | 10 | 7 |
| SE | 21654 | ENSG00000231889 | TRAF3IP2-AS1 | chr6 | + | 111574620 | 111574802 | 111484743 | 111484861 | 111576361 | 111576470 | 9 | 9 | 7 |
| SE | 21655 | ENSG00000231889 | TRAF3IP2-AS1 | chr6 | + | 111574620 | 111574802 | 111565250 | 111565282 | 111576361 | 111576470 | 7 | 8 | 2 |
| SE | 21660 | ENSG00000130449 | ZSWIM6 | chr5 | + | 61472680 | 61473037 | 61332257 | 61332948 | 61490785 | 61490934 | 134 | 101 | 78 |
| SE | 21664 | ENSG00000130449 | ZSWIM6 | chr5 | + | 61530051 | 61530198 | 61525799 | 61525976 | 61538813 | 61538971 | 2 | 2 | 2 |
| SE | 21665 | ENSG00000130449 | ZSWIM6 | chr5 | + | 61531464 | 61531725 | 61530051 | 61530198 | 61538813 | 61538971 | 131 | 122 | 73 |
| SE | 21666 | ENSG00000130449 | ZSWIM6 | chr5 | + | 61531464 | 61531725 | 61530051 | 61530198 | 61541883 | 61541965 | 131 | 124 | 72 |
| SE | 21667 | ENSG00000130449 | ZSWIM6 | chr5 | + | 61535483 | 61535619 | 61531464 | 61531725 | 61538813 | 61538971 | 186 | 111 | 150 |
| SE | 21669 | ENSG00000130449 | ZSWIM6 | chr5 | + | 61538813 | 61538971 | 61530051 | 61530198 | 61541883 | 61541965 | 7 | 3 | 1 |
| SE | 21670 | ENSG00000130449 | ZSWIM6 | chr5 | + | 61538813 | 61538971 | 61531464 | 61531725 | 61541883 | 61541965 | 6 | 5 | 3 |
| SE | 21671 | ENSG00000130449 | ZSWIM6 | chr5 | + | 61538813 | 61538971 | 61535483 | 61535619 | 61539595 | 61539759 | 123 | 65 | 111 |
| SE | 21674 | ENSG00000130449 | ZSWIM6 | chr5 | + | 61539595 | 61539759 | 61538813 | 61538971 | 61541883 | 61541965 | 81 | 63 | 53 |
| SE | 21680 | ENSG00000139433 | GLTP | chr12 | - | 109857525 | 109857659 | 109855618 | 109855769 | 109858682 | 109858741 | 145 | 83 | 102 |
| SE | 21681 | ENSG00000139433 | GLTP | chr12 | - | 109857582 | 109857659 | 109855618 | 109855769 | 109858682 | 109858741 | 23 | 6 | 18 |
| SE | 21683 | ENSG00000139433 | GLTP | chr12 | - | 109858139 | 109858193 | 109857525 | 109857659 | 109858682 | 109858741 | 1 | 0 | 1 |
| SE | 21686 | ENSG00000135341 | MAP3K7 | chr6 | - | 90519257 | 90519319 | 90518446 | 90518562 | 90523677 | 90523783 | 220 | 193 | 176 |
| SE | 21687 | ENSG00000135341 | MAP3K7 | chr6 | - | 90544551 | 90544632 | 90536336 | 90536401 | 90547257 | 90547387 | 10 | 4 | 21 |
| SE | 21689 | ENSG00000135341 | MAP3K7 | chr6 | - | 90560075 | 90560214 | 90556499 | 90556624 | 90561621 | 90561667 | 178 | 144 | 170 |
| SE | 21690 | ENSG00000135341 | MAP3K7 | chr6 | - | 90561621 | 90561667 | 90560075 | 90560214 | 90568557 | 90568623 | 111 | 89 | 109 |
| SE | 21693 | ENSG00000135341 | MAP3K7 | chr6 | - | 90571696 | 90571807 | 90568557 | 90568623 | 90586763 | 90586883 | 125 | 75 | 112 |
| SE | 21694 | ENSG00000133706 | LARS1 | chr5 | - | 146119008 | 146119074 | 146114232 | 146114311 | 146120370 | 146120503 | 3 | 0 | 1 |
| SE | 21695 | ENSG00000133706 | LARS1 | chr5 | - | 146131018 | 146131109 | 146130017 | 146130158 | 146132897 | 146133081 | 249 | 215 | 227 |
| SE | 21696 | ENSG00000133706 | LARS1 | chr5 | - | 146139615 | 146139703 | 146135600 | 146135664 | 146140203 | 146140261 | 0 | 0 | 0 |
| SE | 21697 | ENSG00000133706 | LARS1 | chr5 | - | 146140203 | 146140261 | 146135600 | 146135664 | 146143411 | 146143538 | 43 | 35 | 20 |
| SE | 21698 | ENSG00000133706 | LARS1 | chr5 | - | 146140203 | 146143084 | 146135600 | 146135664 | 146143411 | 146143550 | 97 | 87 | 88 |
| SE | 21699 | ENSG00000133706 | LARS1 | chr5 | - | 146142855 | 146143084 | 146140203 | 146140261 | 146143411 | 146143550 | 42 | 45 | 57 |

| | | | | | | | | | | | | | | | |
|---|---|---|---|---|---|---|---|---|---|---|---|---|---|---|---|
| SE | 21701 | ENSG00000133706 | LARS1 | chr5 | - | 146143411 | 146143550 | 146140203 | 146143084 | 146144266 | 146144349 | 148 | 126 | 152 | [illegible] |
| SE | 21703 | ENSG00000133706 | LARS1 | chr5 | - | 146164309 | 146164471 | 146160373 | 146160486 | 146168127 | 146168265 | 419 | 363 | 330 | [illegible] |
| SE | 21709 | ENSG00000133706 | LARS1 | chr5 | - | 146181011 | 146181187 | 146177546 | 146177665 | 146182487 | 146182520 | 0 | 0 | 0 | [illegible] |
| SE | 21715 | ENSG00000178385 | PLEKHM3 | chr2 | - | 207946366 | 207946512 | 207930925 | 207931119 | 207976650 | 207977450 | 106 | 59 | 102 | [illegible] |
| SE | 21717 | ENSG00000111860 | CEP85L | chr6 | - | 118469071 | 118469303 | 118460771 | 118465568 | 118470536 | 118470644 | 355 | 360 | 439 | [illegible] |
| SE | 21718 | ENSG00000111860 | CEP85L | chr6 | - | 118470536 | 118470644 | 118469071 | 118469303 | 118479870 | 118479921 | 317 | 317 | 349 | [illegible] |
| SE | 21719 | ENSG00000111860 | CEP85L | chr6 | - | 118483705 | 118483858 | 118481778 | 118481933 | 118491685 | 118491865 | 351 | 274 | 351 | [illegible] |
| SE | 21721 | ENSG00000111860 | CEP85L | chr6 | - | 118511297 | 118511415 | 118491430 | 118491865 | 118523801 | 118523920 | 277 | 180 | 312 | [illegible] |
| SE | 21722 | ENSG00000111860 | CEP85L | chr6 | - | 118523801 | 118523920 | 118491430 | 118491865 | 118565528 | 118566316 | 196 | 129 | 193 | [illegible] |
| SE | 21724 | ENSG00000111860 | CEP85L | chr6 | - | 118632452 | 118632611 | 118565034 | 118566316 | 118651196 | 118651591 | 145 | 108 | 163 | [illegible] |
| SE | 21725 | ENSG00000111860 | CEP85L | chr6 | - | 118632452 | 118632611 | 118565034 | 118566316 | 118710035 | 118710068 | 111 | 73 | 132 | [illegible] |
| SE | 21726 | ENSG00000111860 | CEP85L | chr6 | - | 118652699 | 118652808 | 118632452 | 118632611 | 118664410 | 118664521 | 44 | 40 | 78 | [illegible] |
| SE | 21727 | ENSG00000111860 | CEP85L | chr6 | - | 118652699 | 118652808 | 118632452 | 118632611 | 118710035 | 118710075 | 61 | 57 | 106 | [illegible] |
| SE | 21728 | ENSG00000111860 | CEP85L | chr6 | - | 118664410 | 118664521 | 118632452 | 118632611 | 118710035 | 118710068 | 3 | 2 | 5 | [illegible] |
| SE | 21732 | ENSG00000111850 | SMIM8 | chr6 | + | 87391779 | 87391835 | 87375461 | 87375710 | 87392724 | 87392842 | 0 | 0 | 3 | [illegible] |
| SE | 21733 | ENSG00000111850 | SMIM8 | chr6 | + | 87391779 | 87391877 | 87375461 | 87375710 | 87392724 | 87392842 | 0 | 0 | 1 | [illegible] |
| SE | 21734 | ENSG00000188994 | ZNF292 | chr6 | + | 87155190 | 87155361 | 87152832 | 87152869 | 87215902 | 87216057 | 1 | 2 | 0 | [illegible] |
| SE | 21736 | ENSG00000188994 | ZNF292 | chr6 | + | 87214997 | 87215142 | 87213713 | 87213865 | 87215902 | 87216057 | 10 | 3 | 3 | [illegible] |
| SE | 21737 | ENSG00000188994 | ZNF292 | chr6 | + | 87218595 | 87218731 | 87216298 | 87216377 | 87222769 | 87222867 | 229 | 164 | 169 | [illegible] |
| SE | 21738 | ENSG00000188994 | ZNF292 | chr6 | + | 87218595 | 87218731 | 87216298 | 87216377 | 87233324 | 87233527 | 303 | 210 | 227 | [illegible] |
| SE | 21739 | ENSG00000188994 | ZNF292 | chr6 | + | 87233324 | 87233527 | 87218595 | 87218731 | 87243474 | 87243611 | 162 | 114 | 131 | [illegible] |
| SE | 21740 | ENSG00000188994 | ZNF292 | chr6 | + | 87233339 | 87233527 | 87218595 | 87218731 | 87243474 | 87243611 | 73 | 58 | 63 | [illegible] |
| [illegible] | | | | | | | | | | | | | | | |

| | | | | | | | | | | | | | | | | | |
|---|---|---|---|---|---|---|---|---|---|---|---|---|---|---|---|---|---|
| SE | 22255 | ENSG00000130811 | EIF3G | chr19 | - | 10116799 | 10116989 | 10115966 | 10116074 | 10118667 | 10118727 | 586 | 319 | 479 | 461 | 498 | 0 |
| SE | 22257 | ENSG00000130811 | EIF3G | chr19 | - | 10118439 | 10118557 | 10117083 | 10117188 | 10118667 | 10118727 | 4 | 0 | 6 | 0 | 0 | 871 |
| SE | 22265 | ENSG00000137070 | IL11RA | chr9 | + | 34660273 | 34660393 | 34659758 | 34659900 | 34660503 | 34660600 | 46 | 14 | 37 | 27 | 8 | 1 |
| SE | 22266 | ENSG00000137070 | IL11RA | chr9 | + | 34660803 | 34660936 | 34660503 | 34660600 | 34661481 | 34661655 | 22 | 2 | 16 | 25 | 5 | 0 |
| SE | 22267 | ENSG00000137070 | IL11RA | chr9 | + | 34660853 | 34660936 | 34660503 | 34660600 | 34661481 | 34661850 | 25 | 3 | 22 | 25 | 3 | 0 |
| SE | 22268 | ENSG00000181035 | SLC25A42 | chr19 | + | 19101780 | 19101886 | 19095942 | 19096205 | 19104912 | 19104938 | 49 | 9 | 27 | 35 | 35 | 0 |
| SE | 22271 | ENSG00000159082 | SYNJ1 | chr21 | - | 32634860 | 32634881 | 32631777 | 32631801 | 32638907 | 32639125 | 0 | 0 | 1 | 0 | 0 | 28 |
| SE | 22272 | ENSG00000159082 | SYNJ1 | chr21 | - | 32634860 | 32634884 | 32628758 | 32631801 | 32638907 | 32639125 | 0 | 0 | 1 | 0 | 0 | 28 |
| SE | 22273 | ENSG00000159082 | SYNJ1 | chr21 | - | 32642094 | 32642133 | 32641895 | 32641966 | 32645645 | 32645710 | 2 | 0 | 2 | 3 | 7 | 3 |
| SE | 22276 | ENSG00000159082 | SYNJ1 | chr21 | - | 32646392 | 32646602 | 32645645 | 32645789 | 32650183 | 32650346 | 74 | 59 | 52 | 60 | 163 | 0 |
| SE | 22277 | ENSG00000159082 | SYNJ1 | chr21 | - | 32653287 | 32653366 | 32650183 | 32650346 | 32656686 | 32656902 | 63 | 45 | 58 | 68 | 121 | 0 |
| SE | 22279 | ENSG00000159082 | SYNJ1 | chr21 | - | 32676331 | 32676355 | 32673339 | 32673531 | 32678644 | 32678801 | 0 | 3 | 0 | 0 | 0 | 32 |
| SE | 22280 | ENSG00000159082 | SYNJ1 | chr21 | - | 32699837 | 32700105 | 32695056 | 32695282 | 32701960 | 32702047 | 86 | 83 | 60 | 56 | 115 | 5 |
| SE | 22281 | ENSG00000159082 | SYNJ1 | chr21 | - | 32701960 | 32702047 | 32695056 | 32695282 | 32726771 | 32726917 | 17 | 20 | 16 | 20 | 19 | 7 |
| SE | 22339 | ENSG00000166091 | CMTM5 | chr14 | + | 23378668 | 23378869 | 23378348 | 23378501 | 23379030 | 23379123 | 0 | 1 | 0 | 2 | 0 | 26 |
| SE | 22342 | ENSG00000216863 | LY86-AS1 | chr6 | - | 6438575 | 6438732 | 6369001 | 6369155 | 6564294 | 6564388 | 1 | 3 | 0 | 4 | 2 | 0 |
| SE | 22343 | ENSG00000216863 | LY86-AS1 | chr6 | - | 6492892 | 6493138 | 6438575 | 6438732 | 6564294 | 6564388 | 1 | 1 | 1 | 1 | 0 | 0 |
| SE | 22350 | ENSG00000143740 | SNAP47 | chr1 | + | 227731584 | 227731747 | 227728538 | 227728786 | 227758994 | 227759485 | 1 | 0 | 0 | 0 | 1 | 0 |
| SE | 22353 | ENSG00000143740 | SNAP47 | chr1 | + | 227739205 | 227739302 | 227735427 | 227735499 | 227747691 | 227748233 | 0 | 0 | 0 | 0 | 0 | 3 |
| SE | 22356 | ENSG00000143740 | SNAP47 | chr1 | + | 227747691 | 227748233 | 227734995 | 227735499 | 227758994 | 227759485 | 25 | 9 | 28 | 30 | 21 | 1 |
| SE | 22365 | ENSG00000140455 | USP3 | chr15 | + | 63530497 | 63530590 | 63504593 | 63504830 | 63532646 | 63532707 | 1 | 0 | 3 | 3 | 0 | 89 |
| SE | 22369 | ENSG00000140455 | USP3 | chr15 | + | 63533802 | 63533918 | 63532646 | 63532707 | 63553714 | 63553798 | 11 | 6 | 17 | 22 | 16 | 1 |
| SE | 22370 | ENSG00000140455 | USP3 | chr15 | + | 63537024 | 63537156 | 63504592 | 63504830 | 63553714 | 63553733 | 442 | 279 | 394 | 310 | 700 | 2 |
| SE | 22371 | ENSG00000140455 | USP3 | chr15 | + | 63537024 | 63537156 | 63532505 | 63532707 | 63553714 | 63553733 | 603 | 378 | 502 | 424 | 819 | 1 |
| SE | 22372 | ENSG00000140455 | USP3 | chr15 | + | 63537024 | 63537156 | 63533802 | 63533918 | 63553714 | 63553733 | 503 | 311 | 451 | 368 | 825 | 3 |
| SE | 22375 | ENSG00000140455 | USP3 | chr15 | + | 63558105 | 63558188 | 63556666 | 63556748 | 63562894 | 63563008 | 319 | 253 | 322 | 279 | 449 | 2 |
| SE | 22377 | ENSG00000140455 | USP3 | chr15 | + | 63559856 | 63559970 | 63558105 | 63558188 | 63562894 | 63563008 | 538 | 433 | 447 | 451 | 640 | 2 |
| SE | 22379 | ENSG00000140455 | USP3 | chr15 | + | 63562894 | 63563008 | 63559856 | 63559970 | 63570432 | 63570579 | 343 | 225 | 320 | 300 | 441 | 0 |
| SE | 22380 | ENSG00000140455 | USP3 | chr15 | + | 63574045 | 63574152 | 63570432 | 63570579 | 63588304 | 63588396 | 326 | 202 | 265 | 256 | 252 | 0 |
| SE | 22381 | ENSG00000140455 | USP3 | chr15 | + | 63574322 | 63574403 | 63574045 | 63574152 | 63588304 | 63588396 | 603 | 399 | 454 | 423 | 815 | 0 |
| SE | 22382 | ENSG00000166454 | ATMIN | chr16 | + | 81041355 | 81041481 | 81035841 | 81036206 | 81042280 | 81042450 | 181 | 104 | 144 | 157 | 293 | 0 |
| SE | 22383 | ENSG00000166454 | ATMIN | chr16 | + | 81042280 | 81042480 | 81040524 | 81041481 | 81043160 | 81047346 | 349 | 195 | 255 | 327 | 678 | 6 |
| SE | 22384 | ENSG00000284634 | NA | chr12 | + | 9567489 | 9567567 | 9563963 | 9565303 | 9569057 | 9569202 | 0 | 0 | 6 | 4 | 10 | 0 |
| SE | 22385 | ENSG00000284634 | NA | chr12 | + | 9587815 | 9587878 | 9584105 | 9584164 | 9593405 | 9593525 | 0 | 0 | 0 | 1 | 4 | 0 |
| SE | 22386 | ENSG00000284634 | NA | chr12 | + | 9587815 | 9587878 | 9584105 | 9584164 | 9594813 | 9594875 | 0 | 1 | 0 | 1 | 0 | 0 |
| SE | 22387 | ENSG00000284634 | NA | chr12 | + | 9587815 | 9587878 | 9584105 | 9584164 | 9606352 | 9606456 | 0 | 6 | 0 | 1 | 0 | 1 |
| SE | 22388 | ENSG00000284634 | NA | chr12 | + | 9593405 | 9593525 | 9584105 | 9584164 | 9594813 | 9594875 | 0 | 2 | 0 | 9 | 3 | 0 |
| SE | 22389 | ENSG00000284634 | NA | chr12 | + | 9593405 | 9593525 | 9587815 | 9587878 | 9594813 | 9594875 | 0 | 2 | 0 | 8 | 5 | 0 |
| SE | 22390 | ENSG00000284634 | NA | chr12 | + | 9617243 | 9617351 | 9607670 | 9607802 | 9622318 | 9622436 | 11 | 8 | 3 | 16 | 3 | 0 |
| SE | 22393 | ENSG00000284634 | NA | chr12 | + | 9622318 | 9622436 | 9617243 | 9617351 | 9633424 | 9633828 | 8 | 3 | 2 | 6 | 0 | 0 |
| SE | 22394 | ENSG00000284634 | NA | chr12 | + | 9622318 | 9622436 | 9617243 | 9617351 | 9653682 | 9653808 | 7 | 5 | 2 | 9 | 0 | 1 |
| SE | 22395 | ENSG00000284634 | NA | chr12 | + | 9634526 | 9634696 | 9633424 | 9633529 | 9653682 | 9653808 | 3 | 0 | 1 | 1 | 0 | 0 |
| SE | 22396 | ENSG00000284634 | NA | chr12 | + | 9655124 | 9655303 | 9653682 | 9653808 | 9655872 | 9655979 | 10 | 7 | 6 | 2 | 6 | 593 |
| SE | 22397 | ENSG00000284634 | NA | chr12 | + | 9655872 | 9655979 | 9653682 | 9653808 | 9656920 | 9657217 | 734 | 387 | 553 | 529 | 737 | 0 |
| SE | 22399 | ENSG00000151612 | ZNF827 | chr4 | - | 145763089 | 145763117 | 145760774 | 145761598 | 145764987 | 145765165 | 0 | 0 | 2 | 0 | 0 | 0 |
| SE | 22400 | ENSG00000151612 | ZNF827 | chr4 | - | 145763089 | 145763122 | 145757626 | 145761598 | 145763401 | 145763468 | 2 | 2 | 3 | 1 | 0 | 5 |
| SE | 22402 | ENSG00000151612 | ZNF827 | chr4 | - | 145763089 | 145763122 | 145757626 | 145761598 | 145764978 | 145765165 | 0 | 4 | 2 | 0 | 0 | 0 |
| SE | 22403 | ENSG00000151612 | ZNF827 | chr4 | - | 145763089 | 145763122 | 145757626 | 145761598 | 145764987 | 145765165 | 4 | 3 | 3 | 4 | 2 | 0 |
| SE | 22405 | ENSG00000151612 | ZNF827 | chr4 | - | 145763401 | 145763468 | 145757626 | 145761598 | 145764978 | 145765165 | 1 | 1 | 3 | 0 | 12 | 0 |
| SE | 22406 | ENSG00000151612 | ZNF827 | chr4 | - | 145763401 | 145763468 | 145757626 | 145761598 | 145764987 | 145765165 | 2 | 2 | 4 | 2 | 3 | 0 |
| SE | 22407 | ENSG00000151612 | ZNF827 | chr4 | - | 145763401 | 145763468 | 145763089 | 145763122 | 145764974 | 145765165 | 0 | 0 | 0 | 1 | 0 | 0 |
| SE | 22408 | ENSG00000151612 | ZNF827 | chr4 | - | 145763401 | 145763468 | 145763089 | 145763122 | 145764978 | 145765165 | 0 | 0 | 0 | 0 | 0 | 0 |
| SE | 22409 | ENSG00000151612 | ZNF827 | chr4 | - | 145763401 | 145763468 | 145763089 | 145763122 | 145764987 | 145765165 | 2 | 0 | 2 | 0 | 0 | 4 |
| SE | 22410 | ENSG00000151612 | ZNF827 | chr4 | - | 145823421 | 145823525 | 145779373 | 145779511 | 145845955 | 145845992 | 32 | 21 | 26 | 22 | 45 | 0 |
| SE | 22413 | ENSG00000151612 | ZNF827 | chr4 | - | 145885677 | 145886158 | 145870244 | 145870478 | 145902165 | 145902175 | 8 | 6 | 12 | 12 | 9 | 0 |
| SE | 22414 | ENSG00000151612 | ZNF827 | chr4 | - | 145886032 | 145886158 | 145870300 | 145870478 | 145892242 | 145892415 | 21 | 17 | 33 | 24 | 27 | 0 |
| SE | 22415 | ENSG00000151612 | ZNF827 | chr4 | - | 145892242 | 145892409 | 145870244 | 145870478 | 145902165 | 145902737 | 1 | 4 | 7 | 4 | 1 | 0 |
| SE | 22417 | ENSG00000151612 | ZNF827 | chr4 | - | 145892242 | 145892415 | 145870244 | 145870478 | 145902165 | 145902175 | 18 | 10 | 20 | 19 | 15 | 0 |
| SE | 22418 | ENSG00000151612 | ZNF827 | chr4 | - | 145892242 | 145892415 | 145885677 | 145886158 | 145902165 | 145902175 | 39 | 26 | 51 | 43 | 42 | 0 |
| SE | 22434 | ENSG00000079691 | CARMIL1 | chr6 | + | 25528794 | 25528893 | 25520243 | 25520337 | 25537854 | 25537983 | 5 | 19 | 10 | 30 | 65 | 0 |
| SE | 22435 | ENSG00000079691 | CARMIL1 | chr6 | + | 25550909 | 25551085 | 25539946 | 25540078 | 25554008 | 25554096 | 21 | 25 | 17 | 33 | 51 | 1 |
| SE | 22437 | ENSG00000079691 | CARMIL1 | chr6 | + | 25604811 | 25604893 | 25600313 | 25600746 | 25606060 | 25606273 | 1 | 0 | 8 | 2 | 15 | 1 |
| SE | 22438 | ENSG00000079691 | CARMIL1 | chr6 | + | 25606060 | 25606273 | 25600313 | 25600746 | 25610049 | 25610181 | 8 | 2 | 10 | 9 | 0 | 0 |
| SE | 22440 | ENSG00000079691 | CARMIL1 | chr6 | + | 25612736 | 25612886 | 25610049 | 25610181 | 25619446 | 25619693 | 0 | 0 | 0 | 0 | 0 | 8 |
| SE | 22441 | ENSG00000079691 | CARMIL1 | chr6 | + | 25612739 | 25612886 | 25610121 | 25610181 | 25619446 | 25620530 | 0 | 0 | 0 | 1 | 0 | 8 |
| SE | 22454 | ENSG00000151835 | SACS | chr13 | - | 23371077 | 23371165 | 23368401 | 23368487 | 23375118 | 23375269 | 69 | 39 | 56 | 95 | 86 | 0 |
| SE | 22457 | ENSG00000228903 | RASA4CP | chr7 | - | 44036671 | 44036809 | 44034175 | 44034311 | 44038806 | 44038976 | 10 | 17 | 45 | 37 | 37 | 0 |
| SE | 22458 | ENSG00000110013 | SIAE | chr11 | - | 124639709 | 124639867 | 124638541 | 124638737 | 124647364 | 124647498 | 30 | 50 | 14 | 61 | 56 | 0 |
| SE | 22459 | ENSG00000110013 | SIAE | chr11 | - | 124647364 | 124647498 | 124639709 | 124639867 | 124649618 | 124649796 | 8 | 16 | 7 | 26 | 28 | 0 |
| SE | 22460 | ENSG00000110013 | SIAE | chr11 | - | 124648065 | 124648175 | 124647364 | 124647498 | 124649618 | 124649796 | 31 | 36 | 22 | 38 | 31 | 0 |
| SE | 22461 | ENSG00000110013 | SIAE | chr11 | - | 124675570 | 124675677 | 124675257 | 124675402 | 124694282 | 124694521 | 2 | 0 | 1 | 3 | 3 | 0 |
| SE | 22462 | ENSG00000124787 | RPP40 | chr6 | - | 4998715 | 4998841 | 4996221 | 4996420 | 4999808 | 4999904 | 21 | 22 | 38 | 28 | 56 | 3 |
| SE | 22463 | ENSG00000124787 | RPP40 | chr6 | - | 5000562 | 5000631 | 4999808 | 4999904 | 5002100 | 5002245 | 25 | 11 | 41 | 21 | 47 | 0 |
| SE | 22470 | ENSG00000066135 | KDM4A | chr1 | + | 43667771 | 43668019 | 43666953 | 43667091 | 43669099 | 43669299 | 182 | 136 | 158 | 154 | 140 | 0 |
| SE | 22471 | ENSG00000066135 | KDM4A | chr1 | + | 43671504 | 43671875 | 43669099 | 43669299 | 43683683 | 43683804 | 121 | 89 | 147 | 141 | 81 | 0 |
| SE | 22472 | ENSG00000259040 | BLOC1S5-TXNDC5 | chr6 | - | 7888704 | 7888848 | 7885960 | 7886043 | 7889494 | 7889581 | 351 | 148 | 1034 | 470 | 390 | 0 |
| SE | 22473 | ENSG00000259040 | BLOC1S5-TXNDC5 | chr6 | - | 7889494 | 7889581 | 7888704 | 7888848 | 7891620 | 7891736 | 260 | 89 | 720 | 313 | 222 | 0 |
| SE | 22477 | ENSG00000165097 | KDM1B | chr6 | + | 18161326 | 18161454 | 18159882 | 18159982 | 18166266 | 18166378 | 60 | 69 | 33 | 47 | 67 | 0 |
| SE | 22479 | ENSG00000165097 | KDM1B | chr6 | + | 18166266 | 18166378 | 18162834 | 18162924 | 18171362 | 18171479 | 173 | 155 | 128 | 186 | 231 | 0 |
| SE | 22480 | ENSG00000165097 | KDM1B | chr6 | + | 18171362 | 18171479 | 18166266 | 18166378 | 18185771 | 18185810 | 153 | 115 | 104 | 186 | 279 | 0 |
| SE | 22481 | ENSG00000165097 | KDM1B | chr6 | + | 18185771 | 18185810 | 18171362 | 18171479 | 18187791 | 18188002 | 100 | 86 | 70 | 152 | 198 | 0 |
| SE | 22484 | ENSG00000165097 | KDM1B | chr6 | + | 18187791 | 18188002 | 18171362 | 18171479 | 18197056 | 18197233 | 0 | 1 | 0 | 1 | 0 | 4 |
| SE | 22487 | ENSG00000165097 | KDM1B | chr6 | + | 18191196 | 18191381 | 18187791 | 18188002 | 18197056 | 18197233 | 111 | 103 | 68 | 169 | 123 | 0 |
| SE | 22488 | ENSG00000165097 | KDM1B | chr6 | + | 18197056 | 18197233 | 18171362 | 18171479 | 18200438 | 18200576 | 4 | 4 | 5 | 6 | 0 | 0 |
| SE | 22489 | ENSG00000165097 | KDM1B | chr6 | + | 18197586 | 18197661 | 18197056 | 18197233 | 18200438 | 18200576 | 111 | 204 | 103 | 192 | 167 | 0 |
| SE | 22492 | ENSG00000165097 | KDM1B | chr6 | + | 18208128 | 18208206 | 18207397 | 18207529 | 18212487 | 18212604 | 41 | 46 | 36 | 67 | 62 | 1 |
| SE | 22493 | ENSG00000165097 | KDM1B | chr6 | + | 18208131 | 18208206 | 18207397 | 18207529 | 18212487 | 18212604 | 123 | 131 | 97 | 181 | 182 | 1 |
| SE | 22494 | ENSG00000165097 | KDM1B | chr6 | + | 18212487 | 18212604 | 18208128 | 18208206 | 18213655 | 18213781 | 189 | 219 | 140 | 296 | 245 | 0 |
| SE | 22495 | ENSG00000165097 | KDM1B | chr6 | + | 18212487 | 18212604 | 18208128 | 18208206 | 18215006 | 18215129 | 93 | 96 | 74 | 141 | 149 | 0 |
| SE | 22496 | ENSG00000165097 | KDM1B | chr6 | + | 18213655 | 18213781 | 18208128 | 18208206 | 18215006 | 18215129 | 79 | 75 | 29 | 90 | 69 | 0 |
| SE | 22498 | ENSG00000165097 | KDM1B | chr6 | + | 18217732 | 18217880 | 18215006 | 18215129 | 18221908 | 18223848 | 95 | 102 | 37 | 129 | 81 | 0 |
| SE | 22499 | ENSG00000165097 | KDM1B | chr6 | + | 18217732 | 18217885 | 18215006 | 18215129 | 18221908 | 18222043 | 155 | 164 | 62 | 215 | 104 | 0 |
| SE | 22500 | ENSG00000119812 | FAM98A | chr2 | - | 33586561 | 33586678 | 33583669 | 33585697 | 33587239 | 33587320 | 86 | 98 | 103 | 90 | 186 | 2 |
| SE | 22502 | ENSG00000119812 | FAM98A | chr2 | - | 33595488 | 33595637 | 33591922 | 33592214 | 33599168 | 33599284 | 61 | 43 | 47 | 55 | 34 | 0 |
| SE | 22506 | ENSG00000137486 | ARRB1 | chr11 | - | 75267651 | 75267703 | 75260121 | 75266274 | 75268888 | 75268959 | 490 | 307 | 306 | 368 | 324 | 0 |
| SE | 22510 | ENSG00000137486 | ARRB1 | chr11 | - | 75272894 | 75272978 | 75270929 | 75271724 | 75274073 | 75274211 | 171 | 90 | 102 | 143 | 59 | 0 |
| SE | 22512 | ENSG00000137486 | ARRB1 | chr11 | - | 75278608 | 75278744 | 75277363 | 75277448 | 75281074 | 75281142 | 617 | 306 | 276 | 458 | 446 | 1 |
| SE | 22514 | ENSG00000137486 | ARRB1 | chr11 | - | 75290008 | 75290039 | 75287314 | 75287375 | 75351587 | 75351660 | 83 | 37 | 53 | 68 | 42 | 0 |
| SE | 22532 | ENSG00000181991 | MRPS11 | chr15 | + | 88472626 | 88472725 | 88467735 | 88468024 | 88475109 | 88475239 | 119 | 63 | 88 | 69 | 115 | 0 |
| SE | 22533 | ENSG00000181991 | MRPS11 | chr15 | + | 88472626 | 88472725 | 88467735 | 88468024 | 88477871 | 88478205 | 49 | 23 | 37 | 29 | 53 | 0 |
| SE | 22534 | ENSG00000181991 | MRPS11 | chr15 | + | 88472626 | 88472725 | 88467907 | 88469145 | 88475109 | 88475239 | 31 | 22 | 25 | 20 | 36 | 0 |
| SE | 22535 | ENSG00000181991 | MRPS11 | chr15 | + | 88475109 | 88475239 | 88467907 | 88468024 | 88477871 | 88478267 | 3 | 2 | 3 | 5 | 6 | 0 |
| SE | 22536 | ENSG00000181991 | MRPS11 | chr15 | + | 88475109 | 88475239 | 88472506 | 88472725 | 88477871 | 88478267 | 70 | 38 | 53 | 43 | 62 | 0 |
| SE | 22549 | ENSG00000167881 | SRP68 | chr17 | - | 76040902 | 76040978 | 76040418 | 76040474 | 76043828 | 76043958 | 227 | 164 | 256 | 197 | 432 | 0 |
| SE | 22554 | ENSG00000167881 | SRP68 | chr17 | - | 76061109 | 76061219 | 76057402 | 76057543 | 76061491 | 76061574 | 179 | 143 | 170 | 194 | 279 | 0 |
| SE | 22555 | ENSG00000167881 | SRP68 | chr17 | - | 76061109 | 76061219 | 76057402 | 76057543 | 76063975 | 76064171 | 2 | 0 | 2 | 6 | 6 | 0 |
| SE | 22556 | ENSG00000167881 | SRP68 | chr17 | - | 76061109 | 76061219 | 76060307 | 76060390 | 76061491 | 76061574 | 482 | 353 | 448 | 491 | 613 | 0 |
| SE | 22557 | ENSG00000167881 | SRP68 | chr17 | - | 76061491 | 76061574 | 76057402 | 76057543 | 76063975 | 76064171 | 105 | 81 | 147 | 158 | 164 | 0 |
| SE | 22565 | ENSG00000159753 | CARMIL2 | chr16 | + | 67645539 | 67645631 | 67645143 | 67645286 | 67645723 | 67645777 | 2 | 0 | 5 | 6 | 0 | 0 |
| SE | 22566 | ENSG00000159753 | CARMIL2 | chr16 | + | 67651920 | 67652008 | 67651400 | 67651845 | 67652198 | 67652339 | 44 | 31 | 80 | 77 | 37 | 0 |
| SE | 22567 | ENSG00000159753 | CARMIL2 | chr16 | + | 67656193 | 67656299 | 67656030 | 67656067 | 67656423 | 67656645 | 22 | 9 | 24 | 16 | 18 | 0 |
| SE | 22568 | ENSG00000159753 | CARMIL2 | chr16 | + | 67656227 | 67656299 | 67656030 | 67656067 | 67656423 | 67656645 | 49 | 27 | 70 | 49 | 52 | 0 |
| SE | 22569 | ENSG00000159753 | CARMIL2 | chr16 | + | 67656800 | 67656881 | 67656423 | 67656645 | 67657238 | 67657316 | 60 | 27 | 55 | 46 | 53 | 1 |
| SE | 22570 | ENSG00000159753 | CARMIL2 | chr16 | + | 67656800 | 67656881 | 67656423 | 67656645 | 67657405 | 67657569 | 24 | 7 | 9 | 8 | 18 | 0 |
| SE | 22571 | ENSG00000159753 | CARMIL2 | chr16 | + | 67656833 | 67656881 | 67656423 | 67656645 | 67657238 | 67657316 | 18 | 9 | 19 | 24 | 21 | 1 |
| SE | 22573 | ENSG00000159753 | CARMIL2 | chr16 | + | 67657238 | 67657316 | 67656423 | 67656645 | 67657405 | 67657518 | 29 | 11 | 32 | 22 | 10 | 0 |
| SE | 22574 | ENSG00000159753 | CARMIL2 | chr16 | + | 67657238 | 67657316 | 67656800 | 67656881 | 67657405 | 67657518 | 69 | 33 | 84 | 64 | 48 | 0 |
| SE | 22578 | ENSG00000137265 | IRF4 | chr6 | + | 405017 | 405130 | 401423 | 401777 | 407454 | 411443 | 84 | 75 | 182 | 106 | 109 | 0 |
| SE | 22581 | ENSG00000127603 | MACF1 | chr1 | + | 39251845 | 39251941 | 39250013 | 39250103 | 39254297 | 39254375 | 0 | 0 | 0 | 0 | 0 | 217 |
| SE | 22582 | ENSG00000127603 | MACF1 | chr1 | + | 39283188 | 39283301 | 39282207 | 39282374 | 39283408 | 39283515 | 543 | 419 | 554 | 624 | 504 | 0 |
| SE | 22583 | ENSG00000127603 | MACF1 | chr1 | + | 39283408 | 39283515 | 39282207 | 39282374 | 39284065 | 39284185 | 227 | 195 | 215 | 261 | 259 | 0 |
| SE | 22584 | ENSG00000127603 | MACF1 | chr1 | + | 39284065 | 39284185 | 39283408 | 39283515 | 39284332 | 39284428 | 371 | 316 | 347 | 486 | 477 | 0 |
| SE | 22585 | ENSG00000127603 | MACF1 | chr1 | + | 39285603 | 39285758 | 39285296 | 39285390 | 39287285 | 39287562 | 369 | 252 | 372 | 450 | 366 | 2 |
| SE | 22586 | ENSG00000127603 | MACF1 | chr1 | + | 39293457 | 39293619 | 39292765 | 39292843 | 39295045 | 39295150 | 959 | 790 | 762 | 1090 | 1106 | 0 |
| SE | 22589 | ENSG00000127603 | MACF1 | chr1 | + | 39302923 | 39303078 | 39300209 | 39300362 | 39309569 | 39309696 | 701 | 469 | 654 | 857 | 833 | 7 |
| SE | 22590 | ENSG00000127603 | MACF1 | chr1 | + | 39309569 | 39309696 | 39302923 | 39303078 | 39310244 | 39310428 | 405 | 220 | 296 | 433 | 306 | 0 |
| SE | 22592 | ENSG00000127603 | MACF1 | chr1 | + | 39319663 | 39319747 | 39318452 | 39318615 | 39322607 | 39322715 | 578 | 434 | 481 | 574 | 481 | 0 |
| SE | 22593 | ENSG00000127603 | MACF1 | chr1 | + | 39327217 | 39327353 | 39324645 | 39324734 | 39331202 | 39333181 | 507 | 406 | 424 | 529 | 589 | 1 |
| SE | 22595 | ENSG00000127603 | MACF1 | chr1 | + | 39328600 | 39328706 | 39327217 | 39327353 | 39337181 | 39337331 | 8 | 5 | 3 | 9 | 5 | 10 |
| SE | 22598 | ENSG00000127603 | MACF1 | chr1 | + | 39337181 | 39337331 | 39327217 | 39327353 | 39340501 | 39340717 | 273 | 171 | 164 | 289 | 263 | 1 |
| SE | 22599 | ENSG00000127603 | MACF1 | chr1 | + | 39340501 | 39340717 | 39337181 | 39337331 | 39346976 | 39347210 | 267 | 168 | 158 | 278 | 248 | 0 |
| SE | 22600 | ENSG00000127603 | MACF1 | chr1 | + | 39340803 | 39340953 | 39340501 | 39340717 | 39346976 | 39347210 | 688 | 388 | 452 | 654 | 569 | 4 |
| SE | 22602 | ENSG00000127603 | MACF1 | chr1 | + | 39349477 | 39349627 | 39340803 | 39340953 | 39350784 | 39351018 | 179 | 128 | 134 | 189 | 135 | 0 |
| SE | 22603 | ENSG00000127603 | MACF1 | chr1 | + | 39358696 | 39358873 | 39357374 | 39357893 | 39359140 | 39359264 | 468 | 219 | 328 | 400 | 445 | 1 |
| SE | 22608 | ENSG00000127603 | MACF1 | chr1 | + | 39385433 | 39385929 | 39381952 | 39382152 | 39387186 | 39388658 | 809 | 502 | 582 | 725 | 563 | 0 |
| SE | 22609 | ENSG00000127603 | MACF1 | chr1 | + | 39391867 | 39391955 | 39387186 | 39388658 | 39422373 | 39422535 | 2 | 0 | 4 | 0 | 0 | 263 |
| SE | 22611 | ENSG00000127603 | MACF1 | chr1 | + | 39422373 | 39422535 | 39409502 | 39414477 | 39422729 | 39422900 | 510 | 380 | 388 | 449 | 430 | 0 |
| SE | 22612 | ENSG00000127603 | MACF1 | chr1 | + | 39427454 | 39427614 | 39424027 | 39424194 | 39427960 | 39428092 | 313 | 226 | 295 | 379 | 304 | 1 |
| SE | 22613 | ENSG00000127603 | MACF1 | chr1 | + | 39427454 | 39427614 | 39424027 | 39424194 | 39429241 | 39429326 | 152 | 107 | 139 | 181 | 156 | 0 |
| SE | 22614 | ENSG00000127603 | MACF1 | chr1 | + | 39427960 | 39428287 | 39424027 | 39424194 | 39429241 | 39429326 | 224 | 146 | 147 | 181 | 159 | 0 |
| SE | 22615 | ENSG00000127603 | MACF1 | chr1 | + | 39427960 | 39428287 | 39427454 | 39427614 | 39429241 | 39429326 | 387 | 265 | 303 | 379 | 308 | 3 |
| SE | 22616 | ENSG00000127603 | MACF1 | chr1 | + | 39429241 | 39429326 | 39427960 | 39428287 | 39429826 | 39430068 | 426 | 248 | 254 | 333 | 280 | 1 |
| SE | 22619 | ENSG00000127603 | MACF1 | chr1 | + | 39434413 | 39434632 | 39433047 | 39433155 | 39435557 | 39435761 | 723 | 560 | 602 | 858 | 1054 | 0 |
| SE | 22621 | ENSG00000127603 | MACF1 | chr1 | + | 39436481 | 39436490 | 39435557 | 39435761 | 39439273 | 39439500 | 3 | 3 | 0 | 2 | 0 | 0 |
| SE | 22622 | ENSG00000127603 | MACF1 | chr1 | + | 39437776 | 39438008 | 39435557 | 39435761 | 39439273 | 39439500 | 389 | 254 | 285 | 336 | 293 | 0 |
| SE | 22623 | ENSG00000127603 | MACF1 | chr1 | + | 39437776 | 39438008 | 39436481 | 39436490 | 39439273 | 39439500 | 394 | 250 | 281 | 359 | 296 | 0 |
| SE | 22624 | ENSG00000127603 | MACF1 | chr1 | + | 39439273 | 39439500 | 39437776 | 39438008 | 39441002 | 39441125 | 640 | 462 | 452 | 549 | 521 | 0 |
| SE | 22625 | ENSG00000127603 | MACF1 | chr1 | + | 39443445 | 39443574 | 39442713 | 39442911 | 39444661 | 39444835 | 581 | 369 | 438 | 497 | 406 | 0 |
| SE | 22626 | ENSG00000127603 | MACF1 | chr1 | + | 39448032 | 39448152 | 39447691 | 39447898 | 39448593 | 39448763 | 722 | 441 | 538 | 626 | 710 | 0 |
| SE | 22627 | ENSG00000127603 | MACF1 | chr1 | + | 39451051 | 39451211 | 39448593 | 39448763 | 39452155 | 39452350 | 701 | 459 | 504 | 747 | 603 | 0 |
| SE | 22632 | ENSG00000127603 | MACF1 | chr1 | + | 39484600 | 39484730 | 39479797 | 39480009 | 39485537 | 39486067 | 531 | 280 | 465 | 543 | 565 | 0 |
| SE | 22633 | ENSG00000155324 | GRAMD2B | chr5 | + | 126445480 | 126445590 | 126423471 | 126423689 | 126465376 | 126465545 | 0 | 0 | 0 | 0 | 0 | 0 |
| SE | 22634 | ENSG00000155324 | GRAMD2B | chr5 | + | 126445480 | 126445590 | 126423471 | 126423689 | 126465425 | 126465545 | 0 | 0 | 0 | 0 | 0 | 14 |
| SE | 22635 | ENSG00000155324 | GRAMD2B | chr5 | + | 126445480 | 126445611 | 126423408 | 126423689 | 126465425 | 126465545 | 0 | 0 | 0 | 0 | 0 | 14 |
| SE | 22636 | ENSG00000155324 | GRAMD2B | chr5 | + | 126466293 | 126466336 | 126465425 | 126465545 | 126469676 | 126469788 | 1 | 0 | 1 | 0 | 0 | 16 |
| SE | 22637 | ENSG00000155324 | GRAMD2B | chr5 | + | 126466293 | 126466336 | 126465425 | 126465545 | 126472237 | 126472304 | 0 | 0 | 0 | 0 | 0 | 3 |
| SE | 22638 | ENSG00000155324 | GRAMD2B | chr5 | + | 126469676 | 126469788 | 126465151 | 126465545 | 126472237 | 126472304 | 36 | 14 | 34 | 40 | 62 | 3 |
| SE | 22639 | ENSG00000155324 | GRAMD2B | chr5 | + | 126469676 | 126469788 | 126466293 | 126466336 | 126472237 | 126472304 | 21 | 8 | 24 | 28 | 41 | 0 |
| SE | 22640 | ENSG00000155324 | GRAMD2B | chr5 | + | 126477691 | 126477787 | 126473264 | 126473368 | 126480455 | 126480527 | 55 | 33 | 48 | 40 | 86 | 0 |
| SE | 22641 | ENSG00000155324 | GRAMD2B | chr5 | + | 126484398 | 126484524 | 126483462 | 126483574 | 126485685 | 126485773 | 29 | 18 | 17 | 21 | 29 | 0 |
| SE | 22642 | ENSG00000155324 | GRAMD2B | chr5 | + | 126484401 | 126484524 | 126483462 | 126483574 | 126485685 | 126485752 | 37 | 23 | 23 | 27 | 38 | 0 |
| SE | 22643 | ENSG00000250903 | GMDS-DT | chr6 | + | 2263600 | 2263684 | 2248836 | 2248926 | 2329282 | 2329332 | 2 | 8 | 12 | 12 | 5 | 1 |
| SE | 22644 | ENSG00000250903 | GMDS-DT | chr6 | + | 2263600 | 2263684 | 2248836 | 2248926 | 2360543 | 2362108 | 1 | 8 | 12 | 10 | 8 | 0 |
| SE | 22645 | ENSG00000250903 | GMDS-DT | chr6 | + | 2263600 | 2263684 | 2248836 | 2248926 | 2397220 | 2397626 | 1 | 8 | 12 | 10 | 7 | 0 |
| SE | 22646 | ENSG00000250903 | GMDS-DT | chr6 | + | 2263600 | 2263684 | 2248836 | 2248926 | 2398576 | 2398778 | 2 | 9 | 13 | 15 | 5 | 1 |
| SE | 22647 | ENSG00000250903 | GMDS-DT | chr6 | + | 2269697 | 2269799 | 2263600 | 2263684 | 2271815 | 2273417 | 1 | 2 | 8 | 3 | 4 | 0 |
| SE | 22648 | ENSG00000250903 | GMDS-DT | chr6 | + | 2269697 | 2269799 | 2263600 | 2263684 | 2283498 | 2283774 | 0 | 3 | 10 | 3 | 7 | 0 |
| SE | 22649 | ENSG00000250903 | GMDS-DT | chr6 | + | 2269697 | 2269799 | 2263600 | 2263684 | 2397220 | 2397626 | 0 | 2 | 8 | 3 | 4 | 0 |
| SE | 22650 | ENSG00000250903 | GMDS-DT | chr6 | + | 2269697 | 2269799 | 2263600 | 2263684 | 2398576 | 2398778 | 0 | 3 | 8 | 3 | 4 | 1 |
| SE | 22653 | ENSG00000250903 | GMDS-DT | chr6 | + | 2271815 | 2273417 | 2263600 | 2263684 | 2360543 | 2362108 | 0 | 0 | 1 | 0 | 3 | 0 |
| SE | 22654 | ENSG00000250903 | GMDS-DT | chr6 | + | 2271815 | 2273417 | 2263600 | 2263684 | 2397220 | 2397626 | 1 | 2 | 1 | 0 | 4 | 0 |
| SE | 22655 | ENSG00000250903 | GMDS-DT | chr6 | + | 2271815 | 2273417 | 2269697 | 2269799 | 2283498 | 2283774 | 1 | 0 | 0 | 0 | 0 | 0 |
| SE | 22656 | ENSG00000250903 | GMDS-DT | chr6 | + | 2271815 | 2273417 | 2269697 | 2269799 | 2397220 | 2397626 | 2 | 2 | 0 | 0 | 1 | 0 |
| SE | 22661 | ENSG00000250903 | GMDS-DT | chr6 | + | 2329282 | 2329332 | 2248836 | 2248926 | 2398576 | 2398778 | 1 | 0 | 0 | 0 | 0 | 1 |
| SE | 22664 | ENSG00000250903 | GMDS-DT | chr6 | + | 2329282 | 2329404 | 2283516 | 2283763 | 2352338 | 2352702 | 0 | 0 | 0 | 0 | 2 | 0 |
| SE | 22665 | ENSG00000250903 | GMDS-DT | chr6 | + | 2360543 | 2360630 | 2248836 | 2248926 | 2397220 | 2397626 | 0 | 0 | 0 | 0 | 0 | 0 |
| SE | 22666 | ENSG00000250903 | GMDS-DT | chr6 | + | 2360543 | 2360630 | 2248836 | 2248926 | 2398576 | 2398778 | 0 | 0 | 0 | 0 | 0 | 1 |
| SE | 22667 | ENSG00000250903 | GMDS-DT | chr6 | + | 2397220 | 2397626 | 2248836 | 2248926 | 2398576 | 2398778 | 4 | 1 | 4 | 4 | 0 | 1 |
| SE | 22668 | ENSG00000250903 | GMDS-DT | chr6 | + | 2397220 | 2397626 | 2263600 | 2263684 | 2398576 | 2398778 | 4 | 1 | 4 | 3 | 2 | 1 |
| SE | 22669 | ENSG00000250903 | GMDS-DT | chr6 | + | 2397220 | 2397626 | 2269697 | 2269799 | 2398576 | 2398778 | 4 | 1 | 4 | 3 | 0 | 0 |
| SE | 22670 | ENSG00000250903 | GMDS-DT | chr6 | + | 2397220 | 2397626 | 2360543 | 2360630 | 2398576 | 2398778 | 4 | 1 | 4 | 3 | 0 | 0 |
| SE | 22671 | ENSG00000250903 | GMDS-DT | chr6 | + | 2397543 | 2397626 | 2263600 | 2263684 | 2398576 | 2398778 | 3 | 0 | 1 | 0 | 0 | 1 |
| SE | 22675 | ENSG00000250903 | GMDS-DT | chr6 | + | 2453963 | 2454021 | 2398882 | 2399037 | 2480833 | 2481169 | 0 | 0 | 0 | 0 | 0 | 0 |
| SE | 22678 | ENSG00000131844 | MCCC2 | chr5 | + | 71592925 | 71592992 | 71587275 | 71587554 | 71596279 | 71596364 | 91 | 53 | 65 | 77 | 32 | 1 |
| SE | 22681 | ENSG00000131844 | MCCC2 | chr5 | + | 71602505 | 71602633 | 71599658 | 71599760 | 71604355 | 71604468 | 122 | 79 | 118 | 95 | 141 | 0 |
| SE | 22682 | ENSG00000131844 | MCCC2 | chr5 | + | 71602510 | 71602633 | 71599658 | 71599760 | 71604355 | 71604468 | 40 | 33 | 55 | 33 | 47 | 0 |
| SE | 22685 | ENSG00000131844 | MCCC2 | chr5 | + | 71634942 | 71634952 | 71632120 | 71632185 | 71635150 | 71635246 | 2 | 4 | 1 | 0 | 0 | 0 |
| SE | 22686 | ENSG00000131844 | MCCC2 | chr5 | + | 71634942 | 71635042 | 71631538 | 71632185 | 71635150 | 71635246 | 96 | 60 | 105 | 100 | 119 | 0 |
| SE | 22688 | ENSG00000131844 | MCCC2 | chr5 | + | 71636103 | 71636167 | 71635150 | 71635246 | 71639851 | 71640009 | 3 | 3 | 4 | 4 | 6 | 0 |
| SE | 22690 | ENSG00000131844 | MCCC2 | chr5 | + | 71636103 | 71636167 | 71635150 | 71635246 | 71643818 | 71643895 | 2 | 3 | 3 | 2 | 6 | 1 |
| SE | 22693 | ENSG00000131844 | MCCC2 | chr5 | + | 71641002 | 71641075 | 71635150 | 71635246 | 71643818 | 71643895 | 67 | 33 | 57 | 54 | 69 | 1 |
| SE | 22694 | ENSG00000131844 | MCCC2 | chr5 | + | 71641002 | 71641083 | 71635150 | 71635246 | 71643818 | 71643895 | 40 | 14 | 27 | 24 | 34 | 1 |
| SE | 22695 | ENSG00000131844 | MCCC2 | chr5 | + | 71641002 | 71642727 | 71635150 | 71635246 | 71643818 | 71643895 | 43 | 15 | 27 | 24 | 34 | 1 |
| SE | 22696 | ENSG00000131844 | MCCC2 | chr5 | + | 71642663 | 71642727 | 71641002 | 71641075 | 71643770 | 71643895 | 2 | 0 | 0 | 0 | 3 | 2 |
| SE | 22698 | ENSG00000131844 | MCCC2 | chr5 | + | 71643770 | 71643895 | 71640415 | 71641075 | 71649096 | 71649253 | 2 | 4 | 0 | 0 | 1 | 0 |
| SE | 22699 | ENSG00000131844 | MCCC2 | chr5 | + | 71643818 | 71643895 | 71640384 | 71641075 | 71649096 | 71649253 | 10 | 8 | 9 | 16 | 19 | 0 |
| SE | 22700 | ENSG00000131844 | MCCC2 | chr5 | + | 71646210 | 71646277 | 71643042 | 71643895 | 71649096 | 71649253 | 112 | 60 | 122 | 78 | 162 | 0 |
| SE | 22701 | ENSG00000131844 | MCCC2 | chr5 | + | 71649069 | 71649253 | 71646210 | 71646277 | 71650068 | 71650183 | 99 | 74 | 93 | 82 | 110 | 0 |
| SE | 22702 | ENSG00000131844 | MCCC2 | chr5 | + | 71649096 | 71649253 | 71643062 | 71646277 | 71650068 | 71650183 | 193 | 121 | 176 | 150 | 261 | 0 |
| SE | 22709 | ENSG00000164329 | TENT2 | chr5 | + | 79623251 | 79623489 | 79619993 | 79620083 | 79640850 | 79640965 | 459 | 370 | 386 | 416 | 854 | 5 |
| SE | 22710 | ENSG00000164329 | TENT2 | chr5 | + | 79640850 | 79640965 | 79612434 | 79613075 | 79645122 | 79645192 | 1 | 0 | 4 | 1 | 3 | 1 |
| SE | 22711 | ENSG00000164329 | TENT2 | chr5 | + | 79641104 | 79641196 | 79640850 | 79640965 | 79645122 | 79645192 | 164 | 116 | 130 | 148 | 237 | 0 |
| SE | 22714 | ENSG00000164329 | TENT2 | chr5 | + | 79648616 | 79648693 | 79645122 | 79645192 | 79649061 | 79649190 | 313 | 262 | 257 | 300 | 1123 | 0 |
| SE | 22715 | ENSG00000164329 | TENT2 | chr5 | + | 79648616 | 79648693 | 79645122 | 79645192 | 79668891 | 79669028 | 83 | 99 | 76 | 102 | 313 | 3 |
| SE | 22719 | ENSG00000164329 | TENT2 | chr5 | + | 79679578 | 79679670 | 79668720 | 79669028 | 79681981 | 79682061 | 291 | 266 | 314 | 308 | 399 | 0 |
| SE | 22720 | ENSG00000164329 | TENT2 | chr5 | + | 79679578 | 79679670 | 79668720 | 79669028 | 79685198 | 79685233 | 96 | 86 | 113 | 123 | 157 | 1 |
| SE | 22721 | ENSG00000164329 | TENT2 | chr5 | + | 79681981 | 79682061 | 79668720 | 79669028 | 79685198 | 79685233 | 38 | 34 | 50 | 64 | 134 | 1 |
| SE | 22722 | ENSG00000164329 | TENT2 | chr5 | + | 79681981 | 79682061 | 79679578 | 79679670 | 79685198 | 79685233 | 234 | 217 | 252 | 246 | 376 | 0 |
| SE | 22723 | ENSG00000185261 | KIAA0825 | chr5 | - | 94523929 | 94524098 | 94519205 | 94520917 | 94536995 | 94537127 | 25 | 35 | 26 | 34 | 27 | 3 |
| SE | 22724 | ENSG00000185261 | KIAA0825 | chr5 | - | 94536995 | 94537127 | 94523929 | 94524098 | 94582432 | 94582583 | 16 | 27 | 17 | 27 | 36 | 0 |
| SE | 22725 | ENSG00000185261 | KIAA0825 | chr5 | - | 94582432 | 94582583 | 94536995 | 94537127 | 94618499 | 94618597 | 9 | 9 | 8 | 18 | 18 | 3 |
| SE | 22727 | ENSG00000179476 | C14orf28 | chr14 | + | 44903522 | 44903613 | 44903206 | 44903296 | 44904394 | 44904529 | 1 | 0 | 1 | 1 | 7 | 16 |
| SE | 22729 | ENSG00000145908 | ZNF300 | chr5 | - | 150903140 | 150903182 | 150898427 | 150898554 | 150903863 | 150903977 | 16 | 6 | 2 | 5 | 5 | 3 |
| SE | 22730 | ENSG00000145908 | ZNF300 | chr5 | - | 150903328 | 150903383 | 150903140 | 150903182 | 150903863 | 150903977 | 1 | 0 | 3 | 0 | 0 | 12 |
| SE | 22731 | ENSG00000050393 | MCUR1 | chr6 | - | 13788645 | 13788704 | 13787133 | 13787355 | 13791877 | 13791992 | 3 | 6 | 0 | 0 | 3 | 1 |
| SE | 22732 | ENSG00000050393 | MCUR1 | chr6 | - | 13791877 | 13791992 | 13786556 | 13790864 | 13793893 | 13793947 | 92 | 105 | 75 | 257 | 152 | 0 |
| SE | 22735 | ENSG00000050393 | MCUR1 | chr6 | - | 13793893 | 13793947 | 13786556 | 13790864 | 13798832 | 13798904 | 24 | 25 | 18 | 86 | 66 | 0 |
| SE | 22736 | ENSG00000050393 | MCUR1 | chr6 | - | 13793893 | 13793947 | 13791877 | 13791992 | 13798832 | 13798904 | 112 | 126 | 84 | 332 | 200 | 0 |
| SE | 22737 | ENSG00000050393 | MCUR1 | chr6 | - | 13793893 | 13793947 | 13791877 | 13791992 | 13800340 | 13800382 | 37 | 44 | 17 | 81 | 23 | 0 |
| SE | 22738 | ENSG00000050393 | MCUR1 | chr6 | - | 13798832 | 13798904 | 13791877 | 13791992 | 13800340 | 13800382 | 73 | 93 | 44 | 165 | 162 | 0 |
| SE | 22739 | ENSG00000050393 | MCUR1 | chr6 | - | 13798832 | 13798904 | 13793893 | 13793947 | 13800340 | 13800382 | 149 | 177 | 112 | 418 | 337 | 1 |
| SE | 22741 | ENSG00000050393 | MCUR1 | chr6 | - | 13802242 | 13802346 | 13801287 | 13801389 | 13806924 | 13807044 | 104 | 141 | 64 | 230 | 221 | 1 |
| SE | 22742 | ENSG00000050393 | MCUR1 | chr6 | - | 13803729 | 13804315 | 13802242 | 13802346 | 13806924 | 13807044 | 0 | 4 | 0 | 0 | 4 | 62 |
| SE | 22743 | ENSG00000172262 | ZNF131 | chr5 | + | 43079899 | 43079954 | 43068115 | 43068217 | 43099019 | 43099338 | 0 | 2 | 1 | 2 | 1 | 11 |
| SE | 22745 | ENSG00000172262 | ZNF131 | chr5 | + | 43139164 | 43139309 | 43123208 | 43123310 | 43173317 | 43173448 | 84 | 68 | 74 | 97 | 99 | 0 |
| SE | 22746 | ENSG00000172262 | ZNF131 | chr5 | + | 43161248 | 43161931 | 43123208 | 43123310 | 43173317 | 43173448 | 103 | 64 | 79 | 75 | 186 | 0 |
| SE | 22747 | ENSG00000172262 | ZNF131 | chr5 | + | 43161248 | 43161931 | 43139164 | 43139309 | 43173317 | 43173448 | 205 | 136 | 186 | 211 | 335 | 1 |
| SE | 22748 | ENSG00000172262 | ZNF131 | chr5 | + | 43167929 | 43168029 | 43161830 | 43161931 | 43173317 | 43173448 | 0 | 0 | 0 | 1 | 14 | 96 |
| SE | 22749 | ENSG00000172262 | ZNF131 | chr5 | + | 43174446 | 43174494 | 43173317 | 43173448 | 43179828 | 43179950 | 22 | 25 | 13 | 30 | 16 | 0 |
| SE | 22750 | ENSG00000172262 | ZNF131 | chr5 | + | 43179828 | 43179950 | 43173317 | 43173448 | 43191752 | 43192021 | 0 | 0 | 0 | 2 | 0 | 0 |
| SE | 22751 | ENSG00000145730 | PAM | chr5 | + | 102837686 | 102837792 | 102755313 | 102755348 | 102865822 | 102866284 | 2 | 0 | 0 | 0 | 0 | 4 |
| SE | 22752 | ENSG00000145730 | PAM | chr5 | + | 102865822 | 102866284 | 102754782 | 102755348 | 102867272 | 102867309 | 24 | 8 | 11 | 18 | 5 | 0 |
| SE | 22753 | ENSG00000145730 | PAM | chr5 | + | 102867272 | 102867393 | 102754782 | 102755348 | 102901355 | 102901360 | 40 | 36 | 31 | 36 | 27 | 1 |

| | | | | | | | | | | | | | | | | | | |
|---|---|---|---|---|---|---|---|---|---|---|---|---|---|---|---|---|---|---|
| 22255 | 0 | 0 | 0 | 0 | 817 | 677 | 364 | 492 | 853 | 0 | 1 | 0 | 0 | 0 | 298 | 149 | 1 | 1 |
| 22257 | 586 | 864 | 785 | 1194 | 0 | 2 | 7 | 0 | 0 | 1597 | 1211 | 766 | 940 | 1624 | 267 | 149 | 1 | 1 |
| 22265 | 0 | 0 | 0 | 0 | 38 | 12 | 24 | 36 | 31 | 0 | 0 | 0 | 0 | 0 | 269 | 149 | 1 | 1 |
| 22266 | 0 | 0 | 0 | 0 | 31 | 11 | 13 | 19 | 27 | 0 | 0 | 1 | 0 | 0 | 282 | 149 | 1 | 1 |
| 22267 | 0 | 0 | 0 | 0 | 20 | 11 | 12 | 20 | 28 | 0 | 0 | 1 | 0 | 0 | 232 | 149 | 1 | 1 |
| 22268 | 0 | 1 | 2 | 1 | 7 | 47 | 11 | 24 | 77 | 0 | 0 | 0 | 0 | 0 | 255 | 149 | 1 | 1 |
| 22271 | 13 | 10 | 28 | 28 | 0 | 0 | 0 | 0 | 0 | 0 | 21 | 15 | 6 | 17 | 170 | 149 | 1 | 1 |
| 22272 | 13 | 10 | 28 | 28 | 0 | 0 | 0 | 0 | 0 | 0 | 21 | 15 | 6 | 17 | 173 | 149 | 1 | 1 |
| 22273 | 8 | 8 | 11 | 0 | 0 | 0 | 6 | 1 | 0 | 0 | 9 | 7 | 3 | 0 | 188 | 149 | 1 | 1 |
| 22276 | 0 | 1 | 0 | 0 | 32 | 37 | 75 | 24 | 31 | 0 | 0 | 0 | 0 | 0 | 298 | 149 | 1 | 1 |
| 22277 | 2 | 0 | 0 | 0 | 26 | 36 | 46 | 15 | 13 | 0 | 0 | 0 | 0 | 0 | 228 | 149 | 1 | 1 |
| 22279 | 16 | 25 | 42 | 56 | 0 | 0 | 0 | 0 | 0 | 0 | 9 | 14 | 12 | 0 | 173 | 149 | 1 | 1 |
| 22280 | 10 | 3 | 3 | 11 | 39 | 38 | 46 | 20 | 0 | 0 | 8 | 19 | 9 | 0 | 298 | 149 | 1 | 1 |
| 22281 | 0 | 3 | 1 | 2 | 20 | 25 | 39 | 11 | 0 | 1 | 0 | 4 | 5 | 0 | 236 | 149 | 1 | 1 |
| 22339 | 14 | 7 | 46 | 30 | 0 | 0 | 0 | 0 | 18 | 88 | 66 | 32 | 27 | 668 | 298 | 149 | 1 | 1 |
| 22342 | 1 | 0 | 0 | 0 | 0 | 1 | 2 | 0 | 0 | 0 | 0 | 0 | 0 | 0 | 298 | 149 | 1 | 1 |
| 22343 | 1 | 0 | 2 | 0 | 13 | 1 | 2 | 0 | 0 | 0 | 0 | 0 | 0 | 0 | 298 | 149 | 1 | 1 |
| 22350 | 0 | 0 | 0 | 0 | 0 | 0 | 0 | 0 | 0 | 0 | 5 | 0 | 0 | 0 | 298 | 149 | 1 | 1 |
| 22353 | 1 | 3 | 4 | 3 | 0 | 1 | 0 | 0 | 0 | 0 | 4 | 5 | 10 | 12 | 246 | 149 | 1 | 1 |
| 22356 | 0 | 1 | 0 | 0 | 11 | 40 | 21 | 30 | 23 | 0 | 0 | 2 | 0 | 2 | 298 | 149 | 1 | 1 |
| 22365 | 43 | 65 | 58 | 31 | 0 | 3 | 3 | 0 | 0 | 79 | 74 | 96 | 80 | 81 | 242 | 149 | 1 | 1 |
| 22369 | 0 | 0 | 0 | 1 | 16 | 9 | 14 | 6 | 1 | 0 | 0 | 1 | 0 | 0 | 265 | 149 | 1 | 1 |
| 22370 | 3 | 0 | 0 | 0 | 549 | 374 | 493 | 491 | 388 | 0 | 0 | 0 | 0 | 0 | 281 | 149 | 1 | 1 |
| 22371 | 0 | 0 | 0 | 1 | 756 | 527 | 669 | 622 | 545 | 0 | 0 | 1 | 0 | 0 | 281 | 149 | 1 | 1 |
| 22372 | 0 | 0 | 0 | 0 | 571 | 409 | 553 | 554 | 423 | 0 | 0 | 0 | 0 | 0 | 281 | 149 | 1 | 1 |
| 22375 | 4 | 2 | 0 | 0 | 443 | 355 | 463 | 358 | 356 | 0 | 0 | 1 | 0 | 0 | 232 | 149 | 1 | 1 |
| 22377 | 0 | 0 | 0 | 0 | 746 | 500 | 776 | 558 | 563 | 0 | 0 | 0 | 0 | 0 | 263 | 149 | 1 | 1 |
| 22379 | 0 | 0 | 2 | 0 | 483 | 259 | 465 | 398 | 380 | 0 | 0 | 2 | 6 | 0 | 263 | 149 | 1 | 1 |
| 22380 | 0 | 0 | 0 | 0 | 434 | 320 | 467 | 420 | 416 | 0 | 1 | 0 | 0 | 0 | 256 | 149 | 1 | 1 |
| 22381 | 0 | 0 | 0 | 0 | 782 | 502 | 676 | 545 | 664 | 0 | 0 | 0 | 2 | 0 | 230 | 149 | 1 | 1 |
| 22382 | 0 | 0 | 3 | 0 | 156 | 167 | 143 | 150 | 106 | 0 | 0 | 1 | 0 | 0 | 275 | 149 | 1 | 1 |
| 22383 | 0 | 0 | 0 | 0 | 296 | 296 | 241 | 252 | 178 | 0 | 2 | 0 | 2 | 0 | 298 | 149 | 1 | 1 |
| 22384 | 0 | 0 | 1 | 0 | 0 | 0 | 6 | 10 | 0 | 0 | 0 | 4 | 0 | 0 | 227 | 149 | 1 | 1 |
| 22385 | 0 | 0 | 1 | 2 | 0 | 0 | 0 | 1 | 0 | 0 | 0 | 0 | 0 | 0 | 212 | 149 | 1 | 1 |
| 22386 | 3 | 0 | 1 | 0 | 0 | 0 | 0 | 1 | 0 | 0 | 0 | 0 | 0 | 0 | 212 | 149 | 1 | 1 |
| 22387 | 0 | 2 | 0 | 3 | 0 | 0 | 0 | 1 | 0 | 0 | 0 | 0 | 2 | 0 | 212 | 149 | 1 | 1 |
| 22388 | 3 | 0 | 1 | 0 | 0 | 0 | 0 | 5 | 0 | 0 | 0 | 0 | 0 | 0 | 269 | 149 | 1 | 1 |
| 22389 | 1 | 0 | 0 | 0 | 0 | 0 | 0 | 5 | 0 | 0 | 0 | 0 | 0 | 0 | 269 | 149 | 1 | 1 |
| 22390 | 0 | 1 | 0 | 0 | 5 | 10 | 1 | 0 | 7 | 0 | 0 | 0 | 0 | 0 | 257 | 149 | 1 | 1 |
| 22393 | 0 | 0 | 2 | 5 | 0 | 5 | 0 | 0 | 0 | 0 | 0 | 4 | 0 | 0 | 267 | 149 | 1 | 1 |
| 22394 | 0 | 0 | 4 | 4 | 0 | 2 | 0 | 0 | 0 | 0 | 0 | 0 | 0 | 0 | 267 | 149 | 1 | 1 |
| 22395 | 0 | 0 | 1 | 0 | 0 | 0 | 0 | 0 | 0 | 0 | 7 | 0 | 0 | 0 | 298 | 149 | 1 | 1 |
| 22396 | 353 | 438 | 479 | 600 | 0 | 4 | 12 | 0 | 0 | 146 | 305 | 311 | 163 | 149 | 298 | 149 | 1 | 1 |
| 22397 | 0 | 0 | 0 | 0 | 188 | 382 | 313 | 162 | 188 | 0 | 3 | 0 | 0 | 0 | 256 | 149 | 1 | 1 |
| 22399 | 0 | 2 | 3 | 0 | 0 | 1 | 0 | 0 | 0 | 0 | 2 | 0 | 0 | 0 | 177 | 149 | 1 | 1 |
| 22400 | 4 | 3 | 2 | 12 | 0 | 1 | 1 | 0 | 0 | 0 | 1 | 7 | 5 | 0 | 182 | 149 | 1 | 1 |
| 22402 | 2 | 6 | 1 | 2 | 0 | 1 | 0 | 0 | 0 | 0 | 0 | 0 | 0 | 0 | 182 | 149 | 1 | 1 |
| 22403 | 0 | 2 | 3 | 0 | 0 | 1 | 1 | 0 | 0 | 0 | 2 | 0 | 0 | 0 | 182 | 149 | 1 | 1 |
| 22405 | 2 | 6 | 1 | 2 | 0 | 0 | 5 | 0 | 0 | 0 | 0 | 0 | 0 | 0 | 216 | 149 | 1 | 1 |
| 22406 | 0 | 2 | 3 | 0 | 0 | 0 | 9 | 0 | 0 | 0 | 2 | 0 | 0 | 0 | 216 | 149 | 1 | 1 |
| 22407 | 0 | 1 | 0 | 0 | 0 | 0 | 1 | 0 | 0 | 0 | 0 | 0 | 0 | 0 | 216 | 149 | 1 | 1 |
| 22408 | 2 | 0 | 0 | 0 | 0 | 0 | 1 | 0 | 0 | 0 | 0 | 0 | 0 | 0 | 216 | 149 | 1 | 1 |
| 22409 | 1 | 1 | 4 | 2 | 0 | 0 | 3 | 0 | 0 | 0 | 0 | 1 | 0 | 0 | 216 | 149 | 1 | 1 |
| 22410 | 0 | 0 | 0 | 0 | 9 | 24 | 27 | 16 | 3 | 0 | 0 | 4 | 0 | 0 | 253 | 149 | 1 | 1 |
| 22413 | 0 | 0 | 0 | 0 | 31 | 15 | 23 | 15 | 0 | 0 | 3 | 0 | 0 | 0 | 298 | 149 | 1 | 1 |
| 22414 | 0 | 4 | 0 | 1 | 0 | 30 | 17 | 28 | 43 | 0 | 0 | 0 | 0 | 0 | 275 | 149 | 1 | 1 |
| 22415 | 0 | 0 | 0 | 0 | 13 | 4 | 0 | 6 | 0 | 0 | 3 | 0 | 0 | 0 | 298 | 149 | 1 | 1 |
| 22417 | 0 | 0 | 0 | 0 | 0 | 15 | 10 | 12 | 18 | 0 | 3 | 0 | 0 | 0 | 298 | 149 | 1 | 1 |
| 22418 | 1 | 0 | 0 | 0 | 0 | 45 | 29 | 42 | 66 | 0 | 0 | 7 | 0 | 0 | 298 | 149 | 1 | 1 |
| 22434 | 0 | 0 | 0 | 4 | 0 | 71 | 15 | 2 | 17 | 0 | 0 | 0 | 0 | 0 | 248 | 149 | 1 | 1 |
| 22435 | 0 | 0 | 0 | 0 | 18 | 14 | 10 | 2 | 19 | 0 | 0 | 0 | 0 | 0 | 298 | 149 | 1 | 1 |
| 22437 | 0 | 0 | 0 | 0 | 0 | 1 | 0 | 0 | 0 | 0 | 0 | 0 | 0 | 0 | 231 | 149 | 1 | 1 |
| 22438 | 0 | 0 | 0 | 0 | 0 | 5 | 1 | 1 | 0 | 18 | 0 | 0 | 2 | 0 | 298 | 149 | 1 | 1 |
| 22440 | 3 | 5 | 3 | 8 | 0 | 5 | 0 | 0 | 0 | 0 | 0 | 4 | 0 | 8 | 298 | 149 | 1 | 1 |
| 22441 | 3 | 5 | 3 | 8 | 0 | 0 | 0 | 0 | 0 | 0 | 0 | 4 | 0 | 8 | 296 | 149 | 1 | 1 |
| 22454 | 0 | 2 | 0 | 0 | 48 | 34 | 20 | 13 | 15 | 0 | 0 | 0 | 0 | 0 | 237 | 149 | 1 | 1 |
| 22457 | 2 | 0 | 0 | 0 | 14 | 39 | 15 | 34 | 44 | 0 | 0 | 0 | 0 | 0 | 287 | 149 | 1 | 1 |
| 22458 | 0 | 1 | 0 | 2 | 112 | 129 | 28 | 90 | 1513 | 0 | 0 | 0 | 0 | 0 | 298 | 149 | 1 | 1 |
| 22459 | 0 | 0 | 0 | 0 | 64 | 58 | 12 | 22 | 633 | 0 | 0 | 0 | 0 | 2 | 283 | 149 | 1 | 1 |
| 22460 | 0 | 0 | 1 | 0 | 119 | 55 | 53 | 28 | 1097 | 0 | 3 | 0 | 0 | 0 | 259 | 149 | 1 | 1 |
| 22461 | 0 | 0 | 0 | 0 | 14 | 14 | 0 | 3 | 17 | 5 | 2 | 0 | 0 | 5 | 256 | 149 | 1 | 1 |
| 22462 | 4 | 3 | 1 | 6 | 0 | 34 | 2 | 17 | 0 | 0 | 0 | 2 | 0 | 0 | 275 | 149 | 1 | 1 |
| 22463 | 0 | 0 | 0 | 1 | 0 | 18 | 3 | 3 | 0 | 0 | 0 | 0 | 0 | 0 | 218 | 149 | 1 | 1 |
| 22470 | 0 | 0 | 0 | 0 | 350 | 221 | 242 | 257 | 308 | 8 | 0 | 0 | 0 | 0 | 298 | 149 | 1 | 1 |
| 22471 | 0 | 0 | 0 | 0 | 212 | 116 | 109 | 195 | 194 | 0 | 0 | 0 | 0 | 1 | 298 | 149 | 1 | 1 |
| 22472 | 0 | 5 | 0 | 0 | 262 | 921 | 1903 | 168 | 298 | 0 | 0 | 3 | 0 | 0 | 293 | 149 | 1 | 1 |
| 22473 | 0 | 0 | 0 | 0 | 146 | 674 | 1233 | 142 | 242 | 0 | 0 | 5 | 0 | 0 | 236 | 149 | 1 | 1 |
| 22477 | 1 | 0 | 0 | 0 | 87 | 70 | 38 | 177 | 70 | 0 | 0 | 0 | 1 | 0 | 277 | 149 | 1 | 1 |
| 22479 | 0 | 0 | 0 | 1 | 231 | 152 | 116 | 367 | 108 | 0 | 0 | 2 | 0 | 0 | 261 | 149 | 1 | 1 |
| 22480 | 2 | 0 | 0 | 0 | 205 | 105 | 96 | 326 | 72 | 0 | 0 | 0 | 0 | 0 | 266 | 149 | 1 | 1 |
| 22481 | 0 | 0 | 0 | 0 | 104 | 79 | 51 | 232 | 52 | 0 | 0 | 0 | 4 | 0 | 188 | 149 | 1 | 1 |
| 22484 | 4 | 5 | 6 | 0 | 0 | 0 | 0 | 8 | 0 | 8 | 0 | 15 | 22 | 0 | 298 | 149 | 1 | 1 |
| 22487 | 1 | 0 | 1 | 0 | 153 | 80 | 87 | 275 | 93 | 0 | 0 | 0 | 4 | 0 | 298 | 149 | 1 | 1 |
| 22488 | 0 | 0 | 0 | 6 | 8 | 0 | 15 | 23 | 0 | 0 | 0 | 0 | 0 | 0 | 298 | 149 | 1 | 1 |
| 22489 | 0 | 0 | 0 | 0 | 332 | 149 | 139 | 460 | 134 | 0 | 0 | 0 | 1 | 0 | 224 | 149 | 1 | 1 |
| 22492 | 0 | 0 | 0 | 0 | 43 | 39 | 45 | 121 | 63 | 0 | 0 | 0 | 0 | 0 | 227 | 149 | 1 | 1 |
| 22493 | 0 | 0 | 0 | 0 | 188 | 61 | 124 | 245 | 87 | 0 | 0 | 0 | 0 | 0 | 224 | 149 | 1 | 1 |
| 22494 | 0 | 0 | 0 | 0 | 290 | 108 | 181 | 386 | 137 | 0 | 0 | 2 | 0 | 0 | 266 | 149 | 1 | 1 |
| 22495 | 0 | 1 | 1 | 0 | 145 | 44 | 88 | 161 | 82 | 0 | 0 | 2 | 0 | 0 | 266 | 149 | 1 | 1 |
| 22496 | 0 | 1 | 1 | 0 | 151 | 45 | 70 | 180 | 52 | 0 | 0 | 2 | 0 | 0 | 275 | 149 | 1 | 1 |
| 22498 | 0 | 0 | 0 | 0 | 187 | 76 | 101 | 311 | 55 | 0 | 0 | 1 | 0 | 0 | 297 | 149 | 1 | 1 |
| 22499 | 0 | 0 | 0 | 0 | 346 | 134 | 157 | 493 | 96 | 0 | 0 | 1 | 0 | 0 | 298 | 149 | 1 | 1 |
| 22500 | 1 | 0 | 0 | 4 | 55 | 85 | 102 | 79 | 8 | 0 | 3 | 0 | 2 | 0 | 266 | 149 | 1 | 1 |
| 22502 | 0 | 0 | 0 | 0 | 7 | 27 | 49 | 28 | 7 | 0 | 3 | 0 | 0 | 0 | 298 | 149 | 1 | 1 |
| 22506 | 1 | 0 | 0 | 3 | 844 | 358 | 215 | 450 | 1047 | 0 | 0 | 0 | 7 | 0 | 201 | 149 | 1 | 1 |
| 22510 | 0 | 2 | 0 | 0 | 129 | 81 | 53 | 96 | 355 | 0 | 0 | 0 | 0 | 0 | 233 | 149 | 1 | 1 |
| 22512 | 0 | 0 | 0 | 0 | 850 | 388 | 269 | 549 | 1411 | 0 | 0 | 0 | 0 | 5 | 285 | 149 | 1 | 1 |
| 22514 | 0 | 0 | 0 | 0 | 130 | 39 | 23 | 51 | 193 | 0 | 1 | 0 | 0 | 0 | 180 | 149 | 1 | 1 |
| 22532 | 1 | 0 | 1 | 0 | 152 | 85 | 70 | 114 | 181 | 0 | 0 | 0 | 0 | 0 | 248 | 149 | 1 | 1 |
| 22533 | 0 | 0 | 0 | 0 | 26 | 23 | 36 | 45 | 41 | 0 | 2 | 0 | 0 | 0 | 248 | 149 | 1 | 1 |
| 22534 | 0 | 0 | 0 | 0 | 79 | 40 | 27 | 51 | 84 | 0 | 2 | 0 | 0 | 0 | 248 | 149 | 1 | 1 |
| 22535 | 0 | 0 | 0 | 0 | 7 | 7 | 3 | 5 | 6 | 0 | 2 | 0 | 0 | 0 | 279 | 149 | 1 | 1 |
| 22536 | 0 | 0 | 0 | 0 | 93 | 56 | 37 | 69 | 99 | 0 | 0 | 1 | 0 | 0 | 279 | 149 | 1 | 1 |
| 22549 | 2 | 0 | 0 | 1 | 230 | 167 | 229 | 201 | 150 | 0 | 4 | 0 | 2 | 0 | 225 | 149 | 1 | 1 |
| 22554 | 0 | 0 | 0 | 0 | 359 | 270 | 246 | 217 | 223 | 0 | 6 | 0 | 0 | 0 | 259 | 149 | 1 | 1 |
| 22555 | 0 | 0 | 0 | 3 | 11 | 14 | 19 | 30 | 13 | 0 | 0 | 0 | 0 | 0 | 259 | 149 | 1 | 1 |
| 22556 | 0 | 0 | 1 | 0 | 977 | 632 | 617 | 594 | 607 | 0 | 0 | 1 | 2 | 0 | 259 | 149 | 1 | 1 |
| 22557 | 0 | 0 | 0 | 3 | 203 | 167 | 148 | 124 | 160 | 0 | 0 | 0 | 0 | 0 | 232 | 149 | 1 | 1 |
| 22565 | 0 | 0 | 0 | 0 | 0 | 3 | 3 | 0 | 2 | 0 | 3 | 0 | 0 | 2 | 241 | 149 | 1 | 1 |
| 22566 | 0 | 0 | 1 | 0 | 56 | 71 | 22 | 45 | 100 | 0 | 0 | 0 | 0 | 0 | 237 | 149 | 1 | 1 |
| 22567 | 2 | 0 | 0 | 0 | 7 | 22 | 17 | 2 | 23 | 0 | 0 | 0 | 0 | 0 | 255 | 149 | 1 | 1 |
| 22568 | 2 | 0 | 0 | 0 | 39 | 53 | 44 | 24 | 60 | 0 | 0 | 0 | 0 | 0 | 221 | 149 | 1 | 1 |
| 22569 | 0 | 0 | 0 | 0 | 21 | 47 | 29 | 29 | 66 | 0 | 1 | 0 | 0 | 0 | 230 | 149 | 1 | 1 |
| 22570 | 0 | 0 | 0 | 1 | 0 | 26 | 5 | 3 | 16 | 0 | 0 | 0 | 0 | 0 | 230 | 149 | 1 | 1 |
| 22571 | 0 | 0 | 0 | 0 | 8 | 8 | 14 | 16 | 18 | 0 | 1 | 0 | 0 | 0 | 197 | 149 | 1 | 1 |
| 22573 | 0 | 0 | 0 | 1 | 3 | 23 | 23 | 10 | 7 | 0 | 0 | 0 | 0 | 0 | 227 | 149 | 1 | 1 |
| 22574 | 0 | 0 | 0 | 0 | 24 | 48 | 51 | 40 | 57 | 0 | 0 | 1 | 0 | 0 | 227 | 149 | 1 | 1 |
| 22578 | 0 | 0 | 0 | 0 | 97 | 172 | 179 | 109 | 52 | 0 | 2 | 0 | 3 | 0 | 262 | 149 | 1 | 1 |
| 22581 | 205 | 237 | 287 | 304 | 0 | 1 | 0 | 0 | 0 | 133 | 372 | 180 | 124 | 113 | 245 | 149 | 1 | 1 |
| 22582 | 0 | 0 | 1 | 0 | 225 | 679 | 573 | 258 | 298 | 0 | 0 | 0 | 0 | 0 | 262 | 149 | 1 | 1 |
| 22583 | 0 | 0 | 0 | 0 | 90 | 314 | 181 | 98 | 138 | 0 | 0 | 0 | 1 | 0 | 256 | 149 | 1 | 1 |
| 22584 | 0 | 0 | 0 | 0 | 169 | 521 | 380 | 190 | 343 | 0 | 0 | 2 | 0 | 0 | 269 | 149 | 1 | 1 |
| 22585 | 0 | 1 | 0 | 0 | 258 | 564 | 300 | 125 | 186 | 0 | 3 | 0 | 0 | 0 | 298 | 149 | 1 | 1 |
| 22586 | 0 | 0 | 0 | 0 | 408 | 985 | 744 | 398 | 337 | 0 | 0 | 1 | 0 | 0 | 298 | 149 | 1 | 1 |
| 22589 | 2 | 1 | 1 | 0 | 163 | 729 | 550 | 196 | 147 | 0 | 1 | 1 | 8 | 0 | 298 | 149 | 1 | 1 |
| 22590 | 1 | 0 | 2 | 0 | 90 | 424 | 304 | 106 | 99 | 0 | 0 | 3 | 1 | 0 | 276 | 149 | 1 | 1 |
| 22592 | 3 | 0 | 1 | 0 | 186 | 689 | 475 | 198 | 131 | 0 | 2 | 0 | 0 | 0 | 233 | 149 | 1 | 1 |
| 22593 | 0 | 0 | 0 | 0 | 178 | 669 | 361 | 155 | 51 | 0 | 0 | 0 | 0 | 0 | 285 | 149 | 1 | 1 |
| 22595 | 3 | 6 | 11 | 15 | 12 | 9 | 2 | 2 | 0 | 0 | 22 | 35 | 4 | 0 | 255 | 149 | 1 | 1 |
| 22598 | 2 | 1 | 3 | 0 | 33 | 279 | 185 | 36 | 23 | 0 | 3 | 1 | 0 | 0 | 298 | 149 | 1 | 1 |
| 22599 | 0 | 0 | 0 | 2 | 33 | 257 | 150 | 33 | 23 | 0 | 0 | 0 | 0 | 0 | 298 | 149 | 1 | 1 |
| 22600 | 0 | 0 | 0 | 0 | 77 | 723 | 494 | 99 | 81 | 0 | 0 | 0 | 1 | 0 | 298 | 149 | 1 | 1 |
| 22602 | 0 | 0 | 2 | 0 | 41 | 181 | 125 | 43 | 21 | 0 | 0 | 0 | 0 | 0 | 298 | 149 | 1 | 1 |
| 22603 | 0 | 0 | 1 | 0 | 99 | 414 | 263 | 97 | 101 | 0 | 1 | 0 | 0 | 0 | 298 | 149 | 1 | 1 |
| 22608 | 0 | 1 | 0 | 0 | 265 | 900 | 620 | 264 | 96 | 0 | 1 | 0 | 1 | 0 | 298 | 149 | 1 | 1 |
| 22609 | 203 | 220 | 293 | 209 | 0 | 1 | 2 | 0 | 0 | 16 | 224 | 220 | 63 | 3 | 237 | 149 | 1 | 1 |
| 22611 | 0 | 0 | 0 | 0 | 92 | 431 | 355 | 281 | 43 | 0 | 0 | 1 | 0 | 0 | 298 | 149 | 1 | 1 |
| 22612 | 0 | 0 | 0 | 0 | 39 | 313 | 196 | 87 | 62 | 0 | 2 | 0 | 3 | 0 | 298 | 149 | 1 | 1 |
| 22613 | 0 | 0 | 0 | 0 | 27 | 169 | 118 | 37 | 36 | 0 | 0 | 3 | 0 | 0 | 298 | 149 | 1 | 1 |
| 22614 | 0 | 0 | 0 | 0 | 105 | 153 | 150 | 108 | 15 | 0 | 0 | 3 | 0 | 0 | 298 | 149 | 1 | 1 |
| 22615 | 0 | 0 | 0 | 0 | 117 | 295 | 228 | 155 | 41 | 0 | 0 | 0 | 0 | 0 | 298 | 149 | 1 | 1 |
| 22616 | 0 | 0 | 0 | 0 | 130 | 257 | 254 | 133 | 21 | 0 | 0 | 0 | 0 | 0 | 234 | 149 | 1 | 1 |
| 22619 | 0 | 1 | 0 | 0 | 157 | 681 | 779 | 281 | 49 | 6 | 0 | 0 | 0 | 0 | 298 | 149 | 1 | 1 |
| 22621 | 0 | 1 | 2 | 0 | 0 | 0 | 0 | 0 | 0 | 0 | 0 | 2 | 0 | 0 | 158 | 149 | 1 | 1 |
| 22622 | 0 | 1 | 2 | 0 | 142 | 336 | 346 | 166 | 45 | 0 | 0 | 2 | 0 | 0 | 298 | 149 | 1 | 1 |
| 22623 | 1 | 0 | 0 | 0 | 119 | 279 | 275 | 141 | 29 | 0 | 0 | 0 | 0 | 0 | 298 | 149 | 1 | 1 |
| 22624 | 1 | 0 | 0 | 2 | 267 | 552 | 469 | 262 | 55 | 0 | 0 | 1 | 0 | 0 | 298 | 149 | 1 | 1 |
| 22625 | 0 | 0 | 0 | 0 | 143 | 396 | 454 | 141 | 67 | 0 | 0 | 1 | 0 | 0 | 278 | 149 | 1 | 1 |
| 22626 | 0 | 1 | 0 | 0 | 208 | 683 | 535 | 292 | 44 | 0 | 0 | 0 | 0 | 0 | 269 | 149 | 1 | 1 |
| 22627 | 0 | 0 | 0 | 0 | 181 | 663 | 534 | 315 | 52 | 0 | 0 | 0 | 3 | 0 | 298 | 149 | 1 | 1 |
| 22632 | 1 | 0 | 1 | 0 | 70 | 413 | 389 | 198 | 51 | 0 | 5 | 1 | 0 | 0 | 279 | 149 | 1 | 1 |
| 22633 | 0 | 3 | 0 | 4 | 0 | 0 | 1 | 0 | 0 | 0 | 0 | 0 | 0 | 0 | 259 | 149 | 1 | 1 |
| 22634 | 2 | 7 | 10 | 11 | 0 | 0 | 2 | 0 | 0 | 0 | 0 | 11 | 7 | 0 | 259 | 149 | 1 | 1 |
| 22635 | 2 | 7 | 10 | 11 | 0 | 0 | 1 | 0 | 0 | 0 | 0 | 11 | 7 | 0 | 280 | 149 | 1 | 1 |
| 22636 | 6 | 11 | 12 | 21 | 0 | 0 | 0 | 0 | 0 | 0 | 3 | 7 | 1 | 0 | 192 | 149 | 1 | 1 |
| 22637 | 2 | 0 | 2 | 6 | 0 | 0 | 0 | 3 | 0 | 12 | 0 | 1 | 0 | 0 | 192 | 149 | 1 | 1 |
| 22638 | 2 | 0 | 2 | 6 | 10 | 27 | 35 | 19 | 0 | 12 | 0 | 1 | 0 | 0 | 261 | 149 | 1 | 1 |
| 22639 | 0 | 0 | 0 | 0 | 10 | 24 | 28 | 18 | 0 | 0 | 0 | 0 | 3 | 0 | 261 | 149 | 1 | 1 |
| 22640 | 0 | 2 | 0 | 3 | 45 | 71 | 69 | 21 | 0 | 0 | 3 | 4 | 0 | 0 | 245 | 149 | 1 | 1 |
| 22641 | 0 | 0 | 0 | 0 | 5 | 26 | 21 | 1 | 0 | 0 | 2 | 0 | 0 | 0 | 275 | 149 | 1 | 1 |
| 22642 | 0 | 0 | 0 | 0 | 5 | 23 | 25 | 2 | 0 | 0 | 2 | 0 | 0 | 0 | 272 | 149 | 1 | 1 |
| 22643 | 0 | 0 | 0 | 0 | 0 | 4 | 1 | 6 | 2 | 0 | 0 | 0 | 0 | 0 | 233 | 149 | 1 | 1 |
| 22644 | 0 | 0 | 0 | 0 | 0 | 4 | 1 | 9 | 2 | 0 | 0 | 0 | 0 | 3 | 233 | 149 | 1 | 1 |
| 22645 | 0 | 0 | 1 | 0 | 0 | 4 | 1 | 6 | 2 | 0 | 0 | 0 | 0 | 0 | 233 | 149 | 1 | 1 |
| 22646 | 1 | 0 | 0 | 0 | 0 | 10 | 1 | 6 | 2 | 0 | 2 | 2 | 0 | 1 | 233 | 149 | 1 | 1 |
| 22647 | 0 | 1 | 0 | 3 | 0 | 2 | 2 | 2 | 0 | 0 | 0 | 0 | 0 | 0 | 251 | 149 | 1 | 1 |
| 22648 | 0 | 1 | 3 | 0 | 0 | 0 | 2 | 2 | 0 | 0 | 0 | 0 | 0 | 0 | 251 | 149 | 1 | 1 |
| 22649 | 0 | 0 | 0 | 2 | 0 | 0 | 2 | 8 | 0 | 0 | 0 | 0 | 0 | 0 | 251 | 149 | 1 | 1 |
| 22650 | 1 | 1 | 5 | 0 | 0 | 0 | 2 | 2 | 0 | 0 | 6 | 0 | 0 | 0 | 251 | 149 | 1 | 1 |
| 22653 | 0 | 0 | 0 | 3 | 0 | 0 | 0 | 0 | 0 | 0 | 0 | 0 | 3 | 0 | 298 | 149 | 1 | 1 |
| 22654 | 0 | 0 | 0 | 2 | 0 | 1 | 0 | 0 | 0 | 0 | 0 | 0 | 0 | 0 | 298 | 149 | 1 | 1 |
| 22655 | 1 | 2 | 0 | 3 | 0 | 2 | 0 | 0 | 0 | 0 | 0 | 0 | 0 | 0 | 298 | 149 | 1 | 1 |
| 22656 | 0 | 0 | 0 | 0 | 0 | 3 | 0 | 0 | 0 | 0 | 0 | 0 | 6 | 0 | 298 | 149 | 1 | 1 |
| 22661 | 1 | 0 | 0 | 0 | 0 | 0 | 0 | 0 | 0 | 0 | 2 | 2 | 0 | 1 | 199 | 149 | 1 | 1 |
| 22664 | 0 | 0 | 0 | 2 | 0 | 0 | 0 | 0 | 0 | 13 | 0 | 2 | 0 | 0 | 271 | 149 | 1 | 1 |
| 22665 | 0 | 0 | 1 | 0 | 0 | 0 | 0 | 3 | 3 | 0 | 0 | 0 | 0 | 0 | 236 | 149 | 1 | 1 |
| 22666 | 1 | 0 | 0 | 0 | 0 | 0 | 0 | 0 | 3 | 0 | 2 | 2 | 0 | 1 | 236 | 149 | 1 | 1 |
| 22667 | 1 | 0 | 0 | 0 | 0 | 4 | 5 | 0 | 0 | 0 | 2 | 2 | 0 | 1 | 298 | 149 | 1 | 1 |
| 22668 | 1 | 1 | 5 | 0 | 0 | 4 | 5 | 0 | 0 | 0 | 6 | 0 | 0 | 0 | 298 | 149 | 1 | 1 |
| 22669 | 1 | 0 | 0 | 0 | 0 | 4 | 5 | 6 | 0 | 0 | 0 | 0 | 4 | 0 | 298 | 149 | 1 | 1 |
| 22670 | 0 | 0 | 0 | 3 | 0 | 4 | 5 | 3 | 0 | 0 | 0 | 0 | 0 | 0 | 298 | 149 | 1 | 1 |
| 22671 | 1 | 1 | 5 | 0 | 0 | 0 | 3 | 0 | 0 | 0 | 6 | 0 | 0 | 0 | 232 | 149 | 1 | 1 |
| 22675 | 0 | 2 | 0 | 0 | 0 | 0 | 1 | 0 | 4 | 0 | 0 | 0 | 0 | 0 | 207 | 149 | 1 | 1 |
| 22678 | 0 | 2 | 0 | 0 | 80 | 70 | 75 | 56 | 89 | 0 | 0 | 0 | 0 | 0 | 216 | 149 | 1 | 1 |
| 22681 | 1 | 0 | 0 | 0 | 143 | 158 | 103 | 72 | 134 | 0 | 0 | 0 | 0 | 0 | 277 | 149 | 1 | 1 |
| 22682 | 1 | 0 | 0 | 0 | 53 | 49 | 44 | 22 | 63 | 0 | 0 | 0 | 0 | 0 | 272 | 149 | 1 | 1 |
| 22685 | 0 | 1 | 0 | 0 | 0 | 5 | 2 | 0 | 0 | 0 | 0 | 0 | 0 | 0 | 159 | 149 | 1 | 1 |
| 22686 | 0 | 1 | 0 | 0 | 174 | 91 | 102 | 58 | 70 | 0 | 0 | 0 | 0 | 0 | 249 | 149 | 1 | 1 |
| 22688 | 0 | 0 | 0 | 0 | 0 | 9 | 1 | 1 | 7 | 0 | 1 | 0 | 0 | 3 | 213 | 149 | 1 | 1 |
| 22690 | 0 | 0 | 0 | 0 | 0 | 3 | 1 | 1 | 7 | 0 | 0 | 0 | 0 | 0 | 213 | 149 | 1 | 1 |
| 22693 | 0 | 0 | 0 | 0 | 106 | 50 | 43 | 45 | 75 | 0 | 0 | 0 | 0 | 0 | 222 | 149 | 1 | 1 |
| 22694 | 0 | 0 | 0 | 0 | 43 | 20 | 25 | 32 | 18 | 0 | 0 | 0 | 0 | 0 | 230 | 149 | 1 | 1 |
| 22695 | 0 | 0 | 0 | 0 | 43 | 20 | 25 | 35 | 18 | 0 | 0 | 0 | 0 | 0 | 298 | 149 | 1 | 1 |
| 22696 | 3 | 0 | 0 | 1 | 0 | 0 | 0 | 0 | 0 | 0 | 0 | 6 | 0 | 0 | 213 | 149 | 1 | 1 |
| 22698 | 0 | 0 | 0 | 0 | 0 | 0 | 6 | 0 | 0 | 0 | 0 | 3 | 0 | 0 | 274 | 149 | 1 | 1 |
| 22699 | 0 | 0 | 0 | 0 | 15 | 13 | 11 | 6 | 19 | 0 | 0 | 3 | 0 | 0 | 226 | 149 | 1 | 1 |
| 22700 | 2 | 0 | 0 | 0 | 143 | 95 | 75 | 56 | 112 | 0 | 0 | 0 | 0 | 0 | 216 | 149 | 1 | 1 |
| 22701 | 0 | 0 | 0 | 0 | 129 | 85 | 72 | 68 | 53 | 0 | 4 | 0 | 0 | 0 | 298 | 149 | 1 | 1 |
| 22702 | 0 | 0 | 0 | 0 | 228 | 160 | 141 | 114 | 122 | 0 | 4 | 0 | 0 | 0 | 298 | 149 | 1 | 1 |
| 22709 | 0 | 1 | 3 | 0 | 97 | 309 | 291 | 212 | 153 | 0 | 0 | 0 | 0 | 0 | 298 | 149 | 1 | 1 |
| 22710 | 0 | 0 | 1 | 1 | 8 | 9 | 0 | 0 | 0 | 0 | 0 | 0 | 0 | 0 | 264 | 149 | 1 | 1 |
| 22711 | 0 | 0 | 0 | 0 | 34 | 91 | 74 | 29 | 58 | 0 | 4 | 0 | 0 | 0 | 241 | 149 | 1 | 1 |
| 22714 | 1 | 0 | 0 | 0 | 76 | 227 | 137 | 103 | 57 | 0 | 0 | 0 | 0 | 0 | 226 | 149 | 1 | 1 |
| 22715 | 0 | 0 | 1 | 0 | 19 | 66 | 50 | 23 | 19 | 0 | 0 | 0 | 0 | 0 | 226 | 149 | 1 | 1 |
| 22719 | 0 | 0 | 1 | 2 | 99 | 205 | 152 | 90 | 53 | 0 | 0 | 0 | 0 | 0 | 241 | 149 | 1 | 1 |
| 22720 | 0 | 0 | 0 | 0 | 38 | 69 | 57 | 34 | 21 | 0 | 0 | 0 | 0 | 0 | 241 | 149 | 1 | 1 |
| 22721 | 0 | 0 | 0 | 0 | 0 | 24 | 2 | 11 | 0 | 0 | 0 | 0 | 0 | 0 | 229 | 149 | 1 | 1 |
| 22722 | 0 | 0 | 0 | 0 | 61 | 168 | 98 | 67 | 32 | 0 | 4 | 0 | 0 | 0 | 229 | 149 | 1 | 1 |
| 22723 | 1 | 0 | 0 | 0 | 0 | 13 | 13 | 5 | 0 | 0 | 2 | 0 | 0 | 0 | 298 | 149 | 1 | 1 |
| 22724 | 0 | 0 | 1 | 0 | 0 | 7 | 7 | 0 | 8 | 0 | 0 | 0 | 0 | 0 | 281 | 149 | 1 | 1 |
| 22725 | 0 | 0 | 0 | 0 | 0 | 7 | 1 | 0 | 16 | 0 | 0 | 0 | 0 | 0 | 298 | 149 | 1 | 1 |
| 22727 | 6 | 22 | 26 | 35 | 0 | 0 | 0 | 0 | 0 | 11 | 2 | 1 | 3 | 0 | 240 | 149 | 1 | 1 |
| 22729 | 0 | 0 | 0 | 0 | 2 | 0 | 0 | 0 | 0 | 0 | 0 | 0 | 0 | 5 | 191 | 149 | 1 | 1 |
| 22730 | 4 | 2 | 0 | 2 | 0 | 3 | 0 | 0 | 0 | 2 | 0 | 0 | 0 | 0 | 204 | 149 | 1 | 1 |
| 22731 | 0 | 1 | 0 | 0 | 21 | 1 | 2 | 8 | 7 | 0 | 0 | 0 | 0 | 0 | 208 | 149 | 1 | 1 |
| 22732 | 0 | 0 | 0 | 0 | 261 | 139 | 78 | 112 | 732 | 0 | 7 | 0 | 0 | 0 | 264 | 149 | 1 | 1 |
| 22735 | 0 | 0 | 0 | 0 | 95 | 36 | 28 | 114 | 311 | 0 | 0 | 0 | 0 | 6 | 203 | 149 | 1 | 1 |
| 22736 | 0 | 0 | 0 | 1 | 378 | 167 | 106 | 224 | 1022 | 1 | 3 | 0 | 0 | 0 | 203 | 149 | 1 | 1 |
| 22737 | 0 | 0 | 0 | 2 | 86 | 34 | 29 | 26 | 211 | 0 | 2 | 2 | 0 | 0 | 203 | 149 | 1 | 1 |
| 22738 | 0 | 0 | 0 | 2 | 341 | 138 | 124 | 187 | 1082 | 0 | 2 | 2 | 0 | 0 | 221 | 149 | 1 | 1 |
| 22739 | 2 | 1 | 2 | 0 | 632 | 268 | 209 | 385 | 1915 | 0 | 0 | 4 | 0 | 15 | 221 | 149 | 1 | 1 |
| 22741 | 0 | 0 | 0 | 2 | 290 | 166 | 148 | 257 | 1315 | 0 | 0 | 0 | 0 | 0 | 253 | 149 | 1 | 1 |
| 22742 | 80 | 46 | 128 | 111 | 0 | 0 | 0 | 0 | 0 | 196 | 93 | 67 | 106 | 862 | 298 | 149 | 1 | 1 |
| 22743 | 11 | 10 | 2 | 1 | 0 | 0 | 0 | 0 | 0 | 0 | 4 | 3 | 4 | 0 | 204 | 149 | 1 | 1 |
| 22745 | 0 | 0 | 0 | 2 | 16 | 64 | 61 | 10 | 15 | 0 | 0 | 0 | 0 | 0 | 294 | 149 | 1 | 1 |
| 22746 | 0 | 0 | 0 | 2 | 0 | 39 | 42 | 33 | 9 | 0 | 0 | 0 | 0 | 0 | 298 | 149 | 1 | 1 |
| 22747 | 2 | 0 | 0 | 0 | 7 | 113 | 129 | 69 | 21 | 0 | 3 | 0 | 0 | 0 | 298 | 149 | 1 | 1 |
| 22748 | 51 | 72 | 70 | 176 | 0 | 0 | 0 | 0 | 0 | 0 | 31 | 39 | 23 | 8 | 249 | 149 | 1 | 1 |
| 22749 | 0 | 0 | 1 | 0 | 0 | 11 | 15 | 0 | 3 | 0 | 0 | 0 | 0 | 0 | 197 | 149 | 1 | 1 |
| 22750 | 0 | 0 | 0 | 0 | 0 | 1 | 0 | 0 | 0 | 0 | 0 | 0 | 0 | 5 | 271 | 149 | 1 | 1 |
| 22751 | 3 | 3 | 5 | 0 | 0 | 0 | 0 | 0 | 0 | 0 | 4 | 0 | 2 | 6 | 255 | 149 | 1 | 1 |
| 22752 | 0 | 1 | 0 | 0 | 17 | 4 | 4 | 10 | 32 | 0 | 2 | 0 | 0 | 0 | 298 | 149 | 1 | 1 |
| 22753 | 0 | 0 | 0 | 0 | 13 | 21 | 12 | 31 | 0 | 0 | 0 | 0 | 0 | 0 | 270 | 149 | 1 | 1 |

| | | | | | | | | | | | |
|---|---|---|---|---|---|---|---|---|---|---|---|
| 22255 | 1 | 1 | 1 | 1 | 1 | 1 | 0.997 | 1 | 1 | 1 | 0.001 |
| 22257 | 0.003 | 0 | 0.004 | 0 | 0 | 0 | 0.001 | 0.005 | 0 | 0 | 0 |
| 22265 | 0.962 | 1 | 1 | 1 | 1 | 1 | 1 | 1 | 1 | 1 | -0.008 |
| 22266 | 1 | 1 | 1 | 1 | 1 | 1 | 1 | 0.873 | 1 | 1 | 0.025 |
| 22267 | 1 | 1 | 1 | 1 | 1 | 1 | 1 | 0.885 | 1 | 1 | 0.023 |
| 22268 | 1 | 1 | 0.94 | 0.911 | 0.953 | 1 | 1 | 1 | 1 | 1 | -0.039 |
| 22271 | 0 | 0 | 0.081 | 0 | 0 | NA | 0 | 0 | 0 | 0 | 0.016 |
| 22272 | 0 | 0 | 0.079 | 0 | 0 | NA | 0 | 0 | 0 | 0 | 0.016 |
| 22273 | 0.346 | 0 | 0.165 | 0.178 | 1 | NA | 0 | 0.405 | 0.209 | NA | 0.133 |
| 22276 | 1 | 1 | 0.963 | 1 | 1 | 1 | 1 | 1 | 1 | 1 | -0.007 |
| 22277 | 1 | 0.936 | 1 | 1 | 1 | 1 | 1 | 1 | 1 | 1 | -0.013 |
| 22279 | 0 | 0.139 | 0 | 0 | 0 | NA | 0 | 0 | 0 | NA | 0.028 |
| 22280 | 0.896 | 0.806 | 0.909 | 0.903 | 0.839 | 1 | 0.704 | 0.548 | 0.526 | NA | 0.176 |
| 22281 | 0.605 | 1 | 0.771 | 0.927 | 0.857 | 0.927 | 1 | 0.86 | 0.581 | NA | -0.01 |
| 22339 | 0 | 0.034 | 0 | 0.021 | 0 | 0 | 0 | 0 | 0 | 0.013 | 0.008 |
| 22342 | 1 | 0.6 | NA | 1 | 1 | NA | 1 | 1 | NA | NA | -0.1 |
| 22343 | 1 | 0.333 | 1 | 0.2 | NA | 1 | 1 | 1 | NA | NA | -0.367 |
| 22350 | 1 | NA | NA | NA | 1 | NA | 0 | NA | NA | NA | 1 |
| 22353 | 0 | 0 | 0 | 0 | 0 | NA | 0.132 | 0 | 0 | 0 | -0.033 |
| 22356 | 0.926 | 1 | 0.933 | 1 | 1 | 1 | 1 | 0.84 | 1 | 0.852 | 0.033 |
| 22365 | 0.007 | 0 | 0.028 | 0.031 | 0 | 0 | 0.024 | 0.019 | 0 | 0 | 0.005 |
| 22369 | 0.861 | 1 | 1 | 1 | 0.9 | 1 | 1 | 0.887 | 1 | 1 | -0.025 |
| 22370 | 0.992 | 0.98 | 1 | 1 | 1 | 1 | 1 | 1 | 1 | 1 | -0.006 |
| 22371 | 0.997 | 1 | 1 | 1 | 0.998 | 1 | 1 | 0.997 | 1 | 1 | 0 |
| 22372 | 0.989 | 1 | 1 | 1 | 1 | 1 | 1 | 1 | 1 | 1 | -0.002 |
| 22375 | 0.99 | 0.976 | 0.99 | 1 | 1 | 1 | 1 | 0.997 | 1 | 1 | -0.008 |
| 22377 | 0.993 | 1 | 1 | 1 | 1 | 1 | 1 | 1 | 1 | 1 | -0.001 |
| 22379 | 1 | 1 | 1 | 0.988 | 1 | 1 | 1 | 0.992 | 0.974 | 1 | 0.004 |
| 22380 | 1 | 1 | 1 | 1 | 1 | 1 | 0.995 | 1 | 1 | 1 | 0.001 |
| 22381 | 1 | 1 | 1 | 1 | 1 | 1 | 1 | 1 | 0.994 | 1 | 0.001 |
| 22382 | 1 | 1 | 1 | 0.966 | 1 | 1 | 1 | 0.987 | 1 | 1 | -0.004 |
| 22383 | 0.967 | 1 | 1 | 1 | 1 | 1 | 0.987 | 1 | 0.984 | 1 | -0.001 |
| 22384 | NA | NA | 1 | 0.724 | 1 | NA | NA | 0.496 | 1 | NA | 0.16 |
| 22385 | NA | NA | NA | 0.413 | 0.584 | NA | NA | NA | 1 | NA | -0.502 |
| 22386 | NA | 0.19 | NA | 0.413 | NA | NA | NA | NA | 1 | NA | -0.699 |
| 22387 | 0 | 1 | 0 | 1 | 0 | NA | NA | NA | 0.26 | NA | 0.14 |
| 22388 | NA | 0.27 | NA | 0.833 | 1 | NA | NA | NA | 1 | NA | -0.299 |
| 22389 | NA | 0.526 | NA | 1 | 1 | NA | NA | NA | 1 | NA | -0.158 |
| 22390 | 1 | 1 | 0.635 | 1 | 1 | 1 | 1 | 1 | NA | 1 | -0.073 |
| 22393 | 1 | 1 | 1 | 0.626 | 0 | NA | 1 | 0 | NA | NA | 0.225 |
| 22394 | 0.796 | 1 | 1 | 0.557 | 0 | NA | 1 | NA | NA | NA | -0.329 |
| 22395 | 1 | NA | 1 | 0.333 | NA | NA | 0 | NA | NA | NA | 0.778 |
| 22396 | 0.008 | 0.01 | 0.007 | 0.002 | 0.005 | 0 | 0.007 | 0.019 | 0 | 0 | 0.001 |
| 22397 | 1 | 1 | 1 | 1 | 1 | 1 | 0.987 | 1 | 1 | 1 | 0.003 |
| 22399 | NA | NA | 0.457 | 0 | NA | NA | 0.296 | NA | NA | NA | -0.067 |
| 22400 | 0.247 | 0.29 | 0.45 | 0.29 | 0 | NA | 0.45 | 0.105 | 0 | NA | 0.07 |
| 22402 | NA | 0.621 | 0.214 | 0 | 0 | NA | 1 | NA | NA | NA | -0.791 |
| 22403 | 1 | 1 | 0.551 | 0.522 | 1 | NA | 0.29 | 1 | NA | NA | 0.17 |
| 22405 | 1 | 0.256 | 0.256 | 0 | 0.805 | NA | NA | 1 | NA | NA | -0.537 |
| 22406 | 1 | 1 | 0.58 | 0.315 | 1 | NA | 0 | 1 | NA | NA | 0.279 |
| 22407 | NA | NA | 0 | 1 | NA | NA | NA | 1 | NA | NA | -0.5 |
| 22408 | NA | 0 | NA | NA | NA | NA | NA | 1 | NA | NA | -1 |
| 22409 | 0.256 | 0 | 0.58 | 0 | 0 | NA | NA | 0.674 | NA | NA | -0.507 |
| 22410 | 1 | 1 | 1 | 1 | 1 | 1 | 1 | 0.799 | 1 | 1 | 0.04 |
| 22413 | 1 | 1 | 1 | 1 | 1 | 1 | 0.714 | 1 | 1 | NA | 0.072 |
| 22414 | 1 | 1 | 0.817 | 1 | 0.936 | NA | 1 | 1 | 1 | 1 | -0.049 |
| 22415 | 1 | 1 | 1 | 1 | 1 | 1 | 0.4 | NA | 1 | NA | 0.2 |
| 22417 | 1 | 1 | 1 | 1 | 1 | NA | 0.714 | 1 | 1 | 1 | 0.072 |
| 22418 | 1 | 0.929 | 1 | 1 | 1 | NA | 1 | 0.674 | 1 | 1 | 0.067 |
| 22434 | 1 | 1 | 1 | 1 | 0.907 | NA | 1 | 1 | 1 | 1 | -0.019 |
| 22435 | 0.913 | 1 | 1 | 1 | 1 | 1 | 1 | 1 | 1 | 1 | -0.017 |
| 22437 | 0.392 | NA | 1 | 1 | 1 | NA | 1 | NA | NA | NA | -0.152 |
| 22438 | 1 | 1 | 1 | 1 | NA | 0 | 1 | 1 | 0.2 | NA | 0.45 |
| 22440 | 0 | 0 | 0 | 0 | 0 | NA | 1 | 0 | NA | 0 | -0.333 |
| 22441 | 0 | 0 | 0 | 0.144 | 0 | NA | NA | 0 | NA | 0 | 0.029 |
| 22454 | 1 | 1 | 0.946 | 1 | 1 | 1 | 1 | 1 | 1 | 1 | -0.011 |
| 22457 | 1 | 0.815 | 1 | 1 | 1 | 1 | 1 | 1 | 1 | 1 | -0.037 |
| 22458 | 1 | 1 | 0.875 | 1 | 0.933 | 1 | 1 | 1 | 1 | 1 | -0.038 |
| 22459 | 1 | 1 | 1 | 1 | 1 | 1 | 1 | 1 | 1 | 0.994 | 0.001 |
| 22460 | 1 | 1 | 1 | 0.956 | 1 | 1 | 0.913 | 1 | 1 | 1 | 0.009 |
| 22461 | 1 | NA | 1 | 1 | 1 | 0.62 | 0.803 | NA | 1 | 0.664 | 0.228 |
| 22462 | 0.791 | 0.749 | 0.873 | 0.938 | 0.835 | NA | 1 | 0.351 | 1 | NA | 0.054 |
| 22463 | 1 | 1 | 1 | 1 | 0.97 | NA | 1 | 1 | 1 | NA | -0.006 |
| 22470 | 1 | 1 | 1 | 1 | 1 | 0.956 | 1 | 1 | 1 | 1 | 0.009 |
| 22471 | 1 | 1 | 1 | 1 | 1 | 1 | 1 | 1 | 1 | 0.99 | 0.002 |
| 22472 | 1 | 1 | 0.991 | 1 | 1 | 1 | 1 | 0.997 | 1 | 1 | -0.001 |
| 22473 | 1 | 1 | 1 | 1 | 1 | 1 | 1 | 0.994 | 1 | 1 | 0.001 |
| 22477 | 1 | 0.974 | 1 | 1 | 1 | 1 | 1 | 1 | 0.99 | 1 | -0.003 |
| 22479 | 1 | 1 | 1 | 1 | 0.992 | 1 | 1 | 0.971 | 1 | 1 | 0.004 |
| 22480 | 1 | 0.97 | 1 | 1 | 1 | 1 | 1 | 1 | 1 | 1 | -0.006 |
| 22481 | 1 | 1 | 1 | 1 | 1 | 1 | 1 | 1 | 0.979 | 1 | 0.004 |
| 22484 | 0 | 0.111 | 0 | 0.077 | NA | 0 | NA | 0 | 0.154 | NA | -0.004 |
| 22487 | 1 | 0.981 | 1 | 0.988 | 1 | 1 | 1 | 1 | 0.972 | 1 | -0.001 |
| 22488 | 1 | 1 | 1 | 1 | 0 | 1 | NA | 1 | 1 | NA | -0.2 |
| 22489 | 1 | 1 | 1 | 1 | 1 | 1 | 1 | 1 | 0.997 | 1 | 0.001 |
| 22492 | 0.964 | 1 | 1 | 1 | 1 | 1 | 1 | 1 | 1 | 1 | -0.007 |
| 22493 | 0.988 | 1 | 1 | 1 | 1 | 1 | 1 | 1 | 1 | 1 | -0.002 |
| 22494 | 1 | 1 | 1 | 1 | 1 | 1 | 1 | 0.981 | 1 | 1 | 0.004 |
| 22495 | 1 | 1 | 0.976 | 0.987 | 1 | 1 | 1 | 0.961 | 1 | 1 | 0 |
| 22496 | 1 | 1 | 0.94 | 0.98 | 1 | 1 | 1 | 0.95 | 1 | 1 | -0.006 |
| 22498 | 1 | 1 | 1 | 1 | 1 | 1 | 1 | 0.981 | 1 | 1 | 0.004 |
| 22499 | 1 | 1 | 1 | 1 | 1 | 1 | 1 | 0.987 | 1 | 1 | 0.003 |
| 22500 | 0.96 | 0.982 | 1 | 1 | 0.963 | 1 | 0.941 | 1 | 0.957 | 1 | 0.001 |
| 22502 | 1 | 1 | 1 | 1 | 1 | 1 | 0.818 | 1 | 1 | 1 | 0.036 |
| 22506 | 1 | 0.996 | 1 | 1 | 0.988 | 1 | 1 | 1 | 0.979 | 1 | 0.001 |
| 22510 | 1 | 1 | 0.97 | 1 | 1 | 1 | 1 | 1 | 1 | 1 | -0.006 |
| 22512 | 0.997 | 1 | 1 | 1 | 1 | 1 | 1 | 1 | 1 | 0.993 | 0.001 |
| 22514 | 1 | 1 | 1 | 1 | 1 | 1 | 0.97 | 1 | 1 | 1 | 0.006 |
| 22532 | 1 | 0.974 | 1 | 0.976 | 1 | 1 | 1 | 1 | 1 | 1 | -0.01 |
| 22533 | 1 | 1 | 1 | 1 | 1 | 1 | 0.874 | 1 | 1 | 1 | 0.025 |
| 22534 | 1 | 1 | 1 | 1 | 1 | 1 | 0.923 | 1 | 1 | 1 | 0.015 |
| 22535 | 1 | 1 | 1 | 1 | 1 | 1 | 0.651 | 1 | 1 | 1 | 0.07 |
| 22536 | 1 | 1 | 1 | 1 | 1 | 1 | 1 | 0.952 | 1 | 1 | 0.01 |
| 22549 | 1 | 0.982 | 1 | 1 | 0.997 | 1 | 0.965 | 1 | 0.985 | 1 | 0.006 |
| 22554 | 1 | 1 | 1 | 1 | 1 | 1 | 0.963 | 1 | 1 | 1 | 0.007 |
| 22555 | 1 | NA | 1 | 1 | 0.535 | 1 | 1 | 1 | 1 | 1 | -0.116 |
| 22556 | 1 | 1 | 1 | 0.996 | 1 | 1 | 1 | 0.997 | 0.994 | 1 | 0.001 |
| 22557 | 1 | 1 | 1 | 1 | 0.972 | 1 | 1 | 1 | 1 | 1 | -0.006 |
| 22565 | 1 | NA | 1 | 1 | NA | NA | 0.382 | 1 | NA | 0.382 | 0.412 |
| 22566 | 1 | 1 | 1 | 0.98 | 1 | 1 | 1 | 1 | 1 | 1 | -0.004 |
| 22567 | 1 | 0.724 | 1 | 1 | 1 | 1 | 1 | 1 | 1 | 1 | -0.055 |
| 22568 | 1 | 0.901 | 1 | 1 | 1 | 1 | 1 | 1 | 1 | 1 | -0.02 |
| 22569 | 0.975 | 1 | 1 | 1 | 1 | 1 | 0.968 | 1 | 1 | 1 | 0.001 |
| 22570 | 1 | 1 | 1 | 1 | 0.921 | NA | 1 | 1 | 1 | 1 | -0.016 |
| 22571 | 0.932 | 1 | 1 | 1 | 1 | 1 | 0.858 | 1 | 1 | 1 | 0.015 |
| 22573 | 1 | 1 | 1 | 1 | 0.868 | 1 | 1 | 1 | 1 | 1 | -0.026 |
| 22574 | 1 | 1 | 1 | 1 | 1 | 1 | 1 | 0.971 | 1 | 1 | 0.006 |
| 22578 | 1 | 1 | 1 | 1 | 1 | 1 | 0.98 | 1 | 0.954 | 1 | 0.013 |
| 22581 | 0 | 0 | 0 | 0 | 0 | 0 | 0.002 | 0 | 0 | 0 | 0 |
| 22582 | 1 | 1 | 1 | 0.997 | 1 | 1 | 1 | 1 | 1 | 1 | -0.001 |
| 22583 | 1 | 1 | 1 | 1 | 1 | 1 | 1 | 1 | 0.983 | 1 | 0.003 |
| 22584 | 1 | 1 | 1 | 1 | 1 | 1 | 1 | 0.991 | 1 | 1 | 0.002 |
| 22585 | 0.989 | 1 | 0.995 | 1 | 1 | 1 | 0.989 | 1 | 1 | 1 | -0.001 |
| 22586 | 1 | 1 | 1 | 1 | 1 | 1 | 1 | 0.997 | 1 | 1 | 0.001 |
| 22589 | 0.98 | 0.992 | 0.997 | 0.998 | 1 | 1 | 0.997 | 0.996 | 0.925 | 1 | 0.01 |
| 22590 | 1 | 0.992 | 1 | 0.992 | 1 | 1 | 1 | 0.982 | 0.983 | 1 | 0.004 |
| 22592 | 1 | 0.989 | 1 | 0.997 | 1 | 1 | 0.995 | 1 | 1 | 1 | -0.002 |
| 22593 | 0.996 | 1 | 1 | 1 | 1 | 1 | 1 | 1 | 1 | 1 | -0.001 |
| 22595 | 0.319 | 0.493 | 0.226 | 0.323 | 0.163 | 1 | 0.193 | 0.032 | 0.226 | NA | -0.058 |
| 22598 | 0.993 | 0.977 | 0.988 | 0.98 | 1 | 1 | 0.979 | 0.989 | 1 | 1 | -0.006 |
| 22599 | 1 | 1 | 1 | 1 | 0.984 | 1 | 1 | 1 | 1 | 1 | -0.003 |
| 22600 | 0.989 | 1 | 1 | 1 | 1 | 1 | 1 | 1 | 0.98 | 1 | 0.002 |
| 22602 | 1 | 1 | 1 | 0.979 | 1 | 1 | 1 | 1 | 1 | 1 | -0.004 |
| 22603 | 0.996 | 1 | 1 | 0.995 | 1 | 1 | 0.995 | 1 | 1 | 1 | -0.001 |
| 22608 | 1 | 1 | 0.997 | 1 | 1 | 1 | 0.998 | 1 | 0.992 | 1 | 0.001 |
| 22609 | 0.005 | 0 | 0.011 | 0 | 0 | 0 | 0.003 | 0.006 | 0 | 0 | 0.001 |
| 22611 | 1 | 1 | 1 | 1 | 1 | 1 | 1 | 0.994 | 1 | 1 | 0.001 |
| 22612 | 0.994 | 1 | 1 | 1 | 1 | 1 | 0.987 | 1 | 0.935 | 1 | 0.014 |
| 22613 | 1 | 1 | 1 | 1 | 1 | 1 | 1 | 0.952 | 1 | 1 | 0.01 |
| 22614 | 1 | 1 | 1 | 1 | 1 | 1 | 1 | 0.962 | 1 | 1 | 0.008 |
| 22615 | 0.985 | 1 | 1 | 1 | 1 | 1 | 1 | 1 | 1 | 1 | -0.003 |
| 22616 | 0.996 | 1 | 1 | 1 | 1 | 1 | 1 | 1 | 1 | 1 | -0.001 |
| 22619 | 1 | 1 | 0.997 | 1 | 1 | 0.929 | 1 | 1 | 1 | 1 | 0.014 |
| 22621 | 1 | 1 | 0 | 0.485 | NA | NA | NA | 0 | NA | NA | 0.621 |
| 22622 | 1 | 1 | 0.993 | 0.988 | 1 | 1 | 1 | 0.989 | 1 | 1 | -0.002 |
| 22623 | 1 | 0.992 | 1 | 1 | 1 | 1 | 1 | 1 | 1 | 1 | -0.002 |
| 22624 | 1 | 0.996 | 1 | 1 | 0.992 | 1 | 1 | 0.996 | 1 | 1 | -0.002 |
| 22625 | 1 | 1 | 1 | 1 | 1 | 1 | 1 | 0.996 | 1 | 1 | 0.001 |
| 22626 | 1 | 1 | 0.997 | 1 | 1 | 1 | 1 | 1 | 1 | 1 | -0.001 |
| 22627 | 1 | 1 | 1 | 1 | 1 | 1 | 1 | 1 | 0.981 | 1 | 0.004 |
| 22632 | 1 | 0.993 | 1 | 0.997 | 1 | 1 | 0.978 | 0.995 | 1 | 1 | 0.003 |
| 22633 | NA | NA | 0 | NA | 0 | NA | NA | 1 | NA | NA | -1 |
| 22634 | 0 | 0 | 0 | 0 | 0 | NA | NA | 0.095 | 0 | NA | -0.048 |
| 22635 | 0 | 0 | 0 | 0 | 0 | NA | NA | 0.046 | 0 | NA | -0.023 |
| 22636 | 0.046 | 0 | 0.066 | 0 | 0 | NA | 0 | 0 | 0 | NA | 0.022 |
| 22637 | 0 | 0 | NA | 0 | 0 | 0 | NA | 0 | 1 | NA | -0.333 |
| 22638 | 0.873 | 0.8 | 1 | 0.919 | 0.855 | 0.322 | 1 | 0.952 | 1 | NA | 0.071 |
| 22639 | 1 | 1 | 1 | 1 | 1 | 1 | 1 | 1 | 0.774 | NA | 0.056 |
| 22640 | 1 | 1 | 0.936 | 1 | 0.946 | 1 | 0.935 | 0.913 | 1 | NA | 0.014 |
| 22641 | 1 | 1 | 1 | 1 | 1 | 1 | 0.876 | 1 | 1 | NA | 0.031 |
| 22642 | 1 | 1 | 1 | 1 | 1 | 1 | 0.863 | 1 | 1 | NA | 0.034 |
| 22643 | 0.561 | 1 | 1 | 1 | 1 | NA | 1 | 1 | 1 | 1 | -0.088 |
| 22644 | 1 | 1 | 1 | 1 | 1 | NA | 1 | 1 | 1 | 0.299 | 0.175 |
| 22645 | 1 | 1 | 1 | 0.865 | 1 | NA | 1 | 1 | 1 | 1 | -0.027 |
| 22646 | 0.561 | 0.852 | 1 | 1 | 1 | NA | 0.762 | 0.242 | 1 | 0.561 | 0.241 |
| 22647 | 1 | 1 | 0.826 | 1 | 0.442 | NA | 1 | 1 | 1 | NA | -0.146 |
| 22648 | NA | 1 | 0.856 | 0.372 | 1 | NA | NA | 1 | 1 | NA | -0.193 |
| 22649 | NA | 1 | 1 | 1 | 0.543 | NA | NA | 1 | 1 | NA | -0.114 |
| 22650 | 0 | 0.64 | 0.826 | 0.263 | 1 | NA | 0 | 1 | 1 | NA | -0.121 |
| 22653 | NA | NA | 1 | NA | 0.333 | NA | NA | NA | 0 | NA | 0.666 |
| 22654 | 1 | 1 | 1 | NA | 0.5 | NA | 1 | NA | NA | NA | -0.125 |
| 22655 | 1 | 0 | 0 | NA | 0 | NA | 1 | NA | NA | NA | -0.75 |
| 22656 | 1 | 1 | NA | NA | 1 | NA | 1 | NA | 0 | NA | 0.5 |
| 22661 | 0.428 | 0 | NA | NA | NA | NA | 0 | 0 | NA | 0 | 0.214 |
| 22664 | NA | NA | NA | NA | 0.355 | 0 | NA | 0 | NA | NA | 0.355 |
| 22665 | NA | NA | NA | 0 | NA | NA | NA | NA | 1 | 1 | -1 |
| 22666 | 0 | 0 | NA | NA | NA | NA | 0 | 0 | NA | 0.654 | -0.218 |
| 22667 | 0.667 | 0.333 | 1 | 1 | NA | NA | 0.5 | 0.556 | NA | 0 | 0.398 |
| 22668 | 0.667 | 0.333 | 0.667 | 0.231 | 1 | NA | 0.25 | 1 | NA | NA | -0.045 |
| 22669 | 1 | 0.333 | 1 | 1 | NA | NA | 1 | 1 | 0.429 | NA | 0.024 |
| 22670 | 1 | 1 | 1 | 1 | 0 | NA | 1 | 1 | 1 | NA | -0.2 |
| 22671 | 0.658 | 0 | 0.391 | 0 | NA | NA | 0 | 1 | NA | NA | -0.238 |
| 22675 | NA | NA | 0 | NA | NA | NA | NA | 1 | NA | 1 | -1 |
| 22678 | 0.984 | 1 | 0.957 | 1 | 1 | 1 | 1 | 1 | 1 | 1 | -0.012 |
| 22681 | 1 | 0.977 | 1 | 1 | 1 | 1 | 1 | 1 | 1 | 1 | -0.005 |
| 22682 | 1 | 0.948 | 1 | 1 | 1 | 1 | 1 | 1 | 1 | 1 | -0.01 |
| 22685 | 1 | 1 | 0.484 | NA | NA | NA | 1 | 1 | NA | NA | -0.172 |
| 22686 | 1 | 1 | 0.984 | 1 | 1 | 1 | 1 | 1 | 1 | 1 | -0.003 |
| 22688 | 1 | 1 | 1 | 1 | 1 | NA | 0.863 | 1 | 1 | 0.62 | 0.129 |
| 22690 | 0.583 | 1 | 1 | 1 | 1 | NA | 1 | 1 | 1 | 1 | -0.083 |
| 22693 | 0.978 | 1 | 1 | 1 | 1 | 1 | 1 | 1 | 1 | 1 | -0.004 |
| 22694 | 0.963 | 1 | 1 | 1 | 1 | 1 | 1 | 1 | 1 | 1 | -0.007 |
| 22695 | 0.956 | 1 | 1 | 1 | 1 | 1 | 1 | 1 | 1 | 1 | -0.009 |
| 22696 | 0.412 | 0 | NA | NA | 0.677 | NA | NA | 0 | NA | NA | 0.363 |
| 22698 | 1 | 1 | NA | NA | 1 | NA | NA | 0.521 | NA | NA | 0.479 |
| 22699 | 1 | 1 | 1 | 1 | 1 | 1 | 1 | 0.707 | 1 | 1 | 0.059 |
| 22700 | 1 | 0.954 | 1 | 1 | 1 | 1 | 1 | 1 | 1 | 1 | -0.009 |
| 22701 | 1 | 1 | 1 | 1 | 1 | 1 | 0.914 | 1 | 1 | 1 | 0.017 |
| 22702 | 1 | 1 | 1 | 1 | 1 | 1 | 0.952 | 1 | 1 | 1 | 0.01 |
| 22709 | 0.979 | 1 | 0.995 | 0.986 | 1 | 1 | 1 | 1 | 1 | 1 | -0.008 |
| 22710 | 0.361 | NA | 1 | 0.361 | 0.629 | 1 | 1 | NA | NA | NA | -0.412 |
| 22711 | 1 | 1 | 1 | 1 | 1 | 1 | 0.934 | 1 | 1 | 1 | 0.013 |
| 22714 | 1 | 0.994 | 1 | 1 | 1 | 1 | 1 | 1 | 1 | 1 | -0.001 |
| 22715 | 0.948 | 1 | 1 | 0.985 | 1 | 1 | 1 | 1 | 1 | 1 | -0.013 |
| 22719 | 1 | 1 | 1 | 0.995 | 0.992 | 1 | 1 | 1 | 1 | 1 | -0.003 |
| 22720 | 0.983 | 1 | 1 | 1 | 1 | 1 | 1 | 1 | 1 | 1 | -0.003 |
| 22721 | 0.961 | 1 | 1 | 1 | 1 | NA | 1 | 1 | 1 | NA | -0.008 |
| 22722 | 1 | 1 | 1 | 1 | 1 | 1 | 0.965 | 1 | 1 | 1 | 0.007 |
| 22723 | 0.806 | 0.946 | 1 | 1 | 1 | NA | 0.765 | 1 | 1 | NA | 0.029 |
| 22724 | 1 | 1 | 1 | 0.935 | 1 | NA | 1 | 1 | NA | 1 | -0.013 |
| 22725 | 0.6 | 1 | 1 | 1 | 1 | NA | 1 | 1 | NA | 1 | -0.08 |
| 22727 | 0.037 | 0 | 0.027 | 0.023 | 0.11 | 0 | 0 | 0 | 0 | NA | 0.039 |
| 22729 | 0.806 | 1 | 1 | 1 | 1 | 1 | NA | NA | NA | 0 | 0.461 |
| 22730 | 0.057 | 0 | 0.523 | NA | 0 | 0 | 1 | NA | NA | NA | -0.355 |
| 22731 | 0.682 | 1 | 0 | NA | 1 | 1 | 1 | 1 | 1 | 1 | -0.33 |
| 22732 | 1 | 1 | 1 | 1 | 1 | 1 | 0.918 | 1 | 1 | 1 | 0.016 |
| 22735 | 1 | 1 | 1 | 1 | 1 | 1 | 1 | 1 | 1 | 0.974 | 0.005 |
| 22736 | 1 | 1 | 1 | 1 | 0.993 | 0.996 | 0.976 | 1 | 1 | 1 | 0.004 |
| 22737 | 1 | 1 | 1 | 1 | 0.894 | 1 | 0.926 | 0.914 | 1 | 1 | 0.011 |
| 22738 | 1 | 1 | 1 | 1 | 0.982 | 1 | 0.979 | 0.977 | 1 | 1 | 0.005 |
| 22739 | 0.99 | 0.984 | 0.987 | 0.993 | 1 | 1 | 1 | 0.972 | 1 | 0.989 | -0.001 |
| 22741 | 0.984 | 1 | 1 | 1 | 0.985 | 1 | 1 | 1 | 1 | 1 | -0.006 |
| 22742 | 0 | 0.024 | 0 | 0 | 0.018 | 0 | 0 | 0 | 0 | 0 | 0.008 |
| 22743 | 0 | 0.117 | 0.068 | 0.422 | 0.422 | NA | 0 | 0 | 0 | NA | 0.206 |
| 22745 | 1 | 1 | 1 | 1 | 0.962 | 1 | 1 | 1 | 1 | 1 | -0.008 |
| 22746 | 1 | 1 | 1 | 1 | 0.979 | NA | 1 | 1 | 1 | 1 | -0.004 |
| 22747 | 0.99 | 0.971 | 1 | 1 | 1 | 1 | 0.95 | 1 | 1 | 1 | 0.002 |
| 22748 | 0 | 0 | 0 | 0.008 | 0.045 | NA | 0 | 0 | 0 | 0 | 0.011 |
| 22749 | 1 | 1 | 1 | 0.958 | 1 | NA | 1 | 1 | NA | 1 | -0.008 |
| 22750 | NA | NA | NA | 1 | NA | NA | 1 | NA | NA | 0 | 0.5 |
| 22751 | 0.226 | 0 | 0 | 0 | NA | NA | 0 | NA | 0 | 0 | 0.057 |
| 22752 | 1 | 1 | 0.846 | 1 | 1 | 1 | 0.5 | 1 | 1 | 1 | 0.069 |
| 22753 | 0.957 | 1 | 1 | 1 | 1 | 1 | 1 | 1 | 1 | NA | -0.009 |

| | | | | | | | | | | | | | | | | | | | | | | | | | | | | | | | | | | | | | | | | | | | | | | |
|---|---|---|---|---|---|---|---|---|---|---|---|---|---|---|---|---|---|---|---|---|---|---|---|---|---|---|---|---|---|---|---|---|---|---|---|---|---|---|---|---|---|---|---|---|---|---|
| SE | 22754 | ENSG00000145730 | PAM | chr5 | + | 102899935 | 102900002 | 102867272 | 102867393 | 102901355 | 102901413 | 0 | 2 | 1 | 2 | 0 | 46 | 38 | 31 | 44 | 27 | 0 | 1 | 0 | 0 | 0 | 13 | 21 | 14 | 34 | 14 | 216 | 149 | 1 | 1 | 0 | 0.035 | 0.022 | 0.03 | 0 | 0 | 0.032 | 0 | 0 | 0 | 0.011 |
| SE | 22756 | ENSG00000145730 | PAM | chr5 | + | 102924956 | 102925042 | 102913933 | 102914021 | 102926584 | 102926668 | 110 | 75 | 87 | 63 | 32 | 0 | 0 | 0 | 1 | 0 | 58 | 21 | 25 | 46 | 54 | 0 | 0 | 0 | 0 | 0 | 235 | 149 | 1 | 1 | 1 | 1 | 1 | 0.976 | 1 | 1 | 1 | 1 | 1 | 1 | -0.005 |
| SE | 22757 | ENSG00000145730 | PAM | chr5 | + | 102946836 | 102946885 | 102926584 | 102926668 | 102948377 | 102948445 | 77 | 60 | 50 | 56 | 37 | 0 | 0 | 0 | 0 | 0 | 52 | 24 | 15 | 30 | 15 | 0 | 0 | 0 | 3 | 0 | 198 | 149 | 1 | 1 | 1 | 1 | 1 | 1 | 1 | 1 | 1 | 1 | 0.883 | 1 | 0.023 |
| SE | 22759 | ENSG00000145730 | PAM | chr5 | + | 102990271 | 102990401 | 102961157 | 102961229 | 103003032 | 103003149 | 129 | 77 | 72 | 121 | 72 | 0 | 4 | 0 | 0 | 0 | 131 | 66 | 58 | 72 | 111 | 0 | 0 | 0 | 0 | 0 | 279 | 149 | 1 | 1 | 1 | 0.911 | 1 | 1 | 1 | 1 | 1 | 1 | 1 | 1 | -0.018 |
| SE | 22760 | ENSG00000145730 | PAM | chr5 | + | 103005153 | 103005226 | 103003032 | 103003149 | 103006800 | 103006859 | 156 | 68 | 79 | 89 | 83 | 0 | 0 | 0 | 0 | 0 | 100 | 47 | 47 | 86 | 82 | 0 | 0 | 0 | 5 | 0 | 222 | 149 | 1 | 1 | 1 | 1 | 1 | 1 | 1 | 1 | 1 | 1 | 0.92 | 1 | 0.016 |
| SE | 22771 | ENSG00000114904 | NEK4 | chr3 | - | 52737585 | 52737719 | 52708443 | 52711869 | 52739428 | 52739634 | 287 | 334 | 289 | 292 | 422 | 0 | 0 | 0 | 0 | 0 | 453 | 419 | 196 | 636 | 313 | 0 | 0 | 1 | 0 | 0 | 283 | 149 | 1 | 1 | 1 | 1 | 1 | 1 | 1 | 1 | 1 | 0.99 | 1 | 1 | 0.002 |
| SE | 22772 | ENSG00000114904 | NEK4 | chr3 | - | 52739428 | 52739634 | 52737497 | 52737719 | 52743351 | 52743461 | 245 | 296 | 244 | 247 | 358 | 0 | 2 | 0 | 0 | 0 | 438 | 393 | 178 | 568 | 300 | 0 | 0 | 0 | 0 | 0 | 298 | 149 | 1 | 1 | 1 | 0.987 | 1 | 1 | 1 | 1 | 1 | 1 | 1 | 1 | -0.003 |
| SE | 22777 | ENSG00000114904 | NEK4 | chr3 | - | 52747329 | 52747451 | 52746733 | 52746904 | 52749691 | 52749829 | 0 | 2 | 2 | 0 | 0 | 20 | 20 | 11 | 25 | 29 | 0 | 0 | 0 | 0 | 3 | 29 | 13 | 11 | 34 | 25 | 271 | 149 | 1 | 1 | 0 | 0.052 | 0.091 | 0 | 0 | 0 | 0 | 0 | 0 | 0.062 | 0.016 |
| SE | 22780 | ENSG00000114904 | NEK4 | chr3 | - | 52760794 | 52760936 | 52751931 | 52752336 | 52765886 | 52765994 | 57 | 52 | 38 | 47 | 99 | 0 | 0 | 0 | 0 | 0 | 59 | 72 | 35 | 80 | 64 | 0 | 0 | 0 | 2 | 0 | 291 | 149 | 1 | 1 | 1 | 1 | 1 | 1 | 1 | 1 | 1 | 1 | 0.953 | 1 | 0.009 |
| SE | 22781 | ENSG00000114904 | NEK4 | chr3 | - | 52763469 | 52763624 | 52751931 | 52752336 | 52765886 | 52765994 | 119 | 91 | 112 | 129 | 214 | 0 | 0 | 0 | 0 | 0 | 152 | 124 | 56 | 206 | 104 | 0 | 0 | 0 | 2 | 0 | 298 | 149 | 1 | 1 | 1 | 1 | 1 | 1 | 1 | 1 | 1 | 1 | 0.981 | 1 | 0.004 |
| SE | 22782 | ENSG00000114904 | NEK4 | chr3 | - | 52763469 | 52763624 | 52751931 | 52752336 | 52766177 | 52766375 | 0 | 2 | 6 | 0 | 4 | 0 | 0 | 1 | 0 | 0 | 0 | 7 | 2 | 16 | 0 | 0 | 0 | 0 | 0 | 0 | 298 | 149 | 1 | 1 | NA | 1 | 0.75 | NA | 1 | NA | 1 | 1 | 1 | NA | -0.083 |
| SE | 22783 | ENSG00000114904 | NEK4 | chr3 | - | 52763469 | 52763624 | 52760794 | 52760936 | 52765886 | 52765994 | 219 | 185 | 231 | 244 | 487 | 0 | 0 | 0 | 0 | 3 | 248 | 230 | 115 | 361 | 269 | 0 | 7 | 0 | 2 | 0 | 298 | 149 | 1 | 1 | 1 | 1 | 1 | 1 | 0.988 | 1 | 0.943 | 1 | 0.989 | 1 | 0.011 |
| SE | 22784 | ENSG00000114904 | NEK4 | chr3 | - | 52765886 | 52765994 | 52751931 | 52752336 | 52766177 | 52766375 | 37 | 13 | 36 | 28 | 48 | 0 | 0 | 1 | 0 | 0 | 47 | 34 | 14 | 75 | 29 | 0 | 0 | 0 | 0 | 0 | 257 | 149 | 1 | 1 | 1 | 1 | 0.954 | 1 | 1 | 1 | 1 | 1 | 1 | 1 | -0.009 |
| SE | 22788 | ENSG00000138381 | ASNSD1 | chr2 | + | 189663900 | 189663954 | 189661384 | 189661610 | 189665375 | 189665451 | 218 | 210 | 239 | 213 | 377 | 0 | 0 | 0 | 3 | 0 | 49 | 121 | 151 | 72 | 9 | 0 | 0 | 0 | 0 | 0 | 203 | 149 | 1 | 1 | 1 | 1 | 1 | 0.981 | 1 | 1 | 1 | 1 | 1 | 1 | -0.004 |
| SE | 22789 | ENSG00000111962 | UST | chr6 | + | 148941278 | 148941434 | 148886985 | 148887029 | 148953871 | 148953951 | 31 | 18 | 32 | 30 | 16 | 0 | 0 | 0 | 0 | 0 | 0 | 23 | 7 | 11 | 21 | 0 | 0 | 0 | 5 | 0 | 298 | 149 | 1 | 1 | 1 | 1 | 1 | 1 | 1 | NA | 1 | 1 | 0.524 | 1 | 0.119 |
| SE | 22790 | ENSG00000111962 | UST | chr6 | + | 148953871 | 148953951 | 148941278 | 148941434 | 148964409 | 148964563 | 27 | 15 | 30 | 19 | 29 | 1 | 0 | 0 | 1 | 0 | 28 | 53 | 12 | 10 | 37 | 0 | 0 | 0 | 0 | 0 | 229 | 149 | 1 | 1 | 0.946 | 1 | 1 | 0.925 | 1 | 1 | 1 | 1 | 1 | 1 | -0.026 |
| SE | 22793 | ENSG00000131504 | DIAPH1 | chr5 | - | 141518943 | 141518996 | 141515020 | 141517008 | 141524142 | 141524229 | 1 | 0 | 1 | 0 | 0 | 594 | 307 | 420 | 435 | 793 | 0 | 0 | 0 | 0 | 0 | 504 | 389 | 294 | 338 | 874 | 202 | 149 | 1 | 1 | 0.001 | 0 | 0.002 | 0 | 0 | 0 | 0 | 0.003 | 0 | 0 | 0 |
| SE | 22794 | ENSG00000131504 | DIAPH1 | chr5 | - | 141524142 | 141524229 | 141515020 | 141517008 | 141526037 | 141526173 | 1099 | 580 | 753 | 826 | 1461 | 0 | 1 | 0 | 0 | 0 | 1181 | 721 | 599 | 681 | 1768 | 0 | 0 | 0 | 0 | 0 | 236 | 149 | 1 | 1 | 1 | 0.997 | 1 | 1 | 1 | 1 | 1 | 1 | 1 | 1 | -0.001 |
| SE | 22795 | ENSG00000131504 | DIAPH1 | chr5 | - | 141526037 | 141526173 | 141524142 | 141524229 | 141526296 | 141526459 | 2354 | 1606 | 1458 | 2107 | 3013 | 0 | 0 | 0 | 0 | 0 | 4404 | 1955 | 1550 | 2570 | 5331 | 8 | 0 | 0 | 0 | 0 | 285 | 149 | 1 | 1 | 1 | 1 | 1 | 1 | 1 | 0.997 | 1 | 1 | 1 | 1 | 0.001 |
| SE | 22796 | ENSG00000131504 | DIAPH1 | chr5 | - | 141529171 | 141529273 | 141528701 | 141528941 | 141529602 | 141529697 | 985 | 585 | 614 | 775 | 1480 | 0 | 0 | 0 | 0 | 0 | 1361 | 734 | 483 | 724 | 2034 | 0 | 0 | 0 | 5 | 0 | 251 | 149 | 1 | 1 | 1 | 1 | 1 | 1 | 1 | 1 | 1 | 1 | 0.989 | 1 | 0.002 |
| SE | 22797 | ENSG00000131504 | DIAPH1 | chr5 | - | 141529171 | 141529273 | 141528701 | 141528941 | 141534334 | 141534433 | 501 | 312 | 308 | 400 | 743 | 0 | 0 | 0 | 0 | 0 | 675 | 354 | 231 | 374 | 1102 | 0 | 3 | 0 | 8 | 0 | 251 | 149 | 1 | 1 | 1 | 1 | 1 | 1 | 1 | 1 | 0.986 | 1 | 0.965 | 1 | 0.01 |
| SE | 22801 | ENSG00000131504 | DIAPH1 | chr5 | - | 141571427 | 141571436 | 141534334 | 141534433 | 141571925 | 141572040 | 13 | 5 | 8 | 10 | 31 | 0 | 0 | 0 | 0 | 0 | 5 | 4 | 5 | 15 | 14 | 2 | 0 | 1 | 0 | 0 | 158 | 149 | 1 | 1 | 1 | 1 | 1 | 1 | 1 | 0.702 | 1 | 0.825 | 1 | 1 | 0.095 |
| SE | 22802 | ENSG00000131504 | DIAPH1 | chr5 | - | 141580743 | 141580883 | 141579087 | 141579196 | 141582311 | 141582375 | 1849 | 1268 | 1275 | 1687 | 2299 | 2 | 0 | 0 | 0 | 0 | 2837 | 1721 | 1220 | 1955 | 4095 | 0 | 0 | 0 | 0 | 0 | 289 | 149 | 1 | 1 | 0.998 | 1 | 1 | 1 | 1 | 1 | 1 | 1 | 1 | 1 | 0 |
| SE | 22803 | ENSG00000131504 | DIAPH1 | chr5 | - | 141581914 | 141582056 | 141580743 | 141580883 | 141582311 | 141582375 | 6 | 0 | 4 | 0 | 3 | 1009 | 663 | 662 | 810 | 1269 | 0 | 4 | 0 | 0 | 4 | 1479 | 848 | 664 | 995 | 2195 | 291 | 149 | 1 | 1 | 0.003 | 0 | 0.003 | 0 | 0.001 | 0 | 0.002 | 0 | 0 | 0.001 | 0.001 |
| SE | 22804 | ENSG00000131504 | DIAPH1 | chr5 | - | 141584123 | 141584225 | 141583484 | 141583615 | 141587041 | 141587176 | 836 | 433 | 524 | 656 | 915 | 0 | 0 | 1 | 0 | 0 | 1086 | 720 | 498 | 808 | 1566 | 0 | 0 | 0 | 0 | 0 | 251 | 149 | 1 | 1 | 1 | 1 | 0.997 | 1 | 1 | 1 | 1 | 1 | 1 | 1 | -0.001 |
| SE | 22808 | ENSG00000173692 | PSMD1 | chr2 | + | 231062505 | 231062675 | 231062247 | 231062321 | 231066905 | 231067111 | 297 | 221 | 281 | 232 | 451 | 0 | 0 | 0 | 0 | 4 | 188 | 329 | 220 | 325 | 34 | 0 | 0 | 0 | 0 | 0 | 298 | 149 | 1 | 1 | 1 | 1 | 1 | 1 | 0.983 | 1 | 1 | 1 | 1 | 1 | -0.003 |
| SE | 22809 | ENSG00000173692 | PSMD1 | chr2 | + | 231062505 | 231062693 | 231062247 | 231062321 | 231066905 | 231067111 | 174 | 159 | 171 | 142 | 216 | 0 | 0 | 0 | 0 | 4 | 135 | 251 | 176 | 267 | 24 | 0 | 0 | 0 | 0 | 0 | 298 | 149 | 1 | 1 | 1 | 1 | 1 | 1 | 0.964 | 1 | 1 | 1 | 1 | 1 | -0.007 |
| SE | 22810 | ENSG00000173692 | PSMD1 | chr2 | + | 231066905 | 231067111 | 231062505 | 231062675 | 231070024 | 231070106 | 187 | 97 | 179 | 155 | 386 | 0 | 0 | 0 | 0 | 0 | 106 | 137 | 87 | 100 | 11 | 0 | 0 | 0 | 2 | 0 | 298 | 149 | 1 | 1 | 1 | 1 | 1 | 1 | 1 | 1 | 1 | 1 | 0.962 | 1 | 0.008 |
| SE | 22819 | ENSG00000173692 | PSMD1 | chr2 | + | 231152716 | 231152811 | 231146239 | 231146356 | 231153563 | 231153666 | 0 | 0 | 2 | 0 | 0 | 121 | 96 | 135 | 163 | 225 | 0 | 0 | 0 | 0 | 0 | 37 | 95 | 118 | 115 | 19 | 244 | 149 | 1 | 1 | 0 | 0 | 0.009 | 0 | 0 | 0 | 0 | 0 | 0 | 0 | 0.002 |
| SE | 22824 | ENSG00000173692 | PSMD1 | chr2 | + | 231165199 | 231165286 | 231163587 | 231163727 | [illegible] | 231166017 | 86 | 56 | 54 | 63 | 105 | 0 | 0 | 0 | 0 | 0 | 109 | 64 | 62 | 76 | 8 | 0 | 0 | 0 | 2 | 0 | 236 | 149 | 1 | 1 | 1 | 1 | 1 | 1 | 1 | 1 | 1 | 1 | 0.96 | 1 | 0.008 |
| SE | 22825 | ENSG00000173692 | PSMD1 | chr2 | + | 231166380 | 231166491 | 231165870 | 231166017 | 231170565 | 231170721 | 5 | 4 | 0 | 1 | 0 | 146 | 115 | 102 | 138 | 200 | 0 | 0 | 0 | 0 | 0 | 139 | 97 | 97 | 184 | 9 | 260 | 149 | 1 | 1 | 0.019 | 0.02 | 0 | 0.004 | 0 | 0 | 0 | 0 | 0 | 0 | 0.009 |
| SE | 22826 | ENSG00000173692 | PSMD1 | chr2 | + | 231170565 | 231170721 | 231165870 | 231166017 | 231172534 | 231172741 | 187 | 152 | 158 | 203 | 241 | 0 | 0 | 2 | 0 | 0 | 208 | 135 | 124 | 239 | 9 | 0 | 0 | 0 | 0 | 0 | 298 | 149 | 1 | 1 | 1 | 1 | 0.975 | 1 | 1 | 1 | 1 | 1 | 1 | 1 | -0.005 |
| SE | 22827 | ENSG00000173692 | PSMD1 | chr2 | + | 231170578 | 231170721 | 231165870 | 231166017 | 231172534 | 231172556 | 41 | 37 | 56 | 65 | 41 | 0 | 0 | 2 | 0 | 0 | 69 | 38 | 27 | 55 | 0 | 0 | 0 | 0 | 0 | 0 | 292 | 149 | 1 | 1 | 1 | 1 | 0.935 | 1 | 1 | 1 | 1 | 1 | 1 | NA | -0.013 |
| SE | 22828 | ENSG00000177595 | PIDD1 | chr11 | - | 800130 | 800244 | 799814 | 800014 | 800383 | 800451 | 13 | 3 | 2 | 3 | 6 | 1 | 0 | 0 | 0 | 0 | 0 | 6 | 4 | 0 | 12 | 0 | 0 | 0 | 0 | 0 | 263 | 149 | 1 | 1 | 0.88 | 1 | 1 | 1 | 1 | NA | 1 | 1 | NA | 1 | -0.024 |
| SE | 22830 | ENSG00000107968 | MAP3K8 | chr10 | + | 30437175 | 30437406 | 30434183 | 30434378 | 30447781 | 30447949 | 6 | 4 | 6 | 8 | 2 | 3 | 0 | 0 | 0 | 0 | 0 | 6 | 0 | 2 | 3 | 0 | 4 | 0 | 0 | 0 | 298 | 149 | 1 | 1 | 0.5 | 1 | 1 | 1 | 1 | NA | 0.429 | NA | 1 | 1 | 0.09 |
| SE | 22832 | ENSG00000107968 | MAP3K8 | chr10 | + | 30437175 | 30437406 | 30434664 | 30434748 | 30447781 | 30447949 | 2 | 2 | 3 | 4 | 0 | 0 | 0 | 0 | 0 | 0 | 0 | 6 | 8 | 3 | 6 | 0 | 5 | 0 | 0 | 0 | 298 | 149 | 1 | 1 | 1 | 1 | 1 | 1 | NA | NA | 0.375 | 1 | 1 | 1 | 0.156 |
| SE | 22833 | ENSG00000107968 | MAP3K8 | chr10 | + | 30438915 | 30439274 | 30434020 | 30434378 | 30447781 | 30447949 | 134 | 73 | 84 | 97 | 107 | 3 | 0 | 0 | 0 | 0 | 77 | 116 | 149 | 91 | 62 | 0 | 4 | 0 | 0 | 0 | 298 | 149 | 1 | 1 | 0.957 | 1 | 1 | 1 | 1 | 1 | 0.935 | 1 | 1 | 1 | 0.004 |
| SE | 22834 | ENSG00000107968 | MAP3K8 | chr10 | + | 30438915 | 30439274 | 30434664 | 30434748 | 30447781 | 30447949 | 121 | 67 | 73 | 75 | 98 | 0 | 0 | 0 | 0 | 0 | 79 | 108 | 140 | 74 | 32 | 0 | 5 | 0 | 0 | 0 | 298 | 149 | 1 | 1 | 1 | 1 | 1 | 1 | 1 | 1 | 0.915 | 1 | 1 | 1 | 0.017 |
| SE | 22835 | ENSG00000107968 | MAP3K8 | chr10 | + | 30438915 | 30439274 | 30437175 | 30437406 | 30447781 | 30447949 | 192 | 101 | 99 | 136 | 129 | 0 | 1 | 1 | 0 | 0 | 119 | 163 | 194 | 142 | 120 | 0 | 4 | 0 | 2 | 0 | 298 | 149 | 1 | 1 | 1 | 0.981 | 0.98 | 1 | 1 | 1 | 0.953 | 1 | 0.973 | 1 | 0.007 |
| SE | 22837 | ENSG00000107968 | MAP3K8 | chr10 | + | 30459254 | 30459501 | 30458083 | 30458236 | 30460705 | 30461833 | 158 | 47 | 69 | 104 | 97 | 0 | 3 | 0 | 0 | 0 | 113 | 92 | 76 | 81 | 70 | 0 | 0 | 2 | 0 | 0 | 298 | 149 | 1 | 1 | 1 | 0.887 | 1 | 1 | 1 | 1 | 1 | 0.95 | 1 | 1 | -0.013 |
| SE | 22838 | ENSG00000118939 | UCHL3 | chr13 | + | 75549975 | 75549987 | 75549482 | 75549670 | 75560752 | 75560881 | 9 | 5 | 18 | 21 | 31 | 0 | 0 | 1 | 0 | 0 | 9 | 11 | 16 | 7 | 0 | 0 | 0 | 0 | 0 | 0 | 161 | 149 | 1 | 1 | 1 | 1 | 0.943 | 1 | 1 | 1 | 1 | 1 | 1 | NA | -0.011 |
| SE | 22839 | ENSG00000118939 | UCHL3 | chr13 | + | 75549975 | 75549987 | 75549790 | 75549862 | 75560752 | 75560881 | 51 | 24 | 42 | 40 | 41 | 1 | 0 | 1 | 0 | 0 | 23 | 26 | 32 | 13 | 6 | 0 | 1 | 0 | 2 | 0 | 161 | 149 | 1 | 1 | 0.979 | 1 | 0.975 | 1 | 1 | 1 | 0.96 | 1 | 0.857 | 1 | 0.027 |
| SE | 22840 | ENSG00000118939 | UCHL3 | chr13 | + | 75589937 | 75590129 | 75569459 | 75569507 | 75594914 | 75594990 | 0 | 2 | 1 | 2 | 0 | 53 | 58 | 46 | 68 | 122 | 0 | 0 | 1 | 0 | 0 | 18 | 33 | 15 | 47 | 14 | 298 | 149 | 1 | 1 | 0 | 0.017 | 0.011 | 0.014 | 0 | 0 | 0 | 0.032 | 0 | 0 | 0.002 |
| SE | 22841 | ENSG00000118939 | UCHL3 | chr13 | + | 75604768 | 75604827 | 75594914 | 75594990 | 75605728 | 75605930 | 93 | 52 | 85 | 86 | 256 | 0 | 1 | 0 | 0 | 0 | 26 | 54 | 37 | 72 | 9 | 0 | 0 | 0 | 0 | 0 | 208 | 149 | 1 | 1 | 1 | 0.974 | 1 | 1 | 1 | 1 | 1 | 1 | 1 | 1 | -0.005 |
| SE | 22842 | ENSG00000154134 | ROBO3 | chr11 | + | 124877160 | 124877184 | 124876485 | 124876932 | 124877266 | 124877309 | 0 | 0 | 9 | 1 | 2 | 5 | 1 | 15 | 4 | 0 | 0 | 0 | 0 | 0 | 0 | 0 | 0 | 4 | 0 | 0 | 173 | 149 | 1 | 1 | 0 | 0 | 0.341 | 0.177 | 1 | NA | NA | 0 | NA | NA | 0.304 |
| SE | 22844 | ENSG00000154134 | ROBO3 | chr11 | + | 124877160 | 124877309 | 124876821 | 124876932 | 124877518 | 124877796 | 3 | 1 | 3 | 5 | 0 | 0 | 0 | 0 | 0 | 0 | 0 | 0 | 2 | 0 | 0 | 0 | 0 | 0 | 2 | 0 | 298 | 149 | 1 | 1 | 1 | 1 | 1 | 1 | NA | NA | NA | 1 | 0 | NA | 0.5 |
| SE | 22846 | ENSG00000154134 | ROBO3 | chr11 | + | 124877266 | 124877309 | 124876825 | 124876932 | 124877518 | 124877658 | 8 | 2 | 18 | 9 | 0 | 0 | 0 | 0 | 0 | 0 | 0 | 0 | 2 | 0 | 0 | 0 | 0 | 0 | 2 | 0 | 192 | 149 | 1 | 1 | 1 | 1 | 1 | 1 | NA | NA | NA | 1 | 0 | NA | 0.5 |
| SE | 22847 | ENSG00000154134 | ROBO3 | chr11 | + | 124877284 | 124877309 | 124876841 | 124876932 | 124877518 | 124877658 | 0 | 0 | 2 | 1 | 0 | 0 | 0 | 0 | 0 | 0 | 0 | 0 | 2 | 0 | 0 | 0 | 0 | 0 | 2 | 0 | 174 | 149 | 1 | 1 | NA | NA | 1 | 1 | NA | NA | NA | 1 | 0 | NA | 0.5 |
| SE | 22848 | ENSG00000154134 | ROBO3 | chr11 | + | 124877936 | 124878131 | 124877387 | 124877658 | 124878297 | 124878436 | 6 | 1 | 6 | 5 | 5 | 0 | 0 | 0 | 0 | 0 | 0 | 1 | 4 | 14 | 0 | 0 | 0 | 2 | 0 | 0 | 298 | 149 | 1 | 1 | 1 | 1 | 1 | 1 | 1 | NA | 1 | 0.5 | 1 | NA | 0.167 |
| SE | 22850 | ENSG00000154134 | ROBO3 | chr11 | + | 124877936 | 124878131 | 124877518 | 124877752 | 124878297 | 124878436 | 6 | 1 | 1 | 4 | 8 | 1 | 0 | 1 | 0 | 9 | 0 | 1 | 4 | 0 | 0 | 0 | 6 | 1 | 0 | 0 | 298 | 149 | 1 | 1 | 0.75 | 1 | 0.333 | 1 | 0.308 | NA | 0.077 | 0.667 | NA | NA | 0.306 |
| SE | 22854 | ENSG00000154134 | ROBO3 | chr11 | + | 124878297 | 124878436 | 124877518 | 124877752 | 124879189 | 124879341 | 2 | 0 | 1 | 0 | 9 | 2 | 0 | 0 | 0 | 0 | 0 | 6 | 1 | 0 | 0 | 0 | 0 | 0 | 0 | 0 | 288 | 149 | 1 | 1 | 0.341 | NA | 1 | NA | 1 | NA | 1 | 1 | NA | NA | -0.22 |
| SE | 22856 | ENSG00000154134 | ROBO3 | chr11 | + | 124878583 | 124878796 | 124878296 | 124878436 | 124879189 | 124879307 | 25 | 9 | 18 | 14 | 15 | 1 | 0 | 0 | 0 | 0 | 9 | 26 | 11 | 10 | 0 | 0 | 0 | 0 | 0 | 0 | 298 | 149 | 1 | 1 | 0.926 | 1 | 1 | 1 | 1 | 1 | 1 | 1 | 1 | NA | -0.015 |
| SE | 22858 | ENSG00000043143 | JADE2 | chr5 | + | 134552051 | 134552209 | 134537988 | 134538083 | 134559829 | 134559990 | 649 | 359 | 603 | 680 | 533 | 0 | 0 | 3 | 1 | 0 | 408 | 705 | 449 | 284 | 651 | 0 | 0 | 0 | 0 | 0 | 298 | 149 | 1 | 1 | 1 | 1 | 0.99 | 0.997 | 1 | 1 | 1 | 1 | 1 | 1 | -0.003 |
| SE | 22860 | ENSG00000043143 | JADE2 | chr5 | + | 134559829 | 134559990 | 134552051 | 134552209 | 134564493 | 134564610 | 410 | 195 | 379 | 409 | 317 | 0 | 0 | 1 | 0 | 0 | 252 | 429 | 292 | 171 | 417 | 0 | 0 | 0 | 0 | 0 | 298 | 149 | 1 | 1 | 1 | 1 | 0.995 | 1 | 1 | 1 | 1 | 1 | 1 | 1 | -0.001 |
| SE | 22861 | ENSG00000043143 | JADE2 | chr5 | + | 134560745 | 134560957 | 134559829 | 134559990 | 134562199 | 134562367 | 408 | 216 | 416 | 466 | 231 | 0 | 0 | 0 | 0 | 0 | 172 | 348 | 277 | 164 | 337 | 0 | 0 | 4 | 0 | 0 | 298 | 149 | 1 | 1 | 1 | 1 | 1 | 1 | 1 | 1 | 1 | 0.972 | 1 | 1 | 0.006 |
| SE | 22862 | ENSG00000043143 | JADE2 | chr5 | + | 134560745 | 134560957 | 134559829 | 134559990 | 134564493 | 134564610 | 195 | 107 | 205 | 248 | 133 | 0 | 0 | 0 | 0 | 1 | 87 | 171 | 142 | 80 | 184 | 0 | 0 | 0 | 0 | 5 | 298 | 149 | 1 | 1 | 1 | 1 | 1 | 1 | 0.985 | 1 | 1 | 1 | 1 | 0.948 | 0.007 |
| SE | 22863 | ENSG00000043143 | JADE2 | chr5 | + | 134562199 | 134562367 | 134559829 | 134559990 | 134564493 | 134564610 | 167 | 55 | 165 | 151 | 77 | 0 | 0 | 0 | 0 | 1 | 56 | 125 | 83 | 39 | 75 | 0 | 0 | 0 | 0 | 5 | 298 | 149 | 1 | 1 | 1 | 1 | 1 | 1 | 0.975 | 1 | 1 | 1 | 1 | 0.882 | 0.019 |
| SE | 22865 | ENSG00000043143 | JADE2 | chr5 | + | 134564493 | 134564610 | 134559829 | 134559990 | 134573644 | 134573762 | 1 | 4 | 0 | 1 | 1 | 2 | 0 | 0 | 0 | 0 | 0 | 0 | 2 | 0 | 5 | 0 | 0 | 0 | 0 | 0 | 266 | 149 | 1 | 1 | 0.219 | 1 | NA | NA | 1 | NA | NA | 1 | NA | 1 | -0.26 |
| SE | 22866 | ENSG00000043143 | JADE2 | chr5 | + | 134564493 | 134564610 | 134562199 | 134562367 | 134566115 | 134566580 | 265 | 93 | 254 | 233 | 138 | 1 | 0 | 0 | 0 | 0 | 103 | 226 | 140 | 68 | 163 | 0 | 0 | 2 | 0 | 0 | 266 | 149 | 1 | 1 | 0.993 | 1 | 1 | 1 | 1 | 1 | 1 | 0.975 | 1 | 1 | 0.004 |
| SE | 22867 | ENSG00000043143 | JADE2 | chr5 | + | 134566115 | 134566580 | 134564171 | 134564610 | 134573644 | 134573762 | 293 | 131 | 218 | 260 | 188 | 1 | 4 | 0 | 2 | 0 | 130 | 254 | 160 | 92 | 231 | 0 | 0 | 3 | 0 | 0 | 298 | 149 | 1 | 1 | 0.993 | 0.942 | 1 | 0.985 | 1 | 1 | 1 | 0.964 | 1 | 1 | -0.009 |
| SE | 22869 | ENSG00000043143 | JADE2 | chr5 | + | 134573644 | 134573762 | 134566115 | 134566580 | 134578493 | 134579814 | 219 | 125 | 182 | 236 | 248 | 3 | 0 | 0 | 2 | 0 | 127 | 237 | 144 | 111 | 214 | 0 | 0 | 0 | 2 | 0 | 267 | 149 | 1 | 1 | 0.976 | 1 | 1 | 0.985 | 1 | 1 | 1 | 1 | 0.969 | 1 | -0.002 |
| SE | 22871 | ENSG00000043143 | JADE2 | chr5 | + | 134576767 | 134576896 | 134566115 | 134566580 | 134578493 | 134578818 | 80 | 46 | 69 | 78 | 163 | 3 | 0 | 0 | 2 | 0 | 47 | 101 | 35 | 44 | 87 | 0 | 0 | 0 | 2 | 0 | 278 | 149 | 1 | 1 | 0.935 | 1 | 1 | 0.954 | 1 | 1 | 1 | 1 | 0.922 | 1 | -0.007 |
| SE | 22878 | ENSG00000154153 | RETREG1 | chr5 | - | 16481008 | 16481073 | 16478849 | 16478987 | 16483345 | 16483472 | 29 | 14 | 57 | 30 | 90 | 0 | 0 | 0 | 0 | 4 | 0 | 14 | 2 | 5 | 0 | 0 | 0 | 0 | 0 | 0 | 214 | 149 | 1 | 1 | 1 | 1 | 1 | 1 | 0.94 | NA | 1 | 1 | 1 | NA | -0.012 |
| SE | 22879 | ENSG00000154153 | RETREG1 | chr5 | - | 16481008 | 16481093 | 16478033 | 16478987 | 16483345 | 16483472 | 54 | 37 | 88 | 55 | 184 | 0 | 0 | 0 | 0 | 4 | 0 | 27 | 6 | 5 | 0 | 0 | 0 | 0 | 0 | 0 | 234 | 149 | 1 | 1 | 1 | 1 | 1 | 1 | 0.967 | NA | 1 | 1 | 1 | NA | -0.007 |
| SE | 22881 | ENSG00000154153 | RETREG1 | chr5 | - | 16481008 | 16481093 | 16478033 | 16478828 | 16565762 | 16565793 | 31 | 15 | 60 | 35 | 100 | 0 | 1 | 1 | 3 | 0 | 0 | 14 | 2 | 5 | 0 | 0 | 0 | 0 | 0 | 0 | 234 | 149 | 1 | 1 | 1 | 0.905 | 0.974 | 0.881 | 1 | NA | 1 | 1 | 1 | NA | -0.048 |
| SE | 22882 | ENSG00000154153 | RETREG1 | chr5 | - | 16481008 | 16481093 | 16478849 | 16478987 | 16565762 | 16565793 | 41 | 29 | 91 | 47 | 133 | 0 | 1 | 1 | 3 | 0 | 0 | 14 | 6 | 6 | 0 | 0 | 0 | 0 | 0 | 0 | 298 | 149 | 1 | 1 | 1 | 0.935 | 0.978 | 0.887 | 1 | NA | 1 | 1 | 1 | NA | -0.04 |
| SE | 22884 | ENSG00000154153 | RETREG1 | chr5 | - | 16483345 | 16483472 | 16478033 | 16478987 | 16565762 | 16565793 | 8 | 9 | 30 | 11 | 34 | 0 | 1 | 1 | 3 | 0 | 0 | 0 | 4 | 1 | 0 | 0 | 0 | 0 | 0 | 0 | 276 | 149 | 1 | 1 | 1 | 0.829 | 0.942 | 0.664 | 1 | NA | NA | 1 | 1 | NA | -0.113 |
| SE | 22886 | ENSG00000154153 | RETREG1 | chr5 | - | 16483345 | 16483472 | 16481008 | 16481093 | 16565762 | 16565793 | 31 | 31 | 57 | 31 | 115 | 0 | 0 | 0 | 0 | 3 | 0 | 13 | 8 | 1 | 0 | 0 | 0 | 0 | 0 | 0 | 276 | 149 | 1 | 1 | 1 | 1 | 1 | 1 | 0.954 | NA | 1 | 1 | 1 | NA | -0.009 |
| SE | 22890 | ENSG00000140853 | NLRC5 | chr16 | + | 57017073 | 57017188 | 56989484 | 56989617 | 57020700 | 57020769 | 145 | 56 | 163 | 161 | 83 | 1 | 0 | 0 | 0 | 0 | 227 | 301 | 160 | 192 | 424 | 0 | 0 | 0 | 0 | 0 | 264 | 149 | 1 | 1 | 0.988 | 1 | 1 | 1 | 1 | 1 | 1 | 1 | 1 | 1 | -0.002 |
| SE | 22891 | ENSG00000140853 | NLRC5 | chr16 | + | 57022255 | 57022315 | 57020700 | 57021007 | 57023784 | 57023853 | 399 | 190 | 364 | 307 | 223 | 0 | 0 | 0 | 0 | 0 | 615 | 573 | 393 | 562 | 947 | 3 | 0 | 0 | 0 | 0 | 209 | 149 | 1 | 1 | 1 | 1 | 1 | 1 | 1 | 0.993 | 1 | 1 | 1 | 1 | 0.001 |
| SE | 22894 | ENSG00000140853 | NLRC5 | chr16 | + | 57023784 | 57023853 | 57022255 | 57022315 | 57025367 | 57027018 | 384 | 187 | 355 | 323 | 234 | 0 | 0 | 0 | 0 | 2 | 601 | 573 | 359 | 482 | 943 | 0 | 0 | 0 | 0 | 7 | 218 | 149 | 1 | 1 | 1 | 1 | 1 | 1 | 0.988 | 1 | 1 | 1 | 1 | 0.989 | 0 |
| SE | 22897 | ENSG00000140853 | NLRC5 | chr16 | + | 57028301 | 57028385 | 57028071 | 57028155 | 57029772 | 57029856 | 196 | 115 | 213 | 209 | 190 | 0 | 0 | 0 | 0 | 0 | 338 | 271 | 171 | 240 | 436 | 0 | 0 | 0 | 3 | 0 | 233 | 149 | 1 | 1 | 1 | 1 | 1 | 1 | 1 | 1 | 1 | 1 | 0.981 | 1 | 0.004 |
| SE | 22899 | ENSG00000140853 | NLRC5 | chr16 | + | 57033603 | 57033669 | 57031403 | 57031463 | 57034167 | 57034251 | 140 | 67 | 133 | 153 | 87 | 0 | 0 | 1 | 0 | 0 | 421 | 234 | 106 | 166 | 337 | 0 | 0 | 2 | 6 | 0 | 215 | 149 | 1 | 1 | 1 | 1 | 0.989 | 1 | 1 | 1 | 1 | 0.973 | 0.95 | 1 | 0.013 |
| SE | 22900 | ENSG00000140853 | NLRC5 | chr16 | + | 57034511 | 57034946 | 57034167 | 57034251 | 57036099 | 57036183 | 1 | 0 | 0 | 0 | 2 | 80 | 26 | 79 | 74 | 67 | 0 | 2 | 0 | 0 | 0 | 184 | 104 | 59 | 110 | 146 | 298 | 149 | 1 | 1 | 0.006 | 0 | 0 | 0 | 0.015 | 0 | 0.01 | 0 | 0 | 0 | 0.002 |
| SE | 22905 | ENSG00000140853 | NLRC5 | chr16 | + | 57054750 | 57054840 | 57051537 | 57051621 | 57055031 | 57055036 | 197 | 110 | 250 | 200 | 114 | 1 | 1 | 0 | 1 | 5 | 361 | 313 | 206 | 334 | 494 | 8 | 0 | 0 | 0 | 0 | 239 | 149 | 1 | 1 | 0.992 | 0.986 | 1 | 0.992 | 0.934 | 0.966 | 1 | 1 | 1 | 1 | -0.012 |
| SE | 22908 | ENSG00000140853 | NLRC5 | chr16 | + | 57058064 | 57058148 | 57055432 | 57055519 | 57058971 | 57059061 | 69 | 15 | 53 | 58 | 49 | 1 | 0 | 0 | 0 | 0 | 67 | 53 | 44 | 66 | 196 | 0 | 0 | 0 | 0 | 0 | 233 | 149 | 1 | 1 | 0.978 | 1 | 1 | 1 | 1 | 1 | 1 | 1 | 1 | 1 | -0.004 |
| SE | 22911 | ENSG00000140853 | NLRC5 | chr16 | + | 57066533 | 57066614 | 57065211 | 57065298 | 57067386 | 57067470 | 110 | 65 | 114 | 70 | 43 | 0 | 0 | 0 | 0 | 0 | 137 | 146 | 90 | 135 | 259 | 0 | 0 | 1 | 0 | 0 | 230 | 149 | 1 | 1 | 1 | 1 | 1 | 1 | 1 | 1 | 1 | 0.983 | 1 | 1 | 0.003 |
| SE | 22913 | ENSG00000140853 | NLRC5 | chr16 | + | 57069835 | 57069919 | 57067735 | 57067828 | 57070534 | 57070618 | 247 | 96 | 211 | 189 | 101 | 0 | 3 | 0 | 0 | 0 | 238 | 288 | 176 | 353 | 387 | 0 | 6 | 0 | 0 | 5 | 233 | 149 | 1 | 1 | 1 | 0.953 | 1 | 1 | 1 | 1 | 0.968 | 1 | 1 | 0.98 | 0.001 |
| SE | 22915 | ENSG00000140853 | NLRC5 | chr16 | + | 57074599 | 57074683 | 57070534 | 57070618 | 57076818 | 57076902 | 310 | 131 | 271 | 220 | 197 | 0 | 0 | 0 | 0 | 0 | 153 | 339 | 134 | 296 | 303 | 0 | 3 | 0 | 0 | 0 | 233 | 149 | 1 | 1 | 1 | 1 | 1 | 1 | 1 | 1 | 0.986 | 1 | 1 | 1 | 0.003 |
| SE | 22918 | ENSG00000140853 | NLRC5 | chr16 | + | 57077295 | 57077379 | 57076818 | 57076902 | 57077718 | 57077802 | 283 | 107 | 294 | 215 | 218 | 0 | 0 | 0 | 0 | 0 | 263 | 330 | 113 | 212 | 419 | 0 | 6 | 0 | 0 | 0 | 233 | 149 | 1 | 1 | 1 | 1 | 1 | 1 | 1 | 1 | 0.972 | 1 | 1 | 1 | 0.006 |
| SE | 22919 | ENSG00000140853 | NLRC5 | chr16 | + | 57077295 | 57077379 | 57076818 | 57076902 | 57077942 | 57078020 | 192 | 73 | 168 | 146 | 172 | 0 | 0 | 0 | 0 | 0 | 148 | 146 | 78 | 131 | 182 | 0 | 0 | 0 | 6 | 0 | 233 | 149 | 1 | 1 | 1 | 1 | 1 | 1 | 1 | 1 | 1 | 1 | 0.933 | 1 | 0.013 |
| SE | 22920 | ENSG00000140853 | NLRC5 | chr16 | + | 57077718 | 57077802 | 57076818 | 57076902 | 57077942 | 57078020 | 125 | 34 | 114 | 84 | 74 | 0 | 0 | 0 | 0 | 0 | 171 | 87 | 59 | 98 | 263 | 0 | 0 | 0 | 6 | 0 | 233 | 149 | 1 | 1 | 1 | 1 | 1 | 1 | 1 | 1 | 1 | 1 | 0.913 | 1 | 0.017 |
| SE | 22921 | ENSG00000140853 | NLRC5 | chr16 | + | 57077718 | 57077802 | 57077295 | 57077379 | 57077942 | 57078020 | 220 | 68 | 242 | 158 | 122 | 0 | 0 | 2 | 3 | 2 | 286 | 269 | 97 | 179 | 507 | 0 | 0 | 0 | 0 | 0 | 233 | 149 | 1 | 1 | 1 | 1 | 0.987 | 0.971 | 0.975 | 1 | 1 | 1 | 1 | 1 | -0.013 |
| SE | 22922 | ENSG00000140853 | NLRC5 | chr16 | + | 57077942 | 57078020 | 57077718 | 57077802 | 57079049 | 57079133 | 179 | 47 | 164 | 127 | 107 | 0 | 0 | 0 | 0 | 0 | 274 | 159 | 79 | 141 | 397 | 0 | 0 | 0 | 2 | 0 | 227 | 149 | 1 | 1 | 1 | 1 | 1 | 1 | 1 | 1 | 1 | 1 | 0.979 | 1 | 0.004 |
| SE | 22923 | ENSG00000140853 | NLRC5 | chr16 | + | 57081097 | 57081181 | 57079545 | 57079629 | 57081526 | 57081610 | 171 | 59 | 145 | 198 | 78 | 0 | 0 | 0 | 1 | 2 | 181 | 195 | 105 | 147 | 242 | 0 | 2 | 0 | 3 | 0 | 233 | 149 | 1 | 1 | 1 | 1 | 1 | 0.992 | 0.961 | 1 | 0.984 | 1 | 0.969 | 1 | 0 |
| SE | 22925 | ENSG00000145817 | YIPF5 | chr5 | - | 144162217 | 144162399 | 144158161 | 144160559 | 144165431 | 144165604 | 65 | 41 | 63 | 64 | 75 | 0 | 0 | 0 | 1 | 0 | 23 | 67 | 51 | 26 | 24 | 0 | 0 | 0 | 0 | 0 | 298 | 149 | 1 | 1 | 1 | 1 | 1 | 0.97 | 1 | 1 | 1 | 1 | 1 | 1 | -0.006 |
| SE | 22926 | ENSG00000145817 | YIPF5 | chr5 | - | 144164110 | 144164256 | 144162217 | 144162399 | 144165431 | 144165600 | 155 | 78 | 137 | 116 | 80 | 0 | 2 | 2 | 0 | 0 | 24 | 88 | 72 | 63 | 34 | 0 | 0 | 0 | 1 | 0 | 295 | 149 | 1 | 1 | 1 | 0.952 | 0.972 | 1 | 1 | 1 | 1 | 1 | 0.97 | 1 | -0.009 |
| SE | 22928 | ENSG00000145817 | YIPF5 | chr5 | - | 144169845 | 144169965 | 144165431 | 144165604 | 144170534 | 144170640 | 253 | 130 | 227 | 179 | 191 | 0 | 6 | 0 | 2 | 0 | 99 | 144 | 124 | 145 | 143 | 0 | 0 | 0 | 0 | 0 | 269 | 149 | 1 | 1 | 1 | 0.923 | 1 | 0.98 | 1 | 1 | 1 | 1 | 1 | 1 | -0.019 |
| SE | 22932 | ENSG00000105698 | USF2 | chr19 | + | 35278697 | 35278792 | 35271000 | 35271141 | 35278945 | 35279074 | 795 | 668 | 631 | 897 | 877 | 0 | 0 | 1 | 0 | 0 | 1303 | 1004 | 573 | 886 | 2106 | 0 | 0 | 0 | 0 | 0 | 244 | 149 | 1 | 1 | 1 | 1 | 0.997 | 1 | 1 | 1 | 1 | 1 | 1 | 1 | -0.001 |
| SE | 22934 | ENSG00000145723 | GIN1 | chr5 | - | 103097313 | 103097490 | 103096540 | 103096826 | 103097589 | 103097781 | 168 | 76 | 155 | 136 | 340 | 1 | 0 | 0 | 0 | 3 | 85 | 137 | 101 | 26 | 9 | 0 | 0 | 0 | 0 | 0 | 298 | 149 | 1 | 1 | 0.988 | 1 | 1 | 1 | 0.983 | 1 | 1 | 1 | 1 | 1 | -0.006 |
| SE | 22935 | ENSG00000145723 | GIN1 | chr5 | - | 103106715 | 103106909 | 103104540 | 103104846 | 103108568 | 103108714 | 65 | 49 | 32 | 37 | 53 | 3 | 5 | 0 | 3 | 1 | 0 | 25 | 8 | 0 | 0 | 0 | 0 | 0 | 0 | 0 | 298 | 149 | 1 | 1 | 0.915 | 0.831 | 1 | 0.86 | 0.964 | NA | 1 | 1 | NA | NA | -0.086 |
| SE | 22936 | ENSG00000133112 | TPT1 | chr13 | - | 45337466 | 45337553 | 45336898 | 45337388 | 45338659 | 45338776 | 20 | 20 | 39 | 21 | 31 | 41287 | 31083 | 49049 | 43274 | 96359 | 25 | 35 | 29 | 24 | 34 | 37987 | 20785 | 21520 | 23016 | 39942 | 236 | 149 | 1 | 1 | 0 | 0 | 0.001 | 0 | 0 | 0 | 0.001 | 0.001 | 0.001 | 0.001 | -0.001 |
| SE | 22937 | ENSG00000133112 | TPT1 | chr13 | - | 45338659 | 45338776 | 45333470 | 45337388 | 45339496 | 45339533 | 76651 | 57770 | 89929 | 78544 | 190513 | 0 | 0 | 2 | 7 | 0 | 67483 | 37728 | 38914 | 41325 | 72469 | 0 | 0 | 1 | 1 | 0 | 266 | 149 | 1 | 1 | 1 | 1 | 1 | 1 | 1 | 1 | 1 | 1 | 1 | 1 | 0 |
| SE | 22938 | ENSG00000133112 | TPT1 | chr13 | - | 45338659 | 45338776 | 45333470 | 45337388 | 45339993 | 45340258 | 36466 | 28131 | 44128 | 38934 | 90014 | 3 | 0 | 0 | 0 | 0 | 34777 | 18865 | 19425 | 20852 | 35924 | 1 | 8 | 0 | 18 | 0 | 266 | 149 | 1 | 1 | 1 | 1 | 1 | 1 | 1 | 1 | 0.999 | 1 | 0.998 | 1 | 0.001 |
| SE | 22939 | ENSG00000133112 | TPT1 | chr13 | - | 45339496 | 45339602 | 45333470 | 45337388 | 45339993 | 45340184 | 26524 | 16738 | 28640 | 25605 | 41531 | 3 | 0 | 0 | 0 | 0 | 24908 | 15245 | 15945 | 16485 | 29353 | 1 | 8 | 0 | 18 | 0 | 255 | 149 | 1 | 1 | 1 | 1 | 1 | 1 | 1 | 1 | 0.999 | 1 | 0.998 | 1 | 0.001 |
| SE | 22940 | ENSG00000133112 | TPT1 | chr13 | - | 45339496 | 45339602 | 45338184 | 45338776 | 45339993 | 45340184 | 66753 | 46418 | 74478 | 65272 | 142097 | 37 | 37 | 38 | 65 | 80 | 57810 | 34271 | 35475 | 37171 | 66058 | 193 | 144 | 34 | 196 | 131 | 255 | 149 | 1 | 1 | 0.999 | 0.999 | 0.999 | 0.998 | 0.999 | 0.994 | 0.993 | 0.998 | 0.991 | 0.997 | 0.004 |
| SE | 22941 | ENSG00000133112 | TPT1 | chr13 | - | 45339993 | 45340184 | 45333470 | 45337388 | 45340711 | 45340785 | 11209 | 6678 | 11507 | 10905 | 17411 | 0 | 0 | 1 | 0 | 0 | 12226 | 7460 | 7618 | 8553 | 15462 | 0 | 0 | 0 | 0 | 0 | 298 | 149 | 1 | 1 | 1 | 1 | 1 | 1 | 1 | 1 | 1 | 1 | 1 | 1 | 0 |
| SE | 22944 | ENSG00000170234 | PWWP2A | chr5 | - | 160066547 | 160066612 | 160063541 | 160063673 | 160089505 | 160089610 | 0 | 1 | 0 | 0 | 0 | 0 | 0 | 0 | 1 | 2 | 0 | 0 | 0 | 0 | 11 | 0 | 3 | 0 | 0 | 0 | 214 | 149 | 1 | 1 | NA | 1 | NA | 0 | 0 | NA | 0 | NA | NA | 1 | -0.667 |
| SE | 22945 | ENSG00000170234 | PWWP2A | chr5 | - | 160066547 | 160066612 | 160063541 | 160063673 | 160093100 | 160093180 | 0 | 0 | 0 | 1 | 0 | 0 | 0 | 0 | 4 | 1 | 0 | 0 | 0 | 0 | 11 | 0 | 3 | 0 | 0 | 0 | 214 | 149 | 1 | 1 | NA | NA | NA | 0.148 | 0 | NA | 0 | NA | NA | 1 | -0.426 |
| SE | 22949 | ENSG00000170234 | PWWP2A | chr5 | - | 160089505 | 160089610 | 160063541 | 160063673 | 160093100 | 160093180 | 12 | 7 | 11 | 9 | 17 | 0 | 0 | 0 | 4 | 1 | 21 | 3 | 1 | 2 | 15 | 0 | 3 | 0 | 0 | 0 | 254 | 149 | 1 | 1 | 1 | 1 | 1 | 0.569 | 0.909 | 1 | 0.37 | 1 | 1 | 1 | 0.022 |
| SE | 22950 | ENSG00000170234 | PWWP2A | chr5 | - | 160089505 | 160089610 | 160066547 | 160066612 | 160093100 | 160093180 | 12 | 8 | 11 | 8 | 15 | 0 | 0 | 2 | 1 | 0 | 21 | 3 | 1 | 2 | 15 | 0 | 0 | 0 | 0 | 0 | 254 | 149 | 1 | 1 | 1 | 1 | 0.763 | 0.824 | 1 | 1 | 1 | 1 | 1 | 1 | -0.083 |
| SE | 22954 | ENSG00000263798 | NA | chr17 | + | 47978828 | 47978917 | 47977831 | 47977910 | 47980592 | 47980798 | 295 | 218 | 284 | 339 | 176 | 0 | 0 | 0 | 0 | 0 | 574 | 390 | 271 | 536 | 615 | 0 | 0 | 1 | 0 | 0 | 238 | 149 | 1 | 1 | 1 | 1 | 1 | 1 | 1 | 1 | 1 | 0.994 | 1 | 1 | 0.001 |
| SE | 22956 | ENSG00000163468 | CCT3 | chr1 | - | 156324971 | 156325086 | 156320838 | 156321025 | 156333546 | 156333643 | 322 | 236 | 359 | 292 | 342 | 2 | 1 | 0 | 0 | 0 | 370 | 211 | 158 | 183 | 326 | 0 | 0 | 0 | 0 | 0 | 264 | 149 | 1 | 1 | 0.989 | 0.993 | 1 | 1 | 1 | 1 | 1 | 1 | 1 | 1 | -0.004 |
| SE | 22957 | ENSG00000163468 | CCT3 | chr1 | - | 156324971 | 156325089 | 156320838 | 156321025 | 156333546 | 156333622 | 700 | 453 | 703 | 661 | 712 | 2 | 1 | 0 | 0 | 0 | 715 | 515 | 326 | 387 | 762 | 0 | 0 | 0 | 0 | 0 | 267 | 149 | 1 | 1 | 0.995 | 0.996 | 1 | 1 | 1 | 1 | 1 | 1 | 1 | 1 | -0.002 |
| SE | 22958 | ENSG00000163468 | CCT3 | chr1 | - | 156333215 | 156333284 | 156324979 | 156325089 | 156333546 | 156333643 | 0 | 3 | 0 | 0 | 0 | 375 | 217 | 340 | 364 | 359 | 0 | 0 | 0 | 0 | 0 | 345 | 304 | 165 | 203 | 441 | 218 | 149 | 1 | 1 | 0 | 0.009 | 0 | 0 | 0 | 0 | 0 | 0 | 0 | 0 | 0.002 |
| SE | 22959 | ENSG00000163468 | CCT3 | chr1 | - | 156333546 | 156333643 | 156324971 | 156325089 | 156335826 | 156335888 | 281 | 168 | 273 | 274 | 291 | 3 | 0 | 0 | 0 | 0 | 250 | 237 | 130 | 167 | 308 | 0 | 0 | 0 | 0 | 0 | 246 | 149 | 1 | 1 | 0.983 | 1 | 1 | 1 | 1 | 1 | 1 | 1 | 1 | 1 | -0.003 |
| SE | 22960 | ENSG00000163468 | CCT3 | chr1 | - | 156334712 | 156334775 | 156333546 | 156333643 | 156334867 | 156334918 | 504 | 236 | 357 | 411 | 322 | 0 | 0 | 1 | 0 | 0 | 356 | 313 | 183 | 245 | 410 | 0 | 0 | 0 | 0 | 0 | 212 | 149 | 1 | 1 | 1 | 1 | 0.996 | 1 | 1 | 1 | 1 | 1 | 1 | 1 | -0.001 |
| SE | 22961 | ENSG00000163468 | CCT3 | chr1 | - | 156334867 | 156334918 | 156333546 | 156333643 | 156335826 | 156335888 | 161 | 82 | 135 | 123 | 44 | 1 | 2 | 0 | 1 | 0 | 78 | 62 | 41 | 62 | 75 | 0 | 0 | 0 | 0 | 0 | 200 | 149 | 1 | 1 | 0.992 | 0.968 | 1 | 0.989 | 1 | 1 | 1 | 1 | 1 | 1 | -0.01 |
| SE | 22962 | ENSG00000163468 | CCT3 | chr1 | - | 156335826 | 156335888 | 156334867 | 156334918 | 156338153 | 156338214 | 392 | 194 | 311 | 296 | 192 | 0 | 0 | 0 | 1 | 0 | 197 | 221 | 129 | 189 | 185 | 0 | 0 | 0 | 0 | 0 | 211 | 149 | 1 | 1 | 1 | 1 | 1 | 0.995 | 1 | 1 | 1 | 1 | 1 | 1 | -0.001 |
| SE | 22964 | ENSG00000163468 | CCT3 | chr1 | - | 156337078 | 156337240 | 156335826 | 156335888 | 156338153 | 156338247 | 1 | 0 | 0 | 0 | 0 | 246 | 115 | 196 | 190 | 64 | 0 | 0 | 0 | 0 | 0 | 132 | 98 | 81 | 110 | 118 | 298 | 149 | 1 | 1 | 0.002 | 0 | 0 | 0 | 0 | 0 | 0 | 0 | 0 | 0 | 0 |
| SE | 22969 | ENSG00000198612 | COPS8 | chr2 | + | 237089861 | 237089994 | 237088604 | 237088653 | 237093662 | 237094197 | 69 | 26 | 56 | 50 | 60 | 1 | 0 | 0 | 0 | 0 | 26 | 22 | 26 | 21 | 22 | 0 | 0 | 0 | 0 | 0 | 282 | 149 | 1 | 1 | 0.973 | 1 | 1 | 1 | 1 | 1 | 1 | 1 | 1 | 1 | -0.005 |
| SE | 22973 | ENSG00000198612 | COPS8 | chr2 | + | 237095821 | 237095884 | 237094089 | 237094197 | 237096821 | 237096869 | 147 | 77 | 116 | 121 | 170 | 0 | 0 | 0 | 0 | 0 | 133 | 42 | 60 | 73 | 65 | 0 | 0 | 1 | 0 | 0 | 212 | 149 | 1 | 1 | 1 | 1 | 1 | 1 | 1 | 1 | 1 | 0.977 | 1 | 1 | 0.005 |
| SE | 22974 | ENSG00000116459 | ATP5PB | chr1 | + | 111449836 | 111449873 | 111449194 | 111449581 | 111454210 | 111454356 | 256 | 121 | 196 | 205 | 254 | 0 | 0 | 0 | 1 | 0 | 71 | 142 | 164 | 81 | 57 | 0 | 0 | 0 | 0 | 0 | 186 | 149 | 1 | 1 | 1 | 1 | 1 | 0.994 | 1 | 1 | 1 | 1 | 1 | 1 | -0.001 |
| SE | 22976 | ENSG00000116459 | ATP5PB | chr1 | + | 111449836 | 111449873 | 111449194 | 111449581 | 111459456 | 111459636 | 6 | 5 | 3 | 4 | 1 | 0 | 0 | 0 | 0 | 0 | 0 | 2 | 0 | 0 | 0 | 0 | 0 | 1 | 0 | 0 | 186 | 149 | 1 | 1 | 1 | 1 | 1 | 1 | 1 | NA | 1 | 0 | NA | NA | 0.5 |
| SE | 22978 | ENSG00000116459 | ATP5PB | chr1 | + | 111454210 | 111454356 | 111449194 | 111449581 | 111459456 | 111459636 | 0 | 0 | 0 | 1 | 1 | 0 | 0 | 0 | 0 | 0 | 0 | 0 | 0 | 0 | 0 | 0 | 0 | 1 | 0 | 0 | 295 | 149 | 1 | 1 | NA | NA | NA | 1 | 1 | NA | NA | 0 | NA | NA | 1 |
| SE | 22982 | ENSG00000116459 | ATP5PB | chr1 | + | 111454210 | 111454356 | 111449681 | 111449873 | 111459456 | 111459636 | 237 | 111 | 185 | 183 | 219 | 2 | 0 | 0 | 0 | 0 | 71 | 134 | 158 | 74 | 56 | 0 | 0 | 0 | 0 | 0 | 295 | 149 | 1 | 1 | 0.984 | 1 | 1 | 1 | 1 | 1 | 1 | 1 | 1 | 1 | -0.003 |
| SE | 22983 | ENSG00000116459 | ATP5PB | chr1 | + | 111456085 | 111456249 | 111449483 | 111449581 | 111459456 | 111459636 | 0 | 0 | 0 | 0 | 3 | 0 | 0 | 0 | 0 | 0 | 0 | 5 | 1 | 0 | 0 | 0 | 0 | 1 | 0 | 0 | 298 | 149 | 1 | 1 | NA | NA | NA | NA | 1 | NA | 1 | 0.333 | NA | NA | 0.334 |
| SE | 22985 | ENSG00000116459 | ATP5PB | chr1 | + | 111456085 | 111456249 | 111449836 | 111449873 | 111459456 | 111459636 | 11 | 7 | 12 | 14 | 7 | 2 | 0 | 0 | 0 | 0 | 11 | 25 | 31 | 7 | 10 | 0 | 0 | 0 | 0 | 0 | 298 | 149 | 1 | 1 | 0.733 | 1 | 1 | 1 | 1 | 1 | 1 | 1 | 1 | 1 | -0.053 |
| SE | 22986 | ENSG00000116459 | ATP5PB | chr1 | + | 111456085 | 111456249 | 111454210 | 111454356 | 111456629 | 111456748 | 1817 | 1124 | 1435 | 1533 | 2470 | 1 | 0 | 0 | 0 | 0 | 738 | 1079 | 1049 | 953 | 394 | 0 | 0 | 2 | 0 | 0 | 298 | 149 | 1 | 1 | 0.999 | 1 | 1 | 1 | 1 | 1 | 1 | 0.996 | 1 | 1 | 0.001 |
| SE | 22987 | ENSG00000116459 | ATP5PB | chr1 | + | 111456085 | 111456249 | 111454210 | 111454356 | 111459456 | 111459636 | 814 | 487 | 635 | 675 | 932 | 0 | 0 | 0 | 0 | 1 | 299 | 476 | 466 | 418 | 145 | 0 | 0 | 0 | 0 | 0 | 298 | 149 | 1 | 1 | 1 | 1 | 1 | 1 | 0.998 | 1 | 1 | 1 | 1 | 1 | 0 |
| SE | 22989 | ENSG00000116459 | ATP5PB | chr1 | + | 111456226 | 111456249 | 111449836 | 111449873 | 111459456 | 111459636 | 0 | 0 | 0 | 0 | 0 | 2 | 0 | 0 | 0 | 0 | 0 | 5 | 0 | 0 | 0 | 0 | 0 | 0 | 0 | 0 | 172 | 149 | 1 | 1 | 0 | NA | NA | NA | NA | NA | 1 | NA | NA | NA | -1 |
| SE | 22990 | ENSG00000116459 | ATP5PB | chr1 | + | 111456226 | 111456249 | 111454210 | 111454356 | 111456629 | 111456755 | 65 | 42 | 65 | 49 | 119 | 1 | 0 | 0 | 0 | 0 | 13 | 46 | 34 | 45 | 30 | 0 | 0 | 2 | 0 | 0 | 172 | 149 | 1 | 1 | 0.983 | 1 | 1 | 1 | 1 | 1 | 1 | 0.936 | 1 | 1 | 0.009 |
| SE | 22991 | ENSG00000116459 | ATP5PB | chr1 | + | 111456226 | 111456249 | 111454210 | 111454356 | 111459456 | 111459636 | 0 | 0 | 0 | 0 | 0 | 0 | 0 | 0 | 0 | 1 | 0 | 5 | 0 | 0 | 0 | 0 | 0 | 0 | 0 | 0 | 172 | 149 | 1 | 1 | NA | NA | NA | NA | 0 | NA | 1 | NA | NA | NA | -1 |
| SE | 22992 | ENSG00000116459 | ATP5PB | chr1 | + | 111456629 | 111456755 | 111449836 | 111449873 | 111459456 | 111459636 | 935 | 629 | 812 | 858 | 1286 | 2 | 0 | 0 | 0 | 0 | 395 | 585 | 594 | 500 | 188 | 0 | 0 | 0 | 0 | 0 | 275 | 149 | 1 | 1 | 0.996 | 1 | 1 | 1 | 1 | 1 | 1 | 1 | 1 | 1 | -0.001 |
| SE | 22993 | ENSG00000116459 | ATP5PB | chr1 | + | 111456629 | 111456755 | 111449836 | 111449873 | 111460916 | 111461026 | 1 | 1 | 0 | 2 | 2 | 0 | 0 | 0 | 0 | 0 | 0 | 3 | 13 | 2 | 5 | 0 | 4 | 0 | 0 | 0 | 275 | 149 | 1 | 1 | 1 | 1 | NA | 1 | 1 | NA | 0.289 | 1 | 1 | 1 | 0.178 |
| SE | 22994 | ENSG00000116459 | ATP5PB | chr1 | + | 111456629 | 111456755 | 111454210 | 111454356 | 111459456 | 111459636 | 935 | 628 | 812 | 858 | 1286 | 0 | 0 | 0 | 0 | 1 | 395 | 582 | 584 | 498 | 183 | 0 | 0 | 0 | 0 | 0 | 275 | 149 | 1 | 1 | 1 | 1 | 1 | 1 | 0.999 | 1 | 1 | 1 | 1 | 1 | 0 |
| SE | 22995 | ENSG00000116459 | ATP5PB | chr1 | + | 111456629 | 111456755 | 111456085 | 111456249 | 111459456 | 111459636 | 1938 | 1266 | 1618 | 1719 | 2823 | 0 | 0 | 0 | 0 | 3 | 834 | 1185 | 1168 | 1034 | 433 | 0 | 5 | 1 | 0 | 0 | 275 | 149 | 1 | 1 | 1 | 1 | 1 | 1 | 0.998 | 1 | 0.992 | 0.998 | 1 | 1 | 0.002 |
| SE | 22996 | ENSG00000116459 | ATP5PB | chr1 | + | 111459456 | 111459636 | 111449836 | 111449873 | 111460916 | 111461026 | 328 | 196 | 236 | 270 | 552 | 0 | 0 | 0 | 0 | 0 | 162 | 191 | 169 | 150 | 64 | 0 | 4 | 0 | 0 | 0 | 298 | 149 | 1 | 1 | 1 | 1 | 1 | 1 | 1 | 1 | 0.96 | 1 | 1 | 1 | 0.008 |
| SE | 22997 | ENSG00000116459 | ATP5PB | chr1 | + | 111459456 | 111459636 | 111456629 | 111456755 | 111460916 | 111461026 | 1335 | 868 | 1112 | 1181 | 1960 | 0 | 0 | 0 | 2 | 2 | 570 | 822 | 786 | 692 | 276 | 0 | 0 | 1 | 0 | 0 | 298 | 149 | 1 | 1 | 1 | 1 | 1 | 0.997 | 0.998 | 1 | 1 | 0.997 | 1 | 1 | 0 |
| SE | 22998 | ENSG00000205464 | ATP6AP1L | chr5 | + | 82295492 | 82295544 | 82280959 | 82281011 | 82303486 | 82303527 | 6 | 6 | 0 | 4 | 0 | 0 | 0 | 1 | 0 | 0 | 9 | 0 | 1 | 2 | 5 | 0 | 6 | 0 | 2 | 0 | 201 | 149 | 1 | 1 | 1 | 1 | NA | 0.748 | NA | 1 | 0 | 1 | 0.426 | 1 | 0.231 |
| SE | 22999 | ENSG00000205464 | ATP6AP1L | chr5 | + | 82304933 | 82305481 | 82304614 | 82304850 | 82310156 | 82310207 | 3 | 3 | 2 | 0 | 14 | 0 | 0 | 0 | 2 | 1 | 0 | 6 | 0 | 6 | 22 | 0 | 0 | 1 | 0 | 5 | 298 | 149 | 1 | 1 | 1 | 1 | 1 | 0 | 0.875 | NA | 1 | 0 | 1 | 0.688 | 0.103 |
| SE | 23000 | ENSG00000205464 | ATP6AP1L | chr5 | + | 82312610 | 82312851 | 82308858 | 82310207 | 82317997 | 82318300 | 3 | 4 | 5 | 2 | 14 | 1 | 0 | 0 | 0 | 0 | 0 | 6 | 3 | 4 | 13 | 0 | 0 | 0 | 0 | 0 | 298 | 149 | 1 | 1 | 0.6 | 1 | 1 | 1 | 1 | NA | 1 | 1 | 1 | 1 | -0.08 |
| SE | 23007 | ENSG00000206077 | ZDHHC11B | chr5 | - | 716800 | 716865 | 710354 | 712282 | 730433 | 730468 | 2 | 3 | 14 | 13 | 14 | 0 | 0 | 1 | 2 | 0 | 0 | 0 | 0 | 3 | 9 | 0 | 0 | 0 | 0 | 0 | 214 | 149 | 1 | 1 | 1 | 1 | 0.907 | 0.819 | 1 | NA | NA | NA | 1 | 1 | -0.055 |
| SE | 23009 | ENSG00000206077 | ZDHHC11B | chr5 | - | 749251 | 750296 | 748403 | 748559 | 751132 | 751257 | 0 | 0 | 1 | 0 | 2 | 0 | 0 | 1 | 0 | 0 | 0 | 5 | 0 | 0 | 0 | 0 | 0 | 0 | 0 | 0 | 298 | 149 | 1 | 1 | NA | NA | 0.333 | NA | 1 | NA | 1 | NA | NA | NA | -0.334 |
| SE | 23010 | ENSG00000206077 | ZDHHC11B | chr5 | - | 769954 | 770117 | 768834 | 768930 | 784667 | 784729 | 0 | 0 | 0 | 0 | 0 | 1 | 0 | 0 | 0 | 0 | 0 | 0 | 3 | 0 | 0 | 0 | 0 | 0 | 0 | 0 | 298 | 149 | 1 | 1 | 0 | NA | NA | NA | NA | NA | NA | 1 | NA | NA | -1 |
| SE | 23012 | ENSG00000106123 | EPHB6 | chr7 | + | 142861051 | 142861186 | 142855060 | 142855385 | 142862757 | 142862788 | 14 | 14 | 35 | 27 | 27 | 0 | 0 | 0 | 0 | 0 | 67 | 20 | 2 | 31 | 88 | 0 | 0 | 0 | 0 | 3 | 284 | 149 | 1 | 1 | 1 | 1 | 1 | 1 | 1 | 1 | 1 | 1 | 1 | 0.939 | 0.012 |
| SE | 23015 | ENSG00000106123 | EPHB6 | chr7 | + | 142863630 | 142863695 | 142862755 | 142863327 | 142863965 | 142864262 | 25 | 20 | 34 | 19 | 34 | 1 | 0 | 1 | 1 | 1 | 26 | 19 | 11 | 30 | 56 | 0 | 2 | 0 | 2 | 5 | 214 | 149 | 1 | 1 | 0.946 | 1 | 0.959 | 0.93 | 0.959 | 1 | 0.869 | 1 | 0.913 | 0.886 | 0.025 |
| SE | 23016 | ENSG00000106123 | EPHB6 | chr7 | + | 142865959 | 142866316 | 142865474 | 142865630 | 142866480 | 142866605 | 30 | 6 | 16 | 14 | 26 | 0 | 0 | 1 | 0 | 0 | 0 | 9 | 9 | 14 | 25 | 0 | 0 | 0 | 0 | 0 | 298 | 149 | 1 | 1 | 1 | 1 | 0.889 | 1 | 1 | NA | 1 | 1 | 1 | 1 | -0.022 |
| SE | 23021 | ENSG00000183474 | GTF2H2C | chr5 | + | 69562672 | 69562770 | 69560190 | 69560403 | 69565099 | 69565133 | 8 | 12 | 8 | 12 | 6 | 4 | 2 | 1 | 2 | 0 | 0 | 4 | 4 | 15 | 0 | 0 | 0 | 5 | 0 | 0 | 247 | 149 | 1 | 1 | 0.547 | 0.784 | 0.828 | 0.784 | 1 | NA | 1 | 0.326 | 1 | NA | 0.013 |
| SE | 23022 | ENSG00000183474 | GTF2H2C | chr5 | + | 69562672 | 69562770 | 69560208 | 69560242 | 69565099 | 69565133 | 3 | 8 | 7 | 3 | 3 | 0 | 0 | 0 | 0 | 0 | 0 | 2 | 7 | 7 | 0 | 0 | 0 | 0 | 1 | 0 | 247 | 149 | 1 | 1 | 1 | 1 | 1 | 1 | 1 | NA | 1 | 1 | 0.809 | NA | 0.064 |
| SE | 23023 | ENSG00000183474 | GTF2H2C | chr5 | + | 69562672 | 69562770 | 69560258 | 69560601 | 69565099 | 69565133 | 3 | 7 | 6 | 3 | 1 | 0 | 0 | 0 | 0 | 0 | 0 | 2 | 4 | 7 | 0 | 0 | 0 | 0 | 1 | 0 | 247 | 149 | 1 | 1 | 1 | 1 | 1 | 1 | 1 | NA | 1 | 1 | 0.809 | NA | 0.064 |
| SE | 23027 | ENSG00000183474 | GTF2H2C | chr5 | + | 69582365 | 69582429 | 69579756 | 69579864 | 69585593 | 69585596 | 52 | 23 | 72 | 74 | 89 | 0 | 1 | 1 | 0 | 0 | 20 | 23 | 9 | 61 | 24 | 0 | 0 | 0 | 0 | 0 | 213 | 149 | 1 | 1 | 1 | 0.941 | 0.981 | 1 | 1 | 1 | 1 | 1 | 1 | 1 | -0.016 |
| SE | 23030 | ENSG00000152942 | RAD17 | chr5 | + | 69371034 | 69371171 | 69369805 | 69370284 | 69371453 | 69371557 | 23 | 5 | 36 | 32 | 17 | 1 | 0 | 1 | 0 | 0 | 12 | 12 | 3 | 8 | 0 | 0 | 0 | 0 | 0 | 0 | 286 | 149 | 1 | 1 | 0.923 | 1 | 0.949 | 1 | 1 | 1 | 1 | 1 | 1 | NA | -0.026 |
| SE | 23031 | ENSG00000152942 | RAD17 | chr5 | + | 69371034 | 69371171 | 69369805 | 69370284 | 69372033 | 69372217 | 95 | 89 | 114 | 107 | 136 | 0 | 0 | 1 | 3 | 0 | 0 | 104 | 49 | 22 | 17 | 0 | 0 | 0 | 0 | 0 | 286 | 149 | 1 | 1 | 1 | 1 | 0.983 | 0.949 | 1 | NA | 1 | 1 | 1 | 1 | -0.014 |
| SE | 23032 | ENSG00000152942 | RAD17 | chr5 | + | 69371034 | 69371171 | 69369817 | 69369933 | 69372033 | 69372217 | 86 | 97 | 94 | 106 | 124 | 0 | 0 | 2 | 1 | 0 | 0 | 96 | 52 | 16 | 17 | 0 | 0 | 0 | 0 | 0 | 286 | 149 | 1 | 1 | 1 | 1 | 0.961 | 0.982 | 1 | NA | 1 | 1 | 1 | 1 | -0.011 |
| SE | 23036 | ENSG00000152942 | RAD17 | chr5 | + | 69371453 | 69371557 | 69369805 | 69370284 | 69372033 | 69372217 | 15 | 5 | 15 | 13 | 10 | 0 | 0 | 1 | 3 | 0 | 0 | 9 | 10 | 0 | 0 | 0 | 0 | 0 | 0 | 0 | 253 | 149 | 1 | 1 | 1 | 1 | 0.898 | 0.718 | 1 | NA | 1 | 1 | NA | NA | -0.077 |
| SE | 23038 | ENSG00000152942 | RAD17 | chr5 | + | 69371453 | 69371557 | 69371034 | 69371171 | 69373829 | 69374087 | 7 | 2 | 7 | 14 | 3 | 21 | 3 | 4 | 14 | 30 | 12 | 1 | 0 | 0 | 0 | 12 | 9 | 11 | 1 | 0 | 253 | 149 | 1 | 1 | 0.164 | 0.282 | 0.508 | 0.371 | 0.056 | 0.371 | 0.061 | 0 | 0 | NA | 0.168 |
| SE | 23040 | ENSG00000152942 | RAD17 | chr5 | + | 69372033 | 69372217 | 69371453 | 69371557 | 69373829 | 69373882 | 56 | 43 | 61 | 60 | 128 | 0 | 0 | 0 | 3 | 0 | 0 | 17 | 37 | 1 | 0 | 0 | 0 | 0 | 0 | 0 | 298 | 149 | 1 | 1 | 1 | 1 | 1 | 0.909 | 1 | NA | 1 | 1 | 1 | NA | -0.018 |
| SE | 23042 | ENSG00000152942 | RAD17 | chr5 | + | 69381900 | 69382057 | 69374627 | 69374711 | 69384796 | 69384933 | 250 | 195 | 199 | 234 | 454 | 1 | 0 | 0 | 0 | 0 | 36 | 163 | 180 | 75 | 23 | 0 | 0 | 0 | 0 | 0 | 298 | 149 | 1 | 1 | 0.992 | 1 | 1 | 1 | 1 | 1 | 1 | 1 | 1 | 1 | -0.002 |
| SE | 23043 | ENSG00000152942 | RAD17 | chr5 | + | 69384796 | 69384933 | 69381900 | 69382057 | 69386042 | 69386095 | 247 | 202 | 193 | 211 | 364 | 0 | 0 | 3 | 0 | 1 | 55 | 135 | 158 | 25 | 0 | 0 | 4 | 0 | 0 | 0 | 286 | 149 | 1 | 1 | 1 | 1 | 0.971 | 1 | 0.995 | 1 | 0.946 | 1 | 1 | NA | 0.007 |
| SE | 23046 | ENSG00000152942 | RAD17 | chr5 | + | 69396396 | 69396546 | 69393353 | 69393500 | 69400048 | 69400169 | 132 | 100 | 144 | 153 | 168 | 0 | 0 | 1 | 0 | 1 | 19 | 71 | 56 | 30 | 7 | 0 | 0 | 0 | 0 | 0 | 298 | 149 | 1 | 1 | 1 | 1 | 0.986 | 1 | 0.988 | 1 | 1 | 1 | 1 | 1 | -0.005 |
| SE | 23047 | ENSG00000152942 | RAD17 | chr5 | + | 69400048 | 69400169 | 69396396 | 69396546 | 69410492 | 69410550 | 127 | 95 | 131 | 154 | 156 | 0 | 0 | 1 | 0 | 0 | 0 | 66 | 52 | 12 | 0 | 0 | 0 | 0 | 0 | 0 | 270 | 149 | 1 | 1 | 1 | 1 | 0.986 | 1 | 1 | NA | 1 | 1 | 1 | NA | -0.003 |
| SE | 23050 | ENSG00000273841 | TAF9 | chr5 | - | 69366502 | 69366595 | 69364742 | 69365754 | 69369462 | 69369520 | 165 | 117 | 166 | 169 | 345 | 1 | 1 | 2 | 0 | 0 | 76 | 111 | 163 | 145 | 50 | 0 | 3 | 3 | 0 | 0 | 242 | 149 | 1 | 1 | 0.99 | 0.986 | 0.981 | 1 | 1 | 1 | 0.958 | 0.971 | 1 | 1 | 0.006 |
| SE | 23051 | ENSG00000273841 | TAF9 | chr5 | - | 69366502 | 69366595 | 69364742 | 69365754 | 69369656 | 69370013 | 133 | 100 | 129 | 137 | 329 | 0 | 0 | 0 | 0 | 0 | 73 | 89 | 127 | 113 | 40 | 0 | 0 | 0 | 0 | 3 | 242 | 149 | 1 | 1 | 1 | 1 | 1 | 1 | 1 | 1 | 1 | 1 | 1 | 0.891 | 0.022 |
| SE | 23052 | ENSG00000124615 | MOCS1 | chr6 | - | 39909039 | 39909102 | 39904164 | 39907117 | 39909834 | 39909955 | 2 | 0 | 1 | 1 | 2 | 0 | 0 | 0 | 0 | 0 | 0 | 0 | 0 | 0 | 0 | 0 | 0 | 4 | 0 | 0 | 212 | 149 | 1 | 1 | 1 | NA | 1 | 1 | 1 | NA | NA | 0 | NA | NA | 1 |
| SE | 23053 | ENSG00000124615 | MOCS1 | chr6 | - | 39909054 | 39909102 | 39904169 | 39907117 | 39909834 | 39909955 | 1 | 0 | 0 | 1 | 0 | 0 | 0 | 0 | 0 | 0 | 0 | 0 | 0 | 0 | 0 | 0 | 0 | 4 | 0 | 0 | 197 | 149 | 1 | 1 | 1 | NA | NA | 1 | NA | NA | NA | 0 | NA | NA | 1 |
| SE | 23057 | ENSG00000111641 | NOP2 | chr12 | - | 6558015 | 6558137 | 6556870 | 6557642 | 6560097 | 6560326 | 0 | 0 | 0 | 2 | 0 | 73 | 52 | 72 | 60 | 128 | 0 | 0 | 0 | 0 | 0 | 61 | 76 | 51 | 111 | 131 | 271 | 149 | 1 | 1 | 0 | 0 | 0 | 0.018 | 0 | 0 | 0 | 0 | 0 | 0 | 0.004 |
| SE | 23059 | ENSG00000111641 | NOP2 | chr12 | - | 6564149 | 6564248 | 6563890 | 6563946 | 6566528 | 6566617 | 0 | 0 | 2 | 0 | 6 | 0 | 0 | 2 | 4 | 1 | 0 | 1 | 2 | 1 | 0 | 10 | 0 | 0 | 0 | 17 | 248 | 149 | 1 | 1 | NA | NA | 0.375 | 0 | 0.783 | 0 | 1 | 1 | 1 | 0 | -0.214 |
| SE | 23062 | ENSG00000111641 | NOP2 | chr12 | - | 6566100 | 6566336 | 6564149 | 6564248 | 6566528 | 6566559 | 97 | 69 | 83 | 81 | 82 | 0 | 0 | 0 | 0 | 0 | 90 | 56 | 76 | 81 | 142 | 0 | 1 | 0 | 0 | 0 | 298 | 149 | 1 | 1 | 1 | 1 | 1 | 1 | 1 | 1 | 0.966 | 1 | 1 | 1 | 0.007 |
| SE | 23064 | ENSG00000065357 | DGKA | chr12 | + | 55932560 | 55932909 | 55932107 | 55932123 | 55935304 | 55935402 | 0 | 0 | 0 | 0 | 3 | 2 | 0 | 1 | 0 | 0 | 0 | 7 | 0 | 3 | 0 | 0 | 0 | 1 | 0 | 0 | 298 | 149 | 1 | 1 | 0 | NA | 0 | NA | 1 | NA | 1 | 0 | 1 | NA | -0.333 |
| SE | 23065 | ENSG00000065357 | DGKA | chr12 | + | 55932560 | 55932909 | 55932134 | 55932255 | 55935304 | 55935402 | 0 | 0 | 1 | 0 | 3 | 0 | 0 | 1 | 1 | 0 | 0 | 0 | 0 | 3 | 0 | 0 | 0 | 0 | 0 | 0 | 298 | 149 | 1 | 1 | NA | NA | 0.333 | 0 | 1 | NA | NA | NA | 1 | NA | -0.556 |
| SE | 23069 | ENSG00000065357 | DGKA | chr12 | + | 55936652 | 55936856 | 55936422 | 55936567 | 55937016 | 55937032 | 2 | 0 | 0 | 0 | 2 | 269 | 203 | 415 | 296 | 441 | 0 | 0 | 0 | 0 | 0 | 201 | 235 | 154 | 162 | 221 | 298 | 149 | 1 | 1 | 0.004 | 0 | 0 | 0 | 0.002 | 0 | 0 | 0 | 0 | 0 | 0.001 |
| SE | 23070 | ENSG00000065357 | DGKA | chr12 | + | 55937016 | 55937090 | 55936422 | 55936567 | 55937407 | 55937443 | 382 | 285 | 594 | 401 | 593 | 0 | 0 | 0 | 3 | 0 | 315 | 274 | 203 | 239 | 266 | 0 | 0 | 0 | 0 | 0 | 223 | 149 | 1 | 1 | 1 | 1 | 1 | 0.989 | 1 | 1 | 1 | 1 | 1 | 1 | -0.002 |
| SE | 23071 | ENSG00000065357 | DGKA | chr12 | + | 55937407 | 55937543 | 55935335 | 55937090 | 55937977 | 55938025 | 467 | 404 | 683 | 523 | 795 | 0 | 0 | 0 | 0 | 0 | 403 | 273 | 225 | 261 | 277 | 0 | 0 | 0 | 3 | 0 | 285 | 149 | 1 | 1 | 1 | 1 | 1 | 1 | 1 | 1 | 1 | 1 | 0.978 | 1 | 0.004 |
| SE | 23072 | ENSG00000065357 | DGKA | chr12 | + | 55937407 | 55937543 | 55935335 | 55937090 | 55938510 | 55938560 | 226 | 185 | 371 | 241 | 333 | 0 | 1 | 1 | 5 | 0 | 192 | 147 | 124 | 131 | 157 | 0 | 4 | 1 | 0 | 1 | 285 | 149 | 1 | 1 | 1 | 0.99 | 0.995 | 0.962 | 1 | 1 | 0.951 | 0.985 | 1 | 0.988 | 0.005 |
| SE | 23073 | ENSG00000065357 | DGKA | chr12 | + | 55937407 | 55937557 | 55937016 | 55937090 | 55937977 | 55938052 | 233 | 201 | 400 | 253 | 348 | 0 | 0 | 0 | 0 | 0 | 207 | 173 | 130 | 149 | 180 | 0 | 0 | 0 | 3 | 0 | 298 | 149 | 1 | 1 | 1 | 1 | 1 | 1 | 1 | 1 | 1 | 1 | 0.961 | 1 | 0.008 |
| SE | 23075 | ENSG00000065357 | DGKA | chr12 | + | 55937977 | 55938052 | 55937407 | 55937543 | 55938510 | 55938558 | 231 | 211 | 380 | 292 | 498 | 0 | 0 | 0 | 0 | 3 | 216 | 172 | 126 | 158 | 120 | 0 | 0 | 0 | 0 | 0 | 224 | 149 | 1 | 1 | 1 | 1 | 1 | 1 | 0.991 | 1 | 1 | 1 | 1 | 1 | -0.002 |
| SE | 23076 | ENSG00000065357 | DGKA | chr12 | + | 55938510 | 55938560 | 55937977 | 55938052 | 55938914 | 55938957 | 156 | 126 | 264 | 196 | 287 | 0 | 0 | 0 | 0 | 0 | 148 | 126 | 108 | 127 | 113 | 0 | 0 | 3 | 0 | 0 | 199 | 149 | 1 | 1 | 1 | 1 | 1 | 1 | 1 | 1 | 1 | 0.964 | 1 | 1 | 0.007 |
| SE | 23077 | ENSG00000065357 | DGKA | chr12 | + | 55938637 | 55938698 | 55937977 | 55938052 | 55938914 | 55938942 | 55 | 61 | 50 | 73 | 78 | 0 | 0 | 0 | 0 | 0 | 54 | 16 | 24 | 27 | 57 | 0 | 0 | 3 | 0 | 0 | 210 | 149 | 1 | 1 | 1 | 1 | 1 | 1 | 1 | 1 | 1 | 0.85 | 1 | 1 | 0.03 |
| SE | 23078 | ENSG00000065357 | DGKA | chr12 | + | 55938914 | 55938989 | 55938510 | 55938560 | 55939185 | 55939290 | 163 | 147 | 330 | 232 | 295 | 0 | 1 | 1 | 0 | 0 | 142 | 119 | 146 | 174 | 160 | 0 | 0 | 0 | 0 | 0 | 224 | 149 | 1 | 1 | 1 | 0.99 | 0.995 | 1 | 1 | 1 | 1 | 1 | 1 | 1 | -0.003 |
| SE | 23079 | ENSG00000065357 | DGKA | chr12 | + | 55938938 | 55938989 | 55938552 | 55938560 | 55939185 | 55939305 | 57 | 55 | 138 | 115 | 112 | 0 | 1 | 1 | 0 | 0 | 64 | 48 | 59 | 64 | 44 | 0 | 0 | 0 | 0 | 0 | 200 | 149 | 1 | 1 | 1 | 0.976 | 0.99 | 1 | 1 | 1 | 1 | 1 | 1 | 1 | -0.007 |
| SE | 23083 | ENSG00000065357 | DGKA | chr12 | + | 55941997 | 55942037 | 55941509 | 55941584 | 55942173 | 55942263 | 85 | 80 | 125 | 129 | 36 | 0 | 0 | 0 | 2 | 0 | 95 | 100 | 78 | 77 | 130 | 0 | 0 | 0 | 0 | 0 | 189 | 149 | 1 | 1 | 1 | 1 | 1 | 0.981 | 1 | 1 | 1 | 1 | 1 | 1 | -0.004 |
| SE | 23084 | ENSG00000065357 | DGKA | chr12 | + | 55941997 | 55942083 | 55941509 | 55941584 | 55942173 | 55942263 | 568 | 664 | 1028 | 880 | 749 | 0 | 0 | 0 | 2 | 0 | 670 | 531 | 550 | 628 | 703 | 0 | 0 | 0 | 0 | 0 | 235 | 149 | 1 | 1 | 1 | 1 | 1 | 0.996 | 1 | 1 | 1 | 1 | 1 | 1 | -0.001 |
| SE | 23087 | ENSG00000184349 | EFNA5 | chr5 | - | 107387234 | 107387315 | 107376893 | 107381376 | 107387705 | 107387771 | 12 | 5 | 1 | 6 | 7 | 6 | 2 | 1 | 3 | 3 | 0 | 1 | 0 | 0 | 0 | 0 | 0 | 5 | 1 | 0 | 230 | 149 | 1 | 1 | 0.564 | 0.618 | 0.393 | 0.564 | 0.602 | NA | 1 | 0 | 0 | NA | 0.215 |
| SE | 23088 | ENSG00000184349 | EFNA5 | chr5 | - | 107387705 | 107387771 | 107387234 | 107387315 | 107427216 | 107427509 | 19 | 7 | 0 | 7 | 4 | 1 | 0 | 0 | 0 | 0 | 0 | 2 | 2 | 0 | 0 | 0 | 0 | 0 | 0 | 0 | 215 | 149 | 1 | 1 | 0.929 | 1 | NA | 1 | 1 | NA | 1 | 1 | NA | NA | -0.018 |
| SE | 23089 | ENSG00000112996 | MRPS30 | chr5 | + | 44811914 | 44812020 | 44811008 | 44811154 | 44813105 | 44813282 | 213 | 103 | 183 | 205 | 364 | 0 | 0 | 0 | 1 | 0 | 193 | 180 | 126 | 119 | 132 | 0 | 0 | 0 | 0 | 0 | 255 | 149 | 1 | 1 | 1 | 1 | 1 | 0.992 | 1 | 1 | 1 | 1 | 1 | 1 | -0.002 |
| SE | 23090 | ENSG00000112996 | MRPS30 | chr5 | + | 44811914 | 44812020 | 44811008 | 44811154 | 44814912 | 44815066 | 83 | 26 | 62 | 82 | 145 | 0 | 0 | 0 | 0 | 3 | 95 | 84 | 55 | 45 | 47 | 0 | 0 | 0 | 0 | 0 | 255 | 149 | 1 | 1 | 1 | 1 | 1 | 1 | 0.966 | 1 | 1 | 1 | 1 | 1 | -0.007 |
| SE | 23091 | ENSG00000112996 | MRPS30 | chr5 | + | 44813105 | 44813282 | 44811008 | 44811154 | 44814912 | 44815066 | 252 | 139 | 215 | 205 | 452 | 0 | 0 | 0 | 0 | 3 | 210 | 234 | 140 | 141 | 112 | 0 | 0 | 0 | 0 | 0 | 298 | 149 | 1 | 1 | 1 | 1 | 1 | 1 | 0.987 | 1 | 1 | 1 | 1 | 1 | -0.003 |
| SE | 23092 | ENSG00000112996 | MRPS30 | chr5 | + | 44813105 | 44813282 | 44811914 | 44812020 | 44814912 | 44815066 | 387 | 220 | 336 | 335 | 672 | 1 | 3 | 0 | 5 | 1 | 308 | 330 | 212 | 218 | 197 | 0 | 0 | 1 | 0 | 0 | 298 | 149 | 1 | 1 | 0.995 | 0.973 | 1 | 0.971 | 0.997 | 1 | 1 | 0.991 | 1 | 1 | -0.011 |
| SE | 23100 | ENSG00000111912 | NCOA7 | chr6 | + | 125815290 | 125815404 | 125781160 | 125781371 | 125855019 | 125855240 | 115 | 110 | 98 | 128 | 122 | 0 | 0 | 0 | 1 | 0 | 62 | 45 | 119 | 56 | 11 | 0 | 0 | 0 | 0 | 0 | 263 | 149 | 1 | 1 | 1 | 1 | 1 | 0.986 | 1 | 1 | 1 | 1 | 1 | 1 | -0.003 |
| SE | 23104 | ENSG00000111912 | NCOA7 | chr6 | + | 125915332 | 125915480 | 125890641 | 125890810 | 125920942 | 125921068 | 229 | 219 | 184 | 229 | 215 | 1 | 0 | 0 | 1 | 0 | 71 | 157 | 182 | 80 | 19 | 0 | 0 | 0 | 0 | 0 | 297 | 149 | 1 | 1 | 0.991 | 1 | 1 | 0.991 | 1 | 1 | 1 | 1 | 1 | 1 | -0.004 |
| SE | 23105 | ENSG00000111912 | NCOA7 | chr6 | + | 125927662 | 125927758 | 125922681 | 125922834 | 125928173 | 125928247 | 180 | 218 | 152 | 231 | 93 | 0 | 2 | 0 | 0 | 0 | 81 | 180 | 240 | 364 | 37 | 0 | 0 | 0 | 0 | 0 | 245 | 149 | 1 | 1 | 1 | 0.985 | 1 | 1 | 1 | 1 | 1 | 1 | 1 | 1 | -0.003 |
| SE | 23107 | ENSG00000037749 | MFAP3 | chr5 | + | 154049556 | 154050017 | 154038966 | 154039011 | 154052919 | 154052952 | 64 | 46 | 71 | 79 | 99 | 0 | 0 | 2 | 3 | 3 | 0 | 37 | 21 | 20 | 0 | 0 | 0 | 3 | 1 | 0 | 298 | 149 | 1 | 1 | 1 | 1 | 0.947 | 0.929 | 0.943 | NA | 1 | 0.778 | 0.909 | NA | 0.068 |
| SE | 23108 | ENSG00000037749 | MFAP3 | chr5 | + | 154051736 | 154051943 | 154049556 | 154050017 | 154052919 | 154056546 | 0 | 1 | 5 | 1 | 7 | 56 | 43 | 48 | 70 | 93 | 0 | 0 | 0 | 0 | 0 | 0 | 35 | 19 | 16 | 0 | 298 | 149 | 1 | 1 | 0 | 0.011 | 0.05 | 0.007 | 0.036 | NA | 0 | 0 | 0 | NA | 0.021 |
| SE | 23109 | ENSG00000037749 | MFAP3 | chr5 | + | 154051817 | 154051943 | 154049556 | 154050017 | 154052919 | 154057420 | 0 | 1 | 4 | 1 | 3 | 56 | 43 | 48 | 70 | 93 | 0 | 0 | 0 | 0 | 0 | 0 | 35 | 19 | 16 | 0 | 275 | 149 | 1 | 1 | 0 | 0.012 | 0.043 | 0.008 | 0.017 | NA | 0 | 0 | 0 | NA | 0.016 |
| SE | 23110 | ENSG00000037749 | MFAP3 | chr5 | + | 154121717 | 154121897 | 154097022 | 154097271 | 154134844 | 154134916 | 5 | 3 | 0 | 1 | 2 | 0 | 0 | 0 | 0 | 2 | 0 | 0 | 0 | 1 | 0 | 0 | 0 | 0 | 0 | 0 | 298 | 149 | 1 | 1 | 1 | 1 | NA | 1 | 0.333 | NA | NA | NA | 1 | NA | -0.167 |
| SE | 23115 | ENSG00000197375 | SLC22A5 | chr5 | + | 132378163 | 132378235 | 132369972 | 132370365 | 132378377 | 132378481 | 0 | 0 | 0 | 0 | 0 | 6 | 0 | 2 | 1 | 0 | 0 | 0 | 0 | 1 | 0 | 0 | 2 | 0 | 0 | 17 | 221 | 149 | 1 | 1 | 0 | NA | 0 | 0 | NA | NA | 0 | NA | 1 | 0 | -0.333 |
| SE | 23116 | ENSG00000197375 | SLC22A5 | chr5 | + | 132378377 | 132378481 | 132369709 | 132370365 | 132384146 | 132384301 | 11 | 1 | 2 | 5 | 0 | 0 | 0 | 1 | 0 | 0 | 0 | 2 | 0 | 3 | 17 | 0 | 0 | 0 | 0 | 0 | 253 | 149 | 1 | 1 | 1 | 1 | 0.541 | 1 | NA | NA | 1 | NA | 1 | 1 | -0.115 |
| SE | 23119 | ENSG00000197375 | SLC22A5 | chr5 | + | 132390775 | 132390904 | 132388920 | 132389021 | 132392432 | 132392615 | 6 | 3 | 7 | 10 | 10 | 2 | 3 | 0 | 0 | 0 | 6 | 9 | 7 | 0 | 10 | 0 | 3 | 0 | 0 | 0 | 278 | 149 | 1 | 1 | 0.617 | 0.349 | 1 | 1 | 1 | 1 | 0.617 | 1 | NA | 1 | -0.111 |
| SE | 23120 | ENSG00000197375 | SLC22A5 | chr5 | + | 132393675 | 132393811 | 132392432 | 132392615 | 132394184 | 132394272 | 21 | 13 | 25 | 25 | 23 | 0 | 1 | 0 | 0 | 0 | 10 | 18 | 11 | 12 | 10 | 0 | 0 | 0 | 0 | 0 | 285 | 149 | 1 | 1 | 1 | 0.872 | 1 | 1 | 1 | 1 | 1 | 1 | 1 | 1 | -0.026 |
| SE | 23122 | ENSG00000197536 | IRF1-AS1 | chr5 | + | 132449649 | 132449689 | 132419822 | 132419940 | 132486191 | 132488702 | 7 | 5 | 9 | 8 | 1 | 0 | 0 | 1 | 0 | 0 | 5 | 10 | 5 | 3 | 19 | 0 | 0 | 0 | 0 | 0 | 189 | 149 | 1 | 1 | 1 | 1 | 0.876 | 1 | 1 | 1 | 1 | 1 | 1 | 1 | -0.025 |
| SE | 23133 | ENSG00000153982 | GDPD1 | chr17 | + | 59273650 | 59273863 | 59272784 | 59272836 | 59275838 | 59275967 | 5 | 1 | 9 | 9 | 6 | 0 | 0 | 0 | 0 | 4 | 0 | 0 | 2 | 2 | 0 | 0 | 0 | 0 | 0 | 0 | 298 | 149 | 1 | 1 | 1 | 1 | 1 | 1 | 0.429 | NA | NA | 1 | 1 | NA | -0.114 |
| SE | 23134 | ENSG00000079215 | SLC1A3 | chr5 | + | 36607172 | 36607311 | 36606530 | 36606735 | 36608328 | 36608534 | 0 | 0 | 0 | 6 | 0 | 2 | 0 | 1 | 4 | 0 | 0 | 4 | 0 | 0 | 0 | 8 | 10 | 5 | 4 | 0 | 288 | 149 | 1 | 1 | 0 | NA | 0 | 0.437 | NA | 0 | 0.171 | 0 | 0 | NA | 0.103 |
| SE | 23135 | ENSG00000079215 | SLC1A3 | chr5 | + | 36607223 | 36607311 | 36606438 | 36606735 | 36608328 | 36608604 | 0 | 0 | 0 | 3 | 0 | 2 | 0 | 1 | 4 | 0 | 0 | 4 | 0 | 0 | 0 | 8 | 10 | 5 | 4 | 0 | 237 | 149 | 1 | 1 | 0 | NA | 0 | 0.32 | NA | 0 | 0.201 | 0 | 0 | NA | 0.056 |
| SE | 23138 | ENSG00000079215 | SLC1A3 | chr5 | + | 36614936 | 36614979 | 36608328 | 36608604 | 36629449 | 36629587 | 0 | 0 | 0 | 0 | 0 | 3 | 3 | 0 | 1 | 0 | 0 | 0 | 1 | 0 | 0 | 0 | 4 | 4 | 0 | 0 | 192 | 149 | 1 | 1 | NA | 0 | 0 | NA | 0 | NA | NA | 0.162 | 0 | NA | -0.081 |
| SE | 23160 | ENSG00000250999 | NA | chr5 | + | 179662974 | 179663103 | 179658235 | 179658481 | 179664322 | 179664430 | 0 | 1 | 0 | 0 | 0 | 0 | 0 | 0 | 0 | 0 | 0 | 0 | 0 | 0 | 0 | 0 | 0 | 0 | 0 | 1 | 278 | 149 | 1 | 1 | NA | 1 | NA | NA | NA | NA | NA | NA | NA | 0 | 1 |
| SE | 23161 | ENSG00000176783 | RUFY1 | chr5 | + | 179555649 | 179555790 | 179550837 | 179550879 | 179560024 | 179560198 | 1 | 2 | 0 | 0 | 0 | 7 | 4 | 2 | 5 | 5 | 0 | 0 | 0 | 0 | 0 | 14 | 4 | 1 | 10 | 14 | 290 | 149 | 1 | 1 | 0.068 | 0.204 | 0 | 0 | NA | 0 | 0 | 0 | 0 | 0 | 0.068 |
| SE | 23164 | ENSG00000176783 | RUFY1 | chr5 | + | 179577074 | 179577136 | 179569301 | 179569425 | 179580946 | 179581012 | 241 | 353 | 149 | 552 | 435 | 1 | 2 | 0 | 3 | 4 | 1430 | 536 | 272 | 420 | 2516 | 0 | 1 | 5 | 3 | 10 | 211 | 149 | 1 | 1 | 0.994 | 0.992 | 1 | 0.992 | 0.987 | 1 | 0.997 | 0.975 | 0.99 | 0.994 | 0.002 |
| SE | 23165 | ENSG00000176783 | RUFY1 | chr5 | + | 179577074 | 179577136 | 179569301 | 179569425 | 179585795 | 179585865 | 60 | 97 | 46 | 143 | 120 | 2 | 0 | 0 | 0 | 0 | 290 | 112 | 69 | 95 | 567 | 0 | 0 | 2 | 4 | 0 | 211 | 149 | 1 | 1 | 0.955 | 1 | 1 | 1 | 1 | 1 | 1 | 0.961 | 0.944 | 1 | 0.01 |
| SE | 23166 | ENSG00000176783 | RUFY1 | chr5 | + | 179580946 | 179581012 | 179569301 | 179569425 | 179585795 | 179585865 | 145 | 182 | 64 | 275 | 204 | 2 | 0 | 0 | 0 | 0 | 787 | 293 | 156 | 309 | 1494 | 0 | 0 | 2 | 4 | 0 | 215 | 149 | 1 | 1 | 0.98 | 1 | 1 | 1 | 1 | 1 | 1 | 0.982 | 0.982 | 1 | 0.003 |
| SE | 23169 | ENSG00000176783 | RUFY1 | chr5 | + | 179589545 | 179589647 | 179585795 | 179585865 | 179591624 | 179591741 | 400 | 477 | 251 | 775 | 594 | 1 | 0 | 0 | 0 | 0 | 2338 | 728 | 452 | 898 | 4206 | 5 | 0 | 0 | 0 | 0 | 251 | 149 | 1 | 1 | 0.996 | 1 | 1 | 1 | 1 | 0.996 | 1 | 1 | 1 | 1 | 0 |
| SE | 23170 | ENSG00000176783 | RUFY1 | chr5 | + | 179591624 | 179591741 | 179589362 | 179589647 | 179593477 | 179593613 | 480 | 601 | 319 | 890 | 719 | 0 | 0 | 0 | 0 | 1 | 2640 | 716 | 563 | 1028 | 4413 | 0 | 0 | 0 | 0 | 6 | 266 | 149 | 1 | 1 | 1 | 1 | 1 | 1 | 0.998 | 1 | 1 | 1 | 1 | 0.998 | 0 |
| SE | 23171 | ENSG00000176783 | RUFY1 | chr5 | + | 179593477 | 179593645 | 179591624 | 179591741 | 179594865 | 179594898 | 721 | 946 | 482 | 1447 | 1222 | 2 | 1 | 0 | 2 | 0 | 3786 | 1152 | 818 | 1256 | 6373 | 0 | 0 | 0 | 0 | 7 | 298 | 149 | 1 | 1 | 0.994 | 0.998 | 1 | 0.997 | 1 | 1 | 1 | 1 | 1 | 0.998 | -0.002 |
| SE | 23172 | ENSG00000176783 | RUFY1 | chr5 | + | 179596561 | 179596681 | 179594865 | 179594963 | 179598691 | 179598821 | 554 | 760 | 391 | 1311 | 788 | 0 | 0 | 0 | 0 | 2 | 3173 | 958 | 652 | 849 | 6056 | 0 | 0 | 0 | 0 | 0 | 269 | 149 | 1 | 1 | 1 | 1 | 1 | 1 | 0.995 | 1 | 1 | 1 | 1 | 1 | -0.001 |
| SE | 23173 | ENSG00000176783 | RUFY1 | chr5 | + | 179601891 | 179601986 | 179598691 | 179598821 | 179605875 | 179605924 | 129 | 158 | 121 | 359 | 207 | 0 | 0 | 0 | 0 | 0 | 830 | 287 | 167 | 207 | 1337 | 0 | 3 | 0 | 0 | 0 | 244 | 149 | 1 | 1 | 1 | 1 | 1 | 1 | 1 | 1 | 0.983 | 1 | 1 | 1 | 0.003 |
| SE | 23174 | ENSG00000176783 | RUFY1 | chr5 | + | 179607581 | 179607659 | 179605875 | 179605924 | 179609375 | 179610011 | 101 | 123 | 89 | 257 | 181 | 0 | 0 | 0 | 4 | 0 | 547 | 253 | 100 | 137 | 1003 | 0 | 5 | 0 | 0 | 0 | 227 | 149 | 1 | 1 | 1 | 1 | 1 | 0.977 | 1 | 1 | 0.971 | 1 | 1 | 1 | 0.001 |
| SE | 23177 | ENSG00000113240 | CLK4 | chr5 | - | 178608375 | 178608458 | 178605302 | 178605382 | 178612415 | 178612545 | 332 | 268 | 291 | 251 | 614 | 1 | 0 | 0 | 1 | 3 | 59 | 96 | 121 | 31 | 15 | 0 | 0 | 0 | 0 | 0 | 232 | 149 | 1 | 1 | 0.995 | 1 | 1 | 0.994 | 0.992 | 1 | 1 | 1 | 1 | 1 | -0.004 |
| [illegible] | | | | | | | | | | | | | | | | | | | | | | | | | | | | | | | | | | | | | | | | | | | | | | |

| | | | | | | | | | | | | | | | |
|---|---|---|---|---|---|---|---|---|---|---|---|---|---|---|---|
| SE | 23230 | ENSG00000112851 | ERBIN | chr5 | + | 65992709 | 65992907 | 65926555 | 65926806 | 66012048 | 66012127 | 6 | 14 | 7 | [illegible] |
| SE | 23231 | ENSG00000112851 | ERBIN | chr5 | + | 65992709 | 65992907 | 65988632 | 65988682 | 66012048 | 66012127 | 747 | 644 | 564 | [illegible] |
| SE | 23232 | ENSG00000112851 | ERBIN | chr5 | + | 65994746 | 65994864 | 65992709 | 65992907 | 66012048 | 66012127 | 1241 | 1006 | 946 | [illegible] |
| SE | 23233 | ENSG00000112851 | ERBIN | chr5 | + | 66024305 | 66024450 | 66023289 | 66023364 | 66025479 | 66025552 | 845 | 580 | 641 | [illegible] |
| SE | 23234 | ENSG00000112851 | ERBIN | chr5 | + | 66038382 | 66038482 | 66028273 | 66028343 | 66043076 | 66043198 | 1033 | 861 | 746 | [illegible] |
| SE | 23235 | ENSG00000112851 | ERBIN | chr5 | + | 66046352 | 66046538 | 66044136 | 66044310 | 66048666 | 66048781 | 993 | 849 | 886 | [illegible] |
| SE | 23236 | ENSG00000112851 | ERBIN | chr5 | + | 66048666 | 66048781 | 66046352 | 66046538 | 66050782 | 66050966 | 840 | 694 | 926 | [illegible] |
| SE | 23237 | ENSG00000112851 | ERBIN | chr5 | + | 66050782 | 66050966 | 66048666 | 66048781 | 66053405 | 66054951 | 682 | 495 | 833 | [illegible] |
| SE | 23238 | ENSG00000112851 | ERBIN | chr5 | + | 66053405 | 66054951 | 66050782 | 66050966 | 66076315 | 66076408 | 553 | 424 | 516 | [illegible] |
| SE | 23239 | ENSG00000112851 | ERBIN | chr5 | + | 66068876 | 66069020 | 66053405 | 66054951 | 66072168 | 66072291 | 19 | 16 | 10 | [illegible] |
| SE | 23242 | ENSG00000112851 | ERBIN | chr5 | + | 66072168 | 66072291 | 66053405 | 66054951 | 66076315 | 66076408 | 31 | 21 | 31 | [illegible] |
| SE | 23243 | ENSG00000112851 | ERBIN | chr5 | + | 66072168 | 66072291 | 66068876 | 66069020 | 66075023 | 66075230 | 14 | 10 | 19 | [illegible] |
| SE | 23245 | ENSG00000112851 | ERBIN | chr5 | + | 66075023 | 66075230 | 66053405 | 66054951 | 66076874 | 66076949 | 376 | 257 | 279 | [illegible] |
| SE | 23246 | ENSG00000112851 | ERBIN | chr5 | + | 66075023 | 66075230 | 66072168 | 66072291 | 66076315 | 66076408 | 334 | 307 | 313 | [illegible] |
| SE | 23247 | ENSG00000112851 | ERBIN | chr5 | + | 66076315 | 66076408 | 66053405 | 66054951 | 66076874 | 66076949 | 707 | 658 | 477 | [illegible] |
| SE | 23248 | ENSG00000112851 | ERBIN | chr5 | + | 66076315 | 66076408 | 66075023 | 66075230 | 66076874 | 66076949 | 892 | 833 | 653 | [illegible] |
| SE | 23249 | ENSG00000112851 | ERBIN | chr5 | + | 66076874 | 66076949 | 66076109 | 66076408 | 66078422 | 66078546 | 1387 | 1296 | 922 | [illegible] |
| SE | 23251 | ENSG00000108262 | GIT1 | chr17 | - | 29575629 | 29575703 | 29575287 | 29575470 | 29575811 | 29575829 | 78 | 37 | 58 | [illegible] |
| SE | 23255 | ENSG00000108262 | GIT1 | chr17 | - | 29577183 | 29577247 | 29576862 | 29576996 | 29577644 | 29577742 | 21 | 12 | 17 | [illegible] |
| SE | 23256 | ENSG00000108262 | GIT1 | chr17 | - | 29577644 | 29577742 | 29576862 | 29576996 | 29578298 | 29578371 | 30 | 18 | 35 | [illegible] |
| SE | 23260 | ENSG00000145604 | SKP2 | chr5 | + | 36163644 | 36163756 | 36152770 | 36153042 | 36166518 | 36166662 | 46 | 15 | 38 | [illegible] |
| SE | 23261 | ENSG00000145604 | SKP2 | chr5 | + | 36163644 | 36163756 | 36152770 | 36153042 | 36176964 | 36177016 | 23 | 9 | 19 | [illegible] |
| SE | 23262 | ENSG00000145604 | SKP2 | chr5 | + | 36163644 | 36164695 | 36152770 | 36153042 | 36166518 | 36166662 | 24 | 9 | 19 | [illegible] |
| SE | 23263 | ENSG00000145604 | SKP2 | chr5 | + | 36163644 | 36166662 | 36152108 | 36153042 | 36171602 | 36171733 | 23 | 10 | 20 | [illegible] |
| SE | 23264 | ENSG00000145604 | SKP2 | chr5 | + | 36163644 | 36166662 | 36152108 | 36153042 | 36176964 | 36177016 | 23 | 10 | 19 | [illegible] |
| SE | 23265 | ENSG00000145604 | SKP2 | chr5 | + | 36164534 | 36164695 | 36163644 | 36163756 | 36166518 | 36166662 | 1 | 0 | 0 | [illegible] |
| SE | 23266 | ENSG00000145604 | SKP2 | chr5 | + | 36164627 | 36164695 | 36163644 | 36163756 | 36166518 | 36166662 | 1 | 0 | 0 | [illegible] |
| SE | 23267 | ENSG00000145604 | SKP2 | chr5 | + | 36166518 | 36166662 | 36152770 | 36153042 | 36171602 | 36171733 | 6 | 4 | 1 | [illegible] |
| SE | 23268 | ENSG00000145604 | SKP2 | chr5 | + | 36166518 | 36166662 | 36152770 | 36153042 | 36176964 | 36177016 | 6 | 4 | 0 | [illegible] |
| SE | 23269 | ENSG00000145604 | SKP2 | chr5 | + | 36166518 | 36166662 | 36163371 | 36163756 | 36168312 | 36168447 | 48 | 14 | 42 | [illegible] |
| SE | 23272 | ENSG00000145604 | SKP2 | chr5 | + | 36168312 | 36168447 | 36163644 | 36166662 | 36171602 | 36171733 | 26 | 4 | 23 | [illegible] |
| SE | 23273 | ENSG00000145604 | SKP2 | chr5 | + | 36170343 | 36170442 | 36166518 | 36166662 | 36171602 | 36171733 | 53 | 33 | 47 | [illegible] |
| SE | 23274 | ENSG00000145604 | SKP2 | chr5 | + | 36170343 | 36170442 | 36166518 | 36166662 | 36176964 | 36177016 | 0 | 0 | 0 | [illegible] |
| SE | 23277 | ENSG00000145604 | SKP2 | chr5 | + | 36171602 | 36171733 | 36152770 | 36153042 | 36176964 | 36177016 | 97 | 51 | 58 | [illegible] |
| SE | 23280 | ENSG00000145604 | SKP2 | chr5 | + | 36178597 | 36178810 | 36177184 | 36177292 | 36181817 | 36182594 | 6 | 2 | 5 | [illegible] |
| SE | 23282 | ENSG00000145604 | SKP2 | chr5 | + | 36191411 | 36191702 | 36177184 | 36177292 | 36192620 | 36192695 | 0 | 1 | 0 | [illegible] |
| SE | 23283 | ENSG00000145604 | SKP2 | chr5 | + | 36191522 | 36191702 | 36177184 | 36177292 | 36192620 | 36192695 | 0 | 1 | 0 | [illegible] |
| SE | 23286 | ENSG00000164187 | LMBRD2 | chr5 | - | 36122279 | 36122463 | 36117734 | 36117916 | 36122847 | 36122961 | 61 | 51 | 62 | [illegible] |
| SE | 23287 | ENSG00000164187 | LMBRD2 | chr5 | - | 36137273 | 36137441 | 36136308 | 36136519 | 36141106 | 36141202 | 112 | 65 | 119 | [illegible] |
| SE | 23293 | ENSG00000164182 | NDUFAF2 | chr5 | + | 61045523 | 61045604 | 60945182 | 60945382 | 61045839 | 61045978 | 0 | 0 | 0 | [illegible] |
| SE | 23296 | ENSG00000164182 | NDUFAF2 | chr5 | + | 61045839 | 61045978 | 61045523 | 61045604 | 61073124 | 61073214 | 2 | 1 | 2 | [illegible] |
| SE | 23302 | ENSG00000273294 | C1QTNF3-AMACR | chr5 | - | 34023908 | 34024008 | 34020586 | 34020742 | 34028753 | 34028883 | 18 | 6 | 14 | [illegible] |
| SE | 23303 | ENSG00000273294 | C1QTNF3-AMACR | chr5 | - | 34033303 | 34033458 | 34028753 | 34028883 | 34035646 | 34035758 | 32 | 12 | 29 | [illegible] |
| SE | 23312 | ENSG00000119862 | LGALSL | chr2 | + | 64455588 | 64455677 | 64455343 | 64455415 | 64458284 | 64461381 | 27 | 37 | 5 | [illegible] |
| SE | 23314 | ENSG00000119862 | LGALSL | chr2 | + | 64456287 | 64456465 | 64455343 | 64455415 | 64458284 | 64458440 | 79 | 97 | 33 | [illegible] |
| SE | 23316 | ENSG00000168268 | NT5DC2 | chr3 | - | 52524816 | 52524881 | 52524384 | 52524611 | 52524962 | 52525103 | 8 | 1 | 11 | [illegible] |
| SE | 23349 | ENSG00000086189 | DIMT1 | chr5 | - | 62351106 | 62351223 | 62347283 | 62349212 | 62352585 | 62352683 | 0 | 0 | 0 | [illegible] |
| SE | 23350 | ENSG00000086189 | DIMT1 | chr5 | - | 62392170 | 62392234 | 62390875 | 62390982 | 62392925 | 62392990 | 213 | 192 | 222 | [illegible] |
| SE | 23351 | ENSG00000086189 | DIMT1 | chr5 | - | 62398428 | 62398560 | 62394483 | 62394607 | 62398645 | 62398739 | 21 | 10 | 27 | [illegible] |
| SE | 23352 | ENSG00000086189 | DIMT1 | chr5 | - | 62398510 | 62398536 | 62394483 | 62394607 | 62398645 | 62398739 | 18 | 11 | 22 | [illegible] |
| SE | 23353 | ENSG00000086189 | DIMT1 | chr5 | - | 62398510 | 62398560 | 62394483 | 62394607 | 62398645 | 62398739 | 90 | 60 | 88 | [illegible] |
| SE | 23354 | ENSG00000086189 | DIMT1 | chr5 | - | 62399780 | 62399878 | 62398819 | 62398881 | 62402035 | 62402122 | 0 | 0 | 0 | [illegible] |
| SE | 23356 | ENSG00000086189 | DIMT1 | chr5 | - | 62402035 | 62402122 | 62398645 | 62398881 | 62403272 | 62403346 | 62 | 33 | 59 | [illegible] |
| SE | 23358 | ENSG00000147044 | CASK | chrX | - | 41523950 | 41524034 | 41514933 | 41520596 | 41531006 | 41531209 | 53 | 28 | 75 | [illegible] |
| SE | 23362 | ENSG00000147044 | CASK | chrX | - | 41553718 | 41553915 | 41534892 | 41534973 | 41559778 | 41559847 | 16 | 16 | 36 | [illegible] |
| SE | 23363 | ENSG00000147044 | CASK | chrX | - | 41553718 | 41553915 | 41542690 | 41542806 | 41555599 | 41555635 | 41 | 38 | 53 | [illegible] |
| SE | 23364 | ENSG00000147044 | CASK | chrX | - | 41553718 | 41553915 | 41542690 | 41542806 | 41559778 | 41559847 | 47 | 44 | 82 | [illegible] |
| SE | 23365 | ENSG00000147044 | CASK | chrX | - | 41553718 | 41553915 | 41542690 | 41542806 | 41561558 | 41561644 | 31 | 28 | 48 | [illegible] |
| SE | 23366 | ENSG00000147044 | CASK | chrX | - | 41555599 | 41555635 | 41542690 | 41542806 | 41559778 | 41559847 | 6 | 1 | 3 | [illegible] |
| SE | 23367 | ENSG00000147044 | CASK | chrX | - | 41555599 | 41555635 | 41553718 | 41553915 | 41557031 | 41557100 | 4 | 8 | 6 | [illegible] |
| SE | 23370 | ENSG00000147044 | CASK | chrX | - | 41557031 | 41557100 | 41555599 | 41555635 | 41559778 | 41559847 | 5 | 3 | 6 | [illegible] |
| SE | 23371 | ENSG00000147044 | CASK | chrX | - | 41559778 | 41559847 | 41542690 | 41542806 | 41561558 | 41561644 | 43 | 51 | 63 | [illegible] |
| SE | 23372 | ENSG00000147044 | CASK | chrX | - | 41559778 | 41559847 | 41553718 | 41553915 | 41561558 | 41561644 | 59 | 67 | 98 | [illegible] |
| SE | 23373 | ENSG00000147044 | CASK | chrX | - | 41569667 | 41569746 | 41561558 | 41561644 | 41578339 | 41578453 | 92 | 88 | 129 | [illegible] |
| SE | 23378 | ENSG00000147044 | CASK | chrX | - | 41787177 | 41787283 | 41745496 | 41745601 | 41853096 | 41853227 | 63 | 52 | 81 | [illegible] |
| SE | 23379 | ENSG00000147044 | CASK | chrX | - | 41787177 | 41787283 | 41745496 | 41745601 | 41853114 | 41853193 | 142 | 97 | 136 | [illegible] |
| SE | 23380 | ENSG00000147044 | CASK | chrX | - | 41853096 | 41853227 | 41787177 | 41787283 | 41922929 | 41923010 | 28 | 27 | 41 | [illegible] |
| SE | 23381 | ENSG00000147044 | CASK | chrX | - | 41853114 | 41853227 | 41786809 | 41787283 | 41922929 | 41922947 | 106 | 70 | 93 | [illegible] |
| SE | 23384 | ENSG00000153113 | CAST | chr5 | + | 96695835 | 96695907 | 96675538 | 96675601 | 96722638 | 96722698 | 386 | 278 | 302 | [illegible] |
| SE | 23385 | ENSG00000153113 | CAST | chr5 | + | 96712348 | 96712473 | 96695835 | 96695907 | 96722638 | 96722698 | 1 | 0 | 1 | [illegible] |
| SE | 23386 | ENSG00000153113 | CAST | chr5 | + | 96712348 | 96712473 | 96702825 | 96702915 | 96722638 | 96722698 | 0 | 0 | 1 | [illegible] |
| SE | 23387 | ENSG00000153113 | CAST | chr5 | + | 96722638 | 96722698 | 96695784 | 96695907 | 96727488 | 96727530 | 552 | 393 | 407 | [illegible] |
| SE | 23388 | ENSG00000153113 | CAST | chr5 | + | 96722638 | 96722698 | 96695784 | 96695907 | 96729152 | 96729209 | 182 | 122 | 146 | [illegible] |
| SE | 23393 | ENSG00000153113 | CAST | chr5 | + | 96727488 | 96727530 | 96695784 | 96695907 | 96729152 | 96729209 | 144 | 115 | 85 | [illegible] |
| SE | 23394 | ENSG00000153113 | CAST | chr5 | + | 96727488 | 96727530 | 96722638 | 96722698 | 96729152 | 96729209 | 506 | 377 | 334 | [illegible] |
| SE | 23396 | ENSG00000153113 | CAST | chr5 | + | 96727488 | 96727530 | 96722638 | 96722718 | 96729152 | 96729209 | 143 | 116 | 86 | [illegible] |
| SE | 23398 | ENSG00000153113 | CAST | chr5 | + | 96729611 | 96729725 | 96727488 | 96727530 | 96736171 | 96736240 | 32 | 30 | 25 | [illegible] |
| SE | 23400 | ENSG00000153113 | CAST | chr5 | + | 96730779 | 96730860 | 96729589 | 96729725 | 96737848 | 96737947 | 508 | 365 | 387 | [illegible] |
| SE | 23401 | ENSG00000153113 | CAST | chr5 | + | 96736171 | 96736240 | 96729611 | 96729725 | 96737848 | 96737947 | 428 | 277 | 272 | [illegible] |
| SE | 23402 | ENSG00000153113 | CAST | chr5 | + | 96736171 | 96736240 | 96730779 | 96730860 | 96737848 | 96737947 | 1236 | 909 | 754 | [illegible] |
| SE | 23404 | ENSG00000153113 | CAST | chr5 | + | 96741265 | 96741358 | 96740037 | 96740118 | 96741493 | 96741580 | 179 | 90 | 71 | [illegible] |
| SE | 23405 | ENSG00000153113 | CAST | chr5 | + | 96742654 | 96742756 | 96741493 | 96741580 | 96746341 | 96746425 | 645 | 387 | 419 | [illegible] |
| SE | 23406 | ENSG00000153113 | CAST | chr5 | + | 96742654 | 96742756 | 96741493 | 96741580 | 96747344 | 96747392 | 309 | 176 | 173 | [illegible] |
| SE | 23407 | ENSG00000153113 | CAST | chr5 | + | 96746341 | 96746425 | 96741493 | 96741580 | 96747344 | 96747392 | 242 | 146 | 121 | [illegible] |
| SE | 23409 | ENSG00000153113 | CAST | chr5 | + | 96746341 | 96746425 | 96743539 | 96743746 | 96747344 | 96747392 | 244 | 148 | 121 | [illegible] |
| SE | 23411 | ENSG00000153113 | CAST | chr5 | + | 96750586 | 96750682 | 96748517 | 96748613 | 96754059 | 96754161 | 715 | 454 | 491 | [illegible] |
| SE | 23412 | ENSG00000153113 | CAST | chr5 | + | 96754059 | 96754161 | 96750586 | 96750682 | 96754657 | 96754741 | 457 | 326 | 348 | [illegible] |
| SE | 23413 | ENSG00000153113 | CAST | chr5 | + | 96757177 | 96757336 | 96754657 | 96754741 | 96757443 | 96757494 | 0 | 0 | 1 | [illegible] |
| SE | 23414 | ENSG00000153113 | CAST | chr5 | + | 96757443 | 96757494 | 96754657 | 96754741 | 96757582 | 96757654 | 673 | 422 | 395 | [illegible] |
| SE | 23416 | ENSG00000153113 | CAST | chr5 | + | 96762273 | 96762372 | 96757582 | 96757654 | 96765223 | 96765325 | 590 | 380 | 364 | [illegible] |
| SE | 23417 | ENSG00000153113 | CAST | chr5 | + | 96770530 | 96770602 | 96767906 | 96767999 | 96771643 | 96771679 | 556 | 360 | 317 | [illegible] |
| SE | 23419 | ENSG00000160201 | U2AF1 | chr21 | - | 43101365 | 43101432 | 43100452 | 43100519 | 43104314 | 43104402 | 0 | 1 | 0 | [illegible] |
| SE | 23421 | ENSG00000148110 | MFSD14B | chr9 | + | 94440984 | 94441099 | 94438400 | 94438528 | 94446822 | 94446936 | 36 | 18 | 30 | [illegible] |
| SE | 23422 | ENSG00000148110 | MFSD14B | chr9 | + | 94444963 | 94445182 | 94438400 | 94438528 | 94446822 | 94446936 | 71 | 36 | 60 | [illegible] |
| SE | 23424 | ENSG00000148110 | MFSD14B | chr9 | + | 94451455 | 94451527 | 94446822 | 94446936 | 94453957 | 94454070 | 229 | 93 | 182 | [illegible] |
| SE | 23426 | ENSG00000148110 | MFSD14B | chr9 | + | 94453957 | 94454070 | 94451455 | 94451527 | 94458321 | 94458464 | 149 | 61 | 126 | [illegible] |
| SE | 23427 | ENSG00000148110 | MFSD14B | chr9 | + | 94456239 | 94456337 | 94451455 | 94451527 | 94458321 | 94458464 | 79 | 30 | 59 | [illegible] |
| SE | 23433 | ENSG00000172062 | SMN1 | chr5 | + | 70946706 | 70946815 | 70946065 | 70946176 | 70952438 | 70953016 | 0 | 0 | 0 | [illegible] |
| SE | 23440 | ENSG00000123575 | FAM199X | chrX | + | 104186065 | 104186215 | 104175622 | 104175842 | 104186459 | 104186621 | 361 | 201 | 209 | [illegible] |
| SE | 23442 | ENSG00000166888 | STAT6 | chr12 | - | 57099293 | 57099440 | 57099014 | 57099078 | 57099995 | 57100090 | 474 | 272 | 371 | [illegible] |
| SE | 23446 | ENSG00000166888 | STAT6 | chr12 | - | 57104463 | 57104586 | 57102828 | 57102921 | 57104725 | 57104813 | 566 | 253 | 378 | [illegible] |
| SE | 23447 | ENSG00000166888 | STAT6 | chr12 | - | 57106190 | 57106339 | 57105467 | 57105599 | 57106527 | 57106580 | 988 | 596 | 733 | [illegible] |
| SE | 23449 | ENSG00000166888 | STAT6 | chr12 | - | 57106692 | 57106831 | 57106527 | 57106580 | 57107230 | 57107314 | 814 | 469 | 650 | [illegible] |
| SE | 23451 | ENSG00000166888 | STAT6 | chr12 | - | 57107604 | 57107743 | 57107230 | 57107314 | 57111109 | 57111374 | 192 | 109 | 163 | [illegible] |
| SE | 23454 | ENSG00000166888 | STAT6 | chr12 | - | 57108162 | 57108299 | 57107230 | 57107314 | 57111109 | 57111374 | 119 | 65 | 97 | [illegible] |
| SE | 23455 | ENSG00000166888 | STAT6 | chr12 | - | 57108162 | 57108299 | 57107230 | 57107314 | 57111128 | 57111231 | 659 | 341 | 449 | [illegible] |
| SE | 23456 | ENSG00000166888 | STAT6 | chr12 | - | 57108162 | 57108299 | 57107604 | 57107743 | 57111109 | 57111374 | 455 | 204 | 324 | [illegible] |
| SE | 23457 | ENSG00000166888 | STAT6 | chr12 | - | 57108162 | 57108299 | 57107604 | 57107743 | 57111128 | 57111231 | 997 | 480 | 678 | [illegible] |
| SE | 23462 | ENSG00000113282 | CLINT1 | chr5 | - | 157789362 | 157789513 | 157785746 | 157787992 | 157791702 | 157791724 | 401 | 292 | 378 | [illegible] |
| SE | 23463 | ENSG00000113282 | CLINT1 | chr5 | - | 157789362 | 157789567 | 157785742 | 157787992 | 157791702 | 157791995 | 283 | 206 | 272 | [illegible] |
| SE | 23468 | ENSG00000113282 | CLINT1 | chr5 | - | 157809627 | 157809805 | 157805865 | 157806112 | 157813062 | 157813227 | 845 | 658 | 739 | [illegible] |
| SE | 23469 | ENSG00000171530 | TBCA | chr5 | - | 77693265 | 77693352 | 77691165 | 77691498 | 77708241 | 77708347 | 956 | 553 | 862 | [illegible] |
| SE | 23472 | ENSG00000153140 | CETN3 | chr5 | - | 90396492 | 90396564 | 90393709 | 90394107 | 90399357 | 90399549 | 0 | 0 | 2 | [illegible] |
| SE | 23477 | ENSG00000131446 | MGAT1 | chr5 | - | 180795837 | 180795875 | 180792372 | 180793097 | 180803601 | 180803904 | 0 | 0 | 1 | [illegible] |
| SE | 23479 | ENSG00000131446 | MGAT1 | chr5 | - | 180795837 | 180795875 | 180792372 | 180793097 | 180808501 | 180808758 | 0 | 1 | 1 | [illegible] |
| SE | 23480 | ENSG00000131446 | MGAT1 | chr5 | - | 180795837 | 180795875 | 180792372 | 180793097 | 180808647 | 180809076 | 1 | 0 | 1 | [illegible] |
| SE | 23482 | ENSG00000131446 | MGAT1 | chr5 | - | 180808501 | 180808597 | 180793061 | 180793097 | 180815413 | 180815652 | 2 | 3 | 7 | [illegible] |
| SE | 23484 | ENSG00000196670 | ZFP62 | chr5 | - | 180851987 | 180852060 | 180847614 | 180851493 | 180860527 | 180860663 | 11 | 2 | 5 | [illegible] |
| SE | 23485 | ENSG00000196670 | ZFP62 | chr5 | - | 180851987 | 180852060 | 180847614 | 180851493 | 180861218 | 180861276 | 5 | 1 | 3 | [illegible] |
| SE | 23497 | ENSG00000129625 | REEP5 | chr5 | - | 112887014 | 112887183 | 112876384 | 112878835 | 112902379 | 112902518 | 838 | 541 | 614 | [illegible] |
| SE | 23498 | ENSG00000129625 | REEP5 | chr5 | - | 112887014 | 112887183 | 112876384 | 112878835 | 112921162 | 112921256 | 468 | 302 | 333 | [illegible] |
| SE | 23499 | ENSG00000129625 | REEP5 | chr5 | - | 112892449 | 112892817 | 112887014 | 112887183 | 112902379 | 112902518 | 3 | 5 | 0 | [illegible] |
| SE | 23500 | ENSG00000129625 | REEP5 | chr5 | - | 112902379 | 112902518 | 112878084 | 112878835 | 112921162 | 112921256 | 236 | 119 | 195 | [illegible] |
| SE | 23501 | ENSG00000129625 | REEP5 | chr5 | - | 112902379 | 112902518 | 112887014 | 112887183 | 112921162 | 112921256 | 611 | 360 | 473 | [illegible] |
| SE | 23502 | ENSG00000113068 | PFDN1 | chr5 | - | 140281448 | 140281533 | 140245034 | 140246057 | 140300415 | 140300582 | 146 | 135 | 120 | [illegible] |
| SE | 23505 | ENSG00000154822 | PLCL2 | chr3 | + | 17014707 | 17014911 | 17009673 | 17012160 | 17067955 | 17068065 | 210 | 139 | 223 | [illegible] |
| SE | 23506 | ENSG00000154822 | PLCL2 | chr3 | + | 17042873 | 17042949 | 17009673 | 17012160 | 17067955 | 17068065 | 126 | 117 | 114 | [illegible] |
| SE | 23507 | ENSG00000154822 | PLCL2 | chr3 | + | 17042873 | 17042949 | 17014707 | 17014911 | 17067955 | 17068065 | 342 | 301 | 359 | [illegible] |
| SE | 23508 | ENSG00000154822 | PLCL2 | chr3 | + | 17067955 | 17068065 | 17042873 | 17042949 | 17089732 | 17090593 | 368 | 297 | 383 | [illegible] |
| SE | 23514 | ENSG00000100284 | TOM1 | chr22 | + | 35321958 | 35322037 | 35317876 | 35317961 | 35322998 | 35323177 | 47 | 38 | 60 | [illegible] |
| SE | 23519 | ENSG00000100284 | TOM1 | chr22 | + | 35330346 | 35330480 | 35327270 | 35327387 | 35332980 | 35333014 | 154 | 67 | 126 | [illegible] |
| SE | 23520 | ENSG00000100284 | TOM1 | chr22 | + | 35331295 | 35331495 | 35327270 | 35327387 | 35332980 | 35333014 | 3 | 0 | 0 | [illegible] |
| SE | 23521 | ENSG00000100284 | TOM1 | chr22 | + | 35331295 | 35331495 | 35330346 | 35330480 | 35332980 | 35333014 | 3 | 0 | 0 | [illegible] |
| SE | 23524 | ENSG00000100284 | TOM1 | chr22 | + | 35346929 | 35346969 | 35344185 | 35345784 | 35347054 | 35347973 | 52 | 26 | 50 | [illegible] |
| SE | 23525 | ENSG00000100284 | TOM1 | chr22 | + | 35346929 | 35346972 | 35345724 | 35345784 | 35347054 | 35347992 | 28 | 9 | 26 | [illegible] |
| SE | 23527 | ENSG00000104497 | SNX16 | chr8 | - | 81823791 | 81823940 | 81803091 | 81803228 | 81829429 | 81829516 | 54 | 40 | 37 | [illegible] |
| SE | 23528 | ENSG00000104497 | SNX16 | chr8 | - | 81823791 | 81823940 | 81803091 | 81803228 | 81839611 | 81840082 | 20 | 4 | 7 | [illegible] |
| SE | 23529 | ENSG00000104497 | SNX16 | chr8 | - | 81829429 | 81829516 | 81803091 | 81803228 | 81839611 | 81839751 | 20 | 16 | 21 | [illegible] |
| SE | 23531 | ENSG00000104497 | SNX16 | chr8 | - | 81839838 | 81840082 | 81839611 | 81839751 | 81842121 | 81842183 | 26 | 11 | 18 | [illegible] |
| SE | 23538 | ENSG00000163635 | ATXN7 | chr3 | + | 63864897 | 63864999 | 63864498 | 63864688 | 63898398 | 63898497 | 5 | 2 | 4 | [illegible] |
| SE | 23541 | ENSG00000163635 | ATXN7 | chr3 | + | 63873748 | 63873846 | 63864556 | 63864999 | 63898398 | 63898497 | 0 | 0 | 2 | [illegible] |
| SE | 23542 | ENSG00000163635 | ATXN7 | chr3 | + | 63873748 | 63873846 | 63864639 | 63864688 | 63898398 | 63898497 | 0 | 0 | 2 | [illegible] |
| SE | 23544 | ENSG00000163635 | ATXN7 | chr3 | + | 63912587 | 63912923 | 63898398 | 63898497 | 63952378 | 63952483 | 3 | 8 | 10 | [illegible] |
| SE | 23546 | ENSG00000163635 | ATXN7 | chr3 | + | 63952378 | 63952483 | 63913156 | 63913225 | 63979914 | 63980167 | 110 | 78 | 85 | [illegible] |
| SE | 23548 | ENSG00000163635 | ATXN7 | chr3 | + | 63988058 | 63988324 | 63982938 | 63983021 | 63990175 | 63990374 | 329 | 251 | 289 | [illegible] |
| SE | 23549 | ENSG00000163635 | ATXN7 | chr3 | + | 63988058 | 63988324 | 63982938 | 63983021 | 63990737 | 63990859 | 283 | 222 | 250 | [illegible] |
| SE | 23550 | ENSG00000163635 | ATXN7 | chr3 | + | 63990175 | 63990374 | 63982938 | 63983021 | 63990737 | 63990859 | 141 | 84 | 180 | [illegible] |
| SE | 23555 | ENSG00000124532 | MRS2 | chr6 | + | 24405167 | 24405241 | 24402907 | 24403236 | 24408407 | 24408444 | 34 | 13 | 53 | [illegible] |
| SE | 23556 | ENSG00000124532 | MRS2 | chr6 | + | 24408407 | 24408444 | 24405167 | 24405241 | 24412221 | 24412395 | 7 | 4 | 17 | [illegible] |
| SE | 23557 | ENSG00000124532 | MRS2 | chr6 | + | 24409460 | 24409573 | 24408407 | 24408444 | 24412221 | 24412395 | 47 | 46 | 61 | [illegible] |
| SE | 23559 | ENSG00000124532 | MRS2 | chr6 | + | 24415032 | 24415163 | 24412221 | 24412395 | 24416396 | 24416513 | 101 | 52 | 119 | [illegible] |
| SE | 23565 | ENSG00000204574 | ABCF1 | chr6 | + | 30579930 | 30580005 | 30578469 | 30578577 | 30580405 | 30580448 | 167 | 146 | 178 | [illegible] |
| SE | 23567 | ENSG00000204574 | ABCF1 | chr6 | + | 30584191 | 30584331 | 30583804 | 30583890 | 30584417 | 30584566 | 129 | 75 | 127 | [illegible] |
| SE | 23568 | ENSG00000204574 | ABCF1 | chr6 | + | 30590305 | 30590378 | 30590148 | 30590213 | 30590534 | 30590743 | 113 | 55 | 99 | [illegible] |
| SE | 23570 | ENSG00000254996 | ANKHD1-EIF4EBP3 | chr5 | + | 140485120 | 140485248 | 140482579 | 140482667 | 140485588 | 140485732 | 297 | 196 | 248 | [illegible] |
| SE | 23572 | ENSG00000254996 | ANKHD1-EIF4EBP3 | chr5 | + | 140506834 | 140506977 | 140505723 | 140505869 | 140509636 | 140509812 | 344 | 289 | 325 | [illegible] |
| SE | 23573 | ENSG00000131503 | ANKHD1 | chr5 | + | 140507784 | 140507998 | 140506834 | 140506977 | 140509636 | 140509812 | 492 | 348 | 446 | [illegible] |
| SE | 23575 | ENSG00000254996 | ANKHD1-EIF4EBP3 | chr5 | + | 140513362 | 140513479 | 140510018 | 140510181 | 140524065 | 140524240 | 130 | 107 | 124 | [illegible] |
| SE | 23589 | ENSG00000188725 | SMIM15 | chr5 | - | 61162216 | 61162359 | 61161087 | 61161227 | 61162445 | 61162468 | 88 | 28 | 51 | [illegible] |
| SE | 23590 | ENSG00000112941 | TENT4A | chr5 | + | 6739731 | 6739852 | 6738682 | 6738729 | 6746213 | 6746427 | 44 | 50 | 79 | [illegible] |
| SE | 23591 | ENSG00000112941 | TENT4A | chr5 | + | 6742489 | 6742597 | 6739731 | 6739852 | 6746213 | 6746427 | 47 | 34 | 80 | [illegible] |
| SE | 23594 | ENSG00000112941 | TENT4A | chr5 | + | 6751038 | 6751197 | 6750330 | 6750503 | 6754750 | 6755030 | 33 | 34 | 37 | [illegible] |
| SE | 23595 | ENSG00000112941 | TENT4A | chr5 | + | 6752872 | 6753037 | 6751038 | 6751197 | 6754750 | 6755030 | 36 | 34 | 39 | [illegible] |
| SE | 23597 | ENSG00000116991 | SIPA1L2 | chr1 | - | 232404124 | 232404178 | 232403447 | 232403571 | 232415493 | 232415625 | 0 | 0 | 0 | [illegible] |
| SE | 23598 | ENSG00000116991 | SIPA1L2 | chr1 | - | 232419874 | 232419920 | 232415493 | 232415625 | 232425588 | 232425808 | 0 | 0 | 0 | [illegible] |
| SE | 23600 | ENSG00000116991 | SIPA1L2 | chr1 | - | 232439107 | 232439496 | 232432246 | 232432471 | 232441290 | 232441394 | 11 | 11 | 10 | [illegible] |
| SE | 23602 | ENSG00000116991 | SIPA1L2 | chr1 | - | 232471370 | 232471528 | 232464839 | 232465416 | 232479626 | 232479730 | 37 | 26 | 26 | [illegible] |
| SE | 23603 | ENSG00000116991 | SIPA1L2 | chr1 | - | 232479626 | 232479730 | 232471370 | 232471528 | 232483791 | 232483966 | 30 | 29 | 20 | [illegible] |
| SE | 23604 | ENSG00000116991 | SIPA1L2 | chr1 | - | 232493526 | 232493660 | 232490685 | 232491062 | 232513856 | 232515566 | 53 | 33 | 27 | [illegible] |
| SE | 23606 | ENSG00000116991 | SIPA1L2 | chr1 | - | 232533489 | 232533583 | 232513856 | 232515608 | 232629868 | 232630122 | 1 | 1 | 7 | [illegible] |
| SE | 23607 | ENSG00000116991 | SIPA1L2 | chr1 | - | 232574173 | 232574222 | 232513856 | 232515608 | 232629868 | 232630133 | 1 | 2 | 1 | [illegible] |
| SE | 23609 | ENSG00000116991 | SIPA1L2 | chr1 | - | 232574173 | 232574222 | 232533489 | 232533583 | 232629868 | 232630133 | 0 | 0 | 0 | [illegible] |
| SE | 23610 | ENSG00000116991 | SIPA1L2 | chr1 | - | 232608826 | 232608988 | 232574173 | 232574222 | 232629868 | 232630122 | 0 | 0 | 0 | [illegible] |
| SE | 23611 | ENSG00000116991 | SIPA1L2 | chr1 | - | 232608911 | 232608988 | 232574173 | 232574222 | 232629868 | 232630109 | 1 | 8 | 1 | [illegible] |
| SE | 23612 | ENSG00000113558 | SKP1 | chr5 | - | 134160986 | 134161130 | 134157337 | 134158595 | 134167169 | 134167243 | 2334 | 1598 | 1832 | [illegible] |
| SE | 23613 | ENSG00000113558 | SKP1 | chr5 | - | 134167169 | 134167243 | 134160986 | 134161130 | 134173925 | 134174022 | 735 | 449 | 609 | [illegible] |
| SE | 23614 | ENSG00000113558 | SKP1 | chr5 | - | 134170950 | 134171038 | 134167169 | 134167243 | 134173700 | 134173766 | 0 | 0 | 1 | [illegible] |
| SE | 23615 | ENSG00000113558 | SKP1 | chr5 | - | 134170950 | 134171038 | 134167169 | 134167243 | 134173925 | 134174022 | 4 | 1 | 2 | [illegible] |
| SE | 23617 | ENSG00000113558 | SKP1 | chr5 | - | 134173700 | 134173766 | 134170950 | 134171038 | 134173925 | 134174022 | 0 | 1 | 5 | [illegible] |
| SE | 23618 | ENSG00000113558 | SKP1 | chr5 | - | 134174461 | 134174514 | 134173925 | 134174022 | 134176613 | 134176719 | 2 | 1 | 7 | [illegible] |
| SE | 23624 | ENSG00000135074 | ADAM19 | chr5 | - | 157490309 | 157490454 | 157489101 | 157489186 | 157491614 | 157491723 | 184 | 109 | 263 | [illegible] |
| SE | 23627 | ENSG00000135074 | ADAM19 | chr5 | - | 157505668 | 157505808 | 157499572 | 157499662 | 157507055 | 157507140 | 78 | 28 | 62 | [illegible] |
| SE | 23628 | ENSG00000135074 | ADAM19 | chr5 | - | 157507055 | 157507140 | 157505668 | 157505808 | 157509300 | 157509467 | 127 | 45 | 144 | [illegible] |
| SE | 23629 | ENSG00000135074 | ADAM19 | chr5 | - | 157513423 | 157513505 | 157509300 | 157509467 | 157518822 | 157518888 | 35 | 40 | 47 | [illegible] |
| SE | 23630 | ENSG00000135074 | ADAM19 | chr5 | - | 157513433 | 157513505 | 157509300 | 157509467 | 157518822 | 157518888 | 114 | 73 | 122 | [illegible] |
| SE | 23633 | ENSG00000135074 | ADAM19 | chr5 | - | 157530806 | 157530883 | 157519838 | 157520031 | 157537912 | 157537991 | 198 | 118 | 208 | [illegible] |
| SE | 23634 | ENSG00000135074 | ADAM19 | chr5 | - | 157564372 | 157564443 | 157537912 | 157537991 | 157570894 | 157570980 | 141 | 80 | 154 | [illegible] |
| SE | 23635 | ENSG00000113263 | ITK | chr5 | + | 157211286 | 157211368 | 157208888 | 157208993 | 157214190 | 157214319 | 608 | 363 | 726 | [illegible] |
| SE | 23636 | ENSG00000113263 | ITK | chr5 | + | 157211286 | 157211368 | 157208888 | 157208993 | 157217866 | 157217907 | 184 | 105 | 214 | [illegible] |
| SE | 23638 | ENSG00000113263 | ITK | chr5 | + | 157214190 | 157214319 | 157208888 | 157208993 | 157217866 | 157217907 | 175 | 104 | 220 | [illegible] |
| SE | 23639 | ENSG00000113263 | ITK | chr5 | + | 157214190 | 157214319 | 157211286 | 157211368 | 157217866 | 157217907 | 600 | 367 | 748 | [illegible] |
| SE | 23640 | ENSG00000113263 | ITK | chr5 | + | 157222862 | 157223014 | 157217866 | 157217907 | 157228295 | 157228361 | 644 | 414 | 765 | [illegible] |
| SE | 23642 | ENSG00000113263 | ITK | chr5 | + | 157238108 | 157238191 | 157232339 | 157232394 | 157240061 | 157240131 | 684 | 358 | 803 | [illegible] |
| SE | 23643 | ENSG00000113263 | ITK | chr5 | + | 157240061 | 157240195 | 157238108 | 157238191 | 157241645 | 157241720 | 679 | 341 | 796 | [illegible] |
| SE | 23644 | ENSG00000113263 | ITK | chr5 | + | 157240061 | 157240195 | 157238108 | 157238191 | 157243622 | 157243794 | 419 | 196 | 505 | [illegible] |
| SE | 23645 | ENSG00000113263 | ITK | chr5 | + | 157240061 | 157241720 | 157238108 | 157238191 | 157243622 | 157243794 | 686 | 378 | 803 | [illegible] |
| SE | 23646 | ENSG00000113263 | ITK | chr5 | + | 157241645 | 157241720 | 157238108 | 157238191 | 157243622 | 157243794 | 247 | 165 | 279 | [illegible] |
| SE | 23647 | ENSG00000113263 | ITK | chr5 | + | 157241645 | 157241720 | 157240061 | 157240195 | 157243622 | 157243794 | 513 | 315 | 576 | [illegible] |
| SE | 23648 | ENSG00000113263 | ITK | chr5 | + | 157241645 | 157241720 | 157240061 | 157240195 | 157245725 | 157245790 | 156 | 88 | 197 | [illegible] |
| SE | 23649 | ENSG00000113263 | ITK | chr5 | + | 157243622 | 157243794 | 157240061 | 157240195 | 157245725 | 157245790 | 11 | 10 | 8 | [illegible] |
| SE | 23650 | ENSG00000113263 | ITK | chr5 | + | 157243622 | 157243794 | 157240061 | 157241720 | 157245725 | 157245790 | 387 | 252 | 410 | [illegible] |
| SE | 23651 | ENSG00000113263 | ITK | chr5 | + | 157243622 | 157243794 | 157240061 | 157241720 | 157245880 | 157245999 | 377 | 244 | 403 | [illegible] |
| SE | 23653 | ENSG00000113263 | ITK | chr5 | + | 157244261 | 157244478 | 157243622 | 157243794 | 157245880 | 157245999 | 224 | 117 | 332 | [illegible] |
| SE | 23654 | ENSG00000113263 | ITK | chr5 | + | 157245725 | 157245790 | 157240061 | 157241720 | 157245880 | 157245999 | 84 | 63 | 98 | [illegible] |
| SE | 23655 | ENSG00000113263 | ITK | chr5 | + | 157245725 | 157245790 | 157243622 | 157243794 | 157245880 | 157245999 | 93 | 71 | 104 | [illegible] |
| SE | 23656 | ENSG00000113263 | ITK | chr5 | + | 157245725 | 157245790 | 157244261 | 157244478 | 157245880 | 157245999 | 212 | 129 | 271 | [illegible] |
| SE | 23657 | ENSG00000116906 | GNPAT | chr1 | + | 231250960 | 231251143 | 231241211 | 231241456 | 231258221 | 231258309 | 44 | 24 | 39 | [illegible] |
| SE | 23659 | ENSG00000116906 | GNPAT | chr1 | + | 231258221 | 231258309 | 231241215 | 231241456 | 231260506 | 231260683 | 2 | 5 | 2 | [illegible] |
| SE | 23661 | ENSG00000116906 | GNPAT | chr1 | + | 231262722 | 231262852 | 231260506 | 231260683 | 231265292 | 231265420 | 189 | 117 | 197 | [illegible] |
| SE | 23664 | ENSG00000116906 | GNPAT | chr1 | + | 231267679 | 231267903 | 231266276 | 231266407 | 231269862 | 231270056 | 63 | 32 | 56 | [illegible] |
| SE | 23666 | ENSG00000116906 | GNPAT | chr1 | + | 231267679 | 231267911 | 231266276 | 231266407 | 231269862 | 231270056 | 61 | 30 | 56 | [illegible] |
| SE | 23668 | ENSG00000116906 | GNPAT | chr1 | + | 231270757 | 231271000 | 231267679 | 231267903 | 231272311 | 231272391 | 84 | 57 | 66 | [illegible] |
| SE | 23669 | ENSG00000116906 | GNPAT | chr1 | + | 231276134 | 231276196 | 231275404 | 231275498 | 231277498 | 231277530 | 196 | 179 | 157 | [illegible] |
| SE | 23672 | ENSG00000133466 | C1QTNF6 | chr22 | - | 37185870 | 37186046 | 37185217 | 37185455 | 37190864 | 37190928 | 3 | 0 | 0 | [illegible] |
| SE | 23673 | ENSG00000133466 | C1QTNF6 | chr22 | - | 37188162 | 37188476 | 37185870 | 37186046 | 37190864 | 37190928 | 0 | 0 | 0 | [illegible] |
| SE | 23675 | ENSG00000133466 | C1QTNF6 | chr22 | - | 37191671 | 37191737 | 37190864 | 37190928 | 37197602 | 37197748 | 9 | 1 | 6 | [illegible] |
| SE | 23676 | ENSG00000133466 | C1QTNF6 | chr22 | - | 37195380 | 37195458 | 37190864 | 37190928 | 37197602 | 37197748 | 0 | 5 | 1 | [illegible] |
| SE | 23677 | ENSG00000133466 | C1QTNF6 | chr22 | - | 37195380 | 37195458 | 37191671 | 37191737 | 37197602 | 37197748 | 2 | 0 | 3 | [illegible] |
| SE | 23678 | ENSG00000133466 | C1QTNF6 | chr22 | - | 37195380 | 37195512 | 37190864 | 37190928 | 37197602 | 37197841 | 2 | 7 | 5 | [illegible] |
| SE | 23679 | ENSG00000133466 | C1QTNF6 | chr22 | - | 37195380 | 37195512 | 37191671 | 37191737 | 37197602 | 37197841 | 4 | 3 | 8 | [illegible] |
| SE | 23686 | ENSG00000112977 | DAP | chr5 | - | 10748174 | 10748271 | 10683095 | 10683571 | 10761013 | 10761136 | 82 | 50 | 34 | [illegible] |
| SE | 23690 | ENSG00000132842 | AP3B1 | chr5 | - | 78015409 | 78015548 | 78002325 | 78003055 | 78020691 | 78020789 | 178 | 107 | 157 | [illegible] |
| SE | 23691 | ENSG00000132842 | AP3B1 | chr5 | - | 78015409 | 78015548 | 78002325 | 78003055 | 78034360 | 78034445 | 92 | 62 | 82 | [illegible] |
| SE | 23692 | ENSG00000132842 | AP3B1 | chr5 | - | 78020691 | 78020789 | 78002325 | 78003055 | 78034360 | 78034445 | 78 | 66 | 68 | [illegible] |
| SE | 23693 | ENSG00000132842 | AP3B1 | chr5 | - | 78020691 | 78020789 | 78015409 | 78015548 | 78034360 | 78034445 | 158 | 108 | 142 | [illegible] |
| SE | 23694 | ENSG00000132842 | AP3B1 | chr5 | - | 78089392 | 78089499 | 78039042 | 78039274 | 78100952 | 78101025 | 296 | 259 | 212 | [illegible] |
| SE | 23695 | ENSG00000132842 | AP3B1 | chr5 | - | 78141142 | 78141319 | 78129120 | 78129307 | 78156257 | 78156367 | 361 | 278 | 301 | [illegible] |
| SE | 23696 | ENSG00000132842 | AP3B1 | chr5 | - | 78175625 | 78175697 | 78165609 | 78165672 | 78175783 | 78175838 | 292 | 208 | 211 | [illegible] |
| SE | 23697 | ENSG00000132842 | AP3B1 | chr5 | - | 78175783 | 78175838 | 78175625 | 78175697 | 78177338 | 78177436 | 295 | 291 | 201 | [illegible] |
| SE | 23698 | ENSG00000132842 | AP3B1 | chr5 | - | 78177338 | 78177436 | 78175783 | 78175838 | 78181506 | 78181662 | 458 | 423 | 313 | [illegible] |
| SE | 23699 | ENSG00000132842 | AP3B1 | chr5 | - | 78267519 | 78267595 | 78240861 | 78240936 | 78294451 | 78294698 | 133 | 130 | 115 | [illegible] |
| SE | 23700 | ENSG00000164398 | ACSL6 | chr5 | - | 131962534 | 131962624 | 131960519 | 131960621 | 131966415 | 131966532 | 24 | 28 | 54 | [illegible] |
| SE | 23701 | ENSG00000164398 | ACSL6 | chr5 | - | 131962534 | 131962678 | 131960519 | 131960621 | 131966415 | 131966532 | 51 | 61 | 127 | [illegible] |

[illegible]

| | | | | | | | | | | | | | | |
|---|---|---|---|---|---|---|---|---|---|---|---|---|---|---|
| SE | 24196 | ENSG00000104731 | KLHDC4 | chr16 | - | 87755193 | 87755292 | 87748672 | 87748809 | 87761948 | 87762040 | 95 | 49 | 70 [illegible] |
| SE | 24198 | ENSG00000104731 | KLHDC4 | chr16 | - | 87755193 | 87756477 | 87748672 | 87748809 | 87761948 | 87762016 | 156 | 64 | 96 [illegible] |
| SE | 24207 | ENSG00000104731 | KLHDC4 | chr16 | - | 87765065 | 87765340 | 87761948 | 87762150 | 87765791 | 87765961 | 0 | 0 | 1 [illegible] |
| SE | 24210 | ENSG00000113742 | CPEB4 | chr5 | + | 173932449 | 173932500 | 173910522 | 173910604 | 173942883 | 173943049 | 16 | 15 | 14 [illegible] |
| SE | 24214 | ENSG00000113742 | CPEB4 | chr5 | + | 173932449 | 173932500 | 173918215 | 173918254 | 173944966 | 173945092 | 12 | 7 | 6 [illegible] |
| SE | 24218 | ENSG00000113742 | CPEB4 | chr5 | + | 173943025 | 173943049 | 173918215 | 173918254 | 173944966 | 173945140 | 0 | 2 | 1 [illegible] |
| SE | 24222 | ENSG00000237693 | IRGM | chr5 | + | 150877979 | 150878114 | 150848496 | 150848654 | 150879584 | 150879646 | 2 | 5 | 0 [illegible] |
| SE | 24225 | ENSG00000006837 | CDKL3 | chr5 | - | 134308573 | 134308727 | 134308000 | 134308466 | 134312291 | 134312380 | 0 | 3 | 4 [illegible] |
| SE | 24226 | ENSG00000273345 | NA | chr5 | - | 134308573 | 134308727 | 134308137 | 134308466 | 134319357 | 134319497 | 0 | 2 | 3 [illegible] |
| SE | 24227 | ENSG00000273345 | NA | chr5 | - | 134308573 | 134308727 | 134308137 | 134308466 | 134321790 | 134321903 | 0 | 2 | 3 [illegible] |
| SE | 24228 | ENSG00000273345 | NA | chr5 | - | 134312291 | 134312380 | 134308137 | 134308466 | 134319357 | 134319497 | 2 | 1 | 2 [illegible] |
| SE | 24229 | ENSG00000273345 | NA | chr5 | - | 134312291 | 134312380 | 134308137 | 134308466 | 134321790 | 134321903 | 3 | 0 | 0 [illegible] |
| SE | 24230 | ENSG00000006837 | CDKL3 | chr5 | - | 134312291 | 134312380 | 134308573 | 134308727 | 134319357 | 134319497 | 1 | 2 | 3 [illegible] |
| SE | 24232 | ENSG00000273345 | NA | chr5 | - | 134319357 | 134319497 | 134308137 | 134308466 | 134321790 | 134321903 | 3 | 0 | 3 [illegible] |
| SE | 24234 | ENSG00000273345 | NA | chr5 | - | 134319357 | 134319497 | 134312291 | 134312380 | 134321790 | 134321903 | 4 | 1 | 3 [illegible] |
| SE | 24236 | ENSG00000155660 | PDIA4 | chr7 | - | 149005140 | 149005374 | 149003061 | 149004209 | 149005896 | 149006053 | 630 | 302 | 793 [illegible] |
| SE | 24237 | ENSG00000155660 | PDIA4 | chr7 | - | 149005896 | 149006053 | 149005140 | 149005374 | 149008158 | 149008310 | 659 | 322 | 837 [illegible] |
| SE | 24238 | ENSG00000155660 | PDIA4 | chr7 | - | 149005896 | 149006056 | 149005140 | 149005374 | 149008158 | 149008310 | 374 | 190 | 474 [illegible] |
| SE | 24239 | ENSG00000155660 | PDIA4 | chr7 | - | 149020966 | 149021147 | 149018991 | 149019197 | 149028320 | 149028505 | 709 | 397 | 833 [illegible] |
| SE | 24240 | ENSG00000155660 | PDIA4 | chr7 | - | 149027848 | 149027992 | 149020966 | 149021147 | 149028320 | 149028468 | 2 | 0 | 1 [illegible] |
| SE | 24242 | ENSG00000182985 | CADM1 | chr11 | - | 115198405 | 115198438 | 115178643 | 115178775 | 115209573 | 115209657 | 1 | 0 | 2 [illegible] |
| SE | 24252 | ENSG00000196569 | LAMA2 | chr6 | + | 129490981 | 129491079 | 129490682 | 129490795 | 129491900 | 129492077 | 0 | 0 | 0 [illegible] |
| SE | 24268 | ENSG00000115758 | ODC1 | chr2 | - | 10443229 | 10443313 | 10442011 | 10442174 | 10443489 | 10443571 | 225 | 274 | 195 [illegible] |
| SE | 24269 | ENSG00000115758 | ODC1 | chr2 | - | 10444094 | 10444267 | 10443701 | 10443836 | 10444473 | 10444547 | 183 | 171 | 138 [illegible] |
| SE | 24271 | ENSG00000115758 | ODC1 | chr2 | - | 10444473 | 10444547 | 10444094 | 10444267 | 10444930 | 10445049 | 79 | 72 | 55 [illegible] |
| SE | 24273 | ENSG00000115758 | ODC1 | chr2 | - | 10444473 | 10444647 | 10444094 | 10444267 | 10444930 | 10445049 | 255 | 246 | 209 [illegible] |
| SE | 24274 | ENSG00000115758 | ODC1 | chr2 | - | 10445154 | 10445264 | 10444930 | 10445049 | 10448120 | 10448309 | 137 | 134 | 137 [illegible] |
| SE | 24282 | ENSG00000104671 | DCTN6 | chr8 | + | 30175084 | 30175190 | 30164110 | 30164175 | 30177125 | 30177214 | 100 | 89 | 80 [illegible] |
| SE | 24283 | ENSG00000104671 | DCTN6 | chr8 | + | 30175084 | 30175190 | 30165715 | 30165901 | 30177125 | 30177214 | 37 | 46 | 28 [illegible] |
| SE | 24286 | ENSG00000196458 | ZNF605 | chr12 | - | 132932712 | 132932805 | 132921461 | 132927162 | 132933034 | 132933155 | 4 | 6 | 0 [illegible] |
| SE | 24287 | ENSG00000196458 | ZNF605 | chr12 | - | 132948147 | 132948270 | 132944797 | 132945797 | 132956242 | 132956262 | 8 | 0 | 7 [illegible] |
| SE | 24290 | ENSG00000125846 | ZNF133 | chr20 | + | 18289898 | 18290074 | 18288282 | 18288604 | 18305680 | 18305807 | 0 | 2 | 0 [illegible] |
| SE | 24292 | ENSG00000125846 | ZNF133 | chr20 | + | 18297984 | 18298062 | 18288282 | 18288604 | 18305667 | 18305807 | 1 | 0 | 0 [illegible] |
| SE | 24294 | ENSG00000125846 | ZNF133 | chr20 | + | 18298288 | 18298464 | 18297984 | 18298062 | 18305007 | 18305178 | 3 | 2 | 5 [illegible] |
| SE | 24295 | ENSG00000125846 | ZNF133 | chr20 | + | 18298288 | 18298464 | 18297984 | 18298062 | 18305667 | 18305807 | 2 | 1 | 5 [illegible] |
| SE | 24296 | ENSG00000125846 | ZNF133 | chr20 | + | 18305007 | 18305178 | 18288282 | 18288604 | 18305667 | 18305807 | 0 | 1 | 2 [illegible] |
| SE | 24297 | ENSG00000125846 | ZNF133 | chr20 | + | 18305007 | 18305178 | 18288282 | 18288604 | 18305680 | 18305807 | 2 | 0 | 0 [illegible] |
| SE | 24298 | ENSG00000125846 | ZNF133 | chr20 | + | 18305007 | 18305178 | 18288282 | 18288604 | 18306297 | 18306393 | 2 | 0 | 0 [illegible] |
| SE | 24300 | ENSG00000125846 | ZNF133 | chr20 | + | 18305007 | 18305178 | 18289898 | 18290074 | 18305680 | 18305807 | 4 | 0 | 0 [illegible] |
| SE | 24301 | ENSG00000125846 | ZNF133 | chr20 | + | 18305007 | 18305178 | 18297984 | 18298062 | 18305667 | 18305807 | 0 | 1 | 2 [illegible] |
| SE | 24303 | ENSG00000125846 | ZNF133 | chr20 | + | 18305007 | 18305178 | 18298288 | 18298464 | 18305667 | 18305807 | 1 | 2 | 2 [illegible] |
| SE | 24305 | ENSG00000125846 | ZNF133 | chr20 | + | 18305667 | 18305807 | 18305007 | 18305178 | 18306297 | 18306393 | 4 | 3 | 6 [illegible] |
| SE | 24307 | ENSG00000125846 | ZNF133 | chr20 | + | 18305680 | 18305807 | 18305007 | 18305178 | 18306297 | 18306393 | 6 | 2 | 4 [illegible] |
| SE | 24308 | ENSG00000125846 | ZNF133 | chr20 | + | 18306297 | 18306393 | 18288282 | 18288604 | 18315068 | 18315279 | 9 | 2 | 3 [illegible] |
| SE | 24309 | ENSG00000125846 | ZNF133 | chr20 | + | 18306297 | 18306393 | 18288282 | 18288604 | 18315071 | 18316994 | 3 | 4 | 1 [illegible] |
| SE | 24313 | ENSG00000058799 | YIPF1 | chr1 | - | 53866199 | 53866382 | 53860055 | 53860153 | 53866757 | 53866924 | 64 | 28 | 74 [illegible] |
| SE | 24320 | ENSG00000058799 | YIPF1 | chr1 | - | 53886666 | 53886721 | 53883112 | 53883276 | 53889712 | 53889798 | 1 | 0 | 0 [illegible] |
| SE | 24324 | ENSG00000185404 | SP140L | chr2 | + | 230357804 | 230357967 | 230328756 | 230328831 | 230370907 | 230370967 | 110 | 85 | 95 [illegible] |
| SE | 24326 | ENSG00000185404 | SP140L | chr2 | + | 230358963 | 230359132 | 230357804 | 230357967 | 230370907 | 230370967 | 295 | 201 | 318 [illegible] |
| SE | 24327 | ENSG00000185404 | SP140L | chr2 | + | 230361613 | 230361697 | 230357804 | 230357967 | 230370907 | 230370967 | 66 | 61 | 78 [illegible] |
| SE | 24328 | ENSG00000185404 | SP140L | chr2 | + | 230361613 | 230361697 | 230358963 | 230359132 | 230370907 | 230370967 | 235 | 200 | 284 [illegible] |
| SE | 24329 | ENSG00000185404 | SP140L | chr2 | + | 230361613 | 230361697 | 230358963 | 230359132 | 230383509 | 230383575 | 112 | 92 | 120 [illegible] |
| SE | 24332 | ENSG00000185404 | SP140L | chr2 | + | 230389918 | 230390023 | 230388558 | 230388633 | 230393413 | 230393461 | 94 | 86 | 103 [illegible] |
| SE | 24336 | ENSG00000185404 | SP140L | chr2 | + | 230392086 | 230392229 | 230389918 | 230390023 | 230393413 | 230393461 | 114 | 122 | 140 [illegible] |
| SE | 24340 | ENSG00000088808 | PPP1R13B | chr14 | - | 103736002 | 103736202 | 103733194 | 103735195 | 103737693 | 103737860 | 28 | 16 | 33 [illegible] |
| SE | 24341 | ENSG00000088808 | PPP1R13B | chr14 | - | 103736002 | 103736202 | 103733194 | 103735195 | 103738678 | 103738812 | 12 | 8 | 19 [illegible] |
| SE | 24342 | ENSG00000088808 | PPP1R13B | chr14 | - | 103737693 | 103737860 | 103733194 | 103735195 | 103738678 | 103738812 | 16 | 10 | 7 [illegible] |
| SE | 24343 | ENSG00000088808 | PPP1R13B | chr14 | - | 103737693 | 103737860 | 103736002 | 103736202 | 103738678 | 103738812 | 33 | 19 | 21 [illegible] |
| SE | 24349 | ENSG00000088808 | PPP1R13B | chr14 | - | 103784794 | 103784914 | 103778591 | 103778821 | 103797370 | 103797488 | 76 | 33 | 52 [illegible] |
| [illegible] | | | | | | | | | | | | | | |

| | | | | | | | | | | | | | |
|---|---|---|---|---|---|---|---|---|---|---|---|---|---|
| SE | 24689 | ENSG00000118690 | ARMC2 | chr6 | + | 108928087 | 108928233 | 108912334 | 108912558 | 108953032 | 108953351 | 6 | 3 | 2 | [illegible] |
| SE | 24690 | ENSG00000118690 | ARMC2 | chr6 | + | 108936899 | 108936999 | 108928087 | 108928233 | 108953032 | 108953351 | 7 | 5 | 18 | [illegible] |
| SE | 24691 | ENSG00000118690 | ARMC2 | chr6 | + | 108964979 | 108965140 | 108964179 | 108964312 | 108973356 | 108974472 | 8 | 5 | 14 | [illegible] |
| SE | 24692 | ENSG00000204628 | RACK1 | chr5 | - | 181237224 | 181237319 | 181236928 | 181237042 | 181237608 | 181237719 | 15 | 3 | 34 | [illegible] |
| SE | 24693 | ENSG00000204628 | RACK1 | chr5 | - | 181237608 | 181237719 | 181236896 | 181237042 | 181238098 | 181238239 | 4787 | 2125 | 5246 | [illegible] |
| SE | 24695 | ENSG00000204628 | RACK1 | chr5 | - | 181238098 | 181238239 | 181236936 | 181237042 | 181239066 | 181239177 | 3346 | 1586 | 3571 | [illegible] |
| SE | 24696 | ENSG00000204628 | RACK1 | chr5 | - | 181238098 | 181238239 | 181236936 | 181237042 | 181239486 | 181239582 | 0 | 2 | 8 | [illegible] |
| SE | 24697 | ENSG00000204628 | RACK1 | chr5 | - | 181238098 | 181238239 | 181237608 | 181237719 | 181239066 | 181239177 | 6754 | 3235 | 7323 | [illegible] |
| SE | 24698 | ENSG00000204628 | RACK1 | chr5 | - | 181239066 | 181239177 | 181236933 | 181237042 | 181239486 | 181239582 | 1031 | 580 | 1262 | [illegible] |
| SE | 24699 | ENSG00000204628 | RACK1 | chr5 | - | 181239066 | 181239177 | 181238098 | 181238239 | 181239486 | 181239582 | 4431 | 2205 | 4879 | [illegible] |
| SE | 24700 | ENSG00000204628 | RACK1 | chr5 | - | 181240215 | 181240327 | 181239322 | 181239582 | 181241491 | 181241639 | 5 | 0 | 1 | [illegible] |
| SE | 24701 | ENSG00000204628 | RACK1 | chr5 | - | 181240215 | 181240345 | 181239486 | 181239582 | 181241491 | 181241563 | 0 | 1 | 2 | [illegible] |
| SE | 24702 | ENSG00000204628 | RACK1 | chr5 | - | 181241491 | 181241563 | 181239486 | 181239582 | 181242169 | 181242345 | 824 | 465 | 1029 | [illegible] |
| SE | 24703 | ENSG00000204628 | RACK1 | chr5 | - | 181241491 | 181241563 | 181239486 | 181239582 | 181242173 | 181242345 | 825 | 463 | 1028 | [illegible] |
| SE | 24704 | ENSG00000204628 | RACK1 | chr5 | - | 181241491 | 181241639 | 181239037 | 181239582 | 181242169 | 181242345 | 2713 | 1379 | 3207 | [illegible] |
| SE | 24705 | ENSG00000204628 | RACK1 | chr5 | - | 181241491 | 181241639 | 181239037 | 181239582 | 181242173 | 181242206 | 4733 | 2299 | 5263 | [illegible] |
| SE | 24706 | ENSG00000204628 | RACK1 | chr5 | - | 181243253 | 181243405 | 181242249 | 181242345 | 181243691 | 181243904 | 0 | 0 | 1 | [illegible] |
| SE | 24708 | ENSG00000204628 | RACK1 | chr5 | - | 181243285 | 181243405 | 181242173 | 181242345 | 181243691 | 181243904 | 16 | 3 | 5 | [illegible] |
| SE | 24712 | ENSG00000087470 | DNM1L | chr12 | + | 32708152 | 32708224 | 32707366 | 32707413 | 32710928 | 32711015 | 79 | 78 | 56 | [illegible] |
| SE | 24714 | ENSG00000087470 | DNM1L | chr12 | + | 32710928 | 32711015 | 32707366 | 32707413 | 32718642 | 32718763 | 0 | 3 | 1 | [illegible] |
| SE | 24719 | ENSG00000087470 | DNM1L | chr12 | + | 32731853 | 32731943 | 32731355 | 32731511 | 32733714 | 32733807 | 141 | 141 | 104 | [illegible] |
| SE | 24720 | ENSG00000087470 | DNM1L | chr12 | + | 32732537 | 32732610 | 32731853 | 32731943 | 32733714 | 32733807 | 2 | 0 | 0 | [illegible] |
| SE | 24721 | ENSG00000087470 | DNM1L | chr12 | + | 32737864 | 32737942 | 32737104 | 32737161 | 32738263 | 32738296 | 5 | 11 | 14 | [illegible] |
| SE | 24722 | ENSG00000087470 | DNM1L | chr12 | + | 32737864 | 32737942 | 32737104 | 32737161 | 32740063 | 32740240 | 1 | 6 | 4 | [illegible] |
| SE | 24723 | ENSG00000087470 | DNM1L | chr12 | + | 32738263 | 32738296 | 32737104 | 32737161 | 32740063 | 32740240 | 22 | 12 | 15 | [illegible] |
| SE | 24724 | ENSG00000087470 | DNM1L | chr12 | + | 32738263 | 32738296 | 32737864 | 32737942 | 32740063 | 32740240 | 12 | 12 | 21 | [illegible] |
| SE | 24725 | ENSG00000087470 | DNM1L | chr12 | + | 32740063 | 32740240 | 32737104 | 32737161 | 32740408 | 32740467 | 105 | 102 | 79 | [illegible] |
| SE | 24726 | ENSG00000087470 | DNM1L | chr12 | + | 32740408 | 32740518 | 32739844 | 32740240 | 32742588 | 32742684 | 225 | 219 | 169 | [illegible] |
| SE | 24727 | ENSG00000087470 | DNM1L | chr12 | + | 32742588 | 32742748 | 32740153 | 32740518 | 32743353 | 32743581 | 317 | 339 | 260 | [illegible] |
| SE | 24730 | ENSG00000158987 | RAPGEF6 | chr5 | - | 131442348 | 131442537 | 131439580 | 131439715 | 131446482 | 131446703 | 266 | 203 | 416 | [illegible] |
| SE | 24731 | ENSG00000158987 | RAPGEF6 | chr5 | - | 131450022 | 131450046 | 131446482 | 131446703 | 131453053 | 131453177 | 0 | 0 | 0 | [illegible] |
| SE | 24732 | ENSG00000158987 | RAPGEF6 | chr5 | - | 131461704 | 131462088 | 131455800 | 131456012 | 131464040 | 131464281 | 314 | 267 | 391 | [illegible] |
| SE | 24733 | ENSG00000158987 | RAPGEF6 | chr5 | - | 131461704 | 131462088 | 131455800 | 131456012 | 131472586 | 131472744 | 155 | 126 | 178 | [illegible] |
| SE | 24734 | ENSG00000158987 | RAPGEF6 | chr5 | - | 131464040 | 131464281 | 131455800 | 131456012 | 131472586 | 131472744 | 193 | 200 | 275 | [illegible] |
| SE | 24735 | ENSG00000158987 | RAPGEF6 | chr5 | - | 131464040 | 131464281 | 131461704 | 131462088 | 131472586 | 131472744 | 349 | 341 | 487 | [illegible] |
| SE | 24738 | ENSG00000158987 | RAPGEF6 | chr5 | - | 131495552 | 131495660 | 131492581 | 131492785 | 131498442 | 131498518 | 242 | 266 | 383 | [illegible] |
| SE | 24739 | ENSG00000158987 | RAPGEF6 | chr5 | - | 131505363 | 131505522 | 131504625 | 131504778 | 131508070 | 131508157 | 853 | 1096 | 1128 | [illegible] |
| SE | 24743 | ENSG00000158987 | RAPGEF6 | chr5 | - | 131548046 | 131548190 | 131521389 | 131521521 | 131561977 | 131562047 | 206 | 228 | 267 | [illegible] |
| SE | 24744 | ENSG00000158987 | RAPGEF6 | chr5 | - | 131561977 | 131562047 | 131521389 | 131521521 | 131592382 | 131592466 | 35 | 24 | 38 | [illegible] |
| SE | 24761 | ENSG00000250802 | ZBED3-AS1 | chr5 | + | 77098654 | 77098747 | 77087001 | 77087619 | 77105655 | 77105689 | 0 | 0 | 1 | [illegible] |
| SE | 24767 | ENSG00000250802 | ZBED3-AS1 | chr5 | + | 77100235 | 77100391 | 77098654 | 77098747 | 77105655 | 77105689 | 0 | 0 | 0 | [illegible] |
| SE | 24785 | ENSG00000273217 | NA | chr5 | - | 131548046 | 131548190 | 131521389 | 131521521 | 131592382 | 131592466 | 120 | 137 | 169 | [illegible] |
| SE | 24791 | ENSG00000056277 | ZNF280C | chrX | - | 130226758 | 130226905 | 130220348 | 130220480 | 130227681 | 130227782 | 11 | 31 | 12 | [illegible] |
| SE | 24792 | ENSG00000056277 | ZNF280C | chrX | - | 130227681 | 130227782 | 130226758 | 130226905 | 130228976 | 130229134 | 16 | 8 | 10 | [illegible] |
| SE | 24793 | ENSG00000056277 | ZNF280C | chrX | - | 130230509 | 130230727 | 130228976 | 130229134 | 130236213 | 130236320 | 70 | 35 | 40 | [illegible] |
| SE | 24798 | ENSG00000112357 | PEX7 | chr6 | + | 136866626 | 136866733 | 136826313 | 136826469 | 136869889 | 136870003 | 6 | 13 | 10 | [illegible] |
| SE | 24799 | ENSG00000112357 | PEX7 | chr6 | + | 136869889 | 136870003 | 136826313 | 136826469 | 136898141 | 136898241 | 2 | 0 | 1 | [illegible] |
| SE | 24800 | ENSG00000112357 | PEX7 | chr6 | + | 136870731 | 136870839 | 136869889 | 136870003 | 136872197 | 136872253 | 2 | 0 | 0 | [illegible] |
| SE | 24801 | ENSG00000112357 | PEX7 | chr6 | + | 136872197 | 136872253 | 136869889 | 136870003 | 136898141 | 136898241 | 10 | 2 | 11 | [illegible] |
| SE | 24802 | ENSG00000120129 | DUSP1 | chr5 | - | 172769574 | 172769794 | 172768095 | 172769132 | 172770160 | 172770306 | 1695 | 1077 | 1262 | [illegible] |
| SE | 24803 | ENSG00000120129 | DUSP1 | chr5 | - | 172769574 | 172769794 | 172768095 | 172769132 | 172770585 | 172771195 | 1386 | 907 | 1056 | [illegible] |
| SE | 24804 | ENSG00000120129 | DUSP1 | chr5 | - | 172770160 | 172770306 | 172768095 | 172769132 | 172770585 | 172771195 | 231 | 89 | 100 | [illegible] |
| SE | 24805 | ENSG00000120129 | DUSP1 | chr5 | - | 172770160 | 172770306 | 172769574 | 172769794 | 172770585 | 172771195 | 532 | 254 | 304 | [illegible] |
| SE | 24807 | ENSG00000103932 | RPAP1 | chr15 | - | 41522764 | 41522960 | 41522097 | 41522250 | 41523244 | 41523354 | 13 | 2 | 11 | [illegible] |
| SE | 24808 | ENSG00000103932 | RPAP1 | chr15 | - | 41524990 | 41525148 | 41524095 | 41524254 | 41526897 | 41527068 | 9 | 10 | 8 | [illegible] |
| SE | 24809 | ENSG00000103932 | RPAP1 | chr15 | - | 41524990 | 41525165 | 41524095 | 41524254 | 41526897 | 41526970 | 3 | 6 | 4 | [illegible] |
| SE | 24811 | ENSG00000288643 | NA | chr5 | + | 62526141 | 62526257 | 62515387 | 62515501 | 62530708 | 62530785 | 30 | 34 | 37 | [illegible] |
| SE | 24812 | ENSG00000288643 | NA | chr5 | + | 62526141 | 62526257 | 62515387 | 62515501 | 62536701 | 62536781 | 13 | 10 | 19 | [illegible] |
| SE | 24813 | ENSG00000288643 | NA | chr5 | + | 62530708 | 62530785 | 62515387 | 62515501 | 62536701 | 62536781 | 15 | 11 | 20 | [illegible] |
| SE | 24814 | ENSG00000288643 | NA | chr5 | + | 62530708 | 62530785 | 62526141 | 62526257 | 62536701 | 62536781 | 31 | 36 | 33 | [illegible] |
| SE | 24817 | ENSG00000164576 | SAP30L | chr5 | + | 154451090 | 154451213 | 154445996 | 154446805 | 154453401 | 154453500 | 165 | 93 | 94 | [illegible] |
| SE | 24826 | ENSG00000128335 | APOL2 | chr22 | - | 36231339 | 36231466 | 36226208 | 36228280 | 36233152 | 36233241 | 297 | 191 | 254 | [illegible] |
| SE | 24828 | ENSG00000128335 | APOL2 | chr22 | - | 36237060 | 36237185 | 36233401 | 36233455 | 36239440 | 36239527 | 2 | 0 | 1 | [illegible] |
| SE | 24830 | ENSG00000170085 | SIMC1 | chr5 | + | 176270390 | 176270520 | 176238452 | 176238637 | 176289653 | 176289799 | 2 | 1 | 0 | [illegible] |
| SE | 24831 | ENSG00000170085 | SIMC1 | chr5 | + | 176270390 | 176270520 | 176238452 | 176238637 | 176295029 | 176295262 | 2 | 0 | 0 | [illegible] |
| SE | 24832 | ENSG00000170085 | SIMC1 | chr5 | + | 176270390 | 176270520 | 176263416 | 176263502 | 176289653 | 176289799 | 1 | 1 | 2 | [illegible] |
| SE | 24833 | ENSG00000170085 | SIMC1 | chr5 | + | 176289653 | 176290955 | 176238423 | 176238637 | 176295029 | 176295262 | 53 | 35 | 42 | [illegible] |
| SE | 24837 | ENSG00000104320 | NBN | chr8 | - | 89947823 | 89947892 | 89946139 | 89946295 | 89953243 | 89953354 | 227 | 208 | 176 | [illegible] |
| SE | 24838 | ENSG00000104320 | NBN | chr8 | - | 89978219 | 89978323 | 89971172 | 89971290 | 89980733 | 89980893 | 185 | 104 | 151 | [illegible] |
| SE | 24840 | ENSG00000104320 | NBN | chr8 | - | 89980733 | 89980852 | 89978277 | 89978323 | 89981374 | 89981523 | 133 | 54 | 81 | [illegible] |
| SE | 24842 | ENSG00000104320 | NBN | chr8 | - | 89980733 | 89980893 | 89978219 | 89978323 | 89981374 | 89981523 | 297 | 156 | 207 | [illegible] |
| SE | 24843 | ENSG00000104320 | NBN | chr8 | - | 89981374 | 89981523 | 89980733 | 89980852 | 89981952 | 89982002 | 37 | 23 | 38 | [illegible] |
| SE | 24844 | ENSG00000104320 | NBN | chr8 | - | 89981374 | 89981523 | 89980733 | 89980852 | 89982721 | 89982855 | 87 | 48 | 71 | [illegible] |
| SE | 24845 | ENSG00000104320 | NBN | chr8 | - | 89981374 | 89981523 | 89980733 | 89980893 | 89981952 | 89982002 | 196 | 121 | 149 | [illegible] |
| SE | 24846 | ENSG00000104320 | NBN | chr8 | - | 89981374 | 89981523 | 89980733 | 89980893 | 89982721 | 89982855 | 246 | 146 | 182 | [illegible] |
| SE | 24847 | ENSG00000104320 | NBN | chr8 | - | 89981952 | 89982002 | 89980733 | 89980852 | 89982721 | 89982855 | 16 | 9 | 14 | [illegible] |
| SE | 24848 | ENSG00000104320 | NBN | chr8 | - | 89981952 | 89982002 | 89980733 | 89980893 | 89982721 | 89982855 | 16 | 9 | 14 | [illegible] |
| SE | 24859 | ENSG00000283515 | NA | chr19 | + | 58312165 | 58312462 | 58306087 | 58306216 | 58314747 | 58315183 | 166 | 147 | 205 | [illegible] |
| SE | 24860 | ENSG00000122642 | FKBP9 | chr7 | + | 32963543 | 32963702 | 32957403 | 32957794 | 32974616 | 32974644 | 0 | 0 | 0 | [illegible] |
| SE | 24861 | ENSG00000122642 | FKBP9 | chr7 | + | 32974616 | 32974762 | 32957437 | 32957794 | 32975181 | 32975371 | 1 | 1 | 0 | [illegible] |
| SE | 24863 | ENSG00000155846 | PPARGC1B | chr5 | + | 149761431 | 149761612 | 149730302 | 149730420 | 149817612 | 149817746 | 0 | 0 | 0 | [illegible] |
| SE | 24869 | ENSG00000185627 | PSMD13 | chr11 | + | 238997 | 239076 | 236965 | 237144 | 244160 | 244216 | 49 | 28 | 48 | [illegible] |
| SE | 24870 | ENSG00000185627 | PSMD13 | chr11 | + | 244040 | 244075 | 237003 | 237144 | 244160 | 244216 | 18 | 15 | 12 | [illegible] |
| SE | 24872 | ENSG00000185627 | PSMD13 | chr11 | + | 244040 | 244216 | 238997 | 239076 | 244425 | 244469 | 98 | 86 | 109 | [illegible] |
| SE | 24873 | ENSG00000185627 | PSMD13 | chr11 | + | 244160 | 244216 | 238997 | 239076 | 244425 | 244469 | 86 | 74 | 92 | [illegible] |
| SE | 24874 | ENSG00000185627 | PSMD13 | chr11 | + | 248775 | 248855 | 247276 | 247448 | 248931 | 249057 | 481 | 275 | 373 | [illegible] |
| SE | 24875 | ENSG00000185627 | PSMD13 | chr11 | + | 250802 | 250865 | 248606 | 249057 | 252504 | 252827 | 103 | 47 | 70 | [illegible] |
| SE | 24876 | ENSG00000185627 | PSMD13 | chr11 | + | 251545 | 251626 | 250802 | 250865 | 251819 | 251894 | 455 | 263 | 391 | [illegible] |
| SE | 24884 | ENSG00000104951 | IL4I1 | chr19 | - | 49927693 | 49927815 | 49904198 | 49904321 | 49929325 | 49929416 | 21 | 17 | 17 | [illegible] |
| SE | 24885 | ENSG00000113441 | LNPEP | chr5 | + | 96979137 | 96979978 | 96936079 | 96936174 | 96985079 | 96985218 | 476 | 353 | 435 | [illegible] |
| SE | 24886 | ENSG00000113441 | LNPEP | chr5 | + | 96993014 | 96993135 | 96986538 | 96986670 | 96993816 | 96993971 | 648 | 482 | 554 | [illegible] |
| SE | 24887 | ENSG00000113441 | LNPEP | chr5 | + | 96996389 | 96996503 | 96993816 | 96993971 | 96998013 | 96998145 | 273 | 204 | 256 | [illegible] |
| SE | 24889 | ENSG00000113441 | LNPEP | chr5 | + | 97026616 | 97026757 | 97024520 | 97024682 | 97027732 | 97027814 | 618 | 398 | 500 | [illegible] |
| SE | 24890 | ENSG00000198482 | ZNF808 | chr19 | + | 52532907 | 52533009 | 52527667 | 52527711 | 52543265 | 52543347 | 71 | 51 | 41 | [illegible] |
| SE | 24891 | ENSG00000198482 | ZNF808 | chr19 | + | 52543265 | 52543347 | 52532907 | 52533009 | 52553106 | 52553431 | 91 | 68 | 47 | [illegible] |
| SE | 24912 | ENSG00000164221 | CCDC112 | chr5 | - | 115269702 | 115269798 | 115268881 | 115269000 | 115271212 | 115271626 | 61 | 48 | 32 | [illegible] |
| SE | 24913 | ENSG00000164221 | CCDC112 | chr5 | - | 115269702 | 115269798 | 115268881 | 115269000 | 115275215 | 115275606 | 34 | 24 | 15 | [illegible] |
| SE | 24923 | ENSG00000116062 | MSH6 | chr2 | + | 47790926 | 47791123 | 47783081 | 47783493 | 47795893 | 47796036 | 86 | 55 | 78 | [illegible] |
| SE | 24924 | ENSG00000116062 | MSH6 | chr2 | + | 47790926 | 47791123 | 47783081 | 47783493 | 47798610 | 47801155 | 36 | 17 | 30 | [illegible] |
| SE | 24925 | ENSG00000116062 | MSH6 | chr2 | + | 47790926 | 47791123 | 47783150 | 47783231 | 47795893 | 47796036 | 51 | 44 | 49 | [illegible] |
| SE | 24927 | ENSG00000116062 | MSH6 | chr2 | + | 47795893 | 47796063 | 47783127 | 47783493 | 47798610 | 47798627 | 41 | 29 | 31 | [illegible] |
| SE | 24929 | ENSG00000116062 | MSH6 | chr2 | + | 47798610 | 47801155 | 47795893 | 47796063 | 47803419 | 47803628 | 174 | 118 | 125 | [illegible] |
| SE | 24930 | ENSG00000116062 | MSH6 | chr2 | + | 47798614 | 47801155 | 47795893 | 47796063 | 47803419 | 47803685 | 133 | 89 | 94 | [illegible] |
| SE | 24931 | ENSG00000118707 | TGIF2 | chr20 | + | 36576214 | 36576403 | 36573722 | 36573745 | 36578740 | 36578966 | 0 | 0 | 0 | [illegible] |
| SE | 24933 | ENSG00000118707 | TGIF2 | chr20 | + | 36590909 | 36591009 | 36578826 | 36578966 | 36591504 | 36591923 | 42 | 10 | 24 | [illegible] |
| SE | 24935 | ENSG00000072364 | AFF4 | chr5 | - | 132883339 | 132883560 | 132875394 | 132881186 | 132885075 | 132885119 | 179 | 126 | 183 | [illegible] |
| SE | 24936 | ENSG00000072364 | AFF4 | chr5 | - | 132889078 | 132889173 | 132888096 | 132888160 | 132892163 | 132892404 | 120 | 91 | 115 | [illegible] |
| SE | 24938 | ENSG00000072364 | AFF4 | chr5 | - | 132896322 | 132897240 | 132893029 | 132893118 | 132898229 | 132898392 | 436 | 322 | 368 | [illegible] |
| SE | 24940 | ENSG00000072364 | AFF4 | chr5 | - | 132902441 | 132902487 | 132899586 | 132899641 | 132927120 | 132927207 | 12 | 12 | 12 | [illegible] |
| SE | 24943 | ENSG00000072364 | AFF4 | chr5 | - | 132934146 | 132934941 | 132932177 | 132932222 | 132937066 | 132937193 | 280 | 217 | 232 | [illegible] |
| SE | 24949 | ENSG00000163297 | ANTXR2 | chr4 | - | 80035971 | 80036032 | 80033471 | 80033570 | 80054271 | 80054352 | 323 | 289 | 236 | [illegible] |
| SE | 24950 | ENSG00000163297 | ANTXR2 | chr4 | - | 80069435 | 80069507 | 80055931 | 80056013 | 80079352 | 80079600 | 51 | 47 | 30 | [illegible] |
| SE | 24953 | ENSG00000113575 | PPP2CA | chr5 | - | 134200984 | 134201074 | 134199085 | 134199204 | 134201847 | 134202021 | 200 | 145 | 222 | [illegible] |
| SE | 24961 | ENSG00000177169 | ULK1 | chr12 | + | 131915334 | 131915421 | 131915082 | 131915231 | 131915890 | 131916159 | 62 | 34 | 30 | [illegible] |
| SE | 24962 | ENSG00000177169 | ULK1 | chr12 | + | 131917410 | 131917554 | 131916952 | 131917062 | 131918496 | 131918681 | 15 | 3 | 15 | [illegible] |
| SE | 24968 | ENSG00000150712 | MTMR12 | chr5 | - | 32239000 | 32239173 | 32234961 | 32235129 | 32242056 | 32242127 | 194 | 127 | 107 | [illegible] |
| SE | 24970 | ENSG00000150712 | MTMR12 | chr5 | - | 32248001 | 32248126 | 32243520 | 32243599 | 32248771 | 32248878 | 227 | 128 | 150 | [illegible] |
| SE | 24971 | ENSG00000150712 | MTMR12 | chr5 | - | 32255692 | 32255768 | 32248771 | 32248878 | 32263112 | 32263242 | 179 | 103 | 122 | [illegible] |
| SE | 24972 | ENSG00000150712 | MTMR12 | chr5 | - | 32263112 | 32263242 | 32255692 | 32255768 | 32268700 | 32268794 | 273 | 209 | 198 | [illegible] |
| SE | 24974 | ENSG00000150712 | MTMR12 | chr5 | - | 32271832 | 32271905 | 32270816 | 32270947 | 32312757 | 32312924 | 98 | 81 | 81 | [illegible] |
| SE | 24975 | ENSG00000150712 | MTMR12 | chr5 | - | 32273979 | 32274122 | 32270816 | 32270947 | 32312757 | 32312987 | 3 | 2 | 4 | [illegible] |
| SE | 24976 | ENSG00000150712 | MTMR12 | chr5 | - | 32273979 | 32274122 | 32271832 | 32271905 | 32312757 | 32312987 | 94 | 104 | 91 | [illegible] |
| SE | 24980 | ENSG00000196189 | SEMA4A | chr1 | + | 156154549 | 156154717 | 156153330 | 156153476 | 156156413 | 156156574 | 59 | 24 | 28 | [illegible] |
| SE | 24983 | ENSG00000196189 | SEMA4A | chr1 | + | 156162943 | 156163094 | 156161226 | 156161518 | 156172825 | 156173006 | 166 | 150 | 148 | [illegible] |
| SE | 24984 | ENSG00000196189 | SEMA4A | chr1 | + | 156174821 | 156174940 | 156172825 | 156173006 | 156175085 | 156175232 | 78 | 47 | 49 | [illegible] |
| SE | 24996 | ENSG00000240535 | NA | chr5 | - | 55036494 | 55036755 | 55033735 | 55033863 | 55039524 | 55039580 | 0 | 0 | 1 | [illegible] |
| SE | 24998 | ENSG00000145685 | LHFPL2 | chr5 | - | 78632263 | 78632368 | 78564812 | 78564871 | 78648498 | 78648772 | 3 | 2 | 2 | [illegible] |
| SE | 25000 | ENSG00000113249 | HAVCR1 | chr5 | - | 157032853 | 157032870 | 157029739 | 157029841 | 157037246 | 157037361 | 0 | 3 | 0 | [illegible] |
| SE | 25001 | ENSG00000113249 | HAVCR1 | chr5 | - | 157032853 | 157032887 | 157029412 | 157029841 | 157037246 | 157037361 | 6 | 5 | 0 | [illegible] |
| SE | 25004 | ENSG00000166578 | IQCD | chr12 | - | 113197540 | 113197612 | 113195445 | 113195889 | 113200579 | 113200813 | 2 | 0 | 8 | [illegible] |
| SE | 25007 | ENSG00000166578 | IQCD | chr12 | - | 113200579 | 113200813 | 113197540 | 113197612 | 113207437 | 113208259 | 3 | 1 | 0 | [illegible] |
| SE | 25012 | ENSG00000150593 | PDCD4 | chr10 | + | 110887664 | 110887886 | 110884770 | 110885366 | 110889532 | 110889630 | 1488 | 1335 | 1401 | [illegible] |
| SE | 25013 | ENSG00000150593 | PDCD4 | chr10 | + | 110889532 | 110889630 | 110887664 | 110887886 | 110890555 | 110890670 | 941 | 771 | 960 | [illegible] |
| SE | 25014 | ENSG00000150593 | PDCD4 | chr10 | + | 110894090 | 110894198 | 110890555 | 110890670 | 110894411 | 110894522 | 635 | 468 | 586 | [illegible] |
| SE | 25015 | ENSG00000150593 | PDCD4 | chr10 | + | 110894411 | 110894522 | 110894090 | 110894198 | 110895947 | 110896087 | 1108 | 931 | 1113 | [illegible] |
| SE | 25016 | ENSG00000264343 | NOTCH2NLA | chr1 | - | 146189299 | 146189381 | 146164750 | 146165010 | 146228708 | 146229026 | 79 | 55 | 34 | [illegible] |
| SE | 25017 | ENSG00000113272 | THG1L | chr5 | + | 157738621 | 157738734 | 157737909 | 157737994 | 157739320 | 157739698 | 0 | 0 | 1 | [illegible] |
| SE | 25019 | ENSG00000281938 | NA | chr5 | - | 131942634 | 131942695 | 131933512 | 131933641 | 131944664 | 131944752 | 0 | 3 | 0 | [illegible] |
| SE | 25020 | ENSG00000186871 | ERCC6L | chrX | - | 72237830 | 72237948 | 72204656 | 72208698 | 72238843 | 72239027 | 0 | 0 | 0 | [illegible] |
| SE | 25021 | ENSG00000171467 | ZNF318 | chr6 | - | 43340789 | 43340908 | 43336069 | 43340502 | 43342111 | 43342211 | 374 | 239 | 282 | [illegible] |
| SE | 25022 | ENSG00000171467 | ZNF318 | chr6 | - | 43342675 | 43342879 | 43331697 | 43331852 | 43348323 | 43348625 | 94 | 54 | 87 | [illegible] |
| SE | 25025 | ENSG00000148444 | COMMD3 | chr10 | + | 22318271 | 22318306 | 22318104 | 22318168 | 22318652 | 22318713 | 94 | 49 | 81 | [illegible] |
| SE | 25027 | ENSG00000269897 | COMMD3-BMI1 | chr10 | + | 22326889 | 22326986 | 22326430 | 22326561 | 22327949 | 22328058 | 99 | 53 | 68 | [illegible] |
| SE | 25028 | ENSG00000269897 | COMMD3-BMI1 | chr10 | + | 22327594 | 22327650 | 22326889 | 22326986 | 22327949 | 22328058 | 169 | 115 | 147 | [illegible] |
| SE | 25029 | ENSG00000269897 | COMMD3-BMI1 | chr10 | + | 22327741 | 22327792 | 22327594 | 22327650 | 22327949 | 22328058 | 401 | 325 | 375 | [illegible] |
| SE | 25030 | ENSG00000269897 | COMMD3-BMI1 | chr10 | + | 22329047 | 22329128 | 22328599 | 22328698 | 22329212 | 22329542 | 526 | 349 | 359 | [illegible] |
| SE | 25036 | ENSG00000205302 | SNX2 | chr5 | + | 122799691 | 122799855 | 122795265 | 122795383 | 122801868 | 122801935 | 652 | 492 | 447 | [illegible] |
| SE | 25037 | ENSG00000205302 | SNX2 | chr5 | + | 122799694 | 122799855 | 122795265 | 122795383 | 122801868 | 122801935 | 403 | 337 | 289 | [illegible] |
| SE | 25038 | ENSG00000205302 | SNX2 | chr5 | + | 122818817 | 122819023 | 122817274 | 122817373 | 122826049 | 122826193 | 892 | 704 | 576 | [illegible] |
| SE | 25039 | ENSG00000205302 | SNX2 | chr5 | + | 122827378 | 122827459 | 122826049 | 122826193 | 122827574 | 122827646 | 378 | 377 | 248 | [illegible] |
| SE | 25040 | ENSG00000164308 | ERAP2 | chr5 | + | 96883791 | 96883930 | 96879563 | 96880260 | 96886654 | 96886789 | 627 | 970 | 1047 | [illegible] |
| SE | 25041 | ENSG00000164308 | ERAP2 | chr5 | + | 96886654 | 96886789 | 96883791 | 96883930 | 96889184 | 96889305 | 832 | 1306 | 1266 | [illegible] |
| SE | 25042 | ENSG00000164308 | ERAP2 | chr5 | + | 96889184 | 96889305 | 96886654 | 96886789 | 96892298 | 96892453 | 748 | 1200 | 1200 | [illegible] |
| SE | 25044 | ENSG00000164308 | ERAP2 | chr5 | + | 96896372 | 96896504 | 96895245 | 96895359 | 96896731 | 96896863 | 413 | 588 | 914 | [illegible] |
| SE | 25045 | ENSG00000164308 | ERAP2 | chr5 | + | 96896731 | 96896863 | 96896372 | 96896504 | 96900120 | 96900189 | 408 | 613 | 841 | [illegible] |
| SE | 25048 | ENSG00000164308 | ERAP2 | chr5 | + | 96901505 | 96901681 | 96900120 | 96900189 | 96902273 | 96902353 | 446 | 679 | 960 | [illegible] |
| SE | 25049 | ENSG00000164308 | ERAP2 | chr5 | + | 96909579 | 96909764 | 96908960 | 96909117 | 96912636 | 96912798 | 337 | 386 | 638 | [illegible] |
| SE | 25050 | ENSG00000164308 | ERAP2 | chr5 | + | 96909579 | 96909809 | 96908960 | 96909117 | 96912636 | 96912798 | 166 | 160 | 276 | [illegible] |
| SE | 25057 | ENSG00000064652 | SNX24 | chr5 | + | 122946054 | 122946159 | 122936732 | 122936817 | 122999911 | 122999984 | 33 | 30 | 31 | [illegible] |
| SE | 25058 | ENSG00000064652 | SNX24 | chr5 | + | 122965442 | 122965521 | 122946054 | 122946159 | 122999911 | 123000006 | 0 | 0 | 0 | [illegible] |
| SE | 25059 | ENSG00000064652 | SNX24 | chr5 | + | 122983017 | 122983095 | 122946054 | 122946159 | 122998207 | 122998314 | 1 | 0 | 0 | [illegible] |
| SE | 25060 | ENSG00000064652 | SNX24 | chr5 | + | 122983017 | 122983095 | 122946054 | 122946159 | 122999911 | 123000006 | 0 | 0 | 1 | [illegible] |
| SE | 25061 | ENSG00000064652 | SNX24 | chr5 | + | 122999911 | 123000006 | 122983017 | 122983095 | 123001404 | 123001437 | 19 | 23 | 13 | [illegible] |
| SE | 25062 | ENSG00000064652 | SNX24 | chr5 | + | 123001939 | 123002004 | 123001404 | 123001437 | 123007681 | 123007774 | 20 | 16 | 10 | [illegible] |
| SE | 25063 | ENSG00000064652 | SNX24 | chr5 | + | 123001939 | 123002080 | 123001404 | 123001437 | 123007681 | 123008610 | 28 | 24 | 15 | [illegible] |
| SE | 25070 | ENSG00000172239 | PAIP1 | chr5 | - | 43533737 | 43533792 | 43529785 | 43529879 | 43534852 | 43534970 | 326 | 243 | 252 | [illegible] |
| SE | 25071 | ENSG00000172239 | PAIP1 | chr5 | - | 43536818 | 43536944 | 43535533 | 43535640 | 43538923 | 43539035 | 248 | 183 | 195 | [illegible] |
| SE | 25072 | ENSG00000172239 | PAIP1 | chr5 | - | 43538923 | 43539035 | 43536818 | 43536944 | 43543003 | 43543116 | 220 | 170 | 185 | [illegible] |
| SE | 25073 | ENSG00000172239 | PAIP1 | chr5 | - | 43543003 | 43543116 | 43538923 | 43539035 | 43547727 | 43547913 | 337 | 259 | 278 | [illegible] |
| SE | 25074 | ENSG00000172239 | PAIP1 | chr5 | - | 43547727 | 43547913 | 43543003 | 43543116 | 43555829 | 43555961 | 368 | 204 | 290 | [illegible] |
| SE | 25075 | ENSG00000172239 | PAIP1 | chr5 | - | 43555829 | 43555999 | 43547727 | 43547913 | 43556818 | 43557077 | 190 | 80 | 141 | [illegible] |
| SE | 25083 | ENSG00000140743 | CDR2 | chr16 | - | 22364901 | 22365014 | 22345935 | 22347823 | 22374230 | 22374619 | 90 | 60 | 107 | [illegible] |
| SE | 25085 | ENSG00000140743 | CDR2 | chr16 | - | 22422160 | 22422214 | 22364901 | 22365014 | 22429795 | 22429948 | 2 | 7 | 2 | [illegible] |
| SE | 25086 | ENSG00000156206 | CFAP161 | chr15 | + | 81127589 | 81127724 | 81099407 | 81099521 | 81135269 | 81135359 | 25 | 48 | 15 | [illegible] |
| SE | 25090 | ENSG00000156206 | CFAP161 | chr15 | + | 81138050 | 81138135 | 81136515 | 81136748 | 81143661 | 81143820 | 27 | 38 | 18 | [illegible] |
| SE | 25091 | ENSG00000156206 | CFAP161 | chr15 | + | 81147857 | 81147931 | 81143661 | 81143820 | 81148337 | 81149179 | 6 | 5 | 0 | [illegible] |
| SE | 25094 | ENSG00000153310 | CYRIB | chr8 | - | 129846803 | 129846874 | 129839592 | 129842205 | 129849242 | 129849369 | 814 | 476 | 509 | [illegible] |
| SE | 25096 | ENSG00000153310 | CYRIB | chr8 | - | 129850834 | 129850914 | 129846803 | 129846874 | 129852161 | 129852278 | 402 | 266 | 253 | [illegible] |
| SE | 25097 | ENSG00000153310 | CYRIB | chr8 | - | 129855610 | 129855747 | 129854265 | 129854343 | 129862228 | 129862326 | 1365 | 883 | 931 | [illegible] |
| SE | 25100 | ENSG00000153310 | CYRIB | chr8 | - | 129862228 | 129862334 | 129855582 | 129855747 | 129871374 | 129871496 | 1093 | 693 | 800 | [illegible] |
| SE | 25101 | ENSG00000153310 | CYRIB | chr8 | - | 129871374 | 129871496 | 129854265 | 129854343 | 129879388 | 129879471 | 852 | 651 | 508 | [illegible] |
| SE | 25103 | ENSG00000153310 | CYRIB | chr8 | - | 129879388 | 129879471 | 129871374 | 129871496 | 129880375 | 129880458 | 609 | 466 | 375 | [illegible] |
| SE | 25104 | ENSG00000153310 | CYRIB | chr8 | - | 129879388 | 129879471 | 129871374 | 129871496 | 129896748 | 129896854 | 607 | 463 | 377 | [illegible] |
| SE | 25106 | ENSG00000153310 | CYRIB | chr8 | - | 129879388 | 129879471 | 129871374 | 129871496 | 129970942 | 129970972 | 610 | 461 | 376 | [illegible] |
| SE | 25107 | ENSG00000153310 | CYRIB | chr8 | - | 129879596 | 129879789 | 129879388 | 129879471 | 129903311 | 129903350 | 3 | 1 | 0 | [illegible] |
| SE | 25109 | ENSG00000153310 | CYRIB | chr8 | - | 129880375 | 129880458 | 129871374 | 129871496 | 129903311 | 129903350 | 1 | 0 | 0 | [illegible] |
| SE | 25110 | ENSG00000153310 | CYRIB | chr8 | - | 129880375 | 129880458 | 129879388 | 129879471 | 129896671 | 129896854 | 2 | 1 | 1 | [illegible] |
| SE | 25111 | ENSG00000153310 | CYRIB | chr8 | - | 129880375 | 129880458 | 129879388 | 129879471 | 129896748 | 129896854 | 3 | 1 | 1 | [illegible] |
| SE | 25112 | ENSG00000153310 | CYRIB | chr8 | - | 129880375 | 129880458 | 129879388 | 129879471 | 129903311 | 129903350 | 3 | 5 | 1 | [illegible] |
| SE | 25113 | ENSG00000153310 | CYRIB | chr8 | - | 129894486 | 129894564 | 129879388 | 129879471 | 129903311 | 129903350 | 0 | 0 | 4 | [illegible] |
| SE | 25115 | ENSG00000153310 | CYRIB | chr8 | - | 129896671 | 129896854 | 129880375 | 129880458 | 129903311 | 129903350 | 13 | 6 | 5 | [illegible] |
| SE | 25118 | ENSG00000153310 | CYRIB | chr8 | - | 129896748 | 129896854 | 129880375 | 129880458 | 129903311 | 129903350 | 10 | 6 | 5 | [illegible] |
| SE | 25121 | ENSG00000153310 | CYRIB | chr8 | - | 129903311 | 129903350 | 129879388 | 129879471 | 129904498 | 129904585 | 425 | 248 | 272 | [illegible] |
| SE | 25122 | ENSG00000153310 | CYRIB | chr8 | - | 129903311 | 129903350 | 129879388 | 129879471 | 129912359 | 129912417 | 161 | 116 | 115 | [illegible] |
| SE | 25123 | ENSG00000153310 | CYRIB | chr8 | - | 129903311 | 129903350 | 129879388 | 129879471 | 129970942 | 129970995 | 504 | 358 | 311 | [illegible] |
| SE | 25124 | ENSG00000153310 | CYRIB | chr8 | - | 129904498 | 129904585 | 129879388 | 129879471 | 129912359 | 129912417 | 1 | 0 | 1 | [illegible] |
| SE | 25125 | ENSG00000153310 | CYRIB | chr8 | - | 129904498 | 129904585 | 129879388 | 129879471 | 129970942 | 129970972 | 18 | 12 | 9 | [illegible] |
| SE | 25126 | ENSG00000153310 | CYRIB | chr8 | - | 129904498 | 129904585 | 129903311 | 129903350 | 129912359 | 129912417 | 172 | 106 | 109 | [illegible] |
| SE | 25130 | ENSG00000153310 | CYRIB | chr8 | - | 129904498 | 129904585 | 129903311 | 129903350 | 130016369 | 130016451 | 168 | 106 | 110 | [illegible] |
| SE | 25131 | ENSG00000153310 | CYRIB | chr8 | - | 129912359 | 129912417 | 129879388 | 129879471 | 129970942 | 129970995 | 0 | 0 | 0 | [illegible] |
| SE | 25132 | ENSG00000153310 | CYRIB | chr8 | - | 129912359 | 129912417 | 129903311 | 129903350 | 129970942 | 129970995 | 0 | 0 | 0 | [illegible] |
| SE | 25133 | ENSG00000153310 | CYRIB | chr8 | - | 129912359 | 129912417 | 129904498 | 129904585 | 129970942 | 129970995 | 5 | 0 | 0 | [illegible] |
| SE | 25134 | ENSG00000153310 | CYRIB | chr8 | - | 129912469 | 129912515 | 129903311 | 129903350 | 129939607 | 129939872 | 1 | 0 | 3 | [illegible] |
| SE | 25136 | ENSG00000153310 | CYRIB | chr8 | - | 129912469 | 129912515 | 129904498 | 129904585 | 129939607 | 129939872 | 1 | 3 | 1 | [illegible] |
| SE | 25139 | ENSG00000153310 | CYRIB | chr8 | - | 129970942 | 129970995 | 129903311 | 129903350 | 130016369 | 130016405 | 342 | 248 | 204 | [illegible] |
| SE | 25140 | ENSG00000153310 | CYRIB | chr8 | - | 129970942 | 129970995 | 129903311 | 129903350 | 130016606 | 130016637 | 277 | 212 | 174 | [illegible] |
| SE | 25141 | ENSG00000153310 | CYRIB | chr8 | - | 129970942 | 129970995 | 129904498 | 129904585 | 130016369 | 130016405 | 115 | 44 | 55 | [illegible] |
| SE | 25150 | ENSG00000155329 | ZCCHC10 | chr5 | - | 133006758 | 133006920 | 133000131 | 133000173 | 133022840 | 133022906 | 75 | 54 | 47 | [illegible] |
| SE | 25151 | ENSG00000155329 | ZCCHC10 | chr5 | - | 133006758 | 133006920 | 133000131 | 133000173 | 133026496 | 133026549 | 82 | 50 | 46 | [illegible] |
| SE | 25152 | ENSG00000155329 | ZCCHC10 | chr5 | - | 133011338 | 133011455 | 133006758 | 133006920 | 133022840 | 133022906 | 2 | 0 | 0 | [illegible] |
| SE | 25154 | ENSG00000155329 | ZCCHC10 | chr5 | - | 133011338 | 133011455 | 133006758 | 133006920 | 133026496 | 133026549 | 1 | 0 | 0 | [illegible] |
| SE | 25155 | ENSG00000155329 | ZCCHC10 | chr5 | - | 133022840 | 133022906 | 133000131 | 133000173 | 133026496 | 133026549 | 0 | 0 | 0 | [illegible] |
| SE | 25157 | ENSG00000155329 | ZCCHC10 | chr5 | - | 133022840 | 133022906 | 133006758 | 133006920 | 133026496 | 133026549 | 10 | 8 | 7 | [illegible] |
| SE | 25165 | ENSG00000164219 | PGGT1B | chr5 | - | 115221823 | 115222008 | 115216864 | 115216973 | 115236389 | 115236522 | 75 | 88 | 71 | [illegible] |
| SE | 25166 | ENSG00000164219 | PGGT1B | chr5 | - | 115230975 | 115231021 | 115216864 | 115216973 | 115236389 | 115236522 | 78 | 41 | 42 | [illegible] |
| SE | 25168 | ENSG00000164219 | PGGT1B | chr5 | - | 115236389 | 115236522 | 115230975 | 115231021 | 115237857 | 115238009 | 255 | 204 | 174 | [illegible] |
| SE | 25169 | ENSG00000164219 | PGGT1B | chr5 | - | 115253136 | 115253255 | 115241538 | 115241606 | 115262711 | 115262851 | 155 | 89 | 107 | [illegible] |
| SE | 25181 | ENSG00000112759 | SLC29A1 | chr6 | + | 44221585 | 44221685 | 44219635 | 44219782 | 44227262 | 44227342 | 1 | 0 | 0 | [illegible] |
| SE | 25184 | ENSG00000112759 | SLC29A1 | chr6 | + | 44226060 | 44226167 | 44219552 | 44219782 | 44227259 | 44227342 | 3 | 0 | 4 | [illegible] |
| SE | 25185 | ENSG00000112759 | SLC29A1 | chr6 | + | 44226060 | 44226167 | 44219552 | 44219782 | 44227262 | 44227342 | 3 | 0 | 2 | [illegible] |
| SE | 25189 | ENSG00000112759 | SLC29A1 | chr6 | + | 44227259 | 44227342 | 44219627 | 44219782 | 44229389 | 44229471 | 10 | 5 | 7 | [illegible] |
| SE | 25190 | ENSG00000112759 | SLC29A1 | chr6 | + | 44227262 | 44227342 | 44219614 | 44219782 | 44229389 | 44229471 | 17 | 6 | 9 | [illegible] |
| SE | 25192 | ENSG00000214253 | FIS1 | chr7 | - | 101244006 | 101244139 | 101240829 | 101240906 | 101244897 | 101245048 | 573 | 316 | 470 | [illegible] |
| SE | 25195 | ENSG00000113732 | ATP6V0E1 | chr5 | + | 172994774 | 172994822 | 172983770 | 172983964 | 173020237 | 173020367 | 504 | 208 | 371 | [illegible] |
| SE | 25196 | ENSG00000113732 | ATP6V0E1 | chr5 | + | 173020237 | 173020367 | 172994774 | 172994822 | 173034398 | 173034892 | 1305 | 620 | 922 | [illegible] |
| SE | 25198 | ENSG00000160783 | PMF1 | chr1 | + | 156220676 | 156220887 | 156213004 | 156213176 | 156232319 | 156232425 | 0 | 1 | 0 | [illegible] |
| SE | 25199 | ENSG00000160783 | PMF1 | chr1 | + | 156223345 | 156223560 | 156213004 | 156213176 | 156232319 | 156232425 | 0 | 0 | 0 | [illegible] |
| SE | 25200 | ENSG00000160783 | PMF1 | chr1 | + | 156223345 | 156223560 | 156220676 | 156220887 | 156232319 | 156232425 | 0 | 0 | 0 | [illegible] |
| SE | 25204 | ENSG00000160783 | PMF1 | chr1 | + | 156232319 | 156232425 | 156212992 | 156213176 | 156236287 | 156236483 | 127 | 45 | 94 | [illegible] |
| SE | 25206 | ENSG00000160783 | PMF1 | chr1 | + | 156233627 | 156233728 | 156213006 | 156213176 | 156236287 | 156236483 | 156 | 119 | 138 | [illegible] |
| SE | 25209 | ENSG00000138640 | FAM13A | chr4 | - | 88737471 | 88737555 | 88731832 | 88732198 | 88739029 | 88739125 | 14 | 12 | 9 | [illegible] |
| SE | 25210 | ENSG00000138640 | FAM13A | chr4 | - | 88746931 | 88747015 | 88739029 | 88739125 | 88747630 | 88747851 | 56 | 37 | 18 | [illegible] |
| SE | 25211 | ENSG00000138640 | FAM13A | chr4 | - | 88747630 | 88747851 | 88746931 | 88747015 | 88748951 | 88749033 | 69 | 43 | 21 | [illegible] |

| | | | | | | | | | | | | | | | | | | | | | | | | | | | | | | | | | | | | | | | | | | | | | | |
|---|---|---|---|---|---|---|---|---|---|---|---|---|---|---|---|---|---|---|---|---|---|---|---|---|---|---|---|---|---|---|---|---|---|---|---|---|---|---|---|---|---|---|---|---|---|---|
| SE | 25212 | ENSG00000138640 | FAM13A | chr4 | - | 88767982 | 88768059 | 88767552 | 88767595 | 88781164 | 88781351 | 55 | 54 | 26 | 65 | 96 | 0 | 2 | 0 | 1 | 0 | 75 | 32 | 47 | 60 | 29 | 0 | 0 | 0 | 0 | 0 | 226 | 149 | 1 | 1 | 1 | 0.947 | 1 | 0.977 | 1 | 1 | 1 | 1 | 1 | 1 | -0.015 |
| SE | 25213 | ENSG00000138640 | FAM13A | chr4 | - | 88790585 | 88790627 | 88787752 | 88787932 | 88805010 | 88805052 | 39 | 45 | 33 | 39 | 52 | 1 | 1 | 0 | 1 | 5 | 48 | 25 | 43 | 23 | 9 | 0 | 0 | 2 | 1 | 0 | 191 | 149 | 1 | 1 | 0.968 | 0.972 | 1 | 0.968 | 0.89 | 1 | 1 | 0.944 | 0.947 | 1 | -0.019 |
| SE | 25214 | ENSG00000138640 | FAM13A | chr4 | - | 88790585 | 88790627 | 88787752 | 88787932 | 88822975 | 88823231 | 27 | 17 | 14 | 25 | 25 | 0 | 0 | 1 | 2 | 0 | 0 | 5 | 7 | 1 | 4 | 0 | 0 | 2 | 0 | 0 | 191 | 149 | 1 | 1 | 1 | 1 | 0.916 | 0.907 | 1 | NA | 1 | 0.732 | 1 | 1 | 0.032 |
| SE | 25215 | ENSG00000138640 | FAM13A | chr4 | - | 88805010 | 88805052 | 88787752 | 88787932 | 88822975 | 88823201 | 2 | 2 | 3 | 5 | 6 | 0 | 0 | 1 | 2 | 0 | 13 | 0 | 6 | 3 | 0 | 0 | 0 | 2 | 0 | 0 | 191 | 149 | 1 | 1 | 1 | 1 | 0.701 | 0.661 | 1 | 1 | NA | 0.701 | 1 | NA | -0.028 |
| SE | 25217 | ENSG00000138640 | FAM13A | chr4 | - | 88805010 | 88805052 | 88790585 | 88790627 | 88851019 | 88851183 | 6 | 9 | 7 | 4 | 5 | 0 | 0 | 0 | 0 | 0 | 6 | 10 | 17 | 0 | 0 | 0 | 0 | 0 | 0 | 7 | 191 | 149 | 1 | 1 | 1 | 1 | 1 | 1 | 1 | 1 | 1 | 1 | NA | 0 | 0.25 |
| SE | 25219 | ENSG00000138640 | FAM13A | chr4 | - | 88938087 | 88938241 | 88906378 | 88906462 | 89020459 | 89020669 | 5 | 0 | 1 | 2 | 2 | 0 | 0 | 0 | 0 | 0 | 0 | 0 | 0 | 0 | 0 | 0 | 3 | 0 | 0 | 0 | 298 | 149 | 1 | 1 | 1 | NA | 1 | 1 | 1 | NA | 0 | NA | NA | NA | 1 |
| SE | 25220 | ENSG00000138640 | FAM13A | chr4 | - | 88990972 | 88991150 | 88936282 | 88938241 | 89008950 | 89009078 | 0 | 0 | 5 | 0 | 0 | 0 | 0 | 0 | 0 | 0 | 6 | 0 | 0 | 0 | 0 | 0 | 4 | 0 | 0 | 0 | 298 | 149 | 1 | 1 | NA | NA | 1 | NA | NA | 1 | NA | 0 | NA | NA | 0.5 |
| SE | 25222 | ENSG00000138640 | FAM13A | chr4 | - | 89008950 | 89009078 | 88938087 | 88938241 | 89020459 | 89020669 | 0 | 1 | 0 | 0 | 0 | 0 | 0 | 1 | 0 | 0 | 0 | 0 | 4 | 5 | 0 | 0 | 0 | 0 | 0 | 0 | 277 | 149 | 1 | 1 | NA | 1 | 0 | NA | NA | NA | NA | 1 | 1 | NA | -0.5 |
| SE | 25223 | ENSG00000138640 | FAM13A | chr4 | - | 89008950 | 89009078 | 88990972 | 88991150 | 89020459 | 89020669 | 0 | 1 | 5 | 0 | 0 | 1 | 0 | 1 | 0 | 0 | 6 | 0 | 0 | 5 | 0 | 0 | 0 | 0 | 0 | 0 | 277 | 149 | 1 | 1 | 0 | 1 | 0.729 | NA | NA | 1 | NA | NA | 1 | NA | -0.424 |
| SE | 25224 | ENSG00000231256 | CFAP97D1 | chr17 | + | 43783179 | 43783303 | 43781773 | 43781892 | 43783836 | 43783893 | 0 | 0 | 1 | 0 | 0 | 0 | 2 | 0 | 0 | 0 | 5 | 3 | 0 | 1 | 0 | 0 | 0 | 0 | 2 | 0 | 273 | 149 | 1 | 1 | NA | 0 | 1 | NA | NA | 1 | 1 | NA | 0.214 | NA | -0.238 |
| SE | 25228 | ENSG00000205571 | SMN2 | chr5 | + | 70071281 | 70071390 | 70070640 | 70070751 | 70076520 | 70076927 | 0 | 0 | 2 | 6 | 2 | 18 | 26 | 28 | 17 | 13 | 0 | 0 | 0 | 0 | 0 | 16 | 0 | 20 | 4 | 0 | 258 | 149 | 1 | 1 | 0 | 0 | 0.04 | 0.169 | 0.082 | 0 | NA | 0 | 0 | NA | 0.058 |
| SE | 25229 | ENSG00000205571 | SMN2 | chr5 | + | 70071281 | 70071390 | 70070640 | 70070751 | 70077018 | 70077596 | 0 | 1 | 1 | 1 | 0 | 18 | 21 | 14 | 10 | 8 | 0 | 0 | 0 | 0 | 0 | 13 | 0 | 14 | 32 | 0 | 258 | 149 | 1 | 1 | 0 | 0.027 | 0.04 | 0.055 | 0 | 0 | NA | 0 | 0 | NA | 0.024 |
| SE | 25230 | ENSG00000205571 | SMN2 | chr5 | + | 70076520 | 70076574 | 70070640 | 70070751 | 70077018 | 70077160 | 26 | 32 | 30 | 19 | 16 | 18 | 21 | 14 | 10 | 8 | 16 | 0 | 19 | 4 | 0 | 13 | 0 | 14 | 32 | 0 | 203 | 149 | 1 | 1 | 0.515 | 0.528 | 0.611 | 0.582 | 0.595 | 0.475 | NA | 0.499 | 0.084 | NA | 0.214 |
| SE | 25236 | ENSG00000183621 | ZNF438 | chr10 | - | 30848530 | 30850367 | 30835566 | 30835640 | 30876997 | 30877065 | 20 | 10 | 27 | 17 | 35 | 1 | 0 | 0 | 1 | 0 | 15 | 17 | 15 | 3 | 0 | 0 | 0 | 3 | 0 | 0 | 298 | 149 | 1 | 1 | 0.909 | 1 | 1 | 0.895 | 1 | 1 | 1 | 0.714 | 1 | NA | 0.032 |
| SE | 25237 | ENSG00000183621 | ZNF438 | chr10 | - | 30850221 | 30850367 | 30835566 | 30835640 | 30876997 | 30877065 | 20 | 10 | 27 | 17 | 35 | 1 | 0 | 0 | 1 | 0 | 15 | 17 | 17 | 3 | 0 | 0 | 0 | 3 | 0 | 0 | 295 | 149 | 1 | 1 | 0.91 | 1 | 1 | 0.896 | 1 | 1 | 1 | 0.741 | 1 | NA | 0.026 |
| SE | 25238 | ENSG00000183621 | ZNF438 | chr10 | - | 30857679 | 30857778 | 30848530 | 30850367 | 30876997 | 30877065 | 4 | 3 | 10 | 2 | 14 | 20 | 10 | 27 | 17 | 35 | 18 | 0 | 8 | 1 | 0 | 15 | 17 | 13 | 3 | 0 | 248 | 149 | 1 | 1 | 0.107 | 0.153 | 0.182 | 0.066 | 0.194 | 0.419 | 0 | 0.27 | 0.167 | NA | -0.074 |
| SE | 25239 | ENSG00000183621 | ZNF438 | chr10 | - | 30875330 | 30875509 | 30848530 | 30850367 | 30876997 | 30877065 | 0 | 0 | 0 | 0 | 1 | 20 | 10 | 27 | 17 | 35 | 0 | 0 | 0 | 0 | 0 | 15 | 17 | 13 | 3 | 0 | 298 | 149 | 1 | 1 | 0 | 0 | 0 | 0 | 0.014 | 0 | 0 | 0 | 0 | NA | 0.003 |
| SE | 25240 | ENSG00000183621 | ZNF438 | chr10 | - | 30910359 | 30910440 | 30908932 | 30909015 | 30941574 | 30941651 | 6 | 4 | 9 | 3 | 4 | 15 | 10 | 3 | 9 | 15 | 0 | 13 | 1 | 0 | 6 | 0 | 8 | 9 | 2 | 0 | 230 | 149 | 1 | 1 | 0.206 | 0.206 | 0.66 | 0.178 | 0.147 | NA | 0.513 | 0.067 | 0 | 1 | -0.116 |
| SE | 25248 | ENSG00000126456 | IRF3 | chr19 | - | 49660712 | 49660828 | 49659569 | 49659833 | 49662424 | 49662617 | 103 | 30 | 85 | 115 | 106 | 2 | 0 | 0 | 0 | 0 | 97 | 58 | 35 | 74 | 149 | 0 | 0 | 0 | 0 | 0 | 265 | 149 | 1 | 1 | 0.967 | 1 | 1 | 1 | 1 | 1 | 1 | 1 | 1 | 1 | -0.007 |
| SE | 25250 | ENSG00000126456 | IRF3 | chr19 | - | 49661449 | 49662328 | 49660712 | 49660828 | 49662424 | 49662617 | 130 | 56 | 145 | 121 | 112 | 2 | 0 | 0 | 0 | 0 | 148 | 98 | 86 | 88 | 210 | 0 | 0 | 0 | 0 | 0 | 298 | 149 | 1 | 1 | 0.97 | 1 | 1 | 1 | 1 | 1 | 1 | 1 | 1 | 1 | -0.006 |
| SE | 25251 | ENSG00000126456 | IRF3 | chr19 | - | 49661947 | 49662328 | 49659571 | 49659833 | 49662424 | 49662617 | 123 | 53 | 132 | 113 | 90 | 2 | 0 | 0 | 0 | 0 | 148 | 97 | 80 | 76 | 197 | 0 | 0 | 0 | 0 | 0 | 298 | 149 | 1 | 1 | 0.969 | 1 | 1 | 1 | 1 | 1 | 1 | 1 | 1 | 1 | -0.006 |
| SE | 25252 | ENSG00000126456 | IRF3 | chr19 | - | 49661947 | 49662328 | 49660712 | 49660828 | 49662424 | 49662617 | 187 | 70 | 187 | 197 | 178 | 2 | 0 | 0 | 0 | 0 | 212 | 151 | 110 | 102 | 290 | 0 | 0 | 0 | 0 | 0 | 298 | 149 | 1 | 1 | 0.979 | 1 | 1 | 1 | 1 | 1 | 1 | 1 | 1 | 1 | -0.004 |
| SE | 25254 | ENSG00000126456 | IRF3 | chr19 | - | 49663342 | 49663514 | 49663187 | 49663258 | 49664673 | 49664846 | 284 | 108 | 299 | 273 | 367 | 0 | 0 | 0 | 0 | 0 | 148 | 206 | 108 | 141 | 423 | 0 | 0 | 0 | 0 | 4 | 298 | 149 | 1 | 1 | 1 | 1 | 1 | 1 | 1 | 1 | 1 | 1 | 1 | 0.981 | 0.004 |
| SE | 25256 | ENSG00000126456 | IRF3 | chr19 | - | 49663342 | 49663514 | 49663187 | 49663258 | 49665634 | 49665779 | 187 | 67 | 168 | 145 | 263 | 0 | 0 | 1 | 0 | 0 | 111 | 145 | 76 | 75 | 209 | 0 | 0 | 0 | 0 | 0 | 298 | 149 | 1 | 1 | 1 | 1 | 0.988 | 1 | 1 | 1 | 1 | 1 | 1 | 1 | -0.002 |
| SE | 25257 | ENSG00000126456 | IRF3 | chr19 | - | 49664442 | 49664673 | 49663342 | 49663514 | 49665630 | 49665705 | 9 | 8 | 17 | 4 | 3 | 0 | 0 | 0 | 1 | 0 | 0 | 6 | 1 | 10 | 23 | 0 | 0 | 0 | 0 | 0 | 298 | 149 | 1 | 1 | 1 | 1 | 1 | 0.667 | 1 | NA | 1 | 1 | 1 | 1 | -0.067 |
| SE | 25259 | ENSG00000126456 | IRF3 | chr19 | - | 49664552 | 49664673 | 49663187 | 49663258 | 49665634 | 49665847 | 0 | 0 | 3 | 3 | 0 | 0 | 0 | 1 | 0 | 0 | 1 | 0 | 0 | 0 | 0 | 0 | 0 | 0 | 0 | 0 | 270 | 149 | 1 | 1 | NA | NA | 0.623 | 1 | NA | 1 | NA | NA | NA | NA | -0.189 |
| SE | 25260 | ENSG00000126456 | IRF3 | chr19 | - | 49664552 | 49664673 | 49663342 | 49663514 | 49665630 | 49665788 | 14 | 8 | 4 | 3 | 10 | 0 | 0 | 0 | 1 | 0 | 0 | 0 | 0 | 8 | 44 | 0 | 0 | 0 | 0 | 0 | 270 | 149 | 1 | 1 | 1 | 1 | 1 | 0.623 | 1 | 1 | NA | NA | 1 | 1 | -0.075 |
| SE | 25261 | ENSG00000126456 | IRF3 | chr19 | - | 49664552 | 49664673 | 49663342 | 49663514 | 49665634 | 49665847 | 8 | 2 | 5 | 5 | 7 | 0 | 0 | 2 | 0 | 0 | 1 | 0 | 0 | 0 | 24 | 0 | 0 | 0 | 0 | 0 | 270 | 149 | 1 | 1 | 1 | 1 | 0.58 | 1 | 1 | 1 | NA | NA | NA | 1 | -0.084 |
| SE | 25262 | ENSG00000126456 | IRF3 | chr19 | - | 49664586 | 49664673 | 49663420 | 49663514 | 49665634 | 49665754 | 0 | 0 | 3 | 3 | 6 | 0 | 0 | 2 | 0 | 0 | 2 | 0 | 0 | 0 | 0 | 0 | 0 | 0 | 0 | 0 | 236 | 149 | 1 | 1 | NA | NA | 0.486 | 1 | 1 | 1 | NA | NA | NA | NA | -0.171 |
| SE | 25263 | ENSG00000126456 | IRF3 | chr19 | - | 49664673 | 49664846 | 49663187 | 49663514 | 49665630 | 49665705 | 113 | 48 | 154 | 145 | 115 | 0 | 0 | 0 | 1 | 0 | 37 | 71 | 39 | 68 | 221 | 0 | 0 | 0 | 0 | 0 | 298 | 149 | 1 | 1 | 1 | 1 | 1 | 0.986 | 1 | 1 | 1 | 1 | 1 | 1 | -0.003 |
| SE | 25264 | ENSG00000126456 | IRF3 | chr19 | - | 49664673 | 49664846 | 49663187 | 49663514 | 49665634 | 49665847 | 106 | 43 | 133 | 135 | 107 | 0 | 0 | 2 | 0 | 0 | 47 | 70 | 32 | 75 | 214 | 0 | 0 | 0 | 0 | 0 | 298 | 149 | 1 | 1 | 1 | 1 | 0.971 | 1 | 1 | 1 | 1 | 1 | 1 | 1 | -0.006 |
| SE | 25265 | ENSG00000126456 | IRF3 | chr19 | - | 49664673 | 49664950 | 49663342 | 49663514 | 49665634 | 49665779 | 97 | 41 | 133 | 128 | 104 | 0 | 0 | 2 | 0 | 0 | 37 | 61 | 32 | 66 | 214 | 0 | 0 | 0 | 0 | 0 | 298 | 149 | 1 | 1 | 1 | 1 | 0.971 | 1 | 1 | 1 | 1 | 1 | 1 | 1 | -0.006 |
| SE | 25266 | ENSG00000249476 | NA | chr5 | - | 109319758 | 109320635 | 109237119 | 109238118 | 109321393 | 109321477 | 4 | 1 | 8 | 0 | 3 | 0 | 0 | 0 | 2 | 0 | 12 | 11 | 7 | 7 | 0 | 0 | 0 | 0 | 0 | 0 | 298 | 149 | 1 | 1 | 1 | 1 | 1 | 0 | 1 | 1 | 1 | 1 | 1 | NA | -0.2 |
| SE | 25268 | ENSG00000119685 | TTLL5 | chr14 | + | 75683549 | 75683656 | 75681544 | 75681627 | 75690191 | 75690282 | 54 | 48 | 36 | 53 | 87 | 0 | 0 | 0 | 2 | 0 | 38 | 72 | 53 | 68 | 56 | 0 | 0 | 0 | 0 | 0 | 256 | 149 | 1 | 1 | 1 | 1 | 1 | 0.939 | 1 | 1 | 1 | 1 | 1 | 1 | -0.012 |
| SE | 25273 | ENSG00000119685 | TTLL5 | chr14 | + | 75776746 | 75776850 | 75775483 | 75775630 | 75779574 | 75779702 | 97 | 69 | 59 | 101 | 105 | 0 | 0 | 0 | 2 | 0 | 120 | 112 | 72 | 80 | 54 | 0 | 0 | 0 | 0 | 0 | 253 | 149 | 1 | 1 | 1 | 1 | 1 | 0.967 | 1 | 1 | 1 | 1 | 1 | 1 | -0.007 |
| SE | 25274 | ENSG00000119685 | TTLL5 | chr14 | + | 75882684 | 75882902 | 75863666 | 75863862 | 75902141 | 75902224 | 6 | 13 | 20 | 9 | 47 | 0 | 0 | 0 | 0 | 0 | 0 | 23 | 7 | 28 | 3 | 0 | 0 | 4 | 0 | 0 | 298 | 149 | 1 | 1 | 1 | 1 | 1 | 1 | 1 | NA | 1 | 0.467 | 1 | 1 | 0.133 |
| SE | 25275 | ENSG00000119685 | TTLL5 | chr14 | + | 75906554 | 75906677 | 75902141 | 75902224 | 75954423 | 75954828 | 0 | 0 | 0 | 0 | 0 | 4 | 5 | 18 | 9 | 22 | 0 | 0 | 0 | 10 | 0 | 0 | 4 | 7 | 10 | 0 | 272 | 149 | 1 | 1 | 0 | 0 | 0 | 0 | 0 | NA | 0 | 0 | 0.354 | NA | -0.118 |
| SE | 25276 | ENSG00000119685 | TTLL5 | chr14 | + | 75941423 | 75941494 | 75902139 | 75902224 | 75954423 | 75954823 | 0 | 1 | 0 | 0 | 0 | 4 | 5 | 18 | 9 | 22 | 0 | 0 | 0 | 0 | 0 | 0 | 4 | 7 | 10 | 0 | 220 | 149 | 1 | 1 | 0 | 0.119 | 0 | 0 | 0 | NA | 0 | 0 | 0 | NA | 0.024 |
| SE | 25283 | ENSG00000184584 | STING1 | chr5 | - | 139477328 | 139477515 | 139475532 | 139476454 | 139477744 | 139477908 | 122 | 82 | 83 | 70 | 55 | 0 | 0 | 3 | 0 | 0 | 139 | 123 | 79 | 134 | 212 | 0 | 2 | 0 | 0 | 0 | 298 | 149 | 1 | 1 | 1 | 1 | 0.933 | 1 | 1 | 1 | 0.969 | 1 | 1 | 1 | -0.007 |
| SE | 25286 | ENSG00000184584 | STING1 | chr5 | - | 139477744 | 139477908 | 139477328 | 139477515 | 139478269 | 139478508 | 1 | 0 | 2 | 2 | 0 | 95 | 81 | 67 | 89 | 100 | 0 | 0 | 2 | 0 | 0 | 73 | 66 | 61 | 99 | 150 | 298 | 149 | 1 | 1 | 0.005 | 0 | 0.015 | 0.011 | 0 | 0 | 0 | 0.016 | 0 | 0 | 0.003 |
| SE | 25289 | ENSG00000184584 | STING1 | chr5 | - | 139480789 | 139480898 | 139478269 | 139478508 | 139482209 | 139482293 | 85 | 37 | 66 | 45 | 81 | 1 | 0 | 0 | 0 | 0 | 160 | 60 | 41 | 86 | 149 | 0 | 0 | 0 | 0 | 0 | 258 | 149 | 1 | 1 | 0.98 | 1 | 1 | 1 | 1 | 1 | 1 | 1 | 1 | 1 | -0.004 |
| SE | 25290 | ENSG00000184584 | STING1 | chr5 | - | 139481158 | 139481342 | 139480789 | 139480898 | 139482209 | 139482302 | 87 | 37 | 86 | 64 | 35 | 1 | 0 | 0 | 0 | 0 | 142 | 93 | 62 | 71 | 154 | 0 | 0 | 0 | 3 | 0 | 298 | 149 | 1 | 1 | 0.978 | 1 | 1 | 1 | 1 | 1 | 1 | 1 | 0.922 | 1 | 0.011 |
| SE | 25291 | ENSG00000184584 | STING1 | chr5 | - | 139481158 | 139481704 | 139480789 | 139480898 | 139482209 | 139482293 | 224 | 101 | 176 | 148 | 179 | 1 | 0 | 0 | 0 | 0 | 242 | 194 | 112 | 151 | 321 | 0 | 0 | 0 | 3 | 0 | 298 | 149 | 1 | 1 | 0.991 | 1 | 1 | 1 | 1 | 1 | 1 | 1 | 0.962 | 1 | 0.006 |
| SE | 25292 | ENSG00000184584 | STING1 | chr5 | - | 139481477 | 139481550 | 139481158 | 139481342 | 139482209 | 139482302 | 29 | 17 | 11 | 14 | 15 | 1 | 0 | 0 | 0 | 0 | 17 | 34 | 10 | 15 | 56 | 0 | 0 | 0 | 0 | 0 | 222 | 149 | 1 | 1 | 0.951 | 1 | 1 | 1 | 1 | 1 | 1 | 1 | 1 | 1 | -0.01 |
| SE | 25293 | ENSG00000184584 | STING1 | chr5 | - | 139481477 | 139481704 | 139481158 | 139481342 | 139482209 | 139482299 | 173 | 84 | 108 | 105 | 159 | 1 | 0 | 0 | 0 | 0 | 117 | 138 | 64 | 101 | 223 | 0 | 0 | 0 | 0 | 0 | 298 | 149 | 1 | 1 | 0.989 | 1 | 1 | 1 | 1 | 1 | 1 | 1 | 1 | 1 | -0.002 |
| SE | 25297 | ENSG00000134644 | PUM1 | chr1 | - | 30941997 | 30942123 | 30941150 | 30941272 | 30950132 | 30950261 | 212 | 136 | 224 | 249 | 171 | 0 | 0 | 0 | 1 | 0 | 208 | 233 | 137 | 194 | 144 | 0 | 0 | 6 | 0 | 0 | 275 | 149 | 1 | 1 | 1 | 1 | 1 | 0.993 | 1 | 1 | 1 | 0.925 | 1 | 1 | 0.014 |
| SE | 25300 | ENSG00000134644 | PUM1 | chr1 | - | 30945345 | 30945483 | 30941997 | 30942123 | 30950126 | 30950261 | 359 | 192 | 302 | 356 | 450 | 0 | 1 | 1 | 2 | 0 | 296 | 372 | 176 | 200 | 227 | 0 | 0 | 1 | 0 | 0 | 287 | 149 | 1 | 1 | 1 | 0.99 | 0.994 | 0.989 | 1 | 1 | 1 | 0.989 | 1 | 1 | -0.003 |
| SE | 25302 | ENSG00000134644 | PUM1 | chr1 | - | 30953713 | 30953818 | 30952233 | 30952363 | 30964673 | 30964872 | 97 | 64 | 75 | 82 | 66 | 0 | 0 | 1 | 0 | 0 | 71 | 48 | 38 | 16 | 34 | 0 | 0 | 0 | 0 | 0 | 254 | 149 | 1 | 1 | 1 | 1 | 0.978 | 1 | 1 | 1 | 1 | 1 | 1 | 1 | -0.004 |
| SE | 25303 | ENSG00000134644 | PUM1 | chr1 | - | 30953713 | 30953981 | 30952233 | 30952363 | 30964673 | 30964910 | 243 | 127 | 219 | 240 | 203 | 0 | 0 | 1 | 0 | 0 | 174 | 196 | 161 | 115 | 102 | 0 | 0 | 0 | 0 | 0 | 298 | 149 | 1 | 1 | 1 | 1 | 0.991 | 1 | 1 | 1 | 1 | 1 | 1 | 1 | -0.002 |
| SE | 25305 | ENSG00000134644 | PUM1 | chr1 | - | 30967098 | 30967310 | 30966146 | 30966278 | 30968353 | 30968492 | 119 | 41 | 85 | 89 | 45 | 3 | 0 | 0 | 2 | 0 | 43 | 131 | 81 | 117 | 58 | 0 | 0 | 0 | 0 | 0 | 298 | 149 | 1 | 1 | 0.952 | 1 | 1 | 0.957 | 1 | 1 | 1 | 1 | 1 | 1 | -0.018 |
| SE | 25306 | ENSG00000134644 | PUM1 | chr1 | - | 30967166 | 30967310 | 30965981 | 30966278 | 30968353 | 30968492 | 237 | 80 | 175 | 188 | 87 | 3 | 0 | 0 | 2 | 0 | 70 | 183 | 120 | 165 | 143 | 0 | 0 | 0 | 0 | 0 | 293 | 149 | 1 | 1 | 0.976 | 1 | 1 | 0.98 | 1 | 1 | 1 | 1 | 1 | 1 | -0.009 |
| SE | 25307 | ENSG00000134644 | PUM1 | chr1 | - | 30974650 | 30974802 | 30968353 | 30968492 | 30992389 | 30992660 | 8 | 7 | 12 | 23 | 13 | 0 | 0 | 0 | 0 | 0 | 21 | 21 | 7 | 30 | 13 | 0 | 2 | 0 | 0 | 0 | 298 | 149 | 1 | 1 | 1 | 1 | 1 | 1 | 1 | 1 | 0.84 | 1 | 1 | 1 | 0.032 |
| SE | 25311 | ENSG00000134644 | PUM1 | chr1 | - | 30981311 | 30981405 | 30974650 | 30974802 | 30992389 | 30992660 | 90 | 36 | 60 | 78 | 67 | 0 | 0 | 0 | 3 | 0 | 67 | 111 | 74 | 95 | 22 | 0 | 0 | 0 | 5 | 0 | 243 | 149 | 1 | 1 | 1 | 1 | 1 | 0.941 | 1 | 1 | 1 | 1 | 0.921 | 1 | 0.004 |
| SE | 25312 | ENSG00000134644 | PUM1 | chr1 | - | 30992389 | 30992660 | 30981311 | 30981405 | 31006993 | 31007102 | 111 | 62 | 87 | 108 | 110 | 0 | 0 | 2 | 0 | 0 | 109 | 106 | 62 | 62 | 43 | 0 | 0 | 0 | 2 | 0 | 298 | 149 | 1 | 1 | 1 | 1 | 0.956 | 1 | 1 | 1 | 1 | 1 | 0.939 | 1 | 0.003 |
| SE | 25314 | ENSG00000134644 | PUM1 | chr1 | - | 30995053 | 30995220 | 30992389 | 30992660 | 31006993 | 31007102 | 255 | 151 | 218 | 265 | 339 | 1 | 0 | 0 | 0 | 0 | 247 | 189 | 180 | 190 | 153 | 0 | 0 | 3 | 0 | 0 | 298 | 149 | 1 | 1 | 0.992 | 1 | 1 | 1 | 1 | 1 | 1 | 0.968 | 1 | 1 | 0.005 |
| SE | 25315 | ENSG00000134644 | PUM1 | chr1 | - | 30995053 | 30995220 | 30992389 | 30992660 | 31028795 | 31028864 | 266 | 159 | 218 | 271 | 355 | 0 | 0 | 0 | 0 | 0 | 254 | 199 | 183 | 191 | 137 | 0 | 0 | 1 | 0 | 0 | 298 | 149 | 1 | 1 | 1 | 1 | 1 | 1 | 1 | 1 | 1 | 0.989 | 1 | 1 | 0.002 |
| SE | 25316 | ENSG00000134644 | PUM1 | chr1 | - | 31005852 | 31006031 | 30981311 | 30981405 | 31006993 | 31007102 | 191 | 195 | 222 | 241 | 323 | 0 | 0 | 2 | 0 | 0 | 372 | 299 | 259 | 282 | 165 | 0 | 0 | 0 | 2 | 0 | 298 | 149 | 1 | 1 | 1 | 1 | 0.982 | 1 | 1 | 1 | 1 | 1 | 0.986 | 1 | -0.001 |
| SE | 25319 | ENSG00000134644 | PUM1 | chr1 | - | 31006993 | 31007102 | 30992389 | 30992660 | 31028795 | 31028864 | 125 | 72 | 112 | 125 | 115 | 0 | 0 | 0 | 0 | 0 | 193 | 123 | 104 | 108 | 86 | 0 | 0 | 1 | 0 | 0 | 258 | 149 | 1 | 1 | 1 | 1 | 1 | 1 | 1 | 1 | 1 | 0.984 | 1 | 1 | 0.003 |
| SE | 25322 | ENSG00000134644 | PUM1 | chr1 | - | 31055312 | 31055380 | 31028795 | 31028864 | 31059298 | 31059577 | 3 | 3 | 3 | 1 | 8 | 0 | 4 | 5 | 1 | 3 | 12 | 5 | 3 | 3 | 0 | 3 | 3 | 0 | 0 | 0 | 217 | 149 | 1 | 1 | 1 | 0.34 | 0.292 | 0.407 | 0.647 | 0.733 | 0.534 | 1 | 1 | NA | -0.28 |
| SE | 25324 | ENSG00000108828 | VAT1 | chr17 | - | 43017840 | 43017930 | 43016066 | 43016548 | 43018035 | 43018045 | 131 | 55 | 128 | 131 | 203 | 0 | 2 | 0 | 0 | 0 | 248 | 187 | 139 | 402 | 421 | 0 | 0 | 0 | 3 | 0 | 239 | 149 | 1 | 1 | 1 | 0.945 | 1 | 1 | 1 | 1 | 1 | 1 | 0.988 | 1 | -0.009 |
| SE | 25330 | ENSG00000196411 | EPHB4 | chr7 | - | 100806419 | 100806569 | 100805500 | 100805694 | 100807364 | 100807580 | 9 | 3 | 6 | 17 | 18 | 0 | 0 | 0 | 2 | 0 | 0 | 4 | 2 | 0 | 2 | 0 | 0 | 0 | 0 | 0 | 298 | 149 | 1 | 1 | 1 | 1 | 1 | 0.81 | 1 | NA | 1 | 1 | NA | 1 | -0.038 |
| SE | 25331 | ENSG00000196411 | EPHB4 | chr7 | - | 100817191 | 100817357 | 100813918 | 100814021 | 100818519 | 100818644 | 4 | 2 | 16 | 11 | 3 | 0 | 2 | 0 | 0 | 0 | 7 | 3 | 0 | 0 | 0 | 0 | 0 | 0 | 0 | 0 | 298 | 149 | 1 | 1 | 1 | 0.333 | 1 | 1 | 1 | 1 | 1 | NA | NA | NA | -0.133 |
| SE | 25340 | ENSG00000130803 | ZNF317 | chr19 | + | 9155924 | 9156041 | 9140396 | 9140592 | 9156611 | 9156748 | 58 | 32 | 65 | 68 | 30 | 0 | 0 | 1 | 0 | 0 | 114 | 87 | 63 | 65 | 74 | 0 | 1 | 0 | 0 | 3 | 266 | 149 | 1 | 1 | 1 | 1 | 0.973 | 1 | 1 | 1 | 0.98 | 1 | 1 | 0.933 | 0.012 |
| SE | 25341 | ENSG00000130803 | ZNF317 | chr19 | + | 9156611 | 9156748 | 9155924 | 9156041 | 9157267 | 9157394 | 71 | 30 | 72 | 83 | 51 | 0 | 0 | 0 | 0 | 3 | 92 | 92 | 83 | 62 | 89 | 0 | 0 | 0 | 0 | 0 | 286 | 149 | 1 | 1 | 1 | 1 | 1 | 1 | 0.899 | 1 | 1 | 1 | 1 | 1 | -0.02 |
| SE | 25348 | ENSG00000112992 | NNT | chr5 | + | 43612907 | 43613137 | 43609515 | 43609551 | 43624031 | 43624120 | 2 | 8 | 3 | 2 | 2 | 0 | 0 | 0 | 0 | 0 | 0 | 3 | 26 | 1 | 7 | 0 | 0 | 1 | 0 | 0 | 298 | 149 | 1 | 1 | 1 | 1 | 1 | 1 | 1 | NA | 1 | 0.929 | 1 | 1 | 0.018 |
| SE | 25350 | ENSG00000112992 | NNT | chr5 | + | 43615847 | 43616065 | 43612907 | 43613137 | 43624031 | 43624120 | 162 | 70 | 115 | 131 | 153 | 0 | 1 | 0 | 0 | 0 | 85 | 100 | 76 | 37 | 102 | 0 | 0 | 0 | 0 | 0 | 298 | 149 | 1 | 1 | 1 | 0.972 | 1 | 1 | 1 | 1 | 1 | 1 | 1 | 1 | -0.006 |
| SE | 25353 | ENSG00000112992 | NNT | chr5 | + | 43624031 | 43624120 | 43615847 | 43616065 | 43628199 | 43628387 | 95 | 77 | 97 | 88 | 112 | 0 | 0 | 0 | 2 | 0 | 156 | 85 | 84 | 62 | 80 | 0 | 0 | 2 | 0 | 0 | 238 | 149 | 1 | 1 | 1 | 1 | 1 | 0.965 | 1 | 1 | 0.964 | 1 | 1 | 1 | 0 |
| SE | 25354 | ENSG00000112992 | NNT | chr5 | + | 43628199 | 43628387 | 43615847 | 43616065 | 43644191 | 43644325 | 181 | 140 | 158 | 181 | 328 | 0 | 0 | 1 | 0 | 0 | 140 | 143 | 117 | 160 | 116 | 0 | 0 | 0 | 0 | 0 | 298 | 149 | 1 | 1 | 1 | 1 | 0.988 | 1 | 1 | 1 | 1 | 1 | 1 | 1 | -0.002 |
| SE | 25355 | ENSG00000112992 | NNT | chr5 | + | 43644191 | 43644325 | 43628199 | 43628387 | 43644610 | 43644802 | 578 | 423 | 451 | 526 | 888 | 0 | 0 | 0 | 0 | 2 | 495 | 350 | 312 | 343 | 382 | 0 | 0 | 0 | 0 | 0 | 283 | 149 | 1 | 1 | 1 | 1 | 1 | 1 | 0.996 | 1 | 1 | 1 | 1 | 1 | -0.001 |
| SE | 25358 | ENSG00000112992 | NNT | chr5 | + | 43650476 | 43650587 | 43647923 | 43649308 | 43651738 | 43651884 | 135 | 67 | 113 | 113 | 97 | 0 | 0 | 0 | 1 | 0 | 101 | 54 | 57 | 70 | 89 | 0 | 0 | 0 | 0 | 0 | 260 | 149 | 1 | 1 | 1 | 1 | 1 | 0.985 | 1 | 1 | 1 | 1 | 1 | 1 | -0.003 |
| SE | 25360 | ENSG00000112992 | NNT | chr5 | + | 43653017 | 43653213 | 43651738 | 43651884 | 43659170 | 43659350 | 45 | 14 | 30 | 32 | 26 | 0 | 0 | 0 | 1 | 0 | 40 | 21 | 16 | 27 | 12 | 0 | 0 | 0 | 0 | 0 | 298 | 149 | 1 | 1 | 1 | 1 | 1 | 0.941 | 1 | 1 | 1 | 1 | 1 | 1 | -0.012 |
| SE | 25361 | ENSG00000112992 | NNT | chr5 | + | 43700118 | 43700237 | 43677724 | 43677806 | 43702620 | 43702736 | 197 | 120 | 195 | 235 | 374 | 0 | 0 | 0 | 0 | 1 | 265 | 190 | 86 | 107 | 231 | 0 | 2 | 0 | 0 | 0 | 268 | 149 | 1 | 1 | 1 | 1 | 1 | 1 | 0.995 | 1 | 0.981 | 1 | 1 | 1 | 0.003 |
| SE | 25364 | ENSG00000113296 | THBS4 | chr5 | + | 80070642 | 80070750 | 80070305 | 80070410 | 80071020 | 80071180 | 0 | 0 | 0 | 0 | 4 | 0 | 0 | 0 | 0 | 0 | 9 | 9 | 1 | 106 | 19 | 0 | 0 | 0 | 2 | 0 | 257 | 149 | 1 | 1 | NA | NA | NA | NA | 1 | 1 | 1 | 1 | 0.968 | 1 | 0.006 |
| SE | 25371 | ENSG00000092020 | PPP2R3C | chr14 | - | 35091069 | 35091207 | 35087950 | 35088010 | 35096557 | 35096633 | 167 | 105 | 111 | 150 | 253 | 0 | 1 | 0 | 0 | 2 | 5 | 76 | 70 | 41 | 59 | 0 | 0 | 3 | 0 | 0 | 287 | 149 | 1 | 1 | 1 | 0.982 | 1 | 1 | 0.985 | 1 | 1 | 0.924 | 1 | 1 | 0.009 |
| SE | 25373 | ENSG00000092020 | PPP2R3C | chr14 | - | 35107303 | 35107374 | 35099251 | 35099384 | 35108138 | 35108236 | 282 | 178 | 190 | 220 | 251 | 1 | 5 | 0 | 0 | 0 | 107 | 138 | 115 | 67 | 62 | 0 | 1 | 0 | 0 | 0 | 220 | 149 | 1 | 1 | 0.995 | 0.96 | 1 | 1 | 1 | 1 | 0.989 | 1 | 1 | 1 | -0.007 |
| SE | 25375 | ENSG00000092020 | PPP2R3C | chr14 | - | 35108138 | 35108236 | 35099251 | 35099384 | 35110524 | 35110629 | 0 | 0 | 1 | 0 | 0 | 0 | 0 | 0 | 0 | 0 | 0 | 1 | 0 | 0 | 0 | 0 | 0 | 1 | 0 | 1 | 247 | 149 | 1 | 1 | NA | NA | 1 | NA | NA | NA | 1 | 0 | NA | 0 | 0.667 |
| SE | 25376 | ENSG00000092020 | PPP2R3C | chr14 | - | 35109818 | 35109847 | 35108138 | 35108236 | 35110524 | 35110629 | 65 | 43 | 44 | 35 | 55 | 0 | 0 | 1 | 2 | 0 | 1 | 32 | 11 | 10 | 3 | 0 | 0 | 0 | 0 | 0 | 178 | 149 | 1 | 1 | 1 | 1 | 0.974 | 0.936 | 1 | 1 | 1 | 1 | 1 | 1 | -0.018 |
| SE | 25377 | ENSG00000092020 | PPP2R3C | chr14 | - | 35109818 | 35109901 | 35108138 | 35108236 | 35110524 | 35110630 | 117 | 81 | 77 | 81 | 97 | 0 | 0 | 1 | 2 | 0 | 1 | 46 | 40 | 29 | 3 | 0 | 0 | 0 | 0 | 0 | 232 | 149 | 1 | 1 | 1 | 1 | 0.98 | 0.963 | 1 | 1 | 1 | 1 | 1 | 1 | -0.011 |
| SE | 25378 | ENSG00000092020 | PPP2R3C | chr14 | - | 35109818 | 35109931 | 35098662 | 35099384 | 35110524 | 35110629 | 119 | 116 | 91 | 77 | 108 | 0 | 0 | 0 | 0 | 0 | 25 | 106 | 45 | 60 | 17 | 0 | 0 | 1 | 0 | 0 | 262 | 149 | 1 | 1 | 1 | 1 | 1 | 1 | 1 | 1 | 1 | 0.962 | 1 | 0.906 | 0.026 |
| SE | 25379 | ENSG00000092020 | PPP2R3C | chr14 | - | 35109818 | 35109931 | 35107647 | 35108236 | 35110524 | 35110629 | 308 | 241 | 235 | 220 | 281 | 0 | 0 | 1 | 2 | 0 | 64 | 185 | 108 | 100 | 25 | 0 | 0 | 1 | 0 | 0 | 262 | 149 | 1 | 1 | 1 | 1 | 0.993 | 0.984 | 1 | 1 | 1 | 1 | 1 | 1 | -0.005 |
| SE | 25380 | ENSG00000092020 | PPP2R3C | chr14 | - | 35109847 | 35109931 | 35108138 | 35108236 | 35110524 | 35110629 | 52 | 68 | 50 | 33 | 77 | 0 | 0 | 1 | 2 | 0 | 22 | 53 | 21 | 32 | 0 | 0 | 0 | 0 | 0 | 0 | 233 | 149 | 1 | 1 | 1 | 1 | 0.97 | 0.913 | 1 | 1 | 1 | 1 | 1 | NA | -0.023 |
| SE | 25381 | ENSG00000092020 | PPP2R3C | chr14 | - | 35110524 | 35110629 | 35098718 | 35099384 | 35121901 | 35121949 | 4 | 1 | 5 | 3 | 11 | 0 | 0 | 0 | 0 | 0 | 9 | 2 | 5 | 4 | 1 | 0 | 0 | 0 | 1 | 0 | 254 | 149 | 1 | 1 | 1 | 1 | 1 | 1 | 1 | 1 | 1 | 1 | 0.701 | 1 | 0.06 |
| SE | 25382 | ENSG00000092020 | PPP2R3C | chr14 | - | 35110524 | 35110629 | 35108138 | 35108236 | 35116609 | 35116737 | 53 | 59 | 28 | 28 | 64 | 2 | 0 | 0 | 0 | 0 | 0 | 24 | 26 | 31 | 0 | 0 | 0 | 0 | 0 | 0 | 254 | 149 | 1 | 1 | 0.94 | 1 | 1 | 1 | 1 | NA | 1 | 1 | 1 | NA | -0.012 |
| SE | 25383 | ENSG00000092020 | PPP2R3C | chr14 | - | 35116609 | 35116737 | 35109922 | 35110629 | 35121901 | 35121965 | 90 | 122 | 72 | 45 | 126 | 5 | 1 | 7 | 8 | 16 | 0 | 60 | 46 | 54 | 0 | 9 | 2 | 4 | 4 | 0 | 277 | 149 | 1 | 1 | 0.906 | 0.985 | 0.847 | 0.752 | 0.809 | 0 | 0.942 | 0.861 | 0.879 | NA | 0.189 |
| SE | 25386 | ENSG00000113648 | MACROH2A1 | chr5 | - | 135343259 | 135343434 | 135333899 | 135335141 | 135360496 | 135360604 | 814 | 601 | 542 | 694 | 2146 | 0 | 0 | 0 | 0 | 0 | 1325 | 871 | 545 | 877 | 1005 | 0 | 0 | 1 | 3 | 0 | 298 | 149 | 1 | 1 | 1 | 1 | 1 | 1 | 1 | 1 | 1 | 0.996 | 0.993 | 1 | 0.002 |
| SE | 25387 | ENSG00000113648 | MACROH2A1 | chr5 | - | 135345967 | 135346057 | 135333899 | 135335141 | 135360496 | 135360607 | 5 | 4 | 2 | 1 | 21 | 0 | 0 | 0 | 0 | 0 | 7 | 23 | 24 | 32 | 18 | 0 | 0 | 1 | 3 | 0 | 239 | 149 | 1 | 1 | 1 | 1 | 1 | 1 | 1 | 1 | 1 | 0.937 | 0.869 | 1 | 0.039 |
| SE | 25388 | ENSG00000113648 | MACROH2A1 | chr5 | - | 135345967 | 135346057 | 135343259 | 135343434 | 135360496 | 135360607 | 179 | 107 | 133 | 147 | 166 | 0 | 0 | 0 | 0 | 0 | 267 | 219 | 134 | 260 | 273 | 0 | 0 | 4 | 0 | 0 | 239 | 149 | 1 | 1 | 1 | 1 | 1 | 1 | 1 | 1 | 1 | 0.954 | 1 | 1 | 0.009 |
| SE | 25392 | ENSG00000113648 | MACROH2A1 | chr5 | - | 135360496 | 135360607 | 135345967 | 135346057 | 135369405 | 135369603 | 299 | 217 | 189 | 219 | 211 | 0 | 0 | 0 | 0 | 0 | 813 | 430 | 393 | 615 | 653 | 0 | 2 | 4 | 0 | 0 | 260 | 149 | 1 | 1 | 1 | 1 | 1 | 1 | 1 | 1 | 0.992 | 0.983 | 1 | 1 | 0.005 |
| SE | 25393 | ENSG00000113648 | MACROH2A1 | chr5 | - | 135360496 | 135360607 | 135345967 | 135346057 | 135388921 | 135389059 | 0 | 4 | 1 | 1 | 0 | 1 | 0 | 0 | 0 | 0 | 0 | 3 | 6 | 11 | 1 | 0 | 1 | 1 | 11 | 0 | 260 | 149 | 1 | 1 | 0 | 1 | 1 | 1 | NA | NA | 0.632 | 0.775 | 0.364 | 1 | 0.057 |
| SE | 25394 | ENSG00000113648 | MACROH2A1 | chr5 | - | 135360496 | 135360780 | 135345967 | 135346057 | 135369405 | 135369603 | 0 | 4 | 0 | 1 | 0 | 0 | 0 | 0 | 0 | 0 | 0 | 3 | 3 | 11 | 1 | 0 | 2 | 4 | 0 | 0 | 298 | 149 | 1 | 1 | NA | 1 | NA | 1 | NA | NA | 0.429 | 0.273 | 1 | 1 | 0.325 |
| SE | 25395 | ENSG00000113648 | MACROH2A1 | chr5 | - | 135369405 | 135369603 | 135360415 | 135360604 | 135370035 | 135370142 | 1033 | 869 | 748 | 955 | 865 | 0 | 0 | 0 | 0 | 0 | 2937 | 1686 | 1505 | 2338 | 2766 | 11 | 0 | 0 | 16 | 0 | 298 | 149 | 1 | 1 | 1 | 1 | 1 | 1 | 1 | 0.993 | 1 | 1 | 0.986 | 1 | 0.004 |
| SE | 25396 | ENSG00000113648 | MACROH2A1 | chr5 | - | 135369405 | 135369603 | 135360496 | 135360607 | 135370035 | 135370142 | 955 | 747 | 653 | 811 | 713 | 0 | 1 | 0 | 0 | 0 | 2639 | 1539 | 1357 | 2150 | 2483 | 0 | 0 | 2 | 3 | 0 | 298 | 149 | 1 | 1 | 1 | 0.997 | 1 | 1 | 1 | 1 | 1 | 0.997 | 0.997 | 1 | 0.001 |
| SE | 25397 | ENSG00000113648 | MACROH2A1 | chr5 | - | 135370035 | 135370142 | 135360496 | 135360604 | 135388921 | 135389059 | 244 | 192 | 147 | 209 | 219 | 0 | 0 | 0 | 0 | 0 | 855 | 506 | 448 | 654 | 692 | 0 | 0 | 2 | 1 | 0 | 256 | 149 | 1 | 1 | 1 | 1 | 1 | 1 | 1 | 1 | 1 | 0.992 | 0.997 | 1 | 0.002 |
| SE | 25398 | ENSG00000113648 | MACROH2A1 | chr5 | - | 135370035 | 135370142 | 135360496 | 135360607 | 135388921 | 135389059 | 244 | 193 | 147 | 209 | 219 | 0 | 1 | 1 | 2 | 0 | 844 | 506 | 450 | 641 | 692 | 0 | 0 | 3 | 0 | 5 | 256 | 149 | 1 | 1 | 1 | 0.991 | 0.988 | 0.984 | 1 | 1 | 1 | 0.989 | 1 | 0.988 | -0.003 |
| SE | 25399 | ENSG00000113648 | MACROH2A1 | chr5 | - | 135388921 | 135389126 | 135370035 | 135370142 | 135399061 | 135399177 | 410 | 323 | 259 | 394 | 290 | 1 | 0 | 2 | 0 | 0 | 1327 | 887 | 748 | 1088 | 1159 | 0 | 4 | 1 | 4 | 0 | 298 | 149 | 1 | 1 | 0.995 | 1 | 0.985 | 1 | 1 | 1 | 0.991 | 0.997 | 0.993 | 1 | 0 |
| SE | 25400 | ENSG00000113648 | MACROH2A1 | chr5 | - | 135388921 | 135389296 | 135370035 | 135370142 | 135399061 | 135399214 | 329 | 279 | 214 | 311 | 280 | 1 | 0 | 2 | 0 | 0 | 1112 | 707 | 639 | 898 | 955 | 0 | 4 | 1 | 4 | 0 | 298 | 149 | 1 | 1 | 0.994 | 1 | 0.982 | 1 | 1 | 1 | 0.989 | 0.997 | 0.991 | 1 | 0 |
| SE | 25401 | ENSG00000186704 | DTX2P1 | chr7 | + | 76990268 | 76990369 | 76980620 | 76981260 | 77000308 | 77000388 | 15 | 15 | 7 | 18 | 0 | 0 | 0 | 0 | 0 | 0 | 15 | 7 | 3 | 6 | 8 | 0 | 0 | 1 | 0 | 0 | 250 | 149 | 1 | 1 | 1 | 1 | 1 | 1 | NA | | 1 | 0.641 | 1 | 1 | 0.072 |
| SE | 25403 | ENSG00000186704 | DTX2P1 | chr7 | + | 77003288 | 77003449 | 77002162 | 77002321 | 77004217 | 77004308 | 0 | 0 | 1 | 0 | 1 | 19 | 12 | 10 | 9 | 8 | 0 | 0 | 0 | 1 | 1 | 55 | 7 | 5 | 17 | 23 | 298 | 149 | 1 | 1 | 0 | 0 | 0.048 | 0 | 0.059 | 0 | 0 | 0 | 0.029 | 0.021 | 0.011 |
| SE | 25406 | ENSG00000251513 | LIX1-AS1 | chr5 | + | 97183007 | 97183240 | 97103025 | 97103094 | 97431743 | 97431900 | 7 | 7 | 4 | 8 | 9 | 1 | 0 | 0 | 0 | 0 | 0 | 6 | 0 | 2 | 19 | 0 | 0 | 0 | 0 | 0 | 298 | 149 | 1 | 1 | 0.778 | 1 | 1 | 1 | 1 | NA | 1 | NA | 1 | 1 | -0.044 |
| SE | 25407 | ENSG00000251513 | LIX1-AS1 | chr5 | + | 97183504 | 97183669 | 97183007 | 97183240 | 97431743 | 97431900 | 2 | 3 | 0 | 1 | 0 | 0 | 0 | 0 | 0 | 1 | 0 | 20 | 0 | 0 | 0 | 0 | 0 | 0 | 0 | 0 | 298 | 149 | 1 | 1 | 1 | 1 | NA | 1 | 0 | NA | 1 | NA | NA | NA | -0.25 |
| SE | 25408 | ENSG00000182903 | ZNF721 | chr4 | - | 435769 | 435879 | 425814 | 427119 | 437445 | 437511 | 18 | 17 | 16 | 15 | 16 | 0 | 1 | 1 | 0 | 1 | 0 | 11 | 7 | 3 | 0 | 0 | 0 | 0 | 0 | 0 | 259 | 149 | 1 | 1 | 1 | 0.907 | 0.902 | 1 | 0.902 | NA | 1 | 1 | 1 | NA | -0.058 |
| SE | 25410 | ENSG00000182903 | ZNF721 | chr4 | - | 438583 | 438632 | 437445 | 437511 | 472574 | 472701 | 0 | 0 | 1 | 1 | 0 | 11 | 8 | 15 | 12 | 1 | 0 | 0 | 0 | 0 | 0 | 12 | 10 | 1 | 5 | 1 | 198 | 149 | 1 | 1 | 0 | 0 | 0.048 | 0.059 | 0 | 0 | 0 | 0 | 0 | 0 | 0.021 |
| SE | 25411 | ENSG00000182903 | ZNF721 | chr4 | - | 444342 | 444432 | 438583 | 438632 | 472574 | 472701 | 99 | 105 | 103 | 107 | 153 | 0 | 0 | 0 | 3 | 0 | 11 | 27 | 43 | 23 | 27 | 0 | 0 | 0 | 1 | 0 | 239 | 149 | 1 | 1 | 1 | 1 | 1 | 0.957 | 1 | 1 | 1 | 1 | 0.935 | 1 | 0.004 |
| SE | 25415 | ENSG00000020633 | RUNX3 | chr1 | - | 24907258 | 24907417 | 24899510 | 24902666 | 24927573 | 24927730 | 105 | 27 | 51 | 45 | 20 | 0 | 0 | 0 | 0 | 0 | 54 | 41 | 91 | 75 | 129 | 0 | 0 | 0 | 4 | 0 | 298 | 149 | 1 | 1 | 1 | 1 | 1 | 1 | 1 | 1 | 1 | 1 | 0.904 | 1 | 0.019 |
| SE | 25416 | ENSG00000020633 | RUNX3 | chr1 | - | 24919239 | 24919344 | 24899510 | 24902666 | 24927573 | 24927730 | 391 | 109 | 180 | 268 | 142 | 0 | 0 | 0 | 0 | 0 | 191 | 279 | 306 | 116 | 249 | 0 | 0 | 0 | 4 | 0 | 254 | 149 | 1 | 1 | 1 | 1 | 1 | 1 | 1 | 1 | 1 | 1 | 0.944 | 1 | 0.011 |
| SE | 25418 | ENSG00000020633 | RUNX3 | chr1 | - | 24927573 | 24927730 | 24919239 | 24919344 | 24929586 | 24929852 | 740 | 195 | 350 | 515 | 215 | 0 | 0 | 0 | 0 | 0 | 339 | 532 | 558 | 297 | 581 | 0 | 2 | 0 | 0 | 0 | 298 | 149 | 1 | 1 | 1 | 1 | 1 | 1 | 1 | 1 | 0.993 | 1 | 1 | 1 | 0.001 |
| SE | 25424 | ENSG00000247796 | NA | chr5 | + | 53112239 | 53113034 | 53109841 | 53110148 | 53113858 | 53114109 | 0 | 2 | 0 | 0 | 0 | 1 | 0 | 2 | 1 | 4 | 0 | 0 | 1 | 0 | 0 | 0 | 0 | 0 | 0 | 0 | 298 | 149 | 1 | 1 | 0 | 1 | 0 | 0 | 0 | NA | NA | 1 | NA | NA | -0.8 |
| SE | 25433 | ENSG00000288674 | NA | chr1 | + | 226881887 | 226882048 | 226875364 | 226875550 | 226885537 | 226885679 | 23 | 19 | 19 | 20 | 10 | 0 | 0 | 0 | 1 | 0 | 47 | 31 | 12 | 20 | 41 | 0 | 0 | 0 | 0 | 0 | 298 | 149 | 1 | 1 | 1 | 1 | 1 | 0.909 | 1 | 1 | 1 | 1 | 1 | 1 | -0.018 |
| SE | 25434 | ENSG00000288674 | NA | chr1 | + | 226883704 | 226883919 | 226875364 | 226875550 | 226885537 | 226885679 | 6 | 4 | 10 | 15 | 7 | 0 | 0 | 0 | 1 | 0 | 6 | 6 | 0 | 4 | 0 | 0 | 0 | 0 | 0 | 0 | 298 | 149 | 1 | 1 | 1 | 1 | 1 | 0.882 | 1 | 1 | 1 | NA | 1 | NA | -0.024 |
| SE | 25437 | ENSG00000288674 | NA | chr1 | + | 226965670 | 226965737 | 226964999 | 226965410 | 226977448 | 226977523 | 68 | 26 | 73 | 62 | 22 | 1 | 2 | 0 | 0 | 0 | 47 | 95 | 67 | 46 | 98 | 0 | 0 | 0 | 0 | 0 | 216 | 149 | 1 | 1 | 0.979 | 0.9 | 1 | 1 | 1 | 1 | 1 | 1 | 1 | 1 | -0.024 |
| SE | 25442 | ENSG00000011243 | AKAP8L | chr19 | - | 15398629 | 15399410 | 15397713 | 15397855 | 15400793 | 15400864 | 15 | 11 | 14 | 8 | 8 | 0 | 0 | 0 | 0 | 0 | 3 | 0 | 0 | 11 | 6 | 0 | 0 | 2 | 8 | 0 | 298 | 149 | 1 | 1 | 1 | 1 | 1 | 1 | 1 | 1 | NA | 0 | 0.407 | 1 | 0.398 |
| SE | 25447 | ENSG00000011243 | AKAP8L | chr19 | - | 15399831 | 15399984 | 15399301 | 15399410 | 15400294 | 15400358 | 0 | 1 | 0 | 0 | 0 | 101 | 61 | 71 | 99 | 149 | 0 | 0 | 0 | 0 | 0 | 276 | 139 | 112 | 138 | 221 | 298 | 149 | 1 | 1 | 0 | 0.008 | 0 | 0 | 0 | 0 | 0 | 0 | 0 | 0 | 0.002 |
| SE | 25449 | ENSG00000011243 | AKAP8L | chr19 | - | 15400294 | 15400358 | 15397713 | 15399410 | 15400793 | 15400864 | 134 | 93 | 96 | 139 | 181 | 0 | 0 | 0 | 0 | 0 | 344 | 179 | 172 | 226 | 351 | 0 | 0 | 0 | 1 | 0 | 213 | 149 | 1 | 1 | 1 | 1 | 1 | 1 | 1 | 1 | 1 | 1 | 0.994 | 1 | 0.001 |
| SE | 25450 | ENSG00000011243 | AKAP8L | chr19 | - | 15403474 | 15403555 | 15401149 | 15401603 | 15403657 | 15403715 | 57 | 46 | 82 | 80 | 84 | 0 | 0 | 1 | 0 | 0 | 91 | 98 | 33 | 43 | 78 | 0 | 0 | 0 | 0 | 0 | 230 | 149 | 1 | 1 | 1 | 1 | 0.982 | 1 | 1 | 1 | 1 | 1 | 1 | 1 | -0.004 |
| SE | 25451 | ENSG00000011243 | AKAP8L | chr19 | - | 15403474 | 15403555 | 15401149 | 15401603 | 15404009 | 15404042 | 51 | 31 | 61 | 58 | 73 | 0 | 0 | 0 | 0 | 0 | 91 | 64 | 28 | 40 | 73 | 0 | 3 | 0 | 0 | 0 | 230 | 149 | 1 | 1 | 1 | 1 | 1 | 1 | 1 | 1 | 0.933 | 1 | 1 | 1 | 0.013 |
| SE | 25452 | ENSG00000011243 | AKAP8L | chr19 | - | 15403474 | 15403715 | 15401149 | 15401603 | 15404009 | 15404042 | 314 | 227 | 372 | 413 | 406 | 0 | 0 | 0 | 0 | 0 | 557 | 425 | 303 | 340 | 656 | 0 | 3 | 0 | 0 | 0 | 298 | 149 | 1 | 1 | 1 | 1 | 1 | 1 | 1 | 1 | 0.986 | 1 | 1 | 1 | 0.003 |
| SE | 25454 | ENSG00000011243 | AKAP8L | chr19 | - | 15403657 | 15403715 | 15401149 | 15401603 | 15404009 | 15404042 | 51 | 25 | 73 | 77 | 50 | 0 | 0 | 0 | 0 | 0 | 74 | 63 | 67 | 61 | 105 | 0 | 3 | 0 | 0 | 0 | 207 | 149 | 1 | 1 | 1 | 1 | 1 | 1 | 1 | 1 | 0.938 | 1 | 1 | 1 | 0.012 |
| SE | 25458 | ENSG00000217128 | FNIP1 | chr5 | - | 131677702 | 131677872 | 131671504 | 131672924 | 131679028 | 131679175 | 159 | 112 | 153 | 198 | 423 | 0 | 0 | 0 | 0 | 0 | 115 | 112 | 171 | 45 | 62 | 0 | 0 | 1 | 0 | 0 | 298 | 149 | 1 | 1 | 1 | 1 | 1 | 1 | 1 | 1 | 1 | 0.988 | 1 | 1 | 0.002 |
| SE | 25462 | ENSG00000217128 | FNIP1 | chr5 | - | 131719316 | 131719417 | 131718985 | 131719060 | 131730903 | 131731038 | 118 | 79 | 87 | 75 | 142 | 0 | 0 | 0 | 1 | 0 | 6 | 64 | 127 | 37 | 14 | 0 | 0 | 0 | 0 | 0 | 250 | 149 | 1 | 1 | 1 | 1 | 1 | 0.978 | 1 | 1 | 1 | 1 | 1 | 1 | -0.004 |
| SE | 25464 | ENSG00000136830 | NIBAN2 | chr9 | - | 127517825 | 127517941 | 127517111 | 127517216 | 127523678 | 127523846 | 63 | 33 | 44 | 57 | 25 | 0 | 0 | 0 | 0 | 0 | 308 | 98 | 53 | 111 | 231 | 0 | 0 | 2 | 0 | 0 | 265 | 149 | 1 | 1 | 1 | 1 | 1 | 1 | 1 | 1 | 1 | 0.937 | 1 | 1 | 0.013 |
| SE | 25466 | ENSG00000189067 | LITAF | chr16 | - | 11551706 | 11551804 | 11547998 | 11549745 | 11553532 | 11553689 | 4 | 0 | 0 | 6 | 6 | 312 | 98 | 289 | 286 | 172 | 0 | 0 | 0 | 0 | 0 | 193 | 353 | 338 | 223 | 263 | 247 | 149 | 1 | 1 | 0.008 | 0 | 0 | 0.012 | 0.021 | 0 | 0 | 0 | 0 | 0 | 0.008 |
| SE | 25476 | ENSG00000082397 | EPB41L3 | chr18 | - | 5396200 | 5396332 | 5395608 | 5395707 | 5397057 | 5397426 | 173 | 223 | 81 | 199 | 255 | 0 | 0 | 0 | 0 | 0 | 296 | 250 | 102 | 289 | 100 | 5 | 0 | 0 | 0 | 0 | 281 | 149 | 1 | 1 | 1 | 1 | 1 | 1 | 1 | 0.969 | 1 | 1 | 1 | 1 | 0.006 |
| SE | 25477 | ENSG00000082397 | EPB41L3 | chr18 | - | 5398020 | 5398143 | 5397057 | 5397426 | 5400496 | 5400595 | 2 | 0 | 0 | 0 | 0 | 0 | 0 | 2 | 0 | 0 | 0 | 0 | 0 | 1 | 0 | 9 | 0 | 0 | 0 | 0 | 272 | 149 | 1 | 1 | 1 | NA | 0 | NA | NA | 0 | NA | NA | 1 | NA | 0 |
| SE | 25478 | ENSG00000082397 | EPB41L3 | chr18 | - | 5398020 | 5398143 | 5397057 | 5397426 | 5400968 | 5401043 | 2 | 0 | 0 | 0 | 0 | 0 | 4 | 0 | 5 | 5 | 0 | 0 | 0 | 1 | 0 | 4 | 0 | 0 | 4 | 0 | 272 | 149 | 1 | 1 | 1 | 0 | NA | 0 | 0 | 0 | NA | NA | 0.12 | NA | 0.19 |
| SE | 25479 | ENSG00000082397 | EPB41L3 | chr18 | - | 5398020 | 5398143 | 5397057 | 5397426 | 5406776 | 5406968 | 2 | 0 | 0 | 0 | 0 | 22 | 24 | 11 | 20 | 41 | 0 | 0 | 2 | 1 | 0 | 32 | 31 | 12 | 46 | 6 | 272 | 149 | 1 | 1 | 0.047 | 0 | 0 | 0 | 0 | 0 | 0 | 0.084 | 0.012 | 0 | -0.01 |
| SE | 25480 | ENSG00000082397 | EPB41L3 | chr18 | - | 5398020 | 5401043 | 5397057 | 5397426 | 5406776 | 5406968 | 2 | 0 | 0 | 0 | 0 | 22 | 24 | 11 | 20 | 41 | 0 | 0 | 0 | 3 | 0 | 32 | 31 | 12 | 46 | 6 | 298 | 149 | 1 | 1 | 0.043 | 0 | 0 | 0 | 0 | 0 | 0 | 0 | 0.032 | 0 | 0.002 |
| SE | 25483 | ENSG00000082397 | EPB41L3 | chr18 | - | 5400496 | 5400595 | 5398020 | 5398143 | 5406776 | 5406968 | 0 | 0 | 0 | 0 | 1 | 0 | 0 | 0 | 0 | 0 | 0 | 0 | 0 | 0 | 0 | 0 | 0 | 2 | 0 | 0 | 248 | 149 | 1 | 1 | NA | NA | NA | NA | 1 | NA | NA | 0 | NA | NA | 1 |
| SE | 25484 | ENSG00000082397 | EPB41L3 | chr18 | - | 5400968 | 5401043 | 5397057 | 5397426 | 5406764 | 5406968 | 0 | 3 | 0 | 1 | 2 | 0 | 2 | 2 | 1 | 0 | 4 | 0 | 0 | 2 | 0 | 0 | 0 | 0 | 0 | 0 | 224 | 149 | 1 | 1 | NA | 0.499 | 0 | 0.399 | 1 | 1 | NA | NA | 1 | NA | -0.525 |
| SE | 25490 | ENSG00000082397 | EPB41L3 | chr18 | - | 5433468 | 5433556 | 5428312 | 5428465 | 5433902 | 5434121 | 75 | 109 | 43 | 72 | 54 | 0 | 0 | 0 | 0 | 1 | 170 | 43 | 54 | 119 | 47 | 0 | 0 | 1 | 0 | 0 | 237 | 149 | 1 | 1 | 1 | 1 | 1 | 1 | 0.971 | 1 | 1 | 0.971 | 1 | 1 | 0 |
| SE | 25491 | ENSG00000082397 | EPB41L3 | chr18 | - | 5433902 | 5434121 | 5428312 | 5428465 | 5445139 | 5445244 | 0 | 0 | 0 | 0 | 1 | 1 | 0 | 0 | 0 | 0 | 0 | 0 | 1 | 0 | 0 | 0 | 8 | 0 | 0 | 0 | 298 | 149 | 1 | 1 | 1 | 0 | NA | NA | 1 | NA | 0 | 1 | NA | NA | 0 |
| SE | 25498 | ENSG00000134986 | NREP | chr5 | - | 111734669 | 111734742 | 111730516 | 111731046 | 111735429 | 111735507 | 0 | 0 | 0 | 2 | 0 | 34 | 12 | 33 | 29 | 59 | 0 | 3 | 0 | 1 | 0 | 51 | 33 | 21 | 84 | 132 | 222 | 149 | 1 | 1 | 0 | 0 | 0 | 0.044 | 0 | 0 | 0.058 | 0 | 0.008 | 0 | -0.004 |
| SE | 25501 | ENSG00000103540 | CCP110 | chr16 | + | 19524779 | 19525019 | 19523938 | 19524088 | 19525718 | 19525861 | 3 | 0 | 1 | 3 | 3 | 0 | 2 | 2 | 8 | 12 | 0 | 11 | 4 | 0 | 0 | 0 | 0 | 0 | 3 | 0 | 298 | 149 | 1 | 1 | 1 | 0 | 0.2 | 0.158 | 0.111 | NA | 1 | 1 | 0 | NA | -0.373 |
| SE | 25502 | ENSG00000103540 | CCP110 | chr16 | + | 19524779 | 19525224 | 19523941 | 19524088 | 19525718 | 19525799 | 2 | 0 | 1 | 0 | 3 | 0 | 2 | 2 | 8 | 12 | 0 | 11 | 4 | 0 | 0 | 0 | 0 | 0 | 3 | 0 | 298 | 149 | 1 | 1 | 1 | 0 | 0.2 | 0 | 0.111 | NA | 1 | 1 | 0 | NA | -0.404 |
| SE | 25504 | ENSG00000103540 | CCP110 | chr16 | + | 19527866 | 19528022 | 19523856 | 19524088 | 19532415 | 19532544 | 80 | 85 | 44 | 40 | 100 | 0 | 0 | 0 | 0 | 0 | 0 | 47 | 31 | 39 | 25 | 0 | 6 | 0 | 0 | 0 | 298 | 149 | 1 | 1 | 1 | 1 | 1 | 1 | 1 | NA | 0.797 | 1 | 1 | 1 | 0.051 |
| SE | 25505 | ENSG00000103540 | CCP110 | chr16 | + | 19532415 | 19532544 | 19527835 | 19528022 | 19535939 | 19537587 | 117 | 116 | 65 | 65 | 218 | 0 | 0 | 2 | 0 | 0 | 3 | 56 | 70 | 54 | 35 | 0 | 0 | 0 | 0 | 0 | 278 | 149 | 1 | 1 | 1 | 1 | 0.946 | 1 | 1 | 1 | 1 | 1 | 1 | 1 | -0.011 |
| SE | 25508 | ENSG00000103540 | CCP110 | chr16 | + | 19544796 | 19544898 | 19542877 | 19542994 | 19545093 | 19545210 | 79 | 51 | 42 | 59 | 53 | 0 | 0 | 1 | 0 | 0 | 0 | 21 | 5 | 0 | 0 | 0 | 0 | 0 | 0 | 0 | 251 | 149 | 1 | 1 | 1 | 0.968 | 1 | 1 | 1 | NA | 1 | 1 | NA | NA | -0.006 |
| SE | 25509 | ENSG00000103540 | CCP110 | chr16 | + | 19547954 | 19548014 | 19546411 | 19546474 | 19548514 | 19548600 | 30 | 27 | 17 | 15 | 35 | 1 | 0 | 3 | 0 | 0 | 0 | 1 | 23 | 10 | 0 | 0 | 0 | 1 | 0 | 0 | 209 | 149 | 1 | 1 | 0.955 | 1 | 0.802 | 1 | 1 | NA | 1 | 0.943 | 1 | NA | -0.03 |
| SE | 25510 | ENSG00000103540 | CCP110 | chr16 | + | 19547954 | 19548014 | 19546411 | 19546474 | 19551195 | 19553400 | 47 | 51 | 29 | 40 | 47 | 1 | 2 | 2 | 5 | 4 | 0 | 9 | 22 | 10 | 0 | 0 | 0 | 2 | 4 | 0 | 209 | 149 | 1 | 1 | 0.971 | 0.948 | 0.912 | 0.851 | 0.893 | NA | 1 | 0.887 | 0.641 | NA | 0.072 |
| SE | 25511 | ENSG00000103540 | CCP110 | chr16 | + | 19548514 | 19548600 | 19546411 | 19546474 | 19551195 | 19551622 | 10 | 7 | 12 | 8 | 9 | 1 | 2 | 2 | 5 | 4 | 0 | 2 | 17 | 0 | 0 | 0 | 0 | 2 | 4 | 0 | 235 | 149 | 1 | 1 | 0.864 | 0.689 | 0.792 | 0.504 | 0.588 | NA | 1 | 0.843 | 0 | NA | 0.073 |
| SE | 25512 | ENSG00000103540 | CCP110 | chr16 | + | 19548514 | 19548600 | 19546848 | 19548014 | 19551195 | 19551622 | 26 | 14 | 14 | 13 | 26 | 35 | 31 | 17 | 30 | 29 | 0 | 2 | 32 | 0 | 0 | 0 | 8 | 15 | 1 | 0 | 235 | 149 | 1 | 1 | 0.32 | 0.223 | 0.343 | 0.216 | 0.362 | NA | 0.137 | 0.575 | 0 | NA | 0.055 |
| SE | 25513 | ENSG00000142686 | C1orf216 | chr1 | - | 35716500 | 35716675 | 35716197 | 35716326 | 35718706 | 35718894 | 8 | 4 | 0 | 0 | 0 | 1 | 1 | 1 | 2 | 0 | 0 | 0 | 0 | 3 | 0 | 0 | 6 | 0 | 5 | 0 | 298 | 149 | 1 | 1 | 0.8 | 0.667 | 0 | 0 | NA | NA | 0 | NA | 0.231 | NA | 0.251 |
| SE | 25517 | ENSG00000123836 | PFKFB2 | chr1 | + | 207060845 | 207060930 | 207054700 | 207054802 | 207061952 | 207062078 | 0 | 3 | 5 | 0 | 1 | 8 | 17 | 14 | 13 | 48 | 0 | 0 | 1 | 0 | 0 | 5 | 10 | 38 | 9 | 0 | 234 | 149 | 1 | 1 | 0 | 0.101 | 0.185 | 0 | 0.013 | 0 | 0 | 0.016 | 0 | NA | 0.056 |
| SE | 25518 | ENSG00000123836 | PFKFB2 | chr1 | + | 207063142 | 207063209 | 207062546 | 207062716 | 207063346 | 207063421 | 30 | 29 | 35 | 40 | 46 | 0 | 0 | 0 | 0 | 0 | 24 | 27 | 36 | 28 | 9 | 0 | 0 | 2 | 0 | 0 | 216 | 149 | 1 | 1 | 1 | 1 | 1 | 1 | 1 | 1 | 1 | 0.925 | 1 | 1 | 0.015 |
| SE | 25519 | ENSG00000123836 | PFKFB2 | chr1 | + | 207065035 | 207065160 | 207063772 | 207063829 | 207067498 | 207067706 | 100 | 60 | 65 | 50 | 126 | 0 | 0 | 0 | 3 | 0 | 23 | 74 | 118 | 34 | 41 | 0 | 0 | 0 | 0 | 0 | 274 | 149 | 1 | 1 | 1 | 1 | 1 | 0.901 | 1 | 1 | 1 | 1 | 1 | 1 | -0.02 |
| SE | 25520 | ENSG00000123836 | PFKFB2 | chr1 | + | 207070279 | 207070409 | 207069423 | 207069528 | 207071187 | 207071250 | 44 | 49 | 56 | 50 | 91 | 0 | 0 | 2 | 0 | 0 | 0 | 42 | 61 | 53 | 0 | 0 | 0 | 0 | 0 | 0 | 279 | 149 | 1 | 1 | 1 | 1 | 0.937 | 1 | 1 | NA | 1 | 1 | 1 | NA | -0.013 |
| SE | 25527 | ENSG00000131732 | ZCCHC9 | chr5 | + | 81308560 | 81308711 | 81301602 | 81305141 | 81308945 | 81309038 | 79 | 56 | 83 | 61 | 146 | 0 | 0 | 0 | 1 | 0 | 21 | 51 | 45 | 14 | 6 | 0 | 0 | 0 | 0 | 0 | 298 | 149 | 1 | 1 | 1 | 1 | 1 | 0.968 | 1 | 1 | 1 | 1 | 1 | 1 | -0.006 |
| SE | 25528 | ENSG00000203739 | PRDX6-AS1 | chr1 | - | 173440462 | 173440750 | 173417792 | 173418622 | 173441918 | 173442048 | 17 | 3 | 7 | 7 | 10 | 0 | 0 | 0 | 0 | 4 | 20 | 21 | 9 | 8 | 0 | 0 | 0 | 0 | 0 | 0 | 298 | 149 | 1 | 1 | 1 | 1 | 1 | 1 | 0.556 | 1 | 1 | 1 | 1 | NA | -0.089 |
| SE | 25529 | ENSG00000203739 | PRDX6-AS1 | chr1 | - | 173440462 | 173440750 | 173417792 | 173418622 | 173476631 | 173476996 | 5 | 0 | 2 | 2 | 3 | 0 | 0 | 0 | 0 | 4 | 0 | 4 | 1 | 0 | 0 | 0 | 0 | 0 | 0 | 0 | 298 | 149 | 1 | 1 | 1 | NA | 1 | 1 | 0.273 | NA | 1 | 1 | NA | NA | -0.182 |
| SE | 25530 | ENSG00000203739 | PRDX6-AS1 | chr1 | - | 173441918 | 173442048 | 173417792 | 173418622 | 173476631 | 173477171 | 0 | 0 | 0 | 0 | 4 | 0 | 0 | 0 | 0 | 4 | 0 | 0 | 2 | 0 | 0 | 0 | 0 | 0 | 0 | 0 | 279 | 149 | 1 | 1 | NA | NA | NA | NA | 0.348 | NA | NA | 1 | NA | NA | -0.652 |
| SE | 25531 | ENSG00000203739 | PRDX6-AS1 | chr1 | - | 173441918 | 173442048 | 173440462 | 173440750 | 173476631 | 173477171 | 12 | 3 | 3 | 5 | 7 | 0 | 0 | 0 | 0 | 0 | 20 | 14 | 11 | 8 | 0 | 0 | 0 | 1 | 0 | 0 | 279 | 149 | 1 | 1 | 1 | 1 | 1 | 1 | 1 | 1 | 1 | 0.855 | 1 | NA | 0.036 |
| SE | 25534 | ENSG00000203739 | PRDX6-AS1 | chr1 | - | 173441918 | 173442097 | 173440462 | 173440750 | 173476631 | 173476996 | 12 | 3 | 5 | 5 | 7 | 0 | 0 | 0 | 0 | 0 | 20 | 14 | 9 | 8 | 0 | 0 | 0 | 1 | 0 | 0 | 298 | 149 | 1 | 1 | 1 | 1 | 1 | 1 | 1 | 1 | 1 | 0.818 | 1 | NA | 0.045 |
| SE | 25535 | ENSG00000129595 | EPB41L4A | chr5 | - | 112205420 | 112205504 | 112204374 | 112204488 | 112209891 | 112209982 | 14 | 5 | 7 | 9 | 3 | 0 | 0 | 0 | 0 | 0 | 0 | 3 | 6 | 1 | 5 | 0 | 1 | 4 | 0 | 0 | 233 | 149 | 1 | 1 | 1 | 1 | 1 | 1 | 1 | NA | 0.657 | 0.49 | 1 | 1 | 0.213 |
| SE | 25536 | ENSG00000129595 | EPB41L4A | chr5 | - | 112209891 | 112209982 | 112205420 | 112205504 | 112234633 | 112234755 | 7 | 3 | 8 | 15 | 2 | 0 | 0 | 2 | 0 | 0 | 5 | 3 | 5 | 2 | 0 | 0 | 0 | 0 | 0 | 0 | 240 | 149 | 1 | 1 | 1 | 1 | 0.713 | 1 | 1 | 1 | 1 | 1 | 1 | NA | -0.057 |
| SE | 25537 | ENSG00000129595 | EPB41L4A | chr5 | - | 112259228 | 112259292 | 112240718 | 112240810 | 112259890 | 112259979 | 22 | 6 | 9 | 14 | 16 | 2 | 0 | 0 | 0 | 0 | 0 | 0 | 5 | 0 | 0 | 0 | 0 | 0 | 0 | 0 | 213 | 149 | 1 | 1 | 0.885 | 1 | 1 | 1 | 1 | NA | NA | 1 | NA | NA | -0.023 |
| SE | 25541 | ENSG00000151292 | CSNK1G3 | chr5 | + | 123553106 | 123553147 | 123545414 | 123545841 | 123557494 | 123557564 | 388 | 238 | 315 | 362 | 1000 | 1 | 0 | 3 | 3 | 6 | 185 | 307 | 203 | 179 | 427 | 0 | 4 | 5 | 0 | 0 | 190 | 149 | 1 | 1 | 0.997 | 1 | 0.988 | 0.99 | 0.992 | 1 | 0.984 | 0.97 | 1 | 1 | 0.003 |
| SE | 25542 | ENSG00000151292 | CSNK1G3 | chr5 | + | 123553106 | 123553147 | 123545414 | 123545841 | 123573392 | 123573541 | 85 | 47 | 103 | 83 | 332 | 0 | 1 | 0 | 0 | 0 | 39 | 68 | 39 | 52 | 135 | 0 | 0 | 0 | 0 | 0 | 190 | 149 | 1 | 1 | 1 | 0.974 | 1 | 1 | 1 | 1 | 1 | 1 | 1 | 1 | -0.005 |
| SE | 25543 | ENSG00000151292 | CSNK1G3 | chr5 | + | 123557494 | 123557564 | 123545414 | 123545841 | 123573392 | 123573541 | 101 | 86 | 92 | 111 | 244 | 0 | 1 | 0 | 0 | 0 | 38 | 103 | 72 | 22 | 32 | 0 | 0 | 0 | 0 | 0 | 219 | 149 | 1 | 1 | 1 | 0.983 | 1 | 1 | 1 | 1 | 1 | 1 | 1 | 1 | -0.003 |
| SE | 25544 | ENSG00000151292 | CSNK1G3 | chr5 | + | 123557494 | 123557564 | 123552964 | 123553147 | 123573392 | 123573541 | 413 | 282 | 317 | 406 | 918 | 0 | 0 | 2 | 0 | 0 | 197 | 350 | 248 | 159 | 353 | 0 | 0 | 0 | 0 | 0 | 219 | 149 | 1 | 1 | 1 | 1 | 0.991 | 1 | 1 | 1 | 1 | 1 | 1 | 1 | -0.002 |
| SE | 25546 | ENSG00000151292 | CSNK1G3 | chr5 | + | 123588067 | 123588153 | 123573392 | 123573541 | 123588426 | 123588511 | 68 | 50 | 40 | 68 | 97 | 0 | 0 | 0 | 0 | 0 | 23 | 35 | 63 | 4 | 1 | 0 | 0 | 3 | 0 | 0 | 235 | 149 | 1 | 1 | 1 | 1 | 1 | 1 | 1 | 1 | 1 | 0.93 | 1 | 1 | 0.014 |
| SE | 25547 | ENSG00000151292 | CSNK1G3 | chr5 | + | 123595038 | 123595134 | 123591318 | 123591414 | 123604723 | 123604830 | 8 | 1 | 2 | 5 | 21 | 122 | 79 | 111 | 110 | 206 | 0 | 4 | 4 | 2 | 0 | 0 | 32 | 73 | 12 | 2 | 245 | 149 | 1 | 1 | 0.038 | 0.008 | 0.011 | 0.027 | 0.058 | NA | 0.071 | 0.032 | 0.092 | 0 | -0.02 |
| SE | 25548 | ENSG00000151292 | CSNK1G3 | chr5 | + | 123605338 | 123605362 | 123604723 | 123604830 | 123614341 | 123614455 | 3 | 1 | 0 | 5 | 0 | 83 | 38 | 58 | 84 | 118 | 0 | 3 | 4 | 0 | 0 | 4 | 29 | 36 | 2 | 0 | 173 | 149 | 1 | 1 | 0.03 | 0.022 | 0 | 0.049 | 0 | 0 | 0.082 | 0.087 | 0 | NA | -0.022 |
| SE | 25550 | ENSG00000157916 | RER1 | chr1 | + | 2395783 | 2395871 | 2391774 | 2391958 | 2403034 | 2403064 | 4 | 4 | 3 | 6 | 0 | 0 | 0 | 0 | 1 | 0 | 0 | 9 | 7 | 3 | 26 | 0 | 6 | 1 | 0 | 0 | 237 | 149 | 1 | 1 | 1 | 1 | 1 | 0.79 | NA | NA | 0.485 | 0.815 | 1 | 1 | 0.123 |
| SE | 25552 | ENSG00000157916 | RER1 | chr1 | + | 2397115 | 2397220 | 2394004 | 2395871 | 2403034 | 2403064 | 159 | 81 | 135 | 144 | 154 | 0 | 0 | 0 | 0 | 0 | 82 | 102 | 60 | 61 | 150 | 0 | 0 | 0 | 1 | 0 | 254 | 149 | 1 | 1 | 1 | 1 | 1 | 1 | 1 | 1 | 1 | 1 | 0.973 | 1 | 0.005 |
| SE | 25554 | ENSG00000157916 | RER1 | chr1 | + | 2400856 | 2400935 | 2399414 | 2399514 | 2402206 | 2402316 | 563 | 313 | 492 | 475 | 515 | 0 | 0 | 0 | 0 | 0 | 754 | 496 | 321 | 440 | 902 | 0 | 0 | 0 | 0 | 4 | 228 | 149 | 1 | 1 | 1 | 1 | 1 | 1 | 1 | 1 | 1 | 1 | 1 | 0.993 | 0.001 |
| SE | 25555 | ENSG00000248489 | LINC02062 | chr5 | + | 98976770 | 98976900 | 98971854 | 98972076 | 98987443 | 98988012 | 2 | 1 | 0 | 2 | 3 | 0 | 1 | 0 | 0 | 0 | 0 | 6 | 0 | 0 | 0 | 0 | 0 | 0 | 0 | 0 | 279 | 149 | 1 | 1 | 1 | 0.348 | NA | 1 | 1 | NA | 1 | NA | NA | NA | -0.163 |
| SE | 25556 | ENSG00000168394 | TAP1 | chr6 | - | 32847918 | 32848092 | 32847512 | 32847675 | 32848651 | 32848840 | 617 | 252 | 430 | 430 | 265 | 0 | 1 | 1 | 2 | 0 | 703 | 562 | 314 | 512 | 787 | 0 | 0 | 0 | 1 | 0 | 298 | 149 | 1 | 1 | 1 | 0.992 | 0.995 | 0.991 | 1 | 1 | 1 | 1 | 0.996 | 1 | -0.004 |
| SE | 25559 | ENSG00000168394 | TAP1 | chr6 | - | 32852108 | 32852239 | 32850943 | 32851149 | 32852387 | 32852502 | 495 | 181 | 319 | 323 | 307 | 1 | 0 | 0 | 0 | 0 | 482 | 423 | 187 | 484 | 456 | 0 | 0 | 0 | 7 | 0 | 280 | 149 | 1 | 1 | 0.996 | 1 | 1 | 1 | 1 | 1 | 1 | 1 | 0.974 | 1 | 0.004 |
| SE | 25560 | ENSG00000246089 | NA | chr8 | - | 6406527 | 6406708 | 6403550 | 6405120 | 6407081 | 6407142 | 0 | 0 | 0 | 0 | 0 | 0 | 0 | 0 | 4 | 0 | 0 | 0 | 2 | 0 | 0 | 0 | 0 | 0 | 0 | 0 | 298 | 149 | 1 | 1 | NA | NA | NA | 0 | NA | NA | NA | 1 | NA | NA | -1 |
| SE | 25564 | ENSG00000179397 | CATSPERE | chr1 | + | 244477540 | 244477614 | 244457492 | 244457569 | 244477905 | 244477975 | 3 | 3 | 1 | 5 | 7 | 1 | 0 | 3 | 3 | 0 | 2 | 0 | 3 | 2 | 0 | 0 | 0 | 0 | 3 | 0 | 223 | 149 | 1 | 1 | 0.667 | 1 | 0.182 | 0.527 | 1 | 1 | NA | 1 | 0.308 | NA | -0.094 |
| SE | 25565 | ENSG00000179397 | CATSPERE | chr1 | + | 244490446 | 244490471 | 244479716 | 244479784 | 244499001 | 244499079 | 3 | 2 | 2 | 0 | 0 | 4 | 0 | 0 | 1 | 5 | 0 | 0 | 0 | 0 | 0 | 0 | 0 | 0 | 2 | 0 | 174 | 149 | 1 | 1 | 0.391 | 1 | 1 | 0 | 0 | NA | NA | NA | 0 | NA | 0.478 |
| SE | 25566 | ENSG00000179397 | CATSPERE | chr1 | + | 244499001 | 244499079 | 244479716 | 244479784 | 244515276 | 244515327 | 2 | 0 | 0 | 2 | 3 | 1 | 0 | 1 | 0 | 0 | 0 | 0 | 2 | 0 | 0 | 2 | 0 | 3 | 0 | 0 | 227 | 149 | 1 | 1 | 0.568 | NA | 0 | 1 | 1 | 0 | NA | 0 | 1 | NA | 0.309 |
| SE | 25567 | ENSG00000179397 | CATSPERE | chr1 | + | 244499001 | 244499079 | 244479716 | 244479784 | 244518591 | 244518698 | 4 | 0 | 0 | 1 | 0 | 2 | 1 | 0 | 2 | 3 | 0 | 0 | 0 | 0 | 0 | 0 | 0 | 8 | 2 | 0 | 227 | 149 | 1 | 1 | 0.568 | 0 | NA | 0.247 | 0 | NA | NA | 0 | 0 | NA | 0.204 |
| SE | 25570 | ENSG00000179397 | CATSPERE | chr1 | + | 244499001 | 244499079 | 244479716 | 244479784 | 244572329 | 244572772 | 0 | 2 | 0 | 1 | 0 | 2 | 0 | 0 | 0 | 0 | 0 | 0 | 0 | 0 | 0 | 0 | 0 | 1 | 0 | 0 | 227 | 149 | 1 | 1 | 0 | 1 | NA | 1 | NA | NA | NA | 0 | NA | NA | 0.667 |
| SE | 25571 | ENSG00000179397 | CATSPERE | chr1 | + | 244515276 | 244515327 | 244479716 | 244479784 | 244518591 | 244518698 | 2 | 0 | 4 | 1 | 1 | 2 | 1 | 0 | 2 | 3 | 2 | 0 | 8 | 3 | 0 | 0 | 0 | 8 | 2 | 0 | 200 | 149 | 1 | 1 | 0.427 | 0 | 1 | 0.271 | 0.199 | 1 | NA | 0.427 | 0.528 | NA | -0.272 |
| SE | 25572 | ENSG00000179397 | CATSPERE | chr1 | + | 244515276 | 244515327 | 244499001 | 244499079 | 244518591 | 244518698 | 3 | 0 | 3 | 2 | 4 | 4 | 0 | 0 | 0 | 0 | 0 | 0 | 5 | 5 | 0 | 0 | 0 | 0 | 0 | 0 | 200 | 149 | 1 | 1 | 0.358 | NA | 1 | 1 | 1 | NA | NA | 1 | 1 | NA | -0.16 |
| SE | 25573 | ENSG00000179397 | CATSPERE | chr1 | + | 244518591 | 244518698 | 244479716 | 244479784 | 244552321 | 244552465 | 2 | 1 | 1 | 1 | 2 | 1 | 0 | 0 | 0 | 0 | 0 | 0 | 4 | 2 | 0 | 0 | 0 | 0 | 0 | 0 | 256 | 149 | 1 | 1 | 0.538 | 1 | 1 | 1 | 1 | NA | NA | 1 | 1 | NA | -0.092 |
| SE | 25578 | ENSG00000179397 | CATSPERE | chr1 | + | 244552321 | 244552814 | 244518591 | 244518698 | 244560667 | 244561145 | 3 | 7 | 6 | 7 | 16 | 1 | 0 | 1 | 1 | 0 | 0 | 0 | 0 | 1 | 0 | 0 | 0 | 0 | 0 | 0 | 298 | 149 | 1 | 1 | 0.6 | 1 | 0.75 | 0.778 | 1 | NA | NA | NA | 1 | NA | -0.174 |
| SE | 25579 | ENSG00000179397 | CATSPERE | chr1 | + | 244560667 | 244561145 | 244479716 | 244479784 | 244572329 | 244572772 | 4 | 1 | 3 | 0 | 13 | 2 | 0 | 0 | 0 | 0 | 0 | 0 | 0 | 2 | 0 | 0 | 0 | 1 | 0 | 0 | 298 | 149 | 1 | 1 | 0.5 | 1 | 1 | NA | 1 | NA | NA | 0 | 1 | NA | 0.375 |
| SE | 25580 | ENSG00000179397 | CATSPERE | chr1 | + | 244560667 | 244561145 | 244499001 | 244499079 | 244572329 | 244572772 | 6 | 1 | 3 | 0 | 16 | 0 | 2 | 0 | 0 | 0 | 0 | 0 | 0 | 2 | 0 | 0 | 0 | 0 | 0 | 0 | 298 | 149 | 1 | 1 | 1 | 0.2 | 1 | NA | 1 | NA | NA | NA | 1 | NA | -0.2 |
| SE | 25581 | ENSG00000179397 | CATSPERE | chr1 | + | 244581795 | 244581854 | 244572329 | 244572772 | 244588481 | 244588534 | 2 | 1 | 1 | 0 | 0 | 0 | 0 | 0 | 0 | 0 | 0 | 0 | 0 | 0 | 0 | 0 | 8 | 0 | 0 | 0 | 208 | 149 | 1 | 1 | 1 | 1 | 1 | NA | NA | NA | 0 | NA | NA | NA | 1 |
| SE | 25583 | ENSG00000179397 | CATSPERE | chr1 | + | 244605694 | 244605794 | 244593498 | 244593578 | 244610239 | 244610326 | 2 | 8 | 2 | 5 | 0 | 2 | 0 | 1 | 0 | 1 | 0 | 6 | 0 | 14 | 0 | 3 | 0 | 0 | 3 | 0 | 249 | 149 | 1 | 1 | 0.374 | 1 | 0.545 | 1 | 0 | 0 | 1 | NA | 0.736 | NA | 0.005 |
| SE | 25595 | ENSG00000129596 | CDO1 | chr5 | - | 115809621 | 115809697 | 115806348 | 115806518 | 115811160 | 115811315 | 0 | 0 | 0 | 0 | 0 | 3 | 0 | 2 | 0 | 7 | 0 | 1 | 0 | 0 | 0 | 0 | 2 | 4 | 0 | 0 | 225 | 149 | 1 | 1 | 0 | NA | 0 | NA | 0 | NA | 0.249 | 0 | NA | NA | -0.124 |
| SE | 25597 | ENSG00000129596 | CDO1 | chr5 | - | 115813897 | 115813954 | 115813188 | 115813258 | 115816227 | 115816657 | 0 | 0 | 0 | 0 | 1 | 1 | 0 | 0 | 5 | 3 | 0 | 0 | 6 | 0 | 0 | 0 | 2 | 0 | 1 | 0 | 206 | 149 | 1 | 1 | 0 | NA | NA | 0 | 0.194 | NA | 0 | NA | 0.813 | NA | -0.342 |
| SE | 25598 | ENSG00000128872 | TMOD2 | chr15 | + | 51766372 | 51766567 | 51751596 | 51751712 | 51773711 | 51773834 | 14 | 13 | 13 | 7 | 1 | 0 | 0 | 0 | 2 | 0 | 0 | 6 | 9 | 0 | 0 | 0 | 0 | 0 | 0 | 0 | 298 | 149 | 1 | 1 | 1 | 1 | 1 | 0.636 | 1 | NA | | 1 | NA | NA | -0.073 |
| SE | 25599 | ENSG00000128872 | TMOD2 | chr15 | + | 51767070 | 51767152 | 51766372 | 51766567 | 51768261 | 51768418 | 0 | 0 | 0 | 4 | 0 | 29 | 42 | 40 | 36 | 64 | 0 | 0 | 0 | 0 | 0 | 20 | 21 | 12 | 1 | 8 | 231 | 149 | 1 | 1 | 0 | 0 | 0 | 0.067 | 0 | 0 | 0 | 0 | 0 | 0 | 0.013 |
| SE | 25600 | ENSG00000128872 | TMOD2 | chr15 | + | 51768261 | 51768418 | 51766372 | 51766567 | 51773711 | 51773834 | 52 | 81 | 65 | 68 | 131 | 0 | 0 | 2 | 0 | 0 | 20 | 40 | 29 | 6 | 8 | 0 | 0 | 0 | 0 | 0 | 298 | 149 | 1 | 1 | 1 | 1 | 0.942 | 1 | 1 | 1 | 1 | 1 | 1 | 1 | -0.012 |
| SE | 25601 | ENSG00000128872 | TMOD2 | chr15 | + | 51773711 | 51773834 | 51768261 | 51768418 | 51776931 | 51777018 | 53 | 64 | 39 | 56 | 102 | 2 | 1 | 0 | 0 | 0 | 14 | 42 | 28 | 22 | 0 | 0 | 5 | 0 | 0 | 0 | 272 | 149 | 1 | 1 | 0.936 | 0.972 | 1 | 1 | 1 | 1 | 0.821 | 1 | 1 | NA | 0.026 |
| SE | 25602 | ENSG00000128872 | TMOD2 | chr15 | + | 51781043 | 51781174 | 51776931 | 51777018 | 51782720 | 51782828 | 72 | 60 | 51 | 76 | 91 | 1 | 1 | 0 | 0 | 0 | 33 | 53 | 23 | 34 | 0 | 0 | 0 | 0 | 0 | 0 | 280 | 149 | 1 | 1 | 0.975 | 0.97 | 1 | 1 | 1 | 1 | 1 | 1 | 1 | NA | -0.011 |
| SE | 25603 | ENSG00000128872 | TMOD2 | chr15 | + | 51781043 | 51781174 | 51776931 | 51777018 | 51798196 | 51798340 | 33 | 30 | 25 | 27 | 47 | 1 | 0 | 0 | 1 | 1 | 9 | 18 | 13 | 16 | 0 | 0 | 0 | 0 | 0 | 0 | 280 | 149 | 1 | 1 | 0.946 | 1 | 1 | 0.935 | 0.962 | 1 | 1 | 1 | 1 | NA | -0.031 |
| SE | 25604 | ENSG00000128872 | TMOD2 | chr15 | + | 51782720 | 51782828 | 51776931 | 51777018 | 51798196 | 51798340 | 59 | 55 | 39 | 51 | 49 | 1 | 0 | 0 | 1 | 1 | 21 | 11 | 23 | 25 | 0 | 0 | 0 | 0 | 0 | 0 | 257 | 149 | 1 | 1 | 0.972 | 1 | 1 | 0.967 | 0.966 | 1 | 1 | 1 | 1 | NA | -0.019 |
| SE | 25605 | ENSG00000128872 | TMOD2 | chr15 | + | 51782720 | 51782828 | 51781043 | 51781174 | 51798196 | 51798340 | 97 | 85 | 64 | 100 | 90 | 0 | 2 | 1 | 0 | 0 | 45 | 46 | 35 | 43 | 0 | 0 | 0 | 2 | 0 | 0 | 257 | 149 | 1 | 1 | 1 | 0.961 | 0.974 | 1 | 1 | 1 | 1 | 0.91 | 1 | NA | 0.009 |
| SE | 25609 | ENSG00000151640 | DPYSL4 | chr10 | + | 132192282 | 132192520 | 132190746 | 132190835 | 132192657 | 132192832 | 0 | 0 | 0 | 0 | 0 | 3 | 0 | 0 | 0 | 0 | 0 | 7 | 0 | 0 | 0 | 0 | 0 | 0 | 0 | 0 | 298 | 149 | 1 | 1 | 0 | NA | NA | NA | NA | NA | 1 | NA | NA | NA | -1 |
| SE | 25610 | ENSG00000151640 | DPYSL4 | chr10 | + | 132192368 | 132192520 | 132190746 | 132190835 | 132192657 | 132192842 | 0 | 0 | 0 | 0 | 0 | 3 | 0 | 0 | 0 | 0 | 0 | 7 | 0 | 0 | 0 | 0 | 0 | 0 | 0 | 0 | 298 | 149 | 1 | 1 | 0 | NA | NA | NA | NA | NA | 1 | NA | NA | NA | -1 |
| SE | 25612 | ENSG00000151640 | DPYSL4 | chr10 | + | 132202645 | 132202825 | 132201945 | 132202116 | 132203761 | 132203927 | 0 | 1 | 2 | 0 | 0 | 1 | 0 | 0 | 0 | 0 | 0 | 8 | 0 | 0 | 0 | 0 | 0 | 0 | 0 | 0 | 298 | 149 | 1 | 1 | 0 | 1 | 1 | NA | NA | NA | 1 | NA | NA | NA | -0.333 |
| SE | 25613 | ENSG00000113595 | TRIM23 | chr5 | - | 65613726 | 65613855 | 65611669 | 65611881 | 65614097 | 65614219 | 16 | 14 | 14 | 31 | 21 | 17 | 14 | 30 | 20 | 25 | 0 | 16 | 14 | 18 | 5 | 0 | 3 | 19 | 3 | 0 | 278 | 149 | 1 | 1 | 0.335 | 0.349 | 0.2 | 0.454 | 0.31 | NA | 0.741 | 0.283 | 0.763 | 1 | -0.367 |
| SE | 25614 | ENSG00000113595 | TRIM23 | chr5 | - | 65618092 | 65618255 | 65614097 | 65614219 | 65624193 | 65624296 | 91 | 56 | 67 | 55 | 53 | 1 | 0 | 0 | 1 | 0 | 40 | 24 | 37 | 16 | 1 | 0 | 0 | 0 | 0 | 0 | 298 | 149 | 1 | 1 | 0.978 | 1 | 1 | 0.965 | 1 | 1 | 1 | 1 | 1 | 1 | -0.011 |
| SE | 25617 | ENSG00000235172 | LINC01366 | chr5 | + | 170332847 | 170332937 | 170331428 | 170331558 | 170333133 | 170333760 | 2 | 0 | 0 | 1 | 0 | 2 | 0 | 0 | 0 | 0 | 7 | 0 | 2 | 0 | 0 | 0 | 0 | 0 | 0 | 0 | 239 | 149 | 1 | 1 | 0.384 | NA | NA | 1 | NA | 1 | NA | 1 | NA | NA | -0.308 |
| SE | 25620 | ENSG00000106624 | AEBP1 | chr7 | + | 44107808 | 44107931 | 44107628 | 44107700 | 44108006 | 44108084 | 7 | 10 | 25 | 39 | 25 | 0 | 0 | 0 | 1 | 0 | 6 | 19 | 12 | 0 | 16 | 10 | 0 | 1 | 0 | 0 | 272 | 149 | 1 | 1 | 1 | 1 | 1 | 0.955 | 1 | 0.247 | 1 | 0.868 | NA | 1 | 0.212 |
| SE | 25621 | ENSG00000106624 | AEBP1 | chr7 | + | 44109656 | 44110124 | 44109287 | 44109341 | 44110206 | 44110225 | 30 | 5 | 27 | 14 | 15 | 0 | 0 | 0 | 0 | 0 | 0 | 50 | 8 | 3 | 28 | 0 | 0 | 1 | 0 | 10 | 298 | 149 | 1 | 1 | 1 | 1 | 1 | 1 | 1 | NA | 1 | 0.8 | 1 | 0.583 | 0.154 |
| SE | 25623 | ENSG00000106624 | AEBP1 | chr7 | + | 44110724 | 44110809 | 44110206 | 44110346 | 44110912 | 44111057 | 18 | 16 | 18 | 11 | 14 | 0 | 0 | 0 | 0 | 2 | 0 | 17 | 1 | 1 | 0 | 0 | 0 | 0 | 0 | 0 | 234 | 149 | 1 | 1 | 1 | 1 | 1 | 1 | 0.817 | NA | 1 | 1 | 1 | NA | -0.037 |
| SE | 25634 | ENSG00000197506 | SLC28A3 | chr9 | - | 84310411 | 84310587 | 84309628 | 84309714 | 84313358 | 84313454 | 0 | 1 | 4 | 7 | 4 | 0 | 0 | 3 | 3 | 1 | 0 | 2 | 1 | 18 | 7 | 0 | 0 | 0 | 44 | 3 | 298 | 149 | 1 | 1 | NA | 1 | 0.4 | 0.538 | 0.667 | NA | 1 | 1 | 0.17 | 0.538 | -0.026 |
| SE | 25635 | ENSG00000197506 | SLC28A3 | chr9 | - | 84313358 | 84313454 | 84309628 | 84309714 | 84340573 | 84340659 | 0 | 2 | 3 | 4 | 2 | 0 | 0 | 0 | 0 | 0 | 0 | 0 | 0 | 54 | 18 | 0 | 0 | 0 | 1 | 0 | 245 | 149 | 1 | 1 | NA | 1 | 1 | 1 | 1 | NA | NA | NA | 0.97 | 1 | 0.015 |
| SE | 25641 | ENSG00000213123 | DYNLT2B | chr3 | - | 196305655 | 196305773 | 196296005 | 196296069 | 196306942 | 196307012 | 0 | 0 | 1 | 1 | 0 | 12 | 4 | 0 | 2 | 4 | 0 | 0 | 0 | 0 | 0 | 0 | 3 | 0 | 0 | 0 | 267 | 149 | 1 | 1 | 0 | 0 | 1 | 0.218 | 0 | NA | 0 | NA | NA | NA | 0.244 |
| SE | 25642 | ENSG00000213123 | DYNLT2B | chr3 | - | 196316097 | 196316231 | 196306942 | 196307012 | 196318039 | 196318240 | 18 | 8 | 4 | 8 | 10 | 0 | 0 | 0 | 2 | 0 | 0 | 15 | 4 | 8 | 36 | 0 | 0 | 0 | 0 | 0 | 283 | 149 | 1 | 1 | 1 | 1 | 1 | 0.678 | 1 | NA | 1 | 1 | 1 | 1 | -0.064 |
| SE | 25654 | ENSG00000248664 | NA | chr5 | - | 69128033 | 69128108 | 69115234 | 69115402 | 69130342 | 69130545 | 0 | 0 | 0 | 0 | 0 | 1 | 1 | 0 | 0 | 0 | 18 | 0 | 0 | 0 | 1 | 0 | 0 | 0 | 0 | 0 | 224 | 149 | 1 | 1 | 0 | 0 | NA | NA | NA | 1 | NA | NA | NA | 1 | -1 |
| SE | 25661 | ENSG00000164209 | SLC25A46 | chr5 | + | 110743729 | 110743787 | 110741677 | 110742089 | 110746268 | 110746346 | 174 | 149 | 153 | 183 | 154 | 2 | 0 | 1 | 0 | 0 | 14 | 96 | 123 | 52 | 16 | 0 | 0 | 0 | 0 | 0 | 207 | 149 | 1 | 1 | 0.984 | 1 | 0.991 | 1 | 1 | 1 | 1 | 1 | 1 | 1 | -0.005 |
| SE | 25663 | ENSG00000164209 | SLC25A46 | chr5 | + | 110755995 | 110756121 | 110755119 | 110755521 | 110756701 | 110756759 | 1 | 0 | 4 | 3 | 2 | 145 | 105 | 122 | 107 | 205 | 0 | 0 | 0 | 0 | 0 | 15 | 78 | 51 | 60 | 10 | 275 | 149 | 1 | 1 | 0.004 | 0 | 0.017 | 0.015 | 0.005 | 0 | 0 | 0 | 0 | 0 | 0.008 |
| SE | 25664 | ENSG00000164209 | SLC25A46 | chr5 | + | 110756701 | 110756759 | 110753301 | 110755521 | 110761203 | 110761655 | 173 | 126 | 160 | 139 | 255 | 0 | 0 | 0 | 0 | 3 | 21 | 90 | 61 | 72 | 10 | 0 | 2 | 0 | 0 | 0 | 207 | 149 | 1 | 1 | 1 | 1 | 1 | 1 | 0.984 | 1 | 0.97 | 1 | 1 | 1 | 0.003 |
| SE | 25669 | ENSG00000154721 | JAM2 | chr21 | + | 25702169 | 25702269 | 25698676 | 25698879 | 25705978 | 25706086 | 0 | 0 | 8 | 1 | 0 | 0 | 0 | 0 | 0 | 0 | 5 | 0 | 1 | 0 | 0 | 0 | 0 | 0 | 0 | 2 | 249 | 149 | 1 | 1 | NA | NA | 1 | 1 | NA | 1 | NA | 1 | NA | 0 | 0.333 |
| SE | 25670 | ENSG00000154721 | JAM2 | chr21 | + | 25711366 | 25711568 | 25709433 | 25709449 | 25712339 | 25712382 | 0 | 0 | 0 | 0 | 4 | 0 | 2 | 0 | 1 | 5 | 0 | 0 | 0 | 14 | 0 | 5 | 0 | 0 | 0 | 0 | 298 | 149 | 1 | 1 | NA | 0 | NA | 0 | 0.286 | 0 | NA | NA | NA | 1 | -0.405 |
| SE | 25673 | ENSG00000179152 | TCAIM | chr3 | + | 44354738 | 44354811 | 44338451 | 44338834 | 44361364 | 44361518 | 8 | 7 | 1 | 5 | 5 | 0 | 0 | 0 | 0 | 0 | 0 | 7 | 3 | 0 | 0 | 0 | 0 | 1 | 0 | 0 | 222 | 149 | 1 | 1 | 1 | 1 | 1 | 1 | 1 | NA | 1 | 0.668 | NA | NA | 0.166 |
| SE | 25675 | ENSG00000179152 | TCAIM | chr3 | + | 44367455 | 44367708 | 44361364 | 44361518 | 44392854 | 44392977 | 39 | 27 | 48 | 43 | 46 | 0 | 1 | 0 | 0 | 0 | 30 | 26 | 6 | 12 | 3 | 0 | 0 | 0 | 0 | 0 | 298 | 149 | 1 | 1 | 1 | 0.931 | 1 | 1 | 1 | 1 | 1 | 1 | 1 | 1 | -0.014 |
| SE | 25676 | ENSG00000179152 | TCAIM | chr3 | + | 44367531 | 44367708 | 44361364 | 44361518 | 44392854 | 44392977 | 40 | 33 | 48 | 43 | 51 | 0 | 1 | 0 | 0 | 0 | 40 | 29 | 7 | 20 | 3 | 0 | 0 | 0 | 0 | 0 | 298 | 149 | 1 | 1 | 1 | 0.943 | 1 | 1 | 1 | 1 | 1 | 1 | 1 | 1 | -0.011 |
| SE | 25677 | ENSG00000179152 | TCAIM | chr3 | + | 44391207 | 44391331 | 44367531 | 44367708 | 44392854 | 44392977 | 2 | 0 | 1 | 0 | 0 | 31 | 24 | 38 | 33 | 28 | 0 | 0 | 0 | 0 | 0 | 23 | 26 | 3 | 7 | 3 | 273 | 149 | 1 | 1 | 0.034 | 0 | 0.014 | 0 | 0 | 0 | 0 | 0 | 0 | 0 | 0.01 |
| SE | 25678 | ENSG00000179152 | TCAIM | chr3 | + | 44392854 | 44392977 | 44367455 | 44367708 | 44396399 | 44396497 | 46 | 50 | 67 | 56 | 65 | 0 | 0 | 4 | 0 | 0 | 28 | 31 | 10 | 14 | 11 | 0 | 0 | 0 | 0 | 0 | 272 | 149 | 1 | 1 | 1 | 1 | 0.902 | 1 | 1 | 1 | 1 | 1 | 1 | 1 | -0.02 |
| SE | 25679 | ENSG00000179152 | TCAIM | chr3 | + | 44396399 | 44396742 | 44392854 | 44392977 | 44396742 | 44396834 | 35 | 37 | 51 | 43 | 54 | 0 | 2 | 0 | 0 | 0 | 38 | 21 | 11 | 10 | 8 | 0 | 0 | 0 | 0 | 0 | 247 | 149 | 1 | 1 | 1 | 0.918 | 1 | 1 | 1 | 1 | 1 | 1 | 1 | 1 | -0.016 |
| SE | 25680 | ENSG00000179152 | TCAIM | chr3 | + | 44401202 | 44401334 | 44400354 | 44400587 | 44407441 | 44407740 | 72 | 90 | 51 | 111 | 100 | 2 | 1 | 0 | 0 | 1 | 24 | 33 | 28 | 13 | 0 | 0 | 4 | 0 | 0 | 0 | 281 | 149 | 1 | 1 | 0.95 | 0.979 | 1 | 1 | 0.981 | 1 | 0.814 | 1 | 1 | NA | 0.028 |
| SE | 25682 | ENSG00000188295 | ZNF669 | chr1 | - | 247101730 | 247101791 | 247099961 | 247101319 | 247104196 | 247104278 | 11 | 12 | 7 | 9 | 13 | 0 | 0 | 0 | 0 | 0 | 5 | 19 | 4 | 9 | 0 | 0 | 3 | 0 | 2 | 0 | 210 | 149 | 1 | 1 | 1 | 1 | 1 | 1 | 1 | 1 | 0.818 | 1 | 0.761 | NA | 0.105 |
| SE | 25683 | ENSG00000188295 | ZNF669 | chr1 | - | 247101986 | 247102113 | 247101050 | 247101319 | 247104196 | 247104289 | 8 | 10 | 3 | 17 | 2 | 0 | 0 | 0 | 0 | 0 | 0 | 5 | 6 | 3 | 23 | 0 | 3 | 0 | 2 | 0 | 276 | 149 | 1 | 1 | 1 | 1 | 1 | 1 | 1 | NA | 0.474 | 1 | 0.447 | 1 | 0.27 |
| SE | 25684 | ENSG00000188295 | ZNF669 | chr1 | - | 247101986 | 247102113 | 247101730 | 247101791 | 247104196 | 247104289 | 24 | 17 | 14 | 43 | 10 | 0 | 0 | 0 | 2 | 0 | 22 | 24 | 9 | 7 | 20 | 0 | 0 | 0 | 0 | 0 | 276 | 149 | 1 | 1 | 1 | 1 | 1 | 0.921 | 1 | 1 | 1 | 1 | 1 | 1 | -0.016 |
| SE | 25685 | ENSG00000188295 | ZNF669 | chr1 | - | 247101990 | 247102113 | 247101730 | 247101791 | 247104196 | 247104278 | 5 | 8 | 3 | 13 | 2 | 0 | 0 | 0 | 2 | 0 | 0 | 5 | 4 | 2 | 10 | 0 | 0 | 0 | 0 | 0 | 272 | 149 | 1 | 1 | 1 | 1 | 1 | 0.781 | 1 | NA | 1 | 1 | 1 | 1 | -0.044 |
| SE | 25690 | ENSG00000128694 | OSGEPL1 | chr2 | - | 189750549 | 189750656 | 189746659 | 189747168 | 189752652 | 189752724 | 37 | 23 | 24 | 28 | 72 | 0 | 0 | 0 | 1 | 0 | 0 | 12 | 11 | 3 | 4 | 0 | 0 | 0 | 0 | 0 | 256 | 149 | 1 | 1 | 1 | 1 | 1 | 0.942 | 1 | NA | 1 | 1 | 1 | 1 | -0.012 |
| SE | 25710 | ENSG00000247774 | PCED1B-AS1 | chr12 | - | 47209280 | 47209458 | 47206378 | 47206588 | 47216383 | 47216408 | 68 | 51 | 139 | 79 | 170 | 0 | 0 | 0 | 2 | 2 | 39 | 59 | 30 | 29 | 33 | 0 | 0 | 0 | 0 | 0 | 298 | 149 | 1 | 1 | 1 | 1 | 1 | 0.952 | 0.977 | 1 | 1 | 1 | 1 | 1 | -0.014 |
| SE | 25712 | ENSG00000247774 | PCED1B-AS1 | chr12 | - | 47209280 | 47209482 | 47205900 | 47206069 | 47216164 | 47216200 | 46 | 11 | 62 | 48 | 119 | 12 | 2 | 13 | 9 | 13 | 18 | 28 | 36 | 4 | 0 | 0 | 0 | 4 | 2 | 0 | 298 | 149 | 1 | 1 | 0.657 | 0.733 | 0.705 | 0.727 | 0.821 | 1 | 1 | 0.818 | 0.5 | NA | -0.101 |
| SE | 25721 | ENSG00000247774 | PCED1B-AS1 | chr12 | - | 47210639 | 47210761 | 47206510 | 47206588 | 47216383 | 47216408 | 25 | 5 | 35 | 21 | 57 | 0 | 0 | 0 | 2 | 2 | 0 | 8 | 18 | 10 | 0 | 0 | 0 | 0 | 0 | 0 | 271 | 149 | 1 | 1 | 1 | 1 | 1 | 0.852 | 0.94 | NA | 1 | 1 | 1 | NA | -0.042 |
| SE | 25725 | ENSG00000247774 | PCED1B-AS1 | chr12 | - | 47210639 | 47210761 | 47209280 | 47209482 | 47216383 | 47216408 | 30 | 16 | 31 | 33 | 58 | 1 | 0 | 0 | 1 | 0 | 0 | 8 | 11 | 0 | 9 | 0 | 0 | 0 | 0 | 0 | 271 | 149 | 1 | 1 | 0.943 | 1 | 1 | 1 | 0.97 | NA | 1 | 1 | NA | 1 | -0.017 |
| SE | 25728 | ENSG00000247774 | PCED1B-AS1 | chr12 | - | 47216164 | 47216264 | 47209280 | 47209482 | 47216383 | 47216408 | 41 | 8 | 39 | 25 | 45 | 1 | 0 | 0 | 0 | 1 | 28 | 9 | 20 | 0 | 9 | 0 | 0 | 0 | 0 | 0 | 249 | 149 | 1 | 1 | 0.961 | 1 | 1 | 1 | 0.964 | 1 | 1 | 1 | NA | 1 | -0.015 |
| SE | 25729 | ENSG00000196302 | GUSBP15 | chr5 | - | 70518320 | 70518405 | 70516386 | 70516482 | 70522000 | 70522188 | 15 | 15 | 9 | 10 | 9 | 0 | 0 | 1 | 0 | 0 | 0 | 1 | 13 | 4 | 16 | 0 | 0 | 0 | 0 | 0 | 234 | 149 | 1 | 1 | 1 | 1 | 0.851 | 1 | 1 | NA | 1 | 1 | 1 | 1 | -0.03 |
| SE | 25732 | ENSG00000168288 | MMADHC | chr2 | - | 149571084 | 149571171 | 149569636 | 149570168 | 149576436 | 149576542 | 143 | 90 | 138 | 129 | 220 | 1 | 0 | 1 | 0 | 0 | 59 | 99 | 123 | 114 | 27 | 0 | 0 | 0 | 0 | 0 | 236 | 149 | 1 | 1 | 0.989 | 1 | 0.989 | 1 | 1 | 1 | 1 | 1 | 1 | 1 | -0.004 |
| SE | 25733 | ENSG00000168288 | MMADHC | chr2 | - | 149572229 | 149572331 | 149571084 | 149571171 | 149575710 | 149575841 | 1 | 0 | 0 | 0 | 0 | 302 | 147 | 264 | 218 | 743 | 0 | 0 | 0 | 0 | 0 | 75 | 156 | 191 | 134 | 61 | 251 | 149 | 1 | 1 | 0.002 | 0 | 0 | 0 | 0 | 0 | 0 | 0 | 0 | 0 | 0 |
| SE | 25734 | ENSG00000168288 | MMADHC | chr2 | - | 149575710 | 149575841 | 149569636 | 149570168 | 149576436 | 149576542 | 158 | 105 | 155 | 140 | 565 | 1 | 0 | 1 | 0 | 0 | 32 | 107 | 129 | 109 | 25 | 0 | 0 | 0 | 0 | 0 | 280 | 149 | 1 | 1 | 0.988 | 1 | 0.988 | 1 | 1 | 1 | 1 | 1 | 1 | 1 | -0.005 |
| SE | 25735 | ENSG00000168288 | MMADHC | chr2 | - | 149575710 | 149575841 | 149571084 | 149571171 | 149576436 | 149576542 | 459 | 249 | 418 | 354 | 1277 | 0 | 0 | 0 | 1 | 1 | 99 | 260 | 315 | 236 | 86 | 0 | 0 | 0 | 0 | 0 | 280 | 149 | 1 | 1 | 1 | 1 | 1 | 0.995 | 0.999 | 1 | 1 | 1 | 1 | 1 | -0.001 |
| SE | 25737 | ENSG00000168288 | MMADHC | chr2 | - | 149587088 | 149587149 | 149582126 | 149582271 | 149587663 | 149587775 | 194 | 78 | 149 | 149 | 128 | 0 | 0 | 0 | 1 | 0 | 64 | 113 | 95 | 105 | 22 | 0 | 0 | 0 | 0 | 0 | 210 | 149 | 1 | 1 | 1 | 1 | 1 | 0.991 | 1 | 1 | 1 | 1 | 1 | 1 | -0.002 |

| | | | | | | | | | |
|---|---|---|---|---|---|---|---|---|---|
| SE | 25740 | ENSG00000157107 | FCHO2 | chr5 | + | 72990711 | 72990864 | 72990477 | 72990619 |
| SE | 25741 | ENSG00000157107 | FCHO2 | chr5 | + | 73006444 | 73006549 | 72990711 | 72990864 |
| SE | 25742 | ENSG00000157107 | FCHO2 | chr5 | + | 73015625 | 73015724 | 73006444 | 73006549 |
| SE | 25743 | ENSG00000157107 | FCHO2 | chr5 | + | 73063840 | 73063944 | 73058420 | 73058524 |
| SE | 25762 | ENSG00000001084 | GCLC | chr6 | - | 53500431 | 53500513 | 53500199 | 53500350 |
| SE | 25763 | ENSG00000001084 | GCLC | chr6 | - | 53508594 | 53508711 | 53507479 | 53507618 |
| SE | 25764 | ENSG00000001084 | GCLC | chr6 | - | 53516108 | 53516217 | 53514438 | 53514497 |
| SE | 25765 | ENSG00000001084 | GCLC | chr6 | - | 53516108 | 53516222 | 53514438 | 53514497 |
| SE | 25766 | ENSG00000001084 | GCLC | chr6 | - | 53516108 | 53516240 | 53514438 | 53514497 |
| SE | 25767 | ENSG00000001084 | GCLC | chr6 | - | 53522414 | 53522527 | 53520777 | 53520960 |
| SE | 25770 | ENSG00000113391 | FAM172A | chr5 | - | 93824147 | 93824267 | 93784194 | 93784499 |
| SE | 25771 | ENSG00000113391 | FAM172A | chr5 | - | 93881469 | 93881688 | 93824147 | 93824267 |
| SE | 25773 | ENSG00000113391 | FAM172A | chr5 | - | 93964457 | 93964523 | 93881469 | 93881688 |
| SE | 25774 | ENSG00000113391 | FAM172A | chr5 | - | 93964457 | 93964523 | 93958776 | 93958968 |
| SE | 25775 | ENSG00000113391 | FAM172A | chr5 | - | 93964457 | 93964523 | 93958776 | 93958968 |
| SE | 25776 | ENSG00000113391 | FAM172A | chr5 | - | 94050730 | 94050831 | 93958776 | 93958968 |
| SE | 25778 | ENSG00000113391 | FAM172A | chr5 | - | 94053125 | 94053227 | 93964457 | 93964523 |
| SE | 25784 | ENSG00000047621 | C12orf4 | chr12 | - | 4500137 | 4500340 | 4491176 | 4491234 |
| SE | 25785 | ENSG00000047621 | C12orf4 | chr12 | - | 4502310 | 4502482 | 4500137 | 4500340 |
| SE | 25786 | ENSG00000047621 | C12orf4 | chr12 | - | 4517060 | 4517189 | 4505305 | 4505380 |
| SE | 25787 | ENSG00000047621 | C12orf4 | chr12 | - | 4518057 | 4518255 | 4505305 | 4505380 |
| SE | 25788 | ENSG00000047621 | C12orf4 | chr12 | - | 4518773 | 4518880 | 4518057 | 4518255 |

[illegible]

| | | | | | | | | | | | | | | | | | | | | | | | | | | |
|---|---|---|---|---|---|---|---|---|---|---|---|---|---|---|---|---|---|---|---|---|---|---|---|---|---|---|
| SE | 26252 | ENSG00000248099 | INSL3 | chr19 | - | 17820395 | 17820490 | 17816513 | 17817059 | 17821316 | 17821574 | 2 | 0 | 2 | 0 | 0 | 0 | 0 | 0 | 2 | 3 | 0 | 0 | 0 | 0 | 0 |
| SE | 26269 | ENSG00000137449 | CPEB2 | chr4 | + | 15054127 | 15054217 | 15052413 | 15052584 | 15058420 | 15058539 | 59 | 32 | 42 | 90 | 59 | 0 | 0 | 0 | 0 | 0 | 40 | 32 | 54 | 54 | 58 |
| SE | 26270 | ENSG00000137449 | CPEB2 | chr4 | + | 15059186 | 15059301 | 15058420 | 15058539 | 15062078 | 15062260 | 52 | 76 | 53 | 107 | 82 | 1 | 0 | 0 | 0 | 0 | 62 | 72 | 84 | 89 | 65 |
| SE | 26271 | ENSG00000101911 | PRPS2 | chrX | + | 12810021 | 12810146 | 12809233 | 12809332 | 12819506 | 12819680 | 138 | 110 | 184 | 160 | 147 | 2 | 0 | 0 | 0 | 0 | 110 | 168 | 118 | 105 | 46 |
| SE | 26272 | ENSG00000101911 | PRPS2 | chrX | + | 12819506 | 12819680 | 12810021 | 12810146 | 12820643 | 12820803 | 194 | 141 | 220 | 202 | 291 | 1 | 0 | 1 | 0 | 0 | 187 | 248 | 175 | 185 | 15 |
| SE | 26273 | ENSG00000151876 | FBXO4 | chr5 | + | 41927012 | 41927248 | 41925253 | 41925498 | 41928517 | 41928691 | 4 | 9 | 17 | 6 | 15 | 0 | 0 | 0 | 0 | 3 | 0 | 9 | 3 | 0 | 6 |
| SE | 26274 | ENSG00000151876 | FBXO4 | chr5 | + | 41927012 | 41927248 | 41925253 | 41925498 | 41929696 | 41929917 | 41 | 33 | 61 | 20 | 102 | 0 | 0 | 1 | 1 | 1 | 48 | 50 | 12 | 5 | 37 |
| SE | 26275 | ENSG00000151876 | FBXO4 | chr5 | + | 41928517 | 41928691 | 41925298 | 41925498 | 41929696 | 41929917 | 0 | 2 | 1 | 0 | 9 | 0 | 0 | 1 | 1 | 1 | 0 | 5 | 5 | 0 | 0 |
| SE | 26278 | ENSG00000150756 | ATPSCKMT | chr5 | - | 10235210 | 10235261 | 10225506 | 10227647 | 10236477 | 10236615 | 10 | 8 | 10 | 5 | 17 | 8 | 1 | 5 | 4 | 10 | 10 | 0 | 12 | 2 | 0 |
| SE | 26279 | ENSG00000150756 | ATPSCKMT | chr5 | - | 10235210 | 10235261 | 10225506 | 10227647 | 10236521 | 10236615 | 9 | 4 | 5 | 4 | 2 | 0 | 0 | 0 | 0 | 0 | 10 | 0 | 8 | 0 | 0 |
| SE | 26282 | ENSG00000248092 | NNT-AS1 | chr5 | - | 43583466 | 43583566 | 43571593 | 43575935 | 43602777 | 43602965 | 10 | 8 | 14 | 22 | 10 | 1 | 0 | 0 | 0 | 0 | 0 | 6 | 5 | 8 | 0 |
| SE | 26283 | ENSG00000248092 | NNT-AS1 | chr5 | - | 43583466 | 43583566 | 43578268 | 43578579 | 43602777 | 43602965 | 1 | 4 | 5 | 10 | 5 | 0 | 0 | 2 | 0 | 0 | 0 | 0 | 0 | 3 | 0 |
| SE | 26298 | ENSG00000100320 | RBFOX2 | chr22 | - | 35756104 | 35756144 | 35746472 | 35746561 | 35759887 | 35760020 | 4 | 0 | 6 | 2 | 0 | 0 | 0 | 0 | 0 | 0 | 0 | 0 | 0 | 0 | 0 |
| SE | 26304 | ENSG00000113593 | PPWD1 | chr5 | + | 65568917 | 65568994 | 65567512 | 65567615 | 65569636 | 65569732 | 5 | 1 | 1 | 4 | 2 | 4 | 2 | 0 | 2 | 0 | 0 | 0 | 0 | 2 | 0 |
| SE | 26305 | ENSG00000113593 | PPWD1 | chr5 | + | 65571838 | 65572286 | 65563238 | 65563506 | 65576878 | 65576926 | 190 | 169 | 198 | 122 | 216 | 0 | 0 | 0 | 2 | 0 | 19 | 97 | 75 | 19 | 31 |
| SE | 26313 | ENSG00000113300 | CNOT6 | chr5 | + | 180495420 | 180495588 | 180494506 | 180494763 | 180529274 | 180529388 | 0 | 1 | 0 | 1 | 0 | 3 | 0 | 10 | 7 | 7 | 0 | 0 | 0 | 3 | 0 |
| SE | 26314 | ENSG00000113300 | CNOT6 | chr5 | + | 180495651 | 180495755 | 180494411 | 180494763 | 180529274 | 180529388 | 1 | 2 | 4 | 0 | 0 | 3 | 0 | 10 | 7 | 7 | 0 | 4 | 4 | 0 | 0 |
| SE | 26315 | ENSG00000113300 | CNOT6 | chr5 | + | 180529274 | 180529388 | 180494378 | 180494763 | 180549930 | 180550117 | 84 | 73 | 85 | 93 | 115 | 0 | 1 | 0 | 0 | 0 | 90 | 84 | 78 | 64 | 31 |
| SE | 26316 | ENSG00000113300 | CNOT6 | chr5 | + | 180549930 | 180550117 | 180529274 | 180529388 | 180564488 | 180564682 | 90 | 73 | 79 | 92 | 116 | 1 | 0 | 0 | 0 | 0 | 90 | 84 | 80 | 69 | 31 |
| SE | 26318 | ENSG00000113300 | CNOT6 | chr5 | + | 180565819 | 180565977 | 180564674 | 180564743 | 180567087 | 180567242 | 216 | 143 | 175 | 185 | 312 | 0 | 0 | 1 | 4 | 0 | 45 | 119 | 75 | 120 | 30 |
| SE | 26320 | ENSG00000113300 | CNOT6 | chr5 | + | 180567848 | 180568003 | 180567087 | 180567242 | 180571229 | 180571432 | 133 | 88 | 127 | 97 | 164 | 0 | 1 | 0 | 0 | 0 | 28 | 97 | 75 | 116 | 29 |
| SE | 26321 | ENSG00000113300 | CNOT6 | chr5 | + | 180567848 | 180568003 | 180567087 | 180567242 | 180573987 | 180578347 | 130 | 88 | 127 | 97 | 166 | 0 | 0 | 1 | 0 | 0 | 28 | 97 | 75 | 116 | 29 |
| SE | 26322 | ENSG00000113300 | CNOT6 | chr5 | + | 180569109 | 180569340 | 180567087 | 180567242 | 180571229 | 180571432 | 111 | 87 | 94 | 100 | 263 | 0 | 1 | 0 | 0 | 0 | 27 | 75 | 60 | 100 | 29 |
| SE | 26323 | ENSG00000113300 | CNOT6 | chr5 | + | 180569109 | 180569340 | 180567087 | 180567242 | 180573987 | 180578347 | 9 | 4 | 7 | 8 | 23 | 0 | 0 | 1 | 0 | 0 | 4 | 8 | 2 | 3 | 0 |
| SE | 26324 | ENSG00000113300 | CNOT6 | chr5 | + | 180569109 | 180569340 | 180567848 | 180568003 | 180571229 | 180571432 | 201 | 143 | 157 | 169 | 417 | 3 | 0 | 0 | 0 | 0 | 47 | 144 | 93 | 137 | 37 |
| SE | 26325 | ENSG00000113300 | CNOT6 | chr5 | + | 180569109 | 180569340 | 180567848 | 180568003 | 180573987 | 180578347 | 99 | 60 | 70 | 77 | 177 | 0 | 0 | 0 | 0 | 2 | 24 | 77 | 35 | 40 | 8 |
| SE | 26326 | ENSG00000113300 | CNOT6 | chr5 | + | 180571229 | 180571432 | 180567087 | 180567242 | 180573987 | 180578347 | 135 | 88 | 95 | 98 | 364 | 0 | 0 | 1 | 0 | 0 | 41 | 84 | 58 | 71 | 24 |
| SE | 26327 | ENSG00000113300 | CNOT6 | chr5 | + | 180571229 | 180571432 | 180567848 | 180568003 | 180573987 | 180578347 | 138 | 87 | 95 | 98 | 364 | 0 | 0 | 0 | 0 | 2 | 41 | 84 | 58 | 71 | 24 |
| SE | 26332 | ENSG00000139625 | MAP3K12 | chr12 | - | 53483348 | 53483486 | 53482564 | 53483189 | 53483606 | 53483723 | 38 | 35 | 59 | 50 | 37 | 0 | 0 | 0 | 1 | 0 | 18 | 35 | 26 | 20 | 79 |
| SE | 26334 | ENSG00000139625 | MAP3K12 | chr12 | - | 53484256 | 53484365 | 53483910 | 53484020 | 53485055 | 53485214 | 56 | 33 | 67 | 59 | 38 | 0 | 0 | 2 | 2 | 0 | 40 | 93 | 40 | 15 | 56 |
| SE | 26336 | ENSG00000048028 | USP28 | chr11 | - | 113801482 | 113801678 | 113797873 | 113799415 | 113803797 | 113803877 | 306 | 107 | 152 | 223 | 294 | 0 | 0 | 0 | 1 | 0 | 85 | 202 | 137 | 117 | 25 |
| SE | 26337 | ENSG00000048028 | USP28 | chr11 | - | 113803157 | 113803281 | 113797873 | 113799415 | 113803797 | 113803877 | 143 | 41 | 57 | 84 | 109 | 0 | 0 | 0 | 1 | 0 | 32 | 82 | 57 | 31 | 6 |
| SE | 26338 | ENSG00000048028 | USP28 | chr11 | - | 113803157 | 113803281 | 113801482 | 113801678 | 113803797 | 113803877 | 353 | 95 | 143 | 224 | 298 | 0 | 0 | 0 | 1 | 0 | 86 | 259 | 182 | 98 | 42 |
| SE | 26339 | ENSG00000048028 | USP28 | chr11 | - | 113804867 | 113805046 | 113804672 | 113804751 | 113808297 | 113808437 | 254 | 78 | 110 | 177 | 123 | 0 | 0 | 0 | 1 | 0 | 19 | 142 | 69 | 64 | 4 |
| SE | 26341 | ENSG00000048028 | USP28 | chr11 | - | 113808022 | 113808136 | 113806488 | 113806584 | 113808297 | 113808437 | 2 | 2 | 0 | 1 | 2 | 1 | 0 | 4 | 0 | 6 | 0 | 3 | 0 | 0 | 0 |
| SE | 26343 | ENSG00000048028 | USP28 | chr11 | - | 113820291 | 113820358 | 113817830 | 113817837 | 113823604 | 113823700 | 2 | 0 | 1 | 2 | 6 | 38 | 16 | 31 | 19 | 31 | 0 | 3 | 12 | 0 | 0 |
| SE | 26344 | ENSG00000048028 | USP28 | chr11 | - | 113823604 | 113823700 | 113817657 | 113817837 | 113829196 | 113829345 | 36 | 15 | 26 | 21 | 28 | 0 | 0 | 0 | 0 | 0 | 0 | 19 | 9 | 11 | 0 |
| SE | 26345 | ENSG00000048028 | USP28 | chr11 | - | 113827232 | 113827360 | 113823604 | 113823700 | 113829196 | 113829345 | 239 | 79 | 101 | 148 | 104 | 0 | 0 | 0 | 2 | 0 | 129 | 192 | 77 | 107 | 3 |
| SE | 26347 | ENSG00000048028 | USP28 | chr11 | - | 113830866 | 113830943 | 113829196 | 113829345 | 113831919 | 113831989 | 246 | 93 | 119 | 175 | 174 | 2 | 0 | 0 | 1 | 0 | 133 | 153 | 99 | 74 | 23 |
| SE | 26349 | ENSG00000048028 | USP28 | chr11 | - | 113830866 | 113830946 | 113829196 | 113829345 | 113831919 | 113831993 | 190 | 69 | 93 | 129 | 130 | 2 | 0 | 0 | 0 | 0 | 101 | 129 | 83 | 52 | 29 |
| SE | 26351 | ENSG00000048028 | USP28 | chr11 | - | 113831919 | 113831989 | 113829196 | 113829345 | 113833419 | 113833557 | 2 | 0 | 2 | 0 | 0 | 1 | 5 | 1 | 0 | 1 | 0 | 0 | 0 | 0 | 0 |
| SE | 26356 | ENSG00000048028 | USP28 | chr11 | - | 113833419 | 113833557 | 113830882 | 113830943 | 113834248 | 113834335 | 251 | 87 | 97 | 121 | 153 | 0 | 0 | 0 | 1 | 0 | 70 | 128 | 99 | 57 | 85 |
| SE | 26361 | ENSG00000048028 | USP28 | chr11 | - | 113854257 | 113854335 | 113841662 | 113841768 | 113874516 | 113874800 | 9 | 6 | 8 | 5 | 12 | 0 | 0 | 0 | 1 | 0 | 8 | 19 | 3 | 0 | 0 |
| SE | 26362 | ENSG00000048028 | USP28 | chr11 | - | 113854257 | 113854335 | 113841662 | 113841768 | 113875444 | 113875528 | 3 | 0 | 5 | 4 | 2 | 0 | 3 | 2 | 3 | 0 | 0 | 9 | 0 | 3 | 0 |
| SE | 26363 | ENSG00000048028 | USP28 | chr11 | - | 113854257 | 113854335 | 113852500 | 113852633 | 113875444 | 113875532 | 72 | 29 | 21 | 48 | 38 | 0 | 0 | 0 | 0 | 0 | 11 | 46 | 29 | 33 | 5 |
| SE | 26364 | ENSG00000048028 | USP28 | chr11 | - | 113874516 | 113874800 | 113854257 | 113854335 | 113875444 | 113875536 | 10 | 8 | 6 | 6 | 11 | 12 | 7 | 4 | 6 | 0 | 9 | 17 | 6 | 0 | 0 |
| SE | 26366 | ENSG00000048028 | USP28 | chr11 | - | 113874516 | 113874879 | 113854257 | 113854335 | 113875444 | 113875533 | 11 | 8 | 6 | 6 | 11 | 12 | 7 | 4 | 6 | 0 | 9 | 17 | 6 | 0 | 0 |
| SE | 26367 | ENSG00000049883 | PTCD2 | chr5 | + | 72321320 | 72321376 | 72320398 | 72320509 | 72322171 | 72322264 | 0 | 0 | 0 | 0 | 0 | 20 | 9 | 13 | 17 | 0 | 3 | 1 | 0 | 0 | 0 |
| SE | 26368 | ENSG00000049883 | PTCD2 | chr5 | + | 72326611 | 72326741 | 72322171 | 72322264 | 72331257 | 72331375 | 39 | 33 | 30 | 29 | 13 | 1 | 0 | 0 | 2 | 1 | 3 | 12 | 16 | 18 | 0 |
| SE | 26374 | ENSG00000049883 | PTCD2 | chr5 | + | 72335017 | 72335096 | 72331257 | 72331375 | 72335793 | 72335885 | 17 | 8 | 22 | 13 | 21 | 2 | 1 | 1 | 1 | 4 | 0 | 8 | 5 | 8 | 0 |
| SE | 26375 | ENSG00000151881 | TMEM267 | chr5 | - | 43453657 | 43454043 | 43444251 | 43446557 | 43476113 | 43476258 | 30 | 27 | 36 | 56 | 69 | 0 | 0 | 0 | 0 | 7 | 0 | 11 | 19 | 3 | 0 |
| SE | 26376 | ENSG00000151881 | TMEM267 | chr5 | - | 43453657 | 43454043 | 43444251 | 43446557 | 43479889 | 43479987 | 28 | 27 | 36 | 52 | 62 | 4 | 0 | 0 | 0 | 2 | 0 | 11 | 13 | 3 | 0 |
| SE | 26377 | ENSG00000151881 | TMEM267 | chr5 | - | 43453657 | 43454043 | 43444251 | 43446557 | 43483821 | 43483865 | 37 | 29 | 39 | 65 | 76 | 2 | 0 | 0 | 0 | 1 | 10 | 11 | 16 | 3 | 0 |
| SE | 26378 | ENSG00000151881 | TMEM267 | chr5 | - | 43454454 | 43454605 | 43453657 | 43454043 | 43483821 | 43483891 | 0 | 0 | 1 | 0 | 3 | 9 | 2 | 5 | 13 | 14 | 0 | 0 | 1 | 0 | 0 |
| SE | 26380 | ENSG00000151881 | TMEM267 | chr5 | - | 43476113 | 43476258 | 43453657 | 43454043 | 43479889 | 43479997 | 2 | 0 | 2 | 4 | 7 | 0 | 0 | 2 | 0 | 0 | 0 | 0 | 6 | 0 | 0 |
| SE | 26381 | ENSG00000151881 | TMEM267 | chr5 | - | 43479889 | 43479987 | 43445978 | 43446557 | 43483821 | 43483836 | 4 | 0 | 1 | 0 | 2 | 2 | 0 | 0 | 0 | 1 | 0 | 0 | 0 | 0 | 0 |
| SE | 26382 | ENSG00000151881 | TMEM267 | chr5 | - | 43479889 | 43479987 | 43453657 | 43454043 | 43483821 | 43483836 | 0 | 0 | 3 | 0 | 0 | 9 | 2 | 5 | 13 | 14 | 0 | 0 | 0 | 0 | 0 |
| SE | 26383 | ENSG00000056097 | ZFR | chr5 | - | 32379114 | 32379210 | 32364163 | 32364275 | 32385507 | 32385649 | 66 | 54 | 81 | 73 | 67 | 0 | 1 | 0 | 0 | 0 | 45 | 55 | 76 | 43 | 2 |
| SE | 26385 | ENSG00000056097 | ZFR | chr5 | - | 32390274 | 32390437 | 32388468 | 32388674 | 32395158 | 32395304 | 721 | 541 | 660 | 732 | 1112 | 0 | 0 | 0 | 0 | 0 | 443 | 723 | 729 | 489 | 105 |

| | | | | | | | | | | | | | | | |
|---|---|---|---|---|---|---|---|---|---|---|---|---|---|---|---|
| SE | 26730 | ENSG00000125089 | SH3TC1 | chr4 | + | 8204782 | 8205053 | 8199370 | 8199405 | 8205166 | 8205366 | 0 | 0 | 0 | [illegible] |
| SE | 26736 | ENSG00000125089 | SH3TC1 | chr4 | + | 8216956 | 8217167 | 8216110 | 8216257 | 8218244 | 8218347 | 37 | 26 | 20 | [illegible] |
| SE | 26745 | ENSG00000174720 | LARP7 | chr4 | + | 112637643 | 112637787 | 112637125 | 112637239 | 112644667 | 112644871 | 3 | 0 | 0 | [illegible] |
| SE | 26747 | ENSG00000174720 | LARP7 | chr4 | + | 112638617 | 112638673 | 112637192 | 112637239 | 112644667 | 112644871 | 2 | 3 | 1 | [illegible] |
| SE | 26749 | ENSG00000174720 | LARP7 | chr4 | + | 112638617 | 112638673 | 112637569 | 112637603 | 112644667 | 112644871 | 0 | 2 | 1 | [illegible] |
| SE | 26750 | ENSG00000174720 | LARP7 | chr4 | + | 112644424 | 112644488 | 112637125 | 112637239 | 112644667 | 112644871 | 1 | 0 | 1 | [illegible] |
| SE | 26751 | ENSG00000174720 | LARP7 | chr4 | + | 112644424 | 112644488 | 112637569 | 112637603 | 112644667 | 112644871 | 1 | 0 | 0 | [illegible] |
| SE | 26753 | ENSG00000174720 | LARP7 | chr4 | + | 112644424 | 112644488 | 112638617 | 112638673 | 112644667 | 112644871 | 2 | 0 | 0 | [illegible] |
| SE | 26754 | ENSG00000174720 | LARP7 | chr4 | + | 112645550 | 112645662 | 112644667 | 112644871 | 112646350 | 112646451 | 1 | 0 | 0 | [illegible] |
| SE | 26757 | ENSG00000127804 | METTL16 | chr17 | - | 2438108 | 2438198 | 2420730 | 2420904 | 2441489 | 2441559 | 111 | 82 | 128 | [illegible] |
| SE | 26758 | ENSG00000127804 | METTL16 | chr17 | - | 2441489 | 2441559 | 2420730 | 2420904 | 2464207 | 2464350 | 54 | 31 | 46 | [illegible] |
| SE | 26762 | ENSG00000127804 | METTL16 | chr17 | - | 2473523 | 2473664 | 2467760 | 2467876 | 2477685 | 2477885 | 281 | 202 | 277 | [illegible] |
| SE | 26763 | ENSG00000127804 | METTL16 | chr17 | - | 2475240 | 2475299 | 2473523 | 2473664 | 2477685 | 2477885 | 1 | 0 | 0 | [illegible] |
| SE | 26764 | ENSG00000127804 | METTL16 | chr17 | - | 2502203 | 2502331 | 2477685 | 2477885 | 2511758 | 2511835 | 125 | 51 | 109 | [illegible] |
| SE | 26792 | ENSG00000111728 | ST8SIA1 | chr12 | - | 22255279 | 22255389 | 22248825 | 22249098 | 22287148 | 22287194 | 7 | 4 | 11 | [illegible] |
| SE | 26795 | ENSG00000163738 | MTHFD2L | chr4 | + | 74160054 | 74160088 | 74158107 | 74158281 | 74174505 | 74174690 | 6 | 13 | 1 | [illegible] |
| SE | 26797 | ENSG00000163738 | MTHFD2L | chr4 | + | 74181685 | 74181791 | 74180686 | 74180866 | 74199793 | 74199946 | 3 | 4 | 2 | [illegible] |
| SE | 26800 | ENSG00000163281 | GNPDA2 | chr4 | - | 44710952 | 44711137 | 44703245 | 44707926 | 44726473 | 44726523 | 102 | 44 | 53 | [illegible] |
| SE | 26801 | ENSG00000163281 | GNPDA2 | chr4 | - | 44717112 | 44717295 | 44703245 | 44707926 | 44726473 | 44726523 | 2 | 1 | 1 | [illegible] |
| SE | 26805 | ENSG00000163281 | GNPDA2 | chr4 | - | 44718308 | 44718410 | 44717112 | 44717295 | 44726473 | 44726523 | 47 | 19 | 31 | [illegible] |
| SE | 26806 | ENSG00000163281 | GNPDA2 | chr4 | - | 44722083 | 44722242 | 44710952 | 44711137 | 44726473 | 44726545 | 13 | 8 | 20 | [illegible] |
| SE | 26807 | ENSG00000163281 | GNPDA2 | chr4 | - | 44722083 | 44722242 | 44717112 | 44717295 | 44726473 | 44726545 | 13 | 8 | 20 | [illegible] |
| SE | 26808 | ENSG00000163281 | GNPDA2 | chr4 | - | 44722083 | 44722242 | 44718308 | 44718410 | 44726473 | 44726545 | 49 | 41 | 57 | [illegible] |
| SE | 26809 | ENSG00000138760 | SCARB2 | chr4 | - | 76174143 | 76174313 | 76169866 | 76169985 | 76175790 | 76175910 | 481 | 400 | 396 | [illegible] |
| SE | 26811 | ENSG00000138760 | SCARB2 | chr4 | - | 76176436 | 76176528 | 76175790 | 76175910 | 76179516 | 76179695 | 309 | 251 | 237 | [illegible] |
| SE | 26818 | ENSG00000138760 | SCARB2 | chr4 | - | 76180953 | 76181092 | 76175790 | 76175910 | 76195706 | 76195864 | 2 | 1 | 0 | [illegible] |
| SE | 26819 | ENSG00000138760 | SCARB2 | chr4 | - | 76180953 | 76181101 | 76175790 | 76175910 | 76195706 | 76195864 | 93 | 76 | 49 | [illegible] |
| SE | 26820 | ENSG00000138760 | SCARB2 | chr4 | - | 76184656 | 76184780 | 76181013 | 76181101 | 76195706 | 76195864 | 0 | 0 | 4 | [illegible] |
| SE | 26821 | ENSG00000186468 | RPS23 | chr5 | - | 82277692 | 82277852 | 82275877 | 82276518 | 82278319 | 82278341 | 3762 | 2015 | 3912 | [illegible] |
| SE | 26822 | ENSG00000130165 | ELOF1 | chr19 | - | 11554231 | 11554365 | 11553627 | 11554081 | 11559190 | 11559230 | 152 | 39 | 109 | [illegible] |
| SE | 26823 | ENSG00000130165 | ELOF1 | chr19 | - | 11554819 | 11554946 | 11554231 | 11554365 | 11555173 | 11555253 | 0 | 0 | 4 | [illegible] |
| SE | 26825 | ENSG00000130165 | ELOF1 | chr19 | - | 11555173 | 11555253 | 11554231 | 11554365 | 11555344 | 11555467 | 0 | 2 | 2 | [illegible] |
| SE | 26827 | ENSG00000130165 | ELOF1 | chr19 | - | 11555173 | 11555253 | 11554819 | 11554946 | 11559190 | 11559236 | 0 | 0 | 2 | [illegible] |
| SE | 26828 | ENSG00000134153 | EMC7 | chr15 | - | 34088052 | 34088133 | 34084016 | 34084486 | 34090316 | 34090455 | 637 | 333 | 501 | [illegible] |
| SE | 26829 | ENSG00000134153 | EMC7 | chr15 | - | 34088052 | 34088133 | 34084016 | 34084486 | 34095894 | 34096014 | 387 | 194 | 281 | [illegible] |
| SE | 26830 | ENSG00000134153 | EMC7 | chr15 | - | 34090316 | 34090455 | 34084016 | 34084486 | 34095894 | 34096014 | 241 | 129 | 194 | [illegible] |
| SE | 26831 | ENSG00000134153 | EMC7 | chr15 | - | 34090316 | 34090455 | 34088052 | 34088133 | 34095894 | 34096014 | 508 | 271 | 419 | [illegible] |
| SE | 26833 | ENSG00000169245 | CXCL10 | chr4 | - | 76022690 | 76022817 | 76022365 | 76022455 | 76023370 | 76023497 | 27 | 69 | 26 | [illegible] |
| SE | 26839 | ENSG00000172738 | TMEM217 | chr6 | - | 37215940 | 37215966 | 37212570 | 37212996 | 37218425 | 37218696 | 0 | 0 | 0 | [illegible] |
| SE | 26848 | ENSG00000109265 | CRACD | chr4 | + | 56298213 | 56298349 | 56272320 | 56272492 | 56307534 | 56307565 | 4 | 0 | 0 | [illegible] |
| SE | 26851 | ENSG00000137076 | TLN1 | chr9 | - | 35699355 | 35699461 | 35699031 | 35699156 | 35699973 | 35700081 | 1347 | 719 | 725 | [illegible] |
| SE | 26852 | ENSG00000137076 | TLN1 | chr9 | - | 35703774 | 35703900 | 35703558 | 35703676 | 35703990 | 35704174 | 1138 | 649 | 647 | [illegible] |
| SE | 26853 | ENSG00000137076 | TLN1 | chr9 | - | 35704668 | 35704815 | 35704331 | 35704498 | 35705550 | 35705670 | 1485 | 758 | 879 | [illegible] |
| SE | 26855 | ENSG00000137076 | TLN1 | chr9 | - | 35707730 | 35707892 | 35707347 | 35707488 | 35708340 | 35708484 | 2110 | 964 | 1051 | [illegible] |
| SE | 26856 | ENSG00000137076 | TLN1 | chr9 | - | 35710796 | 35710886 | 35710560 | 35710683 | 35710988 | 35711082 | 1007 | 479 | 593 | [illegible] |
| SE | 26857 | ENSG00000137076 | TLN1 | chr9 | - | 35710988 | 35711082 | 35710560 | 35710683 | 35711254 | 35711394 | 748 | 340 | 377 | [illegible] |
| SE | 26858 | ENSG00000137076 | TLN1 | chr9 | - | 35714238 | 35714373 | 35713952 | 35714081 | 35714573 | 35714687 | 1128 | 764 | 574 | [illegible] |
| SE | 26859 | ENSG00000137076 | TLN1 | chr9 | - | 35716389 | 35716556 | 35715058 | 35715187 | 35717145 | 35717440 | 1428 | 642 | 750 | [illegible] |
| SE | 26860 | ENSG00000137076 | TLN1 | chr9 | - | 35717145 | 35717440 | 35715058 | 35715187 | 35717618 | 35717786 | 593 | 237 | 311 | [illegible] |
| SE | 26861 | ENSG00000137076 | TLN1 | chr9 | - | 35722860 | 35722921 | 35722118 | 35722223 | 35723951 | 35724079 | 2343 | 1638 | 1357 | [illegible] |
| SE | 26862 | ENSG00000137076 | TLN1 | chr9 | - | 35722860 | 35722921 | 35722118 | 35722223 | 35724191 | 35724334 | 715 | 503 | 413 | [illegible] |
| SE | 26863 | ENSG00000137076 | TLN1 | chr9 | - | 35723951 | 35724079 | 35722118 | 35722223 | 35724191 | 35724334 | 2714 | 1834 | 1468 | [illegible] |
| SE | 26864 | ENSG00000137076 | TLN1 | chr9 | - | 35723951 | 35724079 | 35722860 | 35722921 | 35724191 | 35724334 | 4338 | 2966 | 2407 | [illegible] |
| SE | 26865 | ENSG00000137076 | TLN1 | chr9 | - | 35724191 | 35724334 | 35722118 | 35722223 | 35724571 | 35724724 | 1938 | 1298 | 1076 | [illegible] |
| SE | 26866 | ENSG00000137076 | TLN1 | chr9 | - | 35724571 | 35724724 | 35722118 | 35722223 | 35725223 | 35725321 | 3 | 0 | 0 | [illegible] |
| SE | 26867 | ENSG00000137076 | TLN1 | chr9 | - | 35724829 | 35724959 | 35724571 | 35724724 | 35725223 | 35725321 | 3803 | 3186 | 1985 | [illegible] |
| SE | 26868 | ENSG00000137076 | TLN1 | chr9 | - | 35725564 | 35725727 | 35725223 | 35725321 | 35732074 | 35732104 | 2123 | 1547 | 1306 | [illegible] |
| SE | 26869 | ENSG00000128059 | PPAT | chr4 | - | 56403979 | 56404118 | 56403288 | 56403401 | 56406494 | 56406701 | 2 | 0 | 2 | [illegible] |
| SE | 26870 | ENSG00000151229 | SLC2A13 | chr12 | - | 39864761 | 39864882 | 39829407 | 39830228 | 39871797 | 39871881 | 14 | 15 | 15 | [illegible] |
| SE | 26871 | ENSG00000103995 | CEP152 | chr15 | - | 48741600 | 48741704 | 48737937 | 48739288 | 48741946 | 48742100 | 9 | 10 | 10 | [illegible] |
| SE | 26872 | ENSG00000103995 | CEP152 | chr15 | - | 48741630 | 48741704 | 48738150 | 48739288 | 48741946 | 48742014 | 3 | 1 | 5 | [illegible] |
| SE | 26873 | ENSG00000103995 | CEP152 | chr15 | - | 48748442 | 48748610 | 48744895 | 48744992 | 48752348 | 48752469 | 28 | 17 | 9 | [illegible] |
| SE | 26874 | ENSG00000103995 | CEP152 | chr15 | - | 48752348 | 48752469 | 48744895 | 48744992 | 48755902 | 48756553 | 10 | 21 | 9 | [illegible] |
| SE | 26875 | ENSG00000103995 | CEP152 | chr15 | - | 48760134 | 48760266 | 48755902 | 48756553 | 48762390 | 48762672 | 16 | 24 | 9 | [illegible] |
| SE | 26877 | ENSG00000184985 | SORCS2 | chr4 | + | 7728349 | 7728462 | 7726779 | 7726903 | 7729586 | 7729712 | 4 | 0 | 3 | [illegible] |
| SE | 26883 | ENSG00000120549 | KIAA1217 | chr10 | + | 24542882 | 24544481 | 24542692 | 24542770 | 24544980 | 24545103 | 0 | 0 | 0 | [illegible] |
| SE | 26885 | ENSG00000138107 | ACTR1A | chr10 | - | 102481836 | 102481898 | 102481131 | 102481172 | 102482000 | 102482175 | 261 | 157 | 196 | [illegible] |
| SE | 26887 | ENSG00000138107 | ACTR1A | chr10 | - | 102482000 | 102482175 | 102481131 | 102481172 | 102484159 | 102484376 | 0 | 0 | 1 | [illegible] |
| SE | 26888 | ENSG00000138107 | ACTR1A | chr10 | - | 102483010 | 102483103 | 102481131 | 102481172 | 102484159 | 102484374 | 74 | 40 | 57 | [illegible] |
| SE | 26889 | ENSG00000138107 | ACTR1A | chr10 | - | 102483010 | 102483103 | 102482000 | 102482175 | 102484159 | 102484374 | 232 | 111 | 163 | [illegible] |
| SE | 26890 | ENSG00000138107 | ACTR1A | chr10 | - | 102484159 | 102484376 | 102483010 | 102483103 | 102488149 | 102488275 | 136 | 60 | 86 | [illegible] |
| SE | 26892 | ENSG00000138107 | ACTR1A | chr10 | - | 102490548 | 102490613 | 102489062 | 102489138 | 102502599 | 102502678 | 142 | 52 | 105 | [illegible] |
| SE | 26893 | ENSG00000160685 | ZBTB7B | chr1 | + | 155002842 | 155002943 | 155002656 | 155002686 | 155014654 | 155015814 | 63 | 39 | 31 | [illegible] |
| SE | 26894 | ENSG00000160685 | ZBTB7B | chr1 | + | 155010257 | 155010397 | 155002629 | 155002943 | 155010915 | 155011015 | 24 | 9 | 21 | [illegible] |
| SE | 26901 | ENSG00000160613 | PCSK7 | chr11 | - | 117207066 | 117207160 | 117206679 | 117206798 | 117207653 | 117207735 | 92 | 29 | 60 | [illegible] |
| SE | 26907 | ENSG00000160613 | PCSK7 | chr11 | - | 117218465 | 117218568 | 117207969 | 117208064 | 117219056 | 117219164 | 69 | 26 | 70 | [illegible] |
| SE | 26909 | ENSG00000160613 | PCSK7 | chr11 | - | 117227156 | 117227322 | 117224699 | 117224755 | 117228215 | 117228350 | 167 | 51 | 122 | [illegible] |
| SE | 26910 | ENSG00000160613 | PCSK7 | chr11 | - | 117230348 | 117230468 | 117229376 | 117229856 | 117232026 | 117232073 | 165 | 75 | 117 | [illegible] |
| SE | 26912 | ENSG00000078967 | UBE2D4 | chr7 | + | 43928013 | 43928111 | 43926437 | 43926556 | 43938430 | 43938494 | 0 | 1 | 0 | [illegible] |
| SE | 26914 | ENSG00000078967 | UBE2D4 | chr7 | + | 43942953 | 43943031 | 43938430 | 43938494 | 43948631 | 43948737 | 5 | 1 | 0 | [illegible] |
| SE | 26915 | ENSG00000078967 | UBE2D4 | chr7 | + | 43947158 | 43947229 | 43942953 | 43943031 | 43948631 | 43948737 | 0 | 0 | 1 | [illegible] |
| SE | 26916 | ENSG00000078967 | UBE2D4 | chr7 | + | 43948631 | 43948737 | 43938430 | 43938494 | 43950598 | 43950692 | 8 | 6 | 2 | [illegible] |
| SE | 26917 | ENSG00000078967 | UBE2D4 | chr7 | + | 43950598 | 43950692 | 43948601 | 43948737 | 43952649 | 43952907 | 14 | 11 | 9 | [illegible] |
| SE | 26918 | ENSG00000138772 | ANXA3 | chr4 | + | 78552184 | 78552345 | 78551826 | 78551859 | 78554435 | 78554488 | 0 | 0 | 0 | [illegible] |
| SE | 26919 | ENSG00000138772 | ANXA3 | chr4 | + | 78571032 | 78571173 | 78554435 | 78554488 | 78573179 | 78573267 | 0 | 0 | 0 | [illegible] |
| SE | 26921 | ENSG00000138772 | ANXA3 | chr4 | + | 78579026 | 78579121 | 78573179 | 78573267 | 78582176 | 78582290 | 7 | 4 | 10 | [illegible] |
| SE | 26922 | ENSG00000138772 | ANXA3 | chr4 | + | 78604276 | 78604399 | 78601509 | 78601568 | 78610055 | 78610363 | 5 | 5 | 20 | [illegible] |
| SE | 26923 | ENSG00000180316 | PNPLA1 | chr6 | + | 36307586 | 36307712 | 36306291 | 36306376 | 36311762 | 36313955 | 4 | 2 | 3 | [illegible] |
| SE | 26924 | ENSG00000161912 | ADCY10P1 | chr6 | + | 41105891 | 41106035 | 41105109 | 41105148 | 41108304 | 41108509 | 1 | 0 | 5 | [illegible] |
| SE | 26926 | ENSG00000161912 | ADCY10P1 | chr6 | + | 41118923 | 41119107 | 41118128 | 41118184 | 41120522 | 41120657 | 18 | 20 | 15 | [illegible] |
| SE | 26927 | ENSG00000161912 | ADCY10P1 | chr6 | + | 41118953 | 41119107 | 41118128 | 41118184 | 41119273 | 41119463 | 1 | 3 | 8 | [illegible] |
| SE | 26928 | ENSG00000161912 | ADCY10P1 | chr6 | + | 41119240 | 41119463 | 41118923 | 41119107 | 41120522 | 41120657 | 0 | 0 | 0 | [illegible] |
| SE | 26930 | ENSG00000161912 | ADCY10P1 | chr6 | + | 41124484 | 41124820 | 41122976 | 41123249 | 41127544 | 41127608 | 7 | 3 | 10 | [illegible] |
| SE | 26931 | ENSG00000161912 | ADCY10P1 | chr6 | + | 41124484 | 41124820 | 41122976 | 41123249 | 41132293 | 41132513 | 7 | 4 | 11 | [illegible] |
| SE | 26932 | ENSG00000161912 | ADCY10P1 | chr6 | + | 41124484 | 41124820 | 41122976 | 41123249 | 41134556 | 41134729 | 6 | 1 | 6 | [illegible] |
| SE | 26936 | ENSG00000161912 | ADCY10P1 | chr6 | + | 41127544 | 41127608 | 41124484 | 41124820 | 41132293 | 41132513 | 2 | 5 | 5 | [illegible] |
| SE | 26937 | ENSG00000161912 | ADCY10P1 | chr6 | + | 41127544 | 41127608 | 41124484 | 41124820 | 41134556 | 41134729 | 0 | 1 | 4 | [illegible] |
| SE | 26938 | ENSG00000161912 | ADCY10P1 | chr6 | + | 41132293 | 41132513 | 41122976 | 41123249 | 41134556 | 41134729 | 5 | 5 | 12 | [illegible] |
| SE | 26939 | ENSG00000161912 | ADCY10P1 | chr6 | + | 41132293 | 41132513 | 41124484 | 41124820 | 41134556 | 41134729 | 3 | 7 | 16 | [illegible] |
| SE | 26940 | ENSG00000161912 | ADCY10P1 | chr6 | + | 41132293 | 41132513 | 41127544 | 41127608 | 41134556 | 41134729 | 4 | 9 | 15 | [illegible] |
| SE | 26941 | ENSG00000161912 | ADCY10P1 | chr6 | + | 41132874 | 41133000 | 41132293 | 41132513 | 41134556 | 41134729 | 0 | 2 | 1 | [illegible] |
| SE | 26943 | ENSG00000161912 | ADCY10P1 | chr6 | + | 41137356 | 41137561 | 41134556 | 41134729 | 41138608 | 41138726 | 0 | 1 | 5 | [illegible] |
| SE | 26944 | ENSG00000161912 | ADCY10P1 | chr6 | + | 41137356 | 41137561 | 41134556 | 41134729 | 41140597 | 41140835 | 0 | 0 | 2 | [illegible] |
| SE | 26945 | ENSG00000161912 | ADCY10P1 | chr6 | + | 41137356 | 41137561 | 41135184 | 41135341 | 41138608 | 41138726 | 0 | 1 | 5 | [illegible] |
| SE | 26946 | ENSG00000161912 | ADCY10P1 | chr6 | + | 41139855 | 41140069 | 41134556 | 41134729 | 41140597 | 41140835 | 21 | 30 | 21 | [illegible] |
| SE | 26950 | ENSG00000144659 | SLC25A38 | chr3 | + | 39391440 | 39391620 | 39390422 | 39390459 | 39391852 | 39392021 | 80 | 32 | 64 | [illegible] |
| SE | 26952 | ENSG00000144659 | SLC25A38 | chr3 | + | 39391852 | 39392021 | 39391440 | 39391620 | 39394409 | 39394418 | 145 | 72 | 154 | [illegible] |
| SE | 26953 | ENSG00000144659 | SLC25A38 | chr3 | + | 39393680 | 39393776 | 39391852 | 39392021 | 39394409 | 39394576 | 0 | 1 | 0 | [illegible] |
| SE | 26955 | ENSG00000172869 | DMXL1 | chr5 | + | 119071554 | 119071656 | 119071204 | 119071257 | 119097978 | 119098104 | 45 | 45 | 43 | [illegible] |
| SE | 26957 | ENSG00000172869 | DMXL1 | chr5 | + | 119101934 | 119102006 | 119097978 | 119098104 | 119110150 | 119110283 | 35 | 19 | 19 | [illegible] |
| SE | 26958 | ENSG00000172869 | DMXL1 | chr5 | + | 119101934 | 119102006 | 119097978 | 119098104 | 119114474 | 119114541 | 35 | 17 | 17 | [illegible] |
| SE | 26959 | ENSG00000172869 | DMXL1 | chr5 | + | 119105179 | 119105258 | 119101934 | 119102006 | 119110150 | 119110283 | 176 | 203 | 149 | [illegible] |
| SE | 26961 | ENSG00000172869 | DMXL1 | chr5 | + | 119110150 | 119110283 | 119097978 | 119098104 | 119114474 | 119114541 | 210 | 225 | 146 | [illegible] |
| SE | 26963 | ENSG00000172869 | DMXL1 | chr5 | + | 119110150 | 119110283 | 119105179 | 119105258 | 119114474 | 119114541 | 334 | 361 | 236 | [illegible] |
| SE | 26964 | ENSG00000172869 | DMXL1 | chr5 | + | 119114474 | 119114541 | 119110150 | 119110283 | 119116157 | 119116336 | 388 | 431 | 273 | [illegible] |
| SE | 26965 | ENSG00000172869 | DMXL1 | chr5 | + | 119118814 | 119119004 | 119116157 | 119116336 | 119120970 | 119121139 | 437 | 394 | 313 | [illegible] |
| SE | 26966 | ENSG00000172869 | DMXL1 | chr5 | + | 119165182 | 119165280 | 119164506 | 119164676 | 119166615 | 119166781 | 64 | 48 | 48 | [illegible] |
| SE | 26967 | ENSG00000172869 | DMXL1 | chr5 | + | 119167602 | 119167864 | 119166615 | 119166781 | 119170189 | 119171280 | 157 | 114 | 141 | [illegible] |
| SE | 26968 | ENSG00000172869 | DMXL1 | chr5 | + | 119171777 | 119171969 | 119170189 | 119171280 | 119175260 | 119175337 | 137 | 139 | 108 | [illegible] |
| SE | 26969 | ENSG00000172869 | DMXL1 | chr5 | + | 119171777 | 119171969 | 119170189 | 119171280 | 119177356 | 119177484 | 89 | 80 | 68 | [illegible] |
| SE | 26970 | ENSG00000172869 | DMXL1 | chr5 | + | 119175260 | 119175337 | 119170189 | 119171280 | 119177356 | 119177484 | 58 | 52 | 46 | [illegible] |
| SE | 26971 | ENSG00000172869 | DMXL1 | chr5 | + | 119175260 | 119175337 | 119171777 | 119171969 | 119177356 | 119177484 | 107 | 111 | 86 | [illegible] |
| SE | 26972 | ENSG00000172869 | DMXL1 | chr5 | + | 119206833 | 119206896 | 119203318 | 119203436 | 119216900 | 119216987 | 10 | 14 | 14 | [illegible] |
| SE | 26974 | ENSG00000172869 | DMXL1 | chr5 | + | 119238988 | 119239080 | 119236313 | 119237414 | 119240418 | 119240471 | 72 | 38 | 66 | [illegible] |
| SE | 26975 | ENSG00000172869 | DMXL1 | chr5 | + | 119244358 | 119244576 | 119240418 | 119240471 | 119246994 | 119249120 | 208 | 165 | 154 | [illegible] |
| SE | 26976 | ENSG00000187091 | PLCD1 | chr3 | - | 38008457 | 38008636 | 38008234 | 38008367 | 38009041 | 38009158 | 42 | 6 | 27 | [illegible] |
| SE | 26977 | ENSG00000187091 | PLCD1 | chr3 | - | 38010360 | 38010562 | 38009903 | 38010275 | 38011213 | 38011445 | 19 | 4 | 20 | [illegible] |
| SE | 26978 | ENSG00000026297 | RNASET2 | chr6 | - | 166931043 | 166931118 | 166922112 | 166929791 | 166934090 | 166934136 | 1020 | 738 | 782 | [illegible] |
| SE | 26979 | ENSG00000026297 | RNASET2 | chr6 | - | 166931043 | 166931118 | 166922112 | 166929791 | 166938894 | 166939008 | 383 | 280 | 300 | [illegible] |
| SE | 26980 | ENSG00000026297 | RNASET2 | chr6 | - | 166934090 | 166934136 | 166922112 | 166929791 | 166938894 | 166939008 | 68 | 73 | 85 | [illegible] |
| SE | 26982 | ENSG00000026297 | RNASET2 | chr6 | - | 166938894 | 166939008 | 166922112 | 166929791 | 166956096 | 166956473 | 1 | 1 | 1 | [illegible] |
| SE | 26983 | ENSG00000026297 | RNASET2 | chr6 | - | 166938894 | 166939008 | 166933364 | 166934136 | 166943018 | 166943089 | 1075 | 767 | 867 | [illegible] |
| SE | 26984 | ENSG00000026297 | RNASET2 | chr6 | - | 166938894 | 166939008 | 166933364 | 166934136 | 166946681 | 166946739 | 524 | 374 | 426 | [illegible] |
| SE | 26985 | ENSG00000026297 | RNASET2 | chr6 | - | 166943018 | 166943089 | 166933364 | 166934136 | 166946681 | 166946739 | 69 | 49 | 48 | [illegible] |
| SE | 26986 | ENSG00000026297 | RNASET2 | chr6 | - | 166943018 | 166943089 | 166938595 | 166939008 | 166946681 | 166946739 | 552 | 376 | 436 | [illegible] |
| SE | 26989 | ENSG00000026297 | RNASET2 | chr6 | - | 166943018 | 166943809 | 166938894 | 166939008 | 166946681 | 166946739 | 248 | 235 | 216 | [illegible] |
| SE | 26990 | ENSG00000026297 | RNASET2 | chr6 | - | 166946681 | 166946739 | 166938894 | 166939008 | 166948569 | 166948625 | 197 | 147 | 164 | [illegible] |
| SE | 26991 | ENSG00000026297 | RNASET2 | chr6 | - | 166946681 | 166946739 | 166943018 | 166943089 | 166952487 | 166952548 | 274 | 150 | 222 | [illegible] |
| SE | 26995 | ENSG00000026297 | RNASET2 | chr6 | - | 166952487 | 166952548 | 166943018 | 166943089 | 166956096 | 166956191 | 144 | 89 | 119 | [illegible] |
| SE | 26996 | ENSG00000026297 | RNASET2 | chr6 | - | 166952487 | 166952548 | 166948565 | 166948625 | 166956096 | 166956191 | 435 | 326 | 332 | [illegible] |
| SE | 26999 | ENSG00000026297 | RNASET2 | chr6 | - | 166955494 | 166955627 | 166952487 | 166952548 | 166956096 | 166956611 | 7 | 4 | 6 | [illegible] |
| SE | 27001 | ENSG00000026297 | RNASET2 | chr6 | - | 166956096 | 166956191 | 166952487 | 166952548 | 166957125 | 166957184 | 181 | 111 | 113 | [illegible] |
| SE | 27005 | ENSG00000137802 | MAPKBP1 | chr15 | + | 41813620 | 41813781 | 41812515 | 41813101 | 41815258 | 41815405 | 27 | 14 | 17 | [illegible] |
| SE | 27007 | ENSG00000137802 | MAPKBP1 | chr15 | + | 41814549 | 41814739 | 41812515 | 41813781 | 41815258 | 41815405 | 62 | 18 | 38 | [illegible] |
| SE | 27009 | ENSG00000137802 | MAPKBP1 | chr15 | + | 41817613 | 41817735 | 41817387 | 41817458 | 41818008 | 41818084 | 20 | 19 | 15 | [illegible] |
| SE | 27010 | ENSG00000118777 | ABCG2 | chr4 | - | 88097452 | 88097607 | 88095519 | 88095609 | 88099323 | 88099448 | 0 | 2 | 0 | [illegible] |
| SE | 27011 | ENSG00000118777 | ABCG2 | chr4 | - | 88211665 | 88211803 | 88139792 | 88140014 | 88230993 | 88231628 | 1 | 0 | 0 | [illegible] |
| SE | 27012 | ENSG00000118762 | PKD2 | chr4 | + | 88043232 | 88043457 | 88038250 | 88038501 | 88046641 | 88046870 | 34 | 12 | 12 | [illegible] |
| SE | 27013 | ENSG00000118762 | PKD2 | chr4 | + | 88046641 | 88046870 | 88038250 | 88038501 | 88051990 | 88052158 | 25 | 14 | 22 | [illegible] |
| SE | 27016 | ENSG00000118762 | PKD2 | chr4 | + | 88061905 | 88062004 | 88057982 | 88058103 | 88065373 | 88065495 | 59 | 45 | 37 | [illegible] |
| SE | 27017 | ENSG00000118762 | PKD2 | chr4 | + | 88067897 | 88068061 | 88065373 | 88065879 | 88074811 | 88074959 | 55 | 36 | 49 | [illegible] |
| SE | 27024 | ENSG00000198646 | NCOA6 | chr20 | - | 34727258 | 34727407 | 34714773 | 34715365 | 34732558 | 34732595 | 99 | 77 | 77 | [illegible] |
| SE | 27025 | ENSG00000198646 | NCOA6 | chr20 | - | 34736689 | 34736758 | 34732558 | 34732595 | 34746806 | 34746928 | 20 | 3 | 18 | [illegible] |
| SE | 27030 | ENSG00000198646 | NCOA6 | chr20 | - | 34757219 | 34758104 | 34746806 | 34746928 | 34758804 | 34758933 | 39 | 25 | 29 | [illegible] |
| SE | 27031 | ENSG00000198646 | NCOA6 | chr20 | - | 34757219 | 34758104 | 34754721 | 34754868 | 34768463 | 34768586 | 50 | 34 | 37 | [illegible] |
| SE | 27033 | ENSG00000198646 | NCOA6 | chr20 | - | 34776292 | 34776448 | 34768463 | 34768586 | 34782120 | 34782404 | 298 | 243 | 222 | [illegible] |
| SE | 27034 | ENSG00000198646 | NCOA6 | chr20 | - | 34792449 | 34792563 | 34782120 | 34782404 | 34825471 | 34825630 | 72 | 59 | 53 | [illegible] |
| SE | 27055 | ENSG00000171497 | PPID | chr4 | - | 158713118 | 158713260 | 158710761 | 158710848 | 158715296 | 158715403 | 259 | 227 | 236 | [illegible] |
| SE | 27056 | ENSG00000171497 | PPID | chr4 | - | 158713118 | 158713260 | 158710761 | 158710848 | 158717011 | 158717055 | 85 | 60 | 75 | [illegible] |
| SE | 27057 | ENSG00000171497 | PPID | chr4 | - | 158715296 | 158715403 | 158713118 | 158713260 | 158715561 | 158715668 | 237 | 201 | 214 | [illegible] |
| SE | 27058 | ENSG00000171497 | PPID | chr4 | - | 158715561 | 158715668 | 158713118 | 158713260 | 158717011 | 158717055 | 5 | 2 | 4 | [illegible] |
| SE | 27059 | ENSG00000171497 | PPID | chr4 | - | 158715561 | 158715684 | 158713118 | 158713260 | 158717011 | 158717200 | 134 | 106 | 113 | [illegible] |
| SE | 27060 | ENSG00000171497 | PPID | chr4 | - | 158719179 | 158719286 | 158717011 | 158717200 | 158721342 | 158721483 | 175 | 121 | 142 | [illegible] |
| SE | 27061 | ENSG00000062194 | GPBP1 | chr5 | + | 57214073 | 57214193 | 57175447 | 57176400 | 57221375 | 57221396 | 315 | 237 | 335 | [illegible] |
| SE | 27063 | ENSG00000062194 | GPBP1 | chr5 | + | 57214073 | 57214193 | 57175447 | 57176400 | 57235965 | 57236032 | 246 | 194 | 244 | [illegible] |
| SE | 27064 | ENSG00000062194 | GPBP1 | chr5 | + | 57221375 | 57221396 | 57175945 | 57176400 | 57230845 | 57230969 | 10 | 10 | 20 | [illegible] |
| SE | 27066 | ENSG00000062194 | GPBP1 | chr5 | + | 57235965 | 57236032 | 57231097 | 57231321 | 57246299 | 57246484 | 307 | 316 | 286 | [illegible] |
| SE | 27068 | ENSG00000062194 | GPBP1 | chr5 | + | 57249408 | 57249576 | 57247074 | 57247215 | 57250953 | 57251012 | 1665 | 1078 | 1271 | [illegible] |
| SE | 27069 | ENSG00000062194 | GPBP1 | chr5 | + | 57249453 | 57249576 | 57247074 | 57247215 | 57250953 | 57251141 | 800 | 535 | 578 | [illegible] |
| SE | 27070 | ENSG00000062194 | GPBP1 | chr5 | + | 57250953 | 57251141 | 57248861 | 57249576 | 57261179 | 57261238 | 1309 | 863 | 996 | [illegible] |
| SE | 27071 | ENSG00000002549 | LAP3 | chr4 | + | 17579823 | 17579939 | 17577197 | 17577567 | 17581759 | 17581814 | 207 | 233 | 180 | [illegible] |
| SE | 27072 | ENSG00000002549 | LAP3 | chr4 | + | 17582287 | 17582393 | 17581759 | 17581814 | 17583482 | 17583642 | 227 | 308 | 189 | [illegible] |
| SE | 27073 | ENSG00000002549 | LAP3 | chr4 | + | 17584971 | 17585136 | 17583430 | 17583642 | 17588818 | 17588977 | 374 | 452 | 362 | [illegible] |
| SE | 27074 | ENSG00000002549 | LAP3 | chr4 | + | 17595409 | 17595534 | 17588818 | 17588977 | 17597045 | 17597134 | 261 | 330 | 256 | [illegible] |
| SE | 27079 | ENSG00000002549 | LAP3 | chr4 | + | 17598455 | 17598558 | 17595409 | 17595534 | 17604587 | 17604667 | 168 | 198 | 166 | [illegible] |
| SE | 27080 | ENSG00000002549 | LAP3 | chr4 | + | 17598455 | 17598558 | 17597045 | 17597134 | 17604587 | 17604667 | 289 | 331 | 273 | [illegible] |
| SE | 27081 | ENSG00000002549 | LAP3 | chr4 | + | 17604587 | 17604667 | 17598455 | 17598558 | 17606828 | 17606884 | 240 | 293 | 235 | [illegible] |
| SE | 27082 | ENSG00000187210 | GCNT1 | chr9 | + | 76460059 | 76460177 | 76459181 | 76459305 | 76500915 | 76501061 | 4 | 5 | 3 | [illegible] |
| SE | 27083 | ENSG00000187210 | GCNT1 | chr9 | + | 76501752 | 76501931 | 76500915 | 76501061 | 76502238 | 76502416 | 1 | 0 | 0 | [illegible] |
| SE | 27085 | ENSG00000187210 | GCNT1 | chr9 | + | 76631396 | 76631495 | 76609248 | 76609342 | 76642193 | 76642286 | 0 | 0 | 0 | [illegible] |
| SE | 27086 | ENSG00000156265 | MAP3K7CL | chr21 | + | 29080551 | 29080708 | 29077470 | 29077713 | 29085071 | 29085143 | 0 | 1 | 1 | [illegible] |
| SE | 27087 | ENSG00000156265 | MAP3K7CL | chr21 | + | 29080551 | 29080708 | 29077470 | 29077713 | 29085812 | 29085917 | 0 | 2 | 3 | [illegible] |
| SE | 27089 | ENSG00000156265 | MAP3K7CL | chr21 | + | 29085071 | 29085143 | 29080551 | 29080708 | 29085812 | 29085917 | 0 | 1 | 0 | [illegible] |
| SE | 27091 | ENSG00000156265 | MAP3K7CL | chr21 | + | 29091667 | 29091761 | 29085812 | 29085917 | 29092438 | 29092581 | 8 | 6 | 4 | [illegible] |
| SE | 27093 | ENSG00000156265 | MAP3K7CL | chr21 | + | 29109039 | 29109183 | 29092253 | 29092581 | 29133305 | 29133414 | 1 | 0 | 4 | [illegible] |
| SE | 27094 | ENSG00000156265 | MAP3K7CL | chr21 | + | 29130794 | 29130923 | 29092438 | 29092581 | 29133305 | 29133414 | 3 | 2 | 8 | [illegible] |
| SE | 27095 | ENSG00000156265 | MAP3K7CL | chr21 | + | 29130794 | 29130923 | 29109039 | 29109183 | 29133305 | 29133414 | 4 | 3 | 9 | [illegible] |
| SE | 27097 | ENSG00000156265 | MAP3K7CL | chr21 | + | 29133305 | 29133414 | 29092438 | 29092581 | 29149188 | 29149250 | 5 | 0 | 1 | [illegible] |
| SE | 27098 | ENSG00000156265 | MAP3K7CL | chr21 | + | 29133305 | 29133414 | 29109039 | 29109183 | 29149188 | 29149250 | 3 | 0 | 2 | [illegible] |
| SE | 27099 | ENSG00000156265 | MAP3K7CL | chr21 | + | 29133305 | 29133414 | 29122054 | 29122213 | 29149188 | 29149250 | 3 | 0 | 1 | [illegible] |
| SE | 27100 | ENSG00000151247 | EIF4E | chr4 | - | 98884921 | 98885061 | 98879275 | 98881142 | 98887078 | 98887167 | 512 | 460 | 490 | [illegible] |
| SE | 27101 | ENSG00000151247 | EIF4E | chr4 | - | 98886453 | 98886546 | 98884921 | 98885061 | 98887078 | 98887192 | 0 | 3 | 0 | [illegible] |
| SE | 27102 | ENSG00000151247 | EIF4E | chr4 | - | 98887888 | 98887952 | 98886350 | 98887192 | 98891236 | 98891332 | 599 | 465 | 461 | [illegible] |
| SE | 27105 | ENSG00000105755 | ETHE1 | chr19 | - | 43511436 | 43511566 | 43508774 | 43508864 | 43526514 | 43526659 | 63 | 29 | 43 | [illegible] |
| SE | 27108 | ENSG00000109475 | RPL34 | chr4 | + | 108621950 | 108622024 | 108620568 | 108620600 | 108622104 | 108622204 | 5729 | 3057 | 7071 | [illegible] |
| SE | 27116 | ENSG00000070756 | PABPC1 | chr8 | - | 100704925 | 100705029 | 100704296 | 100704390 | 100705588 | 100705673 | 4227 | 3244 | 4368 | [illegible] |
| SE | 27117 | ENSG00000070756 | PABPC1 | chr8 | - | 100704925 | 100705056 | 100703137 | 100704390 | 100705588 | 100705673 | 9164 | 6496 | 9053 | [illegible] |
| SE | 27119 | ENSG00000070756 | PABPC1 | chr8 | - | 100706886 | 100706997 | 100706650 | 100706805 | 100709132 | 100709223 | 9492 | 6026 | 9640 | [illegible] |
| SE | 27120 | ENSG00000070756 | PABPC1 | chr8 | - | 100709458 | 100709673 | 100709135 | 100709223 | 100712361 | 100712457 | 2876 | 1909 | 2984 | [illegible] |
| SE | 27121 | ENSG00000070756 | PABPC1 | chr8 | - | 100709458 | 100709731 | 100709132 | 100709223 | 100712361 | 100712404 | 5024 | 3580 | 5187 | [illegible] |
| SE | 27122 | ENSG00000070756 | PABPC1 | chr8 | - | 100712361 | 100712457 | 100709222 | 100709731 | 100712651 | 100712789 | 3594 | 2681 | 3759 | [illegible] |
| SE | 27123 | ENSG00000070756 | PABPC1 | chr8 | - | 100712651 | 100712789 | 100712361 | 100712457 | 100713086 | 100713181 | 13425 | 10504 | 14067 | [illegible] |
| SE | 27124 | ENSG00000070756 | PABPC1 | chr8 | - | 100713086 | 100713181 | 100712651 | 100712789 | 100715461 | 100715496 | 15558 | 12349 | 16013 | [illegible] |
| SE | 27125 | ENSG00000070756 | PABPC1 | chr8 | - | 100713086 | 100713181 | 100712651 | 100712789 | 100715470 | 100715601 | 7130 | 5291 | 7584 | [illegible] |
| SE | 27126 | ENSG00000070756 | PABPC1 | chr8 | - | 100713086 | 100713181 | 100712651 | 100712789 | 100717772 | 100717888 | 6996 | 5164 | 7402 | [illegible] |
| SE | 27127 | ENSG00000070756 | PABPC1 | chr8 | - | 100715461 | 100715601 | 100712651 | 100712789 | 100717772 | 100717888 | 7293 | 5502 | 7495 | [illegible] |
| SE | 27128 | ENSG00000070756 | PABPC1 | chr8 | - | 100715461 | 100715601 | 100712651 | 100713181 | 100717772 | 100717888 | 15912 | 12726 | 16161 | [illegible] |
| SE | 27129 | ENSG00000070756 | PABPC1 | chr8 | - | 100715470 | 100715601 | 100712651 | 100712789 | 100717772 | 100717888 | 6910 | 5177 | 7100 | [illegible] |
| SE | 27130 | ENSG00000070756 | PABPC1 | chr8 | - | 100715470 | 100715601 | 100713086 | 100713181 | 100717772 | 100717888 | 7047 | 5306 | 7288 | [illegible] |
| SE | 27133 | ENSG00000197020 | ZNF100 | chr19 | - | 21744016 | 21744112 | 21726192 | 21727989 | 21744940 | 21745067 | 14 | 29 | 13 | [illegible] |
| SE | 27134 | ENSG00000197020 | ZNF100 | chr19 | - | 21744016 | 21744112 | 21726192 | 21727989 | 21765693 | 21765783 | 11 | 17 | 12 | [illegible] |
| SE | 27135 | ENSG00000197020 | ZNF100 | chr19 | - | 21744016 | 21744115 | 21722770 | 21727989 | 21744940 | 21745067 | 17 | 25 | 18 | [illegible] |
| SE | 27136 | ENSG00000197020 | ZNF100 | chr19 | - | 21744016 | 21744115 | 21722770 | 21727989 | 21765693 | 21765786 | 11 | 17 | 12 | [illegible] |
| SE | 27138 | ENSG00000197020 | ZNF100 | chr19 | - | 21744940 | 21745067 | 21744016 | 21744112 | 21765693 | 21765783 | 14 | 21 | 14 | [illegible] |
| SE | 27139 | ENSG00000197020 | ZNF100 | chr19 | - | 21744940 | 21745067 | 21744016 | 21744115 | 21765693 | 21765783 | 17 | 18 | 19 | [illegible] |
| SE | 27141 | ENSG00000197020 | ZNF100 | chr19 | - | 21765693 | 21765786 | 21722770 | 21727989 | 21767426 | 21767549 | 5 | 8 | 5 | [illegible] |
| SE | 27149 | ENSG00000109534 | GAR1 | chr4 | + | 109816152 | 109816378 | 109815509 | 109815545 | 109817935 | 109818090 | 70 | 85 | 61 | [illegible] |
| SE | 27152 | ENSG00000131507 | NDFIP1 | chr5 | + | 142132211 | 142132342 | 142131807 | 142131895 | 142135729 | 142135817 | 515 | 337 | 545 | [illegible] |
| SE | 27153 | ENSG00000131507 | NDFIP1 | chr5 | + | 142135729 | 142135817 | 142132211 | 142132342 | 142140562 | 142140629 | 241 | 144 | 230 | [illegible] |
| SE | 27155 | ENSG00000131507 | NDFIP1 | chr5 | + | 142137733 | 142137858 | 142135729 | 142135817 | 142144570 | 142144676 | 359 | 267 | 368 | [illegible] |
| SE | 27156 | ENSG00000131507 | NDFIP1 | chr5 | + | 142140562 | 142140629 | 142135729 | 142135817 | 142144570 | 142144676 | 92 | 84 | 102 | [illegible] |
| SE | 27157 | ENSG00000131507 | NDFIP1 | chr5 | + | 142140562 | 142140629 | 142137733 | 142137858 | 142144570 | 142144676 | 665 | 474 | 646 | [illegible] |
| SE | 27158 | ENSG00000131507 | NDFIP1 | chr5 | + | 142140562 | 142140629 | 142137733 | 142137858 | 142151730 | 142154440 | 394 | 271 | 338 | [illegible] |
| SE | 27159 | ENSG00000131507 | NDFIP1 | chr5 | + | 142144570 | 142144676 | 142137733 | 142137858 | 142151730 | 142154440 | 228 | 153 | 244 | [illegible] |
| SE | 27160 | ENSG00000131507 | NDFIP1 | chr5 | + | 142144570 | 142144676 | 142140562 | 142140629 | 142151730 | 142154440 | 513 | 372 | 568 | [illegible] |
| SE | 27163 | ENSG00000159202 | UBE2Z | chr17 | + | 48916075 | 48916187 | 48910689 | 48910880 | 48921159 | 48921272 | 142 | 123 | 145 | [illegible] |
| SE | 27164 | ENSG00000159202 | UBE2Z | chr17 | + | 48922846 | 48922937 | 48921159 | 48921272 | 48926963 | 48927054 | 577 | 449 | 490 | [illegible] |
| SE | 27165 | ENSG00000228252 | COL6A4P2 | chr3 | + | 130231445 | 130231508 | 130228186 | 130228222 | 130233445 | 130233508 | 0 | 2 | 0 | [illegible] |
| SE | 27168 | ENSG00000228252 | COL6A4P2 | chr3 | + | 130250716 | 130250728 | 130248616 | 130248751 | 130250817 | 130251038 | 0 | 7 | 0 | [illegible] |
| SE | 27170 | ENSG00000156500 | PABIR3 | chrX | + | 134814770 | 134814849 | 134807168 | 134807708 | 134829225 | 134829282 | 47 | 39 | 62 | [illegible] |
| SE | 27171 | ENSG00000156500 | PABIR3 | chrX | + | 134831635 | 134831668 | 134829225 | 134829282 | 134832944 | 134833128 | 0 | 0 | 1 | [illegible] |
| SE | 27172 | ENSG00000156500 | PABIR3 | chrX | + | 134831635 | 134831668 | 134829225 | 134829282 | 134838726 | 134839603 | 0 | 0 | 0 | [illegible] |
| SE | 27173 | ENSG00000156500 | PABIR3 | chrX | + | 134832944 | 134833128 | 134829225 | 134829282 | 134838726 | 134839561 | 0 | 2 | 2 | [illegible] |

| | | | | | | | | | | |
|---|---|---|---|---|---|---|---|---|---|---|
| SE | 27175 | ENSG00000156500 | PABIR3 | chrX | + | 134832944 | 134833128 | 134831635 | 134831668 | [illegible] |
| SE | 27176 | ENSG00000156500 | PABIR3 | chrX | + | 134832944 | 134833128 | 134831635 | 134831668 | [illegible] |
| SE | 27178 | ENSG00000156500 | PABIR3 | chrX | + | 134834073 | 134834226 | 134831635 | 134831668 | [illegible] |
| SE | 27179 | ENSG00000156500 | PABIR3 | chrX | + | 134835674 | 134835796 | 134831635 | 134831668 | [illegible] |
| SE | 27185 | ENSG00000156500 | PABIR3 | chrX | + | 134847382 | 134847475 | 134845346 | 134845401 | [illegible] |
| SE | 27187 | ENSG00000156500 | PABIR3 | chrX | + | 134847382 | 134847475 | 134845346 | 134845401 | [illegible] |
| SE | 27188 | ENSG00000156500 | PABIR3 | chrX | + | 134847882 | 134847971 | 134847382 | 134847475 | [illegible] |
| SE | 27189 | ENSG00000156500 | PABIR3 | chrX | + | 134847882 | 134847971 | 134847382 | 134847475 | [illegible] |
| SE | 27190 | ENSG00000156500 | PABIR3 | chrX | + | 134847882 | 134847971 | 134847382 | 134847475 | [illegible] |
| SE | 27191 | ENSG00000156500 | PABIR3 | chrX | + | 134847885 | 134847971 | 134847382 | 134847475 | [illegible] |
| SE | 27193 | ENSG00000156500 | PABIR3 | chrX | + | 134849166 | 134849228 | 134845346 | 134845401 | [illegible] |
| SE | 27194 | ENSG00000156500 | PABIR3 | chrX | + | 134849166 | 134849228 | 134847382 | 134847475 | [illegible] |
| SE | 27195 | ENSG00000156500 | PABIR3 | chrX | + | 134849166 | 134849228 | 134847382 | 134847475 | [illegible] |
| SE | 27196 | ENSG00000156500 | PABIR3 | chrX | + | 134849166 | 134849228 | 134847882 | 134847971 | [illegible] |
| SE | 27198 | ENSG00000156500 | PABIR3 | chrX | + | 134852799 | 134852896 | 134845346 | 134845401 | [illegible] |
| SE | 27199 | ENSG00000156500 | PABIR3 | chrX | + | 134852799 | 134852896 | 134847382 | 134847475 | [illegible] |
| SE | 27201 | ENSG00000156500 | PABIR3 | chrX | + | 134852799 | 134852896 | 134849166 | 134849228 | [illegible] |
| SE | 27206 | ENSG00000111875 | ASF1A | chr6 | + | 118900765 | 118900881 | 118894151 | 118894522 | [illegible] |
| SE | 27207 | ENSG00000111875 | ASF1A | chr6 | + | 118905651 | 118905828 | 118900765 | 118900881 | [illegible] |
| SE | 27210 | ENSG00000147166 | ITGB1BP2 | chrX | + | 71304541 | 71304604 | 71304186 | 71304300 | [illegible] |
| SE | 27216 | ENSG00000114541 | FRMD4B | chr3 | - | 69216262 | 69216349 | 69197714 | 69198774 | [illegible] |
| SE | 27218 | ENSG00000114541 | FRMD4B | chr3 | - | 69287751 | 69287836 | 69250042 | 69250099 | [illegible] |
| SE | 27219 | ENSG00000114541 | FRMD4B | chr3 | - | 69302342 | 69302435 | 69287644 | 69287836 | [illegible] |
| SE | 27223 | ENSG00000163812 | ZDHHC3 | chr3 | - | 44929305 | 44929436 | 44915256 | 44918143 | [illegible] |
| SE | 27230 | ENSG00000163812 | ZDHHC3 | chr3 | - | 44933117 | 44933199 | 44915256 | 44918143 | [illegible] |
| SE | 27231 | ENSG00000163812 | ZDHHC3 | chr3 | - | 44933117 | 44933199 | 44915260 | 44926847 | [illegible] |
| SE | 27232 | ENSG00000163812 | ZDHHC3 | chr3 | - | 44933117 | 44933199 | 44915260 | 44926847 | [illegible] |
| SE | 27233 | ENSG00000163812 | ZDHHC3 | chr3 | - | 44933887 | 44933984 | 44915260 | 44926847 | [illegible] |
| SE | 27234 | ENSG00000163812 | ZDHHC3 | chr3 | - | 44933887 | 44933984 | 44933117 | 44933199 | [illegible] |
| SE | 27235 | ENSG00000163812 | ZDHHC3 | chr3 | - | 44945167 | 44945292 | 44915256 | 44918143 | [illegible] |
| SE | 27236 | ENSG00000163812 | ZDHHC3 | chr3 | - | 44945167 | 44945292 | 44933887 | 44933984 | [illegible] |
| SE | 27237 | ENSG00000163812 | ZDHHC3 | chr3 | - | 44945167 | 44945292 | 44933887 | 44933984 | [illegible] |
| SE | 27238 | ENSG00000163812 | ZDHHC3 | chr3 | - | 44958587 | 44958685 | 44945167 | 44945292 | [illegible] |
| SE | 27239 | ENSG00000163812 | ZDHHC3 | chr3 | - | 44958587 | 44958685 | 44945167 | 44945292 | [illegible] |
| SE | 27240 | ENSG00000163812 | ZDHHC3 | chr3 | - | 44959130 | 44959460 | 44945167 | 44945292 | [illegible] |
| SE | 27246 | ENSG00000073331 | ALPK1 | chr4 | + | 112315800 | 112315852 | 112297342 | 112297469 | [illegible] |
| SE | 27248 | ENSG00000073331 | ALPK1 | chr4 | + | 112382397 | 112382552 | 112377677 | 112377898 | [illegible] |
| SE | 27249 | ENSG00000073331 | ALPK1 | chr4 | + | 112423725 | 112424003 | 112411826 | 112412025 | [illegible] |
| SE | 27250 | ENSG00000073331 | ALPK1 | chr4 | + | 112423943 | 112424003 | 112411826 | 112412025 | [illegible] |
| SE | 27252 | ENSG00000073331 | ALPK1 | chr4 | + | 112429148 | 112429253 | 112427569 | 112427665 | [illegible] |
| SE | 27253 | ENSG00000073331 | ALPK1 | chr4 | + | 112430447 | 112432581 | 112427569 | 112427665 | [illegible] |
| SE | 27254 | ENSG00000073331 | ALPK1 | chr4 | + | 112430447 | 112432581 | 112429148 | 112429253 | [illegible] |
| SE | 27256 | ENSG00000125744 | RTN2 | chr19 | - | 45489345 | 45489553 | 45488759 | 45488986 | [illegible] |
| SE | 27257 | ENSG00000125744 | RTN2 | chr19 | - | 45493159 | 45493378 | 45489345 | 45489553 | [illegible] |
| SE | 27258 | ENSG00000077232 | DNAJC10 | chr2 | + | 182717015 | 182717072 | 182716254 | 182716483 | [illegible] |
| SE | 27259 | ENSG00000077232 | DNAJC10 | chr2 | + | 182723227 | 182723383 | 182722024 | 182722075 | [illegible] |
| SE | 27260 | ENSG00000077232 | DNAJC10 | chr2 | + | 182729847 | 182729941 | 182728575 | 182728994 | [illegible] |
| SE | 27261 | ENSG00000077232 | DNAJC10 | chr2 | + | 182731029 | 182731107 | 182729834 | 182729941 | [illegible] |
| SE | 27262 | ENSG00000077232 | DNAJC10 | chr2 | + | 182732482 | 182732542 | 182731029 | 182731107 | [illegible] |
| SE | 27263 | ENSG00000077232 | DNAJC10 | chr2 | + | 182732498 | 182732542 | 182731029 | 182731107 | [illegible] |
| SE | 27265 | ENSG00000077232 | DNAJC10 | chr2 | + | 182732498 | 182732542 | 182731029 | 182731107 | [illegible] |
| SE | 27268 | ENSG00000077232 | DNAJC10 | chr2 | + | 182736248 | 182736386 | 182731029 | 182731107 | [illegible] |
| SE | 27269 | ENSG00000077232 | DNAJC10 | chr2 | + | 182736248 | 182736386 | 182732482 | 182732542 | [illegible] |
| SE | 27271 | ENSG00000077232 | DNAJC10 | chr2 | + | 182739544 | 182739709 | 182731029 | 182731107 | [illegible] |
| SE | 27273 | ENSG00000077232 | DNAJC10 | chr2 | + | 182742374 | 182742512 | 182741242 | 182741356 | [illegible] |
| SE | 27277 | ENSG00000077232 | DNAJC10 | chr2 | + | 182754702 | 182754805 | 182752071 | 182752188 | [illegible] |
| SE | 27278 | ENSG00000077232 | DNAJC10 | chr2 | + | 182762681 | 182762801 | 182759159 | 182759307 | [illegible] |
| SE | 27286 | ENSG00000064726 | BTBD1 | chr15 | - | 83018706 | 83018853 | 83016422 | 83018225 | [illegible] |
| SE | 27287 | ENSG00000064726 | BTBD1 | chr15 | - | 83020674 | 83020762 | 83016422 | 83018225 | [illegible] |
| SE | 27288 | ENSG00000064726 | BTBD1 | chr15 | - | 83020674 | 83020762 | 83018706 | 83018853 | [illegible] |
| SE | 27289 | ENSG00000064726 | BTBD1 | chr15 | - | 83020674 | 83020762 | 83018706 | 83018853 | [illegible] |
| SE | 27290 | ENSG00000064726 | BTBD1 | chr15 | - | 83028615 | 83028728 | 83018739 | 83018853 | [illegible] |
| SE | 27292 | ENSG00000064726 | BTBD1 | chr15 | - | 83041727 | 83041925 | 83030135 | 83030328 | [illegible] |
| SE | 27294 | ENSG00000136813 | ECPAS | chr9 | - | 111373971 | 111374038 | 111373314 | 111373406 | [illegible] |
| SE | 27295 | ENSG00000136813 | ECPAS | chr9 | - | 111375112 | 111375202 | 111373971 | 111374038 | [illegible] |
| SE | 27296 | ENSG00000136813 | ECPAS | chr9 | - | 111378579 | 111378730 | 111376475 | 111376541 | [illegible] |
| SE | 27297 | ENSG00000136813 | ECPAS | chr9 | - | 111383210 | 111383332 | 111378579 | 111378730 | [illegible] |
| SE | 27298 | ENSG00000136813 | ECPAS | chr9 | - | 111384521 | 111384569 | 111383210 | 111383332 | [illegible] |
| SE | 27299 | ENSG00000136813 | ECPAS | chr9 | - | 111385336 | 111385442 | 111383210 | 111383332 | [illegible] |
| SE | 27300 | ENSG00000136813 | ECPAS | chr9 | - | 111389555 | 111389723 | 111386376 | 111386456 | [illegible] |
| SE | 27301 | ENSG00000136813 | ECPAS | chr9 | - | 111392767 | 111392882 | 111391755 | 111391824 | [illegible] |
| SE | 27302 | ENSG00000136813 | ECPAS | chr9 | - | 111408570 | 111408672 | 111397029 | 111397153 | [illegible] |
| SE | 27304 | ENSG00000136813 | ECPAS | chr9 | - | 111413894 | 111413986 | 111412013 | 111412148 | [illegible] |
| SE | 27305 | ENSG00000136813 | ECPAS | chr9 | - | 111414428 | 111414651 | 111412013 | 111412148 | [illegible] |
| SE | 27307 | ENSG00000136813 | ECPAS | chr9 | - | 111423198 | 111423248 | 111422133 | 111422200 | [illegible] |
| SE | 27309 | ENSG00000136813 | ECPAS | chr9 | - | 111440371 | 111440521 | 111436939 | 111437108 | [illegible] |
| SE | 27316 | ENSG00000166913 | YWHAB | chr20 | + | 44886878 | 44886978 | 44885751 | 44885886 | [illegible] |
| SE | 27317 | ENSG00000166913 | YWHAB | chr20 | + | 44886878 | 44886978 | 44885751 | 44885886 | [illegible] |
| SE | 27318 | ENSG00000166913 | YWHAB | chr20 | + | 44886878 | 44886978 | 44885751 | 44885886 | [illegible] |
| SE | 27324 | ENSG00000166913 | YWHAB | chr20 | + | 44903035 | 44903121 | 44901530 | 44901833 | [illegible] |
| SE | 27325 | ENSG00000166913 | YWHAB | chr20 | + | 44904967 | 44905131 | 44903992 | 44904116 | [illegible] |
| SE | 27327 | ENSG00000196284 | SUPT3H | chr6 | - | 45321795 | 45321891 | 45105921 | 45106006 | [illegible] |
| SE | 27328 | ENSG00000196284 | SUPT3H | chr6 | - | 45322878 | 45322967 | 45105921 | 45106006 | [illegible] |
| SE | 27329 | ENSG00000196284 | SUPT3H | chr6 | - | 45365200 | 45365301 | 45105921 | 45106006 | [illegible] |
| SE | 27331 | ENSG00000170876 | TMEM43 | chr3 | + | 14133738 | 14133809 | 14132865 | 14132935 | [illegible] |
| SE | 27332 | ENSG00000170876 | TMEM43 | chr3 | + | 14135157 | 14135232 | 14134769 | 14134891 | [illegible] |
| SE | 27333 | ENSG00000157869 | RAB28 | chr4 | - | 13369870 | 13369965 | 13361353 | 13361633 | [illegible] |
| SE | 27336 | ENSG00000157869 | RAB28 | chr4 | - | 13381490 | 13381594 | 13376544 | 13376622 | [illegible] |
| SE | 27338 | ENSG00000157869 | RAB28 | chr4 | - | 13460698 | 13460828 | 13381490 | 13381594 | [illegible] |
| SE | 27339 | ENSG00000157869 | RAB28 | chr4 | - | 13474317 | 13474406 | 13381490 | 13381594 | [illegible] |
| SE | 27340 | ENSG00000157869 | RAB28 | chr4 | - | 13474317 | 13474406 | 13460698 | 13460828 | [illegible] |
| SE | 27341 | ENSG00000080493 | SLC4A4 | chr4 | + | 71453494 | 71453669 | 71451187 | 71451301 | [illegible] |
| SE | 27342 | ENSG00000080493 | SLC4A4 | chr4 | + | 71472698 | 71472970 | 71466443 | 71466577 | [illegible] |
| SE | 27346 | ENSG00000167995 | BEST1 | chr11 | + | 61955717 | 61955951 | 61955342 | 61955387 | [illegible] |
| SE | 27347 | ENSG00000167995 | BEST1 | chr11 | + | 61958145 | 61958298 | 61957386 | 61957464 | [illegible] |
| SE | 27348 | ENSG00000167995 | BEST1 | chr11 | + | 61958145 | 61958298 | 61957386 | 61957464 | [illegible] |
| SE | 27349 | ENSG00000167995 | BEST1 | chr11 | + | 61958145 | 61958501 | 61957386 | 61957464 | [illegible] |
| SE | 27351 | ENSG00000167995 | BEST1 | chr11 | + | 61959497 | 61959578 | 61958145 | 61958298 | [illegible] |
| SE | 27352 | ENSG00000167995 | BEST1 | chr11 | + | 61959891 | 61960043 | 61959497 | 61959578 | [illegible] |
| SE | 27353 | ENSG00000167995 | BEST1 | chr11 | + | 61959891 | 61962893 | 61958145 | 61959578 | [illegible] |
| SE | 27354 | ENSG00000167995 | BEST1 | chr11 | + | 61962254 | 61962893 | 61959497 | 61959578 | [illegible] |
| SE | 27361 | ENSG00000232021 | LEF1-AS1 | chr4 | + | 108173418 | 108173519 | 108171907 | 108171979 | [illegible] |
| SE | 27363 | ENSG00000232021 | LEF1-AS1 | chr4 | + | 108175791 | 108175993 | 108173418 | 108173519 | [illegible] |
| SE | 27365 | ENSG00000138670 | RASGEF1B | chr4 | - | 81434638 | 81434734 | 81433839 | 81433963 | [illegible] |
| SE | 27366 | ENSG00000138670 | RASGEF1B | chr4 | - | 81447503 | 81447578 | 81445742 | 81445838 | [illegible] |
| SE | 27367 | ENSG00000138670 | RASGEF1B | chr4 | - | 81456650 | 81456788 | 81448068 | 81448281 | [illegible] |
| SE | 27376 | ENSG00000138738 | PRDM5 | chr4 | - | 120781142 | 120781303 | 120777187 | 120777281 | [illegible] |
| SE | 27377 | ENSG00000138738 | PRDM5 | chr4 | - | 120798266 | 120798424 | 120784997 | 120785091 | [illegible] |
| SE | 27379 | ENSG00000138738 | PRDM5 | chr4 | - | 120816831 | 120816924 | 120816452 | 120816574 | [illegible] |
| SE | 27380 | ENSG00000138758 | SEPTIN11 | chr4 | + | 76973009 | 76973070 | 76972389 | 76972590 | [illegible] |
| SE | 27383 | ENSG00000138758 | SEPTIN11 | chr4 | + | 76995780 | 76995938 | 76974477 | 76974807 | [illegible] |
| SE | 27384 | ENSG00000138758 | SEPTIN11 | chr4 | + | 76995780 | 76995938 | 76987808 | 76987870 | [illegible] |
| SE | 27385 | ENSG00000138758 | SEPTIN11 | chr4 | + | 77005600 | 77005796 | 76996424 | 76996539 | [illegible] |
| SE | 27386 | ENSG00000138758 | SEPTIN11 | chr4 | + | 77011734 | 77011921 | 77005600 | 77005796 | [illegible] |
| SE | 27389 | ENSG00000181982 | CCDC149 | chr4 | - | 24821054 | 24821087 | 24819858 | 24819975 | [illegible] |
| SE | 27391 | ENSG00000181982 | CCDC149 | chr4 | - | 24836435 | 24836508 | 24834947 | 24835032 | [illegible] |
| SE | 27392 | ENSG00000181982 | CCDC149 | chr4 | - | 24894991 | 24895118 | 24876535 | 24876697 | [illegible] |
| SE | 27396 | ENSG00000167216 | KATNAL2 | chr18 | + | 47075276 | 47075368 | 47069481 | 47069600 | [illegible] |
| SE | 27399 | ENSG00000214367 | HAUS3 | chr4 | - | 2129178 | 2129256 | 2128112 | 2128227 | [illegible] |
| SE | 27401 | ENSG00000214367 | HAUS3 | chr4 | - | 2156787 | 2156853 | 2131232 | 2131290 | [illegible] |
| SE | 27402 | ENSG00000214367 | HAUS3 | chr4 | - | 2157857 | 2157911 | 2131232 | 2131290 | [illegible] |
| SE | 27403 | ENSG00000214367 | HAUS3 | chr4 | - | 2157857 | 2157911 | 2156787 | 2156853 | [illegible] |
| SE | 27404 | ENSG00000214367 | HAUS3 | chr4 | - | 2159154 | 2159211 | 2157857 | 2157911 | [illegible] |
| SE | 27406 | ENSG00000214367 | HAUS3 | chr4 | - | 2193203 | 2193316 | 2179307 | 2179465 | [illegible] |
| SE | 27411 | ENSG00000164125 | GASK1B | chr4 | - | 158130785 | 158131012 | 158124473 | 158127614 | [illegible] |
| SE | 27413 | ENSG00000164125 | GASK1B | chr4 | - | 158170465 | 158171599 | 158155610 | 158155825 | [illegible] |
| SE | 27421 | ENSG00000163995 | ABLIM2 | chr4 | - | 8009049 | 8009102 | 8007024 | 8008200 | [illegible] |
| SE | 27425 | ENSG00000163995 | ABLIM2 | chr4 | - | 8029655 | 8029776 | 8009049 | 8009102 | [illegible] |
| SE | 27428 | ENSG00000163945 | UVSSA | chr4 | + | 1353029 | 1353413 | 1351714 | 1351835 | [illegible] |
| SE | 27429 | ENSG00000163945 | UVSSA | chr4 | + | 1364507 | 1364571 | 1355116 | 1355245 | [illegible] |
| SE | 27431 | ENSG00000163945 | UVSSA | chr4 | + | 1380640 | 1380747 | 1380046 | 1380230 | [illegible] |
| SE | 27433 | ENSG00000156194 | PPEF2 | chr4 | - | 75882925 | 75883075 | 75876286 | 75876673 | [illegible] |
| SE | 27436 | ENSG00000136925 | TSTD2 | chr9 | - | 97610345 | 97610451 | 97606142 | 97606261 | [illegible] |
| SE | 27439 | ENSG00000136925 | TSTD2 | chr9 | - | 97617756 | 97617877 | 97611573 | 97611699 | [illegible] |
| SE | 27440 | ENSG00000136925 | TSTD2 | chr9 | - | 97625680 | 97625997 | 97611573 | 97611699 | [illegible] |
| SE | 27441 | ENSG00000136925 | TSTD2 | chr9 | - | 97625680 | 97625997 | 97611573 | 97611699 | [illegible] |
| SE | 27442 | ENSG00000136925 | TSTD2 | chr9 | - | 97625680 | 97625997 | 97617756 | 97617877 | [illegible] |
| SE | 27443 | ENSG00000136925 | TSTD2 | chr9 | - | 97627305 | 97627612 | 97625680 | 97625997 | [illegible] |
| SE | 27444 | ENSG00000136925 | TSTD2 | chr9 | - | 97627397 | 97627612 | 97611573 | 97611699 | [illegible] |
| SE | 27445 | ENSG00000136925 | TSTD2 | chr9 | - | 97627397 | 97627612 | 97625680 | 97625997 | [illegible] |
| SE | 27450 | ENSG00000126522 | ASL | chr7 | + | 66082367 | 66082451 | 66075936 | 66076093 | [illegible] |
| SE | 27451 | ENSG00000126522 | ASL | chr7 | + | 66082879 | 66082936 | 66082367 | 66082451 | [illegible] |
| SE | 27452 | ENSG00000126522 | ASL | chr7 | + | 66083076 | 66083174 | 66082879 | 66082936 | [illegible] |
| SE | 27453 | ENSG00000126522 | ASL | chr7 | + | 66086584 | 66086662 | 66083076 | 66083174 | [illegible] |
| SE | 27456 | ENSG00000126522 | ASL | chr7 | + | 66087728 | 66087791 | 66083076 | 66083174 | [illegible] |
| SE | 27463 | ENSG00000008513 | ST3GAL1 | chr8 | - | 133476277 | 133476600 | 133475718 | 133476074 | [illegible] |
| SE | 27464 | ENSG00000008513 | ST3GAL1 | chr8 | - | 133499134 | 133499189 | 133465893 | 133466090 | [illegible] |
| SE | 27466 | ENSG00000008513 | ST3GAL1 | chr8 | - | 133499134 | 133499189 | 133475718 | 133476600 | [illegible] |
| SE | 27467 | ENSG00000008513 | ST3GAL1 | chr8 | - | 133507198 | 133507321 | 133499134 | 133499189 | [illegible] |
| SE | 27472 | ENSG00000008513 | ST3GAL1 | chr8 | - | 133545773 | 133545926 | 133476028 | 133476600 | [illegible] |
| SE | 27473 | ENSG00000008513 | ST3GAL1 | chr8 | - | 133545773 | 133545926 | 133499134 | 133499189 | [illegible] |
| SE | 27474 | ENSG00000008513 | ST3GAL1 | chr8 | - | 133545773 | 133545926 | 133507198 | 133507321 | [illegible] |
| SE | 27475 | ENSG00000109944 | JHY | chr11 | + | 122924896 | 122925010 | 122903924 | 122904444 | [illegible] |
| SE | 27476 | ENSG00000109944 | JHY | chr11 | + | 122934419 | 122935075 | 122924896 | 122925010 | [illegible] |
| SE | 27477 | ENSG00000109944 | JHY | chr11 | + | 122946497 | 122946792 | 122924896 | 122925010 | [illegible] |
| SE | 27479 | ENSG00000104517 | UBR5 | chr8 | - | 102258315 | 102258424 | 102257631 | 102257717 | [illegible] |
| SE | 27480 | ENSG00000104517 | UBR5 | chr8 | - | 102258984 | 102259129 | 102257631 | 102257717 | [illegible] |
| SE | 27482 | ENSG00000104517 | UBR5 | chr8 | - | 102261914 | 102262070 | 102261145 | 102261259 | [illegible] |
| SE | 27483 | ENSG00000104517 | UBR5 | chr8 | - | 102264473 | 102264558 | 102261914 | 102262070 | [illegible] |
| SE | 27484 | ENSG00000104517 | UBR5 | chr8 | - | 102266946 | 102267007 | 102265099 | 102265278 | [illegible] |
| SE | 27485 | ENSG00000104517 | UBR5 | chr8 | - | 102270038 | 102270183 | 102268977 | 102269108 | [illegible] |
| SE | 27486 | ENSG00000104517 | UBR5 | chr8 | - | 102270038 | 102270183 | 102268977 | 102269108 | [illegible] |
| SE | 27487 | ENSG00000104517 | UBR5 | chr8 | - | 102271133 | 102271245 | 102268977 | 102269108 | [illegible] |
| SE | 27491 | ENSG00000104517 | UBR5 | chr8 | - | 102287432 | 102287564 | 102286359 | 102286617 | [illegible] |
| SE | 27492 | ENSG00000104517 | UBR5 | chr8 | - | 102287432 | 102287564 | 102286359 | 102286617 | [illegible] |
| SE | 27493 | ENSG00000104517 | UBR5 | chr8 | - | 102288154 | 102288266 | 102286359 | 102286617 | [illegible] |
| SE | 27495 | ENSG00000104517 | UBR5 | chr8 | - | 102289452 | 102289576 | 102287432 | 102287564 | [illegible] |
| SE | 27496 | ENSG00000104517 | UBR5 | chr8 | - | 102299451 | 102299496 | 102298866 | 102298971 | [illegible] |
| SE | 27498 | ENSG00000104517 | UBR5 | chr8 | - | 102299451 | 102299547 | 102298847 | 102298971 | [illegible] |
| SE | 27499 | ENSG00000104517 | UBR5 | chr8 | - | 102299999 | 102300151 | 102298847 | 102298971 | [illegible] |
| SE | 27500 | ENSG00000104517 | UBR5 | chr8 | - | 102299999 | 102300151 | 102299451 | 102299547 | [illegible] |
| SE | 27501 | ENSG00000104517 | UBR5 | chr8 | - | 102305084 | 102305295 | 102304042 | 102304169 | [illegible] |
| SE | 27502 | ENSG00000104517 | UBR5 | chr8 | - | 102311298 | 102311496 | 102305084 | 102305295 | [illegible] |
| SE | 27503 | ENSG00000104517 | UBR5 | chr8 | - | 102323310 | 102323495 | 102314743 | 102314853 | [illegible] |
| SE | 27504 | ENSG00000104517 | UBR5 | chr8 | - | 102326545 | 102326663 | 102323310 | 102323495 | [illegible] |
| SE | 27507 | ENSG00000104517 | UBR5 | chr8 | - | 102342472 | 102342666 | 102329282 | 102329399 | [illegible] |
| SE | 27508 | ENSG00000104517 | UBR5 | chr8 | - | 102342472 | 102342684 | 102329282 | 102329399 | [illegible] |
| SE | 27509 | ENSG00000104517 | UBR5 | chr8 | - | 102346902 | 102347092 | 102346235 | 102346395 | [illegible] |
| SE | 27510 | ENSG00000104517 | UBR5 | chr8 | - | 102360588 | 102360665 | 102359651 | 102360190 | [illegible] |
| SE | 27513 | ENSG00000198189 | HSD17B11 | chr4 | - | 87340489 | 87340606 | 87336514 | 87337366 | [illegible] |
| SE | 27515 | ENSG00000198189 | HSD17B11 | chr4 | - | 87374698 | 87374830 | 87372708 | 87372815 | [illegible] |
| SE | 27516 | ENSG00000198189 | HSD17B11 | chr4 | - | 87382254 | 87382362 | 87374698 | 87374830 | [illegible] |
| SE | 27517 | ENSG00000198189 | HSD17B11 | chr4 | - | 87385789 | 87385907 | 87382254 | 87382362 | [illegible] |
| SE | 27519 | ENSG00000131591 | C1orf159 | chr1 | - | 1083914 | 1084011 | 1082915 | 1082987 | [illegible] |
| SE | 27520 | ENSG00000131591 | C1orf159 | chr1 | - | 1083914 | 1084086 | 1081822 | 1082987 | [illegible] |
| SE | 27523 | ENSG00000131591 | C1orf159 | chr1 | - | 1090875 | 1090983 | 1090352 | 1090428 | [illegible] |
| SE | 27524 | ENSG00000131591 | C1orf159 | chr1 | - | 1091277 | 1091374 | 1090352 | 1090428 | [illegible] |
| SE | 27525 | ENSG00000131591 | C1orf159 | chr1 | - | 1091277 | 1091374 | 1090875 | 1090983 | [illegible] |
| SE | 27526 | ENSG00000131591 | C1orf159 | chr1 | - | 1105955 | 1106049 | 1091990 | 1092103 | [illegible] |
| SE | 27529 | ENSG00000284968 | NA | chr4 | - | 86933394 | 86933542 | 86924629 | 86926326 | [illegible] |
| SE | 27536 | ENSG00000197647 | ZNF433 | chr19 | - | 12030139 | 12030198 | 12018165 | 12018301 | [illegible] |
| SE | 27540 | ENSG00000083444 | PLOD1 | chr1 | + | 11944521 | 11944662 | 11934693 | 11934855 | [illegible] |
| SE | 27541 | ENSG00000083444 | PLOD1 | chr1 | + | 11952622 | 11952735 | 11950356 | 11950520 | [illegible] |
| SE | 27544 | ENSG00000083444 | PLOD1 | chr1 | + | 11957841 | 11957943 | 11956916 | 11957014 | [illegible] |
| SE | 27545 | ENSG00000083444 | PLOD1 | chr1 | + | 11964174 | 11964300 | 11963531 | 11963636 | [illegible] |
| SE | 27546 | ENSG00000115956 | PLEK | chr2 | + | 68382541 | 68382633 | 68380722 | 68380904 | [illegible] |
| SE | 27547 | ENSG00000115956 | PLEK | chr2 | + | 68382541 | 68382633 | 68380722 | 68380904 | [illegible] |
| SE | 27548 | ENSG00000115956 | PLEK | chr2 | + | 68382541 | 68382633 | 68380722 | 68380904 | [illegible] |
| SE | 27549 | ENSG00000115956 | PLEK | chr2 | + | 68386501 | 68386686 | 68380722 | 68380904 | [illegible] |
| SE | 27550 | ENSG00000115956 | PLEK | chr2 | + | 68386501 | 68386686 | 68380722 | 68380904 | [illegible] |
| SE | 27551 | ENSG00000115956 | PLEK | chr2 | + | 68386501 | 68386686 | 68382541 | 68382633 | [illegible] |
| SE | 27552 | ENSG00000115956 | PLEK | chr2 | + | 68386501 | 68386686 | 68382541 | 68382633 | [illegible] |
| SE | 27553 | ENSG00000115956 | PLEK | chr2 | + | 68388386 | 68388491 | 68380722 | 68380904 | [illegible] |
| SE | 27554 | ENSG00000115956 | PLEK | chr2 | + | 68388386 | 68388491 | 68382541 | 68382633 | [illegible] |
| SE | 27555 | ENSG00000115956 | PLEK | chr2 | + | 68388386 | 68388491 | 68386501 | 68386686 | [illegible] |
| SE | 27556 | ENSG00000115956 | PLEK | chr2 | + | 68393161 | 68393245 | 68387941 | 68388491 | [illegible] |
| SE | 27577 | ENSG00000109654 | TRIM2 | chr4 | + | 153295312 | 153296036 | 153275892 | 153276130 | [illegible] |
| SE | 27582 | ENSG00000132694 | ARHGEF11 | chr1 | - | 156936815 | 156937005 | 156934839 | 156936058 | [illegible] |
| SE | 27584 | ENSG00000132694 | ARHGEF11 | chr1 | - | 156939148 | 156939285 | 156938417 | 156938513 | [illegible] |
| SE | 27585 | ENSG00000132694 | ARHGEF11 | chr1 | - | 156943934 | 156944102 | 156942689 | 156942780 | [illegible] |
| SE | 27586 | ENSG00000132694 | ARHGEF11 | chr1 | - | 156954891 | 156954921 | 156951572 | 156951699 | [illegible] |
| SE | 27587 | ENSG00000132694 | ARHGEF11 | chr1 | - | 156957791 | 156957815 | 156956419 | 156956564 | [illegible] |
| SE | 27588 | ENSG00000132694 | ARHGEF11 | chr1 | - | 156963519 | 156963594 | 156963202 | 156963304 | [illegible] |
| SE | 27590 | ENSG00000132694 | ARHGEF11 | chr1 | - | 156976982 | 156977054 | 156971696 | 156971816 | [illegible] |
| SE | 27594 | ENSG00000132694 | ARHGEF11 | chr1 | - | 156980436 | 156980486 | 156979228 | 156979286 | [illegible] |
| SE | 27597 | ENSG00000039123 | MTREX | chr5 | + | 55324131 | 55324198 | 55322326 | 55322464 | [illegible] |
| SE | 27598 | ENSG00000039123 | MTREX | chr5 | + | 55328698 | 55328811 | 55327715 | 55327778 | [illegible] |
| SE | 27599 | ENSG00000039123 | MTREX | chr5 | + | 55343330 | 55343455 | 55341680 | 55341771 | [illegible] |
| SE | 27600 | ENSG00000039123 | MTREX | chr5 | + | 55350918 | 55351029 | 55349572 | 55349652 | [illegible] |
| SE | 27601 | ENSG00000039123 | MTREX | chr5 | + | 55358572 | 55358698 | 55353167 | 55353269 | [illegible] |
| SE | 27602 | ENSG00000039123 | MTREX | chr5 | + | 55400232 | 55400421 | 55397415 | 55397526 | [illegible] |
| SE | 27605 | ENSG00000001629 | ANKIB1 | chr7 | + | 92319329 | 92319512 | 92307358 | 92307656 | [illegible] |
| SE | 27607 | ENSG00000001629 | ANKIB1 | chr7 | + | 92327782 | 92327900 | 92307358 | 92307656 | [illegible] |
| SE | 27608 | ENSG00000001629 | ANKIB1 | chr7 | + | 92327782 | 92327900 | 92319329 | 92319512 | [illegible] |
| SE | 27609 | ENSG00000001629 | ANKIB1 | chr7 | + | 92343023 | 92343232 | 92327782 | 92327900 | [illegible] |
| SE | 27611 | ENSG00000001629 | ANKIB1 | chr7 | + | 92350949 | 92351094 | 92344977 | 92345066 | [illegible] |
| SE | 27612 | ENSG00000001629 | ANKIB1 | chr7 | + | 92352475 | 92352642 | 92344977 | 92345066 | [illegible] |
| SE | 27613 | ENSG00000001629 | ANKIB1 | chr7 | + | 92387797 | 92387884 | 92386508 | 92386643 | [illegible] |
| SE | 27614 | ENSG00000001629 | ANKIB1 | chr7 | + | 92389970 | 92390116 | 92387974 | 92388041 | [illegible] |
| SE | 27615 | ENSG00000001629 | ANKIB1 | chr7 | + | 92392240 | 92392292 | 92391165 | 92391344 | [illegible] |
| SE | 27616 | ENSG00000179088 | C12orf42 | chr12 | - | 103267608 | 103267708 | 103263307 | 103263371 | [illegible] |
| SE | 27617 | ENSG00000179088 | C12orf42 | chr12 | - | 103267633 | 103267708 | 103263325 | 103263371 | [illegible] |
| SE | 27618 | ENSG00000179088 | C12orf42 | chr12 | - | 103268857 | 103268962 | 103267852 | 103267886 | [illegible] |

| | | | | | | | | | | | | | | | | | | | | | | | | | | | | | | | | | | | | | | | | | | | | | | |
|---|---|---|---|---|---|---|---|---|---|---|---|---|---|---|---|---|---|---|---|---|---|---|---|---|---|---|---|---|---|---|---|---|---|---|---|---|---|---|---|---|---|---|---|---|---|---|
| SE | 27619 | ENSG00000179088 | C12orf42 | chr12 | - | 103270148 | 103270308 | 103269819 | 103269853 | 103273791 | 103273883 | 5 | 5 | 3 | 6 | 14 | 1 | 0 | 2 | 0 | 1 | 0 | 3 | 0 | 0 | 0 | 0 | 5 | 0 | 0 | 0 | 298 | 149 | 1 | 1 | 0.714 | 1 | 0.429 | 1 | 0.875 | NA | 0.231 | NA | NA | NA | 0.573 |
| SE | 27620 | ENSG00000179088 | C12orf42 | chr12 | - | 103270148 | 103270308 | 103269819 | 103269853 | 103277149 | 103277210 | 5 | 6 | 3 | 6 | 16 | 1 | 0 | 1 | 0 | 2 | 0 | 5 | 0 | 0 | 0 | 0 | 3 | 0 | 0 | 2 | 298 | 149 | 1 | 1 | 0.714 | 1 | 0.6 | 1 | 0.8 | NA | 0.455 | NA | NA | 0 | 0.595 |
| SE | 27621 | ENSG00000179088 | C12orf42 | chr12 | - | 103273791 | 103273883 | 103269806 | 103269853 | 103277149 | 103277210 | 2 | 0 | 2 | 0 | 1 | 1 | 0 | 1 | 0 | 2 | 0 | 5 | 0 | 0 | 0 | 0 | 3 | 0 | 0 | 2 | 241 | 149 | 1 | 1 | 0.553 | NA | 0.553 | NA | 0.236 | NA | 0.507 | NA | NA | 0 | 0.194 |
| SE | 27622 | ENSG00000179088 | C12orf42 | chr12 | - | 103273791 | 103273883 | 103270148 | 103270308 | 103277149 | 103277210 | 1 | 0 | 0 | 1 | 1 | 0 | 1 | 0 | 1 | 3 | 0 | 0 | 0 | 0 | 0 | 0 | 2 | 0 | 0 | 0 | 241 | 149 | 1 | 1 | 1 | 0 | NA | 0.382 | 0.171 | NA | 0 | NA | NA | NA | 0.388 |
| SE | 27623 | ENSG00000179088 | C12orf42 | chr12 | - | 103305973 | 103306345 | 103301942 | 103302559 | 103368886 | 103368998 | 9 | 10 | 7 | 6 | 20 | 3 | 0 | 1 | 7 | 2 | 0 | 15 | 1 | 0 | 0 | 0 | 10 | 2 | 0 | 0 | 298 | 149 | 1 | 1 | 0.6 | 1 | 0.778 | 0.3 | 0.833 | NA | 0.429 | 0.2 | NA | NA | 0.388 |
| SE | 27624 | ENSG00000179088 | C12orf42 | chr12 | - | 103368011 | 103368106 | 103305973 | 103306345 | 103368886 | 103368998 | 0 | 0 | 0 | 0 | 1 | 8 | 10 | 5 | 4 | 19 | 0 | 0 | 0 | 0 | 0 | 0 | 13 | 1 | 0 | 0 | 244 | 149 | 1 | 1 | 0 | 0 | 0 | 0 | 0.031 | NA | 0 | 0 | NA | NA | 0.006 |
| SE | 27625 | ENSG00000179088 | C12orf42 | chr12 | - | 103368886 | 103368998 | 103277149 | 103277210 | 103401606 | 103401675 | 12 | 8 | 16 | 18 | 35 | 1 | 0 | 0 | 0 | 0 | 0 | 25 | 2 | 4 | 6 | 0 | 0 | 0 | 0 | 0 | 261 | 149 | 1 | 1 | 0.873 | 1 | 1 | 1 | 1 | NA | 1 | 1 | 1 | 1 | -0.025 |
| SE | 27626 | ENSG00000179088 | C12orf42 | chr12 | - | 103368886 | 103368998 | 103305973 | 103306345 | 103401606 | 103401675 | 17 | 16 | 20 | 17 | 43 | 0 | 0 | 1 | 0 | 0 | 0 | 35 | 3 | 4 | 4 | 0 | 0 | 0 | 0 | 0 | 261 | 149 | 1 | 1 | 1 | 1 | 0.919 | 1 | 1 | NA | 1 | 1 | 1 | 1 | -0.016 |
| SE | 27629 | ENSG00000179088 | C12orf42 | chr12 | - | 103401606 | 103401675 | 103368886 | 103368998 | 103441467 | 103441591 | 21 | 9 | 23 | 20 | 35 | 1 | 0 | 0 | 0 | 0 | 0 | 57 | 4 | 4 | 4 | 0 | 0 | 0 | 0 | 0 | 218 | 149 | 1 | 1 | 0.935 | 1 | 1 | 1 | 1 | NA | 1 | 1 | 1 | 1 | -0.013 |
| SE | 27633 | ENSG00000156253 | RWDD2B | chr21 | - | 29009594 | 29009692 | 29007899 | 29008621 | 29019210 | 29019340 | 0 | 2 | 8 | 0 | 0 | 12 | 5 | 11 | 4 | 21 | 0 | 0 | 0 | 1 | 0 | 6 | 16 | 2 | 10 | 0 | 247 | 149 | 1 | 1 | 0 | 0.194 | 0.305 | 0 | 0 | 0 | 0 | 0 | 0.057 | NA | 0.086 |
| SE | 27634 | ENSG00000156253 | RWDD2B | chr21 | - | 29009626 | 29009692 | 29008394 | 29008621 | 29019210 | 29019360 | 0 | 2 | 4 | 0 | 0 | 12 | 5 | 11 | 4 | 21 | 0 | 0 | 0 | 1 | 0 | 6 | 16 | 2 | 10 | 0 | 215 | 149 | 1 | 1 | 0 | 0.217 | 0.201 | 0 | 0 | 0 | 0 | 0 | 0.065 | NA | 0.067 |
| SE | 27637 | ENSG00000122008 | POLK | chr5 | + | 75534827 | 75534964 | 75511890 | 75511914 | 75547009 | 75547157 | 7 | 10 | 5 | 22 | 19 | 5 | 17 | 9 | 12 | 12 | 27 | 9 | 5 | 1 | 0 | 4 | 11 | 2 | 0 | 0 | 286 | 149 | 1 | 1 | 0.422 | 0.235 | 0.224 | 0.489 | 0.452 | 0.779 | 0.299 | 0.566 | 1 | NA | -0.297 |
| SE | 27638 | ENSG00000122008 | POLK | chr5 | + | 75534827 | 75534964 | 75511890 | 75511914 | 75552471 | 75552591 | 0 | 2 | 0 | 2 | 0 | 1 | 4 | 2 | 2 | 5 | 0 | 0 | 1 | 0 | 0 | 0 | 0 | 4 | 0 | 0 | 286 | 149 | 1 | 1 | 0 | 0.207 | 0 | 0.343 | 0 | NA | NA | 0.115 | NA | NA | -0.005 |
| SE | 27639 | ENSG00000122008 | POLK | chr5 | + | 75534827 | 75534964 | 75512068 | 75512946 | 75547009 | 75547157 | 6 | 8 | 5 | 20 | 19 | 0 | 2 | 0 | 0 | 0 | 27 | 9 | 5 | 1 | 0 | 0 | 0 | 0 | 0 | 0 | 286 | 149 | 1 | 1 | 1 | 0.676 | 1 | 1 | 1 | 1 | 1 | 1 | 1 | NA | -0.065 |
| SE | 27641 | ENSG00000122008 | POLK | chr5 | + | 75547009 | 75547157 | 75534827 | 75534964 | 75552471 | 75552591 | 67 | 74 | 46 | 69 | 103 | 0 | 1 | 0 | 0 | 0 | 39 | 45 | 38 | 11 | 11 | 0 | 0 | 1 | 0 | 0 | 297 | 149 | 1 | 1 | 1 | 0.974 | 1 | 1 | 1 | 1 | 1 | 0.95 | 1 | 1 | 0.005 |
| SE | 27644 | ENSG00000122008 | POLK | chr5 | + | 75569339 | 75569492 | 75552471 | 75552591 | 75573737 | 75573837 | 205 | 195 | 111 | 182 | 197 | 1 | 0 | 0 | 1 | 0 | 55 | 77 | 60 | 33 | 7 | 0 | 0 | 0 | 0 | 0 | 298 | 149 | 1 | 1 | 0.99 | 1 | 1 | 0.989 | 1 | 1 | 1 | 1 | 1 | 1 | -0.004 |
| SE | 27645 | ENSG00000122008 | POLK | chr5 | + | 75573737 | 75573869 | 75569339 | 75569492 | 75576779 | 75576917 | 177 | 154 | 100 | 140 | 144 | 0 | 2 | 0 | 0 | 0 | 35 | 51 | 22 | 26 | 5 | 0 | 0 | 0 | 0 | 0 | 281 | 149 | 1 | 1 | 1 | 0.976 | 1 | 1 | 1 | 1 | 1 | 1 | 1 | 1 | -0.005 |
| SE | 27646 | ENSG00000122008 | POLK | chr5 | + | 75581208 | 75581448 | 75576779 | 75576933 | 75583292 | 75583417 | 123 | 115 | 83 | 165 | 241 | 0 | 0 | 0 | 0 | 1 | 13 | 17 | 14 | 6 | 0 | 0 | 0 | 0 | 0 | 0 | 298 | 149 | 1 | 1 | 1 | 1 | 1 | 1 | 0.992 | 1 | 1 | 1 | 1 | NA | -0.002 |
| SE | 27647 | ENSG00000122008 | POLK | chr5 | + | 75582037 | 75582123 | 75581208 | 75581448 | 75583292 | 75583417 | 0 | 3 | 0 | 1 | 0 | 56 | 56 | 39 | 84 | 126 | 0 | 0 | 0 | 0 | 0 | 0 | 6 | 4 | 5 | 0 | 235 | 149 | 1 | 1 | 0 | 0.033 | 0 | 0.007 | 0 | NA | 0 | 0 | 0 | NA | 0.008 |
| SE | 27648 | ENSG00000122008 | POLK | chr5 | + | 75584759 | 75584926 | 75583292 | 75583417 | 75587025 | 75587058 | 154 | 201 | 102 | 143 | 201 | 0 | 0 | 0 | 3 | 0 | 15 | 27 | 24 | 10 | 0 | 0 | 0 | 0 | 0 | 0 | 298 | 149 | 1 | 1 | 1 | 1 | 1 | 0.96 | 1 | 1 | 1 | 1 | 1 | NA | -0.008 |
| SE | 27649 | ENSG00000122008 | POLK | chr5 | + | 75590343 | 75590440 | 75587025 | 75587058 | 75593877 | 75594049 | 109 | 133 | 92 | 91 | 133 | 3 | 1 | 0 | 3 | 0 | 0 | 32 | 20 | 16 | 3 | 0 | 0 | 0 | 0 | 0 | 246 | 149 | 1 | 1 | 0.957 | 0.988 | 1 | 0.948 | 1 | NA | 1 | 1 | 1 | 1 | -0.021 |
| SE | 27650 | ENSG00000122008 | POLK | chr5 | + | 75590343 | 75590440 | 75587025 | 75587058 | 75593965 | 75594049 | 40 | 46 | 41 | 33 | 52 | 0 | 4 | 0 | 2 | 9 | 0 | 19 | 8 | 10 | 0 | 0 | 0 | 0 | 0 | 0 | 246 | 149 | 1 | 1 | 1 | 0.874 | 1 | 0.909 | 0.778 | NA | 1 | 1 | 1 | NA | -0.088 |
| SE | 27653 | ENSG00000122008 | POLK | chr5 | + | 75593877 | 75594049 | 75590343 | 75590440 | 75596221 | 75597178 | 183 | 226 | 136 | 160 | 209 | 5 | 8 | 7 | 7 | 11 | 0 | 33 | 35 | 13 | 6 | 0 | 0 | 6 | 5 | 10 | 298 | 149 | 1 | 1 | 0.948 | 0.934 | 0.907 | 0.92 | 0.905 | NA | 1 | 0.745 | 0.565 | 0.231 | 0.288 |
| SE | 27654 | ENSG00000122008 | POLK | chr5 | + | 75593965 | 75594049 | 75587025 | 75587058 | 75596221 | 75597178 | 12 | 24 | 13 | 12 | 37 | 1 | 0 | 3 | 5 | 2 | 0 | 8 | 1 | 0 | 0 | 8 | 0 | 0 | 0 | 0 | 233 | 149 | 1 | 1 | 0.885 | 1 | 0.735 | 0.605 | 0.922 | 0 | 1 | 1 | NA | NA | 0.163 |
| SE | 27655 | ENSG00000122008 | POLK | chr5 | + | 75593965 | 75594049 | 75590343 | 75590440 | 75596221 | 75597178 | 16 | 28 | 17 | 11 | 38 | 5 | 8 | 7 | 7 | 11 | 0 | 8 | 3 | 0 | 0 | 0 | 0 | 6 | 5 | 10 | 233 | 149 | 1 | 1 | 0.672 | 0.691 | 0.608 | 0.501 | 0.688 | NA | 1 | 0.242 | 0 | 0 | 0.322 |
| SE | 27658 | ENSG00000134851 | TMEM165 | chr4 | + | 55403235 | 55403338 | 55398922 | 55399253 | 55411613 | 55411839 | 7 | 0 | 2 | 8 | 2 | 1 | 1 | 0 | 2 | 0 | 0 | 1 | 2 | 0 | 0 | 0 | 0 | 2 | 0 | 0 | 252 | 149 | 1 | 1 | 0.805 | 0 | 1 | 0.703 | 1 | NA | 1 | 0.372 | NA | NA | 0.016 |
| SE | 27659 | ENSG00000134851 | TMEM165 | chr4 | + | 55411613 | 55411839 | 55395956 | 55396396 | 55417802 | 55417985 | 29 | 18 | 15 | 12 | 28 | 0 | 0 | 0 | 0 | 0 | 37 | 22 | 3 | 15 | 16 | 0 | 0 | 1 | 0 | 0 | 298 | 149 | 1 | 1 | 1 | 1 | 1 | 1 | 1 | 1 | 1 | 0.6 | 1 | 1 | 0.08 |
| SE | 27661 | ENSG00000134851 | TMEM165 | chr4 | + | 55417071 | 55417247 | 55411613 | 55411839 | 55417802 | 55417985 | 361 | 226 | 238 | 267 | 337 | 0 | 0 | 0 | 0 | 1 | 463 | 332 | 121 | 235 | 388 | 0 | 0 | 0 | 0 | 0 | 298 | 149 | 1 | 1 | 1 | 1 | 1 | 1 | 0.994 | 1 | 1 | 1 | 1 | 1 | -0.001 |
| SE | 27665 | ENSG00000134851 | TMEM165 | chr4 | + | 55418047 | 55418164 | 55411613 | 55411839 | 55424537 | 55424643 | 16 | 23 | 20 | 13 | 9 | 0 | 0 | 0 | 2 | 0 | 3 | 45 | 68 | 33 | 0 | 12 | 1 | 3 | 0 | 0 | 266 | 149 | 1 | 1 | 1 | 1 | 1 | 0.785 | 1 | 0.123 | 0.962 | 0.927 | 1 | NA | 0.204 |
| SE | 27697 | ENSG00000186431 | FCAR | chr19 | + | 54885234 | 54885525 | 54874247 | 54874323 | 54888006 | 54888220 | 92 | 60 | 45 | 111 | 104 | 0 | 0 | 1 | 0 | 0 | 148 | 132 | 239 | 541 | 300 | 0 | 0 | 0 | 0 | 0 | 298 | 149 | 1 | 1 | 1 | 1 | 0.957 | 1 | 1 | 1 | 1 | 1 | 1 | 1 | -0.009 |
| SE | 27698 | ENSG00000186431 | FCAR | chr19 | + | 54885234 | 54885525 | 54875329 | 54875365 | 54888006 | 54888220 | 145 | 83 | 86 | 140 | 146 | 0 | 0 | 1 | 0 | 0 | 272 | 226 | 369 | 864 | 394 | 0 | 0 | 0 | 0 | 0 | 298 | 149 | 1 | 1 | 1 | 1 | 0.977 | 1 | 1 | 1 | 1 | 1 | 1 | 1 | -0.005 |
| SE | 27704 | ENSG00000164483 | SAMD3 | chr6 | - | 130145339 | 130145422 | 130144314 | 130144804 | 130154824 | 130155025 | 225 | 65 | 132 | 191 | 224 | 1 | 0 | 0 | 0 | 0 | 18 | 62 | 38 | 35 | 7 | 0 | 0 | 0 | 0 | 0 | 232 | 149 | 1 | 1 | 0.993 | 1 | 1 | 1 | 1 | 1 | 1 | 1 | 1 | 1 | -0.001 |
| SE | 27705 | ENSG00000164483 | SAMD3 | chr6 | - | 130146009 | 130146181 | 130144683 | 130144804 | 130154824 | 130154853 | 669 | 275 | 339 | 525 | 709 | 1 | 0 | 0 | 0 | 0 | 0 | 230 | 128 | 46 | 28 | 0 | 0 | 0 | 0 | 0 | 298 | 149 | 1 | 1 | 0.997 | 1 | 1 | 1 | 1 | NA | 1 | 1 | 1 | 1 | -0.001 |
| SE | 27706 | ENSG00000164483 | SAMD3 | chr6 | - | 130146009 | 130146181 | 130145339 | 130145422 | 130154824 | 130154853 | 1349 | 574 | 707 | 1057 | 1317 | 0 | 0 | 0 | 1 | 0 | 10 | 443 | 289 | 146 | 37 | 0 | 0 | 0 | 0 | 0 | 298 | 149 | 1 | 1 | 1 | 1 | 1 | 0.998 | 1 | 1 | 1 | 1 | 1 | 1 | 0 |
| SE | 27707 | ENSG00000164483 | SAMD3 | chr6 | - | 130175840 | 130176008 | 130154824 | 130155025 | 130184102 | 130184187 | 621 | 281 | 317 | 579 | 544 | 0 | 0 | 0 | 1 | 0 | 48 | 312 | 174 | 86 | 102 | 0 | 0 | 0 | 0 | 0 | 298 | 149 | 1 | 1 | 1 | 1 | 1 | 0.997 | 1 | 1 | 1 | 1 | 1 | 1 | -0.001 |
| SE | 27709 | ENSG00000164483 | SAMD3 | chr6 | - | 130184102 | 130184187 | 130175757 | 130176008 | 130184437 | 130184620 | 654 | 232 | 277 | 548 | 499 | 3 | 0 | 0 | 0 | 0 | 65 | 331 | 169 | 105 | 129 | 0 | 2 | 0 | 0 | 0 | 234 | 149 | 1 | 1 | 0.993 | 1 | 1 | 1 | 1 | 1 | 0.991 | 1 | 1 | 1 | 0 |
| SE | 27710 | ENSG00000164483 | SAMD3 | chr6 | - | 130184437 | 130184620 | 130175757 | 130176008 | 130209494 | 130209533 | 60 | 29 | 29 | 66 | 53 | 0 | 0 | 1 | 0 | 0 | 6 | 26 | 26 | 7 | 0 | 0 | 0 | 0 | 0 | 0 | 298 | 149 | 1 | 1 | 1 | 1 | 0.935 | 1 | 1 | 1 | 1 | 1 | 1 | NA | -0.013 |
| SE | 27711 | ENSG00000164483 | SAMD3 | chr6 | - | 130184437 | 130184623 | 130175840 | 130176008 | 130209494 | 130209602 | 241 | 98 | 135 | 242 | 197 | 0 | 0 | 1 | 0 | 0 | 20 | 117 | 72 | 25 | 20 | 0 | 0 | 0 | 0 | 0 | 298 | 149 | 1 | 1 | 1 | 1 | 0.985 | 1 | 1 | 1 | 1 | 1 | 1 | 1 | -0.003 |
| SE | 27712 | ENSG00000164483 | SAMD3 | chr6 | - | 130209494 | 130209602 | 130184437 | 130184623 | 130214336 | 130214526 | 192 | 82 | 111 | 199 | 158 | 5 | 1 | 0 | 0 | 0 | 20 | 106 | 58 | 25 | 20 | 0 | 2 | 1 | 0 | 0 | 257 | 149 | 1 | 1 | 0.957 | 0.979 | 1 | 1 | 1 | 1 | 0.968 | 0.971 | 1 | 1 | -0.001 |
| SE | 27713 | ENSG00000164483 | SAMD3 | chr6 | - | 130209494 | 130209608 | 130184437 | 130184623 | 130214336 | 130214520 | 502 | 244 | 331 | 535 | 397 | 5 | 1 | 0 | 0 | 0 | 70 | 275 | 166 | 87 | 57 | 0 | 2 | 1 | 0 | 0 | 263 | 149 | 1 | 1 | 0.983 | 0.993 | 1 | 1 | 1 | 1 | 0.987 | 0.989 | 1 | 1 | 0 |
| SE | 27718 | ENSG00000083896 | YTHDC1 | chr4 | - | 68316313 | 68316448 | 68310386 | 68314323 | 68318518 | 68318581 | 556 | 445 | 569 | 577 | 1106 | 1 | 0 | 0 | 0 | 0 | 100 | 373 | 602 | 194 | 14 | 0 | 1 | 0 | 5 | 0 | 284 | 149 | 1 | 1 | 0.997 | 1 | 1 | 1 | 1 | 1 | 0.995 | 1 | 0.953 | 1 | 0.01 |
| SE | 27719 | ENSG00000083896 | YTHDC1 | chr4 | - | 68322748 | 68322915 | 68320122 | 68320205 | 68324138 | 68324223 | 550 | 480 | 638 | 603 | 1011 | 0 | 0 | 0 | 0 | 0 | 54 | 391 | 704 | 210 | 34 | 0 | 0 | 2 | 0 | 0 | 298 | 149 | 1 | 1 | 1 | 1 | 1 | 1 | 1 | 1 | 1 | 0.994 | 1 | 1 | 0.001 |
| SE | 27720 | ENSG00000083896 | YTHDC1 | chr4 | - | 68330001 | 68330119 | 68324138 | 68324223 | 68332102 | 68332197 | 300 | 239 | 310 | 330 | 500 | 2 | 0 | 0 | 4 | 1 | 23 | 240 | 391 | 106 | 15 | 0 | 0 | 3 | 0 | 0 | 267 | 149 | 1 | 1 | 0.988 | 1 | 1 | 0.979 | 0.996 | 1 | 1 | 0.986 | 1 | 1 | -0.005 |
| SE | 27721 | ENSG00000083896 | YTHDC1 | chr4 | - | 68330201 | 68330310 | 68330001 | 68330119 | 68332102 | 68332197 | 403 | 288 | 410 | 476 | 659 | 3 | 0 | 0 | 3 | 6 | 31 | 247 | 432 | 97 | 0 | 0 | 8 | 11 | 8 | 0 | 258 | 149 | 1 | 1 | 0.987 | 1 | 1 | 0.989 | 0.984 | 1 | 0.947 | 0.958 | 0.875 | NA | 0.047 |
| SE | 27722 | ENSG00000083896 | YTHDC1 | chr4 | - | 68332793 | 68332847 | 68332102 | 68332197 | 68333307 | 68333397 | 294 | 221 | 231 | 317 | 408 | 17 | 7 | 7 | 15 | 16 | 43 | 149 | 261 | 59 | 0 | 0 | 16 | 9 | 8 | 0 | 203 | 149 | 1 | 1 | 0.927 | 0.959 | 0.96 | 0.939 | 0.949 | 1 | 0.872 | 0.955 | 0.844 | NA | 0.029 |
| SE | 27724 | ENSG00000083896 | YTHDC1 | chr4 | - | 68344048 | 68344080 | 68338282 | 68338384 | 68349725 | 68350045 | 0 | 0 | 0 | 1 | 0 | 138 | 64 | 117 | 122 | 145 | 0 | 0 | 0 | 0 | 0 | 42 | 88 | 165 | 54 | 4 | 181 | 149 | 1 | 1 | 0 | 0 | 0 | 0.007 | 0 | 0 | 0 | 0 | 0 | 0 | 0.001 |
| SE | 27727 | ENSG00000152818 | UTRN | chr6 | + | 144403122 | 144403184 | 144291736 | 144291907 | 144421877 | 144421970 | 574 | 643 | 406 | 650 | 718 | 1 | 0 | 0 | 0 | 0 | 336 | 485 | 406 | 217 | 99 | 0 | 0 | 1 | 0 | 0 | 211 | 149 | 1 | 1 | 0.998 | 1 | 1 | 1 | 1 | 1 | 1 | 0.997 | 1 | 1 | 0 |
| SE | 27728 | ENSG00000152818 | UTRN | chr6 | + | 144403122 | 144403184 | 144344100 | 144344304 | 144421877 | 144421970 | 855 | 885 | 805 | 1130 | 1039 | 1 | 0 | 1 | 2 | 0 | 536 | 1054 | 761 | 256 | 204 | 0 | 0 | 0 | 0 | 0 | 211 | 149 | 1 | 1 | 0.998 | 1 | 0.998 | 0.997 | 1 | 1 | 1 | 1 | 1 | 1 | -0.001 |
| SE | 27729 | ENSG00000152818 | UTRN | chr6 | + | 144423985 | 144424078 | 144423548 | 144423626 | 144426286 | 144426328 | 1253 | 1237 | 1045 | 1468 | 1497 | 0 | 0 | 2 | 0 | 3 | 747 | 1294 | 1099 | 421 | 276 | 0 | 0 | 0 | 0 | 0 | 242 | 149 | 1 | 1 | 1 | 1 | 0.997 | 1 | 0.997 | 1 | 1 | 1 | 1 | 1 | -0.001 |
| SE | 27730 | ENSG00000152818 | UTRN | chr6 | + | 144428777 | 144428893 | 144426286 | 144426459 | 144429580 | 144429741 | 1221 | 876 | 874 | 1112 | 674 | 1 | 0 | 0 | 1 | 0 | 313 | 883 | 641 | 311 | 238 | 0 | 0 | 0 | 0 | 0 | 265 | 149 | 1 | 1 | 0.999 | 1 | 1 | 0.998 | 1 | 1 | 1 | 1 | 1 | 1 | -0.001 |
| SE | 27733 | ENSG00000152818 | UTRN | chr6 | + | 144444280 | 144444382 | 144440351 | 144440471 | 144447210 | 144447318 | 571 | 478 | 452 | 528 | 521 | 1 | 0 | 0 | 0 | 0 | 110 | 489 | 376 | 103 | 7 | 0 | 0 | 0 | 0 | 0 | 251 | 149 | 1 | 1 | 0.997 | 1 | 1 | 1 | 1 | 1 | 1 | 1 | 1 | 1 | -0.001 |
| SE | 27735 | ENSG00000152818 | UTRN | chr6 | + | 144448599 | 144448769 | 144447601 | 144447781 | 144451369 | 144451493 | 1799 | 1660 | 1540 | 1728 | 2283 | 0 | 0 | 0 | 0 | 0 | 205 | 1643 | 1144 | 295 | 54 | 0 | 0 | 1 | 0 | 0 | 298 | 149 | 1 | 1 | 1 | 1 | 1 | 1 | 1 | 1 | 1 | 0.998 | 1 | 1 | 0 |
| SE | 27736 | ENSG00000152818 | UTRN | chr6 | + | 144451369 | 144451493 | 144448599 | 144448769 | 144453781 | 144453869 | 1145 | 1132 | 929 | 1098 | 1103 | 0 | 0 | 0 | 1 | 0 | 189 | 1083 | 799 | 243 | 58 | 0 | 0 | 0 | 0 | 0 | 273 | 149 | 1 | 1 | 1 | 1 | 1 | 0.998 | 1 | 1 | 1 | 1 | 1 | 1 | 0 |
| SE | 27737 | ENSG00000152818 | UTRN | chr6 | + | 144482208 | 144482388 | 144479811 | 144479982 | 144485384 | 144485519 | 1150 | 1033 | 977 | 1247 | 734 | 0 | 0 | 0 | 0 | 0 | 219 | 820 | 766 | 162 | 122 | 0 | 5 | 2 | 0 | 0 | 298 | 149 | 1 | 1 | 1 | 1 | 1 | 1 | 1 | 1 | 0.988 | 0.995 | 1 | 1 | 0.003 |
| SE | 27738 | ENSG00000152818 | UTRN | chr6 | + | 144482208 | 144482388 | 144479811 | 144479982 | 144487547 | 144487697 | 611 | 553 | 539 | 686 | 372 | 0 | 0 | 0 | 3 | 0 | 114 | 450 | 478 | 108 | 61 | 0 | 0 | 0 | 0 | 0 | 298 | 149 | 1 | 1 | 1 | 1 | 1 | 0.991 | 1 | 1 | 1 | 1 | 1 | 1 | -0.002 |
| SE | 27739 | ENSG00000152818 | UTRN | chr6 | + | 144485384 | 144485519 | 144479811 | 144479982 | 144487547 | 144487697 | 540 | 436 | 398 | 526 | 298 | 0 | 0 | 0 | 3 | 0 | 140 | 418 | 338 | 79 | 70 | 0 | 0 | 0 | 0 | 0 | 284 | 149 | 1 | 1 | 1 | 1 | 1 | 0.989 | 1 | 1 | 1 | 1 | 1 | 1 | -0.002 |
| SE | 27740 | ENSG00000152818 | UTRN | chr6 | + | 144485384 | 144485519 | 144482208 | 144482388 | 144487547 | 144487697 | 1079 | 917 | 836 | 1087 | 660 | 0 | 1 | 0 | 0 | 0 | 245 | 783 | 624 | 133 | 131 | 0 | 0 | 0 | 0 | 0 | 284 | 149 | 1 | 1 | 1 | 0.998 | 1 | 1 | 1 | 1 | 1 | 1 | 1 | 1 | 0 |
| SE | 27741 | ENSG00000152818 | UTRN | chr6 | + | 144490070 | 144490199 | 144488672 | 144488834 | 144490928 | 144491102 | 861 | 556 | 595 | 745 | 598 | 0 | 0 | 0 | 2 | 0 | 121 | 522 | 418 | 179 | 174 | 0 | 0 | 2 | 3 | 0 | 278 | 149 | 1 | 1 | 1 | 1 | 1 | 0.995 | 1 | 1 | 1 | 0.991 | 0.97 | 1 | 0.007 |
| SE | 27742 | ENSG00000152818 | UTRN | chr6 | + | 144513908 | 144514037 | 144510943 | 144511123 | 144516228 | 144516387 | 432 | 405 | 364 | 448 | 643 | 1 | 0 | 0 | 0 | 0 | 51 | 343 | 209 | 56 | 18 | 0 | 0 | 0 | 0 | 0 | 278 | 149 | 1 | 1 | 0.996 | 1 | 1 | 1 | 1 | 1 | 1 | 1 | 1 | 1 | -0.001 |
| SE | 27743 | ENSG00000152818 | UTRN | chr6 | + | 144514649 | 144514820 | 144513908 | 144514037 | 144516228 | 144516387 | 663 | 495 | 449 | 610 | 579 | 0 | 0 | 0 | 0 | 0 | 64 | 483 | 267 | 77 | 26 | 0 | 3 | 0 | 0 | 0 | 298 | 149 | 1 | 1 | 1 | 1 | 1 | 1 | 1 | 1 | 0.988 | 1 | 1 | 1 | 0.002 |
| SE | 27744 | ENSG00000152818 | UTRN | chr6 | + | 144516228 | 144516387 | 144514649 | 144514820 | 144516810 | 144516948 | 944 | 748 | 643 | 819 | 899 | 1 | 2 | 0 | 1 | 0 | 118 | 695 | 481 | 111 | 110 | 0 | 1 | 0 | 0 | 0 | 298 | 149 | 1 | 1 | 0.998 | 0.995 | 1 | 0.998 | 1 | 1 | 0.997 | 1 | 1 | 1 | -0.001 |
| SE | 27746 | ENSG00000152818 | UTRN | chr6 | + | 144533084 | 144533260 | 144531051 | 144531202 | 144537581 | 144537717 | 1041 | 883 | 741 | 1068 | 722 | 2 | 4 | 0 | 0 | 0 | 136 | 712 | 474 | 172 | 87 | 0 | 0 | 0 | 0 | 0 | 298 | 149 | 1 | 1 | 0.996 | 0.991 | 1 | 1 | 1 | 1 | 1 | 1 | 1 | 1 | -0.003 |
| SE | 27747 | ENSG00000152818 | UTRN | chr6 | + | 144542794 | 144542870 | 144539293 | 144539443 | 144548639 | 144548854 | 642 | 405 | 453 | 485 | 706 | 0 | 0 | 0 | 0 | 1 | 52 | 330 | 253 | 64 | 96 | 0 | 0 | 0 | 0 | 0 | 225 | 149 | 1 | 1 | 1 | 1 | 1 | 1 | 0.998 | 1 | 1 | 1 | 1 | 1 | 0 |
| SE | 27748 | ENSG00000152818 | UTRN | chr6 | + | 144550964 | 144551082 | 144548639 | 144548854 | 144554687 | 144554893 | 839 | 743 | 658 | 841 | 555 | 0 | 1 | 0 | 0 | 0 | 212 | 630 | 554 | 101 | 122 | 0 | 0 | 0 | 0 | 0 | 267 | 149 | 1 | 1 | 1 | 0.998 | 1 | 1 | 1 | 1 | 1 | 1 | 1 | 1 | 0 |
| SE | 27749 | ENSG00000152818 | UTRN | chr6 | + | 144554687 | 144554893 | 144550964 | 144551082 | 144557156 | 144557311 | 918 | 750 | 697 | 913 | 560 | 5 | 4 | 0 | 4 | 0 | 188 | 618 | 508 | 107 | 104 | 0 | 0 | 0 | 0 | 0 | 298 | 149 | 1 | 1 | 0.989 | 0.989 | 1 | 0.991 | 1 | 1 | 1 | 1 | 1 | 1 | -0.006 |
| SE | 27750 | ENSG00000152818 | UTRN | chr6 | + | 144577098 | 144577288 | 144557156 | 144557311 | 144678405 | 144678578 | 811 | 559 | 547 | 585 | 489 | 0 | 0 | 0 | 2 | 0 | 129 | 418 | 371 | 84 | 64 | 0 | 0 | 0 | 0 | 0 | 298 | 149 | 1 | 1 | 1 | 1 | 1 | 0.993 | 1 | 1 | 1 | 1 | 1 | 1 | -0.001 |
| SE | 27751 | ENSG00000152818 | UTRN | chr6 | + | 144678405 | 144678578 | 144577098 | 144577288 | 144700086 | 144700243 | 812 | 749 | 589 | 738 | 814 | 2 | 5 | 2 | 0 | 5 | 131 | 494 | 347 | 98 | 49 | 0 | 0 | 7 | 0 | 0 | 298 | 149 | 1 | 1 | 0.995 | 0.987 | 0.993 | 1 | 0.988 | 1 | 1 | 0.961 | 1 | 1 | 0 |
| SE | 27752 | ENSG00000152818 | UTRN | chr6 | + | 144678405 | 144678578 | 144583209 | 144583600 | 144700086 | 144700243 | 433 | 502 | 323 | 458 | 575 | 0 | 0 | 0 | 0 | 0 | 55 | 283 | 173 | 63 | 21 | 0 | 0 | 0 | 0 | 3 | 298 | 149 | 1 | 1 | 1 | 1 | 1 | 1 | 1 | 1 | 1 | 1 | 1 | 0.778 | 0.044 |
| SE | 27753 | ENSG00000152818 | UTRN | chr6 | + | 144700086 | 144700243 | 144678405 | 144678578 | 144730356 | 144730486 | 949 | 1106 | 776 | 1083 | 1487 | 0 | 1 | 0 | 0 | 0 | 256 | 665 | 542 | 176 | 66 | 0 | 0 | 0 | 0 | 0 | 298 | 149 | 1 | 1 | 1 | 0.998 | 1 | 1 | 1 | 1 | 1 | 1 | 1 | 1 | 0 |
| SE | 27754 | ENSG00000152818 | UTRN | chr6 | + | 144757928 | 144757989 | 144754719 | 144754798 | 144771906 | 144771968 | 959 | 1018 | 636 | 1035 | 1752 | 0 | 1 | 0 | 0 | 0 | 221 | 571 | 608 | 161 | 129 | 0 | 0 | 0 | 0 | 0 | 210 | 149 | 1 | 1 | 1 | 0.999 | 1 | 1 | 1 | 1 | 1 | 1 | 1 | 1 | 0 |
| SE | 27755 | ENSG00000152818 | UTRN | chr6 | + | 144789193 | 144789279 | 144781921 | 144782123 | 144793833 | 144793991 | 1158 | 1133 | 533 | 1100 | 1264 | 0 | 2 | 1 | 0 | 0 | 182 | 554 | 428 | 176 | 160 | 0 | 0 | 0 | 0 | 0 | 235 | 149 | 1 | 1 | 1 | 0.997 | 0.997 | 1 | 1 | 1 | 1 | 1 | 1 | 1 | -0.001 |
| SE | 27756 | ENSG00000152818 | UTRN | chr6 | + | 144803035 | 144803147 | 144797744 | 144797990 | 144820881 | 144821018 | 597 | 366 | 402 | 519 | 659 | 0 | 1 | 0 | 3 | 3 | 10 | 259 | 191 | 53 | 22 | 0 | 0 | 0 | 0 | 0 | 261 | 149 | 1 | 1 | 1 | 0.995 | 1 | 0.99 | 0.992 | 1 | 1 | 1 | 1 | 1 | -0.005 |
| SE | 27759 | ENSG00000152818 | UTRN | chr6 | + | 144839172 | 144839284 | 144836300 | 144836541 | 144846804 | 144846827 | 282 | 163 | 212 | 234 | 235 | 0 | 2 | 1 | 0 | 0 | 24 | 117 | 104 | 12 | 13 | 0 | 0 | 0 | 0 | 0 | 261 | 149 | 1 | 1 | 1 | 0.979 | 0.992 | 1 | 1 | 1 | 1 | 1 | 1 | 1 | -0.006 |
| SE | 27760 | ENSG00000152818 | UTRN | chr6 | + | 144840739 | 144840832 | 144836300 | 144836541 | 144846804 | 144846822 | 51 | 37 | 50 | 50 | 24 | 0 | 2 | 1 | 0 | 0 | 0 | 23 | 36 | 2 | 0 | 0 | 0 | 0 | 0 | 0 | 242 | 149 | 1 | 1 | 1 | 0.919 | 0.969 | 1 | 1 | NA | 1 | 1 | 1 | NA | -0.022 |
| SE | 27761 | ENSG00000152818 | UTRN | chr6 | + | 144840739 | 144840832 | 144839048 | 144839284 | 144846804 | 144846822 | 265 | 161 | 201 | 247 | 145 | 0 | 0 | 0 | 1 | 0 | 30 | 151 | 124 | 22 | 20 | 0 | 0 | 0 | 0 | 0 | 242 | 149 | 1 | 1 | 1 | 1 | 1 | 0.993 | 1 | 1 | 1 | 1 | 1 | 1 | -0.001 |
| SE | 27762 | ENSG00000152818 | UTRN | chr6 | + | 144846804 | 144846827 | 144840739 | 144840832 | 144850988 | 144851003 | 156 | 88 | 128 | 141 | 48 | 0 | 1 | 3 | 1 | 0 | 8 | 81 | 75 | 14 | 4 | 0 | 0 | 0 | 0 | 0 | 172 | 149 | 1 | 1 | 1 | 0.987 | 0.974 | 0.992 | 1 | 1 | 1 | 1 | 1 | 1 | -0.009 |
| SE | 27763 | ENSG00000137494 | ANKRD42 | chr11 | + | 83206057 | 83206165 | 83198478 | 83198642 | 83210299 | 83210416 | 35 | 34 | 46 | 49 | 34 | 0 | 0 | 0 | 0 | 0 | 44 | 51 | 21 | 12 | 50 | 0 | 0 | 2 | 0 | 0 | 257 | 149 | 1 | 1 | 1 | 1 | 1 | 1 | 1 | 1 | 1 | 0.859 | 1 | 1 | 0.028 |
| SE | 27764 | ENSG00000137494 | ANKRD42 | chr11 | + | 83240758 | 83240934 | 83236403 | 83236509 | 83247942 | 83248971 | 9 | 11 | 4 | 5 | 5 | 0 | 0 | 2 | 0 | 0 | 0 | 12 | 2 | 1 | 12 | 0 | 0 | 0 | 0 | 0 | 298 | 149 | 1 | 1 | 1 | 1 | 0.5 | 1 | 1 | NA | 1 | 1 | 1 | 1 | -0.1 |
| SE | 27765 | ENSG00000137494 | ANKRD42 | chr11 | + | 83245497 | 83245624 | 83240758 | 83240934 | 83247942 | 83248087 | 14 | 13 | 12 | 10 | 19 | 1 | 0 | 1 | 0 | 0 | 2 | 0 | 12 | 0 | 12 | 0 | 0 | 0 | 0 | 0 | 276 | 149 | 1 | 1 | 0.883 | 1 | 0.866 | 1 | 1 | 1 | NA | 1 | NA | 1 | -0.05 |
| SE | 27766 | ENSG00000121064 | SCPEP1 | chr17 | + | 56981081 | 56981230 | 56978130 | 56978235 | 56985377 | 56985427 | 103 | 52 | 64 | 75 | 53 | 0 | 1 | 0 | 0 | 0 | 236 | 231 | 94 | 192 | 296 | 0 | 0 | 0 | 0 | 0 | 298 | 149 | 1 | 1 | 1 | 0.963 | 1 | 1 | 1 | 1 | 1 | 1 | 1 | 1 | -0.007 |
| SE | 27767 | ENSG00000121064 | SCPEP1 | chr17 | + | 56981116 | 56981230 | 56978128 | 56978235 | 56985377 | 56985467 | 65 | 35 | 37 | 42 | 35 | 0 | 1 | 0 | 0 | 0 | 152 | 147 | 51 | 118 | 170 | 0 | 0 | 0 | 0 | 0 | 263 | 149 | 1 | 1 | 1 | 0.952 | 1 | 1 | 1 | 1 | 1 | 1 | 1 | 1 | -0.01 |
| SE | 27768 | ENSG00000121064 | SCPEP1 | chr17 | + | 56985377 | 56985467 | 56981081 | 56981230 | 56987694 | 56987850 | 118 | 67 | 75 | 85 | 109 | 0 | 1 | 0 | 0 | 1 | 288 | 329 | 106 | 201 | 362 | 0 | 0 | 0 | 0 | 0 | 239 | 149 | 1 | 1 | 1 | 0.977 | 1 | 1 | 0.985 | 1 | 1 | 1 | 1 | 1 | -0.008 |
| SE | 27770 | ENSG00000121064 | SCPEP1 | chr17 | + | 56991098 | 56991171 | 56988215 | 56988290 | 56994980 | 56995018 | 297 | 179 | 240 | 256 | 263 | 0 | 0 | 0 | 0 | 0 | 1258 | 984 | 392 | 758 | 1257 | 0 | 5 | 0 | 0 | 10 | 222 | 149 | 1 | 1 | 1 | 1 | 1 | 1 | 1 | 1 | 0.992 | 1 | 1 | 0.988 | 0.004 |
| SE | 27771 | ENSG00000121064 | SCPEP1 | chr17 | + | 56994980 | 56995018 | 56988215 | 56988290 | 56995506 | 56995544 | 115 | 80 | 76 | 68 | 118 | 0 | 0 | 0 | 0 | 0 | 421 | 367 | 122 | 282 | 510 | 0 | 0 | 2 | 0 | 0 | 187 | 149 | 1 | 1 | 1 | 1 | 1 | 1 | 1 | 1 | 1 | 0.98 | 1 | 1 | 0.004 |
| SE | 27772 | ENSG00000121064 | SCPEP1 | chr17 | + | 56995506 | 56995635 | 56994980 | 56995018 | 56996961 | 56996996 | 514 | 331 | 387 | 383 | 440 | 0 | 0 | 1 | 1 | 0 | 2006 | 1519 | 600 | 1125 | 1794 | 0 | 1 | 1 | 0 | 0 | 278 | 149 | 1 | 1 | 1 | 1 | 0.995 | 0.995 | 1 | 1 | 0.999 | 0.997 | 1 | 1 | -0.001 |
| SE | 27775 | ENSG00000121064 | SCPEP1 | chr17 | + | 57002017 | 57002181 | 57000854 | 57000992 | 57006172 | 57006586 | 722 | 630 | 579 | 540 | 694 | 0 | 0 | 0 | 0 | 0 | 2486 | 1742 | 845 | 1697 | 2239 | 0 | 0 | 2 | 0 | 0 | 298 | 149 | 1 | 1 | 1 | 1 | 1 | 1 | 1 | 1 | 1 | 0.995 | 1 | 1 | 0.001 |
| SE | 27777 | ENSG00000177058 | SLC38A9 | chr5 | - | 55627890 | 55627980 | 55625844 | 55626659 | 55633753 | 55633902 | 25 | 30 | 25 | 24 | 45 | 1 | 2 | 0 | 1 | 0 | 6 | 39 | 10 | 9 | 3 | 0 | 0 | 0 | 0 | 0 | 239 | 149 | 1 | 1 | 0.94 | 0.903 | 1 | 0.937 | 1 | 1 | 1 | 1 | 1 | 1 | -0.044 |

SE (continue)

| | | | | | | | | | | | | Inclusion_counts_HC | | | | | Skipping_counts_HC | | | | | Inclusion counts_SLE | | | | | Skipping_counts_SLE | | | | | | | | | Inclusion_level(normalized) | | | | | | | | | | |
|---|---|---|---|---|---|---|---|---|---|---|---|---|---|---|---|---|---|---|---|---|---|---|---|---|---|---|---|---|---|---|---|---|---|---|---|---|---|---|---|---|---|---|---|---|---|---|
| Splicing_event | ID | Gene_ID | Gene_Symbol | Chromosome | Strand | ExonStart_0base | ExonEnd | UpstreamES | UpstreamEE | DownstreamES | DownstreamEE | SRR13214146 | SRR13214147 | SRR13214148 | SRR13214149 | SRR13214150 | SRR13214146 | SRR13214147 | SRR13214148 | SRR13214149 | SRR13214150 | SRR13214141 | SRR13214242 | SRR13214143 | SRR13214144 | SRR13214145 | SRR13214141 | SRR13214242 | SRR13214143 | SRR13214144 | SRR13214145 | IncFormLen | SkipFormLen | p value | FDR | SRR13214146 | SRR13214147 | SRR13214148 | SRR13214149 | SRR13214150 | SRR13214141 | SRR13214242 | SRR13214143 | SRR13214144 | SRR13214145 | Inclevel_diff |
| SE | 27780 | ENSG00000177058 | SLC38A9 | chr5 | - | 55633753 | 55633902 | 55625846 | 55626659 | 55645788 | 55645895 | 1 | 2 | 0 | 1 | 0 | 0 | 0 | 0 | 0 | 0 | 0 | 3 | 0 | 0 | 0 | 10 | 0 | 1 | 0 | 0 | 298 | 149 | 1 | 1 | 1 | 1 | NA | 1 | NA | 0 | 1 | 0 | NA | NA | 0.667 |
| SE | 27781 | ENSG00000177058 | SLC38A9 | chr5 | - | 55633753 | 55633902 | 55627890 | 55627980 | 55645788 | 55645895 | 17 | 16 | 14 | 13 | 15 | 0 | 0 | 0 | 0 | 0 | 0 | 32 | 7 | 7 | 0 | 6 | 2 | 0 | 0 | 0 | 298 | 149 | 1 | 1 | 1 | 1 | 1 | 1 | 1 | 0 | 0.889 | 1 | 1 | NA | 0.278 |
| SE | 27782 | ENSG00000177058 | SLC38A9 | chr5 | - | 55635543 | 55635657 | 55633395 | 55633902 | 55645788 | 55645868 | 103 | 58 | 53 | 90 | 254 | 0 | 0 | 0 | 0 | 0 | 10 | 68 | 44 | 31 | 21 | 0 | 3 | 0 | 0 | 0 | 263 | 149 | 1 | 1 | 1 | 1 | 1 | 1 | 1 | 1 | 0.928 | 1 | 1 | 1 | 0.014 |
| SE | 27783 | ENSG00000177058 | SLC38A9 | chr5 | - | 55645788 | 55645895 | 55633694 | 55635657 | 55649206 | 55649314 | 107 | 66 | 55 | 91 | 293 | 2 | 0 | 0 | 2 | 1 | 41 | 99 | 50 | 44 | 39 | 0 | 0 | 0 | 0 | 0 | 256 | 149 | 1 | 1 | 0.969 | 1 | 1 | 0.964 | 0.994 | 1 | 1 | 1 | 1 | 1 | -0.015 |
| SE | 27790 | ENSG00000177058 | SLC38A9 | chr5 | - | 55664692 | 55664863 | 55656714 | 55656774 | 55697845 | 55697992 | 71 | 66 | 46 | 48 | 126 | 0 | 0 | 0 | 0 | 0 | 27 | 36 | 14 | 19 | 31 | 0 | 0 | 1 | 0 | 0 | 298 | 149 | 1 | 1 | 1 | 1 | 1 | 1 | 1 | 1 | 1 | 0.875 | 1 | 1 | 0.025 |
| SE | 27791 | ENSG00000177058 | SLC38A9 | chr5 | - | 55669227 | 55669321 | 55664559 | 55664863 | 55669556 | 55669620 | 78 | 68 | 43 | 76 | 79 | 1 | 1 | 3 | 0 | 0 | 12 | 48 | 20 | 26 | 4 | 0 | 0 | 0 | 0 | 0 | 243 | 149 | 1 | 1 | 0.98 | 0.977 | 0.898 | 1 | 1 | 1 | 1 | 1 | 1 | 1 | -0.029 |
| SE | 27792 | ENSG00000177058 | SLC38A9 | chr5 | - | 55672562 | 55672695 | 55669757 | 55669879 | 55697845 | 55697992 | 133 | 98 | 93 | 107 | 160 | 2 | 0 | 1 | 1 | 0 | 40 | 90 | 41 | 46 | 4 | 0 | 0 | 0 | 0 | 0 | 282 | 149 | 1 | 1 | 0.972 | 1 | 0.98 | 0.983 | 1 | 1 | 1 | 1 | 1 | 1 | -0.013 |
| SE | 27793 | ENSG00000177058 | SLC38A9 | chr5 | - | 55672562 | 55672822 | 55669757 | 55669879 | 55697845 | 55697994 | 35 | 27 | 25 | 18 | 32 | 2 | 0 | 1 | 1 | 0 | 9 | 22 | 11 | 22 | 1 | 0 | 0 | 0 | 0 | 0 | 298 | 149 | 1 | 1 | 0.897 | 1 | 0.926 | 0.9 | 1 | 1 | 1 | 1 | 1 | 1 | -0.055 |
| SE | 27796 | ENSG00000177058 | SLC38A9 | chr5 | - | 55702738 | 55702894 | 55697845 | 55697992 | 55704042 | 55704140 | 11 | 6 | 6 | 10 | 10 | 0 | 1 | 0 | 0 | 0 | 5 | 22 | 1 | 0 | 16 | 0 | 0 | 0 | 0 | 0 | 298 | 149 | 1 | 1 | 1 | 0.75 | 1 | 1 | 1 | 1 | 1 | 1 | NA | 1 | -0.05 |
| SE | 27799 | ENSG00000177058 | SLC38A9 | chr5 | - | 55704042 | 55704140 | 55697958 | 55697992 | 55704321 | 55704434 | 3 | 1 | 0 | 0 | 2 | 17 | 18 | 8 | 6 | 8 | 0 | 0 | 0 | 0 | 0 | 3 | 6 | 5 | 0 | 0 | 247 | 149 | 1 | 1 | 0.096 | 0.032 | 0 | 0 | 0.131 | 0 | 0 | 0 | NA | NA | 0.052 |
| SE | 27800 | ENSG00000177058 | SLC38A9 | chr5 | - | 55704042 | 55704140 | 55702738 | 55702894 | 55704321 | 55704434 | 5 | 2 | 0 | 0 | 2 | 1 | 4 | 2 | 3 | 0 | 0 | 0 | 0 | 0 | 0 | 0 | 0 | 0 | 3 | 0 | 247 | 149 | 1 | 1 | 0.751 | 0.232 | 0 | 0 | 1 | NA | NA | NA | 0 | NA | 0.397 |
| SE | 27803 | ENSG00000177058 | SLC38A9 | chr5 | - | 55704321 | 55704434 | 55697845 | 55697992 | 55712216 | 55712293 | 15 | 12 | 6 | 4 | 8 | 0 | 0 | 2 | 2 | 0 | 3 | 5 | 3 | 0 | 0 | 0 | 0 | 0 | 0 | 0 | 262 | 149 | 1 | 1 | 1 | 1 | 0.63 | 0.532 | 1 | 1 | 1 | 1 | NA | NA | -0.168 |
| SE | 27804 | ENSG00000177058 | SLC38A9 | chr5 | - | 55704321 | 55704434 | 55702738 | 55702894 | 55711451 | 55711499 | 32 | 21 | 9 | 19 | 32 | 1 | 0 | 0 | 0 | 4 | 0 | 20 | 2 | 27 | 0 | 0 | 0 | 0 | 0 | 0 | 262 | 149 | 1 | 1 | 0.948 | 1 | 1 | 1 | 0.82 | NA | 1 | 1 | 1 | NA | -0.046 |
| SE | 27805 | ENSG00000177058 | SLC38A9 | chr5 | - | 55711351 | 55711499 | 55697845 | 55697992 | 55712216 | 55712291 | 3 | 0 | 1 | 2 | 6 | 0 | 0 | 2 | 2 | 0 | 0 | 0 | 3 | 0 | 0 | 0 | 0 | 0 | 0 | 0 | 297 | 149 | 1 | 1 | 1 | NA | 0.201 | 0.334 | 1 | NA | NA | 1 | NA | NA | -0.366 |
| SE | 27806 | ENSG00000177058 | SLC38A9 | chr5 | - | 55711351 | 55711499 | 55704321 | 55704434 | 55712216 | 55712291 | 3 | 0 | 1 | 2 | 6 | 2 | 0 | 2 | 0 | 1 | 0 | 0 | 3 | 2 | 0 | 0 | 0 | 0 | 0 | 0 | 297 | 149 | 1 | 1 | 0.429 | NA | 0.201 | 1 | 0.751 | NA | NA | 1 | 1 | NA | -0.405 |
| SE | 27807 | ENSG00000177058 | SLC38A9 | chr5 | - | 55711451 | 55711499 | 55697845 | 55697992 | 55712216 | 55712261 | 45 | 28 | 43 | 51 | 43 | 0 | 0 | 2 | 2 | 0 | 17 | 18 | 22 | 7 | 4 | 0 | 0 | 0 | 0 | 0 | 197 | 149 | 1 | 1 | 1 | 1 | 0.942 | 0.951 | 1 | 1 | 1 | 1 | 1 | 1 | -0.021 |
| SE | 27808 | ENSG00000177058 | SLC38A9 | chr5 | - | 55711451 | 55711499 | 55704274 | 55704434 | 55712216 | 55712261 | 34 | 24 | 11 | 23 | 32 | 2 | 0 | 2 | 0 | 1 | 0 | 21 | 7 | 24 | 0 | 0 | 0 | 0 | 0 | 0 | 197 | 149 | 1 | 1 | 0.928 | 1 | 0.806 | 1 | 0.96 | NA | 1 | 1 | 1 | NA | -0.061 |
| SE | 27809 | ENSG00000109814 | UGDH | chr4 | - | 39509759 | 39509907 | 39508565 | 39508660 | 39510352 | 39510550 | 105 | 94 | 141 | 157 | 150 | 1 | 0 | 0 | 0 | 0 | 82 | 99 | 74 | 80 | 43 | 0 | 0 | 0 | 0 | 0 | 297 | 149 | 1 | 1 | 0.981 | 1 | 1 | 1 | 1 | 1 | 1 | 1 | 1 | 1 | -0.004 |
| SE | 27810 | ENSG00000109814 | UGDH | chr4 | - | 39510660 | 39510861 | 39510352 | 39510550 | 39514082 | 39514184 | 79 | 31 | 87 | 81 | 141 | 0 | 2 | 1 | 2 | 0 | 100 | 21 | 49 | 70 | 8 | 0 | 0 | 0 | 2 | 0 | 298 | 149 | 1 | 1 | 1 | 0.886 | 0.978 | 0.953 | 1 | 1 | 1 | 1 | 0.946 | 1 | -0.026 |
| SE | 27811 | ENSG00000109814 | UGDH | chr4 | - | 39514082 | 39514180 | 39510696 | 39510861 | 39521350 | 39521519 | 9 | 0 | 8 | 9 | 27 | 0 | 0 | 0 | 0 | 0 | 19 | 1 | 2 | 7 | 0 | 0 | 0 | 0 | 0 | 1 | 247 | 149 | 1 | 1 | 1 | NA | 1 | 1 | 1 | 1 | 1 | 1 | 1 | 0 | 0.2 |
| SE | 27812 | ENSG00000109814 | UGDH | chr4 | - | 39514082 | 39514184 | 39510660 | 39510861 | 39521350 | 39521519 | 79 | 38 | 61 | 59 | 97 | 0 | 0 | 0 | 0 | 0 | 58 | 23 | 41 | 50 | 5 | 0 | 0 | 0 | 0 | 1 | 251 | 149 | 1 | 1 | 1 | 1 | 1 | 1 | 1 | 1 | 1 | 1 | 1 | 0.748 | 0.05 |
| SE | 27813 | ENSG00000109814 | UGDH | chr4 | - | 39521350 | 39521519 | 39514082 | 39514184 | 39527282 | 39527437 | 113 | 63 | 94 | 77 | 97 | 0 | 0 | 0 | 0 | 2 | 69 | 43 | 52 | 62 | 7 | 0 | 0 | 0 | 0 | 0 | 298 | 149 | 1 | 1 | 1 | 1 | 1 | 1 | 0.96 | 1 | 1 | 1 | 1 | 1 | -0.008 |
| SE | 27815 | ENSG00000109814 | UGDH | chr4 | - | 39521350 | 39521544 | 39514082 | 39514184 | 39527282 | 39527437 | 71 | 39 | 53 | 51 | 70 | 0 | 0 | 0 | 0 | 2 | 47 | 22 | 39 | 44 | 5 | 0 | 0 | 0 | 0 | 0 | 298 | 149 | 1 | 1 | 1 | 1 | 1 | 1 | 0.946 | 1 | 1 | 1 | 1 | 1 | -0.011 |
| SE | 27820 | ENSG00000165521 | EML5 | chr14 | - | 88616141 | 88616242 | 88612430 | 88615854 | 88616725 | 88616879 | 13 | 7 | 12 | 5 | 37 | 5 | 1 | 2 | 2 | 4 | 0 | 0 | 9 | 1 | 0 | 0 | 0 | 0 | 0 | 0 | 250 | 149 | 1 | 1 | 0.608 | 0.807 | 0.781 | 0.598 | 0.846 | NA | NA | 1 | 1 | NA | -0.272 |
| SE | 27821 | ENSG00000165521 | EML5 | chr14 | - | 88622603 | 88622718 | 88621112 | 88621301 | 88624969 | 88625127 | 5 | 13 | 15 | 10 | 7 | 0 | 1 | 3 | 2 | 0 | 0 | 0 | 3 | 0 | 0 | 0 | 0 | 0 | 0 | 0 | 264 | 149 | 1 | 1 | 1 | 0.88 | 0.738 | 0.738 | 1 | NA | NA | 1 | NA | NA | -0.129 |
| SE | 27822 | ENSG00000165521 | EML5 | chr14 | - | 88622603 | 88622718 | 88621112 | 88621301 | 88626837 | 88626900 | 3 | 5 | 12 | 3 | 5 | 0 | 0 | 0 | 0 | 1 | 0 | 0 | 3 | 0 | 0 | 0 | 0 | 0 | 0 | 0 | 264 | 149 | 1 | 1 | 1 | 1 | 1 | 1 | 0.738 | NA | NA | 1 | NA | NA | -0.052 |
| SE | 27824 | ENSG00000165521 | EML5 | chr14 | - | 88624969 | 88625127 | 88622603 | 88622718 | 88626837 | 88626900 | 5 | 13 | 12 | 13 | 5 | 0 | 1 | 0 | 0 | 0 | 0 | 0 | 0 | 5 | 0 | 0 | 0 | 0 | 0 | 0 | 298 | 149 | 1 | 1 | 1 | 0.867 | 1 | 1 | 1 | NA | NA | NA | 1 | NA | -0.027 |
| SE | 27825 | ENSG00000165521 | EML5 | chr14 | - | 88627645 | 88627819 | 88626837 | 88627046 | 88634468 | 88634489 | 11 | 12 | 11 | 14 | 25 | 1 | 0 | 2 | 3 | 0 | 0 | 4 | 1 | 0 | 0 | 0 | 0 | 0 | 0 | 0 | 298 | 149 | 1 | 1 | 0.846 | 1 | 0.733 | 0.7 | 1 | NA | 1 | 1 | NA | NA | -0.144 |
| SE | 27826 | ENSG00000165521 | EML5 | chr14 | - | 88634468 | 88634489 | 88626837 | 88627046 | 88638808 | 88638907 | 1 | 0 | 3 | 7 | 0 | 0 | 0 | 0 | 0 | 0 | 0 | 0 | 0 | 0 | 0 | 11 | 0 | 0 | 0 | 0 | 170 | 149 | 1 | 1 | 1 | NA | 1 | 1 | NA | 0 | NA | NA | NA | NA | 1 |
| SE | 27827 | ENSG00000165521 | EML5 | chr14 | - | 88638808 | 88638907 | 88634468 | 88634489 | 88642892 | 88643022 | 11 | 19 | 16 | 18 | 37 | 0 | 0 | 0 | 0 | 0 | 0 | 8 | 0 | 6 | 0 | 0 | 0 | 1 | 0 | 0 | 248 | 149 | 1 | 1 | 1 | 1 | 1 | 1 | 1 | NA | 1 | 0 | 1 | NA | 0.333 |
| SE | 27829 | ENSG00000165521 | EML5 | chr14 | - | 88649911 | 88649926 | 88644432 | 88644511 | 88657375 | 88657502 | 0 | 0 | 3 | 0 | 0 | 5 | 5 | 13 | 5 | 14 | 0 | 0 | 0 | 0 | 0 | 0 | 0 | 1 | 0 | 0 | 164 | 149 | 1 | 1 | 0 | 0 | 0.173 | 0 | 0 | NA | NA | 0 | NA | NA | 0.035 |
| SE | 27830 | ENSG00000165521 | EML5 | chr14 | - | 88688270 | 88688359 | 88687215 | 88687327 | 88694306 | 88694407 | 12 | 9 | 6 | 11 | 6 | 0 | 0 | 1 | 1 | 0 | 0 | 0 | 1 | 0 | 0 | 0 | 0 | 0 | 0 | 0 | 238 | 149 | 1 | 1 | 1 | 1 | 0.79 | 0.873 | 1 | NA | NA | 1 | NA | NA | -0.067 |
| SE | 27831 | ENSG00000165521 | EML5 | chr14 | - | 88688270 | 88688473 | 88687215 | 88687327 | 88694306 | 88694407 | 19 | 23 | 13 | 10 | 19 | 0 | 0 | 1 | 1 | 0 | 0 | 0 | 5 | 0 | 0 | 0 | 0 | 0 | 0 | 0 | 298 | 149 | 1 | 1 | 1 | 1 | 0.867 | 0.833 | 1 | NA | NA | 1 | NA | NA | -0.06 |
| SE | 27832 | ENSG00000165521 | EML5 | chr14 | - | 88705719 | 88705873 | 88705481 | 88705588 | 88706258 | 88706426 | 1 | 3 | 0 | 0 | 0 | 7 | 2 | 7 | 3 | 18 | 0 | 0 | 0 | 0 | 0 | 0 | 4 | 1 | 0 | 0 | 298 | 149 | 1 | 1 | 0.067 | 0.429 | 0 | 0 | 0 | NA | 0 | 0 | NA | NA | 0.099 |
| SE | 27833 | ENSG00000165521 | EML5 | chr14 | - | 88706258 | 88706426 | 88705481 | 88705588 | 88712270 | 88712483 | 16 | 6 | 13 | 6 | 31 | 3 | 6 | 1 | 5 | 15 | 0 | 6 | 6 | 1 | 0 | 0 | 0 | 1 | 5 | 0 | 298 | 149 | 1 | 1 | 0.727 | 0.333 | 0.867 | 0.375 | 0.508 | NA | 1 | 0.75 | 0.091 | NA | -0.052 |
| SE | 27836 | ENSG00000231769 | NA | chr6 | - | 46122947 | 46123109 | 46102539 | 46102659 | 46129503 | 46129831 | 11 | 4 | 2 | 3 | 1 | 6 | 0 | 1 | 1 | 12 | 0 | 1 | 7 | 0 | 0 | 0 | 4 | 1 | 0 | 0 | 298 | 149 | 1 | 1 | 0.478 | 1 | 0.5 | 0.6 | 0.04 | NA | 0.111 | 0.778 | NA | NA | 0.079 |
| SE | 27837 | ENSG00000231769 | NA | chr6 | - | 46122947 | 46123109 | 46102539 | 46102659 | 46129589 | 46129706 | 9 | 2 | 2 | 0 | 6 | 12 | 1 | 3 | 2 | 7 | 0 | 2 | 4 | 0 | 0 | 0 | 16 | 2 | 0 | 0 | 298 | 149 | 1 | 1 | 0.273 | 0.5 | 0.25 | 0 | 0.3 | NA | 0.059 | 0.5 | NA | NA | -0.015 |
| SE | 27839 | ENSG00000101493 | ZNF516 | chr18 | - | 76371466 | 76371571 | 76361767 | 76362557 | 76378854 | 76380285 | 21 | 20 | 13 | 24 | 2 | 0 | 1 | 0 | 0 | 1 | 33 | 11 | 33 | 22 | 14 | 0 | 0 | 0 | 1 | 0 | 254 | 149 | 1 | 1 | 1 | 0.921 | 1 | 1 | 0.54 | 1 | 1 | 1 | 0.928 | 1 | -0.093 |
| SE | 27840 | ENSG00000101493 | ZNF516 | chr18 | - | 76463027 | 76463141 | 76441244 | 76443211 | 76495143 | 76495242 | 6 | 0 | 2 | 0 | 0 | 1 | 4 | 1 | 2 | 0 | 4 | 2 | 2 | 0 | 0 | 5 | 0 | 8 | 1 | 3 | 263 | 149 | 1 | 1 | 0.773 | 0 | 0.531 | 0 | NA | 0.312 | 1 | 0.124 | 0 | 0 | 0.039 |
| SE | 27845 | ENSG00000272690 | LINC02018 | chr3 | + | 75437055 | 75437275 | 75435365 | 75435413 | 75443507 | 75443646 | 0 | 0 | 1 | 0 | 0 | 0 | 3 | 0 | 0 | 0 | 0 | 1 | 2 | 0 | 0 | 15 | 4 | 0 | 2 | 0 | 298 | 149 | 1 | 1 | NA | 0 | 1 | NA | NA | 0 | 0.111 | 1 | 0 | NA | 0.222 |
| SE | 27854 | ENSG00000272690 | LINC02018 | chr3 | + | 75439784 | 75439846 | 75435316 | 75435413 | 75443507 | 75443646 | 1 | 0 | 0 | 2 | 0 | 0 | 3 | 0 | 0 | 0 | 0 | 0 | 0 | 0 | 0 | 15 | 4 | 0 | 2 | 0 | 211 | 149 | 1 | 1 | 1 | 0 | NA | 1 | NA | 0 | 0 | NA | 0 | NA | 0.667 |
| SE | 27856 | ENSG00000272690 | LINC02018 | chr3 | + | 75439784 | 75439846 | 75437249 | 75437275 | 75443507 | 75443646 | 1 | 0 | 0 | 2 | 0 | 0 | 0 | 1 | 0 | 0 | 0 | 1 | 2 | 0 | 0 | 0 | 0 | 0 | 0 | 0 | 211 | 149 | 1 | 1 | 1 | NA | 0 | 1 | NA | NA | 1 | 1 | NA | NA | -0.333 |
| SE | 27863 | ENSG00000272690 | LINC02018 | chr3 | + | 75451209 | 75451402 | 75443507 | 75443646 | 75537631 | 75538029 | 1 | 0 | 0 | 0 | 7 | 0 | 0 | 0 | 0 | 0 | 8 | 0 | 0 | 0 | 0 | 0 | 0 | 0 | 1 | 0 | 298 | 149 | 1 | 1 | 1 | NA | NA | NA | 1 | 1 | NA | NA | 0 | NA | 0.5 |
| SE | 27875 | ENSG00000213468 | FIRRE | chrX | - | 131711650 | 131711720 | 131691542 | 131692636 | 131749305 | 131749458 | 0 | 0 | 0 | 0 | 0 | 0 | 2 | 1 | 0 | 0 | 0 | 0 | 1 | 0 | 0 | 0 | 0 | 0 | 0 | 0 | 219 | 149 | 1 | 1 | NA | 0 | 0 | NA | NA | NA | NA | 1 | NA | NA | -1 |
| SE | 27876 | ENSG00000213468 | FIRRE | chrX | - | 131711650 | 131711720 | 131709497 | 131709556 | 131749305 | 131749458 | 0 | 0 | 0 | 0 | 2 | 0 | 0 | 0 | 0 | 0 | 0 | 0 | 1 | 0 | 0 | 0 | 0 | 0 | 1 | 0 | 219 | 149 | 1 | 1 | NA | NA | NA | NA | 1 | NA | NA | 1 | 0 | NA | 0.5 |
| SE | 27877 | ENSG00000213468 | FIRRE | chrX | - | 131711650 | 131711744 | 131691557 | 131692636 | 131743095 | 131743248 | 0 | 0 | 0 | 0 | 2 | 0 | 0 | 0 | 0 | 0 | 0 | 0 | 0 | 0 | 0 | 4 | 0 | 0 | 0 | 0 | 243 | 149 | 1 | 1 | NA | NA | NA | NA | 1 | 0 | NA | NA | NA | NA | 1 |
| SE | 27880 | ENSG00000213468 | FIRRE | chrX | - | 131743095 | 131743248 | 131691529 | 131692636 | 131749305 | 131749458 | 0 | 2 | 0 | 4 | 2 | 0 | 2 | 1 | 0 | 0 | 11 | 0 | 1 | 4 | 0 | 0 | 0 | 0 | 0 | 0 | 298 | 149 | 1 | 1 | NA | 0.333 | 0 | 1 | 1 | 1 | NA | 1 | 1 | NA | -0.417 |
| SE | 27882 | ENSG00000213468 | FIRRE | chrX | - | 131756610 | 131756709 | 131755597 | 131755746 | 131768548 | 131768696 | 19 | 27 | 19 | 27 | 36 | 0 | 0 | 0 | 0 | 0 | 36 | 39 | 35 | 30 | 46 | 0 | 1 | 0 | 0 | 0 | 248 | 149 | 1 | 1 | 1 | 1 | 1 | 1 | 1 | 1 | 0.959 | 1 | 1 | 1 | 0.008 |
| SE | 27883 | ENSG00000213468 | FIRRE | chrX | - | 131756642 | 131756709 | 131755597 | 131755746 | 131768548 | 131768696 | 3 | 3 | 2 | 1 | 2 | 0 | 0 | 0 | 0 | 0 | 0 | 9 | 5 | 4 | 7 | 0 | 1 | 0 | 0 | 0 | 216 | 149 | 1 | 1 | 1 | 1 | 1 | 1 | 1 | NA | 0.861 | 1 | 1 | 1 | 0.035 |
| SE | 27885 | ENSG00000213468 | FIRRE | chrX | - | 131783886 | 131784039 | 131777622 | 131777744 | 131794296 | 131794466 | 2 | 3 | 6 | 10 | 2 | 0 | 0 | 1 | 0 | 0 | 5 | 11 | 4 | 11 | 5 | 0 | 0 | 0 | 0 | 0 | 298 | 149 | 1 | 1 | 1 | 1 | 0.75 | 1 | 1 | 1 | 1 | 1 | 1 | 1 | -0.05 |
| SE | 27886 | ENSG00000213468 | FIRRE | chrX | - | 131785156 | 131785258 | 131783886 | 131784039 | 131793993 | 131794115 | 2 | 9 | 10 | 5 | 5 | 0 | 2 | 0 | 0 | 0 | 0 | 25 | 4 | 6 | 0 | 0 | 5 | 0 | 3 | 0 | 251 | 149 | 1 | 1 | 1 | 0.728 | 1 | 1 | 1 | NA | 0.748 | 1 | 0.543 | NA | 0.182 |
| SE | 27887 | ENSG00000213468 | FIRRE | chrX | - | 131785156 | 131785258 | 131783886 | 131784039 | 131794296 | 131794466 | 1 | 9 | 8 | 5 | 0 | 0 | 1 | 0 | 0 | 0 | 0 | 23 | 4 | 6 | 0 | 0 | 0 | 0 | 0 | 0 | 251 | 149 | 1 | 1 | 1 | 0.842 | 1 | 1 | NA | NA | 1 | 1 | 1 | NA | -0.039 |
| SE | 27888 | ENSG00000213468 | FIRRE | chrX | - | 131785156 | 131785258 | 131783886 | 131784039 | 131794323 | 131794466 | 1 | 7 | 7 | 5 | 0 | 0 | 3 | 0 | 0 | 0 | 0 | 17 | 5 | 6 | 0 | 0 | 0 | 0 | 0 | 0 | 251 | 149 | 1 | 1 | 1 | 0.581 | 1 | 1 | NA | NA | 1 | 1 | 1 | NA | -0.105 |
| SE | 27889 | ENSG00000213468 | FIRRE | chrX | - | 131793993 | 131794115 | 131783886 | 131784039 | 131794296 | 131794466 | 0 | 6 | 4 | 0 | 9 | 0 | 1 | 0 | 0 | 0 | 0 | 14 | 0 | 3 | 0 | 0 | 0 | 0 | 0 | 0 | 271 | 149 | 1 | 1 | NA | 0.767 | 1 | NA | 1 | NA | 1 | NA | 1 | NA | -0.078 |
| SE | 27890 | ENSG00000213468 | FIRRE | chrX | - | 131793993 | 131794115 | 131783886 | 131784039 | 131794323 | 131794466 | 0 | 2 | 0 | 0 | 0 | 0 | 3 | 0 | 0 | 0 | 0 | 5 | 0 | 0 | 0 | 0 | 0 | 0 | 0 | 0 | 271 | 149 | 1 | 1 | NA | 0.268 | NA | NA | NA | NA | 1 | NA | NA | NA | -0.732 |
| SE | 27891 | ENSG00000213468 | FIRRE | chrX | - | 131793993 | 131794115 | 131785156 | 131785258 | 131794296 | 131794466 | 1 | 6 | 7 | 0 | 14 | 0 | 2 | 1 | 0 | 0 | 0 | 17 | 0 | 0 | 0 | 0 | 6 | 0 | 0 | 0 | 271 | 149 | 1 | 1 | 1 | 0.623 | 0.794 | NA | 1 | NA | 0.609 | NA | NA | NA | 0.245 |
| SE | 27892 | ENSG00000213468 | FIRRE | chrX | - | 131793993 | 131794115 | 131785156 | 131785258 | 131794323 | 131794466 | 1 | 2 | 1 | 0 | 5 | 0 | 0 | 0 | 0 | 0 | 0 | 2 | 0 | 0 | 0 | 0 | 0 | 1 | 0 | 0 | 271 | 149 | 1 | 1 | 1 | 1 | 1 | NA | 1 | NA | 1 | 0 | NA | NA | 0.5 |
| SE | 27894 | ENSG00000213468 | FIRRE | chrX | - | 131804971 | 131805037 | 131794113 | 131794466 | 131825221 | 131825365 | 3 | 0 | 3 | 0 | 0 | 0 | 2 | 0 | 1 | 1 | 0 | 0 | 0 | 0 | 0 | 0 | 1 | 8 | 4 | 0 | 215 | 149 | 1 | 1 | 1 | 0 | 1 | 0 | 0 | NA | 0 | 0 | 0 | NA | 0.4 |
| SE | 27895 | ENSG00000150977 | RILPL2 | chr12 | - | 123423043 | 123423157 | 123415038 | 123415921 | 123430507 | 123430659 | 320 | 178 | 174 | 263 | 247 | 2 | 1 | 0 | 0 | 0 | 1029 | 403 | 532 | 497 | 858 | 6 | 2 | 9 | 0 | 0 | 263 | 149 | 1 | 1 | 0.989 | 0.99 | 1 | 1 | 1 | 0.99 | 0.991 | 0.971 | 1 | 1 | 0.005 |
| SE | 27903 | ENSG00000118564 | FBXL5 | chr4 | - | 15612265 | 15612414 | 15604380 | 15605799 | 15625251 | 15625977 | 765 | 512 | 532 | 642 | 991 | 0 | 0 | 3 | 0 | 0 | 308 | 574 | 477 | 563 | 91 | 0 | 0 | 1 | 0 | 0 | 298 | 149 | 1 | 1 | 1 | 1 | 0.989 | 1 | 1 | 1 | 1 | 0.996 | 1 | 1 | -0.001 |
| SE | 27904 | ENSG00000118564 | FBXL5 | chr4 | - | 15626872 | 15626955 | 15625251 | 15625977 | 15627884 | 15628033 | 672 | 485 | 439 | 584 | 789 | 0 | 0 | 0 | 2 | 0 | 284 | 535 | 527 | 553 | 124 | 0 | 5 | 0 | 0 | 0 | 232 | 149 | 1 | 1 | 1 | 1 | 1 | 0.995 | 1 | 1 | 0.986 | 1 | 1 | 1 | 0.002 |
| SE | 27908 | ENSG00000118564 | FBXL5 | chr4 | - | 15638507 | 15638694 | 15636372 | 15636676 | 15640787 | 15640883 | 796 | 644 | 608 | 644 | 1590 | 0 | 0 | 0 | 0 | 1 | 458 | 553 | 682 | 666 | 94 | 0 | 3 | 0 | 0 | 0 | 298 | 149 | 1 | 1 | 1 | 1 | 1 | 1 | 0.999 | 1 | 0.989 | 1 | 1 | 1 | 0.002 |
| SE | 27909 | ENSG00000118564 | FBXL5 | chr4 | - | 15640787 | 15640883 | 15638507 | 15638694 | 15644492 | 15644657 | 775 | 652 | 464 | 684 | 1159 | 4 | 0 | 1 | 0 | 3 | 568 | 550 | 671 | 836 | 100 | 0 | 1 | 0 | 0 | 0 | 245 | 149 | 1 | 1 | 0.992 | 1 | 0.996 | 1 | 0.996 | 1 | 0.997 | 1 | 1 | 1 | -0.003 |
| SE | 27912 | ENSG00000118564 | FBXL5 | chr4 | - | 15644492 | 15644657 | 15640787 | 15640883 | 15656228 | 15656342 | 495 | 431 | 274 | 457 | 540 | 0 | 0 | 0 | 0 | 0 | 418 | 377 | 481 | 628 | 74 | 0 | 0 | 2 | 4 | 0 | 298 | 149 | 1 | 1 | 1 | 1 | 1 | 1 | 1 | 1 | 1 | 0.992 | 0.987 | 1 | 0.004 |
| SE | 27913 | ENSG00000118564 | FBXL5 | chr4 | - | 15644492 | 15644657 | 15640787 | 15640883 | 15659726 | 15659922 | 500 | 440 | 279 | 458 | 546 | 1 | 0 | 0 | 0 | 0 | 418 | 391 | 482 | 645 | 74 | 0 | 0 | 0 | 0 | 0 | 298 | 149 | 1 | 1 | 0.996 | 1 | 1 | 1 | 1 | 1 | 1 | 1 | 1 | 1 | -0.001 |
| SE | 27915 | ENSG00000118564 | FBXL5 | chr4 | - | 15644492 | 15644708 | 15640787 | 15640883 | 15656228 | 15656342 | 503 | 437 | 274 | 462 | 547 | 0 | 0 | 0 | 0 | 0 | 428 | 385 | 508 | 665 | 74 | 0 | 0 | 2 | 4 | 0 | 298 | 149 | 1 | 1 | 1 | 1 | 1 | 1 | 1 | 1 | 1 | 0.992 | 0.988 | 1 | 0.004 |
| SE | 27916 | ENSG00000118564 | FBXL5 | chr4 | - | 15644492 | 15644708 | 15640787 | 15640883 | 15659726 | 15659864 | 566 | 485 | 297 | 488 | 610 | 1 | 0 | 0 | 0 | 0 | 446 | 423 | 515 | 674 | 77 | 0 | 0 | 0 | 0 | 0 | 298 | 149 | 1 | 1 | 0.996 | 1 | 1 | 1 | 1 | 1 | 1 | 1 | 1 | 1 | -0.001 |
| SE | 27917 | ENSG00000118564 | FBXL5 | chr4 | - | 15656228 | 15656342 | 15640840 | 15640883 | 15659726 | 15659922 | 2 | 2 | 2 | 2 | 5 | 1 | 0 | 0 | 0 | 0 | 0 | 0 | 16 | 22 | 0 | 0 | 0 | 0 | 0 | 0 | 263 | 149 | 1 | 1 | 0.531 | 1 | 1 | 1 | 1 | NA | NA | 1 | 1 | NA | -0.094 |
| SE | 27918 | ENSG00000118564 | FBXL5 | chr4 | - | 15656228 | 15656342 | 15644492 | 15644657 | 15659726 | 15659922 | 2 | 2 | 2 | 4 | 7 | 5 | 9 | 5 | 3 | 8 | 0 | 0 | 23 | 23 | 0 | 0 | 14 | 10 | 22 | 0 | 263 | 149 | 1 | 1 | 0.185 | 0.112 | 0.185 | 0.43 | 0.331 | NA | 0 | 0.566 | 0.372 | NA | -0.064 |
| SE | 27919 | ENSG00000118564 | FBXL5 | chr4 | - | 15656228 | 15656342 | 15644492 | 15644657 | 15659740 | 15659867 | 4 | 0 | 0 | 2 | 6 | 1 | 1 | 1 | 0 | 6 | 0 | 4 | 20 | 8 | 0 | 0 | 0 | 2 | 9 | 0 | 263 | 149 | 1 | 1 | 0.694 | 0 | 0 | 1 | 0.362 | NA | 1 | 0.85 | 0.335 | NA | -0.317 |
| SE | 27921 | ENSG00000118564 | FBXL5 | chr4 | - | 15656228 | 15656342 | 15644492 | 15644708 | 15659740 | 15659867 | 11 | 4 | 0 | 6 | 8 | 18 | 26 | 0 | 9 | 20 | 10 | 11 | 39 | 38 | 0 | 0 | 11 | 16 | 9 | 0 | 263 | 149 | 1 | 1 | 0.257 | 0.08 | NA | 0.274 | 0.185 | 1 | 0.362 | 0.58 | 0.705 | NA | -0.463 |
| SE | 27922 | ENSG00000138686 | BBS7 | chr4 | - | 121833230 | 121833395 | 121828618 | 121828728 | 121835143 | 121835283 | 35 | 19 | 45 | 43 | 129 | 0 | 1 | 0 | 1 | 0 | 0 | 48 | 10 | 4 | 0 | 0 | 0 | 0 | 0 | 0 | 298 | 149 | 1 | 1 | 1 | 0.905 | 1 | 0.956 | 1 | NA | 1 | 1 | 1 | NA | -0.028 |
| SE | 27923 | ENSG00000138686 | BBS7 | chr4 | - | 121839630 | 121839696 | 121835143 | 121835283 | 121843926 | 121844001 | 56 | 40 | 48 | 60 | 168 | 0 | 0 | 0 | 2 | 0 | 0 | 22 | 9 | 3 | 4 | 0 | 0 | 0 | 0 | 0 | 215 | 149 | 1 | 1 | 1 | 1 | 1 | 0.954 | 1 | NA | 1 | 1 | 1 | 1 | -0.009 |
| SE | 27924 | ENSG00000138686 | BBS7 | chr4 | - | 121858991 | 121859178 | 121855488 | 121855561 | 121861503 | 121861679 | 44 | 33 | 33 | 35 | 87 | 1 | 2 | 1 | 0 | 0 | 26 | 21 | 8 | 1 | 0 | 0 | 0 | 0 | 0 | 0 | 298 | 149 | 1 | 1 | 0.957 | 0.892 | 0.943 | 1 | 1 | 1 | 1 | 1 | 1 | NA | -0.042 |
| SE | 27926 | ENSG00000047315 | POLR2B | chr4 | + | 56985279 | 56985457 | 56978868 | 56979004 | 56986353 | 56986426 | 4 | 0 | 2 | 0 | 1 | 76 | 31 | 71 | 61 | 148 | 0 | 6 | 0 | 0 | 0 | 8 | 62 | 27 | 16 | 8 | 298 | 149 | 1 | 1 | 0.026 | 0 | 0.014 | 0 | 0.003 | 0 | 0.046 | 0 | 0 | 0 | -0.001 |
| SE | 27927 | ENSG00000047315 | POLR2B | chr4 | + | 56990747 | 56990898 | 56986353 | 56986426 | 56994403 | 56994516 | 205 | 110 | 182 | 160 | 433 | 6 | 0 | 0 | 2 | 0 | 20 | 138 | 82 | 78 | 41 | 0 | 4 | 0 | 0 | 0 | 298 | 149 | 1 | 1 | 0.945 | 1 | 1 | 0.976 | 1 | 1 | 0.945 | 1 | 1 | 1 | -0.005 |
| SE | 27928 | ENSG00000047315 | POLR2B | chr4 | + | 56994403 | 56994516 | 56990747 | 56990898 | 56994646 | 56994866 | 327 | 159 | 242 | 232 | 420 | 0 | 0 | 0 | 0 | 0 | 45 | 157 | 108 | 67 | 24 | 0 | 4 | 0 | 0 | 0 | 262 | 149 | 1 | 1 | 1 | 1 | 1 | 1 | 1 | 1 | 0.957 | 1 | 1 | 1 | 0.009 |
| SE | 27929 | ENSG00000047315 | POLR2B | chr4 | + | 56995250 | 56995409 | 56994646 | 56994866 | 56999616 | 56999781 | 605 | 368 | 443 | 428 | 827 | 0 | 1 | 0 | 0 | 0 | 113 | 306 | 192 | 164 | 26 | 0 | 0 | 0 | 0 | 0 | 298 | 149 | 1 | 1 | 1 | 0.995 | 1 | 1 | 1 | 1 | 1 | 1 | 1 | 1 | -0.001 |
| SE | 27930 | ENSG00000047315 | POLR2B | chr4 | + | 57005245 | 57005442 | 56999616 | 56999781 | 57005599 | 57005719 | 731 | 548 | 602 | 571 | 876 | 0 | 0 | 0 | 1 | 0 | 103 | 465 | 466 | 299 | 4 | 0 | 0 | 0 | 0 | 0 | 298 | 149 | 1 | 1 | 1 | 1 | 1 | 0.997 | 1 | 1 | 1 | 1 | 1 | 1 | -0.001 |
| SE | 27931 | ENSG00000047315 | POLR2B | chr4 | + | 57010360 | 57010504 | 57006815 | 57007002 | 57010747 | 57010887 | 520 | 371 | 465 | 458 | 606 | 0 | 0 | 1 | 0 | 0 | 83 | 249 | 369 | 196 | 3 | 0 | 0 | 1 | 0 | 0 | 293 | 149 | 1 | 1 | 1 | 1 | 0.996 | 1 | 1 | 1 | 1 | 0.995 | 1 | 1 | 0 |
| SE | 27932 | ENSG00000047315 | POLR2B | chr4 | + | 57010747 | 57010887 | 57009558 | 57010504 | 57010988 | 57011100 | 1045 | 775 | 899 | 948 | 1028 | 0 | 0 | 0 | 2 | 0 | 188 | 593 | 881 | 440 | 21 | 0 | 0 | 0 | 0 | 0 | 289 | 149 | 1 | 1 | 1 | 1 | 1 | 0.996 | 1 | 1 | 1 | 1 | 1 | 1 | -0.001 |
| SE | 27933 | ENSG00000047315 | POLR2B | chr4 | + | 57015501 | 57015656 | 57010988 | 57011100 | 57017042 | 57017241 | 618 | 466 | 479 | 548 | 535 | 0 | 0 | 0 | 0 | 0 | 94 | 349 | 560 | 290 | 12 | 0 | 0 | 3 | 0 | 0 | 298 | 149 | 1 | 1 | 1 | 1 | 1 | 1 | 1 | 1 | 1 | 0.989 | 1 | 1 | 0.002 |
| SE | 27934 | ENSG00000047315 | POLR2B | chr4 | + | 57017559 | 57017728 | 57017042 | 57017241 | 57020898 | 57020995 | 375 | 210 | 370 | 264 | 646 | 0 | 0 | 0 | 0 | 0 | 16 | 226 | 211 | 124 | 3 | 0 | 0 | 0 | 3 | 0 | 298 | 149 | 1 | 1 | 1 | 1 | 1 | 1 | 1 | 1 | 1 | 1 | 0.954 | 1 | 0.009 |
| SE | 27935 | ENSG00000047315 | POLR2B | chr4 | + | 57022151 | 57022246 | 57020898 | 57020995 | 57023329 | 57023580 | 401 | 329 | 434 | 367 | 511 | 2 | 2 | 0 | 3 | 0 | 35 | 352 | 318 | 213 | 21 | 0 | 4 | 2 | 0 | 0 | 244 | 149 | 1 | 1 | 0.992 | 0.99 | 1 | 0.987 | 1 | 1 | 0.982 | 0.99 | 1 | 1 | -0.001 |
| SE | 27936 | ENSG00000047315 | POLR2B | chr4 | + | 57024004 | 57024112 | 57023490 | 57023751 | 57024885 | 57024962 | 652 | 431 | 561 | 541 | 960 | 0 | 0 | 0 | 0 | 0 | 79 | 421 | 321 | 171 | 25 | 0 | 4 | 1 | 0 | 0 | 257 | 149 | 1 | 1 | 1 | 1 | 1 | 1 | 1 | 1 | 0.984 | 0.995 | 1 | 1 | 0.004 |
| SE | 27938 | ENSG00000084092 | NOA1 | chr4 | - | 56968383 | 56968515 | 56966619 | 56966736 | 56973147 | 56973353 | 226 | 153 | 280 | 245 | 523 | 0 | 0 | 2 | 0 | 1 | 251 | 167 | 128 | 106 | 258 | 0 | 0 | 0 | 0 | 0 | 281 | 149 | 1 | 1 | 1 | 1 | 0.987 | 1 | 0.996 | 1 | 1 | 1 | 1 | 1 | -0.003 |
| SE | 27939 | ENSG00000128040 | SPINK2 | chr4 | - | 56811684 | 56811794 | 56809860 | 56810184 | 56820535 | 56820579 | 7 | 3 | 11 | 5 | 9 | 0 | 1 | 0 | 0 | 0 | 0 | 5 | 1 | 2 | 3 | 0 | 0 | 0 | 0 | 0 | 259 | 149 | 1 | 1 | 1 | 0.633 | 1 | 1 | 1 | NA | 1 | 1 | 1 | 1 | -0.073 |
| SE | 27944 | ENSG00000214944 | ARHGEF28 | chr5 | + | 73897961 | 73898093 | 73894392 | 73894575 | 73901183 | 73901284 | 14 | 0 | 0 | 6 | 4 | 1 | 0 | 0 | 0 | 0 | 0 | 0 | 2 | 1 | 0 | 0 | 0 | 0 | 0 | 0 | 281 | 149 | 1 | 1 | 0.881 | NA | NA | 1 | 1 | NA | NA | 1 | 1 | NA | -0.04 |
| SE | 27945 | ENSG00000214944 | ARHGEF28 | chr5 | + | 73909411 | 73909897 | 73904357 | 73904405 | 73911274 | 73911575 | 8 | 0 | 0 | 0 | 0 | 0 | 0 | 0 | 0 | 1 | 0 | 0 | 0 | 1 | 1 | 0 | 0 | 0 | 0 | 0 | 298 | 149 | 1 | 1 | 1 | NA | NA | NA | 0 | NA | NA | NA | 1 | 1 | -0.5 |
| SE | 27951 | ENSG00000128609 | NDUFA5 | chr7 | - | 123550469 | 123550584 | 123545610 | 123545676 | 123557403 | 123557781 | 134 | 145 | 129 | 174 | 189 | 0 | 0 | 0 | 1 | 1 | 32 | 124 | 152 | 62 | 39 | 0 | 0 | 0 | 0 | 0 | 264 | 149 | 1 | 1 | 1 | 1 | 1 | 0.99 | 0.991 | 1 | 1 | 1 | 1 | 1 | -0.004 |
| SE | 27952 | ENSG00000128609 | NDUFA5 | chr7 | - | 123550469 | 123550586 | 123545610 | 123545676 | 123557403 | 123557448 | 214 | 243 | 236 | 313 | 295 | 0 | 0 | 0 | 1 | 1 | 75 | 203 | 254 | 104 | 60 | 0 | 0 | 0 | 0 | 0 | 266 | 149 | 1 | 1 | 1 | 1 | 1 | 0.994 | 0.994 | 1 | 1 | 1 | 1 | 1 | -0.002 |
| SE | 27956 | ENSG00000128609 | NDUFA5 | chr7 | - | 123557403 | 123557448 | 123545610 | 123550586 | 123557774 | 123557801 | 75 | 94 | 100 | 128 | 101 | 3 | 0 | 1 | 0 | 0 | 32 | 79 | 91 | 40 | 18 | 0 | 0 | 0 | 0 | 0 | 194 | 149 | 1 | 1 | 0.95 | 1 | 0.987 | 1 | 1 | 1 | 1 | 1 | 1 | 1 | -0.013 |
| SE | 27957 | ENSG00000128609 | NDUFA5 | chr7 | - | 123557403 | 123557507 | 123550469 | 123550586 | 123557774 | 123557804 | 34 | 37 | 46 | 47 | 54 | 3 | 0 | 1 | 0 | 0 | 21 | 25 | 38 | 17 | 21 | 0 | 0 | 0 | 0 | 0 | 253 | 149 | 1 | 1 | 0.87 | 1 | 0.964 | 1 | 1 | 1 | 1 | 1 | 1 | 1 | -0.033 |
| SE | 27959 | ENSG00000129422 | MTUS1 | chr8 | - | 17675185 | 17675252 | 17655862 | 17656065 | 17684327 | 17684542 | 24 | 16 | 33 | 18 | 40 | 0 | 2 | 0 | 1 | 0 | 0 | 4 | 2 | 11 | 0 | 2 | 0 | 0 | 0 | 0 | 216 | 149 | 1 | 1 | 1 | 0.847 | 1 | 0.925 | 1 | 0 | 1 | 1 | 1 | NA | 0.204 |
| SE | 27960 | ENSG00000129422 | MTUS1 | chr8 | - | 17684327 | 17684542 | 17655862 | 17656065 | 17697256 | 17697670 | 9 | 12 | 25 | 18 | 34 | 0 | 1 | 0 | 2 | 0 | 2 | 0 | 3 | 5 | 0 | 0 | 0 | 0 | 0 | 0 | 298 | 149 | 1 | 1 | 1 | 0.857 | 1 | 0.818 | 1 | 1 | NA | 1 | 1 | NA | -0.065 |
| SE | 27964 | ENSG00000004468 | CD38 | chr4 | + | 15816510 | 15816640 | 15778274 | 15778647 | 15824880 | 15825016 | 134 | 49 | 178 | 98 | 104 | 0 | 0 | 0 | 0 | 0 | 85 | 222 | 222 | 124 | 59 | 5 | 0 | 0 | 0 | 0 | 279 | 149 | 1 | 1 | 1 | 1 | 1 | 1 | 1 | 0.901 | 1 | 1 | 1 | 1 | 0.02 |
| SE | 27967 | ENSG00000004468 | CD38 | chr4 | + | 15834216 | 15834302 | 15816510 | 15816640 | 15838091 | 15838165 | 65 | 34 | 61 | 27 | 54 | 0 | 0 | 0 | 0 | 2 | 5 | 89 | 83 | 52 | 21 | 0 | 4 | 0 | 0 | 0 | 235 | 149 | 1 | 1 | 1 | 1 | 1 | 1 | 0.945 | 1 | 0.934 | 1 | 1 | 1 | 0.002 |
| SE | 27972 | ENSG00000187583 | PLEKHN1 | chr1 | + | 973499 | 973640 | 973185 | 973326 | 973832 | 974051 | 1 | 0 | 0 | 2 | 0 | 0 | 0 | 0 | 2 | 0 | 0 | 0 | 0 | 2 | 0 | 0 | 0 | 0 | 0 | 0 | 290 | 149 | 1 | 1 | 1 | NA | NA | 0.339 | NA | NA | NA | NA | 1 | NA | -0.331 |
| SE | 27973 | ENSG00000273247 | RAB33B-AS1 | chr4 | - | 139415936 | 139416012 | 139413815 | 139414024 | 139453684 | 139454034 | 3 | 6 | 1 | 2 | 0 | 3 | 2 | 4 | 2 | 2 | 0 | 2 | 4 | 2 | 0 | 0 | 0 | 0 | 3 | 4 | 225 | 149 | 1 | 1 | 0.398 | 0.665 | 0.142 | 0.398 | 0 | NA | 1 | 1 | 0.306 | 0 | -0.256 |
| SE | 27974 | ENSG00000273247 | RAB33B-AS1 | chr4 | - | 139419869 | 139419923 | 139415936 | 139416012 | 139453684 | 139453776 | 0 | 0 | 0 | 1 | 0 | 1 | 0 | 0 | 0 | 0 | 0 | 0 | 0 | 0 | 0 | 0 | 0 | 2 | 0 | 0 | 203 | 149 | 1 | 1 | 0 | NA | NA | 1 | NA | NA | NA | 0 | NA | NA | 0.5 |
| SE | 27976 | ENSG00000147853 | AK3 | chr9 | - | 4722505 | 4722625 | 4719134 | 4719307 | 4740936 | 4741202 | 62 | 37 | 73 | 71 | 51 | 0 | 1 | 0 | 0 | 0 | 11 | 44 | 33 | 59 | 17 | 0 | 0 | 0 | 0 | 0 | 269 | 149 | 1 | 1 | 1 | 0.953 | 1 | 1 | 1 | 1 | 1 | 1 | 1 | 1 | -0.009 |
| SE | 27988 | ENSG00000162980 | ARL5A | chr2 | - | 151791757 | 151791908 | 151789751 | 151789882 | 151803006 | 151803324 | 12 | 3 | 4 | 8 | 0 | 0 | 0 | 2 | 0 | 0 | 0 | 11 | 4 | 8 | 30 | 0 | 3 | 0 | 0 | 0 | 298 | 149 | 1 | 1 | 1 | 1 | 0.5 | 1 | NA | NA | 0.647 | 1 | 1 | 1 | -0.037 |
| SE | 27989 | ENSG00000162980 | ARL5A | chr2 | - | 151810377 | 151810561 | 151806820 | 151806972 | 151812356 | 151812440 | 8 | 4 | 4 | 0 | 2 | 292 | 293 | 200 | 243 | 591 | 0 | 11 | 1 | 2 | 0 | 234 | 218 | 264 | 126 | 74 | 298 | 149 | 1 | 1 | 0.014 | 0.007 | 0.01 | 0 | 0.002 | 0 | 0.025 | 0.002 | 0.008 | 0 | 0 |
| SE | 27991 | ENSG00000162980 | ARL5A | chr2 | - | 151814168 | 151814316 | 151806820 | 151806972 | 151815138 | 151815199 | 321 | 286 | 208 | 260 | 682 | 0 | 1 | 0 | 0 | 0 | 209 | 219 | 226 | 136 | 87 | 0 | 0 | 0 | 0 | 0 | 297 | 149 | 1 | 1 | 1 | 0.993 | 1 | 1 | 1 | 1 | 1 | 1 | 1 | 1 | -0.001 |
| SE | 27994 | ENSG00000137601 | NEK1 | chr4 | - | 169438082 | 169438259 | 169433544 | 169433665 | 169463242 | 169463395 | 42 | 59 | 46 | 47 | 111 | 5 | 1 | 5 | 4 | 3 | 0 | 25 | 21 | 18 | 0 | 0 | 0 | 0 | 0 | 0 | 298 | 149 | 1 | 1 | 0.808 | 0.967 | 0.821 | 0.855 | 0.949 | NA | 1 | 1 | 1 | NA | -0.12 |
| SE | 27995 | ENSG00000137601 | NEK1 | chr4 | - | 169477123 | 169477352 | 169463242 | 169463395 | 169477431 | 169477497 | 86 | 58 | 57 | 45 | 133 | 1 | 0 | 0 | 0 | 1 | 23 | 35 | 26 | 49 | 14 | 0 | 0 | 0 | 0 | 0 | 298 | 149 | 1 | 1 | 0.977 | 1 | 1 | 1 | 0.985 | 1 | 1 | 1 | 1 | 1 | -0.008 |
| SE | 27996 | ENSG00000137601 | NEK1 | chr4 | - | 169477123 | 169477352 | 169463242 | 169463395 | 169479402 | 169479534 | 56 | 25 | 34 | 20 | 64 | 2 | 3 | 2 | 2 | 1 | 0 | 24 | 9 | 17 | 0 | 0 | 0 | 0 | 0 | 0 | 298 | 149 | 1 | 1 | 0.933 | 0.806 | 0.895 | 0.833 | 0.97 | NA | 1 | 1 | 1 | NA | -0.113 |
| SE | 27998 | ENSG00000137601 | NEK1 | chr4 | - | 169477431 | 169477497 | 169477123 | 169477352 | 169479402 | 169479534 | 64 | 49 | 42 | 54 | 114 | 3 | 0 | 0 | 0 | 0 | 48 | 25 | 26 | 50 | 56 | 0 | 0 | 0 | 0 | 0 | 215 | 149 | 1 | 1 | 0.937 | 1 | 1 | 1 | 1 | 1 | 1 | 1 | 1 | 1 | -0.013 |
| SE | 28000 | ENSG00000137601 | NEK1 | chr4 | - | 169507036 | 169507132 | 169479402 | 169479534 | 169507714 | 169507792 | 33 | 25 | 20 | 42 | 30 | 0 | 0 | 0 | 0 | 0 | 17 | 8 | 16 | 32 | 24 | 0 | 2 | 0 | 0 | 0 | 245 | 149 | 1 | 1 | 1 | 1 | 1 | 1 | 1 | 1 | 0.709 | 1 | 1 | 1 | 0.058 |
| SE | 28001 | ENSG00000137601 | NEK1 | chr4 | - | 169508768 | 169508852 | 169508247 | 169508331 | 169537808 | 169537911 | 0 | 0 | 0 | 1 | 0 | 21 | 30 | 15 | 21 | 19 | 0 | 0 | 0 | 0 | 0 | 6 | 15 | 21 | 34 | 25 | 233 | 149 | 1 | 1 | 0 | 0 | 0 | 0.03 | 0 | 0 | 0 | 0 | 0 | 0 | 0.006 |
| SE | 28002 | ENSG00000137601 | NEK1 | chr4 | - | 169537808 | 169537911 | 169508247 | 169508331 | 169555719 | 169555851 | 29 | 42 | 21 | 37 | 31 | 0 | 0 | 1 | 0 | 0 | 6 | 23 | 24 | 40 | 40 | 0 | 0 | 0 | 0 | 0 | 252 | 149 | 1 | 1 | 1 | 1 | 0.925 | 1 | 1 | 1 | 1 | 1 | 1 | 1 | -0.015 |
| SE | 28004 | ENSG00000137601 | NEK1 | chr4 | - | 169561479 | 169561554 | 169555931 | 169556095 | 169561686 | 169561737 | 26 | 31 | 28 | 51 | 54 | 0 | 3 | 0 | 0 | 0 | 0 | 13 | 4 | 2 | 0 | 0 | 0 | 0 | 0 | 0 | 224 | 149 | 1 | 1 | 1 | 0.873 | 1 | 1 | 1 | NA | 1 | 1 | 1 | NA | -0.025 |
| SE | 28005 | ENSG00000137601 | NEK1 | chr4 | - | 169576927 | 169577079 | 169562136 | 169562196 | 169580841 | 169580902 | 68 | 79 | 55 | 61 | 91 | 0 | 0 | 1 | 2 | 1 | 14 | 25 | 38 | 14 | 9 | 0 | 0 | 0 | 0 | 0 | 298 | 149 | 1 | 1 | 1 | 1 | 0.965 | 0.938 | 0.978 | 1 | 1 | 1 | 1 | 1 | -0.024 |
| SE | 28011 | ENSG00000138698 | RAP1GDS1 | chr4 | + | 98343138 | 98343261 | 98293404 | 98293515 | 98352475 | 98352567 | 176 | 149 | 128 | 156 | 137 | 0 | 0 | 0 | 1 | 0 | 184 | 115 | 99 | 128 | 45 | 0 | 1 | 0 | 0 | 0 | 272 | 149 | 1 | 1 | 1 | 1 | 1 | 0.988 | 1 | 1 | 0.984 | 1 | 1 | 1 | 0.001 |
| SE | 28012 | ENSG00000138698 | RAP1GDS1 | chr4 | + | 98343138 | 98343319 | 98293404 | 98293515 | 98352475 | 98352588 | 92 | 72 | 69 | 93 | 76 | 0 | 0 | 0 | 1 | 0 | 70 | 65 | 59 | 50 | 17 | 0 | 1 | 0 | 0 | 0 | 298 | 149 | 1 | 1 | 1 | 1 | 1 | 0.979 | 1 | 1 | 0.97 | 1 | 1 | 1 | 0.002 |
| SE | 28013 | ENSG00000138698 | RAP1GDS1 | chr4 | + | 98352475 | 98352601 | 98343138 | 98343261 | 98379016 | 98379163 | 166 | 128 | 124 | 135 | 116 | 0 | 0 | 3 | 1 | 0 | 158 | 80 | 99 | 132 | 66 | 0 | 0 | 0 | 2 | 0 | 275 | 149 | 1 | 1 | 1 | 1 | 0.957 | 0.987 | 1 | 1 | 1 | 1 | 0.973 | 1 | -0.006 |
| SE | 28018 | ENSG00000138698 | RAP1GDS1 | chr4 | + | 98379016 | 98379163 | 98352475 | 98352601 | 98404476 | 98404602 | 77 | 70 | 57 | 68 | 54 | 0 | 0 | 3 | 1 | 0 | 55 | 36 | 51 | 74 | 38 | 0 | 0 | 1 | 0 | 0 | 296 | 149 | 1 | 1 | 1 | 1 | 0.905 | 0.972 | 1 | 1 | 1 | 0.963 | 1 | 1 | -0.017 |
| SE | 28019 | ENSG00000138698 | RAP1GDS1 | chr4 | + | 98387435 | 98387503 | 98379016 | 98379163 | 98391951 | 98392080 | 0 | 3 | 0 | 2 | 0 | 112 | 68 | 72 | 116 | 131 | 2 | 3 | 1 | 0 | 0 | 55 | 79 | 49 | 97 | 54 | 217 | 149 | 1 | 1 | 0 | 0.029 | 0 | 0.012 | 0 | 0.024 | 0.025 | 0.014 | 0 | 0 | -0.004 |
| SE | 28020 | ENSG00000138698 | RAP1GDS1 | chr4 | + | 98387435 | 98387507 | 98352475 | 98352601 | 98391951 | 98391989 | 0 | 0 | 1 | 0 | 1 | 10 | 5 | 12 | 7 | 16 | 0 | 0 | 0 | 0 | 0 | 0 | 5 | 5 | 1 | 0 | 221 | 149 | 1 | 1 | 0 | 0 | 0.053 | 0 | 0.04 | NA | 0 | 0 | 0 | NA | 0.019 |
| SE | 28021 | ENSG00000138698 | RAP1GDS1 | chr4 | + | 98391951 | 98392080 | 98352475 | 98352601 | 98404476 | 98404586 | 131 | 123 | 108 | 152 | 226 | 0 | 0 | 3 | 1 | 0 | 84 | 110 | 88 | 95 | 57 | 0 | 0 | 1 | 0 | 0 | 278 | 149 | 1 | 1 | 1 | 1 | 0.951 | 0.988 | 1 | 1 | 1 | 0.979 | 1 | 1 | -0.008 |
| SE | 28022 | ENSG00000138698 | RAP1GDS1 | chr4 | + | 98391951 | 98392080 | 98379016 | 98379163 | 98404476 | 98404586 | 233 | 184 | 168 | 260 | 335 | 0 | 0 | 0 | 0 | 0 | 136 | 184 | 132 | 191 | 105 | 0 | 3 | 0 | 0 | 0 | 278 | 149 | 1 | 1 | 1 | 1 | 1 | 1 | 1 | 1 | 0.97 | 1 | 1 | 1 | 0.006 |
| SE | 28023 | ENSG00000138698 | RAP1GDS1 | chr4 | + | 98404476 | 98404602 | 98391951 | 98392080 | 98416744 | 98416888 | 266 | 217 | 174 | 273 | 388 | 2 | 0 | 2 | 0 | 0 | 180 | 233 | 148 | 211 | 184 | 0 | 1 | 0 | 0 | 7 | 275 | 149 | 1 | 1 | 0.986 | 1 | 0.979 | 1 | 1 | 1 | 0.992 | 1 | 1 | 0.934 | 0.008 |
| SE | 28025 | ENSG00000138698 | RAP1GDS1 | chr4 | + | 98421254 | 98421394 | 98420018 | 98420144 | 98433935 | 98434062 | 237 | 172 | 152 | 212 | 244 | 0 | 0 | 0 | 0 | 0 | 162 | 187 | 123 | 140 | 121 | 0 | 0 | 1 | 0 | 0 | 289 | 149 | 1 | 1 | 1 | 1 | 1 | 1 | 1 | 1 | 1 | 0.984 | 1 | 1 | 0.003 |
| SE | 28026 | ENSG00000138698 | RAP1GDS1 | chr4 | + | 98421257 | 98421394 | 98420018 | 98420144 | 98433935 | 98434062 | 122 | 76 | 81 | 103 | 119 | 0 | 0 | 0 | 0 | 0 | 69 | 88 | 57 | 71 | 57 | 0 | 0 | 1 | 0 | 0 | 286 | 149 | 1 | 1 | 1 | 1 | 1 | 1 | 1 | 1 | 1 | 0.967 | 1 | 1 | 0.007 |
| SE | 28027 | ENSG00000138698 | RAP1GDS1 | chr4 | + | 98433935 | 98434062 | 98421254 | 98421394 | 98441989 | 98442117 | 77 | 44 | 59 | 55 | 53 | 1 | 0 | 0 | 0 | 0 | 36 | 52 | 23 | 34 | 26 | 0 | 0 | 0 | 0 | 0 | 276 | 149 | 1 | 1 | 0.977 | 1 | 1 | 1 | 1 | 1 | 1 | 1 | 1 | 1 | -0.005 |
| SE | 28028 | ENSG00000138698 | RAP1GDS1 | chr4 | + | 98436939 | 98437068 | 98433935 | 98434062 | 98441989 | 98442117 | 207 | 96 | 160 | 187 | 385 | 1 | 1 | 2 | 0 | 0 | 115 | 137 | 73 | 131 | 48 | 0 | 0 | 0 | 2 | 0 | 278 | 149 | 1 | 1 | 0.991 | 0.981 | 0.977 | 1 | 1 | 1 | 1 | 1 | 0.972 | 1 | -0.005 |
| SE | 28029 | ENSG00000120519 | SLC10A7 | chr4 | - | 146256825 | 146256936 | 146256490 | 146256520 | 146258691 | 146258837 | 26 | 12 | 25 | 15 | 25 | 25 | 6 | 12 | 16 | 24 | 1 | 30 | 8 | 7 | 0 | 0 | 9 | 21 | 14 | 0 | 260 | 149 | 1 | 1 | 0.373 | 0.534 | 0.544 | 0.349 | 0.374 | 1 | 0.656 | 0.179 | 0.223 | NA | -0.08 |
| SE | 28030 | ENSG00000120519 | SLC10A7 | chr4 | - | 146283191 | 146283265 | 146258691 | 146258837 | 146292928 | 146292980 | 67 | 33 | 45 | 58 | 78 | 0 | 0 | 2 | 0 | 2 | 6 | 30 | 15 | 12 | 21 | 0 | 0 | 0 | 0 | 0 | 223 | 149 | 1 | 1 | 1 | 1 | 0.938 | 1 | 0.963 | 1 | 1 | 1 | 1 | 1 | -0.02 |
| SE | 28031 | ENSG00000120519 | SLC10A7 | chr4 | - | 146305925 | 146306009 | 146293929 | 146294095 | 146325960 | 146325996 | 53 | 20 | 37 | 32 | 58 | 2 | 0 | 2 | 1 | 6 | 0 | 50 | 34 | 20 | 0 | 0 | 0 | 1 | 0 | 0 | 233 | 149 | 1 | 1 | 0.944 | 1 | 0.922 | 0.953 | 0.861 | NA | 1 | 0.956 | 1 | NA | -0.049 |
| SE | 28035 | ENSG00000120519 | SLC10A7 | chr4 | - | 146509912 | 146510049 | 146503848 | 146503924 | 146521617 | 146521717 | 56 | 29 | 55 | 48 | 32 | 0 | 1 | 0 | 0 | 0 | 7 | 40 | 26 | 1 | 8 | 0 | 0 | 0 | 0 | 0 | 286 | 149 | 1 | 1 | 1 | 0.938 | 1 | 1 | 1 | 1 | 1 | 1 | 1 | 1 | -0.012 |
| SE | 28036 | ENSG00000120519 | SLC10A7 | chr4 | - | 146517037 | 146517120 | 146503848 | 146503924 | 146521617 | 146521717 | 69 | 59 | 54 | 62 | 77 | 0 | 1 | 0 | 0 | 0 | 13 | 53 | 44 | 21 | 0 | 0 | 0 | 0 | 0 | 0 | 232 | 149 | 1 | 1 | 1 | 0.974 | 1 | 1 | 1 | 1 | 1 | 1 | 1 | NA | -0.005 |
| SE | 28037 | ENSG00000120519 | SLC10A7 | chr4 | - | 146517037 | 146517120 | 146509912 | 146510049 | 146521617 | 146521717 | 172 | 147 | 152 | 138 | 203 | 0 | 1 | 0 | 0 | 2 | 34 | 114 | 104 | 34 | 13 | 0 | 0 | 0 | 0 | 0 | 232 | 149 | 1 | 1 | 1 | 0.99 | 1 | 1 | 0.985 | 1 | 1 | 1 | 1 | 1 | -0.005 |
| SE | 28038 | ENSG00000049656 | CLPTM1L | chr5 | - | 1320615 | 1320731 | 1317751 | 1318453 | 1321634 | 1321679 | 227 | 88 | 191 | 164 | 244 | 0 | 0 | 0 | 0 | 0 | 174 | 264 | 163 | 131 | 180 | 0 | 2 | 0 | 0 | 0 | 265 | 149 | 1 | 1 | 1 | 1 | 1 | 1 | 1 | 1 | 0.987 | 1 | 1 | 1 | 0.003 |
| SE | 28039 | ENSG00000049656 | CLPTM1L | chr5 | - | 1323786 | 1323869 | 1322833 | 1322911 | 1324762 | 1324813 | 227 | 99 | 204 | 181 | 331 | 0 | 0 | 1 | 0 | 0 | 135 | 245 | 143 | 150 | 212 | 0 | 0 | 1 | 4 | 0 | 232 | 149 | 1 | 1 | 1 | 1 | 0.992 | 1 | 1 | 1 | 1 | 0.989 | 0.96 | 1 | 0.009 |
| SE | 28046 | ENSG00000108961 | RANGRF | chr17 | + | 8289257 | 8289414 | 8288884 | 8289072 | 8289812 | 8290092 | 58 | 13 | 33 | 59 | 53 | 0 | 0 | 0 | 0 | 0 | 94 | 17 | 14 | 24 | 54 | 0 | 0 | 0 | 1 | 0 | 298 | 149 | 1 | 1 | 1 | 1 | 1 | 1 | 1 | 1 | 1 | 1 | 0.923 | 1 | 0.015 |
| SE | 28048 | ENSG00000114738 | MAPKAPK3 | chr3 | + | 50617513 | 50617784 | 50617169 | 50617226 | 50640365 | 50640505 | 188 | 94 | 116 | 152 | 71 | 0 | 0 | 0 | 0 | 0 | 229 | 129 | 97 | 215 | 408 | 0 | 0 | 0 | 0 | 1 | 298 | 149 | 1 | 1 | 1 | 1 | 1 | 1 | 1 | 1 | 1 | 1 | 1 | 0.995 | 0.001 |
| SE | 28049 | ENSG00000114738 | MAPKAPK3 | chr3 | + | 50640365 | 50640505 | 50617389 | 50617784 | 50641706 | 50641771 | 423 | 237 | 280 | 367 | 200 | 2 | 0 | 0 | 0 | 0 | 704 | 349 | 239 | 596 | 1050 | 0 | 0 | 0 | 0 | 0 | 289 | 149 | 1 | 1 | 0.991 | 1 | 1 | 1 | 1 | 1 | 1 | 1 | 1 | 1 | -0.002 |
| SE | 28050 | ENSG00000114738 | MAPKAPK3 | chr3 | + | 50641706 | 50641771 | 50640365 | 50640505 | 50642252 | 50642332 | 279 | 175 | 196 | 246 | 147 | 2 | 1 | 0 | 1 | 0 | 585 | 291 | 173 | 444 | 724 | 2 | 0 | 4 | 7 | 2 | 214 | 149 | 1 | 1 | 0.99 | 0.992 | 1 | 0.994 | 1 | 0.995 | 1 | 0.968 | 0.978 | 0.996 | 0.008 |
| SE | 28053 | ENSG00000114738 | MAPKAPK3 | chr3 | + | 50645709 | 50645785 | 50642252 | 50642332 | 50646739 | 50646825 | 5 | 3 | 7 | 5 | 0 | 1 | 0 | 0 | 0 | 0 | 22 | 11 | 5 | 4 | 0 | 0 | 0 | 0 | 4 | 6 | 225 | 149 | 1 | 1 | 0.768 | 1 | 1 | 1 | NA | 1 | 1 | 1 | 0.398 | 0 | 0.262 |
| SE | 28054 | ENSG00000114738 | MAPKAPK3 | chr3 | + | 50646139 | 50646264 | 50645709 | 50645785 | 50646739 | 50646825 | 551 | 347 | 341 | 406 | 418 | 0 | 0 | 1 | 0 | 0 | 1075 | 457 | 383 | 783 | 1090 | 0 | 0 | 0 | 2 | 0 | 274 | 149 | 1 | 1 | 1 | 1 | 0.995 | 1 | 1 | 1 | 1 | 1 | 0.995 | 1 | 0 |
| SE | 28055 | ENSG00000114738 | MAPKAPK3 | chr3 | + | 50646739 | 50646825 | 50646139 | 50646264 | 50647122 | 50647203 | 442 | 260 | 281 | 335 | 271 | 0 | 0 | 0 | 0 | 0 | 873 | 346 | 319 | 622 | 883 | 0 | 0 | 0 | 7 | 0 | 235 | 149 | 1 | 1 | 1 | 1 | 1 | 1 | 1 | 1 | 1 | 1 | 0.983 | 1 | 0.003 |
| SE | 28058 | ENSG00000282804 | NA | chr6 | - | 24806366 | 24806473 | 24797372 | 24797837 | 24809716 | 24809798 | 419 | 295 | 408 | 351 | 299 | 0 | 1 | 0 | 0 | 0 | 220 | 172 | 132 | 247 | 48 | 0 | 0 | 0 | 0 | 0 | 256 | 149 | 1 | 1 | 1 | 0.994 | 1 | 1 | 1 | 1 | 1 | 1 | 1 | 1 | -0.001 |
| SE | 28062 | ENSG00000121350 | PYROXD1 | chr12 | + | 21441206 | 21441300 | 21440367 | 21440448 | 21445346 | 21445466 | 0 | 0 | 1 | 0 | 0 | 108 | 64 | 78 | 74 | 191 | 0 | 0 | 0 | 0 | 0 | 12 | 73 | 38 | 16 | 16 | 243 | 149 | 1 | 1 | 0 | 0 | 0.008 | 0 | 0 | 0 | 0 | 0 | 0 | 0 | 0.002 |
| SE | 28063 | ENSG00000121350 | PYROXD1 | chr12 | + | 21449562 | 21449691 | 21445346 | 21445466 | 21452080 | 21452154 | 176 | 142 | 129 | 137 | 324 | 0 | 0 | 0 | 1 | 1 | 29 | 125 | 62 | 27 | 3 | 0 | 0 | 0 | 0 | 0 | 278 | 149 | 1 | 1 | 1 | 1 | 1 | 0.987 | 0.994 | 1 | 1 | 1 | 1 | 1 | -0.004 |
| SE | 28065 | ENSG00000121350 | PYROXD1 | chr12 | + | 21452080 | 21452154 | 21445346 | 21445466 | 21455131 | 21455292 | 31 | 24 | 32 | 32 | 45 | 2 | 0 | 5 | 0 | 0 | 12 | 5 | 3 | 2 | 0 | 0 | 0 | 0 | 0 | 0 | 223 | 149 | 1 | 1 | 0.912 | 1 | 0.81 | 1 | 1 | 1 | 1 | 1 | 1 | NA | -0.056 |
| SE | 28066 | ENSG00000121350 | PYROXD1 | chr12 | + | 21452080 | 21452154 | 21449562 | 21449691 | 21455131 | 21455292 | 145 | 120 | 103 | 121 | 246 | 15 | 13 | 15 | 14 | 22 | 34 | 90 | 40 | 16 | 0 | 12 | 13 | 13 | 5 | 0 | 223 | 149 | 1 | 1 | 0.866 | 0.86 | 0.821 | 0.852 | 0.882 | 0.654 | 0.822 | 0.673 | 0.681 | NA | 0.149 |
| SE | 28069 | ENSG00000121350 | PYROXD1 | chr12 | + | 21462007 | 21462125 | 21460902 | 21461154 | 21462739 | 21462746 | 47 | 30 | 26 | 25 | 67 | 1 | 4 | 9 | 19 | 0 | 0 | 14 | 17 | 2 | 0 | 10 | 5 | 9 | 0 | 0 | 267 | 149 | 1 | 1 | 0.963 | 0.807 | 0.617 | 0.423 | 1 | 0 | 0.61 | 0.513 | 1 | NA | 0.231 |
| SE | 28070 | ENSG00000084676 | NCOA1 | chr2 | + | 24554388 | 24554573 | 24491253 | 24491602 | 24564294 | 24564430 | 5 | 0 | 1 | 7 | 5 | 16 | 2 | 10 | 3 | 7 | 13 | 0 | 2 | 4 | 0 | 4 | 7 | 7 | 10 | 0 | 298 | 149 | 1 | 1 | 0.135 | 0 | 0.048 | 0.538 | 0.263 | 0.619 | 0 | 0.125 | 0.167 | NA | -0.031 |
| SE | 28071 | ENSG00000084676 | NCOA1 | chr2 | + | 24554388 | 24554573 | 24492022 | 24492188 | 24556014 | 24556055 | 26 | 20 | 10 | 32 | 22 | 0 | 0 | 0 | 0 | 0 | 28 | 26 | 15 | 53 | 69 | 0 | 3 | 0 | 0 | 0 | 298 | 149 | 1 | 1 | 1 | 1 | 1 | 1 | 1 | 1 | 0.812 | 1 | 1 | 1 | 0.038 |
| SE | 28072 | ENSG00000084676 | NCOA1 | chr2 | + | 24554388 | 24554573 | 24492022 | 24492188 | 24564294 | 24564430 | 4 | 0 | 1 | 7 | 4 | 1 | 0 | 2 | 2 | 0 | 13 | 0 | 2 | 4 | 0 | 0 | 9 | 3 | 2 | 0 | 298 | 149 | 1 | 1 | 0.667 | NA | 0.2 | 0.636 | 1 | 1 | 0 | 0.25 | 0.5 | NA | 0.188 |
| SE | 28075 | ENSG00000084676 | NCOA1 | chr2 | + | 24564294 | 24564430 | 24491253 | 24491602 | 24584475 | 24584484 | 213 | 156 | 164 | 203 | 282 | 6 | 0 | 0 | 0 | 1 | 214 | 256 | 272 | 155 | 174 | 0 | 0 | 0 | 0 | 0 | 285 | 149 | 1 | 1 | 0.949 | 1 | 1 | 1 | 0.993 | 1 | 1 | 1 | 1 | 1 | -0.012 |
| SE | 28076 | ENSG00000084676 | NCOA1 | chr2 | + | 24564294 | 24564430 | 24491253 | 24491602 | 24643965 | 24644055 | 21 | 5 | 14 | 10 | 13 | 2 | 0 | 0 | 0 | 0 | 4 | 16 | 8 | 8 | 12 | 0 | 0 | 0 | 0 | 0 | 285 | 149 | 1 | 1 | 0.846 | 1 | 1 | 1 | 1 | 1 | 1 | 1 | 1 | 1 | -0.031 |
| SE | 28078 | ENSG00000084676 | NCOA1 | chr2 | + | 24584475 | 24584560 | 24491253 | 24491602 | 24643965 | 24644122 | 97 | 53 | 59 | 86 | 137 | 2 | 0 | 0 | 0 | 0 | 103 | 112 | 105 | 59 | 45 | 0 | 0 | 0 | 0 | 0 | 234 | 149 | 1 | 1 | 0.969 | 1 | 1 | 1 | 1 | 1 | 1 | 1 | 1 | 1 | -0.006 |
| SE | 28080 | ENSG00000084676 | NCOA1 | chr2 | + | 24584475 | 24584560 | 24564294 | 24564430 | 24658660 | 24658766 | 103 | 89 | 78 | 111 | 143 | 0 | 0 | 0 | 1 | 0 | 142 | 136 | 164 | 86 | 92 | 0 | 0 | 0 | 0 | 0 | 234 | 149 | 1 | 1 | 1 | 1 | 1 | 0.986 | 1 | 1 | 1 | 1 | 1 | 1 | -0.003 |
| SE | 28082 | ENSG00000084676 | NCOA1 | chr2 | + | 24658660 | 24658766 | 24584475 | 24584560 | 24665748 | 24665915 | 103 | 87 | 84 | 107 | 178 | 0 | 0 | 0 | 0 | 0 | 91 | 115 | 137 | 145 | 90 | 0 | 0 | 1 | 0 | 0 | 255 | 149 | 1 | 1 | 1 | 1 | 1 | 1 | 1 | 1 | 1 | 0.988 | 1 | 1 | 0.002 |
| SE | 28084 | ENSG00000084676 | NCOA1 | chr2 | + | 24665748 | 24665915 | 24658660 | 24658766 | 24682950 | 24683128 | 183 | 142 | 124 | 188 | 255 | 0 | 0 | 0 | 4 | 0 | 166 | 195 | 270 | 226 | 113 | 0 | 1 | 0 | 0 | 0 | 298 | 149 | 1 | 1 | 1 | 1 | 1 | 0.959 | 1 | 1 | 0.99 | 1 | 1 | 1 | -0.006 |
| SE | 28085 | ENSG00000084676 | NCOA1 | chr2 | + | 24673365 | 24673463 | 24658660 | 24658766 | 24682950 | 24683128 | 570 | 402 | 253 | 418 | 593 | 0 | 0 | 0 | 4 | 0 | 489 | 376 | 560 | 479 | 179 | 0 | 1 | 0 | 0 | 0 | 247 | 149 | 1 | 1 | 1 | 1 | 1 | 0.984 | 1 | 1 | 0.996 | 1 | 1 | 1 | -0.002 |
| SE | 28087 | ENSG00000084676 | NCOA1 | chr2 | + | 24682950 | 24683128 | 24673365 | 24673463 | 24691480 | 24691660 | 1102 | 840 | 558 | 903 | 1162 | 0 | 1 | 0 | 0 | 0 | 954 | 784 | 1151 | 955 | 408 | 0 | 0 | 0 | 0 | 0 | 298 | 149 | 1 | 1 | 1 | 0.998 | 1 | 1 | 1 | 1 | 1 | 1 | 1 | 1 | 0 |
| SE | 28088 | ENSG00000084676 | NCOA1 | chr2 | + | 24693251 | 24693347 | 24691480 | 24691660 | 24697657 | 24697798 | 349 | 261 | 249 | 395 | 404 | 0 | 0 | 1 | 0 | 0 | 184 | 345 | 344 | 296 | 28 | 0 | 0 | 0 | 0 | 0 | 245 | 149 | 1 | 1 | 1 | 1 | 0.993 | 1 | 1 | 1 | 1 | 1 | 1 | 1 | -0.001 |
| SE | 28090 | ENSG00000084676 | NCOA1 | chr2 | + | 24705085 | 24705233 | 24697657 | 24697798 | 24706567 | 24707888 | 472 | 272 | 346 | 500 | 552 | 0 | 2 | 0 | 0 | 0 | 264 | 442 | 473 | 387 | 102 | 0 | 2 | 0 | 0 | 0 | 297 | 149 | 1 | 1 | 1 | 0.986 | 1 | 1 | 1 | 1 | 0.991 | 1 | 1 | 1 | -0.001 |
| SE | 28091 | ENSG00000084676 | NCOA1 | chr2 | + | 24706567 | 24707888 | 24705085 | 24705233 | 24710930 | 24711111 | 425 | 260 | 256 | 442 | 560 | 0 | 0 | 0 | 1 | 0 | 219 | 355 | 477 | 410 | 125 | 0 | 0 | 0 | 0 | 0 | 298 | 149 | 1 | 1 | 1 | 1 | 1 | 0.995 | 1 | 1 | 1 | 1 | 1 | 1 | -0.001 |
| SE | 28092 | ENSG00000084676 | NCOA1 | chr2 | + | 24728307 | 24728476 | 24726588 | 24726706 | 24739431 | 24739533 | 231 | 187 | 186 | 205 | 210 | 0 | 0 | 0 | 0 | 0 | 44 | 260 | 257 | 247 | 95 | 0 | 0 | 1 | 0 | 0 | 298 | 149 | 1 | 1 | 1 | 1 | 1 | 1 | 1 | 1 | 1 | 0.992 | 1 | 1 | 0.002 |
| SE | 28094 | ENSG00000084676 | NCOA1 | chr2 | + | 24762686 | 24762776 | 24757972 | 24758156 | 24768078 | 24768135 | 357 | 219 | 184 | 246 | 306 | 0 | 0 | 3 | 0 | 0 | 254 | 230 | 321 | 184 | 118 | 0 | 3 | 1 | 0 | 0 | 239 | 149 | 1 | 1 | 1 | 1 | 0.975 | 1 | 1 | 1 | 0.98 | 0.995 | 1 | 1 | 0 |
| SE | 28095 | ENSG00000084676 | NCOA1 | chr2 | + | 24762686 | 24762776 | 24757972 | 24758156 | 24768220 | 24770702 | 287 | 168 | 131 | 227 | 237 | 0 | 0 | 0 | 2 | 0 | 233 | 187 | 211 | 129 | 52 | 0 | 0 | 3 | 0 | 0 | 239 | 149 | 1 | 1 | 1 | 1 | 1 | 0.986 | 1 | 1 | 1 | 0.978 | 1 | 1 | 0.002 |
| SE | 28096 | ENSG00000084676 | NCOA1 | chr2 | + | 24768078 | 24768135 | 24757972 | 24758156 | 24768220 | 24770702 | 64 | 25 | 38 | 49 | 83 | 0 | 0 | 0 | 2 | 0 | 60 | 19 | 81 | 64 | 7 | 0 | 0 | 3 | 0 | 0 | 206 | 149 | 1 | 1 | 1 | 1 | 1 | 0.947 | 1 | 1 | 1 | 0.951 | 1 | 1 | -0.001 |
| SE | 28102 | ENSG00000121406 | ZNF549 | chr19 | + | 57535143 | 57535270 | 57531069 | 57531108 | 57552384 | 57557542 | 7 | 3 | 3 | 8 | 6 | 1 | 0 | 0 | 0 | 0 | 0 | 6 | 2 | 0 | 0 | 0 | 0 | 0 | 0 | 0 | 276 | 149 | 1 | 1 | 0.791 | 1 | 1 | 1 | 1 | NA | 1 | 1 | NA | NA | -0.042 |
| SE | 28105 | ENSG00000125386 | FAM193A | chr4 | + | 2657802 | 2657880 | 2646684 | 2646832 | 2659557 | 2659670 | 146 | 97 | 106 | 138 | 130 | 2 | 0 | 0 | 0 | 0 | 209 | 184 | 100 | 216 | 179 | 0 | 0 | 0 | 0 | 0 | 227 | 149 | 1 | 1 | 0.98 | 1 | 1 | 1 | 1 | 1 | 1 | 1 | 1 | 1 | -0.004 |
| SE | 28106 | ENSG00000125386 | FAM193A | chr4 | + | 2659811 | 2660054 | 2659557 | 2659670 | 2662837 | 2662991 | 185 | 107 | 132 | 175 | 145 | 0 | 0 | 0 | 1 | 3 | 191 | 193 | 169 | 208 | 383 | 0 | 0 | 0 | 0 | 0 | 298 | 149 | 1 | 1 | 1 | 1 | 1 | 0.989 | 0.96 | 1 | 1 | 1 | 1 | 1 | -0.01 |
| SE | 28107 | ENSG00000125386 | FAM193A | chr4 | + | 2672120 | 2672372 | 2663108 | 2663288 | 2690697 | 2690970 | 104 | 71 | 90 | 121 | 125 | 0 | 0 | 0 | 0 | 0 | 162 | 97 | 108 | 130 | 218 | 0 | 0 | 0 | 0 | 1 | 298 | 149 | 1 | 1 | 1 | 1 | 1 | 1 | 1 | 1 | 1 | 1 | 1 | 0.991 | 0.002 |
| SE | 28109 | ENSG00000125386 | FAM193A | chr4 | + | 2696362 | 2696593 | 2694945 | 2695129 | 2699679 | 2700501 | 194 | 155 | 155 | 215 | 165 | 2 | 0 | 0 | 0 | 0 | 497 | 225 | 193 | 338 | 486 | 0 | 0 | 0 | 0 | 0 | 298 | 149 | 1 | 1 | 0.98 | 1 | 1 | 1 | 1 | 1 | 1 | 1 | 1 | 1 | -0.004 |
| SE | 28113 | ENSG00000075407 | ZNF37A | chr10 | + | 38094930 | 38095224 | 38094333 | 38094490 | 38095537 | 38095559 | 4 | 6 | 12 | 2 | 19 | 2 | 4 | 3 | 1 | 0 | 0 | 9 | 8 | 0 | 5 | 0 | 0 | 1 | 5 | 0 | 298 | 149 | 1 | 1 | 0.5 | 0.429 | 0.667 | 0.5 | 1 | NA | 1 | 0.8 | 0 | 1 | -0.081 |
| SE | 28114 | ENSG00000283930 | PLD5P1 | chr10 | + | 38114754 | 38114881 | 38096573 | 38096632 | 38115194 | 38115290 | 32 | 37 | 21 | 32 | 29 | 0 | 0 | 0 | 0 | 3 | 5 | 33 | 15 | 6 | 24 | 0 | 0 | 0 | 0 | 0 | 276 | 149 | 1 | 1 | 1 | 1 | 1 | 1 | 0.839 | 1 | 1 | 1 | 1 | 1 | -0.032 |
| SE | 28117 | ENSG00000198707 | CEP290 | chr12 | - | 88059897 | 88060020 | 88058847 | 88059020 | 88060829 | 88060994 | 33 | 26 | 19 | 34 | 19 | 0 | 0 | 2 | 0 | 0 | 0 | 22 | 2 | 1 | 0 | 0 | 0 | 0 | 0 | 0 | 272 | 149 | 1 | 1 | 1 | 1 | 0.839 | 1 | 1 | NA | 1 | 1 | 1 | NA | -0.032 |
| SE | 28118 | ENSG00000198707 | CEP290 | chr12 | - | 88060829 | 88060994 | 88059897 | 88060020 | 88062691 | 88062778 | 51 | 52 | 31 | 43 | 70 | 2 | 4 | 2 | 1 | 1 | 16 | 17 | 1 | 1 | 0 | 0 | 0 | 0 | 0 | 0 | 298 | 149 | 1 | 1 | 0.927 | 0.867 | 0.886 | 0.956 | 0.972 | 1 | 1 | 1 | 1 | NA | -0.078 |
| SE | 28119 | ENSG00000198707 | CEP290 | chr12 | - | 88063980 | 88064115 | 88062691 | 88062778 | 88068521 | 88068645 | 96 | 90 | 54 | 75 | 152 | 0 | 1 | 0 | 0 | 0 | 10 | 7 | 2 | 3 | 0 | 0 | 0 | 0 | 0 | 0 | 284 | 149 | 1 | 1 | 1 | 0.979 | 1 | 1 | 1 | 1 | 1 | 1 | 1 | NA | -0.004 |
| SE | 28120 | ENSG00000198707 | CEP290 | chr12 | - | 88068037 | 88068046 | 88063980 | 88064115 | 88068521 | 88068645 | 0 | 1 | 0 | 0 | 0 | 50 | 35 | 21 | 34 | 74 | 0 | 1 | 0 | 0 | 0 | 3 | 1 | 0 | 0 | 0 | 158 | 149 | 1 | 1 | 0 | 0.026 | 0 | 0 | 0 | 0 | 0.485 | NA | NA | NA | -0.237 |
| SE | 28121 | ENSG00000198707 | CEP290 | chr12 | - | 88071780 | 88071926 | 88071293 | 88071449 | 88077696 | 88077918 | 21 | 13 | 16 | 11 | 33 | 0 | 1 | 0 | 0 | 0 | 0 | 0 | 2 | 0 | 0 | 0 | 0 | 0 | 0 | 0 | 295 | 149 | 1 | 1 | 1 | 0.868 | 1 | 1 | 1 | NA | NA | 1 | NA | NA | -0.026 |
| SE | 28122 | ENSG00000198707 | CEP290 | chr12 | - | 88077221 | 88077344 | 88071780 | 88071926 | 88077696 | 88077918 | 13 | 25 | 8 | 17 | 27 | 0 | 4 | 3 | 3 | 8 | 0 | 1 | 0 | 0 | 0 | 0 | 0 | 0 | 0 | 0 | 272 | 149 | 1 | 1 | 1 | 0.774 | 0.594 | 0.756 | 0.649 | NA | 1 | NA | NA | NA | -0.245 |
| SE | 28123 | ENSG00000198707 | CEP290 | chr12 | - | 88079091 | 88079229 | 88077696 | 88077918 | 88080181 | 88080395 | 88 | 77 | 80 | 110 | 196 | 1 | 0 | 1 | 0 | 0 | 0 | 3 | 10 | 0 | 0 | 0 | 0 | 0 | 0 | 0 | 287 | 149 | 1 | 1 | 0.979 | 1 | 0.976 | 1 | 1 | NA | 1 | 1 | NA | NA | -0.009 |
| SE | 28124 | ENSG00000198707 | CEP290 | chr12 | - | 88087779 | 88087944 | 88086390 | 88086498 | 88089031 | 88089487 | 109 | 145 | 115 | 181 | 231 | 0 | 1 | 0 | 0 | 0 | 10 | 10 | 10 | 2 | 15 | 0 | 0 | 0 | 0 | 0 | 298 | 149 | 1 | 1 | 1 | 0.986 | 1 | 1 | 1 | 1 | 1 | 1 | 1 | 1 | -0.003 |
| SE | 28125 | ENSG00000198707 | CEP290 | chr12 | - | 88090727 | 88090839 | 88089031 | 88089487 | 88092680 | 88092832 | 25 | 46 | 33 | 27 | 76 | 0 | 1 | 0 | 0 | 0 | 0 | 8 | 1 | 0 | 7 | 0 | 0 | 0 | 0 | 0 | 261 | 149 | 1 | 1 | 1 | 0.963 | 1 | 1 | 1 | NA | 1 | 1 | NA | 1 | -0.007 |
| SE | 28126 | ENSG00000198707 | CEP290 | chr12 | - | 88092680 | 88092832 | 88089031 | 88090839 | 88093769 | 88093975 | 33 | 49 | 30 | 34 | 74 | 4 | 0 | 1 | 0 | 6 | 0 | 0 | 0 | 0 | 0 | 0 | 3 | 0 | 0 | 0 | 298 | 149 | 1 | 1 | 0.805 | 1 | 0.938 | 1 | 0.86 | NA | 0 | NA | NA | NA | 0.921 |
| SE | 28128 | ENSG00000198707 | CEP290 | chr12 | - | 88109065 | 88109181 | 88106995 | 88107098 | 88111201 | 88111351 | 77 | 84 | 63 | 94 | 66 | 2 | 0 | 0 | 0 | 0 | 0 | 14 | 4 | 7 | 5 | 0 | 0 | 0 | 0 | 0 | 265 | 149 | 1 | 1 | 0.956 | 1 | 1 | 1 | 1 | NA | 1 | 1 | 1 | 1 | -0.009 |
| SE | 28131 | ENSG00000198707 | CEP290 | chr12 | - | 88115477 | 88115570 | 88115097 | 88115182 | 88117032 | 88117145 | 4 | 3 | 4 | 1 | 3 | 70 | 79 | 41 | 75 | 94 | 0 | 0 | 0 | 0 | 0 | 0 | 4 | 1 | 15 | 0 | 242 | 149 | 1 | 1 | 0.034 | 0.023 | 0.057 | 0.008 | 0.019 | NA | 0 | 0 | 0 | NA | 0.028 |
| SE | 28134 | ENSG00000198707 | CEP290 | chr12 | - | 88117032 | 88117145 | 88115097 | 88115182 | 88118482 | 88118570 | 94 | 107 | 64 | 101 | 127 | 0 | 2 | 0 | 0 | 0 | 0 | 8 | 1 | 15 | 0 | 0 | 0 | 0 | 0 | 0 | 262 | 149 | 1 | 1 | 1 | 0.968 | 1 | 1 | 1 | NA | 1 | 1 | 1 | NA | -0.006 |
| SE | 28135 | ENSG00000198707 | CEP290 | chr12 | - | 88119020 | 88119150 | 88118642 | 88118743 | 88120113 | 88120276 | 0 | 2 | 4 | 5 | 5 | 8 | 7 | 8 | 8 | 46 | 0 | 0 | 0 | 0 | 0 | 0 | 3 | 0 | 0 | 3 | 279 | 149 | 1 | 1 | 0 | 0.132 | 0.211 | 0.25 | 0.055 | NA | 0 | NA | NA | 0 | 0.13 |
| SE | 28136 | ENSG00000198707 | CEP290 | chr12 | - | 88120996 | 88121166 | 88118482 | 88120276 | 88125245 | 88125369 | 78 | 97 | 70 | 101 | 86 | 0 | 1 | 0 | 0 | 0 | 4 | 22 | 4 | 0 | 5 | 0 | 0 | 0 | 0 | 0 | 298 | 149 | 1 | 1 | 1 | 0.98 | 1 | 1 | 1 | 1 | 1 | 1 | NA | 1 | -0.004 |
| SE | 28137 | ENSG00000198707 | CEP290 | chr12 | - | 88130544 | 88130565 | 88130267 | 88130420 | 88131164 | 88131218 | 19 | 32 | 15 | 22 | 41 | 0 | 0 | 0 | 0 | 0 | 0 | 4 | 3 | 2 | 0 | 0 | 2 | 0 | 0 | 0 | 170 | 149 | 1 | 1 | 1 | 1 | 1 | 1 | 1 | NA | 0.637 | 1 | 1 | NA | 0.121 |
| SE | 28138 | ENSG00000198707 | CEP290 | chr12 | - | 88130544 | 88130565 | 88130267 | 88130420 | 88136642 | 88136786 | 11 | 17 | 10 | 13 | 3 | 0 | 0 | 0 | 0 | 1 | 0 | 4 | 1 | 2 | 0 | 0 | 0 | 0 | 0 | 0 | 170 | 149 | 1 | 1 | 1 | 1 | 1 | 1 | 0.724 | NA | 1 | 1 | 1 | NA | -0.055 |
| SE | 28139 | ENSG00000198707 | CEP290 | chr12 | - | 88131164 | 88131218 | 88130267 | 88130420 | 88136642 | 88136786 | 4 | 14 | 2 | 13 | 16 | 0 | 0 | 0 | 0 | 1 | 0 | 2 | 0 | 0 | 0 | 0 | 0 | 0 | 0 | 0 | 203 | 149 | 1 | 1 | 1 | 1 | 1 | 1 | 0.922 | NA | 1 | NA | NA | NA | -0.016 |
| SE | 28140 | ENSG00000198707 | CEP290 | chr12 | - | 88131164 | 88131218 | 88130544 | 88130565 | 88136642 | 88136786 | 11 | 31 | 9 | 26 | 54 | 3 | 2 | 2 | 4 | 0 | 0 | 0 | 2 | 0 | 0 | 0 | 0 | 0 | 0 | 0 | 203 | 149 | 1 | 1 | 0.729 | 0.919 | 0.768 | 0.827 | 1 | NA | NA | 1 | NA | NA | -0.151 |
| SE | 28143 | ENSG00000198478 | SH3BGRL2 | chr6 | + | 79673613 | 79673799 | 79631328 | 79631506 | 79696484 | 79696565 | 168 | 237 | 53 | 437 | 169 | 0 | 0 | 0 | 3 | 0 | 140 | 227 | 215 | 267 | 227 | 0 | 0 | 0 | 0 | 0 | 298 | 149 | 1 | 1 | 1 | 1 | 1 | 0.986 | 1 | 1 | 1 | 1 | 1 | 1 | -0.003 |
| SE | 28144 | ENSG00000198478 | SH3BGRL2 | chr6 | + | 79696484 | 79696565 | 79673613 | 79673799 | 79699497 | 79703655 | 239 | 378 | 68 | 576 | 405 | 0 | 0 | 0 | 0 | 2 | 227 | 230 | 213 | 253 | 221 | 0 | 0 | 0 | 5 | 0 | 230 | 149 | 1 | 1 | 1 | 1 | 1 | 1 | 0.992 | 1 | 1 | 1 | 0.97 | 1 | 0.004 |
| SE | 28145 | ENSG00000267364 | NA | chr17 | + | 35316355 | 35316409 | 35313495 | 35313644 | 35321734 | 35321865 | 2 | 0 | 2 | 3 | 0 | 0 | 0 | 6 | 1 | 0 | 0 | 0 | 3 | 3 | 6 | 7 | 0 | 0 | 0 | 0 | 203 | 149 | 1 | 1 | 1 | NA | 0.197 | 0.688 | NA | 0 | NA | 1 | 1 | 1 | -0.122 |
| SE | 28146 | ENSG00000267364 | NA | chr17 | + | 35316355 | 35316409 | 35313495 | 35313644 | 35323147 | 35323343 | 1 | 2 | 1 | 0 | 0 | 10 | 2 | 1 | 5 | 0 | 0 | 0 | 1 | 3 | 0 | 0 | 2 | 6 | 4 | 0 | 203 | 149 | 1 | 1 | 0.068 | 0.423 | 0.423 | 0 | NA | NA | 0 | 0.109 | 0.355 | NA | 0.074 |
| SE | 28147 | ENSG00000267364 | NA | chr17 | + | 35316355 | 35316409 | 35313625 | 35313706 | 35321734 | 35321865 | 3 | 0 | 2 | 3 | 0 | 1 | 5 | 5 | 3 | 7 | 0 | 0 | 4 | 0 | 6 | 0 | 1 | 6 | 2 | 14 | 203 | 149 | 1 | 1 | 0.688 | 0 | 0.227 | 0.423 | 0 | NA | 0 | 0.329 | 0 | 0.239 | 0.126 |
| SE | 28148 | ENSG00000267364 | NA | chr17 | + | 35316355 | 35316409 | 35313625 | 35313706 | 35321736 | 35321865 | 0 | 0 | 1 | 0 | 0 | 0 | 0 | 0 | 0 | 0 | 0 | 0 | 0 | 4 | 0 | 0 | 0 | 1 | 0 | 0 | 203 | 149 | 1 | 1 | NA | NA | 1 | NA | NA | NA | NA | 0 | 1 | NA | 0.5 |
| SE | 28149 | ENSG00000267364 | NA | chr17 | + | 35316355 | 35316409 | 35313625 | 35313706 | 35323147 | 35323343 | 0 | 2 | 3 | 1 | 0 | 16 | 5 | 9 | 20 | 14 | 0 | 0 | 0 | 0 | 0 | 0 | 21 | 8 | 0 | 13 | 203 | 149 | 1 | 1 | 0 | 0.227 | 0.197 | 0.035 | 0 | NA | 0 | 0 | NA | 0 | 0.092 |
| SE | 28152 | ENSG00000267364 | NA | chr17 | + | 35321734 | 35321865 | 35316355 | 35316409 | 35323147 | 35323260 | 14 | 1 | 16 | 5 | 3 | 0 | 2 | 2 | 1 | 0 | 3 | 9 | 8 | 0 | 12 | 0 | 0 | 0 | 0 | 0 | 280 | 149 | 1 | 1 | 1 | 0.21 | 0.81 | 0.727 | 1 | 1 | 1 | 1 | NA | 1 | -0.251 |
| SE | 28153 | ENSG00000267364 | NA | chr17 | + | 35321736 | 35321865 | 35313523 | 35313706 | 35323147 | 35323268 | 11 | 0 | 15 | 4 | 3 | 16 | 5 | 9 | 20 | 14 | 3 | 9 | 5 | 0 | 4 | 0 | 21 | 8 | 0 | 13 | 278 | 149 | 1 | 1 | 0.269 | 0 | 0.472 | 0.097 | 0.103 | 1 | 0.187 | 0.251 | NA | 0.142 | -0.207 |
| SE | 28156 | ENSG00000109189 | USP46 | chr4 | - | 52604500 | 52604584 | 52601856 | 52602054 | 52610540 | 52610617 | 32 | 36 | 29 | 26 | 23 | 0 | 0 | 1 | 0 | 0 | 16 | 16 | 30 | 5 | 4 | 0 | 0 | 0 | 0 | 0 | 233 | 149 | 1 | 1 | 1 | 1 | 0.949 | 1 | 1 | 1 | 1 | 1 | 1 | 1 | -0.01 |
| SE | 28157 | ENSG00000109189 | USP46 | chr4 | - | 52629633 | 52629685 | 52628060 | 52628163 | 52631063 | 52631144 | 1 | 0 | 0 | 4 | 0 | 31 | 12 | 48 | 35 | 49 | 0 | 0 | 0 | 0 | 0 | 0 | 52 | 61 | 34 | 8 | 201 | 149 | 1 | 1 | 0.023 | 0 | 0 | 0.078 | 0 | NA | 0 | 0 | 0 | 0 | 0.02 |
| SE | 28158 | ENSG00000109189 | USP46 | chr4 | - | 52631063 | 52631144 | 52627949 | 52628163 | 52659114 | 52659301 | 31 | 16 | 53 | 39 | 50 | 0 | 1 | 0 | 0 | 0 | 0 | 67 | 69 | 35 | 8 | 0 | 0 | 0 | 0 | 0 | 230 | 149 | 1 | 1 | 1 | 0.912 | 1 | 1 | 1 | NA | 1 | 1 | 1 | 1 | -0.018 |
| SE | 28160 | ENSG00000109189 | USP46 | chr4 | - | 52658173 | 52658281 | 52631063 | 52631144 | 52659114 | 52659275 | 4 | 0 | 0 | 0 | 0 | 9 | 9 | 24 | 24 | 14 | 0 | 1 | 13 | 2 | 0 | 0 | 31 | 36 | 18 | 0 | 257 | 149 | 1 | 1 | 0.205 | 0 | 0 | 0 | 0 | NA | 0.018 | 0.173 | 0.061 | NA | -0.043 |
| SE | 28161 | ENSG00000138796 | HADH | chr4 | + | 108004545 | 108004917 | 107990015 | 107990064 | 108009758 | 108009887 | 0 | 0 | 0 | 0 | 3 | 13 | 6 | 16 | 13 | 14 | 0 | 0 | 0 | 0 | 0 | 27 | 3 | 10 | 11 | 19 | 298 | 149 | 1 | 1 | 0 | 0 | 0 | 0 | 0.097 | 0 | 0 | 0 | 0 | 0 | 0.019 |
| SE | 28162 | ENSG00000138796 | HADH | chr4 | + | [illegible] | [illegible] | [illegible] | [illegible] | [illegible] | [illegible] | [illegible] | [illegible] | [illegible] | [illegible] | [illegible] | [illegible] | [illegible] | [illegible] | [illegible] | [illegible] | [illegible] | [illegible] | [illegible] | [illegible] | [illegible] | [illegible] | [illegible] | [illegible] | [illegible] | [illegible] | [illegible] | [illegible] | [illegible] | [illegible] | [illegible] | [illegible] | [illegible] | [illegible] | [illegible] | [illegible] | [illegible] | [illegible] | [illegible] | [illegible] | [illegible] |

| | | | | | | | | | | | | | | |
|---|---|---|---|---|---|---|---|---|---|---|---|---|---|---|
| SE | 28169 | ENSG00000138796 | HADH | chr4 | + | 108027687 | 108027760 | 108023473 | 108023563 | 108027938 | 108028069 | 22 | 21 | [illegible] |
| SE | 28170 | ENSG00000138796 | HADH | chr4 | + | 108027687 | 108027760 | 108023473 | 108023563 | 108033175 | 108033292 | 64 | 59 | [illegible] |
| SE | 28171 | ENSG00000138796 | HADH | chr4 | + | 108027687 | 108028069 | 108023473 | 108023563 | 108033175 | 108033292 | 23 | 23 | [illegible] |
| SE | 28172 | ENSG00000138796 | HADH | chr4 | + | 108027938 | 108028069 | 108023473 | 108023563 | 108033175 | 108033292 | 2 | 3 | [illegible] |
| SE | 28177 | ENSG00000198915 | RASGEF1A | chr10 | - | 43196475 | 43196548 | 43194534 | 43196268 | 43196975 | 43197099 | 49 | 21 | [illegible] |
| SE | 28180 | ENSG00000170448 | NFXL1 | chr4 | - | 47862845 | 47862915 | 47855058 | 47855163 | 47875126 | 47875293 | 37 | 18 | [illegible] |
| SE | 28181 | ENSG00000170448 | NFXL1 | chr4 | - | 47877011 | 47877191 | 47875126 | 47875293 | 47878524 | 47878665 | 0 | 0 | [illegible] |
| SE | 28183 | ENSG00000170448 | NFXL1 | chr4 | - | 47896522 | 47896647 | 47894179 | 47894302 | 47897966 | 47898081 | 92 | 59 | [illegible] |
| SE | 28184 | ENSG00000164190 | NIPBL | chr5 | + | 36958103 | 36958231 | 36955471 | 36955637 | 36961483 | 36961583 | 606 | 474 | [illegible] |
| SE | 28186 | ENSG00000164190 | NIPBL | chr5 | + | 36970875 | 36971036 | 36962122 | 36962274 | 36971944 | 36972041 | 374 | 277 | [illegible] |
| SE | 28188 | ENSG00000164190 | NIPBL | chr5 | + | 36984675 | 36986301 | 36975775 | 36976402 | 37000816 | 37000888 | 198 | 149 | [illegible] |
| SE | 28189 | ENSG00000164190 | NIPBL | chr5 | + | 36995621 | 36995804 | 36975775 | 36976402 | 37000816 | 37000888 | 12 | 16 | [illegible] |
| SE | 28190 | ENSG00000164190 | NIPBL | chr5 | + | 36995621 | 36995804 | 36984675 | 36986301 | 36996471 | 36997002 | 412 | 373 | [illegible] |
| SE | 28191 | ENSG00000164190 | NIPBL | chr5 | + | 36995621 | 36995804 | 36984675 | 36986301 | 37000372 | 37000570 | 685 | 561 | [illegible] |
| SE | 28194 | ENSG00000164190 | NIPBL | chr5 | + | 37000372 | 37000570 | 36995621 | 36995804 | 37000816 | 37000888 | 447 | 333 | [illegible] |
| SE | 28195 | ENSG00000164190 | NIPBL | chr5 | + | 37008622 | 37008723 | 37008007 | 37008088 | 37010086 | 37010225 | 512 | 469 | [illegible] |
| SE | 28196 | ENSG00000164190 | NIPBL | chr5 | + | 37017018 | 37017162 | 37016037 | 37016170 | 37019310 | 37019400 | 922 | 885 | [illegible] |
| SE | 28197 | ENSG00000164190 | NIPBL | chr5 | + | 37026228 | 37026327 | 37024584 | 37024719 | 37027358 | 37027412 | 339 | 267 | [illegible] |
| SE | 28198 | ENSG00000164190 | NIPBL | chr5 | + | 37036378 | 37036487 | 37027358 | 37027412 | 37038601 | 37038738 | 379 | 259 | [illegible] |
| SE | 28199 | ENSG00000164190 | NIPBL | chr5 | + | 37038601 | 37038738 | 37027358 | 37027412 | 37044346 | 37044487 | 151 | 85 | [illegible] |
| SE | 28200 | ENSG00000164190 | NIPBL | chr5 | + | 37046108 | 37046199 | 37045442 | 37045597 | 37048501 | 37048675 | 497 | 407 | [illegible] |
| SE | 28201 | ENSG00000164190 | NIPBL | chr5 | + | 37049110 | 37049301 | 37048501 | 37048675 | 37051778 | 37051886 | 365 | 305 | [illegible] |
| SE | 28202 | ENSG00000164190 | NIPBL | chr5 | + | 37057185 | 37057332 | 37052365 | 37052566 | 37058890 | 37059165 | 454 | 303 | [illegible] |
| SE | 28203 | ENSG00000164190 | NIPBL | chr5 | + | 37057185 | 37057332 | 37052365 | 37052566 | 37060843 | 37061018 | 231 | 154 | [illegible] |
| SE | 28204 | ENSG00000164190 | NIPBL | chr5 | + | 37057185 | 37057332 | 37052365 | 37052566 | 37063789 | 37063978 | 227 | 149 | [illegible] |
| SE | 28205 | ENSG00000164190 | NIPBL | chr5 | + | 37058890 | 37059165 | 37052365 | 37052566 | 37060843 | 37061018 | 156 | 110 | [illegible] |
| SE | 28207 | ENSG00000164190 | NIPBL | chr5 | + | 37060843 | 37061018 | 37052365 | 37052566 | 37063789 | 37063978 | 296 | 259 | [illegible] |
| SE | 28208 | ENSG00000164190 | NIPBL | chr5 | + | 37060843 | 37061018 | 37057185 | 37057332 | 37063789 | 37063978 | 296 | 260 | [illegible] |
| SE | 28211 | ENSG00000204084 | INPP5B | chr1 | - | 37864311 | 37864423 | 37860696 | 37862430 | 37865760 | 37865888 | 154 | 87 | [illegible] |
| SE | 28212 | ENSG00000204084 | INPP5B | chr1 | - | 37864311 | 37864423 | 37860696 | 37862430 | 37868500 | 37868614 | 95 | 56 | [illegible] |
| SE | 28213 | ENSG00000204084 | INPP5B | chr1 | - | 37868500 | 37868614 | 37860696 | 37862430 | 37872929 | 37873165 | 49 | 16 | [illegible] |
| SE | 28214 | ENSG00000204084 | INPP5B | chr1 | - | 37868500 | 37868614 | 37866458 | 37866543 | 37872929 | 37873165 | 120 | 53 | [illegible] |
| SE | 28216 | ENSG00000204084 | INPP5B | chr1 | - | 37870669 | 37870720 | 37869924 | 37870014 | 37872929 | 37873028 | 0 | 0 | [illegible] |
| SE | 28217 | ENSG00000204084 | INPP5B | chr1 | - | 37873992 | 37874155 | 37872929 | 37873165 | 37875605 | 37875716 | 99 | 80 | [illegible] |
| SE | 28218 | ENSG00000204084 | INPP5B | chr1 | - | 37882806 | 37882918 | 37880084 | 37880194 | 37885637 | 37885825 | 121 | 92 | [illegible] |
| SE | 28219 | ENSG00000204084 | INPP5B | chr1 | - | 37882806 | 37882918 | 37880084 | 37880194 | 37886887 | 37887004 | 65 | 46 | [illegible] |
| SE | 28220 | ENSG00000204084 | INPP5B | chr1 | - | 37885637 | 37885825 | 37880084 | 37880194 | 37886887 | 37887004 | 112 | 53 | [illegible] |
| SE | 28222 | ENSG00000204084 | INPP5B | chr1 | - | 37931672 | 37932053 | 37891357 | 37891454 | 37940687 | 37940798 | 58 | 19 | [illegible] |
| SE | 28223 | ENSG00000204084 | INPP5B | chr1 | - | 37931912 | 37932053 | 37891357 | 37891454 | 37940687 | 37940798 | 23 | 8 | [illegible] |
| SE | 28227 | ENSG00000205937 | RNPS1 | chr16 | - | 2262277 | 2262431 | 2255584 | 2255726 | 2262739 | 2262842 | 582 | 405 | [illegible] |
| SE | 28228 | ENSG00000205937 | RNPS1 | chr16 | - | 2264175 | 2264331 | 2263095 | 2263287 | 2264572 | 2264760 | 748 | 452 | [illegible] |
| SE | 28229 | ENSG00000205937 | RNPS1 | chr16 | - | 2264182 | 2264331 | 2263213 | 2263287 | 2264572 | 2264760 | 228 | 148 | [illegible] |
| SE | 28232 | ENSG00000205937 | RNPS1 | chr16 | - | 2266109 | 2266216 | 2264572 | 2264760 | 2267662 | 2267833 | 4 | 0 | [illegible] |
| SE | 28235 | ENSG00000205937 | RNPS1 | chr16 | - | 2267188 | 2267237 | 2264572 | 2264760 | 2268054 | 2268114 | 0 | 3 | [illegible] |
| SE | 28236 | ENSG00000157326 | DHRS4 | chr14 | + | 23959901 | 23960003 | 23955034 | 23955212 | 23965761 | 23965832 | 46 | 13 | [illegible] |
| SE | 28237 | ENSG00000157326 | DHRS4 | chr14 | + | 23959901 | 23960003 | 23955034 | 23955212 | 23966282 | 23966417 | 17 | 1 | [illegible] |
| SE | 28245 | ENSG00000157326 | DHRS4 | chr14 | + | 23967390 | 23967455 | 23967210 | 23967266 | 23968756 | 23969274 | 1 | 0 | [illegible] |
| SE | 28261 | ENSG00000138443 | ABI2 | chr2 | + | 203351554 | 203351678 | 203328281 | 203328631 | 203355153 | 203355324 | 0 | 0 | [illegible] |
| SE | 28262 | ENSG00000138443 | ABI2 | chr2 | + | 203351554 | 203351678 | 203328281 | 203328631 | 203366876 | 203367044 | 0 | 0 | [illegible] |
| SE | 28264 | ENSG00000138443 | ABI2 | chr2 | + | 203355153 | 203355324 | 203351554 | 203351678 | 203366876 | 203366957 | 0 | 0 | [illegible] |
| SE | 28265 | ENSG00000138443 | ABI2 | chr2 | + | 203366876 | 203367044 | 203328279 | 203328631 | 203376056 | 203376132 | 18 | 13 | [illegible] |
| SE | 28266 | ENSG00000138443 | ABI2 | chr2 | + | 203366876 | 203367044 | 203328279 | 203328631 | 203380207 | 203380306 | 104 | 82 | [illegible] |
| SE | 28269 | ENSG00000138443 | ABI2 | chr2 | + | 203376056 | 203376132 | 203328394 | 203328631 | 203380207 | 203380273 | 3 | 3 | [illegible] |
| SE | 28271 | ENSG00000138443 | ABI2 | chr2 | + | 203380207 | 203380384 | 203366876 | 203367044 | 203391045 | 203391143 | 149 | 122 | [illegible] |
| SE | 28273 | ENSG00000138443 | ABI2 | chr2 | + | 203391045 | 203391143 | 203380207 | 203380384 | 203394699 | 203394846 | 83 | 62 | [illegible] |
| SE | 28276 | ENSG00000138443 | ABI2 | chr2 | + | 203394699 | 203394846 | 203391043 | 203391143 | 203395658 | 203395780 | 33 | 15 | [illegible] |
| SE | 28277 | ENSG00000138443 | ABI2 | chr2 | + | 203395655 | 203395780 | 203391045 | 203391143 | 203402575 | 203402734 | 26 | 12 | [illegible] |
| SE | 28279 | ENSG00000138443 | ABI2 | chr2 | + | 203395658 | 203395780 | 203391045 | 203391143 | 203402575 | 203402734 | 20 | 13 | [illegible] |
| SE | 28289 | ENSG00000171490 | RSL1D1 | chr16 | - | 11846500 | 11846602 | 11841906 | 11842000 | 11846694 | 11846843 | 624 | 458 | [illegible] |
| SE | 28290 | ENSG00000171490 | RSL1D1 | chr16 | - | 11849257 | 11849385 | 11847667 | 11847806 | 11850278 | 11850418 | 0 | 0 | [illegible] |
| SE | 28291 | ENSG00000171490 | RSL1D1 | chr16 | - | 11850278 | 11850413 | 11847667 | 11847806 | 11851407 | 11851580 | 367 | 224 | [illegible] |
| SE | 28292 | ENSG00000171490 | RSL1D1 | chr16 | - | 11850278 | 11850418 | 11847573 | 11847806 | 11851407 | 11851508 | 689 | 420 | [illegible] |
| SE | 28307 | ENSG00000001036 | FUCA2 | chr6 | - | 143501931 | 143502122 | 143496951 | 143497497 | 143503912 | 143504252 | 135 | 99 | [illegible] |
| SE | 28308 | ENSG00000001036 | FUCA2 | chr6 | - | 143502354 | 143502565 | 143501931 | 143502122 | 143503912 | 143504252 | 367 | 196 | [illegible] |
| SE | 28313 | ENSG00000123545 | NDUFAF4 | chr6 | - | 96896743 | 96896847 | 96889314 | 96891391 | 96897665 | 96897874 | 148 | 76 | [illegible] |
| SE | 28316 | ENSG00000088538 | DOCK3 | chr3 | + | 51089242 | 51089284 | 51075355 | 51075440 | 51090229 | 51090384 | 4 | 5 | [illegible] |
| SE | 28317 | ENSG00000088538 | DOCK3 | chr3 | + | 51146548 | 51146630 | 51090229 | 51090384 | 51159243 | 51159304 | 6 | 3 | [illegible] |
| SE | 28318 | ENSG00000088538 | DOCK3 | chr3 | + | 51227282 | 51227445 | 51225648 | 51225773 | 51227981 | 51228088 | 9 | 2 | [illegible] |
| SE | 28319 | ENSG00000088538 | DOCK3 | chr3 | + | 51270814 | 51271007 | 51260155 | 51260326 | 51275078 | 51275206 | 10 | 2 | [illegible] |
| SE | 28323 | ENSG00000139505 | MTMR6 | chr13 | - | 25254384 | 25254434 | 25253763 | 25253964 | 25257195 | 25257321 | 89 | 30 | [illegible] |
| SE | 28324 | ENSG00000139505 | MTMR6 | chr13 | - | 25260573 | 25260687 | 25258620 | 25258692 | 25261667 | 25261808 | 0 | 0 | [illegible] |
| SE | 28325 | ENSG00000139505 | MTMR6 | chr13 | - | 25261667 | 25261802 | 25258559 | 25258692 | 25265818 | 25265947 | 165 | 102 | [illegible] |
| SE | 28326 | ENSG00000141385 | AFG3L2 | chr18 | - | 12337340 | 12337535 | 12328943 | 12329783 | 12340200 | 12340401 | 149 | 132 | [illegible] |
| SE | 28328 | ENSG00000141385 | AFG3L2 | chr18 | - | 12348272 | 12348383 | 12344131 | 12344247 | 12351084 | 12351210 | 113 | 85 | [illegible] |
| SE | 28329 | ENSG00000141385 | AFG3L2 | chr18 | - | 12353004 | 12353158 | 12351305 | 12351413 | 12356693 | 12356831 | 154 | 74 | [illegible] |
| SE | 28331 | ENSG00000141385 | AFG3L2 | chr18 | - | 12363781 | 12363856 | 12359926 | 12360051 | 12366964 | 12367117 | 183 | 86 | [illegible] |
| SE | 28332 | ENSG00000141385 | AFG3L2 | chr18 | - | 12367275 | 12367382 | 12366964 | 12367117 | 12370848 | 12370926 | 294 | 186 | [illegible] |
| SE | 28340 | ENSG00000070831 | CDC42 | chr1 | + | 22078746 | 22078881 | 22078428 | 22078583 | 22081721 | 22081794 | 10 | 13 | [illegible] |
| SE | 28341 | ENSG00000070831 | CDC42 | chr1 | + | 22078746 | 22078881 | 22078428 | 22078583 | 22086438 | 22086548 | 1 | 0 | [illegible] |
| SE | 28342 | ENSG00000070831 | CDC42 | chr1 | + | 22081721 | 22081794 | 22078428 | 22078583 | 22086438 | 22086548 | 3224 | 2129 | [illegible] |
| SE | 28343 | ENSG00000070831 | CDC42 | chr1 | + | 22081721 | 22081794 | 22078705 | 22078881 | 22086438 | 22086548 | 1098 | 678 | [illegible] |
| SE | 28344 | ENSG00000070831 | CDC42 | chr1 | + | 22086668 | 22086866 | 22086438 | 22086548 | 22091427 | 22091576 | 4956 | 3328 | [illegible] |
| SE | 28357 | ENSG00000145907 | G3BP1 | chr5 | + | 151790322 | 151790404 | 151786571 | 151786715 | 151790888 | 151790990 | 394 | 181 | [illegible] |
| SE | 28358 | ENSG00000145907 | G3BP1 | chr5 | + | 151790888 | 151791062 | 151786571 | 151786715 | 151794158 | 151794222 | 247 | 145 | [illegible] |
| SE | 28363 | ENSG00000145907 | G3BP1 | chr5 | + | 151794158 | 151794249 | 151790888 | 151791062 | 151795478 | 151795575 | 518 | 325 | [illegible] |
| SE | 28367 | ENSG00000145907 | G3BP1 | chr5 | + | 151797226 | 151797428 | 151794158 | 151795575 | 151799211 | 151799313 | 878 | 569 | [illegible] |
| SE | 28368 | ENSG00000145907 | G3BP1 | chr5 | + | 151797288 | 151797428 | 151795478 | 151795575 | 151799211 | 151799313 | 461 | 304 | [illegible] |
| SE | 28369 | ENSG00000145907 | G3BP1 | chr5 | + | 151799888 | 151800000 | 151799211 | 151799313 | 151800217 | 151800258 | 822 | 538 | [illegible] |
| SE | 28370 | ENSG00000145907 | G3BP1 | chr5 | + | 151799888 | 151800043 | 151799211 | 151799313 | 151800217 | 151800346 | 525 | 359 | [illegible] |
| SE | 28371 | ENSG00000145907 | G3BP1 | chr5 | + | 151799898 | 151800000 | 151799211 | 151799313 | 151800217 | 151800346 | 250 | 149 | [illegible] |
| SE | 28374 | ENSG00000145907 | G3BP1 | chr5 | + | 151800217 | 151800346 | 151799211 | 151799313 | 151803884 | 151804141 | 0 | 0 | [illegible] |
| SE | 28377 | ENSG00000145907 | G3BP1 | chr5 | + | 151803537 | 151803636 | 151800759 | 151800869 | 151803884 | 151804213 | 0 | 0 | [illegible] |
| SE | 28378 | ENSG00000145907 | G3BP1 | chr5 | + | 151803884 | 151803996 | 151800759 | 151800869 | 151804405 | 151812785 | 197 | 107 | [illegible] |
| SE | 28379 | ENSG00000145907 | G3BP1 | chr5 | + | 151803884 | 151804141 | 151800759 | 151800869 | 151804405 | 151805349 | 251 | 135 | [illegible] |
| SE | 28380 | ENSG00000145907 | G3BP1 | chr5 | + | 151803884 | 151804162 | 151800759 | 151800869 | 151804405 | 151812785 | 251 | 135 | [illegible] |
| SE | 28381 | ENSG00000204842 | ATXN2 | chr12 | - | 111453676 | 111453845 | 111451946 | 111452085 | 111456028 | 111456256 | 1 | 0 | [illegible] |
| SE | 28383 | ENSG00000204842 | ATXN2 | chr12 | - | 111455044 | 111455119 | 111453676 | 111453845 | 111456028 | 111456256 | 0 | 0 | [illegible] |
| SE | 28384 | ENSG00000204842 | ATXN2 | chr12 | - | 111461138 | 111461305 | 111457213 | 111457359 | 111464661 | 111464715 | 0 | 0 | [illegible] |
| SE | 28385 | ENSG00000204842 | ATXN2 | chr12 | - | 111461138 | 111461305 | 111457213 | 111457359 | 111470107 | 111470240 | 0 | 0 | [illegible] |
| SE | 28389 | ENSG00000204842 | ATXN2 | chr12 | - | 111479069 | 111479102 | 111470557 | 111470742 | 111485264 | 111485331 | 1 | 0 | [illegible] |
| SE | 28390 | ENSG00000204842 | ATXN2 | chr12 | - | 111479069 | 111479102 | 111470557 | 111470742 | 111485270 | 111485331 | 0 | 0 | [illegible] |
| SE | 28393 | ENSG00000204842 | ATXN2 | chr12 | - | 111482786 | 111482831 | 111479069 | 111479102 | 111485264 | 111485331 | 0 | 2 | [illegible] |
| SE | 28394 | ENSG00000204842 | ATXN2 | chr12 | - | 111482786 | 111482831 | 111479069 | 111479102 | 111485270 | 111485331 | 0 | 0 | [illegible] |
| SE | 28411 | ENSG00000131495 | NDUFA2 | chr5 | - | 140647255 | 140647362 | 140645284 | 140645678 | 140647482 | 140647630 | 79 | 34 | [illegible] |
| SE | 28414 | ENSG00000148719 | DNAJB12 | chr10 | - | 72344949 | 72345127 | 72343365 | 72343511 | 72354764 | 72354919 | 176 | 132 | [illegible] |
| SE | 28418 | ENSG00000174652 | ZNF266 | chr19 | - | 9417827 | 9417908 | 9415653 | 9415742 | 9418504 | 9418631 | 246 | 142 | [illegible] |
| SE | 28419 | ENSG00000174652 | ZNF266 | chr19 | - | 9418761 | 9418895 | 9418504 | 9418631 | 9419050 | 9419299 | 46 | 24 | [illegible] |
| SE | 28420 | ENSG00000174652 | ZNF266 | chr19 | - | 9418761 | 9419299 | 9418504 | 9418631 | 9420064 | 9420218 | 98 | 23 | [illegible] |
| SE | 28421 | ENSG00000174652 | ZNF266 | chr19 | - | 9418765 | 9418895 | 9418504 | 9418631 | 9419050 | 9419299 | 59 | 37 | [illegible] |
| SE | 28423 | ENSG00000174652 | ZNF266 | chr19 | - | 9419050 | 9419299 | 9418504 | 9418631 | 9420064 | 9420218 | 65 | 13 | [illegible] |
| SE | 28424 | ENSG00000174652 | ZNF266 | chr19 | - | 9419216 | 9419299 | 9418504 | 9418631 | 9420064 | 9420218 | 85 | 41 | [illegible] |
| SE | 28425 | ENSG00000174652 | ZNF266 | chr19 | - | 9433667 | 9433783 | 9420064 | 9420218 | 9434064 | 9434235 | 8 | 10 | [illegible] |
| SE | 28426 | ENSG00000174652 | ZNF266 | chr19 | - | 9433667 | 9433783 | 9420064 | 9420218 | 9434074 | 9434235 | 18 | 10 | [illegible] |
| SE | 28427 | ENSG00000174652 | ZNF266 | chr19 | - | 9433667 | 9433786 | 9420064 | 9420218 | 9434064 | 9434235 | 9 | 8 | [illegible] |
| SE | 28428 | ENSG00000174652 | ZNF266 | chr19 | - | 9433667 | 9433786 | 9420064 | 9420218 | 9434074 | 9434235 | 25 | 24 | [illegible] |
| SE | 28431 | ENSG00000174652 | ZNF266 | chr19 | - | 9434074 | 9434235 | 9420064 | 9420218 | 9435107 | 9435193 | 6 | 4 | [illegible] |
| SE | 28432 | ENSG00000174652 | ZNF266 | chr19 | - | 9434797 | 9434859 | 9420064 | 9420218 | 9435107 | 9435193 | 7 | 2 | [illegible] |
| SE | 28433 | ENSG00000174652 | ZNF266 | chr19 | - | 9434797 | 9434859 | 9434064 | 9434235 | 9435107 | 9435193 | 23 | 7 | [illegible] |
| SE | 28434 | ENSG00000106991 | ENG | chr9 | - | 127818715 | 127818832 | 127818119 | 127818377 | 127819621 | 127819660 | 64 | 23 | [illegible] |
| SE | 28435 | ENSG00000106991 | ENG | chr9 | - | 127827044 | 127827186 | 127826509 | 127826672 | 127829686 | 127829804 | 0 | 0 | [illegible] |
| SE | 28436 | ENSG00000106991 | ENG | chr9 | - | 127829686 | 127829827 | 127826509 | 127826672 | 127843093 | 127843245 | 64 | 30 | [illegible] |
| SE | 28437 | ENSG00000106991 | ENG | chr9 | - | 127843093 | 127843245 | 127829686 | 127829827 | 127854288 | 127854636 | 49 | 30 | [illegible] |
| SE | 28442 | ENSG00000106692 | FKTN | chr9 | + | 105570568 | 105570897 | 105558137 | 105559109 | 105573654 | 105573746 | 1 | 1 | [illegible] |
| SE | 28443 | ENSG00000106692 | FKTN | chr9 | + | 105570750 | 105570897 | 105558137 | 105558165 | 105573654 | 105573746 | 4 | 1 | [illegible] |
| SE | 28444 | ENSG00000106692 | FKTN | chr9 | + | 105573654 | 105573746 | 105558129 | 105558165 | 105574944 | 105575137 | 9 | 19 | [illegible] |
| SE | 28445 | ENSG00000106692 | FKTN | chr9 | + | 105574197 | 105574278 | 105573654 | 105573746 | 105574944 | 105575137 | 0 | 17 | [illegible] |
| SE | 28446 | ENSG00000106692 | FKTN | chr9 | + | 105574197 | 105574332 | 105573654 | 105573746 | 105574944 | 105575018 | 9 | 11 | [illegible] |
| SE | 28447 | ENSG00000106692 | FKTN | chr9 | + | 105598085 | 105598183 | 105596597 | 105596657 | 105598630 | 105598701 | 0 | 1 | [illegible] |
| SE | 28448 | ENSG00000106692 | FKTN | chr9 | + | 105598085 | 105598183 | 105596597 | 105596657 | 105600917 | 105601041 | 0 | 1 | [illegible] |
| SE | 28449 | ENSG00000106692 | FKTN | chr9 | + | 105598085 | 105598183 | 105596597 | 105596657 | 105601144 | 105601348 | 1 | 2 | [illegible] |
| SE | 28450 | ENSG00000106692 | FKTN | chr9 | + | 105598505 | 105598701 | 105598085 | 105598183 | 105601144 | 105601348 | 0 | 0 | [illegible] |
| SE | 28452 | ENSG00000106692 | FKTN | chr9 | + | 105598630 | 105598701 | 105596597 | 105596657 | 105601144 | 105601348 | 0 | 0 | [illegible] |
| SE | 28454 | ENSG00000106692 | FKTN | chr9 | + | 105598630 | 105598701 | 105598085 | 105598183 | 105601144 | 105601348 | 0 | 0 | [illegible] |
| SE | 28455 | ENSG00000106692 | FKTN | chr9 | + | 105600917 | 105601041 | 105596597 | 105596657 | 105601144 | 105601348 | 3 | 5 | [illegible] |
| SE | 28456 | ENSG00000106692 | FKTN | chr9 | + | 105600917 | 105601041 | 105598085 | 105598183 | 105601144 | 105601348 | 3 | 5 | [illegible] |
| SE | 28457 | ENSG00000106692 | FKTN | chr9 | + | 105600917 | 105601041 | 105598630 | 105598701 | 105601144 | 105601348 | 3 | 5 | [illegible] |
| SE | 28458 | ENSG00000106692 | FKTN | chr9 | + | 105603698 | 105603816 | 105599930 | 105601348 | 105604214 | 105604492 | 0 | 0 | [illegible] |
| SE | 28460 | ENSG00000106692 | FKTN | chr9 | + | 105615277 | 105615407 | 105607818 | 105607951 | 105617958 | 105618092 | 25 | 35 | [illegible] |
| SE | 28461 | ENSG00000106692 | FKTN | chr9 | + | 105615277 | 105615407 | 105607818 | 105607951 | 105619933 | 105620061 | 8 | 17 | [illegible] |
| SE | 28462 | ENSG00000106692 | FKTN | chr9 | + | 105615280 | 105615407 | 105607818 | 105607951 | 105617958 | 105618092 | 19 | 20 | [illegible] |
| SE | 28463 | ENSG00000106692 | FKTN | chr9 | + | 105617958 | 105618092 | 105607818 | 105607951 | 105619905 | 105620061 | 2 | 13 | [illegible] |
| SE | 28464 | ENSG00000106692 | FKTN | chr9 | + | 105617958 | 105618092 | 105607818 | 105607951 | 105619933 | 105620061 | 14 | 14 | [illegible] |
| SE | 28465 | ENSG00000106692 | FKTN | chr9 | + | 105617958 | 105618092 | 105615277 | 105615407 | 105619933 | 105620061 | 27 | 31 | [illegible] |
| SE | 28466 | ENSG00000106692 | FKTN | chr9 | + | 105619933 | 105620061 | 105607818 | 105607951 | 105635050 | 105635148 | 9 | 16 | [illegible] |
| SE | 28467 | ENSG00000106692 | FKTN | chr9 | + | 105623049 | 105623221 | 105619905 | 105620061 | 105633355 | 105633474 | 5 | 4 | [illegible] |
| SE | 28468 | ENSG00000106692 | FKTN | chr9 | + | 105623049 | 105623221 | 105619905 | 105620061 | 105635050 | 105637062 | 8 | 4 | [illegible] |
| SE | 28469 | ENSG00000106692 | FKTN | chr9 | + | 105623049 | 105623221 | 105620043 | 105620092 | 105635050 | 105637062 | 3 | 0 | [illegible] |
| SE | 28472 | ENSG00000106692 | FKTN | chr9 | + | 105633355 | 105633474 | 105619933 | 105620061 | 105635050 | 105635148 | 0 | 2 | [illegible] |
| SE | 28473 | ENSG00000106692 | FKTN | chr9 | + | 105633355 | 105633474 | 105623049 | 105623221 | 105635050 | 105635148 | 0 | 1 | [illegible] |
| SE | 28475 | ENSG00000104972 | LILRB1 | chr19 | + | 54631025 | 54631107 | 54630409 | 54630633 | 54631270 | 54631306 | 133 | 35 | [illegible] |
| SE | 28480 | ENSG00000104972 | LILRB1 | chr19 | + | 54636493 | 54636652 | 54635556 | 54635609 | 54636731 | 54637076 | 160 | 85 | [illegible] |
| SE | 28481 | ENSG00000104972 | LILRB1 | chr19 | + | 54636496 | 54636652 | 54635556 | 54635609 | 54636731 | 54637076 | 168 | 86 | [illegible] |
| SE | 28482 | ENSG00000248727 | LINC01948 | chr5 | - | 56464008 | 56464116 | 56451626 | 56457943 | 56467711 | 56467812 | 0 | 2 | [illegible] |
| SE | 28487 | ENSG00000155229 | MMS19 | chr10 | - | 97466503 | 97466585 | 97466058 | 97466159 | 97466775 | 97466901 | 56 | 25 | [illegible] |
| SE | 28488 | ENSG00000155229 | MMS19 | chr10 | - | 97466775 | 97466901 | 97466503 | 97466585 | 97467504 | 97467525 | 76 | 39 | [illegible] |
| SE | 28491 | ENSG00000155229 | MMS19 | chr10 | - | 97470774 | 97470834 | 97470128 | 97470203 | 97476682 | 97476744 | 16 | 9 | [illegible] |
| SE | 28492 | ENSG00000155229 | MMS19 | chr10 | - | 97470774 | 97470861 | 97470128 | 97470203 | 97476682 | 97476744 | 65 | 42 | [illegible] |
| SE | 28495 | ENSG00000155229 | MMS19 | chr10 | - | 97477854 | 97477929 | 97476916 | 97477416 | 97478303 | 97478389 | 209 | 119 | [illegible] |
| SE | 28503 | ENSG00000169062 | UPF3A | chr13 | + | 114286518 | 114286629 | 114281952 | 114282127 | 114291633 | 114291718 | 1 | 1 | [illegible] |
| SE | 28508 | ENSG00000169062 | UPF3A | chr13 | + | 114291633 | 114291792 | 114286518 | 114286629 | 114301730 | 114301867 | 38 | 47 | [illegible] |
| SE | 28509 | ENSG00000169062 | UPF3A | chr13 | + | 114298839 | 114299000 | 114286518 | 114286629 | 114301730 | 114301756 | 76 | 97 | [illegible] |
| SE | 28511 | ENSG00000169062 | UPF3A | chr13 | + | 114298839 | 114299000 | 114291604 | 114291792 | 114304788 | 114305808 | 18 | 23 | [illegible] |
| SE | 28512 | ENSG00000169062 | UPF3A | chr13 | + | 114301730 | 114301867 | 114291633 | 114291792 | 114304788 | 114304949 | 37 | 48 | [illegible] |
| SE | 28513 | ENSG00000169062 | UPF3A | chr13 | + | 114301730 | 114302025 | 114298839 | 114299000 | 114304788 | 114305808 | 143 | 160 | [illegible] |
| SE | 28531 | ENSG00000237651 | C2orf74 | chr2 | + | 61157861 | 61158012 | 61145129 | 61145196 | 61159317 | 61159439 | 1 | 8 | [illegible] |
| SE | 28532 | ENSG00000237651 | C2orf74 | chr2 | + | 61157861 | 61158012 | 61145129 | 61145196 | 61162841 | 61162955 | 1 | 13 | [illegible] |
| SE | 28533 | ENSG00000237651 | C2orf74 | chr2 | + | 61159317 | 61159439 | 61145130 | 61145196 | 61162841 | 61162955 | 4 | 0 | [illegible] |
| SE | 28534 | ENSG00000237651 | C2orf74 | chr2 | + | 61159317 | 61159439 | 61157861 | 61158012 | 61162841 | 61162955 | 2 | 1 | [illegible] |
| SE | 28535 | ENSG00000237651 | C2orf74 | chr2 | + | 61162841 | 61162955 | 61145067 | 61145196 | 61163051 | 61163142 | 61 | 33 | [illegible] |
| SE | 28544 | ENSG00000170088 | TMEM192 | chr4 | - | 165100627 | 165100892 | 165088467 | 165088602 | 165102949 | 165103096 | 91 | 66 | [illegible] |
| SE | 28545 | ENSG00000170088 | TMEM192 | chr4 | - | 165111648 | 165111679 | 165102949 | 165103096 | 165112746 | 165112854 | 3 | 3 | [illegible] |
| SE | 28553 | ENSG00000089091 | DZANK1 | chr20 | - | 18394253 | 18394350 | 18393710 | 18393811 | 18396471 | 18396546 | 9 | 11 | [illegible] |
| SE | 28554 | ENSG00000089091 | DZANK1 | chr20 | - | 18394674 | 18394858 | 18394253 | 18394350 | 18396471 | 18396546 | 0 | 2 | [illegible] |
| SE | 28555 | ENSG00000089091 | DZANK1 | chr20 | - | 18398522 | 18398626 | 18396471 | 18396546 | 18412645 | 18412835 | 10 | 8 | [illegible] |
| SE | 28558 | ENSG00000089091 | DZANK1 | chr20 | - | 18443346 | 18443464 | 18432494 | 18433765 | 18448983 | 18448988 | 8 | 3 | [illegible] |
| SE | 28559 | ENSG00000089091 | DZANK1 | chr20 | - | 18451853 | 18451962 | 18448983 | 18449069 | 18452614 | 18452682 | 1 | 0 | [illegible] |
| SE | 28560 | ENSG00000089091 | DZANK1 | chr20 | - | 18452614 | 18452682 | 18448983 | 18449069 | 18453730 | 18453827 | 5 | 3 | [illegible] |
| SE | 28561 | ENSG00000089091 | DZANK1 | chr20 | - | 18460152 | 18460306 | 18455246 | 18455361 | 18465249 | 18465377 | 11 | 18 | [illegible] |
| SE | 28562 | ENSG00000241258 | CRCP | chr7 | + | 66127703 | 66127740 | 66114603 | 66114970 | 66130743 | 66130842 | 38 | 20 | [illegible] |
| SE | 28563 | ENSG00000241258 | CRCP | chr7 | + | 66127703 | 66127740 | 66114603 | 66114970 | 66145442 | 66145500 | 0 | 4 | [illegible] |
| SE | 28565 | ENSG00000241258 | CRCP | chr7 | + | 66130743 | 66130842 | 66114603 | 66114970 | 66145442 | 66145500 | 2 | 2 | [illegible] |
| SE | 28570 | ENSG00000241258 | CRCP | chr7 | + | 66145442 | 66145500 | 66134279 | 66134374 | 66152207 | 66152511 | 175 | 131 | [illegible] |
| SE | 28573 | ENSG00000119414 | PPP6C | chr9 | - | 125153532 | 125153742 | 125146572 | 125149921 | 125153905 | 125153985 | 622 | 405 | [illegible] |
| SE | 28574 | ENSG00000119414 | PPP6C | chr9 | - | 125153905 | 125153985 | 125146572 | 125149921 | 125158240 | 125158382 | 284 | 165 | [illegible] |
| SE | 28575 | ENSG00000119414 | PPP6C | chr9 | - | 125158240 | 125158382 | 125153905 | 125153985 | 125160840 | 125160906 | 749 | 496 | [illegible] |
| SE | 28579 | ENSG00000119414 | PPP6C | chr9 | - | 125171084 | 125171180 | 125160840 | 125160906 | 125189643 | 125189803 | 513 | 273 | [illegible] |
| SE | 28583 | ENSG00000081087 | OSTM1 | chr6 | - | 108049252 | 108049418 | 108041408 | 108044840 | 108049531 | 108049583 | 161 | 143 | [illegible] |
| SE | 28587 | ENSG00000186352 | ANKRD37 | chr4 | + | 185396831 | 185396950 | 185396044 | 185396424 | 185397149 | 185397255 | 4 | 1 | [illegible] |
| SE | 28589 | ENSG00000105821 | DNAJC2 | chr7 | - | 103326542 | 103326684 | 103324481 | 103324562 | 103327331 | 103327378 | 107 | 109 | [illegible] |
| SE | 28590 | ENSG00000105821 | DNAJC2 | chr7 | - | 103326542 | 103326684 | 103324481 | 103324562 | 103327655 | 103327754 | 175 | 155 | [illegible] |
| SE | 28591 | ENSG00000105821 | DNAJC2 | chr7 | - | 103327331 | 103327378 | 103324481 | 103324562 | 103327655 | 103327754 | 1 | 2 | [illegible] |
| SE | 28593 | ENSG00000105821 | DNAJC2 | chr7 | - | 103337735 | 103337811 | 103327655 | 103327754 | 103341763 | 103341865 | 167 | 155 | [illegible] |
| SE | 28594 | ENSG00000113369 | ARRDC3 | chr5 | - | 91373683 | 91373838 | 91368630 | 91371456 | 91374113 | 91374276 | 463 | 231 | [illegible] |
| SE | 28608 | ENSG00000268173 | NA | chr19 | + | 18175604 | 18175697 | 18175310 | 18175385 | 18177139 | 18177292 | 2848 | 1424 | [illegible] |
| SE | 28609 | ENSG00000268173 | NA | chr19 | + | 18177139 | 18177292 | 18175604 | 18175697 | 18177710 | 18177764 | 3170 | 1651 | [illegible] |
| SE | 28613 | ENSG00000102445 | RUBCNL | chr13 | - | 46345446 | 46345600 | 46344740 | 46344831 | 46349285 | 46349347 | 34 | 37 | [illegible] |
| SE | 28616 | ENSG00000102445 | RUBCNL | chr13 | - | 46356431 | 46356496 | 46350112 | 46350351 | 46359485 | 46359631 | 98 | 54 | [illegible] |
| SE | 28617 | ENSG00000102445 | RUBCNL | chr13 | - | 46359485 | 46359631 | 46356431 | 46356496 | 46362537 | 46362598 | 84 | 54 | [illegible] |
| SE | 28618 | ENSG00000102445 | RUBCNL | chr13 | - | 46361440 | 46361573 | 46359485 | 46359631 | 46362537 | 46362598 | 141 | 118 | [illegible] |
| SE | 28624 | ENSG00000102445 | RUBCNL | chr13 | - | 46371499 | 46371648 | 46368732 | 46368815 | 46371940 | 46372597 | 0 | 1 | [illegible] |
| SE | 28636 | ENSG00000102445 | RUBCNL | chr13 | - | 46377889 | 46378005 | 46371940 | 46372210 | 46387133 | 46387230 | 2 | 4 | [illegible] |
| SE | 28637 | ENSG00000102445 | RUBCNL | chr13 | - | 46377889 | 46378005 | 46371940 | 46372597 | 46378446 | 46378645 | 2 | 6 | [illegible] |
| SE | 28638 | ENSG00000102445 | RUBCNL | chr13 | - | 46377889 | 46378005 | 46371940 | 46372597 | 46385254 | 46385298 | 2 | 6 | [illegible] |
| SE | 28640 | ENSG00000102445 | RUBCNL | chr13 | - | 46377889 | 46378005 | 46371940 | 46372597 | 46387676 | 46387831 | 2 | 6 | [illegible] |
| SE | 28641 | ENSG00000102445 | RUBCNL | chr13 | - | 46377889 | 46378005 | 46376491 | 46376584 | 46386415 | 46386725 | 1 | 1 | [illegible] |
| SE | 28643 | ENSG00000102445 | RUBCNL | chr13 | - | 46385254 | 46385298 | 46377889 | 46378005 | 46386415 | 46386725 | 0 | 0 | [illegible] |
| SE | 28644 | ENSG00000102445 | RUBCNL | chr13 | - | 46386415 | 46386725 | 46377889 | 46378005 | 46387133 | 46387500 | 1 | 1 | [illegible] |
| SE | 28645 | ENSG00000058063 | ATP11B | chr3 | + | 182820259 | 182820376 | 182793502 | 182793786 | 182828119 | 182828209 | 239 | 241 | [illegible] |
| SE | 28646 | ENSG00000058063 | ATP11B | chr3 | + | 182842074 | 182842122 | 182837070 | 182837174 | 182845457 | 182845522 | 363 | 248 | [illegible] |

| | | | | | | | | | | | | | | | | | | |
|---|---|---|---|---|---|---|---|---|---|---|---|---|---|---|---|---|---|---|
| SE | 28647 | ENSG00000058063 | ATP11B | chr3 | + | 182845457 | 182845522 | 182842074 | 182842122 | 182848475 | 182848557 | 247 | 196 | 220 | 336 | 356 | 1 | 0 |
| SE | 28649 | ENSG00000058063 | ATP11B | chr3 | + | 182880878 | 182880947 | 182879495 | 182879649 | 182884752 | 182884898 | 100 | 61 | 72 | 97 | 111 | 2 | 0 |
| SE | 28650 | ENSG00000058063 | ATP11B | chr3 | + | 182880878 | 182880981 | 182879495 | 182879649 | 182884752 | 182884898 | 447 | 240 | 363 | 411 | 539 | 2 | 0 |
| SE | 28651 | ENSG00000058063 | ATP11B | chr3 | + | 182884752 | 182884898 | 182879495 | 182879649 | 182887585 | 182887713 | 2 | 0 | 0 | 4 | 7 | 0 | 0 |
| SE | 28653 | ENSG00000058063 | ATP11B | chr3 | + | 182902480 | 182902566 | 182898606 | 182898772 | 182905793 | 182905919 | 0 | 0 | 0 | 0 | 0 | 0 | 0 |
| SE | 28654 | ENSG00000058063 | ATP11B | chr3 | + | 182902480 | 182902566 | 182898606 | 182898772 | 182913860 | 182913994 | 0 | 0 | 0 | 0 | 0 | 121 | 100 |
| SE | 28655 | ENSG00000058063 | ATP11B | chr3 | + | 182905793 | 182905919 | 182898684 | 182898772 | 182913860 | 182913978 | 0 | 0 | 6 | 1 | 0 | 121 | 100 |
| SE | 28659 | ENSG00000002586 | CD99 | chrX | + | 2723313 | 2723364 | 2722626 | 2722674 | 2726259 | 2726268 | 1487 | 751 | 878 | 1249 | 1432 | 0 | 0 |
| SE | 28660 | ENSG00000002586 | CD99 | chrX | + | 2723313 | 2723364 | 2722626 | 2722674 | 2738199 | 2738256 | 372 | 241 | 212 | 330 | 384 | 0 | 0 |
| SE | 28661 | ENSG00000002586 | CD99 | chrX | + | 2726259 | 2726373 | 2722626 | 2722674 | 2738199 | 2738256 | 700 | 328 | 454 | 614 | 527 | 0 | 0 |
| SE | 28663 | ENSG00000002586 | CD99 | chrX | + | 2726259 | 2726373 | 2723313 | 2723364 | 2738202 | 2738256 | 671 | 300 | 399 | 519 | 630 | 0 | 0 |
| SE | 28664 | ENSG00000002586 | CD99 | chrX | + | 2727292 | 2727343 | 2726259 | 2726373 | 2733356 | 2733667 | 0 | 0 | 0 | 0 | 0 | 13 | 14 |
| SE | 28665 | ENSG00000002586 | CD99 | chrX | + | 2733356 | 2733374 | 2726259 | 2726373 | 2738199 | 2738256 | 0 | 1 | 4 | 2 | 0 | 1247 | 586 |
| SE | 28667 | ENSG00000002586 | CD99 | chrX | + | 2737880 | 2738256 | 2726340 | 2726373 | 2740778 | 2741125 | 639 | 316 | 393 | 490 | 487 | 0 | 0 |
| SE | 28668 | ENSG00000002586 | CD99 | chrX | + | 2738199 | 2738256 | 2726259 | 2726373 | 2740778 | 2741101 | 1871 | 891 | 1168 | 1582 | 1555 | 0 | 0 |
| SE | 28669 | ENSG00000002586 | CD99 | chrX | + | 2738202 | 2738256 | 2726259 | 2726373 | 2740778 | 2740804 | 706 | 349 | 445 | 555 | 570 | 0 | 0 |
| SE | 28670 | ENSG00000140443 | IGF1R | chr15 | + | 98824179 | 98824242 | 98707830 | 98708107 | 98891324 | 98891637 | 0 | 0 | 0 | 0 | 8 | 69 | 32 |
| SE | 28672 | ENSG00000140443 | IGF1R | chr15 | + | 98891324 | 98891637 | 98707561 | 98708107 | 98896756 | 98896849 | 105 | 43 | 184 | 86 | 97 | 0 | 0 |
| SE | 28673 | ENSG00000140443 | IGF1R | chr15 | + | 98908684 | 98908899 | 98899476 | 98899621 | 98911314 | 98911398 | 141 | 68 | 178 | 94 | 161 | 0 | 0 |
| SE | 28674 | ENSG00000140443 | IGF1R | chr15 | + | 98923875 | 98924012 | 98922147 | 98922431 | 98924524 | 98924684 | 104 | 85 | 214 | 92 | 113 | 0 | 0 |
| SE | 28675 | ENSG00000140443 | IGF1R | chr15 | + | 98930234 | 98930305 | 98929557 | 98929660 | 98935315 | 98935366 | 48 | 30 | 61 | 40 | 69 | 0 | 0 |
| SE | 28677 | ENSG00000140443 | IGF1R | chr15 | + | 98948573 | 98948708 | 98942922 | 98943052 | 98957060 | 98964530 | 161 | 141 | 290 | 139 | 314 | 0 | 0 |
| SE | 28678 | ENSG00000185033 | SEMA4B | chr15 | + | 90201457 | 90201735 | 90184962 | 90185081 | 90217438 | 90217554 | 25 | 5 | 10 | 11 | 0 | 0 | 1 |
| SE | 28682 | ENSG00000185033 | SEMA4B | chr15 | + | 90225281 | 90225427 | 90225071 | 90225178 | 90225660 | 90225827 | 9 | 2 | 3 | 9 | 2 | 0 | 0 |
| SE | 28694 | ENSG00000112249 | ASCC3 | chr6 | - | 100642580 | 100642749 | 100638600 | 100638821 | 100644030 | 100644129 | 202 | 176 | 150 | 201 | 371 | 0 | 0 |
| SE | 28695 | ENSG00000112249 | ASCC3 | chr6 | - | 100650537 | 100650714 | 100647225 | 100647451 | 100651559 | 100651646 | 199 | 129 | 144 | 207 | 353 | 0 | 1 |
| SE | 28696 | ENSG00000112249 | ASCC3 | chr6 | - | 100650537 | 100650714 | 100647225 | 100647451 | 100652724 | 100652889 | 73 | 52 | 49 | 81 | 116 | 0 | 0 |
| SE | 28697 | ENSG00000112249 | ASCC3 | chr6 | - | 100651559 | 100651646 | 100647225 | 100647451 | 100652724 | 100652889 | 27 | 25 | 17 | 36 | 78 | 0 | 0 |
| SE | 28698 | ENSG00000112249 | ASCC3 | chr6 | - | 100651559 | 100651646 | 100650537 | 100650714 | 100652724 | 100652889 | 151 | 101 | 112 | 164 | 315 | 0 | 0 |
| SE | 28701 | ENSG00000112249 | ASCC3 | chr6 | - | 100848147 | 100848707 | 100805759 | 100805880 | 100864063 | 100864214 | 517 | 485 | 372 | 538 | 632 | 5 | 0 |
| SE | 28703 | ENSG00000126003 | PLAGL2 | chr20 | - | 32201918 | 32202292 | 32192503 | 32197682 | 32207640 | 32207743 | 74 | 48 | 65 | 104 | 75 | 0 | 0 |
| SE | 28704 | ENSG00000158163 | DZIP1L | chr3 | - | 138071642 | 138071835 | 138068150 | 138068367 | 138077498 | 138077632 | 2 | 0 | 0 | 1 | 0 | 0 | 0 |
| SE | 28707 | ENSG00000164120 | HPGD | chr4 | - | 174493150 | 174493314 | 174490174 | 174492094 | 174495547 | 174495624 | 169 | 103 | 126 | 206 | 223 | 5 | 0 |
| SE | 28708 | ENSG00000164120 | HPGD | chr4 | - | 174495547 | 174495624 | 174491457 | 174492094 | 174508695 | 174508792 | 79 | 37 | 44 | 81 | 51 | 0 | 0 |
| SE | 28709 | ENSG00000164120 | HPGD | chr4 | - | 174508695 | 174508792 | 174495360 | 174495624 | 174517970 | 174518077 | 230 | 123 | 123 | 234 | 203 | 0 | 0 |
| SE | 28715 | ENSG00000168995 | SIGLEC7 | chr19 | + | 51144911 | 51144959 | 51142306 | 51142802 | 51145854 | 51146121 | 31 | 0 | 7 | 16 | 11 | 5 | 1 |
| SE | 28716 | ENSG00000168995 | SIGLEC7 | chr19 | + | 51144911 | 51144959 | 51142306 | 51142802 | 51147220 | 51147317 | 3 | 0 | 1 | 2 | 0 | 0 | 1 |
| SE | 28717 | ENSG00000168995 | SIGLEC7 | chr19 | + | 51145854 | 51146121 | 51142300 | 51142802 | 51147220 | 51147317 | 14 | 9 | 9 | 6 | 4 | 0 | 1 |
| SE | 28718 | ENSG00000168995 | SIGLEC7 | chr19 | + | 51145854 | 51146121 | 51144911 | 51144959 | 51146753 | 51146850 | 105 | 15 | 45 | 58 | 57 | 0 | 0 |
| SE | 28719 | ENSG00000168995 | SIGLEC7 | chr19 | + | 51145854 | 51146121 | 51144911 | 51144959 | 51147220 | 51147317 | 59 | 8 | 21 | 30 | 24 | 0 | 0 |
| SE | 28720 | ENSG00000168995 | SIGLEC7 | chr19 | + | 51146753 | 51146850 | 51144911 | 51144959 | 51147220 | 51147317 | 59 | 13 | 36 | 38 | 20 | 0 | 0 |
| SE | 28724 | ENSG00000010256 | UQCRC1 | chr3 | - | 48601351 | 48601467 | 48600974 | 48601118 | 48603563 | 48603643 | 186 | 83 | 153 | 135 | 115 | 0 | 1 |
| SE | 28727 | ENSG00000010256 | UQCRC1 | chr3 | - | 48609161 | 48609302 | 48605769 | 48605856 | 48609551 | 48609631 | 294 | 125 | 160 | 157 | 102 | 0 | 0 |
| SE | 28736 | ENSG00000182093 | GET1 | chr21 | + | 39408605 | 39408677 | 39406561 | 39406731 | 39410843 | 39410937 | 0 | 1 | 2 | 0 | 0 | 0 | 0 |
| SE | 28740 | ENSG00000147548 | NSD3 | chr8 | - | 38278305 | 38278412 | 38276295 | 38276500 | 38279539 | 38279681 | 221 | 241 | 208 | 208 | 317 | 0 | 3 |
| SE | 28741 | ENSG00000147548 | NSD3 | chr8 | - | 38279539 | 38279681 | 38278305 | 38278412 | 38281499 | 38281583 | 203 | 183 | 171 | 200 | 234 | 1 | 0 |
| SE | 28742 | ENSG00000147548 | NSD3 | chr8 | - | 38289392 | 38289505 | 38288486 | 38288756 | 38290474 | 38290677 | 108 | 112 | 108 | 124 | 228 | 0 | 1 |
| SE | 28744 | ENSG00000147548 | NSD3 | chr8 | - | 38299443 | 38299590 | 38295795 | 38295952 | 38314646 | 38314773 | 60 | 26 | 58 | 42 | 86 | 0 | 0 |
| SE | 28745 | ENSG00000147548 | NSD3 | chr8 | - | 38304586 | 38304757 | 38295795 | 38295952 | 38314646 | 38314773 | 7 | 3 | 6 | 2 | 11 | 0 | 0 |
| SE | 28746 | ENSG00000147548 | NSD3 | chr8 | - | 38304586 | 38304757 | 38299443 | 38299590 | 38305247 | 38305445 | 112 | 70 | 111 | 143 | 173 | 0 | 0 |
| SE | 28747 | ENSG00000147548 | NSD3 | chr8 | - | 38304586 | 38304757 | 38299443 | 38299590 | 38314646 | 38314773 | 66 | 31 | 62 | 75 | 97 | 0 | 0 |
| SE | 28748 | ENSG00000147548 | NSD3 | chr8 | - | 38305247 | 38305445 | 38299443 | 38299590 | 38314646 | 38314773 | 40 | 14 | 27 | 25 | 29 | 0 | 0 |
| SE | 28749 | ENSG00000147548 | NSD3 | chr8 | - | 38305247 | 38305445 | 38304586 | 38304757 | 38314646 | 38314773 | 88 | 56 | 76 | 95 | 113 | 0 | 0 |
| SE | 28751 | ENSG00000147548 | NSD3 | chr8 | - | 38331430 | 38331585 | 38329377 | 38329893 | 38337304 | 38337467 | 731 | 557 | 818 | 808 | 1481 | 0 | 0 |
| SE | 28752 | ENSG00000147548 | NSD3 | chr8 | - | 38337304 | 38337467 | 38331430 | 38331585 | 38338535 | 38338607 | 560 | 393 | 595 | 558 | 986 | 0 | 0 |
| SE | 28753 | ENSG00000147548 | NSD3 | chr8 | - | 38338535 | 38338607 | 38337304 | 38337467 | 38347496 | 38348215 | 321 | 260 | 335 | 342 | 527 | 0 | 0 |
| SE | 28754 | ENSG00000147548 | NSD3 | chr8 | - | 38363872 | 38363900 | 38348095 | 38348215 | 38381589 | 38381731 | 0 | 0 | 0 | 0 | 0 | 0 | 2 |
| SE | 28755 | ENSG00000147548 | NSD3 | chr8 | - | 38363872 | 38363900 | 38348095 | 38348215 | 38381798 | 38382227 | 0 | 0 | 0 | 1 | 0 | 81 | 54 |
| SE | 28758 | ENSG00000151715 | TMEM45B | chr11 | + | 129854609 | 129854816 | 129852474 | 129852660 | 129855707 | 129855892 | 13 | 4 | 10 | 0 | 0 | 0 | 0 |
| SE | 28759 | ENSG00000151715 | TMEM45B | chr11 | + | 129855707 | 129855892 | 129854609 | 129854816 | 129857312 | 129857458 | 10 | 3 | 11 | 2 | 5 | 0 | 1 |
| SE | 28762 | ENSG00000170571 | EMB | chr5 | - | 50399858 | 50399913 | 50396191 | 50399290 | 50403177 | 50403454 | 303 | 160 | 224 | 228 | 809 | 1 | 0 |
| SE | 28763 | ENSG00000170571 | EMB | chr5 | - | 50402285 | 50402319 | 50396191 | 50399290 | 50403177 | 50403454 | 315 | 154 | 177 | 258 | 196 | 1 | 0 |
| SE | 28764 | ENSG00000170571 | EMB | chr5 | - | 50402285 | 50402319 | 50399858 | 50399913 | 50403177 | 50403454 | 1172 | 560 | 764 | 870 | 1264 | 4 | 0 |
| SE | 28765 | ENSG00000170571 | EMB | chr5 | - | 50403177 | 50403454 | 50399858 | 50399913 | 50405724 | 50405852 | 2437 | 1207 | 1511 | 1733 | 3358 | 0 | 2 |
| SE | 28766 | ENSG00000170571 | EMB | chr5 | - | 50405724 | 50405852 | 50403177 | 50403454 | 50410876 | 50410965 | 4136 | 2110 | 2649 | 3114 | 5800 | 4 | 1 |
| SE | 28768 | ENSG00000170571 | EMB | chr5 | - | 50410876 | 50410965 | 50405724 | 50405852 | 50428143 | 50428227 | 1447 | 800 | 908 | 1091 | 1977 | 0 | 0 |
| SE | 28769 | ENSG00000170571 | EMB | chr5 | - | 50411196 | 50411221 | 50405724 | 50405852 | 50428143 | 50428227 | 1 | 0 | 1 | 0 | 0 | 0 | 0 |
| SE | 28770 | ENSG00000170571 | EMB | chr5 | - | 50411196 | 50411221 | 50410876 | 50410965 | 50428143 | 50428227 | 259 | 125 | 181 | 216 | 263 | 3 | 0 |
| SE | 28771 | ENSG00000170571 | EMB | chr5 | - | 50411196 | 50411383 | 50405724 | 50405852 | 50428143 | 50428227 | 375 | 164 | 296 | 381 | 627 | 0 | 0 |
| SE | 28772 | ENSG00000170571 | EMB | chr5 | - | 50411196 | 50411383 | 50410876 | 50410965 | 50428143 | 50428227 | 2033 | 1016 | 1428 | 1737 | 3193 | 3 | 0 |
| SE | 28774 | ENSG00000112293 | GPLD1 | chr6 | - | 24445725 | 24445831 | 24445545 | 24445639 | 24446837 | 24446979 | 2 | 1 | 3 | 1 | 0 | 0 | 0 |
| SE | 28787 | ENSG00000154370 | TRIM11 | chr1 | - | 228400963 | 228401194 | 228397142 | 228397165 | 228402065 | 228402161 | 47 | 26 | 57 | 40 | 9 | 2 | 0 |
| SE | 28788 | ENSG00000154370 | TRIM11 | chr1 | - | 228402065 | 228402161 | 228397142 | 228397165 | 228406153 | 228406187 | 9 | 0 | 7 | 5 | 0 | 0 | 0 |
| SE | 28789 | ENSG00000133048 | CHI3L1 | chr1 | - | 203180469 | 203180652 | 203179760 | 203179877 | 203181161 | 203181285 | 3 | 1 | 0 | 18 | 9 | 1 | 0 |
| SE | 28790 | ENSG00000133048 | CHI3L1 | chr1 | - | 203186315 | 203186345 | 203185183 | 203185385 | 203186598 | 203186704 | 0 | 0 | 2 | 6 | 12 | 0 | 0 |
| SE | 28791 | ENSG00000020426 | MNAT1 | chr14 | + | 60796216 | 60796369 | 60734760 | 60734951 | 60798086 | 60798160 | 149 | 102 | 114 | 123 | 155 | 0 | 0 |
| SE | 28792 | ENSG00000020426 | MNAT1 | chr14 | + | 60796216 | 60796369 | 60734760 | 60734951 | 60808324 | 60808428 | 84 | 42 | 57 | 54 | 42 | 1 | 0 |
| SE | 28793 | ENSG00000020426 | MNAT1 | chr14 | + | 60798086 | 60798160 | 60734760 | 60734951 | 60808324 | 60808428 | 35 | 58 | 36 | 37 | 128 | 1 | 0 |
| SE | 28794 | ENSG00000020426 | MNAT1 | chr14 | + | 60798086 | 60798160 | 60796216 | 60796369 | 60808324 | 60808428 | 102 | 116 | 91 | 105 | 237 | 3 | 2 |
| SE | 28795 | ENSG00000020426 | MNAT1 | chr14 | + | 60818721 | 60818847 | 60811985 | 60812127 | 60879713 | 60879835 | 93 | 99 | 106 | 125 | 156 | 0 | 0 |
| SE | 28797 | ENSG00000116678 | LEPR | chr1 | + | 65605037 | 65605237 | 65601842 | 65601960 | 65608752 | 65608901 | 52 | 60 | 25 | 43 | 65 | 1 | 0 |
| SE | 28798 | ENSG00000116678 | LEPR | chr1 | + | 65622905 | 65622981 | 65621352 | 65621458 | 65633151 | 65635409 | 82 | 91 | 20 | 102 | 121 | 0 | 0 |
| SE | 28799 | ENSG00000124203 | ZNF831 | chr20 | + | 59146230 | 59146374 | 59123380 | 59123505 | 59190983 | 59194757 | 8 | 3 | 8 | 5 | 3 | 1 | 0 |
| SE | 28800 | ENSG00000124203 | ZNF831 | chr20 | + | 59159651 | 59160895 | 59146230 | 59146374 | 59190983 | 59194757 | 10 | 11 | 3 | 9 | 5 | 6 | 0 |
| SE | 28801 | ENSG00000124203 | ZNF831 | chr20 | + | 59206904 | 59207056 | 59195868 | 59196005 | 59252977 | 59253138 | 268 | 116 | 182 | 249 | 178 | 0 | 0 |
| SE | 28802 | ENSG00000124203 | ZNF831 | chr20 | + | 59252977 | 59253138 | 59206904 | 59207056 | 59253897 | 59259113 | 193 | 88 | 134 | 210 | 281 | 1 | 0 |
| SE | 28803 | ENSG00000178163 | ZNF518B | chr4 | - | 10454593 | 10454988 | 10446101 | 10446539 | 10457316 | 10457408 | 18 | 17 | 20 | 18 | 21 | 12 | 5 |
| SE | 28804 | ENSG00000178163 | ZNF518B | chr4 | - | 10454804 | 10454988 | 10439879 | 10446539 | 10457316 | 10457426 | 18 | 22 | 14 | 13 | 19 | 12 | 5 |
| SE | 28805 | ENSG00000196935 | SRGAP1 | chr12 | + | 64091275 | 64091378 | 64086998 | 64087026 | 64094931 | 64094992 | 1 | 1 | 0 | 0 | 5 | 0 | 0 |
| SE | 28806 | ENSG00000196935 | SRGAP1 | chr12 | + | 64091892 | 64091926 | 64086998 | 64087026 | 64094931 | 64094992 | 0 | 0 | 0 | 0 | 0 | 0 | 0 |
| SE | 28874 | ENSG00000196104 | SPOCK3 | chr4 | - | 167062491 | 167062537 | 167000348 | 167000463 | 167233984 | 167234065 | 2 | 0 | 2 | 2 | 3 | 0 | 0 |
| SE | 28882 | ENSG00000215424 | MCM3AP-AS1 | chr21 | + | 46230326 | 46230527 | 46229230 | 46229266 | 46240833 | 46241019 | 1 | 0 | 0 | 0 | 1 | 0 | 0 |
| SE | 28883 | ENSG00000215424 | MCM3AP-AS1 | chr21 | + | 46236223 | 46236326 | 46229220 | 46229266 | 46240833 | 46241019 | 0 | 0 | 4 | 0 | 0 | 0 | 0 |
| SE | 28886 | ENSG00000164597 | COG5 | chr7 | - | 107210525 | 107210605 | 107201371 | 107203630 | 107211098 | 107211225 | 69 | 75 | 100 | 56 | 116 | 0 | 0 |
| SE | 28887 | ENSG00000164597 | COG5 | chr7 | - | 107211098 | 107211225 | 107201371 | 107203630 | 107230614 | 107230691 | 59 | 80 | 61 | 41 | 79 | 0 | 0 |
| SE | 28888 | ENSG00000164597 | COG5 | chr7 | - | 107236449 | 107236687 | 107230614 | 107230691 | 107248395 | 107248499 | 101 | 123 | 107 | 77 | 145 | 0 | 3 |
| SE | 28890 | ENSG00000164597 | COG5 | chr7 | - | 107256731 | 107256794 | 107248395 | 107248499 | 107258272 | 107258383 | 35 | 69 | 81 | 50 | 72 | 0 | 2 |
| SE | 28891 | ENSG00000164597 | COG5 | chr7 | - | 107258272 | 107258383 | 107248395 | 107248499 | 107281299 | 107281399 | 58 | 67 | 72 | 47 | 55 | 1 | 1 |
| SE | 28892 | ENSG00000164597 | COG5 | chr7 | - | 107258272 | 107258383 | 107248395 | 107248499 | 107283570 | 107283732 | 0 | 2 | 1 | 4 | 3 | 0 | 0 |
| SE | 28893 | ENSG00000164597 | COG5 | chr7 | - | 107281299 | 107281399 | 107248395 | 107248499 | 107283570 | 107283732 | 44 | 49 | 65 | 27 | 26 | 0 | 0 |
| SE | 28896 | ENSG00000164597 | COG5 | chr7 | - | 107372594 | 107372760 | 107362307 | 107362420 | 107412501 | 107412632 | 360 | 371 | 379 | 363 | 595 | 1 | 1 |
| SE | 28897 | ENSG00000164597 | COG5 | chr7 | - | 107412501 | 107412632 | 107372594 | 107372760 | 107527236 | 107527357 | 257 | 249 | 271 | 244 | 293 | 0 | 0 |
| SE | 28898 | ENSG00000164597 | COG5 | chr7 | - | 107548277 | 107548332 | 107547997 | 107548180 | 107554284 | 107554342 | 372 | 526 | 352 | 318 | 326 | 0 | 1 |
| SE | 28899 | ENSG00000145495 | MARCHF6 | chr5 | + | 10378758 | 10378832 | 10377797 | 10377894 | 10381799 | 10381943 | 307 | 233 | 269 | 345 | 295 | 0 | 0 |
| SE | 28902 | ENSG00000145495 | MARCHF6 | chr5 | + | 10381799 | 10381943 | 10378758 | 10378832 | 10391541 | 10391731 | 238 | 193 | 210 | 280 | 248 | 1 | 0 |
| SE | 28904 | ENSG00000145495 | MARCHF6 | chr5 | + | 10386993 | 10387066 | 10378758 | 10378832 | 10390331 | 10390491 | 369 | 339 | 378 | 516 | 363 | 2 | 0 |
| SE | 28905 | ENSG00000145495 | MARCHF6 | chr5 | + | 10386993 | 10387066 | 10378758 | 10378832 | 10391541 | 10391731 | 19 | 12 | 7 | 8 | 11 | 1 | 0 |
| SE | 28906 | ENSG00000145495 | MARCHF6 | chr5 | + | 10386993 | 10387066 | 10381799 | 10381943 | 10390331 | 10390491 | 593 | 476 | 602 | 778 | 559 | 9 | 4 |
| SE | 28908 | ENSG00000145495 | MARCHF6 | chr5 | + | 10390331 | 10390500 | 10378758 | 10378832 | 10391541 | 10391574 | 503 | 340 | 424 | 535 | 338 | 1 | 0 |
| SE | 28909 | ENSG00000145495 | MARCHF6 | chr5 | + | 10390331 | 10390500 | 10381799 | 10381943 | 10391541 | 10391574 | 510 | 344 | 419 | 538 | 339 | 4 | 0 |
| SE | 28912 | ENSG00000145495 | MARCHF6 | chr5 | + | 10402532 | 10402607 | 10402383 | 10402452 | 10403406 | 10403541 | 196 | 107 | 195 | 196 | 252 | 0 | 0 |
| SE | 28913 | ENSG00000145495 | MARCHF6 | chr5 | + | 10407101 | 10407202 | 10405557 | 10405677 | 10410138 | 10410276 | 387 | 188 | 415 | 308 | 618 | 0 | 7 |
| SE | 28915 | ENSG00000145495 | MARCHF6 | chr5 | + | 10415472 | 10415669 | 10414432 | 10414502 | 10417269 | 10417404 | 217 | 105 | 208 | 193 | 127 | 1 | 0 |
| SE | 28916 | ENSG00000145495 | MARCHF6 | chr5 | + | 10415487 | 10415669 | 10414432 | 10414502 | 10417269 | 10417404 | 629 | 368 | 589 | 620 | 450 | 1 | 0 |
| SE | 28918 | ENSG00000145495 | MARCHF6 | chr5 | + | 10417269 | 10417404 | 10415472 | 10415669 | 10429892 | 10430028 | 198 | 85 | 187 | 171 | 114 | 0 | 0 |
| SE | 28920 | ENSG00000145495 | MARCHF6 | chr5 | + | 10423734 | 10423824 | 10415487 | 10415669 | 10429892 | 10430028 | 7 | 2 | 6 | 11 | 10 | 0 | 0 |
| SE | 28921 | ENSG00000145495 | MARCHF6 | chr5 | + | 10423734 | 10423824 | 10417269 | 10417404 | 10429892 | 10430028 | 54 | 25 | 74 | 59 | 49 | 0 | 0 |
| SE | 28923 | ENSG00000145495 | MARCHF6 | chr5 | + | 10429892 | 10430028 | 10417269 | 10417404 | 10433593 | 10433684 | 354 | 191 | 373 | 400 | 728 | 0 | 0 |
| SE | 28924 | ENSG00000167635 | ZNF146 | chr19 | + | 36218121 | 36218195 | 36215095 | 36215155 | 36228747 | 36228819 | 37 | 22 | 19 | 13 | 24 | 0 | 1 |
| SE | 28925 | ENSG00000167635 | ZNF146 | chr19 | + | 36218121 | 36218195 | 36215105 | 36215196 | 36228747 | 36228819 | 48 | 30 | 37 | 17 | 33 | 7 | 4 |
| SE | 28926 | ENSG00000167635 | ZNF146 | chr19 | + | 36228747 | 36228819 | 36218121 | 36218195 | 36235658 | 36236059 | 99 | 51 | 66 | 68 | 83 | 0 | 0 |
| SE | 28927 | ENSG00000144959 | NCEH1 | chr3 | - | 172635915 | 172636087 | 172630248 | 172634092 | 172645622 | 172645692 | 107 | 55 | 96 | 84 | 167 | 0 | 0 |
| SE | 28928 | ENSG00000144959 | NCEH1 | chr3 | - | 172647861 | 172648114 | 172645622 | 172645692 | 172710846 | 172711052 | 16 | 5 | 5 | 7 | 6 | 1 | 0 |
| SE | 28930 | ENSG00000112079 | STK38 | chr6 | - | 36496710 | 36496805 | 36493891 | 36495914 | 36497779 | 36497875 | 992 | 722 | 733 | 986 | 1453 | 0 | 0 |
| SE | 28931 | ENSG00000112079 | STK38 | chr6 | - | 36497779 | 36497875 | 36496710 | 36496805 | 36499872 | 36499990 | 401 | 262 | 243 | 336 | 629 | 0 | 0 |
| SE | 28932 | ENSG00000112079 | STK38 | chr6 | - | 36498362 | 36498486 | 36497779 | 36497875 | 36499872 | 36499990 | 1167 | 808 | 843 | 1036 | 1413 | 0 | 0 |
| SE | 28933 | ENSG00000112079 | STK38 | chr6 | - | 36498362 | 36498486 | 36497779 | 36497875 | 36506582 | 36506644 | 802 | 583 | 624 | 737 | 1024 | 4 | 2 |
| SE | 28937 | ENSG00000112079 | STK38 | chr6 | - | 36515337 | 36515492 | 36507499 | 36507602 | 36521733 | 36521817 | 408 | 263 | 280 | 338 | 516 | 0 | 1 |
| SE | 28939 | ENSG00000112079 | STK38 | chr6 | - | 36517716 | 36517840 | 36515337 | 36515492 | 36524340 | 36524463 | 864 | 527 | 577 | 816 | 1459 | 0 | 0 |
| SE | 28940 | ENSG00000112079 | STK38 | chr6 | - | 36521733 | 36521817 | 36515337 | 36515492 | 36524340 | 36524463 | 675 | 431 | 509 | 577 | 782 | 0 | 0 |
| SE | 28941 | ENSG00000112079 | STK38 | chr6 | - | 36521733 | 36521817 | 36517716 | 36517840 | 36524340 | 36524463 | 2195 | 1640 | 1649 | 1998 | 2469 | 0 | 0 |
| SE | 28942 | ENSG00000112079 | STK38 | chr6 | - | 36524340 | 36524463 | 36521733 | 36521817 | 36540071 | 36540207 | 1553 | 1151 | 1164 | 1382 | 1748 | 0 | 3 |
| SE | 28943 | ENSG00000112079 | STK38 | chr6 | - | 36525590 | 36525642 | 36524340 | 36524463 | 36540071 | 36540207 | 1300 | 926 | 935 | 1128 | 1190 | 0 | 0 |
| SE | 28944 | ENSG00000112079 | STK38 | chr6 | - | 36540071 | 36540207 | 36525590 | 36525642 | 36547189 | 36547479 | 766 | 508 | 574 | 702 | 702 | 4 | 0 |
| SE | 28945 | ENSG00000109670 | FBXW7 | chr4 | - | 152346929 | 152347071 | 152337801 | 152337936 | 152411302 | 152411872 | 123 | 103 | 122 | 121 | 92 | 0 | 0 |
| SE | 28946 | ENSG00000109670 | FBXW7 | chr4 | - | 152350041 | 152350124 | 152346811 | 152347071 | 152411302 | 152411872 | 295 | 169 | 200 | 221 | 352 | 1 | 0 |
| SE | 28947 | ENSG00000109670 | FBXW7 | chr4 | - | 152411302 | 152411872 | 152350041 | 152350124 | 152534940 | 152536063 | 227 | 149 | 143 | 207 | 239 | 0 | 0 |
| SE | 28950 | ENSG00000109670 | FBXW7 | chr4 | - | 152451659 | 152451776 | 152412479 | 152412529 | 152534940 | 152535033 | 0 | 0 | 0 | 0 | 0 | 15 | 6 |
| SE | 28951 | ENSG00000109670 | FBXW7 | chr4 | - | 152534940 | 152535033 | 152412479 | 152412529 | 152535126 | 152535191 | 44 | 23 | 33 | 25 | 11 | 0 | 0 |
| SE | 28952 | ENSG00000109670 | FBXW7 | chr4 | - | 152534940 | 152535033 | 152530272 | 152530850 | 152535126 | 152535191 | 50 | 27 | 43 | 56 | 15 | 1 | 0 |
| SE | 28957 | ENSG00000251600 | NA | chr4 | + | 143808313 | 143808370 | 143738974 | 143739117 | 143832559 | 143832615 | 0 | 5 | 0 | 6 | 0 | 0 | 0 |
| SE | 28958 | ENSG00000251600 | NA | chr4 | + | 143808313 | 143808370 | 143738974 | 143739117 | 143859015 | 143859115 | 2 | 9 | 1 | 8 | 3 | 4 | 0 |
| SE | 28959 | ENSG00000251600 | NA | chr4 | + | 143808313 | 143808370 | 143738974 | 143739117 | 143861141 | 143861246 | 0 | 5 | 1 | 5 | 0 | 0 | 1 |
| SE | 28960 | ENSG00000251600 | NA | chr4 | + | 143808313 | 143808370 | 143738974 | 143739117 | 143865970 | 143866076 | 0 | 4 | 0 | 5 | 1 | 0 | 0 |
| SE | 28961 | ENSG00000251600 | NA | chr4 | + | 143832559 | 143832615 | 143738974 | 143739117 | 143859015 | 143859115 | 0 | 5 | 0 | 4 | 12 | 4 | 0 |
| SE | 28965 | ENSG00000251600 | NA | chr4 | + | 143861141 | 143861246 | 143738974 | 143739117 | 143865970 | 143866076 | 22 | 39 | 9 | 66 | 104 | 0 | 0 |
| SE | 28966 | ENSG00000251600 | NA | chr4 | + | 143861141 | 143861246 | 143808313 | 143808370 | 143865970 | 143866076 | 22 | 38 | 10 | 66 | 104 | 1 | 0 |
| SE | 28967 | ENSG00000251600 | NA | chr4 | + | 143937866 | 143937923 | 143865970 | 143866076 | 143967087 | 143967179 | 1 | 1 | 0 | 0 | 0 | 0 | 0 |
| SE | 28968 | ENSG00000251600 | NA | chr4 | + | 143937866 | 143937923 | 143865970 | 143866076 | 143969251 | 143969379 | 2 | 1 | 0 | 0 | 0 | 2 | 6 |
| SE | 28969 | ENSG00000251600 | NA | chr4 | + | 143937866 | 143937923 | 143865970 | 143866076 | 143973109 | 143973251 | 0 | 1 | 1 | 4 | 0 | 0 | 1 |
| SE | 28970 | ENSG00000251600 | NA | chr4 | + | 143937866 | 143937923 | 143865970 | 143866076 | 143978919 | 143978982 | 1 | 1 | 0 | 0 | 0 | 0 | 0 |
| SE | 28971 | ENSG00000251600 | NA | chr4 | + | 143937866 | 143937923 | 143865970 | 143866076 | 143979849 | 143979952 | 0 | 2 | 0 | 0 | 0 | 0 | 0 |
| SE | 28972 | ENSG00000251600 | NA | chr4 | + | 143937866 | 143937923 | 143865970 | 143866076 | 143979854 | 143979952 | 0 | 1 | 0 | 0 | 0 | 0 | 2 |
| SE | 28973 | ENSG00000251600 | NA | chr4 | + | 143937866 | 143937923 | 143912539 | 143912638 | 143967087 | 143967179 | 2 | 12 | 2 | 14 | 14 | 0 | 0 |
| SE | 28974 | ENSG00000251600 | NA | chr4 | + | 143953634 | 143953690 | 143865970 | 143866076 | 143969251 | 143969379 | 1 | 0 | 0 | 0 | 0 | 2 | 6 |
| SE | 28975 | ENSG00000251600 | NA | chr4 | + | 143967087 | 143967179 | 143865970 | 143866076 | 143973109 | 143973251 | 0 | 0 | 0 | 1 | 3 | 0 | 1 |
| SE | 28976 | ENSG00000251600 | NA | chr4 | + | 143967087 | 143967179 | 143865970 | 143866076 | 143979849 | 143979952 | 0 | 0 | 0 | 0 | 3 | 0 | 0 |
| SE | 28978 | ENSG00000251600 | NA | chr4 | + | 143967087 | 143967179 | 143937866 | 143937923 | 143973109 | 143973251 | 1 | 0 | 0 | 1 | 0 | 1 | 0 |
| SE | 28979 | ENSG00000251600 | NA | chr4 | + | 143967087 | 143967179 | 143937866 | 143937923 | 143979849 | 143979952 | 1 | 0 | 0 | 0 | 0 | 2 | 1 |
| SE | 28980 | ENSG00000251600 | NA | chr4 | + | 143969251 | 143969379 | 143865970 | 143866076 | 143979849 | 143979952 | 3 | 6 | 0 | 9 | 9 | 0 | 0 |
| SE | 28981 | ENSG00000251600 | NA | chr4 | + | 143969251 | 143969379 | 143937866 | 143937923 | 143979849 | 143979952 | 3 | 3 | 0 | 0 | 0 | 2 | 1 |
| SE | 28982 | ENSG00000251600 | NA | chr4 | + | 143970424 | 143970534 | 143967087 | 143967179 | 143973109 | 143973251 | 0 | 0 | 0 | 0 | 1 | 0 | 0 |
| SE | 28984 | ENSG00000251600 | NA | chr4 | + | 143973109 | 143973251 | 143865970 | 143866076 | 143978919 | 143979068 | 0 | 1 | 0 | 1 | 5 | 0 | 0 |
| SE | 28985 | ENSG00000251600 | NA | chr4 | + | 143973109 | 143973251 | 143865970 | 143866076 | 143979849 | 143979952 | 0 | 3 | 0 | 5 | 7 | 0 | 0 |
| SE | 28987 | ENSG00000251600 | NA | chr4 | + | 143973109 | 143973251 | 143937866 | 143937923 | 143979849 | 143979952 | 1 | 2 | 1 | 11 | 3 | 2 | 1 |
| SE | 28988 | ENSG00000251600 | NA | chr4 | + | 143973109 | 143973251 | 143967087 | 143967179 | 143979849 | 143979952 | 0 | 2 | 0 | 5 | 2 | 0 | 0 |
| SE | 28989 | ENSG00000251600 | NA | chr4 | + | 143978919 | 143978982 | 143937866 | 143937923 | 143979849 | 143979952 | 2 | 0 | 0 | 0 | 0 | 2 | 1 |
| SE | 28990 | ENSG00000251600 | NA | chr4 | + | 143978919 | 143979068 | 143865970 | 143866076 | 143979849 | 143979952 | 0 | 0 | 0 | 0 | 0 | 0 | 0 |
| SE | 28992 | ENSG00000251600 | NA | chr4 | + | 144095296 | 144095374 | 144089900 | 144090131 | 144103315 | 144103514 | 0 | 0 | 0 | 0 | 0 | 0 | 0 |
| SE | 28993 | ENSG00000016864 | GLT8D1 | chr3 | - | 52695420 | 52695587 | 52695189 | 52695302 | 52695927 | 52696040 | 160 | 115 | 177 | 263 | 449 | 0 | 0 |
| SE | 28996 | ENSG00000016864 | GLT8D1 | chr3 | - | 52703086 | 52703255 | 52700444 | 52700496 | 52705446 | 52705771 | 0 | 0 | 0 | 1 | 0 | 53 | 38 |
| SE | 28998 | ENSG00000016864 | GLT8D1 | chr3 | - | 52703798 | 52703928 | 52700444 | 52700496 | 52705446 | 52705654 | 3 | 2 | 1 | 3 | 1 | 53 | 38 |
| SE | 28999 | ENSG00000016864 | GLT8D1 | chr3 | - | 52704723 | 52704948 | 52703798 | 52703928 | 52705446 | 52705654 | 2 | 0 | 7 | 7 | 6 | 0 | 0 |
| SE | 29001 | ENSG00000016864 | GLT8D1 | chr3 | - | 52704723 | 52704952 | 52700444 | 52700496 | 52705702 | 52705791 | 34 | 23 | 44 | 29 | 27 | 1 | 0 |
| SE | 29005 | ENSG00000123352 | SPATS2 | chr12 | + | 49371227 | 49371290 | 49366904 | 49367587 | 49441729 | 49441816 | 1 | 1 | 2 | 1 | 1 | 0 | 1 |
| SE | 29008 | ENSG00000123352 | SPATS2 | chr12 | + | 49406472 | 49406565 | 49367517 | 49367587 | 49460769 | 49461037 | 0 | 0 | 0 | 0 | 3 | 2 | 1 |
| SE | 29011 | ENSG00000123352 | SPATS2 | chr12 | + | 49441729 | 49441816 | 49367383 | 49367587 | 49460769 | 49461028 | 0 | 1 | 2 | 1 | 1 | 2 | 1 |
| SE | 29014 | ENSG00000123352 | SPATS2 | chr12 | + | 49454224 | 49454385 | 49367495 | 49367587 | 49460769 | 49461037 | 0 | 1 | 0 | 0 | 0 | 2 | 1 |
| SE | 29015 | ENSG00000123352 | SPATS2 | chr12 | + | 49454224 | 49454385 | 49441729 | 49441816 | 49460769 | 49461037 | 0 | 1 | 0 | 1 | 3 | 0 | 1 |
| SE | 29017 | ENSG00000123352 | SPATS2 | chr12 | + | 49494740 | 49495002 | 49490681 | 49490731 | 49496832 | 49496889 | 36 | 21 | 39 | 36 | 114 | 0 | 0 |
| SE | 29019 | ENSG00000123352 | SPATS2 | chr12 | + | 49500069 | 49500205 | 49494740 | 49495002 | 49514554 | 49514613 | 6 | 3 | 9 | 20 | 33 | 1 | 0 |
| SE | 29022 | ENSG00000078814 | MYH7B | chr20 | + | 34986099 | 34986198 | 34985065 | 34985129 | 34986885 | 34986989 | 0 | 0 | 0 | 0 | 0 | 0 | 2 |
| SE | 29023 | ENSG00000164603 | BMT2 | chr7 | - | 112915305 | 112915447 | 112895568 | 112895739 | 112939590 | 112939875 | 59 | 32 | 62 | 63 | 136 | 2 | 0 |
| SE | 29024 | ENSG00000164603 | BMT2 | chr7 | - | 112935186 | 112935264 | 112915305 | 112915447 | 112939590 | 112939618 | 3 | 0 | 0 | 0 | 0 | 23 | 8 |
| SE | 29025 | ENSG00000182150 | ERCC6L2 | chr9 | + | 95897848 | 95897971 | 95880868 | 95881293 | 95907077 | 95907271 | 88 | 78 | 105 | 115 | 129 | 0 | 5 |
| SE | 29026 | ENSG00000182150 | ERCC6L2 | chr9 | + | 95897848 | 95897971 | 95880868 | 95881293 | 95915667 | 95915829 | 66 | 58 | 69 | 77 | 88 | 0 | 1 |
| SE | 29028 | ENSG00000182150 | ERCC6L2 | chr9 | + | 95907077 | 95907271 | 95880868 | 95881293 | 95915667 | 95915829 | 83 | 62 | 101 | 85 | 132 | 0 | 1 |
| SE | 29031 | ENSG00000182150 | ERCC6L2 | chr9 | + | 95916226 | 95916434 | 95915667 | 95915829 | 95921174 | 95921315 | 197 | 201 | 221 | 258 | 320 | 0 | 2 |
| SE | 29033 | ENSG00000182150 | ERCC6L2 | chr9 | + | 95955913 | 95956013 | 95941453 | 95941549 | 95966561 | 95966714 | 109 | 111 | 121 | 127 | 189 | 0 | 2 |
| SE | 29034 | ENSG00000182150 | ERCC6L2 | chr9 | + | 95966561 | 95966714 | 95955913 | 95956013 | 95967458 | 95967625 | 43 | 65 | 65 | 73 | 111 | 0 | 0 |
| SE | 29037 | ENSG00000182150 | ERCC6L2 | chr9 | + | 95967318 | 95967625 | 95966561 | 95966714 | 95970575 | 95970656 | 14 | 9 | 25 | 9 | 10 | 89 | 62 |
| SE | 29039 | ENSG00000182150 | ERCC6L2 | chr9 | + | 95967458 | 95967625 | 95966561 | 95966714 | 95970575 | 95970656 | 14 | 16 | 33 | 13 | 18 | 89 | 62 |
| SE | 29040 | ENSG00000182150 | ERCC6L2 | chr9 | + | 95970575 | 95970656 | 95966561 | 95966714 | 95971932 | 95972373 | 109 | 97 | 121 | 106 | 201 | 2 | 0 |
| SE | 29042 | ENSG00000182150 | ERCC6L2 | chr9 | + | 95970575 | 95973088 | 95966561 | 95966714 | 95978060 | 95978215 | 194 | 192 | 211 | 201 | 366 | 0 | 0 |
| SE | 29043 | ENSG00000182150 | ERCC6L2 | chr9 | + | 95971543 | 95971595 | 95970575 | 95970656 | 95971932 | 95973088 | 2 | 3 | 1 | 8 | 4 | 43 | 56 |
| SE | 29044 | ENSG00000182150 | ERCC6L2 | chr9 | + | 95971932 | 95973088 | 95966561 | 95966714 | 95978060 | 95978215 | 121 | 147 | 154 | 141 | 248 | 0 | 0 |
| SE | 29046 | ENSG00000182150 | ERCC6L2 | chr9 | + | 96004519 | 96004701 | 95978060 | 95978215 | 96012224 | 96013647 | 165 | 194 | 237 | 231 | 440 | 1 | 0 |
| SE | 29047 | ENSG00000182150 | ERCC6L2 | chr9 | + | 96012224 | 96013647 | 96004519 | 96004701 | 96038875 | 96040580 | 56 | 40 | 59 | 47 | 137 | 1 | 2 |
| SE | 29049 | ENSG00000241549 | GUSBP2 | chr6 | - | 26885358 | 26885443 | 26883342 | 26883517 | 26889009 | 26889197 | 13 | 14 | 5 | 0 | 10 | 3 | 0 |
| SE | 29050 | ENSG00000241549 | GUSBP2 | chr6 | - | 26887963 | 26888051 | 26885358 | 26885443 | 26889009 | 26889198 | 2 | 0 | 0 | 1 | 1 | 8 | 10 |
| SE | 29051 | ENSG00000241549 | GUSBP2 | chr6 | - | 26951085 | 26951159 | 26925119 | 26925275 | 26956388 | 26956554 | 1 | 2 | 3 | 0 | 0 | 11 | 7 |
| SE | 29055 | ENSG00000100416 | TRMU | chr22 | + | 46352263 | 46352330 | 46352120 | 46352174 | 46355443 | 46355588 | 27 | 2 | 28 | 26 | 27 | 0 | 0 |
| SE | 29063 | ENSG00000163625 | WDFY3 | chr4 | - | 84677198 | 84677396 | 84669596 | 84672991 | 84678167 | 84678279 | 60 | 37 | 33 | 51 | 73 | 0 | 0 |
| SE | 29064 | ENSG00000163625 | WDFY3 | chr4 | - | 84678167 | 84678279 | 84677198 | 84677396 | 84682373 | 84682470 | 14 | 16 | 12 | 16 | 33 | 1 | 0 |
| SE | 29065 | ENSG00000163625 | WDFY3 | chr4 | - | 84678918 | 84679242 | 84678167 | 84678279 | 84682373 | 84682470 | 58 | 30 | 30 | 40 | 44 | 0 | 0 |
| SE | 29066 | ENSG00000163625 | WDFY3 | chr4 | - | 84683942 | 84684125 | 84682373 | 84682470 | 84688085 | 84688265 | 63 | 39 | 35 | 31 | 49 | 0 | 0 |
| SE | 29070 | ENSG00000163625 | WDFY3 | chr4 | - | 84690505 | 84690664 | 84688085 | 84688265 | 84691630 | 84691785 | 72 | 31 | 38 | 40 | 67 | 0 | 0 |
| SE | 29072 | ENSG00000163625 | WDFY3 | chr4 | - | 84705393 | 84705511 | 84704337 | 84704444 | 84708908 | 84709028 | 78 | 42 | 34 | 72 | 118 | 1 | 1 |
| SE | 29075 | ENSG00000163625 | WDFY3 | chr4 | - | 84756925 | 84757161 | 84755265 | 84755400 | 84765809 | 84766027 | 62 | 31 | 20 | 57 | 65 | 0 | 0 |

| | | | | | | | | | | | | | |
|---|---|---|---|---|---|---|---|---|---|---|---|---|---|
| SE | 29076 | ENSG00000163625 | WDFY3 | chr4 | - | 84765809 | 84766027 | 84756925 | 84757161 | 84766251 | 84766372 | 84 | 65 |
| SE | 29077 | ENSG00000163625 | WDFY3 | chr4 | - | 84796520 | 84796752 | 84794878 | 84794979 | 84797995 | 84798108 | 97 | 55 |
| SE | 29078 | ENSG00000163625 | WDFY3 | chr4 | - | 84808333 | 84808417 | 84803289 | 84803467 | 84809886 | 84810344 | 0 | 0 |
| SE | 29080 | ENSG00000163625 | WDFY3 | chr4 | - | 84826814 | 84826981 | 84821083 | 84821551 | 84829003 | 84829190 | 138 | 122 |
| SE | 29081 | ENSG00000163625 | WDFY3 | chr4 | - | 84831412 | 84831605 | 84829003 | 84829190 | 84836928 | 84837090 | 64 | 33 |
| SE | 29082 | ENSG00000163625 | WDFY3 | chr4 | - | 84836928 | 84837090 | 84831412 | 84831605 | 84841153 | 84841263 | 97 | 45 |
| SE | 29085 | ENSG00000163625 | WDFY3 | chr4 | - | 84879405 | 84879494 | 84860563 | 84860622 | 84932269 | 84932363 | 1 | 2 |
| SE | 29089 | ENSG00000163625 | WDFY3 | chr4 | - | 84932269 | 84932363 | 84860411 | 84860622 | 84966208 | 84966391 | 2 | 3 |
| SE | 29093 | ENSG00000109762 | SNX25 | chr4 | + | 185258847 | 185259064 | 185247293 | 185247378 | 185264437 | 185264498 | 91 | 28 |
| SE | 29094 | ENSG00000109762 | SNX25 | chr4 | + | 185266968 | 185267155 | 185264437 | 185264610 | 185288011 | 185288082 | 145 | 70 |
| SE | 29095 | ENSG00000109762 | SNX25 | chr4 | + | 185288011 | 185288082 | 185266968 | 185267155 | 185310634 | 185310816 | 85 | 40 |
| SE | 29096 | ENSG00000109762 | SNX25 | chr4 | + | 185320732 | 185320864 | 185310634 | 185310816 | 185323527 | 185323800 | 118 | 87 |
| SE | 29099 | ENSG00000109762 | SNX25 | chr4 | + | 185332594 | 185332759 | 185323527 | 185323800 | 185341975 | 185342116 | 66 | 35 |
| SE | 29100 | ENSG00000109762 | SNX25 | chr4 | + | 185339378 | 185339510 | 185323527 | 185323800 | 185341975 | 185342116 | 42 | 32 |
| SE | 29101 | ENSG00000109762 | SNX25 | chr4 | + | 185339378 | 185339510 | 185332594 | 185332759 | 185341975 | 185342116 | 73 | 63 |
| SE | 29102 | ENSG00000109762 | SNX25 | chr4 | + | 185346536 | 185346650 | 185341975 | 185342116 | 185357670 | 185357737 | 20 | 6 |
| SE | 29103 | ENSG00000109762 | SNX25 | chr4 | + | 185351444 | 185351609 | 185346536 | 185346650 | 185353484 | 185353602 | 69 | 50 |
| SE | 29104 | ENSG00000109762 | SNX25 | chr4 | + | 185353484 | 185353602 | 185346536 | 185346650 | 185357670 | 185357737 | 22 | 18 |
| SE | 29111 | ENSG00000061918 | GUCY1B1 | chr4 | + | 155796376 | 155796510 | 155795340 | 155795457 | 155799876 | 155800074 | 30 | 83 |
| SE | 29113 | ENSG00000272146 | ARF4-AS1 | chr3 | + | 57600058 | 57600169 | 57597795 | 57598051 | 57600742 | 57600920 | 4 | 2 |
| SE | 29117 | ENSG00000248672 | LY75-CD302 | chr2 | - | 159777937 | 159777964 | 159771850 | 159772053 | 159780004 | 159780178 | 382 | 354 |
| SE | 29118 | ENSG00000241399 | CD302 | chr2 | - | 159780004 | 159780178 | 159777937 | 159777964 | 159780881 | 159780998 | 517 | 550 |
| SE | 29119 | ENSG00000248672 | LY75-CD302 | chr2 | - | 159783358 | 159783469 | 159780881 | 159780998 | 159806972 | 159807140 | 94 | 99 |
| SE | 29120 | ENSG00000248672 | LY75-CD302 | chr2 | - | 159783358 | 159783469 | 159780881 | 159780998 | 159808448 | 159808571 | 88 | 92 |
| SE | 29121 | ENSG00000248672 | LY75-CD302 | chr2 | - | 159806972 | 159807140 | 159780881 | 159780998 | 159808448 | 159808571 | 169 | 152 |
| SE | 29123 | ENSG00000248672 | LY75-CD302 | chr2 | - | 159810525 | 159810675 | 159808448 | 159808571 | 159815404 | 159815573 | 293 | 225 |
| SE | 29124 | ENSG00000248672 | LY75-CD302 | chr2 | - | 159819725 | 159819920 | 159816805 | 159817032 | 159831669 | 159831786 | 192 | 190 |
| SE | 29125 | ENSG00000248672 | LY75-CD302 | chr2 | - | 159835479 | 159835645 | 159834043 | 159834211 | 159842244 | 159842374 | 222 | 171 |
| SE | 29126 | ENSG00000248672 | LY75-CD302 | chr2 | - | 159840728 | 159840955 | 159835479 | 159835645 | 159842244 | 159842374 | 314 | 267 |
| SE | 29127 | ENSG00000248672 | LY75-CD302 | chr2 | - | 159842244 | 159842374 | 159835479 | 159835645 | 159849979 | 159850140 | 56 | 50 |
| SE | 29129 | ENSG00000248672 | LY75-CD302 | chr2 | - | 159852200 | 159852340 | 159849979 | 159850140 | 159853272 | 159853352 | 132 | 118 |
| SE | 29131 | ENSG00000248672 | LY75-CD302 | chr2 | - | 159878632 | 159878721 | 159878323 | 159878493 | 159879258 | 159879369 | 534 | 666 |
| SE | 29132 | ENSG00000248672 | LY75-CD302 | chr2 | - | 159881082 | 159881240 | 159879258 | 159879369 | 159882123 | 159882315 | 691 | 518 |
| SE | 29134 | ENSG00000248672 | LY75-CD302 | chr2 | - | 159881082 | 159881240 | 159879258 | 159879369 | 159886419 | 159886530 | 350 | 273 |
| SE | 29135 | ENSG00000248672 | LY75-CD302 | chr2 | - | 159881082 | 159881240 | 159879258 | 159879369 | 159890212 | 159890377 | 350 | 273 |
| SE | 29138 | ENSG00000248672 | LY75-CD302 | chr2 | - | 159885152 | 159885293 | 159879258 | 159879369 | 159886419 | 159886530 | 57 | 50 |
| SE | 29139 | ENSG00000248672 | LY75-CD302 | chr2 | - | 159885152 | 159885293 | 159879258 | 159879369 | 159890212 | 159890377 | 5 | 3 |
| SE | 29141 | ENSG00000248672 | LY75-CD302 | chr2 | - | 159885152 | 159885293 | 159881082 | 159881240 | 159890212 | 159890377 | 3 | 1 |
| SE | 29142 | ENSG00000248672 | LY75-CD302 | chr2 | - | 159886419 | 159886530 | 159879258 | 159879369 | 159890212 | 159890377 | 95 | 42 |
| SE | 29143 | ENSG00000248672 | LY75-CD302 | chr2 | - | 159886419 | 159886530 | 159881082 | 159881240 | 159890212 | 159890377 | 95 | 42 |
| SE | 29144 | ENSG00000248672 | LY75-CD302 | chr2 | - | 159886419 | 159886530 | 159885152 | 159885293 | 159890212 | 159890377 | 150 | 89 |
| SE | 29145 | ENSG00000248672 | LY75-CD302 | chr2 | - | 159890212 | 159890377 | 159886419 | 159886530 | 159898687 | 159899059 | 113 | 56 |
| SE | 29147 | ENSG00000285476 | NA | chr5 | + | 176113955 | 176114061 | 176113799 | 176113867 | 176115592 | 176116252 | 1 | 10 |
| SE | 29148 | ENSG00000251623 | FAM153B | chr5 | + | 176119814 | 176120001 | 176119544 | 176119699 | 176120846 | 176120980 | 0 | 2 |
| SE | 29157 | ENSG00000214870 | NA | chr7 | + | 26495808 | 26495881 | 26495018 | 26495199 | 26496113 | 26496349 | 0 | 0 |
| SE | 29159 | ENSG00000144043 | TEX261 | chr2 | - | 70989748 | 70989816 | 70985941 | 70988715 | 70993695 | 70993775 | 2 | 6 |
| SE | 29161 | ENSG00000144043 | TEX261 | chr2 | - | 70993695 | 70993775 | 70991788 | 70991983 | 70994687 | 70994803 | 27 | 15 |
| SE | 29163 | ENSG00000106052 | TAX1BP1 | chr7 | + | 27769675 | 27769834 | 27758030 | 27758133 | 27827737 | 27827820 | 2 | 6 |
| SE | 29164 | ENSG00000106052 | TAX1BP1 | chr7 | + | 27796115 | 27796219 | 27794322 | 27794446 | 27799964 | 27800090 | 828 | 633 |
| SE | 29166 | ENSG00000106052 | TAX1BP1 | chr7 | + | 27816348 | 27816394 | 27799964 | 27800090 | 27816889 | 27817038 | 383 | 312 |
| SE | 29168 | ENSG00000106052 | TAX1BP1 | chr7 | + | 27816348 | 27816520 | 27799964 | 27800090 | 27816889 | 27817038 | 183 | 174 |
| SE | 29170 | ENSG00000106052 | TAX1BP1 | chr7 | + | 27816889 | 27817038 | 27816348 | 27816394 | 27827737 | 27827820 | 612 | 460 |
| SE | 29171 | ENSG00000106052 | TAX1BP1 | chr7 | + | 27827737 | 27827820 | 27816889 | 27817038 | 27828627 | 27828864 | 488 | 348 |
| SE | 29172 | ENSG00000106052 | TAX1BP1 | chr7 | + | 27828627 | 27828783 | 27827800 | 27827820 | 27840720 | 27844564 | 359 | 219 |
| SE | 29175 | ENSG00000227355 | NA | chr9 | + | 121440375 | 121440527 | 121375230 | 121375325 | 121451974 | 121452173 | 0 | 0 |
| SE | 29176 | ENSG00000227355 | NA | chr9 | + | 121440375 | 121440527 | 121375230 | 121375325 | 121462656 | 121462770 | 0 | 0 |
| SE | 29178 | ENSG00000227355 | NA | chr9 | + | 121451974 | 121452173 | 121440375 | 121440527 | 121462656 | 121462770 | 2 | 2 |
| SE | 29181 | ENSG00000129116 | PALLD | chr4 | + | 168877722 | 168878394 | 168832027 | 168832171 | 168890921 | 168891057 | 3 | 0 |
| SE | 29182 | ENSG00000129116 | PALLD | chr4 | + | 168894578 | 168894677 | 168890921 | 168891057 | 168898492 | 168898686 | 57 | 15 |
| SE | 29184 | ENSG00000129116 | PALLD | chr4 | + | 168898492 | 168898714 | 168894578 | 168894677 | 168903756 | 168903835 | 77 | 21 |
| SE | 29185 | ENSG00000129116 | PALLD | chr4 | + | 168913926 | 168914021 | 168903756 | 168903906 | 168915894 | 168916027 | 64 | 16 |
| SE | 29186 | ENSG00000129116 | PALLD | chr4 | + | 168921533 | 168921741 | 168915894 | 168916027 | 168924254 | 168924420 | 107 | 26 |
| SE | 29187 | ENSG00000129116 | PALLD | chr4 | + | 168924254 | 168924420 | 168921533 | 168921741 | 168924944 | 168925078 | 101 | 17 |
| SE | 29194 | ENSG00000196850 | PPTC7 | chr12 | - | 110539821 | 110539945 | 110533244 | 110537095 | 110551788 | 110551968 | 6 | 8 |
| SE | 29196 | ENSG00000120708 | TGFBI | chr5 | + | 136044781 | 136044870 | 136044057 | 136044122 | 136046334 | 136046440 | 0 | 0 |
| SE | 29197 | ENSG00000120708 | TGFBI | chr5 | + | 136046334 | 136046495 | 136044057 | 136044122 | 136046850 | 136047015 | 938 | 430 |
| SE | 29198 | ENSG00000120708 | TGFBI | chr5 | + | 136046334 | 136046556 | 136044057 | 136044122 | 136046850 | 136047015 | 455 | 237 |
| SE | 29200 | ENSG00000120708 | TGFBI | chr5 | + | 136052906 | 136053119 | 136045744 | 136049580 | 136053942 | 136054080 | 853 | 426 |
| SE | 29201 | ENSG00000120708 | TGFBI | chr5 | + | 136053017 | 136053119 | 136049438 | 136049580 | 136053942 | 136054080 | 339 | 172 |
| SE | 29205 | ENSG00000101109 | STK4 | chr20 | + | 44973300 | 44973433 | 44972077 | 44972158 | 44974099 | 44975684 | 47 | 32 |
| SE | 29207 | ENSG00000101109 | STK4 | chr20 | + | 44981828 | 44981943 | 44978442 | 44978571 | 44987131 | 44987296 | 2318 | 2020 |
| SE | 29208 | ENSG00000101109 | STK4 | chr20 | + | 44987131 | 44987296 | 44981828 | 44981943 | 44995089 | 44995257 | 2075 | 1835 |
| SE | 29209 | ENSG00000101109 | STK4 | chr20 | + | 44995089 | 44995257 | 44987131 | 44987296 | 44997168 | 44997306 | 1557 | 1181 |
| SE | 29211 | ENSG00000101109 | STK4 | chr20 | + | 45024972 | 45025130 | 45001166 | 45001353 | 45075017 | 45079948 | 810 | 567 |
| SE | 29213 | ENSG00000181215 | C4orf50 | chr4 | - | 5965023 | 5965145 | 5956701 | 5959626 | 5967413 | 5967462 | 3 | 1 |
| SE | 29214 | ENSG00000181215 | C4orf50 | chr4 | - | 5980173 | 5980338 | 5975898 | 5975955 | 5988346 | 5989647 | 1 | 2 |
| SE | 29215 | ENSG00000181215 | C4orf50 | chr4 | - | 6026202 | 6026278 | 6022703 | 6025013 | 6029715 | 6029781 | 0 | 1 |
| SE | 29217 | ENSG00000164118 | CEP44 | chr4 | + | 174297965 | 174298062 | 174283903 | 174284061 | 174299071 | 174299210 | 82 | 39 |
| SE | 29219 | ENSG00000164118 | CEP44 | chr4 | + | 174297965 | 174298062 | 174283968 | 174284140 | 174299071 | 174299210 | 65 | 29 |
| SE | 29220 | ENSG00000164118 | CEP44 | chr4 | + | 174316529 | 174316567 | 174316165 | 174316290 | 174317334 | 174318665 | 14 | 8 |
| SE | 29222 | ENSG00000023287 | RB1CC1 | chr8 | - | 52630469 | 52630528 | 52628031 | 52628168 | 52634920 | 52634968 | 138 | 86 |
| SE | 29233 | ENSG00000163110 | PDLIM5 | chr4 | + | 94656817 | 94656911 | 94654459 | 94654640 | 94657426 | 94657547 | 1 | 0 |
| SE | 29235 | ENSG00000166004 | CEP295 | chr11 | + | 93683558 | 93683742 | 93679411 | 93679552 | 93687643 | 93687865 | 37 | 21 |
| SE | 29236 | ENSG00000166004 | CEP295 | chr11 | + | 93683963 | 93684128 | 93679411 | 93679552 | 93687643 | 93687865 | 32 | 15 |
| SE | 29237 | ENSG00000166004 | CEP295 | chr11 | + | 93683963 | 93684128 | 93683558 | 93683742 | 93687643 | 93687865 | 78 | 41 |
| SE | 29238 | ENSG00000166004 | CEP295 | chr11 | + | 93691926 | 93692030 | 93691682 | 93691775 | 93695496 | 93695634 | 43 | 39 |
| SE | 29239 | ENSG00000166004 | CEP295 | chr11 | + | 93695496 | 93695634 | 93691926 | 93692030 | 93696319 | 93696417 | 37 | 27 |
| SE | 29240 | ENSG00000166004 | CEP295 | chr11 | + | 93695496 | 93695673 | 93691926 | 93692030 | 93696319 | 93696417 | 63 | 124 |
| SE | 29241 | ENSG00000166004 | CEP295 | chr11 | + | 93696319 | 93696417 | 93695496 | 93695673 | 93696681 | 93696854 | 76 | 132 |
| SE | 29242 | ENSG00000166004 | CEP295 | chr11 | + | 93702459 | 93702637 | 93696681 | 93700186 | 93702775 | 93702919 | 61 | 40 |
| SE | 29243 | ENSG00000166004 | CEP295 | chr11 | + | 93702459 | 93702637 | 93696681 | 93700186 | 93706744 | 93706897 | 54 | 22 |
| SE | 29244 | ENSG00000166004 | CEP295 | chr11 | + | 93702775 | 93702919 | 93696681 | 93700186 | 93706744 | 93706897 | 18 | 25 |
| SE | 29246 | ENSG00000166004 | CEP295 | chr11 | + | 93721953 | 93722050 | 93721311 | 93721412 | 93723040 | 93723289 | 37 | 11 |
| SE | 29247 | ENSG00000166004 | CEP295 | chr11 | + | 93721953 | 93722050 | 93721311 | 93721412 | 93725650 | 93725831 | 33 | 10 |
| SE | 29248 | ENSG00000166004 | CEP295 | chr11 | + | 93723040 | 93723289 | 93721953 | 93722050 | 93725650 | 93725831 | 10 | 2 |
| SE | 29249 | ENSG00000166004 | CEP295 | chr11 | + | 93724253 | 93724375 | 93722501 | 93723289 | 93725650 | 93725831 | 71 | 12 |
| SE | 29250 | ENSG00000166004 | CEP295 | chr11 | + | 93725650 | 93725831 | 93722501 | 93723289 | 93726975 | 93727115 | 16 | 14 |
| SE | 29251 | ENSG00000123213 | NLN | chr5 | + | 65740894 | 65741000 | 65722246 | 65722414 | 65758566 | 65758826 | 1 | 0 |
| SE | 29253 | ENSG00000123213 | NLN | chr5 | + | 65777426 | 65777534 | 65762959 | 65763108 | 65780178 | 65780281 | 53 | 62 |
| SE | 29254 | ENSG00000123213 | NLN | chr5 | + | 65785774 | 65785910 | 65781260 | 65781421 | 65788117 | 65788484 | 45 | 74 |
| SE | 29255 | ENSG00000123213 | NLN | chr5 | + | 65788117 | 65788484 | 65785774 | 65785910 | 65792453 | 65792601 | 49 | 65 |
| SE | 29256 | ENSG00000123213 | NLN | chr5 | + | 65792453 | 65792601 | 65785774 | 65785910 | 65809514 | 65809701 | 3 | 4 |
| SE | 29257 | ENSG00000123213 | NLN | chr5 | + | 65792453 | 65792655 | 65785774 | 65785910 | 65809514 | 65809701 | 24 | 13 |
| SE | 29260 | ENSG00000157800 | SLC37A3 | chr7 | - | 140343411 | 140343563 | 140337283 | 140337349 | 140345215 | 140345263 | 42 | 30 |
| SE | 29261 | ENSG00000157800 | SLC37A3 | chr7 | - | 140343411 | 140343563 | 140337283 | 140337349 | 140345868 | 140345970 | 16 | 7 |
| SE | 29262 | ENSG00000157800 | SLC37A3 | chr7 | - | 140345215 | 140345263 | 140337283 | 140337349 | 140345868 | 140345907 | 20 | 9 |
| SE | 29267 | ENSG00000157800 | SLC37A3 | chr7 | - | 140352061 | 140352146 | 140337283 | 140337349 | 140364407 | 140364491 | 0 | 0 |
| SE | 29270 | ENSG00000157800 | SLC37A3 | chr7 | - | 140355667 | 140355764 | 140352061 | 140352146 | 140364407 | 140364491 | 11 | 2 |
| SE | 29272 | ENSG00000157800 | SLC37A3 | chr7 | - | 140364407 | 140364491 | 140355667 | 140355764 | 140380281 | 140380390 | 0 | 1 |
| SE | 29275 | ENSG00000157800 | SLC37A3 | chr7 | - | 140369589 | 140369682 | 140364407 | 140364491 | 140380281 | 140380390 | 40 | 25 |
| SE | 29276 | ENSG00000157800 | SLC37A3 | chr7 | - | 140379213 | 140379307 | 140369589 | 140369682 | 140380281 | 140380390 | 0 | 0 |
| SE | 29282 | ENSG00000204815 | ODAD4 | chr17 | + | 41955216 | 41955317 | 41949152 | 41949349 | 41961381 | 41961466 | 0 | 0 |
| SE | 29283 | ENSG00000109572 | CLCN3 | chr4 | + | 169635912 | 169636088 | 169620525 | 169621063 | 169680049 | 169680207 | 315 | 333 |
| SE | 29284 | ENSG00000109572 | CLCN3 | chr4 | + | 169663526 | 169663617 | 169635982 | 169636088 | 169680049 | 169680207 | 0 | 1 |
| SE | 29285 | ENSG00000109572 | CLCN3 | chr4 | + | 169663526 | 169663617 | 169660061 | 169660448 | 169680049 | 169680207 | 0 | 0 |
| SE | 29286 | ENSG00000109572 | CLCN3 | chr4 | + | 169687657 | 169687757 | 169680049 | 169680207 | 169689042 | 169689145 | 215 | 355 |
| SE | 29287 | ENSG00000109572 | CLCN3 | chr4 | + | 169690529 | 169690652 | 169689042 | 169689230 | 169692113 | 169692320 | 193 | 228 |
| SE | 29290 | ENSG00000109572 | CLCN3 | chr4 | + | 169703997 | 169704184 | 169697188 | 169697734 | 169713078 | 169713187 | 88 | 95 |
| SE | 29291 | ENSG00000109572 | CLCN3 | chr4 | + | 169706867 | 169707266 | 169697188 | 169697734 | 169713078 | 169713187 | 196 | 231 |
| SE | 29292 | ENSG00000109572 | CLCN3 | chr4 | + | 169706867 | 169707266 | 169703997 | 169704184 | 169713078 | 169713187 | 250 | 275 |
| SE | 29296 | ENSG00000254870 | ATP6V1G2-DDX39B | chr6 | - | 31538762 | 31538855 | 31536499 | 31536683 | 31539146 | 31539274 | 658 | 448 |
| SE | 29305 | ENSG00000285713 | NA | chr4 | - | 145081659 | 145081750 | 145064571 | 145064692 | 145095984 | 145096111 | 77 | 41 |
| SE | 29306 | ENSG00000069869 | NEDD4 | chr15 | - | 55830513 | 55830586 | 55826921 | 55829999 | 55833007 | 55833104 | 62 | 56 |
| SE | 29307 | ENSG00000069869 | NEDD4 | chr15 | - | 55834037 | 55834145 | 55833007 | 55833104 | 55837788 | 55837849 | 31 | 40 |
| SE | 29308 | ENSG00000069869 | NEDD4 | chr15 | - | 55834226 | 55834286 | 55834037 | 55834145 | 55837788 | 55837849 | 55 | 46 |
| SE | 29310 | ENSG00000069869 | NEDD4 | chr15 | - | 55856130 | 55856196 | 55852423 | 55852543 | 55860406 | 55860574 | 24 | 13 |
| SE | 29312 | ENSG00000069869 | NEDD4 | chr15 | - | 55872414 | 55872476 | 55869578 | 55869681 | 55873957 | 55874008 | 58 | 27 |
| SE | 29316 | ENSG00000069869 | NEDD4 | chr15 | - | 55873957 | 55874008 | 55872414 | 55872476 | 55924645 | 55924699 | 47 | 23 |
| SE | 29317 | ENSG00000069869 | NEDD4 | chr15 | - | 55905211 | 55905281 | 55873957 | 55874008 | 55924645 | 55924699 | 0 | 0 |
| SE | 29319 | ENSG00000069869 | NEDD4 | chr15 | - | 55951375 | 55951414 | 55924645 | 55924699 | 55966472 | 55966546 | 11 | 10 |
| SE | 29321 | ENSG00000069869 | NEDD4 | chr15 | - | 55991925 | 55992005 | 55966472 | 55966546 | 55993510 | 55993602 | 1 | 2 |
| SE | 29322 | ENSG00000069869 | NEDD4 | chr15 | - | 55991925 | 55992005 | 55987208 | 55988001 | 55993510 | 55993602 | 0 | 0 |
| SE | 29324 | ENSG00000067082 | KLF6 | chr10 | - | 3783215 | 3783281 | 3781807 | 3782214 | 3784912 | 3785183 | 3 | 5 |
| SE | 29325 | ENSG00000121210 | TMEM131L | chr4 | + | 153473844 | 153473888 | 153467210 | 153467281 | 153550072 | 153550141 | 122 | 87 |
| SE | 29326 | ENSG00000121210 | TMEM131L | chr4 | + | 153556965 | 153557082 | 153555786 | 153555910 | 153558257 | 153558368 | 130 | 107 |
| SE | 29327 | ENSG00000121210 | TMEM131L | chr4 | + | 153556965 | 153557082 | 153555786 | 153555910 | 153580825 | 153580903 | 35 | 37 |
| SE | 29328 | ENSG00000121210 | TMEM131L | chr4 | + | 153558257 | 153558368 | 153555786 | 153555910 | 153580825 | 153580903 | 64 | 47 |
| SE | 29329 | ENSG00000121210 | TMEM131L | chr4 | + | 153558257 | 153558368 | 153556965 | 153557082 | 153580825 | 153580903 | 161 | 119 |
| SE | 29330 | ENSG00000121210 | TMEM131L | chr4 | + | 153558257 | 153558368 | 153556965 | 153557082 | 153581406 | 153581560 | 87 | 66 |
| SE | 29331 | ENSG00000121210 | TMEM131L | chr4 | + | 153580825 | 153580903 | 153556965 | 153557082 | 153581406 | 153581560 | 120 | 81 |
| SE | 29342 | ENSG00000121210 | TMEM131L | chr4 | + | 153602151 | 153602338 | 153598589 | 153598732 | 153602541 | 153602727 | 189 | 152 |
| SE | 29343 | ENSG00000121210 | TMEM131L | chr4 | + | 153612251 | 153612400 | 153603801 | 153604430 | 153620755 | 153620880 | 315 | 432 |
| SE | 29344 | ENSG00000121210 | TMEM131L | chr4 | + | 153627604 | 153627687 | 153626146 | 153626225 | 153632717 | 153632838 | 191 | 129 |
| SE | 29345 | ENSG00000121210 | TMEM131L | chr4 | + | 153627604 | 153627687 | 153626146 | 153626225 | 153634191 | 153634280 | 76 | 44 |
| SE | 29346 | ENSG00000121210 | TMEM131L | chr4 | + | 153632717 | 153632838 | 153626146 | 153626225 | 153634191 | 153634280 | 62 | 45 |
| SE | 29348 | ENSG00000121210 | TMEM131L | chr4 | + | 153633235 | 153633359 | 153632363 | 153632838 | 153634191 | 153634326 | 1 | 0 |
| SE | 29350 | ENSG00000121210 | TMEM131L | chr4 | + | 153635431 | 153635571 | 153634191 | 153634280 | 153636300 | 153636711 | 210 | 157 |
| SE | 29359 | ENSG00000074660 | SCARF1 | chr17 | - | 1643441 | 1643967 | 1639911 | 1640666 | 1644833 | 1644935 | 6 | 0 |
| SE | 29371 | ENSG00000101161 | PRPF6 | chr20 | + | 63994915 | 63995092 | 63993406 | 63993485 | 63995326 | 63995482 | 478 | 330 |
| SE | 29372 | ENSG00000101161 | PRPF6 | chr20 | + | 64010199 | 64010318 | 64001076 | 64001239 | 64016722 | 64016845 | 133 | 63 |
| SE | 29374 | ENSG00000101161 | PRPF6 | chr20 | + | 64016722 | 64016845 | 64011284 | 64011503 | 64022756 | 64022878 | 236 | 116 |
| SE | 29383 | ENSG00000170242 | USP47 | chr11 | + | 11873799 | 11873859 | 11842065 | 11842224 | 11880176 | 11880380 | 17 | 10 |
| SE | 29387 | ENSG00000170242 | USP47 | chr11 | + | 11891967 | 11892106 | 11884466 | 11884580 | 11902714 | 11902860 | 132 | 104 |
| SE | 29388 | ENSG00000170242 | USP47 | chr11 | + | 11897596 | 11897693 | 11884466 | 11884580 | 11902714 | 11902860 | 162 | 128 |
| SE | 29389 | ENSG00000170242 | USP47 | chr11 | + | 11897596 | 11897693 | 11891967 | 11892106 | 11902714 | 11902860 | 342 | 276 |
| SE | 29390 | ENSG00000170242 | USP47 | chr11 | + | 11905398 | 11905548 | 11903262 | 11903342 | 11920155 | 11920251 | 343 | 223 |
| SE | 29392 | ENSG00000170242 | USP47 | chr11 | + | 11950363 | 11950482 | 11949888 | 11950004 | 11952740 | 11952777 | 426 | 305 |
| SE | 29394 | ENSG00000170242 | USP47 | chr11 | + | 11954896 | 11954944 | 11950872 | 11952871 | 11955033 | 11955164 | 152 | 132 |
| SE | 29397 | ENSG00000127419 | TMEM175 | chr4 | + | 947708 | 947892 | 932386 | 932540 | 950420 | 950518 | 11 | 0 |
| SE | 29401 | ENSG00000127419 | TMEM175 | chr4 | + | 948115 | 948154 | 947708 | 947892 | 950420 | 950518 | 13 | 8 |
| SE | 29402 | ENSG00000127419 | TMEM175 | chr4 | + | 948115 | 948615 | 947708 | 947892 | 950420 | 950518 | 17 | 11 |
| SE | 29404 | ENSG00000127419 | TMEM175 | chr4 | + | 948410 | 948615 | 948115 | 948154 | 950423 | 950518 | 9 | 0 |
| SE | 29410 | ENSG00000214174 | AMZ2P1 | chr17 | - | 64973241 | 64973517 | 64972753 | 64972927 | 64974554 | 64975568 | 30 | 30 |
| SE | 29411 | ENSG00000153234 | NR4A2 | chr2 | - | 156326149 | 156326207 | 156325099 | 156326000 | 156326717 | 156326920 | 11 | 9 |
| SE | 29413 | ENSG00000153234 | NR4A2 | chr2 | - | 156326149 | 156326328 | 156324436 | 156326000 | 156326717 | 156326920 | 36 | 25 |
| SE | 29428 | ENSG00000124783 | SSR1 | chr6 | - | 7295087 | 7295102 | 7287569 | 7289931 | 7295391 | 7295485 | 2 | 1 |
| SE | 29430 | ENSG00000124783 | SSR1 | chr6 | - | 7295384 | 7295485 | 7289675 | 7289931 | 7298746 | 7298823 | 0 | 0 |
| SE | 29431 | ENSG00000124783 | SSR1 | chr6 | - | 7295391 | 7295485 | 7281142 | 7289931 | 7297922 | 7298001 | 773 | 479 |
| SE | 29434 | ENSG00000124783 | SSR1 | chr6 | - | 7297922 | 7298001 | 7295055 | 7295485 | 7298746 | 7298823 | 691 | 483 |
| SE | 29435 | ENSG00000124783 | SSR1 | chr6 | - | 7301309 | 7301368 | 7298746 | 7298823 | 7303549 | 7303637 | 196 | 136 |
| SE | 29436 | ENSG00000124783 | SSR1 | chr6 | - | 7301309 | 7301572 | 7298746 | 7298823 | 7303549 | 7303637 | 1148 | 755 |
| SE | 29437 | ENSG00000124783 | SSR1 | chr6 | - | 7303549 | 7303633 | 7301389 | 7301572 | 7309916 | 7310029 | 352 | 233 |
| SE | 29438 | ENSG00000124783 | SSR1 | chr6 | - | 7303549 | 7303637 | 7301309 | 7301572 | 7309916 | 7310029 | 1071 | 756 |
| SE | 29439 | ENSG00000124783 | SSR1 | chr6 | - | 7309916 | 7310029 | 7301309 | 7301572 | 7313041 | 7313174 | 261 | 138 |
| SE | 29451 | ENSG00000130338 | TULP4 | chr6 | + | 158461562 | 158461729 | 158452133 | 158452268 | 158479750 | 158479975 | 65 | 33 |
| SE | 29453 | ENSG00000130338 | TULP4 | chr6 | + | 158494752 | 158494846 | 158493572 | 158493717 | 158498668 | 158498693 | 48 | 54 |
| SE | 29455 | ENSG00000163629 | PTPN13 | chr4 | + | 86686709 | 86686775 | 86672364 | 86672543 | 86689004 | 86689061 | 8 | 13 |
| SE | 29456 | ENSG00000163629 | PTPN13 | chr4 | + | 86693586 | 86693674 | 86689004 | 86689190 | 86701240 | 86701801 | 26 | 25 |
| SE | 29457 | ENSG00000163629 | PTPN13 | chr4 | + | 86693586 | 86693674 | 86689004 | 86689190 | 86716529 | 86716625 | 19 | 12 |
| SE | 29458 | ENSG00000163629 | PTPN13 | chr4 | + | 86701240 | 86701801 | 86689004 | 86689190 | 86716529 | 86716625 | 5 | 8 |
| SE | 29459 | ENSG00000163629 | PTPN13 | chr4 | + | 86701240 | 86701801 | 86693586 | 86693674 | 86716529 | 86716625 | 15 | 24 |
| SE | 29460 | ENSG00000163629 | PTPN13 | chr4 | + | 86741633 | 86741816 | 86735593 | 86735746 | 86744965 | 86745128 | 17 | 24 |
| SE | 29461 | ENSG00000163629 | PTPN13 | chr4 | + | 86753008 | 86753065 | 86744965 | 86745128 | 86758259 | 86758349 | 6 | 5 |
| SE | 29462 | ENSG00000163629 | PTPN13 | chr4 | + | 86753008 | 86753065 | 86751026 | 86751124 | 86758259 | 86758349 | 24 | 7 |
| SE | 29463 | ENSG00000077684 | JADE1 | chr4 | + | 128816366 | 128816463 | 128809843 | 128809877 | 128831732 | 128831810 | 0 | 0 |
| SE | 29464 | ENSG00000077684 | JADE1 | chr4 | + | 128816366 | 128816463 | 128809918 | 128810291 | 128831732 | 128831810 | 0 | 0 |
| SE | 29465 | ENSG00000077684 | JADE1 | chr4 | + | 128816366 | 128816463 | 128810030 | 128810386 | 128831732 | 128831810 | 0 | 0 |
| SE | 29466 | ENSG00000077684 | JADE1 | chr4 | + | 128842952 | 128843038 | 128831732 | 128831810 | 128846374 | 128846532 | 103 | 61 |
| SE | 29467 | ENSG00000077684 | JADE1 | chr4 | + | 128846374 | 128846532 | 128842952 | 128843038 | 128848979 | 128849040 | 237 | 130 |
| SE | 29469 | ENSG00000077684 | JADE1 | chr4 | + | 128855629 | 128855797 | 128852056 | 128852268 | 128861703 | 128862225 | 72 | 55 |
| SE | 29470 | ENSG00000077684 | JADE1 | chr4 | + | 128857337 | 128857454 | 128852056 | 128852268 | 128861703 | 128862225 | 86 | 36 |
| SE | 29471 | ENSG00000077684 | JADE1 | chr4 | + | 128857337 | 128857454 | 128852056 | 128852268 | 128867855 | 128867973 | 0 | 1 |
| SE | 29472 | ENSG00000077684 | JADE1 | chr4 | + | 128857337 | 128857454 | 128855629 | 128855797 | 128861703 | 128862225 | 222 | 103 |
| SE | 29473 | ENSG00000077684 | JADE1 | chr4 | + | 128861703 | 128862225 | 128852056 | 128852268 | 128867855 | 128867973 | 38 | 25 |
| SE | 29474 | ENSG00000077684 | JADE1 | chr4 | + | 128861703 | 128862225 | 128857337 | 128857454 | 128867855 | 128867973 | 146 | 71 |
| SE | 29477 | ENSG00000173020 | GRK2 | chr11 | + | 67279625 | 67279700 | 67279417 | 67279519 | 67279838 | 67279900 | 955 | 522 |
| SE | 29479 | ENSG00000173020 | GRK2 | chr11 | + | 67282270 | 67282365 | 67281821 | 67281952 | 67282434 | 67282477 | 330 | 114 |
| SE | 29481 | ENSG00000173020 | GRK2 | chr11 | + | 67284210 | 67284373 | 67283853 | 67283949 | 67284846 | 67284983 | 574 | 260 |
| SE | 29482 | ENSG00000185684 | EP400P1 | chr12 | + | 132084963 | 132085316 | 132084284 | 132084399 | 132090848 | 132090943 | 9 | 12 |
| SE | 29484 | ENSG00000185684 | EP400P1 | chr12 | + | 132090848 | 132090943 | 132084282 | 132084399 | 132103996 | 132104170 | 6 | 1 |
| SE | 29485 | ENSG00000185684 | EP400P1 | chr12 | + | 132090848 | 132090943 | 132084479 | 132085316 | 132103996 | 132104170 | 10 | 10 |
| SE | 29486 | ENSG00000185684 | EP400P1 | chr12 | + | 132091394 | 132091516 | 132090848 | 132090943 | 132103996 | 132104170 | 0 | 0 |
| SE | 29487 | ENSG00000185684 | EP400P1 | chr12 | + | 132103996 | 132105304 | 132090848 | 132090943 | 132108571 | 132108682 | 27 | 9 |
| SE | 29491 | ENSG00000185684 | EP400P1 | chr12 | + | 132125193 | 132125396 | 132121924 | 132121980 | 132125853 | 132126047 | 32 | 8 |
| SE | 29492 | ENSG00000185684 | EP400P1 | chr12 | + | 132125193 | 132125408 | 132121924 | 132121980 | 132125853 | 132126340 | 20 | 5 |
| SE | 29495 | ENSG00000182957 | SPATA13 | chr13 | + | 24284134 | 24284271 | 24251693 | 24251862 | 24286213 | 24286393 | 344 | 228 |
| SE | 29496 | ENSG00000182957 | SPATA13 | chr13 | + | 24286213 | 24286393 | 24284134 | 24284271 | 24288998 | 24289178 | 216 | 162 |
| SE | 29498 | ENSG00000182957 | SPATA13 | chr13 | + | 24288998 | 24289178 | 24286213 | 24286393 | 24290651 | 24290884 | 297 | 190 |
| SE | 29499 | ENSG00000182957 | SPATA13 | chr13 | + | 24297362 | 24297735 | 24294738 | 24294868 | 24300400 | 24300475 | 267 | 167 |

| | | | | | | | | | | | | | | | | | | | | | | | | | | | | | | | | | |
|---|---|---|---|---|---|---|---|---|---|---|---|---|---|---|---|---|---|---|---|---|---|---|---|---|---|---|---|---|---|---|---|---|---|
| 29076 | 57 | 116 | 119 | 0 | 0 | 0 | 0 | 0 | 104 | 142 | 142 | 169 | 28 | 0 | 0 | 0 | 1 | 0 | 298 | 149 | 1 | 1 | 1 | 1 | 1 | 1 | 1 | 1 | 1 | 1 | 0.988 | 1 | 0.002 |
| 29077 | 54 | 73 | 86 | 1 | 2 | 0 | 0 | 2 | 72 | 58 | 92 | 141 | 28 | 0 | 3 | 2 | 0 | 0 | 298 | 149 | 1 | 1 | 0.98 | 0.932 | 1 | 1 | 0.956 | 1 | 0.906 | 0.958 | 1 | 1 | 0.001 |
| 29078 | 0 | 0 | 0 | 42 | 53 | 29 | 45 | 84 | 0 | 0 | 1 | 1 | 0 | 103 | 57 | 94 | 94 | 18 | 233 | 149 | 1 | 1 | 0 | 0 | 0 | 0 | 0 | 0 | 0 | 0.007 | 0.007 | 0 | -0.003 |
| 29080 | 54 | 95 | 139 | 4 | 0 | 0 | 0 | 0 | 87 | 154 | 199 | 153 | 87 | 0 | 0 | 5 | 0 | 0 | 298 | 149 | 1 | 1 | 0.945 | 1 | 1 | 1 | 1 | 1 | 1 | 0.952 | 1 | 1 | -0.001 |
| 29081 | 32 | 54 | 29 | 0 | 0 | 0 | 0 | 0 | 48 | 92 | 69 | 53 | 38 | 0 | 0 | 0 | 1 | 0 | 298 | 149 | 1 | 1 | 1 | 1 | 1 | 1 | 1 | 1 | 1 | 1 | 0.964 | 1 | 0.007 |
| 29082 | 43 | 64 | 44 | 2 | 0 | 0 | 0 | 0 | 124 | 80 | 84 | 62 | 78 | 0 | 0 | 0 | 0 | 0 | 298 | 149 | 1 | 1 | 0.96 | 1 | 1 | 1 | 1 | 1 | 1 | 1 | 1 | 1 | -0.008 |
| 29085 | 0 | 2 | 5 | 0 | 2 | 0 | 0 | 0 | 9 | 1 | 2 | 2 | 6 | 0 | 1 | 0 | 5 | 0 | 238 | 149 | 1 | 1 | 1 | 0.385 | NA | 1 | 1 | 1 | 0.385 | 1 | 0.2 | 1 | 0.129 |
| 29089 | 0 | 4 | 0 | 1 | 0 | 0 | 0 | 0 | 0 | 11 | 6 | 14 | 3 | 0 | 0 | 0 | 0 | 4 | 243 | 149 | 1 | 1 | 0.551 | 1 | NA | 1 | NA | NA | 1 | 1 | 1 | 0.315 | 0.022 |
| 29093 | 80 | 66 | 142 | 1 | 0 | 0 | 1 | 0 | 47 | 104 | 70 | 31 | 81 | 0 | 0 | 0 | 0 | 0 | 298 | 149 | 1 | 1 | 0.978 | 1 | 1 | 0.971 | 1 | 1 | 1 | 1 | 1 | 1 | -0.01 |
| 29094 | 122 | 127 | 261 | 0 | 2 | 0 | 1 | 2 | 53 | 253 | 109 | 68 | 49 | 0 | 0 | 0 | 0 | 0 | 298 | 149 | 1 | 1 | 1 | 0.946 | 1 | 0.984 | 0.985 | 1 | 1 | 1 | 1 | 1 | -0.017 |
| 29095 | 59 | 73 | 125 | 0 | 0 | 0 | 0 | 0 | 18 | 160 | 58 | 41 | 2 | 0 | 1 | 0 | 0 | 0 | 220 | 149 | 1 | 1 | 1 | 1 | 1 | 1 | 1 | 1 | 0.991 | 1 | 1 | 1 | 0.002 |
| 29096 | 113 | 122 | 185 | 0 | 0 | 1 | 1 | 1 | 87 | 135 | 96 | 50 | 16 | 0 | 0 | 0 | 0 | 0 | 281 | 149 | 1 | 1 | 1 | 1 | 0.984 | 0.985 | 0.99 | 1 | 1 | 1 | 1 | 1 | -0.008 |
| 29099 | 62 | 82 | 138 | 0 | 0 | 0 | 0 | 4 | 22 | 67 | 49 | 20 | 4 | 0 | 0 | 1 | 0 | 0 | 298 | 149 | 1 | 1 | 1 | 1 | 1 | 1 | 0.945 | 1 | 1 | 0.961 | 1 | 1 | -0.003 |
| 29100 | 19 | 31 | 40 | 0 | 0 | 0 | 0 | 4 | 12 | 60 | 38 | 19 | 12 | 0 | 0 | 1 | 0 | 0 | 281 | 149 | 1 | 1 | 1 | 1 | 1 | 1 | 0.841 | 1 | 1 | 0.953 | 1 | 1 | -0.022 |
| 29101 | 57 | 72 | 174 | 0 | 0 | 0 | 0 | 0 | 34 | 82 | 54 | 30 | 6 | 0 | 0 | 1 | 0 | 0 | 281 | 149 | 1 | 1 | 1 | 1 | 1 | 1 | 1 | 1 | 1 | 0.966 | 1 | 1 | 0.007 |
| 29102 | 34 | 28 | 56 | 0 | 0 | 0 | 0 | 0 | 8 | 27 | 21 | 8 | 0 | 0 | 0 | 1 | 0 | 0 | 263 | 149 | 1 | 1 | 1 | 1 | 1 | 1 | 1 | 1 | 1 | 0.922 | 1 | NA | 0.019 |
| 29103 | 95 | 77 | 89 | 11 | 11 | 6 | 6 | 24 | 0 | 87 | 29 | 14 | 7 | 0 | 12 | 23 | 12 | 0 | 298 | 149 | 1 | 1 | 0.758 | 0.694 | 0.888 | 0.865 | 0.65 | NA | 0.784 | 0.387 | 0.368 | 1 | 0.136 |
| 29104 | 23 | 18 | 60 | 1 | 1 | 0 | 2 | 3 | 0 | 27 | 27 | 19 | 0 | 0 | 0 | 2 | 0 | 0 | 267 | 149 | 1 | 1 | 0.925 | 0.909 | 1 | 0.834 | 0.918 | NA | 1 | 0.883 | 1 | NA | -0.044 |
| 29111 | 25 | 130 | 87 | 0 | 0 | 0 | 0 | 0 | 291 | 57 | 69 | 99 | 200 | 0 | 0 | 0 | 0 | 6 | 283 | 149 | 1 | 1 | 1 | 1 | 1 | 1 | 1 | 1 | 1 | 1 | 1 | 0.946 | 0.011 |
| 29113 | 0 | 0 | 0 | 0 | 0 | 2 | 0 | 2 | 0 | 3 | 0 | 0 | 0 | 0 | 1 | 1 | 0 | 0 | 260 | 149 | 1 | 1 | 1 | 1 | 0 | NA | 0 | NA | 0.632 | 0 | NA | NA | 0.184 |

[illegible]

| | | | | | | | | | | | | | | |
|---|---|---|---|---|---|---|---|---|---|---|---|---|---|---|
| SE | 29500 | ENSG00000151458 | ANKRD50 | chr4 | - | 124678675 | 124678905 | 124668983 | 124672534 | 124709999 | 124711274 | 40 | 19 | [illegible] |
| SE | 29502 | ENSG00000109756 | RAPGEF2 | chr4 | + | 159193199 | 159193256 | 159186641 | 159186712 | 159210499 | 159210583 | 55 | 56 | [illegible] |
| SE | 29503 | ENSG00000109756 | RAPGEF2 | chr4 | + | 159210499 | 159210583 | 159193199 | 159193256 | 159238808 | 159238884 | 66 | 96 | [illegible] |
| SE | 29504 | ENSG00000109756 | RAPGEF2 | chr4 | + | 159210499 | 159210583 | 159193199 | 159193256 | 159304341 | 159304473 | 11 | 26 | [illegible] |
| SE | 29505 | ENSG00000109756 | RAPGEF2 | chr4 | + | 159238808 | 159238884 | 159210499 | 159210583 | 159241200 | 159241363 | 76 | 101 | [illegible] |
| SE | 29506 | ENSG00000109756 | RAPGEF2 | chr4 | + | 159241200 | 159241368 | 159210499 | 159210583 | 159304341 | 159304473 | 6 | 6 | [illegible] |
| SE | 29509 | ENSG00000109756 | RAPGEF2 | chr4 | + | 159314590 | 159314768 | 159304341 | 159304473 | 159322346 | 159322363 | 98 | 84 | [illegible] |
| SE | 29510 | ENSG00000109756 | RAPGEF2 | chr4 | + | 159331430 | 159331538 | 159330333 | 159330498 | 159331925 | 159332034 | 73 | 96 | [illegible] |
| SE | 29511 | ENSG00000109756 | RAPGEF2 | chr4 | + | 159331629 | 159331833 | 159331430 | 159331538 | 159331925 | 159332034 | 189 | 194 | [illegible] |
| SE | 29512 | ENSG00000109756 | RAPGEF2 | chr4 | + | 159331925 | 159332034 | 159330333 | 159330498 | 159332450 | 159332697 | 40 | 46 | [illegible] |
| SE | 29513 | ENSG00000109756 | RAPGEF2 | chr4 | + | 159344035 | 159344059 | 159343280 | 159343404 | 159345105 | 159345329 | 20 | 24 | [illegible] |
| SE | 29515 | ENSG00000109756 | RAPGEF2 | chr4 | + | 159351058 | 159351133 | 159350136 | 159350289 | 159352684 | 159352910 | 16 | 20 | [illegible] |
| SE | 29516 | ENSG00000109756 | RAPGEF2 | chr4 | + | 159351058 | 159351181 | 159350227 | 159350289 | 159352684 | 159352910 | 17 | 24 | [illegible] |
| SE | 29517 | ENSG00000109756 | RAPGEF2 | chr4 | + | 159352684 | 159352910 | 159350136 | 159350289 | 159353486 | 159353504 | 81 | 73 | [illegible] |
| SE | 29521 | ENSG00000119242 | CCDC92 | chr12 | - | 123944668 | 123944740 | 123944271 | 123944364 | 123945703 | 123945800 | 0 | 2 | [illegible] |
| SE | 29522 | ENSG00000119242 | CCDC92 | chr12 | - | 123944668 | 123944740 | 123944271 | 123944364 | 123956283 | 123956444 | 0 | 2 | [illegible] |
| SE | 29524 | ENSG00000119242 | CCDC92 | chr12 | - | 123945703 | 123945800 | 123944271 | 123944364 | 123972528 | 123972831 | 0 | 0 | [illegible] |
| SE | 29528 | ENSG00000119242 | CCDC92 | chr12 | - | 123956283 | 123956444 | 123944668 | 123944740 | 123972528 | 123972616 | 0 | 0 | [illegible] |
| SE | 29530 | ENSG00000224565 | LINC01754 | chr20 | - | 47411267 | 47411393 | 47397148 | 47397345 | 47412069 | 47412327 | 3 | 0 | [illegible] |
| SE | 29531 | ENSG00000100650 | SRSF5 | chr14 | + | 69768137 | 69768282 | 69766113 | 69767255 | 69768603 | 69768674 | 2335 | 1623 | [illegible] |
| SE | 29532 | ENSG00000100650 | SRSF5 | chr14 | + | 69768603 | 69768674 | 69768137 | 69768282 | 69768797 | 69768896 | 2510 | 1879 | [illegible] |
| SE | 29533 | ENSG00000100650 | SRSF5 | chr14 | + | 69768603 | 69768674 | 69768137 | 69768282 | 69769181 | 69769251 | 549 | 360 | [illegible] |
| SE | 29534 | ENSG00000100650 | SRSF5 | chr14 | + | 69768612 | 69768674 | 69768137 | 69768282 | 69768797 | 69768896 | 652 | 581 | [illegible] |
| SE | 29535 | ENSG00000100650 | SRSF5 | chr14 | + | 69768612 | 69768674 | 69768137 | 69768282 | 69769181 | 69769251 | 10 | 11 | [illegible] |
| SE | 29536 | ENSG00000100650 | SRSF5 | chr14 | + | 69768797 | 69768896 | 69768137 | 69768282 | 69769181 | 69769251 | 2756 | 2449 | [illegible] |
| [illegible] | [illegible] | [illegible] | [illegible] | [illegible] | [illegible] | [illegible] | [illegible] | [illegible] | [illegible] | [illegible] | [illegible] | [illegible] | [illegible] | [illegible] |

| | | | | | | | | | | | | | | | | | | | | | | | | | | | | | | | | | | | | | | | | | | | | | | |
|---|---|---|---|---|---|---|---|---|---|---|---|---|---|---|---|---|---|---|---|---|---|---|---|---|---|---|---|---|---|---|---|---|---|---|---|---|---|---|---|---|---|---|---|---|---|---|
| SE | 29916 | ENSG00000106070 | GRB10 | chr7 | - | 50710874 | 50710990 | 50703820 | 50703908 | 50755886 | 50756056 | 4 | 0 | 1 | 0 | 0 | 0 | 0 | 0 | 0 | 0 | 0 | 12 | 0 | 0 | 16 | 0 | 0 | 0 | 3 | 0 | 265 | 149 | 1 | 1 | 1 | NA | 1 | NA | NA | NA | 1 | NA | 0 | 1 | 0.333 |
| SE | 29917 | ENSG00000106070 | GRB10 | chr7 | - | 50710874 | 50710990 | 50703820 | 50703908 | 50793223 | 50793432 | 2 | 0 | 1 | 0 | 0 | 1 | 0 | 0 | 0 | 0 | 0 | 12 | 2 | 0 | 18 | 0 | 0 | 0 | 3 | 0 | 265 | 149 | 1 | 1 | 0.529 | NA | 1 | NA | NA | NA | 1 | 1 | 0 | 1 | 0.014 |
| SE | 29918 | ENSG00000106070 | GRB10 | chr7 | - | 50710874 | 50710990 | 50705139 | 50705323 | 50726012 | 50726035 | 0 | 0 | 2 | 0 | 0 | 0 | 0 | 0 | 1 | 0 | 9 | 8 | 0 | 4 | 5 | 0 | 0 | 0 | 0 | 0 | 265 | 149 | 1 | 1 | NA | NA | 1 | 0 | NA | 1 | 1 | NA | 1 | 1 | -0.5 |
| SE | 29920 | ENSG00000106070 | GRB10 | chr7 | - | 50729299 | 50729399 | 50727792 | 50727890 | 50732271 | 50732368 | 0 | 0 | 0 | 0 | 0 | 1 | 0 | 0 | 1 | 0 | 0 | 0 | 0 | 2 | 7 | 1 | 2 | 3 | 3 | 0 | 249 | 149 | 1 | 1 | 0 | NA | NA | 0 | NA | 0 | 0 | 0 | 0.285 | 1 | -0.257 |
| SE | 29921 | ENSG00000106070 | GRB10 | chr7 | - | 50732271 | 50732368 | 50703820 | 50703908 | 50755886 | 50755903 | 0 | 0 | 2 | 1 | 2 | 0 | 0 | 0 | 0 | 0 | 32 | 4 | 0 | 16 | 14 | 0 | 0 | 0 | 3 | 0 | 246 | 149 | 1 | 1 | NA | NA | 1 | 1 | 1 | 1 | 1 | NA | 0.764 | 1 | 0.059 |
| SE | 29922 | ENSG00000106070 | GRB10 | chr7 | - | 50732271 | 50732368 | 50703820 | 50703908 | 50793223 | 50793432 | 0 | 0 | 0 | 0 | 0 | 1 | 0 | 0 | 0 | 0 | 26 | 0 | 0 | 8 | 0 | 0 | 0 | 0 | 3 | 0 | 246 | 149 | 1 | 1 | 0 | NA | NA | NA | NA | 1 | NA | NA | 0.618 | NA | -0.809 |
| SE | 29923 | ENSG00000106070 | GRB10 | chr7 | - | 50732271 | 50732368 | 50710874 | 50710990 | 50755886 | 50755903 | 0 | 0 | 2 | 1 | 2 | 2 | 0 | 0 | 0 | 0 | 6 | 3 | 0 | 5 | 11 | 0 | 0 | 0 | 0 | 0 | 246 | 149 | 1 | 1 | 0 | NA | 1 | 1 | 1 | 1 | 1 | NA | 1 | 1 | -0.25 |
| SE | 29925 | ENSG00000106070 | GRB10 | chr7 | - | 50732271 | 50732368 | 50727792 | 50727890 | 50755886 | 50755903 | 1 | 0 | 2 | 2 | 2 | 0 | 0 | 0 | 3 | 0 | 6 | 5 | 3 | 8 | 11 | 0 | 0 | 0 | 0 | 0 | 246 | 149 | 1 | 1 | 1 | NA | 1 | 0.288 | 1 | 1 | 1 | 1 | 1 | 1 | -0.178 |
| SE | 29926 | ENSG00000106070 | GRB10 | chr7 | - | 50755886 | 50756056 | 50703820 | 50703908 | 50793223 | 50793313 | 0 | 0 | 0 | 3 | 0 | 1 | 0 | 0 | 0 | 0 | 0 | 2 | 0 | 3 | 0 | 0 | 0 | 0 | 3 | 0 | 298 | 149 | 1 | 1 | 0 | NA | NA | 1 | NA | NA | 1 | NA | 0.333 | NA | -0.166 |
| SE | 29927 | ENSG00000106070 | GRB10 | chr7 | - | 50755886 | 50756056 | 50710874 | 50710990 | 50793223 | 50793313 | 2 | 0 | 0 | 3 | 0 | 0 | 0 | 0 | 0 | 0 | 0 | 2 | 0 | 0 | 0 | 0 | 0 | 2 | 0 | 2 | 298 | 149 | 1 | 1 | 1 | NA | NA | 1 | NA | NA | 1 | 0 | NA | 0 | 0.667 |
| SE | 29928 | ENSG00000106070 | GRB10 | chr7 | - | 50755886 | 50756056 | 50732271 | 50732368 | 50793223 | 50793313 | 0 | 0 | 2 | 5 | 2 | 0 | 0 | 0 | 0 | 0 | 6 | 6 | 0 | 8 | 14 | 1 | 0 | 0 | 0 | 0 | 298 | 149 | 1 | 1 | NA | NA | 1 | 1 | 1 | 0.75 | 1 | NA | 1 | 1 | 0.062 |
| SE | 29930 | ENSG00000106070 | GRB10 | chr7 | - | 50780626 | 50780736 | 50755886 | 50756056 | 50793223 | 50793437 | 0 | 0 | 0 | 0 | 0 | 0 | 0 | 0 | 3 | 0 | 0 | 1 | 0 | 0 | 0 | 0 | 2 | 0 | 0 | 0 | 259 | 149 | 1 | 1 | NA | NA | NA | 0 | NA | NA | 0.223 | NA | NA | NA | -0.223 |
| SE | 29931 | ENSG00000248019 | FAM13A-AS1 | chr4 | + | 88722150 | 88722344 | 88711854 | 88712017 | 88723208 | 88723315 | 21 | 16 | 28 | 22 | 39 | 2 | 1 | 0 | 5 | 0 | 45 | 51 | 23 | 19 | 0 | 0 | 0 | 0 | 2 | 0 | 298 | 149 | 1 | 1 | 0.84 | 0.889 | 1 | 0.688 | 1 | 1 | 1 | 1 | 0.826 | NA | -0.073 |
| SE | 29932 | ENSG00000197142 | ACSL5 | chr10 | + | 112398900 | 112399009 | 112394917 | 112395102 | 112404510 | 112404575 | 210 | 125 | 225 | 198 | 208 | 0 | 3 | 1 | 0 | 0 | 331 | 184 | 104 | 175 | 176 | 0 | 0 | 4 | 1 | 0 | 258 | 149 | 1 | 1 | 1 | 0.96 | 0.992 | 1 | 1 | 1 | 1 | 0.938 | 0.99 | 1 | 0.005 |
| SE | 29933 | ENSG00000197142 | ACSL5 | chr10 | + | 112409506 | 112409685 | 112408421 | 112408521 | 112410462 | 112410495 | 236 | 178 | 222 | 211 | 443 | 0 | 0 | 0 | 0 | 1 | 253 | 255 | 113 | 151 | 186 | 0 | 0 | 0 | 0 | 0 | 298 | 149 | 1 | 1 | 1 | 1 | 1 | 1 | 0.996 | 1 | 1 | 1 | 1 | 1 | -0.001 |
| SE | 29934 | ENSG00000197142 | ACSL5 | chr10 | + | 112410583 | 112410635 | 112410462 | 112410495 | 112411455 | 112411529 | 177 | 132 | 163 | 166 | 346 | 0 | 2 | 0 | 0 | 3 | 87 | 162 | 45 | 102 | 108 | 0 | 0 | 0 | 0 | 0 | 201 | 149 | 1 | 1 | 1 | 0.98 | 1 | 1 | 0.988 | 1 | 1 | 1 | 1 | 1 | -0.006 |
| SE | 29937 | ENSG00000197142 | ACSL5 | chr10 | + | 112411455 | 112411529 | 112410583 | 112410635 | 112411901 | 112411979 | 179 | 126 | 144 | 146 | 315 | 0 | 0 | 0 | 2 | 0 | 67 | 151 | 50 | 88 | 110 | 0 | 0 | 0 | 0 | 0 | 223 | 149 | 1 | 1 | 1 | 1 | 1 | 0.98 | 1 | 1 | 1 | 1 | 1 | 1 | -0.004 |
| SE | 29939 | ENSG00000197142 | ACSL5 | chr10 | + | 112417845 | 112417941 | 112416887 | 112417022 | 112421592 | 112421626 | 327 | 276 | 306 | 344 | 439 | 0 | 1 | 0 | 0 | 0 | 459 | 480 | 262 | 380 | 409 | 0 | 0 | 0 | 0 | 0 | 245 | 149 | 1 | 1 | 1 | 0.994 | 1 | 1 | 1 | 1 | 1 | 1 | 1 | 1 | -0.001 |
| SE | 29941 | ENSG00000197142 | ACSL5 | chr10 | + | 112426787 | 112426859 | 112426257 | 112426359 | 112427217 | 112428363 | 365 | 312 | 226 | 308 | 478 | 1 | 0 | 0 | 2 | 7 | 462 | 325 | 207 | 397 | 189 | 10 | 0 | 1 | 0 | 0 | 221 | 149 | 1 | 1 | 0.996 | 1 | 1 | 0.99 | 0.979 | 0.969 | 1 | 0.993 | 1 | 1 | 0.001 |
| SE | 29942 | ENSG00000136051 | WASHC4 | chr12 | + | 105111124 | 105111264 | 105107713 | 105107861 | 105114215 | 105114269 | 813 | 659 | 532 | 691 | 581 | 2 | 5 | 1 | 6 | 0 | 144 | 329 | 201 | 87 | 100 | 0 | 0 | 0 | 0 | 0 | 289 | 149 | 1 | 1 | 0.995 | 0.985 | 0.996 | 0.983 | 1 | 1 | 1 | 1 | 1 | 1 | -0.008 |
| SE | 29943 | ENSG00000136051 | WASHC4 | chr12 | + | 105115183 | 105115229 | 105114361 | 105114427 | 105115660 | 105115728 | 815 | 722 | 528 | 744 | 821 | 1 | 0 | 4 | 1 | 0 | 229 | 365 | 179 | 98 | 87 | 0 | 0 | 0 | 0 | 0 | 195 | 149 | 1 | 1 | 0.998 | 1 | 0.99 | 0.998 | 1 | 1 | 1 | 1 | 1 | 1 | -0.003 |
| SE | 29944 | ENSG00000136051 | WASHC4 | chr12 | + | 105126003 | 105126127 | 105122117 | 105122238 | 105126234 | 105126362 | 593 | 449 | 376 | 518 | 831 | 0 | 2 | 0 | 0 | 0 | 60 | 175 | 83 | 17 | 2 | 0 | 0 | 0 | 0 | 0 | 273 | 149 | 1 | 1 | 1 | 0.992 | 1 | 1 | 1 | 1 | 1 | 1 | 1 | 1 | -0.002 |
| SE | 29945 | ENSG00000136051 | WASHC4 | chr12 | + | 105126234 | 105126362 | 105122117 | 105122238 | 105127128 | 105127289 | 534 | 478 | 377 | 478 | 788 | 4 | 0 | 0 | 0 | 0 | 34 | 110 | 102 | 54 | 15 | 0 | 0 | 1 | 0 | 0 | 277 | 149 | 1 | 1 | 0.986 | 1 | 1 | 1 | 1 | 1 | 1 | 0.982 | 1 | 1 | 0.001 |
| SE | 29946 | ENSG00000136051 | WASHC4 | chr12 | + | 105133769 | 105133896 | 105127128 | 105127289 | 105137885 | 105138011 | 476 | 438 | 352 | 436 | 995 | 0 | 1 | 3 | 3 | 0 | 9 | 117 | 71 | 23 | 25 | 0 | 0 | 0 | 0 | 0 | 276 | 149 | 1 | 1 | 1 | 0.996 | 0.984 | 0.987 | 1 | 1 | 1 | 1 | 1 | 1 | -0.007 |
| SE | 29947 | ENSG00000136051 | WASHC4 | chr12 | + | 105133769 | 105133899 | 105127128 | 105127289 | 105137885 | 105138011 | 226 | 176 | 158 | 200 | 339 | 0 | 1 | 3 | 3 | 0 | 6 | 55 | 41 | 15 | 3 | 0 | 0 | 0 | 0 | 0 | 279 | 149 | 1 | 1 | 1 | 0.989 | 0.966 | 0.973 | 1 | 1 | 1 | 1 | 1 | 1 | -0.014 |
| SE | 29948 | ENSG00000136051 | WASHC4 | chr12 | + | 105137885 | 105138011 | 105133769 | 105133896 | 105140293 | 105140401 | 531 | 573 | 453 | 529 | 1217 | 0 | 1 | 1 | 0 | 0 | 12 | 153 | 64 | 21 | 47 | 0 | 1 | 0 | 0 | 0 | 275 | 149 | 1 | 1 | 1 | 0.997 | 0.996 | 1 | 1 | 1 | 0.988 | 1 | 1 | 1 | 0.001 |
| SE | 29951 | ENSG00000136051 | WASHC4 | chr12 | + | 105143126 | 105143243 | 105141166 | 105141246 | 105144717 | 105144872 | 9 | 4 | 1 | 5 | 5 | 0 | 0 | 1 | 0 | 0 | 15 | 2 | 2 | 2 | 0 | 0 | 0 | 0 | 0 | 0 | 266 | 149 | 1 | 1 | 1 | 1 | 0.359 | 1 | 1 | 1 | 1 | 1 | 1 | NA | -0.128 |
| SE | 29952 | ENSG00000136051 | WASHC4 | chr12 | + | 105143126 | 105143243 | 105142452 | 105142558 | 105144286 | 105144455 | 336 | 268 | 206 | 266 | 470 | 0 | 3 | 0 | 2 | 0 | 14 | 93 | 44 | 26 | 3 | 0 | 0 | 0 | 0 | 0 | 266 | 149 | 1 | 1 | 1 | 0.98 | 1 | 0.987 | 1 | 1 | 1 | 1 | 1 | 1 | -0.007 |
| SE | 29953 | ENSG00000136051 | WASHC4 | chr12 | + | 105144286 | 105144455 | 105143126 | 105143243 | 105144717 | 105144872 | 510 | 440 | 342 | 459 | 663 | 1 | 1 | 0 | 0 | 0 | 29 | 125 | 81 | 48 | 13 | 0 | 1 | 0 | 0 | 0 | 298 | 149 | 1 | 1 | 0.996 | 0.995 | 1 | 1 | 1 | 1 | 0.984 | 1 | 1 | 1 | 0.001 |
| SE | 29954 | ENSG00000136051 | WASHC4 | chr12 | + | 105146451 | 105146526 | 105144717 | 105144872 | 105147041 | 105147146 | 932 | 881 | 621 | 961 | 903 | 0 | 0 | 0 | 0 | 1 | 59 | 316 | 128 | 45 | 18 | 0 | 0 | 0 | 0 | 0 | 224 | 149 | 1 | 1 | 1 | 1 | 1 | 1 | 0.998 | 1 | 1 | 1 | 1 | 1 | 0 |
| SE | 29955 | ENSG00000136051 | WASHC4 | chr12 | + | 105147041 | 105147146 | 105144717 | 105144872 | 105149614 | 105149749 | 260 | 309 | 205 | 281 | 428 | 0 | 1 | 0 | 0 | 0 | 7 | 77 | 43 | 15 | 7 | 0 | 0 | 0 | 0 | 0 | 254 | 149 | 1 | 1 | 1 | 0.995 | 1 | 1 | 1 | 1 | 1 | 1 | 1 | 1 | -0.001 |
| SE | 29957 | ENSG00000136051 | WASHC4 | chr12 | + | 105155109 | 105155208 | 105152345 | 105152451 | 105156725 | 105156792 | 3 | 0 | 1 | 1 | 0 | 212 | 152 | 192 | 201 | 408 | 0 | 0 | 0 | 2 | 0 | 5 | 37 | 7 | 7 | 0 | 248 | 149 | 1 | 1 | 0.008 | 0 | 0.003 | 0.003 | 0 | 0 | 0 | 0 | 0.147 | NA | -0.034 |
| SE | 29959 | ENSG00000136051 | WASHC4 | chr12 | + | 105160000 | 105160148 | 105157235 | 105157322 | 105162748 | 105162802 | 663 | 582 | 535 | 648 | 1101 | 0 | 2 | 0 | 2 | 1 | 63 | 180 | 41 | 47 | 58 | 0 | 0 | 0 | 0 | 0 | 297 | 149 | 1 | 1 | 1 | 0.993 | 1 | 0.994 | 0.998 | 1 | 1 | 1 | 1 | 1 | -0.003 |
| SE | 29960 | ENSG00000136051 | WASHC4 | chr12 | + | 105160000 | 105160148 | 105157235 | 105157322 | 105164110 | 105164307 | 419 | 391 | 341 | 382 | 751 | 0 | 0 | 0 | 2 | 0 | 63 | 92 | 23 | 28 | 45 | 0 | 0 | 0 | 0 | 0 | 297 | 149 | 1 | 1 | 1 | 1 | 1 | 0.99 | 1 | 1 | 1 | 1 | 1 | 1 | -0.002 |
| SE | 29961 | ENSG00000136051 | WASHC4 | chr12 | + | 105162748 | 105162845 | 105160000 | 105160148 | 105164110 | 105164307 | 506 | 445 | 382 | 542 | 689 | 0 | 1 | 0 | 5 | 0 | 6 | 155 | 55 | 27 | 24 | 0 | 0 | 0 | 0 | 0 | 246 | 149 | 1 | 1 | 1 | 0.996 | 1 | 0.985 | 1 | 1 | 1 | 1 | 1 | 1 | -0.004 |
| SE | 29962 | ENSG00000136051 | WASHC4 | chr12 | + | 105164110 | 105164307 | 105162748 | 105162845 | 105164640 | 105164740 | 501 | 458 | 381 | 527 | 758 | 0 | 1 | 0 | 1 | 1 | 27 | 150 | 64 | 23 | 15 | 0 | 0 | 0 | 0 | 0 | 298 | 149 | 1 | 1 | 1 | 0.996 | 1 | 0.996 | 0.997 | 1 | 1 | 1 | 1 | 1 | -0.002 |
| SE | 29963 | ENSG00000136051 | WASHC4 | chr12 | + | 105164110 | 105164307 | 105162748 | 105162845 | 105164672 | 105164729 | 320 | 284 | 222 | 327 | 440 | 0 | 0 | 1 | 0 | 0 | 6 | 82 | 45 | 11 | 11 | 0 | 0 | 0 | 0 | 0 | 298 | 149 | 1 | 1 | 1 | 1 | 0.991 | 1 | 1 | 1 | 1 | 1 | 1 | 1 | -0.002 |
| SE | 29965 | ENSG00000143624 | INTS3 | chr1 | + | 153748688 | 153748755 | 153747278 | 153747363 | 153751094 | 153751239 | 64 | 54 | 47 | 82 | 79 | 2 | 0 | 0 | 3 | 0 | 48 | 39 | 24 | 88 | 105 | 0 | 0 | 0 | 0 | 0 | 216 | 149 | 1 | 1 | 0.957 | 1 | 1 | 0.95 | 1 | 1 | 1 | 1 | 1 | 1 | -0.019 |
| SE | 29966 | ENSG00000143624 | INTS3 | chr1 | + | 153752278 | 153752408 | 153750853 | 153751239 | 153757571 | 153757763 | 32 | 27 | 28 | 24 | 44 | 1 | 0 | 0 | 2 | 0 | 49 | 23 | 20 | 44 | 49 | 0 | 0 | 0 | 0 | 0 | 279 | 149 | 1 | 1 | 0.945 | 1 | 1 | 0.865 | 1 | 1 | 1 | 1 | 1 | 1 | -0.038 |
| SE | 29967 | ENSG00000143624 | INTS3 | chr1 | + | 153753700 | 153753818 | 153752278 | 153752408 | 153754641 | 153754739 | 1 | 0 | 0 | 1 | 0 | 64 | 37 | 60 | 66 | 106 | 0 | 0 | 0 | 7 | 0 | 126 | 77 | 38 | 54 | 92 | 267 | 149 | 1 | 1 | 0.009 | 0 | 0 | 0.008 | 0 | 0 | 0 | 0 | 0.067 | 0 | -0.01 |
| SE | 29968 | ENSG00000143624 | INTS3 | chr1 | + | 153754641 | 153754739 | 153752076 | 153752408 | 153757571 | 153757763 | 111 | 101 | 100 | 146 | 181 | 0 | 0 | 0 | 2 | 0 | 258 | 150 | 93 | 131 | 156 | 0 | 0 | 0 | 0 | 0 | 247 | 149 | 1 | 1 | 1 | 1 | 1 | 0.978 | 1 | 1 | 1 | 1 | 1 | 1 | -0.004 |
| SE | 29971 | ENSG00000143624 | INTS3 | chr1 | + | 153760310 | 153760390 | 153757571 | 153757763 | 153761569 | 153761676 | 0 | 0 | 1 | 0 | 2 | 0 | 0 | 0 | 0 | 0 | 14 | 3 | 0 | 10 | 5 | 0 | 1 | 0 | 0 | 0 | 229 | 149 | 1 | 1 | NA | NA | 1 | NA | 1 | 1 | 0.661 | NA | 1 | 1 | 0.085 |
| SE | 29972 | ENSG00000143624 | INTS3 | chr1 | + | 153760310 | 153760390 | 153757571 | 153757763 | 153762727 | 153762847 | 0 | 0 | 1 | 0 | 2 | 0 | 2 | 0 | 1 | 0 | 14 | 3 | 0 | 11 | 5 | 10 | 0 | 7 | 0 | 0 | 229 | 149 | 1 | 1 | NA | 0 | 1 | 0 | 1 | 0.477 | 1 | 0 | 1 | 1 | -0.195 |
| SE | 29973 | ENSG00000143624 | INTS3 | chr1 | + | 153760826 | 153760918 | 153760310 | 153760390 | 153761569 | 153761676 | 118 | 82 | 75 | 119 | 76 | 0 | 0 | 0 | 0 | 0 | 184 | 130 | 46 | 165 | 184 | 0 | 0 | 1 | 0 | 0 | 241 | 149 | 1 | 1 | 1 | 1 | 1 | 1 | 1 | 1 | 1 | 0.966 | 1 | 1 | 0.007 |
| SE | 29976 | ENSG00000143624 | INTS3 | chr1 | + | 153761569 | 153761676 | 153760310 | 153760390 | 153762727 | 153762847 | 85 | 66 | 63 | 95 | 45 | 1 | 0 | 0 | 0 | 0 | 164 | 117 | 78 | 154 | 182 | 0 | 0 | 0 | 1 | 0 | 256 | 149 | 1 | 1 | 0.98 | 1 | 1 | 1 | 1 | 1 | 1 | 1 | 0.989 | 1 | -0.002 |
| SE | 29978 | ENSG00000143624 | INTS3 | chr1 | + | 153763232 | 153763362 | 153762727 | 153762847 | 153764689 | 153764734 | 79 | 62 | 94 | 91 | 60 | 0 | 0 | 0 | 0 | 0 | 208 | 142 | 72 | 179 | 170 | 0 | 0 | 0 | 2 | 0 | 279 | 149 | 1 | 1 | 1 | 1 | 1 | 1 | 1 | 1 | 1 | 1 | 0.98 | 1 | 0.004 |
| SE | 29979 | ENSG00000143624 | INTS3 | chr1 | + | 153763831 | 153763886 | 153762727 | 153762847 | 153764689 | 153764734 | 1 | 0 | 2 | 0 | 2 | 0 | 0 | 0 | 0 | 0 | 16 | 4 | 8 | 17 | 0 | 0 | 0 | 0 | 2 | 0 | 204 | 149 | 1 | 1 | 1 | NA | 1 | NA | 1 | 1 | 1 | 1 | 0.861 | NA | 0.035 |
| SE | 29980 | ENSG00000143624 | INTS3 | chr1 | + | 153763831 | 153763886 | 153763232 | 153763362 | 153764117 | 153764221 | 123 | 57 | 92 | 122 | 86 | 0 | 0 | 0 | 1 | 0 | 190 | 88 | 52 | 126 | 236 | 0 | 0 | 0 | 2 | 0 | 204 | 149 | 1 | 1 | 1 | 1 | 1 | 0.989 | 1 | 1 | 1 | 1 | 0.979 | 1 | 0.002 |
| SE | 29984 | ENSG00000143624 | INTS3 | chr1 | + | 153764689 | 153764734 | 153763232 | 153763362 | 153764943 | 153765063 | 15 | 12 | 9 | 19 | 7 | 0 | 0 | 0 | 0 | 0 | 17 | 27 | 11 | 36 | 28 | 0 | 1 | 0 | 0 | 0 | 194 | 149 | 1 | 1 | 1 | 1 | 1 | 1 | 1 | 1 | 0.954 | 1 | 1 | 1 | 0.009 |
| SE | 29985 | ENSG00000165475 | CRYL1 | chr13 | - | 20405903 | 20406128 | 20404634 | 20404741 | 20413281 | 20413387 | 0 | 2 | 0 | 0 | 0 | 59 | 42 | 42 | 36 | 64 | 0 | 0 | 0 | 0 | 4 | 50 | 22 | 29 | 39 | 182 | 298 | 149 | 1 | 1 | 0 | 0.023 | 0 | 0 | 0 | 0 | 0 | 0 | 0 | 0.011 | 0.002 |
| SE | 29986 | ENSG00000165475 | CRYL1 | chr13 | - | 20413281 | 20413387 | 20404634 | 20404741 | 20432101 | 20432296 | 109 | 67 | 98 | 87 | 140 | 0 | 0 | 0 | 1 | 0 | 130 | 64 | 47 | 67 | 293 | 0 | 0 | 0 | 0 | 6 | 255 | 149 | 1 | 1 | 1 | 1 | 1 | 0.981 | 1 | 1 | 1 | 1 | 1 | 0.966 | 0.003 |
| SE | 29987 | ENSG00000165475 | CRYL1 | chr13 | - | 20418397 | 20419084 | 20413281 | 20413387 | 20432101 | 20432296 | 0 | 0 | 2 | 0 | 2 | 60 | 30 | 66 | 54 | 85 | 5 | 0 | 1 | 0 | 0 | 83 | 44 | 22 | 39 | 176 | 298 | 149 | 1 | 1 | 0 | 0 | 0.015 | 0 | 0.012 | 0.029 | 0 | 0.022 | 0 | 0 | -0.005 |
| SE | 29989 | ENSG00000165475 | CRYL1 | chr13 | - | 20472965 | 20473074 | 20439592 | 20439754 | 20489369 | 20489496 | 1 | 1 | 2 | 3 | 0 | 74 | 39 | 50 | 51 | 89 | 0 | 0 | 0 | 0 | 0 | 99 | 26 | 18 | 33 | 145 | 258 | 149 | 1 | 1 | 0.008 | 0.015 | 0.023 | 0.033 | 0 | 0 | 0 | 0 | 0 | 0 | 0.016 |
| SE | 29995 | ENSG00000185305 | ARL15 | chr5 | - | 54154579 | 54154639 | 54113132 | 54113410 | 54171783 | 54171928 | 65 | 78 | 49 | 91 | 70 | 0 | 0 | 0 | 0 | 0 | 29 | 131 | 87 | 51 | 28 | 0 | 0 | 1 | 0 | 0 | 209 | 149 | 1 | 1 | 1 | 1 | 1 | 1 | 1 | 1 | 1 | 0.984 | 1 | 1 | 0.003 |
| SE | 29996 | ENSG00000185305 | ARL15 | chr5 | - | 54154579 | 54154639 | 54113132 | 54113410 | 54310431 | 54310573 | 19 | 16 | 11 | 25 | 40 | 0 | 0 | 0 | 0 | 8 | 0 | 18 | 17 | 10 | 11 | 0 | 0 | 1 | 0 | 0 | 209 | 149 | 1 | 1 | 1 | 1 | 1 | 1 | 0.781 | NA | 1 | 0.924 | 1 | 1 | -0.025 |
| SE | 29999 | ENSG00000185305 | ARL15 | chr5 | - | 54171783 | 54171928 | 54154390 | 54154639 | 54310294 | 54310415 | 55 | 84 | 44 | 89 | 34 | 0 | 0 | 0 | 0 | 0 | 52 | 136 | 71 | 43 | 66 | 0 | 5 | 0 | 0 | 0 | 294 | 149 | 1 | 1 | 1 | 1 | 1 | 1 | 1 | 1 | 0.932 | 1 | 1 | 1 | 0.014 |
| SE | 30001 | ENSG00000185305 | ARL15 | chr5 | - | 54307887 | 54307946 | 54154579 | 54154639 | 54310294 | 54310415 | 0 | 1 | 0 | 0 | 0 | 0 | 0 | 0 | 0 | 0 | 0 | 0 | 0 | 0 | 0 | 0 | 5 | 0 | 0 | 0 | 208 | 149 | 1 | 1 | NA | 1 | NA | NA | NA | NA | 0 | NA | NA | NA | 1 |
| SE | 30002 | ENSG00000185305 | ARL15 | chr5 | - | 54307887 | 54307946 | 54154579 | 54154639 | 54310431 | 54310582 | 0 | 0 | 0 | 0 | 0 | 0 | 2 | 0 | 3 | 0 | 12 | 0 | 0 | 0 | 0 | 0 | 0 | 0 | 0 | 5 | 208 | 149 | 1 | 1 | NA | 0 | NA | 0 | NA | 1 | NA | NA | NA | 0 | -0.5 |
| SE | 30006 | ENSG00000151012 | SLC7A11 | chr4 | - | 138182296 | 138182393 | 138180640 | 138180790 | 138183201 | 138183305 | 1 | 0 | 1 | 1 | 0 | 0 | 0 | 0 | 0 | 0 | 0 | 1 | 0 | 0 | 0 | 2 | 0 | 0 | 0 | 0 | 246 | 149 | 1 | 1 | 1 | NA | 1 | 1 | NA | 0 | 1 | NA | NA | NA | 0.5 |
| SE | 30008 | ENSG00000248049 | UBA6-AS1 | chr4 | + | 67716922 | 67717021 | 67701394 | 67701484 | 67720421 | 67721018 | 2 | 0 | 0 | 0 | 0 | 10 | 11 | 10 | 21 | 8 | 0 | 0 | 0 | 0 | 0 | 0 | 14 | 3 | 18 | 35 | 248 | 149 | 1 | 1 | 0.107 | 0 | 0 | 0 | 0 | NA | 0 | 0 | 0 | 0 | 0.021 |
| SE | 30009 | ENSG00000248049 | UBA6-AS1 | chr4 | + | 67720421 | 67720632 | 67701279 | 67701484 | 67721476 | 67722505 | 10 | 11 | 10 | 21 | 8 | 1 | 0 | 7 | 1 | 0 | 0 | 14 | 3 | 18 | 35 | 0 | 0 | 0 | 0 | 0 | 298 | 149 | 1 | 1 | 0.833 | 1 | 0.417 | 0.913 | 1 | NA | 1 | 1 | 1 | 1 | -0.167 |
| SE | 30012 | ENSG00000248049 | UBA6-AS1 | chr4 | + | 67730073 | 67730170 | 67728539 | 67728686 | 67731505 | 67732345 | 38 | 5 | 17 | 4 | 67 | 0 | 0 | 3 | 0 | 1 | 0 | 2 | 7 | 14 | 0 | 0 | 0 | 2 | 0 | 0 | 246 | 149 | 1 | 1 | 1 | 1 | 0.774 | 1 | 0.976 | NA | 1 | 0.679 | 1 | NA | 0.057 |
| SE | 30013 | ENSG00000248049 | UBA6-AS1 | chr4 | + | 67730079 | 67730170 | 67728539 | 67728686 | 67731505 | 67732596 | 15 | 2 | 7 | 0 | 23 | 0 | 0 | 3 | 0 | 1 | 0 | 0 | 2 | 0 | 0 | 0 | 0 | 2 | 0 | 0 | 240 | 149 | 1 | 1 | 1 | 1 | 0.592 | NA | 0.935 | NA | NA | 0.383 | NA | NA | 0.499 |
| SE | 30014 | ENSG00000248049 | UBA6-AS1 | chr4 | + | 67731505 | 67732345 | 67730073 | 67730170 | 67745826 | 67746366 | 20 | 3 | 10 | 0 | 26 | 0 | 1 | 1 | 0 | 0 | 0 | 0 | 4 | 7 | 0 | 0 | 0 | 0 | 0 | 0 | 298 | 149 | 1 | 1 | 1 | 0.6 | 0.833 | NA | 1 | NA | NA | 1 | 1 | NA | -0.142 |
| SE | 30016 | ENSG00000248049 | UBA6-AS1 | chr4 | + | 67753620 | 67753718 | 67751680 | 67751714 | 67754158 | 67754240 | 0 | 11 | 2 | 1 | 2 | 0 | 0 | 0 | 0 | 1 | 0 | 0 | 2 | 0 | 0 | 0 | 0 | 0 | 0 | 0 | 247 | 149 | 1 | 1 | NA | 1 | 1 | 1 | 0.547 | NA | NA | 1 | NA | NA | -0.113 |
| SE | 30017 | ENSG00000248049 | UBA6-AS1 | chr4 | + | 67754158 | 67754240 | 67731505 | 67732345 | 67755188 | 67755255 | 7 | 0 | 0 | 0 | 2 | 0 | 0 | 0 | 0 | 2 | 0 | 0 | 0 | 0 | 7 | 0 | 0 | 0 | 0 | 0 | 231 | 149 | 1 | 1 | 1 | NA | NA | NA | 0.392 | NA | NA | NA | NA | 1 | -0.304 |
| SE | 30032 | ENSG00000144290 | SLC4A10 | chr2 | + | 161904003 | 161904178 | 161900910 | 161901011 | 161905641 | 161905887 | 14 | 22 | 31 | 15 | 17 | 9 | 7 | 8 | 7 | 1 | 0 | 0 | 0 | 0 | 0 | 0 | 2 | 0 | 0 | 0 | 298 | 149 | 1 | 1 | 0.437 | 0.611 | 0.66 | 0.517 | 0.895 | NA | 0 | NA | NA | NA | 0.624 |
| SE | 30033 | ENSG00000144290 | SLC4A10 | chr2 | + | 161904775 | 161904909 | 161900910 | 161901011 | 161905641 | 161905887 | 35 | 19 | 47 | 46 | 39 | 9 | 7 | 8 | 7 | 1 | 0 | 0 | 0 | 0 | 0 | 0 | 2 | 0 | 0 | 0 | 283 | 149 | 1 | 1 | 0.672 | 0.588 | 0.756 | 0.776 | 0.954 | NA | 0 | NA | NA | NA | 0.749 |
| SE | 30035 | ENSG00000144290 | SLC4A10 | chr2 | + | 161947565 | 161947727 | 161942791 | 161942897 | 161949147 | 161949261 | 88 | 73 | 109 | 88 | 176 | 1 | 2 | 4 | 3 | 6 | 0 | 4 | 0 | 0 | 0 | 0 | 0 | 0 | 0 | 0 | 298 | 149 | 1 | 1 | 0.978 | 0.948 | 0.932 | 0.936 | 0.936 | NA | 1 | NA | NA | NA | -0.054 |
| SE | 30036 | ENSG00000144290 | SLC4A10 | chr2 | + | 161950686 | 161950848 | 161949147 | 161949261 | 161958486 | 161958555 | 51 | 34 | 68 | 58 | 102 | 1 | 3 | 1 | 0 | 0 | 0 | 8 | 0 | 0 | 0 | 0 | 0 | 0 | 0 | 0 | 298 | 149 | 1 | 1 | 0.962 | 0.85 | 0.971 | 1 | 1 | NA | 1 | NA | NA | NA | -0.043 |
| SE | 30037 | ENSG00000144290 | SLC4A10 | chr2 | + | 161956988 | 161957240 | 161950686 | 161950848 | 161958486 | 161958555 | 49 | 51 | 112 | 69 | 97 | 5 | 9 | 18 | 2 | 9 | 0 | 0 | 2 | 0 | 0 | 0 | 0 | 0 | 0 | 0 | 298 | 149 | 1 | 1 | 0.831 | 0.739 | 0.757 | 0.945 | 0.843 | NA | NA | 1 | NA | NA | -0.177 |
| SE | 30040 | ENSG00000144290 | SLC4A10 | chr2 | + | 161965050 | 161965173 | 161964134 | 161964308 | 161974248 | 161974316 | 105 | 112 | 219 | 94 | 282 | 0 | 3 | 0 | 0 | 0 | 0 | 0 | 1 | 4 | 0 | 0 | 0 | 0 | 0 | 0 | 272 | 149 | 1 | 1 | 1 | 0.953 | 1 | 1 | 1 | NA | NA | 1 | 1 | NA | -0.009 |
| SE | 30043 | ENSG00000155744 | FAM126B | chr2 | - | 200992886 | 200993003 | 200992280 | 200992368 | 200997444 | 200997540 | 167 | 126 | 101 | 123 | 209 | 0 | 0 | 0 | 0 | 1 | 58 | 157 | 214 | 30 | 36 | 0 | 5 | 1 | 0 | 0 | 266 | 149 | 1 | 1 | 1 | 1 | 1 | 1 | 0.992 | 1 | 0.946 | 0.992 | 1 | 1 | 0.011 |
| SE | 30045 | ENSG00000155744 | FAM126B | chr2 | - | 200997444 | 200997540 | 200992280 | 200992368 | 201008972 | 201009088 | 27 | 20 | 26 | 24 | 40 | 0 | 0 | 0 | 0 | 0 | 0 | 36 | 33 | 12 | 16 | 0 | 0 | 0 | 4 | 0 | 245 | 149 | 1 | 1 | 1 | 1 | 1 | 1 | 1 | NA | 1 | 1 | 0.646 | 1 | 0.089 |
| SE | 30046 | ENSG00000155744 | FAM126B | chr2 | - | 200997444 | 200997540 | 200992886 | 200993003 | 201008972 | 201009088 | 106 | 74 | 68 | 78 | 115 | 0 | 2 | 0 | 0 | 1 | 33 | 96 | 125 | 26 | 35 | 0 | 0 | 0 | 0 | 0 | 245 | 149 | 1 | 1 | 1 | 0.957 | 1 | 1 | 0.986 | 1 | 1 | 1 | 1 | 1 | -0.011 |
| SE | 30051 | ENSG00000155744 | FAM126B | chr2 | - | 201052172 | 201052269 | 201045495 | 201045593 | 201061306 | 201061424 | 0 | 0 | 3 | 0 | 1 | 1 | 2 | 0 | 7 | 0 | 2 | 0 | 11 | 0 | 0 | 0 | 0 | 10 | 0 | 0 | 246 | 149 | 1 | 1 | 0 | 0 | 1 | 0 | 1 | 1 | NA | 0.4 | NA | NA | -0.3 |
| SE | 30052 | ENSG00000155744 | FAM126B | chr2 | - | 201052172 | 201052269 | 201045495 | 201045593 | 201071609 | 201071653 | 0 | 0 | 2 | 0 | 0 | 38 | 18 | 12 | 26 | 9 | 0 | 0 | 2 | 0 | 0 | 5 | 13 | 30 | 10 | 5 | 246 | 149 | 1 | 1 | 0 | 0 | 0.092 | 0 | 0 | 0 | 0 | 0.039 | 0 | 0 | 0.011 |
| SE | 30053 | ENSG00000155744 | FAM126B | chr2 | - | 201061306 | 201061424 | 201023958 | 201024039 | 201071453 | 201071579 | 0 | 0 | 1 | 0 | 2 | 2 | 1 | 0 | 2 | 3 | 0 | 0 | 3 | 0 | 0 | 0 | 0 | 0 | 0 | 0 | 267 | 149 | 1 | 1 | 0 | 0 | 1 | 0 | 0.271 | NA | NA | 1 | NA | NA | -0.746 |
| SE | 30054 | ENSG00000155744 | FAM126B | chr2 | - | 201061306 | 201061424 | 201023958 | 201024039 | 201071609 | 201071671 | 0 | 0 | 0 | 0 | 0 | 3 | 0 | 1 | 1 | 1 | 3 | 0 | 0 | 0 | 0 | 0 | 0 | 0 | 0 | 0 | 267 | 149 | 1 | 1 | 0 | NA | 0 | 0 | 0 | 1 | NA | NA | NA | NA | -1 |
| SE | 30055 | ENSG00000155744 | FAM126B | chr2 | - | 201061306 | 201061424 | 201045495 | 201045593 | 201071453 | 201071579 | 1 | 0 | 1 | 7 | 2 | 15 | 3 | 9 | 24 | 3 | 0 | 0 | 13 | 0 | 0 | 0 | 9 | 11 | 4 | 0 | 267 | 149 | 1 | 1 | 0.036 | 0 | 0.058 | 0.14 | 0.271 | NA | 0 | 0.397 | 0 | NA | -0.031 |
| SE | 30057 | ENSG00000155744 | FAM126B | chr2 | - | 201061306 | 201061424 | 201052172 | 201052269 | 201071609 | 201071671 | 0 | 0 | 2 | 0 | 1 | 0 | 0 | 1 | 0 | 0 | 5 | 0 | 6 | 0 | 0 | 0 | 0 | 0 | 0 | 0 | 267 | 149 | 1 | 1 | NA | NA | 0.527 | NA | 1 | 1 | NA | 1 | NA | NA | -0.236 |
| SE | 30059 | ENSG00000171298 | GAA | chr17 | + | 80101810 | 80101925 | 80101586 | 80101625 | 80104554 | 80105132 | 2 | 0 | 2 | 0 | 4 | 7 | 1 | 3 | 0 | 2 | 0 | 9 | 2 | 7 | 8 | 7 | 11 | 9 | 7 | 18 | 264 | 149 | 1 | 1 | 0.139 | 0 | 0.273 | NA | 0.53 | 0 | 0.316 | 0.111 | 0.361 | 0.201 | 0.038 |
| SE | 30065 | ENSG00000138814 | PPP3CA | chr4 | - | 101098366 | 101098512 | 101093775 | 101093915 | 101099610 | 101099722 | 409 | 299 | 302 | 372 | 806 | 0 | 1 | 0 | 0 | 0 | 161 | 346 | 324 | 349 | 102 | 0 | 3 | 0 | 0 | 0 | 295 | 149 | 1 | 1 | 1 | 0.993 | 1 | 1 | 1 | 1 | 0.983 | 1 | 1 | 1 | 0.002 |
| SE | 30066 | ENSG00000138814 | PPP3CA | chr4 | - | 101108073 | 101108161 | 101099610 | 101099722 | 101108953 | 101109078 | 0 | 1 | 0 | 0 | 0 | 127 | 73 | 88 | 91 | 135 | 0 | 0 | 0 | 0 | 0 | 38 | 38 | 80 | 115 | 17 | 237 | 149 | 1 | 1 | 0 | 0.009 | 0 | 0 | 0 | 0 | 0 | 0 | 0 | 0 | 0.002 |
| SE | 30069 | ENSG00000115289 | PCGF1 | chr2 | - | 74505551 | 74505638 | 74505338 | 74505419 | 74506731 | 74506884 | 9 | 8 | 5 | 4 | 29 | 1 | 0 | 0 | 0 | 0 | 9 | 7 | 6 | 19 | 23 | 0 | 0 | 0 | 0 | 0 | 236 | 149 | 1 | 1 | 0.85 | 1 | 1 | 1 | 1 | 1 | 1 | 1 | 1 | 1 | -0.03 |
| SE | 30073 | ENSG00000115289 | PCGF1 | chr2 | - | 74506180 | 74506252 | 74505951 | 74506057 | 74506731 | 74506884 | 55 | 35 | 30 | 34 | 64 | 0 | 1 | 0 | 1 | 0 | 26 | 35 | 38 | 60 | 58 | 0 | 0 | 0 | 0 | 0 | 221 | 149 | 1 | 1 | 1 | 0.959 | 1 | 0.958 | 1 | 1 | 1 | 1 | 1 | 1 | -0.017 |
| SE | 30074 | ENSG00000164109 | MAD2L1 | chr4 | - | 120061974 | 120062095 | 120060873 | 120060977 | 120065671 | 120065818 | 34 | 32 | 84 | 57 | 79 | 0 | 0 | 0 | 2 | 0 | 14 | 43 | 78 | 36 | 5 | 0 | 0 | 0 | 0 | 0 | 270 | 149 | 1 | 1 | 1 | 1 | 1 | 0.94 | 1 | 1 | 1 | 1 | 1 | 1 | -0.012 |
| SE | 30079 | ENSG00000137713 | PPP2R1B | chr11 | - | 111737447 | 111737569 | 111726907 | 111727057 | 111742052 | 111742144 | 2 | 2 | 3 | 7 | 6 | 2 | 0 | 0 | 0 | 0 | 7 | 0 | 2 | 5 | 6 | 0 | 0 | 0 | 0 | 0 | 271 | 149 | 1 | 1 | 0.355 | 1 | 1 | 1 | 1 | 1 | NA | 1 | 1 | 1 | -0.129 |
| SE | 30081 | ENSG00000137713 | PPP2R1B | chr11 | - | 111759803 | 111759951 | 111755294 | 111755450 | 111764804 | 111764905 | 50 | 18 | 26 | 18 | 114 | 0 | 0 | 0 | 0 | 0 | 0 | 14 | 23 | 25 | 4 | 0 | 0 | 4 | 0 | 0 | 297 | 149 | 1 | 1 | 1 | 1 | 1 | 1 | 1 | NA | 1 | 0.743 | 1 | 1 | 0.064 |
| SE | 30084 | ENSG00000137713 | PPP2R1B | chr11 | - | 111764804 | 111764905 | 111760818 | 111761051 | 111765293 | 111765384 | 159 | 95 | 102 | 196 | 134 | 0 | 0 | 0 | 0 | 0 | 100 | 165 | 146 | 176 | 42 | 0 | 0 | 0 | 3 | 0 | 250 | 149 | 1 | 1 | 1 | 1 | 1 | 1 | 1 | 1 | 1 | 1 | 0.972 | 1 | 0.006 |
| SE | 30085 | ENSG00000137713 | PPP2R1B | chr11 | - | 111765293 | 111765384 | 111760818 | 111761051 | 111766247 | 111766367 | 23 | 14 | 22 | 55 | 29 | 0 | 0 | 0 | 0 | 0 | 34 | 26 | 42 | 58 | 2 | 0 | 2 | 0 | 0 | 0 | 240 | 149 | 1 | 1 | 1 | 1 | 1 | 1 | 1 | 1 | 0.89 | 1 | 1 | 1 | 0.022 |
| SE | 30100 | ENSG00000113522 | RAD50 | chr5 | + | 132579316 | 132579502 | 132575776 | 132575863 | 132587561 | 132587690 | 6 | 2 | 2 | 6 | 7 | 0 | 1 | 5 | 0 | 0 | 0 | 1 | 4 | 0 | 0 | 0 | 0 | 0 | 0 | 0 | 298 | 149 | 1 | 1 | 1 | 0.5 | 0.167 | 1 | 1 | NA | 1 | 1 | NA | NA | -0.267 |
| SE | 30101 | ENSG00000113522 | RAD50 | chr5 | + | 132579316 | 132579502 | 132575776 | 132575928 | 132587561 | 132587690 | 110 | 100 | 86 | 119 | 175 | 0 | 0 | 0 | 1 | 2 | 14 | 51 | 40 | 16 | 2 | 0 | 3 | 1 | 0 | 0 | 298 | 149 | 1 | 1 | 1 | 1 | 1 | 0.983 | 0.978 | 1 | 0.895 | 0.952 | 1 | 1 | 0.023 |
| SE | 30102 | ENSG00000283782 | NA | chr5 | + | 132579316 | 132580066 | 132575776 | 132575928 | 132587561 | 132587690 | 200 | 217 | 155 | 221 | 340 | 0 | 0 | 0 | 1 | 2 | 26 | 99 | 74 | 23 | 14 | 0 | 3 | 1 | 0 | 0 | 298 | 149 | 1 | 1 | 1 | 1 | 1 | 0.991 | 0.988 | 1 | 0.943 | 0.974 | 1 | 1 | 0.012 |
| SE | 30105 | ENSG00000113522 | RAD50 | chr5 | + | 132589630 | 132589837 | 132587923 | 132588089 | 132591223 | 132591406 | 156 | 198 | 158 | 176 | 226 | 0 | 0 | 0 | 3 | 0 | 13 | 83 | 65 | 11 | 9 | 0 | 0 | 0 | 0 | 0 | 298 | 149 | 1 | 1 | 1 | 1 | 1 | 0.967 | 1 | 1 | 1 | 1 | 1 | 1 | -0.007 |
| SE | 30109 | ENSG00000113522 | RAD50 | chr5 | + | 132592763 | 132592965 | 132591876 | 132592034 | 132595103 | 132595201 | 3 | 0 | 2 | 0 | 7 | 1 | 4 | 2 | 3 | 2 | 0 | 1 | 7 | 0 | 0 | 0 | 0 | 5 | 5 | 0 | 298 | 149 | 1 | 1 | 0.6 | 0 | 0.333 | 0 | 0.636 | NA | 1 | 0.412 | 0 | NA | -0.157 |
| SE | 30110 | ENSG00000113522 | RAD50 | chr5 | + | 132594561 | 132594677 | 132591876 | 132592034 | 132594868 | 132595044 | 0 | 0 | 0 | 0 | 4 | 98 | 90 | 90 | 80 | 173 | 0 | 0 | 0 | 0 | 0 | 2 | 41 | 18 | 10 | 8 | 265 | 149 | 1 | 1 | 0 | 0 | 0 | 0 | 0.013 | 0 | 0 | 0 | 0 | 0 | 0.003 |
| SE | 30111 | ENSG00000113522 | RAD50 | chr5 | + | 132594561 | 132594677 | 132591876 | 132592034 | 132595103 | 132595201 | 0 | 0 | 0 | 0 | 4 | 1 | 4 | 2 | 3 | 2 | 0 | 0 | 0 | 0 | 0 | 0 | 0 | 5 | 5 | 0 | 265 | 149 | 1 | 1 | 0 | 0 | 0 | 0 | 0.529 | NA | NA | 0 | 0 | NA | 0.106 |
| SE | 30113 | ENSG00000113522 | RAD50 | chr5 | + | 132595103 | 132595201 | 132591876 | 132592034 | 132595572 | 132595810 | 11 | 9 | 3 | 5 | 5 | 0 | 0 | 7 | 4 | 3 | 0 | 0 | 14 | 8 | 0 | 0 | 0 | 2 | 0 | 0 | 247 | 149 | 1 | 1 | 1 | 1 | 0.205 | 0.43 | 0.501 | NA | NA | 0.809 | 1 | NA | -0.277 |
| SE | 30114 | ENSG00000113522 | RAD50 | chr5 | + | 132595572 | 132595810 | 132591876 | 132592034 | 132603299 | 132603489 | 45 | 40 | 40 | 46 | 65 | 0 | 0 | 0 | 1 | 0 | 9 | 38 | 22 | 2 | 0 | 0 | 0 | 0 | 0 | 0 | 298 | 149 | 1 | 1 | 1 | 1 | 1 | 0.958 | | 1 | 1 | 1 | 1 | NA | -0.008 |
| SE | 30115 | ENSG00000113522 | RAD50 | chr5 | + | 132600061 | 132600138 | 132595572 | 132595810 | 132603299 | 132603405 | 4 | 2 | 4 | 3 | 6 | 45 | 40 | 33 | 42 | 62 | 0 | 0 | 0 | 0 | 0 | 9 | 38 | 20 | 2 | 0 | 226 | 149 | 1 | 1 | 0.055 | 0.032 | 0.074 | 0.045 | 0.06 | 0 | 0 | 0 | 0 | NA | 0.053 |
| SE | 30116 | ENSG00000113522 | RAD50 | chr5 | + | 132642177 | 132642394 | 132640671 | 132640805 | 132643004 | 132646069 | 30 | 4 | 18 | 25 | 36 | 0 | 0 | 0 | 0 | 7 | 0 | 9 | 1 | 0 | 0 | 0 | 0 | 3 | 0 | 0 | 298 | 149 | 1 | 1 | 1 | 1 | 1 | 1 | 0.72 | NA | 1 | 0.143 | NA | NA | 0.372 |
| SE | 30117 | ENSG00000154144 | TBRG1 | chr11 | + | 124624930 | 124625001 | 124622835 | 124623233 | 124625670 | 124625903 | 78 | 81 | 105 | 138 | 153 | 1 | 0 | 0 | 2 | 2 | 108 | 59 | 59 | 102 | 126 | 0 | 0 | 0 | 6 | 0 | 220 | 149 | 1 | 1 | 0.981 | 1 | 1 | 0.979 | 0.981 | 1 | 1 | 1 | 0.92 | 1 | 0.004 |
| SE | 30121 | ENSG00000154144 | TBRG1 | chr11 | + | 124626472 | 124626609 | 124625670 | 124625903 | 124626692 | 124626751 | 4 | 8 | 6 | 19 | 13 | 6 | 3 | 5 | 2 | 5 | 0 | 9 | 11 | 3 | 9 | 0 | 1 | 5 | 4 | 11 | 286 | 149 | 1 | 1 | 0.258 | 0.581 | 0.385 | 0.832 | 0.575 | NA | 0.824 | 0.534 | 0.281 | 0.299 | 0.042 |
| SE | 30126 | ENSG00000154144 | TBRG1 | chr11 | + | 124631239 | 124631417 | 124630744 | 124630855 | 124632092 | 124632452 | 116 | 57 | 90 | 105 | 112 | 0 | 0 | 0 | 0 | 0 | 24 | 97 | 87 | 98 | 51 | 0 | 0 | 0 | 5 | 0 | 298 | 149 | 1 | 1 | 1 | 1 | 1 | 1 | 1 | 1 | 1 | 1 | 0.907 | 1 | 0.019 |
| SE | 30127 | ENSG00000154144 | TBRG1 | chr11 | + | 124631274 | 124631417 | 124630744 | 124630855 | 124632092 | 124632238 | 212 | 121 | 174 | 168 | 205 | 0 | 0 | 0 | 0 | 0 | 72 | 164 | 164 | 141 | 88 | 0 | 0 | 0 | 5 | 0 | 292 | 149 | 1 | 1 | 1 | 1 | 1 | 1 | 1 | 1 | 1 | 1 | 0.935 | 1 | 0.013 |
| SE | 30128 | ENSG00000150961 | SEC24D | chr4 | - | 118732732 | 118732912 | 118731315 | 118731507 | 118738260 | 118738379 | 133 | 81 | 111 | 117 | 172 | 1 | 0 | 0 | 0 | 0 | 75 | 148 | 116 | 88 | 40 | 0 | 0 | 0 | 0 | 0 | 298 | 149 | 1 | 1 | 0.985 | 1 | 1 | 1 | 1 | 1 | 1 | 1 | 1 | 1 | -0.003 |
| SE | 30130 | ENSG00000150961 | SEC24D | chr4 | - | 118739148 | 118739287 | 118738260 | 118738379 | 118740662 | 118740808 | 218 | 143 | 172 | 166 | 159 | 0 | 0 | 0 | 2 | 0 | 181 | 214 | 183 | 141 | 52 | 0 | 4 | 0 | 0 | 0 | 288 | 149 | 1 | 1 | 1 | 1 | 1 | 0.977 | 1 | 1 | 0.965 | 1 | 1 | 1 | 0.002 |
| SE | 30133 | ENSG00000150961 | SEC24D | chr4 | - | 118757720 | 118757845 | 118752696 | 118752888 | 118764801 | 118764917 | 153 | 75 | 120 | 119 | 123 | 1 | 0 | 0 | 0 | 0 | 91 | 117 | 134 | 90 | 49 | 0 | 0 | 0 | 0 | 0 | 274 | 149 | 1 | 1 | 0.988 | 1 | 1 | 1 | 1 | 1 | 1 | 1 | 1 | 1 | -0.002 |
| SE | 30135 | ENSG00000150961 | SEC24D | chr4 | - | 118760390 | 118760607 | 118757720 | 118757845 | 118764801 | 118764917 | 0 | 1 | 0 | 0 | 0 | 113 | 54 | 94 | 81 | 93 | 0 | 0 | 0 | 0 | 0 | 53 | 80 | 91 | 65 | 47 | 298 | 149 | 1 | 1 | 0 | 0.009 | 0 | 0 | 0 | 0 | 0 | 0 | 0 | 0 | 0.002 |
| SE | 30138 | ENSG00000150961 | SEC24D | chr4 | - | 118817263 | 118817412 | 118815450 | 118815726 | 118824619 | 118824749 | 61 | 29 | 42 | 51 | 47 | 0 | 0 | 1 | 0 | 0 | 119 | 86 | 27 | 62 | 23 | 0 | 0 | 0 | 0 | 2 | 298 | 149 | 1 | 1 | 1 | 1 | 0.955 | 1 | 1 | 1 | 1 | 1 | 1 | 0.852 | 0.021 |
| SE | 30139 | ENSG00000150961 | SEC24D | chr4 | - | 118824619 | 118824749 | 118817263 | 118817412 | 118833578 | 118833737 | 51 | 31 | 47 | 63 | 61 | 0 | 0 | 0 | 0 | 0 | 78 | 81 | 51 | 72 | 44 | 0 | 3 | 0 | 0 | 0 | 279 | 149 | 1 | 1 | 1 | 1 | 1 | 1 | 1 | 1 | 0.935 | 1 | 1 | 1 | 0.013 |
| SE | 30142 | ENSG00000198856 | OSTC | chr4 | + | 108655563 | 108655657 | 108650584 | 108650794 | 108667246 | 108667818 | 198 | 82 | 174 | 189 | 191 | 0 | 1 | 0 | 0 | 0 | 142 | 89 | 139 | 67 | 66 | 0 | 0 | 0 | 0 | 0 | 243 | 149 | 1 | 1 | 1 | 0.98 | 1 | 1 | 1 | 1 | 1 | 1 | 1 | 1 | -0.004 |
| SE | 30144 | ENSG00000198856 | OSTC | chr4 | + | 108663232 | 108663306 | 108657449 | 108657647 | 108667246 | 108667395 | 0 | 0 | 0 | 1 | 0 | 241 | 153 | 234 | 209 | 414 | 0 | 0 | 0 | 0 | 0 | 92 | 130 | 146 | 91 | 27 | 223 | 149 | 1 | 1 | 0 | 0 | 0 | 0.003 | 0 | 0 | 0 | 0 | 0 | 0 | 0.001 |
| SE | 30146 | ENSG00000005059 | MCUB | chr4 | + | 109648483 | 109648649 | 109560989 | 109561017 | 109659010 | 109659086 | 0 | 0 | 1 | 0 | 0 | 0 | 0 | 1 | 3 | 1 | 0 | 0 | 0 | 0 | 0 | 9 | 0 | 0 | 0 | 0 | 298 | 149 | 1 | 1 | NA | NA | 0.333 | 0 | 0 | 0 | NA | NA | NA | NA | 0.111 |
| SE | 30147 | ENSG00000005059 | MCUB | chr4 | + | 109659010 | 109659086 | 109560208 | 109560436 | 109660194 | 109660346 | 205 | 153 | 223 | 214 | 267 | 0 | 0 | 1 | 3 | 0 | 176 | 128 | 139 | 134 | 167 | 0 | 0 | 0 | 0 | 0 | 225 | 149 | 1 | 1 | 1 | 1 | 0.993 | 0.979 | 1 | 1 | 1 | 1 | 1 | 1 | -0.006 |
| SE | 30148 | ENSG00000005059 | MCUB | chr4 | + | 109660194 | 109660365 | 109659010 | 109659086 | 109664289 | 109664394 | 683 | 617 | 744 | 757 | 1156 | 0 | 0 | 2 | 1 | 0 | 1049 | 405 | 547 | 529 | 695 | 0 | 0 | 0 | 2 | 0 | 298 | 149 | 1 | 1 | 1 | 1 | 0.995 | 0.997 | 1 | 1 | 1 | 1 | 0.992 | 1 | 0 |
| SE | 30150 | ENSG00000005059 | MCUB | chr4 | + | 109682581 | 109682742 | 109664289 | 109664394 | 109684442 | 109684646 | 783 | 747 | 932 | 887 | 1571 | 0 | 1 | 0 | 0 | 0 | 1255 | 531 | 671 | 665 | 850 | 0 | 0 | 0 | 0 | 0 | 298 | 149 | 1 | 1 | 1 | 0.997 | 1 | 1 | 1 | 1 | 1 | 1 | 1 | 1 | -0.001 |
| SE | 30151 | ENSG00000005059 | MCUB | chr4 | + | 109684442 | 109684646 | 109682581 | 109682742 | 109685250 | 109685367 | 560 | 388 | 597 | 456 | 894 | 0 | 0 | 3 | 0 | 14 | 311 | 289 | 301 | 336 | 442 | 6 | 0 | 2 | 3 | 0 | 298 | 149 | 1 | 1 | 1 | 1 | 0.99 | 1 | 0.97 | 0.963 | 1 | 0.987 | 0.982 | 1 | 0.006 |
| SE | 30153 | ENSG00000177054 | ZDHHC13 | chr11 | + | 19152157 | 19152320 | 19150726 | 19150791 | 19152558 | 19152625 | 101 | 83 | 122 | 86 | 148 | 1 | 0 | 0 | 0 | 0 | 27 | 40 | 46 | 91 | 15 | 0 | 0 | 0 | 0 | 0 | 298 | 149 | 1 | 1 | 0.981 | 1 | 1 | 1 | 1 | 1 | 1 | 1 | 1 | 1 | -0.004 |
| SE | 30154 | ENSG00000177054 | ZDHHC13 | chr11 | + | 19152157 | 19152320 | 19150726 | 19150791 | 19155795 | 19155929 | 50 | 43 | 57 | 49 | 80 | 0 | 0 | 0 | 0 | 0 | 7 | 28 | 16 | 38 | 4 | 0 | 0 | 0 | 1 | 0 | 298 | 149 | 1 | 1 | 1 | 1 | 1 | 1 | 1 | 1 | 1 | 1 | 0.95 | 1 | 0.01 |
| SE | 30155 | ENSG00000177054 | ZDHHC13 | chr11 | + | 19152558 | 19152684 | 19150726 | 19150791 | 19155795 | 19155929 | 34 | 12 | 23 | 17 | 22 | 0 | 0 | 0 | 0 | 0 | 0 | 30 | 7 | 29 | 7 | 0 | 0 | 0 | 1 | 0 | 275 | 149 | 1 | 1 | 1 | 1 | 1 | 1 | 1 | NA | 1 | 1 | 0.94 | 1 | 0.015 |
| SE | 30156 | ENSG00000177054 | ZDHHC13 | chr11 | + | 19152558 | 19152684 | 19152157 | 19152320 | 19155795 | 19155929 | 84 | 52 | 87 | 54 | 90 | 2 | 0 | 0 | 0 | 0 | 20 | 46 | 37 | 82 | 18 | 0 | 4 | 0 | 0 | 0 | 275 | 149 | 1 | 1 | 0.958 | 1 | 1 | 1 | 1 | 1 | 0.862 | 1 | 1 | 1 | 0.019 |
| SE | 30159 | ENSG00000163811 | WDR43 | chr2 | + | 28901986 | 28902124 | 28894666 | 28894923 | 28906459 | 28906581 | 130 | 120 | 143 | 142 | 182 | 0 | 3 | 0 | 1 | 3 | 45 | 111 | 173 | 71 | 10 | 0 | 0 | 2 | 0 | 0 | 287 | 149 | 1 | 1 | 1 | 0.954 | 1 | 0.987 | 0.969 | 1 | 1 | 0.978 | 1 | 1 | -0.014 |
| SE | 30160 | ENSG00000163811 | WDR43 | chr2 | + | 28906459 | 28906581 | 28901986 | 28902124 | 28912589 | 28912710 | 159 | 134 | 188 | 165 | 249 | 0 | 1 | 0 | 0 | 0 | 50 | 132 | 180 | 119 | 25 | 0 | 0 | 0 | 0 | 0 | 271 | 149 | 1 | 1 | 1 | 0.987 | 1 | 1 | 1 | 1 | 1 | 1 | 1 | 1 | -0.003 |
| SE | 30163 | ENSG00000163811 | WDR43 | chr2 | + | 28930057 | 28930149 | 28929578 | 28929710 | 28935520 | 28935607 | 0 | 0 | 0 | 0 | 1 | 82 | 62 | 102 | 70 | 168 | 0 | 2 | 0 | 0 | 0 | 8 | 55 | 79 | 50 | 6 | 241 | 149 | 1 | 1 | 0 | 0 | 0 | 0 | 0.004 | 0 | 0.022 | 0 | 0 | 0 | -0.004 |
| SE | 30164 | ENSG00000163811 | WDR43 | chr2 | + | 28935520 | 28935607 | 28929578 | 28929710 | 28936921 | 28936953 | 109 | 85 | 136 | 109 | 211 | 0 | 0 | 0 | 0 | 0 | 28 | 76 | 105 | 61 | 12 | 0 | 0 | 1 | 3 | 0 | 236 | 149 | 1 | 1 | 1 | 1 | 1 | 1 | 1 | 1 | 1 | 0.985 | 0.928 | 1 | 0.017 |
| SE | 30165 | ENSG00000163811 | WDR43 | chr2 | + | 28936921 | 28936953 | 28929578 | 28929710 | 28941460 | 28941574 | 2 | 0 | 0 | 0 | 0 | 0 | 0 | 0 | 0 | 0 | 0 | 0 | 0 | 8 | 0 | 0 | 0 | 0 | 1 | 0 | 181 | 149 | 1 | 1 | 1 | NA | NA | NA | NA | NA | NA | NA | 0.868 | NA | 0.132 |
| SE | 30169 | ENSG00000166446 | CDYL2 | chr16 | - | 80608091 | 80608235 | 80597906 | 80604546 | 80612625 | 80612836 | 6 | 1 | 14 | 8 | 3 | 0 | 0 | 0 | 0 | 0 | 35 | 16 | 6 | 7 | 0 | 0 | 0 | 1 | 0 | 0 | 293 | 149 | 1 | 1 | 1 | 1 | 1 | 1 | 1 | 1 | 1 | 0.753 | 1 | NA | 0.062 |
| SE | 30170 | ENSG00000166446 | CDYL2 | chr16 | - | 80612625 | 80612836 | 80597906 | 80604546 | 80620762 | 80620935 | 4 | 6 | 7 | 1 | 0 | 0 | 0 | 0 | 0 | 0 | 7 | 10 | 8 | 0 | 2 | 0 | 2 | 7 | 0 | 0 | 298 | 149 | 1 | 1 | 1 | 1 | 1 | 1 | NA | 1 | 0.714 | 0.364 | NA | 1 | 0.231 |
| SE | 30171 | ENSG00000166446 | CDYL2 | chr16 | - | 80633018 | 80633236 | 80620762 | 80620935 | 80684537 | 80685129 | 12 | 28 | 30 | 18 | 21 | 0 | 0 | 0 | 0 | 0 | 34 | 51 | 42 | 50 | 46 | 0 | 4 | 0 | 0 | 0 | 298 | 149 | 1 | 1 | 1 | 1 | 1 | 1 | 1 | 1 | 0.864 | 1 | 1 | 1 | 0.027 |
| SE | 30187 | ENSG00000168397 | ATG4B | chr2 | + | 241651263 | 241651335 | 241651006 | 241651111 | 241653511 | 241653610 | 237 | 154 | 152 | 201 | 311 | 0 | 1 | 1 | 1 | 4 | 333 | 360 | 171 | 234 | 373 | 0 | 0 | 4 | 0 | 4 | 221 | 149 | 1 | 1 | 1 | 0.99 | 0.99 | 0.993 | 0.981 | 1 | 1 | 0.966 | 1 | 0.984 | 0.001 |
| SE | 30188 | ENSG00000168397 | ATG4B | chr2 | + | 241651263 | 241651335 | 241651006 | 241651111 | 241659107 | 241659187 | 109 | 66 | 79 | 89 | 96 | 0 | 0 | 0 | 1 | 0 | 147 | 169 | 89 | 117 | 207 | 0 | 0 | 0 | 0 | 0 | 221 | 149 | 1 | 1 | 1 | 1 | 1 | 0.984 | 1 | 1 | 1 | 1 | 1 | 1 | -0.003 |
| SE | 30189 | ENSG00000168397 | ATG4B | chr2 | + | 241651883 | 241651974 | 241651263 | 241651335 | 241653306 | 241653369 | 1 | 0 | 2 | 1 | 0 | 22 | 13 | 17 | 14 | 28 | 0 | 2 | 0 | 6 | 0 | 5 | 52 | 12 | 31 | 47 | 240 | 149 | 1 | 1 | 0.027 | 0 | 0.068 | 0.042 | 0 | 0 | 0.023 | 0 | 0.107 | 0 | 0.001 |
| SE | 30190 | ENSG00000168397 | ATG4B | chr2 | + | 241651883 | 241651974 | 241651263 | 241651335 | 241653511 | 241653610 | 2 | 0 | 1 | 1 | 0 | 128 | 90 | 75 | 115 | 215 | 0 | 2 | 0 | 6 | 0 | 186 | 192 | 82 | 117 | 166 | 240 | 149 | 1 | 1 | 0.01 | 0 | 0.008 | 0.005 | 0 | 0 | 0.006 | 0 | 0.031 | 0 | -0.003 |
| SE | 30191 | ENSG00000168397 | ATG4B | chr2 | + | 241651883 | 241652012 | 241651263 | 241651335 | 241653306 | 241653333 | 1 | 1 | 1 | 1 | 0 | 22 | 13 | 17 | 14 | 28 | 0 | 2 | 0 | 13 | 0 | 5 | 52 | 12 | 31 | 47 | 278 | 149 | 1 | 1 | 0.024 | 0.04 | 0.031 | 0.037 | 0 | 0 | 0.02 | 0 | 0.184 | 0 | -0.014 |
| SE | 30193 | ENSG00000168397 | ATG4B | chr2 | + | 241653306 | 241653369 | 241651883 | 241651974 | 241653511 | 241653524 | 11 | 15 | 17 | 11 | 3 | 1 | 0 | 0 | 0 | 0 | 19 | 34 | 9 | 21 | 40 | 0 | 0 | 0 | 0 | 0 | 212 | 149 | 1 | 1 | 0.885 | 1 | 1 | 1 | 1 | 1 | 1 | 1 | 1 | 1 | -0.023 |
| SE | 30194 | ENSG00000168397 | ATG4B | chr2 | + | 241653511 | 241653610 | 241651009 | 241651335 | 241654545 | 241654647 | 219 | 132 | 159 | 221 | 355 | 0 | 0 | 0 | 0 | 0 | 288 | 319 | 178 | 188 | 333 | 0 | 0 | 1 | 4 | 0 | 248 | 149 | 1 | 1 | 1 | 1 | 1 | 1 | 1 | 1 | 1 | 0.991 | 0.966 | 1 | 0.009 |
| SE | 30195 | ENSG00000168397 | ATG4B | chr2 | + | 241654545 | 241654647 | 241651009 | 241651335 | 241659107 | 241659772 | 2 | 3 | 1 | 2 | 2 | 0 | 0 | 0 | 0 | 0 | 9 | 22 | 15 | 26 | 21 | 0 | 1 | 0 | 0 | 0 | 251 | 149 | 1 | 1 | 1 | 1 | 1 | 1 | 1 | 1 | 0.929 | 1 | 1 | 1 | 0.014 |
| SE | 30196 | ENSG00000168397 | ATG4B | chr2 | + | 241654545 | 241654647 | 241651009 | 241651335 | 241666644 | 241666838 | 0 | 0 | 2 | 0 | 0 | 0 | 0 | 0 | 0 | 0 | 10 | 0 | 3 | 4 | 0 | 0 | 0 | 1 | 0 | 0 | 251 | 149 | 1 | 1 | NA | NA | 1 | NA | NA | 1 | NA | 0.64 | 1 | NA | 0.12 |
| SE | 30199 | ENSG00000168397 | ATG4B | chr2 | + | 241659107 | 241659187 | 241651263 | 241651335 | 241666644 | 241666769 | 33 | 25 | 48 | 43 | 61 | 0 | 0 | 0 | 0 | 0 | 48 | 40 | 37 | 33 | 93 | 0 | 0 | 1 | 0 | 0 | 229 | 149 | 1 | 1 | 1 | 1 | 1 | 1 | 1 | 1 | 1 | 0.96 | 1 | 1 | 0.008 |
| SE | 30200 | ENSG00000168397 | ATG4B | chr2 | + | 241659107 | 241659187 | 241655270 | 241655343 | 241666644 | 241666769 | 135 | 59 | 121 | 105 | 129 | 0 | 1 | 0 | 0 | 0 | 122 | 109 | 105 | 64 | 248 | 0 | 0 | 0 | 0 | 0 | 229 | 149 | 1 | 1 | 1 | 0.975 | 1 | 1 | 1 | 1 | 1 | 1 | 1 | 1 | -0.005 |
| SE | 30203 | ENSG00000111266 | DUSP16 | chr12 | - | 12554535 | 12554659 | 12521080 | 12521463 | 12560999 | 12561077 | 16 | 17 | 12 | 21 | 30 | 11 | 4 | 18 | 10 | 5 | 8 | 12 | 6 | 5 | 0 | 0 | 2 | 53 | 8 | 0 | 273 | 149 | 1 | 1 | 0.443 | 0.699 | 0.267 | 0.534 | 0.766 | 1 | 0.766 | 0.058 | 0.254 | NA | 0.022 |
| SE | 30211 | ENSG00000132376 | INPP5K | chr17 | - | 1497935 | 1498122 | 1496665 | 1496803 | 1506979 | 1507089 | 199 | 138 | 149 | 223 | 141 | 1 | 0 | 0 | 0 | 0 | 240 | 255 | 165 | 161 | 328 | 0 | 0 | 0 | 0 | 0 | 298 | 149 | 1 | 1 | 0.99 | 1 | 1 | 1 | 1 | 1 | 1 | 1 | 1 | 1 | -0.002 |
| SE | 30212 | ENSG00000132376 | INPP5K | chr17 | - | 1506979 | 1507089 | 1497935 | 1498122 | 1509177 | 1509353 | 70 | 64 | 66 | 92 | 71 | 0 | 0 | 0 | 0 | 0 | 111 | 117 | 60 | 69 | 141 | 0 | 6 | 0 | 0 | 0 | 259 | 149 | 1 | 1 | 1 | 1 | 1 | 1 | 1 | 1 | 0.918 | 1 | 1 | 1 | 0.016 |
| SE | 30213 | ENSG00000132376 | INPP5K | chr17 | - | 1508114 | 1508226 | 1506979 | 1507089 | 1509167 | 1509353 | 106 | 76 | 86 | 117 | 110 | 0 | 0 | 0 | 0 | 0 | 146 | 144 | 71 | 112 | 195 | 0 | 0 | 0 | 4 | 0 | 261 | 149 | 1 | 1 | 1 | 1 | 1 | 1 | 1 | 1 | 1 | 1 | 0.941 | 1 | 0.012 |
| SE | 30216 | ENSG00000132376 | INPP5K | chr17 | - | 1509177 | 1509353 | 1497935 | 1498122 | 1513871 | 1513979 | 1 | 2 | 3 | 1 | 1 | 0 | 0 | 2 | 0 | 0 | 3 | 14 | 0 | 18 | 11 | 0 | 0 | 1 | 0 | 0 | 298 | 149 | 1 | 1 | 1 | 1 | 0.429 | 1 | 1 | 1 | 1 | 0 | 1 | 1 | 0.086 |
| SE | 30227 | ENSG00000132376 | INPP5K | chr17 | - | 1513871 | 1513979 | 1513452 | 1513554 | 1515568 | 1515585 | 7 | 1 | 1 | 5 | 12 | 0 | 0 | 0 | 0 | 0 | 5 | 8 | 5 | 12 | 14 | 0 | 0 | 2 | 0 | 0 | 257 | 149 | 1 | 1 | 1 | 1 | 1 | 1 | 1 | 1 | 1 | 0.592 | 1 | 1 | 0.082 |
| SE | 30228 | ENSG00000132376 | INPP5K | chr17 | - | 1515426 | 1515585 | 1513871 | 1513979 | 1515887 | 1516118 | 1 | 0 | 0 | 1 | 0 | 9 | 0 | 2 | 3 | 11 | 3 | 0 | 1 | 2 | 20 | 0 | 7 | 1 | 0 | 6 | 298 | 149 | 1 | 1 | 0.053 | NA | 0 | 0.143 | 0 | 1 | 0 | 0.333 | 1 | 0.625 | -0.543 |
| SE | 30230 | ENSG00000132376 | INPP5K | chr17 | - | 1515568 | 1515585 | 1513871 | 1513979 | 1515887 | 1516118 | 0 | 0 | 0 | 1 | 0 | 9 | 0 | 2 | 3 | 11 | 0 | 0 | 0 | 0 | 0 | 0 | 7 | 1 | 0 | 6 | 166 | 149 | 1 | 1 | 0 | NA | 0 | 0.23 | 0 | NA | 0 | 0 | NA | 0 | 0.058 |
| SE | 30232 | ENSG00000132376 | INPP5K | chr17 | - | 1515887 | 1516118 | 1515426 | 1515585 | 1516455 | 1516612 | 5 | 3 | 2 | 1 | 1 | 1 | 1 | 0 | 3 | 0 | 0 | 6 | 3 | 1 | 7 | 0 | 7 | 2 | 0 | 0 | 298 | 149 | 1 | 1 | 0.714 | 0.6 | 1 | 0.143 | 1 | NA | 0.3 | 0.429 | 1 | 1 | 0.009 |
| SE | 30234 | ENSG00000005700 | IBTK | chr6 | - | 82191072 | 82191216 | 82181878 | 82182028 | 82191786 | 82191854 | 360 | 208 | 291 | 263 | 569 | 0 | 0 | 0 | 0 | 0 | 16 | 206 | 196 | 40 | 15 | 0 | 0 | 1 | 0 | 0 | 293 | 149 | 1 | 1 | 1 | 1 | 1 | 1 | 1 | 1 | 1 | 0.99 | 1 | 1 | 0.002 |
| SE | 30235 | ENSG00000005700 | IBTK | chr6 | - | 82196297 | 82196401 | 82194478 | 82194642 | 82200140 | 82200253 | 51 | 38 | 31 | 55 | 128 | 2 | 4 | 2 | 2 | 4 | 1 | 40 | 16 | 6 | 0 | 0 | 5 | 0 | 0 | 0 | 253 | 149 | 1 | 1 | 0.938 | 0.848 | 0.901 | 0.942 | 0.95 | 1 | 0.825 | 1 | 1 | NA | -0.04 |
| SE | 30236 | ENSG00000005700 | IBTK | chr6 | - | 82196297 | 82196446 | 82194478 | 82194595 | 82200140 | 82200253 | 86 | 58 | 66 | 83 | 152 | 2 | 0 | 0 | 0 | 2 | 0 | 22 | 17 | 9 | 23 | 0 | 0 | 0 | 0 | 0 | 298 | 149 | 1 | 1 | 0.956 | 1 | 1 | 1 | 0.974 | NA | 1 | 1 | 1 | 1 | -0.014 |
| SE | 30238 | ENSG00000005700 | IBTK | chr6 | - | 82211489 | 82211572 | 82204856 | 82204958 | 82212706 | 82212793 | 57 | 37 | 58 | 41 | 62 | 0 | 0 | 0 | 0 | 4 | 11 | 34 | 27 | 6 | 2 | 0 | 0 | 0 | 0 | 0 | 232 | 149 | 1 | 1 | 1 | 1 | 1 | 1 | 0.909 | 1 | 1 | 1 | 1 | 1 | -0.018 |
| SE | 30239 | ENSG00000005700 | IBTK | chr6 | - | 82211489 | 82211572 | 82211366 | 82211404 | 82212706 | 82212793 | 132 | 92 | 132 | 100 | 224 | 1 | 0 | 2 | 3 | 0 | 51 | 76 | 76 | 23 | 8 | 0 | 0 | 1 | 0 | 0 | 232 | 149 | 1 | 1 | 0.988 | 1 | 0.977 | 0.955 | 1 | 1 | 1 | 0.98 | 1 | 1 | -0.012 |
| SE | 30240 | ENSG00000005700 | IBTK | chr6 | - | 82212706 | 82212793 | 82204856 | 82204958 | 82214226 | 82214829 | 22 | 10 | 22 | 16 | 32 | 0 | 0 | 0 | 1 | 0 | 0 | 11 | 15 | 2 | 0 | 0 | 0 | 0 | 0 | 0 | 236 | 149 | 1 | 1 | 1 | 1 | 1 | 0.91 | 1 | NA | 1 | 1 | 1 | NA | -0.018 |
| SE | 30242 | ENSG00000005700 | IBTK | chr6 | - | 82216075 | 82216250 | 82214226 | 82214829 | 82217959 | 82218137 | 110 | 60 | 98 | 105 | 222 | 0 | 1 | 0 | 1 | 3 | 0 | 36 | 45 | 13 | 28 | 0 | 0 | 11 | 2 | 0 | 298 | 149 | 1 | 1 | 1 | 0.968 | 1 | 0.981 | 0.974 | NA | 1 | 0.672 | 0.765 | 1 | 0.125 |
| SE | 30243 | ENSG00000005700 | IBTK | chr6 | - | 82223439 | 82223620 | 82220589 | 82220713 | 82225476 | 82225647 | 76 | 69 | 88 | 73 | 112 | 1 | 0 | 0 | 0 | 0 | 0 | 54 | 64 | 31 | 4 | 0 | 0 | 0 | 0 | 0 | 298 | 149 | 1 | 1 | 0.974 | 1 | 1 | 1 | 1 | NA | 1 | 1 | 1 | 1 | -0.005 |
| SE | 30244 | ENSG00000005700 | IBTK | chr6 | - | 82224067 | 82224185 | 82223439 | 82223620 | 82225476 | 82225647 | 466 | 410 | 384 | 472 | 443 | 0 | 1 | 0 | 0 | 0 | 144 | 308 | 373 | 69 | 22 | 0 | 0 | 0 | 0 | 0 | 267 | 149 | 1 | 1 | 1 | 0.996 | 1 | 1 | 1 | 1 | 1 | 1 | 1 | 1 | -0.001 |
| SE | 30245 | ENSG00000005700 | IBTK | chr6 | - | 82231717 | 82231842 | 82227163 | 82227302 | 82234158 | 82234255 | 225 | 141 | 222 | 208 | 203 | 0 | 0 | 0 | 0 | 0 | 56 | 114 | 172 | 63 | 12 | 0 | 0 | 0 | 1 | 0 | 274 | 149 | 1 | 1 | 1 | 1 | 1 | 1 | 1 | 1 | 1 | 1 | 0.972 | 1 | 0.006 |
| SE | 30246 | ENSG00000005700 | IBTK | chr6 | - | 82231717 | 82231842 | 82227163 | 82227302 | 82240165 | 82240565 | 93 | 49 | 75 | 72 | 86 | 2 | 0 | 0 | 0 | 1 | 27 | 49 | 69 | 10 | 2 | 0 | 0 | 0 | 0 | 0 | 274 | 149 | 1 | 1 | 0.962 | 1 | 1 | 1 | 0.979 | 1 | 1 | 1 | 1 | 1 | -0.012 |
| SE | 30247 | ENSG00000005700 | IBTK | chr6 | - | 82234158 | 82234255 | 82227163 | 82227302 | 82240165 | 82240565 | 53 | 28 | 60 | 46 | 54 | 2 | 0 | 0 | 0 | 1 | 32 | 15 | 41 | 32 | 6 | 0 | 0 | 0 | 0 | 0 | 246 | 149 | 1 | 1 | 0.941 | 1 | 1 | 1 | 0.97 | 1 | 1 | 1 | 1 | 1 | -0.018 |
| SE | 30248 | ENSG00000005700 | IBTK | chr6 | - | 82234158 | 82234255 | 82227163 | 82227302 | 82247561 | 82247671 | 1 | 0 | 0 | 0 | 0 | 0 | 0 | 0 | 2 | 0 | 0 | 0 | 1 | 0 | 0 | 0 | 0 | 0 | 0 | 0 | 246 | 149 | 1 | 1 | 1 | NA | NA | 0 | NA | NA | NA | 1 | NA | NA | -0.5 |
| SE | 30249 | ENSG00000005700 | IBTK | chr6 | - | 82234158 | 82234255 | 82231717 | 82231842 | 82240165 | 82240565 | 183 | 123 | 206 | 180 | 171 | 1 | 3 | 2 | 1 | 0 | 61 | 80 | 144 | 84 | 16 | 0 | 0 | 0 | 0 | 0 | 246 | 149 | 1 | 1 | 0.991 | 0.961 | 0.984 | 0.991 | 1 | 1 | 1 | 1 | 1 | 1 | -0.015 |
| SE | 30250 | ENSG00000005700 | IBTK | chr6 | - | 82240165 | 82240843 | 82227163 | 82227302 | 82247561 | 82247671 | 32 | 20 | 25 | 27 | 19 | 0 | 0 | 0 | 2 | 0 | 0 | 25 | 11 | 5 | 14 | 0 | 0 | 0 | 0 | 0 | 298 | 149 | 1 | 1 | 1 | 1 | 1 | 0.871 | 1 | NA | 1 | 1 | 1 | 1 | -0.026 |
| SE | 30251 | ENSG00000005700 | IBTK | chr6 | - | 82240165 | 82240843 | 82234158 | 82234255 | 82247561 | 82247671 | 182 | 123 | 197 | 165 | 135 | 2 | 0 | 0 | 0 | 0 | 61 | 97 | 138 | 74 | 27 | 0 | 0 | 1 | 0 | 0 | 298 | 149 | 1 | 1 | 0.978 | 1 | 1 | 1 | 1 | 1 | 1 | 0.986 | 1 | 1 | -0.002 |
| SE | 30252 | ENSG00000059769 | DNAJC25 | chr9 | + | 111642856 | 111643094 | 111631359 | 111631743 | 111647106 | 111647259 | 14 | 15 | 24 | 27 | 30 | 0 | 0 | 1 | 0 | 1 | 12 | 10 | 19 | 6 | 9 | 0 | 0 | 0 | 0 | 0 | 298 | 149 | 1 | 1 | 1 | 1 | 0.923 | 1 | 0.938 | 1 | 1 | 1 | 1 | 1 | -0.028 |
| SE | 30253 | ENSG00000059769 | DNAJC25 | chr9 | + | 111649452 | 111649923 | 111647044 | 111647259 | 111653099 | 111654345 | 57 | 32 | 45 | 38 | 66 | 0 | 2 | 0 | 0 | 0 | 40 | 42 | 78 | 40 | 11 | 0 | 0 | 0 | 0 | 0 | 298 | 149 | 1 | 1 | 1 | 0.889 | 1 | 1 | 1 | 1 | 1 | 1 | 1 | 1 | -0.022 |
| SE | 30254 | ENSG00000153347 | FAM81B | chr5 | + | 95396110 | 95396175 | 95392793 | 95392897 | 95413946 | 95414190 | 1 | 14 | 4 | 11 | 14 | 1 | 1 | 0 | 0 | 0 | 10 | 9 | 8 | 10 | 13 | 0 | 1 | 0 | 0 | 0 | 214 | 149 | 1 | 1 | 0.41 | 0.907 | 1 | 1 | 1 | 1 | 0.862 | 1 | 1 | 1 | -0.109 |
| SE | 30256 | ENSG00000153347 | FAM81B | chr5 | + | 95420283 | 95420402 | 95413946 | 95414190 | 95428602 | 95428732 | 0 | 5 | 2 | 3 | 4 | 0 | 0 | 0 | 0 | 0 | 6 | 7 | 0 | 3 | 5 | 0 | 2 | 0 | 0 | 0 | 268 | 149 | 1 | 1 | NA | 1 | 1 | 1 | 1 | 1 | 0.661 | NA | 1 | 1 | 0.085 |
| SE | 30257 | ENSG00000153347 | FAM81B | chr5 | + | 95420283 | 95420402 | 95413946 | 95414190 | 95446561 | 95446697 | 0 | 6 | 3 | 2 | 9 | 0 | 0 | 0 | 0 | 2 | 6 | 6 | 1 | 6 | 11 | 0 | 1 | 0 | 0 | 0 | 268 | 149 | 1 | 1 | NA | 1 | 1 | 1 | 0.714 | 1 | 0.769 | 1 | 1 | 1 | -0.025 |
| SE | 30258 | ENSG00000153347 | FAM81B | chr5 | + | 95424049 | 95424190 | 95420283 | 95420402 | 95428602 | 95428732 | 0 | 1 | 0 | 0 | 0 | 0 | 0 | 1 | 1 | 0 | 0 | 0 | 0 | 0 | 0 | 0 | 1 | 0 | 0 | 0 | 290 | 149 | 1 | 1 | NA | 1 | 0 | 0 | NA | NA | 0 | NA | NA | NA | 0.333 |
| SE | 30259 | ENSG00000153347 | FAM81B | chr5 | + | 95428602 | 95428732 | 95413946 | 95414190 | 95446561 | 95446689 | 0 | 1 | 0 | 1 | 0 | 0 | 0 | 0 | 0 | 2 | 0 | 7 | 1 | 3 | 0 | 0 | 1 | 0 | 0 | 0 | 279 | 149 | 1 | 1 | NA | 1 | NA | 1 | 0 | NA | 0.789 | 1 | 1 | NA | -0.263 |
| SE | 30260 | ENSG00000153347 | FAM81B | chr5 | + | 95428602 | 95428732 | 95420283 | 95420402 | 95446561 | 95446689 | 0 | 1 | 1 | 2 | 0 | 0 | 1 | 2 | 0 | 5 | 0 | 6 | 1 | 3 | 0 | 0 | 0 | 1 | 3 | 6 | 279 | 149 | 1 | 1 | NA | 0.348 | 0.211 | 1 | 0 | NA | 1 | 0.348 | 0.348 | 0 | -0.034 |
| SE | 30261 | ENSG00000153347 | FAM81B | chr5 | + | 95436799 | 95436906 | 95428602 | 95428732 | 95446561 | 95446689 | 0 | 2 | 1 | 2 | 4 | 0 | 1 | 0 | 1 | 0 | 0 | 0 | 0 | 3 | 0 | 0 | 5 | 1 | 3 | 0 | 256 | 149 | 1 | 1 | NA | 0.538 | 1 | 0.538 | 1 | NA | 0 | 0 | 0.368 | NA | 0.646 |
| SE | 30262 | ENSG00000153347 | FAM81B | chr5 | + | 95436799 | 95436906 | 95428602 | 95428732 | 95448268 | 95448464 | 0 | 2 | 1 | 0 | 0 | 0 | 0 | 0 | 1 | 2 | 0 | 0 | 2 | 2 | 0 | 0 | 2 | 0 | 0 | 0 | 256 | 149 | 1 | 1 | NA | 1 | 1 | 0 | 0 | NA | NA | 1 | 1 | NA | -0.5 |
| SE | 30263 | ENSG00000153347 | FAM81B | chr5 | + | 95446561 | 95446697 | 95428602 | 95428732 | 95448268 | 95448464 | 2 | 2 | 5 | 8 | 7 | 0 | 0 | 0 | 1 | 2 | 0 | 11 | 7 | 13 | 16 | 0 | 0 | 0 | 0 | 0 | 285 | 149 | 1 | 1 | 1 | 1 | 1 | 0.807 | 0.647 | NA | 1 | 1 | 1 | 1 | -0.109 |
| SE | 30264 | ENSG00000153347 | FAM81B | chr5 | + | 95446561 | 95446697 | 95436799 | 95436906 | 95448268 | 95448464 | 2 | 1 | 6 | 9 | 11 | 0 | 0 | 1 | 0 | 0 | 0 | 6 | 6 | 12 | 16 | 0 | 0 | 2 | 1 | 0 | 285 | 149 | 1 | 1 | 1 | 1 | 0.758 | 1 | 1 | NA | 1 | 0.611 | 0.863 | 1 | 0.083 |
| SE | 30266 | ENSG00000178104 | PDE4DIP | chr1 | + | 148866410 | 148866532 | 148863249 | 148863305 | 148868470 | 148868622 | 6 | 0 | 0 | 0 | 0 | 4 | 6 | 0 | 8 | 1 | 8 | 0 | 0 | 0 | 0 | 20 | 1 | 0 | 2 | 2 | 271 | 149 | 1 | 1 | 0.452 | 0 | NA | 0 | 0 | 0.18 | 0 | NA | 0 | 0 | 0.068 |
| SE | 30267 | ENSG00000178104 | PDE4DIP | chr1 | + | 148866413 | 148866505 | 148863249 | 148863305 | 148868470 | 148868618 | 0 | 0 | 0 | 0 | 0 | 4 | 6 | 0 | 8 | 1 | 0 | 0 | 0 | 2 | 0 | 20 | 1 | 0 | 2 | 2 | 241 | 149 | 1 | 1 | 0 | 0 | NA | 0 | 0 | 0 | 0 | NA | 0.382 | 0 | -0.096 |
| SE | 30268 | ENSG00000178104 | PDE4DIP | chr1 | + | 148866413 | 148866532 | 148863249 | 148863305 | 148868470 | 148868609 | 6 | 0 | 0 | 0 | 2 | 4 | 6 | 0 | 8 | 1 | 4 | 0 | 0 | 2 | 0 | 20 | 1 | 0 | 2 | 2 | 268 | 149 | 1 | 1 | 0.455 | 0 | NA | 0 | 0.527 | 0.1 | 0 | NA | 0.357 | 0 | 0.131 |
| SE | 30270 | ENSG00000178104 | PDE4DIP | chr1 | + | 148889783 | 148889894 | 148868470 | 148868622 | 148929196 | 148929273 | 8 | 6 | 3 | 8 | 4 | 0 | 0 | 0 | 0 | 0 | 18 | 0 | 4 | 2 | 0 | 0 | 0 | 1 | 0 | 0 | 260 | 149 | 1 | 1 | 1 | 1 | 1 | 1 | 1 | 1 | NA | 0.696 | 1 | NA | 0.101 |
| SE | 30274 | ENSG00000178104 | PDE4DIP | chr1 | + | 149008443 | 149008495 | 149007200 | 149007436 | 149009567 | 149009791 | 1 | 1 | 5 | 1 | 0 | 0 | 0 | 4 | 0 | 0 | 13 | 0 | 4 | 2 | 0 | 0 | 0 | 1 | 0 | 0 | 201 | 149 | 1 | 1 | 1 | 1 | 0.481 | 1 | NA | 1 | NA | 0.748 | 1 | NA | -0.046 |
| SE | 30280 | ENSG00000236283 | NA | chr2 | + | 164912313 | 164912435 | 164848841 | 164849402 | 164945583 | 164945671 | 1 | 2 | 0 | 0 | 2 | 0 | 0 | 0 | 2 | 0 | 0 | 2 | 2 | 0 | 0 | 0 | 0 | 0 | 0 | 0 | 271 | 149 | 1 | 1 | 1 | 1 | NA | 0 | 1 | NA | 1 | 1 | NA | NA | -0.25 |
| SE | 30283 | ENSG00000236283 | NA | chr2 | + | 164959054 | 164959143 | 164949934 | 164950133 | 164959254 | 164959360 | 0 | 1 | 0 | 1 | 0 | 1 | 2 | 1 | 0 | 0 | 0 | 0 | 0 | 0 | 0 | 0 | 2 | 0 | 0 | 0 | 238 | 149 | 1 | 1 | 0 | 0.238 | 0 | 1 | NA | NA | 0 | NA | NA | NA | 0.309 |
| SE | 30286 | ENSG00000114480 | GBE1 | chr3 | - | 81648855 | 81648991 | 81642570 | 81642990 | 81649795 | 81649878 | 46 | 43 | 55 | 33 | 44 | 0 | 0 | 0 | 1 | 3 | 27 | 45 | 42 | 51 | 29 | 0 | 3 | 0 | 0 | 0 | 285 | 149 | 1 | 1 | 1 | 1 | 1 | 0.945 | 0.885 | 1 | 0.887 | 1 | 1 | 1 | -0.011 |
| SE | 30287 | ENSG00000129810 | SGO1 | chr3 | - | 20174248 | 20175055 | 20171042 | 20171232 | 20176600 | 20176659 | 2 | 4 | 2 | 7 | 6 | 4 | 0 | 2 | 0 | 0 | 0 | 7 | 4 | 11 | 0 | 0 | 3 | 2 | 0 | 0 | 298 | 149 | 1 | 1 | 0.2 | 1 | 0.333 | 1 | 1 | NA | 0.538 | 0.5 | 1 | NA | 0.027 |
| SE | 30288 | ENSG00000129810 | SGO1 | chr3 | - | 20175004 | 20175055 | 20171042 | 20171232 | 20176600 | 20176659 | 4 | 2 | 0 | 0 | 2 | 4 | 0 | 2 | 0 | 0 | 0 | 0 | 0 | 10 | 0 | 0 | 3 | 2 | 0 | 0 | 200 | 149 | 1 | 1 | 0.427 | 1 | 0 | NA | 1 | NA | 0 | 0 | 1 | NA | 0.273 |
| SE | 30290 | ENSG00000148154 | UGCG | chr9 | + | 111929499 | 111929678 | 111926383 | 111926496 | 111932169 | 111932359 | 112 | 123 | 119 | 181 | 125 | 0 | 1 | 0 | 0 | 0 | 27 | 130 | 69 | 337 | 35 | 0 | 2 | 0 | 1 | 0 | 298 | 149 | 1 | 1 | 1 | 0.984 | 1 | 1 | 1 | 1 | 0.97 | 1 | 0.994 | 1 | 0.004 |
| SE | 30291 | ENSG00000148154 | UGCG | chr9 | + | 111931270 | 111931357 | 111929499 | 111929678 | 111932169 | 111932359 | 227 | 219 | 235 | 259 | 307 | 0 | 0 | 0 | 0 | 1 | 37 | 182 | 180 | 788 | 24 | 0 | 4 | 0 | 0 | 0 | 236 | 149 | 1 | 1 | 1 | 1 | 1 | 1 | 0.995 | 1 | 0.966 | 1 | 1 | 1 | 0.006 |
| SE | 30292 | ENSG00000138785 | INTS12 | chr4 | - | 105702885 | 105703023 | 105699849 | 105700014 | 105703647 | 105703809 | 3 | 7 | 3 | 2 | 8 | 6 | 1 | 6 | 11 | 16 | 0 | 6 | 8 | 4 | 24 | 0 | 9 | 9 | 13 | 0 | 287 | 149 | 1 | 1 | 0.206 | 0.784 | 0.206 | 0.086 | 0.206 | NA | 0.257 | 0.316 | 0.138 | 1 | -0.13 |
| SE | 30293 | ENSG00000138785 | INTS12 | chr4 | - | 105702885 | 105703023 | 105699849 | 105700014 | 105707918 | 105708639 | 4 | 1 | 1 | 0 | 2 | 2 | 1 | 6 | 2 | 15 | 0 | 4 | 1 | 2 | 16 | 0 | 7 | 0 | 1 | 6 | 287 | 149 | 1 | 1 | 0.509 | 0.342 | 0.08 | 0 | 0.065 | NA | 0.229 | 1 | 0.509 | 0.581 | -0.381 |
| SE | 30295 | ENSG00000138785 | INTS12 | chr4 | - | 105703647 | 105703809 | 105699849 | 105700014 | 105707918 | 105708639 | 7 | 1 | 6 | 11 | 16 | 2 | 1 | 6 | 2 | 15 | 0 | 9 | 9 | 13 | 0 | 0 | 7 | 0 | 1 | 6 | 298 | 149 | 1 | 1 | 0.636 | 0.333 | 0.333 | 0.733 | 0.348 | NA | 0.391 | 1 | 0.867 | 0 | -0.088 |
| SE | 30296 | ENSG00000138785 | INTS12 | chr4 | - | 105703647 | 105703809 | 105699849 | 105700014 | 105708637 | 105708659 | 6 | 3 | 6 | 11 | 16 | 3 | 1 | 0 | 7 | 5 | 0 | 13 | 9 | 13 | 0 | 12 | 9 | 5 | 8 | 0 | 298 | 149 | 1 | 1 | 0.5 | 0.6 | 1 | 0.44 | 0.615 | 0 | 0.419 | 0.474 | 0.448 | NA | 0.296 |
| SE | 30297 | ENSG00000138785 | INTS12 | chr4 | - | 105703647 | 105703809 | 105702885 | 105703023 | 105707918 | 105708639 | 2 | 7 | 2 | 2 | 6 | 2 | 1 | 0 | 0 | 0 | 0 | 2 | 8 | 2 | 8 | 0 | 0 | 1 | 0 | 0 | 298 | 149 | 1 | 1 | 0.333 | 0.778 | 1 | 1 | 1 | NA | 1 | 0.8 | 1 | 1 | -0.128 |
| SE | 30301 | ENSG00000250412 | KLHL2P1 | chr4 | + | 119344157 | 119344264 | 119334328 | 119334453 | 119354109 | 119354231 | 4 | 1 | 1 | 0 | 1 | 0 | 0 | 0 | 0 | 0 | 0 | 0 | 0 | 0 | 0 | 0 | 2 | 0 | 0 | 0 | 256 | 149 | 1 | 1 | 1 | 1 | 1 | NA | 1 | NA | 0 | NA | NA | NA | 1 |
| SE | 30302 | ENSG00000135299 | ANKRD6 | chr6 | + | 89566833 | 89567096 | 89433151 | 89433375 | 89595915 | 89596014 | 6 | 11 | 7 | 8 | 10 | 3 | 0 | 0 | 1 | 0 | 0 | 3 | 9 | 5 | 0 | 0 | 0 | 0 | 0 | 0 | 298 | 149 | 1 | 1 | 0.5 | 1 | 1 | 0.8 | 1 | NA | 1 | 1 | 1 | NA | -0.14 |
| SE | 30303 | ENSG00000135299 | ANKRD6 | chr6 | + | 89566833 | 89567096 | 89433151 | 89433375 | 89603028 | 89603071 | 1 | 1 | 1 | 0 | 5 | 0 | 0 | 1 | 0 | 0 | 0 | 0 | 2 | 1 | 0 | 0 | 0 | 0 | 0 | 0 | 298 | 149 | 1 | 1 | 1 | 1 | 0.333 | NA | 1 | NA | NA | 1 | 1 | NA | -0.167 |
| SE | 30304 | ENSG00000135299 | ANKRD6 | chr6 | + | 89595915 | 89596014 | 89433203 | 89433375 | 89603028 | 89603049 | 5 | 1 | 1 | 1 | 0 | 0 | 0 | 1 | 0 | 0 | 0 | 7 | 0 | 4 | 0 | 0 | 0 | 0 | 0 | 0 | 248 | 149 | 1 | 1 | 1 | 1 | 0.375 | 1 | NA | NA | 1 | NA | 1 | NA | -0.156 |
| SE | 30306 | ENSG00000135299 | ANKRD6 | chr6 | + | 89606006 | 89606105 | 89602992 | 89603127 | 89612271 | 89612370 | 7 | 8 | 0 | 2 | 0 | 2 | 0 | 0 | 0 | 0 | 0 | 7 | 10 | 5 | 0 | 0 | 0 | 0 | 0 | 0 | 248 | 149 | 1 | 1 | 0.678 | 1 | NA | 1 | NA | NA | 1 | 1 | 1 | NA | -0.107 |
| SE | 30307 | ENSG00000135299 | ANKRD6 | chr6 | + | 89612271 | 89612370 | 89606006 | 89606105 | 89623409 | 89623544 | 2 | 5 | 0 | 2 | 0 | 0 | 0 | 0 | 1 | 0 | 0 | 0 | 2 | 10 | 0 | 0 | 0 | 0 | 0 | 0 | 248 | 149 | 1 | 1 | 1 | 1 | NA | 0.546 | NA | NA | NA | 1 | 1 | NA | -0.151 |
| SE | 30312 | ENSG00000135299 | ANKRD6 | chr6 | + | 89628246 | 89628341 | 89627582 | 89627696 | 89629111 | 89629223 | 0 | 0 | 0 | 0 | 2 | 3 | 4 | 3 | 1 | 4 | 7 | 0 | 0 | 0 | 0 | 0 | 0 | 4 | 10 | 12 | 244 | 149 | 1 | 1 | 0 | 0 | 0 | 0 | 0.234 | 1 | NA | 0 | 0 | 0 | -0.203 |
| SE | 30315 | ENSG00000020577 | SAMD4A | chr14 | + | 54748814 | 54748924 | 54702061 | 54702580 | 54751450 | 54751537 | 6 | 4 | 5 | 13 | 2 | 0 | 0 | 0 | 0 | 1 | 2 | 3 | 2 | 21 | 3 | 0 | 0 | 0 | 0 | 0 | 259 | 149 | 1 | 1 | 1 | 1 | 1 | 1 | 0.535 | 1 | 1 | 1 | 1 | 1 | -0.093 |
| SE | 30316 | ENSG00000020577 | SAMD4A | chr14 | + | 54784339 | 54784381 | 54776410 | 54776540 | 54784536 | 54784620 | 1 | 0 | 0 | 0 | 2 | 4 | 0 | 1 | 2 | 0 | 0 | 0 | 0 | 0 | 0 | 0 | 0 | 0 | 13 | 0 | 191 | 149 | 1 | 1 | 0.163 | NA | 0 | 0 | 1 | NA | NA | NA | 0 | NA | 0.291 |
| SE | 30317 | ENSG00000020577 | SAMD4A | chr14 | + | 54784339 | 54784447 | 54776413 | 54776540 | 54784536 | 54784620 | 4 | 0 | 2 | 3 | 3 | 4 | 0 | 1 | 2 | 0 | 0 | 0 | 0 | 0 | 0 | 0 | 0 | 0 | 13 | 0 | 257 | 149 | 1 | 1 | 0.367 | NA | 0.537 | 0.465 | 1 | NA | NA | NA | 0 | NA | 0.592 |
| SE | 30323 | ENSG00000100298 | APOBEC3H | chr22 | + | 39101917 | 39102042 | 39101236 | 39101504 | 39102511 | 39102571 | 13 | 2 | 4 | 7 | 2 | 0 | 0 | 0 | 0 | 3 | 0 | 21 | 10 | 9 | 11 | 0 | 0 | 0 | 0 | 0 | 274 | 149 | 1 | 1 | 1 | 1 | 1 | 1 | 0.266 | NA | 1 | 1 | 1 | 1 | -0.147 |
| SE | 30324 | ENSG00000100298 | APOBEC3H | chr22 | + | 39102511 | 39102605 | 39101917 | 39102042 | 39103688 | 39103998 | 0 | 2 | 1 | 7 | 0 | 8 | 4 | 5 | 11 | 9 | 0 | 2 | 5 | 0 | 0 | 5 | 29 | 2 | 0 | 16 | 243 | 149 | 1 | 1 | 0 | 0.235 | 0.109 | 0.281 | 0 | 0 | 0.041 | 0.605 | NA | 0 | -0.037 |
| SE | 30326 | ENSG00000164039 | BDH2 | chr4 | - | 103085348 | 103085462 | 103082870 | 103082929 | 103086479 | 103086540 | 88 | 30 | 60 | 70 | 64 | 4 | 2 | 0 | 5 | 2 | 38 | 51 | 38 | 43 | 0 | 0 | 0 | 1 | 0 | 0 | 263 | 149 | 1 | 1 | 0.926 | 0.895 | 1 | 0.888 | 0.948 | 1 | 1 | 0.956 | 1 | NA | -0.058 |

| | | | | | | | | | | | | | | | | | | | | | | | | | | | | | | | | | | | | | | | | | | | | | | |
|---|---|---|---|---|---|---|---|---|---|---|---|---|---|---|---|---|---|---|---|---|---|---|---|---|---|---|---|---|---|---|---|---|---|---|---|---|---|---|---|---|---|---|---|---|---|---|
| SE | 30327 | ENSG00000164039 | BDH2 | chr4 | - | 103091619 | 103091737 | 103091176 | 103091285 | 103092599 | 103092696 | 0 | 0 | 0 | 0 | 0 | 66 | 40 | 51 | 65 | 92 | 0 | 4 | 0 | 0 | 0 | 23 | 54 | 59 | 36 | 12 | 267 | 149 | 1 | 1 | 0 | 0 | 0 | 0 | 0 | 0 | 0.04 | 0 | 0 | 0 | -0.008 |
| SE | 30332 | ENSG00000162430 | SELENON | chr1 | + | 25808579 | 25808789 | 25805141 | 25805275 | 25809025 | 25809150 | 73 | 23 | 61 | 60 | 65 | 0 | 0 | 0 | 0 | 0 | 119 | 119 | 51 | 83 | 220 | 0 | 0 | 2 | 0 | 0 | 298 | 149 | 1 | 1 | 1 | 1 | 1 | 1 | 1 | 1 | 1 | 0.927 | 1 | 1 | 0.015 |
| SE | 30334 | ENSG00000227345 | PARG | chr10 | - | 49869475 | 49869555 | 49865320 | 49865381 | 49879672 | 49879830 | 149 | 108 | 145 | 130 | 125 | 0 | 0 | 0 | 0 | 0 | 89 | 144 | 82 | 122 | 10 | 0 | 4 | 0 | 0 | 0 | 229 | 149 | 1 | 1 | 1 | 1 | 1 | 1 | 1 | 1 | 0.959 | 1 | 1 | 1 | 0.008 |
| SE | 30338 | ENSG00000227345 | PARG | chr10 | - | 49933176 | 49934163 | 49932099 | 49932283 | 49941508 | 49941986 | 123 | 76 | 110 | 112 | 263 | 0 | 0 | 0 | 0 | 0 | 47 | 76 | 42 | 50 | 41 | 0 | 1 | 0 | 0 | 0 | 298 | 149 | 1 | 1 | 1 | 1 | 1 | 1 | 1 | 1 | 0.974 | 1 | 1 | 1 | 0.005 |
| SE | 30348 | ENSG00000108591 | DRG2 | chr17 | + | 18099632 | 18099723 | 18099016 | 18099077 | 18100362 | 18100435 | 57 | 18 | 47 | 66 | 62 | 3 | 0 | 1 | 1 | 0 | 86 | 119 | 23 | 56 | 100 | 0 | 0 | 0 | 0 | 0 | 240 | 149 | 1 | 1 | 0.922 | 1 | 0.967 | 0.976 | 1 | 1 | 1 | 1 | 1 | 1 | -0.027 |
| SE | 30350 | ENSG00000108591 | DRG2 | chr17 | + | 18106432 | 18106470 | 18104622 | 18104681 | 18107153 | 18107959 | 34 | 13 | 16 | 20 | 7 | 2 | 0 | 0 | 1 | 0 | 37 | 23 | 10 | 11 | 17 | 0 | 0 | 0 | 0 | 0 | 187 | 149 | 1 | 1 | 0.931 | 1 | 1 | 0.941 | 1 | 1 | 1 | 1 | 1 | 1 | -0.026 |
| SE | 30351 | ENSG00000108591 | DRG2 | chr17 | + | 18106432 | 18106486 | 18104559 | 18104681 | 18107153 | 18107600 | 98 | 41 | 48 | 63 | 60 | 2 | 0 | 0 | 1 | 0 | 136 | 109 | 33 | 58 | 109 | 0 | 0 | 0 | 0 | 0 | 203 | 149 | 1 | 1 | 0.973 | 1 | 1 | 0.979 | 1 | 1 | 1 | 1 | 1 | 1 | -0.01 |
| SE | 30355 | ENSG00000185090 | MANEAL | chr1 | + | 37796743 | 37796820 | 37795736 | 37795846 | 37799566 | 37800448 | 3 | 1 | 1 | 4 | 0 | 4 | 0 | 0 | 0 | 0 | 0 | 0 | 0 | 0 | 6 | 0 | 0 | 0 | 0 | 0 | 226 | 149 | 1 | 1 | 0.331 | 1 | 1 | 1 | NA | NA | NA | NA | NA | 1 | -0.167 |
| SE | 30365 | ENSG00000248242 | NA | chr4 | - | 104666896 | 104667016 | 104653873 | 104653921 | 104746639 | 104746680 | 0 | 0 | 0 | 0 | 0 | 3 | 6 | 0 | 8 | 17 | 5 | 1 | 0 | 0 | 5 | 33 | 1 | 3 | 6 | 22 | 269 | 149 | 1 | 1 | 0 | 0 | NA | 0 | 0 | 0.077 | 0.356 | 0 | 0 | 0.112 | -0.109 |
| SE | 30366 | ENSG00000248242 | NA | chr4 | - | 104746639 | 104746680 | 104653873 | 104653921 | 104927994 | 104928112 | 3 | 4 | 0 | 5 | 7 | 2 | 1 | 0 | 1 | 4 | 0 | 0 | 1 | 0 | 7 | 3 | 0 | 0 | 0 | 0 | 190 | 149 | 1 | 1 | 0.541 | 0.758 | NA | 0.797 | 0.578 | 0 | NA | 1 | NA | 1 | 0.002 |
| SE | 30367 | ENSG00000248242 | NA | chr4 | - | 104927994 | 104928112 | 104746639 | 104746680 | 104966019 | 104966081 | 0 | 1 | 0 | 2 | 0 | 0 | 0 | 0 | 0 | 0 | 0 | 0 | 0 | 0 | 6 | 0 | 0 | 1 | 0 | 0 | 267 | 149 | 1 | 1 | NA | 1 | NA | 1 | NA | NA | NA | 0 | NA | 1 | 0.5 |
| SE | 30369 | ENSG00000145348 | TBCK | chr4 | - | 106194717 | 106194754 | 106193608 | 106193770 | 106212749 | 106212835 | 98 | 48 | 74 | 94 | 145 | 0 | 2 | 0 | 4 | 0 | 5 | 30 | 24 | 37 | 10 | 0 | 0 | 0 | 0 | 0 | 186 | 149 | 1 | 1 | 1 | 0.951 | 1 | 0.95 | 1 | 1 | 1 | 1 | 1 | 1 | -0.02 |
| SE | 30370 | ENSG00000145348 | TBCK | chr4 | - | 106233587 | 106233650 | 106232937 | 106233064 | 106235268 | 106235367 | 67 | 53 | 51 | 57 | 180 | 0 | 1 | 0 | 0 | 0 | 11 | 22 | 28 | 18 | 19 | 0 | 0 | 0 | 0 | 0 | 212 | 149 | 1 | 1 | 1 | 0.974 | 1 | 1 | 1 | 1 | 1 | 1 | 1 | 1 | -0.005 |
| SE | 30372 | ENSG00000145348 | TBCK | chr4 | - | 106247138 | 106247287 | 106244625 | 106244764 | 106248244 | 106248306 | 124 | 94 | 67 | 97 | 191 | 0 | 0 | 0 | 1 | 2 | 19 | 82 | 45 | 36 | 34 | 0 | 0 | 0 | 2 | 0 | 298 | 149 | 1 | 1 | 1 | 1 | 1 | 0.98 | 0.979 | 1 | 1 | 1 | 0.9 | 1 | 0.012 |
| SE | 30373 | ENSG00000145348 | TBCK | chr4 | - | 106247138 | 106247287 | 106244625 | 106244764 | 106248920 | 106248982 | 90 | 58 | 49 | 69 | 134 | 0 | 0 | 0 | 0 | 5 | 7 | 45 | 27 | 13 | 33 | 0 | 0 | 0 | 0 | 0 | 298 | 149 | 1 | 1 | 1 | 1 | 1 | 1 | 0.931 | 1 | 1 | 1 | 1 | 1 | -0.014 |
| SE | 30374 | ENSG00000145348 | TBCK | chr4 | - | 106247138 | 106247287 | 106244625 | 106244764 | 106250417 | 106250470 | 90 | 57 | 49 | 67 | 136 | 0 | 0 | 0 | 0 | 4 | 7 | 45 | 27 | 13 | 33 | 0 | 0 | 0 | 0 | 0 | 298 | 149 | 1 | 1 | 1 | 1 | 1 | 1 | 0.944 | 1 | 1 | 1 | 1 | 1 | -0.011 |
| SE | 30375 | ENSG00000145348 | TBCK | chr4 | - | 106248244 | 106248306 | 106244625 | 106244764 | 106248920 | 106248982 | 22 | 15 | 9 | 18 | 30 | 0 | 0 | 0 | 0 | 5 | 0 | 14 | 8 | 9 | 0 | 0 | 0 | 0 | 0 | 0 | 211 | 149 | 1 | 1 | 1 | 1 | 1 | 1 | 0.809 | NA | 1 | 1 | 1 | NA | -0.038 |
| SE | 30376 | ENSG00000145348 | TBCK | chr4 | - | 106248244 | 106248306 | 106247138 | 106247287 | 106248920 | 106248982 | 53 | 51 | 27 | 47 | 86 | 0 | 1 | 0 | 2 | 0 | 12 | 51 | 26 | 30 | 1 | 0 | 0 | 0 | 0 | 0 | 211 | 149 | 1 | 1 | 1 | 0.973 | 1 | 0.943 | 1 | 1 | 1 | 1 | 1 | 1 | -0.017 |
| SE | 30377 | ENSG00000145348 | TBCK | chr4 | - | 106248920 | 106248982 | 106244625 | 106244764 | 106250417 | 106250478 | 14 | 13 | 5 | 12 | 25 | 0 | 0 | 0 | 0 | 4 | 6 | 6 | 8 | 6 | 0 | 0 | 0 | 0 | 0 | 0 | 211 | 149 | 1 | 1 | 1 | 1 | 1 | 1 | 0.815 | 1 | 1 | 1 | 1 | NA | -0.037 |
| SE | 30378 | ENSG00000145348 | TBCK | chr4 | - | 106248920 | 106248982 | 106247138 | 106247287 | 106250417 | 106250478 | 14 | 14 | 5 | 14 | 20 | 0 | 0 | 0 | 0 | 2 | 6 | 6 | 8 | 6 | 0 | 0 | 0 | 0 | 0 | 0 | 211 | 149 | 1 | 1 | 1 | 1 | 1 | 1 | 0.876 | 1 | 1 | 1 | 1 | NA | -0.025 |
| SE | 30380 | ENSG00000145348 | TBCK | chr4 | - | 106254946 | 106255047 | 106251865 | 106252007 | 106260436 | 106260510 | 32 | 32 | 50 | 28 | 76 | 26 | 12 | 18 | 35 | 36 | 0 | 32 | 31 | 2 | 9 | 0 | 15 | 12 | 13 | 0 | 250 | 149 | 1 | 1 | 0.423 | 0.614 | 0.623 | 0.323 | 0.557 | NA | 0.56 | 0.606 | 0.084 | 1 | -0.054 |
| SE | 30381 | ENSG00000145348 | TBCK | chr4 | - | 106260436 | 106260510 | 106251865 | 106252007 | 106295093 | 106295166 | 12 | 7 | 5 | 9 | 12 | 0 | 0 | 4 | 1 | 3 | 0 | 1 | 8 | 10 | 0 | 0 | 0 | 0 | 0 | 1 | 223 | 149 | 1 | 1 | 1 | 1 | 0.455 | 0.857 | 0.728 | NA | 1 | 1 | 1 | 0 | 0.058 |
| SE | 30382 | ENSG00000145348 | TBCK | chr4 | - | 106262097 | 106262212 | 106260436 | 106260510 | 106295093 | 106295166 | 116 | 93 | 118 | 121 | 176 | 0 | 0 | 0 | 0 | 1 | 30 | 125 | 70 | 47 | 35 | 0 | 0 | 0 | 0 | 0 | 264 | 149 | 1 | 1 | 1 | 1 | 1 | 1 | 0.99 | 1 | 1 | 1 | 1 | 1 | -0.002 |
| SE | 30383 | ENSG00000145348 | TBCK | chr4 | - | 106309926 | 106309967 | 106308767 | 106308989 | 106315930 | 106316235 | 1 | 0 | 2 | 4 | 8 | 16 | 29 | 36 | 34 | 21 | 0 | 0 | 0 | 0 | 0 | 0 | 22 | 4 | 7 | 0 | 190 | 149 | 1 | 1 | 0.047 | 0 | 0.042 | 0.084 | 0.23 | NA | 0 | 0 | 0 | NA | 0.081 |
| SE | 30388 | ENSG00000120137 | PANK3 | chr5 | - | 168566012 | 168566266 | 168563888 | 168564065 | 168568645 | 168568998 | 276 | 244 | 240 | 246 | 369 | 0 | 1 | 1 | 0 | 0 | 107 | 177 | 226 | 105 | 48 | 0 | 0 | 0 | 0 | 0 | 298 | 149 | 1 | 1 | 1 | 0.992 | 0.992 | 1 | 1 | 1 | 1 | 1 | 1 | 1 | -0.003 |
| SE | 30389 | ENSG00000120137 | PANK3 | chr5 | - | 168568645 | 168568998 | 168566012 | 168566266 | 168579255 | 168579368 | 146 | 118 | 139 | 136 | 216 | 0 | 0 | 0 | 0 | 0 | 30 | 73 | 112 | 38 | 7 | 0 | 0 | 2 | 0 | 0 | 298 | 149 | 1 | 1 | 1 | 1 | 1 | 1 | 1 | 1 | 1 | 0.966 | 1 | 1 | 0.007 |
| SE | 30390 | ENSG00000120137 | PANK3 | chr5 | - | 168578633 | 168578707 | 168568645 | 168568998 | 168579255 | 168579350 | 0 | 0 | 0 | 2 | 0 | 35 | 30 | 34 | 37 | 42 | 0 | 0 | 0 | 0 | 0 | 7 | 15 | 38 | 3 | 1 | 223 | 149 | 1 | 1 | 0 | 0 | 0 | 0.035 | 0 | 0 | 0 | 0 | 0 | 0 | 0.007 |
| SE | 30393 | ENSG00000184271 | POU6F1 | chr12 | - | 51191595 | 51191764 | 51186935 | 51190592 | 51192329 | 51192471 | 16 | 14 | 43 | 19 | 18 | 0 | 0 | 0 | 1 | 0 | 60 | 32 | 11 | 4 | 50 | 0 | 0 | 0 | 0 | 0 | 298 | 149 | 1 | 1 | 1 | 1 | 1 | 0.905 | 1 | 1 | 1 | 1 | 1 | 1 | -0.019 |
| SE | 30395 | ENSG00000184271 | POU6F1 | chr12 | - | 51196798 | 51196927 | 51195969 | 51196173 | 51197769 | 51198023 | 24 | 6 | 42 | 15 | 30 | 0 | 0 | 0 | 0 | 1 | 0 | 20 | 23 | 8 | 15 | 0 | 0 | 0 | 0 | 11 | 278 | 149 | 1 | 1 | 1 | 1 | 1 | 1 | 0.941 | NA | 1 | 1 | 1 | 0.422 | 0.133 |
| SE | 30396 | ENSG00000184271 | POU6F1 | chr12 | - | 51196798 | 51196946 | 51196130 | 51196173 | 51197769 | 51197785 | 10 | 5 | 20 | 7 | 20 | 0 | 0 | 0 | 0 | 1 | 0 | 11 | 12 | 3 | 7 | 0 | 0 | 0 | 0 | 11 | 297 | 149 | 1 | 1 | 1 | 1 | 1 | 1 | 0.909 | NA | 1 | 1 | 1 | 0.242 | 0.171 |
| SE | 30398 | ENSG00000184271 | POU6F1 | chr12 | - | 51199746 | 51199868 | 51198549 | 51198775 | 51204172 | 51204368 | 16 | 7 | 7 | 7 | 6 | 1 | 0 | 0 | 0 | 0 | 23 | 7 | 1 | 6 | 4 | 0 | 0 | 0 | 0 | 0 | 271 | 149 | 1 | 1 | 0.898 | 1 | 1 | 1 | 1 | 1 | 1 | 1 | 1 | 1 | -0.02 |
| SE | 30399 | ENSG00000184271 | POU6F1 | chr12 | - | 51206788 | 51206883 | 51204172 | 51204368 | 51217641 | 51217667 | 17 | 17 | 10 | 7 | 7 | 1 | 0 | 0 | 0 | 6 | 29 | 0 | 8 | 0 | 29 | 0 | 2 | 0 | 0 | 0 | 244 | 149 | 1 | 1 | 0.912 | 1 | 1 | 1 | 0.416 | 1 | 0 | 1 | NA | 1 | 0.116 |
| SE | 30400 | ENSG00000138678 | GPAT3 | chr4 | + | 83588209 | 83588299 | 83587254 | 83587329 | 83590198 | 83590292 | 53 | 41 | 49 | 22 | 93 | 0 | 0 | 0 | 1 | 0 | 139 | 46 | 31 | 43 | 92 | 0 | 0 | 0 | 0 | 0 | 239 | 149 | 1 | 1 | 1 | 1 | 1 | 0.932 | 1 | 1 | 1 | 1 | 1 | 1 | -0.014 |
| SE | 30401 | ENSG00000138678 | GPAT3 | chr4 | + | 83598050 | 83598179 | 83597429 | 83597515 | 83598643 | 83598723 | 228 | 218 | 131 | 142 | 244 | 0 | 2 | 0 | 0 | 1 | 569 | 193 | 175 | 397 | 427 | 0 | 0 | 3 | 4 | 0 | 278 | 149 | 1 | 1 | 1 | 0.983 | 1 | 1 | 0.992 | 1 | 1 | 0.969 | 0.982 | 1 | 0.005 |
| SE | 30405 | ENSG00000153064 | BANK1 | chr4 | + | 101829807 | 101830206 | 101813825 | 101813938 | 101855034 | 101855189 | 330 | 325 | 259 | 437 | 692 | 4 | 1 | 2 | 3 | 0 | 105 | 231 | 69 | 50 | 76 | 0 | 0 | 0 | 0 | 0 | 298 | 149 | 1 | 1 | 0.976 | 0.994 | 0.985 | 0.986 | 1 | 1 | 1 | 1 | 1 | 1 | -0.012 |
| SE | 30406 | ENSG00000153064 | BANK1 | chr4 | + | 102021513 | 102021592 | 101917992 | 101918189 | 102025200 | 102025509 | 321 | 327 | 235 | 461 | 877 | 0 | 0 | 0 | 0 | 2 | 68 | 256 | 85 | 50 | 67 | 0 | 0 | 0 | 0 | 0 | 228 | 149 | 1 | 1 | 1 | 1 | 1 | 1 | 0.997 | 1 | 1 | 1 | 1 | 1 | -0.001 |
| SE | 30407 | ENSG00000153064 | BANK1 | chr4 | + | 102029959 | 102030265 | 102025200 | 102025509 | 102043838 | 102043907 | 197 | 225 | 153 | 304 | 483 | 3 | 3 | 0 | 0 | 0 | 54 | 112 | 24 | 27 | 57 | 0 | 0 | 0 | 0 | 0 | 298 | 149 | 1 | 1 | 0.97 | 0.974 | 1 | 1 | 1 | 1 | 1 | 1 | 1 | 1 | -0.011 |
| SE | 30408 | ENSG00000153064 | BANK1 | chr4 | + | 102060210 | 102060389 | 102043838 | 102043907 | 102063074 | 102063138 | 359 | 327 | 259 | 457 | 741 | 0 | 1 | 0 | 1 | 0 | 20 | 255 | 59 | 36 | 18 | 0 | 0 | 0 | 0 | 0 | 298 | 149 | 1 | 1 | 1 | 0.994 | 1 | 0.996 | 1 | 1 | 1 | 1 | 1 | 1 | -0.002 |
| SE | 30409 | ENSG00000163510 | CWC22 | chr2 | - | 179952470 | 179952598 | 179950824 | 179950926 | 179954204 | 179954357 | 177 | 94 | 146 | 161 | 348 | 0 | 1 | 0 | 2 | 0 | 0 | 23 | 19 | 13 | 0 | 0 | 0 | 0 | 0 | 0 | 277 | 149 | 1 | 1 | 1 | 0.981 | 1 | 0.977 | 1 | NA | 1 | 1 | 1 | NA | -0.008 |
| SE | 30410 | ENSG00000163510 | CWC22 | chr2 | - | 179973633 | 179973802 | 179973192 | 179973246 | 179978189 | 179978318 | 141 | 72 | 117 | 100 | 258 | 0 | 0 | 0 | 0 | 6 | 0 | 19 | 20 | 5 | 3 | 0 | 0 | 0 | 0 | 0 | 298 | 149 | 1 | 1 | 1 | 1 | 1 | 1 | 0.956 | NA | 1 | 1 | 1 | 1 | -0.009 |
| SE | 30412 | ENSG00000064886 | CHI3L2 | chr1 | + | 111230741 | 111230943 | 111229691 | 111229881 | 111231237 | 111231294 | 71 | 8 | 70 | 22 | 80 | 0 | 0 | 0 | 0 | 0 | 36 | 61 | 21 | 20 | 46 | 0 | 4 | 0 | 0 | 0 | 298 | 149 | 1 | 1 | 1 | 1 | 1 | 1 | 1 | 1 | 0.884 | 1 | 1 | 1 | 0.023 |
| SE | 30413 | ENSG00000064886 | CHI3L2 | chr1 | + | 111231237 | 111231294 | 111230741 | 111230943 | 111234906 | 111235000 | 78 | 23 | 121 | 40 | 137 | 0 | 0 | 0 | 0 | 3 | 39 | 109 | 17 | 20 | 80 | 0 | 0 | 0 | 0 | 0 | 206 | 149 | 1 | 1 | 1 | 1 | 1 | 1 | 0.971 | 1 | 1 | 1 | 1 | 1 | -0.006 |
| SE | 30417 | ENSG00000074219 | TEAD2 | chr19 | - | 49343230 | 49343398 | 49342437 | 49342590 | 49347189 | 49347363 | 2 | 0 | 5 | 1 | 1 | 0 | 0 | 0 | 0 | 0 | 0 | 0 | 0 | 0 | 10 | 0 | 0 | 0 | 0 | 13 | 298 | 149 | 1 | 1 | 1 | NA | 1 | 1 | 1 | NA | NA | NA | NA | 0.278 | 0.722 |
| SE | 30418 | ENSG00000074219 | TEAD2 | chr19 | - | 49348956 | 49349059 | 49348702 | 49348845 | 49351300 | 49351365 | 0 | 0 | 2 | 0 | 0 | 5 | 0 | 0 | 0 | 15 | 0 | 0 | 0 | 0 | 0 | 0 | 0 | 2 | 2 | 0 | 252 | 149 | 1 | 1 | 0 | NA | 1 | NA | 0 | NA | NA | 0 | 0 | NA | 0.333 |
| SE | 30420 | ENSG00000074219 | TEAD2 | chr19 | - | 49355958 | 49355970 | 49355302 | 49355419 | 49357251 | 49357314 | 0 | 0 | 0 | 1 | 5 | 4 | 2 | 1 | 3 | 14 | 0 | 0 | 0 | 0 | 0 | 0 | 0 | 2 | 2 | 0 | 161 | 149 | 1 | 1 | 0 | 0 | 0 | 0.236 | 0.248 | NA | NA | 0 | 0 | NA | 0.097 |
| SE | 30424 | ENSG00000103502 | CDIPT | chr16 | - | 29862585 | 29862720 | 29861105 | 29861259 | 29862814 | 29862865 | 65 | 33 | 43 | 43 | 51 | 0 | 1 | 1 | 1 | 2 | 61 | 48 | 21 | 39 | 50 | 0 | 5 | 0 | 0 | 0 | 284 | 149 | 1 | 1 | 1 | 0.945 | 0.958 | 0.958 | 0.93 | 1 | 0.834 | 1 | 1 | 1 | -0.009 |
| SE | 30427 | ENSG00000153006 | SREK1IP1 | chr5 | - | 64741056 | 64741200 | 64727670 | 64728179 | 64754314 | 64754362 | 86 | 64 | 65 | 113 | 118 | 0 | 0 | 0 | 3 | 0 | 56 | 37 | 63 | 49 | 31 | 0 | 0 | 0 | 0 | 0 | 293 | 149 | 1 | 1 | 1 | 1 | 1 | 0.95 | 1 | 1 | 1 | 1 | 1 | 1 | -0.01 |
| SE | 30429 | ENSG00000153006 | SREK1IP1 | chr5 | - | 64754622 | 64754770 | 64754314 | 64754362 | 64765044 | 64765121 | 1 | 0 | 0 | 0 | 0 | 2 | 0 | 7 | 2 | 3 | 0 | 0 | 0 | 0 | 0 | 0 | 2 | 0 | 0 | 0 | 297 | 149 | 1 | 1 | 0.201 | NA | 0 | 0 | 0 | NA | 0 | NA | NA | NA | 0.05 |
| SE | 30432 | ENSG00000250696 | NA | chr4 | + | 69201171 | 69201284 | 69200419 | 69200578 | 69211065 | 69211147 | 1 | 0 | 1 | 0 | 0 | 4 | 2 | 0 | 0 | 2 | 0 | 0 | 0 | 0 | 0 | 0 | 1 | 0 | 0 | 0 | 262 | 149 | 1 | 1 | 0.124 | 0 | 1 | NA | 0 | NA | 0 | NA | NA | NA | 0.281 |
| SE | 30435 | ENSG00000164038 | SLC9B2 | chr4 | - | 103048892 | 103049020 | 103047050 | 103047226 | 103050239 | 103050382 | 9 | 8 | 29 | 18 | 20 | 0 | 0 | 2 | 0 | 0 | 0 | 9 | 16 | 4 | 0 | 0 | 0 | 0 | 0 | 0 | 277 | 149 | 1 | 1 | 1 | 1 | 0.886 | 1 | 1 | NA | 1 | 1 | 1 | NA | -0.023 |
| SE | 30436 | ENSG00000164038 | SLC9B2 | chr4 | - | 103057800 | 103057971 | 103050239 | 103050382 | 103066326 | 103066507 | 31 | 18 | 45 | 40 | 35 | 6 | 2 | 12 | 6 | 5 | 0 | 16 | 13 | 11 | 56 | 0 | 2 | 4 | 0 | 0 | 298 | 149 | 1 | 1 | 0.721 | 0.818 | 0.652 | 0.769 | 0.778 | NA | 0.8 | 0.619 | 1 | 1 | -0.107 |
| SE | 30439 | ENSG00000055955 | ITIH4 | chr3 | - | 52818461 | 52818536 | 52818051 | 52818283 | 52819392 | 52819518 | 14 | 12 | 18 | 23 | 22 | 0 | 0 | 0 | 0 | 0 | 9 | 29 | 34 | 50 | 40 | 0 | 0 | 4 | 0 | 0 | 224 | 149 | 1 | 1 | 1 | 1 | 1 | 1 | 1 | 1 | 1 | 0.85 | 1 | 1 | 0.03 |
| SE | 30451 | ENSG00000086061 | DNAJA1 | chr9 | + | 33026474 | 33026616 | 33025272 | 33025383 | 33026812 | 33026990 | 654 | 598 | 436 | 688 | 794 | 0 | 0 | 0 | 0 | 0 | 600 | 418 | 555 | 735 | 100 | 0 | 0 | 0 | 3 | 0 | 291 | 149 | 1 | 1 | 1 | 1 | 1 | 1 | 1 | 1 | 1 | 1 | 0.992 | 1 | 0.002 |
| SE | 30455 | ENSG00000086061 | DNAJA1 | chr9 | + | 33029884 | 33029989 | 33026812 | 33026990 | 33030439 | 33030667 | 356 | 420 | 362 | 440 | 573 | 0 | 0 | 0 | 0 | 0 | 423 | 268 | 359 | 489 | 70 | 0 | 0 | 0 | 0 | 5 | 254 | 149 | 1 | 1 | 1 | 1 | 1 | 1 | 1 | 1 | 1 | 1 | 1 | 0.891 | 0.022 |
| SE | 30456 | ENSG00000086061 | DNAJA1 | chr9 | + | 33030439 | 33030667 | 33029884 | 33029989 | 33034215 | 33034330 | 643 | 607 | 627 | 781 | 908 | 0 | 0 | 1 | 0 | 0 | 694 | 556 | 586 | 818 | 146 | 0 | 0 | 0 | 0 | 0 | 298 | 149 | 1 | 1 | 1 | 1 | 0.997 | 1 | 1 | 1 | 1 | 1 | 1 | 1 | -0.001 |
| SE | 30458 | ENSG00000086061 | DNAJA1 | chr9 | + | 33037014 | 33037115 | 33036573 | 33036689 | 33038684 | 33038726 | 584 | 508 | 478 | 620 | 627 | 2 | 0 | 1 | 0 | 0 | 306 | 366 | 431 | 516 | 54 | 0 | 0 | 0 | 0 | 0 | 250 | 149 | 1 | 1 | 0.994 | 1 | 0.997 | 1 | 1 | 1 | 1 | 1 | 1 | 1 | -0.002 |
| SE | 30460 | ENSG00000196209 | SIRPB2 | chr20 | - | 1473117 | 1473211 | 1472667 | 1473012 | 1473791 | 1473870 | 13 | 5 | 1 | 11 | 0 | 3 | 1 | 0 | 0 | 0 | 14 | 2 | 7 | 8 | 3 | 0 | 0 | 0 | 0 | 0 | 243 | 149 | 1 | 1 | 0.727 | 0.754 | 1 | 1 | NA | 1 | 1 | 1 | 1 | 1 | -0.13 |
| SE | 30462 | ENSG00000196209 | SIRPB2 | chr20 | - | 1477337 | 1477403 | 1473791 | 1473870 | 1479699 | 1480065 | 6 | 0 | 4 | 5 | 0 | 0 | 0 | 0 | 0 | 0 | 0 | 0 | 1 | 2 | 0 | 0 | 0 | 0 | 2 | 0 | 215 | 149 | 1 | 1 | 1 | NA | 1 | 1 | NA | NA | NA | 1 | 0.409 | NA | 0.295 |
| SE | 30463 | ENSG00000196209 | SIRPB2 | chr20 | - | 1477337 | 1477403 | 1473791 | 1473870 | 1479993 | 1480065 | 6 | 0 | 3 | 5 | 0 | 0 | 0 | 0 | 0 | 0 | 0 | 0 | 0 | 0 | 0 | 0 | 0 | 0 | 3 | 0 | 215 | 149 | 1 | 1 | 1 | NA | 1 | 1 | NA | NA | NA | NA | 0 | NA | 1 |
| SE | 30464 | ENSG00000196209 | SIRPB2 | chr20 | - | 1478265 | 1478607 | 1473791 | 1473870 | 1479699 | 1480065 | 55 | 21 | 10 | 20 | 10 | 0 | 0 | 0 | 0 | 0 | 51 | 16 | 16 | 54 | 3 | 0 | 0 | 0 | 2 | 0 | 298 | 149 | 1 | 1 | 1 | 1 | 1 | 1 | 1 | 1 | 1 | 1 | 0.931 | 1 | 0.014 |
| SE | 30465 | ENSG00000196209 | SIRPB2 | chr20 | - | 1478265 | 1478607 | 1473791 | 1473870 | 1479993 | 1480065 | 10 | 1 | 1 | 5 | 0 | 0 | 0 | 0 | 0 | 0 | 9 | 8 | 2 | 4 | 4 | 0 | 0 | 0 | 3 | 0 | 298 | 149 | 1 | 1 | 1 | 1 | 1 | 1 | NA | 1 | 1 | 1 | 0.4 | 1 | 0.12 |
| SE | 30466 | ENSG00000196209 | SIRPB2 | chr20 | - | 1478265 | 1478607 | 1476804 | 1477403 | 1479699 | 1480065 | 153 | 82 | 31 | 72 | 32 | 1 | 1 | 1 | 0 | 0 | 106 | 51 | 47 | 133 | 71 | 0 | 0 | 1 | 4 | 0 | 298 | 149 | 1 | 1 | 0.987 | 0.976 | 0.939 | 1 | 1 | 1 | 1 | 0.959 | 0.943 | 1 | 0 |
| SE | 30467 | ENSG00000196209 | SIRPB2 | chr20 | - | 1478265 | 1478607 | 1476804 | 1477403 | 1479993 | 1480065 | 108 | 62 | 22 | 57 | 22 | 3 | 0 | 0 | 0 | 0 | 64 | 43 | 33 | 83 | 72 | 0 | 0 | 0 | 0 | 0 | 298 | 149 | 1 | 1 | 0.947 | 1 | 1 | 1 | 1 | 1 | 1 | 1 | 1 | 1 | -0.011 |
| SE | 30470 | ENSG00000196209 | SIRPB2 | chr20 | - | 1479699 | 1480065 | 1477337 | 1477403 | 1491274 | 1491420 | 55 | 26 | 10 | 47 | 2 | 0 | 0 | 0 | 0 | 0 | 13 | 36 | 22 | 71 | 47 | 0 | 4 | 0 | 0 | 0 | 298 | 149 | 1 | 1 | 1 | 1 | 1 | 1 | 1 | 1 | 0.818 | 1 | 1 | 1 | 0.036 |
| SE | 30471 | ENSG00000196209 | SIRPB2 | chr20 | - | 1479993 | 1480065 | 1476804 | 1477403 | 1491274 | 1491383 | 47 | 17 | 9 | 33 | 2 | 0 | 0 | 0 | 0 | 0 | 11 | 34 | 21 | 34 | 19 | 0 | 4 | 0 | 0 | 0 | 221 | 149 | 1 | 1 | 1 | 1 | 1 | 1 | 1 | 1 | 0.851 | 1 | 1 | 1 | 0.03 |
| SE | 30473 | ENSG00000152503 | TRIM36 | chr5 | - | 115141278 | 115141374 | 115137362 | 115137616 | 115144597 | 115144744 | 8 | 12 | 6 | 7 | 10 | 0 | 0 | 0 | 0 | 3 | 8 | 0 | 3 | 0 | 10 | 0 | 0 | 0 | 0 | 0 | 245 | 149 | 1 | 1 | 1 | 1 | 1 | 1 | 0.67 | 1 | NA | 1 | NA | 1 | -0.066 |
| SE | 30479 | ENSG00000014824 | SLC30A9 | chr4 | + | 42035274 | 42035333 | 42023301 | 42023384 | 42038985 | 42039053 | 167 | 155 | 156 | 156 | 256 | 0 | 1 | 0 | 0 | 0 | 53 | 167 | 100 | 89 | 25 | 0 | 0 | 0 | 0 | 0 | 208 | 149 | 1 | 1 | 1 | 0.991 | 1 | 1 | 1 | 1 | 1 | 1 | 1 | 1 | -0.002 |
| SE | 30480 | ENSG00000014824 | SLC30A9 | chr4 | + | 42060190 | 42060246 | 42049376 | 42049479 | 42062985 | 42063121 | 101 | 86 | 129 | 111 | 128 | 0 | 1 | 1 | 0 | 0 | 8 | 46 | 77 | 46 | 6 | 0 | 0 | 0 | 0 | 0 | 205 | 149 | 1 | 1 | 1 | 0.984 | 0.989 | 1 | 1 | 1 | 1 | 1 | 1 | 1 | -0.005 |
| SE | 30481 | ENSG00000014824 | SLC30A9 | chr4 | + | 42070525 | 42070691 | 42067084 | 42067192 | 42075656 | 42075786 | 167 | 111 | 132 | 160 | 214 | 1 | 0 | 1 | 2 | 3 | 3 | 124 | 91 | 54 | 0 | 0 | 0 | 0 | 3 | 0 | 298 | 149 | 1 | 1 | 0.988 | 1 | 0.985 | 0.976 | 0.973 | 1 | 1 | 1 | 0.9 | NA | 0.009 |
| SE | 30482 | ENSG00000014824 | SLC30A9 | chr4 | + | 42070525 | 42070691 | 42067084 | 42067192 | 42078211 | 42078325 | 56 | 38 | 47 | 52 | 97 | 1 | 0 | 0 | 0 | 0 | 0 | 51 | 44 | 17 | 0 | 0 | 0 | 0 | 0 | 0 | 298 | 149 | 1 | 1 | 0.966 | 1 | 1 | 1 | 1 | NA | 1 | 1 | 1 | NA | -0.007 |
| SE | 30483 | ENSG00000014824 | SLC30A9 | chr4 | + | 42070525 | 42070691 | 42067084 | 42067192 | 42086081 | 42086369 | 56 | 32 | 45 | 54 | 99 | 0 | 0 | 0 | 0 | 0 | 0 | 51 | 43 | 17 | 0 | 8 | 0 | 0 | 0 | 0 | 298 | 149 | 1 | 1 | 1 | 1 | 1 | 1 | 1 | 0 | 1 | 1 | 1 | NA | 0.25 |
| SE | 30484 | ENSG00000014824 | SLC30A9 | chr4 | + | 42075656 | 42075786 | 42067084 | 42067192 | 42078211 | 42078325 | 135 | 109 | 127 | 157 | 208 | 1 | 0 | 0 | 0 | 0 | 15 | 91 | 91 | 57 | 5 | 0 | 0 | 0 | 0 | 0 | 279 | 149 | 1 | 1 | 0.986 | 1 | 1 | 1 | 1 | 1 | 1 | 1 | 1 | 1 | -0.003 |
| SE | 30485 | ENSG00000014824 | SLC30A9 | chr4 | + | 42075656 | 42075786 | 42070525 | 42070691 | 42078211 | 42078325 | 245 | 187 | 211 | 263 | 322 | 0 | 6 | 2 | 0 | 0 | 18 | 164 | 139 | 91 | 5 | 0 | 0 | 1 | 0 | 0 | 279 | 149 | 1 | 1 | 1 | 0.943 | 0.983 | 1 | 1 | 1 | 1 | 0.987 | 1 | 1 | -0.012 |
| SE | 30486 | ENSG00000014824 | SLC30A9 | chr4 | + | 42078211 | 42078325 | 42067084 | 42067192 | 42086081 | 42086369 | 55 | 66 | 88 | 102 | 150 | 0 | 0 | 0 | 0 | 0 | 0 | 79 | 69 | 18 | 0 | 8 | 0 | 0 | 0 | 0 | 263 | 149 | 1 | 1 | 1 | 1 | 1 | 1 | 1 | 0 | 1 | 1 | 1 | NA | 0.25 |
| SE | 30487 | ENSG00000014824 | SLC30A9 | chr4 | + | 42078211 | 42078325 | 42070525 | 42070691 | 42086081 | 42086369 | 54 | 72 | 90 | 102 | 150 | 0 | 0 | 0 | 2 | 2 | 0 | 79 | 69 | 18 | 0 | 0 | 0 | 0 | 0 | 0 | 263 | 149 | 1 | 1 | 1 | 1 | 1 | 0.967 | 0.977 | NA | 1 | 1 | 1 | NA | -0.011 |
| SE | 30490 | ENSG00000163138 | PACRGL | chr4 | + | 20700670 | 20700787 | 20700426 | 20700578 | 20701757 | 20701884 | 1 | 0 | 0 | 0 | 0 | 0 | 0 | 1 | 0 | 0 | 0 | 0 | 0 | 3 | 0 | 0 | 0 | 0 | 0 | 0 | 266 | 149 | 1 | 1 | 1 | NA | 0 | NA | NA | NA | NA | NA | 1 | NA | -0.5 |
| SE | 30491 | ENSG00000163138 | PACRGL | chr4 | + | 20701509 | 20701667 | 20700465 | 20700787 | 20701757 | 20701854 | 0 | 1 | 0 | 0 | 3 | 1 | 0 | 0 | 0 | 0 | 0 | 0 | 0 | 4 | 0 | 0 | 0 | 0 | 3 | 0 | 298 | 149 | 1 | 1 | 0 | 1 | NA | NA | 1 | NA | NA | NA | 0.4 | NA | 0.267 |
| SE | 30492 | ENSG00000163138 | PACRGL | chr4 | + | 20701518 | 20701667 | 20700438 | 20700787 | 20701757 | 20701884 | 0 | 1 | 0 | 0 | 3 | 1 | 0 | 0 | 0 | 0 | 0 | 0 | 0 | 4 | 0 | 0 | 0 | 0 | 3 | 0 | 298 | 149 | 1 | 1 | 0 | 1 | NA | NA | 1 | NA | NA | NA | 0.4 | NA | 0.267 |
| SE | 30493 | ENSG00000163138 | PACRGL | chr4 | + | 20701725 | 20701884 | 20700459 | 20700787 | 20702140 | 20702196 | 8 | 4 | 6 | 2 | 2 | 0 | 0 | 0 | 1 | 0 | 0 | 3 | 8 | 8 | 0 | 0 | 0 | 0 | 0 | 0 | 298 | 149 | 1 | 1 | 1 | 1 | 1 | 0.5 | 1 | NA | 1 | 1 | 1 | NA | -0.1 |
| SE | 30494 | ENSG00000163138 | PACRGL | chr4 | + | 20701757 | 20701884 | 20700412 | 20700578 | 20702140 | 20702221 | 8 | 4 | 7 | 2 | 2 | 0 | 0 | 2 | 0 | 0 | 0 | 3 | 8 | 8 | 0 | 0 | 0 | 0 | 0 | 0 | 276 | 149 | 1 | 1 | 1 | 1 | 0.654 | 1 | 1 | NA | 1 | 1 | 1 | NA | -0.069 |
| SE | 30496 | ENSG00000163138 | PACRGL | chr4 | + | 20701757 | 20701884 | 20700437 | 20700706 | 20702140 | 20702221 | 8 | 4 | 6 | 2 | 2 | 0 | 0 | 0 | 0 | 1 | 0 | 3 | 8 | 8 | 0 | 0 | 0 | 0 | 0 | 0 | 276 | 149 | 1 | 1 | 1 | 1 | 1 | 1 | 0.519 | NA | 1 | 1 | 1 | NA | -0.096 |
| SE | 30498 | ENSG00000163138 | PACRGL | chr4 | + | 20701757 | 20701884 | 20700458 | 20700617 | 20702140 | 20702221 | 8 | 6 | 6 | 2 | 2 | 0 | 0 | 1 | 0 | 0 | 0 | 3 | 8 | 8 | 0 | 0 | 0 | 0 | 0 | 0 | 276 | 149 | 1 | 1 | 1 | 1 | 0.764 | 1 | 1 | NA | 1 | 1 | 1 | NA | -0.047 |
| SE | 30501 | ENSG00000163138 | PACRGL | chr4 | + | 20701757 | 20701884 | 20700465 | 20700611 | 20704465 | 20704524 | 1 | 3 | 0 | 0 | 0 | 0 | 0 | 1 | 0 | 5 | 0 | 0 | 0 | 0 | 0 | 0 | 3 | 1 | 0 | 0 | 276 | 149 | 1 | 1 | 1 | 1 | 0 | NA | 0 | NA | 0 | 0 | NA | NA | 0.5 |
| SE | 30502 | ENSG00000163138 | PACRGL | chr4 | + | 20701757 | 20701884 | 20700670 | 20700787 | 20702140 | 20702221 | 9 | 4 | 6 | 2 | 2 | 0 | 0 | 0 | 1 | 0 | 0 | 3 | 8 | 11 | 0 | 0 | 0 | 0 | 0 | 0 | 276 | 149 | 1 | 1 | 1 | 1 | 1 | 0.519 | 1 | NA | 1 | 1 | 1 | NA | -0.096 |
| SE | 30503 | ENSG00000163138 | PACRGL | chr4 | + | 20701757 | 20701884 | 20700670 | 20700787 | 20704465 | 20704524 | 2 | 2 | 0 | 0 | 0 | 4 | 0 | 1 | 0 | 0 | 0 | 0 | 0 | 3 | 0 | 0 | 0 | 0 | 0 | 0 | 276 | 149 | 1 | 1 | 0.213 | 1 | 0 | NA | NA | NA | NA | NA | 1 | NA | -0.596 |
| SE | 30504 | ENSG00000163138 | PACRGL | chr4 | + | 20702140 | 20702221 | 20700418 | 20700617 | 20704465 | 20704524 | 3 | 3 | 6 | 5 | 5 | 4 | 0 | 0 | 1 | 0 | 0 | 3 | 4 | 0 | 0 | 0 | 0 | 0 | 0 | 0 | 230 | 149 | 1 | 1 | 0.327 | 1 | 1 | 0.764 | 1 | NA | 1 | 1 | NA | NA | -0.182 |
| SE | 30505 | ENSG00000163138 | PACRGL | chr4 | + | 20702140 | 20702221 | 20700457 | 20700787 | 20704465 | 20704524 | 3 | 3 | 5 | 6 | 5 | 4 | 0 | 1 | 0 | 0 | 0 | 3 | 4 | 0 | 0 | 0 | 0 | 0 | 0 | 0 | 230 | 149 | 1 | 1 | 0.327 | 1 | 0.764 | 1 | 1 | NA | 1 | 1 | NA | NA | -0.182 |
| SE | 30508 | ENSG00000163138 | PACRGL | chr4 | + | 20702140 | 20702221 | 20701757 | 20701884 | 20702674 | 20702694 | 6 | 4 | 4 | 2 | 1 | 0 | 0 | 0 | 0 | 0 | 0 | 3 | 6 | 2 | 0 | 0 | 0 | 0 | 2 | 0 | 230 | 149 | 1 | 1 | 1 | 1 | 1 | 1 | 1 | NA | 1 | 1 | 0.393 | NA | 0.202 |
| SE | 30509 | ENSG00000163138 | PACRGL | chr4 | + | 20702140 | 20702221 | 20701757 | 20701884 | 20704465 | 20704524 | 11 | 8 | 11 | 7 | 7 | 1 | 2 | 0 | 0 | 0 | 0 | 6 | 12 | 8 | 0 | 0 | 0 | 0 | 0 | 0 | 230 | 149 | 1 | 1 | 0.877 | 0.722 | 1 | 1 | 1 | NA | 1 | 1 | 1 | NA | -0.08 |
| SE | 30511 | ENSG00000163138 | PACRGL | chr4 | + | 20707802 | 20707870 | 20704659 | 20704814 | 20709682 | 20709714 | 17 | 7 | 11 | 13 | 40 | 0 | 3 | 1 | 0 | 0 | 8 | 10 | 13 | 5 | 0 | 0 | 0 | 0 | 0 | 0 | 217 | 149 | 1 | 1 | 1 | 0.616 | 0.883 | 1 | 1 | 1 | 1 | 1 | 1 | NA | -0.1 |
| SE | 30512 | ENSG00000163138 | PACRGL | chr4 | + | 20707802 | 20707870 | 20704659 | 20704814 | 20712787 | 20712922 | 9 | 4 | 2 | 8 | 5 | 0 | 0 | 0 | 2 | 4 | 0 | 0 | 4 | 2 | 0 | 0 | 0 | 0 | 0 | 0 | 217 | 149 | 1 | 1 | 1 | 1 | 1 | 0.733 | 0.462 | NA | NA | 1 | 1 | NA | -0.161 |
| SE | 30513 | ENSG00000163138 | PACRGL | chr4 | + | 20709682 | 20709773 | 20704659 | 20704814 | 20712787 | 20712831 | 0 | 11 | 3 | 2 | 14 | 0 | 0 | 0 | 2 | 4 | 0 | 2 | 0 | 2 | 0 | 0 | 0 | 0 | 0 | 0 | 240 | 149 | 1 | 1 | NA | 1 | 1 | 0.383 | 0.685 | NA | 1 | NA | 1 | NA | -0.233 |
| SE | 30514 | ENSG00000163138 | PACRGL | chr4 | + | 20709682 | 20709773 | 20704659 | 20704814 | 20713431 | 20713521 | 0 | 3 | 1 | 0 | 0 | 4 | 6 | 8 | 16 | 0 | 0 | 0 | 2 | 0 | 5 | 0 | 3 | 0 | 0 | 0 | 240 | 149 | 1 | 1 | 0 | 0.237 | 0.072 | 0 | NA | NA | 0 | 1 | NA | 1 | -0.589 |
| SE | 30515 | ENSG00000163138 | PACRGL | chr4 | + | 20709682 | 20709773 | 20707802 | 20707870 | 20712787 | 20712831 | 7 | 11 | 11 | 14 | 49 | 1 | 0 | 2 | 7 | 0 | 8 | 12 | 9 | 4 | 0 | 0 | 0 | 0 | 0 | 0 | 240 | 149 | 1 | 1 | 0.813 | 1 | 0.773 | 0.554 | 1 | 1 | 1 | 1 | 1 | NA | -0.172 |
| SE | 30516 | ENSG00000163138 | PACRGL | chr4 | + | 20712787 | 20712922 | 20704659 | 20704814 | 20713431 | 20713516 | 27 | 9 | 16 | 16 | 35 | 4 | 6 | 8 | 16 | 0 | 26 | 12 | 2 | 0 | 2 | 0 | 3 | 0 | 0 | 0 | 284 | 149 | 1 | 1 | 0.78 | 0.44 | 0.512 | 0.344 | 1 | 1 | 0.677 | 1 | NA | 1 | -0.304 |
| SE | 30519 | ENSG00000163138 | PACRGL | chr4 | + | 20716106 | 20716248 | 20713431 | 20713539 | 20718815 | 20718976 | 0 | 2 | 0 | 0 | 1 | 3 | 2 | 0 | 0 | 0 | 0 | 0 | 0 | 0 | 5 | 8 | 0 | 0 | 0 | 0 | 291 | 149 | 1 | 1 | 0 | 0.339 | NA | NA | 1 | 0 | NA | NA | NA | 1 | -0.054 |
| SE | 30520 | ENSG00000163138 | PACRGL | chr4 | + | 20716106 | 20716248 | 20713431 | 20713539 | 20724807 | 20724889 | 0 | 0 | 0 | 0 | 3 | 14 | 20 | 13 | 18 | 29 | 0 | 0 | 0 | 0 | 11 | 14 | 10 | 11 | 0 | 4 | 291 | 149 | 1 | 1 | 0 | 0 | 0 | 0 | 0.05 | 0 | 0 | 0 | NA | 0.585 | -0.136 |
| SE | 30521 | ENSG00000163138 | PACRGL | chr4 | + | 20716106 | 20716248 | 20713431 | 20713539 | 20727284 | 20727589 | 0 | 0 | 0 | 0 | 1 | 9 | 7 | 8 | 15 | 22 | 0 | 0 | 0 | 0 | 5 | 17 | 6 | 4 | 0 | 0 | 291 | 149 | 1 | 1 | 0 | 0 | 0 | 0 | 0.023 | 0 | 0 | 0 | NA | 1 | -0.245 |
| SE | 30522 | ENSG00000163138 | PACRGL | chr4 | + | 20718815 | 20718976 | 20713431 | 20713539 | 20724807 | 20724879 | 17 | 2 | 0 | 0 | 0 | 14 | 20 | 13 | 18 | 29 | 16 | 0 | 0 | 0 | 0 | 14 | 10 | 11 | 0 | 4 | 298 | 149 | 1 | 1 | 0.378 | 0.048 | 0 | 0 | 0 | 0.364 | 0 | 0 | NA | 0 | -0.006 |
| SE | 30523 | ENSG00000163138 | PACRGL | chr4 | + | 20718815 | 20718976 | 20713431 | 20713539 | 20727284 | 20732557 | 3 | 4 | 0 | 0 | 0 | 9 | 7 | 8 | 15 | 22 | 8 | 0 | 0 | 0 | 0 | 17 | 6 | 4 | 0 | 0 | 298 | 149 | 1 | 1 | 0.143 | 0.222 | 0 | 0 | 0 | 0.19 | 0 | 0 | NA | NA | 0.01 |
| SE | 30524 | ENSG00000163138 | PACRGL | chr4 | + | 20718815 | 20718976 | 20716106 | 20716248 | 20724807 | 20724879 | 14 | 2 | 0 | 0 | 0 | 0 | 0 | 0 | 0 | 2 | 8 | 0 | 0 | 0 | 0 | 0 | 0 | 0 | 0 | 6 | 298 | 149 | 1 | 1 | 1 | 1 | NA | NA | 0 | 1 | NA | NA | NA | 0 | 0.167 |
| SE | 30527 | ENSG00000163161 | ERCC3 | chr2 | - | 127261227 | 127261346 | 127259295 | 127259448 | 127271335 | 127271453 | 219 | 140 | 204 | 208 | 160 | 2 | 0 | 0 | 0 | 0 | 294 | 281 | 202 | 250 | 209 | 0 | 0 | 0 | 0 | 0 | 268 | 149 | 1 | 1 | 0.984 | 1 | 1 | 1 | 1 | 1 | 1 | 1 | 1 | 1 | -0.003 |
| SE | 30528 | ENSG00000163161 | ERCC3 | chr2 | - | 127271335 | 127271453 | 127260764 | 127261346 | 127272864 | 127272961 | 179 | 118 | 160 | 180 | 156 | 1 | 1 | 0 | 2 | 0 | 298 | 223 | 182 | 202 | 135 | 0 | 0 | 0 | 0 | 0 | 267 | 149 | 1 | 1 | 0.99 | 0.985 | 1 | 0.98 | 1 | 1 | 1 | 1 | 1 | 1 | -0.009 |
| SE | 30529 | ENSG00000163161 | ERCC3 | chr2 | - | 127272864 | 127272961 | 127271335 | 127271453 | 127279172 | 127279261 | 122 | 107 | 138 | 148 | 214 | 0 | 0 | 1 | 0 | 0 | 223 | 171 | 131 | 152 | 60 | 0 | 0 | 0 | 0 | 0 | 246 | 149 | 1 | 1 | 1 | 1 | 0.988 | 1 | 1 | 1 | 1 | 1 | 1 | 1 | -0.002 |
| SE | 30530 | ENSG00000163161 | ERCC3 | chr2 | - | 127280446 | 127280631 | 127279172 | 127279375 | 127286702 | 127286797 | 201 | 95 | 206 | 232 | 178 | 1 | 0 | 1 | 1 | 0 | 369 | 228 | 178 | 288 | 314 | 0 | 0 | 0 | 1 | 0 | 298 | 149 | 1 | 1 | 0.99 | 1 | 0.99 | 0.991 | 1 | 1 | 1 | 1 | 0.993 | 1 | -0.004 |
| SE | 30532 | ENSG00000163161 | ERCC3 | chr2 | - | 127284170 | 127284289 | 127280446 | 127280631 | 127286702 | 127287017 | 3 | 0 | 0 | 1 | 0 | 71 | 20 | 67 | 71 | 27 | 0 | 0 | 0 | 0 | 0 | 98 | 49 | 43 | 92 | 80 | 268 | 149 | 1 | 1 | 0.023 | 0 | 0 | 0.008 | 0 | 0 | 0 | 0 | 0 | 0 | 0.006 |
| SE | 30536 | ENSG00000132182 | NUP210 | chr3 | - | 13351878 | 13351980 | 13343174 | 13343303 | 13352079 | 13352184 | 242 | 80 | 181 | 165 | 69 | 2 | 0 | 0 | 0 | 0 | 121 | 270 | 99 | 123 | 206 | 0 | 0 | 0 | 0 | 0 | 251 | 149 | 1 | 1 | 0.986 | 1 | 1 | 1 | 1 | 1 | 1 | 1 | 1 | 1 | -0.003 |
| SE | 30537 | ENSG00000132182 | NUP210 | chr3 | - | 13365945 | 13366091 | 13360269 | 13360491 | 13371833 | 13372032 | 427 | 148 | 303 | 332 | 342 | 0 | 1 | 0 | 0 | 0 | 300 | 403 | 177 | 207 | 378 | 0 | 0 | 0 | 0 | 0 | 295 | 149 | 1 | 1 | 1 | 0.987 | 1 | 1 | 1 | 1 | 1 | 1 | 1 | 1 | -0.003 |
| SE | 30538 | ENSG00000132182 | NUP210 | chr3 | - | 13365945 | 13366091 | 13360269 | 13360491 | 13373717 | 13373873 | 150 | 41 | 99 | 116 | 91 | 1 | 0 | 0 | 0 | 0 | 121 | 157 | 86 | 93 | 175 | 0 | 0 | 0 | 0 | 0 | 295 | 149 | 1 | 1 | 0.987 | 1 | 1 | 1 | 1 | 1 | 1 | 1 | 1 | 1 | -0.003 |
| SE | 30539 | ENSG00000132182 | NUP210 | chr3 | - | 13371833 | 13372032 | 13360269 | 13360491 | 13373717 | 13373873 | 314 | 103 | 230 | 225 | 265 | 1 | 0 | 0 | 0 | 0 | 156 | 293 | 125 | 130 | 291 | 0 | 0 | 0 | 0 | 0 | 298 | 149 | 1 | 1 | 0.994 | 1 | 1 | 1 | 1 | 1 | 1 | 1 | 1 | 1 | -0.001 |
| SE | 30543 | ENSG00000132182 | NUP210 | chr3 | - | 13373717 | 13373873 | 13371833 | 13372032 | 13375503 | 13375641 | 469 | 159 | 368 | 351 | 372 | 0 | 2 | 0 | 0 | 0 | 265 | 389 | 194 | 175 | 431 | 0 | 0 | 1 | 4 | 0 | 298 | 149 | 1 | 1 | 1 | 0.975 | 1 | 1 | 1 | 1 | 1 | 0.99 | 0.956 | 1 | 0.006 |
| SE | 30544 | ENSG00000132182 | NUP210 | chr3 | - | 13386274 | 13386407 | 13379562 | 13379721 | 13388302 | 13388453 | 510 | 300 | 527 | 483 | 557 | 2 | 0 | 2 | 0 | 0 | 439 | 675 | 330 | 334 | 643 | 0 | 0 | 0 | 4 | 0 | 282 | 149 | 1 | 1 | 0.993 | 1 | 0.993 | 1 | 1 | 1 | 1 | 1 | 0.978 | 1 | 0.002 |
| SE | 30545 | ENSG00000132182 | NUP210 | chr3 | - | 13391210 | 13391307 | 13388302 | 13388453 | 13397356 | 13397488 | 286 | 92 | 233 | 242 | 165 | 0 | 0 | 0 | 1 | 0 | 173 | 277 | 176 | 290 | 341 | 0 | 0 | 0 | 0 | 0 | 246 | 149 | 1 | 1 | 1 | 1 | 1 | 0.993 | 1 | 1 | 1 | 1 | 1 | 1 | -0.001 |
| SE | 30550 | ENSG00000249307 | LINC01088 | chr4 | + | 79306690 | 79306728 | 79211684 | 79211804 | 79308272 | 79308472 | 0 | 25 | 2 | 0 | 0 | 0 | 1 | 3 | 0 | 0 | 4 | 0 | 0 | 9 | 0 | 0 | 0 | 0 | 0 | 0 | 187 | 149 | 1 | 1 | NA | 0.952 | 0.347 | NA | NA | 1 | NA | NA | 1 | NA | -0.351 |
| SE | 30551 | ENSG00000249307 | LINC01088 | chr4 | + | 79307063 | 79307231 | 79306690 | 79306728 | 79308272 | 79308755 | 0 | 0 | 0 | 0 | 0 | 0 | 34 | 0 | 0 | 0 | 0 | 0 | 6 | 0 | 0 | 0 | 0 | 0 | 12 | 0 | 298 | 149 | 1 | 1 | NA | 0 | NA | NA | | NA | NA | 1 | 0 | NA | -0.5 |
| SE | 30553 | ENSG00000170365 | SMAD1 | chr4 | + | 145483073 | 145483176 | 145481870 | 145482038 | 145514437 | 145515013 | 0 | 0 | 0 | 1 | 0 | 2 | 2 | 1 | 0 | 0 | 0 | 0 | 0 | 0 | 0 | 0 | 5 | 2 | 3 | 0 | 252 | 149 | 1 | 1 | 0 | 0 | 0 | 1 | NA | NA | 0 | 0 | 0 | NA | 0.25 |
| SE | 30556 | ENSG00000274070 | CASTOR2 | chr7 | + | 75024439 | 75024534 | 75021873 | 75021956 | 75024633 | 75031528 | 17 | 5 | 12 | 18 | 6 | 1 | 0 | 0 | 0 | 0 | 2 | 15 | 6 | 16 | 56 | 0 | 0 | 0 | 0 | 0 | 244 | 149 | 1 | 1 | 0.912 | 1 | 1 | 1 | 1 | 1 | 1 | 1 | 1 | 1 | -0.018 |
| SE | 30557 | ENSG00000118503 | TNFAIP3 | chr6 | + | 137875687 | 137875835 | 137874844 | 137875035 | 137875995 | 137876166 | 1648 | 1012 | 1853 | 2091 | 4285 | 0 | 0 | 0 | 0 | 0 | 2533 | 2844 | 5308 | 3418 | 2132 | 0 | 2 | 0 | 0 | 0 | 297 | 149 | 1 | 1 | 1 | 1 | 1 | 1 | 1 | 1 | 0.999 | 1 | 1 | 1 | 0 |
| SE | 30558 | ENSG00000118503 | TNFAIP3 | chr6 | + | 137875995 | 137876166 | 137875687 | 137875835 | 137877075 | 137877256 | 1874 | 1262 | 2242 | 2516 | 3391 | 2 | 0 | 1 | 2 | 1 | 3259 | 3379 | 6590 | 4377 | 2811 | 0 | 3 | 11 | 26 | 0 | 298 | 149 | 1 | 1 | 0.998 | 1 | 0.999 | 0.998 | 0.999 | 1 | 0.998 | 0.997 | 0.988 | 1 | 0.002 |
| SE | 30559 | ENSG00000118503 | TNFAIP3 | chr6 | + | 137877075 | 137877256 | 137875837 | 137876166 | 137878431 | 137878525 | 2253 | 1593 | 2868 | 3127 | 3504 | 0 | 0 | 1 | 0 | 0 | 3834 | 4016 | 8074 | 5509 | 3324 | 0 | 0 | 0 | 2 | 0 | 298 | 149 | 1 | 1 | 1 | 1 | 0.999 | 1 | 1 | 1 | 1 | 1 | 0.999 | 1 | 0 |
| SE | 30561 | ENSG00000173838 | MARCHF10 | chr17 | - | 62705538 | 62705581 | 62701313 | 62701758 | 62711230 | 62711344 | 0 | 0 | 1 | 4 | 0 | 0 | 0 | 0 | 0 | 5 | 0 | 0 | 1 | 0 | 0 | 0 | 0 | 0 | 0 | 0 | 192 | 149 | 1 | 1 | NA | NA | 1 | 1 | 0 | NA | NA | 1 | NA | NA | -0.333 |
| SE | 30566 | ENSG00000196396 | PTPN1 | chr20 | + | 50561362 | 50561453 | 50510382 | 50510590 | 50564968 | 50565069 | 711 | 528 | 654 | 822 | 792 | 0 | 0 | 2 | 1 | 0 | 1809 | 1307 | 1245 | 1147 | 1639 | 1 | 0 | 0 | 0 | 0 | 240 | 149 | 1 | 1 | 1 | 1 | 0.995 | 0.998 | 1 | 0.999 | 1 | 1 | 1 | 1 | -0.001 |
| SE | 30567 | ENSG00000196396 | PTPN1 | chr20 | + | 50568379 | 50568478 | 50564968 | 50565069 | 50574516 | 50574654 | 826 | 679 | 802 | 972 | 1143 | 0 | 0 | 3 | 0 | 2 | 2153 | 1549 | 1453 | 1472 | 2211 | 6 | 0 | 0 | 7 | 3 | 248 | 149 | 1 | 1 | 1 | 1 | 0.994 | 1 | 0.997 | 0.995 | 1 | 1 | 0.992 | 0.998 | 0.001 |
| SE | 30569 | ENSG00000196396 | PTPN1 | chr20 | + | 50579702 | 50579926 | 50579167 | 50579329 | 50582691 | 50584760 | 356 | 240 | 372 | 360 | 840 | 0 | 0 | 0 | 0 | 2 | 762 | 583 | 561 | 541 | 559 | 0 | 0 | 2 | 0 | 0 | 298 | 149 | 1 | 1 | 1 | 1 | 1 | 1 | 0.995 | 1 | 1 | 0.993 | 1 | 1 | 0 |
| SE | 30570 | ENSG00000196396 | PTPN1 | chr20 | + | 50581264 | 50581460 | 50579702 | 50579926 | 50582691 | 50584760 | 190 | 57 | 171 | 221 | 211 | 0 | 0 | 0 | 0 | 2 | 244 | 265 | 217 | 211 | 378 | 0 | 0 | 0 | 0 | 0 | 298 | 149 | 1 | 1 | 1 | 1 | 1 | 1 | 0.981 | 1 | 1 | 1 | 1 | 1 | -0.004 |
| SE | 30571 | ENSG00000158195 | WASF2 | chr1 | - | 27409691 | 27410206 | 27404229 | 27408346 | 27412571 | 27412727 | 1234 | 733 | 1082 | 1153 | 1458 | 0 | 0 | 0 | 0 | 1 | 1776 | 1285 | 975 | 1275 | 2331 | 0 | 0 | 0 | 4 | 0 | 298 | 149 | 1 | 1 | 1 | 1 | 1 | 1 | 0.999 | 1 | 1 | 1 | 0.994 | 1 | 0.001 |
| SE | 30572 | ENSG00000158195 | WASF2 | chr1 | - | 27412571 | 27412727 | 27409691 | 27410206 | 27414832 | 27414963 | 1463 | 950 | 1210 | 1387 | 2526 | 0 | 0 | 1 | 0 | 0 | 2287 | 1540 | 1115 | 1726 | 2695 | 14 | 0 | 0 | 2 | 0 | 298 | 149 | 1 | 1 | 1 | 1 | 0.998 | 1 | 1 | 0.988 | 1 | 1 | 0.998 | 1 | 0.002 |
| SE | 30573 | ENSG00000158195 | WASF2 | chr1 | - | 27414832 | 27414963 | 27409691 | 27410206 | 27415984 | 27416102 | 398 | 336 | 309 | 394 | 408 | 0 | 0 | 0 | 0 | 0 | 1120 | 501 | 341 | 644 | 1306 | 0 | 0 | 1 | 0 | 0 | 280 | 149 | 1 | 1 | 1 | 1 | 1 | 1 | 1 | 1 | 1 | 0.995 | 1 | 1 | 0.001 |
| SE | 30574 | ENSG00000158195 | WASF2 | chr1 | - | 27415984 | 27416102 | 27414832 | 27414963 | 27418268 | 27418422 | 856 | 697 | 667 | 878 | 814 | 0 | 0 | 0 | 0 | 0 | 2451 | 1089 | 695 | 1388 | 2661 | 12 | 0 | 0 | 4 | 0 | 267 | 149 | 1 | 1 | 1 | 1 | 1 | 1 | 1 | 0.991 | 1 | 1 | 0.995 | 1 | 0.003 |
| SE | 30575 | ENSG00000288695 | NA | chr4 | + | 55361116 | 55361250 | 55359345 | 55359488 | 55364073 | 55364271 | 0 | 0 | 1 | 0 | 0 | 38 | 9 | 24 | 32 | 14 | 0 | 0 | 2 | 0 | 0 | 19 | 13 | 15 | 7 | 53 | 283 | 149 | 1 | 1 | 0 | 0 | 0.021 | 0 | 0 | 0 | 0 | 0.066 | 0 | 0 | -0.009 |
| SE | 30576 | ENSG00000288695 | NA | chr4 | + | 55364073 | 55364271 | 55359345 | 55359488 | 55367587 | 55367722 | 93 | 30 | 50 | 73 | 58 | 1 | 0 | 1 | 2 | 0 | 67 | 40 | 35 | 21 | 145 | 0 | 0 | 0 | 0 | 0 | 298 | 149 | 1 | 1 | 0.979 | 1 | 0.962 | 0.948 | 1 | 1 | 1 | 1 | 1 | 1 | -0.022 |
| SE | 30577 | ENSG00000288695 | NA | chr4 | + | 55364073 | 55364271 | 55361116 | 55361250 | 55369831 | 55370534 | 4 | 2 | 2 | 1 | 0 | 0 | 0 | 0 | 0 | 0 | 6 | 0 | 0 | 0 | 0 | 0 | 0 | 2 | 0 | 0 | 298 | 149 | 1 | 1 | 1 | 1 | 1 | 1 | NA | 1 | NA | 0 | NA | NA | 0.5 |
| SE | 30583 | ENSG00000166510 | CCDC68 | chr18 | - | 54937956 | 54938097 | 54936832 | 54936958 | 54940996 | 54941083 | 0 | 5 | 0 | 0 | 2 | 0 | 0 | 3 | 0 | 0 | 0 | 4 | 0 | 0 | 0 | 0 | 0 | 0 | 0 | 0 | 290 | 149 | 1 | 1 | NA | 1 | 0 | NA | 1 | NA | 1 | NA | NA | NA | -0.333 |
| SE | 30584 | ENSG00000152784 | PRDM8 | chr4 | + | 80184649 | 80184753 | 80184301 | 80184351 | 80191471 | 80191569 | 1 | 0 | 2 | 0 | 0 | 4 | 0 | 2 | 0 | 0 | 0 | 0 | 0 | 0 | 6 | 0 | 4 | 0 | 0 | 0 | 253 | 149 | 1 | 1 | 0.128 | NA | 0.371 | NA | NA | NA | 0 | NA | NA | 1 | -0.251 |
| SE | 30585 | ENSG00000152784 | PRDM8 | chr4 | + | 80185364 | 80185518 | 80184649 | 80184753 | 80191471 | 80191569 | 1 | 1 | 1 | 1 | 0 | 0 | 0 | 2 | 0 | 0 | 4 | 0 | 0 | 0 | 0 | 0 | 0 | 0 | 0 | 6 | 298 | 149 | 1 | 1 | 1 | 1 | 0.2 | 1 | NA | 1 | NA | NA | NA | 0 | 0.3 |
| SE | 30586 | ENSG00000152784 | PRDM8 | chr4 | + | 80191471 | 80191569 | 80183878 | 80184055 | 80194112 | 80194239 | 6 | 7 | 2 | 5 | 4 | 0 | 0 | 1 | 0 | 0 | 8 | 1 | 7 | 0 | 11 | 0 | 1 | 0 | 0 | 0 | 247 | 149 | 1 | 1 | 1 | 1 | 0.547 | 1 | 1 | 1 | 0.376 | 1 | NA | 1 | 0.065 |
| SE | 30587 | ENSG00000152784 | PRDM8 | chr4 | + | 80191471 | 80191569 | 80183878 | 80184055 | 80196351 | 80196497 | 2 | 0 | 0 | 2 | 7 | 0 | 0 | 0 | 0 | 6 | 4 | 2 | 1 | 5 | 3 | 3 | 5 | 0 | 0 | 0 | 247 | 149 | 1 | 1 | 1 | NA | NA | 1 | 0.413 | 0.446 | 0.194 | 1 | 1 | 1 | 0.076 |
| SE | 30588 | ENSG00000152784 | PRDM8 | chr4 | + | 80191471 | 80191569 | 80184301 | 80184351 | 80194112 | 80194239 | 7 | 5 | 4 | 5 | 2 | 0 | 0 | 0 | 1 | 0 | 8 | 5 | 7 | 0 | 11 | 0 | 3 | 0 | 0 | 0 | 247 | 149 | 1 | 1 | 1 | 1 | 1 | 0.751 | 1 | 1 | 0.501 | 1 | NA | 1 | 0.075 |
| SE | 30589 | ENSG00000152784 | PRDM8 | chr4 | + | 80191471 | 80191569 | 80185269 | 80185518 | 80194112 | 80194239 | 3 | 5 | 3 | 6 | 2 | 1 | 0 | 0 | 0 | 0 | 12 | 1 | 7 | 0 | 11 | 0 | 0 | 0 | 0 | 0 | 247 | 149 | 1 | 1 | 0.644 | 1 | 1 | 1 | 1 | 1 | 1 | 1 | NA | 1 | -0.071 |
| SE | 30590 | ENSG00000152784 | PRDM8 | chr4 | + | 80194112 | 80194239 | 80183878 | 80184055 | 80196351 | 80196598 | 9 | 4 | 3 | 4 | 3 | 0 | 0 | 0 | 0 | 6 | 0 | 2 | 2 | 0 | 0 | 3 | 5 | 0 | 0 | 0 | 276 | 149 | 1 | 1 | 1 | 1 | 1 | 1 | 0.213 | 0 | 0.178 | 1 | NA | NA | 0.45 |
| SE | 30595 | ENSG00000188158 | NHS | chrX | + | 17719343 | 17719406 | 17692334 | 17692468 | 17721440 | 17721633 | 2 | 1 | 3 | 2 | 3 | 0 | 0 | 0 | 0 | 0 | 4 | 0 | 1 | 4 | 0 | 0 | 0 | 2 | 0 | 0 | 212 | 149 | 1 | 1 | 1 | 1 | 1 | 1 | 1 | 1 | NA | 0.26 | 1 | NA | 0.247 |
| SE | 30600 | ENSG00000165533 | TTC8 | chr14 | + | 88841424 | 88841514 | 88841036 | 88841196 | 88843805 | 88843850 | 0 | 2 | 3 | 3 | 0 | 0 | 0 | 0 | 0 | 2 | 0 | 0 | 2 | 5 | 6 | 0 | 0 | 0 | 0 | 0 | 239 | 149 | 1 | 1 | NA | 1 | 1 | 1 | 0 | NA | NA | 1 | 1 | 1 | -0.25 |
| SE | 30605 | ENSG00000165533 | TTC8 | chr14 | + | 88875025 | 88875109 | 88872329 | 88872452 | 88877293 | 88877951 | 12 | 12 | 12 | 6 | 6 | 1 | 0 | 1 | 4 | 0 | 0 | 7 | 15 | 0 | 0 | 0 | 3 | 0 | 3 | 0 | 233 | 149 | 1 | 1 | 0.885 | 1 | 0.885 | 0.49 | 1 | NA | 0.599 | 1 | 0 | NA | 0.319 |
| SE | 30611 | ENSG00000170502 | NUDT9 | chr4 | + | 87454370 | 87454455 | 87451588 | 87451735 | 87457842 | 87457918 | 170 | 141 | 129 | 117 | 163 | 0 | 2 | 0 | 0 | 0 | 141 | 76 | 59 | 98 | 30 | 0 | 0 | 0 | 0 | 0 | 234 | 149 | 1 | 1 | 1 | 0.978 | 1 | 1 | 1 | 1 | 1 | 1 | 1 | 1 | -0.004 |
| SE | 30614 | ENSG00000131042 | LILRB2 | chr19 | - | 54276263 | 54276301 | 54275950 | 54276003 | 54276806 | 54276929 | 138 | 103 | 49 | 132 | 71 | 0 | 0 | 0 | 0 | 0 | 193 | 173 | 108 | 210 | 272 | 0 | 0 | 0 | 0 | 2 | 187 | 149 | 1 | 1 | 1 | 1 | 1 | 1 | 1 | 1 | 1 | 1 | 1 | 0.991 | 0.002 |
| SE | 30616 | ENSG00000131042 | LILRB2 | chr19 | - | 54276806 | 54276929 | 54276380 | 54276456 | 54277549 | 54277597 | 294 | 199 | 124 | 331 | 169 | 0 | 1 | 0 | 0 | 0 | 650 | 503 | 216 | 402 | 684 | 0 | 1 | 0 | 0 | 0 | 272 | 149 | 1 | 1 | 1 | 0.991 | 1 | 1 | 1 | 1 | 0.996 | 1 | 1 | 1 | -0.001 |
| SE | 30624 | ENSG00000168152 | THAP9 | chr4 | + | 82910696 | 82910811 | 82907784 | 82907935 | 82916943 | 82917040 | 1 | 1 | 0 | 2 | 3 | 7 | 4 | 5 | 9 | 28 | 4 | 6 | 0 | 0 | 0 | 0 | 0 | 2 | 0 | 0 | 264 | 149 | 1 | 1 | 0.075 | 0.124 | 0 | 0.111 | 0.057 | 1 | 1 | 0 | NA | NA | -0.593 |
| SE | 30630 | ENSG00000189308 | LIN54 | chr4 | - | 82939888 | 82939962 | 82939538 | 82939736 | 82946257 | 82946474 | 91 | 53 | 63 | 73 | 108 | 0 | 0 | 0 | 0 | 0 | 48 | 61 | 58 | 55 | 11 | 0 | 3 | 0 | 0 | 0 | 223 | 149 | 1 | 1 | 1 | 1 | 1 | 1 | 1 | 1 | 0.931 | 1 | 1 | 1 | 0.014 |
| SE | 30631 | ENSG00000189308 | LIN54 | chr4 | - | 82946257 | 82946474 | 82939888 | 82939962 | 82970326 | 82970465 | 101 | 61 | 68 | 94 | 134 | 0 | 0 | 2 | 2 | 0 | 40 | 68 | 52 | 61 | 17 | 0 | 0 | 0 | 0 | 0 | 298 | 149 | 1 | 1 | 1 | 1 | 0.944 | 0.959 | 1 | 1 | 1 | 1 | 1 | 1 | -0.019 |
| SE | 30636 | ENSG00000189308 | LIN54 | chr4 | - | 82978882 | 82979006 | 82970326 | 82970469 | 82984160 | 82984876 | 73 | 86 | 100 | 84 | 118 | 0 | 0 | 0 | 3 | 0 | 86 | 97 | 173 | 96 | 69 | 0 | 0 | 0 | 0 | 0 | 273 | 149 | 1 | 1 | 1 | 1 | 1 | 0.939 | 1 | 1 | 1 | 1 | 1 | 1 | -0.012 |
| SE | 30642 | ENSG00000101844 | ATG4A | chrX | + | 108138112 | 108138191 | 108137803 | 108137991 | 108150151 | 108150175 | 63 | 38 | 52 | 53 | 98 | 0 | 0 | 0 | 2 | 0 | 66 | 70 | 39 | 45 | 38 | 0 | 0 | 0 | 0 | 0 | 228 | 149 | 1 | 1 | 1 | 1 | 1 | 0.945 | 1 | 1 | 1 | 1 | 1 | 1 | -0.011 |
| SE | 30643 | ENSG00000109272 | PF4V1 | chr4 | + | 73853782 | 73853909 | 73853295 | 73853462 | 73854034 | 73854483 | 94 | 112 | 89 | 702 | 73 | 0 | 0 | 0 | 0 | 0 | 208 | 1497 | 255 | 350 | 6745 | 0 | 0 | 0 | 0 | 1 | 276 | 149 | 1 | 1 | 1 | 1 | 1 | 1 | 1 | 1 | 1 | 1 | 1 | 1 | 0 |
| SE | 30644 | ENSG00000042062 | RIPOR3 | chr20 | - | 50610852 | 50610906 | 50609572 | 50609722 | 50611180 | 50611204 | 1 | 2 | 0 | 1 | 6 | 0 | 0 | 0 | 1 | 0 | 13 | 0 | 0 | 0 | 19 | 0 | 0 | 0 | 0 | 0 | 203 | 149 | 1 | 1 | 1 | 1 | NA | 0.423 | 1 | 1 | NA | NA | NA | 1 | -0.144 |
| SE | 30645 | ENSG00000145734 | BDP1 | chr5 | + | 71461816 | 71461926 | 71458578 | 71458855 | 71464057 | 71464117 | 122 | 139 | 125 | 117 | 88 | 1 | 0 | 1 | 2 | 0 | 44 | 69 | 90 | 98 | 47 | 0 | 0 | 0 | 0 | 0 | 259 | 149 | 1 | 1 | 0.986 | 1 | 0.986 | 0.971 | 1 | 1 | 1 | 1 | 1 | 1 | -0.011 |
| SE | 30646 | ENSG00000145734 | BDP1 | chr5 | + | 71470394 | 71470489 | 71467353 | 71467487 | 71483841 | 71483896 | 179 | 202 | 154 | 161 | 194 | 0 | 0 | 0 | 0 | 0 | 55 | 88 | 81 | 69 | 16 | 4 | 0 | 0 | 0 | 0 | 244 | 149 | 1 | 1 | 1 | 1 | 1 | 1 | 1 | 0.894 | 1 | 1 | 1 | 1 | 0.021 |
| SE | 30647 | ENSG00000145734 | BDP1 | chr5 | + | 71490983 | 71491131 | 71489403 | 71489682 | 71495249 | 71495408 | 247 | 183 | 203 | 218 | 235 | 0 | 0 | 2 | 3 | 0 | 22 | 75 | 62 | 17 | 13 | 0 | 0 | 0 | 2 | 0 | 297 | 149 | 1 | 1 | 1 | 1 | 0.981 | 0.973 | 1 | 1 | 1 | 1 | 0.81 | 1 | 0.029 |
| SE | 30648 | ENSG00000145734 | BDP1 | chr5 | + | 71497269 | 71497426 | 71494464 | 71495408 | 71501561 | 71501653 | 280 | 294 | 306 | 313 | 349 | 0 | 0 | 3 | 0 | 4 | 23 | 122 | 93 | 45 | 71 | 0 | 0 | 2 | 1 | 0 | 298 | 149 | 1 | 1 | 1 | 1 | 0.981 | 1 | 0.978 | 1 | 1 | 0.959 | 0.957 | 1 | 0.009 |
| SE | 30649 | ENSG00000145734 | BDP1 | chr5 | + | 71501561 | 71501653 | 71494464 | 71495408 | 71502598 | 71502791 | 99 | 91 | 95 | 93 | 127 | 0 | 0 | 0 | 1 | 0 | 0 | 39 | 66 | 22 | 11 | 0 | 0 | 0 | 0 | 0 | 241 | 149 | 1 | 1 | 1 | 1 | 1 | 0.983 | 1 | NA | 1 | 1 | 1 | 1 | -0.003 |
| SE | 30650 | ENSG00000145734 | BDP1 | chr5 | + | 71502598 | 71502791 | 71501561 | 71501653 | 71504620 | 71504751 | 224 | 197 | 193 | 161 | 256 | 0 | 0 | 1 | 0 | 0 | 33 | 96 | 99 | 47 | 14 | 0 | 0 | 0 | 0 | 0 | 298 | 149 | 1 | 1 | 1 | 1 | 0.99 | 1 | 1 | 1 | 1 | 1 | 1 | 1 | -0.002 |
| SE | 30651 | ENSG00000145734 | BDP1 | chr5 | + | 71509464 | 71511151 | 71504620 | 71504751 | 71512240 | 71512428 | 302 | 334 | 270 | 329 | 352 | 0 | 2 | 2 | 0 | 0 | 27 | 73 | 114 | 30 | 2 | 0 | 0 | 1 | 0 | 0 | 298 | 149 | 1 | 1 | 1 | 0.988 | 0.985 | 1 | 1 | 1 | 1 | 0.983 | 1 | 1 | -0.002 |
| SE | 30652 | ENSG00000145734 | BDP1 | chr5 | + | 71513184 | 71513407 | 71512240 | 71512428 | 71514943 | 71515122 | 279 | 297 | 224 | 279 | 320 | 2 | 2 | 0 | 0 | 0 | 26 | 105 | 100 | 39 | 16 | 0 | 0 | 0 | 0 | 0 | 298 | 149 | 1 | 1 | 0.986 | 0.987 | 1 | 1 | 1 | 1 | 1 | 1 | 1 | 1 | -0.005 |
| SE | 30653 | ENSG00000145734 | BDP1 | chr5 | + | 71516060 | 71516271 | 71514943 | 71515122 | 71517321 | 71517452 | 380 | 411 | 367 | 383 | 444 | 1 | 0 | 0 | 0 | 0 | 41 | 174 | 114 | 49 | 11 | 0 | 0 | 0 | 0 | 0 | 298 | 149 | 1 | 1 | 0.995 | 1 | 1 | 1 | 1 | 1 | 1 | 1 | 1 | 1 | -0.001 |
| SE | 30654 | ENSG00000145734 | BDP1 | chr5 | + | 71532307 | 71532427 | 71523938 | 71524323 | 71539041 | 71539078 | 113 | 146 | 120 | 146 | 313 | 0 | 0 | 0 | 3 | 0 | 18 | 44 | 23 | 9 | 6 | 0 | 0 | 0 | 0 | 0 | 269 | 149 | 1 | 1 | 1 | 1 | 1 | 0.964 | 1 | 1 | 1 | 1 | 1 | 1 | -0.007 |
| SE | 30655 | ENSG00000145734 | BDP1 | chr5 | + | 71539556 | 71539649 | 71539041 | 71539078 | 71541453 | 71541682 | 140 | 128 | 157 | 205 | 371 | 0 | 0 | 0 | 3 | 0 | 15 | 30 | 17 | 18 | 8 | 0 | 0 | 0 | 0 | 0 | 242 | 149 | 1 | 1 | 1 | 1 | 1 | 0.977 | 1 | 1 | 1 | 1 | 1 | 1 | -0.005 |
| SE | 30657 | ENSG00000145734 | BDP1 | chr5 | + | 71544356 | 71544507 | 71542104 | 71542265 | 71548681 | 71548745 | 90 | 88 | 65 | 51 | 109 | 0 | 0 | 0 | 1 | 3 | 13 | 62 | 17 | 23 | 2 | 0 | 0 | 0 | 0 | 0 | 298 | 149 | 1 | 1 | 1 | 1 | 1 | 0.962 | 0.948 | 1 | 1 | 1 | 1 | 1 | -0.018 |
| SE | 30658 | ENSG00000145734 | BDP1 | chr5 | + | 71545038 | 71545219 | 71544356 | 71544507 | 71548681 | 71548745 | 199 | 253 | 166 | 186 | 359 | 0 | 2 | 0 | 2 | 0 | 10 | 62 | 27 | 29 | 19 | 0 | 0 | 0 | 0 | 0 | 298 | 149 | 1 | 1 | 1 | 0.984 | 1 | 0.979 | 1 | 1 | 1 | 1 | 1 | 1 | -0.007 |
| SE | 30660 | ENSG00000145734 | BDP1 | chr5 | + | 71562273 | 71562520 | 71559981 | 71560237 | 71564753 | 71567820 | 87 | 89 | 65 | 96 | 133 | 0 | 0 | 0 | 1 | 0 | 7 | 22 | 10 | 5 | 12 | 0 | 0 | 0 | 0 | 0 | 298 | 149 | 1 | 1 | 1 | 1 | 1 | 0.98 | 1 | 1 | 1 | 1 | 1 | 1 | -0.004 |
| SE | 30661 | ENSG00000145734 | BDP1 | chr5 | + | 71562774 | 71562895 | 71562273 | 71562520 | 71564753 | 71567820 | 51 | 47 | 36 | 47 | 79 | 32 | 32 | 15 | 20 | 26 | 6 | 14 | 5 | 0 | 2 | 2 | 11 | 1 | 0 | 3 | 270 | 149 | 1 | 1 | 0.468 | 0.448 | 0.57 | 0.565 | 0.626 | 0.623 | 0.413 | 0.734 | NA | 0.269 | 0.026 |
| SE | 30663 | ENSG00000112893 | MAN2A1 | chr5 | + | 109755328 | 109755456 | 109729341 | 109729513 | 109767534 | 109767708 | 256 | 194 | 320 | 228 | 636 | 0 | 0 | 0 | 1 | 0 | 102 | 200 | 143 | 88 | 53 | 0 | 0 | 0 | 0 | 0 | 277 | 149 | 1 | 1 | 1 | 1 | 1 | 0.992 | 1 | 1 | 1 | 1 | 1 | 1 | -0.002 |
| SE | 30666 | ENSG00000112893 | MAN2A1 | chr5 | + | 109789459 | 109789527 | 109788933 | 109789048 | 109817272 | 109817438 | 227 | 160 | 231 | 194 | 340 | 2 | 0 | 0 | 0 | 0 | 47 | 146 | 109 | 48 | 54 | 0 | 0 | 0 | 0 | 0 | 217 | 149 | 1 | 1 | 0.987 | 1 | 1 | 1 | 1 | 1 | 1 | 1 | 1 | 1 | -0.003 |
| SE | 30667 | ENSG00000112893 | MAN2A1 | chr5 | + | 109817272 | 109817438 | 109789459 | 109789527 | 109819668 | 109819887 | 245 | 181 | 218 | 236 | 326 | 0 | 0 | 0 | 2 | 0 | 68 | 171 | 108 | 87 | 76 | 0 | 0 | 0 | 0 | 0 | 298 | 149 | 1 | 1 | 1 | 1 | 1 | 0.983 | 1 | 1 | 1 | 1 | 1 | 1 | -0.003 |
| SE | 30668 | ENSG00000112893 | MAN2A1 | chr5 | + | 109820219 | 109820342 | 109819668 | 109819887 | 109823722 | 109823837 | 417 | 281 | 341 | 327 | 850 | 0 | 2 | 1 | 1 | 0 | 84 | 202 | 194 | 103 | 135 | 0 | 0 | 0 | 0 | 0 | 272 | 149 | 1 | 1 | 1 | 0.987 | 0.995 | 0.994 | 1 | 1 | 1 | 1 | 1 | 1 | -0.005 |
| SE | 30669 | ENSG00000112893 | MAN2A1 | chr5 | + | 109845864 | 109846006 | 109842327 | 109842461 | 109847656 | 109847790 | 188 | 124 | 221 | 179 | 445 | 0 | 1 | 1 | 0 | 0 | 16 | 79 | 122 | 23 | 28 | 0 | 0 | 1 | 0 | 0 | 291 | 149 | 1 | 1 | 1 | 0.984 | 0.991 | 1 | 1 | 1 | 1 | 0.984 | 1 | 1 | -0.002 |
| SE | 30670 | ENSG00000132466 | ANKRD17 | chr4 | - | 73077354 | 73077533 | 73076939 | 73077104 | 73078641 | 73078890 | 187 | 140 | 141 | 203 | 268 | 1 | 0 | 0 | 0 | 0 | 17 | 114 | 68 | 97 | 0 | 0 | 0 | 0 | 0 | 0 | 298 | 149 | 1 | 1 | 0.989 | 1 | 1 | 1 | 1 | 1 | 1 | 1 | 1 | NA | -0.002 |
| SE | 30673 | ENSG00000132466 | ANKRD17 | chr4 | - | 73094078 | 73094228 | 73090666 | 73092300 | 73097116 | 73097272 | 485 | 435 | 393 | 503 | 775 | 0 | 0 | 0 | 0 | 0 | 225 | 444 | 284 | 290 | 13 | 0 | 0 | 1 | 0 | 0 | 298 | 149 | 1 | 1 | 1 | 1 | 1 | 1 | 1 | 1 | 1 | 0.993 | 1 | 1 | 0.001 |
| SE | 30675 | ENSG00000132466 | ANKRD17 | chr4 | - | 73113791 | 73113908 | 73102375 | 73102547 | 73118687 | 73118850 | 115 | 128 | 109 | 148 | 242 | 1 | 0 | 0 | 0 | 6 | 68 | 117 | 79 | 101 | 4 | 0 | 0 | 1 | 0 | 0 | 266 | 149 | 1 | 1 | 0.985 | 1 | 1 | 1 | 0.958 | 1 | 1 | 0.978 | 1 | 1 | -0.007 |
| SE | 30678 | ENSG00000132466 | ANKRD17 | chr4 | - | 73113791 | 73113908 | 73113083 | 73113311 | 73120161 | 73120337 | 9 | 3 | 3 | 2 | 11 | 0 | 0 | 0 | 0 | 0 | 8 | 0 | 4 | 4 | 0 | 0 | 0 | 0 | 3 | 0 | 266 | 149 | 1 | 1 | 1 | 1 | 1 | 1 | 1 | 1 | NA | 1 | 0.428 | NA | 0.191 |
| SE | 30679 | ENSG00000132466 | ANKRD17 | chr4 | - | 73115820 | 73115916 | 73102375 | 73102547 | 73118687 | 73118850 | 126 | 96 | 94 | 88 | 149 | 1 | 0 | 0 | 0 | 6 | 49 | 84 | 86 | 66 | 0 | 0 | 0 | 1 | 0 | 0 | 245 | 149 | 1 | 1 | 0.987 | 1 | 1 | 1 | 0.938 | 1 | 1 | 0.981 | 1 | NA | -0.01 |
| SE | 30680 | ENSG00000132466 | ANKRD17 | chr4 | - | 73115820 | 73115916 | 73102375 | 73102547 | 73120161 | 73120337 | 0 | 3 | 0 | 2 | 4 | 1 | 0 | 0 | 0 | 0 | 10 | 7 | 5 | 11 | 0 | 0 | 8 | 4 | 0 | 0 | 245 | 149 | 1 | 1 | 0 | 1 | NA | 1 | 1 | 1 | 0.347 | 0.432 | 1 | NA | 0.055 |
| SE | 30681 | ENSG00000132466 | ANKRD17 | chr4 | - | 73115820 | 73115916 | 73113083 | 73113311 | 73120161 | 73120337 | 0 | 3 | 0 | 2 | 2 | 0 | 0 | 0 | 0 | 0 | 0 | 7 | 12 | 9 | 0 | 0 | 0 | 0 | 3 | 0 | 245 | 149 | 1 | 1 | NA | 1 | NA | 1 | 1 | NA | 1 | 1 | 0.646 | NA | 0.118 |
| SE | 30682 | ENSG00000132466 | ANKRD17 | chr4 | - | 73115820 | 73115916 | 73113791 | 73113908 | 73118687 | 73118850 | 430 | 378 | 341 | 361 | 569 | 0 | 0 | 0 | 1 | 0 | 148 | 285 | 255 | 287 | 22 | 5 | 1 | 0 | 0 | 0 | 245 | 149 | 1 | 1 | 1 | 1 | 1 | 0.995 | 1 | 0.947 | 0.994 | 1 | 1 | 1 | 0.011 |
| SE | 30687 | ENSG00000132466 | ANKRD17 | chr4 | - | 73118687 | 73118850 | 73115820 | 73115916 | 73120880 | 73121094 | 203 | 166 | 168 | 190 | 241 | 3 | 0 | 0 | 1 | 0 | 65 | 133 | 137 | 136 | 6 | 0 | 3 | 0 | 2 | 0 | 298 | 149 | 1 | 1 | 0.971 | 1 | 1 | 0.99 | 1 | 1 | 0.957 | 1 | 0.971 | 1 | 0.007 |
| SE | 30688 | ENSG00000132466 | ANKRD17 | chr4 | - | 73120161 | 73120337 | 73115820 | 73115916 | 73120880 | 73121094 | 74 | 72 | 83 | 87 | 83 | 3 | 0 | 0 | 1 | 0 | 27 | 109 | 93 | 96 | 15 | 0 | 3 | 0 | 2 | 0 | 298 | 149 | 1 | 1 | 0.925 | 1 | 1 | 0.978 | 1 | 1 | 0.948 | 1 | 0.96 | 1 | -0.001 |
| SE | 30689 | ENSG00000132466 | ANKRD17 | chr4 | - | 73120161 | 73120337 | 73118687 | 73118850 | 73120880 | 73121094 | 150 | 128 | 179 | 164 | 154 | 0 | 0 | 0 | 0 | 0 | 61 | 162 | 128 | 177 | 24 | 0 | 0 | 0 | 4 | 0 | 298 | 149 | 1 | 1 | 1 | 1 | 1 | 1 | 1 | 1 | 1 | 1 | 0.957 | 1 | 0.009 |
| SE | 30690 | ENSG00000132466 | ANKRD17 | chr4 | - | 73120880 | 73121094 | 73120161 | 73120337 | 73124912 | 73125058 | 71 | 69 | 83 | 80 | 81 | 0 | 0 | 0 | 1 | 0 | 22 | 98 | 83 | 85 | 15 | 0 | 0 | 0 | 0 | 0 | 298 | 149 | 1 | 1 | 1 | 1 | 1 | 0.976 | 1 | 1 | 1 | 1 | 1 | 1 | -0.005 |
| SE | 30691 | ENSG00000132466 | ANKRD17 | chr4 | - | 73121616 | 73121759 | 73120880 | 73121094 | 73124912 | 73125058 | 295 | 253 | 319 | 386 | 317 | 0 | 0 | 0 | 0 | 0 | 108 | 270 | 314 | 282 | 94 | 0 | 0 | 0 | 1 | 0 | 292 | 149 | 1 | 1 | 1 | 1 | 1 | 1 | 1 | 1 | 1 | 1 | 0.993 | 1 | 0.001 |
| SE | 30692 | ENSG00000132466 | ANKRD17 | chr4 | - | 73125200 | 73125312 | 73124912 | 73125058 | 73135116 | 73135262 | 191 | 155 | 194 | 223 | 164 | 0 | 0 | 0 | 0 | 0 | 176 | 165 | 180 | 189 | 71 | 7 | 0 | 0 | 0 | 0 | 261 | 149 | 1 | 1 | 1 | 1 | 1 | 1 | 1 | 0.935 | 1 | 1 | 1 | 1 | 0.013 |
| SE | 30695 | ENSG00000132466 | ANKRD17 | chr4 | - | 73144744 | 73144832 | 73142639 | 73142767 | 73146763 | 73146873 | 254 | 168 | 201 | 264 | 367 | 0 | 0 | 0 | 0 | 0 | 36 | 213 | 203 | 126 | 70 | 0 | 0 | 0 | 2 | 0 | 237 | 149 | 1 | 1 | 1 | 1 | 1 | 1 | 1 | 1 | 1 | 1 | 0.975 | 1 | 0.005 |
| SE | 30697 | ENSG00000132466 | ANKRD17 | chr4 | - | 73161191 | 73161348 | 73156018 | 73156166 | 73177379 | 73177533 | 515 | 604 | 615 | 711 | 870 | 0 | 0 | 1 | 4 | 0 | 604 | 828 | 800 | 713 | 368 | 0 | 4 | 0 | 0 | 0 | 298 | 149 | 1 | 1 | 1 | 1 | 0.997 | 0.989 | 1 | 1 | 0.99 | 1 | 1 | 1 | -0.001 |
| SE | 30699 | ENSG00000132466 | ANKRD17 | chr4 | - | 73177379 | 73177533 | 73161191 | 73161348 | 73258275 | 73258322 | 340 | 384 | 438 | 452 | 538 | 1 | 3 | 1 | 0 | 0 | 331 | 541 | 425 | 435 | 222 | 0 | 0 | 2 | 0 | 0 | 298 | 149 | 1 | 1 | 0.994 | 0.985 | 0.995 | 1 | 1 | 1 | 1 | 0.991 | 1 | 1 | -0.003 |
| SE | 30702 | ENSG00000196954 | CASP4 | chr11 | - | 104944747 | 104944851 | 104942865 | 104942973 | 104947082 | 104947192 | 750 | 431 | 480 | 582 | 574 | 0 | 3 | 0 | 0 | 0 | 693 | 687 | 472 | 565 | 498 | 0 | 0 | 3 | 0 | 0 | 253 | 149 | 1 | 1 | 1 | 0.988 | 1 | 1 | 1 | 1 | 1 | 0.989 | 1 | 1 | 0 |
| SE | 30703 | ENSG00000196954 | CASP4 | chr11 | - | 104948532 | 104948676 | 104947082 | 104947192 | 104949542 | 104949777 | 1026 | 555 | 668 | 859 | 901 | 0 | 1 | 0 | 1 | 0 | 1052 | 937 | 636 | 886 | 657 | 0 | 0 | 2 | 0 | 0 | 293 | 149 | 1 | 1 | 1 | 0.996 | 1 | 0.998 | 1 | 1 | 1 | 0.994 | 1 | 1 | 0 |
| SE | 30704 | ENSG00000196954 | CASP4 | chr11 | - | 104950924 | 104951098 | 104949401 | 104949777 | 104951895 | 104952005 | 1098 | 668 | 826 | 979 | 1457 | 0 | 0 | 0 | 2 | 0 | 1356 | 955 | 693 | 1089 | 875 | 0 | 0 | 5 | 0 | 0 | 298 | 149 | 1 | 1 | 1 | 1 | 1 | 0.996 | 1 | 1 | 1 | 0.986 | 1 | 1 | 0.002 |
| SE | 30710 | ENSG00000196954 | CASP4 | chr11 | - | 104954746 | 104955001 | 104949542 | 104949777 | 104968518 | 104968547 | 267 | 142 | 176 | 195 | 429 | 0 | 0 | 0 | 0 | 0 | 232 | 260 | 283 | 303 | 200 | 0 | 0 | 5 | 0 | 0 | 298 | 149 | 1 | 1 | 1 | 1 | 1 | 1 | 1 | 1 | 1 | 0.966 | 1 | 1 | 0.007 |
| SE | 30712 | ENSG00000215158 | NA | chr5 | - | 34175444 | 34175537 | 34164697 | 34169153 | 34175821 | 34175957 | 34 | 25 | 22 | 34 | 38 | 2 | 3 | 2 | 4 | 0 | 0 | 32 | 19 | 36 | 8 | 0 | 0 | 1 | 0 | 8 | 242 | 149 | 1 | 1 | 0.913 | 0.837 | 0.871 | 0.84 | 1 | NA | 1 | 0.921 | 1 | 0.381 | 0.067 |
| SE | 30713 | ENSG00000215158 | NA | chr5 | - | 34179012 | 34179097 | 34175821 | 34175957 | 34182679 | 34182867 | 70 | 18 | 48 | 48 | 32 | 0 | 0 | 0 | 0 | 0 | 9 | 20 | 39 | 22 | 165 | 0 | 2 | 0 | 0 | 0 | 234 | 149 | 1 | 1 | 1 | 1 | 1 | 1 | 1 | 1 | 0.864 | 1 | 1 | 1 | 0.027 |
| SE | 30725 | ENSG00000125375 | DMAC2L | chr14 | + | 50314590 | 50314626 | 50312323 | 50313060 | 50321482 | 50321594 | 2 | 0 | 1 | 1 | 0 | 0 | 0 | 1 | 0 | 0 | 6 | 2 | 0 | 0 | 7 | 0 | 0 | 0 | 0 | 0 | 185 | 149 | 1 | 1 | 1 | NA | 0.446 | 1 | NA | 1 | 1 | NA | NA | 1 | -0.185 |
| SE | 30727 | ENSG00000125375 | DMAC2L | chr14 | + | 50323944 | 50324116 | 50322510 | 50322806 | 50324546 | 50324706 | 0 | 3 | 2 | 3 | 9 | 0 | 0 | 0 | 2 | 0 | 0 | 1 | 0 | 1 | 4 | 0 | 0 | 0 | 0 | 0 | 298 | 149 | 1 | 1 | NA | 1 | 1 | 0.429 | 1 | NA | 1 | NA | 1 | 1 | -0.143 |
| SE | 30728 | ENSG00000125375 | DMAC2L | chr14 | + | 50323944 | 50324116 | 50322510 | 50322806 | 50325608 | 50325645 | 41 | 24 | 34 | 39 | 59 | 0 | 0 | 0 | 0 | 0 | 12 | 26 | 28 | 14 | 71 | 0 | 0 | 2 | 0 | 0 | 298 | 149 | 1 | 1 | 1 | 1 | 1 | 1 | 1 | 1 | 1 | 0.875 | 1 | 1 | 0.025 |
| SE | 30730 | ENSG00000244486 | SCARF2 | chr22 | - | 20431017 | 20431537 | 20430689 | 20430908 | 20431744 | 20431846 | 0 | 0 | 0 | 2 | 0 | 0 | 0 | 2 | 0 | 0 | 0 | 0 | 0 | 2 | 0 | 0 | 0 | 0 | 0 | 6 | 298 | 149 | 1 | 1 | NA | NA | 0 | 1 | NA | NA | NA | NA | 1 | 0 | 0 |
| SE | 30732 | ENSG00000249437 | NAIP | chr5 | - | 70979868 | 70980036 | 70976905 | 70977058 | 70983774 | 70985886 | 1 | 0 | 0 | 0 | 0 | 1 | 4 | 1 | 6 | 0 | 0 | 0 | 1 | 0 | 0 | 0 | 7 | 3 | 5 | 13 | 298 | 149 | 1 | 1 | 0.333 | 0 | 0 | 0 | NA | NA | 0 | 0.143 | 0 | 0 | 0.048 |
| SE | 30733 | ENSG00000249437 | NAIP | chr5 | - | 71011274 | 71011374 | 71003597 | 71003857 | 71011711 | 71011772 | 251 | 229 | 112 | 202 | 249 | 0 | 0 | 0 | 0 | 0 | 431 | 498 | 285 | 486 | 219 | 0 | 2 | 0 | 0 | 0 | 249 | 149 | 1 | 1 | 1 | 1 | 1 | 1 | 1 | 1 | 0.993 | 1 | 1 | 1 | 0.001 |
| SE | 30737 | ENSG00000249437 | NAIP | chr5 | - | 71012347 | 71012918 | 71003597 | 71003857 | 71020658 | 71020710 | 67 | 59 | 39 | 57 | 76 | 0 | 4 | 0 | 0 | 1 | 195 | 205 | 114 | 129 | 44 | 0 | 3 | 0 | 0 | 0 | 298 | 149 | 1 | 1 | 1 | 0.881 | 1 | 1 | 0.974 | 1 | 1 | 0.95 | 1 | 1 | -0.019 |
| SE | 30738 | ENSG00000250220 | NA | chr4 | + | 73269823 | 73269953 | 73259208 | 73259408 | 73302263 | 73302385 | 0 | 0 | 1 | 2 | 0 | 0 | 2 | 0 | 0 | 0 | 0 | 0 | 0 | 3 | 0 | 0 | 0 | 0 | 0 | 0 | 279 | 149 | 1 | 1 | NA | 0 | 1 | 1 | NA | NA | NA | NA | 1 | NA | -0.333 |
| SE | 30739 | ENSG00000250220 | NA | chr4 | + | 73307075 | 73307181 | 73302263 | 73302385 | 73317826 | 73317953 | 0 | 0 | 0 | 0 | 0 | 0 | 0 | 2 | 0 | 17 | 0 | 2 | 0 | 0 | 0 | 0 | 0 | 0 | 0 | 0 | 255 | 149 | 1 | 1 | NA | NA | 0 | NA | 0 | NA | 1 | NA | NA | NA | -1 |
| SE | 30745 | ENSG00000174125 | TLR1 | chr4 | - | 38800856 | 38800948 | 38790847 | 38791231 | 38804753 | 38804787 | 2 | 1 | 3 | 2 | 1 | 0 | 0 | 1 | 0 | 0 | 0 | 0 | 8 | 3 | 0 | 0 | 0 | 0 | 0 | 0 | 241 | 149 | 1 | 1 | 1 | 1 | 0.65 | 1 | 1 | NA | NA | 1 | 1 | NA | -0.07 |
| SE | 30746 | ENSG00000174125 | TLR1 | chr4 | - | 38800856 | 38800948 | 38790847 | 38791231 | 38805111 | 38805193 | 2 | 0 | 3 | 2 | 0 | 0 | 0 | 0 | 0 | 0 | 0 | 7 | 4 | 0 | 0 | 0 | 1 | 0 | 0 | 0 | 241 | 149 | 1 | 1 | 1 | NA | 1 | 1 | NA | NA | 0.812 | 1 | NA | NA | 0.094 |
| SE | 30747 | ENSG00000174125 | TLR1 | chr4 | - | 38800856 | 38800948 | 38796254 | 38798898 | 38804305 | 38804376 | 52 | 10 | 27 | 30 | 83 | 9 | 1 | 7 | 7 | 18 | 0 | 20 | 31 | 13 | 0 | 0 | 3 | 0 | 5 | 0 | 241 | 149 | 1 | 1 | 0.781 | 0.861 | 0.705 | 0.726 | 0.74 | NA | 0.805 | 1 | 0.616 | NA | -0.044 |
| SE | 30748 | ENSG00000174125 | TLR1 | chr4 | - | 38800856 | 38800948 | 38796254 | 38798898 | 38804753 | 38804787 | 25 | 17 | 13 | 18 | 70 | 2 | 30 | 15 | 3 | 25 | 0 | 18 | 16 | 4 | 0 | 0 | 7 | 8 | 1 | 0 | 241 | 149 | 1 | 1 | 0.885 | 0.259 | 0.349 | 0.788 | 0.634 | NA | 0.614 | 0.553 | 0.712 | NA | -0.043 |

| | | | | | | | | | | | | | | | | | | | | | | | |
|---|---|---|---|---|---|---|---|---|---|---|---|---|---|---|---|---|---|---|---|---|---|---|---|
| SE | 30750 | ENSG00000174125 | TLR1 | chr4 | - | 38804305 | 38804382 | 38798560 | 38798898 | 38804753 | 38804787 | 9 | 1 | 7 | 7 | 22 | 2 | 30 | 15 | 3 | 25 | 0 | 3 |
| SE | 30752 | ENSG00000174125 | TLR1 | chr4 | - | 38804305 | 38804382 | 38800471 | 38800948 | 38804753 | 38804787 | 33 | 2 | 24 | 18 | 29 | 9 | 13 | 9 | 8 | 17 | 0 | 4 |
| SE | 30753 | ENSG00000174125 | TLR1 | chr4 | - | 38804305 | 38804382 | 38800471 | 38800948 | 38805111 | 38805533 | 7 | 0 | 13 | 3 | 12 | 0 | 0 | 0 | 0 | 0 | 0 | 2 |
| SE | 30754 | ENSG00000174125 | TLR1 | chr4 | - | 38804753 | 38804787 | 38800856 | 38800948 | 38805111 | 38805193 | 10 | 13 | 8 | 8 | 14 | 0 | 0 | 0 | 0 | 0 | 0 | 0 |
| SE | 30755 | ENSG00000174125 | TLR1 | chr4 | - | 38804753 | 38804787 | 38804305 | 38804382 | 38805111 | 38805193 | 36 | 4 | 26 | 20 | 26 | 0 | 0 | 2 | 0 | 0 | 0 | 5 |
| SE | 30757 | ENSG00000197712 | FAM114A1 | chr4 | + | 38868397 | 38868546 | 38867805 | 38867834 | 38878070 | 38878426 | 0 | 0 | 1 | 0 | 4 | 1 | 0 | 6 | 0 | 0 | 0 | 0 |
| SE | 30759 | ENSG00000197712 | FAM114A1 | chr4 | + | 38891742 | 38891830 | 38878070 | 38878426 | 38905521 | 38905635 | 0 | 0 | 0 | 6 | 14 | 0 | 1 | 0 | 0 | 4 | 15 | 0 |
| SE | 30776 | ENSG00000145331 | TRMT10A | chr4 | - | 99550884 | 99550990 | 99546708 | 99549356 | 99553784 | 99553934 | 32 | 21 | 32 | 17 | 59 | 0 | 0 | 0 | 0 | 10 | 0 | 1 |
| SE | 30778 | ENSG00000137203 | TFAP2A | chr6 | - | 10402491 | 10402610 | 10400447 | 10400589 | 10404507 | 10404594 | 0 | 0 | 1 | 0 | 0 | 0 | 0 | 0 | 0 | 0 | 0 | 0 |
| SE | 30780 | ENSG00000102796 | DHRS12 | chr13 | - | 51769155 | 51769293 | 51767992 | 51768296 | 51773929 | 51774034 | 13 | 19 | 19 | 15 | 10 | 0 | 0 | 1 | 0 | 0 | 0 | 24 |
| SE | 30781 | ENSG00000102796 | DHRS12 | chr13 | - | 51771820 | 51771911 | 51767994 | 51769293 | 51773920 | 51774034 | 5 | 3 | 9 | 7 | 21 | 0 | 0 | 1 | 0 | 2 | 0 | 7 |
| SE | 30783 | ENSG00000102796 | DHRS12 | chr13 | - | 51771820 | 51771911 | 51770783 | 51771495 | 51773929 | 51774034 | 12 | 13 | 14 | 16 | 20 | 0 | 1 | 0 | 0 | 0 | 9 | 10 |
| SE | 30786 | ENSG00000102796 | DHRS12 | chr13 | - | 51790010 | 51790092 | 51773929 | 51774034 | 51791164 | 51791257 | 14 | 17 | 25 | 17 | 52 | 0 | 0 | 0 | 0 | 0 | 25 | 19 |
| SE | 30787 | ENSG00000102796 | DHRS12 | chr13 | - | 51791164 | 51791257 | 51773929 | 51774034 | 51799533 | 51799667 | 6 | 5 | 7 | 11 | 30 | 0 | 0 | 0 | 0 | 0 | 19 | 2 |
| SE | 30795 | ENSG00000064042 | LIMCH1 | chr4 | + | 41680005 | 41680098 | 41676378 | 41676462 | 41680954 | 41681059 | 0 | 0 | 1 | 0 | 0 | 0 | 0 | 0 | 0 | 0 | 2 | 5 |
| SE | 30797 | ENSG00000203778 | FAM229B | chr6 | + | 112097040 | 112097201 | 112087590 | 112087720 | 112099269 | 112099408 | 1 | 1 | 6 | 4 | 12 | 0 | 1 | 0 | 0 | 0 | 0 | 3 |
| SE | 30798 | ENSG00000100412 | ACO2 | chr22 | + | 41469347 | 41469508 | 41469136 | 41469182 | 41507790 | 41508049 | 0 | 0 | 0 | 0 | 0 | 1 | 1 | 4 | 1 | 0 | 0 | 0 |
| SE | 30799 | ENSG00000100412 | ACO2 | chr22 | + | 41477912 | 41478127 | 41469127 | 41469182 | 41499725 | 41499862 | 0 | 0 | 1 | 1 | 0 | 77 | 20 | 54 | 55 | 39 | 0 | 0 |
| SE | 30801 | ENSG00000100412 | ACO2 | chr22 | + | 41499725 | 41499862 | 41469366 | 41469508 | 41507790 | 41508049 | 116 | 59 | 91 | 81 | 145 | 0 | 0 | 0 | 0 | 0 | 103 | 110 |
| SE | 30802 | ENSG00000100412 | ACO2 | chr22 | + | 41502629 | 41502771 | 41499545 | 41499862 | 41507790 | 41508049 | 0 | 0 | 3 | 0 | 0 | 136 | 62 | 108 | 102 | 148 | 0 | 1 |
| SE | 30804 | ENSG00000100412 | ACO2 | chr22 | + | 41507445 | 41507606 | 41499725 | 41499862 | 41507790 | 41508049 | 4 | 3 | 5 | 0 | 0 | 136 | 62 | 108 | 102 | 148 | 0 | 0 |
| SE | 30806 | ENSG00000100412 | ACO2 | chr22 | + | 41515766 | 41515917 | 41515376 | 41515535 | 41517526 | 41517631 | 158 | 77 | 128 | 116 | 96 | 0 | 0 | 0 | 0 | 0 | 129 | 221 |
| SE | 30807 | ENSG00000100412 | ACO2 | chr22 | + | 41515766 | 41515917 | 41515376 | 41515535 | 41517596 | 41517631 | 66 | 22 | 52 | 36 | 28 | 0 | 0 | 0 | 0 | 0 | 57 | 49 |
| SE | 30808 | ENSG00000100412 | ACO2 | chr22 | + | 41515766 | 41515992 | 41515376 | 41515535 | 41517526 | 41517631 | 66 | 22 | 49 | 36 | 28 | 0 | 0 | 0 | 0 | 0 | 57 | 47 |
| SE | 30809 | ENSG00000100412 | ACO2 | chr22 | + | 41517296 | 41517469 | 41515766 | 41515917 | 41517596 | 41517631 | 2 | 0 | 0 | 0 | 0 | 0 | 0 | 3 | 0 | 0 | 0 | 9 |
| SE | 30810 | ENSG00000100412 | ACO2 | chr22 | + | 41517942 | 41518408 | 41517526 | 41517631 | 41520170 | 41520276 | 1 | 0 | 9 | 0 | 0 | 0 | 0 | 0 | 0 | 0 | 0 | 2 |
| SE | 30811 | ENSG00000100412 | ACO2 | chr22 | + | 41517942 | 41518572 | 41517526 | 41517631 | 41520170 | 41520276 | 105 | 56 | 121 | 97 | 99 | 0 | 0 | 0 | 0 | 0 | 189 | 145 |
| SE | 30812 | ENSG00000100412 | ACO2 | chr22 | + | 41517942 | 41518572 | 41517526 | 41517631 | 41522829 | 41522987 | 1 | 1 | 9 | 0 | 1 | 1 | 0 | 0 | 0 | 0 | 0 | 2 |
| SE | 30813 | ENSG00000100412 | ACO2 | chr22 | + | 41518480 | 41518572 | 41517526 | 41517631 | 41520170 | 41520276 | 197 | 125 | 186 | 188 | 203 | 0 | 0 | 0 | 0 | 0 | 340 | 359 |
| SE | 30814 | ENSG00000100412 | ACO2 | chr22 | + | 41518480 | 41518572 | 41517526 | 41517631 | 41522829 | 41522982 | 50 | 26 | 35 | 32 | 40 | 1 | 0 | 0 | 0 | 0 | 38 | 76 |
| SE | 30815 | ENSG00000100412 | ACO2 | chr22 | + | 41518495 | 41518572 | 41517526 | 41517631 | 41520170 | 41520276 | 83 | 46 | 97 | 75 | 86 | 0 | 0 | 0 | 0 | 0 | 154 | 119 |
| SE | 30816 | ENSG00000100412 | ACO2 | chr22 | + | 41518495 | 41518572 | 41517526 | 41517631 | 41522829 | 41522987 | 0 | 1 | 0 | 0 | 1 | 1 | 0 | 0 | 0 | 0 | 0 | 0 |
| SE | 30817 | ENSG00000100412 | ACO2 | chr22 | + | 41520170 | 41520276 | 41516581 | 41518572 | 41522829 | 41522982 | 300 | 187 | 283 | 291 | 361 | 4 | 1 | 0 | 0 | 1 | 556 | 510 |
| SE | 30818 | ENSG00000100412 | ACO2 | chr22 | + | 41520170 | 41520276 | 41517526 | 41517631 | 41522829 | 41522982 | 135 | 86 | 128 | 130 | 191 | 1 | 0 | 0 | 0 | 0 | 252 | 223 |
| SE | 30819 | ENSG00000100412 | ACO2 | chr22 | + | 41522829 | 41522987 | 41520170 | 41520276 | 41523204 | 41523278 | 348 | 195 | 355 | 355 | 378 | 0 | 0 | 1 | 0 | 0 | 635 | 522 |
| SE | 30821 | ENSG00000100412 | ACO2 | chr22 | + | 41527138 | 41527202 | 41526261 | 41526453 | 41527287 | 41527420 | 1 | 0 | 1 | 0 | 0 | 131 | 55 | 114 | 89 | 69 | 0 | 0 |
| SE | 30822 | ENSG00000187621 | TCL6 | chr14 | + | 95654508 | 95654626 | 95652376 | 95652489 | 95662479 | 95662664 | 10 | 2 | 2 | 15 | 14 | 0 | 0 | 1 | 0 | 0 | 0 | 12 |
| SE | 30823 | ENSG00000187621 | TCL6 | chr14 | + | 95662479 | 95662664 | 95654508 | 95654626 | 95663255 | 95663635 | 1 | 0 | 1 | 4 | 1 | 1 | 2 | 1 | 9 | 8 | 0 | 7 |
| SE | 30824 | ENSG00000187621 | TCL6 | chr14 | + | 95662479 | 95662664 | 95654508 | 95654626 | 95669550 | 95669617 | 0 | 0 | 1 | 0 | 1 | 3 | 0 | 0 | 1 | 0 | 0 | 7 |
| SE | 30825 | ENSG00000187621 | TCL6 | chr14 | + | 95663255 | 95663635 | 95654508 | 95654626 | 95669550 | 95669617 | 4 | 2 | 1 | 9 | 8 | 3 | 0 | 0 | 1 | 0 | 0 | 3 |
| SE | 30827 | ENSG00000187621 | TCL6 | chr14 | + | 95668125 | 95669617 | 95663255 | 95665076 | 95670441 | 95670577 | 6 | 5 | 9 | 21 | 5 | 0 | 0 | 0 | 0 | 0 | 0 | 52 |
| SE | 30831 | ENSG00000258315 | C17orf49 | chr17 | + | 7016909 | 7017014 | 7016494 | 7016525 | 7017256 | 7017375 | 34 | 25 | 30 | 14 | 58 | 0 | 1 | 0 | 0 | 2 | 47 | 10 |
| SE | 30853 | ENSG00000109133 | TMEM33 | chr4 | + | 41938601 | 41938696 | 41935128 | 41935529 | 41943746 | 41943814 | 22 | 20 | 22 | 16 | 24 | 0 | 0 | 0 | 0 | 0 | 25 | 23 |
| SE | 30854 | ENSG00000109133 | TMEM33 | chr4 | + | 41938601 | 41938696 | 41935128 | 41935529 | 41944792 | 41944926 | 18 | 22 | 20 | 21 | 19 | 1 | 0 | 0 | 0 | 0 | 25 | 18 |
| SE | 30856 | ENSG00000109133 | TMEM33 | chr4 | + | 41939720 | 41939828 | 41938601 | 41938696 | 41943746 | 41943814 | 6 | 4 | 3 | 5 | 3 | 5 | 4 | 2 | 0 | 9 | 0 | 9 |
| SE | 30858 | ENSG00000109133 | TMEM33 | chr4 | + | 41943746 | 41943814 | 41935128 | 41935529 | 41944792 | 41944926 | 97 | 53 | 46 | 74 | 99 | 1 | 0 | 0 | 0 | 0 | 10 | 62 |
| SE | 30860 | ENSG00000164934 | DCAF13 | chr8 | + | 103420263 | 103420463 | 103414713 | 103415516 | 103420974 | 103421082 | 95 | 57 | 118 | 94 | 131 | 0 | 0 | 1 | 1 | 0 | 3 | 48 |
| SE | 30861 | ENSG00000164934 | DCAF13 | chr8 | + | 103420263 | 103420463 | 103414713 | 103415516 | 103426055 | 103426145 | 57 | 28 | 52 | 40 | 37 | 2 | 0 | 2 | 0 | 0 | 3 | 30 |
| SE | 30862 | ENSG00000164934 | DCAF13 | chr8 | + | 103420974 | 103421082 | 103414713 | 103415516 | 103426055 | 103426145 | 116 | 83 | 120 | 123 | 169 | 2 | 0 | 2 | 0 | 0 | 14 | 75 |
| SE | 30864 | ENSG00000164934 | DCAF13 | chr8 | + | 103430611 | 103430689 | 103427096 | 103427252 | 103432658 | 103432741 | 134 | 94 | 138 | 127 | 205 | 0 | 0 | 3 | 0 | 0 | 20 | 54 |
| SE | 30865 | ENSG00000134049 | IER3IP1 | chr18 | - | 47157435 | 47157537 | 47152833 | 47156232 | 47176186 | 47176364 | 147 | 97 | 152 | 114 | 179 | 0 | 0 | 1 | 1 | 0 | 97 | 95 |
| SE | 30866 | ENSG00000134852 | CLOCK | chr4 | - | 55442431 | 55442634 | 55438281 | 55438407 | 55443686 | 55443896 | 40 | 44 | 54 | 82 | 108 | 2 | 0 | 0 | 0 | 0 | 24 | 37 |
| SE | 30868 | ENSG00000134852 | CLOCK | chr4 | - | 55453053 | 55453129 | 55450090 | 55450232 | 55455896 | 55456003 | 22 | 10 | 6 | 10 | 39 | 0 | 0 | 0 | 0 | 0 | 0 | 5 |
| SE | 30870 | ENSG00000134852 | CLOCK | chr4 | - | 55453676 | 55453824 | 55453053 | 55453129 | 55456217 | 55456300 | 42 | 48 | 47 | 54 | 138 | 0 | 0 | 1 | 0 | 0 | 5 | 14 |
| SE | 30871 | ENSG00000134852 | CLOCK | chr4 | - | 55455896 | 55456003 | 55453053 | 55453129 | 55456217 | 55456300 | 46 | 22 | 31 | 61 | 65 | 0 | 0 | 1 | 0 | 0 | 0 | 24 |
| SE | 30872 | ENSG00000134852 | CLOCK | chr4 | - | 55455896 | 55456003 | 55453676 | 55453824 | 55456217 | 55456300 | 120 | 61 | 81 | 125 | 167 | 0 | 0 | 0 | 0 | 1 | 0 | 47 |
| SE | 30873 | ENSG00000134852 | CLOCK | chr4 | - | 55475962 | 55476054 | 55470716 | 55470806 | 55478814 | 55478963 | 108 | 127 | 71 | 103 | 186 | 1 | 2 | 0 | 0 | 2 | 0 | 71 |
| SE | 30874 | ENSG00000134852 | CLOCK | chr4 | - | 55478814 | 55478963 | 55475962 | 55476054 | 55479639 | 55479699 | 124 | 126 | 77 | 108 | 152 | 2 | 0 | 1 | 2 | 0 | 12 | 60 |
| SE | 30875 | ENSG00000134852 | CLOCK | chr4 | - | 55486392 | 55486549 | 55482738 | 55482828 | 55489373 | 55489465 | 29 | 44 | 25 | 16 | 38 | 55 | 23 | 28 | 41 | 60 | 18 | 35 |
| SE | 30877 | ENSG00000134852 | CLOCK | chr4 | - | 55509911 | 55510065 | 55489373 | 55489465 | 55546781 | 55546909 | 46 | 39 | 22 | 24 | 44 | 4 | 0 | 1 | 1 | 1 | 41 | 21 |
| SE | 30879 | ENSG00000003056 | M6PR | chr12 | - | 8942415 | 8942542 | 8940362 | 8940869 | 8943404 | 8943420 | 596 | 332 | 509 | 549 | 468 | 0 | 0 | 0 | 0 | 0 | 961 | 537 |
| SE | 30882 | ENSG00000003056 | M6PR | chr12 | - | 8946715 | 8946824 | 8946228 | 8946387 | 8949487 | 8949645 | 7 | 1 | 0 | 0 | 5 | 100 | 2 | 4 | 4 | 20 | 0 | 17 |
| SE | 30909 | ENSG00000205726 | ITSN1 | chr21 | + | 33772060 | 33772323 | 33765874 | 33766012 | 33774728 | 33774878 | 11 | 18 | 5 | 11 | 9 | 0 | 0 | 0 | 0 | 0 | 1 | 29 |
| SE | 30912 | ENSG00000205726 | ITSN1 | chr21 | + | 33810974 | 33811222 | 33799807 | 33799929 | 33813912 | 33814072 | 10 | 6 | 12 | 12 | 25 | 0 | 1 | 0 | 0 | 0 | 1 | 8 |
| SE | 30915 | ENSG00000205726 | ITSN1 | chr21 | + | 33826817 | 33826863 | 33819240 | 33819323 | 33829623 | 33829686 | 3 | 1 | 4 | 7 | 0 | 8 | 4 | 2 | 8 | 2 | 0 | 2 |
| SE | 30916 | ENSG00000205726 | ITSN1 | chr21 | + | 33828967 | 33829050 | 33826817 | 33826863 | 33829623 | 33829745 | 0 | 0 | 2 | 0 | 0 | 4 | 1 | 7 | 12 | 2 | 0 | 0 |
| SE | 30918 | ENSG00000205726 | ITSN1 | chr21 | + | 33858685 | 33858792 | 33856735 | 33856857 | 33865150 | 33865334 | 2 | 1 | 0 | 0 | 0 | 0 | 0 | 0 | 1 | 0 | 0 | 2 |
| SE | 30924 | ENSG00000263327 | TAPT1-AS1 | chr4 | + | 16288883 | 16289045 | 16238770 | 16238871 | 16320064 | 16320140 | 0 | 0 | 4 | 0 | 10 | 0 | 1 | 0 | 0 | 0 | 0 | 0 |
| SE | 30929 | ENSG00000288539 | NA | chr5 | - | 52872284 | 52872429 | 52868672 | 52868799 | 52872694 | 52872925 | 0 | 0 | 0 | 0 | 5 | 1 | 0 | 0 | 0 | 0 | 0 | 0 |
| SE | 30930 | ENSG00000100422 | CERK | chr22 | - | 46711085 | 46711149 | 46707842 | 46707988 | 46712167 | 46712293 | 104 | 68 | 141 | 111 | 84 | 0 | 0 | 0 | 0 | 0 | 76 | 140 |
| SE | 30931 | ENSG00000017260 | ATP2C1 | chr3 | + | 130854185 | 130854269 | 130850782 | 130850928 | 130855880 | 130856223 | 0 | 1 | 0 | 0 | 0 | 0 | 1 | 1 | 5 | 0 | 0 | 1 |
| SE | 30932 | ENSG00000017260 | ATP2C1 | chr3 | + | 130894589 | 130894775 | 130894014 | 130894337 | 130930415 | 130930526 | 54 | 27 | 44 | 41 | 60 | 2 | 7 | 1 | 1 | 1 | 0 | 26 |
| SE | 30933 | ENSG00000017260 | ATP2C1 | chr3 | + | 130894707 | 130894775 | 130893990 | 130894337 | 130930415 | 130930526 | 47 | 20 | 37 | 33 | 55 | 2 | 7 | 1 | 1 | 1 | 0 | 30 |
| SE | 30934 | ENSG00000017260 | ATP2C1 | chr3 | + | 130894730 | 130894775 | 130894181 | 130894337 | 130930415 | 130930526 | 30 | 12 | 19 | 15 | 51 | 2 | 7 | 1 | 1 | 1 | 0 | 16 |
| SE | 30936 | ENSG00000017260 | ATP2C1 | chr3 | + | 130937427 | 130937463 | 130934621 | 130934711 | 130940629 | 130940691 | 164 | 160 | 158 | 196 | 257 | 1 | 0 | 0 | 0 | 0 | 190 | 131 |
| SE | 30938 | ENSG00000017260 | ATP2C1 | chr3 | + | 130969291 | 130969396 | 130967329 | 130967419 | 130975331 | 130975488 | 138 | 90 | 132 | 174 | 199 | 0 | 0 | 0 | 1 | 0 | 94 | 107 |
| SE | 30939 | ENSG00000017260 | ATP2C1 | chr3 | + | 130969291 | 130969396 | 130967329 | 130967419 | 130980581 | 130980679 | 52 | 41 | 45 | 71 | 93 | 0 | 0 | 0 | 0 | 0 | 57 | 32 |
| SE | 30940 | ENSG00000017260 | ATP2C1 | chr3 | + | 130975331 | 130975488 | 130967329 | 130967419 | 130980581 | 130980679 | 14 | 19 | 5 | 15 | 5 | 0 | 0 | 0 | 0 | 0 | 15 | 12 |
| SE | 30941 | ENSG00000017260 | ATP2C1 | chr3 | + | 130975331 | 130975488 | 130969291 | 130969396 | 130980581 | 130980679 | 101 | 71 | 94 | 122 | 111 | 1 | 0 | 0 | 0 | 0 | 52 | 86 |
| SE | 30948 | ENSG00000017260 | ATP2C1 | chr3 | + | 130980581 | 130980679 | 130979248 | 130979419 | 130992950 | 130993001 | 338 | 310 | 236 | 336 | 227 | 0 | 1 | 0 | 0 | 0 | 213 | 240 |
| SE | 30949 | ENSG00000017260 | ATP2C1 | chr3 | + | 130997605 | 130997753 | 130996679 | 130996796 | 130998293 | 130998389 | 190 | 213 | 252 | 258 | 269 | 1 | 0 | 1 | 0 | 0 | 122 | 166 |
| SE | 30950 | ENSG00000017260 | ATP2C1 | chr3 | + | 130999517 | 130999659 | 130998293 | 130998389 | 131016151 | 131016684 | 137 | 162 | 115 | 172 | 163 | 0 | 0 | 0 | 0 | 0 | 90 | 87 |
| SE | 30960 | ENSG00000185222 | TCEAL9 | chrX | + | 103357082 | 103357161 | 103356505 | 103356606 | 103357614 | 103358462 | 1 | 1 | 1 | 1 | 0 | 0 | 0 | 1 | 0 | 0 | 14 | 1 |
| SE | 30964 | ENSG00000137807 | KIF23 | chr15 | + | 69439903 | 69440077 | 69438247 | 69438405 | 69444789 | 69445041 | 2 | 5 | 5 | 2 | 2 | 0 | 0 | 1 | 0 | 0 | 0 | 2 |
| SE | 30966 | ENSG00000137807 | KIF23 | chr15 | + | 69440767 | 69441079 | 69440307 | 69440487 | 69444789 | 69445041 | 6 | 4 | 7 | 3 | 13 | 8 | 1 | 5 | 3 | 3 | 0 | 11 |
| SE | 30969 | ENSG00000137807 | KIF23 | chr15 | + | 69446282 | 69446364 | 69446008 | 69446091 | 69446870 | 69446941 | 14 | 10 | 23 | 3 | 13 | 0 | 2 | 0 | 0 | 0 | 9 | 11 |
| SE | 30970 | ENSG00000137807 | KIF23 | chr15 | + | 69446285 | 69446364 | 69446008 | 69446091 | 69446870 | 69446941 | 6 | 3 | 10 | 1 | 7 | 0 | 2 | 0 | 0 | 0 | 0 | 4 |
| SE | 30971 | ENSG00000137807 | KIF23 | chr15 | + | 69446430 | 69446587 | 69446008 | 69446091 | 69446870 | 69446941 | 0 | 0 | 0 | 1 | 1 | 0 | 2 | 0 | 0 | 0 | 0 | 0 |
| SE | 30973 | ENSG00000137269 | LRRC1 | chr6 | + | 53878992 | 53879071 | 53842109 | 53842227 | 53882886 | 53882976 | 13 | 3 | 8 | 8 | 0 | 0 | 0 | 0 | 0 | 2 | 4 | 0 |
| SE | 30975 | ENSG00000145675 | PIK3R1 | chr5 | + | 68273389 | 68273482 | 68226289 | 68227009 | 68273938 | 68274013 | 390 | 190 | 366 | 352 | 353 | 2 | 0 | 0 | 0 | 0 | 40 | 265 |
| SE | 30976 | ENSG00000145675 | PIK3R1 | chr5 | + | 68273389 | 68273482 | 68226289 | 68227009 | 68279601 | 68279733 | 208 | 92 | 171 | 186 | 162 | 0 | 0 | 0 | 0 | 1 | 0 | 134 |
| SE | 30977 | ENSG00000145675 | PIK3R1 | chr5 | + | 68273938 | 68274013 | 68226289 | 68227009 | 68279601 | 68279733 | 135 | 67 | 133 | 176 | 102 | 0 | 0 | 0 | 0 | 1 | 24 | 93 |
| SE | 30979 | ENSG00000145675 | PIK3R1 | chr5 | + | 68292258 | 68292361 | 68280926 | 68281006 | 68293100 | 68293199 | 858 | 556 | 877 | 1079 | 1157 | 2 | 0 | 2 | 0 | 5 | 144 | 691 |
| SE | 30982 | ENSG00000145675 | PIK3R1 | chr5 | + | 68294535 | 68294678 | 68293708 | 68293834 | 68295147 | 68295324 | 982 | 781 | 980 | 1207 | 1773 | 0 | 0 | 0 | 0 | 0 | 113 | 733 |
| SE | 30983 | ENSG00000145675 | PIK3R1 | chr5 | + | 68296170 | 68296341 | 68295419 | 68295488 | 68297411 | 68298583 | 1017 | 703 | 1151 | 1117 | 2055 | 0 | 0 | 0 | 0 | 0 | 79 | 702 |
| SE | 30984 | ENSG00000116237 | ICMT | chr1 | - | 6231901 | 6232119 | 6221192 | 6225262 | 6234404 | 6234511 | 143 | 79 | 113 | 123 | 120 | 1 | 0 | 0 | 0 | 0 | 202 | 189 |
| SE | 30985 | ENSG00000116237 | ICMT | chr1 | - | 6231901 | 6232119 | 6221192 | 6225262 | 6234885 | 6234974 | 144 | 79 | 116 | 126 | 121 | 0 | 0 | 2 | 0 | 0 | 205 | 193 |
| SE | 30986 | ENSG00000116237 | ICMT | chr1 | - | 6233473 | 6233643 | 6231901 | 6232119 | 6234404 | 6234511 | 58 | 18 | 29 | 51 | 34 | 0 | 0 | 0 | 0 | 0 | 49 | 66 |
| SE | 30988 | ENSG00000116237 | ICMT | chr1 | - | 6234404 | 6234511 | 6223490 | 6225262 | 6234885 | 6234974 | 2 | 0 | 0 | 0 | 0 | 0 | 0 | 2 | 0 | 0 | 0 | 3 |
| SE | 30989 | ENSG00000116237 | ICMT | chr1 | - | 6234404 | 6234511 | 6231901 | 6232119 | 6234885 | 6234974 | 1 | 0 | 0 | 0 | 0 | 1 | 0 | 3 | 3 | 1 | 0 | 5 |
| SE | 30990 | ENSG00000116237 | ICMT | chr1 | - | 6234404 | 6234511 | 6233473 | 6233643 | 6234885 | 6234974 | 5 | 1 | 3 | 2 | 0 | 5 | 2 | 2 | 9 | 0 | 8 | 6 |
| SE | 30991 | ENSG00000198792 | TMEM184B | chr22 | - | 38222641 | 38222662 | 38221144 | 38221710 | 38224784 | 38224979 | 0 | 0 | 0 | 0 | 0 | 84 | 10 | 47 | 55 | 25 | 0 | 1 |
| SE | 30992 | ENSG00000198792 | TMEM184B | chr22 | - | 38230668 | 38230744 | 38226598 | 38226870 | 38231243 | 38231334 | 141 | 48 | 100 | 107 | 135 | 1 | 0 | 0 | 0 | 0 | 58 | 105 |
| SE | 30993 | ENSG00000198792 | TMEM184B | chr22 | - | 38245934 | 38246100 | 38231243 | 38231334 | 38247769 | 38248019 | 53 | 12 | 27 | 28 | 31 | 0 | 0 | 0 | 0 | 1 | 0 | 53 |
| SE | 30994 | ENSG00000198792 | TMEM184B | chr22 | - | 38246784 | 38246885 | 38245934 | 38246100 | 38247769 | 38248019 | 0 | 0 | 0 | 0 | 0 | 6 | 3 | 6 | 4 | 0 | 0 | 0 |
| SE | 30995 | ENSG00000198792 | TMEM184B | chr22 | - | 38246805 | 38246885 | 38245934 | 38246100 | 38247769 | 38248019 | 0 | 0 | 0 | 0 | 0 | 6 | 3 | 6 | 4 | 0 | 0 | 0 |
| SE | 30996 | ENSG00000054793 | ATP9A | chr20 | - | 51607526 | 51607584 | 51604816 | 51605020 | 51608517 | 51608626 | 24 | 4 | 1 | 13 | 1 | 2 | 0 | 0 | 0 | 0 | 0 | 12 |
| SE | 30997 | ENSG00000054793 | ATP9A | chr20 | - | 51622073 | 51622172 | 51618953 | 51619043 | 51625191 | 51625362 | 28 | 0 | 0 | 3 | 2 | 0 | 0 | 0 | 0 | 0 | 0 | 9 |
| SE | 30998 | ENSG00000113504 | SLC12A7 | chr5 | - | 1056581 | 1056596 | 1053348 | 1053482 | 1057470 | 1057649 | 2 | 0 | 2 | 2 | 0 | 101 | 38 | 18 | 74 | 29 | 0 | 0 |
| SE | 30999 | ENSG00000113504 | SLC12A7 | chr5 | - | 1085231 | 1085473 | 1065282 | 1065478 | 1093532 | 1093575 | 1 | 0 | 0 | 0 | 0 | 0 | 0 | 0 | 2 | 0 | 0 | 2 |
| SE | 31000 | ENSG00000113504 | SLC12A7 | chr5 | - | 1086902 | 1087033 | 1085231 | 1085473 | 1093532 | 1093575 | 34 | 16 | 9 | 15 | 18 | 0 | 0 | 0 | 0 | 0 | 28 | 16 |
| SE | 31002 | ENSG00000113504 | SLC12A7 | chr5 | - | 1088981 | 1089128 | 1088305 | 1088360 | 1093532 | 1093575 | 118 | 62 | 24 | 67 | 73 | 0 | 0 | 0 | 0 | 0 | 77 | 93 |
| SE | 31004 | ENSG00000163694 | RBM47 | chr4 | - | 40432650 | 40432862 | 40423266 | 40426143 | 40437770 | 40438924 | 87 | 112 | 101 | 104 | 112 | 0 | 0 | 0 | 3 | 0 | 399 | 188 |
| SE | 31005 | ENSG00000163694 | RBM47 | chr4 | - | 40436440 | 40436647 | 40423266 | 40426143 | 40437770 | 40438797 | 13 | 4 | 14 | 10 | 6 | 0 | 0 | 0 | 3 | 0 | 22 | 7 |
| SE | 31007 | ENSG00000163694 | RBM47 | chr4 | - | 40436440 | 40436885 | 40423680 | 40426143 | 40437770 | 40438924 | 2 | 3 | 4 | 0 | 4 | 0 | 0 | 0 | 3 | 0 | 16 | 3 |
| SE | 31011 | ENSG00000163694 | RBM47 | chr4 | - | 40544421 | 40544506 | 40466576 | 40466699 | 40629395 | 40629864 | 21 | 37 | 23 | 33 | 39 | 0 | 0 | 0 | 0 | 0 | 95 | 86 |
| SE | 31017 | ENSG00000085982 | USP40 | chr2 | - | 233488238 | 233488304 | 233485766 | 233485977 | 233489364 | 233489483 | 18 | 11 | 15 | 31 | 39 | 2 | 0 | 0 | 0 | 0 | 8 | 29 |
| SE | 31020 | ENSG00000085982 | USP40 | chr2 | - | 233524491 | 233524562 | 233523169 | 233523489 | 233525477 | 233525562 | 53 | 29 | 50 | 59 | 71 | 0 | 1 | 0 | 0 | 0 | 56 | 64 |
| SE | 31021 | ENSG00000085982 | USP40 | chr2 | - | 233525477 | 233525562 | 233524491 | 233524562 | 233527406 | 233527578 | 22 | 10 | 29 | 28 | 26 | 0 | 0 | 0 | 0 | 0 | 34 | 27 |
| SE | 31022 | ENSG00000085982 | USP40 | chr2 | - | 233533478 | 233533779 | 233529430 | 233529512 | 233540661 | 233540766 | 80 | 56 | 46 | 54 | 161 | 4 | 0 | 0 | 0 | 0 | 51 | 66 |
| SE | 31023 | ENSG00000085982 | USP40 | chr2 | - | 233540661 | 233540766 | 233529430 | 233529512 | 233542267 | 233542363 | 4 | 0 | 0 | 0 | 10 | 0 | 0 | 0 | 0 | 1 | 0 | 0 |
| SE | 31024 | ENSG00000085982 | USP40 | chr2 | - | 233540661 | 233540769 | 233529430 | 233529512 | 233542267 | 233542363 | 47 | 16 | 14 | 36 | 82 | 0 | 0 | 0 | 0 | 1 | 31 | 32 |
| SE | 31026 | ENSG00000038219 | BOD1L1 | chr4 | - | 13588721 | 13588792 | 13587698 | 13587771 | 13590385 | 13590446 | 285 | 288 | 249 | 282 | 352 | 0 | 2 | 0 | 1 | 1 | 15 | 100 |
| SE | 31027 | ENSG00000038219 | BOD1L1 | chr4 | - | 13590385 | 13590446 | 13587698 | 13587771 | 13591922 | 13591966 | 51 | 34 | 36 | 39 | 78 | 0 | 0 | 0 | 0 | 0 | 7 | 8 |
| SE | 31028 | ENSG00000038219 | BOD1L1 | chr4 | - | 13595859 | 13595944 | 13591922 | 13591966 | 13597103 | 13597168 | 139 | 121 | 135 | 152 | 187 | 0 | 0 | 0 | 3 | 0 | 0 | 28 |
| SE | 31029 | ENSG00000038219 | BOD1L1 | chr4 | - | 13597103 | 13597168 | 13595859 | 13595944 | 13598945 | 13605084 | 126 | 114 | 121 | 128 | 164 | 2 | 2 | 0 | 0 | 0 | 12 | 35 |
| SE | 31040 | ENSG00000124459 | ZNF45 | chr19 | - | 43925324 | 43925458 | 43924401 | 43924561 | 43931246 | 43931365 | 20 | 8 | 16 | 13 | 21 | 1 | 0 | 0 | 2 | 0 | 23 | 10 |
| SE | 31041 | ENSG00000124459 | ZNF45 | chr19 | - | 43931246 | 43931365 | 43925324 | 43925458 | 43932603 | 43932690 | 10 | 6 | 11 | 5 | 23 | 5 | 2 | 3 | 4 | 1 | 0 | 6 |
| SE | 31042 | ENSG00000124459 | ZNF45 | chr19 | - | 43931246 | 43931365 | 43929956 | 43930000 | 43932603 | 43932690 | 11 | 10 | 12 | 7 | 38 | 2 | 0 | 0 | 0 | 0 | 0 | 8 |
| SE | 31043 | ENSG00000124459 | ZNF45 | chr19 | - | 43932603 | 43932690 | 43925324 | 43925458 | 43934425 | 43934694 | 21 | 7 | 9 | 14 | 12 | 2 | 0 | 0 | 3 | 2 | 0 | 10 |
| SE | 31044 | ENSG00000124459 | ZNF45 | chr19 | - | 43932603 | 43932690 | 43929956 | 43930000 | 43934425 | 43934694 | 18 | 5 | 6 | 10 | 11 | 1 | 0 | 0 | 0 | 0 | 0 | 10 |
| SE | 31050 | ENSG00000103174 | NAGPA | chr16 | - | 5028879 | 5029008 | 5027979 | 5028185 | 5030384 | 5030493 | 56 | 22 | 49 | 42 | 119 | 0 | 1 | 0 | 0 | 0 | 97 | 45 |
| SE | 31052 | ENSG00000103174 | NAGPA | chr16 | - | 5031744 | 5031884 | 5030384 | 5030493 | 5033272 | 5033358 | 21 | 16 | 21 | 14 | 30 | 0 | 0 | 0 | 1 | 0 | 50 | 23 |
| SE | 31056 | ENSG00000065882 | TBC1D1 | chr4 | + | 38027787 | 38027879 | 38021585 | 38021718 | 38035587 | 38035698 | 347 | 278 | 309 | 295 | 331 | 0 | 0 | 0 | 2 | 0 | 718 | 403 |
| SE | 31057 | ENSG00000065882 | TBC1D1 | chr4 | + | 38035587 | 38035698 | 38027787 | 38027879 | 38044361 | 38044490 | 452 | 321 | 381 | 401 | 434 | 4 | 0 | 4 | 0 | 0 | 1013 | 478 |
| SE | 31058 | ENSG00000065882 | TBC1D1 | chr4 | + | 38044361 | 38044490 | 38035587 | 38035698 | 38045816 | 38045903 | 400 | 288 | 354 | 433 | 629 | 0 | 0 | 0 | 0 | 0 | 726 | 427 |
| SE | 31062 | ENSG00000065882 | TBC1D1 | chr4 | + | 38045816 | 38045903 | 38044361 | 38044490 | 38049617 | 38049804 | 334 | 223 | 275 | 350 | 501 | 0 | 0 | 2 | 1 | 2 | 526 | 376 |
| SE | 31063 | ENSG00000065882 | TBC1D1 | chr4 | + | 38049617 | 38049898 | 38035587 | 38035698 | 38054198 | 38054244 | 116 | 71 | 89 | 103 | 94 | 0 | 0 | 0 | 0 | 0 | 224 | 133 |
| SE | 31065 | ENSG00000065882 | TBC1D1 | chr4 | + | 38051898 | 38052060 | 38049617 | 38049898 | 38053105 | 38053225 | 0 | 0 | 0 | 0 | 0 | 3 | 1 | 2 | 0 | 1 | 0 | 0 |
| SE | 31066 | ENSG00000065882 | TBC1D1 | chr4 | + | 38051898 | 38052060 | 38049617 | 38049898 | 38054198 | 38054338 | 0 | 0 | 0 | 0 | 0 | 114 | 71 | 89 | 103 | 94 | 0 | 0 |
| SE | 31069 | ENSG00000065882 | TBC1D1 | chr4 | + | 38060619 | 38060658 | 38054198 | 38054338 | 38089931 | 38090117 | 1 | 0 | 4 | 0 | 0 | 69 | 32 | 84 | 80 | 55 | 0 | 0 |
| SE | 31070 | ENSG00000065882 | TBC1D1 | chr4 | + | 38089931 | 38090117 | 38054198 | 38054338 | 38095928 | 38096090 | 329 | 166 | 311 | 262 | 353 | 2 | 0 | 0 | 0 | 1 | 728 | 306 |
| SE | 31071 | ENSG00000065882 | TBC1D1 | chr4 | + | 38095928 | 38096090 | 38089931 | 38090117 | 38102998 | 38103157 | 523 | 290 | 459 | 358 | 812 | 0 | 0 | 0 | 0 | 0 | 1079 | 354 |
| SE | 31076 | ENSG00000065882 | TBC1D1 | chr4 | + | 38124961 | 38125131 | 38118032 | 38118192 | 38133083 | 38133257 | 304 | 180 | 252 | 265 | 497 | 0 | 1 | 0 | 4 | 0 | 546 | 373 |
| SE | 31080 | ENSG00000108509 | CAMTA2 | chr17 | - | 4977057 | 4977192 | 4974384 | 4974500 | 4978503 | 4978569 | 67 | 36 | 54 | 72 | 30 | 0 | 0 | 0 | 0 | 0 | 30 | 44 |
| SE | 31082 | ENSG00000108509 | CAMTA2 | chr17 | - | 4981677 | 4981742 | 4981224 | 4981359 | 4982088 | 4982227 | 7 | 15 | 9 | 9 | 13 | 0 | 1 | 0 | 3 | 0 | 26 | 9 |
| SE | 31083 | ENSG00000108509 | CAMTA2 | chr17 | - | 4981677 | 4981742 | 4981224 | 4981359 | 4982756 | 4982892 | 15 | 20 | 9 | 8 | 22 | 0 | 0 | 0 | 0 | 0 | 26 | 9 |
| SE | 31084 | ENSG00000108509 | CAMTA2 | chr17 | - | 4981677 | 4981831 | 4981224 | 4981359 | 4982088 | 4982160 | 64 | 55 | 70 | 70 | 80 | 0 | 1 | 0 | 3 | 0 | 123 | 79 |
| SE | 31085 | ENSG00000108509 | CAMTA2 | chr17 | - | 4981677 | 4981831 | 4981224 | 4981359 | 4982756 | 4982892 | 34 | 34 | 40 | 31 | 26 | 0 | 0 | 0 | 0 | 0 | 38 | 30 |
| SE | 31086 | ENSG00000108509 | CAMTA2 | chr17 | - | 4982088 | 4982160 | 4981224 | 4981359 | 4982756 | 4982892 | 21 | 22 | 15 | 29 | 26 | 0 | 0 | 0 | 0 | 0 | 34 | 15 |
| SE | 31088 | ENSG00000108509 | CAMTA2 | chr17 | - | 4982088 | 4982175 | 4981224 | 4981359 | 4982756 | 4982892 | 6 | 1 | 3 | 3 | 9 | 0 | 0 | 0 | 0 | 0 | 6 | 0 |
| SE | 31090 | ENSG00000108509 | CAMTA2 | chr17 | - | 4982088 | 4982227 | 4981224 | 4981359 | 4982756 | 4982892 | 12 | 15 | 7 | 19 | 19 | 0 | 0 | 0 | 0 | 0 | 12 | 8 |
| SE | 31093 | ENSG00000108509 | CAMTA2 | chr17 | - | 4982975 | 4983043 | 4982756 | 4982892 | 4985879 | 4985983 | 117 | 62 | 69 | 100 | 85 | 0 | 1 | 0 | 0 | 0 | 209 | 117 |
| SE | 31094 | ENSG00000108509 | CAMTA2 | chr17 | - | 4985879 | 4985983 | 4982975 | 4983043 | 4986191 | 4986213 | 77 | 46 | 41 | 66 | 40 | 0 | 0 | 0 | 0 | 0 | 122 | 84 |
| SE | 31097 | ENSG00000099949 | LZTR1 | chr22 | + | 20987503 | 20987583 | 20983026 | 20983089 | 20988009 | 20988113 | 33 | 16 | 25 | 23 | 6 | 1 | 0 | 0 | 0 | 0 | 24 | 30 |
| SE | 31100 | ENSG00000099949 | LZTR1 | chr22 | + | 20995367 | 20995484 | 20994869 | 20995026 | 20995762 | 20995872 | 3 | 0 | 2 | 3 | 2 | 0 | 0 | 0 | 1 | 0 | 4 | 0 |
| SE | 31101 | ENSG00000113649 | TCERG1 | chr5 | + | 146455055 | 146455281 | 146447310 | 146447408 | 146457182 | 146457335 | 76 | 54 | 47 | 53 | 25 | 0 | 0 | 0 | 0 | 1 | 32 | 27 |
| SE | 31102 | ENSG00000113649 | TCERG1 | chr5 | + | 146462017 | 146462096 | 146459214 | 146459337 | 146463550 | 146463793 | 7 | 1 | 4 | 3 | 0 | 14 | 2 | 4 | 10 | 27 | 0 | 0 |
| SE | 31105 | ENSG00000113649 | TCERG1 | chr5 | + | 146483539 | 146483629 | 146481175 | 146482727 | 146492919 | 146492965 | 229 | 177 | 211 | 203 | 276 | 0 | 0 | 0 | 0 | 0 | 159 | 215 |
| SE | 31106 | ENSG00000113649 | TCERG1 | chr5 | + | 146503823 | 146504006 | 146503374 | 146503539 | 146507027 | 146507207 | 89 | 97 | 90 | 74 | 137 | 5 | 20 | 10 | 11 | 11 | 10 | 94 |
| SE | 31107 | ENSG00000113649 | TCERG1 | chr5 | + | 146504363 | 146504494 | 146503823 | 146504006 | 146507027 | 146507177 | 1 | 2 | 0 | 2 | 0 | 40 | 35 | 34 | 33 | 45 | 0 | 1 |
| SE | 31109 | ENSG00000113649 | TCERG1 | chr5 | + | 146507744 | 146507778 | 146507027 | 146507207 | 146507872 | 146507956 | 1 | 4 | 0 | 4 | 1 | 164 | 105 | 121 | 143 | 164 | 0 | 0 |
| SE | 31114 | ENSG00000116133 | DHCR24 | chr1 | - | 54879748 | 54879805 | 54875941 | 54876047 | 54883617 | 54883773 | 1 | 0 | 0 | 1 | 0 | 35 | 5 | 6 | 6 | 20 | 0 | 3 |
| SE | 31116 | ENSG00000198586 | TLK1 | chr2 | - | 170996652 | 170996760 | 170990822 | 170993956 | 171006146 | 171006282 | 203 | 202 | 194 | 320 | 841 | 0 | 0 | 0 | 0 | 0 | 38 | 171 |
| SE | 31117 | ENSG00000198586 | TLK1 | chr2 | - | 170997711 | 170997823 | 170996652 | 170996760 | 171006146 | 171006282 | 513 | 535 | 475 | 682 | 1455 | 5 | 0 | 0 | 3 | 1 | 195 | 551 |
| SE | 31118 | ENSG00000198586 | TLK1 | chr2 | - | 171011372 | 171011454 | 171006971 | 171007063 | 171014850 | 171014948 | 542 | 590 | 458 | 709 | 1183 | 0 | 0 | 1 | 0 | 0 | 220 | 469 |
| SE | 31119 | ENSG00000198586 | TLK1 | chr2 | - | 171014850 | 171014948 | 171011372 | 171011454 | 171028338 | 171028405 | 437 | 511 | 420 | 698 | 986 | 0 | 0 | 0 | 0 | 1 | 290 | 384 |
| SE | 31122 | ENSG00000198586 | TLK1 | chr2 | - | 171028338 | 171028405 | 171014850 | 171014948 | 171046173 | 171046362 | 556 | 656 | 502 | 874 | 1163 | 0 | 0 | 0 | 1 | 4 | 416 | 498 |
| SE | 31123 | ENSG00000198586 | TLK1 | chr2 | - | 171028338 | 171028405 | 171014850 | 171014948 | 171049813 | 171049950 | 197 | 240 | 154 | 300 | 440 | 2 | 2 | 0 | 0 | 0 | 158 | 151 |
| SE | 31124 | ENSG00000198586 | TLK1 | chr2 | - | 171028338 | 171028405 | 171022871 | 171023129 | 171046173 | 171046362 | 304 | 353 | 246 | 430 | 572 | 2 | 0 | 0 | 0 | 0 | 180 | 256 |
| SE | 31125 | ENSG00000198586 | TLK1 | chr2 | - | 171046173 | 171046362 | 171014850 | 171014948 | 171049813 | 171049950 | 517 | 617 | 391 | 830 | 1574 | 2 | 2 | 0 | 0 | 0 | 441 | 540 |
| SE | 31128 | ENSG00000198586 | TLK1 | chr2 | - | 171058150 | 171058197 | 171056470 | 171056566 | 171061080 | 171061156 | 368 | 407 | 377 | 645 | 680 | 6 | 1 | 0 | 0 | 5 | 379 | 484 |
| SE | 31129 | ENSG00000198586 | TLK1 | chr2 | - | 171059923 | 171059986 | 171058150 | 171058197 | 171061080 | 171061156 | 0 | 0 | 1 | 3 | 0 | 266 | 293 | 259 | 439 | 499 | 0 | 0 |
| SE | 31131 | ENSG00000198586 | TLK1 | chr2 | - | 171082780 | 171082852 | 171061080 | 171061156 | 171117738 | 171117857 | 472 | 496 | 396 | 673 | 420 | 0 | 0 | 0 | 7 | 0 | 310 | 545 |
| SE | 31132 | ENSG00000198586 | TLK1 | chr2 | - | 171082780 | 171082852 | 171061080 | 171061156 | 171160289 | 171160495 | 124 | 128 | 98 | 188 | 152 | 0 | 0 | 0 | 0 | 0 | 78 | 179 |
| SE | 31133 | ENSG00000198586 | TLK1 | chr2 | - | 171117738 | 171117857 | 171061080 | 171061156 | 171160289 | 171160495 | 203 | 161 | 184 | 318 | 122 | 0 | 0 | 0 | 0 | 0 | 128 | 184 |
| SE | 31134 | ENSG00000198586 | TLK1 | chr2 | - | 171117738 | 171117857 | 171082780 | 171082852 | 171160289 | 171160495 | 577 | 566 | 509 | 849 | 415 | 2 | 2 | 5 | 5 | 0 | 380 | 585 |
| SE | 31135 | ENSG00000134825 | TMEM258 | chr11 | - | 61789784 | 61789990 | 61789129 | 61789235 | 61790374 | 61790602 | 184 | 88 | 197 | 179 | 91 | 0 | 0 | 1 | 1 | 0 | 248 | 217 |
| SE | 31138 | ENSG00000171476 | HOPX | chr4 | - | 56650682 | 56650812 | 56647988 | 56648797 | 56655856 | 56656012 | 1 | 0 | 0 | 1 | 1 | 124 | 19 | 39 | 42 | 23 | 0 | 0 |
| SE | 31147 | ENSG00000138031 | ADCY3 | chr2 | - | 24831661 | 24831749 | 24830708 | 24830825 | 24834484 | 24834646 | 17 | 12 | 11 | 26 | 18 | 1 | 0 | 0 | 0 | 0 | 66 | 18 |
| SE | 31150 | ENSG00000008311 | AASS | chr7 | - | 122078861 | 122078950 | 122077835 | 122078014 | 122079596 | 122079712 | 14 | 5 | 14 | 15 | 10 | 1 | 0 | 1 | 0 | 0 | 0 | 10 |
| SE | 31153 | ENSG00000008311 | AASS | chr7 | - | 122081499 | 122081595 | 122078861 | 122079712 | 122086011 | 122086115 | 15 | 1 | 9 | 8 | 2 | 1 | 0 | 3 | 0 | 2 | 0 | 5 |
| SE | 31154 | ENSG00000008311 | AASS | chr7 | - | 122082787 | 122082891 | 122081499 | 122081595 | 122086011 | 122086153 | 3 | 0 | 6 | 4 | 0 | 6 | 0 | 2 | 3 | 0 | 0 | 0 |
| SE | 31156 | ENSG00000008311 | AASS | chr7 | - | 122091702 | 122091843 | 122086011 | 122086179 | 122092842 | 122092951 | 20 | 12 | 11 | 7 | 5 | 0 | 0 | 0 | 1 | 0 | 6 | 4 |
| SE | 31160 | ENSG00000162378 | ZYG11B | chr1 | + | 52779852 | 52779993 | 52771019 | 52771774 | 52784876 | 52785053 | 99 | 77 | 52 | 103 | 156 | 1 | 0 | 0 | 1 | 0 | 62 | 104 |
| SE | 31161 | ENSG00000162378 | ZYG11B | chr1 | + | 52790002 | 52790067 | 52784876 | 52785053 | 52796291 | 52796391 | 49 | 34 | 31 | 62 | 64 | 0 | 1 | 0 | 0 | 0 | 26 | 28 |
| SE | 31162 | ENSG00000162378 | ZYG11B | chr1 | + | 52805470 | 52805524 | 52802091 | 52802139 | 52813535 | 52813733 | 4 | 2 | 0 | 4 | 2 | 56 | 45 | 22 | 50 | 72 | 0 | 5 |
| SE | 31167 | ENSG00000239521 | CASTOR3 | chr7 | - | 100211393 | 100211994 | 100200652 | 100202603 | 100213046 | 100213133 | 12 | 5 | 15 | 11 | 11 | 6 | 0 | 0 | 2 | 0 | 0 | 7 |
| SE | 31168 | ENSG00000239521 | CASTOR3 | chr7 | - | 100213046 | 100213133 | 100200652 | 100202603 | 100222596 | 100222720 | 6 | 3 | 2 | 6 | 3 | 2 | 3 | 2 | 3 | 0 | 0 | 13 |
| SE | 31169 | ENSG00000239521 | CASTOR3 | chr7 | - | 100213046 | 100213133 | 100211393 | 100211994 | 100222596 | 100222720 | 9 | 7 | 10 | 15 | 8 | 1 | 0 | 0 | 3 | 0 | 0 | 10 |
| SE | 31171 | ENSG00000239521 | CASTOR3 | chr7 | - | 100223569 | 100223698 | 100222596 | 100222720 | 100223881 | 100223943 | 34 | 19 | 68 | 51 | 25 | 0 | 0 | 0 | 2 | 0 | 57 | 21 |
| SE | 31172 | ENSG00000163104 | SMARCAD1 | chr4 | + | 94213034 | 94213152 | 94208345 | 94208584 | 94226118 | 94226296 | 1 | 0 | 0 | 3 | 3 | 64 | 29 | 46 | 40 | 114 | 0 | 0 |
| SE | 31173 | ENSG00000163104 | SMARCAD1 | chr4 | + | 94226118 | 94226296 | 94208345 | 94208584 | 94233953 | 94234122 | 144 | 91 | 125 | 101 | 235 | 2 | 1 | 0 | 3 | 0 | 9 | 57 |
| SE | 31174 | ENSG00000163104 | SMARCAD1 | chr4 | + | 94233953 | 94234122 | 94226118 | 94226296 | 94236951 | 94237018 | 197 | 178 | 182 | 165 | 304 | 0 | 0 | 1 | 2 | 0 | 19 | 73 |
| SE | 31175 | ENSG00000163104 | SMARCAD1 | chr4 | + | 94240905 | 94241006 | 94236951 | 94237018 | 94249653 | 94249755 | 202 | 213 | 140 | 194 | 380 | 0 | 0 | 0 | 1 | 0 | 30 | 77 |
| SE | 31177 | ENSG00000163104 | SMARCAD1 | chr4 | + | 94250751 | 94250833 | 94249653 | 94249755 | 94252615 | 94253007 | 151 | 125 | 126 | 117 | 286 | 0 | 0 | 0 | 0 | 0 | 0 | 56 |
| SE | 31178 | ENSG00000163104 | SMARCAD1 | chr4 | + | 94273616 | 94273716 | 94270566 | 94270818 | 94274737 | 94274797 | 62 | 36 | 74 | 92 | 95 | 0 | 0 | 2 | 0 | 0 | 0 | 8 |
| SE | 31179 | ENSG00000163104 | SMARCAD1 | chr4 | + | 94277021 | 94277159 | 94276338 | 94276474 | 94278421 | 94278517 | 131 | 90 | 129 | 106 | 192 | 1 | 0 | 2 | 3 | 8 | 0 | 56 |
| SE | 31181 | ENSG00000163104 | SMARCAD1 | chr4 | + | 94284959 | 94285069 | 94283120 | 94283303 | 94289472 | 94289704 | 226 | 201 | 167 | 187 | 382 | 1 | 0 | 1 | 2 | 3 | 8 | 76 |
| SE | 31182 | ENSG00000185000 | DGAT1 | chr8 | - | 144318260 | 144318362 | 144318094 | 144318169 | 144318460 | 144318566 | 82 | 34 | 44 | 65 | 54 | 2 | 0 | 0 | 0 | 0 | 100 | 40 |
| SE | 31183 | ENSG00000185000 | DGAT1 | chr8 | - | 144319027 | 144319068 | 144318834 | 144318920 | 144321320 | 144321408 | 24 | 20 | 19 | 47 | 38 | 0 | 2 | 0 | 0 | 0 | 29 | 9 |
| SE | 31184 | ENSG00000109680 | TBC1D19 | chr4 | + | 26613168 | 26613241 | 26584083 | 26584292 | 26614407 | 26614453 | 30 | 18 | 13 | 11 | 10 | 1 | 0 | 4 | 1 | 0 | 6 | 10 |
| SE | 31185 | ENSG00000109680 | TBC1D19 | chr4 | + | 26614407 | 26614453 | 26584083 | 26584292 | 26620612 | 26620667 | 13 | 12 | 15 | 9 | 15 | 0 | 0 | 1 | 0 | 0 | 6 | 10 |
| SE | 31187 | ENSG00000109680 | TBC1D19 | chr4 | + | 26620612 | 26620688 | 26584083 | 26584292 | 26637210 | 26637285 | 14 | 14 | 16 | 7 | 6 | 0 | 1 | 0 | 0 | 0 | 0 | 1 |
| SE | 31188 | ENSG00000109680 | TBC1D19 | chr4 | + | 26620612 | 26620688 | 26614407 | 26614453 | 26637210 | 26637285 | 43 | 37 | 35 | 18 | 30 | 2 | 0 | 0 | 0 | 0 | 12 | 17 |
| SE | 31189 | ENSG00000109680 | TBC1D19 | chr4 | + | 26717932 | 26718017 | 26688344 | 26688407 | 26720080 | 26720125 | 49 | 38 | 51 | 29 | 50 | 7 | 2 | 1 | 6 | 6 | 15 | 15 |
| SE | 31190 | ENSG00000109680 | TBC1D19 | chr4 | + | 26735454 | 26735487 | 26720080 | 26720125 | 26739863 | 26739973 | 20 | 8 | 11 | 24 | 27 | 1 | 0 | 0 | 0 | 0 | 0 | 6 |
| SE | 31191 | ENSG00000109680 | TBC1D19 | chr4 | + | 26742507 | 26742599 | 26739863 | 26739973 | 26748410 | 26748526 | 24 | 28 | 20 | 35 | 68 | 1 | 0 | 0 | 0 | 0 | 0 | 4 |
| SE | 31192 | ENSG00000109680 | TBC1D19 | chr4 | + | 26753819 | 26753890 | 26748410 | 26748526 | 26754872 | 26754947 | 8 | 3 | 9 | 9 | 25 | 0 | 1 | 0 | 0 | 0 | 0 | 0 |

| | | | | | | | | | | | | | | | | | | | | | | | |
|---|---|---|---|---|---|---|---|---|---|---|---|---|---|---|---|---|---|---|---|---|---|---|---|
| 30750 | 0 | 5 | 0 | 0 | 7 | 8 | 1 | 0 | 226 | 149 | 1 | 1 | 0.748 | 0.022 | 0.235 | 0.606 | 0.367 | NA | 0.22 | 0 | 0.767 | NA | 0.067 |
| 30752 | 23 | 9 | 0 | 0 | 0 | 12 | 5 | 0 | 226 | 149 | 1 | 1 | 0.707 | 0.092 | 0.637 | 0.597 | 0.529 | NA | 1 | 0.558 | 0.543 | NA | -0.188 |
| 30753 | 7 | 0 | 0 | 0 | 7 | 0 | 0 | 0 | 226 | 149 | 1 | 1 | 1 | NA | 1 | 1 | 1 | NA | 0.159 | 1 | NA | NA | 0.42 |
| 30754 | 12 | 5 | 0 | 0 | 7 | 0 | 0 | 0 | 183 | 149 | 1 | 1 | 1 | 1 | 1 | 1 | 1 | NA | 0 | 1 | 1 | NA | 0.333 |
| 30755 | 15 | 17 | 1 | 0 | 0 | 0 | 0 | 0 | 183 | 149 | 1 | 1 | 1 | 1 | 0.914 | 1 | 1 | NA | 1 | 1 | 1 | 1 | -0.017 |
| 30757 | 0 | 2 | 6 | 0 | 0 | 3 | 1 | 3 | 298 | 149 | 1 | 1 | 0 | NA | 0.077 | NA | 1 | NA | NA | 0 | 0.5 | 0.5 | 0.026 |
| 30759 | 5 | 29 | 15 | 0 | 0 | 0 | 0 | 3 | 237 | 149 | 1 | 1 | NA | 0 | NA | 1 | 0.688 | 1 | NA | 1 | 1 | 0.759 | -0.377 |
| 30776 | 3 | 0 | 4 | 0 | 0 | 0 | 0 | 0 | 255 | 149 | 1 | 1 | 1 | 1 | 1 | 1 | 0.775 | NA | 1 | 1 | NA | 1 | -0.045 |
| 30778 | 0 | 0 | 6 | 0 | 0 | 0 | 8 | 0 | 268 | 149 | 1 | 1 | NA | NA | 1 | NA | NA | NA | NA | NA | 0 | 1 | 0.5 |
| 30780 | 26 | 3 | 3 | 0 | 0 | 0 | 0 | 0 | 287 | 149 | 1 | 1 | 1 | 1 | 0.908 | 1 | 1 | NA | 1 | 1 | 1 | 1 | -0.018 |
| 30781 | 6 | 2 | 1 | 0 | 0 | 0 | 0 | 0 | 240 | 149 | 1 | 1 | 1 | 1 | 0.848 | 1 | 0.867 | NA | 1 | 1 | 1 | 1 | -0.057 |
| 30783 | 12 | 5 | 35 | 0 | 0 | 0 | 0 | 0 | 240 | 149 | 1 | 1 | 1 | 0.89 | 1 | 1 | 1 | 1 | 1 | 1 | 1 | 1 | -0.022 |
| 30786 | 19 | 19 | 41 | 0 | 0 | 1 | 2 | 0 | 231 | 149 | 1 | 1 | 1 | 1 | 1 | 1 | 1 | 1 | 1 | 0.925 | 0.86 | 1 | 0.043 |
| 30787 | 2 | 12 | 8 | 0 | 0 | 0 | 1 | 0 | 242 | 149 | 1 | 1 | 1 | 1 | 1 | 1 | 1 | 1 | 1 | 1 | 0.881 | 1 | 0.024 |
| 30795 | 2 | 3 | 0 | 0 | 0 | 1 | 0 | 0 | 242 | 149 | 1 | 1 | NA | NA | 1 | NA | NA | 1 | 1 | 0.552 | 1 | NA | 0.112 |
| 30797 | 5 | 3 | 4 | 0 | 0 | 0 | 0 | 5 | 298 | 149 | 1 | 1 | 1 | 0.333 | 1 | 1 | 1 | NA | 1 | 1 | 1 | 0.286 | 0.045 |
| 30798 | 1 | 0 | 0 | 0 | 4 | 3 | 8 | 0 | 298 | 149 | 1 | 1 | 0 | 0 | 0 | 0 | NA | NA | 0 | 0.143 | 0 | NA | -0.048 |
| 30799 | 0 | 0 | 6 | 59 | 77 | 31 | 81 | 100 | 298 | 149 | 1 | 1 | 0 | 0 | 0.009 | 0.009 | 0 | 0 | 0 | 0 | 0 | 0.029 | -0.002 |
| 30801 | 55 | 90 | 229 | 0 | 0 | 1 | 0 | 0 | 286 | 149 | 1 | 1 | 1 | 1 | 1 | 1 | 1 | 1 | 1 | 0.966 | 1 | 1 | 0.007 |
| 30802 | 0 | 0 | 0 | 103 | 128 | 66 | 94 | 264 | 291 | 149 | 1 | 1 | 0 | 0 | 0.014 | 0 | 0 | 0 | 0.004 | 0 | 0 | 0 | 0.002 |
| 30804 | 0 | 0 | 0 | 103 | 128 | 66 | 94 | 264 | 298 | 149 | 1 | 1 | 0.014 | 0.024 | 0.023 | 0 | 0 | 0 | 0 | 0 | 0 | 0 | 0.012 |
| 30806 | 74 | 99 | 233 | 0 | 1 | 0 | 0 | 0 | 298 | 149 | 1 | 1 | 1 | 1 | 1 | 1 | 1 | 1 | 0.991 | 1 | 1 | 1 | 0.002 |
| 30807 | 22 | 36 | 98 | 0 | 0 | 2 | 0 | 0 | 298 | 149 | 1 | 1 | 1 | 1 | 1 | 1 | 1 | 1 | 1 | 0.846 | 1 | 1 | 0.031 |
| 30808 | 22 | 34 | 98 | 0 | 1 | 0 | 0 | 0 | 298 | 149 | 1 | 1 | 1 | 1 | 1 | 1 | 1 | 1 | 0.959 | 1 | 1 | 1 | 0.008 |
| 30809 | 17 | 10 | 11 | 0 | 2 | 0 | 2 | 0 | 298 | 149 | 1 | 1 | 1 | NA | 0 | NA | NA | NA | 0.692 | 1 | 0.714 | 1 | -0.351 |
| 30810 | 1 | 0 | 0 | 0 | 0 | 2 | 0 | 0 | 298 | 149 | 1 | 1 | 1 | NA | 1 | NA | NA | NA | 1 | 0.2 | NA | NA | 0.4 |
| 30811 | 71 | 105 | 160 | 0 | 0 | 2 | 0 | 0 | 298 | 149 | 1 | 1 | 1 | 1 | 1 | 1 | 1 | 1 | 1 | 0.947 | 1 | 1 | 0.011 |
| 30812 | 2 | 0 | 0 | 0 | 0 | 0 | 0 | 0 | 298 | 149 | 1 | 1 | 0.333 | 1 | 1 | NA | 1 | NA | 1 | 1 | NA | NA | -0.167 |
| 30813 | 145 | 180 | 379 | 0 | 0 | 2 | 0 | 0 | 241 | 149 | 1 | 1 | 1 | 1 | 1 | 1 | 1 | 1 | 1 | 0.978 | 1 | 1 | 0.004 |
| 30814 | 39 | 36 | 121 | 0 | 0 | 0 | 0 | 0 | 241 | 149 | 1 | 1 | 0.969 | 1 | 1 | 1 | 1 | 1 | 1 | 1 | 1 | 1 | -0.006 |
| 30815 | 39 | 95 | 133 | 0 | 0 | 2 | 0 | 0 | 226 | 149 | 1 | 1 | 1 | 1 | 1 | 1 | 1 | 1 | 1 | 0.928 | 1 | 1 | 0.014 |
| 30816 | 1 | 0 | 0 | 0 | 0 | 0 | 0 | 0 | 226 | 149 | 1 | 1 | 0 | 1 | NA | NA | 1 | NA | NA | 1 | NA | NA | -0.333 |
| 30817 | 214 | 254 | 453 | 0 | 0 | 1 | 0 | 5 | 255 | 149 | 1 | 1 | 0.978 | 0.991 | 1 | 1 | 0.995 | 1 | 1 | 0.992 | 1 | 0.981 | -0.002 |
| 30818 | 100 | 107 | 180 | 0 | 0 | 0 | 0 | 0 | 255 | 149 | 1 | 1 | 0.987 | 1 | 1 | 1 | 1 | 1 | 1 | 1 | 1 | 1 | -0.003 |
| 30819 | 285 | 345 | 596 | 0 | 0 | 2 | 0 | 0 | 298 | 149 | 1 | 1 | 1 | 1 | 0.994 | 1 | 1 | 1 | 0.992 | 1 | 1 | 1 | 0 |
| 30821 | 0 | 0 | 0 | 162 | 169 | 91 | 128 | 247 | 213 | 149 | 1 | 1 | 0.005 | 0 | 0.006 | 0 | 0 | 0 | 0 | 0 | 0 | 0 | 0.002 |
| 30822 | 0 | 2 | 35 | 0 | 0 | 0 | 0 | 0 | 267 | 149 | 1 | 1 | 1 | 1 | 0.527 | 1 | 1 | NA | 1 | NA | 1 | 1 | -0.095 |
| 30823 | 0 | 1 | 0 | 0 | 3 | 2 | 1 | 7 | 298 | 149 | 1 | 1 | 0.333 | 0 | 0.333 | 0.182 | 0.059 | NA | 0.538 | 0 | 0.333 | 0 | -0.036 |
| 30824 | 0 | 0 | 0 | 0 | 0 | 0 | 0 | 0 | 298 | 149 | 1 | 1 | 0 | NA | 1 | 0 | 1 | NA | 1 | NA | NA | NA | -0.5 |
| 30825 | 2 | 1 | 7 | 0 | 0 | 0 | 0 | 0 | 298 | 149 | 1 | 1 | 0.4 | 1 | 1 | 0.818 | 1 | NA | 1 | 1 | 1 | 1 | -0.156 |
| 30827 | 4 | 8 | 11 | 0 | 0 | 2 | 0 | 0 | 298 | 149 | 1 | 1 | 1 | 1 | 1 | 1 | 1 | NA | 1 | 0.5 | 1 | 1 | 0.125 |
| 30831 | 18 | 20 | 44 | 0 | 0 | 0 | 0 | 0 | 254 | 149 | 1 | 1 | 1 | 0.936 | 1 | 1 | 0.944 | 1 | 1 | 1 | 1 | 1 | -0.024 |
| 30853 | 41 | 7 | 18 | 0 | 0 | 3 | 0 | 0 | 244 | 149 | 1 | 1 | 1 | 1 | 1 | 1 | 1 | 1 | 1 | 0.893 | 1 | 1 | 0.021 |
| 30854 | 27 | 7 | 15 | 0 | 0 | 0 | 0 | 0 | 244 | 149 | 1 | 1 | 0.917 | 1 | 1 | 1 | 1 | 1 | 1 | 1 | 1 | 1 | -0.017 |
| 30856 | 1 | 0 | 0 | 0 | 5 | 15 | 0 | 3 | 257 | 149 | 1 | 1 | 0.41 | 0.367 | 0.465 | 1 | 0.162 | NA | 0.511 | 0.037 | NA | 0 | 0.298 |
| 30858 | 58 | 60 | 13 | 0 | 0 | 0 | 0 | 0 | 217 | 149 | 1 | 1 | 0.985 | 1 | 1 | 1 | 1 | 1 | 1 | 1 | 1 | 1 | -0.003 |
| 30860 | 89 | 56 | 6 | 0 | 0 | 0 | 0 | 0 | 298 | 149 | 1 | 1 | 1 | 1 | 0.983 | 0.979 | 1 | 1 | 1 | 1 | 1 | 1 | -0.008 |
| 30861 | 66 | 23 | 0 | 0 | 0 | 0 | 0 | 0 | 298 | 149 | 1 | 1 | 0.934 | 1 | 0.929 | 1 | 1 | 1 | 1 | 1 | 1 | NA | -0.027 |
| 30862 | 81 | 68 | 22 | 0 | 0 | 0 | 0 | 0 | 257 | 149 | 1 | 1 | 0.971 | 1 | 0.972 | 1 | 1 | 1 | 1 | 1 | 1 | 1 | -0.011 |
| 30864 | 44 | 28 | 7 | 0 | 0 | 0 | 0 | 0 | 227 | 149 | 1 | 1 | 1 | 1 | 0.968 | 1 | 1 | 1 | 1 | 1 | 1 | 1 | -0.006 |
| 30865 | 74 | 69 | 20 | 0 | 0 | 0 | 0 | 0 | 251 | 149 | 1 | 1 | 1 | 1 | 0.989 | 0.985 | 1 | 1 | 1 | 1 | 1 | 1 | -0.005 |
| 30866 | 29 | 9 | 7 | 0 | 0 | 0 | 0 | 0 | 298 | 149 | 1 | 1 | 0.909 | 1 | 1 | 1 | 1 | 1 | 1 | 1 | 1 | 1 | -0.018 |
| 30868 | 4 | 0 | 7 | 0 | 1 | 0 | 0 | 0 | 225 | 149 | 1 | 1 | 1 | 1 | 1 | 1 | 1 | NA | 0.768 | 1 | NA | 1 | 0.077 |
| 30870 | 11 | 11 | 7 | 0 | 0 | 0 | 0 | 0 | 297 | 149 | 1 | 1 | 1 | 1 | 0.959 | 1 | 1 | 1 | 1 | 1 | 1 | 1 | -0.008 |
| 30871 | 16 | 4 | 8 | 0 | 0 | 0 | 0 | 0 | 256 | 149 | 1 | 1 | 1 | 1 | 0.947 | 1 | 1 | NA | 1 | 1 | 1 | 1 | -0.011 |
| 30872 | 53 | 10 | 14 | 0 | 0 | 0 | 0 | 0 | 256 | 149 | 1 | 1 | 1 | 1 | 1 | 1 | 0.99 | NA | 1 | 1 | 1 | 1 | -0.002 |
| 30873 | 93 | 17 | 25 | 0 | 5 | 8 | 0 | 0 | 241 | 149 | 1 | 1 | 0.985 | 0.975 | 1 | 1 | 0.983 | NA | 0.898 | 0.878 | 1 | 1 | 0.045 |
| 30874 | 63 | 8 | 24 | 0 | 0 | 0 | 0 | 0 | 298 | 149 | 1 | 1 | 0.969 | 1 | 0.975 | 0.964 | 1 | 1 | 1 | 1 | 1 | 1 | -0.018 |
| 30875 | 26 | 10 | 0 | 6 | 23 | 13 | 20 | 0 | 298 | 149 | 1 | 1 | 0.209 | 0.489 | 0.309 | 0.163 | 0.241 | 0.6 | 0.432 | 0.5 | 0.2 | NA | -0.151 |
| 30877 | 23 | 7 | 0 | 0 | 0 | 1 | 9 | 0 | 298 | 149 | 1 | 1 | 0.852 | 1 | 0.917 | 0.923 | 0.957 | 1 | 1 | 0.92 | 0.28 | NA | 0.13 |
| 30879 | 337 | 534 | 841 | 5 | 0 | 0 | 1 | 0 | 276 | 149 | 1 | 1 | 1 | 1 | 1 | 1 | 1 | 0.99 | 1 | 1 | 0.997 | 1 | 0.003 |
| 30882 | 2 | 9 | 18 | 0 | 159 | 10 | 60 | 139 | 258 | 149 | 1 | 1 | 0.039 | 0.224 | 0 | 0 | 0.126 | NA | 0.058 | 0.104 | 0.08 | 0.07 | 0 |
| 30909 | 10 | 18 | 51 | 0 | 2 | 0 | 0 | 0 | 298 | 149 | 1 | 1 | 1 | 1 | 1 | 1 | 1 | 1 | 0.879 | 1 | 1 | 1 | 0.024 |
| 30912 | 11 | 25 | 4 | 0 | 0 | 0 | 0 | 0 | 298 | 149 | 1 | 1 | 1 | 0.75 | 1 | 1 | 1 | 1 | 1 | 1 | 1 | 1 | -0.05 |
| 30915 | 0 | 9 | 0 | 10 | 6 | 1 | 19 | 0 | 195 | 149 | 1 | 1 | 0.223 | 0.16 | 0.604 | 0.401 | 0 | 0 | 0.203 | 0 | 0.266 | NA | 0.16 |
| 30916 | 0 | 0 | 0 | 8 | 2 | 0 | 9 | 0 | 232 | 149 | 1 | 1 | 0 | 0 | 0.155 | 0 | 0 | 0 | 0 | NA | 0 | NA | 0.031 |
| 30918 | 0 | 0 | 0 | 0 | 0 | 0 | 0 | 0 | 256 | 149 | 1 | 1 | 1 | 1 | NA | 0 | NA | NA | 1 | NA | NA | NA | -0.333 |
| 30924 | 2 | 0 | 0 | 0 | 0 | 0 | 0 | 0 | 298 | 149 | 1 | 1 | NA | 0 | 1 | NA | 1 | NA | NA | 1 | NA | NA | -0.333 |
| 30929 | 0 | 0 | 0 | 0 | 0 | 0 | 0 | 11 | 294 | 149 | 1 | 1 | 0 | NA | NA | NA | 1 | NA | NA | NA | NA | 0 | 0.5 |
| 30930 | 101 | 129 | 161 | 5 | 1 | 0 | 0 | 0 | 213 | 149 | 1 | 1 | 1 | 1 | 1 | 1 | 1 | 0.914 | 0.99 | 1 | 1 | 1 | 0.019 |
| 30931 | 0 | 0 | 0 | 0 | 0 | 2 | 4 | 12 | 233 | 149 | 1 | 1 | NA | 0.39 | 0 | 0 | NA | NA | 1 | 0 | 0 | 0 | -0.12 |
| 30932 | 24 | 26 | 15 | 0 | 0 | 6 | 0 | 0 | 298 | 149 | 1 | 1 | 0.931 | 0.659 | 0.957 | 0.953 | 0.968 | NA | 1 | 0.667 | 1 | 1 | -0.023 |
| 30933 | 15 | 32 | 15 | 0 | 0 | 6 | 0 | 0 | 217 | 149 | 1 | 1 | 0.942 | 0.662 | 0.962 | 0.958 | 0.974 | NA | 1 | 0.632 | 1 | 1 | -0.008 |
| 30934 | 8 | 16 | 13 | 0 | 0 | 6 | 0 | 0 | 194 | 149 | 1 | 1 | 0.92 | 0.568 | 0.936 | 0.92 | 0.975 | NA | 1 | 0.506 | 1 | 1 | -0.013 |
| 30936 | 134 | 108 | 156 | 0 | 0 | 0 | 0 | 0 | 185 | 149 | 1 | 1 | 0.992 | 1 | 1 | 1 | 1 | 1 | 1 | 1 | 1 | 1 | -0.002 |
| 30938 | 108 | 97 | 41 | 0 | 2 | 0 | 0 | 0 | 254 | 149 | 1 | 1 | 1 | 1 | 1 | 0.99 | 1 | 1 | 0.969 | 1 | 1 | 1 | 0.004 |
| 30939 | 43 | 41 | 18 | 0 | 0 | 3 | 0 | 0 | 254 | 149 | 1 | 1 | 1 | 1 | 1 | 1 | 1 | 1 | 1 | 0.894 | 1 | 1 | 0.021 |
| 30940 | 46 | 11 | 2 | 0 | 0 | 3 | 0 | 0 | 298 | 149 | 1 | 1 | 1 | 1 | 1 | 1 | 1 | 1 | 1 | 0.885 | 1 | 1 | 0.023 |
| 30941 | 116 | 68 | 25 | 0 | 0 | 0 | 0 | 0 | 298 | 149 | 1 | 1 | 0.981 | 1 | 1 | 1 | 1 | 1 | 1 | 1 | 1 | 1 | -0.004 |
| 30948 | 205 | 254 | 91 | 0 | 0 | 0 | 0 | 0 | 247 | 149 | 1 | 1 | 1 | 0.995 | 1 | 1 | 1 | 1 | 1 | 1 | 1 | 1 | -0.001 |
| 30949 | 219 | 124 | 62 | 0 | 0 | 0 | 0 | 0 | 297 | 149 | 1 | 1 | 0.99 | 1 | 0.992 | 1 | 1 | 1 | 1 | 1 | 1 | 1 | -0.004 |
| 30950 | 116 | 90 | 34 | 0 | 0 | 3 | 0 | 0 | 291 | 149 | 1 | 1 | 1 | 1 | 1 | 1 | 1 | 1 | 1 | 0.952 | 1 | 1 | 0.01 |
| 30960 | 0 | 4 | 0 | 0 | 1 | 0 | 5 | 0 | 228 | 149 | 1 | 1 | 1 | 1 | 0.395 | 1 | NA | 1 | 0.395 | NA | 0.343 | NA | 0.269 |
| 30964 | 3 | 7 | 0 | 0 | 0 | 0 | 0 | 0 | 298 | 149 | 1 | 1 | 1 | 1 | 0.714 | 1 | 1 | NA | 1 | 1 | 1 | NA | -0.057 |
| 30966 | 14 | 1 | 0 | 0 | 10 | 4 | 1 | 0 | 298 | 149 | 1 | 1 | 0.273 | 0.667 | 0.412 | 0.333 | 0.684 | NA | 0.355 | 0.636 | 0.333 | NA | 0.032 |
| 30969 | 8 | 11 | 0 | 0 | 5 | 1 | 0 | 0 | 231 | 149 | 1 | 1 | 1 | 0.763 | 1 | 1 | 1 | 1 | 0.587 | 0.838 | 1 | NA | 0.096 |
| 30970 | 4 | 0 | 0 | 0 | 5 | 1 | 0 | 0 | 228 | 149 | 1 | 1 | 1 | 0.495 | 1 | 1 | 1 | NA | 0.343 | 0.723 | NA | NA | 0.366 |
| 30971 | 0 | 0 | 0 | 0 | 5 | 1 | 0 | 0 | 298 | 149 | 1 | 1 | NA | 0 | NA | 1 | 1 | NA | 0 | 0 | NA | NA | 0.667 |
| 30973 | 0 | 1 | 0 | 0 | 0 | 0 | 0 | 0 | 228 | 149 | 1 | 1 | 1 | 1 | 1 | 1 | 0 | 1 | NA | NA | 1 | NA | -0.2 |
| 30975 | 456 | 81 | 58 | 0 | 0 | 0 | 0 | 0 | 242 | 149 | 1 | 1 | 0.992 | 1 | 1 | 1 | 1 | 1 | 1 | 1 | 1 | 1 | -0.002 |
| 30976 | 202 | 47 | 33 | 0 | 0 | 1 | 0 | 0 | 242 | 149 | 1 | 1 | 1 | 1 | 1 | 1 | 0.99 | NA | 1 | 0.992 | 1 | 1 | 0 |
| 30977 | 185 | 41 | 10 | 0 | 0 | 1 | 0 | 0 | 224 | 149 | 1 | 1 | 1 | 1 | 1 | 1 | 0.985 | 1 | 1 | 0.992 | 1 | 1 | -0.001 |
| 30979 | 2071 | 357 | 31 | 0 | 0 | 3 | 0 | 0 | 252 | 149 | 1 | 1 | 0.996 | 1 | 0.996 | 1 | 0.993 | 1 | 1 | 0.998 | 1 | 1 | -0.003 |
| 30982 | 2106 | 246 | 29 | 0 | 0 | 1 | 0 | 0 | 292 | 149 | 1 | 1 | 1 | 1 | 1 | 1 | 1 | 1 | 1 | 0.999 | 1 | 1 | 0 |
| 30983 | 1623 | 222 | 43 | 0 | 0 | 1 | 0 | 0 | 298 | 149 | 1 | 1 | 1 | 1 | 1 | 1 | 1 | 1 | 1 | 0.999 | 1 | 1 | 0 |
| 30984 | 49 | 173 | 229 | 0 | 0 | 0 | 0 | 0 | 298 | 149 | 1 | 1 | 0.986 | 1 | 1 | 1 | 1 | 1 | 1 | 1 | 1 | 1 | -0.003 |
| 30985 | 49 | 173 | 258 | 0 | 0 | 0 | 0 | 0 | 298 | 149 | 1 | 1 | 1 | 1 | 0.967 | 1 | 1 | 1 | 1 | 1 | 1 | 1 | -0.007 |
| 30986 | 12 | 10 | 54 | 0 | 2 | 0 | 0 | 0 | 298 | 149 | 1 | 1 | 1 | 1 | 1 | 1 | 1 | 1 | 0.943 | 1 | 1 | 1 | 0.011 |
| 30988 | 0 | 0 | 4 | 0 | 0 | 0 | 0 | 0 | 256 | 149 | 1 | 1 | 1 | NA | 0 | NA | NA | NA | 1 | NA | NA | 1 | -0.5 |
| 30989 | 0 | 0 | 4 | 3 | 6 | 0 | 0 | 29 | 256 | 149 | 1 | 1 | 0.368 | NA | 0 | 0 | 0 | 0 | 0.327 | NA | NA | 0.074 | -0.042 |
| 30990 | 2 | 0 | 8 | 12 | 9 | 0 | 4 | 0 | 256 | 149 | 1 | 1 | 0.368 | 0.225 | 0.466 | 0.115 | NA | 0.28 | 0.28 | 1 | 0 | 1 | -0.219 |
| 30991 | 1 | 0 | 0 | 52 | 28 | 23 | 51 | 36 | 170 | 149 | 1 | 1 | 0 | 0 | 0 | 0 | 0 | 0 | 0.03 | 0.037 | 0 | 0 | -0.013 |
| 30992 | 58 | 55 | 107 | 0 | 0 | 0 | 0 | 0 | 225 | 149 | 1 | 1 | 0.989 | 1 | 1 | 1 | 1 | 1 | 1 | 1 | 1 | 1 | -0.002 |
| 30993 | 19 | 13 | 11 | 0 | 0 | 0 | 0 | 0 | 298 | 149 | 1 | 1 | 1 | 1 | 1 | 1 | 0.939 | NA | 1 | 1 | 1 | 1 | -0.012 |
| 30994 | 2 | 0 | 15 | 0 | 14 | 0 | 3 | 0 | 250 | 149 | 1 | 1 | 0 | 0 | 0 | 0 | NA | NA | 0 | 1 | 0 | 1 | -0.5 |
| 30995 | 3 | 0 | 0 | 0 | 14 | 0 | 3 | 0 | 229 | 149 | 1 | 1 | 0 | 0 | 0 | 0 | NA | NA | 0 | 1 | 0 | NA | -0.333 |
| 30996 | 4 | 0 | 13 | 0 | 0 | 0 | 0 | 0 | 207 | 149 | 1 | 1 | 0.896 | 1 | 1 | 1 | 1 | NA | 1 | 1 | NA | 1 | -0.021 |
| 30997 | 1 | 7 | 0 | 0 | 0 | 1 | 0 | 0 | 248 | 149 | 1 | 1 | 1 | NA | NA | 1 | 1 | NA | 1 | 0.375 | 1 | NA | 0.208 |
| 30998 | 0 | 0 | 1 | 72 | 52 | 39 | 31 | 54 | 164 | 149 | 1 | 1 | 0.018 | 0 | 0.092 | 0.024 | 0 | 0 | 0 | 0 | 0 | 0.017 | 0.023 |
| 30999 | 0 | 0 | 0 | 0 | 0 | 1 | 0 | 0 | 298 | 149 | 1 | 1 | 1 | NA | NA | 0 | NA | NA | 1 | 0 | NA | NA | 0 |
| 31000 | 13 | 12 | 26 | 0 | 2 | 0 | 0 | 0 | 280 | 149 | 1 | 1 | 1 | 1 | 1 | 1 | 1 | 1 | 0.81 | 1 | 1 | 1 | 0.038 |
| 31002 | 40 | 43 | 127 | 0 | 1 | 0 | 0 | 0 | 296 | 149 | 1 | 1 | 1 | 1 | 1 | 1 | 1 | 1 | 0.979 | 1 | 1 | 1 | 0.004 |
| 31004 | 184 | 187 | 224 | 0 | 0 | 0 | 0 | 1 | 298 | 149 | 1 | 1 | 1 | 1 | 1 | 0.945 | 1 | 1 | 1 | 1 | 1 | 0.991 | -0.009 |
| 31005 | 13 | 5 | 21 | 0 | 0 | 0 | 0 | 1 | 298 | 149 | 1 | 1 | 1 | 1 | 1 | 0.625 | 1 | 1 | 1 | 1 | 1 | 0.913 | -0.058 |
| 31007 | 6 | 0 | 12 | 0 | 0 | 0 | 0 | 1 | 298 | 149 | 1 | 1 | 1 | 1 | 1 | 0 | 1 | 1 | 1 | 1 | NA | 0.857 | -0.164 |
| 31011 | 52 | 109 | 127 | 5 | 0 | 0 | 0 | 0 | 234 | 149 | 1 | 1 | 1 | 1 | 1 | 1 | 1 | 0.924 | 1 | 1 | 1 | 1 | 0.015 |
| 31017 | 7 | 7 | 40 | 0 | 0 | 0 | 0 | 0 | 215 | 149 | 1 | 1 | 0.862 | 1 | 1 | 1 | 1 | 1 | 1 | 1 | 1 | 1 | -0.028 |
| 31020 | 49 | 47 | 32 | 0 | 0 | 0 | 0 | 0 | 220 | 149 | 1 | 1 | 1 | 0.952 | 1 | 1 | 1 | 1 | 1 | 1 | 1 | 1 | -0.01 |
| 31021 | 42 | 14 | 23 | 0 | 0 | 0 | 2 | 0 | 234 | 149 | 1 | 1 | 1 | 1 | 1 | 1 | 1 | 1 | 1 | 1 | 0.817 | 1 | 0.037 |
| 31022 | 44 | 9 | 1 | 0 | 0 | 2 | 0 | 0 | 298 | 149 | 1 | 1 | 0.909 | 1 | 1 | 1 | 1 | 1 | 1 | 0.917 | 1 | 1 | -0.002 |
| 31023 | 2 | 2 | 0 | 0 | 0 | 1 | 0 | 0 | 254 | 149 | 1 | 1 | 1 | NA | NA | NA | 0.854 | NA | NA | 0.54 | 1 | NA | 0.157 |
| 31024 | 17 | 9 | 1 | 0 | 0 | 1 | 0 | 0 | 257 | 149 | 1 | 1 | 1 | 1 | 1 | 1 | 0.979 | 1 | 1 | 0.908 | 1 | 1 | 0.014 |
| 31026 | 119 | 46 | 35 | 0 | 0 | 0 | 0 | 0 | 220 | 149 | 1 | 1 | 1 | 0.99 | 1 | 0.995 | 0.996 | 1 | 1 | 1 | 1 | 1 | -0.004 |
| 31027 | 13 | 0 | 8 | 0 | 0 | 1 | 0 | 0 | 210 | 149 | 1 | 1 | 1 | 1 | 1 | 1 | 1 | 1 | 1 | 0.902 | NA | 1 | 0.024 |
| 31028 | 60 | 5 | 26 | 0 | 0 | 0 | 0 | 0 | 234 | 149 | 1 | 1 | 1 | 1 | 1 | 0.97 | 1 | NA | 1 | 1 | 1 | 1 | -0.006 |
| 31029 | 59 | 7 | 12 | 0 | 0 | 0 | 0 | 0 | 214 | 149 | 1 | 1 | 0.978 | 0.975 | 1 | 1 | 1 | 1 | 1 | 1 | 1 | 1 | -0.009 |
| 31040 | 3 | 7 | 0 | 0 | 0 | 0 | 0 | 0 | 283 | 149 | 1 | 1 | 0.913 | 1 | 1 | 0.774 | 1 | 1 | 1 | 1 | 1 | NA | -0.063 |
| 31041 | 3 | 7 | 0 | 0 | 0 | 1 | 7 | 0 | 268 | 149 | 1 | 1 | 0.527 | 0.625 | 0.671 | 0.41 | 0.927 | NA | 1 | 0.625 | 0.357 | NA | -0.029 |
| 31042 | 8 | 19 | 0 | 0 | 0 | 1 | 0 | 0 | 268 | 149 | 1 | 1 | 0.754 | 1 | 1 | 1 | 1 | NA | 1 | 0.816 | 1 | NA | 0.012 |
| 31043 | 2 | 11 | 0 | 0 | 12 | 0 | 0 | 0 | 236 | 149 | 1 | 1 | 0.869 | 1 | 1 | 0.747 | 0.791 | NA | 0.345 | 1 | 1 | NA | 0.1 |
| 31044 | 2 | 4 | 0 | 0 | 0 | 0 | 0 | 0 | 236 | 149 | 1 | 1 | 0.919 | 1 | 1 | 1 | 1 | NA | 1 | 1 | 1 | NA | -0.016 |
| 31050 | 30 | 63 | 73 | 0 | 0 | 0 | 0 | 0 | 278 | 149 | 1 | 1 | 1 | 0.922 | 1 | 1 | 1 | 1 | 1 | 1 | 1 | 1 | -0.016 |
| 31052 | 9 | 20 | 31 | 0 | 0 | 0 | 0 | 0 | 289 | 149 | 1 | 1 | 1 | 1 | 1 | 0.878 | 1 | 1 | 1 | 1 | 1 | 1 | -0.024 |
| 31056 | 438 | 475 | 823 | 0 | 0 | 0 | 6 | 0 | 241 | 149 | 1 | 1 | 1 | 1 | 1 | 0.989 | 1 | 1 | 1 | 1 | 0.98 | 1 | 0.002 |
| 31057 | 516 | 556 | 986 | 0 | 4 | 0 | 3 | 0 | 260 | 149 | 1 | 1 | 0.985 | 1 | 0.982 | 1 | 1 | 1 | 0.986 | 1 | 0.991 | 1 | -0.002 |
| 31058 | 404 | 517 | 922 | 0 | 0 | 0 | 0 | 5 | 278 | 149 | 1 | 1 | 1 | 1 | 1 | 1 | 1 | 1 | 1 | 1 | 1 | 0.99 | 0.002 |
| 31062 | 315 | 416 | 824 | 4 | 2 | 4 | 1 | 0 | 236 | 149 | 1 | 1 | 1 | 1 | 0.989 | 0.995 | 0.994 | 0.988 | 0.992 | 0.98 | 0.996 | 1 | 0.004 |
| 31063 | 121 | 135 | 248 | 0 | 4 | 2 | 1 | 0 | 298 | 149 | 1 | 1 | 1 | 1 | 1 | 1 | 1 | 1 | 0.943 | 0.968 | 0.985 | 1 | 0.021 |
| 31065 | 4 | 0 | 0 | 4 | 6 | 0 | 1 | 0 | 298 | 149 | 1 | 1 | 0 | 0 | 0 | NA | 0 | 0 | 0 | 1 | 0 | NA | -0.25 |
| 31066 | 1 | 0 | 0 | 224 | 118 | 116 | 133 | 243 | 298 | 149 | 1 | 1 | 0 | 0 | 0 | 0 | 0 | 0 | 0 | 0.004 | 0 | 0 | -0.001 |
| 31069 | 2 | 0 | 0 | 163 | 129 | 74 | 60 | 155 | 188 | 149 | 1 | 1 | 0.011 | 0 | 0.036 | 0 | 0 | 0 | 0 | 0.021 | 0 | 0 | 0.005 |
| 31070 | 323 | 253 | 475 | 0 | 0 | 0 | 0 | 0 | 298 | 149 | 1 | 1 | 0.988 | 1 | 1 | 1 | 0.994 | 1 | 1 | 1 | 1 | 1 | -0.004 |
| 31071 | 537 | 367 | 633 | 0 | 0 | 3 | 0 | 0 | 298 | 149 | 1 | 1 | 1 | 1 | 1 | 1 | 1 | 1 | 1 | 0.989 | 1 | 1 | 0.002 |
| 31076 | 352 | 389 | 614 | 0 | 0 | 1 | 2 | 6 | 298 | 149 | 1 | 1 | 1 | 0.989 | 1 | 0.971 | 1 | 1 | 1 | 0.994 | 0.99 | 0.981 | -0.001 |
| 31080 | 19 | 54 | 103 | 0 | 0 | 0 | 2 | 0 | 284 | 149 | 1 | 1 | 1 | 1 | 1 | 1 | 1 | 1 | 1 | 1 | 0.934 | 1 | 0.013 |
| 31082 | 3 | 22 | 19 | 0 | 0 | 0 | 0 | 0 | 214 | 149 | 1 | 1 | 1 | 0.913 | 1 | 0.676 | 1 | 1 | 1 | 1 | 1 | 1 | -0.082 |
| 31083 | 7 | 22 | 28 | 0 | 0 | 0 | 1 | 0 | 214 | 149 | 1 | 1 | 1 | 1 | 1 | 1 | 1 | 1 | 1 | 1 | 0.939 | 1 | 0.012 |
| 31084 | 58 | 95 | 208 | 0 | 0 | 0 | 0 | 0 | 298 | 149 | 1 | 1 | 1 | 0.965 | 1 | 0.921 | 1 | 1 | 1 | 1 | 1 | 1 | -0.023 |
| 31085 | 19 | 37 | 76 | 0 | 0 | 0 | 1 | 0 | 298 | 149 | 1 | 1 | 1 | 1 | 1 | 1 | 1 | 1 | 1 | 1 | 0.949 | 1 | 0.01 |
| 31086 | 3 | 17 | 33 | 0 | 0 | 0 | 1 | 0 | 221 | 149 | 1 | 1 | 1 | 1 | 1 | 1 | 1 | 1 | 1 | 1 | 0.92 | 1 | 0.016 |
| 31088 | 3 | 0 | 4 | 0 | 0 | 0 | 1 | 0 | 236 | 149 | 1 | 1 | 1 | 1 | 1 | 1 | 1 | 1 | NA | 1 | 0 | 1 | 0.25 |
| 31090 | 7 | 19 | 22 | 0 | 0 | 0 | 1 | 0 | 288 | 149 | 1 | 1 | 1 | 1 | 1 | 1 | 1 | 1 | 1 | 1 | 0.908 | 1 | 0.018 |
| 31093 | 73 | 145 | 207 | 0 | 0 | 0 | 0 | 0 | 217 | 149 | 1 | 1 | 1 | 0.977 | 1 | 1 | 1 | 1 | 1 | 1 | 1 | 1 | -0.005 |
| 31094 | 55 | 70 | 180 | 3 | 0 | 0 | 0 | 0 | 253 | 149 | 1 | 1 | 1 | 1 | 1 | 1 | 1 | 0.96 | 1 | 1 | 1 | 1 | 0.008 |
| 31097 | 11 | 23 | 33 | 0 | 0 | 0 | 0 | 0 | 229 | 149 | 1 | 1 | 0.955 | 1 | 1 | 1 | 1 | 1 | 1 | 1 | 1 | 1 | -0.009 |
| 31100 | 0 | 0 | 0 | 0 | 0 | 0 | 0 | 3 | 266 | 149 | 1 | 1 | 1 | NA | 1 | 0.627 | 1 | 1 | NA | NA | NA | 0 | 0.407 |
| 31101 | 28 | 42 | 30 | 0 | 0 | 0 | 0 | 0 | 298 | 149 | 1 | 1 | 1 | 1 | 1 | 1 | 0.926 | 1 | 1 | 1 | 1 | 1 | -0.015 |
| 31102 | 5 | 8 | 0 | 0 | 20 | 7 | 13 | 0 | 228 | 149 | 1 | 1 | 0.246 | 0.246 | 0.395 | 0.164 | 0 | NA | 0 | 0.318 | 0.287 | NA | 0.009 |
| 31105 | 187 | 192 | 44 | 0 | 0 | 2 | 0 | 0 | 239 | 149 | 1 | 1 | 1 | 1 | 1 | 1 | 1 | 1 | 1 | 0.983 | 1 | 1 | 0.003 |
| 31106 | 84 | 46 | 0 | 0 | 13 | 35 | 10 | 0 | 298 | 149 | 1 | 1 | 0.899 | 0.708 | 0.818 | 0.771 | 0.862 | 1 | 0.783 | 0.545 | 0.697 | NA | 0.055 |
| 31107 | 2 | 0 | 0 | 10 | 52 | 35 | 19 | 0 | 280 | 149 | 1 | 1 | 0.013 | 0.03 | 0 | 0.031 | 0 | 0 | 0.01 | 0.03 | 0 | NA | 0.005 |
| 31109 | 0 | 6 | 0 | 15 | 151 | 145 | 71 | 19 | 183 | 149 | 1 | 1 | 0.005 | 0.03 | 0 | 0.022 | 0.005 | 0 | 0 | 0 | 0.064 | 0 | 0 |
| 31114 | 1 | 0 | 0 | 0 | 16 | 10 | 5 | 35 | 206 | 149 | 1 | 1 | 0.02 | 0 | 0 | 0.108 | 0 | NA | 0.119 | 0.067 | 0 | 0 | -0.021 |
| 31116 | 187 | 78 | 35 | 0 | 0 | 6 | 0 | 0 | 257 | 149 | 1 | 1 | 1 | 1 | 1 | 1 | 1 | 1 | 1 | 0.948 | 1 | 1 | 0.01 |
| 31117 | 525 | 202 | 142 | 0 | 0 | 5 | 2 | 0 | 261 | 149 | 1 | 1 | 0.983 | 1 | 1 | 0.992 | 0.999 | 1 | 1 | 0.984 | 0.983 | 1 | 0.001 |
| 31118 | 438 | 230 | 263 | 0 | 0 | 0 | 0 | 0 | 231 | 149 | 1 | 1 | 1 | 1 | 0.997 | 1 | 1 | 1 | 1 | 1 | 1 | 1 | -0.001 |
| 31119 | 398 | 227 | 267 | 0 | 0 | 0 | 0 | 0 | 247 | 149 | 1 | 1 | 1 | 1 | 1 | 1 | 0.998 | 1 | 1 | 1 | 1 | 1 | 0 |
| 31122 | 523 | 328 | 338 | 5 | 4 | 0 | 0 | 4 | 216 | 149 | 1 | 1 | 1 | 1 | 1 | 0.998 | 0.995 | 0.983 | 0.988 | 1 | 1 | 0.983 | 0.008 |
| 31123 | 186 | 86 | 121 | 0 | 0 | 0 | 0 | 0 | 216 | 149 | 1 | 1 | 0.985 | 0.988 | 1 | 1 | 1 | 1 | 1 | 1 | 1 | 1 | -0.005 |
| 31124 | 265 | 204 | 167 | 0 | 0 | 3 | 0 | 0 | 216 | 149 | 1 | 1 | 0.991 | 1 | 1 | 1 | 1 | 1 | 1 | 0.984 | 1 | 1 | 0.001 |
| 31125 | 437 | 428 | 427 | 0 | 0 | 0 | 0 | 0 | 298 | 149 | 1 | 1 | 0.992 | 0.994 | 1 | 1 | 1 | 1 | 1 | 1 | 1 | 1 | -0.003 |
| 31128 | 419 | 350 | 580 | 0 | 0 | 2 | 3 | 0 | 196 | 149 | 1 | 1 | 0.979 | 0.997 | 1 | 1 | 0.99 | 1 | 1 | 0.994 | 0.989 | 1 | -0.003 |
| 31129 | 2 | 0 | 0 | 265 | 362 | 317 | 214 | 393 | 212 | 149 | 1 | 1 | 0 | 0 | 0.003 | 0.005 | 0 | 0 | 0 | 0.004 | 0 | 0 | 0.001 |
| 31131 | 448 | 315 | 497 | 0 | 0 | 0 | 0 | 0 | 221 | 149 | 1 | 1 | 1 | 1 | 1 | 0.985 | 1 | 1 | 1 | 1 | 1 | 1 | -0.003 |
| 31132 | 140 | 95 | 156 | 0 | 0 | 4 | 0 | 0 | 221 | 149 | 1 | 1 | 1 | 1 | 1 | 1 | 1 | 1 | 1 | 0.959 | 1 | 1 | 0.008 |
| 31133 | 159 | 109 | 195 | 0 | 0 | 4 | 0 | 0 | 268 | 149 | 1 | 1 | 1 | 1 | 1 | 1 | 1 | 1 | 1 | 0.957 | 1 | 1 | 0.009 |
| 31134 | 499 | 342 | 547 | 0 | 8 | 2 | 0 | 1 | 268 | 149 | 1 | 1 | 0.994 | 0.994 | 0.983 | 0.99 | 1 | 1 | 0.976 | 0.993 | 1 | 0.997 | -0.001 |
| 31135 | 86 | 179 | 166 | 2 | 0 | 0 | 0 | 0 | 298 | 149 | 1 | 1 | 1 | 1 | 0.99 | 0.989 | 1 | 0.984 | 1 | 1 | 1 | 1 | -0.001 |
| 31138 | 0 | 0 | 0 | 1 | 22 | 10 | 12 | 46 | 279 | 149 | 1 | 1 | 0.004 | 0 | 0 | 0.013 | 0.023 | 0 | 0 | 0 | 0 | 0 | 0.008 |
| 31147 | 27 | 24 | 78 | 0 | 0 | 0 | 0 | 0 | 237 | 149 | 1 | 1 | 0.914 | 1 | 1 | 1 | 1 | 1 | 1 | 1 | 1 | 1 | -0.017 |
| 31150 | 13 | 2 | 0 | 0 | 0 | 0 | 0 | 0 | 238 | 149 | 1 | 1 | 0.898 | 1 | 0.898 | 1 | 1 | NA | 1 | 1 | 1 | NA | -0.041 |
| 31153 | 5 | 0 | 0 | 0 | 0 | 0 | 0 | 0 | 245 | 149 | 1 | 1 | 0.901 | 1 | 0.646 | 1 | 0.378 | NA | 1 | 1 | NA | NA | -0.215 |
| 31154 | 0 | 0 | 0 | 0 | 1 | 1 | 0 | 0 | 253 | 149 | 1 | 1 | 0.227 | NA | 0.639 | 0.44 | NA | NA | 0 | 0 | NA | NA | 0.435 |
| 31156 | 4 | 2 | 0 | 0 | 0 | 0 | 0 | 0 | 290 | 149 | 1 | 1 | 1 | 1 | 1 | 0.782 | 1 | 1 | 1 | 1 | 1 | NA | -0.044 |
| 31160 | 54 | 115 | 41 | 0 | 0 | 0 | 0 | 0 | 290 | 149 | 1 | 1 | 0.981 | 1 | 1 | 0.981 | 1 | 1 | 1 | 1 | 1 | 1 | -0.008 |
| 31161 | 39 | 49 | 42 | 0 | 0 | 0 | 0 | 0 | 214 | 149 | 1 | 1 | 1 | 0.959 | 1 | 1 | 1 | 1 | 1 | 1 | 1 | 1 | -0.008 |
| 31162 | 7 | 0 | 0 | 40 | 45 | 44 | 26 | 12 | 203 | 149 | 1 | 1 | 0.05 | 0.032 | 0 | 0.055 | 0.02 | 0 | 0.075 | 0.105 | 0 | 0 | -0.005 |
| 31167 | 3 | 1 | 6 | 0 | 12 | 2 | 3 | 0 | 298 | 149 | 1 | 1 | 0.5 | 1 | 1 | 0.733 | 1 | NA | 0.226 | 0.429 | 0.143 | 1 | 0.397 |
| 31168 | 4 | 3 | 0 | 0 | 3 | 6 | 0 | 0 | 236 | 149 | 1 | 1 | 0.654 | 0.387 | 0.387 | 0.558 | 1 | NA | 0.732 | 0.296 | 1 | NA | -0.079 |
| 31169 | 5 | 0 | 6 | 0 | 0 | 0 | 0 | 14 | 236 | 149 | 1 | 1 | 0.85 | 1 | 1 | 0.759 | 1 | NA | 1 | 1 | NA | 0.213 | 0.184 |
| 31171 | 40 | 39 | 53 | 0 | 0 | 0 | 0 | 0 | 278 | 149 | 1 | 1 | 1 | 1 | 1 | 0.932 | 1 | 1 | 1 | 1 | 1 | 1 | -0.014 |
| 31172 | 0 | 0 | 0 | 0 | 26 | 14 | 0 | 0 | 267 | 149 | 1 | 1 | 0.009 | 0 | 0 | 0.04 | 0.014 | NA | 0 | 0 | NA | NA | 0.013 |
| 31173 | 39 | 1 | 5 | 0 | 0 | 0 | 0 | 0 | 298 | 149 | 1 | 1 | 0.973 | 0.978 | 1 | 0.944 | 1 | 1 | 1 | 1 | 1 | 1 | -0.021 |
| 31174 | 51 | 26 | 5 | 0 | 0 | 0 | 0 | 0 | 298 | 149 | 1 | 1 | 1 | 1 | 0.989 | 0.976 | 1 | 1 | 1 | 1 | 1 | 1 | -0.007 |
| 31175 | 48 | 23 | 16 | 0 | 0 | 0 | 0 | 0 | 250 | 149 | 1 | 1 | 1 | 1 | 1 | 0.991 | 1 | 1 | 1 | 1 | 1 | 1 | -0.002 |
| 31177 | 34 | 29 | 12 | 0 | 0 | 1 | 0 | 0 | 231 | 149 | 1 | 1 | 1 | 1 | 1 | 1 | 1 | NA | 1 | 0.956 | 1 | 1 | 0.011 |
| 31178 | 12 | 3 | 0 | 0 | 0 | 0 | 0 | 0 | 249 | 149 | 1 | 1 | 1 | 1 | 0.957 | 1 | 1 | NA | 1 | 1 | 1 | NA | -0.009 |
| 31179 | 30 | 0 | 0 | 0 | 4 | 0 | 0 | 0 | 287 | 149 | 1 | 1 | 0.986 | 1 | 0.971 | 0.948 | 0.926 | NA | 0.879 | 1 | NA | NA | 0.027 |
| 31181 | 52 | 4 | 0 | 0 | 0 | 2 | 0 | 0 | 259 | 149 | 1 | 1 | 0.992 | 1 | 0.99 | 0.982 | 0.987 | 1 | 1 | 0.937 | 1 | NA | 0.006 |
| 31182 | 58 | 60 | 239 | 0 | 0 | 0 | 0 | 0 | 251 | 149 | 1 | 1 | 0.961 | 1 | 1 | 1 | 1 | 1 | 1 | 1 | 1 | 1 | -0.008 |
| 31183 | 34 | 36 | 103 | 0 | 0 | 0 | 0 | 0 | 190 | 149 | 1 | 1 | 1 | 0.887 | 1 | 1 | 1 | 1 | 1 | 1 | 1 | 1 | -0.023 |
| 31184 | 11 | 0 | 6 | 0 | 5 | 0 | 0 | 0 | 222 | 149 | 1 | 1 | 0.953 | 1 | 0.686 | 0.881 | 1 | 1 | 0.573 | 1 | NA | 1 | 0.011 |
| 31185 | 7 | 0 | 6 | 0 | 0 | 0 | 0 | 0 | 195 | 149 | 1 | 1 | 1 | 1 | 0.92 | 1 | 1 | 1 | 1 | 1 | NA | 1 | -0.016 |
| 31187 | 8 | 0 | 0 | 0 | 0 | 0 | 0 | 0 | 225 | 149 | 1 | 1 | 1 | 0.903 | 1 | 1 | 1 | NA | 1 | 1 | NA | NA | -0.019 |
| 31188 | 18 | 0 | 6 | 0 | 0 | 0 | 0 | 0 | 225 | 149 | 1 | 1 | 0.934 | 1 | 1 | 1 | 1 | 1 | 1 | 1 | NA | 1 | -0.013 |
| 31189 | 30 | 2 | 0 | 0 | 0 | 2 | 0 | 0 | 234 | 149 | 1 | 1 | 0.817 | 0.924 | 0.97 | 0.755 | 0.841 | 1 | 1 | 0.905 | 1 | NA | -0.115 |
| 31190 | 6 | 4 | 0 | 0 | 0 | 0 | 0 | 0 | 182 | 149 | 1 | 1 | 0.942 | 1 | 1 | 1 | 1 | NA | | 1 | 1 | NA | -0.012 |
| 31191 | 12 | 1 | 6 | 0 | 0 | 0 | 0 | 0 | 241 | 149 | 1 | 1 | 0.937 | 1 | 1 | 1 | 1 | NA | 1 | 1 | 1 | 1 | -0.013 |
| 31192 | 1 | 0 | 0 | 0 | 0 | 0 | 0 | 0 | 220 | 149 | 1 | 1 | 1 | 0.67 | 1 | 1 | 1 | NA | NA | 1 | NA | NA | -0.066 |

| | | | | | | | | | | | | | | | | | | | | | | | | | | | | | | | | | | | | | | | | | | | | | | |
|---|---|---|---|---|---|---|---|---|---|---|---|---|---|---|---|---|---|---|---|---|---|---|---|---|---|---|---|---|---|---|---|---|---|---|---|---|---|---|---|---|---|---|---|---|---|---|
| SE | 31202 | ENSG00000025293 | PHF20 | chr20 | + | 35782327 | 35782609 | 35772021 | 35772079 | 35800093 | 35800173 | 0 | 0 | 1 | 2 | 0 | 0 | 0 | 2 | 2 | 0 | 0 | 0 | 0 | 0 | 0 | 23 | 1 | 28 | 11 | 3 | 298 | 149 | 1 | 1 | NA | NA | 0.2 | 0.333 | NA | 0 | 0 | 0 | 0 | 0 | 0.267 |
| SE | 31206 | ENSG00000025293 | PHF20 | chr20 | + | 35801490 | 35801605 | 35772014 | 35772079 | 35842572 | 35842744 | 103 | 58 | 76 | 76 | 61 | 0 | 0 | 1 | 0 | 0 | 98 | 40 | 147 | 62 | 33 | 0 | 0 | 0 | 0 | 0 | 264 | 149 | 1 | 1 | 1 | 1 | 0.977 | 1 | 1 | 1 | 1 | 1 | 1 | 1 | -0.005 |
| SE | 31208 | ENSG00000025293 | PHF20 | chr20 | + | 35842572 | 35842744 | 35801490 | 35801605 | 35847349 | 35847434 | 267 | 158 | 213 | 253 | 243 | 2 | 1 | 0 | 0 | 0 | 202 | 151 | 322 | 178 | 118 | 0 | 0 | 1 | 0 | 0 | 298 | 149 | 1 | 1 | 0.985 | 0.988 | 1 | 1 | 1 | 1 | 1 | 0.994 | 1 | 1 | -0.004 |
| SE | 31210 | ENSG00000025293 | PHF20 | chr20 | + | 35845384 | 35845508 | 35842572 | 35842744 | 35847349 | 35847430 | 4 | 0 | 0 | 0 | 0 | 160 | 97 | 134 | 173 | 180 | 0 | 0 | 1 | 0 | 0 | 110 | 110 | 166 | 112 | 91 | 273 | 149 | 1 | 1 | 0.013 | 0 | 0 | 0 | 0 | 0 | 0 | 0.003 | 0 | 0 | 0.002 |
| SE | 31215 | ENSG00000025293 | PHF20 | chr20 | + | 35858301 | 35858381 | 35842572 | 35842744 | 35863012 | 35863248 | 84 | 59 | 54 | 86 | 162 | 1 | 1 | 0 | 0 | 2 | 87 | 95 | 71 | 56 | 52 | 0 | 1 | 1 | 0 | 4 | 229 | 149 | 1 | 1 | 0.982 | 0.975 | 1 | 1 | 0.981 | 1 | 0.984 | 0.979 | 1 | 0.894 | 0.016 |
| SE | 31217 | ENSG00000025293 | PHF20 | chr20 | + | 35858301 | 35858381 | 35847349 | 35847434 | 35863012 | 35863248 | 208 | 165 | 166 | 221 | 374 | 1 | 0 | 0 | 0 | 0 | 180 | 157 | 160 | 109 | 68 | 0 | 0 | 0 | 4 | 0 | 229 | 149 | 1 | 1 | 0.993 | 1 | 1 | 1 | 1 | 1 | 1 | 1 | 0.947 | 1 | 0.009 |
| SE | 31222 | ENSG00000025293 | PHF20 | chr20 | + | 35869437 | 35869551 | 35858301 | 35858381 | 35870954 | 35871120 | 127 | 72 | 107 | 104 | 166 | 1 | 0 | 0 | 0 | 0 | 92 | 67 | 56 | 69 | 41 | 0 | 0 | 0 | 0 | 0 | 263 | 149 | 1 | 1 | 0.986 | 1 | 1 | 1 | 1 | 1 | 1 | 1 | 1 | 1 | -0.003 |
| SE | 31225 | ENSG00000025293 | PHF20 | chr20 | + | 35913248 | 35913347 | 35899369 | 35899648 | 35917483 | 35917662 | 44 | 52 | 58 | 56 | 142 | 0 | 0 | 1 | 0 | 0 | 38 | 38 | 60 | 23 | 28 | 0 | 0 | 0 | 0 | 0 | 248 | 149 | 1 | 1 | 1 | 1 | 0.972 | 1 | 1 | 1 | 1 | 1 | 1 | 1 | -0.006 |
| SE | 31226 | ENSG00000025293 | PHF20 | chr20 | + | 35914032 | 35914197 | 35913248 | 35913347 | 35917483 | 35917662 | 181 | 135 | 176 | 163 | 238 | 0 | 0 | 0 | 0 | 2 | 133 | 137 | 215 | 124 | 76 | 0 | 0 | 1 | 0 | 0 | 298 | 149 | 1 | 1 | 1 | 1 | 1 | 1 | 0.983 | 1 | 1 | 0.991 | 1 | 1 | -0.002 |
| SE | 31228 | ENSG00000025293 | PHF20 | chr20 | + | 35931248 | 35931444 | 35927779 | 35927879 | 35938696 | 35939108 | 255 | 170 | 211 | 243 | 216 | 1 | 0 | 1 | 0 | 0 | 195 | 105 | 214 | 134 | 92 | 0 | 0 | 0 | 0 | 0 | 298 | 149 | 1 | 1 | 0.992 | 1 | 0.991 | 1 | 1 | 1 | 1 | 1 | 1 | 1 | -0.003 |
| SE | 31229 | ENSG00000121060 | TRIM25 | chr17 | - | 56895342 | 56895441 | 56887908 | 56892229 | 56895520 | 56895604 | 207 | 164 | 150 | 223 | 267 | 0 | 0 | 0 | 0 | 0 | 727 | 299 | 330 | 480 | 295 | 0 | 0 | 0 | 3 | 3 | 248 | 149 | 1 | 1 | 1 | 1 | 1 | 1 | 1 | 1 | 1 | 1 | 0.99 | 0.983 | 0.005 |
| SE | 31230 | ENSG00000121060 | TRIM25 | chr17 | - | 56895520 | 56895604 | 56895342 | 56895441 | 56895925 | 56895952 | 172 | 139 | 153 | 198 | 277 | 0 | 0 | 0 | 2 | 0 | 642 | 253 | 263 | 458 | 262 | 0 | 8 | 0 | 0 | 0 | 233 | 149 | 1 | 1 | 1 | 1 | 1 | 0.984 | 1 | 1 | 0.953 | 1 | 1 | 1 | 0.006 |
| SE | 31234 | ENSG00000121060 | TRIM25 | chr17 | - | 56899114 | 56899180 | 56895769 | 56895952 | 56901418 | 56901578 | 220 | 147 | 186 | 253 | 542 | 0 | 0 | 0 | 0 | 0 | 1173 | 417 | 410 | 687 | 347 | 0 | 0 | 1 | 0 | 0 | 215 | 149 | 1 | 1 | 1 | 1 | 1 | 1 | 1 | 1 | 1 | 0.996 | 1 | 1 | 0.001 |
| SE | 31235 | ENSG00000121060 | TRIM25 | chr17 | - | 56899114 | 56899180 | 56895769 | 56895952 | 56904254 | 56904343 | 48 | 27 | 39 | 61 | 76 | 0 | 0 | 0 | 0 | 0 | 187 | 79 | 71 | 81 | 66 | 0 | 4 | 0 | 0 | 0 | 215 | 149 | 1 | 1 | 1 | 1 | 1 | 1 | 1 | 1 | 0.932 | 1 | 1 | 1 | 0.014 |
| SE | 31236 | ENSG00000121060 | TRIM25 | chr17 | - | 56901418 | 56901578 | 56895769 | 56895952 | 56904254 | 56904343 | 241 | 215 | 200 | 301 | 633 | 0 | 0 | 0 | 0 | 0 | 1322 | 529 | 444 | 774 | 526 | 0 | 4 | 0 | 0 | 0 | 298 | 149 | 1 | 1 | 1 | 1 | 1 | 1 | 1 | 1 | 0.985 | 1 | 1 | 1 | 0.003 |
| SE | 31237 | ENSG00000121060 | TRIM25 | chr17 | - | 56901418 | 56901578 | 56898960 | 56899180 | 56904254 | 56904343 | 425 | 349 | 361 | 509 | 1117 | 0 | 1 | 1 | 2 | 0 | 2347 | 913 | 826 | 1444 | 840 | 5 | 3 | 1 | 6 | 0 | 298 | 149 | 1 | 1 | 1 | 0.994 | 0.994 | 0.992 | 1 | 0.996 | 0.993 | 0.998 | 0.992 | 1 | 0 |
| SE | 31238 | ENSG00000121060 | TRIM25 | chr17 | - | 56901418 | 56901578 | 56898960 | 56899180 | 56908467 | 56908563 | 184 | 135 | 161 | 208 | 484 | 0 | 0 | 0 | 0 | 0 | 1025 | 384 | 383 | 670 | 314 | 0 | 0 | 0 | 0 | 5 | 298 | 149 | 1 | 1 | 1 | 1 | 1 | 1 | 1 | 1 | 1 | 1 | 1 | 0.969 | 0.006 |
| SE | 31239 | ENSG00000121060 | TRIM25 | chr17 | - | 56904254 | 56904488 | 56898960 | 56899180 | 56908467 | 56908563 | 129 | 88 | 93 | 156 | 129 | 0 | 0 | 0 | 0 | 0 | 515 | 202 | 214 | 345 | 249 | 0 | 0 | 0 | 0 | 5 | 298 | 149 | 1 | 1 | 1 | 1 | 1 | 1 | 1 | 1 | 1 | 1 | 1 | 0.961 | 0.008 |
| SE | 31240 | ENSG00000121060 | TRIM25 | chr17 | - | 56904254 | 56904488 | 56901418 | 56901578 | 56908467 | 56908563 | 370 | 302 | 292 | 455 | 762 | 0 | 1 | 0 | 0 | 0 | 1832 | 728 | 656 | 1113 | 775 | 0 | 0 | 0 | 0 | 0 | 298 | 149 | 1 | 1 | 1 | 0.993 | 1 | 1 | 1 | 1 | 1 | 1 | 1 | 1 | -0.001 |
| SE | 31242 | ENSG00000121060 | TRIM25 | chr17 | - | 56912104 | 56912165 | 56908467 | 56908563 | 56912758 | 56913117 | 2 | 0 | 0 | 2 | 4 | 0 | 0 | 0 | 1 | 0 | 2 | 14 | 8 | 4 | 4 | 0 | 3 | 0 | 0 | 0 | 210 | 149 | 1 | 1 | 1 | NA | NA | 0.587 | 1 | 1 | 0.768 | 1 | 1 | 1 | -0.091 |
| SE | 31244 | ENSG00000047365 | ARAP2 | chr4 | - | 36006902 | 36007046 | 35985229 | 35985367 | 36012562 | 36012831 | 16 | 4 | 13 | 17 | 19 | 0 | 0 | 0 | 0 | 0 | 0 | 8 | 5 | 0 | 0 | 0 | 3 | 0 | 0 | 0 | 293 | 149 | 1 | 1 | 1 | 1 | 1 | 1 | 1 | NA | 0.576 | 1 | NA | NA | 0.212 |
| SE | 31245 | ENSG00000047365 | ARAP2 | chr4 | - | 36012562 | 36012831 | 36005364 | 36007046 | 36045978 | 36046034 | 20 | 6 | 16 | 26 | 21 | 0 | 0 | 0 | 1 | 0 | 0 | 8 | 5 | 0 | 0 | 0 | 0 | 0 | 0 | 0 | 298 | 149 | 1 | 1 | 1 | 1 | 1 | 0.929 | 1 | NA | 1 | 1 | NA | NA | -0.014 |
| SE | 31246 | ENSG00000047365 | ARAP2 | chr4 | - | 36015385 | 36015632 | 36012562 | 36012831 | 36015899 | 36015958 | 4 | 0 | 1 | 0 | 4 | 0 | 0 | 0 | 0 | 0 | 0 | 0 | 4 | 0 | 0 | 5 | 6 | 0 | 0 | 0 | 298 | 149 | 1 | 1 | 1 | NA | 1 | NA | 1 | 0 | 0 | 1 | NA | NA | 0.667 |
| SE | 31247 | ENSG00000047365 | ARAP2 | chr4 | - | 36015385 | 36015632 | 36012562 | 36012831 | 36019143 | 36019286 | 3 | 0 | 2 | 1 | 7 | 0 | 4 | 2 | 5 | 0 | 0 | 0 | 0 | 0 | 0 | 3 | 0 | 0 | 0 | 0 | 298 | 149 | 1 | 1 | 1 | 0 | 0.333 | 0.091 | 1 | 0 | NA | NA | NA | NA | 0.485 |
| SE | 31249 | ENSG00000047365 | ARAP2 | chr4 | - | 36015899 | 36015958 | 36012562 | 36012831 | 36019143 | 36019286 | 0 | 0 | 0 | 0 | 0 | 0 | 4 | 2 | 5 | 0 | 0 | 6 | 0 | 0 | 0 | 3 | 0 | 0 | 0 | 0 | 208 | 149 | 1 | 1 | NA | 0 | 0 | 0 | NA | 0 | 1 | NA | NA | NA | -0.5 |
| SE | 31250 | ENSG00000047365 | ARAP2 | chr4 | - | 36015899 | 36015958 | 36015385 | 36015632 | 36019143 | 36019286 | 2 | 0 | 0 | 0 | 0 | 1 | 0 | 1 | 1 | 3 | 0 | 0 | 4 | 0 | 0 | 0 | 0 | 0 | 0 | 0 | 208 | 149 | 1 | 1 | 0.589 | NA | 0 | 0 | 0 | NA | NA | 1 | NA | NA | -0.853 |
| SE | 31251 | ENSG00000047365 | ARAP2 | chr4 | - | 36019143 | 36019286 | 36012562 | 36012831 | 36045978 | 36046076 | 1 | 5 | 3 | 6 | 0 | 4 | 3 | 4 | 10 | 2 | 3 | 7 | 0 | 0 | 0 | 0 | 0 | 0 | 0 | 0 | 292 | 149 | 1 | 1 | 0.113 | 0.46 | 0.277 | 0.234 | 0 | 1 | 1 | NA | NA | NA | -0.783 |
| SE | 31252 | ENSG00000047365 | ARAP2 | chr4 | - | 36019143 | 36019286 | 36015385 | 36015632 | 36045978 | 36046076 | 2 | 1 | 2 | 2 | 3 | 1 | 0 | 0 | 0 | 0 | 0 | 7 | 0 | 0 | 0 | 0 | 0 | 0 | 0 | 0 | 292 | 149 | 1 | 1 | 0.505 | 1 | 1 | 1 | 1 | NA | 1 | NA | NA | NA | -0.099 |
| SE | 31253 | ENSG00000047365 | ARAP2 | chr4 | - | 36046709 | 36046849 | 36045978 | 36046076 | 36052005 | 36052053 | 7 | 8 | 5 | 12 | 2 | 0 | 0 | 0 | 3 | 0 | 3 | 0 | 0 | 0 | 0 | 0 | 0 | 0 | 0 | 0 | 289 | 149 | 1 | 1 | 1 | 1 | 1 | 0.673 | 1 | 1 | NA | NA | NA | NA | -0.065 |
| SE | 31254 | ENSG00000047365 | ARAP2 | chr4 | - | 36052005 | 36052053 | 36046709 | 36046849 | 36057969 | 36058143 | 0 | 1 | 0 | 3 | 0 | 1 | 0 | 1 | 3 | 0 | 0 | 0 | 0 | 0 | 0 | 0 | 0 | 2 | 0 | 0 | 197 | 149 | 1 | 1 | 0 | 1 | 0 | 0.431 | NA | NA | NA | 0 | NA | NA | 0.358 |
| SE | 31256 | ENSG00000047365 | ARAP2 | chr4 | - | 36073688 | 36073823 | 36066003 | 36068278 | 36082250 | 36082286 | 122 | 58 | 65 | 119 | 165 | 2 | 0 | 2 | 1 | 1 | 16 | 47 | 40 | 0 | 1 | 0 | 0 | 3 | 0 | 0 | 284 | 149 | 1 | 1 | 0.97 | 1 | 0.945 | 0.984 | 0.989 | 1 | 1 | 0.875 | NA | 1 | 0.009 |
| SE | 31257 | ENSG00000047365 | ARAP2 | chr4 | - | 36080215 | 36080279 | 36066003 | 36068278 | 36082250 | 36082286 | 112 | 83 | 64 | 104 | 123 | 2 | 0 | 2 | 1 | 1 | 5 | 43 | 24 | 10 | 0 | 0 | 0 | 3 | 0 | 0 | 213 | 149 | 1 | 1 | 0.975 | 1 | 0.957 | 0.986 | 0.989 | 1 | 1 | 0.848 | 1 | NA | 0.019 |
| SE | 31260 | ENSG00000047365 | ARAP2 | chr4 | - | 36121178 | 36121326 | 36117060 | 36117135 | 36124861 | 36124967 | 382 | 319 | 250 | 372 | 278 | 0 | 2 | 1 | 0 | 2 | 36 | 129 | 86 | 40 | 5 | 0 | 0 | 1 | 0 | 0 | 297 | 149 | 1 | 1 | 1 | 0.988 | 0.992 | 1 | 0.986 | 1 | 1 | 0.977 | 1 | 1 | -0.002 |
| SE | 31261 | ENSG00000047365 | ARAP2 | chr4 | - | 36147547 | 36147746 | 36147295 | 36147359 | 36148404 | 36148507 | 521 | 410 | 367 | 455 | 299 | 0 | 0 | 0 | 1 | 0 | 74 | 173 | 145 | 42 | 26 | 0 | 0 | 0 | 0 | 0 | 298 | 149 | 1 | 1 | 1 | 1 | 1 | 0.996 | 1 | 1 | 1 | 1 | 1 | 1 | -0.001 |
| SE | 31262 | ENSG00000047365 | ARAP2 | chr4 | - | 36150899 | 36151044 | 36148404 | 36148507 | 36158729 | 36158864 | 515 | 326 | 355 | 538 | 461 | 0 | 0 | 1 | 0 | 0 | 91 | 226 | 128 | 66 | 34 | 0 | 0 | 0 | 0 | 0 | 294 | 149 | 1 | 1 | 1 | 1 | 0.994 | 1 | 1 | 1 | 1 | 1 | 1 | 1 | -0.001 |
| SE | 31264 | ENSG00000047365 | ARAP2 | chr4 | - | 36160458 | 36160641 | 36159330 | 36159505 | 36161464 | 36161500 | 444 | 335 | 273 | 380 | 514 | 1 | 0 | 0 | 0 | 0 | 40 | 190 | 126 | 49 | 35 | 0 | 0 | 0 | 0 | 0 | 298 | 149 | 1 | 1 | 0.996 | 1 | 1 | 1 | 1 | 1 | 1 | 1 | 1 | 1 | -0.001 |
| SE | 31265 | ENSG00000047365 | ARAP2 | chr4 | - | 36177826 | 36178005 | 36166931 | 36167047 | 36187450 | 36187571 | 717 | 558 | 457 | 649 | 635 | 4 | 4 | 0 | 5 | 4 | 76 | 343 | 202 | 41 | 40 | 0 | 3 | 0 | 0 | 0 | 298 | 149 | 1 | 1 | 0.989 | 0.986 | 1 | 0.985 | 0.988 | 1 | 0.983 | 1 | 1 | 1 | -0.007 |
| SE | 31266 | ENSG00000047365 | ARAP2 | chr4 | - | 36187450 | 36187571 | 36177826 | 36178005 | 36193577 | 36193647 | 523 | 389 | 334 | 497 | 494 | 1 | 0 | 1 | 2 | 1 | 50 | 219 | 132 | 40 | 57 | 0 | 4 | 1 | 0 | 0 | 270 | 149 | 1 | 1 | 0.997 | 1 | 0.995 | 0.993 | 0.996 | 1 | 0.968 | 0.986 | 1 | 1 | 0.005 |
| SE | 31267 | ENSG00000047365 | ARAP2 | chr4 | - | 36187450 | 36187571 | 36177826 | 36178005 | 36210389 | 36210743 | 278 | 231 | 182 | 270 | 286 | 1 | 5 | 1 | 0 | 0 | 27 | 127 | 54 | 10 | 17 | 0 | 0 | 0 | 0 | 0 | 270 | 149 | 1 | 1 | 0.994 | 0.962 | 0.99 | 1 | 1 | 1 | 1 | 1 | 1 | 1 | -0.011 |
| SE | 31268 | ENSG00000047365 | ARAP2 | chr4 | - | 36193577 | 36193647 | 36177826 | 36178005 | 36210389 | 36210743 | 79 | 76 | 34 | 61 | 58 | 1 | 5 | 1 | 0 | 0 | 0 | 27 | 35 | 20 | 8 | 0 | 0 | 0 | 0 | 0 | 219 | 149 | 1 | 1 | 0.982 | 0.912 | 0.959 | 1 | 1 | NA | 1 | 1 | 1 | 1 | -0.029 |
| SE | 31269 | ENSG00000047365 | ARAP2 | chr4 | - | 36193577 | 36193647 | 36177826 | 36178005 | 36228581 | 36229645 | 1 | 0 | 0 | 1 | 0 | 0 | 0 | 0 | 0 | 2 | 0 | 7 | 0 | 0 | 0 | 0 | 0 | 0 | 0 | 0 | 219 | 149 | 1 | 1 | 1 | NA | NA | 1 | 0 | NA | 1 | NA | NA | NA | -0.333 |
| SE | 31270 | ENSG00000047365 | ARAP2 | chr4 | - | 36193577 | 36193647 | 36187450 | 36187571 | 36210389 | 36210743 | 330 | 236 | 187 | 290 | 267 | 2 | 0 | 3 | 4 | 1 | 23 | 119 | 113 | 50 | 43 | 0 | 0 | 0 | 0 | 0 | 219 | 149 | 1 | 1 | 0.991 | 1 | 0.977 | 0.98 | 0.995 | 1 | 1 | 1 | 1 | 1 | -0.011 |
| SE | 31271 | ENSG00000047365 | ARAP2 | chr4 | - | 36210389 | 36210743 | 36193577 | 36193647 | 36212395 | 36212487 | 812 | 543 | 521 | 723 | 564 | 0 | 0 | 2 | 0 | 0 | 120 | 410 | 304 | 153 | 112 | 0 | 0 | 0 | 0 | 0 | 298 | 149 | 1 | 1 | 1 | 1 | 0.992 | 1 | 1 | 1 | 1 | 1 | 1 | 1 | -0.002 |
| SE | 31272 | ENSG00000047365 | ARAP2 | chr4 | - | 36212395 | 36212487 | 36193577 | 36193647 | 36228581 | 36229645 | 0 | 0 | 2 | 0 | 0 | 0 | 0 | 0 | 0 | 0 | 0 | 0 | 0 | 0 | 0 | 0 | 3 | 0 | 0 | 0 | 241 | 149 | 1 | 1 | NA | NA | 1 | NA | NA | NA | 0 | NA | NA | NA | 1 |
| SE | 31274 | ENSG00000047365 | ARAP2 | chr4 | - | 36213242 | 36213319 | 36210389 | 36210743 | 36214421 | 36214480 | 300 | 245 | 221 | 322 | 232 | 0 | 0 | 0 | 2 | 0 | 58 | 241 | 144 | 83 | 21 | 0 | 0 | 0 | 0 | 0 | 226 | 149 | 1 | 1 | 1 | 1 | 1 | 0.991 | 1 | 1 | 1 | 1 | 1 | 1 | -0.002 |
| SE | 31275 | ENSG00000047365 | ARAP2 | chr4 | - | 36213242 | 36213319 | 36212395 | 36212487 | 36228581 | 36229645 | 330 | 192 | 194 | 271 | 240 | 0 | 0 | 2 | 0 | 0 | 50 | 136 | 117 | 66 | 60 | 0 | 0 | 0 | 0 | 0 | 226 | 149 | 1 | 1 | 1 | 1 | 0.985 | 1 | 1 | 1 | 1 | 1 | 1 | 1 | -0.003 |
| SE | 31276 | ENSG00000047365 | ARAP2 | chr4 | - | 36214421 | 36214480 | 36213242 | 36213319 | 36228581 | 36229645 | 1012 | 808 | 607 | 988 | 859 | 1 | 0 | 0 | 1 | 0 | 227 | 552 | 444 | 228 | 104 | 0 | 0 | 0 | 0 | 0 | 208 | 149 | 1 | 1 | 0.999 | 1 | 1 | 0.999 | 1 | 1 | 1 | 1 | 1 | 1 | 0 |
| SE | 31277 | ENSG00000047365 | ARAP2 | chr4 | - | 36228581 | 36229645 | 36214421 | 36214480 | 36244178 | 36244514 | 713 | 558 | 457 | 683 | 569 | 0 | 0 | 0 | 0 | 0 | 129 | 413 | 301 | 167 | 54 | 0 | 4 | 0 | 0 | 0 | 298 | 149 | 1 | 1 | 1 | 1 | 1 | 1 | 1 | 1 | 0.981 | 1 | 1 | 1 | 0.004 |
| SE | 31283 | ENSG00000106066 | CPVL | chr7 | - | 29030576 | 29030759 | 28995234 | 28995882 | 29064060 | 29064234 | 1417 | 1078 | 740 | 1228 | 1596 | 2 | 0 | 0 | 0 | 0 | 3632 | 779 | 572 | 1531 | 2286 | 0 | 0 | 0 | 0 | 0 | 298 | 149 | 1 | 1 | 0.997 | 1 | 1 | 1 | 1 | 1 | 1 | 1 | 1 | 1 | -0.001 |
| SE | 31287 | ENSG00000106066 | CPVL | chr7 | - | 29066022 | 29066121 | 29064060 | 29064234 | 29071772 | 29071904 | 1318 | 935 | 716 | 1032 | 1012 | 0 | 0 | 3 | 0 | 2 | 3035 | 573 | 475 | 1165 | 1496 | 18 | 2 | 0 | 3 | 5 | 248 | 149 | 1 | 1 | 1 | 1 | 0.993 | 1 | 0.997 | 0.99 | 0.994 | 1 | 0.996 | 0.994 | 0.003 |
| SE | 31288 | ENSG00000106066 | CPVL | chr7 | - | 29086483 | 29086550 | 29072300 | 29072423 | 29092622 | 29092702 | 840 | 512 | 489 | 687 | 847 | 0 | 0 | 0 | 0 | 0 | 1726 | 421 | 279 | 663 | 1239 | 0 | 1 | 0 | 0 | 0 | 216 | 149 | 1 | 1 | 1 | 1 | 1 | 1 | 1 | 1 | 0.997 | 1 | 1 | 1 | 0.001 |
| SE | 31289 | ENSG00000106066 | CPVL | chr7 | - | 29092622 | 29092702 | 29072300 | 29072423 | 29112703 | 29112822 | 1 | 0 | 0 | 1 | 0 | 0 | 0 | 1 | 0 | 0 | 0 | 1 | 0 | 0 | 2 | 0 | 0 | 0 | 0 | 0 | 229 | 149 | 1 | 1 | 1 | NA | 0 | 1 | NA | NA | 1 | NA | NA | 1 | -0.333 |
| SE | 31290 | ENSG00000106066 | CPVL | chr7 | - | 29095083 | 29095142 | 29092622 | 29092702 | 29112703 | 29112822 | 263 | 132 | 148 | 214 | 209 | 2 | 0 | 0 | 1 | 0 | 569 | 132 | 118 | 198 | 404 | 0 | 0 | 0 | 0 | 2 | 208 | 149 | 1 | 1 | 0.989 | 1 | 1 | 0.994 | 1 | 1 | 1 | 1 | 1 | 0.993 | -0.002 |
| SE | 31291 | ENSG00000106066 | CPVL | chr7 | - | 29096102 | 29096217 | 29095083 | 29095142 | 29112703 | 29112822 | 1103 | 652 | 660 | 748 | 1272 | 1 | 0 | 0 | 0 | 0 | 1886 | 496 | 319 | 834 | 1744 | 0 | 0 | 0 | 0 | 0 | 264 | 149 | 1 | 1 | 0.998 | 1 | 1 | 1 | 1 | 1 | 1 | 1 | 1 | 1 | 0 |
| SE | 31292 | ENSG00000106066 | CPVL | chr7 | - | 29096102 | 29096217 | 29095083 | 29095142 | 29120892 | 29121071 | 564 | 320 | 326 | 374 | 566 | 0 | 1 | 0 | 0 | 0 | 1029 | 255 | 188 | 434 | 918 | 0 | 1 | 0 | 0 | 0 | 264 | 149 | 1 | 1 | 1 | 0.994 | 1 | 1 | 1 | 1 | 0.993 | 1 | 1 | 1 | 0 |
| SE | 31293 | ENSG00000106066 | CPVL | chr7 | - | 29112703 | 29112822 | 29095083 | 29095142 | 29120892 | 29121071 | 303 | 167 | 202 | 220 | 239 | 0 | 1 | 0 | 0 | 0 | 418 | 141 | 74 | 173 | 426 | 0 | 1 | 0 | 0 | 0 | 268 | 149 | 1 | 1 | 1 | 0.989 | 1 | 1 | 1 | 1 | 0.987 | 1 | 1 | 1 | 0 |
| SE | 31294 | ENSG00000106066 | CPVL | chr7 | - | 29112703 | 29112822 | 29096102 | 29096217 | 29120892 | 29121071 | 840 | 497 | 531 | 594 | 943 | 1 | 2 | 0 | 0 | 0 | 1269 | 382 | 210 | 572 | 1252 | 0 | 0 | 5 | 0 | 0 | 268 | 149 | 1 | 1 | 0.998 | 0.993 | 1 | 1 | 1 | 1 | 1 | 0.959 | 1 | 1 | 0.006 |
| SE | 31296 | ENSG00000204977 | TRIM13 | chr13 | + | 50000994 | 50001108 | 49997666 | 49997763 | 50011934 | 50012327 | 0 | 0 | 0 | 2 | 0 | 36 | 17 | 49 | 38 | 76 | 0 | 6 | 0 | 0 | 0 | 61 | 34 | 56 | 13 | 30 | 263 | 149 | 1 | 1 | 0 | 0 | 0 | 0.029 | 0 | 0 | 0.091 | 0 | 0 | 0 | -0.012 |
| SE | 31298 | ENSG00000186638 | KIF24 | chr9 | - | 34290173 | 34290389 | 34286616 | 34286704 | 34306251 | 34306441 | 2 | 2 | 2 | 1 | 0 | 0 | 0 | 0 | 0 | 0 | 0 | 3 | 0 | 7 | 0 | 0 | 0 | 1 | 0 | 0 | 298 | 149 | 1 | 1 | 1 | 1 | 1 | 1 | NA | NA | 1 | 0 | 1 | NA | 0.333 |
| SE | 31299 | ENSG00000186638 | KIF24 | chr9 | - | 34297016 | 34297114 | 34290173 | 34290389 | 34306251 | 34306441 | 5 | 2 | 4 | 10 | 0 | 0 | 0 | 0 | 0 | 0 | 0 | 7 | 4 | 5 | 0 | 0 | 3 | 0 | 0 | 0 | 247 | 149 | 1 | 1 | 1 | 1 | 1 | 1 | NA | NA | 0.585 | 1 | 1 | NA | 0.138 |
| SE | 31304 | ENSG00000013288 | MAN2B2 | chr4 | + | 6593172 | 6593350 | 6589044 | 6589160 | 6594533 | 6594732 | 165 | 56 | 112 | 122 | 62 | 0 | 0 | 2 | 0 | 0 | 161 | 207 | 93 | 168 | 247 | 0 | 0 | 0 | 0 | 0 | 298 | 149 | 1 | 1 | 1 | 1 | 0.966 | 1 | 1 | 1 | 1 | 1 | 1 | 1 | -0.007 |
| SE | 31305 | ENSG00000013288 | MAN2B2 | chr4 | + | 6597112 | 6597303 | 6594533 | 6594732 | 6600622 | 6600756 | 57 | 15 | 69 | 53 | 13 | 0 | 3 | 0 | 0 | 0 | 126 | 81 | 34 | 85 | 191 | 0 | 0 | 0 | 0 | 0 | 298 | 149 | 1 | 1 | 1 | 0.714 | 1 | 1 | 1 | 1 | 1 | 1 | 1 | 1 | -0.057 |
| SE | 31306 | ENSG00000013288 | MAN2B2 | chr4 | + | 6598197 | 6598354 | 6597112 | 6597303 | 6600622 | 6600756 | 51 | 18 | 46 | 37 | 41 | 0 | 0 | 2 | 1 | 0 | 69 | 40 | 44 | 48 | 98 | 0 | 0 | 0 | 0 | 7 | 298 | 149 | 1 | 1 | 1 | 1 | 0.92 | 0.949 | 1 | 1 | 1 | 1 | 1 | 0.875 | -0.001 |
| SE | 31311 | ENSG00000087266 | SH3BP2 | chr4 | + | 2794162 | 2794262 | 2793084 | 2793138 | 2796373 | 2796646 | 2 | 1 | 0 | 0 | 0 | 2 | 1 | 0 | 1 | 0 | 4 | 0 | 0 | 0 | 0 | 9 | 2 | 0 | 0 | 0 | 249 | 149 | 1 | 1 | 0.374 | 0.374 | NA | 0 | NA | 0.21 | 0 | NA | NA | NA | 0.144 |
| SE | 31312 | ENSG00000087266 | SH3BP2 | chr4 | + | 2809778 | 2809829 | 2793070 | 2793138 | 2820613 | 2820753 | 1 | 0 | 0 | 1 | 0 | 10 | 3 | 4 | 5 | 0 | 0 | 1 | 0 | 0 | 5 | 0 | 8 | 9 | 16 | 25 | 200 | 149 | 1 | 1 | 0.069 | 0 | 0 | 0.13 | NA | NA | 0.085 | 0 | 0 | 0.13 | -0.004 |
| SE | 31313 | ENSG00000087266 | SH3BP2 | chr4 | + | 2809778 | 2809829 | 2799068 | 2799305 | 2820613 | 2820753 | 0 | 0 | 1 | 1 | 0 | 7 | 0 | 5 | 3 | 1 | 0 | 1 | 6 | 0 | 1 | 9 | 1 | 3 | 11 | 10 | 200 | 149 | 1 | 1 | 0 | NA | 0.13 | 0.199 | 0 | 0 | 0.427 | 0.598 | 0 | 0.069 | -0.137 |
| SE | 31314 | ENSG00000087266 | SH3BP2 | chr4 | + | 2814810 | 2814917 | 2812289 | 2812477 | 2820613 | 2820753 | 0 | 0 | 1 | 0 | 0 | 43 | 25 | 19 | 25 | 16 | 0 | 0 | 3 | 0 | 0 | 7 | 37 | 17 | 8 | 14 | 256 | 149 | 1 | 1 | 0 | 0 | 0.03 | 0 | 0 | 0 | 0 | 0.093 | 0 | 0 | -0.013 |
| SE | 31315 | ENSG00000087266 | SH3BP2 | chr4 | + | 2820613 | 2820753 | 2798997 | 2799305 | 2822934 | 2823037 | 110 | 42 | 57 | 74 | 37 | 1 | 0 | 0 | 0 | 0 | 83 | 51 | 68 | 109 | 225 | 0 | 0 | 0 | 0 | 0 | 289 | 149 | 1 | 1 | 0.983 | 1 | 1 | 1 | 1 | 1 | 1 | 1 | 1 | 1 | -0.003 |
| SE | 31321 | ENSG00000109684 | CLNK | chr4 | - | 10525840 | 10525922 | 10520790 | 10520831 | 10528075 | 10528094 | 3 | 1 | 3 | 2 | 2 | 0 | 0 | 0 | 0 | 0 | 0 | 6 | 0 | 0 | 0 | 0 | 0 | 1 | 0 | 0 | 231 | 149 | 1 | 1 | 1 | 1 | 1 | 1 | 1 | NA | 1 | 0 | NA | NA | 0.5 |
| SE | 31322 | ENSG00000143294 | PRCC | chr1 | + | 156786607 | 156787174 | 156782281 | 156782329 | 156791696 | 156791792 | 161 | 81 | 121 | 164 | 166 | 1 | 0 | 0 | 0 | 0 | 213 | 197 | 167 | 161 | 358 | 0 | 0 | 1 | 0 | 0 | 298 | 149 | 1 | 1 | 0.988 | 1 | 1 | 1 | 1 | 1 | 1 | 1 | 1 | 1 | -0.002 |
| SE | 31324 | ENSG00000143294 | PRCC | chr1 | + | 156791696 | 156791792 | 156786607 | 156787174 | 156794664 | 156794808 | 149 | 68 | 121 | 153 | 97 | 2 | 0 | 0 | 1 | 0 | 153 | 143 | 131 | 139 | 249 | 0 | 0 | 2 | 0 | 0 | 245 | 149 | 1 | 1 | 0.978 | 1 | 1 | 0.989 | 1 | 1 | 1 | 0.976 | 1 | 1 | -0.002 |
| SE | 31325 | ENSG00000143294 | PRCC | chr1 | + | 156797275 | 156797341 | 156794664 | 156794808 | 156800373 | 156800497 | 236 | 155 | 204 | 251 | 309 | 0 | 0 | 0 | 1 | 0 | 292 | 287 | 171 | 198 | 385 | 0 | 0 | 0 | 0 | 0 | 215 | 149 | 1 | 1 | 1 | 1 | 1 | 0.994 | 1 | 1 | 1 | 1 | 1 | 1 | -0.001 |
| SE | 31328 | ENSG00000229036 | VDAC1P8 | chr6 | + | 143504522 | 143504597 | 143503930 | 143504142 | 143505339 | 143505457 | 0 | 0 | 0 | 0 | 0 | 0 | 3 | 0 | 0 | 0 | 0 | 4 | 0 | 0 | 0 | 0 | 0 | 0 | 0 | 0 | 224 | 149 | 1 | 1 | NA | 0 | NA | NA | NA | NA | 1 | NA | NA | NA | -1 |
| SE | 31332 | ENSG00000159788 | RGS12 | chr4 | + | 3386415 | 3386437 | 3369878 | 3370321 | 3414071 | 3414241 | 3 | 2 | 5 | 3 | 0 | 0 | 0 | 0 | 0 | 1 | 6 | 1 | 2 | 0 | 0 | 0 | 0 | 0 | 0 | 0 | 171 | 149 | 1 | 1 | 1 | 1 | 1 | 1 | 0 | 1 | 1 | 1 | NA | NA | -0.2 |
| SE | 31335 | ENSG00000157426 | AASDH | chr4 | - | 56342834 | 56342966 | 56338286 | 56338791 | 56343561 | 56343684 | 46 | 20 | 79 | 37 | 141 | 4 | 4 | 7 | 2 | 16 | 0 | 16 | 14 | 0 | 0 | 0 | 2 | 0 | 4 | 0 | 281 | 149 | 1 | 1 | 0.859 | 0.726 | 0.857 | 0.907 | 0.824 | NA | 0.809 | 1 | 0 | NA | 0.232 |
| SE | 31336 | ENSG00000157426 | AASDH | chr4 | - | 56342834 | 56342966 | 56338286 | 56338791 | 56349262 | 56350058 | 27 | 15 | 49 | 21 | 54 | 4 | 0 | 2 | 2 | 3 | 0 | 10 | 11 | 0 | 0 | 0 | 0 | 0 | 0 | 0 | 281 | 149 | 1 | 1 | 0.782 | 1 | 0.929 | 0.848 | 0.905 | NA | 1 | 1 | NA | NA | -0.107 |
| SE | 31337 | ENSG00000157426 | AASDH | chr4 | - | 56343561 | 56343684 | 56338286 | 56338791 | 56349262 | 56350058 | 21 | 8 | 12 | 11 | 20 | 4 | 0 | 2 | 2 | 3 | 0 | 4 | 0 | 4 | 0 | 0 | 0 | 0 | 0 | 0 | 272 | 149 | 1 | 1 | 0.742 | 1 | 0.767 | 0.751 | 0.785 | NA | 1 | NA | 1 | NA | -0.191 |
| SE | 31338 | ENSG00000157426 | AASDH | chr4 | - | 56343561 | 56343684 | 56342834 | 56342966 | 56349262 | 56350058 | 34 | 7 | 33 | 22 | 92 | 0 | 0 | 0 | 1 | 0 | 0 | 8 | 2 | 0 | 0 | 0 | 0 | 0 | 0 | 0 | 272 | 149 | 1 | 1 | 1 | 1 | 1 | 0.923 | 1 | NA | 1 | 1 | NA | NA | -0.015 |
| SE | 31339 | ENSG00000157426 | AASDH | chr4 | - | 56345126 | 56345290 | 56343561 | 56343684 | 56349262 | 56350058 | 77 | 61 | 57 | 71 | 143 | 19 | 4 | 8 | 11 | 10 | 0 | 39 | 8 | 16 | 0 | 0 | 2 | 2 | 0 | 0 | 298 | 149 | 1 | 1 | 0.67 | 0.884 | 0.781 | 0.763 | 0.877 | NA | 0.907 | 0.667 | 1 | NA | -0.063 |
| SE | 31340 | ENSG00000157426 | AASDH | chr4 | - | 56351341 | 56351457 | 56348927 | 56350058 | 56353403 | 56353596 | 62 | 53 | 60 | 62 | 102 | 0 | 0 | 0 | 0 | 3 | 4 | 38 | 7 | 13 | 0 | 0 | 0 | 0 | 0 | 0 | 265 | 149 | 1 | 1 | 1 | 1 | 1 | 1 | 0.95 | 1 | 1 | 1 | 1 | NA | -0.01 |
| SE | 31341 | ENSG00000157426 | AASDH | chr4 | - | 56354038 | 56354211 | 56353403 | 56353596 | 56355181 | 56355423 | 25 | 24 | 41 | 34 | 38 | 0 | 0 | 0 | 0 | 0 | 0 | 7 | 0 | 0 | 0 | 0 | 3 | 0 | 0 | 0 | 298 | 149 | 1 | 1 | 1 | 1 | 1 | 1 | 1 | NA | 0.538 | NA | NA | NA | 0.462 |
| SE | 31342 | ENSG00000157426 | AASDH | chr4 | - | 56354704 | 56354811 | 56354038 | 56354211 | 56355181 | 56355423 | 73 | 63 | 52 | 88 | 123 | 0 | 0 | 3 | 0 | 0 | 9 | 38 | 27 | 32 | 6 | 0 | 0 | 0 | 0 | 0 | 256 | 149 | 1 | 1 | 1 | 1 | 0.91 | 1 | 1 | 1 | 1 | 1 | 1 | 1 | -0.018 |
| SE | 31343 | ENSG00000157426 | AASDH | chr4 | - | 56371450 | 56371643 | 56355181 | 56355423 | 56378147 | 56378464 | 105 | 61 | 88 | 95 | 127 | 14 | 10 | 8 | 3 | 14 | 0 | 49 | 22 | 17 | 3 | 0 | 14 | 19 | 11 | 0 | 298 | 149 | 1 | 1 | 0.789 | 0.753 | 0.846 | 0.941 | 0.819 | NA | 0.636 | 0.367 | 0.436 | 1 | 0.22 |
| SE | 31345 | ENSG00000157426 | AASDH | chr4 | - | 56382476 | 56382597 | 56378147 | 56378464 | 56384069 | 56384341 | 83 | 57 | 88 | 69 | 115 | 1 | 1 | 0 | 0 | 2 | 8 | 34 | 50 | 29 | 16 | 0 | 0 | 0 | 0 | 0 | 270 | 149 | 1 | 1 | 0.979 | 0.969 | 1 | 1 | 0.969 | 1 | 1 | 1 | 1 | 1 | -0.017 |
| SE | 31349 | ENSG00000197653 | DNAH10 | chr12 | + | 123793941 | 123794112 | 123789926 | 123790121 | 123796655 | 123796832 | 0 | 0 | 0 | 3 | 1 | 0 | 1 | 0 | 0 | 0 | 0 | 1 | 3 | 2 | 0 | 0 | 0 | 0 | 0 | 0 | 298 | 149 | 1 | 1 | NA | 0 | NA | 1 | 1 | NA | 1 | 1 | 1 | NA | -0.333 |
| SE | 31350 | ENSG00000197653 | DNAH10 | chr12 | + | 123800215 | 123800388 | 123799245 | 123799371 | 123801280 | 123801432 | 0 | 0 | 1 | 0 | 1 | 0 | 0 | 0 | 1 | 0 | 0 | 0 | 3 | 2 | 0 | 0 | 0 | 0 | 0 | 0 | 298 | 149 | 1 | 1 | NA | NA | 1 | 0 | 1 | NA | NA | 1 | 1 | NA | -0.333 |
| SE | 31354 | ENSG00000124535 | WRNIP1 | chr6 | + | 2779262 | 2779492 | 2765392 | 2766444 | 2783405 | 2783561 | 55 | 28 | 53 | 73 | 55 | 0 | 0 | 0 | 2 | 0 | 46 | 38 | 37 | 21 | 174 | 0 | 0 | 0 | 0 | 0 | 298 | 149 | 1 | 1 | 1 | 1 | 1 | 0.948 | 1 | 1 | 1 | 1 | 1 | 1 | -0.01 |
| SE | 31355 | ENSG00000124535 | WRNIP1 | chr6 | + | 2784323 | 2784403 | 2783405 | 2783561 | 2785006 | 2785744 | 106 | 62 | 109 | 164 | 103 | 1 | 0 | 0 | 0 | 0 | 94 | 125 | 66 | 130 | 524 | 0 | 5 | 0 | 0 | 0 | 229 | 149 | 1 | 1 | 0.986 | 1 | 1 | 1 | 1 | 1 | 0.942 | 1 | 1 | 1 | 0.009 |
| SE | 31356 | ENSG00000005436 | GCFC2 | chr2 | - | 75665928 | 75666053 | 75662704 | 75664783 | 75670137 | 75670284 | 57 | 30 | 33 | 28 | 101 | 2 | 0 | 0 | 0 | 0 | 0 | 8 | 8 | 14 | 7 | 0 | 0 | 0 | 0 | 0 | 274 | 149 | 1 | 1 | 0.939 | 1 | 1 | 1 | 1 | NA | 1 | 1 | 1 | 1 | -0.012 |
| SE | 31357 | ENSG00000005436 | GCFC2 | chr2 | - | 75673443 | 75673520 | 75671949 | 75672016 | 75687826 | 75687892 | 42 | 30 | 30 | 40 | 74 | 0 | 2 | 0 | 0 | 0 | 0 | 11 | 14 | 1 | 6 | 0 | 0 | 0 | 0 | 0 | 226 | 149 | 1 | 1 | 1 | 0.908 | 1 | 1 | 1 | NA | 1 | 1 | 1 | 1 | -0.018 |
| SE | 31358 | ENSG00000005436 | GCFC2 | chr2 | - | 75680192 | 75680314 | 75673443 | 75673520 | 75687826 | 75687977 | 51 | 44 | 32 | 47 | 123 | 1 | 2 | 3 | 2 | 0 | 9 | 20 | 27 | 3 | 0 | 0 | 0 | 0 | 0 | 0 | 271 | 149 | 1 | 1 | 0.966 | 0.924 | 0.854 | 0.928 | 1 | 1 | 1 | 1 | 1 | NA | -0.066 |
| SE | 31359 | ENSG00000005436 | GCFC2 | chr2 | - | 75689025 | 75689225 | 75687826 | 75687977 | 75689968 | 75690081 | 122 | 84 | 111 | 129 | 166 | 0 | 1 | 0 | 0 | 0 | 0 | 30 | 41 | 26 | 17 | 0 | 0 | 0 | 0 | 0 | 298 | 149 | 1 | 1 | 1 | 0.977 | 1 | 1 | 1 | NA | 1 | 1 | 1 | 1 | -0.005 |
| SE | 31360 | ENSG00000005436 | GCFC2 | chr2 | - | 75690637 | 75690719 | 75689968 | 75690081 | 75691976 | 75692100 | 118 | 73 | 91 | 74 | 215 | 0 | 0 | 0 | 3 | 0 | 10 | 45 | 40 | 23 | 17 | 0 | 0 | 0 | 0 | 0 | 231 | 149 | 1 | 1 | 1 | 1 | 1 | 0.941 | 1 | 1 | 1 | 1 | 1 | 1 | -0.012 |
| SE | 31362 | ENSG00000005436 | GCFC2 | chr2 | - | 75694240 | 75694427 | 75691976 | 75692100 | 75701189 | 75701287 | 89 | 72 | 68 | 79 | 191 | 0 | 0 | 0 | 0 | 0 | 9 | 45 | 27 | 4 | 18 | 0 | 0 | 1 | 0 | 0 | 298 | 149 | 1 | 1 | 1 | 1 | 1 | 1 | 1 | 1 | 1 | 0.931 | 1 | 1 | 0.014 |
| SE | 31363 | ENSG00000005436 | GCFC2 | chr2 | - | 75696199 | 75696315 | 75694240 | 75694427 | 75701189 | 75701287 | 85 | 55 | 98 | 97 | 143 | 1 | 2 | 0 | 0 | 1 | 8 | 36 | 46 | 4 | 16 | 0 | 0 | 0 | 0 | 0 | 265 | 149 | 1 | 1 | 0.98 | 0.939 | 1 | 1 | 0.988 | 1 | 1 | 1 | 1 | 1 | -0.019 |
| SE | 31368 | ENSG00000184661 | CDCA2 | chr8 | + | 25472112 | 25472184 | 25469895 | 25469980 | 25479912 | 25480124 | 0 | 0 | 0 | 2 | 0 | 0 | 0 | 1 | 5 | 6 | 0 | 0 | 5 | 0 | 0 | 0 | 0 | 9 | 2 | 0 | 221 | 149 | 1 | 1 | NA | NA | 0 | 0.212 | 0 | NA | NA | 0.272 | 0 | NA | -0.065 |
| SE | 31375 | ENSG00000117525 | F3 | chr1 | - | 94532320 | 94532480 | 94529172 | 94530596 | 94533089 | 94533268 | 1 | 0 | 0 | 2 | 0 | 0 | 0 | 0 | 1 | 0 | 1 | 0 | 0 | 9 | 0 | 0 | 0 | 0 | 1 | 0 | 298 | 149 | 1 | 1 | 1 | NA | NA | 0.5 | NA | 1 | NA | NA | 0.818 | NA | -0.159 |
| SE | 31377 | ENSG00000268758 | ADGRE4P | chr19 | - | 6954603 | 6954699 | 6952499 | 6953318 | 6957043 | 6957224 | 6 | 21 | 13 | 17 | 17 | 0 | 0 | 0 | 0 | 1 | 0 | 0 | 0 | 4 | 0 | 0 | 0 | 0 | 0 | 0 | 245 | 149 | 1 | 1 | 1 | 1 | 1 | 1 | 0.912 | NA | NA | NA | 1 | NA | -0.018 |
| SE | 31378 | ENSG00000268758 | ADGRE4P | chr19 | - | 6957043 | 6957224 | 6954603 | 6954699 | 6960296 | 6960388 | 14 | 19 | 24 | 22 | 12 | 0 | 0 | 0 | 0 | 1 | 0 | 10 | 2 | 6 | 0 | 0 | 0 | 0 | 0 | 0 | 298 | 149 | 1 | 1 | 1 | 1 | 1 | 1 | 0.857 | NA | 1 | 1 | 1 | NA | -0.029 |
| SE | 31380 | ENSG00000268758 | ADGRE4P | chr19 | - | 6972239 | 6972404 | 6971168 | 6971363 | 6974490 | 6974674 | 13 | 45 | 30 | 16 | 42 | 0 | 0 | 0 | 2 | 0 | 9 | 0 | 3 | 0 | 1 | 0 | 0 | 0 | 0 | 0 | 298 | 149 | 1 | 1 | 1 | 1 | 1 | 0.8 | 1 | 1 | NA | 1 | NA | 1 | -0.04 |
| SE | 31381 | ENSG00000268758 | ADGRE4P | chr19 | - | 6977247 | 6977355 | 6974490 | 6974674 | 6981079 | 6981215 | 9 | 25 | 21 | 9 | 23 | 0 | 0 | 0 | 0 | 0 | 3 | 0 | 9 | 0 | 1 | 8 | 0 | 0 | 0 | 0 | 257 | 149 | 1 | 1 | 1 | 1 | 1 | 1 | 1 | 0.179 | NA | 1 | NA | 1 | 0.274 |
| SE | 31384 | ENSG00000151552 | QDPR | chr4 | - | 17504378 | 17504475 | 17501718 | 17501859 | 17509270 | 17509363 | 30 | 24 | 35 | 46 | 44 | 1 | 0 | 1 | 2 | 0 | 30 | 39 | 20 | 32 | 48 | 0 | 0 | 5 | 3 | 0 | 246 | 149 | 1 | 1 | 0.948 | 1 | 0.955 | 0.933 | 1 | 1 | 1 | 0.708 | 0.866 | 1 | 0.052 |
| SE | 31385 | ENSG00000151552 | QDPR | chr4 | - | 17504378 | 17504475 | 17501718 | 17501859 | 17511949 | 17512086 | 17 | 7 | 17 | 17 | 24 | 1 | 0 | 0 | 0 | 0 | 4 | 26 | 4 | 19 | 22 | 0 | 0 | 0 | 0 | 0 | 246 | 149 | 1 | 1 | 0.911 | 1 | 1 | 1 | 1 | 1 | 1 | 1 | 1 | 1 | -0.018 |
| SE | 31386 | ENSG00000151552 | QDPR | chr4 | - | 17509270 | 17509363 | 17501718 | 17501859 | 17511949 | 17511981 | 1 | 0 | 8 | 2 | 1 | 1 | 0 | 0 | 0 | 0 | 0 | 0 | 5 | 0 | 0 | 0 | 0 | 0 | 0 | 0 | 242 | 149 | 1 | 1 | 0.381 | NA | 1 | 1 | 1 | NA | NA | 1 | NA | NA | -0.155 |
| SE | 31387 | ENSG00000151552 | QDPR | chr4 | - | 17509270 | 17509363 | 17504378 | 17504475 | 17511949 | 17511981 | 13 | 17 | 27 | 30 | 21 | 0 | 0 | 1 | 0 | 0 | 26 | 13 | 16 | 13 | 26 | 0 | 0 | 0 | 0 | 0 | 242 | 149 | 1 | 1 | 1 | 1 | 0.943 | 1 | 1 | 1 | 1 | 1 | 1 | 1 | -0.011 |
| SE | 31398 | ENSG00000123933 | MXD4 | chr4 | - | 2261724 | 2261824 | 2257981 | 2258011 | 2261916 | 2262022 | 142 | 32 | 32 | 51 | 12 | 0 | 0 | 1 | 0 | 0 | 79 | 85 | 53 | 33 | 161 | 0 | 0 | 0 | 4 | 0 | 249 | 149 | 1 | 1 | 1 | 1 | 0.95 | 1 | 1 | 1 | 1 | 1 | 0.832 | 1 | 0.024 |
| SE | 31399 | ENSG00000109606 | DHX15 | chr4 | - | 24529600 | 24529770 | 24527474 | 24528041 | 24532863 | 24532951 | 854 | 635 | 659 | 725 | 700 | 0 | 0 | 0 | 0 | 1 | 128 | 328 | 342 | 142 | 51 | 0 | 0 | 0 | 0 | 0 | 298 | 149 | 1 | 1 | 1 | 1 | 1 | 1 | 0.997 | 1 | 1 | 1 | 1 | 1 | -0.001 |
| SE | 31400 | ENSG00000109606 | DHX15 | chr4 | - | 24540839 | 24540948 | 24539708 | 24540299 | 24541872 | 24541899 | 194 | 113 | 177 | 210 | 267 | 1 | 0 | 1 | 0 | 1 | 25 | 81 | 71 | 26 | 10 | 0 | 0 | 0 | 2 | 0 | 258 | 149 | 1 | 1 | 0.991 | 1 | 0.99 | 1 | 0.994 | 1 | 1 | 1 | 0.882 | 1 | 0.019 |
| SE | 31401 | ENSG00000109606 | DHX15 | chr4 | - | 24570653 | 24570847 | 24556250 | 24556410 | 24576242 | 24576678 | 489 | 378 | 498 | 504 | 485 | 0 | 2 | 2 | 0 | 0 | 117 | 263 | 280 | 230 | 132 | 0 | 0 | 0 | 0 | 0 | 298 | 149 | 1 | 1 | 1 | 0.99 | 0.992 | 1 | 1 | 1 | 1 | 1 | 1 | 1 | -0.004 |
| SE | 31402 | ENSG00000109606 | DHX15 | chr4 | - | 24576242 | 24576678 | 24570653 | 24570847 | 24584322 | 24584554 | 473 | 383 | 443 | 486 | 461 | 0 | 0 | 0 | 1 | 0 | 128 | 315 | 320 | 211 | 138 | 0 | 0 | 0 | 0 | 0 | 298 | 149 | 1 | 1 | 1 | 1 | 1 | 0.996 | 1 | 1 | 1 | 1 | 1 | 1 | -0.001 |
| SE | 31405 | ENSG00000155508 | CNOT8 | chr5 | + | 154858638 | 154858801 | 154858516 | 154858543 | 154865191 | 154865385 | 2 | 0 | 0 | 0 | 3 | 1 | 0 | 0 | 0 | 0 | 7 | 0 | 0 | 0 | 0 | 0 | 0 | 3 | 0 | 0 | 298 | 149 | 1 | 1 | 0.5 | NA | NA | NA | 1 | 1 | NA | 0 | NA | NA | 0.25 |
| SE | 31406 | ENSG00000155508 | CNOT8 | chr5 | + | 154858638 | 154858816 | 154858240 | 154858525 | 154865191 | 154865385 | 9 | 3 | 6 | 2 | 2 | 1 | 0 | 0 | 0 | 0 | 0 | 1 | 4 | 0 | 0 | 0 | 0 | 3 | 0 | 0 | 298 | 149 | 1 | 1 | 0.818 | 1 | 1 | 1 | 1 | NA | 1 | 0.4 | NA | NA | 0.264 |
| SE | 31411 | ENSG00000155508 | CNOT8 | chr5 | + | 154859404 | 154859496 | 154858651 | 154858768 | 154859694 | 154859870 | 0 | 2 | 0 | 1 | 0 | 0 | 0 | 0 | 0 | 0 | 0 | 0 | 0 | 0 | 0 | 0 | 0 | 2 | 0 | 0 | 241 | 149 | 1 | 1 | NA | 1 | NA | 1 | NA | NA | NA | 0 | NA | NA | 1 |
| SE | 31412 | ENSG00000155508 | CNOT8 | chr5 | + | 154859404 | 154859496 | 154858651 | 154858768 | 154870660 | 154870822 | 0 | 2 | 0 | 1 | 2 | 1 | 0 | 0 | 4 | 0 | 0 | 0 | 0 | 0 | 0 | 0 | 7 | 5 | 0 | 0 | 241 | 149 | 1 | 1 | 0 | 1 | NA | 0.134 | 1 | NA | 0 | 0 | NA | NA | 0.533 |
| SE | 31413 | ENSG00000155508 | CNOT8 | chr5 | + | 154859404 | 154859496 | 154858651 | 154858768 | 154871729 | 154871874 | 0 | 2 | 0 | 1 | 0 | 0 | 0 | 0 | 1 | 0 | 0 | 0 | 0 | 0 | 0 | 0 | 0 | 0 | 3 | 0 | 241 | 149 | 1 | 1 | NA | 1 | NA | 0.382 | NA | NA | NA | NA | 0 | NA | 0.691 |
| SE | 31414 | ENSG00000155508 | CNOT8 | chr5 | + | 154859694 | 154859873 | 154858624 | 154858768 | 154863206 | 154863395 | 1 | 0 | 0 | 1 | 0 | 105 | 90 | 81 | 89 | 65 | 0 | 0 | 2 | 0 | 0 | 33 | 50 | 54 | 18 | 41 | 298 | 149 | 1 | 1 | 0.005 | 0 | 0 | 0.006 | 0 | 0 | 0 | 0.018 | 0 | 0 | -0.001 |
| SE | 31416 | ENSG00000155508 | CNOT8 | chr5 | + | 154863206 | 154863395 | 154858516 | 154858816 | 154870660 | 154870822 | 9 | 13 | 12 | 8 | 3 | 0 | 0 | 0 | 0 | 0 | 0 | 7 | 9 | 7 | 0 | 0 | 3 | 0 | 0 | 0 | 298 | 149 | 1 | 1 | 1 | 1 | 1 | 1 | 1 | NA | 0.538 | 1 | 1 | NA | 0.154 |
| SE | 31418 | ENSG00000155508 | CNOT8 | chr5 | + | 154863206 | 154863395 | 154858536 | 154858801 | 154870660 | 154870822 | 3 | 2 | 4 | 1 | 0 | 1 | 0 | 0 | 0 | 0 | 6 | 0 | 1 | 0 | 0 | 0 | 0 | 0 | 0 | 0 | 298 | 149 | 1 | 1 | 0.6 | 1 | 1 | 1 | NA | 1 | NA | 1 | NA | NA | -0.1 |
| SE | 31420 | ENSG00000155508 | CNOT8 | chr5 | + | 154863206 | 154863395 | 154858626 | 154858768 | 154867691 | 154867733 | 105 | 90 | 81 | 89 | 65 | 0 | 0 | 0 | 5 | 0 | 33 | 50 | 54 | 24 | 41 | 0 | 0 | 0 | 0 | 0 | 298 | 149 | 1 | 1 | 1 | 1 | 1 | 0.899 | 1 | 1 | 1 | 1 | 1 | 1 | -0.02 |
| SE | 31422 | ENSG00000155508 | CNOT8 | chr5 | + | 154863206 | 154863395 | 154858626 | 154858768 | 154871729 | 154871781 | 105 | 91 | 81 | 91 | 65 | 0 | 0 | 0 | 1 | 0 | 33 | 50 | 54 | 18 | 41 | 0 | 0 | 0 | 3 | 0 | 298 | 149 | 1 | 1 | 1 | 1 | 1 | 0.978 | 1 | 1 | 1 | 1 | 0.75 | 1 | 0.046 |
| SE | 31423 | ENSG00000155508 | CNOT8 | chr5 | + | 154863206 | 154863395 | 154859352 | 154859496 | 154865191 | 154865240 | 217 | 161 | 220 | 202 | 314 | 0 | 0 | 0 | 0 | 6 | 169 | 172 | 157 | 118 | 82 | 0 | 0 | 0 | 2 | 0 | 298 | 149 | 1 | 1 | 1 | 1 | 1 | 1 | 0.963 | 1 | 1 | 1 | 0.967 | 1 | -0.001 |
| SE | 31424 | ENSG00000155508 | CNOT8 | chr5 | + | 154863206 | 154863395 | 154859352 | 154859496 | 154870660 | 154870822 | 4 | 6 | 4 | 12 | 6 | 0 | 0 | 0 | 0 | 2 | 11 | 0 | 2 | 0 | 0 | 0 | 0 | 0 | 0 | 0 | 298 | 149 | 1 | 1 | 1 | 1 | 1 | 1 | 0.6 | 1 | NA | 1 | NA | NA | -0.08 |
| SE | 31425 | ENSG00000155508 | CNOT8 | chr5 | + | 154863206 | 154863395 | 154859352 | 154859496 | 154871729 | 154871781 | 4 | 5 | 4 | 14 | 6 | 0 | 0 | 0 | 0 | 0 | 11 | 0 | 1 | 0 | 0 | 0 | 0 | 3 | 0 | 0 | 298 | 149 | 1 | 1 | 1 | 1 | 1 | 1 | 1 | 1 | 0 | 1 | NA | NA | 0.333 |
| SE | 31428 | ENSG00000155508 | CNOT8 | chr5 | + | 154865191 | 154865385 | 154858516 | 154858816 | 154870660 | 154870704 | 276 | 216 | 251 | 202 | 388 | 0 | 0 | 0 | 0 | 0 | 266 | 238 | 232 | 157 | 148 | 0 | 3 | 0 | 0 | 0 | 298 | 149 | 1 | 1 | 1 | 1 | 1 | 1 | 1 | 1 | 0.975 | 1 | 1 | 1 | 0.005 |
| SE | 31431 | ENSG00000155508 | CNOT8 | chr5 | + | 154865191 | 154865385 | 154858538 | 154858768 | 154871729 | 154871781 | 32 | 13 | 25 | 25 | 10 | 0 | 0 | 0 | 1 | 0 | 14 | 10 | 39 | 19 | 23 | 0 | 0 | 0 | 3 | 0 | 298 | 149 | 1 | 1 | 1 | 1 | 1 | 0.926 | 1 | 1 | 1 | 1 | 0.76 | 1 | 0.033 |
| SE | 31432 | ENSG00000155508 | CNOT8 | chr5 | + | 154865191 | 154865385 | 154858638 | 154858801 | 154870660 | 154870704 | 269 | 215 | 249 | 201 | 386 | 1 | 0 | 0 | 0 | 0 | 273 | 237 | 230 | 157 | 148 | 0 | 0 | 0 | 0 | 0 | 298 | 149 | 1 | 1 | 0.993 | 1 | 1 | 1 | 1 | 1 | 1 | 1 | 1 | 1 | -0.001 |
| SE | 31433 | ENSG00000155508 | CNOT8 | chr5 | + | 154865191 | 154865385 | 154863206 | 154863395 | 154867691 | 154867733 | 224 | 166 | 226 | 205 | 321 | 0 | 0 | 0 | 0 | 0 | 167 | 194 | 181 | 148 | 82 | 0 | 0 | 0 | 6 | 0 | 298 | 149 | 1 | 1 | 1 | 1 | 1 | 1 | 1 | 1 | 1 | 1 | 0.925 | 1 | 0.015 |
| SE | 31434 | ENSG00000155508 | CNOT8 | chr5 | + | 154865191 | 154865385 | 154863206 | 154863395 | 154870660 | 154870704 | 481 | 372 | 465 | 391 | 694 | 0 | 2 | 0 | 0 | 0 | 424 | 409 | 386 | 275 | 230 | 0 | 0 | 1 | 0 | 0 | 298 | 149 | 1 | 1 | 1 | 0.989 | 1 | 1 | 1 | 1 | 1 | 0.995 | 1 | 1 | -0.001 |
| SE | 31435 | ENSG00000155508 | CNOT8 | chr5 | + | 154865191 | 154865385 | 154863206 | 154863395 | 154871729 | 154871781 | 230 | 161 | 226 | 197 | 315 | 0 | 1 | 0 | 2 | 0 | 170 | 174 | 156 | 121 | 82 | 0 | 0 | 0 | 0 | 0 | 298 | 149 | 1 | 1 | 1 | 0.988 | 1 | 0.98 | 1 | 1 | 1 | 1 | 1 | 1 | -0.006 |
| SE | 31436 | ENSG00000155508 | CNOT8 | chr5 | + | 154867691 | 154867792 | 154858626 | 154858768 | 154870660 | 154870822 | 13 | 7 | 7 | 21 | 23 | 1 | 0 | 0 | 4 | 0 | 2 | 17 | 10 | 20 | 0 | 0 | 7 | 5 | 0 | 0 | 250 | 149 | 1 | 1 | 0.886 | 1 | 1 | 0.758 | 1 | 1 | 0.591 | 0.544 | 1 | NA | 0.145 |
| SE | 31437 | ENSG00000155508 | CNOT8 | chr5 | + | 154867691 | 154867792 | 154858626 | 154858768 | 154871729 | 154871874 | 0 | 0 | 0 | 5 | 0 | 0 | 0 | 0 | 1 | 0 | 0 | 0 | 0 | 0 | 0 | 0 | 0 | 0 | 3 | 0 | 250 | 149 | 1 | 1 | NA | NA | NA | 0.749 | NA | NA | NA | NA | 0 | NA | 0.749 |
| SE | 31438 | ENSG00000155508 | CNOT8 | chr5 | + | 154867691 | 154867792 | 154863206 | 154863395 | 154870660 | 154870822 | 13 | 7 | 7 | 16 | 23 | 0 | 2 | 0 | 0 | 0 | 2 | 17 | 10 | 26 | 0 | 0 | 0 | 1 | 0 | 0 | 250 | 149 | 1 | 1 | 1 | 0.676 | 1 | 1 | 1 | 1 | 1 | 0.856 | 1 | NA | -0.029 |
| SE | 31439 | ENSG00000155508 | CNOT8 | chr5 | + | 154867691 | 154867792 | 154863206 | 154863395 | 154871729 | 154871874 | 0 | 0 | 0 | 1 | 0 | 0 | 1 | 0 | 2 | 0 | 0 | 0 | 0 | 6 | 0 | 0 | 0 | 0 | 0 | 0 | 250 | 149 | 1 | 1 | NA | 0 | NA | 0.23 | NA | NA | NA | NA | 1 | NA | -0.885 |
| SE | 31441 | ENSG00000155508 | CNOT8 | chr5 | + | 154867691 | 154867792 | 154865191 | 154865385 | 154871729 | 154871874 | 5 | 7 | 5 | 13 | 11 | 17 | 4 | 10 | 7 | 7 | 2 | 4 | 18 | 27 | 0 | 12 | 2 | 0 | 3 | 0 | 250 | 149 | 1 | 1 | 0.149 | 0.511 | 0.23 | 0.525 | 0.484 | 0.09 | 0.544 | 1 | 0.843 | NA | -0.239 |
| SE | 31442 | ENSG00000155508 | CNOT8 | chr5 | + | 154870660 | 154870822 | 154858638 | 154858768 | 154871729 | 154871831 | 153 | 118 | 121 | 160 | 229 | 0 | 0 | 0 | 1 | 0 | 53 | 143 | 188 | 165 | 105 | 0 | 0 | 0 | 3 | 0 | 298 | 149 | 1 | 1 | 1 | 1 | 1 | 0.988 | 1 | 1 | 1 | 1 | 0.965 | 1 | 0.005 |
| SE | 31443 | ENSG00000155508 | CNOT8 | chr5 | + | 154870660 | 154870822 | 154859404 | 154859496 | 154871729 | 154871831 | 152 | 118 | 121 | 156 | 231 | 0 | 0 | 0 | 0 | 0 | 53 | 136 | 183 | 165 | 105 | 0 | 3 | 0 | 0 | 0 | 298 | 149 | 1 | 1 | 1 | 1 | 1 | 1 | 1 | 1 | 0.958 | 1 | 1 | 1 | 0.008 |
| SE | 31444 | ENSG00000155508 | CNOT8 | chr5 | + | 154870660 | 154870822 | 154863206 | 154863395 | 154871729 | 154871831 | 152 | 120 | 121 | 156 | 229 | 0 | 1 | 0 | 2 | 0 | 53 | 136 | 184 | 165 | 105 | 0 | 0 | 0 | 0 | 0 | 298 | 149 | 1 | 1 | 1 | 0.984 | 1 | 0.975 | 1 | 1 | 1 | 1 | 1 | 1 | -0.008 |
| SE | 31446 | ENSG00000155508 | CNOT8 | chr5 | + | 154870660 | 154870822 | 154867691 | 154867792 | 154871729 | 154871831 | 172 | 131 | 136 | 179 | 257 | 0 | 0 | 0 | 1 | 0 | 62 | 173 | 204 | 194 | 105 | 0 | 0 | 0 | 0 | 0 | 298 | 149 | 1 | 1 | 1 | 1 | 1 | 0.989 | 1 | 1 | 1 | 1 | 1 | 1 | -0.002 |
| SE | 31448 | ENSG00000159496 | RGL4 | chr22 | + | 23697170 | 23697245 | 23696483 | 23696688 | 23697837 | 23697861 | 94 | 32 | 96 | 108 | 64 | 0 | 4 | 0 | 0 | 1 | 90 | 139 | 122 | 706 | 230 | 0 | 0 | 0 | 0 | 0 | 224 | 149 | 1 | 1 | 1 | 0.842 | 1 | 1 | 0.977 | 1 | 1 | 1 | 1 | 1 | -0.036 |
| SE | 31455 | ENSG00000136261 | BZW2 | chr7 | + | 16689796 | 16689906 | 16682779 | 16682845 | 16696914 | 16697047 | 2 | 7 | 4 | 2 | 3 | 0 | 0 | 0 | 0 | 0 | 2 | 0 | 1 | 2 | 0 | 0 | 0 | 0 | 2 | 0 | 259 | 149 | 1 | 1 | 1 | 1 | 1 | 1 | 1 | 1 | NA | 1 | 0.365 | NA | 0.212 |
| SE | 31456 | ENSG00000136261 | BZW2 | chr7 | + | 16694833 | 16695004 | 16682779 | 16682845 | 16696914 | 16696941 | 29 | 38 | 43 | 40 | 35 | 0 | 0 | 0 | 0 | 0 | 31 | 12 | 26 | 8 | 6 | 0 | 0 | 0 | 2 | 0 | 298 | 149 | 1 | 1 | 1 | 1 | 1 | 1 | 1 | 1 | 1 | 1 | 0.667 | 1 | 0.067 |
| SE | 31457 | ENSG00000136261 | BZW2 | chr7 | + | 16694833 | 16695004 | 16689796 | 16689906 | 16696914 | 16696941 | 81 | 80 | 101 | 92 | 119 | 0 | 2 | 0 | 0 | 1 | 74 | 28 | 60 | 35 | 31 | 0 | 0 | 0 | 0 | 0 | 298 | 149 | 1 | 1 | 1 | 0.952 | 1 | 1 | 0.983 | 1 | 1 | 1 | 1 | 1 | -0.013 |
| SE | 31458 | ENSG00000136261 | BZW2 | chr7 | + | 16698047 | 16698186 | 16696914 | 16697061 | 16704546 | 16704662 | 132 | 83 | 169 | 182 | 270 | 0 | 0 | 2 | 1 | 0 | 181 | 86 | 72 | 58 | 84 | 3 | 0 | 0 | 0 | 0 | 288 | 149 | 1 | 1 | 1 | 1 | 0.978 | 0.989 | 1 | 0.969 | 1 | 1 | 1 | 1 | 0 |
| SE | 31461 | ENSG00000178950 | GAK | chr4 | - | 851749 | 851974 | 850935 | 851084 | 859605 | 859722 | 162 | 86 | 107 | 146 | 53 | 1 | 0 | 2 | 0 | 0 | 221 | 167 | 127 | 138 | 197 | 0 | 0 | 0 | 3 | 0 | 298 | 149 | 1 | 1 | 0.988 | 1 | 0.964 | 1 | 1 | 1 | 1 | 1 | 0.958 | 1 | -0.001 |
| SE | 31462 | ENSG00000178950 | GAK | chr4 | - | 859361 | 859493 | 850935 | 851084 | 859605 | 859722 | 0 | 0 | 1 | 1 | 0 | 1 | 0 | 2 | 0 | 0 | 0 | 0 | 0 | 8 | 0 | 0 | 0 | 0 | 3 | 0 | 281 | 149 | 1 | 1 | 0 | NA | 0.21 | 1 | NA | NA | NA | NA | 0.586 | NA | -0.183 |
| SE | 31463 | ENSG00000178950 | GAK | chr4 | - | 859361 | 859493 | 851749 | 851974 | 859605 | 859722 | 4 | 0 | 1 | 3 | 0 | 44 | 19 | 36 | 34 | 13 | 0 | 4 | 0 | 0 | 3 | 68 | 25 | 33 | 36 | 35 | 281 | 149 | 1 | 1 | 0.046 | 0 | 0.015 | 0.045 | 0 | 0 | 0.078 | 0 | 0 | 0.043 | -0.003 |
| SE | 31468 | ENSG00000178950 | GAK | chr4 | - | 888846 | 888970 | 884036 | 884086 | 890531 | 890622 | 169 | 58 | 118 | 138 | 71 | 0 | 0 | 0 | 0 | 0 | 149 | 133 | 89 | 105 | 248 | 0 | 0 | 2 | 0 | 0 | 273 | 149 | 1 | 1 | 1 | 1 | 1 | 1 | 1 | 1 | 1 | 0.96 | 1 | 1 | 0.008 |
| SE | 31469 | ENSG00000178950 | GAK | chr4 | - | 896459 | 896549 | 893873 | 894009 | 898032 | 898158 | 393 | 219 | 355 | 339 | 520 | 2 | 0 | 0 | 0 | 0 | 471 | 481 | 225 | 390 | 640 | 10 | 0 | 0 | 0 | 0 | 239 | 149 | 1 | 1 | 0.992 | 1 | 1 | 1 | 1 | 0.967 | 1 | 1 | 1 | 1 | 0.005 |
| SE | 31470 | ENSG00000178950 | GAK | chr4 | - | 898032 | 898158 | 893942 | 894009 | 904636 | 904779 | 126 | 74 | 106 | 116 | 120 | 0 | 0 | 0 | 0 | 0 | 201 | 136 | 73 | 140 | 217 | 0 | 0 | 0 | 5 | 0 | 275 | 149 | 1 | 1 | 1 | 1 | 1 | 1 | 1 | 1 | 1 | 1 | 0.938 | 1 | 0.012 |
| SE | 31474 | ENSG00000178950 | GAK | chr4 | - | 912734 | 912794 | 912029 | 912174 | 913606 | 913668 | 34 | 11 | 21 | 23 | 13 | 0 | 0 | 1 | 0 | 0 | 39 | 25 | 20 | 33 | 17 | 0 | 0 | 0 | 0 | 0 | 209 | 149 | 1 | 1 | 1 | 1 | 0.937 | 1 | 1 | 1 | 1 | 1 | 1 | 1 | -0.013 |
| SE | 31476 | ENSG00000178950 | GAK | chr4 | - | 913606 | 913668 | 904636 | 904779 | 932042 | 932220 | 68 | 27 | 44 | 41 | 15 | 2 | 0 | 1 | 0 | 0 | 76 | 47 | 26 | 47 | 71 | 0 | 1 | 0 | 0 | 0 | 211 | 149 | 1 | 1 | 0.96 | 1 | 0.969 | 1 | 1 | 1 | 0.971 | 1 | 1 | 1 | -0.008 |
| SE | 31477 | ENSG00000111885 | MAN1A1 | chr6 | - | 119189663 | 119189883 | 119188404 | 119188577 | 119193776 | 119193892 | 876 | 402 | 660 | 687 | 576 | 0 | 0 | 2 | 0 | 0 | 311 | 814 | 975 | 393 | 312 | 0 | 0 | 0 | 0 | 0 | 298 | 149 | 1 | 1 | 1 | 1 | 0.994 | 1 | 1 | 1 | 1 | 1 | 1 | 1 | -0.001 |
| SE | 31478 | ENSG00000111885 | MAN1A1 | chr6 | - | 119201253 | 119201347 | 119193776 | 119193892 | 119204758 | 119204882 | 473 | 210 | 360 | 405 | 378 | 0 | 0 | 0 | 0 | 0 | 241 | 467 | 539 | 227 | 180 | 3 | 0 | 0 | 0 | 0 | 243 | 149 | 1 | 1 | 1 | 1 | 1 | 1 | 1 | 0.98 | 1 | 1 | 1 | 1 | 0.004 |
| SE | 31479 | ENSG00000111885 | MAN1A1 | chr6 | - | 119204758 | 119204882 | 119201253 | 119201347 | 119290682 | 119290763 | 246 | 110 | 185 | 236 | 188 | 0 | 0 | 0 | 0 | 0 | 117 | 263 | 319 | 145 | 112 | 0 | 2 | 2 | 0 | 0 | 273 | 149 | 1 | 1 | 1 | 1 | 1 | 1 | 1 | 1 | 0.986 | 0.989 | 1 | 1 | 0.005 |
| SE | 31483 | ENSG00000125733 | TRIP10 | chr19 | + | 6746028 | 6746196 | 6744799 | 6744994 | 6746451 | 6746561 | 0 | 0 | 0 | 0 | 0 | 8 | 7 | 4 | 16 | 9 | 0 | 0 | 2 | 0 | 0 | 25 | 8 | 17 | 15 | 36 | 298 | 149 | 1 | 1 | 0 | 0 | 0 | 0 | 0 | 0 | 0 | 0.056 | 0 | 0 | -0.011 |
| SE | 31485 | ENSG00000161013 | MGAT4B | chr5 | - | 179798346 | 179798434 | 179798105 | 179798277 | 179798512 | 179798576 | 96 | 44 | 42 | 93 | 49 | 0 | 0 | 0 | 2 | 0 | 160 | 113 | 75 | 126 | 530 | 0 | 0 | 0 | 0 | 0 | 237 | 149 | 1 | 1 | 1 | 1 | 1 | 0.967 | 1 | 1 | 1 | 1 | 1 | 1 | -0.007 |
| SE | 31486 | ENSG00000161013 | MGAT4B | chr5 | - | 179799202 | 179799250 | 179798927 | 179799121 | 179799505 | 179800068 | 28 | 15 | 11 | 23 | 31 | 0 | 0 | 0 | 0 | 0 | 5 | 33 | 9 | 14 | 172 | 0 | 0 | 0 | 1 | 0 | 197 | 149 | 1 | 1 | 1 | 1 | 1 | 1 | 1 | 1 | 1 | 1 | 0.914 | 1 | 0.017 |
| SE | 31488 | ENSG00000161013 | MGAT4B | chr5 | - | 179799202 | 179799307 | 179798927 | 179799121 | 179799505 | 179799636 | 60 | 39 | 50 | 67 | 62 | 0 | 0 | 0 | 0 | 0 | 69 | 83 | 35 | 44 | 412 | 0 | 0 | 0 | 1 | 0 | 254 | 149 | 1 | 1 | 1 | 1 | 1 | 1 | 1 | 1 | 1 | 1 | 0.963 | 1 | 0.007 |
| SE | 31489 | ENSG00000161013 | MGAT4B | chr5 | - | 179799202 | 179799310 | 179798927 | 179799121 | 179799505 | 179799576 | 178 | 111 | 146 | 225 | 181 | 0 | 0 | 0 | 0 | 0 | 228 | 224 | 113 | 136 | 1109 | 0 | 0 | 0 | 1 | 0 | 257 | 149 | 1 | 1 | 1 | 1 | 1 | 1 | 1 | 1 | 1 | 1 | 0.987 | 1 | 0.003 |
| SE | 31492 | ENSG00000161013 | MGAT4B | chr5 | - | 179801333 | 179801467 | 179800906 | 179800953 | 179801553 | 179801692 | 72 | 44 | 58 | 56 | 19 | 0 | 0 | 0 | 0 | 1 | 96 | 60 | 15 | 55 | 393 | 0 | 0 | 0 | 0 | 0 | 283 | 149 | 1 | 1 | 1 | 1 | 1 | 1 | 0.909 | 1 | 1 | 1 | 1 | 1 | -0.018 |
| SE | 31499 | ENSG00000232995 | RGS5 | chr1 | - | 163308543 | 163308631 | 163306232 | 163306329 | 163319181 | 163319291 | 5 | 3 | 9 | 4 | 14 | 1 | 0 | 0 | 3 | 0 | 0 | 6 | 6 | 0 | 0 | 0 | 10 | 0 | 0 | 0 | 237 | 149 | 1 | 1 | 0.759 | 1 | 1 | 0.456 | 1 | NA | 0.274 | 1 | NA | NA | 0.206 |
| SE | 31500 | ENSG00000143248 | RGS5 | chr1 | - | 163308543 | 163308631 | 163306232 | 163306329 | 163321621 | 163321787 | 0 | 2 | 2 | 4 | 11 | 0 | 0 | 0 | 0 | 0 | 0 | 0 | 2 | 0 | 0 | 0 | 0 | 1 | 0 | 0 | 237 | 149 | 1 | 1 | NA | 1 | 1 | 1 | 1 | NA | NA | 0.557 | NA | NA | 0.443 |
| SE | 31501 | ENSG00000232995 | RGS5 | chr1 | - | 163319181 | 163319291 | 163306232 | 163306329 | 163321621 | 163321704 | 17 | 21 | 16 | 10 | 26 | 0 | 0 | 0 | 0 | 0 | 0 | 22 | 3 | 6 | 0 | 0 | 0 | 1 | 0 | 0 | 259 | 149 | 1 | 1 | 1 | 1 | 1 | 1 | 1 | NA | 1 | 0.633 | 1 | NA | 0.122 |
| SE | 31502 | ENSG00000143248 | RGS5 | chr1 | - | 163319181 | 163319291 | 163308543 | 163308631 | 163321621 | 163321772 | 21 | 22 | 23 | 9 | 32 | 0 | 0 | 0 | 2 | 3 | 0 | 20 | 7 | 6 | 0 | 0 | 0 | 0 | 0 | 0 | 259 | 149 | 1 | 1 | 1 | 1 | 1 | 0.721 | 0.86 | NA | 1 | 1 | 1 | NA | -0.084 |
| SE | 31504 | ENSG00000099282 | TSPAN15 | chr10 | + | 69506123 | 69506240 | 69504437 | 69504485 | 69506828 | 69506856 | 0 | 0 | 13 | 9 | 6 | 0 | 0 | 0 | 1 | 0 | 0 | 0 | 2 | 10 | 23 | 0 | 0 | 0 | 0 | 0 | 266 | 149 | 1 | 1 | NA | NA | 1 | 0.834 | 1 | NA | NA | 1 | 1 | 1 | -0.055 |
| SE | 31507 | ENSG00000174227 | PIGG | chr4 | + | 501100 | 501170 | 500395 | 500601 | 515972 | 516185 | 9 | 11 | 9 | 18 | 17 | 1 | 2 | 0 | 0 | 0 | 11 | 13 | 27 | 31 | 37 | 0 | 0 | 0 | 0 | 0 | 219 | 149 | 1 | 1 | 0.86 | 0.789 | 1 | 1 | 1 | 1 | 1 | 1 | 1 | 1 | -0.07 |
| SE | 31508 | ENSG00000174227 | PIGG | chr4 | + | 505717 | 505927 | 500395 | 500601 | 507404 | 507593 | 247 | 203 | 278 | 245 | 291 | 1 | 0 | 0 | 0 | 1 | 629 | 314 | 248 | 370 | 540 | 9 | 0 | 0 | 0 | 0 | 298 | 149 | 1 | 1 | 0.992 | 1 | 1 | 1 | 0.993 | 0.972 | 1 | 1 | 1 | 1 | 0.003 |
| SE | 31509 | ENSG00000174227 | PIGG | chr4 | + | 505717 | 505927 | 501100 | 501170 | 507404 | 507593 | 167 | 129 | 166 | 147 | 201 | 1 | 0 | 1 | 0 | 0 | 505 | 211 | 235 | 279 | 354 | 0 | 0 | 1 | 0 | 0 | 298 | 149 | 1 | 1 | 0.988 | 1 | 0.988 | 1 | 1 | 1 | 1 | 0.992 | 1 | 1 | -0.003 |
| SE | 31510 | ENSG00000174227 | PIGG | chr4 | + | 506760 | 506866 | 505717 | 505927 | 507404 | 507538 | 0 | 0 | 0 | 0 | 0 | 151 | 120 | 147 | 129 | 173 | 0 | 8 | 0 | 0 | 0 | 397 | 190 | 172 | 217 | 293 | 255 | 149 | 1 | 1 | 0 | 0 | 0 | 0 | 0 | 0 | 0.024 | 0 | 0 | 0 | -0.005 |
| SE | 31512 | ENSG00000174227 | PIGG | chr4 | + | 507404 | 507593 | 500395 | 500601 | 515972 | 516185 | 3 | 0 | 2 | 5 | 1 | 1 | 2 | 0 | 0 | 0 | 25 | 0 | 2 | 11 | 0 | 0 | 0 | 0 | 0 | 0 | 298 | 149 | 1 | 1 | 0.6 | 0 | 1 | 1 | 1 | 1 | NA | 1 | 1 | NA | -0.28 |
| SE | 31513 | ENSG00000174227 | PIGG | chr4 | + | 507404 | 507593 | 501100 | 501170 | 515972 | 516185 | 3 | 0 | 3 | 5 | 0 | 0 | 0 | 0 | 2 | 0 | 16 | 0 | 3 | 11 | 0 | 0 | 0 | 0 | 0 | 0 | 298 | 149 | 1 | 1 | 1 | NA | 1 | 0.556 | NA | 1 | NA | 1 | 1 | NA | -0.148 |
| SE | 31514 | ENSG00000174227 | PIGG | chr4 | + | 508828 | 508970 | 500395 | 500601 | 515972 | 516185 | 131 | 86 | 123 | 129 | 109 | 1 | 2 | 0 | 0 | 0 | 180 | 84 | 117 | 193 | 107 | 0 | 0 | 0 | 0 | 0 | 291 | 149 | 1 | 1 | 0.985 | 0.957 | 1 | 1 | 1 | 1 | 1 | 1 | 1 | 1 | -0.012 |
| SE | 31516 | ENSG00000174227 | PIGG | chr4 | + | 508828 | 508970 | 507404 | 507593 | 521055 | 521227 | 90 | 47 | 93 | 66 | 114 | 0 | 0 | 0 | 0 | 0 | 216 | 102 | 105 | 124 | 83 | 0 | 0 | 0 | 1 | 0 | 291 | 149 | 1 | 1 | 1 | 1 | 1 | 1 | 1 | 1 | 1 | 1 | 0.984 | 1 | 0.003 |
| SE | 31517 | ENSG00000174227 | PIGG | chr4 | + | 515972 | 516185 | 507404 | 507593 | 521055 | 521227 | 50 | 34 | 60 | 58 | 48 | 0 | 0 | 0 | 0 | 0 | 81 | 31 | 59 | 76 | 34 | 0 | 0 | 0 | 1 | 0 | 298 | 149 | 1 | 1 | 1 | 1 | 1 | 1 | 1 | 1 | 1 | 1 | 0.974 | 1 | 0.005 |
| SE | 31518 | ENSG00000174227 | PIGG | chr4 | + | 515972 | 516185 | 508828 | 508970 | 521055 | 521227 | 188 | 120 | 184 | 185 | 159 | 0 | 1 | 0 | 0 | 0 | 279 | 118 | 178 | 274 | 141 | 0 | 0 | 0 | 0 | 0 | 298 | 149 | 1 | 1 | 1 | 0.984 | 1 | 1 | 1 | 1 | 1 | 1 | 1 | 1 | -0.003 |
| SE | 31519 | ENSG00000174227 | PIGG | chr4 | + | 515972 | 516185 | 508828 | 508970 | 523458 | 523913 | 140 | 86 | 126 | 134 | 111 | 0 | 0 | 0 | 0 | 0 | 220 | 87 | 121 | 209 | 107 | 0 | 2 | 0 | 0 | 0 | 298 | 149 | 1 | 1 | 1 | 1 | 1 | 1 | 1 | 1 | 0.956 | 1 | 1 | 1 | 0.009 |
| SE | 31523 | ENSG00000174227 | PIGG | chr4 | + | 523458 | 523913 | 515972 | 516185 | 527038 | 527230 | 6 | 2 | 15 | 10 | 7 | 0 | 0 | 0 | 1 | 0 | 19 | 2 | 2 | 13 | 0 | 0 | 0 | 0 | 0 | 0 | 298 | 149 | 1 | 1 | 1 | 1 | 1 | 0.833 | 1 | 1 | 1 | 1 | 1 | NA | -0.033 |
| SE | 31524 | ENSG00000174227 | PIGG | chr4 | + | 523458 | 523913 | 521659 | 521941 | 527038 | 527230 | 20 | 4 | 21 | 14 | 8 | 0 | 0 | 0 | 0 | 0 | 35 | 3 | 10 | 23 | 19 | 0 | 0 | 0 | 2 | 0 | 298 | 149 | 1 | 1 | 1 | 1 | 1 | 1 | 1 | 1 | 1 | 1 | 0.852 | 1 | 0.03 |
| SE | 31525 | ENSG00000174227 | PIGG | chr4 | + | 523458 | 523913 | 521659 | 521941 | 530435 | 530498 | 14 | 2 | 6 | 6 | 1 | 0 | 0 | 0 | 0 | 0 | 22 | 1 | 8 | 10 | 19 | 0 | 0 | 1 | 0 | 0 | 298 | 149 | 1 | 1 | 1 | 1 | 1 | 1 | 1 | 1 | 1 | 0.8 | 1 | 1 | 0.04 |
| SE | 31526 | ENSG00000174227 | PIGG | chr4 | + | 527038 | 527230 | 521659 | 521941 | 530435 | 530725 | 5 | 0 | 3 | 2 | 4 | 0 | 0 | 0 | 0 | 0 | 0 | 4 | 3 | 2 | 0 | 0 | 0 | 1 | 0 | 0 | 298 | 149 | 1 | 1 | 1 | NA | 1 | 1 | 1 | NA | 1 | 0.6 | 1 | NA | 0.133 |
| SE | 31528 | ENSG00000174227 | PIGG | chr4 | + | 530435 | 530745 | 527038 | 527230 | 533817 | 533981 | 11 | 1 | 10 | 4 | 11 | 0 | 0 | 0 | 6 | 0 | 5 | 6 | 6 | 4 | 0 | 0 | 0 | 0 | 0 | 0 | 298 | 149 | 1 | 1 | 1 | 1 | 1 | 0.25 | 1 | 1 | 1 | 1 | 1 | NA | -0.15 |
| SE | 31529 | ENSG00000155903 | RASA2 | chr3 | + | 141529707 | 141529802 | 141516327 | 141516431 | 141540532 | 141540573 | 505 | 335 | 449 | 355 | 722 | 4 | 0 | 0 | 0 | 0 | 96 | 145 | 270 | 95 | 79 | 0 | 0 | 0 | 0 | 0 | 244 | 149 | 1 | 1 | 0.987 | 1 | 1 | 1 | 1 | 1 | 1 | 1 | 1 | 1 | -0.003 |
| SE | 31531 | ENSG00000155903 | RASA2 | chr3 | + | 141553856 | 141553940 | 141540532 | 141540609 | 141555839 | 141555912 | 350 | 220 | 347 | 226 | 502 | 1 | 0 | 0 | 1 | 3 | 71 | 66 | 151 | 76 | 35 | 0 | 0 | 2 | 0 | 0 | 233 | 149 | 1 | 1 | 0.996 | 1 | 1 | 0.993 | 0.991 | 1 | 1 | 0.98 | 1 | 1 | 0 |
| SE | 31532 | ENSG00000155903 | RASA2 | chr3 | + | 141553856 | 141553940 | 141540532 | 141540609 | 141558885 | 141558962 | 214 | 124 | 225 | 125 | 189 | 0 | 0 | 0 | 1 | 0 | 23 | 45 | 72 | 33 | 17 | 0 | 0 | 0 | 0 | 0 | 233 | 149 | 1 | 1 | 1 | 1 | 1 | 0.988 | 1 | 1 | 1 | 1 | 1 | 1 | -0.002 |
| SE | 31533 | ENSG00000155903 | RASA2 | chr3 | + | 141555839 | 141555912 | 141540532 | 141540609 | 141558885 | 141558962 | 76 | 58 | 79 | 66 | 238 | 0 | 0 | 0 | 1 | 0 | 16 | 18 | 34 | 29 | 20 | 0 | 0 | 0 | 0 | 0 | 222 | 149 | 1 | 1 | 1 | 1 | 1 | 0.978 | 1 | 1 | 1 | 1 | 1 | 1 | -0.004 |
| SE | 31534 | ENSG00000155903 | RASA2 | chr3 | + | 141555839 | 141555912 | 141553856 | 141553940 | 141558885 | 141558962 | 220 | 157 | 206 | 168 | 566 | 2 | 1 | 0 | 2 | 0 | 64 | 42 | 116 | 72 | 34 | 0 | 0 | 0 | 0 | 0 | 222 | 149 | 1 | 1 | 0.987 | 0.991 | 1 | 0.983 | 1 | 1 | 1 | 1 | 1 | 1 | -0.008 |
| SE | 31535 | ENSG00000155903 | RASA2 | chr3 | + | 141571405 | 141571554 | 141570911 | 141571068 | 141572608 | 141572723 | 445 | 394 | 406 | 366 | 574 | 0 | 0 | 0 | 1 | 0 | 139 | 207 | 315 | 79 | 59 | 0 | 0 | 0 | 0 | 0 | 298 | 149 | 1 | 1 | 1 | 1 | 1 | 0.995 | 1 | 1 | 1 | 1 | 1 | 1 | -0.001 |
| SE | 31536 | ENSG00000155903 | RASA2 | chr3 | + | 141573943 | 141574067 | 141573146 | 141573221 | 141576999 | 141577106 | 515 | 369 | 405 | 374 | 983 | 0 | 0 | 4 | 0 | 0 | 100 | 223 | 277 | 67 | 43 | 0 | 0 | 0 | 0 | 0 | 273 | 149 | 1 | 1 | 1 | 1 | 0.982 | 1 | 1 | 1 | 1 | 1 | 1 | 1 | -0.004 |
| SE | 31538 | ENSG00000155903 | RASA2 | chr3 | + | 141579436 | 141579489 | 141578523 | 141578754 | 141580367 | 141580445 | 0 | 2 | 2 | 0 | 0 | 0 | 2 | 3 | 4 | 0 | 0 | 0 | 0 | 0 | 0 | 0 | 0 | 2 | 0 | 0 | 202 | 149 | 1 | 1 | NA | 0.425 | 0.33 | 0 | NA | NA | NA | 0 | NA | NA | 0.252 |
| SE | 31539 | ENSG00000155903 | RASA2 | chr3 | + | 141581099 | 141581177 | 141580367 | 141580451 | 141586024 | 141586098 | 358 | 244 | 348 | 255 | 437 | 0 | 0 | 0 | 0 | 0 | 42 | 165 | 183 | 56 | 28 | 0 | 0 | 0 | 2 | 0 | 227 | 149 | 1 | 1 | 1 | 1 | 1 | 1 | 1 | 1 | 1 | 1 | 0.948 | 1 | 0.01 |
| SE | 31540 | ENSG00000155903 | RASA2 | chr3 | + | 141586024 | 141586098 | 141581099 | 141581177 | 141586645 | 141586752 | 458 | 279 | 398 | 325 | 381 | 3 | 0 | 1 | 0 | 3 | 62 | 222 | 224 | 83 | 55 | 0 | 0 | 0 | 0 | 0 | 223 | 149 | 1 | 1 | 0.99 | 1 | 0.996 | 1 | 0.988 | 1 | 1 | 1 | 1 | 1 | -0.005 |
| SE | 31542 | ENSG00000174946 | GPR171 | chr3 | - | 151200908 | 151200997 | 151197831 | 151199439 | 151203103 | 151203216 | 70 | 38 | 53 | 32 | 53 | 19 | 10 | 19 | 12 | 8 | 0 | 24 | 17 | 14 | 0 | 0 | 5 | 5 | 0 | 0 | 238 | 149 | 1 | 1 | 0.698 | 0.704 | 0.636 | 0.625 | 0.806 | NA | 0.75 | 0.68 | 1 | NA | -0.116 |
| SE | 31545 | ENSG00000171608 | PIK3CD | chr1 | + | 9716958 | 9717108 | 9716275 | 9716619 | 9717536 | 9717581 | 203 | 66 | 153 | 198 | 113 | 0 | 0 | 0 | 0 | 0 | 183 | 176 | 98 | 151 | 195 | 0 | 0 | 0 | 0 | 4 | 298 | 149 | 1 | 1 | 1 | 1 | 1 | 1 | 1 | 1 | 1 | 1 | 1 | 0.961 | 0.008 |
| SE | 31547 | ENSG00000171608 | PIK3CD | chr1 | + | 9717063 | 9717108 | 9716439 | 9716619 | 9717536 | 9717626 | 48 | 22 | 53 | 57 | 21 | 0 | 0 | 0 | 0 | 0 | 67 | 60 | 46 | 51 | 64 | 0 | 0 | 0 | 0 | 4 | 194 | 149 | 1 | 1 | 1 | 1 | 1 | 1 | 1 | 1 | 1 | 1 | 1 | 0.925 | 0.015 |
| SE | 31549 | ENSG00000171608 | PIK3CD | chr1 | + | 9717536 | 9717626 | 9716439 | 9716619 | 9718693 | 9718915 | 121 | 66 | 96 | 131 | 40 | 0 | 0 | 0 | 0 | 0 | 135 | 109 | 77 | 114 | 157 | 7 | 0 | 0 | 0 | 0 | 239 | 149 | 1 | 1 | 1 | 1 | 1 | 1 | 1 | 0.923 | 1 | 1 | 1 | 1 | 0.015 |
| SE | 31550 | ENSG00000171608 | PIK3CD | chr1 | + | 9717536 | 9717626 | 9716958 | 9717108 | 9718693 | 9718915 | 209 | 99 | 177 | 229 | 71 | 0 | 0 | 0 | 0 | 0 | 232 | 209 | 128 | 184 | 265 | 0 | 0 | 0 | 2 | 0 | 239 | 149 | 1 | 1 | 1 | 1 | 1 | 1 | 1 | 1 | 1 | 1 | 0.983 | 1 | 0.003 |
| SE | 31554 | ENSG00000163913 | IFT122 | chr3 | + | 129456113 | 129456264 | 129451913 | 129451998 | 129458598 | 129458677 | 2 | 0 | 1 | 0 | 2 | 26 | 19 | 15 | 16 | 15 | 0 | 0 | 0 | 0 | 0 | 50 | 32 | 19 | 40 | 74 | 298 | 149 | 1 | 1 | 0.037 | 0 | 0.032 | 0 | 0.062 | 0 | 0 | 0 | 0 | 0 | 0.026 |
| SE | 31555 | ENSG00000163913 | IFT122 | chr3 | + | 129457921 | 129458015 | 129451913 | 129451998 | 129458598 | 129458677 | 0 | 0 | 0 | 0 | 0 | 26 | 19 | 15 | 16 | 15 | 0 | 3 | 0 | 0 | 2 | 50 | 32 | 19 | 40 | 74 | 243 | 149 | 1 | 1 | 0 | 0 | 0 | 0 | 0 | 0 | 0.054 | 0 | 0 | 0.016 | -0.014 |
| SE | 31559 | ENSG00000163913 | IFT122 | chr3 | + | 129460853 | 129461006 | 129458598 | 129458677 | 129461227 | 129461304 | 0 | 0 | 0 | 0 | 0 | 25 | 15 | 17 | 26 | 9 | 0 | 2 | 2 | 0 | 0 | 100 | 24 | 14 | 41 | 55 | 298 | 149 | 1 | 1 | 0 | 0 | 0 | 0 | 0 | 0 | 0.04 | 0.067 | 0 | 0 | -0.021 |
| SE | 31567 | ENSG00000163913 | IFT122 | chr3 | + | 129499901 | 129500068 | 129495445 | 129495607 | 129502710 | 129502764 | 23 | 7 | 21 | 19 | 23 | 0 | 0 | 0 | 0 | 0 | 78 | 22 | 17 | 29 | 35 | 0 | 0 | 0 | 2 | 0 | 298 | 149 | 1 | 1 | 1 | 1 | 1 | 1 | 1 | 1 | 1 | 1 | 0.879 | 1 | 0.024 |
| SE | 31568 | ENSG00000163913 | IFT122 | chr3 | + | 129504318 | 129504421 | 129502710 | 129502882 | 129506408 | 129506549 | 4 | 5 | 6 | 9 | 4 | 0 | 0 | 0 | 2 | 0 | 0 | 3 | 8 | 19 | 2 | 0 | 2 | 0 | 2 | 0 | 252 | 149 | 1 | 1 | 1 | 1 | 1 | 0.727 | 1 | NA | 0.47 | 1 | 0.849 | 1 | 0.116 |
| SE | 31569 | ENSG00000116668 | SWT1 | chr1 | + | 185180390 | 185180450 | 185174371 | 185175113 | 185184242 | 185184344 | 11 | 12 | 16 | 8 | 15 | 0 | 0 | 0 | 0 | 0 | 0 | 4 | 7 | 0 | 0 | 0 | 0 | 0 | 0 | 0 | 209 | 149 | 1 | 1 | 1 | 0.895 | 1 | 1 | 1 | NA | 1 | 1 | NA | NA | -0.021 |
| SE | 31570 | ENSG00000116668 | SWT1 | chr1 | + | 185181945 | 185182057 | 185180390 | 185180450 | 185184242 | 185184344 | 40 | 26 | 30 | 48 | 50 | 0 | 0 | 0 | 0 | 0 | 0 | 26 | 9 | 6 | 2 | 0 | 0 | 1 | 0 | 0 | 261 | 149 | 1 | 1 | 1 | 1 | 1 | 1 | 1 | NA | 1 | 0.837 | 1 | 1 | 0.041 |
| SE | 31571 | ENSG00000116668 | SWT1 | chr1 | + | 185184242 | 185184344 | 185181945 | 185182057 | 185184742 | 185184931 | 41 | 17 | 15 | 43 | 72 | 0 | 1 | 0 | 0 | 0 | 0 | 33 | 8 | 23 | 9 | 0 | 0 | 0 | 0 | 0 | 251 | 149 | 1 | 1 | 1 | 0.91 | 1 | 1 | 1 | NA | 1 | 1 | 1 | 1 | -0.018 |
| SE | 31573 | ENSG00000116668 | SWT1 | chr1 | + | 185214506 | 185214655 | 185206624 | 185206763 | 185221848 | 185222036 | 115 | 129 | 77 | 106 | 350 | 3 | 0 | 0 | 0 | 0 | 140 | 96 | 58 | 51 | 183 | 0 | 2 | 0 | 0 | 0 | 298 | 149 | 1 | 1 | 0.95 | 1 | 1 | 1 | 1 | 1 | 0.96 | 1 | 1 | 1 | -0.002 |
| SE | 31576 | ENSG00000116668 | SWT1 | chr1 | + | 185221848 | 185222036 | 185214506 | 185214655 | 185231576 | 185231708 | 125 | 148 | 100 | 139 | 349 | 1 | 0 | 0 | 0 | 0 | 224 | 130 | 81 | 84 | 203 | 0 | 0 | 0 | 0 | 0 | 298 | 149 | 1 | 1 | 0.984 | 1 | 1 | 1 | 1 | 1 | 1 | 1 | 1 | 1 | -0.003 |
| SE | 31577 | ENSG00000116668 | SWT1 | chr1 | + | 185276603 | 185276668 | 185271322 | 185271389 | 185290673 | 185291007 | 21 | 32 | 31 | 41 | 18 | 0 | 0 | 1 | 0 | 0 | 0 | 4 | 2 | 2 | 3 | 0 | 0 | 0 | 0 | 0 | 214 | 149 | 1 | 1 | 1 | 1 | 0.956 | 1 | 1 | NA | 1 | 1 | 1 | 1 | -0.009 |
| SE | 31580 | ENSG00000129071 | MBD4 | chr3 | - | 129433097 | 129433247 | 129432175 | 129432606 | 129433849 | 129433984 | 426 | 312 | 441 | 479 | 431 | 0 | 0 | 0 | 2 | 1 | 117 | 263 | 271 | 261 | 82 | 0 | 0 | 0 | 0 | 0 | 298 | 149 | 1 | 1 | 1 | 1 | 1 | 0.992 | 0.995 | 1 | 1 | 1 | 1 | 1 | -0.003 |
| SE | 31581 | ENSG00000129071 | MBD4 | chr3 | - | 129434061 | 129434136 | 129433849 | 129433984 | 129436442 | 129437308 | 203 | 138 | 163 | 197 | 302 | 0 | 0 | 0 | 1 | 0 | 57 | 160 | 129 | 128 | 33 | 0 | 0 | 0 | 0 | 0 | 224 | 149 | 1 | 1 | 1 | 1 | 1 | 0.992 | 1 | 1 | 1 | 1 | 1 | 1 | -0.002 |

| | | | | | | | | | | | |
|---|---|---|---|---|---|---|---|---|---|---|---|
| SE | 31587 | ENSG00000183624 | HMCES | chr3 | + | 129288853 | 129288997 | 129279709 | 129279915 | 129298353 | 129298535 |
| SE | 31590 | ENSG00000183624 | HMCES | chr3 | + | 129298353 | 129298535 | 129290678 | 129290804 | 129301949 | 129302005 |
| SE | 31591 | ENSG00000183624 | HMCES | chr3 | + | 129301949 | 129302142 | 129298353 | 129298535 | 129304588 | 129304751 |
| SE | 31593 | ENSG00000082805 | ERC1 | chr12 | + | 1027747 | 1028572 | 991237 | 991487 | 1083163 | 1083580 |
| SE | 31594 | ENSG00000082805 | ERC1 | chr12 | + | 1028338 | 1028572 | 991262 | 991487 | 1083163 | 1083580 |
| SE | 31595 | ENSG00000082805 | ERC1 | chr12 | + | 1110191 | 1110347 | 1104749 | 1104824 | 1115865 | 1116033 |
| SE | 31601 | ENSG00000082805 | ERC1 | chr12 | + | 1263033 | 1263165 | 1236768 | 1236904 | 1289851 | 1290012 |
| SE | 31607 | ENSG00000181789 | COPG1 | chr3 | + | 129252280 | 129252361 | 129249605 | 129249746 | 129252622 | 129252694 |
| SE | 31608 | ENSG00000181789 | COPG1 | chr3 | + | 129254984 | 129255077 | 129254667 | 129254743 | 129256067 | 129256154 |
| SE | 31609 | ENSG00000181789 | COPG1 | chr3 | + | 129267023 | 129267099 | 129265548 | 129265792 | 129267936 | 129268040 |
| SE | 31612 | ENSG00000181789 | COPG1 | chr3 | + | 129274063 | 129274136 | 129272806 | 129272904 | 129274837 | 129274976 |
| SE | 31614 | ENSG00000168026 | TTC21A | chr3 | + | 39110028 | 39110233 | 39109084 | 39109214 | 39110850 | 39111018 |
| SE | 31617 | ENSG00000168026 | TTC21A | chr3 | + | 39119921 | 39120020 | 39118068 | 39118153 | 39120996 | 39121189 |
| SE | 31618 | ENSG00000168026 | TTC21A | chr3 | + | 39126260 | 39126390 | 39125331 | 39125532 | 39128330 | 39128488 |
| SE | 31619 | ENSG00000168026 | TTC21A | chr3 | + | 39130247 | 39130358 | 39130078 | 39130151 | 39130700 | 39130839 |
| SE | 31620 | ENSG00000168026 | TTC21A | chr3 | + | 39130247 | 39130358 | 39130078 | 39130151 | 39130991 | 39131095 |
| SE | 31621 | ENSG00000168026 | TTC21A | chr3 | + | 39130247 | 39130358 | 39130078 | 39130151 | 39133051 | 39133240 |
| SE | 31622 | ENSG00000168026 | TTC21A | chr3 | + | 39130700 | 39130839 | 39130078 | 39130151 | 39130991 | 39131095 |
| SE | 31623 | ENSG00000168026 | TTC21A | chr3 | + | 39130700 | 39130839 | 39130247 | 39130358 | 39130991 | 39131095 |
| SE | 31625 | ENSG00000168026 | TTC21A | chr3 | + | 39130991 | 39131095 | 39130247 | 39130358 | 39133051 | 39133240 |
| SE | 31630 | ENSG00000111321 | LTBR | chr12 | + | 6387797 | 6387910 | 6386346 | 6386444 | 6388397 | 6388505 |
| SE | 31633 | ENSG00000111321 | LTBR | chr12 | + | 6390111 | 6390340 | 6388799 | 6388825 | 6390659 | 6390735 |
| SE | 31635 | ENSG00000264230 | ANXA8L1 | chr10 | + | 46384205 | 46384285 | 46383455 | 46383546 | 46384773 | 46384833 |
| SE | 31637 | ENSG00000131389 | SLC6A6 | chr3 | + | 14416411 | 14416453 | 14402575 | 14402847 | 14443623 | 14443863 |
| SE | 31641 | ENSG00000131389 | SLC6A6 | chr3 | + | 14445716 | 14445851 | 14443623 | 14443863 | 14447581 | 14447816 |
| SE | 31643 | ENSG00000131389 | SLC6A6 | chr3 | + | 14466515 | 14466650 | 14447581 | 14447816 | 14477204 | 14477342 |
| SE | 31648 | ENSG00000131389 | SLC6A6 | chr3 | + | 14472204 | 14472317 | 14467852 | 14467956 | 14477204 | 14477342 |
| SE | 31649 | ENSG00000131389 | SLC6A6 | chr3 | + | 14472204 | 14472317 | 14467852 | 14467956 | 14478465 | 14478568 |
| SE | 31650 | ENSG00000131389 | SLC6A6 | chr3 | + | 14472204 | 14472317 | 14467852 | 14467956 | 14479084 | 14479185 |
| SE | 31653 | ENSG00000131389 | SLC6A6 | chr3 | + | 14477204 | 14477342 | 14467852 | 14467956 | 14478465 | 14478568 |
| SE | 31654 | ENSG00000131389 | SLC6A6 | chr3 | + | 14477204 | 14477342 | 14467852 | 14467956 | 14479084 | 14479185 |
| SE | 31655 | ENSG00000131389 | SLC6A6 | chr3 | + | 14477204 | 14477342 | 14468087 | 14468212 | 14479084 | 14479185 |
| SE | 31656 | ENSG00000131389 | SLC6A6 | chr3 | + | 14477204 | 14477342 | 14468087 | 14468212 | 14481670 | 14481841 |
| SE | 31657 | ENSG00000131389 | SLC6A6 | chr3 | + | 14477204 | 14477342 | 14472204 | 14472317 | 14478465 | 14478568 |
| SE | 31658 | ENSG00000131389 | SLC6A6 | chr3 | + | 14477204 | 14477342 | 14472204 | 14472317 | 14479084 | 14479185 |
| SE | 31659 | ENSG00000131389 | SLC6A6 | chr3 | + | 14477204 | 14477342 | 14472204 | 14472317 | 14481670 | 14481841 |
| SE | 31661 | ENSG00000131389 | SLC6A6 | chr3 | + | 14478465 | 14478568 | 14472204 | 14472317 | 14479084 | 14479185 |
| SE | 31662 | ENSG00000131389 | SLC6A6 | chr3 | + | 14478465 | 14478568 | 14472204 | 14472317 | 14481670 | 14481841 |
| SE | 31664 | ENSG00000131389 | SLC6A6 | chr3 | + | 14478465 | 14478568 | 14477204 | 14477342 | 14481670 | 14481841 |
| SE | 31665 | ENSG00000131389 | SLC6A6 | chr3 | + | 14479084 | 14479185 | 14468087 | 14468212 | 14481670 | 14481841 |
| SE | 31666 | ENSG00000131389 | SLC6A6 | chr3 | + | 14479084 | 14479185 | 14472204 | 14472317 | 14481670 | 14481841 |
| SE | 31667 | ENSG00000131389 | SLC6A6 | chr3 | + | 14479084 | 14479185 | 14477204 | 14477342 | 14481670 | 14481841 |
| SE | 31668 | ENSG00000131389 | SLC6A6 | chr3 | + | 14479084 | 14479185 | 14478465 | 14478568 | 14481670 | 14481841 |
| SE | 31673 | ENSG00000137171 | KLC4 | chr6 | + | 43071566 | 43071619 | 43071274 | 43071374 | 43071851 | 43071922 |
| SE | 31675 | ENSG00000137171 | KLC4 | chr6 | + | 43073222 | 43073338 | 43072823 | 43072964 | 43073901 | 43073965 |
| SE | 31676 | ENSG00000137171 | KLC4 | chr6 | + | 43073901 | 43073965 | 43073222 | 43073338 | 43074621 | 43074908 |
| SE | 31677 | ENSG00000174776 | WDR49 | chr3 | - | 167505306 | 167505416 | 167500152 | 167500299 | 167522314 | 167522385 |
| SE | 31678 | ENSG00000174776 | WDR49 | chr3 | - | 167527819 | 167528017 | 167522314 | 167522484 | 167529051 | 167529239 |
| SE | 31679 | ENSG00000174776 | WDR49 | chr3 | - | 167529051 | 167529239 | 167522331 | 167522484 | 167531114 | 167531189 |
| SE | 31680 | ENSG00000174776 | WDR49 | chr3 | - | 167554649 | 167554798 | 167536869 | 167537000 | 167560063 | 167560228 |
| SE | 31681 | ENSG00000174776 | WDR49 | chr3 | - | 167554682 | 167554798 | 167536869 | 167537000 | 167560063 | 167560228 |
| SE | 31684 | ENSG00000174776 | WDR49 | chr3 | - | 167566829 | 167566894 | 167560063 | 167560228 | 167575917 | 167576151 |
| SE | 31685 | ENSG00000174776 | WDR49 | chr3 | - | 167566829 | 167566894 | 167560063 | 167560228 | 167602126 | 167602275 |
| SE | 31686 | ENSG00000174776 | WDR49 | chr3 | - | 167566829 | 167566894 | 167560063 | 167560228 | 167653260 | 167653499 |
| SE | 31687 | ENSG00000174776 | WDR49 | chr3 | - | 167575917 | 167576151 | 167536869 | 167537000 | 167602126 | 167602275 |
| SE | 31688 | ENSG00000174776 | WDR49 | chr3 | - | 167575917 | 167576151 | 167560063 | 167560228 | 167602126 | 167602275 |
| SE | 31689 | ENSG00000174776 | WDR49 | chr3 | - | 167575917 | 167576151 | 167560063 | 167560228 | 167653260 | 167653499 |
| SE | 31691 | ENSG00000174776 | WDR49 | chr3 | - | 167575917 | 167576151 | 167566829 | 167566894 | 167653260 | 167653499 |
| SE | 31692 | ENSG00000174776 | WDR49 | chr3 | - | 167620428 | 167620603 | 167604300 | 167604468 | 167626851 | 167627292 |
| SE | 31693 | ENSG00000174776 | WDR49 | chr3 | - | 167621466 | 167621643 | 167620428 | 167620603 | 167626851 | 167627292 |
| SE | 31694 | ENSG00000174776 | WDR49 | chr3 | - | 167626851 | 167627292 | 167566829 | 167566894 | 167653260 | 167653499 |
| SE | 31695 | ENSG00000174776 | WDR49 | chr3 | - | 167626851 | 167627292 | 167604300 | 167604468 | 167653260 | 167653499 |
| SE | 31697 | ENSG00000174776 | WDR49 | chr3 | - | 167633408 | 167633532 | 167560063 | 167560228 | 167653260 | 167653499 |
| SE | 31698 | ENSG00000174776 | WDR49 | chr3 | - | 167633408 | 167633532 | 167604300 | 167604468 | 167653260 | 167653499 |
| SE | 31699 | ENSG00000174776 | WDR49 | chr3 | - | 167633408 | 167633532 | 167627132 | 167627292 | 167653260 | 167653499 |
| SE | 31703 | ENSG00000196428 | TSC22D2 | chr3 | + | 150423048 | 150423120 | 150408334 | 150411308 | 150458375 | 150458679 |
| SE | 31706 | ENSG00000196428 | TSC22D2 | chr3 | + | 150457075 | 150457127 | 150411008 | 150411308 | 150458375 | 150458679 |
| SE | 31707 | ENSG00000196428 | TSC22D2 | chr3 | + | 150457075 | 150457127 | 150423048 | 150423120 | 150458375 | 150458679 |
| SE | 31709 | ENSG00000145012 | LPP | chr3 | + | 188225404 | 188225527 | 188153283 | 188153363 | 188406111 | 188406260 |
| SE | 31711 | ENSG00000145012 | LPP | chr3 | + | 188225404 | 188225527 | 188153874 | 188154021 | 188406111 | 188406260 |
| SE | 31716 | ENSG00000145012 | LPP | chr3 | + | 188341662 | 188341719 | 188153283 | 188153363 | 188406111 | 188406313 |
| SE | 31717 | ENSG00000145012 | LPP | chr3 | + | 188341662 | 188341719 | 188153941 | 188154021 | 188406111 | 188406313 |
| SE | 31720 | ENSG00000145012 | LPP | chr3 | + | 188406111 | 188406313 | 188341662 | 188341719 | 188484591 | 188484595 |
| SE | 31721 | ENSG00000145012 | LPP | chr3 | + | 188524664 | 188524787 | 188484589 | 188484704 | 188609160 | 188609184 |
| SE | 31722 | ENSG00000145012 | LPP | chr3 | + | 188708266 | 188708393 | 188225404 | 188225527 | 188760112 | 188760201 |
| SE | 31724 | ENSG00000145012 | LPP | chr3 | + | 188759262 | 188759500 | 188758738 | 188758768 | 188760112 | 188760282 |
| SE | 31732 | ENSG00000047849 | MAP4 | chr3 | - | 47875684 | 47875900 | 47871916 | 47872100 | 47877416 | 47877523 |
| SE | 31736 | ENSG00000047849 | MAP4 | chr3 | - | 47909037 | 47912421 | 47902949 | 47903000 | 47914816 | 47914939 |
| SE | 31737 | ENSG00000047849 | MAP4 | chr3 | - | 47914816 | 47914939 | 47877416 | 47877523 | 47918718 | 47918841 |
| SE | 31738 | ENSG00000047849 | MAP4 | chr3 | - | 47915950 | 47917174 | 47914816 | 47914939 | 47918718 | 47918841 |
| SE | 31739 | ENSG00000047849 | MAP4 | chr3 | - | 47918718 | 47918841 | 47877416 | 47877523 | 47921764 | 47921878 |
| SE | 31749 | ENSG00000211897 | IGHG3 | chr14 | - | 105770480 | 105770525 | 105770104 | 105770149 | 105770668 | 105770719 |
| SE | 31752 | ENSG00000135535 | CD164 | chr6 | - | 109370410 | 109370467 | 109367820 | 109368325 | 109376073 | 109376112 |
| SE | 31754 | ENSG00000135535 | CD164 | chr6 | - | 109377899 | 109377971 | 109370410 | 109370467 | 109379578 | 109379662 |
| SE | 31755 | ENSG00000144034 | TPRKB | chr2 | - | 73732162 | 73732285 | 73730263 | 73730736 | 73734428 | 73734591 |
| SE | 31757 | ENSG00000144034 | TPRKB | chr2 | - | 73734428 | 73734591 | 73732162 | 73732285 | 73737301 | 73737335 |
| SE | 31758 | ENSG00000144034 | TPRKB | chr2 | - | 73734527 | 73734588 | 73732162 | 73732285 | 73737301 | 73737386 |
| SE | 31759 | ENSG00000144034 | TPRKB | chr2 | - | 73734527 | 73734591 | 73732162 | 73732285 | 73737301 | 73737375 |
| SE | 31761 | ENSG00000206573 | THUMPD3-AS1 | chr3 | - | 9388086 | 9388127 | 9385263 | 9385923 | 9390279 | 9390497 |
| SE | 31762 | ENSG00000206573 | THUMPD3-AS1 | chr3 | - | 9388086 | 9388127 | 9385263 | 9385923 | 9390289 | 9390497 |
| SE | 31769 | ENSG00000206573 | THUMPD3-AS1 | chr3 | - | 9395505 | 9395579 | 9391311 | 9391826 | 9396559 | 9396683 |
| SE | 31770 | ENSG00000206573 | THUMPD3-AS1 | chr3 | - | 9395505 | 9395579 | 9391311 | 9391826 | 9397423 | 9397439 |
| SE | 31784 | ENSG00000204463 | BAG6 | chr6 | - | 31640644 | 31640704 | 31640198 | 31640306 | 31640791 | 31640937 |
| SE | 31785 | ENSG00000204463 | BAG6 | chr6 | - | 31642828 | 31643115 | 31642111 | 31642403 | 31643889 | 31643977 |
| SE | 31787 | ENSG00000204463 | BAG6 | chr6 | - | 31644081 | 31644194 | 31643889 | 31643977 | 31644945 | 31645198 |
| SE | 31796 | ENSG00000204463 | BAG6 | chr6 | - | 31649198 | 31649395 | 31648910 | 31648964 | 31649509 | 31649627 |
| SE | 31797 | ENSG00000204463 | BAG6 | chr6 | - | 31651655 | 31651776 | 31649509 | 31649627 | 31652234 | 31652323 |
| SE | 31798 | ENSG00000152133 | GPATCH11 | chr2 | + | 37089639 | 37089866 | 37088368 | 37088440 | 37090680 | 37090722 |
| SE | 31799 | ENSG00000152133 | GPATCH11 | chr2 | + | 37089639 | 37089866 | 37088368 | 37088440 | 37091915 | 37092036 |
| SE | 31800 | ENSG00000152133 | GPATCH11 | chr2 | + | 37090680 | 37090722 | 37088368 | 37088440 | 37091915 | 37092036 |
| SE | 31801 | ENSG00000152133 | GPATCH11 | chr2 | + | 37090680 | 37090722 | 37089639 | 37089866 | 37091915 | 37092036 |
| SE | 31802 | ENSG00000182185 | RAD51B | chr14 | + | 67825463 | 67825577 | 67823519 | 67823627 | 67835079 | 67835106 |
| SE | 31803 | ENSG00000182185 | RAD51B | chr14 | + | 67835079 | 67835196 | 67825463 | 67825577 | 67865002 | 67865087 |
| SE | 31805 | ENSG00000182185 | RAD51B | chr14 | + | 67865002 | 67865139 | 67835079 | 67835196 | 67885868 | 67885988 |
| SE | 31806 | ENSG00000182185 | RAD51B | chr14 | + | 68411423 | 68411527 | 68291883 | 68291980 | 68468171 | 68468250 |
| SE | 31808 | ENSG00000182185 | RAD51B | chr14 | + | 68468554 | 68468599 | 68468171 | 68468250 | 68497123 | 68497881 |
| SE | 31815 | ENSG00000112701 | SENP6 | chr6 | + | 75621531 | 75621625 | 75601508 | 75602576 | 75622750 | 75622858 |
| SE | 31819 | ENSG00000112701 | SENP6 | chr6 | + | 75633580 | 75633726 | 75623899 | 75623960 | 75634706 | 75634811 |
| SE | 31823 | ENSG00000112701 | SENP6 | chr6 | + | 75663220 | 75663518 | 75659261 | 75659407 | 75666711 | 75666941 |
| SE | 31824 | ENSG00000112701 | SENP6 | chr6 | + | 75675434 | 75675468 | 75670552 | 75670720 | 75675859 | 75676054 |
| SE | 31825 | ENSG00000112701 | SENP6 | chr6 | + | 75695803 | 75695923 | 75678810 | 75678927 | 75697424 | 75697517 |
| SE | 31827 | ENSG00000112701 | SENP6 | chr6 | + | 75711327 | 75711416 | 75709526 | 75709630 | 75713512 | 75713581 |
| SE | 31828 | ENSG00000112701 | SENP6 | chr6 | + | 75713674 | 75713825 | 75713512 | 75713581 | 75715384 | 75715877 |
| SE | 31836 | ENSG00000185278 | ZBTB37 | chr1 | + | 173868924 | 173869120 | 173868354 | 173868405 | 173870196 | 173871148 |
| SE | 31839 | ENSG00000113811 | SELENOK | chr3 | - | 53885825 | 53885904 | 53885300 | 53885561 | 53886850 | 53886934 |
| SE | 31840 | ENSG00000113811 | SELENOK | chr3 | - | 53885825 | 53885912 | 53884416 | 53885561 | 53886850 | 53886934 |
| SE | 31841 | ENSG00000113811 | SELENOK | chr3 | - | 53886850 | 53886934 | 53885825 | 53885912 | 53891769 | 53891859 |
| SE | 31843 | ENSG00000113811 | SELENOK | chr3 | - | 53890925 | 53891060 | 53888394 | 53888483 | 53891769 | 53891885 |
| SE | 31844 | ENSG00000066777 | ARFGEF1 | chr8 | - | 67216589 | 67216662 | 67211482 | 67211615 | 67217781 | 67217920 |
| SE | 31845 | ENSG00000066777 | ARFGEF1 | chr8 | - | 67219430 | 67219560 | 67218002 | 67218138 | 67224902 | 67225033 |
| SE | 31846 | ENSG00000066777 | ARFGEF1 | chr8 | - | 67226022 | 67226183 | 67224902 | 67225033 | 67227136 | 67227309 |
| SE | 31848 | ENSG00000066777 | ARFGEF1 | chr8 | - | 67238342 | 67238493 | 67232854 | 67232945 | 67238734 | 67238754 |
| SE | 31849 | ENSG00000066777 | ARFGEF1 | chr8 | - | 67253450 | 67253622 | 67251298 | 67251450 | 67257731 | 67257816 |
| SE | 31850 | ENSG00000066777 | ARFGEF1 | chr8 | - | 67259814 | 67259926 | 67258084 | 67258290 | 67266005 | 67266207 |
| SE | 31852 | ENSG00000066777 | ARFGEF1 | chr8 | - | 67267090 | 67267230 | 67266875 | 67266984 | 67267342 | 67267442 |
| SE | 31856 | ENSG00000066777 | ARFGEF1 | chr8 | - | 67299208 | 67299355 | 67296430 | 67296610 | 67301223 | 67301380 |
| SE | 31857 | ENSG00000143669 | LYST | chr1 | - | 235686948 | 235687047 | 235677479 | 235677619 | 235693349 | 235693486 |
| SE | 31858 | ENSG00000143669 | LYST | chr1 | - | 235693349 | 235693486 | 235686948 | 235687047 | 235697082 | 235697272 |
| SE | 31859 | ENSG00000143669 | LYST | chr1 | - | 235697082 | 235697272 | 235693349 | 235693486 | 235702746 | 235702977 |
| SE | 31861 | ENSG00000143669 | LYST | chr1 | - | 235720660 | 235720905 | 235716711 | 235716778 | 235724027 | 235724180 |
| SE | 31863 | ENSG00000143669 | LYST | chr1 | - | 235733502 | 235733691 | 235731031 | 235731177 | 235733829 | 235733906 |
| SE | 31865 | ENSG00000143669 | LYST | chr1 | - | 235751209 | 235751362 | 235746335 | 235746527 | 235752004 | 235752171 |
| SE | 31866 | ENSG00000143669 | LYST | chr1 | - | 235751209 | 235751362 | 235746335 | 235746527 | 235753043 | 235753274 |
| SE | 31868 | ENSG00000143669 | LYST | chr1 | - | 235752004 | 235752171 | 235746335 | 235746527 | 235753043 | 235753274 |
| SE | 31869 | ENSG00000143669 | LYST | chr1 | - | 235752004 | 235752171 | 235746335 | 235746527 | 235755477 | 235755647 |
| SE | 31870 | ENSG00000143669 | LYST | chr1 | - | 235752004 | 235752171 | 235751209 | 235751362 | 235753043 | 235753274 |
| SE | 31871 | ENSG00000143669 | LYST | chr1 | - | 235752004 | 235752171 | 235751209 | 235751362 | 235755477 | 235755647 |
| SE | 31873 | ENSG00000143669 | LYST | chr1 | - | 235753043 | 235753274 | 235751209 | 235751362 | 235755477 | 235755647 |
| SE | 31874 | ENSG00000143669 | LYST | chr1 | - | 235753043 | 235753274 | 235752004 | 235752171 | 235755477 | 235755647 |
| SE | 31875 | ENSG00000143669 | LYST | chr1 | - | 235757280 | 235757458 | 235755477 | 235755647 | 235758971 | 235759599 |
| SE | 31876 | ENSG00000143669 | LYST | chr1 | - | 235758971 | 235759599 | 235755477 | 235755647 | 235762719 | 235762851 |
| SE | 31877 | ENSG00000143669 | LYST | chr1 | - | 235762719 | 235762851 | 235758971 | 235759599 | 235766078 | 235766277 |
| SE | 31881 | ENSG00000143669 | LYST | chr1 | - | 235773841 | 235773991 | 235766078 | 235766277 | 235777062 | 235777308 |
| SE | 31882 | ENSG00000143669 | LYST | chr1 | - | 235773841 | 235773991 | 235770159 | 235770297 | 235774912 | 235775086 |
| SE | 31883 | ENSG00000143669 | LYST | chr1 | - | 235774912 | 235775086 | 235766078 | 235766277 | 235777062 | 235777308 |
| SE | 31885 | ENSG00000143669 | LYST | chr1 | - | 235774912 | 235775086 | 235773841 | 235773991 | 235777062 | 235777308 |
| SE | 31886 | ENSG00000143669 | LYST | chr1 | - | 235777062 | 235777308 | 235766078 | 235766277 | 235787199 | 235787373 |
| SE | 31887 | ENSG00000143669 | LYST | chr1 | - | 235777062 | 235777308 | 235774912 | 235775086 | 235780864 | 235781055 |
| SE | 31888 | ENSG00000143669 | LYST | chr1 | - | 235777062 | 235777308 | 235774912 | 235775086 | 235787199 | 235787373 |
| SE | 31889 | ENSG00000143669 | LYST | chr1 | - | 235780864 | 235781055 | 235777062 | 235777308 | 235781926 | 235782087 |
| SE | 31890 | ENSG00000143669 | LYST | chr1 | - | 235781926 | 235782087 | 235777062 | 235777308 | 235787199 | 235787373 |
| SE | 31891 | ENSG00000143669 | LYST | chr1 | - | 235787199 | 235787373 | 235781926 | 235782087 | 235788700 | 235788845 |
| SE | 31892 | ENSG00000143669 | LYST | chr1 | - | 235788700 | 235788845 | 235781926 | 235782087 | 235791698 | 235792125 |
| SE | 31893 | ENSG00000143669 | LYST | chr1 | - | 235791698 | 235792125 | 235788700 | 235788845 | 235793502 | 235793612 |
| SE | 31895 | ENSG00000143669 | LYST | chr1 | - | 235802907 | 235803064 | 235800870 | 235801097 | 235805742 | 235806772 |
| SE | 31896 | ENSG00000143669 | LYST | chr1 | - | 235802907 | 235803064 | 235800870 | 235801097 | 235830225 | 235830424 |
| SE | 31897 | ENSG00000143669 | LYST | chr1 | - | 235804503 | 235804665 | 235802907 | 235803064 | 235805742 | 235806772 |
| SE | 31899 | ENSG00000143669 | LYST | chr1 | - | 235805742 | 235806772 | 235800870 | 235801097 | 235830225 | 235830424 |
| SE | 31901 | ENSG00000143669 | LYST | chr1 | - | 235808454 | 235810534 | 235800870 | 235801097 | 235830225 | 235830424 |
| SE | 31903 | ENSG00000143669 | LYST | chr1 | - | 235812970 | 235813061 | 235808454 | 235810534 | 235830225 | 235830424 |
| SE | 31907 | ENSG00000163932 | PRKCD | chr3 | + | 53165103 | 53165215 | 53161208 | 53161428 | 53173492 | 53173608 |
| SE | 31910 | ENSG00000163932 | PRKCD | chr3 | + | 53173492 | 53173608 | 53161355 | 53161428 | 53178403 | 53178537 |
| SE | 31912 | ENSG00000163932 | PRKCD | chr3 | + | 53173492 | 53173608 | 53168740 | 53168811 | 53178403 | 53178537 |
| SE | 31913 | ENSG00000163932 | PRKCD | chr3 | + | 53173492 | 53173608 | 53168796 | 53168820 | 53178403 | 53178537 |
| SE | 31914 | ENSG00000163932 | PRKCD | chr3 | + | 53173526 | 53173608 | 53161240 | 53161428 | 53178403 | 53178537 |
| SE | 31916 | ENSG00000163932 | PRKCD | chr3 | + | 53173526 | 53173608 | 53168740 | 53168811 | 53178403 | 53178537 |
| SE | 31918 | ENSG00000163932 | PRKCD | chr3 | + | 53184873 | 53184974 | 53183342 | 53183581 | 53185603 | 53185700 |
| SE | 31919 | ENSG00000163932 | PRKCD | chr3 | + | 53185603 | 53185700 | 53184873 | 53184974 | 53185926 | 53186027 |
| SE | 31920 | ENSG00000163932 | PRKCD | chr3 | + | 53185603 | 53185700 | 53184873 | 53184974 | 53186166 | 53186340 |
| SE | 31921 | ENSG00000163932 | PRKCD | chr3 | + | 53185926 | 53186027 | 53184873 | 53184974 | 53186166 | 53186340 |
| SE | 31923 | ENSG00000163932 | PRKCD | chr3 | + | 53186166 | 53186340 | 53185603 | 53185700 | 53186603 | 53186695 |
| SE | 31924 | ENSG00000163932 | PRKCD | chr3 | + | 53186603 | 53186695 | 53186166 | 53186340 | 53188719 | 53188858 |
| SE | 31925 | ENSG00000163932 | PRKCD | chr3 | + | 53187339 | 53187402 | 53186603 | 53186695 | 53188719 | 53188858 |
| SE | 31926 | ENSG00000163932 | PRKCD | chr3 | + | 53189057 | 53189246 | 53188719 | 53188858 | 53189872 | 53190001 |
| SE | 31928 | ENSG00000278259 | MYO19 | chr17 | - | 36501851 | 36501901 | 36501068 | 36501235 | 36503096 | 36503200 |
| SE | 31930 | ENSG00000278259 | MYO19 | chr17 | - | 36510745 | 36510917 | 36509061 | 36509135 | 36511364 | 36511455 |
| SE | 31931 | ENSG00000278259 | MYO19 | chr17 | - | 36511364 | 36511455 | 36510745 | 36510917 | 36513428 | 36513505 |
| SE | 31935 | ENSG00000278259 | MYO19 | chr17 | - | 36527550 | 36527699 | 36525227 | 36525341 | 36532526 | 36532681 |
| SE | 31936 | ENSG00000278259 | MYO19 | chr17 | - | 36528063 | 36528202 | 36527550 | 36527699 | 36532526 | 36532681 |
| SE | 31949 | ENSG00000163703 | CRELD1 | chr3 | + | 9942812 | 9942896 | 9940849 | 9941206 | 9943076 | 9943153 |
| SE | 31953 | ENSG00000163703 | CRELD1 | chr3 | + | 9943993 | 9944095 | 9943076 | 9943515 | 9944364 | 9944886 |
| SE | 31957 | ENSG00000137502 | RAB30 | chr11 | - | 83034171 | 83034272 | 82997223 | 82997324 | 83034726 | 83034782 |
| SE | 31961 | ENSG00000137502 | RAB30 | chr11 | - | 83034726 | 83034782 | 83034171 | 83034272 | 83071690 | 83071896 |
| SE | 31963 | ENSG00000012223 | LTF | chr3 | - | 46437939 | 46438129 | 46435644 | 46436229 | 46439295 | 46439480 |
| SE | 31964 | ENSG00000012223 | LTF | chr3 | - | 46443440 | 46443582 | 46441415 | 46441483 | 46445280 | 46445436 |
| SE | 31965 | ENSG00000012223 | LTF | chr3 | - | 46449853 | 46450028 | 46448862 | 46449017 | 46450494 | 46450673 |
| SE | 31966 | ENSG00000012223 | LTF | chr3 | - | 46450494 | 46450673 | 46448862 | 46449017 | 46454304 | 46454360 |
| SE | 31967 | ENSG00000012223 | LTF | chr3 | - | 46459655 | 46459819 | 46456289 | 46456398 | 46464824 | 46464868 |
| SE | 31973 | ENSG00000168936 | TMEM129 | chr4 | - | 1717515 | 1717675 | 1715951 | 1717428 | 1718151 | 1718626 |
| SE | 31974 | ENSG00000168936 | TMEM129 | chr4 | - | 1717515 | 1717703 | 1716600 | 1717428 | 1718151 | 1718235 |
| SE | 31976 | ENSG00000082293 | COL19A1 | chr6 | + | 69938037 | 69938100 | 69932782 | 69932863 | 69959995 | 69960040 |
| SE | 31978 | ENSG00000082293 | COL19A1 | chr6 | + | 70130181 | 70130223 | 70121879 | 70121942 | 70137684 | 70137747 |
| SE | 31979 | ENSG00000082293 | COL19A1 | chr6 | + | 70156669 | 70156723 | 70156315 | 70156369 | 70161899 | 70161953 |
| SE | 31984 | ENSG00000122970 | IFT81 | chr12 | + | 110143381 | 110143545 | 110136775 | 110136860 | 110146952 | 110147048 |
| SE | 32010 | ENSG00000142798 | HSPG2 | chr1 | - | 21873374 | 21873424 | 21872858 | 21873091 | 21873924 | 21874011 |
| SE | 32012 | ENSG00000100003 | SEC14L2 | chr22 | + | 30397748 | 30397870 | 30397050 | 30397170 | 30399642 | 30399718 |
| SE | 32013 | ENSG00000100003 | SEC14L2 | chr22 | + | 30399642 | 30399718 | 30396994 | 30397170 | 30406341 | 30406385 |
| SE | 32017 | ENSG00000100003 | SEC14L2 | chr22 | + | 30409106 | 30409282 | 30407414 | 30407603 | 30409425 | 30409477 |
| SE | 32018 | ENSG00000100003 | SEC14L2 | chr22 | + | 30409186 | 30409282 | 30407414 | 30407603 | 30409425 | 30409486 |
| SE | 32022 | ENSG00000100003 | SEC14L2 | chr22 | + | 30415947 | 30416087 | 30410595 | 30410679 | 30416233 | 30416403 |
| SE | 32029 | ENSG00000163702 | IL17RC | chr3 | + | 9920924 | 9920969 | 9920490 | 9920602 | 9923880 | 9924020 |
| SE | 32030 | ENSG00000163702 | IL17RC | chr3 | + | 9928579 | 9928630 | 9928304 | 9928486 | 9929851 | 9929880 |
| SE | 32035 | ENSG00000163702 | IL17RC | chr3 | + | 9932819 | 9932858 | 9932607 | 9932703 | 9932952 | 9933593 |
| SE | 32043 | ENSG00000126970 | ZC4H2 | chrX | - | 64919041 | 64919204 | 64915806 | 64917896 | 64920080 | 64920253 |
| SE | 32044 | ENSG00000126970 | ZC4H2 | chrX | - | 64919041 | 64919236 | 64917779 | 64917896 | 64920080 | 64920253 |
| SE | 32045 | ENSG00000081307 | UBA5 | chr3 | + | 132660898 | 132661048 | 132660600 | 132660698 | 132665822 | 132665868 |
| SE | 32046 | ENSG00000081307 | UBA5 | chr3 | + | 132665822 | 132665868 | 132659945 | 132660698 | 132668817 | 132668927 |
| SE | 32047 | ENSG00000081307 | UBA5 | chr3 | + | 132665822 | 132665868 | 132660361 | 132661048 | 132668817 | 132668927 |
| SE | 32048 | ENSG00000081307 | UBA5 | chr3 | + | 132665983 | 132666073 | 132665738 | 132665868 | 132668817 | 132668927 |

| | | | | | | | | | | | | | | | | | | | | | | | | | | | | | | | | | | | |
|---|---|---|---|---|---|---|---|---|---|---|---|---|---|---|---|---|---|---|---|---|---|---|---|---|---|---|---|---|---|---|---|---|---|---|---|
| 31587 | 80 | 18 | 46 | 65 | 71 | 1 | 1 | 1 | 2 | 0 | 52 | 42 | 55 | 21 | 47 | 0 | 0 | 3 | 1 | 4 | 293 | 149 | 1 | 1 | 0.976 | 0.902 | 0.959 | 0.943 | 1 | 1 | 1 | 0.903 | 0.914 | 0.857 | 0.021 |
| 31590 | 355 | 233 | 305 | 375 | 426 | 0 | 0 | 0 | 0 | 0 | 354 | 368 | 325 | 199 | 415 | 0 | 3 | 0 | 6 | 0 | 298 | 149 | 1 | 1 | 1 | 1 | 1 | 1 | 1 | 1 | 0.984 | 1 | 0.943 | 1 | 0.015 |
| 31591 | 356 | 235 | 389 | 448 | 422 | 3 | 0 | 0 | 2 | 0 | 377 | 445 | 384 | 295 | 370 | 0 | 0 | 0 | 0 | 0 | 298 | 149 | 1 | 1 | 0.983 | 1 | 1 | 0.991 | 1 | 1 | 1 | 1 | 1 | 1 | -0.005 |
| 31593 | 88 | 69 | 59 | 77 | 54 | 0 | 3 | 0 | 0 | 0 | 68 | 37 | 93 | 61 | 23 | 0 | 0 | 0 | 0 | 0 | 298 | 149 | 1 | 1 | 1 | 0.92 | 1 | 1 | 1 | 1 | 1 | 1 | 1 | 1 | -0.016 |
| 31594 | 87 | 66 | 55 | 66 | 54 | 0 | 3 | 0 | 0 | 0 | 68 | 37 | 88 | 61 | 23 | 0 | 0 | 0 | 0 | 0 | 298 | 149 | 1 | 1 | 1 | 0.917 | 1 | 1 | 1 | 1 | 1 | 1 | 1 | 1 | -0.017 |
| 31595 | 86 | 98 | 78 | 98 | 82 | 0 | 0 | 1 | 0 | 0 | 55 | 77 | 142 | 111 | 25 | 0 | 0 | 0 | 0 | 0 | 298 | 149 | 1 | 1 | 1 | 1 | 0.975 | 1 | 1 | 1 | 1 | 1 | 1 | 1 | -0.005 |
| 31601 | 91 | 86 | 122 | 107 | 108 | 0 | 4 | 2 | 0 | 0 | 161 | 82 | 118 | 125 | 14 | 0 | 6 | 0 | 0 | 0 | 281 | 149 | 1 | 1 | 1 | 0.919 | 0.97 | 1 | 1 | 1 | 0.879 | 1 | 1 | 1 | 0.002 |
| 31607 | 32 | 20 | 37 | 63 | 53 | 0 | 0 | 2 | 0 | 0 | 66 | 56 | 52 | 70 | 41 | 0 | 0 | 0 | 0 | 0 | 230 | 149 | 1 | 1 | 1 | 1 | 0.923 | 1 | 1 | 1 | 1 | 1 | 1 | 1 | -0.015 |
| 31608 | 134 | 66 | 138 | 153 | 114 | 0 | 0 | 0 | 0 | 0 | 150 | 170 | 113 | 105 | 218 | 0 | 0 | 1 | 1 | 0 | 242 | 149 | 1 | 1 | 1 | 1 | 1 | 1 | 1 | 1 | 1 | 1 | 0.985 | 1 | 0.003 |
| 31609 | 184 | 117 | 172 | 160 | 137 | 0 | 0 | 0 | 0 | 0 | 295 | 211 | 167 | 226 | 245 | 0 | 0 | 0 | 3 | 0 | 225 | 149 | 1 | 1 | 1 | 1 | 1 | 1 | 1 | 1 | 1 | 1 | 0.98 | 1 | 0.004 |
| 31612 | 0 | 0 | 0 | 0 | 0 | 149 | 84 | 128 | 135 | 93 | 8 | 0 | 0 | 0 | 0 | 117 | 148 | 115 | 153 | 235 | 222 | 149 | 1 | 1 | 0 | 0 | 0 | 0 | 0 | 0.044 | 0 | 0 | 0 | 0 | -0.009 |
| 31614 | 5 | 0 | 10 | 0 | 0 | 0 | 2 | 0 | 0 | 0 | 0 | 0 | 5 | 5 | 10 | 0 | 0 | 0 | 0 | 0 | 298 | 149 | 1 | 1 | 1 | 0 | 1 | NA | NA | NA | NA | 1 | 1 | 1 | -0.333 |
| 31617 | 2 | 7 | 6 | 6 | 1 | 0 | 0 | 1 | 0 | 0 | 27 | 0 | 0 | 1 | 15 | 0 | 3 | 0 | 3 | 0 | 248 | 149 | 1 | 1 | 1 | 1 | 0.783 | 1 | 1 | 1 | 0 | NA | 0.167 | 1 | 0.415 |
| 31618 | 1 | 4 | 2 | 0 | 6 | 0 | 0 | 2 | 0 | 0 | 13 | 0 | 2 | 0 | 0 | 0 | 2 | 0 | 0 | 0 | 279 | 149 | 1 | 1 | 1 | 1 | 0.348 | NA | 1 | 1 | NA | 0.348 | NA | NA | 0.163 |
| 31619 | 4 | 0 | 2 | 3 | 2 | 0 | 0 | 1 | 0 | 0 | 0 | 3 | 17 | 1 | 0 | 2 | 0 | 0 | 0 | 0 | 260 | 149 | 1 | 1 | 1 | NA | 0.534 | 1 | 1 | 0 | 1 | 1 | 1 | NA | 0.133 |
| 31620 | 4 | 0 | 2 | 1 | 1 | 0 | 0 | 2 | 1 | 0 | 0 | 2 | 16 | 0 | 0 | 0 | 2 | 0 | 0 | 0 | 260 | 149 | 1 | 1 | 1 | NA | 0.364 | 0.364 | 1 | NA | 0.364 | 1 | NA | NA | 0 |
| 31621 | 4 | 0 | 2 | 1 | 0 | 0 | 0 | 0 | 0 | 0 | 0 | 6 | 16 | 0 | 0 | 9 | 0 | 0 | 0 | 0 | 260 | 149 | 1 | 1 | 1 | NA | 1 | 1 | NA | 0 | 1 | 1 | NA | NA | 0.333 |
| 31622 | 2 | 2 | 7 | 2 | 14 | 0 | 0 | 2 | 1 | 0 | 4 | 15 | 7 | 0 | 0 | 0 | 2 | 0 | 0 | 0 | 288 | 149 | 1 | 1 | 1 | 1 | 0.644 | 0.509 | 1 | 1 | 0.795 | 1 | NA | NA | -0.101 |
| 31623 | 2 | 2 | 6 | 4 | 16 | 0 | 0 | 0 | 0 | 1 | 2 | 16 | 8 | 1 | 0 | 0 | 0 | 0 | 0 | 0 | 288 | 149 | 1 | 1 | 1 | 1 | 1 | 1 | 0.892 | 1 | 1 | 1 | 1 | NA | -0.022 |
| 31625 | 3 | 1 | 4 | 5 | 17 | 0 | 0 | 0 | 0 | 0 | 0 | 0 | 2 | 8 | 10 | 0 | 4 | 0 | 0 | 0 | 253 | 149 | 1 | 1 | 1 | 1 | 1 | 1 | 1 | NA | 0 | 1 | 1 | 1 | 0.25 |
| 31630 | 0 | 0 | 3 | 2 | 0 | 42 | 13 | 16 | 35 | 11 | 0 | 1 | 0 | 0 | 0 | 54 | 67 | 16 | 57 | 110 | 262 | 149 | 1 | 1 | 0 | 0 | 0.096 | 0.031 | 0 | 0 | 0.008 | 0 | 0 | 0 | 0.024 |
| 31633 | 68 | 43 | 33 | 46 | 35 | 2 | 0 | 1 | 0 | 0 | 183 | 57 | 46 | 85 | 77 | 26 | 4 | 0 | 0 | 10 | 298 | 149 | 1 | 1 | 0.944 | 1 | 0.943 | 1 | 1 | 0.779 | 0.877 | 1 | 1 | 0.794 | 0.087 |
| 31635 | 1 | 0 | 1 | 0 | 0 | 1 | 0 | 0 | 0 | 0 | 0 | 0 | 0 | 6 | 0 | 0 | 0 | 0 | 0 | 0 | 229 | 149 | 1 | 1 | 0.394 | NA | 1 | NA | NA | NA | NA | 1 | NA | NA | -0.303 |
| 31637 | 63 | 24 | 35 | 43 | 11 | 0 | 0 | 0 | 1 | 0 | 110 | 89 | 70 | 62 | 75 | 0 | 0 | 0 | 0 | 0 | 191 | 149 | 1 | 1 | 1 | 1 | 1 | 0.971 | 1 | 1 | 0.986 | 1 | 1 | 1 | -0.003 |
| 31641 | 352 | 191 | 255 | 358 | 361 | 0 | 0 | 0 | 0 | 0 | 676 | 576 | 420 | 477 | 865 | 0 | 1 | 0 | 3 | 0 | 284 | 149 | 1 | 1 | 1 | 1 | 1 | 1 | 1 | 1 | 0.997 | 1 | 0.988 | 1 | 0.003 |
| 31643 | 0 | 0 | 0 | 0 | 0 | 0 | 0 | 0 | 1 | 0 | 8 | 8 | 6 | 3 | 0 | 0 | 0 | 0 | 0 | 0 | 284 | 149 | 1 | 1 | NA | NA | NA | 0 | NA | 1 | 1 | 1 | 1 | NA | -1 |
| 31648 | 60 | 39 | 29 | 55 | 48 | 0 | 0 | 1 | 1 | 2 | 112 | 49 | 55 | 40 | 142 | 0 | 6 | 1 | 0 | 0 | 262 | 149 | 1 | 1 | 1 | 1 | 0.943 | 0.969 | 0.932 | 1 | 1 | 0.839 | 0.958 | 1 | 0.009 |
| 31649 | 3 | 7 | 1 | 6 | 0 | 0 | 0 | 1 | 3 | 0 | 0 | 0 | 6 | 5 | 2 | 0 | 1 | 3 | 0 | 0 | 262 | 149 | 1 | 1 | 1 | 1 | 0.363 | 0.532 | NA | NA | 0 | 0.532 | 1 | 1 | 0.091 |
| 31650 | 3 | 7 | 1 | 6 | 0 | 1 | 0 | 2 | 0 | 0 | 0 | 0 | 3 | 5 | 2 | 0 | 5 | 0 | 12 | 0 | 262 | 149 | 1 | 1 | 0.63 | 1 | 0.221 | 1 | NA | NA | 0 | 1 | 0.192 | 1 | 0.165 |
| 31653 | 80 | 59 | 58 | 96 | 77 | 0 | 0 | 1 | 3 | 0 | 141 | 103 | 88 | 77 | 175 | 0 | 1 | 3 | 0 | 0 | 287 | 149 | 1 | 1 | 1 | 1 | 0.968 | 0.943 | 1 | 1 | 0.982 | 0.938 | 1 | 1 | -0.002 |
| 31654 | 0 | 0 | 1 | 1 | 8 | 1 | 0 | 2 | 0 | 0 | 29 | 0 | 14 | 5 | 14 | 0 | 5 | 0 | 12 | 0 | 287 | 149 | 1 | 1 | 0 | NA | 0.206 | 1 | 1 | 1 | 0 | 1 | 0.178 | 1 | -0.084 |
| 31655 | 0 | 0 | 0 | 1 | 6 | 0 | 0 | 0 | 0 | 0 | 29 | 0 | 12 | 10 | 14 | 0 | 0 | 0 | 0 | 8 | 287 | 149 | 1 | 1 | NA | NA | NA | 1 | 1 | 1 | NA | 1 | 1 | 0.476 | 0.131 |
| 31656 | 0 | 1 | 0 | 1 | 3 | 0 | 0 | 0 | 0 | 0 | 0 | 0 | 6 | 7 | 0 | 0 | 0 | 2 | 0 | 0 | 287 | 149 | 1 | 1 | NA | 1 | NA | 1 | 1 | NA | NA | 0.609 | 1 | NA | 0.196 |
| 31657 | 137 | 95 | 93 | 158 | 132 | 0 | 0 | 0 | 0 | 0 | 280 | 155 | 142 | 127 | 352 | 0 | 0 | 4 | 0 | 0 | 287 | 149 | 1 | 1 | 1 | 1 | 1 | 1 | 1 | 1 | 1 | 0.949 | 1 | 1 | 0.01 |
| 31658 | 51 | 36 | 31 | 59 | 50 | 0 | 0 | 0 | 0 | 0 | 157 | 49 | 67 | 42 | 167 | 0 | 0 | 1 | 0 | 0 | 287 | 149 | 1 | 1 | 1 | 1 | 1 | 1 | 1 | 1 | 1 | 0.972 | 1 | 1 | 0.006 |
| 31659 | [illegible] | [illegible] | [illegible] | [illegible] | [illegible] | [illegible] | [illegible] | [illegible] | [illegible] | [illegible] | [illegible] | [illegible] | [illegible] | [illegible] | [illegible] | [illegible] | [illegible] | [illegible] | [illegible] | 0 | 287 | 149 | 1 | 1 | 1 | 1 | 0.801 | 0.968 | 1 | 1 | 1 | 0.969 | 1 | 1 | -0.04 |
| 31661 | [illegible] | [illegible] | [illegible] | [illegible] | [illegible] | [illegible] | [illegible] | [illegible] | [illegible] | [illegible] | [illegible] | [illegible] | [illegible] | [illegible] | [illegible] | [illegible] | [illegible] | [illegible] | [illegible] | 0 | 252 | 149 | 1 | 1 | 1 | 1 | 1 | 1 | 1 | 1 | 1 | 0.981 | 1 | 1 | 0.004 |
| 31662 | [illegible] | [illegible] | [illegible] | [illegible] | [illegible] | [illegible] | [illegible] | [illegible] | [illegible] | [illegible] | [illegible] | [illegible] | [illegible] | [illegible] | [illegible] | [illegible] | [illegible] | [illegible] | [illegible] | 0 | 252 | 149 | 1 | 1 | NA | 1 | 0 | 0.372 | NA | NA | NA | 0.703 | 1 | NA | -0.394 |
| 31664 | [illegible] | [illegible] | [illegible] | [illegible] | [illegible] | [illegible] | [illegible] | [illegible] | [illegible] | [illegible] | [illegible] | [illegible] | [illegible] | [illegible] | [illegible] | [illegible] | [illegible] | [illegible] | [illegible] | 0 | 252 | 149 | 1 | 1 | 1 | 0.95 | 1 | 1 | 0.932 | 1 | 1 | 0.935 | 0.967 | 1 | -0.004 |
| 31665 | [illegible] | [illegible] | [illegible] | [illegible] | [illegible] | [illegible] | [illegible] | [illegible] | [illegible] | [illegible] | [illegible] | [illegible] | [illegible] | [illegible] | [illegible] | [illegible] | [illegible] | [illegible] | [illegible] | 0 | 250 | 149 | 1 | 1 | 1 | 1 | 1 | 1 | 1 | 1 | 1 | 0.973 | 1 | 1 | 0.005 |
| 31666 | [illegible] | [illegible] | [illegible] | [illegible] | [illegible] | [illegible] | [illegible] | [illegible] | [illegible] | [illegible] | [illegible] | [illegible] | [illegible] | [illegible] | [illegible] | [illegible] | [illegible] | [illegible] | [illegible] | 0 | 250 | 149 | 1 | 1 | 1 | 1 | 0.888 | 0.979 | 1 | 1 | 1 | 0.986 | 1 | 1 | -0.024 |

| | | | | | | | | | | | | | | |
|---|---|---|---|---|---|---|---|---|---|---|---|---|---|---|
| SE | 32051 | ENSG00000182568 | SATB1 | chr3 | - | 18351312 | 18351408 | 18347939 | 18349682 | 18351991 | 18352195 | 1 | 0 | [illegible] |
| SE | 32052 | ENSG00000182568 | SATB1 | chr3 | - | 18351991 | 18352195 | 18345376 | 18349682 | 18378169 | 18378325 | 660 | 752 | [illegible] |
| SE | 32053 | ENSG00000182568 | SATB1 | chr3 | - | 18386398 | 18386611 | 18378169 | 18378325 | 18394461 | 18394916 | 540 | 430 | [illegible] |
| SE | 32054 | ENSG00000182568 | SATB1 | chr3 | - | 18415110 | 18415234 | 18397178 | 18397290 | 18416006 | 18416133 | 1083 | 724 | [illegible] |
| SE | 32055 | ENSG00000182568 | SATB1 | chr3 | - | 18415110 | 18415234 | 18410797 | 18411054 | 18416006 | 18416133 | 764 | 443 | [illegible] |
| SE | 32056 | ENSG00000182568 | SATB1 | chr3 | - | 18416006 | 18416133 | 18415110 | 18415234 | 18416901 | 18417078 | 1050 | 628 | [illegible] |
| SE | 32058 | ENSG00000182568 | SATB1 | chr3 | - | 18420756 | 18420991 | 18416900 | 18417078 | 18445517 | 18445588 | 257 | 226 | [illegible] |
| SE | 32060 | ENSG00000182568 | SATB1 | chr3 | - | 18436788 | 18436871 | 18420756 | 18420991 | 18438564 | 18438692 | 2 | 0 | [illegible] |
| SE | 32063 | ENSG00000242086 | MUC20-OT1 | chr3 | + | 195668518 | 195668641 | 195666065 | 195666214 | 195671131 | 195671300 | 114 | 84 | [illegible] |
| SE | 32066 | ENSG00000242086 | MUC20-OT1 | chr3 | + | 195686297 | 195686389 | 195685817 | 195685939 | 195688532 | 195688797 | 0 | 0 | [illegible] |
| SE | 32067 | ENSG00000242086 | MUC20-OT1 | chr3 | + | 195686603 | 195686718 | 195683749 | 195683865 | 195688532 | 195688797 | 0 | 2 | [illegible] |
| SE | 32068 | ENSG00000242086 | MUC20-OT1 | chr3 | + | 195686606 | 195686718 | 195683776 | 195683865 | 195688532 | 195688797 | 0 | 2 | [illegible] |
| SE | 32069 | ENSG00000242086 | MUC20-OT1 | chr3 | + | 195686606 | 195686718 | 195686297 | 195686389 | 195688532 | 195688797 | 0 | 2 | [illegible] |
| SE | 32075 | ENSG00000242086 | MUC20-OT1 | chr3 | + | 195688532 | 195688797 | 195683751 | 195683865 | 195699029 | 195699207 | 62 | 41 | [illegible] |
| SE | 32076 | ENSG00000242086 | MUC20-OT1 | chr3 | + | 195688532 | 195688797 | 195683751 | 195683865 | 195708133 | 195708345 | 54 | 32 | [illegible] |
| SE | 32077 | ENSG00000242086 | MUC20-OT1 | chr3 | + | 195688532 | 195688797 | 195683751 | 195683865 | 195708746 | 195708806 | 54 | 30 | [illegible] |
| SE | 32083 | ENSG00000242086 | MUC20-OT1 | chr3 | + | 195696360 | 195696589 | 195683751 | 195683865 | 195699029 | 195699116 | 46 | 60 | [illegible] |
| SE | 32086 | ENSG00000242086 | MUC20-OT1 | chr3 | + | 195699029 | 195699207 | 195683751 | 195683865 | 195708133 | 195708268 | 5 | 2 | [illegible] |
| SE | 32090 | ENSG00000242086 | MUC20-OT1 | chr3 | + | 195701379 | 195701432 | 195699029 | 195699207 | 195708133 | 195708435 | 0 | 0 | [illegible] |
| SE | 32091 | ENSG00000242086 | MUC20-OT1 | chr3 | + | 195701379 | 195701495 | 195699029 | 195699207 | 195708133 | 195708807 | 5 | 0 | [illegible] |
| SE | 32093 | ENSG00000242086 | MUC20-OT1 | chr3 | + | 195701379 | 195701669 | 195699029 | 195699207 | 195708133 | 195708268 | 11 | 2 | [illegible] |
| SE | 32099 | ENSG00000242086 | MUC20-OT1 | chr3 | + | 195708133 | 195708268 | 195683749 | 195683865 | 195708746 | 195708801 | 4 | 5 | [illegible] |
| SE | 32104 | ENSG00000242086 | MUC20-OT1 | chr3 | + | 195708428 | 195708435 | 195708153 | 195708268 | 195708746 | 195708841 | 0 | 0 | [illegible] |
| SE | 32105 | ENSG00000242086 | MUC20-OT1 | chr3 | + | 195708428 | 195708513 | 195708133 | 195708268 | 195708746 | 195708807 | 1 | 0 | [illegible] |
| SE | 32106 | ENSG00000242086 | MUC20-OT1 | chr3 | + | 195708428 | 195708555 | 195708153 | 195708268 | 195708746 | 195708841 | 0 | 0 | [illegible] |
| SE | 32112 | ENSG00000242086 | MUC20-OT1 | chr3 | + | 195718023 | 195718196 | 195708656 | 195708841 | 195722500 | 195722648 | 0 | 0 | [illegible] |
| SE | 32113 | ENSG00000242086 | MUC20-OT1 | chr3 | + | 195718023 | 195718196 | 195708656 | 195708841 | 195724679 | 195724818 | 0 | 1 | [illegible] |
| SE | 32115 | ENSG00000242086 | MUC20-OT1 | chr3 | + | 195718026 | 195718196 | 195708746 | 195708841 | 195724679 | 195724737 | 0 | 1 | [illegible] |
| SE | 32117 | ENSG00000242086 | MUC20-OT1 | chr3 | + | 195718043 | 195718196 | 195708746 | 195708841 | 195724679 | 195725022 | 0 | 1 | [illegible] |
| SE | 32118 | ENSG00000242086 | MUC20-OT1 | chr3 | + | 195722500 | 195722818 | 195708241 | 195708841 | 195724679 | 195724753 | 1 | 0 | [illegible] |
| SE | 32119 | ENSG00000242086 | MUC20-OT1 | chr3 | + | 195729342 | 195729551 | 195708746 | 195708841 | 195729647 | 195729739 | 0 | 0 | [illegible] |
| SE | 32123 | ENSG00000084463 | WBP11 | chr12 | - | 14794536 | 14794736 | 14793730 | 14793922 | 14794970 | 14795104 | 543 | 344 | [illegible] |
| SE | 32124 | ENSG00000084463 | WBP11 | chr12 | - | 14794536 | 14794736 | 14793730 | 14793922 | 14796806 | 14797003 | 314 | 206 | [illegible] |
| SE | 32125 | ENSG00000084463 | WBP11 | chr12 | - | 14794536 | 14794736 | 14793730 | 14793922 | 14799634 | 14799728 | 313 | 206 | [illegible] |
| SE | 32126 | ENSG00000084463 | WBP11 | chr12 | - | 14794970 | 14795104 | 14793730 | 14793922 | 14796806 | 14797003 | 300 | 236 | [illegible] |
| SE | 32127 | ENSG00000084463 | WBP11 | chr12 | - | 14794970 | 14795104 | 14793730 | 14793922 | 14799634 | 14799728 | 1 | 1 | [illegible] |
| SE | 32132 | ENSG00000084463 | WBP11 | chr12 | - | 14796806 | 14797003 | 14794970 | 14795104 | 14799634 | 14799728 | 357 | 290 | [illegible] |
| SE | 32133 | ENSG00000084463 | WBP11 | chr12 | - | 14800751 | 14800783 | 14799440 | 14799728 | 14801319 | 14801428 | 51 | 44 | [illegible] |
| SE | 32135 | ENSG00000179344 | HLA-DQB1 | chr6 | - | 32660858 | 32660882 | 32659466 | 32660249 | 32661346 | 32661457 | 0 | 0 | [illegible] |
| SE | 32137 | ENSG00000179344 | HLA-DQB1 | chr6 | - | 32664797 | 32665067 | 32661966 | 32662248 | 32666498 | 32666638 | 113 | 423 | [illegible] |
| SE | 32139 | ENSG00000198964 | SGMS1 | chr10 | - | 50311261 | 50311415 | 50305585 | 50307321 | 50343491 | 50344345 | 0 | 0 | [illegible] |
| SE | 32140 | ENSG00000198964 | SGMS1 | chr10 | - | 50327204 | 50327322 | 50311261 | 50311415 | 50343491 | 50344345 | 96 | 55 | [illegible] |
| SE | 32142 | ENSG00000198964 | SGMS1 | chr10 | - | 50341216 | 50341434 | 50340386 | 50340464 | 50343491 | 50344246 | 0 | 0 | [illegible] |
| SE | 32145 | ENSG00000198964 | SGMS1 | chr10 | - | 50460672 | 50460814 | 50433475 | 50433556 | 50519830 | 50519921 | 76 | 62 | [illegible] |
| SE | 32147 | ENSG00000198964 | SGMS1 | chr10 | - | 50519830 | 50519921 | 50466889 | 50466932 | 50590152 | 50590247 | 87 | 82 | [illegible] |
| SE | 32148 | ENSG00000089220 | PEBP1 | chr12 | + | 118138038 | 118138148 | 118136123 | 118136344 | 118139450 | 118139551 | 209 | 99 | [illegible] |
| SE | 32149 | ENSG00000089220 | PEBP1 | chr12 | + | 118139450 | 118139551 | 118138038 | 118138148 | 118144585 | 118145109 | 329 | 212 | [illegible] |
| SE | 32150 | ENSG00000253438 | PCAT1 | chr8 | + | 126589191 | 126589290 | 126557874 | 126557931 | 126617388 | 126617741 | 3 | 3 | [illegible] |
| SE | 32151 | ENSG00000253438 | PCAT1 | chr8 | + | 126589191 | 126589290 | 126557874 | 126557931 | 126713285 | 126713415 | 2 | 1 | [illegible] |
| SE | 32152 | ENSG00000253438 | PCAT1 | chr8 | + | 126592273 | 126592418 | 126589191 | 126589290 | 126713285 | 126713674 | 3 | 0 | [illegible] |
| SE | 32153 | ENSG00000253438 | PCAT1 | chr8 | + | 126707182 | 126707240 | 126589191 | 126589290 | 126713285 | 126713415 | 0 | 0 | [illegible] |
| SE | 32154 | ENSG00000253438 | PCAT1 | chr8 | + | 126708917 | 126709147 | 126707182 | 126707240 | 126713285 | 126713415 | 0 | 0 | [illegible] |
| SE | 32155 | ENSG00000253438 | PCAT1 | chr8 | + | 126713285 | 126713674 | 126589191 | 126589290 | 126877671 | 126877790 | 1 | 1 | [illegible] |
| SE | 32157 | ENSG00000253438 | PCAT1 | chr8 | + | 126878374 | 126878480 | 126877671 | 126877790 | 127006554 | 127006618 | 0 | 0 | [illegible] |
| SE | 32158 | ENSG00000253438 | PCAT1 | chr8 | + | 126926317 | 126926399 | 126877671 | 126877790 | 127006554 | 127006618 | 0 | 0 | [illegible] |
| SE | 32159 | ENSG00000253438 | PCAT1 | chr8 | + | 126938255 | 126938310 | 126877671 | 126877790 | 127006554 | 127006618 | 0 | 1 | [illegible] |
| SE | 32161 | ENSG00000253438 | PCAT1 | chr8 | + | 127013302 | 127013774 | 127006554 | 127006618 | 127339280 | 127339435 | 0 | 0 | [illegible] |
| SE | 32163 | ENSG00000253438 | PCAT1 | chr8 | + | 127322072 | 127322226 | 127006554 | 127006618 | 127339280 | 127339435 | 0 | 0 | [illegible] |
| SE | 32164 | ENSG00000253438 | PCAT1 | chr8 | + | 127408752 | 127409031 | 127398892 | 127399018 | 127414887 | 127415022 | 0 | 0 | [illegible] |
| SE | 32169 | ENSG00000164053 | ATRIP | chr3 | + | 48450036 | 48450170 | 48446732 | 48447092 | 48451728 | 48451899 | 36 | 28 | [illegible] |
| SE | 32173 | ENSG00000164053 | ATRIP | chr3 | + | 48461360 | 48461445 | 48460708 | 48460799 | 48463744 | 48463881 | 0 | 0 | [illegible] |
| SE | 32174 | ENSG00000164053 | ATRIP | chr3 | + | 48464581 | 48464662 | 48464040 | 48464132 | 48464830 | 48465083 | 6 | 1 | [illegible] |
| SE | 32175 | ENSG00000261218 | NA | chr16 | + | 81740259 | 81740379 | 81739073 | 81739436 | 81755869 | 81755966 | 0 | 0 | [illegible] |
| SE | 32176 | ENSG00000261218 | NA | chr16 | + | 81740259 | 81740379 | 81739302 | 81739734 | 81755869 | 81755966 | 0 | 0 | [illegible] |
| SE | 32177 | ENSG00000261218 | NA | chr16 | + | 81740263 | 81740379 | 81739081 | 81739385 | 81755869 | 81755966 | 0 | 0 | [illegible] |
| SE | 32179 | ENSG00000130758 | MAP3K10 | chr19 | + | 40204484 | 40204633 | 40198374 | 40198555 | 40205120 | 40205296 | 16 | 0 | [illegible] |
| SE | 32180 | ENSG00000130758 | MAP3K10 | chr19 | + | 40205120 | 40205296 | 40198396 | 40198555 | 40209102 | 40209219 | 2 | 2 | [illegible] |
| SE | 32182 | ENSG00000113971 | NPHP3 | chr3 | - | 132682702 | 132682818 | 132680608 | 132682090 | 132683398 | 132683524 | 111 | 106 | [illegible] |
| SE | 32183 | ENSG00000113971 | NPHP3 | chr3 | - | 132682702 | 132682818 | 132680608 | 132682090 | 132684553 | 132684794 | 55 | 50 | [illegible] |
| SE | 32184 | ENSG00000113971 | NPHP3 | chr3 | - | 132682702 | 132682818 | 132680608 | 132682090 | 132686259 | 132686387 | 55 | 51 | [illegible] |
| SE | 32187 | ENSG00000113971 | NPHP3 | chr3 | - | 132683398 | 132683524 | 132681627 | 132682090 | 132684553 | 132684685 | 45 | 50 | [illegible] |
| SE | 32188 | ENSG00000113971 | NPHP3 | chr3 | - | 132683398 | 132683524 | 132681627 | 132682090 | 132686259 | 132686387 | 2 | 3 | [illegible] |
| SE | 32190 | ENSG00000113971 | NPHP3 | chr3 | - | 132684553 | 132684794 | 132680608 | 132682090 | 132686259 | 132686387 | 21 | 18 | [illegible] |
| SE | 32193 | ENSG00000113971 | NPHP3 | chr3 | - | 132690527 | 132690650 | 132689073 | 132689263 | 132691191 | 132691286 | 52 | 51 | [illegible] |
| SE | 32194 | ENSG00000113971 | NPHP3 | chr3 | - | 132696730 | 132696813 | 132694826 | 132694965 | 132697259 | 132697362 | 0 | 1 | [illegible] |
| SE | 32195 | ENSG00000113971 | NPHP3 | chr3 | - | 132699352 | 132699450 | 132697259 | 132697362 | 132699917 | 132700061 | 12 | 25 | [illegible] |
| SE | 32197 | ENSG00000113971 | NPHP3 | chr3 | - | 132713125 | 132713286 | 132708100 | 132708257 | 132718993 | 132719144 | 55 | 52 | [illegible] |
| SE | 32198 | ENSG00000113971 | NPHP3 | chr3 | - | 132715084 | 132715218 | 132713125 | 132713286 | 132716756 | 132716813 | 46 | 49 | [illegible] |
| SE | 32199 | ENSG00000113971 | NPHP3 | chr3 | - | 132716756 | 132716909 | 132713125 | 132713286 | 132718993 | 132719144 | 19 | 33 | [illegible] |
| SE | 32200 | ENSG00000113971 | NPHP3 | chr3 | - | 132717954 | 132718117 | 132716756 | 132716909 | 132718993 | 132719144 | 10 | 9 | [illegible] |
| SE | 32202 | ENSG00000108641 | B9D1 | chr17 | - | 19343789 | 19343857 | 19334694 | 19337748 | 19347268 | 19347331 | 1 | 4 | [illegible] |
| SE | 32204 | ENSG00000108641 | B9D1 | chr17 | - | 19371548 | 19371633 | 19360320 | 19360388 | 19377858 | 19377897 | 0 | 0 | [illegible] |
| SE | 32205 | ENSG00000108641 | B9D1 | chr17 | - | 19371548 | 19371633 | 19364435 | 19364601 | 19377858 | 19377897 | 0 | 0 | [illegible] |
| SE | 32206 | ENSG00000108641 | B9D1 | chr17 | - | 19372114 | 19372224 | 19371235 | 19371633 | 19377858 | 19377890 | 0 | 0 | [illegible] |
| SE | 32207 | ENSG00000108641 | B9D1 | chr17 | - | 19372114 | 19372239 | 19371559 | 19371633 | 19377858 | 19378193 | 0 | 0 | [illegible] |
| SE | 32209 | ENSG00000114650 | SCAP | chr3 | - | 47415097 | 47415180 | 47414826 | 47414993 | 47417121 | 47417207 | 41 | 9 | [illegible] |
| SE | 32210 | ENSG00000114650 | SCAP | chr3 | - | 47417303 | 47417823 | 47417121 | 47417207 | 47418133 | 47418249 | 33 | 9 | [illegible] |
| SE | 32211 | ENSG00000114650 | SCAP | chr3 | - | 47417303 | 47417826 | 47417121 | 47417207 | 47418133 | 47418249 | 20 | 6 | [illegible] |
| SE | 32212 | ENSG00000114650 | SCAP | chr3 | - | 47422441 | 47422536 | 47420930 | 47421029 | 47435007 | 47435137 | 41 | 9 | [illegible] |
| SE | 32215 | ENSG00000114650 | SCAP | chr3 | - | 47427156 | 47427262 | 47425996 | 47426169 | 47427446 | 47427667 | 204 | 68 | [illegible] |
| SE | 32216 | ENSG00000114650 | SCAP | chr3 | - | 47427446 | 47427667 | 47425996 | 47426169 | 47435007 | 47435137 | 1 | 3 | [illegible] |
| SE | 32218 | ENSG00000114650 | SCAP | chr3 | - | 47434924 | 47435137 | 47428560 | 47428670 | 47442871 | 47443094 | 47 | 28 | [illegible] |
| SE | 32219 | ENSG00000114650 | SCAP | chr3 | - | 47435007 | 47435137 | 47428512 | 47428670 | 47442871 | 47443091 | 142 | 75 | [illegible] |
| SE | 32227 | ENSG00000145743 | FBXL17 | chr5 | - | 108186116 | 108186247 | 108020798 | 108021001 | 108224120 | 108224228 | 150 | 113 | [illegible] |
| SE | 32228 | ENSG00000145743 | FBXL17 | chr5 | - | 108348398 | 108348530 | 108224120 | 108224228 | 108364737 | 108364767 | 275 | 234 | [illegible] |
| SE | 32229 | ENSG00000145743 | FBXL17 | chr5 | - | 108367830 | 108367953 | 108364737 | 108364995 | 108380698 | 108380923 | 156 | 144 | [illegible] |
| SE | 32230 | ENSG00000156931 | VPS8 | chr3 | + | 184813744 | 184813867 | 184812195 | 184812225 | 184815783 | 184815820 | 0 | 0 | [illegible] |
| SE | 32232 | ENSG00000156931 | VPS8 | chr3 | + | 184824146 | 184824232 | 184812194 | 184812225 | 184824544 | 184824785 | 0 | 0 | [illegible] |
| SE | 32233 | ENSG00000156931 | VPS8 | chr3 | + | 184824930 | 184825048 | 184824544 | 184824785 | 184826162 | 184826231 | 0 | 3 | [illegible] |
| SE | 32234 | ENSG00000156931 | VPS8 | chr3 | + | 184832688 | 184832819 | 184826162 | 184826231 | 184834648 | 184834660 | 123 | 114 | [illegible] |
| SE | 32238 | ENSG00000156931 | VPS8 | chr3 | + | 184843239 | 184843245 | 184839697 | 184839752 | 184849070 | 184849195 | 0 | 0 | [illegible] |
| SE | 32239 | ENSG00000156931 | VPS8 | chr3 | + | 184849935 | 184850022 | 184849028 | 184849195 | 184852499 | 184852538 | 133 | 69 | [illegible] |
| SE | 32240 | ENSG00000156931 | VPS8 | chr3 | + | 184859984 | 184860065 | 184855710 | 184855818 | 184862896 | 184863067 | 287 | 253 | [illegible] |
| SE | 32242 | ENSG00000156931 | VPS8 | chr3 | + | 184882348 | 184882433 | 184870715 | 184870805 | 184886109 | 184886156 | 2 | 2 | [illegible] |
| SE | 32243 | ENSG00000156931 | VPS8 | chr3 | + | 184886109 | 184886156 | 184870715 | 184870805 | 184894702 | 184894925 | 80 | 77 | [illegible] |
| SE | 32244 | ENSG00000156931 | VPS8 | chr3 | + | 184886109 | 184886156 | 184870715 | 184870805 | 184898564 | 184898654 | 15 | 4 | [illegible] |
| SE | 32246 | ENSG00000156931 | VPS8 | chr3 | + | 184894702 | 184894925 | 184870715 | 184870805 | 184898564 | 184898654 | 109 | 78 | [illegible] |
| SE | 32247 | ENSG00000156931 | VPS8 | chr3 | + | 184894702 | 184894925 | 184885968 | 184886156 | 184898564 | 184898654 | 179 | 155 | [illegible] |
| SE | 32248 | ENSG00000156931 | VPS8 | chr3 | + | 184900920 | 184900972 | 184898564 | 184898654 | 184913518 | 184913561 | 71 | 53 | [illegible] |
| SE | 32249 | ENSG00000156931 | VPS8 | chr3 | + | 184914980 | 184915053 | 184913518 | 184913561 | 184915354 | 184915408 | 120 | 90 | [illegible] |
| SE | 32252 | ENSG00000156931 | VPS8 | chr3 | + | 185024335 | 185024389 | 184999695 | 184999861 | 185048478 | 185048559 | 79 | 71 | [illegible] |
| SE | 32255 | ENSG00000144681 | STAC | chr3 | + | 36485762 | 36485925 | 36484976 | 36485058 | 36486133 | 36486249 | 0 | 0 | [illegible] |
| SE | 32256 | ENSG00000144681 | STAC | chr3 | + | 36528847 | 36528985 | 36528695 | 36528747 | 36546190 | 36546488 | 0 | 0 | [illegible] |
| SE | 32257 | ENSG00000203865 | ATP1A1-AS1 | chr1 | - | 116400895 | 116401131 | 116390563 | 116393026 | 116405606 | 116406148 | 0 | 0 | [illegible] |
| SE | 32265 | ENSG00000144645 | OSBPL10 | chr3 | - | 31670796 | 31670983 | 31668641 | 31668824 | 31683633 | 31684114 | 27 | 18 | [illegible] |
| SE | 32267 | ENSG00000144645 | OSBPL10 | chr3 | - | 31797754 | 31797841 | 31747909 | 31748120 | 31830039 | 31830105 | 0 | 0 | [illegible] |
| SE | 32268 | ENSG00000144645 | OSBPL10 | chr3 | - | 31797754 | 31797841 | 31747909 | 31748120 | 31876432 | 31876512 | 0 | 0 | [illegible] |
| SE | 32269 | ENSG00000144645 | OSBPL10 | chr3 | - | 31830039 | 31830231 | 31747909 | 31748120 | 31876432 | 31876512 | 34 | 29 | [illegible] |
| SE | 32270 | ENSG00000144645 | OSBPL10 | chr3 | - | 31879429 | 31879553 | 31876432 | 31876512 | 31879654 | 31879830 | 2 | 1 | [illegible] |
| SE | 32271 | ENSG00000144645 | OSBPL10 | chr3 | - | 31879654 | 31879830 | 31876432 | 31876512 | 31980898 | 31980946 | 31 | 16 | [illegible] |
| SE | 32272 | ENSG00000114745 | GORASP1 | chr3 | - | 39100304 | 39100454 | 39099352 | 39099503 | 39101015 | 39101102 | 19 | 7 | [illegible] |
| SE | 32273 | ENSG00000114745 | GORASP1 | chr3 | - | 39100304 | 39100503 | 39099352 | 39099503 | 39101015 | 39101102 | 20 | 7 | [illegible] |
| SE | 32274 | ENSG00000114745 | GORASP1 | chr3 | - | 39100304 | 39100877 | 39099352 | 39099503 | 39101015 | 39101102 | 31 | 15 | [illegible] |
| SE | 32280 | ENSG00000114745 | GORASP1 | chr3 | - | 39102677 | 39102881 | 39100787 | 39101102 | 39103472 | 39103511 | 17 | 15 | [illegible] |
| SE | 32281 | ENSG00000114745 | GORASP1 | chr3 | - | 39102692 | 39102881 | 39101022 | 39101102 | 39103472 | 39103553 | 11 | 9 | [illegible] |
| SE | 32282 | ENSG00000114745 | GORASP1 | chr3 | - | 39103472 | 39103553 | 39100787 | 39101102 | 39107478 | 39107569 | 6 | 1 | [illegible] |
| SE | 32284 | ENSG00000163964 | PIGX | chr3 | + | 196716857 | 196716921 | 196712670 | 196713124 | 196722414 | 196722556 | 34 | 21 | [illegible] |
| SE | 32285 | ENSG00000163964 | PIGX | chr3 | + | 196717808 | 196717954 | 196716857 | 196716921 | 196721176 | 196721356 | 2 | 2 | [illegible] |
| SE | 32290 | ENSG00000163964 | PIGX | chr3 | + | 196728701 | 196728755 | 196727922 | 196728136 | 196730991 | 196731092 | 1 | 3 | [illegible] |
| SE | 32291 | ENSG00000163964 | PIGX | chr3 | + | 196728701 | 196728755 | 196727922 | 196728136 | 196733758 | 196733902 | 0 | 1 | [illegible] |
| SE | 32292 | ENSG00000163964 | PIGX | chr3 | + | 196730991 | 196731092 | 196727922 | 196728136 | 196733758 | 196733902 | 74 | 35 | [illegible] |
| SE | 32293 | ENSG00000163964 | PIGX | chr3 | + | 196730991 | 196731092 | 196728701 | 196728755 | 196733758 | 196733902 | 45 | 20 | [illegible] |
| SE | 32294 | ENSG00000241553 | ARPC4 | chr3 | + | 9797658 | 9797777 | 9793081 | 9793124 | 9800184 | 9800296 | 799 | 378 | [illegible] |
| SE | 32297 | ENSG00000241553 | ARPC4 | chr3 | + | 9800184 | 9800296 | 9797658 | 9797777 | 9801660 | 9801756 | 1479 | 781 | [illegible] |
| SE | 32298 | ENSG00000241553 | ARPC4 | chr3 | + | 9800184 | 9800296 | 9799878 | 9800067 | 9801660 | 9801756 | 734 | 425 | [illegible] |
| SE | 32301 | ENSG00000241553 | ARPC4 | chr3 | + | 9803842 | 9804013 | 9801660 | 9801756 | 9806209 | 9806417 | 1522 | 858 | [illegible] |
| SE | 32309 | ENSG00000124570 | SERPINB6 | chr6 | - | 2954591 | 2954709 | 2953043 | 2953186 | 2955523 | 2955670 | 574 | 344 | [illegible] |
| SE | 32310 | ENSG00000124570 | SERPINB6 | chr6 | - | 2955523 | 2955670 | 2954591 | 2954709 | 2959167 | 2959342 | 541 | 310 | [illegible] |
| SE | 32311 | ENSG00000124570 | SERPINB6 | chr6 | - | 2956695 | 2956814 | 2955523 | 2955670 | 2959167 | 2959342 | 1 | 0 | [illegible] |
| SE | 32312 | ENSG00000124570 | SERPINB6 | chr6 | - | 2959167 | 2959342 | 2955523 | 2955670 | 2968778 | 2968913 | 212 | 118 | [illegible] |
| SE | 32313 | ENSG00000124570 | SERPINB6 | chr6 | - | 2959167 | 2959342 | 2955523 | 2955670 | 2971116 | 2971206 | 220 | 115 | [illegible] |
| SE | 32317 | ENSG00000124570 | SERPINB6 | chr6 | - | 2960969 | 2961096 | 2959167 | 2959342 | 2971116 | 2971284 | 0 | 0 | [illegible] |
| SE | 32321 | ENSG00000124570 | SERPINB6 | chr6 | - | 2966399 | 2966590 | 2959167 | 2959342 | 2970695 | 2971238 | 5 | 1 | [illegible] |
| SE | 32323 | ENSG00000124570 | SERPINB6 | chr6 | - | 2968778 | 2968913 | 2955523 | 2955670 | 2971116 | 2971276 | 7 | 1 | [illegible] |
| SE | 32326 | ENSG00000124570 | SERPINB6 | chr6 | - | 2968778 | 2969102 | 2959167 | 2959342 | 2970695 | 2970714 | 11 | 7 | [illegible] |
| SE | 32328 | ENSG00000124570 | SERPINB6 | chr6 | - | 2970695 | 2970824 | 2959167 | 2959342 | 2971247 | 2971354 | 0 | 0 | [illegible] |
| SE | 32333 | ENSG00000068745 | IP6K2 | chr3 | - | 48694693 | 48694824 | 48693487 | 48694240 | 48695089 | 48695418 | 0 | 0 | [illegible] |
| SE | 32334 | ENSG00000068745 | IP6K2 | chr3 | - | 48694701 | 48694824 | 48694174 | 48694240 | 48695089 | 48695418 | 0 | 0 | [illegible] |
| SE | 32335 | ENSG00000068745 | IP6K2 | chr3 | - | 48708119 | 48708237 | 48695089 | 48695418 | 48711366 | 48711459 | 5 | 0 | [illegible] |
| SE | 32336 | ENSG00000068745 | IP6K2 | chr3 | - | 48708119 | 48708237 | 48695089 | 48695418 | 48715314 | 48715527 | 3 | 0 | [illegible] |
| SE | 32337 | ENSG00000068745 | IP6K2 | chr3 | - | 48708119 | 48708237 | 48695089 | 48695418 | 48717156 | 48717210 | 4 | 0 | [illegible] |
| SE | 32338 | ENSG00000068745 | IP6K2 | chr3 | - | 48708119 | 48708237 | 48695089 | 48695421 | 48711366 | 48711459 | 2 | 0 | [illegible] |
| SE | 32339 | ENSG00000068745 | IP6K2 | chr3 | - | 48708119 | 48708237 | 48695089 | 48695421 | 48715314 | 48715527 | 0 | 0 | [illegible] |
| SE | 32340 | ENSG00000068745 | IP6K2 | chr3 | - | 48708119 | 48708237 | 48695089 | 48695421 | 48717156 | 48717210 | 1 | 0 | [illegible] |
| SE | 32341 | ENSG00000068745 | IP6K2 | chr3 | - | 48711366 | 48711459 | 48695089 | 48695418 | 48715314 | 48715527 | 0 | 0 | [illegible] |
| SE | 32343 | ENSG00000068745 | IP6K2 | chr3 | - | 48711366 | 48711459 | 48695089 | 48695418 | 48717156 | 48717188 | 0 | 0 | [illegible] |
| SE | 32344 | ENSG00000068745 | IP6K2 | chr3 | - | 48711366 | 48711459 | 48695089 | 48695421 | 48715314 | 48715527 | 0 | 0 | [illegible] |
| SE | 32345 | ENSG00000068745 | IP6K2 | chr3 | - | 48711366 | 48711459 | 48695089 | 48695421 | 48715388 | 48715529 | 0 | 0 | [illegible] |
| SE | 32347 | ENSG00000068745 | IP6K2 | chr3 | - | 48711366 | 48711459 | 48708119 | 48708237 | 48715314 | 48715527 | 1 | 0 | [illegible] |
| SE | 32350 | ENSG00000068745 | IP6K2 | chr3 | - | 48715314 | 48715527 | 48695089 | 48695418 | 48717156 | 48717188 | 9 | 7 | [illegible] |
| SE | 32351 | ENSG00000068745 | IP6K2 | chr3 | - | 48715314 | 48715527 | 48708119 | 48708237 | 48717156 | 48717188 | 4 | 6 | [illegible] |
| SE | 32352 | ENSG00000068745 | IP6K2 | chr3 | - | 48715314 | 48715527 | 48711366 | 48711459 | 48717156 | 48717188 | 4 | 6 | [illegible] |
| SE | 32353 | ENSG00000068745 | IP6K2 | chr3 | - | 48715388 | 48715527 | 48694050 | 48695418 | 48717156 | 48717192 | 5 | 7 | [illegible] |
| SE | 32355 | ENSG00000068745 | IP6K2 | chr3 | - | 48716431 | 48716477 | 48715314 | 48715527 | 48717156 | 48717215 | 1 | 0 | [illegible] |
| SE | 32358 | ENSG00000110844 | PRPF40B | chr12 | + | 49632595 | 49632623 | 49631859 | 49631925 | 49632854 | 49632880 | 4 | 6 | [illegible] |
| SE | 32359 | ENSG00000110844 | PRPF40B | chr12 | + | 49633013 | 49633124 | 49631859 | 49632880 | 49633426 | 49633547 | 7 | 18 | [illegible] |
| SE | 32360 | ENSG00000110844 | PRPF40B | chr12 | + | 49635098 | 49635263 | 49634537 | 49634602 | 49635364 | 49635473 | 0 | 9 | [illegible] |
| SE | 32361 | ENSG00000110844 | PRPF40B | chr12 | + | 49642579 | 49642675 | 49642234 | 49642372 | 49642929 | 49642978 | 6 | 5 | [illegible] |
| SE | 32362 | ENSG00000013441 | CLK1 | chr2 | - | 200854615 | 200854695 | 200853902 | 200853993 | 200855003 | 200855086 | 559 | 420 | [illegible] |
| SE | 32363 | ENSG00000013441 | CLK1 | chr2 | - | 200854615 | 200854695 | 200853902 | 200853993 | 200856681 | 200856811 | 29 | 25 | [illegible] |
| SE | 32364 | ENSG00000013441 | CLK1 | chr2 | - | 200855003 | 200855086 | 200853902 | 200853993 | 200856681 | 200856811 | 233 | 151 | [illegible] |
| SE | 32365 | ENSG00000013441 | CLK1 | chr2 | - | 200855003 | 200855086 | 200854615 | 200854695 | 200856681 | 200856811 | 791 | 566 | [illegible] |
| SE | 32366 | ENSG00000013441 | CLK1 | chr2 | - | 200856681 | 200856811 | 200854615 | 200854695 | 200856890 | 200856985 | 552 | 366 | [illegible] |
| SE | 32367 | ENSG00000013441 | CLK1 | chr2 | - | 200856681 | 200856811 | 200855003 | 200855086 | 200857717 | 200857884 | 452 | 319 | [illegible] |
| SE | 32368 | ENSG00000013441 | CLK1 | chr2 | - | 200856890 | 200856985 | 200856681 | 200856811 | 200857717 | 200857884 | 892 | 592 | [illegible] |
| SE | 32375 | ENSG00000013441 | CLK1 | chr2 | - | 200857972 | 200858089 | 200856890 | 200856985 | 200859679 | 200859746 | 764 | 616 | [illegible] |
| SE | 32376 | ENSG00000013441 | CLK1 | chr2 | - | 200859679 | 200859746 | 200857972 | 200858089 | 200861237 | 200861466 | 561 | 356 | [illegible] |
| SE | 32377 | ENSG00000013441 | CLK1 | chr2 | - | 200859679 | 200859747 | 200857972 | 200858089 | 200861237 | 200861466 | 314 | 213 | [illegible] |
| SE | 32379 | ENSG00000013441 | CLK1 | chr2 | - | 200861237 | 200861466 | 200859679 | 200859746 | 200861701 | 200861862 | 799 | 537 | [illegible] |
| SE | 32380 | ENSG00000013441 | CLK1 | chr2 | - | 200861701 | 200861862 | 200859679 | 200861466 | 200864563 | 200864656 | 700 | 462 | [illegible] |
| SE | 32384 | ENSG00000225697 | SLC26A6 | chr3 | - | 48626217 | 48626354 | 48625722 | 48626000 | 48626630 | 48626685 | 59 | 49 | [illegible] |
| SE | 32385 | ENSG00000225697 | SLC26A6 | chr3 | - | 48628621 | 48628714 | 48628168 | 48628541 | 48629641 | 48629711 | 38 | 31 | [illegible] |
| SE | 32386 | ENSG00000225697 | SLC26A6 | chr3 | - | 48631648 | 48631797 | 48631223 | 48631306 | 48632973 | 48633084 | 0 | 1 | [illegible] |
| SE | 32388 | ENSG00000225697 | SLC26A6 | chr3 | - | 48632244 | 48632337 | 48631879 | 48632044 | 48632973 | 48633084 | 3 | 1 | [illegible] |
| SE | 32389 | ENSG00000225697 | SLC26A6 | chr3 | - | 48632244 | 48632396 | 48631879 | 48632044 | 48632973 | 48633084 | 22 | 3 | [illegible] |
| SE | 32396 | ENSG00000129173 | E2F8 | chr11 | - | 19234743 | 19235058 | 19234359 | 19234521 | 19237313 | 19237470 | 2 | 2 | [illegible] |
| SE | 32401 | ENSG00000169814 | BTD | chr3 | + | 15630001 | 15630120 | 15601744 | 15601894 | 15635423 | 15635688 | 0 | 0 | [illegible] |
| SE | 32403 | ENSG00000169814 | BTD | chr3 | + | 15630001 | 15630120 | 15601750 | 15601770 | 15635423 | 15635688 | 0 | 0 | [illegible] |
| SE | 32406 | ENSG00000169814 | BTD | chr3 | + | 15632590 | 15632866 | 15601751 | 15601770 | 15635423 | 15635429 | 2 | 0 | [illegible] |
| SE | 32407 | ENSG00000169814 | BTD | chr3 | + | 15632590 | 15632866 | 15601763 | 15602310 | 15635423 | 15635429 | 2 | 0 | [illegible] |
| SE | 32409 | ENSG00000159189 | C1QC | chr1 | + | 22644010 | 22644204 | 22643632 | 22643714 | 22647226 | 22647887 | 5 | 5 | [illegible] |
| SE | 32411 | ENSG00000136040 | PLXNC1 | chr12 | + | 94181445 | 94181580 | 94169152 | 94169293 | 94209589 | 94209629 | 178 | 183 | [illegible] |
| SE | 32412 | ENSG00000136040 | PLXNC1 | chr12 | + | 94186372 | 94186473 | 94169152 | 94169293 | 94209589 | 94209629 | 204 | 226 | [illegible] |
| SE | 32413 | ENSG00000136040 | PLXNC1 | chr12 | + | 94186372 | 94186473 | 94181445 | 94181580 | 94209589 | 94209629 | 402 | 458 | [illegible] |
| SE | 32415 | ENSG00000136040 | PLXNC1 | chr12 | + | 94209589 | 94209704 | 94186372 | 94186473 | 94237663 | 94237803 | 274 | 312 | [illegible] |
| SE | 32416 | ENSG00000136040 | PLXNC1 | chr12 | + | 94220015 | 94220163 | 94209589 | 94209704 | 94224227 | 94224315 | 454 | 436 | [illegible] |
| SE | 32417 | ENSG00000136040 | PLXNC1 | chr12 | + | 94224227 | 94224315 | 94220015 | 94220163 | 94226604 | 94226707 | 394 | 388 | [illegible] |
| SE | 32418 | ENSG00000136040 | PLXNC1 | chr12 | + | 94226604 | 94226707 | 94224227 | 94224315 | 94227148 | 94227235 | 430 | 415 | [illegible] |
| SE | 32419 | ENSG00000136040 | PLXNC1 | chr12 | + | 94240484 | 94240664 | 94237663 | 94237803 | 94243937 | 94244025 | 627 | 622 | [illegible] |
| SE | 32422 | ENSG00000136040 | PLXNC1 | chr12 | + | 94297315 | 94297423 | 94294485 | 94294540 | 94300909 | 94301057 | 8 | 8 | [illegible] |
| SE | 32427 | ENSG00000254154 | CRYZL2P-SEC16B | chr1 | - | 177958839 | 177958975 | 177958131 | 177958362 | 177960341 | 177960403 | 0 | 0 | [illegible] |
| SE | 32439 | ENSG00000131370 | SH3BP5 | chr3 | - | 15269712 | 15269877 | 15262157 | 15262289 | 15304102 | 15304231 | 264 | 107 | [illegible] |
| SE | 32440 | ENSG00000131370 | SH3BP5 | chr3 | - | 15271600 | 15271671 | 15269712 | 15269877 | 15273115 | 15273405 | 0 | 0 | [illegible] |
| SE | 32441 | ENSG00000131370 | SH3BP5 | chr3 | - | 15271600 | 15271671 | 15269712 | 15269877 | 15304102 | 15304231 | 1 | 0 | [illegible] |
| SE | 32444 | ENSG00000131370 | SH3BP5 | chr3 | - | 15273115 | 15273405 | 15271600 | 15271671 | 15304102 | 15304231 | 2 | 2 | [illegible] |
| SE | 32445 | ENSG00000131370 | SH3BP5 | chr3 | - | 15304102 | 15304231 | 15273115 | 15273405 | 15330503 | 15330566 | 66 | 46 | [illegible] |
| SE | 32446 | ENSG00000234231 | ANAPC1P4 | chr2 | + | 87705904 | 87705970 | 87704332 | 87704600 | 87707761 | 87707910 | 4 | 0 | [illegible] |
| SE | 32448 | ENSG00000131375 | CAPN7 | chr3 | + | 15210801 | 15210922 | 15206151 | 15206597 | 15212103 | 15212212 | 3 | 1 | [illegible] |
| SE | 32449 | ENSG00000131375 | CAPN7 | chr3 | + | 15212103 | 15212212 | 15206245 | 15206597 | 15217424 | 15217529 | 67 | 35 | [illegible] |

| | | | | | | | | | | | | | | | | | | | | | | | | | | | | | | | | | | | | | | | | | | | | | | |
|---|---|---|---|---|---|---|---|---|---|---|---|---|---|---|---|---|---|---|---|---|---|---|---|---|---|---|---|---|---|---|---|---|---|---|---|---|---|---|---|---|---|---|---|---|---|---|
| SE | 32450 | ENSG00000131375 | CAPN7 | chr3 | + | 15220780 | 15220981 | 15218472 | 15218540 | 15223474 | 15223561 | 269 | 230 | 254 | 222 | 460 | 0 | 0 | 0 | 0 | 2 | 70 | 177 | 242 | 105 | 47 | 0 | 0 | 0 | 0 | 0 | 298 | 149 | 1 | 1 | 1 | 1 | 1 | 1 | 0.991 | 1 | 1 | 1 | 1 | 1 | -0.002 |
| SE | 32451 | ENSG00000131375 | CAPN7 | chr3 | + | 15228973 | 15229059 | 15227838 | 15227965 | 15230441 | 15230535 | 122 | 120 | 125 | 116 | 202 | 2 | 0 | 0 | 0 | 0 | 58 | 69 | 121 | 43 | 20 | 0 | 0 | 0 | 0 | 0 | 235 | 149 | 1 | 1 | 0.975 | 1 | 1 | 1 | 1 | 1 | 1 | 1 | 1 | 1 | -0.005 |
| SE | 32454 | ENSG00000131375 | CAPN7 | chr3 | + | 15240753 | 15240853 | 15240472 | 15240617 | 15245525 | 15245671 | 68 | 78 | 68 | 65 | 119 | 0 | 0 | 0 | 1 | 0 | 0 | 37 | 51 | 29 | 8 | 0 | 0 | 0 | 0 | 0 | 249 | 149 | 1 | 1 | 1 | 1 | 1 | 0.975 | 1 | NA | 1 | 1 | 1 | 1 | -0.005 |
| SE | 32455 | ENSG00000131375 | CAPN7 | chr3 | + | 15240753 | 15240853 | 15240472 | 15240617 | 15246731 | 15246794 | 68 | 78 | 66 | 65 | 119 | 0 | 0 | 0 | 0 | 3 | 0 | 37 | 52 | 29 | 8 | 0 | 2 | 0 | 0 | 0 | 249 | 149 | 1 | 1 | 1 | 1 | 1 | 1 | 0.96 | NA | 0.917 | 1 | 1 | 1 | 0.013 |
| SE | 32457 | ENSG00000131375 | CAPN7 | chr3 | + | 15242177 | 15242253 | 15240472 | 15240617 | 15245525 | 15245671 | 49 | 31 | 51 | 50 | 106 | 0 | 0 | 0 | 1 | 0 | 13 | 27 | 32 | 36 | 16 | 0 | 0 | 0 | 0 | 0 | 225 | 149 | 1 | 1 | 1 | 1 | 1 | 0.971 | 1 | 1 | 1 | 1 | 1 | 1 | -0.006 |
| SE | 32458 | ENSG00000131375 | CAPN7 | chr3 | + | 15242177 | 15242253 | 15240472 | 15240617 | 15246731 | 15246794 | 6 | 9 | 6 | 15 | 22 | 0 | 0 | 0 | 0 | 3 | 0 | 27 | 19 | 0 | 0 | 0 | 2 | 0 | 0 | 0 | 225 | 149 | 1 | 1 | 1 | 1 | 1 | 1 | 0.829 | NA | 0.899 | 1 | NA | NA | 0.016 |
| SE | 32459 | ENSG00000131375 | CAPN7 | chr3 | + | 15242177 | 15242253 | 15240753 | 15240853 | 15245525 | 15245671 | 93 | 84 | 97 | 89 | 223 | 0 | 0 | 2 | 0 | 0 | 13 | 38 | 85 | 45 | 30 | 0 | 0 | 0 | 0 | 0 | 225 | 149 | 1 | 1 | 1 | 1 | 0.97 | 1 | 1 | 1 | 1 | 1 | 1 | 1 | -0.006 |
| SE | 32460 | ENSG00000131375 | CAPN7 | chr3 | + | 15242177 | 15242253 | 15240753 | 15240853 | 15246731 | 15246794 | 23 | 31 | 32 | 38 | 65 | 0 | 0 | 0 | 0 | 0 | 15 | 27 | 72 | 15 | 0 | 0 | 0 | 1 | 0 | 0 | 225 | 149 | 1 | 1 | 1 | 1 | 1 | 1 | 1 | 1 | 1 | 0.979 | 1 | NA | 0.005 |
| SE | 32461 | ENSG00000131375 | CAPN7 | chr3 | + | 15245525 | 15245671 | 15240472 | 15240617 | 15246731 | 15246794 | 162 | 166 | 176 | 145 | 279 | 0 | 0 | 0 | 0 | 3 | 48 | 94 | 86 | 49 | 0 | 0 | 2 | 0 | 0 | 0 | 295 | 149 | 1 | 1 | 1 | 1 | 1 | 1 | 0.979 | 1 | 0.96 | 1 | 1 | NA | 0.006 |
| SE | 32462 | ENSG00000131375 | CAPN7 | chr3 | + | 15245525 | 15245671 | 15240753 | 15240853 | 15246731 | 15246794 | 162 | 166 | 178 | 144 | 279 | 0 | 0 | 0 | 0 | 0 | 48 | 94 | 86 | 49 | 0 | 0 | 0 | 1 | 0 | 0 | 295 | 149 | 1 | 1 | 1 | 1 | 1 | 1 | 1 | 1 | 1 | 0.977 | 1 | NA | 0.006 |
| SE | 32464 | ENSG00000131375 | CAPN7 | chr3 | + | 15246731 | 15246794 | 15242177 | 15242253 | 15247326 | 15247457 | 159 | 134 | 191 | 185 | 155 | 0 | 0 | 2 | 0 | 0 | 81 | 106 | 224 | 105 | 11 | 0 | 0 | 0 | 0 | 0 | 212 | 149 | 1 | 1 | 1 | 1 | 0.985 | 1 | 1 | 1 | 1 | 1 | 1 | 1 | -0.003 |
| SE | 32465 | ENSG00000131375 | CAPN7 | chr3 | + | 15250930 | 15251023 | 15247236 | 15247457 | 15251115 | 15251302 | 137 | 112 | 157 | 110 | 273 | 0 | 1 | 2 | 2 | 1 | 10 | 57 | 88 | 27 | 11 | 0 | 0 | 0 | 0 | 0 | 242 | 149 | 1 | 1 | 1 | 0.986 | 0.98 | 0.971 | 0.994 | 1 | 1 | 1 | 1 | 1 | -0.014 |
| SE | 32466 | ENSG00000131375 | CAPN7 | chr3 | + | 15250974 | 15251023 | 15247326 | 15247457 | 15251115 | 15251246 | 33 | 35 | 42 | 25 | 95 | 0 | 1 | 2 | 2 | 1 | 2 | 11 | 23 | 8 | 0 | 0 | 0 | 0 | 0 | 0 | 198 | 149 | 1 | 1 | 1 | 0.963 | 0.94 | 0.904 | 0.986 | 1 | 1 | 1 | 1 | NA | -0.041 |
| SE | 32467 | ENSG00000203709 | MIR29B2CHG | chr1 | - | 207818961 | 207819146 | 207817238 | 207817414 | 207822203 | 207822702 | 1 | 0 | 3 | 1 | 1 | 1 | 3 | 2 | 0 | 16 | 2 | 4 | 2 | 0 | 8 | 17 | 7 | 0 | 0 | 3 | 298 | 149 | 1 | 1 | 0.333 | 0 | 0.429 | 1 | 0.03 | 0.056 | 0.222 | 1 | NA | 0.571 | -0.104 |
| SE | 32468 | ENSG00000203709 | MIR29B2CHG | chr1 | - | 207818961 | 207819146 | 207817238 | 207817414 | 207867925 | 207868023 | 1 | 1 | 0 | 0 | 5 | 11 | 3 | 17 | 0 | 8 | 2 | 0 | 4 | 0 | 8 | 0 | 6 | 0 | 0 | 0 | 298 | 149 | 1 | 1 | 0.043 | 0.143 | 0 | NA | 0.238 | 1 | 0 | 1 | NA | 1 | -0.644 |
| SE | 32469 | ENSG00000186448 | ZNF197 | chr3 | + | 44630813 | 44630946 | 44629073 | 44629544 | 44631061 | 44631221 | 1 | 6 | 12 | 10 | 0 | 18 | 5 | 3 | 4 | 3 | 0 | 0 | 7 | 5 | 0 | 0 | 3 | 3 | 11 | 11 | 282 | 149 | 1 | 1 | 0.029 | 0.388 | 0.679 | 0.569 | 0 | NA | 0 | 0.552 | 0.194 | 0 | 0.147 |
| SE | 32472 | ENSG00000196378 | ZNF34 | chr8 | - | 144778037 | 144778164 | 144773971 | 144774605 | 144778438 | 144778525 | 2 | 5 | 5 | 11 | 1 | 0 | 0 | 0 | 0 | 0 | 0 | 0 | 8 | 1 | 23 | 4 | 0 | 0 | 2 | 0 | 276 | 149 | 1 | 1 | 1 | 1 | 1 | 1 | 1 | 0 | NA | 1 | 0.213 | 1 | 0.447 |
| SE | 32474 | ENSG00000196345 | ZKSCAN7 | chr3 | + | 44565486 | 44565655 | 44556929 | 44557470 | 44567911 | 44568003 | 15 | 7 | 18 | 13 | 17 | 1 | 0 | 0 | 0 | 0 | 18 | 5 | 8 | 5 | 0 | 0 | 0 | 0 | 0 | 0 | 298 | 149 | 1 | 1 | 0.882 | 1 | 1 | 1 | 1 | 1 | 1 | 1 | 1 | NA | -0.024 |
| SE | 32491 | ENSG00000107140 | TESK1 | chr9 | + | 35606836 | 35606983 | 35606236 | 35606285 | 35607326 | 35607409 | 77 | 40 | 31 | 73 | 68 | 1 | 0 | 0 | 0 | 0 | 143 | 42 | 33 | 40 | 143 | 0 | 0 | 0 | 0 | 0 | 296 | 149 | 1 | 1 | 0.975 | 1 | 1 | 1 | 1 | 1 | 1 | 1 | 1 | 1 | -0.005 |
| SE | 32492 | ENSG00000107140 | TESK1 | chr9 | + | 35606836 | 35606983 | 35606236 | 35606285 | 35607581 | 35607672 | 52 | 16 | 22 | 47 | 36 | 0 | 0 | 0 | 1 | 0 | 76 | 20 | 12 | 27 | 78 | 0 | 0 | 0 | 0 | 0 | 296 | 149 | 1 | 1 | 1 | 1 | 1 | 0.959 | 1 | 1 | 1 | 1 | 1 | 1 | -0.008 |
| SE | 32493 | ENSG00000107140 | TESK1 | chr9 | + | 35607326 | 35607409 | 35606236 | 35606285 | 35607581 | 35607672 | 32 | 14 | 30 | 34 | 48 | 0 | 0 | 0 | 1 | 0 | 9 | 31 | 6 | 29 | 37 | 0 | 0 | 0 | 0 | 0 | 232 | 149 | 1 | 1 | 1 | 1 | 1 | 0.956 | 1 | 1 | 1 | 1 | 1 | 1 | -0.009 |
| SE | 32500 | ENSG00000280571 | NA | chr3 | - | 42786027 | 42786162 | 42758124 | 42758351 | 42794156 | 42794275 | 224 | 178 | 182 | 224 | 120 | 0 | 1 | 0 | 0 | 0 | 147 | 206 | 110 | 113 | 78 | 0 | 0 | 0 | 0 | 0 | 284 | 149 | 1 | 1 | 1 | 0.989 | 1 | 1 | 1 | 1 | 1 | 1 | 1 | 1 | -0.002 |
| SE | 32505 | ENSG00000114857 | NKTR | chr3 | + | 42620015 | 42620043 | 42619663 | 42619708 | 42621428 | 42621516 | 31 | 27 | 19 | 32 | 52 | 17 | 13 | 6 | 13 | 19 | 16 | 13 | 11 | 0 | 38 | 7 | 3 | 5 | 0 | 0 | 177 | 149 | 1 | 1 | 0.606 | 0.636 | 0.727 | 0.674 | 0.697 | 0.658 | 0.785 | 0.649 | NA | 1 | -0.105 |
| SE | 32506 | ENSG00000114857 | NKTR | chr3 | + | 42621428 | 42621516 | 42619663 | 42619708 | 42630545 | 42630575 | 28 | 35 | 24 | 24 | 41 | 0 | 0 | 0 | 0 | 1 | 7 | 5 | 5 | 0 | 0 | 0 | 0 | 0 | 0 | 0 | 237 | 149 | 1 | 1 | 1 | 1 | 1 | 1 | 0.963 | 1 | 1 | 1 | NA | NA | -0.007 |
| SE | 32508 | ENSG00000112486 | CCR6 | chr6 | + | 167123118 | 167123223 | 167111806 | 167112014 | 167136037 | 167136143 | 1 | 1 | 7 | 5 | 1 | 23 | 17 | 37 | 24 | 28 | 0 | 0 | 0 | 12 | 0 | 0 | 8 | 31 | 5 | 0 | 254 | 149 | 1 | 1 | 0.025 | 0.033 | 0.1 | 0.109 | 0.021 | NA | 0 | 0 | 0.585 | NA | -0.137 |
| SE | 32509 | ENSG00000112486 | CCR6 | chr6 | + | 167123138 | 167123223 | 167111806 | 167112014 | 167136037 | 167136143 | 1 | 1 | 5 | 5 | 1 | 23 | 17 | 37 | 24 | 28 | 0 | 0 | 0 | 0 | 0 | 0 | 8 | 31 | 5 | 0 | 234 | 149 | 1 | 1 | 0.027 | 0.036 | 0.079 | 0.117 | 0.022 | NA | 0 | 0 | 0 | NA | 0.056 |
| SE | 32511 | ENSG00000213585 | VDAC1 | chr5 | - | 133992305 | 133992355 | 133991001 | 133991154 | 133992945 | 133993015 | 218 | 147 | 190 | 227 | 200 | 5 | 0 | 0 | 4 | 0 | 466 | 269 | 193 | 394 | 324 | 0 | 0 | 2 | 0 | 0 | 199 | 149 | 1 | 1 | 0.97 | 1 | 1 | 0.977 | 1 | 1 | 1 | 0.986 | 1 | 1 | -0.008 |
| SE | 32513 | ENSG00000080823 | MOK | chr14 | - | 102231706 | 102231821 | 102229213 | 102229657 | 102232534 | 102232671 | 6 | 0 | 0 | 0 | 4 | 0 | 1 | 0 | 0 | 3 | 0 | 0 | 0 | 2 | 0 | 0 | 0 | 0 | 4 | 12 | 264 | 149 | 1 | 1 | 1 | 0 | NA | NA | 0.429 | NA | NA | NA | 0.22 | 0 | 0.366 |
| SE | 32515 | ENSG00000080823 | MOK | chr14 | - | 102232534 | 102232671 | 102229456 | 102229657 | 102233687 | 102233789 | 0 | 2 | 0 | 0 | 3 | 0 | 0 | 0 | 2 | 0 | 0 | 0 | 0 | 6 | 12 | 0 | 0 | 0 | 0 | 0 | 286 | 149 | 1 | 1 | NA | 1 | NA | 0 | 1 | NA | NA | NA | 1 | 1 | -0.333 |
| SE | 32517 | ENSG00000080823 | MOK | chr14 | - | 102232534 | 102232708 | 102229119 | 102229657 | 102233687 | 102233789 | 0 | 1 | 0 | 0 | 3 | 0 | 0 | 0 | 2 | 0 | 0 | 0 | 0 | 4 | 12 | 0 | 0 | 0 | 0 | 0 | 298 | 149 | 1 | 1 | NA | 1 | NA | 0 | 1 | NA | NA | NA | 1 | 1 | -0.333 |
| SE | 32530 | ENSG00000206053 | JPT2 | chr16 | + | 1683508 | 1683592 | 1678276 | 1678356 | 1685438 | 1685587 | 5 | 2 | 2 | 0 | 7 | 11 | 4 | 9 | 14 | 2 | 0 | 1 | 3 | 0 | 0 | 0 | 21 | 4 | 2 | 28 | 233 | 149 | 1 | 1 | 0.225 | 0.242 | 0.124 | 0 | 0.691 | NA | 0.03 | 0.324 | 0 | 0 | 0.168 |
| SE | 32532 | ENSG00000206053 | JPT2 | chr16 | + | 1685438 | 1685587 | 1678275 | 1678356 | 1691842 | 1691985 | 59 | 16 | 48 | 53 | 57 | 0 | 0 | 0 | 0 | 0 | 13 | 45 | 19 | 15 | 96 | 0 | 2 | 0 | 0 | 0 | 298 | 149 | 1 | 1 | 1 | 1 | 1 | 1 | 1 | 1 | 0.918 | 1 | 1 | 1 | 0.016 |
| SE | 32536 | ENSG00000197808 | ZNF461 | chr19 | - | 36656447 | 36656543 | 36638146 | 36640043 | 36658298 | 36658425 | 10 | 2 | 8 | 9 | 18 | 0 | 0 | 0 | 0 | 0 | 0 | 0 | 6 | 2 | 0 | 0 | 0 | 6 | 0 | 0 | 245 | 149 | 1 | 1 | 1 | 1 | 1 | 1 | 1 | NA | NA | 0.378 | 1 | NA | 0.311 |
| SE | 32539 | ENSG00000197808 | ZNF461 | chr19 | - | 36658298 | 36658425 | 36656447 | 36656543 | 36664697 | 36664786 | 35 | 16 | 27 | 33 | 29 | 1 | 0 | 0 | 0 | 0 | 0 | 19 | 28 | 19 | 43 | 0 | 0 | 0 | 0 | 0 | 276 | 149 | 1 | 1 | 0.95 | 1 | 1 | 1 | 1 | NA | 1 | 1 | 1 | 1 | -0.01 |
| SE | 32540 | ENSG00000197808 | ZNF461 | chr19 | - | 36658298 | 36658446 | 36656447 | 36656543 | 36664697 | 36664786 | 18 | 8 | 12 | 17 | 23 | 1 | 0 | 0 | 0 | 0 | 0 | 9 | 14 | 10 | 0 | 0 | 0 | 0 | 0 | 0 | 297 | 149 | 1 | 1 | 0.9 | 1 | 1 | 1 | 1 | NA | 1 | 1 | 1 | NA | -0.02 |
| SE | 32541 | ENSG00000197808 | ZNF461 | chr19 | - | 36658335 | 36658425 | 36656447 | 36656543 | 36664697 | 36664786 | 22 | 12 | 12 | 16 | 9 | 1 | 0 | 0 | 0 | 0 | 0 | 9 | 11 | 11 | 23 | 0 | 0 | 0 | 0 | 0 | 239 | 149 | 1 | 1 | 0.932 | 1 | 1 | 1 | 1 | NA | 1 | 1 | 1 | 1 | -0.014 |
| SE | 32542 | ENSG00000134602 | STK26 | chrX | + | 132023507 | 132023659 | 132023301 | 132023407 | 132054630 | 132054861 | 152 | 84 | 121 | 120 | 42 | 0 | 0 | 4 | 0 | 0 | 19 | 81 | 41 | 57 | 12 | 0 | 0 | 0 | 0 | 0 | 298 | 149 | 1 | 1 | 1 | 1 | 0.938 | 1 | 1 | 1 | 1 | 1 | 1 | 1 | -0.012 |
| SE | 32545 | ENSG00000134602 | STK26 | chrX | + | 132063432 | 132063489 | 132054630 | 132054861 | 132068214 | 132068323 | 339 | 237 | 290 | 405 | 560 | 3 | 0 | 2 | 0 | 0 | 70 | 148 | 105 | 131 | 48 | 0 | 0 | 2 | 0 | 0 | 206 | 149 | 1 | 1 | 0.988 | 1 | 0.991 | 1 | 1 | 1 | 1 | 0.974 | 1 | 1 | 0.001 |
| SE | 32547 | ENSG00000134602 | STK26 | chrX | + | 132072267 | 132072361 | 132071068 | 132071217 | 132072812 | 132072875 | 713 | 358 | 547 | 562 | 987 | 2 | 1 | 2 | 5 | 1 | 118 | 283 | 235 | 149 | 16 | 0 | 0 | 2 | 0 | 0 | 243 | 149 | 1 | 1 | 0.995 | 0.995 | 0.994 | 0.986 | 0.998 | 1 | 1 | 0.986 | 1 | 1 | -0.004 |
| SE | 32550 | ENSG00000211456 | SACM1L | chr3 | + | 45690347 | 45690387 | 45689240 | 45689497 | 45703437 | 45703535 | 15 | 12 | 19 | 8 | 5 | 55 | 24 | 52 | 37 | 16 | 4 | 1 | 18 | 2 | 0 | 7 | 23 | 12 | 6 | 0 | 189 | 149 | 1 | 1 | 0.177 | 0.283 | 0.224 | 0.146 | 0.198 | 0.311 | 0.033 | 0.542 | 0.208 | NA | -0.068 |
| SE | 32552 | ENSG00000211456 | SACM1L | chr3 | + | 45731300 | 45731380 | 45723474 | 45723543 | 45732052 | 45732151 | 415 | 304 | 304 | 382 | 282 | 2 | 2 | 1 | 8 | 0 | 51 | 146 | 153 | 90 | 12 | 0 | 6 | 0 | 0 | 0 | 229 | 149 | 1 | 1 | 0.993 | 0.99 | 0.995 | 0.969 | 1 | 1 | 0.941 | 1 | 1 | 1 | 0.001 |
| SE | 32553 | ENSG00000211456 | SACM1L | chr3 | + | 45732052 | 45732151 | 45723474 | 45723543 | 45735234 | 45735373 | 127 | 85 | 98 | 135 | 131 | 0 | 0 | 0 | 0 | 3 | 16 | 30 | 29 | 21 | 4 | 0 | 0 | 0 | 0 | 0 | 248 | 149 | 1 | 1 | 1 | 1 | 1 | 1 | 0.963 | 1 | 1 | 1 | 1 | 1 | -0.007 |
| SE | 32556 | ENSG00000211456 | SACM1L | chr3 | + | 45739586 | 45739644 | 45738780 | 45738873 | 45743532 | 45743735 | 73 | 54 | 55 | 81 | 60 | 1 | 0 | 3 | 0 | 6 | 0 | 35 | 7 | 20 | 5 | 0 | 1 | 0 | 0 | 0 | 207 | 149 | 1 | 1 | 0.981 | 1 | 0.93 | 1 | 0.878 | NA | 0.962 | 1 | 1 | 1 | -0.033 |
| SE | 32557 | ENSG00000240682 | ISY1 | chr3 | - | 129130549 | 129130636 | 129127414 | 129130188 | 129134073 | 129134195 | 118 | 86 | 129 | 119 | 182 | 0 | 0 | 0 | 2 | 0 | 340 | 118 | 141 | 187 | 237 | 0 | 1 | 0 | 0 | 0 | 236 | 149 | 1 | 1 | 1 | 1 | 1 | 0.974 | 1 | 1 | 0.987 | 1 | 1 | 1 | -0.003 |
| SE | 32559 | ENSG00000240682 | ISY1 | chr3 | - | 129134831 | 129134954 | 129130549 | 129130636 | 129140367 | 129140485 | 138 | 126 | 137 | 162 | 129 | 0 | 0 | 0 | 0 | 2 | 289 | 100 | 193 | 173 | 292 | 0 | 3 | 0 | 0 | 0 | 272 | 149 | 1 | 1 | 1 | 1 | 1 | 1 | 0.972 | 1 | 0.948 | 1 | 1 | 1 | 0.005 |
| SE | 32564 | ENSG00000261796 | ISY1-RAB43 | chr3 | - | 129130087 | 129130188 | 129094985 | 129095169 | 129130549 | 129130636 | 67 | 50 | 87 | 71 | 115 | 0 | 0 | 0 | 0 | 0 | 192 | 62 | 72 | 86 | 85 | 0 | 0 | 4 | 0 | 0 | 250 | 149 | 1 | 1 | 1 | 1 | 1 | 1 | 1 | 1 | 1 | 0.915 | 1 | 1 | 0.017 |
| SE | 32568 | ENSG00000113851 | CRBN | chr3 | - | 3153423 | 3153488 | 3150010 | 3151045 | 3153959 | 3154075 | 187 | 136 | 144 | 192 | 614 | 0 | 1 | 0 | 0 | 0 | 34 | 147 | 85 | 63 | 20 | 0 | 0 | 0 | 0 | 0 | 214 | 149 | 1 | 1 | 1 | 0.99 | 1 | 1 | 1 | 1 | 1 | 1 | 1 | 1 | -0.002 |
| SE | 32572 | ENSG00000113851 | CRBN | chr3 | - | 3156218 | 3156281 | 3154549 | 3154831 | 3167633 | 3167793 | 267 | 232 | 226 | 238 | 637 | 0 | 0 | 0 | 0 | 1 | 30 | 204 | 131 | 58 | 47 | 0 | 0 | 0 | 0 | 0 | 212 | 149 | 1 | 1 | 1 | 1 | 1 | 1 | 0.998 | 1 | 1 | 1 | 1 | 1 | 0 |
| SE | 32573 | ENSG00000113851 | CRBN | chr3 | - | 3167633 | 3167793 | 3153423 | 3156281 | 3172775 | 3172923 | 409 | 339 | 349 | 343 | 687 | 1 | 2 | 5 | 0 | 0 | 116 | 283 | 212 | 124 | 152 | 0 | 3 | 0 | 0 | 0 | 298 | 149 | 1 | 1 | 0.995 | 0.988 | 0.972 | 1 | 1 | 1 | 0.979 | 1 | 1 | 1 | -0.005 |
| SE | 32574 | ENSG00000113851 | CRBN | chr3 | - | 3172775 | 3172925 | 3167429 | 3167793 | 3174058 | 3174121 | 622 | 439 | 459 | 548 | 877 | 0 | 0 | 0 | 0 | 0 | 276 | 418 | 434 | 349 | 379 | 0 | 0 | 1 | 0 | 0 | 298 | 149 | 1 | 1 | 1 | 1 | 1 | 1 | 1 | 1 | 1 | 0.995 | 1 | 1 | 0.001 |
| SE | 32580 | ENSG00000072756 | TRNT1 | chr3 | + | 3127945 | 3128018 | 3126969 | 3126990 | 3129013 | 3129188 | 5 | 9 | 6 | 2 | 9 | 15 | 7 | 11 | 11 | 4 | 0 | 0 | 1 | 0 | 10 | 0 | 0 | 5 | 0 | 3 | 222 | 149 | 1 | 1 | 0.183 | 0.463 | 0.268 | 0.109 | 0.602 | NA | NA | 0.118 | NA | 0.691 | -0.079 |
| SE | 32581 | ENSG00000072756 | TRNT1 | chr3 | + | 3127945 | 3128018 | 3126979 | 3127289 | 3129013 | 3129188 | 5 | 9 | 6 | 2 | 9 | 1 | 0 | 0 | 1 | 0 | 0 | 0 | 1 | 0 | 10 | 0 | 9 | 0 | 0 | 3 | 222 | 149 | 1 | 1 | 0.77 | 1 | 1 | 0.573 | 1 | NA | 0 | 1 | NA | 0.691 | 0.305 |
| SE | 32583 | ENSG00000072756 | TRNT1 | chr3 | + | 3136675 | 3136797 | 3129013 | 3129188 | 3137259 | 3137453 | 2 | 0 | 0 | 0 | 0 | 80 | 93 | 61 | 70 | 56 | 0 | 0 | 0 | 0 | 0 | 31 | 36 | 40 | 14 | 15 | 271 | 149 | 1 | 1 | 0.014 | 0 | 0 | 0 | 0 | 0 | 0 | 0 | 0 | 0 | 0.003 |
| SE | 32585 | ENSG00000072756 | TRNT1 | chr3 | + | 3139588 | 3139710 | 3137259 | 3137453 | 3140509 | 3140728 | 0 | 3 | 0 | 0 | 1 | 58 | 46 | 74 | 80 | 94 | 0 | 0 | 0 | 0 | 0 | 37 | 39 | 33 | 27 | 40 | 271 | 149 | 1 | 1 | 0 | 0.035 | 0 | 0 | 0.006 | 0 | 0 | 0 | 0 | 0 | 0.008 |
| SE | 32586 | ENSG00000072756 | TRNT1 | chr3 | + | 3139588 | 3139710 | 3137259 | 3137453 | 3144583 | 3144710 | 0 | 1 | 0 | 0 | 0 | 7 | 2 | 1 | 0 | 2 | 0 | 0 | 0 | 0 | 0 | 0 | 0 | 3 | 0 | 0 | 271 | 149 | 1 | 1 | 0 | 0.216 | 0 | NA | 0 | NA | NA | 0 | NA | NA | 0.054 |
| SE | 32589 | ENSG00000072756 | TRNT1 | chr3 | + | 3142893 | 3143058 | 3137355 | 3137453 | 3144583 | 3144710 | 0 | 0 | 0 | 0 | 11 | 7 | 2 | 1 | 0 | 2 | 0 | 0 | 0 | 1 | 0 | 0 | 0 | 3 | 0 | 0 | 298 | 149 | 1 | 1 | 0 | 0 | 0 | NA | 0.733 | NA | NA | 0 | 1 | NA | -0.317 |
| SE | 32590 | ENSG00000072756 | TRNT1 | chr3 | + | 3142893 | 3143058 | 3140539 | 3140648 | 3144583 | 3144710 | 0 | 0 | 0 | 0 | 10 | 98 | 74 | 89 | 103 | 122 | 0 | 0 | 0 | 1 | 0 | 61 | 39 | 42 | 27 | 56 | 298 | 149 | 1 | 1 | 0 | 0 | 0 | 0 | 0.039 | 0 | 0 | 0 | 0.018 | 0 | 0.004 |
| SE | 32591 | ENSG00000072756 | TRNT1 | chr3 | + | 3146415 | 3146623 | 3144583 | 3144710 | 3147449 | 3147703 | 30 | 13 | 27 | 21 | 45 | 8 | 10 | 3 | 6 | 19 | 12 | 2 | 0 | 0 | 0 | 0 | 10 | 6 | 7 | 0 | 298 | 149 | 1 | 1 | 0.652 | 0.394 | 0.818 | 0.636 | 0.542 | 1 | 0.091 | 0 | 0 | NA | 0.336 |
| SE | 32592 | ENSG00000072756 | TRNT1 | chr3 | + | 3146429 | 3146563 | 3144583 | 3144710 | 3147449 | 3147703 | 43 | 32 | 27 | 34 | 50 | 8 | 10 | 3 | 6 | 19 | 0 | 9 | 7 | 7 | 8 | 0 | 10 | 6 | 7 | 0 | 283 | 149 | 1 | 1 | 0.739 | 0.628 | 0.826 | 0.749 | 0.581 | NA | 0.322 | 0.381 | 0.345 | 1 | 0.193 |
| SE | 32593 | ENSG00000072756 | TRNT1 | chr3 | + | 3146429 | 3146623 | 3144583 | 3144710 | 3147449 | 3147703 | 68 | 44 | 52 | 49 | 95 | 8 | 10 | 3 | 6 | 19 | 0 | 11 | 7 | 7 | 8 | 0 | 10 | 6 | 7 | 0 | 298 | 149 | 1 | 1 | 0.81 | 0.688 | 0.897 | 0.803 | 0.714 | NA | 0.355 | 0.368 | 0.333 | 1 | 0.268 |
| SE | 32594 | ENSG00000072756 | TRNT1 | chr3 | + | 3149093 | 3149233 | 3147905 | 3148187 | 3150539 | 3150879 | 15 | 12 | 18 | 10 | 37 | 0 | 0 | 0 | 0 | 4 | 0 | 3 | 1 | 3 | 0 | 0 | 0 | 0 | 0 | 0 | 289 | 149 | 1 | 1 | 1 | 1 | 1 | 1 | 0.827 | NA | 1 | 1 | 1 | NA | -0.035 |
| SE | 32599 | ENSG00000091181 | IL5RA | chr3 | - | 3095298 | 3095444 | 3091803 | 3092362 | 3097869 | 3098057 | 29 | 7 | 10 | 6 | 25 | 1 | 3 | 0 | 0 | 0 | 0 | 6 | 7 | 0 | 0 | 0 | 0 | 0 | 0 | 0 | 295 | 149 | 1 | 1 | 0.936 | 0.541 | 1 | 1 | 1 | NA | 1 | 1 | NA | NA | -0.105 |
| SE | 32601 | ENSG00000078618 | NRDC | chr1 | - | 51789567 | 51789657 | 51789190 | 51789433 | 51790532 | 51790649 | 488 | 285 | 372 | 469 | 844 | 0 | 0 | 1 | 0 | 0 | 98 | 452 | 372 | 399 | 98 | 0 | 0 | 0 | 0 | 0 | 239 | 149 | 1 | 1 | 1 | 1 | 0.996 | 1 | 1 | 1 | 1 | 1 | 1 | 1 | -0.001 |
| SE | 32603 | ENSG00000078618 | NRDC | chr1 | - | 51791577 | 51791661 | 51790899 | 51790990 | 51792045 | 51792098 | 402 | 307 | 340 | 397 | 545 | 3 | 0 | 1 | 1 | 2 | 142 | 386 | 355 | 449 | 86 | 0 | 0 | 5 | 3 | 0 | 233 | 149 | 1 | 1 | 0.988 | 1 | 0.995 | 0.996 | 0.994 | 1 | 1 | 0.978 | 0.99 | 1 | 0.001 |
| SE | 32604 | ENSG00000078618 | NRDC | chr1 | - | 51792045 | 51792098 | 51791577 | 51791661 | 51792376 | 51792424 | 294 | 197 | 232 | 274 | 343 | 0 | 0 | 0 | 0 | 6 | 138 | 289 | 262 | 323 | 53 | 0 | 0 | 2 | 7 | 0 | 202 | 149 | 1 | 1 | 1 | 1 | 1 | 1 | 0.977 | 1 | 1 | 0.99 | 0.971 | 1 | 0.003 |
| SE | 32605 | ENSG00000078618 | NRDC | chr1 | - | 51798248 | 51798411 | 51794822 | 51794854 | 51800555 | 51800683 | 211 | 119 | 162 | 217 | 208 | 0 | 0 | 1 | 0 | 6 | 105 | 181 | 156 | 175 | 22 | 0 | 0 | 0 | 0 | 0 | 298 | 149 | 1 | 1 | 1 | 1 | 0.988 | 1 | 0.945 | 1 | 1 | 1 | 1 | 1 | -0.013 |
| SE | 32606 | ENSG00000078618 | NRDC | chr1 | - | 51800555 | 51800683 | 51798248 | 51798411 | 51803813 | 51803964 | 203 | 131 | 187 | 220 | 267 | 1 | 0 | 0 | 0 | 0 | 56 | 150 | 149 | 192 | 31 | 0 | 0 | 0 | 1 | 0 | 277 | 149 | 1 | 1 | 0.991 | 1 | 1 | 1 | 1 | 1 | 1 | 1 | 0.99 | 1 | 0 |
| SE | 32607 | ENSG00000078618 | NRDC | chr1 | - | 51803813 | 51803964 | 51800555 | 51800683 | 51805509 | 51805561 | 309 | 174 | 260 | 327 | 369 | 0 | 0 | 0 | 0 | 6 | 75 | 166 | 207 | 235 | 41 | 0 | 0 | 0 | 0 | 0 | 298 | 149 | 1 | 1 | 1 | 1 | 1 | 1 | 0.969 | 1 | 1 | 1 | 1 | 1 | -0.006 |
| SE | 32609 | ENSG00000078618 | NRDC | chr1 | - | 51811993 | 51812098 | 51810280 | 51810404 | 51814034 | 51814071 | 564 | 429 | 492 | 631 | 943 | 1 | 0 | 0 | 2 | 6 | 297 | 545 | 462 | 392 | 52 | 0 | 0 | 0 | 0 | 0 | 254 | 149 | 1 | 1 | 0.997 | 1 | 1 | 0.995 | 0.989 | 1 | 1 | 1 | 1 | 1 | -0.004 |
| SE | 32610 | ENSG00000078618 | NRDC | chr1 | - | 51819799 | 51819873 | 51818065 | 51818135 | 51821497 | 51821555 | 667 | 514 | 466 | 618 | 1124 | 0 | 0 | 0 | 4 | 0 | 353 | 565 | 596 | 540 | 146 | 0 | 0 | 0 | 5 | 0 | 223 | 149 | 1 | 1 | 1 | 1 | 1 | 0.99 | 1 | 1 | 1 | 1 | 0.986 | 1 | 0.001 |
| SE | 32611 | ENSG00000078618 | NRDC | chr1 | - | 51823663 | 51823786 | 51821497 | 51821555 | 51825286 | 51825382 | 594 | 449 | 425 | 553 | 1119 | 0 | 0 | 0 | 0 | 4 | 226 | 521 | 472 | 420 | 145 | 0 | 0 | 0 | 0 | 0 | 272 | 149 | 1 | 1 | 1 | 1 | 1 | 1 | 0.994 | 1 | 1 | 1 | 1 | 1 | -0.001 |
| SE | 32612 | ENSG00000078618 | NRDC | chr1 | - | 51823663 | 51823786 | 51821497 | 51821555 | 51827795 | 51827869 | 342 | 285 | 267 | 311 | 646 | 0 | 1 | 0 | 0 | 0 | 128 | 334 | 293 | 263 | 88 | 0 | 0 | 0 | 0 | 0 | 272 | 149 | 1 | 1 | 1 | 0.994 | 1 | 1 | 1 | 1 | 1 | 1 | 1 | 1 | -0.001 |
| SE | 32613 | ENSG00000078618 | NRDC | chr1 | - | 51825286 | 51825382 | 51821497 | 51821555 | 51827795 | 51827869 | 78 | 49 | 61 | 68 | 107 | 0 | 1 | 0 | 0 | 0 | 32 | 61 | 55 | 57 | 18 | 0 | 0 | 0 | 0 | 0 | 245 | 149 | 1 | 1 | 1 | 0.968 | 1 | 1 | 1 | 1 | 1 | 1 | 1 | 1 | -0.006 |
| SE | 32615 | ENSG00000078618 | NRDC | chr1 | - | 51827795 | 51827869 | 51825286 | 51825382 | 51834016 | 51834073 | 397 | 251 | 288 | 365 | 604 | 0 | 1 | 0 | 0 | 4 | 233 | 292 | 276 | 295 | 89 | 0 | 0 | 2 | 0 | 0 | 223 | 149 | 1 | 1 | 1 | 0.994 | 1 | 1 | 0.99 | 1 | 1 | 0.989 | 1 | 1 | -0.001 |
| SE | 32616 | ENSG00000078618 | NRDC | chr1 | - | 51834016 | 51834170 | 51825286 | 51825382 | 51836130 | 51836212 | 283 | 225 | 167 | 250 | 279 | 1 | 0 | 0 | 0 | 0 | 211 | 268 | 230 | 303 | 106 | 0 | 0 | 0 | 0 | 0 | 298 | 149 | 1 | 1 | 0.993 | 1 | 1 | 1 | 1 | 1 | 1 | 1 | 1 | 1 | -0.001 |
| SE | 32617 | ENSG00000078618 | NRDC | chr1 | - | 51836130 | 51836212 | 51834016 | 51834170 | 51840225 | 51840514 | 461 | 377 | 259 | 423 | 487 | 0 | 0 | 0 | 0 | 0 | 376 | 431 | 368 | 446 | 183 | 6 | 0 | 0 | 0 | 0 | 231 | 149 | 1 | 1 | 1 | 1 | 1 | 1 | 1 | 0.976 | 1 | 1 | 1 | 1 | 0.005 |
| SE | 32620 | ENSG00000189079 | ARID2 | chr12 | + | 45811417 | 45811551 | 45731216 | 45731314 | 45817669 | 45817888 | 256 | 184 | 222 | 220 | 250 | 0 | 2 | 0 | 0 | 0 | 62 | 164 | 213 | 107 | 38 | 0 | 0 | 1 | 0 | 0 | 283 | 149 | 1 | 1 | 1 | 0.98 | 1 | 1 | 1 | 1 | 1 | 0.991 | 1 | 1 | -0.002 |
| SE | 32621 | ENSG00000189079 | ARID2 | chr12 | + | 45836740 | 45836991 | 45836588 | 45836655 | 45837320 | 45837417 | 362 | 275 | 339 | 336 | 346 | 0 | 5 | 2 | 1 | 0 | 158 | 201 | 350 | 188 | 88 | 0 | 0 | 4 | 0 | 0 | 298 | 149 | 1 | 1 | 1 | 0.965 | 0.988 | 0.994 | 1 | 1 | 1 | 0.978 | 1 | 1 | -0.006 |
| SE | 32624 | ENSG00000189079 | ARID2 | chr12 | + | 45848835 | 45848970 | 45846855 | 45846937 | 45849579 | 45849776 | 333 | 379 | 316 | 403 | 311 | 0 | 0 | 0 | 0 | 0 | 76 | 264 | 314 | 139 | 36 | 0 | 0 | 1 | 0 | 0 | 284 | 149 | 1 | 1 | 1 | 1 | 1 | 1 | 1 | 1 | 1 | 0.994 | 1 | 1 | 0.001 |
| SE | 32625 | ENSG00000189079 | ARID2 | chr12 | + | 45860800 | 45860949 | 45850035 | 45852896 | 45891779 | 45891918 | 154 | 136 | 163 | 177 | 133 | 0 | 0 | 0 | 0 | 0 | 11 | 118 | 140 | 53 | 15 | 0 | 0 | 2 | 0 | 0 | 298 | 149 | 1 | 1 | 1 | 1 | 1 | 1 | 1 | 1 | 1 | 0.972 | 1 | 1 | 0.006 |
| SE | 32626 | ENSG00000189079 | ARID2 | chr12 | + | 45904341 | 45904398 | 45893629 | 45893721 | 45904933 | 45908040 | 0 | 0 | 0 | 0 | 0 | 64 | 36 | 45 | 49 | 107 | 0 | 4 | 4 | 0 | 0 | 0 | 20 | 19 | 9 | 6 | 206 | 149 | 1 | 1 | 0 | 0 | 0 | 0 | 0 | NA | 0.126 | 0.132 | 0 | 0 | -0.065 |
| SE | 32636 | ENSG00000177963 | RIC8A | chr11 | + | 212836 | 212981 | 212596 | 212759 | 213298 | 213418 | 257 | 111 | 190 | 228 | 202 | 0 | 1 | 0 | 0 | 0 | 290 | 259 | 169 | 150 | 361 | 0 | 0 | 0 | 0 | 0 | 294 | 149 | 1 | 1 | 1 | 0.983 | 1 | 1 | 1 | 1 | 1 | 1 | 1 | 1 | -0.003 |
| SE | 32637 | ENSG00000113838 | TBCCD1 | chr3 | - | 186551128 | 186551279 | 186546066 | 186546955 | 186554253 | 186554751 | 79 | 80 | 63 | 88 | 159 | 0 | 0 | 0 | 0 | 0 | 145 | 41 | 34 | 43 | 14 | 0 | 3 | 0 | 0 | 0 | 298 | 149 | 1 | 1 | 1 | 1 | 1 | 1 | 1 | 1 | 0.872 | 1 | 1 | 1 | 0.026 |
| SE | 32638 | ENSG00000113838 | TBCCD1 | chr3 | - | 186556408 | 186556775 | 186551128 | 186551279 | 186558416 | 186558572 | 29 | 16 | 30 | 31 | 33 | 0 | 0 | 0 | 0 | 0 | 10 | 15 | 10 | 14 | 36 | 0 | 0 | 1 | 0 | 0 | 298 | 149 | 1 | 1 | 1 | 1 | 1 | 1 | 1 | 1 | 1 | 0.833 | 1 | 1 | 0.033 |
| SE | 32639 | ENSG00000113838 | TBCCD1 | chr3 | - | 186558416 | 186558572 | 186551128 | 186551279 | 186563993 | 186564372 | 69 | 35 | 55 | 47 | 82 | 3 | 0 | 0 | 0 | 0 | 37 | 28 | 19 | 24 | 28 | 0 | 0 | 0 | 0 | 0 | 298 | 149 | 1 | 1 | 0.92 | 1 | 1 | 1 | 1 | 1 | 1 | 1 | 1 | 1 | -0.016 |
| SE | 32640 | ENSG00000113838 | TBCCD1 | chr3 | - | 186563993 | 186564372 | 186558416 | 186558572 | 186570106 | 186570543 | 69 | 37 | 57 | 47 | 84 | 0 | 0 | 0 | 0 | 0 | 37 | 28 | 18 | 24 | 28 | 0 | 0 | 0 | 0 | 3 | 298 | 149 | 1 | 1 | 1 | 1 | 1 | 1 | 1 | 1 | 1 | 1 | 1 | 0.824 | 0.035 |
| SE | 32642 | ENSG00000196549 | MME | chr3 | + | 155114993 | 155115155 | 155085055 | 155085094 | 155116478 | 155116559 | 13 | 22 | 43 | 39 | 53 | 0 | 0 | 0 | 0 | 0 | 26 | 34 | 192 | 0 | 0 | 0 | 0 | 3 | 0 | 0 | 298 | 149 | 1 | 1 | 1 | 1 | 1 | 1 | 1 | 1 | 1 | 0.97 | NA | NA | 0.01 |
| SE | 32643 | ENSG00000196549 | MME | chr3 | + | 155141990 | 155142127 | 155140190 | 155140292 | 155143442 | 155143571 | 13 | 22 | 33 | 24 | 104 | 0 | 0 | 2 | 0 | 0 | 0 | 6 | 160 | 2 | 0 | 0 | 0 | 2 | 0 | 0 | 286 | 149 | 1 | 1 | 1 | 1 | 0.896 | 1 | 1 | NA | 1 | 0.977 | 1 | NA | -0.013 |
| SE | 32644 | ENSG00000196549 | MME | chr3 | + | 155142236 | 155142330 | 155141990 | 155142127 | 155143442 | 155143571 | 10 | 13 | 20 | 22 | 64 | 2 | 0 | 0 | 2 | 1 | 0 | 3 | 150 | 5 | 0 | 0 | 0 | 2 | 0 | 0 | 243 | 149 | 1 | 1 | 0.754 | 1 | 1 | 0.871 | 0.975 | NA | 1 | 0.979 | 1 | NA | -0.073 |
| SE | 32645 | ENSG00000196549 | MME | chr3 | + | 155147143 | 155147224 | 155144358 | 155144457 | 155148549 | 155148653 | 26 | 30 | 76 | 82 | 176 | 0 | 0 | 0 | 0 | 0 | 0 | 63 | 548 | 18 | 0 | 0 | 0 | 1 | 0 | 0 | 230 | 149 | 1 | 1 | 1 | 1 | 1 | 1 | 1 | NA | 1 | 0.997 | 1 | NA | 0.001 |
| SE | 32646 | ENSG00000196549 | MME | chr3 | + | 155156979 | 155157069 | 155148549 | 155148653 | 155160389 | 155160448 | 0 | 0 | 0 | 0 | 0 | 8 | 5 | 19 | 24 | 54 | 0 | 1 | 0 | 0 | 0 | 0 | 13 | 176 | 2 | 0 | 239 | 149 | 1 | 1 | 0 | 0 | 0 | 0 | 0 | NA | 0.046 | 0 | 0 | NA | -0.015 |
| SE | 32647 | ENSG00000196549 | MME | chr3 | + | 155168491 | 155168625 | 155166901 | 155167021 | 155168731 | 155168797 | 16 | 10 | 28 | 13 | 62 | 0 | 3 | 0 | 2 | 0 | 0 | 1 | 203 | 1 | 0 | 0 | 0 | 4 | 0 | 0 | 283 | 149 | 1 | 1 | 1 | 0.637 | 1 | 0.774 | 1 | NA | 1 | 0.964 | 1 | NA | -0.106 |
| SE | 32649 | ENSG00000150995 | ITPR1 | chr3 | + | 4521023 | 4521094 | 4516475 | 4516583 | 4627762 | 4627878 | 88 | 67 | 106 | 92 | 106 | 0 | 0 | 0 | 0 | 0 | 129 | 161 | 83 | 43 | 60 | 0 | 2 | 0 | 0 | 0 | 220 | 149 | 1 | 1 | 1 | 1 | 1 | 1 | 1 | 1 | 0.982 | 1 | 1 | 1 | 0.004 |
| SE | 32650 | ENSG00000150995 | ITPR1 | chr3 | + | 4639383 | 4639470 | 4627762 | 4627878 | 4642092 | 4642251 | 114 | 82 | 113 | 100 | 102 | 0 | 0 | 0 | 0 | 0 | 137 | 210 | 85 | 97 | 103 | 0 | 2 | 0 | 0 | 0 | 236 | 149 | 1 | 1 | 1 | 1 | 1 | 1 | 1 | 1 | 0.985 | 1 | 1 | 1 | 0.003 |
| SE | 32652 | ENSG00000150995 | ITPR1 | chr3 | + | 4667376 | 4667549 | 4665137 | 4665296 | 4669653 | 4669773 | 176 | 155 | 153 | 195 | 180 | 0 | 0 | 0 | 0 | 0 | 195 | 332 | 143 | 174 | 163 | 0 | 0 | 6 | 0 | 0 | 298 | 149 | 1 | 1 | 1 | 1 | 1 | 1 | 1 | 1 | 1 | 0.923 | 1 | 1 | 0.015 |
| SE | 32653 | ENSG00000150995 | ITPR1 | chr3 | + | 4676613 | 4676801 | 4675067 | 4675221 | 4680552 | 4680691 | 174 | 117 | 134 | 161 | 137 | 0 | 0 | 0 | 0 | 0 | 274 | 235 | 175 | 170 | 89 | 0 | 0 | 0 | 4 | 0 | 298 | 149 | 1 | 1 | 1 | 1 | 1 | 1 | 1 | 1 | 1 | 1 | 0.955 | 1 | 0.009 |
| SE | 32654 | ENSG00000150995 | ITPR1 | chr3 | + | 4676613 | 4676801 | 4675067 | 4675221 | 4681363 | 4681418 | 69 | 56 | 62 | 66 | 65 | 0 | 1 | 0 | 0 | 0 | 117 | 116 | 85 | 70 | 35 | 0 | 0 | 0 | 0 | 0 | 298 | 149 | 1 | 1 | 1 | 0.966 | 1 | 1 | 1 | 1 | 1 | 1 | 1 | 1 | -0.007 |
| SE | 32655 | ENSG00000150995 | ITPR1 | chr3 | + | 4680552 | 4680691 | 4675067 | 4675221 | 4681363 | 4681418 | 77 | 48 | 64 | 59 | 99 | 0 | 1 | 0 | 0 | 0 | 78 | 130 | 54 | 129 | 38 | 0 | 0 | 0 | 0 | 0 | 288 | 149 | 1 | 1 | 1 | 0.961 | 1 | 1 | 1 | 1 | 1 | 1 | 1 | 1 | -0.008 |
| SE | 32657 | ENSG00000150995 | ITPR1 | chr3 | + | 4681363 | 4681418 | 4676613 | 4676801 | 4683385 | 4683551 | 26 | 9 | 20 | 23 | 36 | 0 | 0 | 0 | 0 | 0 | 26 | 34 | 22 | 13 | 26 | 0 | 0 | 1 | 0 | 0 | 204 | 149 | 1 | 1 | 1 | 1 | 1 | 1 | 1 | 1 | 1 | 0.941 | 1 | 1 | 0.012 |
| SE | 32658 | ENSG00000150995 | ITPR1 | chr3 | + | 4688494 | 4688620 | 4685068 | 4685206 | 4691143 | 4691344 | 142 | 69 | 88 | 108 | 123 | 0 | 0 | 0 | 0 | 0 | 106 | 148 | 115 | 99 | 68 | 0 | 6 | 0 | 0 | 0 | 275 | 149 | 1 | 1 | 1 | 1 | 1 | 1 | 1 | 1 | 0.93 | 1 | 1 | 1 | 0.014 |
| SE | 32659 | ENSG00000150995 | ITPR1 | chr3 | + | 4691143 | 4691344 | 4688494 | 4688620 | 4693489 | 4693741 | 161 | 88 | 133 | 161 | 159 | 1 | 0 | 0 | 0 | 5 | 113 | 193 | 141 | 95 | 63 | 0 | 0 | 0 | 0 | 0 | 298 | 149 | 1 | 1 | 0.988 | 1 | 1 | 1 | 0.941 | 1 | 1 | 1 | 1 | 1 | -0.014 |
| SE | 32661 | ENSG00000150995 | ITPR1 | chr3 | + | 4706166 | 4706351 | 4702829 | 4702950 | 4710324 | 4710473 | 173 | 101 | 138 | 156 | 190 | 2 | 0 | 0 | 0 | 0 | 110 | 376 | 191 | 142 | 159 | 0 | 7 | 1 | 0 | 0 | 298 | 149 | 1 | 1 | 0.977 | 1 | 1 | 1 | 1 | 1 | 0.964 | 0.99 | 1 | 1 | 0.005 |
| SE | 32664 | ENSG00000150995 | ITPR1 | chr3 | + | 4725545 | 4725581 | 4711585 | 4711868 | 4727122 | 4727173 | 0 | 0 | 1 | 3 | 2 | 4 | 2 | 1 | 1 | 4 | 9 | 4 | 1 | 0 | 0 | 2 | 0 | 1 | 0 | 0 | 185 | 149 | 1 | 1 | 0 | 0 | 0.446 | 0.707 | 0.287 | 0.784 | 1 | 0.446 | NA | NA | -0.455 |
| SE | 32666 | ENSG00000150995 | ITPR1 | chr3 | + | 4725545 | 4725581 | 4711585 | 4711868 | 4733087 | 4733220 | 0 | 0 | 0 | 3 | 0 | 178 | 95 | 147 | 168 | 153 | 0 | 0 | 0 | 0 | 0 | 113 | 260 | 115 | 186 | 96 | 185 | 149 | 1 | 1 | 0 | 0 | 0 | 0.014 | 0 | 0 | 0 | 0 | 0 | 0 | 0.003 |
| SE | 32671 | ENSG00000150995 | ITPR1 | chr3 | + | 4733087 | 4733220 | 4711600 | 4711868 | 4766529 | 4766710 | 147 | 65 | 100 | 132 | 115 | 0 | 0 | 1 | 0 | 0 | 77 | 167 | 91 | 128 | 60 | 0 | 0 | 5 | 0 | 0 | 282 | 149 | 1 | 1 | 1 | 1 | 0.981 | 1 | 1 | 1 | 1 | 0.906 | 1 | 1 | 0.015 |
| SE | 32674 | ENSG00000150995 | ITPR1 | chr3 | + | 4787946 | 4788139 | 4783815 | 4783920 | 4795064 | 4795187 | 158 | 127 | 147 | 165 | 263 | 0 | 0 | 0 | 0 | 0 | 115 | 168 | 78 | 121 | 28 | 0 | 0 | 3 | 0 | 0 | 298 | 149 | 1 | 1 | 1 | 1 | 1 | 1 | 1 | 1 | 1 | 0.929 | 1 | 1 | 0.014 |
| SE | 32675 | ENSG00000150995 | ITPR1 | chr3 | + | 4787946 | 4788139 | 4783815 | 4783920 | 4800424 | 4800600 | 53 | 53 | 41 | 70 | 99 | 0 | 0 | 0 | 0 | 0 | 46 | 49 | 30 | 71 | 19 | 0 | 0 | 0 | 2 | 0 | 298 | 149 | 1 | 1 | 1 | 1 | 1 | 1 | 1 | 1 | 1 | 1 | 0.947 | 1 | 0.011 |
| SE | 32676 | ENSG00000150995 | ITPR1 | chr3 | + | 4795064 | 4795187 | 4783815 | 4783920 | 4800424 | 4800600 | 89 | 63 | 80 | 83 | 106 | 0 | 0 | 0 | 0 | 0 | 51 | 85 | 33 | 47 | 16 | 0 | 0 | 0 | 2 | 0 | 272 | 149 | 1 | 1 | 1 | 1 | 1 | 1 | 1 | 1 | 1 | 1 | 0.928 | 1 | 0.014 |
| SE | 32680 | ENSG00000150995 | ITPR1 | chr3 | + | 4811264 | 4811460 | 4806102 | 4806267 | 4813141 | 4813234 | 180 | 81 | 124 | 200 | 163 | 0 | 2 | 0 | 1 | 0 | 111 | 198 | 84 | 109 | 34 | 0 | 2 | 0 | 0 | 0 | 298 | 149 | 1 | 1 | 1 | 0.953 | 1 | 0.99 | 1 | 1 | 0.98 | 1 | 1 | 1 | -0.007 |
| SE | 32681 | ENSG00000058056 | USP13 | chr3 | + | 179681877 | 179682003 | 179653031 | 179653393 | 179690240 | 179690301 | 23 | 33 | 26 | 33 | 33 | 0 | 0 | 1 | 0 | 0 | 36 | 19 | 12 | 14 | 8 | 0 | 0 | 1 | 0 | 0 | 275 | 149 | 1 | 1 | 1 | 1 | 0.934 | 1 | 1 | 1 | 1 | 0.867 | 1 | 1 | 0.013 |
| SE | 32685 | ENSG00000058056 | USP13 | chr3 | + | 179690240 | 179690301 | 179653123 | 179653393 | 179701007 | 179701129 | 7 | 11 | 7 | 7 | 18 | 1 | 0 | 0 | 2 | 4 | 0 | 5 | 6 | 2 | 7 | 0 | 0 | 0 | 0 | 0 | 210 | 149 | 1 | 1 | 0.832 | 1 | 1 | 0.713 | 0.761 | NA | 1 | 1 | 1 | 1 | -0.139 |
| SE | 32687 | ENSG00000058056 | USP13 | chr3 | + | 179690240 | 179690301 | 179681877 | 179682003 | 179701007 | 179701129 | 23 | 35 | 24 | 27 | 41 | 0 | 1 | 0 | 0 | 0 | 26 | 16 | 15 | 15 | 11 | 0 | 1 | 2 | 1 | 0 | 210 | 149 | 1 | 1 | 1 | 0.961 | 1 | 1 | 1 | 1 | 0.919 | 0.842 | 0.914 | 1 | 0.057 |
| SE | 32688 | ENSG00000058056 | USP13 | chr3 | + | 179690240 | 179690301 | 179681877 | 179682003 | 179706933 | 179707076 | 9 | 15 | 8 | 8 | 13 | 1 | 0 | 1 | 0 | 0 | 8 | 3 | 6 | 4 | 4 | 0 | 0 | 0 | 0 | 0 | 210 | 149 | 1 | 1 | 0.865 | 1 | 0.85 | 1 | 1 | 1 | 1 | 1 | 1 | 1 | -0.057 |
| SE | 32689 | ENSG00000058056 | USP13 | chr3 | + | 179690243 | 179690301 | 179653123 | 179653393 | 179701007 | 179701129 | 6 | 11 | 6 | 7 | 17 | 1 | 0 | 0 | 2 | 4 | 0 | 5 | 5 | 2 | 7 | 0 | 0 | 0 | 0 | 0 | 207 | 149 | 1 | 1 | 0.812 | 1 | 1 | 0.716 | 0.754 | NA | 1 | 1 | 1 | 1 | -0.144 |
| SE | 32691 | ENSG00000058056 | USP13 | chr3 | + | 179690243 | 179690301 | 179681877 | 179682003 | 179701007 | 179701129 | 6 | 11 | 6 | 7 | 17 | 0 | 1 | 0 | 0 | 0 | 0 | 5 | 5 | 2 | 7 | 0 | 1 | 2 | 1 | 0 | 207 | 149 | 1 | 1 | 1 | 0.888 | 1 | 1 | 1 | NA | 0.783 | 0.643 | 0.59 | 1 | 0.224 |
| SE | 32694 | ENSG00000058056 | USP13 | chr3 | + | 179701007 | 179701129 | 179681877 | 179682003 | 179706933 | 179707076 | 14 | 16 | 17 | 18 | 21 | 1 | 0 | 1 | 0 | 0 | 0 | 20 | 12 | 4 | 7 | 0 | 0 | 0 | 0 | 0 | 271 | 149 | 1 | 1 | 0.885 | 1 | 0.903 | 1 | 1 | NA | 1 | 1 | 1 | 1 | -0.042 |
| SE | 32704 | ENSG00000058056 | USP13 | chr3 | + | 179711668 | 179711779 | 179710713 | 179710839 | 179719939 | 179720034 | 0 | 0 | 0 | 5 | 0 | 0 | 0 | 2 | 3 | 0 | 0 | 0 | 2 | 0 | 0 | 0 | 0 | 0 | 0 | 0 | 260 | 149 | 1 | 1 | NA | NA | 0 | 0.489 | NA | NA | NA | 1 | NA | NA | -0.756 |
| SE | 32707 | ENSG00000058056 | USP13 | chr3 | + | 179719939 | 179720034 | 179708714 | 179708957 | 179730188 | 179730260 | 6 | 1 | 4 | 3 | 3 | 0 | 0 | 0 | 0 | 0 | 20 | 1 | 4 | 5 | 0 | 0 | 0 | 0 | 6 | 0 | 244 | 149 | 1 | 1 | 1 | 1 | 1 | 1 | 1 | 1 | 1 | 1 | 0.337 | NA | 0.166 |
| SE | 32714 | ENSG00000058056 | USP13 | chr3 | + | 179752284 | 179752373 | 179744999 | 179745217 | 179754731 | 179754854 | 19 | 26 | 44 | 29 | 35 | 0 | 4 | 0 | 0 | 0 | 10 | 17 | 27 | 5 | 5 | 0 | 0 | 0 | 0 | 0 | 238 | 149 | 1 | 1 | 1 | 0.803 | 1 | 1 | 1 | 1 | 1 | 1 | 1 | 1 | -0.039 |
| SE | 32719 | ENSG00000058056 | USP13 | chr3 | + | 179757051 | 179757078 | 179754731 | 179754854 | 179761111 | 179761255 | 15 | 13 | 12 | 7 | 7 | 0 | 0 | 2 | 0 | 0 | 12 | 3 | 11 | 3 | 0 | 0 | 0 | 0 | 0 | 0 | 176 | 149 | 1 | 1 | 1 | 1 | 0.836 | 1 | 1 | 1 | 1 | 1 | 1 | NA | -0.033 |
| SE | 32720 | ENSG00000058056 | USP13 | chr3 | + | 179761111 | 179761255 | 179757051 | 179757078 | 179764001 | 179764168 | 28 | 17 | 23 | 16 | 15 | 0 | 0 | 0 | 1 | 0 | 24 | 11 | 21 | 11 | 0 | 0 | 5 | 0 | 0 | 0 | 293 | 149 | 1 | 1 | 1 | 1 | 1 | 0.891 | 1 | 1 | 0.528 | 1 | 1 | NA | 0.096 |
| SE | 32726 | ENSG00000058056 | USP13 | chr3 | + | 179765694 | 179765848 | 179763936 | 179764168 | 179781738 | 179781823 | 27 | 13 | 19 | 26 | 31 | 0 | 0 | 1 | 0 | 0 | 13 | 39 | 12 | 8 | 0 | 0 | 0 | 0 | 0 | 0 | 298 | 149 | 1 | 1 | 1 | 1 | 0.905 | 1 | 1 | 1 | 1 | 1 | 1 | NA | -0.019 |
| SE | 32734 | ENSG00000163349 | HIPK1 | chr1 | + | 113933139 | 113933234 | 113929858 | 113929939 | 113940381 | 113940702 | 1 | 0 | 2 | 2 | 0 | 22 | 14 | 16 | 11 | 0 | 0 | 0 | 0 | 0 | 0 | 16 | 14 | 10 | 9 | 10 | 244 | 149 | 1 | 1 | 0.027 | 0 | 0.071 | 0.1 | NA | 0 | 0 | 0 | 0 | 0 | 0.05 |
| SE | 32735 | ENSG00000163349 | HIPK1 | chr1 | + | 113933139 | 113933234 | 113930504 | 113930624 | 113940381 | 113940702 | 1 | 0 | 1 | 2 | 0 | 0 | 0 | 0 | 0 | 0 | 0 | 0 | 0 | 0 | 0 | 0 | 4 | 0 | 0 | 0 | 244 | 149 | 1 | 1 | 1 | NA | 1 | 1 | NA | NA | 0 | NA | NA | NA | 1 |
| SE | 32736 | ENSG00000163349 | HIPK1 | chr1 | + | 113940381 | 113941459 | 113929709 | 113929939 | 113952765 | 113952889 | 301 | 141 | 213 | 231 | 199 | 0 | 0 | 1 | 0 | 0 | 297 | 251 | 207 | 180 | 83 | 0 | 0 | 0 | 0 | 0 | 298 | 149 | 1 | 1 | 1 | 1 | 0.991 | 1 | 1 | 1 | 1 | 1 | 1 | 1 | -0.002 |
| SE | 32742 | ENSG00000163349 | HIPK1 | chr1 | + | 113963386 | 113963521 | 113962316 | 113962438 | 113966129 | 113966272 | 467 | 266 | 334 | 467 | 324 | 0 | 1 | 2 | 0 | 0 | 365 | 379 | 423 | 434 | 155 | 0 | 0 | 0 | 0 | 0 | 284 | 149 | 1 | 1 | 1 | 0.993 | 0.989 | 1 | 1 | 1 | 1 | 1 | 1 | 1 | -0.004 |
| SE | 32749 | ENSG00000234492 | RPL34-DT | chr4 | - | 108598537 | 108598695 | 108590362 | 108590480 | 108602712 | 108602845 | 1 | 1 | 0 | 6 | 7 | 0 | 0 | 0 | 0 | 0 | 0 | 0 | 4 | 2 | 0 | 0 | 0 | 2 | 0 | 0 | 298 | 149 | 1 | 1 | 1 | 1 | NA | 1 | 1 | NA | NA | 0.5 | 1 | NA | 0.25 |
| SE | 32750 | ENSG00000234492 | RPL34-DT | chr4 | - | 108602712 | 108602845 | 108598537 | 108598695 | 108608522 | 108608650 | 1 | 1 | 0 | 3 | 2 | 0 | 0 | 2 | 0 | 0 | 2 | 0 | 4 | 2 | 0 | 0 | 0 | 0 | 0 | 0 | 282 | 149 | 1 | 1 | 1 | 1 | 0 | 1 | 1 | 1 | NA | 1 | 1 | NA | -0.2 |
| SE | 32759 | ENSG00000168291 | PDHB | chr3 | - | 58433630 | 58433684 | 58431730 | 58431984 | 58433767 | 58433818 | 99 | 90 | 65 | 82 | 27 | 0 | 0 | 0 | 2 | 0 | 141 | 50 | 54 | 60 | 48 | 0 | 0 | 0 | 0 | 0 | 203 | 149 | 1 | 1 | 1 | 1 | 1 | 0.968 | 1 | 1 | 1 | 1 | 1 | 1 | -0.006 |
| SE | 32761 | ENSG00000168297 | PXK | chr3 | + | 58365873 | 58365924 | 58332879 | 58333090 | 58369430 | 58369478 | 47 | 34 | 22 | 50 | 73 | 0 | 0 | 0 | 2 | 0 | 38 | 47 | 29 | 58 | 38 | 0 | 0 | 0 | 0 | 0 | 200 | 149 | 1 | 1 | 1 | 1 | 1 | 0.949 | 1 | 1 | 1 | 1 | 1 | 1 | -0.01 |
| SE | 32762 | ENSG00000168297 | PXK | chr3 | + | 58365873 | 58365924 | 58332879 | 58333090 | 58382513 | 58382678 | 0 | 2 | 5 | 8 | 4 | 5 | 0 | 1 | 2 | 0 | 0 | 4 | 3 | 12 | 6 | 0 | 0 | 0 | 0 | 0 | 200 | 149 | 1 | 1 | 0 | 1 | 0.788 | 0.749 | 1 | NA | 1 | 1 | 1 | 1 | -0.293 |
| SE | 32764 | ENSG00000168297 | PXK | chr3 | + | 58369430 | 58369478 | 58365873 | 58365924 | 58382513 | 58382678 | 60 | 38 | 27 | 55 | 96 | 0 | 1 | 0 | 0 | 3 | 67 | 62 | 44 | 51 | 30 | 0 | 0 | 0 | 0 | 0 | 197 | 149 | 1 | 1 | 1 | 0.966 | 1 | 1 | 0.96 | 1 | 1 | 1 | 1 | 1 | -0.015 |
| SE | 32766 | ENSG00000168297 | PXK | chr3 | + | 58382513 | 58382700 | 58369430 | 58369478 | 58394997 | 58395102 | 61 | 43 | 28 | 53 | 109 | 0 | 0 | 0 | 0 | 0 | 72 | 53 | 46 | 50 | 24 | 0 | 0 | 0 | 3 | 0 | 298 | 149 | 1 | 1 | 1 | 1 | 1 | 1 | 1 | 1 | 1 | 1 | 0.893 | 1 | 0.021 |
| SE | 32767 | ENSG00000168297 | PXK | chr3 | + | 58390581 | 58390659 | 58382513 | 58382700 | 58391146 | 58391220 | 156 | 112 | 94 | 148 | 274 | 3 | 1 | 0 | 0 | 0 | 280 | 188 | 139 | 170 | 161 | 0 | 0 | 1 | 0 | 0 | 227 | 149 | 1 | 1 | 0.972 | 0.987 | 1 | 1 | 1 | 1 | 1 | 0.989 | 1 | 1 | -0.006 |
| SE | 32769 | ENSG00000168297 | PXK | chr3 | + | 58391146 | 58391220 | 58382513 | 58382700 | 58394997 | 58395102 | 3 | 0 | 0 | 0 | 0 | 3 | 4 | 1 | 3 | 11 | 0 | 0 | 1 | 0 | 0 | 0 | 0 | 10 | 4 | 0 | 223 | 149 | 1 | 1 | 0.401 | 0 | 0 | 0 | 0 | NA | NA | 0.063 | 0 | NA | 0.049 |
| SE | 32770 | ENSG00000168297 | PXK | chr3 | + | 58391146 | 58391220 | 58390581 | 58390659 | 58391772 | 58391847 | 172 | 134 | 124 | 164 | 294 | 0 | 0 | 0 | 0 | 0 | 290 | 209 | 130 | 205 | 166 | 0 | 0 | 0 | 5 | 0 | 223 | 149 | 1 | 1 | 1 | 1 | 1 | 1 | 1 | 1 | 1 | 1 | 0.965 | 1 | 0.007 |
| SE | 32773 | ENSG00000168297 | PXK | chr3 | + | 58391772 | 58391847 | 58391146 | 58391220 | 58394997 | 58395102 | 206 | 176 | 133 | 163 | 304 | 1 | 0 | 0 | 0 | 0 | 316 | 247 | 147 | 184 | 220 | 0 | 0 | 0 | 0 | 0 | 224 | 149 | 1 | 1 | 0.993 | 1 | 1 | 1 | 1 | 1 | 1 | 1 | 1 | 1 | -0.001 |
| SE | 32774 | ENSG00000168297 | PXK | chr3 | + | 58397604 | 58397722 | 58397038 | 58397200 | 58399298 | 58399377 | 218 | 149 | 167 | 212 | 314 | 0 | 0 | 0 | 0 | 0 | 188 | 185 | 257 | 244 | 268 | 0 | 0 | 4 | 0 | 0 | 267 | 149 | 1 | 1 | 1 | 1 | 1 | 1 | 1 | 1 | 1 | 0.973 | 1 | 1 | 0.005 |
| SE | 32775 | ENSG00000168297 | PXK | chr3 | + | 58399298 | 58399377 | 58397604 | 58397722 | 58403861 | 58403910 | 153 | 127 | 133 | 171 | 279 | 0 | 0 | 0 | 2 | 0 | 198 | 130 | 203 | 186 | 178 | 0 | 2 | 0 | 0 | 0 | 228 | 149 | 1 | 1 | 1 | 1 | 1 | 0.982 | 1 | 1 | 0.977 | 1 | 1 | 1 | 0.001 |
| SE | 32777 | ENSG00000168297 | PXK | chr3 | + | 58412900 | 58412963 | 58410089 | 58410159 | 58420535 | 58420568 | 33 | 36 | 34 | 42 | 83 | 0 | 2 | 0 | 0 | 0 | 76 | 54 | 50 | 47 | 24 | 0 | 0 | 0 | 0 | 0 | 212 | 149 | 1 | 1 | 1 | 0.927 | 1 | 1 | 1 | 1 | 1 | 1 | 1 | 1 | -0.015 |
| SE | 32779 | ENSG00000168297 | PXK | chr3 | + | 58412903 | 58412963 | 58410089 | 58410159 | 58420535 | 58420568 | 24 | 15 | 19 | 14 | 22 | 0 | 2 | 0 | 0 | 0 | 13 | 40 | 20 | 38 | 16 | 0 | 0 | 0 | 0 | 0 | 209 | 149 | 1 | 1 | 1 | 0.842 | 1 | 1 | 1 | 1 | 1 | 1 | 1 | 1 | -0.032 |
| SE | 32781 | ENSG00000168297 | PXK | chr3 | + | 58420535 | 58420568 | 58412900 | 58412963 | 58423460 | 58423494 | 2 | 5 | 5 | 2 | 7 | 0 | 0 | 0 | 0 | 1 | 0 | 4 | 10 | 10 | 6 | 0 | 3 | 0 | 0 | 0 | 182 | 149 | 1 | 1 | 1 | 1 | 1 | 1 | 0.851 | NA | 0.522 | 1 | 1 | 1 | 0.09 |
| SE | 32784 | ENSG00000168297 | PXK | chr3 | + | 58423460 | 58423494 | 58420535 | 58420568 | 58424751 | 58426019 | 0 | 2 | 0 | 0 | 0 | 2 | 6 | 3 | 5 | 12 | 0 | 8 | 10 | 6 | 0 | 0 | 3 | 5 | 17 | 0 | 183 | 149 | 1 | 1 | 0 | 0.213 | 0 | 0 | 0 | NA | 0.685 | 0.62 | 0.223 | NA | -0.467 |
| SE | 32787 | ENSG00000163686 | ABHD6 | chr3 | + | 58269320 | 58269434 | 58267188 | 58267345 | 58270931 | 58271061 | 45 | 31 | 26 | 38 | 20 | 1 | 0 | 0 | 1 | 0 | 0 | 16 | 14 | 6 | 10 | 0 | 0 | 0 | 0 | 0 | 263 | 149 | 1 | 1 | 0.962 | 1 | 1 | 0.956 | 1 | NA | 1 | 1 | 1 | 1 | -0.016 |
| SE | 32788 | ENSG00000157601 | MX1 | chr21 | + | 41421837 | 41421871 | 41420278 | 41420440 | 41421974 | 41422095 | 0 | 0 | 0 | 0 | 0 | 1 | 0 | 0 | 0 | 0 | 0 | 0 | 0 | 1 | 0 | 0 | 0 | 0 | 0 | 0 | 183 | 149 | 1 | 1 | 0 | NA | NA | NA | NA | NA | NA | NA | 1 | NA | -1 |
| SE | 32790 | ENSG00000157601 | MX1 | chr21 | + | 41427205 | 41427304 | 41425821 | 41426161 | 41427754 | 41427866 | 23 | 37 | 23 | 58 | 7 | 0 | 0 | 1 | 0 | 0 | 107 | 105 | 22 | 137 | 31 | 0 | 0 | 0 | 0 | 0 | 248 | 149 | 1 | 1 | 1 | 1 | 0.933 | 1 | 1 | 1 | 1 | 1 | 1 | 1 | -0.013 |
| SE | 32793 | ENSG00000157601 | MX1 | chr21 | + | 41429456 | 41429559 | 41427754 | 41427866 | 41430532 | 41430608 | 2 | 0 | 0 | 2 | 5 | 72 | 103 | 78 | 179 | 79 | 0 | 1 | 0 | 0 | 0 | 429 | 333 | 80 | 394 | 80 | 252 | 149 | 1 | 1 | 0.016 | 0 | 0 | 0.007 | 0.036 | 0 | 0.002 | 0 | 0 | 0 | 0.011 |
| SE | 32794 | ENSG00000157601 | MX1 | chr21 | + | 41430532 | 41430608 | 41427754 | 41427866 | 41432030 | 41432175 | 39 | 56 | 57 | 106 | 35 | 1 | 0 | 0 | 0 | 0 | 276 | 197 | 49 | 259 | 35 | 0 | 0 | 1 | 1 | 0 | 225 | 149 | 1 | 1 | 0.963 | 1 | 1 | 1 | 1 | 1 | 1 | 0.97 | 0.994 | 1 | 0 |
| SE | 32796 | ENSG00000157601 | MX1 | chr21 | + | 41437014 | 41437152 | 41435836 | 41436029 | 41439693 | 41439848 | 341 | 505 | 353 | 768 | 548 | 0 | 0 | 0 | 3 | 0 | 3402 | 1325 | 655 | 2582 | 696 | 1 | 0 | 11 | 8 | 0 | 287 | 149 | 1 | 1 | 1 | 1 | 1 | 0.993 | 1 | 0.999 | 1 | 0.969 | 0.994 | 1 | 0.006 |
| SE | 32797 | ENSG00000157601 | MX1 | chr21 | + | 41437014 | 41437152 | 41435836 | 41436029 | 41440886 | 41441025 | 136 | 211 | 129 | 347 | 196 | 0 | 0 | 0 | 0 | 0 | 1766 | 607 | 290 | 1307 | 365 | 0 | 0 | 0 | 4 | 0 | 287 | 149 | 1 | 1 | 1 | 1 | 1 | 1 | 1 | 1 | 1 | 1 | 0.994 | 1 | 0.001 |
| SE | 32798 | ENSG00000157601 | MX1 | chr21 | + | 41439693 | 41439848 | 41435836 | 41436029 | 41440886 | 41441025 | 294 | 414 | 286 | 583 | 408 | 0 | 0 | 0 | 0 | 0 | 2313 | 1191 | 451 | 1859 | 547 | 0 | 0 | 0 | 4 | 0 | 298 | 149 | 1 | 1 | 1 | 1 | 1 | 1 | 1 | 1 | 1 | 1 | 0.996 | 1 | 0.001 |
| SE | 32800 | ENSG00000157601 | MX1 | chr21 | + | 41441715 | 41441914 | 41440886 | 41441025 | 41445447 | 41445570 | 252 | 308 | 240 | 486 | 420 | 0 | 0 | 0 | 0 | 0 | 2000 | 912 | 375 | 1654 | 476 | 0 | 0 | 0 | 1 | 0 | 298 | 149 | 1 | 1 | 1 | 1 | 1 | 1 | 1 | 1 | 1 | 1 | 0.999 | 1 | 0 |
| SE | 32801 | ENSG00000157601 | MX1 | chr21 | + | 41441715 | 41443866 | 41440886 | 41441025 | 41445447 | 41445570 | 457 | 674 | 442 | 855 | 662 | 0 | 0 | 0 | 0 | 0 | 3672 | 1582 | 647 | 2851 | 750 | 0 | 0 | 0 | 1 | 0 | 298 | 149 | 1 | 1 | 1 | 1 | 1 | 1 | 1 | 1 | 1 | 1 | 0.999 | 1 | 0 |
| SE | 32804 | ENSG00000157601 | MX1 | chr21 | + | 41449136 | 41449295 | 41445999 | 41446141 | 41451190 | 41451243 | 156 | 312 | 146 | 359 | 363 | 0 | 0 | 0 | 2 | 0 | 1343 | 526 | 173 | 996 | 247 | 0 | 0 | 0 | 0 | 0 | 298 | 149 | 1 | 1 | 1 | 1 | 1 | 0.989 | 1 | 1 | 1 | 1 | 1 | 1 | -0.002 |
| SE | 32807 | ENSG00000157601 | MX1 | chr21 | + | 41451166 | 41451243 | 41449136 | 41449295 | 41452620 | 41452869 | 370 | 545 | 317 | 767 | 690 | 0 | 0 | 0 | 0 | 4 | 3725 | 1450 | 559 | 2427 | 544 | 18 | 0 | 0 | 2 | 0 | 226 | 149 | 1 | 1 | 1 | 1 | 1 | 1 | 0.991 | 0.993 | 1 | 1 | 0.999 | 1 | 0 |
| SE | 32809 | ENSG00000157601 | MX1 | chr21 | + | 41451190 | 41451243 | 41449136 | 41449295 | 41452620 | 41452869 | 106 | 147 | 79 | 232 | 160 | 0 | 0 | 0 | 0 | 4 | 1520 | 523 | 230 | 886 | 203 | 18 | 0 | 0 | 2 | 0 | 202 | 149 | 1 | 1 | 1 | 1 | 1 | 1 | 0.967 | 0.984 | 1 | 1 | 0.997 | 1 | -0.003 |
| SE | 32810 | ENSG00000157601 | MX1 | chr21 | + | 41452620 | 41452869 | 41445999 | 41446141 | 41458527 | 41458758 | 182 | 227 | 149 | 329 | 367 | 1 | 1 | 0 | 0 | 0 | 1080 | 544 | 178 | 905 | 193 | 10 | 0 | 0 | 0 | 0 | 298 | 149 | 1 | 1 | 0.989 | 0.991 | 1 | 1 | 1 | 0.982 | 1 | 1 | 1 | 1 | 0 |
| SE | 32811 | ENSG00000256427 | NA | chr12 | + | 9241869 | 9241962 | 9241343 | 9241459 | 9246468 | 9246548 | 1 | 0 | 3 | 1 | 2 | 1 | 0 | 4 | 1 | 2 | 0 | 0 | 0 | 0 | 2 | 0 | 0 | 2 | 0 | 0 | 242 | 149 | 1 | 1 | 0.381 | NA | 0.316 | 0.381 | 0.381 | NA | NA | 0 | NA | 1 | -0.135 |
| SE | 32812 | ENSG00000256427 | NA | chr12 | + | 9247580 | 9247677 | 9246468 | 9246548 | 9255558 | 9255692 | 9 | 10 | 18 | 14 | 34 | 0 | 0 | 0 | 0 | 0 | 0 | 17 | 6 | 0 | 0 | 0 | 0 | 3 | 0 | 0 | 246 | 149 | 1 | 1 | 1 | 1 | 1 | 1 | 1 | NA | 1 | 0.548 | NA | NA | 0.226 |
| SE | 32813 | ENSG00000233006 | MIR3936HG | chr5 | - | 132352488 | 132352631 | 132349814 | 132349957 | 132360079 | 132360136 | 0 | 0 | 0 | 2 | 0 | 0 | 7 | 3 | 1 | 0 | 0 | 0 | 0 | 0 | 0 | 5 | 2 | 4 | 0 | 0 | 292 | 149 | 1 | 1 | NA | 0 | 0 | 0.505 | NA | 0 | 0 | 0 | NA | NA | 0.168 |
| SE | 32816 | ENSG00000088387 | DOCK9 | chr13 | - | 98804998 | 98805209 | 98800287 | 98800478 | 98807660 | 98807807 | 140 | 102 | 226 | 212 | 456 | 0 | 0 | 1 | 0 | 0 | 16 | 36 | 85 | 42 | 0 | 0 | 0 | 2 | 0 | 0 | 298 | 149 | 1 | 1 | 1 | 1 | 0.991 | 1 | 1 | 1 | 1 | 0.955 | 1 | NA | 0.009 |
| SE | 32817 | ENSG00000088387 | DOCK9 | chr13 | - | 98808639 | 98808675 | 98807660 | 98807807 | 98809351 | 98809465 | 0 | 0 | 2 | 4 | 0 | 33 | 27 | 75 | 52 | 72 | 0 | 0 | 0 | 0 | 0 | 15 | 18 | 25 | 19 | 0 | 185 | 149 | 1 | 1 | 0 | 0 | 0.021 | 0.058 | 0 | 0 | 0 | 0 | 0 | NA | 0.016 |
| SE | 32819 | ENSG00000088387 | DOCK9 | chr13 | - | 98809351 | 98809465 | 98807660 | 98807807 | 98810168 | 98810291 | 54 | 54 | 135 | 91 | 133 | 0 | 0 | 1 | 0 | 0 | 15 | 33 | 41 | 26 | 4 | 0 | 0 | 0 | 0 | 0 | 263 | 149 | 1 | 1 | 1 | 1 | 0.987 | 1 | 1 | 1 | 1 | 1 | 1 | 1 | -0.003 |
| SE | 32822 | ENSG00000088387 | DOCK9 | chr13 | - | 98829306 | 98829522 | 98826829 | 98826887 | 98829642 | 98829756 | 129 | 103 | 169 | 137 | 108 | 0 | 0 | 0 | 1 | 0 | 44 | 38 | 98 | 64 | 19 | 0 | 0 | 0 | 2 | 0 | 298 | 149 | 1 | 1 | 1 | 1 | 1 | 0.986 | 1 | 1 | 1 | 1 | 0.941 | 1 | 0.009 |
| SE | 32823 | ENSG00000088387 | DOCK9 | chr13 | - | 98845328 | 98845372 | 98837493 | 98837609 | 98845923 | 98846060 | 2 | 1 | 10 | 3 | 2 | 37 | 21 | 40 | 42 | 37 | 0 | 1 | 6 | 5 | 0 | 0 | 15 | 27 | 0 | 0 | 193 | 149 | 1 | 1 | 0.04 | 0.035 | 0.162 | 0.052 | 0.04 | NA | 0.049 | 0.146 | 1 | NA | -0.333 |
| SE | 32824 | ENSG00000088387 | DOCK9 | chr13 | - | 98845328 | 98845372 | 98837493 | 98837609 | 98848591 | 98848639 | 2 | 2 | 13 | 0 | 4 | 0 | 1 | 0 | 0 | 0 | 0 | 0 | 0 | 1 | 0 | 11 | 0 | 4 | 3 | 0 | 193 | 149 | 1 | 1 | 1 | 0.607 | 1 | NA | 1 | 0 | NA | 0 | 0.205 | NA | 0.833 |
| SE | 32825 | ENSG00000088387 | DOCK9 | chr13 | - | 98845923 | 98845991 | 98837493 | 98837609 | 98848591 | 98848639 | 18 | 12 | 27 | 32 | 15 | 0 | 1 | 0 | 0 | 0 | 3 | 6 | 19 | 0 | 0 | 11 | 0 | 4 | 3 | 0 | 217 | 149 | 1 | 1 | 1 | 0.892 | 1 | 1 | 1 | 0.158 | 1 | 0.765 | 0 | NA | 0.498 |
| SE | 32827 | ENSG00000088387 | DOCK9 | chr13 | - | 98845923 | 98846060 | 98845328 | 98845372 | 98848591 | 98848639 | 58 | 26 | 48 | 49 | 87 | 1 | 1 | 5 | 0 | 4 | 0 | 17 | 22 | 7 | 4 | 0 | 0 | 0 | 1 | 0 | 286 | 149 | 1 | 1 | 0.968 | 0.931 | 0.833 | 1 | 0.919 | NA | 1 | 1 | 0.785 | 1 | -0.016 |
| SE | 32829 | ENSG00000088387 | DOCK9 | chr13 | - | 98846519 | 98846556 | 98845947 | 98846060 | 98848591 | 98848639 | 0 | 0 | 1 | 0 | 0 | 61 | 26 | 49 | 52 | 86 | 0 | 0 | 0 | 0 | 0 | 0 | 16 | 16 | 2 | 4 | 186 | 149 | 1 | 1 | 0 | 0 | 0.016 | 0 | 0 | NA | 0 | 0 | 0 | 0 | 0.003 |
| SE | 32830 | ENSG00000088387 | DOCK9 | chr13 | - | 98850046 | 98850113 | 98848591 | 98848639 | 98853407 | 98853522 | 173 | 98 | 199 | 188 | 206 | 0 | 0 | 0 | 0 | 0 | 16 | 50 | 70 | 17 | 8 | 0 | 0 | 3 | 0 | 0 | 216 | 149 | 1 | 1 | 1 | 1 | 1 | 1 | 1 | 1 | 1 | 0.942 | 1 | 1 | 0.012 |
| SE | 32832 | ENSG00000088387 | DOCK9 | chr13 | - | 98860404 | 98860522 | 98855897 | 98856031 | 98863018 | 98863132 | 56 | 42 | 92 | 89 | 84 | 1 | 0 | 0 | 0 | 0 | 0 | 26 | 46 | 18 | 7 | 0 | 0 | 0 | 0 | 0 | 267 | 149 | 1 | 1 | 0.969 | 1 | 1 | 1 | 1 | NA | 1 | 1 | 1 | 1 | -0.006 |
| SE | 32833 | ENSG00000088387 | DOCK9 | chr13 | - | 98863369 | 98863548 | 98863018 | 98863132 | 98867424 | 98867536 | 80 | 54 | 128 | 90 | 134 | 0 | 0 | 0 | 0 | 0 | 14 | 8 | 62 | 26 | 10 | 0 | 0 | 2 | 0 | 0 | 298 | 149 | 1 | 1 | 1 | 1 | 1 | 1 | 1 | 1 | 1 | 0.939 | 1 | 1 | 0.012 |
| SE | 32834 | ENSG00000088387 | DOCK9 | chr13 | - | 98879897 | 98879969 | 98868230 | 98868377 | 98880546 | 98880672 | 75 | 61 | 120 | 101 | 144 | 0 | 2 | 0 | 0 | 1 | 10 | 43 | 47 | 23 | 0 | 0 | 0 | 0 | 0 | 0 | 221 | 149 | 1 | 1 | 1 | 0.954 | 1 | 1 | 0.99 | 1 | 1 | 1 | 1 | NA | -0.011 |
| SE | 32835 | ENSG00000088387 | DOCK9 | chr13 | - | 98884970 | 98885092 | 98883812 | 98883899 | 98885707 | 98885831 | 117 | 132 | 168 | 162 | 113 | 0 | 0 | 0 | 3 | 0 | 30 | 33 | 60 | 36 | 8 | 0 | 0 | 0 | 0 | 0 | 271 | 149 | 1 | 1 | 1 | 1 | 1 | 0.967 | 1 | 1 | 1 | 1 | 1 | 1 | -0.007 |
| SE | 32842 | ENSG00000184470 | TXNRD2 | chr22 | - | 19919542 | 19919599 | 19918859 | 19919004 | 19931029 | 19931095 | 44 | 19 | 19 | 28 | 9 | 0 | 0 | 1 | 0 | 0 | 34 | 22 | 18 | 21 | 90 | 0 | 0 | 0 | 0 | 0 | 206 | 149 | 1 | 1 | 1 | 1 | 0.932 | 1 | 1 | 1 | 1 | 1 | 1 | 1 | -0.014 |
| SE | 32847 | ENSG00000144867 | SRPRB | chr3 | + | 133811116 | 133811199 | 133807745 | 133807823 | 133815589 | 133815726 | 117 | 48 | 98 | 85 | 99 | 0 | 0 | 0 | 3 | 0 | 57 | 73 | 87 | 39 | 92 | 0 | 0 | 0 | 0 | 4 | 232 | 149 | 1 | 1 | 1 | 1 | 1 | 0.948 | 1 | 1 | 1 | 1 | 1 | 0.937 | 0.002 |
| SE | 32852 | ENSG00000175105 | ZNF654 | chr3 | + | 88086256 | 88086402 | 88059254 | 88059505 | 88113114 | 88113196 | 73 | 19 | 40 | 54 | 18 | 0 | 0 | 0 | 1 | 0 | 0 | 24 | 30 | 7 | 19 | 0 | 0 | 0 | 0 | 0 | 295 | 149 | 1 | 1 | 1 | 1 | 1 | 0.965 | 1 | NA | 1 | 1 | 1 | 1 | -0.007 |
| SE | 32855 | ENSG00000175105 | ZNF654 | chr3 | + | 88128808 | 88129011 | 88113114 | 88113196 | 88129686 | 88129826 | 39 | 23 | 28 | 42 | 59 | 16 | 2 | 7 | 7 | 7 | 9 | 21 | 19 | 0 | 2 | 0 | 0 | 2 | 0 | 0 | 298 | 149 | 1 | 1 | 0.549 | 0.852 | 0.667 | 0.75 | 0.808 | 1 | 1 | 0.826 | NA | 1 | -0.231 |
| SE | 32856 | ENSG00000175105 | ZNF654 | chr3 | + | 88128808 | 88129011 | 88126133 | 88126269 | 88129686 | 88129826 | 53 | 34 | 42 | 54 | 105 | 0 | 0 | 0 | 1 | 1 | 20 | 26 | 22 | 3 | 2 | 0 | 0 | 0 | 0 | 0 | 298 | 149 | 1 | 1 | 1 | 1 | 1 | 0.964 | 0.981 | 1 | 1 | 1 | 1 | 1 | -0.011 |
| SE | 32858 | ENSG00000163320 | CGGBP1 | chr3 | - | 88058018 | 88058223 | 88051943 | 88055999 | 88140969 | 88141078 | 243 | 167 | 173 | 215 | 193 | 2 | 3 | 0 | 1 | 0 | 186 | 189 | 131 | 137 | 129 | 3 | 0 | 0 | 0 | 2 | 298 | 149 | 1 | 1 | 0.984 | 0.965 | 1 | 0.991 | 1 | 0.969 | 1 | 1 | 1 | 0.97 | 0 |
| SE | 32859 | ENSG00000232912 | RERE-AS1 | chr1 | + | 8428481 | 8428588 | 8424644 | 8424701 | 8429826 | 8429972 | 2 | 1 | 3 | 1 | 0 | 0 | 0 | 2 | 1 | 0 | 0 | 4 | 2 | 0 | 2 | 0 | 0 | 0 | 4 | 2 | 256 | 149 | 1 | 1 | 1 | 1 | 0.466 | 0.368 | NA | NA | 1 | 1 | 0 | 0.368 | 0.117 |
| SE | 32860 | ENSG00000232912 | RERE-AS1 | chr1 | + | 8429826 | 8429972 | 8424644 | 8424701 | 8434569 | 8435013 | 0 | 0 | 3 | 1 | 0 | 0 | 0 | 0 | 0 | 0 | 0 | 0 | 0 | 4 | 2 | 0 | 0 | 0 | 2 | 0 | 295 | 149 | 1 | 1 | NA | NA | 1 | 1 | NA | NA | NA | NA | 0.503 | 1 | 0.248 |
| SE | 32864 | ENSG00000164244 | PRRC1 | chr5 | + | 127524530 | 127524920 | 127523459 | 127523582 | 127526617 | 127526778 | 229 | 116 | 200 | 222 | 406 | 0 | 0 | 0 | 1 | 0 | 63 | 122 | 128 | 109 | 100 | 0 | 0 | 0 | 0 | 0 | 298 | 149 | 1 | 1 | 1 | 1 | 1 | 0.991 | 1 | 1 | 1 | 1 | 1 | 1 | -0.002 |
| SE | 32865 | ENSG00000164244 | PRRC1 | chr5 | + | 127530293 | 127530396 | 127526617 | 127526778 | 127533622 | 127533786 | 98 | 53 | 97 | 104 | 149 | 0 | 1 | 0 | 0 | 0 | 25 | 99 | 102 | 40 | 51 | 0 | 0 | 0 | 0 | 0 | 252 | 149 | 1 | 1 | 1 | 0.969 | 1 | 1 | 1 | 1 | 1 | 1 | 1 | 1 | -0.006 |
| SE | 32866 | ENSG00000164244 | PRRC1 | chr5 | + | 127539039 | 127539143 | 127533622 | 127533786 | 127547818 | 127547921 | 255 | 160 | 224 | 219 | 263 | 0 | 0 | 0 | 0 | 0 | 146 | 216 | 228 | 125 | 54 | 0 | 0 | 1 | 0 | 0 | 253 | 149 | 1 | 1 | 1 | 1 | 1 | 1 | 1 | 1 | 1 | 0.993 | 1 | 1 | 0.001 |
| SE | 32867 | ENSG00000068650 | ATP11A | chr13 | + | 112804956 | 112805046 | 112785134 | 112785257 | 112806212 | 112806293 | 130 | 82 | 100 | 93 | 143 | 0 | 0 | 0 | 0 | 0 | 374 | 190 | 222 | 370 | 260 | 0 | 0 | 1 | 0 | 0 | 239 | 149 | 1 | 1 | 1 | 1 | 1 | 1 | 1 | 1 | 1 | 0.993 | 1 | 1 | 0.001 |
| SE | 32868 | ENSG00000068650 | ATP11A | chr13 | + | 112806212 | 112806293 | 112785134 | 112785257 | 112810618 | 112810699 | 139 | 102 | 95 | 121 | 77 | 0 | 0 | 0 | 0 | 0 | 255 | 157 | 126 | 229 | 283 | 0 | 0 | 0 | 7 | 0 | 230 | 149 | 1 | 1 | 1 | 1 | 1 | 1 | 1 | 1 | 1 | 1 | 0.955 | 1 | 0.009 |
| SE | 32869 | ENSG00000068650 | ATP11A | chr13 | + | 112810618 | 112810726 | 112806212 | 112806293 | 112816082 | 112816211 | 386 | 278 | 257 | 313 | 315 | 0 | 0 | 0 | 0 | 0 | 1026 | 506 | 498 | 833 | 839 | 0 | 0 | 0 | 5 | 0 | 257 | 149 | 1 | 1 | 1 | 1 | 1 | 1 | 1 | 1 | 1 | 1 | 0.99 | 1 | 0.002 |
| SE | 32870 | ENSG00000068650 | ATP11A | chr13 | + | 112819899 | 112819950 | 112819303 | 112819407 | 112823344 | 112823409 | 142 | 80 | 92 | 120 | 86 | 0 | 0 | 0 | 0 | 0 | 116 | 134 | 106 | 204 | 241 | 0 | 4 | 0 | 0 | 0 | 200 | 149 | 1 | 1 | 1 | 1 | 1 | 1 | 1 | 1 | 0.961 | 1 | 1 | 1 | 0.008 |
| SE | 32871 | ENSG00000068650 | ATP11A | chr13 | + | 112824343 | 112824425 | 112823344 | 112823409 | 112825429 | 112825580 | 182 | 124 | 122 | 160 | 139 | 0 | 0 | 0 | 0 | 0 | 349 | 176 | 124 | 349 | 278 | 0 | 0 | 0 | 3 | 0 | 231 | 149 | 1 | 1 | 1 | 1 | 1 | 1 | 1 | 1 | 1 | 1 | 0.987 | 1 | 0.003 |
| SE | 32873 | ENSG00000068650 | ATP11A | chr13 | + | 112826693 | 112826891 | 112824343 | 112824425 | 112831374 | 112831548 | 82 | 73 | 52 | 70 | 94 | 0 | 0 | 0 | 0 | 0 | 226 | 121 | 79 | 172 | 121 | 0 | 0 | 3 | 0 | 0 | 298 | 149 | 1 | 1 | 1 | 1 | 1 | 1 | 1 | 1 | 1 | 0.929 | 1 | 1 | 0.014 |
| SE | 32874 | ENSG00000068650 | ATP11A | chr13 | + | 112842275 | 112842379 | 112836177 | 112836251 | 112851036 | 112851218 | 406 | 396 | 214 | 340 | 360 | 0 | 0 | 0 | 1 | 2 | 1272 | 465 | 354 | 842 | 884 | 0 | 0 | 2 | 1 | 0 | 253 | 149 | 1 | 1 | 1 | 1 | 1 | 0.995 | 0.991 | 1 | 1 | 0.99 | 0.998 | 1 | 0 |
| SE | 32876 | ENSG00000068650 | ATP11A | chr13 | + | 112855910 | 112856085 | 112854278 | 112854530 | 112857817 | 112857920 | 62 | 70 | 61 | 59 | 84 | 0 | 0 | 0 | 0 | 0 | 139 | 137 | 65 | 159 | 141 | 0 | 0 | 3 | 1 | 0 | 298 | 149 | 1 | 1 | 1 | 1 | 1 | 1 | 1 | 1 | 1 | 0.915 | 0.988 | 1 | 0.019 |
| SE | 32880 | ENSG00000068650 | ATP11A | chr13 | + | 112859392 | 112859452 | 112858144 | 112858290 | 112860286 | 112860414 | 140 | 108 | 86 | 99 | 179 | [illegible] | [illegible] | [illegible] | [illegible] | [illegible] | [illegible] | [illegible] | [illegible] | [illegible] | [illegible] | [illegible] | [illegible] | [illegible] | [illegible] | [illegible] | [illegible] | [illegible] | [illegible] | [illegible] | [illegible] | [illegible] | [illegible] | [illegible] | [illegible] | [illegible] | [illegible] | [illegible] | [illegible] | [illegible] | [illegible] |
| SE | 32881 | ENSG00000068650 | ATP11A | chr13 | + | 112860286 | 112860414 | 112859392 | 112859452 | 112862439 | 112862575 | 152 | 148 | 117 | 154 | 165 | [illegible] | [illegible] | [illegible] | [illegible] | [illegible] | [illegible] | [illegible] | [illegible] | [illegible] | [illegible] | [illegible] | [illegible] | [illegible] | [illegible] | [illegible] | [illegible] | [illegible] | [illegible] | [illegible] | [illegible] | [illegible] | [illegible] | [illegible] | [illegible] | [illegible] | [illegible] | [illegible] | [illegible] | [illegible] | [illegible] |
| SE | 32882 | ENSG00000068650 | ATP11A | chr13 | + | 112860286 | 112860414 | 112859392 | 112859452 | 112871734 | 112871800 | 115 | 97 | 80 | 95 | 155 | [illegible] | [illegible] | [illegible] | [illegible] | [illegible] | [illegible] | [illegible] | [illegible] | [illegible] | [illegible] | [illegible] | [illegible] | [illegible] | [illegible] | [illegible] | [illegible] | [illegible] | [illegible] | [illegible] | [illegible] | [illegible] | [illegible] | [illegible] | [illegible] | [illegible] | [illegible] | [illegible] | [illegible] | [illegible] | [illegible] |
| SE | 32883 | ENSG00000068650 | ATP11A | chr13 | + | 112862439 | 112862575 | 112859392 | 112859452 | 112871734 | 112871800 | 82 | 70 | 64 | 70 | 68 | [illegible] | [illegible] | [illegible] | [illegible] | [illegible] | [illegible] | [illegible] | [illegible] | [illegible] | [illegible] | [illegible] | [illegible] | [illegible] | [illegible] | [illegible] | [illegible] | [illegible] | [illegible] | [illegible] | [illegible] | [illegible] | [illegible] | [illegible] | [illegible] | [illegible] | [illegible] | [illegible] | [illegible] | [illegible] | [illegible] |
| SE | 32892 | ENSG00000068650 | ATP11A | chr13 | + | 112880545 | 112880649 | 112878216 | 112878303 | [illegible] | [illegible] | [illegible] | [illegible] | [illegible] | [illegible] | [illegible] | [illegible] | [illegible] | [illegible] | [illegible] | [illegible] | [illegible] | [illegible] | [illegible] | [illegible] | [illegible] | [illegible] | [illegible] | [illegible] | [illegible] | [illegible] | [illegible] | [illegible] | [illegible] | [illegible] | [illegible] | [illegible] | [illegible] | [illegible] | [illegible] | [illegible] | [illegible] | [illegible] | [illegible] | [illegible] | -1 |
| SE | 32893 | ENSG00000068650 | ATP11A | chr13 | + | 112880545 | 112880649 | 112878216 | 112878303 | [illegible] | [illegible] | [illegible] | [illegible] | [illegible] | [illegible] | [illegible] | [illegible] | [illegible] | [illegible] | [illegible] | [illegible] | [illegible] | [illegible] | [illegible] | [illegible] | [illegible] | [illegible] | [illegible] | [illegible] | [illegible] | [illegible] | [illegible] | [illegible] | [illegible] | [illegible] | [illegible] | [illegible] | [illegible] | [illegible] | [illegible] | [illegible] | [illegible] | [illegible] | [illegible] | [illegible] | [illegible] |
| SE | 32894 | ENSG00000068650 | ATP11A | chr13 | + | 112880545 | 112880649 | 112878216 | 112878303 | [illegible] | [illegible] | [illegible] | [illegible] | [illegible] | [illegible] | [illegible] | [illegible] | [illegible] | [illegible] | [illegible] | [illegible] | [illegible] | [illegible] | [illegible] | [illegible] | [illegible] | [illegible] | [illegible] | [illegible] | [illegible] | [illegible] | [illegible] | [illegible] | [illegible] | [illegible] | [illegible] | [illegible] | [illegible] | [illegible] | [illegible] | [illegible] | [illegible] | [illegible] | [illegible] | [illegible] | [illegible] |
| SE | 32895 | ENSG00000075711 | DLG1 | chr3 | - | 197051576 | 197051668 | 197042559 | 197044729 | 197059888 | 197059998 | 374 | 307 | 308 | 380 | 897 | [illegible] | [illegible] | [illegible] | [illegible] | [illegible] | [illegible] | [illegible] | [illegible] | [illegible] | [illegible] | [illegible] | [illegible] | [illegible] | [illegible] | [illegible] | [illegible] | [illegible] | [illegible] | [illegible] | [illegible] | [illegible] | [illegible] | [illegible] | [illegible] | [illegible] | [illegible] | [illegible] | [illegible] | [illegible] | [illegible] |
| SE | 32896 | ENSG00000075711 | DLG1 | chr3 | - | 197059888 | 197059998 | 197051576 | 197051668 | [illegible] | [illegible] | 539 | 448 | 447 | 548 | 1272 | [illegible] | [illegible] | [illegible] | [illegible] | [illegible] | [illegible] | [illegible] | [illegible] | [illegible] | [illegible] | [illegible] | [illegible] | [illegible] | [illegible] | [illegible] | [illegible] | [illegible] | [illegible] | [illegible] | [illegible] | [illegible] | [illegible] | [illegible] | [illegible] | [illegible] | [illegible] | [illegible] | [illegible] | [illegible] | [illegible] |
| SE | 32900 | ENSG00000075711 | DLG1 | chr3 | - | 197069218 | 197069260 | 197066703 | 197066754 | 197075836 | 197075870 | 86 | 36 | 60 | 65 | 130 | [illegible] | [illegible] | [illegible] | [illegible] | [illegible] | [illegible] | [illegible] | [illegible] | [illegible] | [illegible] | [illegible] | [illegible] | [illegible] | [illegible] | [illegible] | [illegible] | [illegible] | [illegible] | [illegible] | [illegible] | [illegible] | [illegible] | [illegible] | [illegible] | [illegible] | [illegible] | [illegible] | [illegible] | [illegible] | [illegible] |
| SE | 32901 | ENSG00000075711 | DLG1 | chr3 | - | 197069218 | 197069260 | 197066703 | 197066754 | 197076585 | 197076682 | 75 | 54 | 66 | 59 | 135 | [illegible] | [illegible] | [illegible] | [illegible] | [illegible] | [illegible] | [illegible] | [illegible] | [illegible] | [illegible] | [illegible] | [illegible] | [illegible] | [illegible] | [illegible] | [illegible] | [illegible] | [illegible] | [illegible] | [illegible] | [illegible] | [illegible] | [illegible] | [illegible] | [illegible] | [illegible] | [illegible] | [illegible] | [illegible] | [illegible] |
| SE | [illegible] | ENSG00000075711 | DLG1 | chr3 | - | [illegible] | [illegible] | [illegible] | [illegible] | [illegible] | [illegible] | [illegible] | [illegible] | [illegible] | [illegible] | [illegible] | [illegible] | [illegible] | [illegible] | [illegible] | [illegible] | [illegible] | [illegible] | [illegible] | [illegible] | [illegible] | [illegible] | [illegible] | [illegible] | [illegible] | [illegible] | [illegible] | [illegible] | [illegible] | [illegible] | [illegible] | [illegible] | [illegible] | [illegible] | [illegible] | [illegible] | [illegible] | [illegible] | [illegible] | [illegible] | [illegible] |

| | | | | | | | | | | | | | | | | | | | | | | | | | | | | | | | | | | | | | | | | | | | | | | |
|---|---|---|---|---|---|---|---|---|---|---|---|---|---|---|---|---|---|---|---|---|---|---|---|---|---|---|---|---|---|---|---|---|---|---|---|---|---|---|---|---|---|---|---|---|---|---|
| SE | 32910 | ENSG00000075711 | DLG1 | chr3 | - | 197081050 | 197081117 | 197076585 | 197076685 | 197085579 | 197085756 | 121 | 85 | 123 | 136 | 227 | 0 | 1 | 0 | 0 | 0 | 55 | 77 | 96 | 77 | 25 | 0 | 0 | 0 | 0 | 0 | 216 | 149 | 1 | 1 | 1 | 0.983 | 1 | 1 | 1 | 1 | 1 | 1 | 1 | 1 | -0.003 |
| SE | 32911 | ENSG00000075711 | DLG1 | chr3 | - | 197085295 | 197085756 | 197081050 | 197081117 | 197090911 | 197091026 | 122 | 93 | 136 | 183 | 198 | 2 | 0 | 0 | 0 | 2 | 31 | 79 | 131 | 48 | 38 | 0 | 0 | 0 | 0 | 0 | 298 | 149 | 1 | 1 | 0.968 | 1 | 1 | 1 | 0.98 | 1 | 1 | 1 | 1 | 1 | -0.01 |
| SE | 32912 | ENSG00000075711 | DLG1 | chr3 | - | 197085579 | 197085756 | 197080477 | 197081117 | 197090911 | 197091026 | 254 | 180 | 275 | 332 | 432 | 2 | 0 | 0 | 0 | 2 | 89 | 175 | 239 | 124 | 60 | 0 | 0 | 0 | 0 | 0 | 298 | 149 | 1 | 1 | 0.984 | 1 | 1 | 1 | 0.991 | 1 | 1 | 1 | 1 | 1 | -0.005 |
| SE | 32913 | ENSG00000075711 | DLG1 | chr3 | - | 197090911 | 197091026 | 197085295 | 197085756 | 197104902 | 197105005 | 290 | 226 | 279 | 355 | 476 | 2 | 0 | 0 | 0 | 1 | 65 | 164 | 301 | 159 | 132 | 0 | 0 | 0 | 1 | 0 | 264 | 149 | 1 | 1 | 1 | 1 | 1 | 1 | 0.996 | 1 | 1 | 1 | 0.989 | 1 | 0.001 |
| SE | 32915 | ENSG00000075711 | DLG1 | chr3 | - | 197104902 | 197105005 | 197090911 | 197091026 | 197119409 | 197119530 | 74 | 67 | 68 | 109 | 125 | 0 | 0 | 0 | 0 | 0 | 10 | 39 | 77 | 31 | 26 | 0 | 0 | 4 | 0 | 0 | 252 | 149 | 1 | 1 | 1 | 1 | 1 | 1 | 1 | 1 | 1 | 0.919 | 1 | 1 | 0.016 |
| SE | 32916 | ENSG00000075711 | DLG1 | chr3 | - | 197104902 | 197105005 | 197090911 | 197091026 | 197130526 | 197130671 | 74 | 67 | 68 | 109 | 125 | 0 | 0 | 0 | 0 | 3 | 0 | 35 | 75 | 31 | 31 | 0 | 0 | 0 | 1 | 0 | 252 | 149 | 1 | 1 | 1 | 1 | 1 | 1 | 0.961 | NA | 1 | 1 | 0.948 | 1 | 0.005 |
| SE | 32917 | ENSG00000075711 | DLG1 | chr3 | - | 197115926 | 197116083 | 197090911 | 197091026 | 197119409 | 197119530 | 152 | 153 | 176 | 182 | 302 | 0 | 0 | 0 | 0 | 0 | 120 | 118 | 173 | 146 | 150 | 0 | 0 | 4 | 0 | 0 | 298 | 149 | 1 | 1 | 1 | 1 | 1 | 1 | 1 | 1 | 1 | 0.956 | 1 | 1 | 0.009 |
| SE | 32918 | ENSG00000075711 | DLG1 | chr3 | - | 197115926 | 197116083 | 197090911 | 197091026 | 197130526 | 197130671 | 0 | 1 | 0 | 1 | 0 | 0 | 0 | 0 | 0 | 3 | 0 | 3 | 10 | 3 | 0 | 0 | 0 | 0 | 1 | 0 | 298 | 149 | 1 | 1 | NA | 1 | NA | 1 | 0 | NA | 1 | 1 | 0.6 | NA | -0.2 |
| SE | 32920 | ENSG00000075711 | DLG1 | chr3 | - | 197115926 | 197116083 | 197104902 | 197105005 | 197130526 | 197130671 | 217 | 178 | 180 | 235 | 365 | 0 | 1 | 0 | 0 | 0 | 57 | 124 | 228 | 141 | 123 | 0 | 0 | 0 | 0 | 5 | 298 | 149 | 1 | 1 | 1 | 0.989 | 1 | 1 | 1 | 1 | 1 | 1 | 1 | 0.925 | 0.013 |
| SE | 32921 | ENSG00000075711 | DLG1 | chr3 | - | 197119409 | 197119530 | 197090911 | 197091026 | 197130526 | 197130671 | 118 | 115 | 159 | 170 | 255 | 0 | 0 | 0 | 0 | 3 | 46 | 96 | 95 | 76 | 27 | 0 | 0 | 0 | 1 | 0 | 270 | 149 | 1 | 1 | 1 | 1 | 1 | 1 | 0.979 | 1 | 1 | 1 | 0.977 | 1 | 0 |
| SE | 32925 | ENSG00000075711 | DLG1 | chr3 | - | 197119409 | 197119530 | 197115926 | 197116083 | 197130526 | 197130671 | 270 | 265 | 334 | 351 | 557 | 0 | 1 | 0 | 1 | 0 | 166 | 211 | 249 | 218 | 177 | 0 | 0 | 1 | 0 | 0 | 270 | 149 | 1 | 1 | 1 | 0.993 | 1 | 0.995 | 1 | 1 | 1 | 0.993 | 1 | 1 | -0.001 |
| SE | 32926 | ENSG00000075711 | DLG1 | chr3 | - | 197130526 | 197130671 | 197090911 | 197091026 | 197136541 | 197136678 | 128 | 97 | 137 | 136 | 256 | 0 | 0 | 0 | 0 | 0 | 29 | 58 | 57 | 43 | 41 | 0 | 3 | 0 | 0 | 0 | 294 | 149 | 1 | 1 | 1 | 1 | 1 | 1 | 1 | 1 | 0.907 | 1 | 1 | 1 | 0.019 |
| SE | 32927 | ENSG00000075711 | DLG1 | chr3 | - | 197130526 | 197130671 | 197119409 | 197119530 | 197136541 | 197136678 | 266 | 227 | 319 | 328 | 555 | 0 | 0 | 0 | 0 | 0 | 83 | 170 | 162 | 141 | 96 | 0 | 0 | 6 | 0 | 0 | 294 | 149 | 1 | 1 | 1 | 1 | 1 | 1 | 1 | 1 | 1 | 0.932 | 1 | 1 | 0.014 |
| SE | 32928 | ENSG00000075711 | DLG1 | chr3 | - | 197141229 | 197141353 | 197140139 | 197140264 | 197142717 | 197142768 | 1 | 1 | 0 | 0 | 0 | 198 | 153 | 196 | 178 | 186 | 0 | 0 | 0 | 0 | 0 | 79 | 89 | 108 | 49 | 57 | 273 | 149 | 1 | 1 | 0.003 | 0.004 | 0 | 0 | 0 | 0 | 0 | 0 | 0 | 0 | 0.001 |
| SE | 32930 | ENSG00000075711 | DLG1 | chr3 | - | 197142717 | 197142768 | 197140139 | 197140264 | 197194424 | 197194589 | 69 | 35 | 58 | 69 | 58 | 0 | 0 | 0 | 3 | 0 | 31 | 38 | 54 | 16 | 15 | 0 | 0 | 0 | 0 | 0 | 200 | 149 | 1 | 1 | 1 | 1 | 1 | 0.945 | 1 | 1 | 1 | 1 | 1 | 1 | -0.011 |
| SE | 32931 | ENSG00000075711 | DLG1 | chr3 | - | 197149742 | 197149796 | 197140139 | 197140264 | 197194424 | 197194589 | 74 | 55 | 72 | 61 | 106 | 0 | 0 | 0 | 3 | 0 | 50 | 61 | 27 | 14 | 14 | 0 | 0 | 0 | 0 | 0 | 203 | 149 | 1 | 1 | 1 | 1 | 1 | 0.937 | 1 | 1 | 1 | 1 | 1 | 1 | -0.013 |
| SE | 32932 | ENSG00000075711 | DLG1 | chr3 | - | 197149742 | 197149796 | 197142717 | 197142768 | 197161639 | 197161738 | 107 | 97 | 136 | 99 | 136 | 0 | 0 | 0 | 0 | 2 | 64 | 63 | 57 | 34 | 30 | 0 | 0 | 0 | 0 | 0 | 203 | 149 | 1 | 1 | 1 | 1 | 1 | 1 | 0.98 | 1 | 1 | 1 | 1 | 1 | -0.004 |
| SE | 32935 | ENSG00000075711 | DLG1 | chr3 | - | 197155668 | 197155763 | 197149742 | 197149796 | 197194424 | 197194589 | 0 | 0 | 0 | 1 | 0 | 159 | 119 | 136 | 144 | 182 | 0 | 0 | 0 | 0 | 0 | 80 | 86 | 77 | 43 | 29 | 244 | 149 | 1 | 1 | 0 | 0 | 0 | 0.004 | 0 | 0 | 0 | 0 | 0 | 0 | 0.001 |
| SE | 32938 | ENSG00000075711 | DLG1 | chr3 | - | 197161639 | 197161738 | 197149742 | 197149796 | 197183574 | 197183808 | 7 | 13 | 16 | 15 | 23 | 1 | 0 | 0 | 1 | 0 | 8 | 16 | 13 | 13 | 2 | 0 | 0 | 2 | 0 | 0 | 248 | 149 | 1 | 1 | 0.808 | 1 | 1 | 0.9 | 1 | 1 | 1 | 0.796 | 1 | 1 | -0.018 |
| SE | 32940 | ENSG00000075711 | DLG1 | chr3 | - | 197194424 | 197194589 | 197142717 | 197142768 | 197282678 | 197282845 | 230 | 196 | 218 | 237 | 569 | 0 | 1 | 0 | 0 | 0 | 78 | 115 | 162 | 80 | 45 | 0 | 0 | 0 | 0 | 0 | 298 | 149 | 1 | 1 | 1 | 0.99 | 1 | 1 | 1 | 1 | 1 | 1 | 1 | 1 | -0.002 |
| SE | 32942 | ENSG00000075711 | DLG1 | chr3 | - | 197296345 | 197296477 | 197282609 | 197282845 | 197297185 | 197297204 | 251 | 228 | 192 | 253 | 513 | 0 | 0 | 0 | 1 | 3 | 110 | 131 | 167 | 69 | 59 | 0 | 0 | 0 | 0 | 0 | 281 | 149 | 1 | 1 | 1 | 1 | 1 | 0.993 | 0.989 | 1 | 1 | 1 | 1 | 1 | -0.004 |
| SE | 32944 | ENSG00000075711 | DLG1 | chr3 | - | 197297352 | 197297426 | 197297185 | 197297235 | 197298535 | 197298612 | 1 | 3 | 3 | 3 | 4 | 4 | 4 | 2 | 1 | 0 | 8 | 0 | 2 | 7 | 11 | 0 | 0 | 2 | 0 | 0 | 223 | 149 | 1 | 1 | 0.143 | 0.334 | 0.501 | 0.667 | 1 | 1 | NA | 0.401 | 1 | 1 | -0.321 |
| SE | 32945 | ENSG00000114127 | XRN1 | chr3 | - | 142313135 | 142313174 | 142312597 | 142312758 | 142318591 | 142318694 | 3 | 3 | 10 | 6 | 8 | 58 | 45 | 67 | 60 | 66 | 0 | 4 | 1 | 9 | 0 | 0 | 36 | 51 | 14 | 0 | 188 | 149 | 1 | 1 | 0.039 | 0.05 | 0.106 | 0.073 | 0.088 | NA | 0.081 | 0.015 | 0.338 | NA | -0.073 |
| SE | 32946 | ENSG00000114127 | XRN1 | chr3 | - | 142318591 | 142318694 | 142312597 | 142312758 | 142329433 | 142329615 | 66 | 59 | 74 | 65 | 128 | 0 | 0 | 0 | 0 | 0 | 0 | 32 | 50 | 11 | 0 | 0 | 0 | 3 | 0 | 0 | 252 | 149 | 1 | 1 | 1 | 1 | 1 | 1 | 1 | NA | 1 | 0.908 | 1 | NA | 0.031 |
| SE | 32947 | ENSG00000114127 | XRN1 | chr3 | - | 142318789 | 142318903 | 142318591 | 142318694 | 142329433 | 142329615 | 248 | 223 | 247 | 214 | 697 | 24 | 22 | 29 | 29 | 79 | 18 | 157 | 93 | 41 | 0 | 0 | 19 | 19 | 1 | 0 | 263 | 149 | 1 | 1 | 0.854 | 0.852 | 0.828 | 0.807 | 0.833 | 1 | 0.824 | 0.735 | 0.959 | NA | -0.045 |
| SE | 32948 | ENSG00000114127 | XRN1 | chr3 | - | 142318789 | 142318903 | 142318591 | 142318694 | 142332374 | 142332424 | 143 | 120 | 144 | 128 | 426 | 2 | 4 | 5 | 6 | 4 | 9 | 94 | 52 | 24 | 0 | 12 | 2 | 0 | 4 | 0 | 263 | 149 | 1 | 1 | 0.976 | 0.944 | 0.942 | 0.924 | 0.984 | 0.298 | 0.964 | 1 | 0.773 | NA | 0.195 |
| SE | 32950 | ENSG00000114127 | XRN1 | chr3 | - | 142329433 | 142329615 | 142318789 | 142318903 | 142332374 | 142332531 | 364 | 339 | 379 | 320 | 864 | 1 | 0 | 0 | 3 | 0 | 14 | 198 | 160 | 48 | 28 | 0 | 0 | 0 | 0 | 0 | 298 | 149 | 1 | 1 | 0.995 | 1 | 1 | 0.982 | 1 | 1 | 1 | 1 | 1 | 1 | -0.005 |
| SE | 32951 | ENSG00000114127 | XRN1 | chr3 | - | 142332966 | 142333089 | 142332374 | 142332531 | 142335447 | 142335505 | 199 | 202 | 142 | 184 | 412 | 0 | 0 | 0 | 1 | 0 | 0 | 85 | 106 | 49 | 18 | 0 | 0 | 0 | 0 | 0 | 272 | 149 | 1 | 1 | 1 | 1 | 1 | 0.99 | 1 | NA | 1 | 1 | 1 | 1 | -0.002 |
| SE | 32952 | ENSG00000114127 | XRN1 | chr3 | - | 142332966 | 142333089 | 142332374 | 142332534 | 142335447 | 142335505 | 339 | 311 | 253 | 318 | 626 | 2 | 0 | 0 | 0 | 0 | 12 | 164 | 153 | 72 | 25 | 0 | 0 | 0 | 0 | 0 | 272 | 149 | 1 | 1 | 0.989 | 1 | 1 | 1 | 1 | 1 | 1 | 1 | 1 | 1 | -0.002 |
| SE | 32953 | ENSG00000114127 | XRN1 | chr3 | - | 142347233 | 142347342 | 142335447 | 142335509 | 142355400 | 142355496 | 308 | 280 | 253 | 322 | 347 | 0 | 0 | 3 | 2 | 0 | 22 | 155 | 174 | 62 | 40 | 0 | 0 | 0 | 0 | 0 | 258 | 149 | 1 | 1 | 1 | 1 | 0.98 | 0.989 | 1 | 1 | 1 | 1 | 1 | 1 | -0.006 |
| SE | 32954 | ENSG00000114127 | XRN1 | chr3 | - | 142370484 | 142370620 | 142365309 | 142365366 | 142371238 | 142371328 | 424 | 413 | 261 | 397 | 438 | 0 | 0 | 1 | 7 | 0 | 91 | 264 | 311 | 58 | 43 | 0 | 0 | 0 | 0 | 0 | 285 | 149 | 1 | 1 | 1 | 1 | 0.993 | 0.967 | 1 | 1 | 1 | 1 | 1 | 1 | -0.008 |
| SE | 32956 | ENSG00000114127 | XRN1 | chr3 | - | 142403868 | 142403989 | 142403673 | 142403772 | 142404906 | 142405073 | 437 | 429 | 304 | 385 | 535 | 0 | 0 | 0 | 0 | 1 | 106 | 217 | 245 | 58 | 55 | 0 | 0 | 0 | 0 | 0 | 270 | 149 | 1 | 1 | 1 | 1 | 1 | 1 | 0.997 | 1 | 1 | 1 | 1 | 1 | -0.001 |
| SE | 32957 | ENSG00000114127 | XRN1 | chr3 | - | 142412543 | 142412663 | 142404906 | 142405076 | 142414134 | 142414291 | 277 | 301 | 242 | 311 | 505 | 4 | 0 | 0 | 0 | 0 | 32 | 154 | 208 | 32 | 31 | 0 | 0 | 0 | 0 | 0 | 269 | 149 | 1 | 1 | 0.975 | 1 | 1 | 1 | 1 | 1 | 1 | 1 | 1 | 1 | -0.005 |
| SE | 32959 | ENSG00000114127 | XRN1 | chr3 | - | 142414134 | 142414291 | 142412543 | 142412663 | 142418503 | 142418609 | 186 | 184 | 161 | 224 | 330 | 0 | 0 | 0 | 0 | 0 | 32 | 110 | 133 | 27 | 25 | 0 | 1 | 0 | 0 | 0 | 298 | 149 | 1 | 1 | 1 | 1 | 1 | 1 | 1 | 1 | 0.982 | 1 | 1 | 1 | 0.004 |
| SE | 32960 | ENSG00000114127 | XRN1 | chr3 | - | 142417139 | 142417229 | 142412543 | 142412663 | 142418503 | 142418609 | 116 | 90 | 70 | 95 | 178 | 0 | 0 | 0 | 0 | 0 | 0 | 54 | 51 | 15 | 3 | 0 | 1 | 0 | 0 | 0 | 239 | 149 | 1 | 1 | 1 | 1 | 1 | 1 | 1 | NA | 0.971 | 1 | 1 | 1 | 0.007 |
| SE | 32962 | ENSG00000114127 | XRN1 | chr3 | - | 142421475 | 142421543 | 142421015 | 142421153 | 142422581 | 142422750 | 1060 | 1117 | 881 | 1148 | 1102 | 4 | 0 | 0 | 0 | 0 | 91 | 909 | 1088 | 296 | 96 | 0 | 0 | 0 | 0 | 0 | 217 | 149 | 1 | 1 | 0.995 | 1 | 1 | 1 | 1 | 1 | 1 | 1 | 1 | 1 | -0.001 |
| SE | 32963 | ENSG00000114127 | XRN1 | chr3 | - | 142425221 | 142425332 | 142423559 | 142423642 | 142425428 | 142425535 | 780 | 746 | 613 | 733 | 954 | 0 | 0 | 0 | 0 | 0 | 132 | 606 | 725 | 272 | 85 | 0 | 7 | 0 | 0 | 0 | 260 | 149 | 1 | 1 | 1 | 1 | 1 | 1 | 1 | 1 | 0.98 | 1 | 1 | 1 | 0.004 |
| SE | 32964 | ENSG00000114127 | XRN1 | chr3 | - | 142425221 | 142425332 | 142423559 | 142423642 | 142426743 | 142426841 | 242 | 217 | 196 | 203 | 308 | 4 | 0 | 1 | 3 | 0 | 12 | 196 | 188 | 69 | 19 | 0 | 0 | 0 | 0 | 0 | 260 | 149 | 1 | 1 | 0.972 | 1 | 0.991 | 0.975 | 1 | 1 | 1 | 1 | 1 | 1 | -0.012 |
| SE | 32965 | ENSG00000114127 | XRN1 | chr3 | - | 142425428 | 142425538 | 142423559 | 142423642 | 142426743 | 142426841 | 385 | 338 | 338 | 338 | 332 | 4 | 0 | 1 | 3 | 0 | 36 | 242 | 276 | 78 | 16 | 0 | 0 | 0 | 0 | 0 | 259 | 149 | 1 | 1 | 0.982 | 1 | 0.995 | 0.985 | 1 | 1 | 1 | 1 | 1 | 1 | -0.008 |
| SE | 32966 | ENSG00000114127 | XRN1 | chr3 | - | 142425428 | 142425538 | 142423559 | 142423642 | 142432660 | 142432893 | 8 | 5 | 1 | 11 | 2 | 16 | 13 | 14 | 21 | 13 | 40 | 13 | 3 | 1 | 0 | 0 | 4 | 5 | 9 | 0 | 259 | 149 | 1 | 1 | 0.223 | 0.181 | 0.039 | 0.232 | 0.081 | 1 | 0.652 | 0.257 | 0.06 | NA | -0.341 |
| SE | 32967 | ENSG00000114127 | XRN1 | chr3 | - | 142425428 | 142425538 | 142425221 | 142425332 | 142426743 | 142426841 | 918 | 860 | 751 | 864 | 976 | 1 | 0 | 0 | 0 | 0 | 156 | 645 | 808 | 278 | 82 | 0 | 0 | 0 | 2 | 0 | 259 | 149 | 1 | 1 | 0.998 | 1 | 1 | 1 | 1 | 1 | 1 | 1 | 0.988 | 1 | 0.002 |
| SE | 32972 | ENSG00000114346 | ECT2 | chr3 | + | 172752149 | 172752436 | 172750762 | 172750857 | 172754508 | 172754660 | 0 | 0 | 0 | 0 | 0 | 9 | 7 | 8 | 8 | 8 | 0 | 0 | 4 | 0 | 0 | 0 | 11 | 2 | 0 | 0 | 298 | 149 | 1 | 1 | 0 | 0 | 0 | 0 | 0 | NA | 0 | 0.5 | NA | NA | -0.25 |
| SE | 32973 | ENSG00000114346 | ECT2 | chr3 | + | 172752149 | 172752472 | 172750684 | 172750857 | 172754508 | 172754585 | 0 | 1 | 0 | 0 | 0 | 9 | 7 | 8 | 8 | 8 | 0 | 1 | 0 | 0 | 0 | 0 | 11 | 2 | 0 | 0 | 298 | 149 | 1 | 1 | 0 | 0.067 | 0 | 0 | 0 | NA | 0.043 | 0 | NA | NA | -0.008 |
| SE | 32974 | ENSG00000114346 | ECT2 | chr3 | + | 172752149 | 172752472 | 172750958 | 172751116 | 172754508 | 172754585 | 0 | 1 | 0 | 0 | 0 | 0 | 0 | 0 | 0 | 5 | 0 | 1 | 0 | 0 | 0 | 0 | 0 | 0 | 0 | 0 | 298 | 149 | 1 | 1 | NA | 1 | NA | NA | 0 | NA | 1 | NA | NA | NA | -0.5 |
| SE | 32975 | ENSG00000114346 | ECT2 | chr3 | + | 172754508 | 172754660 | 172750681 | 172750857 | 172755294 | 172755374 | 31 | 23 | 28 | 25 | 41 | 0 | 0 | 0 | 0 | 0 | 4 | 33 | 13 | 7 | 0 | 0 | 0 | 1 | 0 | 0 | 298 | 149 | 1 | 1 | 1 | 1 | 1 | 1 | 1 | 1 | 1 | 0.867 | 1 | NA | 0.033 |
| SE | 32976 | ENSG00000114346 | ECT2 | chr3 | + | 172755482 | 172755575 | 172755294 | 172755374 | 172756982 | 172757009 | 3 | 4 | 1 | 0 | 3 | 16 | 12 | 12 | 20 | 44 | 0 | 0 | 3 | 0 | 0 | 4 | 24 | 8 | 19 | 0 | 242 | 149 | 1 | 1 | 0.103 | 0.17 | 0.049 | 0 | 0.04 | 0 | 0 | 0.188 | 0 | NA | 0.025 |
| SE | 32977 | ENSG00000114346 | ECT2 | chr3 | + | 172760155 | 172760263 | 172758979 | 172759069 | 172761609 | 172761634 | 18 | 15 | 20 | 23 | 39 | 0 | 0 | 1 | 5 | 1 | 7 | 26 | 10 | 7 | 0 | 0 | 0 | 0 | 0 | 0 | 257 | 149 | 1 | 1 | 1 | 1 | 0.921 | 0.727 | 0.958 | 1 | 1 | 1 | 1 | NA | -0.079 |
| SE | 32978 | ENSG00000114346 | ECT2 | chr3 | + | 172769006 | 172769143 | 172764277 | 172764413 | 172773902 | 172774022 | 16 | 11 | 18 | 29 | 19 | 0 | 0 | 0 | 0 | 0 | 0 | 5 | 11 | 2 | 17 | 0 | 0 | 0 | 5 | 0 | 286 | 149 | 1 | 1 | 1 | 1 | 1 | 1 | 1 | NA | 1 | 1 | 0.172 | 1 | 0.207 |
| SE | 32979 | ENSG00000114346 | ECT2 | chr3 | + | 172818547 | 172818814 | 172816690 | 172816837 | 172820147 | 172820311 | 5 | 0 | 0 | 0 | 0 | 6 | 8 | 11 | 5 | 3 | 0 | 0 | 0 | 0 | 0 | 0 | 9 | 6 | 0 | 0 | 298 | 149 | 1 | 1 | 0.294 | 0 | 0 | 0 | 0 | NA | 0 | 0 | NA | NA | 0.059 |
| SE | 32982 | ENSG00000146192 | FGD2 | chr6 | + | 37010972 | 37011050 | 37008833 | 37009065 | 37013608 | 37013765 | 74 | 49 | 43 | 52 | 56 | 0 | 0 | 2 | 0 | 0 | 167 | 86 | 16 | 26 | 105 | 0 | 0 | 0 | 0 | 0 | 227 | 149 | 1 | 1 | 1 | 1 | 0.934 | 1 | 1 | 1 | 1 | 1 | 1 | 1 | -0.013 |
| SE | 32994 | ENSG00000114316 | USP4 | chr3 | - | 49280743 | 49280847 | 49278169 | 49278451 | 49284465 | 49284584 | 7 | 8 | 5 | 6 | 14 | 0 | 0 | 0 | 0 | 0 | 8 | 3 | 5 | 12 | 5 | 0 | 0 | 0 | 1 | 0 | 253 | 149 | 1 | 1 | 1 | 1 | 1 | 1 | 1 | 1 | 1 | 1 | 0.876 | 1 | 0.025 |
| SE | 32996 | ENSG00000114316 | USP4 | chr3 | - | 49283986 | 49284136 | 49277143 | 49278451 | 49284465 | 49284584 | 148 | 92 | 90 | 94 | 107 | 0 | 0 | 0 | 0 | 0 | 276 | 147 | 100 | 148 | 239 | 0 | 0 | 0 | 1 | 0 | 298 | 149 | 1 | 1 | 1 | 1 | 1 | 1 | 1 | 1 | 1 | 1 | 0.987 | 1 | 0.003 |
| SE | 32997 | ENSG00000114316 | USP4 | chr3 | - | 49283986 | 49284136 | 49280743 | 49280847 | 49284465 | 49284584 | 341 | 199 | 230 | 243 | 322 | 2 | 0 | 3 | 1 | 4 | 523 | 304 | 249 | 342 | 408 | 0 | 0 | 0 | 2 | 0 | 298 | 149 | 1 | 1 | 0.988 | 1 | 0.975 | 0.992 | 0.976 | 1 | 1 | 1 | 0.988 | 1 | -0.011 |
| SE | 32998 | ENSG00000114316 | USP4 | chr3 | - | 49286097 | 49286325 | 49284848 | 49284919 | 49292509 | 49292598 | 435 | 319 | 282 | 399 | 434 | 4 | 0 | 1 | 1 | 0 | 1198 | 460 | 510 | 763 | 1025 | 0 | 4 | 1 | 6 | 0 | 298 | 149 | 1 | 1 | 0.982 | 1 | 0.993 | 0.995 | 1 | 1 | 0.983 | 0.996 | 0.985 | 1 | 0.001 |
| SE | 33000 | ENSG00000114316 | USP4 | chr3 | - | 49294406 | 49294598 | 49292509 | 49292598 | 49302383 | 49302542 | 143 | 129 | 115 | 176 | 133 | 0 | 0 | 0 | 0 | 0 | 493 | 173 | 209 | 274 | 446 | 0 | 0 | 2 | 7 | 0 | 298 | 149 | 1 | 1 | 1 | 1 | 1 | 1 | 1 | 1 | 1 | 0.981 | 0.951 | 1 | 0.014 |
| SE | 33001 | ENSG00000114316 | USP4 | chr3 | - | 49300466 | 49300691 | 49294406 | 49294598 | 49302383 | 49302542 | 71 | 36 | 70 | 66 | 97 | 0 | 0 | 0 | 0 | 0 | 212 | 71 | 68 | 94 | 70 | 0 | 0 | 2 | 0 | 0 | 298 | 149 | 1 | 1 | 1 | 1 | 1 | 1 | 1 | 1 | 1 | 0.944 | 1 | 1 | 0.011 |
| SE | 33003 | ENSG00000114316 | USP4 | chr3 | - | 49300466 | 49300691 | 49298551 | 49298635 | 49302437 | 49302542 | 86 | 57 | 87 | 100 | 82 | 1 | 0 | 0 | 0 | 0 | 119 | 120 | 81 | 112 | 105 | 0 | 0 | 0 | 0 | 0 | 298 | 149 | 1 | 1 | 0.977 | 1 | 1 | 1 | 1 | 1 | 1 | 1 | 1 | 1 | -0.005 |
| SE | 33004 | ENSG00000114316 | USP4 | chr3 | - | 49302383 | 49302542 | 49298551 | 49298635 | 49305714 | 49305888 | 116 | 54 | 97 | 130 | 134 | 0 | 0 | 0 | 1 | 0 | 289 | 133 | 133 | 140 | 216 | 0 | 0 | 1 | 0 | 0 | 298 | 149 | 1 | 1 | 1 | 1 | 1 | 0.985 | 1 | 1 | 1 | 0.985 | 1 | 1 | 0 |
| SE | 33005 | ENSG00000114316 | USP4 | chr3 | - | 49302437 | 49302542 | 49298551 | 49298635 | 49305714 | 49305888 | 73 | 37 | 59 | 69 | 83 | 0 | 0 | 0 | 1 | 0 | 139 | 61 | 47 | 81 | 104 | 0 | 0 | 1 | 0 | 0 | 254 | 149 | 1 | 1 | 1 | 1 | 1 | 0.976 | 1 | 1 | 1 | 0.965 | 1 | 1 | 0.002 |
| SE | 33006 | ENSG00000114316 | USP4 | chr3 | - | 49305714 | 49305888 | 49302383 | 49302542 | 49310619 | 49310737 | 230 | 108 | 183 | 210 | 207 | 1 | 0 | 0 | 0 | 0 | 363 | 260 | 223 | 254 | 299 | 0 | 0 | 0 | 0 | 0 | 298 | 149 | 1 | 1 | 0.991 | 1 | 1 | 1 | 1 | 1 | 1 | 1 | 1 | 1 | -0.002 |
| SE | 33007 | ENSG00000114316 | USP4 | chr3 | - | 49305714 | 49305888 | 49302383 | 49302542 | 49327685 | 49327816 | 117 | 48 | 95 | 120 | 116 | 0 | 0 | 0 | 0 | 0 | 237 | 128 | 102 | 122 | 181 | 0 | 0 | 3 | 0 | 0 | 298 | 149 | 1 | 1 | 1 | 1 | 1 | 1 | 1 | 1 | 1 | 0.944 | 1 | 1 | 0.011 |
| SE | 33008 | ENSG00000114316 | USP4 | chr3 | - | 49310619 | 49310737 | 49305714 | 49305888 | 49311513 | 49311618 | 199 | 132 | 165 | 209 | 152 | 0 | 0 | 1 | 0 | 0 | 225 | 214 | 180 | 221 | 199 | 0 | 0 | 1 | 1 | 0 | 267 | 149 | 1 | 1 | 1 | 1 | 0.989 | 1 | 1 | 1 | 1 | 0.99 | 0.992 | 1 | 0.001 |
| SE | 33015 | ENSG00000114316 | USP4 | chr3 | - | 49324701 | 49324763 | 49305714 | 49305888 | 49327685 | 49327816 | 1 | 2 | 1 | 1 | 0 | 0 | 0 | 1 | 0 | 0 | 9 | 8 | 6 | 9 | 0 | 0 | 3 | 0 | 0 | 0 | 211 | 149 | 1 | 1 | 1 | 1 | 0.414 | 1 | NA | 1 | 0.653 | 1 | 1 | NA | -0.06 |
| SE | 33018 | ENSG00000114316 | USP4 | chr3 | - | 49324893 | 49325039 | 49324696 | 49324763 | 49335468 | 49335596 | 145 | 95 | 127 | 144 | 178 | 0 | 0 | 0 | 0 | 0 | 321 | 235 | 151 | 254 | 354 | 0 | 0 | 0 | 1 | 0 | 295 | 149 | 1 | 1 | 1 | 1 | 1 | 1 | 1 | 1 | 1 | 1 | 0.992 | 1 | 0.002 |
| SE | 33019 | ENSG00000114316 | USP4 | chr3 | - | 49325718 | 49325845 | 49324696 | 49324763 | 49327685 | 49327816 | 425 | 255 | 338 | 400 | 637 | 2 | 0 | 1 | 0 | 0 | 871 | 430 | 328 | 594 | 984 | 1 | 4 | 0 | 1 | 6 | 276 | 149 | 1 | 1 | 0.991 | 1 | 0.995 | 1 | 1 | 0.998 | 0.983 | 1 | 0.997 | 0.989 | 0.004 |
| SE | 33020 | ENSG00000114316 | USP4 | chr3 | - | 49325718 | 49325845 | 49324696 | 49324763 | 49335468 | 49335596 | 0 | 0 | 1 | 1 | 4 | 0 | 0 | 0 | 0 | 0 | 8 | 5 | 7 | 11 | 3 | 0 | 0 | 0 | 1 | 0 | 276 | 149 | 1 | 1 | NA | NA | 1 | 1 | 1 | 1 | 1 | 1 | 0.856 | 1 | 0.029 |
| SE | 33022 | ENSG00000114316 | USP4 | chr3 | - | 49325718 | 49325845 | 49324893 | 49325039 | 49335468 | 49335596 | 239 | 154 | 218 | 242 | 315 | 0 | 0 | 0 | 0 | 0 | 426 | 277 | 230 | 280 | 589 | 0 | 0 | 1 | 0 | 0 | 276 | 149 | 1 | 1 | 1 | 1 | 1 | 1 | 1 | 1 | 1 | 0.992 | 1 | 1 | 0.002 |
| SE | 33023 | ENSG00000114316 | USP4 | chr3 | - | 49327685 | 49327816 | 49324696 | 49324763 | 49335468 | 49335596 | 427 | 272 | 298 | 325 | 391 | 0 | 0 | 0 | 0 | 0 | 689 | 384 | 420 | 523 | 782 | 0 | 0 | 0 | 1 | 0 | 280 | 149 | 1 | 1 | 1 | 1 | 1 | 1 | 1 | 1 | 1 | 1 | 0.996 | 1 | 0.001 |
| SE | 33024 | ENSG00000114316 | USP4 | chr3 | - | 49327685 | 49327816 | 49324893 | 49325039 | 49335468 | 49335596 | 425 | 283 | 297 | 328 | 391 | 0 | 0 | 0 | 0 | 0 | 708 | 389 | 424 | 532 | 783 | 0 | 0 | 1 | 0 | 0 | 280 | 149 | 1 | 1 | 1 | 1 | 1 | 1 | 1 | 1 | 1 | 0.996 | 1 | 1 | 0.001 |
| SE | 33025 | ENSG00000114316 | USP4 | chr3 | - | 49327685 | 49327816 | 49325718 | 49325845 | 49335468 | 49335596 | 891 | 545 | 672 | 761 | 1082 | 0 | 0 | 0 | 0 | 0 | 1600 | 828 | 766 | 1140 | 1826 | 0 | 0 | 2 | 0 | 3 | 280 | 149 | 1 | 1 | 1 | 1 | 1 | 1 | 1 | 1 | 1 | 0.995 | 1 | 0.997 | 0.002 |
| SE | 33028 | ENSG00000183323 | CCDC125 | chr5 | - | 69292187 | 69292362 | 69285336 | 69285467 | 69294792 | 69294900 | 78 | 45 | 76 | 83 | 99 | 1 | 0 | 0 | 0 | 0 | 57 | 83 | 74 | 268 | 19 | 0 | 0 | 0 | 0 | 0 | 298 | 149 | 1 | 1 | 0.975 | 1 | 1 | 1 | 1 | 1 | 1 | 1 | 1 | 1 | -0.005 |
| SE | 33032 | ENSG00000183323 | CCDC125 | chr5 | - | 69303846 | 69303929 | 69300011 | 69300127 | 69306816 | 69306902 | 34 | 28 | 40 | 47 | 79 | 0 | 3 | 0 | 1 | 0 | 55 | 41 | 62 | 119 | 13 | 0 | 0 | 0 | 0 | 0 | 232 | 149 | 1 | 1 | 1 | 0.857 | 1 | 0.968 | 1 | 1 | 1 | 1 | 1 | 1 | -0.035 |
| SE | 33033 | ENSG00000183323 | CCDC125 | chr5 | - | 69303846 | 69303929 | 69300011 | 69300127 | 69307950 | 69308028 | 19 | 5 | 11 | 17 | 48 | 0 | 0 | 0 | 1 | 0 | 39 | 26 | 31 | 60 | 21 | 0 | 0 | 0 | 0 | 0 | 232 | 149 | 1 | 1 | 1 | 1 | 1 | 0.916 | 1 | 1 | 1 | 1 | 1 | 1 | -0.017 |
| SE | 33034 | ENSG00000183323 | CCDC125 | chr5 | - | 69306816 | 69306902 | 69300011 | 69300127 | 69307950 | 69308028 | 39 | 32 | 12 | 29 | 15 | 0 | 0 | 0 | 1 | 0 | 39 | 47 | 48 | 78 | 12 | 0 | 0 | 0 | 0 | 0 | 235 | 149 | 1 | 1 | 1 | 1 | 1 | 0.948 | 1 | 1 | 1 | 1 | 1 | 1 | -0.01 |
| SE | 33040 | ENSG00000188315 | C3orf62 | chr3 | - | 49274048 | 49274140 | 49268595 | 49271445 | 49276426 | 49276841 | 46 | 40 | 30 | 50 | 75 | 0 | 0 | 0 | 2 | 0 | 68 | 41 | 54 | 50 | 68 | 0 | 0 | 0 | 0 | 0 | 241 | 149 | 1 | 1 | 1 | 1 | 1 | 0.939 | 1 | 1 | 1 | 1 | 1 | 1 | -0.012 |
| SE | 33041 | ENSG00000154917 | RAB6B | chr3 | - | 133864583 | 133864642 | 133841609 | 133841663 | 133895396 | 133895595 | 28 | 24 | 4 | 36 | 40 | 0 | 0 | 0 | 1 | 0 | 187 | 52 | 26 | 36 | 422 | 0 | 0 | 0 | 0 | 0 | 208 | 149 | 1 | 1 | 1 | 1 | 1 | 0.963 | 1 | 1 | 1 | 1 | 1 | 1 | -0.007 |
| SE | 33047 | ENSG00000239697 | TNFSF12 | chr17 | + | 7556777 | 7556902 | 7550942 | 7550978 | 7557098 | 7557232 | 144 | 64 | 102 | 81 | 70 | 0 | 0 | 0 | 1 | 0 | 91 | 114 | 60 | 61 | 115 | 0 | 2 | 0 | 0 | 3 | 274 | 149 | 1 | 1 | 1 | 1 | 1 | 0.978 | 1 | 1 | 0.969 | 1 | 1 | 0.954 | 0.011 |
| SE | 33048 | ENSG00000109586 | GALNT7 | chr4 | + | 173247979 | 173248440 | 173168810 | 173168961 | 173292107 | 173292274 | 154 | 101 | 95 | 152 | 156 | 0 | 0 | 1 | 0 | 0 | 33 | 144 | 96 | 91 | 24 | 0 | 0 | 0 | 2 | 1 | 298 | 149 | 1 | 1 | 1 | 1 | 0.979 | 1 | 1 | 1 | 1 | 1 | 0.958 | 0.923 | 0.02 |
| SE | 33050 | ENSG00000109586 | GALNT7 | chr4 | + | 173295763 | 173295843 | 173295395 | 173295526 | 173297794 | 173297977 | 22 | 47 | 12 | 30 | 50 | 0 | 0 | 0 | 2 | 0 | 2 | 42 | 5 | 9 | 0 | 0 | 0 | 0 | 3 | 0 | 229 | 149 | 1 | 1 | 1 | 1 | 1 | 0.907 | 1 | 1 | 1 | 1 | 0.661 | NA | 0.066 |
| SE | 33051 | ENSG00000109586 | GALNT7 | chr4 | + | 173295763 | 173295843 | 173295395 | 173295526 | 173298114 | 173298297 | 33 | 30 | 26 | 35 | 62 | 7 | 2 | 1 | 3 | 3 | 9 | 37 | 8 | 22 | 0 | 0 | 0 | 2 | 3 | 0 | 229 | 149 | 1 | 1 | 0.754 | 0.907 | 0.944 | 0.884 | 0.931 | 1 | 1 | 0.722 | 0.827 | NA | -0.003 |
| SE | 33052 | ENSG00000109586 | GALNT7 | chr4 | + | 173313957 | 173314176 | 173303995 | 173304118 | 173317633 | 173317732 | 214 | 192 | 106 | 183 | 213 | 0 | 1 | 0 | 0 | 0 | 100 | 138 | 105 | 91 | 35 | 0 | 0 | 2 | 0 | 0 | 298 | 149 | 1 | 1 | 1 | 0.99 | 1 | 1 | 1 | 1 | 1 | 0.963 | 1 | 1 | 0.005 |
| SE | 33066 | ENSG00000088726 | TMEM40 | chr3 | - | 12736763 | 12736835 | 12736577 | 12736652 | 12737706 | 12737754 | 128 | 158 | 47 | 295 | 233 | 2 | 0 | 0 | 0 | 0 | 535 | 336 | 175 | 205 | 1744 | 0 | 0 | 1 | 0 | 0 | 221 | 149 | 1 | 1 | 0.977 | 1 | 1 | 1 | 1 | 1 | 1 | 0.992 | 1 | 1 | -0.003 |
| SE | 33067 | ENSG00000088726 | TMEM40 | chr3 | - | 12738135 | 12738168 | 12737517 | 12737754 | 12738552 | 12738588 | 60 | 76 | 23 | 119 | 92 | 0 | 0 | 0 | 0 | 0 | 195 | 140 | 90 | 105 | 728 | 0 | 0 | 0 | 3 | 0 | 182 | 149 | 1 | 1 | 1 | 1 | 1 | 1 | 1 | 1 | 1 | 1 | 0.966 | 1 | 0.007 |
| SE | 33075 | ENSG00000182230 | FAM153B | chr5 | + | 176070966 | 176070997 | 176063708 | 176063764 | 176074635 | 176074829 | 4 | 8 | 2 | 2 | 2 | 0 | 4 | 12 | 6 | 1 | 0 | 8 | 5 | 0 | 0 | 0 | 0 | 1 | 0 | 7 | 180 | 149 | 1 | 1 | 1 | 0.623 | 0.121 | 0.216 | 0.623 | NA | 1 | 0.805 | NA | 0 | -0.085 |
| SE | 33076 | ENSG00000182230 | FAM153B | chr5 | + | 176074635 | 176074829 | 176063708 | 176063764 | 176089462 | 176089521 | 0 | 5 | 13 | 7 | 1 | 0 | 2 | 2 | 0 | 0 | 0 | 0 | 2 | 0 | 7 | 0 | 0 | 1 | 0 | 0 | 298 | 149 | 1 | 1 | NA | 0.556 | 0.765 | 1 | 1 | NA | NA | 0.5 | NA | 1 | 0.08 |
| SE | 33080 | ENSG00000260261 | NA | chr3 | - | 195947334 | 195947479 | 195913077 | 195913770 | 195949241 | 195950116 | 1 | 0 | 0 | 5 | 2 | 2 | 0 | 1 | 0 | 2 | 9 | 0 | 0 | 1 | 0 | 0 | 0 | 0 | 4 | 0 | 294 | 149 | 1 | 1 | 0.202 | NA | 0 | 1 | 0.336 | 1 | NA | NA | 0.112 | NA | -0.172 |
| SE | 33088 | ENSG00000144741 | SLC25A26 | chr3 | + | 66346363 | 66346408 | 66263331 | 66263379 | 66369477 | 66369542 | 10 | 4 | 9 | 8 | 23 | 1 | 0 | 0 | 0 | 0 | 0 | 2 | 5 | 0 | 1 | 0 | 3 | 0 | 0 | 0 | 194 | 149 | 1 | 1 | 0.885 | 1 | 1 | 1 | 1 | NA | 0.339 | 1 | NA | 1 | 0.197 |
| SE | 33091 | ENSG00000144741 | SLC25A26 | chr3 | + | 66362859 | 66362929 | 66263331 | 66263379 | 66369477 | 66369542 | 5 | 7 | 9 | 10 | 19 | 1 | 0 | 0 | 0 | 0 | 0 | 22 | 7 | 13 | 16 | 0 | 3 | 0 | 0 | 0 | 219 | 149 | 1 | 1 | 0.773 | 1 | 1 | 1 | 1 | NA | 0.833 | 1 | 1 | 1 | -0.004 |
| SE | 33092 | ENSG00000144741 | SLC25A26 | chr3 | + | 66362859 | 66362929 | 66346363 | 66346408 | 66369477 | 66369542 | 50 | 16 | 39 | 50 | 100 | 0 | 0 | 1 | 0 | 3 | 25 | 31 | 33 | 39 | 24 | 0 | 0 | 0 | 0 | 0 | 219 | 149 | 1 | 1 | 1 | 1 | 0.964 | 1 | 0.958 | 1 | 1 | 1 | 1 | 1 | -0.016 |
| SE | 33093 | ENSG00000144741 | SLC25A26 | chr3 | + | 66362859 | 66362929 | 66356030 | 66356084 | 66369477 | 66369542 | 9 | 7 | 11 | 10 | 16 | 1 | 0 | 0 | 0 | 0 | 0 | 10 | 7 | 11 | 18 | 0 | 0 | 0 | 0 | 0 | 219 | 149 | 1 | 1 | 0.86 | 1 | 1 | 1 | 1 | NA | 1 | 1 | 1 | 1 | -0.028 |
| SE | 33094 | ENSG00000144741 | SLC25A26 | chr3 | + | 66371201 | 66371413 | 66370528 | 66370602 | 66377689 | 66378056 | 0 | 0 | 2 | 0 | 0 | 21 | 1 | 18 | 17 | 8 | 0 | 0 | 0 | 0 | 0 | 42 | 16 | 19 | 12 | 25 | 298 | 149 | 1 | 1 | 0 | 0 | 0.053 | 0 | 0 | 0 | 0 | 0 | 0 | 0 | 0.011 |
| SE | 33095 | ENSG00000240303 | ACAD11 | chr3 | - | 132559832 | 132559942 | 132558137 | 132559085 | 132561100 | 132561217 | 27 | 41 | 45 | 27 | 7 | 0 | 0 | 0 | 0 | 0 | 17 | 20 | 34 | 10 | 8 | 0 | 2 | 0 | 0 | 0 | 259 | 149 | 1 | 1 | 1 | 1 | 1 | 1 | 1 | 1 | 0.852 | 1 | 1 | 1 | 0.03 |
| SE | 33097 | ENSG00000240303 | ACAD11 | chr3 | - | 132618633 | 132618772 | 132605097 | 132605205 | 132626690 | 132626817 | 26 | 21 | 30 | 42 | 46 | 0 | 0 | 0 | 1 | 0 | 8 | 14 | 21 | 13 | 26 | 0 | 2 | 0 | 0 | 0 | 288 | 149 | 1 | 1 | 1 | 1 | 1 | 0.956 | 1 | 1 | 1 | 0.845 | 1 | 1 | 0.022 |
| SE | 33098 | ENSG00000240303 | ACAD11 | chr3 | - | 132619467 | 132619545 | 132618249 | 132618772 | 132626690 | 132626817 | 162 | 145 | 151 | 188 | 186 | 0 | 0 | 0 | 0 | 2 | 89 | 101 | 180 | 107 | 90 | 0 | 0 | 0 | 0 | 0 | 227 | 149 | 1 | 1 | 1 | 1 | 1 | 1 | 0.984 | 1 | 1 | 1 | 1 | 1 | -0.003 |
| SE | 33099 | ENSG00000240303 | ACAD11 | chr3 | - | 132630436 | 132630558 | 132628339 | 132628446 | 132631340 | 132631479 | 107 | 112 | 121 | 101 | 121 | 1 | 0 | 1 | 0 | 1 | 55 | 83 | 107 | 85 | 33 | 0 | 0 | 0 | 0 | 0 | 271 | 149 | 1 | 1 | 0.983 | 1 | 0.985 | 1 | 0.985 | 1 | 1 | 1 | 1 | 1 | -0.009 |
| SE | 33101 | ENSG00000029993 | HMGB3 | chrX | + | 150986050 | 150986190 | 150985530 | 150985749 | 150987127 | 150987297 | 26 | 13 | 47 | 19 | 47 | 0 | 0 | 0 | 0 | 0 | 101 | 83 | 96 | 119 | 26 | 0 | 0 | 0 | 1 | 0 | 289 | 149 | 1 | 1 | 1 | 1 | 1 | 1 | 1 | 1 | 1 | 1 | 0.984 | 1 | 0.003 |
| SE | 33103 | ENSG00000178035 | IMPDH2 | chr3 | - | 49027709 | 49027916 | 49026518 | 49027047 | 49028198 | 49028322 | 186 | 140 | 216 | 173 | 176 | 0 | 0 | 0 | 0 | 2 | 167 | 108 | 108 | 108 | 182 | 0 | 0 | 0 | 0 | 0 | 298 | 149 | 1 | 1 | 1 | 1 | 1 | 1 | 0.978 | 1 | 1 | 1 | 1 | 1 | -0.004 |
| SE | 33107 | ENSG00000171649 | ZIK1 | chr19 | + | 57584951 | 57584990 | 57584144 | 57584389 | 57588538 | 57588615 | 1 | 0 | 0 | 0 | 0 | 0 | 0 | 0 | 0 | 0 | 0 | 5 | 0 | 0 | 0 | 0 | 4 | 0 | 0 | 0 | 188 | 149 | 1 | 1 | 1 | NA | NA | NA | NA | NA | 0.498 | NA | NA | NA | 0.502 |
| SE | 33109 | ENSG00000171649 | ZIK1 | chr19 | + | 57584951 | 57584990 | 57584144 | 57584612 | 57588538 | 57588615 | 2 | 5 | 0 | 0 | 0 | 3 | 0 | 4 | 0 | 2 | 0 | 2 | 0 | 0 | 0 | 0 | 6 | 0 | 0 | 0 | 188 | 149 | 1 | 1 | 0.346 | 1 | 0 | NA | 0 | NA | 0.209 | NA | NA | NA | 0.128 |
| SE | 33112 | ENSG00000172053 | QARS1 | chr3 | - | 49097109 | 49097287 | 49095932 | 49096079 | 49097991 | 49098117 | 0 | 2 | 2 | 1 | 1 | 49 | 51 | 48 | 62 | 53 | 0 | 0 | 0 | 0 | 0 | 57 | 36 | 30 | 28 | 136 | 298 | 149 | 1 | 1 | 0 | 0.019 | 0.02 | 0.008 | 0.009 | 0 | 0 | 0 | 0 | 0 | 0.011 |
| SE | 33113 | ENSG00000172053 | QARS1 | chr3 | - | 49097133 | 49097287 | 49095932 | 49096079 | 49097991 | 49098117 | 1 | 2 | 2 | 1 | 2 | 49 | 51 | 48 | 62 | 53 | 0 | 0 | 0 | 0 | 0 | 57 | 36 | 30 | 28 | 136 | 298 | 149 | 1 | 1 | 0.01 | 0.019 | 0.02 | 0.008 | 0.019 | 0 | 0 | 0 | 0 | 0 | 0.015 |
| SE | 33115 | ENSG00000172053 | QARS1 | chr3 | - | 49097991 | 49098117 | 49095931 | 49096079 | 49098191 | 49098258 | 120 | 118 | 119 | 158 | 139 | 4 | 0 | 0 | 1 | 1 | 130 | 122 | 47 | 62 | 259 | 0 | 0 | 0 | 5 | 0 | 275 | 149 | 1 | 1 | 0.942 | 1 | 1 | 0.988 | 0.987 | 1 | 1 | 1 | 0.87 | 1 | 0.009 |
| SE | 33120 | ENSG00000172053 | QARS1 | chr3 | - | 49099509 | 49099647 | 49099329 | 49099431 | 49099760 | 49099853 | 234 | 89 | 240 | 164 | 189 | 0 | 0 | 1 | 0 | 0 | 174 | 215 | 115 | 111 | 270 | 0 | 0 | 0 | 0 | 0 | 287 | 149 | 1 | 1 | 1 | 1 | 0.992 | 1 | 1 | 1 | 1 | 1 | 1 | 1 | -0.002 |
| SE | 33122 | ENSG00000172053 | QARS1 | chr3 | - | 49100574 | 49100674 | 49100379 | 49100458 | 49101354 | 49101441 | 656 | 337 | 648 | 577 | 684 | 0 | 0 | 0 | 0 | 0 | 832 | 637 | 497 | 463 | 1116 | 0 | 0 | 0 | 9 | 0 | 249 | 149 | 1 | 1 | 1 | 1 | 1 | 1 | 1 | 1 | 1 | 1 | 0.969 | 1 | 0.006 |
| SE | 33125 | ENSG00000172053 | QARS1 | chr3 | - | 49101619 | 49101705 | 49101354 | 49101441 | 49102204 | 49102265 | 342 | 181 | 372 | 269 | 414 | 0 | 0 | 0 | 0 | 0 | 438 | 349 | 217 | 291 | 581 | 0 | 3 | 2 | 0 | 0 | 235 | 149 | 1 | 1 | 1 | 1 | 1 | 1 | 1 | 1 | 0.987 | 0.986 | 1 | 1 | 0.005 |
| SE | 33126 | ENSG00000172053 | QARS1 | chr3 | - | 49101827 | 49101899 | 49101354 | 49101441 | 49102204 | 49102265 | 107 | 68 | 115 | 96 | 95 | 0 | 0 | 0 | 0 | 0 | 162 | 133 | 114 | 79 | 247 | 0 | 3 | 2 | 0 | 0 | 221 | 149 | 1 | 1 | 1 | 1 | 1 | 1 | 1 | 1 | 0.968 | 0.975 | 1 | 1 | 0.011 |
| SE | 33127 | ENSG00000172053 | QARS1 | chr3 | - | 49101827 | 49101899 | 49101619 | 49101705 | 49102204 | 49102265 | 343 | 169 | 373 | 235 | 359 | 0 | 0 | 2 | 0 | 0 | 361 | 350 | 242 | 260 | 599 | 0 | 0 | 0 | 0 | 0 | 221 | 149 | 1 | 1 | 1 | 1 | 0.992 | 1 | 1 | 1 | 1 | 1 | 1 | 1 | -0.002 |
| SE | 33129 | ENSG00000172053 | QARS1 | chr3 | - | 49102418 | 49102472 | 49102204 | 49102265 | 49103344 | 49103409 | 282 | 187 | 330 | 276 | 238 | 0 | 0 | 0 | 0 | 0 | 449 | 319 | 267 | 226 | 564 | 7 | 0 | 0 | 0 | 5 | 203 | 149 | 1 | 1 | 1 | 1 | 1 | 1 | 1 | 0.979 | 1 | 1 | 1 | 0.988 | 0.007 |
| SE | 33130 | ENSG00000172053 | QARS1 | chr3 | - | 49103344 | 49103409 | 49102418 | 49102472 | 49103630 | 49103706 | 364 | 242 | 448 | 351 | 259 | 0 | 0 | 0 | 0 | 0 | 558 | 402 | 306 | 261 | 685 | 5 | 0 | 0 | 0 | 0 | 214 | 149 | 1 | 1 | 1 | 1 | 1 | 1 | 1 | 0.987 | 1 | 1 | 1 | 1 | 0.003 |
| SE | 33131 | ENSG00000172053 | QARS1 | chr3 | - | 49103344 | 49103431 | 49102418 | 49102472 | 49103630 | 49103706 | 132 | 90 | 169 | 124 | 86 | 0 | 0 | 0 | 0 | 0 | 189 | 185 | 130 | 112 | 204 | 5 | 0 | 0 | 0 | 0 | 236 | 149 | 1 | 1 | 1 | 1 | 1 | 1 | 1 | 0.96 | 1 | 1 | 1 | 1 | 0.008 |
| SE | 33132 | ENSG00000172053 | QARS1 | chr3 | - | 49103344 | 49103466 | 49102418 | 49102472 | 49103630 | 49103706 | 134 | 96 | 179 | 135 | 93 | 0 | 0 | 0 | 0 | 0 | 181 | 185 | 155 | 127 | 220 | 5 | 0 | 0 | 0 | 0 | 271 | 149 | 1 | 1 | 1 | 1 | 1 | 1 | 1 | 0.952 | 1 | 1 | 1 | 1 | 0.01 |
| SE | 33139 | ENSG00000198218 | QRICH1 | chr3 | - | 49044389 | 49044504 | 49033046 | 49033228 | 49047068 | 49047246 | 74 | 50 | 119 | 81 | 139 | 0 | 0 | 0 | 0 | 0 | 119 | 135 | 129 | 148 | 179 | 0 | 0 | 0 | 5 | 0 | 264 | 149 | 1 | 1 | 1 | 1 | 1 | 1 | 1 | 1 | 1 | 1 | 0.944 | 1 | 0.011 |
| SE | 33140 | ENSG00000198218 | QRICH1 | chr3 | - | 49046424 | 49046579 | 49033082 | 49033228 | 49047068 | 49047246 | 215 | 83 | 188 | 181 | 166 | 0 | 0 | 0 | 0 | 0 | 134 | 183 | 151 | 140 | 166 | 0 | 0 | 0 | 5 | 0 | 298 | 149 | 1 | 1 | 1 | 1 | 1 | 1 | 1 | 1 | 1 | 1 | 0.933 | 1 | 0.013 |
| SE | 33142 | ENSG00000198218 | QRICH1 | chr3 | - | 49056861 | 49057890 | 49047068 | 49047246 | 49076708 | 49077038 | 198 | 115 | 151 | 227 | 185 | 0 | 0 | 0 | 1 | 0 | 382 | 247 | 236 | 295 | 293 | 0 | 0 | 0 | 0 | 0 | 298 | 149 | 1 | 1 | 1 | 1 | 1 | 0.991 | 1 | 1 | 1 | 1 | 1 | 1 | -0.002 |
| SE | 33143 | ENSG00000198218 | QRICH1 | chr3 | - | 49092396 | 49092456 | 49076708 | 49077038 | 49093893 | 49094071 | 41 | 22 | 35 | 24 | 31 | 0 | 0 | 0 | 0 | 0 | 83 | 86 | 39 | 92 | 34 | 0 | 0 | 1 | 0 | 0 | 209 | 149 | 1 | 1 | 1 | 1 | 1 | 1 | 1 | 1 | 1 | 0.965 | 1 | 1 | 0.007 |
| SE | 33145 | ENSG00000198218 | QRICH1 | chr3 | - | 49092396 | 49092456 | 49076708 | 49077038 | 49094136 | 49094230 | 40 | 22 | 36 | 24 | 31 | 0 | 1 | 1 | 0 | 0 | 83 | 86 | 39 | 92 | 34 | 0 | 0 | 0 | 0 | 0 | 209 | 149 | 1 | 1 | 1 | 0.94 | 0.962 | 1 | 1 | 1 | 1 | 1 | 1 | 1 | -0.02 |
| SE | 33147 | ENSG00000175161 | CADM2 | chr3 | + | 85802046 | 85802196 | 84959545 | 84959668 | 85883290 | 85883443 | 2 | 1 | 0 | 1 | 0 | 0 | 1 | 0 | 0 | 0 | 0 | 0 | 0 | 0 | 2 | 0 | 0 | 0 | 0 | 0 | 298 | 149 | 1 | 1 | 1 | 0.333 | NA | 1 | NA | NA | NA | NA | NA | 1 | -0.222 |
| SE | 33151 | ENSG00000183726 | TMEM50A | chr1 | + | 25351625 | 25351693 | 25342960 | 25343073 | 25352881 | 25352974 | 870 | 626 | 491 | 797 | 1463 | 0 | 0 | 0 | 1 | 0 | 1126 | 763 | 635 | 816 | 1679 | 0 | 2 | 0 | 0 | 0 | 217 | 149 | 1 | 1 | 1 | 1 | 1 | 0.998 | 1 | 1 | 0.996 | 1 | 1 | 1 | 0 |
| SE | 33152 | ENSG00000183726 | TMEM50A | chr1 | + | 25351625 | 25351693 | 25342960 | 25343073 | 25356792 | 25356853 | 229 | 116 | 125 | 191 | 292 | 0 | 0 | 0 | 0 | 0 | 378 | 212 | 169 | 269 | 466 | 6 | 1 | 0 | 2 | 0 | 217 | 149 | 1 | 1 | 1 | 1 | 1 | 1 | 1 | 0.977 | 0.993 | 1 | 0.989 | 1 | 0.008 |
| SE | 33153 | ENSG00000183726 | TMEM50A | chr1 | + | 25352881 | 25352974 | 25342960 | 25343073 | 25356792 | 25356853 | 885 | 617 | 473 | 790 | 1749 | 0 | 0 | 0 | 0 | 0 | 1365 | 719 | 608 | 895 | 1694 | 6 | 1 | 0 | 2 | 0 | 242 | 149 | 1 | 1 | 1 | 1 | 1 | 1 | 1 | 0.993 | 0.998 | 1 | 0.996 | 1 | 0.003 |
| SE | 33156 | ENSG00000058729 | RIOK2 | chr5 | - | 97166266 | 97166371 | 97165050 | 97165147 | 97167466 | 97167577 | 5 | 10 | 6 | 3 | 0 | 96 | 70 | 95 | 71 | 138 | 0 | 3 | 0 | 2 | 0 | 0 | 42 | 53 | 2 | 0 | 254 | 149 | 1 | 1 | 0.03 | 0.077 | 0.036 | 0.024 | 0 | NA | 0.04 | 0 | 0.37 | NA | -0.103 |
| SE | 33157 | ENSG00000058729 | RIOK2 | chr5 | - | 97168759 | 97168852 | 97166745 | 97167991 | 97171205 | 97171397 | 138 | 108 | 89 | 107 | 174 | 0 | 0 | 0 | 0 | 4 | 0 | 29 | 75 | 12 | 0 | 0 | 0 | 0 | 0 | 0 | 242 | 149 | 1 | 1 | 1 | 1 | 1 | 1 | 0.964 | NA | 1 | 1 | 1 | NA | -0.007 |
| SE | 33158 | ENSG00000058729 | RIOK2 | chr5 | - | 97173174 | 97173263 | 97171205 | 97171397 | 97177115 | 97177291 | 100 | 87 | 83 | 97 | 246 | 0 | 0 | 2 | 1 | 2 | 20 | 17 | 55 | 11 | 6 | 0 | 0 | 0 | 0 | 0 | 238 | 149 | 1 | 1 | 1 | 1 | 0.963 | 0.984 | 0.987 | 1 | 1 | 1 | 1 | 1 | -0.013 |
| SE | 33160 | ENSG00000004766 | VPS50 | chr7 | + | 93239865 | 93239950 | 93232365 | 93232500 | 93252652 | 93252775 | 44 | 42 | 42 | 45 | 12 | 0 | 0 | 0 | 0 | 0 | 0 | 16 | 29 | 6 | 13 | 0 | 0 | 5 | 0 | 0 | 234 | 149 | 1 | 1 | 1 | 1 | 1 | 1 | 1 | NA | 1 | 0.787 | 1 | 1 | 0.053 |
| SE | 33162 | ENSG00000004766 | VPS50 | chr7 | + | 93246101 | 93246129 | 93239865 | 93239934 | 93252652 | 93252775 | 3 | 0 | 2 | 1 | 0 | 68 | 66 | 49 | 66 | 37 | 0 | 0 | 2 | 0 | 0 | 1 | 24 | 20 | 10 | 19 | 177 | 149 | 1 | 1 | 0.036 | 0 | 0.033 | 0.013 | 0 | 0 | 0 | 0.078 | 0 | 0 | 0.001 |
| SE | 33163 | ENSG00000004766 | VPS50 | chr7 | + | 93252652 | 93252775 | 93239865 | 93239934 | 93253859 | 93253931 | 194 | 178 | 141 | 144 | 104 | 2 | 3 | 1 | 0 | 0 | 78 | 79 | 72 | 43 | 44 | 0 | 0 | 0 | 0 | 0 | 272 | 149 | 1 | 1 | 0.982 | 0.97 | 0.987 | 1 | 1 | 1 | 1 | 1 | 1 | 1 | -0.012 |
| SE | 33164 | ENSG00000004766 | VPS50 | chr7 | + | 93252652 | 93252775 | 93239865 | 93239950 | 93253859 | 93253931 | 126 | 112 | 92 | 78 | 70 | 0 | 1 | 0 | 0 | 0 | 77 | 55 | 52 | 33 | 31 | 0 | 0 | 0 | 0 | 0 | 272 | 149 | 1 | 1 | 1 | 0.984 | 1 | 1 | 1 | 1 | 1 | 1 | 1 | 1 | -0.003 |
| SE | 33165 | ENSG00000004766 | VPS50 | chr7 | + | 93272634 | 93272733 | 93271065 | 93271262 | 93276164 | 93276305 | 219 | 166 | 181 | 169 | 372 | 0 | 1 | 0 | 0 | 0 | 47 | 30 | 19 | 16 | 16 | 0 | 0 | 0 | 0 | 0 | 248 | 149 | 1 | 1 | 1 | 0.99 | 1 | 1 | 1 | 1 | 1 | 1 | 1 | 1 | -0.002 |
| SE | 33168 | ENSG00000004766 | VPS50 | chr7 | + | 93323610 | 93323732 | 93311165 | 93311272 | 93334116 | 93334197 | 112 | 47 | 77 | 79 | 219 | 0 | 0 | 1 | 0 | 7 | 0 | 25 | 13 | 0 | 22 | 0 | 0 | 4 | 0 | 0 | 271 | 149 | 1 | 1 | 1 | 1 | 0.977 | 1 | 0.945 | NA | 1 | 0.641 | NA | 1 | 0.104 |
| SE | 33169 | ENSG00000004766 | VPS50 | chr7 | + | 93341426 | 93341575 | 93334116 | 93334197 | 93348710 | 93348798 | 159 | 77 | 84 | 83 | 175 | 0 | 0 | 1 | 0 | 0 | 25 | 34 | 11 | 0 | 1 | 0 | 0 | 0 | 0 | 0 | 298 | 149 | 1 | 1 | 1 | 1 | 0.977 | 1 | 1 | 1 | 1 | 1 | NA | 1 | -0.005 |
| SE | 33182 | ENSG00000145545 | SRD5A1 | chr5 | + | 6651841 | 6652008 | 6633439 | 6633869 | 6656077 | 6656179 | 60 | 37 | 39 | 66 | 38 | 0 | 0 | 0 | 0 | 0 | 319 | 116 | 32 | 84 | 246 | 6 | 1 | 0 | 0 | 0 | 298 | 149 | 1 | 1 | 1 | 1 | 1 | 1 | 1 | 0.964 | 0.983 | 1 | 1 | 1 | 0.011 |
| SE | 33187 | ENSG00000147231 | RADX | chrX | + | 106639526 | 106639687 | 106637759 | 106637924 | 106640551 | 106640721 | 3 | 3 | 5 | 3 | 24 | 0 | 0 | 0 | 0 | 0 | 0 | 2 | 2 | 0 | 0 | 0 | 0 | 4 | 0 | 0 | 298 | 149 | 1 | 1 | 1 | 1 | 1 | 1 | 1 | NA | 1 | 0.2 | NA | NA | 0.4 |
| SE | 33188 | ENSG00000100478 | AP4S1 | chr14 | + | 31032970 | 31033006 | 31025646 | 31025787 | 31066125 | 31066334 | 0 | 0 | 1 | 0 | 0 | 21 | 19 | 34 | 4 | 14 | 0 | 0 | 2 | 2 | 0 | 6 | 20 | 7 | 16 | 18 | 185 | 149 | 1 | 1 | 0 | 0 | 0.023 | 0 | 0 | 0 | 0 | 0.187 | 0.091 | 0 | -0.051 |
| SE | 33189 | ENSG00000100478 | AP4S1 | chr14 | + | 31066125 | 31066334 | 31025637 | 31025787 | 31069842 | 31069929 | 47 | 48 | 69 | 32 | 83 | 0 | 0 | 0 | 2 | 0 | 38 | 55 | 14 | 34 | 57 | 0 | 0 | 0 | 0 | 0 | 298 | 149 | 1 | 1 | 1 | 1 | 1 | 0.889 | 1 | 1 | 1 | 1 | 1 | 1 | -0.022 |
| SE | 33190 | ENSG00000100478 | AP4S1 | chr14 | + | 31072904 | 31072973 | 31069842 | 31069929 | 31080572 | 31080576 | 81 | 56 | 101 | 61 | 177 | 1 | 0 | 0 | 0 | 0 | 75 | 42 | 24 | 28 | 64 | 0 | 0 | 0 | 0 | 0 | 218 | 149 | 1 | 1 | 0.982 | 1 | 1 | 1 | 1 | 1 | 1 | 1 | 1 | 1 | -0.004 |
| SE | 33201 | ENSG00000163930 | BAP1 | chr3 | - | 52407398 | 52407460 | 52407172 | 52407316 | 52407957 | 52408047 | 120 | 68 | 115 | 92 | 88 | 1 | 0 | 0 | 0 | 0 | 126 | 145 | 79 | 111 | 100 | 0 | 0 | 0 | 0 | 0 | 211 | 149 | 1 | 1 | 0.988 | 1 | 1 | 1 | 1 | 1 | 1 | 1 | 1 | 1 | -0.002 |
| SE | 33202 | ENSG00000163930 | BAP1 | chr3 | - | 52408473 | 52408606 | 52407957 | 52408077 | 52409553 | 52409608 | 179 | 107 | 190 | 129 | 267 | 1 | 0 | 0 | 0 | 0 | 202 | 190 | 94 | 157 | 305 | 0 | 0 | 3 | 0 | 0 | 282 | 149 | 1 | 1 | 0.99 | 1 | 1 | 1 | 1 | 1 | 1 | 0.943 | 1 | 1 | 0.009 |
| SE | 33204 | ENSG00000183666 | GUSBP1 | chr5 | + | 21471461 | 21471646 | 21461782 | 21461866 | 21473356 | 21474438 | 0 | 0 | 2 | 0 | 0 | 7 | 2 | 3 | 8 | 1 | 0 | 0 | 0 | 0 | 0 | 12 | 8 | 3 | 0 | 0 | 298 | 149 | 1 | 1 | 0 | 0 | 0.25 | 0 | 0 | 0 | 0 | 0 | NA | NA | 0.05 |
| SE | 33205 | ENSG00000183666 | GUSBP1 | chr5 | + | 21491320 | 21491508 | 21461782 | 21461866 | 21495097 | 21495182 | 115 | 78 | 98 | 85 | 98 | 0 | 0 | 3 | 0 | 0 | 80 | 61 | 37 | 85 | 50 | 0 | 0 | 0 | 2 | 0 | 298 | 149 | 1 | 1 | 1 | 1 | 0.942 | 1 | 1 | 1 | 1 | 1 | 0.955 | 1 | -0.003 |
| SE | 33206 | ENSG00000221978 | CCNL2 | chr1 | - | 1388491 | 1388510 | 1387953 | 1388065 | 1388625 | 1388743 | 3 | 3 | 3 | 2 | 0 | 1 | 1 | 12 | 9 | 4 | 0 | 0 | 0 | 1 | 0 | 0 | 1 | 6 | 3 | 22 | 168 | 149 | 1 | 1 | 0.727 | 0.727 | 0.181 | 0.165 | 0 | NA | 0 | 0 | 0.228 | 0 | 0.303 |
| SE | 33207 | ENSG00000221978 | CCNL2 | chr1 | - | 1388491 | 1388510 | 1387953 | 1388065 | 1390229 | 1390309 | 1 | 0 | 1 | 0 | 0 | 201 | 102 | 222 | 195 | 179 | 0 | 0 | 0 | 0 | 0 | 240 | 137 | 150 | 184 | 190 | 168 | 149 | 1 | 1 | 0.004 | 0 | 0.004 | 0 | 0 | 0 | 0 | 0 | 0 | 0 | 0.002 |
| SE | 33208 | ENSG00000221978 | CCNL2 | chr1 | - | 1388625 | 1388743 | 1387953 | 1388065 | 1388949 | 1389047 | 0 | 1 | 7 | 8 | 0 | 0 | 1 | 1 | 0 | 0 | 0 | 0 | 5 | 3 | 13 | 0 | 0 | 0 | 0 | 0 | 267 | 149 | 1 | 1 | NA | 0.358 | 0.796 | 1 | NA | NA | NA | 1 | 1 | 1 | -0.282 |
| SE | 33210 | ENSG00000221978 | CCNL2 | chr1 | - | 1388625 | 1388743 | 1388491 | 1388510 | 1390229 | 1390371 | 4 | 1 | 6 | 3 | 1 | 1 | 0 | 0 | 0 | 0 | 0 | 0 | 4 | 5 | 13 | 0 | 0 | 0 | 0 | 0 | 267 | 149 | 1 | 1 | 0.691 | 1 | 1 | 1 | 1 | NA | NA | 1 | 1 | 1 | -0.062 |
| SE | 33212 | ENSG00000221978 | CCNL2 | chr1 | - | 1388949 | 1389047 | 1387953 | 1388065 | 1390229 | 1390371 | 0 | 2 | 2 | 0 | 0 | 201 | 102 | 222 | 195 | 179 | 0 | 0 | 0 | 0 | 0 | 240 | 137 | 150 | 184 | 190 | 247 | 149 | 1 | 1 | 0 | 0.012 | 0.005 | 0 | 0 | 0 | 0 | 0 | 0 | 0 | 0.003 |
| SE | 33213 | ENSG00000221978 | CCNL2 | chr1 | - | 1388949 | 1389047 | 1388625 | 1388743 | 1390229 | 1390371 | 0 | 1 | 1 | 1 | 0 | 2 | 0 | 8 | 2 | 5 | 0 | 0 | 0 | 0 | 0 | 0 | 1 | 5 | 4 | 13 | 247 | 149 | 1 | 1 | 0 | 1 | 0.07 | 0.232 | 0 | NA | 0 | 0 | 0 | 0 | 0.26 |
| SE | 33218 | ENSG00000247596 | TWF2 | chr3 | - | 52229660 | 52229782 | 52228611 | 52229201 | 52229919 | 52230070 | 270 | 131 | 220 | 209 | 144 | 0 | 0 | 0 | 0 | 0 | 270 | 222 | 203 | 231 | 374 | 0 | 0 | 0 | 0 | 3 | 271 | 149 | 1 | 1 | 1 | 1 | 1 | 1 | 1 | 1 | 1 | 1 | 1 | 0.986 | 0.003 |
| SE | 33219 | ENSG00000247596 | TWF2 | chr3 | - | 52230869 | 52230995 | 52229919 | 52230070 | 52231126 | 52231231 | 333 | 172 | 261 | 267 | 281 | 0 | 0 | 0 | 0 | 0 | 409 | 441 | 157 | 356 | 609 | 0 | 0 | 0 | 6 | 0 | 275 | 149 | 1 | 1 | 1 | 1 | 1 | 1 | 1 | 1 | 1 | 1 | 0.97 | 1 | 0.006 |
| SE | 33223 | ENSG00000247596 | TWF2 | chr3 | - | 52235028 | 52235106 | 52231443 | 52231539 | 52246678 | 52246788 | 0 | 0 | 2 | 0 | 0 | 0 | 0 | 0 | 0 | 0 | 0 | 1 | 0 | 1 | 0 | 0 | 2 | 0 | 0 | 0 | 227 | 149 | 1 | 1 | NA | NA | 1 | NA | NA | NA | 0.247 | NA | 1 | NA | 0.377 |
| SE | 33226 | ENSG00000023330 | ALAS1 | chr3 | + | 52199209 | 52199440 | 52198085 | 52198255 | 52202506 | 52202734 | 48 | 51 | 70 | 80 | 74 | 0 | 0 | 0 | 0 | 0 | 173 | 81 | 70 | 215 | 269 | 0 | 0 | 0 | 3 | 0 | 298 | 149 | 1 | 1 | 1 | 1 | 1 | 1 | 1 | 1 | 1 | 1 | 0.973 | 1 | 0.005 |
| SE | 33227 | ENSG00000023330 | ALAS1 | chr3 | + | 52199220 | 52199440 | 52198131 | 52198255 | 52202506 | 52202734 | 31 | 36 | 50 | 49 | 71 | 0 | 0 | 0 | 0 | 0 | 151 | 54 | 59 | 137 | 167 | 0 | 0 | 0 | 3 | 0 | 298 | 149 | 1 | 1 | 1 | 1 | 1 | 1 | 1 | 1 | 1 | 1 | 0.958 | 1 | 0.008 |
| SE | 33228 | ENSG00000023330 | ALAS1 | chr3 | + | 52204692 | 52204915 | 52203862 | 52204012 | 52205838 | 52206023 | 172 | 90 | 114 | 116 | 144 | 0 | 0 | 0 | 0 | 0 | 355 | 166 | 123 | 323 | 215 | 8 | 0 | 0 | 0 | 0 | 298 | 149 | 1 | 1 | 1 | 1 | 1 | 1 | 1 | 0.957 | 1 | 1 | 1 | 1 | 0.009 |
| SE | 33229 | ENSG00000023330 | ALAS1 | chr3 | + | 52204692 | 52204915 | 52203862 | 52204012 | 52206571 | 52206751 | 85 | 52 | 54 | 75 | 103 | 0 | 2 | 1 | 1 | 0 | 242 | 78 | 83 | 205 | 151 | 20 | 3 | 2 | 0 | 16 | 298 | 149 | 1 | 1 | 1 | 0.929 | 0.964 | 0.974 | 1 | 0.858 | 0.929 | 0.954 | 1 | 0.825 | 0.06 |
| SE | 33230 | ENSG00000023330 | ALAS1 | chr3 | + | 52205838 | 52206023 | 52203862 | 52204012 | 52206571 | 52206751 | 88 | 46 | 50 | 44 | 83 | 0 | 2 | 1 | 1 | 0 | 88 | 61 | 51 | 91 | 54 | 20 | 3 | 2 | 0 | 16 | 298 | 149 | 1 | 1 | 1 | 0.92 | 0.962 | 0.957 | 1 | 0.688 | 0.91 | 0.927 | 1 | 0.628 | 0.137 |
| SE | 33233 | ENSG00000023330 | ALAS1 | chr3 | + | 52208082 | 52208247 | 52206571 | 52206751 | 52212257 | 52212420 | 47 | 33 | 43 | 51 | 57 | 3 | 0 | 0 | 0 | 1 | 113 | 45 | 32 | 121 | 128 | 0 | 0 | 0 | 0 | 0 | 298 | 149 | 1 | 1 | 0.887 | 1 | 1 | 1 | 0.966 | 1 | 1 | 1 | 1 | 1 | -0.029 |
| SE | 33234 | ENSG00000023330 | ALAS1 | chr3 | + | 52211282 | 52211551 | 52206631 | 52206751 | 52212257 | 52212420 | 45 | 20 | 33 | 41 | 54 | 3 | 0 | 0 | 0 | 1 | 97 | 57 | 41 | 100 | 106 | 0 | 0 | 0 | 0 | 0 | 298 | 149 | 1 | 1 | 0.882 | 1 | 1 | 1 | 0.964 | 1 | 1 | 1 | 1 | 1 | -0.031 |
| SE | 33235 | ENSG00000023330 | ALAS1 | chr3 | + | 52211282 | 52211551 | 52208082 | 52208247 | 52212257 | 52212420 | 112 | 47 | 96 | 106 | 137 | 0 | 0 | 0 | 1 | 0 | 207 | 116 | 92 | 255 | 224 | 0 | 0 | 0 | 0 | 0 | 298 | 149 | 1 | 1 | 1 | 1 | 1 | 0.981 | 1 | 1 | 1 | 1 | 1 | 1 | -0.004 |
| SE | 33236 | ENSG00000187961 | KLHL17 | chr1 | + | 961292 | 961552 | 960583 | 960800 | 961628 | 961750 | 3 | 0 | 0 | 1 | 0 | 0 | 0 | 0 | 0 | 0 | 0 | 0 | 0 | 0 | 4 | 0 | 2 | 0 | 0 | 0 | 298 | 149 | 1 | 1 | 1 | NA | NA | 1 | NA | NA | 0 | NA | NA | 1 | 0.5 |
| SE | 33238 | ENSG00000164087 | POC1A | chr3 | - | 52096568 | 52096712 | 52075225 | 52075985 | 52125112 | 52125181 | 10 | 4 | 3 | 6 | 11 | 1 | 1 | 0 | 0 | 0 | 8 | 0 | 2 | 13 | 11 | 0 | 0 | 0 | 0 | 0 | 293 | 149 | 1 | 1 | 0.836 | 0.67 | 1 | 1 | 1 | 1 | NA | 1 | 1 | 1 | -0.099 |
| SE | 33239 | ENSG00000164087 | POC1A | chr3 | - | 52122378 | 52122477 | 52075285 | 52075985 | 52125112 | 52125181 | 1 | 1 | 3 | 2 | 1 | 1 | 1 | 0 | 0 | 0 | 10 | 10 | 1 | 3 | 14 | 0 | 0 | 0 | 0 | 0 | 248 | 149 | 1 | 1 | 0.375 | 0.375 | 1 | 1 | 1 | 1 | 1 | 1 | 1 | 1 | -0.25 |
| SE | 33241 | ENSG00000164087 | POC1A | chr3 | - | 52138168 | 52138302 | 52125112 | 52125181 | 52145845 | 52145961 | 0 | 2 | 3 | 4 | 9 | 0 | 0 | 0 | 0 | 0 | 21 | 11 | 2 | 18 | 35 | 0 | 0 | 3 | 0 | 0 | 283 | 149 | 1 | 1 | NA | 1 | 1 | 1 | 1 | 1 | 1 | 0.26 | 1 | 1 | 0.148 |
| SE | 33242 | ENSG00000162244 | RPL29 | chr3 | - | 51994772 | 51994857 | 51993627 | 51994126 | 51995041 | 51995106 | 8 | 3 | 8 | 8 | 8 | 3485 | 1812 | 4503 | 3680 | 6478 | 0 | 5 | 3 | 6 | 0 | 4378 | 3427 | 2421 | 2135 | 4888 | 234 | 149 | 1 | 1 | 0.001 | 0.001 | 0.001 | 0.001 | 0.001 | 0 | 0.001 | 0.001 | 0.002 | 0 | 0 |
| SE | 33243 | ENSG00000162244 | RPL29 | chr3 | - | 51995041 | 51995106 | 51993521 | 51994126 | 51995424 | 51995469 | 3486 | 1811 | 4502 | 3680 | 6478 | 0 | 0 | 0 | 0 | 0 | 4398 | 3433 | 2426 | 2137 | 4890 | 12 | 0 | 0 | 0 | 9 | 214 | 149 | 1 | 1 | 1 | 1 | 1 | 1 | 1 | 0.996 | 1 | 1 | 1 | 0.997 | 0.001 |
| SE | 33246 | ENSG00000243989 | ACY1 | chr3 | + | 51985206 | 51985271 | 51984046 | 51984158 | 51985360 | 51985465 | 5 | 1 | 3 | 7 | 2 | 0 | 1 | 0 | 4 | 0 | 11 | 3 | 0 | 1 | 7 | 0 | 0 | 0 | 0 | 0 | 214 | 149 | 1 | 1 | 1 | 0.41 | 1 | 0.549 | 1 | 1 | 1 | NA | 1 | 1 | -0.208 |
| SE | 33247 | ENSG00000243989 | ACY1 | chr3 | + | 51985206 | 51985271 | 51984046 | 51984158 | 51985851 | 51985946 | 1 | 0 | 2 | 0 | 0 | 0 | 0 | 0 | 0 | 5 | 11 | 0 | 0 | 0 | 0 | 0 | 0 | 0 | 0 | 0 | 214 | 149 | 1 | 1 | 1 | NA | 1 | NA | 0 | 1 | NA | NA | NA | NA | -0.333 |
| SE | 33249 | ENSG00000243989 | ACY1 | chr3 | + | 51985360 | 51985465 | 51984046 | 51984158 | 51985851 | 51985897 | 7 | 2 | 11 | 6 | 2 | 0 | 0 | 0 | 0 | 5 | 28 | 4 | 5 | 3 | 0 | 0 | 0 | 0 | 0 | 0 | 254 | 149 | 1 | 1 | 1 | 1 | 1 | 1 | 0.19 | 1 | 1 | 1 | 1 | NA | -0.162 |
| SE | 33251 | ENSG00000243989 | ACY1 | chr3 | + | 51985400 | 51985465 | 51984046 | 51984158 | 51985851 | 51985946 | 4 | 0 | 9 | 2 | 2 | 0 | 0 | 0 | 0 | 5 | 16 | 4 | 5 | 2 | 0 | 0 | 0 | 0 | 0 | 0 | 214 | 149 | 1 | 1 | 1 | NA | 1 | 1 | 0.218 | 1 | 1 | 1 | 1 | NA | -0.196 |
| SE | 33256 | ENSG00000248487 | ABHD14A | chr3 | + | 51978258 | 51978374 | 51977870 | 51978082 | 51980392 | 51980468 | 18 | 5 | 14 | 23 | 13 | 0 | 0 | 0 | 1 | 0 | 29 | 2 | 11 | 8 | 28 | 0 | 0 | 0 | 0 | 0 | 265 | 149 | 1 | 1 | 1 | 1 | 1 | 0.928 | 1 | 1 | 1 | 1 | 1 | 1 | -0.014 |
| SE | 33260 | ENSG00000257335 | MGAM | chr7 | + | 142004389 | 142004488 | 141945820 | 141945997 | 142005528 | 142005657 | 0 | 0 | 0 | 1 | 0 | 0 | 0 | 0 | 0 | 0 | 0 | 6 | 0 | 2 | 0 | 0 | 0 | 0 | 1 | 0 | 248 | 149 | 1 | 1 | NA | NA | NA | 1 | NA | NA | 1 | NA | 0.546 | NA | 0.227 |
| SE | 33261 | ENSG00000257335 | MGAM | chr7 | + | 142004389 | 142004488 | 141995878 | 141995930 | 142005528 | 142005657 | 0 | 0 | 0 | 1 | 0 | 2 | 0 | 0 | 0 | 0 | 0 | 4 | 0 | 2 | 0 | 0 | 0 | 7 | 0 | 0 | 248 | 149 | 1 | 1 | 0 | NA | NA | 1 | NA | NA | 1 | 0 | 1 | NA | -0.167 |
| SE | 33263 | ENSG00000257335 | MGAM | chr7 | + | 142036168 | 142036285 | 142034669 | 142034841 | 142036822 | 142036977 | 34 | 13 | 16 | 19 | 24 | 0 | 0 | 0 | 0 | 0 | 1 | 82 | 132 | 95 | 5 | 0 | 0 | 1 | 0 | 0 | 266 | 149 | 1 | 1 | 1 | 1 | 1 | 1 | 1 | 1 | 1 | 0.987 | 1 | 1 | 0.003 |
| SE | 33264 | ENSG00000257335 | MGAM | chr7 | + | 142047784 | 142047873 | 142040721 | 142040846 | 142050234 | 142050284 | 40 | 3 | 11 | 21 | 24 | 0 | 0 | 0 | 0 | 0 | 53 | 42 | 105 | 99 | 24 | 0 | 0 | 1 | 0 | 0 | 238 | 149 | 1 | 1 | 1 | 1 | 1 | 1 | 1 | 1 | 1 | 0.985 | 1 | 1 | 0.003 |
| SE | 33266 | ENSG00000257335 | MGAM | chr7 | + | 142055999 | 142056096 | 142055557 | 142055726 | 142056829 | 142056942 | 20 | 4 | 14 | 7 | 33 | 1 | 0 | 0 | 0 | 0 | 3 | 38 | 59 | 63 | 30 | 0 | 0 | 0 | 0 | 0 | 246 | 149 | 1 | 1 | 0.924 | 1 | 1 | 1 | 1 | 1 | 1 | 1 | 1 | 1 | -0.015 |
| SE | 33268 | ENSG00000257335 | MGAM | chr7 | + | 142067340 | 142067425 | 142066572 | 142066721 | 142068646 | 142068703 | 0 | 6 | 4 | 2 | 21 | 0 | 0 | 0 | 0 | 0 | 14 | 24 | 31 | 102 | 21 | 0 | 0 | 4 | 1 | 0 | 234 | 149 | 1 | 1 | NA | 1 | 1 | 1 | 1 | 1 | 1 | 0.832 | 0.985 | 1 | 0.037 |
| SE | 33269 | ENSG00000257335 | MGAM | chr7 | + | 142082474 | 142082571 | 142082041 | 142082210 | 142083300 | 142083413 | 0 | 6 | 7 | 15 | 12 | 0 | 0 | 0 | 0 | 0 | 12 | 33 | 117 | 149 | 56 | 0 | 0 | 3 | 0 | 0 | 246 | 149 | 1 | 1 | NA | 1 | 1 | 1 | 1 | 1 | 1 | 0.959 | 1 | 1 | 0.008 |
| SE | 33270 | ENSG00000257335 | MGAM | chr7 | + | 142086654 | 142086717 | 142086217 | 142086328 | 142091912 | 142092047 | 0 | 0 | 6 | 20 | 6 | 0 | 0 | 0 | 0 | 0 | 10 | 39 | 78 | 55 | 8 | 0 | 0 | 1 | 0 | 0 | 212 | 149 | 1 | 1 | NA | NA | 1 | 1 | 1 | 1 | 1 | 0.982 | 1 | 1 | 0.004 |
| SE | 33271 | ENSG00000257335 | MGAM | chr7 | + | 142092520 | 142092608 | 142091912 | 142092047 | 142093411 | 142093550 | 0 | 3 | 6 | 25 | 36 | 0 | 0 | 0 | 0 | 0 | 34 | 36 | 157 | 129 | 12 | 0 | 3 | 1 | 0 | 0 | 237 | 149 | 1 | 1 | NA | 1 | 1 | 1 | 1 | 1 | 0.883 | 0.99 | 1 | 1 | 0.025 |
| SE | 33273 | ENSG00000257335 | MGAM | chr7 | + | 142100801 | 142100890 | 142099612 | 142099737 | 142102629 | 142102679 | 48 | 8 | 22 | 21 | 25 | 0 | 0 | 0 | 0 | 0 | 5 | 76 | 144 | 149 | 42 | 0 | 0 | 1 | 0 | 0 | 238 | 149 | 1 | 1 | 1 | 1 | 1 | 1 | 1 | 1 | 1 | 0.989 | 1 | 1 | 0.002 |
| SE | 33274 | ENSG00000257335 | MGAM | chr7 | + | 142102629 | 142102679 | 142100801 | 142100890 | 142103268 | 142103439 | 35 | 7 | 14 | 12 | 20 | 0 | 0 | 0 | 0 | 0 | 0 | 57 | 84 | 128 | 25 | 0 | 0 | 1 | 0 | 0 | 199 | 149 | 1 | 1 | 1 | 1 | 1 | 1 | 1 | NA | 1 | 0.984 | 1 | 1 | 0.004 |
| SE | 33275 | ENSG00000043093 | DCUN1D1 | chr3 | - | 182947237 | 182947334 | 182938073 | 182945173 | 182961225 | 182961356 | 149 | 97 | 131 | 168 | 329 | 0 | 1 | 0 | 0 | 0 | 14 | 81 | 39 | 24 | 18 | 0 | 0 | 0 | 0 | 0 | 246 | 149 | 1 | 1 | 1 | 0.983 | 1 | 1 | 1 | 1 | 1 | 1 | 1 | 1 | -0.003 |
| SE | 33276 | ENSG00000043093 | DCUN1D1 | chr3 | - | 182947549 | 182947632 | 182947237 | 182947334 | 182961225 | 182961356 | 358 | 202 | 263 | 342 | 486 | 0 | 0 | 0 | 1 | 0 | 40 | 175 | 85 | 46 | 28 | 0 | 0 | 0 | 0 | 0 | 232 | 149 | 1 | 1 | 1 | 1 | 1 | 0.995 | 1 | 1 | 1 | 1 | 1 | 1 | -0.001 |
| SE | 33277 | ENSG00000043093 | DCUN1D1 | chr3 | - | 182963880 | 182964049 | 182961225 | 182961356 | 182965536 | 182965753 | 757 | 618 | 570 | 803 | 1095 | 0 | 0 | 0 | 0 | 1 | 171 | 552 | 400 | 246 | 181 | 0 | 0 | 0 | 0 | 0 | 298 | 149 | 1 | 1 | 1 | 1 | 1 | 1 | 0.998 | 1 | 1 | 1 | 1 | 1 | 0 |
| SE | 33278 | ENSG00000043093 | DCUN1D1 | chr3 | - | 182965536 | 182965753 | 182963880 | 182964049 | 182980486 | 182980538 | 389 | 309 | 284 | 362 | 548 | 3 | 0 | 2 | 0 | 0 | 77 | 253 | 174 | 117 | 107 | 0 | 0 | 0 | 0 | 0 | 298 | 149 | 1 | 1 | 0.985 | 1 | 0.986 | 1 | 1 | 1 | 1 | 1 | 1 | 1 | -0.006 |
| SE | 33279 | ENSG00000043093 | DCUN1D1 | chr3 | - | 182965678 | 182965753 | 182963880 | 182964049 | 182980486 | 182980520 | 2 | 8 | 7 | 13 | 13 | 3 | 0 | 2 | 0 | 0 | 0 | 4 | 7 | 4 | 3 | 0 | 0 | 0 | 0 | 0 | 224 | 149 | 1 | 1 | 0.307 | 1 | 0.7 | 1 | 1 | NA | 1 | 1 | 1 | 1 | -0.199 |
| SE | 33286 | ENSG00000213186 | TRIM59 | chr3 | - | 160447731 | 160447842 | 160438840 | 160439186 | 160448725 | 160448795 | 0 | 2 | 1 | 0 | 8 | 37 | 34 | 41 | 37 | 74 | 0 | 8 | 7 | 0 | 0 | 0 | 18 | 24 | 7 | 10 | 260 | 149 | 1 | 1 | 0 | 0.033 | 0.014 | 0 | 0.058 | NA | 0.203 | 0.143 | 0 | 0 | -0.065 |
| SE | 33287 | ENSG00000213186 | TRIM59 | chr3 | - | 160448725 | 160448795 | 160435502 | 160439186 | 160449471 | 160449725 | 24 | 28 | 29 | 20 | 66 | 0 | 0 | 0 | 1 | 0 | 0 | 16 | 17 | 7 | 10 | 0 | 0 | 0 | 0 | 0 | 219 | 149 | 1 | 1 | 1 | 1 | 1 | 0.932 | 1 | NA | 1 | 1 | 1 | 1 | -0.014 |
| SE | 33288 | ENSG00000213186 | TRIM59 | chr3 | - | 160448725 | 160448795 | 160435502 | 160439186 | 160449683 | 160449760 | 25 | 28 | 32 | 27 | 69 | 1 | 2 | 0 | 1 | 0 | 0 | 12 | 17 | 4 | 10 | 0 | 0 | 0 | 0 | 0 | 219 | 149 | 1 | 1 | 0.944 | 0.905 | 1 | 0.948 | 1 | NA | 1 | 1 | 1 | 1 | -0.041 |
| SE | 33289 | ENSG00000213186 | TRIM59 | chr3 | - | 160448725 | 160448795 | 160435502 | 160439186 | 160449716 | 160449765 | 36 | 35 | 36 | 30 | 71 | 3 | 2 | 9 | 6 | 2 | 0 | 14 | 24 | 4 | 10 | 0 | 7 | 0 | 0 | 3 | 219 | 149 | 1 | 1 | 0.891 | 0.923 | 0.731 | 0.773 | 0.96 | NA | 0.576 | 1 | 1 | 0.694 | 0.038 |
| SE | 33291 | ENSG00000123737 | EXOSC9 | chr4 | + | 121804621 | 121804759 | 121802914 | 121803017 | 121807539 | 121807622 | 174 | 146 | 223 | 208 | 321 | 0 | 0 | 1 | 2 | 0 | 125 | 172 | 208 | 59 | 30 | 0 | 3 | 1 | 0 | 0 | 287 | 149 | 1 | 1 | 1 | 1 | 0.991 | 0.982 | 1 | 1 | 1 | 0.973 | 0.968 | 1 | 0.006 |
| SE | 33292 | ENSG00000123737 | EXOSC9 | chr4 | + | 121807539 | 121807622 | 121802914 | 121803017 | 121811582 | 121811641 | 1 | 0 | 1 | 2 | 0 | 0 | 0 | 2 | 0 | 0 | 0 | 0 | 3 | 1 | 0 | 0 | 0 | 0 | 0 | 0 | 232 | 149 | 1 | 1 | 1 | NA | 0.243 | 1 | NA | NA | NA | 1 | 1 | NA | -0.252 |
| SE | 33293 | ENSG00000123737 | EXOSC9 | chr4 | + | 121807539 | 121807622 | 121804621 | 121804759 | 121809966 | 121810073 | 99 | 90 | 141 | 150 | 199 | 0 | 0 | 0 | 1 | 0 | 106 | 94 | 143 | 41 | 24 | 0 | 3 | 0 | 0 | 0 | 232 | 149 | 1 | 1 | 1 | 1 | 1 | 0.99 | 1 | 1 | 0.953 | 1 | 1 | 1 | 0.007 |
| SE | 33294 | ENSG00000123737 | EXOSC9 | chr4 | + | 121809833 | 121809916 | 121807539 | 121807622 | 121811582 | 121811671 | 3 | 2 | 0 | 6 | 8 | 1 | 0 | 0 | 0 | 0 | 0 | 0 | 1 | 0 | 0 | 0 | 0 | 0 | 0 | 0 | 232 | 149 | 1 | 1 | 0.658 | 1 | NA | 1 | 1 | NA | NA | 1 | NA | NA | -0.086 |
| SE | 33295 | ENSG00000123737 | EXOSC9 | chr4 | + | 121809966 | 121810073 | 121807539 | 121807622 | 121811582 | 121811671 | 31 | 39 | 42 | 60 | 61 | 1 | 0 | 0 | 0 | 0 | 52 | 31 | 64 | 31 | 9 | 0 | 0 | 0 | 0 | 0 | 256 | 149 | 1 | 1 | 0.947 | 1 | 1 | 1 | 1 | 1 | 1 | 1 | 1 | 1 | -0.011 |
| SE | 33296 | ENSG00000123737 | EXOSC9 | chr4 | + | 121809966 | 121810099 | 121804621 | 121804759 | 121813233 | 121813380 | 1 | 0 | 0 | 1 | 0 | 0 | 1 | 0 | 0 | 0 | 0 | 1 | 0 | 0 | 0 | 0 | 0 | 0 | 0 | 0 | 282 | 149 | 1 | 1 | 1 | 0 | NA | 1 | NA | NA | 1 | NA | NA | NA | -0.333 |
| SE | 33297 | ENSG00000123737 | EXOSC9 | chr4 | + | 121809966 | 121810099 | 121807539 | 121807622 | 121811582 | 121811641 | 119 | 113 | 119 | 174 | 152 | 1 | 0 | 0 | 0 | 0 | 90 | 97 | 178 | 121 | 9 | 0 | 0 | 0 | 0 | 0 | 282 | 149 | 1 | 1 | 0.984 | 1 | 1 | 1 | 1 | 1 | 1 | 1 | 1 | 1 | -0.003 |
| SE | 33298 | ENSG00000123737 | EXOSC9 | chr4 | + | 121811582 | 121811671 | 121809966 | 121810099 | 121813233 | 121813293 | 132 | 129 | 125 | 168 | 136 | 1 | 0 | 0 | 0 | 0 | 55 | 104 | 186 | 144 | 1 | 0 | 0 | 0 | 0 | 0 | 238 | 149 | 1 | 1 | 0.988 | 1 | 1 | 1 | 1 | 1 | 1 | 1 | 1 | 1 | -0.002 |
| SE | 33299 | ENSG00000123737 | EXOSC9 | chr4 | + | 121813233 | 121813380 | 121811582 | 121811671 | 121813865 | 121814047 | 264 | 196 | 248 | 261 | 385 | 1 | 1 | 0 | 0 | 0 | 94 | 236 | 348 | 151 | 7 | 0 | 1 | 0 | 0 | 0 | 296 | 149 | 1 | 1 | 0.993 | 0.99 | 1 | 1 | 1 | 1 | 0.992 | 1 | 1 | 1 | -0.002 |
| SE | 33301 | ENSG00000123737 | EXOSC9 | chr4 | + | 121816368 | 121816447 | 121813754 | 121814047 | 121816771 | 121816840 | 159 | 139 | 134 | 132 | 277 | 0 | 1 | 0 | 1 | 1 | 80 | 171 | 223 | 116 | 18 | 0 | 0 | 0 | 6 | 0 | 228 | 149 | 1 | 1 | 1 | 0.989 | 1 | 0.989 | 0.995 | 1 | 1 | 1 | 0.927 | 1 | 0.009 |
| SE | 33302 | ENSG00000125247 | TMTC4 | chr13 | - | 100612397 | 100612510 | 100606357 | 100606427 | 100614315 | 100614430 | 35 | 27 | 13 | 25 | 36 | 0 | 0 | 0 | 0 | 0 | 5 | 11 | 9 | 19 | 0 | 0 | 0 | 1 | 0 | 0 | 262 | 149 | 1 | 1 | 1 | 1 | 1 | 1 | 1 | 1 | 1 | 0.837 | 1 | NA | 0.041 |
| SE | 33303 | ENSG00000125247 | TMTC4 | chr13 | - | 100635678 | 100635779 | 100635064 | 100635195 | 100636531 | 100636632 | 0 | 0 | 2 | 0 | 1 | 21 | 1 | 12 | 7 | 1 | 0 | 0 | 0 | 1 | 0 | 0 | 5 | 12 | 15 | 6 | 250 | 149 | 1 | 1 | 0 | 0 | 0.09 | 0 | 0.373 | NA | 0 | 0 | 0.038 | 0 | 0.083 |
| SE | 33305 | ENSG00000125247 | TMTC4 | chr13 | - | 100662963 | 100663180 | 100642210 | 100642311 | 100664220 | 100664336 | 29 | 12 | 45 | 19 | 12 | 3 | 0 | 0 | 0 | 0 | 23 | 7 | 6 | 10 | 9 | 0 | 0 | 0 | 0 | 0 | 298 | 149 | 1 | 1 | 0.829 | 1 | 1 | 1 | 1 | 1 | 1 | 1 | 1 | 1 | -0.034 |
| SE | 33307 | ENSG00000125247 | TMTC4 | chr13 | - | 100670282 | 100670569 | 100668578 | 100668794 | 100674743 | 100674879 | 6 | 10 | 3 | 7 | 0 | 7 | 0 | 4 | 1 | 0 | 11 | 0 | 6 | 2 | 0 | 17 | 0 | 3 | 1 | 8 | 298 | 149 | 1 | 1 | 0.3 | 1 | 0.273 | 0.778 | NA | 0.244 | NA | 0.5 | 0.5 | 0 | 0.277 |
| SE | 33309 | ENSG00000166128 | RAB8B | chr15 | + | 63223041 | 63223073 | 63189559 | 63189748 | 63223847 | 63223979 | 17 | 4 | 11 | 2 | 12 | 10 | 4 | 1 | 3 | 1 | 0 | 8 | 17 | 7 | 0 | 0 | 7 | 7 | 0 | 0 | 181 | 149 | 1 | 1 | 0.583 | 0.452 | 0.901 | 0.354 | 0.908 | NA | 0.485 | 0.667 | 1 | NA | -0.078 |
| SE | 33312 | ENSG00000166128 | RAB8B | chr15 | + | 63223847 | 63223979 | 63189602 | 63189748 | 63249644 | 63249705 | 10 | 4 | 0 | 2 | 1 | 1 | 3 | 0 | 0 | 4 | 0 | 7 | 8 | 0 | 0 | 0 | 0 | 0 | 0 | 0 | 281 | 149 | 1 | 1 | 0.841 | 0.414 | NA | 1 | 0.117 | NA | 1 | 1 | NA | NA | -0.407 |
| SE | 33313 | ENSG00000166128 | RAB8B | chr15 | + | 63223847 | 63223979 | 63223041 | 63223073 | 63244755 | 63244816 | 44 | 19 | 54 | 29 | 36 | 14 | 12 | 15 | 17 | 49 | 0 | 27 | 52 | 18 | 0 | 0 | 19 | 31 | 7 | 0 | 281 | 149 | 1 | 1 | 0.625 | 0.456 | 0.656 | 0.475 | 0.28 | NA | 0.43 | 0.471 | 0.577 | NA | 0.006 |
| SE | 33314 | ENSG00000166128 | RAB8B | chr15 | + | 63244755 | 63244816 | 63189593 | 63189748 | 63249644 | 63249705 | 1033 | 880 | 609 | 850 | 951 | 1 | 3 | 0 | 0 | 4 | 120 | 747 | 829 | 234 | 116 | 0 | 0 | 0 | 0 | 0 | 210 | 149 | 1 | 1 | 0.999 | 0.995 | 1 | 1 | 0.994 | 1 | 1 | 1 | 1 | 1 | -0.002 |
| SE | 33315 | ENSG00000166128 | RAB8B | chr15 | + | 63244755 | 63244816 | 63189593 | 63189748 | 63255506 | 63255584 | 262 | 173 | 171 | 212 | 199 | 0 | 0 | 0 | 0 | 0 | 35 | 225 | 219 | 53 | 21 | 0 | 0 | 1 | 0 | 0 | 210 | 149 | 1 | 1 | 1 | 1 | 1 | 1 | 1 | 1 | 1 | 0.994 | 1 | 1 | 0.001 |
| SE | 33316 | ENSG00000166128 | RAB8B | chr15 | + | 63244755 | 63244816 | 63223847 | 63223979 | 63249644 | 63249705 | 443 | 368 | 252 | 319 | 577 | 0 | 0 | 0 | 0 | 0 | 47 | 254 | 314 | 92 | 43 | 0 | 0 | 1 | 0 | 0 | 210 | 149 | 1 | 1 | 1 | 1 | 1 | 1 | 1 | 1 | 1 | 0.996 | 1 | 1 | 0.001 |
| SE | 33317 | ENSG00000166128 | RAB8B | chr15 | + | 63249644 | 63249705 | 63189559 | 63189748 | 63255506 | 63255584 | 416 | 322 | 241 | 338 | 481 | 0 | 0 | 0 | 0 | 0 | 46 | 251 | 238 | 104 | 12 | 0 | 0 | 1 | 0 | 0 | 210 | 149 | 1 | 1 | 1 | 1 | 1 | 1 | 1 | 1 | 1 | 0.994 | 1 | 1 | 0.001 |
| SE | 33318 | ENSG00000166128 | RAB8B | chr15 | + | 63249644 | 63249705 | 63244755 | 63244816 | 63255506 | 63255584 | 1224 | 1052 | 717 | 1007 | 1297 | 5 | 3 | 4 | 1 | 3 | 131 | 809 | 911 | 314 | 100 | 0 | 2 | 5 | 4 | 0 | 210 | 149 | 1 | 1 | 0.994 | 0.996 | 0.992 | 0.999 | 0.997 | 1 | 0.997 | 0.992 | 0.982 | 1 | 0.001 |

| | | | | | | | | | | | | | | |
|---|---|---|---|---|---|---|---|---|---|---|---|---|---|---|
| SE | 33319 | ENSG00000166128 | RAB8B | chr15 | + | 63251283 | 63251342 | 63249644 | 63249705 | 63255506 | 63255584 | 1 | 0 | [illegible] |
| SE | 33320 | ENSG00000166128 | RAB8B | chr15 | + | 63255506 | 63255584 | 63249644 | 63249705 | 63256504 | 63256594 | 1909 | 1887 | [illegible] |
| SE | 33321 | ENSG00000166128 | RAB8B | chr15 | + | 63256504 | 63256594 | 63249644 | 63249705 | 63259626 | 63259692 | 516 | 301 | [illegible] |
| SE | 33322 | ENSG00000166128 | RAB8B | chr15 | + | 63262691 | 63262742 | 63259626 | 63259692 | 63263526 | 63263596 | 860 | 513 | [illegible] |
| SE | 33324 | ENSG00000188313 | PLSCR1 | chr3 | - | 146520165 | 146520180 | 146517011 | 146517167 | 146525604 | 146525647 | 0 | 0 | [illegible] |
| SE | 33327 | ENSG00000188313 | PLSCR1 | chr3 | - | 146521543 | 146521705 | 146516902 | 146517167 | 146521832 | 146521981 | 270 | 302 | [illegible] |
| SE | 33330 | ENSG00000188313 | PLSCR1 | chr3 | - | 146521543 | 146522053 | 146517005 | 146517167 | 146533469 | 146533550 | 113 | 122 | [illegible] |
| SE | 33331 | ENSG00000188313 | PLSCR1 | chr3 | - | 146521832 | 146521981 | 146517088 | 146517167 | 146525604 | 146525647 | 6 | 1 | [illegible] |
| SE | 33332 | ENSG00000188313 | PLSCR1 | chr3 | - | 146521832 | 146521981 | 146521543 | 146521705 | 146525604 | 146525647 | 163 | 181 | [illegible] |
| SE | 33336 | ENSG00000188313 | PLSCR1 | chr3 | - | 146521832 | 146522053 | 146521543 | 146521705 | 146525604 | 146525647 | 315 | 325 | [illegible] |
| SE | 33337 | ENSG00000188313 | PLSCR1 | chr3 | - | 146525604 | 146525647 | 146517005 | 146517167 | 146533469 | 146533550 | 16 | 15 | [illegible] |
| SE | 33338 | ENSG00000188313 | PLSCR1 | chr3 | - | 146525604 | 146525647 | 146521543 | 146522053 | 146533469 | 146533550 | 80 | 82 | [illegible] |
| SE | 33340 | ENSG00000188313 | PLSCR1 | chr3 | - | 146536539 | 146536565 | 146533469 | 146533550 | 146544466 | 146544530 | 157 | 124 | [illegible] |
| SE | 33341 | ENSG00000188313 | PLSCR1 | chr3 | - | 146537995 | 146538044 | 146536539 | 146536565 | 146541133 | 146541180 | 6 | 7 | [illegible] |
| SE | 33346 | ENSG00000105373 | NOP53 | chr19 | + | 47750907 | 47751107 | 47750177 | 47750286 | 47751519 | 47751590 | 4607 | 2537 | [illegible] |
| SE | 33357 | ENSG00000187742 | SECISBP2 | chr9 | + | 89319651 | 89319797 | 89318499 | 89318612 | 89325426 | 89325676 | 51 | 30 | [illegible] |
| SE | 33360 | ENSG00000187742 | SECISBP2 | chr9 | + | 89332907 | 89332986 | 89328659 | 89328886 | 89338457 | 89338580 | 54 | 57 | [illegible] |
| SE | 33365 | ENSG00000187742 | SECISBP2 | chr9 | + | 89334521 | 89334730 | 89334024 | 89334155 | 89338457 | 89338580 | 104 | 101 | [illegible] |
| SE | 33367 | ENSG00000187742 | SECISBP2 | chr9 | + | 89334635 | 89334730 | 89334024 | 89334155 | 89338457 | 89338510 | 79 | 73 | [illegible] |
| SE | 33372 | ENSG00000213523 | SRA1 | chr5 | - | 140551060 | 140551169 | 140550067 | 140550911 | 140557146 | 140557446 | 119 | 74 | [illegible] |
| SE | 33373 | ENSG00000213523 | SRA1 | chr5 | - | 140551060 | 140552184 | 140550533 | 140550911 | 140557146 | 140557446 | 152 | 80 | [illegible] |
| SE | 33374 | ENSG00000213523 | SRA1 | chr5 | - | 140551981 | 140552184 | 140537339 | 140537661 | 140557146 | 140557272 | 31 | 8 | [illegible] |
| SE | 33379 | ENSG00000168591 | TMUB2 | chr17 | + | 44187675 | 44187743 | 44187233 | 44187431 | 44187906 | 44188009 | 17 | 6 | [illegible] |
| SE | 33381 | ENSG00000168591 | TMUB2 | chr17 | + | 44187906 | 44188009 | 44187233 | 44187431 | 44188934 | 44189299 | 28 | 6 | [illegible] |
| SE | 33383 | ENSG00000168591 | TMUB2 | chr17 | + | 44187906 | 44188009 | 44187271 | 44187743 | 44190500 | 44191226 | 73 | 22 | [illegible] |
| SE | 33384 | ENSG00000168591 | TMUB2 | chr17 | + | 44188934 | 44189091 | 44187274 | 44187743 | 44190500 | 44191711 | 18 | 11 | [illegible] |
| SE | 33385 | ENSG00000168591 | TMUB2 | chr17 | + | 44188934 | 44189091 | 44187931 | 44188009 | 44190500 | 44191711 | 31 | 7 | [illegible] |
| SE | 33386 | ENSG00000168591 | TMUB2 | chr17 | + | 44188934 | 44189299 | 44187906 | 44188009 | 44190500 | 44191711 | 34 | 7 | [illegible] |
| SE | 33387 | ENSG00000168591 | TMUB2 | chr17 | + | 44188934 | 44189588 | 44187205 | 44187743 | 44190500 | 44191726 | 159 | 70 | [illegible] |
| SE | 33388 | ENSG00000168591 | TMUB2 | chr17 | + | 44189021 | 44189588 | 44187616 | 44187743 | 44190500 | 44191712 | 155 | 69 | [illegible] |
| SE | 33389 | ENSG00000168591 | TMUB2 | chr17 | + | 44189021 | 44189588 | 44187931 | 44188009 | 44190500 | 44191712 | 152 | 69 | [illegible] |
| SE | 33390 | ENSG00000168591 | TMUB2 | chr17 | + | 44189206 | 44189299 | 44188934 | 44189091 | 44189423 | 44189588 | 1 | 1 | [illegible] |
| SE | 33391 | ENSG00000168591 | TMUB2 | chr17 | + | 44189410 | 44189588 | 44188934 | 44189299 | 44190500 | 44191226 | 147 | 61 | [illegible] |
| SE | 33392 | ENSG00000168591 | TMUB2 | chr17 | + | 44189423 | 44189588 | 44188850 | 44189299 | 44190500 | 44190553 | 146 | 61 | [illegible] |
| SE | 33393 | ENSG00000168591 | TMUB2 | chr17 | + | 44189423 | 44189588 | 44188934 | 44189091 | 44190500 | 44190553 | 141 | 60 | [illegible] |
| SE | 33396 | ENSG00000254087 | LYN | chr8 | + | 55941854 | 55941991 | 55879834 | 55880103 | 55947617 | 55947723 | 227 | 196 | [illegible] |
| SE | 33397 | ENSG00000254087 | LYN | chr8 | + | 55946447 | 55946493 | 55879834 | 55880103 | 55947617 | 55947723 | 331 | 248 | [illegible] |
| SE | 33398 | ENSG00000254087 | LYN | chr8 | + | 55946447 | 55946493 | 55941854 | 55941991 | 55947617 | 55947723 | 1069 | 876 | [illegible] |
| SE | 33400 | ENSG00000254087 | LYN | chr8 | + | 55947617 | 55947723 | 55946447 | 55946493 | 55950458 | 55950557 | 2654 | 2419 | [illegible] |
| SE | 33401 | ENSG00000254087 | LYN | chr8 | + | 55950458 | 55950557 | 55947617 | 55947723 | 55950680 | 55950748 | 1983 | 1741 | [illegible] |
| SE | 33402 | ENSG00000254087 | LYN | chr8 | + | 55953831 | 55953984 | 55951965 | 55952115 | 55966714 | 55966897 | 1329 | 1179 | [illegible] |
| SE | 33403 | ENSG00000254087 | LYN | chr8 | + | 55966714 | 55966897 | 55953831 | 55953984 | 55969716 | 55969793 | 635 | 485 | [illegible] |
| SE | 33404 | ENSG00000254087 | LYN | chr8 | + | 55969716 | 55969793 | 55953831 | 55953984 | 55998345 | 55998448 | 180 | 129 | [illegible] |
| SE | 33406 | ENSG00000254087 | LYN | chr8 | + | 55998345 | 55998499 | 55947617 | 55947723 | 56009907 | 56014168 | 1 | 0 | [illegible] |
| SE | 33407 | ENSG00000254087 | LYN | chr8 | + | 55998345 | 55998499 | 55953831 | 55953984 | 56009907 | 56014168 | 1 | 0 | [illegible] |
| SE | 33408 | ENSG00000254087 | LYN | chr8 | + | 55998345 | 55998499 | 55969716 | 55969793 | 56009907 | 56014168 | 420 | 277 | [illegible] |
| SE | 33409 | ENSG00000254087 | LYN | chr8 | + | 55999417 | 55999549 | 55998345 | 55998499 | 56009907 | 56014168 | 1295 | 1023 | [illegible] |
| SE | 33410 | ENSG00000096063 | SRPK1 | chr6 | - | 35857260 | 35857368 | 35842534 | 35842604 | 35869009 | 35869110 | 185 | 102 | [illegible] |
| SE | 33412 | ENSG00000096063 | SRPK1 | chr6 | - | 35869009 | 35869138 | 35856933 | 35857368 | 35869481 | 35869551 | 81 | 52 | [illegible] |
| SE | 33414 | ENSG00000096063 | SRPK1 | chr6 | - | 35872562 | 35872728 | 35870933 | 35870959 | 35874232 | 35874339 | 111 | 139 | [illegible] |
| SE | 33426 | ENSG00000135870 | RC3H1 | chr1 | - | 173951985 | 173952138 | 173947368 | 173947582 | 173961076 | 173961244 | 526 | 562 | [illegible] |
| SE | 33427 | ENSG00000135870 | RC3H1 | chr1 | - | 173964838 | 173965120 | 173963972 | 173964187 | 173970504 | 173970617 | 336 | 185 | [illegible] |
| SE | 33428 | ENSG00000135870 | RC3H1 | chr1 | - | 173970504 | 173970617 | 173964838 | 173965120 | 173972508 | 173972627 | 229 | 96 | [illegible] |
| SE | 33430 | ENSG00000135870 | RC3H1 | chr1 | - | 173978487 | 173978620 | 173972508 | 173972627 | 173980808 | 173981009 | 250 | 143 | [illegible] |
| SE | 33431 | ENSG00000135870 | RC3H1 | chr1 | - | 173983417 | 173983657 | 173982726 | 173982902 | 173984498 | 173984619 | 216 | 127 | [illegible] |
| SE | 33432 | ENSG00000135870 | RC3H1 | chr1 | - | 173992754 | 173993135 | 173984498 | 173984619 | 174022095 | 174022357 | 165 | 91 | [illegible] |
| SE | 33433 | ENSG00000010278 | CD9 | chr12 | + | 6200383 | 6200565 | 6199874 | 6200072 | 6225425 | 6225534 | 20 | 16 | [illegible] |
| SE | 33434 | ENSG00000010278 | CD9 | chr12 | + | 6200421 | 6200565 | 6199714 | 6200072 | 6225425 | 6225534 | 20 | 16 | [illegible] |
| SE | 33437 | ENSG00000010278 | CD9 | chr12 | + | 6232936 | 6233005 | 6232631 | 6232729 | 6233411 | 6233486 | 0 | 0 | [illegible] |
| SE | 33438 | ENSG00000010278 | CD9 | chr12 | + | 6235475 | 6235565 | 6235228 | 6235327 | 6236191 | 6236260 | 84 | 133 | [illegible] |
| SE | 33441 | ENSG00000156976 | EIF4A2 | chr3 | + | 186784428 | 186784477 | 186783576 | 186783639 | 186784563 | 186784696 | 839 | 492 | [illegible] |
| SE | 33442 | ENSG00000156976 | EIF4A2 | chr3 | + | 186784431 | 186784477 | 186783304 | 186783639 | 186784563 | 186784659 | 1500 | 889 | [illegible] |
| SE | 33443 | ENSG00000156976 | EIF4A2 | chr3 | + | 186784563 | 186784696 | 186784202 | 186784477 | 186785882 | 186786044 | 2436 | 1508 | [illegible] |
| SE | 33446 | ENSG00000156976 | EIF4A2 | chr3 | + | 186786501 | 186786645 | 186786163 | 186786273 | 186787126 | 186787264 | 3007 | 2029 | [illegible] |
| SE | 33448 | ENSG00000156976 | EIF4A2 | chr3 | + | 186787478 | 186787584 | 186787126 | 186787264 | 186789124 | 186789140 | 5 | 5 | [illegible] |
| SE | 33450 | ENSG00000156976 | EIF4A2 | chr3 | + | 186787494 | 186787584 | 186787126 | 186787264 | 186789124 | 186789881 | 1043 | 639 | [illegible] |
| SE | 33451 | ENSG00000156976 | EIF4A2 | chr3 | + | 186787802 | 186787882 | 186787313 | 186787584 | 186788309 | 186788416 | 1095 | 790 | [illegible] |
| SE | 33452 | ENSG00000156976 | EIF4A2 | chr3 | + | 186787802 | 186787882 | 186787313 | 186787584 | 186789124 | 186789140 | 1915 | 1103 | [illegible] |
| SE | 33453 | ENSG00000156976 | EIF4A2 | chr3 | + | 186788309 | 186788416 | 186787494 | 186787584 | 186789124 | 186789881 | 10 | 7 | [illegible] |
| SE | 33455 | ENSG00000156976 | EIF4A2 | chr3 | + | 186788309 | 186788420 | 186787313 | 186787584 | 186789124 | 186789291 | 55 | 54 | [illegible] |
| SE | 33457 | ENSG00000165782 | PIP4P1 | chr14 | - | 20459205 | 20459296 | 20457680 | 20458702 | 20459387 | 20459440 | 193 | 90 | [illegible] |
| SE | 33460 | ENSG00000114648 | KLHL18 | chr3 | + | 47319652 | 47319783 | 47282943 | 47283094 | 47322567 | 47322707 | 84 | 30 | [illegible] |
| SE | 33461 | ENSG00000114648 | KLHL18 | chr3 | + | 47333156 | 47333317 | 47329950 | 47330149 | 47334682 | 47334819 | 58 | 44 | [illegible] |
| SE | 33462 | ENSG00000114648 | KLHL18 | chr3 | + | 47340571 | 47340676 | 47336534 | 47336757 | 47342718 | 47342830 | 37 | 14 | [illegible] |
| SE | 33463 | ENSG00000114648 | KLHL18 | chr3 | + | 47340571 | 47340676 | 47336534 | 47336757 | 47343554 | 47344577 | 25 | 7 | [illegible] |
| SE | 33464 | ENSG00000114648 | KLHL18 | chr3 | + | 47342718 | 47342830 | 47336534 | 47336757 | 47343554 | 47344577 | 30 | 15 | [illegible] |
| SE | 33465 | ENSG00000114648 | KLHL18 | chr3 | + | 47342718 | 47342830 | 47340571 | 47340676 | 47343554 | 47344577 | 42 | 22 | [illegible] |
| SE | 33466 | ENSG00000164327 | RICTOR | chr5 | - | 38942832 | 38942971 | 38937919 | 38942378 | 38944445 | 38944569 | 290 | 149 | [illegible] |
| SE | 33467 | ENSG00000164327 | RICTOR | chr5 | - | 38944445 | 38944569 | 38942832 | 38942971 | 38944912 | 38945068 | 164 | 81 | [illegible] |
| SE | 33468 | ENSG00000164327 | RICTOR | chr5 | - | 38946467 | 38946552 | 38945490 | 38945610 | 38947263 | 38947441 | 109 | 74 | [illegible] |
| SE | 33469 | ENSG00000164327 | RICTOR | chr5 | - | 38946467 | 38946552 | 38945490 | 38945724 | 38947263 | 38947441 | 173 | 118 | [illegible] |
| SE | 33472 | ENSG00000164327 | RICTOR | chr5 | - | 38962875 | 38963041 | 38962484 | 38962586 | 38964791 | 38964892 | 330 | 203 | [illegible] |
| SE | 33473 | ENSG00000164327 | RICTOR | chr5 | - | 38966640 | 38966721 | 38964791 | 38964892 | 38967160 | 38967227 | 179 | 146 | [illegible] |
| SE | 33474 | ENSG00000164327 | RICTOR | chr5 | - | 38975536 | 38975604 | 38969686 | 38971959 | 38978582 | 38978648 | 210 | 166 | [illegible] |
| SE | 33477 | ENSG00000164327 | RICTOR | chr5 | - | 38978582 | 38978650 | 38975536 | 38975604 | 38990948 | 38991075 | 46 | 38 | [illegible] |
| SE | 33478 | ENSG00000164327 | RICTOR | chr5 | - | 38981866 | 38982036 | 38975536 | 38975604 | 38990948 | 38991075 | 174 | 133 | [illegible] |
| SE | 33479 | ENSG00000164327 | RICTOR | chr5 | - | 38981866 | 38982036 | 38978582 | 38978650 | 38990948 | 38991075 | 389 | 305 | [illegible] |
| SE | 33480 | ENSG00000164327 | RICTOR | chr5 | - | 38996818 | 38996882 | 38990948 | 38991075 | 39002534 | 39002666 | 276 | 203 | [illegible] |
| SE | 33481 | ENSG00000164327 | RICTOR | chr5 | - | 39021038 | 39021136 | 39003557 | 39003622 | 39074110 | 39074158 | 225 | 164 | [illegible] |
| SE | 33482 | ENSG00000164327 | RICTOR | chr5 | - | 39074110 | 39074158 | 39021038 | 39021136 | 39074328 | 39074389 | 136 | 105 | [illegible] |
| SE | 33483 | ENSG00000125037 | EMC3 | chr3 | - | 9969718 | 9969801 | 9962536 | 9964197 | 9970581 | 9970661 | 415 | 378 | [illegible] |
| SE | 33484 | ENSG00000125037 | EMC3 | chr3 | - | 9973627 | 9973709 | 9970581 | 9970661 | 9974383 | 9974488 | 614 | 613 | [illegible] |
| SE | 33485 | ENSG00000125037 | EMC3 | chr3 | - | 9973627 | 9973709 | 9970581 | 9970661 | 9976956 | 9977050 | 305 | 300 | [illegible] |
| SE | 33486 | ENSG00000125037 | EMC3 | chr3 | - | 9974383 | 9974488 | 9970581 | 9970661 | 9976956 | 9977050 | 198 | 220 | [illegible] |
| SE | 33487 | ENSG00000125037 | EMC3 | chr3 | - | 9974383 | 9974488 | 9973505 | 9973709 | 9976956 | 9977050 | 509 | 535 | [illegible] |
| SE | 33488 | ENSG00000125037 | EMC3 | chr3 | - | 9976956 | 9977050 | 9974383 | 9974488 | 9977388 | 9977446 | 324 | 300 | [illegible] |
| SE | 33494 | ENSG00000164068 | RNF123 | chr3 | + | 49691424 | 49691509 | 49691129 | 49691247 | 49697142 | 49697204 | 29 | 28 | [illegible] |
| SE | 33495 | ENSG00000164068 | RNF123 | chr3 | + | 49697044 | 49697222 | 49691424 | 49691509 | 49697362 | 49697412 | 32 | 17 | [illegible] |
| SE | 33496 | ENSG00000164068 | RNF123 | chr3 | + | 49697142 | 49697222 | 49691424 | 49691509 | 49697362 | 49697457 | 54 | 34 | [illegible] |
| SE | 33497 | ENSG00000164068 | RNF123 | chr3 | + | 49698754 | 49698822 | 49698439 | 49698526 | 49698979 | 49699105 | 37 | 10 | [illegible] |
| SE | 33498 | ENSG00000164068 | RNF123 | chr3 | + | 49713737 | 49713825 | 49712478 | 49713587 | 49713909 | 49713997 | 26 | 12 | [illegible] |
| SE | 33499 | ENSG00000145029 | NICN1 | chr3 | - | 49425882 | 49425996 | 49424948 | 49425438 | 49426251 | 49426407 | 24 | 14 | [illegible] |
| SE | 33500 | ENSG00000120686 | UFM1 | chr13 | + | 38354238 | 38354296 | 38349995 | 38350055 | 38358092 | 38358132 | 344 | 225 | [illegible] |
| SE | 33501 | ENSG00000120686 | UFM1 | chr13 | + | 38358092 | 38358132 | 38354238 | 38354296 | 38359300 | 38359333 | 375 | 255 | [illegible] |
| SE | 33502 | ENSG00000120686 | UFM1 | chr13 | + | 38359300 | 38359333 | 38358092 | 38358132 | 38360710 | 38361125 | 408 | 302 | [illegible] |
| SE | 33512 | ENSG00000134072 | CAMK1 | chr3 | - | 9765758 | 9765890 | 9763138 | 9763213 | 9767666 | 9767781 | 56 | 50 | [illegible] |
| SE | 33516 | ENSG00000175455 | CCDC14 | chr3 | - | 123948690 | 123948785 | 123946802 | 123947319 | 123948895 | 123948933 | 245 | 168 | [illegible] |
| SE | 33517 | ENSG00000175455 | CCDC14 | chr3 | - | 123955842 | 123955965 | 123948895 | 123949132 | 123956045 | 123956115 | 103 | 106 | [illegible] |
| SE | 33518 | ENSG00000175455 | CCDC14 | chr3 | - | 123955842 | 123955965 | 123948895 | 123949132 | 123961143 | 123961381 | 87 | 92 | [illegible] |
| SE | 33519 | ENSG00000175455 | CCDC14 | chr3 | - | 123955842 | 123956115 | 123948895 | 123949132 | 123956354 | 123956427 | 131 | 143 | [illegible] |
| SE | 33521 | ENSG00000175455 | CCDC14 | chr3 | - | 123956045 | 123956115 | 123948895 | 123949132 | 123956354 | 123956427 | 35 | 45 | [illegible] |
| SE | 33522 | ENSG00000175455 | CCDC14 | chr3 | - | 123956354 | 123956427 | 123948895 | 123949132 | 123961143 | 123961225 | 0 | 0 | [illegible] |
| SE | 33524 | ENSG00000175455 | CCDC14 | chr3 | - | 123956739 | 123956795 | 123956354 | 123956427 | 123961143 | 123961209 | 9 | 14 | [illegible] |
| SE | 33528 | ENSG00000082996 | RNF13 | chr3 | + | 149852515 | 149852596 | 149813203 | 149813353 | 149872028 | 149872049 | 261 | 176 | [illegible] |
| SE | 33529 | ENSG00000082996 | RNF13 | chr3 | + | 149852515 | 149852596 | 149813203 | 149813353 | 149895472 | 149895498 | 10 | 1 | [illegible] |
| SE | 33530 | ENSG00000082996 | RNF13 | chr3 | + | 149872028 | 149872154 | 149813214 | 149813353 | 149895472 | 149895498 | 702 | 443 | [illegible] |
| SE | 33531 | ENSG00000082996 | RNF13 | chr3 | + | 149872028 | 149872154 | 149852515 | 149852596 | 149895472 | 149895498 | 1140 | 717 | [illegible] |
| SE | 33532 | ENSG00000082996 | RNF13 | chr3 | + | 149902071 | 149902162 | 149895472 | 149895560 | 149911977 | 149912083 | 713 | 527 | [illegible] |
| SE | 33533 | ENSG00000082996 | RNF13 | chr3 | + | 149902071 | 149902162 | 149895472 | 149895560 | 149960055 | 149960136 | 311 | 222 | [illegible] |
| SE | 33535 | ENSG00000082996 | RNF13 | chr3 | + | 149911977 | 149912083 | 149902071 | 149902162 | 149921133 | 149921184 | 901 | 674 | [illegible] |
| SE | 33536 | ENSG00000082996 | RNF13 | chr3 | + | 149921133 | 149921227 | 149902071 | 149902162 | 149960055 | 149960136 | 224 | 154 | [illegible] |
| SE | 33538 | ENSG00000241404 | EGFL8 | chr6 | + | 32166700 | 32166810 | 32165235 | 32166620 | 32166909 | 32167005 | 10 | 1 | [illegible] |
| SE | 33539 | ENSG00000213533 | STIMATE | chr3 | - | 52842810 | 52842960 | 52836732 | 52840610 | 52843720 | 52843798 | 167 | 174 | [illegible] |
| SE | 33540 | ENSG00000213533 | STIMATE | chr3 | - | 52843720 | 52843798 | 52842810 | 52842960 | 52844828 | 52844941 | 99 | 82 | [illegible] |
| SE | 33541 | ENSG00000213533 | STIMATE | chr3 | - | 52844828 | 52844941 | 52843720 | 52843798 | 52852602 | 52852698 | 47 | 38 | [illegible] |
| SE | 33542 | ENSG00000213533 | STIMATE | chr3 | - | 52849791 | 52849913 | 52844828 | 52844941 | 52852602 | 52852698 | 70 | 32 | [illegible] |
| SE | 33543 | ENSG00000213533 | STIMATE | chr3 | - | 52895877 | 52895955 | 52855395 | 52855444 | 52897290 | 52897543 | 2 | 1 | [illegible] |
| SE | 33544 | ENSG00000164081 | TEX264 | chr3 | + | 51672209 | 51672301 | 51671174 | 51671288 | 51674270 | 51674562 | 2 | 0 | [illegible] |
| SE | 33545 | ENSG00000164081 | TEX264 | chr3 | + | 51672209 | 51672301 | 51671174 | 51671288 | 51684412 | 51684573 | 1 | 0 | [illegible] |
| SE | 33546 | ENSG00000164081 | TEX264 | chr3 | + | 51672340 | 51672392 | 51671174 | 51671288 | 51674270 | 51674562 | 3 | 1 | [illegible] |
| SE | 33547 | ENSG00000164081 | TEX264 | chr3 | + | 51672340 | 51672392 | 51671174 | 51671288 | 51684412 | 51684607 | 1 | 0 | [illegible] |
| SE | 33548 | ENSG00000164081 | TEX264 | chr3 | + | 51672340 | 51672392 | 51671894 | 51672083 | 51674270 | 51674562 | 2 | 0 | [illegible] |
| SE | 33550 | ENSG00000164081 | TEX264 | chr3 | + | 51674270 | 51674562 | 51671254 | 51672301 | 51684412 | 51684467 | 35 | 13 | [illegible] |
| SE | 33552 | ENSG00000164081 | TEX264 | chr3 | + | 51674270 | 51674562 | 51672340 | 51672392 | 51684412 | 51684467 | 39 | 17 | [illegible] |
| SE | 33553 | ENSG00000164081 | TEX264 | chr3 | + | 51681341 | 51681430 | 51674476 | 51674562 | 51684412 | 51684559 | 0 | 0 | [illegible] |
| SE | 33555 | ENSG00000009790 | TRAF3IP3 | chr1 | + | 209758725 | 209758854 | 209756135 | 209756309 | 209759033 | 209759134 | 1 | 0 | [illegible] |
| SE | 33556 | ENSG00000009790 | TRAF3IP3 | chr1 | + | 209759033 | 209759134 | 209756031 | 209756309 | 209759981 | 209760117 | 629 | 439 | [illegible] |
| SE | 33557 | ENSG00000009790 | TRAF3IP3 | chr1 | + | 209759033 | 209759134 | 209756031 | 209756309 | 209762514 | 209762596 | 239 | 155 | [illegible] |
| SE | 33558 | ENSG00000009790 | TRAF3IP3 | chr1 | + | 209759981 | 209760324 | 209756031 | 209756309 | 209762514 | 209762662 | 17 | 16 | [illegible] |
| SE | 33560 | ENSG00000009790 | TRAF3IP3 | chr1 | + | 209759981 | 209760384 | 209756052 | 209756309 | 209762514 | 209762662 | 251 | 157 | [illegible] |
| SE | 33562 | ENSG00000009790 | TRAF3IP3 | chr1 | + | 209759981 | 209760384 | 209759033 | 209759134 | 209762812 | 209762870 | 395 | 290 | [illegible] |
| SE | 33564 | ENSG00000009790 | TRAF3IP3 | chr1 | + | 209762514 | 209762662 | 209759033 | 209759134 | 209762812 | 209762870 | 218 | 137 | [illegible] |
| SE | 33567 | ENSG00000009790 | TRAF3IP3 | chr1 | + | 209763362 | 209763392 | 209763067 | 209763092 | 209763491 | 209763587 | 685 | 509 | [illegible] |
| SE | 33569 | ENSG00000009790 | TRAF3IP3 | chr1 | + | 209777351 | 209777487 | 209775598 | 209775736 | 209778110 | 209778173 | 1393 | 1037 | [illegible] |
| SE | 33570 | ENSG00000009790 | TRAF3IP3 | chr1 | + | 209777351 | 209777487 | 209775598 | 209775736 | 209779314 | 209779364 | 798 | 594 | [illegible] |
| SE | 33571 | ENSG00000009790 | TRAF3IP3 | chr1 | + | 209777351 | 209777487 | 209775598 | 209775736 | 209779430 | 209779655 | 784 | 580 | [illegible] |
| SE | 33572 | ENSG00000009790 | TRAF3IP3 | chr1 | + | 209777351 | 209777487 | 209775598 | 209775736 | 209781344 | 209781458 | 782 | 574 | [illegible] |
| SE | 33573 | ENSG00000009790 | TRAF3IP3 | chr1 | + | 209777351 | 209777487 | 209775598 | 209775736 | 209782055 | 209782313 | 795 | 574 | [illegible] |
| SE | 33575 | ENSG00000009790 | TRAF3IP3 | chr1 | + | 209778110 | 209778173 | 209775598 | 209775736 | 209779314 | 209779364 | 150 | 79 | [illegible] |
| SE | 33576 | ENSG00000009790 | TRAF3IP3 | chr1 | + | 209778110 | 209778173 | 209775598 | 209775736 | 209779430 | 209779655 | 7 | 4 | [illegible] |
| SE | 33578 | ENSG00000009790 | TRAF3IP3 | chr1 | + | 209778110 | 209778173 | 209775598 | 209775736 | 209782055 | 209782307 | 1 | 0 | [illegible] |
| SE | 33581 | ENSG00000009790 | TRAF3IP3 | chr1 | + | 209778110 | 209778173 | 209777351 | 209777487 | 209780469 | 209780606 | 234 | 164 | [illegible] |
| SE | 33582 | ENSG00000009790 | TRAF3IP3 | chr1 | + | 209778110 | 209778173 | 209777351 | 209777487 | 209781344 | 209781458 | 227 | 164 | [illegible] |
| SE | 33584 | ENSG00000009790 | TRAF3IP3 | chr1 | + | 209779314 | 209779374 | 209775598 | 209775736 | 209781344 | 209781458 | 2 | 3 | [illegible] |
| SE | 33585 | ENSG00000009790 | TRAF3IP3 | chr1 | + | 209779314 | 209779374 | 209775598 | 209775736 | 209782055 | 209782307 | 19 | 5 | [illegible] |
| SE | 33587 | ENSG00000009790 | TRAF3IP3 | chr1 | + | 209779314 | 209779374 | 209777351 | 209777487 | 209781344 | 209781458 | 8 | 17 | [illegible] |
| SE | 33590 | ENSG00000009790 | TRAF3IP3 | chr1 | + | 209779314 | 209779374 | 209778110 | 209778173 | 209781344 | 209781458 | 295 | 258 | [illegible] |
| SE | 33591 | ENSG00000009790 | TRAF3IP3 | chr1 | + | 209779314 | 209779374 | 209778110 | 209778173 | 209782055 | 209782307 | 334 | 255 | [illegible] |
| SE | 33592 | ENSG00000009790 | TRAF3IP3 | chr1 | + | 209780469 | 209780606 | 209777351 | 209777487 | 209781344 | 209781458 | 138 | 194 | [illegible] |
| SE | 33593 | ENSG00000009790 | TRAF3IP3 | chr1 | + | 209780469 | 209780606 | 209777351 | 209777487 | 209782055 | 209782307 | 32 | 1 | [illegible] |
| SE | 33594 | ENSG00000009790 | TRAF3IP3 | chr1 | + | 209780469 | 209780606 | 209778110 | 209778173 | 209781344 | 209781458 | 145 | 194 | [illegible] |
| SE | 33595 | ENSG00000009790 | TRAF3IP3 | chr1 | + | 209780469 | 209780606 | 209778110 | 209778173 | 209782055 | 209782307 | 39 | 1 | [illegible] |
| SE | 33598 | ENSG00000009790 | TRAF3IP3 | chr1 | + | 209781344 | 209781458 | 209775598 | 209775736 | 209782055 | 209782307 | 142 | 239 | [illegible] |
| SE | 33609 | ENSG00000034533 | ASTE1 | chr3 | - | 131015181 | 131015256 | 131013874 | 131014387 | 131016143 | 131016339 | 0 | 1 | [illegible] |
| SE | 33610 | ENSG00000034533 | ASTE1 | chr3 | - | 131016989 | 131017049 | 131016143 | 131016339 | 131018505 | 131018716 | 4 | 2 | [illegible] |
| SE | 33611 | ENSG00000034533 | ASTE1 | chr3 | - | 131018505 | 131018716 | 131016143 | 131016339 | 131024004 | 131025331 | 60 | 27 | [illegible] |
| SE | 33612 | ENSG00000034533 | ASTE1 | chr3 | - | 131024004 | 131024238 | 131018504 | 131018716 | 131025163 | 131025331 | 52 | 25 | [illegible] |
| SE | 33613 | ENSG00000034533 | ASTE1 | chr3 | - | 131025473 | 131025594 | 131024004 | 131025331 | 131026187 | 131026609 | 112 | 65 | [illegible] |
| SE | 33614 | ENSG00000034533 | ASTE1 | chr3 | - | 131025473 | 131025594 | 131024004 | 131025331 | 131026506 | 131026825 | 135 | 83 | [illegible] |
| SE | 33616 | ENSG00000173402 | DAG1 | chr3 | + | 49476848 | 49476900 | 49470135 | 49470433 | 49510418 | 49510799 | 0 | 0 | [illegible] |
| SE | 33619 | ENSG00000173402 | DAG1 | chr3 | + | 49476848 | 49476905 | 49470131 | 49470433 | 49510418 | 49510819 | 0 | 1 | [illegible] |
| SE | 33623 | ENSG00000173402 | DAG1 | chr3 | + | 49488625 | 49488745 | 49470276 | 49470433 | 49510418 | 49510765 | 0 | 0 | [illegible] |
| SE | 33631 | ENSG00000181322 | NME9 | chr3 | - | 138324174 | 138324594 | 138319477 | 138319581 | 138324872 | 138324930 | 0 | 0 | [illegible] |
| SE | 33636 | ENSG00000023892 | DEF6 | chr6 | + | 35310458 | 35310644 | 35309669 | 35309810 | 35317890 | 35317999 | 123 | 46 | [illegible] |
| SE | 33638 | ENSG00000023892 | DEF6 | chr6 | + | 35312301 | 35312538 | 35309669 | 35309810 | 35317890 | 35317999 | 0 | 0 | [illegible] |
| SE | 33639 | ENSG00000023892 | DEF6 | chr6 | + | 35312301 | 35312538 | 35310458 | 35310644 | 35317890 | 35317999 | 184 | 98 | [illegible] |
| SE | 33640 | ENSG00000023892 | DEF6 | chr6 | + | 35312625 | 35312772 | 35312301 | 35312538 | 35317890 | 35317999 | 331 | 177 | [illegible] |
| SE | 33641 | ENSG00000010295 | IFFO1 | chr12 | - | 6548064 | 6548157 | 6541511 | 6541642 | 6548424 | 6548545 | 109 | 74 | [illegible] |
| SE | 33642 | ENSG00000010295 | IFFO1 | chr12 | - | 6548064 | 6548160 | 6541511 | 6541642 | 6548424 | 6548545 | 134 | 71 | [illegible] |
| SE | 33643 | ENSG00000010295 | IFFO1 | chr12 | - | 6548424 | 6548545 | 6548064 | 6548157 | 6549475 | 6549484 | 34 | 24 | [illegible] |
| SE | 33645 | ENSG00000010295 | IFFO1 | chr12 | - | 6548667 | 6548825 | 6548064 | 6548545 | 6549475 | 6549484 | 161 | 114 | [illegible] |
| SE | 33646 | ENSG00000010295 | IFFO1 | chr12 | - | 6548667 | 6548849 | 6548424 | 6548545 | 6549475 | 6549484 | 120 | 76 | [illegible] |
| SE | 33647 | ENSG00000010295 | IFFO1 | chr12 | - | 6549475 | 6549484 | 6548064 | 6548160 | 6549755 | 6549896 | 1 | 1 | [illegible] |
| SE | 33650 | ENSG00000010295 | IFFO1 | chr12 | - | 6550940 | 6551001 | 6550694 | 6550790 | 6555256 | 6556042 | 74 | 33 | [illegible] |
| SE | 33652 | ENSG00000176261 | ZBTB8OS | chr1 | - | 32627507 | 32627544 | 32620819 | 32621948 | 32631826 | 32631879 | 140 | 92 | [illegible] |
| SE | 33653 | ENSG00000176261 | ZBTB8OS | chr1 | - | 32627507 | 32627544 | 32620819 | 32621948 | 32633644 | 32633727 | 46 | 40 | [illegible] |
| SE | 33654 | ENSG00000176261 | ZBTB8OS | chr1 | - | 32627507 | 32627544 | 32620819 | 32621948 | 32650432 | 32650525 | 4 | 8 | [illegible] |
| SE | 33656 | ENSG00000176261 | ZBTB8OS | chr1 | - | 32631826 | 32631879 | 32620819 | 32621948 | 32650432 | 32650525 | 0 | 0 | [illegible] |
| SE | 33660 | ENSG00000176261 | ZBTB8OS | chr1 | - | 32631826 | 32631879 | 32627507 | 32627544 | 32650432 | 32650525 | 87 | 46 | [illegible] |
| SE | 33663 | ENSG00000176261 | ZBTB8OS | chr1 | - | 32633261 | 32633354 | 32627507 | 32627544 | 32633644 | 32633727 | 1 | 0 | [illegible] |
| SE | 33664 | ENSG00000176261 | ZBTB8OS | chr1 | - | 32633261 | 32633354 | 32632207 | 32632345 | 32633644 | 32633727 | 2 | 0 | [illegible] |
| SE | 33665 | ENSG00000176261 | ZBTB8OS | chr1 | - | 32633644 | 32633727 | 32620819 | 32621948 | 32650432 | 32650525 | 2 | 1 | [illegible] |
| SE | 33669 | ENSG00000176261 | ZBTB8OS | chr1 | - | 32633644 | 32633727 | 32627507 | 32627544 | 32650432 | 32650525 | 30 | 31 | [illegible] |
| SE | 33670 | ENSG00000176261 | ZBTB8OS | chr1 | - | 32633644 | 32633727 | 32631824 | 32631879 | 32633950 | 32634061 | 234 | 117 | [illegible] |
| SE | 33671 | ENSG00000176261 | ZBTB8OS | chr1 | - | 32633644 | 32633727 | 32631824 | 32631879 | 32634767 | 32634792 | 44 | 28 | [illegible] |
| SE | 33674 | ENSG00000176261 | ZBTB8OS | chr1 | - | 32633950 | 32634072 | 32627507 | 32627544 | 32650401 | 32650525 | 1 | 0 | [illegible] |
| SE | 33675 | ENSG00000176261 | ZBTB8OS | chr1 | - | 32633950 | 32634072 | 32627507 | 32627544 | 32650432 | 32650525 | 0 | 0 | [illegible] |
| SE | 33676 | ENSG00000176261 | ZBTB8OS | chr1 | - | 32633950 | 32634072 | 32631826 | 32631879 | 32634701 | 32634792 | 3 | 7 | [illegible] |
| SE | 33677 | ENSG00000176261 | ZBTB8OS | chr1 | - | 32633950 | 32634072 | 32631826 | 32631879 | 32634767 | 32634792 | 63 | 38 | [illegible] |
| SE | 33678 | ENSG00000176261 | ZBTB8OS | chr1 | - | 32633950 | 32634072 | 32631826 | 32631879 | 32650432 | 32650525 | 0 | 0 | [illegible] |
| SE | 33679 | ENSG00000176261 | ZBTB8OS | chr1 | - | 32633950 | 32634072 | 32633644 | 32633727 | 32634767 | 32634792 | 269 | 132 | [illegible] |
| SE | 33683 | ENSG00000176261 | ZBTB8OS | chr1 | - | 32633950 | 32634110 | 32627507 | 32627544 | 32650401 | 32650525 | 1 | 2 | [illegible] |
| SE | 33684 | ENSG00000176261 | ZBTB8OS | chr1 | - | 32633950 | 32634110 | 32627507 | 32627544 | 32650432 | 32650903 | 3 | 4 | [illegible] |
| SE | 33685 | ENSG00000176261 | ZBTB8OS | chr1 | - | 32633950 | 32634110 | 32631824 | 32631879 | 32634701 | 32634792 | 4 | 1 | [illegible] |
| SE | 33686 | ENSG00000176261 | ZBTB8OS | chr1 | - | 32633950 | 32634110 | 32631824 | 32631879 | 32634767 | 32634792 | 12 | 4 | [illegible] |
| SE | 33687 | ENSG00000176261 | ZBTB8OS | chr1 | - | 32633950 | 32634110 | 32631824 | 32631879 | 32650432 | 32650903 | 3 | 4 | [illegible] |
| SE | 33688 | ENSG00000176261 | ZBTB8OS | chr1 | - | 32633950 | 32634110 | 32633644 | 32633727 | 32634767 | 32634792 | 221 | 103 | [illegible] |
| SE | 33691 | ENSG00000176261 | ZBTB8OS | chr1 | - | 32634701 | 32634792 | 32631826 | 32631879 | 32650432 | 32650525 | 10 | 4 | [illegible] |
| SE | 33692 | ENSG00000176261 | ZBTB8OS | chr1 | - | 32634701 | 32634792 | 32633950 | 32634072 | 32650432 | 32650525 | 16 | 8 | [illegible] |

| | | | | | | | | | | | | | |
|---|---|---|---|---|---|---|---|---|---|---|---|---|---|
| SE | 33694 | ENSG00000176261 | ZBTB8OS | chr1 | - | 32634767 | 32634792 | 32627507 | 32627544 | 32650401 | 32650525 | 0 | 0 |
| SE | 33695 | ENSG00000176261 | ZBTB8OS | chr1 | - | 32634767 | 32634792 | 32627507 | 32627544 | 32650432 | 32650535 | 6 | 4 |
| SE | 33696 | ENSG00000176261 | ZBTB8OS | chr1 | - | 32634767 | 32634792 | 32631826 | 32631879 | 32650432 | 32650535 | 3 | 0 |
| SE | 33697 | ENSG00000176261 | ZBTB8OS | chr1 | - | 32634767 | 32634792 | 32633644 | 32633727 | 32650401 | 32650525 | 0 | 0 |
| SE | 33698 | ENSG00000176261 | ZBTB8OS | chr1 | - | 32634767 | 32634792 | 32633644 | 32633727 | 32650432 | 32650535 | 2 | 0 |
| SE | 33699 | ENSG00000176261 | ZBTB8OS | chr1 | - | 32634767 | 32634792 | 32633950 | 32634072 | 32650401 | 32650525 | 7 | 3 |
| SE | 33702 | ENSG00000176261 | ZBTB8OS | chr1 | - | 32634767 | 32634792 | 32633950 | 32634110 | 32650401 | 32650525 | 0 | 0 |
| SE | 33708 | ENSG00000172046 | USP19 | chr3 | - | 49114173 | 49114284 | 49113991 | 49114093 | 49114762 | 49114873 | 99 | 59 |
| SE | 33709 | ENSG00000172046 | USP19 | chr3 | - | 49116046 | 49116162 | 49115723 | 49115944 | 49116279 | 49116351 | 163 | 84 |
| SE | 33710 | ENSG00000172046 | USP19 | chr3 | - | 49116450 | 49116607 | 49116279 | 49116351 | 49116726 | 49116943 | 138 | 79 |
| SE | 33712 | ENSG00000172046 | USP19 | chr3 | - | 49116726 | 49116943 | 49116450 | 49116607 | 49117656 | 49117711 | 38 | 24 |
| SE | 33714 | ENSG00000172046 | USP19 | chr3 | - | 49117208 | 49117358 | 49116733 | 49116943 | 49117436 | 49117570 | 0 | 0 |
| SE | 33715 | ENSG00000172046 | USP19 | chr3 | - | 49117436 | 49117570 | 49116450 | 49116607 | 49117656 | 49117785 | 25 | 6 |
| SE | 33718 | ENSG00000155975 | VPS37A | chr8 | + | 17265770 | 17265815 | 17246930 | 17247369 | 17265906 | 17265981 | 0 | 0 |
| SE | 33720 | ENSG00000155975 | VPS37A | chr8 | + | 17268257 | 17268372 | 17247122 | 17247369 | 17268855 | 17268956 | 115 | 88 |
| SE | 33721 | ENSG00000155975 | VPS37A | chr8 | + | 17268855 | 17268956 | 17246933 | 17247369 | 17274732 | 17274958 | 129 | 96 |
| SE | 33722 | ENSG00000155975 | VPS37A | chr8 | + | 17271992 | 17272063 | 17268855 | 17268956 | 17274732 | 17274958 | 0 | 0 |
| SE | 33725 | ENSG00000196455 | PIK3R4 | chr3 | - | 130686210 | 130686422 | 130684249 | 130684381 | 130690489 | 130690654 | 56 | 20 |
| SE | 33728 | ENSG00000196455 | PIK3R4 | chr3 | - | 130733547 | 130734130 | 130730307 | 130730442 | 130744485 | 130745264 | 109 | 52 |
| SE | 33729 | ENSG00000196455 | PIK3R4 | chr3 | - | 130735868 | 130736002 | 130733547 | 130734130 | 130744485 | 130745264 | 157 | 123 |
| SE | 33730 | ENSG00000105982 | RNF32 | chr7 | + | 156642976 | 156643087 | 156640783 | 156641004 | 156643800 | 156643892 | 0 | 0 |
| SE | 33731 | ENSG00000105982 | RNF32 | chr7 | + | 156643800 | 156643892 | 156640748 | 156640955 | 156644498 | 156644539 | 2 | 0 |
| SE | 33740 | ENSG00000128513 | POT1 | chr7 | - | 124823796 | 124824074 | 124822385 | 124822551 | 124825251 | 124825357 | 51 | 28 |
| SE | 33741 | ENSG00000128513 | POT1 | chr7 | - | 124823970 | 124824074 | 124822400 | 124822551 | 124825251 | 124825357 | 41 | 21 |
| SE | 33742 | ENSG00000128513 | POT1 | chr7 | - | 124829253 | 124829342 | 124827200 | 124827305 | 124835278 | 124835414 | 61 | 56 |
| SE | 33745 | ENSG00000128513 | POT1 | chr7 | - | 124848539 | 124848659 | 124846941 | 124846998 | 124851871 | 124851951 | 0 | 2 |
| SE | 33746 | ENSG00000128513 | POT1 | chr7 | - | 124851871 | 124851951 | 124846941 | 124846998 | 124852971 | 124853138 | 132 | 84 |
| SE | 33747 | ENSG00000128513 | POT1 | chr7 | - | 124863349 | 124863640 | 124858780 | 124859112 | 124870910 | 124871041 | 122 | 99 |
| SE | 33748 | ENSG00000128513 | POT1 | chr7 | - | 124865128 | 124865299 | 124863349 | 124863640 | 124870910 | 124871041 | 2 | 0 |
| SE | 33750 | ENSG00000128513 | POT1 | chr7 | - | 124870910 | 124871041 | 124863349 | 124863640 | 124892265 | 124892380 | 136 | 115 |
| SE | 33754 | ENSG00000128513 | POT1 | chr7 | - | 124897459 | 124897595 | 124897164 | 124897212 | 124898260 | 124898374 | 1 | 1 |
| SE | 33755 | ENSG00000128513 | POT1 | chr7 | - | 124915573 | 124915646 | 124898260 | 124898374 | 124928814 | 124928996 | 136 | 108 |
| SE | 33756 | ENSG00000128513 | POT1 | chr7 | - | 124928814 | 124928996 | 124915573 | 124915646 | 124929793 | 124929819 | 77 | 62 |
| SE | 33757 | ENSG00000128513 | POT1 | chr7 | - | 124928814 | 124928999 | 124915573 | 124915646 | 124929793 | 124929816 | 88 | 67 |
| SE | 33765 | ENSG00000180376 | CCDC66 | chr3 | + | 56559568 | 56559594 | 56558844 | 56558910 | 56560861 | 56561012 | 13 | 18 |
| SE | 33766 | ENSG00000180376 | CCDC66 | chr3 | + | 56559568 | 56559594 | 56558844 | 56558910 | 56563683 | 56564125 | 128 | 95 |
| SE | 33767 | ENSG00000180376 | CCDC66 | chr3 | + | 56560861 | 56561012 | 56558845 | 56558910 | 56563683 | 56563825 | 9 | 7 |
| SE | 33768 | ENSG00000180376 | CCDC66 | chr3 | + | 56560861 | 56561012 | 56559568 | 56559594 | 56561256 | 56561378 | 13 | 17 |
| SE | 33769 | ENSG00000180376 | CCDC66 | chr3 | + | 56560861 | 56561012 | 56559568 | 56559594 | 56563683 | 56563825 | 22 | 24 |
| SE | 33770 | ENSG00000180376 | CCDC66 | chr3 | + | 56561256 | 56561378 | 56559568 | 56559594 | 56563683 | 56564125 | 6 | 1 |
| SE | 33771 | ENSG00000180376 | CCDC66 | chr3 | + | 56561256 | 56561378 | 56559568 | 56559594 | 56566593 | 56566759 | 1 | 0 |
| SE | 33772 | ENSG00000180376 | CCDC66 | chr3 | + | 56561256 | 56561378 | 56560861 | 56561012 | 56563683 | 56564125 | 5 | 1 |
| SE | 33774 | ENSG00000180376 | CCDC66 | chr3 | + | 56563683 | 56564125 | 56561256 | 56561378 | 56566593 | 56566739 | 280 | 285 |
| SE | 33779 | ENSG00000180376 | CCDC66 | chr3 | + | 56571180 | 56571302 | 56566782 | 56567053 | 56592969 | 56593101 | 79 | 79 |
| SE | 33780 | ENSG00000180376 | CCDC66 | chr3 | + | 56572336 | 56572428 | 56571180 | 56571302 | 56592969 | 56593061 | 8 | 12 |
| SE | 33782 | ENSG00000180376 | CCDC66 | chr3 | + | 56615127 | 56615272 | 56613588 | 56613750 | 56615921 | 56616053 | 72 | 62 |
| SE | 33783 | ENSG00000180376 | CCDC66 | chr3 | + | 56615921 | 56616053 | 56613588 | 56613750 | 56617111 | 56617301 | 22 | 15 |
| SE | 33784 | ENSG00000134109 | EDEM1 | chr3 | + | 5188180 | 5188314 | 5187645 | 5187684 | 5195208 | 5195281 | 29 | 18 |
| SE | 33785 | ENSG00000134109 | EDEM1 | chr3 | + | 5199591 | 5199695 | 5195105 | 5195281 | 5201752 | 5201779 | 243 | 170 |
| SE | 33786 | ENSG00000134109 | EDEM1 | chr3 | + | 5205066 | 5205241 | 5202965 | 5203149 | 5207152 | 5207273 | 193 | 153 |
| SE | 33787 | ENSG00000134109 | EDEM1 | chr3 | + | 5208092 | 5208263 | 5207152 | 5207273 | 5210174 | 5210248 | 330 | 245 |
| SE | 33789 | ENSG00000134109 | EDEM1 | chr3 | + | 5210174 | 5210248 | 5208092 | 5208263 | 5213318 | 5213522 | 61 | 42 |
| SE | 33790 | ENSG00000134109 | EDEM1 | chr3 | + | 5211119 | 5211216 | 5208092 | 5208263 | 5213318 | 5213522 | 153 | 127 |
| SE | 33791 | ENSG00000134109 | EDEM1 | chr3 | + | 5211119 | 5211216 | 5210174 | 5210248 | 5213318 | 5213522 | 327 | 250 |
| SE | 33796 | ENSG00000009830 | POMT2 | chr14 | - | 77279536 | 77279586 | 77278728 | 77278869 | 77279822 | 77279928 | 2 | 0 |
| SE | 33797 | ENSG00000009830 | POMT2 | chr14 | - | 77279536 | 77279622 | 77278728 | 77278869 | 77279822 | 77279928 | 3 | 0 |
| SE | 33798 | ENSG00000009830 | POMT2 | chr14 | - | 77279536 | 77279683 | 77278728 | 77278869 | 77279822 | 77279928 | 2 | 0 |
| SE | 33805 | ENSG00000009830 | POMT2 | chr14 | - | 77291513 | 77291664 | 77291313 | 77291380 | 77296163 | 77296273 | 0 | 0 |
| SE | 33806 | ENSG00000009830 | POMT2 | chr14 | - | 77292769 | 77293290 | 77291313 | 77291380 | 77296163 | 77296273 | 0 | 0 |
| SE | 33808 | ENSG00000009830 | POMT2 | chr14 | - | 77298688 | 77298771 | 77296163 | 77296273 | 77304691 | 77304800 | 5 | 0 |
| SE | 33811 | ENSG00000009830 | POMT2 | chr14 | - | 77299454 | 77299561 | 77298688 | 77298771 | 77302834 | 77302943 | 1 | 2 |
| SE | 33812 | ENSG00000009830 | POMT2 | chr14 | - | 77299454 | 77299561 | 77298688 | 77298771 | 77304691 | 77304800 | 1 | 2 |
| SE | 33817 | ENSG00000009830 | POMT2 | chr14 | - | 77301089 | 77301249 | 77299454 | 77299561 | 77302834 | 77302943 | 3 | 0 |
| SE | 33819 | ENSG00000009830 | POMT2 | chr14 | - | 77302834 | 77302943 | 77298688 | 77298771 | 77304691 | 77304800 | 7 | 2 |
| SE | 33823 | ENSG00000009830 | POMT2 | chr14 | - | 77308965 | 77309027 | 77306336 | 77306441 | 77311948 | 77312033 | 0 | 0 |
| SE | 33824 | ENSG00000009830 | POMT2 | chr14 | - | 77311948 | 77312033 | 77306245 | 77306441 | 77320433 | 77320476 | 3 | 2 |
| SE | 33828 | ENSG00000041802 | LSG1 | chr3 | - | 194670008 | 194670135 | 194666451 | 194666572 | 194672063 | 194672162 | 177 | 113 |
| SE | 33829 | ENSG00000231770 | TMEM44-AS1 | chr3 | + | 194589020 | 194589142 | 194584010 | 194584448 | 194589473 | 194589626 | 0 | 0 |
| SE | 33830 | ENSG00000231770 | TMEM44-AS1 | chr3 | + | 194589024 | 194589142 | 194584013 | 194584448 | 194589473 | 194589626 | 0 | 1 |
| SE | 33831 | ENSG00000231770 | TMEM44-AS1 | chr3 | + | 194589473 | 194589626 | 194589020 | 194589142 | 194589854 | 194590249 | 5 | 2 |
| SE | 33837 | ENSG00000138182 | KIF20B | chr10 | + | 89705293 | 89705441 | 89701589 | 89701680 | 89709166 | 89709253 | 28 | 32 |
| SE | 33838 | ENSG00000138182 | KIF20B | chr10 | + | 89709166 | 89709253 | 89705293 | 89705441 | 89709926 | 89710065 | 11 | 17 |
| SE | 33839 | ENSG00000138182 | KIF20B | chr10 | + | 89709166 | 89709253 | 89705293 | 89705441 | 89710960 | 89711090 | 11 | 17 |
| SE | 33840 | ENSG00000138182 | KIF20B | chr10 | + | 89709344 | 89709461 | 89709166 | 89709253 | 89709926 | 89710065 | 89 | 97 |
| SE | 33841 | ENSG00000138182 | KIF20B | chr10 | + | 89709344 | 89709461 | 89709166 | 89709253 | 89710960 | 89711090 | 33 | 29 |
| SE | 33842 | ENSG00000138182 | KIF20B | chr10 | + | 89709926 | 89710065 | 89705293 | 89705441 | 89710960 | 89711090 | 37 | 50 |
| SE | 33843 | ENSG00000138182 | KIF20B | chr10 | + | 89709926 | 89710065 | 89709166 | 89709253 | 89710960 | 89711090 | 37 | 50 |
| SE | 33844 | ENSG00000138182 | KIF20B | chr10 | + | 89709926 | 89710065 | 89709344 | 89709461 | 89710960 | 89711090 | 92 | 118 |
| SE | 33845 | ENSG00000138182 | KIF20B | chr10 | + | 89718709 | 89718872 | 89717575 | 89717722 | 89719418 | 89719706 | 42 | 43 |
| SE | 33846 | ENSG00000109065 | NAT9 | chr17 | - | 74772482 | 74772576 | 74771959 | 74772395 | 74772895 | 74773039 | 0 | 0 |
| SE | 33847 | ENSG00000109065 | NAT9 | chr17 | - | 74773865 | 74773939 | 74773575 | 74773685 | 74775621 | 74775707 | 1 | 0 |
| SE | 33848 | ENSG00000109065 | NAT9 | chr17 | - | 74773865 | 74774055 | 74773575 | 74773688 | 74775621 | 74775707 | 0 | 0 |
| SE | 33863 | ENSG00000171109 | MFN1 | chr3 | + | 179348844 | 179348963 | 179347708 | 179347810 | 179351899 | 179352035 | 209 | 69 |
| SE | 33864 | ENSG00000171109 | MFN1 | chr3 | + | 179362357 | 179362482 | 179358839 | 179359002 | 179364296 | 179364405 | 196 | 109 |
| SE | 33865 | ENSG00000171109 | MFN1 | chr3 | + | 179364296 | 179364405 | 179362357 | 179362482 | 179365117 | 179365225 | 293 | 191 |
| SE | 33866 | ENSG00000171109 | MFN1 | chr3 | + | 179367438 | 179367592 | 179365117 | 179365225 | 179368035 | 179368103 | 569 | 351 |
| SE | 33867 | ENSG00000171109 | MFN1 | chr3 | + | 179375219 | 179375341 | 179368035 | 179368103 | 179377041 | 179377168 | 264 | 143 |
| SE | 33868 | ENSG00000171109 | MFN1 | chr3 | + | 179377041 | 179377168 | 179375219 | 179375341 | 179377343 | 179377448 | 221 | 144 |
| SE | 33869 | ENSG00000171109 | MFN1 | chr3 | + | 179378340 | 179378814 | 179377343 | 179377448 | 179385568 | 179385721 | 212 | 113 |
| SE | 33870 | ENSG00000132153 | DHX30 | chr3 | + | 47810656 | 47810711 | 47805325 | 47805420 | 47815962 | 47816128 | 43 | 16 |
| SE | 33872 | ENSG00000132153 | DHX30 | chr3 | + | 47818021 | 47818117 | 47810656 | 47810711 | 47827346 | 47827477 | 29 | 17 |
| SE | 33874 | ENSG00000132153 | DHX30 | chr3 | + | 47819235 | 47819264 | 47818021 | 47818117 | 47824766 | 47825004 | 2 | 4 |
| SE | 33876 | ENSG00000132153 | DHX30 | chr3 | + | 47822096 | 47822231 | 47819235 | 47819264 | 47824766 | 47824863 | 0 | 1 |
| SE | 33877 | ENSG00000132153 | DHX30 | chr3 | + | 47827346 | 47827477 | 47810656 | 47810711 | 47840876 | 47841178 | 2 | 3 |
| SE | 33879 | ENSG00000132153 | DHX30 | chr3 | + | 47827346 | 47827477 | 47819226 | 47819264 | 47829023 | 47829134 | 104 | 59 |
| SE | 33880 | ENSG00000132153 | DHX30 | chr3 | + | 47829023 | 47829134 | 47827346 | 47827477 | 47840876 | 47841178 | 195 | 116 |
| SE | 33881 | ENSG00000132153 | DHX30 | chr3 | + | 47841616 | 47841737 | 47840876 | 47841178 | 47843105 | 47843255 | 172 | 84 |
| SE | 33882 | ENSG00000132153 | DHX30 | chr3 | + | 47845699 | 47845852 | 47843105 | 47843255 | 47846164 | 47847001 | 218 | 87 |
| SE | 33885 | ENSG00000132153 | DHX30 | chr3 | + | 47848623 | 47848817 | 47848468 | 47848550 | 47848919 | 47848996 | 53 | 24 |
| SE | 33886 | ENSG00000132153 | DHX30 | chr3 | + | 47849629 | 47849769 | 47849450 | 47849554 | 47849866 | 47850187 | 31 | 13 |
| SE | 33890 | ENSG00000157036 | EXOG | chr3 | + | 38496842 | 38496929 | 38496126 | 38496530 | 38497628 | 38497684 | 1 | 0 |
| SE | 33891 | ENSG00000157036 | EXOG | chr3 | + | 38497628 | 38497684 | 38496271 | 38496530 | 38501354 | 38501494 | 4 | 1 |
| SE | 33892 | ENSG00000157036 | EXOG | chr3 | + | 38497628 | 38497778 | 38496339 | 38496530 | 38500089 | 38500164 | 2 | 3 |
| SE | 33893 | ENSG00000157036 | EXOG | chr3 | + | 38497628 | 38497778 | 38496339 | 38496530 | 38501354 | 38501494 | 9 | 9 |
| SE | 33894 | ENSG00000157036 | EXOG | chr3 | + | 38498766 | 38498836 | 38497628 | 38497684 | 38498925 | 38498989 | 0 | 0 |
| SE | 33895 | ENSG00000157036 | EXOG | chr3 | + | 38498766 | 38498836 | 38497628 | 38497684 | 38501354 | 38501494 | 0 | 0 |
| SE | 33896 | ENSG00000157036 | EXOG | chr3 | + | 38498766 | 38498836 | 38497628 | 38497689 | 38498925 | 38498989 | 0 | 0 |
| SE | 33898 | ENSG00000157036 | EXOG | chr3 | + | 38498766 | 38498836 | 38497628 | 38497778 | 38498925 | 38498989 | 2 | 0 |
| SE | 33900 | ENSG00000157036 | EXOG | chr3 | + | 38498925 | 38498989 | 38497628 | 38497684 | 38501354 | 38501494 | 1 | 0 |
| SE | 33901 | ENSG00000157036 | EXOG | chr3 | + | 38498925 | 38498989 | 38497628 | 38497689 | 38500089 | 38500164 | 0 | 0 |
| SE | 33904 | ENSG00000157036 | EXOG | chr3 | + | 38498925 | 38498989 | 38497628 | 38497778 | 38500089 | 38500164 | 0 | 0 |
| SE | 33905 | ENSG00000157036 | EXOG | chr3 | + | 38498925 | 38498989 | 38497628 | 38497778 | 38501354 | 38501494 | 1 | 0 |
| SE | 33907 | ENSG00000157036 | EXOG | chr3 | + | 38499680 | 38499747 | 38497628 | 38497778 | 38500089 | 38500164 | 1 | 0 |
| SE | 33908 | ENSG00000157036 | EXOG | chr3 | + | 38499680 | 38499747 | 38497628 | 38497778 | 38501354 | 38501494 | 0 | 0 |
| SE | 33909 | ENSG00000157036 | EXOG | chr3 | + | 38499680 | 38499747 | 38498925 | 38498989 | 38500089 | 38500164 | 1 | 0 |
| SE | 33910 | ENSG00000157036 | EXOG | chr3 | + | 38499680 | 38499747 | 38498925 | 38498989 | 38501354 | 38501494 | 0 | 0 |
| SE | 33912 | ENSG00000157036 | EXOG | chr3 | + | 38500089 | 38500164 | 38496737 | 38497689 | 38501354 | 38501494 | 3 | 1 |
| SE | 33915 | ENSG00000157036 | EXOG | chr3 | + | 38500089 | 38500164 | 38498925 | 38498989 | 38501354 | 38501494 | 3 | 2 |
| SE | 33916 | ENSG00000157036 | EXOG | chr3 | + | 38500089 | 38500164 | 38499680 | 38499747 | 38501354 | 38501494 | 4 | 1 |
| SE | 33917 | ENSG00000157036 | EXOG | chr3 | + | 38501354 | 38501494 | 38496344 | 38496530 | 38503614 | 38503691 | 41 | 44 |
| SE | 33918 | ENSG00000157036 | EXOG | chr3 | + | 38513749 | 38513868 | 38506853 | 38506968 | 38523900 | 38524403 | 0 | 0 |
| SE | 33919 | ENSG00000157036 | EXOG | chr3 | + | 38519261 | 38519377 | 38506853 | 38506968 | 38523900 | 38524060 | 1 | 0 |
| SE | 33920 | ENSG00000215256 | DHRS4-AS1 | chr14 | - | 23953308 | 23953419 | 23938761 | 23941158 | 23953773 | 23954033 | 2 | 2 |
| SE | 33926 | ENSG00000160796 | NBEAL2 | chr3 | + | 46991405 | 46991688 | 46991218 | 46991304 | 46991839 | 46991946 | 290 | 102 |
| SE | 33928 | ENSG00000160796 | NBEAL2 | chr3 | + | 46994454 | 46994553 | 46993936 | 46994020 | 46995031 | 46995633 | 78 | 25 |
| SE | 33929 | ENSG00000160796 | NBEAL2 | chr3 | + | 46995931 | 46996051 | 46995713 | 46995846 | 46996270 | 46996592 | 192 | 70 |
| SE | 33930 | ENSG00000160796 | NBEAL2 | chr3 | + | 46996750 | 46996833 | 46996270 | 46996592 | 46996953 | 46997046 | 132 | 44 |
| SE | 33932 | ENSG00000160796 | NBEAL2 | chr3 | + | 46998958 | 46999117 | 46998716 | 46998879 | 46999314 | 46999474 | 60 | 13 |
| SE | 33933 | ENSG00000160796 | NBEAL2 | chr3 | + | 47001919 | 47002288 | 47001688 | 47001826 | 47002370 | 47002520 | 165 | 72 |
| SE | 33934 | ENSG00000160796 | NBEAL2 | chr3 | + | 47003173 | 47003309 | 47002956 | 47003081 | 47003815 | 47003976 | 170 | 54 |
| SE | 33936 | ENSG00000160796 | NBEAL2 | chr3 | + | 47005945 | 47006063 | 47005737 | 47005847 | 47006164 | 47006262 | 250 | 108 |
| SE | 33940 | ENSG00000160796 | NBEAL2 | chr3 | + | 47007240 | 47007350 | 47007065 | 47007155 | 47007524 | 47007697 | 193 | 105 |
| SE | 33942 | ENSG00000160796 | NBEAL2 | chr3 | + | 47008069 | 47008186 | 47007815 | 47007910 | 47008282 | 47008441 | 152 | 65 |
| SE | 33948 | ENSG00000140451 | PIF1 | chr15 | - | 64821366 | 64821520 | 64821166 | 64821281 | 64822265 | 64822391 | 1 | 0 |
| SE | 33950 | ENSG00000109790 | KLHL5 | chr4 | + | 39063571 | 39063797 | 39062939 | 39063035 | 39075964 | 39076147 | 0 | 1 |
| SE | 33951 | ENSG00000109790 | KLHL5 | chr4 | + | 39075964 | 39076147 | 39045059 | 39045096 | 39081102 | 39081182 | 82 | 92 |
| SE | 33952 | ENSG00000109790 | KLHL5 | chr4 | + | 39075964 | 39076147 | 39062231 | 39063035 | 39081102 | 39081182 | 104 | 143 |
| SE | 33953 | ENSG00000109790 | KLHL5 | chr4 | + | 39081962 | 39082159 | 39081102 | 39081239 | 39086514 | 39086727 | 252 | 316 |
| SE | 33954 | ENSG00000109790 | KLHL5 | chr4 | + | 39103286 | 39103511 | 39096691 | 39096878 | 39107568 | 39107731 | 206 | 363 |
| SE | 33959 | ENSG00000165195 | PIGA | chrX | - | 15326949 | 15327255 | 15325913 | 15326046 | 15331215 | 15331992 | 0 | 0 |
| SE | 33962 | ENSG00000165195 | PIGA | chrX | - | 15331215 | 15331992 | 15325913 | 15326046 | 15335500 | 15335523 | 23 | 31 |
| SE | 33963 | ENSG00000165195 | PIGA | chrX | - | 15331589 | 15331992 | 15325913 | 15326046 | 15335500 | 15335503 | 10 | 14 |
| SE | 33964 | ENSG00000165195 | PIGA | chrX | - | 15331819 | 15331992 | 15325913 | 15326046 | 15335500 | 15335544 | 8 | 13 |
| SE | 33965 | ENSG00000165195 | PIGA | chrX | - | 15331873 | 15331992 | 15325606 | 15326046 | 15335500 | 15335523 | 0 | 0 |
| SE | 33974 | ENSG00000079102 | RUNX1T1 | chr8 | - | 92062545 | 92062709 | 92017225 | 92017431 | 92095027 | 92095235 | 0 | 0 |
| SE | 33988 | ENSG00000114742 | WDR48 | chr3 | + | 39063049 | 39063190 | 39052012 | 39052073 | 39065810 | 39065889 | 272 | 146 |
| SE | 33991 | ENSG00000114742 | WDR48 | chr3 | + | 39066726 | 39066875 | 39066547 | 39066630 | 39068770 | 39068820 | 230 | 137 |
| SE | 33992 | ENSG00000114742 | WDR48 | chr3 | + | 39066745 | 39066875 | 39066547 | 39066630 | 39068770 | 39068859 | 409 | 265 |
| SE | 33993 | ENSG00000114742 | WDR48 | chr3 | + | 39074725 | 39074950 | 39069642 | 39069744 | 39077138 | 39077213 | 279 | 167 |
| SE | 33994 | ENSG00000114742 | WDR48 | chr3 | + | 39077138 | 39077213 | 39074725 | 39074950 | 39078136 | 39078239 | 314 | 224 |
| SE | 33995 | ENSG00000114742 | WDR48 | chr3 | + | 39078136 | 39078239 | 39077138 | 39077213 | 39079710 | 39079808 | 437 | 317 |
| SE | 33997 | ENSG00000114742 | WDR48 | chr3 | + | 39079710 | 39079808 | 39077138 | 39077213 | 39084154 | 39084187 | 79 | 57 |
| SE | 33998 | ENSG00000114742 | WDR48 | chr3 | + | 39079710 | 39079808 | 39078010 | 39078239 | 39084154 | 39084187 | 398 | 315 |
| SE | 33999 | ENSG00000114742 | WDR48 | chr3 | + | 39084644 | 39084741 | 39084154 | 39084262 | 39085514 | 39085610 | 205 | 126 |
| SE | 34003 | ENSG00000060971 | ACAA1 | chr3 | - | 38126161 | 38126700 | 38125825 | 38125881 | 38129289 | 38129388 | 83 | 45 |
| SE | 34005 | ENSG00000060971 | ACAA1 | chr3 | - | 38126509 | 38126700 | 38125825 | 38125881 | 38129289 | 38129388 | 62 | 24 |
| SE | 34006 | ENSG00000060971 | ACAA1 | chr3 | - | 38127785 | 38127866 | 38125825 | 38125881 | 38129289 | 38129388 | 115 | 104 |
| SE | 34008 | ENSG00000060971 | ACAA1 | chr3 | - | 38127785 | 38127866 | 38125825 | 38126700 | 38131595 | 38131638 | 198 | 113 |
| SE | 34009 | ENSG00000060971 | ACAA1 | chr3 | - | 38129289 | 38129388 | 38125825 | 38126700 | 38131595 | 38131638 | 48 | 49 |
| SE | 34010 | ENSG00000060971 | ACAA1 | chr3 | - | 38129289 | 38129388 | 38127630 | 38127866 | 38131595 | 38131638 | 224 | 184 |
| SE | 34011 | ENSG00000060971 | ACAA1 | chr3 | - | 38129289 | 38129388 | 38127630 | 38127866 | 38131925 | 38132005 | 176 | 126 |
| SE | 34012 | ENSG00000060971 | ACAA1 | chr3 | - | 38131595 | 38131638 | 38127785 | 38127866 | 38131925 | 38131946 | 19 | 15 |
| SE | 34019 | ENSG00000173137 | ADCK5 | chr8 | + | 144377392 | 144377490 | 144374041 | 144374107 | 144379386 | 144379490 | 0 | 0 |
| SE | 34026 | ENSG00000114859 | CLCN2 | chr3 | - | 184352012 | 184352117 | 184346934 | 184347021 | 184352292 | 184352331 | 5 | 9 |
| SE | 34027 | ENSG00000114859 | CLCN2 | chr3 | - | 184352292 | 184352331 | 184352012 | 184352117 | 184352442 | 184352496 | 4 | 5 |
| SE | 34032 | ENSG00000114859 | CLCN2 | chr3 | - | 184357619 | 184357698 | 184357361 | 184357487 | 184357961 | 184358095 | 0 | 0 |
| SE | 34033 | ENSG00000114859 | CLCN2 | chr3 | - | 184357778 | 184357856 | 184357619 | 184357698 | 184357961 | 184358095 | 5 | 0 |
| SE | 34045 | ENSG00000114867 | EIF4G1 | chr3 | + | 184315488 | 184315545 | 184314501 | 184314674 | 184316131 | 184316218 | 8 | 0 |
| SE | 34047 | ENSG00000114867 | EIF4G1 | chr3 | + | 184315488 | 184315545 | 184314501 | 184314674 | 184317746 | 184317816 | 8 | 0 |
| SE | 34048 | ENSG00000114867 | EIF4G1 | chr3 | + | 184315762 | 184315856 | 184314494 | 184314674 | 184316131 | 184316218 | 54 | 23 |
| SE | 34051 | ENSG00000114867 | EIF4G1 | chr3 | + | 184315762 | 184315856 | 184315165 | 184315545 | 184316131 | 184316218 | 109 | 48 |
| SE | 34052 | ENSG00000114867 | EIF4G1 | chr3 | + | 184315762 | 184315856 | 184315165 | 184315545 | 184317320 | 184317431 | 52 | 24 |
| SE | 34057 | ENSG00000114867 | EIF4G1 | chr3 | + | 184316131 | 184316218 | 184314494 | 184314674 | 184319688 | 184319801 | 0 | 0 |
| SE | 34058 | ENSG00000114867 | EIF4G1 | chr3 | + | 184316131 | 184316218 | 184315165 | 184315545 | 184317320 | 184317431 | 59 | 44 |
| SE | 34059 | ENSG00000114867 | EIF4G1 | chr3 | + | 184316131 | 184316218 | 184315165 | 184315545 | 184317716 | 184317816 | 0 | 0 |
| SE | 34060 | ENSG00000114867 | EIF4G1 | chr3 | + | 184316131 | 184316218 | 184315407 | 184315856 | 184317320 | 184317431 | 119 | 68 |
| SE | 34063 | ENSG00000114867 | EIF4G1 | chr3 | + | 184317320 | 184317431 | 184315762 | 184315856 | 184317716 | 184317816 | 3 | 0 |
| SE | 34067 | ENSG00000114867 | EIF4G1 | chr3 | + | 184317320 | 184317497 | 184314541 | 184314674 | 184317749 | 184317816 | 14 | 4 |
| SE | 34069 | ENSG00000114867 | EIF4G1 | chr3 | + | 184317320 | 184317497 | 184315163 | 184315545 | 184317746 | 184317816 | 4 | 2 |
| SE | 34072 | ENSG00000114867 | EIF4G1 | chr3 | + | 184317320 | 184317497 | 184316071 | 184316218 | 184317746 | 184317816 | 115 | 72 |
| SE | 34079 | ENSG00000114867 | EIF4G1 | chr3 | + | 184322361 | 184322450 | 184321281 | 184322103 | 184322543 | 184322730 | 575 | 478 |
| SE | 34081 | ENSG00000114867 | EIF4G1 | chr3 | + | 184323082 | 184323218 | 184322820 | 184322954 | 184323407 | 184323593 | 390 | 270 |
| SE | 34083 | ENSG00000114867 | EIF4G1 | chr3 | + | 184323082 | 184323241 | 184322820 | 184322954 | 184323407 | 184323593 | 627 | 406 |
| SE | 34084 | ENSG00000114867 | EIF4G1 | chr3 | + | 184324877 | 184325114 | 184324200 | 184324347 | 184325268 | 184325299 | 1719 | 1163 |
| SE | 34085 | ENSG00000114867 | EIF4G1 | chr3 | + | 184325268 | 184325373 | 184324877 | 184325114 | 184325479 | 184325639 | 1097 | 708 |
| SE | 34087 | ENSG00000114867 | EIF4G1 | chr3 | + | 184325850 | 184325951 | 184325298 | 184325639 | 184326880 | 184326908 | 276 | 185 |
| SE | 34089 | ENSG00000114867 | EIF4G1 | chr3 | + | 184326526 | 184326629 | 184325298 | 184325639 | 184326880 | 184326908 | 267 | 156 |
| SE | 34090 | ENSG00000114867 | EIF4G1 | chr3 | + | 184326526 | 184326629 | 184325850 | 184325951 | 184326880 | 184326908 | 545 | 330 |
| SE | 34091 | ENSG00000114867 | EIF4G1 | chr3 | + | 184326880 | 184326983 | 184326526 | 184326629 | 184327215 | 184327303 | 574 | 356 |
| SE | 34092 | ENSG00000114867 | EIF4G1 | chr3 | + | 184327829 | 184328002 | 184327585 | 184327704 | 184328630 | 184328756 | 373 | 148 |
| SE | 34093 | ENSG00000114867 | EIF4G1 | chr3 | + | 184328630 | 184328756 | 184327585 | 184327704 | 184331265 | 184331364 | 0 | 0 |
| SE | 34095 | ENSG00000135083 | CCNJL | chr5 | - | 160254128 | 160254335 | 160251652 | 160253798 | 160255548 | 160255708 | 0 | 0 |
| SE | 34100 | ENSG00000135083 | CCNJL | chr5 | - | 160294727 | 160294963 | 160280524 | 160280738 | 160311857 | 160311972 | 0 | 0 |
| SE | 34101 | ENSG00000135083 | CCNJL | chr5 | - | 160311857 | 160311972 | 160280524 | 160280738 | 160312744 | 160312928 | 2 | 3 |
| SE | 34102 | ENSG00000176894 | PXMP2 | chr12 | + | 132695883 | 132696046 | 132690262 | 132690376 | 132701249 | 132701369 | 16 | 9 |
| SE | 34103 | ENSG00000176894 | PXMP2 | chr12 | + | 132695883 | 132696046 | 132690262 | 132690376 | 132704618 | 132704898 | 8 | 9 |
| SE | 34104 | ENSG00000176894 | PXMP2 | chr12 | + | 132701249 | 132701369 | 132690262 | 132690376 | 132704618 | 132704900 | 2 | 1 |
| SE | 34108 | ENSG00000129235 | TXNDC17 | chr17 | + | 6641752 | 6641834 | 6641026 | 6641227 | 6642248 | 6642324 | 31 | 12 |
| SE | 34109 | ENSG00000129235 | TXNDC17 | chr17 | + | 6641752 | 6641932 | 6641033 | 6641227 | 6642248 | 6642324 | 7 | 3 |
| SE | 34111 | ENSG00000075785 | RAB7A | chr3 | + | 128788913 | 128789070 | 128726162 | 128726359 | 128795359 | 128795420 | 1 | 0 |
| SE | 34112 | ENSG00000075785 | RAB7A | chr3 | + | 128795359 | 128795420 | 128726135 | 128726359 | 128797942 | 128798036 | 595 | 386 |
| SE | 34114 | ENSG00000075785 | RAB7A | chr3 | + | 128806371 | 128806449 | 128797942 | 128798069 | 128807542 | 128807671 | 337 | 249 |
| SE | 34115 | ENSG00000075785 | RAB7A | chr3 | + | 128806371 | 128806590 | 128797942 | 128798069 | 128807542 | 128807594 | 1262 | 949 |
| SE | 34116 | ENSG00000075785 | RAB7A | chr3 | + | 128806371 | 128807671 | 128797942 | 128798069 | 128813326 | 128814796 | 1115 | 807 |
| SE | 34117 | ENSG00000075785 | RAB7A | chr3 | + | 128807542 | 128807671 | 128797942 | 128798069 | 128813326 | 128813541 | 477 | 313 |
| SE | 34118 | ENSG00000075785 | RAB7A | chr3 | + | 128811145 | 128811257 | 128807542 | 128807671 | 128813326 | 128814762 | 0 | 0 |
| SE | 34120 | ENSG00000116095 | PLEKHA3 | chr2 | + | 178499210 | 178499254 | 178495495 | 178495660 | 178501060 | 178501176 | 179 | 130 |
| SE | 34148 | ENSG00000181458 | TMEM45A | chr3 | + | 100519076 | 100519316 | 100492650 | 100492928 | 100519461 | 100519608 | 4 | 0 |
| SE | 34149 | ENSG00000181458 | TMEM45A | chr3 | + | 100519461 | 100519608 | 100492650 | 100492928 | 100555208 | 100555401 | 2 | 0 |

| | | | | | | | | | | | | | | |
|---|---|---|---|---|---|---|---|---|---|---|---|---|---|---|
| SE | 34150 | ENSG00000181458 | TMEM45A | chr3 | + | 100519532 | 100519608 | 100492659 | 100492928 | 100555208 | 100555401 | 0 | 0 | [illegible] |
| SE | 34151 | ENSG00000181458 | TMEM45A | chr3 | + | 100555208 | 100555401 | 100519461 | 100519608 | 100558404 | 100558527 | 3 | 0 | [illegible] |
| SE | 34153 | ENSG00000181458 | TMEM45A | chr3 | + | 100570608 | 100574659 | 100568923 | 100568967 | 100576924 | 100577444 | 0 | 0 | [illegible] |
| SE | 34155 | ENSG00000102144 | PGK1 | chrX | + | 78113743 | 78113899 | 78109866 | 78109917 | 78114015 | 78114160 | 1261 | 745 | [illegible] |
| SE | 34156 | ENSG00000102144 | PGK1 | chrX | + | 78118050 | 78118170 | 78117311 | 78117415 | 78122834 | 78122903 | 1184 | 732 | [illegible] |
| SE | 34161 | ENSG00000198585 | NUDT16 | chr3 | + | 131382045 | 131382315 | 131381773 | 131381942 | 131383161 | 131383272 | 19 | 6 | [illegible] |
| SE | 34167 | ENSG00000163660 | CCNL1 | chr3 | - | 157151485 | 157151591 | 157150281 | 157150381 | 157151728 | 157151930 | 5 | 2 | [illegible] |
| SE | 34173 | ENSG00000204427 | ABHD16A | chr6 | - | 31688248 | 31688305 | 31688040 | 31688103 | 31688722 | 31688786 | 82 | 53 | [illegible] |
| SE | 34174 | ENSG00000204427 | ABHD16A | chr6 | - | 31689014 | 31689119 | 31688722 | 31688786 | 31689580 | 31689704 | 73 | 37 | [illegible] |
| SE | 34175 | ENSG00000204427 | ABHD16A | chr6 | - | 31690077 | 31690127 | 31689580 | 31689704 | 31690538 | 31690602 | 13 | 7 | [illegible] |
| SE | 34176 | ENSG00000204427 | ABHD16A | chr6 | - | 31696947 | 31697033 | 31693358 | 31693432 | 31700941 | 31701028 | 102 | 62 | [illegible] |
| SE | 34178 | ENSG00000204427 | ABHD16A | chr6 | - | 31702073 | 31702130 | 31700941 | 31701028 | 31702611 | 31702984 | 6 | 1 | [illegible] |
| SE | 34179 | ENSG00000204427 | ABHD16A | chr6 | - | 31702073 | 31702130 | 31700941 | 31701028 | 31703149 | 31703311 | 4 | 1 | [illegible] |
| SE | 34185 | ENSG00000175193 | PARL | chr3 | - | 183856431 | 183856551 | 183844230 | 183844326 | 183862752 | 183862801 | 3 | 1 | [illegible] |
| SE | 34186 | ENSG00000175193 | PARL | chr3 | - | 183862696 | 183862801 | 183844230 | 183844326 | 183866624 | 183866765 | 113 | 94 | [illegible] |
| SE | 34187 | ENSG00000175193 | PARL | chr3 | - | 183862752 | 183862801 | 183844230 | 183844326 | 183866624 | 183866765 | 273 | 185 | [illegible] |
| SE | 34188 | ENSG00000175193 | PARL | chr3 | - | 183866624 | 183866765 | 183862606 | 183862801 | 183867864 | 183868060 | 488 | 363 | [illegible] |
| SE | 34189 | ENSG00000175193 | PARL | chr3 | - | 183867864 | 183868060 | 183866624 | 183866765 | 183884721 | 183884727 | 316 | 212 | [illegible] |
| SE | 34191 | ENSG00000154767 | XPC | chr3 | - | 14148561 | 14148731 | 14147466 | 14148001 | 14148813 | 14148928 | 129 | 81 | [illegible] |
| SE | 34193 | ENSG00000154767 | XPC | chr3 | - | 14159740 | 14159830 | 14158010 | 14158892 | 14164812 | 14164933 | 168 | 130 | [illegible] |
| [illegible] | | | | | | | | | | | | | | |

| | | | | | | | | | | | | | | |
|---|---|---|---|---|---|---|---|---|---|---|---|---|---|---|
| SE | 34756 | ENSG00000236548 | RNF217-AS1 | chr6 | - | 124941389 | 124941524 | 124935007 | 124935117 | 124947930 | 124948098 | 0 | 1 | [illegible] |
| SE | 34758 | ENSG00000236548 | RNF217-AS1 | chr6 | - | 124941389 | 124941604 | 124935007 | 124935117 | 124947930 | 124948098 | 0 | 0 | [illegible] |
| SE | 34759 | ENSG00000236548 | RNF217-AS1 | chr6 | - | 124945298 | 124945439 | 124935007 | 124935117 | 124947930 | 124948098 | 0 | 2 | [illegible] |
| SE | 34760 | ENSG00000236548 | RNF217-AS1 | chr6 | - | 124945298 | 124945439 | 124941389 | 124941524 | 124947930 | 124948098 | 0 | 2 | [illegible] |
| SE | 34764 | ENSG00000112149 | CD83 | chr6 | + | 14133648 | 14133755 | 14117949 | 14118065 | 14135107 | 14136906 | 38 | 31 | [illegible] |
| SE | 34769 | ENSG00000100034 | PPM1F | chr22 | - | 21925568 | 21925662 | 21919424 | 21923471 | 21931147 | 21931248 | 138 | 55 | [illegible] |
| SE | 34770 | ENSG00000100034 | PPM1F | chr22 | - | 21939531 | 21939680 | 21934023 | 21934226 | 21945842 | 21946108 | 323 | 172 | [illegible] |
| SE | 34771 | ENSG00000100034 | PPM1F | chr22 | - | 21939531 | 21939680 | 21934023 | 21934226 | 21952791 | 21952831 | 188 | 96 | [illegible] |
| SE | 34772 | ENSG00000100034 | PPM1F | chr22 | - | 21945842 | 21946108 | 21934023 | 21934226 | 21952791 | 21952831 | 56 | 22 | [illegible] |
| SE | 34774 | ENSG00000075790 | BCAP29 | chr7 | + | 107580645 | 107580864 | 107580271 | 107580301 | 107583881 | 107583982 | 53 | 45 | [illegible] |
| SE | 34775 | ENSG00000075790 | BCAP29 | chr7 | + | 107580758 | 107580864 | 107580245 | 107580301 | 107583881 | 107583982 | 67 | 45 | [illegible] |
| SE | 34776 | ENSG00000075790 | BCAP29 | chr7 | + | 107593954 | 107594105 | 107583881 | 107583982 | 107595866 | 107595921 | 338 | 198 | [illegible] |
| SE | 34778 | ENSG00000075790 | BCAP29 | chr7 | + | 107595866 | 107596002 | 107583881 | 107583982 | 107600396 | 107600505 | 203 | 168 | [illegible] |
| SE | 34779 | ENSG00000075790 | BCAP29 | chr7 | + | 107595866 | 107596002 | 107593954 | 107594105 | 107600396 | 107600505 | 377 | 297 | [illegible] |
| SE | 34780 | ENSG00000075790 | BCAP29 | chr7 | + | 107595868 | 107596002 | 107593954 | 107594105 | 107600396 | 107600505 | 200 | 163 | [illegible] |
| SE | 34782 | ENSG00000075790 | BCAP29 | chr7 | + | 107613653 | 107613732 | 107613331 | 107613432 | 107616260 | 107616389 | 9 | 1 | [illegible] |
| SE | 34788 | ENSG00000075790 | BCAP29 | chr7 | + | 107616260 | 107616389 | 107613653 | 107613732 | 107618327 | 107618635 | 3 | 4 | [illegible] |
| SE | 34789 | ENSG00000075790 | BCAP29 | chr7 | + | 107616260 | 107616389 | 107613653 | 107613732 | 107621629 | 107621953 | 1 | 1 | [illegible] |
| SE | 34791 | ENSG00000100292 | HMOX1 | chr22 | + | 35389863 | 35389963 | 35386684 | 35387176 | 35393467 | 35394207 | 173 | 69 | [illegible] |
| SE | 34793 | ENSG00000136152 | COG3 | chr13 | + | 45492158 | 45492250 | 45491411 | 45491538 | 45493346 | 45493486 | 134 | 122 | [illegible] |
| SE | 34794 | ENSG00000136152 | COG3 | chr13 | + | 45493346 | 45493486 | 45492158 | 45492250 | 45496151 | 45496312 | 252 | 205 | [illegible] |
| SE | 34795 | ENSG00000136152 | COG3 | chr13 | + | 45493346 | 45493486 | 45492158 | 45492250 | 45503243 | 45503349 | 94 | 92 | [illegible] |
| SE | 34796 | ENSG00000136152 | COG3 | chr13 | + | 45496151 | 45496312 | 45492158 | 45492250 | 45503243 | 45503349 | 78 | 39 | [illegible] |
| SE | 34798 | ENSG00000136152 | COG3 | chr13 | + | 45511764 | 45511854 | 45509691 | 45509816 | 45516142 | 45516263 | 159 | 107 | [illegible] |
| SE | 34799 | ENSG00000136152 | COG3 | chr13 | + | 45524975 | 45525051 | 45518959 | 45519094 | 45529790 | 45529918 | 78 | 75 | [illegible] |
| [illegible] | | | | | | | | | | | | | | |

| | | | | | | | | | | | | | | | | | | | | | | | | | | | | | | | | | | | | | | | | | | | | | | |
|---|---|---|---|---|---|---|---|---|---|---|---|---|---|---|---|---|---|---|---|---|---|---|---|---|---|---|---|---|---|---|---|---|---|---|---|---|---|---|---|---|---|---|---|---|---|---|
| SE | 35123 | ENSG00000266338 | NBPF15 | chr1 | - | 144459365 | 144459483 | 144456536 | 144456805 | 144460842 | 144460961 | 20 | 13 | 18 | 7 | 8 | 1 | 0 | 1 | 0 | 0 | 11 | 24 | 16 | 5 | 4 | 0 | 0 | 0 | 0 | 0 | 267 | 149 | 1 | 1 | 0.918 | 1 | 0.909 | 1 | 1 | 1 | 1 | 1 | 1 | 1 | -0.035 |
| SE | 35126 | ENSG00000266338 | NBPF15 | chr1 | - | 144460842 | 144460961 | 144459365 | 144459483 | 144461380 | 144461674 | 5 | 7 | 5 | 6 | 7 | 0 | 0 | 2 | 0 | 0 | 11 | 16 | 5 | 1 | 4 | 0 | 0 | 0 | 0 | 0 | 268 | 149 | 1 | 1 | 1 | 1 | 0.582 | 1 | 1 | 1 | 1 | 1 | 1 | 1 | -0.084 |
| SE | 35127 | ENSG00000244405 | ETV5 | chr3 | - | 186084168 | 186084280 | 186081123 | 186081175 | 186105304 | 186105355 | 0 | 1 | 0 | 0 | 0 | 1 | 0 | 0 | 2 | 4 | 0 | 0 | 0 | 0 | 0 | 0 | 4 | 2 | 4 | 7 | 261 | 149 | 1 | 1 | 0 | 1 | NA | 0 | 0 | NA | 0 | 0 | 0 | 0 | 0.25 |
| SE | 35129 | ENSG00000288520 | NA | chr6 | - | 154167919 | 154168113 | 154159827 | 154160040 | 154246590 | 154246760 | 147 | 110 | 216 | 160 | 378 | 0 | 0 | 0 | 0 | 0 | 21 | 85 | 136 | 80 | 0 | 0 | 0 | 0 | 1 | 0 | 298 | 149 | 1 | 1 | 1 | 1 | 1 | 1 | 1 | 1 | 1 | 1 | 0.976 | NA | 0.006 |
| SE | 35130 | ENSG00000288520 | NA | chr6 | - | 154221256 | 154221328 | 154214217 | 154214276 | 154223169 | 154223243 | 265 | 272 | 381 | 300 | 627 | 4 | 0 | 2 | 0 | 0 | 86 | 99 | 168 | 111 | 41 | 0 | 5 | 0 | 0 | 0 | 221 | 149 | 1 | 1 | 0.978 | 1 | 0.992 | 1 | 1 | 1 | 0.93 | 1 | 1 | 1 | 0.008 |
| SE | 35131 | ENSG00000288520 | NA | chr6 | - | 154221256 | 154221328 | 154214217 | 154214276 | 154246590 | 154246760 | 113 | 138 | 163 | 146 | 266 | 0 | 0 | 0 | 4 | 1 | 40 | 39 | 73 | 47 | 18 | 0 | 0 | 0 | 0 | 0 | 221 | 149 | 1 | 1 | 1 | 1 | 1 | 0.961 | 0.994 | 1 | 1 | 1 | 1 | 1 | -0.009 |
| SE | 35132 | ENSG00000288520 | NA | chr6 | - | 154223169 | 154223243 | 154214217 | 154214276 | 154246590 | 154246760 | 80 | 70 | 116 | 65 | 182 | 0 | 0 | 0 | 4 | 1 | 15 | 28 | 44 | 22 | 4 | 0 | 0 | 0 | 0 | 0 | 223 | 149 | 1 | 1 | 1 | 1 | 1 | 0.916 | 0.992 | 1 | 1 | 1 | 1 | 1 | -0.018 |
| SE | 35133 | ENSG00000288520 | NA | chr6 | - | 154223169 | 154223243 | 154221256 | 154221328 | 154246590 | 154246760 | 220 | 190 | 341 | 213 | 550 | 0 | 0 | 0 | 0 | 1 | 61 | 90 | 132 | 89 | 28 | 0 | 0 | 0 | 0 | 0 | 223 | 149 | 1 | 1 | 1 | 1 | 1 | 1 | 0.997 | 1 | 1 | 1 | 1 | 1 | -0.001 |
| SE | 35134 | ENSG00000288520 | NA | chr6 | - | 154246590 | 154246760 | 154223169 | 154223243 | 154265911 | 154265964 | 186 | 159 | 277 | 174 | 431 | 0 | 0 | 0 | 0 | 3 | 47 | 74 | 114 | 81 | 17 | 0 | 0 | 0 | 0 | 0 | 298 | 149 | 1 | 1 | 1 | 1 | 1 | 1 | 0.986 | 1 | 1 | 1 | 1 | 1 | -0.003 |
| SE | 35136 | ENSG00000007923 | DNAJC11 | chr1 | - | 6636116 | 6636246 | 6634167 | 6635700 | 6637197 | 6637340 | 75 | 34 | 47 | 83 | 57 | 0 | 0 | 0 | 0 | 0 | 158 | 88 | 49 | 98 | 89 | 0 | 0 | 0 | 2 | 0 | 279 | 149 | 1 | 1 | 1 | 1 | 1 | 1 | 1 | 1 | 1 | 1 | 0.963 | 1 | 0.007 |
| SE | 35137 | ENSG00000007923 | DNAJC11 | chr1 | - | 6639901 | 6640057 | 6637446 | 6638364 | 6644557 | 6644674 | 101 | 52 | 73 | 96 | 96 | 1 | 0 | 0 | 2 | 0 | 131 | 108 | 60 | 136 | 118 | 0 | 0 | 1 | 4 | 0 | 298 | 149 | 1 | 1 | 0.981 | 1 | 1 | 0.96 | 1 | 1 | 1 | 0.968 | 0.944 | 1 | 0.006 |
| SE | 35139 | ENSG00000007923 | DNAJC11 | chr1 | - | 6651528 | 6651602 | 6645788 | 6645978 | 6652828 | 6652951 | 71 | 54 | 74 | 91 | 144 | 0 | 0 | 1 | 0 | 0 | 143 | 88 | 32 | 56 | 136 | 0 | 0 | 0 | 0 | 0 | 223 | 149 | 1 | 1 | 1 | 1 | 0.98 | 1 | 1 | 1 | 1 | 1 | 1 | 1 | -0.004 |
| SE | 35146 | ENSG00000107758 | PPP3CB | chr10 | - | 73448663 | 73448690 | 73446491 | 73446576 | 73454411 | 73454489 | 1 | 0 | 0 | 0 | 0 | 179 | 149 | 156 | 157 | 261 | 0 | 0 | 0 | 0 | 0 | 180 | 153 | 92 | 149 | 129 | 176 | 149 | 1 | 1 | 0.005 | 0 | 0 | 0 | 0 | 0 | 0 | 0 | 0 | 0 | 0.001 |
| SE | 35147 | ENSG00000107758 | PPP3CB | chr10 | - | 73454411 | 73454489 | 73446491 | 73446576 | 73467552 | 73467619 | 318 | 275 | 242 | 258 | 471 | 0 | 0 | 0 | 0 | 2 | 328 | 306 | 196 | 264 | 293 | 0 | 0 | 0 | 0 | 0 | 227 | 149 | 1 | 1 | 1 | 1 | 1 | 1 | 0.994 | 1 | 1 | 1 | 1 | 1 | -0.001 |
| SE | 35149 | ENSG00000107758 | PPP3CB | chr10 | - | 73474918 | 73475030 | 73471467 | 73471613 | 73478498 | 73478623 | 581 | 354 | 407 | 517 | 1052 | 2 | 0 | 1 | 2 | 0 | 440 | 447 | 273 | 364 | 421 | 0 | 0 | 0 | 0 | 0 | 261 | 149 | 1 | 1 | 0.994 | 1 | 0.996 | 0.993 | 1 | 1 | 1 | 1 | 1 | 1 | -0.003 |
| SE | 35150 | ENSG00000107758 | PPP3CB | chr10 | - | 73478498 | 73478623 | 73474918 | 73475030 | 73479316 | 73479517 | 556 | 375 | 474 | 534 | 959 | 1 | 0 | 0 | 0 | 0 | 527 | 462 | 310 | 417 | 449 | 0 | 0 | 0 | 0 | 0 | 274 | 149 | 1 | 1 | 0.997 | 1 | 1 | 1 | 1 | 1 | 1 | 1 | 1 | 1 | -0.001 |
| SE | 35153 | ENSG00000158411 | MITD1 | chr2 | - | 99169549 | 99169610 | 99169262 | 99169470 | 99170536 | 99170652 | 383 | 318 | 234 | 280 | 419 | 0 | 0 | 0 | 0 | 0 | 314 | 165 | 200 | 218 | 223 | 0 | 0 | 2 | 0 | 0 | 210 | 149 | 1 | 1 | 1 | 1 | 1 | 1 | 1 | 1 | 1 | 0.986 | 1 | 1 | 0.003 |
| SE | 35156 | ENSG00000188001 | TPRG1 | chr3 | + | 189121661 | 189121861 | 189121285 | 189121371 | 189123578 | 189123700 | 15 | 20 | 5 | 10 | 18 | 0 | 0 | 1 | 0 | 0 | 0 | 17 | 13 | 10 | 8 | 0 | 3 | 0 | 0 | 3 | 298 | 149 | 1 | 1 | 1 | 1 | 0.714 | 1 | 1 | NA | 0.739 | 1 | 1 | 0.571 | 0.115 |
| SE | 35160 | ENSG00000188001 | TPRG1 | chr3 | + | 189147583 | 189147647 | 189132398 | 189132697 | 189150655 | 189150805 | 32 | 17 | 6 | 10 | 26 | 1 | 0 | 2 | 0 | 0 | 0 | 20 | 3 | 1 | 0 | 0 | 0 | 0 | 0 | 11 | 213 | 149 | 1 | 1 | 0.957 | 1 | 0.677 | 1 | 1 | NA | 1 | 1 | 1 | 0 | 0.177 |
| SE | 35161 | ENSG00000188001 | TPRG1 | chr3 | + | 189147583 | 189147647 | 189132398 | 189132697 | 189207375 | 189207594 | 28 | 12 | 6 | 16 | 19 | 0 | 0 | 0 | 0 | 0 | 0 | 6 | 1 | 1 | 0 | 0 | 5 | 0 | 0 | 0 | 213 | 149 | 1 | 1 | 1 | 1 | 1 | 1 | 1 | NA | 0.456 | 1 | 1 | NA | 0.181 |
| SE | 35162 | ENSG00000188001 | TPRG1 | chr3 | + | 189147583 | 189147647 | 189132398 | 189132697 | 189215291 | 189215383 | 22 | 11 | 5 | 9 | 9 | 0 | 0 | 3 | 1 | 0 | 0 | 4 | 1 | 1 | 0 | 0 | 0 | 0 | 0 | 0 | 213 | 149 | 1 | 1 | 1 | 1 | 0.538 | 0.863 | 1 | NA | 1 | 1 | 1 | NA | -0.12 |
| SE | 35163 | ENSG00000188001 | TPRG1 | chr3 | + | 189150655 | 189150805 | 189147583 | 189147647 | 189150903 | 189151136 | 11 | 7 | 1 | 1 | 17 | 3 | 0 | 0 | 0 | 0 | 0 | 16 | 2 | 0 | 0 | 0 | 0 | 0 | 0 | 0 | 298 | 149 | 1 | 1 | 0.647 | 1 | 1 | 1 | 1 | NA | 1 | 1 | NA | NA | -0.071 |
| SE | 35164 | ENSG00000188001 | TPRG1 | chr3 | + | 189150655 | 189150872 | 189147583 | 189147647 | 189207375 | 189207594 | 10 | 7 | 1 | 1 | 17 | 6 | 2 | 1 | 7 | 10 | 0 | 16 | 2 | 0 | 0 | 0 | 2 | 0 | 0 | 0 | 298 | 149 | 1 | 1 | 0.455 | 0.636 | 0.333 | 0.067 | 0.459 | NA | 0.8 | 1 | NA | NA | -0.51 |
| SE | 35165 | ENSG00000188001 | TPRG1 | chr3 | + | 189207375 | 189207594 | 189132398 | 189132697 | 189215291 | 189215383 | 8 | 1 | 5 | 7 | 8 | 0 | 0 | 3 | 1 | 0 | 0 | 15 | 4 | 1 | 9 | 0 | 0 | 0 | 0 | 0 | 298 | 149 | 1 | 1 | 1 | 1 | 0.455 | 0.778 | 1 | NA | 1 | 1 | 1 | 1 | -0.153 |
| SE | 35166 | ENSG00000188001 | TPRG1 | chr3 | + | 189207375 | 189207594 | 189147583 | 189147647 | 189215291 | 189215383 | 14 | 3 | 6 | 14 | 18 | 0 | 1 | 0 | 0 | 0 | 0 | 12 | 4 | 1 | 9 | 0 | 0 | 0 | 0 | 0 | 298 | 149 | 1 | 1 | 1 | 0.6 | 1 | 1 | 1 | NA | 1 | 1 | 1 | 1 | -0.08 |
| SE | 35167 | ENSG00000188001 | TPRG1 | chr3 | + | 189217876 | 189218003 | 189215291 | 189215383 | 189219567 | 189219662 | 0 | 3 | 2 | 4 | 0 | 2 | 0 | 0 | 0 | 0 | 0 | 7 | 8 | 1 | 0 | 0 | 0 | 0 | 0 | 0 | 276 | 149 | 1 | 1 | 0 | 1 | 1 | 1 | NA | NA | 1 | 1 | 1 | NA | -0.25 |
| SE | 35169 | ENSG00000101596 | SMCHD1 | chr18 | + | 2666156 | 2666232 | 2655725 | 2656261 | 2666869 | 2667031 | 343 | 336 | 449 | 449 | 637 | 3 | 0 | 0 | 2 | 0 | 126 | 402 | 399 | 188 | 140 | 0 | 2 | 4 | 2 | 0 | 225 | 149 | 1 | 1 | 0.987 | 1 | 1 | 0.993 | 1 | 1 | 0.993 | 0.985 | 0.984 | 1 | 0.004 |
| SE | 35170 | ENSG00000101596 | SMCHD1 | chr18 | + | 2666869 | 2667031 | 2655725 | 2656261 | 2688393 | 2688508 | 4 | 2 | 4 | 7 | 1 | 0 | 0 | 0 | 0 | 0 | 17 | 15 | 53 | 3 | 7 | 0 | 0 | 0 | 0 | 1 | 298 | 149 | 1 | 1 | 1 | 1 | 1 | 1 | 1 | 1 | 1 | 1 | 1 | 0.778 | 0.044 |
| SE | 35173 | ENSG00000101596 | SMCHD1 | chr18 | + | 2688627 | 2688747 | 2688393 | 2688508 | 2694526 | 2694693 | 1050 | 1178 | 1132 | 1432 | 1416 | 0 | 0 | 0 | 2 | 0 | 779 | 1420 | 2133 | 790 | 268 | 0 | 0 | 0 | 0 | 0 | 269 | 149 | 1 | 1 | 1 | 1 | 1 | 0.997 | 1 | 1 | 1 | 1 | 1 | 1 | -0.001 |
| SE | 35175 | ENSG00000101596 | SMCHD1 | chr18 | + | 2697830 | 2698041 | 2697031 | 2697122 | 2700538 | 2700659 | 1032 | 976 | 1043 | 1253 | 1316 | 0 | 0 | 0 | 0 | 0 | 286 | 946 | 1174 | 421 | 128 | 0 | 0 | 0 | 1 | 0 | 298 | 149 | 1 | 1 | 1 | 1 | 1 | 1 | 1 | 1 | 1 | 1 | 0.995 | 1 | 0.001 |
| SE | 35176 | ENSG00000101596 | SMCHD1 | chr18 | + | 2700734 | 2700918 | 2700538 | 2700659 | 2703691 | 2703886 | 1167 | 1100 | 1024 | 1209 | 1395 | 2 | 5 | 0 | 4 | 0 | 327 | 787 | 1149 | 369 | 117 | 0 | 2 | 0 | 6 | 0 | 298 | 149 | 1 | 1 | 0.997 | 0.991 | 1 | 0.993 | 1 | 1 | 0.995 | 1 | 0.969 | 1 | 0.003 |
| SE | 35177 | ENSG00000101596 | SMCHD1 | chr18 | + | 2703691 | 2703886 | 2700734 | 2700918 | 2705693 | 2705807 | 890 | 1100 | 903 | 996 | 991 | 0 | 1 | 0 | 2 | 0 | 202 | 613 | 1134 | 353 | 67 | 0 | 0 | 1 | 0 | 0 | 298 | 149 | 1 | 1 | 1 | 0.998 | 1 | 0.996 | 1 | 1 | 1 | 0.998 | 1 | 1 | -0.001 |
| SE | 35178 | ENSG00000101596 | SMCHD1 | chr18 | + | 2718157 | 2718235 | 2707806 | 2707920 | 2718314 | 2718434 | 742 | 720 | 744 | 842 | 928 | 0 | 3 | 4 | 1 | 4 | 102 | 313 | 643 | 129 | 59 | 0 | 1 | 2 | 0 | 2 | 227 | 149 | 1 | 1 | 1 | 0.994 | 0.992 | 0.998 | 0.993 | 1 | 0.995 | 0.995 | 1 | 0.951 | 0.007 |
| SE | 35179 | ENSG00000101596 | SMCHD1 | chr18 | + | 2718314 | 2718434 | 2707806 | 2707920 | 2724898 | 2724995 | 55 | 74 | 80 | 83 | 49 | 0 | 0 | 0 | 0 | 1 | 14 | 91 | 201 | 26 | 10 | 0 | 0 | 0 | 0 | 0 | 269 | 149 | 1 | 1 | 1 | 1 | 1 | 1 | 0.964 | 1 | 1 | 1 | 1 | 1 | -0.007 |
| SE | 35180 | ENSG00000101596 | SMCHD1 | chr18 | + | 2718314 | 2718434 | 2718157 | 2718235 | 2722518 | 2722550 | 886 | 858 | 916 | 923 | 1269 | 0 | 0 | 1 | 0 | 0 | 120 | 394 | 682 | 129 | 45 | 0 | 0 | 0 | 0 | 0 | 269 | 149 | 1 | 1 | 1 | 1 | 0.998 | 1 | 1 | 1 | 1 | 1 | 1 | 1 | 0 |
| SE | 35182 | ENSG00000101596 | SMCHD1 | chr18 | + | 2722518 | 2722663 | 2718314 | 2718434 | 2726451 | 2726524 | 418 | 357 | 400 | 395 | 667 | 0 | 0 | 1 | 0 | 1 | 31 | 169 | 252 | 44 | 12 | 0 | 0 | 2 | 0 | 0 | 294 | 149 | 1 | 1 | 1 | 1 | 0.995 | 1 | 0.997 | 1 | 1 | 0.985 | 1 | 1 | 0.001 |
| SE | 35183 | ENSG00000101596 | SMCHD1 | chr18 | + | 2724898 | 2724995 | 2718314 | 2718434 | 2726451 | 2726524 | 235 | 235 | 278 | 296 | 548 | 0 | 0 | 1 | 0 | 1 | 40 | 187 | 377 | 60 | 8 | 0 | 0 | 2 | 0 | 0 | 246 | 149 | 1 | 1 | 1 | 1 | 0.994 | 1 | 0.997 | 1 | 1 | 0.991 | 1 | 1 | 0 |
| SE | 35184 | ENSG00000101596 | SMCHD1 | chr18 | + | 2724898 | 2724995 | 2722518 | 2722663 | 2726451 | 2726524 | 728 | 683 | 747 | 778 | 1398 | 0 | 0 | 0 | 0 | 0 | 90 | 365 | 513 | 79 | 30 | 0 | 3 | 1 | 0 | 0 | 246 | 149 | 1 | 1 | 1 | 1 | 1 | 1 | 1 | 1 | 0.987 | 0.997 | 1 | 1 | 0.003 |
| SE | 35185 | ENSG00000101596 | SMCHD1 | chr18 | + | 2729274 | 2729409 | 2728413 | 2728596 | 2732264 | 2732492 | 892 | 896 | 911 | 1052 | 1168 | 0 | 0 | 0 | 0 | 0 | 101 | 433 | 769 | 153 | 72 | 0 | 0 | 2 | 0 | 0 | 284 | 149 | 1 | 1 | 1 | 1 | 1 | 1 | 1 | 1 | 1 | 0.995 | 1 | 1 | 0.001 |
| SE | 35186 | ENSG00000101596 | SMCHD1 | chr18 | + | 2729274 | 2729409 | 2728413 | 2728596 | 2738396 | 2738545 | 414 | 439 | 449 | 489 | 683 | 0 | 0 | 2 | 0 | 0 | 55 | 211 | 385 | 89 | 28 | 0 | 0 | 0 | 0 | 0 | 284 | 149 | 1 | 1 | 1 | 1 | 0.992 | 1 | 1 | 1 | 1 | 1 | 1 | 1 | -0.002 |
| SE | 35187 | ENSG00000101596 | SMCHD1 | chr18 | + | 2732264 | 2732492 | 2728413 | 2728596 | 2738396 | 2738545 | 729 | 793 | 695 | 962 | 1135 | 0 | 0 | 2 | 0 | 0 | 105 | 429 | 607 | 193 | 163 | 0 | 0 | 0 | 0 | 0 | 298 | 149 | 1 | 1 | 1 | 1 | 0.994 | 1 | 1 | 1 | 1 | 1 | 1 | 1 | -0.001 |
| SE | 35188 | ENSG00000101596 | SMCHD1 | chr18 | + | 2732264 | 2732492 | 2729274 | 2729409 | 2738396 | 2738545 | 1209 | 1252 | 1160 | 1530 | 1620 | 0 | 1 | 1 | 0 | 0 | 151 | 651 | 994 | 257 | 207 | 0 | 0 | 0 | 0 | 0 | 298 | 149 | 1 | 1 | 1 | 0.998 | 0.998 | 1 | 1 | 1 | 1 | 1 | 1 | 1 | -0.001 |
| SE | 35190 | ENSG00000101596 | SMCHD1 | chr18 | + | 2743760 | 2743928 | 2740702 | 2740821 | 2747521 | 2747647 | 1334 | 1364 | 1276 | 1821 | 1803 | 3 | 2 | 0 | 1 | 1 | 234 | 925 | 978 | 326 | 164 | 0 | 6 | 0 | 0 | 0 | 298 | 149 | 1 | 1 | 0.996 | 0.997 | 1 | 0.999 | 0.999 | 1 | 0.987 | 1 | 1 | 1 | 0.001 |
| SE | 35192 | ENSG00000101596 | SMCHD1 | chr18 | + | 2751277 | 2751393 | 2750349 | 2750507 | 2752487 | 2752552 | 1284 | 1395 | 1259 | 1698 | 1131 | 1 | 0 | 0 | 2 | 3 | 281 | 864 | 1083 | 237 | 140 | 0 | 6 | 2 | 0 | 0 | 265 | 149 | 1 | 1 | 0.999 | 1 | 1 | 0.998 | 0.995 | 1 | 0.988 | 0.997 | 1 | 1 | 0.001 |
| SE | 35194 | ENSG00000101596 | SMCHD1 | chr18 | + | 2762104 | 2762236 | 2760400 | 2760739 | 2763636 | 2763789 | 731 | 660 | 720 | 763 | 989 | 2 | 1 | 0 | 0 | 0 | 97 | 331 | 470 | 58 | 92 | 0 | 0 | 0 | 0 | 0 | 281 | 149 | 1 | 1 | 0.995 | 0.997 | 1 | 1 | 1 | 1 | 1 | 1 | 1 | 1 | -0.002 |
| SE | 35196 | ENSG00000101596 | SMCHD1 | chr18 | + | 2764068 | 2764106 | 2763875 | 2763969 | 2769693 | 2769820 | 0 | 0 | 0 | 1 | 0 | 0 | 0 | 1 | 4 | 1 | 0 | 0 | 1 | 0 | 0 | 0 | 0 | 3 | 6 | 0 | 187 | 149 | 1 | 1 | NA | NA | 0 | 0.166 | 0 | NA | NA | 0.21 | 0 | NA | -0.05 |
| SE | 35197 | ENSG00000101596 | SMCHD1 | chr18 | + | 2771532 | 2771618 | 2769988 | 2770108 | 2772249 | 2772372 | 708 | 818 | 768 | 832 | 1545 | 0 | 0 | 0 | 0 | 0 | 159 | 386 | 496 | 135 | 118 | 0 | 0 | 0 | 3 | 0 | 235 | 149 | 1 | 1 | 1 | 1 | 1 | 1 | 1 | 1 | 1 | 1 | 0.966 | 1 | 0.007 |
| SE | 35199 | ENSG00000101596 | SMCHD1 | chr18 | + | 2777805 | 2777915 | 2772249 | 2772372 | 2778168 | 2778239 | 485 | 491 | 554 | 608 | 700 | 0 | 0 | 0 | 1 | 0 | 46 | 266 | 288 | 63 | 57 | 0 | 0 | 0 | 0 | 0 | 259 | 149 | 1 | 1 | 1 | 1 | 1 | 0.997 | 1 | 1 | 1 | 1 | 1 | 1 | -0.001 |
| SE | 35202 | ENSG00000101596 | SMCHD1 | chr18 | + | 2796406 | 2796521 | 2795948 | 2796107 | 2802527 | 2802804 | 431 | 417 | 433 | 487 | 523 | 1 | 0 | 0 | 0 | 0 | 14 | 198 | 298 | 34 | 2 | 0 | 0 | 0 | 0 | 0 | 264 | 149 | 1 | 1 | 0.996 | 1 | 1 | 1 | 1 | 1 | 1 | 1 | 1 | 1 | -0.001 |
| SE | 35204 | ENSG00000102362 | SYTL4 | chrX | - | 100703092 | 100703185 | 100701927 | 100702107 | 100704810 | 100704886 | 2 | 5 | 1 | 10 | 4 | 0 | 2 | 0 | 0 | 0 | 18 | 9 | 11 | 10 | 48 | 0 | 0 | 3 | 0 | 0 | 242 | 149 | 1 | 1 | 1 | 0.606 | 1 | 1 | 1 | 1 | 1 | 0.693 | 1 | 1 | -0.017 |
| SE | 35206 | ENSG00000102362 | SYTL4 | chrX | - | 100704810 | 100704886 | 100701927 | 100702107 | 100731417 | 100731510 | 0 | 2 | 0 | 0 | 0 | 0 | 0 | 0 | 2 | 0 | 0 | 0 | 4 | 0 | 0 | 0 | 0 | 0 | 0 | 5 | 225 | 149 | 1 | 1 | NA | 1 | NA | 0 | NA | NA | NA | 1 | NA | 0 | 0 |
| SE | 35207 | ENSG00000102362 | SYTL4 | chrX | - | 100704810 | 100704886 | 100701927 | 100702107 | 100732003 | 100732123 | 0 | 3 | 0 | 0 | 0 | 3 | 3 | 1 | 6 | 0 | 0 | 0 | 3 | 0 | 1 | 6 | 2 | 4 | 7 | 37 | 225 | 149 | 1 | 1 | 0 | 0.398 | 0 | 0 | NA | 0 | 0 | 0.332 | 0 | 0.018 | 0.029 |
| SE | 35208 | ENSG00000102362 | SYTL4 | chrX | - | 100704810 | 100704886 | 100703092 | 100703185 | 100732003 | 100732123 | 2 | 4 | 0 | 8 | 1 | 0 | 0 | 0 | 6 | 0 | 8 | 0 | 10 | 0 | 17 | 0 | 0 | 5 | 0 | 2 | 225 | 149 | 1 | 1 | 1 | 1 | NA | 0.469 | 1 | 1 | NA | 0.57 | NA | 0.849 | 0.061 |
| SE | 35209 | ENSG00000224914 | LINC00863 | chr10 | + | 87344436 | 87344495 | 87341684 | 87341720 | 87354619 | 87354897 | 11 | 1 | 1 | 2 | 13 | 0 | 0 | 2 | 0 | 0 | 9 | 13 | 1 | 7 | 6 | 0 | 0 | 1 | 0 | 0 | 208 | 149 | 1 | 1 | 1 | 1 | 0.264 | 1 | 1 | 1 | 1 | 0.417 | 1 | 1 | -0.031 |
| SE | 35210 | ENSG00000224914 | LINC00863 | chr10 | + | 87354619 | 87354897 | 87344436 | 87344495 | 87355090 | 87355425 | 17 | 5 | 3 | 9 | 32 | 0 | 0 | 0 | 0 | 1 | 25 | 26 | 6 | 21 | 23 | 0 | 0 | 0 | 0 | 0 | 298 | 149 | 1 | 1 | 1 | 1 | 1 | 1 | 0.941 | 1 | 1 | 1 | 1 | 1 | -0.012 |
| SE | 35213 | ENSG00000185798 | WDR53 | chr3 | - | 196560995 | 196561491 | 196554176 | 196554807 | 196566876 | 196567307 | 73 | 46 | 65 | 45 | 71 | 0 | 2 | 0 | 0 | 0 | 70 | 58 | 37 | 48 | 15 | 0 | 0 | 2 | 0 | 0 | 298 | 149 | 1 | 1 | 1 | 0.92 | 1 | 1 | 1 | 1 | 1 | 0.902 | 1 | 1 | 0.004 |
| SE | 35215 | ENSG00000185798 | WDR53 | chr3 | - | 196560995 | 196561491 | 196554176 | 196554807 | 196568518 | 196568554 | 45 | 42 | 51 | 38 | 37 | 0 | 1 | 0 | 0 | 0 | 63 | 39 | 33 | 37 | 15 | 0 | 0 | 0 | 0 | 0 | 298 | 149 | 1 | 1 | 1 | 0.955 | 1 | 1 | 1 | 1 | 1 | 1 | 1 | 1 | -0.009 |
| SE | 35216 | ENSG00000185798 | WDR53 | chr3 | - | 196566876 | 196567307 | 196554176 | 196554807 | 196568518 | 196568554 | 4 | 8 | 4 | 3 | 0 | 0 | 1 | 0 | 0 | 0 | 0 | 15 | 4 | 2 | 0 | 0 | 0 | 0 | 0 | 0 | 298 | 149 | 1 | 1 | 1 | 0.8 | 1 | 1 | NA | NA | 1 | 1 | 1 | NA | -0.05 |
| SE | 35217 | ENSG00000185798 | WDR53 | chr3 | - | 196566876 | 196567307 | 196560995 | 196561491 | 196568518 | 196568554 | 33 | 10 | 18 | 10 | 34 | 1 | 0 | 0 | 0 | 0 | 7 | 34 | 6 | 13 | 0 | 0 | 0 | 0 | 0 | 0 | 298 | 149 | 1 | 1 | 0.943 | 1 | 1 | 1 | 1 | 1 | 1 | 1 | 1 | NA | -0.011 |
| SE | 35228 | ENSG00000074935 | TUBE1 | chr6 | - | 112072757 | 112072898 | 112071901 | 112072076 | 112074709 | 112074850 | 80 | 64 | 102 | 90 | 173 | 2 | 0 | 4 | 3 | 4 | 21 | 23 | 46 | 18 | 0 | 0 | 0 | 0 | 2 | 0 | 290 | 149 | 1 | 1 | 0.954 | 1 | 0.929 | 0.939 | 0.957 | 1 | 1 | 1 | 0.822 | NA | 0 |
| SE | 35229 | ENSG00000074935 | TUBE1 | chr6 | - | 112075936 | 112076112 | 112074709 | 112074850 | 112076321 | 112076509 | 35 | 23 | 38 | 47 | 93 | 0 | 0 | 0 | 0 | 1 | 0 | 1 | 3 | 8 | 0 | 0 | 0 | 0 | 1 | 0 | 298 | 149 | 1 | 1 | 1 | 1 | 1 | 1 | 0.979 | NA | 1 | 1 | 0.8 | NA | 0.062 |
| SE | 35230 | ENSG00000074935 | TUBE1 | chr6 | - | 112075936 | 112076171 | 112074709 | 112074850 | 112076321 | 112076509 | 23 | 20 | 24 | 30 | 69 | 0 | 0 | 0 | 0 | 1 | 0 | 2 | 2 | 3 | 0 | 0 | 0 | 0 | 1 | 0 | 298 | 149 | 1 | 1 | 1 | 1 | 1 | 1 | 0.972 | NA | 1 | 1 | 0.6 | NA | 0.128 |
| SE | 35231 | ENSG00000074935 | TUBE1 | chr6 | - | 112079632 | 112079754 | 112076321 | 112076509 | 112081091 | 112081203 | 63 | 49 | 73 | 88 | 80 | 1 | 0 | 2 | 2 | 0 | 0 | 49 | 25 | 20 | 12 | 0 | 0 | 1 | 0 | 0 | 271 | 149 | 1 | 1 | 0.972 | 1 | 0.953 | 0.96 | 1 | NA | 1 | 0.932 | 1 | 1 | -0.006 |
| SE | 35232 | ENSG00000074935 | TUBE1 | chr6 | - | 112084188 | 112084246 | 112081091 | 112081207 | 112085674 | 112085701 | 15 | 12 | 32 | 18 | 19 | 0 | 0 | 0 | 0 | 0 | 0 | 2 | 2 | 2 | 6 | 0 | 0 | 1 | 0 | 0 | 207 | 149 | 1 | 1 | 1 | 1 | 1 | 1 | 1 | NA | 1 | 0.59 | 1 | 1 | 0.103 |
| SE | 35233 | ENSG00000074935 | TUBE1 | chr6 | - | 112084188 | 112084246 | 112081091 | 112081207 | 112086555 | 112086608 | 45 | 42 | 82 | 63 | 55 | 1 | 2 | 2 | 0 | 0 | 6 | 31 | 31 | 22 | 12 | 0 | 0 | 2 | 0 | 0 | 207 | 149 | 1 | 1 | 0.97 | 0.938 | 0.967 | 1 | 1 | 1 | 1 | 0.918 | 1 | 1 | -0.009 |
| SE | 35234 | ENSG00000182899 | RPL35A | chr3 | + | 197951158 | 197951311 | 197950935 | 197950978 | 197954002 | 197954120 | 2419 | 1150 | 2571 | 2355 | 3107 | 0 | 0 | 0 | 0 | 0 | 1362 | 1313 | 976 | 1081 | 2062 | 0 | 0 | 1 | 0 | 0 | 298 | 149 | 1 | 1 | 1 | 1 | 1 | 1 | 1 | 1 | 1 | 0.998 | 1 | 1 | 0 |
| SE | 35235 | ENSG00000182899 | RPL35A | chr3 | + | 197951158 | 197951328 | 197950935 | 197950978 | 197954002 | 197954147 | 956 | 403 | 1021 | 949 | 958 | 0 | 0 | 0 | 0 | 0 | 624 | 567 | 411 | 428 | 943 | 0 | 0 | 1 | 0 | 0 | 298 | 149 | 1 | 1 | 1 | 1 | 1 | 1 | 1 | 1 | 1 | 0.995 | 1 | 1 | 0.001 |
| SE | 35237 | ENSG00000182899 | RPL35A | chr3 | + | 197953487 | 197953660 | 197951158 | 197951311 | 197954002 | 197954147 | 0 | 0 | 4 | 1 | 0 | 1464 | 748 | 1552 | 1415 | 2156 | 0 | 0 | 0 | 4 | 0 | 738 | 755 | 570 | 656 | 1122 | 298 | 149 | 1 | 1 | 0 | 0 | 0.001 | 0 | 0 | 0 | 0 | 0 | 0.003 | 0 | 0 |
| SE | 35264 | ENSG00000103064 | SLC7A6 | chr16 | + | 68266592 | 68266721 | 68264515 | 68264576 | 68274690 | 68274948 | 35 | 48 | 59 | 53 | 39 | 0 | 0 | 0 | 0 | 0 | 72 | 10 | 41 | 29 | 28 | 0 | 1 | 0 | 0 | 0 | 278 | 149 | 1 | 1 | 1 | 1 | 1 | 1 | 1 | 1 | 0.843 | 1 | 1 | 1 | 0.031 |
| SE | 35268 | ENSG00000103064 | SLC7A6 | chr16 | + | 68274690 | 68275249 | 68266590 | 68266721 | 68287745 | 68287871 | 65 | 66 | 98 | 83 | 54 | 0 | 0 | 0 | 0 | 0 | 104 | 35 | 63 | 56 | 37 | 0 | 1 | 0 | 0 | 0 | 298 | 149 | 1 | 1 | 1 | 1 | 1 | 1 | 1 | 1 | 0.946 | 1 | 1 | 1 | 0.011 |
| SE | 35270 | ENSG00000103064 | SLC7A6 | chr16 | + | 68287745 | 68287871 | 68274690 | 68275249 | 68290395 | 68290540 | 88 | 57 | 160 | 121 | 119 | 1 | 0 | 0 | 0 | 0 | 37 | 47 | 41 | 87 | 29 | 0 | 0 | 0 | 0 | 0 | 275 | 149 | 1 | 1 | 0.979 | 1 | 1 | 1 | 1 | 1 | 1 | 1 | 1 | 1 | -0.004 |
| SE | 35271 | ENSG00000103064 | SLC7A6 | chr16 | + | 68287745 | 68287871 | 68274690 | 68275249 | 68291208 | 68291332 | 32 | 23 | 39 | 31 | 15 | 1 | 0 | 0 | 0 | 0 | 38 | 31 | 22 | 31 | 17 | 0 | 0 | 0 | 0 | 0 | 275 | 149 | 1 | 1 | 0.945 | 1 | 1 | 1 | 1 | 1 | 1 | 1 | 1 | 1 | -0.011 |
| SE | 35272 | ENSG00000103064 | SLC7A6 | chr16 | + | 68290395 | 68290540 | 68274690 | 68275249 | 68291208 | 68291332 | 21 | 17 | 29 | 23 | 52 | 1 | 0 | 0 | 0 | 0 | 18 | 28 | 17 | 9 | 5 | 0 | 0 | 0 | 0 | 0 | 294 | 149 | 1 | 1 | 0.914 | 1 | 1 | 1 | 1 | 1 | 1 | 1 | 1 | 1 | -0.017 |
| SE | 35273 | ENSG00000103064 | SLC7A6 | chr16 | + | 68290395 | 68290540 | 68287745 | 68287871 | 68291208 | 68291332 | 81 | 53 | 162 | 120 | 159 | 2 | 2 | 1 | 1 | 0 | 23 | 51 | 38 | 72 | 19 | 6 | 6 | 2 | 4 | 2 | 294 | 149 | 1 | 1 | 0.954 | 0.931 | 0.988 | 0.984 | 1 | 0.66 | 0.812 | 0.906 | 0.901 | 0.828 | 0.15 |
| SE | 35276 | ENSG00000103064 | SLC7A6 | chr16 | + | 68291208 | 68291332 | 68290395 | 68290540 | 68291557 | 68291578 | 74 | 45 | 113 | 80 | 154 | 0 | 0 | 0 | 0 | 0 | 31 | 78 | 51 | 33 | 49 | 0 | 3 | 0 | 0 | 0 | 273 | 149 | 1 | 1 | 1 | 1 | 1 | 1 | 1 | 1 | 0.934 | 1 | 1 | 1 | 0.013 |
| SE | 35282 | ENSG00000173950 | XXYLT1 | chr3 | - | 195183475 | 195183629 | 195180393 | 195180515 | 195226708 | 195226856 | 1 | 0 | 0 | 0 | 0 | 1 | 0 | 4 | 5 | 0 | 0 | 12 | 4 | 0 | 0 | 4 | 0 | 5 | 0 | 0 | 298 | 149 | 1 | 1 | 0.333 | NA | 0 | 0 | NA | 0 | 1 | 0.286 | NA | NA | -0.318 |
| SE | 35286 | ENSG00000173950 | XXYLT1 | chr3 | - | 195270088 | 195270205 | 195226723 | 195226856 | 195270554 | 195270861 | 0 | 0 | 0 | 0 | 0 | 5 | 4 | 2 | 6 | 0 | 4 | 0 | 0 | 0 | 0 | 18 | 12 | 17 | 15 | 0 | 266 | 149 | 1 | 1 | 0 | 0 | 0 | 0 | NA | 0.111 | 0 | 0 | 0 | NA | -0.028 |
| SE | 35289 | ENSG00000145016 | RUBCN | chr3 | - | 197677479 | 197677541 | 197676216 | 197677038 | 197681128 | 197681367 | 84 | 62 | 76 | 96 | 31 | 0 | 0 | 0 | 0 | 0 | 152 | 117 | 71 | 191 | 97 | 0 | 0 | 0 | 2 | 0 | 211 | 149 | 1 | 1 | 1 | 1 | 1 | 1 | 1 | 1 | 1 | 1 | 0.985 | 1 | 0.003 |
| SE | 35292 | ENSG00000145016 | RUBCN | chr3 | - | 197696953 | 197697049 | 197695865 | 197695981 | 197700612 | 197701146 | 111 | 93 | 69 | 126 | 123 | 0 | 0 | 2 | 1 | 1 | 227 | 145 | 97 | 211 | 172 | 0 | 0 | 2 | 0 | 0 | 245 | 149 | 1 | 1 | 1 | 1 | 0.955 | 0.987 | 0.987 | 1 | 1 | 0.967 | 1 | 1 | -0.008 |
| SE | 35293 | ENSG00000145016 | RUBCN | chr3 | - | 197699186 | 197699231 | 197695865 | 197695981 | 197700612 | 197701146 | 0 | 0 | 0 | 2 | 1 | 0 | 0 | 2 | 1 | 1 | 0 | 0 | 0 | 0 | 0 | 0 | 0 | 2 | 0 | 0 | 194 | 149 | 1 | 1 | NA | NA | 0 | 0.606 | 0.434 | NA | NA | 0 | NA | NA | 0.347 |
| SE | 35295 | ENSG00000145016 | RUBCN | chr3 | - | 197701707 | 197701864 | 197700459 | 197701146 | 197703547 | 197703654 | 82 | 42 | 58 | 75 | 133 | 2 | 0 | 0 | 0 | 0 | 239 | 123 | 105 | 155 | 282 | 0 | 0 | 0 | 0 | 4 | 298 | 149 | 1 | 1 | 0.953 | 1 | 1 | 1 | 1 | 1 | 1 | 1 | 1 | 0.972 | -0.004 |
| SE | 35297 | ENSG00000145016 | RUBCN | chr3 | - | 197717976 | 197718130 | 197705091 | 197705175 | 197749268 | 197749699 | 45 | 21 | 25 | 30 | 22 | 0 | 0 | 0 | 0 | 0 | 8 | 31 | 11 | 65 | 40 | 0 | 0 | 1 | 0 | 0 | 298 | 149 | 1 | 1 | 1 | 1 | 1 | 1 | 1 | 1 | 1 | 0.846 | 1 | 1 | 0.031 |
| SE | 35300 | ENSG00000145107 | TM4SF19 | chr3 | - | 196326954 | 196327032 | 196324270 | 196324440 | 196327389 | 196327591 | 7 | 1 | 3 | 3 | 0 | 2 | 0 | 0 | 0 | 3 | 0 | 0 | 4 | 0 | 0 | 0 | 0 | 1 | 0 | 0 | 227 | 149 | 1 | 1 | 0.697 | 1 | 1 | 1 | 0 | NA | NA | 1 | NA | NA | -0.261 |
| SE | 35301 | ENSG00000163961 | RNF168 | chr3 | - | 196483769 | 196483891 | 196475230 | 196475312 | 196487398 | 196487578 | 181 | 158 | 185 | 215 | 256 | 0 | 1 | 0 | 0 | 4 | 23 | 137 | 188 | 136 | 8 | 0 | 0 | 0 | 0 | 0 | 271 | 149 | 1 | 1 | 1 | 0.989 | 1 | 1 | 0.972 | 1 | 1 | 1 | 1 | 1 | -0.008 |
| SE | 35302 | ENSG00000163961 | RNF168 | chr3 | - | 196487398 | 196487578 | 196475230 | 196475312 | 196502872 | 196503677 | 5 | 3 | 5 | 1 | 7 | 0 | 0 | 0 | 0 | 0 | 16 | 4 | 0 | 7 | 0 | 0 | 1 | 0 | 0 | 0 | 298 | 149 | 1 | 1 | 1 | 1 | 1 | 1 | 1 | 1 | 0.667 | NA | 1 | NA | 0.111 |
| SE | 35306 | ENSG00000165288 | BRWD3 | chrX | - | 80707426 | 80707503 | 80704677 | 80704846 | 80716156 | 80716250 | 28 | 32 | 37 | 40 | 99 | 1 | 0 | 0 | 0 | 1 | 0 | 7 | 22 | 14 | 0 | 0 | 0 | 3 | 0 | 0 | 226 | 149 | 1 | 1 | 0.949 | 1 | 1 | 1 | 0.985 | NA | 1 | 0.829 | 1 | NA | 0.044 |
| SE | 35307 | ENSG00000165288 | BRWD3 | chrX | - | 80709427 | 80709577 | 80704677 | 80704846 | 80716156 | 80716250 | 149 | 147 | 117 | 167 | 302 | 1 | 0 | 0 | 0 | 1 | 117 | 111 | 75 | 54 | 34 | 0 | 0 | 3 | 0 | 0 | 298 | 149 | 1 | 1 | 0.987 | 1 | 1 | 1 | 0.993 | 1 | 1 | 0.926 | 1 | 1 | 0.011 |
| SE | 35309 | ENSG00000165288 | BRWD3 | chrX | - | 80723747 | 80723876 | 80722561 | 80722787 | 80724932 | 80725067 | 183 | 139 | 148 | 192 | 402 | 1 | 0 | 0 | 0 | 0 | 55 | 106 | 126 | 23 | 38 | 0 | 0 | 0 | 0 | 0 | 278 | 149 | 1 | 1 | 0.99 | 1 | 1 | 1 | 1 | 1 | 1 | 1 | 1 | 1 | -0.002 |
| SE | 35311 | ENSG00000165288 | BRWD3 | chrX | - | 80791853 | 80791952 | 80745568 | 80745729 | 80808538 | 80808598 | 87 | 74 | 85 | 104 | 32 | 2 | 0 | 0 | 1 | 0 | 79 | 42 | 56 | 40 | 55 | 0 | 0 | 0 | 0 | 0 | 248 | 149 | 1 | 1 | 0.963 | 1 | 1 | 0.984 | 1 | 1 | 1 | 1 | 1 | 1 | -0.011 |
| SE | 35313 | ENSG00000165288 | BRWD3 | chrX | - | 80808538 | 80808598 | 80791853 | 80791952 | 80809012 | 80809042 | 41 | 17 | 16 | 24 | 10 | 2 | 0 | 0 | 0 | 0 | 4 | 4 | 12 | 16 | 15 | 0 | 0 | 0 | 0 | 0 | 209 | 149 | 1 | 1 | 0.936 | 1 | 1 | 1 | 1 | 1 | 1 | 1 | 1 | 1 | -0.013 |
| SE | 35314 | ENSG00000165288 | BRWD3 | chrX | - | 80809012 | 80809042 | 80808538 | 80808598 | 80809245 | 80809304 | 65 | 28 | 24 | 36 | 14 | 0 | 0 | 0 | 2 | 0 | 4 | 25 | 19 | 21 | 19 | 0 | 0 | 0 | 0 | 0 | 179 | 149 | 1 | 1 | 1 | 1 | 1 | 0.937 | 1 | 1 | 1 | 1 | 1 | 1 | -0.013 |
| SE | 35315 | ENSG00000165288 | BRWD3 | chrX | - | 80809245 | 80809304 | 80809012 | 80809042 | 80809440 | 80809683 | 68 | 24 | 22 | 35 | 13 | 1 | 0 | 0 | 0 | 0 | 4 | 11 | 17 | 14 | 22 | 0 | 0 | 0 | 0 | 0 | 208 | 149 | 1 | 1 | 0.98 | 1 | 1 | 1 | 1 | 1 | 1 | 1 | 1 | 1 | -0.004 |
| SE | 35316 | ENSG00000105717 | PBX4 | chr19 | - | 19563508 | 19563615 | 19561706 | 19562117 | 19564932 | 19565089 | 20 | 2 | 25 | 12 | 24 | 0 | 0 | 0 | 0 | 0 | 28 | 33 | 16 | 8 | 88 | 0 | 0 | 1 | 0 | 0 | 256 | 149 | 1 | 1 | 1 | 1 | 1 | 1 | 1 | 1 | 1 | 0.903 | 1 | 1 | 0.019 |
| SE | 35338 | ENSG00000184203 | PPP1R2 | chr3 | - | 195519017 | 195519185 | 195514427 | 195516942 | 195523691 | 195523786 | 935 | 848 | 1010 | 1089 | 2305 | 1 | 0 | 0 | 0 | 0 | 568 | 651 | 969 | 502 | 269 | 0 | 0 | 0 | 1 | 0 | 298 | 149 | 1 | 1 | 0.998 | 1 | 1 | 1 | 1 | 1 | 1 | 1 | 0.996 | 1 | 0 |
| SE | 35340 | ENSG00000184203 | PPP1R2 | chr3 | - | 195524818 | 195524896 | 195523691 | 195523786 | 195529793 | 195529861 | 472 | 434 | 612 | 627 | 1089 | 3 | 0 | 3 | 1 | 0 | 418 | 339 | 584 | 345 | 154 | 0 | 0 | 0 | 0 | 0 | 227 | 149 | 1 | 1 | 0.99 | 1 | 0.993 | 0.998 | 1 | 1 | 1 | 1 | 1 | 1 | -0.004 |
| SE | 35343 | ENSG00000114473 | IQCG | chr3 | - | 197892627 | 197892759 | 197891445 | 197891547 | 197912674 | 197912749 | 52 | 36 | 39 | 38 | 30 | 1 | 0 | 0 | 0 | 0 | 39 | 79 | 45 | 22 | 22 | 0 | 0 | 0 | 0 | 0 | 281 | 149 | 1 | 1 | 0.965 | 1 | 1 | 1 | 1 | 1 | 1 | 1 | 1 | 1 | -0.007 |
| SE | 35344 | ENSG00000114473 | IQCG | chr3 | - | 197912484 | 197912536 | 197892627 | 197892759 | 197912674 | 197912952 | 0 | 0 | 2 | 0 | 0 | 14 | 12 | 17 | 11 | 10 | 0 | 0 | 0 | 0 | 0 | 15 | 23 | 19 | 4 | 0 | 201 | 149 | 1 | 1 | 0 | 0 | 0.08 | 0 | 0 | 0 | 0 | 0 | 0 | NA | 0.016 |
| SE | 35345 | ENSG00000114473 | IQCG | chr3 | - | 197945619 | 197945686 | 197943756 | 197944051 | 197959528 | 197959719 | 46 | 33 | 47 | 37 | 42 | 2 | 0 | 0 | 0 | 1 | 21 | 29 | 24 | 21 | 4 | 0 | 0 | 0 | 0 | 0 | 216 | 149 | 1 | 1 | 0.941 | 1 | 1 | 1 | 0.967 | 1 | 1 | 1 | 1 | 1 | -0.018 |
| SE | 35347 | ENSG00000164073 | MFSD8 | chr4 | - | 127921117 | 127921198 | 127919290 | 127920836 | 127921523 | 127921771 | 6 | 1 | 1 | 4 | 11 | 10 | 1 | 9 | 11 | 17 | 0 | 4 | 3 | 0 | 0 | 0 | 10 | 3 | 7 | 0 | 230 | 149 | 1 | 1 | 0.28 | 0.393 | 0.067 | 0.191 | 0.295 | NA | 0.206 | 0.393 | 0 | NA | 0.046 |
| SE | 35348 | ENSG00000164073 | MFSD8 | chr4 | - | 127921523 | 127921771 | 127917798 | 127920836 | 127930682 | 127930709 | 52 | 12 | 39 | 47 | 98 | 1 | 0 | 0 | 0 | 0 | 4 | 17 | 22 | 27 | 21 | 0 | 0 | 0 | 0 | 0 | 298 | 149 | 1 | 1 | 0.963 | 1 | 1 | 1 | 1 | 1 | 1 | 1 | 1 | 1 | -0.007 |
| SE | 35349 | ENSG00000164073 | MFSD8 | chr4 | - | 127921523 | 127921963 | 127920556 | 127920836 | 127930682 | 127930817 | 38 | 17 | 45 | 29 | 65 | 1 | 0 | 0 | 0 | 0 | 38 | 27 | 13 | 19 | 7 | 0 | 0 | 0 | 0 | 0 | 298 | 149 | 1 | 1 | 0.95 | 1 | 1 | 1 | 1 | 1 | 1 | 1 | 1 | 1 | -0.01 |
| SE | 35352 | ENSG00000164073 | MFSD8 | chr4 | - | 127930682 | 127930817 | 127917924 | 127921771 | 127932984 | 127933093 | 117 | 44 | 84 | 92 | 167 | 0 | 0 | 0 | 0 | 2 | 13 | 18 | 53 | 54 | 27 | 0 | 0 | 0 | 0 | 0 | 284 | 149 | 1 | 1 | 1 | 1 | 1 | 1 | 0.978 | 1 | 1 | 1 | 1 | 1 | -0.004 |
| SE | 35353 | ENSG00000164073 | MFSD8 | chr4 | - | 127930682 | 127930817 | 127921523 | 127921963 | 127932984 | 127933093 | 111 | 52 | 94 | 79 | 144 | 0 | 0 | 2 | 2 | 0 | 47 | 32 | 46 | 58 | 13 | 0 | 0 | 0 | 0 | 0 | 284 | 149 | 1 | 1 | 1 | 1 | 0.961 | 0.954 | 1 | 1 | 1 | 1 | 1 | 1 | -0.017 |
| SE | 35355 | ENSG00000164073 | MFSD8 | chr4 | - | 127938782 | 127938836 | 127932421 | 127933093 | 127942044 | 127942158 | 50 | 37 | 45 | 46 | 83 | 1 | 0 | 0 | 0 | 0 | 15 | 14 | 28 | 30 | 5 | 0 | 0 | 0 | 0 | 0 | 203 | 149 | 1 | 1 | 0.973 | 1 | 1 | 1 | 1 | 1 | 1 | 1 | 1 | 1 | -0.005 |
| SE | 35356 | ENSG00000164073 | MFSD8 | chr4 | - | 127938782 | 127938838 | 127932318 | 127933093 | 127939852 | 127939886 | 103 | 75 | 91 | 91 | 189 | 8 | 1 | 1 | 1 | 8 | 47 | 40 | 47 | 51 | 19 | 0 | 0 | 3 | 0 | 1 | 205 | 149 | 1 | 1 | 0.903 | 0.982 | 0.985 | 0.985 | 0.945 | 1 | 1 | 0.919 | 1 | 0.932 | -0.01 |
| SE | 35357 | ENSG00000164073 | MFSD8 | chr4 | - | 127938782 | 127938838 | 127932318 | 127933093 | 127942044 | 127942047 | 53 | 40 | 50 | 52 | 83 | 1 | 0 | 0 | 0 | 0 | 15 | 22 | 33 | 30 | 5 | 0 | 0 | 0 | 0 | 0 | 205 | 149 | 1 | 1 | 0.975 | 1 | 1 | 1 | 1 | 1 | 1 | 1 | 1 | 1 | -0.005 |
| SE | 35358 | ENSG00000164073 | MFSD8 | chr4 | - | 127938782 | 127938838 | 127932318 | 127933093 | 127943751 | 127943992 | 50 | 37 | 46 | 48 | 84 | 0 | 0 | 0 | 1 | 0 | 15 | 16 | 34 | 30 | 5 | 0 | 0 | 0 | 0 | 0 | 205 | 149 | 1 | 1 | 1 | 1 | 1 | 0.972 | 1 | 1 | 1 | 1 | 1 | 1 | -0.006 |
| SE | 35360 | ENSG00000164073 | MFSD8 | chr4 | - | 127939852 | 127939997 | 127932984 | 127933093 | 127942044 | 127942158 | 35 | 26 | 28 | 30 | 66 | 1 | 0 | 0 | 0 | 0 | 0 | 14 | 11 | 5 | 1 | 0 | 0 | 0 | 0 | 0 | 294 | 149 | 1 | 1 | 0.947 | 1 | 1 | 1 | 1 | NA | 1 | 1 | 1 | 1 | -0.011 |
| SE | 35361 | ENSG00000164073 | MFSD8 | chr4 | - | 127939852 | 127939997 | 127932984 | 127933093 | 127943751 | 127943992 | 8 | 4 | 2 | 3 | 8 | 0 | 0 | 0 | 1 | 0 | 0 | 0 | 3 | 0 | 1 | 0 | 0 | 0 | 0 | 0 | 294 | 149 | 1 | 1 | 1 | 1 | 1 | 0.603 | 1 | NA | NA | 1 | NA | 1 | -0.079 |
| SE | 35366 | ENSG00000164073 | MFSD8 | chr4 | - | 127942044 | 127942158 | 127932984 | 127933093 | 127943751 | 127943992 | 11 | 14 | 21 | 14 | 12 | 0 | 0 | 0 | 1 | 0 | 6 | 16 | 6 | 3 | 0 | 0 | 0 | 0 | 0 | 0 | 263 | 149 | 1 | 1 | 1 | 1 | 1 | 0.888 | 1 | 1 | 1 | 1 | 1 | NA | -0.022 |
| SE | 35367 | ENSG00000164073 | MFSD8 | chr4 | - | 127942044 | 127942158 | 127938782 | 127938838 | 127943751 | 127943992 | 13 | 17 | 25 | 19 | 12 | 0 | 0 | 0 | 1 | 1 | 6 | 22 | 11 | 3 | 0 | 0 | 0 | 6 | 0 | 0 | 263 | 149 | 1 | 1 | 1 | 1 | 1 | 0.915 | 0.872 | 1 | 1 | 0.509 | 1 | NA | 0.08 |
| SE | 35368 | ENSG00000164073 | MFSD8 | chr4 | - | 127942044 | 127942158 | 127939778 | 127939997 | 127943751 | 127943992 | 38 | 38 | 46 | 43 | 70 | 1 | 3 | 1 | 2 | 0 | 6 | 30 | 14 | 8 | 0 | 0 | 0 | 0 | 0 | 0 | 263 | 149 | 1 | 1 | 0.956 | 0.878 | 0.963 | 0.924 | 1 | 1 | 1 | 1 | 1 | NA | -0.056 |
| SE | 35372 | ENSG00000164073 | MFSD8 | chr4 | - | 127957500 | 127957590 | 127949803 | 127949847 | 127965071 | 127965143 | 19 | 12 | 20 | 18 | 13 | 0 | 0 | 1 | 0 | 0 | 5 | 13 | 18 | 9 | 0 | 0 | 0 | 0 | 0 | 0 | 239 | 149 | 1 | 1 | 1 | 1 | 0.926 | 1 | 1 | 1 | 1 | 1 | 1 | NA | -0.015 |
| SE | 35373 | ENSG00000164073 | MFSD8 | chr4 | - | 127957500 | 127957592 | 127949803 | 127949847 | 127965071 | 127965074 | 50 | 31 | 57 | 37 | 44 | 0 | 0 | 1 | 0 | 0 | 5 | 28 | 36 | 20 | 20 | 0 | 0 | 0 | 0 | 0 | 241 | 149 | 1 | 1 | 1 | 1 | 0.972 | 1 | 1 | 1 | 1 | 1 | 1 | 1 | -0.006 |
| SE | 35374 | ENSG00000164073 | MFSD8 | chr4 | - | 127965071 | 127965208 | 127957500 | 127957592 | 127965896 | 127965949 | 32 | 19 | 41 | 20 | 28 | 0 | 0 | 6 | 0 | 1 | 0 | 15 | 19 | 14 | 20 | 0 | 0 | 3 | 0 | 0 | 286 | 149 | 1 | 1 | 1 | 1 | 0.781 | 1 | 0.936 | NA | 1 | 0.767 | 1 | 1 | 0.002 |
| SE | 35375 | ENSG00000164073 | MFSD8 | chr4 | - | 127965071 | 127965222 | 127957500 | 127957592 | 127965896 | 127965938 | 32 | 19 | 41 | 20 | 28 | 0 | 0 | 6 | 0 | 1 | 0 | 15 | 19 | 15 | 20 | 0 | 0 | 3 | 0 | 0 | 298 | 149 | 1 | 1 | 1 | 1 | 0.774 | 1 | 0.933 | NA | 1 | 0.76 | 1 | 1 | 0.001 |
| SE | 35376 | ENSG00000164073 | MFSD8 | chr4 | - | 127965071 | 127965482 | 127957500 | 127957590 | 127965896 | 127965995 | 0 | 0 | 2 | 0 | 0 | 0 | 1 | 0 | 0 | 0 | 0 | 0 | 1 | 0 | 0 | 0 | 0 | 0 | 0 | 0 | 298 | 149 | 1 | 1 | NA | 0 | 1 | NA | NA | NA | NA | 1 | NA | NA | -0.5 |
| SE | 35382 | ENSG00000140262 | TCF12 | chr15 | + | 57091788 | 57091891 | 56921025 | 56921098 | 57166401 | 57166428 | 166 | 134 | 170 | 175 | 449 | 0 | 1 | 0 | 0 | 0 | 102 | 91 | 150 | 88 | 46 | 0 | 0 | 0 | 0 | 0 | 252 | 149 | 1 | 1 | 1 | 0.988 | 1 | 1 | 1 | 1 | 1 | 1 | 1 | 1 | -0.002 |
| SE | 35383 | ENSG00000140262 | TCF12 | chr15 | + | 57129782 | 57129972 | 57091788 | 57091891 | 57166401 | 57166466 | 0 | 0 | 0 | 0 | 0 | 192 | 145 | 190 | 194 | 500 | 0 | 0 | 4 | 2 | 0 | 102 | 111 | 169 | 101 | 56 | 298 | 149 | 1 | 1 | 0 | 0 | 0 | 0 | 0 | 0 | 0 | 0.012 | 0.01 | 0 | -0.004 |
| SE | 35385 | ENSG00000140262 | TCF12 | chr15 | + | 57251349 | 57251423 | 57243471 | 57243550 | 57253261 | 57253468 | 102 | 80 | 84 | 106 | 127 | 0 | 0 | 1 | 1 | 0 | 13 | 105 | 60 | 18 | 43 | 0 | 0 | 0 | 0 | 0 | 223 | 149 | 1 | 1 | 1 | 1 | 0.982 | 0.986 | 1 | 1 | 1 | 1 | 1 | 1 | -0.006 |
| SE | 35386 | ENSG00000140262 | TCF12 | chr15 | + | 57252420 | 57252492 | 57251168 | 57251423 | 57253261 | 57253468 | 2 | 3 | 4 | 4 | 7 | 36 | 27 | 39 | 52 | 50 | 0 | 7 | 1 | 0 | 0 | 0 | 45 | 33 | 7 | 27 | 221 | 149 | 1 | 1 | 0.036 | 0.07 | 0.065 | 0.049 | 0.086 | NA | 0.095 | 0.02 | 0 | 0 | 0.032 |
| SE | 35387 | ENSG00000140262 | TCF12 | chr15 | + | 57282444 | 57282598 | 57273029 | 57273262 | 57286156 | 57286251 | 143 | 126 | 132 | 134 | 268 | 0 | 0 | 0 | 0 | 0 | 15 | 139 | 121 | 63 | 9 | 0 | 2 | 1 | 0 | 0 | 298 | 149 | 1 | 1 | 1 | 1 | 1 | 1 | 1 | 1 | 0.972 | 0.984 | 1 | 1 | 0.009 |
| SE | 35388 | ENSG00000161267 | BDH1 | chr3 | - | 197532411 | 197532522 | 197522639 | 197522781 | 197533488 | 197533561 | 53 | 33 | 76 | 52 | 45 | 0 | 0 | 1 | 0 | 0 | 40 | 48 | 17 | 67 | 57 | 0 | 0 | 0 | 0 | 0 | 260 | 149 | 1 | 1 | 1 | 1 | 0.978 | 1 | 1 | 1 | 1 | 1 | 1 | 1 | -0.004 |
| SE | 35393 | ENSG00000161267 | BDH1 | chr3 | - | 197554561 | 197554713 | 197546360 | 197546486 | 197555780 | 197555968 | 2 | 0 | 0 | 0 | 0 | 3 | 2 | 3 | 5 | 2 | 6 | 5 | 0 | 0 | 0 | 0 | 19 | 0 | 0 | 0 | 298 | 149 | 1 | 1 | 0.25 | 0 | 0 | 0 | 0 | 1 | 0.116 | NA | NA | NA | -0.508 |
| SE | 35397 | ENSG00000122068 | FYTTD1 | chr3 | + | 197768438 | 197768587 | 197756425 | 197756557 | 197770131 | 197770161 | 1046 | 731 | 786 | 977 | 1129 | 0 | 0 | 0 | 0 | 4 | 246 | 644 | 700 | 378 | 90 | 0 | 0 | 0 | 0 | 0 | 298 | 149 | 1 | 1 | 1 | 1 | 1 | 1 | 0.993 | 1 | 1 | 1 | 1 | 1 | -0.001 |
| SE | 35399 | ENSG00000122068 | FYTTD1 | chr3 | + | 197773402 | 197773499 | 197768438 | 197768587 | 197774148 | 197774210 | 230 | 167 | 156 | 242 | 335 | 1 | 0 | 0 | 0 | 0 | 57 | 155 | 188 | 131 | 29 | 0 | 0 | 0 | 0 | 0 | 246 | 149 | 1 | 1 | 0.993 | 1 | 1 | 1 | 1 | 1 | 1 | 1 | 1 | 1 | -0.001 |
| SE | 35403 | ENSG00000115539 | PDCL3 | chr2 | + | 100571589 | 100571798 | 100569577 | 100569721 | 100576353 | 100576739 | 245 | 109 | 254 | 211 | 244 | 0 | 0 | 0 | 0 | 0 | 167 | 180 | 182 | 208 | 47 | 4 | 0 | 0 | 0 | 0 | 298 | 149 | 1 | 1 | 1 | 1 | 1 | 1 | 1 | 0.954 | 1 | 1 | 1 | 1 | 0.009 |
| SE | 35405 | ENSG00000174891 | RSRC1 | chr3 | + | 158122102 | 158122298 | 158110067 | 158110223 | 158203071 | 158203245 | 13 | 7 | 12 | 12 | 12 | 0 | 0 | 1 | 0 | 0 | 0 | 28 | 6 | 4 | 0 | 0 | 0 | 0 | 0 | 0 | 298 | 149 | 1 | 1 | 1 | 1 | 0.857 | 1 | 1 | NA | 1 | 1 | 1 | NA | -0.029 |
| SE | 35407 | ENSG00000174891 | RSRC1 | chr3 | + | 158132085 | 158132263 | 158123865 | 158123991 | 158203071 | 158203245 | 4 | 0 | 1 | 0 | 3 | 75 | 66 | 86 | 92 | 111 | 0 | 0 | 0 | 0 | 0 | 0 | 47 | 65 | 47 | 14 | 298 | 149 | 1 | 1 | 0.026 | 0 | 0.006 | 0 | 0.013 | NA | 0 | 0 | 0 | 0 | 0.009 |
| SE | 35409 | ENSG00000174891 | RSRC1 | chr3 | + | 158203071 | 158203269 | 158123865 | 158123991 | 158298038 | 158298075 | 75 | 66 | 86 | 92 | 111 | 3 | 3 | 0 | 0 | 3 | 0 | 47 | 65 | 47 | 14 | 0 | 0 | 5 | 0 | 0 | 298 | 149 | 1 | 1 | 0.926 | 0.917 | 1 | 1 | 0.949 | NA | 1 | 0.867 | 1 | 1 | -0.008 |
| SE | 35410 | ENSG00000174891 | RSRC1 | chr3 | + | 158225656 | 158225733 | 158203071 | 158203245 | 158231169 | 158231278 | 3 | 6 | 4 | 0 | 5 | 1 | 0 | 0 | 0 | 0 | 0 | 0 | 1 | 1 | 3 | 0 | 3 | 0 | 0 | 0 | 226 | 149 | 1 | 1 | 0.664 | 1 | 1 | NA | 1 | NA | 0 | 1 | 1 | 1 | 0.166 |
| SE | 35412 | ENSG00000174891 | RSRC1 | chr3 | + | 158231169 | 158231278 | 158203071 | 158203245 | 158298038 | 158298075 | 1 | 2 | 0 | 0 | 0 | 73 | 70 | 58 | 94 | 146 | 0 | 3 | 0 | 0 | 0 | 27 | 41 | 30 | 7 | 0 | 258 | 149 | 1 | 1 | 0.008 | 0.016 | 0 | 0 | 0 | 0 | 0.041 | 0 | 0 | NA | -0.005 |
| SE | 35413 | ENSG00000174891 | RSRC1 | chr3 | + | 158231169 | 158231278 | 158225656 | 158225733 | 158298038 | 158298075 | 0 | 3 | 0 | 0 | 0 | 6 | 1 | 7 | 4 | 14 | 0 | 0 | 0 | 0 | 0 | 25 | 0 | 4 | 0 | 0 | 258 | 149 | 1 | 1 | 0 | 0.634 | 0 | 0 | 0 | 0 | NA | 0 | NA | NA | 0.127 |
| SE | 35415 | ENSG00000174891 | RSRC1 | chr3 | + | 158354856 | 158354908 | 158298038 | 158298075 | 158460934 | 158460962 | 102 | 98 | 84 | 119 | 207 | 0 | 1 | 0 | 0 | 0 | 23 | 49 | 43 | 33 | 0 | 0 | 0 | 0 | 0 | 0 | 201 | 149 | 1 | 1 | 1 | 0.986 | 1 | 1 | 1 | 1 | 1 | 1 | 1 | NA | -0.003 |
| SE | 35422 | ENSG00000186001 | LRCH3 | chr3 | + | 197817175 | 197817302 | 197814907 | 197815052 | 197820324 | 197820430 | 498 | 389 | 370 | 445 | 794 | 0 | 1 | 3 | 0 | 0 | 686 | 449 | 404 | 436 | 454 | 0 | 0 | 0 | 0 | 0 | 276 | 149 | 1 | 1 | 1 | 0.995 | 0.985 | 1 | 1 | 1 | 1 | 1 | 1 | 1 | -0.004 |
| SE | 35425 | ENSG00000186001 | LRCH3 | chr3 | + | 197829563 | 197829673 | 197826877 | 197827014 | 197830769 | 197830863 | 248 | 207 | 264 | 271 | 442 | 0 | 0 | 0 | 0 | 2 | 262 | 255 | 139 | 260 | 257 | 0 | 0 | 0 | 0 | 0 | 259 | 149 | 1 | 1 | 1 | 1 | 1 | 1 | 0.992 | 1 | 1 | 1 | 1 | 1 | -0.002 |
| SE | 35426 | ENSG00000186001 | LRCH3 | chr3 | + | 197835673 | 197835738 | 197832196 | 197832317 | 197839320 | 197839397 | 133 | 94 | 110 | 112 | 138 | 0 | 2 | 0 | 0 | 0 | 163 | 185 | 73 | 89 | 205 | 0 | 4 | 0 | 2 | 0 | 214 | 149 | 1 | 1 | 1 | 0.97 | 1 | 1 | 1 | 1 | 0.97 | 1 | 0.969 | 1 | 0.006 |
| SE | 35427 | ENSG00000186001 | LRCH3 | chr3 | + | 197835673 | 197835822 | 197832196 | 197832317 | 197839320 | 197839397 | 312 | 245 | 286 | 305 | 403 | 0 | 2 | 0 | 0 | 0 | 381 | 391 | 231 | 295 | 377 | 0 | 4 | 0 | 2 | 0 | 298 | 149 | 1 | 1 | 1 | 0.984 | 1 | 1 | 1 | 1 | 0.98 | 1 | 0.987 | 1 | 0.003 |
| SE | 35433 | ENSG00000186001 | LRCH3 | chr3 | + | 197865422 | 197865471 | 197858817 | 197858905 | 197866111 | 197866179 | 88 | 53 | 79 | 70 | 115 | 0 | 0 | 0 | 0 | 5 | 93 | 41 | 58 | 72 | 68 | 0 | 0 | 0 | 0 | 0 | 198 | 149 | 1 | 1 | 1 | 1 | 1 | 1 | 0.945 | 1 | 1 | 1 | 1 | 1 | -0.011 |
| SE | 35434 | ENSG00000186001 | LRCH3 | chr3 | + | 197865422 | 197865471 | 197858817 | 197858905 | 197870159 | 197870278 | 12 | 9 | 21 | 26 | 30 | 0 | 1 | 0 | 0 | 3 | 0 | 5 | 13 | 14 | 19 | 0 | 0 | 0 | 0 | 0 | 198 | 149 | 1 | 1 | 1 | 0.871 | 1 | 1 | 0.883 | NA | 1 | 1 | 1 | 1 | -0.049 |
| SE | 35435 | ENSG00000186001 | LRCH3 | chr3 | + | 197866111 | 197866219 | 197858820 | 197858905 | 197870159 | 197870176 | 46 | 20 | 29 | 40 | 50 | 0 | 1 | 0 | 0 | 3 | 31 | 20 | 18 | 24 | 61 | 0 | 0 | 0 | 0 | 0 | 257 | 149 | 1 | 1 | 1 | 0.921 | 1 | 1 | 0.906 | 1 | 1 | 1 | 1 | 1 | -0.035 |
| SE | 35437 | ENSG00000186001 | LRCH3 | chr3 | + | 197870159 | 197870278 | 197866111 | 197866219 | 197871324 | 197871462 | 232 | 124 | 185 | 222 | 348 | 1 | 0 | 0 | 2 | 0 | 190 | 124 | 108 | 120 | 181 | 0 | 0 | 0 | 2 | 0 | 268 | 149 | 1 | 1 | 0.992 | 1 | 1 | 0.984 | 1 | 1 | 1 | 1 | 0.971 | 1 | 0.001 |
| SE | 35438 | ENSG00000186001 | LRCH3 | chr3 | + | 197871324 | 197871462 | 197870159 | 197870278 | 197875697 | 197875760 | 239 | 124 | 184 | 228 | 345 | 1 | 0 | 0 | 0 | 0 | 170 | 123 | 139 | 99 | 173 | 0 | 0 | 0 | 0 | 0 | 287 | 149 | 1 | 1 | 0.992 | 1 | 1 | 1 | 1 | 1 | 1 | 1 | 1 | 1 | -0.002 |
| SE | 35439 | ENSG00000186001 | LRCH3 | chr3 | + | 197875697 | 197875775 | 197871324 | 197871462 | 197883540 | 197883772 | 167 | 79 | 121 | 151 | 239 | 0 | 0 | 0 | 0 | 0 | 81 | 91 | 126 | 74 | 110 | 0 | 0 | 0 | 0 | 1 | 227 | 149 | 1 | 1 | 1 | 1 | 1 | 1 | 1 | 1 | 1 | 1 | 1 | 0.986 | 0.003 |
| SE | 35440 | ENSG00000186001 | LRCH3 | chr3 | + | 197880511 | 197880757 | 197875697 | 197875775 | 197883540 | 197888436 | 1 | 0 | 0 | 0 | 0 | 122 | 58 | 87 | 107 | 175 | 0 | 0 | 1 | 0 | 0 | 70 | 76 | 90 | 59 | 68 | 298 | 149 | 1 | 1 | 0.004 | 0 | 0 | 0 | 0 | 0 | 0 | 0.006 | 0 | 0 | 0 |
| SE | 35448 | ENSG00000111642 | CHD4 | chr12 | - | 6578845 | 6578917 | 6577784 | 6578546 | 6581043 | 6581173 | 356 | 276 | 328 | 441 | 462 | 0 | 0 | 0 | 0 | 0 | 1090 | 520 | 442 | 698 | 876 | 0 | 0 | 0 | 5 | 0 | 221 | 149 | 1 | 1 | 1 | 1 | 1 | 1 | 1 | 1 | 1 | 1 | 0.989 | 1 | 0.002 |
| SE | 35449 | ENSG00000111642 | CHD4 | chr12 | - | 6580581 | 6580679 | 6578845 | 6578917 | 6581043 | 6581173 | 0 | 5 | 3 | 1 | 3 | 320 | 254 | 303 | 402 | 418 | 0 | 1 | 4 | 3 | 0 | 1002 | 488 | 397 | 640 | 830 | 247 | 149 | 1 | 1 | 0 | 0.012 | 0.006 | 0.001 | 0.004 | 0 | 0.001 | 0.006 | 0.003 | 0 | 0.003 |
| SE | 35450 | ENSG00000111642 | CHD4 | chr12 | - | 6580581 | 6580699 | 6578845 | 6578917 | 6581043 | 6581173 | 0 | 1 | 1 | 1 | 0 | 320 | 254 | 303 | 402 | 418 | 0 | 1 | 0 | 3 | 0 | 1002 | 488 | 397 | 640 | 830 | 267 | 149 | 1 | 1 | 0 | 0.002 | 0.002 | 0.001 | 0 | 0 | 0.001 | 0 | 0.003 | 0 | 0 |
| SE | 35451 | ENSG00000111642 | CHD4 | chr12 | - | 6581648 | 6581814 | 6581290 | 6581388 | 6582136 | 6582281 | 671 | 522 | 773 | 881 | 734 | 0 | 0 | 0 | 0 | 0 | 1753 | 1041 | 897 | 1357 | 1556 | 0 | 7 | 0 | 2 | 0 | 298 | 149 | 1 | 1 | 1 | 1 | 1 | 1 | 1 | 1 | 0.987 | 1 | 0.997 | 1 | 0.003 |
| SE | 35457 | ENSG00000111642 | CHD4 | chr12 | - | 6590072 | 6590213 | 6588297 | 6588422 | 6591465 | 6591583 | 1 | 0 | 0 | 0 | 6 | 310 | 184 | 316 | 300 | 241 | 0 | 0 | 0 | 0 | 0 | 451 | 389 | 231 | 280 | 404 | 290 | 149 | 1 | 1 | 0.002 | 0 | 0 | 0 | 0.013 | 0 | 0 | 0 | 0 | 0 | 0.003 |
| SE | 35459 | ENSG00000168566 | SNRNP48 | chr6 | + | 7599672 | 7599680 | 7595026 | 7595101 | 7601335 | 7601524 | 13 | 5 | 13 | 16 | 17 | 36 | 16 | 12 | 30 | 73 | 0 | 5 | 2 | 6 | 0 | 6 | 30 | 17 | 12 | 0 | 157 | 149 | 1 | 1 | 0.255 | 0.229 | 0.507 | 0.336 | 0.181 | 0 | 0.137 | 0.1 | 0.322 | NA | 0.162 |
| SE | 35461 | ENSG00000161217 | PCYT1A | chr3 | - | 196239546 | 196239735 | 196234367 | 196238894 | 196247366 | 196247518 | 217 | 137 | 149 | 135 | 263 | 0 | 0 | 0 | 0 | 0 | 228 | 161 | 177 | 139 | 239 | 0 | 4 | 0 | 0 | 0 | 298 | 149 | 1 | 1 | 1 | 1 | 1 | 1 | 1 | 1 | 0.953 | 1 | 1 | 1 | 0.009 |
| SE | 35463 | ENSG00000161217 | PCYT1A | chr3 | - | 196267279 | 196267365 | 196257787 | 196257887 | 196270414 | 196270541 | 1 | 1 | 3 | 4 | 1 | 105 | 45 | 91 | 93 | 121 | 0 | 0 | 3 | 0 | 0 | 77 | 100 | 63 | 71 | 92 | 235 | 149 | 1 | 1 | 0.006 | 0.014 | 0.02 | 0.027 | 0.005 | 0 | 0 | 0.029 | 0 | 0 | 0.009 |
| SE | 35464 | ENSG00000161217 | PCYT1A | chr3 | - | 196270414 | 196270541 | 196257787 | 196257887 | 196287614 | 196287663 | 130 | 48 | 108 | 131 | 135 | 0 | 0 | 0 | 0 | 0 | 112 | 120 | 76 | 85 | 122 | 0 | 0 | 0 | 5 | 0 | 276 | 149 | 1 | 1 | 1 | 1 | 1 | 1 | 1 | 1 | 1 | 1 | 0.902 | 1 | 0.02 |
| SE | 35465 | ENSG00000161217 | PCYT1A | chr3 | - | 196283474 | 196283538 | 196270414 | 196270541 | 196287531 | 196287726 | 6 | 0 | 1 | 2 | 2 | 5 | 0 | 5 | 1 | 0 | 0 | 3 | 0 | 7 | 2 | 0 | 6 | 8 | 0 | 8 | 213 | 149 | 1 | 1 | 0.456 | NA | 0.123 | 0.583 | 1 | NA | 0.259 | 0 | 1 | 0.149 | 0.189 |
| SE | 35467 | ENSG00000175110 | MRPS22 | chr3 | + | 139346850 | 139347044 | 139344038 | 139344198 | 139348159 | 139348324 | 172 | 92 | 123 | 105 | 151 | 0 | 0 | 0 | 0 | 0 | 140 | 126 | 156 | 141 | 18 | 4 | 0 | 0 | 0 | 0 | 298 | 149 | 1 | 1 | 1 | 1 | 1 | 1 | 1 | 0.946 | 1 | 1 | 1 | 1 | 0.011 |
| SE | 35468 | ENSG00000175110 | MRPS22 | chr3 | + | 139346877 | 139347044 | 139343955 | 139344198 | 139348159 | 139348324 | 279 | 160 | 186 | 191 | 215 | 0 | 0 | 0 | 0 | 0 | 211 | 226 | 240 | 249 | 61 | 4 | 0 | 0 | 0 | 0 | 298 | 149 | 1 | 1 | 1 | 1 | 1 | 1 | 1 | 0.963 | 1 | 1 | 1 | 1 | 0.007 |
| SE | 35469 | ENSG00000175110 | MRPS22 | chr3 | + | 139346880 | 139347044 | 139344019 | 139344198 | 139348159 | 139348324 | 178 | 98 | 138 | 123 | 159 | 0 | 0 | 0 | 0 | 0 | 155 | 143 | 158 | 151 | 19 | 4 | 0 | 0 | 0 | 0 | 298 | 149 | 1 | 1 | 1 | 1 | 1 | 1 | 1 | 0.951 | 1 | 1 | 1 | 1 | 0.01 |
| SE | 35470 | ENSG00000175110 | MRPS22 | chr3 | + | 139349327 | 139349434 | 139348159 | 139348324 | 139350178 | 139350322 | 1 | 1 | 0 | 1 | 0 | 79 | 66 | 76 | 82 | 62 | 0 | 0 | 0 | 7 | 0 | 141 | 80 | 85 | 99 | 20 | 256 | 149 | 1 | 1 | 0.007 | 0.009 | 0 | 0.007 | 0 | 0 | 0 | 0 | 0.04 | 0 | -0.003 |
| SE | 35471 | ENSG00000175110 | MRPS22 | chr3 | + | 139350178 | 139351060 | 139348159 | 139348324 | 139352646 | 139352792 | 154 | 111 | 140 | 135 | 146 | 0 | 2 | 0 | 0 | 1 | 184 | 151 | 147 | 196 | 77 | 0 | 0 | 0 | 0 | 0 | 298 | 149 | 1 | 1 | 1 | 0.965 | 1 | 1 | 0.986 | 1 | 1 | 1 | 1 | 1 | -0.01 |
| SE | 35472 | ENSG00000175110 | MRPS22 | chr3 | + | 139352646 | 139352792 | 139350178 | 139351060 | 139355681 | 139355790 | 358 | 241 | 291 | 263 | 477 | 0 | 0 | 1 | 0 | 0 | 360 | 285 | 358 | 407 | 179 | 0 | 0 | 0 | 0 | 0 | 295 | 149 | 1 | 1 | 1 | 1 | 0.993 | 1 | 1 | 1 | 1 | 1 | 1 | 1 | -0.001 |
| SE | 35473 | ENSG00000198075 | SULT1C4 | chr2 | + | 108382384 | 108382482 | 108381761 | 108381887 | 108383092 | 108383219 | 0 | 0 | 2 | 0 | 0 | 0 | 0 | 0 | 0 | 0 | 0 | 0 | 0 | 0 | 0 | 2 | 0 | 0 | 0 | 0 | 247 | 149 | 1 | 1 | NA | NA | 1 | NA | NA | 0 | NA | NA | NA | NA | 1 |
| SE | 35479 | ENSG00000172830 | SSH3 | chr11 | + | 67309396 | 67309543 | 67308411 | 67308458 | 67309767 | 67309968 | 14 | 1 | 18 | 6 | 7 | 0 | 0 | 1 | 0 | 0 | 12 | 24 | 8 | 24 | 25 | 0 | 0 | 0 | 0 | 0 | 296 | 149 | 1 | 1 | 1 | 1 | 0.901 | 1 | 1 | 1 | 1 | 1 | 1 | 1 | -0.02 |
| SE | 35481 | ENSG00000019169 | MARCO | chr2 | + | 118971498 | 118971534 | 118970113 | 118970338 | 118974332 | 118974440 | 9 | 25 | 19 | 8 | 3 | 0 | 1 | 0 | 0 | 0 | 97 | 17 | 7 | 2 | 80 | 0 | 0 | 0 | 0 | 0 | 185 | 149 | 1 | 1 | 1 | 0.953 | 1 | 1 | 1 | 1 | 1 | 1 | 1 | 1 | -0.009 |
| SE | 35484 | ENSG00000247193 | NA | chr4 | + | 36272946 | 36273048 | 36256517 | 36256661 | 36273234 | 36274220 | 2 | 0 | 0 | 0 | 0 | 0 | 0 | 0 | 0 | 2 | 0 | 0 | 2 | 0 | 0 | 0 | 0 | 0 | 0 | 4 | 251 | 149 | 1 | 1 | 1 | NA | NA | NA | 0 | NA | NA | 1 | NA | 0 | 0 |
| SE | 35486 | ENSG00000282870 | FRG1DP | chr20 | + | 29094346 | 29094461 | 29091593 | 29091719 | 29096727 | 29096832 | 0 | 0 | 0 | 0 | 0 | 0 | 0 | 0 | 0 | 2 | 0 | 0 | 0 | 3 | 0 | 0 | 0 | 0 | 0 | 0 | 264 | 149 | 1 | 1 | NA | NA | NA | NA | 0 | NA | NA | NA | 1 | NA | -1 |
| SE | 35487 | ENSG00000282870 | FRG1DP | chr20 | + | 29096727 | 29096832 | 29094346 | 29094461 | 29100368 | 29100460 | 0 | 1 | 0 | 0 | 0 | 0 | 1 | 0 | 0 | 0 | 0 | 0 | 0 | 0 | 0 | 0 | 0 | 0 | 3 | 0 | 254 | 149 | 1 | 1 | NA | 0.37 | NA | NA | NA | NA | NA | NA | 0 | NA | 0.37 |
| SE | 35488 | ENSG00000051382 | PIK3CB | chr3 | - | 138656141 | 138656274 | 138652697 | 138655526 | 138657689 | 138657835 | 229 | 182 | 148 | 214 | 254 | 0 | 0 | 0 | 0 | 1 | 143 | 159 | 128 | 180 | 134 | 0 | 0 | 2 | 2 | 0 | 282 | 149 | 1 | 1 | 1 | 1 | 1 | 1 | 0.993 | 1 | 1 | 0.971 | 0.979 | 1 | 0.009 |
| SE | 35489 | ENSG00000051382 | PIK3CB | chr3 | - | 138657689 | 138657835 | 138656141 | 138656274 | 138663905 | 138664029 | 201 | 156 | 132 | 173 | 175 | 0 | 0 | 0 | 0 | 0 | 129 | 191 | 131 | 151 | 126 | 0 | 2 | 0 | 0 | 0 | 295 | 149 | 1 | 1 | 1 | 1 | 1 | 1 | 1 | 1 | 0.98 | 1 | 1 | 1 | 0.004 |
| SE | 35491 | ENSG00000051382 | PIK3CB | chr3 | - | 138681966 | 138682045 | 138665035 | 138665203 | 138683677 | 138683787 | 97 | 64 | 113 | 106 | 178 | 0 | 0 | 0 | 0 | 0 | 62 | 154 | 99 | 135 | 209 | 0 | 2 | 0 | 0 | 0 | 228 | 149 | 1 | 1 | 1 | 1 | 1 | 1 | 1 | 1 | 0.981 | 1 | 1 | 1 | 0.004 |
| SE | 35492 | ENSG00000051382 | PIK3CB | chr3 | - | 138681966 | 138682060 | 138665035 | 138665203 | 138683677 | 138683787 | 54 | 32 | 42 | 39 | 71 | 0 | 0 | 0 | 0 | 0 | 35 | 58 | 41 | 59 | 68 | 0 | 2 | 0 | 0 | 0 | 243 | 149 | 1 | 1 | 1 | 1 | 1 | 1 | 1 | 1 | 0.947 | 1 | 1 | 1 | 0.011 |
| SE | 35493 | ENSG00000051382 | PIK3CB | chr3 | - | 138684624 | 138684803 | 138683677 | 138683787 | 138688874 | 138688974 | 143 | 133 | 176 | 199 | 210 | 0 | 0 | 0 | 1 | 0 | 75 | 242 | 183 | 226 | 274 | 0 | 0 | 0 | 0 | 0 | 298 | 149 | 1 | 1 | 1 | 1 | 1 | 0.99 | 1 | 1 | 1 | 1 | 1 | 1 | -0.002 |
| SE | 35499 | ENSG00000051382 | PIK3CB | chr3 | - | 138690999 | 138691143 | 138688874 | 138688974 | 138694785 | 138694907 | 181 | 159 | 175 | 242 | 202 | 2 | 0 | 0 | 0 | 0 | 85 | 220 | 197 | 282 | 276 | 0 | 0 | 0 | 0 | 0 | 293 | 149 | 1 | 1 | 0.979 | 1 | 1 | 1 | 1 | 1 | 1 | 1 | 1 | 1 | -0.004 |
| SE | 35500 | ENSG00000051382 | PIK3CB | chr3 | - | 138694785 | 138694907 | 138684624 | 138684803 | 138698906 | 138699095 | 107 | 85 | 95 | 125 | 181 | 0 | 2 | 4 | 0 | 0 | 105 | 130 | 115 | 105 | 184 | 0 | 5 | 2 | 0 | 0 | 271 | 149 | 1 | 1 | 1 | 0.959 | 0.929 | 1 | 1 | 1 | 0.935 | 0.969 | 1 | 1 | -0.003 |
| SE | 35504 | ENSG00000051382 | PIK3CB | chr3 | - | 138712207 | 138712304 | 138707158 | 138707289 | 138714467 | 138714486 | 158 | 158 | 161 | 151 | 350 | 0 | 0 | 0 | 0 | 0 | 120 | 122 | 112 | 200 | 122 | 0 | 0 | 1 | 0 | 0 | 246 | 149 | 1 | 1 | 1 | 1 | 1 | 1 | 1 | 1 | 1 | 0.985 | 1 | 1 | 0.003 |
| SE | 35505 | ENSG00000051382 | PIK3CB | chr3 | - | 138734633 | 138734804 | 138733360 | 138733438 | 138737706 | 138737886 | 293 | 276 | 348 | 341 | 493 | 0 | 0 | 0 | 1 | 5 | 151 | 339 | 351 | 480 | 340 | 4 | 0 | 1 | 0 | 0 | 298 | 149 | 1 | 1 | 1 | 1 | 1 | 0.994 | 0.98 | 0.95 | 1 | 0.994 | 1 | 1 | 0.006 |
| SE | 35506 | ENSG00000051382 | PIK3CB | chr3 | - | 138737706 | 138737886 | 138734633 | 138734804 | 138742557 | 138742755 | 323 | 270 | 346 | 299 | 469 | 0 | 1 | 0 | 1 | 0 | 300 | 426 | 439 | 550 | 511 | 0 | 0 | 1 | 0 | 0 | 298 | 149 | 1 | 1 | 1 | 0.993 | 1 | 0.993 | 1 | 1 | 1 | 1 | 1 | 1 | -0.003 |
| SE | 35507 | ENSG00000051382 | PIK3CB | chr3 | - | 138742557 | 138742755 | 138734633 | 138734804 | 138755753 | 138755979 | 0 | 2 | 2 | 2 | 0 | 0 | 0 | 0 | 0 | 0 | 0 | 0 | 0 | 0 | 5 | 0 | 1 | 0 | 0 | 0 | 298 | 149 | 1 | 1 | NA | 1 | 1 | 1 | NA | NA | 0 | NA | NA | 1 | 0.5 |
| SE | 35508 | ENSG00000051382 | PIK3CB | chr3 | - | 138742557 | 138742781 | 138734633 | 138734804 | 138755753 | 138755979 | 97 | 132 | 131 | 140 | 206 | 0 | 0 | 0 | 0 | 0 | 137 | 198 | 150 | 258 | 220 | 0 | 1 | 0 | 0 | 0 | 298 | 149 | 1 | 1 | 1 | 1 | 1 | 1 | 1 | 1 | 0.99 | 1 | 1 | 1 | 0.002 |
| SE | 35510 | ENSG00000051382 | PIK3CB | chr3 | - | 138796462 | 138796567 | 138759172 | 138759359 | 138834694 | 138834833 | 44 | 37 | 49 | 69 | 93 | 2 | 0 | 1 | 2 | 0 | 20 | 64 | 42 | 93 | 41 | 0 | 0 | 0 | 0 | 17 | 254 | 149 | 1 | 1 | 0.928 | 1 | 0.966 | 0.953 | 1 | 1 | 1 | 1 | 1 | 0.586 | 0.052 |
| SE | 35515 | ENSG00000197980 | LEKR1 | chr3 | + | 156927428 | 156927604 | 156920574 | 156920694 | 156942528 | 156942714 | 2 | 0 | 2 | 2 | 1 | 0 | 1 | 0 | 0 | 1 | 0 | 6 | 2 | 3 | 0 | 0 | 0 | 0 | 0 | 0 | 298 | 149 | 1 | 1 | 1 | 0 | 1 | 1 | 0.333 | NA | 1 | 1 | 1 | NA | -0.333 |
| SE | 35516 | ENSG00000197980 | LEKR1 | chr3 | + | 156942528 | 156942714 | 156829285 | 156829377 | 156979193 | 156979275 | 0 | 2 | 0 | 0 | 0 | 0 | 0 | 0 | 0 | 0 | 0 | 6 | 0 | 0 | 0 | 0 | 0 | 1 | 0 | 0 | 298 | 149 | 1 | 1 | NA | 1 | NA | NA | NA | NA | 1 | 0 | NA | NA | 0.5 |
| SE | 35517 | ENSG00000197980 | LEKR1 | chr3 | + | 156979818 | 156980028 | 156979193 | 156979275 | 156992652 | 156992730 | 0 | 0 | 0 | 6 | 0 | 0 | 0 | 0 | 0 | 0 | 0 | 0 | 0 | 0 | 0 | 0 | 0 | 0 | 1 | 0 | 298 | 149 | 1 | 1 | NA | NA | NA | 1 | NA | NA | NA | NA | 0 | NA | 1 |
| SE | 35520 | ENSG00000166323 | C11orf65 | chr11 | - | 108405428 | 108405559 | 108393207 | 108393378 | 108406762 | 108406963 | 4 | 7 | 6 | 4 | 6 | 2 | 1 | 1 | 0 | 0 | 0 | 19 | 7 | 0 | 0 | 0 | 0 | 0 | 0 | 0 | 280 | 149 | 1 | 1 | 0.516 | 0.788 | 0.761 | 1 | 1 | NA | 1 | 1 | NA | NA | -0.187 |
| SE | 35521 | ENSG00000166323 | C11orf65 | chr11 | - | 108405428 | 108405559 | 108393207 | 108393378 | 108461478 | 108461568 | 4 | 5 | 3 | 2 | 7 | 0 | 0 | 0 | 1 | 0 | 0 | 8 | 2 | 0 | 0 | 0 | 0 | 0 | 0 | 0 | 280 | 149 | 1 | 1 | 1 | 1 | 1 | 0.516 | 1 | NA | 1 | 1 | NA | NA | -0.097 |
| SE | 35522 | ENSG00000166323 | C11orf65 | chr11 | - | 108406762 | 108406963 | 108393207 | 108393378 | 108461478 | 108461568 | 4 | 1 | 2 | 0 | 0 | 0 | 0 | 0 | 1 | 0 | 0 | 0 | 3 | 0 | 0 | 0 | 0 | 0 | 0 | 0 | 298 | 149 | 1 | 1 | 1 | 1 | 1 | 0 | NA | NA | NA | 1 | NA | NA | -0.25 |
| SE | 35523 | ENSG00000166323 | C11orf65 | chr11 | - | 108406762 | 108406963 | 108405428 | 108405559 | 108461478 | 108461568 | 2 | 2 | 4 | 2 | 0 | 0 | 0 | 0 | 0 | 1 | 0 | 11 | 8 | 0 | 0 | 0 | 0 | 0 | 0 | 0 | 298 | 149 | 1 | 1 | 1 | 1 | 1 | 1 | 0 | NA | 1 | 1 | NA | NA | -0.2 |
| SE | 35524 | ENSG00000166323 | C11orf65 | chr11 | - | 108407095 | 108407149 | 108406762 | 108406963 | 108431745 | 108431838 | 0 | 0 | 0 | 0 | 0 | 0 | 0 | 0 | 2 | 0 | 0 | 4 | 2 | 0 | 0 | 0 | 0 | 0 | 0 | 0 | 203 | 149 | 1 | 1 | NA | NA | NA | 0 | NA | NA | 1 | 1 | NA | NA | -1 |
| SE | 35525 | ENSG00000166323 | C11orf65 | chr11 | - | 108431745 | 108431838 | 108406762 | 108406963 | 108461478 | 108461568 | 0 | 0 | 0 | 2 | 0 | 2 | 0 | 1 | 0 | 0 | 0 | 0 | 2 | 0 | 0 | 0 | 0 | 3 | 0 | 0 | 242 | 149 | 1 | 1 | 0 | NA | 0 | 1 | NA | NA | NA | 0.291 | NA | NA | 0.042 |
| SE | 35527 | ENSG00000175066 | GK5 | chr3 | - | 142170324 | 142170458 | 142157526 | 142165770 | 142171418 | 142171478 | 346 | 191 | 151 | 254 | 109 | 1 | 1 | 0 | 0 | 0 | 113 | 158 | 236 | 129 | 73 | 0 | 0 | 3 | 4 | 0 | 283 | 149 | 1 | 1 | 0.995 | 0.99 | 1 | 1 | 1 | 1 | 1 | 0.969 | 1 | 1 | 0.003 |
| SE | 35529 | ENSG00000175066 | GK5 | chr3 | - | 142181460 | 142181565 | 142177481 | 142177576 | 142182922 | 142183049 | 223 | 86 | 121 | 163 | 99 | 0 | 0 | 2 | 0 | 0 | 121 | 110 | 166 | 88 | 31 | 0 | 1 | 1 | 0 | 0 | 254 | 149 | 1 | 1 | 1 | 1 | 0.973 | 1 | 1 | 1 | 0.985 | 0.99 | 1 | 1 | 0 |
| SE | 35532 | ENSG00000175066 | GK5 | chr3 | - | 142187703 | 142187779 | 142186451 | 142186513 | 142198801 | 142198933 | 145 | 40 | 55 | 91 | 125 | 0 | 0 | 0 | 1 | 0 | 55 | 61 | 106 | 44 | 12 | 0 | 0 | 0 | 0 | 0 | 225 | 149 | 1 | 1 | 1 | 1 | 1 | 0.984 | 1 | 1 | 1 | 1 | 1 | 1 | -0.003 |
| SE | 35533 | ENSG00000175066 | GK5 | chr3 | - | 142198801 | 142198933 | 142187703 | 142187779 | 142204694 | 142204788 | 238 | 86 | 90 | 151 | 195 | 0 | 1 | 1 | 0 | 1 | 51 | 96 | 138 | 84 | 22 | 0 | 0 | 0 | 0 | 0 | 281 | 149 | 1 | 1 | 1 | 0.979 | 0.979 | 1 | 0.99 | 1 | 1 | 1 | 1 | 1 | -0.01 |
| SE | 35534 | ENSG00000175066 | GK5 | chr3 | - | 142215598 | 142215692 | 142213525 | 142213601 | 142225308 | 142225416 | 99 | 60 | 49 | 74 | 82 | 0 | 1 | 0 | 0 | 0 | 2 | 64 | 93 | 28 | 3 | 0 | 0 | 0 | 0 | 0 | 243 | 149 | 1 | 1 | 1 | 0.974 | 1 | 1 | 1 | 1 | 1 | 1 | 1 | 1 | -0.005 |
| SE | 35544 | ENSG00000173905 | GOLIM4 | chr3 | - | 168010742 | 168010823 | 168008688 | 168010418 | 168024525 | 168024594 | 19 | 32 | 5 | 14 | 23 | 0 | 1 | 0 | 0 | 0 | 0 | 0 | 31 | 20 | 0 | 0 | 0 | 0 | 0 | 0 | 230 | 149 | 1 | 1 | 1 | 0.954 | 1 | 1 | 1 | NA | NA | 1 | 1 | NA | -0.009 |
| SE | 35545 | ENSG00000173905 | GOLIM4 | chr3 | - | 168040785 | 168040869 | 168036835 | 168036994 | 168041391 | 168041474 | 3 | 2 | 5 | 0 | 4 | 19 | 27 | 17 | 22 | 8 | 48 | 5 | 3 | 0 | 0 | 15 | 21 | 22 | 33 | 0 | 233 | 149 | 1 | 1 | 0.092 | 0.045 | 0.158 | 0 | 0.242 | 0.672 | 0.132 | 0.08 | 0 | NA | -0.114 |
| SE | 35546 | ENSG00000173905 | GOLIM4 | chr3 | - | 168041391 | 168041474 | 168036835 | 168036994 | 168043378 | 168043529 | 59 | 65 | 49 | 52 | 69 | 3 | 2 | 2 | 3 | 6 | 48 | 51 | 55 | 67 | 0 | 0 | 0 | 0 | 0 | 0 | 232 | 149 | 1 | 1 | 0.927 | 0.954 | 0.94 | 0.918 | 0.881 | 1 | 1 | 1 | 1 | NA | -0.076 |
| SE | 35547 | ENSG00000071794 | HLTF | chr3 | - | 149039580 | 149039693 | 149039048 | 149039229 | 149040030 | 149040156 | 75 | 55 | 68 | 39 | 127 | 0 | 1 | 0 | 0 | 0 | 0 | 26 | 4 | 0 | 6 | 0 | 0 | 0 | 0 | 0 | 262 | 149 | 1 | 1 | 1 | 0.969 | 1 | 1 | 1 | NA | 1 | 1 | NA | 1 | -0.006 |
| SE | 35548 | ENSG00000071794 | HLTF | chr3 | - | 149041489 | 149041668 | 149040030 | 149040156 | 149042165 | 149042290 | 80 | 38 | 60 | 49 | 98 | 2 | 1 | 1 | 3 | 0 | 0 | 19 | 22 | 0 | 5 | 0 | 0 | 0 | 0 | 0 | 298 | 149 | 1 | 1 | 0.952 | 0.95 | 0.968 | 0.891 | 1 | NA | 1 | 1 | NA | 1 | -0.048 |
| SE | 35549 | ENSG00000071794 | HLTF | chr3 | - | 149041489 | 149041680 | 149040030 | 149040156 | 149042165 | 149042290 | 61 | 23 | 39 | 26 | 49 | 2 | 1 | 1 | 3 | 0 | 0 | 8 | 8 | 0 | 5 | 0 | 0 | 0 | 0 | 0 | 298 | 149 | 1 | 1 | 0.938 | 0.92 | 0.951 | 0.812 | 1 | NA | 1 | 1 | NA | 1 | -0.076 |
| SE | 35550 | ENSG00000071794 | HLTF | chr3 | - | 149046079 | 149046259 | 149042165 | 149042290 | 149048027 | 149048163 | 55 | 38 | 47 | 48 | 88 | 0 | 2 | 0 | 0 | 0 | 13 | 42 | 9 | 0 | 0 | 0 | 0 | 0 | 0 | 0 | 298 | 149 | 1 | 1 | 1 | 0.905 | 1 | 1 | 1 | 1 | 1 | 1 | NA | NA | -0.019 |
| SE | 35551 | ENSG00000071794 | HLTF | chr3 | - | 149052044 | 149052144 | 149050269 | 149050375 | 149055302 | 149055400 | 0 | 0 | 1 | 5 | 0 | 47 | 29 | 28 | 42 | 39 | 0 | 0 | 0 | 0 | 0 | 0 | 30 | 18 | 2 | 4 | 249 | 149 | 1 | 1 | 0 | 0 | 0.021 | 0.067 | 0 | NA | 0 | 0 | 0 | 0 | 0.018 |
| SE | 35552 | ENSG00000071794 | HLTF | chr3 | - | 149055302 | 149055400 | 149050231 | 149050375 | 149059717 | 149059804 | 114 | 83 | 84 | 146 | 132 | 0 | 2 | 0 | 0 | 0 | 20 | 57 | 42 | 4 | 13 | 0 | 0 | 0 | 0 | 0 | 247 | 149 | 1 | 1 | 1 | 0.962 | 1 | 1 | 1 | 1 | 1 | 1 | 1 | 1 | -0.008 |
| SE | 35554 | ENSG00000182606 | TRAK1 | chr3 | + | 42184684 | 42184801 | 42176813 | 42176890 | 42188044 | 42188145 | 195 | 175 | 163 | 196 | 227 | 0 | 0 | 1 | 0 | 0 | 642 | 300 | 195 | 353 | 435 | 0 | 0 | 0 | 0 | 0 | 266 | 149 | 1 | 1 | 1 | 1 | 0.989 | 1 | 1 | 1 | 1 | 1 | 1 | 1 | -0.002 |
| SE | 35555 | ENSG00000182606 | TRAK1 | chr3 | + | 42188044 | 42188145 | 42184684 | 42184801 | 42189015 | 42189124 | 149 | 84 | 103 | 155 | 152 | 0 | 0 | 0 | 0 | 0 | 261 | 160 | 105 | 164 | 225 | 0 | 0 | 2 | 2 | 0 | 250 | 149 | 1 | 1 | 1 | 1 | 1 | 1 | 1 | 1 | 1 | 0.969 | 0.98 | 1 | 0.01 |
| SE | 35556 | ENSG00000182606 | TRAK1 | chr3 | + | 42189015 | 42189124 | 42188044 | 42188145 | 42191557 | 42191636 | 245 | 139 | 163 | 260 | 241 | 0 | 3 | 0 | 0 | 1 | 397 | 253 | 167 | 281 | 382 | 0 | 0 | 0 | 0 | 0 | 258 | 149 | 1 | 1 | 1 | 0.964 | 1 | 1 | 0.993 | 1 | 1 | 1 | 1 | 1 | -0.009 |
| SE | 35557 | ENSG00000182606 | TRAK1 | chr3 | + | 42191557 | 42191636 | 42189015 | 42189124 | 42193074 | 42193205 | 243 | 159 | 162 | 265 | 270 | 0 | 0 | 0 | 0 | 0 | 402 | 282 | 176 | 322 | 329 | 0 | 0 | 2 | 0 | 0 | 228 | 149 | 1 | 1 | 1 | 1 | 1 | 1 | 1 | 1 | 1 | 0.983 | 1 | 1 | 0.003 |
| SE | 35560 | ENSG00000182606 | TRAK1 | chr3 | + | 42200817 | 42201054 | 42199176 | 42199253 | 42202435 | 42202752 | 124 | 66 | 74 | 119 | 148 | 0 | 0 | 0 | 0 | 0 | 240 | 151 | 85 | 114 | 205 | 0 | 3 | 0 | 0 | 0 | 298 | 149 | 1 | 1 | 1 | 1 | 1 | 1 | 1 | 1 | 0.962 | 1 | 1 | 1 | 0.008 |
| SE | 35561 | ENSG00000182606 | TRAK1 | chr3 | + | 42220498 | 42220591 | 42219493 | 42219596 | 42222941 | 42225148 | 0 | 0 | 0 | 0 | 0 | 16 | 19 | 21 | 22 | 17 | 0 | 0 | 0 | 0 | 3 | 26 | 43 | 13 | 12 | 32 | 242 | 149 | 1 | 1 | 0 | 0 | 0 | 0 | 0 | 0 | 0 | 0 | 0 | 0.055 | -0.011 |

| | | | | | | | | | | | | | | | | | | | | | | | | | | | | | | | | | | | | | | | | | | | | | | |
|---|---|---|---|---|---|---|---|---|---|---|---|---|---|---|---|---|---|---|---|---|---|---|---|---|---|---|---|---|---|---|---|---|---|---|---|---|---|---|---|---|---|---|---|---|---|---|
| SE | 35562 | ENSG00000120685 | PROSER1 | chr13 | - | 39012690 | 39014476 | 39012082 | 39012233 | 39017499 | 39017544 | 92 | 63 | 72 | 89 | 113 | 3 | 0 | 0 | 0 | 0 | 129 | 95 | 78 | 104 | 34 | 0 | 0 | 0 | 0 | 0 | 298 | 149 | 1 | 1 | 0.939 | 1 | 1 | 1 | 1 | 1 | 1 | 1 | 1 | 1 | -0.012 |
| SE | 35566 | ENSG00000120685 | PROSER1 | chr13 | - | 39023051 | 39023130 | 39017499 | 39017544 | 39026276 | 39026387 | 1 | 1 | 0 | 3 | 1 | 1 | 0 | 0 | 1 | 0 | 0 | 5 | 0 | 4 | 0 | 11 | 0 | 0 | 0 | 0 | 228 | 149 | 1 | 1 | 0.395 | 1 | NA | 0.662 | 1 | 0 | 1 | NA | 1 | NA | 0.098 |
| SE | 35567 | ENSG00000120685 | PROSER1 | chr13 | - | 39023051 | 39023130 | 39022325 | 39022412 | 39024472 | 39024556 | 145 | 112 | 95 | 126 | 367 | 0 | 0 | 1 | 0 | 0 | 169 | 106 | 123 | 116 | 71 | 0 | 0 | 0 | 0 | 0 | 228 | 149 | 1 | 1 | 1 | 1 | 0.984 | 1 | 1 | 1 | 1 | 1 | 1 | 1 | -0.003 |
| SE | 35569 | ENSG00000120685 | PROSER1 | chr13 | - | 39024472 | 39024556 | 39023051 | 39023130 | 39026276 | 39026387 | 160 | 114 | 143 | 128 | 425 | 0 | 0 | 0 | 2 | 0 | 139 | 149 | 139 | 94 | 114 | 0 | 0 | 0 | 0 | 0 | 233 | 149 | 1 | 1 | 1 | 1 | 1 | 0.976 | 1 | 1 | 1 | 1 | 1 | 1 | -0.005 |
| SE | 35570 | ENSG00000120685 | PROSER1 | chr13 | - | 39028226 | 39028320 | 39026276 | 39026387 | 39029280 | 39029375 | 160 | 84 | 150 | 121 | 397 | 0 | 0 | 0 | 0 | 1 | 131 | 158 | 108 | 81 | 86 | 0 | 0 | 0 | 0 | 0 | 243 | 149 | 1 | 1 | 1 | 1 | 1 | 1 | 0.996 | 1 | 1 | 1 | 1 | 1 | -0.001 |
| SE | 35577 | ENSG00000095951 | HIVEP1 | chr6 | + | 12020247 | 12020418 | 12015525 | 12015668 | 12089183 | 12089237 | 0 | 0 | 1 | 2 | 0 | 56 | 59 | 69 | 70 | 145 | 0 | 0 | 0 | 0 | 0 | 43 | 41 | 63 | 26 | 14 | 298 | 149 | 1 | 1 | 0 | 0 | 0.007 | 0.014 | 0 | 0 | 0 | 0 | 0 | 0 | 0.004 |
| SE | 35578 | ENSG00000095951 | HIVEP1 | chr6 | + | 12089183 | 12089237 | 12015525 | 12015668 | 12119889 | 12119934 | 87 | 95 | 119 | 97 | 277 | 2 | 1 | 0 | 0 | 0 | 51 | 55 | 69 | 41 | 22 | 0 | 0 | 0 | 0 | 0 | 203 | 149 | 1 | 1 | 0.97 | 0.986 | 1 | 1 | 1 | 1 | 1 | 1 | 1 | 1 | -0.009 |
| SE | 35579 | ENSG00000095951 | HIVEP1 | chr6 | + | 12119889 | 12125870 | 12089183 | 12089237 | 12129758 | 12129892 | 123 | 163 | 164 | 146 | 335 | 0 | 0 | 0 | 0 | 0 | 36 | 112 | 122 | 80 | 37 | 0 | 0 | 0 | 3 | 0 | 298 | 149 | 1 | 1 | 1 | 1 | 1 | 1 | 1 | 1 | 1 | 1 | 0.93 | 1 | 0.014 |
| SE | 35580 | ENSG00000095951 | HIVEP1 | chr6 | + | 12129758 | 12129892 | 12119889 | 12125870 | 12130766 | 12130942 | 120 | 180 | 164 | 156 | 174 | 4 | 7 | 11 | 4 | 6 | 0 | 122 | 162 | 138 | 30 | 0 | 4 | 7 | 1 | 0 | 283 | 149 | 1 | 1 | 0.94 | 0.931 | 0.887 | 0.954 | 0.939 | NA | 0.941 | 0.924 | 0.986 | 1 | -0.033 |
| SE | 35581 | ENSG00000095951 | HIVEP1 | chr6 | + | 12130766 | 12130942 | 12125745 | 12125870 | 12135790 | 12135892 | 56 | 80 | 74 | 57 | 119 | 0 | 1 | 1 | 7 | 1 | 2 | 18 | 69 | 34 | 0 | 0 | 0 | 3 | 0 | 0 | 298 | 149 | 1 | 1 | 1 | 0.976 | 0.974 | 0.803 | 0.983 | 1 | 1 | 0.92 | 1 | NA | -0.033 |
| SE | 35582 | ENSG00000095951 | HIVEP1 | chr6 | + | 12135790 | 12135892 | 12130766 | 12130942 | 12161438 | 12161891 | 81 | 111 | 99 | 81 | 152 | 2 | 2 | 0 | 0 | 0 | 16 | 29 | 100 | 47 | 0 | 0 | 8 | 0 | 0 | 0 | 251 | 149 | 1 | 1 | 0.96 | 0.971 | 1 | 1 | 1 | 1 | 0.683 | 1 | 1 | NA | 0.065 |
| SE | 35583 | ENSG00000114331 | ACAP2 | chr3 | - | 195285795 | 195285857 | 195274744 | 195279428 | 195291705 | 195291815 | 197 | 157 | 182 | 188 | 290 | 0 | 0 | 0 | 1 | 0 | 17 | 102 | 109 | 59 | 9 | 0 | 0 | 0 | 0 | 0 | 211 | 149 | 1 | 1 | 1 | 1 | 1 | 0.993 | 1 | 1 | 1 | 1 | 1 | 1 | -0.001 |
| SE | 35584 | ENSG00000114331 | ACAP2 | chr3 | - | 195289120 | 195289231 | 195285327 | 195285857 | 195291705 | 195291815 | 531 | 374 | 412 | 549 | 604 | 0 | 0 | 2 | 4 | 0 | 103 | 318 | 416 | 174 | 6 | 0 | 0 | 0 | 0 | 0 | 260 | 149 | 1 | 1 | 1 | 1 | 0.992 | 0.987 | 1 | 1 | 1 | 1 | 1 | 1 | -0.004 |
| SE | 35585 | ENSG00000114331 | ACAP2 | chr3 | - | 195291705 | 195291815 | 195289120 | 195289231 | 195292264 | 195292452 | 536 | 403 | 421 | 566 | 502 | 1 | 2 | 2 | 4 | 1 | 121 | 366 | 446 | 206 | 9 | 0 | 0 | 1 | 0 | 0 | 259 | 149 | 1 | 1 | 0.997 | 0.991 | 0.992 | 0.988 | 0.997 | 1 | 1 | 0.996 | 1 | 1 | -0.006 |
| SE | 35586 | ENSG00000114331 | ACAP2 | chr3 | - | 195295195 | 195295300 | 195294718 | 195294811 | 195295707 | 195295892 | 6 | 2 | 1 | 1 | 9 | 482 | 378 | 324 | 476 | 627 | 0 | 7 | 7 | 0 | 0 | 203 | 254 | 378 | 184 | 19 | 254 | 149 | 1 | 1 | 0.007 | 0.003 | 0.002 | 0.001 | 0.008 | 0 | 0.016 | 0.011 | 0 | 0 | -0.001 |
| SE | 35587 | ENSG00000114331 | ACAP2 | chr3 | - | 195301574 | 195301644 | 195297189 | 195297281 | 195301965 | 195302115 | 983 | 879 | 740 | 990 | 1011 | 1 | 0 | 1 | 0 | 0 | 396 | 566 | 824 | 651 | 136 | 0 | 0 | 0 | 0 | 0 | 219 | 149 | 1 | 1 | 0.999 | 1 | 0.998 | 1 | 1 | 1 | 1 | 1 | 1 | 1 | -0.001 |
| SE | 35588 | ENSG00000114331 | ACAP2 | chr3 | - | 195301574 | 195301644 | 195297189 | 195297281 | 195306510 | 195306616 | 363 | 332 | 316 | 383 | 385 | 1 | 0 | 0 | 0 | 0 | 223 | 272 | 388 | 257 | 42 | 0 | 1 | 0 | 0 | 0 | 219 | 149 | 1 | 1 | 0.996 | 1 | 1 | 1 | 1 | 1 | 0.995 | 1 | 1 | 1 | 0 |
| SE | 35589 | ENSG00000114331 | ACAP2 | chr3 | - | 195301965 | 195302174 | 195297189 | 195297281 | 195306510 | 195306616 | 335 | 314 | 271 | 301 | 399 | 1 | 0 | 0 | 0 | 0 | 113 | 203 | 258 | 140 | 48 | 0 | 1 | 0 | 0 | 0 | 298 | 149 | 1 | 1 | 0.994 | 1 | 1 | 1 | 1 | 1 | 0.99 | 1 | 1 | 1 | 0.001 |
| SE | 35590 | ENSG00000114331 | ACAP2 | chr3 | - | 195301965 | 195302174 | 195301574 | 195301644 | 195306510 | 195306616 | 955 | 861 | 694 | 910 | 1025 | 1 | 0 | 0 | 0 | 0 | 292 | 497 | 694 | 534 | 142 | 6 | 0 | 0 | 0 | 0 | 298 | 149 | 1 | 1 | 0.998 | 1 | 1 | 1 | 1 | 0.961 | 1 | 1 | 1 | 1 | 0.007 |
| SE | 35592 | ENSG00000114331 | ACAP2 | chr3 | - | 195320700 | 195320813 | 195308785 | 195308837 | 195326884 | 195326959 | 381 | 303 | 297 | 409 | 654 | 1 | 1 | 3 | 1 | 0 | 183 | 197 | 252 | 180 | 97 | 0 | 1 | 1 | 0 | 0 | 262 | 149 | 1 | 1 | 0.995 | 0.994 | 0.983 | 0.996 | 1 | 1 | 0.991 | 0.993 | 1 | 1 | -0.003 |
| SE | 35594 | ENSG00000114331 | ACAP2 | chr3 | - | 195333027 | 195333123 | 195326331 | 195326959 | 195336931 | 195336965 | 673 | 573 | 501 | 633 | 1561 | 1 | 0 | 0 | 0 | 0 | 213 | 284 | 473 | 329 | 162 | 0 | 0 | 0 | 2 | 0 | 245 | 149 | 1 | 1 | 0.998 | 1 | 1 | 1 | 1 | 1 | 1 | 1 | 0.99 | 1 | 0.002 |
| SE | 35595 | ENSG00000114331 | ACAP2 | chr3 | - | 195342470 | 195342654 | 195335898 | 195336976 | 195345258 | 195345317 | 818 | 626 | 629 | 754 | 1768 | 0 | 0 | 0 | 0 | 2 | 306 | 379 | 491 | 349 | 186 | 0 | 0 | 0 | 0 | 0 | 298 | 149 | 1 | 1 | 1 | 1 | 1 | 1 | 0.998 | 1 | 1 | 1 | 1 | 1 | 0 |
| SE | 35597 | ENSG00000114331 | ACAP2 | chr3 | - | 195392089 | 195392147 | 195381902 | 195382022 | 195410706 | 195411009 | 139 | 83 | 99 | 120 | 255 | 0 | 0 | 0 | 0 | 0 | 87 | 79 | 107 | 124 | 58 | 0 | 2 | 0 | 0 | 0 | 207 | 149 | 1 | 1 | 1 | 1 | 1 | 1 | 1 | 1 | 0.966 | 1 | 1 | 1 | 0.007 |
| SE | 35598 | ENSG00000114331 | ACAP2 | chr3 | - | 195392089 | 195392147 | 195381902 | 195382022 | 195442794 | 195442853 | 166 | 103 | 126 | 151 | 256 | 1 | 0 | 0 | 0 | 1 | 55 | 85 | 118 | 109 | 54 | 0 | 0 | 0 | 0 | 0 | 207 | 149 | 1 | 1 | 0.992 | 1 | 1 | 1 | 0.995 | 1 | 1 | 1 | 1 | 1 | -0.003 |
| SE | 35599 | ENSG00000114331 | ACAP2 | chr3 | - | 195410706 | 195411009 | 195381902 | 195382022 | 195442794 | 195442905 | 1 | 1 | 0 | 0 | 0 | 1 | 0 | 0 | 0 | 1 | 0 | 2 | 0 | 0 | 0 | 0 | 0 | 0 | 0 | 0 | 298 | 149 | 1 | 1 | 0.333 | 1 | NA | NA | 0 | NA | 1 | NA | NA | NA | -0.556 |
| SE | 35603 | ENSG00000114331 | ACAP2 | chr3 | - | 195412845 | 195412916 | 195410706 | 195411009 | 195442794 | 195442905 | 4 | 1 | 2 | 3 | 0 | 1 | 1 | 0 | 0 | 0 | 11 | 0 | 0 | 15 | 8 | 0 | 0 | 0 | 0 | 0 | 220 | 149 | 1 | 1 | 0.73 | 0.404 | 1 | 1 | NA | 1 | NA | NA | 1 | 1 | -0.217 |
| SE | 35604 | ENSG00000285942 | NA | chr15 | - | 42530914 | 42531069 | 42528326 | 42528501 | 42545076 | 42545262 | 4 | 0 | 0 | 0 | 4 | 0 | 0 | 0 | 0 | 1 | 23 | 0 | 0 | 0 | 0 | 0 | 0 | 0 | 0 | 0 | 298 | 149 | 1 | 1 | 1 | NA | NA | NA | 0.667 | 1 | NA | NA | NA | NA | -0.166 |
| SE | 35617 | ENSG00000054967 | RELT | chr11 | + | 73392210 | 73392468 | 73391143 | 73391223 | 73393836 | 73393917 | 80 | 29 | 52 | 65 | 21 | 0 | 0 | 0 | 2 | 0 | 111 | 39 | 39 | 92 | 161 | 0 | 0 | 0 | 0 | 0 | 298 | 149 | 1 | 1 | 1 | 1 | 1 | 0.942 | 1 | 1 | 1 | 1 | 1 | 1 | -0.012 |
| SE | 35618 | ENSG00000054967 | RELT | chr11 | + | 73394476 | 73394734 | 73394235 | 73394317 | 73395086 | 73395285 | 62 | 41 | 52 | 79 | 21 | 0 | 0 | 0 | 0 | 0 | 195 | 101 | 96 | 122 | 164 | 0 | 0 | 0 | 1 | 0 | 298 | 149 | 1 | 1 | 1 | 1 | 1 | 1 | 1 | 1 | 1 | 1 | 0.984 | 1 | 0.003 |
| SE | 35627 | ENSG00000205981 | DNAJC19 | chr3 | - | 180988177 | 180988229 | 180987179 | 180988096 | 180989599 | 180989703 | 72 | 43 | 85 | 91 | 30 | 3 | 1 | 2 | 7 | 3 | 0 | 48 | 38 | 38 | 74 | 0 | 0 | 0 | 0 | 0 | 201 | 149 | 1 | 1 | 0.947 | 0.97 | 0.969 | 0.906 | 0.881 | NA | 1 | 1 | 1 | 1 | -0.065 |
| SE | 35630 | ENSG00000138246 | DNAJC13 | chr3 | + | 132450646 | 132450847 | 132447897 | 132447939 | 132453297 | 132453504 | 405 | 367 | 267 | 356 | 554 | 1 | 0 | 0 | 1 | 7 | 13 | 392 | 192 | 92 | 29 | 0 | 0 | 0 | 0 | 0 | 298 | 149 | 1 | 1 | 0.995 | 1 | 1 | 0.994 | 0.975 | 1 | 1 | 1 | 1 | 1 | -0.007 |
| SE | 35632 | ENSG00000138246 | DNAJC13 | chr3 | + | 132457268 | 132457368 | 132456662 | 132456832 | 132460249 | 132460357 | 203 | 164 | 152 | 210 | 269 | 0 | 0 | 2 | 0 | 0 | 10 | 157 | 100 | 29 | 0 | 0 | 0 | 0 | 0 | 0 | 249 | 149 | 1 | 1 | 1 | 1 | 0.978 | 1 | 1 | 1 | 1 | 1 | 1 | NA | -0.004 |
| SE | 35633 | ENSG00000138246 | DNAJC13 | chr3 | + | 132461049 | 132461205 | 132460249 | 132460357 | 132462466 | 132462523 | 185 | 118 | 139 | 224 | 193 | 0 | 0 | 0 | 2 | 0 | 28 | 157 | 82 | 37 | 7 | 0 | 2 | 1 | 0 | 0 | 298 | 149 | 1 | 1 | 1 | 1 | 1 | 0.982 | 1 | 1 | 0.975 | 0.976 | 1 | 1 | 0.006 |
| SE | 35634 | ENSG00000138246 | DNAJC13 | chr3 | + | 132462466 | 132462523 | 132461049 | 132461205 | 132463695 | 132463817 | 124 | 93 | 120 | 184 | 130 | 0 | 0 | 0 | 0 | 0 | 27 | 135 | 92 | 37 | 11 | 0 | 0 | 3 | 0 | 0 | 206 | 149 | 1 | 1 | 1 | 1 | 1 | 1 | 1 | 1 | 1 | 0.957 | 1 | 1 | 0.009 |
| SE | 35635 | ENSG00000138246 | DNAJC13 | chr3 | + | 132462466 | 132462523 | 132461049 | 132461205 | 132465994 | 132466070 | 35 | 24 | 30 | 41 | 14 | 0 | 0 | 0 | 0 | 0 | 0 | 52 | 27 | 24 | 0 | 0 | 0 | 2 | 0 | 0 | 206 | 149 | 1 | 1 | 1 | 1 | 1 | 1 | 1 | NA | 1 | 0.907 | 1 | NA | 0.031 |
| SE | 35636 | ENSG00000138246 | DNAJC13 | chr3 | + | 132463695 | 132463817 | 132461049 | 132461205 | 132465994 | 132466070 | 58 | 60 | 48 | 69 | 69 | 0 | 0 | 0 | 0 | 0 | 0 | 70 | 60 | 11 | 0 | 0 | 0 | 2 | 0 | 0 | 271 | 149 | 1 | 1 | 1 | 1 | 1 | 1 | 1 | NA | 1 | 0.943 | 1 | NA | 0.019 |
| SE | 35637 | ENSG00000138246 | DNAJC13 | chr3 | + | 132463695 | 132463817 | 132462466 | 132462523 | 132465994 | 132466070 | 148 | 141 | 137 | 218 | 185 | 2 | 0 | 0 | 0 | 0 | 27 | 151 | 124 | 24 | 11 | 0 | 0 | 0 | 0 | 0 | 271 | 149 | 1 | 1 | 0.976 | 1 | 1 | 1 | 1 | 1 | 1 | 1 | 1 | 1 | -0.005 |
| SE | 35640 | ENSG00000138246 | DNAJC13 | chr3 | + | 132466298 | 132466394 | 132463695 | 132463817 | 132467169 | 132467313 | 114 | 93 | 77 | 108 | 113 | 0 | 0 | 1 | 0 | 0 | 0 | 81 | 99 | 32 | 8 | 0 | 4 | 0 | 0 | 0 | 245 | 149 | 1 | 1 | 1 | 1 | 0.979 | 1 | 1 | NA | 0.925 | 1 | 1 | 1 | 0.015 |
| SE | 35644 | ENSG00000138246 | DNAJC13 | chr3 | + | 132495087 | 132495166 | 132494143 | 132494259 | 132496527 | 132496663 | 222 | 170 | 128 | 165 | 371 | 10 | 20 | 6 | 10 | 10 | 0 | 145 | 93 | 46 | 9 | 0 | 6 | 4 | 0 | 0 | 228 | 149 | 1 | 1 | 0.936 | 0.847 | 0.933 | 0.915 | 0.96 | NA | 0.94 | 0.938 | 1 | 1 | -0.051 |
| SE | 35645 | ENSG00000138246 | DNAJC13 | chr3 | + | 132499733 | 132499808 | 132499125 | 132499310 | 132500793 | 132500913 | 148 | 119 | 98 | 123 | 190 | 2 | 1 | 1 | 3 | 0 | 0 | 98 | 64 | 26 | 0 | 0 | 0 | 0 | 0 | 0 | 224 | 149 | 1 | 1 | 0.98 | 0.988 | 0.985 | 0.965 | 1 | NA | 1 | 1 | 1 | NA | -0.016 |
| SE | 35646 | ENSG00000138246 | DNAJC13 | chr3 | + | 132502288 | 132502468 | 132500793 | 132500913 | 132503213 | 132503381 | 192 | 147 | 171 | 182 | 228 | 0 | 0 | 0 | 1 | 0 | 22 | 184 | 63 | 31 | 0 | 0 | 0 | 0 | 0 | 0 | 298 | 149 | 1 | 1 | 1 | 1 | 1 | 0.989 | 1 | 1 | 1 | 1 | 1 | NA | -0.002 |
| SE | 35647 | ENSG00000138246 | DNAJC13 | chr3 | + | 132511066 | 132511244 | 132507236 | 132507353 | 132513007 | 132513099 | 194 | 152 | 152 | 191 | 352 | 2 | 1 | 0 | 0 | 0 | 1 | 207 | 122 | 24 | 0 | 0 | 0 | 0 | 0 | 0 | 298 | 149 | 1 | 1 | 0.98 | 0.987 | 1 | 1 | 1 | 1 | 1 | 1 | 1 | NA | -0.007 |
| SE | 35648 | ENSG00000138246 | DNAJC13 | chr3 | + | 132513007 | 132513099 | 132507236 | 132507353 | 132514570 | 132514670 | 79 | 41 | 58 | 43 | 78 | 0 | 0 | 0 | 1 | 0 | 8 | 41 | 35 | 6 | 0 | 0 | 0 | 0 | 0 | 0 | 241 | 149 | 1 | 1 | 1 | 1 | 1 | 0.964 | 1 | 1 | 1 | 1 | 1 | NA | -0.007 |
| SE | 35649 | ENSG00000138246 | DNAJC13 | chr3 | + | 132514570 | 132514670 | 132513007 | 132513099 | 132516421 | 132516496 | 268 | 198 | 217 | 214 | 461 | 0 | 0 | 0 | 1 | 0 | 30 | 212 | 118 | 32 | 0 | 0 | 0 | 0 | 0 | 0 | 249 | 149 | 1 | 1 | 1 | 1 | 1 | 0.992 | 1 | 1 | 1 | 1 | 1 | NA | -0.002 |
| SE | 35654 | ENSG00000161204 | ABCF3 | chr3 | + | 184187396 | 184187761 | 184186506 | 184186875 | 184187860 | 184187901 | 95 | 37 | 63 | 66 | 45 | 0 | 0 | 0 | 0 | 2 | 95 | 49 | 61 | 65 | 75 | 0 | 0 | 0 | 0 | 0 | 298 | 149 | 1 | 1 | 1 | 1 | 1 | 1 | 0.918 | 1 | 1 | 1 | 1 | 1 | -0.016 |
| SE | 35655 | ENSG00000161204 | ABCF3 | chr3 | + | 184187663 | 184187761 | 184186795 | 184186875 | 184187860 | 184187958 | 46 | 24 | 29 | 33 | 19 | 0 | 0 | 0 | 0 | 2 | 66 | 39 | 31 | 25 | 41 | 0 | 0 | 0 | 0 | 0 | 247 | 149 | 1 | 1 | 1 | 1 | 1 | 1 | 0.851 | 1 | 1 | 1 | 1 | 1 | -0.03 |
| SE | 35657 | ENSG00000161204 | ABCF3 | chr3 | + | 184189387 | 184189443 | 184189087 | 184189277 | 184189556 | 184189725 | 82 | 27 | 81 | 48 | 78 | 0 | 0 | 0 | 0 | 0 | 91 | 81 | 87 | 72 | 74 | 0 | 1 | 0 | 0 | 0 | 205 | 149 | 1 | 1 | 1 | 1 | 1 | 1 | 1 | 1 | 0.983 | 1 | 1 | 1 | 0.003 |
| SE | 35661 | ENSG00000010404 | IDS | chrX | - | 149496345 | 149496516 | 149490313 | 149490440 | 149498106 | 149498266 | 718 | 374 | 583 | 639 | 738 | 0 | 0 | 0 | 0 | 0 | 465 | 561 | 664 | 498 | 661 | 0 | 0 | 0 | 5 | 0 | 298 | 149 | 1 | 1 | 1 | 1 | 1 | 1 | 1 | 1 | 1 | 1 | 0.98 | 1 | 0.004 |
| SE | 35666 | ENSG00000010404 | IDS | chrX | - | 149504156 | 149504293 | 149500948 | 149501037 | 149505034 | 149505177 | 86 | 30 | 74 | 81 | 47 | 2 | 0 | 1 | 0 | 0 | 87 | 120 | 64 | 51 | 106 | 0 | 0 | 0 | 0 | 0 | 286 | 149 | 1 | 1 | 0.957 | 1 | 0.975 | 1 | 1 | 1 | 1 | 1 | 1 | 1 | -0.014 |
| SE | 35670 | ENSG00000120949 | TNFRSF8 | chr1 | + | 12126182 | 12126236 | 12125898 | 12126052 | 12135587 | 12135613 | 19 | 10 | 4 | 11 | 2 | 0 | 0 | 0 | 1 | 0 | 0 | 0 | 1 | 5 | 6 | 0 | 0 | 0 | 0 | 4 | 203 | 149 | 1 | 1 | 1 | 1 | 1 | 0.89 | 1 | NA | NA | 1 | 1 | 0.524 | 0.137 |
| SE | 35673 | ENSG00000132591 | ERAL1 | chr17 | + | 28856504 | 28856582 | 28856263 | 28856391 | 28857938 | 28857985 | 41 | 23 | 52 | 42 | 60 | 0 | 0 | 0 | 0 | 0 | 32 | 59 | 40 | 69 | 45 | 0 | 0 | 0 | 1 | 0 | 227 | 149 | 1 | 1 | 1 | 1 | 1 | 1 | 1 | 1 | 1 | 1 | 0.978 | 1 | 0.004 |
| SE | 35674 | ENSG00000132591 | ERAL1 | chr17 | + | 28856504 | 28856685 | 28856263 | 28856391 | 28857938 | 28857985 | 18 | 13 | 31 | 17 | 23 | 0 | 0 | 0 | 0 | 0 | 45 | 28 | 30 | 25 | 21 | 0 | 0 | 0 | 1 | 0 | 298 | 149 | 1 | 1 | 1 | 1 | 1 | 1 | 1 | 1 | 1 | 1 | 0.926 | 1 | 0.015 |
| SE | 35677 | ENSG00000132591 | ERAL1 | chr17 | + | 28858575 | 28858633 | 28858373 | 28858486 | 28858963 | 28859113 | 13 | 6 | 8 | 4 | 3 | 0 | 0 | 0 | 0 | 2 | 2 | 13 | 9 | 20 | 3 | 0 | 0 | 0 | 0 | 0 | 207 | 149 | 1 | 1 | 1 | 1 | 1 | 1 | 0.519 | 1 | 1 | 1 | 1 | 1 | -0.096 |
| SE | 35678 | ENSG00000132591 | ERAL1 | chr17 | + | 28858575 | 28858824 | 28858037 | 28858486 | 28858963 | 28859113 | 179 | 85 | 201 | 141 | 228 | 0 | 0 | 0 | 0 | 2 | 169 | 255 | 140 | 175 | 307 | 0 | 0 | 0 | 0 | 0 | 298 | 149 | 1 | 1 | 1 | 1 | 1 | 1 | 0.983 | 1 | 1 | 1 | 1 | 1 | -0.003 |
| SE | 35681 | ENSG00000171643 | S100Z | chr5 | + | 76870165 | 76870284 | 76850000 | 76850155 | 76875303 | 76875500 | 27 | 20 | 20 | 10 | 70 | 0 | 1 | 2 | 0 | 0 | 23 | 39 | 11 | 22 | 55 | 0 | 0 | 0 | 0 | 0 | 268 | 149 | 1 | 1 | 1 | 0.917 | 0.848 | 1 | 1 | 1 | 1 | 1 | 1 | 1 | -0.047 |
| SE | 35682 | ENSG00000171643 | S100Z | chr5 | + | 76877673 | 76877834 | 76875300 | 76875500 | 76920716 | 76921026 | 50 | 40 | 47 | 31 | 40 | 0 | 0 | 0 | 1 | 0 | 152 | 46 | 27 | 62 | 73 | 0 | 0 | 1 | 0 | 0 | 298 | 149 | 1 | 1 | 1 | 1 | 1 | 0.939 | 1 | 1 | 1 | 0.931 | 1 | 1 | 0.002 |
| SE | 35684 | ENSG00000143554 | SLC27A3 | chr1 | + | 153778686 | 153778885 | 153778462 | 153778553 | 153779113 | 153779211 | 116 | 58 | 69 | 92 | 111 | 1 | 0 | 0 | 1 | 0 | 312 | 144 | 47 | 100 | 194 | 0 | 0 | 0 | 0 | 0 | 298 | 149 | 1 | 1 | 0.983 | 1 | 1 | 1 | 1 | 1 | 1 | 1 | 1 | 1 | -0.003 |
| SE | 35685 | ENSG00000143554 | SLC27A3 | chr1 | + | 153778757 | 153778885 | 153778511 | 153778553 | 153779113 | 153779211 | 90 | 37 | 51 | 66 | 87 | 1 | 0 | 0 | 0 | 0 | 190 | 105 | 28 | 72 | 123 | 0 | 0 | 0 | 0 | 0 | 277 | 149 | 1 | 1 | 0.98 | 1 | 1 | 1 | 1 | 1 | 1 | 1 | 1 | 1 | -0.004 |
| SE | 35686 | ENSG00000143554 | SLC27A3 | chr1 | + | 153778845 | 153778885 | 153778462 | 153778553 | 153779113 | 153779211 | 47 | 19 | 22 | 22 | 39 | 1 | 0 | 0 | 0 | 0 | 61 | 35 | 13 | 24 | 54 | 0 | 0 | 0 | 0 | 0 | 189 | 149 | 1 | 1 | 0.974 | 1 | 1 | 1 | 1 | 1 | 1 | 1 | 1 | 1 | -0.005 |
| SE | 35687 | ENSG00000175556 | LONRF3 | chrX | + | 118978344 | 118978463 | 118974613 | 118975597 | 118989407 | 118989672 | 10 | 6 | 28 | 24 | 0 | 0 | 0 | 0 | 0 | 1 | 0 | 24 | 15 | 11 | 0 | 13 | 0 | 2 | 12 | 0 | 268 | 149 | 1 | 1 | 1 | 1 | 1 | 1 | 0 | 0 | 1 | 0.807 | 0.338 | NA | 0.264 |
| SE | 35688 | ENSG00000175556 | LONRF3 | chrX | + | 118978344 | 118978463 | 118974613 | 118975597 | 119006120 | 119006235 | 6 | 1 | 8 | 9 | 0 | 0 | 0 | 0 | 0 | 0 | 0 | 8 | 6 | 9 | 0 | 0 | 6 | 0 | 0 | 0 | 268 | 149 | 1 | 1 | 1 | 1 | 1 | 1 | NA | NA | 0.426 | 1 | 1 | NA | 0.191 |
| SE | 35690 | ENSG00000175556 | LONRF3 | chrX | + | 118982820 | 118982943 | 118978344 | 118978463 | 119006120 | 119006235 | 7 | 5 | 16 | 5 | 3 | 0 | 0 | 0 | 0 | 0 | 27 | 10 | 6 | 23 | 14 | 0 | 0 | 1 | 0 | 0 | 272 | 149 | 1 | 1 | 1 | 1 | 1 | 1 | 1 | 1 | 1 | 0.767 | 1 | 1 | 0.047 |
| SE | 35692 | ENSG00000175556 | LONRF3 | chrX | + | 118989407 | 118989672 | 118974613 | 118975597 | 119006120 | 119006235 | 0 | 0 | 0 | 0 | 1 | 0 | 0 | 0 | 0 | 0 | 13 | 2 | 2 | 12 | 0 | 0 | 6 | 0 | 0 | 0 | 298 | 149 | 1 | 1 | NA | NA | NA | NA | 1 | 1 | 0.143 | 1 | 1 | NA | 0.214 |
| SE | 35693 | ENSG00000175556 | LONRF3 | chrX | + | 118989407 | 118989672 | 118978344 | 118978463 | 118990469 | 118990560 | 24 | 11 | 46 | 30 | 7 | 0 | 0 | 0 | 0 | 0 | 0 | 36 | 28 | 28 | 5 | 0 | 0 | 0 | 5 | 0 | 298 | 149 | 1 | 1 | 1 | 1 | 1 | 1 | 1 | NA | 1 | 1 | 0.737 | 1 | 0.066 |
| SE | 35694 | ENSG00000175556 | LONRF3 | chrX | + | 118989407 | 118989672 | 118978344 | 118978463 | 119006120 | 119006235 | 4 | 5 | 20 | 15 | 0 | 0 | 0 | 0 | 0 | 0 | 0 | 18 | 10 | 2 | 0 | 0 | 0 | 1 | 0 | 0 | 298 | 149 | 1 | 1 | 1 | 1 | 1 | 1 | NA | NA | 1 | 0.833 | 1 | NA | 0.056 |
| SE | 35695 | ENSG00000175556 | LONRF3 | chrX | + | 118989407 | 118989672 | 118982820 | 118982943 | 119006120 | 119006235 | 11 | 4 | 13 | 3 | 4 | 0 | 0 | 0 | 0 | 0 | 27 | 10 | 16 | 20 | 12 | 0 | 0 | 0 | 1 | 0 | 298 | 149 | 1 | 1 | 1 | 1 | 1 | 1 | 1 | 1 | 1 | 1 | 0.909 | 1 | 0.018 |
| SE | 35696 | ENSG00000175556 | LONRF3 | chrX | + | 118990469 | 118990560 | 118978344 | 118978463 | 119006120 | 119006235 | 8 | 1 | 17 | 12 | 4 | 0 | 0 | 0 | 0 | 0 | 0 | 6 | 2 | 13 | 0 | 0 | 0 | 1 | 0 | 0 | 240 | 149 | 1 | 1 | 1 | 1 | 1 | 1 | 1 | NA | 1 | 0.554 | 1 | NA | 0.149 |
| SE | 35697 | ENSG00000175556 | LONRF3 | chrX | + | 118990469 | 118990560 | 118989407 | 118989672 | 119006120 | 119006235 | 26 | 6 | 43 | 26 | 10 | 0 | 0 | 0 | 0 | 0 | 0 | 26 | 19 | 33 | 5 | 0 | 2 | 0 | 0 | 0 | 240 | 149 | 1 | 1 | 1 | 1 | 1 | 1 | 1 | NA | 0.89 | 1 | 1 | 1 | 0.027 |
| SE | 35698 | ENSG00000175556 | LONRF3 | chrX | + | 119006120 | 119006235 | 118974613 | 118975597 | 119009125 | 119009247 | 12 | 7 | 15 | 9 | 3 | 0 | 0 | 0 | 0 | 0 | 0 | 2 | 6 | 8 | 0 | 0 | 0 | 0 | 2 | 2 | 264 | 149 | 1 | 1 | 1 | 1 | 1 | 1 | 1 | NA | 1 | 1 | 0.693 | 0 | 0.327 |
| SE | 35699 | ENSG00000175556 | LONRF3 | chrX | + | 119006120 | 119006235 | 118990469 | 118990560 | 119009125 | 119009247 | 25 | 10 | 38 | 29 | 10 | 0 | 0 | 0 | 0 | 0 | 0 | 13 | 20 | 28 | 0 | 0 | 0 | 1 | 1 | 0 | 264 | 149 | 1 | 1 | 1 | 1 | 1 | 1 | 1 | NA | 1 | 0.919 | 0.94 | NA | 0.047 |
| SE | 35700 | ENSG00000175556 | LONRF3 | chrX | + | 119011814 | 119011973 | 119009125 | 119009247 | 119014206 | 119014356 | 11 | 8 | 21 | 8 | 0 | 1 | 0 | 0 | 0 | 0 | 0 | 5 | 23 | 32 | 0 | 0 | 0 | 0 | 3 | 0 | 298 | 149 | 1 | 1 | 0.846 | 1 | 1 | 1 | NA | NA | 1 | 1 | 0.842 | NA | 0.014 |
| SE | 35703 | ENSG00000175556 | LONRF3 | chrX | + | 119014206 | 119014356 | 119009125 | 119009247 | 119017534 | 119017983 | 13 | 6 | 17 | 13 | 1 | 0 | 0 | 0 | 0 | 0 | 14 | 13 | 13 | 35 | 0 | 0 | 0 | 0 | 2 | 0 | 298 | 149 | 1 | 1 | 1 | 1 | 1 | 1 | 1 | 1 | 1 | 1 | 0.897 | NA | 0.026 |
| SE | 35706 | ENSG00000100151 | PICK1 | chr22 | + | 38070837 | 38070891 | 38069032 | 38069122 | 38071681 | 38071728 | 19 | 8 | 24 | 25 | 11 | 1 | 0 | 1 | 0 | 0 | 23 | 50 | 28 | 40 | 94 | 0 | 0 | 0 | 0 | 0 | 203 | 149 | 1 | 1 | 0.933 | 1 | 0.946 | 1 | 1 | 1 | 1 | 1 | 1 | 1 | -0.024 |
| SE | 35707 | ENSG00000259456 | ADNP-AS1 | chr20 | + | 50933719 | 50933827 | 50931374 | 50931449 | 50944356 | 50945134 | 5 | 0 | 6 | 1 | 2 | 0 | 0 | 1 | 0 | 0 | 12 | 0 | 2 | 0 | 4 | 0 | 3 | 1 | 0 | 0 | 257 | 149 | 1 | 1 | 1 | NA | 0.777 | 1 | 1 | 1 | 0 | 0.537 | NA | 1 | 0.31 |
| SE | 35708 | ENSG00000259456 | ADNP-AS1 | chr20 | + | 50933723 | 50933827 | 50931371 | 50931449 | 50944356 | 50945148 | 8 | 2 | 1 | 4 | 9 | 0 | 0 | 1 | 0 | 0 | 24 | 0 | 4 | 0 | 4 | 0 | 3 | 1 | 0 | 0 | 253 | 149 | 1 | 1 | 1 | 1 | 0.371 | 1 | 1 | 1 | 0 | 0.702 | NA | 1 | 0.199 |
| SE | 35718 | ENSG00000185507 | IRF7 | chr11 | - | 614173 | 614399 | 613784 | 613865 | 614475 | 614534 | 21 | 11 | 18 | 24 | 0 | 0 | 0 | 0 | 1 | 0 | 7 | 31 | 0 | 8 | 12 | 0 | 3 | 2 | 3 | 0 | 298 | 149 | 1 | 1 | 1 | 1 | 1 | 0.923 | NA | 1 | 0.838 | 0 | 0.571 | 1 | 0.299 |
| SE | 35722 | ENSG00000130787 | HIP1R | chr12 | + | 122855979 | 122856163 | 122855830 | 122855903 | 122856255 | 122856344 | 160 | 83 | 141 | 130 | 51 | 0 | 0 | 0 | 0 | 0 | 146 | 242 | 76 | 87 | 231 | 0 | 0 | 0 | 3 | 0 | 298 | 149 | 1 | 1 | 1 | 1 | 1 | 1 | 1 | 1 | 1 | 0.927 | 1 | 1 | 0.015 |
| SE | 35724 | ENSG00000130787 | HIP1R | chr12 | + | 122861307 | 122861514 | 122861130 | 122861192 | 122861705 | 122862084 | 105 | 49 | 121 | 108 | 91 | 0 | 0 | 0 | 1 | 0 | 151 | 237 | 77 | 50 | 192 | 0 | 0 | 0 | 0 | 0 | 298 | 149 | 1 | 1 | 1 | 1 | 1 | 0.982 | 1 | 1 | 1 | 1 | 1 | 1 | -0.004 |
| SE | 35725 | ENSG00000087269 | NOP14 | chr4 | - | 2944072 | 2944226 | 2942191 | 2942351 | 2945127 | 2945229 | 96 | 37 | 84 | 51 | 24 | 0 | 0 | 0 | 0 | 0 | 49 | 61 | 34 | 29 | 54 | 3 | 0 | 0 | 0 | 0 | 298 | 149 | 1 | 1 | 1 | 1 | 1 | 1 | 1 | 0.891 | 1 | 1 | 1 | 1 | 0.022 |
| SE | 35726 | ENSG00000087269 | NOP14 | chr4 | - | 2949933 | 2950213 | 2948277 | 2948408 | 2951113 | 2951245 | 275 | 187 | 221 | 225 | 271 | 0 | 0 | 0 | 2 | 0 | 553 | 281 | 219 | 338 | 457 | 10 | 0 | 0 | 0 | 0 | 298 | 149 | 1 | 1 | 1 | 1 | 1 | 0.983 | 1 | 0.965 | 1 | 1 | 1 | 1 | 0.004 |
| SE | 35727 | ENSG00000087269 | NOP14 | chr4 | - | 2951113 | 2951245 | 2949933 | 2950213 | 2952274 | 2952397 | 311 | 257 | 280 | 291 | 375 | 0 | 0 | 0 | 0 | 0 | 694 | 375 | 298 | 374 | 555 | 0 | 0 | 0 | 0 | 4 | 281 | 149 | 1 | 1 | 1 | 1 | 1 | 1 | 1 | 1 | 1 | 1 | 1 | 0.987 | 0.003 |
| SE | 35729 | ENSG00000163840 | DTX3L | chr3 | + | 122565858 | 122566070 | 122564337 | 122564613 | 122568488 | 122570024 | 416 | 301 | 310 | 393 | 622 | 0 | 1 | 0 | 2 | 4 | 388 | 355 | 189 | 660 | 78 | 2 | 3 | 0 | 0 | 0 | 298 | 149 | 1 | 1 | 1 | 0.993 | 1 | 0.99 | 0.987 | 0.99 | 0.983 | 1 | 1 | 1 | -0.001 |
| SE | 35730 | ENSG00000163840 | DTX3L | chr3 | + | 122565858 | 122566070 | 122564337 | 122564613 | 122570454 | 122570672 | 123 | 67 | 95 | 119 | 81 | 0 | 0 | 2 | 0 | 0 | 134 | 96 | 48 | 218 | 44 | 0 | 2 | 0 | 1 | 0 | 298 | 149 | 1 | 1 | 1 | 1 | 0.96 | 1 | 1 | 1 | 0.96 | 1 | 0.991 | 1 | 0.002 |
| SE | 35731 | ENSG00000163840 | DTX3L | chr3 | + | 122568488 | 122570024 | 122564337 | 122564613 | 122570454 | 122570672 | 261 | 258 | 239 | 314 | 409 | 0 | 0 | 2 | 0 | 0 | 365 | 248 | 155 | 558 | 45 | 0 | 2 | 0 | 1 | 0 | 298 | 149 | 1 | 1 | 1 | 1 | 0.984 | 1 | 1 | 1 | 0.984 | 1 | 0.996 | 1 | 0.001 |
| SE | 35732 | ENSG00000163840 | DTX3L | chr3 | + | 122568488 | 122570024 | 122565858 | 122566070 | 122570454 | 122570672 | 554 | 493 | 454 | 587 | 946 | 0 | 2 | 0 | 1 | 0 | 617 | 504 | 297 | 1000 | 79 | 0 | 0 | 1 | 0 | 0 | 298 | 149 | 1 | 1 | 1 | 0.992 | 1 | 0.997 | 1 | 1 | 1 | 0.993 | 1 | 1 | -0.001 |
| SE | 35733 | ENSG00000163840 | DTX3L | chr3 | + | 122568488 | 122570024 | 122565858 | 122566070 | 122571677 | 122575203 | 294 | 236 | 215 | 277 | 541 | 0 | 0 | 0 | 0 | 0 | 261 | 261 | 142 | 442 | 34 | 0 | 0 | 1 | 4 | 0 | 298 | 149 | 1 | 1 | 1 | 1 | 1 | 1 | 1 | 1 | 1 | 0.986 | 0.982 | 1 | 0.006 |
| SE | 35734 | ENSG00000163840 | DTX3L | chr3 | + | 122570454 | 122570672 | 122565858 | 122566070 | 122571677 | 122571747 | 176 | 178 | 195 | 261 | 226 | 0 | 0 | 0 | 0 | 0 | 165 | 114 | 107 | 393 | 58 | 0 | 0 | 1 | 4 | 0 | 298 | 149 | 1 | 1 | 1 | 1 | 1 | 1 | 1 | 1 | 1 | 0.982 | 0.98 | 1 | 0.008 |
| SE | 35735 | ENSG00000163840 | DTX3L | chr3 | + | 122570454 | 122570672 | 122568488 | 122570024 | 122571677 | 122571747 | 437 | 432 | 434 | 571 | 631 | 1 | 0 | 0 | 2 | 0 | 528 | 353 | 261 | 947 | 103 | 7 | 2 | 0 | 0 | 0 | 298 | 149 | 1 | 1 | 0.995 | 1 | 1 | 0.993 | 1 | 0.974 | 0.989 | 1 | 1 | 1 | 0.005 |
| SE | 35738 | ENSG00000162736 | NCSTN | chr1 | + | 160343703 | 160343802 | 160343390 | 160343481 | 160344721 | 160344826 | 3 | 0 | 0 | 1 | 5 | 164 | 111 | 104 | 138 | 124 | 18 | 5 | 0 | 0 | 10 | 593 | 392 | 218 | 381 | 662 | 248 | 149 | 1 | 1 | 0.011 | 0 | 0 | 0.004 | 0.024 | 0.018 | 0.008 | 0 | 0 | 0.009 | 0.001 |
| SE | 35740 | ENSG00000162736 | NCSTN | chr1 | + | 160344475 | 160344614 | 160343315 | 160343481 | 160344721 | 160344826 | 0 | 0 | 0 | 0 | 0 | 164 | 111 | 104 | 138 | 124 | 0 | 0 | 0 | 1 | 0 | 593 | 392 | 218 | 381 | 662 | 288 | 149 | 1 | 1 | 0 | 0 | 0 | 0 | 0 | 0 | 0 | 0 | 0.001 | 0 | 0 |
| SE | 35741 | ENSG00000162736 | NCSTN | chr1 | + | 160348998 | 160349122 | 160344721 | 160344826 | 160349548 | 160349668 | 322 | 210 | 218 | 269 | 311 | 0 | 0 | 0 | 0 | 0 | 940 | 623 | 356 | 596 | 980 | 1 | 3 | 0 | 0 | 0 | 273 | 149 | 1 | 1 | 1 | 1 | 1 | 1 | 1 | 0.998 | 0.991 | 1 | 1 | 1 | 0.002 |
| SE | 35742 | ENSG00000162736 | NCSTN | chr1 | + | 160348998 | 160349122 | 160344721 | 160344826 | 160351695 | 160351805 | 195 | 137 | 121 | 160 | 146 | 0 | 0 | 0 | 0 | 0 | 778 | 452 | 273 | 460 | 755 | 0 | 0 | 0 | 0 | 2 | 273 | 149 | 1 | 1 | 1 | 1 | 1 | 1 | 1 | 1 | 1 | 1 | 1 | 0.995 | 0.001 |
| SE | 35749 | ENSG00000162736 | NCSTN | chr1 | + | 160351695 | 160351805 | 160351221 | 160351372 | 160352886 | 160352991 | 190 | 114 | 133 | 153 | 288 | 1 | 0 | 0 | 0 | 0 | 355 | 242 | 145 | 237 | 360 | 0 | 0 | 0 | 0 | 0 | 259 | 149 | 1 | 1 | 0.991 | 1 | 1 | 1 | 1 | 1 | 1 | 1 | 1 | 1 | -0.002 |
| SE | 35751 | ENSG00000162736 | NCSTN | chr1 | + | 160352053 | 160352206 | 160351695 | 160351805 | 160352886 | 160352991 | 302 | 237 | 254 | 258 | 199 | 0 | 0 | 0 | 0 | 0 | 575 | 467 | 272 | 362 | 674 | 0 | 3 | 0 | 0 | 0 | 298 | 149 | 1 | 1 | 1 | 1 | 1 | 1 | 1 | 1 | 0.987 | 1 | 1 | 1 | 0.003 |
| SE | 35752 | ENSG00000162736 | NCSTN | chr1 | + | 160352125 | 160352206 | 160351695 | 160351805 | 160352886 | 160352991 | 32 | 27 | 28 | 46 | 25 | 0 | 0 | 0 | 0 | 0 | 34 | 34 | 27 | 29 | 91 | 0 | 3 | 0 | 0 | 0 | 230 | 149 | 1 | 1 | 1 | 1 | 1 | 1 | 1 | 1 | 0.88 | 1 | 1 | 1 | 0.024 |
| SE | 35758 | ENSG00000162736 | NCSTN | chr1 | + | 160355862 | 160355958 | 160354117 | 160354290 | 160356599 | 160356754 | 0 | 0 | 0 | 0 | 0 | 0 | 1 | 1 | 2 | 1 | 0 | 5 | 2 | 0 | 11 | 17 | 2 | 0 | 19 | 8 | 245 | 149 | 1 | 1 | NA | 0 | 0 | 0 | 0 | 0 | 0.603 | 1 | 0 | 0.455 | -0.412 |
| SE | 35759 | ENSG00000162736 | NCSTN | chr1 | + | 160355862 | 160355958 | 160355654 | 160355757 | 160356259 | 160356347 | 280 | 148 | 221 | 208 | 461 | 0 | 1 | 0 | 0 | 0 | 347 | 461 | 249 | 367 | 397 | 0 | 0 | 0 | 0 | 0 | 245 | 149 | 1 | 1 | 1 | 0.989 | 1 | 1 | 1 | 1 | 1 | 1 | 1 | 1 | -0.002 |
| SE | 35760 | ENSG00000162736 | NCSTN | chr1 | + | 160356259 | 160356347 | 160354117 | 160354290 | 160356599 | 160356754 | 96 | 62 | 64 | 84 | 79 | 0 | 1 | 1 | 2 | 1 | 224 | 150 | 86 | 133 | 249 | 17 | 2 | 0 | 19 | 8 | 237 | 149 | 1 | 1 | 1 | 0.975 | 0.976 | 0.964 | 0.98 | 0.892 | 0.979 | 1 | 0.815 | 0.951 | 0.052 |
| SE | 35761 | ENSG00000162736 | NCSTN | chr1 | + | 160356259 | 160356347 | 160355862 | 160355958 | 160356599 | 160356754 | 293 | 156 | 209 | 230 | 392 | 0 | 0 | 0 | 0 | 0 | 465 | 456 | 250 | 381 | 520 | 0 | 2 | 0 | 0 | 4 | 237 | 149 | 1 | 1 | 1 | 1 | 1 | 1 | 1 | 1 | 0.993 | 1 | 1 | 0.988 | 0.004 |
| SE | 35769 | ENSG00000188687 | SLC4A5 | chr2 | - | 74232468 | 74232647 | 74231235 | 74231308 | 74233401 | 74233563 | 4 | 0 | 9 | 9 | 7 | 2 | 0 | 1 | 0 | 3 | 0 | 0 | 0 | 4 | 5 | 0 | 0 | 0 | 0 | 0 | 298 | 149 | 1 | 1 | 0.5 | NA | 0.818 | 1 | 0.538 | NA | NA | NA | 1 | 1 | -0.286 |
| SE | 35770 | ENSG00000188687 | SLC4A5 | chr2 | - | 74235100 | 74235214 | 74233401 | 74233563 | 74239334 | 74239535 | 5 | 4 | 5 | 6 | 3 | 0 | 0 | 0 | 1 | 0 | 0 | 0 | 0 | 5 | 0 | 0 | 0 | 0 | 0 | 0 | 263 | 149 | 1 | 1 | 1 | 1 | 1 | 0.773 | 1 | NA | NA | NA | 1 | NA | -0.045 |
| SE | 35771 | ENSG00000188687 | SLC4A5 | chr2 | - | 74241993 | 74242052 | 74239334 | 74239535 | 74247035 | 74247307 | 4 | 3 | 8 | 4 | 8 | 0 | 0 | 2 | 0 | 0 | 0 | 4 | 3 | 4 | 0 | 0 | 0 | 0 | 0 | 0 | 208 | 149 | 1 | 1 | 1 | 1 | 0.741 | 1 | 1 | NA | 1 | 1 | 1 | NA | -0.052 |
| [illegible] | | | | | | | | | | | | | | | | | | | | | | | | | | | | | | | | | | | | | | | | | | | | | | |

| | | | | | | | | | | | | | | |
|---|---|---|---|---|---|---|---|---|---|---|---|---|---|---|
| SE | 36007 | ENSG00000258653 | NA | chr14 | + | 73909340 | 73909430 | 73904901 | 73905083 | 73923394 | 73923522 | 26 | 24 | [illegible] |
| SE | 36008 | ENSG00000258653 | NA | chr14 | + | 73920979 | 73921105 | 73904901 | 73905083 | 73923394 | 73923522 | 27 | 9 | [illegible] |
| SE | 36009 | ENSG00000258653 | NA | chr14 | + | 73920979 | 73921105 | 73909340 | 73909430 | 73923394 | 73923522 | 79 | 48 | [illegible] |
| SE | 36015 | ENSG00000164463 | CREBRF | chr5 | + | 173086500 | 173086626 | 173080584 | 173080784 | 173090314 | 173091401 | 673 | 486 | [illegible] |
| SE | 36016 | ENSG00000164463 | CREBRF | chr5 | + | 173090314 | 173091401 | 173086500 | 173086626 | 173108623 | 173108818 | 613 | 430 | [illegible] |
| SE | 36017 | ENSG00000164463 | CREBRF | chr5 | + | 173108623 | 173108818 | 173090314 | 173091401 | 173110521 | 173110711 | 517 | 362 | [illegible] |
| SE | 36018 | ENSG00000164463 | CREBRF | chr5 | + | 173112305 | 173112379 | 173108623 | 173110711 | 173123079 | 173123202 | 531 | 414 | [illegible] |
| SE | 36019 | ENSG00000164463 | CREBRF | chr5 | + | 173123079 | 173123202 | 173112305 | 173112379 | 173133629 | 173139284 | 420 | 310 | [illegible] |
| SE | 36021 | ENSG00000177694 | NAALADL2 | chr3 | + | 174536620 | 174536683 | 174523412 | 174523548 | 174550568 | 174550637 | 0 | 5 | [illegible] |
| SE | 36024 | ENSG00000177694 | NAALADL2 | chr3 | + | 175460044 | 175460175 | 175447228 | 175447372 | 175463400 | 175463493 | 0 | 0 | [illegible] |
| SE | 36026 | ENSG00000145819 | ARHGAP26 | chr5 | + | 142875109 | 142875171 | 142873399 | 142873495 | 142879373 | 142879445 | 286 | 259 | [illegible] |
| SE | 36027 | ENSG00000145819 | ARHGAP26 | chr5 | + | 142885297 | 142885399 | 142879373 | 142879445 | 142894237 | 142894335 | 487 | 536 | [illegible] |
| SE | 36028 | ENSG00000145819 | ARHGAP26 | chr5 | + | 142894237 | 142894348 | 142885297 | 142885399 | 142901934 | 142902039 | 501 | 549 | [illegible] |
| SE | 36030 | ENSG00000145819 | ARHGAP26 | chr5 | + | 142901934 | 142902039 | 142894237 | 142894348 | 142907703 | 142907804 | 167 | 163 | [illegible] |
| SE | 36031 | ENSG00000145819 | ARHGAP26 | chr5 | + | 142903539 | 142903669 | 142894237 | 142894348 | 142907703 | 142907804 | 383 | 275 | [illegible] |
| SE | 36032 | ENSG00000145819 | ARHGAP26 | chr5 | + | 142903539 | 142903669 | 142901934 | 142902039 | 142907703 | 142907804 | 524 | 394 | [illegible] |
| SE | 36033 | ENSG00000145819 | ARHGAP26 | chr5 | + | 142907703 | 142907804 | 142903539 | 142903669 | 142913198 | 142913293 | 498 | 373 | [illegible] |
| SE | 36035 | ENSG00000145819 | ARHGAP26 | chr5 | + | 142932046 | 142932125 | 142913198 | 142913293 | 143014079 | 143014116 | 294 | 270 | [illegible] |
| SE | 36036 | ENSG00000145819 | ARHGAP26 | chr5 | + | 143014079 | 143014116 | 142932046 | 142932125 | 143037195 | 143037223 | 209 | 159 | [illegible] |
| SE | 36037 | ENSG00000145819 | ARHGAP26 | chr5 | + | 143037195 | 143037261 | 143014079 | 143014116 | 143041524 | 143041714 | 51 | 66 | [illegible] |
| SE | 36039 | ENSG00000145819 | ARHGAP26 | chr5 | + | 143041815 | 143041890 | 143014079 | 143014116 | 143120987 | 143121147 | 2 | 2 | [illegible] |
| SE | 36040 | ENSG00000145819 | ARHGAP26 | chr5 | + | 143166026 | 143166097 | 143147230 | 143147381 | 143172958 | 143173032 | 1 | 0 | [illegible] |
| SE | 36042 | ENSG00000145819 | ARHGAP26 | chr5 | + | 143172958 | 143173032 | 143147230 | 143147381 | 143207197 | 143207473 | 0 | 1 | [illegible] |
| SE | 36044 | ENSG00000145819 | ARHGAP26 | chr5 | + | 143172958 | 143173032 | 143147230 | 143147381 | 143213996 | 143214088 | 0 | 1 | [illegible] |
| SE | 36045 | ENSG00000145819 | ARHGAP26 | chr5 | + | 143172958 | 143173032 | 143166026 | 143166097 | 143207197 | 143207473 | 0 | 0 | [illegible] |
| SE | 36047 | ENSG00000145819 | ARHGAP26 | chr5 | + | 143207197 | 143207473 | 143147230 | 143147381 | 143213996 | 143214088 | 30 | 21 | [illegible] |
| SE | 36049 | ENSG00000145819 | ARHGAP26 | chr5 | + | 143207197 | 143207473 | 143172958 | 143173032 | 143213996 | 143214088 | 0 | 0 | [illegible] |
| SE | 36050 | ENSG00000145819 | ARHGAP26 | chr5 | + | 143207382 | 143207473 | 143147230 | 143147381 | 143213996 | 143214088 | 0 | 0 | [illegible] |
| SE | 36051 | ENSG00000145819 | ARHGAP26 | chr5 | + | 143207382 | 143207473 | 143172958 | 143173032 | 143213996 | 143214088 | 0 | 0 | [illegible] |
| [illegible] | | | | | | | | | | | | | | |

| | | | | | | | | | | |
|---|---|---|---|---|---|---|---|---|---|---|
| SE | 36398 | ENSG00000276291 | FRG1HP | chr9 | + | 41009793 | 41009843 | 41008880 | 41009006 | 41011616 |
| SE | 36399 | ENSG00000276291 | FRG1HP | chr9 | + | 41011616 | 41011730 | 41008880 | 41009006 | 41032833 |
| SE | 36407 | ENSG00000116191 | RALGPS2 | chr1 | + | 178883465 | 178883533 | 178878901 | 178878992 | 178885075 |
| SE | 36409 | ENSG00000116191 | RALGPS2 | chr1 | + | 178897663 | 178897756 | 178893837 | 178894024 | 178902105 |
| SE | 36410 | ENSG00000116191 | RALGPS2 | chr1 | + | 178902105 | 178902211 | 178897663 | 178897756 | 178906775 |
| SE | 36411 | ENSG00000116191 | RALGPS2 | chr1 | + | 178902105 | 178902211 | 178897663 | 178897756 | 178916329 |
| SE | 36412 | ENSG00000116191 | RALGPS2 | chr1 | + | 178906775 | 178906867 | 178897663 | 178897756 | 178916329 |
| SE | 36413 | ENSG00000116191 | RALGPS2 | chr1 | + | 178906775 | 178906867 | 178902105 | 178902211 | 178916329 |
| SE | 36414 | ENSG00000179262 | RAD23A | chr19 | + | 12948176 | 12948358 | 12947847 | 12948009 | 12948496 |
| SE | 36415 | ENSG00000179262 | RAD23A | chr19 | + | 12948176 | 12948358 | 12947847 | 12948009 | 12948685 |
| SE | 36416 | ENSG00000179262 | RAD23A | chr19 | + | 12948496 | 12948552 | 12947847 | 12948009 | 12948685 |
| SE | 36419 | ENSG00000179262 | RAD23A | chr19 | + | 12949080 | 12949159 | 12948685 | 12948813 | 12949274 |
| SE | 36420 | ENSG00000179262 | RAD23A | chr19 | + | 12949080 | 12949159 | 12948685 | 12948813 | 12949277 |
| SE | 36421 | ENSG00000179262 | RAD23A | chr19 | + | 12949274 | 12949408 | 12948688 | 12948813 | 12952688 |
| SE | 36422 | ENSG00000179262 | RAD23A | chr19 | + | 12949277 | 12949408 | 12948685 | 12948813 | 12952688 |
| SE | 36428 | ENSG00000121858 | TNFSF10 | chr3 | - | 172509216 | 172509321 | 172505507 | 172506919 | 172523252 |
| SE | 36430 | ENSG00000121858 | TNFSF10 | chr3 | - | 172511616 | 172511659 | 172505507 | 172506919 | 172523252 |
| SE | 36433 | ENSG00000121858 | TNFSF10 | chr3 | - | 172514860 | 172514998 | 172505661 | 172506919 | 172523252 |
| SE | 36442 | ENSG00000100462 | PRMT5 | chr14 | - | 22923050 | 22923160 | 22922741 | 22922835 | 22924007 |
| SE | 36444 | ENSG00000100462 | PRMT5 | chr14 | - | 22927525 | 22927660 | 22924878 | 22925040 | 22928125 |
| SE | 36468 | ENSG00000162302 | RPS6KA4 | chr11 | + | 64359377 | 64359403 | 64359181 | 64359290 | 64360162 |
| SE | 36471 | ENSG00000162302 | RPS6KA4 | chr11 | + | 64370224 | 64370384 | 64369698 | 64369893 | 64370562 |
| SE | 36478 | ENSG00000124596 | OARD1 | chr6 | - | 41070075 | 41070134 | 41052264 | 41052363 | 41071131 |
| SE | 36480 | ENSG00000124596 | OARD1 | chr6 | - | 41070075 | 41070134 | 41066787 | 41067437 | 41071595 |
| SE | 36481 | ENSG00000124596 | OARD1 | chr6 | - | 41070075 | 41070134 | 41068840 | 41068953 | 41071131 |
| SE | 36483 | ENSG00000124596 | OARD1 | chr6 | - | 41071131 | 41071276 | 41069037 | 41070134 | 41071595 |
| SE | 36496 | ENSG00000163626 | COX18 | chr4 | - | 73068028 | 73068129 | 73065249 | 73065413 | 73069316 |
| SE | 36497 | ENSG00000163626 | COX18 | chr4 | - | 73068028 | 73068133 | 73065249 | 73065413 | 73069316 |
| SE | 36498 | ENSG00000168405 | CMAHP | chr6 | - | 25084479 | 25084576 | 25081108 | 25081879 | 25086783 |
| SE | 36499 | ENSG00000168405 | CMAHP | chr6 | - | 25084479 | 25084576 | 25081108 | 25081879 | 25091895 |
| SE | 36501 | ENSG00000168405 | CMAHP | chr6 | - | 25086783 | 25086899 | 25081108 | 25081879 | 25091895 |
| SE | 36503 | ENSG00000168405 | CMAHP | chr6 | - | 25086783 | 25086899 | 25084479 | 25084576 | 25091895 |
| SE | 36506 | ENSG00000168405 | CMAHP | chr6 | - | 25091895 | 25092012 | 25086616 | 25086899 | 25097060 |
| SE | 36507 | ENSG00000168405 | CMAHP | chr6 | - | 25093513 | 25093688 | 25086783 | 25086899 | 25097060 |
| SE | 36508 | ENSG00000168405 | CMAHP | chr6 | - | 25093513 | 25093688 | 25091895 | 25092012 | 25097060 |
| SE | 36509 | ENSG00000168405 | CMAHP | chr6 | - | 25106358 | 25106494 | 25097060 | 25097201 | 25108575 |
| SE | 36510 | ENSG00000168405 | CMAHP | chr6 | - | 25109955 | 25110103 | 25109499 | 25109581 | 25113591 |
| SE | 36511 | ENSG00000168405 | CMAHP | chr6 | - | 25109955 | 25110103 | 25109499 | 25109581 | 25114990 |
| SE | 36515 | ENSG00000168405 | CMAHP | chr6 | - | 25166289 | 25166349 | 25137619 | 25137823 | 25369945 |
| SE | 36518 | ENSG00000168405 | CMAHP | chr6 | - | 25260347 | 25260576 | 25255188 | 25255306 | 25369945 |
| SE | 36519 | ENSG00000168405 | CMAHP | chr6 | - | 25389118 | 25389167 | 25369945 | 25370079 | 25420301 |
| SE | 36520 | ENSG00000168405 | CMAHP | chr6 | - | 25389118 | 25389167 | 25369945 | 25370079 | 25452036 |
| SE | 36522 | ENSG00000168405 | CMAHP | chr6 | - | 25420301 | 25420383 | 25369945 | 25370079 | 25452036 |
| SE | 36523 | ENSG00000168405 | CMAHP | chr6 | - | 25420301 | 25420383 | 25389118 | 25389167 | 25452036 |
| SE | 36525 | ENSG00000169230 | PRELID1 | chr5 | + | 177304292 | 177304430 | 177303854 | 177304077 | 177304624 |
| SE | 36526 | ENSG00000169230 | PRELID1 | chr5 | + | 177305870 | 177305984 | 177304575 | 177304850 | 177306097 |
| SE | 36528 | ENSG00000169230 | PRELID1 | chr5 | + | 177306097 | 177306176 | 177305870 | 177305984 | 177306421 |
| SE | 36571 | ENSG00000147996 | CBWD5 | chr9 | + | 65677127 | 65677225 | 65675836 | 65676130 | 65679165 |
| SE | 36572 | ENSG00000147996 | CBWD5 | chr9 | + | 65677127 | 65677229 | 65675940 | 65676130 | 65679165 |
| SE | 36579 | ENSG00000147996 | CBWD5 | chr9 | + | 65694904 | 65695044 | 65693385 | 65693473 | 65698514 |
| SE | 36581 | ENSG00000147996 | CBWD5 | chr9 | + | 65700250 | 65700337 | 65698514 | 65698561 | 65702969 |
| SE | 36582 | ENSG00000147996 | CBWD5 | chr9 | + | 65702969 | 65703014 | 65700250 | 65700337 | 65708901 |
| SE | 36583 | ENSG00000147996 | CBWD5 | chr9 | + | 65702969 | 65703014 | 65700250 | 65700337 | 65720040 |
| SE | 36584 | ENSG00000147996 | CBWD5 | chr9 | + | 65704330 | 65704414 | 65702969 | 65703014 | 65708901 |
| SE | 36585 | ENSG00000147996 | CBWD5 | chr9 | + | 65704330 | 65704547 | 65702969 | 65703014 | 65708901 |
| SE | 36586 | ENSG00000147996 | CBWD5 | chr9 | + | 65707012 | 65707152 | 65702969 | 65703014 | 65708901 |
| SE | 36587 | ENSG00000147996 | CBWD5 | chr9 | + | 65708901 | 65708958 | 65700286 | 65700337 | 65720040 |
| SE | 36588 | ENSG00000147996 | CBWD5 | chr9 | + | 65708901 | 65708958 | 65702969 | 65703014 | 65720040 |
| SE | 36589 | ENSG00000147996 | CBWD5 | chr9 | + | 65731629 | 65731698 | 65719384 | 65720092 | 65731801 |
| SE | 36590 | ENSG00000147996 | CBWD5 | chr9 | + | 65732946 | 65733122 | 65731801 | 65731867 | 65733509 |
| SE | 36591 | ENSG00000147996 | CBWD5 | chr9 | + | 65732992 | 65733122 | 65731801 | 65731867 | 65733509 |
| SE | 36595 | ENSG00000169180 | XPO6 | chr16 | - | 28112903 | 28113050 | 28107521 | 28107677 | 28121669 |
| SE | 36596 | ENSG00000169180 | XPO6 | chr16 | - | 28117317 | 28117462 | 28112903 | 28113050 | 28121669 |
| SE | 36597 | ENSG00000169180 | XPO6 | chr16 | - | 28121669 | 28121762 | 28117036 | 28117462 | 28125688 |
| SE | 36599 | ENSG00000169180 | XPO6 | chr16 | - | 28132333 | 28132403 | 28125688 | 28125848 | 28133840 |
| SE | 36600 | ENSG00000169180 | XPO6 | chr16 | - | 28133840 | 28133933 | 28125688 | 28125848 | 28146093 |
| SE | 36601 | ENSG00000169180 | XPO6 | chr16 | - | 28133840 | 28133933 | 28132333 | 28132403 | 28135215 |
| SE | 36602 | ENSG00000169180 | XPO6 | chr16 | - | 28135215 | 28135324 | 28133840 | 28133933 | 28146093 |
| SE | 36603 | ENSG00000169180 | XPO6 | chr16 | - | 28146093 | 28146203 | 28135215 | 28135324 | 28152658 |
| SE | 36606 | ENSG00000169180 | XPO6 | chr16 | - | 28152658 | 28152785 | 28146090 | 28146203 | 28155701 |
| SE | 36607 | ENSG00000169180 | XPO6 | chr16 | - | 28152658 | 28152785 | 28146090 | 28146203 | 28155945 |
| SE | 36609 | ENSG00000169180 | XPO6 | chr16 | - | 28152929 | 28152964 | 28152658 | 28152785 | 28155701 |
| SE | 36610 | ENSG00000169180 | XPO6 | chr16 | - | 28152929 | 28152964 | 28152658 | 28152785 | 28155945 |
| SE | 36612 | ENSG00000169180 | XPO6 | chr16 | - | 28154587 | 28154612 | 28152929 | 28152964 | 28155701 |
| SE | 36613 | ENSG00000169180 | XPO6 | chr16 | - | 28155701 | 28155771 | 28146093 | 28146203 | 28155945 |
| SE | 36615 | ENSG00000169180 | XPO6 | chr16 | - | 28155701 | 28155771 | 28152658 | 28152785 | 28155945 |
| SE | 36616 | ENSG00000169180 | XPO6 | chr16 | - | 28155701 | 28155771 | 28152658 | 28152785 | 28156073 |
| SE | 36617 | ENSG00000169180 | XPO6 | chr16 | - | 28155701 | 28155771 | 28152929 | 28152964 | 28155945 |
| SE | 36618 | ENSG00000169180 | XPO6 | chr16 | - | 28155701 | 28155771 | 28152929 | 28152964 | 28156073 |
| SE | 36619 | ENSG00000169180 | XPO6 | chr16 | - | 28166507 | 28166585 | 28156073 | 28156527 | 28169749 |
| SE | 36620 | ENSG00000169180 | XPO6 | chr16 | - | 28166507 | 28166585 | 28156073 | 28156527 | 28175897 |
| SE | 36621 | ENSG00000169180 | XPO6 | chr16 | - | 28169749 | 28169909 | 28166507 | 28166585 | 28175897 |
| SE | 36625 | ENSG00000169180 | XPO6 | chr16 | - | 28180940 | 28181031 | 28177219 | 28177332 | 28208986 |
| SE | 36630 | ENSG00000287860 | NA | chr7 | - | 6987224 | 6987410 | 6961764 | 6961922 | 7104393 |
| SE | 36631 | ENSG00000287860 | NA | chr7 | - | 7000243 | 7000376 | 6987224 | 6987410 | 7104393 |
| SE | 36638 | ENSG00000274265 | NA | chr1 | + | 149177157 | 149177212 | 149175769 | 149176250 | 149180295 |
| SE | 36639 | ENSG00000274265 | NA | chr1 | + | 149177157 | 149177212 | 149175769 | 149176250 | 149190635 |
| SE | 36642 | ENSG00000274265 | NA | chr1 | + | 149180295 | 149180547 | 149175943 | 149176250 | 149190635 |
| SE | 36644 | ENSG00000274265 | NA | chr1 | + | 149180295 | 149180547 | 149177157 | 149177212 | 149190635 |
| SE | 36645 | ENSG00000274265 | NA | chr1 | + | 149180295 | 149180547 | 149177157 | 149177212 | 149250564 |
| SE | 36647 | ENSG00000274265 | NA | chr1 | + | 149190635 | 149190795 | 149176021 | 149176250 | 149259501 |
| SE | 36648 | ENSG00000274265 | NA | chr1 | + | 149190635 | 149190795 | 149177157 | 149177212 | 149250564 |
| SE | 36649 | ENSG00000274265 | NA | chr1 | + | 149190635 | 149190795 | 149177157 | 149177212 | 149259501 |
| SE | 36650 | ENSG00000274265 | NA | chr1 | + | 149190635 | 149190795 | 149180295 | 149180547 | 149250564 |
| SE | 36661 | ENSG00000180329 | CCDC43 | chr17 | - | 44680584 | 44680643 | 44677439 | 44679043 | 44682002 |
| SE | 36662 | ENSG00000180329 | CCDC43 | chr17 | - | 44680584 | 44680652 | 44677482 | 44679043 | 44682002 |
| SE | 36663 | ENSG00000180329 | CCDC43 | chr17 | - | 44683871 | 44683959 | 44681730 | 44682138 | 44689549 |
| SE | 36668 | ENSG00000180530 | NRIP1 | chr21 | - | 14991200 | 14991306 | 14968154 | 14968526 | 15043494 |
| SE | 36671 | ENSG00000180530 | NRIP1 | chr21 | - | 15014343 | 15014466 | 14961234 | 14968526 | 15064744 |
| SE | 36673 | ENSG00000180530 | NRIP1 | chr21 | - | 15043494 | 15043574 | 14961234 | 14968526 | 15064744 |
| SE | 36675 | ENSG00000214922 | HLA-F-AS1 | chr6 | - | 29728364 | 29728495 | 29726600 | 29727139 | 29737965 |
| SE | 36676 | ENSG00000214922 | HLA-F-AS1 | chr6 | - | 29728364 | 29728495 | 29726600 | 29727139 | 29748899 |
| SE | 36677 | ENSG00000214922 | HLA-F-AS1 | chr6 | - | 29738550 | 29738886 | 29738334 | 29738455 | 29745472 |
| SE | 36678 | ENSG00000214922 | HLA-F-AS1 | chr6 | - | 29738550 | 29738886 | 29738334 | 29738455 | 29747147 |
| SE | 36679 | ENSG00000214922 | HLA-F-AS1 | chr6 | - | 29738550 | 29738886 | 29738334 | 29738455 | 29748899 |
| SE | 36680 | ENSG00000214922 | HLA-F-AS1 | chr6 | - | 29745472 | 29745727 | 29726600 | 29727139 | 29748899 |
| SE | 36682 | ENSG00000214922 | HLA-F-AS1 | chr6 | - | 29745472 | 29745727 | 29738334 | 29738455 | 29748899 |
| SE | 36683 | ENSG00000214922 | HLA-F-AS1 | chr6 | - | 29745472 | 29745727 | 29738550 | 29738886 | 29747147 |
| SE | 36684 | ENSG00000214922 | HLA-F-AS1 | chr6 | - | 29745472 | 29745727 | 29738550 | 29738886 | 29748899 |
| SE | 36685 | ENSG00000214922 | HLA-F-AS1 | chr6 | - | 29747147 | 29747425 | 29738334 | 29738455 | 29748899 |
| SE | 36692 | ENSG00000158062 | UBXN11 | chr1 | - | 26296918 | 26296995 | 26294191 | 26294331 | 26297961 |
| SE | 36693 | ENSG00000158062 | UBXN11 | chr1 | - | 26297426 | 26297481 | 26294204 | 26294331 | 26297961 |
| SE | 36694 | ENSG00000158062 | UBXN11 | chr1 | - | 26297426 | 26297481 | 26296918 | 26296995 | 26297961 |
| SE | 36697 | ENSG00000158062 | UBXN11 | chr1 | - | 26301693 | 26301722 | 26297961 | 26298062 | 26302812 |
| SE | 36698 | ENSG00000158062 | UBXN11 | chr1 | - | 26301693 | 26301722 | 26297961 | 26298062 | 26306591 |
| SE | 36700 | ENSG00000158062 | UBXN11 | chr1 | - | 26302812 | 26302918 | 26301693 | 26301722 | 26303042 |
| SE | 36702 | ENSG00000158062 | UBXN11 | chr1 | - | 26303042 | 26303167 | 26301693 | 26301722 | 26306591 |
| SE | 36721 | ENSG00000246082 | NUDT16L2P | chr3 | + | 131362171 | 131362441 | 131361897 | 131362068 | 131363355 |
| SE | 36724 | ENSG00000180116 | C12orf40 | chr12 | + | 39692048 | 39692112 | 39684866 | 39684927 | 39716755 |
| SE | 36728 | ENSG00000085265 | FCN1 | chr9 | - | 134911132 | 134911267 | 134903231 | 134910045 | 134912485 |
| SE | 36729 | ENSG00000085265 | FCN1 | chr9 | - | 134912485 | 134912615 | 134911132 | 134911267 | 134913580 |
| SE | 36730 | ENSG00000085265 | FCN1 | chr9 | - | 134913015 | 134913143 | 134912485 | 134912615 | 134913580 |
| SE | 36731 | ENSG00000085265 | FCN1 | chr9 | - | 134913580 | 134913613 | 134913015 | 134913143 | 134914384 |
| SE | 36732 | ENSG00000085265 | FCN1 | chr9 | - | 134914384 | 134914420 | 134913580 | 134913613 | 134914755 |
| SE | 36733 | ENSG00000085265 | FCN1 | chr9 | - | 134914384 | 134914420 | 134913580 | 134913613 | 134916347 |
| SE | 36734 | ENSG00000085265 | FCN1 | chr9 | - | 134914755 | 134914809 | 134913580 | 134913613 | 134916347 |
| SE | 36735 | ENSG00000085265 | FCN1 | chr9 | - | 134914755 | 134914809 | 134914384 | 134914420 | 134916347 |
| SE | 36736 | ENSG00000085265 | FCN1 | chr9 | - | 134916347 | 134916461 | 134914755 | 134914809 | 134917768 |
| SE | 36737 | ENSG00000159685 | CHCHD6 | chr3 | + | 126730560 | 126730630 | 126727077 | 126727186 | 126733077 |
| SE | 36738 | ENSG00000159685 | CHCHD6 | chr3 | + | 126780339 | 126780430 | 126733077 | 126733222 | 126852646 |
| SE | 36739 | ENSG00000159685 | CHCHD6 | chr3 | + | 126953024 | 126953184 | 126914679 | 126914750 | 126957415 |
| SE | 36740 | ENSG00000173193 | PARP14 | chr3 | + | 122685184 | 122685318 | 122680838 | 122681070 | 122687079 |
| SE | 36741 | ENSG00000173193 | PARP14 | chr3 | + | 122692300 | 122692543 | 122687079 | 122687113 | 122695425 |
| SE | 36742 | ENSG00000173193 | PARP14 | chr3 | + | 122695425 | 122695662 | 122692300 | 122692543 | 122699389 |
| SE | 36744 | ENSG00000173193 | PARP14 | chr3 | + | 122699389 | 122701635 | 122695425 | 122695662 | 122704526 |
| SE | 36745 | ENSG00000173193 | PARP14 | chr3 | + | 122703741 | 122703978 | 122695425 | 122695662 | 122704526 |
| SE | 36746 | ENSG00000173193 | PARP14 | chr3 | + | 122703741 | 122703978 | 122699389 | 122701635 | 122704526 |
| SE | 36748 | ENSG00000173193 | PARP14 | chr3 | + | 122704526 | 122704748 | 122699389 | 122701635 | 122708189 |
| SE | 36750 | ENSG00000173193 | PARP14 | chr3 | + | 122708189 | 122708268 | 122699389 | 122701635 | 122713423 |
| SE | 36751 | ENSG00000173193 | PARP14 | chr3 | + | 122718358 | 122718958 | 122718070 | 122718277 | 122720254 |
| SE | 36758 | ENSG00000163539 | CLASP2 | chr3 | - | 33501651 | 33501768 | 33496244 | 33498717 | 33516022 |
| SE | 36759 | ENSG00000163539 | CLASP2 | chr3 | - | 33510557 | 33510764 | 33496244 | 33498717 | 33516022 |
| SE | 36761 | ENSG00000163539 | CLASP2 | chr3 | - | 33514579 | 33514679 | 33510557 | 33510764 | 33516022 |
| SE | 36762 | ENSG00000163539 | CLASP2 | chr3 | - | 33535232 | 33535461 | 33516980 | 33517174 | 33538788 |
| SE | 36763 | ENSG00000163539 | CLASP2 | chr3 | - | 33551251 | 33551395 | 33544697 | 33544841 | 33559306 |
| SE | 36764 | ENSG00000163539 | CLASP2 | chr3 | - | 33559124 | 33559175 | 33551251 | 33551395 | 33559306 |
| SE | 36766 | ENSG00000163539 | CLASP2 | chr3 | - | 33566731 | 33566734 | 33560807 | 33560971 | 33570726 |
| SE | 36767 | ENSG00000163539 | CLASP2 | chr3 | - | 33570726 | 33570790 | 33560807 | 33560971 | 33573109 |
| SE | 36769 | ENSG00000163539 | CLASP2 | chr3 | - | 33577201 | 33577264 | 33576168 | 33576275 | 33581820 |
| SE | 36770 | ENSG00000163539 | CLASP2 | chr3 | - | 33594950 | 33594968 | 33592394 | 33592496 | 33602951 |
| SE | 36771 | ENSG00000163539 | CLASP2 | chr3 | - | 33596710 | 33596734 | 33594950 | 33594968 | 33602951 |
| SE | 36772 | ENSG00000163539 | CLASP2 | chr3 | - | 33604153 | 33604209 | 33602442 | 33603125 | 33606590 |
| SE | 36773 | ENSG00000163539 | CLASP2 | chr3 | - | 33612000 | 33612071 | 33608566 | 33608626 | 33619602 |
| SE | 36774 | ENSG00000163539 | CLASP2 | chr3 | - | 33644756 | 33644903 | 33632291 | 33632371 | 33663444 |
| SE | 36780 | ENSG00000163539 | CLASP2 | chr3 | - | 33684358 | 33684456 | 33663444 | 33663515 | 33688276 |
| SE | 36781 | ENSG00000163539 | CLASP2 | chr3 | - | 33687059 | 33687135 | 33663444 | 33663515 | 33688276 |
| SE | 36782 | ENSG00000163539 | CLASP2 | chr3 | - | 33687059 | 33687135 | 33684358 | 33684456 | 33688276 |
| SE | 36783 | ENSG00000163539 | CLASP2 | chr3 | - | 33688276 | 33688368 | 33684358 | 33684456 | 33689828 |
| SE | 36784 | ENSG00000240207 | NA | chr3 | + | 158733950 | 158734037 | 158732262 | 158732528 | 158740388 |
| SE | 36786 | ENSG00000240207 | NA | chr3 | + | 158742029 | 158742144 | 158740388 | 158740466 | 158742371 |
| SE | 36787 | ENSG00000240207 | NA | chr3 | + | 158742029 | 158742144 | 158740388 | 158740466 | 158783878 |
| SE | 36789 | ENSG00000058262 | SEC61A1 | chr3 | + | 128052834 | 128052902 | 128052436 | 128052559 | 128055515 |
| SE | 36790 | ENSG00000058262 | SEC61A1 | chr3 | + | 128055515 | 128055581 | 128052488 | 128052902 | 128056708 |
| SE | 36791 | ENSG00000058262 | SEC61A1 | chr3 | + | 128055672 | 128055751 | 128055515 | 128055581 | 128056708 |
| SE | 36793 | ENSG00000058262 | SEC61A1 | chr3 | + | 128056708 | 128056840 | 128055672 | 128055751 | 128060101 |
| SE | 36794 | ENSG00000058262 | SEC61A1 | chr3 | + | 128064876 | 128065037 | 128060507 | 128060661 | 128066953 |
| SE | 36795 | ENSG00000058262 | SEC61A1 | chr3 | + | 128065020 | 128065256 | 128060507 | 128060661 | 128066953 |
| SE | 36797 | ENSG00000058262 | SEC61A1 | chr3 | + | 128066953 | 128067151 | 128064876 | 128065037 | 128067420 |
| SE | 36830 | ENSG00000168038 | ULK4 | chr3 | - | 41463086 | 41463253 | 41455496 | 41455595 | 41566024 |
| SE | 36831 | ENSG00000168038 | ULK4 | chr3 | - | 41566024 | 41566130 | 41463086 | 41463253 | 41615668 |
| SE | 36833 | ENSG00000168038 | ULK4 | chr3 | - | 41705253 | 41705305 | 41681752 | 41681804 | 41715236 |
| SE | 36834 | ENSG00000168038 | ULK4 | chr3 | - | 41717727 | 41717861 | 41715446 | 41715568 | 41754360 |
| SE | 36837 | ENSG00000168038 | ULK4 | chr3 | - | 41819422 | 41819506 | 41800131 | 41800293 | 41835863 |
| SE | 36838 | ENSG00000168038 | ULK4 | chr3 | - | 41835863 | 41835971 | 41819422 | 41819506 | 41883873 |
| SE | 36840 | ENSG00000168038 | ULK4 | chr3 | - | 41911316 | 41911386 | 41907844 | 41907941 | 41911540 |
| SE | 36850 | ENSG00000134504 | KCTD1 | chr18 | - | 26459619 | 26459925 | 26454909 | 26455901 | 26476514 |
| SE | 36851 | ENSG00000134504 | KCTD1 | chr18 | - | 26476514 | 26476659 | 26454909 | 26455901 | 26501071 |
| SE | 36861 | ENSG00000175792 | RUVBL1 | chr3 | - | 128101558 | 128101648 | 128100594 | 128100744 | 128104772 |
| SE | 36862 | ENSG00000175792 | RUVBL1 | chr3 | - | 128104772 | 128104924 | 128101558 | 128101648 | 128112887 |
| SE | 36863 | ENSG00000175792 | RUVBL1 | chr3 | - | 128112887 | 128113020 | 128104772 | 128104924 | 128119327 |
| SE | 36864 | ENSG00000175792 | RUVBL1 | chr3 | - | 128119327 | 128119414 | 128112887 | 128113020 | 128123583 |
| SE | 36865 | ENSG00000133422 | MORC2 | chr22 | - | 30927529 | 30927803 | 30926712 | 30926871 | 30928018 |
| SE | 36868 | ENSG00000133422 | MORC2 | chr22 | - | 30939958 | 30940041 | 30939620 | 30939706 | 30940757 |
| SE | 36870 | ENSG00000133422 | MORC2 | chr22 | - | 30963224 | 30963334 | 30958640 | 30958694 | 30967065 |
| SE | 36882 | ENSG00000122779 | TRIM24 | chr7 | + | 138576372 | 138576445 | 138573506 | 138573642 | 138577419 |
| SE | 36883 | ENSG00000122779 | TRIM24 | chr7 | + | 138580561 | 138580694 | 138579203 | 138579532 | 138581696 |
| SE | 36885 | ENSG00000109943 | CRTAM | chr11 | + | 122850067 | 122850214 | 122838499 | 122838592 | 122853942 |
| SE | 36895 | ENSG00000261040 | WFDC21P | chr17 | - | 60085931 | 60085985 | 60085102 | 60085203 | 60088196 |
| SE | 36896 | ENSG00000261040 | WFDC21P | chr17 | - | 60085931 | 60086079 | 60085102 | 60085203 | 60088196 |
| SE | 36897 | ENSG00000261040 | WFDC21P | chr17 | - | 60085931 | 60086126 | 60085054 | 60085203 | 60088196 |
| SE | 36901 | ENSG00000196126 | HLA-DRB1 | chr6 | - | 32580246 | 32580270 | 32578768 | 32579104 | 32580745 |
| SE | 36902 | ENSG00000196126 | HLA-DRB1 | chr6 | - | 32580745 | 32580856 | 32580246 | 32580270 | 32581556 |
| SE | 36903 | ENSG00000196126 | HLA-DRB1 | chr6 | - | 32581556 | 32581838 | 32580745 | 32580856 | 32584108 |
| SE | 36904 | ENSG00000184106 | TREML3P | chr6 | - | 41215761 | 41215819 | 41209598 | 41209730 | 41217126 |
| SE | 36905 | ENSG00000184106 | TREML3P | chr6 | - | 41215761 | 41215819 | 41209598 | 41209730 | 41217652 |
| SE | 36906 | ENSG00000184106 | TREML3P | chr6 | - | 41217126 | 41217177 | 41209598 | 41209730 | 41217652 |
| SE | 36907 | ENSG00000184106 | TREML3P | chr6 | - | 41217126 | 41217177 | 41215761 | 41215819 | 41217652 |
| SE | 36908 | ENSG00000158234 | FAIM | chr3 | + | 138610603 | 138611037 | 138608605 | 138608937 | 138621406 |
| SE | 36909 | ENSG00000158234 | FAIM | chr3 | + | 138619710 | 138619770 | 138608699 | 138608937 | 138621406 |
| SE | 36917 | ENSG00000135622 | SEMA4F | chr2 | + | 74662731 | 74662825 | 74657852 | 74657951 | 74673456 |
| SE | 36922 | ENSG00000135622 | SEMA4F | chr2 | + | 74675748 | 74675909 | 74675524 | 74675634 | 74679275 |
| SE | 36925 | ENSG00000164944 | VIRMA | chr8 | - | 94499373 | 94499506 | 94496327 | 94496480 | 94506499 |
| SE | 36926 | ENSG00000164944 | VIRMA | chr8 | - | 94510416 | 94510652 | 94509687 | 94509940 | 94511184 |
| SE | 36927 | ENSG00000164944 | VIRMA | chr8 | - | 94514868 | 94514951 | 94511995 | 94512089 | 94517787 |
| SE | 36928 | ENSG00000164944 | VIRMA | chr8 | - | 94517787 | 94517942 | 94514868 | 94514951 | 94518984 |
| SE | 36929 | ENSG00000164944 | VIRMA | chr8 | - | 94518984 | 94519476 | 94517787 | 94517942 | 94526222 |
| SE | 36930 | ENSG00000164944 | VIRMA | chr8 | - | 94530962 | 94531085 | 94529069 | 94529342 | 94534838 |
| SE | 36932 | ENSG00000143512 | HHIPL2 | chr1 | - | 222538886 | 222539062 | 222538728 | 222538774 | 222540009 |
| SE | 36936 | ENSG00000127329 | PTPRB | chr12 | - | 70594466 | 70594724 | 70591733 | 70592545 | 70596048 |

| | | | | | | | | | | | | | | | | | | | | | | | | | | | | | | | | | | | | | | | | | | | | | | |
|---|---|---|---|---|---|---|---|---|---|---|---|---|---|---|---|---|---|---|---|---|---|---|---|---|---|---|---|---|---|---|---|---|---|---|---|---|---|---|---|---|---|---|---|---|---|---|
| SE | 36944 | ENSG00000138459 | SLC35A5 | chr3 | + | 112573888 | 112573956 | 112570438 | 112570670 | 112580545 | 112580559 | 106 | 88 | 61 | 75 | 142 | 1 | 0 | 0 | 0 | 0 | 19 | 42 | 28 | 41 | 8 | 0 | 0 | 0 | 0 | 0 | 217 | 149 | 1 | 1 | 0.986 | 1 | 1 | 1 | 1 | 1 | 1 | 1 | 1 | 1 | -0.003 |
| SE | 36945 | ENSG00000138459 | SLC35A5 | chr3 | + | 112581178 | 112581326 | 112580545 | 112580706 | 112582670 | 112584156 | 40 | 31 | 30 | 31 | 47 | 0 | 0 | 0 | 0 | 0 | 0 | 33 | 14 | 6 | 5 | 0 | 4 | 0 | 0 | 0 | 297 | 149 | 1 | 1 | 1 | 1 | 1 | 1 | 1 | NA | 0.805 | 1 | 1 | 1 | 0.049 |
| SE | 36946 | ENSG00000138459 | SLC35A5 | chr3 | + | 112581244 | 112581326 | 112580545 | 112580706 | 112582670 | 112582730 | 23 | 22 | 20 | 23 | 39 | 0 | 0 | 0 | 0 | 0 | 0 | 18 | 13 | 2 | 0 | 0 | 4 | 0 | 0 | 0 | 231 | 149 | 1 | 1 | 1 | 1 | 1 | 1 | 1 | NA | 0.744 | 1 | 1 | NA | 0.085 |
| SE | 36947 | ENSG00000006704 | GTF2IRD1 | chr7 | + | 74544754 | 74544802 | 74539878 | 74539968 | 74545743 | 74545809 | 2 | 0 | 2 | 0 | 0 | 0 | 0 | 0 | 0 | 0 | 0 | 0 | 4 | 0 | 0 | 11 | 0 | 0 | 0 | 0 | 197 | 149 | 1 | 1 | 1 | NA | 1 | NA | NA | 0 | NA | 1 | NA | NA | 0.5 |
| SE | 36953 | ENSG00000178222 | RNF212 | chr4 | - | 1081418 | 1081467 | 1079642 | 1079688 | 1081566 | 1081619 | 0 | 3 | 0 | 2 | 10 | 0 | 0 | 1 | 0 | 0 | 0 | 2 | 0 | 0 | 0 | 0 | 0 | 1 | 0 | 0 | 198 | 149 | 1 | 1 | NA | 1 | 0 | 1 | 1 | NA | 1 | 0 | NA | NA | 0.25 |
| SE | 36954 | ENSG00000178222 | RNF212 | chr4 | - | 1081566 | 1081619 | 1079642 | 1079688 | 1085895 | 1085954 | 0 | 0 | 3 | 0 | 5 | 0 | 1 | 0 | 0 | 0 | 0 | 0 | 1 | 0 | 0 | 0 | 0 | 0 | 0 | 0 | 202 | 149 | 1 | 1 | NA | 0 | 1 | NA | 1 | NA | NA | 1 | NA | NA | -0.333 |
| SE | 36955 | ENSG00000178222 | RNF212 | chr4 | - | 1099715 | 1099927 | 1096764 | 1096839 | 1108342 | 1108404 | 0 | 1 | 0 | 0 | 3 | 0 | 0 | 2 | 2 | 4 | 0 | 4 | 0 | 5 | 0 | 6 | 0 | 0 | 0 | 4 | 298 | 149 | 1 | 1 | NA | 1 | 0 | 0 | 0.273 | 0 | 1 | NA | 1 | 0 | -0.182 |
| SE | 36956 | ENSG00000178222 | RNF212 | chr4 | - | 1099715 | 1099927 | 1096764 | 1096839 | 1113355 | 1113541 | 0 | 2 | 0 | 0 | 3 | 0 | 0 | 0 | 1 | 0 | 0 | 2 | 0 | 5 | 0 | 0 | 0 | 0 | 0 | 0 | 298 | 149 | 1 | 1 | NA | 1 | NA | 0 | 1 | NA | 1 | NA | 1 | NA | -0.333 |
| SE | 36957 | ENSG00000178222 | RNF212 | chr4 | - | 1108342 | 1108404 | 1096462 | 1096839 | 1113355 | 1113526 | 0 | 0 | 2 | 2 | 0 | 0 | 0 | 0 | 1 | 0 | 6 | 0 | 0 | 4 | 0 | 0 | 0 | 0 | 0 | 0 | 211 | 149 | 1 | 1 | NA | NA | 1 | 0.585 | NA | 1 | NA | NA | NA | 1 | -0.208 |
| SE | 36958 | ENSG00000178222 | RNF212 | chr4 | - | 1108342 | 1108404 | 1099715 | 1099927 | 1113355 | 1113526 | 0 | 0 | 0 | 0 | 0 | 0 | 1 | 0 | 0 | 0 | 0 | 2 | 0 | 0 | 0 | 0 | 0 | 0 | 0 | 0 | 211 | 149 | 1 | 1 | NA | 0 | NA | NA | NA | NA | 1 | NA | NA | NA | -1 |
| SE | 36960 | ENSG00000174564 | IL20RB | chr3 | + | 136989440 | 136989565 | 136982159 | 136982350 | 136995413 | 136995556 | 2 | 0 | 0 | 0 | 4 | 0 | 0 | 0 | 0 | 0 | 0 | 10 | 4 | 0 | 9 | 0 | 0 | 4 | 0 | 0 | 274 | 149 | 1 | 1 | 1 | NA | NA | NA | 1 | NA | 1 | 0.352 | NA | 1 | 0.216 |
| SE | 36962 | ENSG00000174564 | IL20RB | chr3 | + | 136995413 | 136995556 | 136982159 | 136982350 | 137010112 | 137011079 | 1 | 4 | 0 | 1 | 4 | 0 | 1 | 0 | 0 | 0 | 0 | 0 | 4 | 0 | 0 | 0 | 0 | 1 | 3 | 0 | 292 | 149 | 1 | 1 | 1 | 1 | 0 | 1 | 1 | NA | NA | 0.671 | 0 | NA | 0.465 |
| SE | 36968 | ENSG00000153914 | SREK1 | chr5 | + | 66170584 | 66170947 | 66163582 | 66170170 | 66174945 | 66175041 | 114 | 154 | 121 | 130 | 157 | 2 | 6 | 4 | 2 | 5 | 11 | 33 | 40 | 12 | 0 | 10 | 0 | 0 | 0 | 0 | 298 | 149 | 1 | 1 | 0.966 | 0.928 | 0.938 | 0.97 | 0.94 | 0.355 | 1 | 1 | 1 | NA | 0.11 |
| SE | 36973 | ENSG00000114125 | RNF7 | chr3 | + | 141743508 | 141743556 | 141738296 | 141738509 | 141745158 | 141745517 | 41 | 15 | 27 | 40 | 31 | 0 | 0 | 0 | 2 | 0 | 43 | 56 | 50 | 42 | 48 | 0 | 0 | 0 | 0 | 0 | 197 | 149 | 1 | 1 | 1 | 1 | 1 | 0.938 | 1 | 1 | 1 | 1 | 1 | 1 | -0.012 |
| SE | 36976 | ENSG00000137824 | RMDN3 | chr15 | - | 40738500 | 40738576 | 40737964 | 40738042 | 40740132 | 40740193 | 33 | 21 | 34 | 23 | 23 | 0 | 0 | 0 | 0 | 0 | 43 | 29 | 27 | 20 | 21 | 0 | 0 | 0 | 2 | 0 | 225 | 149 | 1 | 1 | 1 | 1 | 1 | 1 | 1 | 1 | 1 | 1 | 0.869 | 1 | 0.026 |
| SE | 36998 | ENSG00000233452 | STXBP5-AS1 | chr6 | - | 146850824 | 146851331 | 146843113 | 146844712 | 146911551 | 146911665 | 11 | 6 | 2 | 13 | 7 | 1 | 0 | 0 | 0 | 0 | 11 | 6 | 8 | 18 | 55 | 0 | 0 | 0 | 0 | 0 | 298 | 149 | 1 | 1 | 0.846 | 1 | 1 | 1 | 1 | 1 | 1 | 1 | 1 | 1 | -0.031 |
| SE | 37002 | ENSG00000233452 | STXBP5-AS1 | chr6 | - | 146860617 | 146860743 | 146850824 | 146851331 | 146911551 | 146911663 | 0 | 1 | 0 | 4 | 5 | 0 | 1 | 0 | 13 | 4 | 0 | 4 | 8 | 0 | 20 | 11 | 6 | 1 | 10 | 43 | 275 | 149 | 1 | 1 | NA | 0.351 | NA | 0.143 | 0.404 | 0 | 0.265 | 0.813 | 0 | 0.201 | 0.044 |
| SE | 37003 | ENSG00000233452 | STXBP5-AS1 | chr6 | - | 146860617 | 146860743 | 146850824 | 146851331 | 146972044 | 146972070 | 0 | 2 | 0 | 3 | 2 | 0 | 2 | 0 | 0 | 0 | 0 | 2 | 2 | 0 | 13 | 0 | 0 | 2 | 4 | 0 | 275 | 149 | 1 | 1 | NA | 0.351 | NA | 1 | 1 | NA | 1 | 0.351 | 0 | 1 | 0.196 |
| SE | 37007 | ENSG00000233452 | STXBP5-AS1 | chr6 | - | 146864383 | 146864602 | 146860617 | 146860743 | 146911551 | 146911665 | 1 | 0 | 2 | 0 | 0 | 0 | 0 | 0 | 2 | 3 | 0 | 0 | 0 | 0 | 0 | 0 | 2 | 6 | 0 | 7 | 298 | 149 | 1 | 1 | 1 | NA | 1 | 0 | 0 | NA | 0 | 0 | NA | 0 | 0.5 |
| SE | 37010 | ENSG00000233452 | STXBP5-AS1 | chr6 | - | 146911551 | 146911663 | 146851117 | 146851331 | 146972044 | 146972070 | 0 | 1 | 0 | 10 | 4 | 0 | 2 | 0 | 0 | 0 | 11 | 6 | 1 | 9 | 43 | 0 | 0 | 2 | 4 | 0 | 261 | 149 | 1 | 1 | NA | 0.222 | NA | 1 | 1 | 1 | 1 | 0.222 | 0.562 | 1 | -0.016 |
| SE | 37013 | ENSG00000233452 | STXBP5-AS1 | chr6 | - | 146911551 | 146911663 | 146860617 | 146860743 | 146972044 | 146972070 | 0 | 0 | 0 | 2 | 3 | 0 | 1 | 0 | 1 | 0 | 0 | 0 | 5 | 0 | 7 | 0 | 0 | 0 | 0 | 0 | 261 | 149 | 1 | 1 | NA | 0 | NA | 0.533 | 1 | NA | NA | 1 | NA | 1 | -0.489 |
| SE | 37017 | ENSG00000233452 | STXBP5-AS1 | chr6 | - | 146911551 | 146911665 | 146850820 | 146851331 | 146972044 | 146972070 | 1 | 2 | 0 | 12 | 4 | 0 | 2 | 0 | 0 | 0 | 11 | 6 | 1 | 9 | 43 | 0 | 0 | 2 | 4 | 0 | 263 | 149 | 1 | 1 | 1 | 0.362 | NA | 1 | 1 | 1 | 1 | 0.221 | 0.56 | 1 | 0.084 |
| SE | 37020 | ENSG00000233452 | STXBP5-AS1 | chr6 | - | 146911551 | 146911665 | 146860617 | 146860743 | 146915675 | 146915885 | 0 | 2 | 4 | 6 | 8 | 0 | 0 | 0 | 0 | 0 | 36 | 9 | 10 | 7 | 47 | 0 | 0 | 0 | 0 | 7 | 263 | 149 | 1 | 1 | NA | 1 | 1 | 1 | 1 | 1 | 1 | 1 | 1 | 0.792 | 0.042 |
| SE | 37021 | ENSG00000233452 | STXBP5-AS1 | chr6 | - | 146911551 | 146911665 | 146860617 | 146860743 | 146972044 | 146972070 | 1 | 1 | 0 | 2 | 3 | 0 | 1 | 0 | 1 | 0 | 0 | 0 | 6 | 0 | 7 | 0 | 0 | 0 | 0 | 0 | 263 | 149 | 1 | 1 | 1 | 0.362 | NA | 0.531 | 1 | NA | NA | 1 | NA | 1 | -0.277 |
| SE | 37025 | ENSG00000233452 | STXBP5-AS1 | chr6 | - | 146962390 | 146962566 | 146911551 | 146911665 | 146972044 | 146972070 | 0 | 0 | 1 | 0 | 0 | 1 | 1 | 0 | 2 | 0 | 0 | 0 | 0 | 0 | 6 | 0 | 0 | 1 | 0 | 0 | 298 | 149 | 1 | 1 | 0 | 0 | 1 | 0 | NA | NA | NA | 0 | NA | 1 | -0.25 |
| SE | 37033 | ENSG00000233452 | STXBP5-AS1 | chr6 | - | 146972044 | 146972070 | 146911551 | 146911665 | 147001440 | 147001543 | 1 | 1 | 1 | 5 | 0 | 0 | 0 | 0 | 1 | 0 | 0 | 4 | 1 | 0 | 0 | 0 | 0 | 0 | 0 | 0 | 175 | 149 | 1 | 1 | 1 | 1 | 1 | 0.81 | NA | NA | 1 | 1 | NA | NA | -0.047 |
| SE | 37042 | ENSG00000233452 | STXBP5-AS1 | chr6 | - | 147001440 | 147001543 | 146972044 | 146972070 | 147074804 | 147074847 | 2 | 7 | 3 | 5 | 7 | 0 | 0 | 0 | 1 | 0 | 0 | 8 | 5 | 6 | 12 | 0 | 0 | 0 | 0 | 0 | 252 | 149 | 1 | 1 | 1 | 1 | 1 | 0.747 | 1 | NA | 1 | 1 | 1 | 1 | -0.051 |
| SE | 37044 | ENSG00000233452 | STXBP5-AS1 | chr6 | - | 147061543 | 147061670 | 147001440 | 147001543 | 147074804 | 147074847 | 0 | 0 | 0 | 2 | 0 | 1 | 3 | 2 | 5 | 7 | 0 | 0 | 0 | 0 | 0 | 0 | 8 | 3 | 6 | 6 | 276 | 149 | 1 | 1 | 0 | 0 | 0 | 0.178 | 0 | NA | 0 | 0 | 0 | 0 | 0.036 |
| SE | 37045 | ENSG00000233452 | STXBP5-AS1 | chr6 | - | 147073243 | 147073357 | 147001440 | 147001543 | 147074804 | 147074847 | 2 | 0 | 0 | 0 | 0 | 1 | 3 | 2 | 5 | 7 | 0 | 0 | 0 | 0 | 0 | 0 | 8 | 3 | 6 | 6 | 263 | 149 | 1 | 1 | 0.531 | 0 | 0 | 0 | 0 | NA | 0 | 0 | 0 | 0 | 0.106 |
| SE | 37050 | ENSG00000233452 | STXBP5-AS1 | chr6 | - | 147108471 | 147108610 | 147074804 | 147074847 | 147119416 | 147119477 | 4 | 0 | 1 | 2 | 10 | 1 | 3 | 1 | 6 | 4 | 3 | 8 | 0 | 0 | 0 | 0 | 0 | 3 | 0 | 0 | 288 | 149 | 1 | 1 | 0.674 | 0 | 0.341 | 0.147 | 0.564 | 1 | 1 | 0 | NA | NA | -0.321 |
| SE | 37051 | ENSG00000233452 | STXBP5-AS1 | chr6 | - | 147108471 | 147108610 | 147074804 | 147074847 | 147173592 | 147173647 | 1 | 0 | 1 | 0 | 0 | 0 | 0 | 0 | 0 | 5 | 0 | 4 | 0 | 0 | 5 | 0 | 0 | 0 | 6 | 6 | 288 | 149 | 1 | 1 | 1 | NA | 1 | NA | 0 | NA | 1 | NA | 0 | 0.301 | 0.233 |
| SE | 37052 | ENSG00000233452 | STXBP5-AS1 | chr6 | - | 147119416 | 147119477 | 147074804 | 147074847 | 147173645 | 147173647 | 3 | 5 | 1 | 5 | 13 | 0 | 0 | 0 | 0 | 5 | 0 | 1 | 3 | 0 | 0 | 0 | 0 | 0 | 6 | 6 | 210 | 149 | 1 | 1 | 1 | 1 | 1 | 1 | 0.648 | NA | 1 | 1 | 0 | 0 | 0.43 |
| SE | 37053 | ENSG00000233452 | STXBP5-AS1 | chr6 | - | 147119416 | 147119477 | 147108471 | 147108610 | 147173592 | 147173647 | 5 | 2 | 0 | 1 | 14 | 0 | 0 | 0 | 0 | 0 | 3 | 5 | 0 | 0 | 0 | 0 | 0 | 0 | 0 | 5 | 210 | 149 | 1 | 1 | 1 | 1 | NA | 1 | 1 | 1 | 1 | NA | NA | 0 | 0.333 |
| SE | 37054 | ENSG00000233452 | STXBP5-AS1 | chr6 | - | 147170012 | 147170083 | 147119416 | 147119477 | 147173592 | 147173647 | 0 | 3 | 0 | 3 | 10 | 2 | 3 | 1 | 4 | 18 | 0 | 0 | 1 | 0 | 0 | 3 | 5 | 1 | 0 | 0 | 220 | 149 | 1 | 1 | 0 | 0.404 | 0 | 0.337 | 0.273 | 0 | 0 | 0.404 | NA | NA | 0.068 |
| SE | 37064 | ENSG00000069535 | MAOB | chrX | - | 43775174 | 43775272 | 43769306 | 43769418 | 43778681 | 43778739 | 4 | 6 | 0 | 8 | 7 | 0 | 0 | 0 | 0 | 0 | 37 | 41 | 15 | 10 | 382 | 0 | 0 | 0 | 0 | 9 | 247 | 149 | 1 | 1 | 1 | 1 | NA | 1 | 1 | 1 | 1 | 1 | 1 | 0.962 | 0.008 |
| SE | 37072 | ENSG00000005812 | FBXL3 | chr13 | - | 77015408 | 77015580 | 77005259 | 77007788 | 77018599 | 77018686 | 405 | 233 | 328 | 319 | 385 | 0 | 0 | 0 | 1 | 0 | 92 | 205 | 275 | 84 | 39 | 0 | 0 | 1 | 0 | 0 | 298 | 149 | 1 | 1 | 1 | 1 | 1 | 0.994 | 1 | 1 | 1 | 0.993 | 1 | 1 | 0 |
| SE | 37085 | ENSG00000127481 | UBR4 | chr1 | - | 19110741 | 19110832 | 19110379 | 19110464 | 19112523 | 19112867 | 117 | 82 | 121 | 147 | 192 | 0 | 0 | 0 | 0 | 0 | 283 | 218 | 138 | 192 | 204 | 0 | 0 | 3 | 0 | 1 | 240 | 149 | 1 | 1 | 1 | 1 | 1 | 1 | 1 | 1 | 1 | 0.966 | 1 | 0.992 | 0.008 |
| SE | 37086 | ENSG00000127481 | UBR4 | chr1 | - | 19113698 | 19113827 | 19112523 | 19112867 | 19113944 | 19114070 | 328 | 219 | 343 | 354 | 412 | 0 | 0 | 0 | 0 | 0 | 513 | 453 | 293 | 475 | 452 | 0 | 0 | 0 | 3 | 0 | 278 | 149 | 1 | 1 | 1 | 1 | 1 | 1 | 1 | 1 | 1 | 1 | 0.988 | 1 | 0.002 |
| SE | 37088 | ENSG00000127481 | UBR4 | chr1 | - | 19118871 | 19118957 | 19117220 | 19117414 | 19119556 | 19119701 | 149 | 111 | 173 | 196 | 321 | 0 | 0 | 0 | 0 | 0 | 368 | 240 | 216 | 350 | 307 | 0 | 0 | 0 | 1 | 0 | 235 | 149 | 1 | 1 | 1 | 1 | 1 | 1 | 1 | 1 | 1 | 1 | 0.996 | 1 | 0.001 |
| SE | 37089 | ENSG00000127481 | UBR4 | chr1 | - | 19120179 | 19120348 | 19119556 | 19119701 | 19121188 | 19121434 | 195 | 120 | 186 | 209 | 375 | 0 | 0 | 0 | 0 | 0 | 445 | 188 | 196 | 313 | 292 | 0 | 0 | 2 | 0 | 0 | 298 | 149 | 1 | 1 | 1 | 1 | 1 | 1 | 1 | 1 | 1 | 0.98 | 1 | 1 | 0.004 |
| SE | 37090 | ENSG00000127481 | UBR4 | chr1 | - | 19125602 | 19125699 | 19124476 | 19124690 | 19126445 | 19126655 | 4 | 0 | 0 | 0 | 0 | 148 | 120 | 147 | 169 | 272 | 0 | 0 | 2 | 0 | 0 | 299 | 206 | 202 | 168 | 314 | 246 | 149 | 1 | 1 | 0.016 | 0 | 0 | 0 | 0 | 0 | 0 | 0.006 | 0 | 0 | 0.002 |
| SE | 37091 | ENSG00000127481 | UBR4 | chr1 | - | 19126445 | 19126655 | 19125540 | 19124690 | 19127622 | 19127701 | 253 | 182 | 255 | 300 | 449 | 0 | 1 | 0 | 0 | 0 | 467 | 362 | 281 | 292 | 469 | 0 | 0 | 1 | 2 | 0 | 298 | 149 | 1 | 1 | 1 | 0.989 | 1 | 1 | 1 | 1 | 1 | 0.993 | 0.986 | 1 | 0.002 |
| SE | 37092 | ENSG00000127481 | UBR4 | chr1 | - | 19128210 | 19128318 | 19127622 | 19127739 | 19128977 | 19129074 | 177 | 100 | 260 | 229 | 178 | 0 | 0 | 0 | 0 | 0 | 336 | 206 | 155 | 208 | 188 | 0 | 4 | 0 | 1 | 0 | 257 | 149 | 1 | 1 | 1 | 1 | 1 | 1 | 1 | 1 | 0.968 | 1 | 0.992 | 1 | 0.008 |
| SE | 37093 | ENSG00000127481 | UBR4 | chr1 | - | 19128977 | 19129074 | 19128210 | 19128318 | 19138006 | 19138181 | 147 | 89 | 239 | 203 | 142 | 0 | 0 | 0 | 0 | 0 | 260 | 185 | 145 | 174 | 171 | 0 | 0 | 1 | 0 | 0 | 246 | 149 | 1 | 1 | 1 | 1 | 1 | 1 | 1 | 1 | 1 | 0.989 | 1 | 1 | 0.002 |
| SE | 37094 | ENSG00000127481 | UBR4 | chr1 | - | 19140787 | 19140892 | 19139082 | 19139220 | 19141346 | 19141524 | 144 | 58 | 71 | 117 | 67 | 0 | 1 | 0 | 0 | 0 | 131 | 123 | 115 | 172 | 200 | 0 | 0 | 0 | 0 | 0 | 254 | 149 | 1 | 1 | 1 | 0.971 | 1 | 1 | 1 | 1 | 1 | 1 | 1 | 1 | -0.006 |
| SE | 37095 | ENSG00000127481 | UBR4 | chr1 | - | 19140787 | 19140892 | 19139082 | 19139220 | 19143979 | 19144091 | 40 | 19 | 14 | 35 | 18 | 0 | 0 | 0 | 0 | 0 | 41 | 44 | 37 | 61 | 51 | 0 | 0 | 0 | 3 | 0 | 254 | 149 | 1 | 1 | 1 | 1 | 1 | 1 | 1 | 1 | 1 | 1 | 0.923 | 1 | 0.015 |
| SE | 37097 | ENSG00000127481 | UBR4 | chr1 | - | 19149762 | 19149795 | 19148562 | 19148626 | 19150576 | 19150793 | 2 | 1 | 0 | 0 | 0 | 69 | 45 | 88 | 87 | 55 | 0 | 0 | 0 | 0 | 0 | 97 | 122 | 105 | 135 | 163 | 182 | 149 | 1 | 1 | 0.023 | 0.018 | 0 | 0 | 0 | 0 | 0 | 0 | 0 | 0 | 0.008 |
| SE | 37103 | ENSG00000127481 | UBR4 | chr1 | - | 19174318 | 19174447 | 19173438 | 19173621 | 19174953 | 19175033 | 317 | 217 | 281 | 332 | 513 | 0 | 0 | 0 | 0 | 0 | 416 | 548 | 326 | 501 | 461 | 0 | 0 | 0 | 3 | 0 | 278 | 149 | 1 | 1 | 1 | 1 | 1 | 1 | 1 | 1 | 1 | 1 | 0.989 | 1 | 0.002 |
| SE | 37104 | ENSG00000127481 | UBR4 | chr1 | - | 19174318 | 19174447 | 19173438 | 19173621 | 19176591 | 19176727 | 177 | 123 | 161 | 172 | 276 | 1 | 0 | 0 | 0 | 0 | 250 | 277 | 172 | 250 | 238 | 0 | 0 | 0 | 3 | 0 | 278 | 149 | 1 | 1 | 0.99 | 1 | 1 | 1 | 1 | 1 | 1 | 1 | 0.978 | 1 | 0.002 |
| SE | 37105 | ENSG00000127481 | UBR4 | chr1 | - | 19174953 | 19175033 | 19173438 | 19173621 | 19176591 | 19176727 | 138 | 62 | 132 | 135 | 141 | 1 | 0 | 0 | 0 | 0 | 170 | 236 | 143 | 206 | 168 | 0 | 0 | 0 | 3 | 0 | 229 | 149 | 1 | 1 | 0.989 | 1 | 1 | 1 | 1 | 1 | 1 | 1 | 0.978 | 1 | 0.002 |
| SE | 37106 | ENSG00000127481 | UBR4 | chr1 | - | 19174953 | 19175033 | 19174318 | 19174447 | 19176591 | 19176727 | 277 | 157 | 254 | 295 | 375 | 0 | 2 | 0 | 0 | 0 | 327 | 507 | 298 | 457 | 391 | 0 | 0 | 1 | 0 | 0 | 229 | 149 | 1 | 1 | 1 | 0.981 | 1 | 1 | 1 | 1 | 1 | 0.995 | 1 | 1 | -0.003 |
| SE | 37107 | ENSG00000127481 | UBR4 | chr1 | - | 19185098 | 19185286 | 19184015 | 19184175 | 19186539 | 19186657 | 268 | 148 | 227 | 287 | 274 | 0 | 0 | 0 | 0 | 0 | 282 | 385 | 253 | 252 | 438 | 0 | 4 | 0 | 0 | 0 | 298 | 149 | 1 | 1 | 1 | 1 | 1 | 1 | 1 | 1 | 0.98 | 1 | 1 | 1 | 0.004 |
| SE | 37109 | ENSG00000127481 | UBR4 | chr1 | - | 19192480 | 19192540 | 19187440 | 19187540 | 19193432 | 19193557 | 11 | 4 | 11 | 10 | 6 | 1 | 0 | 0 | 0 | 0 | 24 | 7 | 36 | 44 | 13 | 0 | 0 | 3 | 0 | 0 | 209 | 149 | 1 | 1 | 0.887 | 1 | 1 | 1 | 1 | 1 | 1 | 0.895 | 1 | 1 | -0.002 |
| SE | 37111 | ENSG00000127481 | UBR4 | chr1 | - | 19199650 | 19199754 | 19198798 | 19198928 | 19201717 | 19201815 | 378 | 217 | 277 | 358 | 344 | 0 | 0 | 0 | 2 | 4 | 433 | 515 | 328 | 365 | 551 | 6 | 0 | 1 | 2 | 0 | 253 | 149 | 1 | 1 | 1 | 1 | 1 | 0.991 | 0.981 | 0.977 | 1 | 0.995 | 0.991 | 1 | 0.002 |
| SE | 37112 | ENSG00000127481 | UBR4 | chr1 | - | 19201717 | 19201815 | 19199650 | 19199754 | 19210072 | 19210266 | 263 | 148 | 187 | 248 | 174 | 1 | 0 | 0 | 0 | 0 | 317 | 305 | 221 | 223 | 348 | 0 | 0 | 0 | 0 | 0 | 247 | 149 | 1 | 1 | 0.994 | 1 | 1 | 1 | 1 | 1 | 1 | 1 | 1 | 1 | -0.001 |
| SE | 37116 | ENSG00000086475 | SEPHS1 | chr10 | - | 13322834 | 13323047 | 13317427 | 13319356 | 13328350 | 13328450 | 129 | 77 | 149 | 121 | 100 | 0 | 0 | 0 | 1 | 0 | 112 | 148 | 152 | 164 | 159 | 0 | 0 | 0 | 2 | 0 | 298 | 149 | 1 | 1 | 1 | 1 | 1 | 0.984 | 1 | 1 | 1 | 1 | 0.976 | 1 | 0.002 |
| SE | 37117 | ENSG00000086475 | SEPHS1 | chr10 | - | 13333816 | 13333971 | 13329697 | 13329788 | 13336242 | 13336350 | 143 | 149 | 196 | 116 | 244 | 0 | 1 | 0 | 0 | 0 | 263 | 120 | 164 | 191 | 158 | 0 | 0 | 0 | 0 | 0 | 298 | 149 | 1 | 1 | 1 | 0.987 | 1 | 1 | 1 | 1 | 1 | 1 | 1 | 1 | -0.003 |
| SE | 37119 | ENSG00000261553 | NA | chr13 | + | 75594914 | 75594990 | 75569459 | 75569507 | 75621762 | 75621868 | 29 | 32 | 25 | 33 | 51 | 2 | 0 | 0 | 0 | 0 | 0 | 13 | 6 | 20 | 4 | 0 | 0 | 0 | 0 | 0 | 225 | 149 | 1 | 1 | 0.906 | 1 | 1 | 1 | 1 | NA | 1 | 1 | 1 | 1 | -0.019 |
| SE | 37120 | ENSG00000261553 | NA | chr13 | + | 75623270 | 75623320 | 75621762 | 75621868 | 75713181 | 75713252 | 17 | 3 | 5 | 19 | 11 | 2 | 0 | 0 | 0 | 0 | 32 | 0 | 4 | 3 | 3 | 0 | 0 | 0 | 0 | 21 | 199 | 149 | 1 | 1 | 0.864 | 1 | 1 | 1 | 1 | 1 | NA | 1 | 1 | 0.097 | 0.199 |
| SE | 37121 | ENSG00000144848 | ATG3 | chr3 | - | 112550191 | 112550262 | 112548532 | 112548640 | 112553279 | 112553329 | 453 | 413 | 453 | 470 | 689 | 0 | 1 | 0 | 5 | 0 | 932 | 331 | 396 | 623 | 434 | 0 | 0 | 0 | 4 | 0 | 220 | 149 | 1 | 1 | 1 | 0.996 | 1 | 0.985 | 1 | 1 | 1 | 1 | 0.991 | 1 | -0.002 |
| SE | 37122 | ENSG00000144848 | ATG3 | chr3 | - | 112550191 | 112550262 | 112548532 | 112548640 | 112558375 | 112558417 | 202 | 211 | 215 | 222 | 377 | 0 | 0 | 0 | 0 | 0 | 432 | 108 | 197 | 321 | 225 | 0 | 2 | 0 | 0 | 0 | 220 | 149 | 1 | 1 | 1 | 1 | 1 | 1 | 1 | 1 | 0.973 | 1 | 1 | 1 | 0.005 |
| SE | 37123 | ENSG00000144848 | ATG3 | chr3 | - | 112553279 | 112553329 | 112548532 | 112548640 | 112558375 | 112558417 | 10 | 22 | 14 | 26 | 19 | 0 | 0 | 0 | 0 | 0 | 27 | 25 | 43 | 18 | 2 | 0 | 2 | 0 | 0 | 0 | 199 | 149 | 1 | 1 | 1 | 1 | 1 | 1 | 1 | 1 | 0.903 | 1 | 1 | 1 | 0.019 |
| SE | 37127 | ENSG00000172139 | SLC9C1 | chr3 | - | 112278728 | 112278857 | 112277694 | 112277860 | 112280682 | 112280783 | 1 | 0 | 3 | 4 | 2 | 0 | 0 | 0 | 0 | 0 | 0 | 1 | 0 | 0 | 2 | 0 | 1 | 0 | 0 | 0 | 278 | 149 | 1 | 1 | 1 | NA | 1 | 1 | 1 | NA | 0.349 | NA | NA | 1 | 0.326 |
| SE | 37129 | ENSG00000121578 | B4GALT4 | chr3 | - | 119218649 | 119218772 | 119211739 | 119216344 | 119224057 | 119224245 | 76 | 39 | 64 | 26 | 69 | 2 | 0 | 0 | 5 | 6 | 39 | 53 | 64 | 43 | 0 | 0 | 11 | 3 | 2 | 0 | 272 | 149 | 1 | 1 | 0.954 | 1 | 1 | 0.74 | 0.863 | 1 | 0.725 | 0.921 | 0.922 | NA | 0.019 |
| SE | 37130 | ENSG00000121578 | B4GALT4 | chr3 | - | 119218649 | 119218772 | 119211739 | 119216344 | 119226808 | 119227041 | 50 | 18 | 42 | 16 | 48 | 1 | 0 | 0 | 0 | 0 | 26 | 26 | 24 | 27 | 0 | 0 | 0 | 0 | 0 | 0 | 272 | 149 | 1 | 1 | 0.965 | 1 | 1 | 1 | 1 | 1 | 1 | 1 | 1 | NA | -0.007 |
| SE | 37131 | ENSG00000121578 | B4GALT4 | chr3 | - | 119218711 | 119218772 | 119216238 | 119216344 | 119224057 | 119224245 | 26 | 25 | 22 | 8 | 28 | 2 | 0 | 0 | 5 | 6 | 8 | 14 | 67 | 25 | 0 | 0 | 11 | 3 | 2 | 0 | 210 | 149 | 1 | 1 | 0.902 | 1 | 1 | 0.532 | 0.768 | 1 | 0.475 | 0.941 | 0.899 | NA | 0.012 |
| SE | 37132 | ENSG00000121578 | B4GALT4 | chr3 | - | 119218711 | 119218772 | 119216238 | 119216344 | 119226808 | 119227041 | 4 | 6 | 4 | 1 | 3 | 1 | 0 | 0 | 0 | 0 | 0 | 3 | 18 | 8 | 0 | 0 | 0 | 0 | 0 | 0 | 210 | 149 | 1 | 1 | 0.739 | 1 | 1 | 1 | 1 | NA | 1 | 1 | 1 | NA | -0.052 |
| SE | 37133 | ENSG00000121578 | B4GALT4 | chr3 | - | 119224057 | 119224245 | 119211739 | 119216344 | 119226808 | 119227041 | 49 | 28 | 36 | 31 | 42 | 1 | 0 | 0 | 0 | 0 | 4 | 48 | 63 | 55 | 18 | 0 | 0 | 0 | 0 | 0 | 298 | 149 | 1 | 1 | 0.961 | 1 | 1 | 1 | 1 | 1 | 1 | 1 | 1 | 1 | -0.008 |
| SE | 37134 | ENSG00000121578 | B4GALT4 | chr3 | - | 119224057 | 119224245 | 119218649 | 119218772 | 119226808 | 119227041 | 81 | 57 | 68 | 36 | 67 | 2 | 0 | 1 | 0 | 0 | 17 | 64 | 124 | 76 | 18 | 0 | 0 | 3 | 0 | 0 | 298 | 149 | 1 | 1 | 0.953 | 1 | 0.971 | 1 | 1 | 1 | 1 | 0.954 | 1 | 1 | -0.006 |
| SE | 37138 | ENSG00000121578 | B4GALT4 | chr3 | - | 119235144 | 119235227 | 119226808 | 119227041 | 119236852 | 119237070 | 0 | 6 | 0 | 4 | 0 | 4 | 0 | 2 | 0 | 0 | 7 | 17 | 16 | 4 | 0 | 0 | 0 | 3 | 0 | 0 | 232 | 149 | 1 | 1 | 0 | 1 | 0 | 1 | NA | 1 | 1 | 0.774 | 1 | NA | -0.444 |
| SE | 37142 | ENSG00000121578 | B4GALT4 | chr3 | - | 119236852 | 119237070 | 119229846 | 119230244 | 119240849 | 119240878 | 48 | 16 | 36 | 25 | 14 | 0 | 0 | 0 | 0 | 0 | 8 | 67 | 23 | 18 | 16 | 0 | 0 | 3 | 0 | 0 | 298 | 149 | 1 | 1 | 1 | 1 | 1 | 1 | 1 | 1 | 0.918 | 1 | 1 | 1 | 0.016 |
| SE | 37143 | ENSG00000121578 | B4GALT4 | chr3 | - | 119240315 | 119240530 | 119236852 | 119237070 | 119240849 | 119240907 | 1 | 0 | 0 | 0 | 0 | 4 | 6 | 10 | 5 | 0 | 0 | 9 | 4 | 0 | 0 | 4 | 9 | 9 | 4 | 6 | 298 | 149 | 1 | 1 | 0.111 | 0 | 0 | 0 | NA | 0 | 0.333 | 0.182 | 0 | 0 | -0.075 |
| SE | 37145 | ENSG00000114520 | SNX4 | chr3 | - | 125457265 | 125457365 | 125453809 | 125453955 | 125460770 | 125460801 | 183 | 145 | 113 | 138 | 174 | 0 | 0 | 0 | 4 | 0 | 14 | 83 | 78 | 59 | 21 | 0 | 0 | 0 | 0 | 0 | 249 | 149 | 1 | 1 | 1 | 1 | 1 | 0.954 | 1 | 1 | 1 | 1 | 1 | 1 | -0.009 |
| SE | 37146 | ENSG00000114520 | SNX4 | chr3 | - | 125480246 | 125480319 | 125476694 | 125476756 | 125497340 | 125497388 | 56 | 36 | 25 | 45 | 39 | 1 | 0 | 0 | 0 | 0 | 13 | 24 | 15 | 8 | 13 | 0 | 0 | 0 | 0 | 0 | 222 | 149 | 1 | 1 | 0.974 | 1 | 1 | 1 | 1 | 1 | 1 | 1 | 1 | 1 | -0.005 |
| SE | 37148 | ENSG00000114520 | SNX4 | chr3 | - | 125498058 | 125498194 | 125497983 | 125497983 | 125504622 | 125504744 | 220 | 146 | 152 | 159 | 299 | 0 | 0 | 0 | 0 | 0 | 92 | 180 | 119 | 60 | 26 | 0 | 0 | 0 | 2 | 0 | 285 | 149 | 1 | 1 | 1 | 1 | 1 | 1 | 1 | 1 | 1 | 1 | 0.94 | 1 | 0.012 |
| SE | 37149 | ENSG00000114520 | SNX4 | chr3 | - | 125504622 | 125504744 | 125498058 | 125498194 | 125520031 | 125520191 | 105 | 66 | 64 | 65 | 150 | 2 | 0 | 0 | 0 | 1 | 13 | 65 | 59 | 16 | 18 | 0 | 0 | 2 | 0 | 0 | 271 | 149 | 1 | 1 | 0.967 | 1 | 1 | 1 | 0.988 | 1 | 1 | 0.942 | 1 | 1 | 0.003 |
| SE | 37150 | ENSG00000151726 | ACSL1 | chr4 | - | 184763166 | 184763255 | 184762406 | 184762523 | 184764852 | 184764925 | 175 | 113 | 117 | 156 | 178 | 0 | 0 | 0 | 0 | 0 | 437 | 506 | 1139 | 607 | 250 | 1 | 0 | 0 | 0 | 0 | 238 | 149 | 1 | 1 | 1 | 1 | 1 | 1 | 1 | 0.996 | 1 | 1 | 1 | 1 | 0.001 |
| SE | 37151 | ENSG00000151726 | ACSL1 | chr4 | - | 184764852 | 184764925 | 184763166 | 184763255 | 184765890 | 184765986 | 216 | 126 | 145 | 176 | 136 | 0 | 0 | 0 | 0 | 0 | 545 | 603 | 1354 | 744 | 308 | 0 | 0 | 1 | 0 | 0 | 222 | 149 | 1 | 1 | 1 | 1 | 1 | 1 | 1 | 1 | 1 | 0.999 | 1 | 1 | 0 |
| SE | 37152 | ENSG00000151726 | ACSL1 | chr4 | - | 184765890 | 184765986 | 184764852 | 184764925 | 184766621 | 184766702 | 162 | 102 | 118 | 137 | 120 | 0 | 0 | 0 | 0 | 0 | 390 | 421 | 958 | 534 | 227 | 0 | 0 | 1 | 0 | 0 | 245 | 149 | 1 | 1 | 1 | 1 | 1 | 1 | 1 | 1 | 1 | 0.998 | 1 | 1 | 0 |
| SE | 37153 | ENSG00000151726 | ACSL1 | chr4 | - | 184770254 | 184770332 | 184768315 | 184768450 | 184773080 | 184773154 | 6 | 6 | 7 | 7 | 13 | 0 | 0 | 0 | 0 | 0 | 28 | 44 | 12 | 36 | 0 | 0 | 2 | 0 | 4 | 0 | 227 | 149 | 1 | 1 | 1 | 1 | 1 | 1 | 1 | 1 | 0.935 | 1 | 0.855 | NA | 0.052 |
| SE | 37154 | ENSG00000151726 | ACSL1 | chr4 | - | 184770398 | 184770476 | 184768029 | 184768450 | 184773080 | 184773154 | 154 | 94 | 92 | 106 | 237 | 0 | 0 | 0 | 0 | 0 | 259 | 252 | 533 | 289 | 110 | 0 | 2 | 0 | 4 | 0 | 227 | 149 | 1 | 1 | 1 | 1 | 1 | 1 | 1 | 1 | 0.988 | 1 | 0.979 | 1 | 0.007 |
| SE | 37160 | ENSG00000151726 | ACSL1 | chr4 | - | 184788378 | 184788513 | 184783926 | 184783991 | 184788616 | 184788731 | 0 | 0 | 1 | 0 | 0 | 90 | 64 | 68 | 65 | 75 | 0 | 2 | 3 | 4 | 0 | 266 | 328 | 864 | 472 | 141 | 284 | 149 | 1 | 1 | 0 | 0 | 0.008 | 0 | 0 | 0 | 0.003 | 0.002 | 0.004 | 0 | 0 |
| SE | 37161 | ENSG00000151726 | ACSL1 | chr4 | - | 184788616 | 184788731 | 184783926 | 184783991 | 184825915 | 184826119 | 81 | 54 | 53 | 60 | 73 | 0 | 0 | 0 | 0 | 0 | 219 | 296 | 785 | 436 | 141 | 0 | 0 | 2 | 0 | 0 | 264 | 149 | 1 | 1 | 1 | 1 | 1 | 1 | 1 | 1 | 1 | 0.996 | 1 | 1 | 0.001 |
| SE | 37185 | ENSG00000114446 | IFT57 | chr3 | - | 108163662 | 108163729 | 108160811 | 108162655 | 108167792 | 108167864 | 17 | 13 | 26 | 16 | 33 | 0 | 0 | 0 | 0 | 7 | 8 | 0 | 7 | 1 | 0 | 0 | 0 | 0 | 0 | 0 | 216 | 149 | 1 | 1 | 1 | 1 | 1 | 1 | 0.765 | 1 | NA | 1 | 1 | NA | -0.047 |
| SE | 37187 | ENSG00000114446 | IFT57 | chr3 | - | 108166853 | 108166985 | 108165430 | 108165493 | 108167792 | 108167864 | 218 | 160 | 247 | 187 | 284 | 0 | 0 | 0 | 0 | 4 | 108 | 165 | 146 | 100 | 12 | 0 | 1 | 0 | 0 | 0 | 281 | 149 | 1 | 1 | 1 | 1 | 1 | 1 | 0.974 | 1 | 0.989 | 1 | 1 | 1 | -0.003 |
| SE | 37188 | ENSG00000114446 | IFT57 | chr3 | - | 108219409 | 108219572 | 108218534 | 108218633 | 108222110 | 108222360 | 125 | 75 | 104 | 104 | 122 | 0 | 1 | 1 | 0 | 4 | 37 | 76 | 70 | 46 | 29 | 0 | 0 | 0 | 0 | 0 | 298 | 149 | 1 | 1 | 1 | 0.974 | 0.981 | 1 | 0.938 | 1 | 1 | 1 | 1 | 1 | -0.021 |
| SE | 37194 | ENSG00000033327 | GAB2 | chr11 | - | 78318142 | 78318219 | 78280600 | 78280901 | 78417645 | 78417822 | 0 | 0 | 0 | 0 | 0 | 27 | 6 | 8 | 26 | 25 | 0 | 1 | 0 | 0 | 0 | 44 | 35 | 47 | 34 | 71 | 226 | 149 | 1 | 1 | 0 | 0 | 0 | 0 | 0 | 0 | 0.018 | 0 | 0 | 0 | -0.004 |
| SE | 37196 | ENSG00000033327 | GAB2 | chr11 | - | 78364015 | 78364103 | 78318142 | 78318219 | 78417645 | 78417779 | 1 | 1 | 0 | 0 | 1 | 0 | 0 | 0 | 0 | 0 | 0 | 0 | 6 | 0 | 0 | 0 | 1 | 0 | 0 | 0 | 237 | 149 | 1 | 1 | 1 | 1 | NA | NA | 1 | NA | 0 | 1 | NA | NA | 0.5 |
| SE | 37197 | ENSG00000149480 | MTA2 | chr11 | - | 62595263 | 62595492 | 62594980 | 62595070 | 62595751 | 62595891 | 542 | 224 | 436 | 439 | 472 | 1 | 0 | 0 | 0 | 0 | 517 | 567 | 262 | 467 | 652 | 3 | 0 | 0 | 0 | 0 | 298 | 149 | 1 | 1 | 0.996 | 1 | 1 | 1 | 1 | 0.989 | 1 | 1 | 1 | 1 | 0.001 |
| SE | 37200 | ENSG00000149480 | MTA2 | chr11 | - | 62597315 | 62597415 | 62596278 | 62596337 | 62597609 | 62597712 | 122 | 81 | 130 | 123 | 233 | 0 | 0 | 0 | 0 | 0 | 265 | 128 | 99 | 138 | 262 | 0 | 0 | 0 | 3 | 0 | 249 | 149 | 1 | 1 | 1 | 1 | 1 | 1 | 1 | 1 | 1 | 1 | 0.965 | 1 | 0.007 |
| SE | 37201 | ENSG00000149480 | MTA2 | chr11 | - | 62598023 | 62598141 | 62597609 | 62597712 | 62598521 | 62598639 | 187 | 107 | 168 | 175 | 205 | 0 | 0 | 0 | 0 | 2 | 247 | 210 | 128 | 253 | 316 | 0 | 0 | 0 | 0 | 0 | 267 | 149 | 1 | 1 | 1 | 1 | 1 | 1 | 0.983 | 1 | 1 | 1 | 1 | 1 | -0.003 |
| SE | 37203 | ENSG00000149480 | MTA2 | chr11 | - | 62598521 | 62598639 | 62598108 | 62598390 | 62600165 | 62600259 | 507 | 298 | 411 | 472 | 457 | 2 | 0 | 0 | 0 | 0 | 770 | 618 | 304 | 670 | 718 | 0 | 0 | 0 | 0 | 0 | 267 | 149 | 1 | 1 | 0.993 | 1 | 1 | 1 | 1 | 1 | 1 | 1 | 1 | 1 | -0.001 |
| SE | 37207 | ENSG00000163611 | SPICE1 | chr3 | - | 113450335 | 113450516 | 113448037 | 113448140 | 113453465 | 113453950 | 115 | 131 | 110 | 115 | 96 | 2 | 7 | 3 | 2 | 0 | 21 | 47 | 71 | 31 | 0 | 12 | 0 | 5 | 0 | 0 | 298 | 149 | 1 | 1 | 0.966 | 0.903 | 0.948 | 0.966 | 1 | 0.467 | 1 | 0.877 | 1 | NA | 0.121 |
| SE | 37208 | ENSG00000163611 | SPICE1 | chr3 | - | 113465652 | 113465784 | 113460616 | 113460764 | 113468138 | 113468404 | 75 | 82 | 81 | 93 | 95 | 0 | 0 | 0 | 0 | 0 | 21 | 39 | 75 | 30 | 18 | 0 | 3 | 0 | 0 | 0 | 281 | 149 | 1 | 1 | 1 | 1 | 1 | 1 | 1 | 1 | 0.873 | 1 | 1 | 1 | 0.025 |
| SE | 37209 | ENSG00000153551 | CMTM7 | chr3 | + | 32441839 | 32442013 | 32391697 | 32392065 | 32449453 | 32449552 | 62 | 32 | 58 | 43 | 28 | 0 | 0 | 0 | 0 | 0 | 116 | 65 | 31 | 45 | 74 | 0 | 0 | 0 | 0 | 5 | 298 | 149 | 1 | 1 | 1 | 1 | 1 | 1 | 1 | 1 | 1 | 1 | 1 | 0.881 | 0.024 |
| SE | 37210 | ENSG00000153551 | CMTM7 | chr3 | + | 32441839 | 32442013 | 32391697 | 32392065 | 32452391 | 32452473 | 23 | 2 | 19 | 12 | 7 | 0 | 0 | 0 | 0 | 2 | 26 | 19 | 14 | 11 | 45 | 0 | 0 | 0 | 0 | 0 | 298 | 149 | 1 | 1 | 1 | 1 | 1 | 1 | 0.636 | 1 | 1 | 1 | 1 | 1 | -0.073 |
| SE | 37211 | ENSG00000153551 | CMTM7 | chr3 | + | 32449453 | 32449552 | 32391852 | 32392065 | 32452391 | 32452473 | 40 | 25 | 59 | 66 | 34 | 0 | 0 | 0 | 0 | 2 | 77 | 66 | 46 | 64 | 104 | 0 | 0 | 0 | 0 | 0 | 248 | 149 | 1 | 1 | 1 | 1 | 1 | 1 | 0.911 | 1 | 1 | 1 | 1 | 1 | -0.018 |
| SE | 37213 | ENSG00000153551 | CMTM7 | chr3 | + | 32449453 | 32449552 | 32441839 | 32442013 | 32454240 | 32454358 | 35 | 25 | 35 | 22 | 23 | 0 | 0 | 0 | 2 | 0 | 89 | 45 | 16 | 31 | 39 | 0 | 0 | 0 | 0 | 0 | 248 | 149 | 1 | 1 | 1 | 1 | 1 | 0.869 | 1 | 1 | 1 | 1 | 1 | 1 | -0.026 |
| SE | 37215 | ENSG00000153551 | CMTM7 | chr3 | + | 32452391 | 32452473 | 32441839 | 32442013 | 32454240 | 32454542 | 70 | 20 | 91 | 60 | 101 | 0 | 0 | 0 | 2 | 0 | 50 | 121 | 63 | 78 | 225 | 0 | 0 | 0 | 0 | 0 | 231 | 149 | 1 | 1 | 1 | 1 | 1 | 0.951 | 1 | 1 | 1 | 1 | 1 | 1 | -0.01 |
| SE | 37216 | ENSG00000153551 | CMTM7 | chr3 | + | 32452391 | 32452473 | 32449453 | 32449552 | 32454240 | 32454542 | 111 | 47 | 153 | 130 | 128 | 0 | 0 | 0 | 0 | 0 | 133 | 188 | 109 | 145 | 303 | 0 | 4 | 2 | 0 | 0 | 231 | 149 | 1 | 1 | 1 | 1 | 1 | 1 | 1 | 1 | 0.968 | 0.972 | 1 | 1 | 0.012 |
| SE | 37222 | ENSG00000187164 | SHTN1 | chr10 | - | 116915374 | 116915484 | 116911789 | 116911843 | 116921433 | 116921516 | 245 | 116 | 106 | 124 | 324 | 0 | 2 | 0 | 0 | 0 | 72 | 143 | 196 | 189 | 21 | 0 | 1 | 0 | 0 | 0 | 259 | 149 | 1 | 1 | 1 | 0.971 | 1 | 1 | 1 | 1 | 0.988 | 1 | 1 | 1 | -0.003 |
| SE | 37223 | ENSG00000187164 | SHTN1 | chr10 | - | 116929848 | 116930002 | 116927791 | 116927891 | 116940465 | 116940612 | 246 | 228 | 122 | 88 | 596 | 0 | 0 | 0 | 0 | 1 | 147 | 215 | 244 | 339 | 80 | 3 | 0 | 3 | 0 | 0 | 298 | 149 | 1 | 1 | 1 | 1 | 1 | 1 | 0.997 | 0.961 | 1 | 0.976 | 1 | 1 | 0.012 |
| SE | 37224 | ENSG00000187164 | SHTN1 | chr10 | - | 116979255 | 116979308 | 116968651 | 116968712 | 117005021 | 117005187 | 45 | 41 | 18 | 26 | 42 | 0 | 0 | 0 | 1 | 0 | 19 | 26 | 36 | 34 | 9 | 0 | 0 | 0 | 0 | 0 | 202 | 149 | 1 | 1 | 1 | 1 | 1 | 0.95 | 1 | 1 | 1 | 1 | 1 | 1 | -0.01 |
| SE | 37228 | ENSG00000284707 | NA | chr7 | - | 97869777 | 97869813 | 97868907 | 97869179 | 97872350 | 97872449 | 28 | 11 | 10 | 35 | 20 | 1 | 0 | 1 | 0 | 0 | 13 | 10 | 10 | 12 | 7 | 0 | 0 | 0 | 0 | 0 | 185 | 149 | 1 | 1 | 0.958 | 1 | 0.89 | 1 | 1 | 1 | 1 | 1 | 1 | 1 | -0.03 |
| SE | 37232 | ENSG00000284707 | NA | chr7 | - | 97945507 | 97945615 | 97936946 | 97937160 | 97972134 | 97972299 | 8 | 4 | 8 | 11 | 14 | 4 | 2 | 8 | 0 | 0 | 0 | 5 | 11 | 8 | 0 | 0 | 0 | 0 | 5 | 0 | 257 | 149 | 1 | 1 | 0.537 | 0.537 | 0.367 | 1 | 1 | NA | 1 | 1 | 0.481 | NA | -0.139 |
| SE | 37234 | ENSG00000150457 | LATS2 | chr13 | - | 20979690 | 20979797 | 20973035 | 20975364 | 20983223 | 20983806 | 48 | 21 | 15 | 39 | 29 | 0 | 0 | 0 | 2 | 0 | 16 | 8 | 26 | 25 | 8 | 0 | 0 | 3 | 0 | 0 | 256 | 149 | 1 | 1 | 1 | 1 | 1 | 0.919 | 1 | 1 | 1 | 0.835 | 1 | 1 | 0.017 |
| SE | 37235 | ENSG00000150457 | LATS2 | chr13 | - | 20981465 | 20981648 | 20973035 | 20975364 | 20983223 | 20983806 | 68 | 42 | 33 | 98 | 70 | 0 | 0 | 0 | 2 | 0 | 234 | 46 | 83 | 146 | 113 | 0 | 0 | 3 | 0 | 0 | 298 | 149 | 1 | 1 | 1 | 1 | 1 | 0.961 | 1 | 1 | 1 | 0.933 | 1 | 1 | 0.006 |
| SE | 37236 | ENSG00000150457 | LATS2 | chr13 | - | 20981465 | 20981648 | 20979690 | 20979797 | 20983223 | 20983806 | 83 | 53 | 46 | 124 | 81 | 0 | 0 | 0 | 0 | 1 | 258 | 53 | 98 | 168 | 163 | 0 | 0 | 0 | 0 | 0 | 298 | 149 | 1 | 1 | 1 | 1 | 1 | 1 | 0.976 | 1 | 1 | 1 | 1 | 1 | -0.005 |
| SE | 37237 | ENSG00000189283 | FHIT | chr3 | - | 59886219 | 59886287 | 59749701 | 59752321 | 59922345 | 59922414 | 0 | 0 | 1 | 0 | 0 | 20 | 6 | 27 | 36 | 90 | 0 | 0 | 0 | 0 | 0 | 12 | 3 | 24 | 8 | 7 | 217 | 149 | 1 | 1 | 0 | 0 | 0.025 | 0 | 0 | 0 | 0 | 0 | 0 | 0 | 0.005 |
| SE | 37238 | ENSG00000189283 | FHIT | chr3 | - | 59922345 | 59922414 | 59751406 | 59752321 | 60080790 | 60080840 | 7 | 2 | 14 | 18 | 31 | 0 | 0 | 0 | 0 | 3 | 6 | 3 | 11 | 3 | 0 | 0 | 0 | 0 | 0 | 0 | 218 | 149 | 1 | 1 | 1 | 1 | 1 | 1 | 0.876 | 1 | 1 | 1 | 1 | NA | -0.025 |
| SE | 37239 | ENSG00000189283 | FHIT | chr3 | - | 60011370 | 60011400 | 59922345 | 59922414 | 60014006 | 60014075 | 19 | 14 | 31 | 15 | 37 | 0 | 0 | 0 | 0 | 0 | 6 | 4 | 36 | 19 | 15 | 0 | 0 | 0 | 3 | 0 | 179 | 149 | 1 | 1 | 1 | 1 | 1 | 1 | 1 | 1 | 1 | 1 | 0.841 | 1 | 0.032 |
| SE | 37240 | ENSG00000189283 | FHIT | chr3 | - | 60011370 | 60011400 | 59922345 | 59922414 | 60080790 | 60080840 | 3 | 1 | 4 | 3 | 12 | 3 | 0 | 0 | 6 | 1 | 6 | 0 | 6 | 5 | 0 | 0 | 0 | 2 | 0 | 0 | 179 | 149 | 1 | 1 | 0.454 | 1 | 1 | 0.294 | 0.909 | 1 | NA | 0.714 | 1 | NA | -0.173 |
| SE | 37242 | ENSG00000189283 | FHIT | chr3 | - | 60014006 | 60014075 | 59922345 | 59922414 | 60080790 | 60080840 | 0 | 0 | 0 | 0 | 3 | 3 | 0 | 0 | 6 | 1 | 0 | 0 | 0 | 0 | 0 | 0 | 0 | 2 | 0 | 0 | 218 | 149 | 1 | 1 | 0 | NA | NA | 0 | 0.672 | NA | NA | 0 | NA | NA | 0.224 |
| SE | 37243 | ENSG00000189283 | FHIT | chr3 | - | 60014006 | 60014075 | 60011370 | 60011400 | 60080790 | 60080840 | 8 | 8 | 12 | 6 | 14 | 1 | 0 | 0 | 0 | 0 | 0 | 0 | 13 | 5 | 7 | 0 | 0 | 0 | 0 | 0 | 218 | 149 | 1 | 1 | 0.845 | 1 | 1 | 1 | 1 | NA | NA | 1 | 1 | 1 | -0.031 |
| SE | 37244 | ENSG00000189283 | FHIT | chr3 | - | 60077369 | 60077526 | 59922345 | 59922414 | 60080790 | 60080840 | 4 | 0 | 2 | 1 | 1 | 3 | 0 | 0 | 6 | 1 | 0 | 0 | 0 | 2 | 0 | 0 | 0 | 2 | 0 | 0 | 298 | 149 | 1 | 1 | 0.4 | NA | 1 | 0.077 | 0.333 | NA | NA | 0 | 1 | NA | -0.048 |
| SE | 37245 | ENSG00000189283 | FHIT | chr3 | - | 60077369 | 60077526 | 60014006 | 60014075 | 60080790 | 60080840 | 4 | 0 | 2 | 1 | 1 | 0 | 0 | 0 | 0 | 3 | 0 | 0 | 0 | 2 | 0 | 0 | 0 | 0 | 0 | 0 | 298 | 149 | 1 | 1 | 1 | NA | 1 | 1 | 0.143 | NA | NA | NA | 1 | NA | -0.214 |
| SE | 37246 | ENSG00000189283 | FHIT | chr3 | - | 60765523 | 60765563 | 60536859 | 60536979 | 60821918 | 60822011 | 0 | 0 | 0 | 1 | 0 | 5 | 7 | 4 | 3 | 10 | 0 | 0 | 0 | 0 | 0 | 12 | 10 | 1 | 8 | 15 | 189 | 149 | 1 | 1 | 0 | 0 | 0 | 0.208 | 0 | 0 | 0 | 0 | 0 | 0 | 0.042 |
| SE | 37247 | ENSG00000189283 | FHIT | chr3 | - | 61166972 | 61167042 | 61042046 | 61042099 | 61200616 | 61200665 | 1 | 2 | 0 | 0 | 0 | 3 | 11 | 2 | 4 | 5 | 0 | 0 | 0 | 0 | 0 | 21 | 15 | 12 | 0 | 4 | 219 | 149 | 1 | 1 | 0.185 | 0.11 | 0 | 0 | 0 | 0 | 0 | 0 | NA | 0 | 0.059 |
| SE | 37252 | ENSG00000178694 | NSUN3 | chr3 | + | 94094139 | 94094294 | 94084106 | 94084450 | 94095032 | 94095094 | 102 | 78 | 51 | 107 | 129 | 0 | 0 | 0 | 1 | 0 | 28 | 86 | 63 | 36 | 20 | 0 | 0 | 0 | 0 | 0 | 298 | 149 | 1 | 1 | 1 | 1 | 1 | 0.982 | 1 | 1 | 1 | 1 | 1 | 1 | -0.004 |
| SE | 37254 | ENSG00000266086 | NA | chr17 | - | 58006342 | 58006527 | 58004922 | 58005973 | 58006943 | 58007246 | 477 | 225 | 457 | 434 | 229 | 1 | 0 | 0 | 0 | 0 | 277 | 270 | 224 | 237 | 104 | 0 | 0 | 0 | 0 | 0 | 298 | 149 | 1 | 1 | 0.996 | 1 | 1 | 1 | 1 | 1 | 1 | 1 | 1 | 1 | -0.001 |
| SE | 37261 | ENSG00000099330 | OCEL1 | chr19 | + | 17228255 | 17228309 | 17227839 | 17228005 | 17228802 | 17228826 | 37 | 32 | 33 | 32 | 63 | 1 | 0 | 0 | 0 | 0 | 21 | 27 | 28 | 37 | 97 | 0 | 0 | 0 | 0 | 0 | 203 | 149 | 1 | 1 | 0.964 | 1 | 1 | 1 | 1 | 1 | 1 | 1 | 1 | 1 | -0.007 |
| SE | 37263 | ENSG00000122707 | RECK | chr9 | + | 36056979 | 36057013 | 36052264 | 36052323 | 36058826 | 36058901 | 1 | 0 | 2 | 0 | 1 | 17 | 2 | 16 | 11 | 14 | 0 | 0 | 0 | 0 | 0 | 0 | 5 | 2 | 4 | 0 | 183 | 149 | 1 | 1 | 0.046 | 0 | 0.092 | 0 | 0.055 | NA | 0 | 0 | 0 | NA | 0.039 |
| SE | 37264 | ENSG00000122707 | RECK | chr9 | + | 36060118 | 36060155 | 36058826 | 36058901 | 36065576 | 36065624 | 6 | 0 | 2 | 0 | 7 | 0 | 1 | 2 | 0 | 0 | 0 | 0 | 1 | 1 | 0 | 0 | 0 | 0 | 0 | 0 | 186 | 149 | 1 | 1 | 1 | 0 | 0.445 | NA | 1 | NA | NA | 1 | 1 | NA | -0.389 |
| SE | 37265 | ENSG00000122707 | RECK | chr9 | + | 36063794 | 36063880 | 36060118 | 36060155 | 36065576 | 36065624 | 26 | 16 | 39 | 43 | 90 | 2 | 0 | 0 | 0 | 0 | 10 | 12 | 10 | 3 | 0 | 0 | 0 | 0 | 0 | 0 | 235 | 149 | 1 | 1 | 0.892 | 1 | 1 | 1 | 1 | 1 | 1 | 1 | 1 | NA | -0.022 |
| SE | 37266 | ENSG00000122707 | RECK | chr9 | + | 36066768 | 36066870 | 36065576 | 36065624 | 36080604 | 36080638 | 1 | 0 | 0 | 0 | 2 | 22 | 21 | 24 | 28 | 62 | 0 | 0 | 0 | 0 | 0 | 18 | 15 | 10 | 4 | 2 | 251 | 149 | 1 | 1 | 0.026 | 0 | 0 | 0 | 0.019 | 0 | 0 | 0 | 0 | 0 | 0.009 |
| SE | 37267 | ENSG00000122707 | RECK | chr9 | + | 36087693 | 36087961 | 36083364 | 36083562 | 36091163 | 36091343 | 71 | 45 | 60 | 74 | 95 | 0 | 2 | 1 | 0 | 0 | 22 | 32 | 35 | 23 | 0 | 0 | 0 | 0 | 0 | 0 | 298 | 149 | 1 | 1 | 1 | 0.918 | 0.968 | 1 | 1 | 1 | 1 | 1 | 1 | NA | -0.023 |
| SE | 37268 | ENSG00000122707 | RECK | chr9 | + | 36112304 | 36112476 | 36109956 | 36110079 | 36116984 | 36117177 | 16 | 16 | 16 | 13 | 25 | 1 | 1 | 0 | 0 | 0 | 0 | 4 | 1 | 7 | 0 | 0 | 2 | 1 | 4 | 0 | 298 | 149 | 1 | 1 | 0.889 | 0.889 | 1 | 1 | 1 | NA | 0.5 | 0.333 | 0.467 | NA | 0.522 |
| SE | 37269 | ENSG00000122707 | RECK | chr9 | + | 36118756 | 36118967 | 36116984 | 36117177 | 36120662 | 36120736 | 11 | 1 | 13 | 11 | 22 | 0 | 0 | 0 | 0 | 4 | 0 | 3 | 1 | 4 | 0 | 0 | 0 | 0 | 0 | 0 | 298 | 149 | 1 | 1 | 1 | 1 | 1 | 1 | 0.733 | NA | 1 | 1 | 1 | NA | -0.053 |
| SE | 37271 | ENSG00000174738 | NR1D2 | chr3 | + | 23954536 | 23954803 | 23945285 | 23945594 | 23959670 | 23959815 | 23 | 15 | 18 | 5 | 15 | 0 | 0 | 0 | 3 | 0 | 9 | 10 | 11 | 25 | 0 | 0 | 0 | 0 | 0 | 0 | 298 | 149 | 1 | 1 | 1 | 1 | 1 | 0.455 | 1 | 1 | 1 | 1 | 1 | NA | -0.109 |
| SE | 37273 | ENSG00000174738 | NR1D2 | chr3 | + | 23956036 | 23956125 | 23954536 | 23954803 | 23961976 | 23962334 | 83 | 48 | 51 | 64 | 91 | 1 | 0 | 0 | 1 | 0 | 22 | 15 | 46 | 25 | 4 | 0 | 0 | 0 | 0 | 0 | 238 | 149 | 1 | 1 | 0.981 | 1 | 1 | 0.976 | 1 | 1 | 1 | 1 | 1 | 1 | -0.009 |
| SE | 37274 | ENSG00000174738 | NR1D2 | chr3 | + | 23959670 | 23959815 | 23954536 | 23954803 | 23961976 | 23962605 | 182 | 107 | 152 | 159 | 394 | 1 | 0 | 0 | 1 | 0 | 86 | 74 | 126 | 70 | 38 | 0 | 0 | 0 | 0 | 0 | 294 | 149 | 1 | 1 | 0.989 | 1 | 1 | 0.988 | 1 | 1 | 1 | 1 | 1 | 1 | -0.005 |
| SE | 37275 | ENSG00000174738 | NR1D2 | chr3 | + | 23959670 | 23959815 | 23956036 | 23956125 | 23961976 | 23962605 | 273 | 161 | 225 | 232 | 540 | 0 | 1 | 0 | 0 | 0 | 85 | 112 | 183 | 90 | 47 | 0 | 0 | 3 | 0 | 0 | 294 | 149 | 1 | 1 | 1 | 0.988 | 1 | 1 | 1 | 1 | 1 | 0.969 | 1 | 1 | 0.004 |
| SE | 37276 | ENSG00000174738 | NR1D2 | chr3 | + | 23959670 | 23959924 | 23954625 | 23954803 | 23961976 | 23962334 | 9 | 3 | 10 | 2 | 15 | 1 | 0 | 0 | 1 | 0 | 9 | 10 | 7 | 19 | 0 | 0 | 0 | 0 | 0 | 0 | 298 | 149 | 1 | 1 | 0.818 | 1 | 1 | 0.5 | 1 | 1 | 1 | 1 | 1 | NA | -0.136 |
| SE | 37277 | ENSG00000174738 | NR1D2 | chr3 | + | 23959670 | 23959924 | 23956036 | 23956125 | 23961976 | 23962334 | 100 | 57 | 83 | 75 | 162 | 0 | 1 | 0 | 0 | 0 | 8 | 48 | 64 | 39 | 9 | 0 | 0 | 3 | 0 | 0 | 298 | 149 | 1 | 1 | 1 | 0.966 | 1 | 1 | 1 | 1 | 1 | 0.914 | 1 | 1 | 0.01 |
| SE | 37278 | ENSG00000174738 | NR1D2 | chr3 | + | 23961976 | 23962605 | 23956036 | 23956125 | 23964976 | 23965162 | 155 | 75 | 127 | 144 | 157 | 1 | 0 | 0 | 0 | 0 | 68 | 64 | 157 | 34 | 21 | 0 | 0 | 0 | 0 | 0 | 298 | 149 | 1 | 1 | 0.987 | 1 | 1 | 1 | 1 | 1 | 1 | 1 | 1 | 1 | -0.003 |
| SE | 37279 | ENSG00000174738 | NR1D2 | chr3 | + | 23961976 | 23962605 | 23959670 | 23959815 | 23967812 | 23968023 | 178 | 106 | 142 | 159 | 385 | 0 | 0 | 0 | 0 | 1 | 77 | 64 | 121 | 53 | 38 | 0 | 0 | 0 | 0 | 0 | 298 | 149 | 1 | 1 | 1 | 1 | 1 | 1 | 0.995 | 1 | 1 | 1 | 1 | 1 | -0.001 |
| SE | 37282 | ENSG00000174738 | NR1D2 | chr3 | + | 23964976 | 23965162 | 23961976 | 23962605 | 23967812 | 23968023 | 311 | 209 | 251 | 293 | 271 | 0 | 2 | 0 | 1 | 0 | 145 | 154 | 273 | 59 | 48 | 0 | 0 | 0 | 0 | 0 | 298 | 149 | 1 | 1 | 1 | 0.981 | 1 | 0.993 | 1 | 1 | 1 | 1 | 1 | 1 | -0.005 |
| SE | 37283 | ENSG00000174738 | NR1D2 | chr3 | + | 23964976 | 23965162 | 23961976 | 23962605 | 23977222 | 23977543 | 159 | 76 | 133 | 150 | 169 | 0 | 0 | 0 | 0 | 0 | 68 | 66 | 169 | 54 | 21 | 0 | 2 | 3 | 0 | 0 | 298 | 149 | 1 | 1 | 1 | 1 | 1 | 1 | 1 | 1 | 0.943 | 0.966 | 1 | 1 | 0.018 |
| SE | 37284 | ENSG00000174738 | NR1D2 | chr3 | + | 23964976 | 23965162 | 23963223 | 23963330 | 23977222 | 23977543 | 29 | 18 | 32 | 24 | 38 | 0 | 0 | 0 | 0 | 0 | 5 | 44 | 47 | 51 | 0 | 0 | 5 | 0 | 0 | 0 | 298 | 149 | 1 | 1 | 1 | 1 | 1 | 1 | 1 | 1 | 0.815 | 1 | 1 | NA | 0.046 |
| SE | 37289 | ENSG00000151576 | QTRT2 | chr3 | + | 114056998 | 114057106 | 114056736 | 114056864 | 114070625 | 114070838 | 9 | 9 | 13 | 15 | 32 | 10 | 2 | 0 | 0 | 0 | 0 | 1 | 0 | 2 | 3 | 0 | 0 | 0 | 0 | 0 | 257 | 149 | 1 | 1 | 0.343 | 0.723 | 1 | 1 | 1 | NA | 1 | NA | 1 | 1 | -0.187 |
| SE | 37291 | ENSG00000151576 | QTRT2 | chr3 | + | 114065236 | 114065457 | 114056998 | 114057106 | 114066227 | 114066283 | 70 | 52 | 64 | 62 | 165 | 0 | 0 | 0 | 1 | 0 | 41 | 38 | 54 | 62 | 20 | 0 | 0 | 0 | 0 | 0 | 298 | 149 | 1 | 1 | 1 | 1 | 1 | 0.969 | 1 | 1 | 1 | 1 | 1 | 1 | -0.006 |
| SE | 37292 | ENSG00000151576 | QTRT2 | chr3 | + | 114065236 | 114065457 | 114056998 | 114057106 | 114070625 | 114070751 | 17 | 30 | 14 | 19 | 40 | 1 | 0 | 0 | 0 | 0 | 11 | 11 | 1 | 10 | 0 | 0 | 0 | 0 | 0 | 0 | 298 | 149 | 1 | 1 | 0.895 | 1 | 1 | 1 | 1 | 1 | 1 | 1 | 1 | NA | -0.021 |
| SE | 37293 | ENSG00000151576 | QTRT2 | chr3 | + | 114066227 | 114066283 | 114056998 | 114057106 | 114070625 | 114070806 | 3 | 0 | 2 | 0 | 0 | 1 | 0 | 0 | 0 | 0 | 0 | 5 | 0 | 15 | 0 | 0 | 0 | 0 | 0 | 0 | 205 | 149 | 1 | 1 | 0.686 | NA | 1 | NA | NA | NA | 1 | NA | 1 | NA | -0.157 |
| SE | 37294 | ENSG00000151576 | QTRT2 | chr3 | + | 114066227 | 114066283 | 114065236 | 114065457 | 114070625 | 114070806 | 26 | 17 | 27 | 32 | 78 | 0 | 0 | 0 | 0 | 1 | 30 | 25 | 35 | 49 | 10 | 0 | 0 | 0 | 0 | 0 | 205 | 149 | 1 | 1 | 1 | 1 | 1 | 1 | 0.983 | 1 | 1 | 1 | 1 | 1 | -0.003 |
| SE | 37299 | ENSG00000163814 | CDCP1 | chr3 | - | 45089053 | 45089141 | 45082276 | 45086067 | 45091172 | 45091538 | 1 | 3 | 0 | 0 | 0 | 0 | 2 | 0 | 0 | 0 | 31 | 12 | 0 | 4 | 10 | 0 | 0 | 0 | 0 | 0 | 237 | 149 | 1 | 1 | 1 | 0.485 | NA | NA | NA | 1 | 1 | NA | 1 | 1 | -0.258 |
| SE | 37300 | ENSG00000168028 | RPSA | chr3 | + | 39408605 | 39408724 | 39407602 | 39407786 | 39410753 | 39410849 | 8728 | 4399 | 10268 | 8214 | 19333 | 8 | 4 | 21 | 16 | 41 | 4608 | 5724 | 4637 | 3657 | 7913 | 21 | 7 | 13 | 15 | 3 | 268 | 149 | 1 | 1 | 0.998 | 0.998 | 0.996 | 0.997 | 0.996 | 0.992 | 0.998 | 0.995 | 0.993 | 0.999 | 0.002 |
| SE | 37301 | ENSG00000168028 | RPSA | chr3 | + | 39408605 | 39408724 | 39407602 | 39407786 | 39411648 | 39411777 | 3326 | 1729 | 3907 | 3063 | 8097 | 1 | 0 | 0 | 0 | 2 | 1771 | 2112 | 1783 | 1399 | 3139 | 0 | 0 | 0 | 0 | 0 | 268 | 149 | 1 | 1 | 0.999 | 1 | 1 | 1 | 1 | 1 | 1 | 1 | 1 | 1 | 0 |
| SE | 37303 | ENSG00000168028 | RPSA | chr3 | + | 39410753 | 39410999 | 39407602 | 39407786 | 39411648 | 39411777 | 1985 | 986 | 2605 | 2137 | 2576 | 1 | 0 | 0 | 0 | 2 | 1408 | 1408 | 1208 | 836 | 2068 | 0 | 0 | 0 | 0 | 0 | 298 | 149 | 1 | 1 | 0.999 | 1 | 1 | 1 | 0.998 | 1 | 1 | 1 | 1 | 1 | -0.001 |
| SE | 37304 | ENSG00000168028 | RPSA | chr3 | + | 39410753 | 39410999 | 39407602 | 39407786 | 39411895 | 39412059 | 8 | 4 | 21 | 16 | 41 | 0 | 0 | 1 | 0 | 0 | 21 | 7 | 17 | 15 | 3 | 0 | 0 | 0 | 0 | 0 | 298 | 149 | 1 | 1 | 1 | 1 | 0.913 | 1 | 1 | 1 | 1 | 1 | 1 | 1 | -0.017 |
| SE | 37305 | ENSG00000168028 | RPSA | chr3 | + | 39410753 | 39410999 | 39408203 | 39408724 | 39411648 | 39411777 | 7414 | 3671 | 9001 | 7297 | 13830 | 0 | 0 | 1 | 0 | 0 | 4265 | 5042 | 4072 | 3101 | 6865 | 0 | 2 | 0 | 0 | 0 | 298 | 149 | 1 | 1 | 1 | 1 | 1 | 1 | 1 | 1 | 0.999 | 1 | 1 | 1 | 0 |
| SE | 37306 | ENSG00000168028 | RPSA | chr3 | + | 39411648 | 39411777 | 39407602 | 39407786 | 39411895 | 39412059 | 1410 | 550 | 1527 | 1259 | 1208 | 0 | 0 | 1 | 0 | 0 | 468 | 815 | 644 | 498 | 1170 | 0 | 0 | 0 | 0 | 0 | 278 | 149 | 1 | 1 | 1 | 1 | 0.999 | 1 | 1 | 1 | 1 | 1 | 1 | 1 | 0 |
| SE | 37307 | ENSG00000168028 | RPSA | chr3 | + | 39411648 | 39411777 | 39410753 | 39410999 | 39411895 | 39412059 | 3386 | 1532 | 4111 | 3381 | 3741 | 0 | 0 | 0 | 0 | 0 | 1855 | 2221 | 1839 | 1319 | 3235 | 0 | 0 | 4 | 0 | 0 | 278 | 149 | 1 | 1 | 1 | 1 | 1 | 1 | 1 | 1 | 1 | 0.996 | 1 | 1 | 0.001 |
| SE | 37308 | ENSG00000163606 | CD200R1 | chr3 | - | 112924489 | 112924535 | 112921204 | 112923799 | 112925084 | 112925193 | 24 | 23 | 49 | 35 | 59 | 0 | 0 | 1 | 1 | 0 | 0 | 7 | 0 | 2 | 3 | 0 | 0 | 0 | 0 | 0 | 195 | 149 | 1 | 1 | 1 | 1 | 0.974 | 0.964 | 1 | NA | 1 | NA | 1 | 1 | -0.012 |
| SE | 37309 | ENSG00000163606 | CD200R1 | chr3 | - | 112924489 | 112924535 | 112921204 | 112923799 | 112928815 | 112929064 | 16 | 6 | 18 | 17 | 32 | 1 | 2 | 1 | 1 | 8 | 0 | 4 | 0 | 0 | 3 | 0 | 0 | 0 | 0 | 0 | 195 | 149 | 1 | 1 | 0.924 | 0.696 | 0.932 | 0.929 | 0.753 | NA | 1 | NA | NA | NA | -0.153 |
| SE | 37310 | ENSG00000163606 | CD200R1 | chr3 | - | 112925084 | 112925193 | 112921204 | 112923799 | 112928815 | 112929064 | 12 | 13 | 21 | 22 | 18 | 1 | 2 | 1 | 1 | 8 | 0 | 0 | 4 | 0 | 3 | 0 | 0 | 0 | 0 | 0 | 258 | 149 | 1 | 1 | 0.874 | 0.79 | 0.924 | 0.927 | 0.565 | NA | NA | 1 | NA | 1 | -0.184 |
| SE | 37311 | ENSG00000163606 | CD200R1 | chr3 | - | 112925084 | 112925193 | 112924489 | 112924535 | 112928815 | 112929064 | 28 | 32 | 58 | 43 | 51 | 8 | 2 | 7 | 4 | 7 | 0 | 3 | 4 | 2 | 6 | 0 | 0 | 0 | 0 | 0 | 258 | 149 | 1 | 1 | 0.669 | 0.902 | 0.827 | 0.861 | 0.808 | NA | 1 | 1 | 1 | 1 | -0.187 |
| SE | 37312 | ENSG00000163606 | CD200R1 | chr3 | - | 112931105 | 112931171 | 112928917 | 112929507 | 112947855 | 112947924 | 95 | 58 | 117 | 89 | 176 | 1 | 0 | 1 | 0 | 0 | 8 | 30 | 3 | 8 | 0 | 0 | 0 | 0 | 0 | 0 | 215 | 149 | 1 | 1 | 0.985 | 1 | 0.988 | 1 | 1 | 1 | 1 | 1 | 1 | NA | -0.005 |
| SE | 37313 | ENSG00000163606 | CD200R1 | chr3 | - | 112931105 | 112931171 | 112928917 | 112929507 | 112974790 | 112974912 | 51 | 34 | 71 | 44 | 90 | 0 | 0 | 0 | 0 | 0 | 0 | 25 | 3 | 2 | 0 | 0 | 2 | 0 | 1 | 0 | 215 | 149 | 1 | 1 | 1 | 1 | 1 | 1 | 1 | NA | 0.897 | 1 | 0.581 | NA | 0.174 |
| SE | 37314 | ENSG00000163606 | CD200R1 | chr3 | - | 112947855 | 112947924 | 112928917 | 112929507 | 112974790 | 112974912 | 51 | 13 | 41 | 29 | 60 | 0 | 0 | 0 | 0 | 0 | 0 | 18 | 0 | 0 | 0 | 0 | 2 | 0 | 1 | 0 | 218 | 149 | 1 | 1 | 1 | 1 | 1 | 1 | 1 | NA | 0.86 | NA | 0 | NA | 0.57 |
| SE | 37315 | ENSG00000163606 | CD200R1 | chr3 | - | 112947855 | 112947924 | 112931105 | 112931171 | 112974790 | 112974912 | 96 | 46 | 105 | 79 | 159 | 3 | 9 | 20 | 5 | 13 | 8 | 26 | 2 | 6 | 0 | 0 | 3 | 2 | 0 | 0 | 218 | 149 | 1 | 1 | 0.956 | 0.777 | 0.782 | 0.915 | 0.893 | 1 | 0.856 | 0.406 | 1 | NA | 0.049 |
| SE | 37316 | ENSG00000004534 | RBM6 | chr3 | + | 49962575 | 49962685 | 49940006 | 49940225 | 50048244 | 50048319 | 73 | 46 | 61 | 66 | 37 | 0 | 0 | 0 | 0 | 0 | 41 | 60 | 38 | 46 | 59 | 0 | 2 | 0 | 0 | 0 | 259 | 149 | 1 | 1 | 1 | 1 | 1 | 1 | 1 | 1 | 0.945 | 1 | 1 | 1 | 0.011 |
| SE | 37323 | ENSG00000004534 | RBM6 | chr3 | + | 50048244 | 50048319 | 49999439 | 49999513 | 50054334 | 50054395 | 308 | 269 | 270 | 291 | 636 | 0 | 0 | 0 | 0 | 0 | 202 | 291 | 193 | 217 | 115 | 0 | 0 | 0 | 0 | 5 | 224 | 149 | 1 | 1 | 1 | 1 | 1 | 1 | 1 | 1 | 1 | 1 | 1 | 0.939 | 0.012 |
| SE | 37324 | ENSG00000004534 | RBM6 | chr3 | + | 50054334 | 50054395 | 50048244 | 50048319 | 50057727 | 50057806 | 380 | 326 | 320 | 352 | 708 | 0 | 2 | 0 | 3 | 0 | 264 | 349 | 276 | 320 | 146 | 0 | 4 | 2 | 4 | 0 | 210 | 149 | 1 | 1 | 1 | 0.991 | 1 | 0.988 | 1 | 1 | 0.984 | 0.99 | 0.983 | 1 | 0.004 |
| SE | 37325 | ENSG00000004534 | RBM6 | chr3 | + | 50059631 | 50059746 | 50058401 | 50058562 | 50060955 | 50060998 | 185 | 118 | 153 | 160 | 270 | 0 | 0 | 0 | 1 | 0 | 139 | 158 | 104 | 144 | 105 | 0 | 1 | 3 | 0 | 0 | 264 | 149 | 1 | 1 | 1 | 0.985 | 1 | 0.989 | 1 | 1 | 0.989 | 0.951 | 1 | 1 | 0.007 |
| SE | 37326 | ENSG00000004534 | RBM6 | chr3 | + | 50059648 | 50059746 | 50058401 | 50058562 | 50060955 | 50060998 | 428 | 311 | 360 | 373 | 504 | 0 | 1 | 0 | 1 | 0 | 339 | 320 | 216 | 298 | 188 | 0 | 1 | 3 | 0 | 0 | 247 | 149 | 1 | 1 | 1 | 0.995 | 1 | 0.996 | 1 | 1 | 0.995 | 0.977 | 1 | 1 | 0.004 |
| SE | 37327 | ENSG00000004534 | RBM6 | chr3 | + | 50060800 | 50060835 | 50059697 | 50059746 | 50060955 | 50060998 | 1 | 4 | 3 | 4 | 7 | 322 | 219 | 262 | 273 | 358 | 0 | 0 | 0 | 0 | 0 | 257 | 248 | 163 | 222 | 156 | 184 | 149 | 1 | 1 | 0.003 | 0.015 | 0.009 | 0.012 | 0.016 | 0 | 0 | 0 | 0 | 0 | 0.011 |
| SE | 37328 | ENSG00000004534 | RBM6 | chr3 | + | 50060955 | 50060998 | 50059342 | 50059746 | 50061139 | 50061172 | 342 | 233 | 281 | 280 | 380 | 0 | 1 | 0 | 0 | 0 | 283 | 251 | 169 | 222 | 168 | 0 | 0 | 0 | 0 | 0 | 192 | 149 | 1 | 1 | 1 | 0.994 | 1 | 1 | 1 | 1 | 1 | 1 | 1 | 1 | -0.001 |
| SE | 37329 | ENSG00000004534 | RBM6 | chr3 | + | 50065030 | 50065126 | 50061461 | 50062108 | 50066241 | 50066502 | 285 | 279 | 296 | 320 | 388 | 0 | 4 | 0 | 0 | 0 | 298 | 375 | 278 | 414 | 206 | 0 | 0 | 4 | 3 | 0 | 245 | 149 | 1 | 1 | 1 | 0.977 | 1 | 1 | 1 | 1 | 1 | 0.977 | 0.988 | 1 | 0.002 |
| SE | 37330 | ENSG00000004534 | RBM6 | chr3 | + | 50065578 | 50065632 | 50065030 | 50065126 | 50066241 | 50066442 | 0 | 2 | 2 | 2 | 0 | 175 | 177 | 183 | 186 | 250 | 0 | 0 | 0 | 0 | 0 | 229 | 228 | 162 | 253 | 158 | 203 | 149 | 1 | 1 | 0 | 0.008 | 0.008 | 0.008 | 0 | 0 | 0 | 0 | 0 | 0 | 0.005 |
| SE | 37331 | ENSG00000004534 | RBM6 | chr3 | + | 50068689 | 50068764 | 50066241 | 50066502 | 50075200 | 50075330 | 153 | 134 | 172 | 141 | 182 | 0 | 0 | 1 | 0 | 0 | 165 | 178 | 186 | 219 | 78 | 0 | 0 | 0 | 0 | 0 | 224 | 149 | 1 | 1 | 1 | 1 | 0.991 | 1 | 1 | 1 | 1 | 1 | 1 | 1 | -0.002 |
| SE | 37332 | ENSG00000004534 | RBM6 | chr3 | + | 50070454 | 50070552 | 50068689 | 50068764 | 50075200 | 50075330 | 264 | 232 | 220 | 264 | 233 | 0 | 0 | 2 | 0 | 0 | 233 | 294 | 251 | 316 | 96 | 0 | 0 | 3 | 0 | 0 | 247 | 149 | 1 | 1 | 1 | 1 | 0.985 | 1 | 1 | 1 | 1 | 0.981 | 1 | 1 | 0.001 |
| SE | 37335 | ENSG00000241163 | LINC00877 | chr3 | - | 72055860 | 72055962 | 72035299 | 72036520 | 72059686 | 72059830 | 21 | 43 | 4 | 25 | 21 | 0 | 0 | 1 | 0 | 0 | 18 | 6 | 4 | 28 | 16 | 0 | 0 | 0 | 0 | 0 | 251 | 149 | 1 | 1 | 1 | 1 | 0.704 | 1 | 1 | 1 | 1 | 1 | 1 | 1 | -0.059 |
| SE | 37336 | ENSG00000241163 | LINC00877 | chr3 | - | 72060509 | 72060611 | 72059437 | 72060109 | 72060802 | 72061094 | 7 | 24 | 8 | 13 | 15 | 2 | 0 | 0 | 5 | 0 | 0 | 0 | 8 | 4 | 0 | 0 | 0 | 3 | 6 | 0 | 251 | 149 | 1 | 1 | 0.675 | 1 | 1 | 0.607 | 1 | NA | NA | 0.613 | 0.284 | NA | 0.408 |
| SE | 37337 | ENSG00000241163 | LINC00877 | chr3 | - | 72060509 | 72061094 | 72059686 | 72060109 | 72099903 | 72100010 | 11 | 22 | 12 | 21 | 14 | 2 | 0 | 0 | 2 | 0 | 19 | 0 | 11 | 12 | 0 | 0 | 0 | 0 | 0 | 0 | 298 | 149 | 1 | 1 | 0.733 | 1 | 1 | 0.84 | 1 | 1 | NA | 1 | 1 | NA | -0.085 |
| SE | 37338 | ENSG00000241163 | LINC00877 | chr3 | - | 72060802 | 72061094 | 72059686 | 72060109 | 72099903 | 72100010 | 8 | 7 | 5 | 19 | 1 | 2 | 0 | 0 | 2 | 0 | 19 | 0 | 8 | 14 | 0 | 0 | 0 | 0 | 0 | 0 | 298 | 149 | 1 | 1 | 0.667 | 1 | 1 | 0.826 | 1 | 1 | NA | 1 | 1 | NA | -0.101 |
| SE | 37345 | ENSG00000075826 | SEC31B | chr10 | - | 100489251 | 100489398 | 100488857 | 100488974 | 100489702 | 100489761 | 8 | 5 | 14 | 18 | 18 | 1 | 0 | 0 | 0 | 0 | 10 | 34 | 7 | 9 | 22 | 0 | 0 | 0 | 0 | 0 | 296 | 149 | 1 | 1 | 0.801 | 1 | 1 | 1 | 1 | 1 | 1 | 1 | 1 | 1 | -0.04 |
| SE | 37346 | ENSG00000075826 | SEC31B | chr10 | - | 100489702 | 100489761 | 100488857 | 100489398 | 100490705 | 100490883 | 9 | 1 | 4 | 10 | 12 | 0 | 0 | 0 | 0 | 1 | 0 | 6 | 6 | 1 | 35 | 0 | 0 | 2 | 0 | 0 | 208 | 149 | 1 | 1 | 1 | 1 | 1 | 1 | 0.896 | NA | 1 | 0.682 | 1 | 1 | 0.059 |
| SE | 37348 | ENSG00000075826 | SEC31B | chr10 | - | 100490705 | 100490883 | 100489702 | 100489761 | 100495384 | 100495546 | 21 | 19 | 33 | 35 | 15 | 0 | 1 | 0 | 0 | 0 | 15 | 40 | 34 | 19 | 77 | 0 | 0 | 0 | 3 | 0 | 298 | 149 | 1 | 1 | 1 | 0.905 | 1 | 1 | 1 | 1 | 1 | 1 | 0.76 | 1 | 0.029 |
| SE | 37356 | ENSG00000173706 | HEG1 | chr3 | - | 124971042 | 124971148 | 124970552 | 124970801 | 124971993 | 124972156 | 2 | 0 | 2 | 0 | 0 | 0 | 0 | 0 | 0 | 0 | 0 | 5 | 0 | 0 | 0 | 0 | 4 | 0 | 0 | 0 | 255 | 149 | 1 | 1 | 1 | NA | 1 | NA | NA | NA | 0.422 | NA | NA | NA | 0.578 |
| SE | 37357 | ENSG00000173706 | HEG1 | chr3 | - | 124971042 | 124971148 | 124970552 | 124970801 | 124973730 | 124973907 | 1 | 0 | 2 | 0 | 0 | 64 | 49 | 50 | 123 | 78 | 0 | 5 | 0 | 0 | 0 | 52 | 71 | 63 | 71 | 49 | 255 | 149 | 1 | 1 | 0.009 | 0 | 0.023 | 0 | 0 | 0 | 0.04 | 0 | 0 | 0 | -0.002 |
| SE | 37358 | ENSG00000173706 | HEG1 | chr3 | - | 124973730 | 124973905 | 124965709 | 124970801 | 124977858 | 124977946 | 115 | 77 | 89 | 188 | 120 | 1 | 0 | 0 | 0 | 0 | 91 | 89 | 78 | 95 | 61 | 0 | 0 | 0 | 0 | 0 | 298 | 149 | 1 | 1 | 0.983 | 1 | 1 | 1 | 1 | 1 | 1 | 1 | 1 | 1 | -0.003 |
| SE | 37359 | ENSG00000173706 | HEG1 | chr3 | - | 124977858 | 124977946 | 124973730 | 124973905 | 124990786 | 124990824 | 76 | 44 | 57 | 92 | 64 | 0 | 0 | 0 | 0 | 1 | 78 | 25 | 22 | 34 | 17 | 0 | 4 | 0 | 0 | 0 | 237 | 149 | 1 | 1 | 1 | 1 | 1 | 1 | 0.976 | 1 | 0.797 | 1 | 1 | 1 | 0.036 |
| SE | 37360 | ENSG00000173706 | HEG1 | chr3 | - | 125001851 | 125002012 | 124997688 | 124997823 | 125002256 | 125002315 | 67 | 29 | 42 | 48 | 65 | 1 | 0 | 1 | 0 | 2 | 24 | 36 | 24 | 27 | 12 | 3 | 4 | 0 | 1 | 4 | 298 | 149 | 1 | 1 | 0.971 | 1 | 0.955 | 1 | 0.942 | 0.8 | 0.818 | 1 | 0.931 | 0.6 | 0.144 |
| SE | 37361 | ENSG00000173706 | HEG1 | chr3 | - | 125010438 | 125010555 | 125009704 | 125009824 | 125012622 | 125013990 | 104 | 29 | 31 | 79 | 28 | 1 | 0 | 0 | 0 | 1 | 39 | 95 | 64 | 76 | 39 | 0 | 0 | 0 | 0 | 0 | 266 | 149 | 1 | 1 | 0.983 | 1 | 1 | 1 | 0.94 | 1 | 1 | 1 | 1 | 1 | -0.015 |
| SE | 37363 | ENSG00000182378 | PLCXD1 | chrX | + | 281481 | 281684 | 276321 | 276394 | 284166 | 284314 | 3 | 0 | 1 | 3 | 4 | 0 | 0 | 2 | 0 | 0 | 0 | 14 | 0 | 5 | 19 | 0 | 5 | 0 | 0 | 10 | 298 | 149 | 1 | 1 | 1 | NA | 0.2 | 1 | 1 | NA | 0.583 | NA | 1 | 0.487 | 0.11 |
| SE | 37364 | ENSG00000166927 | MS4A7 | chr11 | + | 60383128 | 60383288 | 60378484 | 60378664 | 60386716 | 60386773 | 237 | 192 | 96 | 180 | 174 | 1 | 0 | 0 | 0 | 0 | 88 | 114 | 28 | 88 | 20 | 0 | 0 | 0 | 0 | 0 | 298 | 149 | 1 | 1 | 0.992 | 1 | 1 | 1 | 1 | 1 | 1 | 1 | 1 | 1 | -0.002 |
| SE | 37365 | ENSG00000166927 | MS4A7 | chr11 | + | 60383128 | 60383288 | 60378484 | 60378664 | 60393786 | 60393813 | 143 | 134 | 70 | 105 | 93 | 1 | 1 | 0 | 0 | 0 | 61 | 81 | 10 | 62 | 8 | 0 | 0 | 0 | 0 | 0 | 298 | 149 | 1 | 1 | 0.986 | 0.985 | 1 | 1 | 1 | 1 | 1 | 1 | 1 | 1 | -0.006 |
| SE | 37366 | ENSG00000166927 | MS4A7 | chr11 | + | 60383128 | 60383288 | 60378484 | 60378664 | 60394945 | 60395254 | 142 | 134 | 70 | 107 | 95 | 0 | 1 | 0 | 0 | 0 | 61 | 79 | 10 | 66 | 8 | 0 | 0 | 0 | 0 | 0 | 298 | 149 | 1 | 1 | 1 | 0.985 | 1 | 1 | 1 | 1 | 1 | 1 | 1 | 1 | -0.003 |
| SE | 37367 | ENSG00000166927 | MS4A7 | chr11 | + | 60383179 | 60383288 | 60378563 | 60378664 | 60386716 | 60386774 | 59 | 40 | 20 | 50 | 46 | 1 | 0 | 0 | 0 | 0 | 22 | 12 | 13 | 16 | 12 | 0 | 0 | 0 | 0 | 0 | 258 | 149 | 1 | 1 | 0.971 | 1 | 1 | 1 | 1 | 1 | 1 | 1 | 1 | 1 | -0.006 |
| SE | 37370 | ENSG00000166927 | MS4A7 | chr11 | + | 60385087 | 60385222 | 60383128 | 60383288 | 60393786 | 60395948 | 16 | 35 | 16 | 12 | 32 | 1 | 0 | 0 | 0 | 0 | 16 | 22 | 3 | 3 | 5 | 0 | 2 | 0 | 0 | 0 | 284 | 149 | 1 | 1 | 0.894 | 1 | 1 | 1 | 1 | 1 | 0.852 | 1 | 1 | 1 | 0.008 |
| SE | 37374 | ENSG00000166927 | MS4A7 | chr11 | + | 60386716 | 60386773 | 60383128 | 60383288 | 60393786 | 60394522 | 56 | 35 | 12 | 46 | 50 | 1 | 0 | 0 | 0 | 0 | 14 | 25 | 9 | 13 | 7 | 0 | 2 | 0 | 0 | 0 | 206 | 149 | 1 | 1 | 0.976 | 1 | 1 | 1 | 1 | 1 | 0.9 | 1 | 1 | 1 | 0.015 |
| SE | 37375 | ENSG00000166927 | MS4A7 | chr11 | + | 60386716 | 60386773 | 60385087 | 60385222 | 60389389 | 60389596 | 105 | 140 | 66 | 75 | 145 | 0 | 0 | 0 | 1 | 0 | 65 | 44 | 13 | 19 | 35 | 0 | 0 | 0 | 0 | 0 | 206 | 149 | 1 | 1 | 1 | 1 | 1 | 0.982 | 1 | 1 | 1 | 1 | 1 | 1 | -0.004 |
| SE | 37376 | ENSG00000166927 | MS4A7 | chr11 | + | 60386716 | 60386773 | 60385087 | 60385222 | 60393786 | 60394522 | 2 | 3 | 4 | 0 | 12 | 1 | 0 | 0 | 0 | 0 | 4 | 10 | 1 | 0 | 4 | 0 | 0 | 0 | 0 | 0 | 206 | 149 | 1 | 1 | 0.591 | 1 | 1 | NA | 1 | 1 | 1 | 1 | NA | 1 | -0.102 |
| SE | 37379 | ENSG00000166927 | MS4A7 | chr11 | + | 60389389 | 60389596 | 60385087 | 60385222 | 60394945 | 60395254 | 3 | 4 | 3 | 4 | 4 | 0 | 5 | 0 | 0 | 0 | 0 | 1 | 5 | 2 | 0 | 0 | 0 | 0 | 0 | 0 | 298 | 149 | 1 | 1 | 1 | 0.286 | 1 | 1 | 1 | NA | 1 | 1 | 1 | NA | -0.143 |
| SE | 37380 | ENSG00000166927 | MS4A7 | chr11 | + | 60392684 | 60392786 | 60383128 | 60383288 | 60393786 | 60393813 | 40 | 37 | 13 | 31 | 22 | 1 | 0 | 0 | 0 | 0 | 3 | 12 | 9 | 5 | 0 | 0 | 2 | 0 | 0 | 0 | 251 | 149 | 1 | 1 | 0.96 | 1 | 1 | 1 | 1 | 1 | 0.781 | 1 | 1 | NA | 0.047 |
| SE | 37381 | ENSG00000166927 | MS4A7 | chr11 | + | 60392684 | 60392786 | 60383128 | 60383288 | 60394945 | 60395254 | 5 | 1 | 2 | 3 | 5 | 0 | 0 | 0 | 2 | 2 | 0 | 0 | 1 | 4 | 0 | 0 | 0 | 4 | 0 | 0 | 251 | 149 | 1 | 1 | 1 | 1 | 1 | 0.471 | 0.597 | NA | NA | 1 | 0.372 | NA | 0.128 |
| SE | 37390 | ENSG00000010327 | STAB1 | chr3 | + | 52502631 | 52502727 | 52502158 | 52502228 | 52502998 | 52503109 | 53 | 21 | 62 | 39 | 27 | 0 | 0 | 0 | 0 | 0 | 114 | 113 | 52 | 82 | 210 | 0 | 0 | 1 | 0 | 0 | 245 | 149 | 1 | 1 | 1 | 1 | 1 | 1 | 1 | 1 | 1 | 0.969 | 1 | 1 | 0.006 |
| SE | 37391 | ENSG00000010327 | STAB1 | chr3 | + | 52504027 | 52504155 | 52503771 | 52503902 | 52504460 | 52504549 | 70 | 26 | 48 | 44 | 14 | 0 | 0 | 0 | 0 | 0 | 147 | 109 | 48 | 76 | 268 | 5 | 0 | 0 | 0 | 0 | 277 | 149 | 1 | 1 | 1 | 1 | 1 | 1 | 1 | 0.941 | 1 | 1 | 1 | 1 | 0.012 |
| SE | 37392 | ENSG00000010327 | STAB1 | chr3 | + | 52504460 | 52504549 | 52504027 | 52504155 | 52504738 | 52504876 | 84 | 38 | 57 | 50 | 20 | 0 | 0 | 1 | 0 | 0 | 234 | 170 | 86 | 119 | 366 | 0 | 0 | 0 | 0 | 0 | 238 | 149 | 1 | 1 | 1 | 0.96 | 1 | 1 | 1 | 1 | 1 | 1 | 1 | 1 | -0.008 |
| SE | 37395 | ENSG00000010327 | STAB1 | chr3 | + | 52514700 | 52514829 | 52514364 | 52514496 | 52514988 | 52515045 | 46 | 17 | 34 | 34 | 4 | 0 | 0 | 0 | 0 | 0 | 116 | 67 | 50 | 101 | 182 | 0 | 0 | 1 | 0 | 0 | 278 | 149 | 1 | 1 | 1 | 1 | 1 | 1 | 1 | 1 | 1 | 0.964 | 1 | 1 | 0.007 |
| SE | 37396 | ENSG00000010327 | STAB1 | chr3 | + | 52516042 | 52516238 | 52515422 | 52515506 | 52516355 | 52516451 | 92 | 61 | 68 | 87 | 9 | 0 | 0 | 0 | 3 | 1 | 203 | 195 | 122 | 159 | 353 | 0 | 0 | 3 | 4 | 0 | 298 | 149 | 1 | 1 | 1 | 1 | 1 | 0.935 | 0.818 | 1 | 1 | 0.953 | 0.952 | 1 | -0.03 |
| SE | 37397 | ENSG00000010327 | STAB1 | chr3 | + | 52521360 | 52521510 | 52520641 | 52521005 | 52521595 | 52521700 | 28 | 21 | 20 | 23 | 0 | 2 | 2 | 0 | 0 | 0 | 55 | 34 | 21 | 35 | 31 | 0 | 0 | 0 | 0 | 0 | 298 | 149 | 1 | 1 | 0.875 | 0.84 | 1 | 1 | NA | 1 | 1 | 1 | 1 | 1 | -0.071 |
| SE | 37399 | ENSG00000143862 | ARL8A | chr1 | - | 202135149 | 202135220 | 202133403 | 202134516 | 202135458 | 202135526 | 263 | 164 | 159 | 252 | 531 | 0 | 0 | 0 | 0 | 0 | 728 | 298 | 403 | 486 | 563 | 0 | 0 | 0 | 1 | 0 | 220 | 149 | 1 | 1 | 1 | 1 | 1 | 1 | 1 | 1 | 1 | 1 | 0.997 | 1 | 0.001 |
| SE | 37401 | ENSG00000143862 | ARL8A | chr1 | - | 202135706 | 202135800 | 202135458 | 202135526 | 202138367 | 202138448 | 117 | 70 | 67 | 111 | 208 | 0 | 0 | 0 | 2 | 0 | 314 | 129 | 187 | 175 | 268 | 0 | 0 | 0 | 0 | 0 | 243 | 149 | 1 | 1 | 1 | 1 | 1 | 0.971 | 1 | 1 | 1 | 1 | 1 | 1 | -0.006 |
| SE | 37402 | ENSG00000143862 | ARL8A | chr1 | - | 202137964 | 202138038 | 202135706 | 202135800 | 202138367 | 202138448 | 283 | 136 | 175 | 253 | 278 | 0 | 0 | 0 | 0 | 2 | 485 | 363 | 311 | 391 | 481 | 0 | 0 | 0 | 0 | 6 | 223 | 149 | 1 | 1 | 1 | 1 | 1 | 1 | 0.989 | 1 | 1 | 1 | 1 | 0.982 | 0.001 |
| SE | 37403 | ENSG00000143862 | ARL8A | chr1 | - | 202138367 | 202138448 | 202135706 | 202135800 | 202144449 | 202144737 | 60 | 16 | 42 | 35 | 37 | 0 | 0 | 0 | 1 | 0 | 63 | 76 | 41 | 85 | 105 | 0 | 3 | 0 | 0 | 0 | 230 | 149 | 1 | 1 | 1 | 1 | 1 | 0.958 | 1 | 1 | 0.943 | 1 | 1 | 1 | 0.003 |
| SE | 37404 | ENSG00000116489 | CAPZA1 | chr1 | + | 112646754 | 112646794 | 112619831 | 112619883 | 112647209 | 112647273 | 0 | 9 | 2 | 0 | 2 | 648 | 457 | 539 | 552 | 925 | 0 | 4 | 1 | 0 | 8 | 107 | 748 | 370 | 240 | 95 | 189 | 149 | 1 | 1 | 0 | 0.015 | 0.003 | 0 | 0.002 | 0 | 0.004 | 0.002 | 0 | 0.062 | -0.01 |
| SE | 37405 | ENSG00000116489 | CAPZA1 | chr1 | + | 112647209 | 112647273 | 112619830 | 112619883 | 112649417 | 112649469 | 971 | 704 | 780 | 856 | 1390 | 0 | 0 | 0 | 0 | 1 | 199 | 1146 | 550 | 403 | 135 | 0 | 0 | 0 | 0 | 0 | 213 | 149 | 1 | 1 | 1 | 1 | 1 | 1 | 0.999 | 1 | 1 | 1 | 1 | 1 | 0 |
| SE | 37407 | ENSG00000116489 | CAPZA1 | chr1 | + | 112654464 | 112654671 | 112653597 | 112653661 | 112659021 | 112659101 | 3102 | 2105 | 2584 | 2683 | 5295 | 2 | 0 | 0 | 1 | 0 | 474 | 2638 | 1433 | 1065 | 669 | 0 | 0 | 0 | 0 | 0 | 298 | 149 | 1 | 1 | 0.999 | 1 | 1 | 0.999 | 1 | 1 | 1 | 1 | 1 | 1 | 0 |

| | | | | | | | | | | | | | | | | | | | | | | | | | | | | | | | | | | | | | | | | | | | | | | |
|---|---|---|---|---|---|---|---|---|---|---|---|---|---|---|---|---|---|---|---|---|---|---|---|---|---|---|---|---|---|---|---|---|---|---|---|---|---|---|---|---|---|---|---|---|---|---|
| SE | 37408 | ENSG00000116489 | CAPZA1 | chr1 | + | 112659021 | 112659101 | 112654464 | 112654671 | 112659700 | 112659779 | 2298 | 1495 | 1717 | 1921 | 3632 | 1 | 0 | 1 | 0 | 0 | 320 | 1804 | 1044 | 754 | 484 | 0 | 0 | 0 | 0 | 0 | 229 | 149 | 1 | 1 | 0.999 | 1 | 0.999 | 1 | 1 | 1 | 1 | 1 | 1 | 1 | 0 |
| SE | 37409 | ENSG00000116489 | CAPZA1 | chr1 | + | 112667073 | 112667145 | 112659328 | 112659779 | 112669542 | 112669605 | 1735 | 1107 | 1137 | 1334 | 2482 | 1 | 0 | 0 | 0 | 0 | 208 | 1250 | 748 | 454 | 115 | 0 | 0 | 0 | 0 | 0 | 221 | 149 | 1 | 1 | 0.999 | 1 | 1 | 1 | 1 | 1 | 1 | 1 | 1 | 1 | 0 |
| SE | 37426 | ENSG00000114850 | SSR3 | chr3 | - | 156548865 | 156549003 | 156544307 | 156544439 | 156553654 | 156553781 | 186 | 93 | 207 | 157 | 254 | 1 | 0 | 1 | 0 | 0 | 182 | 122 | 180 | 108 | 59 | 0 | 0 | 0 | 0 | 0 | 287 | 149 | 1 | 1 | 0.99 | 1 | 0.991 | 1 | 1 | 1 | 1 | 1 | 1 | 1 | -0.004 |
| SE | 37427 | ENSG00000114850 | SSR3 | chr3 | - | 156548904 | 156549003 | 156543953 | 156544439 | 156553654 | 156553781 | 551 | 383 | 669 | 499 | 1043 | 1 | 0 | 1 | 0 | 0 | 573 | 320 | 628 | 347 | 228 | 0 | 0 | 0 | 0 | 0 | 248 | 149 | 1 | 1 | 0.997 | 1 | 0.998 | 1 | 1 | 1 | 1 | 1 | 1 | 1 | -0.001 |
| SE | 37428 | ENSG00000114850 | SSR3 | chr3 | - | 156553654 | 156553781 | 156548865 | 156549003 | 156554956 | 156555117 | 359 | 191 | 419 | 322 | 411 | 1 | 0 | 0 | 0 | 0 | 314 | 236 | 342 | 274 | 90 | 0 | 0 | 0 | 7 | 0 | 276 | 149 | 1 | 1 | 1 | 1 | 1 | 1 | 1 | 1 | 1 | 1 | 0.955 | 1 | 0.009 |
| SE | 37434 | ENSG00000153157 | SYCP2L | chr6 | + | 10902676 | 10902762 | 10898823 | 10898848 | 10906019 | 10906054 | 8 | 3 | 7 | 2 | 0 | 0 | 0 | 0 | 1 | 0 | 0 | 0 | 4 | 0 | 0 | 0 | 0 | 0 | 0 | 0 | 235 | 149 | 1 | 1 | 1 | 1 | 1 | 0.559 | NA | NA | NA | 1 | NA | NA | -0.11 |
| SE | 37435 | ENSG00000153157 | SYCP2L | chr6 | + | 10902874 | 10902963 | 10902676 | 10902762 | 10906019 | 10906054 | 3 | 0 | 1 | 9 | 3 | 0 | 2 | 0 | 0 | 0 | 10 | 0 | 0 | 0 | 0 | 0 | 0 | 4 | 0 | 0 | 238 | 149 | 1 | 1 | 1 | 0 | 1 | 1 | 1 | 1 | NA | 0 | NA | NA | 0.3 |
| SE | 37438 | ENSG00000153157 | SYCP2L | chr6 | + | 10942458 | 10942529 | 10935057 | 10935187 | 10942676 | 10942746 | 4 | 2 | 0 | 7 | 8 | 0 | 0 | 0 | 1 | 0 | 7 | 0 | 0 | 0 | 0 | 0 | 0 | 0 | 0 | 0 | 220 | 149 | 1 | 1 | 1 | 1 | NA | 0.826 | 1 | 1 | NA | NA | NA | NA | -0.043 |
| SE | 37447 | ENSG00000189180 | ZNF33A | chr10 | + | 38012297 | 38012350 | 38010649 | 38010783 | 38016870 | 38017015 | 91 | 66 | 98 | 92 | 83 | 0 | 1 | 0 | 2 | 0 | 29 | 34 | 56 | 47 | 55 | 0 | 0 | 0 | 0 | 0 | 202 | 149 | 1 | 1 | 1 | 0.98 | 1 | 0.971 | 1 | 1 | 1 | 1 | 1 | 1 | -0.01 |
| SE | 37451 | ENSG00000158427 | TMSB15B | chrX | + | 103964227 | 103964294 | 103962674 | 103962723 | 103964505 | 103964622 | 1 | 0 | 1 | 0 | 0 | 0 | 2 | 0 | 0 | 0 | 0 | 0 | 4 | 0 | 0 | 0 | 0 | 0 | 0 | 0 | 216 | 149 | 1 | 1 | 1 | 0 | 1 | NA | NA | NA | NA | 1 | NA | NA | -0.333 |
| SE | 37452 | ENSG00000163681 | SLMAP | chr3 | + | 57820612 | 57820705 | 57757119 | 57757849 | 57831382 | 57831530 | 1 | 1 | 0 | 0 | 0 | 84 | 76 | 101 | 98 | 105 | 0 | 0 | 0 | 0 | 0 | 49 | 68 | 62 | 62 | 52 | 242 | 149 | 1 | 1 | 0.007 | 0.008 | 0 | 0 | 0 | 0 | 0 | 0 | 0 | 0 | 0.003 |
| SE | 37453 | ENSG00000163681 | SLMAP | chr3 | + | 57841298 | 57841371 | 57831382 | 57831530 | 57847196 | 57847233 | 127 | 92 | 86 | 84 | 140 | 0 | 0 | 1 | 0 | 0 | 49 | 86 | 76 | 55 | 27 | 0 | 0 | 0 | 0 | 0 | 222 | 149 | 1 | 1 | 1 | 1 | 0.983 | 1 | 1 | 1 | 1 | 1 | 1 | 1 | -0.003 |
| SE | 37455 | ENSG00000163681 | SLMAP | chr3 | + | 57860698 | 57860839 | 57858087 | 57858159 | 57861948 | 57862086 | 274 | 334 | 253 | 346 | 515 | 0 | 0 | 0 | 0 | 2 | 358 | 392 | 471 | 367 | 156 | 0 | 0 | 0 | 6 | 0 | 290 | 149 | 1 | 1 | 1 | 1 | 1 | 1 | 0.992 | 1 | 1 | 1 | 0.969 | 1 | 0.005 |
| SE | 37459 | ENSG00000163681 | SLMAP | chr3 | + | 57864806 | 57864857 | 57864547 | 57864716 | 57890045 | 57891206 | 17 | 21 | 25 | 38 | 34 | 0 | 0 | 2 | 0 | 0 | 1 | 20 | 20 | 29 | 5 | 0 | 0 | 0 | 0 | 0 | 200 | 149 | 1 | 1 | 1 | 1 | 0.903 | 1 | 1 | 1 | 1 | 1 | 1 | 1 | -0.019 |
| SE | 37461 | ENSG00000163681 | SLMAP | chr3 | + | 57865241 | 57865292 | 57864547 | 57864716 | 57871635 | 57871698 | 6 | 8 | 13 | 7 | 19 | 7 | 11 | 7 | 8 | 11 | 0 | 2 | 31 | 8 | 5 | 0 | 7 | 8 | 0 | 0 | 200 | 149 | 1 | 1 | 0.39 | 0.351 | 0.58 | 0.395 | 0.563 | NA | 0.176 | 0.743 | 1 | 1 | -0.274 |
| SE | 37463 | ENSG00000163681 | SLMAP | chr3 | + | 57865241 | 57865292 | 57864547 | 57864716 | 57890045 | 57891206 | 0 | 3 | 1 | 1 | 3 | 0 | 0 | 2 | 0 | 0 | 0 | 0 | 14 | 0 | 5 | 0 | 0 | 0 | 0 | 0 | 200 | 149 | 1 | 1 | NA | 1 | 0.271 | 1 | 1 | NA | NA | 1 | NA | 1 | -0.182 |
| SE | 37466 | ENSG00000163681 | SLMAP | chr3 | + | 57865241 | 57865292 | 57864806 | 57864857 | 57890045 | 57891206 | 24 | 28 | 25 | 24 | 23 | 0 | 0 | 2 | 2 | 0 | 5 | 31 | 41 | 18 | 11 | 0 | 0 | 0 | 0 | 0 | 200 | 149 | 1 | 1 | 1 | 1 | 0.903 | 0.899 | 1 | 1 | 1 | 1 | 1 | 1 | -0.04 |
| SE | 37467 | ENSG00000163681 | SLMAP | chr3 | + | 57871635 | 57871698 | 57864547 | 57864716 | 57890040 | 57890100 | 17 | 30 | 25 | 37 | 36 | 38 | 40 | 10 | 50 | 45 | 0 | 33 | 32 | 8 | 0 | 7 | 8 | 3 | 4 | 0 | 212 | 149 | 1 | 1 | 0.239 | 0.345 | 0.637 | 0.342 | 0.36 | 0 | 0.744 | 0.882 | 0.584 | NA | -0.168 |
| SE | 37468 | ENSG00000163681 | SLMAP | chr3 | + | 57871635 | 57871698 | 57864547 | 57864716 | 57890045 | 57891206 | 4 | 4 | 2 | 2 | 6 | 0 | 0 | 2 | 0 | 0 | 0 | 2 | 5 | 0 | 0 | 0 | 0 | 0 | 0 | 0 | 212 | 149 | 1 | 1 | 1 | 1 | 0.413 | 1 | 1 | NA | 1 | 1 | NA | NA | -0.117 |
| SE | 37472 | ENSG00000163681 | SLMAP | chr3 | + | 57871635 | 57871698 | 57864806 | 57864857 | 57890045 | 57891206 | 26 | 22 | 23 | 34 | 34 | 0 | 0 | 2 | 2 | 0 | 1 | 17 | 20 | 22 | 12 | 0 | 0 | 0 | 0 | 0 | 212 | 149 | 1 | 1 | 1 | 1 | 0.89 | 0.923 | 1 | 1 | 1 | 1 | 1 | 1 | -0.037 |
| SE | 37473 | ENSG00000163681 | SLMAP | chr3 | + | 57871635 | 57871698 | 57864806 | 57864857 | 57896510 | 57896591 | 26 | 22 | 23 | 34 | 34 | 0 | 2 | 0 | 0 | 0 | 1 | 17 | 19 | 22 | 12 | 0 | 0 | 0 | 0 | 0 | 212 | 149 | 1 | 1 | 1 | 0.885 | 1 | 1 | 1 | 1 | 1 | 1 | 1 | 1 | -0.023 |
| SE | 37475 | ENSG00000163681 | SLMAP | chr3 | + | 57871635 | 57871698 | 57865241 | 57865292 | 57890045 | 57891206 | 19 | 16 | 16 | 17 | 16 | 0 | 2 | 0 | 0 | 0 | 7 | 20 | 17 | 14 | 6 | 0 | 0 | 0 | 0 | 0 | 212 | 149 | 1 | 1 | 1 | 0.849 | 1 | 1 | 1 | 1 | 1 | 1 | 1 | 1 | -0.03 |
| SE | 37476 | ENSG00000163681 | SLMAP | chr3 | + | 57883960 | 57884026 | 57864806 | 57864857 | 57890040 | 57890100 | 0 | 1 | 0 | 0 | 0 | 68 | 26 | 27 | 41 | 47 | 0 | 0 | 0 | 0 | 0 | 0 | 17 | 10 | 19 | 8 | 215 | 149 | 1 | 1 | 0 | 0.026 | 0 | 0 | 0 | NA | 0 | 0 | 0 | 0 | 0.005 |
| SE | 37477 | ENSG00000163681 | SLMAP | chr3 | + | 57883960 | 57884026 | 57871635 | 57871698 | 57890040 | 57890100 | 0 | 1 | 0 | 2 | 0 | 27 | 44 | 35 | 48 | 56 | 0 | 0 | 2 | 0 | 0 | 0 | 57 | 44 | 13 | 0 | 215 | 149 | 1 | 1 | 0 | 0.016 | 0 | 0.028 | 0 | NA | 0 | 0.031 | 0 | NA | -0.002 |
| SE | 37478 | ENSG00000163681 | SLMAP | chr3 | + | 57890040 | 57890100 | 57864547 | 57864716 | 57896246 | 57896371 | 18 | 15 | 5 | 29 | 24 | 0 | 1 | 0 | 0 | 0 | 0 | 19 | 15 | 9 | 5 | 0 | 0 | 0 | 0 | 0 | 209 | 149 | 1 | 1 | 1 | 0.914 | 1 | 1 | 1 | NA | 1 | 1 | 1 | 1 | -0.017 |
| SE | 37480 | ENSG00000163681 | SLMAP | chr3 | + | 57890040 | 57890100 | 57864806 | 57864857 | 57896510 | 57896590 | 93 | 52 | 43 | 61 | 75 | 0 | 2 | 0 | 0 | 0 | 8 | 18 | 23 | 36 | 23 | 0 | 0 | 0 | 0 | 0 | 209 | 149 | 1 | 1 | 1 | 0.949 | 1 | 1 | 1 | 1 | 1 | 1 | 1 | 1 | -0.01 |
| SE | 37481 | ENSG00000163681 | SLMAP | chr3 | + | 57890040 | 57890100 | 57871635 | 57871698 | 57896510 | 57896590 | 51 | 68 | 53 | 68 | 84 | 0 | 0 | 0 | 1 | 0 | 8 | 47 | 45 | 28 | 15 | 0 | 0 | 0 | 0 | 0 | 209 | 149 | 1 | 1 | 1 | 1 | 1 | 0.98 | 1 | 1 | 1 | 1 | 1 | 1 | -0.004 |
| SE | 37487 | ENSG00000163681 | SLMAP | chr3 | + | 57912380 | 57912701 | 57909075 | 57909150 | 57916905 | 57917077 | 109 | 127 | 123 | 86 | 182 | 0 | 0 | 0 | 3 | 1 | 0 | 71 | 56 | 61 | 7 | 0 | 0 | 0 | 0 | 0 | 298 | 149 | 1 | 1 | 1 | 1 | 1 | 0.935 | 0.989 | NA | 1 | 1 | 1 | 1 | -0.015 |
| SE | 37488 | ENSG00000163681 | SLMAP | chr3 | + | 57912380 | 57912701 | 57909075 | 57909150 | 57922888 | 57923023 | 109 | 127 | 123 | 86 | 182 | 0 | 0 | 0 | 2 | 0 | 0 | 73 | 52 | 61 | 7 | 0 | 0 | 0 | 0 | 0 | 298 | 149 | 1 | 1 | 1 | 1 | 1 | 0.956 | 1 | NA | 1 | 1 | 1 | 1 | -0.009 |
| SE | 37489 | ENSG00000163681 | SLMAP | chr3 | + | 57913157 | 57913275 | 57912294 | 57912701 | 57916905 | 57916971 | 128 | 125 | 113 | 130 | 311 | 0 | 0 | 0 | 0 | 0 | 26 | 126 | 91 | 93 | 8 | 0 | 1 | 4 | 0 | 0 | 267 | 149 | 1 | 1 | 1 | 1 | 1 | 1 | 1 | 1 | 0.986 | 0.927 | 1 | 1 | 0.017 |
| SE | 37490 | ENSG00000163681 | SLMAP | chr3 | + | 57913157 | 57913275 | 57912294 | 57912701 | 57922888 | 57923023 | 45 | 56 | 33 | 43 | 125 | 0 | 0 | 0 | 0 | 0 | 0 | 38 | 28 | 39 | 3 | 0 | 3 | 0 | 0 | 0 | 267 | 149 | 1 | 1 | 1 | 1 | 1 | 1 | 1 | NA | 0.876 | 1 | 1 | 1 | 0.031 |
| SE | 37491 | ENSG00000163681 | SLMAP | chr3 | + | 57916905 | 57917077 | 57909075 | 57909150 | 57922888 | 57923023 | 105 | 109 | 99 | 126 | 235 | 0 | 0 | 0 | 2 | 0 | 8 | 79 | 75 | 28 | 0 | 0 | 0 | 0 | 0 | 0 | 298 | 149 | 1 | 1 | 1 | 1 | 1 | 0.969 | 1 | 1 | 1 | 1 | 1 | NA | -0.006 |
| SE | 37492 | ENSG00000163681 | SLMAP | chr3 | + | 57916905 | 57917077 | 57912380 | 57912701 | 57922888 | 57923023 | 105 | 109 | 99 | 123 | 234 | 0 | 0 | 0 | 0 | 0 | 8 | 80 | 79 | 28 | 0 | 0 | 3 | 0 | 0 | 0 | 298 | 149 | 1 | 1 | 1 | 1 | 1 | 1 | 1 | 1 | 0.93 | 1 | 1 | NA | 0.017 |
| SE | 37493 | ENSG00000163681 | SLMAP | chr3 | + | 57916905 | 57917077 | 57913157 | 57913275 | 57922888 | 57923023 | 191 | 179 | 184 | 215 | 421 | 0 | 0 | 0 | 1 | 0 | 34 | 167 | 143 | 88 | 5 | 0 | 0 | 0 | 0 | 0 | 298 | 149 | 1 | 1 | 1 | 1 | 1 | 0.991 | 1 | 1 | 1 | 1 | 1 | 1 | -0.002 |
| SE | 37502 | ENSG00000165115 | KIF27 | chr9 | - | 83859155 | 83859371 | 83853628 | 83853835 | 83867683 | 83867860 | 12 | 15 | 20 | 13 | 17 | 0 | 0 | 2 | 2 | 0 | 0 | 11 | 10 | 0 | 0 | 0 | 0 | 0 | 0 | 0 | 298 | 149 | 1 | 1 | 1 | 1 | 0.833 | 0.765 | 1 | NA | 1 | 1 | NA | NA | -0.08 |
| SE | 37503 | ENSG00000165115 | KIF27 | chr9 | - | 83859155 | 83859371 | 83853628 | 83853835 | 83870518 | 83870632 | 6 | 9 | 8 | 10 | 15 | 0 | 2 | 2 | 1 | 0 | 0 | 6 | 10 | 0 | 0 | 0 | 8 | 2 | 0 | 0 | 298 | 149 | 1 | 1 | 1 | 0.692 | 0.667 | 0.833 | 1 | NA | 0.273 | 0.714 | NA | NA | 0.345 |
| SE | 37504 | ENSG00000165115 | KIF27 | chr9 | - | 83867683 | 83867860 | 83853628 | 83853835 | 83870518 | 83870632 | 6 | 9 | 13 | 6 | 2 | 0 | 2 | 2 | 1 | 0 | 0 | 0 | 5 | 2 | 0 | 0 | 8 | 2 | 0 | 0 | 298 | 149 | 1 | 1 | 1 | 0.692 | 0.765 | 0.75 | 1 | NA | 0 | 0.556 | 1 | NA | 0.323 |
| SE | 37505 | ENSG00000165115 | KIF27 | chr9 | - | 83867683 | 83867860 | 83859155 | 83859371 | 83870518 | 83870632 | 12 | 19 | 23 | 13 | 9 | 0 | 4 | 0 | 6 | 5 | 0 | 5 | 5 | 2 | 0 | 0 | 0 | 0 | 0 | 0 | 298 | 149 | 1 | 1 | 1 | 0.704 | 1 | 0.52 | 0.474 | NA | 1 | 1 | 1 | NA | -0.26 |
| SE | 37507 | ENSG00000165115 | KIF27 | chr9 | - | 83880296 | 83880494 | 83870518 | 83870632 | 83883812 | 83884018 | 14 | 15 | 10 | 25 | 3 | 0 | 1 | 1 | 0 | 0 | 0 | 5 | 5 | 2 | 0 | 0 | 0 | 0 | 0 | 0 | 298 | 149 | 1 | 1 | 1 | 0.882 | 0.833 | 1 | 1 | NA | 1 | 1 | 1 | NA | -0.057 |
| SE | 37508 | ENSG00000165115 | KIF27 | chr9 | - | 83883812 | 83884018 | 83870518 | 83870632 | 83887040 | 83887196 | 2 | 6 | 5 | 3 | 6 | 1 | 0 | 2 | 4 | 0 | 0 | 3 | 7 | 0 | 0 | 0 | 0 | 0 | 0 | 0 | 298 | 149 | 1 | 1 | 0.5 | 1 | 0.556 | 0.273 | 1 | NA | 1 | 1 | NA | NA | -0.334 |
| SE | 37509 | ENSG00000165115 | KIF27 | chr9 | - | 83903059 | 83904018 | 83899660 | 83899804 | 83908451 | 83908652 | 43 | 21 | 14 | 45 | 20 | 6 | 8 | 11 | 8 | 1 | 11 | 10 | 75 | 5 | 0 | 0 | 3 | 11 | 0 | 0 | 298 | 149 | 1 | 1 | 0.782 | 0.568 | 0.389 | 0.738 | 0.909 | 1 | 0.625 | 0.773 | 1 | NA | -0.172 |
| SE | 37510 | ENSG00000165115 | KIF27 | chr9 | - | 83908451 | 83908652 | 83899660 | 83899804 | 83915293 | 83915678 | 23 | 22 | 25 | 20 | 29 | 0 | 0 | 1 | 0 | 0 | 0 | 14 | 20 | 0 | 0 | 0 | 0 | 0 | 0 | 0 | 298 | 149 | 1 | 1 | 1 | 1 | 0.926 | 1 | 1 | NA | 1 | 1 | NA | NA | -0.015 |
| SE | 37514 | ENSG00000284959 | NA | chr14 | - | 81095433 | 81095566 | 81094214 | 81094324 | 81165192 | 81165381 | 0 | 6 | 0 | 0 | 0 | 0 | 0 | 0 | 0 | 8 | 0 | 0 | 0 | 2 | 9 | 0 | 0 | 0 | 0 | 0 | 282 | 149 | 1 | 1 | NA | 1 | NA | NA | 0 | NA | NA | NA | 1 | 1 | -0.5 |
| SE | 37516 | ENSG00000284959 | NA | chr14 | - | 81099398 | 81099526 | 81095433 | 81095566 | 81165192 | 81165381 | 2 | 11 | 2 | 7 | 9 | 0 | 0 | 0 | 0 | 0 | 1 | 0 | 0 | 3 | 1 | 0 | 0 | 0 | 2 | 0 | 277 | 149 | 1 | 1 | 1 | 1 | 1 | 1 | 1 | 1 | NA | NA | 0.447 | 1 | 0.184 |
| SE | 37521 | ENSG00000130313 | PGLS | chr19 | + | 17517287 | 17517389 | 17515741 | 17516280 | 17517709 | 17517835 | 180 | 73 | 96 | 96 | 96 | 0 | 0 | 2 | 0 | 0 | 243 | 165 | 91 | 186 | 349 | 0 | 0 | 0 | 0 | 0 | 251 | 149 | 1 | 1 | 1 | 1 | 0.966 | 1 | 1 | 1 | 1 | 1 | 1 | 1 | -0.007 |
| SE | 37522 | ENSG00000130313 | PGLS | chr19 | + | 17517709 | 17517850 | 17517287 | 17517389 | 17520943 | 17521030 | 262 | 95 | 174 | 188 | 126 | 1 | 0 | 0 | 0 | 0 | 392 | 201 | 148 | 236 | 474 | 0 | 0 | 0 | 0 | 0 | 290 | 149 | 1 | 1 | 0.993 | 1 | 1 | 1 | 1 | 1 | 1 | 1 | 1 | 1 | -0.001 |
| SE | 37524 | ENSG00000206531 | CD200R1L | chr3 | - | 112837941 | 112838010 | 112829318 | 112829384 | 112845678 | 112845950 | 0 | 0 | 2 | 1 | 5 | 0 | 0 | 0 | 4 | 1 | 6 | 0 | 0 | 0 | 0 | 0 | 0 | 0 | 0 | 0 | 218 | 149 | 1 | 1 | NA | NA | 1 | 0.146 | 0.774 | 1 | NA | NA | NA | NA | -0.36 |
| SE | 37526 | ENSG00000105281 | SLC1A5 | chr19 | - | 46778674 | 46778908 | 46777210 | 46777405 | 46782382 | 46782549 | 133 | 49 | 83 | 66 | 125 | 0 | 0 | 1 | 4 | 0 | 173 | 157 | 40 | 87 | 115 | 0 | 0 | 0 | 1 | 5 | 298 | 149 | 1 | 1 | 1 | 1 | 0.976 | 0.892 | 1 | 1 | 1 | 1 | 0.978 | 0.92 | -0.006 |
| SE | 37530 | ENSG00000167920 | TMEM99 | chr17 | + | 40828056 | 40828296 | 40819121 | 40819208 | 40833114 | 40833199 | 2 | 0 | 0 | 0 | 0 | 9 | 9 | 8 | 14 | 7 | 0 | 0 | 0 | 0 | 0 | 6 | 5 | 0 | 1 | 0 | 298 | 149 | 1 | 1 | 0.1 | 0 | 0 | 0 | 0 | 0 | 0 | NA | 0 | NA | 0.02 |
| SE | 37531 | ENSG00000167920 | TMEM99 | chr17 | + | 40828103 | 40828296 | 40819144 | 40819208 | 40833114 | 40833199 | 2 | 0 | 0 | 0 | 0 | 9 | 9 | 8 | 14 | 7 | 0 | 0 | 0 | 0 | 0 | 6 | 5 | 0 | 1 | 0 | 298 | 149 | 1 | 1 | 0.1 | 0 | 0 | 0 | 0 | 0 | 0 | NA | 0 | NA | 0.02 |
| SE | 37533 | ENSG00000130270 | ATP8B3 | chr19 | - | 1806627 | 1806689 | 1806096 | 1806169 | 1807167 | 1807266 | 4 | 5 | 2 | 7 | 0 | 0 | 0 | 0 | 4 | 0 | 0 | 20 | 4 | 20 | 22 | 0 | 0 | 0 | 0 | 0 | 211 | 149 | 1 | 1 | 1 | 1 | 1 | 0.553 | NA | NA | 1 | 1 | 1 | 1 | -0.112 |
| SE | 37544 | ENSG00000108064 | TFAM | chr10 | + | 58386649 | 58386716 | 58386219 | 58386338 | 58388189 | 58388260 | 0 | 0 | 2 | 0 | 3 | 147 | 91 | 115 | 136 | 184 | 0 | 0 | 0 | 0 | 0 | 29 | 120 | 117 | 93 | 6 | 216 | 149 | 1 | 1 | 0 | 0 | 0.012 | 0 | 0.011 | 0 | 0 | 0 | 0 | 0 | 0.005 |
| SE | 37545 | ENSG00000108064 | TFAM | chr10 | + | 58390764 | 58390860 | 58388669 | 58388819 | 58394357 | 58394414 | 307 | 204 | 260 | 258 | 560 | 15 | 25 | 21 | 42 | 34 | 80 | 129 | 150 | 86 | 0 | 17 | 12 | 32 | 35 | 0 | 245 | 149 | 1 | 1 | 0.926 | 0.832 | 0.883 | 0.789 | 0.909 | 0.741 | 0.867 | 0.74 | 0.599 | NA | 0.131 |
| SE | 37578 | ENSG00000082074 | FYB1 | chr5 | - | 39118873 | 39119036 | 39110355 | 39110389 | 39122335 | 39122402 | 681 | 594 | 660 | 723 | 1845 | 1 | 0 | 0 | 0 | 3 | 118 | 412 | 233 | 244 | 60 | 0 | 0 | 0 | 0 | 0 | 298 | 149 | 1 | 1 | 0.997 | 1 | 1 | 1 | 0.997 | 1 | 1 | 1 | 1 | 1 | -0.001 |
| SE | 37579 | ENSG00000082074 | FYB1 | chr5 | - | 39119534 | 39119634 | 39118873 | 39119036 | 39122335 | 39122402 | 1415 | 1551 | 1414 | 1584 | 2377 | 0 | 0 | 3 | 1 | 0 | 521 | 1047 | 886 | 756 | 271 | 0 | 2 | 0 | 0 | 0 | 249 | 149 | 1 | 1 | 1 | 1 | 0.996 | 0.999 | 1 | 1 | 0.997 | 1 | 1 | 1 | 0 |
| SE | 37580 | ENSG00000082074 | FYB1 | chr5 | - | 39124252 | 39124278 | 39122335 | 39122402 | 39130589 | 39130612 | 292 | 446 | 296 | 399 | 394 | 0 | 0 | 0 | 1 | 0 | 166 | 325 | 375 | 270 | 72 | [illegible] | [illegible] | [illegible] | [illegible] | [illegible] | 175 | 149 | [illegible] | [illegible] | [illegible] | [illegible] | [illegible] | [illegible] | [illegible] | [illegible] | [illegible] | [illegible] | [illegible] | [illegible] | -0.001 |
| SE | 37583 | ENSG00000082074 | FYB1 | chr5 | - | 39127740 | 39127807 | 39124252 | 39124278 | 39130589 | 39130612 | 1208 | 1133 | 1348 | 1369 | 2687 | 2 | 1 | 1 | 1 | 0 | 393 | 923 | 655 | 628 | 273 | 0 | 0 | 0 | 0 | 0 | 216 | 149 | 1 | 1 | 0.998 | 0.999 | 0.999 | 0.999 | 1 | 1 | 1 | 1 | 1 | 1 | -0.001 |
| SE | 37584 | ENSG00000082074 | FYB1 | chr5 | - | 39130589 | 39130612 | 39127740 | 39127807 | 39134207 | 39134349 | 1599 | 1791 | 1451 | 1799 | 2960 | 0 | 0 | 1 | 0 | 0 | 843 | 1317 | 1249 | 1287 | 449 | 0 | 0 | 1 | 0 | 0 | 172 | 149 | 1 | 1 | 1 | 1 | 0.999 | 1 | 1 | 1 | 1 | 0.999 | 1 | 1 | 0 |
| SE | 37586 | ENSG00000082074 | FYB1 | chr5 | - | 39141094 | 39141141 | 39138656 | 39138691 | 39153447 | 39153604 | 88 | 143 | 125 | 152 | 101 | 0 | 0 | 0 | 1 | 0 | 141 | 59 | 111 | 76 | 150 | 0 | 0 | 0 | 0 | 0 | 196 | 149 | 1 | 1 | 1 | 1 | 1 | 0.991 | 1 | 1 | 1 | 1 | 1 | 1 | -0.002 |
| SE | 37588 | ENSG00000082074 | FYB1 | chr5 | - | 39153447 | 39153604 | 39141094 | 39141141 | 39270568 | 39270649 | 611 | 901 | 843 | 917 | 809 | 1 | 0 | 0 | 1 | 0 | 505 | 460 | 648 | 580 | 512 | 0 | 0 | 0 | 0 | 0 | 298 | 149 | 1 | 1 | 0.997 | 1 | 1 | 0.998 | 1 | 1 | 1 | 1 | 1 | 1 | -0.001 |
| SE | 37589 | ENSG00000082074 | FYB1 | chr5 | - | 39201825 | 39202987 | 39141094 | 39141141 | 39270568 | 39270649 | 109 | 82 | 62 | 126 | 101 | 1 | 0 | 0 | 1 | 0 | 162 | 123 | 77 | 90 | 159 | 0 | 0 | 0 | 0 | 0 | 298 | 149 | 1 | 1 | 0.982 | 1 | 1 | 0.984 | 1 | 1 | 1 | 1 | 1 | 1 | -0.007 |
| SE | 37590 | ENSG00000082074 | FYB1 | chr5 | - | 39201825 | 39202987 | 39153447 | 39153604 | 39219442 | 39219563 | 709 | 824 | 733 | 940 | 883 | 2 | 0 | 1 | 1 | 2 | 861 | 482 | 654 | 789 | 498 | 0 | 3 | 1 | 8 | 0 | 298 | 149 | 1 | 1 | 0.994 | 1 | 0.997 | 0.998 | 0.995 | 1 | 0.988 | 0.997 | 0.98 | 1 | 0.004 |
| SE | 37591 | ENSG00000082074 | FYB1 | chr5 | - | 39201825 | 39202987 | 39153447 | 39153604 | 39270568 | 39270649 | 668 | 791 | 713 | 913 | 883 | 0 | 0 | 1 | 1 | 0 | 764 | 495 | 616 | 683 | 591 | 0 | 0 | 3 | 0 | 0 | 298 | 149 | 1 | 1 | 1 | 1 | 0.997 | 0.998 | 1 | 1 | 1 | 0.99 | 1 | 1 | 0.001 |
| SE | 37598 | ENSG00000082074 | FYB1 | chr5 | - | 39211057 | 39211091 | 39153447 | 39153604 | 39212677 | 39213017 | 0 | 0 | 2 | 0 | 0 | 0 | 0 | 0 | 0 | 12 | 0 | 0 | 0 | 4 | 0 | 0 | 1 | 0 | 0 | 0 | 183 | 149 | 1 | 1 | NA | NA | 1 | NA | 0 | NA | 0 | NA | 1 | NA | 0 |
| SE | 37599 | ENSG00000082074 | FYB1 | chr5 | - | 39211057 | 39211091 | 39153447 | 39153604 | 39219442 | 39219563 | 0 | 0 | 0 | 0 | 2 | 2 | 0 | 1 | 1 | 2 | 0 | 0 | 0 | 5 | 0 | 0 | 3 | 1 | 8 | 0 | 183 | 149 | 1 | 1 | 0 | NA | 0 | 0 | 0.449 | NA | 0 | 0 | 0.337 | NA | 0 |
| SE | 37600 | ENSG00000082074 | FYB1 | chr5 | - | 39211057 | 39211091 | 39153447 | 39153604 | 39250670 | 39250806 | 0 | 0 | 0 | 0 | 0 | 3 | 0 | 0 | 2 | 1 | 0 | 0 | 0 | 4 | 0 | 0 | 2 | 1 | 0 | 0 | 183 | 149 | 1 | 1 | 0 | NA | NA | 0 | 0 | NA | 0 | 0 | 1 | NA | -0.333 |
| SE | 37601 | ENSG00000082074 | FYB1 | chr5 | - | 39211057 | 39211091 | 39153447 | 39153604 | 39270568 | 39270828 | 0 | 0 | 0 | 0 | 0 | 0 | 0 | 1 | 1 | 0 | 0 | 0 | 0 | 4 | 0 | 0 | 0 | 3 | 0 | 0 | 183 | 149 | 1 | 1 | NA | NA | 0 | 0 | NA | NA | NA | 0 | 1 | NA | -0.5 |
| SE | 37602 | ENSG00000082074 | FYB1 | chr5 | - | 39211057 | 39211091 | 39153447 | 39153604 | 39274402 | 39274528 | 0 | 0 | 0 | 0 | 0 | 0 | 2 | 0 | 2 | 0 | 0 | 0 | 0 | 4 | 0 | 0 | 0 | 0 | 0 | 0 | 183 | 149 | 1 | 1 | NA | 0 | NA | 0 | NA | NA | NA | NA | 1 | NA | -1 |
| SE | 37607 | ENSG00000082074 | FYB1 | chr5 | - | 39211057 | 39211091 | 39202893 | 39202987 | 39274402 | 39274528 | 1 | 1 | 7 | 2 | 11 | 12 | 24 | 17 | 9 | 17 | 0 | 9 | 4 | 8 | 0 | 0 | 16 | 16 | 34 | 0 | 183 | 149 | 1 | 1 | 0.064 | 0.033 | 0.251 | 0.153 | 0.345 | NA | 0.314 | 0.169 | 0.161 | NA | -0.045 |
| SE | 37609 | ENSG00000082074 | FYB1 | chr5 | - | 39211057 | 39211091 | 39208863 | 39208964 | 39219442 | 39219563 | 0 | 0 | 0 | 0 | 4 | 2 | 0 | 0 | 0 | 0 | 0 | 0 | 1 | 3 | 1 | 3 | 6 | 1 | 0 | 0 | 183 | 149 | 1 | 1 | 0 | NA | NA | NA | 1 | 0 | 0 | 0.449 | 1 | 1 | 0.01 |
| SE | 37610 | ENSG00000082074 | FYB1 | chr5 | - | 39211057 | 39211091 | 39208863 | 39208964 | 39250670 | 39250806 | 0 | 0 | 0 | 0 | 4 | 3 | 0 | 0 | 0 | 0 | 0 | 5 | 1 | 1 | 1 | 0 | 0 | 2 | 0 | 0 | 183 | 149 | 1 | 1 | 0 | NA | NA | NA | 1 | NA | 1 | 0.289 | 1 | 1 | -0.322 |
| SE | 37611 | ENSG00000082074 | FYB1 | chr5 | - | 39211057 | 39211091 | 39208863 | 39208964 | 39270568 | 39270828 | 0 | 0 | 0 | 0 | 4 | 5 | 2 | 0 | 1 | 1 | 0 | 2 | 1 | 1 | 1 | 0 | 3 | 3 | 0 | 3 | 183 | 149 | 1 | 1 | 0 | 0 | NA | 0 | 0.765 | NA | 0.352 | 0.213 | 1 | 0.213 | -0.253 |
| SE | 37612 | ENSG00000082074 | FYB1 | chr5 | - | 39211057 | 39211091 | 39208863 | 39208964 | 39274402 | 39274528 | 0 | 0 | 0 | 0 | 4 | 2 | 0 | 0 | 1 | 1 | 0 | 0 | 1 | 1 | 1 | 0 | 0 | 0 | 0 | 0 | 183 | 149 | 1 | 1 | 0 | NA | NA | 0 | 0.765 | NA | NA | 1 | 1 | 1 | -0.745 |
| SE | 37613 | ENSG00000082074 | FYB1 | chr5 | - | 39212677 | 39213017 | 39153447 | 39153604 | 39219442 | 39219563 | 11 | 10 | 13 | 15 | 13 | 2 | 0 | 1 | 1 | 2 | 0 | 16 | 15 | 6 | 24 | 0 | 3 | 1 | 8 | 0 | 298 | 149 | 1 | 1 | 0.733 | 1 | 0.867 | 0.882 | 0.765 | NA | 0.727 | 0.882 | 0.273 | 1 | 0.129 |
| SE | 37614 | ENSG00000082074 | FYB1 | chr5 | - | 39212677 | 39213017 | 39153447 | 39153604 | 39250670 | 39250806 | 2 | 0 | 1 | 4 | 12 | 3 | 0 | 0 | 2 | 1 | 8 | 2 | 1 | 8 | 4 | 0 | 2 | 1 | 0 | 0 | 298 | 149 | 1 | 1 | 0.25 | NA | 1 | 0.5 | 0.857 | 1 | 0.333 | 0.333 | 1 | 1 | -0.081 |
| SE | 37615 | ENSG00000082074 | FYB1 | chr5 | - | 39212677 | 39213017 | 39153447 | 39153604 | 39270568 | 39270828 | 3 | 0 | 0 | 3 | 14 | 0 | 0 | 1 | 1 | 0 | 0 | 3 | 5 | 4 | 0 | 0 | 0 | 3 | 0 | 0 | 298 | 149 | 1 | 1 | 1 | NA | 0 | 0.6 | 1 | NA | 1 | 0.455 | 1 | NA | -0.168 |
| SE | 37616 | ENSG00000082074 | FYB1 | chr5 | - | 39212677 | 39213017 | 39153447 | 39153604 | 39274402 | 39274528 | 2 | 3 | 0 | 2 | 18 | 0 | 2 | 0 | 2 | 0 | 0 | 1 | 0 | 0 | 0 | 0 | 0 | 0 | 0 | 0 | 298 | 149 | 1 | 1 | 1 | 0.429 | NA | 0.333 | 1 | NA | 1 | NA | NA | NA | -0.309 |
| SE | 37624 | ENSG00000082074 | FYB1 | chr5 | - | 39212677 | 39213017 | 39208863 | 39208964 | 39274402 | 39274528 | 4 | 8 | 1 | 5 | 8 | 2 | 0 | 0 | 1 | 1 | 25 | 4 | 3 | 2 | 10 | 0 | 0 | 0 | 0 | 0 | 298 | 149 | 1 | 1 | 0.5 | 1 | 1 | 0.714 | 0.8 | 1 | 1 | 1 | 1 | 1 | -0.197 |
| SE | 37627 | ENSG00000082074 | FYB1 | chr5 | - | 39212677 | 39213017 | 39211057 | 39211091 | 39270568 | 39270828 | 4 | 4 | 2 | 4 | 7 | 5 | 3 | 4 | 1 | 0 | 0 | 6 | 5 | 4 | 0 | 0 | 14 | 2 | 0 | 6 | 298 | 149 | 1 | 1 | 0.286 | 0.4 | 0.2 | 0.667 | 1 | NA | 0.176 | 0.556 | 1 | 0 | 0.078 |
| SE | 37628 | ENSG00000082074 | FYB1 | chr5 | - | 39212677 | 39213017 | 39211057 | 39211091 | 39274402 | 39274528 | 3 | 7 | 2 | 3 | 11 | 0 | 1 | 1 | 0 | 5 | 0 | 4 | 0 | 0 | 0 | 0 | 0 | 0 | 0 | 0 | 298 | 149 | 1 | 1 | 1 | 0.778 | 0.5 | 1 | 0.524 | NA | 1 | NA | NA | NA | -0.24 |
| SE | 37629 | ENSG00000082074 | FYB1 | chr5 | - | 39250670 | 39250806 | 39153447 | 39153604 | 39270568 | 39270644 | 34 | 2 | 2 | 6 | 4 | 0 | 0 | 1 | 1 | 0 | 7 | 24 | 9 | 4 | 12 | 0 | 0 | 3 | 0 | 0 | 285 | 149 | 1 | 1 | 1 | 1 | 0.511 | 0.758 | 1 | 1 | 1 | 0.611 | 1 | 1 | -0.068 |
| SE | 37630 | ENSG00000082074 | FYB1 | chr5 | - | 39250670 | 39250806 | 39153447 | 39153604 | 39274402 | 39274528 | 4 | 2 | 0 | 2 | 1 | 0 | 2 | 0 | 2 | 0 | 0 | 2 | 1 | 1 | 0 | 0 | 0 | 0 | 0 | 0 | 285 | 149 | 1 | 1 | 1 | 0.343 | NA | 0.343 | 1 | NA | 1 | 1 | 1 | NA | -0.329 |
| SE | 37634 | ENSG00000082074 | FYB1 | chr5 | - | 39250670 | 39250806 | 39208863 | 39208964 | 39274402 | 39274528 | 4 | 2 | 0 | 1 | 0 | 2 | 0 | 0 | 1 | 1 | 0 | 0 | 2 | 1 | 0 | 0 | 0 | 0 | 0 | 0 | 285 | 149 | 1 | 1 | 0.511 | 1 | NA | 0.343 | 0 | NA | NA | 1 | 1 | NA | -0.536 |
| SE | 37636 | ENSG00000082074 | FYB1 | chr5 | - | 39250670 | 39250806 | 39211057 | 39211091 | 39274402 | 39274528 | 1 | 2 | 0 | 1 | 0 | 0 | 1 | 1 | 0 | 5 | 0 | 8 | 3 | 3 | 0 | 0 | 0 | 0 | 0 | 0 | 285 | 149 | 1 | 1 | 1 | 0.511 | 0 | 1 | 0 | NA | 1 | 1 | 1 | NA | -0.498 |
| SE | 37639 | ENSG00000082074 | FYB1 | chr5 | - | 39270568 | 39270828 | 39208863 | 39208964 | 39274402 | 39274472 | 5 | 6 | 0 | 1 | 1 | 2 | 0 | 0 | 1 | 1 | 12 | 3 | 3 | 6 | 3 | 0 | 0 | 0 | 0 | 0 | 298 | 149 | 1 | 1 | 0.556 | 1 | NA | 0.333 | 0.333 | 1 | 1 | 1 | 1 | 1 | -0.445 |
| SE | 37641 | ENSG00000082074 | FYB1 | chr5 | - | 39270568 | 39270828 | 39212677 | 39213017 | 39274402 | 39274472 | 3 | 4 | 0 | 3 | 2 | 2 | 3 | 0 | 2 | 6 | 12 | 2 | 5 | 10 | 0 | 0 | 0 | 0 | 0 | 0 | 298 | 149 | 1 | 1 | 0.429 | 0.4 | NA | 0.429 | 0.143 | 1 | 1 | 1 | 1 | NA | -0.65 |
| SE | 37642 | ENSG00000136631 | VPS45 | chr1 | + | 150076915 | 150076984 | 150076232 | 150076312 | 150077093 | 150077231 | 129 | 84 | 115 | 144 | 246 | 0 | 0 | 0 | 0 | 1 | 152 | 115 | 49 | 68 | 27 | 0 | 0 | 0 | 2 | 0 | 218 | 149 | 1 | 1 | 1 | 1 | 1 | 1 | 0.994 | 1 | 1 | 1 | 0.959 | 1 | 0.007 |
| SE | 37644 | ENSG00000136631 | VPS45 | chr1 | + | 150081341 | 150081476 | 150077188 | 150077779 | 150081883 | 150081997 | 223 | 200 | 173 | 221 | 182 | 0 | 0 | 1 | 0 | 0 | 177 | 167 | 111 | 111 | 52 | 0 | 0 | 0 | 0 | 0 | 284 | 149 | 1 | 1 | 1 | 1 | 0.989 | 1 | 1 | 1 | 1 | 1 | 1 | 1 | -0.002 |
| SE | 37645 | ENSG00000136631 | VPS45 | chr1 | + | 150081406 | 150081476 | 150077668 | 150077779 | 150081883 | 150081926 | 43 | 33 | 23 | 35 | 32 | 0 | 0 | 1 | 0 | 0 | 29 | 29 | 16 | 11 | 11 | 0 | 0 | 0 | 0 | 0 | 219 | 149 | 1 | 1 | 1 | 1 | 0.94 | 1 | 1 | 1 | 1 | 1 | 1 | 1 | -0.012 |
| SE | 37646 | ENSG00000136631 | VPS45 | chr1 | + | 150081883 | 150081997 | 150077188 | 150077779 | 150082715 | 150082883 | 68 | 55 | 34 | 83 | 95 | 0 | 0 | 0 | 0 | 5 | 53 | 40 | 19 | 30 | 17 | 0 | 0 | 0 | 0 | 0 | 263 | 149 | 1 | 1 | 1 | 1 | 1 | 1 | 0.915 | 1 | 1 | 1 | 1 | 1 | -0.017 |
| SE | 37647 | ENSG00000136631 | VPS45 | chr1 | + | 150091936 | 150092095 | 150082715 | 150082883 | 150092301 | 150092409 | 188 | 153 | 129 | 217 | 294 | 0 | 0 | 0 | 1 | 0 | 182 | 207 | 44 | 83 | 19 | 0 | 0 | 0 | 0 | 0 | 298 | 149 | 1 | 1 | 1 | 1 | 1 | 0.991 | 1 | 1 | 1 | 1 | 1 | 1 | -0.002 |
| SE | 37649 | ENSG00000136631 | VPS45 | chr1 | + | 150142837 | 150142931 | 150110495 | 150110627 | 150144708 | 150145288 | 5 | 0 | 0 | 0 | 0 | 88 | 56 | 50 | 52 | 57 | 0 | 0 | 0 | 0 | 0 | 17 | 36 | 20 | 52 | 28 | 243 | 149 | 1 | 1 | 0.034 | 0 | 0 | 0 | 0 | 0 | 0 | 0 | 0 | 0 | 0.007 |
| SE | 37651 | ENSG00000136933 | RABEPK | chr9 | + | 125203007 | 125203066 | 125200541 | 125200906 | 125213369 | 125213522 | 18 | 8 | 13 | 11 | 7 | 0 | 0 | 0 | 0 | 0 | 0 | 17 | 2 | 12 | 14 | 0 | 11 | 0 | 0 | 0 | 208 | 149 | 1 | 1 | 1 | 1 | 1 | 1 | 1 | NA | 0.525 | 1 | 1 | 1 | 0.119 |
| SE | 37653 | ENSG00000136933 | RABEPK | chr9 | + | 125207563 | 125207721 | 125203007 | 125203066 | 125213369 | 125213522 | 73 | 18 | 50 | 37 | 78 | 0 | 0 | 0 | 2 | 1 | 75 | 21 | 20 | 13 | 78 | 0 | 0 | 0 | 3 | 0 | 298 | 149 | 1 | 1 | 1 | 1 | 1 | 0.902 | 0.975 | 1 | 1 | 1 | 0.684 | 1 | 0.039 |
| SE | 37656 | ENSG00000136933 | RABEPK | chr9 | + | 125220538 | 125220700 | 125207563 | 125207721 | 125227909 | 125228059 | 43 | 17 | 24 | 34 | 34 | 1 | 0 | 0 | 1 | 1 | 10 | 43 | 20 | 19 | 28 | 0 | 0 | 0 | 0 | 0 | 298 | 149 | 1 | 1 | 0.956 | 1 | 1 | 0.944 | 0.944 | 1 | 1 | 1 | 1 | 1 | -0.031 |
| SE | 37657 | ENSG00000185267 | CDNF | chr10 | - | 14825478 | 14825620 | 14819244 | 14820158 | 14828144 | 14828272 | 17 | 10 | 20 | 12 | 43 | 0 | 0 | 1 | 0 | 0 | 10 | 15 | 5 | 8 | 3 | 0 | 0 | 0 | 0 | 0 | 291 | 149 | 1 | 1 | 1 | 1 | 0.911 | 1 | 1 | 1 | 1 | 1 | 1 | 1 | -0.018 |
| SE | 37658 | ENSG00000185267 | CDNF | chr10 | - | 14825493 | 14825620 | 14819884 | 14820158 | 14828144 | 14828205 | 5 | 3 | 11 | 4 | 18 | 0 | 0 | 1 | 0 | 0 | 0 | 7 | 3 | 4 | 0 | 0 | 0 | 0 | 0 | 0 | 276 | 149 | 1 | 1 | 1 | 1 | 0.856 | 1 | 1 | NA | 1 | 1 | 1 | NA | -0.029 |
| SE | 37659 | ENSG00000185267 | CDNF | chr10 | - | 14834216 | 14834333 | 14828144 | 14828272 | 14836250 | 14836294 | 0 | 0 | 2 | 0 | 0 | 0 | 0 | 2 | 0 | 3 | 0 | 0 | 0 | 0 | 0 | 0 | 0 | 2 | 0 | 0 | 266 | 149 | 1 | 1 | NA | NA | 0.359 | NA | 0 | NA | NA | 0 | NA | NA | 0.179 |
| SE | 37660 | ENSG00000185267 | CDNF | chr10 | - | 14836250 | 14836294 | 14828144 | 14828272 | 14837831 | 14837922 | 0 | 0 | 0 | 0 | 3 | 1 | 0 | 3 | 1 | 2 | 0 | 0 | 0 | 0 | 0 | 0 | 4 | 0 | 0 | 0 | 193 | 149 | 1 | 1 | 0 | NA | 0 | 0 | 0.537 | NA | 0 | NA | NA | NA | 0.134 |
| SE | 37661 | ENSG00000168615 | ADAM9 | chr8 | + | 39011657 | 39011716 | 39007885 | 39007983 | 39013964 | 39014043 | 13 | 13 | 22 | 23 | 56 | 2 | 0 | 0 | 0 | 0 | 21 | 31 | 28 | 10 | 16 | 0 | 0 | 0 | 0 | 0 | 208 | 149 | 1 | 1 | 0.823 | 1 | 1 | 1 | 1 | 1 | 1 | 1 | 1 | 1 | -0.035 |
| SE | 37663 | ENSG00000168615 | ADAM9 | chr8 | + | 39041945 | 39042117 | 39026676 | 39026810 | 39046001 | 39046058 | 92 | 108 | 57 | 90 | 129 | 0 | 0 | 0 | 0 | 0 | 31 | 121 | 115 | 67 | 73 | 0 | 0 | 3 | 0 | 0 | 298 | 149 | 1 | 1 | 1 | 1 | 1 | 1 | 1 | 1 | 1 | 0.95 | 1 | 1 | 0.01 |
| SE | 37665 | ENSG00000168615 | ADAM9 | chr8 | + | 39046001 | 39046058 | 39026676 | 39026810 | 39054480 | 39054573 | 0 | 2 | 1 | 2 | 1 | 0 | 0 | 2 | 1 | 0 | 0 | 4 | 3 | 0 | 0 | 0 | 14 | 0 | 0 | 0 | 206 | 149 | 1 | 1 | NA | 1 | 0.266 | 0.591 | 1 | NA | 0.171 | 1 | NA | NA | 0.129 |
| SE | 37667 | ENSG00000168615 | ADAM9 | chr8 | + | 39046953 | 39047046 | 39041945 | 39042117 | 39054480 | 39054573 | 0 | 0 | 0 | 0 | 0 | 101 | 75 | 44 | 64 | 97 | 0 | 0 | 4 | 0 | 0 | 24 | 123 | 64 | 60 | 21 | 242 | 149 | 1 | 1 | 0 | 0 | 0 | 0 | 0 | 0 | 0 | 0 | 0.039 | 0 | -0.008 |
| SE | 37668 | ENSG00000168615 | ADAM9 | chr8 | + | 39071297 | 39071403 | 39055576 | 39055772 | 39076146 | 39076275 | 64 | 58 | 43 | 71 | 57 | 1 | 0 | 0 | 0 | 0 | 12 | 106 | 89 | 50 | 16 | 0 | 0 | 0 | 0 | 0 | 255 | 149 | 1 | 1 | 0.974 | 1 | 1 | 1 | 1 | 1 | 1 | 1 | 1 | 1 | -0.005 |
| SE | 37670 | ENSG00000168615 | ADAM9 | chr8 | + | 39071297 | 39071413 | 39055576 | 39055772 | 39076146 | 39076275 | 63 | 58 | 43 | 67 | 57 | 1 | 0 | 0 | 0 | 0 | 12 | 106 | 85 | 47 | 16 | 0 | 0 | 0 | 0 | 0 | 265 | 149 | 1 | 1 | 0.973 | 1 | 1 | 1 | 1 | 1 | 1 | 1 | 1 | 1 | -0.005 |
| SE | 37672 | ENSG00000168615 | ADAM9 | chr8 | + | 39076146 | 39076275 | 39055576 | 39055772 | 39077227 | 39077411 | 5 | 4 | 2 | 5 | 0 | 0 | 4 | 6 | 4 | 1 | 7 | 0 | 3 | 9 | 0 | 0 | 2 | 15 | 3 | 0 | 278 | 149 | 1 | 1 | 1 | 0.349 | 0.152 | 0.401 | 0 | 1 | 0 | 0.097 | 0.617 | NA | -0.048 |
| SE | 37675 | ENSG00000168615 | ADAM9 | chr8 | + | 39082640 | 39082721 | 39077227 | 39077411 | 39090046 | 39090188 | 40 | 22 | 11 | 35 | 20 | 0 | 0 | 0 | 0 | 1 | 18 | 73 | 41 | 51 | 2 | 0 | 1 | 3 | 0 | 0 | 230 | 149 | 1 | 1 | 1 | 1 | 1 | 1 | 0.928 | 1 | 0.979 | 0.899 | 1 | 1 | 0.01 |
| SE | 37677 | ENSG00000168615 | ADAM9 | chr8 | + | 39082967 | 39083073 | 39082640 | 39082721 | 39090135 | 39090188 | 65 | 55 | 27 | 74 | 45 | 0 | 0 | 0 | 0 | 2 | 72 | 95 | 97 | 54 | 32 | 0 | 4 | 0 | 0 | 0 | 255 | 149 | 1 | 1 | 1 | 1 | 1 | 1 | 0.929 | 1 | 0.933 | 1 | 1 | 1 | -0.001 |
| SE | 37680 | ENSG00000168615 | ADAM9 | chr8 | + | 39090046 | 39091346 | 39082967 | 39083073 | 39101862 | 39101930 | 110 | 96 | 68 | 107 | 209 | 0 | 0 | 0 | 0 | 0 | 34 | 171 | 156 | 113 | 40 | 0 | 2 | 3 | 0 | 0 | 298 | 149 | 1 | 1 | 1 | 1 | 1 | 1 | 1 | 1 | 0.977 | 0.963 | 1 | 1 | 0.012 |
| SE | 37682 | ENSG00000168615 | ADAM9 | chr8 | + | 39091258 | 39091346 | 39082967 | 39083073 | 39101862 | 39101930 | 58 | 46 | 28 | 51 | 119 | 0 | 0 | 0 | 0 | 0 | 14 | 80 | 70 | 49 | 14 | 0 | 2 | 3 | 0 | 0 | 237 | 149 | 1 | 1 | 1 | 1 | 1 | 1 | 1 | 1 | 0.962 | 0.936 | 1 | 1 | 0.02 |
| SE | 37683 | ENSG00000168615 | ADAM9 | chr8 | + | 39091671 | 39091764 | 39091258 | 39091346 | 39101862 | 39101930 | 0 | 0 | 0 | 0 | 0 | 75 | 76 | 47 | 70 | 181 | 0 | 0 | 2 | 2 | 0 | 14 | 116 | 106 | 76 | 20 | 242 | 149 | 1 | 1 | 0 | 0 | 0 | 0 | 0 | 0 | 0 | 0.011 | 0.016 | 0 | -0.005 |
| SE | 37686 | ENSG00000147912 | FBXO10 | chr9 | - | 37515903 | 37516085 | 37510891 | 37512721 | 37518124 | 37518438 | 11 | 10 | 29 | 13 | 9 | 0 | 0 | 0 | 0 | 0 | 0 | 10 | 18 | 10 | 7 | 0 | 0 | 1 | 0 | 0 | 298 | 149 | 1 | 1 | 1 | 1 | 1 | 1 | 1 | NA | 1 | 0.9 | 1 | 1 | 0.025 |
| SE | 37687 | ENSG00000147912 | FBXO10 | chr9 | - | 37525101 | 37525172 | 37522824 | 37522977 | 37529123 | 37529260 | 18 | 11 | 28 | 22 | 13 | 0 | 0 | 0 | 1 | 0 | 51 | 65 | 14 | 29 | 59 | 0 | 0 | 0 | 1 | 0 | 220 | 149 | 1 | 1 | 1 | 1 | 1 | 0.937 | 1 | 1 | 1 | 1 | 0.952 | 1 | -0.003 |
| SE | 37690 | ENSG00000197385 | ZNF860 | chr3 | + | 31986247 | 31986409 | 31981770 | 31981902 | 31988659 | 31988925 | 1 | 2 | 0 | 0 | 1 | 5 | 5 | 3 | 5 | 3 | 0 | 1 | 0 | 0 | 0 | 0 | 2 | 0 | 0 | 0 | 298 | 149 | 1 | 1 | 0.091 | 0.167 | 0 | 0 | 0.143 | NA | 0.2 | NA | NA | NA | -0.12 |
| SE | 37691 | ENSG00000163507 | CIP2A | chr3 | - | 108553647 | 108553730 | 108552233 | 108552373 | 108554375 | 108554489 | 24 | 30 | 51 | 32 | 57 | 0 | 0 | 0 | 0 | 3 | 0 | 22 | 14 | 7 | 0 | 0 | 0 | 0 | 0 | 0 | 232 | 149 | 1 | 1 | 1 | 1 | 1 | 1 | 0.924 | NA | 1 | 1 | 1 | NA | -0.015 |
| SE | 37692 | ENSG00000163507 | CIP2A | chr3 | - | 108557217 | 108557414 | 108554375 | 108554489 | 108559756 | 108559867 | 28 | 34 | 54 | 34 | 61 | 0 | 0 | 3 | 0 | 0 | 9 | 21 | 31 | 8 | 0 | 0 | 0 | 0 | 0 | 0 | 298 | 149 | 1 | 1 | 1 | 1 | 0.9 | 1 | 1 | 1 | 1 | 1 | 1 | NA | -0.02 |
| SE | 37693 | ENSG00000163507 | CIP2A | chr3 | - | 108563121 | 108563244 | 108560734 | 108560841 | 108565354 | 108565454 | 8 | 7 | 18 | 16 | 36 | 0 | 0 | 0 | 1 | 0 | 0 | 0 | 7 | 0 | 0 | 0 | 0 | 0 | 0 | 0 | 272 | 149 | 1 | 1 | 1 | 1 | 1 | 0.898 | 1 | NA | NA | 1 | 1 | NA | -0.02 |
| SE | 37694 | ENSG00000163507 | CIP2A | chr3 | - | 108563125 | 108563244 | 108560648 | 108560841 | 108565354 | 108565454 | 18 | 14 | 40 | 29 | 72 | 0 | 0 | 0 | 1 | 0 | 0 | 12 | 2 | 11 | 2 | 0 | 0 | 0 | 0 | 0 | 268 | 149 | 1 | 1 | 1 | 1 | 1 | 0.942 | 1 | NA | 1 | 1 | 1 | 1 | -0.012 |
| SE | 37695 | ENSG00000163507 | CIP2A | chr3 | - | 108582107 | 108582138 | 108581414 | 108581511 | 108582976 | 108583083 | 3 | 6 | 5 | 1 | 0 | 0 | 0 | 1 | 1 | 0 | 0 | 2 | 3 | 3 | 4 | 0 | 0 | 0 | 0 | 0 | 180 | 149 | 1 | 1 | 1 | 1 | 0.805 | 0.453 | NA | NA | 1 | 1 | 1 | 1 | -0.185 |
| SE | 37696 | ENSG00000163507 | CIP2A | chr3 | - | 108582107 | 108582202 | 108581414 | 108581511 | 108582976 | 108583083 | 18 | 16 | 32 | 12 | 9 | 0 | 0 | 1 | 1 | 0 | 9 | 22 | 25 | 13 | 15 | 0 | 0 | 0 | 0 | 0 | 244 | 149 | 1 | 1 | 1 | 1 | 0.951 | 0.88 | | 1 | 1 | 1 | 1 | 1 | -0.034 |
| SE | 37697 | ENSG00000163507 | CIP2A | chr3 | - | 108582976 | 108583083 | 108582107 | 108582138 | 108585064 | 108585212 | 0 | 0 | 13 | 4 | 2 | 0 | 1 | 0 | 0 | 0 | 0 | 0 | 0 | 0 | 0 | 0 | 0 | 0 | 1 | 0 | 256 | 149 | 1 | 1 | NA | 0 | 1 | 1 | 1 | NA | NA | NA | 0 | NA | 0.75 |
| SE | 37699 | ENSG00000163507 | CIP2A | chr3 | - | 108585665 | 108585749 | 108585064 | 108585212 | 108589273 | 108589456 | 1 | 0 | 0 | 0 | 0 | 2 | 1 | 2 | 5 | 2 | 0 | 0 | 0 | 0 | 0 | 0 | 14 | 1 | 4 | 0 | 233 | 149 | 1 | 1 | 0.242 | 0 | 0 | 0 | 0 | NA | 0 | 0 | 0 | NA | 0.048 |
| SE | 37700 | ENSG00000285162 | NA | chr7 | + | 29354624 | 29354663 | 29146848 | 29146960 | 29367931 | 29367987 | 12 | 14 | 10 | 13 | 6 | 0 | 2 | 0 | 0 | 0 | 15 | 11 | 8 | 3 | 0 | 0 | 0 | 0 | 0 | 0 | 188 | 149 | 1 | 1 | 1 | 0.847 | 1 | 1 | 1 | 1 | 1 | 1 | 1 | NA | -0.031 |
| SE | 37702 | ENSG00000285162 | NA | chr7 | + | 29393678 | 29393710 | 29367931 | 29367987 | 29398372 | 29398486 | 70 | 74 | 51 | 75 | 54 | 0 | 0 | 0 | 4 | 0 | 24 | 43 | 50 | 31 | 0 | 0 | 0 | 0 | 0 | 0 | 181 | 149 | 1 | 1 | 1 | 1 | 1 | 0.939 | 1 | 1 | 1 | 1 | 1 | NA | -0.012 |
| SE | 37703 | ENSG00000285162 | NA | chr7 | + | 29398372 | 29398486 | 29393678 | 29393710 | 29400542 | 29400828 | 88 | 109 | 82 | 103 | 93 | 1 | 0 | 0 | 0 | 0 | 42 | 46 | 74 | 51 | 5 | 0 | 0 | 0 | 0 | 0 | 263 | 149 | 1 | 1 | 0.98 | 1 | 1 | 1 | 1 | 1 | 1 | 1 | 1 | 1 | -0.004 |
| SE | 37704 | ENSG00000106069 | CHN2 | chr7 | + | 29480278 | 29480356 | 29400542 | 29400828 | 29495951 | 29496036 | 47 | 47 | 36 | 75 | 106 | 0 | 0 | 2 | 0 | 0 | 63 | 13 | 26 | 28 | 7 | 0 | 0 | 4 | 0 | 0 | 227 | 149 | 1 | 1 | 1 | 1 | 0.922 | 1 | 1 | 1 | 1 | 0.81 | 1 | 1 | 0.022 |
| SE | 37705 | ENSG00000106069 | CHN2 | chr7 | + | 29507227 | 29507365 | 29504743 | 29504821 | 29509300 | 29509406 | 122 | 100 | 64 | 91 | 239 | 0 | 1 | 0 | 0 | 5 | 29 | 41 | 45 | 53 | 23 | 0 | 0 | 4 | 0 | 0 | 287 | 149 | 1 | 1 | 1 | 0.981 | 1 | 1 | 0.961 | 1 | 1 | 0.854 | 1 | 1 | 0.018 |
| SE | 37718 | ENSG00000181450 | ZNF678 | chr1 | + | 227582513 | 227582606 | 227563542 | 227563724 | 227646543 | 227646670 | 20 | 22 | 22 | 28 | 12 | 14 | 5 | 9 | 15 | 10 | 0 | 28 | 14 | 7 | 6 | 0 | 11 | 3 | 3 | 3 | 242 | 149 | 1 | 1 | 0.468 | 0.73 | 0.601 | 0.535 | 0.425 | NA | 0.61 | 0.742 | 0.59 | 0.552 | -0.072 |
| SE | 37719 | ENSG00000181450 | ZNF678 | chr1 | + | 227582513 | 227582606 | 227563542 | 227563724 | 227654335 | 227654353 | 2 | 2 | 4 | 5 | 8 | 0 | 3 | 0 | 4 | 0 | 0 | 24 | 1 | 2 | 0 | 0 | 3 | 1 | 0 | 1 | 242 | 149 | 1 | 1 | 1 | 0.291 | 1 | 0.435 | 1 | NA | 0.831 | 0.381 | 1 | 0 | 0.192 |
| SE | 37720 | ENSG00000181450 | ZNF678 | chr1 | + | 227646543 | 227646670 | 227563555 | 227563724 | 227654335 | 227654768 | 33 | 17 | 25 | 40 | 25 | 0 | 3 | 0 | 4 | 0 | 0 | 16 | 16 | 4 | 6 | 0 | 3 | 1 | 0 | 1 | 276 | 149 | 1 | 1 | 1 | 0.754 | 1 | 0.844 | 1 | NA | 0.742 | 0.896 | 1 | 0.764 | 0.069 |
| SE | 37721 | ENSG00000181450 | ZNF678 | chr1 | + | 227646543 | 227646670 | 227582513 | 227582606 | 227650955 | 227651076 | 42 | 38 | 41 | 62 | 32 | 1 | 0 | 0 | 0 | 0 | 0 | 22 | 15 | 13 | 19 | 0 | 0 | 0 | 0 | 0 | 276 | 149 | 1 | 1 | 0.958 | 1 | 1 | 1 | 1 | NA | 1 | 1 | 1 | 1 | -0.008 |
| SE | 37722 | ENSG00000181450 | ZNF678 | chr1 | + | 227646543 | 227646670 | 227582513 | 227582606 | 227654335 | 227654768 | 35 | 31 | 37 | 50 | 26 | 0 | 1 | 1 | 1 | 8 | 0 | 12 | 26 | 6 | 9 | 0 | 10 | 0 | 0 | 0 | 276 | 149 | 1 | 1 | 1 | 0.944 | 0.952 | 0.964 | 0.637 | NA | 0.393 | 1 | 1 | 1 | 0.051 |
| SE | 37723 | ENSG00000181450 | ZNF678 | chr1 | + | 227650955 | 227651076 | 227582513 | 227582606 | 227654335 | 227654353 | 29 | 17 | 23 | 29 | 30 | 0 | 1 | 1 | 1 | 8 | 0 | 16 | 9 | 0 | 6 | 0 | 10 | 0 | 0 | 0 | 270 | 149 | 1 | 1 | 1 | 0.904 | 0.927 | 0.941 | 0.674 | NA | 0.469 | 1 | NA | 1 | 0.066 |
| SE | 37724 | ENSG00000181450 | ZNF678 | chr1 | + | 227650955 | 227651076 | 227646543 | 227646670 | 227654335 | 227654353 | 58 | 36 | 46 | 71 | 51 | 24 | 12 | 18 | 28 | 20 | 0 | 31 | 11 | 8 | 19 | 0 | 5 | 13 | 1 | 3 | 270 | 149 | 1 | 1 | 0.571 | 0.623 | 0.585 | 0.583 | 0.585 | NA | 0.774 | 0.318 | 0.815 | 0.778 | -0.082 |
| SE | 37726 | ENSG00000174500 | GCSAM | chr3 | - | 112125225 | 112125254 | 112120838 | 112123772 | 112126986 | 112127033 | 43 | 38 | 73 | 38 | 129 | 1 | 4 | 0 | 0 | 7 | 10 | 19 | 17 | 7 | 0 | 0 | 0 | 0 | 0 | 0 | 178 | 149 | 1 | 1 | 0.973 | 0.888 | 1 | 1 | 0.939 | 1 | 1 | 1 | 1 | NA | -0.04 |
| SE | 37727 | ENSG00000174500 | GCSAM | chr3 | - | 112128016 | 112128061 | 112126986 | 112127033 | 112130444 | 112130513 | 53 | 38 | 66 | 39 | 63 | 0 | 0 | 1 | 0 | 1 | 26 | 23 | 10 | 13 | 0 | 0 | 0 | 0 | 0 | 0 | 194 | 149 | 1 | 1 | 1 | 1 | 0.981 | 1 | 0.98 | 1 | 1 | 1 | 1 | NA | -0.008 |
| SE | 37728 | ENSG00000174500 | GCSAM | chr3 | - | 112128016 | 112128061 | 112126986 | 112127033 | 112133091 | 112133225 | 13 | 11 | 17 | 13 | 33 | 0 | 2 | 0 | 0 | 0 | 0 | 9 | 1 | 0 | 0 | 0 | 0 | 0 | 0 | 0 | 194 | 149 | 1 | 1 | 1 | 0.809 | 1 | 1 | 1 | NA | 1 | 1 | NA | NA | -0.038 |
| SE | 37729 | ENSG00000174500 | GCSAM | chr3 | - | 112128016 | 112128228 | 112126986 | 112127033 | 112130444 | 112130513 | 25 | 26 | 31 | 29 | 51 | 0 | 0 | 1 | 0 | 1 | 7 | 15 | 11 | 0 | 4 | 0 | 0 | 0 | 0 | 0 | 298 | 149 | 1 | 1 | 1 | 1 | 0.939 | 1 | 0.962 | 1 | 1 | 1 | NA | 1 | -0.02 |
| SE | 37730 | ENSG00000174500 | GCSAM | chr3 | - | 112130444 | 112130513 | 112126986 | 112127033 | 112133091 | 112133166 | 11 | 8 | 16 | 3 | 5 | 0 | 2 | 0 | 0 | 0 | 14 | 5 | 0 | 11 | 4 | 0 | 0 | 0 | 0 | 0 | 218 | 149 | 1 | 1 | 1 | 0.732 | 1 | 1 | 1 | 1 | 1 | NA | 1 | 1 | -0.054 |
| SE | 37731 | ENSG00000174500 | GCSAM | chr3 | - | 112130444 | 112130513 | 112128016 | 112128061 | 112133091 | 112133166 | 45 | 32 | 56 | 27 | 33 | 0 | 0 | 0 | 1 | 3 | 40 | 19 | 9 | 24 | 4 | 0 | 0 | 0 | 0 | 0 | 218 | 149 | 1 | 1 | 1 | 1 | 1 | 0.949 | 0.883 | 1 | 1 | 1 | 1 | 1 | -0.034 |
| SE | 37732 | ENSG00000174500 | GCSAM | chr3 | - | 112130444 | 112130519 | 112128016 | 112128061 | 112133091 | 112133254 | 24 | 19 | 38 | 12 | 26 | 0 | 0 | 0 | 1 | 3 | 10 | 8 | 9 | 3 | 0 | 0 | 0 | 0 | 0 | 0 | 224 | 149 | 1 | 1 | 1 | 1 | 1 | 0.889 | 0.852 | 1 | 1 | 1 | 1 | NA | -0.052 |
| SE | 37733 | ENSG00000174500 | GCSAM | chr3 | - | 112132636 | 112132910 | 112130444 | 112130513 | 112133091 | 112133225 | 10 | 4 | 18 | 7 | 18 | 30 | 21 | 25 | 19 | 8 | 0 | 10 | 0 | 0 | 0 | 37 | 16 | 0 | 21 | 8 | 298 | 149 | 1 | 1 | 0.143 | 0.087 | 0.265 | 0.156 | 0.529 | 0 | 0.238 | NA | 0 | 0 | 0.177 |
| SE | 37734 | ENSG00000174500 | GCSAM | chr3 | - | 112132636 | 112132910 | 112130444 | 112130519 | 112133091 | 112133225 | 9 | 3 | 10 | 5 | 7 | 7 | 4 | 7 | 0 | 0 | 0 | 5 | 1 | 0 | 0 | 0 | 0 | 0 | 0 | 0 | 298 | 149 | 1 | 1 | 0.391 | 0.273 | 0.417 | 1 | 1 | NA | 1 | 1 | NA | NA | -0.384 |
| SE | 37736 | ENSG00000144827 | ABHD10 | chr3 | + | 111986263 | 111986375 | 111979009 | 111979203 | 111986913 | 111987051 | 55 | 30 | 55 | 48 | 93 | 2 | 0 | 1 | 0 | 0 | 0 | 44 | 44 | 22 | 10 | 0 | 0 | 1 | 0 | 0 | 261 | 149 | 1 | 1 | 0.94 | 1 | 0.969 | 1 | 1 | NA | 1 | 0.962 | 1 | 1 | -0.009 |
| SE | 37744 | ENSG00000186469 | GNG2 | chr14 | + | 51873881 | 51874082 | 51860733 | 51860790 | 51877616 | 51877657 | 0 | 0 | 0 | 6 | 0 | 459 | 139 | 224 | 269 | 168 | 0 | 0 | 0 | 0 | 0 | 69 | 194 | 198 | 155 | 67 | 298 | 149 | 1 | 1 | 0 | 0 | 0 | 0.011 | 0 | 0 | 0 | 0 | 0 | 0 | 0.002 |
| SE | 37746 | ENSG00000186469 | GNG2 | chr14 | + | 51877616 | 51877657 | 51847125 | 51847378 | 51950649 | 51950765 | 206 | 69 | 94 | 124 | 112 | 0 | 0 | 2 | 0 | 0 | 18 | 103 | 122 | 42 | 17 | 0 | 0 | 0 | 0 | 1 | 190 | 149 | 1 | 1 | 1 | 1 | 0.974 | 1 | 1 | 1 | 1 | 1 | 1 | 0.93 | 0.009 |
| SE | 37748 | ENSG00000186469 | GNG2 | chr14 | + | 51913008 | 51913128 | 51877616 | 51877657 | 51939780 | 51940399 | 2 | 3 | 3 | 1 | 0 | 0 | 0 | 0 | 0 | 0 | 0 | 1 | 1 | 2 | 0 | 0 | 0 | 0 | 10 | 0 | 269 | 149 | 1 | 1 | 1 | 1 | 1 | 1 | NA | NA | 1 | 1 | 0.1 | NA | 0.3 |
| SE | 37751 | ENSG00000186469 | GNG2 | chr14 | + | 51939780 | 51940091 | 51877616 | 51877657 | 51950649 | 51950765 | 0 | 1 | 2 | 0 | 0 | 603 | 182 | 288 | 363 | 271 | 0 | 0 | 0 | 10 | 0 | 74 | 275 | 299 | 145 | 66 | 298 | 149 | 1 | 1 | 0 | 0.003 | 0.003 | 0 | 0 | 0 | 0 | 0 | 0.033 | 0 | -0.005 |
| SE | 37752 | ENSG00000186469 | GNG2 | chr14 | + | 51950649 | 51950765 | 51877591 | 51877657 | 51966558 | 51966752 | 1290 | 380 | 652 | 781 | 650 | 0 | 0 | 1 | 0 | 0 | 230 | 587 | 685 | 437 | 161 | 0 | 0 | 0 | 0 | 0 | 265 | 149 | 1 | 1 | 1 | 1 | 0.997 | 1 | 1 | 1 | 1 | 1 | 1 | 1 | -0.001 |
| SE | 37753 | ENSG00000186469 | GNG2 | chr14 | + | 51951886 | 51952132 | 51950649 | 51950765 | 51966558 | 51967040 | 0 | 0 | 0 | 1 | 0 | 862 | 261 | 450 | 534 | 493 | 0 | 0 | 0 | 0 | 0 | 174 | 429 | 512 | 330 | 112 | 298 | 149 | 1 | 1 | 0 | 0 | 0 | 0.001 | 0 | 0 | 0 | 0 | 0 | 0 | 0 |
| SE | 37754 | ENSG00000243696 | NA | chr3 | - | 52831407 | 52831478 | 52830552 | 52830670 | 52833113 | 52833247 | 11 | 4 | 4 | 1 | 2 | 15 | 4 | 4 | 5 | 2 | 0 | 12 | 15 | 24 | 16 | 0 | 16 | 17 | 86 | 10 | 220 | 149 | 1 | 1 | 0.332 | 0.404 | 0.404 | 0.119 | 0.404 | NA | 0.337 | 0.374 | 0.159 | 0.52 | -0.015 |
| SE | 37762 | ENSG00000170954 | ZNF415 | chr19 | - | 53115209 | 53115307 | 53107878 | 53109908 | 53115703 | 53115811 | 0 | 2 | 0 | 0 | 2 | 2 | 1 | 0 | 0 | 10 | 0 | 2 | 0 | 0 | 0 | 0 | 0 | 0 | 0 | 0 | 247 | 149 | 1 | 1 | 0 | 0.547 | NA | NA | 0.108 | NA | 1 | NA | NA | NA | -0.782 |
| SE | 37763 | ENSG00000170954 | ZNF415 | chr19 | - | 53115209 | 53115307 | 53107878 | 53109908 | 53116312 | 53116433 | 0 | 2 | 1 | 0 | 3 | 2 | 12 | 4 | 10 | 1 | 0 | 4 | 0 | 0 | 0 | 0 | 1 | 0 | 0 | 0 | 247 | 149 | 1 | 1 | 0 | 0.091 | 0.131 | 0 | 0.644 | NA | 0.707 | NA | NA | NA | -0.534 |
| SE | 37765 | ENSG00000170954 | ZNF415 | chr19 | - | 53115703 | 53115924 | 53107878 | 53109908 | 53116312 | 53116433 | 2 | 1 | 0 | 2 | 9 | 2 | 12 | 4 | 10 | 1 | 0 | 0 | 0 | 0 | 0 | 0 | 1 | 0 | 0 | 0 | 298 | 149 | 1 | 1 | 0.333 | 0.04 | 0 | 0.091 | 0.818 | NA | 0 | NA | NA | NA | 0.256 |
| SE | 37767 | ENSG00000170954 | ZNF415 | chr19 | - | 53116312 | 53116433 | 53107878 | 53109908 | 53122661 | 53122743 | 4 | 16 | 6 | 11 | 2 | 0 | 0 | 1 | 0 | 0 | 0 | 1 | 0 | 0 | 0 | 0 | 0 | 0 | 0 | 0 | 270 | 149 | 1 | 1 | 1 | 1 | 0.768 | 1 | 1 | NA | 1 | NA | NA | NA | -0.046 |
| SE | 37774 | ENSG00000139719 | VPS33A | chr12 | - | 122232799 | 122232968 | 122229563 | 122232427 | 122235785 | 122235923 | 299 | 125 | 197 | 241 | 345 | 0 | 1 | 0 | 1 | 1 | 221 | 126 | 110 | 175 | 208 | 0 | 0 | 1 | 0 | 0 | 298 | 149 | 1 | 1 | 1 | 0.984 | 1 | 0.992 | 0.994 | 1 | 1 | 0.982 | 1 | 1 | -0.002 |
| SE | 37775 | ENSG00000139719 | VPS33A | chr12 | - | 122235785 | 122235923 | 122232799 | 122232968 | 122238586 | 122238724 | 229 | 129 | 191 | 184 | 384 | 1 | 0 | 0 | 0 | 0 | 244 | 139 | 96 | 160 | 179 | 0 | 0 | 0 | 0 | 0 | 287 | 149 | 1 | 1 | 0.992 | 1 | 1 | 1 | 1 | 1 | 1 | 1 | 1 | 1 | -0.002 |
| SE | 37778 | ENSG00000139719 | VPS33A | chr12 | - | 122242381 | 122242508 | 122239877 | 122239945 | 122244568 | 122244762 | 155 | 96 | 169 | 151 | 184 | 0 | 0 | 1 | 1 | 0 | 185 | 102 | 105 | 175 | 156 | 0 | 0 | 0 | 0 | 0 | 276 | 149 | 1 | 1 | 1 | 1 | 0.989 | 0.988 | 1 | 1 | 1 | 1 | 1 | 1 | -0.005 |
| SE | 37782 | ENSG00000139719 | VPS33A | chr12 | - | 122261260 | 122261447 | 122250982 | 122251099 | 122263571 | 122263699 | 257 | 141 | 226 | 272 | 354 | 0 | 0 | 0 | 0 | 1 | 397 | 305 | 189 | 311 | 252 | 0 | 0 | 1 | 1 | 0 | 298 | 149 | 1 | 1 | 1 | 1 | 1 | 1 | 0.994 | 1 | 1 | 0.99 | 0.994 | 1 | 0.002 |
| SE | 37783 | ENSG00000139719 | VPS33A | chr12 | - | 122264711 | 122264804 | 122264133 | 122264199 | 122266306 | 122266336 | 0 | 0 | 0 | 2 | 2 | 98 | 41 | 98 | 72 | 29 | 0 | 0 | 0 | 0 | 0 | 141 | 114 | 47 | 64 | 63 | 242 | 149 | 1 | 1 | 0 | 0 | 0 | 0.017 | 0.041 | 0 | 0 | 0 | 0 | 0 | 0.012 |
| SE | 37786 | ENSG00000015479 | MATR3 | chr5 | + | 139307238 | 139308327 | 139293741 | 139293772 | 139314674 | 139314736 | 540 | 404 | 459 | 583 | 779 | 3 | 1 | 6 | 4 | 1 | 23 | 250 | 273 | 86 | 23 | 0 | 1 | 1 | 0 | 0 | 298 | 149 | 1 | 1 | 0.989 | 0.995 | 0.975 | 0.986 | 0.997 | 1 | 0.992 | 0.993 | 1 | 1 | -0.009 |
| SE | 37788 | ENSG00000015479 | MATR3 | chr5 | + | 139307240 | 139308327 | 139293717 | 139293772 | 139314674 | 139314736 | 539 | 398 | 453 | 582 | 779 | 3 | 1 | 6 | 4 | 1 | 23 | 247 | 273 | 80 | 23 | 0 | 1 | 1 | 0 | 0 | 298 | 149 | 1 | 1 | 0.989 | 0.995 | 0.974 | 0.986 | 0.997 | 1 | 0.992 | 0.993 | 1 | 1 | -0.009 |
| SE | 37790 | ENSG00000015479 | MATR3 | chr5 | + | 139314674 | 139314736 | 139293679 | 139293805 | 139315696 | 139315738 | 340 | 242 | 276 | 329 | 279 | 0 | 0 | 4 | 0 | 0 | 30 | 103 | 142 | 59 | 23 | 0 | 0 | 0 | 0 | 0 | 211 | 149 | 1 | 1 | 1 | 1 | 0.98 | 1 | 1 | 1 | 1 | 1 | 1 | 1 | -0.004 |
| SE | 37831 | ENSG00000114126 | TFDP2 | chr3 | - | 141963811 | 141963963 | 141959673 | 141959840 | 141970072 | 141970141 | 356 | 142 | 223 | 253 | 521 | 0 | 0 | 0 | 0 | 0 | 143 | 173 | 213 | 217 | 125 | 0 | 1 | 0 | 0 | 0 | 298 | 149 | 1 | 1 | 1 | 1 | 1 | 1 | 1 | 1 | 0.989 | 1 | 1 | 1 | 0.002 |
| SE | 37832 | ENSG00000114126 | TFDP2 | chr3 | - | 141963811 | 141963963 | 141959673 | 141959840 | 141974047 | 141974191 | 161 | 68 | 113 | 109 | 282 | 2 | 0 | 0 | 0 | 0 | 55 | 81 | 99 | 90 | 67 | 0 | 0 | 0 | 0 | 0 | 298 | 149 | 1 | 1 | 0.976 | 1 | 1 | 1 | 1 | 1 | 1 | 1 | 1 | 1 | -0.005 |
| SE | 37833 | ENSG00000114126 | TFDP2 | chr3 | - | 141970072 | 141970141 | 141959673 | 141959840 | 141974047 | 141974191 | 89 | 48 | 62 | 53 | 117 | 2 | 0 | 0 | 0 | 0 | 40 | 32 | 57 | 58 | 57 | 0 | 0 | 0 | 0 | 0 | 218 | 149 | 1 | 1 | 0.968 | 1 | 1 | 1 | 1 | 1 | 1 | 1 | 1 | 1 | -0.006 |
| SE | 37834 | ENSG00000114126 | TFDP2 | chr3 | - | 141970072 | 141970141 | 141963811 | 141963963 | 141974047 | 141974191 | 282 | 119 | 170 | 195 | 352 | 0 | 0 | 0 | 0 | 2 | 128 | 122 | 171 | 185 | 115 | 0 | 1 | 0 | 0 | 0 | 218 | 149 | 1 | 1 | 1 | 1 | 1 | 1 | 0.992 | 1 | 0.988 | 1 | 1 | 1 | 0.001 |
| SE | 37835 | ENSG00000114126 | TFDP2 | chr3 | - | 141993537 | 141993585 | 141978519 | 141978682 | 141995019 | 141995141 | 277 | 115 | 167 | 178 | 261 | 0 | 0 | 0 | 0 | 0 | 151 | 131 | 108 | 168 | 107 | 0 | 0 | 2 | 0 | 3 | 197 | 149 | 1 | 1 | 1 | 1 | 1 | 1 | 1 | 1 | 1 | 0.976 | 1 | 0.964 | 0.012 |
| SE | 37839 | ENSG00000114126 | TFDP2 | chr3 | - | 141995019 | 141995141 | 141978519 | 141978682 | 142054114 | 142054149 | 1 | 0 | 0 | 0 | 4 | 0 | 2 | 0 | 0 | 0 | 2 | 4 | 2 | 6 | 0 | 0 | 0 | 0 | 0 | 0 | 271 | 149 | 1 | 1 | 1 | 0 | NA | NA | 1 | 1 | 1 | 1 | 1 | NA | -0.333 |
| SE | 37845 | ENSG00000114126 | TFDP2 | chr3 | - | 142005440 | 142005544 | 141978536 | 141978682 | 142054114 | 142054149 | 50 | 17 | 38 | 24 | 25 | 0 | 2 | 0 | 0 | 0 | 23 | 24 | 21 | 16 | 28 | 0 | 0 | 0 | 0 | 0 | 253 | 149 | 1 | 1 | 1 | 0.833 | 1 | 1 | 1 | 1 | 1 | 1 | 1 | 1 | -0.033 |
| SE | 37846 | ENSG00000114126 | TFDP2 | chr3 | - | 142005440 | 142005544 | 141978536 | 141978682 | 142093060 | 142093127 | 93 | 39 | 52 | 61 | 73 | 2 | 0 | 0 | 0 | 0 | 42 | 67 | 51 | 51 | 24 | 0 | 0 | 0 | 0 | 0 | 253 | 149 | 1 | 1 | 0.965 | 1 | 1 | 1 | 1 | 1 | 1 | 1 | 1 | 1 | -0.007 |
| SE | 37850 | ENSG00000114126 | TFDP2 | chr3 | - | 142054114 | 142054149 | 141978519 | 141978682 | 142093060 | 142093127 | 9 | 7 | 7 | 3 | 19 | 2 | 0 | 0 | 0 | 0 | 2 | 0 | 5 | 1 | 0 | 0 | 0 | 0 | 0 | 0 | 184 | 149 | 1 | 1 | 0.785 | 1 | 1 | 1 | 1 | 1 | NA | 1 | 1 | NA | -0.043 |
| SE | 37851 | ENSG00000114126 | TFDP2 | chr3 | - | 142054114 | 142054149 | 141995019 | 141995141 | 142093060 | 142093127 | 11 | 5 | 7 | 4 | 23 | 1 | 4 | 0 | 3 | 6 | 4 | 4 | 5 | 7 | 0 | 0 | 0 | 4 | 0 | 0 | 184 | 149 | 1 | 1 | 0.899 | 0.503 | 1 | 0.519 | 0.756 | 1 | 1 | 0.503 | 1 | NA | -0.14 |
| SE | 37854 | ENSG00000114126 | TFDP2 | chr3 | - | 142093060 | 142093127 | 142005440 | 142005544 | 142101734 | 142101757 | 131 | 52 | 80 | 98 | 121 | 1 | 0 | 0 | 1 | 0 | 51 | 86 | 75 | 75 | 41 | 0 | 0 | 0 | 4 | 0 | 216 | 149 | 1 | 1 | 0.989 | 1 | 1 | 0.985 | 1 | 1 | 1 | 1 | 0.928 | 1 | 0.009 |
| SE | 37858 | ENSG00000114573 | ATP6V1A | chr3 | + | 113786231 | 113786383 | 113778740 | 113778835 | 113788712 | 113788875 | 73 | 67 | 64 | 75 | 110 | 1 | 0 | 0 | 0 | 0 | 142 | 78 | 57 | 87 | 72 | 0 | 0 | 0 | 0 | 0 | 298 | 149 | 1 | 1 | 0.973 | 1 | 1 | 1 | 1 | 1 | 1 | 1 | 1 | 1 | -0.005 |
| SE | 37859 | ENSG00000114573 | ATP6V1A | chr3 | + | 113794871 | 113794994 | 113789731 | 113789840 | 113795089 | 113795204 | 92 | 67 | 88 | 96 | 99 | 0 | 0 | 0 | 0 | 0 | 73 | 113 | 75 | 91 | 20 | 0 | 0 | 3 | 0 | 0 | 272 | 149 | 1 | 1 | 1 | 1 | 1 | 1 | 1 | 1 | 1 | 0.932 | 1 | 1 | 0.014 |
| SE | 37861 | ENSG00000135776 | ABCB10 | chr1 | - | 229525935 | 229526116 | 229521591 | 229521635 | 229530198 | 229530408 | 44 | 21 | 28 | 30 | 58 | 0 | 0 | 1 | 0 | 0 | 7 | 30 | 15 | 48 | 6 | 0 | 0 | 0 | 0 | 0 | 298 | 149 | 1 | 1 | 1 | 1 | 0.933 | 1 | 1 | 1 | 1 | 1 | 1 | 1 | -0.013 |
| SE | 37863 | ENSG00000135776 | ABCB10 | chr1 | - | 229542236 | 229542371 | 229540605 | 229540752 | 229547498 | 229547701 | 156 | 71 | 133 | 144 | 90 | 0 | 0 | 0 | 0 | 0 | 121 | 172 | 108 | 127 | 70 | 0 | 0 | 1 | 0 | 0 | 284 | 149 | 1 | 1 | 1 | 1 | 1 | 1 | 1 | 1 | 1 | 0.983 | 1 | 1 | 0.003 |
| SE | 37864 | ENSG00000135776 | ABCB10 | chr1 | - | 229547498 | 229547701 | 229542236 | 229542371 | 229549233 | 229549434 | 95 | 38 | 89 | 87 | 47 | 1 | 0 | 2 | 0 | 0 | 48 | 79 | 53 | 77 | 28 | 0 | 0 | 0 | 0 | 0 | 298 | 149 | 1 | 1 | 0.979 | 1 | 0.957 | 1 | 1 | 1 | 1 | 1 | 1 | 1 | -0.013 |
| SE | 37866 | ENSG00000100100 | PIK3IP1 | chr22 | - | 31289498 | 31289699 | 31281612 | 31283288 | 31289932 | 31290726 | 32 | 29 | 38 | 32 | 52 | 0 | 0 | 1 | 0 | 0 | 3 | 63 | 59 | 26 | 50 | 0 | 0 | 0 | 0 | 0 | 298 | 149 | 1 | 1 | 1 | 1 | 0.95 | 1 | 1 | 1 | 1 | 1 | 1 | 1 | -0.01 |
| SE | 37867 | ENSG00000100100 | PIK3IP1 | chr22 | - | 31290964 | 31291084 | 31289498 | 31289699 | 31292274 | 31292479 | 313 | 144 | 432 | 231 | 112 | 0 | 0 | 0 | 0 | 0 | 158 | 264 | 394 | 102 | 353 | 0 | 6 | 0 | 0 | 0 | 269 | 149 | 1 | 1 | 1 | 1 | 1 | 1 | 1 | 1 | 0.961 | 1 | 1 | 1 | 0.008 |
| SE | 37868 | ENSG00000100100 | PIK3IP1 | chr22 | - | 31291179 | 31291296 | 31290964 | 31291084 | 31292274 | 31292479 | 718 | 385 | 1059 | 674 | 363 | 0 | 0 | 1 | 0 | 0 | 240 | 671 | 683 | 261 | 527 | 0 | 0 | 0 | 0 | 0 | 266 | 149 | 1 | 1 | 1 | 1 | 0.998 | 1 | 1 | 1 | 1 | 1 | 1 | 1 | 0 |
| SE | 37870 | ENSG00000196187 | TMEM63A | chr1 | - | 225848896 | 225849012 | 225848491 | 225848554 | 225849911 | 225850079 | 203 | 122 | 244 | 158 | 133 | 0 | 0 | 1 | 0 | 1 | 87 | 159 | 60 | 192 | 400 | 0 | 0 | 0 | 5 | 0 | 265 | 149 | 1 | 1 | 1 | 1 | 0.993 | 1 | 0.987 | 1 | 1 | 1 | 0.956 | 1 | 0.005 |
| SE | 37871 | ENSG00000196187 | TMEM63A | chr1 | - | 225852663 | 225852769 | 225849911 | 225850079 | 225853628 | 225853791 | 225 | 99 | 253 | 194 | 165 | 0 | 0 | 1 | 0 | 0 | 83 | 154 | 94 | 115 | 375 | 0 | 0 | 0 | 0 | 0 | 255 | 149 | 1 | 1 | 1 | 1 | 0.993 | 1 | 1 | 1 | 1 | 1 | 1 | 1 | -0.001 |
| SE | 37874 | ENSG00000196187 | TMEM63A | chr1 | - | 225853628 | 225853791 | 225852663 | 225852769 | 225855877 | 225855940 | 253 | 118 | 325 | 257 | 223 | 0 | 0 | 0 | 0 | 0 | 73 | 175 | 121 | 135 | 408 | 0 | 3 | 0 | 0 | 3 | 298 | 149 | 1 | 1 | 1 | 1 | 1 | 1 | 1 | 1 | 0.967 | 1 | 1 | 0.986 | 0.009 |
| SE | 37875 | ENSG00000196187 | TMEM63A | chr1 | - | 225855877 | 225855940 | 225853628 | 225853791 | 225856651 | 225856738 | 186 | 82 | 277 | 206 | 169 | 0 | 0 | 2 | 0 | 0 | 59 | 89 | 96 | 108 | 323 | 0 | 0 | 0 | 6 | 0 | 212 | 149 | 1 | 1 | 1 | 1 | 0.99 | 1 | 1 | 1 | 1 | 1 | 0.927 | 1 | 0.013 |
| SE | 37876 | ENSG00000196187 | TMEM63A | chr1 | - | 225856651 | 225856738 | 225855877 | 225855940 | 225856910 | 225857017 | 125 | 78 | 197 | 170 | 120 | 0 | 1 | 0 | 1 | 0 | 48 | 82 | 90 | 84 | 284 | 0 | 0 | 0 | 0 | 0 | 236 | 149 | 1 | 1 | 1 | 0.98 | 1 | 0.991 | 1 | 1 | 1 | 1 | 1 | 1 | -0.006 |
| SE | 37878 | ENSG00000196187 | TMEM63A | chr1 | - | 225856910 | 225857017 | 225855877 | 225855940 | 225859195 | 225859349 | 31 | 33 | 54 | 52 | 72 | 0 | 1 | 0 | 0 | 0 | 25 | 33 | 27 | 46 | 152 | 0 | 0 | 3 | 2 | 0 | 256 | 149 | 1 | 1 | 1 | 0.951 | 1 | 1 | 1 | 1 | 1 | 0.84 | 0.93 | 1 | 0.036 |
| SE | 37879 | ENSG00000196187 | TMEM63A | chr1 | - | 225856910 | 225857017 | 225856651 | 225856738 | 225859195 | 225859349 | 120 | 85 | 195 | 173 | 108 | 0 | 0 | 0 | 1 | 0 | 55 | 76 | 73 | 99 | 333 | 0 | 0 | 0 | 0 | 0 | 256 | 149 | 1 | 1 | 1 | 1 | 1 | 0.99 | 1 | 1 | 1 | 1 | 1 | 1 | -0.002 |
| SE | 37880 | ENSG00000196187 | TMEM63A | chr1 | - | 225859195 | 225859349 | 225856910 | 225857017 | 225860859 | 225860997 | 158 | 130 | 230 | 231 | 349 | 0 | 0 | 1 | 0 | 0 | 118 | 130 | 93 | 173 | 434 | 0 | 0 | 0 | 0 | 0 | 298 | 149 | 1 | 1 | 1 | 1 | 0.991 | 1 | 1 | 1 | 1 | 1 | 1 | 1 | -0.002 |
| SE | 37881 | ENSG00000196187 | TMEM63A | chr1 | - | 225859195 | 225859349 | 225856910 | 225857017 | 225862217 | 225862351 | 38 | 41 | 62 | 73 | 79 | 0 | 0 | 0 | 0 | 0 | 38 | 43 | 41 | 70 | 179 | 0 | 0 | 0 | 3 | 0 | 298 | 149 | 1 | 1 | 1 | 1 | 1 | 1 | 1 | 1 | 1 | 1 | 0.921 | 1 | 0.016 |
| SE | 37882 | ENSG00000196187 | TMEM63A | chr1 | - | 225860859 | 225860997 | 225856910 | 225857017 | 225862217 | 225862351 | 164 | 140 | 199 | 191 | 306 | 0 | 0 | 0 | 0 | 0 | 180 | 99 | 66 | 195 | 255 | 0 | 0 | 0 | 3 | 0 | 287 | 149 | 1 | 1 | 1 | 1 | 1 | 1 | 1 | 1 | 1 | 1 | 0.971 | 1 | 0.006 |
| SE | 37884 | ENSG00000196187 | TMEM63A | chr1 | - | 225866132 | 225866208 | 225865896 | 225866050 | 225866573 | 225866748 | 0 | 0 | 1 | 0 | 0 | 0 | 1 | 3 | 2 | 0 | 0 | 0 | 1 | 0 | 0 | 0 | 6 | 3 | 0 | 0 | 225 | 149 | 1 | 1 | NA | 0 | 0.181 | 0 | NA | NA | 0 | 0.181 | NA | NA | -0.03 |
| SE | 37885 | ENSG00000196187 | TMEM63A | chr1 | - | 225866132 | 225866208 | 225865896 | 225866050 | 225867111 | 225867163 | 1 | 0 | 0 | 0 | 0 | 0 | 0 | 0 | 1 | 0 | 0 | 0 | 1 | 0 | 0 | 0 | 0 | 0 | 0 | 6 | 225 | 149 | 1 | 1 | 1 | NA | NA | 0 | NA | NA | NA | 1 | NA | 0 | 0 |
| SE | 37886 | ENSG00000196187 | TMEM63A | chr1 | - | 225866573 | 225866682 | 225865896 | 225865967 | 225867111 | 225867163 | 180 | 109 | 195 | 217 | 240 | 3 | 0 | 1 | 1 | 0 | 143 | 119 | 83 | 140 | 513 | 0 | 0 | 0 | 6 | 0 | 258 | 149 | 1 | 1 | 0.972 | 1 | 0.991 | 0.992 | 1 | 1 | 1 | 1 | 0.931 | 1 | 0.005 |
| SE | 37887 | ENSG00000196187 | TMEM63A | chr1 | - | 225866573 | 225866682 | 225865896 | 225865967 | 225867887 | 225868030 | 105 | 73 | 115 | 126 | 152 | 1 | 0 | 0 | 0 | 0 | 88 | 53 | 49 | 61 | 295 | 4 | 0 | 0 | 0 | 0 | 258 | 149 | 1 | 1 | 0.984 | 1 | 1 | 1 | 1 | 0.927 | 1 | 1 | 1 | 1 | 0.011 |
| SE | 37889 | ENSG00000196187 | TMEM63A | chr1 | - | 225867111 | 225867163 | 225865896 | 225865967 | 225871075 | 225871113 | 3 | 0 | 0 | 0 | 0 | 0 | 0 | 0 | 0 | 0 | 0 | 0 | 0 | 0 | 0 | 4 | 0 | 0 | 0 | 0 | 201 | 149 | 1 | 1 | 1 | NA | NA | NA | NA | 0 | NA | NA | NA | NA | 1 |
| SE | 37892 | ENSG00000196187 | TMEM63A | chr1 | - | 225867887 | 225868030 | 225865896 | 225865967 | 225871075 | 225871113 | 96 | 58 | 131 | 99 | 98 | 0 | 0 | 0 | 0 | 0 | 127 | 58 | 30 | 85 | 181 | 4 | 0 | 0 | 0 | 0 | 292 | 149 | 1 | 1 | 1 | 1 | 1 | 1 | 1 | 0.942 | 1 | 1 | 1 | 1 | 0.012 |
| SE | 37893 | ENSG00000196187 | TMEM63A | chr1 | - | 225867887 | 225868030 | 225867111 | 225867163 | 225871075 | 225871113 | 206 | 112 | 257 | 237 | 214 | 0 | 0 | 0 | 0 | 0 | 210 | 135 | 78 | 201 | 375 | 0 | 0 | 0 | 1 | 0 | 292 | 149 | 1 | 1 | 1 | 1 | 1 | 1 | 1 | 1 | 1 | 1 | 0.99 | 1 | 0.002 |
| SE | 37896 | ENSG00000184014 | DENND5A | chr11 | - | 9140188 | 9140248 | 9138848 | 9139854 | 9141973 | 9142108 | 4 | 1 | 7 | 1 | 2 | 5 | 0 | 3 | 0 | 5 | 5 | 0 | 7 | 0 | 12 | 6 | 0 | 5 | 9 | 0 | 209 | 149 | 1 | 1 | 0.363 | 1 | 0.625 | 1 | 0.222 | 0.373 | NA | 0.5 | 0 | 1 | 0.174 |
| SE | 37901 | ENSG00000184014 | DENND5A | chr11 | - | 9142721 | 9144278 | 9141939 | 9142108 | 9144994 | 9145113 | 270 | 209 | 248 | 263 | 354 | 0 | 0 | 0 | 0 | 0 | 527 | 297 | 251 | 460 | 360 | 0 | 0 | 0 | 3 | 0 | 298 | 149 | 1 | 1 | 1 | 1 | 1 | 1 | 1 | 1 | 1 | 1 | 0.987 | 1 | 0.003 |
| SE | 37904 | ENSG00000184014 | DENND5A | chr11 | - | 9144096 | 9144278 | 9141939 | 9142108 | 9144994 | 9145113 | 104 | 91 | 94 | 93 | 112 | 0 | 0 | 0 | 0 | 0 | 190 | 143 | 89 | 185 | 106 | 0 | 0 | 0 | 3 | 0 | 298 | 149 | 1 | 1 | 1 | 1 | 1 | 1 | 1 | 1 | 1 | 1 | 0.969 | 1 | 0.006 |
| SE | 37906 | ENSG00000184014 | DENND5A | chr11 | - | 9145669 | 9145815 | 9144096 | 9145113 | 9150080 | 9150209 | 84 | 51 | 56 | 57 | 67 | 0 | 0 | 0 | 0 | 0 | 130 | 90 | 52 | 154 | 86 | 0 | 0 | 0 | 5 | 0 | 295 | 149 | 1 | 1 | 1 | 1 | 1 | 1 | 1 | 1 | 1 | 1 | 0.94 | 1 | 0.012 |
| SE | 37907 | ENSG00000184014 | DENND5A | chr11 | - | 9147029 | 9147151 | 9144096 | 9145113 | 9150080 | 9150209 | 111 | 83 | 90 | 86 | 234 | 0 | 0 | 0 | 0 | 0 | 238 | 85 | 93 | 165 | 114 | 0 | 0 | 0 | 5 | 0 | 271 | 149 | 1 | 1 | 1 | 1 | 1 | 1 | 1 | 1 | 1 | 1 | 0.948 | 1 | 0.01 |
| SE | 37910 | ENSG00000184014 | DENND5A | chr11 | - | 9150679 | 9150764 | 9150080 | 9150209 | 9152357 | 9152442 | 238 | 151 | 252 | 234 | 434 | 0 | 0 | 0 | 0 | 0 | 418 | 246 | 200 | 298 | 291 | 5 | 0 | 1 | 4 | 0 | 234 | 149 | 1 | 1 | 1 | 1 | 1 | 1 | 1 | 0.982 | 1 | 0.992 | 0.979 | 1 | 0.009 |
| SE | 37911 | ENSG00000184014 | DENND5A | chr11 | - | 9152357 | 9152442 | 9150679 | 9150764 | 9160712 | 9160865 | 221 | 209 | 268 | 242 | 414 | 0 | 0 | 0 | 0 | 0 | 527 | 274 | 262 | 337 | 285 | 0 | 1 | 0 | 0 | 0 | 234 | 149 | 1 | 1 | 1 | 1 | 1 | 1 | 1 | 1 | 0.994 | 1 | 1 | 1 | 0.001 |
| SE | 37912 | ENSG00000184014 | DENND5A | chr11 | - | 9158416 | 9158540 | 9152357 | 9152442 | 9160712 | 9160865 | 5 | 0 | 0 | 0 | 1 | 155 | 163 | 172 | 170 | 256 | 0 | 0 | 0 | 3 | 0 | 365 | 203 | 179 | 231 | 214 | 273 | 149 | 1 | 1 | 0.017 | 0 | 0 | 0 | 0.002 | 0 | 0 | 0 | 0.007 | 0 | 0.002 |
| SE | 37913 | ENSG00000184014 | DENND5A | chr11 | - | 9160257 | 9160299 | 9152357 | 9152442 | 9160712 | 9160865 | 1 | 0 | 0 | 0 | 2 | 155 | 163 | 172 | 170 | 256 | 0 | 1 | 1 | 0 | 0 | 365 | 203 | 179 | 231 | 214 | 191 | 149 | 1 | 1 | 0.005 | 0 | 0 | 0 | 0.006 | 0 | 0.004 | 0.004 | 0 | 0 | 0.001 |
| SE | 37915 | ENSG00000184014 | DENND5A | chr11 | - | 9165835 | 9165967 | 9160712 | 9160865 | 9169855 | 9169949 | 243 | 243 | 239 | 280 | 284 | 0 | 0 | 0 | 0 | 0 | 458 | 310 | 245 | 415 | 337 | 0 | 0 | 1 | 0 | 0 | 281 | 149 | 1 | 1 | 1 | 1 | 1 | 1 | 1 | 1 | 1 | 0.992 | 1 | 1 | 0.002 |

| | | | | | | | | | | | | | | | | | | | | | | | | | | | | | | | | | | | | | | | | | | | | | | |
|---|---|---|---|---|---|---|---|---|---|---|---|---|---|---|---|---|---|---|---|---|---|---|---|---|---|---|---|---|---|---|---|---|---|---|---|---|---|---|---|---|---|---|---|---|---|---|
| SE | 37916 | ENSG00000184014 | DENND5A | chr11 | - | 9170626 | 9170777 | 9169855 | 9169949 | 9176771 | 9176902 | 204 | 195 | 184 | 236 | 347 | 0 | 1 | 0 | 1 | 0 | 430 | 293 | 265 | 354 | 320 | 0 | 0 | 0 | 0 | 0 | 298 | 149 | 1 | 1 | 1 | 0.99 | 1 | 0.992 | 1 | 1 | 1 | 1 | 1 | 1 | -0.004 |
| SE | 37921 | ENSG00000184014 | DENND5A | chr11 | - | 9203659 | 9204317 | 9193493 | 9193681 | 9206672 | 9206752 | 300 | 243 | 259 | 254 | 531 | 0 | 0 | 0 | 0 | 0 | 645 | 228 | 366 | 515 | 369 | 0 | 0 | 2 | 0 | 0 | 298 | 149 | 1 | 1 | 1 | 1 | 1 | 1 | 1 | 1 | 1 | 0.989 | 1 | 1 | 0.002 |
| SE | 37922 | ENSG00000184014 | DENND5A | chr11 | - | 9207172 | 9207208 | 9206672 | 9206782 | 9207560 | 9207622 | 0 | 0 | 0 | 0 | 0 | 55 | 28 | 58 | 55 | 35 | 0 | 2 | 0 | 0 | 0 | 52 | 33 | 52 | 59 | 89 | 185 | 149 | 1 | 1 | 0 | 0 | 0 | 0 | 0 | 0 | 0.047 | 0 | 0 | 0 | -0.009 |
| SE | 37925 | ENSG00000067113 | PLPP1 | chr5 | - | 55475298 | 55475450 | 55467868 | 55468149 | 55490959 | 55491114 | 11 | 4 | 8 | 11 | 22 | 0 | 0 | 4 | 0 | 3 | 0 | 1 | 0 | 6 | 1 | 0 | 0 | 0 | 0 | 0 | 298 | 149 | 1 | 1 | 1 | 1 | 0.5 | 1 | 0.786 | NA | 1 | NA | 1 | 1 | -0.143 |
| SE | 37928 | ENSG00000067113 | PLPP1 | chr5 | - | 55490959 | 55491114 | 55475298 | 55475450 | 55534571 | 55534951 | 0 | 0 | 1 | 2 | 3 | 1 | 1 | 3 | 3 | 6 | 0 | 1 | 0 | 0 | 0 | 0 | 0 | 0 | 0 | 0 | 298 | 149 | 1 | 1 | 0 | 0 | 0.143 | 0.25 | 0.2 | NA | 1 | NA | NA | NA | -0.881 |
| SE | 37932 | ENSG00000169908 | TM4SF1 | chr3 | - | 149375442 | 149375588 | 149371686 | 149371867 | 149375679 | 149375769 | 4 | 1 | 0 | 0 | 0 | 0 | 1 | 0 | 0 | 0 | 0 | 8 | 0 | 0 | 19 | 0 | 0 | 0 | 0 | 8 | 295 | 149 | 1 | 1 | 1 | 0.336 | NA | NA | NA | NA | 1 | NA | NA | 0.545 | -0.104 |
| SE | 37934 | ENSG00000148843 | PDCD11 | chr10 | + | 103432128 | 103432234 | 103427328 | 103427391 | 103433947 | 103434037 | 81 | 54 | 92 | 63 | 69 | 1 | 0 | 0 | 0 | 0 | 60 | 42 | 31 | 43 | 93 | 0 | 0 | 2 | 3 | 0 | 255 | 149 | 1 | 1 | 0.979 | 1 | 1 | 1 | 1 | 1 | 1 | 0.901 | 0.893 | 1 | 0.037 |
| SE | 37936 | ENSG00000148843 | PDCD11 | chr10 | + | 103438014 | 103438071 | 103434797 | 103434975 | 103439745 | 103439868 | 22 | 8 | 27 | 15 | 8 | 0 | 0 | 0 | 0 | 2 | 14 | 4 | 3 | 20 | 37 | 0 | 0 | 0 | 0 | 0 | 206 | 149 | 1 | 1 | 1 | 1 | 1 | 1 | 0.743 | 1 | 1 | 1 | 1 | 1 | -0.051 |
| SE | 37937 | ENSG00000148843 | PDCD11 | chr10 | + | 103438685 | 103438808 | 103438014 | 103438071 | 103439745 | 103439868 | 112 | 64 | 111 | 105 | 125 | 1 | 0 | 0 | 0 | 0 | 193 | 120 | 80 | 78 | 173 | 0 | 0 | 0 | 4 | 0 | 272 | 149 | 1 | 1 | 0.984 | 1 | 1 | 1 | 1 | 1 | 1 | 1 | 1 | 1 | -0.003 |
| SE | 37939 | ENSG00000138964 | PARVG | chr22 | + | 44181741 | 44181917 | 44172955 | 44173191 | 44185807 | 44185872 | 0 | 0 | 0 | 0 | 0 | 1 | 0 | 0 | 0 | 0 | 0 | 3 | 0 | 0 | 0 | 4 | 2 | 0 | 4 | 0 | 298 | 149 | 1 | 1 | 0 | NA | NA | NA | NA | 0 | 0.429 | NA | 0 | NA | -0.143 |
| SE | 37941 | ENSG00000138964 | PARVG | chr22 | + | 44181741 | 44181917 | 44181335 | 44181370 | 44183317 | 44183408 | 423 | 240 | 233 | 382 | 270 | 0 | 0 | 0 | 0 | 0 | 833 | 422 | 252 | 448 | 906 | 0 | 0 | 0 | 1 | 0 | 298 | 149 | 1 | 1 | 1 | 1 | 1 | 1 | 1 | 1 | 1 | 1 | 0.996 | 1 | 0.001 |
| SE | 37944 | ENSG00000138964 | PARVG | chr22 | + | 44183317 | 44183408 | 44172983 | 44173191 | 44190550 | 44190666 | 0 | 1 | 1 | 0 | 0 | 1 | 0 | 0 | 0 | 0 | 0 | 0 | 0 | 10 | 10 | 0 | 0 | 0 | 0 | 0 | 240 | 149 | 1 | 1 | 0 | 1 | 1 | NA | NA | NA | NA | NA | 1 | 1 | -0.333 |
| SE | 37945 | ENSG00000138964 | PARVG | chr22 | + | 44183317 | 44183408 | 44181738 | 44181917 | 44185807 | 44185872 | 482 | 275 | 309 | 452 | 294 | 0 | 0 | 0 | 0 | 0 | 1016 | 520 | 296 | 525 | 1116 | 0 | 3 | 0 | 0 | 0 | 240 | 149 | 1 | 1 | 1 | 1 | 1 | 1 | 1 | 1 | 0.991 | 1 | 1 | 1 | 0.002 |
| SE | 37946 | ENSG00000138964 | PARVG | chr22 | + | 44190550 | 44190666 | 44185525 | 44189254 | 44192048 | 44192104 | 609 | 355 | 419 | 614 | 388 | 1 | 0 | 2 | 1 | 7 | 904 | 674 | 369 | 873 | 1213 | 0 | 0 | 2 | 0 | 4 | 265 | 149 | 1 | 1 | 0.997 | 1 | 0.992 | 0.997 | 0.969 | 1 | 1 | 0.99 | 1 | 0.994 | -0.006 |
| SE | 37947 | ENSG00000138964 | PARVG | chr22 | + | 44193800 | 44193823 | 44192048 | 44192104 | 44196154 | 44196213 | 372 | 228 | 264 | 416 | 283 | 0 | 0 | 0 | 0 | 0 | 589 | 433 | 253 | 507 | 649 | 0 | 0 | 0 | 0 | 10 | 172 | 149 | 1 | 1 | 1 | 1 | 1 | 1 | 1 | 1 | 1 | 1 | 1 | 0.983 | 0.003 |
| SE | 37948 | ENSG00000138964 | PARVG | chr22 | + | 44196346 | 44196415 | 44196154 | 44196213 | 44198620 | 44198717 | 473 | 284 | 329 | 414 | 490 | 0 | 0 | 0 | 0 | 2 | 730 | 409 | 241 | 503 | 689 | 0 | 0 | 0 | 0 | 0 | 218 | 149 | 1 | 1 | 1 | 1 | 1 | 1 | 0.994 | 1 | 1 | 1 | 1 | 1 | -0.001 |
| SE | 37949 | ENSG00000138964 | PARVG | chr22 | + | 44205756 | 44205829 | 44198620 | 44198722 | 44206316 | 44206622 | 570 | 334 | 343 | 482 | 674 | 1 | 0 | 2 | 1 | 1 | 972 | 587 | 314 | 643 | 923 | 2 | 1 | 0 | 8 | 0 | 222 | 149 | 1 | 1 | 0.997 | 1 | 0.991 | 0.997 | 0.998 | 0.997 | 0.997 | 1 | 0.982 | 1 | 0.001 |
| SE | 37961 | ENSG00000233396 | LINC01719 | chr1 | + | 146057693 | 146057988 | 146052625 | 146052948 | 146058312 | 146058461 | 0 | 0 | 0 | 1 | 1 | 0 | 0 | 0 | 0 | 2 | 0 | 2 | 0 | 0 | 0 | 4 | 0 | 0 | 0 | 0 | 298 | 149 | 1 | 1 | NA | NA | NA | 1 | 0.2 | NA | 1 | NA | NA | NA | -0.4 |
| SE | 37968 | ENSG00000160883 | HK3 | chr5 | - | 176883769 | 176883869 | 176881943 | 176882127 | 176884038 | 176884134 | 393 | 220 | 179 | 205 | 84 | 0 | 0 | 5 | 2 | 0 | 923 | 510 | 290 | 1055 | 1299 | 8 | 0 | 0 | 4 | 0 | 249 | 149 | 1 | 1 | 1 | 1 | 0.955 | 0.984 | 1 | 0.986 | 1 | 1 | 0.994 | 1 | -0.008 |
| SE | 37969 | ENSG00000160883 | HK3 | chr5 | - | 176888331 | 176888565 | 176887450 | 176887746 | 176888708 | 176888864 | 52 | 32 | 46 | 37 | 19 | 0 | 0 | 0 | 0 | 0 | 97 | 106 | 35 | 108 | 129 | 0 | 2 | 1 | 0 | 0 | 298 | 149 | 1 | 1 | 1 | 1 | 1 | 1 | 1 | 1 | 0.964 | 0.946 | 1 | 1 | 0.018 |
| SE | 37970 | ENSG00000160883 | HK3 | chr5 | - | 176889644 | 176889744 | 176889380 | 176889564 | 176890634 | 176890730 | 364 | 226 | 226 | 267 | 144 | 0 | 0 | 0 | 0 | 0 | 1373 | 691 | 252 | 1063 | 1363 | 0 | 0 | 0 | 5 | 0 | 249 | 149 | 1 | 1 | 1 | 1 | 1 | 1 | 1 | 1 | 1 | 1 | 0.992 | 1 | 0.002 |
| SE | 37971 | ENSG00000160883 | HK3 | chr5 | - | 176890634 | 176890730 | 176889644 | 176889744 | 176891387 | 176891391 | 233 | 142 | 165 | 176 | 95 | 0 | 0 | 3 | 0 | 0 | 889 | 457 | 159 | 739 | 983 | 0 | 1 | 0 | 5 | 0 | 245 | 149 | 1 | 1 | 1 | 1 | 1 | 0.973 | 1 | 1 | 0.996 | 1 | 0.989 | 1 | -0.002 |
| SE | 37972 | ENSG00000160883 | HK3 | chr5 | - | 176890821 | 176890941 | 176890634 | 176890730 | 176891036 | 176891191 | 262 | 140 | 119 | 183 | 126 | 1 | 0 | 0 | 4 | 0 | 812 | 444 | 139 | 720 | 980 | 7 | 0 | 0 | 4 | 0 | 269 | 149 | 1 | 1 | 0.993 | 1 | 1 | 0.962 | 1 | 0.985 | 1 | 1 | 0.99 | 1 | -0.004 |
| SE | 37973 | ENSG00000160883 | HK3 | chr5 | - | 176890821 | 176890941 | 176890634 | 176890730 | 176891387 | 176891391 | 92 | 55 | 35 | 67 | 37 | 1 | 0 | 0 | 0 | 0 | 263 | 156 | 61 | 309 | 394 | 0 | 0 | 0 | 0 | 0 | 269 | 149 | 1 | 1 | 0.981 | 1 | 1 | 1 | 1 | 1 | 1 | 1 | 1 | 1 | -0.004 |
| SE | 37974 | ENSG00000160883 | HK3 | chr5 | - | 176891036 | 176891191 | 176890634 | 176890730 | 176891387 | 176891550 | 82 | 38 | 42 | 61 | 34 | 1 | 0 | 0 | 0 | 0 | 232 | 178 | 46 | 197 | 196 | 0 | 0 | 0 | 0 | 0 | 298 | 149 | 1 | 1 | 0.976 | 1 | 1 | 1 | 1 | 1 | 1 | 1 | 1 | 1 | -0.005 |
| SE | 37975 | ENSG00000160883 | HK3 | chr5 | - | 176891036 | 176891191 | 176890821 | 176890941 | 176891387 | 176891550 | 251 | 123 | 127 | 173 | 123 | 0 | 0 | 1 | 0 | 0 | 774 | 469 | 124 | 604 | 782 | 0 | 3 | 0 | 0 | 0 | 298 | 149 | 1 | 1 | 1 | 1 | 0.984 | 1 | 1 | 1 | 0.987 | 1 | 1 | 1 | -0.001 |
| SE | 37976 | ENSG00000160883 | HK3 | chr5 | - | 176891036 | 176891279 | 176890634 | 176890730 | 176891387 | 176891391 | 10 | 1 | 2 | 10 | 4 | 1 | 0 | 0 | 0 | 0 | 7 | 1 | 4 | 10 | 4 | 0 | 0 | 0 | 0 | 0 | 298 | 149 | 1 | 1 | 0.833 | 1 | 1 | 1 | 1 | 1 | 1 | 1 | 1 | 1 | -0.033 |
| SE | 37977 | ENSG00000160883 | HK3 | chr5 | - | 176891036 | 176891279 | 176890821 | 176890941 | 176891387 | 176891391 | 179 | 86 | 87 | 122 | 93 | 0 | 0 | 1 | 0 | 0 | 549 | 292 | 82 | 417 | 590 | 0 | 3 | 0 | 0 | 0 | 298 | 149 | 1 | 1 | 1 | 1 | 0.978 | 1 | 1 | 1 | 0.98 | 1 | 1 | 1 | 0 |
| SE | 37982 | ENSG00000114670 | NEK11 | chr3 | + | 131027929 | 131028002 | 131026876 | 131027006 | 131029612 | 131029878 | 0 | 0 | 0 | 3 | 4 | 1 | 0 | 0 | 0 | 0 | 0 | 0 | 0 | 3 | 0 | 0 | 0 | 0 | 0 | 0 | 222 | 149 | 1 | 1 | 0 | NA | NA | 1 | 1 | NA | NA | NA | 1 | NA | -0.333 |
| SE | 37984 | ENSG00000114670 | NEK11 | chr3 | + | 131109802 | 131109921 | 131029612 | 131029878 | 131132744 | 131132809 | 1 | 3 | 1 | 2 | 6 | 0 | 0 | 0 | 0 | 0 | 2 | 0 | 0 | 7 | 10 | 0 | 0 | 1 | 0 | 0 | 268 | 149 | 1 | 1 | 1 | 1 | 1 | 1 | 1 | 1 | NA | 0 | 1 | 1 | 0.25 |
| SE | 37985 | ENSG00000114670 | NEK11 | chr3 | + | 131133245 | 131133355 | 131132744 | 131132809 | 131133829 | 131133956 | 1 | 3 | 0 | 0 | 0 | 0 | 1 | 1 | 4 | 15 | 7 | 0 | 0 | 0 | 0 | 0 | 0 | 4 | 14 | 6 | 259 | 149 | 1 | 1 | 1 | 0.633 | 0 | 0 | 0 | 1 | NA | 0 | 0 | 0 | 0.077 |
| SE | 37986 | ENSG00000114670 | NEK11 | chr3 | + | 131133276 | 131133355 | 131132744 | 131132809 | 131133829 | 131133956 | 0 | 0 | 0 | 0 | 0 | 0 | 1 | 1 | 4 | 15 | 1 | 0 | 0 | 0 | 0 | 0 | 0 | 4 | 14 | 6 | 228 | 149 | 1 | 1 | NA | 0 | 0 | 0 | 0 | 1 | NA | 0 | 0 | 0 | -0.25 |
| SE | 37987 | ENSG00000114670 | NEK11 | chr3 | + | 131162407 | 131162527 | 131154869 | 131155121 | 131165425 | 131165519 | 7 | 4 | 7 | 9 | 17 | 0 | 0 | 0 | 2 | 0 | 0 | 4 | 0 | 2 | 0 | 0 | 0 | 0 | 0 | 0 | 269 | 149 | 1 | 1 | 1 | 1 | 1 | 0.714 | 1 | NA | 1 | NA | 1 | NA | -0.057 |
| SE | 37988 | ENSG00000114670 | NEK11 | chr3 | + | 131168829 | 131168937 | 131165276 | 131165519 | 131170772 | 131170887 | 4 | 9 | 5 | 11 | 11 | 0 | 0 | 0 | 0 | 0 | 0 | 4 | 5 | 6 | 0 | 0 | 0 | 0 | 1 | 0 | 257 | 149 | 1 | 1 | 1 | 1 | 1 | 1 | 1 | NA | 1 | 1 | 0.777 | NA | 0.074 |
| SE | 37989 | ENSG00000114670 | NEK11 | chr3 | + | 131273477 | 131273574 | 131243435 | 131243496 | 131349556 | 131349776 | 0 | 1 | 1 | 4 | 8 | 0 | 0 | 0 | 1 | 0 | 0 | 8 | 11 | 0 | 0 | 0 | 3 | 0 | 0 | 0 | 246 | 149 | 1 | 1 | NA | 1 | 1 | 0.708 | 1 | NA | 0.618 | 1 | NA | NA | 0.118 |
| SE | 37993 | ENSG00000095906 | NUBP2 | chr16 | + | 1786536 | 1786655 | 1782931 | 1783036 | 1787676 | 1787831 | 6 | 4 | 14 | 1 | 0 | 0 | 0 | 0 | 1 | 2 | 0 | 17 | 10 | 0 | 4 | 0 | 0 | 1 | 0 | 0 | 268 | 149 | 1 | 1 | 1 | 1 | 1 | 0.357 | 0 | NA | 1 | 0.848 | NA | 1 | -0.278 |
| SE | 37994 | ENSG00000095906 | NUBP2 | chr16 | + | 1786536 | 1786955 | 1782995 | 1783036 | 1787676 | 1787831 | 19 | 11 | 27 | 34 | 9 | 0 | 0 | 0 | 1 | 2 | 21 | 29 | 3 | 0 | 25 | 0 | 0 | 1 | 0 | 0 | 298 | 149 | 1 | 1 | 1 | 1 | 1 | 0.944 | 0.692 | 1 | 1 | 0.6 | NA | 1 | 0.027 |
| SE | 37995 | ENSG00000095906 | NUBP2 | chr16 | + | 1786737 | 1786955 | 1786536 | 1786655 | 1787676 | 1787740 | 18 | 9 | 19 | 33 | 9 | 5 | 2 | 6 | 0 | 0 | 21 | 22 | 3 | 0 | 21 | 0 | 10 | 12 | 0 | 0 | 298 | 149 | 1 | 1 | 0.643 | 0.692 | 0.613 | 1 | 1 | 1 | 0.524 | 0.111 | NA | 1 | 0.131 |
| SE | 37997 | ENSG00000197386 | HTT | chr4 | + | 3062475 | 3062692 | 3041421 | 3042245 | 3071235 | 3071396 | 2 | 0 | 0 | 0 | 0 | 1 | 0 | 0 | 0 | 0 | 0 | 5 | 0 | 0 | 0 | 0 | 0 | 1 | 0 | 0 | 298 | 149 | 1 | 1 | 0.5 | NA | NA | NA | NA | NA | 1 | 0 | NA | NA | 0 |
| SE | 37998 | ENSG00000197386 | HTT | chr4 | + | 3071235 | 3071396 | 3041421 | 3042245 | 3086938 | 3087022 | 6 | 0 | 0 | 0 | 0 | 3 | 0 | 0 | 0 | 0 | 0 | 0 | 1 | 0 | 0 | 0 | 0 | 0 | 0 | 0 | 298 | 149 | 1 | 1 | 0.5 | NA | NA | NA | NA | NA | NA | 1 | NA | NA | -0.5 |
| SE | 37999 | ENSG00000197386 | HTT | chr4 | + | 3086938 | 3087022 | 3074680 | 3075088 | 3099273 | 3099394 | 145 | 99 | 104 | 146 | 208 | 0 | 0 | 0 | 0 | 0 | 273 | 149 | 113 | 177 | 344 | 0 | 0 | 0 | 0 | 2 | 233 | 149 | 1 | 1 | 1 | 1 | 1 | 1 | 1 | 1 | 1 | 1 | 1 | 0.991 | 0.002 |
| SE | 38000 | ENSG00000197386 | HTT | chr4 | + | 3099273 | 3099394 | 3086938 | 3087022 | 3103823 | 3103883 | 275 | 193 | 179 | 226 | 344 | 0 | 2 | 1 | 0 | 0 | 479 | 241 | 246 | 330 | 507 | 0 | 0 | 0 | 0 | 5 | 270 | 149 | 1 | 1 | 1 | 0.982 | 0.99 | 1 | 1 | 1 | 1 | 1 | 1 | 0.982 | -0.002 |
| SE | 38002 | ENSG00000197386 | HTT | chr4 | + | 3103823 | 3103883 | 3086938 | 3087022 | 3105356 | 3105436 | 129 | 79 | 90 | 113 | 145 | 0 | 0 | 0 | 0 | 0 | 233 | 171 | 119 | 103 | 358 | 8 | 0 | 0 | 0 | 0 | 209 | 149 | 1 | 1 | 1 | 1 | 1 | 1 | 1 | 0.954 | 1 | 1 | 1 | 1 | 0.009 |
| SE | 38003 | ENSG00000197386 | HTT | chr4 | + | 3103823 | 3103883 | 3099273 | 3099394 | 3105356 | 3105436 | 267 | 185 | 179 | 216 | 319 | 0 | 0 | 0 | 0 | 1 | 472 | 310 | 275 | 284 | 594 | 0 | 2 | 4 | 3 | 2 | 209 | 149 | 1 | 1 | 1 | 1 | 1 | 1 | 0.996 | 1 | 0.991 | 0.98 | 0.985 | 0.995 | 0.009 |
| SE | 38004 | ENSG00000197386 | HTT | chr4 | + | 3115303 | 3115445 | 3107284 | 3107423 | 3116084 | 3116263 | 178 | 103 | 128 | 157 | 183 | 2 | 0 | 0 | 1 | 0 | 330 | 202 | 120 | 211 | 447 | 0 | 0 | 0 | 7 | 0 | 291 | 149 | 1 | 1 | 0.979 | 1 | 1 | 0.988 | 1 | 1 | 1 | 1 | 0.939 | 1 | 0.006 |
| SE | 38005 | ENSG00000197386 | HTT | chr4 | + | 3122888 | 3122936 | 3121227 | 3121432 | 3125548 | 3125629 | 107 | 60 | 80 | 108 | 102 | 0 | 0 | 1 | 0 | 0 | 161 | 158 | 97 | 141 | 361 | 0 | 0 | 3 | 0 | 0 | 197 | 149 | 1 | 1 | 1 | 1 | 0.984 | 1 | 1 | 1 | 1 | 0.961 | 1 | 1 | 0.005 |
| SE | 38008 | ENSG00000197386 | HTT | chr4 | + | 3135903 | 3135967 | 3134400 | 3134540 | 3136225 | 3136326 | 156 | 114 | 107 | 100 | 160 | 0 | 0 | 0 | 1 | 0 | 263 | 179 | 131 | 187 | 362 | 0 | 0 | 0 | 0 | 0 | 213 | 149 | 1 | 1 | 1 | 1 | 1 | 0.986 | 1 | 1 | 1 | 1 | 1 | 1 | -0.003 |
| SE | 38009 | ENSG00000197386 | HTT | chr4 | + | 3142765 | 3142886 | 3140509 | 3140656 | 3145151 | 3145228 | 182 | 152 | 182 | 194 | 266 | 0 | 0 | 0 | 0 | 0 | 381 | 241 | 151 | 283 | 392 | 0 | 1 | 0 | 0 | 0 | 270 | 149 | 1 | 1 | 1 | 1 | 1 | 1 | 1 | 1 | 0.993 | 1 | 1 | 1 | 0.001 |
| SE | 38011 | ENSG00000197386 | HTT | chr4 | + | 3154292 | 3154419 | 3148004 | 3148207 | 3160281 | 3160392 | 49 | 21 | 37 | 37 | 56 | 0 | 0 | 4 | 0 | 0 | 118 | 66 | 17 | 59 | 89 | 0 | 0 | 0 | 0 | 0 | 276 | 149 | 1 | 1 | 1 | 1 | 0.833 | 1 | 1 | 1 | 1 | 1 | 1 | 1 | -0.033 |
| SE | 38012 | ENSG00000197386 | HTT | chr4 | + | 3157071 | 3157199 | 3148004 | 3148207 | 3160281 | 3160392 | 131 | 94 | 94 | 119 | 210 | 0 | 0 | 4 | 0 | 0 | 231 | 145 | 112 | 126 | 405 | 0 | 0 | 0 | 0 | 0 | 277 | 149 | 1 | 1 | 1 | 1 | 0.927 | 1 | 1 | 1 | 1 | 1 | 1 | 1 | -0.015 |
| SE | 38013 | ENSG00000197386 | HTT | chr4 | + | 3157071 | 3157199 | 3154292 | 3154419 | 3160281 | 3160392 | 228 | 168 | 192 | 230 | 335 | 0 | 0 | 0 | 0 | 0 | 403 | 262 | 204 | 240 | 621 | 9 | 0 | 0 | 0 | 0 | 277 | 149 | 1 | 1 | 1 | 1 | 1 | 1 | 1 | 0.96 | 1 | 1 | 1 | 1 | 0.008 |
| SE | 38014 | ENSG00000197386 | HTT | chr4 | + | 3160281 | 3160392 | 3157071 | 3157199 | 3172319 | 3172397 | 306 | 256 | 261 | 295 | 637 | 0 | 0 | 0 | 0 | 0 | 568 | 408 | 324 | 340 | 1091 | 0 | 5 | 0 | 0 | 0 | 260 | 149 | 1 | 1 | 1 | 1 | 1 | 1 | 1 | 1 | 0.979 | 1 | 1 | 1 | 0.004 |
| SE | 38015 | ENSG00000197386 | HTT | chr4 | + | 3174945 | 3175107 | 3174720 | 3174799 | 3177331 | 3177387 | 237 | 200 | 217 | 255 | 201 | 0 | 0 | 0 | 0 | 0 | 287 | 245 | 187 | 278 | 502 | 0 | 0 | 0 | 5 | 0 | 298 | 149 | 1 | 1 | 1 | 1 | 1 | 1 | 1 | 1 | 1 | 1 | 0.965 | 1 | 0.007 |
| SE | 38016 | ENSG00000197386 | HTT | chr4 | + | 3182353 | 3182470 | 3180514 | 3180651 | 3186596 | 3186719 | 135 | 74 | 101 | 119 | 219 | 0 | 0 | 1 | 0 | 0 | 188 | 136 | 94 | 130 | 242 | 0 | 1 | 0 | 0 | 0 | 266 | 149 | 1 | 1 | 1 | 1 | 0.983 | 1 | 1 | 1 | 0.987 | 1 | 1 | 1 | -0.001 |
| SE | 38019 | ENSG00000197386 | HTT | chr4 | + | 3207280 | 3207357 | 3206806 | 3206983 | 3208772 | 3208911 | 126 | 59 | 89 | 137 | 130 | 1 | 0 | 0 | 0 | 0 | 155 | 139 | 90 | 174 | 339 | 0 | 0 | 0 | 5 | 0 | 226 | 149 | 1 | 1 | 0.988 | 1 | 1 | 1 | 1 | 1 | 1 | 1 | 0.958 | 1 | 0.006 |
| SE | 38020 | ENSG00000197386 | HTT | chr4 | + | 3217764 | 3217952 | 3215109 | 3215211 | 3220181 | 3220308 | 137 | 58 | 96 | 142 | 162 | 0 | 0 | 0 | 0 | 0 | 242 | 164 | 97 | 96 | 289 | 0 | 0 | 2 | 5 | 0 | 298 | 149 | 1 | 1 | 1 | 1 | 1 | 1 | 1 | 1 | 1 | 0.96 | 0.906 | 1 | 0.027 |
| SE | 38023 | ENSG00000197386 | HTT | chr4 | + | 3236148 | 3236254 | 3235564 | 3235778 | 3238446 | 3238609 | 177 | 104 | 139 | 201 | 168 | 0 | 0 | 0 | 0 | 0 | 144 | 250 | 91 | 109 | 501 | 5 | 0 | 2 | 0 | 0 | 255 | 149 | 1 | 1 | 1 | 1 | 1 | 1 | 1 | 0.944 | 1 | 0.964 | 1 | 1 | 0.018 |
| SE | 38028 | ENSG00000104432 | IL7 | chr8 | - | 78721019 | 78721135 | 78719008 | 78719110 | 78721347 | 78721440 | 0 | 3 | 1 | 0 | 0 | 1 | 0 | 0 | 0 | 0 | 6 | 0 | 0 | 0 | 4 | 0 | 0 | 0 | 0 | 0 | 265 | 149 | 1 | 1 | 0 | 1 | 1 | NA | NA | 1 | NA | NA | NA | 1 | -0.333 |
| SE | 38029 | ENSG00000104432 | IL7 | chr8 | - | 78736473 | 78736527 | 78732771 | 78733832 | 78738503 | 78738570 | 25 | 17 | 6 | 7 | 6 | 2 | 0 | 0 | 1 | 0 | 0 | 11 | 0 | 10 | 0 | 0 | 0 | 0 | 0 | 0 | 203 | 149 | 1 | 1 | 0.902 | 1 | 1 | 0.837 | 1 | NA | 1 | NA | 1 | NA | -0.052 |
| SE | 38030 | ENSG00000104432 | IL7 | chr8 | - | 78736473 | 78736527 | 78732771 | 78733832 | 78740001 | 78740082 | 20 | 10 | 4 | 4 | 8 | 0 | 1 | 0 | 0 | 0 | 0 | 6 | 0 | 8 | 0 | 0 | 0 | 0 | 0 | 0 | 203 | 149 | 1 | 1 | 1 | 0.88 | 1 | 1 | 1 | NA | 1 | NA | 1 | NA | -0.024 |
| SE | 38031 | ENSG00000104432 | IL7 | chr8 | - | 78738503 | 78738635 | 78732773 | 78733832 | 78740001 | 78740082 | 9 | 5 | 8 | 4 | 8 | 0 | 1 | 0 | 0 | 0 | 0 | 8 | 0 | 14 | 11 | 0 | 0 | 0 | 0 | 0 | 281 | 149 | 1 | 1 | 1 | 0.726 | 1 | 1 | 1 | NA | 1 | NA | 1 | 1 | -0.055 |
| SE | 38032 | ENSG00000104432 | IL7 | chr8 | - | 78738503 | 78738635 | 78736473 | 78736527 | 78740001 | 78740082 | 19 | 13 | 10 | 6 | 9 | 5 | 0 | 0 | 0 | 3 | 0 | 13 | 0 | 21 | 11 | 0 | 0 | 0 | 0 | 0 | 281 | 149 | 1 | 1 | 0.668 | 1 | 1 | 1 | 0.614 | NA | 1 | NA | 1 | 1 | -0.144 |
| SE | 38034 | ENSG00000104432 | IL7 | chr8 | - | 78746893 | 78747096 | 78740001 | 78740082 | 78798071 | 78798208 | 2 | 0 | 0 | 1 | 0 | 5 | 4 | 5 | 3 | 7 | 0 | 0 | 0 | 0 | 0 | 0 | 3 | 0 | 0 | 0 | 298 | 149 | 1 | 1 | 0.167 | 0 | 0 | 0.143 | 0 | NA | 0 | NA | NA | NA | 0.062 |
| SE | 38038 | ENSG00000215908 | CROCCP2 | chr1 | - | 16621945 | 16622115 | 16619844 | 16620024 | 16623869 | 16623997 | 0 | 0 | 2 | 1 | 0 | 2 | 1 | 0 | 1 | 0 | 0 | 7 | 0 | 0 | 0 | 0 | 0 | 0 | 0 | 0 | 298 | 149 | 1 | 1 | 0 | 0 | 1 | 0.333 | NA | NA | 1 | NA | NA | NA | -0.667 |
| SE | 38040 | ENSG00000215908 | CROCCP2 | chr1 | - | 16621945 | 16622115 | 16619844 | 16620245 | 16625774 | 16626005 | 0 | 0 | 0 | 6 | 0 | 0 | 0 | 0 | 1 | 0 | 1 | 7 | 0 | 0 | 0 | 0 | 0 | 0 | 0 | 0 | 298 | 149 | 1 | 1 | NA | NA | NA | 0.75 | NA | 1 | 1 | NA | NA | NA | -0.25 |
| SE | 38042 | ENSG00000215908 | CROCCP2 | chr1 | - | 16623869 | 16623997 | 16621945 | 16622115 | 16625774 | 16626005 | 0 | 1 | 2 | 3 | 0 | 0 | 0 | 0 | 3 | 0 | 0 | 0 | 0 | 0 | 0 | 1 | 7 | 0 | 0 | 0 | 277 | 149 | 1 | 1 | NA | 1 | 1 | 0.35 | NA | 0 | 0 | NA | NA | NA | 0.783 |
| SE | 38044 | ENSG00000215908 | CROCCP2 | chr1 | - | 16626352 | 16626499 | 16625374 | 16626005 | 16627144 | 16627289 | 75 | 13 | 74 | 107 | 21 | 0 | 0 | 2 | 2 | 0 | 54 | 71 | 84 | 20 | 127 | 0 | 0 | 15 | 3 | 0 | 296 | 149 | 1 | 1 | 1 | 1 | 0.949 | 0.964 | 1 | 1 | 1 | 0.738 | 0.77 | 1 | 0.081 |
| SE | 38047 | ENSG00000215908 | CROCCP2 | chr1 | - | 16633094 | 16633346 | 16632274 | 16632371 | 16642766 | 16642850 | 16 | 11 | 31 | 34 | 12 | 0 | 0 | 0 | 1 | 0 | 52 | 30 | 32 | 25 | 68 | 0 | 0 | 0 | 0 | 0 | 298 | 149 | 1 | 1 | 1 | 1 | 1 | 0.944 | 1 | 1 | 1 | 1 | 1 | 1 | -0.011 |
| SE | 38048 | ENSG00000215908 | CROCCP2 | chr1 | - | 16635026 | 16635168 | 16632274 | 16632371 | 16642766 | 16642850 | 15 | 9 | 8 | 28 | 0 | 0 | 0 | 0 | 1 | 0 | 7 | 3 | 14 | 14 | 0 | 0 | 0 | 0 | 0 | 0 | 291 | 149 | 1 | 1 | 1 | 1 | 1 | 0.935 | NA | 1 | 1 | 1 | 1 | NA | -0.016 |
| SE | 38049 | ENSG00000215908 | CROCCP2 | chr1 | - | 16635026 | 16635168 | 16633094 | 16633346 | 16642766 | 16642850 | 35 | 14 | 49 | 86 | 16 | 1 | 1 | 0 | 0 | 0 | 75 | 30 | 47 | 33 | 42 | 0 | 0 | 0 | 0 | 0 | 291 | 149 | 1 | 1 | 0.947 | 0.878 | 1 | 1 | 1 | 1 | 1 | 1 | 1 | 1 | -0.035 |
| SE | 38050 | ENSG00000215908 | CROCCP2 | chr1 | - | 16635026 | 16635168 | 16633094 | 16633346 | 16643027 | 16643145 | 24 | 6 | 46 | 59 | 16 | 0 | 0 | 0 | 1 | 0 | 57 | 31 | 35 | 19 | 41 | 0 | 4 | 0 | 0 | 0 | 291 | 149 | 1 | 1 | 1 | 1 | 1 | 0.968 | 1 | 1 | 0.799 | 1 | 1 | 1 | 0.034 |
| SE | 38051 | ENSG00000215908 | CROCCP2 | chr1 | - | 16635026 | 16635168 | 16633094 | 16633346 | 16644645 | 16644686 | 23 | 6 | 50 | 58 | 17 | 0 | 0 | 0 | 0 | 0 | 57 | 27 | 33 | 19 | 41 | 0 | 2 | 0 | 0 | 0 | 291 | 149 | 1 | 1 | 1 | 1 | 1 | 1 | 1 | 1 | 0.874 | 1 | 1 | 1 | 0.025 |
| SE | 38054 | ENSG00000215908 | CROCCP2 | chr1 | - | 16643027 | 16643145 | 16633094 | 16633346 | 16644645 | 16644672 | 9 | 2 | 15 | 14 | 7 | 0 | 0 | 0 | 0 | 0 | 8 | 10 | 10 | 2 | 1 | 0 | 2 | 0 | 0 | 0 | 267 | 149 | 1 | 1 | 1 | 1 | 1 | 1 | 1 | 1 | 0.736 | 1 | 1 | 1 | 0.053 |
| SE | 38056 | ENSG00000188389 | PDCD1 | chr2 | - | 241852197 | 241852353 | 241851948 | 241851983 | 241852620 | 241852980 | 25 | 7 | 8 | 15 | 0 | 3 | 0 | 3 | 0 | 0 | 19 | 39 | 9 | 7 | 13 | 0 | 1 | 1 | 0 | 0 | 298 | 149 | 1 | 1 | 0.806 | 1 | 0.571 | 1 | NA | 1 | 0.951 | 0.818 | 1 | 1 | -0.11 |
| SE | 38059 | ENSG00000270249 | NA | chr7 | - | 102566996 | 102567083 | 102541500 | 102541662 | 102569834 | 102569924 | 40 | 58 | 66 | 110 | 124 | 0 | 1 | 0 | 0 | 4 | 259 | 119 | 137 | 131 | 69 | 0 | 0 | 0 | 0 | 0 | 236 | 149 | 1 | 1 | 1 | 0.973 | 1 | 1 | 0.951 | 1 | 1 | 1 | 1 | 1 | -0.015 |
| SE | 38070 | ENSG00000105856 | HBP1 | chr7 | + | 107173456 | 107173591 | 107168999 | 107169027 | 107174447 | 107174527 | 1 | 0 | 0 | 3 | 0 | 0 | 0 | 1 | 0 | 6 | 8 | 5 | 6 | 0 | 0 | 0 | 0 | 0 | 0 | 0 | 284 | 149 | 1 | 1 | 1 | NA | NA | 0.611 | NA | 1 | 1 | 1 | 1 | NA | -0.195 |
| SE | 38073 | ENSG00000105856 | HBP1 | chr7 | + | 107174447 | 107174745 | 107169002 | 107169185 | 107179878 | 107180062 | 4 | 2 | 8 | 5 | 10 | 5 | 8 | 19 | 8 | 15 | 0 | 14 | 8 | 8 | 0 | 0 | 1 | 11 | 1 | 0 | 298 | 149 | 1 | 1 | 0.286 | 0.111 | 0.174 | 0.238 | 0.25 | NA | 0.875 | 0.267 | 0.8 | NA | -0.436 |
| SE | 38075 | ENSG00000105856 | HBP1 | chr7 | + | 107179816 | 107180062 | 107168999 | 107169027 | 107182372 | 107182576 | 427 | 245 | 413 | 384 | 818 | 0 | 0 | 1 | 0 | 0 | 104 | 363 | 510 | 157 | 81 | 0 | 0 | 0 | 0 | 0 | 298 | 149 | 1 | 1 | 1 | 1 | 0.995 | 1 | 1 | 1 | 1 | 1 | 1 | 1 | -0.001 |
| SE | 38076 | ENSG00000105856 | HBP1 | chr7 | + | 107179878 | 107180062 | 107168960 | 107169027 | 107182372 | 107182415 | 486 | 276 | 458 | 431 | 898 | 0 | 0 | 1 | 0 | 0 | 115 | 403 | 577 | 157 | 91 | 0 | 0 | 0 | 0 | 0 | 298 | 149 | 1 | 1 | 1 | 1 | 0.996 | 1 | 1 | 1 | 1 | 1 | 1 | 1 | -0.001 |
| SE | 38077 | ENSG00000105856 | HBP1 | chr7 | + | 107182372 | 107182601 | 107179877 | 107180062 | 107185800 | 107185942 | 939 | 527 | 847 | 812 | 1723 | 0 | 0 | 0 | 0 | 0 | 272 | 719 | 1117 | 378 | 177 | 6 | 0 | 0 | 0 | 0 | 298 | 149 | 1 | 1 | 1 | 1 | 1 | 1 | 1 | 0.958 | 1 | 1 | 1 | 1 | 0.008 |
| SE | 38079 | ENSG00000105856 | HBP1 | chr7 | + | 107190172 | 107190317 | 107186568 | 107186681 | 107195833 | 107196151 | 162 | 88 | 143 | 172 | 219 | 0 | 0 | 0 | 0 | 0 | 102 | 136 | 249 | 133 | 13 | 0 | 3 | 2 | 0 | 0 | 294 | 149 | 1 | 1 | 1 | 1 | 1 | 1 | 1 | 1 | 0.958 | 0.984 | 1 | 1 | 0.012 |
| SE | 38080 | ENSG00000105856 | HBP1 | chr7 | + | 107190172 | 107190317 | 107186568 | 107186681 | 107200159 | 107200301 | 21 | 10 | 13 | 23 | 53 | 0 | 1 | 0 | 0 | 0 | 44 | 56 | 68 | 31 | 6 | 0 | 0 | 5 | 0 | 0 | 294 | 149 | 1 | 1 | 1 | 0.835 | 1 | 1 | 1 | 1 | 1 | 0.873 | 1 | 1 | -0.008 |
| SE | 38081 | ENSG00000105856 | HBP1 | chr7 | + | 107195833 | 107196151 | 107186568 | 107186681 | 107200159 | 107200300 | 215 | 130 | 170 | 172 | 247 | 0 | 1 | 0 | 0 | 0 | 9 | 132 | 268 | 79 | 14 | 0 | 0 | 5 | 0 | 0 | 298 | 149 | 1 | 1 | 1 | 0.985 | 1 | 1 | 1 | 1 | 1 | 0.964 | 1 | 1 | 0.004 |
| SE | 38082 | ENSG00000105856 | HBP1 | chr7 | + | 107195833 | 107196151 | 107190172 | 107190317 | 107200159 | 107200300 | 359 | 208 | 302 | 322 | 418 | 2 | 0 | 1 | 1 | 5 | 67 | 214 | 449 | 181 | 21 | 0 | 5 | 1 | 0 | 0 | 298 | 149 | 1 | 1 | 0.989 | 1 | 0.993 | 0.994 | 0.977 | 1 | 0.955 | 0.996 | 1 | 1 | 0 |
| SE | 38084 | ENSG00000160584 | SIK3 | chr11 | - | 116861273 | 116861383 | 116859264 | 116859604 | 116861840 | 116861926 | 33 | 17 | 28 | 54 | 53 | 0 | 0 | 0 | 0 | 0 | 151 | 45 | 47 | 52 | 56 | 0 | 4 | 0 | 0 | 0 | 259 | 149 | 1 | 1 | 1 | 1 | 1 | 1 | 1 | 1 | 0.866 | 1 | 1 | 1 | 0.027 |
| SE | 38088 | ENSG00000160584 | SIK3 | chr11 | - | 116875373 | 116875451 | 116875157 | 116875267 | 116875865 | 116876009 | 142 | 66 | 106 | 85 | 149 | 0 | 0 | 1 | 0 | 0 | 151 | 120 | 84 | 107 | 154 | 0 | 0 | 0 | 0 | 0 | 227 | 149 | 1 | 1 | 1 | 1 | 0.986 | 1 | 1 | 1 | 1 | 1 | 1 | 1 | -0.003 |
| SE | 38093 | ENSG00000160584 | SIK3 | chr11 | - | 116927218 | 116927380 | 116897192 | 116897317 | 116954043 | 116954107 | 198 | 126 | 238 | 234 | 301 | 1 | 0 | 1 | 0 | 0 | 192 | 225 | 196 | 258 | 187 | 0 | 3 | 0 | 0 | 0 | 298 | 149 | 1 | 1 | 0.99 | 1 | 0.992 | 1 | 1 | 1 | 0.974 | 1 | 1 | 1 | 0.002 |
| SE | 38094 | ENSG00000160584 | SIK3 | chr11 | - | 116950132 | 116950256 | 116927218 | 116927380 | 116954043 | 116954107 | 0 | 0 | 0 | 0 | 0 | 104 | 72 | 116 | 127 | 125 | 3 | 2 | 0 | 0 | 3 | 141 | 120 | 123 | 162 | 104 | 273 | 149 | 1 | 1 | 0 | 0 | 0 | 0 | 0 | 0.011 | 0.009 | 0 | 0 | 0.015 | -0.007 |
| SE | 38097 | ENSG00000100258 | LMF2 | chr22 | - | 50505228 | 50505334 | 50504801 | 50504984 | 50505402 | 50505537 | 12 | 5 | 13 | 12 | 5 | 1 | 0 | 0 | 0 | 0 | 5 | 12 | 2 | 8 | 20 | 0 | 0 | 0 | 0 | 0 | 255 | 149 | 1 | 1 | 0.875 | 1 | 1 | 1 | 1 | 1 | 1 | 1 | 1 | 1 | -0.025 |
| SE | 38098 | ENSG00000100258 | LMF2 | chr22 | - | 50506637 | 50506666 | 50506284 | 50506502 | 50506781 | 50507035 | 15 | 5 | 26 | 14 | 9 | 2 | 0 | 0 | 0 | 0 | 11 | 17 | 7 | 30 | 35 | 0 | 0 | 0 | 0 | 0 | 178 | 149 | 1 | 1 | 0.863 | 1 | 1 | 1 | 1 | 1 | 1 | 1 | 1 | 1 | -0.027 |
| SE | 38101 | ENSG00000143811 | PYCR2 | chr1 | - | 225922203 | 225922383 | 225921857 | 225922079 | 225923700 | 225923771 | 104 | 48 | 91 | 104 | 28 | 0 | 0 | 0 | 0 | 0 | 64 | 50 | 39 | 58 | 554 | 0 | 1 | 0 | 0 | 0 | 298 | 149 | 1 | 1 | 1 | 1 | 1 | 1 | 1 | 1 | 0.962 | 1 | 1 | 1 | 0.008 |
| SE | 38102 | ENSG00000143811 | PYCR2 | chr1 | - | 225923700 | 225923771 | 225922203 | 225922383 | 225924043 | 225924250 | 42 | 30 | 56 | 56 | 23 | 1 | 0 | 3 | 1 | 0 | 8 | 16 | 17 | 18 | 348 | 0 | 0 | 0 | 0 | 0 | 220 | 149 | 1 | 1 | 0.966 | 1 | 0.927 | 0.974 | 1 | 1 | 1 | 1 | 1 | 1 | -0.027 |
| SE | 38104 | ENSG00000110871 | COQ5 | chr12 | - | 120510016 | 120510123 | 120503968 | 120504081 | 120516566 | 120516788 | 61 | 26 | 44 | 37 | 137 | 0 | 0 | 0 | 2 | 0 | 32 | 45 | 10 | 48 | 19 | 0 | 0 | 0 | 0 | 0 | 256 | 149 | 1 | 1 | 1 | 1 | 1 | 0.915 | 1 | 1 | 1 | 1 | 1 | 1 | -0.017 |
| SE | 38105 | ENSG00000110871 | COQ5 | chr12 | - | 120516566 | 120516788 | 120510016 | 120510123 | 120522213 | 120522363 | 109 | 76 | 116 | 102 | 283 | 1 | 0 | 0 | 2 | 0 | 130 | 116 | 28 | 105 | 86 | 0 | 0 | 0 | 0 | 0 | 298 | 149 | 1 | 1 | 0.982 | 1 | 1 | 0.962 | 1 | 1 | 1 | 1 | 1 | 1 | -0.011 |
| SE | 38109 | ENSG00000161904 | LEMD2 | chr6 | - | 33781076 | 33781153 | 33780001 | 33780179 | 33784351 | 33784427 | 183 | 95 | 129 | 125 | 165 | 1 | 0 | 0 | 0 | 0 | 322 | 201 | 226 | 188 | 371 | 0 | 0 | 0 | 0 | 0 | 226 | 149 | 1 | 1 | 0.992 | 1 | 1 | 1 | 1 | 1 | 1 | 1 | 1 | 1 | -0.002 |
| SE | 38119 | ENSG00000132394 | EEFSEC | chr3 | + | 128358216 | 128358373 | 128341232 | 128341889 | 128408068 | 128408646 | 63 | 32 | 58 | 43 | 39 | 0 | 0 | 1 | 0 | 0 | 40 | 36 | 53 | 38 | 68 | 0 | 0 | 0 | 3 | 0 | 298 | 149 | 1 | 1 | 1 | 1 | 0.967 | 1 | 1 | 1 | 1 | 1 | 0.864 | 1 | 0.021 |
| SE | 38124 | ENSG00000064961 | HMG20B | chr19 | + | 3575501 | 3575660 | 3574369 | 3574586 | 3576260 | 3576307 | 207 | 194 | 176 | 230 | 214 | 0 | 0 | 0 | 1 | 0 | 411 | 252 | 134 | 194 | 1067 | 0 | 0 | 1 | 0 | 0 | 298 | 149 | 1 | 1 | 1 | 1 | 1 | 0.991 | 1 | 1 | 1 | 0.985 | 1 | 1 | 0.001 |
| SE | 38125 | ENSG00000064961 | HMG20B | chr19 | + | 3575501 | 3575660 | 3574369 | 3574586 | 3576552 | 3576625 | 1 | 0 | 0 | 1 | 0 | 0 | 0 | 0 | 0 | 0 | 5 | 0 | 0 | 1 | 0 | 0 | 4 | 0 | 0 | 0 | 298 | 149 | 1 | 1 | 1 | NA | NA | 1 | NA | 1 | 0 | NA | 1 | NA | 0.333 |
| SE | 38126 | ENSG00000064961 | HMG20B | chr19 | + | 3575539 | 3575660 | 3573691 | 3574586 | 3576260 | 3576307 | 249 | 225 | 211 | 284 | 226 | 0 | 0 | 0 | 1 | 0 | 490 | 302 | 169 | 257 | 1236 | 0 | 1 | 0 | 0 | 0 | 270 | 149 | 1 | 1 | 1 | 1 | 1 | 0.994 | 1 | 1 | 1 | 0.989 | 1 | 1 | 0.001 |
| SE | 38127 | ENSG00000064961 | HMG20B | chr19 | + | 3575539 | 3575660 | 3573691 | 3574586 | 3576552 | 3576602 | 29 | 26 | 31 | 42 | 9 | 0 | 0 | 0 | 0 | 0 | 55 | 46 | 34 | 44 | 108 | 0 | 4 | 0 | 0 | 0 | 270 | 149 | 1 | 1 | 1 | 1 | 1 | 1 | 1 | 1 | 0.864 | 1 | 1 | 1 | 0.027 |
| SE | 38128 | ENSG00000064961 | HMG20B | chr19 | + | 3576260 | 3576307 | 3573691 | 3574586 | 3576552 | 3576602 | 47 | 34 | 21 | 51 | 39 | 0 | 0 | 0 | 0 | 0 | 95 | 41 | 28 | 51 | 188 | 0 | 4 | 0 | 0 | 0 | 196 | 149 | 1 | 1 | 1 | 1 | 1 | 1 | 1 | 1 | 0.886 | 1 | 1 | 1 | 0.023 |
| SE | 38129 | ENSG00000064961 | HMG20B | chr19 | + | 3576260 | 3576307 | 3575501 | 3575660 | 3576552 | 3576602 | 268 | 232 | 200 | 295 | 246 | 1 | 0 | 0 | 3 | 0 | 522 | 298 | 160 | 261 | 1313 | 0 | 0 | 0 | 0 | 0 | 196 | 149 | 1 | 1 | 0.995 | 1 | 1 | 0.987 | 1 | 1 | 1 | 1 | 1 | 1 | -0.004 |
| SE | 38130 | ENSG00000064961 | HMG20B | chr19 | + | 3576891 | 3577107 | 3576351 | 3576625 | 3577980 | 3578113 | 370 | 298 | 262 | 397 | 375 | 1 | 0 | 0 | 0 | 0 | 788 | 421 | 219 | 383 | 1709 | 0 | 0 | 0 | 0 | 0 | 298 | 149 | 1 | 1 | 0.995 | 1 | 1 | 1 | 1 | 1 | 1 | 1 | 1 | 1 | -0.001 |
| SE | 38132 | ENSG00000113048 | MRPS27 | chr5 | - | 72223682 | 72223850 | 72219402 | 72221148 | 72226056 | 72226199 | 277 | 194 | 241 | 240 | 477 | 1 | 1 | 0 | 0 | 2 | 239 | 181 | 177 | 173 | 332 | 0 | 0 | 4 | 0 | 0 | 298 | 149 | 1 | 1 | 0.993 | 0.99 | 1 | 1 | 0.992 | 1 | 1 | 0.957 | 1 | 1 | 0.004 |
| SE | 38134 | ENSG00000113048 | MRPS27 | chr5 | - | 72228265 | 72228368 | 72226056 | 72226199 | 72232442 | 72232558 | 204 | 147 | 186 | 202 | 444 | 0 | 0 | 2 | 0 | 0 | 339 | 237 | 142 | 199 | 360 | 0 | 0 | 3 | 0 | 0 | 252 | 149 | 1 | 1 | 1 | 1 | 0.982 | 1 | 1 | 1 | 1 | 0.966 | 1 | 1 | 0.003 |
| SE | 38135 | ENSG00000113048 | MRPS27 | chr5 | - | 72241461 | 72241688 | 72238074 | 72238128 | 72295530 | 72295589 | 0 | 0 | 1 | 1 | 0 | 108 | 42 | 62 | 85 | 153 | 0 | 0 | 3 | 0 | 0 | 70 | 59 | 49 | 49 | 89 | 298 | 149 | 1 | 1 | 0 | 0 | 0.008 | 0.006 | 0 | 0 | 0 | 0.03 | 0 | 0 | -0.003 |
| SE | 38136 | ENSG00000113048 | MRPS27 | chr5 | - | 72241646 | 72241688 | 72238013 | 72238128 | 72295530 | 72295589 | 0 | 1 | 0 | 1 | 1 | 108 | 42 | 62 | 85 | 153 | 0 | 5 | 1 | 0 | 0 | 70 | 59 | 49 | 49 | 89 | 191 | 149 | 1 | 1 | 0 | 0.018 | 0 | 0.009 | 0.005 | 0 | 0.062 | 0.016 | 0 | 0 | -0.009 |
| SE | 38143 | ENSG00000198087 | CD2AP | chr6 | + | 47544605 | 47544706 | 47533601 | 47533755 | 47554645 | 47554766 | 148 | 160 | 145 | 197 | 183 | 0 | 0 | 0 | 0 | 0 | 29 | 158 | 117 | 117 | 48 | 0 | 0 | 1 | 0 | 0 | 250 | 149 | 1 | 1 | 1 | 1 | 1 | 1 | 1 | 1 | 1 | 0.986 | 1 | 1 | 0.003 |
| SE | 38144 | ENSG00000198087 | CD2AP | chr6 | + | 47554645 | 47554766 | 47544605 | 47544706 | 47574063 | 47574251 | 229 | 255 | 241 | 338 | 397 | 0 | 0 | 1 | 0 | 0 | 67 | 254 | 194 | 127 | 93 | 0 | 0 | 0 | 0 | 0 | 270 | 149 | 1 | 1 | 1 | 1 | 0.993 | 1 | 1 | 1 | 1 | 1 | 1 | 1 | -0.001 |
| SE | 38145 | ENSG00000198087 | CD2AP | chr6 | + | 47580863 | 47580900 | 47579384 | 47579489 | 47582002 | 47582065 | 223 | 204 | 211 | 258 | 779 | 0 | 0 | 0 | 2 | 0 | 47 | 108 | 124 | 60 | 15 | 0 | 0 | 0 | 0 | 0 | 186 | 149 | 1 | 1 | 1 | 1 | 1 | 0.99 | 1 | 1 | 1 | 1 | 1 | 1 | -0.002 |
| SE | 38147 | ENSG00000138750 | NUP54 | chr4 | - | 76124648 | 76124756 | 76117663 | 76118194 | 76130655 | 76130701 | 132 | 86 | 142 | 120 | 239 | 0 | 0 | 1 | 2 | 0 | 47 | 97 | 70 | 23 | 22 | 0 | 0 | 0 | 0 | 0 | 257 | 149 | 1 | 1 | 1 | 1 | 0.988 | 0.972 | 1 | 1 | 1 | 1 | 1 | 1 | -0.008 |
| SE | 38148 | ENSG00000138750 | NUP54 | chr4 | - | 76131229 | 76131284 | 76130655 | 76130749 | 76132522 | 76132719 | 108 | 79 | 154 | 112 | 356 | 0 | 0 | 1 | 0 | 0 | 26 | 48 | 73 | 35 | 9 | 0 | 0 | 0 | 0 | 0 | 204 | 149 | 1 | 1 | 1 | 1 | 0.991 | 1 | 1 | 1 | 1 | 1 | 1 | 1 | -0.002 |
| SE | 38149 | ENSG00000138750 | NUP54 | chr4 | - | 76132522 | 76132719 | 76130655 | 76130749 | 76136185 | 76136412 | 1 | 0 | 1 | 0 | 0 | 0 | 0 | 0 | 0 | 0 | 0 | 2 | 0 | 0 | 0 | 0 | 0 | 0 | 4 | 0 | 298 | 149 | 1 | 1 | 1 | NA | 1 | NA | NA | NA | 1 | NA | 0 | NA | 0.5 |
| SE | 38150 | ENSG00000138750 | NUP54 | chr4 | - | 76132522 | 76132719 | 76131229 | 76131284 | 76134174 | 76134362 | 210 | 148 | 228 | 179 | 744 | 1 | 0 | 0 | 0 | 0 | 43 | 83 | 104 | 37 | 9 | 0 | 0 | 0 | 0 | 0 | 298 | 149 | 1 | 1 | 0.991 | 1 | 1 | 1 | 1 | 1 | 1 | 1 | 1 | 1 | -0.002 |
| SE | 38151 | ENSG00000138750 | NUP54 | chr4 | - | 76134174 | 76134362 | 76132522 | 76132719 | 76136185 | 76136412 | 281 | 201 | 218 | 238 | 687 | 1 | 0 | 0 | 0 | 0 | 221 | 137 | 162 | 111 | 147 | 0 | 2 | 0 | 0 | 0 | 298 | 149 | 1 | 1 | 0.993 | 1 | 1 | 1 | 1 | 1 | 0.972 | 1 | 1 | 1 | 0.004 |
| SE | 38152 | ENSG00000138750 | NUP54 | chr4 | - | 76134174 | 76134362 | 76132522 | 76132719 | 76144148 | 76144292 | 120 | 85 | 106 | 80 | 431 | 0 | 0 | 0 | 0 | 0 | 22 | 39 | 49 | 11 | 0 | 0 | 0 | 2 | 0 | 0 | 298 | 149 | 1 | 1 | 1 | 1 | 1 | 1 | 1 | 1 | 1 | 1 | 0.733 | NA | 0.067 |
| SE | 38153 | ENSG00000138750 | NUP54 | chr4 | - | 76136185 | 76136412 | 76132522 | 76132719 | 76144148 | 76144292 | 15 | 4 | 11 | 11 | 9 | 0 | 0 | 0 | 0 | 0 | 25 | 15 | 5 | 7 | 12 | 0 | 0 | 2 | 0 | 0 | 298 | 149 | 1 | 1 | 1 | 1 | 1 | 1 | 1 | 1 | 1 | 1 | 0.636 | 1 | 0.073 |
| SE | 38164 | ENSG00000137074 | APTX | chr9 | - | 32966675 | 32966773 | 32924896 | 32925028 | 32985970 | 32986030 | 0 | 0 | 0 | 0 | 0 | 1 | 0 | 0 | 2 | 3 | 0 | 1 | 0 | 0 | 8 | 0 | 0 | 0 | 0 | 0 | 247 | 149 | 1 | 1 | 0 | NA | NA | 0 | 0 | NA | 1 | NA | NA | | -1 |
| SE | 38165 | ENSG00000137074 | APTX | chr9 | - | 32966675 | 32966773 | 32947549 | 32947618 | 32985970 | 32986030 | 1 | 0 | 0 | 4 | 0 | 0 | 0 | 0 | 4 | 0 | 0 | 3 | 3 | 0 | 0 | 0 | 0 | 0 | 0 | 0 | 247 | 149 | 1 | 1 | 1 | NA | 1 | NA | 0.311 | NA | 1 | NA | NA | | -0.23 |
| SE | 38166 | ENSG00000137074 | APTX | chr9 | - | 32973431 | 32973652 | 32966675 | 32966773 | 32974457 | 32974561 | 20 | 17 | 30 | 20 | 31 | 2 | 0 | 0 | 0 | 3 | 73 | 22 | 18 | 25 | 6 | 0 | 4 | 0 | 0 | 0 | 298 | 149 | 1 | 1 | 0.833 | 1 | 1 | 0.838 | 1 | 0.733 | 1 | 1 | 1 | | -0.012 |
| SE | 38167 | ENSG00000137074 | APTX | chr9 | - | 32974457 | 32974561 | 32966675 | 32966773 | 32985970 | 32986030 | 4 | 2 | 3 | 2 | 8 | 0 | 0 | 0 | 0 | 0 | 0 | 9 | 8 | 0 | 5 | 0 | 1 | 0 | 0 | 4 | 253 | 149 | 1 | 1 | 1 | 1 | 1 | 1 | 1 | NA | 0.841 | 1 | NA | 0.424 | 0.245 |
| SE | 38168 | ENSG00000137074 | APTX | chr9 | - | 32980065 | 32980460 | 32974457 | 32974561 | 32984630 | 32984857 | 0 | 2 | 0 | 0 | 0 | 14 | 9 | 11 | 19 | 29 | 0 | 0 | 0 | 0 | 0 | 35 | 11 | 9 | 17 | 3 | 298 | 149 | 1 | 1 | 0 | 0.1 | 0 | 0 | 0 | 0 | 0 | 0 | 0 | 0 | 0.02 |
| SE | 38170 | ENSG00000137074 | APTX | chr9 | - | 32984630 | 32984857 | 32947549 | 32947618 | 32985970 | 32986030 | 63 | 29 | 32 | 32 | 71 | 0 | 0 | 0 | 0 | 4 | 27 | 60 | 40 | 24 | 14 | 0 | 0 | 0 | 0 | 0 | 298 | 149 | 1 | 1 | 1 | 1 | 1 | 1 | 0.899 | 1 | 1 | 1 | 1 | 1 | -0.02 |
| SE | 38172 | ENSG00000137074 | APTX | chr9 | - | 32985970 | 32986030 | 32966675 | 32966773 | 32989758 | 32989895 | 0 | 0 | 0 | 1 | 0 | 0 | 0 | 0 | 0 | 0 | 0 | 1 | 0 | 0 | 4 | 0 | 0 | 1 | 0 | 0 | 209 | 149 | 1 | 1 | NA | NA | NA | 1 | NA | NA | 1 | 0 | NA | 1 | 0.333 |
| SE | 38174 | ENSG00000137074 | APTX | chr9 | - | 32987543 | 32987729 | 32985970 | 32986030 | 32988082 | 32988129 | 54 | 27 | 29 | 43 | 101 | 0 | 0 | 0 | 0 | 1 | 57 | 61 | 47 | 25 | 18 | 0 | 0 | 0 | 0 | 0 | 298 | 149 | 1 | 1 | 1 | 1 | 1 | 1 | 0.981 | 1 | 1 | 1 | 1 | 1 | -0.004 |
| SE | 38175 | ENSG00000137074 | APTX | chr9 | - | 32987543 | 32987731 | 32985970 | 32986030 | 32988082 | 32988124 | 68 | 31 | 43 | 46 | 114 | 0 | 0 | 0 | 0 | 1 | 65 | 67 | 50 | 47 | 27 | 0 | 0 | 0 | 0 | 0 | 298 | 149 | 1 | 1 | 1 | 1 | 1 | 1 | 0.983 | 1 | 1 | 1 | 1 | 1 | -0.003 |
| SE | 38176 | ENSG00000137074 | APTX | chr9 | - | 32987543 | 32987731 | 32985970 | 32986030 | 32989758 | 32989895 | 58 | 30 | 29 | 48 | 110 | 0 | 0 | 0 | 1 | 0 | 57 | 70 | 59 | 40 | 30 | 0 | 0 | 0 | 0 | 0 | 298 | 149 | 1 | 1 | 1 | 1 | 1 | 0.96 | 1 | 1 | 1 | 1 | 1 | 1 | -0.008 |
| SE | 38177 | ENSG00000137074 | APTX | chr9 | - | 32987543 | 32987842 | 32985970 | 32986030 | 32988082 | 32988540 | 54 | 27 | 29 | 43 | 102 | 0 | 0 | 0 | 0 | 1 | 57 | 64 | 47 | 25 | 18 | 0 | 0 | 0 | 0 | 0 | 298 | 149 | 1 | 1 | 1 | 1 | 1 | 1 | 0.981 | 1 | 1 | 1 | 1 | 1 | -0.004 |
| SE | 38178 | ENSG00000137074 | APTX | chr9 | - | 32987543 | 32987846 | 32985535 | 32986030 | 32988082 | 32988124 | 54 | 27 | 29 | 43 | 101 | 0 | 0 | 0 | 0 | 1 | 57 | 61 | 47 | 25 | 18 | 0 | 0 | 0 | 0 | 0 | 298 | 149 | 1 | 1 | 1 | 1 | 1 | 1 | 0.981 | 1 | 1 | 1 | 1 | 1 | -0.004 |
| SE | 38179 | ENSG00000137074 | APTX | chr9 | - | 32987543 | 32987846 | 32985535 | 32986030 | 32989758 | 32989895 | 56 | 27 | 29 | 43 | 101 | 0 | 0 | 0 | 1 | 0 | 57 | 61 | 51 | 32 | 18 | 0 | 0 | 0 | 0 | 0 | 298 | 149 | 1 | 1 | 1 | 1 | 1 | 0.956 | 1 | 1 | 1 | 1 | 1 | 1 | -0.009 |
| SE | 38180 | ENSG00000137074 | APTX | chr9 | - | 32987543 | 32988129 | 32985970 | 32986030 | 32989758 | 32989895 | 59 | 35 | 32 | 48 | 102 | 0 | 0 | 0 | 1 | 0 | 60 | 64 | 57 | 27 | 27 | 0 | 0 | 0 | 0 | 0 | 298 | 149 | 1 | 1 | 1 | 1 | 1 | 0.96 | 1 | 1 | 1 | 1 | 1 | 1 | -0.008 |
| SE | 38181 | ENSG00000137074 | APTX | chr9 | - | 32987759 | 32987846 | 32985970 | 32986030 | 32988082 | 32988129 | 1 | 0 | 0 | 0 | 0 | 0 | 0 | 0 | 0 | 1 | 0 | 8 | 0 | 0 | 0 | 0 | 0 | 0 | 0 | 0 | 236 | 149 | 1 | 1 | 1 | NA | NA | NA | 0 | NA | 1 | NA | NA | NA | -0.5 |
| SE | 38182 | ENSG00000137074 | APTX | chr9 | - | 32988082 | 32988124 | 32985970 | 32986030 | 32989758 | 32989895 | 6 | 4 | 2 | 1 | 6 | 0 | 0 | 0 | 1 | 0 | 2 | 6 | 6 | 10 | 2 | 0 | 0 | 0 | 0 | 0 | 191 | 149 | 1 | 1 | 1 | 1 | 1 | 0.438 | 1 | 1 | 1 | 1 | 1 | 1 | -0.112 |
| SE | 38185 | ENSG00000137074 | APTX | chr9 | - | 32988082 | 32988129 | 32985535 | 32986030 | 32989758 | 32989895 | 5 | 5 | 3 | 4 | 1 | 0 | 0 | 0 | 1 | 0 | 3 | 3 | 10 | 2 | 9 | 0 | 0 | 0 | 0 | 0 | 196 | 149 | 1 | 1 | 1 | 1 | 1 | 0.753 | 1 | 1 | 1 | 1 | 1 | 1 | -0.049 |
| SE | 38189 | ENSG00000137074 | APTX | chr9 | - | 32988091 | 32988124 | 32987543 | 32987731 | 32989758 | 32989895 | 5 | 4 | 0 | 2 | 5 | 4 | 3 | 0 | 5 | 9 | 2 | 7 | 6 | 10 | 2 | 0 | 9 | 12 | 15 | 12 | 182 | 149 | 1 | 1 | 0.506 | 0.522 | NA | 0.247 | 0.313 | 1 | 0.389 | 0.29 | 0.353 | 0.12 | -0.033 |
| SE | 38192 | ENSG00000137074 | APTX | chr9 | - | 32989758 | 32989895 | 32987543 | 32988129 | 33001566 | 33001579 | 29 | 19 | 25 | 35 | 14 | 0 | 1 | 0 | 0 | 0 | 64 | 25 | 37 | 40 | 39 | 0 | 0 | 0 | 0 | 0 | 286 | 149 | 1 | 1 | 1 | 0.908 | 1 | 1 | 1 | 1 | 1 | 1 | 1 | 1 | -0.018 |
| SE | 38195 | ENSG00000151287 | TEX30 | chr13 | - | 102767272 | 102767478 | 102765887 | 102766580 | 102768259 | 102768311 | 71 | 29 | 46 | 40 | 51 | 4 | 4 | 6 | 8 | 5 | 36 | 11 | 46 | 10 | 0 | 12 | 0 | 0 | 0 | 0 | 298 | 149 | 1 | 1 | 0.899 | 0.784 | 0.793 | 0.714 | 0.836 | 0.6 | 1 | 1 | 1 | NA | -0.095 |
| SE | 38196 | ENSG00000151287 | TEX30 | chr13 | - | 102769310 | 102769541 | 102768259 | 102768311 | 102770011 | 102770086 | 45 | 38 | 50 | 53 | 25 | 0 | 0 | 2 | 0 | 0 | 25 | 5 | 39 | 3 | 0 | 0 | 0 | 0 | 0 | 0 | 298 | 149 | 1 | 1 | 1 | 1 | 0.926 | 1 | 1 | 1 | 1 | 1 | 1 | NA | -0.015 |
| SE | 38197 | ENSG00000151287 | TEX30 | chr13 | - | 102769310 | 102769541 | 102768259 | 102768311 | 102773681 | 102773755 | 34 | 23 | 34 | 31 | 19 | 1 | 0 | 0 | 0 | 0 | 12 | 5 | 32 | 1 | 0 | 0 | 0 | 0 | 0 | 0 | 298 | 149 | 1 | 1 | 0.944 | 1 | 1 | 1 | 1 | 1 | 1 | 1 | 1 | NA | -0.011 |
| SE | 38199 | ENSG00000151287 | TEX30 | chr13 | - | 102770011 | 102770086 | 102769310 | 102769541 | 102773681 | 102773786 | 17 | 15 | 16 | 25 | 5 | 2 | 0 | 0 | 0 | 0 | 13 | 0 | 8 | 2 | 0 | 0 | 0 | 3 | 0 | 0 | 224 | 149 | 1 | 1 | 0.85 | 1 | 1 | 1 | 1 | 1 | NA | 0.639 | 1 | NA | 0.09 |
| SE | 38212 | ENSG00000121897 | LIAS | chr4 | + | 39465284 | 39465342 | 39463524 | 39463605 | 39467517 | 39467646 | 14 | 16 | 31 | 18 | 21 | 4 | 0 | 0 | 1 | 0 | 13 | 18 | 11 | 8 | 0 | 0 | 0 | 0 | 2 | 0 | 207 | 149 | 1 | 1 | 0.716 | 1 | 1 | 0.928 | 1 | 1 | 1 | 1 | 0.742 | NA | -0.007 |
| SE | 38220 | ENSG00000113108 | APBB3 | chr5 | - | 140560312 | 140560504 | 140558267 | 140558821 | 140561017 | 140561101 | 12 | 9 | 17 | 10 | 8 | 0 | 0 | 0 | 0 | 0 | 18 | 11 | 11 | 8 | 19 | 0 | 0 | 3 | 0 | 0 | 298 | 149 | 1 | 1 | 1 | 1 | 1 | 1 | 1 | 1 | 1 | 0.647 | 1 | 1 | 0.071 |
| SE | 38221 | ENSG00000113108 | APBB3 | chr5 | - | 140560638 | 140560754 | 140560312 | 140560504 | 140561017 | 140561101 | 22 | 14 | 34 | 17 | 21 | 0 | 0 | 1 | 0 | 0 | 28 | 25 | 8 | 39 | 48 | 0 | 0 | 0 | 0 | 0 | 265 | 149 | 1 | 1 | 1 | 1 | 0.95 | 1 | 1 | 1 | 1 | 1 | 1 | 1 | -0.01 |
| SE | 38225 | ENSG00000113108 | APBB3 | chr5 | - | 140561843 | 140561995 | 140561586 | 140561701 | 140562099 | 140562227 | 5 | 1 | 5 | 5 | 5 | 0 | 0 | 2 | 0 | 1 | 0 | 4 | 3 | 2 | 0 | 0 | 0 | 0 | 0 | 0 | 298 | 149 | 1 | 1 | 1 | 1 | 0.556 | 1 | 0.714 | NA | 1 | | 1 | NA | -0.146 |
| SE | 38230 | ENSG00000121753 | ADGRB2 | chr1 | - | 31736572 | 31736723 | 31736320 | 31736390 | 31737428 | 31737531 | 1 | 0 | 0 | 0 | 0 | 0 | 0 | 0 | 0 | 0 | 0 | 0 | 0 | 0 | 0 | 0 | 0 | 1 | 0 | 1 | 298 | 149 | 1 | 1 | 1 | NA | NA | NA | NA | NA | NA | 0 | NA | 0 | 1 |
| SE | 38231 | ENSG00000121753 | ADGRB2 | chr1 | - | 31741372 | 31741479 | 31740346 | 31740541 | 31741623 | 31741725 | 1 | 0 | 0 | 0 | 0 | 1 | 0 | 0 | 0 | 0 | 0 | 0 | 0 | 0 | 0 | 0 | 0 | 0 | 0 | 2 | 256 | 149 | 1 | 1 | 0.368 | NA | NA | NA | NA | NA | NA | NA | NA | 0 | 0.368 |
| SE | 38236 | ENSG00000183604 | SMG1P5 | chr16 | - | 30268401 | 30268561 | 30267552 | 30267672 | 30268935 | 30269083 | 14 | 6 | 12 | 13 | 5 | 0 | 1 | 0 | 0 | 0 | 1 | 18 | 9 | 17 | 0 | 2 | 5 | 0 | 2 | 0 | 298 | 149 | 1 | 1 | 1 | 0.75 | 1 | 1 | 1 | 0.2 | 0.643 | 1 | 0.81 | NA | 0.287 |
| SE | 38237 | ENSG00000183604 | SMG1P5 | chr16 | - | 30277596 | 30277793 | 30277263 | 30277428 | 30278180 | 30278380 | 34 | 29 | 29 | 35 | 32 | 0 | 0 | 0 | 0 | 0 | 44 | 68 | 30 | 32 | 19 | 0 | 0 | 0 | 1 | 0 | 298 | 149 | 1 | 1 | 1 | 1 | 1 | 1 | 1 | 1 | 1 | 1 | 0.941 | 1 | 0.012 |
| SE | 38239 | ENSG00000183604 | SMG1P5 | chr16 | - | 30294114 | 30294187 | 30293098 | 30293196 | 30294565 | 30294691 | 11 | 5 | 13 | 10 | 18 | 0 | 0 | 0 | 0 | 2 | 8 | 14 | 6 | 18 | 0 | 0 | 0 | 0 | 0 | 0 | 222 | 149 | 1 | 1 | 1 | 1 | 1 | 1 | 0.858 | 1 | 1 | 1 | 1 | NA | -0.028 |
| SE | 38241 | ENSG00000183604 | SMG1P5 | chr16 | - | 30305054 | 30305210 | 30301221 | 30301358 | 30305789 | 30305953 | 41 | 19 | 28 | 19 | 33 | 2 | 0 | 0 | 0 | 0 | 58 | 36 | 33 | 52 | 34 | 0 | 0 | 0 | 4 | 0 | 298 | 149 | 1 | 1 | 0.911 | 1 | 1 | 1 | 1 | 1 | 1 | 1 | 0.867 | 1 | 0.009 |
| SE | 38242 | ENSG00000178922 | HYI | chr1 | - | 43452902 | 43453004 | 43452204 | 43452319 | 43453412 | 43453497 | 1 | 0 | 0 | 9 | 0 | 0 | 0 | 1 | 1 | 6 | 0 | 0 | 2 | 5 | 0 | 0 | 0 | 0 | 0 | 0 | 251 | 149 | 1 | 1 | 1 | NA | 0 | 0.842 | 0 | NA | NA | 1 | 1 | NA | -0.539 |
| SE | 38260 | ENSG00000204745 | NA | chr2 | + | 87130102 | 87130194 | 87128546 | 87128815 | 87131968 | 87132117 | 2 | 0 | 0 | 0 | 0 | 0 | 0 | 0 | 0 | 0 | 0 | 0 | 4 | 0 | 0 | 0 | 0 | 8 | 5 | 0 | 241 | 149 | 1 | 1 | 1 | NA | NA | NA | NA | NA | NA | 0.236 | 0 | NA | 0.882 |
| SE | 38261 | ENSG00000204745 | NA | chr2 | + | 87138386 | 87138457 | 87138255 | 87138292 | 87140826 | 87141005 | 0 | 0 | 1 | 0 | 3 | 0 | 0 | 0 | 0 | 0 | 0 | 3 | 6 | 1 | 0 | 0 | 0 | 2 | 0 | 3 | 220 | 149 | 1 | 1 | NA | NA | 1 | NA | 1 | NA | 1 | 0.67 | 1 | 0 | 0.333 |
| SE | 38262 | ENSG00000204745 | NA | chr2 | + | 87185531 | 87185618 | 87182958 | 87183097 | 87187146 | 87187242 | 25 | 21 | 34 | 16 | 28 | 0 | 0 | 0 | 0 | 3 | 10 | 8 | 12 | 17 | 0 | 12 | 0 | 0 | 0 | 0 | 236 | 149 | 1 | 1 | 1 | 1 | 1 | 1 | 0.855 | 0.345 | 1 | 1 | 1 | NA | 0.135 |
| SE | 38263 | ENSG00000134684 | YARS1 | chr1 | - | 32780084 | 32780278 | 32779381 | 32779523 | 32781047 | 32781145 | 217 | 186 | 201 | 211 | 230 | 0 | 0 | 0 | 0 | 0 | 495 | 238 | 247 | 359 | 315 | 0 | 0 | 3 | 0 | 0 | 298 | 149 | 1 | 1 | 1 | 1 | 1 | 1 | 1 | 1 | 1 | 1 | 0.984 | 1 | 0.003 |
| SE | 38264 | ENSG00000134684 | YARS1 | chr1 | - | 32782403 | 32782539 | 32781047 | 32781145 | 32786361 | 32786447 | 473 | 313 | 426 | 529 | 617 | 0 | 0 | 0 | 3 | 0 | 879 | 517 | 435 | 579 | 705 | 0 | 0 | 0 | 0 | 0 | 285 | 149 | 1 | 1 | 1 | 1 | 1 | 0.989 | 1 | 1 | 1 | 1 | 1 | 1 | -0.002 |
| SE | 38265 | ENSG00000134684 | YARS1 | chr1 | - | 32786361 | 32786447 | 32782403 | 32782539 | 32791161 | 32791254 | 157 | 105 | 136 | 183 | 289 | 0 | 1 | 0 | 3 | 0 | 257 | 150 | 111 | 178 | 245 | 0 | 0 | 0 | 0 | 0 | 235 | 149 | 1 | 1 | 1 | 0.985 | 1 | 0.975 | 1 | 1 | 1 | 1 | 1 | 1 | -0.008 |
| SE | 38266 | ENSG00000134684 | YARS1 | chr1 | - | 32786361 | 32787075 | 32782403 | 32782539 | 32791161 | 32791254 | 283 | 184 | 209 | 313 | 446 | 0 | 1 | 0 | 3 | 0 | 428 | 302 | 242 | 352 | 353 | 0 | 0 | 0 | 0 | 0 | 298 | 149 | 1 | 1 | 1 | 0.989 | 1 | 0.981 | 1 | 1 | 1 | 1 | 1 | 1 | -0.006 |
| SE | 38268 | ENSG00000134684 | YARS1 | chr1 | - | 32786939 | 32787075 | 32786361 | 32786447 | 32797762 | 32797843 | 166 | 86 | 122 | 174 | 312 | 0 | 1 | 0 | 0 | 0 | 252 | 193 | 125 | 202 | 187 | 0 | 0 | 0 | 0 | 0 | 285 | 149 | 1 | 1 | 1 | 0.978 | 1 | 1 | 1 | 1 | 1 | 1 | 1 | 1 | -0.004 |
| SE | 38269 | ENSG00000134684 | YARS1 | chr1 | - | 32791161 | 32791254 | 32786361 | 32786447 | 32797762 | 32797843 | 156 | 82 | 97 | 96 | 146 | 0 | 1 | 0 | 0 | 0 | 274 | 212 | 137 | 207 | 202 | 0 | 0 | 0 | 0 | 0 | 242 | 149 | 1 | 1 | 1 | 0.981 | 1 | 1 | 1 | 1 | 1 | 1 | 1 | 1 | -0.004 |
| SE | 38270 | ENSG00000134684 | YARS1 | chr1 | - | 32791161 | 32791254 | 32786361 | 32787075 | 32797591 | 32797638 | 95 | 42 | 44 | 88 | 106 | 0 | 0 | 0 | 0 | 0 | 103 | 118 | 71 | 82 | 49 | 0 | 0 | 0 | 2 | 0 | 242 | 149 | 1 | 1 | 1 | 1 | 1 | 1 | 1 | 1 | 1 | 1 | 0.962 | 1 | 0.008 |
| SE | 38271 | ENSG00000134684 | YARS1 | chr1 | - | 32791161 | 32791254 | 32786361 | 32787075 | 32797762 | 32797843 | 273 | 160 | 163 | 218 | 305 | 1 | 0 | 0 | 0 | 0 | 429 | 363 | 250 | 368 | 307 | 0 | 0 | 0 | 0 | 0 | 242 | 149 | 1 | 1 | 0.994 | 1 | 1 | 1 | 1 | 1 | 1 | 1 | 1 | 1 | -0.001 |
| SE | 38272 | ENSG00000134684 | YARS1 | chr1 | - | 32797591 | 32797638 | 32786939 | 32787075 | 32797762 | 32797843 | 4 | 0 | 3 | 2 | 0 | 1 | 0 | 0 | 0 | 0 | 0 | 4 | 4 | 12 | 0 | 0 | 0 | 0 | 4 | 0 | 196 | 149 | 1 | 1 | 0.753 | NA | 1 | 1 | NA | NA | 1 | 1 | 1 | NA | -0.082 |
| SE | 38274 | ENSG00000134684 | YARS1 | chr1 | - | 32797762 | 32797843 | 32790652 | 32791254 | 32806481 | 32806580 | 213 | 133 | 153 | 167 | 248 | 0 | 0 | 1 | 3 | 1 | 386 | 285 | 214 | 344 | 312 | 0 | 0 | 0 | 4 | 0 | 230 | 149 | 1 | 1 | 1 | 1 | 0.99 | 0.973 | 0.994 | 1 | 1 | 1 | 0.982 | 1 | -0.005 |
| SE | 38275 | ENSG00000134684 | YARS1 | chr1 | - | 32804544 | 32804883 | 32797762 | 32797843 | 32806481 | 32806611 | 0 | 0 | 0 | 3 | 0 | 125 | 59 | 86 | 88 | 85 | 0 | 0 | 0 | 4 | 0 | 214 | 169 | 99 | 184 | 186 | 298 | 149 | 1 | 1 | 0 | 0 | 0 | 0.017 | 0 | 0 | 0 | 0 | 0.011 | 0 | 0.001 |
| SE | 38276 | ENSG00000134684 | YARS1 | chr1 | - | 32806481 | 32806611 | 32797159 | 32797843 | 32810590 | 32810766 | 481 | 224 | 348 | 365 | 348 | 0 | 0 | 0 | 0 | 0 | 591 | 612 | 376 | 519 | 564 | 0 | 1 | 0 | 0 | 9 | 279 | 149 | 1 | 1 | 1 | 1 | 1 | 1 | 1 | 1 | 0.997 | 1 | 1 | 0.971 | 0.006 |
| SE | 38277 | ENSG00000134684 | YARS1 | chr1 | - | 32806481 | 32806611 | 32797159 | 32797843 | 32817187 | 32817357 | 113 | 47 | 80 | 89 | 79 | 0 | 0 | 3 | 0 | 0 | 209 | 159 | 97 | 176 | 171 | 0 | 0 | 0 | 0 | 0 | 279 | 149 | 1 | 1 | 1 | 1 | 0.934 | 1 | 1 | 1 | 1 | 1 | 1 | 1 | -0.013 |
| SE | 38278 | ENSG00000134684 | YARS1 | chr1 | - | 32810590 | 32810766 | 32797762 | 32797843 | 32817187 | 32817380 | 6 | 0 | 3 | 1 | 0 | 0 | 0 | 3 | 0 | 0 | 10 | 6 | 2 | 9 | 9 | 0 | 0 | 0 | 0 | 0 | 298 | 149 | 1 | 1 | 1 | NA | 0.333 | 1 | NA | 1 | 1 | 1 | 1 | 1 | -0.222 |
| SE | 38279 | ENSG00000134684 | YARS1 | chr1 | - | 32810590 | 32810766 | 32806481 | 32806611 | 32810910 | 32811057 | 542 | 270 | 400 | 416 | 378 | 0 | 0 | 0 | 0 | 0 | 570 | 675 | 428 | 481 | 560 | 0 | 0 | 0 | 8 | 0 | 298 | 149 | 1 | 1 | 1 | 1 | 1 | 1 | 1 | 1 | 1 | 1 | 0.968 | 1 | 0.006 |
| SE | 38284 | ENSG00000164330 | EBF1 | chr5 | - | 158712153 | 158712333 | 158707978 | 158708173 | 158712969 | 158713147 | 24 | 17 | 25 | 48 | 72 | 0 | 0 | 0 | 0 | 2 | 27 | 86 | 18 | 15 | 106 | 0 | 0 | 0 | 0 | 0 | 298 | 149 | 1 | 1 | 1 | 1 | 1 | 1 | 0.947 | 1 | 1 | 1 | 1 | 1 | -0.011 |
| SE | 38285 | ENSG00000164330 | EBF1 | chr5 | - | 158796344 | 158796475 | 158777412 | 158777539 | 158823175 | 158823317 | 28 | 29 | 39 | 42 | 65 | 0 | 0 | 1 | 0 | 0 | 57 | 83 | 7 | 30 | 14 | 0 | 0 | 0 | 0 | 0 | 280 | 149 | 1 | 1 | 1 | 1 | 0.954 | 1 | 1 | 1 | 1 | 1 | 1 | 1 | -0.009 |
| SE | 38286 | ENSG00000164330 | EBF1 | chr5 | - | 158823175 | 158823317 | 158796344 | 158796475 | 158840028 | 158840110 | 26 | 33 | 29 | 41 | 71 | 0 | 0 | 0 | 0 | 1 | 27 | 59 | 10 | 30 | 31 | 0 | 0 | 0 | 0 | 0 | 291 | 149 | 1 | 1 | 1 | 1 | 1 | 1 | 0.973 | 1 | 1 | 1 | 1 | 1 | -0.005 |
| SE | 38287 | ENSG00000164330 | EBF1 | chr5 | - | 158823199 | 158823317 | 158796344 | 158796475 | 158840028 | 158840110 | 19 | 20 | 13 | 28 | 42 | 0 | 0 | 0 | 0 | 1 | 19 | 27 | 5 | 17 | 14 | 0 | 0 | 0 | 0 | 0 | 267 | 149 | 1 | 1 | 1 | 1 | 1 | 1 | 0.959 | 1 | 1 | 1 | 1 | 1 | -0.008 |
| SE | 38289 | ENSG00000164330 | EBF1 | chr5 | - | 159096342 | 159096406 | 159095619 | 159095675 | 159096973 | 159097130 | 44 | 26 | 41 | 63 | 62 | 0 | 0 | 0 | 0 | 0 | 44 | 121 | 21 | 19 | 90 | 0 | 0 | 0 | 0 | 5 | 213 | 149 | 1 | 1 | 1 | 1 | 1 | 1 | 1 | 1 | 1 | 1 | 1 | 0.926 | 0.015 |
| SE | 38298 | ENSG00000204954 | C12orf73 | chr12 | - | 103953413 | 103953534 | 103950201 | 103951630 | 103956630 | 103956748 | 5 | 0 | 1 | 5 | 2 | 4 | 0 | 1 | 0 | 0 | 14 | 12 | 1 | 0 | 2 | 0 | 0 | 0 | 0 | 7 | 270 | 149 | 1 | 1 | 0.408 | NA | 0.356 | 1 | 1 | 1 | 1 | 1 | NA | 0.136 | -0.093 |
| SE | 38299 | ENSG00000204954 | C12orf73 | chr12 | - | 103954874 | 103954968 | 103953413 | 103953534 | 103956630 | 103956748 | 0 | 1 | 0 | 2 | 5 | 4 | 0 | 1 | 0 | 0 | 0 | 4 | 8 | 0 | 0 | 7 | 4 | 1 | 0 | 2 | 243 | 149 | 1 | 1 | 0 | 1 | 0 | 1 | 1 | 0 | 0.38 | 0.831 | NA | 0 | 0.297 |
| SE | 38304 | ENSG00000278535 | DHRS11 | chr17 | + | 36594970 | 36595180 | 36591878 | 36592156 | 36598162 | 36598257 | 9 | 0 | 7 | 3 | 3 | 0 | 0 | 0 | 0 | 1 | 0 | 0 | 5 | 0 | 0 | 0 | 0 | 0 | 0 | 0 | 298 | 149 | 1 | 1 | 1 | NA | 1 | 1 | 0.6 | NA | NA | 1 | NA | NA | -0.1 |
| SE | 38305 | ENSG00000278535 | DHRS11 | chr17 | + | 36595026 | 36595180 | 36591886 | 36592156 | 36598162 | 36598257 | 5 | 0 | 4 | 3 | 3 | 0 | 0 | 0 | 0 | 1 | 0 | 0 | 7 | 0 | 0 | 0 | 0 | 0 | 0 | 0 | 298 | 149 | 1 | 1 | 1 | NA | 1 | 1 | 0.6 | NA | NA | 1 | NA | NA | -0.1 |
| SE | 38306 | ENSG00000278535 | DHRS11 | chr17 | + | 36599670 | 36599763 | 36598920 | 36599050 | 36599971 | 36599985 | 3 | 3 | 6 | 1 | 2 | 0 | 0 | 1 | 1 | 0 | 2 | 3 | 0 | 0 | 0 | 0 | 0 | 0 | 0 | 0 | 242 | 149 | 1 | 1 | 1 | 1 | 0.787 | 0.381 | 1 | 1 | 1 | NA | NA | NA | -0.166 |
| SE | 38312 | ENSG00000138172 | CALHM2 | chr10 | - | 103451878 | 103452036 | 103449698 | 103449934 | 103452155 | 103452336 | 3 | 0 | 6 | 2 | 0 | 0 | 0 | 3 | 0 | 0 | 0 | 0 | 0 | 0 | 0 | 0 | 2 | 0 | 0 | 0 | 298 | 149 | 1 | 1 | 1 | NA | 0.5 | 1 | NA | NA | 0 | NA | NA | NA | 0.833 |
| SE | 38314 | ENSG00000138172 | CALHM2 | chr10 | - | 103451878 | 103452036 | 103451082 | 103451285 | 103452155 | 103452336 | 3 | 0 | 6 | 2 | 0 | 24 | 13 | 10 | 14 | 6 | 11 | 0 | 0 | 0 | 0 | 6 | 13 | 0 | 0 | 2 | 298 | 149 | 1 | 1 | 0.059 | 0 | 0.231 | 0.067 | 0 | 0.478 | 0 | NA | NA | 0 | -0.088 |
| SE | 38315 | ENSG00000143742 | SRP9 | chr1 | + | 225783299 | 225783368 | 225777812 | 225778012 | 225789239 | 225790464 | 721 | 331 | 566 | 547 | 483 | 0 | 0 | 0 | 0 | 0 | 184 | 505 | 342 | 323 | 98 | 0 | 5 | 0 | 0 | 0 | 218 | 149 | 1 | 1 | 1 | 1 | 1 | 1 | 1 | 1 | 0.986 | 1 | 1 | 1 | 0.003 |
| SE | 38320 | ENSG00000143742 | SRP9 | chr1 | + | 225789033 | 225789141 | 225783299 | 225783368 | 225789239 | 225789524 | 13 | 3 | 7 | 7 | 5 | 602 | 284 | 456 | 447 | 372 | 0 | 6 | 9 | 8 | 0 | 142 | 421 | 286 | 278 | 78 | 257 | 149 | 1 | 1 | 0.012 | 0.006 | 0.009 | 0.009 | 0.008 | 0 | 0.008 | 0.018 | 0.016 | 0 | 0 |
| SE | 38321 | ENSG00000143742 | SRP9 | chr1 | + | 225789033 | 225789141 | 225786861 | 225786985 | 225789239 | 225789524 | 7 | 4 | 6 | 6 | 5 | 47 | 22 | 44 | 48 | 48 | 0 | 6 | 8 | 5 | 0 | 0 | 51 | 44 | 21 | 11 | 257 | 149 | 1 | 1 | 0.079 | 0.095 | 0.073 | 0.068 | 0.057 | NA | 0.064 | 0.095 | 0.121 | 0 | 0.004 |
| SE | 38323 | ENSG00000162777 | DENND2D | chr1 | - | 111189211 | 111189253 | 111188701 | 111188786 | 111192139 | 111192317 | 629 | 354 | 456 | 389 | 492 | 1 | 0 | 1 | 0 | 0 | 85 | 188 | 106 | 187 | 219 | 0 | 0 | 0 | 0 | 0 | 191 | 149 | 1 | 1 | 0.998 | 1 | 0.997 | 1 | 1 | 1 | 1 | 1 | 1 | 1 | -0.001 |
| SE | 38325 | ENSG00000162777 | DENND2D | chr1 | - | 111192139 | 111192317 | 111188701 | 111188786 | 111195915 | 111196056 | 2 | 2 | 4 | 1 | 1 | 0 | 0 | 0 | 0 | 0 | 0 | 0 | 0 | 7 | 0 | 0 | 0 | 0 | 7 | 0 | 298 | 149 | 1 | 1 | 1 | 1 | 1 | 1 | 1 | NA | NA | NA | 0.333 | NA | 0.667 |
| SE | 38326 | ENSG00000162777 | DENND2D | chr1 | - | 111192139 | 111192317 | 111189211 | 111189253 | 111194577 | 111194726 | 846 | 451 | 613 | 510 | 1020 | 1 | 0 | 0 | 0 | 0 | 110 | 293 | 178 | 222 | 296 | 0 | 0 | 0 | 0 | 0 | 298 | 149 | 1 | 1 | 0.998 | 1 | 1 | 1 | 1 | 1 | 1 | 1 | 1 | 1 | 0 |
| SE | 38327 | ENSG00000162777 | DENND2D | chr1 | - | 111192139 | 111192317 | 111189211 | 111189253 | 111195915 | 111196056 | 365 | 188 | 241 | 222 | 212 | 2 | 0 | 0 | 0 | 0 | 29 | 91 | 33 | 110 | 116 | 0 | 0 | 0 | 0 | 0 | 298 | 149 | 1 | 1 | 0.989 | 1 | 1 | 1 | 1 | 1 | 1 | 1 | 1 | 1 | -0.002 |
| SE | 38328 | ENSG00000162777 | DENND2D | chr1 | - | 111194577 | 111194726 | 111189211 | 111189253 | 111195915 | 111196056 | 372 | 168 | 288 | 235 | 309 | 2 | 0 | 0 | 0 | 0 | 29 | 134 | 63 | 84 | 143 | 0 | 0 | 0 | 0 | 0 | 298 | 149 | 1 | 1 | 0.989 | 1 | 1 | 1 | 1 | 1 | 1 | 1 | 1 | 1 | -0.002 |
| SE | 38329 | ENSG00000162777 | DENND2D | chr1 | - | 111194577 | 111194726 | 111192139 | 111192317 | 111195915 | 111196056 | 853 | 433 | 663 | 524 | 1118 | 1 | 2 | 3 | 1 | 1 | 110 | 336 | 208 | 203 | 323 | 0 | 0 | 0 | 7 | 0 | 298 | 149 | 1 | 1 | 0.998 | 0.991 | 0.991 | 0.996 | 0.998 | 1 | 1 | 1 | 0.935 | 1 | 0.008 |
| SE | 38332 | ENSG00000162777 | DENND2D | chr1 | - | 111197919 | 111197989 | 111195913 | 111196056 | 111198629 | 111198742 | 146 | 87 | 105 | 103 | 89 | 3 | 0 | 2 | 2 | 0 | 20 | 44 | 21 | 23 | 15 | 0 | 1 | 0 | 0 | 0 | 219 | 149 | 1 | 1 | 0.971 | 1 | 0.973 | 0.972 | 1 | 1 | 1 | 0.935 | 1 | 1 | -0.004 |
| SE | 38333 | ENSG00000162777 | DENND2D | chr1 | - | 111197919 | 111197989 | 111197175 | 111197253 | 111198629 | 111198742 | 315 | 173 | 261 | 227 | 192 | 0 | 0 | 1 | 0 | 0 | 70 | 119 | 34 | 53 | 119 | 0 | 1 | 0 | 0 | 0 | 219 | 149 | 1 | 1 | 1 | 1 | 0.994 | 1 | 1 | 1 | 0.988 | 1 | 1 | 1 | 0.001 |
| SE | 38335 | ENSG00000162777 | DENND2D | chr1 | - | 111197919 | 111197989 | 111197175 | 111197253 | 111200392 | 111200646 | 106 | 37 | 75 | 58 | 23 | 1 | 0 | 0 | 0 | 0 | 15 | 28 | 5 | 26 | 66 | 0 | 0 | 0 | 0 | 0 | 219 | 149 | 1 | 1 | 0.986 | 1 | 1 | 1 | 1 | 1 | 1 | 1 | 1 | 1 | -0.003 |
| SE | 38336 | ENSG00000162777 | DENND2D | chr1 | - | 111198629 | 111198742 | 111195915 | 111197989 | 111199622 | 111199798 | 397 | 232 | 362 | 312 | 366 | 1 | 0 | 0 | 0 | 2 | 108 | 154 | 72 | 77 | 113 | 0 | 0 | 0 | 0 | 0 | 262 | 149 | 1 | 1 | 0.996 | 1 | 1 | 1 | 0.99 | 1 | 1 | 1 | 1 | 1 | -0.003 |
| SE | 38337 | ENSG00000162777 | DENND2D | chr1 | - | 111198629 | 111198742 | 111195915 | 111197989 | 111200392 | 111200546 | 179 | 103 | 164 | 156 | 149 | 0 | 0 | 1 | 0 | 0 | 64 | 92 | 38 | 34 | 47 | 0 | 0 | 0 | 0 | 0 | 262 | 149 | 1 | 1 | 1 | 1 | 0.989 | 1 | 1 | 1 | 1 | 1 | 1 | 1 | -0.002 |
| SE | 38339 | ENSG00000162777 | DENND2D | chr1 | - | 111198629 | 111198742 | 111197175 | 111197253 | 111200392 | 111200546 | 0 | 1 | 1 | 0 | 4 | 1 | 0 | 0 | 0 | 0 | 0 | 1 | 0 | 2 | 0 | 0 | 0 | 0 | 0 | 0 | 262 | 149 | 1 | 1 | 0 | 1 | 1 | NA | 1 | NA | 1 | NA | 1 | NA | -0.25 |
| SE | 38340 | ENSG00000162777 | DENND2D | chr1 | - | 111199622 | 111199798 | 111195915 | 111197989 | 111200392 | 111200546 | 249 | 98 | 159 | 164 | 160 | 0 | 0 | 1 | 0 | 0 | 72 | 106 | 28 | 51 | 90 | 0 | 0 | 0 | 0 | 0 | 298 | 149 | 1 | 1 | 1 | 1 | 0.988 | 1 | 1 | 1 | 1 | 1 | 1 | 1 | -0.002 |
| SE | 38341 | ENSG00000162777 | DENND2D | chr1 | - | 111199622 | 111199798 | 111197175 | 111197253 | 111200392 | 111200546 | 248 | 98 | 159 | 164 | 158 | 1 | 0 | 0 | 0 | 0 | 72 | 114 | 28 | 51 | 90 | 0 | 0 | 0 | 0 | 0 | 298 | 149 | 1 | 1 | 0.992 | 1 | 1 | 1 | 1 | 1 | 1 | 1 | 1 | 1 | -0.002 |
| SE | 38343 | ENSG00000162777 | DENND2D | chr1 | - | 111199622 | 111199798 | 111198629 | 111198742 | 111200392 | 111200546 | 468 | 230 | 362 | 321 | 380 | 0 | 1 | 1 | 0 | 4 | 116 | 168 | 62 | 96 | 156 | 0 | 0 | 0 | 2 | 0 | 298 | 149 | 1 | 1 | 1 | 0.991 | 0.995 | 1 | 0.979 | 1 | 1 | 1 | 0.96 | 1 | 0.001 |
| SE | 38344 | ENSG00000162777 | DENND2D | chr1 | - | 111202394 | 111202445 | 111199622 | 111199798 | 111204145 | 111204535 | 2 | 0 | 0 | 0 | 3 | 5 | 0 | 3 | 2 | 0 | 0 | 0 | 0 | 0 | 0 | 0 | 2 | 0 | 0 | 0 | 200 | 149 | 1 | 1 | 0.23 | NA | 0 | 0 | 1 | NA | 0 | NA | NA | | 0.307 |
| SE | 38346 | ENSG00000119411 | BSPRY | chr9 | + | 113368258 | 113368383 | 113362368 | 113362394 | 113369615 | 113371222 | 1 | 0 | 1 | 0 | 0 | 0 | 0 | 0 | 0 | 0 | 0 | 0 | 6 | 3 | 0 | 0 | 2 | 0 | 0 | 0 | 274 | 149 | 1 | 1 | 1 | NA | 1 | NA | NA | NA | 0 | 1 | 1 | NA | 0.333 |
| SE | 38350 | ENSG00000150337 | FCGR1A | chr1 | + | 149783169 | 149783190 | 149782693 | 149782774 | 149788365 | 149788617 | 0 | 0 | 0 | 0 | 0 | 2 | 0 | 0 | 0 | 4 | 0 | 3 | 0 | 0 | 3 | 6 | 2 | 2 | 0 | 0 | 170 | 149 | 1 | 1 | 0 | NA | 0 | NA | 0 | 0 | 0.568 | 0 | NA | 1 | -0.392 |
| SE | 38351 | ENSG00000150337 | FCGR1A | chr1 | + | 149784002 | 149784257 | 149783169 | 149783190 | 149788365 | 149788617 | 77 | 45 | 44 | 52 | 89 | 0 | 0 | 0 | 0 | 0 | 359 | 102 | 23 | 185 | 210 | 0 | 3 | 0 | 0 | 3 | 298 | 149 | 1 | 1 | 1 | 1 | 1 | 1 | 1 | 1 | 0.944 | 1 | 1 | 0.972 | 0.017 |
| SE | 38357 | ENSG00000163002 | NUP35 | chr2 | + | 183130994 | 183131352 | 183130417 | 183130545 | 183133565 | 183133623 | 1 | 0 | 1 | 0 | 0 | 23 | 22 | 33 | 38 | 37 | 0 | 0 | 4 | 0 | 0 | 0 | 17 | 0 | 1 | 7 | 298 | 149 | 1 | 1 | 0.021 | 0 | 0.015 | 0 | 0 | NA | NA | 0.105 | 0 | 0 | -0.028 |
| SE | 38359 | ENSG00000163002 | NUP35 | chr2 | + | 183157443 | 183157513 | 183133565 | 183133623 | 183158282 | 183158411 | 22 | 10 | 23 | 24 | 17 | 0 | 0 | 1 | 0 | 1 | 8 | 6 | 6 | 9 | 0 | 0 | 0 | 3 | 0 | 0 | 219 | 149 | 1 | 1 | 1 | 1 | 0.94 | 1 | 0.92 | 1 | 1 | 0.576 | 1 | NA | 0.078 |
| SE | 38361 | ENSG00000186074 | CD300LF | chr17 | - | 74696194 | 74696217 | 74695724 | 74695859 | 74703034 | 74703098 | 4 | 2 | 2 | 3 | 0 | 0 | 0 | 0 | 0 | 0 | 7 | 0 | 0 | 3 | 0 | 0 | 0 | 1 | 1 | 0 | 172 | 149 | 1 | 1 | 1 | 1 | 1 | 1 | NA | 1 | NA | 0 | 0.722 | NA | 0.426 |
| SE | 38362 | ENSG00000186074 | CD300LF | chr17 | - | 74698368 | 74698481 | 74695724 | 74695859 | 74703034 | 74703098 | 63 | 26 | 22 | 42 | 69 | 0 | 0 | 0 | 0 | 0 | 24 | 17 | 21 | 35 | 63 | 0 | 0 | 1 | 1 | 0 | 262 | 149 | 1 | 1 | 1 | 1 | 1 | 1 | 1 | 1 | 1 | 0.923 | 0.952 | 1 | 0.025 |
| SE | 38363 | ENSG00000186074 | CD300LF | chr17 | - | 74698368 | 74698481 | 74696194 | 74696217 | 74703034 | 74703098 | 110 | 44 | 45 | 71 | 99 | 1 | 1 | 0 | 0 | 0 | 55 | 20 | 37 | 44 | 128 | 0 | 0 | 0 | 0 | 0 | 262 | 149 | 1 | 1 | 0.984 | 0.962 | 1 | 1 | 1 | 1 | 1 | 1 | 1 | 1 | -0.011 |
| SE | 38365 | ENSG00000186074 | CD300LF | chr17 | - | 74703034 | 74703098 | 74695724 | 74695859 | 74704477 | 74704816 | 100 | 43 | 67 | 62 | 98 | 0 | 0 | 0 | 0 | 0 | 137 | 35 | 34 | 78 | 193 | 0 | 0 | 0 | 2 | 0 | 213 | 149 | 1 | 1 | 1 | 1 | 1 | 1 | 1 | 1 | 1 | 1 | 0.965 | 1 | 0.007 |

| | | | | | | | | | | | | | | | | | | | | | | | | | | | | | | | | | | | | | | | | | | | | | | |
|---|---|---|---|---|---|---|---|---|---|---|---|---|---|---|---|---|---|---|---|---|---|---|---|---|---|---|---|---|---|---|---|---|---|---|---|---|---|---|---|---|---|---|---|---|---|---|
| SE | 38372 | ENSG00000112365 | ZBTB24 | chr6 | - | 109467652 | 109467734 | 109462593 | 109466574 | 109475398 | 109475482 | 49 | 33 | 56 | 33 | 44 | 1 | 2 | 0 | 1 | 0 | 10 | 32 | 30 | 10 | 6 | 0 | 0 | 4 | 4 | 0 | 231 | 149 | 1 | 1 | 0.969 | 0.914 | 1 | 0.955 | 1 | 1 | 1 | 0.829 | 0.617 | 1 | 0.078 |
| SE | 38374 | ENSG00000112365 | ZBTB24 | chr6 | - | 109475398 | 109475482 | 109462593 | 109466574 | 109481074 | 109482054 | 0 | 2 | 0 | 1 | 0 | 0 | 0 | 0 | 0 | 0 | 0 | 0 | 6 | 7 | 0 | 0 | 0 | 0 | 0 | 10 | 233 | 149 | 1 | 1 | NA | 1 | NA | 1 | NA | NA | NA | 1 | 1 | 0 | 0.333 |
| SE | 38375 | ENSG00000112365 | ZBTB24 | chr6 | - | 109475398 | 109475482 | 109467652 | 109467734 | 109481074 | 109482054 | 11 | 14 | 25 | 9 | 19 | 0 | 0 | 0 | 0 | 1 | 0 | 11 | 12 | 7 | 6 | 0 | 0 | 0 | 0 | 0 | 233 | 149 | 1 | 1 | 1 | 1 | 1 | 1 | 0.924 | NA | 1 | 1 | 1 | 1 | -0.015 |
| SE | 38376 | ENSG00000112365 | ZBTB24 | chr6 | - | 109476174 | 109476258 | 109475398 | 109475482 | 109481074 | 109482054 | 18 | 15 | 19 | 26 | 15 | 0 | 0 | 0 | 0 | 0 | 5 | 31 | 20 | 19 | 0 | 0 | 0 | 2 | 3 | 0 | 233 | 149 | 1 | 1 | 1 | 1 | 1 | 1 | 1 | 1 | 1 | 0.865 | 0.802 | NA | 0.083 |
| SE | 38379 | ENSG00000138430 | OLA1 | chr2 | - | 174078967 | 174079090 | 174072446 | 174075527 | 174081151 | 174081248 | 361 | 263 | 335 | 353 | 589 | 2 | 1 | 0 | 0 | 3 | 158 | 196 | 271 | 309 | 40 | 4 | 0 | 0 | 0 | 0 | 272 | 149 | 1 | 1 | 0.99 | 0.993 | 1 | 1 | 0.991 | 0.956 | 1 | 1 | 1 | 1 | 0.004 |
| SE | 38380 | ENSG00000138430 | OLA1 | chr2 | - | 174078967 | 174079090 | 174072446 | 174075527 | 174123179 | 174123277 | 131 | 104 | 142 | 145 | 212 | 2 | 0 | 0 | 1 | 4 | 117 | 78 | 129 | 120 | 26 | 0 | 1 | 3 | 0 | 0 | 272 | 149 | 1 | 1 | 0.973 | 1 | 1 | 0.988 | 0.967 | 1 | 0.977 | 0.959 | 1 | 1 | -0.002 |
| SE | 38381 | ENSG00000138430 | OLA1 | chr2 | - | 174081151 | 174081248 | 174072446 | 174075527 | 174123179 | 174123277 | 0 | 1 | 0 | 0 | 0 | 2 | 0 | 0 | 1 | 4 | 0 | 0 | 0 | 0 | 0 | 0 | 1 | 3 | 0 | 0 | 246 | 149 | 1 | 1 | 0 | 1 | NA | 0 | 0 | NA | 0 | 0 | NA | NA | 0.25 |
| SE | 38382 | ENSG00000138430 | OLA1 | chr2 | - | 174081151 | 174081248 | 174078967 | 174079090 | 174123179 | 174123277 | 109 | 81 | 70 | 101 | 171 | 2 | 0 | 0 | 2 | 0 | 19 | 52 | 57 | 77 | 14 | 0 | 0 | 0 | 5 | 0 | 246 | 149 | 1 | 1 | 1 | 1 | 1 | 0.968 | 1 | 1 | 1 | 1 | 0.903 | 1 | 0.013 |
| SE | 38383 | ENSG00000138430 | OLA1 | chr2 | - | 174081923 | 174082064 | 174080907 | 174081248 | 174123179 | 174123277 | 548 | 338 | 489 | 492 | 762 | 0 | 0 | 0 | 0 | 0 | 282 | 337 | 478 | 549 | 83 | 0 | 0 | 0 | 0 | 4 | 290 | 149 | 1 | 1 | 1 | 1 | 1 | 1 | 1 | 1 | 1 | 1 | 1 | 0.914 | 0.017 |
| SE | 38384 | ENSG00000138430 | OLA1 | chr2 | - | 174141824 | 174142000 | 174123594 | 174123675 | 174223032 | 174223160 | 461 | 324 | 377 | 396 | 518 | 0 | 0 | 0 | 0 | 3 | 188 | 291 | 378 | 386 | 137 | 0 | 0 | 0 | 0 | 0 | 298 | 149 | 1 | 1 | 1 | 1 | 1 | 1 | 0.989 | 1 | 1 | 1 | 1 | 1 | -0.002 |
| SE | 38392 | ENSG00000143458 | GABPB2 | chr1 | + | 151073038 | 151073157 | 151070577 | 151070934 | 151088157 | 151088297 | 8 | 4 | 7 | 5 | 3 | 6 | 0 | 5 | 2 | 6 | 0 | 0 | 1 | 5 | 0 | 0 | 0 | 0 | 0 | 0 | 268 | 149 | 1 | 1 | 0.426 | 1 | 0.438 | 0.582 | 0.218 | NA | NA | 1 | 1 | NA | -0.467 |
| SE | 38394 | ENSG00000143458 | GABPB2 | chr1 | + | 151088157 | 151088297 | 151070803 | 151070934 | 151093191 | 151093386 | 7 | 2 | 7 | 4 | 6 | 0 | 0 | 0 | 0 | 1 | 0 | 0 | 1 | 0 | 11 | 0 | 0 | 0 | 0 | 0 | 289 | 149 | 1 | 1 | 1 | 1 | 1 | 1 | 0.756 | NA | NA | 1 | NA | 1 | -0.049 |
| SE | 38396 | ENSG00000143458 | GABPB2 | chr1 | + | 151088189 | 151088297 | 151070760 | 151070934 | 151093191 | 151093386 | 32 | 34 | 22 | 17 | 25 | 0 | 0 | 0 | 0 | 1 | 0 | 16 | 3 | 13 | 14 | 0 | 0 | 0 | 0 | 0 | 257 | 149 | 1 | 1 | 1 | 1 | 1 | 1 | 0.935 | NA | 1 | 1 | 1 | 1 | -0.013 |
| SE | 38397 | ENSG00000143458 | GABPB2 | chr1 | + | 151088189 | 151088297 | 151070760 | 151070934 | 151097851 | 151097915 | 31 | 32 | 20 | 15 | 33 | 0 | 0 | 1 | 0 | 0 | 0 | 16 | 2 | 13 | 3 | 0 | 0 | 0 | 0 | 0 | 257 | 149 | 1 | 1 | 1 | 1 | 0.921 | 1 | 1 | NA | 1 | 1 | 1 | 1 | -0.016 |
| SE | 38399 | ENSG00000143458 | GABPB2 | chr1 | + | 151088469 | 151088517 | 151088157 | 151088297 | 151093191 | 151093386 | 0 | 0 | 10 | 2 | 0 | 1 | 2 | 2 | 2 | 0 | 0 | 0 | 0 | 0 | 5 | 0 | 0 | 1 | 0 | 11 | 197 | 149 | 1 | 1 | 0 | 0 | 0.791 | 0.431 | NA | NA | NA | 0 | NA | 0.256 | 0.177 |
| SE | 38403 | ENSG00000143458 | GABPB2 | chr1 | + | 151093191 | 151093386 | 151070577 | 151070934 | 151097851 | 151097915 | 125 | 74 | 119 | 126 | 225 | 1 | 0 | 1 | 0 | 0 | 90 | 63 | 72 | 88 | 59 | 0 | 0 | 0 | 0 | 0 | 298 | 149 | 1 | 1 | 1 | 1 | 0.983 | 1 | 1 | 1 | 1 | 1 | 1 | 1 | -0.003 |
| SE | 38405 | ENSG00000143458 | GABPB2 | chr1 | + | 151093191 | 151093386 | 151090405 | 151090573 | 151097851 | 151097915 | 176 | 97 | 182 | 182 | 288 | 0 | 1 | 0 | 2 | 1 | 172 | 88 | 100 | 124 | 76 | 0 | 0 | 2 | 4 | 0 | 298 | 149 | 1 | 1 | 1 | 0.98 | 1 | 0.978 | 0.993 | 1 | 1 | 0.962 | 0.939 | 1 | 0.01 |
| SE | 38406 | ENSG00000143458 | GABPB2 | chr1 | + | 151097851 | 151098002 | 151070760 | 151070934 | 151107036 | 151107222 | 52 | 39 | 55 | 59 | 55 | 0 | 1 | 0 | 0 | 0 | 62 | 39 | 53 | 72 | 66 | 0 | 0 | 0 | 0 | 0 | 298 | 149 | 1 | 1 | 1 | 0.951 | 1 | 1 | 1 | 1 | 1 | 1 | 1 | 1 | -0.01 |
| SE | 38409 | ENSG00000132549 | VPS13B | chr8 | + | 99038422 | 99038566 | 99013759 | 99013935 | 99096311 | 99096432 | 369 | 379 | 372 | 364 | 141 | 0 | 0 | 0 | 0 | 0 | 48 | 241 | 218 | 73 | 85 | 0 | 0 | 1 | 0 | 0 | 293 | 149 | 1 | 1 | 1 | 1 | 1 | 1 | 1 | 1 | 1 | 0.991 | 1 | 1 | 0.002 |
| SE | 38411 | ENSG00000132549 | VPS13B | chr8 | + | 99111097 | 99111279 | 99102952 | 99103120 | 99115699 | 99115874 | 476 | 464 | 308 | 370 | 708 | 0 | 0 | 0 | 0 | 0 | 70 | 332 | 304 | 66 | 53 | 0 | 0 | 0 | 5 | 0 | 298 | 149 | 1 | 1 | 1 | 1 | 1 | 1 | 1 | 1 | 1 | 1 | 0.868 | 1 | 0.026 |
| SE | 38414 | ENSG00000132549 | VPS13B | chr8 | + | 99115699 | 99115874 | 99111097 | 99111279 | 99121176 | 99121445 | 637 | 660 | 428 | 503 | 835 | 0 | 1 | 0 | 1 | 0 | 45 | 440 | 435 | 130 | 42 | 0 | 0 | 0 | 0 | 0 | 298 | 149 | 1 | 1 | 1 | 0.997 | 1 | 0.996 | 1 | 1 | 1 | 1 | 1 | 1 | -0.001 |
| SE | 38415 | ENSG00000132549 | VPS13B | chr8 | + | 99134631 | 99134727 | 99121176 | 99121445 | 99135014 | 99135137 | 505 | 527 | 431 | 476 | 368 | 1 | 0 | 0 | 0 | 0 | 36 | 327 | 277 | 113 | 38 | 0 | 0 | 0 | 0 | 0 | 245 | 149 | 1 | 1 | 0.997 | 1 | 1 | 1 | 1 | 1 | 1 | 1 | 1 | 1 | -0.001 |
| SE | 38416 | ENSG00000132549 | VPS13B | chr8 | + | 99135014 | 99135137 | 99121176 | 99121445 | 99135595 | 99135733 | 212 | 203 | 174 | 199 | 178 | 0 | 0 | 0 | 2 | 0 | 8 | 124 | 121 | 45 | 23 | 0 | 0 | 0 | 0 | 0 | 272 | 149 | 1 | 1 | 1 | 1 | 1 | 0.982 | 1 | 1 | 1 | 1 | 1 | 1 | -0.004 |
| SE | 38420 | ENSG00000132549 | VPS13B | chr8 | + | 99170038 | 99170163 | 99147840 | 99148010 | 99192875 | 99193057 | 105 | 87 | 89 | 104 | 73 | 0 | 1 | 0 | 0 | 0 | 0 | 65 | 59 | 17 | 8 | 0 | 0 | 2 | 0 | 0 | 274 | 149 | 1 | 1 | 1 | 0.979 | 1 | 1 | 1 | NA | 1 | 0.941 | 1 | 1 | 0.011 |
| SE | 38422 | ENSG00000132549 | VPS13B | chr8 | + | 99274197 | 99274332 | 99192875 | 99193057 | 99275080 | 99275254 | 390 | 304 | 319 | 410 | 517 | 0 | 0 | 2 | 1 | 0 | 71 | 263 | 217 | 90 | 59 | 0 | 0 | 0 | 0 | 0 | 284 | 149 | 1 | 1 | 1 | 1 | 0.988 | 0.995 | 1 | 1 | 1 | 1 | 1 | 1 | -0.003 |
| SE | 38424 | ENSG00000132549 | VPS13B | chr8 | + | 99429569 | 99429647 | 99391556 | 99391704 | 99431536 | 99431664 | 1 | 1 | 0 | 0 | 0 | 110 | 94 | 104 | 67 | 81 | 0 | 0 | 0 | 0 | 0 | 10 | 72 | 49 | 2 | 14 | 227 | 149 | 1 | 1 | 0.006 | 0.007 | 0 | 0 | 0 | 0 | 0 | 0 | 0 | 0 | 0.003 |
| SE | 38425 | ENSG00000132549 | VPS13B | chr8 | + | 99431520 | 99431664 | 99391556 | 99391704 | 99442400 | 99442411 | 249 | 248 | 218 | 235 | 296 | 0 | 0 | 1 | 4 | 0 | 72 | 198 | 155 | 66 | 101 | 0 | 0 | 0 | 0 | 0 | 293 | 149 | 1 | 1 | 1 | 1 | 0.991 | 0.968 | 1 | 1 | 1 | 1 | 1 | 1 | -0.008 |
| SE | 38426 | ENSG00000132549 | VPS13B | chr8 | + | 99431536 | 99431664 | 99391556 | 99391704 | 99442400 | 99442635 | 347 | 333 | 315 | 293 | 366 | 0 | 0 | 1 | 4 | 0 | 82 | 254 | 202 | 68 | 115 | 0 | 0 | 0 | 0 | 0 | 277 | 149 | 1 | 1 | 1 | 1 | 0.994 | 0.975 | 1 | 1 | 1 | 1 | 1 | 1 | -0.006 |
| SE | 38427 | ENSG00000132549 | VPS13B | chr8 | + | 99442400 | 99442635 | 99391556 | 99391704 | 99467413 | 99467634 | 81 | 71 | 68 | 60 | 84 | 0 | 1 | 0 | 0 | 0 | 33 | 100 | 61 | 12 | 36 | 0 | 0 | 0 | 0 | 0 | 298 | 149 | 1 | 1 | 1 | 0.973 | 1 | 1 | 1 | 1 | 1 | 1 | 1 | 1 | -0.005 |
| SE | 38428 | ENSG00000132549 | VPS13B | chr8 | + | 99444119 | 99444261 | 99442400 | 99442635 | 99467413 | 99467634 | 0 | 0 | 3 | 0 | 0 | 81 | 71 | 67 | 56 | 84 | 0 | 4 | 0 | 0 | 0 | 33 | 100 | 61 | 12 | 36 | 291 | 149 | 1 | 1 | 0 | 0 | 0.022 | 0 | 0 | 0 | 0.02 | 0 | 0 | 0 | 0 |
| SE | 38430 | ENSG00000132549 | VPS13B | chr8 | + | 99502835 | 99502950 | 99501686 | 99501858 | 99511103 | 99511512 | 142 | 109 | 92 | 148 | 139 | 0 | 2 | 0 | 0 | 2 | 26 | 70 | 79 | 30 | 31 | 0 | 0 | 0 | 0 | 0 | 264 | 149 | 1 | 1 | 1 | 0.969 | 1 | 1 | 0.975 | 1 | 1 | 1 | 1 | 1 | -0.011 |
| SE | 38433 | ENSG00000132549 | VPS13B | chr8 | + | 99507769 | 99507911 | 99501686 | 99501858 | 99511103 | 99511340 | 18 | 14 | 11 | 10 | 8 | 0 | 2 | 0 | 0 | 2 | 11 | 9 | 6 | 2 | 14 | 0 | 0 | 0 | 0 | 0 | 291 | 149 | 1 | 1 | 1 | 0.782 | 1 | 1 | 0.672 | 1 | 1 | 1 | 1 | 1 | -0.109 |
| SE | 38436 | ENSG00000132549 | VPS13B | chr8 | + | 99511103 | 99511512 | 99501686 | 99501858 | 99520898 | 99521010 | 147 | 160 | 118 | 198 | 144 | 0 | 0 | 0 | 0 | 0 | 68 | 111 | 113 | 15 | 23 | 0 | 4 | 0 | 0 | 0 | 298 | 149 | 1 | 1 | 1 | 1 | 1 | 1 | 1 | 1 | 0.933 | 1 | 1 | 1 | 0.013 |
| SE | 38438 | ENSG00000132549 | VPS13B | chr8 | + | 99641810 | 99642498 | 99577489 | 99577633 | 99661353 | 99661491 | 182 | 118 | 135 | 139 | 206 | 0 | 0 | 0 | 0 | 3 | 18 | 131 | 115 | 35 | 0 | 0 | 0 | 0 | 0 | 0 | 298 | 149 | 1 | 1 | 1 | 1 | 1 | 1 | 0.972 | 1 | 1 | 1 | 1 | NA | -0.006 |
| SE | 38439 | ENSG00000132549 | VPS13B | chr8 | + | 99776774 | 99776956 | 99766773 | 99766970 | 99778681 | 99779031 | 103 | 50 | 80 | 88 | 143 | 0 | 0 | 0 | 0 | 0 | 3 | 83 | 63 | 25 | 13 | 0 | 0 | 3 | 0 | 0 | 298 | 149 | 1 | 1 | 1 | 1 | 1 | 1 | 1 | 1 | 1 | 0.913 | 1 | 1 | 0.017 |
| SE | 38440 | ENSG00000132549 | VPS13B | chr8 | + | 99819411 | 99819582 | 99818712 | 99818888 | 99819920 | 99820122 | 200 | 148 | 141 | 198 | 221 | 1 | 0 | 0 | 0 | 1 | 60 | 157 | 140 | 81 | 11 | 0 | 0 | 0 | 0 | 0 | 298 | 149 | 1 | 1 | 0.99 | 1 | 1 | 1 | 0.991 | 1 | 1 | 1 | 1 | 1 | -0.004 |
| SE | 38441 | ENSG00000132549 | VPS13B | chr8 | + | 99819411 | 99819582 | 99818712 | 99818888 | 99821293 | 99821482 | 93 | 51 | 54 | 99 | 155 | 0 | 1 | 0 | 0 | 0 | 27 | 86 | 90 | 26 | 0 | 0 | 0 | 0 | 0 | 0 | 298 | 149 | 1 | 1 | 1 | 0.962 | 1 | 1 | 1 | 1 | 1 | 1 | 1 | NA | -0.008 |
| SE | 38442 | ENSG00000132549 | VPS13B | chr8 | + | 99819920 | 99820122 | 99818712 | 99818888 | 99821293 | 99821482 | 125 | 108 | 114 | 116 | 84 | 0 | 1 | 0 | 0 | 0 | 37 | 133 | 94 | 41 | 15 | 0 | 0 | 0 | 0 | 0 | 298 | 149 | 1 | 1 | 1 | 0.982 | 1 | 1 | 1 | 1 | 1 | 1 | 1 | 1 | -0.004 |
| SE | 38443 | ENSG00000132549 | VPS13B | chr8 | + | 99819920 | 99820122 | 99819411 | 99819582 | 99821293 | 99821482 | 231 | 205 | 201 | 216 | 149 | 0 | 0 | 0 | 1 | 0 | 70 | 207 | 145 | 96 | 26 | 0 | 3 | 1 | 0 | 0 | 298 | 149 | 1 | 1 | 1 | 1 | 1 | 0.991 | 1 | 1 | 0.972 | 0.986 | 1 | 1 | 0.007 |
| SE | 38444 | ENSG00000132549 | VPS13B | chr8 | + | 99832368 | 99832652 | 99823831 | 99823978 | 99835196 | 99835324 | 92 | 51 | 90 | 73 | 96 | 0 | 0 | 1 | 0 | 0 | 11 | 63 | 58 | 22 | 12 | 0 | 0 | 0 | 0 | 0 | 298 | 149 | 1 | 1 | 1 | 1 | 0.978 | 1 | 1 | 1 | 1 | 1 | 1 | 1 | -0.004 |
| SE | 38446 | ENSG00000132549 | VPS13B | chr8 | + | 99868288 | 99868465 | 99861775 | 99861946 | 99870784 | 99870887 | 40 | 25 | 45 | 40 | 35 | 0 | 0 | 0 | 0 | 0 | 10 | 27 | 20 | 12 | 0 | 0 | 2 | 0 | 7 | 0 | 298 | 149 | 1 | 1 | 1 | 1 | 1 | 1 | 1 | 1 | 0.871 | 1 | 0.462 | NA | 0.167 |
| SE | 38447 | ENSG00000132549 | VPS13B | chr8 | + | 99870784 | 99870887 | 99868288 | 99868465 | 99871447 | 99871697 | 24 | 17 | 32 | 28 | 32 | 0 | 2 | 0 | 0 | 0 | 0 | 25 | 17 | 6 | 4 | 0 | 0 | 0 | 0 | 0 | 252 | 149 | 1 | 1 | 1 | 0.834 | 1 | 1 | 1 | NA | 1 | 1 | 1 | 1 | -0.033 |
| SE | 38451 | ENSG00000116871 | MAP7D1 | chr1 | + | 36176198 | 36176581 | 36173363 | 36173478 | 36176696 | 36176842 | 114 | 58 | 73 | 66 | 12 | 0 | 0 | 0 | 1 | 1 | 43 | 77 | 45 | 75 | 171 | 0 | 0 | 0 | 4 | 0 | 298 | 149 | 1 | 1 | 1 | 1 | 1 | 0.971 | 0.857 | 1 | 1 | 1 | 0.904 | 1 | -0.015 |
| SE | 38453 | ENSG00000147251 | DOCK11 | chrX | + | 118543510 | 118543593 | 118542925 | 118543015 | 118545322 | 118545392 | 1172 | 867 | 786 | 1027 | 1099 | 0 | 0 | 0 | 0 | 0 | 164 | 789 | 588 | 175 | 202 | 0 | 2 | 1 | 0 | 5 | 232 | 149 | 1 | 1 | 1 | 1 | 1 | 1 | 1 | 1 | 0.996 | 0.997 | 1 | 0.963 | 0.009 |
| SE | 38454 | ENSG00000147251 | DOCK11 | chrX | + | 118545322 | 118545392 | 118543510 | 118543593 | 118546020 | 118546116 | 1096 | 831 | 773 | 991 | 1430 | 0 | 0 | 0 | 0 | 0 | 139 | 773 | 599 | 147 | 212 | 0 | 6 | 2 | 0 | 0 | 219 | 149 | 1 | 1 | 1 | 1 | 1 | 1 | 1 | 1 | 0.989 | 0.995 | 1 | 1 | 0.003 |
| SE | 38455 | ENSG00000147251 | DOCK11 | chrX | + | 118546020 | 118546116 | 118543510 | 118543593 | 118561382 | 118561517 | 214 | 181 | 168 | 202 | 263 | 0 | 0 | 0 | 0 | 0 | 55 | 180 | 120 | 60 | 28 | 0 | 3 | 0 | 1 | 0 | 245 | 149 | 1 | 1 | 1 | 1 | 1 | 1 | 1 | 1 | 0.973 | 1 | 0.973 | 1 | 0.011 |
| SE | 38456 | ENSG00000147251 | DOCK11 | chrX | + | 118561382 | 118561517 | 118546020 | 118546116 | 118566004 | 118566182 | 1073 | 845 | 736 | 1014 | 1353 | 1 | 0 | 0 | 0 | 0 | 229 | 794 | 625 | 219 | 100 | 0 | 0 | 0 | 0 | 0 | 284 | 149 | 1 | 1 | 0.998 | 1 | 1 | 1 | 1 | 1 | 1 | 1 | 1 | 1 | 0 |
| SE | 38457 | ENSG00000147251 | DOCK11 | chrX | + | 118566004 | 118566182 | 118561382 | 118561517 | 118566573 | 118566653 | 1116 | 869 | 739 | 1146 | 1543 | 3 | 0 | 1 | 0 | 0 | 206 | 846 | 773 | 208 | 68 | 0 | 0 | 6 | 0 | 0 | 298 | 149 | 1 | 1 | 0.995 | 1 | 0.997 | 1 | 1 | 1 | 1 | 0.985 | 1 | 1 | 0.001 |
| SE | 38458 | ENSG00000147251 | DOCK11 | chrX | + | 118584734 | 118584857 | 118580096 | 118580179 | 118588136 | 118588321 | 213 | 154 | 173 | 196 | 314 | 0 | 0 | 0 | 4 | 0 | 23 | 147 | 100 | 42 | 19 | 0 | 0 | 0 | 0 | 0 | 272 | 149 | 1 | 1 | 1 | 1 | 1 | 0.964 | 1 | 1 | 1 | 1 | 1 | 1 | -0.007 |
| SE | 38459 | ENSG00000147251 | DOCK11 | chrX | + | 118585040 | 118585117 | 118584734 | 118584857 | 118588136 | 118588321 | 730 | 512 | 560 | 608 | 1065 | 0 | 1 | 0 | 0 | 0 | 159 | 451 | 334 | 134 | 82 | 0 | 0 | 0 | 0 | 0 | 226 | 149 | 1 | 1 | 1 | 0.997 | 1 | 1 | 1 | 1 | 1 | 1 | 1 | 1 | -0.001 |
| SE | 38461 | ENSG00000147251 | DOCK11 | chrX | + | 118605237 | 118605356 | 118599138 | 118599228 | 118608071 | 118608141 | 628 | 473 | 460 | 546 | 578 | 0 | 3 | 0 | 2 | 1 | 35 | 387 | 231 | 57 | 67 | 0 | 0 | 0 | 0 | 0 | 268 | 149 | 1 | 1 | 1 | 0.989 | 1 | 0.993 | 0.997 | 1 | 1 | 1 | 1 | 1 | -0.004 |
| SE | 38463 | ENSG00000147251 | DOCK11 | chrX | + | 118618549 | 118618728 | 118615599 | 118615711 | 118624538 | 118624655 | 414 | 354 | 337 | 475 | 657 | 2 | 0 | 0 | 0 | 0 | 20 | 253 | 145 | 54 | 52 | 0 | 0 | 0 | 0 | 0 | 298 | 149 | 1 | 1 | 0.99 | 1 | 1 | 1 | 1 | 1 | 1 | 1 | 1 | 1 | -0.002 |
| SE | 38465 | ENSG00000147251 | DOCK11 | chrX | + | 118636345 | 118636412 | 118630378 | 118630490 | 118638079 | 118638127 | 903 | 804 | 675 | 803 | 1002 | 0 | 0 | 0 | 1 | 0 | 115 | 606 | 597 | 115 | 77 | 0 | 0 | 0 | 0 | 0 | 216 | 149 | 1 | 1 | 1 | 1 | 1 | 0.998 | 1 | 1 | 1 | 1 | 1 | 1 | 0 |
| SE | 38466 | ENSG00000147251 | DOCK11 | chrX | + | 118636345 | 118636412 | 118630378 | 118630490 | 118641189 | 118641305 | 204 | 158 | 147 | 165 | 232 | 0 | 1 | 0 | 1 | 4 | 21 | 104 | 148 | 11 | 17 | 0 | 0 | 0 | 0 | 0 | 216 | 149 | 1 | 1 | 1 | 0.991 | 1 | 0.991 | 0.976 | 1 | 1 | 1 | 1 | 1 | -0.008 |
| SE | 38468 | ENSG00000147251 | DOCK11 | chrX | + | 118638079 | 118638127 | 118636345 | 118636412 | 118641189 | 118641305 | 558 | 532 | 399 | 499 | 453 | 1 | 2 | 0 | 3 | 7 | 89 | 383 | 375 | 90 | 36 | 0 | 7 | 0 | 0 | 0 | 197 | 149 | 1 | 1 | 0.998 | 0.995 | 1 | 0.992 | 0.98 | 1 | 0.976 | 1 | 1 | 1 | -0.002 |
| SE | 38469 | ENSG00000147251 | DOCK11 | chrX | + | 118639434 | 118639577 | 118638079 | 118638127 | 118641189 | 118641305 | 790 | 557 | 670 | 678 | 1452 | 3 | 0 | 0 | 1 | 0 | 49 | 527 | 402 | 91 | 48 | 0 | 3 | 4 | 0 | 0 | 292 | 149 | 1 | 1 | 0.993 | 1 | 1 | 0.997 | 1 | 1 | 0.989 | 0.981 | 1 | 1 | 0.004 |
| SE | 38471 | ENSG00000147251 | DOCK11 | chrX | + | 118641189 | 118641305 | 118638079 | 118638127 | 118648944 | 118649127 | 28 | 13 | 15 | 27 | 2 | 0 | 0 | 0 | 0 | 0 | 0 | 16 | 12 | 2 | 1 | 0 | 6 | 0 | 0 | 0 | 265 | 149 | 1 | 1 | 1 | 1 | 1 | 1 | 1 | NA | 0.6 | 1 | 1 | 1 | 0.1 |
| SE | 38472 | ENSG00000147251 | DOCK11 | chrX | + | 118641189 | 118641305 | 118639434 | 118639577 | 118648944 | 118649127 | 514 | 325 | 399 | 453 | 760 | 0 | 0 | 2 | 0 | 0 | 34 | 311 | 276 | 65 | 25 | 0 | 0 | 0 | 0 | 0 | 265 | 149 | 1 | 1 | 1 | 1 | 0.991 | 1 | 1 | 1 | 1 | 1 | 1 | 1 | -0.002 |
| SE | 38475 | ENSG00000147251 | DOCK11 | chrX | + | 118654601 | 118654817 | 118651963 | 118652077 | 118654903 | 118654961 | 538 | 413 | 436 | 531 | 816 | 0 | 1 | 0 | 0 | 0 | 29 | 311 | 194 | 39 | 42 | 0 | 0 | 0 | 0 | 0 | 298 | 149 | 1 | 1 | 1 | 0.995 | 1 | 1 | 1 | 1 | 1 | 1 | 1 | 1 | -0.001 |
| SE | 38476 | ENSG00000147251 | DOCK11 | chrX | + | 118654903 | 118654961 | 118651963 | 118652077 | 118662685 | 118662792 | 75 | 49 | 64 | 56 | 97 | 1 | 3 | 2 | 6 | 1 | 5 | 33 | 59 | 13 | 0 | 0 | 5 | 3 | 7 | 0 | 207 | 149 | 1 | 1 | 0.982 | 0.922 | 0.958 | 0.87 | 0.986 | 1 | 0.826 | 0.934 | 0.572 | NA | 0.111 |
| SE | 38479 | ENSG00000147251 | DOCK11 | chrX | + | 118685519 | 118685537 | 118683102 | 118683217 | 118685687 | 118686147 | 0 | 1 | 0 | 3 | 3 | 325 | 146 | 164 | 220 | 336 | 0 | 2 | 4 | 0 | 0 | 18 | 89 | 96 | 12 | 0 | 167 | 149 | 1 | 1 | 0 | 0.006 | 0 | 0.012 | 0.008 | 0 | 0.02 | 0.036 | 0 | NA | -0.009 |
| SE | 38480 | ENSG00000173930 | SLCO4C1 | chr5 | - | 102270623 | 102270806 | 102263683 | 102263780 | 102291342 | 102291606 | 224 | 92 | 81 | 116 | 116 | 4 | 0 | 0 | 0 | 0 | 52 | 129 | 91 | 95 | 23 | 0 | 0 | 0 | 0 | 0 | 298 | 149 | 1 | 1 | 0.966 | 1 | 1 | 1 | 1 | 1 | 1 | 1 | 1 | 1 | -0.007 |
| SE | 38481 | ENSG00000173930 | SLCO4C1 | chr5 | - | 102291342 | 102291606 | 102270623 | 102270806 | 102295907 | 102296284 | 179 | 80 | 75 | 110 | 93 | 3 | 0 | 0 | 1 | 0 | 47 | 116 | 78 | 85 | 23 | 0 | 5 | 0 | 0 | 0 | 298 | 149 | 1 | 1 | 0.968 | 1 | 1 | 0.982 | 1 | 1 | 0.921 | 1 | 1 | 1 | 0.006 |
| SE | 38482 | ENSG00000198793 | MTOR | chr1 | - | 11111513 | 11111753 | 11109648 | 11109729 | 11112851 | 11112917 | 0 | 0 | 0 | 2 | 0 | 74 | 29 | 70 | 77 | 58 | 0 | 0 | 0 | 0 | 0 | 93 | 50 | 27 | 80 | 66 | 298 | 149 | 1 | 1 | 0 | 0 | 0 | 0.013 | 0 | 0 | 0 | 0 | 0 | 0 | 0.003 |
| SE | 38483 | ENSG00000198793 | MTOR | chr1 | - | 11115395 | 11115468 | 11114812 | 11114887 | 11121245 | 11121368 | 42 | 24 | 52 | 40 | 24 | 0 | 0 | 0 | 0 | 0 | 111 | 46 | 22 | 70 | 57 | 0 | 0 | 1 | 0 | 0 | 222 | 149 | 1 | 1 | 1 | 1 | 1 | 1 | 1 | 1 | 1 | 0.937 | 1 | 1 | 0.013 |
| SE | 38485 | ENSG00000198793 | MTOR | chr1 | - | 11124497 | 11124633 | 11121978 | 11122126 | 11126621 | 11126796 | 181 | 92 | 164 | 231 | 165 | 1 | 0 | 0 | 0 | 0 | 104 | 178 | 111 | 99 | 128 | 0 | 0 | 0 | 0 | 0 | 285 | 149 | 1 | 1 | 0.99 | 1 | 1 | 1 | 1 | 1 | 1 | 1 | 1 | 1 | -0.002 |
| SE | 38486 | ENSG00000198793 | MTOR | chr1 | - | 11126621 | 11126796 | 11124497 | 11124633 | 11127009 | 11127144 | 138 | 82 | 136 | 173 | 162 | 0 | 0 | 0 | 1 | 0 | 119 | 116 | 68 | 116 | 85 | 0 | 2 | 0 | 0 | 0 | 298 | 149 | 1 | 1 | 1 | 1 | 1 | 0.989 | 1 | 1 | 0.967 | 1 | 1 | 1 | 0.004 |
| SE | 38488 | ENSG00000198793 | MTOR | chr1 | - | 11144967 | 11145045 | 11144647 | 11144755 | 11146675 | 11146791 | 58 | 29 | 39 | 67 | 31 | 0 | 0 | 0 | 0 | 0 | 49 | 83 | 30 | 55 | 46 | 0 | 1 | 0 | 0 | 0 | 227 | 149 | 1 | 1 | 1 | 1 | 1 | 1 | 1 | 1 | 0.982 | 1 | 1 | 1 | 0.004 |
| SE | 38489 | ENSG00000198793 | MTOR | chr1 | - | 11146675 | 11146791 | 11144967 | 11145045 | 11150125 | 11150226 | 72 | 50 | 53 | 69 | 25 | 0 | 0 | 1 | 0 | 0 | 38 | 85 | 48 | 77 | 54 | 0 | 0 | 0 | 0 | 0 | 265 | 149 | 1 | 1 | 1 | 1 | 0.968 | 1 | 1 | 1 | 1 | 1 | 1 | 1 | -0.006 |
| SE | 38493 | ENSG00000198793 | MTOR | chr1 | - | 11199257 | 11199403 | 11167441 | 11167517 | 11199540 | 11199703 | 97 | 97 | 87 | 102 | 138 | 0 | 1 | 0 | 0 | 0 | 141 | 129 | 136 | 120 | 126 | 1 | 0 | 0 | 0 | 0 | 295 | 149 | 1 | 1 | 1 | 0.98 | 1 | 1 | 1 | 0.986 | 1 | 1 | 1 | 1 | -0.001 |
| SE | 38494 | ENSG00000198793 | MTOR | chr1 | - | 11204560 | 11204703 | 11199540 | 11199703 | 11209311 | 11209458 | 125 | 78 | 104 | 96 | 81 | 0 | 0 | 0 | 0 | 0 | 96 | 148 | 95 | 193 | 158 | 0 | 0 | 4 | 0 | 0 | 292 | 149 | 1 | 1 | 1 | 1 | 1 | 1 | 1 | 1 | 1 | 0.924 | 1 | 1 | 0.015 |
| SE | 38495 | ENSG00000198793 | MTOR | chr1 | - | 11216147 | 11216234 | 11213398 | 11213566 | 11228667 | 11228918 | 123 | 50 | 112 | 90 | 110 | 0 | 0 | 0 | 0 | 0 | 163 | 95 | 79 | 62 | 166 | 0 | 0 | 3 | 0 | 0 | 236 | 149 | 1 | 1 | 1 | 1 | 1 | 1 | 1 | 1 | 1 | 0.943 | 1 | 1 | 0.011 |
| SE | 38496 | ENSG00000198793 | MTOR | chr1 | - | 11230924 | 11231054 | 11228667 | 11228918 | 11232435 | 11232528 | 42 | 28 | 34 | 68 | 79 | 1 | 0 | 0 | 0 | 3 | 120 | 70 | 25 | 27 | 67 | 0 | 0 | 0 | 0 | 0 | 279 | 149 | 1 | 1 | 0.957 | 1 | 1 | 1 | 0.934 | 1 | 1 | 1 | 1 | 1 | -0.022 |
| SE | 38498 | ENSG00000198793 | MTOR | chr1 | - | 11238401 | 11238617 | 11237842 | 11238048 | 11240302 | 11240547 | 114 | 52 | 74 | 105 | 81 | 0 | 0 | 0 | 0 | 0 | 57 | 107 | 63 | 77 | 84 | 0 | 0 | 0 | 3 | 0 | 298 | 149 | 1 | 1 | 1 | 1 | 1 | 1 | 1 | 1 | 1 | 1 | 0.928 | 1 | 0.014 |
| SE | 38499 | ENSG00000198793 | MTOR | chr1 | - | 11241552 | 11241681 | 11240302 | 11240547 | 11243113 | 11243300 | 99 | 62 | 77 | 83 | 154 | 0 | 0 | 0 | 0 | 0 | 43 | 93 | 71 | 63 | 130 | 0 | 0 | 0 | 3 | 0 | 278 | 149 | 1 | 1 | 1 | 1 | 1 | 1 | 1 | 1 | 1 | 1 | 0.918 | 1 | 0.016 |
| SE | 38501 | ENSG00000198793 | MTOR | chr1 | - | 11253838 | 11253973 | 11247818 | 11248094 | 11255991 | 11256192 | 144 | 79 | 130 | 129 | 124 | 0 | 0 | 0 | 1 | 0 | 257 | 200 | 162 | 126 | 167 | 0 | 0 | 0 | 0 | 0 | 284 | 149 | 1 | 1 | 1 | 1 | 1 | 0.985 | 1 | 1 | 1 | 1 | 1 | 1 | -0.003 |
| SE | 38503 | ENSG00000117298 | ECE1 | chr1 | - | 21219958 | 21220131 | 21217280 | 21217978 | 21221746 | 21221842 | 120 | 57 | 115 | 94 | 102 | 0 | 0 | 2 | 0 | 0 | 164 | 155 | 115 | 153 | 304 | 0 | 0 | 0 | 0 | 0 | 298 | 149 | 1 | 1 | 1 | 1 | 0.966 | 1 | 1 | 1 | 1 | 1 | 1 | 1 | -0.007 |
| SE | 38505 | ENSG00000117298 | ECE1 | chr1 | - | 21221746 | 21221842 | 21217246 | 21220131 | 21227158 | 21227226 | 91 | 39 | 75 | 67 | 75 | 1 | 0 | 0 | 0 | 0 | 137 | 99 | 99 | 110 | 256 | 3 | 0 | 0 | 0 | 0 | 245 | 149 | 1 | 1 | 0.982 | 1 | 1 | 1 | 1 | 0.965 | 1 | 1 | 1 | 1 | 0.003 |
| SE | 38506 | ENSG00000117298 | ECE1 | chr1 | - | 21225249 | 21225440 | 21219893 | 21220131 | 21227158 | 21227226 | 180 | 136 | 180 | 206 | 148 | 1 | 0 | 0 | 0 | 0 | 498 | 307 | 279 | 367 | 808 | 3 | 0 | 0 | 0 | 0 | 298 | 149 | 1 | 1 | 0.989 | 1 | 1 | 1 | 1 | 0.988 | 1 | 1 | 1 | 1 | 0 |
| SE | 38508 | ENSG00000117298 | ECE1 | chr1 | - | 21227158 | 21227226 | 21225249 | 21225440 | 21227930 | 21228041 | 211 | 159 | 238 | 295 | 174 | 2 | 0 | 0 | 0 | 0 | 697 | 390 | 360 | 491 | 1115 | 3 | 0 | 0 | 0 | 0 | 217 | 149 | 1 | 1 | 0.986 | 1 | 1 | 1 | 1 | 0.994 | 1 | 1 | 1 | 1 | -0.002 |
| SE | 38509 | ENSG00000117298 | ECE1 | chr1 | - | 21227158 | 21227226 | 21225249 | 21225440 | 21233557 | 21233661 | 67 | 52 | 63 | 89 | 59 | 2 | 0 | 0 | 0 | 0 | 166 | 87 | 90 | 133 | 295 | 0 | 0 | 0 | 0 | 0 | 217 | 149 | 1 | 1 | 0.958 | 1 | 1 | 1 | 1 | 1 | 1 | 1 | 1 | 1 | -0.008 |
| SE | 38510 | ENSG00000117298 | ECE1 | chr1 | - | 21227930 | 21228041 | 21225249 | 21225440 | 21233557 | 21233661 | 170 | 120 | 162 | 145 | 171 | 2 | 0 | 0 | 0 | 0 | 409 | 271 | 250 | 247 | 593 | 0 | 0 | 0 | 0 | 0 | 260 | 149 | 1 | 1 | 0.98 | 1 | 1 | 1 | 1 | 1 | 1 | 1 | 1 | 1 | -0.004 |
| SE | 38511 | ENSG00000117298 | ECE1 | chr1 | - | 21227930 | 21228041 | 21227158 | 21227226 | 21233557 | 21233661 | 319 | 229 | 339 | 356 | 288 | 0 | 1 | 0 | 0 | 0 | 938 | 586 | 533 | 627 | 1438 | 0 | 0 | 0 | 0 | 0 | 260 | 149 | 1 | 1 | 1 | 0.992 | 1 | 1 | 1 | 1 | 1 | 1 | 1 | 1 | -0.002 |
| SE | 38514 | ENSG00000117298 | ECE1 | chr1 | - | 21244988 | 21245103 | 21238133 | 21238244 | 21247220 | 21247363 | 216 | 154 | 257 | 261 | 279 | 0 | 0 | 0 | 0 | 0 | 583 | 523 | 372 | 445 | 726 | 0 | 6 | 0 | 0 | 0 | 264 | 149 | 1 | 1 | 1 | 1 | 1 | 1 | 1 | 1 | 0.98 | 1 | 1 | 1 | 0.004 |
| SE | 38516 | ENSG00000117298 | ECE1 | chr1 | - | 21255946 | 21256138 | 21247220 | 21247363 | 21257524 | 21257590 | 289 | 234 | 273 | 305 | 372 | 0 | 0 | 0 | 0 | 0 | 986 | 555 | 387 | 526 | 1092 | 0 | 2 | 0 | 5 | 0 | 298 | 149 | 1 | 1 | 1 | 1 | 1 | 1 | 1 | 1 | 0.993 | 1 | 0.981 | 1 | 0.005 |
| SE | 38517 | ENSG00000117298 | ECE1 | chr1 | - | 21258692 | 21258839 | 21257524 | 21257590 | 21260270 | 21260392 | 272 | 198 | 266 | 313 | 364 | 0 | 0 | 0 | 1 | 0 | 835 | 586 | 409 | 584 | 1187 | 0 | 0 | 0 | 0 | 0 | 296 | 149 | 1 | 1 | 1 | 1 | 1 | 0.994 | 1 | 1 | 1 | 1 | 1 | 1 | -0.001 |
| SE | 38519 | ENSG00000117298 | ECE1 | chr1 | - | 21279190 | 21279332 | 21272698 | 21272911 | 21290069 | 21290156 | 67 | 38 | 80 | 74 | 21 | 0 | 0 | 0 | 0 | 0 | 68 | 114 | 73 | 148 | 228 | 0 | 0 | 1 | 0 | 0 | 291 | 149 | 1 | 1 | 1 | 1 | 1 | 1 | 1 | 1 | 1 | 0.974 | 1 | 1 | 0.005 |
| SE | 38520 | ENSG00000130024 | PHF10 | chr6 | - | 169705132 | 169705321 | 169703901 | 169704088 | 169705615 | 169705724 | 356 | 267 | 338 | 379 | 302 | 3 | 0 | 0 | 0 | 1 | 190 | 219 | 214 | 217 | 174 | 0 | 6 | 0 | 0 | 0 | 298 | 149 | 1 | 1 | 0.983 | 1 | 1 | 1 | 0.993 | 1 | 0.948 | 1 | 1 | 1 | 0.006 |
| SE | 38523 | ENSG00000130024 | PHF10 | chr6 | - | 169717822 | 169717900 | 169715954 | 169716088 | 169718787 | 169718918 | 324 | 330 | 366 | 319 | 846 | 1 | 2 | 3 | 2 | 8 | 265 | 296 | 299 | 195 | 144 | 0 | 9 | 1 | 0 | 0 | 227 | 149 | 1 | 1 | 0.995 | 0.991 | 0.988 | 0.991 | 0.986 | 1 | 0.956 | 0.995 | 1 | 1 | 0 |
| SE | 38524 | ENSG00000130024 | PHF10 | chr6 | - | 169717822 | 169717900 | 169715954 | 169716088 | 169721004 | 169721111 | 258 | 275 | 319 | 263 | 727 | 0 | 0 | 0 | 0 | 0 | 218 | 246 | 256 | 164 | 131 | 0 | 0 | 0 | 1 | 0 | 227 | 149 | 1 | 1 | 1 | 1 | 1 | 1 | 1 | 1 | 1 | 1 | 0.991 | 1 | 0.002 |
| SE | 38525 | ENSG00000130024 | PHF10 | chr6 | - | 169717822 | 169717900 | 169715954 | 169716088 | 169723844 | 169723931 | 258 | 275 | 318 | 263 | 727 | 0 | 0 | 1 | 0 | 0 | 218 | 246 | 258 | 164 | 131 | 0 | 0 | 0 | 0 | 0 | 227 | 149 | 1 | 1 | 1 | 1 | 0.995 | 1 | 1 | 1 | 1 | 1 | 1 | 1 | -0.001 |
| SE | 38526 | ENSG00000130024 | PHF10 | chr6 | - | 169717822 | 169717906 | 169715954 | 169716088 | 169718787 | 169718918 | 458 | 462 | 551 | 454 | 1290 | 1 | 2 | 3 | 2 | 8 | 424 | 513 | 476 | 269 | 256 | 0 | 9 | 1 | 0 | 0 | 233 | 149 | 1 | 1 | 0.997 | 0.993 | 0.992 | 0.993 | 0.99 | 1 | 0.973 | 0.997 | 1 | 1 | -0.001 |
| SE | 38527 | ENSG00000130024 | PHF10 | chr6 | - | 169717822 | 169717906 | 169715954 | 169716088 | 169721004 | 169721111 | 261 | 277 | 325 | 265 | 733 | 0 | 0 | 0 | 0 | 0 | 218 | 253 | 256 | 169 | 131 | 0 | 0 | 0 | 1 | 0 | 233 | 149 | 1 | 1 | 1 | 1 | 1 | 1 | 1 | 1 | 1 | 1 | 0.991 | 1 | 0.002 |
| SE | 38528 | ENSG00000130024 | PHF10 | chr6 | - | 169718787 | 169718918 | 169715954 | 169716088 | 169721004 | 169721111 | 121 | 105 | 127 | 133 | 221 | 0 | 0 | 0 | 0 | 0 | 71 | 139 | 83 | 47 | 43 | 0 | 0 | 0 | 1 | 0 | 280 | 149 | 1 | 1 | 1 | 1 | 1 | 1 | 1 | 1 | 1 | 1 | 0.962 | 1 | 0.008 |
| SE | 38529 | ENSG00000130024 | PHF10 | chr6 | - | 169718787 | 169718918 | 169717822 | 169717900 | 169721004 | 169721111 | 185 | 158 | 172 | 187 | 332 | 0 | 0 | 1 | 0 | 0 | 118 | 180 | 124 | 78 | 56 | 0 | 0 | 0 | 0 | 0 | 280 | 149 | 1 | 1 | 1 | 1 | 0.989 | 1 | 1 | 1 | 1 | 1 | 1 | 1 | -0.002 |
| SE | 38530 | ENSG00000130024 | PHF10 | chr6 | - | 169718787 | 169718918 | 169717822 | 169717906 | 169721004 | 169721111 | 320 | 292 | 347 | 329 | 773 | 0 | 0 | 0 | 1 | 0 | 277 | 391 | 295 | 156 | 168 | 0 | 0 | 0 | 4 | 0 | 280 | 149 | 1 | 1 | 1 | 1 | 1 | 0.994 | 1 | 1 | 1 | 1 | 0.954 | 1 | 0.008 |
| SE | 38531 | ENSG00000130024 | PHF10 | chr6 | - | 169721004 | 169721111 | 169715954 | 169716088 | 169723844 | 169723931 | 4 | 1 | 9 | 1 | 0 | 0 | 0 | 1 | 0 | 0 | 5 | 5 | 2 | 1 | 0 | 0 | 0 | 0 | 0 | 0 | 256 | 149 | 1 | 1 | 1 | 1 | 0.84 | 1 | NA | 1 | 1 | 1 | 1 | NA | -0.04 |
| SE | 38532 | ENSG00000130024 | PHF10 | chr6 | - | 169721004 | 169721111 | 169717822 | 169717900 | 169723844 | 169723931 | 4 | 1 | 10 | 1 | 0 | 0 | 0 | 0 | 0 | 0 | 5 | 5 | 2 | 0 | 0 | 0 | 0 | 2 | 0 | 0 | 256 | 149 | 1 | 1 | 1 | 1 | 1 | 1 | NA | 1 | 1 | 0.368 | NA | NA | 0.211 |
| SE | 38540 | ENSG00000274049 | INO80B-WBP1 | chr2 | + | 74457333 | 74457538 | 74456102 | 74456272 | 74459642 | 74459745 | 71 | 51 | 78 | 71 | 73 | 0 | 0 | 1 | 0 | 0 | 35 | 36 | 36 | 34 | 57 | 1 | 0 | 0 | 0 | 0 | 298 | 149 | 1 | 1 | 1 | 1 | 0.975 | 1 | 1 | 0.946 | 1 | 1 | 1 | 1 | 0.006 |
| SE | 38541 | ENSG00000274049 | INO80B-WBP1 | chr2 | + | 74457333 | 74457860 | 74456102 | 74456272 | 74459642 | 74459745 | 71 | 51 | 78 | 71 | 73 | 0 | 0 | 1 | 0 | 0 | 35 | 36 | 36 | 34 | 57 | 1 | 0 | 0 | 0 | 0 | 298 | 149 | 1 | 1 | 1 | 1 | 0.975 | 1 | 1 | 0.946 | 1 | 1 | 1 | 1 | 0.006 |
| SE | 38543 | ENSG00000180098 | TRNAU1AP | chr1 | + | 28560632 | 28560732 | 28553086 | 28553737 | 28561112 | 28561260 | 50 | 20 | 49 | 40 | 35 | 3 | 0 | 0 | 0 | 0 | 59 | 28 | 18 | 44 | 57 | 0 | 0 | 0 | 0 | 0 | 249 | 149 | 1 | 1 | 0.909 | 1 | 1 | 1 | 1 | 1 | 1 | 1 | 1 | 1 | -0.018 |
| SE | 38544 | ENSG00000180098 | TRNAU1AP | chr1 | + | 28560632 | 28561398 | 28553639 | 28553737 | 28564702 | 28564834 | 57 | 23 | 61 | 52 | 51 | 0 | 0 | 0 | 0 | 0 | 67 | 34 | 29 | 60 | 90 | 0 | 0 | 0 | 2 | 0 | 298 | 149 | 1 | 1 | 1 | 1 | 1 | 1 | 1 | 1 | 1 | 1 | 0.938 | 1 | 0.012 |
| SE | 38546 | ENSG00000180098 | TRNAU1AP | chr1 | + | 28567293 | 28567413 | 28564702 | 28564834 | 28571175 | 28571338 | 154 | 66 | 107 | 105 | 112 | 0 | 0 | 0 | 3 | 0 | 99 | 74 | 63 | 112 | 260 | 0 | 0 | 0 | 0 | 0 | 269 | 149 | 1 | 1 | 1 | 1 | 1 | 0.951 | 1 | 1 | 1 | 1 | 1 | 1 | -0.01 |
| SE | 38547 | ENSG00000180098 | TRNAU1AP | chr1 | + | 28567293 | 28567413 | 28564702 | 28564834 | 28571866 | 28571900 | 76 | 35 | 53 | 63 | 34 | 0 | 0 | 1 | 0 | 0 | 52 | 32 | 44 | 46 | 126 | 0 | 0 | 0 | 0 | 0 | 269 | 149 | 1 | 1 | 1 | 1 | 0.967 | 1 | 1 | 1 | 1 | 1 | 1 | 1 | -0.007 |
| SE | 38548 | ENSG00000180098 | TRNAU1AP | chr1 | + | 28571175 | 28571338 | 28564702 | 28564834 | 28571866 | 28571900 | 46 | 17 | 39 | 21 | 43 | 0 | 0 | 1 | 0 | 0 | 52 | 31 | 16 | 33 | 65 | 0 | 0 | 0 | 0 | 0 | 298 | 149 | 1 | 1 | 1 | 1 | 0.951 | 1 | 1 | 1 | 1 | 1 | 1 | 1 | -0.01 |
| SE | 38550 | ENSG00000180098 | TRNAU1AP | chr1 | + | 28571866 | 28571900 | 28571175 | 28571338 | 28577499 | 28577744 | 47 | 15 | 38 | 22 | 46 | 1 | 0 | 0 | 0 | 0 | 35 | 32 | 22 | 42 | 68 | 0 | 0 | 0 | 0 | 0 | 183 | 149 | 1 | 1 | 0.975 | 1 | 1 | 1 | 1 | 1 | 1 | 1 | 1 | 1 | -0.005 |
| SE | 38552 | ENSG00000165801 | ARHGEF40 | chr14 | + | 21073044 | 21073178 | 21070367 | 21070399 | 21076359 | 21076456 | 3 | 0 | 1 | 2 | 0 | 0 | 0 | 0 | 0 | 0 | 0 | 0 | 0 | 0 | 6 | 0 | 0 | 0 | 1 | 0 | 283 | 149 | 1 | 1 | 1 | NA | 1 | 1 | NA | NA | NA | NA | 0 | 1 | 0.5 |
| SE | 38553 | ENSG00000165801 | ARHGEF40 | chr14 | + | 21073044 | 21073242 | 21070272 | 21070399 | 21075331 | 21075499 | 3 | 0 | 1 | 2 | 0 | 0 | 0 | 0 | 0 | 0 | 0 | 0 | 0 | 4 | 6 | 0 | 0 | 0 | 0 | 2 | 298 | 149 | 1 | 1 | 1 | NA | 1 | 1 | NA | NA | NA | NA | 1 | 0.6 | 0.2 |
| SE | 38554 | ENSG00000165801 | ARHGEF40 | chr14 | + | 21073931 | 21074991 | 21073044 | 21073242 | 21075331 | 21075499 | 14 | 8 | 7 | 17 | 9 | 0 | 0 | 0 | 1 | 0 | 16 | 10 | 7 | 17 | 24 | 0 | 0 | 0 | 0 | 0 | 298 | 149 | 1 | 1 | 1 | 1 | 1 | 0.895 | 1 | 1 | 1 | 1 | 1 | 1 | -0.021 |
| SE | 38555 | ENSG00000165801 | ARHGEF40 | chr14 | + | 21073931 | 21075180 | 21073044 | 21073242 | 21075331 | 21075499 | 149 | 98 | 71 | 137 | 93 | 0 | 0 | 0 | 1 | 0 | 513 | 120 | 121 | 278 | 597 | 0 | 0 | 0 | 0 | 0 | 298 | 149 | 1 | 1 | 1 | 1 | 1 | 0.986 | 1 | 1 | 1 | 1 | 1 | 1 | -0.003 |
| SE | 38556 | ENSG00000165801 | ARHGEF40 | chr14 | + | 21075331 | 21075499 | 21070290 | 21070399 | 21076359 | 21076456 | 4 | 2 | 1 | 4 | 0 | 0 | 0 | 0 | 0 | 0 | 0 | 6 | 0 | 3 | 4 | 0 | 0 | 0 | 1 | 0 | 298 | 149 | 1 | 1 | 1 | 1 | 1 | 1 | NA | NA | 1 | NA | 0.6 | 1 | 0.133 |
| SE | 38559 | ENSG00000165801 | ARHGEF40 | chr14 | + | 21076773 | 21076890 | 21076535 | 21076643 | 21078176 | 21078207 | 88 | 29 | 42 | 67 | 90 | 0 | 0 | 0 | 0 | 0 | 301 | 59 | 54 | 205 | 330 | 0 | 0 | 0 | 3 | 2 | 266 | 149 | 1 | 1 | 1 | 1 | 1 | 1 | 1 | 1 | 1 | 1 | 0.975 | 0.989 | 0.007 |
| SE | 38560 | ENSG00000165801 | ARHGEF40 | chr14 | + | 21078372 | 21078488 | 21078176 | 21078272 | 21078883 | 21079010 | 50 | 27 | 29 | 39 | 50 | 1 | 0 | 0 | 0 | 0 | 194 | 53 | 48 | 102 | 243 | 0 | 4 | 0 | 0 | 0 | 265 | 149 | 1 | 1 | 0.966 | 1 | 1 | 1 | 1 | 1 | 0.882 | 1 | 1 | 1 | 0.017 |
| SE | 38563 | ENSG00000198925 | ATG9A | chr2 | - | 219222049 | 219222167 | 219221079 | 219221302 | 219222271 | 219222450 | 42 | 16 | 24 | 28 | 8 | 0 | 0 | 0 | 0 | 0 | 4 | 16 | 17 | 25 | 68 | 0 | 0 | 2 | 0 | 0 | 267 | 149 | 1 | 1 | 1 | 1 | 1 | 1 | 1 | 1 | 1 | 0.826 | 1 | 1 | 0.035 |
| SE | 38565 | ENSG00000198925 | ATG9A | chr2 | - | 219227921 | 219228053 | 219227769 | 219227813 | 219229009 | 219229342 | 28 | 14 | 27 | 19 | 14 | 0 | 0 | 0 | 2 | 0 | 16 | 8 | 5 | 8 | 15 | 0 | 0 | 0 | 0 | 0 | 281 | 149 | 1 | 1 | 1 | 1 | 1 | 0.834 | 1 | 1 | 1 | 1 | 1 | 1 | -0.033 |
| SE | 38570 | ENSG00000198925 | ATG9A | chr2 | - | 219228433 | 219228482 | 219227921 | 219228053 | 219229009 | 219229325 | 7 | 3 | 5 | 3 | 1 | 0 | 1 | 1 | 4 | 0 | 0 | 8 | 0 | 1 | 13 | 0 | 0 | 0 | 0 | 0 | 198 | 149 | 1 | 1 | 1 | 0.693 | 0.79 | 0.361 | 1 | NA | 1 | NA | 1 | 1 | -0.231 |
| SE | 38571 | ENSG00000198925 | ATG9A | chr2 | - | 219228433 | 219228482 | 219227921 | 219228053 | 219229534 | 219229639 | 7 | 3 | 6 | 3 | 1 | 5 | 4 | 0 | 2 | 0 | 0 | 10 | 0 | 3 | 13 | 0 | 1 | 3 | 0 | 0 | 198 | 149 | 1 | 1 | 0.513 | 0.361 | 1 | 0.53 | 1 | NA | 0.883 | 0 | 1 | 1 | -0.04 |
| SE | 38572 | ENSG00000198925 | ATG9A | chr2 | - | 219228433 | 219228485 | 219227769 | 219227813 | 219229009 | 219229174 | 0 | 3 | 0 | 1 | 0 | 0 | 0 | 0 | 2 | 0 | 1 | 2 | 2 | 0 | 3 | 0 | 0 | 0 | 0 | 0 | 201 | 149 | 1 | 1 | NA | 1 | NA | 0.27 | NA | 1 | 1 | 1 | NA | 1 | -0.365 |
| SE | 38574 | ENSG00000198925 | ATG9A | chr2 | - | 219228433 | 219228485 | 219227921 | 219228053 | 219229009 | 219229174 | 9 | 8 | 7 | 3 | 2 | 0 | 1 | 1 | 4 | 0 | 0 | 14 | 2 | 1 | 24 | 0 | 0 | 0 | 0 | 0 | 201 | 149 | 1 | 1 | 1 | 0.856 | 0.838 | 0.357 | 1 | NA | 1 | 1 | 1 | 1 | -0.19 |
| SE | 38575 | ENSG00000198925 | ATG9A | chr2 | - | 219228433 | 219228485 | 219227921 | 219228053 | 219229534 | 219229594 | 9 | 4 | 8 | 2 | 1 | 5 | 4 | 0 | 2 | 0 | 0 | 8 | 4 | 2 | 22 | 0 | 1 | 3 | 0 | 0 | 201 | 149 | 1 | 1 | 0.572 | 0.426 | 1 | 0.426 | 1 | NA | 0.856 | 0.497 | 1 | 1 | -0.153 |
| SE | 38576 | ENSG00000198925 | ATG9A | chr2 | - | 219228433 | 219228485 | 219227921 | 219228079 | 219229534 | 219229594 | 0 | 0 | 0 | 0 | 0 | 0 | 0 | 0 | 1 | 0 | 0 | 5 | 0 | 0 | 0 | 0 | 0 | 0 | 1 | 9 | 201 | 149 | 1 | 1 | NA | NA | NA | 0 | NA | NA | 1 | NA | NA | NA | -1 |
| SE | 38578 | ENSG00000198925 | ATG9A | chr2 | - | 219229009 | 219229174 | 219228433 | 219228485 | 219229534 | 219229618 | 3 | 8 | 4 | 3 | 1 | 2 | 1 | 3 | 1 | 0 | 0 | 8 | 4 | 0 | 11 | 0 | 0 | 8 | 1 | 9 | 298 | 149 | 1 | 1 | 0.429 | 0.8 | 0.4 | 0.6 | 1 | NA | 1 | 0.2 | 0 | 0.379 | 0.251 |
| SE | 38580 | ENSG00000144746 | ARL6IP5 | chr3 | + | 69089811 | 69089960 | 69085041 | 69085223 | 69101838 | 69101894 | 2 | 2 | 1 | 4 | 14 | 614 | 408 | 381 | 559 | 488 | 0 | 3 | 1 | 0 | 15 | 1030 | 701 | 445 | 616 | 907 | 298 | 149 | 1 | 1 | 0.002 | 0.002 | 0.001 | 0.004 | 0.014 | 0 | 0.002 | 0.001 | 0 | 0.008 | 0.002 |
| SE | 38581 | ENSG00000144746 | ARL6IP5 | chr3 | + | 69099971 | 69100069 | 69085014 | 69085223 | 69101838 | 69102056 | 2 | 0 | 1 | 0 | 0 | 614 | 408 | 381 | 559 | 488 | 0 | 0 | 0 | 0 | 4 | 1030 | 701 | 445 | 616 | 907 | 247 | 149 | 1 | 1 | 0.002 | 0 | 0.002 | 0 | 0 | 0 | 0 | 0 | 0 | 0.003 | 0 |
| SE | 38582 | ENSG00000144746 | ARL6IP5 | chr3 | + | 69101838 | 69102056 | 69084936 | 69085223 | 69104463 | 69104489 | 1435 | 1101 | 970 | 1407 | 1277 | 4 | 0 | 0 | 3 | 0 | 2785 | 1853 | 1151 | 1642 | 2463 | 0 | 0 | 8 | 4 | 2 | 298 | 149 | 1 | 1 | 0.994 | 1 | 1 | 0.996 | 1 | 1 | 1 | 0.986 | 0.995 | 0.998 | 0.002 |
| SE | 38583 | ENSG00000196878 | LAMB3 | chr1 | - | 209622981 | 209623179 | 209622535 | 209622680 | 209623504 | 209623725 | 0 | 0 | 0 | 1 | 0 | 0 | 0 | 0 | 1 | 0 | 0 | 0 | 0 | 0 | 0 | 0 | 0 | 1 | 0 | 0 | 298 | 149 | 1 | 1 | NA | NA | NA | 0.333 | NA | NA | NA | 0 | NA | NA | 0.333 |
| SE | 38584 | ENSG00000196878 | LAMB3 | chr1 | - | 209634446 | 209634638 | 209633069 | 209633133 | 209637907 | 209637981 | 0 | 0 | 3 | 1 | 0 | 0 | 0 | 0 | 0 | 0 | 1 | 3 | 0 | 0 | 0 | 0 | 0 | 0 | 4 | 0 | 298 | 149 | 1 | 1 | NA | NA | 1 | 1 | NA | 1 | 1 | NA | 0 | NA | 0.333 |
| SE | 38585 | ENSG00000196878 | LAMB3 | chr1 | - | 209638533 | 209638648 | 209637907 | 209637981 | 209649963 | 209650118 | 0 | 0 | 3 | 3 | 0 | 0 | 0 | 0 | 0 | 0 | 0 | 0 | 4 | 5 | 0 | 0 | 0 | 0 | 2 | 0 | 264 | 149 | 1 | 1 | NA | NA | 1 | NA | NA | NA | NA | 1 | 0.585 | NA | 0.208 |
| SE | 38587 | ENSG00000132879 | FBXO44 | chr1 | + | 11655669 | 11655699 | 11654909 | 11654952 | 11655805 | 11656100 | 3 | 0 | 1 | 9 | 0 | 4 | 1 | 11 | 2 | 0 | 0 | 0 | 0 | 1 | 0 | 12 | 0 | 3 | 1 | 0 | 179 | 149 | 1 | 1 | 0.384 | 0 | 0.07 | 0.789 | NA | 0 | NA | 0 | 0.454 | NA | 0.159 |
| SE | 38590 | ENSG00000167085 | PHB | chr17 | - | 49411677 | 49411839 | 49409330 | 49409473 | 49413176 | 49413291 | 501 | 230 | 456 | 457 | 806 | 0 | 0 | 0 | 4 | 4 | 424 | 298 | 238 | 289 | 385 | 0 | 0 | 3 | 1 | 0 | 298 | 149 | 1 | 1 | 1 | 1 | 1 | 0.983 | 0.99 | 1 | 1 | 1 | 1 | 1 | -0.005 |
| SE | 38591 | ENSG00000167085 | PHB | chr17 | - | 49413176 | 49413291 | 49411677 | 49411839 | 49414834 | 49414858 | 266 | 124 | 228 | 230 | 413 | 1 | 0 | 0 | 0 | 0 | 189 | 143 | 125 | 140 | 235 | 0 | 0 | 0 | 0 | 0 | 264 | 149 | 1 | 1 | 0.993 | 1 | 1 | 1 | 1 | 1 | 1 | 1 | 1 | 1 | -0.001 |
| SE | 38592 | ENSG00000167085 | PHB | chr17 | - | 49414241 | 49414333 | 49413176 | 49413291 | 49414739 | 49414858 | 7 | 0 | 1 | 1 | 1 | 1 | 0 | 1 | 1 | 3 | 0 | 3 | 2 | 0 | 0 | 0 | 0 | 1 | 0 | 0 | 241 | 149 | 1 | 1 | 0.812 | NA | 0.382 | 0.382 | 0.171 | NA | 1 | 0.553 | NA | NA | -0.34 |
| SE | 38594 | ENSG00000169905 | TOR1AIP2 | chr1 | - | 179847534 | 179847636 | 179839966 | 179846828 | 179850844 | 179851363 | 84 | 116 | 81 | 100 | 127 | 1 | 0 | 0 | 0 | 0 | 36 | 60 | 88 | 58 | 3 | 0 | 0 | 0 | 0 | 0 | 251 | 149 | 1 | 1 | 0.98 | 1 | 1 | 1 | 1 | 1 | 1 | 1 | 1 | 1 | -0.004 |
| SE | 38596 | ENSG00000169905 | TOR1AIP2 | chr1 | - | 179877238 | 179877373 | 179852631 | 179852811 | 179877692 | 179877772 | 34 | 13 | 22 | 14 | 37 | 0 | 0 | 0 | 0 | 0 | 2 | 24 | 30 | 20 | 0 | 0 | 4 | 0 | 0 | 0 | 284 | 149 | 1 | 1 | 1 | 1 | 1 | 1 | 1 | 1 | 0.759 | 1 | 1 | NA | 0.06 |
| SE | 38599 | ENSG00000122406 | RPL5 | chr1 | + | 92833388 | 92833458 | 92832039 | 92832117 | 92836189 | 92836354 | 0 | 0 | 1 | 0 | 0 | 1 | 0 | 0 | 0 | 0 | 0 | 0 | 0 | 0 | 0 | 0 | 1 | 0 | 0 | 0 | 219 | 149 | 1 | 1 | 0 | NA | 1 | NA | NA | NA | 0 | NA | NA | NA | 0.5 |
| SE | 38600 | ENSG00000122406 | RPL5 | chr1 | + | 92833544 | 92833660 | 92832012 | 92832117 | 92836189 | 92836354 | 10 | 3 | 6 | 13 | 0 | 1 | 0 | 0 | 0 | 0 | 8 | 5 | 1 | 0 | 4 | 0 | 1 | 0 | 0 | 0 | 265 | 149 | 1 | 1 | 0.849 | 1 | 1 | 1 | NA | 1 | 0.738 | 1 | NA | 1 | 0.028 |
| SE | 38601 | ENSG00000122406 | RPL5 | chr1 | + | 92833544 | 92833660 | 92833388 | 92833458 | 92836189 | 92836354 | 801 | 417 | 972 | 709 | 617 | 0 | 0 | 1 | 0 | 0 | 290 | 377 | 489 | 290 | 254 | 0 | 0 | 0 | 0 | 0 | 265 | 149 | 1 | 1 | 1 | 1 | 0.998 | 1 | 1 | 1 | 1 | 1 | 1 | 1 | 0 |
| SE | 38602 | ENSG00000122406 | RPL5 | chr1 | + | 92833795 | 92833949 | 92833544 | 92833660 | 92834778 | 92834913 | 4 | 5 | 5 | 7 | 13 | 4605 | 3046 | 5725 | 4507 | 7994 | 0 | 0 | 4 | 6 | 0 | 3213 | 2347 | 2614 | 1780 | 2189 | 298 | 149 | 1 | 1 | 0 | 0.001 | 0 | 0.001 | 0.001 | 0 | 0 | 0.001 | 0.002 | 0 | 0 |
| SE | 38603 | ENSG00000122406 | RPL5 | chr1 | + | 92834778 | 92834913 | 92832761 | 92833660 | 92836189 | 92836271 | 10223 | 6593 | 12532 | 10176 | 16338 | 1 | 0 | 1 | 0 | 0 | 6490 | 5177 | 5680 | 3903 | 4576 | 0 | 0 | 0 | 0 | 0 | 284 | 149 | 1 | 1 | 1 | 1 | 1 | 1 | 1 | 1 | 1 | 1 | 1 | 1 | 0 |
| SE | 38604 | ENSG00000122406 | RPL5 | chr1 | + | 92836189 | 92836392 | 92834778 | 92834913 | 92837455 | 92837469 | 9705 | 5905 | 12010 | 9816 | 15943 | 0 | 0 | 0 | 0 | 1 | 5320 | 4849 | 5207 | 3686 | 4198 | 0 | 0 | 0 | 3 | 0 | 298 | 149 | 1 | 1 | 1 | 1 | 1 | 1 | 1 | 1 | 1 | 1 | 0.998 | 1 | 0 |
| SE | 38606 | ENSG00000122406 | RPL5 | chr1 | + | 92840550 | 92840639 | 92834778 | 92834913 | 92841765 | 92841835 | 74 | 52 | 100 | 66 | 130 | 0 | 0 | 0 | 0 | 0 | 44 | 27 | 47 | 61 | 73 | 0 | 0 | 4 | 0 | 0 | 238 | 149 | 1 | 1 | 1 | 1 | 1 | 1 | 1 | 1 | 1 | 0.88 | 1 | 1 | 0.024 |
| SE | 38608 | ENSG00000204178 | MACO1 | chr1 | + | 25448807 | 25448934 | 25446761 | 25446903 | 25454258 | 25454382 | 95 | 74 | 84 | 85 | 96 | 0 | 0 | 0 | 0 | 0 | 60 | 71 | 76 | 76 | 80 | 0 | 1 | 0 | 0 | 0 | 276 | 149 | 1 | 1 | 1 | 1 | 1 | 1 | 1 | 1 | 0.975 | 1 | 1 | 1 | 0.005 |
| SE | 38611 | ENSG00000204178 | MACO1 | chr1 | + | 25458390 | 25458892 | 25456652 | 25456831 | 25484115 | 25484274 | 231 | 152 | 233 | 232 | 404 | 0 | 0 | 0 | 1 | 0 | 126 | 216 | 226 | 200 | 50 | 0 | 0 | 0 | 0 | 0 | 298 | 149 | 1 | 1 | 1 | 1 | 1 | 0.991 | 1 | 1 | 1 | 1 | 1 | 1 | -0.002 |
| SE | 38612 | ENSG00000204178 | MACO1 | chr1 | + | 25484115 | 25484274 | 25458390 | 25458892 | 25485612 | 25485795 | 206 | 121 | 173 | 204 | 271 | 0 | 0 | 0 | 0 | 0 | 114 | 195 | 208 | 147 | 54 | 0 | 0 | 0 | 6 | 0 | 298 | 149 | 1 | 1 | 1 | 1 | 1 | 1 | 1 | 1 | 1 | 1 | 0.925 | 1 | 0.015 |
| SE | 38614 | ENSG00000204178 | MACO1 | chr1 | + | 25491409 | 25491584 | 25489172 | 25489293 | 25498263 | 25499230 | 214 | 189 | 263 | 248 | 447 | 0 | 0 | 0 | 0 | 0 | 162 | 171 | 213 | 226 | 40 | 0 | 2 | 0 | 0 | 0 | 298 | 149 | 1 | 1 | 1 | 1 | 1 | 1 | 1 | 1 | 0.977 | 1 | 1 | 1 | 0.005 |
| SE | 38624 | ENSG00000112096 | SOD2 | chr6 | - | 159684853 | 159685033 | 159669068 | 159682638 | 159688125 | 159688242 | 1946 | 2090 | 1389 | 2682 | 1386 | 11 | 5 | 1 | 4 | 1 | 8955 | 2625 | 4701 | 5651 | 14962 | 9 | 0 | 12 | 10 | 9 | 298 | 149 | 1 | 1 | 0.989 | 0.995 | 0.999 | 0.997 | 0.999 | 0.998 | 1 | 0.995 | 0.996 | 0.999 | -0.002 |
| SE | 38630 | ENSG00000115325 | DOK1 | chr2 | + | 74555155 | 74555453 | 74554749 | 74554859 | 74556307 | 74557114 | 8 | 0 | 2 | 2 | 2 | 0 | 0 | 1 | 0 | 0 | 4 | 7 | 4 | 0 | 8 | 0 | 0 | 0 | 0 | 0 | 298 | 149 | 1 | 1 | 1 | NA | 0.5 | 1 | 1 | 1 | 1 | 1 | NA | 1 | -0.125 |
| SE | 38631 | ENSG00000115325 | DOK1 | chr2 | + | 74555574 | 74555668 | 74555131 | 74555453 | 74555893 | 74555926 | 65 | 14 | 37 | 53 | 28 | 0 | 0 | 0 | 0 | 0 | 37 | 62 | 32 | 35 | 102 | 0 | 0 | 0 | 3 | 0 | 243 | 149 | 1 | 1 | 1 | 1 | 1 | 1 | 1 | 1 | 1 | 1 | 0.877 | 1 | 0.025 |
| SE | 38636 | ENSG00000165078 | CPA6 | chr8 | - | 67517922 | 67518047 | 67511540 | 67511655 | 67624175 | 67624251 | 0 | 0 | 0 | 3 | 2 | 2 | 0 | 0 | 0 | 0 | 0 | 0 | 4 | 2 | 0 | 0 | 0 | 0 | 0 | 7 | 274 | 149 | 1 | 1 | 0 | NA | NA | 1 | 1 | NA | NA | 1 | 1 | 0 | 0 |
| SE | 38638 | ENSG00000156875 | MFSD14A | chr1 | + | 100058665 | 100058728 | 100049908 | 100050004 | 100059877 | 100060005 | 190 | 105 | 121 | 160 | 148 | 1 | 0 | 0 | 0 | 1 | 205 | 171 | 163 | 182 | 97 | 0 | 0 | 0 | 0 | 0 | 212 | 149 | 1 | 1 | 0.993 | 1 | 1 | 1 | 0.99 | 1 | 1 | 1 | 1 | 1 | -0.003 |
| SE | 38642 | ENSG00000283761 | NA | chr1 | + | 100069613 | 100069685 | 100068472 | 100068586 | 100077164 | 100077269 | 104 | 41 | 88 | 102 | 104 | 0 | 0 | 0 | 4 | 0 | 61 | 72 | 68 | 41 | 8 | 0 | 0 | 0 | 0 | 0 | 221 | 149 | 1 | 1 | 1 | 1 | 1 | 0.945 | 1 | 1 | 1 | 1 | 1 | 1 | -0.011 |
| SE | 38643 | ENSG00000156875 | MFSD14A | chr1 | + | 100077164 | 100077277 | 100067976 | 100068195 | 100078450 | 100078548 | 130 | 92 | 106 | 139 | 357 | 0 | 0 | 0 | 0 | 0 | 103 | 127 | 205 | 145 | 19 | 0 | 2 | 0 | 0 | 0 | 262 | 149 | 1 | 1 | 1 | 1 | 1 | 1 | 1 | 1 | 0.973 | 1 | 1 | 1 | 0.005 |
| SE | 38644 | ENSG00000156875 | MFSD14A | chr1 | + | 100078450 | 100078548 | 100077164 | 100077277 | 100080495 | 100080659 | 256 | 157 | 216 | 258 | 575 | 0 | 1 | 0 | 0 | 0 | 185 | 266 | 356 | 267 | 47 | 0 | 0 | 0 | 0 | 0 | 247 | 149 | 1 | 1 | 1 | 0.99 | 1 | 1 | 1 | 1 | 1 | 1 | 1 | 1 | -0.002 |
| SE | 38646 | ENSG00000073417 | PDE8A | chr15 | + | 85075861 | 85075918 | 85067013 | 85067204 | 85083555 | 85083644 | 22 | 6 | 13 | 15 | 8 | 0 | 3 | 1 | 0 | 0 | 12 | 3 | 7 | 20 | 9 | 0 | 1 | 0 | 0 | 0 | 206 | 149 | 1 | 1 | 1 | 0.591 | 0.904 | 1 | 1 | 1 | 0.685 | 1 | 1 | 1 | -0.038 |
| SE | 38647 | ENSG00000073417 | PDE8A | chr15 | + | 85076732 | 85076787 | 85075861 | 85075918 | 85083555 | 85083644 | 56 | 46 | 55 | 56 | 79 | 2 | 0 | 0 | 0 | 0 | 16 | 33 | 50 | 83 | 28 | 0 | 0 | 0 | 0 | 0 | 204 | 149 | 1 | 1 | 0.953 | 1 | 1 | 1 | 1 | 1 | 1 | 1 | 1 | 1 | -0.009 |
| SE | 38648 | ENSG00000073417 | PDE8A | chr15 | + | 85083555 | 85083644 | 85076732 | 85076787 | 85089337 | 85089416 | 67 | 45 | 64 | 60 | 100 | 0 | 1 | 0 | 0 | 1 | 18 | 47 | 30 | 71 | 44 | 0 | 3 | 0 | 0 | 0 | 238 | 149 | 1 | 1 | 1 | 0.966 | 1 | 1 | 0.984 | 1 | 0.907 | 1 | 1 | 1 | 0.009 |
| SE | 38649 | ENSG00000073417 | PDE8A | chr15 | + | 85089337 | 85089416 | 85083555 | 85083644 | 85090717 | 85090915 | 21 | 4 | 11 | 18 | 40 | 2 | 0 | 0 | 0 | 0 | 0 | 8 | 12 | 6 | 28 | 0 | 0 | 0 | 1 | 0 | 228 | 149 | 1 | 1 | 0.873 | 1 | 1 | 1 | 1 | NA | 1 | 1 | 0.797 | 1 | 0.025 |
| SE | 38651 | ENSG00000073417 | PDE8A | chr15 | + | 85089337 | 85089416 | 85083555 | 85083644 | 85097947 | 85098036 | 25 | 5 | 11 | 17 | 44 | 4 | 3 | 2 | 2 | 0 | 0 | 9 | 7 | 10 | 36 | 0 | 8 | 5 | 7 | 0 | 228 | 149 | 1 | 1 | 0.803 | 0.521 | 0.782 | 0.847 | 1 | NA | 0.424 | 0.478 | 0.483 | 1 | 0.194 |
| SE | 38657 | ENSG00000073417 | PDE8A | chr15 | + | 85115983 | 85116119 | 85115438 | 85115487 | 85120796 | 85121014 | 102 | 99 | 117 | 142 | 219 | 0 | 0 | 0 | 0 | 0 | 90 | 85 | 80 | 65 | 32 | 0 | 0 | 0 | 0 | 2 | 285 | 149 | 1 | 1 | 1 | 1 | 1 | 1 | 1 | 1 | 1 | 1 | 1 | 0.893 | 0.021 |
| SE | 38658 | ENSG00000073417 | PDE8A | chr15 | + | 85115983 | 85116119 | 85115438 | 85115487 | 85126206 | 85126374 | 101 | 99 | 119 | 141 | 219 | 0 | 0 | 0 | 0 | 0 | 90 | 84 | 79 | 65 | 32 | 0 | 0 | 1 | 0 | 0 | 285 | 149 | 1 | 1 | 1 | 1 | 1 | 1 | 1 | 1 | 1 | 0.976 | 1 | 1 | 0.005 |
| SE | 38659 | ENSG00000073417 | PDE8A | chr15 | + | 85117640 | 85117839 | 85115983 | 85116119 | 85120796 | 85121014 | 73 | 49 | 58 | 67 | 129 | 1 | 1 | 2 | 1 | 0 | 63 | 51 | 52 | 53 | 8 | 0 | 2 | 1 | 0 | 0 | 298 | 149 | 1 | 1 | 0.973 | 0.961 | 0.935 | 0.971 | 1 | 1 | 0.927 | 0.963 | 1 | 1 | -0.01 |
| SE | 38660 | ENSG00000073417 | PDE8A | chr15 | + | 85117640 | 85117839 | 85115983 | 85116119 | 85126206 | 85126374 | 33 | 25 | 30 | 40 | 93 | 0 | 1 | 4 | 0 | 0 | 12 | 15 | 21 | 29 | 0 | 0 | 1 | 0 | 0 | 0 | 298 | 149 | 1 | 1 | 1 | 0.926 | 0.789 | 1 | 1 | 1 | 0.882 | 1 | 1 | NA | -0.028 |
| SE | 38661 | ENSG00000073417 | PDE8A | chr15 | + | 85120796 | 85121014 | 85115438 | 85115487 | 85126206 | 85126374 | 2 | 0 | 0 | 0 | 0 | 0 | 0 | 0 | 0 | 0 | 0 | 0 | 0 | 0 | 2 | 0 | 0 | 1 | 0 | 0 | 298 | 149 | 1 | 1 | 1 | NA | NA | NA | NA | NA | NA | 0 | NA | 1 | 0.5 |
| SE | 38662 | ENSG00000073417 | PDE8A | chr15 | + | 85120796 | 85121014 | 85115983 | 85116119 | 85123060 | 85123193 | 38 | 42 | 40 | 58 | 25 | 1 | 0 | 0 | 0 | 0 | 57 | 37 | 23 | 51 | 41 | 0 | 0 | 0 | 4 | 0 | 298 | 149 | 1 | 1 | 0.95 | 1 | 1 | 1 | 1 | 1 | 1 | 1 | 0.864 | 1 | 0.017 |
| SE | 38663 | ENSG00000073417 | PDE8A | chr15 | + | 85120796 | 85121014 | 85115983 | 85116119 | 85126206 | 85126374 | 3 | 1 | 2 | 1 | 0 | 0 | 1 | 4 | 0 | 0 | 0 | 2 | 1 | 0 | 0 | 0 | 1 | 0 | 0 | 0 | 298 | 149 | 1 | 1 | 1 | 0.333 | 0.2 | 1 | NA | NA | 0.5 | 1 | NA | NA | -0.117 |
| SE | 38664 | ENSG00000073417 | PDE8A | chr15 | + | 85120796 | 85121014 | 85117640 | 85117839 | 85126206 | 85126374 | 42 | 24 | 29 | 27 | 36 | 0 | 0 | 1 | 0 | 0 | 51 | 36 | 31 | 24 | 8 | 0 | 0 | 0 | 0 | 0 | 298 | 149 | 1 | 1 | 1 | 1 | 0.935 | 1 | 1 | 1 | 1 | 1 | 1 | 1 | -0.013 |
| SE | 38665 | ENSG00000073417 | PDE8A | chr15 | + | 85123060 | 85123193 | 85115983 | 85116119 | 85126206 | 85126374 | 24 | 11 | 21 | 28 | 35 | 0 | 1 | 4 | 0 | 0 | 0 | 17 | 17 | 40 | 7 | 0 | 1 | 0 | 0 | 0 | 282 | 149 | 1 | 1 | 1 | 0.853 | 0.735 | 1 | 1 | NA | 0.9 | 1 | 1 | 1 | -0.057 |
| SE | 38666 | ENSG00000073417 | PDE8A | chr15 | + | 85123060 | 85123193 | 85120190 | 85121014 | 85126206 | 85126374 | 59 | 49 | 59 | 82 | 50 | 2 | 0 | 0 | 0 | 0 | 51 | 50 | 38 | 72 | 38 | 0 | 0 | 0 | 0 | 0 | 282 | 149 | 1 | 1 | 0.94 | 1 | 1 | 1 | 1 | 1 | 1 | 1 | 1 | 1 | -0.012 |
| SE | 38671 | ENSG00000143365 | RORC | chr1 | - | 151813238 | 151813346 | 151812946 | 151813057 | 151813487 | 151813620 | 27 | 12 | 22 | 19 | 19 | 0 | 0 | 1 | 0 | 0 | 0 | 0 | 0 | 0 | 2 | 0 | 0 | 0 | 0 | 9 | 257 | 149 | 1 | 1 | 1 | 1 | 0.927 | 1 | 1 | NA | NA | NA | NA | 0.114 | 0.871 |
| SE | 38673 | ENSG00000143365 | RORC | chr1 | - | 151813487 | 151813620 | 151812946 | 151813057 | 151814573 | 151814695 | 6 | 1 | 8 | 4 | 0 | 2 | 0 | 0 | 0 | 0 | 0 | 0 | 0 | 0 | 9 | 0 | 0 | 0 | 0 | 0 | 282 | 149 | 1 | 1 | 0.613 | 1 | 1 | 1 | NA | NA | NA | NA | NA | 1 | -0.097 |
| SE | 38675 | ENSG00000143365 | RORC | chr1 | - | 151813487 | 151813702 | 151812946 | 151813057 | 151814573 | 151814695 | 0 | 0 | 1 | 1 | 0 | 2 | 0 | 0 | 0 | 0 | 0 | 0 | 0 | 0 | 9 | 0 | 0 | 0 | 0 | 0 | 298 | 149 | 1 | 1 | 0 | NA | 1 | 1 | NA | NA | NA | NA | NA | 1 | -0.333 |
| SE | 38682 | ENSG00000167258 | CDK12 | chr17 | + | 39490556 | 39490733 | 39470878 | 39471763 | 39492750 | 39492890 | 496 | 346 | 379 | 369 | 417 | 5 | 2 | 3 | 0 | 0 | 794 | 532 | 543 | 796 | 505 | 0 | 0 | 2 | 4 | 8 | 298 | 149 | 1 | 1 | 0.98 | 0.989 | 0.984 | 1 | 1 | 1 | 1 | 0.993 | 0.99 | 0.969 | 0 |
| SE | 38683 | ENSG00000167258 | CDK12 | chr17 | + | 39494523 | 39494694 | 39492750 | 39492890 | 39501249 | 39501439 | 473 | 400 | 492 | 497 | 667 | 1 | 0 | 2 | 3 | 0 | 568 | 627 | 557 | 706 | 334 | 8 | 3 | 4 | 0 | 0 | 298 | 149 | 1 | 1 | 0.996 | 1 | 0.992 | 0.988 | 1 | 0.973 | 0.991 | 0.986 | 1 | 1 | 0.005 |
| SE | 38684 | ENSG00000167258 | CDK12 | chr17 | + | 39501249 | 39501439 | 39494523 | 39494694 | 39509704 | 39509761 | 521 | 415 | 562 | 529 | 731 | 0 | 0 | 0 | 2 | 0 | 378 | 591 | 563 | 633 | 241 | 0 | 0 | 1 | 0 | 0 | 298 | 149 | 1 | 1 | 1 | 1 | 1 | 0.992 | 1 | 1 | 1 | 0.996 | 1 | 1 | -0.001 |
| SE | 38685 | ENSG00000167258 | CDK12 | chr17 | + | 39509704 | 39509761 | 39501249 | 39501439 | 39511528 | 39511630 | 320 | 233 | 331 | 344 | 455 | 1 | 0 | 0 | 0 | 0 | 182 | 342 | 324 | 371 | 144 | 0 | 1 | 0 | 0 | 0 | 206 | 149 | 1 | 1 | 0.996 | 1 | 1 | 1 | 1 | 1 | 0.996 | 1 | 1 | 1 | 0 |
| SE | 38687 | ENSG00000167258 | CDK12 | chr17 | + | 39519955 | 39520087 | 39517439 | 39517556 | 39524673 | 39524885 | 228 | 187 | 272 | 299 | 363 | 4 | 0 | 0 | 0 | 0 | 239 | 244 | 377 | 277 | 128 | 0 | 0 | 0 | 5 | 0 | 281 | 149 | 1 | 1 | 0.968 | 1 | 1 | 1 | 1 | 1 | 1 | 1 | 0.967 | 1 | 0 |
| SE | 38688 | ENSG00000167258 | CDK12 | chr17 | + | 39547767 | 39547830 | 39546616 | 39546758 | 39551037 | 39551141 | 11 | 5 | 2 | 14 | 4 | 0 | 0 | 2 | 1 | 1 | 0 | 9 | 3 | 19 | 5 | 0 | 0 | 0 | 0 | 0 | 212 | 149 | 1 | 1 | 1 | 1 | 0.413 | 0.908 | 0.738 | NA | 1 | 1 | 1 | 1 | -0.188 |
| SE | 38690 | ENSG00000167258 | CDK12 | chr17 | + | 39551037 | 39551142 | 39547820 | 39547830 | 39564759 | 39564830 | 2 | 8 | 3 | 11 | 3 | 0 | 0 | 0 | 0 | 0 | 0 | 5 | 3 | 21 | 0 | 0 | 0 | 0 | 2 | 0 | 254 | 149 | 1 | 1 | 1 | 1 | 1 | 1 | 1 | NA | 1 | 1 | 0.86 | NA | 0.047 |
| SE | 38691 | ENSG00000167258 | CDK12 | chr17 | + | 39556285 | 39556413 | 39547820 | 39547830 | 39564759 | 39564830 | 16 | 14 | 16 | 29 | 29 | 0 | 0 | 0 | 0 | 0 | 7 | 23 | 15 | 23 | 39 | 0 | 0 | 0 | 2 | 0 | 277 | 149 | 1 | 1 | 1 | 1 | 1 | 1 | 1 | 1 | 1 | 1 | 0.861 | 1 | 0.028 |
| SE | 38692 | ENSG00000167258 | CDK12 | chr17 | + | 39556285 | 39556413 | 39551037 | 39551142 | 39564759 | 39564830 | 39 | 31 | 35 | 47 | 66 | 0 | 3 | 1 | 3 | 3 | 0 | 43 | 25 | 38 | 68 | 0 | 1 | 0 | 11 | 0 | 277 | 149 | 1 | 1 | 1 | 0.848 | 0.95 | 0.894 | 0.922 | NA | 0.959 | 1 | 0.65 | 1 | 0.021 |
| SE | 38695 | ENSG00000100379 | KCTD17 | chr22 | + | 37060822 | 37060922 | 37059312 | 37059438 | 37061538 | 37061564 | 12 | 8 | 7 | 5 | 6 | 0 | 0 | 0 | 0 | 0 | 35 | 9 | 2 | 17 | 33 | 0 | 1 | 0 | 3 | 0 | 249 | 149 | 1 | 1 | 1 | 1 | 1 | 1 | 1 | 1 | 0.843 | 1 | 0.772 | 1 | 0.077 |
| SE | 38696 | ENSG00000100379 | KCTD17 | chr22 | + | 37060822 | 37060922 | 37059312 | 37059438 | 37062524 | 37063390 | 11 | 8 | 8 | 6 | 6 | 0 | 0 | 1 | 0 | 0 | 35 | 7 | 2 | 17 | 27 | 0 | 0 | 0 | 0 | 0 | 249 | 149 | 1 | 1 | 1 | 1 | 0.827 | 1 | 1 | 1 | 1 | 1 | 1 | 1 | -0.035 |
| SE | 38697 | ENSG00000100379 | KCTD17 | chr22 | + | 37061103 | 37061175 | 37059312 | 37059438 | 37061538 | 37061629 | 0 | 2 | 0 | 0 | 0 | 0 | 0 | 0 | 0 | 0 | 0 | 0 | 0 | 0 | 11 | 0 | 1 | 0 | 3 | 0 | 221 | 149 | 1 | 1 | NA | 1 | NA | NA | NA | NA | 0 | NA | 0 | 1 | 0.667 |
| SE | 38698 | ENSG00000100379 | KCTD17 | chr22 | + | 37061103 | 37061175 | 37060822 | 37060922 | 37061538 | 37061629 | 0 | 4 | 0 | 0 | 0 | 2 | 0 | 0 | 0 | 0 | 0 | 0 | 0 | 0 | 0 | 0 | 2 | 0 | 0 | 6 | 221 | 149 | 1 | 1 | 0 | 1 | NA | NA | NA | NA | 0 | NA | NA | 0 | 0.5 |
| SE | 38699 | ENSG00000100379 | KCTD17 | chr22 | + | 37061538 | 37061629 | 37059312 | 37059438 | 37062524 | 37063374 | 2 | 1 | 0 | 2 | 2 | 0 | 0 | 1 | 0 | 0 | 0 | 5 | 0 | 3 | 0 | 0 | 0 | 0 | 0 | 0 | 240 | 149 | 1 | 1 | 1 | 1 | 0 | 1 | 1 | NA | 1 | NA | 1 | NA | -0.2 |
| SE | 38700 | ENSG00000100379 | KCTD17 | chr22 | + | 37061538 | 37061629 | 37060822 | 37060922 | 37062524 | 37063374 | 4 | 1 | 0 | 2 | 2 | 1 | 0 | 1 | 1 | 0 | 0 | 6 | 0 | 0 | 6 | 0 | 0 | 0 | 0 | 0 | 240 | 149 | 1 | 1 | 0.713 | 1 | 0 | 0.554 | 1 | NA | 1 | NA | NA | 1 | -0.347 |
| SE | 38701 | ENSG00000116690 | PRG4 | chr1 | + | 186300090 | 186300213 | 186296845 | 186296951 | 186301591 | 186301711 | 0 | 1 | 0 | 0 | 0 | 0 | 0 | 0 | 0 | 0 | 0 | 0 | 0 | 0 | 0 | 0 | 0 | 1 | 0 | 0 | 272 | 149 | 1 | 1 | NA | 1 | NA | NA | NA | NA | NA | 0 | NA | NA | 1 |
| SE | 38707 | ENSG00000228794 | LINC01128 | chr1 | + | 829002 | 829104 | 827597 | 827775 | 841199 | 841373 | 6 | 4 | 32 | 15 | 2 | 1 | 0 | 1 | 0 | 0 | 3 | 14 | 0 | 4 | 7 | 0 | 0 | 0 | 0 | 0 | 251 | 149 | 1 | 1 | 0.781 | 1 | 0.95 | 1 | 1 | 1 | 1 | NA | 1 | 1 | -0.054 |
| SE | 38709 | ENSG00000228794 | LINC01128 | chr1 | + | 841199 | 841373 | 829002 | 829104 | 851926 | 852110 | 5 | 4 | 26 | 13 | 1 | 1 | 2 | 6 | 7 | 0 | 3 | 14 | 0 | 0 | 7 | 0 | 1 | 0 | 0 | 0 | 298 | 149 | 1 | 1 | 0.714 | 0.5 | 0.684 | 0.481 | 1 | 1 | 0.875 | NA | NA | 1 | -0.283 |
| SE | 38711 | ENSG00000228794 | LINC01128 | chr1 | + | 847653 | 847806 | 829002 | 829104 | 851926 | 852094 | 11 | 2 | 14 | 3 | 13 | 1 | 2 | 6 | 7 | 0 | 3 | 3 | 8 | 4 | 0 | 0 | 1 | 0 | 0 | 0 | 298 | 149 | 1 | 1 | 0.846 | 0.333 | 0.538 | 0.176 | 1 | 1 | 0.6 | 1 | 1 | NA | -0.321 |
| SE | 38714 | ENSG00000228794 | LINC01128 | chr1 | + | 849483 | 849602 | 847653 | 847806 | 850180 | 850351 | 9 | 0 | 7 | 3 | 0 | 0 | 0 | 0 | 0 | 0 | 3 | 5 | 1 | 0 | 0 | 0 | 0 | 0 | 0 | 7 | 268 | 149 | 1 | 1 | 1 | NA | 1 | 1 | NA | NA | 1 | 1 | NA | 0 | 0.333 |
| SE | 38715 | ENSG00000228794 | LINC01128 | chr1 | + | 849483 | 849602 | 847653 | 847806 | 851926 | 852110 | 9 | 0 | 6 | 6 | 2 | 8 | 0 | 7 | 1 | 6 | 0 | 3 | 1 | 0 | 0 | 3 | 0 | 4 | 0 | 0 | 268 | 149 | 1 | 1 | 0.385 | NA | 0.323 | 0.769 | 0.156 | 0 | 1 | 0.122 | NA | NA | 0.034 |
| SE | 38716 | ENSG00000228794 | LINC01128 | chr1 | + | 850177 | 850351 | 849483 | 849602 | 851926 | 852110 | 3 | 2 | 3 | 0 | 4 | 0 | 0 | 2 | 3 | 2 | 0 | 2 | 0 | 10 | 7 | 0 | 0 | 0 | 0 | 0 | 298 | 149 | 1 | 1 | 1 | 1 | 0.429 | 0 | 0.5 | NA | 1 | NA | 1 | 1 | -0.414 |
| SE | 38718 | ENSG00000228794 | LINC01128 | chr1 | + | 850180 | 850351 | 849483 | 849602 | 851926 | 852014 | 0 | 1 | 5 | 0 | 4 | 0 | 0 | 2 | 3 | 2 | 0 | 4 | 0 | 5 | 7 | 0 | 0 | 0 | 0 | 0 | 298 | 149 | 1 | 1 | NA | 1 | 0.556 | 0 | 0.5 | NA | 1 | NA | 1 | 1 | -0.486 |
| SE | 38719 | ENSG00000228794 | LINC01128 | chr1 | + | 851926 | 852014 | 829002 | 829104 | 852670 | 852766 | 1 | 1 | 5 | 3 | 0 | 0 | 1 | 1 | 1 | 0 | 0 | 1 | 0 | 0 | 0 | 0 | 0 | 0 | 0 | 0 | 237 | 149 | 1 | 1 | 1 | 0.386 | 0.759 | 0.654 | NA | NA | 1 | NA | NA | NA | -0.3 |
| SE | 38720 | ENSG00000228794 | LINC01128 | chr1 | + | 851926 | 852110 | 829002 | 829104 | 852670 | 852766 | 18 | 10 | 24 | 22 | 37 | 0 | 1 | 1 | 1 | 0 | 13 | 5 | 2 | 2 | 0 | 0 | 0 | 0 | 0 | 0 | 298 | 149 | 1 | 1 | 1 | 0.833 | 0.923 | 0.917 | 1 | 1 | 1 | 1 | 1 | NA | -0.065 |
| SE | 38721 | ENSG00000228794 | LINC01128 | chr1 | + | 851926 | 852110 | 847653 | 847806 | 853390 | 853497 | 18 | 1 | 11 | 2 | 12 | 0 | 0 | 0 | 1 | 0 | 3 | 7 | 13 | 0 | 2 | 0 | 0 | 0 | 0 | 0 | 298 | 149 | 1 | 1 | 1 | 1 | 1 | 0.5 | 1 | 1 | 1 | 1 | NA | 1 | -0.1 |
| SE | 38722 | ENSG00000228794 | LINC01128 | chr1 | + | 851926 | 852766 | 847653 | 847806 | 853390 | 853602 | 12 | 6 | 27 | 10 | 23 | 0 | 0 | 0 | 1 | 0 | 3 | 8 | 4 | 3 | 0 | 0 | 0 | 0 | 0 | 0 | 298 | 149 | 1 | 1 | 1 | 1 | 1 | 0.833 | 1 | 1 | 1 | 1 | 1 | NA | -0.033 |
| SE | 38724 | ENSG00000159140 | SON | chr21 | + | 33549475 | 33555391 | 33546212 | 33546379 | 33557155 | 33557316 | 864 | 542 | 751 | 952 | 1405 | 1 | 0 | 0 | 0 | 0 | 674 | 730 | 725 | 1001 | 477 | 0 | 3 | 0 | 0 | 0 | 298 | 149 | 1 | 1 | 0.998 | 1 | 1 | 1 | 1 | 1 | 0.992 | 1 | 1 | 1 | 0.001 |
| SE | 38725 | ENSG00000159140 | SON | chr21 | + | 33557155 | 33557316 | 33549475 | 33555391 | 33557545 | 33559056 | 646 | 411 | 609 | 783 | 1086 | 0 | 0 | 0 | 0 | 0 | 629 | 603 | 620 | 860 | 325 | 0 | 2 | 0 | 2 | 0 | 298 | 149 | 1 | 1 | 1 | 1 | 1 | 1 | 1 | 1 | 0.993 | 1 | 0.995 | 1 | 0.002 |
| SE | 38726 | ENSG00000159140 | SON | chr21 | + | 33567156 | 33567267 | 33559586 | 33559775 | 33568970 | 33569087 | 1415 | 1129 | 1192 | 1387 | 1967 | 0 | 3 | 0 | 0 | 0 | 1427 | 1093 | 848 | 1190 | 540 | 0 | 0 | 0 | 0 | 0 | 260 | 149 | 1 | 1 | 1 | 0.995 | 1 | 1 | 1 | 1 | 1 | 1 | 1 | 1 | -0.001 |
| SE | 38729 | ENSG00000159140 | SON | chr21 | + | 33573307 | 33573455 | 33568970 | 33569087 | 33575595 | 33575667 | 1235 | 963 | 1017 | 1189 | 1717 | 0 | 0 | 0 | 3 | 1 | 1232 | 1008 | 789 | 1060 | 508 | 0 | 0 | 0 | 0 | 0 | 297 | 149 | 1 | 1 | 1 | 1 | 1 | 0.995 | 0.999 | 1 | 1 | 1 | 1 | 1 | -0.001 |
| SE | 38744 | ENSG00000086062 | B4GALT1 | chr9 | - | 33120418 | 33120606 | 33115990 | 33116113 | 33135188 | 33135424 | 1032 | 417 | 758 | 808 | 869 | 1 | 0 | 5 | 0 | 4 | 1250 | 1282 | 1184 | 1216 | 1848 | 0 | 7 | 5 | 9 | 1 | 298 | 149 | 1 | 1 | 0.998 | 1 | 0.987 | 1 | 0.991 | 1 | 0.989 | 0.992 | 0.985 | 0.999 | 0.002 |
| SE | 38759 | ENSG00000148399 | DPH7 | chr9 | - | 137574751 | 137574843 | 137564892 | 137565154 | 137576079 | 137576167 | 4 | 5 | 6 | 5 | 10 | 0 | 0 | 0 | 0 | 0 | 0 | 2 | 1 | 10 | 17 | 0 | 0 | 1 | 0 | 0 | 241 | 149 | 1 | 1 | 1 | 1 | 1 | 1 | 1 | NA | 1 | 0.382 | 1 | 1 | 0.154 |
| SE | 38760 | ENSG00000148399 | DPH7 | chr9 | - | 137574751 | 137574846 | 137565084 | 137565154 | 137576079 | 137576167 | 0 | 0 | 0 | 1 | 0 | 0 | 0 | 0 | 0 | 0 | 0 | 3 | 0 | 1 | 6 | 0 | 0 | 1 | 0 | 0 | 244 | 149 | 1 | 1 | NA | NA | NA | 1 | NA | NA | 1 | 0 | 1 | 1 | 0.25 |
| SE | 38761 | ENSG00000148399 | DPH7 | chr9 | - | 137575708 | 137575892 | 137574751 | 137574843 | 137576079 | 137576167 | 8 | 8 | 14 | 5 | 6 | 7 | 13 | 14 | 15 | 15 | 0 | 14 | 1 | 0 | 7 | 0 | 10 | 4 | 10 | 11 | 298 | 149 | 1 | 1 | 0.364 | 0.235 | 0.333 | 0.143 | 0.167 | NA | 0.412 | 0.111 | 0 | 0.241 | 0.057 |
| SE | 38767 | ENSG00000148399 | DPH7 | chr9 | - | 137577576 | 137577603 | 137576308 | 137576402 | 137578624 | 137578717 | 2 | 0 | 2 | 1 | 0 | 2 | 0 | 0 | 0 | 0 | 0 | 4 | 3 | 0 | 5 | 0 | 3 | 2 | 0 | 0 | 176 | 149 | 1 | 1 | 0.458 | NA | 1 | 1 | NA | NA | 0.53 | 0.559 | NA | 1 | 0.123 |
| SE | 38768 | ENSG00000181804 | SLC9A9 | chr3 | - | 143382059 | 143382114 | 143363483 | 143363563 | 143467036 | 143467190 | 45 | 40 | 26 | 39 | 11 | 0 | 0 | 1 | 0 | 0 | 29 | 36 | 39 | 54 | 18 | 0 | 0 | 0 | 0 | 0 | 204 | 149 | 1 | 1 | 1 | 1 | 0.95 | 1 | 1 | 1 | 1 | 1 | 1 | 1 | -0.01 |
| SE | 38769 | ENSG00000181804 | SLC9A9 | chr3 | - | 143652254 | 143652360 | 143552361 | 143552450 | 143693191 | 143693307 | 25 | 10 | 16 | 34 | 47 | 1 | 0 | 0 | 0 | 0 | 12 | 24 | 24 | 29 | 21 | 0 | 1 | 0 | 0 | 0 | 255 | 149 | 1 | 1 | 0.936 | 1 | 1 | 1 | 1 | 1 | 0.933 | 1 | 1 | 1 | 0.001 |

| | | | | | | | | | | | | | | |
|---|---|---|---|---|---|---|---|---|---|---|---|---|---|---|
| SE | 38770 | ENSG00000181804 | SLC9A9 | chr3 | - | 143652254 | 143652360 | 143578584 | 143578723 | 143693191 | 143693307 | 55 | 52 | [illegible] |
| SE | 38771 | ENSG00000181804 | SLC9A9 | chr3 | - | 143693191 | 143693307 | 143652254 | 143652360 | 143795000 | 143795077 | 67 | 40 | [illegible] |
| SE | 38772 | ENSG00000181804 | SLC9A9 | chr3 | - | 143796825 | 143796903 | 143795000 | 143795077 | 143832018 | 143832221 | 158 | 129 | [illegible] |
| SE | 38773 | ENSG00000181804 | SLC9A9 | chr3 | - | 143832018 | 143832221 | 143796825 | 143796903 | 143848147 | 143848458 | 276 | 240 | [illegible] |
| SE | 38775 | ENSG00000162601 | MYSM1 | chr1 | - | 58663449 | 58663522 | 58661405 | 58661511 | 58665498 | 58665631 | 3 | 9 | [illegible] |
| SE | 38776 | ENSG00000162601 | MYSM1 | chr1 | - | 58665498 | 58665631 | 58661405 | 58661511 | 58667037 | 58667226 | 166 | 137 | [illegible] |
| SE | 38777 | ENSG00000162601 | MYSM1 | chr1 | - | 58667037 | 58667226 | 58661405 | 58661511 | 58667846 | 58667921 | 69 | 65 | [illegible] |
| SE | 38778 | ENSG00000162601 | MYSM1 | chr1 | - | 58668429 | 58668566 | 58667846 | 58667921 | 58668983 | 58669038 | 39 | 31 | [illegible] |
| SE | 38779 | ENSG00000162601 | MYSM1 | chr1 | - | 58668631 | 58668682 | 58667846 | 58667921 | 58668983 | 58669038 | 91 | 79 | [illegible] |
| SE | 38783 | ENSG00000162601 | MYSM1 | chr1 | - | 58689293 | 58689367 | 58689037 | 58689116 | 58690225 | 58690249 | 6 | 11 | [illegible] |
| SE | 38790 | ENSG00000111961 | SASH1 | chr6 | + | 148471416 | 148471503 | 148468544 | 148468585 | 148474109 | 148474222 | 12 | 54 | [illegible] |
| SE | 38799 | ENSG00000072422 | RHOBTB1 | chr10 | - | 60872042 | 60872290 | 60871471 | 60871651 | 60874953 | 60875043 | 9 | 19 | [illegible] |
| SE | 38800 | ENSG00000072422 | RHOBTB1 | chr10 | - | 60872184 | 60872290 | 60869437 | 60871651 | 60874953 | 60875042 | 25 | 41 | [illegible] |
| SE | 38801 | ENSG00000072422 | RHOBTB1 | chr10 | - | 60877907 | 60878058 | 60874953 | 60875042 | 60886111 | 60886230 | 16 | 20 | [illegible] |
| SE | 38802 | ENSG00000072422 | RHOBTB1 | chr10 | - | 60886111 | 60886230 | 60877907 | 60878058 | 60888211 | 60889185 | 30 | 33 | [illegible] |
| SE | 38803 | ENSG00000072422 | RHOBTB1 | chr10 | - | 60888211 | 60889185 | 60886111 | 60886230 | 60910886 | 60910990 | 25 | 29 | [illegible] |
| SE | 38804 | ENSG00000072422 | RHOBTB1 | chr10 | - | 60892809 | 60892995 | 60888211 | 60889185 | 60910886 | 60910990 | 28 | 18 | [illegible] |
| SE | 38806 | ENSG00000072422 | RHOBTB1 | chr10 | - | 60941803 | 60941854 | 60911350 | 60911552 | 60943970 | 60944185 | 4 | 2 | [illegible] |
| SE | 38808 | ENSG00000132749 | TESMIN | chr11 | - | 68742317 | 68742394 | 68738461 | 68738788 | 68747207 | 68747366 | 0 | 0 | [illegible] |
| SE | 38810 | ENSG00000132749 | TESMIN | chr11 | - | 68744990 | 68745111 | 68742317 | 68742394 | 68747207 | 68747366 | 2 | 4 | [illegible] |
| SE | 38811 | ENSG00000006625 | GGCT | chr7 | - | 30498802 | 30498938 | 30496620 | 30497235 | 30500535 | 30500680 | 72 | 49 | [illegible] |
| SE | 38813 | ENSG00000229180 | NA | chr7 | - | 66542502 | 66542642 | 66541241 | 66541468 | 66545600 | 66545686 | 0 | 0 | [illegible] |
| SE | 38816 | ENSG00000229180 | NA | chr7 | - | 66573384 | 66573550 | 66553484 | 66554837 | 66592256 | 66592386 | 41 | 56 | [illegible] |
| SE | 38817 | ENSG00000229180 | NA | chr7 | - | 66576754 | 66576929 | 66573384 | 66573550 | 66588511 | 66588580 | 32 | 19 | [illegible] |
| SE | 38819 | ENSG00000229180 | NA | chr7 | - | 66576754 | 66576932 | 66573384 | 66573550 | 66588511 | 66588580 | 34 | 19 | [illegible] |
| SE | 38821 | ENSG00000229180 | NA | chr7 | - | 66578528 | 66578727 | 66576754 | 66576929 | 66588511 | 66588580 | 0 | 0 | [illegible] |
| SE | 38822 | ENSG00000229180 | NA | chr7 | - | 66578528 | 66578727 | 66576754 | 66576932 | 66588511 | 66588580 | 0 | 0 | [illegible] |
| SE | 38823 | ENSG00000229180 | NA | chr7 | - | 66588511 | 66588580 | 66573384 | 66573550 | 66592256 | 66592386 | 0 | 0 | [illegible] |
| SE | 38824 | ENSG00000229180 | NA | chr7 | - | 66588511 | 66588580 | 66576754 | 66576929 | 66592256 | 66592386 | 0 | 1 | [illegible] |
| SE | 38830 | ENSG00000122674 | CCZ1 | chr7 | + | 5911860 | 5911922 | 5910034 | 5910116 | 5912842 | 5912954 | 45 | 73 | [illegible] |
| SE | 38832 | ENSG00000122674 | CCZ1 | chr7 | + | 5918866 | 5918900 | 5912842 | 5912954 | 5919848 | 5919966 | 87 | 60 | [illegible] |
| SE | 38833 | ENSG00000122674 | CCZ1 | chr7 | + | 5918866 | 5918900 | 5912842 | 5912954 | 5923386 | 5923545 | 9 | 4 | [illegible] |
| SE | 38834 | ENSG00000122674 | CCZ1 | chr7 | + | 5919848 | 5919966 | 5912842 | 5912954 | 5923386 | 5923545 | 110 | 68 | [illegible] |
| SE | 38835 | ENSG00000122674 | CCZ1 | chr7 | + | 5919848 | 5919966 | 5918866 | 5918900 | 5923386 | 5923545 | 189 | 121 | [illegible] |
| SE | 38836 | ENSG00000173366 | NA | chr3 | - | 52225526 | 52225635 | 52221080 | 52224312 | 52229660 | 52229773 | 14 | 5 | [illegible] |
| SE | 38839 | ENSG00000164307 | ERAP1 | chr5 | - | 96793399 | 96793513 | 96792060 | 96792192 | 96797174 | 96797309 | 573 | 329 | [illegible] |
| SE | 38840 | ENSG00000164307 | ERAP1 | chr5 | - | 96793802 | 96793957 | 96793399 | 96793513 | 96795041 | 96795162 | 435 | 196 | [illegible] |
| SE | 38841 | ENSG00000164307 | ERAP1 | chr5 | - | 96795041 | 96795162 | 96793399 | 96793513 | 96797174 | 96797309 | 302 | 174 | [illegible] |
| SE | 38842 | ENSG00000164307 | ERAP1 | chr5 | - | 96797174 | 96797309 | 96795041 | 96795162 | 96800861 | 96801000 | 697 | 348 | [illegible] |
| SE | 38843 | ENSG00000164307 | ERAP1 | chr5 | - | 96800861 | 96801000 | 96795041 | 96795162 | 96803402 | 96803943 | 131 | 66 | [illegible] |
| SE | 38845 | ENSG00000164307 | ERAP1 | chr5 | - | 96803402 | 96803827 | 96797246 | 96797309 | 96807859 | 96807922 | 2 | 1 | [illegible] |
| SE | 38846 | ENSG00000164307 | ERAP1 | chr5 | - | 96804820 | 96804940 | 96803539 | 96803939 | 96807859 | 96807922 | 2 | 2 | [illegible] |
| SE | 38847 | ENSG00000145386 | CCNA2 | chr4 | - | 121818799 | 121818913 | 121818043 | 121818177 | 121819371 | 121819579 | 17 | 25 | [illegible] |
| SE | 38852 | ENSG00000117054 | ACADM | chr1 | + | 75726825 | 75726947 | 75724708 | 75724817 | 75728400 | 75728488 | 0 | 1 | [illegible] |
| SE | 38862 | ENSG00000117054 | ACADM | chr1 | + | 75733050 | 75733149 | 75732852 | 75732922 | 75733527 | 75733628 | 0 | 0 | [illegible] |
| SE | 38863 | ENSG00000117054 | ACADM | chr1 | + | 75733050 | 75733149 | 75732852 | 75732922 | 75734790 | 75734871 | 0 | 0 | [illegible] |
| SE | 38866 | ENSG00000117054 | ACADM | chr1 | + | 75734790 | 75734869 | 75732852 | 75732922 | 75739979 | 75740110 | 4 | 1 | [illegible] |
| SE | 38867 | ENSG00000117054 | ACADM | chr1 | + | 75734790 | 75734871 | 75732852 | 75732922 | 75739979 | 75740110 | 129 | 69 | [illegible] |
| SE | 38868 | ENSG00000117054 | ACADM | chr1 | + | 75734790 | 75734875 | 75732852 | 75732922 | 75739979 | 75740110 | 5 | 2 | [illegible] |
| SE | 38870 | ENSG00000117054 | ACADM | chr1 | + | 75742167 | 75744391 | 75739979 | 75740110 | 75745834 | 75745914 | 0 | 1 | [illegible] |
| SE | 38877 | ENSG00000117054 | ACADM | chr1 | + | 75750450 | 75750546 | 75749418 | 75749559 | 75761121 | 75761370 | 253 | 259 | [illegible] |
| SE | 38878 | ENSG00000117054 | ACADM | chr1 | + | 75750450 | 75750891 | 75749418 | 75749559 | 75761121 | 75761370 | 66 | 77 | [illegible] |
| SE | 38879 | ENSG00000117054 | ACADM | chr1 | + | 75750769 | 75750891 | 75750450 | 75750546 | 75761121 | 75761370 | 0 | 1 | [illegible] |
| SE | 38881 | ENSG00000117054 | ACADM | chr1 | + | 75761121 | 75761370 | 75750450 | 75750546 | 75762691 | 75762809 | 253 | 217 | [illegible] |
| SE | 38882 | ENSG00000137818 | RPLP1 | chr15 | + | 69453646 | 69453721 | 69452817 | 69453020 | 69455169 | 69455287 | 5367 | 2495 | [illegible] |
| SE | 38886 | ENSG00000137818 | RPLP1 | chr15 | + | 69455169 | 69455287 | 69453018 | 69453721 | 69455427 | 69455546 | 8057 | 3968 | [illegible] |
| SE | 38889 | ENSG00000111843 | TMEM14C | chr6 | + | 10724960 | 10725037 | 10723124 | 10723241 | 10725906 | 10726008 | 51 | 19 | [illegible] |
| SE | 38890 | ENSG00000111843 | TMEM14C | chr6 | + | 10725906 | 10726008 | 10723108 | 10723241 | 10728639 | 10728727 | 134 | 44 | [illegible] |
| SE | 38891 | ENSG00000111843 | TMEM14C | chr6 | + | 10728639 | 10728727 | 10725906 | 10726008 | 10730614 | 10730763 | 243 | 89 | [illegible] |
| SE | 38894 | ENSG00000151466 | SCLT1 | chr4 | - | 128874414 | 128874466 | 128870848 | 128871543 | 128888678 | 128888774 | 5 | 2 | [illegible] |
| SE | 38896 | ENSG00000151466 | SCLT1 | chr4 | - | 128891058 | 128891137 | 128888678 | 128888774 | 128936654 | 128936704 | 38 | 37 | [illegible] |
| SE | 38899 | ENSG00000151466 | SCLT1 | chr4 | - | 128948495 | 128948570 | 128946006 | 128946152 | 128952768 | 128952840 | 136 | 155 | [illegible] |
| SE | 38902 | ENSG00000151466 | SCLT1 | chr4 | - | 128965226 | 128965318 | 128959599 | 128959777 | 128997873 | 128997939 | 8 | 16 | [illegible] |
| SE | 38903 | ENSG00000151466 | SCLT1 | chr4 | - | 128970377 | 128970468 | 128965226 | 128965318 | 128997873 | 128997939 | 24 | 28 | [illegible] |
| SE | 38904 | ENSG00000151466 | SCLT1 | chr4 | - | 128992166 | 128992237 | 128959599 | 128959777 | 128997873 | 128997939 | 106 | 96 | [illegible] |
| SE | 38905 | ENSG00000151466 | SCLT1 | chr4 | - | 128992166 | 128992237 | 128970377 | 128970468 | 128997873 | 128997939 | 215 | 229 | [illegible] |
| SE | 38906 | ENSG00000151466 | SCLT1 | chr4 | - | 129003740 | 129003876 | 128999671 | 128999794 | 129039040 | 129039096 | 561 | 377 | [illegible] |
| SE | 38908 | ENSG00000151466 | SCLT1 | chr4 | - | 129043394 | 129043467 | 129039040 | 129039096 | 129043992 | 129044051 | 149 | 170 | [illegible] |
| SE | 38909 | ENSG00000122873 | CISD1 | chr10 | + | 58276117 | 58276212 | 58269177 | 58269304 | 58277116 | 58277322 | 0 | 0 | [illegible] |
| SE | 38919 | ENSG00000067066 | SP100 | chr2 | + | 230442936 | 230443099 | 230417590 | 230417665 | 230444177 | 230444346 | 1118 | 1023 | [illegible] |
| SE | 38921 | ENSG00000067066 | SP100 | chr2 | + | 230446818 | 230446902 | 230444177 | 230444346 | 230449087 | 230449150 | 588 | 659 | [illegible] |
| SE | 38922 | ENSG00000067066 | SP100 | chr2 | + | 230446818 | 230446902 | 230444177 | 230444346 | 230450171 | 230450255 | 236 | 307 | [illegible] |
| SE | 38924 | ENSG00000067066 | SP100 | chr2 | + | 230449087 | 230449150 | 230446818 | 230446902 | 230449313 | 230449380 | 122 | 139 | [illegible] |
| SE | 38925 | ENSG00000067066 | SP100 | chr2 | + | 230449087 | 230449150 | 230446818 | 230446902 | 230449560 | 230449710 | 415 | 375 | [illegible] |
| SE | 38926 | ENSG00000067066 | SP100 | chr2 | + | 230449087 | 230449150 | 230446818 | 230446902 | 230450171 | 230450255 | 115 | 132 | [illegible] |
| SE | 38927 | ENSG00000067066 | SP100 | chr2 | + | 230449087 | 230449150 | 230446818 | 230446902 | 230461261 | 230461414 | 127 | 138 | [illegible] |
| SE | 38928 | ENSG00000067066 | SP100 | chr2 | + | 230449560 | 230449710 | 230446818 | 230446902 | 230450171 | 230450255 | 463 | 394 | [illegible] |
| SE | 38929 | ENSG00000067066 | SP100 | chr2 | + | 230449560 | 230449710 | 230446818 | 230446902 | 230461261 | 230461414 | 3 | 4 | [illegible] |
| SE | 38930 | ENSG00000067066 | SP100 | chr2 | + | 230449560 | 230449710 | 230449087 | 230449150 | 230450171 | 230450255 | 767 | 639 | [illegible] |
| SE | 38932 | ENSG00000067066 | SP100 | chr2 | + | 230449560 | 230449710 | 230449087 | 230449150 | 230462434 | 230462518 | 305 | 245 | [illegible] |
| SE | 38933 | ENSG00000067066 | SP100 | chr2 | + | 230450171 | 230450255 | 230446818 | 230446902 | 230461261 | 230461328 | 284 | 201 | [illegible] |
| SE | 38935 | ENSG00000067066 | SP100 | chr2 | + | 230450171 | 230450255 | 230449087 | 230449150 | 230462434 | 230462518 | 5 | 4 | [illegible] |
| SE | 38936 | ENSG00000067066 | SP100 | chr2 | + | 230450171 | 230450255 | 230449374 | 230449710 | 230461261 | 230461328 | 716 | 562 | [illegible] |
| SE | 38937 | ENSG00000067066 | SP100 | chr2 | + | 230450171 | 230450255 | 230449374 | 230449710 | 230462434 | 230462518 | 157 | 123 | [illegible] |
| SE | 38938 | ENSG00000067066 | SP100 | chr2 | + | 230461261 | 230461414 | 230449087 | 230449150 | 230462434 | 230462518 | 546 | 539 | [illegible] |
| SE | 38939 | ENSG00000067066 | SP100 | chr2 | + | 230461261 | 230461414 | 230449560 | 230449710 | 230462434 | 230462518 | 537 | 537 | [illegible] |
| SE | 38944 | ENSG00000067066 | SP100 | chr2 | + | 230469042 | 230469096 | 230466300 | 230466354 | 230473323 | 230473440 | 0 | 3 | [illegible] |
| SE | 38945 | ENSG00000067066 | SP100 | chr2 | + | 230469042 | 230469096 | 230467119 | 230467215 | 230470014 | 230470098 | 461 | 338 | [illegible] |
| SE | 38946 | ENSG00000067066 | SP100 | chr2 | + | 230470014 | 230470098 | 230469042 | 230469096 | 230473323 | 230473440 | 270 | 222 | [illegible] |
| SE | 38947 | ENSG00000067066 | SP100 | chr2 | + | 230470014 | 230470098 | 230469042 | 230469096 | 230473383 | 230473440 | 282 | 235 | [illegible] |
| SE | 38948 | ENSG00000067066 | SP100 | chr2 | + | 230493791 | 230493911 | 230474393 | 230474447 | 230494415 | 230494460 | 0 | 0 | [illegible] |
| SE | 38950 | ENSG00000067066 | SP100 | chr2 | + | 230503065 | 230503110 | 230498460 | 230498535 | 230504185 | 230504290 | 190 | 241 | [illegible] |
| SE | 38951 | ENSG00000067066 | SP100 | chr2 | + | 230504185 | 230504290 | 230503065 | 230503110 | 230506302 | 230506445 | 302 | 318 | [illegible] |
| SE | 38952 | ENSG00000067066 | SP100 | chr2 | + | 230504185 | 230504290 | 230503065 | 230503110 | 230507992 | 230508031 | 121 | 151 | [illegible] |
| SE | 38953 | ENSG00000067066 | SP100 | chr2 | + | 230506302 | 230506445 | 230503065 | 230503110 | 230507992 | 230508031 | 120 | 107 | [illegible] |
| SE | 38954 | ENSG00000067066 | SP100 | chr2 | + | 230506302 | 230506445 | 230504185 | 230504290 | 230507992 | 230508031 | 300 | 275 | [illegible] |
| SE | 38955 | ENSG00000067066 | SP100 | chr2 | + | 230507992 | 230508031 | 230506302 | 230506445 | 230511124 | 230511166 | 134 | 133 | [illegible] |
| SE | 38956 | ENSG00000067066 | SP100 | chr2 | + | 230511124 | 230511166 | 230507992 | 230508031 | 230539266 | 230539382 | 126 | 117 | [illegible] |
| SE | 38957 | ENSG00000067066 | SP100 | chr2 | + | 230541300 | 230541372 | 230540875 | 230540996 | 230541891 | 230542035 | 208 | 208 | [illegible] |
| SE | 38958 | ENSG00000067066 | SP100 | chr2 | + | 230541891 | 230542035 | 230541300 | 230541372 | 230542835 | 230543122 | 310 | 297 | [illegible] |
| SE | 38960 | ENSG00000101457 | DNTTIP1 | chr20 | + | 45801998 | 45802055 | 45801401 | 45801458 | 45803332 | 45803378 | 22 | 12 | [illegible] |
| SE | 38962 | ENSG00000101457 | DNTTIP1 | chr20 | + | 45809113 | 45809185 | 45805305 | 45805366 | 45810884 | 45810940 | 95 | 47 | [illegible] |
| SE | 38965 | ENSG00000116406 | EDEM3 | chr1 | - | 184693365 | 184693432 | 184690502 | 184692145 | 184694043 | 184694472 | 3 | 3 | [illegible] |
| SE | 38968 | ENSG00000116406 | EDEM3 | chr1 | - | 184726254 | 184726389 | 184723750 | 184723856 | 184732836 | 184732990 | 495 | 405 | [illegible] |
| SE | 38969 | ENSG00000116406 | EDEM3 | chr1 | - | 184734530 | 184734643 | 184732782 | 184732990 | 184737024 | 184737064 | 223 | 181 | [illegible] |
| SE | 38970 | ENSG00000116406 | EDEM3 | chr1 | - | 184749546 | 184749592 | 184737610 | 184737711 | 184754488 | 184754858 | 93 | 56 | [illegible] |
| SE | 38971 | ENSG00000198818 | SFT2D1 | chr6 | - | 166322856 | 166322886 | 166319727 | 166320256 | 166328275 | 166328357 | 69 | 43 | [illegible] |
| SE | 38972 | ENSG00000198818 | SFT2D1 | chr6 | - | 166322856 | 166324595 | 166319859 | 166320256 | 166328275 | 166328357 | 81 | 55 | [illegible] |
| SE | 38973 | ENSG00000198818 | SFT2D1 | chr6 | - | 166324536 | 166324595 | 166322856 | 166322886 | 166328275 | 166328357 | 34 | 19 | [illegible] |
| SE | 38974 | ENSG00000198818 | SFT2D1 | chr6 | - | 166326131 | 166326167 | 166322856 | 166324595 | 166328275 | 166328357 | 102 | 78 | [illegible] |
| SE | 38976 | ENSG00000160298 | C21orf58 | chr21 | - | 46301996 | 46302154 | 46300429 | 46300794 | 46302484 | 46302576 | 9 | 4 | [illegible] |
| SE | 38977 | ENSG00000160298 | C21orf58 | chr21 | - | 46318784 | 46318961 | 46318011 | 46318220 | 46320636 | 46320825 | 3 | 2 | [illegible] |
| SE | 38978 | ENSG00000160298 | C21orf58 | chr21 | - | 46318784 | 46318961 | 46318011 | 46318220 | 46322638 | 46323869 | 3 | 1 | [illegible] |
| SE | 38981 | ENSG00000004399 | PLXND1 | chr3 | - | 129561645 | 129561709 | 129560688 | 129560723 | 129561799 | 129561903 | 199 | 120 | [illegible] |
| SE | 38982 | ENSG00000004399 | PLXND1 | chr3 | - | 129567491 | 129567604 | 129566526 | 129566631 | 129567697 | 129567805 | 116 | 37 | [illegible] |
| SE | 38983 | ENSG00000004399 | PLXND1 | chr3 | - | 129570785 | 129570935 | 129569188 | 129569957 | 129571039 | 129571204 | 173 | 64 | [illegible] |
| SE | 38985 | ENSG00000004399 | PLXND1 | chr3 | - | 129575765 | 129575855 | 129575468 | 129575562 | 129578328 | 129578433 | 80 | 29 | [illegible] |
| SE | 38986 | ENSG00000004399 | PLXND1 | chr3 | - | 129584384 | 129584562 | 129584124 | 129584233 | 129585951 | 129586081 | 144 | 72 | [illegible] |
| SE | 38987 | ENSG00000157216 | SSBP3 | chr1 | - | 54241473 | 54241509 | 54240904 | 54240959 | 54242163 | 54242212 | 270 | 96 | [illegible] |
| SE | 38993 | ENSG00000157216 | SSBP3 | chr1 | - | 54257126 | 54257186 | 54251793 | 54251860 | 54401860 | 54401945 | 50 | 13 | [illegible] |
| SE | 38996 | ENSG00000157216 | SSBP3 | chr1 | - | 54281437 | 54281527 | 54242163 | 54243299 | 54401860 | 54401945 | 451 | 141 | [illegible] |
| SE | 38997 | ENSG00000157216 | SSBP3 | chr1 | - | 54281437 | 54281527 | 54251793 | 54251860 | 54401860 | 54401945 | 458 | 142 | [illegible] |
| SE | 38998 | ENSG00000157216 | SSBP3 | chr1 | - | 54281437 | 54281527 | 54257126 | 54257186 | 54401860 | 54401945 | 706 | 231 | [illegible] |
| SE | 38999 | ENSG00000157216 | SSBP3 | chr1 | - | 54404857 | 54404930 | 54404575 | 54404637 | 54413355 | 54413479 | 188 | 79 | [illegible] |
| SE | 39000 | ENSG00000003400 | CASP10 | chr2 | + | 201185770 | 201186124 | 201182897 | 201183308 | 201187705 | 201187799 | 211 | 113 | [illegible] |
| SE | 39004 | ENSG00000003400 | CASP10 | chr2 | + | 201187705 | 201187799 | 201185770 | 201186124 | 201200442 | 201200728 | 57 | 21 | [illegible] |
| SE | 39006 | ENSG00000003400 | CASP10 | chr2 | + | 201192983 | 201193119 | 201187679 | 201187799 | 201203729 | 201203766 | 88 | 60 | [illegible] |
| SE | 39007 | ENSG00000003400 | CASP10 | chr2 | + | 201192983 | 201193119 | 201187679 | 201187799 | 201205881 | 201205973 | 87 | 59 | [illegible] |
| SE | 39009 | ENSG00000003400 | CASP10 | chr2 | + | 201195841 | 201195948 | 201187705 | 201187799 | 201200442 | 201200728 | 18 | 5 | [illegible] |
| SE | 39010 | ENSG00000003400 | CASP10 | chr2 | + | 201195841 | 201195948 | 201187705 | 201187799 | 201203729 | 201203766 | 96 | 42 | [illegible] |
| SE | 39011 | ENSG00000003400 | CASP10 | chr2 | + | 201195841 | 201195948 | 201187705 | 201187799 | 201208074 | 201208183 | 21 | 5 | [illegible] |
| SE | 39016 | ENSG00000003400 | CASP10 | chr2 | + | 201203729 | 201203766 | 201187705 | 201187799 | 201208074 | 201208160 | 1 | 0 | [illegible] |
| SE | 39017 | ENSG00000003400 | CASP10 | chr2 | + | 201203729 | 201203766 | 201192983 | 201193119 | 201205881 | 201205973 | 13 | 18 | [illegible] |
| SE | 39018 | ENSG00000003400 | CASP10 | chr2 | + | 201203729 | 201203766 | 201192983 | 201193119 | 201208074 | 201208160 | 1 | 0 | [illegible] |
| SE | 39021 | ENSG00000003400 | CASP10 | chr2 | + | 201203729 | 201203766 | 201195841 | 201195948 | 201209069 | 201209562 | 49 | 27 | [illegible] |
| SE | 39026 | ENSG00000003400 | CASP10 | chr2 | + | 201205881 | 201205973 | 201203729 | 201203766 | 201209069 | 201209562 | 64 | 34 | [illegible] |
| SE | 39028 | ENSG00000003400 | CASP10 | chr2 | + | 201208074 | 201208183 | 201195841 | 201195948 | 201209069 | 201209562 | 67 | 39 | [illegible] |
| SE | 39029 | ENSG00000003400 | CASP10 | chr2 | + | 201208074 | 201208183 | 201203729 | 201203766 | 201209069 | 201209562 | 61 | 36 | [illegible] |
| SE | 39030 | ENSG00000003400 | CASP10 | chr2 | + | 201208074 | 201208183 | 201205881 | 201205973 | 201209069 | 201209562 | 162 | 80 | [illegible] |
| SE | 39039 | ENSG00000005448 | WDR54 | chr2 | + | 74422119 | 74422375 | 74421758 | 74421816 | 74422869 | 74422932 | 11 | 3 | [illegible] |
| SE | 39040 | ENSG00000005448 | WDR54 | chr2 | + | 74422152 | 74422375 | 74421729 | 74421816 | 74422869 | 74422932 | 14 | 4 | [illegible] |
| SE | 39041 | ENSG00000005448 | WDR54 | chr2 | + | 74423477 | 74423531 | 74423190 | 74423385 | 74423854 | 74423982 | 23 | 9 | [illegible] |
| SE | 39042 | ENSG00000005448 | WDR54 | chr2 | + | 74424874 | 74424975 | 74423318 | 74423982 | 74425074 | 74425237 | 28 | 20 | [illegible] |
| SE | 39043 | ENSG00000118985 | ELL2 | chr5 | - | 95889085 | 95889130 | 95885097 | 95888987 | 95891102 | 95891274 | 74 | 36 | [illegible] |
| SE | 39045 | ENSG00000118985 | ELL2 | chr5 | - | 95913770 | 95913934 | 95900955 | 95901080 | 95943001 | 95943049 | 1 | 0 | [illegible] |
| SE | 39047 | ENSG00000118985 | ELL2 | chr5 | - | 95919423 | 95919545 | 95913325 | 95913934 | 95932631 | 95932713 | 56 | 40 | [illegible] |
| SE | 39049 | ENSG00000118985 | ELL2 | chr5 | - | 95932631 | 95932713 | 95913819 | 95913934 | 95943001 | 95943049 | 2 | 1 | [illegible] |
| SE | 39053 | ENSG00000274588 | DGKK | chrX | - | 50422445 | 50422526 | 50420402 | 50420507 | 50424247 | 50424358 | 16 | 12 | [illegible] |
| SE | 39056 | ENSG00000172992 | DCAKD | chr17 | - | 45033934 | 45034042 | 45032043 | 45032121 | 45034186 | 45034390 | 3 | 2 | [illegible] |
| SE | 39059 | ENSG00000196172 | ZNF681 | chr19 | - | 23754822 | 23754897 | 23744963 | 23745323 | 23758746 | 23758837 | 4 | 5 | [illegible] |
| SE | 39060 | ENSG00000196172 | ZNF681 | chr19 | - | 23754822 | 23754918 | 23739194 | 23745323 | 23758746 | 23758840 | 7 | 5 | [illegible] |
| SE | 39062 | ENSG00000196172 | ZNF681 | chr19 | - | 23755424 | 23755551 | 23754822 | 23754918 | 23758746 | 23758891 | 2 | 2 | [illegible] |
| SE | 39064 | ENSG00000172175 | MALT1 | chr18 | + | 58700440 | 58700591 | 58696365 | 58696487 | 58709975 | 58710072 | 165 | 107 | [illegible] |
| SE | 39066 | ENSG00000172175 | MALT1 | chr18 | + | 58700440 | 58700591 | 58696365 | 58696487 | 58715934 | 58715967 | 158 | 98 | [illegible] |
| SE | 39067 | ENSG00000172175 | MALT1 | chr18 | + | 58709377 | 58709556 | 58696365 | 58696487 | 58709975 | 58710072 | 155 | 113 | [illegible] |
| SE | 39068 | ENSG00000172175 | MALT1 | chr18 | + | 58709377 | 58709556 | 58696365 | 58696487 | 58714082 | 58714109 | 1 | 1 | [illegible] |
| SE | 39073 | ENSG00000172175 | MALT1 | chr18 | + | 58709975 | 58710072 | 58709377 | 58709556 | 58714082 | 58714109 | 244 | 180 | [illegible] |
| SE | 39075 | ENSG00000172175 | MALT1 | chr18 | + | 58714082 | 58714109 | 58696365 | 58696487 | 58715934 | 58715967 | 67 | 55 | [illegible] |
| SE | 39076 | ENSG00000172175 | MALT1 | chr18 | + | 58714082 | 58714109 | 58700440 | 58700591 | 58715934 | 58715967 | 70 | 59 | [illegible] |
| SE | 39077 | ENSG00000172175 | MALT1 | chr18 | + | 58720379 | 58720422 | 58715934 | 58715967 | 58723047 | 58723251 | 0 | 0 | [illegible] |
| SE | 39080 | ENSG00000172175 | MALT1 | chr18 | + | 58741864 | 58742014 | 58735201 | 58735329 | 58744337 | 58744495 | 831 | 809 | [illegible] |
| SE | 39081 | ENSG00000172175 | MALT1 | chr18 | + | 58744337 | 58744495 | 58741430 | 58742014 | 58745665 | 58745791 | 567 | 501 | [illegible] |
| SE | 39082 | ENSG00000173171 | MTX1 | chr1 | + | 155212126 | 155212219 | 155210547 | 155210627 | 155212384 | 155212567 | 56 | 31 | [illegible] |
| SE | 39086 | ENSG00000171729 | TMEM51 | chr1 | + | 15214894 | 15215431 | 15152531 | 15152784 | 15219325 | 15219346 | 0 | 1 | [illegible] |
| SE | 39087 | ENSG00000171729 | TMEM51 | chr1 | + | 15214894 | 15215431 | 15210489 | 15210562 | 15219325 | 15219346 | 0 | 2 | [illegible] |
| SE | 39089 | ENSG00000109320 | NFKB1 | chr4 | + | 102510879 | 102510949 | 102501404 | 102501788 | 102529835 | 102529914 | 0 | 0 | [illegible] |
| SE | 39090 | ENSG00000109320 | NFKB1 | chr4 | + | 102510879 | 102510949 | 102503374 | 102503490 | 102525511 | 102525557 | 3 | 3 | [illegible] |
| SE | 39092 | ENSG00000109320 | NFKB1 | chr4 | + | 102525511 | 102525557 | 102510879 | 102510949 | 102529835 | 102529914 | 49 | 51 | [illegible] |
| SE | 39093 | ENSG00000109320 | NFKB1 | chr4 | + | 102529835 | 102529914 | 102525511 | 102525557 | 102533844 | 102533885 | 103 | 60 | [illegible] |
| SE | 39099 | ENSG00000109320 | NFKB1 | chr4 | + | 102537857 | 102537956 | 102533844 | 102533885 | 102566986 | 102567135 | 293 | 247 | [illegible] |
| SE | 39100 | ENSG00000109320 | NFKB1 | chr4 | + | 102566986 | 102567135 | 102537857 | 102537956 | 102576875 | 102576902 | 507 | 409 | [illegible] |
| SE | 39103 | ENSG00000109320 | NFKB1 | chr4 | + | 102596830 | 102596872 | 102596137 | 102596332 | 102597519 | 102597661 | 0 | 1 | [illegible] |
| SE | 39104 | ENSG00000109320 | NFKB1 | chr4 | + | 102606495 | 102606697 | 102600894 | 102601009 | 102607149 | 102607319 | 156 | 108 | [illegible] |
| SE | 39105 | ENSG00000109320 | NFKB1 | chr4 | + | 102612043 | 102612110 | 102610574 | 102610699 | 102612433 | 102612606 | 186 | 157 | [illegible] |
| SE | 39106 | ENSG00000117228 | GBP1 | chr1 | - | 89055112 | 89056228 | 89054681 | 89054875 | 89056853 | 89057134 | 412 | 521 | [illegible] |
| SE | 39107 | ENSG00000116120 | FARSB | chr2 | - | 222619644 | 222619737 | 222613810 | 222613928 | 222623649 | 222623730 | 89 | 71 | [illegible] |
| SE | 39109 | ENSG00000116120 | FARSB | chr2 | - | 222640861 | 222640931 | 222639579 | 222639695 | 222642850 | 222643005 | 102 | 108 | [illegible] |
| SE | 39110 | ENSG00000159692 | CTBP1 | chr4 | - | 1213477 | 1213605 | 1212912 | 1213030 | 1214342 | 1214462 | 316 | 109 | [illegible] |
| SE | 39111 | ENSG00000159692 | CTBP1 | chr4 | - | 1225359 | 1225566 | 1215990 | 1216205 | 1228198 | 1228343 | 407 | 134 | [illegible] |
| SE | 39112 | ENSG00000159692 | CTBP1 | chr4 | - | 1238182 | 1238337 | 1228198 | 1228343 | 1241324 | 1241463 | 1077 | 391 | [illegible] |
| SE | 39113 | ENSG00000159692 | CTBP1 | chr4 | - | 1238182 | 1238337 | 1228198 | 1228343 | 1248915 | 1249137 | 605 | 191 | [illegible] |
| SE | 39114 | ENSG00000159692 | CTBP1 | chr4 | - | 1238182 | 1238337 | 1228198 | 1228343 | 1249648 | 1249953 | 599 | 189 | [illegible] |
| SE | 39115 | ENSG00000159692 | CTBP1 | chr4 | - | 1241324 | 1241519 | 1228198 | 1228343 | 1248915 | 1248976 | 15 | 9 | [illegible] |
| SE | 39116 | ENSG00000159692 | CTBP1 | chr4 | - | 1241324 | 1241519 | 1228198 | 1228343 | 1249648 | 1249931 | 2 | 0 | [illegible] |
| SE | 39118 | ENSG00000159692 | CTBP1 | chr4 | - | 1241324 | 1241519 | 1238182 | 1238337 | 1249648 | 1249931 | 480 | 202 | [illegible] |
| SE | 39122 | ENSG00000099985 | OSM | chr22 | - | 30265001 | 30265144 | 30262828 | 30264464 | 30266765 | 30266851 | 10 | 6 | [illegible] |
| SE | 39123 | ENSG00000134352 | IL6ST | chr5 | - | 55947492 | 55947589 | 55942669 | 55942751 | 55951463 | 55951604 | 227 | 177 | [illegible] |
| SE | 39126 | ENSG00000134352 | IL6ST | chr5 | - | 55951463 | 55951604 | 55942669 | 55942751 | 55954809 | 55954992 | 0 | 4 | [illegible] |
| SE | 39128 | ENSG00000134352 | IL6ST | chr5 | - | 55951928 | 55952075 | 55951463 | 55951604 | 55952249 | 55952351 | 433 | 540 | [illegible] |
| SE | 39129 | ENSG00000134352 | IL6ST | chr5 | - | 55951928 | 55952075 | 55951463 | 55951604 | 55954809 | 55954992 | 237 | 320 | [illegible] |
| SE | 39130 | ENSG00000134352 | IL6ST | chr5 | - | 55952249 | 55952351 | 55951463 | 55951604 | 55954809 | 55954992 | 124 | 121 | [illegible] |
| SE | 39131 | ENSG00000134352 | IL6ST | chr5 | - | 55952249 | 55952351 | 55951928 | 55952075 | 55954809 | 55954992 | 308 | 327 | [illegible] |
| SE | 39133 | ENSG00000134352 | IL6ST | chr5 | - | 55957208 | 55957291 | 55955979 | 55956235 | 55960401 | 55960480 | 447 | 594 | [illegible] |
| SE | 39135 | ENSG00000134352 | IL6ST | chr5 | - | 55960401 | 55960561 | 55957208 | 55957291 | 55963351 | 55963506 | 460 | 609 | [illegible] |
| SE | 39136 | ENSG00000134352 | IL6ST | chr5 | - | 55963351 | 55963506 | 55960401 | 55960561 | 55968275 | 55968396 | 246 | 324 | [illegible] |
| SE | 39141 | ENSG00000134352 | IL6ST | chr5 | - | 55968275 | 55968396 | 55963351 | 55963506 | 55976214 | 55976278 | 5 | 13 | [illegible] |
| SE | 39145 | ENSG00000134352 | IL6ST | chr5 | - | 55969549 | 55969855 | 55963351 | 55963506 | 55976214 | 55976278 | 273 | 317 | [illegible] |
| SE | 39147 | ENSG00000134352 | IL6ST | chr5 | - | 55969549 | 55969855 | 55968275 | 55968396 | 55976214 | 55976278 | 581 | 702 | [illegible] |
| SE | 39148 | ENSG00000134352 | IL6ST | chr5 | - | 55969759 | 55969855 | 55963351 | 55963506 | 55976214 | 55976278 | 211 | 245 | [illegible] |
| SE | 39150 | ENSG00000134352 | IL6ST | chr5 | - | 55969759 | 55969855 | 55968275 | 55968396 | 55976214 | 55976278 | 213 | 245 | [illegible] |
| SE | 39151 | ENSG00000134352 | IL6ST | chr5 | - | 55976214 | 55976293 | 55969549 | 55969855 | 55982723 | 55982800 | 395 | 431 | [illegible] |
| SE | 39152 | ENSG00000134352 | IL6ST | chr5 | - | 55982723 | 55982811 | 55976214 | 55976293 | 55994783 | 55994792 | 293 | 355 | [illegible] |
| SE | 39153 | ENSG00000165240 | ATP7A | chrX | + | 77922998 | 77923042 | 77910738 | 77910835 | 77962622 | 77962814 | 3 | 0 | [illegible] |
| SE | 39154 | ENSG00000165240 | ATP7A | chrX | + | 77922998 | 77923042 | 77910738 | 77910835 | 77971620 | 77971761 | 5 | 2 | [illegible] |
| SE | 39155 | ENSG00000165240 | ATP7A | chrX | + | 77922998 | 77923052 | 77910739 | 77910835 | 77962622 | 77962814 | 2 | 0 | [illegible] |
| SE | 39156 | ENSG00000165240 | ATP7A | chrX | + | 77922998 | 77923052 | 77910739 | 77910835 | 77971620 | 77971761 | 2 | 2 | [illegible] |
| SE | 39158 | ENSG00000165240 | ATP7A | chrX | + | 77962622 | 77962814 | 77922998 | 77923042 | 77971620 | 77971761 | 9 | 17 | [illegible] |
| SE | 39159 | ENSG00000165240 | ATP7A | chrX | + | 77962622 | 77962814 | 77922998 | 77923052 | 77971620 | 77971761 | 8 | 17 | [illegible] |
| SE | 39160 | ENSG00000165240 | ATP7A | chrX | + | 77974744 | 77974808 | 77971620 | 77971761 | 77975510 | 77975726 | 0 | 1 | [illegible] |

| | | | | | | | | | | | | | | |
|---|---|---|---|---|---|---|---|---|---|---|---|---|---|---|
| SE | 39161 | ENSG00000165240 | ATP7A | chrX | + | 77974744 | 77974808 | 77971620 | 77971761 | 77988241 | 77988731 | 0 | 1 | [illegible] |
| SE | 39162 | ENSG00000165240 | ATP7A | chrX | + | 77989232 | 77989958 | 77988241 | 77988731 | 77998477 | 77998684 | 96 | 80 | [illegible] |
| SE | 39163 | ENSG00000165240 | ATP7A | chrX | + | 77998477 | 77998684 | 77989232 | 77989958 | 78003072 | 78003236 | 91 | 89 | [illegible] |
| SE | 39164 | ENSG00000165240 | ATP7A | chrX | + | 77998477 | 77998684 | 77989232 | 77989958 | 78009101 | 78009263 | 41 | 45 | [illegible] |
| SE | 39165 | ENSG00000165240 | ATP7A | chrX | + | 78003072 | 78003236 | 77989232 | 77989958 | 78009101 | 78009263 | 58 | 64 | [illegible] |
| SE | 39168 | ENSG00000165240 | ATP7A | chrX | + | 78011448 | 78011674 | 78009101 | 78009263 | 78012878 | 78013112 | 27 | 28 | [illegible] |
| SE | 39171 | ENSG00000165240 | ATP7A | chrX | + | 78020944 | 78021079 | 78020243 | 78020398 | 78031399 | 78031582 | 24 | 15 | [illegible] |
| SE | 39172 | ENSG00000165240 | ATP7A | chrX | + | 78029249 | 78029444 | 78020944 | 78021079 | 78031399 | 78031582 | 42 | 30 | [illegible] |
| SE | 39173 | ENSG00000165240 | ATP7A | chrX | + | 78031399 | 78031582 | 78020243 | 78020398 | 78033604 | 78033821 | 92 | 82 | [illegible] |
| SE | 39174 | ENSG00000165240 | ATP7A | chrX | + | 78031399 | 78031582 | 78029249 | 78029444 | 78033604 | 78033821 | 121 | 104 | [illegible] |
| SE | 39175 | ENSG00000165240 | ATP7A | chrX | + | 78033604 | 78033821 | 78031399 | 78031582 | 78038835 | 78038982 | 142 | 136 | [illegible] |
| SE | 39179 | ENSG00000121775 | TMEM39B | chr1 | + | 32076762 | 32076846 | 32075602 | 32075822 | 32091674 | 32091744 | 0 | 0 | [illegible] |
| SE | 39182 | ENSG00000121775 | TMEM39B | chr1 | + | 32094783 | 32094971 | 32091674 | 32092011 | 32100441 | 32100562 | 38 | 20 | [illegible] |
| SE | 39183 | ENSG00000121775 | TMEM39B | chr1 | + | 32095659 | 32095762 | 32094783 | 32094971 | 32100441 | 32100562 | 0 | 1 | [illegible] |
| SE | 39184 | ENSG00000121775 | TMEM39B | chr1 | + | 32100441 | 32100562 | 32094783 | 32094971 | 32102430 | 32102863 | 28 | 15 | [illegible] |
| SE | 39199 | ENSG00000235652 | FBXO30-DT | chr6 | + | 145881578 | 145881648 | 145828360 | 145828456 | 145883204 | 145883661 | 1 | 0 | [illegible] |
| SE | 39201 | ENSG00000134489 | HRH4 | chr18 | + | 24468787 | 24468951 | 24460636 | 24460921 | 24476746 | 24479961 | 25 | 14 | [illegible] |
| SE | 39202 | ENSG00000286104 | NA | chr19 | + | 58586154 | 58586356 | 58575444 | 58575588 | 58598966 | 58599575 | 13 | 0 | [illegible] |
| SE | 39203 | ENSG00000286104 | NA | chr19 | + | 58591497 | 58591624 | 58586154 | 58586356 | 58598966 | 58599801 | 0 | 4 | [illegible] |
| SE | 39204 | ENSG00000150753 | CCT5 | chr5 | + | 10254144 | 10254205 | 10250215 | 10250445 | 10258110 | 10258303 | 193 | 138 | [illegible] |
| SE | 39206 | ENSG00000150753 | CCT5 | chr5 | + | 10254673 | 10254838 | 10254144 | 10254205 | 10258110 | 10258303 | 387 | 199 | [illegible] |
| SE | 39208 | ENSG00000150753 | CCT5 | chr5 | + | 10254765 | 10254838 | 10254144 | 10254205 | 10258110 | 10258303 | 1 | 1 | [illegible] |
| SE | 39209 | ENSG00000150753 | CCT5 | chr5 | + | 10254795 | 10254838 | 10250242 | 10250445 | 10258110 | 10258247 | 1 | 0 | [illegible] |
| SE | 39210 | ENSG00000150753 | CCT5 | chr5 | + | 10255954 | 10256153 | 10254144 | 10254205 | 10258110 | 10258247 | 236 | 184 | [illegible] |
| SE | 39211 | ENSG00000150753 | CCT5 | chr5 | + | 10255954 | 10256153 | 10254673 | 10254838 | 10258110 | 10258247 | 612 | 424 | [illegible] |
| SE | 39212 | ENSG00000150753 | CCT5 | chr5 | + | 10258110 | 10258303 | 10250318 | 10250445 | 10258385 | 10258535 | 309 | 226 | [illegible] |
| SE | 39213 | ENSG00000150753 | CCT5 | chr5 | + | 10260791 | 10260911 | 10258385 | 10258535 | 10261559 | 10261745 | 704 | 537 | [illegible] |
| SE | 39215 | ENSG00000150753 | CCT5 | chr5 | + | 10260791 | 10260911 | 10258385 | 10258535 | 10264655 | 10264902 | 337 | 276 | [illegible] |
| SE | 39217 | ENSG00000150753 | CCT5 | chr5 | + | 10261559 | 10261745 | 10258385 | 10258535 | 10263133 | 10263314 | 1 | 0 | [illegible] |
| SE | 39218 | ENSG00000150753 | CCT5 | chr5 | + | 10261559 | 10261745 | 10258385 | 10258535 | 10264655 | 10264902 | 0 | 1 | [illegible] |
| SE | 39219 | ENSG00000150753 | CCT5 | chr5 | + | 10261559 | 10261745 | 10260791 | 10260911 | 10262480 | 10262618 | 607 | 442 | [illegible] |
| SE | 39220 | ENSG00000150753 | CCT5 | chr5 | + | 10261559 | 10261745 | 10260791 | 10260911 | 10264655 | 10264902 | 367 | 264 | [illegible] |
| SE | 39222 | ENSG00000150753 | CCT5 | chr5 | + | 10262480 | 10262618 | 10258385 | 10258535 | 10264655 | 10264902 | 1 | 2 | [illegible] |
| SE | 39223 | ENSG00000150753 | CCT5 | chr5 | + | 10262480 | 10262618 | 10260791 | 10260911 | 10264655 | 10264902 | 1 | 2 | [illegible] |
| SE | 39224 | ENSG00000150753 | CCT5 | chr5 | + | 10262480 | 10262618 | 10261559 | 10261745 | 10263133 | 10263314 | 651 | 472 | [illegible] |
| SE | 39225 | ENSG00000150753 | CCT5 | chr5 | + | 10262480 | 10262618 | 10261559 | 10261745 | 10264655 | 10264902 | 227 | 171 | [illegible] |
| SE | 39226 | ENSG00000150753 | CCT5 | chr5 | + | 10263133 | 10263314 | 10258385 | 10258535 | 10264655 | 10264902 | 510 | 410 | [illegible] |
| SE | 39227 | ENSG00000150753 | CCT5 | chr5 | + | 10263133 | 10263314 | 10261559 | 10261745 | 10264655 | 10264902 | 511 | 409 | [illegible] |
| SE | 39228 | ENSG00000150753 | CCT5 | chr5 | + | 10263133 | 10263314 | 10261974 | 10262618 | 10264655 | 10264902 | 935 | 713 | [illegible] |
| SE | 39229 | ENSG00000197497 | ZNF665 | chr19 | - | 53183951 | 53184080 | 53182883 | 53182943 | 53193311 | 53193374 | 0 | 0 | [illegible] |
| SE | 39230 | ENSG00000197497 | ZNF665 | chr19 | - | 53190185 | 53190320 | 53182883 | 53182943 | 53193311 | 53193327 | 0 | 0 | [illegible] |
| SE | 39238 | ENSG00000197948 | FCHSD1 | chr5 | - | 141645770 | 141645932 | 141645019 | 141645148 | 141646086 | 141646191 | 157 | 55 | [illegible] |
| SE | 39239 | ENSG00000197948 | FCHSD1 | chr5 | - | 141646602 | 141646722 | 141646086 | 141646191 | 141647134 | 141647230 | 84 | 44 | [illegible] |
| SE | 39240 | ENSG00000197948 | FCHSD1 | chr5 | - | 141647134 | 141647230 | 141646086 | 141646722 | 141647397 | 141647483 | 61 | 33 | [illegible] |
| SE | 39244 | ENSG00000197948 | FCHSD1 | chr5 | - | 141650358 | 141650404 | 141649886 | 141649954 | 141651019 | 141651117 | 94 | 42 | [illegible] |
| SE | 39245 | ENSG00000144935 | TRPC1 | chr3 | + | 142736378 | 142736533 | 142724033 | 142724731 | 142748257 | 142748460 | 3 | 3 | [illegible] |
| SE | 39246 | ENSG00000144935 | TRPC1 | chr3 | + | 142743484 | 142743586 | 142736378 | 142736533 | 142748257 | 142748460 | 5 | 2 | [illegible] |
| SE | 39247 | ENSG00000144935 | TRPC1 | chr3 | + | 142791018 | 142791158 | 142784703 | 142785040 | 142802168 | 142802344 | 8 | 6 | [illegible] |
| SE | 39248 | ENSG00000144935 | TRPC1 | chr3 | + | 142792823 | 142792967 | 142791018 | 142791158 | 142802168 | 142802344 | 9 | 11 | [illegible] |
| SE | 39251 | ENSG00000113810 | SMC4 | chr3 | + | 160400821 | 160400965 | 160399638 | 160399697 | 160401914 | 160401923 | 7 | 18 | [illegible] |
| SE | 39255 | ENSG00000113810 | SMC4 | chr3 | + | 160401914 | 160402093 | 160399668 | 160399749 | 160402675 | 160402867 | 99 | 58 | [illegible] |
| SE | 39256 | ENSG00000113810 | SMC4 | chr3 | + | 160401914 | 160402093 | 160399668 | 160399749 | 160404327 | 160404336 | 5 | 6 | [illegible] |
| SE | 39258 | ENSG00000113810 | SMC4 | chr3 | + | 160402675 | 160402867 | 160399642 | 160399749 | 160404327 | 160404336 | 185 | 164 | [illegible] |
| SE | 39260 | ENSG00000113810 | SMC4 | chr3 | + | 160402675 | 160402867 | 160401914 | 160402093 | 160404377 | 160404504 | 105 | 67 | [illegible] |
| SE | 39262 | ENSG00000113810 | SMC4 | chr3 | + | 160412325 | 160412453 | 160411919 | 160411977 | 160413472 | 160413484 | 59 | 62 | [illegible] |
| SE | 39265 | ENSG00000113810 | SMC4 | chr3 | + | 160419357 | 160419543 | 160417722 | 160417956 | 160420739 | 160420901 | 213 | 179 | [illegible] |
| SE | 39266 | ENSG00000113810 | SMC4 | chr3 | + | 160431031 | 160431205 | 160430598 | 160430743 | 160431642 | 160431825 | 289 | 248 | [illegible] |
| SE | 39267 | ENSG00000113810 | SMC4 | chr3 | + | 160432282 | 160432515 | 160431642 | 160431825 | 160433025 | 160433209 | 217 | 221 | [illegible] |
| SE | 39268 | ENSG00000081870 | HSPB11 | chr1 | - | 53928371 | 53928440 | 53923532 | 53923947 | 53930038 | 53930145 | 344 | 274 | [illegible] |
| SE | 39269 | ENSG00000081870 | HSPB11 | chr1 | - | 53928371 | 53928440 | 53923532 | 53923947 | 53939984 | 53940097 | 29 | 30 | [illegible] |
| SE | 39270 | ENSG00000081870 | HSPB11 | chr1 | - | 53930038 | 53930145 | 53923532 | 53923947 | 53939984 | 53940097 | 136 | 121 | [illegible] |
| SE | 39271 | ENSG00000081870 | HSPB11 | chr1 | - | 53930038 | 53930145 | 53928371 | 53928440 | 53939984 | 53940097 | 448 | 362 | [illegible] |
| SE | 39272 | ENSG00000081870 | HSPB11 | chr1 | - | 53939984 | 53940097 | 53930038 | 53930145 | 53945554 | 53945572 | 183 | 184 | [illegible] |
| SE | 39275 | ENSG00000128050 | PAICS | chr4 | + | 56446356 | 56446377 | 56441662 | 56441860 | 56446694 | 56446873 | 0 | 0 | [illegible] |
| SE | 39276 | ENSG00000128050 | PAICS | chr4 | + | 56451871 | 56452052 | 56450618 | 56450702 | 56459371 | 56459499 | 58 | 36 | [illegible] |
| SE | 39278 | ENSG00000115318 | LOXL3 | chr2 | - | 74533881 | 74533993 | 74532257 | 74533679 | 74534099 | 74534236 | 57 | 32 | [illegible] |
| SE | 39281 | ENSG00000130695 | CEP85 | chr1 | + | 26244165 | 26244318 | 26239761 | 26239838 | 26255170 | 26255865 | 43 | 41 | [illegible] |
| SE | 39282 | ENSG00000130695 | CEP85 | chr1 | + | 26255170 | 26255865 | 26244165 | 26244318 | 26257596 | 26257730 | 60 | 46 | [illegible] |
| SE | 39283 | ENSG00000130695 | CEP85 | chr1 | + | 26258142 | 26258260 | 26257596 | 26257730 | 26268482 | 26268635 | 24 | 10 | [illegible] |
| SE | 39285 | ENSG00000130695 | CEP85 | chr1 | + | 26268482 | 26268635 | 26258142 | 26258260 | 26269459 | 26269614 | 30 | 29 | [illegible] |
| SE | 39286 | ENSG00000130695 | CEP85 | chr1 | + | 26272020 | 26272071 | 26271013 | 26271107 | 26274963 | 26275071 | 38 | 44 | [illegible] |
| SE | 39287 | ENSG00000130695 | CEP85 | chr1 | + | 26274963 | 26275071 | 26271464 | 26272071 | 26276534 | 26276760 | 65 | 43 | [illegible] |
| SE | 39288 | ENSG00000141293 | SKAP1 | chr17 | - | 48136474 | 48136560 | 48133445 | 48133816 | 48137228 | 48137334 | 0 | 0 | [illegible] |
| SE | 39290 | ENSG00000141293 | SKAP1 | chr17 | - | 48162468 | 48162566 | 48137228 | 48137337 | 48170608 | 48170659 | 369 | 203 | [illegible] |
| SE | 39291 | ENSG00000141293 | SKAP1 | chr17 | - | 48162468 | 48162569 | 48136937 | 48137337 | 48170608 | 48170659 | 476 | 268 | [illegible] |
| SE | 39292 | ENSG00000141293 | SKAP1 | chr17 | - | 48170608 | 48170659 | 48162468 | 48162566 | 48180053 | 48180248 | 167 | 106 | [illegible] |
| SE | 39294 | ENSG00000141293 | SKAP1 | chr17 | - | 48180053 | 48180248 | 48162468 | 48162569 | 48345904 | 48346005 | 2 | 0 | [illegible] |
| SE | 39295 | ENSG00000141293 | SKAP1 | chr17 | - | 48182393 | 48182457 | 48180053 | 48180248 | 48187842 | 48187926 | 94 | 61 | [illegible] |
| SE | 39296 | ENSG00000141293 | SKAP1 | chr17 | - | 48182393 | 48182457 | 48180053 | 48180248 | 48345904 | 48345965 | 75 | 37 | [illegible] |
| SE | 39298 | ENSG00000141293 | SKAP1 | chr17 | - | 48184722 | 48184847 | 48182393 | 48182457 | 48189422 | 48189500 | 295 | 216 | [illegible] |
| SE | 39300 | ENSG00000141293 | SKAP1 | chr17 | - | 48187842 | 48187926 | 48182393 | 48182457 | 48189422 | 48189500 | 143 | 96 | [illegible] |
| SE | 39301 | ENSG00000141293 | SKAP1 | chr17 | - | 48187842 | 48187926 | 48184722 | 48184847 | 48189422 | 48189500 | 248 | 169 | [illegible] |
| SE | 39303 | ENSG00000141293 | SKAP1 | chr17 | - | 48189422 | 48189500 | 48184722 | 48184847 | 48345904 | 48346005 | 153 | 57 | [illegible] |
| SE | 39305 | ENSG00000141293 | SKAP1 | chr17 | - | 48248886 | 48249029 | 48189422 | 48189500 | 48345904 | 48345965 | 0 | 1 | [illegible] |
| SE | 39307 | ENSG00000141293 | SKAP1 | chr17 | - | 48396679 | 48396785 | 48363788 | 48363814 | 48430074 | 48430214 | 325 | 151 | [illegible] |
| SE | 39309 | ENSG00000116514 | RNF19B | chr1 | - | 32938396 | 32938528 | 32936444 | 32937259 | 32944018 | 32944159 | 131 | 104 | [illegible] |
| SE | 39311 | ENSG00000116514 | RNF19B | chr1 | - | 32942251 | 32942459 | 32936444 | 32937259 | 32944018 | 32944159 | 118 | 67 | [illegible] |
| SE | 39312 | ENSG00000116514 | RNF19B | chr1 | - | 32942251 | 32942459 | 32938396 | 32938528 | 32944018 | 32944159 | 246 | 144 | [illegible] |
| SE | 39313 | ENSG00000116514 | RNF19B | chr1 | - | 32944018 | 32944159 | 32942251 | 32942459 | 32945513 | 32945628 | 364 | 237 | [illegible] |
| SE | 39314 | ENSG00000116514 | RNF19B | chr1 | - | 32945513 | 32945628 | 32944018 | 32944159 | 32946401 | 32946564 | 450 | 348 | [illegible] |
| SE | 39315 | ENSG00000116514 | RNF19B | chr1 | - | 32945513 | 32945628 | 32944018 | 32944159 | 32948221 | 32948363 | 223 | 164 | [illegible] |
| SE | 39317 | ENSG00000116514 | RNF19B | chr1 | - | 32946401 | 32946564 | 32945513 | 32945628 | 32948221 | 32948363 | 382 | 314 | [illegible] |
| SE | 39318 | ENSG00000116514 | RNF19B | chr1 | - | 32949568 | 32949774 | 32948221 | 32948363 | 32964050 | 32964685 | 136 | 94 | [illegible] |
| SE | 39319 | ENSG00000116514 | RNF19B | chr1 | - | 32949568 | 32949774 | 32948221 | 32948366 | 32964050 | 32964685 | 50 | 45 | [illegible] |
| SE | 39322 | ENSG00000145388 | METTL14 | chr4 | + | 118697181 | 118697323 | 118694435 | 118694526 | 118700549 | 118700642 | 195 | 210 | [illegible] |
| SE | 39323 | ENSG00000145388 | METTL14 | chr4 | + | 118700549 | 118700642 | 118697181 | 118697323 | 118703934 | 118704051 | 187 | 160 | [illegible] |
| SE | 39324 | ENSG00000145388 | METTL14 | chr4 | + | 118705610 | 118705821 | 118703934 | 118704051 | 118709997 | 118715430 | 217 | 168 | [illegible] |
| SE | 39325 | ENSG00000159588 | CCDC17 | chr1 | - | 45621876 | 45621995 | 45621557 | 45621735 | 45622240 | 45622348 | 0 | 1 | [illegible] |
| SE | 39328 | ENSG00000159588 | CCDC17 | chr1 | - | 45623216 | 45623439 | 45622954 | 45623117 | 45623735 | 45624057 | 0 | 2 | [illegible] |
| SE | 39329 | ENSG00000159588 | CCDC17 | chr1 | - | 45623556 | 45623652 | 45622954 | 45623117 | 45623735 | 45624053 | 1 | 0 | [illegible] |
| SE | 39331 | ENSG00000288684 | NA | chr9 | - | 32551364 | 32551469 | 32550773 | 32550968 | 32552433 | 32552568 | 0 | 0 | [illegible] |
| SE | 39333 | ENSG00000144559 | TAMM41 | chr3 | - | 11807832 | 11807895 | 11790441 | 11790581 | 11809516 | 11809682 | 6 | 4 | [illegible] |
| SE | 39334 | ENSG00000144559 | TAMM41 | chr3 | - | 11808334 | 11808583 | 11807832 | 11807895 | 11809516 | 11810059 | 2 | 0 | [illegible] |
| SE | 39337 | ENSG00000144559 | TAMM41 | chr3 | - | 11826717 | 11826805 | 11809516 | 11809682 | 11829713 | 11829910 | 3 | 1 | [illegible] |
| SE | 39341 | ENSG00000144559 | TAMM41 | chr3 | - | 11844890 | 11845096 | 11844028 | 11844211 | 11846501 | 11846885 | 15 | 0 | [illegible] |
| SE | 39346 | ENSG00000158864 | NDUFS2 | chr1 | + | 161208004 | 161208127 | 161206406 | 161206597 | 161209192 | 161209313 | 0 | 0 | [illegible] |
| SE | 39348 | ENSG00000158864 | NDUFS2 | chr1 | + | 161209842 | 161209931 | 161209482 | 161209595 | 161210110 | 161210188 | 109 | 78 | [illegible] |
| SE | 39349 | ENSG00000158864 | NDUFS2 | chr1 | + | 161209856 | 161209931 | 161209482 | 161209595 | 161210110 | 161210188 | 286 | 181 | [illegible] |
| SE | 39353 | ENSG00000158864 | NDUFS2 | chr1 | + | 161210303 | 161210389 | 161209482 | 161210188 | 161210590 | 161210696 | 342 | 234 | [illegible] |
| SE | 39355 | ENSG00000158864 | NDUFS2 | chr1 | + | 161212350 | 161212480 | 161209482 | 161210710 | 161213379 | 161213475 | 294 | 166 | [illegible] |
| SE | 39356 | ENSG00000158864 | NDUFS2 | chr1 | + | 161212587 | 161213182 | 161212350 | 161212480 | 161213379 | 161213475 | 0 | 0 | [illegible] |
| SE | 39358 | ENSG00000158864 | NDUFS2 | chr1 | + | 161212591 | 161212654 | 161212350 | 161212480 | 161213379 | 161213475 | 2 | 0 | [illegible] |
| SE | 39359 | ENSG00000158864 | NDUFS2 | chr1 | + | 161212591 | 161213182 | 161212350 | 161212480 | 161213379 | 161213475 | 1 | 0 | [illegible] |
| SE | 39360 | ENSG00000158864 | NDUFS2 | chr1 | + | 161213086 | 161213182 | 161212350 | 161212480 | 161213379 | 161213475 | 0 | 0 | [illegible] |
| SE | 39361 | ENSG00000158864 | NDUFS2 | chr1 | + | 161213089 | 161213182 | 161212350 | 161212480 | 161213379 | 161213475 | 0 | 0 | [illegible] |
| SE | 39362 | ENSG00000158864 | NDUFS2 | chr1 | + | 161213089 | 161213182 | 161212591 | 161212654 | 161213379 | 161213475 | 0 | 0 | [illegible] |
| SE | 39364 | ENSG00000285159 | NA | chr6 | + | 158857061 | 158859109 | 158853587 | 158853744 | 158860490 | 158860697 | 5 | 1 | [illegible] |
| SE | 39366 | ENSG00000185745 | IFIT1 | chr10 | + | 89393106 | 89393341 | 89392545 | 89392717 | 89402280 | 89406486 | 0 | 3 | [illegible] |
| SE | 39367 | ENSG00000174437 | ATP2A2 | chr12 | + | 110282603 | 110282621 | 110281115 | 110281907 | 110282712 | 110282795 | 136 | 93 | [illegible] |
| SE | 39369 | ENSG00000174437 | ATP2A2 | chr12 | + | 110322991 | 110323072 | 110296598 | 110296737 | 110326389 | 110326475 | 497 | 416 | [illegible] |
| SE | 39370 | ENSG00000174437 | ATP2A2 | chr12 | + | 110343231 | 110343434 | 110342227 | 110342448 | 110344885 | 110344971 | 294 | 154 | [illegible] |
| SE | 39371 | ENSG00000174437 | ATP2A2 | chr12 | + | 110345248 | 110345382 | 110344885 | 110344971 | 110346000 | 110346118 | 284 | 149 | [illegible] |
| SE | 39372 | ENSG00000174437 | ATP2A2 | chr12 | + | 110346000 | 110346118 | 110345248 | 110345382 | 110346200 | 110346321 | 212 | 109 | [illegible] |
| SE | 39374 | ENSG00000130052 | STARD8 | chrX | + | 68715293 | 68715375 | 68712913 | 68712985 | 68716367 | 68716431 | 21 | 16 | [illegible] |
| SE | 39377 | ENSG00000220804 | LINC01881 | chr2 | + | 242094886 | 242095027 | 242088632 | 242088801 | 242114638 | 242114741 | 79 | 68 | [illegible] |
| SE | 39378 | ENSG00000220804 | LINC01881 | chr2 | + | 242114638 | 242114741 | 242094886 | 242095027 | 242116182 | 242116232 | 93 | 97 | [illegible] |
| SE | 39383 | ENSG00000220804 | LINC01881 | chr2 | + | 242122286 | 242122455 | 242118964 | 242119057 | 242130654 | 242130812 | 117 | 109 | [illegible] |
| SE | 39385 | ENSG00000220804 | LINC01881 | chr2 | + | 242130654 | 242130812 | 242118964 | 242119057 | 242138783 | 242138888 | 9 | 5 | [illegible] |
| SE | 39391 | ENSG00000220804 | LINC01881 | chr2 | + | 242138783 | 242138888 | 242122286 | 242122455 | 242175168 | 242175694 | 8 | 10 | [illegible] |
| SE | 39393 | ENSG00000220804 | LINC01881 | chr2 | + | 242138783 | 242138888 | 242130654 | 242130866 | 242159348 | 242159498 | 14 | 16 | [illegible] |
| SE | 39395 | ENSG00000220804 | LINC01881 | chr2 | + | 242159348 | 242159498 | 242138783 | 242138888 | 242159959 | 242160153 | 44 | 45 | [illegible] |
| SE | 39396 | ENSG00000160712 | IL6R | chr1 | + | 154434518 | 154434700 | 154430482 | 154430606 | 154434989 | 154435156 | 364 | 227 | [illegible] |
| SE | 39397 | ENSG00000160712 | IL6R | chr1 | + | 154435968 | 154436110 | 154434989 | 154435156 | 154437434 | 154437544 | 207 | 156 | [illegible] |
| SE | 39402 | ENSG00000160712 | IL6R | chr1 | + | 154449910 | 154449980 | 154448124 | 154448171 | 154454487 | 154454581 | 231 | 173 | [illegible] |
| SE | 39408 | ENSG00000240230 | COX19 | chr7 | - | 919967 | 920050 | 905059 | 905172 | 926552 | 926644 | 152 | 50 | [illegible] |
| SE | 39409 | ENSG00000240230 | COX19 | chr7 | - | 973180 | 973292 | 964851 | 969456 | 975427 | 975549 | 230 | 98 | [illegible] |
| SE | 39411 | ENSG00000137393 | RNF144B | chr6 | + | 18399498 | 18399699 | 18387349 | 18387630 | 18427580 | 18427685 | 73 | 77 | [illegible] |
| SE | 39413 | ENSG00000137393 | RNF144B | chr6 | + | 18427580 | 18427685 | 18399498 | 18399699 | 18457154 | 18457359 | 32 | 35 | [illegible] |
| SE | 39414 | ENSG00000137393 | RNF144B | chr6 | + | 18427580 | 18427685 | 18399498 | 18399699 | 18459606 | 18459751 | 33 | 31 | [illegible] |
| SE | 39415 | ENSG00000137393 | RNF144B | chr6 | + | 18427580 | 18427685 | 18399498 | 18399699 | 18463290 | 18463380 | 29 | 31 | [illegible] |
| SE | 39416 | ENSG00000137393 | RNF144B | chr6 | + | 18439683 | 18439744 | 18399498 | 18399699 | 18457154 | 18457359 | 50 | 44 | [illegible] |
| SE | 39417 | ENSG00000137393 | RNF144B | chr6 | + | 18439683 | 18439744 | 18399498 | 18399699 | 18459606 | 18459751 | 2 | 1 | [illegible] |
| SE | 39418 | ENSG00000137393 | RNF144B | chr6 | + | 18439683 | 18439744 | 18399498 | 18399699 | 18463290 | 18463380 | 0 | 1 | [illegible] |
| SE | 39421 | ENSG00000137393 | RNF144B | chr6 | + | 18439683 | 18439744 | 18427580 | 18427685 | 18463290 | 18463380 | 12 | 19 | [illegible] |
| SE | 39422 | ENSG00000137393 | RNF144B | chr6 | + | 18457154 | 18457359 | 18399498 | 18399699 | 18459606 | 18459751 | 101 | 125 | [illegible] |
| SE | 39423 | ENSG00000137393 | RNF144B | chr6 | + | 18457154 | 18457359 | 18399498 | 18399699 | 18463290 | 18463380 | 0 | 1 | [illegible] |
| SE | 39425 | ENSG00000137393 | RNF144B | chr6 | + | 18457154 | 18457359 | 18427580 | 18427685 | 18463290 | 18463380 | 4 | 5 | [illegible] |
| SE | 39428 | ENSG00000137393 | RNF144B | chr6 | + | 18459606 | 18459751 | 18399498 | 18399699 | 18463290 | 18463380 | 39 | 99 | [illegible] |
| SE | 39429 | ENSG00000137393 | RNF144B | chr6 | + | 18459606 | 18459751 | 18427580 | 18427685 | 18463290 | 18463380 | 44 | 99 | [illegible] |
| SE | 39432 | ENSG00000055917 | PUM2 | chr2 | - | 20253821 | 20254014 | 20248690 | 20251716 | 20254862 | 20254984 | 1821 | 1413 | [illegible] |
| SE | 39433 | ENSG00000055917 | PUM2 | chr2 | - | 20253821 | 20254014 | 20248690 | 20251716 | 20255215 | 20255341 | 1053 | 804 | [illegible] |
| SE | 39434 | ENSG00000055917 | PUM2 | chr2 | - | 20254862 | 20254984 | 20248690 | 20251716 | 20255215 | 20255341 | 250 | 172 | [illegible] |
| SE | 39435 | ENSG00000055917 | PUM2 | chr2 | - | 20254862 | 20254984 | 20253821 | 20254014 | 20255215 | 20255341 | 995 | 769 | [illegible] |
| SE | 39436 | ENSG00000055917 | PUM2 | chr2 | - | 20258242 | 20258371 | 20256032 | 20256170 | 20260336 | 20260466 | 808 | 492 | [illegible] |
| SE | 39437 | ENSG00000055917 | PUM2 | chr2 | - | 20263192 | 20263460 | 20260336 | 20260466 | 20278582 | 20278819 | 380 | 287 | [illegible] |
| SE | 39441 | ENSG00000055917 | PUM2 | chr2 | - | 20297552 | 20297678 | 20294375 | 20294518 | 20307977 | 20308071 | 225 | 130 | [illegible] |
| SE | 39450 | ENSG00000055917 | PUM2 | chr2 | - | 20332966 | 20333063 | 20331716 | 20331937 | 20350596 | 20350702 | 19 | 13 | [illegible] |
| SE | 39451 | ENSG00000055917 | PUM2 | chr2 | - | 20332966 | 20333063 | 20331716 | 20331937 | 20350809 | 20350834 | 15 | 10 | [illegible] |
| SE | 39452 | ENSG00000213024 | NUP62 | chr19 | - | 49911104 | 49911343 | 49909854 | 49909884 | 49927693 | 49927815 | 0 | 1 | [illegible] |
| SE | 39453 | ENSG00000213024 | NUP62 | chr19 | - | 49911104 | 49911366 | 49909580 | 49909884 | 49927693 | 49927815 | 0 | 1 | [illegible] |
| SE | 39454 | ENSG00000213024 | NUP62 | chr19 | - | 49911104 | 49911366 | 49909580 | 49909884 | 49929325 | 49929504 | 0 | 1 | [illegible] |
| SE | 39455 | ENSG00000213024 | NUP62 | chr19 | - | 49911104 | 49911370 | 49909736 | 49909884 | 49929325 | 49929475 | 0 | 1 | [illegible] |
| SE | 39460 | ENSG00000213024 | NUP62 | chr19 | - | 49918507 | 49918621 | 49909394 | 49909884 | 49929349 | 49929409 | 0 | 3 | [illegible] |
| SE | 39461 | ENSG00000213024 | NUP62 | chr19 | - | 49918507 | 49918621 | 49909394 | 49909884 | 49929416 | 49929504 | 0 | 3 | [illegible] |
| SE | 39466 | ENSG00000213024 | NUP62 | chr19 | - | 49927693 | 49927815 | 49918507 | 49918621 | 49929325 | 49929429 | 21 | 20 | [illegible] |
| SE | 39473 | ENSG00000213024 | NUP62 | chr19 | - | 49927693 | 49927848 | 49918507 | 49918621 | 49929325 | 49929435 | 17 | 18 | [illegible] |
| SE | 39478 | ENSG00000142794 | NBPF3 | chr1 | + | 21466086 | 21466156 | 21444947 | 21445219 | 21466995 | 21467309 | 13 | 1 | [illegible] |
| SE | 39479 | ENSG00000142794 | NBPF3 | chr1 | + | 21468687 | 21468897 | 21444947 | 21445219 | 21470631 | 21470734 | 22 | 22 | [illegible] |
| SE | 39480 | ENSG00000142794 | NBPF3 | chr1 | + | 21474899 | 21474951 | 21473376 | 21473585 | 21478143 | 21478196 | 22 | 25 | [illegible] |
| SE | 39481 | ENSG00000011028 | MRC2 | chr17 | + | 62671997 | 62672152 | 62671648 | 62671837 | 62674062 | 62674170 | 0 | 1 | [illegible] |
| SE | 39484 | ENSG00000158793 | NIT1 | chr1 | + | 161118785 | 161118881 | 161118104 | 161118178 | 161119133 | 161119307 | 30 | 11 | [illegible] |
| SE | 39485 | ENSG00000158793 | NIT1 | chr1 | + | 161118785 | 161118881 | 161118113 | 161118209 | 161119133 | 161119307 | 21 | 6 | [illegible] |
| SE | 39486 | ENSG00000158793 | NIT1 | chr1 | + | 161118785 | 161118881 | 161118116 | 161118551 | 161119133 | 161119307 | 21 | 6 | [illegible] |
| SE | 39487 | ENSG00000158793 | NIT1 | chr1 | + | 161118785 | 161118881 | 161118139 | 161118239 | 161119133 | 161119307 | 22 | 6 | [illegible] |
| SE | 39488 | ENSG00000158793 | NIT1 | chr1 | + | 161118803 | 161118881 | 161118085 | 161118178 | 161119133 | 161119388 | 19 | 5 | [illegible] |
| SE | 39490 | ENSG00000010322 | NISCH | chr3 | + | 52455605 | 52455734 | 52455117 | 52455153 | 52457842 | 52457926 | 34 | 42 | [illegible] |
| SE | 39491 | ENSG00000010322 | NISCH | chr3 | + | 52458661 | 52458844 | 52457842 | 52457926 | 52470858 | 52470907 | 134 | 79 | [illegible] |
| SE | 39495 | ENSG00000010322 | NISCH | chr3 | + | 52480183 | 52480295 | 52479748 | 52479862 | 52485777 | 52485827 | 123 | 69 | [illegible] |
| SE | 39498 | ENSG00000010322 | NISCH | chr3 | + | 52484512 | 52484637 | 52480183 | 52480295 | 52485777 | 52485827 | 128 | 41 | [illegible] |
| SE | 39499 | ENSG00000010322 | NISCH | chr3 | + | 52486469 | 52486552 | 52485784 | 52485827 | 52487195 | 52487653 | 0 | 0 | [illegible] |
| SE | 39501 | ENSG00000010322 | NISCH | chr3 | + | 52489335 | 52489678 | 52487195 | 52488605 | 52491351 | 52491513 | 37 | 18 | [illegible] |
| SE | 39503 | ENSG00000010322 | NISCH | chr3 | + | 52490074 | 52490231 | 52489335 | 52489678 | 52491351 | 52491513 | 43 | 17 | [illegible] |
| SE | 39504 | ENSG00000010322 | NISCH | chr3 | + | 52490704 | 52490833 | 52487195 | 52488605 | 52491351 | 52491513 | 54 | 18 | [illegible] |
| SE | 39505 | ENSG00000010322 | NISCH | chr3 | + | 52490704 | 52490833 | 52489335 | 52489678 | 52491351 | 52491513 | 57 | 19 | [illegible] |
| SE | 39507 | ENSG00000141179 | PCTP | chr17 | + | 55771105 | 55771201 | 55767334 | 55767452 | 55773655 | 55773895 | 23 | 26 | [illegible] |
| SE | 39513 | ENSG00000118260 | CREB1 | chr2 | + | 207567462 | 207567563 | 207560225 | 207560372 | 207570178 | 207570288 | 365 | 252 | [illegible] |
| SE | 39514 | ENSG00000118260 | CREB1 | chr2 | + | 207570178 | 207570321 | 207560225 | 207560372 | 207577504 | 207577631 | 0 | 2 | [illegible] |
| SE | 39516 | ENSG00000118260 | CREB1 | chr2 | + | 207576630 | 207576693 | 207575271 | 207575454 | 207577109 | 207577224 | 5 | 9 | [illegible] |
| SE | 39519 | ENSG00000186517 | ARHGAP30 | chr1 | - | 161049423 | 161049689 | 161046945 | 161049334 | 161051313 | 161051715 | 612 | 285 | [illegible] |
| SE | 39520 | ENSG00000186517 | ARHGAP30 | chr1 | - | 161051313 | 161051715 | 161049423 | 161049689 | 161052285 | 161052363 | 873 | 373 | [illegible] |
| SE | 39521 | ENSG00000186517 | ARHGAP30 | chr1 | - | 161052439 | 161052543 | 161052285 | 161052363 | 161052625 | 161052797 | 974 | 551 | [illegible] |
| SE | 39525 | ENSG00000186517 | ARHGAP30 | chr1 | - | 161053257 | 161053385 | 161052625 | 161052797 | 161054365 | 161054473 | 709 | 318 | [illegible] |
| SE | 39533 | ENSG00000143515 | ATP8B2 | chr1 | + | 154328104 | 154328172 | 154325524 | 154325702 | 154330395 | 154330454 | 40 | 12 | [illegible] |
| SE | 39534 | ENSG00000143515 | ATP8B2 | chr1 | + | 154331443 | 154331505 | 154331047 | 154331146 | 154331605 | 154331678 | 219 | 121 | [illegible] |
| SE | 39535 | ENSG00000143515 | ATP8B2 | chr1 | + | 154331605 | 154331678 | 154331443 | 154331505 | 154331953 | 154332024 | 196 | 125 | [illegible] |
| SE | 39536 | ENSG00000143515 | ATP8B2 | chr1 | + | 154332617 | 154332697 | 154331953 | 154332024 | 154334106 | 154334265 | 185 | 102 | [illegible] |
| SE | 39539 | ENSG00000143515 | ATP8B2 | chr1 | + | 154342479 | 154342523 | 154340853 | 154341062 | 154342795 | 154342961 | 180 | 71 | [illegible] |
| SE | 39540 | ENSG00000143515 | ATP8B2 | chr1 | + | 154343112 | 154343301 | 154342795 | 154342961 | 154343452 | 154343568 | 495 | 246 | [illegible] |
| SE | 39541 | ENSG00000143515 | ATP8B2 | chr1 | + | 154345799 | 154345883 | 154345321 | 154345545 | 154346230 | 154346476 | 183 | 100 | [illegible] |
| SE | 39542 | ENSG00000174547 | MRPL11 | chr11 | - | 66436801 | 66436909 | 66435833 | 66436112 | 66437103 | 66437222 | 0 | 1 | [illegible] |
| SE | 39543 | ENSG00000174547 | MRPL11 | chr11 | - | 66436815 | 66436909 | 66435837 | 66436112 | 66437103 | 66437263 | 0 | 1 | [illegible] |
| SE | 39554 | ENSG00000144554 | FANCD2 | chr3 | + | 10032831 | 10032972 | 10028619 | 10028721 | 10034468 | 10034513 | 78 | 67 | [illegible] |
| SE | 39555 | ENSG00000144554 | FANCD2 | chr3 | + | 10034468 | 10034536 | 10028619 | 10028721 | 10035172 | 10035233 | 1 | 0 | [illegible] |
| SE | 39556 | ENSG00000144554 | FANCD2 | chr3 | + | 10034468 | 10034536 | 10032831 | 10032972 | 10034694 | 10034798 | 41 | 35 | [illegible] |
| SE | 39557 | ENSG00000144554 | FANCD2 | chr3 | + | 10034694 | 10034798 | 10034468 | 10034536 | 10035172 | 10035233 | 56 | 44 | [illegible] |
| SE | 39558 | ENSG00000144554 | FANCD2 | chr3 | + | 10036286 | 10036339 | 10035172 | 10035233 | 10039278 | 10039357 | 48 | 38 | [illegible] |
| SE | 39559 | ENSG00000144554 | FANCD2 | chr3 | + | 10039278 | 10039357 | 10036286 | 10036339 | 10039720 | 10039845 | 49 | 40 | [illegible] |
| SE | 39560 | ENSG00000144554 | FANCD2 | chr3 | + | 10041622 | 10041710 | 10039720 | 10039845 | 10042558 | 10042663 | 11 | 15 | [illegible] |
| SE | 39561 | ENSG00000144554 | FANCD2 | chr3 | + | 10043049 | 10043150 | 10042558 | 10042663 | 10043483 | 10043592 | 17 | 16 | [illegible] |

| | | | | | | | | | | | | | | |
|---|---|---|---|---|---|---|---|---|---|---|---|---|---|---|
| SE | 39562 | ENSG00000144554 | FANCD2 | chr3 | + | 10049373 | 10049505 | 10043828 | 10043864 | 10052386 | 10052497 | 24 | 18 | [illegible] |
| SE | 39563 | ENSG00000144554 | FANCD2 | chr3 | + | 10052386 | 10052497 | 10049373 | 10049505 | 10060293 | 10060403 | 42 | 25 | [illegible] |
| SE | 39565 | ENSG00000144554 | FANCD2 | chr3 | + | 10067208 | 10067317 | 10065863 | 10065979 | 10072870 | 10072964 | 45 | 30 | [illegible] |
| SE | 39566 | ENSG00000144554 | FANCD2 | chr3 | + | 10072870 | 10072981 | 10067208 | 10067317 | 10073252 | 10073362 | 25 | 28 | [illegible] |
| SE | 39569 | ENSG00000144554 | FANCD2 | chr3 | + | 10087133 | 10087264 | 10085811 | 10085922 | 10088448 | 10088542 | 36 | 36 | [illegible] |
| SE | 39570 | ENSG00000144554 | FANCD2 | chr3 | + | 10090291 | 10090385 | 10088827 | 10088950 | 10092180 | 10092252 | 10 | 13 | [illegible] |
| SE | 39571 | ENSG00000144554 | FANCD2 | chr3 | + | 10093284 | 10093323 | 10092180 | 10092252 | 10094288 | 10094363 | 21 | 15 | [illegible] |
| SE | 39572 | ENSG00000144554 | FANCD2 | chr3 | + | 10094288 | 10094363 | 10093284 | 10093323 | 10095199 | 10095274 | 37 | 20 | [illegible] |
| SE | 39573 | ENSG00000106330 | MOSPD3 | chr7 | + | 100613193 | 100613269 | 100612539 | 100612996 | 100613476 | 100613706 | 26 | 1 | [illegible] |
| SE | 39574 | ENSG00000140307 | GTF2A2 | chr15 | - | 59642135 | 59642262 | 59638061 | 59639157 | 59650668 | 59650764 | 419 | 289 | [illegible] |
| SE | 39575 | ENSG00000140307 | GTF2A2 | chr15 | - | 59642135 | 59642262 | 59638061 | 59639157 | 59652205 | 59652326 | 203 | 112 | [illegible] |
| SE | 39576 | ENSG00000140307 | GTF2A2 | chr15 | - | 59650668 | 59650764 | 59642135 | 59642262 | 59652205 | 59652326 | 193 | 159 | [illegible] |
| SE | 39578 | ENSG00000140307 | GTF2A2 | chr15 | - | 59650668 | 59650773 | 59638865 | 59639157 | 59652205 | 59652326 | 168 | 106 | [illegible] |
| SE | 39579 | ENSG00000140307 | GTF2A2 | chr15 | - | 59650668 | 59650773 | 59642135 | 59642262 | 59652205 | 59652326 | 380 | 282 | [illegible] |
| SE | 39583 | ENSG00000140307 | GTF2A2 | chr15 | - | 59652205 | 59652326 | 59650668 | 59650773 | 59657405 | 59657515 | 315 | 196 | [illegible] |
| SE | 39585 | ENSG00000140307 | GTF2A2 | chr15 | - | 59656814 | 59657033 | 59652205 | 59652326 | 59657405 | 59657514 | 2 | 0 | [illegible] |
| SE | 39586 | ENSG00000197568 | HHLA3 | chr1 | + | 70366145 | 70366328 | 70354810 | 70355111 | 70366443 | 70366537 | 4 | 2 | [illegible] |
| SE | 39587 | ENSG00000197568 | HHLA3 | chr1 | + | 70366145 | 70366328 | 70354854 | 70355107 | 70366443 | 70366537 | 4 | 4 | [illegible] |
| SE | 39591 | ENSG00000186148 | NA | chr2 | + | 109964687 | 109964802 | 109947363 | 109947845 | 109968069 | 109968198 | 0 | 0 | [illegible] |
| SE | 39592 | ENSG00000135951 | TSGA10 | chr2 | - | 99020279 | 99020482 | 99018535 | 99018640 | 99035229 | 99035439 | 4 | 4 | [illegible] |
| SE | 39593 | ENSG00000135951 | TSGA10 | chr2 | - | 99020279 | 99020482 | 99018535 | 99018640 | 99064938 | 99065124 | 2 | 3 | [illegible] |
| SE | 39594 | ENSG00000135951 | TSGA10 | chr2 | - | 99035229 | 99035439 | 99018535 | 99018640 | 99064938 | 99065124 | 2 | 5 | [illegible] |
| SE | 39595 | ENSG00000135951 | TSGA10 | chr2 | - | 99035229 | 99035439 | 99020279 | 99020482 | 99064938 | 99065124 | 4 | 4 | [illegible] |
| SE | 39596 | ENSG00000135951 | TSGA10 | chr2 | - | 99073017 | 99073073 | 99071705 | 99071874 | 99078658 | 99078813 | 13 | 6 | [illegible] |
| SE | 39598 | ENSG00000135951 | TSGA10 | chr2 | - | 99103966 | 99104118 | 99071705 | 99071874 | 99105358 | 99105436 | 10 | 16 | [illegible] |
| SE | 39599 | ENSG00000135951 | TSGA10 | chr2 | - | 99117543 | 99117759 | 99110849 | 99110915 | 99127047 | 99127146 | 53 | 53 | [illegible] |
| SE | 39600 | ENSG00000135951 | TSGA10 | chr2 | - | 99117543 | 99117759 | 99110849 | 99110915 | 99141427 | 99141572 | 35 | 39 | [illegible] |
| SE | 39601 | ENSG00000135951 | TSGA10 | chr2 | - | 99117543 | 99117759 | 99110849 | 99110915 | 99154692 | 99154962 | 35 | 39 | [illegible] |
| SE | 39602 | ENSG00000135951 | TSGA10 | chr2 | - | 99118550 | 99118686 | 99117543 | 99117759 | 99118776 | 99118801 | 5 | 9 | [illegible] |
| SE | 39603 | ENSG00000135951 | TSGA10 | chr2 | - | 99118550 | 99118686 | 99117543 | 99117759 | 99127047 | 99127176 | 6 | 6 | [illegible] |
| SE | 39604 | ENSG00000135951 | TSGA10 | chr2 | - | 99118776 | 99118801 | 99117566 | 99117759 | 99127047 | 99127176 | 3 | 0 | [illegible] |
| SE | 39606 | ENSG00000135951 | TSGA10 | chr2 | - | 99127047 | 99127176 | 99110849 | 99110915 | 99141427 | 99141572 | 2 | 8 | [illegible] |
| SE | 39608 | ENSG00000135951 | TSGA10 | chr2 | - | 99127047 | 99127176 | 99117543 | 99117759 | 99141427 | 99141572 | 20 | 19 | [illegible] |
| SE | 39612 | ENSG00000164142 | FAM160A1 | chr4 | + | 151454700 | 151454808 | 151409175 | 151409465 | 151482523 | 151482648 | 6 | 2 | [illegible] |
| SE | 39613 | ENSG00000164142 | FAM160A1 | chr4 | + | 151454700 | 151454808 | 151409175 | 151409465 | 151566137 | 151566364 | 4 | 2 | [illegible] |
| SE | 39614 | ENSG00000164142 | FAM160A1 | chr4 | + | 151482523 | 151482648 | 151409351 | 151409465 | 151566137 | 151566258 | 6 | 2 | [illegible] |
| SE | 39615 | ENSG00000164142 | FAM160A1 | chr4 | + | 151482523 | 151482648 | 151454700 | 151454808 | 151566137 | 151566258 | 10 | 4 | [illegible] |
| SE | 39617 | ENSG00000164142 | FAM160A1 | chr4 | + | 151656231 | 151656410 | 151649458 | 151650592 | 151656759 | 151656898 | 0 | 7 | [illegible] |
| SE | 39624 | ENSG00000091483 | FH | chr1 | - | 241502442 | 241502570 | 241498737 | 241500590 | 241504041 | 241504245 | 202 | 79 | [illegible] |
| SE | 39625 | ENSG00000091483 | FH | chr1 | - | 241508602 | 241508760 | 241506002 | 241506168 | 241511966 | 241512143 | 32 | 13 | [illegible] |
| SE | 39626 | ENSG00000091483 | FH | chr1 | - | 241508602 | 241508785 | 241506002 | 241506168 | 241511966 | 241512143 | 109 | 59 | [illegible] |
| SE | 39627 | ENSG00000091483 | FH | chr1 | - | 241519059 | 241519088 | 241517181 | 241517376 | 241519590 | 241519722 | 0 | 1 | [illegible] |
| SE | 39629 | ENSG00000090989 | EXOC1 | chr4 | + | 55870677 | 55870905 | 55868335 | 55868523 | 55871100 | 55871233 | 141 | 79 | [illegible] |
| SE | 39630 | ENSG00000090989 | EXOC1 | chr4 | + | 55871848 | 55871958 | 55871100 | 55871233 | 55877916 | 55878066 | 77 | 50 | [illegible] |
| SE | 39631 | ENSG00000090989 | EXOC1 | chr4 | + | 55883822 | 55883928 | 55877916 | 55878066 | 55890222 | 55890386 | 139 | 134 | [illegible] |
| SE | 39632 | ENSG00000090989 | EXOC1 | chr4 | + | 55888887 | 55888932 | 55882726 | 55883928 | 55890222 | 55890386 | 3 | 0 | [illegible] |
| SE | 39633 | ENSG00000090989 | EXOC1 | chr4 | + | 55890222 | 55890386 | 55883767 | 55883928 | 55891314 | 55891422 | 204 | 216 | [illegible] |
| SE | 39636 | ENSG00000090989 | EXOC1 | chr4 | + | 55892614 | 55892711 | 55891314 | 55891422 | 55893551 | 55893641 | 15 | 12 | [illegible] |
| SE | 39637 | ENSG00000090989 | EXOC1 | chr4 | + | 55892634 | 55892711 | 55891314 | 55891422 | 55893551 | 55893627 | 81 | 70 | [illegible] |
| SE | 39638 | ENSG00000090989 | EXOC1 | chr4 | + | 55893551 | 55893780 | 55892634 | 55892711 | 55896716 | 55896900 | 112 | 68 | [illegible] |
| SE | 39639 | ENSG00000090989 | EXOC1 | chr4 | + | 55899684 | 55899884 | 55896716 | 55896900 | 55902343 | 55902538 | 187 | 193 | [illegible] |
| SE | 39640 | ENSG00000090989 | EXOC1 | chr4 | + | 55900558 | 55900692 | 55899684 | 55899884 | 55902343 | 55902538 | 4 | 3 | [illegible] |
| SE | 39642 | ENSG00000197774 | EME2 | chr16 | + | 1775040 | 1775132 | 1774259 | 1774352 | 1775314 | 1775408 | 0 | 0 | [illegible] |
| SE | 39643 | ENSG00000197774 | EME2 | chr16 | + | 1776745 | 1777405 | 1776067 | 1776339 | 1778454 | 1781708 | 1 | 0 | [illegible] |
| SE | 39644 | ENSG00000148290 | SURF1 | chr9 | - | 133352693 | 133352730 | 133352445 | 133352608 | 133353748 | 133353940 | 23 | 19 | [illegible] |
| SE | 39645 | ENSG00000148290 | SURF1 | chr9 | - | 133352693 | 133352766 | 133352445 | 133352608 | 133353748 | 133353940 | 204 | 88 | [illegible] |
| SE | 39646 | ENSG00000148290 | SURF1 | chr9 | - | 133353748 | 133353940 | 133352693 | 133352766 | 133354658 | 133354741 | 184 | 67 | [illegible] |
| SE | 39647 | ENSG00000148290 | SURF1 | chr9 | - | 133356268 | 133356320 | 133354823 | 133354957 | 133356399 | 133356487 | 12 | 3 | [illegible] |
| SE | 39648 | ENSG00000176171 | BNIP3 | chr10 | - | 131970863 | 131970970 | 131968227 | 131970787 | 131973033 | 131973118 | 56 | 45 | [illegible] |
| SE | 39649 | ENSG00000176171 | BNIP3 | chr10 | - | 131973792 | 131973943 | 131973033 | 131973118 | 131981760 | 131981923 | 67 | 39 | [illegible] |
| SE | 39650 | ENSG00000127946 | HIP1 | chr7 | - | 75546938 | 75547032 | 75545087 | 75545188 | 75547754 | 75547813 | 17 | 35 | [illegible] |
| SE | 39652 | ENSG00000127946 | HIP1 | chr7 | - | 75559731 | 75559915 | 75558166 | 75558255 | 75561328 | 75561401 | 49 | 67 | [illegible] |
| SE | 39653 | ENSG00000127946 | HIP1 | chr7 | - | 75573760 | 75573901 | 75568198 | 75568256 | 75581236 | 75581298 | 79 | 68 | [illegible] |
| SE | 39656 | ENSG00000173545 | ZNF622 | chr5 | - | 16463107 | 16463270 | 16458516 | 16458629 | 16463481 | 16463742 | 239 | 135 | [illegible] |
| SE | 39657 | ENSG00000181555 | SETD2 | chr3 | - | 47042560 | 47042700 | 47037664 | 47037777 | 47056820 | 47057490 | 145 | 130 | [illegible] |
| SE | 39658 | ENSG00000181555 | SETD2 | chr3 | - | 47046486 | 47046621 | 47042560 | 47042700 | 47056820 | 47057490 | 195 | 162 | [illegible] |
| SE | 39659 | ENSG00000181555 | SETD2 | chr3 | - | 47056820 | 47057490 | 47046486 | 47046621 | 47062162 | 47062346 | 139 | 128 | [illegible] |
| SE | 39660 | ENSG00000181555 | SETD2 | chr3 | - | 47062162 | 47062346 | 47056820 | 47057490 | 47067069 | 47067118 | 179 | 180 | [illegible] |
| SE | 39664 | ENSG00000181555 | SETD2 | chr3 | - | 47083719 | 47084382 | 47066991 | 47067118 | 47086194 | 47086314 | 414 | 349 | [illegible] |
| SE | 39666 | ENSG00000181555 | SETD2 | chr3 | - | 47086194 | 47086314 | 47066991 | 47067118 | 47097954 | 47097961 | 4 | 4 | [illegible] |
| SE | 39667 | ENSG00000181555 | SETD2 | chr3 | - | 47086194 | 47086314 | 47083719 | 47084382 | 47088112 | 47088247 | 309 | 236 | [illegible] |
| SE | 39668 | ENSG00000181555 | SETD2 | chr3 | - | 47086194 | 47086314 | 47083719 | 47084382 | 47097954 | 47097961 | 182 | 146 | [illegible] |
| SE | 39669 | ENSG00000181555 | SETD2 | chr3 | - | 47088112 | 47088247 | 47083719 | 47084382 | 47097954 | 47098081 | 54 | 64 | [illegible] |
| SE | 39671 | ENSG00000181555 | SETD2 | chr3 | - | 47088112 | 47088249 | 47083719 | 47084382 | 47097954 | 47097961 | 0 | 1 | [illegible] |
| SE | 39675 | ENSG00000181555 | SETD2 | chr3 | - | 47101457 | 47101555 | 47097954 | 47098081 | 47105996 | 47106120 | 192 | 180 | [illegible] |
| SE | 39676 | ENSG00000181555 | SETD2 | chr3 | - | 47103345 | 47103423 | 47097954 | 47098081 | 47105996 | 47106120 | 137 | 143 | [illegible] |
| SE | 39677 | ENSG00000181555 | SETD2 | chr3 | - | 47103345 | 47103423 | 47097954 | 47098081 | 47113875 | 47114004 | 6 | 7 | [illegible] |
| SE | 39678 | ENSG00000181555 | SETD2 | chr3 | - | 47103345 | 47103423 | 47101457 | 47101555 | 47105996 | 47106120 | 405 | 377 | [illegible] |
| SE | 39679 | ENSG00000181555 | SETD2 | chr3 | - | 47105996 | 47106120 | 47097954 | 47098081 | 47113875 | 47114004 | 160 | 93 | [illegible] |
| SE | 39687 | ENSG00000224699 | LAMTOR5-AS1 | chr1 | + | 110413320 | 110413410 | 110412172 | 110412293 | 110416008 | 110416186 | 1 | 0 | [illegible] |
| SE | 39702 | ENSG00000198734 | F5 | chr1 | - | 169525900 | 169526017 | 169524836 | 169524908 | 169529607 | 169529818 | 39 | 27 | [illegible] |
| SE | 39704 | ENSG00000198734 | F5 | chr1 | - | 169529607 | 169529818 | 169527914 | 169528094 | 169530785 | 169531022 | 76 | 35 | [illegible] |
| SE | 39705 | ENSG00000198734 | F5 | chr1 | - | 169540293 | 169543114 | 169536505 | 169536680 | 169544295 | 169544508 | 79 | 49 | [illegible] |
| SE | 39706 | ENSG00000198734 | F5 | chr1 | - | 169550639 | 169550739 | 169549800 | 169550015 | 169552556 | 169552734 | 159 | 90 | [illegible] |
| SE | 39707 | ENSG00000198734 | F5 | chr1 | - | 169550639 | 169550739 | 169549800 | 169550015 | 169555181 | 169555347 | 65 | 48 | [illegible] |
| SE | 39708 | ENSG00000198734 | F5 | chr1 | - | 169552556 | 169552734 | 169549800 | 169550015 | 169555181 | 169555347 | 62 | 41 | [illegible] |
| SE | 39709 | ENSG00000198734 | F5 | chr1 | - | 169552556 | 169552734 | 169550639 | 169550739 | 169555181 | 169555347 | 160 | 84 | [illegible] |
| SE | 39710 | ENSG00000198734 | F5 | chr1 | - | 169555181 | 169555347 | 169552556 | 169552734 | 169556645 | 169556867 | 98 | 71 | [illegible] |
| SE | 39711 | ENSG00000198734 | F5 | chr1 | - | 169582430 | 169582522 | 169572220 | 169572343 | 169586228 | 169586481 | 38 | 19 | [illegible] |
| SE | 39712 | ENSG00000132716 | DCAF8 | chr1 | - | 160220458 | 160220532 | 160218848 | 160218968 | 160221086 | 160221210 | 3 | 5 | [illegible] |
| SE | 39715 | ENSG00000132716 | DCAF8 | chr1 | - | 160224441 | 160224549 | 160222650 | 160222781 | 160225590 | 160225663 | 357 | 261 | [illegible] |
| SE | 39717 | ENSG00000132716 | DCAF8 | chr1 | - | 160225590 | 160225663 | 160222650 | 160222781 | 160231296 | 160231407 | 68 | 56 | [illegible] |
| SE | 39719 | ENSG00000132716 | DCAF8 | chr1 | - | 160231296 | 160231407 | 160222650 | 160222781 | 160237134 | 160237229 | 229 | 144 | [illegible] |
| SE | 39720 | ENSG00000132716 | DCAF8 | chr1 | - | 160231296 | 160231407 | 160225590 | 160225663 | 160237134 | 160237229 | 508 | 388 | [illegible] |
| SE | 39723 | ENSG00000132716 | DCAF8 | chr1 | - | 160243959 | 160244034 | 160239221 | 160240370 | 160262275 | 160262448 | 208 | 181 | [illegible] |
| SE | 39733 | ENSG00000232706 | NUTM2HP | chr10 | + | 50682928 | 50683065 | 50681827 | 50681956 | 50683510 | 50683891 | 0 | 0 | [illegible] |
| SE | 39737 | ENSG00000132581 | SDF2 | chr17 | - | 28651485 | 28651577 | 28648924 | 28649276 | 28662045 | 28662189 | 8 | 4 | [illegible] |
| SE | 39738 | ENSG00000132581 | SDF2 | chr17 | - | 28655286 | 28655483 | 28648345 | 28649276 | 28661725 | 28661819 | 162 | 96 | [illegible] |
| SE | 39739 | ENSG00000132581 | SDF2 | chr17 | - | 28655286 | 28655483 | 28648345 | 28649276 | 28662045 | 28662132 | 146 | 78 | [illegible] |
| SE | 39740 | ENSG00000132581 | SDF2 | chr17 | - | 28655286 | 28655483 | 28651485 | 28651577 | 28662045 | 28662132 | 39 | 19 | [illegible] |
| SE | 39741 | ENSG00000240038 | AMY2B | chr1 | + | 103565434 | 103565594 | 103554699 | 103555109 | 103571556 | 103571770 | 173 | 144 | [illegible] |
| SE | 39744 | ENSG00000116478 | HDAC1 | chr1 | + | 32302620 | 32302733 | 32292082 | 32292218 | 32316664 | 32316782 | 454 | 285 | [illegible] |
| SE | 39745 | ENSG00000116478 | HDAC1 | chr1 | + | 32316664 | 32316782 | 32302620 | 32302733 | 32324478 | 32324553 | 975 | 636 | [illegible] |
| SE | 39746 | ENSG00000116478 | HDAC1 | chr1 | + | 32324478 | 32324553 | 32316664 | 32316782 | 32326938 | 32327077 | 778 | 480 | [illegible] |
| SE | 39748 | ENSG00000116478 | HDAC1 | chr1 | + | 32329067 | 32329160 | 32327188 | 32327677 | 32330577 | 32330686 | 175 | 76 | [illegible] |
| SE | 39751 | ENSG00000149554 | CHEK1 | chr11 | + | 125627606 | 125627830 | 125626315 | 125626833 | 125629231 | 125629296 | 12 | 14 | [illegible] |
| SE | 39754 | ENSG00000149554 | CHEK1 | chr11 | + | 125656205 | 125656335 | 125655224 | 125655347 | 125675927 | 125676041 | 0 | 2 | [illegible] |
| SE | 39757 | ENSG00000182866 | LCK | chr1 | + | 32274324 | 32274434 | 32251238 | 32251371 | 32274736 | 32274818 | 394 | 171 | [illegible] |
| SE | 39759 | ENSG00000182866 | LCK | chr1 | + | 32279580 | 32279747 | 32276606 | 32276786 | 32279840 | 32279994 | 356 | 193 | [illegible] |
| SE | 39760 | ENSG00000182866 | LCK | chr1 | + | 32279670 | 32279747 | 32276606 | 32276786 | 32279840 | 32279909 | 887 | 421 | [illegible] |
| SE | 39761 | ENSG00000182866 | LCK | chr1 | + | 32280078 | 32280210 | 32279670 | 32279994 | 32285513 | 32286160 | 1220 | 621 | [illegible] |
| SE | 39764 | ENSG00000132185 | FCRLA | chr1 | + | 161710474 | 161710492 | 161706971 | 161707343 | 161711933 | 161712218 | 0 | 0 | [illegible] |
| SE | 39765 | ENSG00000132185 | FCRLA | chr1 | + | 161710759 | 161710912 | 161706971 | 161707343 | 161711933 | 161712218 | 37 | 13 | [illegible] |
| SE | 39766 | ENSG00000132185 | FCRLA | chr1 | + | 161710759 | 161710912 | 161706971 | 161707343 | 161713084 | 161713705 | 36 | 13 | [illegible] |
| SE | 39769 | ENSG00000132185 | FCRLA | chr1 | + | 161711207 | 161711474 | 161710759 | 161710912 | 161713084 | 161713452 | 50 | 12 | [illegible] |
| SE | 39770 | ENSG00000132185 | FCRLA | chr1 | + | 161711207 | 161712218 | 161710288 | 161710912 | 161713084 | 161714352 | 122 | 24 | [illegible] |
| SE | 39771 | ENSG00000132185 | FCRLA | chr1 | + | 161711933 | 161712218 | 161706971 | 161707343 | 161713084 | 161713468 | 73 | 12 | [illegible] |
| SE | 39772 | ENSG00000132185 | FCRLA | chr1 | + | 161711933 | 161712218 | 161710759 | 161710912 | 161713084 | 161713468 | 73 | 12 | [illegible] |
| SE | 39773 | ENSG00000132185 | FCRLA | chr1 | + | 161711933 | 161712218 | 161711207 | 161711474 | 161713084 | 161713468 | 104 | 35 | [illegible] |
| SE | 39776 | ENSG00000072274 | TFRC | chr3 | - | 196069454 | 196069568 | 196068031 | 196068130 | 196072002 | 196072152 | 52 | 36 | [illegible] |
| SE | 39778 | ENSG00000072274 | TFRC | chr3 | - | 196072002 | 196072152 | 196069454 | 196069568 | 196073929 | 196074125 | 119 | 86 | [illegible] |
| SE | 39779 | ENSG00000072274 | TFRC | chr3 | - | 196073929 | 196074125 | 196072002 | 196072152 | 196075158 | 196075360 | 274 | 211 | [illegible] |
| SE | 39782 | ENSG00000072274 | TFRC | chr3 | - | 196075158 | 196075360 | 196073929 | 196074125 | 196077063 | 196077122 | 269 | 250 | [illegible] |
| SE | 39783 | ENSG00000072274 | TFRC | chr3 | - | 196077063 | 196077122 | 196075158 | 196075360 | 196082042 | 196082090 | 105 | 115 | [illegible] |
| SE | 39785 | ENSG00000152102 | FAM168B | chr2 | - | 131052890 | 131053015 | 131047875 | 131052452 | 131055271 | 131055449 | 144 | 45 | [illegible] |
| SE | 39787 | ENSG00000106459 | NRF1 | chr7 | + | 129630077 | 129630168 | 129611738 | 129611824 | 129657342 | 129657574 | 2 | 2 | [illegible] |
| SE | 39788 | ENSG00000106459 | NRF1 | chr7 | + | 129641724 | 129641822 | 129611746 | 129611824 | 129657342 | 129657574 | 0 | 0 | [illegible] |
| SE | 39790 | ENSG00000106459 | NRF1 | chr7 | + | 129709074 | 129709233 | 129690405 | 129690546 | 129710373 | 129710571 | 175 | 72 | [illegible] |
| SE | 39796 | ENSG00000116299 | ELAPOR1 | chr1 | + | 109173687 | 109173837 | 109173473 | 109173579 | 109185044 | 109185133 | 38 | 17 | [illegible] |
| SE | 39797 | ENSG00000116299 | ELAPOR1 | chr1 | + | 109185044 | 109185133 | 109173687 | 109173837 | 109188176 | 109188354 | 21 | 14 | [illegible] |
| SE | 39799 | ENSG00000117385 | P3H1 | chr1 | - | 42751640 | 42751740 | 42750185 | 42750336 | 42752273 | 42752369 | 2 | 0 | [illegible] |
| SE | 39801 | ENSG00000117385 | P3H1 | chr1 | - | 42756400 | 42756455 | 42755547 | 42755637 | 42757782 | 42757922 | 1 | 2 | [illegible] |
| SE | 39802 | ENSG00000117385 | P3H1 | chr1 | - | 42759200 | 42759390 | 42758851 | 42758983 | 42762322 | 42762475 | 31 | 13 | [illegible] |
| SE | 39803 | ENSG00000117385 | P3H1 | chr1 | - | 42759200 | 42759393 | 42758851 | 42758983 | 42762322 | 42762403 | 22 | 10 | [illegible] |
| SE | 39812 | ENSG00000144711 | IQSEC1 | chr3 | - | 12902772 | 12902822 | 12897219 | 12899434 | 12908348 | 12908525 | 16 | 15 | [illegible] |
| SE | 39813 | ENSG00000144711 | IQSEC1 | chr3 | - | 12908348 | 12908525 | 12900576 | 12901522 | 12911628 | 12911728 | 380 | 200 | [illegible] |
| SE | 39816 | ENSG00000144711 | IQSEC1 | chr3 | - | 12915103 | 12915133 | 12913427 | 12913553 | 12920429 | 12920596 | 38 | 14 | [illegible] |
| SE | 39817 | ENSG00000144711 | IQSEC1 | chr3 | - | 12915593 | 12915733 | 12913427 | 12913553 | 12920429 | 12920596 | 172 | 86 | [illegible] |
| SE | 39818 | ENSG00000144711 | IQSEC1 | chr3 | - | 12915593 | 12915733 | 12915103 | 12915133 | 12920429 | 12920596 | 571 | 330 | [illegible] |
| SE | 39821 | ENSG00000144711 | IQSEC1 | chr3 | - | 13047477 | 13047522 | 12941570 | 12941865 | 13072991 | 13073429 | 6 | 0 | [illegible] |
| SE | 39822 | ENSG00000215267 | AKR1C7P | chr10 | - | 5277127 | 5277294 | 5275172 | 5275237 | 5282775 | 5282898 | 10 | 5 | [illegible] |
| SE | 39824 | ENSG00000253214 | NA | chr8 | + | 81157613 | 81157803 | 81154264 | 81154406 | 81158581 | 81158710 | 0 | 3 | [illegible] |
| SE | 39833 | ENSG00000085465 | OVGP1 | chr1 | - | 111421275 | 111421461 | 111419609 | 111419726 | 111421564 | 111421673 | 7 | 10 | [illegible] |
| SE | 39834 | ENSG00000085465 | OVGP1 | chr1 | - | 111421275 | 111421673 | 111419609 | 111419726 | 111422926 | 111423051 | 4 | 9 | [illegible] |
| SE | 39835 | ENSG00000085465 | OVGP1 | chr1 | - | 111421564 | 111421673 | 111419609 | 111419726 | 111422926 | 111423051 | 0 | 4 | [illegible] |
| SE | 39836 | ENSG00000080854 | IGSF9B | chr11 | - | 133929670 | 133929782 | 133926915 | 133927091 | 133930983 | 133931134 | 2 | 1 | [illegible] |
| SE | 39837 | ENSG00000080854 | IGSF9B | chr11 | - | 133935616 | 133935762 | 133932070 | 133932213 | 133937375 | 133937493 | 3 | 1 | [illegible] |
| SE | 39838 | ENSG00000080854 | IGSF9B | chr11 | - | 133936052 | 133936194 | 133935616 | 133935762 | 133937375 | 133937493 | 4 | 2 | [illegible] |
| SE | 39841 | ENSG00000143452 | HORMAD1 | chr1 | - | 150708255 | 150708407 | 150706552 | 150706809 | 150708893 | 150708961 | 36 | 27 | [illegible] |
| SE | 39843 | ENSG00000143452 | HORMAD1 | chr1 | - | 150711832 | 150711853 | 150711544 | 150711571 | 150714084 | 150714121 | 19 | 10 | [illegible] |
| SE | 39844 | ENSG00000143452 | HORMAD1 | chr1 | - | 150711832 | 150711853 | 150711544 | 150711571 | 150717137 | 150717282 | 1 | 2 | [illegible] |
| SE | 39845 | ENSG00000143452 | HORMAD1 | chr1 | - | 150714084 | 150714121 | 150711544 | 150711571 | 150717137 | 150717282 | 0 | 1 | [illegible] |
| SE | 39847 | ENSG00000143452 | HORMAD1 | chr1 | - | 150714084 | 150714121 | 150711832 | 150711853 | 150717137 | 150717282 | 14 | 6 | [illegible] |
| SE | 39848 | ENSG00000143452 | HORMAD1 | chr1 | - | 150714614 | 150714678 | 150713727 | 150714121 | 150717137 | 150717282 | 5 | 4 | [illegible] |
| SE | 39849 | ENSG00000143452 | HORMAD1 | chr1 | - | 150714614 | 150714678 | 150713727 | 150714121 | 150719472 | 150719538 | 4 | 2 | [illegible] |
| SE | 39850 | ENSG00000143452 | HORMAD1 | chr1 | - | 150717137 | 150717282 | 150711544 | 150711571 | 150719472 | 150719538 | 4 | 8 | [illegible] |
| SE | 39851 | ENSG00000143452 | HORMAD1 | chr1 | - | 150717137 | 150717282 | 150714084 | 150714121 | 150719472 | 150719538 | 13 | 15 | [illegible] |
| SE | 39852 | ENSG00000143452 | HORMAD1 | chr1 | - | 150717137 | 150717282 | 150714614 | 150714678 | 150719472 | 150719538 | 4 | 10 | [illegible] |
| SE | 39853 | ENSG00000143452 | HORMAD1 | chr1 | - | 150717137 | 150717307 | 150714614 | 150714678 | 150719472 | 150719538 | 1 | 2 | [illegible] |
| SE | 39858 | ENSG00000169239 | CA5B | chrX | + | 15749970 | 15750165 | 15738269 | 15738352 | 15764577 | 15764665 | 218 | 144 | [illegible] |
| SE | 39862 | ENSG00000143457 | GOLPH3L | chr1 | - | 150661813 | 150661928 | 150646229 | 150648748 | 150694655 | 150694850 | 40 | 31 | [illegible] |
| SE | 39863 | ENSG00000143457 | GOLPH3L | chr1 | - | 150663631 | 150663763 | 150646229 | 150648748 | 150694655 | 150694850 | 63 | 49 | [illegible] |
| SE | 39875 | ENSG00000196821 | ILRUN | chr6 | - | 34606554 | 34606904 | 34587287 | 34590600 | 34646600 | 34646798 | 502 | 296 | [illegible] |
| SE | 39876 | ENSG00000196821 | ILRUN | chr6 | - | 34606554 | 34606904 | 34587287 | 34590600 | 34654624 | 34654779 | 307 | 183 | [illegible] |
| SE | 39877 | ENSG00000196821 | ILRUN | chr6 | - | 34646600 | 34646798 | 34587287 | 34590600 | 34654624 | 34654779 | 320 | 151 | [illegible] |
| SE | 39888 | ENSG00000143420 | ENSA | chr1 | - | 150625641 | 150625802 | 150622843 | 150622859 | 150627466 | 150627592 | 190 | 94 | [illegible] |
| SE | 39889 | ENSG00000143420 | ENSA | chr1 | - | 150625641 | 150625808 | 150622124 | 150622859 | 150627466 | 150627592 | 657 | 368 | [illegible] |
| SE | 39890 | ENSG00000143420 | ENSA | chr1 | - | 150626478 | 150626526 | 150625554 | 150625808 | 150627311 | 150627592 | 6 | 3 | [illegible] |
| SE | 39892 | ENSG00000143420 | ENSA | chr1 | - | 150627311 | 150627592 | 150625464 | 150625808 | 150629413 | 150629502 | 218 | 128 | [illegible] |
| SE | 39893 | ENSG00000143420 | ENSA | chr1 | - | 150627397 | 150627592 | 150623621 | 150625808 | 150629413 | 150629577 | 217 | 128 | [illegible] |
| SE | 39894 | ENSG00000143420 | ENSA | chr1 | - | 150627466 | 150627592 | 150623280 | 150625808 | 150629413 | 150629470 | 636 | 375 | [illegible] |
| SE | 39895 | ENSG00000271723 | MROH7-TTC4 | chr1 | + | 54659114 | 54659189 | 54652852 | 54654157 | 54665166 | 54665240 | 0 | 0 | [illegible] |
| SE | 39898 | ENSG00000271723 | MROH7-TTC4 | chr1 | + | 54717491 | 54717653 | 54716599 | 54716717 | 54721162 | 54721240 | 140 | 91 | [illegible] |
| SE | 39903 | ENSG00000271723 | MROH7-TTC4 | chr1 | + | 54731654 | 54731700 | 54731485 | 54731549 | 54733628 | 54733710 | 16 | 10 | [illegible] |
| SE | 39904 | ENSG00000271723 | MROH7-TTC4 | chr1 | + | 54731654 | 54731700 | 54731485 | 54731549 | 54737581 | 54737664 | 0 | 0 | [illegible] |
| SE | 39906 | ENSG00000271723 | MROH7-TTC4 | chr1 | + | 54733628 | 54733710 | 54731485 | 54731549 | 54737581 | 54737664 | 19 | 11 | [illegible] |
| SE | 39907 | ENSG00000271723 | MROH7-TTC4 | chr1 | + | 54733628 | 54733710 | 54731485 | 54731700 | 54737581 | 54737664 | 53 | 51 | [illegible] |
| SE | 39921 | ENSG00000204580 | DDR1 | chr6 | + | 30885624 | 30885680 | 30884352 | 30884710 | 30888687 | 30888814 | 2 | 0 | [illegible] |
| SE | 39924 | ENSG00000204580 | DDR1 | chr6 | + | 30892295 | 30892542 | 30892001 | 30892188 | 30893067 | 30893163 | 3 | 3 | [illegible] |
| SE | 39925 | ENSG00000204580 | DDR1 | chr6 | + | 30895403 | 30895514 | 30894505 | 30894671 | 30896620 | 30896865 | 2 | 2 | [illegible] |
| SE | 39926 | ENSG00000204580 | DDR1 | chr6 | + | 30895403 | 30895514 | 30894505 | 30894671 | 30896730 | 30896865 | 2 | 1 | [illegible] |
| SE | 39927 | ENSG00000204580 | DDR1 | chr6 | + | 30896620 | 30896865 | 30894505 | 30894671 | 30897013 | 30897141 | 9 | 3 | [illegible] |
| SE | 39928 | ENSG00000204580 | DDR1 | chr6 | + | 30896730 | 30896865 | 30894587 | 30894671 | 30897013 | 30897141 | 8 | 3 | [illegible] |
| SE | 39930 | ENSG00000206530 | CFAP44 | chr3 | - | 113401583 | 113401739 | 113401239 | 113401287 | 113403851 | 113404016 | 22 | 54 | [illegible] |
| SE | 39931 | ENSG00000285943 | NA | chr3 | - | 113427186 | 113427339 | 113416524 | 113416627 | 113433564 | 113433669 | 19 | 17 | [illegible] |
| SE | 39942 | ENSG00000132535 | DLG4 | chr17 | - | 7192944 | 7193117 | 7191892 | 7192002 | 7193482 | 7193584 | 56 | 52 | [illegible] |
| SE | 39943 | ENSG00000132535 | DLG4 | chr17 | - | 7203192 | 7203329 | 7202717 | 7203047 | 7203423 | 7203593 | 72 | 34 | [illegible] |
| SE | 39944 | ENSG00000132535 | DLG4 | chr17 | - | 7208074 | 7208239 | 7204198 | 7204252 | 7217117 | 7217206 | 3 | 1 | [illegible] |
| SE | 39945 | ENSG00000132535 | DLG4 | chr17 | - | 7208173 | 7208239 | 7204198 | 7204252 | 7217117 | 7217188 | 8 | 3 | [illegible] |
| SE | 39946 | ENSG00000164253 | WDR41 | chr5 | - | 77426943 | 77427034 | 77425969 | 77426191 | 77432892 | 77432955 | 2 | 0 | [illegible] |
| SE | 39948 | ENSG00000164253 | WDR41 | chr5 | - | 77437335 | 77437424 | 77436260 | 77436394 | 77440812 | 77440997 | 61 | 27 | [illegible] |
| SE | 39949 | ENSG00000164253 | WDR41 | chr5 | - | 77438239 | 77438361 | 77436359 | 77436394 | 77440812 | 77440997 | 94 | 43 | [illegible] |
| SE | 39950 | ENSG00000164253 | WDR41 | chr5 | - | 77438239 | 77438361 | 77437335 | 77437424 | 77440812 | 77440997 | 193 | 85 | [illegible] |
| SE | 39951 | ENSG00000164253 | WDR41 | chr5 | - | 77449759 | 77449860 | 77440812 | 77440997 | 77453816 | 77453928 | 86 | 46 | [illegible] |
| SE | 39953 | ENSG00000164253 | WDR41 | chr5 | - | 77451290 | 77451353 | 77440812 | 77440997 | 77453816 | 77453910 | 39 | 20 | [illegible] |
| SE | 39959 | ENSG00000164253 | WDR41 | chr5 | - | 77464760 | 77464809 | 77463092 | 77463226 | 77489456 | 77489557 | 95 | 54 | [illegible] |
| SE | 39961 | ENSG00000164253 | WDR41 | chr5 | - | 77480080 | 77480189 | 77464760 | 77464809 | 77489456 | 77489572 | 6 | 0 | [illegible] |
| SE | 39962 | ENSG00000164253 | WDR41 | chr5 | - | 77480080 | 77480189 | 77464760 | 77464809 | 77492169 | 77492220 | 3 | 0 | [illegible] |
| SE | 39963 | ENSG00000164253 | WDR41 | chr5 | - | 77489456 | 77489557 | 77464760 | 77464809 | 77492169 | 77492487 | 66 | 41 | [illegible] |
| SE | 39965 | ENSG00000164253 | WDR41 | chr5 | - | 77489456 | 77489572 | 77464760 | 77464809 | 77492169 | 77492195 | 109 | 51 | [illegible] |
| SE | 39966 | ENSG00000164253 | WDR41 | chr5 | - | 77489456 | 77489572 | 77480080 | 77480189 | 77492169 | 77492195 | 31 | 3 | [illegible] |
| SE | 39967 | ENSG00000114767 | RRP9 | chr3 | - | 51934471 | 51934551 | 51933707 | 51933781 | 51934630 | 51934776 | 22 | 11 | [illegible] |
| SE | 39970 | ENSG00000135537 | AFG1L | chr6 | + | 108346987 | 108347039 | 108323824 | 108324048 | 108355653 | 108355755 | 17 | 22 | [illegible] |
| SE | 39971 | ENSG00000135537 | AFG1L | chr6 | + | 108346987 | 108347039 | 108323824 | 108324048 | 108356689 | 108356751 | 8 | 12 | [illegible] |
| SE | 39972 | ENSG00000135537 | AFG1L | chr6 | + | 108355653 | 108355755 | 108323824 | 108324048 | 108356689 | 108356751 | 7 | 3 | [illegible] |
| SE | 39974 | ENSG00000135537 | AFG1L | chr6 | + | 108401995 | 108402054 | 108366232 | 108366332 | 108447213 | 108447296 | 11 | 8 | [illegible] |
| SE | 39975 | ENSG00000135537 | AFG1L | chr6 | + | 108447213 | 108447296 | 108401995 | 108402054 | 108476864 | 108476935 | 11 | 7 | [illegible] |
| SE | 39976 | ENSG00000135537 | AFG1L | chr6 | + | 108510211 | 108510352 | 108477191 | 108477292 | 108519696 | 108519810 | 20 | 12 | [illegible] |
| SE | 39979 | ENSG00000080822 | CLDND1 | chr3 | - | 98517051 | 98517189 | 98515479 | 98516879 | 98518884 | 98518995 | 253 | 143 | [illegible] |

| | | | | | | | | | | | |
|---|---|---|---|---|---|---|---|---|---|---|---|
| SE | 39980 | ENSG00000080822 | CLDND1 | chr3 | - | 98517051 | 98517189 | 98515482 | 98515832 | 98518884 | 98518995 | 190 | 100 |
| SE | 39981 | ENSG00000080822 | CLDND1 | chr3 | - | 98517157 | 98517189 | 98516776 | 98516879 | 98518884 | 98518995 | 9 | 4 |
| SE | 39983 | ENSG00000080822 | CLDND1 | chr3 | - | 98521132 | 98521442 | 98518734 | 98518995 | 98522541 | 98522754 | 117 | 86 |
| SE | 39984 | ENSG00000080822 | CLDND1 | chr3 | - | 98521132 | 98521442 | 98518734 | 98518995 | 98522848 | 98522875 | 423 | 260 |
| SE | 39986 | ENSG00000080822 | CLDND1 | chr3 | - | 98521652 | 98521696 | 98521132 | 98521437 | 98522848 | 98522955 | 0 | 0 |
| SE | 39987 | ENSG00000080822 | CLDND1 | chr3 | - | 98521652 | 98521696 | 98521132 | 98521442 | 98522541 | 98522606 | 2 | 0 |
| SE | 39988 | ENSG00000080822 | CLDND1 | chr3 | - | 98521652 | 98521696 | 98521132 | 98521442 | 98522848 | 98522955 | 2 | 0 |
| SE | 39989 | ENSG00000080822 | CLDND1 | chr3 | - | 98521652 | 98521703 | 98521132 | 98521437 | 98522848 | 98522875 | 0 | 0 |
| SE | 39990 | ENSG00000080822 | CLDND1 | chr3 | - | 98521652 | 98521703 | 98521132 | 98521442 | 98522541 | 98522605 | 2 | 0 |
| SE | 39991 | ENSG00000080822 | CLDND1 | chr3 | - | 98521652 | 98521703 | 98521132 | 98521442 | 98522848 | 98522875 | 2 | 0 |
| SE | 39992 | ENSG00000080822 | CLDND1 | chr3 | - | 98521652 | 98521718 | 98521132 | 98521437 | 98522848 | 98522922 | 0 | 0 |
| SE | 39993 | ENSG00000080822 | CLDND1 | chr3 | - | 98521652 | 98521718 | 98521132 | 98521442 | 98522541 | 98522602 | 2 | 0 |
| SE | 39994 | ENSG00000080822 | CLDND1 | chr3 | - | 98521652 | 98521718 | 98521132 | 98521442 | 98522848 | 98522922 | 2 | 0 |
| SE | 39995 | ENSG00000107566 | ERLIN1 | chr10 | - | 100156144 | 100156234 | 100154859 | 100154939 | 100164003 | 100164095 | 92 | 67 |
| SE | 39997 | ENSG00000107566 | ERLIN1 | chr10 | - | 100164003 | 100164095 | 100156144 | 100156234 | 100174207 | 100174281 | 27 | 39 |
| SE | 39998 | ENSG00000107566 | ERLIN1 | chr10 | - | 100167347 | 100167406 | 100156144 | 100156234 | 100174207 | 100174281 | 52 | 36 |
| SE | 40000 | ENSG00000107566 | ERLIN1 | chr10 | - | 100174207 | 100174281 | 100156144 | 100156234 | 100175944 | 100176070 | 26 | 24 |
| SE | 40001 | ENSG00000107566 | ERLIN1 | chr10 | - | 100175944 | 100176070 | 100174207 | 100174281 | 100178132 | 100178194 | 145 | 97 |
| SE | 40003 | ENSG00000107566 | ERLIN1 | chr10 | - | 100179200 | 100179247 | 100175944 | 100176070 | 100183755 | 100183837 | 11 | 2 |
| SE | 40004 | ENSG00000107566 | ERLIN1 | chr10 | - | 100179200 | 100179247 | 100175944 | 100176070 | 100185513 | 100185637 | 1 | 0 |
| SE | 40005 | ENSG00000107566 | ERLIN1 | chr10 | - | 100183755 | 100183837 | 100175944 | 100176070 | 100185513 | 100185637 | 18 | 11 |
| SE | 40006 | ENSG00000107566 | ERLIN1 | chr10 | - | 100183755 | 100183837 | 100179200 | 100179247 | 100185513 | 100185637 | 51 | 22 |
| SE | 40007 | ENSG00000107566 | ERLIN1 | chr10 | - | 100185513 | 100185637 | 100179200 | 100179247 | 100185961 | 100186033 | 4 | 2 |
| SE | 40008 | ENSG00000203747 | FCGR3A | chr1 | - | 161544700 | 161544958 | 161541758 | 161543199 | 161548420 | 161548675 | 2685 | 1027 |
| SE | 40010 | ENSG00000121289 | CEP89 | chr19 | - | 32899856 | 32899998 | 32887751 | 32887841 | 32901244 | 32901412 | 30 | 26 |
| SE | 40015 | ENSG00000077097 | TOP2B | chr3 | - | 25606042 | 25606122 | 25601099 | 25601225 | 25607170 | 25607375 | 361 | 319 |
| SE | 40016 | ENSG00000077097 | TOP2B | chr3 | - | 25607170 | 25607375 | 25601099 | 25601225 | 25609182 | 25609344 | 158 | 98 |
| SE | 40017 | ENSG00000077097 | TOP2B | chr3 | - | 25609567 | 25609712 | 25609182 | 25609344 | 25612514 | 25612709 | 588 | 453 |
| SE | 40019 | ENSG00000077097 | TOP2B | chr3 | - | 25618653 | 25618849 | 25615430 | 25615586 | 25619861 | 25620062 | 536 | 508 |
| SE | 40020 | ENSG00000077097 | TOP2B | chr3 | - | 25619861 | 25620062 | 25618653 | 25618849 | 25620681 | 25620816 | 964 | 873 |
| SE | 40021 | ENSG00000077097 | TOP2B | chr3 | - | 25626771 | 25626864 | 25626559 | 25626674 | 25627186 | 25627296 | 889 | 722 |
| SE | 40022 | ENSG00000077097 | TOP2B | chr3 | - | 25632445 | 25632583 | 25630800 | 25630939 | 25632692 | 25632794 | 546 | 442 |
| SE | 40023 | ENSG00000077097 | TOP2B | chr3 | - | 25637214 | 25637312 | 25635935 | 25636148 | 25638164 | 25638310 | 751 | 703 |
| SE | 40024 | ENSG00000077097 | TOP2B | chr3 | - | 25638164 | 25638310 | 25637214 | 25637312 | 25642321 | 25642385 | 910 | 765 |
| SE | 40026 | ENSG00000117266 | CDK18 | chr1 | + | 205525138 | 205525195 | 205524231 | 205524357 | 205526064 | 205526179 | 8 | 5 |
| SE | 40037 | ENSG00000143190 | POU2F1 | chr1 | + | 167241406 | 167241610 | 167220875 | 167220958 | 167303473 | 167303574 | 4 | 3 |
| SE | 40040 | ENSG00000143190 | POU2F1 | chr1 | + | 167303473 | 167303574 | 167220889 | 167220958 | 167319013 | 167319056 | 1 | 0 |
| SE | 40042 | ENSG00000143190 | POU2F1 | chr1 | + | 167303473 | 167303574 | 167241406 | 167241610 | 167332469 | 167332535 | 2 | 7 |
| SE | 40043 | ENSG00000143190 | POU2F1 | chr1 | + | 167319013 | 167319056 | 167220890 | 167220958 | 167332469 | 167332535 | 0 | 0 |
| SE | 40045 | ENSG00000143190 | POU2F1 | chr1 | + | 167332469 | 167332535 | 167303473 | 167303574 | 167365466 | 167365567 | 19 | 23 |
| SE | 40049 | ENSG00000143190 | POU2F1 | chr1 | + | 167376552 | 167376587 | 167376028 | 167376155 | 167383856 | 167383951 | 0 | 0 |
| SE | 40052 | ENSG00000204186 | ZDBF2 | chr2 | + | 206280549 | 206280593 | 206279539 | 206279592 | 206281800 | 206281909 | 18 | 16 |
| SE | 40055 | ENSG00000204186 | ZDBF2 | chr2 | + | 206304716 | 206306875 | 206297245 | 206297373 | 206307046 | 206307161 | 17 | 12 |
| SE | 40057 | ENSG00000125347 | IRF1 | chr5 | - | 132484361 | 132484497 | 132472148 | 132472178 | 132485666 | 132485716 | 624 | 330 |
| SE | 40062 | ENSG00000125347 | IRF1 | chr5 | - | 132484361 | 132484497 | 132481608 | 132484075 | 132486521 | 132486686 | 131 | 71 |
| SE | 40068 | ENSG00000125347 | IRF1 | chr5 | - | 132485666 | 132485716 | 132481608 | 132484075 | 132486521 | 132486686 | 1 | 1 |
| SE | 40070 | ENSG00000125347 | IRF1 | chr5 | - | 132485666 | 132485716 | 132484361 | 132484497 | 132486250 | 132486348 | 1178 | 764 |
| SE | 40074 | ENSG00000125347 | IRF1 | chr5 | - | 132486250 | 132486373 | 132483761 | 132484075 | 132486521 | 132486686 | 5 | 3 |
| SE | 40088 | ENSG00000125347 | IRF1 | chr5 | - | 132489581 | 132489740 | 132489391 | 132489483 | 132490544 | 132490773 | 0 | 1 |
| SE | 40091 | ENSG00000162368 | CMPK1 | chr1 | + | 47368468 | 47368615 | 47333789 | 47334116 | 47372954 | 47373107 | 1778 | 1709 |
| SE | 40093 | ENSG00000242114 | MTFP1 | chr22 | + | 30426716 | 30426844 | 30425622 | 30425946 | 30428461 | 30429053 | 10 | 3 |
| SE | 40094 | ENSG00000242114 | MTFP1 | chr22 | + | 30427121 | 30427403 | 30426716 | 30426844 | 30428461 | 30428643 | 9 | 0 |
| SE | 40095 | ENSG00000242114 | MTFP1 | chr22 | + | 30427170 | 30427403 | 30426716 | 30426844 | 30428461 | 30429054 | 10 | 1 |
| SE | 40096 | ENSG00000242114 | MTFP1 | chr22 | + | 30427239 | 30427403 | 30426716 | 30426844 | 30428461 | 30428951 | 11 | 0 |
| SE | 40097 | ENSG00000072840 | EVC | chr4 | + | 5745203 | 5745341 | 5741715 | 5741814 | 5748147 | 5748306 | 10 | 0 |
| SE | 40102 | ENSG00000095321 | CRAT | chr9 | - | 129108407 | 129108478 | 129107813 | 129108077 | 129109178 | 129109277 | 0 | 0 |
| SE | 40106 | ENSG00000095321 | CRAT | chr9 | - | 129108762 | 129108903 | 129108407 | 129108478 | 129109178 | 129109277 | 0 | 1 |
| SE | 40107 | ENSG00000095321 | CRAT | chr9 | - | 129108898 | 129109277 | 129107813 | 129108077 | 129110482 | 129110747 | 6 | 3 |
| SE | 40108 | ENSG00000095321 | CRAT | chr9 | - | 129109102 | 129109277 | 129107813 | 129108077 | 129110482 | 129110710 | 4 | 3 |
| SE | 40110 | ENSG00000095321 | CRAT | chr9 | - | 129109178 | 129109277 | 129107813 | 129108077 | 129110482 | 129110545 | 11 | 6 |
| SE | 40115 | ENSG00000103342 | GSPT1 | chr16 | - | 11875760 | 11875929 | 11868127 | 11873171 | 11877406 | 11877580 | 244 | 184 |
| SE | 40117 | ENSG00000103342 | GSPT1 | chr16 | - | 11877406 | 11877580 | 11876085 | 11876175 | 11883014 | 11883095 | 351 | 232 |
| SE | 40118 | ENSG00000103342 | GSPT1 | chr16 | - | 11883014 | 11883095 | 11877406 | 11877580 | 11885180 | 11885274 | 350 | 208 |
| SE | 40119 | ENSG00000103342 | GSPT1 | chr16 | - | 11886470 | 11886611 | 11885180 | 11885274 | 11886776 | 11886904 | 737 | 519 |
| SE | 40120 | ENSG00000103342 | GSPT1 | chr16 | - | 11886470 | 11886611 | 11885180 | 11885274 | 11891061 | 11891139 | 291 | 183 |
| SE | 40121 | ENSG00000103342 | GSPT1 | chr16 | - | 11886776 | 11886931 | 11885180 | 11885274 | 11891061 | 11891139 | 5 | 2 |
| SE | 40122 | ENSG00000103342 | GSPT1 | chr16 | - | 11886776 | 11886931 | 11886470 | 11886611 | 11887569 | 11887750 | 876 | 729 |
| SE | 40123 | ENSG00000103342 | GSPT1 | chr16 | - | 11886776 | 11886931 | 11886470 | 11886611 | 11891061 | 11891139 | 451 | 338 |
| SE | 40124 | ENSG00000103342 | GSPT1 | chr16 | - | 11887569 | 11887750 | 11886470 | 11886611 | 11891061 | 11891139 | 283 | 266 |
| SE | 40126 | ENSG00000103342 | GSPT1 | chr16 | - | 11894953 | 11894987 | 11891061 | 11891139 | 11896557 | 11896782 | 320 | 300 |
| SE | 40129 | ENSG00000103342 | GSPT1 | chr16 | - | 11897839 | 11897881 | 11896557 | 11896785 | 11897993 | 11898035 | 183 | 109 |
| SE | 40135 | ENSG00000245060 | LINC00847 | chr5 | + | 180831735 | 180831806 | 180830987 | 180831229 | 180834311 | 180835646 | 5 | 1 |
| SE | 40137 | ENSG00000245060 | LINC00847 | chr5 | + | 180831758 | 180831806 | 180831226 | 180831229 | 180834311 | 180835705 | 3 | 1 |
| SE | 40139 | ENSG00000245060 | LINC00847 | chr5 | + | 180832672 | 180832860 | 180831023 | 180831229 | 180834311 | 180835002 | 3 | 1 |
| SE | 40140 | ENSG00000245060 | LINC00847 | chr5 | + | 180832672 | 180832860 | 180831648 | 180831806 | 180834311 | 180835002 | 1 | 1 |
| SE | 40141 | ENSG00000245060 | LINC00847 | chr5 | + | 180832744 | 180832860 | 180831722 | 180831806 | 180834311 | 180835705 | 1 | 1 |
| SE | 40147 | ENSG00000146122 | DAAM2 | chr6 | + | 39888678 | 39888763 | 39887404 | 39887592 | 39891340 | 39891361 | 0 | 0 |
| SE | 40151 | ENSG00000138382 | METTL5 | chr2 | - | 169812456 | 169812506 | 169811756 | 169811858 | 169815476 | 169815528 | 88 | 62 |
| SE | 40152 | ENSG00000138382 | METTL5 | chr2 | - | 169812456 | 169812524 | 169811756 | 169811858 | 169815476 | 169815528 | 3 | 0 |
| SE | 40155 | ENSG00000138382 | METTL5 | chr2 | - | 169819560 | 169819643 | 169811813 | 169812506 | 169821942 | 169822057 | 3 | 0 |
| SE | 40157 | ENSG00000138382 | METTL5 | chr2 | - | 169821091 | 169821273 | 169812456 | 169812506 | 169821942 | 169822057 | 88 | 75 |
| SE | 40159 | ENSG00000138382 | METTL5 | chr2 | - | 169821091 | 169821273 | 169819560 | 169819643 | 169821942 | 169822057 | 214 | 180 |
| SE | 40163 | ENSG00000138382 | METTL5 | chr2 | - | 169821942 | 169822057 | 169821091 | 169821273 | 169824488 | 169824558 | 191 | 157 |
| SE | 40167 | ENSG00000185291 | IL3RA | chrX | + | 1348430 | 1348545 | 1345315 | 1345434 | 1352099 | 1352232 | 97 | 120 |
| SE | 40168 | ENSG00000185291 | IL3RA | chrX | + | 1358860 | 1358887 | 1356220 | 1356336 | 1365137 | 1365252 | 79 | 44 |
| SE | 40169 | ENSG00000115306 | SPTBN1 | chr2 | + | 54618077 | 54618193 | 54617607 | 54617688 | 54621399 | 54621512 | 532 | 335 |
| SE | 40173 | ENSG00000115306 | SPTBN1 | chr2 | + | 54624803 | 54624962 | 54623478 | 54623596 | 54625931 | 54626234 | 433 | 307 |
| SE | 40175 | ENSG00000115306 | SPTBN1 | chr2 | + | 54653608 | 54653853 | 54649614 | 54649989 | 54655069 | 54655208 | 304 | 213 |
| SE | 40176 | ENSG00000115306 | SPTBN1 | chr2 | + | 54655913 | 54655998 | 54655069 | 54655208 | 54657849 | 54658046 | 447 | 321 |
| SE | 40177 | ENSG00000115306 | SPTBN1 | chr2 | + | 54655913 | 54655998 | 54655069 | 54655208 | 54659153 | 54659266 | 122 | 98 |
| SE | 40178 | ENSG00000115306 | SPTBN1 | chr2 | + | 54657849 | 54658046 | 54655069 | 54655208 | 54659153 | 54659266 | 400 | 265 |
| SE | 40179 | ENSG00000115306 | SPTBN1 | chr2 | + | 54657849 | 54658046 | 54655913 | 54655998 | 54659153 | 54659266 | 729 | 496 |
| SE | 40180 | ENSG00000115306 | SPTBN1 | chr2 | + | 54659153 | 54659266 | 54657849 | 54658046 | 54659935 | 54659999 | 596 | 388 |
| SE | 40181 | ENSG00000114544 | SLC41A3 | chr3 | - | 126022785 | 126022932 | 126016730 | 126016875 | 126026334 | 126026479 | 42 | 34 |
| SE | 40189 | ENSG00000231607 | DLEU2 | chr13 | - | 49988585 | 49988621 | 49976332 | 49976454 | 49993792 | 49993961 | 1 | 0 |
| SE | 40190 | ENSG00000231607 | DLEU2 | chr13 | - | 49988585 | 49988621 | 49982551 | 49983754 | 49993792 | 49993961 | 0 | 1 |
| SE | 40191 | ENSG00000231607 | DLEU2 | chr13 | - | 49993792 | 49993961 | 49975946 | 49976454 | 49997073 | 49997136 | 5 | 5 |
| SE | 40195 | ENSG00000231607 | DLEU2 | chr13 | - | 50045117 | 50045232 | 50044639 | 50044768 | 50049583 | 50049662 | 0 | 0 |
| SE | 40204 | ENSG00000267575 | NA | chr19 | + | 27794681 | 27794869 | 27794017 | 27794122 | 27797753 | 27798315 | 1 | 0 |
| SE | 40205 | ENSG00000267575 | NA | chr19 | + | 27797753 | 27798176 | 27793430 | 27793650 | 27805604 | 27805811 | 15 | 13 |
| SE | 40210 | ENSG00000144589 | STK11IP | chr2 | + | 219609096 | 219609213 | 219608582 | 219608788 | 219609362 | 219609540 | 16 | 13 |
| SE | 40211 | ENSG00000144589 | STK11IP | chr2 | + | 219614160 | 219614242 | 219613751 | 219613930 | 219614475 | 219614546 | 21 | 7 |
| SE | 40212 | ENSG00000167315 | ACAA2 | chr18 | - | 49785196 | 49785351 | 49782163 | 49783931 | 49787290 | 49787361 | 423 | 270 |
| SE | 40213 | ENSG00000167315 | ACAA2 | chr18 | - | 49795764 | 49795881 | 49794279 | 49794427 | 49797465 | 49797594 | 416 | 191 |
| SE | 40214 | ENSG00000167315 | ACAA2 | chr18 | - | 49797465 | 49797594 | 49794279 | 49794427 | 49802686 | 49802853 | 74 | 23 |
| SE | 40221 | ENSG00000084693 | AGBL5 | chr2 | + | 27052332 | 27052543 | 27051667 | 27051793 | 27052912 | 27053156 | 0 | 0 |
| SE | 40223 | ENSG00000084693 | AGBL5 | chr2 | + | 27069047 | 27069094 | 27068631 | 27068744 | 27069572 | 27069706 | 2 | 4 |
| SE | 40224 | ENSG00000084693 | AGBL5 | chr2 | + | 27069572 | 27069706 | 27069047 | 27069094 | 27070091 | 27070618 | 10 | 8 |
| SE | 40228 | ENSG00000132768 | DPH2 | chr1 | + | 43970965 | 43971189 | 43970595 | 43970708 | 43972157 | 43972334 | 2 | 0 |
| SE | 40229 | ENSG00000132768 | DPH2 | chr1 | + | 43970965 | 43971189 | 43970595 | 43970817 | 43972157 | 43972334 | 2 | 1 |
| SE | 40231 | ENSG00000132768 | DPH2 | chr1 | + | 43971386 | 43972070 | 43970595 | 43970708 | 43972157 | 43972334 | 14 | 2 |
| SE | 40234 | ENSG00000117408 | IPO13 | chr1 | + | 43949416 | 43950153 | 43946949 | 43947684 | 43956319 | 43956460 | 36 | 9 |
| SE | 40235 | ENSG00000117408 | IPO13 | chr1 | + | 43950085 | 43950153 | 43947569 | 43947684 | 43956319 | 43956460 | 11 | 1 |
| SE | 40236 | ENSG00000117408 | IPO13 | chr1 | + | 43957194 | 43957315 | 43956809 | 43956976 | 43957401 | 43957549 | 28 | 9 |
| SE | 40238 | ENSG00000117408 | IPO13 | chr1 | + | 43960248 | 43960329 | 43958745 | 43958889 | 43960875 | 43961013 | 65 | 20 |
| SE | 40239 | ENSG00000117408 | IPO13 | chr1 | + | 43967314 | 43967496 | 43966929 | 43967019 | 43967585 | 43967752 | 25 | 17 |
| SE | 40240 | ENSG00000197746 | PSAP | chr10 | - | 71818616 | 71818724 | 71816297 | 71817476 | 71819030 | 71819111 | 9051 | 5670 |
| SE | 40241 | ENSG00000197746 | PSAP | chr10 | - | 71818616 | 71818724 | 71816297 | 71817476 | 71819464 | 71819622 | 4657 | 3071 |
| SE | 40242 | ENSG00000197746 | PSAP | chr10 | - | 71819030 | 71819111 | 71816297 | 71817476 | 71819464 | 71819622 | 1974 | 1520 |
| SE | 40243 | ENSG00000197746 | PSAP | chr10 | - | 71819030 | 71819111 | 71818616 | 71818724 | 71819464 | 71819622 | 6327 | 4096 |
| SE | 40244 | ENSG00000197746 | PSAP | chr10 | - | 71819464 | 71819622 | 71819030 | 71819111 | 71819713 | 71819783 | 12001 | 8057 |
| SE | 40245 | ENSG00000197746 | PSAP | chr10 | - | 71821875 | 71822007 | 71820239 | 71820335 | 71825836 | 71825893 | 9559 | 6669 |
| SE | 40246 | ENSG00000197746 | PSAP | chr10 | - | 71821875 | 71822016 | 71820239 | 71820335 | 71825836 | 71825893 | 4078 | 2746 |
| SE | 40247 | ENSG00000197746 | PSAP | chr10 | - | 71823887 | 71823896 | 71821875 | 71822007 | 71825836 | 71825893 | 13 | 10 |
| SE | 40248 | ENSG00000197746 | PSAP | chr10 | - | 71825836 | 71825893 | 71821875 | 71822007 | 71828013 | 71828134 | 7532 | 5428 |
| SE | 40249 | ENSG00000197746 | PSAP | chr10 | - | 71828013 | 71828157 | 71825836 | 71825893 | 71828876 | 71829077 | 8337 | 5344 |
| SE | 40250 | ENSG00000197746 | PSAP | chr10 | - | 71828876 | 71829077 | 71825836 | 71825893 | 71831125 | 71831251 | 3181 | 1886 |
| SE | 40251 | ENSG00000170581 | STAT2 | chr12 | - | 56343303 | 56343531 | 56342617 | 56342869 | 56343824 | 56344135 | 173 | 137 |
| SE | 40252 | ENSG00000170581 | STAT2 | chr12 | - | 56343824 | 56344135 | 56341596 | 56343531 | 56346145 | 56346203 | 533 | 371 |
| SE | 40253 | ENSG00000170581 | STAT2 | chr12 | - | 56348751 | 56348804 | 56348528 | 56348623 | 56348923 | 56349029 | 141 | 80 |
| SE | 40254 | ENSG00000170581 | STAT2 | chr12 | - | 56348751 | 56348804 | 56348528 | 56348623 | 56349344 | 56349509 | 39 | 28 |
| SE | 40256 | ENSG00000170581 | STAT2 | chr12 | - | 56348751 | 56348851 | 56348528 | 56348623 | 56349344 | 56349509 | 42 | 33 |
| SE | 40258 | ENSG00000170581 | STAT2 | chr12 | - | 56348923 | 56349059 | 56348528 | 56348623 | 56349344 | 56349509 | 1 | 0 |
| SE | 40259 | ENSG00000170581 | STAT2 | chr12 | - | 56348923 | 56349059 | 56348528 | 56348623 | 56349425 | 56349509 | 5 | 1 |
| SE | 40261 | ENSG00000170581 | STAT2 | chr12 | - | 56348923 | 56349059 | 56348751 | 56348804 | 56349425 | 56349509 | 100 | 50 |
| SE | 40266 | ENSG00000170581 | STAT2 | chr12 | - | 56349344 | 56349509 | 56348751 | 56348804 | 56350096 | 56350190 | 1 | 0 |
| SE | 40268 | ENSG00000170581 | STAT2 | chr12 | - | 56349588 | 56349636 | 56349344 | 56349509 | 56350096 | 56350190 | 305 | 178 |
| SE | 40270 | ENSG00000170581 | STAT2 | chr12 | - | 56350096 | 56350190 | 56349344 | 56349509 | 56350411 | 56350432 | 149 | 66 |
| SE | 40272 | ENSG00000170581 | STAT2 | chr12 | - | 56350411 | 56350432 | 56350096 | 56350190 | 56350828 | 56350888 | 292 | 154 |
| SE | 40276 | ENSG00000170581 | STAT2 | chr12 | - | 56354777 | 56354863 | 56354418 | 56354614 | 56355275 | 56355351 | 446 | 273 |
| SE | 40277 | ENSG00000170581 | STAT2 | chr12 | - | 56355707 | 56355791 | 56355442 | 56355532 | 56356131 | 56356285 | 94 | 48 |
| SE | 40278 | ENSG00000170581 | STAT2 | chr12 | - | 56355707 | 56355803 | 56355442 | 56355532 | 56356131 | 56356218 | 243 | 118 |
| SE | 40286 | ENSG00000271383 | NBPF19 | chr1 | + | 149478879 | 149479094 | 149477944 | 149478047 | 149480141 | 149480214 | 73 | 72 |
| SE | 40288 | ENSG00000271383 | NBPF19 | chr1 | + | 149482174 | 149482226 | 149480141 | 149480214 | 149487331 | 149487383 | 2 | 1 |
| SE | 40290 | ENSG00000271383 | NBPF19 | chr1 | + | 149487331 | 149487383 | 149486129 | 149486293 | 149488012 | 149488185 | 74 | 52 |
| SE | 40292 | ENSG00000089682 | RBM41 | chrX | - | 107088734 | 107088839 | 107069309 | 107069402 | 107113396 | 107113468 | 12 | 27 |
| SE | 40294 | ENSG00000089682 | RBM41 | chrX | - | 107113396 | 107113468 | 107069309 | 107069402 | 107115351 | 107115556 | 21 | 26 |
| SE | 40301 | ENSG00000234663 | LINC01934 | chr2 | + | 181195346 | 181195472 | 181123931 | 181123980 | 181208032 | 181208262 | 4 | 1 |
| SE | 40304 | ENSG00000234663 | LINC01934 | chr2 | + | 181208032 | 181208262 | 181173723 | 181173865 | 181226470 | 181226539 | 1 | 8 |
| SE | 40305 | ENSG00000234663 | LINC01934 | chr2 | + | 181208032 | 181208262 | 181195346 | 181195472 | 181226470 | 181226539 | 1 | 5 |
| SE | 40307 | ENSG00000234663 | LINC01934 | chr2 | + | 181225003 | 181225047 | 181208032 | 181208262 | 181226470 | 181226539 | 2 | 0 |
| SE | 40308 | ENSG00000234663 | LINC01934 | chr2 | + | 181226470 | 181226535 | 181123834 | 181123980 | 181238250 | 181238450 | 22 | 10 |
| SE | 40309 | ENSG00000234663 | LINC01934 | chr2 | + | 181226470 | 181226535 | 181123834 | 181123980 | 181262828 | 181262977 | 23 | 7 |
| SE | 40310 | ENSG00000234663 | LINC01934 | chr2 | + | 181226470 | 181226535 | 181123834 | 181123980 | 181354207 | 181354343 | 14 | 7 |
| SE | 40311 | ENSG00000234663 | LINC01934 | chr2 | + | 181238250 | 181238450 | 181123834 | 181123980 | 181262828 | 181262977 | 2 | 0 |
| SE | 40312 | ENSG00000234663 | LINC01934 | chr2 | + | 181238250 | 181238450 | 181123834 | 181123980 | 181354207 | 181354343 | 1 | 1 |
| SE | 40313 | ENSG00000234663 | LINC01934 | chr2 | + | 181238250 | 181238450 | 181226470 | 181226535 | 181262828 | 181262977 | 18 | 10 |
| SE | 40314 | ENSG00000234663 | LINC01934 | chr2 | + | 181238250 | 181238450 | 181226470 | 181226535 | 181287974 | 181288074 | 17 | 10 |
| SE | 40315 | ENSG00000234663 | LINC01934 | chr2 | + | 181238250 | 181238450 | 181226470 | 181226535 | 181322660 | 181322730 | 17 | 10 |
| SE | 40316 | ENSG00000234663 | LINC01934 | chr2 | + | 181238250 | 181238450 | 181226470 | 181226535 | 181354207 | 181354343 | 17 | 11 |
| SE | 40317 | ENSG00000234663 | LINC01934 | chr2 | + | 181238250 | 181238450 | 181226470 | 181226535 | 181362686 | 181362902 | 19 | 13 |
| SE | 40318 | ENSG00000234663 | LINC01934 | chr2 | + | 181238250 | 181238450 | 181226470 | 181226539 | 181262828 | 181262977 | 4 | 0 |
| SE | 40319 | ENSG00000234663 | LINC01934 | chr2 | + | 181238250 | 181238450 | 181226470 | 181226539 | 181322660 | 181322730 | 3 | 0 |
| SE | 40320 | ENSG00000234663 | LINC01934 | chr2 | + | 181238250 | 181238450 | 181226470 | 181226539 | 181354207 | 181354343 | 3 | 1 |
| SE | 40321 | ENSG00000234663 | LINC01934 | chr2 | + | 181238250 | 181238450 | 181226470 | 181226539 | 181362686 | 181362902 | 5 | 3 |
| SE | 40322 | ENSG00000234663 | LINC01934 | chr2 | + | 181256222 | 181256292 | 181226470 | 181226535 | 181262828 | 181262977 | 6 | 0 |
| SE | 40323 | ENSG00000234663 | LINC01934 | chr2 | + | 181256222 | 181256292 | 181226470 | 181226535 | 181322660 | 181322730 | 5 | 0 |
| SE | 40324 | ENSG00000234663 | LINC01934 | chr2 | + | 181256222 | 181256292 | 181226470 | 181226535 | 181354207 | 181354343 | 5 | 0 |
| SE | 40325 | ENSG00000234663 | LINC01934 | chr2 | + | 181256222 | 181256292 | 181226470 | 181226539 | 181262828 | 181262977 | 2 | 0 |
| SE | 40331 | ENSG00000234663 | LINC01934 | chr2 | + | 181262828 | 181262977 | 181123836 | 181123980 | 181354207 | 181354343 | 3 | 3 |
| SE | 40332 | ENSG00000234663 | LINC01934 | chr2 | + | 181262828 | 181262977 | 181226470 | 181226535 | 181322660 | 181322730 | 14 | 8 |
| SE | 40333 | ENSG00000234663 | LINC01934 | chr2 | + | 181262828 | 181262977 | 181226470 | 181226535 | 181341045 | 181341129 | 14 | 8 |
| SE | 40334 | ENSG00000234663 | LINC01934 | chr2 | + | 181262828 | 181262977 | 181226470 | 181226535 | 181354207 | 181354343 | 17 | 9 |
| SE | 40335 | ENSG00000234663 | LINC01934 | chr2 | + | 181262828 | 181262977 | 181226470 | 181226535 | 181362686 | 181362918 | 16 | 8 |
| SE | 40336 | ENSG00000234663 | LINC01934 | chr2 | + | 181262828 | 181262977 | 181226470 | 181226539 | 181322660 | 181322730 | 1 | 3 |
| SE | 40337 | ENSG00000234663 | LINC01934 | chr2 | + | 181262828 | 181262977 | 181226470 | 181226539 | 181354207 | 181354343 | 4 | 4 |
| SE | 40338 | ENSG00000234663 | LINC01934 | chr2 | + | 181262828 | 181262977 | 181226470 | 181226539 | 181362686 | 181362918 | 3 | 3 |
| SE | 40339 | ENSG00000234663 | LINC01934 | chr2 | + | 181262828 | 181262977 | 181238250 | 181238450 | 181322660 | 181322730 | 1 | 0 |
| SE | 40340 | ENSG00000234663 | LINC01934 | chr2 | + | 181262828 | 181262977 | 181238250 | 181238450 | 181354207 | 181354343 | 4 | 1 |
| SE | 40341 | ENSG00000234663 | LINC01934 | chr2 | + | 181262828 | 181262977 | 181238250 | 181238450 | 181362686 | 181362918 | 3 | 0 |
| SE | 40342 | ENSG00000234663 | LINC01934 | chr2 | + | 181262828 | 181262977 | 181256222 | 181256292 | 181322660 | 181322730 | 2 | 0 |
| SE | 40343 | ENSG00000234663 | LINC01934 | chr2 | + | 181262828 | 181262977 | 181256222 | 181256292 | 181354207 | 181354343 | 5 | 1 |
| SE | 40345 | ENSG00000234663 | LINC01934 | chr2 | + | 181287974 | 181288074 | 181226470 | 181226535 | 181322660 | 181322730 | 1 | 0 |
| SE | 40346 | ENSG00000234663 | LINC01934 | chr2 | + | 181287974 | 181288074 | 181226470 | 181226535 | 181362686 | 181362918 | 1 | 0 |
| SE | 40349 | ENSG00000234663 | LINC01934 | chr2 | + | 181322660 | 181322730 | 181226470 | 181226535 | 181341045 | 181341129 | 0 | 1 |
| SE | 40350 | ENSG00000234663 | LINC01934 | chr2 | + | 181322660 | 181322730 | 181226470 | 181226535 | 181354207 | 181354343 | 5 | 1 |
| SE | 40351 | ENSG00000234663 | LINC01934 | chr2 | + | 181322660 | 181322730 | 181226470 | 181226535 | 181362686 | 181362918 | 0 | 2 |
| SE | 40352 | ENSG00000234663 | LINC01934 | chr2 | + | 181322660 | 181322730 | 181226470 | 181226539 | 181354207 | 181354343 | 8 | 1 |
| SE | 40353 | ENSG00000234663 | LINC01934 | chr2 | + | 181322660 | 181322730 | 181226470 | 181226539 | 181362686 | 181362918 | 3 | 2 |
| SE | 40354 | ENSG00000234663 | LINC01934 | chr2 | + | 181322660 | 181322730 | 181238250 | 181238450 | 181354207 | 181354343 | 4 | 0 |
| SE | 40355 | ENSG00000234663 | LINC01934 | chr2 | + | 181322660 | 181322730 | 181238250 | 181238450 | 181362686 | 181362918 | 0 | 1 |
| SE | 40356 | ENSG00000234663 | LINC01934 | chr2 | + | 181322660 | 181322730 | 181256222 | 181256292 | 181354207 | 181354343 | 5 | 0 |
| SE | 40358 | ENSG00000234663 | LINC01934 | chr2 | + | 181322660 | 181322730 | 181262828 | 181262977 | 181354207 | 181354343 | 4 | 0 |
| SE | 40359 | ENSG00000234663 | LINC01934 | chr2 | + | 181322660 | 181322730 | 181262828 | 181262977 | 181362686 | 181362918 | 0 | 1 |
| SE | 40360 | ENSG00000234663 | LINC01934 | chr2 | + | 181322660 | 181322730 | 181287974 | 181288074 | 181362686 | 181362918 | 0 | 1 |
| SE | 40361 | ENSG00000234663 | LINC01934 | chr2 | + | 181341045 | 181341129 | 181226470 | 181226535 | 181354207 | 181354343 | 0 | 5 |
| SE | 40362 | ENSG00000234663 | LINC01934 | chr2 | + | 181341045 | 181341129 | 181262828 | 181262977 | 181354207 | 181354343 | 0 | 0 |
| SE | 40363 | ENSG00000234663 | LINC01934 | chr2 | + | 181341045 | 181341129 | 181322660 | 181322730 | 181354207 | 181354343 | 0 | 0 |
| SE | 40367 | ENSG00000234663 | LINC01934 | chr2 | + | 181354207 | 181354343 | 181262828 | 181262977 | 181362686 | 181362902 | 49 | 34 |
| SE | 40369 | ENSG00000234663 | LINC01934 | chr2 | + | 181362686 | 181362918 | 181226470 | 181226535 | 181398378 | 181399751 | 21 | 20 |
| SE | 40370 | ENSG00000234663 | LINC01934 | chr2 | + | 181363297 | 181363337 | 181362686 | 181362918 | 181398378 | 181399751 | 0 | 2 |
| SE | 40371 | ENSG00000163719 | MTMR14 | chr3 | + | 9650327 | 9650529 | 9649518 | 9649742 | 9653620 | 9653769 | 3 | 1 |
| SE | 40372 | ENSG00000163719 | MTMR14 | chr3 | + | 9650327 | 9650529 | 9649518 | 9649742 | 9662266 | 9662375 | 1 | 2 |
| SE | 40374 | ENSG00000163719 | MTMR14 | chr3 | + | 9653620 | 9653769 | 9649432 | 9649742 | 9669431 | 9669492 | 95 | 46 |
| SE | 40375 | ENSG00000163719 | MTMR14 | chr3 | + | 9653620 | 9653769 | 9650327 | 9650529 | 9662266 | 9662285 | 149 | 82 |
| SE | 40376 | ENSG00000163719 | MTMR14 | chr3 | + | 9662266 | 9662285 | 9649514 | 9649742 | 9669431 | 9669492 | 0 | 0 |
| SE | 40378 | ENSG00000163719 | MTMR14 | chr3 | + | 9662266 | 9662375 | 9649432 | 9649742 | 9669431 | 9669492 | 3 | 2 |
| SE | 40386 | ENSG00000163719 | MTMR14 | chr3 | + | 9672684 | 9672758 | 9671047 | 9671170 | 9677316 | 9677387 | 430 | 237 |
| SE | 40387 | ENSG00000163719 | MTMR14 | chr3 | + | 9672684 | 9672758 | 9671047 | 9671170 | 9677983 | 9678058 | 143 | 75 |

| | | | | | | | | | | | | | | | | | | | | | | | | | | | | | | | | | | | | | | | | | | | | | | |
|---|---|---|---|---|---|---|---|---|---|---|---|---|---|---|---|---|---|---|---|---|---|---|---|---|---|---|---|---|---|---|---|---|---|---|---|---|---|---|---|---|---|---|---|---|---|---|
| SE | 40388 | ENSG00000163719 | MTMR14 | chr3 | + | 9677316 | 9677387 | 9671047 | 9671170 | 9677983 | 9678058 | 49 | 14 | 40 | 41 | 104 | 0 | 0 | 0 | 0 | 0 | 47 | 24 | 28 | 41 | 68 | 0 | 0 | 0 | 4 | 0 | 220 | 149 | 1 | 1 | 1 | 1 | 1 | 1 | 1 | 1 | 1 | 1 | 0.874 | 1 | 0.025 |
| SE | 40389 | ENSG00000163719 | MTMR14 | chr3 | + | 9677316 | 9677387 | 9672684 | 9672758 | 9677983 | 9678058 | 310 | 163 | 213 | 239 | 389 | 0 | 0 | 0 | 0 | 0 | 516 | 333 | 252 | 365 | 515 | 0 | 4 | 0 | 0 | 0 | 220 | 149 | 1 | 1 | 1 | 1 | 1 | 1 | 1 | 1 | 0.983 | 1 | 1 | 1 | 0.003 |
| SE | 40400 | ENSG00000008952 | SEC62 | chr3 | + | 169969286 | 169969383 | 169966813 | 169966858 | 169975607 | 169975716 | 5 | 4 | 6 | 12 | 6 | 286 | 219 | 282 | 303 | 326 | 0 | 0 | 20 | 5 | 0 | 15 | 198 | 189 | 40 | 0 | 246 | 149 | 1 | 1 | 0.01 | 0.011 | 0.013 | 0.023 | 0.011 | 0 | 0 | 0.06 | 0.07 | NA | -0.019 |
| SE | 40401 | ENSG00000008952 | SEC62 | chr3 | + | 169975607 | 169975716 | 169966800 | 169966858 | 169976945 | 169977051 | 738 | 626 | 691 | 771 | 1011 | 0 | 1 | 0 | 0 | 0 | 74 | 546 | 496 | 110 | 3 | 0 | 0 | 0 | 0 | 0 | 258 | 149 | 1 | 1 | 1 | 0.997 | 1 | 1 | 1 | 1 | 1 | 1 | 1 | 1 | -0.001 |
| SE | 40402 | ENSG00000008952 | SEC62 | chr3 | + | 169985804 | 169985865 | 169983160 | 169983253 | 169988239 | 169988359 | 542 | 363 | 443 | 520 | 1034 | 0 | 0 | 1 | 0 | 0 | 93 | 268 | 248 | 122 | 74 | 0 | 0 | 0 | 2 | 0 | 210 | 149 | 1 | 1 | 1 | 1 | 0.997 | 1 | 1 | 1 | 1 | 1 | 0.977 | 1 | 0.004 |
| SE | 40405 | ENSG00000008952 | SEC62 | chr3 | + | 169988239 | 169988359 | 169985804 | 169985865 | 169992593 | 169992721 | 662 | 455 | 632 | 654 | 1242 | 1 | 1 | 0 | 0 | 0 | 95 | 316 | 322 | 122 | 61 | 0 | 0 | 0 | 0 | 0 | 269 | 149 | 1 | 1 | 0.997 | 0.996 | 1 | 1 | 1 | 1 | 1 | 1 | 1 | 1 | -0.001 |
| SE | 40406 | ENSG00000078304 | PPP2R5C | chr14 | + | 101786017 | 101786183 | 101762904 | 101762970 | 101836168 | 101836451 | 309 | 152 | 102 | 213 | 270 | 1 | 2 | 0 | 0 | 0 | 74 | 99 | 216 | 140 | 41 | 0 | 0 | 0 | 0 | 0 | 298 | 149 | 1 | 1 | 0.994 | 0.974 | 1 | 1 | 1 | 1 | 1 | 1 | 1 | 1 | -0.006 |
| SE | 40408 | ENSG00000078304 | PPP2R5C | chr14 | + | 101818991 | 101819085 | 101762904 | 101762970 | 101856685 | 101856885 | 2 | 1 | 4 | 3 | 10 | 36 | 9 | 12 | 11 | 18 | 0 | 6 | 2 | 0 | 0 | 16 | 5 | 13 | 10 | 0 | 243 | 149 | 1 | 1 | 0.033 | 0.064 | 0.17 | 0.143 | 0.254 | 0 | 0.424 | 0.086 | 0 | NA | 0.005 |
| SE | 40409 | ENSG00000078304 | PPP2R5C | chr14 | + | 101818991 | 101819085 | 101786120 | 101786183 | 101824013 | 101824229 | 0 | 1 | 1 | 0 | 12 | 6 | 5 | 4 | 12 | 4 | 0 | 0 | 0 | 0 | 0 | 0 | 13 | 15 | 7 | 10 | 243 | 149 | 1 | 1 | 0 | 0.109 | 0.133 | 0 | 0.648 | NA | 0 | 0 | 0 | 0 | 0.178 |
| SE | 40411 | ENSG00000078304 | PPP2R5C | chr14 | + | 101818991 | 101819085 | 101809904 | 101810036 | 101824013 | 101824229 | 1 | 0 | 1 | 1 | 9 | 2 | 5 | 4 | 4 | 10 | 0 | 0 | 2 | 0 | 0 | 0 | 4 | 0 | 2 | 2 | 243 | 149 | 1 | 1 | 0.235 | 0 | 0.133 | 0.133 | 0.356 | NA | 0 | 1 | 0 | 0 | -0.079 |
| SE | 40413 | ENSG00000078304 | PPP2R5C | chr14 | + | 101824013 | 101824169 | 101809836 | 101810036 | 101831700 | 101831804 | 9 | 5 | 4 | 5 | 11 | 1 | 0 | 9 | 2 | 3 | 0 | 6 | 3 | 7 | 2 | 0 | 3 | 8 | 0 | 0 | 298 | 149 | 1 | 1 | 0.818 | 1 | 0.182 | 0.556 | 0.647 | NA | 0.5 | 0.158 | 1 | 1 | -0.024 |
| SE | 40416 | ENSG00000078304 | PPP2R5C | chr14 | + | 101824013 | 101824169 | 101809836 | 101810036 | 101882160 | 101882205 | 3 | 5 | 4 | 4 | 10 | 2 | 0 | 2 | 0 | 0 | 0 | 4 | 0 | 2 | 2 | 0 | 0 | 0 | 0 | 0 | 298 | 149 | 1 | 1 | 0.429 | 1 | 0.5 | 1 | 1 | NA | 1 | NA | 1 | 1 | -0.214 |
| SE | 40418 | ENSG00000078304 | PPP2R5C | chr14 | + | 101831700 | 101831804 | 101786017 | 101786183 | 101836168 | 101836432 | 13 | 8 | 7 | 6 | 21 | 24 | 14 | 8 | 7 | 15 | 0 | 1 | 9 | 2 | 0 | 0 | 14 | 14 | 8 | 2 | 253 | 149 | 1 | 1 | 0.242 | 0.252 | 0.34 | 0.335 | 0.452 | NA | 0.04 | 0.275 | 0.128 | 0 | 0.213 |
| SE | 40422 | ENSG00000078304 | PPP2R5C | chr14 | + | 101831700 | 101831804 | 101824013 | 101824169 | 101836168 | 101836432 | 12 | 1 | 6 | 4 | 17 | 3 | 1 | 7 | 6 | 8 | 0 | 3 | 9 | 7 | 0 | 0 | 8 | 2 | 4 | 7 | 253 | 149 | 1 | 1 | 0.702 | 0.371 | 0.335 | 0.282 | 0.556 | NA | 0.181 | 0.726 | 0.508 | 0 | 0.095 |
| SE | 40423 | ENSG00000078304 | PPP2R5C | chr14 | + | 101831700 | 101831804 | 101824013 | 101824169 | 101856685 | 101856885 | 11 | 2 | 3 | 7 | 4 | 10 | 4 | 7 | 11 | 20 | 0 | 2 | 3 | 5 | 0 | 0 | 10 | 9 | 0 | 0 | 253 | 149 | 1 | 1 | 0.393 | 0.227 | 0.202 | 0.273 | 0.105 | NA | 0.105 | 0.164 | 1 | NA | -0.183 |
| SE | 40424 | ENSG00000078304 | PPP2R5C | chr14 | + | 101856685 | 101856885 | 101786017 | 101786183 | 101882160 | 101882205 | 892 | 362 | 507 | 625 | 876 | 0 | 0 | 0 | 0 | 0 | 185 | 401 | 342 | 284 | 64 | 0 | 0 | 1 | 0 | 0 | 298 | 149 | 1 | 1 | 1 | 1 | 1 | 1 | 1 | 1 | 1 | 0.994 | 1 | 1 | 0.001 |
| SE | 40425 | ENSG00000078304 | PPP2R5C | chr14 | + | 101856685 | 101856885 | 101809802 | 101810036 | 101882160 | 101882205 | 1009 | 454 | 664 | 780 | 915 | 2 | 0 | 2 | 0 | 0 | 304 | 570 | 445 | 385 | 63 | 0 | 0 | 0 | 0 | 0 | 298 | 149 | 1 | 1 | 0.996 | 1 | 0.994 | 1 | 1 | 1 | 1 | 1 | 1 | 1 | -0.002 |
| SE | 40426 | ENSG00000078304 | PPP2R5C | chr14 | + | 101856685 | 101856885 | 101824013 | 101824169 | 101882160 | 101882205 | 680 | 275 | 407 | 501 | 704 | 1 | 0 | 0 | 0 | 0 | 151 | 326 | 247 | 212 | 41 | 0 | 0 | 0 | 0 | 0 | 298 | 149 | 1 | 1 | 0.997 | 1 | 1 | 1 | 1 | 1 | 1 | 1 | 1 | 1 | -0.001 |
| SE | 40427 | ENSG00000078304 | PPP2R5C | chr14 | + | 101882160 | 101882271 | 101856685 | 101856885 | 101883256 | 101883349 | 1065 | 539 | 681 | 834 | 1164 | 0 | 0 | 0 | 0 | 0 | 247 | 630 | 440 | 399 | 98 | 0 | 0 | 0 | 2 | 0 | 260 | 149 | 1 | 1 | 1 | 1 | 1 | 1 | 1 | 1 | 1 | 1 | 0.991 | 1 | 0.002 |
| SE | 40428 | ENSG00000078304 | PPP2R5C | chr14 | + | 101882160 | 101882271 | 101856685 | 101856885 | 101883431 | 101883562 | 573 | 244 | 355 | 433 | 578 | 0 | 0 | 0 | 1 | 0 | 124 | 276 | 188 | 173 | 37 | 0 | 0 | 0 | 0 | 0 | 260 | 149 | 1 | 1 | 1 | 1 | 1 | 0.996 | 1 | 1 | 1 | 1 | 1 | 1 | -0.001 |
| SE | 40429 | ENSG00000078304 | PPP2R5C | chr14 | + | 101883256 | 101883349 | 101856685 | 101856885 | 101883431 | 101883562 | 232 | 169 | 235 | 278 | 639 | 0 | 0 | 0 | 1 | 0 | 63 | 146 | 151 | 100 | 30 | 0 | 0 | 0 | 0 | 0 | 242 | 149 | 1 | 1 | 1 | 1 | 1 | 0.994 | 1 | 1 | 1 | 1 | 1 | 1 | -0.001 |
| SE | 40430 | ENSG00000078304 | PPP2R5C | chr14 | + | 101883256 | 101883349 | 101882160 | 101882271 | 101883431 | 101883562 | 741 | 468 | 578 | 687 | 1234 | 4 | 1 | 0 | 0 | 3 | 187 | 506 | 412 | 325 | 91 | 0 | 3 | 1 | 0 | 0 | 242 | 149 | 1 | 1 | 0.991 | 0.997 | 1 | 1 | 0.996 | 1 | 0.99 | 0.996 | 1 | 1 | 0 |
| SE | 40431 | ENSG00000078304 | PPP2R5C | chr14 | + | 101883431 | 101883562 | 101856685 | 101856885 | 101894506 | 101894560 | 0 | 0 | 0 | 1 | 0 | 0 | 0 | 0 | 1 | 0 | 0 | 0 | 2 | 0 | 0 | 0 | 0 | 1 | 1 | 0 | 280 | 149 | 1 | 1 | NA | NA | NA | 0.347 | NA | NA | NA | 0.516 | 0 | NA | 0.089 |
| SE | 40432 | ENSG00000078304 | PPP2R5C | chr14 | + | 101883431 | 101883562 | 101856685 | 101856885 | 101901718 | 101901744 | 0 | 0 | 0 | 1 | 0 | 0 | 0 | 1 | 0 | 0 | 0 | 0 | 5 | 0 | 0 | 0 | 6 | 0 | 0 | 0 | 280 | 149 | 1 | 1 | NA | NA | 0 | 1 | NA | NA | 0 | NA | 1 | NA | 0 |
| SE | 40434 | ENSG00000078304 | PPP2R5C | chr14 | + | 101889981 | 101890192 | 101883431 | 101883562 | 101892999 | 101893108 | 2 | 1 | 0 | 0 | 0 | 3 | 3 | 6 | 6 | 6 | 0 | 2 | 3 | 0 | 0 | 8 | 8 | 7 | 0 | 0 | 298 | 149 | 1 | 1 | 0.25 | 0.143 | 0 | 0 | 0 | 0 | 0.111 | 0.176 | NA | NA | -0.017 |
| SE | 40436 | ENSG00000078304 | PPP2R5C | chr14 | + | 101890236 | 101890296 | 101883431 | 101883562 | 101894506 | 101894517 | 275 | 188 | 195 | 291 | 348 | 0 | 0 | 0 | 0 | 0 | 75 | 192 | 202 | 133 | 26 | 0 | 0 | 2 | 0 | 0 | 209 | 149 | 1 | 1 | 1 | 1 | 1 | 1 | 1 | 1 | 1 | 0.986 | 1 | 1 | 0.003 |
| SE | 40437 | ENSG00000078304 | PPP2R5C | chr14 | + | 101890236 | 101890296 | 101883431 | 101883562 | 101901718 | 101901889 | 274 | 187 | 195 | 285 | 338 | 0 | 0 | 0 | 0 | 0 | 75 | 189 | 201 | 133 | 27 | 0 | 0 | 0 | 5 | 0 | 209 | 149 | 1 | 1 | 1 | 1 | 1 | 1 | 1 | 1 | 1 | 1 | 0.95 | 1 | 0.01 |
| SE | 40438 | ENSG00000078304 | PPP2R5C | chr14 | + | 101892999 | 101893108 | 101883431 | 101883562 | 101894506 | 101894517 | 612 | 306 | 461 | 475 | 1245 | 0 | 0 | 0 | 0 | 0 | 118 | 466 | 314 | 250 | 52 | 0 | 0 | 2 | 0 | 0 | 258 | 149 | 1 | 1 | 1 | 1 | 1 | 1 | 1 | 1 | 1 | 0.989 | 1 | 1 | 0.002 |
| SE | 40439 | ENSG00000078304 | PPP2R5C | chr14 | + | 101892999 | 101893108 | 101890236 | 101890296 | 101894506 | 101894517 | 1307 | 615 | 942 | 1028 | 2693 | 2 | 1 | 0 | 6 | 10 | 268 | 937 | 669 | 494 | 106 | 0 | 3 | 4 | 0 | 0 | 258 | 149 | 1 | 1 | 0.997 | 0.997 | 1 | 0.99 | 0.994 | 1 | 0.994 | 0.99 | 1 | 1 | -0.001 |
| SE | 40440 | ENSG00000078304 | PPP2R5C | chr14 | + | 101894506 | 101894560 | 101856685 | 101856885 | 101901718 | 101901744 | 294 | 180 | 271 | 275 | 920 | 0 | 0 | 1 | 0 | 0 | 43 | 186 | 207 | 106 | 23 | 0 | 6 | 0 | 0 | 0 | 203 | 149 | 1 | 1 | 1 | 1 | 0.995 | 1 | 1 | 1 | 0.958 | 1 | 1 | 1 | 0.007 |
| SE | 40441 | ENSG00000078304 | PPP2R5C | chr14 | + | 101894506 | 101894560 | 101883431 | 101883562 | 101901718 | 101901744 | 294 | 180 | 271 | 275 | 920 | 0 | 0 | 0 | 0 | 0 | 43 | 186 | 208 | 105 | 23 | 0 | 0 | 0 | 5 | 0 | 203 | 149 | 1 | 1 | 1 | 1 | 1 | 1 | 1 | 1 | 1 | 1 | 0.939 | 1 | 0.012 |
| SE | 40442 | ENSG00000078304 | PPP2R5C | chr14 | + | 101894506 | 101894560 | 101890236 | 101890296 | 101901718 | 101901744 | 296 | 181 | 271 | 281 | 930 | 0 | 0 | 0 | 0 | 0 | 43 | 189 | 210 | 105 | 23 | 0 | 0 | 0 | 0 | 1 | 203 | 149 | 1 | 1 | 1 | 1 | 1 | 1 | 1 | 1 | 1 | 1 | 1 | 0.944 | 0.011 |
| SE | 40443 | ENSG00000078304 | PPP2R5C | chr14 | + | 101901718 | 101901889 | 101894506 | 101894560 | 101906401 | 101906529 | 1513 | 812 | 1148 | 1284 | 3211 | 0 | 0 | 0 | 4 | 0 | 468 | 1175 | 1087 | 810 | 254 | 4 | 0 | 0 | 0 | 0 | 298 | 149 | 1 | 1 | 1 | 1 | 1 | 0.994 | 1 | 0.983 | 1 | 1 | 1 | 1 | 0.002 |
| SE | 40444 | ENSG00000078304 | PPP2R5C | chr14 | + | 101906401 | 101906529 | 101901718 | 101901889 | 101909588 | 101909690 | 1118 | 626 | 809 | 955 | 1710 | 6 | 0 | 2 | 2 | 7 | 525 | 912 | 970 | 836 | 262 | 0 | 1 | 0 | 0 | 0 | 277 | 149 | 1 | 1 | 0.99 | 1 | 0.995 | 0.996 | 0.992 | 1 | 0.998 | 1 | 1 | 1 | -0.005 |
| SE | 40447 | ENSG00000078304 | PPP2R5C | chr14 | + | 101909588 | 101909690 | 101906401 | 101906529 | 101912400 | 101912473 | 986 | 591 | 703 | 788 | 1143 | 0 | 0 | 2 | 2 | 0 | 385 | 624 | 752 | 691 | 157 | 0 | 0 | 0 | 8 | 0 | 251 | 149 | 1 | 1 | 1 | 1 | 0.995 | 0.996 | 1 | 1 | 1 | 1 | 0.981 | 1 | 0.002 |
| SE | 40448 | ENSG00000078304 | PPP2R5C | chr14 | + | 101912400 | 101912473 | 101909588 | 101909690 | 101917830 | 101917947 | 627 | 398 | 450 | 513 | 748 | 1 | 1 | 0 | 0 | 0 | 251 | 366 | 374 | 362 | 126 | 0 | 0 | 0 | 0 | 0 | 222 | 149 | 1 | 1 | 0.998 | 0.996 | 1 | 1 | 1 | 1 | 1 | 1 | 1 | 1 | -0.001 |
| SE | 40454 | ENSG00000135801 | TAF5L | chr1 | - | 229607773 | 229607835 | 229606833 | 229606876 | 229610105 | 229610210 | 2 | 1 | 0 | 6 | 5 | 0 | 1 | 0 | 0 | 0 | 0 | 19 | 8 | 8 | 0 | 8 | 0 | 0 | 0 | 0 | 211 | 149 | 1 | 1 | 1 | 0.414 | NA | 1 | 1 | 0 | 1 | 1 | 1 | NA | 0.103 |
| SE | 40455 | ENSG00000135801 | TAF5L | chr1 | - | 229610105 | 229610210 | 229599193 | 229602919 | 229614340 | 229614485 | 65 | 44 | 41 | 62 | 117 | 0 | 0 | 0 | 0 | 0 | 21 | 91 | 51 | 78 | 67 | 0 | 0 | 1 | 1 | 0 | 254 | 149 | 1 | 1 | 1 | 1 | 1 | 1 | 1 | 1 | 1 | 0.968 | 0.979 | 1 | 0.011 |
| SE | 40458 | ENSG00000184432 | COPB2 | chr3 | - | 139378040 | 139378126 | 139375597 | 139375614 | 139379046 | 139379173 | 160 | 89 | 121 | 116 | 173 | 0 | 1 | 0 | 0 | 0 | 46 | 90 | 86 | 68 | 0 | 0 | 0 | 1 | 0 | 0 | 235 | 149 | 1 | 1 | 1 | 0.983 | 1 | 1 | 1 | 1 | 1 | 0.982 | 1 | NA | 0.001 |
| SE | 40459 | ENSG00000184432 | COPB2 | chr3 | - | 139378040 | 139378189 | 139375467 | 139375614 | 139379046 | 139379173 | 488 | 270 | 359 | 397 | 559 | 0 | 1 | 0 | 0 | 0 | 152 | 316 | 232 | 199 | 40 | 0 | 0 | 1 | 0 | 0 | 298 | 149 | 1 | 1 | 1 | 0.993 | 1 | 1 | 1 | 1 | 1 | 0.991 | 1 | 1 | 0 |
| SE | 40460 | ENSG00000184432 | COPB2 | chr3 | - | 139379771 | 139379907 | 139379379 | 139379466 | 139383297 | 139383435 | 6 | 0 | 5 | 1 | 7 | 238 | 106 | 158 | 156 | 204 | 0 | 4 | 4 | 0 | 0 | 69 | 148 | 97 | 129 | 26 | 285 | 149 | 1 | 1 | 0.013 | 0 | 0.016 | 0.003 | 0.018 | 0 | 0.014 | 0.021 | 0 | 0 | 0.003 |
| SE | 40461 | ENSG00000184432 | COPB2 | chr3 | - | 139385003 | 139385200 | 139383297 | 139383435 | 139389468 | 139389648 | 9 | 4 | 19 | 8 | 6 | 6 | 0 | 0 | 2 | 0 | 0 | 4 | 16 | 8 | 0 | 0 | 0 | 0 | 0 | 0 | 298 | 149 | 1 | 1 | 0.429 | 1 | 1 | 0.667 | 1 | NA | 1 | 1 | 1 | NA | -0.181 |
| SE | 40464 | ENSG00000131697 | NPHP4 | chr1 | - | 5864337 | 5864517 | 5863889 | 5864033 | 5866372 | 5866458 | 7 | 2 | 7 | 3 | 3 | 0 | 0 | 0 | 1 | 0 | 5 | 5 | 3 | 7 | 29 | 0 | 0 | 0 | 0 | 0 | 298 | 149 | 1 | 1 | 1 | 1 | 1 | 0.6 | 1 | 1 | 1 | 1 | 1 | 1 | -0.08 |
| SE | 40465 | ENSG00000131697 | NPHP4 | chr1 | - | 5865101 | 5865273 | 5864337 | 5864517 | 5866372 | 5866458 | 7 | 9 | 9 | 1 | 1 | 0 | 0 | 3 | 1 | 0 | 11 | 11 | 4 | 8 | 27 | 0 | 0 | 0 | 0 | 0 | 298 | 149 | 1 | 1 | 1 | 1 | 0.6 | 1 | 1 | 1 | 1 | 1 | 1 | 1 | -0.08 |
| SE | 40467 | ENSG00000131697 | NPHP4 | chr1 | - | 5887285 | 5887466 | 5880113 | 5880239 | 5904616 | 5904804 | 5 | 0 | 7 | 6 | 2 | 0 | 0 | 1 | 0 | 0 | 0 | 5 | 0 | 0 | 0 | 0 | 0 | 0 | 0 | 0 | 298 | 149 | 1 | 1 | 1 | NA | 0.778 | 1 | 1 | NA | 1 | NA | NA | NA | -0.055 |
| SE | 40469 | ENSG00000131697 | NPHP4 | chr1 | - | 5890867 | 5891028 | 5887285 | 5887466 | 5904616 | 5904804 | 14 | 3 | 4 | 15 | 5 | 0 | 0 | 0 | 0 | 0 | 0 | 0 | 2 | 8 | 4 | 0 | 2 | 0 | 0 | 0 | 298 | 149 | 1 | 1 | 1 | 1 | 1 | 1 | 1 | NA | 0 | 1 | 1 | 1 | 0.25 |
| SE | 40470 | ENSG00000131697 | NPHP4 | chr1 | - | 5952699 | 5952836 | 5948069 | 5948251 | 5961793 | 5961949 | 7 | 0 | 6 | 4 | 0 | 0 | 0 | 0 | 1 | 0 | 0 | 12 | 21 | 5 | 10 | 0 | 0 | 0 | 5 | 0 | 286 | 149 | 1 | 1 | 1 | NA | 1 | 0.676 | NA | NA | 1 | 1 | 0.343 | 1 | 0.056 |
| SE | 40471 | ENSG00000131697 | NPHP4 | chr1 | - | 5969086 | 5969259 | 5967298 | 5967363 | 5978269 | 5978413 | 36 | 18 | 29 | 51 | 40 | 0 | 0 | 0 | 0 | 0 | 53 | 37 | 40 | 48 | 52 | 0 | 2 | 0 | 0 | 0 | 298 | 149 | 1 | 1 | 1 | 1 | 1 | 1 | 1 | 1 | 0.902 | 1 | 1 | 1 | 0.02 |
| SE | 40475 | ENSG00000164088 | PPM1M | chr3 | + | 52248636 | 52248700 | 52248152 | 52248237 | 52248955 | 52249063 | 104 | 50 | 78 | 66 | 93 | 0 | 0 | 0 | 1 | 0 | 81 | 126 | 58 | 92 | 210 | 0 | 0 | 0 | 0 | 0 | 213 | 149 | 1 | 1 | 1 | 1 | 1 | 0.979 | 1 | 1 | 1 | 1 | 1 | 1 | -0.004 |
| SE | 40477 | ENSG00000285708 | NA | chr3 | - | 70965889 | 70966056 | 70959225 | 70959391 | 70970735 | 70970805 | 690 | 530 | 916 | 913 | 1016 | 0 | 0 | 1 | 1 | 1 | 347 | 800 | 800 | 526 | 218 | 0 | 0 | 0 | 1 | 0 | 298 | 149 | 1 | 1 | 1 | 1 | 0.998 | 0.998 | 0.998 | 1 | 1 | 1 | 0.996 | 1 | 0 |
| SE | 40478 | ENSG00000285708 | NA | chr3 | - | 70965889 | 70966056 | 70959225 | 70959391 | 70976940 | 70977042 | 297 | 249 | 393 | 382 | 470 | 0 | 0 | 0 | 0 | 2 | 102 | 316 | 340 | 275 | 82 | 0 | 0 | 2 | 0 | 0 | 298 | 149 | 1 | 1 | 1 | 1 | 1 | 1 | 0.992 | 1 | 1 | 0.988 | 1 | 1 | 0.001 |
| SE | 40479 | ENSG00000285708 | NA | chr3 | - | 70970735 | 70970805 | 70959225 | 70959391 | 70976940 | 70977042 | 2 | 2 | 7 | 3 | 1 | 0 | 0 | 0 | 0 | 2 | 1 | 0 | 1 | 3 | 0 | 0 | 0 | 2 | 0 | 0 | 219 | 149 | 1 | 1 | 1 | 1 | 1 | 1 | 0.254 | 1 | NA | 0.254 | 1 | NA | 0.099 |
| SE | 40480 | ENSG00000285708 | NA | chr3 | - | 70970735 | 70970805 | 70965889 | 70966056 | 70972554 | 70972676 | 587 | 517 | 824 | 878 | 810 | 5 | 0 | 0 | 1 | 2 | 403 | 772 | 808 | 453 | 234 | 0 | 5 | 0 | 0 | 0 | 219 | 149 | 1 | 1 | 0.988 | 1 | 1 | 0.998 | 0.996 | 1 | 0.991 | 1 | 1 | 1 | -0.002 |
| SE | 40483 | ENSG00000114861 | FOXP1 | chr3 | - | 70972554 | 70972676 | 70968443 | 70970805 | 70976940 | 70977042 | 901 | 762 | 1214 | 1228 | 1021 | 3 | 2 | 6 | 3 | 1 | 571 | 1036 | 1193 | 807 | 417 | 1 | 0 | 9 | 2 | 0 | 271 | 149 | 1 | 1 | 0.994 | 0.995 | 0.991 | 0.996 | 0.998 | 0.997 | 1 | 0.986 | 0.996 | 1 | -0.001 |
| SE | 40485 | ENSG00000285708 | NA | chr3 | - | 71000971 | 71001059 | 70987993 | 70988077 | 71041327 | 71041532 | 262 | 169 | 307 | 233 | 566 | 0 | 0 | 0 | 0 | 0 | 88 | 229 | 252 | 215 | 35 | 0 | 0 | 1 | 0 | 0 | 237 | 149 | 1 | 1 | 1 | 1 | 1 | 1 | 1 | 1 | 1 | 0.994 | 1 | 1 | 0.001 |
| SE | 40486 | ENSG00000285708 | NA | chr3 | - | 71015548 | 71015653 | 70987993 | 70988077 | 71041327 | 71041532 | 105 | 90 | 151 | 137 | 168 | 0 | 0 | 0 | 0 | 0 | 90 | 109 | 137 | 93 | 24 | 0 | 0 | 1 | 0 | 0 | 254 | 149 | 1 | 1 | 1 | 1 | 1 | 1 | 1 | 1 | 1 | 0.988 | 1 | 1 | 0.002 |
| SE | 40488 | ENSG00000285708 | NA | chr3 | - | 71041327 | 71041532 | 71015548 | 71015653 | 71052536 | 71052626 | 284 | 238 | 372 | 321 | 516 | 0 | 0 | 0 | 1 | 0 | 106 | 300 | 339 | 212 | 80 | 0 | 0 | 0 | 0 | 0 | 298 | 149 | 1 | 1 | 1 | 1 | 1 | 0.994 | 1 | 1 | 1 | 1 | 1 | 1 | -0.001 |
| SE | 40489 | ENSG00000285708 | NA | chr3 | - | 71046941 | 71047095 | 71041327 | 71041532 | 71052536 | 71052626 | 254 | 166 | 362 | 301 | 261 | 0 | 0 | 2 | 0 | 1 | 146 | 244 | 278 | 180 | 52 | 0 | 0 | 0 | 0 | 0 | 298 | 149 | 1 | 1 | 1 | 1 | 0.989 | 1 | 0.992 | 1 | 1 | 1 | 1 | 1 | -0.004 |
| SE | 40490 | ENSG00000285708 | NA | chr3 | - | 71112535 | 71112637 | 70977827 | 70978029 | 71299819 | 71299880 | 0 | 0 | 2 | 0 | 4 | 0 | 0 | 0 | 0 | 1 | 0 | 1 | 0 | 0 | 0 | 0 | 0 | 0 | 0 | 0 | 251 | 149 | 1 | 1 | NA | NA | 1 | NA | 0.704 | NA | 1 | NA | NA | NA | -0.148 |
| SE | 40491 | ENSG00000285708 | NA | chr3 | - | 71112535 | 71112637 | 71015548 | 71015653 | 71299819 | 71299880 | 0 | 0 | 0 | 0 | 4 | 0 | 1 | 0 | 0 | 0 | 0 | 5 | 0 | 0 | 0 | 0 | 0 | 0 | 0 | 0 | 251 | 149 | 1 | 1 | NA | 0 | NA | NA | 1 | NA | 1 | NA | NA | NA | -0.5 |
| SE | 40492 | ENSG00000285708 | NA | chr3 | - | 71112535 | 71112637 | 71015548 | 71015653 | 71690009 | 71690165 | 0 | 0 | 0 | 0 | 0 | 1 | 0 | 0 | 0 | 0 | 0 | 8 | 0 | 0 | 0 | 0 | 8 | 0 | 0 | 0 | 251 | 149 | 1 | 1 | 0 | NA | NA | NA | NA | NA | 0.372 | NA | NA | NA | -0.372 |
| SE | 40493 | ENSG00000285708 | NA | chr3 | - | 71112535 | 71112637 | 71053635 | 71053773 | 71299819 | 71299880 | 68 | 46 | 91 | 84 | 100 | 3 | 0 | 0 | 0 | 0 | 7 | 85 | 66 | 27 | 3 | 0 | 0 | 0 | 0 | 0 | 251 | 149 | 1 | 1 | 0.931 | 1 | 1 | 1 | 1 | 1 | 1 | 1 | 1 | 1 | -0.014 |
| SE | 40494 | ENSG00000114861 | FOXP1 | chr3 | - | 71198201 | 71198392 | 71112535 | 71112637 | 71299819 | 71299880 | 179 | 176 | 260 | 162 | 272 | 0 | 0 | 0 | 0 | 4 | 97 | 115 | 176 | 102 | 111 | 0 | 1 | 0 | 0 | 0 | 298 | 149 | 1 | 1 | 1 | 1 | 1 | 1 | 0.971 | 1 | 0.983 | 1 | 1 | 1 | -0.002 |
| SE | 40495 | ENSG00000285708 | NA | chr3 | - | 71299819 | 71299880 | 71015548 | 71015653 | 71359149 | 71359244 | 32 | 20 | 32 | 30 | 40 | 0 | 0 | 0 | 0 | 1 | 16 | 33 | 43 | 14 | 44 | 0 | 0 | 0 | 0 | 0 | 210 | 149 | 1 | 1 | 1 | 1 | 1 | 1 | 0.966 | 1 | 1 | 1 | 1 | 1 | -0.007 |
| SE | 40496 | ENSG00000285708 | NA | chr3 | - | 71299819 | 71299880 | 71112535 | 71112637 | 71493425 | 71493555 | 5 | 2 | 1 | 1 | 0 | 0 | 0 | 2 | 0 | 0 | 0 | 3 | 3 | 1 | 0 | 0 | 4 | 0 | 0 | 0 | 210 | 149 | 1 | 1 | 1 | 1 | 0.262 | 1 | NA | NA | 0.347 | 1 | 1 | NA | 0.033 |
| SE | 40497 | ENSG00000285708 | NA | chr3 | - | 71299819 | 71299880 | 71198201 | 71198392 | 71359149 | 71359244 | 134 | 121 | 183 | 110 | 194 | 0 | 0 | 0 | 2 | 0 | 80 | 106 | 154 | 89 | 112 | 0 | 0 | 3 | 1 | 0 | 210 | 149 | 1 | 1 | 1 | 1 | 1 | 0.975 | 1 | 1 | 1 | 0.973 | 0.984 | 1 | 0.004 |
| SE | 40499 | ENSG00000285708 | NA | chr3 | - | 71335159 | 71335281 | 71299819 | 71299880 | 71493425 | 71493555 | 1 | 8 | 5 | 3 | 3 | 7 | 3 | 1 | 4 | 3 | 0 | 1 | 1 | 0 | 0 | 0 | 3 | 10 | 1 | 0 | 271 | 149 | 1 | 1 | 0.073 | 0.595 | 0.733 | 0.292 | 0.355 | NA | 0.155 | 0.052 | 0 | NA | 0.341 |
| SE | 40500 | ENSG00000285708 | NA | chr3 | - | 71359149 | 71359244 | 71299819 | 71299880 | 71467250 | 71467315 | 58 | 53 | 84 | 57 | 58 | 0 | 0 | 1 | 0 | 0 | 42 | 66 | 47 | 39 | 69 | 0 | 0 | 0 | 0 | 0 | 244 | 149 | 1 | 1 | 1 | 1 | 0.981 | 1 | 1 | 1 | 1 | 1 | 1 | 1 | -0.004 |
| SE | 40502 | ENSG00000285708 | NA | chr3 | - | 71359149 | 71359244 | 71335159 | 71335281 | 71493425 | 71493555 | 71 | 110 | 100 | 81 | 98 | 0 | 0 | 1 | 0 | 0 | 129 | 105 | 99 | 88 | 57 | 0 | 0 | 0 | 0 | 0 | 244 | 149 | 1 | 1 | 1 | 1 | 0.984 | 1 | 1 | 1 | 1 | 1 | 1 | 1 | -0.003 |
| SE | 40503 | ENSG00000285708 | NA | chr3 | - | 71467250 | 71467315 | 71299819 | 71299880 | 71493425 | 71493555 | 2 | 1 | 3 | 0 | 0 | 7 | 3 | 1 | 4 | 3 | 0 | 3 | 2 | 0 | 0 | 0 | 3 | 10 | 1 | 0 | 214 | 149 | 1 | 1 | 0.166 | 0.188 | 0.676 | 0 | 0 | NA | 0.41 | 0.122 | 0 | NA | 0.029 |
| SE | 40505 | ENSG00000285708 | NA | chr3 | - | 71493425 | 71493555 | 71112535 | 71112637 | 71690009 | 71690165 | 10 | 13 | 17 | 17 | 27 | 0 | 0 | 0 | 0 | 0 | 0 | 27 | 30 | 21 | 16 | 0 | 4 | 0 | 0 | 0 | 279 | 149 | 1 | 1 | 1 | 1 | 1 | 1 | 1 | NA | 0.783 | 1 | 1 | 1 | 0.054 |
| SE | 40506 | ENSG00000285708 | NA | chr3 | - | 71690009 | 71690165 | 71493425 | 71493555 | 71710411 | 71710484 | 12 | 17 | 15 | 17 | 27 | 0 | 0 | 0 | 3 | 0 | 0 | 29 | 39 | 22 | 22 | 0 | 0 | 4 | 0 | 0 | 298 | 149 | 1 | 1 | 1 | 1 | 1 | 0.739 | 1 | NA | 1 | 0.83 | 1 | 1 | -0.01 |
| SE | 40508 | ENSG00000285708 | NA | chr3 | - | 71693874 | 71693941 | 71690009 | 71690165 | 71710411 | 71710484 | 166 | 180 | 159 | 178 | 465 | 1 | 2 | 0 | 0 | 0 | 537 | 286 | 323 | 438 | 239 | 0 | 0 | 1 | 0 | 6 | 216 | 149 | 1 | 1 | 0.991 | 0.984 | 1 | 1 | 1 | 1 | 1 | 0.996 | 1 | 0.965 | 0.003 |
| SE | 40509 | ENSG00000285708 | NA | chr3 | - | 71693874 | 71693941 | 71690009 | 71690165 | 71725191 | 71725418 | 161 | 167 | 152 | 168 | 446 | 1 | 1 | 0 | 0 | 0 | 517 | 260 | 307 | 411 | 227 | 0 | 0 | 3 | 0 | 0 | 216 | 149 | 1 | 1 | 0.991 | 0.991 | 1 | 1 | 1 | 1 | 1 | 0.986 | 1 | 1 | -0.001 |
| SE | 40510 | ENSG00000285708 | NA | chr3 | - | 71693874 | 71693941 | 71690009 | 71690165 | 71728475 | 71728623 | 161 | 167 | 152 | 168 | 446 | 0 | 0 | 0 | 0 | 0 | 507 | 260 | 313 | 411 | 227 | 0 | 0 | 2 | 0 | 0 | 216 | 149 | 1 | 1 | 1 | 1 | 1 | 1 | 1 | 1 | 1 | 0.991 | 1 | 1 | 0.002 |
| SE | 40512 | ENSG00000285708 | NA | chr3 | - | 71699613 | 71699708 | 71690009 | 71690165 | 71725191 | 71725418 | 0 | 0 | 0 | 0 | 0 | 1 | 1 | 0 | 0 | 0 | 0 | 0 | 0 | 0 | 12 | 0 | 0 | 3 | 0 | 0 | 244 | 149 | 1 | 1 | 0 | 0 | NA | NA | NA | NA | NA | 0 | NA | 1 | -0.5 |
| SE | 40515 | ENSG00000285708 | NA | chr3 | - | 71699613 | 71699708 | 71693874 | 71693941 | 71725191 | 71725418 | 4 | 0 | 2 | 1 | 0 | 0 | 0 | 0 | 0 | 0 | 0 | 0 | 0 | 0 | 6 | 10 | 0 | 0 | 0 | 0 | 244 | 149 | 1 | 1 | 1 | NA | 1 | 1 | NA | 0 | NA | NA | NA | 1 | 0.5 |
| SE | 40516 | ENSG00000285708 | NA | chr3 | - | 71699613 | 71699708 | 71693874 | 71693941 | 71728475 | 71728623 | 4 | 0 | 2 | 1 | 0 | 0 | 0 | 0 | 0 | 0 | 0 | 0 | 8 | 0 | 6 | 0 | 0 | 6 | 0 | 0 | 244 | 149 | 1 | 1 | 1 | NA | 1 | 1 | NA | NA | NA | 0.449 | NA | 1 | 0.275 |
| SE | 40517 | ENSG00000285708 | NA | chr3 | - | 71710411 | 71710484 | 71690009 | 71690165 | 71725191 | 71725418 | 2 | 3 | 3 | 3 | 0 | 1 | 1 | 0 | 0 | 0 | 0 | 2 | 1 | 0 | 6 | 0 | 0 | 3 | 0 | 0 | 222 | 149 | 1 | 1 | 0.573 | 0.668 | 1 | 1 | NA | NA | 1 | 0.183 | NA | 1 | 0.083 |
| SE | 40518 | ENSG00000285708 | NA | chr3 | - | 71710411 | 71710484 | 71690009 | 71690165 | 71728475 | 71728623 | 4 | 4 | 0 | 0 | 1 | 0 | 0 | 0 | 0 | 0 | 0 | 1 | 7 | 1 | 10 | 0 | 0 | 2 | 0 | 0 | 222 | 149 | 1 | 1 | 1 | 1 | NA | NA | 1 | NA | 1 | 0.701 | 1 | 1 | 0.075 |
| SE | 40521 | ENSG00000285708 | NA | chr3 | - | 71710411 | 71710484 | 71699613 | 71699708 | 71725191 | 71725418 | 80 | 102 | 70 | 103 | 127 | 0 | 0 | 1 | 0 | 0 | 248 | 140 | 129 | 196 | 121 | 0 | 3 | 0 | 0 | 0 | 222 | 149 | 1 | 1 | 1 | 1 | 0.979 | 1 | 1 | 1 | 0.969 | 1 | 1 | 1 | 0.002 |
| SE | 40525 | ENSG00000162627 | SNX7 | chr1 | + | 98701816 | 98701903 | 98698705 | 98698905 | 98738236 | 98738389 | 0 | 0 | 0 | 1 | 1 | 0 | 1 | 0 | 0 | 0 | 0 | 0 | 4 | 2 | 0 | 0 | 0 | 0 | 0 | 0 | 236 | 149 | 1 | 1 | NA | 0 | NA | 1 | 1 | NA | NA | 1 | 1 | NA | -0.333 |
| SE | 40529 | ENSG00000111726 | CMAS | chr12 | + | 22060831 | 22060926 | 22058566 | 22058700 | 22065120 | 22065668 | 93 | 66 | 84 | 87 | 60 | 0 | 0 | 3 | 0 | 0 | 126 | 115 | 114 | 110 | 58 | 0 | 0 | 0 | 0 | 0 | 244 | 149 | 1 | 1 | 1 | 1 | 0.945 | 1 | 1 | 1 | 1 | 1 | 1 | 1 | -0.011 |
| SE | 40531 | ENSG00000111726 | CMAS | chr12 | + | 22062280 | 22062434 | 22060831 | 22060926 | 22065120 | 22065668 | 50 | 25 | 38 | 21 | 39 | 0 | 0 | 1 | 0 | 0 | 22 | 17 | 13 | 22 | 51 | 0 | 2 | 0 | 0 | 0 | 298 | 149 | 1 | 1 | 1 | 1 | 0.95 | 1 | 1 | 1 | 0.81 | 1 | 1 | 1 | 0.028 |
| SE | 40532 | ENSG00000008128 | CDK11A | chr1 | - | 1712263 | 1712400 | 1709478 | 1709640 | 1716345 | 1716478 | 33 | 77 | 32 | 38 | 16 | 0 | 0 | 0 | 0 | 0 | 78 | 6 | 35 | 65 | 93 | 4 | 0 | 0 | 0 | 0 | 286 | 149 | 1 | 1 | 1 | 1 | 1 | 1 | 1 | 0.91 | 1 | 1 | 1 | 1 | 0.018 |
| SE | 40534 | ENSG00000008128 | CDK11A | chr1 | - | 1716345 | 1716478 | 1712263 | 1712400 | 1719327 | 1719455 | 36 | 54 | 18 | 34 | 28 | 0 | 0 | 0 | 2 | 0 | 131 | 14 | 55 | 99 | 126 | 0 | 0 | 0 | 0 | 0 | 282 | 149 | 1 | 1 | 1 | 1 | 1 | 0.9 | 1 | 1 | 1 | 1 | 1 | 1 | -0.02 |
| SE | 40536 | ENSG00000008128 | CDK11A | chr1 | - | 1722707 | 1722831 | 1722587 | 1722634 | 1724257 | 1724324 | 40 | 26 | 22 | 46 | 28 | 0 | 0 | 0 | 1 | 0 | 98 | 31 | 66 | 76 | 48 | 5 | 0 | 0 | 0 | 0 | 273 | 149 | 1 | 1 | 1 | 1 | 1 | 0.962 | 1 | 0.915 | 1 | 1 | 1 | 1 | 0.009 |
| SE | 40537 | ENSG00000122644 | ARL4A | chr7 | + | 12687634 | 12687728 | 12686855 | 12687043 | 12688165 | 12689179 | 13 | 9 | 5 | 5 | 4 | 9 | 8 | 21 | 26 | 18 | 7 | 19 | 33 | 9 | 0 | 10 | 7 | 19 | 11 | 0 | 243 | 149 | 1 | 1 | 0.47 | 0.408 | 0.127 | 0.105 | 0.12 | 0.3 | 0.625 | 0.516 | 0.334 | NA | -0.198 |
| SE | 40538 | ENSG00000122644 | ARL4A | chr7 | + | 12687634 | 12687728 | 12686855 | 12687043 | 12688168 | 12688914 | 4 | 2 | 4 | 0 | 0 | 10 | 12 | 17 | 3 | 10 | 0 | 2 | 6 | 0 | 0 | 8 | 6 | 18 | 2 | 0 | 243 | 149 | 1 | 1 | 0.197 | 0.093 | 0.126 | 0 | 0 | 0 | 0.17 | 0.17 | 0 | NA | -0.002 |
| SE | 40544 | ENSG00000116977 | LGALS8 | chr1 | + | 236523969 | 236524061 | 236518423 | 236518500 | 236525967 | 236526115 | 144 | 119 | 108 | 151 | 58 | 0 | 1 | 0 | 0 | 0 | 91 | 275 | 157 | 152 | 143 | 0 | 0 | 0 | 0 | 0 | 241 | 149 | 1 | 1 | 1 | 0.987 | 1 | 1 | | 1 | 1 | 1 | 1 | 1 | -0.003 |
| SE | 40545 | ENSG00000116977 | LGALS8 | chr1 | + | 236525423 | 236525545 | 236518300 | 236518500 | 236525967 | 236526115 | 1 | 1 | 2 | 0 | 5 | 0 | 1 | 0 | 0 | 0 | 0 | 0 | 5 | 0 | 0 | 0 | 0 | 0 | 0 | 0 | 271 | 149 | 1 | 1 | 1 | 0.355 | 1 | NA | 1 | NA | NA | 1 | NA | NA | -0.161 |
| SE | 40550 | ENSG00000116977 | LGALS8 | chr1 | + | 236541653 | 236541710 | 236540246 | 236540683 | 236542760 | 236542787 | 350 | 311 | 392 | 379 | 730 | 0 | 0 | 0 | 0 | 0 | 146 | 419 | 305 | 166 | 78 | 2 | 0 | 0 | 0 | 0 | 206 | 149 | 1 | 1 | 1 | 1 | 1 | 1 | 1 | 0.981 | 1 | 1 | 1 | 1 | 0.004 |
| SE | 40552 | ENSG00000116977 | LGALS8 | chr1 | + | 236544749 | 236544915 | 236543026 | 236543648 | 236548011 | 236548161 | 557 | 545 | 568 | 534 | 811 | 0 | 0 | 0 | 0 | 0 | 143 | 654 | 485 | 402 | 62 | 0 | 1 | 0 | 0 | 0 | 298 | 149 | 1 | 1 | 1 | 1 | 1 | 1 | 1 | 1 | 0.997 | 1 | 1 | 1 | 0.001 |
| SE | 40553 | ENSG00000121236 | TRIM6 | chr11 | + | 5603245 | 5603735 | 5596108 | 5596181 | 5604533 | 5604629 | 0 | 0 | 0 | 3 | 0 | 0 | 0 | 0 | 0 | 0 | 11 | 4 | 2 | 5 | 0 | 0 | 0 | 0 | 3 | 0 | 298 | 149 | 1 | 1 | NA | NA | NA | 1 | NA | 1 | 1 | 1 | 0.455 | NA | 0.136 |
| SE | 40554 | ENSG00000121236 | TRIM6 | chr11 | + | 5603245 | 5603735 | 5596636 | 5596914 | 5604533 | 5604629 | 1 | 0 | 0 | 3 | 0 | 0 | 0 | 0 | 1 | 0 | 11 | 4 | 2 | 2 | 0 | 0 | 0 | 0 | 3 | 0 | 298 | 149 | 1 | 1 | 1 | NA | NA | 0.6 | NA | 1 | 1 | 1 | 1 | NA | -0.2 |
| SE | 40555 | ENSG00000121236 | TRIM6 | chr11 | + | 5603607 | 5603735 | 5596112 | 5596181 | 5604533 | 5604629 | 0 | 0 | 0 | 3 | 0 | 0 | 0 | 0 | 0 | 0 | 11 | 4 | 2 | 2 | 0 | 0 | 0 | 0 | 3 | 0 | 277 | 149 | 1 | 1 | NA | NA | NA | 1 | NA | 1 | 1 | 1 | 0.264 | NA | 0.184 |
| SE | 40556 | ENSG00000121236 | TRIM6 | chr11 | + | 5603607 | 5603735 | 5596745 | 5596914 | 5604533 | 5604629 | 0 | 0 | 0 | 3 | 0 | 0 | 0 | 0 | 1 | 0 | 11 | 4 | 2 | 2 | 0 | 0 | 0 | 0 | 0 | 0 | 277 | 149 | 1 | 1 | NA | NA | NA | 0.617 | NA | 1 | 1 | 1 | 1 | NA | -0.383 |
| SE | 40559 | ENSG00000086548 | CEACAM6 | chr19 | + | 41766182 | 41766299 | 41761968 | 41762223 | 41770801 | 41772211 | 4 | 0 | 2 | 13 | 0 | 0 | 0 | 0 | 0 | 0 | 141 | 18 | 19 | 884 | 83 | 10 | 0 | 0 | 17 | 0 | 266 | 149 | 1 | 1 | 1 | NA | 1 | 1 | NA | 0.888 | 1 | 1 | 0.967 | 1 | 0.029 |
| SE | 40562 | ENSG00000069998 | HDHD5 | chr22 | - | 17149541 | 17149745 | 17148447 | 17148560 | 17158456 | 17159168 | 32 | 9 | 19 | 30 | 47 | 1 | 0 | 1 | 1 | 0 | 104 | 41 | 11 | 30 | 98 | 0 | 0 | 0 | 0 | 0 | 298 | 149 | 1 | 1 | 0.941 | 1 | 0.905 | 0.938 | 1 | 1 | 1 | 1 | 1 | 1 | -0.043 |
| SE | 40566 | ENSG00000171680 | PLEKHG5 | chr1 | - | 6495515 | 6495589 | 6477528 | 6477658 | 6496492 | 6496569 | 0 | 0 | 1 | 0 | 0 | 1 | 0 | 4 | 1 | 0 | 0 | 0 | 0 | 0 | 0 | 0 | 0 | 5 | 0 | 0 | 223 | 149 | 1 | 1 | 0 | NA | 0.143 | 0 | NA | NA | NA | 0 | NA | NA | 0.048 |
| SE | 40567 | ENSG00000082438 | COBLL1 | chr2 | - | 164703103 | 164703220 | 164700521 | 164700756 | 164704443 | 164704518 | 0 | 1 | 12 | 2 | 7 | 20 | 10 | 19 | 30 | 56 | 0 | 0 | 2 | 0 | 0 | 0 | 14 | 5 | 3 | 0 | 266 | 149 | 1 | 1 | 0 | 0.053 | 0.261 | 0.036 | 0.065 | NA | 0 | 0.183 | 0 | NA | 0.022 |
| SE | 40569 | ENSG00000082438 | COBLL1 | chr2 | - | 164727105 | 164727144 | 164722424 | 164722522 | 164727968 | 164728197 | 2 | 9 | 1 | 5 | 7 | 31 | 17 | 16 | 33 | 73 | 0 | 0 | 0 | 0 | 0 | 0 | 36 | 16 | 9 | 3 | 188 | 149 | 1 | 1 | 0.049 | 0.296 | 0.047 | 0.107 | 0.071 | NA | 0 | 0 | 0 | 0 | 0.114 |
| SE | 40572 | ENSG00000082438 | COBLL1 | chr2 | - | 164779654 | 164779785 | 164743686 | 164743875 | 164841155 | 164841243 | 2 | 0 | 0 | 0 | 0 | 0 | 0 | 1 | 0 | 0 | 0 | 0 | 0 | 0 | 0 | 0 | 10 | 0 | 0 | 0 | 280 | 149 | 1 | 1 | 1 | NA | 0 | NA | NA | NA | 0 | NA | NA | NA | 0.5 |
| SE | 40575 | ENSG00000163875 | MEAF6 | chr1 | - | 37495884 | 37495918 | 37489992 | 37494107 | 37501803 | 37501996 | 396 | 261 | 303 | 347 | 792 | 0 | 0 | 0 | 1 | 0 | 269 | 288 | 204 | 313 | 352 | 0 | 0 | 0 | 0 | 0 | 183 | 149 | 1 | 1 | 1 | 1 | 1 | 0.996 | 1 | 1 | 1 | 1 | 1 | 1 | -0.001 |
| SE | 40579 | ENSG00000163875 | MEAF6 | chr1 | - | 37496706 | 37496736 | 37495884 | 37495918 | 37501803 | 37501996 | 1 | 2 | 5 | 2 | 8 | 287 | 190 | 231 | 258 | 629 | 0 | 4 | 0 | 0 | 0 | 185 | 221 | 137 | 221 | 290 | 179 | 149 | 1 | 1 | 0.003 | 0.009 | 0.018 | 0.006 | 0.01 | 0 | 0.015 | 0 | 0 | 0 | 0.006 |
| SE | 40581 | ENSG00000163875 | MEAF6 | chr1 | - | 37501803 | 37501996 | 37495884 | 37495918 | 37509277 | 37509323 | 572 | 347 | 473 | 535 | 1043 | 0 | 0 | 0 | 0 | 0 | 406 | 428 | 289 | 456 | 605 | 8 | 0 | 0 | 0 | 0 | 298 | 149 | 1 | 1 | 1 | 1 | 1 | 1 | 1 | 0.962 | 1 | 1 | 1 | 1 | 0.008 |
| SE | 40582 | ENSG00000163875 | MEAF6 | chr1 | - | 37509277 | 37509323 | 37501414 | 37501996 | 37509454 | 37509542 | 375 | 211 | 308 | 365 | 570 | 0 | 0 | 0 | 0 | 1 | 326 | 296 | 210 | 343 | 372 | 0 | 0 | 0 | 0 | 0 | 195 | 149 | 1 | 1 | 1 | 1 | 1 | 1 | 0.998 | 1 | 1 | 1 | 1 | 1 | 0 |
| SE | 40583 | ENSG00000163875 | MEAF6 | chr1 | - | 37509454 | 37509542 | 37501414 | 37501996 | 37513422 | 37513520 | 128 | 67 | 100 | 113 | 112 | 0 | 0 | 1 | 0 | 0 | 148 | 81 | 48 | 87 | 107 | 0 | 0 | 0 | 0 | 0 | 237 | 149 | 1 | 1 | 1 | 1 | 0.984 | 1 | 1 | 1 | 1 | 1 | 1 | 1 | -0.003 |
| SE | 40585 | ENSG00000085433 | WDR47 | chr1 | - | 108981732 | 108981864 | 108974535 | 108974754 | 108982608 | 108982779 | 64 | 51 | 40 | 56 | 45 | 1 | 0 | 0 | 3 | 0 | 9 | 40 | 47 | 54 | 14 | 0 | 0 | 3 | 0 | 0 | 281 | 149 | 1 | 1 | 0.971 | 1 | 1 | 0.908 | 1 | 1 | 1 | 0.893 | 1 | 1 | -0.003 |
| SE | 40586 | ENSG00000085433 | WDR47 | chr1 | - | 108981732 | 108981864 | 108974535 | 108974754 | 108983281 | 108983451 | 30 | 31 | 18 | 24 | 25 | 0 | 0 | 0 | 4 | 0 | 4 | 36 | 47 | 28 | 7 | 0 | 0 | 0 | 4 | 0 | 281 | 149 | 1 | 1 | 1 | 1 | 1 | 0.761 | 1 | 1 | 1 | 1 | 0.788 | 1 | -0.005 |
| SE | 40587 | ENSG00000085433 | WDR47 | chr1 | - | 108982608 | 108982779 | 108974535 | 108974754 | 108983281 | 108983451 | 81 | 50 | 38 | 81 | 69 | 0 | 0 | 0 | 4 | 0 | 24 | 70 | 47 | 53 | 10 | 0 | 0 | 0 | 4 | 0 | 298 | 149 | 1 | 1 | 1 | 1 | 1 | 0.91 | 1 | 1 | 1 | 1 | 0.869 | 1 | 0.008 |
| SE | 40589 | ENSG00000085433 | WDR47 | chr1 | - | 108982608 | 108982785 | 108974535 | 108974754 | 108983281 | 108983451 | 11 | 3 | 2 | 6 | 1 | 0 | 0 | 0 | 4 | 0 | 0 | 0 | 3 | 0 | 0 | 0 | 0 | 0 | 4 | 0 | 298 | 149 | 1 | 1 | 1 | 1 | 1 | 0.429 | 1 | NA | NA | 1 | 0 | NA | 0.386 |
| SE | 40592 | ENSG00000085433 | WDR47 | chr1 | - | 108991253 | 108991329 | 108983281 | 108983451 | 108995579 | 108995837 | 28 | 10 | 12 | 28 | 33 | 5 | 5 | 4 | 10 | 4 | 0 | 20 | 8 | 14 | 0 | 12 | 26 | 5 | 18 | 0 | 225 | 149 | 1 | 1 | 0.788 | 0.57 | 0.665 | 0.65 | 0.845 | 0 | 0.337 | 0.514 | 0.34 | NA | 0.406 |
| SE | 40593 | ENSG00000085433 | WDR47 | chr1 | - | 109017517 | 109017601 | 109013819 | 109013925 | 109023354 | 109023521 | 81 | 63 | 80 | 78 | 148 | 0 | 0 | 0 | 0 | 0 | 25 | 71 | 27 | 46 | 21 | 0 | 3 | 0 | 0 | 0 | 233 | 149 | 1 | 1 | 1 | 1 | 1 | 1 | 1 | 1 | 0.938 | 1 | 1 | 1 | 0.012 |
| SE | 40602 | ENSG00000162613 | FUBP1 | chr1 | - | 77945505 | 77945642 | 77944054 | 77944269 | 77949154 | 77949300 | 9 | 1 | 11 | 13 | 25 | 0 | 0 | 0 | 2 | 0 | 0 | 7 | 14 | 11 | 18 | 0 | 2 | 0 | 0 | 0 | 286 | 149 | 1 | 1 | 1 | 1 | 1 | 0.772 | 1 | NA | 0.646 | 1 | 1 | 1 | 0.043 |
| SE | 40603 | ENSG00000162613 | FUBP1 | chr1 | - | 77946529 | 77946576 | 77944054 | 77944269 | 77949154 | 77949267 | 1 | 0 | 4 | 1 | 1 | 0 | 0 | 0 | 2 | 0 | 11 | 1 | 5 | 2 | 0 | 0 | 2 | 0 | 0 | 0 | 196 | 149 | 1 | 1 | 1 | NA | 1 | 0.275 | 1 | 1 | 0.275 | 1 | 1 | NA | 0 |
| SE | 40605 | ENSG00000162613 | FUBP1 | chr1 | - | 77947452 | 77947547 | 77946529 | 77946576 | 77949154 | 77949300 | 5 | 0 | 3 | 1 | 10 | 4 | 5 | 13 | 10 | 7 | 0 | 3 | 0 | 0 | 0 | 21 | 6 | 12 | 6 | 0 | 244 | 149 | 1 | 1 | 0.433 | 0 | 0.124 | 0.058 | 0.466 | 0 | 0.234 | 0 | 0 | NA | 0.158 |
| SE | 40606 | ENSG00000162613 | FUBP1 | chr1 | - | 77947452 | 77947552 | 77946481 | 77946576 | 77949154 | 77949267 | 5 | 0 | 3 | 0 | 0 | 4 | 5 | 13 | 10 | 7 | 0 | 6 | 0 | 3 | 0 | 21 | 6 | 12 | 6 | 0 | 249 | 149 | 1 | 1 | 0.428 | 0 | 0.121 | 0 | 0 | 0 | 0.374 | 0 | 0.23 | NA | -0.041 |
| SE | 40607 | ENSG00000162613 | FUBP1 | chr1 | - | 77949154 | 77949300 | 77944054 | 77948774 | 77955254 | 77955282 | 485 | 356 | 495 | 580 | 231 | 1 | 0 | 5 | 0 | 0 | 559 | 284 | 323 | 435 | 99 | 0 | 1 | 7 | 0 | 0 | 295 | 149 | 1 | 1 | 0.996 | 1 | 0.98 | 1 | 1 | 1 | 0.993 | 0.959 | 1 | 1 | 0.005 |
| SE | 40608 | ENSG00000162613 | FUBP1 | chr1 | - | 77949154 | 77949300 | 77945505 | 77945642 | 77955254 | 77955282 | 252 | 183 | 285 | 330 | 152 | 0 | 0 | 0 | 0 | 0 | 292 | 148 | 186 | 240 | 82 | 0 | 3 | 0 | 0 | 0 | 295 | 149 | 1 | 1 | 1 | 1 | 1 | 1 | 1 | 1 | 0.961 | 1 | 1 | 1 | 0.008 |
| SE | 40609 | ENSG00000162613 | FUBP1 | chr1 | - | 77964645 | 77964747 | 77964253 | 77964356 | 77964869 | 77964968 | 1017 | 1005 | 1023 | 1201 | 1686 | 0 | 0 | 0 | 0 | 1 | 1515 | 1250 | 1062 | 1156 | 318 | 0 | 0 | 0 | 0 | 0 | 251 | 149 | 1 | 1 | 1 | 1 | 1 | 1 | 0.999 | 1 | 1 | 1 | 1 | 1 | 0 |
| SE | 40610 | ENSG00000162613 | FUBP1 | chr1 | - | 77966693 | 77966751 | 77965068 | 77965231 | 77966883 | 77966955 | 2235 | 2405 | 1939 | 2160 | 2442 | 0 | 0 | 0 | 0 | 0 | 3086 | 2059 | 2194 | 2464 | 546 | 0 | 0 | 4 | 0 | 0 | 207 | 149 | 1 | 1 | 1 | 1 | 1 | 1 | 1 | 1 | 1 | 0.997 | 1 | 1 | 0.001 |
| SE | 40611 | ENSG00000162613 | FUBP1 | chr1 | - | 77966883 | 77966955 | 77966693 | 77966751 | 77967048 | 77967098 | 1411 | 1443 | 1294 | 1436 | 1667 | 0 | 0 | 0 | 0 | 0 | 1775 | 1277 | 1320 | 1474 | 365 | 0 | 0 | 0 | 6 | 0 | 221 | 149 | 1 | 1 | 1 | 1 | 1 | 1 | 1 | 1 | 1 | 1 | 0.994 | 1 | 0.001 |
| SE | 40612 | ENSG00000162613 | FUBP1 | chr1 | - | 77967626 | 77967666 | 77967048 | 77967098 | 77968164 | 77968203 | 238 | 231 | 273 | 328 | 295 | 0 | 0 | 0 | 0 | 0 | 308 | 260 | 276 | 273 | 81 | 3 | 0 | 0 | 0 | 0 | 189 | 149 | 1 | 1 | 1 | 1 | 1 | 1 | 1 | 0.988 | 1 | 1 | 1 | 1 | 0.002 |
| SE | 40614 | ENSG00000213390 | ARHGAP19 | chr10 | - | 97229146 | 97229225 | 97222172 | 97226132 | 97229763 | 97229874 | 141 | 83 | 98 | 121 | 270 | 0 | 0 | 0 | 0 | 0 | 366 | 154 | 153 | 264 | 36 | 0 | 0 | 2 | 3 | 0 | 228 | 149 | 1 | 1 | 1 | 1 | 1 | 1 | 1 | 1 | 1 | 0.98 | 0.983 | 1 | 0.007 |
| SE | 40615 | ENSG00000213390 | ARHGAP19 | chr10 | - | 97243967 | 97244159 | 97235216 | 97235315 | 97246271 | 97246337 | 164 | 71 | 123 | 122 | 152 | 0 | 0 | 2 | 2 | 0 | 258 | 142 | 168 | 225 | 43 | 0 | 0 | 0 | 0 | 0 | 298 | 149 | 1 | 1 | 1 | 1 | 0.969 | 0.968 | 1 | 1 | 1 | 1 | 1 | 1 | -0.013 |
| SE | 40616 | ENSG00000213390 | ARHGAP19 | chr10 | - | 97246271 | 97246337 | 97243967 | 97244159 | 97256317 | 97256404 | 93 | 47 | 79 | 64 | 111 | 0 | 0 | 1 | 0 | 2 | 111 | 58 | 75 | 117 | 12 | 0 | 0 | 2 | 0 | 0 | 215 | 149 | 1 | 1 | 1 | 1 | 0.982 | 1 | 0.975 | 1 | 1 | 0.963 | 1 | 1 | -0.001 |
| SE | 40621 | ENSG00000213390 | ARHGAP19 | chr10 | - | 97259401 | 97259628 | 97246271 | 97246337 | 97263419 | 97263629 | 78 | 68 | 63 | 79 | 157 | 3 | 0 | 0 | 0 | 0 | 83 | 88 | 127 | 150 | 43 | 0 | 0 | 0 | 0 | 0 | 298 | 149 | 1 | 1 | 0.929 | 1 | 1 | 1 | 1 | 1 | 1 | 1 | 1 | 1 | -0.014 |
| SE | 40622 | ENSG00000213390 | ARHGAP19 | chr10 | - | 97259401 | 97259628 | 97246271 | 97246337 | 97292571 | 97292623 | 1 | 4 | 2 | 5 | 7 | 0 | 0 | 0 | 0 | 0 | 8 | 7 | 29 | 34 | 0 | 10 | 0 | 0 | 0 | 0 | 298 | 149 | 1 | 1 | 1 | 1 | 1 | 1 | 1 | 0.286 | 1 | 1 | 1 | NA | 0.178 |
| SE | 40623 | ENSG00000213390 | ARHGAP19 | chr10 | - | 97263419 | 97263629 | 97259401 | 97259628 | 97264825 | 97264906 | 135 | 102 | 119 | 162 | 236 | 0 | 0 | 0 | 0 | 0 | 143 | 161 | 177 | 254 | 93 | 0 | 0 | 1 | 0 | 0 | 298 | 149 | 1 | 1 | 1 | 1 | 1 | 1 | 1 | 1 | 1 | 0.989 | 1 | 1 | 0.002 |
| SE | 40627 | ENSG00000143344 | RGL1 | chr1 | + | 183806374 | 183806485 | 183742125 | 183742289 | 183847565 | 183847774 | 4 | 11 | 7 | 8 | 7 | 1 | 1 | 1 | 1 | 0 | 51 | 0 | 9 | 9 | 0 | 0 | 2 | 0 | 0 | 0 | 260 | 149 | 1 | 1 | 0.696 | 0.863 | 0.8 | 0.821 | 1 | 1 | 0 | 1 | 1 | NA | 0.086 |
| SE | 40628 | ENSG00000143344 | RGL1 | chr1 | + | 183806374 | 183806485 | 183805119 | 183805324 | 183847565 | 183847774 | 6 | 14 | 5 | 10 | 1 | 1 | 0 | 0 | 0 | 0 | 113 | 0 | 6 | 19 | 3 | 0 | 0 | 0 | 3 | 0 | 260 | 149 | 1 | 1 | 1 | 1 | 1 | 1 | 1 | 1 | NA | 1 | 0.784 | 1 | 0.054 |
| SE | 40630 | ENSG00000143344 | RGL1 | chr1 | + | 183900149 | 183900236 | 183897807 | 183897897 | 183902567 | 183902600 | 13 | 7 | 6 | 14 | 12 | 0 | 0 | 0 | 0 | 0 | 102 | 0 | 8 | 41 | 18 | 0 | 0 | 1 | 3 | 0 | 236 | 149 | 1 | 1 | 1 | 1 | 1 | 1 | 1 | 1 | NA | 0.835 | 0.896 | 1 | 0.067 |
| SE | 40631 | ENSG00000143344 | RGL1 | chr1 | + | 183902567 | 183902600 | 183897807 | 183897897 | 183904849 | 183904971 | 1 | 0 | 1 | 2 | 1 | 0 | 0 | 0 | 0 | 0 | 0 | 0 | 1 | 3 | 0 | 0 | 0 | 0 | 1 | 0 | 182 | 149 | 1 | 1 | 1 | NA | 1 | 1 | 1 | NA | NA | 1 | 0.711 | NA | 0.145 |
| SE | 40634 | ENSG00000134248 | LAMTOR5 | chr1 | - | 110403918 | 110404036 | 110401248 | 110401583 | 110406317 | 110406379 | 546 | 249 | 428 | 517 | 506 | 2 | 0 | 1 | 0 | 2 | 436 | 456 | 460 | 625 | 528 | 0 | 0 | 0 | 4 | 0 | 267 | 149 | 1 | 1 | 0.993 | 1 | 0.996 | 1 | 0.993 | 1 | 1 | 1 | 0.989 | 1 | -0.001 |
| SE | 40635 | ENSG00000134248 | LAMTOR5 | chr1 | - | 110406317 | 110406379 | 110403608 | 110404036 | 110407032 | 110407128 | 113 | 54 | 95 | 110 | 199 | 0 | 0 | 0 | 1 | 0 | 134 | 130 | 142 | 172 | 193 | 0 | 0 | 0 | 0 | 0 | 211 | 149 | 1 | 1 | 1 | 1 | 1 | 0.987 | 1 | 1 | 1 | 1 | 1 | 1 | -0.003 |
| SE | 40638 | ENSG00000143322 | ABL2 | chr1 | - | 179114877 | 179115030 | 179112308 | 179112398 | 179118586 | 179118764 | 17 | 9 | 20 | 14 | 38 | 0 | 0 | 0 | 0 | 1 | 13 | 20 | 34 | 34 | 11 | 0 | 0 | 0 | 0 | 0 | 298 | 149 | 1 | 1 | 1 | 1 | 1 | 1 | 0.95 | 1 | 1 | 1 | 1 | 1 | -0.01 |
| SE | 40639 | ENSG00000143322 | ABL2 | chr1 | - | 179117331 | 179117516 | 179112308 | 179112398 | 179118586 | 179118764 | 24 | 25 | 46 | 47 | 20 | 0 | 0 | 0 | 0 | 1 | 26 | 20 | 48 | 25 | 0 | 0 | 0 | 0 | 0 | 0 | 298 | 149 | 1 | 1 | 1 | 1 | 1 | 1 | 0.909 | 1 | 1 | 1 | 1 | NA | -0.018 |
| SE | 40641 | ENSG00000143322 | ABL2 | chr1 | - | 179120189 | 179120274 | 179118586 | 179118764 | 179121594 | 179121867 | 43 | 33 | 48 | 61 | 30 | 0 | 0 | 1 | 0 | 0 | 37 | 37 | 50 | 37 | 16 | 0 | 0 | 0 | 0 | 0 | 234 | 149 | 1 | 1 | 1 | 1 | 0.968 | 1 | 1 | 1 | 1 | 1 | 1 | 1 | -0.006 |
| SE | 40642 | ENSG00000143322 | ABL2 | chr1 | - | 179126376 | 179126672 | 179121594 | 179121867 | 179131310 | 179131481 | 44 | 19 | 42 | 30 | 54 | 0 | 0 | 1 | 0 | 0 | 34 | 27 | 20 | 31 | 12 | 0 | 0 | 0 | 0 | 0 | 298 | 149 | 1 | 1 | 1 | 1 | 0.955 | 1 | 1 | 1 | 1 | 1 | 1 | 1 | -0.009 |
| SE | 40643 | ENSG00000143322 | ABL2 | chr1 | - | 179133311 | 179133374 | 179131310 | 179131481 | 179142932 | 179143056 | 0 | 2 | 0 | 1 | 0 | 2 | 0 | 0 | 0 | 0 | 4 | 5 | 0 | 3 | 0 | 0 | 0 | 0 | 14 | 0 | 212 | 149 | 1 | 1 | 0 | 1 | NA | 1 | NA | 1 | 1 | NA | 0.131 | NA | -0.044 |
| SE | 40644 | ENSG00000143322 | ABL2 | chr1 | - | 179133311 | 179133374 | 179131310 | 179131481 | 179229240 | 179229397 | 1 | 3 | 1 | 2 | 0 | 1 | 0 | 0 | 0 | 0 | 4 | 7 | 0 | 4 | 4 | 0 | 0 | 3 | 7 | 0 | 212 | 149 | 1 | 1 | 0.413 | 1 | 1 | 1 | NA | 1 | 1 | 0 | 0.287 | 1 | 0.196 |
| SE | 40645 | ENSG00000162994 | CLHC1 | chr2 | - | 55180509 | 55180712 | 55177601 | 55177781 | 55181569 | 55181744 | 7 | 5 | 6 | 7 | 0 | 0 | 0 | 0 | 1 | 0 | 12 | 6 | 2 | 0 | 0 | 0 | 0 | 0 | 2 | 0 | 298 | 149 | 1 | 1 | 1 | 1 | 1 | 0.778 | NA | 1 | 1 | 1 | 0 | NA | 0.195 |
| SE | 40646 | ENSG00000162994 | CLHC1 | chr2 | - | 55180509 | 55180712 | 55177601 | 55177781 | 55206269 | 55206376 | 3 | 1 | 4 | 4 | 0 | 0 | 0 | 0 | 0 | 0 | 0 | 2 | 3 | 0 | 0 | 5 | 0 | 0 | 0 | 0 | 298 | 149 | 1 | 1 | 1 | 1 | 1 | 1 | NA | 0 | 1 | 1 | NA | NA | 0.333 |
| SE | 40647 | ENSG00000162994 | CLHC1 | chr2 | - | 55181569 | 55181744 | 55177601 | 55177781 | 55206269 | 55206376 | 2 | 3 | 0 | 7 | 0 | 0 | 0 | 0 | 0 | 0 | 12 | 5 | 2 | 4 | 0 | 5 | 0 | 0 | 0 | 0 | 298 | 149 | 1 | 1 | 1 | 1 | NA | 1 | NA | 0.545 | 1 | 1 | 1 | NA | 0.114 |
| SE | 40648 | ENSG00000162994 | CLHC1 | chr2 | - | 55181569 | 55181744 | 55180509 | 55180712 | 55206269 | 55206376 | 6 | 8 | 2 | 9 | 0 | 0 | 1 | 0 | 1 | 0 | 24 | 9 | 4 | 2 | 0 | 0 | 0 | 3 | 0 | 0 | 298 | 149 | 1 | 1 | 1 | 0.8 | 1 | 0.818 | NA | 1 | 1 | 0.4 | 1 | NA | 0.054 |
| SE | 40649 | ENSG00000162994 | CLHC1 | chr2 | - | 55181569 | 55181744 | 55180509 | 55180712 | 55222234 | 55222493 | 4 | 5 | 2 | 3 | 0 | 0 | 0 | 0 | 2 | 0 | 12 | 4 | 2 | 0 | 0 | 0 | 0 | 0 | 0 | 0 | 298 | 149 | 1 | 1 | 1 | 1 | 1 | 0.429 | NA | 1 | 1 | 1 | NA | NA | -0.143 |
| SE | 40651 | ENSG00000162994 | CLHC1 | chr2 | - | 55206817 | 55206940 | 55206269 | 55206376 | 55208625 | 55208710 | 1 | 6 | 0 | 7 | 0 | 1 | 5 | 1 | 4 | 3 | 0 | 0 | 3 | 0 | 0 | 0 | 0 | 0 | 0 | 0 | 272 | 149 | 1 | 1 | 0.354 | 0.397 | 0 | 0.489 | 0 | NA | NA | 1 | NA | NA | -0.752 |
| SE | 40652 | ENSG00000162994 | CLHC1 | chr2 | - | 55206817 | 55206940 | 55206269 | 55206376 | 55209403 | 55209516 | 2 | 1 | 0 | 6 | 0 | 0 | 0 | 1 | 1 | 0 | 0 | 0 | 3 | 0 | 0 | 0 | 5 | 0 | 0 | 0 | 272 | 149 | 1 | 1 | 1 | 1 | 0 | 0.767 | NA | NA | 0 | 1 | NA | NA | 0.192 |
| SE | 40653 | ENSG00000162994 | CLHC1 | chr2 | - | 55206817 | 55206940 | 55206269 | 55206376 | 55209629 | 55209831 | 1 | 1 | 0 | 6 | 5 | 0 | 1 | 0 | 3 | 0 | 0 | 2 | 3 | 0 | 0 | 0 | 0 | 0 | 0 | 0 | 272 | 149 | 1 | 1 | 1 | 0.354 | NA | 0.523 | 1 | NA | 1 | 1 | NA | NA | -0.281 |
| SE | 40654 | ENSG00000162994 | CLHC1 | chr2 | - | 55208625 | 55208710 | 55206269 | 55206376 | 55209403 | 55209516 | 4 | 24 | 2 | 17 | 11 | 0 | 0 | 1 | 1 | 0 | 5 | 14 | 5 | 0 | 0 | 0 | 5 | 0 | 0 | 0 | 234 | 149 | 1 | 1 | 1 | 1 | 0.56 | 0.915 | 1 | 1 | 0.641 | 1 | NA | NA | 0.015 |
| SE | 40655 | ENSG00000162994 | CLHC1 | chr2 | - | 55208625 | 55208710 | 55206269 | 55206376 | 55209629 | 55209831 | 1 | 5 | 2 | 2 | 0 | 0 | 1 | 0 | 3 | 0 | 0 | 5 | 1 | 0 | 0 | 0 | 0 | 0 | 0 | 0 | 234 | 149 | 1 | 1 | 1 | 0.761 | 1 | 0.298 | NA | NA | 1 | 1 | NA | NA | -0.235 |
| SE | 40656 | ENSG00000162994 | CLHC1 | chr2 | - | 55208625 | 55208710 | 55206817 | 55206940 | 55209403 | 55209516 | 4 | 24 | 1 | 14 | 8 | 1 | 0 | 0 | 0 | 0 | 5 | 14 | 5 | 0 | 0 | 0 | 0 | 0 | 0 | 0 | 234 | 149 | 1 | 1 | 0.718 | 1 | 1 | 1 | 1 | 1 | 1 | 1 | NA | NA | -0.056 |
| SE | 40657 | ENSG00000162994 | CLHC1 | chr2 | - | 55208625 | 55208710 | 55206817 | 55206940 | 55209629 | 55209831 | 0 | 4 | 1 | 1 | 0 | 0 | 0 | 0 | 0 | 5 | 0 | 5 | 1 | 0 | 0 | 0 | 2 | 0 | 0 | 0 | 234 | 149 | 1 | 1 | NA | 1 | 1 | 1 | 0 | NA | 0.614 | 1 | NA | NA | -0.057 |
| SE | 40658 | ENSG00000162994 | CLHC1 | chr2 | - | 55209403 | 55209516 | 55206269 | 55206376 | 55209629 | 55209652 | 3 | 4 | 3 | 14 | 8 | 0 | 1 | 0 | 3 | 0 | 0 | 7 | 2 | 6 | 8 | 0 | 0 | 0 | 0 | 0 | 262 | 149 | 1 | 1 | 1 | 0.695 | 1 | 0.726 | 1 | NA | 1 | 1 | 1 | 1 | -0.116 |
| SE | 40659 | ENSG00000162994 | CLHC1 | chr2 | - | 55209403 | 55209516 | 55206817 | 55206940 | 55209629 | 55209652 | 4 | 4 | 2 | 13 | 8 | 0 | 0 | 0 | 0 | 5 | 0 | 2 | 2 | 6 | 8 | 0 | 2 | 0 | 0 | 0 | 262 | 149 | 1 | 1 | 1 | 1 | 1 | 1 | 0.476 | NA | 0.363 | 1 | 1 | 1 | 0.054 |
| SE | 40661 | ENSG00000162994 | CLHC1 | chr2 | - | 55209629 | 55209831 | 55206269 | 55206376 | 55232235 | 55232354 | 0 | 1 | 0 | 3 | 0 | 0 | 0 | 0 | 0 | 0 | 0 | 0 | 1 | 0 | 0 | 0 | 0 | 1 | 0 | 0 | 298 | 149 | 1 | 1 | NA | 1 | NA | 1 | NA | NA | NA | 0.333 | NA | NA | 0.667 |
| SE | 40662 | ENSG00000162994 | CLHC1 | chr2 | - | 55212672 | 55212806 | 55209629 | 55209831 | 55217810 | 55217998 | 2 | 16 | 6 | 31 | 16 | 0 | 0 | 0 | 0 | 0 | 0 | 17 | 3 | 7 | 8 | 0 | 0 | 3 | 0 | 0 | 283 | 149 | 1 | 1 | 1 | 1 | 1 | 1 | 1 | NA | 1 | 0.345 | 1 | 1 | 0.164 |
| SE | 40663 | ENSG00000162994 | CLHC1 | chr2 | - | 55212672 | 55212806 | 55209629 | 55209831 | 55228031 | 55228204 | 2 | 12 | 4 | 21 | 16 | 0 | 0 | 4 | 0 | 0 | 0 | 15 | 3 | 6 | 8 | 0 | 0 | 0 | 0 | 0 | 283 | 149 | 1 | 1 | 1 | 1 | 0.345 | 1 | 1 | NA | 1 | 1 | 1 | 1 | -0.131 |
| SE | 40664 | ENSG00000162994 | CLHC1 | chr2 | - | 55217810 | 55217998 | 55209629 | 55209831 | 55222234 | 55222493 | 0 | 9 | 0 | 3 | 4 | 3 | 0 | 3 | 4 | 4 | 0 | 0 | 9 | 0 | 0 | 0 | 0 | 0 | 0 | 11 | 298 | 149 | 1 | 1 | 0 | 1 | 0 | 0.273 | 0.333 | NA | NA | 1 | NA | 0 | -0.179 |
| SE | 40666 | ENSG00000162994 | CLHC1 | chr2 | - | 55222234 | 55222493 | 55209629 | 55209831 | 55228031 | 55228204 | 3 | 0 | 3 | 10 | 8 | 0 | 0 | 4 | 0 | 0 | 0 | 2 | 1 | 0 | 11 | 0 | 0 | 0 | 0 | 0 | 298 | 149 | 1 | 1 | 1 | NA | 0.273 | 1 | 1 | NA | 1 | 1 | NA | 1 | -0.182 |
| SE | 40667 | ENSG00000162994 | CLHC1 | chr2 | - | 55222234 | 55222493 | 55209629 | 55209831 | 55232235 | 55232274 | 3 | 1 | 7 | 6 | 10 | 0 | 0 | 0 | 0 | 0 | 0 | 13 | 0 | 4 | 15 | 0 | 0 | 1 | 0 | 0 | 298 | 149 | 1 | 1 | 1 | 1 | 1 | 1 | 1 | NA | 1 | 0 | 1 | 1 | 0.25 |
| SE | 40668 | ENSG00000162994 | CLHC1 | chr2 | - | 55222234 | 55222493 | 55218344 | 55218600 | 55232222 | 55232293 | 6 | 5 | 7 | 9 | 28 | 0 | 0 | 0 | 1 | 0 | 0 | 19 | 5 | 5 | 0 | 0 | 0 | 0 | 0 | 0 | 298 | 149 | 1 | 1 | 1 | 1 | 1 | 0.818 | 1 | NA | 1 | 1 | 1 | NA | -0.036 |
| SE | 40670 | ENSG00000162994 | CLHC1 | chr2 | - | 55228031 | 55228204 | 55209629 | 55209831 | 55232235 | 55232354 | 0 | 0 | 5 | 0 | 0 | 0 | 0 | 0 | 0 | 0 | 0 | 3 | 0 | 0 | 0 | 0 | 0 | 1 | 0 | 0 | 298 | 149 | 1 | 1 | NA | NA | 1 | NA | NA | NA | 1 | 0 | NA | NA | 0.5 |
| SE | 40671 | ENSG00000162994 | CLHC1 | chr2 | - | 55228031 | 55228204 | 55222234 | 55222493 | 55232222 | 55232293 | 0 | 2 | 0 | 6 | 6 | 0 | 1 | 3 | 0 | 0 | 0 | 2 | 1 | 0 | 0 | 0 | 0 | 0 | 0 | 0 | 298 | 149 | 1 | 1 | NA | 0.5 | 0 | 1 | 1 | NA | 1 | 1 | NA | NA | -0.375 |
| SE | 40672 | ENSG00000162994 | CLHC1 | chr2 | - | 55228031 | 55228204 | 55222234 | 55222493 | 55232235 | 55232354 | 0 | 0 | 1 | 6 | 4 | 0 | 1 | 4 | 2 | 6 | 0 | 5 | 1 | 0 | 0 | 0 | 13 | 0 | 4 | 4 | 298 | 149 | 1 | 1 | NA | 0 | 0.111 | 0.6 | 0.25 | NA | 0.161 | 1 | 0 | 0 | -0.05 |
| SE | 40677 | ENSG00000112294 | ALDH5A1 | chr6 | + | 24511829 | 24511868 | 24504868 | 24504985 | 24515166 | 24515310 | 2 | 5 | 5 | 4 | 8 | 4 | 0 | 13 | 6 | 3 | 0 | 0 | 0 | 2 | 0 | 18 | 0 | 0 | 0 | 0 | 188 | 149 | 1 | 1 | 0.284 | 1 | 0.234 | 0.346 | 0.679 | 0 | NA | NA | 1 | NA | 0.009 |
| SE | 40680 | ENSG00000112294 | ALDH5A1 | chr6 | + | 24527996 | 24528166 | 24522637 | 24522925 | 24532118 | 24532177 | 38 | 32 | 35 | 39 | 80 | 0 | 0 | 0 | 1 | 0 | 0 | 47 | 26 | 11 | 7 | 0 | 0 | 0 | 0 | 0 | 298 | 149 | 1 | 1 | 1 | 1 | 1 | 0.951 | 1 | NA | 1 | 1 | 1 | 1 | -0.01 |
| SE | 40687 | ENSG00000076641 | PAG1 | chr8 | - | 80993102 | 80993307 | 80991478 | 80991530 | 81029995 | 81030089 | 326 | 293 | 321 | 314 | 257 | 0 | 0 | 0 | 1 | 0 | 52 | 379 | 377 | 275 | 75 | 0 | 0 | 5 | 0 | 0 | 298 | 149 | 1 | 1 | 1 | 1 | 1 | 0.994 | 1 | 1 | 1 | 0.974 | 1 | 1 | 0.004 |
| SE | 40688 | ENSG00000076641 | PAG1 | chr8 | - | 80993102 | 80993307 | 80991478 | 80991530 | 81070111 | 81070170 | 178 | 160 | 180 | 176 | 107 | 0 | 0 | 0 | 1 | 0 | 45 | 232 | 253 | 164 | 43 | 0 | 0 | 0 | 1 | 0 | 298 | 149 | 1 | 1 | 1 | 1 | 1 | 1 | 1 | 1 | 1 | 1 | 0.988 | 1 | 0.002 |
| SE | 40689 | ENSG00000076641 | PAG1 | chr8 | - | 81029995 | 81030089 | 80991478 | 80991530 | 81070111 | 81070170 | 22 | 22 | 31 | 21 | 19 | 0 | 0 | 0 | 0 | 0 | 10 | 24 | 39 | 31 | 26 | 0 | 0 | 0 | 1 | 0 | 243 | 149 | 1 | 1 | 1 | 1 | 1 | 1 | 1 | 1 | 1 | 1 | 0.95 | 1 | 0.01 |
| SE | 40690 | ENSG00000076641 | PAG1 | chr8 | - | 81029995 | 81030089 | 80993102 | 80993307 | 81070111 | 81070170 | 167 | 153 | 168 | 152 | 165 | 1 | 0 | 1 | 0 | 0 | 17 | 172 | 153 | 140 | 49 | 0 | 5 | 0 | 0 | 0 | 243 | 149 | 1 | 1 | 0.99 | 1 | 0.99 | 1 | 1 | 1 | 0.955 | 1 | 1 | 1 | 0.005 |
| SE | 40694 | ENSG00000131188 | PRR7 | chr5 | + | 177453955 | 177454040 | 177446813 | 177446960 | 177454828 | 177455494 | 0 | 0 | 0 | 0 | 0 | 0 | 0 | 0 | 1 | 0 | 0 | 0 | 0 | 0 | 3 | 0 | 0 | 0 | 0 | 0 | 234 | 149 | 1 | 1 | NA | NA | NA | 0 | NA | NA | NA | NA | NA | 1 | -1 |
| SE | 40700 | ENSG00000117448 | AKR1A1 | chr1 | + | 45561788 | 45561878 | 45552435 | 45552563 | 45566568 | 45566688 | 158 | 105 | 98 | 135 | 99 | 0 | 0 | 0 | 3 | 0 | 483 | 154 | 95 | 146 | 230 | 0 | 0 | 0 | 0 | 0 | 239 | 149 | 1 | 1 | 1 | 1 | 1 | 0.966 | 1 | 1 | 1 | 1 | 1 | 1 | -0.007 |
| SE | 40703 | ENSG00000233785 | NA | chrX | - | 23782567 | 23782648 | 23772991 | 23773510 | 23782884 | 23782956 | 0 | 4 | 0 | 0 | 0 | 0 | 0 | 0 | 1 | 0 | 0 | 0 | 0 | 1 | 0 | 0 | 0 | 0 | 0 | 0 | 230 | 149 | 1 | 1 | NA | 1 | NA | 0 | NA | NA | NA | NA | 1 | NA | -0.5 |
| SE | 40704 | ENSG00000004487 | KDM1A | chr1 | + | 23030468 | 23030634 | 23019442 | 23019947 | 23044426 | 23044486 | 84 | 48 | 63 | 83 | 96 | 0 | 0 | 1 | 0 | 0 | 122 | 59 | 35 | 56 | 97 | 0 | 0 | 0 | 0 | 0 | 298 | 149 | 1 | 1 | 1 | 1 | 0.969 | 1 | 1 | 1 | 1 | 1 | 1 | 1 | -0.006 |
| SE | 40706 | ENSG00000004487 | KDM1A | chr1 | + | 23053760 | 23053839 | 23050386 | 23050520 | 23055068 | 23055161 | 164 | 114 | 161 | 177 | 219 | 2 | 0 | 0 | 0 | 2 | 121 | 117 | 134 | 128 | 198 | 0 | 0 | 0 | 5 | 0 | 228 | 149 | 1 | 1 | 0.982 | 1 | 1 | 1 | 0.986 | 1 | 1 | 1 | 0.944 | 1 | 0.005 |
| SE | 40708 | ENSG00000004487 | KDM1A | chr1 | + | 23068538 | 23068681 | 23059072 | 23059167 | 23069060 | 23069151 | 451 | 315 | 425 | 402 | 746 | 0 | 1 | 0 | 0 | 0 | 527 | 449 | 319 | 391 | 453 | 0 | 0 | 0 | 0 | 0 | 292 | 149 | 1 | 1 | 1 | 0.994 | 1 | 1 | 1 | 1 | 1 | 1 | 1 | 1 | -0.001 |
| SE | 40709 | ENSG00000004487 | KDM1A | chr1 | + | 23069060 | 23069151 | 23068538 | 23068681 | 23071224 | 23071359 | 521 | 389 | 453 | 462 | 775 | 0 | 1 | 0 | 0 | 0 | 636 | 525 | 367 | 467 | 585 | 0 | 0 | 0 | 0 | 0 | 240 | 149 | 1 | 1 | 1 | 0.996 | 1 | 1 | 1 | 1 | 1 | 1 | 1 | 1 | -0.001 |
| SE | 40710 | ENSG00000004487 | KDM1A | chr1 | + | 23079552 | 23079667 | 23078965 | 23079177 | 23081445 | 23081573 | 147 | 105 | 122 | 154 | 87 | 0 | 0 | 0 | 0 | 0 | 102 | 113 | 90 | 69 | 125 | 0 | 0 | 1 | 0 | 0 | 264 | 149 | 1 | 1 | 1 | 1 | 1 | 1 | 1 | 1 | 1 | 0.981 | 1 | 1 | 0.004 |
| SE | 40711 | ENSG00000135932 | CAB39 | chr2 | + | 230790871 | 230791036 | 230759958 | 230760115 | 230810262 | 230810304 | 310 | 201 | 218 | 253 | 363 | 0 | 0 | 0 | 0 | 0 | 94 | 174 | 191 | 128 | 78 | 0 | 0 | 0 | 2 | 0 | 298 | 149 | 1 | 1 | 1 | 1 | 1 | 1 | 1 | 1 | 1 | 1 | 0.97 | 1 | 0.006 |
| SE | 40712 | ENSG00000135932 | CAB39 | chr2 | + | 230793212 | 230793331 | 230790871 | 230791036 | 230798728 | 230798897 | 579 | 330 | 449 | 592 | 956 | 0 | 0 | 0 | 0 | 5 | 175 | 401 | 401 | 197 | 104 | 0 | 0 | 0 | 0 | 0 | 268 | 149 | 1 | 1 | 1 | 1 | 1 | 1 | 0.991 | 1 | 1 | 1 | 1 | 1 | -0.002 |
| SE | 40713 | ENSG00000135932 | CAB39 | chr2 | + | 230793212 | 230793331 | 230790871 | 230791036 | 230810262 | 230810304 | 316 | 159 | 242 | 311 | 440 | 0 | 4 | 0 | 0 | 0 | 66 | 209 | 219 | 102 | 37 | 0 | 0 | 0 | 0 | 0 | 268 | 149 | 1 | 1 | 1 | 0.957 | 1 | 1 | 1 | 1 | 1 | 1 | 1 | 1 | -0.009 |
| SE | 40714 | ENSG00000135932 | CAB39 | chr2 | + | 230798728 | 230798897 | 230790871 | 230791036 | 230810262 | 230810304 | 604 | 402 | 436 | 514 | 1072 | 0 | 4 | 0 | 0 | 0 | 107 | 393 | 350 | 228 | 58 | 0 | 0 | 0 | 0 | 0 | 298 | 149 | 1 | 1 | 1 | 0.98 | 1 | 1 | 1 | 1 | 1 | 1 | 1 | 1 | -0.004 |
| SE | 40715 | ENSG00000135932 | CAB39 | chr2 | + | 230798728 | 230798897 | 230793212 | 230793331 | 230810262 | 230810304 | 867 | 574 | 643 | 796 | 1585 | 0 | 0 | 0 | 1 | 0 | 216 | 590 | 535 | 323 | 125 | 0 | 0 | 1 | 0 | 0 | 298 | 149 | 1 | 1 | 1 | 1 | 1 | 0.997 | 1 | 1 | 1 | 0.996 | 1 | 1 | 0 |
| SE | 40717 | ENSG00000135932 | CAB39 | chr2 | + | 230810262 | 230810322 | 230798728 | 230798897 | 230817753 | 230817897 | 276 | 181 | 200 | 226 | 537 | 0 | 0 | 0 | 0 | 0 | 59 | 168 | 163 | 104 | 11 | 0 | 1 | 0 | 0 | 0 | 209 | 149 | 1 | 1 | 1 | 1 | 1 | 1 | 1 | 1 | 0.992 | 1 | 1 | 1 | 0.002 |
| SE | 40718 | ENSG00000135932 | CAB39 | chr2 | + | 230814048 | 230814114 | 230798728 | 230798897 | 230817753 | 230817897 | 238 | 177 | 155 | 170 | 536 | 0 | 0 | 0 | 0 | 0 | 114 | 135 | 215 | 73 | 37 | 0 | 1 | 0 | 0 | 0 | 215 | 149 | 1 | 1 | 1 | 1 | 1 | 1 | 1 | 1 | 0.989 | 1 | 1 | 1 | 0.002 |
| SE | 40719 | ENSG00000135932 | CAB39 | chr2 | + | 230814048 | 230814114 | 230810262 | 230810322 | 230817753 | 230817897 | 999 | 677 | 670 | 742 | 2001 | 2 | 7 | 2 | 5 | 2 | 204 | 573 | 630 | 365 | 146 | 0 | 0 | 2 | 2 | 0 | 215 | 149 | 1 | 1 | 0.997 | 0.985 | 0.996 | 0.99 | 0.999 | 1 | 1 | 0.995 | 0.992 | 1 | -0.004 |
| SE | 40720 | ENSG00000135932 | CAB39 | chr2 | + | 230817753 | 230817897 | 230814048 | 230814114 | 230818515 | 230818704 | 1813 | 1265 | 1274 | 1391 | 3585 | 0 | 1 | 0 | 0 | 0 | 384 | 1122 | 1308 | 726 | 251 | 0 | 4 | 0 | 0 | 0 | 293 | 149 | 1 | 1 | 1 | 0.998 | 1 | 1 | 1 | 1 | 0.993 | 1 | 1 | 1 | 0.001 |
| SE | 40723 | ENSG00000225855 | RUSC1-AS1 | chr1 | - | 155319802 | 155319932 | 155318264 | 155318396 | 155321624 | 155321695 | 0 | 1 | 1 | 0 | 0 | 0 | 0 | 0 | 0 | 0 | 0 | 0 | 0 | 0 | 0 | 0 | 0 | 0 | 0 | 9 | 279 | 149 | 1 | 1 | NA | 1 | 1 | NA | NA | NA | NA | NA | NA | 0 | 1 |
| SE | 40725 | ENSG00000225855 | RUSC1-AS1 | chr1 | - | 155322242 | 155322437 | 155321624 | 155321695 | 155324077 | 155324168 | 5 | 4 | 1 | 1 | 1 | 0 | 0 | 0 | 0 | 2 | 16 | 4 | 1 | 1 | 10 | 0 | 0 | 0 | 0 | 0 | 298 | 149 | 1 | 1 | 1 | 1 | 1 | 1 | 0.2 | 1 | 1 | 1 | 1 | 1 | -0.16 |
| SE | 40733 | ENSG00000184898 | RBM43 | chr2 | - | 151252754 | 151252855 | 151247939 | 151251664 | 151255532 | 151255743 | 109 | 50 | 103 | 79 | 142 | 0 | 0 | 0 | 2 | 0 | 41 | 45 | 44 | 64 | 11 | 0 | 0 | 0 | 0 | 0 | 250 | 149 | 1 | 1 | 1 | 1 | 1 | 0.959 | 1 | 1 | 1 | 1 | 1 | 1 | -0.008 |
| SE | 40735 | ENSG00000073792 | IGF2BP2 | chr3 | - | 185652093 | 185652168 | 185649402 | 185649534 | 185657285 | 185657402 | 33 | 47 | 24 | 53 | 50 | 0 | 0 | 0 | 1 | 0 | 163 | 41 | 34 | 116 | 226 | 0 | 0 | 0 | 0 | 0 | 224 | 149 | 1 | 1 | 1 | 1 | 1 | 0.972 | 1 | 1 | 1 | 1 | 1 | 1 | -0.006 |
| SE | 40737 | ENSG00000073792 | IGF2BP2 | chr3 | - | 185692698 | 185692762 | 185689354 | 185689627 | 185696611 | 185696663 | 31 | 59 | 14 | 56 | 43 | 0 | 0 | 0 | 0 | 0 | 201 | 62 | 29 | 115 | 226 | 0 | 0 | 1 | 0 | 0 | 213 | 149 | 1 | 1 | 1 | 1 | 1 | 1 | 1 | 1 | 1 | 0.953 | 1 | 1 | 0.009 |
| SE | 40738 | ENSG00000073792 | IGF2BP2 | chr3 | - | 185692698 | 185692780 | 185689354 | 185689627 | 185696611 | 185696663 | 8 | 19 | 1 | 14 | 13 | 0 | 0 | 0 | 0 | 0 | 64 | 14 | 14 | 37 | 46 | 0 | 0 | 1 | 0 | 0 | 231 | 149 | 1 | 1 | 1 | 1 | 1 | 1 | 1 | 1 | 1 | 0.9 | 1 | 1 | 0.02 |
| SE | 40739 | ENSG00000073792 | IGF2BP2 | chr3 | - | 185693176 | 185693228 | 185692698 | 185692762 | 185696611 | 185696663 | 0 | 1 | 0 | 1 | 0 | 23 | 40 | 13 | 43 | 32 | 0 | 0 | 0 | 6 | 0 | 137 | 48 | 17 | 83 | 180 | 201 | 149 | 1 | 1 | 0 | 0.018 | 0 | 0.017 | 0 | 0 | 0 | 0 | 0.051 | 0 | -0.003 |
| SE | 40744 | ENSG00000163870 | TPRA1 | chr3 | - | 127575748 | 127575809 | 127575402 | 127575505 | 127575939 | 127576050 | 34 | 9 | 17 | 18 | 12 | 1 | 0 | 2 | 2 | 0 | 0 | 4 | 3 | 1 | 32 | 0 | 0 | 4 | 0 | 2 | 210 | 149 | 1 | 1 | 0.96 | 1 | 0.858 | 0.865 | 1 | NA | 1 | 0.347 | 1 | 0.919 | 0.12 |
| SE | 40745 | ENSG00000163870 | TPRA1 | chr3 | - | 127575939 | 127576050 | 127575748 | 127575809 | 127576616 | 127576696 | 43 | 20 | 29 | 26 | 17 | 2 | 0 | 0 | 0 | 0 | 41 | 8 | 5 | 5 | 31 | 0 | 0 | 0 | 0 | 0 | 260 | 149 | 1 | 1 | 0.925 | 1 | 1 | 1 | 1 | 1 | 1 | 1 | 1 | 1 | -0.015 |
| SE | 40746 | ENSG00000163870 | TPRA1 | chr3 | - | 127576820 | 127576893 | 127576615 | 127576696 | 127576989 | 127577076 | 33 | 19 | 25 | 30 | 13 | 2 | 0 | 0 | 1 | 0 | 48 | 15 | 16 | 5 | 45 | 0 | 1 | 0 | 5 | 0 | 222 | 149 | 1 | 1 | 0.917 | 1 | 1 | 0.953 | 1 | 1 | 0.91 | 1 | 0.402 | 1 | 0.112 |
| SE | 40747 | ENSG00000163870 | TPRA1 | chr3 | - | 127576989 | 127577076 | 127576615 | 127576696 | 127579739 | 127579872 | 6 | 1 | 6 | 3 | 1 | 0 | 0 | 0 | 1 | 0 | 6 | 2 | 1 | 6 | 15 | 0 | 0 | 0 | 0 | 0 | 236 | 149 | 1 | 1 | 1 | 1 | 1 | 0.654 | 1 | 1 | 1 | 1 | 1 | 1 | -0.069 |
| SE | 40748 | ENSG00000163870 | TPRA1 | chr3 | - | 127588069 | 127588160 | 127580021 | 127580163 | 127590409 | 127590699 | 0 | 0 | 0 | 2 | 0 | 21 | 9 | 6 | 10 | 8 | 0 | 0 | 0 | 0 | 0 | 1 | 6 | 5 | 1 | 2 | 240 | 149 | 1 | 1 | 0 | 0 | 0 | 0.11 | 0 | 0 | 0 | 0 | 0 | 0 | 0.022 |
| SE | 40749 | ENSG00000163870 | TPRA1 | chr3 | - | 127590409 | 127590592 | 127580021 | 127580163 | 127590882 | 127591088 | 21 | 9 | 6 | 10 | 8 | 2 | 0 | 0 | 0 | 0 | 1 | 6 | 5 | 1 | 2 | 0 | 0 | 0 | 0 | 0 | 298 | 149 | 1 | 1 | 0.84 | 1 | 1 | 1 | 1 | 1 | 1 | 1 | 1 | 1 | -0.032 |
| SE | 40758 | ENSG00000114026 | OGG1 | chr3 | + | 9750944 | 9751192 | 9749943 | 9750423 | 9751769 | 9751949 | 95 | 46 | 81 | 76 | 64 | 0 | 0 | 0 | 0 | 0 | 76 | 113 | 43 | 40 | 162 | 0 | 0 | 0 | 2 | 0 | 298 | 149 | 1 | 1 | 1 | 1 | 1 | 1 | 1 | 1 | 1 | 1 | 0.909 | 1 | 0.018 |
| SE | 40760 | ENSG00000114026 | OGG1 | chr3 | + | 9754703 | 9754885 | 9751769 | 9751949 | 9756470 | 9756621 | 18 | 9 | 16 | 28 | 10 | 0 | 0 | 0 | 0 | 0 | 18 | 32 | 23 | 21 | 67 | 0 | 0 | 0 | 1 | 0 | 298 | 149 | 1 | 1 | 1 | 1 | 1 | 1 | 1 | 1 | 1 | 1 | 0.913 | 1 | 0.017 |
| SE | 40764 | ENSG00000114026 | OGG1 | chr3 | + | 9754703 | 9754885 | 9751769 | 9751949 | 9787727 | 9788219 | 4 | 3 | 7 | 15 | 4 | 0 | 2 | 0 | 0 | 0 | 6 | 9 | 16 | 12 | 36 | 0 | 0 | 0 | 0 | 0 | 298 | 149 | 1 | 1 | 1 | 0.429 | 1 | 1 | 1 | 1 | 1 | 1 | 1 | 1 | -0.114 |
| SE | 40766 | ENSG00000114026 | OGG1 | chr3 | + | 9756470 | 9756621 | 9751769 | 9751949 | 9787727 | 9788219 | 0 | 0 | 0 | 0 | 0 | 0 | 2 | 0 | 0 | 0 | 0 | 0 | 1 | 1 | 0 | 0 | 0 | 0 | 0 | 0 | 298 | 149 | 1 | 1 | NA | 0 | NA | NA | NA | NA | NA | 1 | 1 | NA | -1 |
| SE | 40768 | ENSG00000114026 | OGG1 | chr3 | + | 9756470 | 9756621 | 9754703 | 9754885 | 9787727 | 9788219 | 14 | 6 | 9 | 14 | 6 | 0 | 0 | 0 | 1 | 0 | 12 | 23 | 8 | 9 | 31 | 0 | 0 | 0 | 0 | 0 | 298 | 149 | 1 | 1 | 1 | 1 | 1 | 0.875 | 1 | 1 | 1 | 1 | 1 | 1 | -0.025 |
| SE | 40770 | ENSG00000114026 | OGG1 | chr3 | + | 9756766 | 9756816 | 9754703 | 9754885 | 9781512 | 9781600 | 0 | 0 | 0 | 5 | 0 | 0 | 0 | 0 | 0 | 0 | 12 | 0 | 0 | 4 | 0 | 0 | 0 | 4 | 0 | 0 | 199 | 149 | 1 | 1 | NA | NA | NA | 1 | NA | 1 | NA | 0 | 1 | NA | 0.333 |
| SE | 40771 | ENSG00000114026 | OGG1 | chr3 | + | 9756766 | 9756816 | 9754703 | 9754885 | 9787727 | 9788219 | 0 | 0 | 0 | 5 | 0 | 0 | 0 | 0 | 1 | 0 | 12 | 0 | 0 | 4 | 0 | 0 | 0 | 0 | 0 | 0 | 199 | 149 | 1 | 1 | NA | NA | NA | 0.789 | NA | 1 | NA | NA | 1 | NA | -0.211 |

| | | | | | | | | | | | | | | | | | | | | | | | | | | | | | | | | | | | | | | | | | | | | | | |
|---|---|---|---|---|---|---|---|---|---|---|---|---|---|---|---|---|---|---|---|---|---|---|---|---|---|---|---|---|---|---|---|---|---|---|---|---|---|---|---|---|---|---|---|---|---|---|
| SE | 40772 | ENSG00000114026 | OGG1 | chr3 | + | 9756766 | 9756816 | 9756470 | 9756621 | 9787727 | 9788219 | 2 | 0 | 0 | 1 | 0 | 0 | 0 | 0 | 0 | 0 | 0 | 0 | 0 | 1 | 0 | 0 | 0 | 1 | 0 | 0 | 199 | 149 | 1 | 1 | 1 | NA | NA | 1 | NA | NA | NA | 0 | 1 | NA | 0.5 |
| SE | 40773 | ENSG00000114026 | OGG1 | chr3 | + | 9759186 | 9759286 | 9756766 | 9756816 | 9765808 | 9766737 | 0 | 0 | 0 | 0 | 4 | 4 | 0 | 0 | 3 | 0 | 0 | 0 | 0 | 0 | 0 | 0 | 0 | 1 | 0 | 8 | 249 | 149 | 1 | 1 | 0 | NA | NA | 0 | 1 | NA | NA | 0 | NA | 0 | 0.333 |
| SE | 40774 | ENSG00000114026 | OGG1 | chr3 | + | 9759233 | 9759286 | 9756766 | 9756816 | 9765808 | 9766540 | 0 | 0 | 1 | 1 | 4 | 4 | 0 | 0 | 3 | 0 | 0 | 6 | 0 | 0 | 0 | 0 | 0 | 1 | 0 | 8 | 202 | 149 | 1 | 1 | 0 | NA | 1 | 0.197 | 1 | NA | 1 | 0 | NA | 0 | 0.216 |
| SE | 40775 | ENSG00000114026 | OGG1 | chr3 | + | 9759233 | 9759286 | 9756766 | 9756816 | 9787727 | 9788219 | 0 | 1 | 1 | 1 | 0 | 0 | 0 | 0 | 0 | 0 | 0 | 3 | 0 | 0 | 0 | 0 | 0 | 0 | 1 | 0 | 202 | 149 | 1 | 1 | NA | 1 | 1 | 1 | NA | NA | 1 | NA | 0 | NA | 0.5 |
| SE | 40776 | ENSG00000114026 | OGG1 | chr3 | + | 9781512 | 9781600 | 9751769 | 9751949 | 9787727 | 9788219 | 0 | 0 | 0 | 0 | 0 | 0 | 2 | 0 | 0 | 0 | 0 | 0 | 7 | 0 | 0 | 0 | 0 | 0 | 0 | 0 | 237 | 149 | 1 | 1 | NA | 0 | NA | NA | NA | NA | NA | 1 | NA | NA | -1 |
| SE | 40777 | ENSG00000114026 | OGG1 | chr3 | + | 9781512 | 9781600 | 9754703 | 9754885 | 9787727 | 9788219 | 0 | 0 | 0 | 0 | 0 | 0 | 0 | 0 | 1 | 0 | 0 | 0 | 9 | 0 | 0 | 0 | 0 | 0 | 0 | 0 | 237 | 149 | 1 | 1 | NA | NA | NA | 0 | NA | NA | NA | 1 | NA | NA | -1 |
| SE | 40779 | ENSG00000103160 | HSDL1 | chr16 | - | 84130336 | 84130431 | 84129985 | 84130171 | 84131101 | 84131327 | 44 | 39 | 44 | 95 | 74 | 0 | 0 | 0 | 2 | 0 | 72 | 111 | 46 | 50 | 59 | 0 | 0 | 0 | 0 | 0 | 244 | 149 | 1 | 1 | 1 | 1 | 1 | 0.967 | 1 | 1 | 1 | 1 | 1 | 1 | -0.007 |
| SE | 40780 | ENSG00000103160 | HSDL1 | chr16 | - | 84135543 | 84135593 | 84131101 | 84131327 | 84145079 | 84145156 | 29 | 23 | 14 | 41 | 19 | 0 | 0 | 0 | 0 | 3 | 23 | 48 | 22 | 19 | 13 | 0 | 0 | 0 | 0 | 0 | 199 | 149 | 1 | 1 | 1 | 1 | 1 | 1 | 0.826 | 1 | 1 | 1 | 1 | 1 | -0.035 |
| SE | 40781 | ENSG00000103160 | HSDL1 | chr16 | - | 84135543 | 84135605 | 84130960 | 84131327 | 84145079 | 84145169 | 44 | 43 | 26 | 61 | 22 | 0 | 0 | 0 | 0 | 3 | 27 | 66 | 32 | 32 | 29 | 0 | 0 | 0 | 0 | 0 | 211 | 149 | 1 | 1 | 1 | 1 | 1 | 1 | 0.838 | 1 | 1 | 1 | 1 | 1 | -0.032 |
| SE | 40782 | ENSG00000103160 | HSDL1 | chr16 | - | 84139222 | 84139306 | 84135543 | 84135605 | 84145079 | 84145175 | 0 | 0 | 0 | 0 | 0 | 9 | 13 | 12 | 16 | 3 | 0 | 2 | 0 | 3 | 0 | 4 | 13 | 9 | 9 | 8 | 233 | 149 | 1 | 1 | 0 | 0 | 0 | 0 | 0 | 0 | 0.09 | 0 | 0.176 | 0 | -0.053 |
| SE | 40783 | ENSG00000154269 | ENPP3 | chr6 | + | 131638446 | 131638576 | 131637303 | 131637462 | 131641454 | 131641530 | 1 | 0 | 0 | 0 | 1 | 8 | 17 | 17 | 7 | 18 | 0 | 0 | 0 | 0 | 0 | 10 | 0 | 0 | 4 | 0 | 279 | 149 | 1 | 1 | 0.063 | 0 | 0 | 0 | 0.029 | 0 | NA | NA | 0 | NA | 0.018 |
| SE | 40784 | ENSG00000154269 | ENPP3 | chr6 | + | 131638450 | 131638576 | 131637234 | 131637462 | 131641454 | 131641530 | 1 | 0 | 0 | 0 | 1 | 8 | 17 | 17 | 7 | 18 | 0 | 0 | 0 | 0 | 0 | 10 | 0 | 0 | 4 | 0 | 275 | 149 | 1 | 1 | 0.063 | 0 | 0 | 0 | 0.029 | 0 | NA | NA | 0 | NA | 0.018 |
| SE | 40787 | ENSG00000154269 | ENPP3 | chr6 | + | 131675079 | 131675189 | 131674161 | 131674281 | 131676735 | 131676797 | 9 | 2 | 11 | 3 | 23 | 5 | 0 | 0 | 0 | 0 | 0 | 0 | 2 | 1 | 0 | 0 | 0 | 0 | 0 | 0 | 259 | 149 | 1 | 1 | 0.509 | 1 | 1 | 1 | 1 | NA | NA | 1 | 1 | NA | -0.098 |
| SE | 40788 | ENSG00000154269 | ENPP3 | chr6 | + | 131733587 | 131733723 | 131726045 | 131726200 | 131737354 | 131737432 | 9 | 11 | 10 | 2 | 41 | 3 | 0 | 0 | 0 | 1 | 0 | 0 | 2 | 0 | 0 | 0 | 0 | 0 | 0 | 0 | 285 | 149 | 1 | 1 | 0.611 | 1 | 1 | 1 | 0.955 | NA | NA | 1 | NA | NA | -0.087 |
| SE | 40795 | ENSG00000244682 | FCGR2C | chr1 | + | 161583720 | 161583814 | 161581338 | 161581549 | 161588421 | 161588442 | 12 | 17 | 3 | 10 | 1 | 102 | 54 | 60 | 79 | 23 | 0 | 1 | 0 | 4 | 10 | 25 | 83 | 0 | 24 | 108 | 243 | 149 | 1 | 1 | 0.067 | 0.162 | 0.03 | 0.072 | 0.026 | 0 | 0.007 | NA | 0.093 | 0.054 | 0.033 |
| SE | 40796 | ENSG00000244682 | FCGR2C | chr1 | + | 161583720 | 161583814 | 161583089 | 161583193 | 161588421 | 161588442 | 24 | 18 | 2 | 17 | 17 | 243 | 54 | 0 | 66 | 92 | 0 | 0 | 2 | 3 | 10 | 75 | 0 | 41 | 0 | 73 | 243 | 149 | 1 | 1 | 0.057 | 0.17 | 1 | 0.136 | 0.102 | 0 | NA | 0.029 | 1 | 0.077 | 0.017 |
| SE | 40798 | ENSG00000244682 | FCGR2C | chr1 | + | 161590080 | 161590218 | 161589561 | 161589819 | 161591143 | 161591291 | 21 | 8 | 3 | 13 | 17 | 6 | 1 | 0 | 2 | 0 | 0 | 6 | 0 | 0 | 0 | 0 | 0 | 0 | 9 | 11 | 287 | 149 | 1 | 1 | 0.645 | 0.806 | 1 | 0.771 | 1 | NA | 1 | NA | 0 | 0 | 0.511 |
| SE | 40800 | ENSG00000244682 | FCGR2C | chr1 | + | 161599629 | 161599792 | 161592149 | 161592242 | 161601239 | 161601364 | 45 | 30 | 8 | 12 | 7 | 0 | 0 | 0 | 0 | 0 | 0 | 1 | 3 | 0 | 8 | 0 | 0 | 0 | 0 | 12 | 298 | 149 | 1 | 1 | 1 | 1 | 1 | 1 | 1 | NA | 1 | 1 | NA | 0.25 | 0.25 |
| SE | 40801 | ENSG00000145335 | SNCA | chr4 | - | 89729193 | 89729277 | 89724098 | 89726660 | 89822245 | 89822388 | 103 | 116 | 39 | 247 | 200 | 0 | 0 | 0 | 1 | 0 | 177 | 194 | 121 | 235 | 1222 | 0 | 1 | 0 | 0 | 0 | 233 | 149 | 1 | 1 | 1 | 1 | 1 | 0.994 | 1 | 1 | 0.992 | 1 | 1 | 1 | 0 |
| SE | 40805 | ENSG00000055332 | EIF2AK2 | chr2 | - | 37107473 | 37107527 | 37099209 | 37107395 | 37109193 | 37109295 | 355 | 449 | 286 | 615 | 564 | 1 | 3 | 0 | 0 | 0 | 441 | 613 | 688 | 1427 | 77 | 0 | 8 | 0 | 9 | 0 | 203 | 149 | 1 | 1 | 0.996 | 0.991 | 1 | 1 | 1 | 1 | 0.983 | 1 | 0.991 | 1 | 0.003 |
| SE | 40808 | ENSG00000055332 | EIF2AK2 | chr2 | - | 37122505 | 37122664 | 37119958 | 37120139 | 37135483 | 37135546 | 518 | 727 | 415 | 760 | 625 | 0 | 0 | 0 | 0 | 0 | 1074 | 1066 | 1295 | 2505 | 73 | 0 | 0 | 0 | 2 | 0 | 298 | 149 | 1 | 1 | 1 | 1 | 1 | 1 | 1 | 1 | 1 | 1 | 0.998 | 1 | 0 |
| SE | 40809 | ENSG00000055332 | EIF2AK2 | chr2 | - | 37126288 | 37126411 | 37119986 | 37120139 | 37135483 | 37135546 | 46 | 79 | 50 | 83 | 81 | 0 | 0 | 0 | 0 | 0 | 49 | 122 | 140 | 200 | 0 | 0 | 0 | 0 | 2 | 0 | 272 | 149 | 1 | 1 | 1 | 1 | 1 | 1 | 1 | 1 | 1 | 1 | 0.982 | NA | 0.004 |
| SE | 40813 | ENSG00000055332 | EIF2AK2 | chr2 | - | 37141552 | 37141701 | 37139630 | 37139757 | 37146852 | 37146961 | 385 | 504 | 353 | 555 | 492 | 0 | 0 | 2 | 0 | 0 | 806 | 927 | 918 | 1921 | 85 | 4 | 2 | 1 | 7 | 0 | 298 | 149 | 1 | 1 | 1 | 1 | 0.989 | 1 | 1 | 0.99 | 0.996 | 0.998 | 0.993 | 1 | 0.002 |
| SE | 40814 | ENSG00000055332 | EIF2AK2 | chr2 | - | 37141552 | 37141701 | 37139630 | 37139757 | 37147687 | 37147806 | 152 | 195 | 124 | 196 | 195 | 0 | 2 | 0 | 0 | 0 | 188 | 319 | 311 | 555 | 16 | 8 | 0 | 0 | 0 | 0 | 298 | 149 | 1 | 1 | 1 | 0.98 | 1 | 1 | 1 | 0.922 | 1 | 1 | 1 | 1 | 0.012 |
| SE | 40815 | ENSG00000055332 | EIF2AK2 | chr2 | - | 37146852 | 37146973 | 37139630 | 37139757 | 37147687 | 37147806 | 191 | 306 | 165 | 294 | 286 | 0 | 2 | 0 | 0 | 0 | 385 | 427 | 491 | 1046 | 47 | 8 | 0 | 0 | 0 | 0 | 270 | 149 | 1 | 1 | 1 | 0.988 | 1 | 1 | 1 | 0.964 | 1 | 1 | 1 | 1 | 0.005 |
| SE | 40817 | ENSG00000055332 | EIF2AK2 | chr2 | - | 37148856 | 37148915 | 37147687 | 37147822 | 37156907 | 37157044 | 39 | 47 | 30 | 59 | 54 | 0 | 0 | 0 | 0 | 0 | 118 | 97 | 91 | 226 | 15 | 0 | 0 | 0 | 1 | 0 | 208 | 149 | 1 | 1 | 1 | 1 | 1 | 1 | 1 | 1 | 1 | 1 | 0.994 | 1 | 0.001 |
| SE | 40818 | ENSG00000055332 | EIF2AK2 | chr2 | - | 37148856 | 37148939 | 37147687 | 37147822 | 37156907 | 37157192 | 52 | 64 | 36 | 76 | 77 | 0 | 0 | 0 | 0 | 0 | 123 | 125 | 128 | 308 | 15 | 0 | 0 | 0 | 1 | 0 | 232 | 149 | 1 | 1 | 1 | 1 | 1 | 1 | 1 | 1 | 1 | 1 | 0.995 | 1 | 0.001 |
| SE | 40819 | ENSG00000055332 | EIF2AK2 | chr2 | - | 37148856 | 37149023 | 37147687 | 37147822 | 37156907 | 37156935 | 67 | 105 | 56 | 108 | 118 | 0 | 0 | 0 | 0 | 0 | 177 | 185 | 188 | 448 | 23 | 0 | 0 | 0 | 1 | 0 | 298 | 149 | 1 | 1 | 1 | 1 | 1 | 1 | 1 | 1 | 1 | 1 | 0.996 | 1 | 0.001 |
| SE | 40820 | ENSG00000055332 | EIF2AK2 | chr2 | - | 37148856 | 37149257 | 37147687 | 37147822 | 37156907 | 37157047 | 66 | 105 | 56 | 108 | 118 | 0 | 0 | 0 | 0 | 0 | 180 | 178 | 188 | 448 | 23 | 0 | 0 | 0 | 1 | 0 | 298 | 149 | 1 | 1 | 1 | 1 | 1 | 1 | 1 | 1 | 1 | 1 | 0.996 | 1 | 0.001 |
| SE | 40825 | ENSG00000078900 | TP73 | chr1 | + | 3731462 | 3731556 | 3730926 | 3731065 | 3732746 | 3733079 | 0 | 0 | 1 | 0 | 0 | 1 | 0 | 0 | 0 | 0 | 0 | 0 | 7 | 0 | 0 | 0 | 0 | 0 | 0 | 0 | 243 | 149 | 1 | 1 | 0 | NA | 1 | NA | NA | NA | NA | 1 | NA | NA | -0.5 |
| SE | 40826 | ENSG00000144591 | GMPPA | chr2 | + | 219499955 | 219500015 | 219498864 | 219498938 | 219500120 | 219500218 | 24 | 16 | 25 | 22 | 7 | 0 | 0 | 1 | 0 | 0 | 19 | 26 | 30 | 34 | 44 | 0 | 0 | 0 | 0 | 0 | 209 | 149 | 1 | 1 | 1 | 1 | 0.947 | 1 | 1 | 1 | 1 | 1 | 1 | 1 | -0.011 |
| SE | 40827 | ENSG00000144591 | GMPPA | chr2 | + | 219499960 | 219500015 | 219498864 | 219498938 | 219500120 | 219500218 | 10 | 9 | 12 | 13 | 8 | 0 | 0 | 1 | 0 | 0 | 14 | 14 | 17 | 11 | 20 | 0 | 0 | 0 | 0 | 0 | 204 | 149 | 1 | 1 | 1 | 1 | 0.898 | 1 | 1 | 1 | 1 | 1 | 1 | 1 | -0.02 |
| SE | 40828 | ENSG00000144591 | GMPPA | chr2 | + | 219500120 | 219500218 | 219498864 | 219498938 | 219501475 | 219501579 | 47 | 19 | 55 | 51 | 62 | 0 | 0 | 1 | 0 | 0 | 51 | 71 | 39 | 42 | 114 | 0 | 0 | 0 | 0 | 4 | 247 | 149 | 1 | 1 | 1 | 1 | 0.971 | 1 | 1 | 1 | 1 | 1 | 1 | 0.945 | 0.005 |
| SE | 40829 | ENSG00000144591 | GMPPA | chr2 | + | 219500177 | 219500218 | 219498890 | 219498938 | 219501475 | 219501579 | 17 | 7 | 23 | 21 | 28 | 0 | 0 | 1 | 0 | 0 | 10 | 30 | 26 | 11 | 38 | 0 | 0 | 0 | 0 | 4 | 190 | 149 | 1 | 1 | 1 | 1 | 0.947 | 1 | 1 | 1 | 1 | 1 | 1 | 0.882 | 0.013 |
| SE | 40830 | ENSG00000144591 | GMPPA | chr2 | + | 219501850 | 219502037 | 219501475 | 219501579 | 219502381 | 219502441 | 110 | 31 | 133 | 106 | 100 | 1 | 0 | 2 | 0 | 0 | 136 | 112 | 131 | 122 | 218 | 1 | 1 | 0 | 0 | 0 | 298 | 149 | 1 | 1 | 0.982 | 1 | 0.971 | 1 | 1 | 0.986 | 0.982 | 1 | 1 | 1 | -0.003 |
| SE | 40831 | ENSG00000144591 | GMPPA | chr2 | + | 219501911 | 219502037 | 219501475 | 219501579 | 219502381 | 219502441 | 65 | 19 | 71 | 66 | 51 | 1 | 0 | 2 | 0 | 0 | 77 | 81 | 68 | 57 | 145 | 1 | 1 | 0 | 0 | 0 | 275 | 149 | 1 | 1 | 0.972 | 1 | 0.951 | 1 | 1 | 0.977 | 0.978 | 1 | 1 | 1 | -0.006 |
| SE | 40834 | ENSG00000144591 | GMPPA | chr2 | + | 219505694 | 219505761 | 219505457 | 219505555 | 219505979 | 219506072 | 10 | 5 | 3 | 3 | 2 | 0 | 0 | 0 | 1 | 0 | 15 | 18 | 5 | 7 | 14 | 0 | 0 | 0 | 0 | 0 | 216 | 149 | 1 | 1 | 1 | 1 | 1 | 0.674 | 1 | 1 | 1 | 1 | 1 | 1 | -0.065 |
| SE | 40835 | ENSG00000144591 | GMPPA | chr2 | + | 219505714 | 219505761 | 219505457 | 219505555 | 219505979 | 219506072 | 21 | 14 | 15 | 20 | 7 | 0 | 0 | 0 | 1 | 0 | 52 | 42 | 17 | 28 | 32 | 0 | 0 | 0 | 0 | 0 | 196 | 149 | 1 | 1 | 1 | 1 | 1 | 0.938 | 1 | 1 | 1 | 1 | 1 | 1 | -0.012 |
| SE | 40836 | ENSG00000126016 | AMOT | chrX | - | 112822254 | 112823188 | 112815357 | 112815877 | 112825071 | 112825220 | 10 | 7 | 13 | 14 | 14 | 0 | 0 | 0 | 0 | 0 | 13 | 2 | 0 | 4 | 0 | 0 | 0 | 3 | 0 | 0 | 298 | 149 | 1 | 1 | 1 | 1 | 1 | 1 | 1 | 1 | 1 | 0 | 1 | NA | 0.25 |
| SE | 40837 | ENSG00000126016 | AMOT | chrX | - | 112822254 | 112823188 | 112815357 | 112815877 | 112832293 | 112832370 | 8 | 5 | 10 | 10 | 13 | 2 | 0 | 6 | 4 | 3 | 8 | 0 | 0 | 1 | 0 | 0 | 0 | 0 | 0 | 0 | 298 | 149 | 1 | 1 | 0.667 | 1 | 0.455 | 0.556 | 0.684 | 1 | NA | NA | 1 | NA | -0.328 |
| SE | 40838 | ENSG00000126016 | AMOT | chrX | - | 112825071 | 112825220 | 112815357 | 112815877 | 112832293 | 112832370 | 1 | 2 | 4 | 0 | 0 | 2 | 0 | 6 | 4 | 3 | 0 | 0 | 6 | 3 | 0 | 0 | 0 | 0 | 0 | 0 | 298 | 149 | 1 | 1 | 0.2 | 1 | 0.25 | 0 | 0 | NA | NA | 1 | 1 | NA | -0.71 |
| SE | 40839 | ENSG00000126016 | AMOT | chrX | - | 112825071 | 112825220 | 112822254 | 112823188 | 112832293 | 112832370 | 4 | 4 | 7 | 4 | 1 | 1 | 0 | 0 | 0 | 0 | 5 | 2 | 3 | 6 | 0 | 0 | 0 | 0 | 0 | 0 | 298 | 149 | 1 | 1 | 0.667 | 1 | 1 | 1 | 1 | 1 | 1 | 1 | 1 | NA | -0.067 |
| SE | 40841 | ENSG00000117020 | AKT3 | chr1 | - | 243504347 | 243505334 | 243499762 | 243499786 | 243512323 | 243512426 | 60 | 60 | 54 | 56 | 112 | 0 | 4 | 0 | 2 | 4 | 0 | 17 | 11 | 11 | 8 | 0 | 0 | 0 | 0 | 0 | 298 | 149 | 1 | 1 | 1 | 0.882 | 1 | 0.933 | 0.933 | NA | 1 | 1 | 1 | 1 | -0.05 |
| SE | 40842 | ENSG00000117020 | AKT3 | chr1 | - | 243504387 | 243505334 | 243499772 | 243499786 | 243512323 | 243512426 | 60 | 60 | 54 | 56 | 112 | 0 | 4 | 0 | 2 | 4 | 0 | 17 | 11 | 11 | 8 | 0 | 0 | 0 | 0 | 0 | 298 | 149 | 1 | 1 | 1 | 0.882 | 1 | 0.933 | 0.933 | NA | 1 | 1 | 1 | 1 | -0.05 |
| SE | 40843 | ENSG00000117020 | AKT3 | chr1 | - | 243545509 | 243545597 | 243512323 | 243512426 | 243563719 | 243563848 | 128 | 128 | 118 | 131 | 251 | 0 | 0 | 0 | 1 | 0 | 24 | 43 | 56 | 42 | 0 | 0 | 0 | 0 | 0 | 0 | 237 | 149 | 1 | 1 | 1 | 1 | 1 | 0.988 | 1 | 1 | 1 | 1 | 1 | NA | -0.002 |
| SE | 40844 | ENSG00000117020 | AKT3 | chr1 | - | 243552728 | 243552943 | 243545509 | 243545597 | 243563719 | 243563848 | 330 | 276 | 263 | 331 | 395 | 1 | 0 | 1 | 0 | 0 | 123 | 150 | 123 | 112 | 33 | 0 | 0 | 0 | 0 | 0 | 298 | 149 | 1 | 1 | 0.994 | 1 | 0.992 | 1 | 1 | 1 | 1 | 1 | 1 | 1 | -0.003 |
| SE | 40846 | ENSG00000117020 | AKT3 | chr1 | - | 243572925 | 243573048 | 243563719 | 243563848 | 243613670 | 243613739 | 355 | 193 | 293 | 310 | 384 | 0 | 0 | 0 | 1 | 0 | 107 | 133 | 95 | 74 | 67 | 0 | 0 | 0 | 0 | 0 | 272 | 149 | 1 | 1 | 1 | 1 | 1 | 0.994 | 1 | 1 | 1 | 1 | 1 | 1 | -0.001 |
| SE | 40847 | ENSG00000117020 | AKT3 | chr1 | - | 243572925 | 243573048 | 243563719 | 243563848 | 243615095 | 243615161 | 192 | 117 | 142 | 172 | 151 | 0 | 0 | 0 | 0 | 4 | 81 | 79 | 70 | 53 | 26 | 0 | 0 | 0 | 0 | 0 | 272 | 149 | 1 | 1 | 1 | 1 | 1 | 1 | 0.954 | 1 | 1 | 1 | 1 | 1 | -0.009 |
| SE | 40848 | ENSG00000117020 | AKT3 | chr1 | - | 243613670 | 243613739 | 243563719 | 243563848 | 243615095 | 243615161 | 660 | 537 | 519 | 559 | 422 | 0 | 0 | 0 | 0 | 4 | 418 | 271 | 315 | 273 | 348 | 0 | 0 | 0 | 0 | 0 | 218 | 149 | 1 | 1 | 1 | 1 | 1 | 1 | 0.986 | 1 | 1 | 1 | 1 | 1 | -0.003 |
| SE | 40850 | ENSG00000117020 | AKT3 | chr1 | - | 243615095 | 243615161 | 243572925 | 243573048 | 243637610 | 243637742 | 306 | 233 | 226 | 271 | 326 | 0 | 0 | 0 | 0 | 0 | 266 | 155 | 121 | 139 | 108 | 0 | 6 | 0 | 0 | 0 | 215 | 149 | 1 | 1 | 1 | 1 | 1 | 1 | 1 | 1 | 0.947 | 1 | 1 | 1 | 0.011 |
| SE | 40851 | ENSG00000117020 | AKT3 | chr1 | - | 243615095 | 243615161 | 243572925 | 243573048 | 243843124 | 243843170 | 4 | 3 | 0 | 0 | 0 | 0 | 0 | 0 | 1 | 0 | 8 | 1 | 6 | 4 | 0 | 0 | 0 | 0 | 0 | 0 | 215 | 149 | 1 | 1 | 1 | 1 | NA | 0 | NA | 1 | 1 | 1 | 1 | NA | -0.333 |
| SE | 40852 | ENSG00000117020 | AKT3 | chr1 | - | 243645892 | 243646037 | 243637610 | 243637742 | 243664771 | 243664812 | 1139 | 958 | 841 | 961 | 1148 | 0 | 0 | 1 | 0 | 1 | 877 | 456 | 544 | 456 | 631 | 0 | 0 | 0 | 0 | 0 | 294 | 149 | 1 | 1 | 1 | 1 | 0.998 | 1 | 0.998 | 1 | 1 | 1 | 1 | 1 | -0.001 |
| SE | 40853 | ENSG00000117020 | AKT3 | chr1 | - | 243664771 | 243664883 | 243645878 | 243646037 | 243695590 | 243695716 | 1071 | 815 | 781 | 956 | 1845 | 1 | 0 | 1 | 9 | 0 | 626 | 409 | 439 | 399 | 604 | 0 | 2 | 1 | 4 | 0 | 261 | 149 | 1 | 1 | 0.998 | 1 | 0.998 | 0.984 | 1 | 1 | 0.992 | 0.996 | 0.983 | 1 | 0.002 |
| SE | 40855 | ENSG00000117020 | AKT3 | chr1 | - | 243695590 | 243695716 | 243664771 | 243664883 | 243739522 | 243739613 | 679 | 476 | 523 | 643 | 1362 | 0 | 0 | 1 | 0 | 0 | 356 | 248 | 274 | 227 | 420 | 0 | 0 | 0 | 0 | 0 | 275 | 149 | 1 | 1 | 1 | 1 | 0.996 | 1 | 1 | 1 | 1 | 1 | 1 | 1 | -0.001 |
| SE | 40856 | ENSG00000117020 | AKT3 | chr1 | - | 243695590 | 243695716 | 243664771 | 243664883 | 243843124 | 243843127 | 1178 | 822 | 949 | 1138 | 2631 | 1 | 0 | 1 | 0 | 0 | 623 | 460 | 453 | 388 | 716 | 0 | 0 | 0 | 0 | 5 | 275 | 149 | 1 | 1 | 0.998 | 1 | 0.998 | 1 | 1 | 1 | 1 | 1 | 1 | 0.987 | 0.002 |
| SE | 40858 | ENSG00000117020 | AKT3 | chr1 | - | 243734957 | 243735116 | 243695590 | 243695716 | 243843124 | 243843282 | 3 | 2 | 2 | 1 | 8 | 505 | 348 | 433 | 498 | 1286 | 0 | 0 | 0 | 0 | 0 | 284 | 218 | 183 | 164 | 301 | 298 | 149 | 1 | 1 | 0.003 | 0.003 | 0.002 | 0.001 | 0.003 | 0 | 0 | 0 | 0 | 0 | 0.002 |
| SE | 40859 | ENSG00000117020 | AKT3 | chr1 | - | 243739522 | 243739613 | 243664771 | 243664883 | 243843124 | 243843282 | 0 | 0 | 1 | 0 | 0 | 1 | 0 | 1 | 0 | 0 | 0 | 0 | 0 | 0 | 0 | 0 | 0 | 0 | 0 | 5 | 240 | 149 | 1 | 1 | 0 | NA | 0.383 | NA | NA | NA | NA | NA | NA | 0 | 0.192 |
| SE | 40862 | ENSG00000117020 | AKT3 | chr1 | - | 243739522 | 243739613 | 243735663 | 243735747 | 243843124 | 243843282 | 1 | 1 | 0 | 0 | 1 | 1 | 0 | 1 | 5 | 1 | 0 | 0 | 0 | 0 | 0 | 0 | 0 | 2 | 0 | 0 | 240 | 149 | 1 | 1 | 0.383 | 1 | 0 | 0 | 0.383 | NA | NA | 0 | NA | NA | 0.353 |
| SE | 40863 | ENSG00000117020 | AKT3 | chr1 | - | 243843124 | 243843282 | 243695590 | 243695716 | 243850039 | 243850041 | 509 | 351 | 437 | 499 | 1286 | 0 | 0 | 0 | 0 | 3 | 284 | 218 | 184 | 164 | 305 | 0 | 0 | 0 | 0 | 0 | 298 | 149 | 1 | 1 | 1 | 1 | 1 | 1 | 0.995 | 1 | 1 | 1 | 1 | 1 | -0.001 |
| SE | 40867 | ENSG00000167196 | FBXO22 | chr15 | + | 75914109 | 75914205 | 75913202 | 75913290 | 75917229 | 75917394 | 85 | 62 | 83 | 63 | 56 | 1 | 0 | 0 | 1 | 0 | 69 | 44 | 35 | 46 | 14 | 0 | 0 | 0 | 0 | 0 | 245 | 149 | 1 | 1 | 0.981 | 1 | 1 | 0.975 | 1 | 1 | 1 | 1 | 1 | 1 | -0.009 |
| SE | 40868 | ENSG00000167196 | FBXO22 | chr15 | + | 75917229 | 75917394 | 75914109 | 75914205 | 75929883 | 75930049 | 136 | 111 | 142 | 100 | 107 | 1 | 0 | 0 | 0 | 0 | 92 | 67 | 70 | 76 | 37 | 0 | 0 | 0 | 0 | 0 | 298 | 149 | 1 | 1 | 0.986 | 1 | 1 | 1 | 1 | 1 | 1 | 1 | 1 | 1 | -0.003 |
| SE | 40869 | ENSG00000167196 | FBXO22 | chr15 | + | 75929883 | 75930049 | 75917229 | 75917394 | 75932684 | 75932798 | 100 | 72 | 111 | 74 | 79 | 0 | 0 | 1 | 0 | 0 | 89 | 56 | 50 | 55 | 53 | 0 | 0 | 0 | 0 | 0 | 298 | 149 | 1 | 1 | 1 | 1 | 0.982 | 1 | 1 | 1 | 1 | 1 | 1 | 1 | -0.004 |
| SE | 40871 | ENSG00000227191 | TRGC2 | chr7 | - | 38242427 | 38242475 | 38239579 | 38240166 | 38245182 | 38245230 | 285 | 165 | 203 | 378 | 687 | 1 | 0 | 0 | 3 | 2 | 115 | 302 | 133 | 176 | 166 | 0 | 0 | 0 | 0 | 0 | 197 | 149 | 1 | 1 | 0.995 | 1 | 1 | 0.99 | 0.996 | 1 | 1 | 1 | 1 | 1 | -0.004 |
| SE | 40873 | ENSG00000173198 | CYSLTR1 | chrX | - | 78283453 | 78283540 | 78271467 | 78273773 | 78327304 | 78327611 | 145 | 89 | 85 | 113 | 202 | 3 | 4 | 1 | 1 | 0 | 19 | 50 | 8 | 15 | 0 | 0 | 1 | 0 | 0 | 0 | 236 | 149 | 1 | 1 | 0.968 | 0.934 | 0.982 | 0.986 | 1 | 1 | 0.969 | 1 | 1 | NA | -0.018 |
| SE | 40875 | ENSG00000116983 | HPCAL4 | chr1 | - | 39691064 | 39691170 | 39684441 | 39684611 | 39691305 | 39691485 | 11 | 1 | 2 | 1 | 0 | 0 | 0 | 1 | 2 | 0 | 0 | 0 | 0 | 4 | 0 | 0 | 1 | 3 | 0 | 0 | 255 | 149 | 1 | 1 | 1 | 1 | 0.539 | 0.226 | NA | NA | 0 | 0 | 1 | NA | 0.358 |
| SE | 40877 | ENSG00000067829 | IDH3G | chrX | - | 153786211 | 153786272 | 153785767 | 153785973 | 153786354 | 153786449 | 138 | 79 | 127 | 127 | 214 | 1 | 0 | 0 | 0 | 0 | 210 | 158 | 92 | 102 | 237 | 0 | 6 | 0 | 1 | 0 | 210 | 149 | 1 | 1 | 0.99 | 1 | 1 | 1 | 1 | 1 | 0.949 | 1 | 0.986 | 1 | 0.011 |
| SE | 40879 | ENSG00000067829 | IDH3G | chrX | - | 153787050 | 153787153 | 153785766 | 153786449 | 153787463 | 153787553 | 100 | 52 | 94 | 106 | 92 | 1 | 0 | 0 | 0 | 0 | 86 | 178 | 90 | 94 | 267 | 0 | 0 | 0 | 1 | 0 | 252 | 149 | 1 | 1 | 0.983 | 1 | 1 | 1 | 1 | 1 | 1 | 1 | 0.982 | 1 | 0 |
| SE | 40880 | ENSG00000067829 | IDH3G | chrX | - | 153787463 | 153787597 | 153787050 | 153787153 | 153787822 | 153787955 | 176 | 97 | 169 | 188 | 152 | 1 | 0 | 0 | 0 | 0 | 244 | 277 | 147 | 202 | 394 | 0 | 0 | 0 | 0 | 0 | 283 | 149 | 1 | 1 | 0.989 | 1 | 1 | 1 | 1 | 1 | 1 | 1 | 1 | 1 | -0.002 |
| SE | 40881 | ENSG00000067829 | IDH3G | chrX | - | 153787463 | 153787597 | 153787050 | 153787153 | 153788074 | 153788135 | 90 | 42 | 83 | 85 | 81 | 0 | 0 | 0 | 0 | 0 | 117 | 113 | 78 | 91 | 214 | 0 | 0 | 0 | 0 | 2 | 283 | 149 | 1 | 1 | 1 | 1 | 1 | 1 | 1 | 1 | 1 | 1 | 1 | 0.983 | 0.003 |
| SE | 40882 | ENSG00000067829 | IDH3G | chrX | - | 153787822 | 153787955 | 153787050 | 153787153 | 153788074 | 153788135 | 71 | 33 | 61 | 55 | 40 | 0 | 0 | 0 | 0 | 0 | 75 | 115 | 46 | 84 | 209 | 0 | 0 | 0 | 0 | 2 | 282 | 149 | 1 | 1 | 1 | 1 | 1 | 1 | 1 | 1 | 1 | 1 | 1 | 0.982 | 0.004 |
| SE | 40883 | ENSG00000067829 | IDH3G | chrX | - | 153787822 | 153787955 | 153787390 | 153787597 | 153788074 | 153788135 | 157 | 88 | 149 | 158 | 111 | 0 | 0 | 0 | 0 | 0 | 202 | 284 | 116 | 198 | 389 | 0 | 0 | 0 | 3 | 0 | 282 | 149 | 1 | 1 | 1 | 1 | 1 | 1 | 1 | 1 | 1 | 1 | 0.972 | 1 | 0.006 |
| SE | 40886 | ENSG00000067829 | IDH3G | chrX | - | 153790563 | 153790575 | 153789837 | 153790292 | 153790809 | 153790851 | 44 | 17 | 46 | 56 | 49 | 1 | 0 | 0 | 0 | 0 | 47 | 69 | 40 | 104 | 156 | 0 | 0 | 0 | 0 | 0 | 161 | 149 | 1 | 1 | 0.976 | 1 | 1 | 1 | 1 | 1 | 1 | 1 | 1 | 1 | -0.005 |
| SE | 40887 | ENSG00000067829 | IDH3G | chrX | - | 153790809 | 153790851 | 153790563 | 153790575 | 153794245 | 153794325 | 51 | 11 | 50 | 53 | 41 | 0 | 1 | 0 | 1 | 1 | 75 | 75 | 38 | 82 | 141 | 0 | 0 | 0 | 4 | 0 | 191 | 149 | 1 | 1 | 1 | 0.896 | 1 | 0.976 | 0.97 | 1 | 1 | 1 | 0.941 | 1 | -0.02 |
| SE | 40888 | ENSG00000067829 | IDH3G | chrX | - | 153790809 | 153790971 | 153790563 | 153790575 | 153794245 | 153794356 | 16 | 8 | 10 | 14 | 19 | 0 | 1 | 0 | 1 | 1 | 29 | 16 | 13 | 53 | 58 | 0 | 0 | 0 | 4 | 0 | 298 | 149 | 1 | 1 | 1 | 0.8 | 1 | 0.875 | 0.905 | 1 | 1 | 1 | 0.869 | 1 | -0.058 |
| SE | 40890 | ENSG00000197050 | ZNF420 | chr19 | + | 37080344 | 37080388 | 37078417 | 37078570 | 37089038 | 37089127 | 12 | 14 | 7 | 24 | 11 | 1 | 0 | 0 | 0 | 0 | 0 | 0 | 4 | 2 | 0 | 0 | 0 | 0 | 0 | 0 | 193 | 149 | 1 | 1 | 0.903 | 1 | 1 | 1 | 1 | NA | NA | 1 | 1 | NA | -0.019 |
| SE | 40891 | ENSG00000197050 | ZNF420 | chr19 | + | 37080344 | 37080388 | 37078417 | 37078570 | 37127127 | 37127139 | 2 | 11 | 0 | 5 | 4 | 0 | 0 | 0 | 4 | 0 | 0 | 0 | 0 | 2 | 0 | 0 | 1 | 0 | 0 | 0 | 193 | 149 | 1 | 1 | 1 | 1 | NA | 0.491 | 1 | NA | 0 | NA | 1 | NA | 0.373 |
| SE | 40893 | ENSG00000197050 | ZNF420 | chr19 | + | 37089038 | 37089127 | 37080344 | 37080388 | 37127127 | 37127139 | 5 | 1 | 4 | 15 | 4 | 0 | 4 | 0 | 1 | 3 | 0 | 0 | 2 | 0 | 0 | 0 | 0 | 0 | 0 | 0 | 238 | 149 | 1 | 1 | 1 | 0.135 | 1 | 0.904 | 0.455 | NA | NA | 1 | NA | NA | -0.301 |
| SE | 40894 | ENSG00000197050 | ZNF420 | chr19 | + | 37090994 | 37091121 | 37089038 | 37089127 | 37127127 | 37127255 | 55 | 65 | 42 | 50 | 74 | 0 | 0 | 1 | 0 | 0 | 0 | 0 | 2 | 15 | 0 | 0 | 0 | 0 | 0 | 0 | 276 | 149 | 1 | 1 | 1 | 1 | 0.958 | 1 | 1 | NA | NA | 1 | NA | NA | -0.008 |
| SE | 40895 | ENSG00000197050 | ZNF420 | chr19 | + | 37096030 | 37096115 | 37090994 | 37091121 | 37111599 | 37111687 | 0 | 1 | 1 | 0 | 2 | 2 | 5 | 7 | 1 | 1 | 0 | 0 | 0 | 0 | 0 | 0 | 11 | 0 | 0 | 0 | 234 | 149 | 1 | 1 | 0 | 0.113 | 0.083 | 0 | 0.56 | NA | 0 | NA | NA | NA | 0.151 |
| SE | 40896 | ENSG00000197050 | ZNF420 | chr19 | + | 37096030 | 37096115 | 37090994 | 37091121 | 37127127 | 37130368 | 0 | 2 | 1 | 0 | 2 | 43 | 40 | 31 | 33 | 64 | 0 | 0 | 0 | 0 | 0 | 0 | 0 | 8 | 0 | 0 | 234 | 149 | 1 | 1 | 0 | 0.031 | 0.02 | 0 | 0.02 | NA | NA | 0 | NA | NA | 0.014 |
| SE | 40897 | ENSG00000197050 | ZNF420 | chr19 | + | 37111599 | 37111711 | 37090994 | 37091121 | 37127127 | 37127139 | 7 | 6 | 8 | 1 | 2 | 43 | 40 | 31 | 33 | 64 | 0 | 20 | 0 | 3 | 0 | 0 | 0 | 8 | 0 | 0 | 261 | 149 | 1 | 1 | 0.085 | 0.079 | 0.128 | 0.017 | 0.018 | NA | 1 | 0 | 1 | NA | -0.601 |
| SE | 40898 | ENSG00000155287 | SLC25A28 | chr10 | - | 99612542 | 99612599 | 99610521 | 99611366 | 99613695 | 99613924 | 63 | 28 | 74 | 77 | 168 | 1 | 0 | 0 | 0 | 0 | 129 | 58 | 55 | 73 | 110 | 0 | 0 | 0 | 0 | 0 | 206 | 149 | 1 | 1 | 0.979 | 1 | 1 | 1 | 1 | 1 | 1 | 1 | 1 | 1 | -0.004 |
| SE | 40899 | ENSG00000155287 | SLC25A28 | chr10 | - | 99612542 | 99612599 | 99610521 | 99611366 | 99620044 | 99620147 | 30 | 10 | 21 | 22 | 38 | 0 | 0 | 1 | 0 | 0 | 54 | 31 | 19 | 51 | 69 | 0 | 0 | 0 | 0 | 0 | 206 | 149 | 1 | 1 | 1 | 1 | 0.938 | 1 | 1 | 1 | 1 | 1 | 1 | 1 | -0.012 |
| SE | 40900 | ENSG00000155287 | SLC25A28 | chr10 | - | 99613695 | 99613924 | 99610521 | 99611366 | 99620044 | 99620439 | 18 | 5 | 14 | 9 | 7 | 0 | 0 | 1 | 0 | 0 | 23 | 20 | 7 | 21 | 26 | 0 | 0 | 0 | 0 | 0 | 298 | 149 | 1 | 1 | 1 | 1 | 0.875 | 1 | 1 | 1 | 1 | 1 | 1 | 1 | -0.025 |
| SE | 40902 | ENSG00000151883 | PARP8 | chr5 | + | 50667084 | 50667186 | 50665898 | 50666069 | 50668070 | 50668125 | 990 | 641 | 677 | 745 | 682 | 0 | 0 | 0 | 1 | 0 | 106 | 478 | 585 | 281 | 70 | 0 | 0 | 0 | 0 | 0 | 251 | 149 | 1 | 1 | 1 | 1 | 1 | 0.998 | 1 | 1 | 1 | 1 | 1 | 1 | 0 |
| SE | 40905 | ENSG00000151883 | PARP8 | chr5 | + | 50668070 | 50668125 | 50666636 | 50667186 | 50750150 | 50750188 | 1414 | 913 | 933 | 1034 | 1111 | 4 | 5 | 8 | 8 | 3 | 163 | 804 | 835 | 402 | 130 | 0 | 7 | 10 | 0 | 0 | 204 | 149 | 1 | 1 | 0.996 | 0.993 | 0.988 | 0.99 | 0.996 | 1 | 0.988 | 0.984 | 1 | 1 | -0.002 |
| SE | 40907 | ENSG00000151883 | PARP8 | chr5 | + | 50738979 | 50739086 | 50668070 | 50668125 | 50744735 | 50744828 | 0 | 0 | 0 | 0 | 3 | 48 | 10 | 40 | 49 | 34 | 0 | 0 | 0 | 0 | 0 | 10 | 19 | 12 | 36 | 7 | 256 | 149 | 1 | 1 | 0 | 0 | 0 | 0 | 0.049 | 0 | 0 | 0 | 0 | 0 | 0.01 |
| SE | 40909 | ENSG00000151883 | PARP8 | chr5 | + | 50741830 | 50741951 | 50668070 | 50668125 | 50750150 | 50750188 | 2 | 0 | 1 | 3 | 2 | 1179 | 793 | 797 | 892 | 870 | 0 | 12 | 0 | 4 | 0 | 154 | 684 | 717 | 344 | 124 | 270 | 149 | 1 | 1 | 0.001 | 0 | 0.001 | 0.002 | 0.001 | 0 | 0.01 | 0 | 0.006 | 0 | -0.002 |
| SE | 40910 | ENSG00000151883 | PARP8 | chr5 | + | 50741830 | 50741951 | 50738979 | 50739086 | 50744735 | 50744828 | 0 | 2 | 4 | 7 | 2 | 0 | 0 | 0 | 0 | 3 | 0 | 16 | 2 | 0 | 0 | 0 | 0 | 0 | 0 | 0 | 270 | 149 | 1 | 1 | NA | 1 | 1 | 1 | 0.269 | NA | 1 | 1 | NA | NA | -0.183 |
| SE | 40913 | ENSG00000151883 | PARP8 | chr5 | + | 50759642 | 50759732 | 50750150 | 50750188 | 50760291 | 50760362 | 3307 | 2199 | 2183 | 2511 | 2305 | 1 | 0 | 0 | 3 | 0 | 554 | 1973 | 2075 | 855 | 371 | 0 | 0 | 0 | 1 | 0 | 239 | 149 | 1 | 1 | 1 | 1 | 1 | 0.998 | 1 | 1 | 1 | 1 | 0.998 | 1 | 0 |
| SE | 40918 | ENSG00000151883 | PARP8 | chr5 | + | 50763147 | 50763242 | 50761820 | 50761898 | 50788522 | 50788589 | 397 | 281 | 341 | 202 | 432 | 1 | 0 | 0 | 1 | 0 | 62 | 177 | 198 | 97 | 24 | 0 | 0 | 0 | 0 | 0 | 244 | 149 | 1 | 1 | 0.996 | 1 | 1 | 0.992 | 1 | 1 | 1 | 1 | 1 | 1 | -0.002 |
| SE | 40921 | ENSG00000151883 | PARP8 | chr5 | + | 50797137 | 50797233 | 50796981 | 50797032 | 50815431 | 50815524 | 585 | 348 | 423 | 455 | 521 | 0 | 0 | 0 | 0 | 4 | 42 | 329 | 327 | 100 | 54 | 0 | 0 | 0 | 0 | 0 | 245 | 149 | 1 | 1 | 1 | 1 | 1 | 1 | 0.988 | 1 | 1 | 1 | 1 | 1 | -0.002 |
| SE | 40923 | ENSG00000151883 | PARP8 | chr5 | + | 50822334 | 50822400 | 50815431 | 50815524 | 50826754 | 50826803 | 11 | 2 | 7 | 11 | 17 | 1 | 0 | 0 | 0 | 0 | 0 | 13 | 22 | 3 | 0 | 0 | 0 | 0 | 0 | 0 | 215 | 149 | 1 | 1 | 0.884 | 1 | 1 | 1 | 1 | NA | 1 | 1 | 1 | NA | -0.023 |
| SE | 40924 | ENSG00000151883 | PARP8 | chr5 | + | 50824907 | 50824975 | 50822334 | 50822400 | 50826754 | 50826803 | 903 | 630 | 679 | 748 | 1274 | 3 | 2 | 4 | 2 | 15 | 61 | 455 | 509 | 171 | 79 | 0 | 4 | 5 | 0 | 0 | 217 | 149 | 1 | 1 | 0.995 | 0.995 | 0.991 | 0.996 | 0.983 | 1 | 0.987 | 0.986 | 1 | 1 | -0.003 |
| SE | 40925 | ENSG00000151883 | PARP8 | chr5 | + | 50828311 | 50828384 | 50827943 | 50828056 | 50829891 | 50829961 | 757 | 417 | 566 | 608 | 882 | 0 | 0 | 1 | 3 | 0 | 107 | 381 | 368 | 118 | 88 | 0 | 0 | 0 | 0 | 0 | 222 | 149 | 1 | 1 | 1 | 1 | 0.997 | 0.993 | 1 | 1 | 1 | 1 | 1 | 1 | -0.002 |
| SE | 40926 | ENSG00000151883 | PARP8 | chr5 | + | 50832780 | 50832854 | 50829891 | 50829961 | 50833978 | 50834048 | 875 | 498 | 629 | 761 | 960 | 1 | 1 | 0 | 0 | 0 | 153 | 434 | 506 | 173 | 114 | 0 | 0 | 0 | 0 | 0 | 223 | 149 | 1 | 1 | 0.998 | 0.997 | 1 | 1 | 1 | 1 | 1 | 1 | 1 | 1 | -0.001 |
| SE | 40927 | ENSG00000151883 | PARP8 | chr5 | + | 50833978 | 50834048 | 50829891 | 50829961 | 50841965 | 50842068 | 1 | 1 | 0 | 1 | 0 | 0 | 0 | 0 | 0 | 2 | 0 | 0 | 1 | 1 | 0 | 0 | 0 | 0 | 0 | 0 | 219 | 149 | 1 | 1 | 1 | 1 | NA | 1 | 0 | NA | NA | 1 | 1 | NA | -0.25 |
| SE | 40928 | ENSG00000151883 | PARP8 | chr5 | + | 50834841 | 50835015 | 50833978 | 50834048 | 50841965 | 50842342 | 366 | 229 | 347 | 357 | 337 | 0 | 0 | 0 | 1 | 0 | 25 | 200 | 252 | 76 | 26 | 0 | 0 | 2 | 4 | 0 | 298 | 149 | 1 | 1 | 1 | 1 | 1 | 0.994 | 1 | 1 | 1 | 0.984 | 0.905 | 1 | 0.021 |
| SE | 40929 | ENSG00000151883 | PARP8 | chr5 | + | 50834930 | 50835015 | 50833978 | 50834048 | 50841965 | 50842068 | 780 | 486 | 665 | 730 | 684 | 0 | 0 | 0 | 1 | 0 | 110 | 402 | 484 | 157 | 82 | 0 | 0 | 2 | 4 | 0 | 234 | 149 | 1 | 1 | 1 | 1 | 1 | 0.998 | 1 | 1 | 1 | 0.994 | 0.962 | 1 | 0.008 |
| SE | 40930 | ENSG00000204219 | TCEA3 | chr1 | - | 23419076 | 23419139 | 23417903 | 23418009 | 23424564 | 23424618 | 11 | 3 | 15 | 13 | 3 | 0 | 0 | 0 | 2 | 1 | 0 | 0 | 1 | 4 | 10 | 0 | 1 | 0 | 0 | 0 | 212 | 149 | 1 | 1 | 1 | 1 | 1 | 0.82 | 0.678 | NA | 0 | 1 | 1 | 1 | 0.15 |
| SE | 40931 | ENSG00000113966 | ARL6 | chr3 | + | 97765990 | 97766089 | 97764520 | 97764977 | 97768080 | 97768230 | 0 | 0 | 0 | 0 | 4 | 1 | 1 | 3 | 3 | 3 | 0 | 0 | 0 | 0 | 0 | 0 | 4 | 1 | 0 | 0 | 248 | 149 | 1 | 1 | 0 | 0 | 0 | 0 | 0.445 | NA | 0 | 0 | NA | NA | 0.089 |
| SE | 40933 | ENSG00000113966 | ARL6 | chr3 | + | 97791770 | 97791826 | 97787989 | 97788119 | 97798023 | 97798203 | 7 | 5 | 2 | 5 | 16 | 0 | 0 | 1 | 0 | 0 | 0 | 0 | 1 | 0 | 0 | 0 | 0 | 0 | 0 | 0 | 205 | 149 | 1 | 1 | 1 | 1 | 0.592 | 1 | 1 | NA | NA | 1 | NA | NA | -0.082 |
| SE | 40937 | ENSG00000168710 | AHCYL1 | chr1 | + | 110019547 | 110019626 | 110018551 | 110018650 | 110020730 | 110020851 | 86 | 23 | 46 | 86 | 124 | 0 | 0 | 0 | 0 | 0 | 112 | 68 | 20 | 66 | 120 | 0 | 3 | 0 | 0 | 0 | 228 | 149 | 1 | 1 | 1 | 1 | 1 | 1 | 1 | 1 | 0.937 | 1 | 1 | 1 | 0.013 |
| SE | 40938 | ENSG00000168710 | AHCYL1 | chr1 | + | 110019547 | 110019626 | 110018551 | 110018650 | 110021673 | 110022045 | 4 | 3 | 2 | 4 | 0 | 0 | 0 | 0 | 0 | 0 | 5 | 8 | 6 | 14 | 0 | 0 | 0 | 0 | 1 | 0 | 228 | 149 | 1 | 1 | 1 | 1 | 1 | 1 | NA | 1 | 1 | 1 | 0.901 | NA | 0.025 |
| SE | 40939 | ENSG00000168710 | AHCYL1 | chr1 | + | 110020730 | 110020851 | 110018551 | 110018650 | 110021673 | 110022045 | 124 | 73 | 69 | 85 | 245 | 0 | 0 | 0 | 0 | 0 | 206 | 95 | 49 | 60 | 173 | 0 | 0 | 0 | 1 | 0 | 270 | 149 | 1 | 1 | 1 | 1 | 1 | 1 | 1 | 1 | 1 | 1 | 0.971 | 1 | 0.006 |
| SE | 40941 | ENSG00000204287 | HLA-DRA | chr6 | + | 32442447 | 32442693 | 32439877 | 32440032 | 32443184 | 32443466 | 7995 | 6071 | 7104 | 7905 | 16161 | 0 | 0 | 3 | 0 | 0 | 26184 | 7181 | 3758 | 7768 | 20205 | 0 | 0 | 0 | 0 | 0 | 298 | 149 | 1 | 1 | 1 | 1 | 0.999 | 1 | 1 | 1 | 1 | 1 | 1 | 1 | 0 |
| SE | 40943 | ENSG00000164776 | PHKG1 | chr7 | - | 56086969 | 56087024 | 56083649 | 56083715 | 56087597 | 56087776 | 1 | 1 | 4 | 2 | 0 | 1 | 0 | 0 | 1 | 0 | 0 | 1 | 6 | 1 | 0 | 0 | 0 | 0 | 0 | 0 | 204 | 149 | 1 | 1 | 0.422 | 1 | 1 | 0.594 | NA | NA | 1 | 1 | 1 | NA | -0.246 |
| SE | 40948 | ENSG00000123473 | STIL | chr1 | - | 47260288 | 47260539 | 47250138 | 47251922 | 47262902 | 47263062 | 7 | 4 | 6 | 4 | 8 | 0 | 0 | 4 | 0 | 0 | 0 | 2 | 10 | 17 | 0 | 0 | 4 | 0 | 0 | 0 | 298 | 149 | 1 | 1 | 1 | 1 | 0.429 | 1 | 1 | NA | 0.2 | 1 | 1 | NA | 0.152 |
| SE | 40949 | ENSG00000123473 | STIL | chr1 | - | 47280240 | 47280363 | 47272075 | 47272241 | 47280945 | 47281209 | 3 | 2 | 3 | 4 | 2 | 1 | 0 | 0 | 0 | 0 | 7 | 9 | 7 | 18 | 0 | 0 | 0 | 1 | 0 | 0 | 272 | 149 | 1 | 1 | 0.622 | 1 | 1 | 1 | 1 | 1 | 1 | 0.793 | 1 | NA | -0.024 |
| SE | 40950 | ENSG00000123473 | STIL | chr1 | - | 47293457 | 47293544 | 47289434 | 47289585 | 47295764 | 47295848 | 11 | 1 | 3 | 13 | 9 | 0 | 0 | 0 | 1 | 0 | 0 | 15 | 4 | 8 | 10 | 0 | 0 | 0 | 0 | 0 | 236 | 149 | 1 | 1 | 1 | 1 | 1 | 0.891 | 1 | NA | 1 | 1 | 1 | 1 | -0.022 |
| SE | 40951 | ENSG00000123473 | STIL | chr1 | - | 47300109 | 47300152 | 47299904 | 47299968 | 47301560 | 47301748 | 0 | 1 | 0 | 1 | 4 | 0 | 0 | 0 | 1 | 0 | 9 | 0 | 7 | 6 | 0 | 0 | 2 | 0 | 0 | 0 | 192 | 149 | 1 | 1 | NA | 1 | NA | 1 | 1 | 1 | 0 | 1 | 1 | NA | 0.25 |
| SE | 40952 | ENSG00000123473 | STIL | chr1 | - | 47312947 | 47313021 | 47310275 | 47310362 | 47314035 | 47314147 | 0 | 1 | 0 | 0 | 0 | 0 | 0 | 0 | 3 | 0 | 0 | 0 | 0 | 0 | 0 | 0 | 1 | 0 | 7 | 0 | 223 | 149 | 1 | 1 | NA | 1 | NA | 0 | NA | NA | 0 | NA | 0 | NA | 0.5 |
| SE | 40953 | ENSG00000123473 | STIL | chr1 | - | 47312947 | 47313156 | 47310275 | 47310362 | 47314035 | 47314147 | 1 | 0 | 1 | 0 | 0 | 0 | 0 | 0 | 3 | 0 | 0 | 0 | 0 | 0 | 0 | 0 | 1 | 0 | 7 | 0 | 298 | 149 | 1 | 1 | 1 | NA | 1 | 0 | NA | NA | 0 | NA | 0 | NA | 0.667 |
| SE | 40959 | ENSG00000263001 | GTF2I | chr7 | + | 74689123 | 74689227 | 74657717 | 74658068 | 74690972 | 74691111 | 145 | 81 | 92 | 145 | 78 | 0 | 0 | 0 | 1 | 0 | 55 | 111 | 46 | 99 | 110 | 0 | 0 | 0 | 0 | 0 | 253 | 149 | 1 | 1 | 1 | 1 | 1 | 0.988 | 1 | 1 | 1 | 1 | 1 | 1 | -0.002 |
| SE | 40960 | ENSG00000263001 | GTF2I | chr7 | + | 74690972 | 74691111 | 74689123 | 74689227 | 74698960 | 74699041 | 333 | 200 | 269 | 323 | 369 | 1 | 0 | 0 | 1 | 0 | 173 | 261 | 136 | 212 | 133 | 0 | 0 | 0 | 0 | 0 | 288 | 149 | 1 | 1 | 0.994 | 1 | 1 | 0.994 | 1 | 1 | 1 | 1 | 1 | 1 | -0.002 |
| SE | 40962 | ENSG00000263001 | GTF2I | chr7 | + | 74700605 | 74700634 | 74700246 | 74700430 | 74705163 | 74705218 | 360 | 240 | 270 | 311 | 389 | 0 | 0 | 0 | 2 | 0 | 314 | 346 | 230 | 289 | 175 | 4 | 0 | 1 | 0 | 0 | 178 | 149 | 1 | 1 | 1 | 1 | 1 | 0.992 | 1 | 0.985 | 1 | 0.995 | 1 | 1 | 0.002 |
| SE | 40963 | ENSG00000263001 | GTF2I | chr7 | + | 74706389 | 74706433 | 74705163 | 74705218 | 74711031 | 74711109 | 574 | 460 | 517 | 535 | 759 | 1 | 0 | 0 | 0 | 0 | 667 | 686 | 411 | 546 | 414 | 0 | 0 | 0 | 0 | 0 | 193 | 149 | 1 | 1 | 0.998 | 1 | 1 | 1 | 1 | 1 | 1 | 1 | 1 | 1 | 0 |
| SE | 40964 | ENSG00000263001 | GTF2I | chr7 | + | 74711031 | 74711109 | 74706389 | 74706433 | 74716893 | 74716950 | 571 | 504 | 527 | 540 | 819 | 1 | 0 | 0 | 0 | 0 | 796 | 705 | 397 | 554 | 411 | 0 | 0 | 1 | 0 | 0 | 227 | 149 | 1 | 1 | 0.997 | 1 | 1 | 1 | 1 | 1 | 1 | 0.996 | 1 | 1 | 0 |
| SE | 40967 | ENSG00000263001 | GTF2I | chr7 | + | 74730228 | 74730294 | 74728785 | 74728896 | 74732478 | 74732662 | 294 | 169 | 296 | 270 | 472 | 0 | 1 | 0 | 0 | 0 | 189 | 249 | 158 | 186 | 134 | 0 | 0 | 0 | 0 | 0 | 215 | 149 | 1 | 1 | 1 | 0.992 | 1 | 1 | 1 | 1 | 1 | 1 | 1 | 1 | -0.002 |
| SE | 40968 | ENSG00000263001 | GTF2I | chr7 | + | 74733922 | 74733981 | 74732478 | 74732662 | 74735460 | 74735513 | 600 | 403 | 545 | 494 | 765 | 0 | 0 | 1 | 0 | 0 | 408 | 579 | 292 | 427 | 307 | 0 | 0 | 0 | 0 | 0 | 208 | 149 | 1 | 1 | 1 | 1 | 0.997 | 1 | 1 | 1 | 1 | 1 | 1 | 1 | -0.001 |
| SE | 40969 | ENSG00000263001 | GTF2I | chr7 | + | 74735460 | 74735532 | 74733595 | 74733981 | 74736499 | 74736683 | 346 | 220 | 282 | 283 | 403 | 0 | 0 | 0 | 0 | 0 | 255 | 341 | 159 | 227 | 164 | 0 | 0 | 3 | 0 | 0 | 221 | 149 | 1 | 1 | 1 | 1 | 1 | 1 | 1 | 1 | 1 | 0.973 | 1 | 1 | 0.005 |
| SE | 40970 | ENSG00000263001 | GTF2I | chr7 | + | 74736499 | 74736683 | 74735460 | 74735532 | 74738044 | 74738103 | 225 | 160 | 204 | 197 | 280 | 0 | 0 | 0 | 1 | 0 | 180 | 247 | 134 | 150 | 130 | 0 | 3 | 0 | 0 | 0 | 298 | 149 | 1 | 1 | 1 | 1 | 1 | 0.99 | 1 | 1 | 0.976 | 1 | 1 | 1 | 0.003 |
| SE | 40971 | ENSG00000263001 | GTF2I | chr7 | + | 74744138 | 74744250 | 74743448 | 74743520 | 74744757 | 74744941 | 22 | 22 | 32 | 22 | 44 | 3 | 3 | 0 | 5 | 6 | 12 | 74 | 91 | 35 | 8 | 0 | 0 | 2 | 2 | 0 | 261 | 149 | 1 | 1 | 0.807 | 0.807 | 1 | 0.715 | 0.807 | 1 | 1 | 0.963 | 0.909 | 1 | -0.147 |
| SE | 40972 | ENSG00000263001 | GTF2I | chr7 | + | 74752089 | 74752170 | 74751360 | 74751416 | 74753093 | 74753177 | 80 | 79 | 96 | 86 | 138 | 0 | 0 | 0 | 0 | 0 | 101 | 97 | 59 | 83 | 75 | 0 | 0 | 1 | 0 | 0 | 230 | 149 | 1 | 1 | 1 | 1 | 1 | 1 | 1 | 1 | 1 | 0.975 | 1 | 1 | 0.005 |
| SE | 40973 | ENSG00000263001 | GTF2I | chr7 | + | 74755508 | 74755537 | 74753845 | 74754029 | 74756819 | 74756861 | 175 | 140 | 148 | 181 | 248 | 0 | 0 | 0 | 1 | 0 | 176 | 189 | 84 | 144 | 98 | 0 | 0 | 0 | 0 | 0 | 178 | 149 | 1 | 1 | 1 | 1 | 1 | 0.993 | 1 | 1 | 1 | 1 | 1 | 1 | -0.001 |
| SE | 40974 | ENSG00000263001 | GTF2I | chr7 | + | 74757958 | 74758000 | 74755239 | 74756861 | 74758779 | 74758855 | 102 | 70 | 71 | 120 | 194 | 0 | 0 | 0 | 0 | 1 | 75 | 86 | 38 | 50 | 44 | 0 | 0 | 0 | 0 | 0 | 191 | 149 | 1 | 1 | 1 | 1 | 1 | 1 | 0.993 | 1 | 1 | 1 | 1 | 1 | -0.001 |
| SE | 40978 | ENSG00000011485 | PPP5C | chr19 | + | 46387092 | 46387235 | 46384376 | 46384909 | 46387365 | 46387453 | 271 | 158 | 239 | 273 | 318 | 0 | 0 | 0 | 1 | 0 | 350 | 309 | 132 | 194 | 334 | 0 | 0 | 0 | 0 | 0 | 292 | 149 | 1 | 1 | 1 | 1 | 1 | 0.993 | 1 | 1 | 1 | 1 | 1 | 1 | -0.001 |
| SE | 40979 | ENSG00000186205 | MTARC1 | chr1 | + | 220798151 | 220798275 | 220797873 | 220798014 | 220798831 | 220798905 | 2 | 0 | 0 | 1 | 1 | 0 | 0 | 0 | 0 | 0 | 0 | 0 | 1 | 0 | 0 | 0 | 2 | 0 | 0 | 0 | 273 | 149 | 1 | 1 | 1 | NA | NA | 1 | 1 | NA | 0 | 1 | NA | NA | 0.5 |
| SE | 40980 | ENSG00000186205 | MTARC1 | chr1 | + | 220798151 | 220798275 | 220797873 | 220798014 | 220805051 | 220805113 | 2 | 1 | 1 | 0 | 0 | 38 | 21 | 32 | 26 | 16 | 0 | 0 | 2 | 1 | 3 | 87 | 152 | 105 | 148 | 180 | 273 | 149 | 1 | 1 | 0.028 | 0.025 | 0.017 | 0 | 0 | 0 | 0 | 0.01 | 0.004 | 0.009 | 0.009 |
| SE | 40981 | ENSG00000186205 | MTARC1 | chr1 | + | 220798151 | 220798275 | 220797873 | 220798065 | 220805051 | 220805113 | 1 | 1 | 1 | 6 | 0 | 0 | 3 | 0 | 1 | 0 | 32 | 12 | 7 | 1 | 6 | 0 | 14 | 5 | 6 | 1 | 273 | 149 | 1 | 1 | 1 | 0.154 | 1 | 0.766 | NA | 1 | 0.319 | 0.433 | 0.083 | 0.766 | 0.21 |
| SE | 40982 | ENSG00000155393 | HEATR3 | chr16 | + | 50066366 | 50066539 | 50065969 | 50066269 | 50068779 | 50068867 | 10 | 5 | 1 | 9 | 0 | 2 | 4 | 0 | 2 | 0 | 0 | 6 | 9 | 8 | 0 | 0 | 0 | 0 | 4 | 0 | 298 | 149 | 1 | 1 | 0.714 | 0.385 | 1 | 0.692 | NA | NA | 1 | 1 | 0.5 | NA | -0.136 |
| SE | 40983 | ENSG00000155393 | HEATR3 | chr16 | + | 50069244 | 50069289 | 50068779 | 50068867 | 50070177 | 50070238 | 0 | 0 | 0 | 2 | 1 | 39 | 17 | 28 | 36 | 30 | 0 | 0 | 0 | 0 | 0 | 1 | 15 | 16 | 44 | 19 | 194 | 149 | 1 | 1 | 0 | 0 | 0 | 0.041 | 0.025 | 0 | 0 | 0 | 0 | 0 | 0.013 |
| SE | 40984 | ENSG00000155393 | HEATR3 | chr16 | + | 50070177 | 50070290 | 50068779 | 50068867 | 50075570 | 50075711 | 37 | 14 | 24 | 43 | 38 | 0 | 1 | 0 | 0 | 0 | 8 | 23 | 25 | 51 | 18 | 0 | 0 | 1 | 0 | 0 | 262 | 149 | 1 | 1 | 1 | 0.888 | 1 | 1 | 1 | 1 | 1 | 0.934 | 1 | 1 | -0.009 |
| SE | 40986 | ENSG00000155393 | HEATR3 | chr16 | + | 50084107 | 50084291 | 50083936 | 50084027 | 50084568 | 50084620 | 71 | 31 | 82 | 71 | 174 | 0 | 0 | 0 | 1 | 0 | 23 | 54 | 39 | 68 | 5 | 0 | 0 | 0 | 0 | 0 | 298 | 149 | 1 | 1 | 1 | 1 | 1 | 0.973 | 1 | 1 | 1 | 1 | 1 | 1 | -0.005 |
| SE | 40987 | ENSG00000155393 | HEATR3 | chr16 | + | 50084133 | 50084291 | 50083936 | 50084027 | 50084568 | 50084651 | 127 | 57 | 122 | 130 | 356 | 0 | 0 | 0 | 1 | 0 | 27 | 64 | 70 | 121 | 34 | 0 | 0 | 0 | 0 | 0 | 298 | 149 | 1 | 1 | 1 | 1 | 1 | 0.985 | 1 | 1 | 1 | 1 | 1 | 1 | -0.003 |
| SE | 40989 | ENSG00000155393 | HEATR3 | chr16 | + | 50100229 | 50100373 | 50094704 | 50094793 | 50102258 | 50102435 | 148 | 96 | 112 | 142 | 187 | 0 | 1 | 0 | 0 | 0 | 98 | 91 | 137 | 123 | 139 | 0 | 0 | 0 | 0 | 0 | 293 | 149 | 1 | 1 | 1 | 0.98 | 1 | 1 | 1 | 1 | 1 | 1 | 1 | 1 | -0.004 |
| SE | 40990 | ENSG00000156136 | DCK | chr4 | + | 71022366 | 71022560 | 70998066 | 70998182 | 71023558 | 71023706 | 757 | 493 | 647 | 811 | 1405 | 0 | 0 | 0 | 0 | 0 | 377 | 535 | 395 | 494 | 100 | 0 | 0 | 2 | 0 | 0 | 298 | 149 | 1 | 1 | 1 | 1 | 1 | 1 | 1 | 1 | 1 | 0.99 | 1 | 1 | 0.002 |
| SE | 40991 | ENSG00000156136 | DCK | chr4 | + | 71022366 | 71022560 | 70998066 | 70998182 | 71025815 | 71025931 | 225 | 152 | 211 | 282 | 542 | 0 | 0 | 0 | 0 | 0 | 90 | 217 | 105 | 199 | 38 | 0 | 0 | 4 | 0 | 0 | 298 | 149 | 1 | 1 | 1 | 1 | 1 | 1 | 1 | 1 | 1 | 0.929 | 1 | 1 | 0.014 |
| SE | 40992 | ENSG00000156136 | DCK | chr4 | + | 71023558 | 71023706 | 70998066 | 70998182 | 71025815 | 71025931 | 663 | 479 | 614 | 773 | 1409 | 0 | 0 | 0 | 0 | 0 | 289 | 501 | 332 | 408 | 120 | 0 | 0 | 4 | 0 | 0 | 297 | 149 | 1 | 1 | 1 | 1 | 1 | 1 | 1 | 1 | 1 | 0.977 | 1 | 1 | 0.005 |
| SE | 40993 | ENSG00000156136 | DCK | chr4 | + | 71023558 | 71023706 | 71022366 | 71022560 | 71025815 | 71025931 | 1197 | 820 | 1052 | 1302 | 2273 | 2 | 0 | 2 | 0 | 1 | 576 | 820 | 620 | 707 | 182 | 0 | 1 | 0 | 4 | 0 | 297 | 149 | 1 | 1 | 0.997 | 1 | 0.996 | 1 | 0.999 | 1 | 0.998 | 1 | 0.989 | 1 | 0.001 |
| SE | 40994 | ENSG00000156136 | DCK | chr4 | + | 71025815 | 71025931 | 71023558 | 71023706 | 71026664 | 71026755 | 850 | 579 | 773 | 918 | 1767 | 4 | 0 | 0 | 0 | 0 | 343 | 593 | 373 | 464 | 130 | 0 | 2 | 0 | 0 | 0 | 265 | 149 | 1 | 1 | 0.992 | 1 | 1 | 1 | 1 | 1 | 1 | 0.991 | 1 | 1 | 0 |
| SE | 40995 | ENSG00000156136 | DCK | chr4 | + | 71028622 | 71028719 | 71026664 | 71026755 | 71029351 | 71030914 | 0 | 3 | 1 | 7 | 4 | 203 | 114 | 200 | 201 | 467 | 0 | 0 | 0 | 0 | 0 | 45 | 106 | 53 | 69 | 59 | 246 | 149 | 1 | 1 | 0 | 0.016 | 0.003 | 0.021 | 0.005 | 0 | 0 | 0 | 0 | 0 | 0.009 |
| SE | 40996 | ENSG00000075240 | GRAMD4 | chr22 | + | 46663837 | 46663863 | 46663039 | 46663172 | 46664025 | 46664117 | 29 | 12 | 20 | 27 | 39 | 0 | 0 | 0 | 2 | 0 | 69 | 18 | 25 | 32 | 70 | 0 | 0 | 0 | 0 | 0 | 175 | 149 | 1 | 1 | 1 | 1 | 1 | 0.92 | 1 | 1 | 1 | 1 | 1 | 1 | -0.016 |
| SE | 40998 | ENSG00000075240 | GRAMD4 | chr22 | + | 46676599 | 46676668 | 46675467 | 46675552 | 46677146 | 46679683 | 31 | 33 | 36 | 35 | 22 | 2 | 1 | 1 | 0 | 0 | 86 | 63 | 16 | 54 | 56 | 16 | 0 | 3 | 0 | 0 | 218 | 149 | 1 | 1 | 0.914 | 0.958 | 0.961 | 1 | 1 | 0.786 | 1 | 0.785 | 1 | 1 | 0.052 |
| SE | 41008 | ENSG00000164091 | WDR82 | chr3 | - | 52258535 | 52258678 | 52254433 | 52257519 | 52259196 | 52259266 | 1087 | 670 | 1065 | 1105 | 1484 | 0 | 0 | 0 | 3 | 1 | 768 | 752 | 730 | 657 | 825 | 0 | 0 | 0 | 0 | 0 | 292 | 149 | 1 | 1 | 1 | 1 | 1 | 0.995 | 0.999 | 1 | 1 | 1 | 1 | 1 | -0.001 |
| SE | 41014 | ENSG00000164091 | WDR82 | chr3 | - | 52270711 | 52270809 | 52266951 | 52267018 | 52278200 | 52278649 | 542 | 240 | 534 | 511 | 345 | 4 | 2 | 3 | 3 | 0 | 344 | 348 | 288 | 275 | 384 | 0 | 2 | 0 | 0 | 5 | 247 | 149 | 1 | 1 | 0.988 | 0.986 | 0.991 | 0.99 | 1 | 1 | 0.991 | 1 | 1 | 0.979 | -0.003 |
| SE | 41019 | ENSG00000137166 | FOXP4 | chr6 | + | 41577985 | 41578081 | 41565744 | 41565964 | 41584768 | 41584818 | 5 | 0 | 0 | 9 | 3 | 0 | 0 | 1 | 0 | 0 | 0 | 0 | 5 | 12 | 24 | 0 | 0 | 1 | 4 | 0 | 245 | 149 | 1 | 1 | 1 | NA | 0 | 1 | 1 | NA | NA | 0.753 | 0.646 | 1 | -0.05 |
| SE | 41020 | ENSG00000137166 | FOXP4 | chr6 | + | 41591220 | 41591322 | 41590270 | 41590347 | 41594869 | 41594991 | 45 | 16 | 11 | 24 | 17 | 0 | 0 | 0 | 2 | 0 | 30 | 44 | 23 | 15 | 108 | 0 | 0 | 0 | 0 | 0 | 251 | 149 | 1 | 1 | 1 | 1 | 1 | 0.877 | 1 | 1 | 1 | 1 | 1 | 1 | -0.025 |
| SE | 41023 | ENSG00000064687 | ABCA7 | chr19 | + | 1044576 | 1044744 | 1043724 | 1043841 | 1045001 | 1045231 | 95 | 72 | 91 | 106 | 32 | 1 | 0 | 4 | 0 | 0 | 56 | 101 | 36 | 42 | 136 | 0 | 0 | 0 | 0 | 0 | 298 | 149 | 1 | 1 | 0.979 | 1 | 0.919 | 1 | 1 | 1 | 1 | 1 | 1 | 1 | -0.02 |
| SE | 41029 | ENSG00000064687 | ABCA7 | chr19 | + | 1056906 | 1057084 | 1056329 | 1056499 | 1057914 | 1058059 | 31 | 8 | 38 | 40 | 21 | 0 | 0 | 0 | 0 | 0 | 19 | 17 | 26 | 25 | 34 | 0 | 0 | 0 | 2 | 0 | 298 | 149 | 1 | 1 | 1 | 1 | 1 | 1 | 1 | 1 | 1 | 1 | 0.862 | 1 | 0.028 |
| SE | 41031 | ENSG00000064687 | ABCA7 | chr19 | + | 1056906 | 1057429 | 1056329 | 1056499 | 1057914 | 1058059 | 64 | 19 | 67 | 70 | 28 | 0 | 0 | 0 | 0 | 0 | 37 | 23 | 50 | 57 | 58 | 0 | 0 | 0 | 2 | 0 | 298 | 149 | 1 | 1 | 1 | 1 | 1 | 1 | 1 | 1 | 1 | 1 | 0.934 | 1 | 0.013 |
| SE | 41033 | ENSG00000064687 | ABCA7 | chr19 | + | 1057313 | 1057429 | 1056329 | 1056499 | 1057914 | 1058059 | 31 | 11 | 27 | 29 | 5 | 0 | 0 | 0 | 0 | 0 | 18 | 15 | 25 | 40 | 24 | 0 | 0 | 0 | 2 | 0 | 265 | 149 | 1 | 1 | 1 | 1 | 1 | 1 | 1 | 1 | 1 | 1 | 0.918 | 1 | 0.016 |
| SE | 41034 | ENSG00000064687 | ABCA7 | chr19 | + | 1057313 | 1057429 | 1056329 | 1056499 | 1058145 | 1058269 | 2 | 1 | 3 | 5 | 4 | 1 | 1 | 0 | 0 | 0 | 0 | 15 | 6 | 8 | 0 | 0 | 0 | 0 | 0 | 0 | 265 | 149 | 1 | 1 | 0.529 | 0.36 | 1 | 1 | 1 | NA | 1 | 1 | 1 | NA | -0.222 |
| SE | 41035 | ENSG00000064687 | ABCA7 | chr19 | + | 1057313 | 1057429 | 1056329 | 1056499 | 1058617 | 1058732 | 0 | 1 | 2 | 2 | 0 | 0 | 0 | 1 | 0 | 0 | 0 | 8 | 4 | 8 | 0 | 0 | 5 | 1 | 0 | 4 | 265 | 149 | 1 | 1 | NA | 1 | 0.529 | 1 | NA | NA | 0.474 | 0.692 | 1 | 0 | 0.301 |
| SE | 41036 | ENSG00000064687 | ABCA7 | chr19 | + | 1057313 | 1057429 | 1056906 | 1057084 | 1057914 | 1058059 | 73 | 17 | 70 | 70 | 19 | 0 | 1 | 0 | 0 | 0 | 50 | 26 | 45 | 68 | 134 | 0 | 0 | 0 | 0 | 0 | 265 | 149 | 1 | 1 | 1 | 0.905 | 1 | 1 | 1 | 1 | 1 | 1 | 1 | 1 | -0.019 |
| SE | 41037 | ENSG00000064687 | ABCA7 | chr19 | + | 1057313 | 1057429 | 1056906 | 1057084 | 1058617 | 1058732 | 24 | 3 | 26 | 30 | 10 | 0 | 0 | 0 | 2 | 0 | 6 | 12 | 17 | 25 | 61 | 0 | 0 | 0 | 0 | 0 | 265 | 149 | 1 | 1 | 1 | 1 | 1 | 0.894 | 1 | 1 | 1 | 1 | 1 | 1 | -0.021 |
| SE | 41038 | ENSG00000064687 | ABCA7 | chr19 | + | 1057914 | 1058059 | 1056329 | 1056499 | 1058145 | 1058269 | 54 | 31 | 65 | 46 | 31 | 1 | 1 | 0 | 0 | 0 | 65 | 53 | 46 | 52 | 68 | 0 | 0 | 0 | 0 | 0 | 294 | 149 | 1 | 1 | 0.965 | 0.94 | 1 | 1 | 1 | 1 | 1 | 1 | 1 | 1 | -0.019 |
| SE | 41042 | ENSG00000064687 | ABCA7 | chr19 | + | 1058145 | 1058269 | 1057313 | 1057429 | 1058617 | 1058732 | 27 | 10 | 33 | 24 | 27 | 0 | 0 | 2 | 0 | 0 | 20 | 55 | 18 | 28 | 27 | 0 | 0 | 0 | 0 | 0 | 273 | 149 | 1 | 1 | 1 | 1 | 0.9 | 1 | 1 | 1 | 1 | 1 | 1 | 1 | -0.02 |
| SE | 41043 | ENSG00000064687 | ABCA7 | chr19 | + | 1058617 | 1058747 | 1058145 | 1058269 | 1058819 | 1058940 | 71 | 35 | 83 | 89 | 36 | 0 | 0 | 1 | 0 | 0 | 91 | 131 | 58 | 67 | 197 | 0 | 0 | 0 | 0 | 0 | 279 | 149 | 1 | 1 | 1 | 1 | 0.978 | 1 | 1 | 1 | 1 | 1 | 1 | 1 | -0.004 |
| SE | 41044 | ENSG00000064687 | ABCA7 | chr19 | + | 1062171 | 1062313 | 1061781 | 1061888 | 1063543 | 1063678 | 78 | 30 | 54 | 85 | 18 | 0 | 0 | 0 | 1 | 0 | 6 | 34 | 23 | 38 | 132 | 0 | 0 | 0 | 0 | 0 | 291 | 149 | 1 | 1 | 1 | 1 | 1 | 0.978 | 1 | 1 | 1 | 1 | 1 | 1 | -0.004 |
| SE | 41046 | ENSG00000112144 | CILK1 | chr6 | - | 53016082 | 53016250 | 53013640 | 53013982 | 53018329 | 53018501 | 97 | 65 | 94 | 114 | 225 | 0 | 0 | 0 | 2 | 0 | 25 | 61 | 63 | 74 | 9 | 0 | 0 | 0 | 0 | 0 | 298 | 149 | 1 | 1 | 1 | 1 | 1 | 0.966 | 1 | 1 | 1 | 1 | 1 | 1 | -0.007 |
| SE | 41047 | ENSG00000112144 | CILK1 | chr6 | - | 53018329 | 53018501 | 53016082 | 53016250 | 53019226 | 53019359 | 76 | 55 | 80 | 102 | 151 | 0 | 0 | 0 | 0 | 4 | 26 | 45 | 31 | 53 | 36 | 0 | 0 | 0 | 0 | 0 | 298 | 149 | 1 | 1 | 1 | 1 | 1 | 1 | 0.95 | 1 | 1 | 1 | 1 | 1 | -0.01 |
| SE | 41048 | ENSG00000112144 | CILK1 | chr6 | - | 53019226 | 53019359 | 53018329 | 53018501 | 53031064 | 53031144 | 111 | 81 | 101 | 116 | 169 | 0 | 0 | 0 | 2 | 0 | 23 | 52 | 49 | 44 | 59 | 0 | 0 | 0 | 0 | 0 | 282 | 149 | 1 | 1 | 1 | 1 | 1 | 0.968 | 1 | 1 | 1 | 1 | 1 | 1 | -0.006 |
| SE | 41049 | ENSG00000112144 | CILK1 | chr6 | - | 53031064 | 53031144 | 53019226 | 53019359 | 53032532 | 53032654 | 111 | 72 | 84 | 105 | 178 | 0 | 0 | 0 | 6 | 0 | 7 | 45 | 47 | 43 | 42 | 0 | 0 | 0 | 0 | 0 | 229 | 149 | 1 | 1 | 1 | 1 | 1 | 0.919 | 1 | 1 | 1 | 1 | 1 | 1 | -0.016 |
| SE | 41050 | ENSG00000112144 | CILK1 | chr6 | - | 53037938 | 53037993 | 53032532 | 53032654 | 53041135 | 53041408 | 57 | 46 | 63 | 65 | 54 | 3 | 1 | 2 | 0 | 0 | 0 | 30 | 24 | 55 | 14 | 0 | 7 | 0 | 0 | 0 | 204 | 149 | 1 | 1 | 0.933 | 0.971 | 0.958 | 1 | 1 | NA | 0.758 | 1 | 1 | 1 | 0.033 |
| SE | 41051 | ENSG00000112144 | CILK1 | chr6 | - | 53060935 | 53061046 | 53041135 | 53041408 | 53061595 | 53061802 | 0 | 1 | 1 | 0 | 0 | 8 | 6 | 9 | 11 | 2 | 7 | 0 | 2 | 0 | 0 | 6 | 12 | 1 | 4 | 0 | 260 | 149 | 1 | 1 | 0 | 0.087 | 0.06 | 0 | 0 | 0.401 | 0 | 0.534 | 0 | NA | -0.204 |
| SE | 41053 | ENSG00000108797 | CNTNAP1 | chr17 | + | 42688875 | 42689047 | 42688461 | 42688611 | 42689520 | 42689627 | 6 | 2 | 2 | 9 | 0 | 0 | 0 | 0 | 1 | 0 | 0 | 6 | 5 | 12 | 48 | 0 | 1 | 1 | 0 | 3 | 298 | 149 | 1 | 1 | 1 | 1 | 1 | 0.818 | NA | NA | 1 | 0.714 | 1 | 0.889 | 0.054 |
| SE | 41060 | ENSG00000160741 | CRTC2 | chr1 | - | 153949114 | 153949384 | 153948457 | 153948644 | 153951259 | 153951604 | 72 | 27 | 68 | 80 | 57 | 0 | 0 | 0 | 0 | 0 | 146 | 114 | 102 | 119 | 299 | 0 | 0 | 0 | 2 | 5 | 298 | 149 | 1 | 1 | 1 | 1 | 1 | 1 | 1 | 1 | 1 | 1 | 0.967 | 0.968 | 0.013 |
| SE | 41061 | ENSG00000160741 | CRTC2 | chr1 | - | 153949114 | 153949384 | 153948457 | 153948644 | 153952017 | 153952262 | 66 | 26 | 62 | 71 | 53 | 0 | 0 | 0 | 0 | 0 | 105 | 103 | 79 | 106 | 221 | 0 | 0 | 0 | 2 | 0 | 298 | 149 | 1 | 1 | 1 | 1 | 1 | 1 | 1 | 1 | 1 | 1 | 0.964 | 1 | 0.007 |
| SE | 41062 | ENSG00000160741 | CRTC2 | chr1 | - | 153951259 | 153951666 | 153948457 | 153948644 | 153952017 | 153952262 | 12 | 8 | 4 | 19 | 5 | 0 | 0 | 0 | 0 | 0 | 89 | 24 | 27 | 44 | 61 | 0 | 0 | 0 | 2 | 0 | 298 | 149 | 1 | 1 | 1 | 1 | 1 | 1 | 1 | 1 | 1 | 1 | 0.917 | 1 | 0.017 |
| SE | 41064 | ENSG00000160741 | CRTC2 | chr1 | - | 153952017 | 153952262 | 153951259 | 153951666 | 153952396 | 153952446 | 141 | 55 | 106 | 133 | 170 | 2 | 0 | 1 | 0 | 2 | 338 | 217 | 194 | 232 | 321 | 0 | 0 | 0 | 0 | 0 | 298 | 149 | 1 | 1 | 0.972 | 1 | 0.981 | 1 | 0.977 | 1 | 1 | 1 | 1 | 1 | -0.014 |
| SE | 41066 | ENSG00000160741 | CRTC2 | chr1 | - | 153953537 | 153953606 | 153953265 | 153953369 | 153954254 | 153954316 | 157 | 90 | 95 | 150 | 113 | 0 | 0 | 1 | 0 | 0 | 210 | 222 | 147 | 134 | 277 | 0 | 0 | 0 | 0 | 0 | 218 | 149 | 1 | 1 | 1 | 1 | 1 | 0.99 | 1 | 1 | 1 | 1 | 1 | 1 | -0.002 |
| SE | 41070 | ENSG00000115183 | TANC1 | chr2 | + | 159163282 | 159163546 | 159150369 | 159150556 | 159169249 | 159169372 | 0 | 0 | 3 | 1 | 0 | 1 | 0 | 0 | 1 | 0 | 0 | 0 | 1 | 0 | 13 | 6 | 0 | 0 | 0 | 0 | 298 | 149 | 1 | 1 | 0 | NA | 1 | 0.333 | NA | 0 | NA | 1 | NA | 1 | -0.222 |
| SE | 41071 | ENSG00000115183 | TANC1 | chr2 | + | 159227818 | 159227965 | 159225687 | 159225779 | 159228795 | 159228896 | 2 | 4 | 4 | 0 | 3 | 1 | 0 | 0 | 0 | 0 | 0 | 5 | 4 | 0 | 0 | 0 | 0 | 2 | 0 | 0 | 296 | 149 | 1 | 1 | 0.502 | 1 | 1 | NA | 1 | NA | 1 | 0.502 | NA | NA | 0.124 |
| SE | 41072 | ENSG00000239305 | RNF103 | chr2 | - | 86612158 | 86612274 | 86603397 | 86605418 | 86620329 | 86620469 | 69 | 70 | 76 | 72 | 78 | 3 | 2 | 0 | 2 | 1 | 28 | 36 | 132 | 27 | 0 | 21 | 0 | 2 | 0 | 0 | 265 | 149 | 1 | 1 | 0.928 | 0.952 | 1 | 0.953 | 0.978 | 0.428 | 1 | 0.974 | 1 | NA | 0.112 |
| SE | 41077 | ENSG00000092841 | MYL6 | chr12 | + | 56159586 | 56159622 | 56158683 | 56158711 | 56160024 | 56160148 | 4 | 3 | 5 | 7 | 3 | 0 | 0 | 0 | 0 | 0 | 21 | 4 | 4 | 16 | 14 | 0 | 0 | 0 | 2 | 0 | 185 | 149 | 1 | 1 | 1 | 1 | 1 | 1 | 1 | 1 | 1 | 1 | 0.866 | 1 | 0.027 |
| SE | 41079 | ENSG00000092841 | MYL6 | chr12 | + | 56159586 | 56160148 | 56158683 | 56158711 | 56160242 | 56160280 | 3349 | 2433 | 2052 | 3632 | 3480 | 0 | 0 | 0 | 0 | 0 | 7813 | 3614 | 2260 | 4367 | 14342 | 18 | 0 | 0 | 0 | 4 | 298 | 149 | 1 | 1 | 1 | 1 | 1 | 1 | 1 | 0.995 | 1 | 1 | 1 | 0.999 | 0.001 |
| SE | 41081 | ENSG00000092841 | MYL6 | chr12 | + | 56159974 | 56160148 | 56158357 | 56158711 | 56160242 | 56160320 | 3070 | 2227 | 1863 | 3255 | 3190 | 0 | 0 | 0 | 0 | 0 | 6848 | 3324 | 2022 | 3981 | 11987 | 18 | 0 | 0 | 0 | 4 | 298 | 149 | 1 | 1 | 1 | 1 | 1 | 1 | 1 | 0.995 | 1 | 1 | 1 | 0.999 | 0.001 |
| SE | 41082 | ENSG00000092841 | MYL6 | chr12 | + | 56160024 | 56160148 | 56158683 | 56158711 | 56160242 | 56160320 | 2284 | 1482 | 1377 | 2300 | 2341 | 0 | 0 | 0 | 0 | 0 | 4112 | 2090 | 1300 | 2387 | 7282 | 18 | 0 | 0 | 0 | 4 | 273 | 149 | 1 | 1 | 1 | 1 | 1 | 1 | 1 | 0.992 | 1 | 1 | 1 | 0.999 | 0.002 |
| SE | 41085 | ENSG00000165480 | SKA3 | chr13 | - | 21155692 | 21155811 | 21153594 | 21155150 | 21157921 | 21158125 | 4 | 14 | 5 | 3 | 12 | 0 | 0 | 1 | 2 | 0 | 39 | 6 | 5 | 27 | 0 | 6 | 0 | 0 | 0 | 0 | 268 | 149 | 1 | 1 | 1 | 1 | 0.735 | 0.455 | 1 | 0.783 | 1 | 1 | 1 | NA | -0.108 |
| SE | 41086 | ENSG00000165480 | SKA3 | chr13 | - | 21161789 | 21161875 | 21159901 | 21159987 | 21172338 | 21172504 | 6 | 12 | 5 | 5 | 23 | 0 | 0 | 0 | 0 | 6 | 15 | 9 | 9 | 17 | 0 | 0 | 0 | 0 | 0 | 0 | 235 | 149 | 1 | 1 | 1 | 1 | 1 | 1 | 0.708 | 1 | 1 | 1 | 1 | NA | -0.058 |
| SE | 41087 | ENSG00000165480 | SKA3 | chr13 | - | 21167987 | 21168399 | 21161789 | 21161875 | 21172338 | 21172504 | 38 | 43 | 28 | 45 | 79 | 0 | 0 | 0 | 0 | 1 | 103 | 38 | 40 | 26 | 80 | 0 | 0 | 0 | 0 | 0 | 298 | 149 | 1 | 1 | 1 | 1 | 1 | 1 | 0.975 | 1 | 1 | 1 | 1 | 1 | -0.005 |
| SE | 41095 | ENSG00000143761 | ARF1 | chr1 | + | 228097077 | 228097262 | 228082707 | 228082765 | 228097341 | 228097452 | 812 | 449 | 642 | 860 | 996 | 0 | 0 | 0 | 0 | 0 | 2227 | 1381 | 953 | 1482 | 3813 | 0 | 0 | 0 | 0 | 12 | 298 | 149 | 1 | 1 | 1 | 1 | 1 | 1 | 1 | 1 | 1 | 1 | 1 | 0.994 | 0.001 |
| SE | 41096 | ENSG00000122483 | CCDC18 | chr1 | + | 93183359 | 93183495 | 93180715 | 93180852 | 93186344 | 93186400 | 7 | 7 | 7 | 2 | 1 | 2 | 0 | 0 | 0 | 0 | 2 | 5 | 3 | 0 | 5 | 0 | 0 | 0 | 0 | 0 | 285 | 149 | 1 | 1 | 0.647 | 1 | 1 | 1 | 1 | 1 | 1 | 1 | NA | 1 | -0.071 |
| SE | 41098 | ENSG00000122483 | CCDC18 | chr1 | + | 93205509 | 93205631 | 93201891 | 93201988 | 93207106 | 93207398 | 14 | 18 | 16 | 9 | 19 | 7 | 3 | 2 | 0 | 3 | 0 | 10 | 3 | 13 | 19 | 0 | 0 | 10 | 0 | 0 | 271 | 149 | 1 | 1 | 0.524 | 0.767 | 0.815 | 1 | 0.777 | NA | 1 | 0.142 | 1 | 1 | -0.009 |
| SE | 41099 | ENSG00000122483 | CCDC18 | chr1 | + | 93210801 | 93210926 | 93207106 | 93207398 | 93212100 | 93212261 | 40 | 34 | 41 | 48 | 63 | 0 | 2 | 1 | 1 | 0 | 5 | 20 | 22 | 9 | 38 | 16 | 0 | 0 | 0 | 0 | 274 | 149 | 1 | 1 | 1 | 0.902 | 0.957 | 0.963 | 1 | 0.145 | 1 | 1 | 1 | 1 | 0.135 |
| SE | 41100 | ENSG00000122483 | CCDC18 | chr1 | + | 93210801 | 93210926 | 93207106 | 93207398 | 93214742 | 93214966 | 13 | 8 | 15 | 16 | 21 | 0 | 0 | 1 | 0 | 0 | 0 | 8 | 6 | 0 | 16 | 0 | 0 | 0 | 0 | 0 | 274 | 149 | 1 | 1 | 1 | 1 | 0.891 | 1 | 1 | NA | 1 | 1 | NA | 1 | -0.022 |
| SE | 41101 | ENSG00000122483 | CCDC18 | chr1 | + | 93210801 | 93210926 | 93207106 | 93207398 | 93214745 | 93214966 | 13 | 8 | 15 | 16 | 21 | 0 | 0 | 0 | 0 | 0 | 0 | 8 | 6 | 0 | 16 | 0 | 0 | 2 | 0 | 0 | 274 | 149 | 1 | 1 | 1 | 1 | 1 | 1 | 1 | NA | 1 | 0.62 | NA | 1 | 0.127 |
| SE | 41102 | ENSG00000122483 | CCDC18 | chr1 | + | 93212100 | 93212261 | 93207106 | 93207398 | 93214742 | 93214966 | 25 | 24 | 18 | 26 | 35 | 0 | 0 | 1 | 0 | 0 | 59 | 12 | 8 | 8 | 21 | 0 | 0 | 0 | 0 | 0 | 298 | 149 | 1 | 1 | 1 | 1 | 0.9 | 1 | 1 | 1 | 1 | 1 | 1 | 1 | -0.02 |
| SE | 41103 | ENSG00000122483 | CCDC18 | chr1 | + | 93212100 | 93212261 | 93207106 | 93207398 | 93214745 | 93214966 | 11 | 12 | 5 | 17 | 18 | 0 | 0 | 0 | 0 | 0 | 21 | 8 | 12 | 11 | 19 | 0 | 0 | 2 | 0 | 0 | 298 | 149 | 1 | 1 | 1 | 1 | 1 | 1 | 1 | 1 | 1 | 0.75 | 1 | 1 | 0.05 |
| SE | 41104 | ENSG00000122483 | CCDC18 | chr1 | + | 93212100 | 93212261 | 93210801 | 93210926 | 93214742 | 93214966 | 53 | 50 | 43 | 57 | 77 | 0 | 0 | 0 | 1 | 0 | 50 | 24 | 27 | 17 | 43 | 0 | 0 | 2 | 0 | 0 | 298 | 149 | 1 | 1 | 1 | 1 | 1 | 0.966 | 1 | 1 | 1 | 0.871 | 1 | 1 | 0.019 |
| SE | 41105 | ENSG00000122483 | CCDC18 | chr1 | + | 93212100 | 93212261 | 93210801 | 93210926 | 93214745 | 93214966 | 39 | 38 | 30 | 48 | 60 | 1 | 0 | 0 | 2 | 0 | 12 | 20 | 31 | 20 | 41 | 0 | 0 | 2 | 0 | 0 | 298 | 149 | 1 | 1 | 0.951 | 1 | 1 | 0.923 | 1 | 1 | 1 | 0.886 | 1 | 1 | -0.002 |
| SE | 41106 | ENSG00000122483 | CCDC18 | chr1 | + | 93217737 | 93217869 | 93216635 | 93216746 | 93221608 | 93221743 | 35 | 43 | 31 | 52 | 72 | 10 | 4 | 2 | 3 | 11 | 93 | 35 | 32 | 50 | 43 | 0 | 1 | 2 | 0 | 2 | 281 | 149 | 1 | 1 | 0.65 | 0.851 | 0.892 | 0.902 | 0.776 | 1 | 0.949 | 0.895 | 1 | 0.919 | -0.138 |
| SE | 41108 | ENSG00000122483 | CCDC18 | chr1 | + | 93232425 | 93232593 | 93226332 | 93226449 | 93236247 | 93236390 | 45 | 84 | 47 | 50 | 124 | 0 | 1 | 0 | 0 | 0 | 13 | 25 | 30 | 15 | 40 | 0 | 0 | 0 | 0 | 0 | 298 | 149 | 1 | 1 | 1 | 0.977 | 1 | 1 | 1 | 1 | 1 | 1 | 1 | 1 | -0.005 |
| SE | 41111 | ENSG00000122483 | CCDC18 | chr1 | + | 93246104 | 93246204 | 93239682 | 93239896 | 93246837 | 93246892 | 37 | 43 | 15 | 28 | 35 | 1 | 0 | 0 | 0 | 0 | 0 | 11 | 12 | 1 | 4 | 0 | 8 | 0 | 0 | 6 | 249 | 149 | 1 | 1 | 0.957 | 1 | 1 | 1 | 1 | NA | 0.451 | 1 | 1 | 0.285 | 0.307 |
| SE | 41112 | ENSG00000122483 | CCDC18 | chr1 | + | 93246837 | 93246954 | 93239682 | 93239896 | 93256334 | 93256538 | 2 | 1 | 0 | 4 | 0 | 0 | 0 | 2 | 0 | 0 | 0 | 8 | 0 | 0 | 0 | 0 | 0 | 0 | 0 | 0 | 266 | 149 | 1 | 1 | 1 | 1 | 0 | 1 | NA | NA | 1 | NA | NA | NA | -0.25 |
| SE | 41113 | ENSG00000122483 | CCDC18 | chr1 | + | 93254470 | 93254614 | 93246837 | 93246954 | 93256334 | 93256538 | 44 | 28 | 29 | 37 | 32 | 1 | 1 | 0 | 4 | 0 | 0 | 8 | 12 | 4 | 13 | 0 | 0 | 0 | 0 | 0 | 293 | 149 | 1 | 1 | 0.957 | 0.934 | 1 | 0.825 | 1 | NA | 1 | 1 | 1 | 1 | -0.057 |
| SE | 41114 | ENSG00000122483 | CCDC18 | chr1 | + | 93270346 | 93270814 | 93264700 | 93264901 | 93278462 | 93278477 | 7 | 4 | 13 | 3 | 4 | 0 | 2 | 4 | 1 | 2 | 14 | 0 | 0 | 0 | 0 | 0 | 0 | 0 | 0 | 0 | 298 | 149 | 1 | 1 | 1 | 0.5 | 0.619 | 0.6 | 0.5 | 1 | NA | NA | NA | NA | -0.356 |
| SE | 41116 | ENSG00000143578 | CREB3L4 | chr1 | + | 153973379 | 153973464 | 153973194 | 153973263 | 153973619 | 153973716 | 23 | 19 | 20 | 23 | 22 | 0 | 0 | 0 | 1 | 0 | 16 | 29 | 24 | 52 | 33 | 0 | 0 | 0 | 0 | 0 | 234 | 149 | 1 | 1 | 1 | 1 | 1 | 0.936 | 1 | 1 | 1 | 1 | 1 | 1 | -0.013 |
| SE | 41121 | ENSG00000115523 | GNLY | chr2 | + | 85697505 | 85697677 | 85695957 | 85696056 | 85698397 | 85698447 | 6657 | 1138 | 1858 | 2221 | 5597 | 0 | 3 | 0 | 0 | 0 | 1710 | 657 | 1425 | 627 | 1120 | 0 | 0 | 0 | 0 | 0 | 298 | 149 | 1 | 1 | 1 | 0.995 | 1 | 1 | 1 | 1 | 1 | 1 | 1 | 1 | -0.001 |
| SE | 41124 | ENSG00000115523 | GNLY | chr2 | + | 85698397 | 85698447 | 85697505 | 85697677 | 85698563 | 85698747 | 18 | 4 | 4 | 8 | 12 | 6160 | 1039 | 1789 | 1955 | 6427 | 0 | 4 | 5 | 2 | 5 | 1244 | 393 | 1178 | 647 | 873 | 199 | 149 | 1 | 1 | 0.002 | 0.003 | 0.002 | 0.003 | 0.001 | 0 | 0.008 | 0.003 | 0.002 | 0.004 | -0.001 |
| SE | 41126 | ENSG00000124406 | ATP8A1 | chr4 | - | 42443564 | 42443672 | 42423616 | 42423705 | 42444577 | 42444634 | 205 | 156 | 246 | 239 | 172 | 0 | 0 | 0 | 0 | 3 | 12 | 121 | 122 | 47 | 70 | 0 | 0 | 1 | 0 | 0 | 257 | 149 | 1 | 1 | 1 | 1 | 1 | 1 | 0.971 | 1 | 1 | 0.986 | 1 | 1 | -0.003 |
| SE | 41127 | ENSG00000124406 | ATP8A1 | chr4 | - | 42451980 | 42452059 | 42446582 | 42446644 | 42455296 | 42455419 | 269 | 242 | 298 | 310 | 254 | 0 | 0 | 1 | 0 | 2 | 13 | 134 | 120 | 79 | 53 | 0 | 2 | 3 | 0 | 0 | 228 | 149 | 1 | 1 | 1 | 1 | 0.995 | 1 | 0.988 | 1 | 0.978 | 0.963 | 1 | 1 | 0.008 |
| SE | 41129 | ENSG00000124406 | ATP8A1 | chr4 | - | 42522159 | 42522299 | 42507015 | 42507154 | 42524762 | 42524847 | 680 | 538 | 563 | 774 | 987 | 0 | 0 | 0 | 0 | 0 | 765 | 636 | 472 | 422 | 1316 | 0 | 0 | 0 | 5 | 0 | 289 | 149 | 1 | 1 | 1 | 1 | 1 | 1 | 1 | 1 | 1 | 1 | 0.978 | 1 | 0.004 |
| SE | 41130 | ENSG00000124406 | ATP8A1 | chr4 | - | 42543916 | 42543986 | 42524762 | 42524847 | 42549012 | 42549062 | 255 | 193 | 241 | 246 | 338 | 2 | 0 | 0 | 0 | 0 | 156 | 147 | 134 | 87 | 175 | 0 | 0 | 0 | 0 | 0 | 219 | 149 | 1 | 1 | 0.989 | 1 | 1 | 1 | 1 | 1 | 1 | 1 | 1 | 1 | -0.002 |
| SE | 41131 | ENSG00000124406 | ATP8A1 | chr4 | - | 42549012 | 42549062 | 42543916 | 42543986 | 42552504 | 42552610 | 55 | 44 | 45 | 49 | 87 | 0 | 0 | 0 | 0 | 1 | 15 | 49 | 29 | 24 | 3 | 0 | 0 | 0 | 0 | 0 | 199 | 149 | 1 | 1 | 1 | 1 | 1 | 1 | 0.985 | 1 | 1 | 1 | 1 | 1 | -0.003 |
| SE | 41133 | ENSG00000124406 | ATP8A1 | chr4 | - | 42552504 | 42552610 | 42543916 | 42543986 | 42574618 | 42574707 | 3 | 2 | 0 | 3 | 9 | 0 | 1 | 0 | 0 | 0 | 0 | 0 | 2 | 0 | 0 | 0 | 0 | 0 | 0 | 0 | 255 | 149 | 1 | 1 | 1 | 0.539 | NA | 1 | 1 | NA | NA | 1 | NA | NA | -0.115 |
| SE | 41136 | ENSG00000124406 | ATP8A1 | chr4 | - | 42574618 | 42574707 | 42555967 | 42556040 | 42575621 | 42575699 | 158 | 124 | 150 | 186 | 246 | 0 | 0 | 0 | 0 | 0 | 37 | 126 | 115 | 57 | 38 | 0 | 0 | 2 | 0 | 0 | 238 | 149 | 1 | 1 | 1 | 1 | 1 | 1 | 1 | 1 | 1 | 0.973 | 1 | 1 | 0.005 |
| SE | 41137 | ENSG00000124406 | ATP8A1 | chr4 | - | 42579812 | 42579978 | 42578259 | 42578387 | 42581620 | 42581732 | 535 | 331 | 447 | 427 | 728 | 0 | 0 | 0 | 1 | 0 | 198 | 335 | 180 | 141 | 149 | 0 | 0 | 0 | 0 | 0 | 298 | 149 | 1 | 1 | 1 | 1 | 1 | 0.995 | 1 | 1 | 1 | 1 | 1 | 1 | -0.001 |

| | | | | | | | | | | | | | | |
|---|---|---|---|---|---|---|---|---|---|---|---|---|---|---|
| SE | 41141 | ENSG00000124406 | ATP8A1 | chr4 | - | 42625613 | 42625713 | 42624535 | 42624634 | 42626994 | 42627109 | 319 | 256 | [illegible] |
| SE | 41143 | ENSG00000009780 | FAM76A | chr1 | + | 27732602 | 27732657 | 27727471 | 27727536 | 27734030 | 27734183 | 44 | 28 | [illegible] |
| SE | 41144 | ENSG00000009780 | FAM76A | chr1 | + | 27734030 | 27734183 | 27732602 | 27732657 | 27744653 | 27744811 | 94 | 50 | [illegible] |
| SE | 41153 | ENSG00000013573 | DDX11 | chr12 | + | 31101832 | 31101982 | 31100998 | 31101130 | 31102242 | 31102311 | 5 | 5 | [illegible] |
| SE | 41154 | ENSG00000013573 | DDX11 | chr12 | + | 31103316 | 31103395 | 31102935 | 31103020 | 31103576 | 31103731 | 4 | 4 | [illegible] |
| SE | 41155 | ENSG00000013573 | DDX11 | chr12 | + | 31103321 | 31103395 | 31102935 | 31103020 | 31103576 | 31103731 | 3 | 4 | [illegible] |
| SE | 41159 | ENSG00000163491 | NEK10 | chr3 | - | 27290616 | 27290754 | 27287697 | 27287743 | 27291261 | 27291390 | 1 | 2 | [illegible] |
| SE | 41160 | ENSG00000163491 | NEK10 | chr3 | - | 27307858 | 27307945 | 27304746 | 27304971 | 27308925 | 27309005 | 4 | 4 | [illegible] |
| SE | 41196 | ENSG00000146416 | AIG1 | chr6 | + | 143136834 | 143136990 | 143060853 | 143061066 | 143165081 | 143165183 | 29 | 11 | [illegible] |
| SE | 41197 | ENSG00000146416 | AIG1 | chr6 | + | 143186669 | 143186815 | 143164943 | 143165183 | 143187628 | 143187814 | 2 | 0 | [illegible] |
| SE | 41199 | ENSG00000146416 | AIG1 | chr6 | + | 143334077 | 143334095 | 143333281 | 143333445 | 143335050 | 143335530 | 0 | 4 | [illegible] |
| SE | 41202 | ENSG00000128789 | PSMG2 | chr18 | + | 12720509 | 12720683 | 12718516 | 12718635 | 12724498 | 12724520 | 395 | 224 | [illegible] |
| SE | 41203 | ENSG00000118200 | CAMSAP2 | chr1 | + | 200807375 | 200807537 | 200760838 | 200761098 | 200815560 | 200815644 | 12 | 19 | [illegible] |
| SE | 41205 | ENSG00000118200 | CAMSAP2 | chr1 | + | 200847209 | 200847292 | 200844781 | 200844869 | 200847639 | 200847709 | 5 | 11 | [illegible] |
| SE | 41206 | ENSG00000118200 | CAMSAP2 | chr1 | + | 200847257 | 200847292 | 200844781 | 200844869 | 200847639 | 200847709 | 1 | 5 | [illegible] |
| SE | 41207 | ENSG00000118200 | CAMSAP2 | chr1 | + | 200857305 | 200857424 | 200856009 | 200856125 | 200857753 | 200858119 | 19 | 25 | [illegible] |
| SE | 41214 | ENSG00000124713 | GNMT | chr6 | + | 42963071 | 42963214 | 42962761 | 42962878 | 42963327 | 42963449 | 7 | 4 | [illegible] |
| SE | 41215 | ENSG00000135828 | RNASEL | chr1 | - | 182581224 | 182581357 | 182576255 | 182576389 | 182582052 | 182582258 | 165 | 129 | [illegible] |
| SE | 41216 | ENSG00000135828 | RNASEL | chr1 | - | 182584080 | 182584166 | 182582052 | 182582258 | 182585326 | 182586970 | 50 | 21 | [illegible] |
| SE | 41217 | ENSG00000176890 | TYMS | chr18 | + | 669071 | 669173 | 662145 | 662320 | 670691 | 670867 | 22 | 31 | [illegible] |
| SE | 41218 | ENSG00000176890 | TYMS | chr18 | + | 670691 | 670867 | 669071 | 669173 | 671379 | 671451 | 36 | 29 | [illegible] |
| SE | 41219 | ENSG00000157181 | ODR4 | chr1 | + | 186379766 | 186379884 | 186375837 | 186375974 | 186383021 | 186383156 | 109 | 122 | [illegible] |
| SE | 41221 | ENSG00000157181 | ODR4 | chr1 | + | 186385987 | 186386083 | 186383021 | 186383156 | 186390710 | 186390851 | 28 | 25 | [illegible] |
| SE | 41222 | ENSG00000157181 | ODR4 | chr1 | + | 186393946 | 186394015 | 186391695 | 186391791 | 186398312 | 186398441 | 71 | 64 | [illegible] |
| SE | 41225 | ENSG00000153815 | CMIP | chr16 | + | 81660883 | 81660946 | 81657774 | 81657816 | 81664268 | 81664349 | 302 | 270 | [illegible] |
| SE | 41226 | ENSG00000153815 | CMIP | chr16 | + | 81660883 | 81660946 | 81657774 | 81657816 | 81670141 | 81670245 | 224 | 211 | [illegible] |
| SE | 41229 | ENSG00000153815 | CMIP | chr16 | + | 81671965 | 81672070 | 81670141 | 81670245 | 81678274 | 81678628 | 302 | 212 | [illegible] |
| SE | 41230 | ENSG00000153815 | CMIP | chr16 | + | 81696559 | 81696667 | 81693438 | 81693487 | 81699684 | 81699801 | 169 | 93 | [illegible] |
| SE | 41231 | ENSG00000153815 | CMIP | chr16 | + | 81699684 | 81699801 | 81696559 | 81696667 | 81701659 | 81701800 | 303 | 212 | [illegible] |
| SE | 41234 | ENSG00000119782 | FKBP1B | chr2 | + | 24054103 | 24054192 | 24053901 | 24053949 | 24054472 | 24054540 | 0 | 0 | [illegible] |
| SE | 41237 | ENSG00000119782 | FKBP1B | chr2 | + | 24054472 | 24054540 | 24054103 | 24054192 | 24060813 | 24060926 | 0 | 0 | [illegible] |
| SE | 41239 | ENSG00000128563 | PRKRIP1 | chr7 | + | 102365973 | 102366076 | 102363894 | 102364411 | 102375822 | 102375949 | 36 | 17 | [illegible] |
| SE | 41240 | ENSG00000239969 | NA | chr7 | + | 102366017 | 102366076 | 102364297 | 102364411 | 102375822 | 102375949 | 12 | 11 | [illegible] |
| SE | 41245 | ENSG00000119640 | ACYP1 | chr14 | - | 75061683 | 75061762 | 75053242 | 75053659 | 75063469 | 75063561 | 10 | 7 | [illegible] |
| SE | 41246 | ENSG00000119640 | ACYP1 | chr14 | - | 75063469 | 75063561 | 75061683 | 75061762 | 75063953 | 75064007 | 9 | 5 | [illegible] |
| SE | 41249 | ENSG00000117505 | DR1 | chr1 | + | 93353907 | 93354071 | 93345906 | 93346865 | 93360492 | 93361168 | 1659 | 1349 | [illegible] |
| SE | 41251 | ENSG00000162819 | BROX | chr1 | + | 222715144 | 222715269 | 222712563 | 222712942 | 222718924 | 222719031 | 2 | 4 | [illegible] |
| SE | 41252 | ENSG00000162819 | BROX | chr1 | + | 222715144 | 222715269 | 222713380 | 222713416 | 222715683 | 222715800 | 1 | 2 | [illegible] |
| SE | 41253 | ENSG00000162819 | BROX | chr1 | + | 222715144 | 222715269 | 222713380 | 222713416 | 222718924 | 222719031 | 3 | 0 | [illegible] |
| SE | 41257 | ENSG00000162819 | BROX | chr1 | + | 222717857 | 222718094 | 222715683 | 222715800 | 222718924 | 222719031 | 0 | 1 | [illegible] |
| SE | 41258 | ENSG00000162819 | BROX | chr1 | + | 222722418 | 222722514 | 222719262 | 222719359 | 222724091 | 222724164 | 323 | 245 | [illegible] |
| SE | 41259 | ENSG00000162819 | BROX | chr1 | + | 222722418 | 222722514 | 222719262 | 222719359 | 222724099 | 222724164 | 148 | 121 | [illegible] |
| SE | 41260 | ENSG00000162819 | BROX | chr1 | + | 222725449 | 222725555 | 222724091 | 222724164 | 222727167 | 222727209 | 269 | 194 | [illegible] |
| SE | 41261 | ENSG00000162819 | BROX | chr1 | + | 222729619 | 222729701 | 222728742 | 222728828 | 222730026 | 222730177 | 347 | 264 | [illegible] |
| SE | 41264 | ENSG00000164506 | STXBP5 | chr6 | + | 147291093 | 147291172 | 147278080 | 147278204 | 147310083 | 147310238 | 180 | 120 | [illegible] |
| SE | 41266 | ENSG00000164506 | STXBP5 | chr6 | + | 147324958 | 147325084 | 147316228 | 147316407 | 147327124 | 147327276 | 179 | 167 | [illegible] |
| SE | 41267 | ENSG00000164506 | STXBP5 | chr6 | + | 147324958 | 147325084 | 147316228 | 147316407 | 147334156 | 147334222 | 80 | 69 | [illegible] |
| SE | 41268 | ENSG00000164506 | STXBP5 | chr6 | + | 147327124 | 147327276 | 147316228 | 147316407 | 147334156 | 147334222 | 85 | 62 | [illegible] |
| SE | 41269 | ENSG00000164506 | STXBP5 | chr6 | + | 147327124 | 147327276 | 147324958 | 147325084 | 147334156 | 147334222 | 184 | 160 | [illegible] |
| SE | 41271 | ENSG00000164506 | STXBP5 | chr6 | + | 147339178 | 147339238 | 147334156 | 147334222 | 147353322 | 147353373 | 12 | 16 | [illegible] |
| SE | 41272 | ENSG00000164506 | STXBP5 | chr6 | + | 147339336 | 147339384 | 147339178 | 147339238 | 147353322 | 147353373 | 15 | 9 | [illegible] |
| SE | 41274 | ENSG00000164506 | STXBP5 | chr6 | + | 147364000 | 147364166 | 147363334 | 147363704 | 147373730 | 147373842 | 252 | 260 | [illegible] |
| SE | 41275 | ENSG00000164506 | STXBP5 | chr6 | + | 147373730 | 147373842 | 147364000 | 147364166 | 147382777 | 147382998 | 251 | 195 | [illegible] |
| SE | 41276 | ENSG00000124614 | RPS10 | chr6 | - | 34417797 | 34417897 | 34417453 | 34417547 | 34418368 | 34418424 | 2 | 0 | [illegible] |
| SE | 41277 | ENSG00000124614 | RPS10 | chr6 | - | 34421729 | 34421807 | 34417453 | 34418424 | 34424668 | 34424840 | 6536 | 3259 | [illegible] |
| SE | 41278 | ENSG00000149781 | FERMT3 | chr11 | + | 64219523 | 64219658 | 64219250 | 64219358 | 64219739 | 64219789 | 469 | 212 | [illegible] |
| SE | 41279 | ENSG00000149781 | FERMT3 | chr11 | + | 64221015 | 64221140 | 64220435 | 64220669 | 64223047 | 64223189 | 654 | 373 | [illegible] |
| SE | 41280 | ENSG00000143498 | TAF1A | chr1 | - | 222563172 | 222563296 | 222561363 | 222561518 | 222564058 | 222564125 | 72 | 82 | [illegible] |
| SE | 41281 | ENSG00000143498 | TAF1A | chr1 | - | 222570534 | 222570665 | 222569509 | 222569668 | 222577444 | 222577643 | 53 | 40 | [illegible] |
| SE | 41282 | ENSG00000143498 | TAF1A | chr1 | - | 222579758 | 222579825 | 222577444 | 222577643 | 222584127 | 222584297 | 13 | 12 | [illegible] |
| SE | 41283 | ENSG00000143498 | TAF1A | chr1 | - | 222579758 | 222579872 | 222577444 | 222577643 | 222584127 | 222584297 | 40 | 32 | [illegible] |
| SE | 41284 | ENSG00000143498 | TAF1A | chr1 | - | 222579758 | 222579872 | 222577444 | 222577643 | 222588442 | 222588565 | 24 | 15 | [illegible] |
| SE | 41285 | ENSG00000143498 | TAF1A | chr1 | - | 222579758 | 222579890 | 222577444 | 222577643 | 222584127 | 222584297 | 27 | 18 | [illegible] |
| SE | 41286 | ENSG00000143498 | TAF1A | chr1 | - | 222579758 | 222579890 | 222577444 | 222577643 | 222588442 | 222588565 | 27 | 15 | [illegible] |
| SE | 41287 | ENSG00000143498 | TAF1A | chr1 | - | 222584127 | 222584297 | 222577444 | 222577643 | 222588442 | 222588565 | 11 | 23 | [illegible] |
| SE | 41288 | ENSG00000143498 | TAF1A | chr1 | - | 222584127 | 222584297 | 222579758 | 222579872 | 222588442 | 222588565 | 24 | 34 | [illegible] |
| SE | 41289 | ENSG00000143498 | TAF1A | chr1 | - | 222584127 | 222584297 | 222579758 | 222579890 | 222588442 | 222588565 | 11 | 20 | [illegible] |
| SE | 41290 | ENSG00000058668 | ATP2B4 | chr1 | + | 203682741 | 203683398 | 203626786 | 203627219 | 203698156 | 203698354 | 429 | 225 | [illegible] |
| SE | 41293 | ENSG00000058668 | ATP2B4 | chr1 | + | 203707861 | 203708104 | 203703651 | 203703813 | 203709300 | 203709542 | 258 | 121 | [illegible] |
| SE | 41294 | ENSG00000058668 | ATP2B4 | chr1 | + | 203707861 | 203708104 | 203703651 | 203703813 | 203710876 | 203711108 | 1 | 10 | [illegible] |
| SE | 41295 | ENSG00000058668 | ATP2B4 | chr1 | + | 203709300 | 203709542 | 203703651 | 203703813 | 203710876 | 203711108 | 316 | 142 | [illegible] |
| SE | 41296 | ENSG00000058668 | ATP2B4 | chr1 | + | 203709300 | 203709542 | 203707861 | 203708104 | 203710876 | 203711108 | 572 | 253 | [illegible] |
| SE | 41300 | ENSG00000058668 | ATP2B4 | chr1 | + | 203733222 | 203733400 | 203727394 | 203727571 | 203739545 | 203739644 | 2 | 1 | [illegible] |
| SE | 41301 | ENSG00000276997 | NA | chr1 | - | 222480302 | 222480426 | 222479367 | 222479558 | 222486056 | 222486121 | 0 | 0 | [illegible] |
| SE | 41304 | ENSG00000269893 | SNHG8 | chr4 | + | 118279106 | 118279137 | 118278754 | 118278792 | 118279388 | 118279431 | 51 | 15 | [illegible] |
| SE | 41306 | ENSG00000127948 | POR | chr7 | + | 75970986 | 75971092 | 75954075 | 75954180 | 75972412 | 75972461 | 2 | 1 | [illegible] |
| SE | 41311 | ENSG00000137411 | VARS2 | chr6 | + | 30915745 | 30915867 | 30915354 | 30915455 | 30915980 | 30916047 | 91 | 55 | [illegible] |
| SE | 41314 | ENSG00000137411 | VARS2 | chr6 | + | 30922449 | 30922554 | 30922115 | 30922241 | 30922705 | 30922774 | 16 | 2 | [illegible] |
| SE | 41316 | ENSG00000137411 | VARS2 | chr6 | + | 30925273 | 30925385 | 30923352 | 30924560 | 30925543 | 30925719 | 10 | 8 | [illegible] |
| SE | 41317 | ENSG00000134108 | ARL8B | chr3 | + | 5128459 | 5128584 | 5122248 | 5122588 | 5170502 | 5170583 | 0 | 1 | [illegible] |
| SE | 41326 | ENSG00000130826 | DKC1 | chrX | + | 154765443 | 154765530 | 154764898 | 154764966 | 154765906 | 154765998 | 129 | 106 | [illegible] |
| SE | 41327 | ENSG00000130826 | DKC1 | chrX | + | 154765906 | 154765998 | 154765443 | 154765530 | 154766215 | 154766400 | 160 | 136 | [illegible] |
| SE | 41328 | ENSG00000130826 | DKC1 | chrX | + | 154766996 | 154767061 | 154766215 | 154766400 | 154767255 | 154767382 | 258 | 162 | [illegible] |
| SE | 41329 | ENSG00000130826 | DKC1 | chrX | + | 154768301 | 154768312 | 154767255 | 154767382 | 154769166 | 154769291 | 0 | 1 | [illegible] |
| SE | 41330 | ENSG00000130826 | DKC1 | chrX | + | 154768301 | 154768432 | 154767255 | 154767382 | 154769166 | 154769210 | 148 | 83 | [illegible] |
| SE | 41332 | ENSG00000130826 | DKC1 | chrX | + | 154770758 | 154770879 | 154769166 | 154769310 | 154773130 | 154773249 | 194 | 110 | [illegible] |
| SE | 41333 | ENSG00000130826 | DKC1 | chrX | + | 154774785 | 154774844 | 154774601 | 154774705 | 154775194 | 154775273 | 0 | 0 | [illegible] |
| SE | 41336 | ENSG00000071282 | LMCD1 | chr3 | + | 8537184 | 8537440 | 8532736 | 8532825 | 8548567 | 8548863 | 0 | 0 | [illegible] |
| SE | 41337 | ENSG00000145287 | PLAC8 | chr4 | - | 83094677 | 83094791 | 83090047 | 83090971 | 83104895 | 83105020 | 2506 | 1549 | [illegible] |
| SE | 41338 | ENSG00000145287 | PLAC8 | chr4 | - | 83104895 | 83105020 | 83092860 | 83094791 | 83107803 | 83107950 | 2319 | 1250 | [illegible] |
| SE | 41341 | ENSG00000145287 | PLAC8 | chr4 | - | 83107803 | 83107950 | 83104895 | 83105020 | 83109727 | 83109843 | 1249 | 671 | [illegible] |
| SE | 41343 | ENSG00000145287 | PLAC8 | chr4 | - | 83134534 | 83134634 | 83117883 | 83117971 | 83136958 | 83137075 | 0 | 0 | [illegible] |
| SE | 41346 | ENSG00000113621 | TXNDC15 | chr5 | + | 134893491 | 134893655 | 134887694 | 134888182 | 134896293 | 134896424 | 253 | 112 | [illegible] |
| SE | 41352 | ENSG00000117118 | SDHB | chr1 | - | 17027748 | 17027865 | 17023972 | 17024074 | 17028599 | 17028736 | 657 | 358 | [illegible] |
| SE | 41353 | ENSG00000117118 | SDHB | chr1 | - | 17028599 | 17028736 | 17023972 | 17024074 | 17033059 | 17033145 | 294 | 165 | [illegible] |
| SE | 41357 | ENSG00000162374 | ELAVL4 | chr1 | + | 50177088 | 50177192 | 50144956 | 50145197 | 50193764 | 50193918 | 0 | 2 | [illegible] |
| SE | 41360 | ENSG00000144401 | METTL21A | chr2 | - | 207620640 | 207620694 | 207581694 | 207582160 | 207621805 | 207621917 | 1 | 0 | [illegible] |
| SE | 41362 | ENSG00000144401 | METTL21A | chr2 | - | 207621805 | 207621917 | 207581793 | 207581946 | 207624228 | 207624404 | 25 | 14 | [illegible] |
| SE | 41367 | ENSG00000143228 | NUF2 | chr1 | + | 163328457 | 163328530 | 163328227 | 163328304 | 163328845 | 163328907 | 0 | 0 | [illegible] |
| SE | 41372 | ENSG00000114796 | KLHL24 | chr3 | + | 183636220 | 183636311 | 183635622 | 183635747 | 183643479 | 183643542 | 14 | 28 | [illegible] |
| SE | 41373 | ENSG00000114796 | KLHL24 | chr3 | + | 183636220 | 183636311 | 183635622 | 183635747 | 183650295 | 183650575 | 0 | 11 | [illegible] |
| SE | 41375 | ENSG00000114796 | KLHL24 | chr3 | + | 183636220 | 183636311 | 183635622 | 183635793 | 183650295 | 183650575 | 1 | 0 | [illegible] |
| SE | 41378 | ENSG00000114796 | KLHL24 | chr3 | + | 183643479 | 183643542 | 183635619 | 183635747 | 183650295 | 183650590 | 97 | 78 | [illegible] |
| SE | 41379 | ENSG00000114796 | KLHL24 | chr3 | + | 183643479 | 183643542 | 183635622 | 183635782 | 183650295 | 183650590 | 94 | 77 | [illegible] |
| SE | 41380 | ENSG00000114796 | KLHL24 | chr3 | + | 183643479 | 183643542 | 183636220 | 183636311 | 183650295 | 183650590 | 107 | 95 | [illegible] |
| SE | 41381 | ENSG00000114796 | KLHL24 | chr3 | + | 183644085 | 183644167 | 183643479 | 183643542 | 183650295 | 183650538 | 0 | 2 | [illegible] |
| SE | 41382 | ENSG00000114796 | KLHL24 | chr3 | + | 183650295 | 183651276 | 183643479 | 183643542 | 183663457 | 183663642 | 210 | 188 | [illegible] |
| SE | 41383 | ENSG00000114796 | KLHL24 | chr3 | + | 183663457 | 183663642 | 183650295 | 183651276 | 183664920 | 183665039 | 165 | 112 | [illegible] |
| SE | 41384 | ENSG00000114796 | KLHL24 | chr3 | + | 183663457 | 183663642 | 183650295 | 183651276 | 183671033 | 183671123 | 60 | 49 | [illegible] |
| SE | 41385 | ENSG00000114796 | KLHL24 | chr3 | + | 183664920 | 183665039 | 183650295 | 183651276 | 183671033 | 183671123 | 50 | 37 | [illegible] |
| SE | 41388 | ENSG00000114796 | KLHL24 | chr3 | + | 183672295 | 183672484 | 183671033 | 183671222 | 183679085 | 183684477 | 263 | 183 | [illegible] |
| SE | 41389 | ENSG00000254635 | WAC-AS1 | chr10 | - | 28525867 | 28526008 | 28523220 | 28524611 | 28532441 | 28533066 | 4 | 1 | [illegible] |
| SE | 41390 | ENSG00000254635 | WAC-AS1 | chr10 | - | 28525947 | 28526008 | 28522651 | 28524611 | 28532441 | 28532743 | 7 | 6 | [illegible] |
| SE | 41392 | ENSG00000117569 | PTBP2 | chr1 | + | 96769702 | 96769875 | 96751424 | 96751500 | 96770707 | 96770851 | 205 | 150 | [illegible] |
| SE | 41393 | ENSG00000117569 | PTBP2 | chr1 | + | 96804784 | 96804939 | 96785058 | 96785254 | 96806418 | 96806452 | 2 | 0 | [illegible] |
| SE | 41394 | ENSG00000117569 | PTBP2 | chr1 | + | 96804799 | 96804939 | 96784947 | 96785254 | 96806418 | 96806452 | 2 | 0 | [illegible] |
| SE | 41395 | ENSG00000117569 | PTBP2 | chr1 | + | 96806418 | 96806452 | 96804784 | 96804939 | 96806865 | 96806958 | 1 | 0 | [illegible] |
| SE | 41397 | ENSG00000068120 | COASY | chr17 | + | 42562295 | 42562487 | 42562150 | 42562219 | 42562611 | 42562919 | 3 | 0 | [illegible] |
| SE | 41398 | ENSG00000068120 | COASY | chr17 | + | 42562355 | 42562487 | 42562150 | 42562219 | 42562611 | 42563322 | 3 | 0 | [illegible] |
| SE | 41402 | ENSG00000054392 | HHAT | chr1 | + | 210348932 | 210349066 | 210328901 | 210329104 | 210362851 | 210362919 | 12 | 12 | [illegible] |
| SE | 41403 | ENSG00000054392 | HHAT | chr1 | + | 210362851 | 210362919 | 210348932 | 210349066 | 210387467 | 210387581 | 6 | 10 | [illegible] |
| SE | 41404 | ENSG00000054392 | HHAT | chr1 | + | 210400467 | 210400662 | 210387467 | 210387581 | 210404463 | 210404679 | 2 | 2 | [illegible] |
| SE | 41406 | ENSG00000054392 | HHAT | chr1 | + | 210418153 | 210418325 | 210404463 | 210404679 | 210464504 | 210464655 | 1 | 0 | [illegible] |
| SE | 41410 | ENSG00000050327 | ARHGEF5 | chr7 | + | 144375513 | 144375675 | 144374731 | 144374889 | 144377119 | 144377190 | 20 | 3 | [illegible] |
| SE | 41412 | ENSG00000013016 | EHD3 | chr2 | + | 31261548 | 31261713 | 31260509 | 31260922 | 31266176 | 31269451 | 82 | 88 | [illegible] |
| SE | 41423 | ENSG00000166928 | MS4A14 | chr11 | + | 60404545 | 60404666 | 60402911 | 60403061 | 60415436 | 60417563 | 6 | 4 | [illegible] |
| SE | 41424 | ENSG00000166928 | MS4A14 | chr11 | + | 60405888 | 60405987 | 60402911 | 60403061 | 60415436 | 60417685 | 0 | 1 | [illegible] |
| SE | 41435 | ENSG00000119285 | HEATR1 | chr1 | - | 236554597 | 236554752 | 236553580 | 236553739 | 236555295 | 236555464 | 260 | 231 | [illegible] |
| SE | 41436 | ENSG00000119285 | HEATR1 | chr1 | - | 236574660 | 236574903 | 236574201 | 236574333 | 236576218 | 236576377 | 216 | 168 | [illegible] |
| SE | 41437 | ENSG00000119285 | HEATR1 | chr1 | - | 236583012 | 236583196 | 236582735 | 236582872 | 236585024 | 236585216 | 141 | 93 | [illegible] |
| SE | 41438 | ENSG00000119285 | HEATR1 | chr1 | - | 236586240 | 236586452 | 236585819 | 236585941 | 236587401 | 236587490 | 137 | 205 | [illegible] |
| SE | 41439 | ENSG00000119285 | HEATR1 | chr1 | - | 236587401 | 236587490 | 236585819 | 236585941 | 236590846 | 236590954 | 3 | 0 | [illegible] |
| SE | 41440 | ENSG00000119285 | HEATR1 | chr1 | - | 236587401 | 236587490 | 236586240 | 236586452 | 236587947 | 236588043 | 97 | 130 | [illegible] |
| SE | 41441 | ENSG00000119285 | HEATR1 | chr1 | - | 236587947 | 236588043 | 236587401 | 236587490 | 236590846 | 236590954 | 108 | 125 | [illegible] |
| SE | 41442 | ENSG00000119285 | HEATR1 | chr1 | - | 236596835 | 236596976 | 236595771 | 236596044 | 236597877 | 236597979 | 110 | 66 | [illegible] |
| SE | 41445 | ENSG00000137776 | SLTM | chr15 | - | 58883625 | 58883786 | 58879049 | 58880107 | 58886974 | 58887007 | 173 | 120 | [illegible] |
| SE | 41446 | ENSG00000137776 | SLTM | chr15 | - | 58888384 | 58888555 | 58887225 | 58887540 | 58889429 | 58889554 | 436 | 469 | [illegible] |
| SE | 41447 | ENSG00000137776 | SLTM | chr15 | - | 58888384 | 58888573 | 58887456 | 58887540 | 58889429 | 58889490 | 240 | 233 | [illegible] |
| SE | 41448 | ENSG00000137776 | SLTM | chr15 | - | 58890280 | 58890461 | 58889429 | 58889554 | 58892896 | 58893060 | 72 | 118 | [illegible] |
| SE | 41452 | ENSG00000137776 | SLTM | chr15 | - | 58912562 | 58912610 | 58901259 | 58901287 | 58913498 | 58913696 | 135 | 159 | [illegible] |
| SE | 41458 | ENSG00000137776 | SLTM | chr15 | - | 58921445 | 58921584 | 58916934 | 58917020 | 58932355 | 58932443 | 0 | 5 | [illegible] |
| SE | 41462 | ENSG00000106780 | MEGF9 | chr9 | - | 120659373 | 120659575 | 120622615 | 120622755 | 120713757 | 120714470 | 242 | 149 | [illegible] |
| SE | 41464 | ENSG00000204152 | TIMM23B | chr10 | + | 49943234 | 49943358 | 49942048 | 49942300 | 49952125 | 49952219 | 6 | 0 | [illegible] |
| SE | 41466 | ENSG00000204152 | TIMM23B | chr10 | + | 49945031 | 49945090 | 49943234 | 49943358 | 49952125 | 49952219 | 48 | 28 | [illegible] |
| SE | 41467 | ENSG00000204152 | TIMM23B | chr10 | + | 49952125 | 49952219 | 49942080 | 49942300 | 49958369 | 49958480 | 2 | 4 | [illegible] |
| SE | 41470 | ENSG00000204152 | TIMM23B | chr10 | + | 49952448 | 49952533 | 49952125 | 49952219 | 49958369 | 49958480 | 39 | 32 | [illegible] |
| SE | 41471 | ENSG00000204152 | TIMM23B | chr10 | + | 49955001 | 49955060 | 49952125 | 49952219 | 49958369 | 49958480 | 32 | 13 | [illegible] |
| SE | 41472 | ENSG00000204152 | TIMM23B | chr10 | + | 49955001 | 49955060 | 49952448 | 49952533 | 49958369 | 49958480 | 141 | 79 | [illegible] |
| SE | 41474 | ENSG00000177989 | ODF3B | chr22 | - | 50531569 | 50531702 | 50531190 | 50531295 | 50531968 | 50532184 | 12 | 1 | [illegible] |
| SE | 41475 | ENSG00000177989 | ODF3B | chr22 | - | 50531569 | 50531773 | 50531190 | 50531295 | 50531968 | 50532077 | 14 | 3 | [illegible] |
| SE | 41484 | ENSG00000270629 | NBPF14 | chr1 | - | 148583140 | 148583192 | 148577982 | 148578034 | 148585140 | 148585213 | 1 | 0 | [illegible] |
| SE | 41486 | ENSG00000270629 | NBPF14 | chr1 | - | 148584470 | 148584676 | 148583140 | 148583192 | 148587300 | 148587403 | 28 | 34 | [illegible] |
| SE | 41488 | ENSG00000270629 | NBPF14 | chr1 | - | 148585140 | 148585213 | 148584470 | 148584676 | 148587300 | 148587403 | 46 | 45 | [illegible] |
| SE | 41490 | ENSG00000270629 | NBPF14 | chr1 | - | 148587300 | 148587403 | 148585140 | 148585213 | 148589183 | 148589364 | 27 | 35 | [illegible] |
| SE | 41493 | ENSG00000270629 | NBPF14 | chr1 | - | 148598273 | 148598343 | 148595542 | 148595752 | 148598612 | 148598695 | 3 | 1 | [illegible] |
| SE | 41494 | ENSG00000126107 | HECTD3 | chr1 | - | 45004247 | 45004377 | 45004059 | 45004134 | 45004599 | 45004806 | 92 | 38 | [illegible] |
| SE | 41495 | ENSG00000126107 | HECTD3 | chr1 | - | 45005793 | 45005883 | 45004599 | 45004806 | 45005996 | 45006116 | 85 | 41 | [illegible] |
| SE | 41501 | ENSG00000136877 | FPGS | chr9 | + | 127807208 | 127807286 | 127806496 | 127807087 | 127807420 | 127807482 | 68 | 35 | [illegible] |
| SE | 41505 | ENSG00000138674 | SEC31A | chr4 | - | 82821036 | 82821108 | 82818508 | 82819253 | 82824554 | 82824674 | 380 | 241 | [illegible] |
| SE | 41506 | ENSG00000138674 | SEC31A | chr4 | - | 82828999 | 82829058 | 82827368 | 82827632 | 82830936 | 82830975 | 68 | 41 | [illegible] |
| SE | 41508 | ENSG00000138674 | SEC31A | chr4 | - | 82830936 | 82830975 | 82827368 | 82827632 | 82842139 | 82842481 | 0 | 1 | [illegible] |
| SE | 41510 | ENSG00000138674 | SEC31A | chr4 | - | 82830936 | 82830975 | 82828999 | 82829058 | 82842184 | 82842481 | 2 | 0 | [illegible] |
| SE | 41511 | ENSG00000138674 | SEC31A | chr4 | - | 82830936 | 82830975 | 82828999 | 82829058 | 82844385 | 82844460 | 0 | 0 | [illegible] |
| SE | 41513 | ENSG00000138674 | SEC31A | chr4 | - | 82842139 | 82842481 | 82830936 | 82830975 | 82844385 | 82844460 | 44 | 34 | [illegible] |
| SE | 41514 | ENSG00000138674 | SEC31A | chr4 | - | 82842184 | 82842481 | 82828999 | 82829058 | 82844385 | 82844509 | 62 | 35 | [illegible] |
| SE | 41515 | ENSG00000138674 | SEC31A | chr4 | - | 82842184 | 82842481 | 82830936 | 82830975 | 82844385 | 82844509 | 46 | 31 | [illegible] |
| SE | 41517 | ENSG00000138674 | SEC31A | chr4 | - | 82848803 | 82848977 | 82843684 | 82844509 | 82851430 | 82851604 | 310 | 187 | [illegible] |
| SE | 41522 | ENSG00000138674 | SEC31A | chr4 | - | 82870324 | 82870424 | 82867154 | 82867316 | 82871943 | 82872086 | 159 | 99 | [illegible] |
| SE | 41523 | ENSG00000138674 | SEC31A | chr4 | - | 82871943 | 82872086 | 82867154 | 82867316 | 82878729 | 82878928 | 0 | 2 | [illegible] |
| SE | 41524 | ENSG00000138674 | SEC31A | chr4 | - | 82871943 | 82872086 | 82870324 | 82870424 | 82874610 | 82874751 | 209 | 120 | [illegible] |
| SE | 41525 | ENSG00000138674 | SEC31A | chr4 | - | 82874610 | 82874751 | 82871932 | 82872086 | 82878729 | 82878928 | 137 | 81 | [illegible] |
| SE | 41526 | ENSG00000138674 | SEC31A | chr4 | - | 82875726 | 82875822 | 82874610 | 82874751 | 82878729 | 82878928 | 574 | 409 | [illegible] |
| SE | 41527 | ENSG00000138674 | SEC31A | chr4 | - | 82878729 | 82878918 | 82875784 | 82875822 | 82880798 | 82880922 | 372 | 273 | [illegible] |
| SE | 41528 | ENSG00000138674 | SEC31A | chr4 | - | 82878729 | 82878928 | 82875726 | 82875822 | 82880798 | 82880922 | 697 | 533 | [illegible] |
| SE | 41529 | ENSG00000138674 | SEC31A | chr4 | - | 82878857 | 82878928 | 82875726 | 82875822 | 82880798 | 82880922 | 174 | 121 | [illegible] |
| SE | 41531 | ENSG00000138674 | SEC31A | chr4 | - | 82880798 | 82880922 | 82878729 | 82878928 | 82881857 | 82881937 | 392 | 301 | [illegible] |
| SE | 41532 | ENSG00000138674 | SEC31A | chr4 | - | 82880798 | 82880922 | 82878729 | 82878928 | 82891087 | 82891178 | 253 | 204 | [illegible] |
| SE | 41533 | ENSG00000138674 | SEC31A | chr4 | - | 82881857 | 82881937 | 82880798 | 82880922 | 82891087 | 82891178 | 120 | 87 | [illegible] |
| SE | 41534 | ENSG00000138674 | SEC31A | chr4 | - | 82881857 | 82881940 | 82878729 | 82878928 | 82891087 | 82891120 | 19 | 13 | [illegible] |
| SE | 41535 | ENSG00000138674 | SEC31A | chr4 | - | 82881857 | 82881940 | 82880798 | 82880922 | 82891087 | 82891120 | 161 | 113 | [illegible] |
| SE | 41536 | ENSG00000138674 | SEC31A | chr4 | - | 82897988 | 82898062 | 82881856 | 82881940 | 82900076 | 82900223 | 9 | 10 | [illegible] |
| SE | 41537 | ENSG00000138674 | SEC31A | chr4 | - | 82897988 | 82898062 | 82881857 | 82881937 | 82899712 | 82899737 | 6 | 12 | [illegible] |
| SE | 41538 | ENSG00000071127 | WDR1 | chr4 | - | 10075688 | 10077186 | 10075148 | 10075484 | 10077303 | 10077448 | 2 | 1 | [illegible] |
| SE | 41539 | ENSG00000071127 | WDR1 | chr4 | - | 10087706 | 10087935 | 10084442 | 10084530 | 10088292 | 10088373 | 830 | 654 | [illegible] |
| SE | 41540 | ENSG00000071127 | WDR1 | chr4 | - | 10087706 | 10087940 | 10084423 | 10084530 | 10088292 | 10088373 | 1878 | 1290 | [illegible] |
| SE | 41541 | ENSG00000071127 | WDR1 | chr4 | - | 10088292 | 10088373 | 10087705 | 10087940 | 10088663 | 10088741 | 1436 | 1000 | [illegible] |
| SE | 41542 | ENSG00000071127 | WDR1 | chr4 | - | 10088663 | 10088741 | 10088286 | 10088373 | 10097710 | 10097891 | 1068 | 802 | [illegible] |
| SE | 41544 | ENSG00000071127 | WDR1 | chr4 | - | 10103895 | 10103986 | 10098989 | 10099139 | 10116112 | 10116234 | 1760 | 1555 | [illegible] |
| SE | 41545 | ENSG00000071127 | WDR1 | chr4 | - | 10108217 | 10108395 | 10103895 | 10103986 | 10116112 | 10116234 | 5 | 0 | [illegible] |
| SE | 41546 | ENSG00000198682 | PAPSS2 | chr10 | + | 87721755 | 87721770 | 87715731 | 87715843 | 87727283 | 87727489 | 0 | 0 | [illegible] |
| SE | 41547 | ENSG00000142945 | KIF2C | chr1 | + | 44753131 | 44753254 | 44750441 | 44750564 | 44753732 | 44753833 | 1 | 2 | [illegible] |
| SE | 41548 | ENSG00000015171 | ZMYND11 | chr10 | + | 197910 | 198005 | 179993 | 180128 | 209888 | 210048 | 1 | 0 | [illegible] |
| SE | 41549 | ENSG00000015171 | ZMYND11 | chr10 | + | 218454 | 218499 | 209888 | 210048 | 221194 | 221356 | 1 | 1 | [illegible] |
| SE | 41551 | ENSG00000015171 | ZMYND11 | chr10 | + | 237584 | 237677 | 236837 | 236915 | 239437 | 239525 | 181 | 88 | [illegible] |
| SE | 41552 | ENSG00000015171 | ZMYND11 | chr10 | + | 239437 | 239525 | 236837 | 236915 | 240055 | 240111 | 35 | 21 | [illegible] |
| SE | 41553 | ENSG00000015171 | ZMYND11 | chr10 | + | 239437 | 239525 | 236837 | 236915 | 240058 | 240111 | 1 | 0 | [illegible] |
| SE | 41554 | ENSG00000015171 | ZMYND11 | chr10 | + | 240892 | 240970 | 239961 | 240111 | 242020 | 242139 | 184 | 121 | [illegible] |
| SE | 41556 | ENSG00000015171 | ZMYND11 | chr10 | + | 247397 | 247466 | 246765 | 246973 | 248335 | 248531 | 294 | 179 | [illegible] |
| SE | 41558 | ENSG00000126106 | TMEM53 | chr1 | - | 44654939 | 44655209 | 44645359 | 44645464 | 44660173 | 44660295 | 2 | 0 | [illegible] |
| SE | 41559 | ENSG00000126106 | TMEM53 | chr1 | - | 44660173 | 44660295 | 44645359 | 44645464 | 44674330 | 44674402 | 1 | 0 | [illegible] |
| SE | 41562 | ENSG00000184831 | APOO | chrX | - | 23833606 | 23833612 | 23833356 | 23833510 | 23840312 | 23840377 | 1 | 1 | [illegible] |
| SE | 41563 | ENSG00000184831 | APOO | chrX | - | 23840312 | 23840377 | 23833362 | 23833510 | 23856301 | 23856382 | 5 | 7 | [illegible] |
| SE | 41564 | ENSG00000184831 | APOO | chrX | - | 23856301 | 23856382 | 23840026 | 23840377 | 23858641 | 23858733 | 25 | 39 | [illegible] |
| SE | 41565 | ENSG00000184831 | APOO | chrX | - | 23868592 | 23868688 | 23858641 | 23858733 | 23878914 | 23879007 | 9 | 22 | [illegible] |
| SE | 41566 | ENSG00000184831 | APOO | chrX | - | 23874402 | 23874457 | 23868592 | 23868688 | 23878914 | 23879034 | 10 | 19 | [illegible] |
| SE | 41567 | ENSG00000184831 | APOO | chrX | - | 23880844 | 23880952 | 23878914 | 23879034 | 23907693 | 23907903 | 6 | 13 | [illegible] |
| SE | 41570 | ENSG00000084072 | PPIE | chr1 | + | 39741894 | 39741921 | 39741365 | 39741409 | 39743215 | 39743297 | 161 | 103 | [illegible] |
| SE | 41571 | ENSG00000084072 | PPIE | chr1 | + | 39741894 | 39741921 | 39741365 | 39741409 | 39743823 | 39743924 | 23 | 17 | [illegible] |
| SE | 41572 | ENSG00000084072 | PPIE | chr1 | + | 39741894 | 39742031 | 39741365 | 39741409 | 39743215 | 39743297 | 28 | 21 | [illegible] |

| | | | | | | | | | | | | | | | | | | | | | | | | | | | | | | | | | | | | | | | | | | | | | | |
|---|---|---|---|---|---|---|---|---|---|---|---|---|---|---|---|---|---|---|---|---|---|---|---|---|---|---|---|---|---|---|---|---|---|---|---|---|---|---|---|---|---|---|---|---|---|---|
| SE | 41573 | ENSG00000084072 | PPIE | chr1 | + | 39741894 | 39743297 | 39741365 | 39741409 | 39743823 | 39743924 | 173 | 122 | 170 | 151 | 250 | 0 | 2 | 0 | 0 | 0 | 216 | 178 | 169 | 173 | 267 | 0 | 0 | 0 | 0 | 0 | 298 | 149 | 1 | 1 | 1 | 0.968 | 1 | 1 | 1 | 1 | 1 | 1 | 1 | 1 | -0.006 |
| SE | 41574 | ENSG00000084072 | PPIE | chr1 | + | 39743215 | 39743297 | 39741365 | 39741409 | 39743823 | 39743924 | 142 | 95 | 119 | 120 | 197 | 0 | 2 | 0 | 0 | 0 | 168 | 109 | 129 | 141 | 229 | 0 | 0 | 0 | 0 | 0 | 231 | 149 | 1 | 1 | 1 | 0.968 | 1 | 1 | 1 | 1 | 1 | 1 | 1 | 1 | -0.006 |
| SE | 41575 | ENSG00000084072 | PPIE | chr1 | + | 39743215 | 39743297 | 39741894 | 39741921 | 39743823 | 39743924 | 281 | 183 | 191 | 237 | 432 | 0 | 0 | 1 | 0 | 0 | 306 | 189 | 221 | 271 | 392 | 0 | 3 | 0 | 0 | 0 | 231 | 149 | 1 | 1 | 1 | 1 | 0.992 | 1 | 1 | 1 | 0.976 | 1 | 1 | 1 | 0.003 |
| SE | 41579 | ENSG00000084072 | PPIE | chr1 | + | 39748902 | 39749088 | 39745374 | 39745498 | 39752909 | 39753052 | 241 | 115 | 237 | 232 | 295 | 0 | 0 | 0 | 1 | 0 | 247 | 130 | 111 | 144 | 361 | 0 | 0 | 0 | 0 | 0 | 298 | 149 | 1 | 1 | 1 | 1 | 1 | 0.991 | 1 | 1 | 1 | 1 | 1 | 1 | -0.002 |
| SE | 41581 | ENSG00000084072 | PPIE | chr1 | + | 39762317 | 39762407 | 39761198 | 39761373 | 39762516 | 39762651 | 3 | 0 | 1 | 0 | 0 | 0 | 0 | 0 | 4 | 0 | 11 | 5 | 3 | 0 | 0 | 0 | 0 | 1 | 0 | 0 | 239 | 149 | 1 | 1 | 1 | NA | 1 | 0 | NA | 1 | 1 | 0.652 | NA | NA | -0.217 |
| SE | 41583 | ENSG00000118308 | IRAG2 | chr12 | + | 25052793 | 25052956 | 25052234 | 25052349 | 25061591 | 25061653 | 106 | 99 | 91 | 105 | 159 | 0 | 1 | 0 | 0 | 0 | 43 | 83 | 73 | 42 | 25 | 0 | 0 | 0 | 0 | 0 | 298 | 149 | 1 | 1 | 1 | 0.98 | 1 | 1 | 1 | 1 | 1 | 1 | 1 | 1 | -0.004 |
| SE | 41586 | ENSG00000118308 | IRAG2 | chr12 | + | 25062819 | 25062900 | 25052270 | 25052956 | 25063722 | 25063816 | 122 | 104 | 82 | 120 | 213 | 1 | 0 | 0 | 0 | 0 | 81 | 74 | 85 | 53 | 47 | 0 | 0 | 0 | 0 | 0 | 230 | 149 | 1 | 1 | 0.988 | 1 | 1 | 1 | 1 | 1 | 1 | 1 | 1 | 1 | -0.002 |
| SE | 41587 | ENSG00000118308 | IRAG2 | chr12 | + | 25062819 | 25062900 | 25052270 | 25052956 | 25066364 | 25066409 | 92 | 71 | 59 | 91 | 131 | 0 | 0 | 0 | 2 | 0 | 51 | 62 | 54 | 36 | 20 | 0 | 0 | 0 | 0 | 0 | 230 | 149 | 1 | 1 | 1 | 1 | 1 | 0.967 | 1 | 1 | 1 | 1 | 1 | 1 | -0.007 |
| SE | 41588 | ENSG00000118308 | IRAG2 | chr12 | + | 25063719 | 25063816 | 25052739 | 25052956 | 25066364 | 25066409 | 108 | 95 | 120 | 135 | 230 | 0 | 0 | 0 | 2 | 0 | 52 | 106 | 87 | 47 | 32 | 0 | 0 | 0 | 0 | 0 | 246 | 149 | 1 | 1 | 1 | 1 | 1 | 0.976 | 1 | 1 | 1 | 1 | 1 | 1 | -0.005 |
| SE | 41589 | ENSG00000118308 | IRAG2 | chr12 | + | 25063719 | 25063816 | 25052739 | 25052956 | 25069349 | 25069431 | 17 | 14 | 14 | 17 | 8 | 0 | 0 | 0 | 1 | 1 | 0 | 7 | 8 | 0 | 0 | 0 | 0 | 0 | 0 | 0 | 246 | 149 | 1 | 1 | 1 | 1 | 1 | 0.911 | 0.829 | NA | 1 | 1 | NA | NA | -0.052 |
| SE | 41590 | ENSG00000118308 | IRAG2 | chr12 | + | 25063719 | 25063816 | 25052739 | 25052956 | 25079243 | 25079290 | 10 | 10 | 9 | 12 | 2 | 0 | 2 | 0 | 5 | 2 | 0 | 2 | 1 | 0 | 0 | 0 | 0 | 0 | 0 | 0 | 246 | 149 | 1 | 1 | 1 | 0.752 | 1 | 0.592 | 0.377 | NA | 1 | 1 | NA | NA | -0.256 |
| SE | 41591 | ENSG00000118308 | IRAG2 | chr12 | + | 25063719 | 25063816 | 25062819 | 25062900 | 25066364 | 25066409 | 348 | 284 | 308 | 357 | 593 | 1 | 0 | 0 | 0 | 0 | 163 | 288 | 243 | 172 | 70 | 0 | 0 | 0 | 0 | 0 | 246 | 149 | 1 | 1 | 0.995 | 1 | 1 | 1 | 1 | 1 | 1 | 1 | 1 | 1 | -0.001 |
| SE | 41592 | ENSG00000118308 | IRAG2 | chr12 | + | 25063722 | 25063816 | 25052641 | 25052956 | 25066364 | 25066512 | 96 | 85 | 109 | 125 | 218 | 0 | 0 | 0 | 2 | 0 | 52 | 99 | 81 | 44 | 32 | 0 | 0 | 0 | 0 | 0 | 243 | 149 | 1 | 1 | 1 | 1 | 1 | 0.975 | 1 | 1 | 1 | 1 | 1 | 1 | -0.005 |
| SE | 41593 | ENSG00000118308 | IRAG2 | chr12 | + | 25063722 | 25063816 | 25052641 | 25052956 | 25069349 | 25069431 | 5 | 4 | 5 | 7 | 6 | 0 | 0 | 0 | 1 | 1 | 0 | 5 | 7 | 0 | 0 | 0 | 0 | 0 | 0 | 0 | 243 | 149 | 1 | 1 | 1 | 1 | 1 | 0.811 | 0.786 | NA | 1 | 1 | NA | NA | -0.081 |
| SE | 41595 | ENSG00000118308 | IRAG2 | chr12 | + | 25063722 | 25063816 | 25062819 | 25062900 | 25066364 | 25066512 | 147 | 128 | 145 | 167 | 322 | 1 | 0 | 0 | 0 | 0 | 95 | 131 | 129 | 79 | 63 | 0 | 0 | 0 | 0 | 0 | 243 | 149 | 1 | 1 | 0.989 | 1 | 1 | 1 | 1 | 1 | 1 | 1 | 1 | 1 | -0.002 |
| SE | 41596 | ENSG00000118308 | IRAG2 | chr12 | + | 25066364 | 25066512 | 25052270 | 25052956 | 25069349 | 25069431 | 146 | 104 | 100 | 142 | 144 | 0 | 0 | 0 | 1 | 1 | 78 | 70 | 124 | 79 | 45 | 0 | 0 | 0 | 0 | 0 | 297 | 149 | 1 | 1 | 1 | 1 | 1 | 0.986 | 0.986 | 1 | 1 | 1 | 1 | 1 | -0.006 |
| SE | 41597 | ENSG00000118308 | IRAG2 | chr12 | + | 25066364 | 25066512 | 25052270 | 25052956 | 25079243 | 25079290 | 2 | 1 | 0 | 2 | 6 | 0 | 2 | 0 | 5 | 2 | 0 | 7 | 4 | 0 | 0 | 0 | 0 | 0 | 0 | 0 | 297 | 149 | 1 | 1 | 1 | 0.201 | NA | 0.167 | 0.601 | NA | 1 | 1 | NA | NA | -0.508 |
| SE | 41599 | ENSG00000118308 | IRAG2 | chr12 | + | 25066364 | 25066512 | 25063719 | 25063816 | 25079243 | 25079290 | 146 | 133 | 160 | 159 | 330 | 0 | 0 | 0 | 2 | 0 | 74 | 135 | 134 | 68 | 40 | 0 | 0 | 0 | 0 | 0 | 297 | 149 | 1 | 1 | 1 | 1 | 1 | 0.976 | 1 | 1 | 1 | 1 | 1 | 1 | -0.005 |
| SE | 41601 | ENSG00000118308 | IRAG2 | chr12 | + | 25069349 | 25069431 | 25063719 | 25063816 | 25079243 | 25079290 | 72 | 96 | 96 | 104 | 148 | 0 | 0 | 0 | 2 | 0 | 11 | 75 | 65 | 49 | 11 | 0 | 0 | 0 | 0 | 0 | 231 | 149 | 1 | 1 | 1 | 1 | 1 | 0.971 | 1 | 1 | 1 | 1 | 1 | 1 | -0.006 |
| SE | 41603 | ENSG00000118308 | IRAG2 | chr12 | + | 25088099 | 25088157 | 25083422 | 25083493 | 25089613 | 25089677 | 280 | 197 | 294 | 313 | 469 | 1 | 3 | 0 | 1 | 0 | 92 | 213 | 308 | 129 | 48 | 0 | 0 | 0 | 0 | 0 | 207 | 149 | 1 | 1 | 0.995 | 0.979 | 1 | 0.996 | 1 | 1 | 1 | 1 | 1 | 1 | -0.006 |
| [illegible] | | | | | | | | | | | | | | | | | | | | | | | | | | | | | | | | | | | | | | | | | | | | | | |

| | | | | | | | | | | | | | | | | | | | | | |
|---|---|---|---|---|---|---|---|---|---|---|---|---|---|---|---|---|---|---|---|---|---|
| SE | 42073 | ENSG00000136636 | KCTD3 | chr1 | + | 215579908 | 215579999 | 215578999 | 215579137 | 215586494 | 215586685 | 32 | 22 | 24 | 29 | 36 | 0 | 0 | 0 | 1 | 0 |
| SE | 42074 | ENSG00000136636 | KCTD3 | chr1 | + | 215618087 | 215618154 | 215611824 | 215611921 | 215618885 | 215618951 | 0 | 0 | 0 | 0 | 2 | 15 | 14 | 11 | 10 | 28 |
| SE | 42076 | ENSG00000266094 | RASSF5 | chr1 | + | 206584386 | 206584684 | 206583119 | 206583379 | 206586825 | 206586842 | 812 | 412 | 577 | 651 | 965 | 0 | 2 | 0 | 1 | 0 |
| SE | 42080 | ENSG00000026751 | SLAMF7 | chr1 | + | 160748193 | 160748514 | 160739056 | 160739356 | 160751344 | 160751448 | 183 | 71 | 136 | 136 | 71 | 0 | 0 | 2 | 0 | 0 |
| SE | 42082 | ENSG00000026751 | SLAMF7 | chr1 | + | 160749820 | 160750093 | 160739056 | 160739356 | 160751344 | 160751448 | 26 | 8 | 8 | 27 | 15 | 0 | 0 | 2 | 0 | 0 |
| SE | 42084 | ENSG00000026751 | SLAMF7 | chr1 | + | 160749820 | 160750093 | 160748193 | 160748514 | 160750303 | 160750423 | 242 | 145 | 219 | 194 | 151 | 0 | 0 | 0 | 0 | 3 |
| SE | 42086 | ENSG00000026751 | SLAMF7 | chr1 | + | 160749820 | 160750093 | 160748193 | 160748514 | 160752185 | 160752246 | 105 | 64 | 91 | 86 | 60 | 0 | 0 | 1 | 5 | 0 |
| SE | 42089 | ENSG00000026751 | SLAMF7 | chr1 | + | 160750303 | 160750423 | 160748193 | 160748514 | 160752185 | 160752248 | 26 | 13 | 17 | 39 | 30 | 0 | 0 | 1 | 5 | 0 |
| SE | 42092 | ENSG00000026751 | SLAMF7 | chr1 | + | 160750303 | 160750423 | 160749820 | 160750093 | 160753105 | 160754814 | 135 | 74 | 126 | 105 | 84 | 2 | 0 | 0 | 0 | 0 |
| SE | 42093 | ENSG00000026751 | SLAMF7 | chr1 | + | 160750303 | 160750423 | 160749820 | 160750132 | 160752185 | 160752248 | 29 | 13 | 20 | 40 | 28 | 1 | 0 | 0 | 0 | 0 |
| SE | 42095 | ENSG00000026751 | SLAMF7 | chr1 | + | 160751344 | 160751448 | 160748193 | 160748514 | 160752185 | 160752248 | 150 | 86 | 119 | 123 | 169 | 0 | 0 | 1 | 5 | 0 |
| SE | 42098 | ENSG00000026751 | SLAMF7 | chr1 | + | 160752185 | 160752248 | 160749820 | 160750093 | 160753105 | 160754814 | 173 | 99 | 137 | 176 | 171 | 2 | 0 | 0 | 0 | 0 |
| SE | 42101 | ENSG00000130413 | STK33 | chr11 | - | 8440677 | 8440753 | 8436026 | 8436139 | 8452821 | 8452906 | 6 | 5 | 12 | 4 | 11 | 2 | 0 | 0 | 1 | 2 |
| SE | 42112 | ENSG00000163956 | LRPAP1 | chr4 | - | 3516386 | 3516718 | 3516115 | 3516198 | 3518033 | 3518192 | 1 | 0 | 0 | 1 | 0 | 170 | 84 | 158 | 167 | 260 |
| SE | 42113 | ENSG00000163956 | LRPAP1 | chr4 | - | 3518033 | 3518192 | 3515881 | 3516198 | 3518870 | 3518991 | 544 | 292 | 474 | 514 | 662 | 0 | 0 | 0 | 0 | 0 |
| SE | 42117 | ENSG00000083812 | ZNF324 | chr19 | + | 58469179 | 58469306 | 58467044 | 58467183 | 58469727 | 58469844 | 39 | 12 | 27 | 43 | 13 | 1 | 0 | 0 | 0 | 0 |
| SE | 42120 | ENSG00000106400 | ZNHIT1 | chr7 | + | 101222603 | 101222774 | 101218164 | 101218217 | 101223476 | 101223556 | 297 | 179 | 350 | 332 | 434 | 0 | 0 | 0 | 1 | 0 |
| SE | 42122 | ENSG00000162804 | SNED1 | chr2 | + | 241087391 | 241087475 | 241082276 | 241082364 | 241088364 | 241088402 | 11 | 3 | 11 | 5 | 1 | 0 | 0 | 0 | 0 | 2 |
| SE | 42124 | ENSG00000117450 | PRDX1 | chr1 | - | 45518937 | 45519054 | 45515653 | 45515807 | 45521828 | 45521884 | 450 | 300 | 428 | 478 | 501 | 0 | 0 | 1 | 0 | 0 |
| SE | 42126 | ENSG00000198952 | SMG5 | chr1 | - | 156251402 | 156251477 | 156250857 | 156250996 | 156252413 | 156252504 | 93 | 53 | 81 | 78 | 77 | 1 | 0 | 0 | 0 | 4 |
| SE | 42127 | ENSG00000198952 | SMG5 | chr1 | - | 156259004 | 156259163 | 156253448 | 156253508 | 156260450 | 156260626 | 101 | 27 | 70 | 74 | 44 | 2 | 0 | 1 | 1 | 0 |
| SE | 42128 | ENSG00000198952 | SMG5 | chr1 | - | 156261332 | 156261408 | 156260450 | 156260626 | 156263394 | 156263570 | 72 | 32 | 74 | 93 | 38 | 0 | 0 | 0 | 0 | 0 |
| SE | 42132 | ENSG00000198952 | SMG5 | chr1 | - | 156273360 | 156273450 | 156272319 | 156272398 | 156274596 | 156274686 | 176 | 118 | 179 | 220 | 181 | 1 | 0 | 0 | 0 | 0 |
| SE | 42133 | ENSG00000198952 | SMG5 | chr1 | - | 156274596 | 156274686 | 156273360 | 156273450 | 156277084 | 156277241 | 162 | 88 | 153 | 193 | 172 | 0 | 0 | 0 | 0 | 0 |
| SE | 42143 | ENSG00000005249 | PRKAR2B | chr7 | + | 107128211 | 107128295 | 107121951 | 107122004 | 107140846 | 107140953 | 551 | 1035 | 165 | 1543 | 836 | 2 | 7 | 2 | 8 | 0 |
| SE | 42144 | ENSG00000005249 | PRKAR2B | chr7 | + | 107156983 | 107157049 | 107153176 | 107153251 | 107159448 | 107161811 | 182 | 336 | 55 | 519 | 696 | 0 | 0 | 0 | 1 | 0 |
| SE | 42145 | ENSG00000005249 | PRKAR2B | chr7 | + | 107157185 | 107157324 | 107156983 | 107157049 | 107159448 | 107161811 | 676 | 1393 | 297 | 1973 | 3626 | 3 | 1 | 4 | 8 | 2 |
| SE | 42147 | ENSG00000163472 | TMEM79 | chr1 | + | 156285183 | 156285267 | 156282934 | 156283065 | 156286259 | 156286473 | 9 | 4 | 2 | 3 | 4 | 0 | 0 | 1 | 0 | 0 |
| SE | 42148 | ENSG00000163472 | TMEM79 | chr1 | + | 156285183 | 156285983 | 156282947 | 156283065 | 156286259 | 156286473 | 15 | 5 | 6 | 7 | 8 | 0 | 0 | 1 | 0 | 0 |
| SE | 42149 | ENSG00000163472 | TMEM79 | chr1 | + | 156285183 | 156285983 | 156282947 | 156283065 | 156291384 | 156292443 | 14 | 5 | 4 | 7 | 12 | 1 | 0 | 0 | 0 | 3 |
| SE | 42151 | ENSG00000163472 | TMEM79 | chr1 | + | 156286259 | 156286473 | 156282934 | 156283065 | 156291384 | 156291525 | 4 | 0 | 7 | 2 | 0 | 1 | 0 | 1 | 0 | 3 |
| SE | 42152 | ENSG00000163472 | TMEM79 | chr1 | + | 156286259 | 156286473 | 156285183 | 156285983 | 156291384 | 156291525 | 5 | 0 | 8 | 2 | 0 | 0 | 0 | 0 | 0 | 4 |
| SE | 42157 | ENSG00000197343 | ZNF655 | chr7 | + | 99562348 | 99562494 | 99561887 | 99561973 | 99571687 | 99571717 | 27 | 10 | 35 | 25 | 44 | 1 | 1 | 6 | 3 | 6 |
| SE | 42160 | ENSG00000197343 | ZNF655 | chr7 | + | 99562367 | 99562494 | 99560532 | 99560695 | 99572244 | 99576453 | 9 | 6 | 7 | 7 | 9 | 14 | 5 | 8 | 2 | 12 |
| SE | 42161 | ENSG00000197343 | ZNF655 | chr7 | + | 99562367 | 99562494 | 99561887 | 99561973 | 99571259 | 99571571 | 87 | 51 | 61 | 63 | 78 | 0 | 0 | 0 | 0 | 0 |
| SE | 42165 | ENSG00000197343 | ZNF655 | chr7 | + | 99571259 | 99571379 | 99560532 | 99560695 | 99572244 | 99572318 | 3 | 0 | 0 | 0 | 0 | 14 | 5 | 8 | 2 | 12 |
| SE | 42166 | ENSG00000197343 | ZNF655 | chr7 | + | 99571687 | 99571792 | 99560532 | 99560695 | 99572244 | 99572318 | 75 | 46 | 53 | 60 | 108 | 14 | 5 | 8 | 2 | 12 |
| SE | 42167 | ENSG00000197343 | ZNF655 | chr7 | + | 99571687 | 99571792 | 99571259 | 99571379 | 99572244 | 99572318 | 67 | 44 | 49 | 57 | 103 | 3 | 0 | 0 | 0 | 0 |
| SE | 42168 | ENSG00000138780 | GSTCD | chr4 | + | 105823230 | 105823275 | 105822953 | 105823069 | 105825671 | 105825800 | 4 | 8 | 12 | 8 | 26 | 0 | 1 | 0 | 0 | 0 |
| SE | 42169 | ENSG00000138780 | GSTCD | chr4 | + | 105834460 | 105834594 | 105825671 | 105825800 | 105837858 | 105837889 | 16 | 4 | 16 | 7 | 33 | 0 | 0 | 1 | 0 | 0 |
| SE | 42172 | ENSG00000255733 | IFNG-AS1 | chr12 | + | 67995940 | 67996073 | 67989444 | 67989579 | 68006089 | 68006146 | 67 | 128 | 134 | 113 | 128 | 7 | 7 | 1 | 2 | 2 |
| SE | 42173 | ENSG00000255733 | IFNG-AS1 | chr12 | + | 67995940 | 67996073 | 67989444 | 67989579 | 68019861 | 68019898 | 42 | 61 | 66 | 56 | 72 | 0 | 0 | 1 | 0 | 0 |
| SE | 42174 | ENSG00000255733 | IFNG-AS1 | chr12 | + | 68006089 | 68006146 | 67989444 | 67989579 | 68019861 | 68019898 | 18 | 41 | 23 | 31 | 13 | 0 | 0 | 1 | 0 | 0 |
| SE | 42177 | ENSG00000255733 | IFNG-AS1 | chr12 | + | 68026650 | 68026708 | 68019861 | 68019898 | 68034783 | 68037094 | 0 | 0 | 0 | 0 | 1 | 4 | 6 | 5 | 0 | 7 |
| SE | 42178 | ENSG00000134318 | ROCK2 | chr2 | - | 11207725 | 11207910 | 11202051 | 11202121 | 11208286 | 11208447 | 237 | 310 | 177 | 282 | 294 | 0 | 0 | 0 | 0 | 0 |
| SE | 42179 | ENSG00000134318 | ROCK2 | chr2 | - | 11215509 | 11215645 | 11215320 | 11215412 | 11216157 | 11216206 | 79 | 113 | 66 | 135 | 168 | 0 | 0 | 0 | 0 | 1 |
| SE | 42183 | ENSG00000134318 | ROCK2 | chr2 | - | 11286538 | 11286639 | 11249660 | 11249798 | 11287654 | 11287736 | 116 | 91 | 45 | 80 | 46 | 2 | 0 | 0 | 0 | 0 |
| SE | 42184 | ENSG00000163818 | LZTFL1 | chr3 | - | 45827355 | 45827459 | 45823315 | 45826332 | 45828438 | 45828615 | 97 | 37 | 107 | 86 | 167 | 1 | 1 | 1 | 6 | 10 |
| SE | 42186 | ENSG00000163818 | LZTFL1 | chr3 | - | 45837926 | 45838047 | 45835589 | 45835784 | 45841988 | 45842116 | 53 | 21 | 72 | 40 | 127 | 2 | 3 | 6 | 5 | 9 |
| SE | 42187 | ENSG00000163818 | LZTFL1 | chr3 | - | 45837926 | 45838051 | 45834237 | 45835784 | 45841988 | 45842119 | 66 | 26 | 87 | 56 | 134 | 2 | 3 | 6 | 5 | 9 |
| SE | 42188 | ENSG00000163818 | LZTFL1 | chr3 | - | 45837926 | 45838051 | 45834237 | 45835784 | 45854985 | 45855074 | 58 | 23 | 74 | 40 | 131 | 1 | 0 | 0 | 1 | 0 |
| SE | 42189 | ENSG00000163818 | LZTFL1 | chr3 | - | 45854985 | 45855074 | 45837926 | 45838051 | 45913119 | 45913177 | 7 | 2 | 6 | 0 | 3 | 0 | 0 | 0 | 0 | 2 |
| SE | 42190 | ENSG00000163818 | LZTFL1 | chr3 | - | 45858939 | 45859016 | 45854985 | 45855074 | 45913119 | 45913177 | 0 | 0 | 2 | 0 | 0 | 6 | 1 | 2 | 0 | 3 |
| SE | 42192 | ENSG00000164241 | C5orf63 | chr5 | - | 127056396 | 127056506 | 127052612 | 127052669 | 127058881 | 127059002 | 8 | 10 | 9 | 23 | 25 | 15 | 12 | 23 | 12 | 15 |
| SE | 42203 | ENSG00000132155 | RAF1 | chr3 | - | 12584846 | 12584981 | 12583600 | 12584657 | 12585121 | 12585253 | 295 | 184 | 259 | 267 | 309 | 0 | 0 | 0 | 0 | 0 |
| SE | 42204 | ENSG00000132155 | RAF1 | chr3 | - | 12590797 | 12590974 | 12586846 | 12587637 | 12591707 | 12591792 | 253 | 165 | 284 | 247 | 305 | 0 | 0 | 1 | 0 | 0 |
| SE | 42205 | ENSG00000132155 | RAF1 | chr3 | - | 12591707 | 12591792 | 12586846 | 12587637 | 12599690 | 12599755 | 108 | 55 | 93 | 101 | 147 | 0 | 0 | 0 | 0 | 0 |
| SE | 42209 | ENSG00000132155 | RAF1 | chr3 | - | 12604135 | 12604289 | 12600198 | 12600415 | 12608765 | 12608923 | 402 | 275 | 412 | 353 | 554 | 1 | 0 | 0 | 0 | 0 |
| SE | 42212 | ENSG00000116266 | STXBP3 | chr1 | + | 108779285 | 108779410 | 108776332 | 108776423 | 108782421 | 108782517 | 376 | 293 | 250 | 356 | 695 | 0 | 0 | 3 | 2 | 0 |
| SE | 42214 | ENSG00000116266 | STXBP3 | chr1 | + | 108800219 | 108800305 | 108798144 | 108798237 | 108807400 | 108807481 | 350 | 314 | 253 | 379 | 390 | 2 | 0 | 0 | 0 | 0 |
| SE | 42217 | ENSG00000123612 | ACVR1C | chr2 | - | 157556092 | 157556332 | 157550161 | 157550392 | 157587186 | 157587417 | 8 | 4 | 7 | 4 | 9 | 1 | 1 | 1 | 2 | 0 |
| SE | 42232 | ENSG00000116663 | FBXO6 | chr1 | + | 11671265 | 11671392 | 11668655 | 11668944 | 11671927 | 11672023 | 99 | 59 | 76 | 80 | 172 | 0 | 1 | 0 | 0 | 0 |
| SE | 42235 | ENSG00000159214 | CCDC24 | chr1 | + | 43992522 | 43992639 | 43991801 | 43992387 | 43993803 | 43993964 | 1 | 1 | 0 | 0 | 0 | 0 | 0 | 0 | 0 | 0 |
| SE | 42236 | ENSG00000159214 | CCDC24 | chr1 | + | 43995107 | 43995162 | 43993803 | 43993964 | 43995600 | 43995670 | 3 | 4 | 9 | 3 | 10 | 0 | 0 | 0 | 0 | 0 |
| SE | 42242 | ENSG00000258465 | NA | chr1 | - | 160243959 | 160244034 | 160239696 | 160240370 | 160263264 | 160263462 | 203 | 177 | 157 | 209 | 144 | 2 | 0 | 0 | 2 | 0 |
| SE | 42243 | ENSG00000258465 | NA | chr1 | - | 160243959 | 160244034 | 160239696 | 160240370 | 160279800 | 160279845 | 203 | 177 | 156 | 205 | 148 | 3 | 0 | 0 | 0 | 0 |
| SE | 42244 | ENSG00000258465 | NA | chr1 | - | 160263264 | 160263462 | 160240141 | 160240370 | 160279800 | 160279845 | 2 | 0 | 1 | 2 | 0 | 3 | 0 | 0 | 0 | 0 |
| SE | 42246 | ENSG00000258465 | NA | chr1 | - | 160263264 | 160263462 | 160243959 | 160244034 | 160279800 | 160279845 | 0 | 0 | 2 | 4 | 0 | 0 | 0 | 0 | 0 | 4 |
| SE | 42247 | ENSG00000258465 | NA | chr1 | - | 160278130 | 160278191 | 160243959 | 160244034 | 160279800 | 160279845 | 1 | 5 | 0 | 0 | 0 | 0 | 0 | 0 | 0 | 4 |
| SE | 42248 | ENSG00000258465 | NA | chr1 | - | 160278130 | 160278191 | 160263264 | 160263462 | 160279800 | 160279845 | 2 | 5 | 0 | 0 | 0 | 0 | 0 | 0 | 1 | 0 |
| SE | 42255 | ENSG00000116001 | TIA1 | chr2 | - | 70215370 | 70215494 | 70214348 | 70214491 | 70216207 | 70216292 | 264 | 160 | 268 | 286 | 374 | 0 | 0 | 3 | 0 | 0 |
| SE | 42256 | ENSG00000116001 | TIA1 | chr2 | - | 70215370 | 70215494 | 70214348 | 70214494 | 70216207 | 70216292 | 397 | 278 | 355 | 401 | 554 | 0 | 0 | 0 | 0 | 0 |
| SE | 42257 | ENSG00000116001 | TIA1 | chr2 | - | 70216207 | 70216292 | 70215370 | 70215494 | 70216403 | 70216499 | 418 | 256 | 360 | 408 | 580 | 2 | 0 | 0 | 0 | 2 |
| SE | 42258 | ENSG00000116001 | TIA1 | chr2 | - | 70216752 | 70216994 | 70216403 | 70216499 | 70224553 | 70224629 | 179 | 149 | 193 | 194 | 246 | 0 | 0 | 0 | 1 | 1 |
| SE | 42259 | ENSG00000116001 | TIA1 | chr2 | - | 70216885 | 70216994 | 70216403 | 70216499 | 70224553 | 70224592 | 237 | 205 | 267 | 293 | 364 | 0 | 0 | 0 | 1 | 1 |
| SE | 42265 | ENSG00000116001 | TIA1 | chr2 | - | 70229058 | 70229091 | 70228343 | 70228462 | 70229263 | 70229318 | 77 | 45 | 55 | 80 | 120 | 0 | 0 | 0 | 0 | 2 |
| SE | 42266 | ENSG00000116001 | TIA1 | chr2 | - | 70236072 | 70236175 | 70230755 | 70230854 | 70248404 | 70248622 | 44 | 27 | 35 | 50 | 47 | 0 | 3 | 0 | 2 | 0 |
| SE | 42267 | ENSG00000116001 | TIA1 | chr2 | - | 70236078 | 70236175 | 70230755 | 70230854 | 70248404 | 70248454 | 276 | 206 | 224 | 261 | 353 | 0 | 3 | 0 | 2 | 0 |
| SE | 42269 | ENSG00000160752 | FDPS | chr1 | + | 155309752 | 155309965 | 155308958 | 155309076 | 155310042 | 155310205 | 7 | 0 | 7 | 7 | 0 | 3 | 0 | 1 | 1 | 0 |
| SE | 42272 | ENSG00000160752 | FDPS | chr1 | + | 155309788 | 155309965 | 155308915 | 155309076 | 155310042 | 155310205 | 7 | 1 | 7 | 7 | 0 | 3 | 0 | 1 | 1 | 0 |
| SE | 42273 | ENSG00000160752 | FDPS | chr1 | + | 155309788 | 155309965 | 155308915 | 155309076 | 155310063 | 155310205 | 1 | 1 | 0 | 0 | 1 | 3 | 0 | 2 | 3 | 0 |
| SE | 42277 | ENSG00000160752 | FDPS | chr1 | + | 155312254 | 155312395 | 155310042 | 155310205 | 155317940 | 155317982 | 212 | 89 | 198 | 179 | 102 | 0 | 0 | 0 | 0 | 0 |
| SE | 42286 | ENSG00000156398 | SFXN2 | chr10 | + | 102715222 | 102715328 | 102714635 | 102714681 | 102726611 | 102726797 | 0 | 0 | 0 | 0 | 0 | 5 | 6 | 6 | 0 | 0 |
| SE | 42290 | ENSG00000156398 | SFXN2 | chr10 | + | 102716613 | 102716711 | 102714537 | 102714681 | 102726611 | 102726777 | 0 | 0 | 1 | 0 | 0 | 5 | 6 | 6 | 0 | 0 |
| SE | 42292 | ENSG00000156398 | SFXN2 | chr10 | + | 102717702 | 102717780 | 102714635 | 102714681 | 102726611 | 102726657 | 0 | 0 | 4 | 2 | 0 | 5 | 6 | 6 | 0 | 0 |
| SE | 42309 | ENSG00000185842 | DNAH14 | chr1 | + | 224960152 | 224960302 | 224954958 | 224955098 | 224964478 | 224964609 | 25 | 18 | 6 | 23 | 44 | 0 | 0 | 0 | 1 | 0 |
| SE | 42310 | ENSG00000185842 | DNAH14 | chr1 | + | 224974090 | 224974153 | 224968758 | 224968874 | 225002782 | 225002927 | 21 | 9 | 5 | 17 | 42 | 0 | 0 | 0 | 0 | 0 |
| SE | 42311 | ENSG00000185842 | DNAH14 | chr1 | + | 225002782 | 225002927 | 224974090 | 224974153 | 225007412 | 225007544 | 13 | 8 | 2 | 23 | 31 | 3 | 0 | 0 | 0 | 0 |
| SE | 42312 | ENSG00000185842 | DNAH14 | chr1 | + | 225023614 | 225023865 | 225007412 | 225007544 | 225038693 | 225038823 | 0 | 4 | 5 | 4 | 6 | 1 | 0 | 0 | 1 | 0 |
| SE | 42314 | ENSG00000185842 | DNAH14 | chr1 | + | 225043885 | 225043983 | 225043743 | 225043789 | 225050209 | 225050376 | 7 | 1 | 0 | 1 | 6 | 1 | 0 | 1 | 0 | 0 |
| SE | 42315 | ENSG00000185842 | DNAH14 | chr1 | + | 225043885 | 225043983 | 225043743 | 225043789 | 225051450 | 225051795 | 9 | 6 | 0 | 1 | 3 | 0 | 0 | 0 | 1 | 1 |
| SE | 42316 | ENSG00000185842 | DNAH14 | chr1 | + | 225050209 | 225050376 | 225043743 | 225043789 | 225051450 | 225051684 | 3 | 1 | 2 | 0 | 1 | 0 | 0 | 0 | 1 | 1 |
| SE | 42317 | ENSG00000185842 | DNAH14 | chr1 | + | 225050209 | 225050376 | 225043885 | 225043983 | 225051450 | 225051684 | 2 | 2 | 1 | 0 | 4 | 2 | 6 | 0 | 0 | 0 |
| SE | 42318 | ENSG00000185842 | DNAH14 | chr1 | + | 225080378 | 225080748 | 225079206 | 225079548 | 225082548 | 225082739 | 0 | 0 | 0 | 3 | 6 | 0 | 0 | 0 | 1 | 0 |
| SE | 42319 | ENSG00000185842 | DNAH14 | chr1 | + | 225100712 | 225100884 | 225097117 | 225097239 | 225117683 | 225117788 | 1 | 1 | 0 | 0 | 0 | 1 | 1 | 0 | 0 | 0 |
| SE | 42325 | ENSG00000185842 | DNAH14 | chr1 | + | 225240592 | 225240822 | 225185290 | 225185425 | 225252300 | 225252417 | 1 | 0 | 0 | 0 | 0 | 3 | 0 | 0 | 0 | 0 |
| SE | 42327 | ENSG00000185842 | DNAH14 | chr1 | + | 225252300 | 225252417 | 225240592 | 225240822 | 225257959 | 225258118 | 0 | 0 | 0 | 0 | 1 | 1 | 0 | 0 | 0 | 0 |
| SE | 42328 | ENSG00000185842 | DNAH14 | chr1 | + | 225257959 | 225258118 | 225252300 | 225252417 | 225259120 | 225259253 | 0 | 0 | 0 | 0 | 1 | 4 | 0 | 0 | 0 | 0 |
| SE | 42336 | ENSG00000215271 | HOMEZ | chr14 | - | 23285291 | 23285611 | 23284990 | 23285028 | 23285912 | 23286117 | 0 | 0 | 0 | 0 | 2 | 2 | 0 | 0 | 0 | 0 |
| SE | 42338 | ENSG00000070759 | TESK2 | chr1 | - | 45345876 | 45345994 | 45343882 | 45345558 | 45346978 | 45347062 | 17 | 10 | 13 | 5 | 12 | 0 | 0 | 0 | 0 | 0 |
| SE | 42342 | ENSG00000070759 | TESK2 | chr1 | - | 45347608 | 45347693 | 45346978 | 45347062 | 45347917 | 45348000 | 11 | 11 | 23 | 22 | 24 | 0 | 0 | 1 | 0 | 0 |
| SE | 42343 | ENSG00000070759 | TESK2 | chr1 | - | 45355302 | 45355449 | 45347917 | 45348000 | 45385911 | 45385960 | 17 | 11 | 20 | 24 | 21 | 1 | 0 | 0 | 0 | 0 |
| SE | 42347 | ENSG00000132773 | TOE1 | chr1 | + | 45341072 | 45341215 | 45340169 | 45340304 | 45341302 | 45341343 | 16 | 10 | 17 | 11 | 24 | 0 | 2 | 0 | 0 | 0 |
| SE | 42348 | ENSG00000132773 | TOE1 | chr1 | + | 45341072 | 45341215 | 45340220 | 45340542 | 45341302 | 45341343 | 14 | 8 | 15 | 9 | 23 | 0 | 0 | 0 | 0 | 0 |
| SE | 42350 | ENSG00000132773 | TOE1 | chr1 | + | 45341948 | 45342107 | 45341472 | 45341569 | 45342383 | 45342643 | 62 | 27 | 45 | 54 | 95 | 0 | 0 | 0 | 0 | 0 |
| SE | 42351 | ENSG00000132773 | TOE1 | chr1 | + | 45342383 | 45342643 | 45341948 | 45342107 | 45342842 | 45343002 | 41 | 19 | 33 | 48 | 76 | 0 | 0 | 0 | 0 | 0 |
| SE | 42352 | ENSG00000198502 | HLA-DRB5 | chr6 | - | 32518053 | 32518077 | 32517352 | 32517752 | 32518555 | 32518666 | 0 | 674 | 766 | 2 | 0 | 0 | 0 | 1 | 0 | 0 |
| SE | 42353 | ENSG00000198502 | HLA-DRB5 | chr6 | - | 32519369 | 32519651 | 32518555 | 32518666 | 32521904 | 32522174 | 117 | 927 | 980 | 642 | 88 | 0 | 2 | 1 | 0 | 0 |
| SE | 42355 | ENSG00000180957 | PITPNB | chr22 | - | 27897129 | 27897137 | 27896551 | 27896626 | 27897800 | 27897892 | 223 | 146 | 187 | 203 | 286 | 1 | 3 | 0 | 0 | 0 |
| SE | 42356 | ENSG00000180957 | PITPNB | chr22 | - | 27897129 | 27897137 | 27896551 | 27896626 | 27910963 | 27911109 | 15 | 8 | 10 | 15 | 21 | 4 | 2 | 6 | 6 | 8 |
| SE | 42358 | ENSG00000180957 | PITPNB | chr22 | - | 27897800 | 27897892 | 27896899 | 27897137 | 27910963 | 27911109 | 300 | 206 | 258 | 272 | 424 | 0 | 0 | 0 | 0 | 0 |
| SE | 42362 | ENSG00000048740 | CELF2 | chr10 | + | 11217424 | 11217507 | 11165483 | 11165682 | 11257737 | 11257872 | 705 | 510 | 580 | 638 | 689 | 1 | 2 | 5 | 2 | 0 |
| SE | 42363 | ENSG00000048740 | CELF2 | chr10 | + | 11249152 | 11249201 | 11165483 | 11165682 | 11257737 | 11257872 | 282 | 230 | 283 | 297 | 259 | 1 | 2 | 5 | 2 | 0 |
| SE | 42365 | ENSG00000048740 | CELF2 | chr10 | + | 11257737 | 11257872 | 11165483 | 11165682 | 11314138 | 11314258 | 1 | 2 | 5 | 3 | 0 | 0 | 0 | 0 | 0 | 0 |
| SE | 42366 | ENSG00000048740 | CELF2 | chr10 | + | 11257737 | 11257872 | 11217424 | 11217507 | 11270665 | 11270824 | 196 | 120 | 151 | 148 | 220 | 0 | 0 | 0 | 0 | 0 |
| SE | 42370 | ENSG00000048740 | CELF2 | chr10 | + | 11288417 | 11288552 | 11257737 | 11257872 | 11314138 | 11314258 | 57 | 44 | 68 | 86 | 23 | 0 | 0 | 0 | 1 | 0 |
| SE | 42371 | ENSG00000048740 | CELF2 | chr10 | + | 11314138 | 11314258 | 11165483 | 11165682 | 11321206 | 11321386 | 159 | 82 | 154 | 153 | 103 | 1 | 0 | 0 | 0 | 0 |
| SE | 42372 | ENSG00000048740 | CELF2 | chr10 | + | 11314138 | 11314270 | 11165483 | 11165682 | 11321206 | 11321386 | 66 | 40 | 66 | 69 | 31 | 1 | 0 | 0 | 0 | 0 |
| SE | 42378 | ENSG00000048740 | CELF2 | chr10 | + | 11321206 | 11321386 | 11314138 | 11314270 | 11328925 | 11329097 | 75 | 42 | 70 | 74 | 31 | 1 | 0 | 0 | 0 | 3 |
| SE | 42379 | ENSG00000048740 | CELF2 | chr10 | + | 11325835 | 11325979 | 11314138 | 11314270 | 11328925 | 11329053 | 416 | 227 | 340 | 332 | 462 | 1 | 0 | 0 | 0 | 3 |
| SE | 42380 | ENSG00000048740 | CELF2 | chr10 | + | 11325835 | 11325979 | 11321188 | 11321386 | 11328925 | 11329053 | 795 | 394 | 664 | 663 | 655 | 6 | 1 | 3 | 0 | 0 |
| SE | 42381 | ENSG00000048740 | CELF2 | chr10 | + | 11325835 | 11325979 | 11321188 | 11321386 | 11330455 | 11330639 | 383 | 169 | 331 | 333 | 193 | 0 | 0 | 0 | 1 | 0 |
| SE | 42382 | ENSG00000048740 | CELF2 | chr10 | + | 11328925 | 11329097 | 11321206 | 11321386 | 11330455 | 11330639 | 90 | 43 | 77 | 104 | 43 | 0 | 0 | 0 | 1 | 0 |
| SE | 42383 | ENSG00000048740 | CELF2 | chr10 | + | 11328925 | 11329097 | 11325835 | 11325979 | 11330455 | 11330639 | 500 | 267 | 414 | 436 | 507 | 4 | 1 | 5 | 2 | 0 |
| SE | 42384 | ENSG00000196743 | GM2A | chr5 | + | 151259754 | 151259916 | 151253184 | 151253297 | 151266730 | 151266880 | 313 | 97 | 141 | 132 | 195 | 0 | 0 | 1 | 0 | 0 |
| SE | 42391 | ENSG00000113194 | FAF2 | chr5 | + | 176453317 | 176453702 | 176453138 | 176453228 | 176479187 | 176479256 | 1 | 2 | 0 | 3 | 0 | 0 | 0 | 0 | 0 | 0 |
| SE | 42393 | ENSG00000113194 | FAF2 | chr5 | + | 176488950 | 176489027 | 176486354 | 176486489 | 176492193 | 176492332 | 179 | 92 | 131 | 175 | 85 | 1 | 0 | 0 | 0 | 3 |
| SE | 42394 | ENSG00000113194 | FAF2 | chr5 | + | 176488950 | 176489027 | 176486354 | 176486489 | 176493998 | 176494084 | 51 | 27 | 48 | 46 | 28 | 0 | 0 | 0 | 0 | 2 |
| SE | 42395 | ENSG00000113194 | FAF2 | chr5 | + | 176492193 | 176492332 | 176486354 | 176486489 | 176493998 | 176494084 | 49 | 20 | 33 | 37 | 54 | 0 | 0 | 0 | 0 | 2 |
| SE | 42397 | ENSG00000113194 | FAF2 | chr5 | + | 176493998 | 176494084 | 176488950 | 176489027 | 176496485 | 176496663 | 1 | 0 | 0 | 1 | 2 | 0 | 0 | 0 | 0 | 2 |
| SE | 42398 | ENSG00000113194 | FAF2 | chr5 | + | 176494183 | 176494275 | 176493998 | 176494084 | 176496485 | 176496663 | 113 | 92 | 112 | 133 | 167 | 1 | 0 | 0 | 0 | 0 |
| SE | 42399 | ENSG00000113194 | FAF2 | chr5 | + | 176496485 | 176496663 | 176493998 | 176494275 | 176498913 | 176498977 | 224 | 207 | 214 | 229 | 283 | 1 | 0 | 3 | 0 | 0 |
| SE | 42401 | ENSG00000143476 | DTL | chr1 | + | 212064916 | 212065029 | 212062883 | 212062949 | 212066811 | 212066885 | 4 | 14 | 18 | 12 | 8 | 0 | 0 | 1 | 0 | 0 |
| SE | 42402 | ENSG00000143476 | DTL | chr1 | + | 212068223 | 212068327 | 212066811 | 212066885 | 212068598 | 212068703 | 3 | 14 | 8 | 10 | 20 | 0 | 0 | 4 | 0 | 0 |
| SE | 42405 | ENSG00000104381 | GDAP1 | chr8 | + | 74351251 | 74351466 | 74350433 | 74350578 | 74360136 | 74360310 | 7 | 2 | 7 | 4 | 15 | 2 | 0 | 2 | 1 | 0 |
| SE | 42406 | ENSG00000104381 | GDAP1 | chr8 | + | 74351273 | 74351321 | 74350394 | 74350578 | 74360136 | 74360149 | 13 | 3 | 1 | 9 | 14 | 2 | 0 | 2 | 1 | 0 |
| SE | 42408 | ENSG00000104381 | GDAP1 | chr8 | + | 74351273 | 74351418 | 74350395 | 74350578 | 74360136 | 74360310 | 5 | 4 | 3 | 8 | 3 | 2 | 0 | 2 | 1 | 0 |
| SE | 42409 | ENSG00000104381 | GDAP1 | chr8 | + | 74351273 | 74351466 | 74350394 | 74350578 | 74360136 | 74360310 | 12 | 6 | 10 | 11 | 18 | 2 | 0 | 2 | 1 | 0 |
| SE | 42411 | ENSG00000104381 | GDAP1 | chr8 | + | 74351453 | 74351466 | 74351273 | 74351321 | 74360136 | 74360310 | 0 | 0 | 0 | 0 | 0 | 10 | 2 | 1 | 6 | 14 |
| SE | 42424 | ENSG00000151131 | C12orf45 | chr12 | + | 104991711 | 104991848 | 104988166 | 104988253 | 104994477 | 104994649 | 59 | 31 | 61 | 50 | 57 | 0 | 1 | 0 | 0 | 0 |
| SE | 42426 | ENSG00000163946 | TASOR | chr3 | - | 56624827 | 56625006 | 56624478 | 56624643 | 56627036 | 56627145 | 297 | 204 | 261 | 292 | 809 | 1 | 0 | 0 | 0 | 2 |
| SE | 42427 | ENSG00000163946 | TASOR | chr3 | - | 56627581 | 56627741 | 56627036 | 56627145 | 56628491 | 56628614 | 568 | 410 | 473 | 500 | 1528 | 0 | 0 | 0 | 0 | 2 |
| SE | 42428 | ENSG00000163946 | TASOR | chr3 | - | 56633043 | 56633843 | 56628491 | 56628614 | 56638705 | 56638765 | 195 | 170 | 142 | 192 | 440 | 0 | 1 | 0 | 2 | 1 |
| SE | 42429 | ENSG00000163946 | TASOR | chr3 | - | 56633043 | 56633966 | 56628491 | 56628614 | 56638705 | 56638765 | 331 | 320 | 267 | 340 | 683 | 0 | 1 | 0 | 2 | 1 |
| SE | 42430 | ENSG00000163946 | TASOR | chr3 | - | 56638705 | 56638765 | 56633043 | 56633843 | 56639985 | 56640130 | 245 | 230 | 208 | 239 | 454 | 2 | 0 | 0 | 0 | 0 |
| SE | 42431 | ENSG00000163946 | TASOR | chr3 | - | 56638705 | 56638765 | 56633043 | 56633966 | 56639985 | 56640130 | 381 | 379 | 332 | 385 | 699 | 2 | 1 | 0 | 2 | 9 |
| SE | 42432 | ENSG00000163946 | TASOR | chr3 | - | 56639985 | 56640130 | 56638705 | 56638765 | 56641348 | 56641752 | 703 | 662 | 526 | 688 | 1049 | 0 | 0 | 0 | 2 | 0 |
| SE | 42433 | ENSG00000163946 | TASOR | chr3 | - | 56641348 | 56641752 | 56639985 | 56640130 | 56660730 | 56660834 | 357 | 311 | 246 | 352 | 428 | 0 | 0 | 0 | 0 | 0 |
| SE | 42434 | ENSG00000163946 | TASOR | chr3 | - | 56646521 | 56647223 | 56641348 | 56641752 | 56660730 | 56660834 | 233 | 206 | 195 | 244 | 282 | 0 | 0 | 0 | 0 | 1 |
| SE | 42435 | ENSG00000163946 | TASOR | chr3 | - | 56648821 | 56648893 | 56646521 | 56647223 | 56648984 | 56649057 | 706 | 619 | 442 | 627 | 774 | 0 | 0 | 2 | 0 | 3 |
| SE | 42437 | ENSG00000163946 | TASOR | chr3 | - | 56660730 | 56660834 | 56646521 | 56647223 | 56660913 | 56661017 | 247 | 206 | 148 | 228 | 316 | 1 | 0 | 0 | 0 | 0 |
| SE | 42439 | ENSG00000163946 | TASOR | chr3 | - | 56662384 | 56662490 | 56660913 | 56661017 | 56666259 | 56666384 | 232 | 127 | 156 | 206 | 199 | 0 | 0 | 0 | 3 | 5 |
| SE | 42440 | ENSG00000163946 | TASOR | chr3 | - | 56662384 | 56662490 | 56660913 | 56661017 | 56668396 | 56668558 | 232 | 125 | 156 | 206 | 201 | 0 | 2 | 0 | 0 | 2 |
| SE | 42441 | ENSG00000163946 | TASOR | chr3 | - | 56662384 | 56662490 | 56660913 | 56661017 | 56669699 | 56669791 | 232 | 125 | 157 | 206 | 199 | 0 | 0 | 1 | 0 | 0 |
| SE | 42443 | ENSG00000163946 | TASOR | chr3 | - | 56663540 | 56663572 | 56660968 | 56661017 | 56668396 | 56668489 | 16 | 23 | 8 | 22 | 24 | 0 | 2 | 0 | 0 | 2 |
| SE | 42444 | ENSG00000163946 | TASOR | chr3 | - | 56663540 | 56663572 | 56660968 | 56661017 | 56669699 | 56669791 | 2 | 9 | 2 | 9 | 2 | 0 | 0 | 1 | 0 | 0 |
| SE | 42445 | ENSG00000163946 | TASOR | chr3 | - | 56663540 | 56663572 | 56662384 | 56662490 | 56666259 | 56666384 | 299 | 146 | 169 | 246 | 338 | 0 | 2 | 0 | 0 | 0 |
| SE | 42446 | ENSG00000163946 | TASOR | chr3 | - | 56663540 | 56663572 | 56662384 | 56662490 | 56668396 | 56668489 | 54 | 44 | 29 | 49 | 73 | 0 | 0 | 0 | 0 | 2 |
| SE | 42447 | ENSG00000163946 | TASOR | chr3 | - | 56663540 | 56663572 | 56662384 | 56662490 | 56669699 | 56669791 | 31 | 15 | 13 | 15 | 18 | 0 | 0 | 1 | 0 | 0 |
| SE | 42448 | ENSG00000163946 | TASOR | chr3 | - | 56666259 | 56666384 | 56660913 | 56661017 | 56668396 | 56668489 | 138 | 69 | 108 | 129 | 216 | 0 | 2 | 0 | 0 | 2 |
| SE | 42450 | ENSG00000163946 | TASOR | chr3 | - | 56666259 | 56666384 | 56662384 | 56662490 | 56668396 | 56668489 | 138 | 71 | 108 | 126 | 211 | 0 | 2 | 0 | 0 | 2 |
| SE | 42454 | ENSG00000163946 | TASOR | chr3 | - | 56668396 | 56668558 | 56660913 | 56661017 | 56669699 | 56669791 | 101 | 75 | 130 | 140 | 224 | 0 | 0 | 1 | 0 | 0 |
| SE | 42455 | ENSG00000163946 | TASOR | chr3 | - | 56668396 | 56668558 | 56662384 | 56662490 | 56669699 | 56669791 | 101 | 73 | 130 | 140 | 224 | 0 | 0 | 1 | 0 | 0 |
| SE | 42457 | ENSG00000163946 | TASOR | chr3 | - | 56668396 | 56668558 | 56666259 | 56666384 | 56669699 | 56669791 | 260 | 155 | 257 | 297 | 513 | 0 | 0 | 0 | 0 | 1 |
| SE | 42459 | ENSG00000163946 | TASOR | chr3 | - | 56670072 | 56670145 | 56669699 | 56669791 | 56673579 | 56673725 | 261 | 253 | 219 | 300 | 274 | 0 | 2 | 0 | 0 | 0 |
| SE | 42460 | ENSG00000163946 | TASOR | chr3 | - | 56671599 | 56671692 | 56669699 | 56669791 | 56673579 | 56673725 | 454 | 365 | 371 | 387 | 616 | 0 | 2 | 0 | 0 | 0 |
| SE | 42461 | ENSG00000163946 | TASOR | chr3 | - | 56671599 | 56671692 | 56670072 | 56670145 | 56673579 | 56673725 | 1385 | 1236 | 1084 | 1151 | 1655 | 0 | 2 | 1 | 1 | 0 |
| SE | 42462 | ENSG00000163946 | TASOR | chr3 | - | 56673579 | 56673725 | 56671599 | 56671692 | 56682675 | 56683017 | 910 | 772 | 727 | 790 | 1120 | 0 | 0 | 2 | 0 | 0 |
| SE | 42476 | ENSG00000173083 | HPSE | chr4 | - | 83306202 | 83306317 | 83302149 | 83302268 | 83309401 | 83309495 | 94 | 88 | 73 | 77 | 80 | 0 | 0 | 0 | 0 | 0 |
| SE | 42477 | ENSG00000173083 | HPSE | chr4 | - | 83308844 | 83308951 | 83302149 | 83302268 | 83309401 | 83309495 | 80 | 42 | 45 | 57 | 47 | 0 | 0 | 0 | 0 | 0 |
| SE | 42478 | ENSG00000173083 | HPSE | chr4 | - | 83308844 | 83308951 | 83306202 | 83306317 | 83309401 | 83309495 | 134 | 91 | 106 | 116 | 84 | 0 | 0 | 0 | 1 | 0 |
| SE | 42479 | ENSG00000173083 | HPSE | chr4 | - | 83310721 | 83310890 | 83310030 | 83310078 | 83313113 | 83313287 | 161 | 74 | 122 | 151 | 149 | 1 | 0 | 0 | 0 | 0 |
| SE | 42480 | ENSG00000173083 | HPSE | chr4 | - | 83310721 | 83310890 | 83310030 | 83310078 | 83319343 | 83319469 | 87 | 42 | 58 | 77 | 73 | 0 | 0 | 0 | 0 | 0 |
| SE | 42481 | ENSG00000173083 | HPSE | chr4 | - | 83313113 | 83313287 | 83310030 | 83310078 | 83319343 | 83319469 | 75 | 21 | 40 | 59 | 48 | 0 | 0 | 0 | 0 | 0 |
| SE | 42485 | ENSG00000251503 | CENPS-CORT | chr1 | + | 10440346 | 10440413 | 10434656 | 10434690 | 10451376 | 10451659 | 12 | 1 | 7 | 7 | 11 | 3 | 0 | 0 | 0 | 0 |
| SE | 42486 | ENSG00000251503 | CENPS-CORT | chr1 | + | 10442264 | 10442441 | 10440346 | 10440413 | 10451376 | 10451411 | 18 | 10 | 14 | 13 | 26 | 1 | 0 | 0 | 1 | 1 |
| SE | 42495 | ENSG00000162687 | KCNT2 | chr1 | - | 196468993 | 196469068 | 196467702 | 196467786 | 196479178 | 196479238 | 0 | 0 | 0 | 7 | 11 | 0 | 0 | 0 | 0 | 0 |
| SE | 42498 | ENSG00000143630 | HCN3 | chr1 | + | 155284538 | 155284757 | 155283973 | 155284135 | 155285164 | 155285311 | 2 | 0 | 0 | 0 | 1 | 0 | 1 | 0 | 0 | 0 |
| SE | 42500 | ENSG00000228878 | SEPTIN7-DT | chr7 | - | 35755732 | 35755817 | 35751907 | 35755195 | 35758178 | 35758273 | 1 | 1 | 0 | 0 | 0 | 7 | 2 | 1 | 2 | 6 |
| SE | 42507 | ENSG00000176444 | CLK2 | chr1 | - | 155266728 | 155266895 | 155264644 | 155264774 | 155268009 | 155268126 | 66 | 45 | 79 | 61 | 75 | 2 | 0 | 0 | 2 | 0 |
| SE | 42511 | ENSG00000172071 | EIF2AK3 | chr2 | - | 88593016 | 88593114 | 88590817 | 88591052 | 88593271 | 88593405 | 0 | 0 | 2 | 5 | 0 | 166 | 142 | 139 | 157 | 198 |
| SE | 42512 | ENSG00000172071 | EIF2AK3 | chr2 | - | 88593271 | 88593405 | 88590817 | 88591052 | 88595468 | 88595498 | 246 | 189 | 204 | 220 | 266 | 0 | 0 | 0 | 0 | 0 |
| SE | 42514 | ENSG00000172071 | EIF2AK3 | chr2 | - | 88595468 | 88595663 | 88593271 | 88593405 | 88613723 | 88613853 | 112 | 72 | 93 | 85 | 108 | 0 | 1 | 0 | 0 | 0 |
| SE | 42518 | ENSG00000133256 | PDE6B | chr4 | + | 665254 | 665329 | 664880 | 664944 | 666530 | 666614 | 32 | 30 | 44 | 34 | 27 | 0 | 0 | 0 | 0 | 1 |
| SE | 42521 | ENSG00000133256 | PDE6B | chr4 | + | 667855 | 668006 | 666530 | 666614 | 670048 | 670782 | 16 | 12 | 16 | 22 | 18 | 1 | 0 | 0 | 0 | 2 |
| SE | 42525 | ENSG00000169231 | THBS3 | chr1 | - | 155203512 | 155203539 | 155203222 | 155203305 | 155204854 | 155204957 | 6 | 7 | 8 | 5 | 5 | 2 | 0 | 0 | 0 | 1 |
| SE | 42546 | ENSG00000169241 | SLC50A1 | chr1 | + | 155136827 | 155136951 | 155135811 | 155135972 | 155137978 | 155138098 | 1 | 1 | 0 | 5 | 0 | 0 | 0 | 0 | 1 | 0 |
| SE | 42551 | ENSG00000169241 | SLC50A1 | chr1 | + | 155137560 | 155137722 | 155135811 | 155135972 | 155137978 | 155138098 | 50 | 22 | 46 | 52 | 47 | 0 | 0 | 0 | 1 | 0 |
| SE | 42556 | ENSG00000169241 | SLC50A1 | chr1 | + | 155137978 | 155138098 | 155136298 | 155136376 | 155138179 | 155138368 | 66 | 59 | 61 | 87 | 39 | 0 | 0 | 0 | 0 | 0 |
| SE | 42559 | ENSG00000198624 | CCDC69 | chr5 | - | 151186022 | 151186124 | 151184343 | 151184441 | 151187385 | 151187459 | 319 | 172 | 301 | 277 | 361 | 0 | 1 | 0 | 0 | 0 |
| SE | 42560 | ENSG00000198624 | CCDC69 | chr5 | - | 151187385 | 151187459 | 151186022 | 151186124 | 151198996 | 151199084 | 609 | 335 | 582 | 553 | 690 | 0 | 0 | 3 | 4 | 1 |
| SE | 42561 | ENSG00000198624 | CCDC69 | chr5 | - | 151198996 | 151199084 | 151187385 | 151187459 | 151201581 | 151201688 | 665 | 339 | 597 | 557 | 667 | 3 | 0 | 3 | 0 | 1 |
| SE | 42562 | ENSG00000198624 | CCDC69 | chr5 | - | 151205399 | 151205475 | 151201581 | 151201688 | 151223922 | 151224092 | 379 | 201 | 368 | 351 | 315 | 1 | 0 | 0 | 2 | 0 |
| SE | 42564 | ENSG00000143952 | VPS54 | chr2 | - | 63913824 | 63913925 | 63913222 | 63913310 | 63914181 | 63914287 | 5 | 3 | 4 | 2 | 11 | 33 | 22 | 26 | 37 | 89 |
| SE | 42565 | ENSG00000143952 | VPS54 | chr2 | - | 63933672 | 63934013 | 63921205 | 63921335 | 63942464 | 63942561 | 138 | 148 | 110 | 143 | 255 | 0 | 0 | 2 | 1 | 1 |
| SE | 42566 | ENSG00000143952 | VPS54 | chr2 | - | 63942464 | 63942561 | 63933672 | 63934013 | 63944599 | 63944655 | 181 | 183 | 150 | 176 | 350 | 0 | 0 | 0 | 2 | 0 |
| SE | 42567 | ENSG00000143952 | VPS54 | chr2 | - | 63942464 | 63942561 | 63933672 | 63934013 | 63947382 | 63947490 | 60 | 81 | 50 | 72 | 147 | 1 | 0 | 0 | 0 | 0 |
| SE | 42568 | ENSG00000143952 | VPS54 | chr2 | - | 63944599 | 63944655 | 63933672 | 63934013 | 63947382 | 63947490 | 41 | 41 | 25 | 42 | 59 | 1 | 0 | 0 | 0 | 0 |
| SE | 42569 | ENSG00000143952 | VPS54 | chr2 | - | 63944599 | 63944655 | 63942464 | 63942561 | 63947382 | 63947490 | 168 | 144 | 127 | 149 | 249 | 0 | 1 | 0 | 0 | 0 |
| SE | 42571 | ENSG00000143952 | VPS54 | chr2 | - | 63965834 | 63965966 | 63962057 | 63962443 | 63968956 | 63968991 | 159 | 115 | 100 | 95 | 199 | 0 | 0 | 0 | 0 | 0 |
| SE | 42574 | ENSG00000158042 | MRPL17 | chr11 | - | 6682746 | 6682815 | 6680384 | 6682402 | 6683122 | 6683340 | 50 | 18 | 26 | 34 | 48 | 0 | 1 | 0 | 0 | 0 |
| SE | 42577 | ENSG00000198380 | GFPT1 | chr2 | - | 69328270 | 69328438 | 69326913 | 69327075 | 69329296 | 69329424 | 129 | 75 | 81 | 164 | 234 | 0 | 0 | 2 | 0 | 4 |
| SE | 42578 | ENSG00000198380 | GFPT1 | chr2 | - | 69329683 | 69329798 | 69329296 | 69329424 | 69337897 | 69338055 | 201 | 178 | 170 | 228 | 202 | 1 | 0 | 0 | 0 | 0 |
| SE | 42579 | ENSG00000198380 | GFPT1 | chr2 | - | 69354258 | 69354312 | 69350077 | 69350183 | 69354488 | 69354568 | 0 | 0 | 0 | 5 | 3 | 87 | 64 | 75 | 105 | 117 |
| SE | 42580 | ENSG00000198380 | GFPT1 | chr2 | - | 69354488 | 69354568 | 69354258 | 69354312 | 69356495 | 69356557 | 34 | 18 | 53 | 57 | 28 | 0 | 0 | 0 | 0 | 2 |
| SE | 42581 | ENSG00000198380 | GFPT1 | chr2 | - | 69359267 | 69359326 | 69358328 | 69358463 | 69363544 | 69363670 | 160 | 106 | 115 | 165 | 232 | 0 | 0 | 0 | 1 | 0 |
| SE | 42583 | ENSG00000198380 | GFPT1 | chr2 | - | 69371309 | 69371472 | 69370000 | 69370108 | 69374005 | 69374113 | 0 | 0 | 0 | 0 | 6 | 49 | 44 | 59 | 67 | 61 |
| SE | 42585 | ENSG00000078140 | UBE2K | chr4 | + | 39737419 | 39737513 | 39698108 | 39698390 | 39745751 | 39745810 | 137 | 92 | 80 | 111 | 203 | 0 | 0 | 1 | 3 | 0 |
| SE | 42586 | ENSG00000078140 | UBE2K | chr4 | + | 39737419 | 39737513 | 39698108 | 39698390 | 39755656 | 39755739 | 46 | 29 | 19 | 47 | 31 | 0 | 1 | 0 | 0 | 0 |

| | | | | | | | | | | | | | | | | | | | | | | | | |
|---|---|---|---|---|---|---|---|---|---|---|---|---|---|---|---|---|---|---|---|---|---|---|---|---|
| 42073 | 14 | 31 | 40 | 37 | 1 | 0 | 0 | 0 | 0 | 0 | 240 | 149 | 1 | 1 | 1 | 1 | 1 | 0.947 | 1 | 1 | 1 | 1 | 1 | 1 | -0.011 |
| 42074 | 0 | 0 | 0 | 0 | 0 | 15 | 7 | 5 | 4 | 0 | 216 | 149 | 1 | 1 | 0 | 0 | 0 | 0 | 0.047 | 0 | 0 | 0 | 0 | NA | 0.009 |
| 42076 | 625 | 510 | 590 | 728 | 797 | 0 | 3 | 2 | 8 | 4 | 298 | 149 | 1 | 1 | 1 | 0.99 | 1 | 0.997 | 1 | 1 | 0.988 | 0.993 | 0.978 | 0.99 | 0.008 |
| 42080 | 115 | 229 | 102 | 122 | 117 | 0 | 0 | 0 | 0 | 0 | 298 | 149 | 1 | 1 | 1 | 1 | 0.971 | 1 | 1 | 1 | 1 | 1 | 1 | 1 | -0.006 |
| 42082 | 21 | 30 | 9 | 2 | 3 | 0 | 0 | 0 | 0 | 0 | 298 | 149 | 1 | 1 | 1 | 1 | 0.667 | 1 | 1 | 1 | 1 | 1 | 1 | 1 | -0.067 |
| 42084 | 247 | 217 | 174 | 135 | 155 | 0 | 4 | 0 | 0 | 0 | 298 | 149 | 1 | 1 | 1 | 1 | 1 | 1 | 0.962 | 1 | 0.964 | 1 | 1 | 1 | 0 |
| 42086 | 85 | 70 | 53 | 59 | 56 | 0 | 0 | 2 | 2 | 0 | 298 | 149 | 1 | 1 | 1 | 1 | 0.978 | 0.896 | 1 | 1 | 1 | 0.93 | 0.937 | 1 | 0.001 |
| 42089 | 22 | 31 | 47 | 36 | 19 | 0 | 0 | 2 | 2 | 0 | 269 | 149 | 1 | 1 | 1 | 1 | 0.904 | 0.812 | 1 | 1 | 1 | 0.929 | 0.909 | 1 | -0.024 |
| 42092 | 134 | 146 | 117 | 76 | 122 | 0 | 0 | 0 | 0 | 0 | 269 | 149 | 1 | 1 | 0.974 | 1 | 1 | 1 | 1 | 1 | 1 | 1 | 1 | 1 | -0.005 |
| 42093 | 22 | 30 | 47 | 40 | 30 | 0 | 0 | 0 | 0 | 0 | 269 | 149 | 1 | 1 | 0.941 | 1 | 1 | 1 | 1 | 1 | 1 | 1 | 1 | 1 | -0.012 |
| 42095 | 114 | 159 | 84 | 98 | 137 | 0 | 0 | 2 | 2 | 0 | 253 | 149 | 1 | 1 | 1 | 1 | 0.986 | 0.935 | 1 | 1 | 1 | 0.961 | 0.967 | 1 | -0.001 |
| 42098 | 130 | 154 | 123 | 106 | 151 | 0 | 0 | 0 | 0 | 0 | 212 | 149 | 1 | 1 | 0.984 | 1 | 1 | 1 | 1 | 1 | 1 | 1 | 1 | 1 | -0.003 |
| 42101 | 3 | 7 | 2 | 6 | 4 | 0 | 0 | 0 | 1 | 0 | 225 | 149 | 1 | 1 | 0.665 | 1 | 1 | 0.726 | 0.785 | 1 | 1 | 1 | 0.799 | 1 | -0.125 |
| 42112 | 0 | 0 | 0 | 0 | 0 | 267 | 248 | 152 | 220 | 346 | 298 | 149 | 1 | 1 | 0.003 | 0 | 0 | 0.003 | 0 | 0 | 0 | 0 | 0 | 0 | 0.001 |
| 42113 | 1032 | 804 | 501 | 695 | 1020 | 0 | 0 | 0 | 4 | 0 | 298 | 149 | 1 | 1 | 1 | 1 | 1 | 1 | 1 | 1 | 1 | 1 | 0.989 | 1 | 0.002 |
| 42117 | 39 | 34 | 24 | 23 | 52 | 0 | 0 | 0 | 0 | 0 | 276 | 149 | 1 | 1 | 0.955 | 1 | 1 | 1 | 1 | 1 | 1 | 1 | 1 | 1 | -0.009 |
| 42120 | 380 | 369 | 167 | 237 | 430 | 0 | 0 | 0 | 3 | 0 | 298 | 149 | 1 | 1 | 1 | 1 | 1 | 0.994 | 1 | 1 | 1 | 1 | 0.975 | 1 | 0.004 |
| 42122 | 0 | 0 | 6 | 2 | 5 | 0 | 0 | 0 | 0 | 0 | 233 | 149 | 1 | 1 | 1 | 1 | 1 | 1 | 0.242 | NA | NA | 1 | 1 | 1 | -0.152 |
| 42124 | 914 | 542 | 272 | 482 | 661 | 0 | 0 | 0 | 0 | 0 | 266 | 149 | 1 | 1 | 1 | 1 | 0.996 | 1 | 1 | 1 | 1 | 1 | 1 | 1 | -0.001 |
| 42126 | 43 | 103 | 43 | 48 | 84 | 0 | 0 | 0 | 0 | 0 | 224 | 149 | 1 | 1 | 0.984 | 1 | 1 | 1 | 0.928 | 1 | 1 | 1 | 1 | 1 | -0.018 |
| 42127 | 101 | 116 | 48 | 86 | 78 | 0 | 0 | 0 | 0 | 12 | 298 | 149 | 1 | 1 | 0.962 | 1 | 0.972 | 0.974 | 1 | 1 | 1 | 1 | 1 | 0.765 | 0.029 |
| 42128 | 44 | 96 | 62 | 88 | 82 | 0 | 0 | 0 | 4 | 0 | 225 | 149 | 1 | 1 | 1 | 1 | 1 | 1 | 1 | 1 | 1 | 1 | 0.936 | 1 | 0.013 |
| 42132 | 203 | 206 | 163 | 235 | 232 | 0 | 0 | 0 | 2 | 0 | 239 | 149 | 1 | 1 | 0.991 | 1 | 1 | 1 | 1 | 1 | 1 | 1 | 0.987 | 1 | 0.001 |
| 42133 | 239 | 194 | 166 | 248 | 258 | 0 | 0 | 0 | 2 | 0 | 239 | 149 | 1 | 1 | 1 | 1 | 1 | 1 | 1 | 1 | 1 | 1 | 0.987 | 1 | 0.003 |
| 42143 | 1870 | 1153 | 1088 | 1379 | 1918 | 9 | 2 | 0 | 5 | 9 | 233 | 149 | 1 | 1 | 0.994 | 0.99 | 0.981 | 0.992 | 1 | 0.993 | 0.997 | 1 | 0.994 | 0.993 | -0.004 |
| 42144 | 460 | 232 | 378 | 362 | 493 | 0 | 0 | 0 | 0 | 0 | 215 | 149 | 1 | 1 | 1 | 1 | 1 | 0.997 | 1 | 1 | 1 | 1 | 1 | 1 | -0.001 |
| 42145 | 1565 | 1019 | 920 | 1262 | 1522 | 0 | 0 | 4 | 6 | 0 | 288 | 149 | 1 | 1 | 0.991 | 0.999 | 0.975 | 0.992 | 0.999 | 1 | 1 | 0.992 | 0.991 | 1 | -0.005 |
| 42147 | 0 | 8 | 2 | 0 | 7 | 0 | 0 | 0 | 0 | 0 | 233 | 149 | 1 | 1 | 1 | 1 | 0.561 | 1 | 1 | NA | 1 | 1 | NA | 1 | -0.088 |
| 42148 | 1 | 8 | 3 | 12 | 21 | 0 | 0 | 0 | 0 | 0 | 298 | 149 | 1 | 1 | 1 | 1 | 0.75 | 1 | 1 | 1 | 1 | 1 | 1 | 1 | -0.05 |
| 42149 | 1 | 8 | 3 | 0 | 21 | 0 | 0 | 0 | 0 | 0 | 298 | 149 | 1 | 1 | 0.875 | 1 | 0.667 | 1 | 0.667 | 1 | 1 | 1 | NA | 1 | -0.158 |
| 42151 | 0 | 5 | 0 | 10 | 0 | 0 | 0 | 0 | 0 | 0 | 298 | 149 | 1 | 1 | 0.667 | NA | 0.778 | 1 | 0 | NA | 1 | NA | 1 | NA | -0.389 |
| 42152 | 0 | 5 | 0 | 22 | 0 | 0 | 0 | 0 | 0 | 0 | 298 | 149 | 1 | 1 | 1 | NA | 1 | 1 | 0 | NA | 1 | NA | 1 | NA | -0.25 |
| 42157 | 0 | 7 | 20 | 24 | 6 | 0 | 0 | 1 | 0 | 0 | 295 | 149 | 1 | 1 | 0.932 | 0.835 | 0.747 | 0.808 | 0.787 | NA | 1 | 0.91 | 1 | 1 | -0.156 |
| 42160 | 0 | 2 | 3 | 7 | 17 | 0 | 3 | 7 | 2 | 0 | 276 | 149 | 1 | 1 | 0.258 | 0.393 | 0.321 | 0.654 | 0.288 | NA | 0.265 | 0.188 | 0.654 | 1 | -0.144 |
| 42161 | 72 | 66 | 33 | 28 | 32 | 0 | 1 | 0 | 0 | 0 | 276 | 149 | 1 | 1 | 1 | 1 | 1 | 1 | 1 | 1 | 0.973 | 1 | 1 | 1 | 0.005 |
| 42165 | 0 | 0 | 0 | 0 | 0 | 0 | 3 | 7 | 2 | 0 | 269 | 149 | 1 | 1 | 0.106 | 0 | 0 | 0 | 0 | NA | 0 | 0 | 0 | NA | 0.021 |
| 42166 | 10 | 34 | 21 | 14 | 0 | 0 | 3 | 7 | 2 | 0 | 254 | 149 | 1 | 1 | 0.759 | 0.844 | 0.795 | 0.946 | 0.841 | 1 | 0.869 | 0.638 | 0.804 | NA | 0.009 |
| 42167 | 12 | 34 | 21 | 14 | 0 | 0 | 0 | 0 | 0 | 0 | 254 | 149 | 1 | 1 | 0.929 | 1 | 1 | 1 | 1 | 1 | 1 | 1 | 1 | NA | -0.014 |
| 42168 | 6 | 8 | 2 | 2 | 0 | 0 | 0 | 0 | 0 | 0 | 194 | 149 | 1 | 1 | 1 | 0.86 | 1 | 1 | 1 | 1 | 1 | 1 | 1 | NA | -0.028 |
| 42169 | 6 | 6 | 4 | 34 | 8 | 0 | 0 | 0 | 0 | 0 | 283 | 149 | 1 | 1 | 1 | 1 | 0.894 | 1 | 1 | 1 | 1 | 1 | 1 | 1 | -0.021 |
| 42172 | 13 | 33 | 31 | 30 | 0 | 0 | 0 | 3 | 0 | 0 | 282 | 149 | 1 | 1 | 0.835 | 0.906 | 0.986 | 0.968 | 0.971 | 1 | 1 | 0.845 | 1 | NA | -0.028 |
| 42173 | 0 | 16 | 26 | 12 | 1 | 0 | 0 | 0 | 0 | 0 | 282 | 149 | 1 | 1 | 1 | 1 | 0.972 | 1 | 1 | NA | 1 | 1 | 1 | 1 | -0.006 |
| 42174 | 1 | 9 | 5 | 6 | 1 | 0 | 0 | 0 | 0 | 0 | 206 | 149 | 1 | 1 | 1 | 1 | 0.943 | 1 | 1 | 1 | 1 | 1 | 1 | 1 | -0.011 |
| 42177 | 0 | 6 | 0 | 0 | 0 | 0 | 0 | 6 | 0 | 1 | 207 | 149 | 1 | 1 | 0 | 0 | 0 | NA | 0.093 | NA | 1 | 0 | NA | 0 | -0.31 |
| 42178 | 65 | 89 | 138 | 19 | 59 | 0 | 0 | 3 | 0 | 0 | 298 | 149 | 1 | 1 | 1 | 1 | 1 | 1 | 1 | 1 | 1 | 0.958 | 1 | 1 | 0.008 |
| 42179 | 17 | 58 | 60 | 18 | 35 | 0 | 0 | 0 | 0 | 0 | 285 | 149 | 1 | 1 | 1 | 1 | 1 | 1 | 0.989 | 1 | 1 | 1 | 1 | 1 | -0.002 |
| 42183 | 18 | 47 | 42 | 34 | 40 | 0 | 0 | 0 | 0 | 0 | 250 | 149 | 1 | 1 | 0.972 | 1 | 1 | 1 | 1 | 1 | 1 | 1 | 1 | 1 | -0.006 |
| 42184 | 19 | 38 | 21 | 36 | 0 | 0 | 0 | 0 | 8 | 0 | 253 | 149 | 1 | 1 | 0.983 | 0.956 | 0.984 | 0.894 | 0.908 | 1 | 1 | 1 | 0.726 | NA | 0.014 |
| 42186 | 0 | 28 | 16 | 8 | 0 | 0 | 0 | 0 | 0 | 0 | 270 | 149 | 1 | 1 | 0.936 | 0.794 | 0.869 | 0.815 | 0.886 | NA | 1 | 1 | 1 | NA | -0.14 |
| 42187 | 0 | 38 | 24 | 14 | 0 | 0 | 0 | 0 | 0 | 0 | 274 | 149 | 1 | 1 | 0.947 | 0.825 | 0.887 | 0.859 | 0.89 | NA | | 1 | 1 | NA | -0.118 |
| 42188 | 0 | 28 | 19 | 8 | 0 | 0 | 0 | 0 | 0 | 0 | 274 | 149 | 1 | 1 | 0.969 | 1 | 1 | 0.956 | 1 | NA | 1 | 1 | 1 | NA | -0.015 |
| 42189 | 0 | 0 | 3 | 0 | 0 | 0 | 0 | 0 | 0 | 0 | 238 | 149 | 1 | 1 | 1 | 1 | 1 | NA | 0.484 | NA | NA | 1 | NA | NA | -0.129 |
| 42190 | 0 | 0 | 4 | 0 | 0 | 0 | 0 | 0 | 0 | 0 | 226 | 149 | 1 | 1 | 0 | 0 | 0.397 | NA | 0 | NA | NA | 1 | NA | NA | -0.901 |
| 42192 | 0 | 3 | 15 | 4 | 0 | 0 | 2 | 0 | 10 | 0 | 259 | 149 | 1 | 1 | 0.235 | 0.324 | 0.184 | 0.524 | 0.489 | NA | 0.463 | | 0.187 | NA | -0.199 |
| 42203 | 302 | 341 | 419 | 441 | 358 | 0 | 2 | 0 | 0 | 0 | 284 | 149 | 1 | 1 | 1 | 1 | 1 | 1 | 1 | 1 | 0.989 | 1 | 1 | 1 | 0.002 |
| 42204 | 303 | 224 | 357 | 509 | 276 | 0 | 0 | 0 | 1 | 0 | 298 | 149 | 1 | 1 | 1 | 1 | 0.993 | 1 | 1 | 1 | 1 | 1 | 0.996 | 1 | -0.001 |
| 42205 | 171 | 88 | 125 | 163 | 173 | 0 | 0 | 0 | 4 | 0 | 234 | 149 | 1 | 1 | 1 | 1 | 1 | 1 | 1 | 1 | 1 | 1 | 0.963 | 1 | 0.007 |
| 42209 | 507 | 490 | 576 | 712 | 807 | 0 | 0 | 0 | 0 | 0 | 298 | 149 | 1 | 1 | 0.995 | 1 | 1 | 1 | 1 | 1 | 1 | 1 | 1 | 1 | -0.001 |
| 42212 | 47 | 182 | 131 | 30 | 14 | 0 | 0 | 1 | 2 | 0 | 274 | 149 | 1 | 1 | 1 | 1 | 0.978 | 0.99 | 1 | 1 | 1 | 0.986 | 0.891 | 1 | 0.018 |
| 42214 | 29 | 184 | 145 | 55 | 37 | 0 | 0 | 0 | 2 | 0 | 235 | 149 | 1 | 1 | 0.991 | 1 | 1 | 1 | 1 | 1 | 1 | 1 | 0.946 | 1 | 0.009 |
| 42217 | 5 | 0 | 2 | 4 | 0 | 0 | 0 | 0 | 0 | 0 | 298 | 149 | 1 | 1 | 0.8 | 0.667 | 0.778 | 0.5 | 1 | 1 | NA | 1 | 1 | NA | -0.251 |
| 42232 | 151 | 121 | 68 | 136 | 158 | 6 | 0 | 0 | 1 | 0 | 276 | 149 | 1 | 1 | 1 | 0.97 | 1 | 1 | 1 | 0.931 | 1 | 1 | 0.987 | 1 | 0.01 |
| 42235 | 0 | 4 | 0 | 0 | 0 | 0 | 3 | 0 | 0 | 0 | 266 | 149 | 1 | 1 | 1 | 1 | NA | NA | NA | NA | 0.428 | NA | NA | NA | 0.572 |
| 42236 | 0 | 0 | 1 | 3 | 0 | 0 | 0 | 0 | 6 | 0 | 204 | 149 | 1 | 1 | 1 | 1 | 1 | 1 | 1 | NA | NA | 1 | 0.268 | NA | 0.366 |
| 42242 | 528 | 256 | 253 | 347 | 203 | 0 | 0 | 0 | 0 | 0 | 224 | 149 | 1 | 1 | 0.985 | 1 | 1 | 0.986 | 1 | 1 | 1 | 1 | 1 | 1 | -0.006 |
| 42243 | 528 | 255 | 244 | 347 | 200 | 0 | 0 | 0 | 0 | 0 | 224 | 149 | 1 | 1 | 0.978 | 1 | 1 | 1 | 1 | 1 | 1 | 1 | 1 | 1 | -0.004 |
| 42244 | 0 | 6 | 5 | 0 | 0 | 0 | 0 | 0 | 0 | 0 | 298 | 149 | 1 | 1 | 0.25 | NA | 1 | 1 | NA | NA | 1 | 1 | NA | NA | -0.25 |
| 42246 | 0 | 8 | 14 | 4 | 12 | 0 | 1 | 0 | 4 | 9 | 298 | 149 | 1 | 1 | NA | NA | 1 | 1 | 0 | NA | 0.8 | 1 | 0.333 | 0.4 | 0.033 |
| 42247 | 0 | 0 | 0 | 0 | 0 | 0 | 1 | 0 | 4 | 9 | 210 | 149 | 1 | 1 | 1 | 1 | NA | NA | 0 | NA | 0 | NA | 0 | 0 | 0.667 |
| 42248 | 0 | 0 | 0 | 0 | 0 | 0 | 6 | 5 | 0 | 0 | 210 | 149 | 1 | 1 | 1 | 1 | 0 | NA | NA | NA | 0 | 0 | NA | NA | 0.667 |
| 42255 | 42 | 216 | 115 | 84 | 51 | 0 | 0 | 0 | 0 | 0 | 273 | 149 | 1 | 1 | 1 | 1 | 0.98 | 1 | 1 | 1 | 1 | 1 | 1 | 1 | -0.004 |
| 42256 | 46 | 314 | 145 | 106 | 77 | 0 | 2 | 0 | 0 | 0 | 273 | 149 | 1 | 1 | 1 | 1 | 1 | 1 | 1 | 1 | 0.988 | 1 | 1 | 1 | 0.002 |
| 42257 | 56 | 348 | 121 | 113 | 87 | 0 | 0 | 5 | 0 | 0 | 234 | 149 | 1 | 1 | 0.993 | 1 | 1 | 1 | 0.995 | 1 | 1 | 0.939 | 1 | 1 | 0.01 |
| 42258 | 34 | 94 | 66 | 64 | 28 | 0 | 0 | 0 | 0 | 0 | 298 | 149 | 1 | 1 | 1 | 1 | 1 | 0.99 | 0.992 | 1 | 1 | 1 | 1 | 1 | -0.004 |
| 42259 | 65 | 193 | 94 | 93 | 57 | 0 | 0 | 0 | 0 | 0 | 258 | 149 | 1 | 1 | 1 | 1 | 1 | 0.994 | 0.995 | 1 | 1 | 1 | 1 | 1 | -0.002 |
| 42265 | 24 | 60 | 41 | 38 | 46 | 0 | 0 | 0 | 0 | 0 | 182 | 149 | 1 | 1 | 1 | 1 | 1 | 1 | 0.98 | 1 | 1 | 1 | 1 | 1 | -0.004 |
| 42266 | 0 | 8 | 8 | 15 | 31 | 0 | 0 | 0 | 0 | 0 | 252 | 149 | 1 | 1 | 1 | 0.842 | 1 | 0.937 | 1 | NA | 1 | 1 | 1 | 1 | -0.044 |
| 42267 | 39 | 117 | 108 | 83 | 68 | 0 | 0 | 0 | 0 | 0 | 246 | 149 | 1 | 1 | 1 | 0.977 | 1 | 0.988 | 1 | 1 | 1 | 1 | 1 | 1 | -0.007 |
| 42269 | 0 | 4 | 0 | 0 | 0 | 12 | 0 | 0 | 6 | 0 | 298 | 149 | 1 | 1 | 0.538 | NA | 0.778 | 0.778 | NA | 0 | 1 | NA | 0 | NA | 0.365 |
| 42272 | 0 | 4 | 0 | 0 | 0 | 12 | 0 | 0 | 6 | 0 | 298 | 149 | 1 | 1 | 0.538 | 1 | 0.778 | 0.778 | NA | 0 | 1 | NA | 0 | NA | 0.44 |
| 42273 | 0 | 0 | 0 | 0 | 0 | 0 | 0 | 0 | 10 | 0 | 298 | 149 | 1 | 1 | 0.143 | 1 | 0 | 0 | 1 | NA | NA | NA | 0 | NA | 0.429 |
| 42277 | 365 | 247 | 94 | 196 | 221 | 0 | 0 | 0 | 6 | 0 | 290 | 149 | 1 | 1 | 1 | 1 | 1 | 0 | 1 | 1 | 1 | | 0.944 | 1 | 0.011 |
| 42286 | 0 | 0 | 5 | 0 | 0 | 0 | 0 | 3 | 0 | 0 | 255 | 149 | 1 | 1 | 0 | 0 | 0 | NA | NA | NA | NA | 0.493 | NA | NA | -0.493 |
| 42290 | 0 | 0 | 0 | 0 | 0 | 0 | 0 | 3 | 0 | 0 | 247 | 149 | 1 | 1 | 0 | 0 | 0.091 | NA | NA | NA | NA | 0 | NA | NA | 0.03 |
| 42292 | 0 | 0 | 0 | 0 | 0 | 0 | 0 | 3 | 0 | 0 | 227 | 149 | 1 | 1 | 0 | 0 | 0.304 | 1 | NA | NA | NA | 0 | NA | NA | 0.326 |
| 42309 | 87 | 22 | 24 | 48 | 101 | 0 | 0 | 0 | 0 | 0 | 298 | 149 | 1 | 1 | 1 | 1 | 1 | 0.92 | 1 | 1 | 1 | 1 | 1 | 1 | -0.016 |
| 42310 | 52 | 17 | 5 | 10 | 41 | 0 | 0 | 2 | 0 | 0 | 212 | 149 | 1 | 1 | 1 | 1 | 1 | 1 | 1 | 1 | 1 | 0.637 | 1 | 1 | 0.073 |
| 42311 | 49 | 8 | 7 | 4 | 4 | 2 | 0 | 0 | 1 | 0 | 294 | 149 | 1 | 1 | 0.687 | 1 | 1 | 1 | 1 | 0.925 | 1 | 1 | 0.67 | 1 | 0.018 |
| 42312 | 0 | 5 | 5 | 4 | 13 | 0 | 0 | 0 | 0 | 0 | 298 | 149 | 1 | 1 | 0 | 1 | 1 | 0.667 | 1 | NA | 1 | 1 | 1 | 1 | -0.267 |
| 42314 | 7 | 0 | 0 | 4 | 0 | 0 | 0 | 0 | 0 | 0 | 247 | 149 | 1 | 1 | 0.809 | 1 | 0 | 1 | 1 | 1 | NA | NA | 1 | NA | -0.238 |
| 42315 | 7 | 0 | 1 | 5 | 0 | 0 | 0 | 0 | 0 | 0 | 247 | 149 | 1 | 1 | 1 | 1 | NA | 0.376 | 0.644 | 1 | NA | 1 | 1 | NA | -0.245 |
| 42316 | 0 | 0 | 0 | 4 | 0 | 0 | 0 | 0 | 0 | 0 | 298 | 149 | 1 | 1 | 1 | 1 | 1 | 0 | 0.333 | NA | NA | NA | 1 | NA | -0.333 |
| 42317 | 0 | 0 | 0 | 6 | 0 | 0 | 0 | 1 | 3 | 0 | 298 | 149 | 1 | 1 | 0.333 | 0.143 | 1 | NA | 1 | NA | NA | 0 | 0.5 | NA | 0.369 |
| 42318 | 0 | 3 | 5 | 0 | 0 | 0 | 0 | 0 | 0 | 0 | 298 | 149 | 1 | 1 | NA | NA | NA | 0.6 | 1 | NA | 1 | 1 | NA | NA | -0.2 |
| 42319 | 0 | 0 | 0 | 0 | 0 | 0 | 0 | 2 | 1 | 2 | 298 | 149 | 1 | 1 | 0.333 | 0.333 | NA | NA | NA | | NA | 0 | 0 | 0 | 0.333 |
| 42325 | 9 | 0 | 2 | 0 | 0 | 0 | 0 | 0 | 0 | 0 | 298 | 149 | 1 | 1 | 0.143 | NA | NA | NA | NA | 1 | NA | 1 | NA | NA | -0.857 |
| 42327 | 18 | 0 | 4 | 0 | 0 | 0 | 0 | 0 | 0 | 0 | 266 | 149 | 1 | 1 | 0 | NA | NA | NA | 1 | 1 | NA | 1 | NA | NA | -0.5 |
| 42328 | 9 | 0 | 2 | 0 | 0 | 0 | 0 | 0 | 0 | 0 | 298 | 149 | 1 | 1 | 0 | NA | NA | NA | 1 | 1 | NA | 1 | NA | NA | -0.5 |
| 42336 | 0 | 0 | 2 | 2 | 4 | 0 | 0 | 0 | 0 | 0 | 298 | 149 | 1 | 1 | 0 | NA | NA | NA | 1 | NA | NA | 1 | 1 | 1 | -0.5 |
| 42338 | 2 | 5 | 0 | 9 | 25 | 14 | 0 | 0 | 0 | 0 | 267 | 149 | 1 | 1 | 1 | 1 | 1 | 1 | 1 | 0.074 | 1 | NA | 1 | 1 | 0.232 |
| 42342 | 24 | 29 | 21 | 14 | 40 | 0 | 0 | 0 | 0 | 0 | 234 | 149 | 1 | 1 | 1 | 1 | 0.936 | 1 | 1 | 1 | 1 | 1 | 1 | 1 | -0.013 |
| 42343 | 26 | 8 | 18 | 20 | 49 | 0 | 0 | 0 | 0 | 0 | 296 | 149 | 1 | 1 | 0.895 | 1 | 1 | 1 | 1 | 1 | 1 | 1 | 1 | 1 | -0.021 |
| 42347 | 28 | 15 | 4 | 19 | 33 | 0 | 0 | 0 | 2 | 0 | 292 | 149 | 1 | 1 | 1 | 0.718 | 1 | 1 | 1 | 1 | 1 | 1 | 0.829 | 1 | -0.022 |
| 42348 | 18 | 10 | 4 | 11 | 24 | 0 | 0 | 0 | 1 | 0 | 292 | 149 | 1 | 1 | 1 | 1 | 1 | 1 | 1 | 1 | 1 | 1 | 0.849 | 1 | 0.03 |
| 42350 | 58 | 79 | 53 | 72 | 92 | 0 | 0 | 0 | 1 | 0 | 298 | 149 | 1 | 1 | 1 | 1 | 1 | 1 | 1 | 1 | 1 | 1 | 0.973 | 1 | 0.005 |
| 42351 | 59 | 69 | 43 | 57 | 73 | 0 | 0 | 0 | 3 | 0 | 298 | 149 | 1 | 1 | 1 | 1 | 1 | 1 | 1 | 1 | 1 | 1 | 0.905 | 1 | 0.019 |
| 42352 | 0 | 0 | 418 | 979 | 2408 | 0 | 0 | 1 | 1 | 0 | 173 | 149 | 1 | 1 | NA | 1 | 0.998 | 1 | NA | NA | NA | 0.997 | 0.999 | 1 | 0.001 |
| 42353 | 125 | 139 | 223 | 1602 | 4241 | 0 | 0 | 1 | 1 | 3 | 298 | 149 | 1 | 1 | 1 | 0.996 | 0.998 | 1 | 1 | 1 | 1 | 0.991 | 0.999 | 0.999 | 0.001 |
| 42355 | 96 | 108 | 67 | 126 | 74 | 0 | 5 | 2 | 0 | 0 | 157 | 149 | 1 | 1 | 0.995 | 0.979 | 1 | 1 | 1 | 1 | 0.953 | 0.97 | 1 | 1 | 0.01 |
| 42356 | 0 | 6 | 3 | 1 | 0 | 0 | 3 | 8 | 5 | 5 | 157 | 149 | 1 | 1 | 0.781 | 0.792 | 0.613 | 0.703 | 0.714 | NA | 0.655 | 0.262 | 0.16 | 0 | 0.451 |
| 42358 | 120 | 138 | 100 | 170 | 94 | 0 | 2 | 0 | 0 | 0 | 241 | 149 | 1 | 1 | 1 | 1 | 1 | 1 | 1 | 1 | 0.977 | 1 | 1 | 1 | 0.005 |
| 42362 | 371 | 723 | 844 | 485 | 300 | 0 | 0 | 4 | 0 | 5 | 232 | 149 | 1 | 1 | 0.998 | 0.994 | 0.987 | 0.995 | 1 | 1 | 1 | 0.993 | 1 | 0.975 | 0.001 |
| 42363 | 277 | 381 | 370 | 237 | 262 | 0 | 0 | 4 | 0 | 5 | 198 | 149 | 1 | 1 | 0.995 | 0.989 | 0.977 | 0.991 | 1 | 1 | 1 | 0.986 | 1 | 0.975 | -0.002 |
| 42365 | 0 | 0 | 5 | 0 | 5 | 0 | 0 | 1 | 0 | 0 | 284 | 149 | 1 | 1 | 1 | 1 | 1 | 1 | NA | NA | NA | 0.724 | NA | 1 | 0.138 |
| 42366 | 86 | 207 | 215 | 83 | 68 | 0 | 1 | 0 | 0 | 0 | 284 | 149 | 1 | 1 | 1 | 1 | 1 | 1 | 1 | 1 | 0.991 | 1 | 1 | 1 | 0.002 |
| 42370 | 103 | 96 | 109 | 53 | 74 | 0 | 0 | 1 | 0 | 0 | 284 | 149 | 1 | 1 | 1 | 1 | 1 | 0.978 | 1 | 1 | 1 | 0.983 | 1 | 1 | -0.001 |
| 42371 | 58 | 127 | 152 | 74 | 80 | 0 | 0 | 0 | 0 | 0 | 269 | 149 | 1 | 1 | 0.989 | 1 | 1 | 1 | 1 | 1 | 1 | 1 | 1 | 1 | -0.002 |
| 42372 | 20 | 96 | 47 | 70 | 61 | 0 | 0 | 0 | 0 | 0 | 281 | 149 | 1 | 1 | 0.972 | 1 | 1 | 1 | 1 | 1 | 1 | 1 | 1 | 1 | -0.006 |
| 42378 | 33 | 100 | 52 | 73 | 61 | 0 | 0 | 0 | 0 | 0 | 298 | 149 | 1 | 1 | 0.974 | 1 | 1 | 1 | 0.838 | 1 | 1 | 1 | 1 | 1 | -0.038 |
| 42379 | 172 | 334 | 408 | 178 | 212 | 0 | 0 | 0 | 0 | 0 | 293 | 149 | 1 | 1 | 0.995 | 1 | 1 | 1 | 0.987 | 1 | 1 | 1 | 1 | 1 | -0.004 |
| 42380 | 272 | 606 | 727 | 396 | 488 | 0 | 4 | 3 | 3 | 0 | 293 | 149 | 1 | 1 | 0.985 | 0.995 | 0.991 | 1 | 1 | 1 | 0.987 | 0.992 | 0.985 | 1 | 0.001 |
| 42381 | 100 | 276 | 322 | 222 | 276 | 0 | 0 | 0 | 0 | 0 | 293 | 149 | 1 | 1 | 1 | 1 | 1 | 0.994 | 1 | 1 | 1 | 1 | 1 | 1 | -0.001 |
| 42382 | 37 | 67 | 44 | 34 | 5 | 0 | 0 | 0 | 0 | 0 | 298 | 149 | 1 | 1 | 1 | 1 | 1 | 0.981 | 1 | 1 | 1 | 1 | 1 | 1 | -0.004 |
| 42383 | 209 | 399 | 449 | 207 | 217 | 0 | 1 | 0 | 0 | 0 | 298 | 149 | 1 | 1 | 0.984 | 0.993 | 0.976 | 0.991 | 1 | 1 | 0.995 | 1 | 1 | 1 | -0.01 |
| 42384 | 236 | 348 | 154 | 227 | 437 | 0 | 0 | 0 | 0 | 0 | 298 | 149 | 1 | 1 | 1 | 1 | 0.986 | 1 | 1 | 1 | 1 | 1 | 1 | 1 | -0.003 |
| 42391 | 0 | 2 | 0 | 0 | 2 | 0 | 1 | 0 | 0 | 0 | 298 | 149 | 1 | 1 | 1 | 1 | NA | 1 | NA | NA | 0.5 | NA | NA | 1 | 0.25 |
| 42393 | 109 | 119 | 186 | 101 | 82 | 0 | 0 | 0 | 0 | 0 | 226 | 149 | 1 | 1 | 0.992 | | 1 | 1 | 0.949 | 1 | 1 | 1 | 1 | 1 | -0.012 |
| 42394 | 36 | 20 | 47 | 48 | 16 | 0 | 0 | 0 | 0 | 0 | 226 | 149 | 1 | 1 | 1 | 1 | 1 | 1 | 0.902 | 1 | 1 | 1 | 1 | 1 | -0.02 |
| 42395 | 26 | 34 | 53 | 47 | 31 | 0 | 0 | 0 | 0 | 0 | 288 | 149 | 1 | 1 | 1 | 1 | 1 | 1 | 0.933 | 1 | 1 | 1 | 1 | 1 | -0.013 |
| 42397 | 0 | 2 | 1 | 0 | 0 | 0 | 0 | 0 | 0 | 0 | 235 | 149 | 1 | 1 | 1 | NA | NA | 1 | 0.388 | NA | 1 | 1 | NA | NA | -0.204 |
| 42398 | 115 | 147 | 243 | 185 | 98 | 0 | 2 | 0 | 0 | 0 | 241 | 149 | 1 | 1 | 0.986 | 1 | 1 | 1 | 1 | 1 | 0.978 | 1 | 1 | 1 | 0.002 |
| 42399 | 264 | 317 | 475 | 406 | 249 | 0 | 0 | 5 | 0 | 0 | 298 | 149 | 1 | 1 | 0.991 | 1 | 0.973 | 1 | 1 | 1 | 1 | 0.979 | 1 | 1 | -0.003 |
| 42401 | 29 | 24 | 24 | 11 | 0 | 0 | 0 | 0 | 0 | 0 | 262 | 149 | 1 | 1 | 1 | 1 | 0.911 | 1 | 1 | 1 | 1 | 1 | 1 | NA | -0.018 |
| 42402 | 0 | 6 | 31 | 27 | 0 | 0 | 0 | 0 | 0 | 0 | 253 | 149 | 1 | 1 | 1 | 1 | 0.541 | 1 | 1 | NA | 1 | 1 | 1 | NA | -0.092 |
| 42405 | 0 | 19 | 3 | 7 | 5 | 0 | 0 | 1 | 6 | 0 | 298 | 149 | 1 | 1 | 0.636 | 1 | 0.636 | 0.667 | 1 | NA | 1 | 0.6 | 0.368 | 1 | 0.046 |
| 42406 | 0 | 19 | 13 | 0 | 4 | 0 | 0 | 1 | 6 | 0 | 197 | 149 | 1 | 1 | 0.831 | 1 | 0.274 | 0.872 | 1 | NA | 1 | 0.908 | 0 | 1 | 0.068 |
| 42408 | 0 | 16 | 17 | 0 | 0 | 0 | 0 | 1 | 6 | 0 | 294 | 149 | 1 | 1 | 0.559 | 1 | 0.432 | 0.802 | 1 | NA | 1 | 0.896 | 0 | NA | 0.127 |
| 42409 | 0 | 35 | 20 | 7 | 5 | 0 | 0 | 1 | 6 | 0 | 298 | 149 | 1 | 1 | 0.75 | 1 | 0.714 | 0.846 | 1 | NA | 1 | 0.909 | 0.368 | 1 | 0.043 |
| 42411 | 0 | 0 | 0 | 4 | 0 | 0 | 13 | 1 | 0 | 4 | 162 | 149 | 1 | 1 | 0 | 0 | 0 | 0 | 0 | NA | 0 | 0 | 1 | 0 | -0.25 |
| 42424 | 60 | 18 | 21 | 12 | 38 | 0 | 0 | 0 | 0 | 0 | 286 | 149 | 1 | 1 | 1 | 0.942 | 1 | 1 | 1 | 1 | 1 | 1 | 1 | 1 | -0.012 |
| 42426 | 24 | 187 | 147 | 21 | 32 | 0 | 0 | 2 | 0 | 0 | 298 | 149 | 1 | 1 | 0.993 | 1 | 1 | 1 | 0.995 | 1 | 1 | 0.974 | 1 | 1 | 0.003 |
| 42427 | 123 | 245 | 200 | 58 | 143 | 0 | 0 | 0 | 0 | 0 | 298 | 149 | 1 | 1 | 1 | 1 | 1 | 1 | 0.997 | 1 | 1 | 1 | 1 | 1 | -0.001 |
| 42428 | 6 | 117 | 83 | 16 | 5 | 0 | 0 | 0 | 0 | 0 | 298 | 149 | 1 | 1 | 1 | 0.988 | 1 | 0.98 | 0.995 | 1 | 1 | 1 | 1 | 1 | -0.007 |
| 42429 | 4 | 202 | 133 | 24 | 20 | 0 | 0 | 0 | 0 | 0 | 298 | 149 | 1 | 1 | 1 | 0.994 | 1 | 0.988 | 0.997 | 1 | 1 | 1 | 1 | 1 | -0.004 |
| 42430 | 15 | 179 | 145 | 43 | 0 | 0 | 0 | 0 | 0 | 0 | 209 | 149 | 1 | 1 | 0.989 | 1 | 1 | 1 | 1 | 1 | 1 | 1 | 1 | NA | -0.002 |
| 42431 | 13 | 265 | 191 | 51 | 15 | 0 | 0 | 4 | 0 | 0 | 209 | 149 | 1 | 1 | 0.993 | 0.996 | 1 | 0.993 | 0.982 | 1 | 1 | 0.971 | 1 | 1 | -0.001 |
| 42432 | 76 | 458 | 340 | 119 | 16 | 0 | 0 | 0 | 0 | 0 | 294 | 149 | 1 | 1 | 1 | 1 | 1 | 0.994 | 1 | 1 | 1 | 1 | 1 | 1 | -0.001 |
| 42433 | 44 | 217 | 177 | 64 | 16 | 0 | 2 | 0 | 0 | 0 | 298 | 149 | 1 | 1 | 1 | 1 | 1 | 1 | 1 | 1 | 0.982 | 1 | 1 | 1 | 0.004 |
| 42434 | 16 | 105 | 106 | 37 | 8 | 0 | 0 | 0 | 0 | 0 | 298 | 149 | 1 | 1 | 1 | 1 | 1 | 1 | 0.993 | 1 | 1 | 1 | 1 | 1 | -0.001 |
| 42435 | 115 | 336 | 291 | 108 | 52 | 0 | 0 | 0 | 0 | 0 | 221 | 149 | 1 | 1 | 1 | 1 | 0.993 | 1 | 0.994 | 1 | 1 | 1 | 1 | 1 | -0.003 |
| 42437 | 51 | 185 | 124 | 46 | 57 | 0 | 0 | 0 | 0 | 0 | 253 | 149 | 1 | 1 | 0.993 | 1 | 1 | 1 | 1 | 1 | 1 | 1 | 1 | 1 | -0.001 |
| 42439 | 56 | 140 | 75 | 49 | 49 | 0 | 0 | 0 | 0 | 0 | 255 | 149 | 1 | 1 | 1 | 1 | 1 | 0.976 | 0.959 | 1 | 1 | 1 | 1 | 1 | -0.013 |
| 42440 | 56 | 140 | 75 | 49 | 49 | 0 | 0 | 0 | 0 | 0 | 255 | 149 | 1 | 1 | 1 | 0.973 | 1 | 1 | 0.983 | 1 | 1 | 1 | 1 | 1 | -0.009 |
| 42441 | 56 | 140 | 75 | 49 | 49 | 0 | 0 | 0 | 0 | 0 | 255 | 149 | 1 | 1 | 1 | 1 | 0.989 | 1 | 1 | 1 | 1 | 1 | 1 | 1 | -0.002 |
| 42443 | 1 | 12 | 15 | 14 | 8 | 0 | 0 | 0 | 0 | 0 | 181 | 149 | 1 | 1 | 1 | 0.904 | 1 | 1 | 0.908 | 1 | 1 | 1 | 1 | 1 | -0.038 |
| 42444 | 0 | 3 | 3 | 4 | 0 | 0 | 0 | 0 | 0 | 0 | 181 | 149 | 1 | 1 | 1 | 1 | 0.622 | 1 | 1 | NA | 1 | 1 | 1 | NA | -0.076 |
| 42445 | 27 | 120 | 87 | 54 | 57 | 0 | 0 | 0 | 0 | 0 | 181 | 149 | 1 | 1 | 1 | 0.984 | 1 | 1 | 1 | 1 | 1 | 1 | 1 | 1 | -0.003 |
| 42446 | 16 | 19 | 19 | 10 | 17 | 0 | 0 | 0 | 0 | 0 | 181 | 149 | 1 | 1 | 1 | 1 | 1 | 1 | 0.968 | 1 | 1 | 1 | 1 | 1 | -0.006 |
| 42447 | 0 | 9 | 6 | 0 | 0 | 0 | 0 | 0 | 0 | 0 | 181 | 149 | 1 | 1 | 1 | 1 | 0.915 | 1 | 1 | NA | 1 | 1 | NA | NA | -0.017 |
| 42448 | 13 | 66 | 54 | 24 | 22 | 0 | 0 | 0 | 0 | 0 | 274 | 149 | 1 | 1 | 1 | 0.949 | 1 | 1 | 0.983 | 1 | 1 | 1 | 1 | 1 | -0.014 |
| 42450 | 13 | 66 | 54 | 24 | 22 | 0 | 0 | 0 | 0 | 0 | 274 | 149 | 1 | 1 | 1 | 1 | 1 | 1 | 0.983 | 1 | 1 | 1 | 1 | 1 | -0.003 |
| 42454 | 27 | 84 | 43 | 18 | 59 | 0 | 0 | 0 | 0 | 0 | 298 | 149 | 1 | 1 | 1 | 1 | 0.985 | 1 | 1 | 1 | 1 | 1 | 1 | 1 | -0.003 |
| 42455 | 27 | 84 | 43 | 18 | 59 | 0 | 0 | 0 | 0 | 0 | 298 | 149 | 1 | 1 | 1 | 1 | 0.985 | 1 | 1 | 1 | 1 | 1 | 1 | 1 | -0.003 |
| 42457 | 45 | 154 | 103 | 60 | 92 | 0 | 0 | 0 | 0 | 0 | 298 | 149 | 1 | 1 | 1 | 1 | 1 | 1 | 0.996 | 1 | 1 | 1 | 1 | 1 | -0.001 |
| 42459 | 121 | 222 | 152 | 63 | 162 | 0 | 0 | 0 | 0 | 0 | 222 | 149 | 1 | 1 | 1 | 0.988 | 1 | 1 | 1 | 1 | 1 | 1 | 1 | 1 | -0.002 |
| 42460 | 347 | 439 | 324 | 152 | 235 | 0 | 0 | 0 | 0 | 0 | 242 | 149 | 1 | 1 | 1 | 0.991 | 1 | 1 | 1 | 1 | 1 | 1 | 1 | 1 | -0.002 |
| 42461 | 1247 | 1179 | 965 | 557 | 734 | 0 | 0 | 1 | 0 | 0 | 242 | 149 | 1 | 1 | 1 | 0.997 | 0.999 | 0.999 | 1 | 1 | 1 | 0.998 | 1 | 1 | -0.001 |
| 42462 | 614 | 744 | 565 | 306 | 370 | 0 | 0 | 6 | 0 | 0 | 295 | 149 | 1 | 1 | 1 | 1 | 0.995 | 1 | 1 | 1 | 1 | 0.979 | 1 | 1 | 0.003 |
| 42476 | 411 | 117 | 46 | 156 | 1159 | 6 | 0 | 0 | 0 | 0 | 264 | 149 | 1 | 1 | 1 | 1 | 1 | 1 | 1 | 0.975 | 1 | 1 | 1 | 1 | 0.005 |
| 42477 | 250 | 54 | 26 | 98 | 765 | 6 | 0 | 0 | 0 | 0 | 256 | 149 | 1 | 1 | 1 | 1 | 1 | 1 | 1 | 0.96 | 1 | 1 | 1 | 1 | 0.008 |
| 42478 | 467 | 135 | 63 | 217 | 1584 | 0 | 0 | 0 | 0 | 0 | 256 | 149 | 1 | 1 | 1 | 1 | 1 | 0.985 | 1 | 1 | 1 | 1 | 1 | 1 | -0.003 |
| 42479 | 450 | 129 | 80 | 219 | 1308 | 0 | 0 | 0 | 1 | 0 | 298 | 149 | 1 | 1 | 0.988 | 1 | 1 | 1 | 1 | 1 | 1 | 1 | 0.991 | 1 | -0.001 |
| 42480 | 201 | 64 | 42 | 130 | 721 | 0 | 0 | 0 | 0 | 2 | 298 | 149 | 1 | 1 | 1 | 1 | 1 | 1 | 1 | 1 | 1 | 1 | 1 | 0.994 | 0.001 |
| 42481 | 109 | 69 | 17 | 56 | 435 | 0 | 0 | 0 | 0 | 2 | 298 | 149 | 1 | 1 | 1 | 1 | 1 | 1 | 1 | 1 | 1 | 1 | 1 | 0.991 | 0.002 |
| 42485 | 22 | 5 | 9 | 2 | 6 | 0 | 0 | 0 | 0 | 0 | 216 | 149 | 1 | 1 | 0.734 | 1 | 1 | 1 | 1 | 1 | 1 | 1 | 1 | 1 | -0.053 |
| 42486 | 20 | 16 | 18 | 9 | 22 | 0 | 0 | 0 | 0 | 0 | 298 | 149 | 1 | 1 | 0.9 | 1 | 1 | 0.867 | 0.929 | 1 | 1 | 1 | 1 | 1 | -0.061 |
| 42495 | 0 | 9 | 2 | 2 | 7 | 0 | 3 | 0 | 0 | 13 | 224 | 149 | 1 | 1 | NA | NA | NA | 1 | 1 | NA | 0.666 | 1 | 1 | 0.264 | 0.268 |
| 42498 | 0 | 0 | 0 | 1 | 12 | 0 | 0 | 0 | 0 | 0 | 298 | 149 | 1 | 1 | 1 | 0 | NA | NA | 1 | NA | NA | NA | 1 | 1 | -0.333 |
| 42500 | 0 | 6 | 2 | 0 | 0 | 1 | 1 | 0 | 0 | 5 | 234 | 149 | 1 | 1 | 0.083 | 0.241 | 0 | 0 | 0 | 0 | 0.793 | 1 | NA | 0 | -0.383 |
| 42507 | 98 | 96 | 30 | 48 | 88 | 5 | 0 | 1 | 0 | 7 | 298 | 149 | 1 | 1 | 0.943 | 1 | 1 | 0.938 | 1 | 0.907 | 1 | 0.938 | 1 | 0.863 | 0.035 |
| 42511 | 0 | 0 | 2 | 0 | 2 | 167 | 122 | 327 | 102 | 12 | 247 | 149 | 1 | 1 | 0 | 0 | 0.009 | 0.019 | 0 | 0 | 0 | 0.004 | 0 | 0.091 | -0.013 |
| 42512 | 170 | 166 | 435 | 117 | 12 | 0 | 1 | 0 | 0 | 0 | 283 | 149 | 1 | 1 | 1 | 1 | 1 | 1 | 1 | 1 | 0.989 | 1 | 1 | 1 | 0.002 |
| 42514 | 25 | 55 | 150 | 23 | 0 | 0 | 0 | 3 | 0 | 0 | 298 | 149 | 1 | 1 | 1 | 0.973 | 1 | 1 | 1 | 1 | 1 | 0.962 | 1 | NA | 0.004 |
| 42518 | 42 | 55 | 41 | 22 | 53 | 0 | 0 | 0 | 0 | 0 | 224 | 149 | 1 | 1 | 1 | 1 | 1 | 1 | 0.947 | 1 | 1 | 1 | 1 | 1 | -0.011 |
| 42521 | 10 | 27 | 18 | 10 | 20 | 2 | 0 | 0 | 0 | 0 | 298 | 149 | 1 | 1 | 0.889 | 1 | 1 | 1 | 0.818 | 0.714 | 1 | 1 | 1 | 1 | -0.001 |
| 42525 | 13 | 1 | 3 | 12 | 29 | 0 | 0 | 0 | 0 | 0 | 176 | 149 | 1 | 1 | 0.717 | 1 | 1 | 1 | 0.809 | 1 | 1 | 1 | 1 | 1 | -0.095 |
| 42546 | 7 | 4 | 1 | 2 | 0 | 0 | 3 | 0 | 0 | 0 | 273 | 149 | 1 | 1 | 1 | 1 | NA | 0.732 | NA | 1 | 0.421 | 1 | 1 | NA | 0.055 |
| 42551 | 42 | 72 | 38 | 23 | 153 | 0 | 3 | 0 | 0 | 0 | 298 | 149 | 1 | 1 | 1 | 1 | 1 | 0.963 | 1 | 1 | 0.923 | 1 | 1 | 1 | 0.008 |
| 42556 | 155 | 128 | 81 | 104 | 379 | 0 | 0 | 0 | 0 | 8 | 269 | 149 | 1 | 1 | 1 | 1 | 1 | 1 | 1 | 1 | 1 | 1 | 1 | 0.963 | 0.007 |
| 42559 | 505 | 476 | 357 | 375 | 556 | 0 | 0 | 0 | 0 | 0 | 251 | 149 | 1 | 1 | 1 | 0.99 | 1 | 1 | 1 | 1 | 1 | 1 | 1 | 1 | -0.002 |
| 42560 | 1031 | 888 | 747 | 666 | 1138 | 0 | 3 | 0 | 7 | 0 | 223 | 149 | 1 | 1 | 1 | 1 | 0.992 | 0.989 | 0.998 | 1 | 0.995 | 1 | 0.985 | 1 | 0 |
| 42561 | 993 | 903 | 787 | 716 | 1192 | 0 | 0 | 0 | 0 | 6 | 237 | 149 | 1 | 1 | 0.993 | 1 | 0.992 | 1 | 0.998 | 1 | 1 | 1 | 1 | 0.992 | -0.002 |
| 42562 | 598 | 484 | 429 | 336 | 721 | 0 | 0 | 1 | 0 | 0 | 225 | 149 | 1 | 1 | 0.996 | 1 | 1 | 0.991 | 1 | 1 | 1 | 0.996 | 1 | 1 | -0.002 |
| 42564 | 0 | 14 | 1 | 0 | 11 | 0 | 19 | 14 | 6 | 0 | 250 | 149 | 1 | 1 | 0.083 | 0.075 | 0.084 | 0.031 | 0.069 | NA | 0.305 | 0.041 | 0 | 1 | -0.268 |
| 42565 | 23 | 103 | 69 | 67 | 36 | 0 | 0 | 0 | 0 | 0 | 298 | 149 | 1 | 1 | 1 | 1 | 0.965 | 0.986 | 0.992 | 1 | 1 | 1 | 1 | 1 | -0.011 |
| 42566 | 32 | 157 | 120 | 98 | 36 | 0 | 0 | 3 | 0 | 0 | 246 | 149 | 1 | 1 | 1 | 1 | 1 | 0.982 | 1 | 1 | 1 | 0.96 | 1 | 1 | 0.004 |
| 42567 | 12 | 74 | 36 | 35 | 22 | 0 | 0 | 0 | 0 | 0 | 246 | 149 | 1 | 1 | 0.973 | 1 | 1 | 1 | 1 | 1 | 1 | 1 | 1 | 1 | -0.005 |
| 42568 | 2 | 34 | 12 | 6 | 0 | 0 | 0 | 0 | 0 | 0 | 205 | 149 | 1 | 1 | 0.968 | 1 | 1 | 1 | 1 | 1 | 1 | 1 | 1 | NA | -0.006 |
| 42569 | 22 | 123 | 88 | 72 | 20 | 0 | 0 | 0 | 0 | 0 | 205 | 149 | 1 | 1 | 1 | 0.991 | 1 | 1 | 1 | 1 | 1 | 1 | 1 | 1 | -0.002 |
| 42571 | 51 | 47 | 40 | 47 | 39 | 0 | 0 | 1 | 0 | 0 | 281 | 149 | 1 | 1 | 1 | 1 | 1 | 1 | 1 | 1 | 1 | 0.955 | 1 | 1 | 0.009 |
| 42574 | 17 | 40 | 15 | 34 | 23 | 0 | 0 | 0 | 0 | 0 | 218 | 149 | 1 | 1 | 1 | 0.925 | 1 | 1 | 1 | 1 | 1 | 1 | 1 | 1 | -0.015 |
| 42577 | 16 | 121 | 157 | 71 | 0 | 0 | 0 | 0 | 0 | 0 | 298 | 149 | 1 | 1 | 1 | 1 | 0.953 | 1 | 0.967 | 1 | 1 | 1 | 1 | NA | -0.016 |
| 42578 | 76 | 280 | 340 | 111 | 40 | 0 | 0 | 0 | 0 | 0 | 264 | 149 | 1 | 1 | 0.991 | 1 | 1 | 1 | 1 | 1 | 1 | 1 | 1 | 1 | -0.002 |
| 42579 | 0 | 0 | 0 | 1 | 0 | 16 | 80 | 121 | 97 | 69 | 203 | 149 | 1 | 1 | 0 | 0 | 0 | 0.034 | 0.018 | 0 | 0 | 0 | 0.008 | 0 | 0.009 |
| 42580 | 29 | 53 | 78 | 52 | 20 | 0 | 0 | 0 | 0 | 0 | 229 | 149 | 1 | 1 | 1 | 1 | 1 | 1 | 0.901 | 1 | 1 | 1 | 1 | 1 | -0.02 |
| 42581 | 158 | 170 | 174 | 160 | 81 | 0 | 0 | 0 | 0 | 0 | 208 | 149 | 1 | 1 | 1 | 1 | 1 | 0.992 | 1 | 1 | 1 | 1 | 1 | 1 | -0.002 |
| 42583 | 0 | 0 | 0 | 0 | 0 | 69 | 67 | 78 | 50 | 24 | 298 | 149 | 1 | 1 | 0 | 0 | 0 | 0 | 0.047 | 0 | 0 | 0 | 0 | 0 | 0.009 |
| 42585 | 67 | 100 | 86 | 51 | 38 | 0 | 0 | 0 | 1 | 0 | 243 | 149 | 1 | 1 | 1 | 1 | 0.98 | 0.958 | 1 | 1 | 1 | 1 | 0.969 | 1 | -0.006 |
| 42586 | 20 | 32 | 26 | 14 | 7 | 0 | 0 | 0 | 0 | 0 | 243 | 149 | 1 | 1 | 1 | 0.947 | 1 | 1 | 1 | 1 | 1 | 1 | 1 | 1 | -0.011 |

| | | | | | | | | | | | | | | |
|---|---|---|---|---|---|---|---|---|---|---|---|---|---|---|
| SE | 42587 | ENSG00000078140 | UBE2K | chr4 | + | 39745751 | 39745810 | 39698138 | 39698390 | 39755656 | 39755739 | 44 | 24 | [illegible] |
| SE | 42589 | ENSG00000078140 | UBE2K | chr4 | + | 39745751 | 39745810 | 39737419 | 39737513 | 39755656 | 39755739 | 140 | 93 | [illegible] |
| SE | 42590 | ENSG00000078140 | UBE2K | chr4 | + | 39755656 | 39755739 | 39698162 | 39698390 | 39774833 | 39774933 | 154 | 85 | [illegible] |
| SE | 42591 | ENSG00000078140 | UBE2K | chr4 | + | 39755656 | 39755739 | 39745751 | 39745810 | 39774833 | 39774933 | 250 | 136 | [illegible] |
| SE | 42594 | ENSG00000078140 | UBE2K | chr4 | + | 39774833 | 39774933 | 39755656 | 39755739 | 39777681 | 39777810 | 294 | 160 | [illegible] |
| SE | 42595 | ENSG00000078140 | UBE2K | chr4 | + | 39774833 | 39774933 | 39755656 | 39755739 | 39778359 | 39779983 | 120 | 70 | [illegible] |
| SE | 42596 | ENSG00000078140 | UBE2K | chr4 | + | 39777681 | 39777810 | 39755656 | 39755739 | 39778359 | 39778384 | 180 | 146 | [illegible] |
| SE | 42600 | ENSG00000116353 | MECR | chr1 | - | 29201373 | 29201518 | 29200515 | 29200589 | 29201942 | 29202045 | 0 | 1 | [illegible] |
| SE | 42601 | ENSG00000116353 | MECR | chr1 | - | 29201942 | 29202045 | 29200323 | 29200589 | 29206761 | 29206905 | 12 | 9 | [illegible] |
| SE | 42602 | ENSG00000116353 | MECR | chr1 | - | 29203130 | 29203233 | 29201942 | 29202045 | 29206761 | 29206828 | 18 | 11 | [illegible] |
| SE | 42603 | ENSG00000116353 | MECR | chr1 | - | 29220837 | 29220921 | 29216587 | 29216685 | 29230730 | 29230897 | 1 | 3 | [illegible] |
| SE | 42604 | ENSG00000116353 | MECR | chr1 | - | 29220837 | 29220921 | 29216587 | 29216812 | 29230730 | 29230897 | 1 | 7 | [illegible] |
| SE | 42607 | ENSG00000116353 | MECR | chr1 | - | 29223139 | 29223260 | 29220716 | 29220921 | 29230730 | 29230901 | 0 | 0 | [illegible] |
| SE | 42608 | ENSG00000116353 | MECR | chr1 | - | 29228880 | 29229054 | 29216600 | 29216812 | 29230730 | 29230942 | 0 | 0 | [illegible] |
| SE | 42610 | ENSG00000116353 | MECR | chr1 | - | 29230354 | 29230406 | 29223139 | 29223260 | 29230730 | 29230891 | 0 | 0 | [illegible] |
| SE | 42612 | ENSG00000168434 | COG7 | chr16 | - | 23416966 | 23417121 | 23413447 | 23413564 | 23418699 | 23418827 | 57 | 26 | [illegible] |
| SE | 42613 | ENSG00000168434 | COG7 | chr16 | - | 23416966 | 23417121 | 23413447 | 23413564 | 23424748 | 23424947 | 32 | 18 | [illegible] |
| SE | 42614 | ENSG00000168434 | COG7 | chr16 | - | 23418699 | 23418827 | 23413447 | 23413564 | 23424748 | 23424947 | 6 | 4 | [illegible] |
| SE | 42616 | ENSG00000168434 | COG7 | chr16 | - | 23442476 | 23442645 | 23434635 | 23434718 | 23445047 | 23445164 | 59 | 43 | [illegible] |
| SE | 42617 | ENSG00000224259 | LINC01133 | chr1 | + | 159965498 | 159965645 | 159961217 | 159961405 | 159976643 | 159977112 | 0 | 2 | [illegible] |
| SE | 42619 | ENSG00000126088 | UROD | chr1 | + | 45013135 | 45013354 | 45012856 | 45013019 | 45013574 | 45013791 | 66 | 52 | [illegible] |
| SE | 42620 | ENSG00000126088 | UROD | chr1 | + | 45013291 | 45013354 | 45012803 | 45013019 | 45013574 | 45013791 | 2 | 1 | [illegible] |
| SE | 42623 | ENSG00000126088 | UROD | chr1 | + | 45013908 | 45014070 | 45013539 | 45013791 | 45014438 | 45014567 | 51 | 16 | [illegible] |
| SE | 42631 | ENSG00000204311 | PJVK | chr2 | + | 178459192 | 178459269 | 178458509 | 178458627 | 178460347 | 178460432 | 0 | 0 | [illegible] |
| SE | 42633 | ENSG00000204311 | PJVK | chr2 | + | 178460347 | 178460446 | 178456009 | 178456151 | 178460981 | 178461204 | 2 | 2 | [illegible] |
| SE | 42634 | ENSG00000204311 | PJVK | chr2 | + | 178460347 | 178460446 | 178458509 | 178458627 | 178460981 | 178461204 | 4 | 4 | [illegible] |
| SE | 42637 | ENSG00000008130 | NADK | chr1 | - | 1753566 | 1753649 | 1751231 | 1753060 | 1754283 | 1754383 | 70 | 44 | [illegible] |
| SE | 42638 | ENSG00000008130 | NADK | chr1 | - | 1754050 | 1754208 | 1751231 | 1753060 | 1754283 | 1754383 | 223 | 109 | [illegible] |
| SE | 42641 | ENSG00000008130 | NADK | chr1 | - | 1756257 | 1756343 | 1754543 | 1754698 | 1756502 | 1756608 | 89 | 40 | [illegible] |
| SE | 42645 | ENSG00000008130 | NADK | chr1 | - | 1757180 | 1757310 | 1756003 | 1756343 | 1761951 | 1762035 | 325 | 171 | [illegible] |
| SE | 42646 | ENSG00000008130 | NADK | chr1 | - | 1761951 | 1762035 | 1756770 | 1757310 | 1765227 | 1765446 | 897 | 571 | [illegible] |
| SE | 42647 | ENSG00000008130 | NADK | chr1 | - | 1761951 | 1762035 | 1756770 | 1757310 | 1779904 | 1779962 | 69 | 23 | [illegible] |
| SE | 42648 | ENSG00000008130 | NADK | chr1 | - | 1765227 | 1765446 | 1756770 | 1757310 | 1779904 | 1779962 | 535 | 356 | [illegible] |
| SE | 42650 | ENSG00000213085 | CFAP45 | chr1 | - | 159876555 | 159876749 | 159872943 | 159873168 | 159877348 | 159877462 | 0 | 3 | [illegible] |
| SE | 42651 | ENSG00000213085 | CFAP45 | chr1 | - | 159876555 | 159876789 | 159872943 | 159873168 | 159877348 | 159877462 | 1 | 1 | [illegible] |
| SE | 42652 | ENSG00000213085 | CFAP45 | chr1 | - | 159886510 | 159886689 | 159884435 | 159884565 | 159887840 | 159888011 | 0 | 6 | [illegible] |
| SE | 42663 | ENSG00000088854 | C20orf194 | chr20 | - | 3304439 | 3304543 | 3298028 | 3298175 | 3315032 | 3315120 | 35 | 39 | [illegible] |
| SE | 42666 | ENSG00000088854 | C20orf194 | chr20 | - | 3324891 | 3324968 | 3322651 | 3322696 | 3326196 | 3326284 | 37 | 26 | [illegible] |
| SE | 42668 | ENSG00000088854 | C20orf194 | chr20 | - | 3348524 | 3348623 | 3343675 | 3343731 | 3359515 | 3359593 | 28 | 31 | [illegible] |
| SE | 42672 | ENSG00000135521 | LTV1 | chr6 | + | 143850130 | 143850218 | 143846050 | 143846224 | 143857302 | 143857444 | 217 | 218 | [illegible] |
| SE | 42676 | ENSG00000116711 | PLA2G4A | chr1 | + | 186854285 | 186854387 | 186828948 | 186829035 | 186870434 | 186870516 | 17 | 16 | [illegible] |
| SE | 42677 | ENSG00000116711 | PLA2G4A | chr1 | + | 186906964 | 186907002 | 186894097 | 186894211 | 186911247 | 186911389 | 42 | 66 | [illegible] |
| SE | 42678 | ENSG00000116711 | PLA2G4A | chr1 | + | 186911247 | 186911389 | 186894097 | 186894211 | 186932762 | 186932899 | 12 | 15 | [illegible] |
| SE | 42679 | ENSG00000116711 | PLA2G4A | chr1 | + | 186932762 | 186932899 | 186911247 | 186911389 | 186939007 | 186939230 | 16 | 24 | [illegible] |
| SE | 42680 | ENSG00000116711 | PLA2G4A | chr1 | + | 186939007 | 186939230 | 186911247 | 186911389 | 186939979 | 186940094 | 14 | 27 | [illegible] |
| SE | 42687 | ENSG00000153561 | RMND5A | chr2 | + | 86740926 | 86741069 | 86720290 | 86720809 | 86751895 | 86752030 | 173 | 136 | [illegible] |
| SE | 42688 | ENSG00000153561 | RMND5A | chr2 | + | 86751895 | 86752030 | 86740926 | 86741069 | 86753457 | 86753545 | 368 | 277 | [illegible] |
| SE | 42689 | ENSG00000153561 | RMND5A | chr2 | + | 86753457 | 86753558 | 86740926 | 86741069 | 86765026 | 86765193 | 179 | 122 | [illegible] |
| SE | 42690 | ENSG00000153561 | RMND5A | chr2 | + | 86753457 | 86753558 | 86751895 | 86752030 | 86765858 | 86766024 | 118 | 97 | [illegible] |
| SE | 42692 | ENSG00000153561 | RMND5A | chr2 | + | 86765858 | 86766024 | 86753457 | 86753558 | 86770022 | 86770125 | 358 | 249 | [illegible] |
| SE | 42693 | ENSG00000153561 | RMND5A | chr2 | + | 86770022 | 86770125 | 86765858 | 86766024 | 86771557 | 86771712 | 522 | 319 | [illegible] |
| SE | 42694 | ENSG00000153561 | RMND5A | chr2 | + | 86771557 | 86771712 | 86765858 | 86766024 | 86773347 | 86773425 | 149 | 143 | [illegible] |
| SE | 42695 | ENSG00000179387 | ELMOD2 | chr4 | + | 140525419 | 140525570 | 140524167 | 140524280 | 140527465 | 140527494 | 117 | 100 | [illegible] |
| SE | 42696 | ENSG00000179387 | ELMOD2 | chr4 | + | 140537411 | 140537541 | 140535416 | 140535830 | 140542573 | 140542642 | 51 | 40 | [illegible] |
| SE | 42697 | ENSG00000179387 | ELMOD2 | chr4 | + | 140540167 | 140540301 | 140537411 | 140537541 | 140542573 | 140542642 | 104 | 62 | [illegible] |
| SE | 42701 | ENSG00000168256 | NKIRAS2 | chr17 | + | 42022375 | 42022640 | 42021563 | 42021671 | 42023653 | 42024220 | 53 | 46 | [illegible] |
| SE | 42702 | ENSG00000168256 | NKIRAS2 | chr17 | + | 42022398 | 42022472 | 42021563 | 42021671 | 42023653 | 42024833 | 59 | 36 | [illegible] |
| SE | 42703 | ENSG00000168256 | NKIRAS2 | chr17 | + | 42022398 | 42022640 | 42021544 | 42021671 | 42023653 | 42023841 | 140 | 89 | [illegible] |
| SE | 42707 | ENSG00000066557 | LRRC40 | chr1 | - | 70159329 | 70159438 | 70155688 | 70155796 | 70173464 | 70173510 | 90 | 68 | [illegible] |
| SE | 42708 | ENSG00000066557 | LRRC40 | chr1 | - | 70173464 | 70173510 | 70159329 | 70159438 | 70173621 | 70173709 | 88 | 67 | [illegible] |
| SE | 42709 | ENSG00000066557 | LRRC40 | chr1 | - | 70173621 | 70173709 | 70173464 | 70173510 | 70175809 | 70175982 | 101 | 90 | [illegible] |
| SE | 42711 | ENSG00000066557 | LRRC40 | chr1 | - | 70187264 | 70187338 | 70184784 | 70184914 | 70189091 | 70189273 | 83 | 29 | [illegible] |
| SE | 42712 | ENSG00000066557 | LRRC40 | chr1 | - | 70187264 | 70187338 | 70184784 | 70184914 | 70205389 | 70205579 | 13 | 7 | [illegible] |
| SE | 42713 | ENSG00000066557 | LRRC40 | chr1 | - | 70189091 | 70189273 | 70184784 | 70184914 | 70205389 | 70205579 | 53 | 32 | [illegible] |
| SE | 42714 | ENSG00000066557 | LRRC40 | chr1 | - | 70189091 | 70189273 | 70187264 | 70187338 | 70205389 | 70205579 | 116 | 54 | [illegible] |
| SE | 42717 | ENSG00000182473 | EXOC7 | chr17 | - | 76084005 | 76084139 | 76081015 | 76083750 | 76084247 | 76084289 | 475 | 300 | [illegible] |
| SE | 42730 | ENSG00000221994 | ZNF630 | chrX | - | 48046202 | 48046310 | 48002594 | 48002653 | 48060449 | 48060545 | 4 | 0 | [illegible] |
| SE | 42732 | ENSG00000221994 | ZNF630 | chrX | - | 48060818 | 48060903 | 48060449 | 48060545 | 48066871 | 48067061 | 3 | 0 | [illegible] |
| SE | 42733 | ENSG00000221994 | ZNF630 | chrX | - | 48060818 | 48060945 | 48060449 | 48060545 | 48066871 | 48067061 | 12 | 4 | [illegible] |
| SE | 42739 | ENSG00000133226 | SRRM1 | chr1 | + | 24648023 | 24648166 | 24646666 | 24646789 | 24648858 | 24648887 | 1 | 4 | [illegible] |
| SE | 42741 | ENSG00000133226 | SRRM1 | chr1 | + | 24650564 | 24650627 | 24649987 | 24650086 | 24651408 | 24651612 | 0 | 0 | [illegible] |
| SE | 42743 | ENSG00000133226 | SRRM1 | chr1 | + | 24661309 | 24661396 | 24660718 | 24660799 | 24662659 | 24662804 | 237 | 197 | [illegible] |
| SE | 42744 | ENSG00000133226 | SRRM1 | chr1 | + | 24661312 | 24661396 | 24660718 | 24660799 | 24662659 | 24662804 | 99 | 71 | [illegible] |
| SE | 42746 | ENSG00000133226 | SRRM1 | chr1 | + | 24670119 | 24670315 | 24669122 | 24669587 | 24671385 | 24671595 | 277 | 174 | [illegible] |
| SE | 42747 | ENSG00000119965 | C10orf88 | chr10 | - | 122937704 | 122938159 | 122930900 | 122932661 | 122948648 | 122948755 | 49 | 42 | [illegible] |
| SE | 42748 | ENSG00000119965 | C10orf88 | chr10 | - | 122948648 | 122948755 | 122937704 | 122938159 | 122951953 | 122952007 | 22 | 17 | [illegible] |
| SE | 42749 | ENSG00000119965 | C10orf88 | chr10 | - | 122948648 | 122948855 | 122937704 | 122938159 | 122951953 | 122952026 | 78 | 58 | [illegible] |
| SE | 42750 | ENSG00000134371 | CDC73 | chr1 | + | 193125111 | 193125217 | 193121982 | 193122331 | 193130173 | 193130243 | 189 | 123 | [illegible] |
| SE | 42751 | ENSG00000134371 | CDC73 | chr1 | + | 193127847 | 193128138 | 193125111 | 193125217 | 193130173 | 193130243 | 0 | 0 | [illegible] |
| SE | 42752 | ENSG00000134371 | CDC73 | chr1 | + | 193141072 | 193141142 | 193138084 | 193138173 | 193141849 | 193142066 | 0 | 4 | [illegible] |
| SE | 42754 | ENSG00000134371 | CDC73 | chr1 | + | 193147866 | 193147965 | 193141849 | 193142066 | 193150303 | 193150381 | 199 | 170 | [illegible] |
| SE | 42755 | ENSG00000134371 | CDC73 | chr1 | + | 193161122 | 193161190 | 193152379 | 193152444 | 193203794 | 193203852 | 0 | 0 | [illegible] |
| SE | 42757 | ENSG00000134371 | CDC73 | chr1 | + | 193237707 | 193237854 | 193236255 | 193236356 | 193249729 | 193249871 | 0 | 0 | [illegible] |
| SE | 42760 | ENSG00000141577 | CEP131 | chr17 | - | 81192735 | 81192843 | 81192475 | 81192593 | 81193925 | 81194127 | 19 | 5 | [illegible] |
| SE | 42761 | ENSG00000143376 | SNX27 | chr1 | + | 151638887 | 151639119 | 151612028 | 151612512 | 151658234 | 151658427 | 158 | 113 | [illegible] |
| SE | 42762 | ENSG00000143376 | SNX27 | chr1 | + | 151638887 | 151639119 | 151612063 | 151612381 | 151658234 | 151658427 | 127 | 102 | [illegible] |
| SE | 42764 | ENSG00000143376 | SNX27 | chr1 | + | 151665932 | 151666011 | 151662165 | 151662270 | 151668471 | 151668635 | 266 | 149 | [illegible] |
| SE | 42765 | ENSG00000143376 | SNX27 | chr1 | + | 151668471 | 151668635 | 151662165 | 151662270 | 151683355 | 151683445 | 276 | 169 | [illegible] |
| SE | 42766 | ENSG00000143376 | SNX27 | chr1 | + | 151683355 | 151683445 | 151668471 | 151668635 | 151692434 | 151692584 | 460 | 300 | [illegible] |
| SE | 42767 | ENSG00000143376 | SNX27 | chr1 | + | 151693423 | 151693483 | 151692910 | 151693039 | 151694369 | 151694485 | 173 | 112 | [illegible] |
| SE | 42768 | ENSG00000001460 | STPG1 | chr1 | - | 24360850 | 24361041 | 24356998 | 24358619 | 24373701 | 24373810 | 4 | 6 | [illegible] |
| SE | 42769 | ENSG00000001460 | STPG1 | chr1 | - | 24369673 | 24369839 | 24360850 | 24361041 | 24373701 | 24373810 | 3 | 9 | [illegible] |
| SE | 42770 | ENSG00000001460 | STPG1 | chr1 | - | 24373701 | 24373810 | 24360850 | 24361041 | 24379652 | 24379823 | 7 | 8 | [illegible] |
| SE | 42771 | ENSG00000001460 | STPG1 | chr1 | - | 24391560 | 24391679 | 24383866 | 24384003 | 24401318 | 24401331 | 10 | 13 | [illegible] |
| SE | 42772 | ENSG00000001460 | STPG1 | chr1 | - | 24391923 | 24392067 | 24391560 | 24391679 | 24401318 | 24401456 | 0 | 0 | [illegible] |
| SE | 42774 | ENSG00000137462 | TLR2 | chr4 | + | 153687916 | 153687988 | 153684264 | 153684360 | 153702891 | 153705702 | 3 | 4 | [illegible] |
| SE | 42777 | ENSG00000137462 | TLR2 | chr4 | + | 153702226 | 153702510 | 153687916 | 153687988 | 153702891 | 153702962 | 3 | 10 | [illegible] |
| SE | 42791 | ENSG00000139842 | CUL4A | chr13 | + | 113233896 | 113233986 | 113233152 | 113233339 | 113235062 | 113235145 | 324 | 199 | [illegible] |
| SE | 42792 | ENSG00000139842 | CUL4A | chr13 | + | 113235062 | 113235145 | 113233896 | 113233986 | 113236822 | 113236890 | 392 | 223 | [illegible] |
| SE | 42798 | ENSG00000137409 | MTCH1 | chr6 | - | 36976497 | 36976598 | 36975657 | 36975717 | 36977198 | 36977250 | 1 | 0 | [illegible] |
| SE | 42799 | ENSG00000137409 | MTCH1 | chr6 | - | 36977198 | 36977250 | 36975657 | 36975717 | 36977633 | 36977673 | 248 | 151 | [illegible] |
| SE | 42800 | ENSG00000137409 | MTCH1 | chr6 | - | 36977633 | 36977691 | 36977198 | 36977250 | 36978077 | 36978155 | 254 | 150 | [illegible] |
| SE | 42802 | ENSG00000143450 | OAZ3 | chr1 | + | 151768057 | 151768233 | 151767224 | 151767299 | 151770171 | 151770257 | 2 | 0 | [illegible] |
| SE | 42803 | ENSG00000143450 | OAZ3 | chr1 | + | 151768144 | 151768233 | 151767247 | 151767299 | 151770171 | 151770257 | 1 | 0 | [illegible] |
| SE | 42806 | ENSG00000051341 | POLQ | chr3 | - | 121471989 | 121472164 | 121468304 | 121468431 | 121476539 | 121476733 | 1 | 6 | [illegible] |
| SE | 42807 | ENSG00000051341 | POLQ | chr3 | - | 121473349 | 121473487 | 121471989 | 121472164 | 121476539 | 121476733 | 1 | 8 | [illegible] |
| SE | 42808 | ENSG00000051341 | POLQ | chr3 | - | 121522002 | 121522149 | 121519870 | 121520083 | 121532989 | 121533209 | 1 | 1 | [illegible] |
| SE | 42809 | ENSG00000051341 | POLQ | chr3 | - | 121529644 | 121529792 | 121522002 | 121522149 | 121532989 | 121533209 | 9 | 2 | [illegible] |
| SE | 42810 | ENSG00000051341 | POLQ | chr3 | - | 121537099 | 121537208 | 121532989 | 121533209 | 121539432 | 121539589 | 0 | 2 | [illegible] |
| SE | 42812 | ENSG00000205578 | POM121B | chr7 | + | 73295724 | 73295901 | 73295321 | 73295424 | 73296748 | 73298437 | 1 | 0 | [illegible] |
| SE | 42816 | ENSG00000140575 | IQGAP1 | chr15 | + | 90429588 | 90429666 | 90426109 | 90426266 | 90433718 | 90433795 | 667 | 454 | [illegible] |
| SE | 42817 | ENSG00000140575 | IQGAP1 | chr15 | + | 90431220 | 90431252 | 90429588 | 90429666 | 90433718 | 90433795 | 2 | 0 | [illegible] |
| SE | 42818 | ENSG00000140575 | IQGAP1 | chr15 | + | 90439331 | 90439399 | 90433718 | 90433795 | 90440501 | 90440528 | 1563 | 1083 | [illegible] |
| SE | 42819 | ENSG00000140575 | IQGAP1 | chr15 | + | 90440501 | 90440615 | 90433718 | 90433795 | 90441505 | 90441684 | 1362 | 895 | [illegible] |
| SE | 42821 | ENSG00000140575 | IQGAP1 | chr15 | + | 90443393 | 90443478 | 90440501 | 90440615 | 90448572 | 90448736 | 280 | 178 | [illegible] |
| SE | 42822 | ENSG00000140575 | IQGAP1 | chr15 | + | 90466268 | 90466436 | 90466000 | 90466091 | 90467449 | 90467592 | 1950 | 1396 | [illegible] |
| SE | 42824 | ENSG00000140575 | IQGAP1 | chr15 | + | 90466268 | 90466436 | 90466000 | 90466091 | 90473714 | 90473798 | 1034 | 714 | [illegible] |
| SE | 42826 | ENSG00000140575 | IQGAP1 | chr15 | + | 90467449 | 90467592 | 90466000 | 90466091 | 90473714 | 90473798 | 3 | 2 | [illegible] |
| SE | 42828 | ENSG00000140575 | IQGAP1 | chr15 | + | 90467449 | 90467592 | 90466268 | 90466436 | 90473714 | 90473798 | 918 | 684 | [illegible] |
| SE | 42829 | ENSG00000140575 | IQGAP1 | chr15 | + | 90472839 | 90473010 | 90466000 | 90466091 | 90473714 | 90473798 | 782 | 513 | [illegible] |
| SE | 42830 | ENSG00000140575 | IQGAP1 | chr15 | + | 90472839 | 90473010 | 90466268 | 90466436 | 90473714 | 90473798 | 782 | 514 | [illegible] |
| SE | 42832 | ENSG00000140575 | IQGAP1 | chr15 | + | 90474063 | 90474133 | 90473895 | 90473967 | 90474484 | 90474693 | 1878 | 1323 | [illegible] |
| SE | 42833 | ENSG00000140575 | IQGAP1 | chr15 | + | 90474484 | 90474693 | 90474063 | 90474133 | 90476662 | 90476818 | 2590 | 2012 | [illegible] |
| SE | 42834 | ENSG00000140575 | IQGAP1 | chr15 | + | 90477664 | 90477889 | 90477066 | 90477230 | 90481959 | 90482100 | 1450 | 1071 | [illegible] |
| SE | 42835 | ENSG00000140575 | IQGAP1 | chr15 | + | 90481959 | 90482100 | 90477664 | 90477889 | 90483360 | 90483593 | 750 | 595 | [illegible] |
| SE | 42836 | ENSG00000140575 | IQGAP1 | chr15 | + | 90482196 | 90482281 | 90481959 | 90482100 | 90483360 | 90483593 | 1468 | 1015 | [illegible] |
| SE | 42838 | ENSG00000140575 | IQGAP1 | chr15 | + | 90494712 | 90494835 | 90492544 | 90492711 | 90497231 | 90497340 | 2139 | 1232 | [illegible] |
| SE | 42844 | ENSG00000123728 | RAP2C | chrX | - | 132216995 | 132217156 | 132214133 | 132214446 | 132217277 | 132217367 | 302 | 209 | [illegible] |
| SE | 42845 | ENSG00000123728 | RAP2C | chrX | - | 132217277 | 132217367 | 132216995 | 132217156 | 132218231 | 132218332 | 16 | 10 | [illegible] |
| SE | 42846 | ENSG00000123728 | RAP2C | chrX | - | 132217915 | 132218012 | 132216995 | 132217806 | 132218231 | 132218341 | 2 | 1 | [illegible] |
| SE | 42847 | ENSG00000123728 | RAP2C | chrX | - | 132218231 | 132218341 | 132216995 | 132217806 | 132219369 | 132219447 | 0 | 1 | [illegible] |
| SE | 42848 | ENSG00000143552 | NUP210L | chr1 | - | 154117724 | 154117880 | 154104011 | 154104210 | 154118670 | 154118808 | 3 | 0 | [illegible] |
| SE | 42849 | ENSG00000152495 | CAMK4 | chr5 | + | 111344023 | 111344102 | 111224381 | 111224644 | 111374849 | 111374912 | 82 | 132 | [illegible] |
| SE | 42850 | ENSG00000152495 | CAMK4 | chr5 | + | 111374849 | 111374912 | 111344023 | 111344102 | 111376859 | 111376942 | 115 | 176 | [illegible] |
| SE | 42852 | ENSG00000152495 | CAMK4 | chr5 | + | 111376859 | 111376942 | 111344023 | 111344102 | 111394709 | 111394749 | 115 | 122 | [illegible] |
| SE | 42853 | ENSG00000152495 | CAMK4 | chr5 | + | 111376859 | 111376942 | 111344023 | 111344102 | 111446685 | 111446776 | 1 | 0 | [illegible] |
| SE | 42855 | ENSG00000152495 | CAMK4 | chr5 | + | 111394709 | 111394782 | 111344023 | 111344102 | 111446685 | 111446776 | 49 | 41 | [illegible] |
| SE | 42856 | ENSG00000152495 | CAMK4 | chr5 | + | 111394709 | 111394782 | 111376859 | 111376942 | 111446685 | 111446776 | 205 | 221 | [illegible] |
| SE | 42857 | ENSG00000152495 | CAMK4 | chr5 | + | 111446685 | 111446776 | 111394709 | 111394782 | 111449128 | 111449193 | 199 | 200 | [illegible] |
| SE | 42858 | ENSG00000152495 | CAMK4 | chr5 | + | 111449128 | 111449203 | 111446685 | 111446776 | 111473310 | 111473324 | 151 | 134 | [illegible] |
| SE | 42859 | ENSG00000152495 | CAMK4 | chr5 | + | 111473310 | 111473386 | 111449128 | 111449203 | 111478380 | 111478507 | 146 | 132 | [illegible] |
| SE | 42862 | ENSG00000187010 | RHD | chr1 | + | 25306595 | 25306729 | 25303321 | 25303459 | 25316999 | 25317079 | 1 | 0 | [illegible] |
| SE | 42863 | ENSG00000187010 | RHD | chr1 | + | 25306595 | 25306729 | 25303321 | 25303459 | 25328897 | 25329211 | 0 | 0 | [illegible] |
| SE | 42864 | ENSG00000187010 | RHD | chr1 | + | 25307646 | 25307816 | 25306595 | 25306729 | 25316999 | 25317079 | 0 | 0 | [illegible] |
| SE | 42872 | ENSG00000151748 | SAV1 | chr14 | - | 50640749 | 50640893 | 50633579 | 50635384 | 50644743 | 50645014 | 114 | 128 | [illegible] |
| SE | 42875 | ENSG00000125637 | PSD4 | chr2 | + | 113182345 | 113183512 | 113173970 | 113174054 | 113184956 | 113185073 | 308 | 230 | [illegible] |
| SE | 42879 | ENSG00000125637 | PSD4 | chr2 | + | 113185364 | 113186255 | 113184956 | 113185073 | 113193257 | 113193370 | 42 | 73 | [illegible] |
| SE | 42880 | ENSG00000125637 | PSD4 | chr2 | + | 113185661 | 113186255 | 113185364 | 113185440 | 113193047 | 113193128 | 2 | 0 | [illegible] |
| SE | 42881 | ENSG00000125637 | PSD4 | chr2 | + | 113185876 | 113186255 | 113185364 | 113185440 | 113193047 | 113193128 | 168 | 122 | [illegible] |
| SE | 42882 | ENSG00000125637 | PSD4 | chr2 | + | 113185960 | 113186255 | 113185364 | 113185440 | 113193047 | 113193128 | 2 | 0 | [illegible] |
| SE | 42884 | ENSG00000125637 | PSD4 | chr2 | + | 113192379 | 113192589 | 113185364 | 113186255 | 113193257 | 113193370 | 173 | 128 | [illegible] |
| SE | 42885 | ENSG00000125637 | PSD4 | chr2 | + | 113193047 | 113193128 | 113184956 | 113185073 | 113193257 | 113193370 | 191 | 160 | [illegible] |
| SE | 42886 | ENSG00000125637 | PSD4 | chr2 | + | 113193047 | 113193128 | 113185364 | 113186255 | 113193257 | 113193370 | 193 | 160 | [illegible] |
| SE | 42887 | ENSG00000125637 | PSD4 | chr2 | + | 113193047 | 113193128 | 113192379 | 113192589 | 113193257 | 113193370 | 412 | 369 | [illegible] |
| SE | 42888 | ENSG00000125637 | PSD4 | chr2 | + | 113193257 | 113193370 | 113193047 | 113193128 | 113193591 | 113193650 | 407 | 344 | [illegible] |
| SE | 42909 | ENSG00000164037 | SLC9B1 | chr4 | - | 102905513 | 102905650 | 102885047 | 102885328 | 102906535 | 102906644 | 1 | 3 | [illegible] |
| SE | 42910 | ENSG00000164037 | SLC9B1 | chr4 | - | 102905513 | 102905650 | 102900927 | 102901332 | 102906535 | 102906644 | 3 | 5 | [illegible] |
| SE | 42911 | ENSG00000164037 | SLC9B1 | chr4 | - | 102910438 | 102910588 | 102906276 | 102906644 | 102911430 | 102911537 | 20 | 6 | [illegible] |
| SE | 42912 | ENSG00000164037 | SLC9B1 | chr4 | - | 102912007 | 102912129 | 102911430 | 102911537 | 102932123 | 102932299 | 0 | 1 | [illegible] |
| SE | 42914 | ENSG00000164037 | SLC9B1 | chr4 | - | 102945192 | 102945320 | 102932123 | 102932299 | 102949256 | 102949427 | 4 | 4 | [illegible] |
| SE | 42917 | ENSG00000164037 | SLC9B1 | chr4 | - | 102946646 | 102946789 | 102945192 | 102945320 | 102949256 | 102949427 | 5 | 0 | [illegible] |
| SE | 42919 | ENSG00000164037 | SLC9B1 | chr4 | - | 102949256 | 102949427 | 102932123 | 102932299 | 102991642 | 102991712 | 2 | 0 | [illegible] |
| SE | 42921 | ENSG00000164037 | SLC9B1 | chr4 | - | 102989799 | 102989941 | 102932123 | 102932299 | 102991642 | 102991712 | 2 | 0 | [illegible] |
| SE | 42926 | ENSG00000164112 | SMIM43 | chr4 | - | 121764130 | 121764245 | 121763763 | 121763872 | 121764816 | 121765170 | 0 | 0 | [illegible] |
| SE | 42927 | ENSG00000114861 | FOXP1 | chr3 | - | 70972010 | 70972180 | 70970735 | 70970805 | 70972554 | 70972676 | 3 | 0 | [illegible] |
| SE | 42928 | ENSG00000114861 | FOXP1 | chr3 | - | 70972010 | 70972180 | 70970735 | 70970805 | 70976940 | 70977042 | 2 | 0 | [illegible] |
| SE | 42929 | ENSG00000114861 | FOXP1 | chr3 | - | 70972010 | 70972676 | 70970735 | 70970805 | 70976940 | 70977042 | 537 | 423 | [illegible] |
| SE | 42931 | ENSG00000114861 | FOXP1 | chr3 | - | 71041327 | 71041532 | 71015548 | 71015653 | 71064823 | 71064926 | 284 | 238 | [illegible] |
| SE | 42932 | ENSG00000114861 | FOXP1 | chr3 | - | 71046941 | 71047089 | 71041327 | 71041532 | 71052536 | 71052626 | 114 | 66 | [illegible] |
| SE | 42933 | ENSG00000114861 | FOXP1 | chr3 | - | 71055112 | 71055433 | 71053717 | 71053773 | 71112535 | 71112637 | 0 | 0 | [illegible] |
| SE | 42935 | ENSG00000114861 | FOXP1 | chr3 | - | 71296432 | 71296445 | 71053635 | 71053773 | 71299819 | 71299880 | 0 | 0 | [illegible] |
| SE | 42936 | ENSG00000114861 | FOXP1 | chr3 | - | 71296432 | 71296445 | 71198201 | 71198392 | 71299819 | 71299880 | 0 | 0 | [illegible] |
| SE | 42938 | ENSG00000114861 | FOXP1 | chr3 | - | 71493425 | 71493555 | 71356488 | 71359244 | 71581548 | 71581697 | 103 | 128 | [illegible] |
| SE | 42940 | ENSG00000163156 | SCNM1 | chr1 | + | 151166470 | 151166541 | 151156663 | 151156722 | 151166933 | 151167022 | 112 | 97 | [illegible] |
| SE | 42941 | ENSG00000163156 | SCNM1 | chr1 | + | 151168143 | 151168338 | 151166470 | 151166541 | 151168985 | 151169111 | 50 | 31 | [illegible] |
| SE | 42942 | ENSG00000163156 | SCNM1 | chr1 | + | 151168143 | 151168338 | 151167325 | 151167414 | 151168985 | 151169111 | 103 | 85 | [illegible] |
| SE | 42951 | ENSG00000196110 | ZNF699 | chr19 | - | 9305071 | 9305124 | 9302377 | 9302504 | 9309349 | 9309838 | 48 | 39 | [illegible] |
| SE | 42957 | ENSG00000121940 | CLCC1 | chr1 | - | 108940044 | 108940142 | 108939635 | 108939782 | 108941404 | 108941494 | 124 | 102 | [illegible] |
| SE | 42958 | ENSG00000121940 | CLCC1 | chr1 | - | 108940044 | 108940142 | 108939635 | 108939782 | 108943474 | 108943615 | 65 | 51 | [illegible] |
| SE | 42959 | ENSG00000121940 | CLCC1 | chr1 | - | 108940044 | 108940142 | 108939635 | 108939782 | 108943835 | 108943882 | 62 | 49 | [illegible] |
| SE | 42962 | ENSG00000121940 | CLCC1 | chr1 | - | 108941404 | 108941498 | 108939635 | 108939782 | 108943474 | 108943615 | 18 | 14 | [illegible] |
| SE | 42967 | ENSG00000121940 | CLCC1 | chr1 | - | 108943474 | 108943615 | 108939635 | 108939782 | 108943835 | 108943882 | 39 | 12 | [illegible] |
| SE | 42969 | ENSG00000121940 | CLCC1 | chr1 | - | 108943474 | 108943615 | 108940044 | 108940142 | 108943835 | 108943882 | 42 | 14 | [illegible] |
| SE | 42970 | ENSG00000121940 | CLCC1 | chr1 | - | 108943474 | 108943615 | 108941404 | 108941498 | 108943835 | 108943882 | 69 | 43 | [illegible] |
| SE | 42973 | ENSG00000121940 | CLCC1 | chr1 | - | 108943835 | 108943907 | 108941404 | 108941498 | 108947610 | 108947718 | 1 | 0 | [illegible] |
| SE | 42979 | ENSG00000121940 | CLCC1 | chr1 | - | 108949819 | 108949921 | 108947610 | 108947718 | 108962308 | 108962469 | 41 | 37 | [illegible] |
| SE | 42980 | ENSG00000121940 | CLCC1 | chr1 | - | 108950308 | 108950355 | 108949819 | 108949921 | 108962308 | 108963453 | 71 | 50 | [illegible] |
| SE | 42981 | ENSG00000121940 | CLCC1 | chr1 | - | 108950308 | 108950394 | 108949819 | 108949921 | 108962308 | 108962469 | 79 | 60 | [illegible] |
| SE | 42982 | ENSG00000121940 | CLCC1 | chr1 | - | 108950308 | 108950438 | 108949819 | 108949921 | 108962308 | 108962469 | 111 | 69 | [illegible] |
| SE | 42983 | ENSG00000121940 | CLCC1 | chr1 | - | 108950308 | 108950438 | 108949819 | 108949921 | 108963360 | 108963425 | 111 | 69 | [illegible] |
| SE | 42984 | ENSG00000121940 | CLCC1 | chr1 | - | 108950308 | 108950448 | 108949819 | 108949921 | 108962308 | 108962464 | 140 | 94 | [illegible] |
| SE | 42985 | ENSG00000121940 | CLCC1 | chr1 | - | 108950308 | 108950448 | 108949819 | 108949921 | 108963360 | 108963457 | 129 | 74 | [illegible] |
| SE | 42987 | ENSG00000121940 | CLCC1 | chr1 | - | 108962308 | 108962464 | 108950308 | 108950448 | 108963342 | 108963484 | 25 | 23 | [illegible] |
| SE | 42989 | ENSG00000121940 | CLCC1 | chr1 | - | 108962308 | 108962469 | 108949819 | 108949921 | 108963360 | 108963413 | 6 | 5 | [illegible] |
| SE | 42990 | ENSG00000121940 | CLCC1 | chr1 | - | 108962308 | 108962469 | 108950308 | 108950438 | 108963360 | 108963413 | 8 | 6 | [illegible] |
| SE | 42991 | ENSG00000121940 | CLCC1 | chr1 | - | 108962308 | 108962469 | 108950308 | 108950438 | 108963380 | 108963475 | 2 | 1 | [illegible] |
| SE | 42992 | ENSG00000121940 | CLCC1 | chr1 | - | 108962308 | 108962469 | 108950308 | 108950448 | 108963342 | 108963484 | 25 | 23 | [illegible] |
| SE | 42994 | ENSG00000121940 | CLCC1 | chr1 | - | 108962308 | 108962469 | 108950308 | 108950448 | 108963380 | 108963475 | 25 | 23 | [illegible] |
| SE | 42995 | ENSG00000065911 | MTHFD2 | chr2 | + | 74201381 | 74201479 | 74187382 | 74187457 | 74205704 | 74205889 | 3 | 2 | [illegible] |
| SE | 42997 | ENSG00000065911 | MTHFD2 | chr2 | + | 74201381 | 74201479 | 74198612 | 74198742 | 74207703 | 74207826 | 5 | 2 | [illegible] |
| SE | 43001 | ENSG00000065911 | MTHFD2 | chr2 | + | 74205704 | 74205889 | 74201381 | 74201479 | 74207703 | 74207826 | 113 | 72 | [illegible] |
| SE | 43002 | ENSG00000065911 | MTHFD2 | chr2 | + | 74205981 | 74206088 | 74205704 | 74205889 | 74207703 | 74207826 | 7 | 0 | [illegible] |
| SE | 43008 | ENSG00000076258 | FMO4 | chr1 | + | 171319817 | 171319957 | 171316185 | 171316327 | 171323003 | 171323192 | 20 | 19 | [illegible] |
| SE | 43009 | ENSG00000076258 | FMO4 | chr1 | + | 171319817 | 171319957 | 171316185 | 171316327 | 171324137 | 171324300 | 15 | 14 | [illegible] |
| SE | 43010 | ENSG00000076258 | FMO4 | chr1 | + | 171319817 | 171319957 | 171316185 | 171316327 | 171337355 | 171337423 | 14 | 14 | [illegible] |
| SE | 43011 | ENSG00000076258 | FMO4 | chr1 | + | 171323003 | 171323192 | 171316185 | 171316327 | 171324137 | 171324300 | 8 | 8 | [illegible] |
| SE | 43014 | ENSG00000076258 | FMO4 | chr1 | + | 171323003 | 171323192 | 171319817 | 171319957 | 171324137 | 171324300 | 14 | 13 | [illegible] |
| SE | 43015 | ENSG00000076258 | FMO4 | chr1 | + | 171323003 | 171323192 | 171319817 | 171319957 | 171337355 | 171337423 | 6 | 5 | [illegible] |
| SE | 43016 | ENSG00000076258 | FMO4 | chr1 | + | 171324137 | 171324300 | 171316185 | 171316327 | 171331639 | 171331782 | 6 | 5 | [illegible] |

| | | | | | | | | | | | | |
|---|---|---|---|---|---|---|---|---|---|---|---|---|
| SE | 43019 | ENSG00000076258 | FMO4 | chr1 | + | 171324137 | 171324300 | 171323003 | 171323192 | 171331639 | 171331782 | 14 | 14 | [illegible] |
| SE | 43021 | ENSG00000076258 | FMO4 | chr1 | + | 171331639 | 171331782 | 171324137 | 171324300 | 171334410 | 171334763 | 6 | 6 | [illegible] |
| SE | 43022 | ENSG00000076258 | FMO4 | chr1 | + | 171332708 | 171332908 | 171331639 | 171331782 | 171334410 | 171334763 | 26 | 14 | [illegible] |
| SE | 43023 | ENSG00000076258 | FMO4 | chr1 | + | 171334410 | 171334763 | 171324137 | 171324300 | 171337355 | 171337425 | 9 | 14 | [illegible] |
| SE | 43024 | ENSG00000144120 | TMEM177 | chr2 | + | 119680831 | 119681123 | 119679202 | 119679276 | 119685657 | 119686507 | 1 | 1 | [illegible] |
| SE | 43025 | ENSG00000117395 | EBNA1BP2 | chr1 | - | 43166825 | 43166919 | 43164642 | 43164805 | 43167159 | 43167235 | 92 | 75 | [illegible] |
| SE | 43026 | ENSG00000117395 | EBNA1BP2 | chr1 | - | 43168938 | 43169028 | 43167159 | 43167235 | 43170755 | 43170879 | 137 | 114 | [illegible] |
| SE | 43027 | ENSG00000117395 | EBNA1BP2 | chr1 | - | 43170755 | 43170879 | 43168938 | 43169028 | 43171478 | 43171651 | 174 | 151 | [illegible] |
| SE | 43029 | ENSG00000111674 | ENO2 | chr12 | + | 6917580 | 6917714 | 6917037 | 6917107 | 6917939 | 6918162 | 74 | 50 | [illegible] |
| SE | 43034 | ENSG00000079308 | TNS1 | chr2 | - | 217813214 | 217813304 | 217812367 | 217812445 | 217813684 | 217813816 | 7 | 5 | [illegible] |
| SE | 43035 | ENSG00000079308 | TNS1 | chr2 | - | 217813214 | 217813307 | 217812367 | 217812445 | 217813684 | 217813816 | 8 | 6 | [illegible] |
| SE | 43040 | ENSG00000079308 | TNS1 | chr2 | - | 217893438 | 217893561 | 217892947 | 217893012 | 217895005 | 217895056 | 7 | 0 | [illegible] |
| SE | 43041 | ENSG00000079308 | TNS1 | chr2 | - | 217900462 | 217900512 | 217897794 | 217897969 | 217906334 | 217906385 | 3 | 1 | [illegible] |
| SE | 43042 | ENSG00000079308 | TNS1 | chr2 | - | 217901709 | 217901823 | 217900462 | 217900512 | 217906334 | 217906385 | 0 | 0 | [illegible] |
| SE | 43043 | ENSG00000079308 | TNS1 | chr2 | - | 217920194 | 217920236 | 217907209 | 217907251 | 217943864 | 217944005 | 3 | 1 | [illegible] |
| SE | 43051 | ENSG00000182287 | AP1S2 | chrX | - | 15828191 | 15828200 | 15825805 | 15827372 | 15845378 | 15845516 | 0 | 1 | [illegible] |
| SE | 43053 | ENSG00000182287 | AP1S2 | chrX | - | 15831451 | 15831568 | 15828191 | 15828200 | 15845378 | 15845516 | 3 | 3 | [illegible] |
| SE | 43056 | ENSG00000182287 | AP1S2 | chrX | - | 15833443 | 15833524 | 15831289 | 15833007 | 15845378 | 15845516 | 17 | 11 | [illegible] |
| SE | 43059 | ENSG00000016391 | CHDH | chr3 | - | 53819531 | 53819674 | 53818937 | 53819040 | 53820473 | 53820608 | 1 | 0 | [illegible] |
| SE | 43060 | ENSG00000101474 | APMAP | chr20 | - | 24964374 | 24964398 | 24962924 | 24964022 | 24968891 | 24969084 | 1 | 0 | [illegible] |
| SE | 43062 | ENSG00000101474 | APMAP | chr20 | - | 24971459 | 24971576 | 24970196 | 24970371 | 24973644 | 24973737 | 1010 | 449 | [illegible] |
| SE | 43063 | ENSG00000101474 | APMAP | chr20 | - | 24978766 | 24978882 | 24973644 | 24973737 | 24983902 | 24984019 | 533 | 177 | [illegible] |
| SE | 43064 | ENSG00000101474 | APMAP | chr20 | - | 24978766 | 24978882 | 24973644 | 24973737 | 24992593 | 24992642 | 305 | 109 | [illegible] |
| SE | 43067 | ENSG00000271810 | NA | chr1 | - | 112702998 | 112703119 | 112702613 | 112702693 | 112703643 | 112703806 | 698 | 395 | [illegible] |
| SE | 43069 | ENSG00000271810 | NA | chr1 | - | 112711007 | 112711071 | 112710743 | 112710851 | 112711265 | 112711355 | 1 | 5 | [illegible] |
| SE | 43070 | ENSG00000178917 | ZNF852 | chr3 | - | 44502541 | 44502668 | 44499301 | 44500634 | 44510552 | 44510636 | 47 | 15 | [illegible] |
| SE | 43075 | ENSG00000185986 | SDHAP3 | chr5 | - | 1579276 | 1579484 | 1576178 | 1576309 | 1584983 | 1585133 | 20 | 16 | [illegible] |
| SE | 43076 | ENSG00000185986 | SDHAP3 | chr5 | - | 1584983 | 1585133 | 1576178 | 1576309 | 1589246 | 1589365 | 9 | 15 | [illegible] |
| SE | 43077 | ENSG00000213160 | KLHL23 | chr2 | + | 169697360 | 169697531 | 169695214 | 169695247 | 169735012 | 169736227 | 8 | 6 | [illegible] |
| SE | 43078 | ENSG00000213160 | KLHL23 | chr2 | + | 169697892 | 169697982 | 169697360 | 169697531 | 169741384 | 169741537 | 0 | 0 | [illegible] |
| SE | 43079 | ENSG00000213160 | KLHL23 | chr2 | + | 169735012 | 169736227 | 169697360 | 169697531 | 169741384 | 169741537 | 1 | 3 | [illegible] |
| SE | 43080 | ENSG00000213160 | KLHL23 | chr2 | + | 169741384 | 169741537 | 169735012 | 169736227 | 169749421 | 169749682 | 4 | 3 | [illegible] |
| SE | 43084 | ENSG00000239900 | ADSL | chr22 | + | 40353072 | 40353117 | 40349831 | 40350035 | 40354247 | 40354327 | 108 | 62 | [illegible] |
| SE | 43089 | ENSG00000239900 | ADSL | chr22 | + | 40354247 | 40354327 | 40349831 | 40353117 | 40358863 | 40358912 | 130 | 91 | [illegible] |
| SE | 43090 | ENSG00000239900 | ADSL | chr22 | + | 40361272 | 40363071 | 40360401 | 40360492 | 40364275 | 40364365 | 108 | 81 | [illegible] |
| SE | 43097 | ENSG00000177728 | TMEM94 | chr17 | + | 75489575 | 75489662 | 75489265 | 75489368 | 75490233 | 75490350 | 52 | 21 | [illegible] |
| SE | 43099 | ENSG00000177728 | TMEM94 | chr17 | + | 75491690 | 75491852 | 75491376 | 75491455 | 75492473 | 75492789 | 8 | 3 | [illegible] |
| SE | 43100 | ENSG00000177728 | TMEM94 | chr17 | + | 75491690 | 75491900 | 75491302 | 75491455 | 75492473 | 75492762 | 25 | 17 | [illegible] |
| SE | 43101 | ENSG00000177728 | TMEM94 | chr17 | + | 75492238 | 75492337 | 75491690 | 75491852 | 75492473 | 75492742 | 1 | 0 | [illegible] |
| SE | 43103 | ENSG00000177728 | TMEM94 | chr17 | + | 75495283 | 75495399 | 75494887 | 75495034 | 75495543 | 75495643 | 26 | 8 | [illegible] |
| SE | 43105 | ENSG00000177728 | TMEM94 | chr17 | + | 75497112 | 75497198 | 75496727 | 75496807 | 75497780 | 75497862 | 27 | 11 | [illegible] |
| SE | 43108 | ENSG00000119048 | UBE2B | chr5 | + | 134376668 | 134376694 | 134374382 | 134374463 | 134388324 | 134388413 | 9 | 0 | [illegible] |
| SE | 43109 | ENSG00000119048 | UBE2B | chr5 | + | 134376668 | 134376694 | 134374382 | 134374463 | 134390224 | 134390321 | 5 | 0 | [illegible] |
| SE | 43110 | ENSG00000119048 | UBE2B | chr5 | + | 134380718 | 134380808 | 134374382 | 134374463 | 134388324 | 134388413 | 498 | 328 | [illegible] |
| SE | 43111 | ENSG00000119048 | UBE2B | chr5 | + | 134380718 | 134380808 | 134374382 | 134374463 | 134390224 | 134390321 | 1 | 3 | [illegible] |
| SE | 43112 | ENSG00000119048 | UBE2B | chr5 | + | 134380718 | 134380808 | 134376668 | 134376694 | 134388324 | 134388413 | 945 | 608 | [illegible] |
| SE | 43113 | ENSG00000119048 | UBE2B | chr5 | + | 134380718 | 134380808 | 134376668 | 134376694 | 134390224 | 134390321 | 137 | 85 | [illegible] |
| SE | 43114 | ENSG00000119048 | UBE2B | chr5 | + | 134388324 | 134388413 | 134374382 | 134374463 | 134390224 | 134390298 | 285 | 187 | [illegible] |
| SE | 43115 | ENSG00000119048 | UBE2B | chr5 | + | 134388324 | 134388413 | 134376668 | 134376694 | 134390224 | 134390298 | 289 | 187 | [illegible] |
| SE | 43116 | ENSG00000119048 | UBE2B | chr5 | + | 134388324 | 134388413 | 134380718 | 134380808 | 134390224 | 134390298 | 1125 | 740 | [illegible] |
| SE | 43117 | ENSG00000119048 | UBE2B | chr5 | + | 134388351 | 134388413 | 134380718 | 134380808 | 134390224 | 134390573 | 53 | 37 | [illegible] |
| SE | 43118 | ENSG00000119048 | UBE2B | chr5 | + | 134388976 | 134389052 | 134388324 | 134388413 | 134390224 | 134390321 | 5 | 4 | [illegible] |
| SE | 43123 | ENSG00000038274 | MAT2B | chr5 | + | 163517560 | 163517674 | 163516517 | 163516711 | 163518192 | 163519149 | 979 | 551 | [illegible] |
| SE | 43124 | ENSG00000136878 | USP20 | chr9 | + | 129852539 | 129852636 | 129849799 | 129849924 | 129856306 | 129856360 | 77 | 37 | [illegible] |
| SE | 43125 | ENSG00000136878 | USP20 | chr9 | + | 129860936 | 129861033 | 129858466 | 129858598 | 129861542 | 129861612 | 144 | 63 | [illegible] |
| SE | 43132 | ENSG00000186470 | BTN3A2 | chr6 | + | 26368449 | 26368608 | 26368177 | 26368267 | 26368720 | 26368800 | 1 | 0 | [illegible] |
| SE | 43134 | ENSG00000186470 | BTN3A2 | chr6 | + | 26370321 | 26370603 | 26368564 | 26368912 | 26372896 | 26373087 | 840 | 507 | [illegible] |
| SE | 43138 | ENSG00000186470 | BTN3A2 | chr6 | + | 26373275 | 26373296 | 26372896 | 26373097 | 26373386 | 26373413 | 328 | 213 | [illegible] |
| SE | 43140 | ENSG00000135392 | DNAJC14 | chr12 | - | 55823069 | 55823189 | 55822571 | 55822732 | 55823401 | 55823508 | 131 | 71 | [illegible] |
| SE | 43141 | ENSG00000135392 | DNAJC14 | chr12 | - | 55829488 | 55829636 | 55827251 | 55828714 | 55830695 | 55830781 | 33 | 12 | [illegible] |
| SE | 43144 | ENSG00000113161 | HMGCR | chr5 | + | 75356342 | 75356448 | 75355364 | 75355522 | 75358656 | 75358827 | 151 | 92 | [illegible] |
| SE | 43146 | ENSG00000113161 | HMGCR | chr5 | + | 75359169 | 75359310 | 75356342 | 75356448 | 75359984 | 75360139 | 2 | 8 | [illegible] |
| SE | 43150 | ENSG00000198590 | C3orf35 | chr3 | + | 37399476 | 37399659 | 37381061 | 37383371 | 37411137 | 37411309 | 4 | 0 | [illegible] |
| SE | 43154 | ENSG00000122335 | SERAC1 | chr6 | - | 158117726 | 158117821 | 158116184 | 158116282 | 158119028 | 158119120 | 33 | 14 | [illegible] |
| SE | 43155 | ENSG00000122335 | SERAC1 | chr6 | - | 158128107 | 158128270 | 158120424 | 158120575 | 158130372 | 158130486 | 54 | 60 | [illegible] |
| SE | 43156 | ENSG00000122335 | SERAC1 | chr6 | - | 158128107 | 158128270 | 158120424 | 158120575 | 158143055 | 158143184 | 32 | 33 | [illegible] |
| SE | 43157 | ENSG00000122335 | SERAC1 | chr6 | - | 158130372 | 158130486 | 158120424 | 158120575 | 158143055 | 158143184 | 6 | 11 | [illegible] |
| SE | 43161 | ENSG00000122335 | SERAC1 | chr6 | - | 158148864 | 158148954 | 158145919 | 158146913 | 158150452 | 158150563 | 20 | 23 | [illegible] |
| SE | 43169 | ENSG00000171262 | FAM98B | chr15 | + | 38464031 | 38464177 | 38454126 | 38454232 | 38465268 | 38465403 | 274 | 179 | [illegible] |
| SE | 43170 | ENSG00000116044 | NFE2L2 | chr2 | - | 177283599 | 177283804 | 177234004 | 177234271 | 177287610 | 177287777 | 3 | 0 | [illegible] |
| SE | 43171 | ENSG00000116044 | NFE2L2 | chr2 | - | 177283599 | 177283804 | 177234004 | 177234271 | 177310963 | 177311035 | 2 | 0 | [illegible] |
| SE | 43174 | ENSG00000183814 | LIN9 | chr1 | - | 226233343 | 226233523 | 226232712 | 226233193 | 226238970 | 226239096 | 50 | 38 | [illegible] |
| SE | 43175 | ENSG00000183814 | LIN9 | chr1 | - | 226250838 | 226250919 | 226238970 | 226239096 | 226277774 | 226277932 | 6 | 11 | [illegible] |
| SE | 43176 | ENSG00000183814 | LIN9 | chr1 | - | 226265532 | 226265634 | 226250838 | 226250919 | 226266212 | 226266332 | 15 | 21 | [illegible] |
| SE | 43177 | ENSG00000183814 | LIN9 | chr1 | - | 226266212 | 226266332 | 226250838 | 226250919 | 226277774 | 226277932 | 2 | 0 | [illegible] |
| SE | 43178 | ENSG00000183814 | LIN9 | chr1 | - | 226267956 | 226268090 | 226266212 | 226266332 | 226277774 | 226277932 | 51 | 80 | [illegible] |
| SE | 43179 | ENSG00000183814 | LIN9 | chr1 | - | 226286332 | 226286458 | 226277774 | 226277932 | 226287663 | 226287797 | 31 | 41 | [illegible] |
| SE | 43180 | ENSG00000183814 | LIN9 | chr1 | - | 226295841 | 226295946 | 226287663 | 226287797 | 226297718 | 226297813 | 33 | 22 | [illegible] |
| SE | 43181 | ENSG00000204252 | HLA-DOA | chr6 | - | 33007079 | 33007215 | 33004181 | 33006841 | 33007310 | 33007592 | 44 | 26 | [illegible] |
| SE | 43186 | ENSG00000169188 | APEX2 | chrX | + | 55002961 | 55003108 | 55002250 | 55002431 | 55003798 | 55003868 | 69 | 39 | [illegible] |
| SE | 43187 | ENSG00000169188 | APEX2 | chrX | + | 55003798 | 55003868 | 55001545 | 55001629 | 55006517 | 55006757 | 14 | 10 | [illegible] |
| SE | 43190 | ENSG00000171853 | TRAPPC12 | chr2 | + | 3387619 | 3388670 | 3379693 | 3379876 | 3401776 | 3401893 | 71 | 44 | [illegible] |
| SE | 43192 | ENSG00000171853 | TRAPPC12 | chr2 | + | 3421880 | 3421994 | 3401776 | 3401893 | 3424524 | 3424663 | 268 | 159 | [illegible] |
| SE | 43193 | ENSG00000171853 | TRAPPC12 | chr2 | + | 3424524 | 3424663 | 3401776 | 3401893 | 3457620 | 3457693 | 5 | 1 | [illegible] |
| SE | 43195 | ENSG00000171853 | TRAPPC12 | chr2 | + | 3443778 | 3443891 | 3424524 | 3424663 | 3477694 | 3477795 | 31 | 26 | [illegible] |
| SE | 43198 | ENSG00000171853 | TRAPPC12 | chr2 | + | 3462857 | 3463071 | 3460300 | 3460336 | 3465596 | 3465695 | 0 | 0 | [illegible] |
| SE | 43199 | ENSG00000171853 | TRAPPC12 | chr2 | + | 3465596 | 3465695 | 3459930 | 3460336 | 3477694 | 3477795 | 128 | 135 | [illegible] |
| SE | 43200 | ENSG00000171853 | TRAPPC12 | chr2 | + | 3466253 | 3466393 | 3465596 | 3465695 | 3477694 | 3478101 | 2 | 0 | [illegible] |
| SE | 43201 | ENSG00000171853 | TRAPPC12 | chr2 | + | 3466253 | 3466393 | 3465596 | 3465695 | 3478845 | 3478871 | 2 | 0 | [illegible] |
| SE | 43202 | ENSG00000171853 | TRAPPC12 | chr2 | + | 3477691 | 3477795 | 3465596 | 3465695 | 3478845 | 3478933 | 102 | 91 | [illegible] |
| SE | 43203 | ENSG00000171853 | TRAPPC12 | chr2 | + | 3477694 | 3477795 | 3465596 | 3465695 | 3478845 | 3478913 | 180 | 174 | [illegible] |
| SE | 43204 | ENSG00000218739 | CEBPZOS | chr2 | + | 37197253 | 37197350 | 37196507 | 37196520 | 37198493 | 37198672 | 0 | 1 | [illegible] |
| SE | 43205 | ENSG00000218739 | CEBPZOS | chr2 | + | 37197253 | 37197350 | 37196507 | 37196520 | 37199703 | 37199819 | 0 | 0 | [illegible] |
| SE | 43207 | ENSG00000218739 | CEBPZOS | chr2 | + | 37198493 | 37198672 | 37197253 | 37197350 | 37199703 | 37199819 | 11 | 15 | [illegible] |
| SE | 43208 | ENSG00000218739 | CEBPZOS | chr2 | + | 37199703 | 37199819 | 37196487 | 37196520 | 37201047 | 37201092 | 234 | 154 | [illegible] |
| SE | 43209 | ENSG00000218739 | CEBPZOS | chr2 | + | 37199703 | 37199819 | 37196487 | 37196520 | 37201641 | 37201726 | 0 | 3 | [illegible] |
| SE | 43210 | ENSG00000218739 | CEBPZOS | chr2 | + | 37201047 | 37201092 | 37196487 | 37196520 | 37201641 | 37201726 | 114 | 55 | [illegible] |
| SE | 43211 | ENSG00000218739 | CEBPZOS | chr2 | + | 37201047 | 37201092 | 37199703 | 37199819 | 37201641 | 37201726 | 350 | 215 | [illegible] |
| SE | 43212 | ENSG00000218739 | CEBPZOS | chr2 | + | 37201641 | 37201726 | 37201047 | 37201092 | 37203249 | 37203495 | 103 | 82 | [illegible] |
| SE | 43213 | ENSG00000181666 | ZNF875 | chr19 | + | 37322184 | 37322232 | 37317886 | 37318186 | 37323532 | 37323570 | 18 | 4 | [illegible] |
| SE | 43214 | ENSG00000181666 | ZNF875 | chr19 | + | 37324795 | 37324936 | 37324207 | 37324265 | 37344609 | 37344797 | 7 | 1 | [illegible] |
| SE | 43215 | ENSG00000181666 | ZNF875 | chr19 | + | 37324795 | 37324936 | 37324207 | 37324265 | 37347189 | 37347196 | 8 | 0 | [illegible] |
| SE | 43216 | ENSG00000181666 | ZNF875 | chr19 | + | 37324795 | 37324936 | 37324207 | 37324265 | 37347776 | 37347872 | 7 | 0 | [illegible] |
| SE | 43219 | ENSG00000181666 | ZNF875 | chr19 | + | 37335168 | 37335257 | 37324795 | 37324936 | 37344609 | 37344774 | 0 | 0 | [illegible] |
| SE | 43220 | ENSG00000181666 | ZNF875 | chr19 | + | 37335168 | 37335257 | 37324795 | 37324936 | 37347189 | 37347316 | 1 | 0 | [illegible] |
| SE | 43224 | ENSG00000181666 | ZNF875 | chr19 | + | 37344609 | 37344774 | 37335168 | 37335257 | 37346422 | 37346675 | 0 | 0 | [illegible] |
| SE | 43225 | ENSG00000181666 | ZNF875 | chr19 | + | 37344609 | 37344774 | 37335168 | 37335257 | 37347189 | 37347316 | 0 | 0 | [illegible] |
| SE | 43226 | ENSG00000181666 | ZNF875 | chr19 | + | 37344933 | 37345021 | 37324207 | 37324265 | 37347189 | 37347258 | 0 | 0 | [illegible] |
| SE | 43227 | ENSG00000181666 | ZNF875 | chr19 | + | 37344933 | 37345021 | 37324207 | 37324265 | 37347776 | 37347872 | 0 | 0 | [illegible] |
| SE | 43228 | ENSG00000181666 | ZNF875 | chr19 | + | 37344933 | 37345021 | 37335168 | 37335257 | 37346422 | 37346675 | 0 | 3 | [illegible] |
| SE | 43229 | ENSG00000181666 | ZNF875 | chr19 | + | 37344933 | 37345021 | 37335168 | 37335257 | 37347189 | 37347258 | 0 | 0 | [illegible] |
| SE | 43232 | ENSG00000181666 | ZNF875 | chr19 | + | 37346422 | 37346492 | 37335168 | 37335257 | 37347189 | 37347316 | 0 | 4 | [illegible] |
| SE | 43234 | ENSG00000181666 | ZNF875 | chr19 | + | 37346887 | 37347052 | 37335168 | 37335257 | 37347189 | 37347316 | 0 | 0 | [illegible] |
| SE | 43236 | ENSG00000181666 | ZNF875 | chr19 | + | 37347189 | 37347316 | 37324795 | 37324936 | 37347776 | 37347810 | 15 | 15 | [illegible] |
| SE | 43238 | ENSG00000181666 | ZNF875 | chr19 | + | 37347189 | 37347316 | 37335168 | 37335257 | 37347776 | 37347810 | 16 | 18 | [illegible] |
| SE | 43240 | ENSG00000181666 | ZNF875 | chr19 | + | 37347231 | 37347316 | 37335168 | 37335257 | 37347776 | 37347872 | 11 | 13 | [illegible] |
| SE | 43243 | ENSG00000181666 | ZNF875 | chr19 | + | 37357961 | 37358039 | 37347562 | 37347872 | 37362105 | 37362174 | 0 | 0 | [illegible] |
| SE | 43245 | ENSG00000181666 | ZNF875 | chr19 | + | 37358540 | 37358661 | 37347776 | 37347872 | 37362108 | 37362274 | 1 | 0 | [illegible] |
| SE | 43246 | ENSG00000181666 | ZNF875 | chr19 | + | 37358540 | 37358661 | 37357961 | 37358039 | 37362108 | 37362274 | 1 | 0 | [illegible] |
| SE | 43248 | ENSG00000181666 | ZNF875 | chr19 | + | 37359425 | 37359546 | 37347776 | 37347872 | 37362105 | 37363864 | 1 | 0 | [illegible] |
| SE | 43250 | ENSG00000181666 | ZNF875 | chr19 | + | 37359425 | 37359546 | 37358546 | 37358661 | 37362108 | 37362146 | 0 | 0 | [illegible] |
| SE | 43253 | ENSG00000145982 | FARS2 | chr6 | + | 5272425 | 5272489 | 5261325 | 5261660 | 5368549 | 5368712 | 0 | 1 | [illegible] |
| SE | 43254 | ENSG00000145982 | FARS2 | chr6 | + | 5368549 | 5369182 | 5261498 | 5261660 | 5404541 | 5404701 | 74 | 25 | [illegible] |
| SE | 43255 | ENSG00000145982 | FARS2 | chr6 | + | 5368549 | 5369182 | 5261498 | 5261660 | 5431040 | 5431172 | 41 | 14 | [illegible] |
| SE | 43256 | ENSG00000145982 | FARS2 | chr6 | + | 5404541 | 5404701 | 5261498 | 5261660 | 5431040 | 5431172 | 50 | 19 | [illegible] |
| SE | 43258 | ENSG00000139372 | TDG | chr12 | + | 103967853 | 103967971 | 103965821 | 103966060 | 103972959 | 103973033 | 0 | 0 | [illegible] |
| SE | 43259 | ENSG00000139372 | TDG | chr12 | + | 103967853 | 103967971 | 103965821 | 103966060 | 103976917 | 103977060 | 0 | 0 | [illegible] |
| SE | 43260 | ENSG00000139372 | TDG | chr12 | + | 103967853 | 103967971 | 103965821 | 103966060 | 103979830 | 103980072 | 0 | 0 | [illegible] |
| SE | 43261 | ENSG00000139372 | TDG | chr12 | + | 103972959 | 103973033 | 103965844 | 103966060 | 103976917 | 103977060 | 14 | 0 | [illegible] |
| SE | 43262 | ENSG00000139372 | TDG | chr12 | + | 103972959 | 103973033 | 103967853 | 103967971 | 103976917 | 103977060 | 7 | 0 | [illegible] |
| SE | 43264 | ENSG00000139372 | TDG | chr12 | + | 103976917 | 103977060 | 103967853 | 103967971 | 103979830 | 103980072 | 58 | 44 | [illegible] |
| SE | 43265 | ENSG00000139372 | TDG | chr12 | + | 103983135 | 103983218 | 103982798 | 103982934 | 103983294 | 103983389 | 260 | 137 | [illegible] |
| SE | 43266 | ENSG00000139372 | TDG | chr12 | + | 103983294 | 103983389 | 103983135 | 103983218 | 103984748 | 103984920 | 220 | 105 | [illegible] |
| SE | 43267 | ENSG00000111325 | OGFOD2 | chr12 | + | 122975810 | 122975867 | 122974579 | 122975121 | 122976653 | 122976767 | 3 | 3 | [illegible] |
| SE | 43268 | ENSG00000111325 | OGFOD2 | chr12 | + | 122976252 | 122976401 | 122975810 | 122975867 | 122976653 | 122976767 | 0 | 0 | [illegible] |
| SE | 43271 | ENSG00000111325 | OGFOD2 | chr12 | + | 122976870 | 122976970 | 122975810 | 122975867 | 122977980 | 122978056 | 2 | 1 | [illegible] |
| SE | 43272 | ENSG00000111325 | OGFOD2 | chr12 | + | 122977980 | 122978057 | 122976870 | 122976970 | 122978359 | 122978547 | 0 | 1 | [illegible] |
| SE | 43273 | ENSG00000111325 | OGFOD2 | chr12 | + | 122977980 | 122978091 | 122976870 | 122976970 | 122978359 | 122978569 | 2 | 2 | [illegible] |
| SE | 43274 | ENSG00000111325 | OGFOD2 | chr12 | + | 122977980 | 122978091 | 122976870 | 122976970 | 122978441 | 122978569 | 2 | 1 | [illegible] |
| SE | 43275 | ENSG00000111325 | OGFOD2 | chr12 | + | 122977980 | 122978199 | 122976870 | 122976970 | 122978359 | 122978569 | 2 | 1 | [illegible] |
| SE | 43276 | ENSG00000187815 | ZFP69 | chr1 | + | 40489528 | 40489624 | 40489087 | 40489214 | 40494920 | 40496288 | 65 | 45 | [illegible] |
| SE | 43277 | ENSG00000245954 | LINC02273 | chr4 | + | 152101483 | 152101580 | 152100817 | 152101218 | 152103626 | 152104705 | 48 | 10 | [illegible] |
| SE | 43278 | ENSG00000167550 | RHEBL1 | chr12 | - | 49066955 | 49067035 | 49066618 | 49066701 | 49069034 | 49069106 | 0 | 0 | [illegible] |
| SE | 43279 | ENSG00000167550 | RHEBL1 | chr12 | - | 49066967 | 49067035 | 49066618 | 49066701 | 49069034 | 49069106 | 7 | 0 | [illegible] |
| SE | 43280 | ENSG00000167962 | ZNF598 | chr16 | - | 1998964 | 1999121 | 1998701 | 1998852 | 1999495 | 2000256 | 37 | 21 | [illegible] |
| SE | 43281 | ENSG00000167962 | ZNF598 | chr16 | - | 2000390 | 2000481 | 1999495 | 2000256 | 2001003 | 2001191 | 24 | 13 | [illegible] |
| SE | 43282 | ENSG00000167962 | ZNF598 | chr16 | - | 2000408 | 2000481 | 1999495 | 2000256 | 2001003 | 2001191 | 15 | 4 | [illegible] |
| SE | 43283 | ENSG00000167962 | ZNF598 | chr16 | - | 2002990 | 2003120 | 2002205 | 2002732 | 2003615 | 2003728 | 39 | 19 | [illegible] |
| SE | 43290 | ENSG00000134698 | AGO4 | chr1 | + | 35816881 | 35817047 | 35808015 | 35808435 | 35822861 | 35822982 | 134 | 47 | [illegible] |
| SE | 43291 | ENSG00000134698 | AGO4 | chr1 | + | 35826747 | 35826835 | 35825925 | 35826060 | 35831426 | 35831574 | 345 | 261 | [illegible] |
| SE | 43292 | ENSG00000134698 | AGO4 | chr1 | + | 35831426 | 35831574 | 35826747 | 35826835 | 35831811 | 35831931 | 412 | 340 | [illegible] |
| SE | 43293 | ENSG00000134698 | AGO4 | chr1 | + | 35832056 | 35832185 | 35831811 | 35831931 | 35832436 | 35832570 | 340 | 270 | [illegible] |
| SE | 43294 | ENSG00000134698 | AGO4 | chr1 | + | 35832056 | 35832185 | 35831811 | 35831931 | 35833989 | 35834174 | 201 | 168 | [illegible] |
| SE | 43297 | ENSG00000134698 | AGO4 | chr1 | + | 35832436 | 35832570 | 35832056 | 35832185 | 35835833 | 35835993 | 143 | 104 | [illegible] |
| SE | 43298 | ENSG00000134698 | AGO4 | chr1 | + | 35833989 | 35834174 | 35832056 | 35832185 | 35835833 | 35835993 | 166 | 96 | [illegible] |
| SE | 43299 | ENSG00000134698 | AGO4 | chr1 | + | 35833989 | 35834174 | 35832436 | 35832570 | 35835833 | 35835993 | 285 | 137 | [illegible] |
| SE | 43300 | ENSG00000134698 | AGO4 | chr1 | + | 35850156 | 35850258 | 35841615 | 35841750 | 35850853 | 35851053 | 236 | 158 | [illegible] |
| SE | 43302 | ENSG00000009413 | REV3L | chr6 | - | 111308013 | 111308245 | 111307360 | 111307570 | 111309852 | 111310099 | 0 | 1 | [illegible] |
| SE | 43304 | ENSG00000009413 | REV3L | chr6 | - | 111311068 | 111311259 | 111309852 | 111310099 | 111315266 | 111315381 | 52 | 33 | [illegible] |
| SE | 43305 | ENSG00000009413 | REV3L | chr6 | - | 111313351 | 111313489 | 111309852 | 111310099 | 111315266 | 111315381 | 54 | 22 | [illegible] |
| SE | 43306 | ENSG00000009413 | REV3L | chr6 | - | 111313351 | 111313489 | 111311068 | 111311259 | 111315266 | 111315381 | 133 | 84 | [illegible] |
| SE | 43307 | ENSG00000009413 | REV3L | chr6 | - | 111322568 | 111322678 | 111315266 | 111315381 | 111329531 | 111329738 | 66 | 50 | [illegible] |
| SE | 43308 | ENSG00000009413 | REV3L | chr6 | - | 111331675 | 111331784 | 111329531 | 111329738 | 111333122 | 111333367 | 92 | 60 | [illegible] |
| SE | 43309 | ENSG00000009413 | REV3L | chr6 | - | 111335468 | 111335610 | 111333122 | 111333367 | 111343924 | 111344043 | 95 | 63 | [illegible] |
| SE | 43310 | ENSG00000009413 | REV3L | chr6 | - | 111358821 | 111359014 | 111357013 | 111357125 | 111363852 | 111363978 | 83 | 72 | [illegible] |
| SE | 43311 | ENSG00000009413 | REV3L | chr6 | - | 111365264 | 111365344 | 111363852 | 111363978 | 111367114 | 111367358 | 155 | 134 | [illegible] |
| SE | 43313 | ENSG00000009413 | REV3L | chr6 | - | 111379981 | 111380219 | 111377700 | 111377843 | 111387764 | 111387913 | 55 | 45 | [illegible] |
| SE | 43315 | ENSG00000009413 | REV3L | chr6 | - | 111388000 | 111388085 | 111387745 | 111387913 | 111389105 | 111389210 | 109 | 85 | [illegible] |
| SE | 43319 | ENSG00000135597 | REPS1 | chr6 | - | 138908667 | 138908816 | 138904865 | 138905132 | 138911275 | 138911371 | 183 | 131 | [illegible] |
| SE | 43321 | ENSG00000135597 | REPS1 | chr6 | - | 138912674 | 138912950 | 138911275 | 138911371 | 138914696 | 138914761 | 75 | 47 | [illegible] |
| SE | 43322 | ENSG00000135597 | REPS1 | chr6 | - | 138912764 | 138912950 | 138911245 | 138911371 | 138914696 | 138914761 | 187 | 122 | [illegible] |
| SE | 43329 | ENSG00000135597 | REPS1 | chr6 | - | 138917554 | 138917646 | 138914696 | 138914761 | 138920214 | 138920316 | 5 | 0 | [illegible] |
| SE | 43334 | ENSG00000135597 | REPS1 | chr6 | - | 138941334 | 138941489 | 138926400 | 138930098 | 138943512 | 138943576 | 152 | 156 | [illegible] |
| SE | 43335 | ENSG00000135597 | REPS1 | chr6 | - | 138941334 | 138941489 | 138926400 | 138930098 | 138945218 | 138945372 | 102 | 87 | [illegible] |
| SE | 43336 | ENSG00000135597 | REPS1 | chr6 | - | 138943512 | 138943576 | 138941334 | 138941489 | 138943852 | 138943926 | 114 | 93 | [illegible] |
| SE | 43337 | ENSG00000135597 | REPS1 | chr6 | - | 138947789 | 138947913 | 138945500 | 138945697 | 138987529 | 138987640 | 164 | 108 | [illegible] |
| SE | 43338 | ENSG00000135597 | REPS1 | chr6 | - | 138951041 | 138951160 | 138947881 | 138947913 | 138987529 | 138987731 | 0 | 0 | [illegible] |
| SE | 43339 | ENSG00000173221 | GLRX | chr5 | - | 95816506 | 95816626 | 95813848 | 95814389 | 95822455 | 95822714 | 828 | 487 | [illegible] |
| SE | 43348 | ENSG00000075188 | NUP37 | chr12 | - | 102075000 | 102075094 | 102073102 | 102074467 | 102076796 | 102076847 | 56 | 22 | [illegible] |
| SE | 43351 | ENSG00000075188 | NUP37 | chr12 | - | 102115802 | 102115864 | 102112174 | 102112232 | 102118362 | 102118583 | 0 | 0 | [illegible] |
| SE | 43353 | ENSG00000113119 | TMCO6 | chr5 | + | 140641869 | 140642053 | 140641305 | 140641780 | 140642314 | 140642413 | 15 | 3 | [illegible] |
| SE | 43355 | ENSG00000113119 | TMCO6 | chr5 | + | 140643563 | 140643675 | 140642906 | 140643041 | 140643779 | 140643966 | 24 | 2 | [illegible] |
| SE | 43356 | ENSG00000113119 | TMCO6 | chr5 | + | 140644099 | 140644188 | 140643779 | 140643966 | 140644572 | 140645311 | 2 | 1 | [illegible] |
| SE | 43357 | ENSG00000113119 | TMCO6 | chr5 | + | 140644099 | 140644194 | 140643779 | 140643966 | 140644572 | 140644740 | 5 | 3 | [illegible] |
| SE | 43361 | ENSG00000090863 | GLG1 | chr16 | - | 74453170 | 74453334 | 74447426 | 74452131 | 74456648 | 74456755 | 403 | 243 | [illegible] |
| SE | 43362 | ENSG00000090863 | GLG1 | chr16 | - | 74456648 | 74456755 | 74447439 | 74453334 | 74457873 | 74457994 | 608 | 469 | [illegible] |
| SE | 43363 | ENSG00000090863 | GLG1 | chr16 | - | 74459681 | 74459789 | 74457873 | 74457994 | 74462093 | 74462195 | 287 | 198 | [illegible] |
| SE | 43365 | ENSG00000090863 | GLG1 | chr16 | - | 74467755 | 74467848 | 74463355 | 74463479 | 74469984 | 74470073 | 1 | 1 | [illegible] |
| SE | 43367 | ENSG00000090863 | GLG1 | chr16 | - | 74472525 | 74472634 | 74472348 | 74472411 | 74474545 | 74474632 | 0 | 2 | [illegible] |
| SE | 43368 | ENSG00000090863 | GLG1 | chr16 | - | 74474545 | 74474632 | 74472348 | 74472411 | 74477395 | 74477533 | 750 | 495 | [illegible] |
| SE | 43369 | ENSG00000090863 | GLG1 | chr16 | - | 74492956 | 74493140 | 74491000 | 74491215 | 74494759 | 74494831 | 1227 | 772 | [illegible] |
| SE | 43370 | ENSG00000090863 | GLG1 | chr16 | - | 74494759 | 74494831 | 74492956 | 74493140 | 74496440 | 74496644 | 923 | 554 | [illegible] |
| SE | 43371 | ENSG00000090863 | GLG1 | chr16 | - | 74496440 | 74496644 | 74494759 | 74494831 | 74503530 | 74503746 | 1024 | 667 | [illegible] |
| SE | 43372 | ENSG00000090863 | GLG1 | chr16 | - | 74532120 | 74532153 | 74508838 | 74508925 | 74606656 | 74606827 | 209 | 144 | [illegible] |
| SE | 43373 | ENSG00000092010 | PSME1 | chr14 | + | 24137142 | 24137205 | 24136897 | 24137017 | 24137320 | 24137431 | 1618 | 863 | [illegible] |
| SE | 43374 | ENSG00000092010 | PSME1 | chr14 | + | 24138048 | 24138117 | 24137699 | 24137797 | 24138195 | 24138263 | 1267 | 826 | [illegible] |
| SE | 43375 | ENSG00000092010 | PSME1 | chr14 | + | 24138473 | 24138560 | 24138344 | 24138399 | 24138735 | 24138787 | 1461 | 931 | [illegible] |
| SE | 43376 | ENSG00000092010 | PSME1 | chr14 | + | 24138473 | 24138560 | 24138344 | 24138403 | 24138735 | 24138787 | 229 | 143 | [illegible] |
| SE | 43377 | ENSG00000092010 | PSME1 | chr14 | + | 24138473 | 24138568 | 24138344 | 24138399 | 24138735 | 24138967 | 913 | 627 | [illegible] |
| SE | 43382 | ENSG00000198791 | CNOT7 | chr8 | - | 17232426 | 17232537 | 17224965 | 17230848 | 17234715 | 17234860 | 586 | 384 | [illegible] |
| SE | 43383 | ENSG00000198791 | CNOT7 | chr8 | - | 17232426 | 17232537 | 17224965 | 17230848 | 17242991 | 17243185 | 330 | 206 | [illegible] |
| SE | 43384 | ENSG00000198791 | CNOT7 | chr8 | - | 17234715 | 17234860 | 17230567 | 17230848 | 17242991 | 17243185 | 5 | 4 | [illegible] |
| SE | 43385 | ENSG00000198791 | CNOT7 | chr8 | - | 17234715 | 17234860 | 17231569 | 17232537 | 17237211 | 17237373 | 481 | 295 | [illegible] |
| SE | 43386 | ENSG00000198791 | CNOT7 | chr8 | - | 17234715 | 17234860 | 17231569 | 17232537 | 17242991 | 17243185 | 269 | 185 | [illegible] |
| SE | 43387 | ENSG00000198791 | CNOT7 | chr8 | - | 17237211 | 17237373 | 17231569 | 17232537 | 17242991 | 17243185 | 853 | 695 | [illegible] |
| SE | 43388 | ENSG00000198791 | CNOT7 | chr8 | - | 17237211 | 17237373 | 17234592 | 17234860 | 17242991 | 17243185 | 1066 | 808 | [illegible] |
| SE | 43389 | ENSG00000198791 | CNOT7 | chr8 | - | 17242991 | 17243185 | 17237211 | 17237373 | 17245035 | 17245091 | 1582 | 1298 | [illegible] |
| SE | 43391 | ENSG00000198791 | CNOT7 | chr8 | - | 17243441 | 17243622 | 17242993 | 17243185 | 17245035 | 17245247 | 7 | 5 | [illegible] |
| SE | 43392 | ENSG00000198791 | CNOT7 | chr8 | - | 17245035 | 17245247 | 17242991 | 17243185 | 17246674 | 17246855 | 825 | 668 | [illegible] |

| | | | | | | | | | | | | | | | | | | | | | | | | | | | | | | | | | | | | | | | | | | | | | | |
|---|---|---|---|---|---|---|---|---|---|---|---|---|---|---|---|---|---|---|---|---|---|---|---|---|---|---|---|---|---|---|---|---|---|---|---|---|---|---|---|---|---|---|---|---|---|---|
| SE | 43393 | ENSG00000198791 | CNOT7 | chr8 | - | 17245856 | 17245931 | 17245035 | 17245247 | 17246522 | 17246609 | 4 | 4 | 7 | 7 | 6 | 0 | 2 | 1 | 0 | 0 | 0 | 9 | 8 | 8 | 0 | 0 | 0 | 1 | 0 | 0 | 224 | 149 | 1 | 1 | 1 | 0.571 | 0.823 | 1 | 1 | NA | 1 | 0.842 | 1 | NA | -0.069 |
| SE | 43395 | ENSG00000213071 | LPAL2 | chr6 | - | 160477129 | 160477288 | 160466808 | 160467017 | 160478507 | 160478688 | 0 | 4 | 2 | 4 | 2 | 11 | 0 | 3 | 3 | 10 | 0 | 0 | 1 | 0 | 0 | 0 | 2 | 1 | 0 | 0 | 298 | 149 | 1 | 1 | 0 | 1 | 0.25 | 0.4 | 0.091 | NA | 0 | 0.333 | NA | NA | 0.182 |
| SE | 43396 | ENSG00000213071 | LPAL2 | chr6 | - | 160491362 | 160491520 | 160487250 | 160487432 | 160492780 | 160492962 | 7 | 2 | 0 | 1 | 7 | 4 | 3 | 0 | 4 | 0 | 0 | 0 | 0 | 1 | 0 | 0 | 0 | 0 | 0 | 0 | 298 | 149 | 1 | 1 | 0.467 | 0.25 | NA | 0.111 | 1 | NA | NA | NA | 1 | NA | -0.543 |
| SE | 43397 | ENSG00000088682 | COQ9 | chr16 | + | 57456503 | 57456626 | 57452800 | 57452936 | 57458245 | 57458350 | 37 | 18 | 30 | 48 | 37 | 0 | 1 | 1 | 1 | 0 | 5 | 38 | 17 | 20 | 38 | 0 | 0 | 2 | 0 | 0 | 272 | 149 | 1 | 1 | 1 | 0.908 | 0.943 | 0.963 | 1 | 1 | 1 | 0.823 | 1 | 1 | -0.002 |
| SE | 43398 | ENSG00000088682 | COQ9 | chr16 | + | 57456503 | 57456646 | 57452800 | 57452936 | 57458245 | 57458350 | 37 | 19 | 32 | 50 | 37 | 0 | 1 | 1 | 1 | 0 | 5 | 40 | 17 | 24 | 38 | 0 | 0 | 2 | 0 | 0 | 292 | 149 | 1 | 1 | 1 | 0.907 | 0.942 | 0.962 | 1 | 1 | 1 | 0.813 | 1 | 1 | 0 |
| SE | 43399 | ENSG00000088682 | COQ9 | chr16 | + | 57456930 | 57457015 | 57456503 | 57456646 | 57458245 | 57458316 | 54 | 26 | 55 | 74 | 45 | 0 | 1 | 0 | 0 | 0 | 41 | 81 | 39 | 26 | 81 | 0 | 0 | 0 | 0 | 0 | 234 | 149 | 1 | 1 | 1 | 0.943 | 1 | 1 | 1 | 1 | 1 | 1 | 1 | 1 | -0.011 |
| SE | 43400 | ENSG00000088682 | COQ9 | chr16 | + | 57458245 | 57458350 | 57452800 | 57452936 | 57459564 | 57459597 | 56 | 31 | 35 | 55 | 27 | 1 | 0 | 0 | 1 | 0 | 28 | 65 | 38 | 45 | 95 | 0 | 4 | 0 | 0 | 0 | 254 | 149 | 1 | 1 | 0.97 | 1 | 1 | 0.97 | 1 | 1 | 0.905 | 1 | 1 | 1 | 0.007 |
| SE | 43401 | ENSG00000088682 | COQ9 | chr16 | + | 57458245 | 57458350 | 57456930 | 57457015 | 57459564 | 57459597 | 90 | 46 | 61 | 96 | 50 | 0 | 0 | 1 | 0 | 0 | 45 | 117 | 57 | 66 | 124 | 0 | 0 | 0 | 0 | 0 | 254 | 149 | 1 | 1 | 1 | 1 | 0.973 | 1 | 1 | 1 | 1 | 1 | 1 | 1 | -0.005 |
| SE | 43409 | ENSG00000160789 | LMNA | chr1 | + | 156138487 | 156138607 | 156137653 | 156137743 | 156139769 | 156139894 | 16 | 8 | 8 | 25 | 13 | 0 | 0 | 0 | 0 | 0 | 16 | 19 | 15 | 5 | 197 | 0 | 0 | 1 | 0 | 0 | 269 | 149 | 1 | 1 | 1 | 1 | 1 | 1 | 1 | 1 | 1 | 0.893 | 1 | 1 | 0.021 |
| SE | 43412 | ENSG00000132849 | PATJ | chr1 | + | 61797286 | 61797428 | 61795466 | 61795558 | 61801622 | 61801769 | 67 | 62 | 87 | 57 | 143 | 0 | 0 | 0 | 3 | 0 | 55 | 65 | 59 | 59 | 36 | 0 | 0 | 0 | 0 | 0 | 291 | 149 | 1 | 1 | 1 | 1 | 1 | 0.907 | 1 | 1 | 1 | 1 | 1 | 1 | -0.019 |
| SE | 43413 | ENSG00000132849 | PATJ | chr1 | + | 61808473 | 61808530 | 61805447 | 61805524 | 61822944 | 61823079 | 53 | 52 | 78 | 79 | 118 | 0 | 1 | 2 | 0 | 0 | 75 | 25 | 49 | 32 | 53 | 0 | 0 | 0 | 0 | 0 | 206 | 149 | 1 | 1 | 1 | 0.974 | 0.966 | 1 | 1 | 1 | 1 | 1 | 1 | 1 | -0.012 |
| SE | 43415 | ENSG00000132849 | PATJ | chr1 | + | 61833653 | 61833785 | 61827421 | 61827583 | 61856029 | 61856239 | 59 | 94 | 117 | 105 | 131 | 0 | 1 | 1 | 2 | 0 | 35 | 34 | 107 | 43 | 0 | 0 | 0 | 0 | 2 | 0 | 281 | 149 | 1 | 1 | 1 | 0.98 | 0.984 | 0.965 | 1 | 1 | 1 | 1 | 0.919 | NA | 0.006 |
| SE | 43416 | ENSG00000132849 | PATJ | chr1 | + | 61861550 | 61861667 | 61856029 | 61856239 | 61864237 | 61864633 | 95 | 188 | 166 | 149 | 214 | 0 | 2 | 0 | 0 | 0 | 89 | 96 | 186 | 108 | 17 | 0 | 0 | 0 | 0 | 0 | 266 | 149 | 1 | 1 | 1 | 0.981 | 1 | 1 | 1 | 1 | 1 | 1 | 1 | 1 | -0.004 |
| SE | 43417 | ENSG00000132849 | PATJ | chr1 | + | 61899582 | 61899654 | 61884236 | 61884408 | 61901281 | 61901459 | 152 | 179 | 175 | 162 | 164 | 2 | 0 | 1 | 0 | 0 | 75 | 61 | 198 | 81 | 29 | 0 | 0 | 0 | 0 | 0 | 221 | 149 | 1 | 1 | 0.981 | 1 | 0.992 | 1 | 1 | 1 | 1 | 1 | 1 | 1 | -0.005 |
| SE | 43418 | ENSG00000132849 | PATJ | chr1 | + | 61899582 | 61899654 | 61884236 | 61884408 | 61908371 | 61908480 | 49 | 55 | 48 | 49 | 37 | 1 | 0 | 1 | 2 | 0 | 21 | 23 | 57 | 36 | 10 | 0 | 0 | 0 | 1 | 0 | 221 | 149 | 1 | 1 | 0.971 | 1 | 0.97 | 0.943 | 1 | 1 | 1 | 1 | 0.96 | 1 | -0.015 |
| SE | 43419 | ENSG00000132849 | PATJ | chr1 | + | 61901281 | 61901459 | 61884236 | 61884408 | 61908371 | 61908480 | 64 | 72 | 95 | 108 | 132 | 1 | 0 | 1 | 2 | 0 | 36 | 69 | 70 | 53 | 0 | 0 | 0 | 0 | 1 | 0 | 298 | 149 | 1 | 1 | 0.97 | 1 | 0.979 | 0.964 | 1 | 1 | 1 | 1 | 0.964 | NA | -0.008 |
| SE | 43420 | ENSG00000132849 | PATJ | chr1 | + | 61901281 | 61901459 | 61899582 | 61899654 | 61908371 | 61908480 | 166 | 196 | 226 | 223 | 260 | 0 | 0 | 0 | 0 | 1 | 90 | 108 | 211 | 98 | 19 | 0 | 0 | 0 | 0 | 0 | 298 | 149 | 1 | 1 | 1 | 1 | 1 | 1 | 0.992 | 1 | 1 | 1 | 1 | 1 | -0.002 |
| SE | 43421 | ENSG00000132849 | PATJ | chr1 | + | 62017855 | 62017947 | 61990167 | 61990364 | 62037976 | 62038049 | 44 | 48 | 52 | 67 | 93 | 0 | 0 | 4 | 0 | 0 | 0 | 32 | 50 | 12 | 3 | 0 | 8 | 3 | 0 | 0 | 241 | 149 | 1 | 1 | 1 | 1 | 0.889 | 1 | 1 | NA | 0.712 | 0.912 | 1 | 1 | 0.072 |
| SE | 43422 | ENSG00000132849 | PATJ | chr1 | + | 62050965 | 62051058 | 62037976 | 62038049 | 62079449 | 62079567 | 56 | 65 | 73 | 69 | 111 | 0 | 0 | 0 | 2 | 0 | 30 | 43 | 81 | 20 | 6 | 0 | 3 | 0 | 0 | 0 | 242 | 149 | 1 | 1 | 1 | 1 | 1 | 0.955 | 1 | 1 | 0.898 | 1 | 1 | 1 | 0.011 |
| SE | 43423 | ENSG00000132849 | PATJ | chr1 | + | 62100318 | 62100408 | 62084514 | 62084648 | 62108436 | 62108520 | 3 | 0 | 0 | 1 | 0 | 21 | 21 | 40 | 20 | 32 | 0 | 0 | 2 | 0 | 0 | 4 | 1 | 38 | 21 | 0 | 239 | 149 | 1 | 1 | 0.082 | 0 | 0 | 0.03 | 0 | 0 | 0 | 0.032 | 0 | NA | 0.014 |
| SE | 43425 | ENSG00000132849 | PATJ | chr1 | + | 62114052 | 62114246 | 62108436 | 62108520 | 62117131 | 62117218 | 28 | 29 | 39 | 32 | 31 | 0 | 0 | 0 | 0 | 2 | 7 | 12 | 33 | 28 | 20 | 0 | 0 | 0 | 0 | 0 | 298 | 149 | 1 | 1 | 1 | 1 | 1 | 1 | 0.886 | 1 | 1 | 1 | 1 | 1 | -0.023 |
| SE | 43426 | ENSG00000132849 | PATJ | chr1 | + | 62116531 | 62116679 | 62114052 | 62114246 | 62117131 | 62117218 | 64 | 39 | 86 | 41 | 125 | 0 | 0 | 4 | 5 | 6 | 0 | 15 | 43 | 23 | 8 | 0 | 0 | 0 | 0 | 0 | 297 | 149 | 1 | 1 | 1 | 1 | 0.915 | 0.804 | 0.913 | NA | 1 | 1 | 1 | 1 | -0.074 |
| SE | 43429 | ENSG00000137274 | BPHL | chr6 | + | 3119284 | 3119512 | 3118375 | 3118847 | 3123656 | 3123760 | 1 | 0 | 1 | 0 | 5 | 0 | 0 | 3 | 3 | 0 | 0 | 5 | 2 | 0 | 0 | 0 | 0 | 0 | 0 | 7 | 298 | 149 | 1 | 1 | 1 | NA | 0.143 | 0 | 1 | NA | 1 | 1 | NA | 0 | -0.131 |
| SE | 43430 | ENSG00000137274 | BPHL | chr6 | + | 3137361 | 3137422 | 3129044 | 3129198 | 3140385 | 3140509 | 7 | 5 | 9 | 7 | 16 | 0 | 0 | 0 | 0 | 0 | 0 | 8 | 0 | 5 | 12 | 0 | 0 | 2 | 0 | 0 | 210 | 149 | 1 | 1 | 1 | 1 | 1 | 1 | 1 | NA | 1 | 0 | 1 | 1 | 0.25 |
| SE | 43431 | ENSG00000137274 | BPHL | chr6 | + | 3137361 | 3137493 | 3129044 | 3129198 | 3140385 | 3140509 | 19 | 7 | 23 | 12 | 66 | 0 | 0 | 0 | 0 | 0 | 0 | 20 | 4 | 28 | 30 | 0 | 0 | 2 | 0 | 0 | 281 | 149 | 1 | 1 | 1 | 1 | 1 | 1 | 1 | NA | 1 | 0.515 | 1 | 1 | 0.121 |
| SE | 43432 | ENSG00000137274 | BPHL | chr6 | + | 3138040 | 3138179 | 3137361 | 3137422 | 3140385 | 3140509 | 0 | 0 | 0 | 2 | 0 | 0 | 0 | 0 | 1 | 0 | 0 | 0 | 1 | 0 | 0 | 0 | 0 | 0 | 0 | 0 | 288 | 149 | 1 | 1 | NA | NA | NA | 0.509 | NA | NA | NA | 1 | NA | NA | -0.491 |
| SE | 43436 | ENSG00000099622 | CIRBP | chr19 | + | 1270866 | 1271036 | 1269265 | 1269410 | 1271328 | 1271467 | 21 | 5 | 27 | 25 | 18 | 0 | 2 | 0 | 0 | 0 | 45 | 41 | 44 | 36 | 78 | 0 | 0 | 0 | 0 | 0 | 298 | 149 | 1 | 1 | 1 | 0.556 | 1 | 1 | 1 | 1 | 1 | 1 | 1 | 1 | -0.089 |
| SE | 43437 | ENSG00000099622 | CIRBP | chr19 | + | 1270927 | 1271036 | 1269267 | 1269410 | 1271328 | 1271467 | 26 | 16 | 21 | 10 | 9 | 0 | 2 | 0 | 0 | 0 | 4 | 13 | 14 | 5 | 19 | 0 | 0 | 0 | 0 | 0 | 258 | 149 | 1 | 1 | 1 | 0.822 | 1 | 1 | 1 | 1 | 1 | 1 | 1 | 1 | -0.036 |
| SE | 43438 | ENSG00000099622 | CIRBP | chr19 | + | 1271120 | 1271246 | 1270927 | 1271036 | 1271328 | 1271467 | 488 | 213 | 409 | 382 | 551 | 1 | 0 | 7 | 2 | 4 | 395 | 405 | 492 | 327 | 677 | 0 | 1 | 5 | 0 | 3 | 275 | 149 | 1 | 1 | 0.996 | 1 | 0.969 | 0.99 | 0.987 | 1 | 0.995 | 0.982 | 1 | 0.992 | -0.005 |
| SE | 43439 | ENSG00000099622 | CIRBP | chr19 | + | 1271139 | 1271246 | 1270866 | 1271036 | 1271328 | 1271360 | 792 | 347 | 626 | 604 | 725 | 1 | 0 | 7 | 2 | 4 | 769 | 816 | 872 | 605 | 1190 | 0 | 1 | 5 | 0 | 3 | 256 | 149 | 1 | 1 | 0.998 | 1 | 0.981 | 0.994 | 0.991 | 1 | 0.998 | 0.99 | 1 | 0.996 | -0.004 |
| SE | 43443 | ENSG00000099622 | CIRBP | chr19 | + | 1271550 | 1271632 | 1270927 | 1271467 | 1271980 | 1271995 | 675 | 331 | 690 | 604 | 672 | 0 | 0 | 1 | 0 | 0 | 636 | 681 | 695 | 564 | 1160 | 0 | 0 | 1 | 3 | 4 | 231 | 149 | 1 | 1 | 1 | 1 | 0.998 | 1 | 1 | 1 | 1 | 0.998 | 0.992 | 0.995 | 0.003 |
| SE | 43444 | ENSG00000099622 | CIRBP | chr19 | + | 1271550 | 1271632 | 1271139 | 1271365 | 1271980 | 1271995 | 293 | 144 | 314 | 245 | 284 | 0 | 0 | 0 | 0 | 0 | 216 | 319 | 243 | 145 | 371 | 0 | 0 | 0 | 2 | 0 | 231 | 149 | 1 | 1 | 1 | 1 | 1 | 1 | 1 | 1 | 1 | 1 | 0.979 | 1 | 0.004 |
| SE | 43445 | ENSG00000099622 | CIRBP | chr19 | + | 1271980 | 1272051 | 1271139 | 1271365 | 1272426 | 1272552 | 212 | 116 | 193 | 192 | 181 | 0 | 0 | 0 | 0 | 3 | 228 | 330 | 192 | 150 | 286 | 0 | 0 | 0 | 0 | 0 | 220 | 149 | 1 | 1 | 1 | 1 | 1 | 1 | 0.976 | 1 | 1 | 1 | 1 | 1 | -0.005 |
| SE | 43449 | ENSG00000204055 | NA | chr9 | + | 129258793 | 129258882 | 129258353 | 129258384 | 129259000 | 129259865 | 2 | 0 | 0 | 0 | 1 | 0 | 0 | 0 | 0 | 4 | 0 | 4 | 0 | 0 | 0 | 0 | 0 | 0 | 0 | 0 | 238 | 149 | 1 | 1 | 1 | NA | NA | NA | 0.135 | NA | 1 | NA | NA | NA | -0.432 |
| SE | 43451 | ENSG00000095261 | PSMD5 | chr9 | - | 120826764 | 120826907 | 120824493 | 120824685 | 120829098 | 120829208 | 95 | 77 | 114 | 98 | 172 | 0 | 0 | 0 | 1 | 0 | 65 | 102 | 45 | 64 | 45 | 0 | 0 | 2 | 0 | 0 | 292 | 149 | 1 | 1 | 1 | 1 | 1 | 0.98 | 1 | 1 | 1 | 0.92 | 1 | 1 | 0.012 |
| [illegible] | | | | | | | | | | | | | | | | | | | | | | | | | | | | | | | | | | | | | | | | | | | | | | |

| | | | | | | | | | | | | | | | | | | | |
|---|---|---|---|---|---|---|---|---|---|---|---|---|---|---|---|---|---|---|---|
| SE | 43891 | ENSG00000179335 | CLK3 | chr15 | + | 74626988 | 74627066 | 74625852 | 74625968 | 74627351 | 74627446 | 1 | 1 | 1 | 1 | 0 | 258 | 139 | 245 |
| SE | 43892 | ENSG00000179335 | CLK3 | chr15 | + | 74627351 | 74627446 | 74623619 | 74625018 | 74627538 | 74627668 | 273 | 182 | 237 | 294 | 418 | 0 | 0 | 3 |
| SE | 43893 | ENSG00000179335 | CLK3 | chr15 | + | 74627351 | 74627446 | 74625801 | 74625968 | 74627538 | 74627668 | 520 | 305 | 466 | 562 | 701 | 0 | 0 | 1 |
| SE | 43896 | ENSG00000197782 | ZNF780A | chr19 | - | 40089259 | 40089414 | 40084744 | 40084798 | 40090165 | 40090235 | 12 | 2 | 10 | 0 | 7 | 20 | 19 | 18 |
| SE | 43898 | ENSG00000184117 | NIPSNAP1 | chr22 | - | 29564237 | 29564371 | 29561791 | 29561862 | 29569192 | 29569287 | 2 | 0 | 0 | 0 | 3 | 58 | 33 | 71 |
| SE | 43899 | ENSG00000184117 | NIPSNAP1 | chr22 | - | 29569192 | 29569287 | 29561791 | 29561862 | 29570161 | 29570207 | 98 | 53 | 126 | 98 | 154 | 0 | 0 | 1 |
| SE | 43900 | ENSG00000184117 | NIPSNAP1 | chr22 | - | 29580098 | 29580155 | 29570404 | 29570532 | 29580984 | 29581074 | 0 | 0 | 2 | 1 | 0 | 5 | 3 | 12 |
| SE | 43903 | ENSG00000112118 | MCM3 | chr6 | - | 52272300 | 52272451 | 52269085 | 52269226 | 52273229 | 52273356 | 297 | 252 | 306 | 333 | 494 | 0 | 0 | 0 |
| SE | 43904 | ENSG00000112118 | MCM3 | chr6 | - | 52276267 | 52276476 | 52273741 | 52273916 | 52277066 | 52277198 | 108 | 47 | 110 | 76 | 51 | 1 | 0 | 0 |
| SE | 43906 | ENSG00000112118 | MCM3 | chr6 | - | 52278741 | 52278850 | 52277534 | 52277688 | 52279360 | 52279452 | 172 | 71 | 147 | 138 | 119 | 0 | 0 | 0 |
| SE | 43910 | ENSG00000112972 | HMGCS1 | chr5 | - | 43295751 | 43295917 | 43294690 | 43294861 | 43297001 | 43297166 | 148 | 91 | 141 | 133 | 173 | 0 | 0 | 0 |
| SE | 43912 | ENSG00000112972 | HMGCS1 | chr5 | - | 43307765 | 43307824 | 43298445 | 43298975 | 43313355 | 43313412 | 5 | 2 | 10 | 2 | 7 | 5 | 3 | 8 |
| SE | 43913 | ENSG00000165029 | ABCA1 | chr9 | - | 104785395 | 104785639 | 104781005 | 104784455 | 104786297 | 104786390 | 33 | 39 | 37 | 47 | 46 | 0 | 0 | 0 |
| SE | 43915 | ENSG00000165029 | ABCA1 | chr9 | - | 104788425 | 104788567 | 104787919 | 104788054 | 104790921 | 104791028 | 48 | 73 | 66 | 84 | 76 | 0 | 0 | 0 |
| SE | 43916 | ENSG00000165029 | ABCA1 | chr9 | - | 104790921 | 104791028 | 104788425 | 104788567 | 104791935 | 104791998 | 48 | 55 | 60 | 76 | 57 | 0 | 0 | 0 |
| SE | 43917 | ENSG00000165029 | ABCA1 | chr9 | - | 104800509 | 104800584 | 104799818 | 104799988 | 104802053 | 104802159 | 46 | 41 | 55 | 66 | 105 | 0 | 0 | 0 |
| SE | 43918 | ENSG00000165029 | ABCA1 | chr9 | - | 104802053 | 104802159 | 104800509 | 104800584 | 104803283 | 104803316 | 31 | 49 | 42 | 70 | 81 | 0 | 0 | 0 |
| SE | 43919 | ENSG00000165029 | ABCA1 | chr9 | - | 104810799 | 104810924 | 104809465 | 104809564 | 104812573 | 104812722 | 17 | 20 | 9 | 12 | 21 | 1 | 0 | 0 |
| SE | 43920 | ENSG00000165029 | ABCA1 | chr9 | - | 104812573 | 104812722 | 104809465 | 104809564 | 104814117 | 104814231 | 8 | 9 | 11 | 9 | 16 | 0 | 0 | 0 |
| SE | 43921 | ENSG00000165029 | ABCA1 | chr9 | - | 104819585 | 104819723 | 104818662 | 104818883 | 104819926 | 104820069 | 11 | 7 | 3 | 18 | 14 | 0 | 1 | 0 |
| SE | 43922 | ENSG00000165029 | ABCA1 | chr9 | - | 104822495 | 104822667 | 104821374 | 104821506 | 104824464 | 104824578 | 22 | 18 | 13 | 22 | 11 | 0 | 1 | 0 |
| SE | 43923 | ENSG00000165029 | ABCA1 | chr9 | - | 104825682 | 104825887 | 104824464 | 104824578 | 104826947 | 104827169 | 11 | 12 | 22 | 17 | 17 | 0 | 1 | 0 |
| SE | 43925 | ENSG00000165029 | ABCA1 | chr9 | - | 104903613 | 104903771 | 104889101 | 104889195 | 104927934 | 104928139 | 15 | 18 | 15 | 17 | 10 | 1 | 0 | 0 |
| SE | 43926 | ENSG00000232077 | LINC01031 | chr1 | + | 193349132 | 193349275 | 193304744 | 193304938 | 193351103 | 193351239 | 0 | 0 | 1 | 0 | 0 | 0 | 0 | 0 |
| SE | 43930 | ENSG00000231304 | SGO1-AS1 | chr3 | + | 20177216 | 20177250 | 20174734 | 20174818 | 20194349 | 20194411 | 0 | 0 | 0 | 1 | 3 | 0 | 1 | 0 |
| SE | 43935 | ENSG00000077782 | FGFR1 | chr8 | - | 38420256 | 38420350 | 38419532 | 38419735 | 38421796 | 38423175 | 0 | 1 | 0 | 0 | 0 | 3 | 1 | 3 |
| SE | 43940 | ENSG00000077782 | FGFR1 | chr8 | - | 38423024 | 38423175 | 38421796 | 38421941 | 38424508 | 38424699 | 2 | 0 | 0 | 0 | 0 | 7 | 5 | 6 |
| SE | 43944 | ENSG00000077782 | FGFR1 | chr8 | - | 38429681 | 38429948 | 38427920 | 38428435 | 38457355 | 38457446 | 4 | 1 | 0 | 0 | 0 | 4 | 0 | 0 |
| SE | 43954 | ENSG00000115165 | CYTIP | chr2 | - | 157418522 | 157418589 | 157414618 | 157416143 | 157427350 | 157427420 | 1215 | 878 | 1172 | 1174 | 1688 | 0 | 1 | 0 |
| SE | 43955 | ENSG00000115165 | CYTIP | chr2 | - | 157430859 | 157430962 | 157430558 | 157430652 | 157434697 | 157434747 | 922 | 674 | 842 | 924 | 927 | 0 | 1 | 0 |
| SE | 43957 | ENSG00000115165 | CYTIP | chr2 | - | 157434369 | 157434424 | 157430859 | 157430962 | 157443846 | 157444006 | 163 | 113 | 138 | 152 | 267 | 0 | 0 | 1 |
| SE | 43958 | ENSG00000115165 | CYTIP | chr2 | - | 157434697 | 157434747 | 157430859 | 157430962 | 157443846 | 157444006 | 87 | 41 | 93 | 84 | 35 | 0 | 0 | 1 |
| SE | 43959 | ENSG00000115165 | CYTIP | chr2 | - | 157434697 | 157434747 | 157434160 | 157434424 | 157443846 | 157444006 | 1434 | 967 | 1341 | 1432 | 1384 | 1 | 0 | 0 |
| SE | 43960 | ENSG00000115165 | CYTIP | chr2 | - | 157438882 | 157438963 | 157434697 | 157434747 | 157439275 | 157439414 | 1 | 0 | 1 | 0 | 0 | 24 | 19 | 25 |
| SE | 43961 | ENSG00000115165 | CYTIP | chr2 | - | 157438882 | 157438963 | 157434697 | 157434747 | 157443846 | 157444098 | 1 | 0 | 2 | 0 | 0 | 1053 | 647 | 992 |
| SE | 43964 | ENSG00000171295 | ZNF440 | chr19 | + | 11830273 | 11830409 | 11814343 | 11814450 | 11830616 | 11830677 | 20 | 27 | 12 | 17 | 15 | 0 | 0 | 0 |
| SE | 43965 | ENSG00000171295 | ZNF440 | chr19 | + | 11830282 | 11830409 | 11814272 | 11814450 | 11830616 | 11830677 | 20 | 26 | 12 | 20 | 15 | 0 | 0 | 0 |
| SE | 43971 | ENSG00000108443 | RPS6KB1 | chr17 | + | 59930357 | 59930432 | 59930116 | 59930174 | 59930799 | 59931052 | 1 | 1 | 0 | 6 | 4 | 2 | 1 | 3 |
| SE | 43974 | ENSG00000108443 | RPS6KB1 | chr17 | + | 59930799 | 59931052 | 59930357 | 59930432 | 59931621 | 59931722 | 2 | 1 | 1 | 9 | 1 | 4 | 5 | 8 |
| SE | 43975 | ENSG00000108443 | RPS6KB1 | chr17 | + | 59931621 | 59931722 | 59930116 | 59930174 | 59934169 | 59934260 | 296 | 276 | 270 | 306 | 397 | 1 | 0 | 1 |
| SE | 43978 | ENSG00000108443 | RPS6KB1 | chr17 | + | 59940835 | 59940943 | 59936214 | 59936541 | 59945405 | 59945518 | 139 | 83 | 96 | 144 | 212 | 0 | 0 | 0 |
| SE | 43980 | ENSG00000114626 | ABTB1 | chr3 | + | 127677464 | 127677563 | 127677167 | 127677286 | 127677675 | 127677843 | 87 | 13 | 84 | 60 | 64 | 0 | 0 | 1 |
| SE | 43981 | ENSG00000114626 | ABTB1 | chr3 | + | 127677675 | 127677843 | 127677464 | 127677563 | 127679984 | 127680185 | 102 | 17 | 84 | 69 | 36 | 1 | 0 | 0 |
| SE | 43983 | ENSG00000114554 | PLXNA1 | chr3 | + | 126991383 | 126991566 | 126988520 | 126989787 | 127003329 | 127003470 | 10 | 13 | 8 | 21 | 27 | 0 | 0 | 0 |
| SE | 43990 | ENSG00000115540 | MOB4 | chr2 | + | 197548335 | 197548415 | 197540350 | 197540437 | 197550274 | 197550386 | 195 | 81 | 153 | 202 | 272 | 0 | 1 | 0 |
| SE | 43993 | ENSG00000160271 | RALGDS | chr9 | - | 133102475 | 133102571 | 133097719 | 133102139 | 133102778 | 133102900 | 303 | 125 | 210 | 242 | 214 | 0 | 0 | 0 |
| SE | 43994 | ENSG00000160271 | RALGDS | chr9 | - | 133103229 | 133103262 | 133102778 | 133102900 | 133103746 | 133103833 | 192 | 106 | 137 | 190 | 182 | 0 | 0 | 0 |
| SE | 43995 | ENSG00000160271 | RALGDS | chr9 | - | 133103746 | 133103833 | 133103229 | 133103262 | 133104262 | 133104331 | 226 | 116 | 167 | 218 | 223 | 2 | 0 | 0 |
| SE | 43997 | ENSG00000160271 | RALGDS | chr9 | - | 133107084 | 133107300 | 133106644 | 133106748 | 133107987 | 133108018 | 128 | 54 | 133 | 169 | 65 | 2 | 0 | 0 |

| | | | | | | | | | | | | | | |
|---|---|---|---|---|---|---|---|---|---|---|---|---|---|---|
| SE | 44334 | ENSG00000115524 | SF3B1 | chr2 | - | 197416740 | 197416911 | 197409769 | 197410007 | 197418508 | 197418588 | 1469 | 1204 | [illegible] |
| SE | 44337 | ENSG00000115524 | SF3B1 | chr2 | - | 197418895 | 197420542 | 197418508 | 197418588 | 197421028 | 197421133 | 2192 | 1735 | [illegible] |
| SE | 44338 | ENSG00000115524 | SF3B1 | chr2 | - | 197420427 | 197420542 | 197418508 | 197418588 | 197421028 | 197421133 | 2428 | 1837 | [illegible] |
| SE | 44339 | ENSG00000115524 | SF3B1 | chr2 | - | 197423807 | 197423974 | 197421028 | 197421133 | 197434971 | 197435027 | 3688 | 2636 | [illegible] |
| SE | 44340 | ENSG00000115524 | SF3B1 | chr2 | - | 197423893 | 197423974 | 197421028 | 197421133 | 197434971 | 197435016 | 7 | 3 | [illegible] |
| SE | 44343 | ENSG00000163517 | HDAC11 | chr3 | + | 13496735 | 13496852 | 13483463 | 13483564 | 13500712 | 13500789 | 5 | 1 | [illegible] |
| SE | 44345 | ENSG00000163517 | HDAC11 | chr3 | + | 13500712 | 13500789 | 13483463 | 13483564 | 13501870 | 13501933 | 4 | 0 | [illegible] |
| SE | 44351 | ENSG00000163517 | HDAC11 | chr3 | + | 13502883 | 13502980 | 13501870 | 13501933 | 13504093 | 13504113 | 8 | 1 | [illegible] |
| SE | 44382 | ENSG00000072210 | ALDH3A2 | chr17 | + | 19657720 | 19657862 | 19656487 | 19656574 | 19661126 | 19661268 | 61 | 35 | [illegible] |
| SE | 44383 | ENSG00000072210 | ALDH3A2 | chr17 | + | 19657744 | 19657862 | 19654905 | 19656574 | 19661126 | 19661268 | 166 | 93 | [illegible] |
| SE | 44386 | ENSG00000072210 | ALDH3A2 | chr17 | + | 19671720 | 19671956 | 19663332 | 19663499 | 19675557 | 19675644 | 1 | 0 | [illegible] |
| SE | 44391 | ENSG00000072210 | ALDH3A2 | chr17 | + | 19673150 | 19673275 | 19671720 | 19671956 | 19675557 | 19675741 | 5 | 1 | [illegible] |
| SE | 44396 | ENSG00000134419 | RPS15A | chr16 | - | 18784737 | 18784823 | 18781294 | 18783102 | 18788062 | 18788105 | 10483 | 5033 | [illegible] |
| SE | 44397 | ENSG00000134419 | RPS15A | chr16 | - | 18786239 | 18786336 | 18782967 | 18784823 | 18788062 | 18788142 | 54 | 17 | [illegible] |
| SE | 44399 | ENSG00000134419 | RPS15A | chr16 | - | 18788062 | 18788142 | 18784020 | 18784823 | 18788980 | 18789118 | 16550 | 7503 | [illegible] |
| SE | 44405 | ENSG00000163346 | PBXIP1 | chr1 | - | 154953670 | 154953757 | 154951470 | 154951535 | 154956068 | 154956099 | 2 | 0 | [illegible] |
| SE | 44407 | ENSG00000163346 | PBXIP1 | chr1 | - | 154953670 | 154953757 | 154951794 | 154951919 | 154956068 | 154956099 | 1 | 10 | [illegible] |
| SE | 44409 | ENSG00000228058 | LINC01736 | chr1 | + | 230005957 | 230006098 | 230002716 | 230002993 | 230006772 | 230007195 | 1 | 0 | [illegible] |
| SE | 44410 | ENSG00000175322 | ZNF519 | chr18 | - | 14080188 | 14080266 | 14078199 | 14078298 | 14084876 | 14085076 | 8 | 1 | [illegible] |
| SE | 44411 | ENSG00000175322 | ZNF519 | chr18 | - | 14084196 | 14084347 | 14080188 | 14080266 | 14084876 | 14085076 | 0 | 0 | [illegible] |
| SE | 44412 | ENSG00000175322 | ZNF519 | chr18 | - | 14123113 | 14123207 | 14122162 | 14122603 | 14124349 | 14124476 | 0 | 0 | [illegible] |
| SE | 44414 | ENSG00000175322 | ZNF519 | chr18 | - | 14124349 | 14124476 | 14099937 | 14106409 | 14132274 | 14132428 | 18 | 4 | [illegible] |
| SE | 44415 | ENSG00000056050 | HPF1 | chr4 | - | 169748743 | 169748842 | 169731703 | 169731876 | 169753675 | 169753835 | 1 | 2 | [illegible] |
| SE | 44418 | ENSG00000056050 | HPF1 | chr4 | - | 169750535 | 169750725 | 169741956 | 169742107 | 169753675 | 169753835 | 216 | 112 | [illegible] |
| SE | 44420 | ENSG00000204435 | CSNK2B | chr6 | + | 31667867 | 31667970 | 31666820 | 31666903 | 31668538 | 31668654 | 485 | 308 | [illegible] |
| SE | 44421 | ENSG00000204435 | CSNK2B | chr6 | + | 31668512 | 31668654 | 31667867 | 31667970 | 31669096 | 31669172 | 321 | 167 | [illegible] |
| SE | 44422 | ENSG00000204435 | CSNK2B | chr6 | + | 31668538 | 31668654 | 31667867 | 31667970 | 31669096 | 31669172 | 670 | 394 | [illegible] |
| SE | 44423 | ENSG00000204435 | CSNK2B | chr6 | + | 31668538 | 31668692 | 31667867 | 31667970 | 31669096 | 31669172 | 287 | 185 | [illegible] |
| SE | 44424 | ENSG00000204435 | CSNK2B | chr6 | + | 31669096 | 31669172 | 31666820 | 31668654 | 31669327 | 31669508 | 148 | 90 | [illegible] |
| SE | 44425 | ENSG00000204435 | CSNK2B | chr6 | + | 31669318 | 31669508 | 31669096 | 31669172 | 31669835 | 31670066 | 573 | 312 | [illegible] |
| SE | 44427 | ENSG00000164011 | ZNF691 | chr1 | + | 42849290 | 42849413 | 42846572 | 42846657 | 42849564 | 42849742 | 7 | 12 | [illegible] |
| SE | 44428 | ENSG00000164011 | ZNF691 | chr1 | + | 42849290 | 42849413 | 42846572 | 42846657 | 42850949 | 42852477 | 4 | 1 | [illegible] |
| SE | 44429 | ENSG00000164011 | ZNF691 | chr1 | + | 42849564 | 42849742 | 42846572 | 42846657 | 42850949 | 42852473 | 1 | 5 | [illegible] |
| SE | 44430 | ENSG00000164011 | ZNF691 | chr1 | + | 42849564 | 42849742 | 42849290 | 42849413 | 42850949 | 42852473 | 6 | 16 | [illegible] |
| SE | 44431 | ENSG00000164011 | ZNF691 | chr1 | + | 42850658 | 42850743 | 42849564 | 42849742 | 42850949 | 42852477 | 1 | 1 | [illegible] |
| SE | 44455 | ENSG00000145703 | IQGAP2 | chr5 | + | 76461569 | 76461669 | 76403284 | 76403591 | 76562395 | 76562552 | 403 | 233 | [illegible] |
| SE | 44456 | ENSG00000145703 | IQGAP2 | chr5 | + | 76562395 | 76562552 | 76461569 | 76461669 | 76570579 | 76570657 | 1100 | 756 | [illegible] |
| SE | 44457 | ENSG00000145703 | IQGAP2 | chr5 | + | 76570579 | 76570657 | 76562395 | 76562552 | 76575692 | 76575769 | 1117 | 801 | [illegible] |
| SE | 44458 | ENSG00000145703 | IQGAP2 | chr5 | + | 76575692 | 76575769 | 76562395 | 76562552 | 76588905 | 76588973 | 417 | 333 | [illegible] |
| SE | 44459 | ENSG00000145703 | IQGAP2 | chr5 | + | 76588905 | 76588973 | 76575692 | 76575769 | 76592837 | 76592925 | 369 | 255 | [illegible] |
| SE | 44460 | ENSG00000145703 | IQGAP2 | chr5 | + | 76589614 | 76589728 | 76588905 | 76588973 | 76592837 | 76592925 | 539 | 404 | [illegible] |
| SE | 44461 | ENSG00000145703 | IQGAP2 | chr5 | + | 76590407 | 76590586 | 76589614 | 76589728 | 76592837 | 76592925 | 723 | 583 | [illegible] |
| SE | 44462 | ENSG00000145703 | IQGAP2 | chr5 | + | 76606178 | 76606303 | 76600811 | 76600972 | 76611019 | 76611183 | 1230 | 732 | [illegible] |
| SE | 44463 | ENSG00000145703 | IQGAP2 | chr5 | + | 76606178 | 76606303 | 76600811 | 76600972 | 76627409 | 76627500 | 541 | 311 | [illegible] |
| SE | 44464 | ENSG00000145703 | IQGAP2 | chr5 | + | 76611019 | 76611183 | 76600811 | 76600972 | 76627409 | 76627500 | 225 | 132 | [illegible] |
| SE | 44465 | ENSG00000145703 | IQGAP2 | chr5 | + | 76611019 | 76611183 | 76606178 | 76606303 | 76627409 | 76627500 | 917 | 562 | [illegible] |
| SE | 44467 | ENSG00000145703 | IQGAP2 | chr5 | + | 76618513 | 76618600 | 76611019 | 76611183 | 76627409 | 76627496 | 1 | 0 | [illegible] |
| SE | 44471 | ENSG00000145703 | IQGAP2 | chr5 | + | 76654199 | 76654271 | 76652749 | 76652833 | 76658458 | 76658667 | 55 | 41 | [illegible] |
| SE | 44472 | ENSG00000145703 | IQGAP2 | chr5 | + | 76654933 | 76655003 | 76654199 | 76654271 | 76658458 | 76658667 | 422 | 316 | [illegible] |
| SE | 44473 | ENSG00000145703 | IQGAP2 | chr5 | + | 76668680 | 76668844 | 76665025 | 76665175 | 76671758 | 76671784 | 1125 | 754 | [illegible] |
| SE | 44474 | ENSG00000145703 | IQGAP2 | chr5 | + | 76673951 | 76674036 | 76673448 | 76673589 | 76674476 | 76674709 | 566 | 451 | [illegible] |
| SE | 44475 | ENSG00000145703 | IQGAP2 | chr5 | + | 76674476 | 76674709 | 76673448 | 76673589 | 76677217 | 76677350 | 578 | 433 | [illegible] |
| SE | 44476 | ENSG00000145703 | IQGAP2 | chr5 | + | 76677217 | 76677350 | 76674476 | 76674709 | 76683114 | 76683217 | 896 | 613 | [illegible] |
| SE | 44477 | ENSG00000145703 | IQGAP2 | chr5 | + | 76683114 | 76683217 | 76674521 | 76674709 | 76693354 | 76693442 | 6 | 17 | [illegible] |
| SE | 44479 | ENSG00000145703 | IQGAP2 | chr5 | + | 76697986 | 76698147 | 76695453 | 76695666 | 76701075 | 76701213 | 567 | 375 | [illegible] |
| SE | 44480 | ENSG00000145703 | IQGAP2 | chr5 | + | 76701075 | 76701213 | 76697986 | 76698147 | 76702481 | 76702590 | 318 | 206 | [illegible] |
| SE | 44483 | ENSG00000090266 | NDUFB2 | chr7 | + | 140700869 | 140700959 | 140696991 | 140697074 | 140702865 | 140703010 | 0 | 0 | [illegible] |
| SE | 44509 | ENSG00000187134 | AKR1C1 | chr10 | + | 4965913 | 4966081 | 4963252 | 4963528 | 4966926 | 4967043 | 4 | 0 | [illegible] |
| SE | 44510 | ENSG00000187134 | AKR1C1 | chr10 | + | 4966009 | 4966081 | 4963448 | 4963528 | 4966926 | 4967043 | 0 | 0 | [illegible] |
| SE | 44511 | ENSG00000285215 | NA | chr1 | + | 120972785 | 120972910 | 120969118 | 120969354 | 120973107 | 120973246 | 13 | 2 | [illegible] |
| SE | 44512 | ENSG00000285215 | NA | chr1 | + | 120972785 | 120972910 | 120969118 | 120969354 | 120973346 | 120973561 | 3 | 0 | [illegible] |
| SE | 44514 | ENSG00000285215 | NA | chr1 | + | 120973107 | 120973246 | 120972785 | 120972910 | 120973346 | 120973561 | 23 | 8 | [illegible] |
| SE | 44517 | ENSG00000135778 | NTPCR | chr1 | + | 232956346 | 232956443 | 232950604 | 232950744 | 232969908 | 232970118 | 25 | 47 | [illegible] |
| SE | 44521 | ENSG00000101940 | WDR13 | chrX | + | 48600318 | 48600626 | 48599352 | 48599462 | 48601783 | 48601964 | 55 | 25 | [illegible] |
| SE | 44523 | ENSG00000119402 | FBXW2 | chr9 | - | 120776930 | 120777021 | 120776092 | 120776226 | 120787768 | 120788278 | 10 | 3 | [illegible] |
| SE | 44524 | ENSG00000119402 | FBXW2 | chr9 | - | 120778350 | 120778545 | 120776092 | 120776226 | 120787768 | 120788186 | 468 | 264 | [illegible] |
| SE | 44526 | ENSG00000122188 | LAX1 | chr1 | + | 203772067 | 203772147 | 203771366 | 203771477 | 203773874 | 203774722 | 108 | 59 | [illegible] |
| SE | 44528 | ENSG00000076321 | KLHL20 | chr1 | + | 173716002 | 173716066 | 173714980 | 173715078 | 173733712 | 173733828 | 40 | 39 | [illegible] |
| SE | 44531 | ENSG00000076321 | KLHL20 | chr1 | + | 173755922 | 173756038 | 173753212 | 173753307 | 173756975 | 173757159 | 48 | 42 | [illegible] |
| SE | 44532 | ENSG00000076321 | KLHL20 | chr1 | + | 173766145 | 173766289 | 173756975 | 173757159 | 173774304 | 173774438 | 88 | 60 | [illegible] |
| SE | 44533 | ENSG00000076321 | KLHL20 | chr1 | + | 173766145 | 173766289 | 173756975 | 173757159 | 173782123 | 173782230 | 60 | 41 | [illegible] |
| SE | 44534 | ENSG00000076321 | KLHL20 | chr1 | + | 173774304 | 173774438 | 173756975 | 173757159 | 173782123 | 173782230 | 0 | 1 | [illegible] |
| SE | 44535 | ENSG00000076321 | KLHL20 | chr1 | + | 173774304 | 173774438 | 173766145 | 173766289 | 173775633 | 173775842 | 53 | 56 | [illegible] |
| SE | 44536 | ENSG00000076321 | KLHL20 | chr1 | + | 173774304 | 173774438 | 173766145 | 173766289 | 173782123 | 173782230 | 29 | 19 | [illegible] |
| SE | 44538 | ENSG00000076321 | KLHL20 | chr1 | + | 173775633 | 173775842 | 173774304 | 173774438 | 173782123 | 173782230 | 51 | 64 | [illegible] |
| SE | 44539 | ENSG00000076321 | KLHL20 | chr1 | + | 173782123 | 173782230 | 173775633 | 173775842 | 173785162 | 173786692 | 42 | 28 | [illegible] |
| SE | 44540 | ENSG00000166526 | ZNF3 | chr7 | - | 100075134 | 100075261 | 100064032 | 100064912 | 100075541 | 100075630 | 35 | 9 | [illegible] |
| SE | 44544 | ENSG00000166526 | ZNF3 | chr7 | - | 100079535 | 100079572 | 100077302 | 100077433 | 100081634 | 100081709 | 1 | 2 | [illegible] |
| SE | 44545 | ENSG00000166526 | ZNF3 | chr7 | - | 100079535 | 100079656 | 100075541 | 100075630 | 100081634 | 100081703 | 10 | 5 | [illegible] |
| SE | 44546 | ENSG00000166526 | ZNF3 | chr7 | - | 100079535 | 100079656 | 100075541 | 100075630 | 100082300 | 100082548 | 0 | 0 | [illegible] |
| SE | 44549 | ENSG00000142676 | RPL11 | chr1 | + | 23693806 | 23693913 | 23692305 | 23692759 | 23694659 | 23694791 | 6419 | 4296 | [illegible] |
| SE | 44550 | ENSG00000142676 | RPL11 | chr1 | + | 23694659 | 23694791 | 23693806 | 23693913 | 23695797 | 23695908 | 11162 | 7852 | [illegible] |
| SE | 44552 | ENSG00000183020 | AP2A2 | chr11 | + | 970168 | 970311 | 959436 | 959505 | 972061 | 972191 | 198 | 157 | [illegible] |
| SE | 44553 | ENSG00000183020 | AP2A2 | chr11 | + | 970168 | 970311 | 959436 | 959505 | 977094 | 977120 | 73 | 60 | [illegible] |
| SE | 44555 | ENSG00000183020 | AP2A2 | chr11 | + | 972061 | 972255 | 970168 | 970311 | 977094 | 977120 | 163 | 118 | [illegible] |
| SE | 44556 | ENSG00000183020 | AP2A2 | chr11 | + | 977094 | 977224 | 972061 | 972255 | 981197 | 981299 | 75 | 43 | [illegible] |
| SE | 44557 | ENSG00000183020 | AP2A2 | chr11 | + | 993283 | 993381 | 992502 | 992685 | 993753 | 993985 | 109 | 57 | [illegible] |
| SE | 44559 | ENSG00000026508 | CD44 | chr11 | + | 35180273 | 35180407 | 35176574 | 35176740 | 35186831 | 35186900 | 1670 | 996 | [illegible] |
| SE | 44562 | ENSG00000026508 | CD44 | chr11 | + | 35198120 | 35198246 | 35189834 | 35190065 | 35201081 | 35201195 | 7 | 2 | [illegible] |
| SE | 44566 | ENSG00000026508 | CD44 | chr11 | + | 35198147 | 35198246 | 35190009 | 35190065 | 35201081 | 35201195 | 0 | 0 | [illegible] |
| SE | 44567 | ENSG00000026508 | CD44 | chr11 | + | 35201081 | 35201195 | 35197856 | 35198246 | 35201667 | 35201731 | 1 | 0 | [illegible] |
| SE | 44568 | ENSG00000026508 | CD44 | chr11 | + | 35201081 | 35201195 | 35197856 | 35198246 | 35201670 | 35201753 | 1 | 0 | [illegible] |
| SE | 44569 | ENSG00000026508 | CD44 | chr11 | + | 35201667 | 35201787 | 35201081 | 35201195 | 35204511 | 35204640 | 3 | 3 | [illegible] |
| SE | 44570 | ENSG00000026508 | CD44 | chr11 | + | 35201670 | 35201787 | 35198120 | 35198246 | 35204511 | 35204640 | 7 | 3 | [illegible] |
| SE | 44571 | ENSG00000026508 | CD44 | chr11 | + | 35201670 | 35201787 | 35198120 | 35198246 | 35214851 | 35214914 | 4 | 0 | [illegible] |
| SE | 44572 | ENSG00000026508 | CD44 | chr11 | + | 35201670 | 35201787 | 35201081 | 35201195 | 35204511 | 35204640 | 3 | 3 | [illegible] |
| SE | 44573 | ENSG00000026508 | CD44 | chr11 | + | 35204511 | 35204640 | 35189834 | 35190065 | 35206111 | 35206243 | 23 | 17 | [illegible] |
| SE | 44576 | ENSG00000026508 | CD44 | chr11 | + | 35204511 | 35204640 | 35198120 | 35198246 | 35208104 | 35208206 | 5 | 4 | [illegible] |
| SE | 44577 | ENSG00000026508 | CD44 | chr11 | + | 35204511 | 35204640 | 35198120 | 35198246 | 35214851 | 35214914 | 2 | 5 | [illegible] |
| SE | 44578 | ENSG00000026508 | CD44 | chr11 | + | 35204511 | 35204640 | 35201667 | 35201787 | 35214851 | 35214914 | 5 | 7 | [illegible] |
| SE | 44583 | ENSG00000026508 | CD44 | chr11 | + | 35208104 | 35208206 | 35189834 | 35189972 | 35214851 | 35214914 | 0 | 3 | [illegible] |
| SE | 44588 | ENSG00000026508 | CD44 | chr11 | + | 35208104 | 35208206 | 35204511 | 35204640 | 35214851 | 35214914 | 4 | 5 | [illegible] |
| SE | 44595 | ENSG00000026508 | CD44 | chr11 | + | 35211245 | 35211449 | 35201620 | 35201787 | 35214851 | 35214861 | 171 | 95 | [illegible] |
| SE | 44598 | ENSG00000026508 | CD44 | chr11 | + | 35214851 | 35214914 | 35189834 | 35190065 | 35219315 | 35219387 | 1330 | 911 | [illegible] |
| SE | 44599 | ENSG00000026508 | CD44 | chr11 | + | 35214851 | 35214914 | 35189834 | 35190065 | 35229128 | 35229164 | 219 | 172 | [illegible] |
| SE | 44600 | ENSG00000026508 | CD44 | chr11 | + | 35219315 | 35219387 | 35214665 | 35214914 | 35221653 | 35221673 | 2652 | 1715 | [illegible] |
| SE | 44601 | ENSG00000026508 | CD44 | chr11 | + | 35221653 | 35221732 | 35214665 | 35214914 | 35229128 | 35229164 | 1660 | 1000 | [illegible] |
| SE | 44604 | ENSG00000026025 | VIM | chr10 | + | 17236293 | 17236379 | 17235845 | 17235889 | 17237229 | 17237593 | 8929 | 5542 | [illegible] |
| SE | 44605 | ENSG00000157978 | LDLRAP1 | chr1 | + | 25562643 | 25562716 | 25557152 | 25557267 | 25563069 | 25563153 | 103 | 109 | [illegible] |
| SE | 44606 | ENSG00000157978 | LDLRAP1 | chr1 | + | 25563069 | 25563153 | 25557152 | 25557267 | 25563660 | 25563791 | 121 | 57 | [illegible] |
| SE | 44607 | ENSG00000164050 | PLXNB1 | chr3 | - | 48413669 | 48413818 | 48413068 | 48413169 | 48413894 | 48414071 | 2 | 0 | [illegible] |
| SE | 44611 | ENSG00000167065 | DUSP18 | chr22 | - | 30663012 | 30663260 | 30652050 | 30652297 | 30663403 | 30663506 | 10 | 6 | [illegible] |
| SE | 44615 | ENSG00000167065 | DUSP18 | chr22 | - | 30666806 | 30666938 | 30652095 | 30652297 | 30667461 | 30667887 | 1 | 0 | [illegible] |
| SE | 44616 | ENSG00000099338 | CATSPERG | chr19 | + | 38337739 | 38337861 | 38337592 | 38337646 | 38343579 | 38343724 | 0 | 0 | [illegible] |
| SE | 44618 | ENSG00000099338 | CATSPERG | chr19 | + | 38346449 | 38346605 | 38344295 | 38344368 | 38358277 | 38358328 | 8 | 9 | [illegible] |
| SE | 44635 | ENSG00000133398 | MED10 | chr5 | - | 6374323 | 6374426 | 6371873 | 6372601 | 6377165 | 6377249 | 341 | 289 | [illegible] |
| SE | 44636 | ENSG00000183137 | CEP57L1 | chr6 | + | 109100191 | 109100286 | 109095152 | 109095575 | 109106034 | 109106090 | 3 | 2 | [illegible] |
| SE | 44637 | ENSG00000183137 | CEP57L1 | chr6 | + | 109100191 | 109100286 | 109095152 | 109095575 | 109145218 | 109145307 | 1 | 0 | [illegible] |
| SE | 44638 | ENSG00000183137 | CEP57L1 | chr6 | + | 109106034 | 109106090 | 109095109 | 109095575 | 109140420 | 109140675 | 1 | 0 | [illegible] |
| SE | 44639 | ENSG00000183137 | CEP57L1 | chr6 | + | 109106034 | 109106090 | 109095109 | 109095575 | 109145218 | 109145381 | 1 | 1 | [illegible] |
| SE | 44640 | ENSG00000183137 | CEP57L1 | chr6 | + | 109106034 | 109106090 | 109095109 | 109095575 | 109146757 | 109146937 | 0 | 0 | [illegible] |
| SE | 44642 | ENSG00000183137 | CEP57L1 | chr6 | + | 109106034 | 109106090 | 109095187 | 109095251 | 109145218 | 109145381 | 1 | 1 | [illegible] |
| SE | 44643 | ENSG00000183137 | CEP57L1 | chr6 | + | 109106034 | 109106090 | 109100191 | 109100286 | 109145218 | 109145381 | 3 | 3 | [illegible] |
| SE | 44644 | ENSG00000183137 | CEP57L1 | chr6 | + | 109124990 | 109125031 | 109095509 | 109095575 | 109145218 | 109145381 | 1 | 0 | [illegible] |
| SE | 44648 | ENSG00000183137 | CEP57L1 | chr6 | + | 109140420 | 109140675 | 109095162 | 109095251 | 109145218 | 109145381 | 9 | 1 | [illegible] |
| SE | 44649 | ENSG00000183137 | CEP57L1 | chr6 | + | 109140420 | 109140675 | 109095181 | 109095575 | 109145218 | 109145381 | 11 | 1 | [illegible] |
| SE | 44650 | ENSG00000183137 | CEP57L1 | chr6 | + | 109140420 | 109140675 | 109095181 | 109095575 | 109146757 | 109146937 | 5 | 0 | [illegible] |
| SE | 44651 | ENSG00000183137 | CEP57L1 | chr6 | + | 109140420 | 109140675 | 109106034 | 109106090 | 109145218 | 109145381 | 10 | 1 | [illegible] |
| SE | 44652 | ENSG00000183137 | CEP57L1 | chr6 | + | 109140420 | 109140675 | 109106034 | 109106090 | 109146757 | 109146937 | 4 | 0 | [illegible] |
| SE | 44653 | ENSG00000183137 | CEP57L1 | chr6 | + | 109145218 | 109145381 | 109095157 | 109095575 | 109146757 | 109146933 | 38 | 30 | [illegible] |
| SE | 44654 | ENSG00000183137 | CEP57L1 | chr6 | + | 109145218 | 109145381 | 109106034 | 109106090 | 109146757 | 109146933 | 32 | 27 | [illegible] |
| SE | 44656 | ENSG00000183137 | CEP57L1 | chr6 | + | 109145218 | 109145381 | 109140420 | 109140675 | 109146757 | 109146933 | 40 | 25 | [illegible] |
| SE | 44658 | ENSG00000183137 | CEP57L1 | chr6 | + | 109155790 | 109155877 | 109155229 | 109155307 | 109159024 | 109159102 | 30 | 44 | [illegible] |
| SE | 44659 | ENSG00000183137 | CEP57L1 | chr6 | + | 109158599 | 109158718 | 109155790 | 109155877 | 109159024 | 109159102 | 0 | 4 | [illegible] |
| SE | 44660 | ENSG00000164331 | ANKRA2 | chr5 | - | 73554860 | 73554986 | 73554321 | 73554388 | 73555487 | 73555585 | 49 | 17 | [illegible] |
| SE | 44661 | ENSG00000164331 | ANKRA2 | chr5 | - | 73555487 | 73555585 | 73554860 | 73554986 | 73557574 | 73557640 | 104 | 57 | [illegible] |
| SE | 44666 | ENSG00000072501 | SMC1A | chrX | - | 53382505 | 53382660 | 53382231 | 53382383 | 53383096 | 53383253 | 337 | 205 | [illegible] |
| SE | 44667 | ENSG00000072501 | SMC1A | chrX | - | 53394777 | 53394888 | 53383096 | 53383253 | 53396226 | 53396378 | 440 | 298 | [illegible] |
| SE | 44668 | ENSG00000072501 | SMC1A | chrX | - | 53399588 | 53399730 | 53396471 | 53396617 | 53403565 | 53403672 | 521 | 399 | [illegible] |
| SE | 44669 | ENSG00000072501 | SMC1A | chrX | - | 53399588 | 53399730 | 53396471 | 53396617 | 53405011 | 53405149 | 260 | 189 | [illegible] |
| SE | 44670 | ENSG00000072501 | SMC1A | chrX | - | 53405011 | 53405149 | 53403776 | 53403893 | 53405244 | 53405391 | 177 | 109 | [illegible] |
| SE | 44671 | ENSG00000072501 | SMC1A | chrX | - | 53405770 | 53405956 | 53405492 | 53405672 | 53409061 | 53409269 | 277 | 213 | [illegible] |
| SE | 44673 | ENSG00000072501 | SMC1A | chrX | - | 53414757 | 53414870 | 53411994 | 53412253 | 53422491 | 53422654 | 0 | 3 | [illegible] |
| SE | 44675 | ENSG00000072501 | SMC1A | chrX | - | 53414980 | 53415169 | 53414757 | 53414870 | 53422491 | 53422654 | 193 | 110 | [illegible] |
| SE | 44677 | ENSG00000189195 | BTBD8 | chr1 | + | 92107883 | 92108001 | 92102472 | 92102669 | 92129686 | 92129776 | 8 | 12 | [illegible] |
| SE | 44678 | ENSG00000189195 | BTBD8 | chr1 | + | 92141114 | 92141211 | 92139349 | 92139430 | 92147179 | 92147268 | 5 | 1 | [illegible] |
| SE | 44679 | ENSG00000189195 | BTBD8 | chr1 | + | 92175062 | 92175249 | 92171398 | 92171460 | 92176828 | 92177546 | 0 | 1 | [illegible] |
| SE | 44684 | ENSG00000185344 | ATP6V0A2 | chr12 | + | 123747606 | 123747725 | 123744881 | 123744972 | 123748574 | 123748785 | 59 | 50 | [illegible] |
| SE | 44688 | ENSG00000159592 | GPBP1L1 | chr1 | - | 45630481 | 45630606 | 45629575 | 45629678 | 45640209 | 45640403 | 160 | 125 | [illegible] |
| SE | 44689 | ENSG00000159592 | GPBP1L1 | chr1 | - | 45633488 | 45633647 | 45630481 | 45630606 | 45634095 | 45634236 | 408 | 329 | [illegible] |
| SE | 44693 | ENSG00000159592 | GPBP1L1 | chr1 | - | 45642426 | 45642499 | 45640209 | 45640403 | 45654542 | 45654829 | 241 | 125 | [illegible] |
| SE | 44694 | ENSG00000159592 | GPBP1L1 | chr1 | - | 45655189 | 45655319 | 45654542 | 45654829 | 45659027 | 45659142 | 488 | 311 | [illegible] |
| SE | 44695 | ENSG00000159592 | GPBP1L1 | chr1 | - | 45660183 | 45661225 | 45659027 | 45659142 | 45686411 | 45686536 | 590 | 469 | [illegible] |
| SE | 44714 | ENSG00000143772 | ITPKB | chr1 | - | 226637678 | 226637750 | 226631689 | 226634886 | 226639556 | 226639658 | 323 | 205 | [illegible] |
| SE | 44718 | ENSG00000135999 | EPC2 | chr2 | + | 148690213 | 148690373 | 148644750 | 148645170 | 148743621 | 148743767 | 168 | 104 | [illegible] |
| SE | 44719 | ENSG00000135999 | EPC2 | chr2 | + | 148743621 | 148743767 | 148690213 | 148690373 | 148753926 | 148754020 | 381 | 306 | [illegible] |
| SE | 44720 | ENSG00000135999 | EPC2 | chr2 | + | 148753926 | 148754133 | 148743621 | 148743767 | 148761781 | 148761809 | 512 | 460 | [illegible] |
| SE | 44722 | ENSG00000135999 | EPC2 | chr2 | + | 148762669 | 148762802 | 148753926 | 148754133 | 148764954 | 148765146 | 115 | 134 | [illegible] |
| SE | 44723 | ENSG00000135999 | EPC2 | chr2 | + | 148762669 | 148762802 | 148753926 | 148754133 | 148769150 | 148769240 | 1 | 1 | [illegible] |
| SE | 44724 | ENSG00000135999 | EPC2 | chr2 | + | 148764954 | 148765146 | 148753926 | 148754133 | 148769150 | 148769240 | 163 | 111 | [illegible] |
| SE | 44725 | ENSG00000135999 | EPC2 | chr2 | + | 148764954 | 148765146 | 148762669 | 148762802 | 148769150 | 148769240 | 334 | 280 | [illegible] |
| SE | 44726 | ENSG00000135999 | EPC2 | chr2 | + | 148781643 | 148781780 | 148771043 | 148771387 | 148783596 | 148783756 | 363 | 305 | [illegible] |
| SE | 44727 | ENSG00000135999 | EPC2 | chr2 | + | 148783596 | 148783756 | 148781643 | 148781780 | 148784667 | 148785001 | 279 | 193 | [illegible] |
| SE | 44728 | ENSG00000138069 | RAB1A | chr2 | - | 65088938 | 65089070 | 65086853 | 65088690 | 65090982 | 65091078 | 722 | 584 | [illegible] |
| SE | 44732 | ENSG00000138069 | RAB1A | chr2 | - | 65090982 | 65091078 | 65088938 | 65089070 | 65104733 | 65104806 | 268 | 184 | [illegible] |
| SE | 44734 | ENSG00000138069 | RAB1A | chr2 | - | 65097970 | 65098066 | 65088938 | 65089070 | 65104733 | 65104806 | 158 | 112 | [illegible] |
| SE | 44735 | ENSG00000138069 | RAB1A | chr2 | - | 65097970 | 65098066 | 65090982 | 65091078 | 65104733 | 65104806 | 437 | 330 | [illegible] |
| SE | 44739 | ENSG00000198265 | HELZ | chr17 | - | 67148568 | 67148714 | 67145742 | 67145890 | 67149866 | 67149985 | 371 | 265 | [illegible] |
| SE | 44740 | ENSG00000198265 | HELZ | chr17 | - | 67149866 | 67149985 | 67148568 | 67148714 | 67151045 | 67151224 | 259 | 147 | [illegible] |
| SE | 44741 | ENSG00000198265 | HELZ | chr17 | - | 67160260 | 67160362 | 67151045 | 67151224 | 67160896 | 67161076 | 221 | 167 | [illegible] |
| SE | 44742 | ENSG00000198265 | HELZ | chr17 | - | 67160260 | 67160365 | 67151045 | 67151224 | 67160896 | 67161076 | 133 | 90 | [illegible] |
| SE | 44743 | ENSG00000198265 | HELZ | chr17 | - | 67160896 | 67161076 | 67151045 | 67151224 | 67166477 | 67166608 | 203 | 147 | [illegible] |
| SE | 44746 | ENSG00000198265 | HELZ | chr17 | - | 67188318 | 67188616 | 67178658 | 67178926 | 67189588 | 67189696 | 588 | 417 | [illegible] |
| SE | 44747 | ENSG00000198265 | HELZ | chr17 | - | 67189588 | 67189696 | 67178658 | 67178926 | 67201128 | 67201185 | 0 | 0 | [illegible] |
| SE | 44751 | ENSG00000198265 | HELZ | chr17 | - | 67203318 | 67203443 | 67200612 | 67201185 | 67215898 | 67215935 | 616 | 447 | [illegible] |
| SE | 44752 | ENSG00000198265 | HELZ | chr17 | - | 67215898 | 67215935 | 67203318 | 67203443 | 67218594 | 67218822 | 359 | 281 | [illegible] |
| SE | 44754 | ENSG00000175137 | SH3BP5L | chr1 | - | 248814710 | 248814877 | 248814448 | 248814610 | 248816533 | 248816662 | 1 | 0 | [illegible] |
| SE | 44755 | ENSG00000175137 | SH3BP5L | chr1 | - | 248820402 | 248820912 | 248816821 | 248816884 | 248821146 | 248821261 | 3 | 0 | [illegible] |
| SE | 44757 | ENSG00000175137 | SH3BP5L | chr1 | - | 248821146 | 248821261 | 248816821 | 248816884 | 248824752 | 248825635 | 1 | 0 | [illegible] |
| SE | 44758 | ENSG00000175137 | SH3BP5L | chr1 | - | 248821146 | 248821261 | 248820402 | 248820912 | 248824752 | 248825635 | 1 | 0 | [illegible] |
| SE | 44759 | ENSG00000171595 | DNAI2 | chr17 | + | 74286976 | 74287098 | 74285039 | 74285201 | 74289593 | 74289736 | 3 | 1 | [illegible] |
| SE | 44760 | ENSG00000171595 | DNAI2 | chr17 | + | 74291019 | 74291133 | 74289593 | 74289736 | 74299717 | 74299857 | 2 | 0 | [illegible] |
| SE | 44770 | ENSG00000145833 | DDX46 | chr5 | + | 134777573 | 134777725 | 134773695 | 134773861 | 134781132 | 134781246 | 237 | 189 | [illegible] |
| SE | 44772 | ENSG00000145833 | DDX46 | chr5 | + | 134782944 | 134783065 | 134781920 | 134782086 | 134784365 | 134784541 | 406 | 321 | [illegible] |
| SE | 44773 | ENSG00000145833 | DDX46 | chr5 | + | 134782944 | 134783065 | 134781920 | 134782086 | 134785464 | 134785586 | 233 | 175 | [illegible] |
| SE | 44775 | ENSG00000145833 | DDX46 | chr5 | + | 134784365 | 134784541 | 134782050 | 134782086 | 134788512 | 134788591 | 2 | 1 | [illegible] |
| SE | 44776 | ENSG00000145833 | DDX46 | chr5 | + | 134784365 | 134784541 | 134782944 | 134783065 | 134785464 | 134785586 | 306 | 299 | [illegible] |
| SE | 44777 | ENSG00000145833 | DDX46 | chr5 | + | 134785464 | 134785586 | 134781920 | 134782086 | 134788512 | 134788591 | 205 | 159 | [illegible] |
| SE | 44778 | ENSG00000145833 | DDX46 | chr5 | + | 134785464 | 134785586 | 134784365 | 134784541 | 134788512 | 134788591 | 328 | 310 | [illegible] |
| SE | 44779 | ENSG00000145833 | DDX46 | chr5 | + | 134789215 | 134789264 | 134788512 | 134788591 | 134790469 | 134790552 | 2 | 0 | [illegible] |
| SE | 44784 | ENSG00000134717 | BTF3L4 | chr1 | + | 52071149 | 52071327 | 52064824 | 52064938 | 52083339 | 52083500 | 2 | 2 | [illegible] |
| SE | 44785 | ENSG00000134717 | BTF3L4 | chr1 | + | 52071149 | 52071327 | 52064824 | 52064938 | 52086111 | 52086610 | 2 | 2 | [illegible] |
| SE | 44787 | ENSG00000138074 | SLC5A6 | chr2 | - | 27201349 | 27201453 | 27200997 | 27201113 | 27201665 | 27201847 | 12 | 3 | [illegible] |
| SE | 44788 | ENSG00000138074 | SLC5A6 | chr2 | - | 27206482 | 27206534 | 27206025 | 27206093 | 27206876 | 27206942 | 5 | 1 | [illegible] |
| SE | 44790 | ENSG00000138074 | SLC5A6 | chr2 | - | 27208613 | 27208686 | 27207257 | 27207790 | 27211466 | 27211533 | 0 | 0 | [illegible] |
| SE | 44791 | ENSG00000138074 | SLC5A6 | chr2 | - | 27212019 | 27212146 | 27211466 | 27211533 | 27212628 | 27212777 | 8 | 5 | [illegible] |
| SE | 44793 | ENSG00000054282 | SDCCAG8 | chr1 | + | 243274542 | 243274656 | 243256033 | 243256240 | 243286271 | 243286397 | 63 | 54 | [illegible] |
| SE | 44794 | ENSG00000054282 | SDCCAG8 | chr1 | + | 243286271 | 243286397 | 243256033 | 243256240 | 243316754 | 243316893 | 0 | 0 | [illegible] |
| SE | 44797 | ENSG00000054282 | SDCCAG8 | chr1 | + | 243304712 | 243304777 | 243286382 | 243286397 | 243316754 | 243316893 | 4 | 2 | [illegible] |
| SE | 44798 | ENSG00000054282 | SDCCAG8 | chr1 | + | 243304712 | 243304777 | 243286382 | 243286397 | 243330539 | 243330692 | 4 | 2 | [illegible] |
| SE | 44800 | ENSG00000054282 | SDCCAG8 | chr1 | + | 243307988 | 243308177 | 243304712 | 243304777 | 243330539 | 243330692 | 24 | 16 | [illegible] |
| SE | 44801 | ENSG00000054282 | SDCCAG8 | chr1 | + | 243316754 | 243316893 | 243286271 | 243286397 | 243330539 | 243330692 | 39 | 46 | [illegible] |
| SE | 44802 | ENSG00000054282 | SDCCAG8 | chr1 | + | 243316754 | 243316893 | 243304712 | 243304777 | 243330539 | 243330692 | 39 | 46 | [illegible] |
| SE | 44804 | ENSG00000054282 | SDCCAG8 | chr1 | + | 243415701 | 243415829 | 243378720 | 243378863 | 243417967 | 243418076 | 43 | 46 | [illegible] |
| SE | 44806 | ENSG00000054282 | SDCCAG8 | chr1 | + | 243417967 | 243418076 | 243378720 | 243378863 | 243426426 | 243426558 | 18 | 30 | [illegible] |
| SE | 44807 | ENSG00000054282 | SDCCAG8 | chr1 | + | 243417967 | 243418076 | 243415701 | 243415829 | 243426426 | 243426558 | 36 | 60 | [illegible] |
| SE | 44808 | ENSG00000054282 | SDCCAG8 | chr1 | + | 243489013 | 243489140 | 243426426 | 243426558 | 243499755 | 243500089 | 13 | 12 | [illegible] |
| SE | 44809 | ENSG00000177873 | ZNF619 | chr3 | + | 40477917 | 40478003 | 40477112 | 40477357 | 40482587 | 40482704 | 3 | 4 | [illegible] |
| SE | 44810 | ENSG00000177873 | ZNF619 | chr3 | + | 40481862 | 40482016 | 40477769 | 40478003 | 40482587 | 40482704 | 13 | 7 | [illegible] |

| | | | | | | | | | | | | | |
|---|---|---|---|---|---|---|---|---|---|---|---|---|---|
| SE | 44811 | ENSG00000177873 | ZNF619 | chr3 | + | 40481862 | 40482016 | 40477769 | 40478003 | 40486805 | 40488259 | 3 | 2 |
| SE | 44814 | ENSG00000177873 | ZNF619 | chr3 | + | 40482587 | 40482704 | 40477917 | 40478003 | 40486805 | 40488259 | 16 | 10 |
| SE | 44815 | ENSG00000177873 | ZNF619 | chr3 | + | 40482587 | 40482704 | 40481862 | 40482016 | 40486805 | 40488259 | 26 | 15 |
| SE | 44817 | ENSG00000177873 | ZNF619 | chr3 | + | 40483673 | 40483723 | 40482587 | 40482704 | 40486805 | 40488386 | 0 | 0 |
| SE | 44845 | ENSG00000143493 | INTS7 | chr1 | - | 211944783 | 211944969 | 211940398 | 211942111 | 211952568 | 211952701 | 40 | 18 |
| SE | 44846 | ENSG00000143493 | INTS7 | chr1 | - | 211946606 | 211946645 | 211944783 | 211944969 | 211952568 | 211952701 | 1 | 3 |
| SE | 44847 | ENSG00000143493 | INTS7 | chr1 | - | 211946606 | 211946705 | 211944783 | 211944969 | 211952568 | 211952701 | 37 | 15 |
| SE | 44848 | ENSG00000143493 | INTS7 | chr1 | - | 211952568 | 211952701 | 211946606 | 211946705 | 211966429 | 211966467 | 54 | 20 |
| SE | 44849 | ENSG00000143493 | INTS7 | chr1 | - | 211952610 | 211952701 | 211946606 | 211946705 | 211966429 | 211966498 | 15 | 4 |
| SE | 44850 | ENSG00000143493 | INTS7 | chr1 | - | 211975165 | 211975372 | 211968512 | 211968603 | 211978271 | 211978511 | 0 | 1 |
| SE | 44851 | ENSG00000143493 | INTS7 | chr1 | - | 211975165 | 211975372 | 211968512 | 211968707 | 211978271 | 211978511 | 59 | 40 |
| SE | 44852 | ENSG00000143493 | INTS7 | chr1 | - | 211976581 | 211976719 | 211975165 | 211975372 | 211978271 | 211978511 | 62 | 29 |
| SE | 44854 | ENSG00000143493 | INTS7 | chr1 | - | 212011374 | 212011421 | 212007249 | 212007449 | 212016885 | 212017023 | 94 | 99 |
| SE | 44855 | ENSG00000143493 | INTS7 | chr1 | - | 212016885 | 212017023 | 212011374 | 212011421 | 212020121 | 212020268 | 97 | 104 |
| SE | 44860 | ENSG00000167384 | ZNF180 | chr19 | - | 44484360 | 44484435 | 44479282 | 44479409 | 44497283 | 44497372 | 22 | 21 |
| SE | 44861 | ENSG00000167384 | ZNF180 | chr19 | - | 44485035 | 44485157 | 44484360 | 44484435 | 44497283 | 44497377 | 0 | 0 |
| SE | 44862 | ENSG00000167384 | ZNF180 | chr19 | - | 44497283 | 44497372 | 44484360 | 44484435 | 44500274 | 44500522 | 15 | 7 |
| SE | 44863 | ENSG00000167384 | ZNF180 | chr19 | - | 44497283 | 44497377 | 44484360 | 44484435 | 44500274 | 44500495 | 15 | 10 |
| SE | 44864 | ENSG00000257815 | PRANCR | chr12 | - | 70221699 | 70221901 | 70219130 | 70219597 | 70222158 | 70222377 | 4 | 2 |
| SE | 44865 | ENSG00000257815 | PRANCR | chr12 | - | 70221699 | 70221901 | 70219130 | 70219597 | 70225210 | 70225390 | 4 | 0 |
| SE | 44866 | ENSG00000257815 | PRANCR | chr12 | - | 70222158 | 70222377 | 70219130 | 70219597 | 70225210 | 70225390 | 0 | 0 |
| SE | 44867 | ENSG00000257815 | PRANCR | chr12 | - | 70222158 | 70222377 | 70221699 | 70221901 | 70225210 | 70225390 | 1 | 2 |
| SE | 44870 | ENSG00000257815 | PRANCR | chr12 | - | 70242180 | 70242229 | 70225210 | 70225390 | 70242796 | 70242893 | 0 | 0 |
| SE | 44871 | ENSG00000257815 | PRANCR | chr12 | - | 70243065 | 70243204 | 70242796 | 70242893 | 70243311 | 70243349 | 3 | 2 |
| SE | 44872 | ENSG00000134744 | TUT4 | chr1 | - | 52436754 | 52436963 | 52435364 | 52435465 | 52438219 | 52438335 | 119 | 61 |
| SE | 44873 | ENSG00000134744 | TUT4 | chr1 | - | 52436754 | 52436975 | 52435364 | 52435465 | 52438219 | 52438335 | 93 | 38 |
| SE | 44874 | ENSG00000134744 | TUT4 | chr1 | - | 52436754 | 52436978 | 52435364 | 52435465 | 52438219 | 52438335 | 133 | 60 |
| SE | 44875 | ENSG00000134744 | TUT4 | chr1 | - | 52438219 | 52438335 | 52435364 | 52435465 | 52445786 | 52445869 | 125 | 105 |
| SE | 44877 | ENSG00000134744 | TUT4 | chr1 | - | 52465069 | 52465173 | 52446589 | 52446667 | 52468180 | 52468267 | 133 | 94 |
| SE | 44880 | ENSG00000134744 | TUT4 | chr1 | - | 52465069 | 52465173 | 52461711 | 52461769 | 52471951 | 52472102 | 83 | 58 |
| SE | 44881 | ENSG00000134744 | TUT4 | chr1 | - | 52468180 | 52468267 | 52446589 | 52446667 | 52471951 | 52472102 | 96 | 103 |
| SE | 44882 | ENSG00000134744 | TUT4 | chr1 | - | 52468180 | 52468267 | 52461711 | 52461769 | 52471951 | 52472102 | 97 | 103 |
| SE | 44883 | ENSG00000134744 | TUT4 | chr1 | - | 52468180 | 52468267 | 52465069 | 52465173 | 52471951 | 52472102 | 269 | 234 |
| SE | 44885 | ENSG00000134744 | TUT4 | chr1 | - | 52477707 | 52477882 | 52474831 | 52475535 | 52493610 | 52493662 | 169 | 154 |
| SE | 44889 | ENSG00000134744 | TUT4 | chr1 | - | 52481803 | 52481923 | 52477707 | 52477882 | 52490731 | 52490801 | 27 | 21 |
| SE | 44890 | ENSG00000134744 | TUT4 | chr1 | - | 52481803 | 52481923 | 52481422 | 52481635 | 52488908 | 52489035 | 137 | 93 |
| SE | 44892 | ENSG00000134744 | TUT4 | chr1 | - | 52488908 | 52489035 | 52481803 | 52481923 | 52490731 | 52490801 | 140 | 121 |
| SE | 44893 | ENSG00000134744 | TUT4 | chr1 | - | 52490731 | 52490801 | 52477707 | 52477882 | 52493610 | 52493662 | 79 | 53 |
| SE | 44894 | ENSG00000134744 | TUT4 | chr1 | - | 52493610 | 52493662 | 52490731 | 52490801 | 52495426 | 52495515 | 171 | 131 |
| SE | 44895 | ENSG00000134744 | TUT4 | chr1 | - | 52495426 | 52495515 | 52493610 | 52493662 | 52509595 | 52509712 | 91 | 61 |
| SE | 44896 | ENSG00000134744 | TUT4 | chr1 | - | 52497005 | 52497183 | 52495426 | 52495515 | 52509595 | 52509712 | 290 | 208 |
| SE | 44897 | ENSG00000134744 | TUT4 | chr1 | - | 52509595 | 52509712 | 52497005 | 52497183 | 52515890 | 52516054 | 580 | 555 |
| SE | 44898 | ENSG00000134744 | TUT4 | chr1 | - | 52509595 | 52509712 | 52497005 | 52497183 | 52552930 | 52553069 | 70 | 44 |
| SE | 44899 | ENSG00000134744 | TUT4 | chr1 | - | 52515890 | 52516054 | 52497005 | 52497183 | 52552930 | 52553090 | 14 | 3 |
| SE | 44900 | ENSG00000134744 | TUT4 | chr1 | - | 52515890 | 52516054 | 52509595 | 52509712 | 52552930 | 52553090 | 524 | 514 |
| SE | 44902 | ENSG00000134744 | TUT4 | chr1 | - | 52529767 | 52529885 | 52526019 | 52526373 | 52534986 | 52535122 | 4 | 3 |
| SE | 44903 | ENSG00000134744 | TUT4 | chr1 | - | 52534986 | 52535122 | 52515890 | 52516054 | 52552930 | 52553085 | 0 | 0 |
| SE | 44905 | ENSG00000134744 | TUT4 | chr1 | - | 52534986 | 52535122 | 52526019 | 52526373 | 52552930 | 52553085 | 0 | 0 |
| SE | 44906 | ENSG00000134744 | TUT4 | chr1 | - | 52538523 | 52538643 | 52526019 | 52526373 | 52552930 | 52553085 | 0 | 0 |
| SE | 44908 | ENSG00000162889 | MAPKAPK2 | chr1 | + | 206729971 | 206730098 | 206729395 | 206729475 | 206730687 | 206730763 | 604 | 286 |
| SE | 44909 | ENSG00000162889 | MAPKAPK2 | chr1 | + | 206729971 | 206730098 | 206729395 | 206729475 | 206731137 | 206731262 | 232 | 111 |
| SE | 44910 | ENSG00000162889 | MAPKAPK2 | chr1 | + | 206730687 | 206730763 | 206729395 | 206729475 | 206731137 | 206731262 | 102 | 57 |
| SE | 44912 | ENSG00000162889 | MAPKAPK2 | chr1 | + | 206731639 | 206731725 | 206731137 | 206731262 | 206731838 | 206731919 | 582 | 352 |
| SE | 44913 | ENSG00000162889 | MAPKAPK2 | chr1 | + | 206731838 | 206731919 | 206731639 | 206731725 | 206732574 | 206734281 | 571 | 333 |
| SE | 44914 | ENSG00000245146 | MALINC1 | chr5 | - | 140106295 | 140106461 | 140105586 | 140105762 | 140108472 | 140108594 | 4 | 2 |
| SE | 44915 | ENSG00000245146 | MALINC1 | chr5 | - | 140106295 | 140106490 | 140105586 | 140105762 | 140108472 | 140108603 | 3 | 3 |
| SE | 44916 | ENSG00000162739 | SLAMF6 | chr1 | - | 160490574 | 160490685 | 160490197 | 160490236 | 160491124 | 160491388 | 749 | 310 |
| SE | 44917 | ENSG00000162739 | SLAMF6 | chr1 | - | 160491124 | 160491388 | 160490574 | 160490685 | 160496060 | 160496393 | 795 | 287 |
| SE | 44920 | ENSG00000249896 | LINC02495 | chr4 | + | 6228016 | 6228139 | 6226925 | 6227003 | 6232859 | 6233936 | 1 | 0 |
| SE | 44921 | ENSG00000112312 | GMNN | chr6 | + | 24784443 | 24784554 | 24784086 | 24784169 | 24785637 | 24785707 | 15 | 22 |
| SE | 44924 | ENSG00000182095 | TNRC18 | chr7 | - | 5359397 | 5359569 | 5356915 | 5357276 | 5361593 | 5361722 | 375 | 263 |
| SE | 44925 | ENSG00000182095 | TNRC18 | chr7 | - | 5359397 | 5359569 | 5356915 | 5357276 | 5361896 | 5362027 | 238 | 175 |
| SE | 44926 | ENSG00000182095 | TNRC18 | chr7 | - | 5361593 | 5361722 | 5356915 | 5357276 | 5361896 | 5362033 | 137 | 80 |
| SE | 44929 | ENSG00000182095 | TNRC18 | chr7 | - | 5376846 | 5376993 | 5376033 | 5376224 | 5377370 | 5377576 | 86 | 9 |
| SE | 44930 | ENSG00000182095 | TNRC18 | chr7 | - | 5390484 | 5390628 | 5387671 | 5389336 | 5394439 | 5394595 | 28 | 7 |
| SE | 44937 | ENSG00000138138 | ATAD1 | chr10 | - | 87770951 | 87771041 | 87767672 | 87767723 | 87776320 | 87776427 | 311 | 233 |
| SE | 44938 | ENSG00000138138 | ATAD1 | chr10 | - | 87776320 | 87776427 | 87770951 | 87771041 | 87784469 | 87784670 | 275 | 175 |
| SE | 44940 | ENSG00000138138 | ATAD1 | chr10 | - | 87784469 | 87784670 | 87776320 | 87776427 | 87790309 | 87790430 | 371 | 212 |
| SE | 44947 | ENSG00000101096 | NFATC2 | chr20 | - | 51435187 | 51435314 | 51386956 | 51391451 | 51435705 | 51435761 | 108 | 57 |
| SE | 44948 | ENSG00000101096 | NFATC2 | chr20 | - | 51454547 | 51454688 | 51435705 | 51435761 | 51473979 | 51474152 | 483 | 213 |
| SE | 44949 | ENSG00000101096 | NFATC2 | chr20 | - | 51473979 | 51474152 | 51454547 | 51454688 | 51475457 | 51475660 | 680 | 288 |
| SE | 44950 | ENSG00000117682 | DHDDS | chr1 | + | 26442707 | 26442873 | 26438167 | 26438284 | 26446315 | 26446382 | 58 | 47 |
| SE | 44951 | ENSG00000117682 | DHDDS | chr1 | + | 26442730 | 26442873 | 26438167 | 26438284 | 26446315 | 26446359 | 121 | 85 |
| SE | 44952 | ENSG00000117682 | DHDDS | chr1 | + | 26442730 | 26442873 | 26438167 | 26438284 | 26447558 | 26447660 | 43 | 32 |
| SE | 44953 | ENSG00000117682 | DHDDS | chr1 | + | 26446315 | 26446432 | 26438167 | 26438284 | 26447558 | 26447615 | 36 | 23 |
| SE | 44954 | ENSG00000117682 | DHDDS | chr1 | + | 26446315 | 26446432 | 26442707 | 26442873 | 26447558 | 26447615 | 115 | 75 |
| SE | 44955 | ENSG00000117682 | DHDDS | chr1 | + | 26446315 | 26446432 | 26442707 | 26442873 | 26457790 | 26457905 | 81 | 51 |
| SE | 44956 | ENSG00000117682 | DHDDS | chr1 | + | 26447558 | 26447660 | 26442730 | 26442873 | 26457790 | 26457905 | 13 | 5 |
| SE | 44957 | ENSG00000117682 | DHDDS | chr1 | + | 26447558 | 26447660 | 26442730 | 26442873 | 26460036 | 26460144 | 0 | 0 |
| SE | 44959 | ENSG00000117682 | DHDDS | chr1 | + | 26457790 | 26457905 | 26442707 | 26442873 | 26460036 | 26460144 | 15 | 4 |
| SE | 44960 | ENSG00000117682 | DHDDS | chr1 | + | 26457790 | 26457905 | 26447558 | 26447660 | 26460036 | 26460144 | 38 | 8 |
| SE | 44964 | ENSG00000132680 | KHDC4 | chr1 | - | 155921860 | 155921918 | 155921384 | 155921527 | 155923626 | 155923687 | 50 | 49 |
| SE | 44965 | ENSG00000132680 | KHDC4 | chr1 | - | 155925631 | 155925843 | 155923626 | 155923687 | 155929295 | 155929375 | 84 | 54 |
| SE | 44967 | ENSG00000132680 | KHDC4 | chr1 | - | 155926675 | 155926839 | 155925631 | 155925843 | 155929295 | 155929375 | 105 | 68 |
| SE | 44968 | ENSG00000132680 | KHDC4 | chr1 | - | 155927103 | 155927156 | 155925631 | 155925843 | 155929295 | 155929375 | 27 | 25 |
| SE | 44969 | ENSG00000132680 | KHDC4 | chr1 | - | 155927103 | 155927156 | 155926470 | 155926839 | 155929295 | 155929375 | 93 | 66 |
| SE | 44970 | ENSG00000132680 | KHDC4 | chr1 | - | 155929711 | 155929840 | 155929295 | 155929375 | 155933632 | 155933849 | 134 | 106 |
| SE | 44975 | ENSG00000138798 | EGF | chr4 | + | 109940945 | 109941145 | 109912882 | 109913462 | 109943841 | 109944069 | 18 | 21 |
| SE | 44976 | ENSG00000138798 | EGF | chr4 | + | 109943253 | 109943435 | 109940775 | 109941145 | 109943841 | 109943972 | 39 | 56 |
| SE | 44978 | ENSG00000138798 | EGF | chr4 | + | 109960866 | 109960989 | 109945072 | 109945275 | 109961862 | 109961985 | 8 | 19 |
| SE | 44979 | ENSG00000138798 | EGF | chr4 | + | 109968970 | 109969119 | 109964400 | 109964537 | 109974702 | 109974807 | 54 | 45 |
| SE | 44981 | ENSG00000138798 | EGF | chr4 | + | 109994732 | 109994880 | 109993246 | 109993369 | 109999678 | 109999846 | 20 | 18 |
| SE | 44982 | ENSG00000138798 | EGF | chr4 | + | 110004504 | 110004622 | 109999678 | 109999846 | 110011201 | 110011689 | 4 | 1 |
| SE | 44985 | ENSG00000244560 | NA | chr7 | + | 149289547 | 149289726 | 149288430 | 149289457 | 149289937 | 149290038 | 8 | 4 |
| SE | 44987 | ENSG00000244560 | NA | chr7 | + | 149291555 | 149291745 | 149289937 | 149290038 | 149292210 | 149292308 | 1 | 0 |
| SE | 44989 | ENSG00000260404 | NA | chr4 | + | 118597093 | 118597225 | 118595911 | 118596022 | 118605583 | 118605819 | 28 | 18 |
| SE | 44990 | ENSG00000260404 | NA | chr4 | + | 118605583 | 118605819 | 118597093 | 118597225 | 118608657 | 118608722 | 19 | 23 |
| SE | 44991 | ENSG00000275131 | PDE4DIPP2 | chr1 | - | 120491581 | 120491628 | 120489859 | 120489916 | 120493157 | 120493301 | 3 | 8 |
| SE | 44992 | ENSG00000275131 | PDE4DIPP2 | chr1 | - | 120491934 | 120492073 | 120491581 | 120491628 | 120493157 | 120493301 | 10 | 11 |
| SE | 44993 | ENSG00000275131 | PDE4DIPP2 | chr1 | - | 120493157 | 120493301 | 120489859 | 120489916 | 120494233 | 120494460 | 18 | 11 |
| SE | 44994 | ENSG00000275131 | PDE4DIPP2 | chr1 | - | 120494233 | 120494460 | 120493157 | 120493301 | 120495029 | 120495146 | 22 | 19 |
| SE | 44995 | ENSG00000275131 | PDE4DIPP2 | chr1 | - | 120495029 | 120495146 | 120494233 | 120494460 | 120497181 | 120497421 | 6 | 9 |
| SE | 44996 | ENSG00000275131 | PDE4DIPP2 | chr1 | - | 120495029 | 120495146 | 120494233 | 120494460 | 120501109 | 120501234 | 4 | 9 |
| SE | 44997 | ENSG00000275131 | PDE4DIPP2 | chr1 | - | 120497181 | 120497421 | 120494233 | 120494460 | 120501109 | 120501234 | 2 | 0 |
| SE | 44998 | ENSG00000275131 | PDE4DIPP2 | chr1 | - | 120497181 | 120497421 | 120495029 | 120495146 | 120501109 | 120501234 | 2 | 0 |
| SE | 44999 | ENSG00000275131 | PDE4DIPP2 | chr1 | - | 120505710 | 120505962 | 120504381 | 120504513 | 120511665 | 120511818 | 16 | 11 |
| SE | 45000 | ENSG00000275131 | PDE4DIPP2 | chr1 | - | 120505710 | 120505962 | 120504381 | 120504513 | 120512469 | 120512693 | 10 | 4 |
| SE | 45001 | ENSG00000275131 | PDE4DIPP2 | chr1 | - | 120509484 | 120509670 | 120505710 | 120505962 | 120511665 | 120511818 | 0 | 0 |
| SE | 45002 | ENSG00000275131 | PDE4DIPP2 | chr1 | - | 120511665 | 120511818 | 120504381 | 120504513 | 120512469 | 120512693 | 25 | 35 |
| SE | 45003 | ENSG00000275131 | PDE4DIPP2 | chr1 | - | 120511665 | 120511818 | 120505710 | 120505962 | 120512469 | 120512693 | 31 | 42 |
| SE | 45004 | ENSG00000275131 | PDE4DIPP2 | chr1 | - | 120513765 | 120513817 | 120512469 | 120512693 | 120514824 | 120515060 | 19 | 20 |
| SE | 45005 | ENSG00000275131 | PDE4DIPP2 | chr1 | - | 120520233 | 120520669 | 120519467 | 120519821 | 120523893 | 120524126 | 38 | 46 |
| SE | 45008 | ENSG00000275131 | PDE4DIPP2 | chr1 | - | 120531877 | 120532089 | 120528386 | 120530384 | 120536310 | 120536339 | 95 | 123 |
| SE | 45009 | ENSG00000275131 | PDE4DIPP2 | chr1 | - | 120536310 | 120536402 | 120531877 | 120532089 | 120538322 | 120538404 | 35 | 46 |
| SE | 45012 | ENSG00000162623 | TYW3 | chr1 | + | 74748750 | 74748822 | 74738689 | 74738788 | 74752291 | 74752394 | 195 | 123 |
| SE | 45013 | ENSG00000162623 | TYW3 | chr1 | + | 74752291 | 74752425 | 74738689 | 74738788 | 74763893 | 74764001 | 13 | 13 |
| SE | 45017 | ENSG00000158019 | BABAM2 | chr2 | + | 28237201 | 28237301 | 28129270 | 28129380 | 28241322 | 28241393 | 156 | 133 |
| SE | 45019 | ENSG00000158019 | BABAM2 | chr2 | + | 28241322 | 28241393 | 28129270 | 28129380 | 28298337 | 28298491 | 2 | 1 |
| SE | 45020 | ENSG00000158019 | BABAM2 | chr2 | + | 28241322 | 28241393 | 28237201 | 28237301 | 28298337 | 28298491 | 21 | 23 |
| SE | 45021 | ENSG00000158019 | BABAM2 | chr2 | + | 28244779 | 28244862 | 28241322 | 28241393 | 28298337 | 28298491 | 253 | 171 |
| SE | 45029 | ENSG00000184500 | PROS1 | chr3 | - | 93928701 | 93928797 | 93927249 | 93927407 | 93973673 | 93973933 | 1 | 0 |
| SE | 45033 | ENSG00000134369 | NAV1 | chr1 | + | 201790553 | 201790577 | 201789739 | 201789792 | 201790688 | 201790766 | 4 | 0 |
| SE | 45036 | ENSG00000134369 | NAV1 | chr1 | + | 201810522 | 201810758 | 201809945 | 201810105 | 201811602 | 201811757 | 4 | 5 |
| SE | 45037 | ENSG00000134369 | NAV1 | chr1 | + | 201812464 | 201812661 | 201811902 | 201811974 | 201813139 | 201813258 | 8 | 9 |
| SE | 45038 | ENSG00000007341 | ST7L | chr1 | - | 112539158 | 112539275 | 112525875 | 112526111 | 112540778 | 112540872 | 1 | 2 |
| SE | 45039 | ENSG00000007341 | ST7L | chr1 | - | 112540778 | 112540872 | 112523517 | 112526111 | 112541950 | 112541975 | 3 | 5 |
| SE | 45040 | ENSG00000007341 | ST7L | chr1 | - | 112541950 | 112542090 | 112523513 | 112526111 | 112550600 | 112550693 | 16 | 1 |
| SE | 45043 | ENSG00000007341 | ST7L | chr1 | - | 112550600 | 112550693 | 112541789 | 112542090 | 112555867 | 112556018 | 13 | 1 |
| SE | 45044 | ENSG00000007341 | ST7L | chr1 | - | 112550600 | 112550693 | 112541789 | 112542090 | 112576985 | 112577088 | 3 | 1 |
| SE | 45045 | ENSG00000007341 | ST7L | chr1 | - | 112555867 | 112556018 | 112541950 | 112542090 | 112576985 | 112577088 | 9 | 8 |
| SE | 45046 | ENSG00000007341 | ST7L | chr1 | - | 112555867 | 112556018 | 112550600 | 112550693 | 112576985 | 112577088 | 20 | 4 |
| SE | 45047 | ENSG00000007341 | ST7L | chr1 | - | 112571274 | 112571351 | 112555867 | 112556018 | 112576985 | 112577088 | 0 | 0 |
| SE | 45048 | ENSG00000007341 | ST7L | chr1 | - | 112576985 | 112577088 | 112555867 | 112556018 | 112578344 | 112578417 | 18 | 15 |
| SE | 45049 | ENSG00000007341 | ST7L | chr1 | - | 112576985 | 112577088 | 112555867 | 112556018 | 112581991 | 112582106 | 7 | 1 |
| SE | 45050 | ENSG00000007341 | ST7L | chr1 | - | 112578344 | 112578417 | 112555867 | 112556018 | 112581991 | 112582106 | 7 | 8 |
| SE | 45052 | ENSG00000007341 | ST7L | chr1 | - | 112578344 | 112578417 | 112555867 | 112556018 | 112597970 | 112598086 | 0 | 0 |
| SE | 45053 | ENSG00000007341 | ST7L | chr1 | - | 112578344 | 112578417 | 112576985 | 112577088 | 112581991 | 112582106 | 17 | 25 |
| SE | 45055 | ENSG00000007341 | ST7L | chr1 | - | 112581991 | 112582106 | 112578344 | 112578417 | 112582374 | 112582472 | 40 | 38 |
| SE | 45057 | ENSG00000007341 | ST7L | chr1 | - | 112582374 | 112582472 | 112578344 | 112578417 | 112591524 | 112591603 | 11 | 8 |
| SE | 45058 | ENSG00000007341 | ST7L | chr1 | - | 112582374 | 112582472 | 112578344 | 112578417 | 112597970 | 112598086 | 1 | 1 |
| SE | 45059 | ENSG00000007341 | ST7L | chr1 | - | 112582374 | 112582472 | 112581991 | 112582106 | 112583971 | 112584126 | 68 | 47 |
| SE | 45061 | ENSG00000007341 | ST7L | chr1 | - | 112583971 | 112584126 | 112582374 | 112582472 | 112597970 | 112598086 | 51 | 27 |
| SE | 45062 | ENSG00000007341 | ST7L | chr1 | - | 112591524 | 112591603 | 112578344 | 112578417 | 112597970 | 112598086 | 31 | 16 |
| SE | 45063 | ENSG00000007341 | ST7L | chr1 | - | 112591524 | 112591603 | 112582374 | 112582472 | 112597970 | 112598086 | 41 | 22 |
| SE | 45065 | ENSG00000007341 | ST7L | chr1 | - | 112591524 | 112591603 | 112583971 | 112584126 | 112600793 | 112600848 | 35 | 17 |
| SE | 45066 | ENSG00000007341 | ST7L | chr1 | - | 112597970 | 112598086 | 112583971 | 112584126 | 112600793 | 112600848 | 13 | 19 |
| SE | 45067 | ENSG00000007341 | ST7L | chr1 | - | 112597970 | 112598086 | 112583971 | 112584126 | 112610840 | 112611003 | 6 | 1 |
| SE | 45068 | ENSG00000007341 | ST7L | chr1 | - | 112597970 | 112598086 | 112591481 | 112591603 | 112600793 | 112600848 | 60 | 51 |
| SE | 45069 | ENSG00000007341 | ST7L | chr1 | - | 112600793 | 112600848 | 112583974 | 112584126 | 112610840 | 112610906 | 18 | 16 |
| SE | 45071 | ENSG00000007341 | ST7L | chr1 | - | 112616812 | 112616844 | 112610840 | 112611003 | 112618246 | 112618369 | 13 | 9 |
| SE | 45072 | ENSG00000007341 | ST7L | chr1 | - | 112616812 | 112616844 | 112610840 | 112611003 | 112618908 | 112619189 | 13 | 11 |
| SE | 45073 | ENSG00000007341 | ST7L | chr1 | - | 112616812 | 112616895 | 112610458 | 112611003 | 112618246 | 112618369 | 29 | 24 |
| SE | 45074 | ENSG00000007341 | ST7L | chr1 | - | 112616812 | 112616895 | 112610458 | 112611003 | 112618908 | 112618941 | 44 | 26 |
| SE | 45076 | ENSG00000007341 | ST7L | chr1 | - | 112618246 | 112618369 | 112610840 | 112611003 | 112618908 | 112619189 | 0 | 0 |
| SE | 45077 | ENSG00000007341 | ST7L | chr1 | - | 112618246 | 112618369 | 112616812 | 112616844 | 112618908 | 112619189 | 3 | 0 |
| SE | 45078 | ENSG00000007341 | ST7L | chr1 | - | 112618246 | 112618369 | 112616812 | 112616844 | 112619196 | 112619664 | 4 | 0 |
| SE | 45079 | ENSG00000007341 | ST7L | chr1 | - | 112618246 | 112618369 | 112616812 | 112616895 | 112618908 | 112619189 | 1 | 3 |
| SE | 45082 | ENSG00000007341 | ST7L | chr1 | - | 112618908 | 112619139 | 112616812 | 112616895 | 112619909 | 112620825 | 16 | 5 |
| SE | 45085 | ENSG00000100181 | TPTEP1 | chr22 | + | 16614076 | 16614178 | 16601910 | 16602215 | 16638578 | 16638740 | 1 | 6 |
| SE | 45086 | ENSG00000100181 | TPTEP1 | chr22 | + | 16622840 | 16622897 | 16602043 | 16602215 | 16637039 | 16637090 | 2 | 1 |
| SE | 45088 | ENSG00000100181 | TPTEP1 | chr22 | + | 16637039 | 16637090 | 16601910 | 16602215 | 16638578 | 16638740 | 6 | 10 |
| SE | 45092 | ENSG00000100181 | TPTEP1 | chr22 | + | 16647604 | 16647785 | 16638578 | 16638740 | 16648526 | 16648648 | 2 | 3 |
| SE | 45093 | ENSG00000100181 | TPTEP1 | chr22 | + | 16647662 | 16647785 | 16638578 | 16638740 | 16648526 | 16648830 | 8 | 17 |
| SE | 45094 | ENSG00000100181 | TPTEP1 | chr22 | + | 16647662 | 16647785 | 16647167 | 16647257 | 16648526 | 16648830 | 2 | 4 |
| SE | 45099 | ENSG00000158887 | MPZ | chr1 | - | 161306707 | 161306921 | 161306328 | 161306464 | 161307257 | 161307424 | 3 | 4 |
| SE | 45100 | ENSG00000158887 | MPZ | chr1 | - | 161306788 | 161306921 | 161306328 | 161306464 | 161307257 | 161307424 | 1 | 0 |
| SE | 45102 | ENSG00000227373 | RABGAP1L-DT | chr1 | - | 174158886 | 174158990 | 174155626 | 174155697 | 174159916 | 174160150 | 0 | 0 |
| SE | 45104 | ENSG00000088035 | ALG6 | chr1 | + | 63396512 | 63396597 | 63370770 | 63371059 | 63402253 | 63402273 | 149 | 46 |
| SE | 45105 | ENSG00000088035 | ALG6 | chr1 | + | 63411145 | 63411155 | 63407061 | 63407126 | 63411681 | 63411806 | 5 | 3 |
| SE | 45106 | ENSG00000088035 | ALG6 | chr1 | + | 63411145 | 63411331 | 63407061 | 63407126 | 63411681 | 63411806 | 32 | 19 |
| SE | 45107 | ENSG00000088035 | ALG6 | chr1 | + | 63411681 | 63411806 | 63411145 | 63411155 | 63411925 | 63412061 | 0 | 5 |
| SE | 45109 | ENSG00000088035 | ALG6 | chr1 | + | 63411925 | 63412061 | 63411145 | 63411155 | 63414060 | 63414146 | 56 | 19 |
| SE | 45110 | ENSG00000088035 | ALG6 | chr1 | + | 63428732 | 63428801 | 63419369 | 63419440 | 63428927 | 63429075 | 51 | 32 |
| SE | 45113 | ENSG00000133731 | IMPA1 | chr8 | - | 81673840 | 81673949 | 81660515 | 81660667 | 81679125 | 81679230 | 3 | 1 |
| SE | 45114 | ENSG00000133731 | IMPA1 | chr8 | - | 81674671 | 81674831 | 81673895 | 81673949 | 81676233 | 81676279 | 2 | 1 |
| SE | 45115 | ENSG00000133731 | IMPA1 | chr8 | - | 81676233 | 81676265 | 81673840 | 81673949 | 81679125 | 81679230 | 6 | 4 |
| SE | 45116 | ENSG00000133731 | IMPA1 | chr8 | - | 81676233 | 81676279 | 81673840 | 81673949 | 81679125 | 81679230 | 39 | 15 |
| SE | 45117 | ENSG00000133731 | IMPA1 | chr8 | - | 81678582 | 81678689 | 81676233 | 81676265 | 81679125 | 81679230 | 0 | 1 |
| SE | 45119 | ENSG00000133731 | IMPA1 | chr8 | - | 81685762 | 81685963 | 81681497 | 81681584 | 81686251 | 81686283 | 2 | 3 |
| SE | 45121 | ENSG00000119203 | CPSF3 | chr2 | + | 9432510 | 9432688 | 9430751 | 9430880 | 9433870 | 9433960 | 211 | 188 |
| SE | 45122 | ENSG00000119203 | CPSF3 | chr2 | + | 9436210 | 9436361 | 9433870 | 9433960 | 9440490 | 9440666 | 155 | 161 |
| SE | 45123 | ENSG00000119203 | CPSF3 | chr2 | + | 9452912 | 9453021 | 9448197 | 9448350 | 9455658 | 9455757 | 100 | 101 |
| SE | 45124 | ENSG00000197928 | ZNF677 | chr19 | - | 53251535 | 53251605 | 53243332 | 53243897 | 53253085 | 53253156 | 34 | 37 |
| SE | 45125 | ENSG00000230124 | ACBD6 | chr1 | - | 180281305 | 180281386 | 180274650 | 180275412 | 180314691 | 180314722 | 2 | 3 |
| SE | 45126 | ENSG00000230124 | ACBD6 | chr1 | - | 180288269 | 180288517 | 180281305 | 180281386 | 180314691 | 180314722 | 64 | 52 |
| SE | 45128 | ENSG00000230124 | ACBD6 | chr1 | - | 180326410 | 180326562 | 180314691 | 180314722 | 180397515 | 180397605 | 0 | 0 |
| SE | 45129 | ENSG00000230124 | ACBD6 | chr1 | - | 180397515 | 180397605 | 180314691 | 180314722 | 180413365 | 180413388 | 128 | 113 |
| SE | 45130 | ENSG00000230124 | ACBD6 | chr1 | - | 180413365 | 180413471 | 180397515 | 180397605 | 180430179 | 180430262 | 123 | 111 |
| SE | 45131 | ENSG00000230124 | ACBD6 | chr1 | - | 180492268 | 180492365 | 180430179 | 180430262 | 180495460 | 180495525 | 137 | 104 |
| SE | 45132 | ENSG00000230124 | ACBD6 | chr1 | - | 180495460 | 180495525 | 180492268 | 180492365 | 180502044 | 180502577 | 166 | 97 |
| SE | 45142 | ENSG00000107099 | DOCK8 | chr9 | + | 271626 | 271729 | 215170 | 215416 | 286460 | 286636 | 69 | 82 |
| SE | 45143 | ENSG00000107099 | DOCK8 | chr9 | + | 276993 | 277086 | 271626 | 271729 | 280790 | 280905 | 3 | 0 |
| SE | 45145 | ENSG00000107099 | DOCK8 | chr9 | + | 276993 | 277086 | 273083 | 273100 | 280790 | 280905 | 0 | 0 |
| SE | 45151 | ENSG00000107099 | DOCK8 | chr9 | + | 284374 | 284567 | 273083 | 273100 | 286460 | 286636 | 0 | 0 |
| SE | 45153 | ENSG00000107099 | DOCK8 | chr9 | + | 304580 | 304704 | 289509 | 289581 | 311953 | 312166 | 1552 | 1283 |
| SE | 45154 | ENSG00000107099 | DOCK8 | chr9 | + | 311953 | 312166 | 289509 | 289581 | 317042 | 317128 | 453 | 300 |
| SE | 45156 | ENSG00000107099 | DOCK8 | chr9 | + | 325670 | 325737 | 317042 | 317128 | 328021 | 328171 | 837 | 548 |
| SE | 45158 | ENSG00000107099 | DOCK8 | chr9 | + | 334224 | 334384 | 332397 | 332478 | 336581 | 336718 | 855 | 691 |
| SE | 45160 | ENSG00000107099 | DOCK8 | chr9 | + | 339005 | 339099 | 336581 | 336718 | 340158 | 340321 | 1877 | 1420 |
| SE | 45161 | ENSG00000107099 | DOCK8 | chr9 | + | 365590 | 366552 | 340158 | 340321 | 368017 | 368135 | 3 | 3 |
| SE | 45162 | ENSG00000107099 | DOCK8 | chr9 | + | 369221 | 369447 | 368017 | 368135 | 370229 | 370300 | 3 | 0 |
| SE | 45163 | ENSG00000107099 | DOCK8 | chr9 | + | 371427 | 371566 | 368017 | 370300 | 372184 | 372286 | 1344 | 937 |
| SE | 45164 | ENSG00000107099 | DOCK8 | chr9 | + | 372184 | 372286 | 369663 | 371566 | 376209 | 376305 | 882 | 552 |
| SE | 45165 | ENSG00000107099 | DOCK8 | chr9 | + | 376976 | 377211 | 376209 | 376305 | 382512 | 382685 | 202 | 97 |
| SE | 45168 | ENSG00000107099 | DOCK8 | chr9 | + | 404917 | 405073 | 399145 | 399259 | 406929 | 407069 | 977 | 658 |
| SE | 45169 | ENSG00000107099 | DOCK8 | chr9 | + | 406929 | 407069 | 399145 | 399259 | 414781 | 414951 | 298 | 154 |
| SE | 45170 | ENSG00000107099 | DOCK8 | chr9 | + | 414781 | 414951 | 406929 | 407069 | 418067 | 418207 | 653 | 359 |
| SE | 45171 | ENSG00000107099 | DOCK8 | chr9 | + | 420400 | 420583 | 418067 | 418207 | 420948 | 421078 | 1729 | 1568 |
| SE | 45173 | ENSG00000107099 | DOCK8 | chr9 | + | 426884 | 426981 | 422047 | 422135 | 429701 | 429854 | 224 | 162 |

| | | | | | | | | | | | | |
|---|---|---|---|---|---|---|---|---|---|---|---|---|
| SE | 45174 | ENSG00000107099 | DOCK8 | chr9 | + | 426884 | 426981 | 422047 | 422135 | 432165 | 432324 | [illegible] |
| SE | 45175 | ENSG00000107099 | DOCK8 | chr9 | + | 428361 | 428496 | 426884 | 426981 | 429701 | 429854 | [illegible] |
| SE | 45176 | ENSG00000107099 | DOCK8 | chr9 | + | 428361 | 428496 | 426884 | 426981 | 432165 | 432324 | [illegible] |
| SE | 45177 | ENSG00000107099 | DOCK8 | chr9 | + | 429701 | 429854 | 422047 | 422135 | 432165 | 432324 | [illegible] |
| SE | 45178 | ENSG00000107099 | DOCK8 | chr9 | + | 429701 | 429854 | 426884 | 426981 | 432165 | 432324 | [illegible] |
| SE | 45179 | ENSG00000107099 | DOCK8 | chr9 | + | 429701 | 429854 | 428361 | 428496 | 432165 | 432324 | [illegible] |
| SE | 45182 | ENSG00000107099 | DOCK8 | chr9 | + | 439244 | 439388 | 434782 | 434975 | 441285 | 441417 | [illegible] |
| SE | 45184 | ENSG00000107099 | DOCK8 | chr9 | + | 443426 | 443516 | 441874 | 442009 | 446369 | 446606 | [illegible] |
| SE | 45186 | ENSG00000141384 | TAF4B | chr18 | + | 26295194 | 26295314 | 26293425 | 26293531 | 26315228 | 26315398 | [illegible] |
| SE | 45188 | ENSG00000141384 | TAF4B | chr18 | + | 26321069 | 26321200 | 26293425 | 26293531 | 26327014 | 26327140 | [illegible] |
| SE | 45189 | ENSG00000141384 | TAF4B | chr18 | + | 26327014 | 26327140 | 26321069 | 26321200 | 26357689 | 26357794 | [illegible] |
| SE | 45195 | ENSG00000170584 | NUDCD2 | chr5 | - | 163456928 | 163457005 | 163453922 | 163454050 | 163457561 | 163457610 | [illegible] |
| SE | 45196 | ENSG00000170584 | NUDCD2 | chr5 | - | 163456928 | 163457080 | 163446525 | 163454050 | 163457561 | 163457610 | [illegible] |
| SE | 45197 | ENSG00000170584 | NUDCD2 | chr5 | - | 163457561 | 163457610 | 163456928 | 163457080 | 163459861 | 163460055 | [illegible] |
| SE | 45199 | ENSG00000121774 | KHDRBS1 | chr1 | + | 32036909 | 32037043 | 32033187 | 32033334 | 32037834 | 32038036 | [illegible] |
| SE | 45200 | ENSG00000121774 | KHDRBS1 | chr1 | + | 32036909 | 32037043 | 32033187 | 32033334 | 32038551 | 32038619 | [illegible] |
| SE | 45201 | ENSG00000121774 | KHDRBS1 | chr1 | + | 32036909 | 32037043 | 32033187 | 32033334 | 32042526 | 32042624 | [illegible] |
| SE | 45202 | ENSG00000121774 | KHDRBS1 | chr1 | + | 32037834 | 32038036 | 32033187 | 32033334 | 32038551 | 32038619 | [illegible] |
| SE | 45203 | ENSG00000121774 | KHDRBS1 | chr1 | + | 32037834 | 32038036 | 32036909 | 32037043 | 32038551 | 32038619 | [illegible] |
| SE | 45204 | ENSG00000121774 | KHDRBS1 | chr1 | + | 32037834 | 32038119 | 32033187 | 32033334 | 32038551 | 32038619 | [illegible] |
| SE | 45205 | ENSG00000121774 | KHDRBS1 | chr1 | + | 32037834 | 32038119 | 32036909 | 32037043 | 32038551 | 32038619 | [illegible] |
| SE | 45206 | ENSG00000121774 | KHDRBS1 | chr1 | + | 32038551 | 32038619 | 32033187 | 32033334 | 32039514 | 32039573 | [illegible] |
| SE | 45208 | ENSG00000121774 | KHDRBS1 | chr1 | + | 32038551 | 32038619 | 32036909 | 32037043 | 32042526 | 32042624 | [illegible] |
| SE | 45210 | ENSG00000121774 | KHDRBS1 | chr1 | + | 32039514 | 32039573 | 32038551 | 32038619 | 32042526 | 32042624 | [illegible] |
| SE | 45222 | ENSG00000035687 | ADSS2 | chr1 | - | 244415980 | 244416078 | 244411286 | 244411436 | 244417627 | 244417752 | [illegible] |
| SE | 45224 | ENSG00000035687 | ADSS2 | chr1 | - | 244432544 | 244432595 | 244424320 | 244424387 | 244436824 | 244436893 | [illegible] |
| SE | 45226 | ENSG00000204599 | TRIM39 | chr6 | + | 30327646 | 30327696 | 30326956 | 30326995 | 30328888 | 30329041 | [illegible] |
| SE | 45227 | ENSG00000204599 | TRIM39 | chr6 | + | 30327651 | 30327696 | 30326834 | 30326995 | 30328888 | 30329041 | [illegible] |
| SE | 45228 | ENSG00000204599 | TRIM39 | chr6 | + | 30327651 | 30327696 | 30326834 | 30326995 | 30328908 | 30329041 | [illegible] |
| SE | 45229 | ENSG00000204599 | TRIM39 | chr6 | + | 30329310 | 30329770 | 30328888 | 30329041 | 30330780 | 30330876 | [illegible] |
| SE | 45232 | ENSG00000025800 | KPNA6 | chr1 | + | 32160614 | 32160703 | 32159399 | 32159531 | 32161946 | 32162046 | [illegible] |
| SE | 45233 | ENSG00000025800 | KPNA6 | chr1 | + | 32160618 | 32160703 | 32159399 | 32159531 | 32161946 | 32162046 | [illegible] |
| SE | 45234 | ENSG00000025800 | KPNA6 | chr1 | + | 32163234 | 32163313 | 32162360 | 32162524 | 32166104 | 32166230 | [illegible] |
| SE | 45235 | ENSG00000025800 | KPNA6 | chr1 | + | 32167168 | 32167296 | 32166104 | 32166230 | 32169881 | 32170060 | [illegible] |
| SE | 45238 | ENSG00000029639 | TFB1M | chr6 | - | 155260272 | 155260400 | 155256133 | 155258082 | 155285157 | 155285277 | [illegible] |
| SE | 45239 | ENSG00000029639 | TFB1M | chr6 | - | 155260272 | 155260568 | 155257976 | 155258082 | 155285157 | 155285199 | [illegible] |
| SE | 45240 | ENSG00000029639 | TFB1M | chr6 | - | 155313137 | 155313244 | 155311187 | 155311339 | 155314295 | 155314493 | [illegible] |
| SE | 45244 | ENSG00000104419 | NDRG1 | chr8 | - | 133247874 | 133247926 | 133246615 | 133246663 | 133248714 | 133248771 | [illegible] |
| SE | 45252 | ENSG00000104419 | NDRG1 | chr8 | - | 133264546 | 133264652 | 133262045 | 133262167 | 133280231 | 133280267 | [illegible] |
| SE | 45258 | ENSG00000104419 | NDRG1 | chr8 | - | 133284248 | 133284329 | 133280231 | 133280267 | 133297133 | 133297239 | [illegible] |
| SE | 45260 | ENSG00000104419 | NDRG1 | chr8 | - | 133288259 | 133288431 | 133284248 | 133284329 | 133297133 | 133297252 | [illegible] |
| SE | 45262 | ENSG00000164403 | SHROOM1 | chr5 | - | 132823849 | 132824419 | 132823622 | 132823764 | 132824614 | 132824821 | [illegible] |
| SE | 45263 | ENSG00000164403 | SHROOM1 | chr5 | - | 132824614 | 132824821 | 132823849 | 132824419 | 132825017 | 132825073 | [illegible] |
| SE | 45264 | ENSG00000274810 | NPHP3-ACAD11 | chr3 | - | 132621131 | 132621292 | 132619467 | 132619545 | 132626690 | 132626817 | [illegible] |
| SE | 45267 | ENSG00000105808 | RASA4 | chr7 | - | 102590158 | 102590296 | 102587634 | 102587770 | 102592284 | 102592323 | [illegible] |
| SE | 45268 | ENSG00000105808 | RASA4 | chr7 | - | 102602323 | 102602405 | 102600256 | 102600388 | 102605857 | 102605987 | [illegible] |
| SE | 45269 | ENSG00000105808 | RASA4 | chr7 | - | 102605857 | 102605987 | 102602323 | 102602405 | 102608760 | 102608822 | [illegible] |
| SE | 45270 | ENSG00000205236 | NA | chr7 | - | 102624188 | 102624351 | 102611085 | 102611142 | 102638713 | 102638834 | [illegible] |
| SE | 45271 | ENSG00000205236 | NA | chr7 | - | 102640203 | 102640414 | 102639122 | 102639199 | 102640611 | 102640773 | [illegible] |
| SE | 45294 | ENSG00000145850 | TIMD4 | chr5 | - | 156948415 | 156948499 | 156926262 | 156926312 | 156949650 | 156949731 | [illegible] |
| SE | 45295 | ENSG00000145850 | TIMD4 | chr5 | - | 156949650 | 156949731 | 156926262 | 156926312 | 156951511 | 156951790 | [illegible] |
| SE | 45298 | ENSG00000286192 | NA | chr7 | - | 2069193 | 2069338 | 1898199 | 1898390 | 2213211 | 2213273 | [illegible] |
| SE | 45299 | ENSG00000286192 | NA | chr7 | - | 2149095 | 2149238 | 2069193 | 2069338 | 2213211 | 2213273 | [illegible] |
| SE | 45300 | ENSG00000286192 | NA | chr7 | - | 2200261 | 2200557 | 2149095 | 2149238 | 2213211 | 2213273 | [illegible] |
| SE | 45302 | ENSG00000286192 | NA | chr7 | - | 2229983 | 2230143 | 2222574 | 2222754 | 2230548 | 2230727 | [illegible] |
| SE | 45304 | ENSG00000163159 | VPS72 | chr1 | - | 151178000 | 151178145 | 151176303 | 151177031 | 151184316 | 151184493 | [illegible] |
| SE | 45305 | ENSG00000163159 | VPS72 | chr1 | - | 151178000 | 151178145 | 151176461 | 151177064 | 151184316 | 151184493 | [illegible] |
| SE | 45306 | ENSG00000163159 | VPS72 | chr1 | - | 151185505 | 151185620 | 151184316 | 151184493 | 151185797 | 151185950 | [illegible] |
| SE | 45307 | ENSG00000163568 | AIM2 | chr1 | - | 159068567 | 159068701 | 159065909 | 159066329 | 159076632 | 159076766 | [illegible] |
| SE | 45308 | ENSG00000163568 | AIM2 | chr1 | - | 159073237 | 159073519 | 159068567 | 159068701 | 159076632 | 159076766 | [illegible] |
| SE | 45316 | ENSG00000116030 | SUMO1 | chr2 | - | 202207867 | 202208002 | 202206649 | 202207321 | 202210734 | 202210806 | [illegible] |
| SE | 45317 | ENSG00000116030 | SUMO1 | chr2 | - | 202210734 | 202210783 | 202206869 | 202207321 | 202214356 | 202214434 | [illegible] |
| SE | 45318 | ENSG00000116030 | SUMO1 | chr2 | - | 202210734 | 202210806 | 202206181 | 202207321 | 202214356 | 202214429 | [illegible] |
| SE | 45320 | ENSG00000116030 | SUMO1 | chr2 | - | 202231542 | 202231732 | 202220031 | 202220106 | 202238439 | 202238566 | [illegible] |
| SE | 45321 | ENSG00000116030 | SUMO1 | chr2 | - | 202233577 | 202233670 | 202220031 | 202220106 | 202238439 | 202238586 | [illegible] |
| SE | 45323 | ENSG00000144320 | LNPK | chr2 | - | 175938312 | 175938383 | 175937343 | 175937514 | 175939551 | 175939657 | [illegible] |
| SE | 45324 | ENSG00000144320 | LNPK | chr2 | - | 175940987 | 175941080 | 175939551 | 175939657 | 175947479 | 175947692 | [illegible] |
| SE | 45325 | ENSG00000144320 | LNPK | chr2 | - | 175979809 | 175979868 | 175970763 | 175970804 | 175992230 | 175992418 | [illegible] |
| SE | 45327 | ENSG00000144320 | LNPK | chr2 | - | 175994133 | 175994280 | 175993180 | 175993223 | 175995552 | 175995646 | [illegible] |
| SE | 45329 | ENSG00000144320 | LNPK | chr2 | - | 175995557 | 175995646 | 175979809 | 175979868 | 176002159 | 176002267 | [illegible] |
| SE | 45330 | ENSG00000179010 | MRFAP1 | chr4 | + | 6640827 | 6641258 | 6640685 | 6640755 | 6641729 | 6641791 | [illegible] |
| SE | 45332 | ENSG00000180879 | SSR4 | chrX | + | 153795927 | 153796126 | 153794675 | 153794754 | 153796433 | 153796552 | [illegible] |
| SE | 45333 | ENSG00000180879 | SSR4 | chrX | + | 153797457 | 153797532 | 153794658 | 153796552 | 153797724 | 153797814 | [illegible] |
| SE | 45334 | ENSG00000180879 | SSR4 | chrX | + | 153797724 | 153797814 | 153796433 | 153796552 | 153798070 | 153798136 | [illegible] |
| SE | 45346 | ENSG00000260271 | NA | chr6 | + | 90355763 | 90355818 | 90346656 | 90346790 | 90356379 | 90356578 | [illegible] |
| SE | 45356 | ENSG00000115267 | IFIH1 | chr2 | - | 162277414 | 162277693 | 162276686 | 162276946 | 162279995 | 162280112 | [illegible] |
| SE | 45359 | ENSG00000115267 | IFIH1 | chr2 | - | 162288134 | 162288355 | 162282365 | 162282576 | 162293563 | 162293668 | [illegible] |
| SE | 45360 | ENSG00000115267 | IFIH1 | chr2 | - | 162293563 | 162293668 | 162288134 | 162288355 | 162306708 | 162306855 | [illegible] |
| SE | 45361 | ENSG00000115267 | IFIH1 | chr2 | - | 162306708 | 162306855 | 162293563 | 162293668 | 162310764 | 162310933 | [illegible] |
| SE | 45362 | ENSG00000143164 | DCAF6 | chr1 | + | 167951799 | 167951861 | 167936558 | 167937008 | 167966628 | 167966721 | [illegible] |
| SE | 45363 | ENSG00000143164 | DCAF6 | chr1 | + | 167966628 | 167966721 | 167951799 | 167951861 | 167974829 | 167975015 | [illegible] |
| SE | 45364 | ENSG00000143164 | DCAF6 | chr1 | + | 167987494 | 167987608 | 167974829 | 167975015 | 167991203 | 167991339 | [illegible] |
| SE | 45365 | ENSG00000143164 | DCAF6 | chr1 | + | 167993225 | 167993440 | 167991203 | 167991339 | 168002481 | 168002575 | [illegible] |
| SE | 45369 | ENSG00000143164 | DCAF6 | chr1 | + | 168019544 | 168019667 | 168004532 | 168004793 | 168043024 | 168043140 | [illegible] |
| SE | 45373 | ENSG00000143164 | DCAF6 | chr1 | + | 168038370 | 168038488 | 168019544 | 168019667 | 168043024 | 168043140 | [illegible] |
| SE | 45377 | ENSG00000143164 | DCAF6 | chr1 | + | 168065589 | 168065746 | 168063620 | 168063759 | 168068357 | 168068463 | [illegible] |
| SE | 45378 | ENSG00000143164 | DCAF6 | chr1 | + | 168066376 | 168066465 | 168063620 | 168063759 | 168068357 | 168068463 | [illegible] |
| SE | 45379 | ENSG00000143164 | DCAF6 | chr1 | + | 168066376 | 168066465 | 168065589 | 168065746 | 168068357 | 168068463 | [illegible] |
| SE | 45380 | ENSG00000143164 | DCAF6 | chr1 | + | 168066382 | 168066465 | 168065589 | 168065746 | 168068357 | 168068463 | [illegible] |
| SE | 45381 | ENSG00000143164 | DCAF6 | chr1 | + | 168068357 | 168068463 | 168066376 | 168066465 | 168075370 | 168075786 | [illegible] |
| SE | 45383 | ENSG00000144468 | RHBDD1 | chr2 | + | 226837547 | 226837770 | 226836055 | 226836087 | 226838072 | 226838154 | [illegible] |
| SE | 45384 | ENSG00000144468 | RHBDD1 | chr2 | + | 226837547 | 226837770 | 226836055 | 226836087 | 226839408 | 226839626 | [illegible] |
| SE | 45386 | ENSG00000144468 | RHBDD1 | chr2 | + | 226837547 | 226837770 | 226836081 | 226836199 | 226864603 | 226864856 | [illegible] |
| SE | 45387 | ENSG00000144468 | RHBDD1 | chr2 | + | 226838072 | 226838154 | 226836033 | 226836087 | 226839408 | 226839627 | [illegible] |
| SE | 45388 | ENSG00000144468 | RHBDD1 | chr2 | + | 226838072 | 226838154 | 226836033 | 226836087 | 226864603 | 226865065 | [illegible] |
| SE | 45389 | ENSG00000144468 | RHBDD1 | chr2 | + | 226838072 | 226838154 | 226837523 | 226837838 | 226839408 | 226839627 | [illegible] |
| SE | 45390 | ENSG00000144468 | RHBDD1 | chr2 | + | 226838072 | 226838154 | 226837523 | 226837838 | 226864603 | 226865065 | [illegible] |
| SE | 45391 | ENSG00000144468 | RHBDD1 | chr2 | + | 226838072 | 226838154 | 226837547 | 226837770 | 226839408 | 226839627 | [illegible] |
| SE | 45396 | ENSG00000144468 | RHBDD1 | chr2 | + | 226867185 | 226867318 | 226864603 | 226865126 | 226906792 | 226906881 | [illegible] |
| SE | 45397 | ENSG00000144468 | RHBDD1 | chr2 | + | 226906792 | 226906881 | 226867185 | 226867318 | 226914207 | 226914351 | [illegible] |
| SE | 45400 | ENSG00000144468 | RHBDD1 | chr2 | + | 226914207 | 226914351 | 226908821 | 226908878 | 226995430 | 226995755 | [illegible] |
| SE | 45401 | ENSG00000156463 | SH3RF2 | chr5 | + | 146049074 | 146049245 | 146047771 | 146047863 | 146055980 | 146056213 | [illegible] |
| SE | 45404 | ENSG00000180385 | EMC3-AS1 | chr3 | + | 9987814 | 9987905 | 9987214 | 9987264 | 9988711 | 9988747 | [illegible] |
| SE | 45405 | ENSG00000180385 | EMC3-AS1 | chr3 | + | 9987905 | 9988007 | 9986892 | 9987264 | 9988711 | 9988747 | [illegible] |
| SE | 45407 | ENSG00000180385 | EMC3-AS1 | chr3 | + | 9988711 | 9988747 | 9986892 | 9987264 | 10002716 | 10002765 | [illegible] |
| SE | 45408 | ENSG00000070190 | DAPP1 | chr4 | + | 99835622 | 99835745 | 99816826 | 99817014 | 99840288 | 99840422 | [illegible] |
| SE | 45409 | ENSG00000070190 | DAPP1 | chr4 | + | 99840288 | 99840422 | 99835622 | 99835745 | 99853217 | 99853348 | [illegible] |
| SE | 45410 | ENSG00000070190 | DAPP1 | chr4 | + | 99853217 | 99853348 | 99840288 | 99840422 | 99861577 | 99861625 | [illegible] |
| SE | 45413 | ENSG00000115446 | UNC50 | chr2 | + | 98616206 | 98616346 | 98610774 | 98610895 | 98616431 | 98616533 | [illegible] |
| SE | 45414 | ENSG00000115446 | UNC50 | chr2 | + | 98616431 | 98616533 | 98610774 | 98610895 | 98618167 | 98618515 | [illegible] |
| SE | 45415 | ENSG00000115446 | UNC50 | chr2 | + | 98616431 | 98616533 | 98616205 | 98616346 | 98618167 | 98618515 | [illegible] |
| SE | 45425 | ENSG00000176542 | USF3 | chr3 | - | 113664312 | 113664409 | 113648384 | 113661425 | 113670120 | 113670203 | [illegible] |
| SE | 45426 | ENSG00000176542 | USF3 | chr3 | - | 113670120 | 113670203 | 113664312 | 113664409 | 113673347 | 113673376 | [illegible] |
| SE | 45428 | ENSG00000176542 | USF3 | chr3 | - | 113677281 | 113677397 | 113674831 | 113674896 | 113696369 | 113696642 | [illegible] |
| SE | 45430 | ENSG00000169683 | LRRC45 | chr17 | + | 82029542 | 82029635 | 82029018 | 82029185 | 82030064 | 82030238 | [illegible] |
| SE | 45431 | ENSG00000162877 | PM20D1 | chr1 | - | 205832597 | 205832766 | 205830279 | 205830379 | 205840251 | 205840323 | [illegible] |
| SE | 45432 | ENSG00000162877 | PM20D1 | chr1 | - | 205840251 | 205840323 | 205832597 | 205832766 | 205841810 | 205841879 | [illegible] |
| SE | 45433 | ENSG00000162877 | PM20D1 | chr1 | - | 205842153 | 205842215 | 205841810 | 205841889 | 205842675 | 205842751 | [illegible] |
| SE | 45434 | ENSG00000153989 | NUS1 | chr6 | + | 117703604 | 117703704 | 117694030 | 117694180 | 117706924 | 117710727 | [illegible] |
| SE | 45435 | ENSG00000283196 | LSP1P5 | chr2 | - | 89580557 | 89580635 | 89542841 | 89543836 | 89581005 | 89581140 | [illegible] |
| SE | 45439 | ENSG00000170791 | CHCHD7 | chr8 | + | 56216388 | 56216506 | 56214597 | 56214667 | 56217330 | 56217689 | [illegible] |
| SE | 45440 | ENSG00000170791 | CHCHD7 | chr8 | + | 56216388 | 56216531 | 56211788 | 56214667 | 56217330 | 56217465 | [illegible] |
| SE | 45441 | ENSG00000170791 | CHCHD7 | chr8 | + | 56216432 | 56216506 | 56214597 | 56214667 | 56217330 | 56217713 | [illegible] |
| SE | 45442 | ENSG00000170791 | CHCHD7 | chr8 | + | 56216432 | 56216531 | 56214444 | 56214667 | 56217330 | 56217590 | [illegible] |
| SE | 45443 | ENSG00000170791 | CHCHD7 | chr8 | + | 56216679 | 56216754 | 56216388 | 56216506 | 56217330 | 56217412 | [illegible] |
| SE | 45445 | ENSG00000100991 | TRPC4AP | chr20 | - | 35006434 | 35006575 | 35005694 | 35005803 | 35007549 | 35007640 | [illegible] |
| SE | 45446 | ENSG00000100991 | TRPC4AP | chr20 | - | 35007549 | 35007640 | 35006434 | 35006575 | 35008663 | 35008747 | [illegible] |
| SE | 45447 | ENSG00000100991 | TRPC4AP | chr20 | - | 35013007 | 35013066 | 35010186 | 35010288 | 35016007 | 35016139 | [illegible] |
| SE | 45450 | ENSG00000100991 | TRPC4AP | chr20 | - | 35035122 | 35035308 | 35021189 | 35021332 | 35044504 | 35044712 | [illegible] |
| SE | 45452 | ENSG00000100991 | TRPC4AP | chr20 | - | 35044504 | 35044712 | 35035122 | 35035308 | 35054975 | 35055031 | [illegible] |
| SE | 45453 | ENSG00000100991 | TRPC4AP | chr20 | - | 35049865 | 35049994 | 35035122 | 35035308 | 35054975 | 35055031 | [illegible] |
| SE | 45454 | ENSG00000100991 | TRPC4AP | chr20 | - | 35049865 | 35049994 | 35044504 | 35044712 | 35054975 | 35055031 | [illegible] |
| SE | 45455 | ENSG00000100991 | TRPC4AP | chr20 | - | 35054975 | 35055031 | 35049865 | 35049994 | 35057513 | 35057571 | [illegible] |
| SE | 45456 | ENSG00000100991 | TRPC4AP | chr20 | - | 35054975 | 35055031 | 35049865 | 35049994 | 35069295 | 35069412 | [illegible] |
| SE | 45457 | ENSG00000100991 | TRPC4AP | chr20 | - | 35057513 | 35057571 | 35049865 | 35049994 | 35069295 | 35069412 | [illegible] |
| SE | 45458 | ENSG00000100991 | TRPC4AP | chr20 | - | 35057513 | 35057571 | 35054975 | 35055031 | 35069295 | 35069412 | [illegible] |
| SE | 45459 | ENSG00000100991 | TRPC4AP | chr20 | - | 35057513 | 35057571 | 35054975 | 35055031 | 35078045 | 35078174 | [illegible] |
| SE | 45460 | ENSG00000100991 | TRPC4AP | chr20 | - | 35069295 | 35069412 | 35054975 | 35055031 | 35078045 | 35078174 | [illegible] |
| SE | 45461 | ENSG00000100991 | TRPC4AP | chr20 | - | 35069295 | 35069412 | 35057513 | 35057571 | 35078045 | 35078174 | [illegible] |
| SE | 45463 | ENSG00000163666 | HESX1 | chr3 | - | 57198752 | 57198952 | 57198390 | 57198492 | 57199761 | 57199938 | [illegible] |
| SE | 45464 | ENSG00000163666 | HESX1 | chr3 | - | 57211334 | 57211502 | 57199761 | 57200028 | 57226295 | 57226336 | [illegible] |
| SE | 45465 | ENSG00000143149 | ALDH9A1 | chr1 | - | 165665017 | 165665130 | 165662215 | 165663144 | 165667308 | 165667450 | [illegible] |
| SE | 45467 | ENSG00000143149 | ALDH9A1 | chr1 | - | 165695251 | 165695397 | 165682824 | 165683110 | 165698377 | 165698513 | [illegible] |
| SE | 45468 | ENSG00000204524 | ZNF805 | chr19 | + | 57243922 | 57244049 | 57240631 | 57240921 | 57248604 | 57248700 | [illegible] |
| SE | 45469 | ENSG00000204524 | ZNF805 | chr19 | + | 57248604 | 57248700 | 57243922 | 57244049 | 57253072 | 57255135 | [illegible] |
| SE | 45490 | ENSG00000183665 | TRMT12 | chr8 | + | 124452151 | 124452342 | 124450916 | 124451257 | 124452451 | 124452741 | [illegible] |
| SE | 45491 | ENSG00000111786 | SRSF9 | chr12 | - | 120464732 | 120464822 | 120463949 | 120464122 | 120465626 | 120465787 | [illegible] |
| SE | 45493 | ENSG00000183831 | ANKRD45 | chr1 | - | 173627064 | 173627159 | 173624786 | 173624925 | 173646845 | 173647013 | [illegible] |
| SE | 45494 | ENSG00000168894 | RNF181 | chr2 | + | 85596518 | 85596649 | 85595724 | 85595849 | 85597103 | 85597178 | [illegible] |
| SE | 45501 | ENSG00000168894 | RNF181 | chr2 | + | 85597103 | 85597178 | 85596821 | 85596931 | 85597444 | 85597604 | [illegible] |
| SE | 45503 | ENSG00000169994 | MYO7B | chr2 | + | 127627183 | 127627310 | 127626974 | 127627092 | 127628371 | 127628535 | [illegible] |
| SE | 45504 | ENSG00000169994 | MYO7B | chr2 | + | 127634175 | 127634289 | 127633257 | 127633363 | 127634595 | 127634683 | [illegible] |
| SE | 45505 | ENSG00000204475 | NCR3 | chr6 | - | 31589525 | 31589633 | 31588894 | 31589176 | 31589781 | 31589899 | [illegible] |
| SE | 45506 | ENSG00000204475 | NCR3 | chr6 | - | 31589974 | 31590126 | 31589781 | 31589899 | 31592678 | 31592749 | [illegible] |
| SE | 45507 | ENSG00000163960 | UBXN7 | chr3 | - | 196361843 | 196361923 | 196347661 | 196356846 | 196362293 | 196362687 | [illegible] |
| SE | 45508 | ENSG00000163960 | UBXN7 | chr3 | - | 196368027 | 196368155 | 196362293 | 196362687 | 196369420 | 196369511 | [illegible] |
| SE | 45509 | ENSG00000163960 | UBXN7 | chr3 | - | 196368027 | 196368155 | 196362293 | 196362687 | 196371895 | 196372042 | [illegible] |
| SE | 45510 | ENSG00000163960 | UBXN7 | chr3 | - | 196369420 | 196369511 | 196362293 | 196362687 | 196371895 | 196372042 | [illegible] |
| SE | 45515 | ENSG00000163960 | UBXN7 | chr3 | - | 196402951 | 196403019 | 196393553 | 196393619 | 196432326 | 196432427 | [illegible] |
| SE | 45523 | ENSG00000186231 | KLHL32 | chr6 | + | 97085125 | 97085341 | 97064627 | 97064726 | 97113782 | 97114509 | [illegible] |
| SE | 45529 | ENSG00000197134 | ZNF257 | chr19 | + | 22060507 | 22060628 | 22052451 | 22052635 | 22072808 | 22072935 | [illegible] |
| SE | 45530 | ENSG00000197134 | ZNF257 | chr19 | + | 22072808 | 22072935 | 22052429 | 22052635 | 22073468 | 22073564 | [illegible] |
| SE | 45531 | ENSG00000197134 | ZNF257 | chr19 | + | 22072808 | 22072935 | 22052429 | 22052635 | 22087976 | 22088405 | [illegible] |
| SE | 45532 | ENSG00000197134 | ZNF257 | chr19 | + | 22073468 | 22073564 | 22052429 | 22052635 | 22087976 | 22091103 | [illegible] |
| SE | 45533 | ENSG00000197134 | ZNF257 | chr19 | + | 22073468 | 22073564 | 22072808 | 22072935 | 22087976 | 22091103 | [illegible] |
| SE | 45541 | ENSG00000139370 | SLC15A4 | chr12 | - | 128799258 | 128799417 | 128793193 | 128794356 | 128800853 | 128801009 | [illegible] |
| SE | 45542 | ENSG00000139370 | SLC15A4 | chr12 | - | 128800853 | 128801009 | 128793193 | 128794356 | 128808787 | 128808956 | [illegible] |
| SE | 45543 | ENSG00000139370 | SLC15A4 | chr12 | - | 128808787 | 128808956 | 128800806 | 128801009 | 128809395 | 128809473 | [illegible] |
| SE | 45548 | ENSG00000082146 | STRADB | chr2 | + | 201457378 | 201457450 | 201454745 | 201454852 | 201458783 | 201458823 | [illegible] |
| SE | 45549 | ENSG00000082146 | STRADB | chr2 | + | 201469952 | 201470052 | 201458783 | 201458864 | 201472954 | 201473076 | [illegible] |
| SE | 45550 | ENSG00000082146 | STRADB | chr2 | + | 201474646 | 201474755 | 201472856 | 201473076 | 201475618 | 201475742 | [illegible] |
| SE | 45551 | ENSG00000082146 | STRADB | chr2 | + | 201478356 | 201478601 | 201478086 | 201478191 | 201479488 | 201479531 | [illegible] |
| SE | 45552 | ENSG00000082146 | STRADB | chr2 | + | 201478562 | 201478601 | 201478149 | 201478191 | 201479488 | 201479531 | [illegible] |
| SE | 45564 | ENSG00000117335 | CD46 | chr1 | + | 207757539 | 207757642 | 207757013 | 207757202 | 207759638 | 207759724 | [illegible] |
| SE | 45565 | ENSG00000117335 | CD46 | chr1 | + | 207761248 | 207761446 | 207759638 | 207759724 | 207767012 | 207767195 | [illegible] |
| SE | 45566 | ENSG00000117335 | CD46 | chr1 | + | 207767012 | 207767195 | 207759638 | 207759724 | 207767778 | 207767823 | [illegible] |
| SE | 45569 | ENSG00000117335 | CD46 | chr1 | + | 207767778 | 207767823 | 207767012 | 207767195 | 207783291 | 207783330 | [illegible] |
| SE | 45570 | ENSG00000117335 | CD46 | chr1 | + | 207767778 | 207767823 | 207767012 | 207767195 | 207785070 | 207785106 | [illegible] |
| SE | 45571 | ENSG00000117335 | CD46 | chr1 | + | 207770320 | 207770362 | 207767012 | 207767195 | 207783291 | 207783330 | [illegible] |
| SE | 45573 | ENSG00000117335 | CD46 | chr1 | + | 207783291 | 207783330 | 207767012 | 207767195 | 207785070 | 207785106 | [illegible] |
| SE | 45574 | ENSG00000117335 | CD46 | chr1 | + | 207783291 | 207783330 | 207767778 | 207767823 | 207785070 | 207785106 | [illegible] |
| SE | 45575 | ENSG00000117335 | CD46 | chr1 | + | 207785070 | 207785106 | 207767778 | 207767823 | 207785618 | 207785682 | [illegible] |
| SE | 45580 | ENSG00000285253 | NA | chr15 | - | 56654384 | 56654503 | 56652487 | 56653599 | 56658423 | 56658486 | [illegible] |
| SE | 45581 | ENSG00000285253 | NA | chr15 | - | 56917146 | 56917212 | 56890013 | 56890167 | 56917539 | 56918571 | [illegible] |
| SE | 45582 | ENSG00000021355 | SERPINB1 | chr6 | - | 2838548 | 2838686 | 2837881 | 2837999 | 2840418 | 2840485 | [illegible] |
| SE | 45585 | ENSG00000091127 | PUS7 | chr7 | - | 105465312 | 105465414 | 105462620 | 105462750 | 105468336 | 105468463 | [illegible] |
| SE | 45586 | ENSG00000091127 | PUS7 | chr7 | - | 105468336 | 105468463 | 105462620 | 105462750 | 105470760 | 105470848 | [illegible] |
| SE | 45587 | ENSG00000091127 | PUS7 | chr7 | - | 105468336 | 105468463 | 105465312 | 105465414 | 105470687 | 105470848 | [illegible] |
| SE | 45588 | ENSG00000091127 | PUS7 | chr7 | - | 105468336 | 105468463 | 105465312 | 105465414 | 105472131 | 105472193 | [illegible] |
| SE | 45589 | ENSG00000091127 | PUS7 | chr7 | - | 105470687 | 105470848 | 105465312 | 105465414 | 105472131 | 105472193 | [illegible] |
| SE | 45590 | ENSG00000091127 | PUS7 | chr7 | - | 105470687 | 105470848 | 105468336 | 105468463 | 105472131 | 105472193 | [illegible] |
| SE | 45591 | ENSG00000091127 | PUS7 | chr7 | - | 105470687 | 105470848 | 105468336 | 105468463 | 105481051 | 105481177 | [illegible] |
| SE | 45592 | ENSG00000091127 | PUS7 | chr7 | - | 105470760 | 105470848 | 105468336 | 105468463 | 105472131 | 105472193 | [illegible] |
| SE | 45593 | ENSG00000091127 | PUS7 | chr7 | - | 105470760 | 105470848 | 105468336 | 105468463 | 105481051 | 105481177 | [illegible] |
| SE | 45594 | ENSG00000091127 | PUS7 | chr7 | - | 105472131 | 105472193 | 105468336 | 105468463 | 105481051 | 105481177 | [illegible] |
| SE | 45595 | ENSG00000091127 | PUS7 | chr7 | - | 105472131 | 105472193 | 105470687 | 105470848 | 105481051 | 105481177 | [illegible] |
| SE | 45596 | ENSG00000091127 | PUS7 | chr7 | - | 105502419 | 105502564 | 105495141 | 105495253 | 105505954 | 105506056 | [illegible] |
| SE | 45598 | ENSG00000091127 | PUS7 | chr7 | - | 105506188 | 105506273 | 105505954 | 105506056 | 105508114 | 105508135 | [illegible] |
| SE | 45599 | ENSG00000049246 | PER3 | chr1 | + | 7786720 | 7786836 | 7785440 | 7785586 | 7788044 | 7788246 | [illegible] |
| SE | 45600 | ENSG00000049246 | PER3 | chr1 | + | 7787197 | 7787246 | 7786720 | 7786836 | 7788044 | 7788338 | [illegible] |
| SE | 45601 | ENSG00000049246 | PER3 | chr1 | + | 7808892 | 7808998 | 7803691 | 7803848 | 7809892 | 7810021 | [illegible] |
| SE | 45605 | ENSG00000164023 | SGMS2 | chr4 | + | 107895309 | 107896008 | 107858471 | 107858553 | 107899574 | 107899692 | [illegible] |
| SE | 45606 | ENSG00000164023 | SGMS2 | chr4 | + | 107908564 | 107908731 | 107903232 | 107903386 | 107910349 | 107910953 | [illegible] |
| SE | 45615 | ENSG00000132781 | MUTYH | chr1 | - | 45331660 | 45331849 | 45331419 | 45331556 | 45332022 | 45332086 | [illegible] |
| SE | 45616 | ENSG00000288208 | NA | chr1 | - | 45332390 | 45332488 | 45332165 | 45332310 | 45332573 | 45332687 | [illegible] |
| SE | 45617 | ENSG00000288208 | NA | chr1 | - | 45332573 | 45332687 | 45332165 | 45332310 | 45332762 | 45332834 | [illegible] |
| SE | 45618 | ENSG00000288208 | NA | chr1 | - | 45332917 | 45332959 | 45332762 | 45332834 | 45333096 | 45333170 | [illegible] |
| SE | 45619 | ENSG00000132781 | MUTYH | chr1 | - | 45332917 | 45332959 | 45332165 | 45332834 | 45333284 | 45333324 | [illegible] |
| SE | 45622 | ENSG00000132781 | MUTYH | chr1 | - | 45333284 | 45333324 | 45332762 | 45332834 | 45334390 | 45334511 | [illegible] |

| | | | | | | | | | | | | | |
|---|---|---|---|---|---|---|---|---|---|---|---|---|---|
| SE | 45624 | ENSG00000288208 | NA | chr1 | - | 45355302 | 45355449 | 45334390 | 45334511 | 45385911 | 45385960 | 14 | 4 |
| SE | 45627 | ENSG00000081721 | DUSP12 | chr1 | + | 161752367 | 161752464 | 161751865 | 161751984 | 161753074 | 161753105 | 148 | 107 |
| SE | 45629 | ENSG00000183019 | MCEMP1 | chr19 | + | 7678349 | 7678400 | 7677928 | 7678241 | 7678490 | 7678604 | 136 | 78 |
| SE | 45637 | ENSG00000118246 | FASTKD2 | chr2 | + | 206774224 | 206774397 | 206772180 | 206772320 | 206787936 | 206788155 | 48 | 45 |
| SE | 45638 | ENSG00000118246 | FASTKD2 | chr2 | + | 206786732 | 206786899 | 206774224 | 206774397 | 206787936 | 206788155 | 155 | 107 |
| SE | 45656 | ENSG00000110651 | CD81 | chr11 | + | 2386554 | 2386639 | 2384662 | 2384682 | 2390411 | 2390526 | 4 | 0 |
| SE | 45658 | ENSG00000110651 | CD81 | chr11 | + | 2395868 | 2395970 | 2395415 | 2395520 | 2396627 | 2396714 | 518 | 169 |
| SE | 45660 | ENSG00000163608 | NEPRO | chr3 | - | 113010642 | 113010704 | 113008170 | 113008430 | 113011044 | 113011165 | 153 | 128 |
| SE | 45661 | ENSG00000163608 | NEPRO | chr3 | - | 113011044 | 113011406 | 113008170 | 113008430 | 113013948 | 113014032 | 1 | 1 |
| SE | 45662 | ENSG00000163608 | NEPRO | chr3 | - | 113012733 | 113013403 | 113011044 | 113011406 | 113013952 | 113014032 | 56 | 47 |
| SE | 45664 | ENSG00000163608 | NEPRO | chr3 | - | 113017447 | 113017602 | 113013948 | 113014032 | 113019541 | 113019644 | 40 | 22 |
| SE | 45668 | ENSG00000109323 | MANBA | chr4 | - | 102633250 | 102633473 | 102630769 | 102632281 | 102634787 | 102635045 | 3 | 0 |
| SE | 45670 | ENSG00000109323 | MANBA | chr4 | - | 102639712 | 102639857 | 102634787 | 102635045 | 102650536 | 102650701 | 69 | 66 |
| SE | 45672 | ENSG00000109323 | MANBA | chr4 | - | 102658858 | 102659028 | 102657681 | 102657900 | 102664684 | 102664852 | 0 | 2 |
| SE | 45673 | ENSG00000109323 | MANBA | chr4 | - | 102664684 | 102664852 | 102650536 | 102650701 | 102689573 | 102689684 | 2 | 3 |
| SE | 45675 | ENSG00000109323 | MANBA | chr4 | - | 102668962 | 102669049 | 102664684 | 102664852 | 102670915 | 102671011 | 78 | 86 |
| SE | 45677 | ENSG00000109323 | MANBA | chr4 | - | 102671892 | 102672054 | 102671280 | 102671398 | 102673918 | 102674070 | 0 | 0 |
| SE | 45679 | ENSG00000109323 | MANBA | chr4 | - | 102672008 | 102672146 | 102671280 | 102671398 | 102673918 | 102674070 | 0 | 1 |
| SE | 45684 | ENSG00000109323 | MANBA | chr4 | - | 102714437 | 102714561 | 102690595 | 102690771 | 102723861 | 102723967 | 87 | 51 |
| SE | 45686 | ENSG00000109323 | MANBA | chr4 | - | 102722870 | 102723041 | 102690595 | 102690771 | 102723861 | 102723921 | 238 | 116 |

[illegible]

| | | | | | | | | | | | | | | | | | | | | | | | | | | | | | | | | | | | | | | | | | | | | | | |
|---|---|---|---|---|---|---|---|---|---|---|---|---|---|---|---|---|---|---|---|---|---|---|---|---|---|---|---|---|---|---|---|---|---|---|---|---|---|---|---|---|---|---|---|---|---|---|
| SE | 46070 | ENSG00000068097 | HEATR6 | chr17 | - | 60059889 | 60060096 | 60059421 | 60059521 | 60069710 | 60069848 | 53 | 28 | [illegible] | [illegible] | [illegible] | [illegible] | [illegible] | [illegible] | [illegible] | [illegible] | 28 | 30 | 32 | 20 | 38 | 0 | 0 | 1 | 0 | 0 | 298 | 149 | 1 | 1 | 0.964 | 1 | 1 | 1 | 0.974 | 1 | 1 | 0.941 | 1 | 1 | -0.001 |
| SE | 46071 | ENSG00000068097 | HEATR6 | chr17 | - | 60066208 | 60066386 | 60059889 | 60060096 | 60067433 | 60067732 | 49 | 27 | [illegible] | [illegible] | [illegible] | [illegible] | [illegible] | [illegible] | [illegible] | [illegible] | 27 | 26 | 27 | 13 | 36 | 0 | 0 | 0 | 1 | 0 | 298 | 149 | 1 | 1 | 1 | 1 | 0.938 | 1 | 1 | 1 | | 1 | 0.867 | 1 | 0.014 |
| SE | 46072 | ENSG00000068097 | HEATR6 | chr17 | - | 60067433 | 60067732 | 60059889 | 60060096 | 60069710 | 60069848 | 39 | 12 | [illegible] | [illegible] | [illegible] | [illegible] | [illegible] | [illegible] | [illegible] | [illegible] | 0 | 24 | 4 | 25 | 30 | 0 | 3 | 0 | 1 | 0 | 298 | 149 | 1 | 1 | 0.83 | 1 | 0.875 | 1 | 1 | NA | 0.8 | 1 | 0.926 | 1 | 0.01 |
| SE | 46073 | ENSG00000068097 | HEATR6 | chr17 | - | 60072214 | 60072329 | 60070705 | 60070807 | 60073163 | 60073279 | 89 | 27 | 48 | 64 | 62 | 0 | 0 | 0 | 0 | 0 | 22 | 33 | 35 | 41 | 69 | 0 | 0 | 0 | 1 | 0 | 264 | 149 | 1 | 1 | 1 | 1 | 1 | 1 | 1 | 1 | 1 | 1 | 0.959 | 1 | 0.008 |
| SE | 46075 | ENSG00000068097 | HEATR6 | chr17 | - | 60073163 | 60073279 | 60072214 | 60072329 | 60076129 | 60076237 | 20 | 8 | 11 | 27 | 25 | 0 | 0 | 0 | 0 | 0 | 12 | 3 | 12 | 15 | 25 | 0 | 1 | 0 | 0 | 0 | 265 | 149 | 1 | 1 | 1 | 1 | 1 | 1 | 1 | | 0.628 | 1 | 1 | 1 | 0.074 |
| SE | 46076 | ENSG00000068097 | HEATR6 | chr17 | - | 60073745 | 60073886 | 60073163 | 60073279 | 60076129 | 60076237 | 51 | 21 | 20 | 27 | 12 | 3 | 0 | 2 | 7 | 5 | 0 | 27 | 15 | 24 | 29 | 0 | 0 | 4 | 0 | 0 | 290 | 149 | 1 | 1 | 0.897 | 1 | 0.837 | 0.665 | 0.552 | NA | 1 | 0.658 | 1 | 1 | -0.124 |
| SE | 46078 | ENSG00000068097 | HEATR6 | chr17 | - | 60073961 | 60074159 | 60073235 | 60073279 | 60076129 | 60076237 | 7 | 5 | 2 | 7 | 2 | 3 | 0 | 2 | 7 | 5 | 9 | 3 | 2 | 5 | 0 | 0 | 0 | 4 | 0 | 0 | 298 | 149 | 1 | 1 | 0.538 | 1 | 0.333 | 0.333 | 0.167 | 1 | 1 | 0.2 | 1 | NA | -0.326 |
| SE | 46080 | ENSG00000068097 | HEATR6 | chr17 | - | 60075704 | 60075833 | 60073745 | 60073886 | 60076129 | 60076237 | 0 | 0 | 0 | 2 | 0 | 18 | 3 | 3 | 5 | 2 | 0 | 0 | 0 | 0 | 3 | 0 | 13 | 2 | 2 | 6 | 278 | 149 | 1 | 1 | 0 | 0 | 0 | 0.177 | 0 | NA | 0 | 0 | 0 | 0.211 | -0.017 |
| SE | 46081 | ENSG00000068097 | HEATR6 | chr17 | - | 60076129 | 60076237 | 60072214 | 60072329 | 60078695 | 60078912 | 6 | 4 | 3 | 11 | 2 | 0 | 0 | 0 | 0 | 0 | 9 | 14 | 7 | 7 | 4 | 0 | 0 | 1 | 0 | 0 | 257 | 149 | 1 | 1 | 1 | 1 | 1 | 1 | 1 | 1 | 1 | 0.802 | 1 | 1 | 0.04 |
| SE | 46083 | ENSG00000148468 | FAM171A1 | chr10 | - | 15215995 | 15216110 | 15211642 | 15214601 | 15220943 | 15221060 | 5 | 8 | 3 | 6 | 16 | 1 | 0 | 2 | 0 | 0 | 0 | 0 | 7 | 0 | 5 | 0 | 0 | 0 | 0 | 8 | 264 | 149 | 1 | 1 | 0.738 | 1 | 0.458 | 1 | 1 | NA | NA | 1 | NA | 0.261 | 0.209 |
| SE | 46084 | ENSG00000148468 | FAM171A1 | chr10 | - | 15248638 | 15248815 | 15220943 | 15221060 | 15254720 | 15254879 | 11 | 1 | 8 | 6 | 5 | 0 | 0 | 0 | 0 | 4 | 0 | 20 | 1 | 0 | 9 | 0 | 0 | 0 | 0 | 0 | 298 | 149 | 1 | 1 | 1 | 1 | 1 | 1 | 0.385 | NA | 1 | 1 | NA | 1 | -0.123 |
| SE | 46089 | ENSG00000146007 | ZMAT2 | chr5 | + | 140703917 | 140703991 | 140702005 | 140702129 | 140704425 | 140704502 | 379 | 232 | 376 | 632 | 691 | 0 | 0 | 0 | 0 | 0 | 1591 | 1003 | 887 | 1054 | 982 | 0 | 1 | 0 | 0 | 0 | 223 | 149 | 1 | 1 | 1 | 1 | 1 | 1 | 1 | 1 | 0.999 | 1 | 1 | 1 | 0 |
| SE | 46090 | ENSG00000146007 | ZMAT2 | chr5 | + | 140703917 | 140704004 | 140702005 | 140702129 | 140704425 | 140704571 | 140 | 73 | 149 | 277 | 145 | 0 | 0 | 0 | 0 | 0 | 622 | 382 | 379 | 407 | 400 | 0 | 1 | 0 | 0 | 0 | 236 | 149 | 1 | 1 | 1 | 1 | 1 | 1 | 1 | 1 | 0.996 | 1 | 1 | 1 | 0.001 |
| SE | 46092 | ENSG00000166912 | MTMR10 | chr15 | - | 30948301 | 30948471 | 30947053 | 30947300 | 30953561 | 30953631 | 77 | 101 | 100 | 110 | 201 | 0 | 0 | 0 | 2 | 0 | 143 | 138 | 147 | 149 | 40 | 0 | 0 | 0 | 0 | 0 | 298 | 149 | 1 | 1 | 1 | 1 | 1 | 0.965 | 1 | 1 | 1 | 1 | 1 | 1 | -0.007 |
| SE | 46094 | ENSG00000166912 | MTMR10 | chr15 | - | 30954762 | 30954893 | 30953561 | 30953631 | 30958862 | 30958951 | 283 | 272 | 266 | 352 | 352 | 0 | 0 | 0 | 0 | 0 | 479 | 447 | 426 | 428 | 156 | 6 | 0 | 0 | 0 | 0 | 280 | 149 | 1 | 1 | 1 | 1 | 1 | 1 | 1 | 0.977 | 1 | 1 | 1 | 1 | 0.005 |
| SE | 46096 | ENSG00000166912 | MTMR10 | chr15 | - | 30959033 | 30959121 | 30958862 | 30958951 | 30960880 | 30960933 | 206 | 200 | 244 | 300 | 411 | 0 | 2 | 0 | 3 | 2 | 289 | 357 | 402 | 404 | 171 | 0 | 0 | 0 | 4 | 0 | 237 | 149 | 1 | 1 | 1 | 0.984 | 1 | 0.984 | 0.992 | 1 | 1 | 1 | 0.984 | 1 | -0.005 |
| SE | 46100 | ENSG00000166912 | MTMR10 | chr15 | - | 30967919 | 30968010 | 30965874 | 30966040 | 30974313 | 30974456 | 97 | 99 | 101 | 128 | 236 | 0 | 0 | 0 | 1 | 0 | 201 | 183 | 178 | 177 | 73 | 0 | 0 | 0 | 0 | 0 | 240 | 149 | 1 | 1 | 1 | 1 | 1 | 0.988 | 1 | 1 | 1 | 1 | 1 | 1 | -0.002 |
| SE | 46102 | ENSG00000166912 | MTMR10 | chr15 | - | 30990776 | 30990837 | 30976818 | 30976955 | 30991446 | 30991578 | 206 | 225 | 163 | 217 | 228 | 1 | 0 | 0 | 1 | 0 | 214 | 258 | 277 | 221 | 119 | 0 | 0 | 0 | 0 | 0 | 210 | 149 | 1 | 1 | 0.993 | 1 | 1 | 0.994 | 1 | 1 | 1 | 1 | 1 | 1 | -0.003 |
| SE | 46104 | ENSG00000106714 | CNTNAP3 | chr9 | - | 39085735 | 39085823 | 39078689 | 39078920 | 39086715 | 39086849 | 0 | 0 | 0 | 0 | 2 | 0 | 0 | 0 | 0 | 0 | 0 | 0 | 0 | 3 | 0 | 0 | 0 | 8 | 0 | 0 | 237 | 149 | 1 | 1 | NA | NA | NA | NA | 1 | NA | NA | 0 | 1 | NA | 0.5 |
| SE | 46106 | ENSG00000106714 | CNTNAP3 | chr9 | - | 39102496 | 39102715 | 39099910 | 39100150 | 39103743 | 39103914 | 0 | 0 | 0 | 1 | 0 | 0 | 0 | 0 | 0 | 0 | 0 | 0 | 2 | 0 | 0 | 0 | 0 | 4 | 0 | 0 | 298 | 149 | 1 | 1 | NA | NA | NA | 1 | NA | NA | NA | 0.2 | NA | NA | 0.8 |
| SE | 46107 | ENSG00000106714 | CNTNAP3 | chr9 | - | 39106484 | 39106607 | 39103690 | 39103914 | 39109159 | 39109287 | 0 | 0 | 0 | 2 | 0 | 0 | 0 | 0 | 1 | 0 | 0 | 0 | 0 | 0 | 0 | 0 | 0 | 9 | 4 | 0 | 272 | 149 | 1 | 1 | NA | NA | NA | 0.523 | NA | NA | NA | 0 | 0 | NA | 0.523 |
| SE | 46110 | ENSG00000106714 | CNTNAP3 | chr9 | - | 39144239 | 39144346 | 39140518 | 39140638 | 39149805 | 39149977 | 0 | 0 | 0 | 0 | 0 | 0 | 0 | 0 | 2 | 0 | 0 | 0 | 3 | 3 | 0 | 0 | 0 | 3 | 0 | 0 | 256 | 149 | 1 | 1 | NA | NA | NA | 0 | NA | NA | NA | 0.368 | 1 | NA | -0.684 |
| SE | 46111 | ENSG00000106714 | CNTNAP3 | chr9 | - | 39149805 | 39149977 | 39140518 | 39140638 | 39165932 | 39166076 | 1 | 1 | 0 | 3 | 3 | 0 | 0 | 0 | 2 | 0 | 0 | 0 | 16 | 0 | 0 | 0 | 0 | 11 | 16 | 6 | 298 | 149 | 1 | 1 | 1 | 1 | NA | 0.429 | 1 | NA | NA | 0.421 | 0 | 0 | 0.717 |
| SE | 46114 | ENSG00000147889 | CDKN2A | chr9 | - | 21993881 | 21994053 | 21970901 | 21971208 | 21994138 | 21994411 | 0 | 1 | 0 | 0 | 0 | 0 | 0 | 5 | 0 | 0 | 0 | 0 | 0 | 0 | 0 | 0 | 0 | 0 | 4 | 3 | 298 | 149 | 1 | 1 | NA | 1 | 0 | NA | NA | NA | NA | NA | 0 | 0 | 0.5 |
| SE | 46117 | ENSG00000181315 | ZNF322 | chr6 | - | 26658557 | 26658646 | 26643658 | 26643728 | 26659440 | 26659685 | 42 | 19 | 14 | 36 | 70 | 1 | 2 | 1 | 5 | 2 | 0 | 24 | 8 | 14 | 0 | 0 | 1 | 3 | 0 | 0 | 238 | 149 | 1 | 1 | 0.963 | 0.856 | 0.898 | 0.818 | 0.956 | NA | 0.938 | 0.625 | 1 | NA | 0.044 |
| SE | 46143 | ENSG00000272888 | CHASERR | chr15 | + | 92885514 | 92885584 | 92882706 | 92883186 | 92886168 | 92886222 | 409 | 384 | 483 | 494 | 1012 | 0 | 0 | 0 | 0 | 2 | 322 | 473 | 662 | 371 | 326 | 0 | 0 | 0 | 0 | 0 | 219 | 149 | 1 | 1 | 1 | 1 | 1 | 1 | 0.997 | 1 | 1 | 1 | 1 | 1 | -0.001 |
| SE | 46144 | ENSG00000272888 | CHASERR | chr15 | + | 92892003 | 92892072 | 92886168 | 92886254 | 92897723 | 92897975 | 920 | 669 | 929 | 933 | 2446 | 0 | 1 | 0 | 0 | 0 | 1028 | 801 | 1430 | 923 | 810 | 0 | 1 | 0 | 0 | 0 | 218 | 149 | 1 | 1 | 1 | 0.998 | 1 | 1 | 1 | 1 | 0.998 | 1 | 1 | 1 | 0 |
| SE | 46145 | ENSG00000126070 | AGO3 | chr1 | + | 35966954 | 35967075 | 35945691 | 35945863 | 35972023 | 35972232 | 121 | 89 | 124 | 129 | 201 | 0 | 1 | 1 | 0 | 0 | 60 | 119 | 169 | 99 | 112 | 0 | 0 | 2 | 0 | 0 | 270 | 149 | 1 | 1 | 1 | 0.98 | 0.986 | 1 | 1 | 1 | 1 | 0.979 | 1 | 1 | -0.003 |
| SE | 46146 | ENSG00000126070 | AGO3 | chr1 | + | 35973374 | 35973511 | 35972023 | 35972232 | 36004340 | 36004475 | 116 | 103 | 157 | 138 | 221 | 0 | 0 | 3 | 1 | 0 | 68 | 184 | 178 | 97 | 98 | 0 | 0 | 0 | 0 | 0 | 286 | 149 | 1 | 1 | 1 | 1 | 0.965 | 0.986 | 1 | 1 | 1 | 1 | 1 | 1 | -0.01 |
| SE | 46149 | ENSG00000126070 | AGO3 | chr1 | + | 36004340 | 36004475 | 35973374 | 35973511 | 36008689 | 36008777 | 88 | 72 | 72 | 85 | 151 | 0 | 0 | 0 | 0 | 2 | 58 | 64 | 48 | 50 | 9 | 0 | 0 | 0 | 0 | 0 | 284 | 149 | 1 | 1 | 1 | 1 | 1 | 1 | 0.975 | 1 | 1 | 1 | 1 | 1 | -0.005 |
| SE | 46151 | ENSG00000126070 | AGO3 | chr1 | + | 36013629 | 36013752 | 36009474 | 36009594 | 36013914 | 36014048 | 210 | 160 | 121 | 240 | 176 | 2 | 0 | 0 | 0 | 0 | 168 | 123 | 112 | 164 | 104 | 0 | 0 | 2 | 0 | 0 | 272 | 149 | 1 | 1 | 0.983 | 1 | 1 | 1 | 1 | 1 | 1 | 0.968 | 1 | 1 | 0.003 |
| SE | 46152 | ENSG00000126070 | AGO3 | chr1 | + | 36013914 | 36014048 | 36013629 | 36013752 | 36027113 | 36027298 | 165 | 128 | 125 | 180 | 169 | 0 | 0 | 1 | 0 | 0 | 159 | 89 | 105 | 115 | 70 | 0 | 0 | 0 | 0 | 0 | 283 | 149 | 1 | 1 | 1 | 1 | 0.985 | 1 | 1 | 1 | 1 | 1 | 1 | 1 | -0.003 |
| SE | 46153 | ENSG00000126070 | AGO3 | chr1 | + | 36027113 | 36027298 | 36013914 | 36014048 | 36034173 | 36034333 | 92 | 68 | 102 | 100 | 97 | 0 | 2 | 0 | 0 | 0 | 95 | 68 | 65 | 38 | 37 | 0 | 1 | 3 | 0 | 0 | 298 | 149 | 1 | 1 | 1 | 0.944 | 1 | 1 | 1 | 1 | 0.971 | 0.915 | 1 | 1 | 0.012 |
| SE | 46154 | ENSG00000126070 | AGO3 | chr1 | + | 36034173 | 36034333 | 36027113 | 36027298 | 36036176 | 36036267 | 69 | 80 | 70 | 78 | 90 | 1 | 0 | 0 | 0 | 0 | 56 | 64 | 55 | 31 | 33 | 0 | 0 | 0 | 0 | 0 | 298 | 149 | 1 | 1 | 0.972 | 1 | 1 | 1 | 1 | 1 | 1 | 1 | 1 | 1 | -0.006 |
| SE | 46155 | ENSG00000126070 | AGO3 | chr1 | + | 36036176 | 36036267 | 36034173 | 36034333 | 36039789 | 36039984 | 73 | 73 | 52 | 58 | 103 | 0 | 0 | 0 | 1 | 0 | 52 | 61 | 54 | 22 | 19 | 0 | 0 | 0 | 0 | 0 | 240 | 149 | 1 | 1 | 1 | 1 | 1 | 0.973 | 1 | 1 | 1 | 1 | 1 | 1 | -0.005 |
| SE | 46156 | ENSG00000126070 | AGO3 | chr1 | + | 36043446 | 36043548 | 36040306 | 36040441 | 36054945 | 36055145 | 79 | 63 | 45 | 76 | 113 | 1 | 0 | 0 | 0 | 0 | 54 | 107 | 55 | 80 | 30 | 0 | 0 | 0 | 0 | 0 | 251 | 149 | 1 | 1 | 0.979 | 1 | 1 | 1 | 1 | 1 | 1 | 1 | 1 | 1 | -0.004 |
| SE | 46157 | ENSG00000126070 | AGO3 | chr1 | + | 36054945 | 36055145 | 36043446 | 36043548 | 36055636 | 36056239 | 60 | 75 | 56 | 67 | 125 | 1 | 0 | 0 | 2 | 2 | 26 | 88 | 44 | 67 | 18 | 0 | 0 | 0 | 0 | 0 | 298 | 149 | 1 | 1 | 0.968 | 1 | 1 | 0.944 | 0.969 | 1 | 1 | 1 | 1 | 1 | -0.024 |
| SE | 46161 | ENSG00000213593 | TMX2 | chr11 | + | 57737835 | 57738026 | 57737607 | 57737668 | 57738973 | 57739039 | 15 | 16 | 21 | 32 | 24 | 0 | 0 | 0 | 0 | 0 | 26 | 32 | 27 | 34 | 48 | 0 | 0 | 0 | 2 | 0 | 298 | 149 | 1 | 1 | 1 | 1 | 1 | 1 | 1 | 1 | 1 | 1 | 0.895 | 1 | 0.021 |
| SE | 46163 | ENSG00000213593 | TMX2 | chr11 | + | 57737912 | 57738026 | 57737607 | 57737668 | 57738973 | 57739039 | 46 | 18 | 25 | 26 | 44 | 0 | 0 | 0 | 0 | 0 | 50 | 45 | 17 | 39 | 54 | 0 | 0 | 0 | 2 | 0 | 263 | 149 | 1 | 1 | 1 | 1 | 1 | 1 | 1 | 1 | 1 | 1 | 0.917 | 1 | 0.017 |
| SE | 46165 | ENSG00000213593 | TMX2 | chr11 | + | 57738353 | 57738430 | 57737835 | 57738026 | 57738663 | 57738770 | 148 | 99 | 131 | 140 | 188 | 1 | 0 | 3 | 0 | 0 | 174 | 99 | 95 | 179 | 152 | 0 | 0 | 0 | 0 | 4 | 226 | 149 | 1 | 1 | 0.99 | 1 | 0.966 | 1 | 1 | 1 | 1 | 1 | 1 | 0.962 | -0.001 |
| SE | 46166 | ENSG00000213593 | TMX2 | chr11 | + | 57738663 | 57738770 | 57737607 | 57737668 | 57738973 | 57739039 | 88 | 63 | 91 | 97 | 116 | 0 | 0 | 0 | 0 | 0 | 135 | 114 | 62 | 116 | 120 | 0 | 0 | 0 | 2 | 0 | 256 | 149 | 1 | 1 | 1 | 1 | 1 | 1 | 1 | 1 | 1 | 1 | 0.971 | 1 | 0.006 |
| SE | 46171 | ENSG00000213593 | TMX2 | chr11 | + | 57738973 | 57739039 | 57738663 | 57738770 | 57739130 | 57739212 | 234 | 145 | 186 | 210 | 303 | 1 | 0 | 0 | 0 | 0 | 278 | 210 | 160 | 250 | 266 | 0 | 0 | 0 | 0 | 0 | 215 | 149 | 1 | 1 | 0.994 | 1 | 1 | 1 | 1 | 1 | 1 | 1 | 1 | 1 | -0.001 |
| SE | 46172 | ENSG00000236438 | FAM157A | chr3 | + | 198157122 | 198157669 | 198153286 | 198153409 | 198167675 | 198167871 | 11 | 8 | 4 | 9 | 0 | 5 | 0 | 0 | 0 | 0 | 53 | 8 | 12 | 51 | 1 | 0 | 0 | 0 | 9 | 0 | 298 | 149 | 1 | 1 | 0.524 | 1 | 1 | 1 | NA | 1 | 1 | 1 | 0.739 | 1 | -0.067 |
| SE | 46174 | ENSG00000100280 | AP1B1 | chr22 | - | 29339753 | 29339774 | 29338989 | 29339133 | 29340655 | 29340857 | 0 | 0 | 3 | 0 | 0 | 40 | 14 | 30 | 30 | 4 | 0 | 6 | 0 | 0 | 0 | 9 | 84 | 30 | 38 | 153 | 170 | 149 | 1 | 1 | 0 | 0 | 0.081 | 0 | 0 | 0 | 0.059 | 0 | 0 | 0 | 0.004 |
| SE | 46176 | ENSG00000100280 | AP1B1 | chr22 | - | 29350034 | 29350150 | 29349217 | 29349383 | 29354649 | 29354871 | 88 | 29 | 46 | 63 | 82 | 0 | 0 | 0 | 0 | 0 | 146 | 65 | 25 | 52 | 186 | 0 | 0 | 0 | 0 | 4 | 265 | 149 | 1 | 1 | 1 | 1 | 1 | 1 | 1 | 1 | 1 | 1 | 1 | 0.963 | 0.007 |
| SE | 46177 | ENSG00000100280 | AP1B1 | chr22 | - | 29351170 | 29351266 | 29350034 | 29350150 | 29354649 | 29354871 | 119 | 68 | 62 | 80 | 98 | 0 | 0 | 0 | 0 | 0 | 176 | 115 | 46 | 86 | 319 | 0 | 0 | 1 | 0 | 0 | 245 | 149 | 1 | 1 | 1 | 1 | 1 | 1 | 1 | 1 | 1 | 0.965 | 1 | 1 | 0.007 |
| SE | 46178 | ENSG00000100280 | AP1B1 | chr22 | - | 29351704 | 29351825 | 29351170 | 29351266 | 29354649 | 29354871 | 339 | 133 | 203 | 220 | 251 | 1 | 0 | 0 | 0 | 0 | 567 | 394 | 134 | 215 | 949 | 0 | 0 | 3 | 0 | 0 | 270 | 149 | 1 | 1 | 0.995 | 1 | 1 | 1 | 1 | 1 | 1 | 0.961 | 1 | 1 | 0.007 |
| SE | 46179 | ENSG00000100280 | AP1B1 | chr22 | - | 29356425 | 29356616 | 29354649 | 29354871 | 29358725 | 29358971 | 100 | 32 | 63 | 59 | 50 | 0 | 0 | 0 | 1 | 0 | 146 | 141 | 59 | 95 | 441 | 0 | 0 | 0 | 1 | 0 | 298 | 149 | 1 | 1 | 1 | 1 | 1 | 0.967 | 1 | 1 | 1 | 1 | 0.979 | 1 | -0.002 |
| SE | 46180 | ENSG00000100280 | AP1B1 | chr22 | - | 29363000 | 29363106 | 29359672 | 29359959 | 29367206 | 29367267 | 340 | 158 | 224 | 265 | 401 | 0 | 0 | 0 | 0 | 0 | 522 | 405 | 185 | 350 | 1097 | 0 | 0 | 0 | 0 | 5 | 255 | 149 | 1 | 1 | 1 | 1 | 1 | 1 | 1 | 1 | 1 | 1 | 1 | 0.992 | 0.002 |
| SE | 46181 | ENSG00000100280 | AP1B1 | chr22 | - | 29367206 | 29367267 | 29363000 | 29363106 | 29388423 | 29388529 | 110 | 58 | 63 | 75 | 124 | 0 | 0 | 0 | 0 | 2 | 168 | 133 | 85 | 132 | 423 | 0 | 0 | 0 | 0 | 0 | 210 | 149 | 1 | 1 | 1 | 1 | 1 | 1 | 0.978 | 1 | 1 | 1 | 1 | 1 | -0.004 |
| SE | 46182 | ENSG00000100280 | AP1B1 | chr22 | - | 29367206 | 29367270 | 29363000 | 29363106 | 29388423 | 29388570 | 236 | 96 | 151 | 187 | 151 | 0 | 0 | 0 | 0 | 2 | 316 | 265 | 136 | 234 | 759 | 0 | 0 | 0 | 0 | 0 | 213 | 149 | 1 | 1 | 1 | 1 | 1 | 1 | 0.981 | 1 | 1 | 1 | 1 | 1 | -0.004 |
| SE | 46191 | ENSG00000185158 | LRRC37B | chr17 | + | 32027768 | 32027840 | 32024710 | 32024782 | 32030655 | 32030727 | 89 | 76 | 97 | 92 | 147 | 0 | 2 | 0 | 1 | 0 | 159 | 91 | 97 | 107 | 57 | 0 | 0 | 0 | 0 | 0 | 221 | 149 | 1 | 1 | 1 | 0.962 | 1 | 0.984 | 1 | 1 | 1 | 1 | 1 | 1 | -0.011 |
| SE | 46195 | ENSG00000185158 | LRRC37B | chr17 | + | 32031377 | 32031458 | 32030655 | 32030727 | 32034909 | 32034981 | 149 | 149 | 186 | 178 | 335 | 0 | 0 | 0 | 0 | 0 | 423 | 162 | 222 | 257 | 151 | 0 | 0 | 0 | 1 | 0 | 230 | 149 | 1 | 1 | 1 | 1 | 1 | 1 | 1 | 1 | 1 | 1 | 0.994 | 1 | 0.001 |
| SE | 46198 | ENSG00000185158 | LRRC37B | chr17 | + | 32049101 | 32049394 | 32047166 | 32047901 | 32050002 | 32050006 | 66 | 73 | 80 | 90 | 85 | 0 | 0 | 0 | 0 | 0 | 101 | 63 | 90 | 156 | 49 | 0 | 0 | 1 | 0 | 0 | 298 | 149 | 1 | 1 | 1 | 1 | 1 | 1 | 1 | 1 | 1 | 0.978 | 1 | 1 | 0.004 |
| SE | 46199 | ENSG00000185158 | LRRC37B | chr17 | + | 32050002 | 32050107 | 32049101 | 32049394 | 32053265 | 32053360 | 40 | 31 | 28 | 45 | 38 | 0 | 0 | 0 | 0 | 0 | 30 | 17 | 27 | 34 | 21 | 4 | 0 | 0 | 0 | 0 | 254 | 149 | 1 | 1 | 1 | 1 | 1 | 1 | 1 | 0.815 | 1 | 1 | 1 | 1 | 0.037 |
| SE | 46201 | ENSG00000117533 | VAMP4 | chr1 | - | 171719170 | 171719221 | 171710713 | 171710814 | 171728523 | 171728570 | 260 | 197 | 207 | 209 | 305 | 0 | 0 | 0 | 0 | 0 | 244 | 154 | 137 | 138 | 226 | 3 | 0 | 0 | 0 | 0 | 200 | 149 | 1 | 1 | 1 | 1 | 1 | 1 | 1 | 0.984 | 1 | 1 | 1 | 1 | 0.003 |
| SE | 46202 | ENSG00000117533 | VAMP4 | chr1 | - | 171728523 | 171728570 | 171719170 | 171719221 | 171738348 | 171738463 | 317 | 236 | 239 | 234 | 343 | 3 | 0 | 1 | 0 | 0 | 276 | 215 | 188 | 189 | 304 | 0 | 2 | 0 | 0 | 0 | 196 | 149 | 1 | 1 | 0.988 | 1 | 0.995 | 1 | 1 | 1 | 0.988 | 1 | 1 | 1 | -0.001 |
| SE | 46203 | ENSG00000114439 | BBX | chr3 | + | 107638501 | 107638545 | 107526326 | 107526398 | 107641790 | 107641864 | 1 | 0 | 0 | 0 | 0 | 0 | 0 | 4 | 4 | 0 | 0 | 0 | 0 | 0 | 0 | 5 | 0 | 8 | 0 | 0 | 193 | 149 | 1 | 1 | 1 | NA | 0 | 0 | NA | 0 | NA | 0 | NA | NA | 0.333 |
| SE | 46204 | ENSG00000114439 | BBX | chr3 | + | 107638501 | 107638545 | 107526326 | 107526398 | 107645835 | 107645909 | 0 | 0 | 0 | 1 | 0 | 111 | 78 | 89 | 83 | 64 | 0 | 0 | 0 | 0 | 0 | 30 | 81 | 69 | 41 | 40 | 193 | 149 | 1 | 1 | 0 | 0 | 0 | 0.009 | 0 | 0 | 0 | 0 | 0 | 0 | 0.002 |
| SE | 46206 | ENSG00000114439 | BBX | chr3 | + | 107641790 | 107641864 | 107599312 | 107599588 | 107645835 | 107645909 | 2 | 8 | 5 | 5 | 5 | 3 | 7 | 1 | 2 | 5 | 7 | 14 | 11 | 0 | 0 | 6 | 0 | 0 | 0 | 5 | 223 | 149 | 1 | 1 | 0.308 | 0.433 | 0.77 | 0.626 | 0.401 | 0.438 | 1 | 1 | NA | 0 | -0.102 |
| SE | 46207 | ENSG00000114439 | BBX | chr3 | + | 107641790 | 107641864 | 107638501 | 107638545 | 107645835 | 107645909 | 3 | 7 | 5 | 5 | 5 | 0 | 0 | 0 | 2 | 0 | 7 | 14 | 14 | 0 | 0 | 0 | 0 | 0 | 0 | 0 | 223 | 149 | 1 | 1 | 1 | 1 | 1 | 0.626 | 1 | 1 | 1 | 1 | NA | NA | -0.075 |
| SE | 46213 | ENSG00000114439 | BBX | chr3 | + | 107677426 | 107677470 | 107645835 | 107645909 | 107710451 | 107710622 | 0 | 0 | 0 | 0 | 0 | 104 | 91 | 89 | 82 | 101 | 0 | 3 | 0 | 0 | 0 | 66 | 88 | 93 | 39 | 23 | 193 | 149 | 1 | 1 | 0 | 0 | 0 | 0 | 0 | 0 | 0.026 | 0 | 0 | 0 | -0.005 |
| SE | 46215 | ENSG00000114439 | BBX | chr3 | + | 107711274 | 107711364 | 107710451 | 107710622 | 107716606 | 107716712 | 0 | 0 | 0 | 1 | 3 | 346 | 257 | 326 | 355 | 649 | 0 | 0 | 0 | 0 | 0 | 184 | 292 | 374 | 161 | 248 | 239 | 149 | 1 | 1 | 0 | 0 | 0 | 0.002 | 0.003 | 0 | 0 | 0 | 0 | 0 | 0.001 |
| SE | 46217 | ENSG00000114439 | BBX | chr3 | + | 107778370 | 107778519 | 107774718 | 107774857 | 107791239 | 107791299 | 248 | 221 | 255 | 279 | 583 | 0 | 1 | 0 | 0 | 0 | 77 | 280 | 283 | 61 | 37 | 0 | 0 | 0 | 0 | 0 | 298 | 149 | 1 | 1 | 1 | 0.991 | 1 | 1 | 1 | 1 | 1 | 1 | 1 | 1 | -0.002 |
| SE | 46219 | ENSG00000114439 | BBX | chr3 | + | 107791239 | 107791299 | 107778370 | 107778519 | 107798522 | 107798659 | 133 | 114 | 134 | 109 | 324 | 0 | 3 | 0 | 4 | 0 | 44 | 161 | 157 | 23 | 11 | 0 | 0 | 3 | 0 | 0 | 209 | 149 | 1 | 1 | 1 | 0.964 | 1 | 0.951 | 1 | 1 | 1 | 0.974 | 1 | 1 | -0.012 |
| SE | 46221 | ENSG00000232995 | RGS5 | chr1 | - | 163305202 | 163305313 | 163300662 | 163300691 | 163306232 | 163306329 | 4 | 3 | 3 | 7 | 9 | 0 | 0 | 0 | 0 | 3 | 0 | 8 | 4 | 0 | 0 | 0 | 0 | 0 | 0 | 0 | 260 | 149 | 1 | 1 | 1 | 1 | 1 | 1 | 0.632 | NA | 1 | 1 | NA | NA | -0.074 |
| SE | 46222 | ENSG00000232995 | RGS5 | chr1 | - | 163306232 | 163306329 | 163248274 | 163248706 | 163319181 | 163319291 | 0 | 2 | 6 | 4 | 4 | 0 | 1 | 0 | 0 | 0 | 0 | 6 | 1 | 0 | 0 | 0 | 0 | 0 | 0 | 0 | 246 | 149 | 1 | 1 | NA | 0.548 | 1 | 1 | 1 | NA | 1 | 1 | NA | NA | -0.113 |
| SE | 46228 | ENSG00000023909 | GCLM | chr1 | - | 93897838 | 93897898 | 93896617 | 93896820 | 93901584 | 93901669 | 88 | 53 | 61 | 74 | 89 | 0 | 0 | 0 | 1 | 0 | 79 | 42 | 37 | 69 | 149 | 0 | 0 | 0 | 0 | 0 | 209 | 149 | 1 | 1 | 1 | 1 | 1 | 0.981 | | 1 | 1 | 1 | 1 | 1 | -0.004 |
| SE | 46229 | ENSG00000023909 | GCLM | chr1 | - | 93904522 | 93904588 | 93901584 | 93901669 | 93909037 | 93909430 | 39 | 20 | 18 | 34 | 18 | 2 | 1 | 0 | 0 | 0 | 19 | 9 | 10 | 30 | 21 | 0 | 0 | 0 | 0 | 0 | 215 | 149 | 1 | 1 | 0.931 | 0.933 | 1 | | 1 | 1 | 1 | 1 | 1 | 1 | -0.027 |
| SE | 46233 | ENSG00000251131 | NA | chr5 | + | 43023918 | 43024097 | 43019235 | 43019316 | 43024198 | 43024607 | 2 | 3 | 5 | 2 | 0 | 0 | 0 | 0 | 0 | 6 | 13 | 0 | 0 | 1 | 0 | 0 | 0 | 0 | 0 | 0 | 298 | 149 | 1 | 1 | 1 | 1 | 1 | 1 | 0 | 1 | NA | NA | 1 | NA | -0.2 |
| SE | 46234 | ENSG00000251131 | NA | chr5 | + | 43023918 | 43024109 | 43019235 | 43019316 | 43024198 | 43024588 | 2 | 1 | 3 | 0 | 3 | 0 | 0 | 0 | 0 | 6 | 10 | 0 | 2 | 1 | 0 | 0 | 0 | 0 | 0 | 0 | 298 | 149 | 1 | 1 | 1 | 1 | 1 | NA | 0.2 | 1 | NA | 1 | 1 | NA | -0.2 |
| SE | 46235 | ENSG00000251131 | NA | chr5 | + | 43023918 | 43024109 | 43019310 | 43020098 | 43024198 | 43024588 | 2 | 0 | 2 | 0 | 3 | 0 | 0 | 3 | 0 | 0 | 0 | 0 | 2 | 1 | 0 | 0 | 0 | 0 | 0 | 0 | 298 | 149 | 1 | 1 | 1 | NA | 0.25 | NA | 1 | NA | NA | 1 | 1 | NA | -0.25 |
| SE | 46236 | ENSG00000139645 | ANKRD52 | chr12 | - | 56243784 | 56243876 | 56237806 | 56243392 | 56244050 | 56244133 | 96 | 38 | 80 | 71 | 166 | 2 | 0 | 0 | 2 | 0 | 117 | 92 | 50 | 72 | 68 | 0 | 0 | 1 | 0 | 0 | 241 | 149 | 1 | 1 | 0.967 | 1 | | 0.956 | 1 | 1 | 1 | 0.969 | 1 | 1 | -0.009 |
| SE | 46239 | ENSG00000139645 | ANKRD52 | chr12 | - | 56253721 | 56253800 | 56253287 | 56253402 | 56254066 | 56254279 | 49 | 25 | 43 | 31 | 23 | 0 | 1 | 0 | 0 | 0 | 16 | 76 | 31 | 56 | 73 | 0 | 0 | 0 | 0 | 0 | 228 | 149 | 1 | 1 | 1 | 0.942 | 1 | 1 | 1 | 1 | 1 | 1 | 1 | 1 | -0.012 |
| SE | 46240 | ENSG00000139645 | ANKRD52 | chr12 | - | 56254577 | 56254720 | 56254066 | 56254279 | 56254864 | 56254952 | 39 | 20 | 47 | 42 | 18 | 0 | 0 | 0 | 0 | 0 | 10 | 40 | 23 | 35 | 47 | 0 | 0 | 2 | 0 | 0 | 292 | 149 | 1 | 1 | 1 | 1 | 1 | 1 | 1 | 1 | 1 | 0.854 | 1 | 1 | 0.029 |
| SE | 46242 | ENSG00000139645 | ANKRD52 | chr12 | - | 56255783 | 56255984 | 56254864 | 56254952 | 56257014 | 56257068 | 25 | 6 | 21 | 18 | 8 | 0 | 0 | 1 | 0 | 0 | 44 | 11 | 15 | 6 | 4 | 0 | 0 | 0 | 0 | 0 | 298 | 149 | 1 | 1 | 1 | 1 | 0.913 | 1 | 1 | 1 | 1 | 1 | 1 | 1 | -0.017 |
| SE | 46244 | ENSG00000198301 | SDAD1 | chr4 | - | 75957517 | 75957708 | 75957324 | 75957409 | 75957846 | 75957941 | 328 | 218 | 308 | 235 | 579 | 1 | 0 | 0 | 0 | 0 | 7 | 92 | 59 | 17 | 20 | 0 | 0 | 0 | 0 | 0 | 298 | 149 | 1 | 1 | 0.994 | 1 | 1 | 1 | 1 | 1 | 1 | 1 | 1 | 1 | -0.001 |
| SE | 46245 | ENSG00000198301 | SDAD1 | chr4 | - | 75960065 | 75960192 | 75957846 | 75957941 | 75961027 | 75961104 | 384 | 306 | 343 | 346 | 617 | 3 | 0 | 0 | 0 | 0 | 27 | 154 | 151 | 24 | 30 | 0 | 0 | 0 | 0 | 0 | 276 | 149 | 1 | 1 | 0.986 | 1 | 1 | 1 | 1 | 1 | 1 | 1 | 1 | 1 | -0.003 |
| SE | 46248 | ENSG00000198301 | SDAD1 | chr4 | - | 75977645 | 75977756 | 75975923 | 75975995 | 75981371 | 75981470 | 178 | 100 | 153 | 135 | 284 | 4 | 6 | 3 | 9 | 5 | 10 | 90 | 57 | 31 | 0 | 10 | 0 | 5 | 0 | 0 | 260 | 149 | 1 | 1 | 0.962 | 0.905 | 0.967 | 0.896 | 0.97 | 0.364 | 1 | 0.867 | 1 | NA | 0.132 |
| SE | 46252 | ENSG00000135148 | TRAFD1 | chr12 | + | 112133139 | 112133320 | 112130510 | 112130569 | 112134737 | 112134873 | 0 | 0 | 0 | 1 | 0 | 293 | 173 | 229 | 228 | 156 | 0 | 0 | 2 | 0 | 0 | 442 | 363 | 261 | 300 | 493 | 298 | 149 | 1 | 1 | 0 | 0 | 0 | 0.002 | 0 | 0 | 0 | 0.004 | 0 | 0 | 0 |
| SE | 46253 | ENSG00000135148 | TRAFD1 | chr12 | + | 112140118 | 112140205 | 112135012 | 112135066 | 112140818 | 112141224 | 0 | 0 | 0 | 5 | 0 | 307 | 173 | 198 | 202 | 309 | 0 | 0 | 0 | 9 | 5 | 468 | 330 | 217 | 397 | 486 | 236 | 149 | 1 | 1 | 0 | 0 | 0 | 0.015 | 0 | 0 | 0 | 0 | 0.014 | 0.006 | -0.001 |
| SE | 46255 | ENSG00000135148 | TRAFD1 | chr12 | + | 112145585 | 112145662 | 112141952 | 112142295 | 112148073 | 112148213 | 198 | 131 | 140 | 137 | 220 | 0 | 0 | 0 | 0 | 0 | 185 | 251 | 120 | 234 | 267 | 0 | 0 | 0 | 0 | 7 | 226 | 149 | 1 | 1 | 1 | 1 | 1 | 1 | 1 | 1 | 1 | 1 | 1 | 0.962 | 0.008 |
| SE | 46256 | ENSG00000135148 | TRAFD1 | chr12 | + | 112148073 | 112148304 | 112145585 | 112145662 | 112149750 | 112149808 | 365 | 206 | 240 | 230 | 355 | 0 | 0 | 0 | 3 | 0 | 457 | 432 | 206 | 423 | 516 | 0 | 0 | 0 | 0 | 0 | 298 | 149 | 1 | 1 | 1 | 1 | 1 | 0.975 | 1 | 1 | 1 | 1 | 1 | 1 | -0.005 |
| SE | 46259 | ENSG00000162851 | TFB2M | chr1 | - | 246544520 | 246544681 | 246540560 | 246541202 | 246548544 | 246548607 | 95 | 98 | 105 | 117 | 201 | 0 | 1 | 0 | 0 | 0 | 118 | 118 | 87 | 99 | 35 | 0 | 0 | 0 | 0 | 0 | 298 | 149 | 1 | 1 | 1 | 0.98 | 1 | 1 | 1 | 1 | 1 | 1 | 1 | 1 | -0.004 |
| SE | 46260 | ENSG00000162851 | TFB2M | chr1 | - | 246548544 | 246548607 | 246544520 | 246544681 | 246551212 | 246551302 | 85 | 74 | 87 | 88 | 192 | 0 | 0 | 0 | 2 | 0 | 70 | 91 | 74 | 69 | 24 | 0 | 4 | 0 | 0 | 0 | 212 | 149 | 1 | 1 | 1 | 1 | 1 | 0.969 | 1 | 1 | 0.941 | 1 | 1 | 1 | 0.006 |
| SE | 46261 | ENSG00000162851 | TFB2M | chr1 | - | 246548544 | 246548607 | 246544520 | 246544681 | 246556572 | 246556721 | 30 | 29 | 29 | 40 | 44 | 1 | 0 | 0 | 0 | 0 | 44 | 38 | 33 | 35 | 11 | 0 | 0 | 0 | 0 | 0 | 212 | 149 | 1 | 1 | 0.955 | 1 | 1 | 1 | 1 | 1 | 1 | 1 | 1 | 1 | -0.009 |
| SE | 46262 | ENSG00000162851 | TFB2M | chr1 | - | 246551212 | 246551302 | 246544520 | 246544681 | 246556572 | 246556721 | 33 | 13 | 22 | 31 | 55 | 1 | 0 | 0 | 0 | 0 | 0 | 30 | 28 | 24 | 5 | 0 | 0 | 0 | 0 | 0 | 239 | 149 | 1 | 1 | 0.954 | 1 | 1 | 1 | 1 | NA | 1 | 1 | 1 | 1 | -0.009 |
| SE | 46263 | ENSG00000162851 | TFB2M | chr1 | - | 246551212 | 246551302 | 246548544 | 246548607 | 246556572 | 246556721 | 86 | 63 | 81 | 80 | 202 | 0 | 0 | 0 | 3 | 0 | 26 | 79 | 68 | 58 | 18 | 0 | 0 | 0 | 0 | 0 | 239 | 149 | 1 | 1 | 1 | 1 | 1 | 0.943 | 1 | 1 | 1 | 1 | 1 | 1 | -0.011 |
| SE | 46264 | ENSG00000162851 | TFB2M | chr1 | - | 246556572 | 246556721 | 246551212 | 246551302 | 246564345 | 246564434 | 66 | 34 | 63 | 59 | 116 | 0 | 0 | 0 | 1 | 0 | 15 | 46 | 54 | 38 | 13 | 0 | 0 | 0 | 0 | 0 | 298 | 149 | 1 | 1 | 1 | 1 | 1 | 0.967 | 1 | 1 | 1 | 1 | 1 | 1 | -0.007 |
| SE | 46266 | ENSG00000162851 | TFB2M | chr1 | - | 246564345 | 246564434 | 246557380 | 246557534 | 246565825 | 246566261 | 78 | 41 | 62 | 69 | 43 | 0 | 0 | 0 | 1 | 0 | 19 | 51 | 46 | 54 | 13 | 0 | 0 | 0 | 0 | 0 | 238 | 149 | 1 | 1 | 1 | 1 | 1 | 0.977 | 1 | 1 | 1 | 1 | 1 | 1 | -0.005 |
| SE | 46267 | ENSG00000231752 | EMBP1 | chr1 | + | 121538322 | 121538420 | 121519131 | 121519459 | 121556509 | 121556674 | 8 | 3 | 6 | 4 | 1 | 2 | 0 | 0 | 0 | 0 | 3 | 0 | 1 | 4 | 1 | 0 | 0 | 0 | 0 | 0 | 247 | 149 | 1 | 1 | 0.707 | 1 | 1 | 1 | 1 | 1 | NA | 1 | 1 | 1 | -0.059 |
| SE | 46270 | ENSG00000107625 | DDX50 | chr10 | + | 68910306 | 68910382 | 68906710 | 68907007 | 68911067 | 68911246 | 165 | 120 | 171 | 150 | 312 | 2 | 0 | 0 | 0 | 0 | 15 | 155 | 113 | 82 | 5 | 0 | 0 | 0 | 0 | 0 | 225 | 149 | 1 | 1 | 0.982 | 1 | 1 | 1 | 1 | 1 | 1 | 1 | 1 | 1 | -0.004 |
| SE | 46271 | ENSG00000107625 | DDX50 | chr10 | + | 68911067 | 68911246 | 68910306 | 68910382 | 68913161 | 68913230 | 189 | 156 | 211 | 198 | 261 | 0 | 2 | 0 | 0 | 0 | 38 | 195 | 128 | 163 | 28 | 0 | 0 | 0 | 0 | 0 | 298 | 149 | 1 | 1 | 1 | 0.975 | 1 | 1 | 1 | 1 | 1 | 1 | 1 | 1 | -0.005 |
| SE | 46272 | ENSG00000107625 | DDX50 | chr10 | + | 68914058 | 68914204 | 68913390 | 68913576 | 68919831 | 68919981 | 239 | 152 | 203 | 189 | 285 | 1 | 1 | 0 | 3 | 1 | 64 | 101 | 111 | 115 | 3 | 0 | 0 | 1 | 0 | 0 | 295 | 149 | 1 | 1 | 0.992 | 0.987 | 1 | 0.97 | 0.993 | 1 | 1 | 0.982 | 1 | 1 | -0.008 |
| SE | 46273 | ENSG00000107625 | DDX50 | chr10 | + | 68934198 | 68934360 | 68919831 | 68919981 | 68934798 | 68934918 | 172 | 73 | 161 | 179 | 478 | 0 | 2 | 1 | 1 | 8 | 12 | 83 | 71 | 40 | 0 | 0 | 0 | 3 | 0 | 0 | 298 | 149 | 1 | 1 | 1 | 0.948 | 0.988 | 0.989 | 0.968 | 1 | 1 | 0.922 | 1 | NA | -0.002 |
| SE | 46274 | ENSG00000107625 | DDX50 | chr10 | + | 68936005 | 68936079 | 68919853 | 68919981 | 68941059 | 68941194 | 0 | 0 | 0 | 0 | 1 | 1 | 0 | 0 | 0 | 0 | 0 | 0 | 3 | 0 | 0 | 0 | 0 | 0 | 0 | 0 | 223 | 149 | 1 | 1 | 0 | NA | NA | NA | 1 | NA | NA | 1 | NA | NA | -0.5 |
| SE | 46275 | ENSG00000107625 | DDX50 | chr10 | + | 68936005 | 68936079 | 68934798 | 68934918 | 68936935 | 68937095 | 79 | 43 | 73 | 82 | 182 | 0 | 0 | 0 | 1 | 0 | 18 | 36 | 35 | 20 | 20 | 0 | 0 | 0 | 0 | 0 | 223 | 149 | 1 | 1 | 1 | 1 | 1 | 0.982 | 1 | 1 | 1 | 1 | 1 | 1 | -0.004 |
| SE | 46276 | ENSG00000107625 | DDX50 | chr10 | + | 68936005 | 68936079 | 68934798 | 68934918 | 68941059 | 68941194 | 24 | 7 | 16 | 24 | 42 | 0 | 0 | 0 | 0 | 2 | 0 | 12 | 11 | 2 | 0 | 0 | 0 | 1 | 0 | 0 | 223 | 149 | 1 | 1 | 1 | 1 | 1 | 1 | 0.933 | NA | 1 | 0.88 | 1 | NA | 0.027 |
| SE | 46277 | ENSG00000107625 | DDX50 | chr10 | + | 68936005 | 68936079 | 68934798 | 68934918 | 68943212 | 68943257 | 23 | 7 | 16 | 29 | 40 | 0 | 0 | 1 | 0 | 0 | 0 | 12 | 8 | 2 | 0 | 0 | 0 | 0 | 0 | 0 | 223 | 149 | 1 | 1 | 1 | 1 | 0.914 | 1 | 1 | NA | 1 | 1 | 1 | NA | -0.017 |
| SE | 46278 | ENSG00000107625 | DDX50 | chr10 | + | 68936935 | 68937095 | 68934798 | 68934918 | 68941059 | 68941194 | 74 | 78 | 66 | 76 | 125 | 0 | 0 | 0 | 0 | 2 | 13 | 49 | 56 | 35 | 14 | 0 | 0 | 1 | 0 | 0 | 298 | 149 | 1 | 1 | 1 | 1 | 1 | 1 | 0.969 | 1 | 1 | 0.966 | 1 | 1 | 0.001 |
| SE | 46280 | ENSG00000107625 | DDX50 | chr10 | + | 68936935 | 68937095 | 68936005 | 68936079 | 68941059 | 68941194 | 131 | 114 | 123 | 133 | 267 | 2 | 0 | 0 | 0 | 2 | 31 | 73 | 83 | 53 | 34 | 0 | 0 | 3 | 0 | 0 | 298 | 149 | 1 | 1 | 0.97 | 1 | 1 | 1 | 0.985 | 1 | 1 | 0.933 | 1 | 1 | 0.004 |
| SE | 46282 | ENSG00000107625 | DDX50 | chr10 | + | 68941059 | 68941194 | 68934798 | 68934918 | 68943212 | 68943257 | 116 | 70 | 143 | 122 | 216 | 0 | 0 | 1 | 0 | 0 | 33 | 89 | 90 | 97 | 5 | 0 | 0 | 0 | 0 | 0 | 284 | 149 | 1 | 1 | 1 | 1 | 0.987 | 1 | 1 | 1 | 1 | 1 | 1 | 1 | -0.003 |
| SE | 46283 | ENSG00000107625 | DDX50 | chr10 | + | 68941059 | 68941194 | 68936005 | 68936079 | 68943212 | 68943257 | 118 | 70 | 143 | 122 | 216 | 1 | 0 | 0 | 5 | 0 | 33 | 89 | 92 | 97 | 5 | 0 | 0 | 0 | 0 | 0 | 284 | 149 | 1 | 1 | 0.984 | 1 | 1 | 0.928 | 1 | 1 | 1 | 1 | 1 | 1 | -0.018 |
| SE | 46284 | ENSG00000107625 | DDX50 | chr10 | + | 68941059 | 68941194 | 68936935 | 68937095 | 68943212 | 68943257 | 190 | 148 | 209 | 197 | 339 | 0 | 0 | 0 | 1 | 0 | 46 | 138 | 145 | 132 | 19 | 0 | 0 | 0 | 0 | 0 | 284 | 149 | 1 | 1 | 1 | 1 | 1 | 0.99 | 1 | 1 | 1 | 1 | 1 | 1 | -0.002 |
| SE | 46286 | ENSG00000151240 | DIP2C | chr10 | - | 422823 | 423033 | 419064 | 419199 | 440870 | 440996 | 4 | 8 | 17 | 7 | 4 | 0 | 0 | 0 | 0 | 0 | 0 | 2 | 9 | 0 | 0 | 0 | 0 | 6 | 0 | 0 | 298 | 149 | 1 | 1 | 1 | 1 | 1 | 1 | 1 | NA | 1 | 0.429 | NA | NA | 0.285 |
| SE | 46288 | ENSG00000197147 | LRRC8B | chr1 | + | 89526961 | 89526986 | 89524835 | 89525022 | 89568246 | 89568307 | 0 | 0 | 0 | 0 | 3 | 3 | 2 | 0 | 0 | 0 | 0 | 0 | 0 | 0 | 0 | 12 | 7 | 3 | 0 | 0 | 174 | 149 | 1 | 1 | 0 | 0 | NA | NA | 1 | 0 | 0 | 0 | NA | NA | 0.333 |
| SE | 46289 | ENSG00000197147 | LRRC8B | chr1 | + | 89549914 | 89549971 | 89526961 | 89526986 | 89568246 | 89568307 | 0 | 7 | 0 | 2 | 2 | 0 | 0 | 0 | 0 | 3 | 7 | 13 | 1 | 2 | 4 | 0 | 0 | 0 | 0 | 0 | 206 | 149 | 1 | 1 | NA | 1 | NA | 1 | 0.325 | 1 | 1 | 1 | 1 | 1 | -0.225 |
| SE | 46291 | ENSG00000197147 | LRRC8B | chr1 | + | 89572473 | 89572612 | 89568438 | 89568493 | 89579590 | 89579688 | 0 | 0 | 2 | 1 | 0 | 21 | 27 | 21 | 20 | 26 | 0 | 0 | 0 | 0 | 0 | 73 | 63 | 36 | 55 | 84 | 288 | 149 | 1 | 1 | 0 | 0 | 0.047 | 0.025 | 0 | 0 | 0 | 0 | 0 | 0 | 0.014 |
| SE | 46292 | ENSG00000197147 | LRRC8B | chr1 | + | 89588312 | 89588368 | 89582624 | 89584789 | 89592770 | 89593366 | 0 | 1 | 0 | 2 | 0 | 10 | 25 | 24 | 21 | 56 | 0 | 0 | 0 | 0 | 0 | 0 | 32 | 26 | 24 | 11 | 205 | 149 | 1 | 1 | 0 | 0.028 | 0 | 0.065 | 0 | NA | 0 | 0 | 0 | 0 | 0.019 |
| SE | 46295 | ENSG00000116830 | TTF2 | chr1 | + | 117073660 | 117073727 | 117062386 | 117062473 | 117074869 | 117075444 | 33 | 55 | 54 | 42 | 66 | 0 | 0 | 0 | 0 | 0 | 145 | 120 | 126 | 110 | 68 | 0 | 3 | 4 | 0 | 0 | 216 | 149 | 1 | 1 | 1 | 1 | 1 | 1 | 1 | 1 | 0.965 | 0.956 | 1 | 1 | 0.016 |
| SE | 46296 | ENSG00000116830 | TTF2 | chr1 | + | 117079567 | 117079649 | 117077915 | 117078043 | 117081827 | 117081947 | 46 | 58 | 50 | 51 | 71 | 0 | 0 | 0 | 0 | 0 | 38 | 35 | 64 | 57 | 34 | 0 | 0 | 2 | 0 | 0 | 231 | 149 | 1 | 1 | 1 | 1 | 1 | 1 | 1 | 1 | 1 | 0.954 | 1 | 1 | 0.009 |
| SE | 46298 | ENSG00000116830 | TTF2 | chr1 | + | 117086416 | 117086522 | 117081827 | 117081947 | 117088800 | 117088982 | 40 | 21 | 33 | 41 | 33 | 0 | 0 | 0 | 0 | 2 | 39 | 12 | 12 | 54 | 5 | 0 | 0 | 0 | 0 | 0 | 255 | 149 | 1 | 1 | 1 | 1 | 1 | 1 | 0.906 | 1 | 1 | 1 | 1 | 1 | -0.019 |
| SE | 46299 | ENSG00000116830 | TTF2 | chr1 | + | 117090054 | 117090208 | 117088800 | 117088982 | 117090531 | 117090623 | 75 | 59 | 47 | 69 | 54 | 0 | 0 | 0 | 0 | 0 | 53 | 93 | 30 | 68 | 28 | 0 | 0 | 2 | 0 | 0 | 298 | 149 | 1 | 1 | 1 | 1 | 1 | 1 | 1 | 1 | 1 | 0.882 | 1 | 1 | 0.024 |
| SE | 46300 | ENSG00000116830 | TTF2 | chr1 | + | 117091816 | 117091950 | 117091327 | 117091410 | 117092730 | 117092901 | 37 | 26 | 31 | 36 | 7 | 0 | 0 | 0 | 1 | 0 | 35 | 18 | 36 | 31 | 32 | 0 | 1 | 0 | 0 | 0 | 283 | 149 | 1 | 1 | 1 | 1 | 1 | 0.95 | 1 | 1 | 0.905 | 1 | 1 | 1 | 0.009 |
| SE | 46303 | ENSG00000116830 | TTF2 | chr1 | + | 117097350 | 117097433 | 117096148 | 117096299 | 117098832 | 117098907 | 20 | 36 | 23 | 35 | 38 | 2 | 0 | 1 | 0 | 0 | 35 | 22 | 12 | 30 | 4 | 0 | 0 | 0 | 0 | 0 | 232 | 149 | 1 | 1 | 0.865 | 1 | 0.937 | 1 | 1 | 1 | 1 | 1 | 1 | 1 | -0.04 |
| SE | 46306 | ENSG00000093183 | SEC22C | chr3 | - | 42555929 | 42555995 | 42547968 | 42553448 | 42557577 | 42557696 | 21 | 6 | 24 | 24 | 18 | 2 | 0 | 0 | 0 | 0 | 4 | 21 | 21 | 17 | 18 | 0 | 0 | 0 | 0 | 0 | 215 | 149 | 1 | 1 | 0.879 | 1 | 1 | 1 | 1 | 1 | 1 | 1 | 1 | 1 | -0.024 |
| SE | 46308 | ENSG00000093183 | SEC22C | chr3 | - | 42557577 | 42557696 | 42555929 | 42555995 | 42561116 | 42561296 | 59 | 18 | 43 | 67 | 52 | 0 | 0 | 1 | 0 | 0 | 29 | 55 | 39 | 35 | 32 | 0 | 0 | 0 | 0 | 0 | 268 | 149 | 1 | 1 | 1 | 1 | 0.96 | 1 | 1 | 1 | 1 | 1 | 1 | 1 | -0.008 |
| SE | 46313 | ENSG00000093183 | SEC22C | chr3 | - | 42565775 | 42565883 | 42563764 | 42563980 | 42568864 | 42569073 | 0 | 0 | 0 | 0 | 0 | 7 | 3 | 7 | 3 | 12 | 0 | 1 | 0 | 0 | 0 | 7 | 7 | 8 | 0 | 1 | 257 | 149 | 1 | 1 | 0 | 0 | 0 | 0 | 0 | 0 | 0.076 | 0 | NA | 0 | -0.019 |
| SE | 46314 | ENSG00000093183 | SEC22C | chr3 | - | 42566743 | 42566836 | 42563522 | 42563686 | 42568864 | 42569073 | 1 | 0 | 0 | 0 | 0 | 24 | 22 | 38 | 28 | 22 | 0 | 0 | 5 | 6 | 1 | 0 | 40 | 20 | 12 | 21 | 242 | 149 | 1 | 1 | 0.025 | 0 | 0 | 0 | 0 | NA | 0 | 0.133 | 0.235 | 0.028 | -0.094 |
| SE | 46315 | ENSG00000093183 | SEC22C | chr3 | - | 42566743 | 42566836 | 42563764 | 42563980 | 42568864 | 42569073 | 1 | 0 | 0 | 0 | 0 | 7 | 3 | 7 | 3 | 12 | 0 | 0 | 2 | 0 | 1 | 7 | 7 | 8 | 0 | 1 | 242 | 149 | 1 | 1 | 0.081 | 0 | 0 | 0 | 0 | 0 | 0 | 0.133 | NA | 0.381 | -0.112 |
| SE | 46316 | ENSG00000093183 | SEC22C | chr3 | - | 42579476 | 42579593 | 42568864 | 42569073 | 42581845 | 42582023 | 0 | 0 | 2 | 0 | 0 | 10 | 6 | 13 | 11 | 1 | 0 | 0 | 0 | 0 | 0 | 12 | 14 | 13 | 6 | 3 | 266 | 149 | 1 | 1 | 0 | 0 | 0.079 | 0 | 0 | 0 | 0 | 0 | 0 | 0 | 0.016 |
| SE | 46318 | ENSG00000093183 | SEC22C | chr3 | - | 42581311 | 42581365 | 42580442 | 42580505 | 42581845 | 42581943 | 0 | 0 | 0 | 0 | 0 | 0 | 0 | 1 | 3 | 2 | 0 | 0 | 0 | 0 | 4 | 0 | 0 | 0 | 1 | 0 | 203 | 149 | 1 | 1 | NA | NA | 0 | 0 | 0 | NA | NA | NA | 0 | 1 | -0.5 |
| SE | 46320 | ENSG00000095564 | BTAF1 | chr10 | + | 91939951 | 91940066 | 91935651 | 91935780 | 91942421 | 91942568 | 332 | 291 | 271 | 312 | 315 | 0 | 0 | 1 | 0 | 0 | 170 | 134 | 211 | 108 | 66 | 0 | 0 | 0 | 0 | 0 | 264 | 149 | 1 | 1 | 1 | 1 | 0.994 | 1 | 1 | 1 | 1 | 1 | 1 | 1 | -0.001 |
| SE | 46322 | ENSG00000095564 | BTAF1 | chr10 | + | 91942421 | 91942568 | 91935651 | 91935780 | 91951402 | 91951566 | 269 | 280 | 297 | 286 | 294 | 1 | 0 | 0 | 0 | 0 | 136 | 161 | 168 | 57 | 45 | 0 | 0 | 0 | 0 | 3 | 296 | 149 | 1 | 1 | 0.993 | 1 | 1 | 1 | 1 | 1 | 1 | 1 | 1 | 0.883 | 0.022 |
| SE | 46323 | ENSG00000095564 | BTAF1 | chr10 | + | 91942421 | 91942568 | 91939951 | 91940066 | 91951402 | 91951566 | 481 | 494 | 482 | 514 | 554 | 5 | 2 | 4 | 0 | 0 | 234 | 258 | 314 | 146 | 107 | 0 | 1 | 2 | 0 | 0 | 296 | 149 | 1 | 1 | 0.98 | 0.992 | 0.984 | 1 | 1 | 1 | 0.992 | 0.988 | 1 | 1 | -0.005 |
| SE | 46325 | ENSG00000095564 | BTAF1 | chr10 | + | 91953736 | 91953873 | 91951402 | 91951566 | 91959064 | 91959154 | 204 | 222 | 217 | 252 | 215 | 1 | 0 | 0 | 0 | 0 | 143 | 175 | 179 | 75 | 54 | 0 | 0 | 0 | 0 | 0 | 286 | 149 | 1 | 1 | 0.991 | 1 | 1 | 1 | 1 | 1 | 1 | 1 | 1 | 1 | -0.002 |
| SE | 46326 | ENSG00000095564 | BTAF1 | chr10 | + | 91956527 | 91956657 | 91951402 | 91951566 | 91959064 | 91959154 | 9 | 8 | 6 | 3 | 5 | 1 | 0 | 0 | 0 | 0 | 0 | 8 | 8 | 0 | 0 | 0 | 0 | 0 | 0 | 0 | 279 | 149 | 1 | 1 | 0.828 | 1 | 1 | 1 | 1 | NA | 1 | 1 | NA | NA | -0.034 |
| SE | 46328 | ENSG00000095564 | BTAF1 | chr10 | + | 91956527 | 91956657 | 91953736 | 91953873 | 91959064 | 91959154 | 226 | 253 | 272 | 294 | 206 | 0 | 0 | 0 | 5 | 0 | 63 | 228 | 210 | 120 | 40 | 0 | 1 | 5 | 0 | 0 | 279 | 149 | 1 | 1 | 1 | 1 | 1 | 0.969 | 1 | 1 | 0.992 | 0.957 | 1 | 1 | 0.004 |
| SE | 46330 | ENSG00000095564 | BTAF1 | chr10 | + | 91957224 | 91957293 | 91956527 | 91956657 | 91959064 | 91959154 | 684 | 583 | 619 | 667 | 777 | 3 | 5 | 5 | 2 | 4 | 188 | 360 | 526 | 154 | 46 | 0 | 0 | 0 | 0 | 0 | 218 | 149 | 1 | 1 | 0.994 | 0.988 | 0.988 | 0.996 | 0.993 | 1 | 1 | 1 | 1 | 1 | -0.008 |
| SE | 46331 | ENSG00000095564 | BTAF1 | chr10 | + | 91964076 | 91964201 | 91962537 | 91962678 | 91966636 | 91966757 | 547 | 445 | 563 | 616 | 724 | 1 | 0 | 0 | 0 | 0 | 66 | 235 | 403 | 173 | 37 | 0 | 0 | 0 | 0 | 0 | 274 | 149 | 1 | 1 | 0.997 | 1 | 1 | 1 | 1 | 1 | 1 | 1 | 1 | 1 | -0.001 |
| SE | 46334 | ENSG00000095564 | BTAF1 | chr10 | + | 91966636 | 91966757 | 91964076 | 91964201 | 91980453 | 91980558 | 412 | 331 | 463 | 482 | 459 | 0 | 7 | 0 | 0 | 0 | 72 | 171 | 313 | 108 | 32 | 0 | 2 | 0 | 0 | 0 | 270 | 149 | 1 | 1 | 1 | 0.963 | 1 | 1 | 1 | 1 | 0.979 | 1 | 1 | 1 | -0.003 |
| SE | 46337 | ENSG00000095564 | BTAF1 | chr10 | + | 91994534 | 91994644 | 91993693 | 91993847 | 91996368 | 91996570 | 221 | 198 | 258 | 189 | 192 | 0 | 0 | 0 | 0 | 2 | 44 | 83 | 116 | 58 | 29 | 0 | 0 | 0 | 0 | 0 | 259 | 149 | 1 | 1 | 1 | 1 | 1 | 1 | 0.982 | 1 | 1 | 1 | 1 | 1 | -0.004 |
| SE | 46339 | ENSG00000095564 | BTAF1 | chr10 | + | 92008828 | 92008950 | 92008122 | 92008275 | 92009040 | 92009208 | 496 | 352 | 431 | 395 | 795 | 0 | 0 | 0 | 2 | 0 | 86 | 163 | 245 | 70 | 24 | 0 | 0 | 0 | 0 | 0 | 271 | 149 | 1 | 1 | 1 | 1 | 1 | 0.991 | 1 | 1 | 1 | 1 | 1 | 1 | -0.002 |
| SE | 46340 | ENSG00000095564 | BTAF1 | chr10 | + | 92009040 | 92009208 | 92008122 | 92008275 | 92011072 | 92011150 | 162 | 131 | 156 | 149 | 212 | 1 | 0 | 0 | 1 | 0 | 30 | 66 | 69 | 14 | 16 | 0 | 0 | 0 | 0 | 0 | 298 | 149 | 1 | 1 | 0.988 | 1 | 1 | 0.987 | 1 | 1 | 1 | 1 | 1 | 1 | -0.005 |
| SE | 46341 | ENSG00000095564 | BTAF1 | chr10 | + | 92011072 | 92011150 | 92009040 | 92009208 | 92011285 | 92011415 | 236 | 202 | 214 | 224 | 351 | 0 | 0 | 0 | 0 | 1 | 30 | 80 | 104 | 22 | 25 | 0 | 0 | 0 | 0 | 0 | 227 | 149 | 1 | 1 | 1 | 1 | 1 | 1 | 0.996 | 1 | 1 | 1 | 1 | 1 | -0.001 |
| SE | 46342 | ENSG00000095564 | BTAF1 | chr10 | + | 92011285 | 92011415 | 92009040 | 92009208 | 92013666 | 92013808 | 137 | 100 | 124 | 149 | 218 | 2 | 0 | 0 | 0 | 0 | 8 | 39 | 59 | 9 | 5 | 0 | 0 | 0 | 0 | 0 | 279 | 149 | 1 | 1 | 0.973 | 1 | 1 | 1 | 1 | 1 | 1 | 1 | 1 | 1 | -0.005 |
| SE | 46343 | ENSG00000095564 | BTAF1 | chr10 | + | 92016339 | 92016465 | 92013898 | 92014029 | 92018782 | 92018935 | 459 | 361 | 466 | 445 | 684 | 0 | 1 | 0 | 0 | 0 | 10 | 163 | 171 | 87 | 68 | 0 | 0 | 0 | 0 | 0 | 275 | 149 | 1 | 1 | 1 | 0.995 | 1 | 1 | 1 | 1 | 1 | 1 | 1 | 1 | -0.001 |
| SE | 46344 | ENSG00000095564 | BTAF1 | chr10 | + | 92024755 | 92024967 | 92018782 | 92018935 | 92026591 | 92026751 | 283 | 249 | 338 | 416 | 328 | 0 | 0 | 1 | 0 | 0 | 22 | 110 | 167 | 39 | 21 | 0 | 0 | 0 | 1 | 0 | 298 | 149 | 1 | 1 | 1 | 1 | 0.994 | 1 | 1 | 1 | 1 | 1 | 0.951 | 1 | 0.009 |
| SE | 46346 | ENSG00000028310 | BRD9 | chr5 | - | 868606 | 869119 | 865413 | 865581 | 869244 | 869404 | 2 | 3 | 3 | 3 | 0 | 11 | 1 | 1 | 1 | 1 | 0 | 3 | 0 | 2 | 17 | 0 | 5 | 1 | 7 | 4 | 298 | 149 | 1 | 1 | 0.083 | 0.6 | 0.6 | 0.6 | 0 | NA | 0.231 | 0 | 0.125 | 0.68 | 0.118 |
| SE | 46348 | ENSG00000028310 | BRD9 | chr5 | - | 869037 | 869119 | 868606 | 868804 | 869244 | 869394 | 1 | 0 | 1 | 2 | 0 | 1 | 0 | 0 | 2 | 0 | 0 | 8 | 0 | 2 | 0 | 0 | 19 | 0 | 1 | 0 | 231 | 149 | 1 | 1 | 0.392 | NA | 1 | 0.392 | NA | NA | 0.214 | NA | 0.563 | NA | 0.206 |
| SE | 46349 | ENSG00000028310 | BRD9 | chr5 | - | 869037 | 869119 | 868606 | 868804 | 870472 | 870553 | 0 | 0 | 0 | 2 | 0 | 0 | 0 | 0 | 0 | 0 | 0 | 4 | 0 | 0 | 0 | 0 | 4 | 0 | 0 | 0 | 231 | 149 | 1 | 1 | NA | NA | NA | 1 | NA | NA | 0.392 | NA | NA | NA | 0.608 |
| SE | 46351 | ENSG00000028310 | BRD9 | chr5 | - | 869244 | 869394 | 868748 | 868804 | 870472 | 870575 | 6 | 4 | 12 | 7 | 0 | 0 | 0 | 0 | 0 | 0 | 12 | 28 | 8 | 7 | 4 | 0 | 4 | 0 | 0 | 0 | 298 | 149 | 1 | 1 | 1 | 1 | 1 | 1 | NA | 1 | 0.778 | 1 | 1 | 1 | 0.044 |
| SE | 46352 | ENSG00000028310 | BRD9 | chr5 | - | 869244 | 869394 | 869037 | 869119 | 870472 | 870575 | 6 | 4 | 13 | 6 | 0 | 0 | 0 | 0 | 2 | 0 | 12 | 13 | 8 | 8 | 4 | 0 | 0 | 0 | 0 | 0 | 298 | 149 | 1 | 1 | 1 | 1 | 1 | 0.6 | NA | 1 | 1 | 1 | 1 | 1 | -0.1 |
| SE | 46354 | ENSG00000028310 | BRD9 | chr5 | - | 869244 | 869404 | 868606 | 869119 | 870472 | 870553 | 4 | 0 | 1 | 6 | 6 | 0 | 0 | 0 | 2 | 0 | 0 | 6 | 4 | 9 | 3 | 0 | 0 | 0 | 0 | 0 | 298 | 149 | 1 | 1 | 1 | NA | 1 | 0.6 | 1 | NA | 1 | 1 | 1 | 1 | -0.1 |
| SE | 46355 | ENSG00000028310 | BRD9 | chr5 | - | 871525 | 871564 | 870287 | 870575 | 876100 | 876203 | 53 | 29 | 74 | 63 | 42 | 0 | 0 | 0 | 1 | 0 | 36 | 60 | 49 | 70 | 156 | 0 | 0 | 0 | 0 | 0 | 188 | 149 | 1 | 1 | 1 | 1 | 1 | 0.98 | 1 | 1 | 1 | 1 | 1 | 1 | -0.004 |
| SE | 46359 | ENSG00000028310 | BRD9 | chr5 | - | 883937 | 884070 | 879793 | 879889 | 886591 | 886707 | 32 | 25 | 35 | 25 | 44 | 0 | 0 | 0 | 0 | 0 | 44 | 20 | 15 | 17 | 64 | 0 | 0 | 1 | 0 | 0 | 282 | 149 | 1 | 1 | 1 | 1 | 1 | 1 | 1 | 1 | 1 | 0.888 | 1 | 1 | 0.022 |
| SE | 46360 | ENSG00000028310 | BRD9 | chr5 | - | 883937 | 884070 | 881106 | 881182 | 887360 | 887471 | 14 | 10 | 12 | 8 | 13 | 0 | 0 | 0 | 0 | 0 | 19 | 14 | 12 | 1 | 27 | 0 | 0 | 2 | 0 | 0 | 282 | 149 | 1 | 1 | 1 | 1 | 1 | 1 | 1 | 1 | 1 | 0.76 | 1 | 1 | 0.048 |
| SE | 46361 | ENSG00000028310 | BRD9 | chr5 | - | 886591 | 886707 | 882955 | 884070 | 887360 | 887471 | 77 | 44 | 76 | 70 | 94 | 1 | 0 | 0 | 0 | 0 | 134 | 75 | 57 | 86 | 140 | 0 | 2 | 0 | 0 | 0 | 265 | 149 | 1 | 1 | 0.977 | 1 | 1 | 1 | 1 | 1 | 0.955 | 1 | 1 | 1 | 0.004 |
| SE | 46362 | ENSG00000028310 | BRD9 | chr5 | - | 886591 | 886767 | 883937 | 884070 | 887360 | 887471 | 33 | 23 | 36 | 29 | 53 | 1 | 0 | 0 | 0 | 0 | 67 | 21 | 20 | 28 | 95 | 0 | 2 | 0 | 0 | 0 | 298 | 149 | 1 | 1 | 0.943 | 1 | 1 | 1 | 1 | 1 | 0.84 | 1 | 1 | 1 | 0.021 |
| SE | 46370 | ENSG00000163406 | SLC15A2 | chr3 | + | 121922219 | 121922302 | 121915615 | 121915693 | 121922774 | 121922861 | 11 | 7 | 8 | 34 | 0 | 0 | 0 | 0 | 1 | 0 | 6 | 11 | 1 | 5 | 22 | 0 | 0 | 0 | 0 | 11 | 232 | 149 | 1 | 1 | 1 | 1 | 1 | 0.956 | NA | 1 | 1 | 1 | 1 | 0.562 | 0.077 |
| SE | 46371 | ENSG00000163406 | SLC15A2 | chr3 | + | 121924944 | 121925033 | 121924350 | 121924383 | 121927757 | 121927839 | 12 | 10 | 7 | 11 | 20 | 0 | 0 | 2 | 2 | 0 | 7 | 1 | 2 | 0 | 18 | 0 | 0 | 0 | 0 | 0 | 238 | 149 | 1 | 1 | 1 | 1 | 0.687 | 0.775 | 1 | 1 | 1 | 1 | NA | 1 | -0.108 |
| SE | 46372 | ENSG00000163406 | SLC15A2 | chr3 | + | 121940383 | 121940488 | 121939190 | 121939495 | 121940830 | 121940982 | 16 | 4 | 6 | 10 | 6 | 0 | 0 | 0 | 0 | 0 | 6 | 13 | 8 | 5 | 13 | 0 | 0 | 2 | 0 | 0 | 254 | 149 | 1 | 1 | 1 | 1 | 1 | 1 | 1 | 1 | 1 | 0.701 | 1 | 1 | 0.06 |
| SE | 46374 | ENSG00000101986 | ABCD1 | chrX | + | 153740573 | 153740719 | 153740091 | 153740237 | 153742986 | 153743071 | 69 | 49 | 35 | 51 | 38 | 0 | 0 | 0 | 0 | 0 | 162 | 86 | 43 | 90 | 142 | 0 | 0 | 0 | 3 | 0 | 295 | 149 | 1 | 1 | 1 | 1 | 1 | 1 | 1 | 1 | 1 | 1 | 0.938 | 1 | 0.012 |
| SE | 46377 | ENSG00000160049 | DFFA | chr1 | - | 10463430 | 10463620 | 10462467 | 10463209 | 10469176 | 10469338 | 144 | 116 | 121 | 144 | 112 | 0 | 2 | 0 | 0 | 0 | 278 | 136 | 77 | 148 | 259 | 0 | 0 | 0 | 0 | 0 | 298 | 149 | 1 | 1 | 1 | 0.967 | 1 | 1 | 1 | 1 | 1 | 1 | 1 | 1 | -0.007 |
| SE | 46378 | ENSG00000160049 | DFFA | chr1 | - | 10467189 | 10467332 | 10462987 | 10463209 | 10469176 | 10469338 | 78 | 43 | 72 | 57 | 46 | 0 | 2 | 0 | 0 | 0 | 98 | 65 | 61 | 54 | 81 | 0 | 0 | 0 | 0 | 0 | 292 | 149 | 1 | 1 | 1 | 0.916 | 1 | 1 | 1 | 1 | 1 | 1 | 1 | 1 | -0.017 |
| SE | 46379 | ENSG00000160049 | DFFA | chr1 | - | 10467189 | 10467332 | 10463430 | 10463620 | 10469176 | 10469338 | 187 | 138 | 180 | 184 | 193 | 1 | 0 | 0 | 0 | 0 | 323 | 162 | 121 | 140 | 323 | 0 | 0 | 0 | 0 | 0 | 292 | 149 | 1 | 1 | 0.99 | 1 | 1 | 1 | 1 | 1 | 1 | 1 | 1 | 1 | -0.002 |
| SE | 46380 | ENSG00000160049 | DFFA | chr1 | - | 10469176 | 10469338 | 10467189 | 10467332 | 10472322 | 10472479 | 134 | 72 | 141 | 116 | 71 | 0 | 0 | 1 | 0 | 0 | 175 | 93 | 85 | 64 | 132 | 0 | 0 | 0 | 0 | 0 | 298 | 149 | 1 | 1 | 1 | 1 | 0.986 | 1 | 1 | 1 | 1 | 1 | 1 | 1 | -0.003 |
| SE | 46381 | ENSG00000136628 | EPRS1 | chr1 | - | 219982771 | 219982844 | 219981377 | 219981457 | 219983188 | 219983398 | 207 | 123 | 193 | 178 | 210 | 1 | 0 | 0 | 0 | 0 | 8 | 86 | 92 | 42 | 0 | 0 | 0 | 0 | 0 | 0 | 222 | 149 | 1 | 1 | 0.993 | 1 | 1 | 1 | 1 | 1 | 1 | 1 | 1 | NA | -0.001 |
| SE | 46382 | ENSG00000136628 | EPRS1 | chr1 | - | 219996982 | 219997342 | 219988589 | 219988823 | 220001137 | 220001255 | 352 | 236 | 404 | 338 | 645 | 1 | 2 | 1 | 0 | 1 | 27 | 241 | 158 | 72 | 16 | 0 | 0 | 0 | 0 | 0 | 298 | 149 | 1 | 1 | 0.994 | 0.983 | 0.995 | 1 | 0.997 | 1 | 1 | 1 | 1 | 1 | -0.006 |
| SE | 46383 | ENSG00000136628 | EPRS1 | chr1 | - | 220001137 | 220001255 | 219996982 | 219997342 | 220005247 | 220005360 | 358 | 301 | 402 | 369 | 778 | 1 | 0 | 0 | 0 | 0 | 14 | 257 | 185 | 111 | 8 | 0 | 0 | 0 | 0 | 0 | 267 | 149 | 1 | 1 | 0.995 | 1 | 1 | 1 | 1 | 1 | 1 | 1 | 1 | 1 | -0.001 |
| SE | 46384 | ENSG00000136628 | EPRS1 | chr1 | - | 220006105 | 220006313 | 220005247 | 220005360 | 220010945 | 220011056 | 316 | 226 | 274 | 307 | 579 | 1 | 0 | 0 | 1 | 0 | 15 | 236 | 152 | 85 | 6 | 0 | 0 | 0 | 5 | 0 | 298 | 149 | 1 | 1 | 0.994 | 1 | 1 | 0.994 | 1 | 1 | 1 | 1 | 0.895 | 1 | 0.019 |
| SE | 46385 | ENSG00000136628 | EPRS1 | chr1 | - | 220007201 | 220007338 | 220006105 | 220006313 | 220010945 | 220011056 | 573 | 490 | 655 | 575 | 828 | 0 | 0 | 0 | 0 | 1 | 18 | 424 | 367 | 233 | 34 | 0 | 0 | 0 | 0 | 0 | 286 | 149 | 1 | 1 | 1 | 1 | 1 | 1 | 0.998 | 1 | 1 | 1 | 1 | 1 | 0 |
| SE | 46386 | ENSG00000136628 | EPRS1 | chr1 | - | 220018125 | 220018146 | 220010945 | 220011056 | 220018448 | 220018508 | 0 | 0 | 0 | 0 | 1 | 266 | 255 | 316 | 295 | 439 | 0 | 0 | 0 | 0 | 0 | 8 | 229 | 211 | 113 | 0 | 170 | 149 | 1 | 1 | 0 | 0 | 0 | 0 | 0.002 | 0 | 0 | 0 | 0 | NA | 0 |
| SE | 46389 | ENSG00000136628 | EPRS1 | chr1 | - | 220025131 | 220025258 | 220024263 | 220024456 | 220030385 | 220030480 | 331 | 241 | 339 | 357 | 559 | 15 | 6 | 11 | 9 | 17 | 52 | 160 | 182 | 79 | 0 | 0 | 21 | 7 | 3 | 0 | 276 | 149 | 1 | 1 | 0.923 | 0.956 | 0.943 | 0.955 | 0.947 | 1 | 0.804 | 0.933 | 0.934 | NA | 0.027 |
| SE | 46391 | ENSG00000136628 | EPRS1 | chr1 | - | 220032386 | 220032526 | 220030385 | 220030480 | 220033501 | 220033658 | 302 | 159 | 263 | 265 | 478 | 1 | 0 | 0 | 0 | 0 | 41 | 131 | 122 | 83 | 22 | 0 | 0 | 0 | 0 | 0 | 289 | 149 | 1 | 1 | 0.994 | 1 | 1 | 1 | 1 | 1 | 1 | 1 | 1 | 1 | -0.001 |
| SE | 46392 | ENSG00000136628 | EPRS1 | chr1 | - | 220033501 | 220033658 | 220030385 | 220030480 | 220034913 | 220035013 | 250 | 117 | 182 | 194 | 154 | 0 | 0 | 0 | 0 | 0 | 0 | 77 | 63 | 86 | 1 | 0 | 0 | 1 | 0 | 0 | 298 | 149 | 1 | 1 | 1 | 1 | 1 | 1 | 1 | NA | 1 | 0.969 | 1 | 1 | 0.008 |
| SE | 46393 | ENSG00000178028 | DMAP1 | chr1 | + | 44214349 | 44214441 | 44213470 | 44213858 | 44214702 | 44214898 | 73 | 59 | 83 | 51 | 85 | 0 | 0 | 0 | 2 | 0 | 41 | 117 | 43 | 65 | 76 | 0 | 0 | 0 | 0 | 0 | 241 | 149 | 1 | 1 | 1 | 1 | 1 | 0.94 | 1 | 1 | 1 | 1 | 1 | 1 | -0.012 |
| SE | 46395 | ENSG00000248932 | NA | chr3 | + | 139419556 | 139419680 | 139389760 | 139389933 | 139423015 | 139423114 | 6 | 8 | 3 | 3 | 5 | 0 | 0 | 2 | 3 | 0 | 5 | 6 | 3 | 0 | 0 | 0 | 0 | 0 | 0 | 0 | 273 | 149 | 1 | 1 | 1 | 1 | 0.45 | 0.353 | 1 | 1 | 1 | 1 | NA | NA | -0.239 |
| SE | 46396 | ENSG00000248932 | NA | chr3 | + | 139419556 | 139419680 | 139389784 | 139390247 | 139423015 | 139423114 | 7 | 6 | 4 | 2 | 5 | 0 | 0 | 1 | 1 | 0 | 5 | 8 | 1 | 3 | 0 | 0 | 4 | 0 | 0 | 0 | 273 | 149 | 1 | 1 | 1 | 1 | 0.686 | 0.522 | 1 | 1 | 0.522 | 1 | 1 | NA | -0.039 |
| SE | 46397 | ENSG00000248932 | NA | chr3 | + | 139419556 | 139419680 | 139389875 | 139390429 | 139423015 | 139423114 | 9 | 4 | 4 | 2 | 3 | 0 | 0 | 1 | 1 | 0 | 13 | 6 | 1 | 0 | 0 | 0 | 0 | 0 | 0 | 0 | 273 | 149 | 1 | 1 | 1 | 1 | 1 | 0.522 | 1 | 1 | 1 | 1 | NA | NA | -0.096 |
| SE | 46400 | ENSG00000248932 | NA | chr3 | + | 139459930 | 139459985 | 139423015 | 139423114 | 139561652 | 139561748 | 0 | 0 | 0 | 1 | 0 | 0 | 0 | 4 | 1 | 3 | 0 | 3 | 1 | 0 | 0 | 0 | 0 | 0 | 0 | 0 | 204 | 149 | 1 | 1 | NA | NA | 0 | 0.422 | 0 | NA | 1 | 1 | NA | NA | -0.859 |
| SE | 46406 | ENSG00000168724 | DNAJC21 | chr5 | + | 34936143 | 34936266 | 34935709 | 34935833 | 34937329 | 34937630 | 103 | 98 | 86 | 135 | 145 | 0 | 0 | 0 | 0 | 0 | 61 | 88 | 117 | 89 | 35 | 0 | 4 | 0 | 0 | 0 | 272 | 149 | 1 | 1 | 1 | 1 | 1 | 1 | 1 | 1 | 0.923 | 1 | 1 | 1 | 0.015 |
| SE | 46407 | ENSG00000168724 | DNAJC21 | chr5 | + | 34941095 | 34941183 | 34938857 | 34939009 | 34944866 | 34945025 | 136 | 108 | 112 | 166 | 213 | 2 | 0 | 1 | 1 | 0 | 104 | 64 | 147 | 108 | 46 | 0 | 0 | 0 | 6 | 0 | 237 | 149 | 1 | 1 | 0.977 | 1 | 0.986 | 0.991 | 1 | 1 | 1 | 1 | 0.919 | 1 | 0.007 |
| SE | 46408 | ENSG00000168724 | DNAJC21 | chr5 | + | 34945721 | 34945803 | 34944866 | 34945025 | 34949542 | 34949677 | 4 | 1 | 8 | 9 | 2 | 0 | 2 | 0 | 0 | 0 | 8 | 5 | 1 | 5 | 0 | 0 | 0 | 0 | 0 | 0 | 231 | 149 | 1 | 1 | 1 | 0.244 | 1 | 1 | 1 | 1 | 1 | 1 | 1 | NA | -0.151 |
| SE | 46409 | ENSG00000168724 | DNAJC21 | chr5 | + | 34945760 | 34945803 | 34944866 | 34945025 | 34949542 | 34949677 | 54 | 33 | 77 | 70 | 94 | 0 | 2 | 0 | 0 | 0 | 4 | 21 | 41 | 74 | 0 | 0 | 0 | 0 | 0 | 0 | 192 | 149 | 1 | 1 | 1 | 0.928 | 1 | 1 | 1 | 1 | 1 | 1 | 1 | NA | -0.014 |
| SE | 46413 | ENSG00000164010 | ERMAP | chr1 | + | 42824547 | 42824752 | 42817123 | 42817253 | 42825622 | 42825738 | 0 | 0 | 0 | 0 | 15 | 7 | 7 | 1 | 6 | 0 | 0 | 0 | 0 | 0 | 0 | 15 | 6 | 0 | 0 | 9 | 298 | 149 | 1 | 1 | 0 | 0 | 0 | 0 | 1 | 0 | 0 | NA | NA | 0 | 0.2 |
| SE | 46414 | ENSG00000164010 | ERMAP | chr1 | + | 42824547 | 42824752 | 42817533 | 42817815 | 42825622 | 42825738 | 0 | 0 | 0 | 0 | 18 | 0 | 0 | 0 | 0 | 0 | 0 | 0 | 0 | 0 | 0 | 0 | 0 | 0 | 0 | 2 | 298 | 149 | 1 | 1 | NA | NA | NA | NA | 1 | NA | NA | NA | NA | 0 | 1 |
| SE | 46415 | ENSG00000164010 | ERMAP | chr1 | + | 42835037 | 42835154 | 42829914 | 42831115 | 42837157 | 42837190 | 16 | 6 | 3 | 8 | 12 | 1 | 0 | 0 | 0 | 0 | 78 | 14 | 19 | 19 | 43 | 0 | 0 | 0 | 0 | 0 | 266 | 149 | 1 | 1 | 0.9 | 1 | 1 | 1 | 1 | 1 | 1 | 1 | 1 | 1 | -0.02 |
| SE | 46418 | ENSG00000164010 | ERMAP | chr1 | + | 42840032 | 42840053 | 42838900 | 42838921 | 42840151 | 42840178 | 33 | 25 | 31 | 27 | 34 | 0 | 0 | 0 | 0 | 0 | 76 | 27 | 36 | 28 | 38 | 0 | 0 | 0 | 3 | 0 | 170 | 149 | 1 | 1 | 1 | 1 | 1 | 1 | 1 | 1 | 1 | 1 | 0.891 | 1 | 0.022 |
| SE | 46419 | ENSG00000118655 | DCLRE1B | chr1 | + | 113906995 | 113907161 | 113905212 | 113905775 | 113908008 | 113908191 | 50 | 6 | 20 | 20 | 30 | 0 | 0 | 1 | 1 | 2 | 35 | 34 | 18 | 32 | 21 | 3 | 0 | 0 | 0 | 0 | 298 | 149 | 1 | 1 | 1 | 1 | 0.909 | 0.909 | 0.882 | 0.854 | 1 | 1 | 1 | 1 | -0.031 |
| SE | 46420 | ENSG00000118655 | DCLRE1B | chr1 | + | 113906995 | 113907161 | 113905212 | 113905775 | 113911130 | 113912381 | 19 | 7 | 8 | 6 | 3 | 0 | 0 | 0 | 0 | 0 | 7 | 13 | 4 | 9 | 0 | 10 | 0 | 0 | 0 | 0 | 298 | 149 | 1 | 1 | 1 | 1 | 1 | 1 | 1 | 0.259 | 1 | 1 | 1 | NA | 0.185 |
| SE | 46424 | ENSG00000287725 | NA | chr11 | + | 69078891 | 69079020 | 69078733 | 69078793 | 69079833 | 69079883 | 15 | 10 | 12 | 12 | 7 | 0 | 0 | 0 | 0 | 0 | 19 | 20 | 15 | 10 | 28 | 0 | 0 | 0 | 1 | 0 | 278 | 149 | 1 | 1 | 1 | 1 | 1 | 1 | 1 | 1 | 1 | 1 | 0.843 | 1 | 0.031 |
| SE | 46428 | ENSG00000272047 | GTF2H5 | chr6 | + | 158170469 | 158170538 | 158168349 | 158168395 | 158191976 | 158199344 | 81 | 44 | 68 | 53 | 71 | 2 | 1 | 0 | 2 | 0 | 44 | 37 | 22 | 61 | 14 | 0 | 9 | 0 | 3 | 7 | 218 | 149 | 1 | 1 | 0.965 | 0.968 | 1 | 0.948 | 1 | 1 | 0.738 | 1 | 0.933 | 0.578 | 0.126 |
| SE | 46429 | ENSG00000272047 | GTF2H5 | chr6 | + | 158187039 | 158187121 | 158170469 | 158170538 | 158191976 | 158192881 | 1 | 2 | 1 | 0 | 2 | 86 | 44 | 69 | 57 | 71 | 0 | 0 | 0 | 0 | 0 | 45 | 39 | 23 | 61 | 17 | 231 | 149 | 1 | 1 | 0.007 | 0.028 | 0.009 | 0 | 0.018 | 0 | 0 | 0 | 0 | 0 | 0.012 |
| SE | 46432 | ENSG00000183475 | ASB7 | chr15 | + | 100629436 | 100630042 | 100612165 | 100612427 | 100648322 | 100651701 | 146 | 105 | 154 | 145 | 230 | 1 | 0 | 0 | 0 | 0 | 96 | 122 | 190 | 92 | 33 | 0 | 2 | 0 | 4 | 0 | 298 | 149 | 1 | 1 | 0.986 | 1 | 1 | 1 | 1 | 1 | 0.968 | 1 | 0.92 | 1 | 0.02 |
| SE | 46434 | ENSG00000150627 | WDR17 | chr4 | + | 176115795 | 176115979 | 176111574 | 176111703 | 176119866 | 176120097 | 9 | 3 | 2 | 4 | 4 | 1 | 0 | 0 | 0 | 0 | 0 | 0 | 0 | 9 | 0 | 0 | 0 | 0 | 0 | 0 | 298 | 149 | 1 | 1 | 0.818 | 1 | 1 | 1 | 1 | NA | NA | NA | 1 | NA | -0.036 |
| SE | 46436 | ENSG00000150627 | WDR17 | chr4 | + | 176145994 | 176146159 | 176141982 | 176142069 | 176148132 | 176148335 | 4 | 10 | 5 | 0 | 6 | 2 | 0 | 0 | 1 | 1 | 0 | 0 | 3 | 0 | 0 | 0 | 0 | 0 | 0 | 0 | 298 | 149 | 1 | 1 | 0.5 | 1 | 1 | 0 | 0.75 | NA | NA | 1 | NA | NA | -0.35 |
| SE | 46451 | ENSG00000115415 | STAT1 | chr2 | - | 190970497 | 190970717 | 190944702 | 190945657 | 190974829 | 190974932 | 723 | 888 | 841 | 827 | 1030 | 0 | 0 | 2 | 0 | 0 | 1176 | 817 | 546 | 1652 | 137 | 1 | 0 | 1 | 2 | 0 | 298 | 149 | 1 | 1 | 1 | 1 | 0.995 | 1 | 1 | 0.998 | 1 | 0.996 | 0.998 | 1 | 0.001 |
| SE | 46452 | ENSG00000115415 | STAT1 | chr2 | - | 190975309 | 190975388 | 190974829 | 190974932 | 190975811 | 190975887 | 3 | 13 | 8 | 9 | 5 | 509 | 634 | 654 | 581 | 926 | 15 | 6 | 11 | 10 | 0 | 843 | 526 | 407 | 1165 | 82 | 228 | 149 | 1 | 1 | 0.004 | 0.013 | 0.008 | 0.01 | 0.004 | 0.011 | 0.007 | 0.017 | 0.006 | 0 | 0 |
| SE | 46453 | ENSG00000115415 | STAT1 | chr2 | - | 190975309 | 190975434 | 190974829 | 190974932 | 190975811 | 190975887 | 9 | 8 | 9 | 12 | 1 | 509 | 634 | 654 | 581 | 926 | 4 | 0 | 12 | 5 | 0 | 843 | 526 | 407 | 1165 | 82 | 274 | 149 | 1 | 1 | 0.01 | 0.007 | 0.007 | 0.011 | 0.001 | 0.003 | 0 | 0.016 | 0.002 | 0 | 0.003 |
| SE | 46454 | ENSG00000115415 | STAT1 | chr2 | - | 190978527 | 190978606 | 190976839 | 190977025 | 190978855 | 190979001 | 0 | 1 | 6 | 3 | 5 | 323 | 234 | 430 | 375 | 402 | 4 | 0 | 0 | 1 | 0 | 485 | 360 | 161 | 612 | 106 | 228 | 149 | 1 | 1 | 0 | 0.003 | 0.009 | 0.005 | 0.008 | 0.005 | 0 | 0 | 0.001 | 0 | 0.004 |
| SE | 46456 | ENSG00000115415 | STAT1 | chr2 | - | 190983641 | 190983740 | 190982187 | 190982518 | 190984309 | 190984393 | 480 | 341 | 584 | 545 | 489 | 0 | 0 | 2 | 1 | 1 | 720 | 468 | 261 | 761 | 222 | 0 | 0 | 0 | 4 | 0 | 248 | 149 | 1 | 1 | 1 | 1 | 0.994 | 0.997 | 0.997 | 1 | 1 | 1 | 0.991 | 1 | -0.001 |
| SE | 46457 | ENSG00000115415 | STAT1 | chr2 | - | 190983641 | 190983758 | 190982382 | 190982518 | 190984309 | 190984393 | 124 | 89 | 170 | 160 | 94 | 0 | 0 | 2 | 1 | 1 | 158 | 119 | 75 | 177 | 62 | 0 | 0 | 0 | 4 | 0 | 266 | 149 | 1 | 1 | 1 | 1 | 0.979 | 0.989 | 0.981 | 1 | 1 | 1 | 0.961 | 1 | -0.002 |
| SE | 46458 | ENSG00000115415 | STAT1 | chr2 | - | 190989614 | 190989674 | 190987038 | 190987068 | 190991227 | 190991320 | 756 | 750 | 867 | 791 | 838 | 0 | 0 | 0 | 4 | 0 | 1306 | 738 | 449 | 1437 | 587 | 0 | 0 | 0 | 0 | 0 | 209 | 149 | 1 | 1 | 1 | 1 | 1 | 0.993 | 1 | 1 | 1 | 1 | 1 | 1 | -0.001 |
| SE | 46459 | ENSG00000115415 | STAT1 | chr2 | - | 190995060 | 190995208 | 190991227 | 190991320 | 190997855 | 190998007 | 568 | 589 | 655 | 604 | 661 | 0 | 2 | 1 | 0 | 0 | 945 | 523 | 340 | 1177 | 428 | 0 | 4 | 2 | 7 | 0 | 297 | 149 | 1 | 1 | 1 | 0.993 | 0.997 | 1 | 1 | 1 | 0.985 | 0.988 | 0.988 | 1 | 0.006 |
| SE | 46460 | ENSG00000115415 | STAT1 | chr2 | - | 190995060 | 190995219 | 190991227 | 190991320 | 190997855 | 190998007 | 1476 | 1643 | 1787 | 1699 | 2268 | 0 | 2 | 1 | 0 | 0 | 3647 | 1786 | 1095 | 3601 | 1120 | 0 | 4 | 2 | 7 | 0 | 298 | 149 | 1 | 1 | 1 | 0.998 | 0.999 | 1 | 1 | 1 | 0.996 | 0.996 | 0.996 | 1 | 0.002 |
| SE | 46467 | ENSG00000185864 | NPIPB4 | chr16 | - | 21840802 | 21841011 | 21839249 | 21839310 | 21843385 | 21843490 | 2 | 0 | 2 | 0 | 0 | 2 | 13 | 45 | 18 | 84 | 0 | 0 | 1 | 1 | 0 | 29 | 18 | 43 | 72 | 44 | 298 | 149 | 1 | 1 | 0.333 | 0 | 0.022 | 0 | 0 | 0 | 0 | 0.011 | 0.007 | 0 | 0.067 |
| SE | 46468 | ENSG00000185864 | NPIPB4 | chr16 | - | 21851769 | 21851855 | 21847412 | 21847541 | 21857258 | 21857416 | 0 | 0 | 4 | 17 | 13 | 0 | 2 | 16 | 1 | 3 | 0 | 8 | 6 | 13 | 19 | 0 | 30 | 7 | 2 | 6 | 235 | 149 | 1 | 1 | NA | 0 | 0.137 | 0.915 | 0.733 | NA | 0.145 | 0.352 | 0.805 | 0.668 | -0.046 |
| SE | 46469 | ENSG00000185864 | NPIPB4 | chr16 | - | 21851769 | 21852046 | 21847481 | 21847541 | 21857258 | 21857416 | 4 | 2 | 4 | 20 | 13 | 0 | 2 | 16 | 1 | 3 | 0 | 10 | 13 | 13 | 19 | 0 | 30 | 7 | 2 | 6 | 298 | 149 | 1 | 1 | 1 | 0.333 | 0.111 | 0.909 | 0.684 | NA | 0.143 | 0.481 | 0.765 | 0.613 | 0.107 |
| SE | 46470 | ENSG00000185864 | NPIPB4 | chr16 | - | 21851914 | 21852046 | 21847412 | 21847541 | 21857258 | 21857345 | 5 | 2 | 4 | 7 | 9 | 0 | 2 | 16 | 1 | 3 | 0 | 13 | 23 | 3 | 8 | 0 | 30 | 7 | 2 | 6 | 281 | 149 | 1 | 1 | 1 | 0.347 | 0.117 | 0.788 | 0.614 | NA | 0.187 | 0.635 | 0.443 | 0.414 | 0.153 |
| SE | 46471 | ENSG00000185864 | NPIPB4 | chr16 | - | 21854083 | 21854421 | 21851948 | 21852046 | 21857258 | 21857416 | 0 | 1 | 0 | 3 | 0 | 4 | 2 | 0 | 1 | 0 | 0 | 1 | 13 | 2 | 3 | 0 | 0 | 7 | 0 | 0 | 298 | 149 | 1 | 1 | 0 | 0.2 | NA | 0.6 | NA | NA | 1 | 0.481 | 1 | 1 | -0.604 |
| SE | 46474 | ENSG00000154518 | ATP5MC3 | chr2 | - | 175180097 | 175180178 | 175178185 | 175179250 | 175181354 | 175181466 | 434 | 188 | 378 | 305 | 340 | 2 | 0 | 0 | 0 | 0 | 557 | 380 | 263 | 392 | 363 | 0 | 0 | 0 | 0 | 0 | 230 | 149 | 1 | 1 | 0.993 | 1 | 1 | 1 | 1 | 1 | 1 | 1 | 1 | 1 | -0.001 |
| SE | 46478 | ENSG00000255154 | HTD2 | chr3 | + | 58310506 | 58310591 | 58306271 | 58306417 | 58316514 | 58316591 | 25 | 23 | 8 | 25 | 18 | 0 | 0 | 0 | 0 | 0 | 0 | 13 | 11 | 10 | 21 | 0 | 0 | 0 | 1 | 0 | 234 | 149 | 1 | 1 | 1 | 1 | 1 | 1 | 1 | NA | 1 | 1 | 0.864 | 1 | 0.034 |
| SE | 46479 | ENSG00000255154 | HTD2 | chr3 | + | 58315700 | 58315819 | 58310506 | 58310591 | 58316514 | 58316580 | 0 | 0 | 2 | 0 | 0 | 21 | 20 | 12 | 24 | 21 | 0 | 0 | 0 | 0 | 0 | 13 | 29 | 13 | 14 | 33 | 268 | 149 | 1 | 1 | 0 | 0 | 0.085 | 0 | 0 | 0 | 0 | 0 | 0 | 0 | 0.017 |
| SE | 46481 | ENSG00000255154 | HTD2 | chr3 | + | 58316514 | 58316591 | 58310506 | 58310591 | 58316914 | 58316993 | 52 | 27 | 22 | 29 | 45 | 2 | 0 | 0 | 0 | 0 | 13 | 50 | 27 | 41 | 69 | 0 | 0 | 0 | 0 | 0 | 226 | 149 | 1 | 1 | 0.945 | 1 | 1 | 1 | 1 | 1 | 1 | 1 | 1 | 1 | -0.011 |
| SE | 46482 | ENSG00000255154 | HTD2 | chr3 | + | 58316914 | 58316993 | 58316497 | 58316591 | 58317439 | 58317775 | 65 | 27 | 30 | 34 | 43 | 0 | 0 | 0 | 0 | 1 | 4 | 32 | 32 | 35 | 75 | 0 | 0 | 0 | 0 | 0 | 228 | 149 | 1 | 1 | 1 | 1 | 1 | 1 | 0.966 | 1 | 1 | 1 | 1 | 1 | -0.007 |
| SE | 46483 | ENSG00000065978 | YBX1 | chr1 | + | 42683402 | 42683466 | 42682417 | 42682731 | 42693489 | 42693523 | 1047 | 801 | 863 | 996 | 1221 | 0 | 0 | 0 | 1 | 0 | 2466 | 1113 | 1092 | 1226 | 2217 | 0 | 1 | 0 | 0 | 0 | 213 | 149 | 1 | 1 | 1 | 1 | 1 | 0.999 | 1 | 1 | 0.999 | 1 | 1 | 1 | 0 |
| SE | 46485 | ENSG00000065978 | YBX1 | chr1 | + | 42696641 | 42696944 | 42696198 | 42696288 | 42697179 | 42697211 | 1689 | 1209 | 1492 | 1675 | 2092 | 0 | 0 | 0 | 0 | 6 | 4402 | 1639 | 1795 | 2239 | 3900 | 5 | 0 | 0 | 12 | 0 | 298 | 149 | 1 | 1 | 1 | 1 | 1 | 1 | 0.994 | 0.998 | 1 | 1 | 0.989 | 1 | 0.001 |
| SE | 46486 | ENSG00000065978 | YBX1 | chr1 | + | 42697179 | 42697262 | 42696198 | 42696288 | 42700780 | 42701046 | 51 | 47 | 37 | 76 | 43 | 0 | 0 | 0 | 0 | 0 | 211 | 58 | 90 | 187 | 197 | 0 | 1 | 0 | 0 | 0 | 232 | 149 | 1 | 1 | 1 | 1 | 1 | 1 | 1 | 1 | 0.974 | 1 | 1 | 1 | 0.005 |

| | | | | | | | | | | | | | | | | | | | | | | | | | | | | | | | | | | | | | | | | | | | | | | |
|---|---|---|---|---|---|---|---|---|---|---|---|---|---|---|---|---|---|---|---|---|---|---|---|---|---|---|---|---|---|---|---|---|---|---|---|---|---|---|---|---|---|---|---|---|---|---|
| SE | 46495 | ENSG00000109184 | DCUN1D4 | chr4 | + | 51860542 | 51860663 | 51843102 | 51843267 | 51863436 | 51863507 | 2 | 1 | 0 | 2 | 0 | 12 | 2 | 9 | 10 | 2 | 0 | 0 | 0 | 0 | 2 | 0 | 5 | 2 | 2 | 4 | 270 | 149 | 1 | 1 | 0.084 | 0.216 | 0 | 0.099 | 0 | NA | 0 | 0 | 0 | 0.216 | 0.026 |
| SE | 46496 | ENSG00000109184 | DCUN1D4 | chr4 | + | 51860542 | 51860663 | 51848204 | 51848367 | 51863436 | 51863507 | 2 | 0 | 1 | 2 | 0 | 3 | 8 | 10 | 20 | 32 | 0 | 0 | 0 | 0 | 2 | 8 | 0 | 0 | 14 | 0 | 270 | 149 | 1 | 1 | 0.269 | 0 | 0.052 | 0.052 | 0 | 0 | NA | NA | 0 | 1 | -0.259 |
| SE | 46497 | ENSG00000109184 | DCUN1D4 | chr4 | + | 51863669 | 51863709 | 51863436 | 51863507 | 51874270 | 51874367 | 67 | 40 | 51 | 74 | 89 | 1 | 0 | 0 | 5 | 0 | 43 | 30 | 49 | 47 | 32 | 0 | 0 | 0 | 0 | 0 | 189 | 149 | 1 | 1 | 0.981 | 1 | 1 | 0.921 | 1 | 1 | 1 | 1 | 1 | 1 | -0.02 |
| SE | 46498 | ENSG00000109184 | DCUN1D4 | chr4 | + | 51886567 | 51886638 | 51877455 | 51877854 | 51899269 | 51899378 | 114 | 80 | 64 | 70 | 149 | 0 | 0 | 0 | 1 | 0 | 79 | 39 | 38 | 34 | 126 | 0 | 0 | 0 | 0 | 0 | 220 | 149 | 1 | 1 | 1 | 1 | 1 | 0.979 | 1 | 1 | 1 | 1 | 1 | 1 | -0.004 |
| SE | 46499 | ENSG00000109184 | DCUN1D4 | chr4 | + | 51886567 | 51887244 | 51877762 | 51877854 | 51899269 | 51899378 | 115 | 79 | 66 | 72 | 149 | 0 | 0 | 0 | 1 | 0 | 79 | 34 | 33 | 32 | 126 | 0 | 0 | 0 | 0 | 0 | 298 | 149 | 1 | 1 | 1 | 1 | 1 | 0.973 | 1 | 1 | 1 | 1 | 1 | 1 | -0.005 |
| SE | 46502 | ENSG00000109184 | DCUN1D4 | chr4 | + | 51891759 | 51891851 | 51886567 | 51887244 | 51899269 | 51899378 | 51 | 58 | 51 | 48 | 96 | 0 | 0 | 1 | 0 | 0 | 0 | 6 | 11 | 15 | 17 | 0 | 0 | 0 | 0 | 0 | 241 | 149 | 1 | 1 | 1 | 1 | 0.969 | 1 | 1 | NA | 1 | 1 | 1 | 1 | -0.006 |
| SE | 46503 | ENSG00000109184 | DCUN1D4 | chr4 | + | 51899269 | 51899378 | 51891759 | 51891851 | 51911069 | 51911114 | 105 | 102 | 77 | 83 | 199 | 0 | 0 | 0 | 0 | 1 | 0 | 19 | 22 | 22 | 6 | 0 | 10 | 0 | 0 | 0 | 258 | 149 | 1 | 1 | 1 | 1 | 1 | 1 | 0.991 | NA | 0.523 | 1 | 1 | 1 | 0.117 |
| SE | 46504 | ENSG00000109184 | DCUN1D4 | chr4 | + | 51905188 | 51905320 | 51899336 | 51899378 | 51911069 | 51911174 | 3 | 4 | 5 | 0 | 0 | 54 | 48 | 42 | 45 | 96 | 0 | 0 | 1 | 0 | 0 | 0 | 10 | 15 | 5 | 0 | 281 | 149 | 1 | 1 | 0.029 | 0.042 | 0.059 | 0 | 0 | NA | 0 | 0.034 | 0 | NA | 0.015 |
| SE | 46505 | ENSG00000109184 | DCUN1D4 | chr4 | + | 51908919 | 51908975 | 51899269 | 51899378 | 51911069 | 51911174 | 4 | 5 | 6 | 6 | 2 | 54 | 48 | 42 | 45 | 96 | 0 | 1 | 0 | 0 | 0 | 0 | 10 | 15 | 5 | 0 | 205 | 149 | 1 | 1 | 0.051 | 0.07 | 0.094 | 0.088 | 0.015 | NA | 0.068 | 0 | 0 | NA | 0.041 |
| SE | 46506 | ENSG00000109184 | DCUN1D4 | chr4 | + | 51911069 | 51911174 | 51899269 | 51899378 | 51913289 | 51913392 | 101 | 81 | 76 | 79 | 144 | 13 | 10 | 7 | 9 | 13 | 0 | 35 | 30 | 10 | 0 | 0 | 0 | 4 | 3 | 0 | 254 | 149 | 1 | 1 | 0.82 | 0.826 | 0.864 | 0.837 | 0.867 | NA | 1 | 0.815 | 0.662 | NA | 0.017 |
| SE | 46508 | ENSG00000177311 | ZBTB38 | chr3 | + | 141386871 | 141386937 | 141381424 | 141381487 | 141403926 | 141404031 | 6 | 2 | 4 | 12 | 8 | 0 | 0 | 0 | 0 | 0 | 0 | 3 | 2 | 0 | 0 | 0 | 1 | 0 | 0 | 0 | 215 | 149 | 1 | 1 | 1 | 1 | 1 | 1 | 1 | NA | 0.675 | 1 | NA | NA | 0.162 |
| SE | 46509 | ENSG00000177311 | ZBTB38 | chr3 | + | 141387440 | 141387887 | 141386878 | 141386937 | 141403926 | 141404016 | 0 | 0 | 4 | 3 | 2 | 6 | 4 | 8 | 5 | 11 | 0 | 0 | 0 | 0 | 0 | 0 | 4 | 2 | 0 | 0 | 298 | 149 | 1 | 1 | 0 | 0 | 0.2 | 0.231 | 0.083 | NA | 0 | 0 | NA | NA | 0.103 |
| SE | 46510 | ENSG00000177311 | ZBTB38 | chr3 | + | 141396258 | 141396503 | 141386871 | 141386937 | 141403926 | 141404031 | 1 | 2 | 2 | 2 | 11 | 6 | 4 | 8 | 5 | 11 | 0 | 8 | 8 | 0 | 0 | 0 | 4 | 2 | 0 | 0 | 298 | 149 | 1 | 1 | 0.077 | 0.2 | 0.111 | 0.167 | 0.333 | NA | 0.5 | 0.667 | NA | NA | -0.406 |
| SE | 46511 | ENSG00000177311 | ZBTB38 | chr3 | + | 141427633 | 141427821 | 141403926 | 141404031 | 141438242 | 141438365 | 2 | 1 | 0 | 0 | 0 | 8 | 2 | 7 | 4 | 5 | 0 | 4 | 0 | 0 | 0 | 0 | 3 | 4 | 0 | 0 | 298 | 149 | 1 | 1 | 0.111 | 0.2 | 0 | 0 | 0 | NA | 0.4 | 0 | NA | NA | -0.138 |
| SE | 46512 | ENSG00000177311 | ZBTB38 | chr3 | + | 141427633 | 141427821 | 141403926 | 141404031 | 141442388 | 141443020 | 3 | 2 | 0 | 0 | 0 | 113 | 61 | 66 | 76 | 115 | 0 | 4 | 2 | 0 | 0 | 16 | 94 | 107 | 8 | 0 | 298 | 149 | 1 | 1 | 0.013 | 0.016 | 0 | 0 | 0 | 0 | 0.021 | 0.009 | 0 | NA | -0.002 |
| SE | 46514 | ENSG00000177311 | ZBTB38 | chr3 | + | 141438242 | 141438365 | 141403926 | 141404031 | 141442388 | 141442670 | 36 | 11 | 27 | 18 | 23 | 113 | 61 | 66 | 76 | 115 | 0 | 22 | 26 | 3 | 0 | 16 | 94 | 107 | 8 | 0 | 272 | 149 | 1 | 1 | 0.149 | 0.09 | 0.183 | 0.115 | 0.099 | 0 | 0.114 | 0.117 | 0.17 | NA | 0.027 |
| SE | 46515 | ENSG00000177311 | ZBTB38 | chr3 | + | 141438242 | 141438365 | 141427633 | 141427821 | 141442388 | 141442670 | 28 | 9 | 20 | 14 | 18 | 1 | 1 | 0 | 0 | 0 | 0 | 19 | 22 | 3 | 0 | 0 | 0 | 2 | 0 | 0 | 272 | 149 | 1 | 1 | 0.939 | 0.831 | 1 | 1 | 1 | NA | 1 | 0.858 | 1 | NA | 0.001 |
| SE | 46516 | ENSG00000177311 | ZBTB38 | chr3 | + | 141438267 | 141438365 | 141403926 | 141404031 | 141442388 | 141442713 | 42 | 8 | 35 | 22 | 28 | 113 | 61 | 66 | 76 | 115 | 0 | 29 | 26 | 2 | 0 | 16 | 94 | 107 | 8 | 0 | 247 | 149 | 1 | 1 | 0.183 | 0.073 | 0.242 | 0.149 | 0.128 | 0 | 0.157 | 0.128 | 0.131 | NA | 0.051 |
| SE | 46520 | ENSG00000223745 | CCDC18-AS1 | chr1 | - | 93269682 | 93269758 | 93264637 | 93264772 | 93273473 | 93273583 | 4 | 3 | 5 | 6 | 3 | 0 | 1 | 1 | 1 | 1 | 12 | 0 | 0 | 0 | 0 | 0 | 2 | 2 | 0 | 0 | 225 | 149 | 1 | 1 | 1 | 0.665 | 0.768 | 0.799 | 0.665 | 1 | 0 | 0 | NA | NA | 0.446 |
| SE | 46521 | ENSG00000223745 | CCDC18-AS1 | chr1 | - | 93269682 | 93269758 | 93264637 | 93264772 | 93305387 | 93305470 | 4 | 8 | 5 | 3 | 3 | 13 | 23 | 32 | 38 | 29 | 12 | 0 | 0 | 0 | 0 | 21 | 16 | 22 | 27 | 0 | 225 | 149 | 1 | 1 | 0.169 | 0.187 | 0.094 | 0.05 | 0.064 | 0.275 | 0 | 0 | NA | NA | 0.044 |
| SE | 46522 | ENSG00000223745 | CCDC18-AS1 | chr1 | - | 93290564 | 93290686 | 93264668 | 93264772 | 93305387 | 93305470 | 3 | 0 | 1 | 1 | 1 | 13 | 23 | 32 | 38 | 29 | 0 | 0 | 0 | 0 | 0 | 21 | 16 | 22 | 27 | 0 | 271 | 149 | 1 | 1 | 0.113 | 0 | 0.017 | 0.014 | 0.019 | 0 | 0 | 0 | 0 | NA | 0.033 |
| SE | 46523 | ENSG00000223745 | CCDC18-AS1 | chr1 | - | 93305387 | 93305470 | 93264639 | 93264772 | 93311489 | 93311608 | 11 | 19 | 29 | 35 | 28 | 0 | 4 | 0 | 1 | 0 | 23 | 20 | 17 | 17 | 0 | 0 | 0 | 0 | 0 | 0 | 232 | 149 | 1 | 1 | 1 | 0.753 | 1 | 0.957 | 1 | 1 | 1 | 1 | 1 | NA | -0.058 |
| SE | 46524 | ENSG00000223745 | CCDC18-AS1 | chr1 | - | 93305387 | 93305470 | 93264639 | 93264772 | 93318167 | 93318228 | 5 | 15 | 22 | 32 | 25 | 0 | 2 | 1 | 0 | 0 | 21 | 15 | 19 | 20 | 0 | 0 | 4 | 2 | 2 | 0 | 232 | 149 | 1 | 1 | 1 | 0.828 | 0.934 | 1 | 1 | 1 | 0.707 | 0.859 | 0.865 | NA | 0.095 |
| SE | 46526 | ENSG00000223745 | CCDC18-AS1 | chr1 | - | 93305387 | 93305470 | 93264639 | 93264772 | 93337376 | 93337469 | 7 | 15 | 20 | 30 | 28 | 0 | 0 | 2 | 0 | 1 | 21 | 22 | 16 | 13 | 0 | 0 | 0 | 0 | 0 | 0 | 232 | 149 | 1 | 1 | 1 | 1 | 0.865 | 1 | 0.947 | 1 | 1 | 1 | 1 | NA | -0.038 |
| SE | 46527 | ENSG00000223745 | CCDC18-AS1 | chr1 | - | 93309539 | 93309634 | 93305387 | 93305470 | 93309714 | 93309770 | 0 | 1 | 0 | 0 | 1 | 13 | 2 | 5 | 5 | 3 | 0 | 0 | 0 | 0 | 0 | 6 | 6 | 7 | 6 | 0 | 244 | 149 | 1 | 1 | 0 | 0.234 | 0 | 0 | 0.169 | 0 | 0 | 0 | 0 | NA | 0.081 |
| SE | 46528 | ENSG00000223745 | CCDC18-AS1 | chr1 | - | 93309714 | 93309790 | 93305387 | 93305470 | 93318167 | 93318228 | 7 | 0 | 1 | 2 | 1 | 0 | 0 | 2 | 2 | 0 | 3 | 6 | 7 | 4 | 0 | 0 | 0 | 3 | 7 | 0 | 225 | 149 | 1 | 1 | 1 | NA | 0.249 | 0.398 | 1 | 1 | 1 | 0.607 | 0.275 | NA | -0.059 |
| SE | 46530 | ENSG00000223745 | CCDC18-AS1 | chr1 | - | 93309714 | 93309790 | 93305387 | 93305470 | 93339356 | 93339536 | 9 | 0 | 1 | 2 | 1 | 0 | 0 | 1 | 0 | 0 | 3 | 6 | 7 | 4 | 0 | 0 | 0 | 0 | 0 | 0 | 225 | 149 | 1 | 1 | 1 | NA | 0.398 | 1 | 1 | 1 | 1 | 1 | 1 | NA | -0.15 |
| SE | 46531 | ENSG00000223745 | CCDC18-AS1 | chr1 | - | 93311489 | 93311608 | 93264704 | 93264772 | 93324634 | 93324729 | 19 | 26 | 10 | 20 | 3 | 2 | 0 | 8 | 7 | 3 | 0 | 3 | 8 | 12 | 6 | 0 | 0 | 0 | 0 | 0 | 268 | 149 | 1 | 1 | 0.841 | 1 | 0.41 | 0.614 | 0.357 | NA | 1 | 1 | 1 | 1 | -0.356 |
| SE | 46534 | ENSG00000223745 | CCDC18-AS1 | chr1 | - | 93315148 | 93315269 | 93305387 | 93305470 | 93318167 | 93318228 | 0 | 1 | 0 | 1 | 0 | 0 | 0 | 2 | 2 | 0 | 0 | 0 | 0 | 1 | 0 | 0 | 0 | 3 | 7 | 0 | 270 | 149 | 1 | 1 | NA | 1 | 0 | 0.216 | NA | NA | NA | 0 | 0.073 | NA | 0.369 |
| SE | 46536 | ENSG00000223745 | CCDC18-AS1 | chr1 | - | 93318167 | 93318228 | 93264644 | 93264772 | 93324634 | 93324691 | 0 | 2 | 3 | 1 | 0 | 2 | 0 | 8 | 7 | 3 | 0 | 4 | 2 | 2 | 0 | 0 | 0 | 0 | 0 | 0 | 210 | 149 | 1 | 1 | 0 | 1 | 0.21 | 0.092 | 0 | NA | 1 | 1 | 1 | NA | -0.74 |
| SE | 46538 | ENSG00000223745 | CCDC18-AS1 | chr1 | - | 93318167 | 93318228 | 93305387 | 93305470 | 93339356 | 93339536 | 0 | 0 | 1 | 0 | 0 | 0 | 0 | 1 | 0 | 0 | 0 | 0 | 0 | 7 | 0 | 0 | 0 | 0 | 0 | 0 | 210 | 149 | 1 | 1 | NA | NA | 0.415 | NA | NA | NA | NA | NA | 1 | NA | -0.585 |
| SE | 46539 | ENSG00000223745 | CCDC18-AS1 | chr1 | - | 93318167 | 93318228 | 93309714 | 93309790 | 93324634 | 93324691 | 0 | 1 | 2 | 1 | 0 | 3 | 0 | 5 | 3 | 2 | 0 | 0 | 0 | 0 | 0 | 12 | 5 | 2 | 0 | 4 | 210 | 149 | 1 | 1 | 0 | 1 | 0.221 | 0.191 | 0 | 0 | 0 | 0 | NA | 0 | 0.282 |
| SE | 46542 | ENSG00000223745 | CCDC18-AS1 | chr1 | - | 93318167 | 93318228 | 93310127 | 93311608 | 93339356 | 93339536 | 0 | 1 | 0 | 0 | 1 | 0 | 5 | 2 | 1 | 0 | 0 | 0 | 0 | 0 | 0 | 0 | 0 | 1 | 0 | 0 | 210 | 149 | 1 | 1 | NA | 0.124 | 0 | 0 | 1 | NA | NA | 0 | NA | NA | 0.281 |
| SE | 46543 | ENSG00000223745 | CCDC18-AS1 | chr1 | - | 93318167 | 93318228 | 93315148 | 93315269 | 93324634 | 93324691 | 0 | 1 | 2 | 1 | 0 | 3 | 5 | 2 | 0 | 0 | 0 | 0 | 0 | 0 | 0 | 0 | 7 | 0 | 0 | 0 | 210 | 149 | 1 | 1 | 0 | 0.124 | 0.415 | 1 | NA | NA | 0 | NA | NA | NA | 0.385 |
| SE | 46545 | ENSG00000223745 | CCDC18-AS1 | chr1 | - | 93324634 | 93324729 | 93305110 | 93305470 | 93337376 | 93337392 | 8 | 20 | 12 | 15 | 7 | 3 | 0 | 0 | 1 | 3 | 0 | 5 | 6 | 3 | 2 | 0 | 7 | 0 | 0 | 0 | 244 | 149 | 1 | 1 | 0.62 | 1 | 1 | 0.902 | 0.588 | NA | 0.304 | 1 | 1 | 1 | -0.004 |
| SE | 46546 | ENSG00000223745 | CCDC18-AS1 | chr1 | - | 93325768 | 93325895 | 93324634 | 93324729 | 93337376 | 93337392 | 73 | 52 | 96 | 81 | 30 | 3 | 0 | 0 | 0 | 0 | 40 | 33 | 35 | 48 | 13 | 0 | 0 | 0 | 0 | 0 | 276 | 149 | 1 | 1 | 0.929 | 1 | 1 | 1 | 1 | 1 | 1 | 1 | 1 | 1 | -0.014 |
| SE | 46548 | ENSG00000223745 | CCDC18-AS1 | chr1 | - | 93326601 | 93326662 | 93325768 | 93325895 | 93338229 | 93339274 | 2 | 9 | 3 | 4 | 0 | 0 | 1 | 0 | 0 | 0 | 0 | 4 | 2 | 0 | 0 | 0 | 0 | 0 | 0 | 0 | 210 | 149 | 1 | 1 | 1 | 0.865 | 1 | 1 | NA | NA | 1 | 1 | NA | NA | -0.034 |
| SE | 46549 | ENSG00000223745 | CCDC18-AS1 | chr1 | - | 93326601 | 93326662 | 93325768 | 93325895 | 93339356 | 93339536 | 2 | 8 | 2 | 4 | 0 | 2 | 0 | 0 | 1 | 0 | 0 | 4 | 2 | 0 | 0 | 0 | 1 | 0 | 0 | 0 | 210 | 149 | 1 | 1 | 0.415 | 1 | 1 | 0.739 | NA | NA | 0.739 | 1 | NA | NA | -0.081 |
| SE | 46550 | ENSG00000223745 | CCDC18-AS1 | chr1 | - | 93337376 | 93337512 | 93325768 | 93325895 | 93338229 | 93338268 | 33 | 51 | 60 | 41 | 27 | 0 | 1 | 0 | 0 | 0 | 19 | 28 | 15 | 20 | 3 | 0 | 0 | 0 | 0 | 0 | 285 | 149 | 1 | 1 | 1 | 0.964 | 1 | 1 | 1 | 1 | 1 | 1 | 1 | 1 | -0.007 |
| SE | 46552 | ENSG00000223745 | CCDC18-AS1 | chr1 | - | 93338229 | 93338268 | 93325768 | 93325895 | 93338373 | 93338531 | 1 | 4 | 3 | 0 | 0 | 0 | 0 | 2 | 0 | 0 | 0 | 0 | 0 | 5 | 0 | 0 | 0 | 0 | 0 | 0 | 188 | 149 | 1 | 1 | 1 | 1 | 0.543 | NA | NA | NA | NA | NA | 1 | NA | -0.152 |
| SE | 46553 | ENSG00000223745 | CCDC18-AS1 | chr1 | - | 93338229 | 93339274 | 93325768 | 93325895 | 93339356 | 93339536 | 51 | 78 | 71 | 78 | 26 | 2 | 0 | 0 | 1 | 0 | 42 | 60 | 69 | 53 | 36 | 0 | 1 | 0 | 0 | 0 | 298 | 149 | 1 | 1 | 0.927 | 1 | 1 | 0.975 | 1 | 1 | 0.968 | 1 | 1 | 1 | -0.013 |
| SE | 46556 | ENSG00000223745 | CCDC18-AS1 | chr1 | - | 93339648 | 93339679 | 93338130 | 93339536 | 93340457 | 93340472 | 84 | 80 | 47 | 63 | 33 | 0 | 0 | 2 | 0 | 0 | 33 | 38 | 44 | 27 | 20 | 0 | 0 | 0 | 0 | 0 | 180 | 149 | 1 | 1 | 1 | 1 | 0.951 | 1 | 1 | 1 | 1 | 1 | 1 | 1 | -0.01 |
| SE | 46557 | ENSG00000223745 | CCDC18-AS1 | chr1 | - | 93339648 | 93339679 | 93338130 | 93339536 | 93345723 | 93345799 | 13 | 9 | 10 | 17 | 4 | 1 | 0 | 0 | 0 | 0 | 12 | 12 | 6 | 5 | 6 | 0 | 0 | 0 | 0 | 0 | 180 | 149 | 1 | 1 | 0.915 | 1 | 1 | 1 | 1 | 1 | 1 | 1 | 1 | 1 | -0.017 |
| SE | 46558 | ENSG00000223745 | CCDC18-AS1 | chr1 | - | 93340457 | 93340510 | 93338130 | 93339536 | 93345723 | 93345799 | 0 | 0 | 7 | 3 | 4 | 1 | 0 | 0 | 0 | 0 | 0 | 0 | 1 | 1 | 0 | 0 | 0 | 0 | 0 | 0 | 202 | 149 | 1 | 1 | 0 | NA | 1 | 1 | 1 | NA | NA | 1 | 1 | NA | -0.25 |
| SE | 46565 | ENSG00000138386 | NAB1 | chr2 | + | 190655976 | 190656159 | 190653730 | 190653780 | 190659157 | 190659995 | 4 | 1 | 9 | 13 | 2 | 2 | 2 | 1 | 1 | 2 | 7 | 0 | 12 | 10 | 0 | 0 | 2 | 3 | 0 | 0 | 298 | 149 | 1 | 1 | 0.5 | 0.2 | 0.818 | 0.867 | 0.333 | 1 | 0 | 0.667 | 1 | NA | -0.123 |
| SE | 46567 | ENSG00000138386 | NAB1 | chr2 | + | 190673100 | 190673152 | 190670325 | 190670459 | 190683737 | 190683827 | 107 | 65 | 93 | 110 | 192 | 0 | 0 | 1 | 0 | 0 | 64 | 90 | 100 | 73 | 40 | 0 | 0 | 0 | 0 | 0 | 201 | 149 | 1 | 1 | 1 | 1 | 0.986 | 1 | 1 | 1 | 1 | 1 | 1 | 1 | -0.003 |
| SE | 46568 | ENSG00000138386 | NAB1 | chr2 | + | 190673100 | 190673152 | 190670325 | 190670459 | 190685475 | 190685638 | 40 | 34 | 40 | 41 | 108 | 0 | 0 | 0 | 0 | 0 | 6 | 32 | 34 | 34 | 34 | 0 | 0 | 1 | 0 | 0 | 201 | 149 | 1 | 1 | 1 | 1 | 1 | 1 | 1 | 1 | 1 | 0.962 | 1 | 1 | 0.008 |
| SE | 46569 | ENSG00000138386 | NAB1 | chr2 | + | 190683737 | 190683827 | 190670325 | 190670459 | 190685475 | 190685638 | 41 | 31 | 18 | 46 | 60 | 0 | 0 | 0 | 0 | 0 | 30 | 37 | 20 | 19 | 32 | 0 | 0 | 1 | 0 | 0 | 239 | 149 | 1 | 1 | 1 | 1 | 1 | 1 | 1 | 1 | 1 | 0.926 | 1 | 1 | 0.015 |
| SE | 46571 | ENSG00000138386 | NAB1 | chr2 | + | 190683737 | 190683827 | 190673100 | 190673152 | 190685478 | 190685638 | 45 | 13 | 23 | 36 | 68 | 0 | 2 | 0 | 0 | 1 | 56 | 40 | 42 | 21 | 0 | 0 | 0 | 4 | 0 | 0 | 239 | 149 | 1 | 1 | 1 | 0.802 | 1 | 1 | 0.977 | 1 | 1 | 0.867 | 1 | NA | -0.011 |
| SE | 46574 | ENSG00000188738 | FSIP2 | chr2 | + | 185747312 | 185747423 | 185746668 | 185746810 | 185753721 | 185753842 | 0 | 2 | 1 | 0 | 0 | 0 | 1 | 0 | 0 | 0 | 10 | 0 | 0 | 0 | 0 | 0 | 0 | 0 | 0 | 0 | 260 | 149 | 1 | 1 | NA | 0.534 | 1 | NA | NA | 1 | NA | NA | NA | NA | -0.233 |
| SE | 46575 | ENSG00000188738 | FSIP2 | chr2 | + | 185753721 | 185753848 | 185747312 | 185747423 | 185760987 | 185761103 | 0 | 1 | 0 | 0 | 0 | 0 | 1 | 0 | 0 | 0 | 10 | 0 | 0 | 0 | 0 | 0 | 0 | 0 | 0 | 0 | 276 | 149 | 1 | 1 | NA | 0.351 | NA | NA | NA | 1 | NA | NA | NA | NA | -0.649 |
| SE | 46578 | ENSG00000058272 | PPP1R12A | chr12 | - | 79779320 | 79779351 | 79778549 | 79778600 | 79781814 | 79781862 | 0 | 2 | 1 | 0 | 0 | 392 | 279 | 267 | 390 | 666 | 0 | 0 | 0 | 0 | 0 | 21 | 189 | 233 | 45 | 12 | 180 | 149 | 1 | 1 | 0 | 0.006 | 0.003 | 0 | 0 | 0 | 0 | 0 | 0 | 0 | 0.002 |
| SE | 46579 | ENSG00000058272 | PPP1R12A | chr12 | - | 79779338 | 79779351 | 79778549 | 79778600 | 79781814 | 79781862 | 0 | 1 | 0 | 1 | 0 | 392 | 279 | 267 | 390 | 666 | 0 | 0 | 0 | 0 | 0 | 21 | 189 | 233 | 45 | 12 | 162 | 149 | 1 | 1 | 0 | 0.003 | 0 | 0.002 | 0 | 0 | 0 | 0 | 0 | 0 | 0.001 |
| SE | 46580 | ENSG00000058272 | PPP1R12A | chr12 | - | 79782570 | 79782595 | 79781814 | 79781862 | 79786373 | 79786478 | 0 | 1 | 0 | 1 | 0 | 441 | 327 | 307 | 416 | 780 | 0 | 0 | 1 | 0 | 0 | 21 | 178 | 252 | 62 | 6 | 174 | 149 | 1 | 1 | 0 | 0.003 | 0 | 0.002 | 0 | 0 | 0 | 0.003 | 0 | 0 | 0 |
| SE | 46581 | ENSG00000058272 | PPP1R12A | chr12 | - | 79786373 | 79786478 | 79781814 | 79781862 | 79786776 | 79788783 | 324 | 260 | 242 | 335 | 606 | 0 | 3 | 0 | 0 | 0 | 21 | 158 | 207 | 54 | 6 | 0 | 0 | 0 | 0 | 0 | 254 | 149 | 1 | 1 | 1 | 0.981 | 1 | 1 | 1 | 1 | 1 | 1 | 1 | 1 | -0.004 |
| SE | 46583 | ENSG00000058272 | PPP1R12A | chr12 | - | 79786776 | 79788783 | 79781814 | 79781862 | 79790466 | 79790483 | 555 | 501 | 369 | 505 | 975 | 0 | 0 | 0 | 0 | 0 | 14 | 316 | 317 | 144 | 10 | 0 | 0 | 1 | 0 | 0 | 298 | 149 | 1 | 1 | 1 | 1 | 1 | 1 | 1 | 1 | 1 | 0.994 | 1 | 1 | 0.001 |
| SE | 46584 | ENSG00000058272 | PPP1R12A | chr12 | - | 79788647 | 79788783 | 79781814 | 79781862 | 79790466 | 79790483 | 565 | 501 | 370 | 516 | 991 | 0 | 0 | 0 | 0 | 0 | 14 | 318 | 318 | 145 | 10 | 0 | 0 | 1 | 0 | 0 | 285 | 149 | 1 | 1 | 1 | 1 | 1 | 1 | 1 | 1 | 1 | 0.994 | 1 | 1 | 0.001 |
| SE | 46586 | ENSG00000058272 | PPP1R12A | chr12 | - | 79790466 | 79790483 | 79786776 | 79788783 | 79793862 | 79793928 | 592 | 517 | 388 | 532 | 1023 | 0 | 3 | 0 | 3 | 1 | 15 | 321 | 344 | 153 | 10 | 0 | 5 | 1 | 0 | 0 | 166 | 149 | 1 | 1 | 1 | 0.994 | 1 | 0.994 | 0.999 | 1 | 0.983 | 0.997 | 1 | 1 | 0.001 |
| SE | 46587 | ENSG00000058272 | PPP1R12A | chr12 | - | 79793862 | 79793928 | 79790466 | 79790483 | 79795637 | 79795759 | 676 | 615 | 465 | 649 | 1201 | 0 | 0 | 0 | 0 | 0 | 39 | 450 | 415 | 176 | 25 | 0 | 0 | 0 | 1 | 0 | 215 | 149 | 1 | 1 | 1 | 1 | 1 | 1 | 1 | 1 | 1 | 1 | 0.992 | 1 | 0.002 |
| SE | 46588 | ENSG00000058272 | PPP1R12A | chr12 | - | 79805591 | 79805768 | 79798493 | 79798584 | 79806165 | 79806297 | 49 | 42 | 24 | 46 | 50 | 52 | 40 | 52 | 64 | 93 | 0 | 16 | 22 | 12 | 11 | 0 | 50 | 49 | 10 | 0 | 298 | 149 | 1 | 1 | 0.32 | 0.344 | 0.188 | 0.264 | 0.212 | NA | 0.138 | 0.183 | 0.375 | 1 | -0.158 |
| SE | 46589 | ENSG00000058272 | PPP1R12A | chr12 | - | 79805591 | 79805768 | 79798493 | 79798584 | 79807225 | 79807330 | 40 | 28 | 21 | 35 | 59 | 8 | 15 | 7 | 18 | 40 | 0 | 9 | 19 | 10 | 6 | 0 | 4 | 4 | 1 | 9 | 298 | 149 | 1 | 1 | 0.714 | 0.483 | 0.6 | 0.493 | 0.424 | NA | 0.529 | 0.704 | 0.833 | 0.25 | -0.036 |
| SE | 46590 | ENSG00000058272 | PPP1R12A | chr12 | - | 79806165 | 79806297 | 79805591 | 79805768 | 79807225 | 79807330 | 49 | 52 | 19 | 45 | 31 | 20 | 11 | 10 | 9 | 32 | 0 | 14 | 14 | 4 | 8 | 0 | 0 | 7 | 0 | 3 | 281 | 149 | 1 | 1 | 0.565 | 0.715 | 0.502 | 0.726 | 0.339 | NA | 1 | 0.515 | 1 | 0.586 | -0.206 |
| SE | 46595 | ENSG00000058272 | PPP1R12A | chr12 | - | 79828319 | 79828464 | 79822115 | 79822190 | 79832331 | 79832491 | 730 | 562 | 540 | 676 | 646 | 6 | 0 | 0 | 0 | 2 | 141 | 443 | 456 | 243 | 85 | 0 | 3 | 0 | 0 | 0 | 294 | 149 | 1 | 1 | 0.984 | 1 | 1 | 1 | 0.994 | 1 | 0.987 | 1 | 1 | 1 | -0.002 |
| SE | 46596 | ENSG00000058272 | PPP1R12A | chr12 | - | 79848787 | 79848931 | 79845301 | 79845420 | 79872807 | 79872872 | 1 | 0 | 0 | 3 | 0 | 431 | 277 | 299 | 392 | 274 | 0 | 0 | 0 | 7 | 0 | 208 | 265 | 291 | 130 | 165 | 293 | 149 | 1 | 1 | 0.001 | 0 | 0 | 0.004 | 0 | 0 | 0 | 0 | 0.027 | 0 | -0.004 |
| SE | 46598 | ENSG00000273189 | NA | chr19 | + | 49478908 | 49479047 | 49476422 | 49476566 | 49480551 | 49480620 | 41 | 27 | 70 | 49 | 29 | 0 | 0 | 0 | 3 | 0 | 29 | 12 | 25 | 10 | 30 | 0 | 0 | 0 | 0 | 4 | 288 | 149 | 1 | 1 | 1 | 1 | 1 | 0.894 | 1 | 1 | 1 | 1 | 1 | 0.795 | 0.02 |
| SE | 46601 | ENSG00000141556 | TBCD | chr17 | + | 82756164 | 82756215 | 82752041 | 82752377 | 82763964 | 82764062 | 34 | 14 | 23 | 22 | 16 | 1 | 0 | 0 | 0 | 0 | 8 | 2 | 9 | 12 | 63 | 0 | 0 | 0 | 0 | 0 | 200 | 149 | 1 | 1 | 0.962 | 1 | 1 | 1 | 1 | 1 | 1 | 1 | 1 | 1 | -0.008 |
| SE | 46607 | ENSG00000141556 | TBCD | chr17 | + | 82814839 | 82814934 | 82809707 | 82809782 | 82870223 | 82870380 | 25 | 17 | 29 | 44 | 8 | 0 | 0 | 0 | 0 | 0 | 16 | 28 | 9 | 16 | 62 | 0 | 0 | 1 | 0 | 0 | 244 | 149 | 1 | 1 | 1 | 1 | 1 | 1 | 1 | 1 | 1 | 0.846 | 1 | 1 | 0.031 |
| SE | 46609 | ENSG00000141556 | TBCD | chr17 | + | 82889667 | 82889697 | 82884144 | 82884202 | 82893546 | 82893632 | 39 | 20 | 43 | 56 | 43 | 1 | 0 | 0 | 0 | 0 | 33 | 60 | 23 | 22 | 49 | 0 | 0 | 0 | 0 | 0 | 179 | 149 | 1 | 1 | 0.97 | 1 | 1 | 1 | 1 | 1 | 1 | 1 | 1 | 1 | -0.006 |
| SE | 46611 | ENSG00000141556 | TBCD | chr17 | + | 82893546 | 82893632 | 82884144 | 82884202 | 82900650 | 82900731 | 38 | 17 | 28 | 54 | 59 | 0 | 0 | 0 | 0 | 1 | 28 | 25 | 14 | 27 | 85 | 0 | 0 | 0 | 0 | 0 | 235 | 149 | 1 | 1 | 1 | 1 | 1 | 1 | 0.974 | 1 | 1 | 1 | 1 | 1 | -0.005 |
| SE | 46613 | ENSG00000141556 | TBCD | chr17 | + | 82893546 | 82893632 | 82889667 | 82889697 | 82905935 | 82906053 | 18 | 9 | 13 | 21 | 18 | 0 | 0 | 0 | 0 | 0 | 0 | 20 | 9 | 10 | 12 | 0 | 0 | 3 | 0 | 0 | 235 | 149 | 1 | 1 | 1 | 1 | 1 | 1 | 1 | NA | 1 | 0.655 | 1 | 1 | 0.086 |
| SE | 46615 | ENSG00000141556 | TBCD | chr17 | + | 82895700 | 82895967 | 82893546 | 82893632 | 82903404 | 82903478 | 0 | 0 | 4 | 2 | 0 | 0 | 0 | 1 | 0 | 0 | 24 | 0 | 0 | 0 | 0 | 3 | 3 | 2 | 6 | 6 | 298 | 149 | 1 | 1 | NA | NA | 0.667 | 1 | NA | 0.8 | 0 | 0 | 0 | 0 | 0.673 |
| SE | 46623 | ENSG00000141556 | TBCD | chr17 | + | 82903404 | 82903478 | 82900541 | 82900731 | 82905935 | 82906053 | 91 | 33 | 94 | 113 | 135 | 0 | 1 | 0 | 2 | 2 | 106 | 117 | 40 | 70 | 173 | 0 | 0 | 0 | 0 | 0 | 223 | 149 | 1 | 1 | 1 | 0.957 | 1 | 0.974 | 0.978 | 1 | 1 | 1 | 1 | 1 | -0.018 |
| SE | 46624 | ENSG00000141556 | TBCD | chr17 | + | 82903404 | 82903478 | 82900541 | 82900731 | 82911757 | 82911789 | 31 | 14 | 29 | 38 | 61 | 0 | 0 | 0 | 2 | 2 | 42 | 32 | 14 | 18 | 82 | 0 | 0 | 0 | 0 | 0 | 223 | 149 | 1 | 1 | 1 | 1 | 1 | 0.927 | 0.953 | 1 | 1 | 1 | 1 | 1 | -0.024 |
| SE | 46628 | ENSG00000141556 | TBCD | chr17 | + | 82907760 | 82907821 | 82905935 | 82906053 | 82909284 | 82909307 | 77 | 24 | 69 | 69 | 51 | 0 | 0 | 0 | 0 | 4 | 57 | 63 | 50 | 41 | 107 | 0 | 0 | 0 | 0 | 0 | 210 | 149 | 1 | 1 | 1 | 1 | 1 | 1 | 0.9 | 1 | 1 | 1 | 1 | 1 | -0.02 |
| SE | 46629 | ENSG00000141556 | TBCD | chr17 | + | 82908296 | 82908437 | 82907760 | 82907821 | 82909284 | 82909307 | 0 | 0 | 0 | 1 | 0 | 56 | 22 | 46 | 41 | 20 | 0 | 0 | 0 | 0 | 0 | 37 | 40 | 41 | 27 | 79 | 290 | 149 | 1 | 1 | 0 | 0 | 0 | 0.012 | 0 | 0 | 0 | 0 | 0 | 0 | 0.002 |
| SE | 46640 | ENSG00000251136 | NA | chr8 | - | 89722014 | 89722156 | 89717427 | 89717728 | 89724628 | 89724966 | 3 | 0 | 0 | 6 | 14 | 0 | 0 | 0 | 1 | 0 | 0 | 8 | 1 | 0 | 0 | 0 | 0 | 0 | 0 | 0 | 291 | 149 | 1 | 1 | 1 | NA | NA | 0.754 | 1 | NA | 1 | 1 | NA | NA | -0.082 |
| SE | 46641 | ENSG00000251136 | NA | chr8 | - | 89724628 | 89724966 | 89717427 | 89717728 | 89757044 | 89757572 | 0 | 0 | 0 | 1 | 0 | 0 | 0 | 0 | 1 | 0 | 0 | 0 | 0 | 5 | 0 | 0 | 0 | 0 | 0 | 0 | 298 | 149 | 1 | 1 | NA | NA | NA | 0.333 | NA | NA | NA | NA | 1 | NA | -0.667 |
| SE | 46644 | ENSG00000162614 | NEXN | chr1 | + | 77925187 | 77925229 | 77918124 | 77918273 | 77926413 | 77926611 | 3 | 11 | 0 | 17 | 3 | 0 | 5 | 0 | 3 | 0 | 44 | 9 | 26 | 6 | 6 | 13 | 0 | 5 | 4 | 0 | 191 | 149 | 1 | 1 | 1 | 0.632 | NA | 0.816 | 1 | 0.725 | 1 | 0.802 | 0.539 | 1 | 0.049 |
| SE | 46645 | ENSG00000162614 | NEXN | chr1 | + | 77933281 | 77933479 | 77929315 | 77929504 | 77935822 | 77935919 | 28 | 97 | 18 | 64 | 74 | 1 | 0 | 0 | 0 | 0 | 71 | 42 | 58 | 68 | 47 | 0 | 0 | 0 | 0 | 3 | 298 | 149 | 1 | 1 | 0.933 | 1 | 1 | 1 | 1 | | 1 | 1 | 1 | 0.887 | 0.009 |
| SE | 46646 | ENSG00000162614 | NEXN | chr1 | + | 77933281 | 77933479 | 77929315 | 77929504 | 77942022 | 77942208 | 21 | 63 | 12 | 41 | 35 | 0 | 0 | 0 | 0 | 0 | 52 | 31 | 30 | 44 | 41 | 0 | 0 | 0 | 4 | 0 | 298 | 149 | 1 | 1 | 1 | 1 | 1 | 1 | 1 | 1 | 1 | 1 | 0.846 | 1 | 0.031 |
| SE | 46647 | ENSG00000162614 | NEXN | chr1 | + | 77935822 | 77936044 | 77929315 | 77929504 | 77942022 | 77942208 | 14 | 41 | 12 | 57 | 61 | 0 | 0 | 0 | 0 | 0 | 0 | 40 | 48 | 27 | 7 | 0 | 0 | 0 | 4 | 0 | 298 | 149 | 1 | 1 | 1 | 1 | 1 | 1 | 1 | NA | 1 | 1 | 0.771 | 1 | 0.057 |
| SE | 46650 | ENSG00000126749 | EMG1 | chr12 | + | 6975089 | 6975148 | 6974551 | 6974693 | 6975695 | 6975977 | 90 | 29 | 94 | 72 | 100 | 0 | 0 | 1 | 0 | 0 | 19 | 48 | 23 | 31 | 32 | 0 | 0 | 0 | 0 | 0 | 208 | 149 | 1 | 1 | 1 | 1 | 0.985 | 1 | 1 | 1 | 1 | 1 | 1 | 1 | -0.003 |
| SE | 46652 | ENSG00000126749 | EMG1 | chr12 | + | 6975228 | 6975378 | 6975089 | 6975148 | 6983001 | 6983206 | 75 | 41 | 88 | 64 | 115 | 0 | 1 | 1 | 1 | 0 | 45 | 41 | 19 | 32 | 22 | 0 | 0 | 1 | 0 | 0 | 298 | 149 | 1 | 1 | 1 | 0.953 | 0.978 | 0.97 | 1 | 1 | 1 | 0.905 | 1 | 1 | -0.001 |
| SE | 46653 | ENSG00000126749 | EMG1 | chr12 | + | 6975228 | 6975378 | 6975089 | 6975148 | 6987781 | 6987838 | 73 | 32 | 80 | 58 | 109 | 0 | 0 | 0 | 1 | 0 | 45 | 42 | 19 | 32 | 22 | 0 | 0 | 0 | 0 | 0 | 298 | 149 | 1 | 1 | 1 | 1 | 1 | 0.967 | 1 | 1 | 1 | 1 | 1 | 1 | -0.007 |
| SE | 46654 | ENSG00000126749 | EMG1 | chr12 | + | 6983001 | 6983206 | 6975089 | 6975148 | 6987781 | 6987838 | 11 | 20 | 15 | 12 | 9 | 0 | 0 | 0 | 1 | 0 | 24 | 11 | 4 | 8 | 1 | 0 | 0 | 0 | 0 | 0 | 298 | 149 | 1 | 1 | 1 | 1 | 1 | 0.857 | 1 | 1 | 1 | 1 | 1 | 1 | -0.029 |
| SE | 46657 | ENSG00000134222 | PSRC1 | chr1 | - | 109281136 | 109281251 | 109280776 | 109281046 | 109281618 | 109282060 | 18 | 2 | 8 | 8 | 18 | 3 | 0 | 0 | 0 | 1 | 14 | 13 | 9 | 7 | 28 | 0 | 0 | 0 | 0 | 0 | 264 | 149 | 1 | 1 | 0.772 | 1 | 1 | 1 | 0.91 | 1 | 1 | 1 | 1 | 1 | -0.064 |
| SE | 46668 | ENSG00000198740 | ZNF652 | chr17 | - | 49298105 | 49298924 | 49289207 | 49291695 | 49311311 | 49311456 | 107 | 71 | 72 | 126 | 258 | 0 | 0 | 2 | 0 | 0 | 85 | 92 | 111 | 83 | 25 | 0 | 0 | 0 | 0 | 0 | 298 | 149 | 1 | 1 | 1 | 1 | 0.947 | 1 | 1 | 1 | 1 | 1 | 1 | 1 | -0.011 |
| SE | 46669 | ENSG00000198740 | ZNF652 | chr17 | - | 49311311 | 49311456 | 49289205 | 49289924 | 49311926 | 49312042 | 287 | 169 | 232 | 287 | 524 | 0 | 0 | 0 | 3 | 0 | 146 | 208 | 216 | 196 | 113 | 0 | 0 | 1 | 0 | 0 | 294 | 149 | 1 | 1 | 1 | 1 | 1 | 0.98 | 1 | 1 | 1 | 0.991 | 1 | 1 | -0.002 |
| SE | 46672 | ENSG00000198740 | ZNF652 | chr17 | - | 49311926 | 49312042 | 49311311 | 49311456 | 49312697 | 49312845 | 347 | 205 | 313 | 308 | 594 | 0 | 0 | 1 | 0 | 9 | 139 | 235 | 210 | 202 | 138 | 0 | 2 | 3 | 2 | 0 | 265 | 149 | 1 | 1 | 1 | 1 | 0.994 | 1 | 0.974 | 1 | 0.985 | 0.975 | 0.983 | 1 | 0.005 |
| SE | 46673 | ENSG00000198740 | ZNF652 | chr17 | - | 49312697 | 49312845 | 49289205 | 49289924 | 49316825 | 49317983 | 73 | 47 | 69 | 75 | 117 | 1 | 0 | 0 | 0 | 0 | 12 | 40 | 32 | 23 | 1 | 0 | 0 | 0 | 0 | 0 | 297 | 149 | 1 | 1 | 0.973 | 1 | 1 | 1 | 1 | 1 | 1 | 1 | 1 | 1 | -0.005 |
| SE | 46676 | ENSG00000117090 | SLAMF1 | chr1 | - | 160619775 | 160619849 | 160612487 | 160612580 | 160624095 | 160624185 | 34 | 23 | 26 | 26 | 14 | 8 | 5 | 3 | 3 | 4 | 0 | 13 | 10 | 0 | 5 | 0 | 0 | 5 | 4 | 0 | 223 | 149 | 1 | 1 | 0.74 | 0.755 | 0.853 | 0.853 | 0.7 | NA | 1 | 0.572 | 0 | 1 | 0.137 |
| SE | 46677 | ENSG00000117090 | SLAMF1 | chr1 | - | 160619775 | 160619849 | 160612487 | 160612580 | 160634612 | 160634897 | 14 | 16 | 11 | 11 | 9 | 0 | 6 | 3 | 1 | 1 | 0 | 8 | 10 | 0 | 5 | 0 | 0 | 1 | 2 | 0 | 223 | 149 | 1 | 1 | 1 | 0.641 | 0.71 | 0.88 | 0.857 | NA | 1 | 0.87 | 0 | 1 | 0.1 |
| SE | 46678 | ENSG00000117090 | SLAMF1 | chr1 | - | 160623502 | 160623585 | 160619775 | 160619849 | 160624095 | 160624185 | 1 | 1 | 8 | 1 | 0 | 20 | 8 | 15 | 17 | 6 | 0 | 7 | 0 | 0 | 5 | 0 | 5 | 0 | 0 | 0 | 232 | 149 | 1 | 1 | 0.031 | 0.074 | 0.255 | 0.036 | 0 | NA | 0.473 | NA | NA | 1 | -0.657 |
| SE | 46679 | ENSG00000117090 | SLAMF1 | chr1 | - | 160624095 | 160624185 | 160612487 | 160612580 | 160634612 | 160634897 | 16 | 9 | 16 | 5 | 7 | 0 | 6 | 3 | 1 | 1 | 0 | 0 | 6 | 4 | 0 | 0 | 0 | 1 | 2 | 0 | 239 | 149 | 1 | 1 | 1 | 0.483 | 0.769 | 0.757 | 0.814 | NA | NA | 0.789 | 0.555 | NA | 0.093 |
| SE | 46680 | ENSG00000117090 | SLAMF1 | chr1 | - | 160624095 | 160624185 | 160619775 | 160619849 | 160634612 | 160634897 | 26 | 12 | 28 | 19 | 9 | 0 | 1 | 0 | 0 | 1 | 0 | 5 | 1 | 0 | 0 | 0 | 0 | 0 | 0 | 0 | 239 | 149 | 1 | 1 | 1 | 0.882 | 1 | 1 | 0.849 | NA | 1 | 1 | NA | NA | -0.054 |
| SE | 46681 | ENSG00000117090 | SLAMF1 | chr1 | - | 160634612 | 160634897 | 160619775 | 160619849 | 160637190 | 160637529 | 75 | 52 | 64 | 56 | 60 | 0 | 0 | 1 | 0 | 0 | 16 | 17 | 17 | 3 | 22 | 0 | 0 | 0 | 0 | 0 | 298 | 149 | 1 | 1 | 1 | 1 | 0.97 | 1 | 1 | 1 | 1 | 1 | 1 | 1 | -0.006 |
| SE | 46683 | ENSG00000143786 | CNIH3 | chr1 | + | 224520999 | 224521327 | 224515861 | 224515876 | 224536935 | 224536981 | 3 | 1 | 0 | 2 | 1 | 0 | 0 | 1 | 0 | 0 | 0 | 8 | 0 | 4 | 0 | 0 | 0 | 0 | 0 | 0 | 298 | 149 | 1 | 1 | 1 | 1 | 0 | 1 | 1 | NA | 1 | NA | 1 | NA | -0.2 |
| SE | 46693 | ENSG00000171103 | TRMT61B | chr2 | - | 28850327 | 28850405 | 28849820 | 28850242 | 28869578 | 28870295 | 22 | 25 | 18 | 19 | 54 | 0 | 1 | 0 | 0 | 0 | 0 | 25 | 21 | 9 | 2 | 0 | 0 | 0 | 0 | 0 | 227 | 149 | 1 | 1 | 1 | 0.943 | 1 | 1 | 1 | NA | | 1 | 1 | 1 | -0.011 |
| SE | 46695 | ENSG00000171103 | TRMT61B | chr2 | - | 28851071 | 28851298 | 28850327 | 28850405 | 28861117 | 28861308 | 14 | 31 | 17 | 22 | 64 | 1 | 0 | 0 | 0 | 0 | 0 | 17 | 10 | 2 | 13 | 0 | 0 | 0 | 0 | 0 | 298 | 149 | 1 | 1 | 0.875 | 1 | 1 | 1 | 1 | NA | 1 | 1 | 1 | 1 | -0.025 |
| SE | 46696 | ENSG00000171103 | TRMT61B | chr2 | - | 28852407 | 28852499 | 28850327 | 28850405 | 28861117 | 28861308 | 16 | 5 | 9 | 11 | 33 | 1 | 0 | 0 | 0 | 0 | 0 | 0 | 0 | 8 | 7 | 0 | 0 | 0 | 0 | 0 | 241 | 149 | 1 | 1 | 0.908 | 1 | 1 | 1 | 1 | NA | NA | NA | 1 | 1 | -0.018 |
| SE | 46698 | ENSG00000171103 | TRMT61B | chr2 | - | 28852407 | 28852499 | 28851071 | 28851298 | 28861117 | 28861308 | 29 | 20 | 20 | 18 | 98 | 0 | 0 | 0 | 0 | 0 | 21 | 22 | 11 | 22 | 31 | 0 | 0 | 0 | 1 | 0 | 241 | 149 | 1 | 1 | 1 | 1 | 1 | 1 | 1 | 1 | 1 | 1 | 0.932 | 1 | 0.014 |
| SE | 46699 | ENSG00000171103 | TRMT61B | chr2 | - | 28861117 | 28861308 | 28850327 | 28850405 | 28869578 | 28870295 | 3 | 2 | 6 | 3 | 0 | 0 | 0 | 4 | 0 | 0 | 0 | 6 | 13 | 8 | 4 | 0 | 0 | 0 | 0 | 0 | 298 | 149 | 1 | 1 | 1 | 1 | 0.429 | 1 | NA | NA | 1 | 1 | 1 | 1 | -0.143 |
| SE | 46700 | ENSG00000171103 | TRMT61B | chr2 | - | 28861117 | 28861308 | 28852407 | 28852499 | 28865016 | 28865119 | 107 | 92 | 87 | 92 | 238 | 1 | 0 | 1 | 0 | 4 | 37 | 47 | 30 | 52 | 76 | 0 | 0 | 0 | 0 | 0 | 298 | 149 | 1 | 1 | 0.982 | 1 | 0.978 | | 0.967 | 1 | 1 | 1 | 1 | 1 | -0.015 |
| SE | 46709 | ENSG00000162714 | ZNF496 | chr1 | - | 247310323 | 247310456 | 247309698 | 247309806 | 247322697 | 247322805 | 37 | 16 | 28 | 13 | 69 | 0 | 0 | 0 | 1 | 0 | 14 | 48 | 20 | 16 | 20 | 0 | 0 | 0 | 0 | 0 | 282 | 149 | 1 | 1 | 1 | 1 | 1 | 0.873 | 1 | 1 | 1 | 1 | 1 | 1 | -0.025 |
| SE | 46710 | ENSG00000162714 | ZNF496 | chr1 | - | 247310323 | 247310456 | 247309698 | 247309806 | 247323153 | 247323230 | 55 | 16 | 38 | 20 | 68 | 0 | 0 | 0 | 0 | 1 | 18 | 55 | 37 | 19 | 39 | 0 | 0 | 0 | 0 | 0 | 282 | 149 | 1 | 1 | 1 | 1 | 1 | 1 | 0.973 | 1 | 1 | 1 | 1 | 1 | -0.005 |
| SE | 46711 | ENSG00000162714 | ZNF496 | chr1 | - | 247322697 | 247322805 | 247309698 | 247309806 | 247323153 | 247323230 | 0 | 0 | 2 | 2 | 0 | 0 | 0 | 0 | 0 | 1 | 0 | 5 | 1 | 0 | 0 | 0 | 0 | 0 | 0 | 0 | 257 | 149 | 1 | 1 | NA | NA | 1 | 1 | 0 | NA | 1 | 1 | NA | NA | -0.333 |
| SE | 46714 | ENSG00000055070 | SZRD1 | chr1 | + | 16387054 | 16387125 | 16367234 | 16367308 | 16393230 | 16393482 | 0 | 0 | 0 | 0 | 0 | 46 | 27 | 44 | 47 | 29 | 0 | 2 | 0 | 0 | 0 | 89 | 33 | 24 | 69 | 112 | 220 | 149 | 1 | 1 | 0 | 0 | 0 | 0 | 0 | 0 | 0.039 | 0 | 0 | 0 | -0.008 |
| SE | 46716 | ENSG00000055070 | SZRD1 | chr1 | + | 16390549 | 16390626 | 16367242 | 16367308 | 16391374 | 16391424 | 0 | 1 | 0 | 0 | 0 | 46 | 39 | 48 | 56 | 50 | 0 | 1 | 0 | 2 | 0 | 36 | 93 | 40 | 65 | 111 | 226 | 149 | 1 | 1 | 0 | 0.017 | 0 | 0 | 0 | 0 | 0.007 | 0 | 0.02 | 0 | -0.002 |
| SE | 46719 | ENSG00000055070 | SZRD1 | chr1 | + | 16391374 | 16391424 | 16387054 | 16387125 | 16393230 | 16393482 | 18 | 13 | 16 | 17 | 11 | 0 | 0 | 0 | 2 | 0 | 33 | 18 | 10 | 25 | 20 | 0 | 2 | 0 | 0 | 0 | 199 | 149 | 1 | 1 | 1 | 1 | 1 | 0.864 | 1 | 1 | 0.871 | 1 | 1 | 1 | -0.001 |
| SE | 46724 | ENSG00000249673 | NOP14-AS1 | chr4 | + | 2936493 | 2936601 | 2935634 | 2935735 | 2937751 | 2937844 | 0 | 0 | 1 | 3 | 0 | 1 | 2 | 7 | 1 | 0 | 0 | 14 | 4 | 5 | 8 | 0 | 7 | 6 | 0 | 0 | 257 | 149 | 1 | 1 | 0 | 0 | 0.076 | 0.635 | NA | NA | 0.537 | 0.279 | 1 | 1 | -0.526 |
| SE | 46726 | ENSG00000249673 | NOP14-AS1 | chr4 | + | 2937974 | 2938142 | 2935139 | 2937844 | 2938692 | 2938792 | 27 | 11 | 37 | 18 | 21 | 1 | 0 | 0 | 0 | 0 | 14 | 52 | 27 | 29 | 39 | 0 | 0 | 0 | 0 | 0 | 298 | 149 | 1 | 1 | 0.931 | 1 | 1 | 1 | 1 | 1 | 1 | 1 | 1 | 1 | -0.014 |
| SE | 46727 | ENSG00000249673 | NOP14-AS1 | chr4 | + | 2937974 | 2938142 | 2935139 | 2937844 | 2940301 | 2940351 | 12 | 8 | 12 | 10 | 10 | 0 | 0 | 0 | 0 | 3 | 14 | 35 | 17 | 11 | 22 | 0 | 0 | 0 | 0 | 0 | 298 | 149 | 1 | 1 | 1 | 1 | 1 | 1 | 0.625 | 1 | 1 | 1 | 1 | 1 | -0.075 |
| SE | 46728 | ENSG00000249673 | NOP14-AS1 | chr4 | + | 2938692 | 2938792 | 2934909 | 2938142 | 2940301 | 2940351 | 17 | 5 | 23 | 10 | 12 | 2 | 0 | 0 | 1 | 0 | 0 | 19 | 13 | 14 | 13 | 0 | 0 | 0 | 0 | 0 | 249 | 149 | 1 | 1 | 0.836 | 1 | 1 | 0.857 | 1 | NA | 1 | 1 | 1 | 1 | -0.061 |
| SE | 46729 | ENSG00000249673 | NOP14-AS1 | chr4 | + | 2938692 | 2938792 | 2935139 | 2937844 | 2940301 | 2940351 | 4 | 2 | 4 | 2 | 3 | 0 | 0 | 0 | 0 | 3 | 0 | 2 | 8 | 0 | 6 | 0 | 0 | 0 | 0 | 0 | 249 | 149 | 1 | 1 | 1 | 1 | 1 | 1 | 0.374 | NA | 1 | 1 | NA | 1 | -0.125 |
| SE | 46730 | ENSG00000249673 | NOP14-AS1 | chr4 | + | 2938692 | 2938792 | 2935139 | 2937844 | 2946872 | 2947120 | 2 | 2 | 5 | 6 | 5 | 1 | 0 | 0 | 0 | 0 | 0 | 3 | 4 | 1 | 0 | 3 | 0 | 0 | 0 | 0 | 249 | 149 | 1 | 1 | 0.545 | 1 | 1 | 1 | 1 | 0 | 1 | 1 | 1 | NA | 0.159 |
| SE | 46732 | ENSG00000163125 | RPRD2 | chr1 | + | 150408905 | 150409052 | 150364707 | 150364919 | 150417595 | 150417725 | 0 | 2 | 0 | 0 | 0 | 96 | 38 | 57 | 76 | 95 | 0 | 0 | 0 | 0 | 0 | 32 | 50 | 46 | 32 | 7 | 296 | 149 | 1 | 1 | 0 | 0.026 | 0 | 0 | 0 | 0 | 0 | 0 | 0 | 0 | 0.005 |
| SE | 46733 | ENSG00000163125 | RPRD2 | chr1 | + | 150440922 | 150441023 | 150417595 | 150417725 | 150443230 | 150443283 | 224 | 157 | 212 | 285 | 335 | 0 | 0 | 0 | 0 | 0 | 180 | 278 | 144 | 203 | 118 | 0 | 0 | 2 | 0 | 0 | 250 | 149 | 1 | 1 | 1 | 1 | 1 | 1 | 1 | 1 | 1 | 0.977 | 1 | 1 | 0.005 |
| SE | 46736 | ENSG00000163125 | RPRD2 | chr1 | + | 150457287 | 150457570 | 150446225 | 150446401 | 150460059 | 150460317 | 306 | 173 | 310 | 356 | 731 | 3 | 0 | 0 | 0 | 0 | 244 | 340 | 254 | 240 | 119 | 0 | 0 | 0 | 0 | 0 | 298 | 149 | 1 | 1 | 0.981 | 1 | 1 | 1 | 1 | 1 | 1 | 1 | 1 | 1 | -0.004 |
| SE | 46737 | ENSG00000163125 | RPRD2 | chr1 | + | 150460059 | 150460317 | 150457287 | 150457570 | 150464526 | 150464727 | 328 | 188 | 313 | 335 | 803 | 2 | 0 | 0 | 2 | 5 | 228 | 354 | 217 | 230 | 152 | 6 | 0 | 0 | 0 | 0 | 298 | 149 | 1 | 1 | 0.988 | 1 | 1 | 0.988 | 0.988 | 0.95 | 1 | 1 | 1 | 1 | 0.003 |
| SE | 46743 | ENSG00000150456 | EEF1AKMT1 | chr13 | - | 20731840 | 20732121 | 20728730 | 20729216 | 20757454 | 20757617 | 14 | 8 | 19 | 16 | 16 | 0 | 1 | 2 | 3 | 3 | 0 | 16 | 2 | 0 | 7 | 0 | 0 | 0 | 0 | 0 | 298 | 149 | 1 | 1 | 1 | 0.8 | 0.826 | 0.727 | 0.727 | NA | 1 | 1 | NA | 1 | -0.184 |
| SE | 46744 | ENSG00000150456 | EEF1AKMT1 | chr13 | - | 20737722 | 20737805 | 20731840 | 20732121 | 20757454 | 20757617 | 17 | 2 | 15 | 11 | 11 | 5 | 2 | 8 | 5 | 1 | 12 | 19 | 7 | 0 | 6 | 0 | 3 | 1 | 0 | 0 | 232 | 149 | 1 | 1 | 0.686 | 0.391 | 0.546 | 0.586 | 0.876 | 1 | 0.803 | 0.818 | NA | 1 | -0.288 |
| SE | 46759 | ENSG00000108179 | PPIF | chr10 | + | 79349075 | 79349106 | 79347468 | 79347743 | 79349664 | 79349753 | 45 | 27 | 43 | 45 | 22 | 0 | 0 | 0 | 0 | 0 | 91 | 74 | 90 | 166 | 150 | 0 | 3 | 0 | 1 | 0 | 180 | 149 | 1 | 1 | 1 | 1 | 1 | 1 | 1 | 1 | 0.953 | 1 | 0.993 | 1 | 0.011 |
| SE | 46760 | ENSG00000108179 | PPIF | chr10 | + | 79351486 | 79351583 | 79349664 | 79349753 | 79353706 | 79355334 | 32 | 22 | 32 | 37 | 27 | 1 | 0 | 1 | 1 | 0 | 86 | 64 | 64 | 116 | 143 | 0 | 1 | 0 | 0 | 0 | 246 | 149 | 1 | 1 | 0.951 | 1 | 0.951 | 0.957 | 1 | 1 | 0.975 | 1 | 1 | 1 | -0.023 |
| SE | 46761 | ENSG00000108179 | PPIF | chr10 | + | 79352316 | 79352392 | 79351486 | 79351583 | 79353644 | 79353706 | 28 | 16 | 21 | 21 | 17 | 0 | 0 | 0 | 0 | 0 | 47 | 8 | 12 | 77 | 81 | 0 | 0 | 0 | 1 | 0 | 225 | 149 | 1 | 1 | 1 | 1 | 1 | 1 | 1 | 1 | 1 | 1 | 0.981 | 1 | 0.004 |
| SE | 46768 | ENSG00000117115 | PADI2 | chr1 | - | 17092407 | 17092533 | 17086520 | 17086699 | 17093566 | 17093684 | 62 | 23 | 51 | 59 | 95 | 1 | 0 | 0 | 0 | 2 | 199 | 119 | 92 | 308 | 288 | 0 | 0 | 0 | 0 | 0 | 275 | 149 | 1 | 1 | 0.971 | 1 | 1 | 1 | 0.963 | 1 | 1 | 1 | 1 | 1 | -0.013 |
| SE | 46769 | ENSG00000117115 | PADI2 | chr1 | - | 17102986 | 17103059 | 17095908 | 17095970 | 17104877 | 17105061 | 22 | 7 | 8 | 18 | 9 | 0 | 0 | 0 | 0 | 0 | 47 | 54 | 51 | 86 | 88 | 0 | 0 | 0 | 4 | 0 | 222 | 149 | 1 | 1 | 1 | 1 | 1 | 1 | 1 | 1 | 1 | 1 | 0.935 | 1 | 0.013 |
| SE | 46770 | ENSG00000225492 | GBP1P1 | chr1 | + | 89410299 | 89410508 | 89407678 | 89407758 | 89420190 | 89420318 | 12 | 0 | 5 | 9 | 1 | 0 | 0 | 0 | 0 | 0 | 27 | 5 | 2 | 13 | 0 | 0 | 0 | 1 | 2 | 0 | 298 | 149 | 1 | 1 | 1 | NA | 1 | 1 | 1 | 1 | 1 | 0.5 | 0.765 | NA | 0.184 |
| SE | 46771 | ENSG00000167257 | RNF214 | chr11 | + | 117234266 | 117234379 | 117232624 | 117232726 | 117238600 | 117238646 | 28 | 24 | 27 | 28 | 29 | 0 | 0 | 0 | 1 | 0 | 40 | 23 | 20 | 32 | 12 | 0 | 0 | 0 | 2 | 0 | 262 | 149 | 1 | 1 | 1 | 1 | 1 | 0.941 | 1 | 1 | 1 | 1 | 0.901 | 1 | 0.008 |
| SE | 46772 | ENSG00000167257 | RNF214 | chr11 | + | 117244444 | 117244585 | 117238600 | 117239860 | 117246808 | 117246840 | 136 | 143 | 149 | 147 | 240 | 0 | 1 | 0 | 0 | 0 | 453 | 155 | 146 | 239 | 185 | 0 | 0 | 0 | 0 | 0 | 290 | 149 | 1 | 1 | 1 | 0.987 | 1 | 1 | 1 | 1 | 1 | 1 | 1 | 1 | -0.003 |
| SE | 46773 | ENSG00000167257 | RNF214 | chr11 | + | 117246808 | 117246948 | 117244444 | 117244585 | 117279907 | 117280004 | 139 | 110 | 123 | 116 | 252 | 0 | 0 | 5 | 0 | 0 | 344 | 152 | 94 | 201 | 190 | 0 | 4 | 0 | 0 | 0 | 289 | 149 | 1 | 1 | 1 | 1 | 0.927 | 1 | 1 | 1 | 0.951 | 1 | 1 | 1 | -0.005 |
| SE | 46774 | ENSG00000167257 | RNF214 | chr11 | + | 117281599 | 117281698 | 117281313 | 117281404 | 117281893 | 117281911 | 120 | 81 | 99 | 108 | 372 | 0 | 0 | 0 | 0 | 3 | 86 | 96 | 44 | 65 | 115 | 1 | 0 | 0 | 3 | 0 | 248 | 149 | 1 | 1 | 1 | 1 | 1 | 1 | 0.987 | 0.981 | 1 | 1 | 0.929 | 1 | 0.015 |
| SE | 46776 | ENSG00000163798 | SLC4A1AP | chr2 | + | 27665099 | 27665295 | 27663470 | 27664577 | 27667267 | 27667390 | 151 | 115 | 125 | 137 | 167 | 0 | 0 | 0 | 0 | 0 | 187 | 120 | 143 | 164 | 66 | 0 | 3 | 2 | 0 | 0 | 298 | 149 | 1 | 1 | 1 | 1 | 1 | 1 | 1 | 1 | 0.952 | 0.973 | 1 | 1 | 0.015 |
| SE | 46780 | ENSG00000163798 | SLC4A1AP | chr2 | + | 27693684 | 27693754 | 27688699 | 27688767 | 27694633 | 27694969 | 111 | 74 | 102 | 128 | 124 | 0 | 0 | 0 | 0 | 0 | 120 | 110 | 80 | 83 | 0 | 0 | 1 | 0 | 0 | 0 | 219 | 149 | 1 | 1 | 1 | 1 | 1 | 1 | 1 | 1 | 0.987 | 1 | 1 | NA | 0.003 |
| SE | 46781 | ENSG00000096060 | FKBP5 | chr6 | - | 35576993 | 35577233 | 35573584 | 35575942 | 35580035 | 35580221 | 1319 | 988 | 1004 | 755 | 1362 | 0 | 0 | 9 | 0 | 0 | 529 | 2596 | 2553 | 703 | 247 | 3 | 3 | 9 | 0 | 0 | 298 | 149 | 1 | 1 | 1 | 1 | 0.982 | 1 | 1 | 0.989 | 0.998 | 0.993 | 1 | 1 | 0 |
| SE | 46782 | ENSG00000096060 | FKBP5 | chr6 | - | 35576993 | 35577233 | 35573584 | 35575942 | 35587033 | 35587117 | 398 | 356 | 367 | 258 | 430 | 0 | 1 | 1 | 1 | 0 | 216 | 1037 | 1003 | 267 | 114 | 0 | 5 | 0 | 0 | 6 | 298 | 149 | 1 | 1 | 1 | 0.994 | 0.995 | 0.992 | 1 | 1 | 0.99 | 1 | 1 | 0.905 | 0.017 |
| SE | 46784 | ENSG00000096060 | FKBP5 | chr6 | - | 35580035 | 35580221 | 35576993 | 35577233 | 35587033 | 35587117 | 1425 | 962 | 968 | 759 | 1357 | 8 | 7 | 5 | 2 | 6 | 472 | 2494 | 2456 | 685 | 213 | 2 | 6 | 2 | 0 | 0 | 298 | 149 | 1 | 1 | 0.989 | 0.986 | 0.99 | 0.995 | 0.991 | 0.992 | 0.995 | 0.998 | 1 | 1 | -0.007 |
| SE | 46785 | ENSG00000096060 | FKBP5 | chr6 | - | 35580035 | 35580221 | 35576993 | 35577233 | 35591129 | 35591220 | 930 | 641 | 646 | 499 | 939 | 0 | 0 | 0 | 0 | 0 | 315 | 1565 | 1556 | 436 | 133 | 0 | 0 | 1 | 0 | 0 | 298 | 149 | 1 | 1 | 1 | 1 | 1 | 1 | 1 | 1 | 1 | 0.999 | 1 | 1 | 0 |
| SE | 46786 | ENSG00000096060 | FKBP5 | chr6 | - | 35587033 | 35587117 | 35576993 | 35577233 | 35591129 | 35591220 | 578 | 469 | 378 | 307 | 449 | 0 | 0 | 0 | 0 | 0 | 278 | 1030 | 1115 | 267 | 142 | 0 | 0 | 1 | 0 | 0 | 233 | 149 | 1 | 1 | 1 | 1 | 1 | 1 | 1 | 1 | 1 | 0.999 | 1 | 1 | 0 |
| SE | 46787 | ENSG00000096060 | FKBP5 | chr6 | - | 35587033 | 35587117 | 35580035 | 35580221 | 35591129 | 35591220 | 1063 | 783 | 696 | 563 | 860 | 1 | 2 | 4 | 0 | 1 | 433 | 1937 | 2015 | 516 | 222 | 0 | 0 | 4 | 0 | 0 | 233 | 149 | 1 | 1 | 0.999 | 0.996 | 0.991 | 1 | 0.998 | 1 | 1 | 0.997 | 1 | 1 | -0.003 |
| SE | 46788 | ENSG00000096060 | FKBP5 | chr6 | - | 35591129 | 35591220 | 35587033 | 35587117 | 35597247 | 35597404 | 1292 | 1013 | 878 | 713 | 875 | 4 | 2 | 2 | 2 | 0 | 627 | 2344 | 2478 | 679 | 285 | 0 | 11 | 4 | 0 | 0 | 240 | 149 | 1 | 1 | 0.995 | 0.997 | 0.996 | 0.996 | 1 | 1 | 0.992 | 0.997 | 1 | 1 | -0.001 |
| SE | 46789 | ENSG00000096060 | FKBP5 | chr6 | - | 35597247 | 35597404 | 35591129 | 35591220 | 35619095 | 35619210 | 1398 | 1108 | 1035 | 750 | 1226 | 0 | 0 | 2 | 2 | 0 | 664 | 2566 | 2729 | 729 | 260 | 0 | 4 | 3 | 0 | 0 | 298 | 149 | 1 | 1 | 1 | 1 | 1 | 0.995 | 1 | 1 | 0.997 | 0.998 | 1 | 1 | 0 |
| SE | 46790 | ENSG00000096060 | FKBP5 | chr6 | - | 35619095 | 35619210 | 35597247 | 35597404 | 35637013 | 35637158 | 617 | 497 | 487 | 333 | 572 | 0 | 0 | 0 | 0 | 0 | 247 | 1148 | 1249 | 332 | 93 | 0 | 1 | 1 | 0 | 0 | 264 | 149 | 1 | 1 | 1 | 1 | 1 | 1 | 1 | 1 | 0.998 | 0.999 | 1 | 1 | 0.001 |
| SE | 46791 | ENSG00000096060 | FKBP5 | chr6 | - | 35620131 | 35620274 | 35619095 | 35619210 | 35637013 | 35637158 | 3890 | 3355 | 2915 | 2367 | 4196 | 17 | 15 | 11 | 7 | 8 | 3463 | 8860 | 9746 | 2930 | 1675 | 10 | 15 | 42 | 7 | 0 | 292 | 149 | 1 | 1 | 0.992 | 0.991 | 0.993 | 0.994 | 0.996 | 0.994 | 0.997 | 0.992 | 0.995 | 1 | -0.002 |
| SE | 46792 | ENSG00000096060 | FKBP5 | chr6 | - | 35637013 | 35637158 | 35597247 | 35597404 | 35642719 | 35642843 | 942 | 700 | 721 | 530 | 1165 | 0 | 1 | 0 | 0 | 0 | 663 | 2093 | 2075 | 565 | 259 | 0 | 0 | 0 | 0 | 0 | 294 | 149 | 1 | 1 | 1 | 0.997 | 1 | 1 | 1 | 1 | 1 | 1 | 1 | 1 | -0.001 |
| SE | 46793 | ENSG00000096060 | FKBP5 | chr6 | - | 35637013 | 35637158 | 35620131 | 35620274 | 35642719 | 35642843 | 2828 | 2366 | 2107 | 1670 | 3325 | 0 | 0 | 0 | 0 | 0 | 2312 | 6038 | 6335 | 1918 | 1003 | 0 | 6 | 3 | 0 | 0 | 294 | 149 | 1 | 1 | 1 | 1 | 1 | 1 | 1 | 1 | 0.998 | 0.999 | 1 | 1 | 0.001 |
| SE | 46794 | ENSG00000145860 | RNF145 | chr5 | - | 159158839 | 159159035 | 159157823 | 159158426 | 159161265 | 159161360 | 133 | 181 | 189 | 207 | 280 | 0 | 2 | 0 | 3 | 0 | 218 | 117 | 86 | 135 | 40 | 0 | 0 | 0 | 0 | 0 | 298 | 149 | 1 | 1 | 1 | 0.978 | 1 | 0.972 | 1 | 1 | 1 | 1 | 1 | 1 | -0.01 |
| SE | 46795 | ENSG00000145860 | RNF145 | chr5 | - | 159168872 | 159169055 | 159162931 | 159163079 | 159169678 | 159169819 | 64 | 41 | 46 | 62 | 59 | 0 | 0 | 0 | 2 | 0 | 59 | 32 | 20 | 39 | 11 | 0 | 0 | 0 | 0 | 0 | 298 | 149 | 1 | 1 | 1 | 1 | 1 | 0.939 | 1 | 1 | 1 | 1 | 1 | 1 | -0.012 |
| SE | 46796 | ENSG00000145860 | RNF145 | chr5 | - | 159168991 | 159169055 | 159162931 | 159163079 | 159169678 | 159169819 | 29 | 26 | 18 | 23 | 12 | 0 | 0 | 0 | 2 | 0 | 25 | 9 | 15 | 15 | 0 | 0 | 0 | 0 | 0 | 0 | 213 | 149 | 1 | 1 | 1 | 1 | 1 | 0.889 | 1 | 1 | 1 | 1 | 1 | NA | -0.022 |
| SE | 46797 | ENSG00000145860 | RNF145 | chr5 | - | 159173982 | 159174158 | 159169678 | 159169819 | 159176631 | 159176867 | 41 | 51 | 78 | 57 | 57 | 1 | 2 | 1 | 3 | 3 | 39 | 13 | 2 | 17 | 0 | 9 | 0 | 4 | 0 | 0 | 298 | 149 | 1 | 1 | 0.953 | 0.927 | 0.975 | 0.905 | 0.905 | 0.684 | 1 | 0.2 | 1 | NA | 0.212 |
| SE | 46798 | ENSG00000145860 | RNF145 | chr5 | - | 159176631 | 159176867 | 159169678 | 159169819 | 159181959 | 159182051 | 44 | 23 | 34 | 28 | 57 | 0 | 0 | 1 | 0 | 0 | 31 | 12 | 13 | 0 | 7 | 0 | 0 | 0 | 0 | 0 | 298 | 149 | 1 | 1 | 1 | 1 | 0.944 | 1 | 1 | 1 | 1 | 1 | NA | 1 | -0.011 |
| SE | 46799 | ENSG00000145860 | RNF145 | chr5 | - | 159176631 | 159176867 | 159169678 | 159169819 | 159194715 | 159194824 | 1 | 2 | 1 | 3 | 3 | 0 | 1 | 0 | 1 | 0 | 9 | 0 | 4 | 0 | 5 | 0 | 0 | 0 | 0 | 0 | 298 | 149 | 1 | 1 | 1 | 0.5 | 1 | 0.6 | 1 | 1 | NA | 1 | NA | 1 | -0.18 |
| SE | 46800 | ENSG00000145860 | RNF145 | chr5 | - | 159181959 | 159182051 | 159169678 | 159169819 | 159194715 | 159194824 | 69 | 29 | 64 | 89 | 40 | 0 | 1 | 0 | 1 | 0 | 58 | 80 | 100 | 46 | 94 | 0 | 0 | 0 | 0 | 0 | 241 | 149 | 1 | 1 | 1 | 0.947 | 1 | 0.982 | 1 | 1 | 1 | 1 | 1 | 1 | -0.014 |
| SE | 46801 | ENSG00000145860 | RNF145 | chr5 | - | 159181959 | 159182051 | 159176631 | 159176867 | 159194715 | 159194824 | 112 | 48 | 96 | 113 | 92 | 0 | 0 | 0 | 0 | 0 | 80 | 91 | 109 | 46 | 101 | 0 | 0 | 0 | 0 | 5 | 241 | 149 | 1 | 1 | 1 | 1 | 1 | 1 | 1 | 1 | 1 | 1 | 1 | 0.926 | 0.015 |
| SE | 46802 | ENSG00000145860 | RNF145 | chr5 | - | 159203433 | 159203656 | 159194715 | 159194824 | 159209230 | 159209510 | 99 | 48 | 60 | 118 | 42 | 0 | 0 | 0 | 0 | 0 | 60 | 88 | 100 | 82 | 99 | 0 | 0 | 2 | 0 | 0 | 298 | 149 | 1 | 1 | 1 | 1 | 1 | 1 | 1 | 1 | 1 | 0.962 | 1 | 1 | 0.008 |
| SE | 46814 | ENSG00000120280 | TASL | chrX | - | 30576751 | 30576866 | 30558808 | 30560356 | 30577637 | 30577766 | 148 | 146 | 94 | 83 | 188 | 0 | 0 | 0 | 0 | 0 | 79 | 50 | 54 | 122 | 48 | 5 | 0 | 0 | 0 | 0 | 264 | 149 | 1 | 1 | 1 | 1 | 1 | 1 | 1 | 0.899 | 1 | 1 | 1 | 1 | 0.02 |
| SE | 46816 | ENSG00000197780 | TAF13 | chr1 | - | 109066134 | 109066232 | 109064139 | 109064693 | 109075920 | 109075971 | 105 | 79 | 58 | 116 | 101 | 0 | 0 | 0 | 0 | 0 | 89 | 60 | 124 | 102 | 18 | 0 | 0 | 0 | 0 | 2 | 247 | 149 | 1 | 1 | 1 | 1 | 1 | 1 | 1 | 1 | 1 | 1 | 1 | 0.844 | 0.031 |
| SE | 46817 | ENSG00000197780 | TAF13 | chr1 | - | 109074986 | 109075065 | 109064139 | 109064693 | 109075920 | 109075971 | 1 | 3 | 3 | 5 | 3 | 0 | 0 | 0 | 0 | 0 | 0 | 0 | 5 | 1 | 0 | 0 | 0 | 0 | 0 | 2 | 228 | 149 | 1 | 1 | 1 | 1 | 1 | 1 | 1 | NA | NA | 1 | 1 | 0 | 0.333 |
| SE | 46823 | ENSG00000028137 | TNFRSF1B | chr1 | + | 12192347 | 12192524 | 12191837 | 12191923 | 12192862 | 12193008 | 287 | 135 | 191 | 221 | 121 | 0 | 1 | 0 | 0 | 0 | 319 | 339 | 262 | 219 | 760 | 0 | 0 | 4 | 0 | 0 | 298 | 149 | 1 | 1 | 1 | 0.985 | 1 | 1 | 1 | 1 | 1 | 0.97 | 1 | 1 | 0.003 |
| SE | 46824 | ENSG00000028137 | TNFRSF1B | chr1 | + | 12192430 | 12192524 | 12191773 | 12191923 | 12192862 | 12193098 | 614 | 252 | 330 | 417 | 212 | 0 | 1 | 0 | 0 | 0 | 549 | 563 | 415 | 385 | 1115 | 0 | 0 | 4 | 0 | 0 | 243 | 149 | 1 | 1 | 1 | 0.994 | 1 | 1 | 1 | 1 | 1 | 0.985 | 1 | 1 | 0.002 |
| SE | 46825 | ENSG00000028137 | TNFRSF1B | chr1 | + | 12193954 | 12194032 | 12192862 | 12193098 | 12194583 | 12194618 | 947 | 445 | 454 | 680 | 520 | 0 | 0 | 0 | 0 | 0 | 1007 | 635 | 530 | 570 | 1488 | 0 | 0 | 0 | 2 | 3 | 227 | 149 | 1 | 1 | 1 | 1 | 1 | 1 | 1 | 1 | 1 | 1 | 0.995 | 0.997 | 0.002 |
| SE | 46826 | ENSG00000028137 | TNFRSF1B | chr1 | + | 12201966 | 12202171 | 12194583 | 12194618 | 12206739 | 12209220 | 1119 | 511 | 552 | 771 | 794 | 2 | 5 | 4 | 0 | 2 | 935 | 671 | 524 | 557 | 1466 | 5 | 0 | 1 | 4 | 0 | 298 | 149 | 1 | 1 | 0.996 | 0.981 | 0.986 | 1 | 0.995 | 0.989 | 1 | 0.996 | 0.986 | 1 | -0.003 |
| SE | 46831 | ENSG00000155330 | C16orf87 | chr16 | - | 46824385 | 46824482 | 46809602 | 46809785 | 46831083 | 46831162 | 63 | 40 | 51 | 67 | 81 | 0 | 0 | 0 | 0 | 0 | 9 | 55 | 88 | 38 | 27 | 0 | 0 | 1 | 0 | 0 | 246 | 149 | 1 | 1 | 1 | 1 | 1 | 1 | 1 | 1 | 1 | 0.982 | 1 | 1 | 0.004 |
| SE | 46833 | ENSG00000066827 | ZFAT | chr8 | - | 134588245 | 134588395 | 134583831 | 134584005 | 134600435 | 134600668 | 59 | 48 | 60 | 69 | 98 | 0 | 0 | 0 | 2 | 0 | 71 | 37 | 25 | 43 | 57 | 0 | 0 | 0 | 0 | 0 | 298 | 149 | 1 | 1 | 1 | 1 | 1 | 0.945 | 1 | 1 | 1 | 1 | 1 | 1 | -0.011 |
| SE | 46834 | ENSG00000066827 | ZFAT | chr8 | - | 134590267 | 134590355 | 134588245 | 134588395 | 134600435 | 134600668 | 32 | 28 | 36 | 50 | 90 | 0 | 0 | 1 | 0 | 0 | 62 | 35 | 7 | 35 | 37 | 0 | 0 | 0 | 2 | 0 | 237 | 149 | 1 | 1 | 1 | 1 | 0.958 | 1 | 1 | 1 | 1 | 1 | 0.917 | 1 | 0.008 |
| SE | 46835 | ENSG00000066827 | ZFAT | chr8 | - | 134599729 | 134599862 | 134590267 | 134590355 | 134600435 | 134600668 | 6 | 0 | 2 | 0 | 0 | 24 | 13 | 25 | 32 | 38 | 0 | 0 | 7 | 2 | 0 | 45 | 25 | 3 | 14 | 34 | 282 | 149 | 1 | 1 | 0.117 | 0 | 0.041 | 0 | 0 | 0 | 0 | 0.552 | 0.07 | 0 | -0.093 |
| SE | 46839 | ENSG00000066827 | ZFAT | chr8 | - | 134610469 | 134610655 | 134608728 | 134608879 | 134637460 | 134637593 | 53 | 72 | 42 | 63 | 101 | 4 | 1 | 4 | 0 | 1 | 79 | 65 | 48 | 51 | 46 | 0 | 1 | 5 | 4 | 0 | 298 | 149 | 1 | 1 | 0.869 | 0.973 | 0.84 | 1 | 0.981 | 1 | 0.97 | 0.828 | 0.864 | 1 | 0 |
| SE | 46841 | ENSG00000066827 | ZFAT | chr8 | - | 134674729 | 134674888 | 134657560 | 134657737 | 134712844 | 134713038 | 2 | 1 | 2 | 5 | 2 | 13 | 8 | 8 | 16 | 17 | 0 | 8 | 2 | 0 | 3 | 0 | 7 | 15 | 10 | 23 | 298 | 149 | 1 | 1 | 0.071 | 0.059 | 0.111 | 0.135 | 0.056 | NA | 0.364 | 0.062 | 0 | 0.061 | -0.035 |
| SE | 46843 | ENSG00000155463 | OXA1L | chr14 | + | 22767317 | 22767409 | 22766689 | 22766764 | 22767957 | 22768171 | 44 | 20 | 44 | 41 | 12 | 1 | 1 | 0 | 1 | 0 | 96 | 55 | 58 | 36 | 128 | 8 | 0 | 0 | 0 | 0 | 241 | 149 | 1 | 1 | 0.965 | 0.925 | 1 | 0.962 | 1 | 0.881 | 1 | 1 | 1 | 1 | -0.006 |
| SE | 46844 | ENSG00000155463 | OXA1L | chr14 | + | 22767957 | 22768171 | 22766521 | 22766764 | 22769790 | 22769934 | 394 | 215 | 359 | 371 | 196 | 0 | 0 | 0 | 0 | 0 | 568 | 494 | 331 | 354 | 707 | 0 | 8 | 0 | 0 | 0 | 298 | 149 | 1 | 1 | 1 | 1 | 1 | 1 | 1 | 1 | 0.969 | 1 | 1 | 1 | 0.006 |
| SE | 46849 | ENSG00000162894 | FCMR | chr1 | - | 206909724 | 206909868 | 206903316 | 206905147 | 206911729 | 206911952 | 14 | 1 | 7 | 13 | 5 | 0 | 0 | 1 | 0 | 0 | 1 | 4 | 3 | 0 | 14 | 0 | 0 | 0 | 0 | 0 | 293 | 149 | 1 | 1 | 1 | 1 | 0.781 | 1 | 1 | 1 | 1 | 1 | NA | 1 | -0.044 |
| SE | 46851 | ENSG00000162894 | FCMR | chr1 | - | 206909724 | 206909868 | 206909461 | 206909520 | 206910209 | 206910340 | 735 | 288 | 579 | 489 | 491 | 0 | 1 | 0 | 0 | 0 | 213 | 408 | 214 | 130 | 252 | 0 | 0 | 0 | 0 | 0 | 293 | 149 | 1 | 1 | 1 | 0.993 | 1 | 1 | 1 | 1 | 1 | 1 | 1 | 1 | -0.001 |
| SE | 46853 | ENSG00000162894 | FCMR | chr1 | - | 206910209 | 206910340 | 206909724 | 206909868 | 206921817 | 206921941 | 319 | 133 | 264 | 225 | 261 | 1 | 0 | 1 | 0 | 0 | 90 | 208 | 88 | 46 | 106 | 0 | 0 | 0 | 0 | 0 | 280 | 149 | 1 | 1 | 0.994 | 1 | 0.993 | 1 | 1 | 1 | 1 | 1 | 1 | 1 | -0.003 |
| SE | 46854 | ENSG00000162894 | FCMR | chr1 | - | 206911729 | 206911952 | 206903316 | 206905147 | 206912928 | 206913042 | 715 | 354 | 896 | 654 | 2022 | 0 | 4 | 0 | 1 | 0 | 357 | 708 | 303 | 246 | 554 | 0 | 0 | 0 | 0 | 0 | 298 | 149 | 1 | 1 | 1 | 0.978 | 1 | 0.997 | 1 | 1 | 1 | 1 | 1 | 1 | -0.005 |
| SE | 46855 | ENSG00000162894 | FCMR | chr1 | - | 206911729 | 206911952 | 206903316 | 206905147 | 206921817 | 206921867 | 1 | 2 | 10 | 5 | 4 | 76 | 25 | 53 | 39 | 27 | 0 | 0 | 0 | 11 | 0 | 0 | 43 | 16 | 2 | 15 | 298 | 149 | 1 | 1 | 0.007 | 0.038 | 0.086 | 0.06 | 0.069 | NA | 0 | 0 | 0.733 | 0 | -0.131 |
| SE | 46856 | ENSG00000162894 | FCMR | chr1 | - | 206911729 | 206911952 | 206909724 | 206909868 | 206921817 | 206921867 | 9 | 3 | 13 | 12 | 4 | 1 | 0 | 1 | 0 | 0 | 0 | 1 | 1 | 11 | 0 | 0 | 0 | 0 | 0 | 0 | 298 | 149 | 1 | 1 | 0.818 | 1 | 0.867 | 1 | 1 | NA | 1 | 1 | 1 | NA | -0.063 |
| SE | 46857 | ENSG00000162894 | FCMR | chr1 | - | 206911729 | 206911952 | 206910209 | 206910340 | 206912928 | 206913042 | 1065 | 476 | 1213 | 891 | 2213 | 1 | 0 | 1 | 0 | 0 | 450 | 968 | 408 | 331 | 731 | 0 | 0 | 0 | 0 | 0 | 298 | 149 | 1 | 1 | 0.998 | 1 | 0.998 | 1 | 1 | 1 | 1 | 1 | 1 | 1 | -0.001 |
| SE | 46858 | ENSG00000162894 | FCMR | chr1 | - | 206911729 | 206911952 | 206910209 | 206910340 | 206913758 | 206914094 | 352 | 122 | 318 | 237 | 194 | 0 | 0 | 2 | 0 | 0 | 93 | 260 | 105 | 85 | 177 | 0 | 0 | 0 | 0 | 0 | 298 | 149 | 1 | 1 | 1 | 1 | 0.988 | 1 | 1 | 1 | 1 | 1 | 1 | 1 | -0.002 |
| SE | 46861 | ENSG00000162894 | FCMR | chr1 | - | 206912928 | 206913042 | 206910209 | 206910340 | 206913758 | 206914094 | 381 | 191 | 485 | 375 | 785 | 0 | 0 | 2 | 0 | 0 | 195 | 385 | 163 | 73 | 220 | 0 | 0 | 0 | 0 | 0 | 263 | 149 | 1 | 1 | 1 | 1 | 0.993 | 1 | 1 | 1 | 1 | 1 | 1 | 1 | -0.001 |
| SE | 46863 | ENSG00000162894 | FCMR | chr1 | - | 206912928 | 206913042 | 206911729 | 206911952 | 206913758 | 206914094 | 1055 | 530 | 1336 | 1006 | 2717 | 2 | 0 | 0 | 0 | 3 | 543 | 1068 | 449 | 306 | 744 | 0 | 0 | 0 | 0 | 0 | 263 | 149 | 1 | 1 | 0.997 | 1 | 1 | 1 | 0.998 | 1 | 1 | 1 | 1 | 1 | -0.001 |
| SE | 46864 | ENSG00000162894 | FCMR | chr1 | - | 206912928 | 206913042 | 206911729 | 206911952 | 206913861 | 206914094 | 606 | 326 | 770 | 536 | 1820 | 1 | 0 | 1 | 1 | 0 | 291 | 617 | 268 | 218 | 512 | 0 | 0 | 0 | 0 | 0 | 263 | 149 | 1 | 1 | 0.997 | 1 | 0.998 | 0.997 | 1 | 1 | 1 | 1 | 1 | 1 | -0.002 |
| SE | 46865 | ENSG00000162894 | FCMR | chr1 | - | 206912928 | 206913042 | 206911729 | 206911952 | 206921817 | 206921998 | 593 | 305 | 749 | 522 | 1727 | 1 | 2 | 9 | 5 | 4 | 307 | 608 | 270 | 215 | 535 | 0 | 0 | 0 | 11 | 0 | 263 | 149 | 1 | 1 | 0.997 | 0.989 | 0.979 | 0.983 | 0.996 | 1 | 1 | 1 | 0.917 | 1 | 0.005 |
| SE | 46866 | ENSG00000162894 | FCMR | chr1 | - | 206913758 | 206914094 | 206910209 | 206910340 | 206921817 | 206921867 | 1245 | 612 | 1261 | 1041 | 1267 | 4 | 0 | 0 | 2 | 5 | 420 | 946 | 397 | 387 | 516 | 0 | 1 | 0 | 0 | 0 | 298 | 149 | 1 | 1 | 0.994 | 1 | 1 | 0.996 | 0.992 | 1 | 0.998 | 1 | 1 | 1 | -0.003 |
| SE | 46871 | ENSG00000162894 | FCMR | chr1 | - | 206921340 | 206921433 | 206905018 | 206905147 | 206921817 | 206921997 | 18 | 5 | 11 | 7 | 11 | 76 | 25 | 53 | 39 | 27 | 0 | 5 | 8 | 2 | 5 | 0 | 43 | 16 | 2 | 15 | 242 | 149 | 1 | 1 | 0.127 | 0.11 | 0.113 | 0.1 | 0.201 | NA | 0.067 | 0.235 | 0.381 | 0.17 | -0.083 |
| SE | 46874 | ENSG00000055609 | KMT2C | chr7 | - | 152139185 | 152139259 | 152134921 | 152138904 | 152139674 | 152139791 | 370 | 311 | 244 | 343 | 452 | 0 | 0 | 2 | 0 | 0 | 111 | 321 | 360 | 194 | 46 | 0 | 0 | 2 | 0 | 0 | 223 | 149 | 1 | 1 | 1 | 1 | 0.988 | 1 | 1 | 1 | 1 | 0.992 | 1 | 1 | -0.001 |
| SE | 46875 | ENSG00000055609 | KMT2C | chr7 | - | 152139674 | 152139791 | 152138979 | 152139259 | 152144712 | 152144881 | 502 | 430 | 400 | 530 | 675 | 0 | 0 | 0 | 0 | 0 | 178 | 501 | 466 | 250 | 69 | 0 | 0 | 0 | 4 | 0 | 266 | 149 | 1 | 1 | 1 | 1 | 1 | 1 | 1 | 1 | 1 | 1 | 0.972 | 1 | 0.006 |
| SE | 46876 | ENSG00000055609 | KMT2C | chr7 | - | 152144712 | 152144881 | 152139674 | 152139791 | 152145152 | 152145295 | 386 | 304 | 342 | 434 | 502 | 0 | 1 | 0 | 0 | 0 | 130 | 408 | 307 | 176 | 56 | 0 | 0 | 0 | 0 | 0 | 298 | 149 | 1 | 1 | 1 | 0.993 | 1 | 1 | 1 | 1 | 1 | 1 | 1 | 1 | -0.001 |
| SE | 46877 | ENSG00000055609 | KMT2C | chr7 | - | 152144712 | 152144881 | 152139674 | 152139791 | 152145164 | 152145295 | 305 | 237 | 273 | 360 | 447 | 0 | 1 | 0 | 1 | 0 | 119 | 324 | 217 | 119 | 47 | 5 | 0 | 0 | 0 | 0 | 298 | 149 | 1 | 1 | 1 | 0.992 | 1 | 0.994 | 1 | 0.922 | 1 | 1 | 1 | 1 | 0.013 |
| SE | 46881 | ENSG00000055609 | KMT2C | chr7 | - | 152157760 | 152157922 | 152156204 | 152156271 | 152158862 | 152159072 | 25 | 23 | 32 | 39 | 66 | 1 | 0 | 0 | 0 | 0 | 12 | 31 | 20 | 9 | 11 | 0 | 0 | 0 | 0 | 0 | 298 | 149 | 1 | 1 | 0.926 | 1 | 1 | 1 | 1 | 1 | 1 | 1 | 1 | 1 | -0.015 |
| SE | 46882 | ENSG00000055609 | KMT2C | chr7 | - | 152157760 | 152157922 | 152156204 | 152156274 | 152158862 | 152159072 | 42 | 29 | 36 | 47 | 86 | 1 | 2 | 0 | 1 | 1 | 12 | 37 | 48 | 34 | 20 | 0 | 0 | 1 | 1 | 0 | 298 | 149 | 1 | 1 | 0.955 | 0.879 | 1 | 0.959 | 0.977 | 1 | 1 | 0.96 | 0.944 | 1 | -0.027 |
| SE | 46883 | ENSG00000055609 | KMT2C | chr7 | - | 152157760 | 152157922 | 152156204 | 152156343 | 152158862 | 152159072 | 35 | 25 | 30 | 50 | 76 | 0 | 0 | 0 | 1 | 0 | 12 | 35 | 21 | 23 | 9 | 0 | 0 | 2 | 0 | 0 | 298 | 149 | 1 | 1 | 1 | 1 | 1 | 0.962 | 1 | 1 | 1 | 0.84 | 1 | 1 | 0.024 |
| SE | 46885 | ENSG00000055609 | KMT2C | chr7 | - | 152157760 | 152157925 | 152156204 | 152156274 | 152158862 | 152159072 | 70 | 53 | 47 | 80 | 137 | 1 | 2 | 0 | 1 | 1 | 0 | 53 | 65 | 36 | 18 | 0 | 0 | 1 | 1 | 0 | 298 | 149 | 1 | 1 | 0.972 | 0.93 | 1 | 0.976 | 0.986 | NA | 1 | 0.97 | 0.947 | 1 | -0.006 |
| SE | 46886 | ENSG00000055609 | KMT2C | chr7 | - | 152157760 | 152157925 | 152156204 | 152156343 | 152158862 | 152159072 | 63 | 49 | 41 | 83 | 127 | 0 | 0 | 0 | 1 | 0 | 0 | 51 | 38 | 25 | 7 | 0 | 0 | 2 | 0 | 0 | 298 | 149 | 1 | 1 | 1 | 1 | 1 | 0.976 | 1 | NA | 1 | 0.905 | 1 | 1 | 0.019 |
| SE | 46889 | ENSG00000055609 | KMT2C | chr7 | - | 152160550 | 152160601 | 152158862 | 152159072 | 152162116 | 152163826 | 5 | 4 | 5 | 3 | 4 | 303 | 323 | 339 | 351 | 347 | 0 | 0 | 0 | 0 | 0 | 137 | 285 | 249 | 219 | 59 | 200 | 149 | 1 | 1 | 0.012 | 0.009 | 0.011 | 0.006 | 0.009 | 0 | 0 | 0 | 0 | 0 | 0.009 |
| SE | 46893 | ENSG00000055609 | KMT2C | chr7 | - | 152195521 | 152195569 | 152194439 | 152194568 | 152195906 | 152196011 | 1 | 4 | 2 | 4 | 8 | 248 | 209 | 216 | 259 | 592 | 0 | 4 | 0 | 0 | 0 | 74 | 238 | 212 | 110 | 74 | 197 | 149 | 1 | 1 | 0.003 | 0.014 | 0.007 | 0.012 | 0.01 | 0 | 0.013 | 0 | 0 | 0 | 0.007 |
| SE | 46895 | ENSG00000055609 | KMT2C | chr7 | - | 152205105 | 152205225 | 152202859 | 152203064 | 152207299 | 152207425 | 220 | 272 | 195 | 276 | 301 | 0 | 0 | 0 | 0 | 1 | 75 | 196 | 109 | 121 | 51 | 0 | 0 | 0 | 0 | 0 | 269 | 149 | 1 | 1 | 1 | 1 | 1 | 1 | 0.994 | 1 | 1 | 1 | 1 | 1 | -0.001 |
| SE | 46896 | ENSG00000055609 | KMT2C | chr7 | - | 152220522 | 152220735 | 152207285 | 152207428 | 152222000 | 152222066 | 432 | 456 | 425 | 623 | 590 | 0 | 0 | 0 | 1 | 0 | 88 | 479 | 467 | 317 | 54 | 0 | 0 | 0 | 0 | 0 | 298 | 149 | 1 | 1 | 1 | 1 | 1 | 0.997 | 1 | 1 | 1 | 1 | 1 | 1 | -0.001 |
| SE | 46898 | ENSG00000055609 | KMT2C | chr7 | - | 152222572 | 152222682 | 152222000 | 152222066 | 152229922 | 152230027 | 135 | 99 | 130 | 197 | 189 | 0 | 0 | 0 | 0 | 0 | 65 | 109 | 158 | 77 | 36 | 0 | 1 | 0 | 0 | 0 | 259 | 149 | 1 | 1 | 1 | 1 | 1 | 1 | 1 | 1 | 0.984 | 1 | 1 | 1 | 0.003 |
| SE | 46899 | ENSG00000055609 | KMT2C | chr7 | - | 152224014 | 152224179 | 152222572 | 152222682 | 152224434 | 152224616 | 324 | 250 | 325 | 412 | 409 | 1 | 0 | 1 | 0 | 0 | 222 | 317 | 261 | 163 | 142 | 0 | 0 | 0 | 2 | 0 | 298 | 149 | 1 | 1 | 0.994 | 1 | 0.994 | 1 | 1 | 1 | 1 | 1 | 0.976 | 1 | 0.002 |
| SE | 46902 | ENSG00000055609 | KMT2C | chr7 | - | 152224434 | 152224616 | 152222572 | 152222682 | 152230219 | 152230321 | 2 | 1 | 1 | 1 | 5 | 0 | 0 | 0 | 0 | 0 | 0 | 8 | 3 | 5 | 0 | 0 | 0 | 2 | 0 | 0 | 298 | 149 | 1 | 1 | 1 | 1 | 1 | 1 | 1 | NA | 1 | 0.429 | 1 | NA | 0.19 |
| SE | 46904 | ENSG00000055609 | KMT2C | chr7 | - | 152229922 | 152230027 | 152222572 | 152222682 | 152230219 | 152230321 | 95 | 77 | 80 | 92 | 80 | 0 | 0 | 0 | 0 | 0 | 39 | 57 | 82 | 73 | 24 | 0 | 0 | 2 | 0 | 0 | 254 | 149 | 1 | 1 | 1 | 1 | 1 | 1 | 1 | 1 | 1 | 0.96 | 1 | 1 | 0.008 |
| SE | 46907 | ENSG00000055609 | KMT2C | chr7 | - | 152247901 | 152248620 | 152235816 | 152235933 | 152251938 | 152252090 | 4 | 7 | 0 | 4 | 9 | 0 | 0 | 1 | 0 | 0 | 0 | 12 | 14 | 11 | 1 | 0 | 0 | 0 | 0 | 0 | 298 | 149 | 1 | 1 | 1 | 1 | 0 | 1 | 1 | NA | 1 | 1 | 1 | 1 | -0.2 |
| SE | 46909 | ENSG00000055609 | KMT2C | chr7 | - | 152250852 | 152250966 | 152249875 | 152249953 | 152251938 | 152252090 | 345 | 284 | 277 | 320 | 474 | 0 | 1 | 0 | 0 | 0 | 57 | 244 | 317 | 160 | 111 | 0 | 0 | 0 | 0 | 0 | 263 | 149 | 1 | 1 | 1 | 0.994 | 1 | 1 | 1 | 1 | 1 | 1 | 1 | 1 | -0.001 |
| SE | 46912 | ENSG00000055609 | KMT2C | chr7 | - | 152273704 | 152273867 | 152265037 | 152265209 | 152286299 | 152286376 | 223 | 207 | 183 | 303 | 175 | 0 | 2 | 0 | 0 | 0 | 148 | 262 | 228 | 173 | 164 | 0 | 0 | 0 | 0 | 0 | 298 | 149 | 1 | 1 | 1 | 0.981 | 1 | 1 | 1 | 1 | 1 | 1 | 1 | 1 | -0.004 |
| SE | 46913 | ENSG00000055609 | KMT2C | chr7 | - | 152286299 | 152286376 | 152273704 | 152273867 | 152309965 | 152310075 | 8 | 10 | 7 | 31 | 7 | 218 | 217 | 194 | 264 | 225 | 18 | 11 | 13 | 5 | 0 | 182 | 175 | 203 | 166 | 64 | 226 | 149 | 1 | 1 | 0.024 | 0.029 | 0.023 | 0.072 | 0.02 | 0.061 | 0.04 | 0.041 | 0.019 | 0 | 0.001 |
| SE | 46914 | ENSG00000055609 | KMT2C | chr7 | - | 152311797 | 152311946 | 152309965 | 152310075 | 152315137 | 152315338 | 834 | 757 | 598 | 1025 | 1045 | 0 | 0 | 2 | 1 | 0 | 862 | 843 | 653 | 765 | 1021 | 0 | 0 | 8 | 0 | 0 | 298 | 149 | 1 | 1 | 1 | 1 | 0.993 | 0.998 | 1 | 1 | 1 | 0.976 | 1 | 1 | 0.003 |
| SE | 46916 | ENSG00000055609 | KMT2C | chr7 | - | 152344186 | 152344255 | 152330600 | 152330742 | 152358586 | 152358675 | 2 | 2 | 0 | 0 | 0 | 12 | 12 | 4 | 5 | 21 | 0 | 5 | 2 | 0 | 5 | 0 | 4 | 10 | 2 | 0 | 218 | 149 | 1 | 1 | 0.102 | 0.102 | 0 | 0 | 0 | NA | 0.461 | 0.12 | 0 | 1 | -0.354 |
| SE | 46917 | ENSG00000055609 | KMT2C | chr7 | - | 152358586 | 152358675 | 152330171 | 152330739 | 152435625 | 152435706 | 203 | 161 | 186 | 187 | 206 | 0 | 0 | 0 | 2 | 0 | 90 | 164 | 130 | 129 | 111 | 0 | 0 | 0 | 0 | 0 | 238 | 149 | 1 | 1 | 1 | 1 | 1 | 0.983 | 1 | 1 | 1 | 1 | 1 | 1 | -0.003 |
| SE | 46921 | ENSG00000141027 | NCOR1 | chr17 | - | 16048844 | 16048988 | 16046899 | 16047093 | 16057513 | 16057723 | 305 | 228 | 321 | 365 | 339 | 0 | 0 | 1 | 0 | 0 | 189 | 230 | 188 | 243 | 153 | 0 | 0 | 2 | 4 | 0 | 293 | 149 | 1 | 1 | 1 | 1 | 0.994 | 1 | 1 | 1 | 1 | 0.98 | 0.969 | 1 | 0.009 |
| SE | 46922 | ENSG00000141027 | NCOR1 | chr17 | - | 16058470 | 16058599 | 16057906 | 16058064 | 16061400 | 16061894 | 349 | 222 | 297 | 357 | 371 | 0 | 0 | 2 | 0 | 0 | 169 | 333 | 199 | 309 | 81 | 0 | 0 | 1 | 0 | 0 | 278 | 149 | 1 | 1 | 1 | 1 | 0.988 | 1 | 1 | 1 | 1 | 0.991 | 1 | 1 | -0.001 |

| | | | | | | | | | | | | | | |
|---|---|---|---|---|---|---|---|---|---|---|---|---|---|---|
| SE | 46923 | ENSG00000141027 | NCOR1 | chr17 | - | 16061400 | 16061894 | 16058470 | 16058599 | 16062104 | 16062270 | 281 | 167 | [illegible] |
| SE | 46924 | ENSG00000141027 | NCOR1 | chr17 | - | 16061757 | 16061894 | 16058470 | 16058599 | 16062104 | 16062270 | 116 | 66 | [illegible] |
| SE | 46925 | ENSG00000141027 | NCOR1 | chr17 | - | 16064067 | 16064187 | 16062104 | 16062270 | 16064869 | 16065019 | 255 | 167 | [illegible] |
| SE | 46926 | ENSG00000141027 | NCOR1 | chr17 | - | 16064869 | 16065019 | 16064067 | 16064187 | 16065484 | 16065691 | 406 | 269 | [illegible] |
| SE | 46927 | ENSG00000141027 | NCOR1 | chr17 | - | 16067893 | 16068115 | 16065484 | 16065691 | 16070164 | 16070525 | 6 | 4 | [illegible] |
| SE | 46930 | ENSG00000141027 | NCOR1 | chr17 | - | 16072144 | 16072228 | 16071408 | 16071665 | 16073428 | 16073569 | 423 | 322 | [illegible] |
| SE | 46931 | ENSG00000141027 | NCOR1 | chr17 | - | 16072144 | 16072228 | 16071408 | 16071665 | 16075533 | 16075702 | 170 | 145 | [illegible] |
| SE | 46932 | ENSG00000141027 | NCOR1 | chr17 | - | 16073428 | 16073569 | 16071408 | 16071665 | 16075533 | 16075702 | 210 | 150 | [illegible] |
| SE | 46933 | ENSG00000141027 | NCOR1 | chr17 | - | 16073428 | 16073569 | 16072144 | 16072228 | 16075533 | 16075702 | 471 | 331 | [illegible] |
| SE | 46936 | ENSG00000141027 | NCOR1 | chr17 | - | 16086281 | 16086442 | 16080606 | 16080727 | 16091862 | 16092058 | 519 | 362 | [illegible] |
| SE | 46938 | ENSG00000141027 | NCOR1 | chr17 | - | 16091862 | 16092058 | 16086281 | 16086442 | 16108785 | 16108912 | 241 | 171 | [illegible] |
| SE | 46939 | ENSG00000141027 | NCOR1 | chr17 | - | 16098366 | 16098496 | 16091563 | 16092058 | 16108785 | 16108912 | 190 | 142 | [illegible] |
| SE | 46940 | ENSG00000141027 | NCOR1 | chr17 | - | 16101249 | 16101757 | 16098366 | 16098496 | 16108785 | 16108912 | 270 | 191 | [illegible] |
| SE | 46941 | ENSG00000141027 | NCOR1 | chr17 | - | 16101249 | 16101805 | 16098366 | 16098496 | 16108785 | 16108912 | 655 | 468 | [illegible] |
| SE | 46942 | ENSG00000141027 | NCOR1 | chr17 | - | 16117887 | 16118027 | 16108785 | 16108912 | 16119422 | 16119485 | 555 | 418 | [illegible] |
| SE | 46945 | ENSG00000141027 | NCOR1 | chr17 | - | 16143605 | 16143696 | 16139007 | 16139186 | 16146375 | 16146548 | 517 | 401 | [illegible] |
| SE | 46946 | ENSG00000141027 | NCOR1 | chr17 | - | 16146375 | 16146548 | 16143605 | 16143696 | 16149450 | 16149517 | 524 | 399 | [illegible] |
| SE | 46948 | ENSG00000141027 | NCOR1 | chr17 | - | 16158759 | 16158873 | 16153338 | 16153395 | 16164978 | 16165161 | 469 | 362 | [illegible] |
| SE | 46949 | ENSG00000141027 | NCOR1 | chr17 | - | 16164978 | 16165161 | 16158759 | 16158873 | 16171802 | 16171995 | 419 | 301 | [illegible] |
| SE | 46950 | ENSG00000141027 | NCOR1 | chr17 | - | 16164978 | 16165161 | 16158759 | 16158873 | 16186553 | 16186687 | 218 | 163 | [illegible] |
| SE | 46951 | ENSG00000141027 | NCOR1 | chr17 | - | 16171802 | 16171995 | 16158759 | 16158873 | 16186553 | 16186687 | 284 | 212 | [illegible] |
| SE | 46952 | ENSG00000141027 | NCOR1 | chr17 | - | 16171802 | 16171995 | 16164870 | 16165161 | 16186553 | 16186687 | 487 | 346 | [illegible] |
| SE | 46954 | ENSG00000141027 | NCOR1 | chr17 | - | 16171802 | 16172020 | 16158759 | 16158873 | 16186553 | 16186687 | 4 | 8 | [illegible] |
| SE | 46955 | ENSG00000141027 | NCOR1 | chr17 | - | 16171802 | 16172020 | 16164978 | 16165161 | 16186553 | 16186687 | 207 | 142 | [illegible] |
| SE | 46957 | ENSG00000141027 | NCOR1 | chr17 | - | 16186553 | 16186687 | 16171802 | 16171995 | 16194461 | 16194613 | 558 | 425 | [illegible] |
| SE | 46958 | ENSG00000141027 | NCOR1 | chr17 | - | 16194461 | 16194639 | 16186553 | 16186687 | 16215361 | 16215529 | 336 | 248 | [illegible] |
| SE | 46976 | ENSG00000023697 | DERA | chr12 | + | 15916116 | 15916316 | 15911328 | 15911414 | 15956935 | 15957033 | 2 | 1 | [illegible] |
| SE | 46980 | ENSG00000092295 | TGM1 | chr14 | - | 24259074 | 24259249 | 24258534 | 24258673 | 24259703 | 24259811 | 3 | 1 | [illegible] |
| SE | 46992 | ENSG00000163462 | TRIM46 | chr1 | + | 155174600 | 155174641 | 155173786 | 155174029 | 155175385 | 155175647 | 0 | 0 | [illegible] |
| SE | 46994 | ENSG00000163462 | TRIM46 | chr1 | + | 155178808 | 155178846 | 155178491 | 155178613 | 155179631 | 155179934 | 0 | 0 | [illegible] |
| SE | 46999 | ENSG00000142949 | PTPRF | chr1 | + | 43602070 | 43602097 | 43598719 | 43598913 | 43603415 | 43603533 | 0 | 0 | [illegible] |
| SE | 47002 | ENSG00000122417 | ODF2L | chr1 | - | 86355344 | 86355383 | 86354773 | 86354859 | 86356443 | 86356602 | 7 | 10 | [illegible] |
| SE | 47003 | ENSG00000122417 | ODF2L | chr1 | - | 86355344 | 86355388 | 86354773 | 86354859 | 86356443 | 86356544 | 13 | 12 | [illegible] |
| SE | 47005 | ENSG00000122417 | ODF2L | chr1 | - | 86356443 | 86356602 | 86354773 | 86354859 | 86358786 | 86358891 | 56 | 68 | [illegible] |
| SE | 47007 | ENSG00000122417 | ODF2L | chr1 | - | 86371017 | 86371153 | 86360425 | 86360536 | 86372430 | 86372540 | 93 | 108 | [illegible] |
| SE | 47008 | ENSG00000122417 | ODF2L | chr1 | - | 86371017 | 86371153 | 86360425 | 86360536 | 86376232 | 86376418 | 30 | 54 | [illegible] |
| SE | 47009 | ENSG00000122417 | ODF2L | chr1 | - | 86372430 | 86372540 | 86360425 | 86360536 | 86376232 | 86376418 | 41 | 46 | [illegible] |
| SE | 47010 | ENSG00000122417 | ODF2L | chr1 | - | 86372430 | 86372540 | 86371017 | 86371153 | 86376232 | 86376418 | 102 | 99 | [illegible] |
| SE | 47011 | ENSG00000122417 | ODF2L | chr1 | - | 86382241 | 86382358 | 86376232 | 86376418 | 86382930 | 86383002 | 94 | 64 | [illegible] |
| SE | 47013 | ENSG00000122417 | ODF2L | chr1 | - | 86386914 | 86387079 | 86385457 | 86385590 | 86396032 | 86396262 | 33 | 32 | [illegible] |
| SE | 47014 | ENSG00000122417 | ODF2L | chr1 | - | 86386914 | 86387086 | 86385457 | 86385590 | 86396032 | 86396280 | 42 | 36 | [illegible] |
| SE | 47015 | ENSG00000186301 | MST1P2 | chr1 | + | 16646786 | 16646901 | 16646596 | 16646709 | 16646979 | 16647116 | 0 | 0 | [illegible] |
| SE | 47017 | ENSG00000175309 | PHYKPL | chr5 | - | 178214795 | 178214885 | 178211889 | 178211970 | 178229939 | 178230320 | 33 | 17 | [illegible] |
| SE | 47019 | ENSG00000175309 | PHYKPL | chr5 | - | 178215275 | 178215430 | 178214795 | 178214885 | 178222851 | 178222934 | 64 | 28 | [illegible] |
| SE | 47023 | ENSG00000175309 | PHYKPL | chr5 | - | 178222354 | 178222580 | 178214795 | 178214885 | 178222851 | 178222934 | 51 | 22 | [illegible] |
| SE | 47024 | ENSG00000175309 | PHYKPL | chr5 | - | 178222354 | 178222580 | 178215275 | 178215430 | 178222851 | 178222934 | 77 | 32 | [illegible] |
| SE | 47025 | ENSG00000175309 | PHYKPL | chr5 | - | 178222851 | 178222934 | 178214795 | 178214885 | 178229939 | 178230073 | 5 | 0 | [illegible] |
| SE | 47026 | ENSG00000175309 | PHYKPL | chr5 | - | 178222851 | 178222934 | 178215275 | 178215430 | 178229939 | 178230073 | 2 | 0 | [illegible] |
| SE | 47030 | ENSG00000175309 | PHYKPL | chr5 | - | 178224615 | 178224729 | 178222851 | 178222934 | 178229939 | 178230073 | 9 | 0 | [illegible] |
| SE | 47034 | ENSG00000175309 | PHYKPL | chr5 | - | 178225354 | 178225429 | 178224447 | 178224564 | 178229939 | 178230049 | 53 | 26 | [illegible] |
| SE | 47036 | ENSG00000175309 | PHYKPL | chr5 | - | 178229939 | 178230099 | 178211889 | 178211970 | 178232491 | 178232541 | 2 | 0 | [illegible] |
| SE | 47038 | ENSG00000175309 | PHYKPL | chr5 | - | 178229939 | 178230099 | 178225354 | 178225429 | 178231668 | 178231829 | 81 | 40 | [illegible] |
| SE | 47039 | ENSG00000175309 | PHYKPL | chr5 | - | 178229939 | 178230099 | 178225354 | 178225429 | 178232491 | 178232541 | 83 | 36 | [illegible] |
| SE | 47043 | ENSG00000175309 | PHYKPL | chr5 | - | 178231668 | 178231829 | 178229939 | 178230099 | 178232491 | 178232630 | 2 | 4 | [illegible] |
| SE | 47044 | ENSG00000175309 | PHYKPL | chr5 | - | 178231668 | 178231866 | 178225354 | 178225429 | 178232491 | 178232768 | 1 | 0 | [illegible] |
| SE | 47050 | ENSG00000126067 | PSMB2 | chr1 | - | 35605232 | 35605282 | 35599540 | 35603374 | 35609245 | 35609408 | 250 | 91 | [illegible] |
| SE | 47051 | ENSG00000126067 | PSMB2 | chr1 | - | 35609245 | 35609408 | 35605232 | 35605282 | 35631273 | 35631344 | 380 | 139 | [illegible] |
| SE | 47052 | ENSG00000126067 | PSMB2 | chr1 | - | 35636309 | 35636432 | 35631273 | 35631344 | 35641341 | 35641526 | 426 | 198 | [illegible] |
| SE | 47053 | ENSG00000087301 | TXNDC16 | chr14 | - | 52455323 | 52455462 | 52440563 | 52440724 | 52470036 | 52470173 | 19 | 13 | [illegible] |
| SE | 47054 | ENSG00000087301 | TXNDC16 | chr14 | - | 52457089 | 52457174 | 52455323 | 52455462 | 52470036 | 52470173 | 34 | 23 | [illegible] |
| SE | 47055 | ENSG00000087301 | TXNDC16 | chr14 | - | 52462811 | 52462970 | 52457089 | 52457174 | 52470036 | 52470173 | 33 | 15 | [illegible] |
| SE | 47056 | ENSG00000087301 | TXNDC16 | chr14 | - | 52488362 | 52488486 | 52482821 | 52482965 | 52490390 | 52490451 | 41 | 50 | [illegible] |
| SE | 47057 | ENSG00000087301 | TXNDC16 | chr14 | - | 52490390 | 52490451 | 52488362 | 52488486 | 52490838 | 52491005 | 24 | 28 | [illegible] |
| SE | 47058 | ENSG00000087301 | TXNDC16 | chr14 | - | 52511239 | 52511390 | 52490838 | 52491005 | 52514879 | 52514970 | 46 | 45 | [illegible] |
| SE | 47059 | ENSG00000087301 | TXNDC16 | chr14 | - | 52514879 | 52514970 | 52511239 | 52511390 | 52519171 | 52519293 | 54 | 24 | [illegible] |
| SE | 47060 | ENSG00000087301 | TXNDC16 | chr14 | - | 52542370 | 52542453 | 52537598 | 52537672 | 52543397 | 52543630 | 73 | 60 | [illegible] |
| SE | 47062 | ENSG00000134463 | ECHDC3 | chr10 | + | 11755407 | 11755608 | 11749494 | 11749592 | 11763223 | 11763351 | 8 | 1 | [illegible] |
| SE | 47064 | ENSG00000196419 | XRCC6 | chr22 | + | 41621989 | 41622086 | 41621162 | 41621345 | 41636112 | 41636251 | 0 | 8 | [illegible] |
| SE | 47066 | ENSG00000196419 | XRCC6 | chr22 | + | 41636112 | 41636251 | 41628117 | 41628230 | 41636515 | 41636770 | 3588 | 2455 | [illegible] |
| SE | 47067 | ENSG00000196419 | XRCC6 | chr22 | + | 41636235 | 41636251 | 41628117 | 41628230 | 41636515 | 41636770 | 35 | 19 | [illegible] |
| SE | 47068 | ENSG00000196419 | XRCC6 | chr22 | + | 41646895 | 41647082 | 41621349 | 41622086 | 41650722 | 41650891 | 377 | 217 | [illegible] |
| SE | 47069 | ENSG00000196419 | XRCC6 | chr22 | + | 41646895 | 41647082 | 41637607 | 41637791 | 41650722 | 41650891 | 945 | 514 | [illegible] |
| SE | 47070 | ENSG00000196419 | XRCC6 | chr22 | + | 41653528 | 41653690 | 41650722 | 41650891 | 41656902 | 41657032 | 1133 | 556 | [illegible] |
| SE | 47071 | ENSG00000196419 | XRCC6 | chr22 | + | 41656902 | 41657032 | 41653528 | 41653690 | 41658251 | 41658352 | 1168 | 563 | [illegible] |
| SE | 47072 | ENSG00000196419 | XRCC6 | chr22 | + | 41656902 | 41657032 | 41653528 | 41653690 | 41661330 | 41661444 | 574 | 270 | [illegible] |
| SE | 47073 | ENSG00000196419 | XRCC6 | chr22 | + | 41658251 | 41658352 | 41653528 | 41653690 | 41661330 | 41661444 | 248 | 141 | [illegible] |
| SE | 47074 | ENSG00000196419 | XRCC6 | chr22 | + | 41658251 | 41658352 | 41656902 | 41657032 | 41661330 | 41661444 | 842 | 434 | [illegible] |
| SE | 47075 | ENSG00000196419 | XRCC6 | chr22 | + | 41658251 | 41658352 | 41656902 | 41657032 | 41663621 | 41664034 | 498 | 230 | [illegible] |
| SE | 47076 | ENSG00000196419 | XRCC6 | chr22 | + | 41661330 | 41661444 | 41656902 | 41657032 | 41663621 | 41664034 | 188 | 114 | [illegible] |
| SE | 47077 | ENSG00000196419 | XRCC6 | chr22 | + | 41661330 | 41661444 | 41658251 | 41658352 | 41663621 | 41664034 | 538 | 320 | [illegible] |
| SE | 47079 | ENSG00000095787 | WAC | chr10 | + | 28533997 | 28534034 | 28533117 | 28533620 | 28535561 | 28535757 | 155 | 124 | [illegible] |
| SE | 47080 | ENSG00000095787 | WAC | chr10 | + | 28533997 | 28534034 | 28533500 | 28533582 | 28535561 | 28535757 | 65 | 55 | [illegible] |
| SE | 47081 | ENSG00000095787 | WAC | chr10 | + | 28533997 | 28534182 | 28532772 | 28533007 | 28535561 | 28535726 | 24 | 15 | [illegible] |
| SE | 47082 | ENSG00000095787 | WAC | chr10 | + | 28533997 | 28534182 | 28533370 | 28533620 | 28535561 | 28535726 | 10 | 7 | [illegible] |
| SE | 47084 | ENSG00000095787 | WAC | chr10 | + | 28535909 | 28536059 | 28535561 | 28535757 | 28537583 | 28537815 | 6 | 16 | [illegible] |
| SE | 47086 | ENSG00000095787 | WAC | chr10 | + | 28535909 | 28536059 | 28535561 | 28535757 | 28583405 | 28583505 | 4 | 15 | [illegible] |
| SE | 47087 | ENSG00000095787 | WAC | chr10 | + | 28537583 | 28537815 | 28535561 | 28535757 | 28583398 | 28583505 | 0 | 2 | [illegible] |
| SE | 47089 | ENSG00000095787 | WAC | chr10 | + | 28537949 | 28538066 | 28537583 | 28537815 | 28583398 | 28583505 | 1 | 0 | [illegible] |
| SE | 47092 | ENSG00000095787 | WAC | chr10 | + | 28595732 | 28596041 | 28590719 | 28590832 | 28610698 | 28610710 | 571 | 434 | [illegible] |
| SE | 47093 | ENSG00000095787 | WAC | chr10 | + | 28608185 | 28608431 | 28590719 | 28590832 | 28610698 | 28610710 | 905 | 681 | [illegible] |
| SE | 47094 | ENSG00000095787 | WAC | chr10 | + | 28608185 | 28608431 | 28595732 | 28596041 | 28610698 | 28610710 | 1066 | 896 | [illegible] |
| SE | 47095 | ENSG00000095787 | WAC | chr10 | + | 28608185 | 28608431 | 28595732 | 28596041 | 28610707 | 28611922 | 350 | 315 | [illegible] |
| SE | 47096 | ENSG00000095787 | WAC | chr10 | + | 28610698 | 28610821 | 28608185 | 28608431 | 28611773 | 28611922 | 1399 | 1124 | [illegible] |
| SE | 47097 | ENSG00000095787 | WAC | chr10 | + | 28610710 | 28610821 | 28608185 | 28608431 | 28611773 | 28611922 | 651 | 512 | [illegible] |
| SE | 47098 | ENSG00000095787 | WAC | chr10 | + | 28611254 | 28611481 | 28610698 | 28610821 | 28611773 | 28611922 | 9 | 8 | [illegible] |
| SE | 47099 | ENSG00000095787 | WAC | chr10 | + | 28614566 | 28614685 | 28611773 | 28611922 | 28616172 | 28616362 | 658 | 410 | [illegible] |
| SE | 47100 | ENSG00000095787 | WAC | chr10 | + | 28614566 | 28614685 | 28611773 | 28611922 | 28617656 | 28617784 | 345 | 225 | [illegible] |
| SE | 47101 | ENSG00000095787 | WAC | chr10 | + | 28614566 | 28616362 | 28610707 | 28611922 | 28617656 | 28617784 | 909 | 623 | [illegible] |
| SE | 47102 | ENSG00000095787 | WAC | chr10 | + | 28615327 | 28615573 | 28614566 | 28614685 | 28616172 | 28616362 | 0 | 0 | [illegible] |
| SE | 47103 | ENSG00000095787 | WAC | chr10 | + | 28616172 | 28616362 | 28611773 | 28611922 | 28617656 | 28617784 | 565 | 402 | [illegible] |
| SE | 47104 | ENSG00000095787 | WAC | chr10 | + | 28616172 | 28616362 | 28614566 | 28614685 | 28617656 | 28617784 | 881 | 594 | [illegible] |
| SE | 47105 | ENSG00000095787 | WAC | chr10 | + | 28616175 | 28616362 | 28614566 | 28614685 | 28617656 | 28617784 | 678 | 468 | [illegible] |
| SE | 47109 | ENSG00000174839 | DENND6A | chr3 | - | 57633264 | 57633354 | 57630924 | 57630978 | 57634557 | 57634622 | 372 | 203 | [illegible] |
| SE | 47110 | ENSG00000174839 | DENND6A | chr3 | - | 57634557 | 57634622 | 57633264 | 57633354 | 57634703 | 57634769 | 227 | 140 | [illegible] |
| SE | 47115 | ENSG00000144824 | PHLDB2 | chr3 | + | 111884063 | 111885412 | 111845820 | 111845935 | 111913318 | 111913702 | 21 | 11 | [illegible] |
| SE | 47116 | ENSG00000144824 | PHLDB2 | chr3 | + | 111932268 | 111932397 | 111920281 | 111920419 | 111939474 | 111939630 | 27 | 8 | [illegible] |
| SE | 47117 | ENSG00000144824 | PHLDB2 | chr3 | + | 111945267 | 111945357 | 111940534 | 111940645 | 111948931 | 111949075 | 50 | 14 | [illegible] |
| SE | 47120 | ENSG00000144824 | PHLDB2 | chr3 | + | 111958274 | 111958295 | 111953929 | 111954029 | 111962107 | 111962312 | 0 | 0 | [illegible] |
| SE | 47122 | ENSG00000144824 | PHLDB2 | chr3 | + | 111967677 | 111967824 | 111966612 | 111966703 | 111969689 | 111969849 | 40 | 17 | [illegible] |
| SE | 47125 | ENSG00000115750 | TAF1B | chr2 | + | 9868275 | 9868429 | 9854325 | 9854421 | 9882705 | 9882805 | 108 | 92 | [illegible] |
| SE | 47126 | ENSG00000115750 | TAF1B | chr2 | + | 9875864 | 9876018 | 9868275 | 9868429 | 9882705 | 9882805 | 166 | 115 | [illegible] |
| SE | 47132 | ENSG00000164124 | TMEM144 | chr4 | + | 158203642 | 158203708 | 158201603 | 158201698 | 158211452 | 158211574 | 0 | 0 | [illegible] |
| SE | 47133 | ENSG00000164124 | TMEM144 | chr4 | + | 158211452 | 158211574 | 158201603 | 158201698 | 158212607 | 158212776 | 3 | 1 | [illegible] |
| SE | 47137 | ENSG00000164124 | TMEM144 | chr4 | + | 158212060 | 158212220 | 158211452 | 158211574 | 158212607 | 158212698 | 4 | 6 | [illegible] |
| SE | 47138 | ENSG00000164124 | TMEM144 | chr4 | + | 158232900 | 158232982 | 158219309 | 158219390 | 158235437 | 158235443 | 15 | 13 | [illegible] |
| SE | 47139 | ENSG00000164124 | TMEM144 | chr4 | + | 158235437 | 158235505 | 158232900 | 158232982 | 158237524 | 158237643 | 19 | 11 | [illegible] |
| SE | 47152 | ENSG00000138617 | PARP16 | chr15 | - | 65248116 | 65248228 | 65234459 | 65234823 | 65259463 | 65259542 | 0 | 1 | [illegible] |
| SE | 47153 | ENSG00000138617 | PARP16 | chr15 | - | 65263148 | 65263320 | 65260884 | 65261026 | 65286252 | 65286843 | 40 | 14 | [illegible] |
| SE | 47156 | ENSG00000138617 | PARP16 | chr15 | - | 65270934 | 65271072 | 65263148 | 65263320 | 65286252 | 65286366 | 12 | 7 | [illegible] |
| SE | 47157 | ENSG00000138617 | PARP16 | chr15 | - | 65270934 | 65271072 | 65266561 | 65266768 | 65286252 | 65286366 | 36 | 14 | [illegible] |
| SE | 47159 | ENSG00000243701 | DUBR | chr3 | + | 107245137 | 107245218 | 107240717 | 107241087 | 107246007 | 107247033 | 0 | 0 | [illegible] |
| SE | 47161 | ENSG00000243701 | DUBR | chr3 | + | 107246007 | 107246052 | 107240674 | 107241087 | 107266819 | 107266967 | 2 | 0 | [illegible] |
| SE | 47162 | ENSG00000243701 | DUBR | chr3 | + | 107246007 | 107246052 | 107240674 | 107241087 | 107285319 | 107285664 | 2 | 0 | [illegible] |
| SE | 47166 | ENSG00000243701 | DUBR | chr3 | + | 107246007 | 107248831 | 107240663 | 107241087 | 107291761 | 107292018 | 4 | 1 | [illegible] |
| SE | 47170 | ENSG00000243701 | DUBR | chr3 | + | 107248577 | 107248831 | 107240714 | 107241087 | 107285319 | 107285531 | 0 | 0 | [illegible] |
| SE | 47172 | ENSG00000243701 | DUBR | chr3 | + | 107285319 | 107285531 | 107240695 | 107241087 | 107291761 | 107291915 | 0 | 2 | [illegible] |
| SE | 47174 | ENSG00000243701 | DUBR | chr3 | + | 107285319 | 107286177 | 107240717 | 107241087 | 107291761 | 107291930 | 0 | 2 | [illegible] |
| SE | 47185 | ENSG00000242808 | SOX2-OT | chr3 | + | 181030569 | 181030689 | 181003774 | 181003933 | 181056696 | 181056786 | 0 | 0 | [illegible] |
| SE | 47187 | ENSG00000242808 | SOX2-OT | chr3 | + | 181056696 | 181056786 | 180989769 | 180989840 | 181088876 | 181088917 | 0 | 0 | [illegible] |
| SE | 47189 | ENSG00000242808 | SOX2-OT | chr3 | + | 181056696 | 181056786 | 181030569 | 181030689 | 181088876 | 181088917 | 0 | 0 | [illegible] |
| SE | 47230 | ENSG00000185104 | FAF1 | chr1 | - | 50451792 | 50452193 | 50441275 | 50441523 | 50475463 | 50475679 | 0 | 0 | [illegible] |
| SE | 47231 | ENSG00000185104 | FAF1 | chr1 | - | 50583651 | 50583715 | 50582617 | 50582699 | 50584684 | 50584811 | 133 | 87 | [illegible] |
| SE | 47232 | ENSG00000185104 | FAF1 | chr1 | - | 50596120 | 50596216 | 50584684 | 50584811 | 50655441 | 50655528 | 112 | 63 | [illegible] |
| SE | 47233 | ENSG00000185104 | FAF1 | chr1 | - | 50744683 | 50744775 | 50738862 | 50738954 | 50787999 | 50788205 | 173 | 146 | [illegible] |
| SE | 47235 | ENSG00000182134 | TDRKH | chr1 | - | 151778684 | 151779006 | 151776438 | 151776599 | 151779950 | 151780128 | 22 | 18 | [illegible] |
| SE | 47236 | ENSG00000182134 | TDRKH | chr1 | - | 151779102 | 151779242 | 151778632 | 151779006 | 151779950 | 151780128 | 68 | 55 | [illegible] |
| SE | 47239 | ENSG00000117500 | TMED5 | chr1 | - | 93156299 | 93156483 | 93149741 | 93154888 | 93158814 | 93158885 | 355 | 252 | [illegible] |
| SE | 47240 | ENSG00000117500 | TMED5 | chr1 | - | 93156299 | 93156483 | 93149741 | 93154888 | 93160128 | 93160226 | 962 | 686 | [illegible] |
| SE | 47241 | ENSG00000117500 | TMED5 | chr1 | - | 93158814 | 93158885 | 93151857 | 93154888 | 93160128 | 93160226 | 12 | 14 | [illegible] |
| SE | 47243 | ENSG00000115239 | ASB3 | chr2 | - | 53700270 | 53700528 | 53693883 | 53694014 | 53714383 | 53714581 | 76 | 55 | [illegible] |
| SE | 47244 | ENSG00000115239 | ASB3 | chr2 | - | 53714383 | 53714581 | 53700270 | 53700528 | 53716565 | 53716743 | 143 | 91 | [illegible] |
| SE | 47245 | ENSG00000115239 | ASB3 | chr2 | - | 53714383 | 53714581 | 53700270 | 53700528 | 53728711 | 53728847 | 28 | 19 | [illegible] |
| SE | 47246 | ENSG00000115239 | ASB3 | chr2 | - | 53716565 | 53716743 | 53700270 | 53700528 | 53728711 | 53728847 | 137 | 72 | [illegible] |
| SE | 47247 | ENSG00000115239 | ASB3 | chr2 | - | 53716565 | 53716743 | 53714383 | 53714581 | 53728711 | 53728847 | 254 | 144 | [illegible] |
| SE | 47250 | ENSG00000115239 | ASB3 | chr2 | - | 53748280 | 53748367 | 53729457 | 53729570 | 53750782 | 53750941 | 2 | 0 | [illegible] |
| SE | 47255 | ENSG00000185475 | TMEM179B | chr11 | + | 62789186 | 62789210 | 62787414 | 62787527 | 62789291 | 62789426 | 2 | 5 | [illegible] |
| SE | 47256 | ENSG00000185475 | TMEM179B | chr11 | + | 62789600 | 62789679 | 62789291 | 62789426 | 62789885 | 62790400 | 123 | 53 | [illegible] |
| SE | 47257 | ENSG00000133116 | KL | chr13 | + | 33055046 | 33055315 | 33053766 | 33054277 | 33060678 | 33061780 | 5 | 0 | [illegible] |
| SE | 47258 | ENSG00000133116 | KL | chr13 | + | 33055046 | 33055365 | 33053766 | 33054277 | 33060678 | 33061780 | 3 | 0 | [illegible] |
| SE | 47261 | ENSG00000113360 | DROSHA | chr5 | - | 31409249 | 31409332 | 31409055 | 31409159 | 31410745 | 31410887 | 91 | 83 | [illegible] |
| SE | 47262 | ENSG00000113360 | DROSHA | chr5 | - | 31410745 | 31410887 | 31406852 | 31406945 | 31421271 | 31421377 | 69 | 38 | [illegible] |
| SE | 47263 | ENSG00000113360 | DROSHA | chr5 | - | 31410745 | 31410887 | 31409249 | 31409332 | 31421271 | 31421377 | 99 | 65 | [illegible] |
| SE | 47270 | ENSG00000113360 | DROSHA | chr5 | - | 31467938 | 31468063 | 31466181 | 31466281 | 31472062 | 31472232 | 124 | 91 | [illegible] |
| SE | 47271 | ENSG00000113360 | DROSHA | chr5 | - | 31467938 | 31468063 | 31466181 | 31466281 | 31483553 | 31483628 | 74 | 59 | [illegible] |
| SE | 47272 | ENSG00000113360 | DROSHA | chr5 | - | 31472062 | 31472232 | 31466181 | 31466281 | 31483553 | 31483628 | 69 | 46 | [illegible] |
| SE | 47274 | ENSG00000113360 | DROSHA | chr5 | - | 31493206 | 31493293 | 31486490 | 31486562 | 31495285 | 31495372 | 128 | 77 | [illegible] |
| SE | 47275 | ENSG00000113360 | DROSHA | chr5 | - | 31495285 | 31495372 | 31486490 | 31486562 | 31504554 | 31504635 | 60 | 37 | [illegible] |
| SE | 47276 | ENSG00000113360 | DROSHA | chr5 | - | 31508620 | 31508775 | 31504554 | 31504635 | 31511034 | 31511176 | 195 | 117 | [illegible] |
| SE | 47282 | ENSG00000113360 | DROSHA | chr5 | - | 31531449 | 31531525 | 31530797 | 31530924 | 31532113 | 31532196 | 19 | 19 | [illegible] |
| SE | 47285 | ENSG00000121957 | GPSM2 | chr1 | + | 108882579 | 108882625 | 108876981 | 108877228 | 108885274 | 108885578 | 0 | 4 | [illegible] |
| SE | 47293 | ENSG00000121957 | GPSM2 | chr1 | + | 108918612 | 108919150 | 108914337 | 108914408 | 108922416 | 108922576 | 3 | 7 | [illegible] |
| SE | 47295 | ENSG00000121957 | GPSM2 | chr1 | + | 108923999 | 108924214 | 108922416 | 108922576 | 108929700 | 108929811 | 23 | 24 | [illegible] |
| SE | 47301 | ENSG00000245552 | NA | chr11 | + | 95204805 | 95204912 | 95157627 | 95157803 | 95208554 | 95209111 | 0 | 3 | [illegible] |
| SE | 47302 | ENSG00000245552 | NA | chr11 | + | 95232589 | 95232674 | 95157627 | 95157803 | 95232879 | 95232919 | 11 | 4 | [illegible] |
| SE | 47303 | ENSG00000245552 | NA | chr11 | + | 95232879 | 95232919 | 95232589 | 95232674 | 95233453 | 95233929 | 13 | 7 | [illegible] |
| SE | 47304 | ENSG00000073282 | TP63 | chr3 | + | 189808271 | 189808526 | 189738641 | 189738774 | 189864231 | 189864418 | 0 | 2 | [illegible] |
| SE | 47308 | ENSG00000073282 | TP63 | chr3 | + | 189890788 | 189890882 | 189889339 | 189889484 | 189894205 | 189894502 | 0 | 0 | [illegible] |
| SE | 47315 | ENSG00000133106 | EPSTI1 | chr13 | - | 42895008 | 42895108 | 42889210 | 42889242 | 42900309 | 42900383 | 74 | 61 | [illegible] |
| SE | 47316 | ENSG00000133106 | EPSTI1 | chr13 | - | 42917540 | 42917624 | 42900309 | 42900383 | 42926335 | 42926429 | 128 | 182 | [illegible] |
| SE | 47319 | ENSG00000133106 | EPSTI1 | chr13 | - | 42963254 | 42963338 | 42953947 | 42954021 | 42969093 | 42969177 | 70 | 86 | [illegible] |
| SE | 47320 | ENSG00000133106 | EPSTI1 | chr13 | - | 42964065 | 42964139 | 42963254 | 42963338 | 42969093 | 42969177 | 297 | 395 | [illegible] |
| SE | 47323 | ENSG00000133106 | EPSTI1 | chr13 | - | 42991098 | 42991182 | 42970611 | 42970670 | 42991977 | 42992180 | 0 | 0 | [illegible] |
| SE | 47324 | ENSG00000166669 | ATF7IP2 | chr16 | + | 10387143 | 10387290 | 10386054 | 10386122 | 10387377 | 10387484 | 107 | 89 | [illegible] |
| SE | 47326 | ENSG00000166669 | ATF7IP2 | chr16 | + | 10387377 | 10387494 | 10386054 | 10386122 | 10414573 | 10414612 | 8 | 5 | [illegible] |
| SE | 47327 | ENSG00000166669 | ATF7IP2 | chr16 | + | 10387377 | 10387494 | 10386054 | 10386122 | 10428867 | 10429016 | 0 | 2 | [illegible] |
| SE | 47328 | ENSG00000166669 | ATF7IP2 | chr16 | + | 10414573 | 10414612 | 10386057 | 10386122 | 10419580 | 10419620 | 0 | 1 | [illegible] |
| SE | 47330 | ENSG00000166669 | ATF7IP2 | chr16 | + | 10414573 | 10414612 | 10387377 | 10387494 | 10428867 | 10429004 | 1 | 0 | [illegible] |
| SE | 47331 | ENSG00000166669 | ATF7IP2 | chr16 | + | 10419580 | 10419623 | 10386054 | 10386122 | 10428867 | 10429004 | 7 | 0 | [illegible] |
| SE | 47332 | ENSG00000166669 | ATF7IP2 | chr16 | + | 10419580 | 10419623 | 10414573 | 10414612 | 10428867 | 10429004 | 9 | 1 | [illegible] |
| SE | 47334 | ENSG00000166669 | ATF7IP2 | chr16 | + | 10428867 | 10429016 | 10419580 | 10419623 | 10430610 | 10430672 | 86 | 48 | [illegible] |
| SE | 47336 | ENSG00000166669 | ATF7IP2 | chr16 | + | 10438100 | 10438235 | 10433524 | 10433649 | 10440363 | 10440462 | 75 | 99 | [illegible] |
| SE | 47338 | ENSG00000166669 | ATF7IP2 | chr16 | + | 10457371 | 10457529 | 10440363 | 10440462 | 10473478 | 10473534 | 14 | 8 | [illegible] |
| SE | 47342 | ENSG00000166669 | ATF7IP2 | chr16 | + | 10480878 | 10480964 | 10473478 | 10473534 | 10481835 | 10482118 | 14 | 24 | [illegible] |
| SE | 47343 | ENSG00000164136 | IL15 | chr4 | + | 141656185 | 141656307 | 141636582 | 141636748 | 141688821 | 141689035 | 4 | 7 | [illegible] |
| SE | 47344 | ENSG00000164136 | IL15 | chr4 | + | 141656185 | 141656307 | 141636582 | 141636748 | 141719365 | 141719476 | 2 | 0 | [illegible] |
| SE | 47345 | ENSG00000164136 | IL15 | chr4 | + | 141656185 | 141656307 | 141636601 | 141637224 | 141688821 | 141689035 | 10 | 5 | [illegible] |
| SE | 47346 | ENSG00000164136 | IL15 | chr4 | + | 141688821 | 141689035 | 141636725 | 141636748 | 141719365 | 141719476 | 0 | 1 | [illegible] |
| SE | 47347 | ENSG00000164136 | IL15 | chr4 | + | 141688821 | 141689035 | 141656185 | 141656307 | 141719365 | 141719476 | 6 | 9 | [illegible] |
| SE | 47348 | ENSG00000164136 | IL15 | chr4 | + | 141688821 | 141689035 | 141656185 | 141656307 | 141719377 | 141719476 | 8 | 12 | [illegible] |
| SE | 47349 | ENSG00000164136 | IL15 | chr4 | + | 141719365 | 141719476 | 141656185 | 141656307 | 141720468 | 141720566 | 5 | 11 | [illegible] |
| SE | 47350 | ENSG00000164136 | IL15 | chr4 | + | 141719377 | 141719476 | 141656185 | 141656307 | 141720468 | 141720566 | 1 | 11 | [illegible] |
| SE | 47355 | ENSG00000156535 | CD109 | chr6 | + | 73697399 | 73697572 | 73695784 | 73696289 | 73723250 | 73723279 | 8 | 7 | [illegible] |
| SE | 47357 | ENSG00000156535 | CD109 | chr6 | + | 73762383 | 73762480 | 73758943 | 73759028 | 73762740 | 73762882 | 15 | 16 | [illegible] |
| SE | 47358 | ENSG00000156535 | CD109 | chr6 | + | 73792625 | 73792802 | 73788467 | 73788612 | 73803219 | 73803301 | 13 | 11 | [illegible] |
| SE | 47359 | ENSG00000156535 | CD109 | chr6 | + | 73806843 | 73807072 | 73803219 | 73803301 | 73808082 | 73808248 | 11 | 9 | [illegible] |
| SE | 47361 | ENSG00000286112 | NA | chr9 | - | 128824770 | 128824869 | 128824071 | 128824174 | 128824976 | 128825049 | 34 | 9 | [illegible] |
| SE | 47362 | ENSG00000286112 | NA | chr9 | - | 128826021 | 128826152 | 128824976 | 128825049 | 128826383 | 128826433 | 69 | 37 | [illegible] |
| SE | 47363 | ENSG00000286112 | NA | chr9 | - | 128832290 | 128832475 | 128829744 | 128830005 | 128833226 | 128833643 | 1 | 0 | [illegible] |
| SE | 47365 | ENSG00000286112 | NA | chr9 | - | 128833739 | 128833826 | 128833226 | 128833643 | 128835322 | 128835402 | 17 | 5 | [illegible] |
| SE | 47366 | ENSG00000171097 | KYAT1 | chr9 | - | 128838217 | 128838367 | 128838050 | 128838137 | 128842653 | 128842801 | 12 | 4 | [illegible] |
| SE | 47367 | ENSG00000286112 | NA | chr9 | - | 128846679 | 128846856 | 128845352 | 128845411 | 128881896 | 128881931 | 0 | 4 | [illegible] |
| SE | 47376 | ENSG00000144426 | NBEAL1 | chr2 | + | 203045635 | 203045778 | 203041764 | 203041856 | 203049813 | 203049975 | 0 | 0 | [illegible] |
| SE | 47377 | ENSG00000144426 | NBEAL1 | chr2 | + | 203056426 | 203056508 | 203049813 | 203049975 | 203057325 | 203057353 | 14 | 26 | [illegible] |
| SE | 47381 | ENSG00000144426 | NBEAL1 | chr2 | + | 203099628 | 203099712 | 203084462 | 203084569 | 203107419 | 203107518 | 15 | 21 | [illegible] |
| SE | 47382 | ENSG00000144426 | NBEAL1 | chr2 | + | 203115984 | 203116070 | 203113014 | 203113318 | 203122253 | 203122343 | 29 | 32 | [illegible] |
| SE | 47383 | ENSG00000144426 | NBEAL1 | chr2 | + | 203138161 | 203138315 | 203136598 | 203136774 | 203138619 | 203138748 | 57 | 37 | [illegible] |
| SE | 47384 | ENSG00000144426 | NBEAL1 | chr2 | + | 203151464 | 203151589 | 203148990 | 203149148 | 203157698 | 203157825 | 40 | 35 | [illegible] |
| SE | 47385 | ENSG00000144426 | NBEAL1 | chr2 | + | 203157698 | 203157825 | 203151464 | 203151589 | 203166148 | 203166297 | 48 | 37 | [illegible] |
| SE | 47387 | ENSG00000144426 | NBEAL1 | chr2 | + | 203183278 | 203183388 | 203180381 | 203180512 | 203188471 | 203188589 | 69 | 88 | [illegible] |

| | | | | | | | | | | | | | | |
|---|---|---|---|---|---|---|---|---|---|---|---|---|---|---|
| SE | 47388 | ENSG00000144426 | NBEAL1 | chr2 | + | 203193794 | 203193911 | 203190291 | 203190389 | 203197301 | 203197391 | 13 | 10 | [illegible] |
| SE | 47389 | ENSG00000144426 | NBEAL1 | chr2 | + | 203193794 | 203193911 | 203190291 | 203190389 | 203199337 | 203199447 | 8 | 2 | [illegible] |
| SE | 47390 | ENSG00000144426 | NBEAL1 | chr2 | + | 203197301 | 203197391 | 203190291 | 203190389 | 203199337 | 203199447 | 19 | 10 | [illegible] |
| SE | 47391 | ENSG00000144426 | NBEAL1 | chr2 | + | 203197301 | 203197391 | 203193025 | 203193911 | 203199337 | 203199447 | 24 | 17 | [illegible] |
| SE | 47392 | ENSG00000144426 | NBEAL1 | chr2 | + | 203201542 | 203201715 | 203199337 | 203199447 | 203202686 | 203202781 | 22 | 20 | [illegible] |
| SE | 47393 | ENSG00000144426 | NBEAL1 | chr2 | + | 203213517 | 203213653 | 203210957 | 203211106 | 203217252 | 203217350 | 3 | 9 | [illegible] |
| SE | 47394 | ENSG00000240288 | GHRLOS | chr3 | + | 10287413 | 10287481 | 10285753 | 10285853 | 10287944 | 10288102 | 20 | 6 | [illegible] |
| SE | 47396 | ENSG00000173372 | C1QA | chr1 | + | 22637609 | 22637779 | 22636627 | 22636702 | 22638832 | 22639331 | 14 | 15 | [illegible] |
| SE | 47397 | ENSG00000104131 | EIF3J | chr15 | + | 44557488 | 44557650 | 44554552 | 44554667 | 44560248 | 44560322 | 850 | 841 | [illegible] |
| SE | 47403 | ENSG00000156804 | FBXO32 | chr8 | - | 123506391 | 123506574 | 123504603 | 123504747 | 123531897 | 123531990 | 77 | 50 | [illegible] |
| SE | 47404 | ENSG00000156804 | FBXO32 | chr8 | - | 123513197 | 123513382 | 123506391 | 123506574 | 123514239 | 123514333 | 169 | 106 | [illegible] |
| SE | 47406 | ENSG00000156804 | FBXO32 | chr8 | - | 123514051 | 123514333 | 123513273 | 123513382 | 123531897 | 123531990 | 58 | 29 | [illegible] |
| SE | 47408 | ENSG00000156804 | FBXO32 | chr8 | - | 123514239 | 123514333 | 123513197 | 123513382 | 123531897 | 123531990 | 156 | 98 | [illegible] |
| SE | 47409 | ENSG00000156804 | FBXO32 | chr8 | - | 123534701 | 123534814 | 123533190 | 123533240 | 123540898 | 123541189 | 156 | 127 | [illegible] |
| SE | 47410 | ENSG00000161973 | CCDC42 | chr17 | - | 8735389 | 8735611 | 8735095 | 8735254 | 8741473 | 8741671 | 0 | 0 | [illegible] |
| SE | 47415 | ENSG00000198171 | DDRGK1 | chr20 | - | 3191764 | 3191821 | 3191189 | 3191238 | 3194829 | 3194868 | 152 | 72 | [illegible] |
| SE | 47418 | ENSG00000115155 | OTOF | chr2 | - | 26467098 | 26467233 | 26466713 | 26466851 | 26467364 | 26467501 | 4 | 12 | [illegible] |
| SE | 47419 | ENSG00000115155 | OTOF | chr2 | - | 26477415 | 26477506 | 26477171 | 26477288 | 26477648 | 26477749 | 4 | 4 | [illegible] |
| SE | 47422 | ENSG00000162366 | PDZK1IP1 | chr1 | - | 47187318 | 47187427 | 47185001 | 47185097 | 47189865 | 47190012 | 14 | 46 | [illegible] |
| SE | 47423 | ENSG00000163867 | ZMYM6 | chr1 | - | 35006950 | 35007098 | 35005131 | 35005272 | 35008751 | 35008792 | 443 | 463 | [illegible] |
| SE | 47424 | ENSG00000163867 | ZMYM6 | chr1 | - | 35011889 | 35012005 | 35010546 | 35011036 | 35012430 | 35012456 | 165 | 145 | [illegible] |
| SE | 47425 | ENSG00000163867 | ZMYM6 | chr1 | - | 35012430 | 35012581 | 35011889 | 35012005 | 35014696 | 35014888 | 125 | 87 | [illegible] |
| SE | 47427 | ENSG00000163867 | ZMYM6 | chr1 | - | 35020382 | 35020467 | 35014987 | 35015162 | 35030546 | 35030713 | 71 | 69 | [illegible] |
| SE | 47439 | ENSG00000198178 | CLEC4C | chr12 | - | 7730796 | 7730912 | 7729382 | 7729740 | 7741420 | 7741531 | 25 | 24 | [illegible] |
| SE | 47441 | ENSG00000198178 | CLEC4C | chr12 | - | 7741420 | 7741531 | 7737428 | 7737574 | 7746330 | 7746423 | 132 | 110 | [illegible] |
| SE | 47442 | ENSG00000198178 | CLEC4C | chr12 | - | 7741420 | 7741531 | 7737428 | 7737574 | 7747317 | 7747481 | 71 | 67 | [illegible] |
| SE | 47443 | ENSG00000198178 | CLEC4C | chr12 | - | 7746330 | 7746423 | 7737428 | 7737574 | 7747317 | 7747339 | 10 | 7 | [illegible] |
| SE | 47450 | ENSG00000156873 | PHKG2 | chr16 | + | 30751548 | 30751603 | 30751105 | 30751217 | 30753231 | 30753297 | 43 | 13 | [illegible] |
| SE | 47451 | ENSG00000156873 | PHKG2 | chr16 | + | 30751548 | 30751603 | 30751105 | 30751281 | 30753231 | 30753297 | 76 | 25 | [illegible] |
| SE | 47453 | ENSG00000156873 | PHKG2 | chr16 | + | 30756366 | 30756520 | 30753393 | 30753557 | 30756589 | 30756715 | 22 | 14 | [illegible] |
| SE | 47454 | ENSG00000083635 | NUFIP1 | chr13 | - | 44949721 | 44949838 | 44943441 | 44943674 | 44959380 | 44959574 | 64 | 49 | [illegible] |
| SE | 47455 | ENSG00000083635 | NUFIP1 | chr13 | - | 44949721 | 44949838 | 44943441 | 44943674 | 44965843 | 44965936 | 32 | 24 | [illegible] |
| SE | 47456 | ENSG00000083635 | NUFIP1 | chr13 | - | 44959380 | 44959574 | 44943441 | 44943674 | 44965843 | 44965936 | 45 | 42 | [illegible] |
| SE | 47458 | ENSG00000083635 | NUFIP1 | chr13 | - | 44965843 | 44965936 | 44959380 | 44959574 | 44979189 | 44979266 | 74 | 73 | [illegible] |
| SE | 47460 | ENSG00000083635 | NUFIP1 | chr13 | - | 44979889 | 44979952 | 44965843 | 44965936 | 44980721 | 44980820 | 11 | 10 | [illegible] |
| SE | 47464 | ENSG00000122420 | PTGFR | chr1 | + | 78497873 | 78497944 | 78492671 | 78493541 | 78536405 | 78539749 | 0 | 0 | [illegible] |
| SE | 47469 | ENSG00000264364 | DYNLL2 | chr17 | + | 58087081 | 58087222 | 58083418 | 58083683 | 58089141 | 58095542 | 269 | 137 | [illegible] |
| SE | 47470 | ENSG00000183023 | SLC8A1 | chr2 | - | 40160764 | 40160864 | 40139400 | 40139676 | 40164853 | 40164984 | 161 | 100 | [illegible] |
| SE | 47473 | ENSG00000183023 | SLC8A1 | chr2 | - | 40174824 | 40174842 | 40164853 | 40164984 | 40428472 | 40430271 | 11 | 27 | [illegible] |
| SE | 47476 | ENSG00000183023 | SLC8A1 | chr2 | - | 40177751 | 40177855 | 40175260 | 40175281 | 40178386 | 40178493 | 2 | 1 | [illegible] |
| SE | 47483 | ENSG00000143363 | PRUNE1 | chr1 | + | 151017811 | 151017904 | 151008419 | 151008671 | 151027232 | 151027327 | 14 | 28 | [illegible] |
| SE | 47486 | ENSG00000143363 | PRUNE1 | chr1 | + | 151018466 | 151018669 | 151017811 | 151017904 | 151027232 | 151027327 | 21 | 15 | [illegible] |
| SE | 47488 | ENSG00000143363 | PRUNE1 | chr1 | + | 151018556 | 151018669 | 151017811 | 151017904 | 151025514 | 151025673 | 6 | 3 | [illegible] |
| SE | 47489 | ENSG00000143363 | PRUNE1 | chr1 | + | 151018556 | 151018669 | 151017811 | 151017904 | 151027232 | 151027327 | 6 | 3 | [illegible] |
| SE | 47490 | ENSG00000143363 | PRUNE1 | chr1 | + | 151024610 | 151024795 | 151008476 | 151008671 | 151025514 | 151025673 | 106 | 89 | [illegible] |
| SE | 47491 | ENSG00000143363 | PRUNE1 | chr1 | + | 151024610 | 151024795 | 151008476 | 151008671 | 151027232 | 151027327 | 1 | 8 | [illegible] |
| SE | 47493 | ENSG00000143363 | PRUNE1 | chr1 | + | 151024610 | 151024795 | 151017811 | 151017904 | 151027232 | 151027327 | 25 | 24 | [illegible] |
| SE | 47494 | ENSG00000143363 | PRUNE1 | chr1 | + | 151024610 | 151024795 | 151018466 | 151018669 | 151025514 | 151025673 | 159 | 133 | [illegible] |
| SE | 47496 | ENSG00000143363 | PRUNE1 | chr1 | + | 151025514 | 151025673 | 151008477 | 151008671 | 151027232 | 151027327 | 69 | 74 | [illegible] |
| SE | 47499 | ENSG00000143363 | PRUNE1 | chr1 | + | 151025514 | 151025673 | 151018466 | 151018669 | 151033805 | 151033837 | 0 | 0 | [illegible] |
| SE | 47501 | ENSG00000143363 | PRUNE1 | chr1 | + | 151027232 | 151027327 | 151018466 | 151018669 | 151033805 | 151034113 | 4 | 5 | [illegible] |
| SE | 47505 | ENSG00000135845 | PIGC | chr1 | - | 172376553 | 172376636 | 172370188 | 172370311 | 172386780 | 172386957 | 3 | 4 | [illegible] |
| SE | 47506 | ENSG00000135845 | PIGC | chr1 | - | 172376553 | 172376636 | 172370188 | 172370311 | 172399818 | 172399916 | 5 | 4 | [illegible] |
| SE | 47507 | ENSG00000135845 | PIGC | chr1 | - | 172378130 | 172378258 | 172376553 | 172376636 | 172399818 | 172399916 | 2 | 0 | [illegible] |
| SE | 47508 | ENSG00000135845 | PIGC | chr1 | - | 172386780 | 172386957 | 172370188 | 172370311 | 172399818 | 172399916 | 0 | 1 | [illegible] |
| SE | 47509 | ENSG00000135845 | PIGC | chr1 | - | 172386780 | 172386957 | 172376553 | 172376636 | 172399818 | 172399916 | 0 | 0 | [illegible] |
| SE | 47510 | ENSG00000135845 | PIGC | chr1 | - | 172386780 | 172386957 | 172376553 | 172376636 | 172415293 | 172415376 | 0 | 0 | [illegible] |
| SE | 47511 | ENSG00000135845 | PIGC | chr1 | - | 172397163 | 172397272 | 172393133 | 172394410 | 172399818 | 172399916 | 7 | 7 | [illegible] |
| SE | 47512 | ENSG00000135845 | PIGC | chr1 | - | 172399818 | 172399916 | 172376553 | 172376636 | 172415293 | 172415326 | 2 | 0 | [illegible] |
| SE | 47513 | ENSG00000135845 | PIGC | chr1 | - | 172399818 | 172399916 | 172386780 | 172386957 | 172415293 | 172415326 | 0 | 0 | [illegible] |
| SE | 47514 | ENSG00000135845 | PIGC | chr1 | - | 172399818 | 172399916 | 172393133 | 172394410 | 172415293 | 172415326 | 1 | 8 | [illegible] |
| SE | 47515 | ENSG00000135845 | PIGC | chr1 | - | 172399818 | 172399916 | 172393133 | 172394410 | 172443987 | 172444083 | 1 | 8 | [illegible] |
| SE | 47516 | ENSG00000135845 | PIGC | chr1 | - | 172415293 | 172415374 | 172393133 | 172394410 | 172443987 | 172444083 | 4 | 0 | [illegible] |
| SE | 47517 | ENSG00000135845 | PIGC | chr1 | - | 172415293 | 172415374 | 172399818 | 172399916 | 172443987 | 172444083 | 2 | 0 | [illegible] |
| SE | 47518 | ENSG00000121766 | ZCCHC17 | chr1 | + | 31310043 | 31310164 | 31296981 | 31297075 | 31318166 | 31318236 | 30 | 24 | [illegible] |
| SE | 47519 | ENSG00000121766 | ZCCHC17 | chr1 | + | 31310043 | 31310164 | 31296981 | 31297075 | 31319108 | 31319166 | 187 | 126 | [illegible] |
| SE | 47520 | ENSG00000121766 | ZCCHC17 | chr1 | + | 31310043 | 31310164 | 31297007 | 31297213 | 31319108 | 31319166 | 139 | 102 | [illegible] |
| SE | 47524 | ENSG00000121766 | ZCCHC17 | chr1 | + | 31319108 | 31319166 | 31296981 | 31297075 | 31338956 | 31339048 | 0 | 0 | [illegible] |
| SE | 47525 | ENSG00000121766 | ZCCHC17 | chr1 | + | 31319108 | 31319166 | 31310043 | 31310164 | 31337174 | 31337275 | 231 | 154 | [illegible] |
| SE | 47527 | ENSG00000121766 | ZCCHC17 | chr1 | + | 31337174 | 31337275 | 31319108 | 31319166 | 31346639 | 31346740 | 180 | 124 | [illegible] |
| SE | 47528 | ENSG00000121766 | ZCCHC17 | chr1 | + | 31338956 | 31339048 | 31319108 | 31319166 | 31346639 | 31346740 | 50 | 36 | [illegible] |
| SE | 47529 | ENSG00000121766 | ZCCHC17 | chr1 | + | 31338956 | 31339048 | 31319108 | 31319166 | 31364031 | 31364953 | 0 | 1 | [illegible] |
| SE | 47530 | ENSG00000121766 | ZCCHC17 | chr1 | + | 31338956 | 31339048 | 31337174 | 31337275 | 31346639 | 31346740 | 167 | 99 | [illegible] |
| SE | 47531 | ENSG00000121766 | ZCCHC17 | chr1 | + | 31346639 | 31346740 | 31319108 | 31319166 | 31364031 | 31364953 | 0 | 0 | [illegible] |
| SE | 47532 | ENSG00000121766 | ZCCHC17 | chr1 | + | 31346639 | 31346740 | 31338956 | 31339048 | 31348828 | 31348968 | 180 | 105 | [illegible] |
| SE | 47534 | ENSG00000121766 | ZCCHC17 | chr1 | + | 31346639 | 31346818 | 31338956 | 31339048 | 31348828 | 31348974 | 102 | 56 | [illegible] |
| SE | 47538 | ENSG00000066322 | ELOVL1 | chr1 | - | 43364927 | 43365008 | 43364737 | 43364794 | 43365185 | 43365268 | 164 | 60 | [illegible] |
| SE | 47539 | ENSG00000066322 | ELOVL1 | chr1 | - | 43364927 | 43365097 | 43364737 | 43364794 | 43365185 | 43365376 | 114 | 44 | [illegible] |
| SE | 47543 | ENSG00000144357 | UBR3 | chr2 | + | 169878524 | 169878574 | 169877493 | 169877637 | 169891164 | 169891231 | 141 | 112 | [illegible] |
| SE | 47544 | ENSG00000144357 | UBR3 | chr2 | + | 169896506 | 169896735 | 169895180 | 169895311 | 169905113 | 169905293 | 312 | 276 | [illegible] |
| SE | 47545 | ENSG00000144357 | UBR3 | chr2 | + | 169905113 | 169905293 | 169896506 | 169896735 | 169906030 | 169906164 | 323 | 322 | [illegible] |
| SE | 47546 | ENSG00000144357 | UBR3 | chr2 | + | 169946287 | 169946392 | 169942492 | 169942634 | 169947541 | 169947715 | 124 | 99 | [illegible] |
| SE | 47547 | ENSG00000144357 | UBR3 | chr2 | + | 170013840 | 170013972 | 170008839 | 170008940 | 170014814 | 170014865 | 0 | 1 | [illegible] |
| SE | 47550 | ENSG00000144357 | UBR3 | chr2 | + | 170014814 | 170014865 | 170013840 | 170013972 | 170015280 | 170015366 | 4 | 2 | [illegible] |
| SE | 47551 | ENSG00000144357 | UBR3 | chr2 | + | 170016832 | 170016919 | 170015280 | 170015366 | 170029345 | 170029448 | 0 | 0 | [illegible] |
| SE | 47552 | ENSG00000144357 | UBR3 | chr2 | + | 170040881 | 170040985 | 170029345 | 170029448 | 170055459 | 170055584 | 356 | 330 | [illegible] |
| SE | 47553 | ENSG00000144357 | UBR3 | chr2 | + | 170055459 | 170055584 | 170040881 | 170040985 | 170061078 | 170061186 | 245 | 214 | [illegible] |
| SE | 47554 | ENSG00000144357 | UBR3 | chr2 | + | 170073403 | 170073607 | 170061317 | 170061443 | 170079813 | 170080023 | 73 | 38 | [illegible] |
| SE | 47555 | ENSG00000144357 | UBR3 | chr2 | + | 170073403 | 170073607 | 170061317 | 170061443 | 170081725 | 170084131 | 4 | 0 | [illegible] |
| SE | 47556 | ENSG00000144357 | UBR3 | chr2 | + | 170073427 | 170073607 | 170061317 | 170061443 | 170079813 | 170080023 | 142 | 86 | [illegible] |
| SE | 47557 | ENSG00000144357 | UBR3 | chr2 | + | 170073427 | 170073607 | 170061317 | 170061443 | 170081725 | 170082259 | 73 | 48 | [illegible] |
| SE | 47558 | ENSG00000144357 | UBR3 | chr2 | + | 170079813 | 170080023 | 170073403 | 170073607 | 170080544 | 170080684 | 125 | 74 | [illegible] |
| SE | 47559 | ENSG00000144357 | UBR3 | chr2 | + | 170080544 | 170080684 | 170073403 | 170073607 | 170081725 | 170082259 | 73 | 64 | [illegible] |
| SE | 47568 | ENSG00000111880 | RNGTT | chr6 | - | 88678352 | 88678419 | 88614271 | 88614395 | 88769773 | 88769874 | 110 | 54 | [illegible] |
| SE | 47569 | ENSG00000111880 | RNGTT | chr6 | - | 88801563 | 88801632 | 88769773 | 88769874 | 88844356 | 88844521 | 94 | 86 | [illegible] |
| SE | 47570 | ENSG00000111880 | RNGTT | chr6 | - | 88849754 | 88849826 | 88844356 | 88844521 | 88853628 | 88853764 | 201 | 208 | [illegible] |
| SE | 47571 | ENSG00000111880 | RNGTT | chr6 | - | 88891805 | 88891915 | 88890494 | 88890596 | 88904714 | 88904955 | 137 | 116 | [illegible] |
| SE | 47572 | ENSG00000111880 | RNGTT | chr6 | - | 88941070 | 88941180 | 88929163 | 88929267 | 88963345 | 88963409 | 94 | 66 | [illegible] |
| SE | 47576 | ENSG00000157445 | CACNA2D3 | chr3 | + | 54984607 | 54984670 | 54969799 | 54969844 | 54987682 | 54987753 | 78 | 58 | [illegible] |
| SE | 47577 | ENSG00000157445 | CACNA2D3 | chr3 | + | 54984607 | 54984670 | 54969799 | 54969844 | 55004762 | 55004838 | 23 | 18 | [illegible] |
| SE | 47578 | ENSG00000157445 | CACNA2D3 | chr3 | + | 54987682 | 54987753 | 54969799 | 54969844 | 55004762 | 55004838 | 19 | 9 | [illegible] |
| SE | 47579 | ENSG00000157445 | CACNA2D3 | chr3 | + | 54987682 | 54987753 | 54984607 | 54984670 | 55004762 | 55004838 | 81 | 49 | [illegible] |
| SE | 47581 | ENSG00000008869 | HEATR5B | chr2 | - | 36988645 | 36988859 | 36980894 | 36981794 | 36990647 | 36990799 | 172 | 69 | [illegible] |
| SE | 47584 | ENSG00000008869 | HEATR5B | chr2 | - | 37005631 | 37005759 | 37003541 | 37003686 | 37007049 | 37007304 | 208 | 107 | [illegible] |
| SE | 47586 | ENSG00000008869 | HEATR5B | chr2 | - | 37013840 | 37014020 | 37008610 | 37008848 | 37019808 | 37019877 | 236 | 118 | [illegible] |
| SE | 47587 | ENSG00000008869 | HEATR5B | chr2 | - | 37019808 | 37019877 | 37013840 | 37014020 | 37027922 | 37028174 | 50 | 22 | [illegible] |
| SE | 47589 | ENSG00000008869 | HEATR5B | chr2 | - | 37027922 | 37028174 | 37020654 | 37020836 | 37032628 | 37032773 | 86 | 41 | [illegible] |
| SE | 47590 | ENSG00000008869 | HEATR5B | chr2 | - | 37028680 | 37028920 | 37027922 | 37028174 | 37032628 | 37032773 | 180 | 111 | [illegible] |
| SE | 47591 | ENSG00000008869 | HEATR5B | chr2 | - | 37041132 | 37041292 | 37040328 | 37040518 | 37049652 | 37049843 | 154 | 52 | [illegible] |
| SE | 47592 | ENSG00000008869 | HEATR5B | chr2 | - | 37065754 | 37065910 | 37064739 | 37064990 | 37068680 | 37068930 | 195 | 164 | [illegible] |
| SE | 47593 | ENSG00000008869 | HEATR5B | chr2 | - | 37070229 | 37070387 | 37068680 | 37068930 | 37072109 | 37072281 | 652 | 495 | [illegible] |
| SE | 47594 | ENSG00000008869 | HEATR5B | chr2 | - | 37070229 | 37070387 | 37068680 | 37068930 | 37075484 | 37075634 | 138 | 82 | [illegible] |
| SE | 47595 | ENSG00000008869 | HEATR5B | chr2 | - | 37072109 | 37072281 | 37068680 | 37068930 | 37075484 | 37075634 | 318 | 224 | [illegible] |
| SE | 47597 | ENSG00000008869 | HEATR5B | chr2 | - | 37079119 | 37079331 | 37076910 | 37077019 | 37083288 | 37083436 | 757 | 601 | [illegible] |
| SE | 47600 | ENSG00000147403 | RPL10 | chrX | + | 154400463 | 154400626 | 154399802 | 154399941 | 154403262 | 154403497 | 7308 | 3453 | [illegible] |
| SE | 47601 | ENSG00000163743 | RCHY1 | chr4 | - | 75491582 | 75491637 | 75490580 | 75490701 | 75491723 | 75491782 | 22 | 32 | [illegible] |
| SE | 47602 | ENSG00000163743 | RCHY1 | chr4 | - | 75491610 | 75491637 | 75490580 | 75490701 | 75491723 | 75491782 | 147 | 100 | [illegible] |
| SE | 47603 | ENSG00000163743 | RCHY1 | chr4 | - | 75494100 | 75494179 | 75491888 | 75491933 | 75508819 | 75508935 | 296 | 191 | [illegible] |
| SE | 47604 | ENSG00000163743 | RCHY1 | chr4 | - | 75494100 | 75494179 | 75491888 | 75491933 | 75508823 | 75508935 | 125 | 90 | [illegible] |
| SE | 47605 | ENSG00000163743 | RCHY1 | chr4 | - | 75509176 | 75509296 | 75508819 | 75508935 | 75514196 | 75514286 | 203 | 107 | [illegible] |
| SE | 47619 | ENSG00000213672 | NCKIPSD | chr3 | - | 48681286 | 48681780 | 48680058 | 48680229 | 48682044 | 48682135 | 47 | 25 | [illegible] |
| SE | 47620 | ENSG00000110619 | CARS1 | chr11 | - | 3002540 | 3002600 | 3001961 | 3002053 | 3005365 | 3005433 | 35 | 37 | [illegible] |
| SE | 47621 | ENSG00000110619 | CARS1 | chr11 | - | 3002540 | 3002600 | 3001969 | 3002174 | 3005365 | 3005433 | 12 | 20 | [illegible] |
| SE | 47622 | ENSG00000110619 | CARS1 | chr11 | - | 3015780 | 3015849 | 3012194 | 3012276 | 3017105 | 3017295 | 111 | 78 | [illegible] |
| SE | 47624 | ENSG00000110619 | CARS1 | chr11 | - | 3018407 | 3018511 | 3017105 | 3017295 | 3018619 | 3018749 | 45 | 13 | [illegible] |
| SE | 47625 | ENSG00000110619 | CARS1 | chr11 | - | 3026675 | 3026797 | 3020219 | 3020332 | 3028995 | 3029084 | 102 | 64 | [illegible] |
| SE | 47633 | ENSG00000157349 | DDX19B | chr16 | + | 70315968 | 70316104 | 70299141 | 70299354 | 70324584 | 70324687 | 13 | 4 | [illegible] |
| SE | 47635 | ENSG00000260537 | NA | chr16 | + | 70324584 | 70324687 | 70317495 | 70317588 | 70325573 | 70325688 | 148 | 110 | [illegible] |
| SE | 47636 | ENSG00000288701 | PRRC2B | chr9 | + | 131436619 | 131436722 | 131432616 | 131432794 | 131438988 | 131439061 | 344 | 202 | [illegible] |
| SE | 47638 | ENSG00000288701 | PRRC2B | chr9 | + | 131447661 | 131447804 | 131447084 | 131447206 | 131455075 | 131455166 | 538 | 335 | [illegible] |
| SE | 47639 | ENSG00000288701 | PRRC2B | chr9 | + | 131455075 | 131455166 | 131447661 | 131447804 | 131459163 | 131459356 | 526 | 369 | [illegible] |
| SE | 47641 | ENSG00000288701 | PRRC2B | chr9 | + | 131467562 | 131467753 | 131464762 | 131465078 | 131470787 | 131470791 | 346 | 200 | [illegible] |
| SE | 47642 | ENSG00000288701 | PRRC2B | chr9 | + | 131473507 | 131473724 | 131470787 | 131470983 | 131477743 | 131477949 | 129 | 75 | [illegible] |
| SE | 47647 | ENSG00000288701 | PRRC2B | chr9 | + | 131479251 | 131479393 | 131478473 | 131478619 | 131481725 | 131481808 | 365 | 213 | [illegible] |
| SE | 47651 | ENSG00000288701 | PRRC2B | chr9 | + | 131486084 | 131486182 | 131484947 | 131485140 | 131487166 | 131487294 | 126 | 50 | [illegible] |
| SE | 47655 | ENSG00000172197 | MBOAT1 | chr6 | - | 20144219 | 20144315 | 20131143 | 20131199 | 20151184 | 20151262 | 50 | 60 | [illegible] |
| SE | 47658 | ENSG00000255284 | NA | chr11 | + | 778112 | 778218 | 777607 | 777695 | 783529 | 783671 | 0 | 0 | [illegible] |
| SE | 47659 | ENSG00000273088 | NA | chr1 | - | 155169790 | 155169857 | 155169408 | 155169560 | 155172564 | 155172892 | 198 | 66 | [illegible] |
| SE | 47660 | ENSG00000007968 | E2F2 | chr1 | - | 23524382 | 23524488 | 23521836 | 23522056 | 23530541 | 23531233 | 5 | 0 | [illegible] |
| SE | 47663 | ENSG00000163479 | SSR2 | chr1 | - | 156011809 | 156011887 | 156009047 | 156009650 | 156014960 | 156015069 | 902 | 385 | [illegible] |
| SE | 47664 | ENSG00000163479 | SSR2 | chr1 | - | 156011809 | 156012565 | 156009061 | 156009650 | 156014960 | 156015069 | 385 | 170 | [illegible] |
| SE | 47665 | ENSG00000163479 | SSR2 | chr1 | - | 156012384 | 156012565 | 156011809 | 156011887 | 156014960 | 156015069 | 2 | 3 | [illegible] |
| SE | 47666 | ENSG00000163479 | SSR2 | chr1 | - | 156012394 | 156012565 | 156011809 | 156011887 | 156014960 | 156015069 | 0 | 1 | [illegible] |
| SE | 47670 | ENSG00000163479 | SSR2 | chr1 | - | 156018131 | 156018368 | 156011809 | 156011887 | 156020012 | 156020167 | 422 | 198 | [illegible] |
| SE | 47671 | ENSG00000163479 | SSR2 | chr1 | - | 156018251 | 156018368 | 156011809 | 156011887 | 156020012 | 156020167 | 418 | 196 | [illegible] |
| SE | 47673 | ENSG00000163479 | SSR2 | chr1 | - | 156020012 | 156020167 | 156018131 | 156018368 | 156020887 | 156020912 | 1512 | 611 | [illegible] |
| SE | 47676 | ENSG00000226891 | LINC01359 | chr1 | - | 64989585 | 64989668 | 64989434 | 64989494 | 64991297 | 64991441 | 0 | 0 | [illegible] |
| SE | 47681 | ENSG00000162869 | PPP1R21 | chr2 | + | 48454555 | 48454741 | 48451007 | 48451076 | 48458125 | 48458227 | 196 | 159 | [illegible] |
| SE | 47682 | ENSG00000162869 | PPP1R21 | chr2 | + | 48454594 | 48454741 | 48451007 | 48451076 | 48458125 | 48458227 | 258 | 220 | [illegible] |
| SE | 47683 | ENSG00000162869 | PPP1R21 | chr2 | + | 48471086 | 48471188 | 48465492 | 48465642 | 48474682 | 48474819 | 63 | 51 | [illegible] |
| SE | 47685 | ENSG00000162869 | PPP1R21 | chr2 | + | 48474682 | 48474819 | 48465492 | 48465642 | 48479923 | 48480016 | 62 | 34 | [illegible] |
| SE | 47686 | ENSG00000162869 | PPP1R21 | chr2 | + | 48474682 | 48474819 | 48471278 | 48471367 | 48479923 | 48480016 | 121 | 101 | [illegible] |
| SE | 47687 | ENSG00000162869 | PPP1R21 | chr2 | + | 48479483 | 48479556 | 48474682 | 48474819 | 48479923 | 48480016 | 2 | 2 | [illegible] |
| SE | 47688 | ENSG00000162869 | PPP1R21 | chr2 | + | 48491017 | 48491170 | 48486630 | 48486758 | 48495678 | 48495771 | 146 | 136 | [illegible] |
| SE | 47689 | ENSG00000162869 | PPP1R21 | chr2 | + | 48495678 | 48495771 | 48491017 | 48491170 | 48498492 | 48498501 | 110 | 74 | [illegible] |
| SE | 47690 | ENSG00000162869 | PPP1R21 | chr2 | + | 48505563 | 48505596 | 48498492 | 48498735 | 48507268 | 48507385 | 144 | 98 | [illegible] |
| SE | 47693 | ENSG00000171148 | TADA3 | chr3 | - | 9784027 | 9784213 | 9779966 | 9780549 | 9787005 | 9787109 | 91 | 54 | [illegible] |
| SE | 47694 | ENSG00000171148 | TADA3 | chr3 | - | 9785315 | 9785425 | 9779966 | 9780549 | 9787005 | 9787109 | 182 | 83 | [illegible] |
| SE | 47696 | ENSG00000143947 | RPS27A | chr2 | + | 55233362 | 55233417 | 55232677 | 55232872 | 55234118 | 55234204 | 5398 | 2658 | [illegible] |
| SE | 47697 | ENSG00000143947 | RPS27A | chr2 | + | 55233362 | 55233417 | 55232677 | 55232872 | 55234830 | 55234959 | 19 | 6 | [illegible] |
| SE | 47698 | ENSG00000143947 | RPS27A | chr2 | + | 55234118 | 55234204 | 55232677 | 55232872 | 55234830 | 55234959 | 6204 | 3553 | [illegible] |
| SE | 47699 | ENSG00000143947 | RPS27A | chr2 | + | 55234118 | 55234204 | 55233362 | 55233417 | 55234830 | 55234959 | 11642 | 6216 | [illegible] |
| SE | 47702 | ENSG00000170624 | SGCD | chr5 | + | 156329533 | 156329579 | 156123855 | 156124019 | 156344488 | 156344677 | 3 | 0 | [illegible] |
| SE | 47705 | ENSG00000170624 | SGCD | chr5 | + | 156594931 | 156595051 | 156589230 | 156589318 | 156647463 | 156647536 | 12 | 2 | [illegible] |
| SE | 47707 | ENSG00000234807 | LINC01135 | chr1 | + | 58893544 | 58893717 | 58785132 | 58785305 | 58899046 | 58901101 | 2 | 0 | [illegible] |
| SE | 47719 | ENSG00000153560 | UBP1 | chr3 | - | 33402800 | 33402904 | 33400961 | 33401016 | 33409235 | 33409346 | 323 | 294 | [illegible] |
| SE | 47721 | ENSG00000153560 | UBP1 | chr3 | - | 33409235 | 33409346 | 33402800 | 33402904 | 33409448 | 33409587 | 458 | 385 | [illegible] |
| SE | 47723 | ENSG00000153560 | UBP1 | chr3 | - | 33425589 | 33425741 | 33416757 | 33416834 | 33439735 | 33439856 | 447 | 272 | [illegible] |
| SE | 47724 | ENSG00000153560 | UBP1 | chr3 | - | 33439735 | 33439925 | 33425589 | 33425741 | 33440333 | 33440510 | 138 | 70 | [illegible] |
| SE | 47727 | ENSG00000145687 | SSBP2 | chr5 | - | 81440557 | 81440636 | 81437429 | 81437458 | 81442652 | 81442723 | 141 | 127 | [illegible] |
| SE | 47729 | ENSG00000145687 | SSBP2 | chr5 | - | 81440557 | 81440636 | 81437429 | 81437458 | 81448789 | 81448825 | 41 | 31 | [illegible] |
| SE | 47731 | ENSG00000145687 | SSBP2 | chr5 | - | 81442652 | 81442723 | 81437429 | 81437458 | 81448789 | 81448825 | 2 | 1 | [illegible] |
| SE | 47732 | ENSG00000145687 | SSBP2 | chr5 | - | 81442652 | 81442723 | 81440537 | 81440636 | 81446867 | 81446922 | 110 | 101 | [illegible] |
| SE | 47733 | ENSG00000145687 | SSBP2 | chr5 | - | 81442652 | 81442723 | 81440537 | 81440636 | 81448789 | 81448825 | 72 | 76 | [illegible] |
| SE | 47734 | ENSG00000145687 | SSBP2 | chr5 | - | 81446867 | 81446922 | 81437429 | 81437458 | 81448789 | 81448825 | 11 | 17 | [illegible] |
| SE | 47738 | ENSG00000145687 | SSBP2 | chr5 | - | 81447909 | 81448063 | 81446867 | 81446922 | 81448789 | 81448825 | 0 | 0 | [illegible] |
| SE | 47739 | ENSG00000145687 | SSBP2 | chr5 | - | 81448789 | 81448825 | 81440537 | 81440636 | 81461054 | 81461103 | 6 | 3 | [illegible] |
| SE | 47740 | ENSG00000145687 | SSBP2 | chr5 | - | 81448789 | 81448825 | 81446867 | 81446922 | 81461054 | 81461103 | 41 | 38 | [illegible] |
| SE | 47745 | ENSG00000145687 | SSBP2 | chr5 | - | 81615472 | 81615557 | 81513627 | 81513717 | 81636556 | 81636618 | 94 | 104 | [illegible] |
| SE | 47746 | ENSG00000145687 | SSBP2 | chr5 | - | 81636556 | 81636618 | 81615055 | 81615557 | 81650266 | 81650339 | 91 | 100 | [illegible] |
| SE | 47747 | ENSG00000145687 | SSBP2 | chr5 | - | 81650266 | 81650339 | 81615055 | 81615557 | 81750980 | 81751016 | 5 | 10 | [illegible] |
| SE | 47748 | ENSG00000145687 | SSBP2 | chr5 | - | 81650266 | 81650339 | 81615055 | 81615557 | 81751667 | 81751797 | 0 | 0 | [illegible] |
| SE | 47750 | ENSG00000203761 | MSTO2P | chr1 | + | 155747761 | 155747896 | 155747562 | 155747680 | 155748399 | 155748531 | 16 | 4 | [illegible] |
| SE | 47751 | ENSG00000203761 | MSTO2P | chr1 | + | 155747976 | 155748127 | 155747761 | 155747896 | 155748399 | 155748531 | 0 | 0 | [illegible] |
| SE | 47752 | ENSG00000100029 | PES1 | chr22 | - | 30579136 | 30579303 | 30578836 | 30578998 | 30579750 | 30579923 | 193 | 119 | [illegible] |
| SE | 47753 | ENSG00000100029 | PES1 | chr22 | - | 30579750 | 30579935 | 30579136 | 30579303 | 30580052 | 30580178 | 184 | 125 | [illegible] |
| SE | 47755 | ENSG00000100029 | PES1 | chr22 | - | 30588020 | 30588174 | 30586997 | 30587395 | 30589190 | 30589270 | 127 | 90 | [illegible] |
| SE | 47758 | ENSG00000080200 | CRYBG3 | chr3 | + | 97843194 | 97843261 | 97822010 | 97822355 | 97864216 | 97864647 | 28 | 27 | [illegible] |
| SE | 47759 | ENSG00000080200 | CRYBG3 | chr3 | + | 97888340 | 97888455 | 97886630 | 97886767 | 97889354 | 97889390 | 21 | 41 | [illegible] |
| SE | 47761 | ENSG00000080200 | CRYBG3 | chr3 | + | 97895958 | 97896085 | 97889354 | 97889390 | 97898882 | 97899025 | 34 | 43 | [illegible] |
| SE | 47762 | ENSG00000080200 | CRYBG3 | chr3 | + | 97898882 | 97899025 | 97895958 | 97896085 | 97899136 | 97899263 | 105 | 107 | [illegible] |
| SE | 47765 | ENSG00000154358 | OBSCN | chr1 | + | 228219323 | 228219599 | 228217012 | 228217288 | 228224458 | 228224734 | 26 | 1 | [illegible] |
| SE | 47767 | ENSG00000154358 | OBSCN | chr1 | + | 228219323 | 228219599 | 228217012 | 228217288 | 228249964 | 228250240 | 16 | 1 | [illegible] |
| SE | 47769 | ENSG00000154358 | OBSCN | chr1 | + | 228224458 | 228224734 | 228219323 | 228219599 | 228243182 | 228243458 | 18 | 5 | [illegible] |
| SE | 47770 | ENSG00000154358 | OBSCN | chr1 | + | 228245412 | 228245688 | 228244295 | 228244571 | 228246527 | 228246642 | 32 | 37 | [illegible] |
| SE | 47771 | ENSG00000154358 | OBSCN | chr1 | + | 228245412 | 228245688 | 228244295 | 228244571 | 228249964 | 228250240 | 22 | 19 | [illegible] |
| SE | 47772 | ENSG00000154358 | OBSCN | chr1 | + | 228246527 | 228246803 | 228244295 | 228244571 | 228249964 | 228250240 | 10 | 14 | [illegible] |
| SE | 47774 | ENSG00000154358 | OBSCN | chr1 | + | 228249964 | 228250240 | 228246527 | 228246803 | 228256650 | 228256926 | 19 | 15 | [illegible] |
| SE | 47779 | ENSG00000154358 | OBSCN | chr1 | + | 228276929 | 228277042 | 228276449 | 228276716 | 228277159 | 228277313 | 18 | 4 | [illegible] |
| SE | 47780 | ENSG00000154358 | OBSCN | chr1 | + | 228277504 | 228277666 | 228277159 | 228277313 | 228277753 | 228277858 | 43 | 39 | [illegible] |
| SE | 47781 | ENSG00000154358 | OBSCN | chr1 | + | 228283017 | 228283284 | 228281932 | 228282205 | 228283502 | 228283769 | 41 | 23 | [illegible] |
| SE | 47782 | ENSG00000154358 | OBSCN | chr1 | + | 228286077 | 228286350 | 228283502 | 228283769 | 228286772 | 228287039 | 29 | 17 | [illegible] |
| SE | 47785 | ENSG00000154358 | OBSCN | chr1 | + | 228294779 | 228295043 | 228294151 | 228294415 | 228298453 | 228298717 | 26 | 39 | [illegible] |
| SE | 47789 | ENSG00000154358 | OBSCN | chr1 | + | 228303674 | 228303938 | 228298453 | 228298717 | 228304302 | 228304566 | 17 | 12 | [illegible] |
| SE | 47790 | ENSG00000154358 | OBSCN | chr1 | + | 228303674 | 228303938 | 228298453 | 228298717 | 228305122 | 228305386 | 0 | 5 | [illegible] |
| SE | 47791 | ENSG00000154358 | OBSCN | chr1 | + | 228303674 | 228303938 | 228298453 | 228298717 | 228306371 | 228306638 | 0 | 5 | [illegible] |
| SE | 47792 | ENSG00000154358 | OBSCN | chr1 | + | 228304302 | 228304566 | 228298453 | 228298717 | 228305122 | 228305386 | 15 | 5 | [illegible] |
| SE | 47794 | ENSG00000154358 | OBSCN | chr1 | + | 228304302 | 228304566 | 228303674 | 228303938 | 228305122 | 228305386 | 32 | 11 | [illegible] |
| SE | 47795 | ENSG00000154358 | OBSCN | chr1 | + | 228304302 | 228304566 | 228303674 | 228303938 | 228306371 | 228306638 | 17 | 7 | [illegible] |

| | | | | | | | | | | | | | | | | | | | | | | | | | | | | | | | | | | | | | | | | | | | | | | |
|---|---|---|---|---|---|---|---|---|---|---|---|---|---|---|---|---|---|---|---|---|---|---|---|---|---|---|---|---|---|---|---|---|---|---|---|---|---|---|---|---|---|---|---|---|---|---|
| SE | 47796 | ENSG00000154358 | OBSCN | chr1 | + | 228305122 | 228305386 | 228298453 | 228298717 | 228306371 | 228306638 | 10 | 11 | 11 | 7 | 6 | 1 | 0 | 0 | 0 | 0 | 0 | 0 | 4 | 4 | 2 | 0 | 0 | 0 | 0 | 0 | 298 | 149 | 1 | 1 | 0.833 | 1 | 1 | 1 | 1 | NA | NA | 1 | 1 | 1 | -0.033 |
| SE | 47797 | ENSG00000154358 | OBSCN | chr1 | + | 228305122 | 228305386 | 228303674 | 228303938 | 228306371 | 228306638 | 9 | 11 | 15 | 7 | 6 | 0 | 0 | 0 | 0 | 2 | 0 | 0 | 4 | 6 | 2 | 0 | 0 | 2 | 0 | 0 | 298 | 149 | 1 | 1 | 1 | 1 | 1 | 1 | 0.6 | NA | NA | 0.5 | 1 | 1 | 0.087 |
| SE | 47798 | ENSG00000154358 | OBSCN | chr1 | + | 228305122 | 228305386 | 228304302 | 228304566 | 228306371 | 228306638 | 24 | 15 | 40 | 20 | 25 | 0 | 0 | 1 | 0 | 0 | 0 | 0 | 15 | 16 | 11 | 0 | 0 | 0 | 0 | 0 | 298 | 149 | 1 | 1 | 1 | 1 | 0.952 | 1 | 1 | NA | NA | 1 | 1 | 1 | -0.01 |
| SE | 47799 | ENSG00000154358 | OBSCN | chr1 | + | 228305161 | 228305386 | 228304302 | 228304566 | 228306371 | 228306638 | 9 | 11 | 11 | 7 | 6 | 0 | 0 | 1 | 0 | 0 | 0 | 0 | 4 | 4 | 2 | 0 | 0 | 0 | 0 | 0 | 298 | 149 | 1 | 1 | 1 | 1 | 0.846 | 1 | 1 | NA | NA | 1 | 1 | 1 | -0.031 |
| SE | 47800 | ENSG00000154358 | OBSCN | chr1 | + | 228308110 | 228308383 | 228307258 | 228307531 | 228309098 | 228309295 | 11 | 2 | 12 | 5 | 9 | 3 | 2 | 7 | 7 | 0 | 0 | 9 | 0 | 3 | 3 | 0 | 0 | 0 | 0 | 0 | 298 | 149 | 1 | 1 | 0.647 | 0.333 | 0.462 | 0.263 | 1 | NA | 1 | NA | 1 | 1 | -0.459 |
| SE | 47802 | ENSG00000154358 | OBSCN | chr1 | + | 228350840 | 228350943 | 228349889 | 228350057 | 228351320 | 228351450 | 14 | 11 | 25 | 11 | 8 | 0 | 0 | 2 | 0 | 0 | 0 | 9 | 3 | 1 | 0 | 0 | 0 | 0 | 0 | 0 | 252 | 149 | 1 | 1 | 1 | 1 | 0.881 | 1 | 1 | NA | 1 | 1 | 1 | NA | -0.024 |
| SE | 47803 | ENSG00000154358 | OBSCN | chr1 | + | 228356150 | 228356193 | 228352950 | 228353062 | 228359596 | 228361196 | 11 | 4 | 21 | 12 | 16 | 0 | 0 | 0 | 0 | 0 | 0 | 6 | 0 | 0 | 0 | 0 | 4 | 0 | 0 | 0 | 192 | 149 | 1 | 1 | 1 | 1 | 1 | 1 | 1 | NA | 0.538 | NA | NA | NA | 0.462 |
| SE | 47804 | ENSG00000154358 | OBSCN | chr1 | + | 228356150 | 228356344 | 228353006 | 228353062 | 228359596 | 228361250 | 4 | 0 | 10 | 14 | 3 | 0 | 0 | 0 | 0 | 0 | 0 | 0 | 0 | 0 | 0 | 0 | 4 | 0 | 0 | 0 | 298 | 149 | 1 | 1 | 1 | NA | 1 | 1 | 1 | NA | 0 | NA | NA | NA | 1 |
| SE | 47806 | ENSG00000014138 | POLA2 | chr11 | + | 65266581 | 65266706 | 65261931 | 65262371 | 65267476 | 65267568 | 81 | 53 | 92 | 80 | 165 | 0 | 0 | 0 | 0 | 0 | 194 | 86 | 80 | 100 | 161 | 0 | 0 | 0 | 1 | 0 | 274 | 149 | 1 | 1 | 1 | 1 | 1 | 1 | 1 | 1 | 1 | 1 | 0.982 | 1 | 0.004 |
| SE | 47807 | ENSG00000014138 | POLA2 | chr11 | + | 65278729 | 65278923 | 65275891 | 65275998 | 65279537 | 65279626 | 56 | 40 | 70 | 69 | 71 | 0 | 0 | 0 | 1 | 0 | 96 | 63 | 51 | 70 | 64 | 0 | 0 | 0 | 0 | 0 | 298 | 149 | 1 | 1 | 1 | 1 | 1 | 0.972 | 1 | 1 | 1 | 1 | 1 | 1 | -0.006 |
| SE | 47825 | ENSG00000187554 | TLR5 | chr1 | - | 223134681 | 223134864 | 223132474 | 223132639 | 223137177 | 223137263 | 37 | 17 | 16 | 26 | 13 | 0 | 0 | 1 | 0 | 0 | 42 | 48 | 11 | 39 | 21 | 0 | 0 | 0 | 0 | 0 | 298 | 149 | 1 | 1 | 1 | 1 | 0.889 | 1 | 1 | 1 | 1 | 1 | 1 | 1 | -0.022 |
| SE | 47826 | ENSG00000187554 | TLR5 | chr1 | - | 223134681 | 223134864 | 223132474 | 223132639 | 223141647 | 223141763 | 21 | 11 | 6 | 6 | 11 | 1 | 0 | 0 | 0 | 0 | 41 | 31 | 9 | 22 | 11 | 0 | 0 | 0 | 0 | 0 | 298 | 149 | 1 | 1 | 0.913 | 1 | 1 | 1 | 1 | 1 | 1 | 1 | 1 | 1 | -0.017 |
| SE | 47830 | ENSG00000187554 | TLR5 | chr1 | - | 223137177 | 223137263 | 223134681 | 223134864 | 223141647 | 223141763 | 17 | 6 | 8 | 14 | 2 | 2 | 2 | 0 | 0 | 0 | 1 | 14 | 2 | 17 | 10 | 0 | 0 | 1 | 0 | 0 | 235 | 149 | 1 | 1 | 0.843 | 0.655 | 1 | 1 | 1 | 1 | 1 | 0.559 | 1 | 1 | -0.012 |
| SE | 47832 | ENSG00000187554 | TLR5 | chr1 | - | 223141647 | 223141763 | 223132474 | 223132639 | 223143195 | 223143246 | 4 | 0 | 0 | 3 | 1 | 0 | 1 | 3 | 1 | 2 | 0 | 2 | 0 | 1 | 0 | 0 | 0 | 0 | 0 | 0 | 265 | 149 | 1 | 1 | 1 | 0 | 0 | 0.628 | 0.219 | NA | 1 | NA | 1 | NA | -0.631 |
| SE | 47833 | ENSG00000187554 | TLR5 | chr1 | - | 223141647 | 223141763 | 223134681 | 223134864 | 223143195 | 223143246 | 5 | 2 | 0 | 3 | 1 | 1 | 0 | 1 | 0 | 1 | 0 | 2 | 1 | 1 | 0 | 0 | 0 | 0 | 4 | 0 | 265 | 149 | 1 | 1 | 0.738 | 1 | 0 | 1 | 0.36 | NA | 1 | 1 | 0.123 | NA | -0.088 |
| SE | 47834 | ENSG00000187554 | TLR5 | chr1 | - | 223141647 | 223141763 | 223137177 | 223137263 | 223143195 | 223143246 | 9 | 1 | 2 | 6 | 1 | 5 | 5 | 3 | 7 | 0 | 0 | 6 | 0 | 8 | 0 | 3 | 3 | 1 | 0 | 0 | 265 | 149 | 1 | 1 | 0.503 | 0.101 | 0.273 | 0.325 | 1 | 0 | 0.529 | 0 | 1 | NA | 0.058 |
| SE | 47836 | ENSG00000116675 | DNAJC6 | chr1 | + | 65364634 | 65364785 | 65254461 | 65254649 | 65365884 | 65365934 | 29 | 41 | 11 | 45 | 49 | 0 | 0 | 0 | 0 | 0 | 156 | 67 | 26 | 85 | 156 | 0 | 0 | 2 | 0 | 0 | 298 | 149 | 1 | 1 | 1 | 1 | 1 | 1 | 1 | 1 | 1 | 0.867 | 1 | 1 | 0.027 |
| SE | 47837 | ENSG00000164405 | UQCRQ | chr5 | + | 132866868 | 132867035 | 132866641 | 132866687 | 132867487 | 132867687 | 149 | 78 | 123 | 103 | 41 | 1 | 0 | 2 | 0 | 0 | 156 | 74 | 35 | 85 | 155 | 0 | 0 | 0 | 0 | 0 | 298 | 149 | 1 | 1 | 0.987 | 1 | 0.969 | 1 | 1 | 1 | 1 | 1 | 1 | 1 | -0.009 |
| SE | 47841 | ENSG00000143013 | LMO4 | chr1 | + | 87331226 | 87331309 | 87329098 | 87329244 | 87332012 | 87332251 | 1 | 0 | 0 | 0 | 0 | 12 | 6 | 11 | 9 | 4 | 0 | 0 | 0 | 0 | 0 | 2 | 11 | 14 | 3 | 21 | 232 | 149 | 1 | 1 | 0.051 | 0 | 0 | 0 | 0 | 0 | 0 | 0 | 0 | 0 | 0.01 |
| SE | 47860 | ENSG00000138768 | USO1 | chr4 | + | 75757496 | 75757573 | 75752539 | 75752604 | 75770438 | 75770539 | 359 | 213 | 275 | 310 | 372 | 1 | 2 | 0 | 0 | 0 | 75 | 140 | 96 | 45 | 46 | 0 | 0 | 4 | 0 | 0 | 226 | 149 | 1 | 1 | 0.996 | 0.986 | 1 | 1 | 1 | 1 | 1 | 0.941 | 1 | 1 | 0.008 |
| SE | 47861 | ENSG00000138768 | USO1 | chr4 | + | 75760593 | 75760605 | 75757496 | 75757573 | 75770438 | 75770539 | 1 | 0 | 1 | 1 | 4 | 232 | 151 | 177 | 205 | 248 | 0 | 0 | 0 | 0 | 0 | 50 | 97 | 74 | 36 | 16 | 161 | 149 | 1 | 1 | 0.004 | 0 | 0.005 | 0.004 | 0.015 | 0 | 0 | 0 | 0 | 0 | 0.006 |
| SE | 47862 | ENSG00000138768 | USO1 | chr4 | + | 75790642 | 75790797 | 75790149 | 75790238 | 75799621 | 75799732 | 78 | 43 | 65 | 84 | 101 | 1 | 0 | 1 | 1 | 2 | 0 | 14 | 10 | 5 | 0 | 0 | 0 | 4 | 0 | 0 | 298 | 149 | 1 | 1 | 0.975 | 1 | 0.97 | 0.977 | 0.962 | NA | 1 | 0.556 | 1 | NA | 0.125 |
| SE | 47863 | ENSG00000138768 | USO1 | chr4 | + | 75793689 | 75793901 | 75790642 | 75790797 | 75799621 | 75799732 | 266 | 145 | 214 | 262 | 211 | 0 | 0 | 1 | 2 | 0 | 0 | 81 | 43 | 5 | 0 | 0 | 0 | 0 | 0 | 0 | 298 | 149 | 1 | 1 | 1 | 1 | 0.991 | 0.985 | 1 | NA | 1 | 1 | 1 | NA | -0.005 |
| [illegible] | | | | | | | | | | | | | | | | | | | | | | | | | | | | | | | | | | | | | | | | | | | | | | |

| | | | | | | | | | | | | |
|---|---|---|---|---|---|---|---|---|---|---|---|---|
| SE | 48233 | ENSG00000138821 | SLC39A8 | chr4 | - | 102263160 | 102263193 | 102259454 | 102259514 | 102267489 | 102267674 | 5 | 4 | [illegible] |
| SE | 48237 | ENSG00000138821 | SLC39A8 | chr4 | - | 102344443 | 102344915 | 102315667 | 102315830 | 102345344 | 102345461 | 18 | 5 | [illegible] |
| SE | 48240 | ENSG00000112414 | ADGRG6 | chr6 | + | 142392947 | 142393000 | 142390257 | 142390343 | 142393895 | 142393958 | 6 | 0 | [illegible] |
| SE | 48241 | ENSG00000112414 | ADGRG6 | chr6 | + | 142405342 | 142405391 | 142403801 | 142403973 | 142405687 | 142405828 | 1 | 0 | [illegible] |
| SE | 48245 | ENSG00000116729 | WLS | chr1 | - | 68193954 | 68194227 | 68159122 | 68159247 | 68232193 | 68232523 | 88 | 56 | [illegible] |
| SE | 48247 | ENSG00000214595 | EML6 | chr2 | + | 54844046 | 54844248 | 54829341 | 54829477 | 54847485 | 54847623 | 6 | 5 | [illegible] |
| SE | 48250 | ENSG00000214595 | EML6 | chr2 | + | 54900696 | 54900896 | 54899640 | 54899782 | 54903043 | 54903196 | 0 | 0 | [illegible] |
| SE | 48251 | ENSG00000214595 | EML6 | chr2 | + | 54901940 | 54902141 | 54899640 | 54899782 | 54903043 | 54903196 | 0 | 0 | [illegible] |
| SE | 48253 | ENSG00000214595 | EML6 | chr2 | + | 54902424 | 54902541 | 54899640 | 54899782 | 54903043 | 54903196 | 0 | 0 | [illegible] |
| SE | 48255 | ENSG00000214595 | EML6 | chr2 | + | 54952593 | 54952692 | 54950649 | 54950779 | 54953982 | 54954156 | 2 | 5 | [illegible] |
| SE | 48257 | ENSG00000214595 | EML6 | chr2 | + | 54966722 | 54966880 | 54964570 | 54964733 | 54966999 | 54967231 | 4 | 1 | [illegible] |
| SE | 48261 | ENSG00000111602 | TIMELESS | chr12 | - | 56428548 | 56428652 | 56428235 | 56428405 | 56428882 | 56429100 | 18 | 28 | [illegible] |
| SE | 48262 | ENSG00000111602 | TIMELESS | chr12 | - | 56430104 | 56430281 | 56428882 | 56429100 | 56430880 | 56430968 | 41 | 14 | [illegible] |
| SE | 48264 | ENSG00000069424 | KCNAB2 | chr1 | + | 5995862 | 5996084 | 5992670 | 5992788 | 6040516 | 6040645 | 0 | 1 | [illegible] |
| SE | 48265 | ENSG00000069424 | KCNAB2 | chr1 | + | 6018634 | 6018835 | 5992665 | 5992788 | 6040516 | 6040645 | 0 | 0 | [illegible] |
| SE | 48268 | ENSG00000069424 | KCNAB2 | chr1 | + | 6040516 | 6040645 | 6026012 | 6026445 | 6072754 | 6072798 | 460 | 226 | [illegible] |
| SE | 48270 | ENSG00000069424 | KCNAB2 | chr1 | + | 6051510 | 6051754 | 6041830 | 6041872 | 6072754 | 6072798 | 2 | 0 | [illegible] |
| SE | 48273 | ENSG00000069424 | KCNAB2 | chr1 | + | 6082194 | 6082274 | 6073414 | 6073770 | 6085203 | 6085248 | 592 | 300 | [illegible] |
| SE | 48275 | ENSG00000069424 | KCNAB2 | chr1 | + | 6085739 | 6085793 | 6085203 | 6085248 | 6087466 | 6087511 | 1 | 0 | [illegible] |
| SE | 48277 | ENSG00000069424 | KCNAB2 | chr1 | + | 6095938 | 6096204 | 6095529 | 6095624 | 6096635 | 6096756 | 7 | 0 | [illegible] |
| SE | 48280 | ENSG00000113532 | ST8SIA4 | chr5 | - | 100856102 | 100856396 | 100806932 | 100812129 | 100895653 | 100895785 | 131 | 105 | [illegible] |
| SE | 48281 | ENSG00000113532 | ST8SIA4 | chr5 | - | 100886342 | 100886600 | 100806932 | 100812129 | 100895653 | 100895785 | 191 | 144 | [illegible] |
| SE | 48282 | ENSG00000113532 | ST8SIA4 | chr5 | - | 100886342 | 100886600 | 100856102 | 100856396 | 100895653 | 100895785 | 376 | 259 | [illegible] |
| SE | 48283 | ENSG00000113532 | ST8SIA4 | chr5 | - | 100895653 | 100895785 | 100885310 | 100886600 | 100902842 | 100903214 | 233 | 168 | [illegible] |
| SE | 48284 | ENSG00000113532 | ST8SIA4 | chr5 | - | 100900314 | 100900463 | 100895661 | 100895785 | 100902842 | 100903224 | 0 | 0 | [illegible] |
| SE | 48288 | ENSG00000229474 | PATL2 | chr15 | - | 44669509 | 44669563 | 44669279 | 44669413 | 44669776 | 44669874 | 87 | 21 | [illegible] |
| SE | 48291 | ENSG00000229474 | PATL2 | chr15 | - | 44672387 | 44672456 | 44669966 | 44670087 | 44673234 | 44673377 | 24 | 2 | [illegible] |
| SE | 48292 | ENSG00000229474 | PATL2 | chr15 | - | 44672387 | 44672456 | 44669966 | 44670087 | 44676474 | 44676565 | 3 | 0 | [illegible] |
| SE | 48293 | ENSG00000229474 | PATL2 | chr15 | - | 44673234 | 44673377 | 44669966 | 44670087 | 44675485 | 44675619 | 0 | 0 | [illegible] |
| SE | 48294 | ENSG00000229474 | PATL2 | chr15 | - | 44673234 | 44673377 | 44672387 | 44672456 | 44674149 | 44674230 | 88 | 27 | [illegible] |
| SE | 48295 | ENSG00000229474 | PATL2 | chr15 | - | 44674149 | 44674230 | 44672387 | 44672456 | 44676474 | 44676824 | 5 | 2 | [illegible] |
| SE | 48296 | ENSG00000229474 | PATL2 | chr15 | - | 44674149 | 44674230 | 44673234 | 44673377 | 44675485 | 44675619 | 69 | 15 | [illegible] |
| SE | 48297 | ENSG00000229474 | PATL2 | chr15 | - | 44674292 | 44674411 | 44673234 | 44673377 | 44675485 | 44675691 | 0 | 0 | [illegible] |
| SE | 48301 | ENSG00000229474 | PATL2 | chr15 | - | 44676474 | 44676565 | 44669966 | 44670087 | 44710095 | 44710214 | 10 | 9 | [illegible] |
| SE | 48311 | ENSG00000174501 | ANKRD36C | chr2 | - | 95859860 | 95860074 | 95857308 | 95857492 | 95876438 | 95876509 | 124 | 133 | [illegible] |
| SE | 48312 | ENSG00000174501 | ANKRD36C | chr2 | - | 95867439 | 95867581 | 95859860 | 95860074 | 95876438 | 95876509 | 119 | 175 | [illegible] |
| SE | 48313 | ENSG00000174501 | ANKRD36C | chr2 | - | 95880426 | 95880499 | 95876438 | 95876509 | 95880594 | 95880623 | 60 | 64 | [illegible] |
| SE | 48314 | ENSG00000174501 | ANKRD36C | chr2 | - | 95891664 | 95891737 | 95889965 | 95889994 | 95891831 | 95891860 | 74 | 88 | [illegible] |
| SE | 48316 | ENSG00000174501 | ANKRD36C | chr2 | - | 95905440 | 95905503 | 95904923 | 95904952 | 95906630 | 95906703 | 0 | 1 | [illegible] |
| SE | 48318 | ENSG00000174501 | ANKRD36C | chr2 | - | 95951347 | 95951408 | 95950748 | 95950779 | 95953938 | 95954005 | 168 | 156 | [illegible] |
| SE | 48319 | ENSG00000174501 | ANKRD36C | chr2 | - | 95953938 | 95954005 | 95951347 | 95951408 | 95958590 | 95958663 | 35 | 37 | [illegible] |
| SE | 48321 | ENSG00000174501 | ANKRD36C | chr2 | - | 95978121 | 95978189 | 95962518 | 95962547 | 95980647 | 95980785 | 266 | 217 | [illegible] |
| SE | 48325 | ENSG00000120798 | NR2C1 | chr12 | - | 95028386 | 95028509 | 95025149 | 95025255 | 95049067 | 95049233 | 90 | 57 | [illegible] |
| SE | 48328 | ENSG00000120798 | NR2C1 | chr12 | - | 95028386 | 95028524 | 95025149 | 95025255 | 95049067 | 95049233 | 104 | 63 | [illegible] |
| SE | 48329 | ENSG00000120798 | NR2C1 | chr12 | - | 95031348 | 95031488 | 95025149 | 95025255 | 95040475 | 95040597 | 36 | 20 | [illegible] |
| SE | 48330 | ENSG00000120798 | NR2C1 | chr12 | - | 95031348 | 95031488 | 95025149 | 95025255 | 95049067 | 95049492 | 32 | 9 | [illegible] |
| SE | 48331 | ENSG00000120798 | NR2C1 | chr12 | - | 95031348 | 95031488 | 95028386 | 95028509 | 95040475 | 95040597 | 38 | 22 | [illegible] |
| SE | 48332 | ENSG00000120798 | NR2C1 | chr12 | - | 95031348 | 95031488 | 95028386 | 95028509 | 95049067 | 95049492 | 34 | 11 | [illegible] |
| SE | 48333 | ENSG00000120798 | NR2C1 | chr12 | - | 95031348 | 95031488 | 95028386 | 95028524 | 95040475 | 95040597 | 102 | 50 | [illegible] |
| SE | 48334 | ENSG00000120798 | NR2C1 | chr12 | - | 95031348 | 95031488 | 95028386 | 95028524 | 95049067 | 95049492 | 98 | 38 | [illegible] |
| SE | 48335 | ENSG00000120798 | NR2C1 | chr12 | - | 95040475 | 95040597 | 95028386 | 95028524 | 95049067 | 95049233 | 29 | 16 | [illegible] |
| SE | 48336 | ENSG00000120798 | NR2C1 | chr12 | - | 95040475 | 95040597 | 95028457 | 95028509 | 95049067 | 95049233 | 29 | 13 | [illegible] |
| SE | 48337 | ENSG00000120798 | NR2C1 | chr12 | - | 95040475 | 95040597 | 95031348 | 95031488 | 95049067 | 95049233 | 58 | 27 | [illegible] |
| SE | 48338 | ENSG00000120798 | NR2C1 | chr12 | - | 95040475 | 95040597 | 95031348 | 95031488 | 95051761 | 95051869 | 17 | 8 | [illegible] |
| SE | 48339 | ENSG00000120798 | NR2C1 | chr12 | - | 95049067 | 95049233 | 95040475 | 95040597 | 95051761 | 95051943 | 118 | 103 | [illegible] |
| SE | 48340 | ENSG00000120798 | NR2C1 | chr12 | - | 95051761 | 95051943 | 95049067 | 95049233 | 95057552 | 95057643 | 189 | 169 | [illegible] |
| SE | 48346 | ENSG00000157193 | LRP8 | chr1 | - | 53246923 | 53247056 | 53245546 | 53245981 | 53250689 | 53250862 | 10 | 10 | [illegible] |
| SE | 48347 | ENSG00000157193 | LRP8 | chr1 | - | 53246955 | 53247056 | 53245544 | 53245981 | 53250689 | 53250862 | 6 | 5 | [illegible] |
| SE | 48349 | ENSG00000157193 | LRP8 | chr1 | - | 53249379 | 53249440 | 53246709 | 53247056 | 53250689 | 53250862 | 0 | 2 | [illegible] |
| SE | 48350 | ENSG00000157193 | LRP8 | chr1 | - | 53249379 | 53249556 | 53242363 | 53247056 | 53250689 | 53250862 | 1 | 6 | [illegible] |
| SE | 48353 | ENSG00000157193 | LRP8 | chr1 | - | 53258008 | 53258074 | 53257239 | 53257464 | 53258318 | 53258471 | 1 | 1 | [illegible] |
| SE | 48355 | ENSG00000157193 | LRP8 | chr1 | - | 53262445 | 53262564 | 53262067 | 53262207 | 53264168 | 53264396 | 14 | 8 | [illegible] |
| SE | 48356 | ENSG00000157193 | LRP8 | chr1 | - | 53264168 | 53264396 | 53262445 | 53262564 | 53266472 | 53266589 | 26 | 5 | [illegible] |
| SE | 48358 | ENSG00000157193 | LRP8 | chr1 | - | 53271027 | 53271153 | 53266472 | 53266699 | 53271226 | 53271250 | 2 | 4 | [illegible] |
| SE | 48359 | ENSG00000157193 | LRP8 | chr1 | - | 53271226 | 53271346 | 53271027 | 53271153 | 53275630 | 53275753 | 5 | 4 | [illegible] |
| SE | 48360 | ENSG00000157193 | LRP8 | chr1 | - | 53271226 | 53271415 | 53271027 | 53271153 | 53275630 | 53275753 | 3 | 3 | [illegible] |
| SE | 48362 | ENSG00000157193 | LRP8 | chr1 | - | 53275630 | 53275753 | 53271226 | 53271346 | 53280586 | 53280715 | 2 | 1 | [illegible] |
| SE | 48363 | ENSG00000157193 | LRP8 | chr1 | - | 53275630 | 53275753 | 53271226 | 53271346 | 53289566 | 53289689 | 1 | 1 | [illegible] |
| SE | 48364 | ENSG00000157193 | LRP8 | chr1 | - | 53276691 | 53277078 | 53271226 | 53275753 | 53280586 | 53280715 | 1 | 0 | [illegible] |
| SE | 48365 | ENSG00000157193 | LRP8 | chr1 | - | 53280586 | 53280715 | 53271226 | 53271346 | 53289566 | 53289689 | 4 | 4 | [illegible] |
| SE | 48366 | ENSG00000157193 | LRP8 | chr1 | - | 53280586 | 53280715 | 53275630 | 53275753 | 53289566 | 53289689 | 5 | 1 | [illegible] |
| SE | 48368 | ENSG00000133302 | SLF1 | chr5 | + | 94643272 | 94643435 | 94630502 | 94630743 | 94649453 | 94649597 | 113 | 63 | [illegible] |
| SE | 48370 | ENSG00000133302 | SLF1 | chr5 | + | 94662297 | 94662351 | 94654629 | 94654752 | 94663749 | 94663908 | 46 | 45 | [illegible] |
| SE | 48371 | ENSG00000133302 | SLF1 | chr5 | + | 94665860 | 94666024 | 94663749 | 94663908 | 94670150 | 94670279 | 22 | 17 | [illegible] |
| SE | 48372 | ENSG00000133302 | SLF1 | chr5 | + | 94678807 | 94678955 | 94670842 | 94671008 | 94686572 | 94686718 | 31 | 9 | [illegible] |
| SE | 48373 | ENSG00000134215 | VAV3 | chr1 | - | 107574046 | 107574198 | 107571160 | 107573372 | 107596211 | 107596341 | 245 | 178 | [illegible] |
| SE | 48375 | ENSG00000134215 | VAV3 | chr1 | - | 107596211 | 107596341 | 107574046 | 107574198 | 107602396 | 107602484 | 368 | 311 | [illegible] |
| SE | 48376 | ENSG00000134215 | VAV3 | chr1 | - | 107609423 | 107609539 | 107603096 | 107603163 | 107609930 | 107609965 | 3 | 0 | [illegible] |
| SE | 48377 | ENSG00000134215 | VAV3 | chr1 | - | 107642618 | 107642755 | 107617566 | 107617632 | 107683487 | 107683533 | 72 | 56 | [illegible] |
| SE | 48379 | ENSG00000134215 | VAV3 | chr1 | - | 107683487 | 107683533 | 107642618 | 107642755 | 107688380 | 107688406 | 85 | 48 | [illegible] |
| SE | 48380 | ENSG00000134215 | VAV3 | chr1 | - | 107683487 | 107683533 | 107642618 | 107642755 | 107704549 | 107704650 | 22 | 10 | [illegible] |
| SE | 48381 | ENSG00000134215 | VAV3 | chr1 | - | 107683487 | 107683533 | 107668838 | 107669472 | 107688380 | 107688406 | 33 | 13 | [illegible] |
| SE | 48382 | ENSG00000134215 | VAV3 | chr1 | - | 107688380 | 107688406 | 107683487 | 107683533 | 107704549 | 107704650 | 130 | 92 | [illegible] |
| SE | 48383 | ENSG00000134215 | VAV3 | chr1 | - | 107704549 | 107704650 | 107688380 | 107688406 | 107704959 | 107705061 | 223 | 152 | [illegible] |
| SE | 48384 | ENSG00000134215 | VAV3 | chr1 | - | 107755426 | 107755513 | 107751116 | 107751202 | 107757260 | 107757329 | 245 | 157 | [illegible] |
| SE | 48385 | ENSG00000134215 | VAV3 | chr1 | - | 107757260 | 107757329 | 107755426 | 107755513 | 107760783 | 107760879 | 290 | 160 | [illegible] |
| SE | 48386 | ENSG00000134215 | VAV3 | chr1 | - | 107760783 | 107760879 | 107757260 | 107757329 | 107765075 | 107765175 | 363 | 222 | [illegible] |
| SE | 48389 | ENSG00000134215 | VAV3 | chr1 | - | 107874900 | 107875017 | 107779433 | 107779492 | 107964665 | 107964853 | 137 | 85 | [illegible] |
| SE | 48390 | ENSG00000162542 | TMCO4 | chr1 | - | 19737371 | 19737456 | 19700767 | 19700885 | 19746455 | 19746599 | 14 | 9 | [illegible] |
| SE | 48392 | ENSG00000162542 | TMCO4 | chr1 | - | 19740776 | 19740941 | 19737371 | 19737456 | 19746455 | 19746599 | 1 | 1 | [illegible] |
| SE | 48393 | ENSG00000162542 | TMCO4 | chr1 | - | 19745531 | 19745651 | 19740776 | 19740941 | 19746455 | 19746599 | 73 | 45 | [illegible] |
| SE | 48394 | ENSG00000162542 | TMCO4 | chr1 | - | 19755633 | 19755766 | 19747162 | 19747260 | 19771307 | 19771427 | 61 | 24 | [illegible] |
| SE | 48395 | ENSG00000162542 | TMCO4 | chr1 | - | 19770541 | 19770569 | 19755633 | 19755766 | 19771307 | 19771427 | 20 | 6 | [illegible] |
| SE | 48396 | ENSG00000162542 | TMCO4 | chr1 | - | 19771307 | 19771482 | 19755633 | 19755766 | 19780579 | 19780764 | 28 | 6 | [illegible] |
| SE | 48397 | ENSG00000162542 | TMCO4 | chr1 | - | 19798070 | 19798215 | 19787025 | 19787117 | 19799854 | 19799940 | 1 | 0 | [illegible] |
| SE | 48399 | ENSG00000162542 | TMCO4 | chr1 | - | 19798136 | 19798215 | 19787025 | 19787117 | 19799854 | 19799916 | 10 | 2 | [illegible] |
| SE | 48401 | ENSG00000183690 | EFHC2 | chrX | - | 44163921 | 44164027 | 44147871 | 44148896 | 44176291 | 44176384 | 34 | 31 | [illegible] |
| SE | 48402 | ENSG00000183690 | EFHC2 | chrX | - | 44235304 | 44235447 | 44232480 | 44232677 | 44242120 | 44242289 | 29 | 50 | [illegible] |
| SE | 48404 | ENSG00000079335 | CDC14A | chr1 | + | 100377545 | 100377621 | 100353761 | 100353852 | 100390731 | 100390824 | 152 | 116 | [illegible] |
| SE | 48405 | ENSG00000079335 | CDC14A | chr1 | + | 100383465 | 100383640 | 100377545 | 100377621 | 100390731 | 100390824 | 0 | 0 | [illegible] |
| SE | 48408 | ENSG00000079335 | CDC14A | chr1 | + | 100455404 | 100455492 | 100442933 | 100442996 | 100467955 | 100468094 | 59 | 51 | [illegible] |
| SE | 48409 | ENSG00000079335 | CDC14A | chr1 | + | 100462650 | 100462881 | 100442933 | 100442996 | 100467955 | 100468094 | 304 | 246 | [illegible] |
| SE | 48410 | ENSG00000079335 | CDC14A | chr1 | + | 100462650 | 100462881 | 100455404 | 100455492 | 100467955 | 100468094 | 412 | 346 | [illegible] |
| SE | 48413 | ENSG00000079335 | CDC14A | chr1 | + | 100498928 | 100499262 | 100498084 | 100498207 | 100518250 | 100520277 | 221 | 120 | [illegible] |
| SE | 48414 | ENSG00000079335 | CDC14A | chr1 | + | 100504813 | 100504937 | 100498928 | 100499262 | 100518250 | 100518253 | 3 | 1 | [illegible] |
| SE | 48415 | ENSG00000134057 | CCNB1 | chr5 | + | 69174250 | 69174409 | 69171269 | 69171452 | 69174876 | 69175113 | 13 | 5 | [illegible] |
| SE | 48418 | ENSG00000086015 | MAST2 | chr1 | + | 46000958 | 46000979 | 45997723 | 45997799 | 46002804 | 46002883 | 0 | 0 | [illegible] |
| SE | 48420 | ENSG00000086015 | MAST2 | chr1 | + | 46006240 | 46006395 | 46002804 | 46002883 | 46008295 | 46008371 | 45 | 34 | [illegible] |
| SE | 48422 | ENSG00000086015 | MAST2 | chr1 | + | 46019595 | 46019697 | 46010729 | 46010939 | 46021949 | 46022082 | 37 | 8 | [illegible] |
| SE | 48423 | ENSG00000086015 | MAST2 | chr1 | + | 46025676 | 46025815 | 46023771 | 46023980 | 46027730 | 46027863 | 37 | 7 | [illegible] |
| SE | 48425 | ENSG00000086015 | MAST2 | chr1 | + | 46030128 | 46030238 | 46029830 | 46029953 | 46030606 | 46030759 | 21 | 24 | [illegible] |
| SE | 48426 | ENSG00000134827 | TCN1 | chr11 | - | 59861526 | 59861679 | 59859181 | 59859267 | 59862581 | 59862635 | 29 | 4 | [illegible] |
| SE | 48427 | ENSG00000134827 | TCN1 | chr11 | - | 59861526 | 59861682 | 59859076 | 59859267 | 59862581 | 59862722 | 56 | 6 | [illegible] |
| SE | 48428 | ENSG00000134827 | TCN1 | chr11 | - | 59862581 | 59862722 | 59859076 | 59859267 | 59866391 | 59866471 | 0 | 0 | [illegible] |
| SE | 48433 | ENSG00000116171 | SCP2 | chr1 | + | 52954739 | 52954804 | 52950754 | 52950886 | 52961502 | 52961629 | 66 | 62 | [illegible] |
| SE | 48435 | ENSG00000116171 | SCP2 | chr1 | + | 52976682 | 52976769 | 52974768 | 52974832 | 52978216 | 52978367 | 48 | 44 | [illegible] |
| SE | 48436 | ENSG00000116171 | SCP2 | chr1 | + | 52978216 | 52978367 | 52976682 | 52976769 | 52980395 | 52980453 | 85 | 79 | [illegible] |
| SE | 48437 | ENSG00000116171 | SCP2 | chr1 | + | 52993257 | 52996000 | 52988028 | 52988136 | 53014889 | 53015043 | 1 | 0 | [illegible] |
| SE | 48438 | ENSG00000116171 | SCP2 | chr1 | + | 53014889 | 53015043 | 52988028 | 52988136 | 53027968 | 53028071 | 528 | 275 | [illegible] |
| SE | 48439 | ENSG00000116171 | SCP2 | chr1 | + | 53027968 | 53028071 | 53014889 | 53015043 | 53038916 | 53039046 | 685 | 334 | [illegible] |
| SE | 48440 | ENSG00000116171 | SCP2 | chr1 | + | 53027977 | 53028071 | 53014927 | 53015043 | 53038916 | 53039046 | 215 | 110 | [illegible] |
| SE | 48443 | ENSG00000104643 | MTMR9 | chr8 | + | 11306189 | 11306407 | 11304840 | 11305014 | 11309526 | 11309688 | 56 | 75 | [illegible] |
| SE | 48444 | ENSG00000104643 | MTMR9 | chr8 | + | 11314922 | 11315064 | 11309526 | 11309688 | 11316672 | 11316893 | 42 | 38 | [illegible] |
| SE | 48445 | ENSG00000104643 | MTMR9 | chr8 | + | 11316672 | 11316893 | 11309526 | 11309688 | 11319686 | 11319838 | 31 | 14 | [illegible] |
| SE | 48446 | ENSG00000104643 | MTMR9 | chr8 | + | 11319686 | 11319838 | 11316672 | 11316893 | 11322624 | 11323057 | 69 | 38 | [illegible] |
| SE | 48448 | ENSG00000251129 | LINC02506 | chr4 | + | 32146052 | 32146267 | 31997375 | 31997822 | 32155313 | 32155406 | 1 | 0 | [illegible] |
| SE | 48450 | ENSG00000251129 | LINC02506 | chr4 | + | 32157955 | 32158081 | 32155313 | 32155406 | 32225023 | 32225121 | 0 | 0 | [illegible] |
| SE | 48460 | ENSG00000131238 | PPT1 | chr1 | - | 40092044 | 40092172 | 40091328 | 40091399 | 40092397 | 40092500 | 833 | 352 | [illegible] |
| SE | 48463 | ENSG00000177030 | DEAF1 | chr11 | - | 678693 | 678822 | 674535 | 674783 | 679687 | 679816 | 64 | 31 | [illegible] |
| SE | 48465 | ENSG00000177030 | DEAF1 | chr11 | - | 679977 | 680074 | 679687 | 679816 | 680962 | 681045 | 2 | 0 | [illegible] |
| SE | 48466 | ENSG00000177030 | DEAF1 | chr11 | - | 680962 | 681089 | 679687 | 679816 | 684897 | 684963 | 107 | 39 | [illegible] |
| SE | 48467 | ENSG00000177030 | DEAF1 | chr11 | - | 686857 | 686997 | 683575 | 684963 | 687910 | 687936 | 204 | 73 | [illegible] |
| SE | 48468 | ENSG00000114395 | CYB561D2 | chr3 | + | 50352008 | 50352046 | 50350857 | 50351560 | 50353240 | 50354032 | 22 | 6 | [illegible] |
| SE | 48469 | ENSG00000259494 | MRPL46 | chr15 | - | 88460152 | 88460382 | 88459477 | 88459863 | 88464702 | 88464813 | 0 | 0 | [illegible] |
| SE | 48473 | ENSG00000143543 | JTB | chr1 | - | 153976975 | 153977013 | 153976692 | 153976775 | 153977338 | 153977394 | 56 | 16 | [illegible] |
| SE | 48475 | ENSG00000117461 | PIK3R3 | chr1 | - | 46045917 | 46046088 | 46040139 | 46043871 | 46046550 | 46046625 | 21 | 17 | [illegible] |
| SE | 48476 | ENSG00000117461 | PIK3R3 | chr1 | - | 46055794 | 46055971 | 46046550 | 46046625 | 46061928 | 46062071 | 37 | 14 | [illegible] |
| SE | 48478 | ENSG00000117461 | PIK3R3 | chr1 | - | 46080641 | 46080750 | 46077514 | 46077613 | 46131846 | 46131956 | 28 | 11 | [illegible] |
| SE | 48479 | ENSG00000117461 | PIK3R3 | chr1 | - | 46131846 | 46131956 | 46080641 | 46080750 | 46132783 | 46132913 | 14 | 4 | [illegible] |
| SE | 48480 | ENSG00000204519 | ZNF551 | chr19 | + | 57685261 | 57685385 | 57681968 | 57682122 | 57686480 | 57688028 | 7 | 2 | [illegible] |
| SE | 48481 | ENSG00000204519 | ZNF551 | chr19 | + | 57685261 | 57685385 | 57681976 | 57682244 | 57686480 | 57688028 | 7 | 4 | [illegible] |
| SE | 48482 | ENSG00000079313 | REXO1 | chr19 | - | 1817706 | 1817780 | 1817178 | 1817329 | 1818481 | 1818595 | 41 | 13 | [illegible] |
| SE | 48483 | ENSG00000079313 | REXO1 | chr19 | - | 1825838 | 1825943 | 1823571 | 1823785 | 1826877 | 1828631 | 69 | 19 | [illegible] |
| SE | 48484 | ENSG00000079313 | REXO1 | chr19 | - | 1830705 | 1830866 | 1827647 | 1828631 | 1848201 | 1848446 | 0 | 0 | [illegible] |
| SE | 48492 | ENSG00000138095 | LRPPRC | chr2 | - | 43894544 | 43894629 | 43889733 | 43889876 | 43899218 | 43899334 | 299 | 200 | [illegible] |
| SE | 48493 | ENSG00000138095 | LRPPRC | chr2 | - | 43894544 | 43896708 | 43889733 | 43889876 | 43899218 | 43899334 | 312 | 234 | [illegible] |
| SE | 48495 | ENSG00000138095 | LRPPRC | chr2 | - | 43895367 | 43895566 | 43894544 | 43894629 | 43896633 | 43896708 | 5 | 0 | [illegible] |
| SE | 48496 | ENSG00000138095 | LRPPRC | chr2 | - | 43895367 | 43895566 | 43894544 | 43894629 | 43899218 | 43899334 | 2 | 0 | [illegible] |
| SE | 48499 | ENSG00000138095 | LRPPRC | chr2 | - | 43896633 | 43896655 | 43894544 | 43894629 | 43899218 | 43899334 | 61 | 34 | [illegible] |
| SE | 48500 | ENSG00000138095 | LRPPRC | chr2 | - | 43896633 | 43896708 | 43889733 | 43889876 | 43899218 | 43899334 | 32 | 40 | [illegible] |
| SE | 48502 | ENSG00000138095 | LRPPRC | chr2 | - | 43896633 | 43896708 | 43895367 | 43895566 | 43899218 | 43899334 | 35 | 40 | [illegible] |
| SE | 48504 | ENSG00000138095 | LRPPRC | chr2 | - | 43912431 | 43912558 | 43905691 | 43905780 | 43918024 | 43918133 | 409 | 276 | [illegible] |
| SE | 48507 | ENSG00000138095 | LRPPRC | chr2 | - | 43918255 | 43918398 | 43912431 | 43912558 | 43925066 | 43925157 | 329 | 235 | [illegible] |
| SE | 48508 | ENSG00000138095 | LRPPRC | chr2 | - | 43919402 | 43919468 | 43918255 | 43918398 | 43925066 | 43925157 | 0 | 0 | [illegible] |
| SE | 48512 | ENSG00000138095 | LRPPRC | chr2 | - | 43945331 | 43945417 | 43943686 | 43943894 | 43946112 | 43946243 | 255 | 177 | [illegible] |
| SE | 48513 | ENSG00000138095 | LRPPRC | chr2 | - | 43947730 | 43947775 | 43947256 | 43947370 | 43948121 | 43948199 | 155 | 124 | [illegible] |
| SE | 48514 | ENSG00000138095 | LRPPRC | chr2 | - | 43957725 | 43957957 | 43957384 | 43957451 | 43960540 | 43960634 | 0 | 1 | [illegible] |
| SE | 48515 | ENSG00000138095 | LRPPRC | chr2 | - | 43974149 | 43974295 | 43973794 | 43973900 | 43974613 | 43974758 | 479 | 350 | [illegible] |
| SE | 48516 | ENSG00000138095 | LRPPRC | chr2 | - | 43974610 | 43974758 | 43974149 | 43974295 | 43975090 | 43975217 | 130 | 77 | [illegible] |
| SE | 48517 | ENSG00000138095 | LRPPRC | chr2 | - | 43974613 | 43974758 | 43974149 | 43974295 | 43975090 | 43975217 | 424 | 281 | [illegible] |
| SE | 48518 | ENSG00000138095 | LRPPRC | chr2 | - | 43976142 | 43976229 | 43975090 | 43975217 | 43976993 | 43977052 | 343 | 228 | [illegible] |
| SE | 48523 | ENSG00000251349 | MSANTD3-TMEFF1 | chr9 | + | 100572517 | 100572676 | 100561396 | 100561520 | 100576515 | 100577636 | 32 | 15 | [illegible] |
| SE | 48527 | ENSG00000111481 | COPZ1 | chr12 | + | 54342205 | 54342287 | 54340371 | 54340615 | 54345459 | 54345515 | 198 | 109 | [illegible] |
| SE | 48529 | ENSG00000111481 | COPZ1 | chr12 | + | 54343224 | 54343316 | 54342205 | 54342287 | 54347766 | 54347844 | 162 | 115 | [illegible] |
| SE | 48530 | ENSG00000111481 | COPZ1 | chr12 | + | 54345459 | 54345515 | 54342205 | 54342287 | 54347766 | 54347840 | 198 | 115 | [illegible] |
| SE | 48531 | ENSG00000111481 | COPZ1 | chr12 | + | 54345459 | 54345515 | 54343224 | 54343316 | 54347766 | 54347840 | 844 | 516 | [illegible] |
| SE | 48532 | ENSG00000111481 | COPZ1 | chr12 | + | 54346611 | 54346674 | 54345459 | 54345515 | 54347766 | 54347844 | 6 | 0 | [illegible] |
| SE | 48536 | ENSG00000186716 | BCR | chr22 | + | 23290338 | 23290413 | 23289516 | 23289621 | 23292540 | 23292638 | 214 | 131 | [illegible] |
| SE | 48537 | ENSG00000186716 | BCR | chr22 | + | 23292540 | 23292638 | 23290338 | 23290413 | 23295023 | 23295155 | 239 | 161 | [illegible] |
| SE | 48538 | ENSG00000186716 | BCR | chr22 | + | 23295023 | 23295155 | 23292540 | 23292638 | 23309423 | 23309483 | 238 | 171 | [illegible] |
| SE | 48539 | ENSG00000186716 | BCR | chr22 | + | 23310323 | 23310433 | 23309423 | 23309483 | 23311696 | 23311836 | 169 | 141 | [illegible] |
| SE | 48540 | ENSG00000186716 | BCR | chr22 | + | 23313967 | 23314073 | 23312886 | 23313021 | 23314551 | 23314714 | 102 | 43 | [illegible] |
| SE | 48543 | ENSG00000126091 | ST3GAL3 | chr1 | + | 43792101 | 43792149 | 43736232 | 43736425 | 43814890 | 43814933 | 16 | 7 | [illegible] |
| SE | 48549 | ENSG00000126091 | ST3GAL3 | chr1 | + | 43819113 | 43819236 | 43814890 | 43814933 | 43824730 | 43824892 | 0 | 0 | [illegible] |
| SE | 48553 | ENSG00000126091 | ST3GAL3 | chr1 | + | 43831654 | 43831794 | 43824831 | 43824892 | 43838218 | 43838261 | 0 | 0 | [illegible] |
| SE | 48556 | ENSG00000126091 | ST3GAL3 | chr1 | + | 43858134 | 43858308 | 43838218 | 43838311 | 43879334 | 43879507 | 0 | 2 | [illegible] |
| SE | 48559 | ENSG00000126091 | ST3GAL3 | chr1 | + | 43879026 | 43879231 | 43838218 | 43838311 | 43879334 | 43879507 | 0 | 0 | [illegible] |
| SE | 48561 | ENSG00000126091 | ST3GAL3 | chr1 | + | 43879334 | 43879507 | 43858134 | 43858308 | 43894382 | 43894477 | 1 | 4 | [illegible] |
| SE | 48569 | ENSG00000126091 | ST3GAL3 | chr1 | + | 43899540 | 43899727 | 43899167 | 43899289 | 43930131 | 43930221 | 0 | 2 | [illegible] |
| SE | 48570 | ENSG00000126091 | ST3GAL3 | chr1 | + | 43900748 | 43900833 | 43899540 | 43899727 | 43920403 | 43920550 | 1 | 1 | [illegible] |
| SE | 48580 | ENSG00000150681 | RGS18 | chr1 | + | 192159219 | 192159321 | 192158461 | 192158756 | 192160377 | 192160439 | 1334 | 2902 | [illegible] |
| SE | 48581 | ENSG00000150681 | RGS18 | chr1 | + | 192159219 | 192159321 | 192158461 | 192158756 | 192161452 | 192161487 | 483 | 1178 | [illegible] |
| SE | 48582 | ENSG00000150681 | RGS18 | chr1 | + | 192159219 | 192159321 | 192158461 | 192158756 | 192181291 | 192181454 | 488 | 1192 | [illegible] |
| SE | 48583 | ENSG00000150681 | RGS18 | chr1 | + | 192160377 | 192160439 | 192158461 | 192158756 | 192161452 | 192161487 | 19 | 41 | [illegible] |
| SE | 48584 | ENSG00000150681 | RGS18 | chr1 | + | 192160377 | 192160439 | 192158461 | 192158756 | 192181291 | 192181458 | 258 | 529 | [illegible] |
| SE | 48585 | ENSG00000150681 | RGS18 | chr1 | + | 192160377 | 192160439 | 192159219 | 192159321 | 192161452 | 192161487 | 582 | 1109 | [illegible] |
| SE | 48586 | ENSG00000150681 | RGS18 | chr1 | + | 192160377 | 192160439 | 192159219 | 192159321 | 192181291 | 192181458 | 1113 | 2231 | [illegible] |
| SE | 48587 | ENSG00000150681 | RGS18 | chr1 | + | 192161452 | 192161487 | 192158513 | 192158756 | 192181291 | 192181454 | 3 | 8 | [illegible] |
| SE | 48588 | ENSG00000150681 | RGS18 | chr1 | + | 192161452 | 192161487 | 192159219 | 192159321 | 192181291 | 192181454 | 3 | 9 | [illegible] |
| SE | 48589 | ENSG00000150681 | RGS18 | chr1 | + | 192161452 | 192161487 | 192160377 | 192160439 | 192163279 | 192163656 | 16 | 8 | [illegible] |
| SE | 48591 | ENSG00000131374 | TBC1D5 | chr3 | - | 17166766 | 17166928 | 17157161 | 17161256 | 17167748 | 17167828 | 239 | 162 | [illegible] |
| SE | 48593 | ENSG00000131374 | TBC1D5 | chr3 | - | 17258505 | 17258591 | 17238162 | 17238419 | 17291894 | 17292001 | 221 | 130 | [illegible] |
| SE | 48594 | ENSG00000131374 | TBC1D5 | chr3 | - | 17291894 | 17292001 | 17258505 | 17258591 | 17307991 | 17308134 | 306 | 208 | [illegible] |
| SE | 48596 | ENSG00000131374 | TBC1D5 | chr3 | - | 17372074 | 17372247 | 17307991 | 17308134 | 17374470 | 17374540 | 658 | 388 | [illegible] |
| SE | 48597 | ENSG00000131374 | TBC1D5 | chr3 | - | 17374470 | 17374540 | 17372074 | 17372247 | 17374628 | 17374679 | 549 | 335 | [illegible] |
| SE | 48598 | ENSG00000131374 | TBC1D5 | chr3 | - | 17376524 | 17376613 | 17374628 | 17374679 | 17383912 | 17384015 | 324 | 191 | [illegible] |
| SE | 48602 | ENSG00000131374 | TBC1D5 | chr3 | - | 17508473 | 17508605 | 17428449 | 17428519 | 17622647 | 17622765 | 184 | 204 | [illegible] |
| SE | 48603 | ENSG00000131374 | TBC1D5 | chr3 | - | 17508473 | 17508605 | 17428449 | 17428519 | 17623848 | 17623913 | 248 | 272 | [illegible] |
| SE | 48604 | ENSG00000131374 | TBC1D5 | chr3 | - | 17508473 | 17508605 | 17428449 | 17428519 | 17742480 | 17742604 | 167 | 195 | [illegible] |
| SE | 48606 | ENSG00000131374 | TBC1D5 | chr3 | - | 17518901 | 17518995 | 17508473 | 17508605 | 17576413 | 17576549 | 23 | 22 | [illegible] |
| SE | 48611 | ENSG00000131374 | TBC1D5 | chr3 | - | 17547015 | 17547076 | 17508473 | 17508605 | 17562018 | 17562142 | 0 | 0 | [illegible] |
| SE | 48612 | ENSG00000131374 | TBC1D5 | chr3 | - | 17547015 | 17547076 | 17508473 | 17508605 | 17586352 | 17586557 | 0 | 0 | [illegible] |
| SE | 48613 | ENSG00000131374 | TBC1D5 | chr3 | - | 17547015 | 17547076 | 17508473 | 17508605 | 17623848 | 17623916 | 0 | 0 | [illegible] |
| SE | 48615 | ENSG00000131374 | TBC1D5 | chr3 | - | 17562018 | 17562142 | 17508473 | 17508605 | 17576413 | 17576549 | 34 | 10 | [illegible] |
| SE | 48622 | ENSG00000131374 | TBC1D5 | chr3 | - | 17562018 | 17562142 | 17518901 | 17518995 | 17622647 | 17622765 | 7 | 3 | [illegible] |
| SE | 48624 | ENSG00000131374 | TBC1D5 | chr3 | - | 17562018 | 17562142 | 17518901 | 17518995 | 17742480 | 17742498 | 1 | 1 | [illegible] |
| SE | 48625 | ENSG00000131374 | TBC1D5 | chr3 | - | 17562018 | 17562142 | 17547015 | 17547076 | 17586352 | 17586557 | 22 | 15 | [illegible] |
| SE | 48627 | ENSG00000131374 | TBC1D5 | chr3 | - | 17575202 | 17575310 | 17508548 | 17508605 | 17576413 | 17576549 | 2 | 0 | [illegible] |
| SE | 48628 | ENSG00000131374 | TBC1D5 | chr3 | - | 17575202 | 17575310 | 17508548 | 17508605 | 17623848 | 17623913 | 1 | 0 | [illegible] |

| | | | | | | | | | | | | | | | | | | | | | | | | |
|---|---|---|---|---|---|---|---|---|---|---|---|---|---|---|---|---|---|---|---|---|---|---|---|---|
| SE | 48629 | ENSG00000131374 | TBC1D5 | chr3 | - | 17575202 | 17575310 | 17508548 | 17508605 | 17742480 | 17742748 | 0 | 0 | 1 | 0 | 0 | 6 | 9 | 5 | 5 | 13 | 0 | 0 | [illegible] |
| SE | 48630 | ENSG00000131374 | TBC1D5 | chr3 | - | 17575202 | 17575310 | 17562018 | 17562142 | 17576413 | 17576549 | 2 | 0 | 5 | 4 | 11 | 1 | 0 | 0 | 9 | 4 | 14 | 1 | [illegible] |
| SE | 48631 | ENSG00000131374 | TBC1D5 | chr3 | - | 17575202 | 17575310 | 17562018 | 17562142 | 17623848 | 17623913 | 1 | 0 | 0 | 0 | 0 | 7 | 5 | 17 | 8 | 12 | 0 | 0 | [illegible] |
| SE | 48632 | ENSG00000131374 | TBC1D5 | chr3 | - | 17575202 | 17575310 | 17562018 | 17562142 | 17742480 | 17742748 | 0 | 0 | 1 | 0 | 0 | 1 | 1 | 3 | 2 | 0 | 0 | 0 | [illegible] |
| SE | 48633 | ENSG00000131374 | TBC1D5 | chr3 | - | 17576413 | 17576549 | 17508473 | 17508605 | 17586352 | 17586466 | 4 | 0 | 9 | 6 | 14 | 14 | 5 | 10 | 9 | 19 | 10 | 1 | [illegible] |
| SE | 48634 | ENSG00000131374 | TBC1D5 | chr3 | - | 17576413 | 17576549 | 17508473 | 17508605 | 17622647 | 17622765 | 4 | 2 | 1 | 3 | 0 | 23 | 18 | 9 | 22 | 43 | 4 | 0 | [illegible] |
| SE | 48636 | ENSG00000131374 | TBC1D5 | chr3 | - | 17576413 | 17576549 | 17518901 | 17518995 | 17586352 | 17586466 | 2 | 0 | 9 | 6 | 14 | 1 | 2 | 3 | 0 | 6 | 10 | 1 | [illegible] |
| SE | 48637 | ENSG00000131374 | TBC1D5 | chr3 | - | 17576413 | 17576549 | 17518901 | 17518995 | 17622647 | 17622765 | 2 | 2 | 1 | 3 | 0 | 3 | 1 | 2 | 4 | 7 | 4 | 0 | [illegible] |
| SE | 48638 | ENSG00000131374 | TBC1D5 | chr3 | - | 17576413 | 17576549 | 17518901 | 17518995 | 17623848 | 17623913 | 0 | 0 | 0 | 0 | 1 | 13 | 9 | 5 | 9 | 9 | 0 | 9 | [illegible] |
| SE | 48641 | ENSG00000131374 | TBC1D5 | chr3 | - | 17576413 | 17576549 | 17562018 | 17562142 | 17623848 | 17623913 | 1 | 0 | 0 | 9 | 5 | 7 | 5 | 17 | 8 | 12 | 0 | 11 | [illegible] |
| SE | 48642 | ENSG00000131374 | TBC1D5 | chr3 | - | 17576413 | 17576549 | 17575202 | 17575310 | 17623848 | 17623913 | 2 | 0 | 5 | 4 | 11 | 1 | 0 | 0 | 0 | 0 | 25 | 10 | [illegible] |
| SE | 48648 | ENSG00000131374 | TBC1D5 | chr3 | - | 17586352 | 17586557 | 17518901 | 17518995 | 17742480 | 17742612 | 1 | 4 | 2 | 0 | 6 | 4 | 0 | 0 | 3 | 1 | 8 | 9 | [illegible] |
| SE | 48655 | ENSG00000131374 | TBC1D5 | chr3 | - | 17622647 | 17622765 | 17428449 | 17428519 | 17623848 | 17623913 | 41 | 22 | 30 | 36 | 85 | 0 | 0 | 0 | 0 | 1 | 21 | 13 | [illegible] |
| SE | 48662 | ENSG00000131374 | TBC1D5 | chr3 | - | 17623848 | 17623913 | 17428449 | 17428519 | 17742480 | 17742564 | 10 | 12 | 11 | 7 | 5 | 0 | 0 | 0 | 0 | 0 | 14 | 14 | [illegible] |
| SE | 48663 | ENSG00000131374 | TBC1D5 | chr3 | - | 17623848 | 17623913 | 17508473 | 17508605 | 17662989 | 17663110 | 52 | 67 | 32 | 44 | 75 | 0 | 2 | 0 | 2 | 0 | 48 | 55 | [illegible] |
| SE | 48665 | ENSG00000131374 | TBC1D5 | chr3 | - | 17623848 | 17623913 | 17518901 | 17518995 | 17646960 | 17647056 | 10 | 8 | 3 | 6 | 9 | 1 | 0 | 0 | 0 | 0 | 0 | 0 | [illegible] |
| SE | 48668 | ENSG00000131374 | TBC1D5 | chr3 | - | 17623848 | 17623913 | 17575202 | 17575310 | 17742480 | 17742564 | 11 | 12 | 11 | 7 | 4 | 0 | 0 | 1 | 0 | 0 | 14 | 14 | [illegible] |
| SE | 48669 | ENSG00000131374 | TBC1D5 | chr3 | - | 17623848 | 17623913 | 17586351 | 17586466 | 17742480 | 17742564 | 15 | 13 | 11 | 7 | 4 | 0 | 1 | 0 | 0 | 0 | 14 | 14 | [illegible] |
| SE | 48670 | ENSG00000131374 | TBC1D5 | chr3 | - | 17623848 | 17623913 | 17586352 | 17586557 | 17742480 | 17742564 | 14 | 14 | 15 | 11 | 12 | 0 | 2 | 0 | 0 | 0 | 14 | 14 | [illegible] |
| SE | 48671 | ENSG00000131374 | TBC1D5 | chr3 | - | 17644040 | 17644120 | 17623848 | 17623913 | 17646960 | 17647056 | 1 | 0 | 2 | 0 | 0 | 4 | 5 | 1 | 1 | 20 | 0 | 0 | [illegible] |
| SE | 48672 | ENSG00000131374 | TBC1D5 | chr3 | - | 17646960 | 17647056 | 17518901 | 17518995 | 17742480 | 17742594 | 1 | 1 | 0 | 0 | 0 | 4 | 0 | 0 | 3 | 1 | 0 | 0 | [illegible] |
| SE | 48673 | ENSG00000131374 | TBC1D5 | chr3 | - | 17646960 | 17647056 | 17623848 | 17623913 | 17672544 | 17672675 | 3 | 5 | 2 | 1 | 16 | 0 | 2 | 3 | 4 | 1 | 0 | 0 | [illegible] |
| SE | 48675 | ENSG00000131374 | TBC1D5 | chr3 | - | 17662989 | 17663110 | 17508473 | 17508605 | 17742480 | 17742748 | 0 | 3 | 2 | 2 | 0 | 6 | 9 | 5 | 5 | 13 | 0 | 6 | [illegible] |
| SE | 48679 | ENSG00000038002 | AGA | chr4 | - | 177436297 | 177436351 | 177434439 | 177434481 | 177437404 | 177437519 | 45 | 24 | 36 | 39 | 33 | 1 | 0 | 1 | 0 | 3 | 27 | 28 | [illegible] |
| SE | 48680 | ENSG00000038002 | AGA | chr4 | - | 177438744 | 177438857 | 177437067 | 177437519 | 177440272 | 177440426 | 86 | 56 | 46 | 60 | 59 | 0 | 1 | 0 | 0 | 0 | 39 | 18 | [illegible] |
| SE | 48681 | ENSG00000038002 | AGA | chr4 | - | 177439575 | 177439688 | 177438744 | 177438857 | 177440272 | 177440426 | 218 | 106 | 126 | 142 | 231 | 0 | 2 | 1 | 0 | 0 | 133 | 117 | [illegible] |
| SE | 48682 | ENSG00000105499 | PLA2G4C | chr19 | - | 48085058 | 48085112 | 48077770 | 48077824 | 48088685 | 48088712 | 1 | 19 | 5 | 14 | 9 | 0 | 0 | 0 | 1 | 0 | 27 | 20 | [illegible] |
| SE | 48683 | ENSG00000105499 | PLA2G4C | chr19 | - | 48085058 | 48085112 | 48077770 | 48077824 | 48090363 | 48090409 | 2 | 8 | 2 | 6 | 3 | 0 | 0 | 0 | 0 | 0 | 16 | 16 | [illegible] |
| SE | 48684 | ENSG00000105499 | PLA2G4C | chr19 | - | 48088685 | 48088712 | 48077770 | 48077824 | 48090363 | 48090417 | 0 | 0 | 1 | 1 | 1 | 0 | 0 | 0 | 0 | 0 | 0 | 1 | [illegible] |
| SE | 48685 | ENSG00000105499 | PLA2G4C | chr19 | - | 48088685 | 48088712 | 48085058 | 48085112 | 48090363 | 48090417 | 2 | 16 | 5 | 10 | 7 | 1 | 1 | 0 | 1 | 0 | 26 | 9 | [illegible] |
| SE | 48688 | ENSG00000105499 | PLA2G4C | chr19 | - | 48104587 | 48104724 | 48099670 | 48099860 | 48105332 | 48105413 | 10 | 36 | 1 | 32 | 45 | 1 | 1 | 0 | 1 | 1 | 42 | 54 | [illegible] |
| SE | 48691 | ENSG00000095380 | NANS | chr9 | + | 98060781 | 98060997 | 98056731 | 98056940 | 98075569 | 98075732 | 69 | 47 | 57 | 70 | 24 | 0 | 0 | 0 | 0 | 0 | 112 | 62 | [illegible] |
| SE | 48693 | ENSG00000160991 | ORAI2 | chr7 | + | 102436201 | 102436333 | 102433529 | 102433661 | 102438943 | 102439181 | 31 | 6 | 29 | 26 | 33 | 2 | 0 | 0 | 0 | 0 | 0 | 7 | [illegible] |
| SE | 48694 | ENSG00000160991 | ORAI2 | chr7 | + | 102436224 | 102436333 | 102433548 | 102433661 | 102438943 | 102439047 | 42 | 8 | 35 | 36 | 36 | 2 | 0 | 0 | 0 | 0 | 8 | 5 | [illegible] |
| SE | 48695 | ENSG00000160991 | ORAI2 | chr7 | + | 102436229 | 102436333 | 102433569 | 102433661 | 102438943 | 102439181 | 24 | 3 | 26 | 25 | 30 | 2 | 0 | 0 | 0 | 0 | 0 | 5 | [illegible] |
| SE | 48706 | ENSG00000143401 | ANP32E | chr1 | - | 150223185 | 150223240 | 150220391 | 150220758 | 150226607 | 150226795 | 82 | 77 | 78 | 110 | 297 | 0 | 0 | 2 | 0 | 0 | 27 | 128 | [illegible] |
| SE | 48707 | ENSG00000143401 | ANP32E | chr1 | - | 150223185 | 150223294 | 150220377 | 150220761 | 150226607 | 150226730 | 44 | 52 | 61 | 58 | 121 | 0 | 2 | 0 | 1 | 1 | 0 | 42 | [illegible] |
| SE | 48708 | ENSG00000143401 | ANP32E | chr1 | - | 150226607 | 150226795 | 150223185 | 150223240 | 150229071 | 150229114 | 525 | 558 | 743 | 813 | 1946 | 0 | 2 | 0 | 0 | 0 | 140 | 917 | [illegible] |
| SE | 48710 | ENSG00000143401 | ANP32E | chr1 | - | 150229071 | 150229114 | 150226607 | 150226795 | 150235732 | 150235829 | 5 | 29 | 46 | 6 | 135 | 0 | 0 | 0 | 0 | 0 | 7 | 42 | [illegible] |
| SE | 48711 | ENSG00000143401 | ANP32E | chr1 | - | 150229071 | 150229144 | 150226607 | 150226795 | 150235732 | 150235983 | 47 | 80 | 115 | 92 | 371 | 0 | 0 | 0 | 0 | 0 | 13 | 129 | [illegible] |
| SE | 48714 | ENSG00000143401 | ANP32E | chr1 | - | 150229071 | 150229237 | 150226607 | 150226795 | 150235732 | 150236022 | 361 | 412 | 561 | 601 | 1405 | 0 | 0 | 0 | 0 | 0 | 101 | 695 | [illegible] |
| SE | 48715 | ENSG00000143401 | ANP32E | chr1 | - | 150230570 | 150230693 | 150226607 | 150226795 | 150235732 | 150236064 | 7 | 4 | 7 | 8 | 11 | 0 | 0 | 0 | 0 | 0 | 29 | 29 | [illegible] |
| SE | 48716 | ENSG00000143401 | ANP32E | chr1 | - | 150230570 | 150230693 | 150229071 | 150229237 | 150231776 | 150231926 | 1114 | 1040 | 1424 | 1340 | 2843 | 1 | 4 | 0 | 2 | 8 | 240 | 1296 | [illegible] |
| SE | 48717 | ENSG00000143401 | ANP32E | chr1 | - | 150230570 | 150230693 | 150229071 | 150229237 | 150235732 | 150236064 | 586 | 536 | 735 | 730 | 1458 | 0 | 1 | 0 | 0 | 0 | 104 | 707 | [illegible] |
| SE | 48718 | ENSG00000143401 | ANP32E | chr1 | - | 150231776 | 150231926 | 150229071 | 150229237 | 150235732 | 150235955 | 122 | 114 | 172 | 130 | 255 | 0 | 1 | 0 | 0 | 0 | 10 | 135 | [illegible] |
| SE | 48720 | ENSG00000106610 | STAG3L4 | chr7 | + | 67305986 | 67306083 | 67302620 | 67302885 | 67308576 | 67308652 | 19 | 7 | 10 | 11 | 5 | 0 | 0 | 0 | 0 | 0 | 19 | 0 | [illegible] |
| SE | 48721 | ENSG00000106610 | STAG3L4 | chr7 | + | 67305986 | 67306083 | 67302620 | 67302885 | 67308934 | 67309139 | 21 | 7 | 10 | 6 | 6 | 0 | 0 | 0 | 2 | 0 | 3 | 2 | [illegible] |
| SE | 48724 | ENSG00000106610 | STAG3L4 | chr7 | + | 67308576 | 67308652 | 67305986 | 67306083 | 67309563 | 67309715 | 5 | 2 | 3 | 7 | 2 | 0 | 1 | 1 | 0 | 2 | 16 | 4 | [illegible] |
| SE | 48725 | ENSG00000106610 | STAG3L4 | chr7 | + | 67308576 | 67309139 | 67305986 | 67306083 | 67309563 | 67309715 | 57 | 34 | 45 | 41 | 38 | 0 | 1 | 1 | 0 | 2 | 125 | 48 | [illegible] |
| SE | 48726 | ENSG00000106610 | STAG3L4 | chr7 | + | 67308934 | 67309139 | 67305986 | 67306083 | 67309563 | 67309715 | 60 | 34 | 45 | 35 | 41 | 0 | 1 | 1 | 0 | 2 | 105 | 46 | [illegible] |
| SE | 48727 | ENSG00000106610 | STAG3L4 | chr7 | + | 67308934 | 67309139 | 67308576 | 67308652 | 67309563 | 67309715 | 52 | 31 | 41 | 34 | 38 | 0 | 0 | 0 | 1 | 0 | 105 | 44 | [illegible] |
| SE | 48729 | ENSG00000076650 | GPATCH1 | chr19 | + | 33090779 | 33090865 | 33088133 | 33088268 | 33093358 | 33093519 | 149 | 74 | 141 | 115 | 187 | 0 | 0 | 0 | 1 | 0 | 23 | 63 | [illegible] |
| SE | 48732 | ENSG00000076650 | GPATCH1 | chr19 | + | 33101494 | 33101574 | 33100172 | 33100274 | 33106694 | 33106899 | 8 | 4 | 10 | 9 | 17 | 0 | 0 | 0 | 1 | 0 | 27 | 14 | [illegible] |
| SE | 48733 | ENSG00000076650 | GPATCH1 | chr19 | + | 33106694 | 33106899 | 33101494 | 33101574 | 33109716 | 33109877 | 32 | 17 | 35 | 35 | 61 | 2 | 0 | 0 | 0 | 0 | 53 | 25 | [illegible] |
| SE | 48734 | ENSG00000076650 | GPATCH1 | chr19 | + | 33111723 | 33111902 | 33109716 | 33110016 | 33112485 | 33112613 | 24 | 27 | 51 | 40 | 31 | 1 | 0 | 0 | 0 | 0 | 0 | 26 | [illegible] |
| SE | 48753 | ENSG00000116473 | RAP1A | chr1 | + | 111703335 | 111703476 | 111697440 | 111697497 | 111709148 | 111709264 | 778 | 638 | 552 | 702 | 1137 | 0 | 0 | 0 | 1 | 0 | 189 | 534 | [illegible] |
| SE | 48754 | ENSG00000116473 | RAP1A | chr1 | + | 111704342 | 111704486 | 111703335 | 111703476 | 111709148 | 111709264 | 1557 | 1359 | 1128 | 1591 | 2588 | 0 | 0 | 0 | 0 | 3 | 412 | 1089 | [illegible] |
| SE | 48758 | ENSG00000186432 | KPNA4 | chr3 | - | 160521778 | 160521910 | 160515451 | 160515580 | 160525799 | 160525844 | 786 | 561 | 723 | 833 | 1508 | 0 | 2 | 0 | 0 | 2 | 245 | 695 | [illegible] |
| SE | 48759 | ENSG00000186432 | KPNA4 | chr3 | - | 160525020 | 160525132 | 160521778 | 160521910 | 160525799 | 160525844 | 0 | 0 | 1 | 0 | 2 | 339 | 239 | 337 | 386 | 731 | 0 | 3 | [illegible] |
| SE | 48760 | ENSG00000186432 | KPNA4 | chr3 | - | 160530854 | 160530940 | 160527952 | 160528039 | 160531461 | 160531557 | 291 | 193 | 242 | 328 | 669 | 0 | 0 | 0 | 0 | 0 | 46 | 218 | [illegible] |
| SE | 48761 | ENSG00000186432 | KPNA4 | chr3 | - | 160535807 | 160535897 | 160535660 | 160535690 | 160536795 | 160536840 | 256 | 213 | 215 | 263 | 504 | 0 | 0 | 0 | 0 | 0 | 72 | 177 | [illegible] |
| SE | 48764 | ENSG00000187024 | PTRH1 | chr9 | - | 127714378 | 127714424 | 127713509 | 127714282 | 127714602 | 127714702 | 10 | 1 | 0 | 2 | 12 | 1 | 0 | 0 | 0 | 0 | 0 | 1 | [illegible] |
| SE | 48766 | ENSG00000142534 | RPS11 | chr19 | + | 49497172 | 49497325 | 49496433 | 49496471 | 49497519 | 49497595 | 7469 | 4934 | 7242 | 7000 | 6871 | 2 | 0 | 3 | 0 | 1 | 12067 | 7620 | [illegible] |
| SE | 48768 | ENSG00000142534 | RPS11 | chr19 | + | 49497193 | 49497325 | 49496364 | 49496471 | 49497519 | 49497595 | 13117 | 8405 | 12965 | 12252 | 12011 | 2 | 0 | 3 | 0 | 1 | 19925 | 13010 | [illegible] |
| SE | 48769 | ENSG00000142534 | RPS11 | chr19 | + | 49497916 | 49498046 | 49497193 | 49497595 | 49499511 | 49499688 | 9499 | 5382 | 9815 | 8676 | 9881 | 0 | 0 | 1 | 0 | 1 | 12160 | 8925 | [illegible] |
| SE | 48771 | ENSG00000116161 | CACYBP | chr1 | + | 175006744 | 175006841 | 175004613 | 175004833 | 175007097 | 175007197 | 311 | 209 | 270 | 346 | 478 | 0 | 0 | 3 | 1 | 0 | 434 | 164 | [illegible] |
| SE | 48774 | ENSG00000058804 | NDC1 | chr1 | - | 53807655 | 53807791 | 53806424 | 53806517 | 53809694 | 53809746 | 129 | 62 | 108 | 123 | 296 | 0 | 0 | 0 | 0 | 0 | 179 | 75 | [illegible] |
| SE | 48776 | ENSG00000058804 | NDC1 | chr1 | - | 53835499 | 53835620 | 53832489 | 53832591 | 53838204 | 53838296 | 32 | 15 | 32 | 20 | 14 | 3 | 1 | 0 | 1 | 2 | 0 | 0 | [illegible] |
| SE | 48777 | ENSG00000153922 | CHD1 | chr5 | - | 98858179 | 98858390 | 98853984 | 98856725 | 98858963 | 98859015 | 472 | 442 | 413 | 463 | 715 | 0 | 1 | 0 | 3 | 3 | 47 | 164 | [illegible] |
| SE | 48778 | ENSG00000153922 | CHD1 | chr5 | - | 98858963 | 98859015 | 98858179 | 98858390 | 98859971 | 98860068 | 484 | 418 | 420 | 452 | 775 | 0 | 0 | 0 | 0 | 3 | 53 | 151 | [illegible] |
| SE | 48779 | ENSG00000153922 | CHD1 | chr5 | - | 98859971 | 98860068 | 98858963 | 98859015 | 98863407 | 98863586 | 436 | 367 | 364 | 407 | 720 | 0 | 0 | 1 | 1 | 0 | 32 | 126 | [illegible] |
| SE | 48781 | ENSG00000153922 | CHD1 | chr5 | - | 98868494 | 98868635 | 98863407 | 98863586 | 98869753 | 98869882 | 892 | 829 | 678 | 943 | 1693 | 0 | 0 | 2 | 0 | 0 | 65 | 300 | [illegible] |
| SE | 48783 | ENSG00000153922 | CHD1 | chr5 | - | 98872050 | 98872201 | 98870686 | 98870803 | 98872416 | 98872555 | 730 | 615 | 616 | 648 | 1195 | 0 | 1 | 0 | 0 | 4 | 16 | 204 | [illegible] |
| SE | 48784 | ENSG00000153922 | CHD1 | chr5 | - | 98872416 | 98872555 | 98872050 | 98872201 | 98873592 | 98873723 | 678 | 571 | 610 | 632 | 1025 | 0 | 0 | 0 | 0 | 3 | 31 | 168 | [illegible] |
| SE | 48785 | ENSG00000153922 | CHD1 | chr5 | - | 98873592 | 98873723 | 98872416 | 98872555 | 98875071 | 98875113 | 617 | 504 | 510 | 597 | 851 | 0 | 0 | 1 | 0 | 3 | 38 | 179 | [illegible] |
| SE | 48786 | ENSG00000153922 | CHD1 | chr5 | - | 98875071 | 98875113 | 98872416 | 98872555 | 98876397 | 98876558 | 45 | 38 | 46 | 26 | 60 | 0 | 0 | 0 | 0 | 0 | 0 | 5 | [illegible] |
| SE | 48789 | ENSG00000153922 | CHD1 | chr5 | - | 98881974 | 98882123 | 98881246 | 98881375 | 98885577 | 98885649 | 320 | 280 | 255 | 328 | 348 | 0 | 0 | 2 | 1 | 0 | 28 | 93 | [illegible] |
| SE | 48790 | ENSG00000153922 | CHD1 | chr5 | - | 98883087 | 98883237 | 98881246 | 98881375 | 98885577 | 98885649 | 288 | 260 | 232 | 301 | 379 | 0 | 0 | 2 | 1 | 0 | 16 | 110 | [illegible] |
| SE | 48791 | ENSG00000153922 | CHD1 | chr5 | - | 98883087 | 98883237 | 98881974 | 98882123 | 98885577 | 98885649 | 664 | 595 | 576 | 699 | 890 | 0 | 4 | 0 | 5 | 1 | 76 | 301 | [illegible] |
| SE | 48792 | ENSG00000153922 | CHD1 | chr5 | - | 98888087 | 98888240 | 98885577 | 98885649 | 98889075 | 98889238 | 484 | 392 | 435 | 480 | 673 | 0 | 0 | 0 | 1 | 0 | 37 | 187 | [illegible] |
| SE | 48793 | ENSG00000153922 | CHD1 | chr5 | - | 98892524 | 98892713 | 98889075 | 98889238 | 98893415 | 98893606 | 445 | 416 | 396 | 541 | 497 | 2 | 0 | 0 | 0 | 0 | 111 | 303 | [illegible] |
| SE | 48794 | ENSG00000153922 | CHD1 | chr5 | - | 98896225 | 98896442 | 98894596 | 98894686 | 98897192 | 98897320 | 575 | 405 | 490 | 510 | 455 | 1 | 0 | 0 | 0 | 0 | 19 | 148 | [illegible] |
| SE | 48795 | ENSG00000153922 | CHD1 | chr5 | - | 98898663 | 98898764 | 98898255 | 98898434 | 98899479 | 98899705 | 364 | 324 | 332 | 366 | 417 | 1 | 0 | 0 | 0 | 0 | 8 | 135 | [illegible] |
| SE | 48801 | ENSG00000064763 | FAR2 | chr12 | + | 29293299 | 29293475 | 29270411 | 29270638 | 29297020 | 29297078 | 65 | 59 | 30 | 71 | 54 | 0 | 0 | 1 | 0 | 0 | 36 | 80 | [illegible] |
| SE | 48802 | ENSG00000064763 | FAR2 | chr12 | + | 29293299 | 29293475 | 29270411 | 29270638 | 29307657 | 29307835 | 22 | 31 | 11 | 33 | 14 | 0 | 0 | 1 | 0 | 0 | 18 | 28 | [illegible] |
| SE | 48803 | ENSG00000064763 | FAR2 | chr12 | + | 29297020 | 29297200 | 29293299 | 29293475 | 29307657 | 29307835 | 70 | 45 | 31 | 61 | 80 | 2 | 0 | 2 | 0 | 0 | 69 | 74 | [illegible] |
| SE | 48804 | ENSG00000064763 | FAR2 | chr12 | + | 29332844 | 29332908 | 29332687 | 29332727 | 29333631 | 29334072 | 1 | 1 | 0 | 1 | 1 | 17 | 22 | 23 | 29 | 28 | 7 | 2 | [illegible] |
| SE | 48811 | ENSG00000038945 | MSR1 | chr8 | - | 16120417 | 16120606 | 16107880 | 16110218 | 16143557 | 16143611 | 10 | 4 | 0 | 14 | 9 | 0 | 0 | 0 | 0 | 0 | 12 | 7 | [illegible] |
| SE | 48812 | ENSG00000038945 | MSR1 | chr8 | - | 16168457 | 16168870 | 16164064 | 16164251 | 16175186 | 16175300 | 63 | 62 | 7 | 130 | 112 | 1 | 0 | 0 | 1 | 0 | 20 | 114 | [illegible] |
| SE | 48815 | ENSG00000038945 | MSR1 | chr8 | - | 16175186 | 16175300 | 16168457 | 16168870 | 16177885 | 16177988 | 34 | 24 | 6 | 70 | 51 | 0 | 0 | 0 | 2 | 0 | 0 | 70 | [illegible] |
| SE | 48817 | ENSG00000163599 | CTLA4 | chr2 | + | 203870585 | 203870648 | 203867942 | 203868051 | 203872707 | 203872812 | 7 | 4 | 8 | 16 | 13 | 0 | 0 | 1 | 0 | 2 | 7 | 3 | [illegible] |
| SE | 48818 | ENSG00000163599 | CTLA4 | chr2 | + | 203870585 | 203870933 | 203867770 | 203868051 | 203872707 | 203872775 | 24 | 6 | 19 | 29 | 28 | 0 | 0 | 1 | 0 | 2 | 27 | 48 | [illegible] |
| SE | 48821 | ENSG00000247950 | SEC24B-AS1 | chr4 | - | 109405649 | 109405773 | 109396193 | 109396279 | 109414257 | 109414433 | 0 | 0 | 1 | 0 | 0 | 1 | 0 | 0 | 1 | 0 | 0 | 0 | [illegible] |
| SE | 48822 | ENSG00000247950 | SEC24B-AS1 | chr4 | - | 109414257 | 109414433 | 109396193 | 109396279 | 109430579 | 109430760 | 1 | 0 | 1 | 1 | 0 | 0 | 0 | 1 | 0 | 0 | 7 | 0 | [illegible] |
| SE | 48827 | ENSG00000247950 | SEC24B-AS1 | chr4 | - | 109430579 | 109430780 | 109414257 | 109414433 | 109431691 | 109431746 | 0 | 1 | 1 | 3 | 1 | 1 | 0 | 0 | 0 | 0 | 8 | 5 | [illegible] |
| SE | 48828 | ENSG00000247950 | SEC24B-AS1 | chr4 | - | 109431940 | 109432104 | 109431691 | 109431746 | 109433733 | 109433817 | 2 | 1 | 0 | 2 | 4 | 0 | 0 | 0 | 0 | 1 | 8 | 5 | [illegible] |
| SE | 48829 | ENSG00000228327 | NA | chr1 | - | 744727 | 744825 | 744194 | 744391 | 746694 | 746818 | 0 | 2 | 0 | 0 | 5 | 0 | 0 | 1 | 0 | 0 | 0 | 13 | [illegible] |
| SE | 48830 | ENSG00000228327 | NA | chr1 | - | 769496 | 769712 | 768547 | 768613 | 772975 | 773107 | 19 | 4 | 38 | 6 | 14 | 1 | 0 | 3 | 0 | 0 | 65 | 8 | [illegible] |
| SE | 48832 | ENSG00000132463 | GRSF1 | chr4 | - | 70824293 | 70824368 | 70815782 | 70820861 | 70825295 | 70825431 | 247 | 119 | 217 | 210 | 460 | 0 | 0 | 0 | 0 | 2 | 142 | 179 | [illegible] |
| SE | 48833 | ENSG00000132463 | GRSF1 | chr4 | - | 70825295 | 70825431 | 70815782 | 70820861 | 70826123 | 70826245 | 137 | 63 | 136 | 141 | 235 | 0 | 0 | 0 | 3 | 0 | 52 | 94 | [illegible] |
| SE | 48834 | ENSG00000132463 | GRSF1 | chr4 | - | 70826123 | 70826245 | 70825295 | 70825431 | 70827851 | 70828036 | 327 | 199 | 291 | 329 | 651 | 0 | 1 | 3 | 6 | 0 | 196 | 223 | [illegible] |
| SE | 48836 | ENSG00000132463 | GRSF1 | chr4 | - | 70832306 | 70832450 | 70827851 | 70828036 | 70836157 | 70836314 | 6 | 0 | 4 | 2 | 3 | 0 | 0 | 1 | 0 | 0 | 6 | 4 | [illegible] |
| SE | 48837 | ENSG00000132463 | GRSF1 | chr4 | - | 70833117 | 70833273 | 70832306 | 70832450 | 70836157 | 70836314 | 301 | 249 | 283 | 298 | 273 | 3 | 0 | 0 | 0 | 0 | 461 | 322 | [illegible] |
| SE | 48842 | ENSG00000065135 | GNAI3 | chr1 | + | 109573895 | 109574037 | 109573736 | 109573779 | 109579203 | 109579361 | 642 | 449 | 414 | 644 | 662 | 3 | 1 | 0 | 0 | 0 | 181 | 412 | [illegible] |
| SE | 48843 | ENSG00000065135 | GNAI3 | chr1 | + | 109579203 | 109579361 | 109573895 | 109574037 | 109582436 | 109582565 | 637 | 429 | 430 | 622 | 779 | 0 | 3 | 1 | 1 | 6 | 126 | 364 | [illegible] |
| SE | 48844 | ENSG00000065135 | GNAI3 | chr1 | + | 109586215 | 109586345 | 109582436 | 109582565 | 109586728 | 109586882 | 606 | 503 | 449 | 602 | 853 | 0 | 0 | 1 | 0 | 0 | 132 | 424 | [illegible] |
| SE | 48845 | ENSG00000102580 | DNAJC3 | chr13 | + | 95723241 | 95723366 | 95709226 | 95709337 | 95757643 | 95757796 | 77 | 25 | 45 | 51 | 58 | 0 | 0 | 0 | 0 | 0 | 6 | 71 | [illegible] |
| SE | 48846 | ENSG00000102580 | DNAJC3 | chr13 | + | 95725177 | 95725252 | 95723241 | 95723366 | 95757643 | 95757796 | 207 | 137 | 154 | 166 | 340 | 0 | 0 | 0 | 0 | 0 | 65 | 193 | [illegible] |
| SE | 48847 | ENSG00000102580 | DNAJC3 | chr13 | + | 95757643 | 95757796 | 95725177 | 95725252 | 95760039 | 95760221 | 379 | 345 | 337 | 378 | 777 | 2 | 0 | 0 | 0 | 0 | 232 | 397 | [illegible] |
| SE | 48848 | ENSG00000102580 | DNAJC3 | chr13 | + | 95785938 | 95786071 | 95763832 | 95763953 | 95787006 | 95787155 | 314 | 275 | 254 | 351 | 551 | 0 | 0 | 3 | 0 | 2 | 82 | 213 | [illegible] |
| SE | 48850 | ENSG00000163563 | MNDA | chr1 | + | 158842133 | 158842418 | 158831350 | 158831557 | 158843278 | 158843415 | 4375 | 4111 | 2540 | 3799 | 9996 | 2 | 3 | 2 | 1 | 3 | 1417 | 3718 | [illegible] |
| SE | 48851 | ENSG00000163563 | MNDA | chr1 | + | 158843278 | 158843415 | 158831350 | 158831557 | 158845586 | 158846003 | 2 | 13 | 2 | 1 | 5 | 0 | 0 | 0 | 0 | 0 | 0 | 0 | [illegible] |
| SE | 48854 | ENSG00000165661 | QSOX2 | chr9 | - | 136221795 | 136221941 | 136219029 | 136219164 | 136223762 | 136223853 | 92 | 30 | 64 | 78 | 56 | 1 | 0 | 0 | 0 | 0 | 93 | 33 | [illegible] |
| SE | 48856 | ENSG00000165661 | QSOX2 | chr9 | - | 136224006 | 136224112 | 136219029 | 136219164 | 136226773 | 136226874 | 1 | 1 | 2 | 5 | 1 | 0 | 0 | 0 | 0 | 0 | 0 | 0 | [illegible] |
| SE | 48860 | ENSG00000066279 | ASPM | chr1 | - | 197100430 | 197105185 | 197095997 | 197096164 | 197117788 | 197117983 | 6 | 4 | 18 | 6 | 6 | 0 | 5 | 5 | 2 | 0 | 5 | 2 | [illegible] |
| SE | 48861 | ENSG00000184677 | ZBTB40 | chr1 | + | 22452658 | 22452946 | 22451850 | 22452004 | 22489879 | 22490645 | 0 | 2 | 2 | 2 | 2 | 26 | 33 | 50 | 36 | 16 | 0 | 4 | [illegible] |
| SE | 48865 | ENSG00000184677 | ZBTB40 | chr1 | + | 22508529 | 22508731 | 22508000 | 22508137 | 22509099 | 22509233 | 158 | 148 | 118 | 175 | 200 | 1 | 0 | 0 | 0 | 0 | 120 | 71 | [illegible] |
| SE | 48867 | ENSG00000184677 | ZBTB40 | chr1 | + | 22517299 | 22517464 | 22512923 | 22513130 | 22520060 | 22520275 | 86 | 55 | 92 | 80 | 40 | 0 | 0 | 0 | 0 | 0 | 112 | 82 | [illegible] |
| SE | 48874 | ENSG00000143514 | TP53BP2 | chr1 | - | 223799898 | 223800047 | 223795814 | 223796590 | 223800699 | 223800726 | 99 | 86 | 83 | 82 | 117 | 0 | 1 | 0 | 0 | 0 | 50 | 160 | [illegible] |
| SE | 48876 | ENSG00000143514 | TP53BP2 | chr1 | - | 223806845 | 223806947 | 223804173 | 223804348 | 223810430 | 223810513 | 185 | 189 | 180 | 180 | 293 | 0 | 0 | 0 | 1 | 0 | 200 | 215 | [illegible] |
| SE | 48877 | ENSG00000143514 | TP53BP2 | chr1 | - | 223810430 | 223810513 | 223806845 | 223806947 | 223814239 | 223814328 | 187 | 168 | 148 | 190 | 272 | 0 | 2 | 0 | 0 | 2 | 184 | 161 | [illegible] |
| SE | 48878 | ENSG00000143514 | TP53BP2 | chr1 | - | 223814239 | 223814353 | 223806845 | 223806947 | 223821219 | 223821367 | 78 | 48 | 54 | 65 | 70 | 0 | 2 | 0 | 0 | 0 | 70 | 93 | [illegible] |
| SE | 48883 | ENSG00000109906 | ZBTB16 | chr11 | + | 114121817 | 114121914 | 114064411 | 114064568 | 114156336 | 114156434 | 3 | 0 | 0 | 1 | 0 | 58 | 10 | 22 | 35 | 6 | 0 | 0 | [illegible] |
| SE | 48885 | ENSG00000109906 | ZBTB16 | chr11 | + | 114156336 | 114156434 | 114062225 | 114064568 | 114186951 | 114187038 | 99 | 29 | 40 | 57 | 17 | 0 | 0 | 0 | 0 | 0 | 23 | 63 | [illegible] |
| SE | 48886 | ENSG00000109906 | ZBTB16 | chr11 | + | 114186951 | 114187038 | 114156336 | 114156434 | 114242166 | 114242337 | 125 | 35 | 52 | 58 | 17 | 2 | 0 | 0 | 0 | 0 | 23 | 90 | [illegible] |
| SE | 48888 | ENSG00000152990 | ADGRA3 | chr4 | - | 22413600 | 22413760 | 22413181 | 22413390 | 22420885 | 22421089 | 7 | 4 | 9 | 3 | 4 | 1 | 0 | 0 | 0 | 0 | 13 | 1 | [illegible] |
| SE | 48889 | ENSG00000152990 | ADGRA3 | chr4 | - | 22413600 | 22413814 | 22413181 | 22413390 | 22420885 | 22421089 | 7 | 6 | 12 | 6 | 6 | 1 | 0 | 0 | 0 | 0 | 13 | 2 | [illegible] |
| SE | 48895 | ENSG00000115107 | STEAP3 | chr2 | + | 119239362 | 119239430 | 119223830 | 119223888 | 119245488 | 119245988 | 0 | 0 | 0 | 0 | 0 | 0 | 0 | 0 | 2 | 0 | 0 | 0 | [illegible] |
| SE | 48896 | ENSG00000115107 | STEAP3 | chr2 | + | 119239362 | 119239430 | 119230619 | 119231034 | 119245488 | 119245988 | 2 | 0 | 0 | 0 | 0 | 3 | 1 | 0 | 0 | 2 | 0 | 0 | [illegible] |
| SE | 48901 | ENSG00000158773 | USF1 | chr1 | - | 161041323 | 161041411 | 161040813 | 161040872 | 161041650 | 161041721 | 152 | 90 | 138 | 145 | 49 | 0 | 0 | 0 | 0 | 0 | 261 | 273 | [illegible] |
| SE | 48905 | ENSG00000158773 | USF1 | chr1 | - | 161042115 | 161042217 | 161041323 | 161041411 | 161042554 | 161042670 | 92 | 50 | 87 | 98 | 54 | 0 | 0 | 0 | 0 | 0 | 222 | 156 | [illegible] |
| SE | 48906 | ENSG00000158773 | USF1 | chr1 | - | 161042115 | 161042217 | 161041325 | 161041846 | 161042554 | 161042670 | 251 | 148 | 235 | 221 | 138 | 0 | 0 | 0 | 0 | 0 | 679 | 423 | [illegible] |
| SE | 48911 | ENSG00000204165 | CXorf65 | chrX | - | 71106339 | 71106426 | 71105999 | 71106137 | 71106531 | 71106740 | 7 | 6 | 15 | 15 | 7 | 0 | 0 | 0 | 0 | 0 | 13 | 19 | [illegible] |
| SE | 48912 | ENSG00000204165 | CXorf65 | chrX | - | 71106358 | 71106426 | 71105999 | 71106137 | 71106531 | 71106788 | 3 | 11 | 10 | 14 | 4 | 0 | 0 | 0 | 0 | 0 | 10 | 16 | [illegible] |
| SE | 48915 | ENSG00000011295 | TTC19 | chr17 | + | 16026539 | 16026702 | 16023672 | 16025171 | 16027373 | 16027535 | 204 | 88 | 152 | 138 | 171 | 0 | 0 | 0 | 2 | 0 | 73 | 139 | [illegible] |
| SE | 48916 | ENSG00000198408 | OGA | chr10 | - | 101787363 | 101787523 | 101784442 | 101786587 | 101790895 | 101791088 | 724 | 710 | 647 | 760 | 674 | 2 | 2 | 0 | 0 | 0 | 304 | 674 | [illegible] |
| SE | 48918 | ENSG00000198408 | OGA | chr10 | - | 101791353 | 101791439 | 101790895 | 101791088 | 101792838 | 101792901 | 273 | 254 | 325 | 304 | 487 | 0 | 0 | 0 | 0 | 0 | 113 | 251 | [illegible] |
| SE | 48919 | ENSG00000198408 | OGA | chr10 | - | 101793685 | 101793744 | 101792838 | 101792901 | 101797979 | 101798017 | 4 | 0 | 2 | 2 | 13 | 1 | 0 | 0 | 0 | 0 | 0 | 5 | [illegible] |
| SE | 48924 | ENSG00000198408 | OGA | chr10 | - | 101800241 | 101800400 | 101798841 | 101799455 | 101806044 | 101806143 | 915 | 982 | 843 | 924 | 1324 | 0 | 0 | 0 | 0 | 0 | 853 | 914 | [illegible] |
| SE | 48925 | ENSG00000198408 | OGA | chr10 | - | 101803734 | 101804019 | 101798841 | 101799455 | 101806044 | 101806143 | 593 | 447 | 537 | 602 | 838 | 0 | 0 | 0 | 0 | 0 | 405 | 670 | [illegible] |
| SE | 48928 | ENSG00000198408 | OGA | chr10 | - | 101807729 | 101807901 | 101806044 | 101806143 | 101810183 | 101810314 | 531 | 424 | 478 | 546 | 1096 | 0 | 0 | 0 | 0 | 2 | 333 | 538 | [illegible] |
| SE | 48930 | ENSG00000198408 | OGA | chr10 | - | 101813554 | 101813606 | 101813029 | 101813127 | 101817823 | 101817866 | 99 | 89 | 93 | 95 | 105 | 0 | 0 | 2 | 1 | 0 | 52 | 77 | [illegible] |
| SE | 48932 | ENSG00000225873 | C3orf86 | chr3 | - | 44422852 | 44422937 | 44421129 | 44421813 | 44428930 | 44429097 | 1 | 2 | 0 | 0 | 0 | 2 | 1 | 0 | 3 | 0 | 6 | 4 | [illegible] |
| SE | 48933 | ENSG00000225873 | C3orf86 | chr3 | - | 44422852 | 44422937 | 44421129 | 44421813 | 44439796 | 44439946 | 2 | 1 | 0 | 0 | 0 | 2 | 4 | 0 | 8 | 16 | 12 | 8 | [illegible] |
| SE | 48934 | ENSG00000225873 | C3orf86 | chr3 | - | 44428930 | 44429097 | 44421141 | 44421813 | 44439796 | 44439946 | 2 | 1 | 0 | 3 | 0 | 2 | 4 | 0 | 8 | 16 | 0 | 0 | [illegible] |
| SE | 48935 | ENSG00000225873 | C3orf86 | chr3 | - | 44428930 | 44429097 | 44422852 | 44422937 | 44439796 | 44439946 | 0 | 1 | 0 | 0 | 0 | 1 | 0 | 0 | 0 | 0 | 0 | 0 | [illegible] |
| SE | 48937 | ENSG00000117174 | ZNHIT6 | chr1 | - | 85657846 | 85657971 | 85649416 | 85654098 | 85677235 | 85677313 | 61 | 56 | 109 | 68 | 115 | 0 | 1 | 0 | 4 | 1 | 0 | 25 | [illegible] |
| SE | 48938 | ENSG00000117174 | ZNHIT6 | chr1 | - | 85706441 | 85706507 | 85706248 | 85706355 | 85707628 | 85707936 | 184 | 153 | 169 | 178 | 131 | 5 | 0 | 0 | 2 | 0 | 47 | 97 | [illegible] |
| SE | 48942 | ENSG00000057468 | MSH4 | chr1 | + | 75876935 | 75877000 | 75867513 | 75867588 | 75878148 | 75878318 | 0 | 0 | 0 | 1 | 0 | 0 | 0 | 0 | 1 | 0 | 0 | 0 | [illegible] |
| SE | 48943 | ENSG00000205534 | SMG1P2 | chr16 | - | 29528416 | 29528576 | 29527567 | 29527687 | 29528950 | 29529098 | 3 | 4 | 3 | 5 | 5 | 0 | 0 | 0 | 0 | 0 | 2 | 5 | [illegible] |
| SE | 48944 | ENSG00000088832 | FKBP1A | chr20 | - | 1371021 | 1371147 | 1369533 | 1370072 | 1372075 | 1372240 | 8 | 1 | 7 | 6 | 4 | 1002 | 521 | 469 | 809 | 1045 | 21 | 0 | [illegible] |
| SE | 48945 | ENSG00000088832 | FKBP1A | chr20 | - | 1371119 | 1371147 | 1368984 | 1370072 | 1372075 | 1372240 | 1 | 0 | 1 | 0 | 0 | 1002 | 521 | 469 | 809 | 1045 | 0 | 0 | [illegible] |
| SE | 48946 | ENSG00000088832 | FKBP1A | chr20 | - | 1372075 | 1372240 | 1368976 | 1370072 | 1375490 | 1375603 | 1522 | 745 | 789 | 1185 | 1635 | 5 | 0 | 0 | 0 | 0 | 2500 | 1011 | [illegible] |
| SE | 48947 | ENSG00000088832 | FKBP1A | chr20 | - | 1372075 | 1372240 | 1368976 | 1370072 | 1392833 | 1392881 | 1006 | 522 | 472 | 809 | 1045 | 2 | 0 | 0 | 0 | 1 | 1801 | 730 | [illegible] |
| SE | 48949 | ENSG00000088832 | FKBP1A | chr20 | - | 1375490 | 1375603 | 1368999 | 1372240 | 1392833 | 1392881 | 657 | 291 | 398 | 492 | 622 | 4 | 1 | 3 | 0 | 0 | 915 | 384 | [illegible] |
| SE | 48951 | ENSG00000088832 | FKBP1A | chr20 | - | 1377429 | 1377511 | 1375490 | 1375603 | 1392833 | 1392881 | 1 | 0 | 0 | 1 | 0 | 180 | 89 | 98 | 149 | 60 | 0 | 6 | [illegible] |
| SE | 48994 | ENSG00000159079 | CFAP298 | chr21 | - | 32603160 | 32603292 | 32601666 | 32601973 | 32604124 | 32604200 | 136 | 89 | 141 | 153 | 102 | 1 | 0 | 0 | 0 | 0 | 155 | 135 | [illegible] |
| SE | 48995 | ENSG00000159079 | CFAP298 | chr21 | - | 32609837 | 32609954 | 32607648 | 32607716 | 32612104 | 32612555 | 37 | 22 | 30 | 44 | 19 | 0 | 2 | 0 | 0 | 0 | 73 | 29 | [illegible] |
| SE | 48996 | ENSG00000182983 | ZNF662 | chr3 | + | 42912016 | 42912069 | 42908801 | 42908909 | 42913200 | 42913303 | 0 | 1 | 0 | 0 | 2 | 1 | 1 | 5 | 4 | 2 | 0 | 0 | [illegible] |
| SE | 48997 | ENSG00000182983 | ZNF662 | chr3 | + | 42913200 | 42913302 | 42908792 | 42908909 | 42914326 | 42919334 | 1 | 2 | 6 | 4 | 2 | 2 | 3 | 2 | 3 | 0 | 0 | 0 | [illegible] |
| SE | 49002 | ENSG00000143416 | SELENBP1 | chr1 | - | 151369003 | 151369189 | 151368198 | 151368319 | 151369712 | 151369769 | 0 | 0 | 1 | 0 | 2 | 1 | 0 | 0 | 0 | 3 | 0 | 3 | [illegible] |
| SE | 49003 | ENSG00000143416 | SELENBP1 | chr1 | - | 151369003 | 151369554 | 151368198 | 151368319 | 151369712 | 151369769 | 0 | 1 | 1 | 1 | 4 | 1 | 0 | 0 | 0 | 3 | 0 | 3 | [illegible] |
| SE | 49007 | ENSG00000152147 | GEMIN6 | chr2 | + | 38779257 | 38779325 | 38778971 | 38779118 | 38781516 | 38781965 | 2 | 1 | 0 | 4 | 0 | 54 | 37 | 67 | 38 | 75 | 0 | 0 | [illegible] |
| SE | 49016 | ENSG00000164509 | IL31RA | chr5 | + | 55859508 | 55859599 | 55853313 | 55853567 | 55872269 | 55872451 | 2 | 5 | 2 | 3 | 1 | 0 | 0 | 0 | 0 | 0 | 64 | 5 | [illegible] |
| SE | 49017 | ENSG00000164509 | IL31RA | chr5 | + | 55866610 | 55866754 | 55859508 | 55859599 | 55868790 | 55868908 | 0 | 1 | 0 | 0 | 4 | 1 | 1 | 1 | 2 | 0 | 8 | 0 | [illegible] |
| SE | 49018 | ENSG00000164509 | IL31RA | chr5 | + | 55868790 | 55868908 | 55859508 | 55859599 | 55872269 | 55872451 | 4 | 0 | 5 | 4 | 0 | 1 | 0 | 2 | 1 | 0 | 26 | 0 | [illegible] |
| SE | 49019 | ENSG00000164509 | IL31RA | chr5 | + | 55899915 | 55900132 | 55896349 | 55896429 | 55906105 | 55906288 | 4 | 0 | 2 | 5 | 5 | 0 | 0 | 1 | 0 | 0 | 14 | 0 | [illegible] |
| SE | 49020 | ENSG00000164509 | IL31RA | chr5 | + | 55913476 | 55913570 | 55910531 | 55910672 | 55914846 | 55914928 | 4 | 0 | 2 | 0 | 5 | 0 | 1 | 0 | 0 | 0 | 3 | 4 | [illegible] |
| SE | 49021 | ENSG00000164509 | IL31RA | chr5 | + | 55913476 | 55913570 | 55910531 | 55910672 | 55916643 | 55916834 | 1 | 0 | 0 | 0 | 3 | 0 | 0 | 0 | 0 | 0 | 0 | 4 | [illegible] |
| SE | 49022 | ENSG00000164509 | IL31RA | chr5 | + | 55914846 | 55914928 | 55910531 | 55910672 | 55916643 | 55916834 | 0 | 2 | 0 | 2 | 0 | 0 | 0 | 0 | 0 | 0 | 0 | 0 | [illegible] |
| SE | 49023 | ENSG00000164509 | IL31RA | chr5 | + | 55914846 | 55914928 | 55913476 | 55913570 | 55916643 | 55916834 | 4 | 1 | 2 | 2 | 3 | 1 | 0 | 0 | 0 | 1 | 3 | 0 | [illegible] |
| SE | 49025 | ENSG00000134453 | RBM17 | chr10 | + | 6101270 | 6101387 | 6096909 | 6097188 | 6104930 | 6105097 | 415 | 335 | 396 | 428 | 584 | 0 | 0 | 0 | 0 | 0 | 524 | 463 | [illegible] |
| SE | 49026 | ENSG00000134453 | RBM17 | chr10 | + | 6104930 | 6105097 | 6101270 | 6101387 | 6106140 | 6106238 | 378 | 287 | 323 | 427 | 451 | 0 | 0 | 1 | 0 | 0 | 390 | 337 | [illegible] |
| SE | 49027 | ENSG00000134453 | RBM17 | chr10 | + | 6113507 | 6113581 | 6112209 | 6112361 | 6114048 | 6114147 | 129 | 121 | 128 | 135 | 155 | 0 | 0 | 0 | 0 | 0 | 161 | 148 | [illegible] |
| SE | 49028 | ENSG00000178498 | DTX3 | chr12 | + | 57605343 | 57605731 | 57604638 | 57604858 | 57606189 | 57606252 | 20 | 21 | 27 | 31 | 19 | 0 | 1 | 2 | 1 | 0 | 12 | 18 | [illegible] |
| SE | 49029 | ENSG00000178498 | DTX3 | chr12 | + | 57605570 | 57605731 | 57604823 | 57604858 | 57605988 | 57606252 | 2 | 0 | 1 | 0 | 0 | 0 | 0 | 0 | 1 | 0 | 0 | 0 | [illegible] |
| SE | 49030 | ENSG00000178498 | DTX3 | chr12 | + | 57605570 | 57605731 | 57604823 | 57604858 | 57606189 | 57606252 | 22 | 20 | 25 | 28 | 19 | 0 | 1 | 2 | 1 | 0 | 12 | 17 | [illegible] |
| SE | 49031 | ENSG00000178498 | DTX3 | chr12 | + | 57605988 | 57606029 | 57604622 | 57604858 | 57606189 | 57606252 | 1 | 0 | 0 | 0 | 0 | 0 | 1 | 2 | 1 | 0 | 17 | 3 | [illegible] |
| SE | 49036 | ENSG00000165119 | HNRNPK | chr9 | - | 83970161 | 83970331 | 83968082 | 83969440 | 83970736 | 83970819 | 3039 | 1984 | 2687 | 3221 | 3388 | 0 | 0 | 0 | 0 | 0 | 4146 | 3213 | [illegible] |
| SE | 49037 | ENSG00000165119 | HNRNPK | chr9 | - | 83970161 | 83970331 | 83968196 | 83969380 | 83970736 | 83970819 | 2989 | 1850 | 2620 | 3146 | 3278 | 0 | 0 | 0 | 1 | 0 | 3944 | 3141 | [illegible] |
| SE | 49038 | ENSG00000165119 | HNRNPK | chr9 | - | 83970736 | 83970819 | 83969495 | 83970331 | 83970896 | 83970912 | 3679 | 2194 | 3275 | 3767 | 3579 | 0 | 0 | 0 | 0 | 0 | 4903 | 3887 | [illegible] |
| SE | 49039 | ENSG00000165119 | HNRNPK | chr9 | - | 83970896 | 83970912 | 83969495 | 83970331 | 83971272 | 83971294 | 102 | 89 | 125 | 130 | 57 | 0 | 0 | 0 | 0 | 0 | 286 | 194 | [illegible] |
| SE | 49040 | ENSG00000165119 | HNRNPK | chr9 | - | 83971272 | 83971356 | 83970896 | 83970912 | 83971671 | 83971726 | 2156 | 1290 | 2008 | 2233 | 1768 | 1 | 0 | 0 | 0 | 0 | 2923 | 2385 | [illegible] |
| SE | 49041 | ENSG00000165119 | HNRNPK | chr9 | - | 83971671 | 83971726 | 83971272 | 83971356 | 83971881 | 83972189 | 1803 | 1189 | 1787 | 2008 | 1332 | 4 | 2 | 11 | 3 | 3 | 2486 | 2087 | [illegible] |
| SE | 49042 | ENSG00000165119 | HNRNPK | chr9 | - | 83971881 | 83972189 | 83971272 | 83971356 | 83973285 | 83973399 | 4 | 6 | 13 | 14 | 7 | 1 | 0 | 0 | 0 | 0 | 21 | 15 | [illegible] |
| SE | 49043 | ENSG00000165119 | HNRNPK | chr9 | - | 83972843 | 83972972 | 83971881 | 83972189 | 83973285 | 83973399 | 3604 | 2266 | 3087 | 3597 | 5485 | 0 | 4 | 2 | 11 | 4 | 3266 | 2645 | [illegible] |
| SE | 49046 | ENSG00000165119 | HNRNPK | chr9 | - | 83976994 | 83977051 | 83975461 | 83975505 | 83977688 | 83977786 | 2318 | 1677 | 2131 | 2468 | 4723 | 7 | 1 | 1 | 0 | 2 | 2658 | 2066 | [illegible] |
| SE | 49047 | ENSG00000165119 | HNRNPK | chr9 | - | 83978372 | 83978452 | 83978194 | 83978279 | 83980152 | 83980190 | 1083 | 616 | 766 | 962 | 733 | 2 | 0 | 0 | 0 | 0 | 1223 | 1028 | [illegible] |
| SE | 49048 | ENSG00000165119 | HNRNPK | chr9 | - | 83978372 | 83978452 | 83978194 | 83978279 | 83980502 | 83980595 | 1433 | 811 | 1091 | 1369 | 965 | 0 | 1 | 0 | 0 | 0 | 1670 | 1464 | [illegible] |
| SE | 49049 | ENSG00000144445 | KANSL1L | chr2 | - | 210026379 | 210026402 | 210025103 | 210025216 | 210027295 | 210027350 | 1 | 0 | 0 | 0 | 0 | 15 | 20 | 20 | 17 | 42 | 0 | 0 | [illegible] |
| SE | 49050 | ENSG00000144445 | KANSL1L | chr2 | - | 210029802 | 210029918 | 210028844 | 210028969 | 210040419 | 210040527 | 21 | 14 | 6 | 12 | 40 | 0 | 0 | 2 | 0 | 2 | 5 | 6 | [illegible] |
| SE | 49051 | ENSG00000144445 | KANSL1L | chr2 | - | 210031420 | 210031546 | 210029802 | 210029918 | 210040419 | 210040527 | 23 | 8 | 12 | 6 | 9 | 7 | 1 | 0 | 0 | 7 | 0 | 14 | [illegible] |
| SE | 49052 | ENSG00000144445 | KANSL1L | chr2 | - | 210040419 | 210040527 | 210031420 | 210031546 | 210043938 | 210044104 | 18 | 5 | 8 | 3 | 18 | 0 | 1 | 0 | 0 | 0 | 0 | 9 | [illegible] |
| SE | 49053 | ENSG00000144445 | KANSL1L | chr2 | - | 210075551 | 210075756 | 210043432 | 210044104 | 210098085 | 210098207 | 24 | 6 | 10 | 6 | 19 | 0 | 0 | 0 | 2 | 0 | 15 | 12 | [illegible] |
| SE | 49055 | ENSG00000010270 | STARD3NL | chr7 | + | 38179089 | 38179145 | 38178350 | 38178420 | 38201713 | 38201966 | 0 | 0 | 0 | 0 | 0 | 1 | 0 | 2 | 0 | 0 | 0 | 0 | [illegible] |
| SE | 49056 | ENSG00000010270 | STARD3NL | chr7 | + | 38179089 | 38179145 | 38178350 | 38178420 | 38202069 | 38202197 | 0 | 0 | 0 | 0 | 0 | 2 | 0 | 0 | 0 | 0 | 0 | 0 | [illegible] |
| SE | 49058 | ENSG00000010270 | STARD3NL | chr7 | + | 38198210 | 38198365 | 38178394 | 38178420 | 38202069 | 38202197 | 0 | 1 | 0 | 0 | 0 | 2 | 0 | 0 | 0 | 0 | 0 | 0 | [illegible] |
| SE | 49060 | ENSG00000010270 | STARD3NL | chr7 | + | 38201713 | 38201966 | 38178389 | 38178420 | 38202069 | 38202197 | 1 | 1 | 2 | 0 | 2 | 2 | 0 | 0 | 0 | 0 | 0 | 0 | [illegible] |
| SE | 49062 | ENSG00000010270 | STARD3NL | chr7 | + | 38201713 | 38201966 | 38179089 | 38179145 | 38202069 | 38202197 | 0 | 1 | 2 | 0 | 2 | 0 | 0 | 0 | 0 | 0 | 0 | 0 | [illegible] |
| SE | 49065 | ENSG00000010270 | STARD3NL | chr7 | + | 38202069 | 38202197 | 38179089 | 38179145 | 38207446 | 38207729 | 4 | 1 | 0 | 0 | 2 | 13 | 6 | 0 | 8 | 20 | 0 | 0 | [illegible] |
| SE | 49066 | ENSG00000010270 | STARD3NL | chr7 | + | 38202069 | 38202197 | 38198210 | 38198365 | 38207446 | 38207729 | 4 | 2 | 0 | 0 | 2 | 0 | 0 | 0 | 1 | 4 | 0 | 0 | [illegible] |
| SE | 49067 | ENSG00000010270 | STARD3NL | chr7 | + | 38202069 | 38202197 | 38201713 | 38201966 | 38207446 | 38207729 | 4 | 2 | 0 | 0 | 4 | 1 | 0 | 3 | 2 | 4 | 0 | 0 | [illegible] |
| SE | 49074 | ENSG00000188739 | RBM34 | chr1 | - | 235135651 | 235135770 | 235131182 | 235131997 | 235136033 | 235136073 | 262 | 234 | 240 | 323 | 321 | 1 | 0 | 0 | 0 | 0 | 94 | 128 | [illegible] |
| SE | 49075 | ENSG00000188739 | RBM34 | chr1 | - | 235136033 | 235136073 | 235135648 | 235135770 | 235137876 | 235137940 | 132 | 116 | 115 | 164 | 190 | 0 | 0 | 0 | 0 | 0 | 116 | 67 | [illegible] |
| SE | 49078 | ENSG00000188739 | RBM34 | chr1 | - | 235154880 | 235155112 | 235152487 | 235152765 | 235160510 | 235160647 | 301 | 231 | 239 | 305 | 396 | 0 | 0 | 0 | 0 | 0 | 214 | 248 | [illegible] |
| SE | 49079 | ENSG00000188739 | RBM34 | chr1 | - | 235154880 | 235155112 | 235152487 | 235152765 | 235160892 | 235161067 | 148 | 146 | 128 | 155 | 245 | 0 | 1 | 0 | 0 | 0 | 81 | 118 | [illegible] |
| SE | 49081 | ENSG00000188739 | RBM34 | chr1 | - | 235160510 | 235160647 | 235152487 | 235152765 | 235160892 | 235161067 | 130 | 54 | 103 | 123 | 147 | 0 | 1 | 0 | 0 | 0 | 75 | 114 | [illegible] |
| SE | 49083 | ENSG00000188739 | RBM34 | chr1 | - | 235160510 | 235160647 | 235160194 | 235160248 | 235160892 | 235161067 | 130 | 56 | 103 | 126 | 149 | 0 | 0 | 0 | 1 | 0 | 75 | 114 | [illegible] |
| SE | 49084 | ENSG00000072736 | NFATC3 | chr16 | + | 68121986 | 68123121 | 68084750 | 68085180 | 68126447 | 68126610 | 1462 | 678 | 1165 | 1427 | 1929 | 0 | 0 | 0 | 1 | 0 | 837 | 1105 | [illegible] |

| | | | | | | | | | | | | | |
|---|---|---|---|---|---|---|---|---|---|---|---|---|---|
| SE | 49086 | ENSG00000072736 | NFATC3 | chr16 | + | 68121986 | 68123121 | 68085343 | 68085784 | 68126447 | 68126610 | 1483 | 683 |
| SE | 49087 | ENSG00000072736 | NFATC3 | chr16 | + | 68121986 | 68123121 | 68085343 | 68085784 | 68157868 | 68158068 | 48 | 13 |
| SE | 49089 | ENSG00000072736 | NFATC3 | chr16 | + | 68126447 | 68126610 | 68085445 | 68085784 | 68157868 | 68158068 | 380 | 230 |
| SE | 49090 | ENSG00000072736 | NFATC3 | chr16 | + | 68126447 | 68126610 | 68121986 | 68123121 | 68157868 | 68158068 | 1815 | 900 |
| SE | 49095 | ENSG00000072736 | NFATC3 | chr16 | + | 68190767 | 68191775 | 68183239 | 68183366 | 68226349 | 68226471 | 474 | 292 |
| SE | 49096 | ENSG00000072736 | NFATC3 | chr16 | + | 68214328 | 68214432 | 68190767 | 68191775 | 68217710 | 68218090 | 20 | 14 |
| SE | 49097 | ENSG00000072736 | NFATC3 | chr16 | + | 68214328 | 68214432 | 68190767 | 68191775 | 68221195 | 68221284 | 22 | 12 |
| SE | 49108 | ENSG00000177943 | MAMDC4 | chr9 | + | 136859208 | 136859317 | 136859004 | 136859132 | 136859885 | 136860064 | 10 | 4 |
| SE | 49109 | ENSG00000103254 | ANTKMT | chr16 | + | 722083 | 722134 | 721800 | 721941 | 722308 | 722590 | 3 | 1 |
| SE | 49116 | ENSG00000153363 | LINC00467 | chr1 | + | 211390563 | 211390670 | 211382771 | 211382925 | 211391615 | 211391697 | 0 | 0 |
| SE | 49117 | ENSG00000153363 | LINC00467 | chr1 | + | 211390563 | 211390670 | 211384441 | 211384534 | 211391615 | 211391697 | 0 | 0 |
| SE | 49119 | ENSG00000153363 | LINC00467 | chr1 | + | 211396766 | 211396831 | 211391826 | 211391976 | 211426210 | 211426283 | 22 | 10 |
| SE | 49120 | ENSG00000153363 | LINC00467 | chr1 | + | 211396766 | 211396841 | 211391826 | 211391976 | 211426210 | 211426283 | 17 | 7 |
| SE | 49121 | ENSG00000153363 | LINC00467 | chr1 | + | 211399376 | 211399437 | 211396766 | 211396831 | 211426210 | 211426283 | 0 | 0 |
| SE | 49123 | ENSG00000153363 | LINC00467 | chr1 | + | 211422538 | 211422670 | 211396766 | 211396831 | 211426210 | 211426283 | 0 | 1 |
| SE | 49126 | ENSG00000000457 | SCYL3 | chr1 | - | 169855795 | 169855957 | 169854269 | 169854964 | 169859040 | 169859212 | 20 | 10 |
| SE | 49127 | ENSG00000000457 | SCYL3 | chr1 | - | 169864368 | 169864508 | 169862612 | 169862797 | 169866895 | 169866973 | 68 | 34 |
| SE | 49128 | ENSG00000000457 | SCYL3 | chr1 | - | 169870254 | 169870357 | 169868927 | 169869039 | 169873695 | 169873752 | 78 | 49 |
| SE | 49129 | ENSG00000000457 | SCYL3 | chr1 | - | 169870254 | 169870357 | 169868927 | 169869128 | 169873695 | 169873752 | 14 | 17 |
| SE | 49130 | ENSG00000000457 | SCYL3 | chr1 | - | 169875977 | 169876091 | 169873695 | 169873752 | 169878633 | 169878819 | 63 | 58 |
| SE | 49131 | ENSG00000156011 | PSD3 | chr8 | - | 18774826 | 18774938 | 18765448 | 18765538 | 18799294 | 18799353 | 0 | 1 |
| SE | 49139 | ENSG00000163528 | CHCHD4 | chr3 | - | 14119144 | 14119313 | 14116425 | 14116524 | 14121916 | 14122086 | 1 | 0 |
| SE | 49141 | ENSG00000174574 | AKIRIN1 | chr1 | + | 38998170 | 38998311 | 38991275 | 38991600 | 39000971 | 39001106 | 265 | 192 |
| SE | 49143 | ENSG00000174574 | AKIRIN1 | chr1 | + | 39000971 | 39001106 | 38991287 | 38991600 | 39003346 | 39003418 | 184 | 117 |
| SE | 49149 | ENSG00000077044 | DGKD | chr2 | + | 233388256 | 233388367 | 233354493 | 233354674 | 233390402 | 233390483 | 214 | 116 |
| SE | 49151 | ENSG00000077044 | DGKD | chr2 | + | 233434379 | 233434484 | 233390402 | 233390483 | 233434768 | 233434899 | 421 | 253 |
| SE | 49153 | ENSG00000077044 | DGKD | chr2 | + | 233437376 | 233437479 | 233436315 | 233436441 | 233441886 | 233441995 | 174 | 88 |
| SE | 49157 | ENSG00000077044 | DGKD | chr2 | + | 233454356 | 233454464 | 233451963 | 233452060 | 233454762 | 233454873 | 2 | 0 |
| SE | 49158 | ENSG00000077044 | DGKD | chr2 | + | 233457220 | 233457328 | 233456898 | 233456995 | 233458283 | 233458397 | 321 | 175 |
| [illegible] | | | | | | | | | | | | | |

| | | | | | | | | | | | | | | | | | | | | | | | |
|---|---|---|---|---|---|---|---|---|---|---|---|---|---|---|---|---|---|---|---|---|---|---|---|
| SE | 49628 | ENSG00000178927 | CYBC1 | chr17 | - | 82449169 | 82449292 | 82445250 | 82445960 | 82450699 | 82450717 | 15 | 4 | 6 | 8 | 0 | 0 | 0 | 2 | 0 | 0 | 0 | 7 |
| SE | 49629 | ENSG00000178927 | CYBC1 | chr17 | - | 82449169 | 82449292 | 82446622 | 82446696 | 82450699 | 82450717 | 11 | 10 | 6 | 12 | 1 | 1 | 0 | 0 | 2 | 0 | 0 | 7 |
| SE | 49631 | ENSG00000178927 | CYBC1 | chr17 | - | 82449173 | 82449292 | 82447579 | 82447621 | 82450699 | 82450743 | 11 | 4 | 6 | 8 | 0 | 1 | 0 | 0 | 3 | 1 | 0 | 7 |
| SE | 49633 | ENSG00000178927 | CYBC1 | chr17 | - | 82449430 | 82449480 | 82449169 | 82449292 | 82450479 | 82450709 | 9 | 5 | 3 | 11 | 3 | 3 | 0 | 1 | 2 | 1 | 13 | 5 |
| SE | 49642 | ENSG00000081760 | AACS | chr12 | + | 125073875 | 125073979 | 125065433 | 125065717 | 125076490 | 125076611 | 40 | 30 | 54 | 38 | 45 | 0 | 0 | 0 | 0 | 0 | 72 | 43 |
| SE | 49644 | ENSG00000081760 | AACS | chr12 | + | 125102678 | 125102793 | 125091425 | 125091523 | 125102999 | 125103081 | 21 | 19 | 35 | 16 | 47 | 0 | 0 | 1 | 0 | 0 | 30 | 5 |
| SE | 49645 | ENSG00000081760 | AACS | chr12 | + | 125107120 | 125107268 | 125102999 | 125103081 | 125114476 | 125114557 | 19 | 23 | 23 | 12 | 40 | 0 | 0 | 0 | 0 | 1 | 38 | 4 |
| SE | 49649 | ENSG00000136895 | GARNL3 | chr9 | + | 127264951 | 127265021 | 127243078 | 127243253 | 127291167 | 127291242 | 1 | 2 | 2 | 4 | 1 | 0 | 0 | 0 | 0 | 0 | 0 | 0 |
| SE | 49650 | ENSG00000136895 | GARNL3 | chr9 | + | 127332273 | 127332349 | 127325068 | 127325095 | 127333022 | 127333086 | 2 | 1 | 0 | 1 | 3 | 0 | 0 | 0 | 1 | 0 | 9 | 0 |
| SE | 49651 | ENSG00000136895 | GARNL3 | chr9 | + | 127335229 | 127335333 | 127333022 | 127333121 | 127336127 | 127336222 | 12 | 4 | 0 | 8 | 12 | 0 | 0 | 0 | 4 | 0 | 3 | 4 |
| SE | 49657 | ENSG00000136895 | GARNL3 | chr9 | + | 127383437 | 127383545 | 127361715 | 127365366 | 127385026 | 127385145 | 1 | 0 | 0 | 0 | 0 | 0 | 0 | 0 | 0 | 0 | 0 | 0 |
| SE | 49659 | ENSG00000183458 | PKD1P3 | chr16 | + | 14919862 | 14920083 | 14919244 | 14919428 | 14920271 | 14920387 | 10 | 1 | 6 | 5 | 2 | 0 | 0 | 0 | 0 | 0 | 16 | 15 |
| SE | 49660 | ENSG00000183426 | NPIPA1 | chr16 | + | 14925100 | 14925189 | 14924504 | 14924638 | 14925732 | 14925876 | 5 | 4 | 4 | 1 | 2 | 6 | 3 | 9 | 4 | 2 | 0 | 2 |
| SE | 49661 | ENSG00000183458 | PKD1P3 | chr16 | + | 14933366 | 14933493 | 14933065 | 14933276 | 14935159 | 14935276 | 7 | 3 | 10 | 3 | 2 | 0 | 0 | 1 | 3 | 0 | 0 | 16 |
| SE | 49662 | ENSG00000132122 | SPATA6 | chr1 | - | 48305786 | 48305878 | 48295372 | 48298893 | 48355669 | 48355769 | 31 | 22 | 26 | 16 | 41 | 0 | 0 | 1 | 0 | 0 | 0 | 27 |
| SE | 49663 | ENSG00000132122 | SPATA6 | chr1 | - | 48305786 | 48305878 | 48295372 | 48298893 | 48355717 | 48355769 | 24 | 18 | 16 | 10 | 23 | 0 | 2 | 0 | 0 | 4 | 0 | 8 |
| SE | 49664 | ENSG00000132122 | SPATA6 | chr1 | - | 48393254 | 48393300 | 48385308 | 48385349 | 48395266 | 48395354 | 0 | 0 | 3 | 0 | 0 | 20 | 17 | 8 | 10 | 12 | 0 | 0 |
| SE | 49670 | ENSG00000238057 | ZEB2-AS1 | chr2 | + | 144518751 | 144518861 | 144518202 | 144518575 | 144519958 | 144520110 | 0 | 6 | 2 | 0 | 2 | 2 | 0 | 0 | 0 | 0 | 0 | 0 |
| SE | 49688 | ENSG00000263956 | NBPF11 | chr1 | - | 148122728 | 148122801 | 148122054 | 148122266 | 148123852 | 148124067 | 55 | 43 | 49 | 41 | 54 | 0 | 0 | 0 | 0 | 0 | 34 | 43 |
| SE | 49690 | ENSG00000263956 | NBPF11 | chr1 | - | 148126828 | 148127038 | 148124898 | 148125001 | 148135671 | 148135813 | 19 | 7 | 29 | 14 | 47 | 0 | 0 | 3 | 0 | 0 | 9 | 2 |
| SE | 49693 | ENSG00000263956 | NBPF11 | chr1 | - | 148135123 | 148135247 | 148126828 | 148127038 | 148135671 | 148135813 | 0 | 0 | 0 | 0 | 4 | 9 | 3 | 13 | 8 | 30 | 0 | 0 |
| SE | 49694 | ENSG00000263956 | NBPF11 | chr1 | - | 148135123 | 148135247 | 148127140 | 148127288 | 148135671 | 148135813 | 0 | 0 | 0 | 0 | 0 | 3 | 2 | 0 | 1 | 0 | 0 | 0 |
| SE | 49695 | ENSG00000263956 | NBPF11 | chr1 | - | 148135671 | 148135813 | 148127140 | 148127288 | 148143414 | 148143686 | 4 | 2 | 0 | 2 | 6 | 0 | 0 | 0 | 0 | 0 | 0 | 1 |
| SE | 49696 | ENSG00000263956 | NBPF11 | chr1 | - | 148137710 | 148137809 | 148135671 | 148135813 | 148143414 | 148143686 | 17 | 12 | 13 | 27 | 23 | 1 | 0 | 0 | 1 | 6 | 4 | 20 |
| SE | 49700 | ENSG00000188732 | FAM221A | chr7 | + | 23692182 | 23692253 | 23691389 | 23691596 | 23698191 | 23698299 | 0 | 0 | 0 | 2 | 0 | 2 | 2 | 1 | 3 | 1 | 0 | 0 |
| SE | 49701 | ENSG00000188732 | FAM221A | chr7 | + | 23692182 | 23692253 | 23691389 | 23691596 | 23700785 | 23700868 | 0 | 0 | 0 | 1 | 0 | 0 | 0 | 2 | 0 | 2 | 0 | 0 |
| SE | 49702 | ENSG00000188732 | FAM221A | chr7 | + | 23692182 | 23694913 | 23691389 | 23691596 | 23698191 | 23698299 | 7 | 3 | 1 | 3 | 5 | 2 | 2 | 1 | 3 | 1 | 0 | 0 |
| SE | 49703 | ENSG00000188732 | FAM221A | chr7 | + | 23698191 | 23698299 | 23691389 | 23691596 | 23700785 | 23700868 | 8 | 4 | 7 | 16 | 4 | 0 | 0 | 2 | 0 | 2 | 0 | 18 |
| SE | 49705 | ENSG00000085840 | ORC1 | chr1 | - | 52383829 | 52383937 | 52383419 | 52383569 | 52384549 | 52384721 | 1 | 1 | 4 | 7 | 0 | 0 | 0 | 0 | 0 | 0 | 16 | 3 |
| SE | 49709 | ENSG00000145741 | BTF3 | chr5 | + | 73499133 | 73499202 | 73498407 | 73498558 | 73502487 | 73502601 | 2655 | 1573 | 2429 | 2148 | 3704 | 1 | 0 | 0 | 1 | 0 | 1390 | 1447 |
| SE | 49711 | ENSG00000145741 | BTF3 | chr5 | + | 73502915 | 73503117 | 73502363 | 73502601 | 73504346 | 73504403 | 5322 | 3185 | 4818 | 4514 | 7547 | 0 | 0 | 0 | 0 | 4 | 3568 | 3209 |
| SE | 49714 | ENSG00000101444 | AHCY | chr20 | - | 34285439 | 34285634 | 34280267 | 34281165 | 34290331 | 34290449 | 57 | 31 | 64 | 69 | 66 | 0 | 0 | 0 | 0 | 0 | 76 | 74 |
| SE | 49715 | ENSG00000146834 | MEPCE | chr7 | + | 100430614 | 100431689 | 100428816 | 100429040 | 100432918 | 100433137 | 48 | 27 | 57 | 34 | 38 | 0 | 0 | 0 | 0 | 0 | 77 | 50 |
| SE | 49716 | ENSG00000171735 | CAMTA1 | chr1 | + | 6806982 | 6807088 | 6785517 | 6785575 | 6825091 | 6825210 | 1 | 0 | 0 | 0 | 1 | 14 | 9 | 17 | 18 | 15 | 0 | 0 |
| SE | 49718 | ENSG00000171735 | CAMTA1 | chr1 | + | 6820180 | 6820250 | 6806987 | 6807088 | 6825091 | 6825210 | 51 | 34 | 29 | 35 | 45 | 1 | 0 | 0 | 0 | 0 | 68 | 57 |
| SE | 49719 | ENSG00000171735 | CAMTA1 | chr1 | + | 6825091 | 6825210 | 6785453 | 6785575 | 6871756 | 6871840 | 158 | 98 | 103 | 159 | 229 | 0 | 0 | 0 | 0 | 4 | 137 | 152 |
| SE | 49721 | ENSG00000171735 | CAMTA1 | chr1 | + | 6871756 | 6871840 | 6825091 | 6825210 | 6886239 | 6886346 | 118 | 74 | 72 | 110 | 189 | 0 | 0 | 0 | 1 | 0 | 141 | 114 |
| SE | 49723 | ENSG00000171735 | CAMTA1 | chr1 | + | 6886239 | 6886346 | 6825109 | 6825210 | 6887653 | 6887842 | 1 | 0 | 8 | 2 | 3 | 9 | 1 | 4 | 0 | 3 | 0 | 0 |
| SE | 49727 | ENSG00000154832 | CXXC1 | chr18 | - | 50284719 | 50284839 | 50284377 | 50284562 | 50285001 | 50285247 | 156 | 79 | 87 | 127 | 128 | 0 | 0 | 0 | 0 | 0 | 153 | 122 |
| SE | 49728 | ENSG00000154832 | CXXC1 | chr18 | - | 50284731 | 50284839 | 50284377 | 50284562 | 50285001 | 50285043 | 353 | 188 | 246 | 294 | 282 | 0 | 0 | 0 | 0 | 0 | 504 | 317 |
| SE | 49729 | ENSG00000154832 | CXXC1 | chr18 | - | 50285001 | 50285247 | 50284719 | 50284839 | 50285748 | 50285928 | 249 | 132 | 166 | 204 | 191 | 1 | 4 | 0 | 2 | 0 | 358 | 206 |
| SE | 49743 | ENSG00000111331 | OAS3 | chr12 | + | 112963312 | 112963457 | 112962651 | 112962902 | 112964234 | 112964408 | 53 | 84 | 70 | 115 | 72 | 0 | 0 | 0 | 0 | 0 | 512 | 179 |
| SE | 49744 | ENSG00000111331 | OAS3 | chr12 | + | 112963312 | 112963457 | 112962651 | 112962902 | 112967935 | 112968174 | 35 | 58 | 50 | 75 | 46 | 0 | 0 | 0 | 0 | 0 | 359 | 114 |
| SE | 49745 | ENSG00000111331 | OAS3 | chr12 | + | 112964234 | 112964408 | 112962651 | 112962902 | 112967935 | 112968174 | 2 | 0 | 0 | 0 | 0 | 0 | 0 | 0 | 0 | 0 | 28 | 0 |
| SE | 49746 | ENSG00000111331 | OAS3 | chr12 | + | 112964234 | 112964408 | 112963312 | 112963457 | 112967935 | 112968174 | 21 | 27 | 21 | 42 | 26 | 0 | 0 | 0 | 0 | 0 | 181 | 68 |
| SE | 49747 | ENSG00000111331 | OAS3 | chr12 | + | 112964238 | 112964408 | 112963312 | 112963457 | 112967935 | 112968174 | 2 | 0 | 0 | 0 | 0 | 0 | 0 | 0 | 0 | 0 | 28 | 0 |
| SE | 49751 | ENSG00000111331 | OAS3 | chr12 | + | 112965896 | 112966029 | 112964234 | 112964408 | 112967935 | 112968174 | 4 | 5 | 0 | 1 | 9 | 2 | 0 | 0 | 0 | 0 | 34 | 4 |
| SE | 49752 | ENSG00000111331 | OAS3 | chr12 | + | 112966222 | 112966349 | 112965743 | 112966029 | 112967417 | 112967593 | 2 | 0 | 0 | 2 | 0 | 41 | 67 | 34 | 48 | 33 | 0 | 0 |
| SE | 49753 | ENSG00000111331 | OAS3 | chr12 | + | 112967417 | 112967583 | 112965743 | 112966029 | 112967935 | 112968174 | 41 | 67 | 34 | 48 | 33 | 0 | 0 | 0 | 0 | 0 | 338 | 78 |
| SE | 49755 | ENSG00000111331 | OAS3 | chr12 | + | 112967417 | 112967593 | 112965743 | 112966029 | 112967935 | 112968174 | 87 | 127 | 67 | 100 | 50 | 0 | 0 | 0 | 0 | 0 | 581 | 142 |
| SE | 49759 | ENSG00000077549 | CAPZB | chr1 | - | 19344357 | 19344434 | 19338774 | 19339617 | 19345186 | 19345252 | 2074 | 1158 | 1449 | 1614 | 2846 | 0 | 2 | 4 | 0 | 0 | 2926 | 1665 |
| SE | 49760 | ENSG00000077549 | CAPZB | chr1 | - | 19344357 | 19345252 | 19339136 | 19339617 | 19378539 | 19378653 | 998 | 534 | 629 | 733 | 1597 | 0 | 0 | 0 | 0 | 0 | 1427 | 805 |
| SE | 49761 | ENSG00000077549 | CAPZB | chr1 | - | 19345186 | 19345252 | 19338775 | 19339617 | 19378539 | 19378653 | 0 | 2 | 0 | 2 | 0 | 0 | 0 | 0 | 0 | 0 | 0 | 0 |
| SE | 49762 | ENSG00000077549 | CAPZB | chr1 | - | 19356634 | 19356751 | 19344357 | 19345252 | 19357421 | 19357563 | 2105 | 1183 | 1724 | 1808 | 2388 | 12 | 0 | 1 | 8 | 7 | 3158 | 2091 |
| SE | 49763 | ENSG00000077549 | CAPZB | chr1 | - | 19356634 | 19356751 | 19344357 | 19345252 | 19378539 | 19378653 | 972 | 541 | 824 | 867 | 990 | 0 | 0 | 1 | 2 | 0 | 1450 | 985 |
| SE | 49764 | ENSG00000077549 | CAPZB | chr1 | - | 19357421 | 19357563 | 19344357 | 19345252 | 19378539 | 19378653 | 756 | 348 | 601 | 572 | 784 | 0 | 0 | 1 | 2 | 0 | 901 | 838 |
| SE | 49765 | ENSG00000077549 | CAPZB | chr1 | - | 19357421 | 19357563 | 19356634 | 19356751 | 19378539 | 19378653 | 1879 | 991 | 1502 | 1507 | 2180 | 1 | 0 | 1 | 1 | 4 | 2609 | 1953 |
| SE | 49766 | ENSG00000077549 | CAPZB | chr1 | - | 19385504 | 19385626 | 19378539 | 19378653 | 19419660 | 19419750 | 1458 | 777 | 1117 | 1209 | 1359 | 3 | 1 | 5 | 0 | 0 | 2132 | 1658 |
| SE | 49767 | ENSG00000077549 | CAPZB | chr1 | - | 19412235 | 19417164 | 19385504 | 19385626 | 19419660 | 19419750 | 2 | 0 | 0 | 0 | 0 | 698 | 366 | 523 | 618 | 502 | 0 | 0 |
| SE | 49770 | ENSG00000213626 | LBH | chr2 | + | 30256701 | 30256803 | 30234404 | 30234507 | 30257432 | 30257548 | 1 | 0 | 0 | 1 | 2 | 482 | 219 | 623 | 499 | 325 | 0 | 0 |
| SE | 49773 | ENSG00000146215 | CRIP3 | chr6 | - | 43307611 | 43307743 | 43306445 | 43306517 | 43307838 | 43307876 | 3 | 0 | 2 | 6 | 3 | 0 | 0 | 0 | 1 | 0 | 11 | 6 |
| SE | 49774 | ENSG00000146215 | CRIP3 | chr6 | - | 43307838 | 43307896 | 43306445 | 43306517 | 43308314 | 43308409 | 0 | 0 | 0 | 1 | 1 | 1 | 0 | 0 | 0 | 0 | 0 | 1 |
| SE | 49790 | ENSG00000116337 | AMPD2 | chr1 | + | 109629326 | 109629490 | 109628082 | 109629235 | 109630232 | 109630406 | 69 | 49 | 46 | 77 | 37 | 0 | 0 | 0 | 1 | 0 | 64 | 67 |
| SE | 49791 | ENSG00000116337 | AMPD2 | chr1 | + | 109629326 | 109629916 | 109629108 | 109629235 | 109630232 | 109630406 | 125 | 64 | 71 | 101 | 55 | 0 | 0 | 0 | 1 | 0 | 110 | 95 |
| SE | 49795 | ENSG00000015153 | YAF2 | chr12 | - | 42172139 | 42172209 | 42161612 | 42161765 | 42199135 | 42199235 | 0 | 1 | 1 | 0 | 0 | 3 | 1 | 0 | 5 | 0 | 0 | 0 |
| SE | 49796 | ENSG00000015153 | YAF2 | chr12 | - | 42172139 | 42172209 | 42161612 | 42161765 | 42235613 | 42235755 | 2 | 1 | 1 | 0 | 0 | 0 | 3 | 0 | 3 | 1 | 0 | 0 |
| SE | 49797 | ENSG00000015153 | YAF2 | chr12 | - | 42172139 | 42172209 | 42161612 | 42161765 | 42237598 | 42237724 | 0 | 2 | 1 | 0 | 0 | 16 | 9 | 13 | 22 | 16 | 0 | 0 |
| SE | 49798 | ENSG00000015153 | YAF2 | chr12 | - | 42199135 | 42199235 | 42161612 | 42161765 | 42210354 | 42210454 | 3 | 2 | 0 | 6 | 0 | 3 | 6 | 4 | 4 | 3 | 0 | 0 |
| SE | 49799 | ENSG00000015153 | YAF2 | chr12 | - | 42199135 | 42199235 | 42161612 | 42161765 | 42210547 | 42210619 | 1 | 0 | 0 | 2 | 0 | 1 | 0 | 0 | 0 | 2 | 0 | 0 |
| SE | 49800 | ENSG00000015153 | YAF2 | chr12 | - | 42199135 | 42199235 | 42161612 | 42161765 | 42237598 | 42237724 | 1 | 0 | 2 | 2 | 0 | 16 | 9 | 13 | 22 | 16 | 0 | 0 |
| SE | 49801 | ENSG00000015153 | YAF2 | chr12 | - | 42199135 | 42199235 | 42161612 | 42161765 | 42238154 | 42238180 | 2 | 0 | 0 | 2 | 0 | 0 | 0 | 0 | 2 | 0 | 0 | 0 |
| SE | 49803 | ENSG00000015153 | YAF2 | chr12 | - | 42210354 | 42210454 | 42161612 | 42161765 | 42210547 | 42210619 | 7 | 9 | 5 | 12 | 5 | 1 | 0 | 0 | 0 | 2 | 0 | 23 |
| SE | 49805 | ENSG00000015153 | YAF2 | chr12 | - | 42210547 | 42210619 | 42161612 | 42161765 | 42235613 | 42235755 | 1 | 0 | 0 | 0 | 0 | 0 | 3 | 0 | 3 | 1 | 0 | 2 |
| SE | 49806 | ENSG00000015153 | YAF2 | chr12 | - | 42210547 | 42210619 | 42161612 | 42161765 | 42237598 | 42237724 | 2 | 0 | 0 | 0 | 0 | 16 | 9 | 13 | 22 | 16 | 0 | 2 |
| SE | 49807 | ENSG00000015153 | YAF2 | chr12 | - | 42210547 | 42210619 | 42161612 | 42161765 | 42238154 | 42238303 | 1 | 0 | 0 | 0 | 0 | 0 | 0 | 0 | 2 | 0 | 0 | 2 |
| SE | 49808 | ENSG00000015153 | YAF2 | chr12 | - | 42210547 | 42210619 | 42199135 | 42199235 | 42237598 | 42237724 | 1 | 0 | 0 | 0 | 0 | 0 | 0 | 2 | 0 | 0 | 0 | 0 |
| SE | 49810 | ENSG00000015153 | YAF2 | chr12 | - | 42212443 | 42212484 | 42210547 | 42210619 | 42237598 | 42237724 | 0 | 1 | 0 | 0 | 0 | 1 | 0 | 0 | 4 | 2 | 0 | 0 |
| SE | 49812 | ENSG00000015153 | YAF2 | chr12 | - | 42235613 | 42235755 | 42161612 | 42161765 | 42237598 | 42237724 | 0 | 3 | 2 | 3 | 1 | 16 | 9 | 13 | 22 | 16 | 0 | 0 |
| SE | 49814 | ENSG00000015153 | YAF2 | chr12 | - | 42235613 | 42235755 | 42210547 | 42210619 | 42237598 | 42237724 | 0 | 0 | 2 | 0 | 0 | 1 | 0 | 0 | 4 | 2 | 0 | 0 |
| SE | 49815 | ENSG00000015153 | YAF2 | chr12 | - | 42237598 | 42237724 | 42161612 | 42161765 | 42238154 | 42238180 | 39 | 27 | 33 | 42 | 38 | 0 | 0 | 0 | 2 | 0 | 2 | 33 |
| SE | 49816 | ENSG00000015153 | YAF2 | chr12 | - | 42237598 | 42237724 | 42199135 | 42199235 | 42238154 | 42238180 | 23 | 18 | 23 | 20 | 22 | 1 | 0 | 0 | 0 | 0 | 2 | 21 |
| SE | 49821 | ENSG00000006837 | CDKL3 | chr5 | - | 134350248 | 134350427 | 134321790 | 134321903 | 134359896 | 134360091 | 0 | 2 | 4 | 0 | 1 | 0 | 1 | 0 | 0 | 0 | 12 | 0 |
| SE | 49824 | ENSG00000006837 | CDKL3 | chr5 | - | 134359896 | 134360091 | 134350248 | 134350427 | 134370997 | 134371041 | 1 | 2 | 5 | 0 | 1 | 3 | 0 | 0 | 1 | 0 | 12 | 0 |
| SE | 49825 | ENSG00000204149 | AGAP6 | chr10 | + | 49988120 | 49988215 | 49987164 | 49987234 | 49988750 | 49988938 | 2 | 0 | 1 | 0 | 4 | 4 | 2 | 3 | 1 | 5 | 0 | 0 |
| SE | 49826 | ENSG00000204149 | AGAP6 | chr10 | + | 49989307 | 49989376 | 49988367 | 49988938 | 49991675 | 49991744 | 43 | 27 | 23 | 32 | 35 | 0 | 0 | 0 | 0 | 0 | 19 | 12 |
| SE | 49827 | ENSG00000204149 | AGAP6 | chr10 | + | 49994394 | 49994429 | 49991376 | 49991744 | 50001995 | 50002096 | 61 | 46 | 57 | 58 | 107 | 0 | 0 | 1 | 0 | 0 | 18 | 28 |
| SE | 49828 | ENSG00000204149 | AGAP6 | chr10 | + | 50001602 | 50001746 | 49994394 | 49994429 | 50001995 | 50002096 | 0 | 0 | 2 | 0 | 0 | 54 | 40 | 49 | 48 | 99 | 0 | 3 |
| SE | 49829 | ENSG00000204149 | AGAP6 | chr10 | + | 50001995 | 50002096 | 49994394 | 49994429 | 50004684 | 50004720 | 106 | 78 | 83 | 90 | 170 | 2 | 0 | 1 | 0 | 3 | 76 | 49 |
| SE | 49830 | ENSG00000204149 | AGAP6 | chr10 | + | 50004684 | 50004720 | 50001995 | 50002096 | 50008024 | 50008076 | 94 | 79 | 88 | 91 | 177 | 0 | 0 | 0 | 0 | 0 | 82 | 64 |
| SE | 49831 | ENSG00000204149 | AGAP6 | chr10 | + | 50004684 | 50004720 | 50001995 | 50002096 | 50008710 | 50009348 | 15 | 16 | 26 | 21 | 50 | 0 | 0 | 0 | 1 | 0 | 19 | 14 |
| SE | 49832 | ENSG00000204149 | AGAP6 | chr10 | + | 50008024 | 50008076 | 50001995 | 50002096 | 50008710 | 50010407 | 38 | 52 | 51 | 61 | 107 | 0 | 0 | 0 | 1 | 0 | 60 | 24 |
| SE | 49833 | ENSG00000204149 | AGAP6 | chr10 | + | 50008024 | 50008076 | 50004463 | 50004720 | 50008710 | 50010407 | 109 | 113 | 107 | 119 | 230 | 0 | 0 | 0 | 0 | 0 | 107 | 75 |
| SE | 49851 | ENSG00000226383 | LINC01876 | chr2 | - | 156227966 | 156228004 | 156205381 | 156205438 | 156254777 | 156254918 | 0 | 4 | 0 | 0 | 0 | 1 | 0 | 0 | 0 | 0 | 0 | 2 |
| SE | 49857 | ENSG00000226383 | LINC01876 | chr2 | - | 156246609 | 156246750 | 156227966 | 156228004 | 156254777 | 156254931 | 0 | 0 | 2 | 1 | 0 | 0 | 4 | 0 | 0 | 0 | 0 | 0 |
| SE | 49859 | ENSG00000226383 | LINC01876 | chr2 | - | 156252035 | 156252128 | 156227966 | 156228004 | 156254777 | 156254931 | 0 | 0 | 0 | 1 | 0 | 0 | 4 | 0 | 0 | 0 | 0 | 0 |
| SE | 49863 | ENSG00000151136 | BTBD11 | chr12 | + | 107581151 | 107581304 | 107543934 | 107544139 | 107610189 | 107610320 | 9 | 5 | 13 | 21 | 17 | 0 | 0 | 0 | 0 | 0 | 19 | 0 |
| SE | 49864 | ENSG00000151136 | BTBD11 | chr12 | + | 107617088 | 107617182 | 107615060 | 107615162 | 107618157 | 107618378 | 8 | 9 | 12 | 8 | 2 | 0 | 1 | 0 | 0 | 0 | 10 | 0 |
| SE | 49865 | ENSG00000151136 | BTBD11 | chr12 | + | 107617300 | 107617436 | 107617088 | 107617182 | 107618157 | 107618378 | 35 | 89 | 98 | 59 | 45 | 1 | 2 | 0 | 1 | 0 | 114 | 45 |
| SE | 49870 | ENSG00000109180 | OCIAD1 | chr4 | + | 48831226 | 48831249 | 48831062 | 48831141 | 48832619 | 48832682 | 3 | 0 | 4 | 16 | 7 | 0 | 0 | 0 | 4 | 0 | 0 | 0 |
| SE | 49871 | ENSG00000109180 | OCIAD1 | chr4 | + | 48831226 | 48831398 | 48831043 | 48831141 | 48832619 | 48832682 | 0 | 1 | 5 | 4 | 6 | 0 | 0 | 0 | 4 | 0 | 0 | 1 |
| SE | 49872 | ENSG00000109180 | OCIAD1 | chr4 | + | 48831226 | 48831433 | 48831062 | 48831141 | 48832619 | 48832682 | 0 | 0 | 4 | 0 | 3 | 0 | 0 | 0 | 4 | 0 | 0 | 1 |
| SE | 49875 | ENSG00000109180 | OCIAD1 | chr4 | + | 48832619 | 48832682 | 48831064 | 48831249 | 48833400 | 48833429 | 214 | 139 | 186 | 190 | 220 | 0 | 0 | 0 | 3 | 0 | 246 | 189 |
| SE | 49877 | ENSG00000109180 | OCIAD1 | chr4 | + | 48832619 | 48832761 | 48831222 | 48831249 | 48833400 | 48833481 | 21 | 10 | 10 | 17 | 18 | 0 | 0 | 0 | 3 | 0 | 38 | 32 |
| SE | 49880 | ENSG00000109180 | OCIAD1 | chr4 | + | 48837159 | 48837252 | 48833400 | 48833481 | 48842635 | 48842689 | 0 | 0 | 1 | 0 | 0 | 453 | 352 | 367 | 404 | 661 | 0 | 0 |
| SE | 49881 | ENSG00000109180 | OCIAD1 | chr4 | + | 48849946 | 48850082 | 48848398 | 48848446 | 48851805 | 48851899 | 1001 | 745 | 855 | 945 | 1899 | 0 | 0 | 0 | 0 | 0 | 1016 | 700 |
| SE | 49884 | ENSG00000109180 | OCIAD1 | chr4 | + | 48851805 | 48851975 | 48849946 | 48850082 | 48857212 | 48857252 | 824 | 532 | 696 | 789 | 1821 | 0 | 0 | 0 | 0 | 2 | 681 | 604 |
| SE | 49885 | ENSG00000109180 | OCIAD1 | chr4 | + | 48857212 | 48857365 | 48851805 | 48851975 | 48860724 | 48861200 | 666 | 462 | 613 | 642 | 1458 | 3 | 0 | 7 | 6 | 3 | 622 | 508 |
| SE | 49886 | ENSG00000109180 | OCIAD1 | chr4 | + | 48857212 | 48857365 | 48852191 | 48852504 | 48860724 | 48861200 | 296 | 249 | 323 | 292 | 572 | 0 | 3 | 0 | 0 | 0 | 335 | 231 |
| SE | 49892 | ENSG00000163600 | ICOS | chr2 | + | 203957798 | 203957883 | 203956658 | 203956765 | 203959585 | 203961577 | 106 | 63 | 128 | 178 | 278 | 0 | 1 | 0 | 0 | 0 | 3 | 95 |
| SE | 49896 | ENSG00000141068 | KSR1 | chr17 | + | 27601359 | 27601401 | 27597267 | 27597436 | 27603833 | 27603888 | 18 | 4 | 4 | 9 | 11 | 3 | 2 | 5 | 3 | 0 | 0 | 11 |
| SE | 49906 | ENSG00000108061 | SHOC2 | chr10 | + | 111009712 | 111009830 | 111009247 | 111009385 | 111011609 | 111011670 | 674 | 559 | 504 | 784 | 1601 | 4 | 0 | 1 | 5 | 3 | 21 | 453 |
| SE | 49907 | ENSG00000138756 | BMP2K | chr4 | + | 78826036 | 78826155 | 78776341 | 78776721 | 78833581 | 78833687 | 191 | 215 | 111 | 132 | 128 | 0 | 0 | 0 | 0 | 0 | 230 | 178 |
| SE | 49908 | ENSG00000138756 | BMP2K | chr4 | + | 78844927 | 78845049 | 78842384 | 78842527 | 78847187 | 78847269 | 188 | 180 | 116 | 184 | 472 | 0 | 1 | 0 | 0 | 0 | 192 | 194 |
| SE | 49909 | ENSG00000138756 | BMP2K | chr4 | + | 78844927 | 78845049 | 78842384 | 78842527 | 78850923 | 78851056 | 104 | 96 | 57 | 115 | 190 | 0 | 0 | 0 | 1 | 0 | 119 | 76 |
| SE | 49910 | ENSG00000138756 | BMP2K | chr4 | + | 78847187 | 78847269 | 78842384 | 78842527 | 78850923 | 78851056 | 64 | 78 | 52 | 78 | 136 | 0 | 0 | 0 | 1 | 0 | 98 | 63 |
| SE | 49913 | ENSG00000138756 | BMP2K | chr4 | + | 78878733 | 78878891 | 78872613 | 78872798 | 78910609 | 78916365 | 90 | 79 | 41 | 49 | 79 | 0 | 0 | 0 | 0 | 0 | 62 | 79 |
| SE | 49915 | ENSG00000144455 | SUMF1 | chr3 | - | 3779240 | 3779322 | 3777724 | 3777803 | 3788913 | 3789074 | 6 | 3 | 6 | 7 | 7 | 0 | 0 | 0 | 0 | 0 | 0 | 10 |
| SE | 49916 | ENSG00000144455 | SUMF1 | chr3 | - | 3782502 | 3782701 | 3779240 | 3779322 | 3788913 | 3789074 | 19 | 19 | 33 | 11 | 77 | 0 | 0 | 2 | 1 | 0 | 56 | 25 |
| SE | 49917 | ENSG00000144455 | SUMF1 | chr3 | - | 3822733 | 3822897 | 3788913 | 3789074 | 3827241 | 3827509 | 13 | 8 | 11 | 6 | 18 | 0 | 1 | 0 | 0 | 0 | 0 | 13 |
| SE | 49919 | ENSG00000144455 | SUMF1 | chr3 | - | 4376329 | 4376389 | 4361145 | 4362254 | 4410864 | 4410978 | 54 | 25 | 35 | 60 | 88 | 0 | 1 | 2 | 1 | 0 | 100 | 46 |
| SE | 49921 | ENSG00000144455 | SUMF1 | chr3 | - | 4449265 | 4449340 | 4410864 | 4410978 | 4452875 | 4453049 | 54 | 29 | 55 | 42 | 20 | 0 | 2 | 0 | 1 | 0 | 83 | 83 |
| SE | 49927 | ENSG00000133657 | ATP13A3 | chr3 | - | 194412198 | 194412288 | 194402676 | 194406116 | 194413758 | 194413839 | 127 | 53 | 78 | 100 | 161 | 20 | 7 | 12 | 20 | 33 | 0 | 7 |
| SE | 49928 | ENSG00000133657 | ATP13A3 | chr3 | - | 194413758 | 194413839 | 194402671 | 194406116 | 194419878 | 194419967 | 63 | 48 | 40 | 60 | 96 | 2 | 0 | 0 | 1 | 3 | 0 | 9 |
| SE | 49929 | ENSG00000133657 | ATP13A3 | chr3 | - | 194413758 | 194413839 | 194402671 | 194406116 | 194425341 | 194425529 | 15 | 5 | 15 | 12 | 28 | 0 | 2 | 0 | 3 | 2 | 0 | 1 |
| SE | 49931 | ENSG00000133657 | ATP13A3 | chr3 | - | 194419878 | 194419967 | 194402671 | 194406116 | 194425341 | 194425529 | 34 | 23 | 25 | 30 | 68 | 0 | 2 | 0 | 3 | 2 | 0 | 6 |
| SE | 49932 | ENSG00000133657 | ATP13A3 | chr3 | - | 194419878 | 194419967 | 194413758 | 194413839 | 194425341 | 194425529 | 121 | 85 | 80 | 115 | 170 | 3 | 1 | 5 | 0 | 0 | 0 | 17 |
| SE | 49935 | ENSG00000133657 | ATP13A3 | chr3 | - | 194427074 | 194427252 | 194425341 | 194425529 | 194428844 | 194428917 | 201 | 138 | 154 | 156 | 286 | 1 | 2 | 0 | 0 | 0 | 0 | 71 |
| SE | 49936 | ENSG00000133657 | ATP13A3 | chr3 | - | 194428844 | 194428917 | 194425341 | 194425529 | 194429677 | 194429774 | 48 | 22 | 35 | 42 | 26 | 0 | 0 | 0 | 0 | 2 | 7 | 29 |
| SE | 49937 | ENSG00000133657 | ATP13A3 | chr3 | - | 194437555 | 194437573 | 194437310 | 194437464 | 194438855 | 194438972 | 194 | 88 | 127 | 132 | 234 | 0 | 0 | 0 | 3 | 0 | 36 | 74 |
| SE | 49940 | ENSG00000133657 | ATP13A3 | chr3 | - | 194455892 | 194455962 | 194454257 | 194454392 | 194459470 | 194459541 | 53 | 22 | 44 | 37 | 46 | 1 | 0 | 0 | 0 | 0 | 0 | 18 |
| SE | 49942 | ENSG00000037474 | NSUN2 | chr5 | - | 6602460 | 6602500 | 6599238 | 6600232 | 6604137 | 6604276 | 123 | 95 | 102 | 121 | 206 | 1 | 1 | 0 | 1 | 0 | 134 | 121 |
| SE | 49945 | ENSG00000037474 | NSUN2 | chr5 | - | 6607199 | 6607384 | 6606819 | 6606912 | 6610954 | 6611085 | 168 | 114 | 111 | 153 | 75 | 0 | 0 | 0 | 0 | 0 | 164 | 71 |
| SE | 49946 | ENSG00000037474 | NSUN2 | chr5 | - | 6609825 | 6609922 | 6606819 | 6606912 | 6610954 | 6611085 | 46 | 37 | 68 | 75 | 45 | 0 | 0 | 0 | 0 | 0 | 77 | 69 |
| SE | 49947 | ENSG00000037474 | NSUN2 | chr5 | - | 6609825 | 6609922 | 6607199 | 6607384 | 6610954 | 6611085 | 135 | 113 | 128 | 171 | 123 | 1 | 0 | 1 | 0 | 0 | 171 | 93 |
| SE | 49948 | ENSG00000037474 | NSUN2 | chr5 | - | 6611724 | 6611798 | 6610954 | 6611085 | 6616726 | 6616857 | 169 | 178 | 193 | 213 | 133 | 0 | 0 | 0 | 0 | 0 | 284 | 139 |
| SE | 49949 | ENSG00000037474 | NSUN2 | chr5 | - | 6621602 | 6621744 | 6620115 | 6620298 | 6623213 | 6623285 | 0 | 0 | 1 | 0 | 1 | 4 | 6 | 3 | 5 | 2 | 9 | 0 |
| SE | 49955 | ENSG00000171793 | CTPS1 | chr1 | + | 40999829 | 41001117 | 40997393 | 40997526 | 41002159 | 41002254 | 27 | 24 | 15 | 33 | 7 | 1 | 0 | 0 | 0 | 0 | 52 | 37 |
| SE | 49956 | ENSG00000171793 | CTPS1 | chr1 | + | 41000434 | 41000566 | 40995916 | 40997526 | 41001028 | 41001117 | 0 | 0 | 0 | 0 | 0 | 27 | 45 | 37 | 33 | 38 | 0 | 0 |
| SE | 49957 | ENSG00000171793 | CTPS1 | chr1 | + | 41000972 | 41001117 | 40997393 | 40997526 | 41002159 | 41002254 | 27 | 22 | 15 | 28 | 7 | 1 | 0 | 0 | 0 | 0 | 32 | 30 |
| SE | 49958 | ENSG00000171793 | CTPS1 | chr1 | + | 41001028 | 41001117 | 40993852 | 40997526 | 41002159 | 41002254 | 50 | 62 | 49 | 51 | 45 | 1 | 0 | 0 | 0 | 0 | 67 | 43 |
| SE | 49960 | ENSG00000171793 | CTPS1 | chr1 | + | 41009444 | 41009589 | 41008658 | 41008890 | 41010160 | 41010254 | 81 | 39 | 85 | 72 | 85 | 0 | 0 | 0 | 0 | 0 | 157 | 48 |
| SE | 49961 | ENSG00000171793 | CTPS1 | chr1 | + | 41010160 | 41010254 | 41009444 | 41009589 | 41011657 | 41011754 | 29 | 21 | 39 | 27 | 34 | 0 | 0 | 0 | 0 | 0 | 25 | 36 |
| SE | 49962 | ENSG00000162976 | SLC66A3 | chr2 | + | 11164203 | 11164261 | 11160624 | 11160694 | 11171924 | 11172045 | 80 | 37 | 50 | 42 | 64 | 0 | 0 | 0 | 1 | 0 | 33 | 41 |
| SE | 49972 | ENSG00000151690 | MFSD6 | chr2 | + | 190469757 | 190469855 | 190435894 | 190437561 | 190488656 | 190488818 | 57 | 64 | 24 | 94 | 78 | 0 | 0 | 0 | 5 | 0 | 36 | 77 |
| SE | 49974 | ENSG00000151690 | MFSD6 | chr2 | + | 190500014 | 190500276 | 190497648 | 190497719 | 190509162 | 190509205 | 127 | 87 | 86 | 86 | 236 | 0 | 0 | 0 | 0 | 0 | 132 | 100 |
| SE | 49976 | ENSG00000215305 | VPS16 | chr20 | + | 2859718 | 2859807 | 2840702 | 2840827 | 2860448 | 2860593 | 8 | 1 | 5 | 7 | 0 | 0 | 0 | 1 | 0 | 0 | 4 | 3 |
| SE | 49977 | ENSG00000215305 | VPS16 | chr20 | + | 2860053 | 2860151 | 2859718 | 2859807 | 2860448 | 2860593 | 4 | 6 | 4 | 14 | 0 | 0 | 1 | 0 | 1 | 0 | 28 | 14 |
| SE | 49978 | ENSG00000215305 | VPS16 | chr20 | + | 2860238 | 2860367 | 2860053 | 2860151 | 2860448 | 2860593 | 55 | 21 | 39 | 63 | 31 | 0 | 1 | 1 | 0 | 0 | 31 | 80 |
| SE | 49979 | ENSG00000215305 | VPS16 | chr20 | + | 2860448 | 2860593 | 2840702 | 2840827 | 2860747 | 2860863 | 43 | 10 | 18 | 39 | 8 | 0 | 0 | 0 | 0 | 0 | 44 | 43 |
| SE | 49980 | ENSG00000215305 | VPS16 | chr20 | + | 2865398 | 2865495 | 2865147 | 2865317 | 2866211 | 2866315 | 174 | 59 | 92 | 209 | 94 | 0 | 0 | 0 | 0 | 0 | 251 | 146 |
| SE | 49982 | ENSG00000065308 | TRAM2 | chr6 | - | 52516005 | 52516122 | 52509527 | 52509586 | 52516627 | 52516737 | 41 | 17 | 43 | 47 | 31 | 0 | 0 | 1 | 0 | 0 | 83 | 34 |
| SE | 49983 | ENSG00000065308 | TRAM2 | chr6 | - | 52516627 | 52516737 | 52516005 | 52516122 | 52535782 | 52535846 | 27 | 13 | 29 | 32 | 20 | 0 | 0 | 0 | 0 | 0 | 55 | 30 |
| SE | 49984 | ENSG00000065308 | TRAM2 | chr6 | - | 52535782 | 52535846 | 52516627 | 52516737 | 52576795 | 52577060 | 6 | 2 | 5 | 12 | 6 | 1 | 1 | 0 | 0 | 0 | 28 | 13 |
| SE | 49985 | ENSG00000159164 | SV2A | chr1 | - | 149907699 | 149907833 | 149906649 | 149906856 | 149908041 | 149908206 | 1 | 4 | 1 | 7 | 4 | 0 | 0 | 0 | 0 | 0 | 0 | 20 |
| SE | 49986 | ENSG00000159164 | SV2A | chr1 | - | 149909460 | 149909571 | 149909191 | 149909280 | 149909800 | 149909890 | 1 | 7 | 0 | 3 | 12 | 0 | 0 | 0 | 0 | 0 | 0 | 3 |
| SE | 49987 | ENSG00000159164 | SV2A | chr1 | - | 149910569 | 149910703 | 149909800 | 149909890 | 149910825 | 149910977 | 2 | 6 | 0 | 6 | 17 | 0 | 0 | 0 | 0 | 0 | 0 | 8 |
| SE | 49988 | ENSG00000241878 | PISD | chr22 | - | 31620552 | 31620599 | 31618976 | 31619836 | 31620995 | 31621142 | 3 | 2 | 3 | 8 | 13 | 1 | 0 | 0 | 0 | 0 | 0 | 7 |
| SE | 49989 | ENSG00000241878 | PISD | chr22 | - | 31620552 | 31620713 | 31618490 | 31619836 | 31620995 | 31621142 | 139 | 62 | 97 | 96 | 102 | 1 | 0 | 0 | 0 | 0 | 221 | 134 |
| SE | 49997 | ENSG00000241878 | PISD | chr22 | - | 31628002 | 31628210 | 31625762 | 31626016 | 31628852 | 31628981 | 2 | 1 | 2 | 2 | 4 | 2 | 0 | 1 | 1 | 0 | 1 | 0 |
| SE | 50003 | ENSG00000241878 | PISD | chr22 | - | 31628852 | 31628981 | 31628002 | 31628210 | 31638365 | 31638445 | 3 | 0 | 1 | 1 | 4 | 0 | 0 | 0 | 1 | 0 | 1 | 0 |
| SE | 50007 | ENSG00000175087 | PDIK1L | chr1 | + | 26112459 | 26112650 | 26111836 | 26111915 | 26114291 | 26114593 | 9 | 0 | 8 | 2 | 1 | 6 | 2 | 9 | 8 | 5 | 0 | 9 |
| SE | 50008 | ENSG00000175087 | PDIK1L | chr1 | + | 26114291 | 26114593 | 26111164 | 26111420 | 26121836 | 26122124 | 36 | 23 | 43 | 31 | 84 | 0 | 0 | 0 | 0 | 1 | 0 | 28 |
| SE | 50009 | ENSG00000175087 | PDIK1L | chr1 | + | 26114291 | 26114593 | 26111823 | 26111915 | 26121836 | 26122124 | 42 | 25 | 52 | 38 | 89 | 0 | 0 | 0 | 1 | 0 | 0 | 37 |
| SE | 50010 | ENSG00000162298 | SYVN1 | chr11 | - | 65131131 | 65131197 | 65130919 | 65131036 | 65131273 | 65131373 | 131 | 53 | 89 | 121 | 151 | 0 | 0 | 0 | 0 | 0 | 36 | 142 |
| SE | 50013 | ENSG00000162298 | SYVN1 | chr11 | - | 65132247 | 65132351 | 65131273 | 65131596 | 65132921 | 65133074 | 52 | 5 | 40 | 41 | 18 | 1 | 1 | 0 | 0 | 0 | 49 | 77 |
| SE | 50014 | ENSG00000162298 | SYVN1 | chr11 | - | 65132247 | 65132780 | 65131469 | 65131596 | 65132921 | 65133074 | 76 | 16 | 62 | 62 | 35 | 1 | 1 | 0 | 0 | 0 | 52 | 114 |
| SE | 50015 | ENSG00000162298 | SYVN1 | chr11 | - | 65132731 | 65132780 | 65131469 | 65131596 | 65132921 | 65133074 | 22 | 14 | 14 | 18 | 19 | 1 | 1 | 0 | 0 | 0 | 0 | 35 |
| SE | 50017 | ENSG00000107902 | LHPP | chr10 | + | 124496960 | 124497024 | 124488421 | 124488575 | 124498035 | 124498128 | 98 | 66 | 123 | 92 | 73 | 0 | 0 | 0 | 1 | 0 | 183 | 77 |
| SE | 50018 | ENSG00000107902 | LHPP | chr10 | + | 124517179 | 124517271 | 124498035 | 124498128 | 124613263 | 124614141 | 54 | 32 | 66 | 51 | 58 | 2 | 1 | 1 | 0 | 0 | 95 | 30 |
| SE | 50019 | ENSG00000157873 | TNFRSF14 | chr1 | + | 2557725 | 2557834 | 2556370 | 2556733 | 2558342 | 2558468 | 153 | 63 | 113 | 127 | 55 | 1 | 0 | 0 | 0 | 0 | 104 | 139 |
| SE | 50025 | ENSG00000165359 | INTS6L | chrX | + | 135569336 | 135569431 | 135556167 | 135556300 | 135572814 | 135573033 | 70 | 57 | 56 | 98 | 148 | 0 | 0 | 0 | 0 | 3 | 76 | 96 |
| SE | 50027 | ENSG00000165359 | INTS6L | chrX | + | 135575083 | 135575226 | 135573938 | 135574062 | 135577192 | 135577295 | 319 | 297 | 277 | 291 | 289 | 0 | 0 | 1 | 0 | 0 | 31 | 170 |
| SE | 50030 | ENSG00000165359 | INTS6L | chrX | + | 135581527 | 135581564 | 135581049 | 135581143 | 135582240 | 135582510 | 26 | 25 | 38 | 39 | 33 | 1 | 1 | 0 | 0 | 0 | 7 | 14 |
| SE | 50031 | ENSG00000165359 | INTS6L | chrX | + | 135581527 | 135581631 | 135581049 | 135581143 | 135582240 | 135582502 | 87 | 64 | 88 | 94 | 111 | 1 | 1 | 0 | 0 | 0 | 18 | 28 |
| SE | 50033 | ENSG00000254876 | STRA6LP | chr9 | + | 97251540 | 97251557 | 97250791 | 97250843 | 97276568 | 97276697 | 3 | 7 | 8 | 5 | 1 | 0 | 1 | 0 | 0 | 0 | 13 | 15 |
| SE | 50034 | ENSG00000254876 | STRA6LP | chr9 | + | 97281515 | 97281669 | 97279361 | 97279473 | 97287659 | 97287735 | 9 | 11 | 20 | 11 | 11 | 1 | 0 | 3 | 0 | 0 | 0 | 11 |
| SE | 50035 | ENSG00000254876 | STRA6LP | chr9 | + | 97292555 | 97292675 | 97291370 | 97291487 | 97293946 | 97294110 | 3 | 0 | 3 | 5 | 0 | 3 | 6 | 4 | 2 | 9 | 0 | 0 |
| SE | 50039 | ENSG00000154222 | CC2D1B | chr1 | - | 52355397 | 52355449 | 52354839 | 52354939 | 52355607 | 52355666 | 91 | 58 | 81 | 90 | 94 | 0 | 0 | 0 | 2 | 0 | 75 | 63 |
| SE | 50042 | ENSG00000154222 | CC2D1B | chr1 | - | 52358685 | 52358758 | 52357780 | 52358461 | 52359026 | 52359157 | 76 | 29 | 78 | 43 | 17 | 0 | 0 | 0 | 2 | 0 | 59 | 44 |
| SE | 50043 | ENSG00000154222 | CC2D1B | chr1 | - | 52359704 | 52359820 | 52359458 | 52359534 | 52360073 | 52360233 | 13 | 13 | 15 | 12 | 15 | 0 | 0 | 1 | 0 | 0 | 0 | 2 |
| SE | 50044 | ENSG00000154222 | CC2D1B | chr1 | - | 52359704 | 52359836 | 52359458 | 52359534 | 52360073 | 52360233 | 17 | 16 | 25 | 23 | 18 | 0 | 0 | 1 | 0 | 0 | 0 | 14 |
| SE | 50046 | ENSG00000137628 | DDX60 | chr4 | - | 168244599 | 168244717 | 168237690 | 168237795 | 168246417 | 168246618 | 1 | 2 | 0 | 0 | 0 | 75 | 51 | 82 | 85 | 75 | 0 | 0 |
| SE | 50047 | ENSG00000137628 | DDX60 | chr4 | - | 168248187 | 168248292 | 168246417 | 168246618 | 168250953 | 168251106 | 171 | 204 | 137 | 187 | 215 | 0 | 0 | 0 | 0 | 1 | 19 | 123 |
| SE | 50048 | ENSG00000137628 | DDX60 | chr4 | - | 168250953 | 168251106 | 168248187 | 168248292 | 168252508 | 168252656 | 269 | 326 | 209 | 330 | 274 | 3 | 0 | 1 | 0 | 0 | 26 | 274 |
| SE | 50049 | ENSG00000137628 | DDX60 | chr4 | - | 168255710 | 168255869 | 168252508 | 168252656 | 168260864 | 168260989 | 225 | 243 | 209 | 325 | 376 | 0 | 2 | 0 | 0 | 0 | 21 | 185 |
| SE | 50050 | ENSG00000137628 | DDX60 | chr4 | - | 168261999 | 168262128 | 168260864 | 168260989 | 168262682 | 168262787 | 203 | 230 | 187 | 255 | 279 | 0 | 0 | 0 | 2 | 0 | 13 | 145 |
| SE | 50051 | ENSG00000137628 | DDX60 | chr4 | - | 168265464 | 168265614 | 168262706 | 168262787 | 168267581 | 168267691 | 0 | 1 | 0 | 3 | 0 | 77 | 94 | 96 | 131 | 125 | 0 | 0 |
| SE | 50053 | ENSG00000137628 | DDX60 | chr4 | - | 168276014 | 168276181 | 168275344 | 168275503 | 168280334 | 168280590 | 446 | 629 | 438 | 569 | 626 | 0 | 0 | 1 | 0 | 0 | 191 | 532 |
| SE | 50055 | ENSG00000137628 | DDX60 | chr4 | - | 168287047 | 168287203 | 168285392 | 168285498 | 168291747 | 168291906 | 42 | 58 | 53 | 55 | 106 | 0 | 0 | 1 | 0 | 0 | 11 | 29 |
| SE | 50056 | ENSG00000137628 | DDX60 | chr4 | - | 168288173 | 168288315 | 168285392 | 168285498 | 168291747 | 168291906 | 52 | 53 | 61 | 85 | 136 | 0 | 0 | 1 | 0 | 0 | 10 | 51 |
| SE | 50057 | ENSG00000137628 | DDX60 | chr4 | - | 168288173 | 168288315 | 168287047 | 168287203 | 168291747 | 168291906 | 139 | 127 | 139 | 209 | 385 | 2 | 2 | 0 | 0 | 0 | 38 | 123 |
| SE | 50062 | ENSG00000102763 | VWA8 | chr13 | - | 41699070 | 41699270 | 41692861 | 41692972 | 41701391 | 41701530 | 58 | 53 | 79 | 67 | 90 | 0 | 0 | 0 | 0 | 0 | 2 | 85 |
| SE | 50063 | ENSG00000102763 | VWA8 | chr13 | - | 41719590 | 41719742 | 41703302 | 41703411 | 41721369 | 41721575 | 95 | 86 | 107 | 127 | 125 | 0 | 0 | 1 | 4 | 0 | 27 | 124 |
| SE | 50064 | ENSG00000102763 | VWA8 | chr13 | - | 41727193 | 41727313 | 41721369 | 41721575 | 41732079 | 41732155 | 49 | 43 | 48 | 67 | 164 | 0 | 0 | 0 | 2 | 0 | 18 | 33 |
| SE | 50065 | ENSG00000102763 | VWA8 | chr13 | - | 41729541 | 41729677 | 41727193 | 41727313 | 41732079 | 41732155 | 213 | 202 | 216 | 176 | 347 | 0 | 2 | 0 | 2 | 0 | 51 | 139 |
| SE | 50066 | ENSG00000102763 | VWA8 | chr13 | - | 41732079 | 41732155 | 41727193 | 41727313 | 41761127 | 41761204 | 25 | 19 | 31 | 27 | 43 | 0 | 0 | 1 | 0 | 0 | 1 | 9 |
| SE | 50067 | ENSG00000102763 | VWA8 | chr13 | - | 41777984 | 41778056 | 41761127 | 41761204 | 41787436 | 41787543 | 41 | 26 | 40 | 26 | 54 | 0 | 0 | 2 | 0 | 0 | 0 | 15 |
| SE | 50069 | ENSG00000102763 | VWA8 | chr13 | - | 41883386 | 41883491 | 41868345 | 41868477 | 41885919 | 41886028 | 94 | 82 | 104 | 119 | 177 | 0 | 0 | 0 | 0 | 0 | 62 | 73 |
| SE | 50071 | ENSG00000102763 | VWA8 | chr13 | - | 41891419 | 41891587 | 41887196 | 41887361 | 41907585 | 41907696 | 69 | 78 | 74 | 130 | 157 | 0 | 2 | 0 | 0 | 0 | 135 | 48 |
| SE | 50072 | ENSG00000053254 | FOXN3 | chr14 | - | 89180700 | 89180806 | 89156171 | 89162969 | 89190386 | 89190449 | 138 | 66 | 102 | 96 | 84 | 0 | 0 | 0 | 0 | 0 | 59 | 98 |
| SE | 50073 | ENSG00000053254 | FOXN3 | chr14 | - | 89180700 | 89180806 | 89156171 | 89162969 | 89280949 | 89281014 | 188 | 96 | 169 | 156 | 129 | 0 | 0 | 0 | 2 | 0 | 78 | 126 |
| SE | 50074 | ENSG00000053254 | FOXN3 | chr14 | - | 89190383 | 89190449 | 89180700 | 89180806 | 89223116 | 89223342 | 31 | 14 | 35 | 28 | 16 | 0 | 0 | 0 | 1 | 0 | 64 | 35 |
| SE | 50076 | ENSG00000053254 | FOXN3 | chr14 | - | 89190383 | 89190449 | 89180700 | 89180806 | 89350671 | 89350808 | 33 | 14 | 35 | 28 | 16 | 0 | 0 | 1 | 0 | 0 | 64 | 35 |
| SE | 50077 | ENSG00000053254 | FOXN3 | chr14 | - | 89190386 | 89190449 | 89162877 | 89162969 | 89280949 | 89281014 | 125 | 83 | 131 | 144 | 269 | 0 | 0 | 0 | 2 | 0 | 132 | 143 |
| SE | 50079 | ENSG00000053254 | FOXN3 | chr14 | - | 89190386 | 89190449 | 89180700 | 89180806 | 89350671 | 89350808 | 8 | 6 | 8 | 15 | 13 | 0 | 0 | 1 | 0 | 0 | 2 | 7 |
| SE | 50080 | ENSG00000053254 | FOXN3 | chr14 | - | 89280949 | 89281014 | 89180700 | 89180806 | 89350671 | 89350791 | 224 | 138 | 196 | 221 | 240 | 0 | 0 | 1 | 0 | 0 | 163 | 165 |
| SE | 50081 | ENSG00000053254 | FOXN3 | chr14 | - | 89280949 | 89281014 | 89190381 | 89190449 | 89350671 | 89350791 | 324 | 204 | 290 | 325 | 504 | 2 | 0 | 0 | 0 | 0 | 316 | 303 |
| SE | 50082 | ENSG00000053254 | FOXN3 | chr14 | - | 89284427 | 89284522 | 89280949 | 89281014 | 89350671 | 89350808 | 0 | 1 | 0 | 0 | 1 | 327 | 220 | 319 | 329 | 502 | 0 | 0 |
| SE | 50083 | ENSG00000053254 | FOXN3 | chr14 | - | 89333113 | 89333259 | 89280949 | 89281014 | 89350671 | 89350808 | 2 | 0 | 3 | 0 | 0 | 327 | 220 | 319 | 329 | 502 | 0 | 6 |

| | | | | | | | | | | | | | |
|---|---|---|---|---|---|---|---|---|---|---|---|---|---|
| SE | 50084 | ENSG00000053254 | FOXN3 | chr14 | - | 89350671 | 89350808 | 89280949 | 89281014 | 89411933 | 89412017 | 599 | 483 |
| SE | 50085 | ENSG00000053254 | FOXN3 | chr14 | - | 89411933 | 89412490 | 89350671 | 89350808 | 89486568 | 89486671 | 341 | 315 |
| SE | 50092 | ENSG00000092969 | TGFB2 | chr1 | + | 218363333 | 218363417 | 218346234 | 218347047 | 218405168 | 218405332 | 0 | 0 |
| SE | 50096 | ENSG00000115828 | QPCT | chr2 | + | 37367231 | 37367408 | 37359579 | 37359858 | 37369684 | 37369702 | 124 | 63 |
| SE | 50097 | ENSG00000115828 | QPCT | chr2 | + | 37372355 | 37372472 | 37369684 | 37369784 | 37372681 | 37372742 | 147 | 80 |
| SE | 50099 | ENSG00000163322 | ABRAXAS1 | chr4 | - | 83467453 | 83467538 | 83463493 | 83463608 | 83469031 | 83469151 | 184 | 199 |
| SE | 50101 | ENSG00000163322 | ABRAXAS1 | chr4 | - | 83469031 | 83469151 | 83467453 | 83467538 | 83472221 | 83472288 | 52 | 64 |
| SE | 50102 | ENSG00000163322 | ABRAXAS1 | chr4 | - | 83469031 | 83469151 | 83467453 | 83467538 | 83476642 | 83476679 | 55 | 66 |
| SE | 50103 | ENSG00000163322 | ABRAXAS1 | chr4 | - | 83470202 | 83470396 | 83467453 | 83467538 | 83472221 | 83472288 | 145 | 124 |
| SE | 50104 | ENSG00000163322 | ABRAXAS1 | chr4 | - | 83470202 | 83470396 | 83467453 | 83467538 | 83476642 | 83476679 | 4 | 6 |
| SE | 50105 | ENSG00000163322 | ABRAXAS1 | chr4 | - | 83470202 | 83470396 | 83469031 | 83469151 | 83472221 | 83472288 | 224 | 195 |
| SE | 50106 | ENSG00000163322 | ABRAXAS1 | chr4 | - | 83470202 | 83470396 | 83469031 | 83469151 | 83476642 | 83476679 | 83 | 77 |
| SE | 50107 | ENSG00000163322 | ABRAXAS1 | chr4 | - | 83472221 | 83472288 | 83467453 | 83467538 | 83476642 | 83476679 | 44 | 41 |
| SE | 50111 | ENSG00000163322 | ABRAXAS1 | chr4 | - | 83476642 | 83476679 | 83469031 | 83469151 | 83482153 | 83482244 | 12 | 8 |
| SE | 50112 | ENSG00000163322 | ABRAXAS1 | chr4 | - | 83476642 | 83476679 | 83472221 | 83472288 | 83482153 | 83482244 | 159 | 129 |
| SE | 50114 | ENSG00000117480 | FAAH | chr1 | + | 46409100 | 46409198 | 46408458 | 46408584 | 46410397 | 46410497 | 3 | 1 |
| SE | 50115 | ENSG00000117480 | FAAH | chr1 | + | 46411611 | 46411651 | 46410813 | 46410854 | 46412142 | 46412251 | 24 | 4 |
| SE | 50116 | ENSG00000132704 | FCRL2 | chr1 | - | 157746870 | 157746899 | 157745732 | 157746774 | 157748552 | 157748618 | 59 | 44 |
| SE | 50117 | ENSG00000132704 | FCRL2 | chr1 | - | 157746870 | 157746899 | 157745732 | 157746774 | 157748874 | 157748960 | 17 | 4 |
| SE | 50118 | ENSG00000132704 | FCRL2 | chr1 | - | 157748552 | 157748618 | 157745732 | 157746774 | 157748874 | 157748960 | 21 | 22 |
| SE | 50120 | ENSG00000132704 | FCRL2 | chr1 | - | 157748874 | 157748960 | 157745732 | 157746774 | 157749649 | 157749677 | 37 | 22 |
| SE | 50121 | ENSG00000132704 | FCRL2 | chr1 | - | 157766854 | 157766971 | 157749649 | 157749677 | 157768413 | 157768701 | 43 | 23 |
| SE | 50123 | ENSG00000132704 | FCRL2 | chr1 | - | 157767230 | 157767509 | 157766854 | 157766971 | 157768413 | 157768701 | 56 | 21 |
| SE | 50125 | ENSG00000132704 | FCRL2 | chr1 | - | 157768413 | 157768701 | 157749649 | 157749677 | 157770408 | 157770666 | 4 | 0 |
| SE | 50126 | ENSG00000132704 | FCRL2 | chr1 | - | 157768413 | 157768701 | 157766854 | 157766971 | 157770408 | 157770666 | 6 | 0 |
| SE | 50127 | ENSG00000132704 | FCRL2 | chr1 | - | 157768413 | 157768701 | 157767230 | 157767509 | 157770408 | 157770666 | 7 | 1 |
| SE | 50134 | ENSG00000245213 | NA | chr4 | - | 173166123 | 173166299 | 173164977 | 173165132 | 173166644 | 173166790 | 2 | 1 |
| SE | 50135 | ENSG00000245213 | NA | chr4 | - | 173166644 | 173166790 | 173143516 | 173145511 | 173167961 | 173168054 | 6 | 4 |
| SE | 50136 | ENSG00000132300 | PTCD3 | chr2 | + | 86107117 | 86107200 | 86106222 | 86106351 | 86108349 | 86108402 | 0 | 0 |
| SE | 50137 | ENSG00000132300 | PTCD3 | chr2 | + | 86108349 | 86108402 | 86106222 | 86106351 | 86108499 | 86108536 | 302 | 224 |
| SE | 50138 | ENSG00000132300 | PTCD3 | chr2 | + | 86110977 | 86111158 | 86108499 | 86108536 | 86116529 | 86116598 | 107 | 79 |
| SE | 50139 | ENSG00000132300 | PTCD3 | chr2 | + | 86111112 | 86111158 | 86108499 | 86108536 | 86116529 | 86116598 | 457 | 353 |
| SE | 50140 | ENSG00000132300 | PTCD3 | chr2 | + | 86118920 | 86119044 | 86117054 | 86117159 | 86119543 | 86119614 | 245 | 157 |
| SE | 50142 | ENSG00000132300 | PTCD3 | chr2 | + | 86118920 | 86119044 | 86117054 | 86117159 | 86123700 | 86123762 | 201 | 123 |
| SE | 50145 | ENSG00000132300 | PTCD3 | chr2 | + | 86121478 | 86121594 | 86117054 | 86117159 | 86123700 | 86123733 | 161 | 76 |
| SE | 50146 | ENSG00000132300 | PTCD3 | chr2 | + | 86121478 | 86121594 | 86118920 | 86119044 | 86123700 | 86123733 | 402 | 268 |
| SE | 50147 | ENSG00000132300 | PTCD3 | chr2 | + | 86123700 | 86123762 | 86118920 | 86119044 | 86124994 | 86125082 | 222 | 134 |
| SE | 50148 | ENSG00000132300 | PTCD3 | chr2 | + | 86124994 | 86125082 | 86123700 | 86123762 | 86125454 | 86125515 | 469 | 269 |
| SE | 50149 | ENSG00000132300 | PTCD3 | chr2 | + | 86124994 | 86125082 | 86123700 | 86123762 | 86125794 | 86125880 | 123 | 72 |
| SE | 50150 | ENSG00000132300 | PTCD3 | chr2 | + | 86125454 | 86125515 | 86123700 | 86123762 | 86125794 | 86125880 | 65 | 53 |
| SE | 50152 | ENSG00000132300 | PTCD3 | chr2 | + | 86125454 | 86125515 | 86124994 | 86125082 | 86127160 | 86127437 | 221 | 136 |
| SE | 50153 | ENSG00000132300 | PTCD3 | chr2 | + | 86125794 | 86125880 | 86124994 | 86125082 | 86127160 | 86127305 | 119 | 78 |
| SE | 50155 | ENSG00000132300 | PTCD3 | chr2 | + | 86127940 | 86127991 | 86127160 | 86127305 | 86130647 | 86130737 | 127 | 104 |
| SE | 50156 | ENSG00000132300 | PTCD3 | chr2 | + | 86130647 | 86130737 | 86127871 | 86127991 | 86131077 | 86131106 | 284 | 244 |
| SE | 50158 | ENSG00000132300 | PTCD3 | chr2 | + | 86134838 | 86134987 | 86134291 | 86134377 | 86136981 | 86137140 | 120 | 93 |
| SE | 50159 | ENSG00000132300 | PTCD3 | chr2 | + | 86136520 | 86136562 | 86134291 | 86134377 | 86136981 | 86137140 | 50 | 26 |
| SE | 50160 | ENSG00000132300 | PTCD3 | chr2 | + | 86136520 | 86136562 | 86134838 | 86134987 | 86136981 | 86137140 | 264 | 171 |
| SE | 50165 | ENSG00000025708 | TYMP | chr22 | - | 50527164 | 50527283 | 50526575 | 50526738 | 50529903 | 50529990 | 230 | 187 |
| SE | 50171 | ENSG00000025708 | TYMP | chr22 | - | 50529135 | 50529338 | 50527587 | 50527717 | 50529495 | 50529719 | 318 | 271 |
| SE | 50172 | ENSG00000025708 | TYMP | chr22 | - | 50529135 | 50529338 | 50527587 | 50527717 | 50529903 | 50529990 | 1 | 1 |
| SE | 50174 | ENSG00000025708 | TYMP | chr22 | - | 50529495 | 50529719 | 50529135 | 50529338 | 50529903 | 50529990 | 527 | 448 |
| SE | 50176 | ENSG00000025708 | TYMP | chr22 | - | 50529495 | 50529731 | 50529135 | 50529338 | 50529903 | 50530032 | 371 | 303 |
| SE | 50177 | ENSG00000025708 | TYMP | chr22 | - | 50529495 | 50529747 | 50529140 | 50529338 | 50529903 | 50530012 | 325 | 271 |
| SE | 50178 | ENSG00000025708 | TYMP | chr22 | - | 50529495 | 50529754 | 50527587 | 50527717 | 50529903 | 50530012 | 2 | 4 |
| SE | 50179 | ENSG00000025708 | TYMP | chr22 | - | 50529495 | 50529754 | 50529135 | 50529338 | 50529903 | 50530012 | 319 | 273 |
| SE | 50183 | ENSG00000185808 | PIGP | chr21 | - | 37069551 | 37069624 | 37067261 | 37067380 | 37072433 | 37072537 | 133 | 115 |
| SE | 50190 | ENSG00000156482 | RPL30 | chr8 | - | 98044826 | 98045088 | 98042644 | 98042775 | 98045346 | 98045399 | 3777 | 1665 |
| SE | 50191 | ENSG00000156482 | RPL30 | chr8 | - | 98044942 | 98045088 | 98024850 | 98025738 | 98045346 | 98045399 | 3731 | 1648 |
| SE | 50192 | ENSG00000156482 | RPL30 | chr8 | - | 98044942 | 98045088 | 98042644 | 98042775 | 98045346 | 98045399 | 10274 | 4900 |
| SE | 50196 | ENSG00000178467 | P4HTM | chr3 | + | 48990832 | 48990914 | 48989888 | 48990610 | 49001437 | 49001562 | 26 | 5 |
| SE | 50198 | ENSG00000178467 | P4HTM | chr3 | + | 49004860 | 49005867 | 49002499 | 49002596 | 49006686 | 49007115 | 1 | 0 |
| SE | 50199 | ENSG00000178467 | P4HTM | chr3 | + | 49006063 | 49006187 | 49004860 | 49005867 | 49006686 | 49007115 | 74 | 24 |
| SE | 50200 | ENSG00000171502 | COL24A1 | chr1 | - | 85734748 | 85734964 | 85729232 | 85730692 | 85737395 | 85737505 | 7 | 11 |
| SE | 50201 | ENSG00000171502 | COL24A1 | chr1 | - | 85736145 | 85736559 | 85734788 | 85734964 | 85737395 | 85737505 | 0 | 0 |
| SE | 50202 | ENSG00000171502 | COL24A1 | chr1 | - | 85960757 | 85960880 | 85911379 | 85911433 | 85961248 | 85961293 | 0 | 0 |
| SE | 50204 | ENSG00000148187 | MRRF | chr9 | + | 122265461 | 122265780 | 122264886 | 122264938 | 122270863 | 122271075 | 12 | 5 |
| SE | 50206 | ENSG00000148187 | MRRF | chr9 | + | 122265461 | 122265780 | 122264886 | 122264938 | 122280442 | 122280598 | 8 | 2 |
| SE | 50208 | ENSG00000148187 | MRRF | chr9 | + | 122270863 | 122271075 | 122265461 | 122265780 | 122280442 | 122280598 | 23 | 16 |
| SE | 50210 | ENSG00000148187 | MRRF | chr9 | + | 122270952 | 122271075 | 122265461 | 122265780 | 122280442 | 122280598 | 19 | 10 |
| SE | 50211 | ENSG00000148187 | MRRF | chr9 | + | 122280442 | 122280598 | 122270807 | 122271075 | 122285168 | 122285287 | 124 | 94 |
| SE | 50212 | ENSG00000148187 | MRRF | chr9 | + | 122285168 | 122285287 | 122280442 | 122280598 | 122291748 | 122291840 | 169 | 144 |
| SE | 50213 | ENSG00000148187 | MRRF | chr9 | + | 122285779 | 122285946 | 122285168 | 122285287 | 122286038 | 122286166 | 43 | 31 |
| SE | 50215 | ENSG00000148187 | MRRF | chr9 | + | 122286038 | 122286166 | 122285168 | 122285287 | 122313226 | 122313330 | 2 | 4 |
| SE | 50216 | ENSG00000148187 | MRRF | chr9 | + | 122291748 | 122291840 | 122285168 | 122285287 | 122313226 | 122313330 | 103 | 103 |
| SE | 50217 | ENSG00000148187 | MRRF | chr9 | + | 122291748 | 122291840 | 122286038 | 122286166 | 122313226 | 122313330 | 43 | 55 |
| SE | 50221 | ENSG00000134330 | IAH1 | chr2 | + | 9475986 | 9476039 | 9474550 | 9474647 | 9478253 | 9478370 | 2 | 0 |
| SE | 50224 | ENSG00000134330 | IAH1 | chr2 | + | 9478221 | 9478370 | 9474536 | 9474647 | 9481285 | 9481447 | 182 | 80 |
| SE | 50225 | ENSG00000134330 | IAH1 | chr2 | + | 9478221 | 9478370 | 9475673 | 9476039 | 9481285 | 9481447 | 247 | 114 |
| SE | 50226 | ENSG00000134330 | IAH1 | chr2 | + | 9478253 | 9478370 | 9474538 | 9474647 | 9481285 | 9481447 | 168 | 70 |
| SE | 50227 | ENSG00000134330 | IAH1 | chr2 | + | 9478253 | 9478370 | 9475986 | 9476039 | 9481285 | 9481447 | 166 | 70 |
| SE | 50228 | ENSG00000134330 | IAH1 | chr2 | + | 9484431 | 9484550 | 9481283 | 9481447 | 9488146 | 9488314 | 120 | 56 |
| SE | 50230 | ENSG00000176018 | LYSMD3 | chr5 | - | 90520363 | 90520461 | 90518290 | 90519425 | 90525034 | 90525300 | 3 | 1 |
| SE | 50232 | ENSG00000176018 | LYSMD3 | chr5 | - | 90526918 | 90527002 | 90525034 | 90525300 | 90529447 | 90529561 | 1 | 0 |
| SE | 50234 | ENSG00000130940 | CASZ1 | chr1 | - | 10655648 | 10655813 | 10654418 | 10654591 | 10658507 | 10658576 | 24 | 4 |
| SE | 50235 | ENSG00000130940 | CASZ1 | chr1 | - | 10656645 | 10656736 | 10654418 | 10654591 | 10658507 | 10658576 | 16 | 13 |
| SE | 50236 | ENSG00000130940 | CASZ1 | chr1 | - | 10656645 | 10656736 | 10654418 | 10654591 | 10659701 | 10660536 | 3 | 0 |
| SE | 50237 | ENSG00000130940 | CASZ1 | chr1 | - | 10656645 | 10656736 | 10655648 | 10655813 | 10658507 | 10658576 | 45 | 16 |
| SE | 50238 | ENSG00000130940 | CASZ1 | chr1 | - | 10658507 | 10658576 | 10654418 | 10654591 | 10659701 | 10660536 | 1 | 0 |
| SE | 50239 | ENSG00000130940 | CASZ1 | chr1 | - | 10658507 | 10658576 | 10656645 | 10656736 | 10659701 | 10660536 | 29 | 15 |
| SE | 50240 | ENSG00000130940 | CASZ1 | chr1 | - | 10659701 | 10660536 | 10654418 | 10654591 | 10665082 | 10665571 | 14 | 3 |
| SE | 50242 | ENSG00000001630 | CYP51A1 | chr7 | - | 92131773 | 92131872 | 92128879 | 92129056 | 92134665 | 92134803 | 25 | 10 |
| SE | 50244 | ENSG00000126012 | KDM5C | chrX | - | 53195230 | 53195410 | 53194930 | 53195068 | 53195915 | 53196054 | 577 | 262 |
| SE | 50246 | ENSG00000126012 | KDM5C | chrX | - | 53198763 | 53198888 | 53198489 | 53198637 | 53198976 | 53199158 | 94 | 53 |
| SE | 50247 | ENSG00000126012 | KDM5C | chrX | - | 53201549 | 53201744 | 53198976 | 53199158 | 53201853 | 53201973 | 200 | 85 |
| SE | 50249 | ENSG00000126012 | KDM5C | chrX | - | 53215794 | 53215976 | 53214688 | 53214847 | 53216073 | 53216197 | 421 | 237 |
| SE | 50251 | ENSG00000126012 | KDM5C | chrX | - | 53218275 | 53218398 | 53217795 | 53217966 | 53224739 | 53225176 | 255 | 165 |
| SE | 50252 | ENSG00000126012 | KDM5C | chrX | - | 53218563 | 53218712 | 53218314 | 53218398 | 53220838 | 53220916 | 0 | 0 |
| SE | 50253 | ENSG00000126012 | KDM5C | chrX | - | 53220838 | 53220916 | 53217795 | 53217966 | 53224739 | 53225176 | 91 | 29 |
| SE | 50254 | ENSG00000126012 | KDM5C | chrX | - | 53220838 | 53220916 | 53218275 | 53218398 | 53224739 | 53225176 | 190 | 82 |
| SE | 50264 | ENSG00000143167 | GPA33 | chr1 | - | 167063581 | 167063737 | 167055729 | 167055849 | 167068921 | 167069138 | 56 | 52 |
| SE | 50265 | ENSG00000143167 | GPA33 | chr1 | - | 167068921 | 167069138 | 167063581 | 167063737 | 167073384 | 167073539 | 45 | 45 |
| SE | 50266 | ENSG00000143167 | GPA33 | chr1 | - | 167073384 | 167073539 | 167068921 | 167069138 | 167090244 | 167090377 | 37 | 29 |
| SE | 50267 | ENSG00000143167 | GPA33 | chr1 | - | 167082244 | 167082311 | 167073384 | 167073539 | 167090244 | 167090326 | 4 | 0 |
| SE | 50268 | ENSG00000275557 | NA | chr1 | + | 149611654 | 149611848 | 149608461 | 149608516 | 149612363 | 149612402 | 0 | 0 |
| SE | 50270 | ENSG00000134909 | ARHGAP32 | chr11 | - | 128986523 | 128986668 | 128985878 | 128986085 | 128988022 | 128988125 | 22 | 10 |
| SE | 50271 | ENSG00000134909 | ARHGAP32 | chr11 | - | 129040927 | 129041009 | 128998318 | 128998468 | 129062279 | 129062357 | 27 | 26 |
| SE | 50273 | ENSG00000134909 | ARHGAP32 | chr11 | - | 129093620 | 129093707 | 129066730 | 129066868 | 129123445 | 129123530 | 17 | 34 |
| SE | 50277 | ENSG00000131115 | ZNF227 | chr19 | + | 44228445 | 44228572 | 44217790 | 44217852 | 44229732 | 44229792 | 31 | 26 |
| SE | 50278 | ENSG00000131115 | ZNF227 | chr19 | + | 44228486 | 44228572 | 44217790 | 44217852 | 44229732 | 44229816 | 13 | 12 |
| SE | 50279 | ENSG00000131115 | ZNF227 | chr19 | + | 44229732 | 44229816 | 44228445 | 44228572 | 44234701 | 44234835 | 25 | 33 |
| SE | 50280 | ENSG00000131115 | ZNF227 | chr19 | + | 44229732 | 44229816 | 44228445 | 44228633 | 44234701 | 44234835 | 8 | 16 |
| SE | 50281 | ENSG00000179632 | MAF1 | chr8 | + | 144105868 | 144105997 | 144105639 | 144105766 | 144106342 | 144106464 | 224 | 87 |
| SE | 50283 | ENSG00000179632 | MAF1 | chr8 | + | 144106554 | 144106674 | 144106342 | 144106464 | 144106834 | 144106963 | 546 | 232 |
| SE | 50284 | ENSG00000179632 | MAF1 | chr8 | + | 144106834 | 144106963 | 144106342 | 144106674 | 144107087 | 144107217 | 548 | 241 |
| SE | 50285 | ENSG00000063978 | RNF4 | chr4 | + | 2470986 | 2471109 | 2469138 | 2469229 | 2490336 | 2490502 | 3 | 0 |
| SE | 50287 | ENSG00000063978 | RNF4 | chr4 | + | 2470986 | 2471109 | 2469180 | 2469258 | 2497006 | 2497103 | 5 | 1 |
| SE | 50288 | ENSG00000063978 | RNF4 | chr4 | + | 2490336 | 2490411 | 2469105 | 2469258 | 2497006 | 2497121 | 67 | 18 |
| SE | 50289 | ENSG00000063978 | RNF4 | chr4 | + | 2490336 | 2490502 | 2469067 | 2469258 | 2497006 | 2497012 | 270 | 130 |
| SE | 50291 | ENSG00000063978 | RNF4 | chr4 | + | 2494563 | 2494752 | 2490336 | 2490502 | 2497006 | 2497079 | 3 | 0 |
| SE | 50292 | ENSG00000063978 | RNF4 | chr4 | + | 2494566 | 2494752 | 2490336 | 2490502 | 2497006 | 2497079 | 2 | 0 |
| SE | 50293 | ENSG00000063978 | RNF4 | chr4 | + | 2497006 | 2497121 | 2490336 | 2490502 | 2500658 | 2500698 | 317 | 191 |
| SE | 50298 | ENSG00000100591 | AHSA1 | chr14 | + | 77462159 | 77462242 | 77459615 | 77459806 | 77464597 | 77464686 | 62 | 38 |
| SE | 50299 | ENSG00000100591 | AHSA1 | chr14 | + | 77462159 | 77462242 | 77459615 | 77459806 | 77468082 | 77468184 | 57 | 34 |
| SE | 50302 | ENSG00000100591 | AHSA1 | chr14 | + | 77462641 | 77462759 | 77462159 | 77462242 | 77468082 | 77468184 | 105 | 71 |
| SE | 50303 | ENSG00000100591 | AHSA1 | chr14 | + | 77464597 | 77464686 | 77459615 | 77459806 | 77468082 | 77468184 | 2 | 2 |
| SE | 50305 | ENSG00000100591 | AHSA1 | chr14 | + | 77464597 | 77464686 | 77462159 | 77462242 | 77468082 | 77468184 | 7 | 3 |
| SE | 50306 | ENSG00000100591 | AHSA1 | chr14 | + | 77464597 | 77464686 | 77462641 | 77462759 | 77465538 | 77465667 | 373 | 234 |
| SE | 50307 | ENSG00000100591 | AHSA1 | chr14 | + | 77464597 | 77464686 | 77462641 | 77462759 | 77468082 | 77468184 | 79 | 52 |
| SE | 50308 | ENSG00000100591 | AHSA1 | chr14 | + | 77465538 | 77465667 | 77462159 | 77462242 | 77468082 | 77468184 | 180 | 102 |
| SE | 50309 | ENSG00000100591 | AHSA1 | chr14 | + | 77465538 | 77465667 | 77462641 | 77462759 | 77468082 | 77468184 | 180 | 102 |
| SE | 50311 | ENSG00000100591 | AHSA1 | chr14 | + | 77465538 | 77465667 | 77464582 | 77464686 | 77468082 | 77468184 | 475 | 285 |
| SE | 50313 | ENSG00000100591 | AHSA1 | chr14 | + | 77468082 | 77468184 | 77465538 | 77465667 | 77469076 | 77469470 | 119 | 76 |
| SE | 50314 | ENSG00000100591 | AHSA1 | chr14 | + | 77468456 | 77468508 | 77468082 | 77468184 | 77469076 | 77469247 | 312 | 198 |
| SE | 50315 | ENSG00000148335 | NTMT1 | chr9 | + | 129632649 | 129632801 | 129626174 | 129626295 | 129634053 | 129634306 | 7 | 5 |
| SE | 50319 | ENSG00000236304 | NA | chr11 | + | 76659607 | 76659694 | 76657055 | 76657156 | 76662628 | 76663866 | 133 | 288 |
| SE | 50322 | ENSG00000060339 | CCAR1 | chr10 | + | 68737844 | 68737889 | 68736875 | 68737048 | 68742375 | 68742408 | 21 | 16 |
| SE | 50323 | ENSG00000060339 | CCAR1 | chr10 | + | 68740628 | 68740661 | 68736875 | 68737048 | 68742375 | 68742408 | 25 | 19 |
| SE | 50324 | ENSG00000060339 | CCAR1 | chr10 | + | 68740628 | 68740661 | 68736875 | 68737048 | 68747160 | 68747275 | 6 | 2 |
| SE | 50326 | ENSG00000060339 | CCAR1 | chr10 | + | 68742375 | 68742569 | 68736875 | 68737048 | 68747160 | 68747275 | 124 | 135 |
| SE | 50327 | ENSG00000060339 | CCAR1 | chr10 | + | 68742375 | 68742569 | 68740628 | 68740661 | 68747160 | 68747275 | 193 | 185 |
| SE | 50328 | ENSG00000060339 | CCAR1 | chr10 | + | 68742427 | 68742569 | 68736875 | 68737048 | 68747160 | 68747275 | 122 | 131 |
| SE | 50329 | ENSG00000060339 | CCAR1 | chr10 | + | 68754713 | 68754827 | 68753851 | 68754077 | 68755369 | 68755536 | 220 | 148 |
| SE | 50331 | ENSG00000060339 | CCAR1 | chr10 | + | 68756525 | 68756632 | 68755475 | 68755536 | 68757293 | 68757377 | 2 | 0 |
| SE | 50332 | ENSG00000060339 | CCAR1 | chr10 | + | 68761006 | 68761192 | 68757293 | 68757377 | 68765887 | 68765927 | 172 | 232 |
| SE | 50333 | ENSG00000060339 | CCAR1 | chr10 | + | 68765887 | 68766079 | 68757293 | 68757377 | 68771205 | 68771301 | 157 | 201 |
| SE | 50334 | ENSG00000060339 | CCAR1 | chr10 | + | 68771205 | 68771445 | 68765887 | 68766079 | 68772987 | 68773099 | 229 | 268 |
| SE | 50335 | ENSG00000060339 | CCAR1 | chr10 | + | 68786135 | 68786218 | 68772987 | 68773099 | 68786545 | 68786692 | 146 | 156 |
| SE | 50336 | ENSG00000060339 | CCAR1 | chr10 | + | 68786545 | 68786692 | 68772987 | 68773099 | 68787926 | 68788047 | 269 | 245 |
| SE | 50337 | ENSG00000060339 | CCAR1 | chr10 | + | 68789709 | 68789915 | 68788142 | 68788328 | 68791206 | 68791266 | 157 | 183 |
| SE | 50338 | ENSG00000227124 | ZNF717 | chr3 | - | 75741275 | 75741368 | 75735867 | 75739345 | 75741609 | 75741736 | 41 | 13 |
| SE | 50342 | ENSG00000227124 | ZNF717 | chr3 | - | 75784914 | 75785002 | 75783305 | 75783364 | 75785383 | 75785583 | 1 | 0 |
| SE | 50343 | ENSG00000259343 | TMC3-AS1 | chr15 | + | 81332344 | 81332541 | 81324337 | 81324561 | 81341561 | 81341708 | 6 | 1 |
| SE | 50346 | ENSG00000259343 | TMC3-AS1 | chr15 | + | 81358335 | 81358503 | 81341561 | 81341708 | 81361918 | 81362036 | 9 | 3 |
| SE | 50354 | ENSG00000003393 | ALS2 | chr2 | - | 201716589 | 201716687 | 201715671 | 201715839 | 201718076 | 201718210 | 0 | 0 |
| SE | 50357 | ENSG00000003393 | ALS2 | chr2 | - | 201719985 | 201720128 | 201718076 | 201718210 | 201723042 | 201723120 | 4 | 1 |
| SE | 50362 | ENSG00000003393 | ALS2 | chr2 | - | 201738978 | 201739068 | 201738669 | 201738735 | 201741673 | 201741854 | 1 | 0 |
| SE | 50368 | ENSG00000003393 | ALS2 | chr2 | - | 201744257 | 201744429 | 201740609 | 201741854 | 201746565 | 201746748 | 30 | 13 |
| SE | 50369 | ENSG00000003393 | ALS2 | chr2 | - | 201754502 | 201754623 | 201753145 | 201753242 | 201757401 | 201757759 | 25 | 10 |
| SE | 50370 | ENSG00000003393 | ALS2 | chr2 | - | 201754502 | 201754671 | 201753145 | 201753242 | 201757401 | 201757759 | 47 | 28 |
| SE | 50376 | ENSG00000058866 | DGKG | chr3 | - | 186161602 | 186161663 | 186147200 | 186150188 | 186164897 | 186164967 | 16 | 16 |
| SE | 50377 | ENSG00000058866 | DGKG | chr3 | - | 186164897 | 186165018 | 186158182 | 186161663 | 186188201 | 186188379 | 23 | 36 |
| SE | 50379 | ENSG00000058866 | DGKG | chr3 | - | 186251758 | 186251919 | 186242503 | 186242568 | 186253092 | 186253182 | 31 | 20 |
| SE | 50382 | ENSG00000058866 | DGKG | chr3 | - | 186261698 | 186261778 | 186257853 | 186257939 | 186267684 | 186267777 | 2 | 0 |
| SE | 50383 | ENSG00000058866 | DGKG | chr3 | - | 186265246 | 186265306 | 186257853 | 186257939 | 186267684 | 186267777 | 1 | 1 |
| SE | 50389 | ENSG00000058866 | DGKG | chr3 | - | 186288709 | 186288880 | 186280669 | 186280744 | 186297420 | 186297483 | 19 | 19 |
| SE | 50391 | ENSG00000181847 | TIGIT | chr3 | + | 114293306 | 114294122 | 114291101 | 114291296 | 114299596 | 114299703 | 0 | 0 |
| SE | 50392 | ENSG00000181847 | TIGIT | chr3 | + | 114293999 | 114294122 | 114291101 | 114291296 | 114299596 | 114299703 | 4 | 0 |
| SE | 50393 | ENSG00000181847 | TIGIT | chr3 | + | 114295544 | 114295874 | 114293306 | 114294122 | 114299596 | 114299703 | 132 | 44 |
| SE | 50394 | ENSG00000185219 | ZNF445 | chr3 | - | 44451313 | 44451482 | 44450867 | 44450962 | 44455120 | 44455696 | 16 | 4 |
| SE | 50395 | ENSG00000185219 | ZNF445 | chr3 | - | 44458243 | 44458364 | 44455120 | 44455696 | 44477590 | 44477670 | 10 | 7 |
| SE | 50398 | ENSG00000272758 | WDR5B-DT | chr3 | + | 122425606 | 122425658 | 122416224 | 122416250 | 122432466 | 122432521 | 0 | 0 |
| SE | 50402 | ENSG00000109536 | FRG1 | chr4 | + | 189952161 | 189952287 | 189943201 | 189943272 | 189955036 | 189955150 | 103 | 77 |
| SE | 50403 | ENSG00000109536 | FRG1 | chr4 | + | 189953067 | 189953125 | 189952161 | 189952287 | 189955036 | 189955150 | 176 | 131 |
| SE | 50406 | ENSG00000109536 | FRG1 | chr4 | + | 189961428 | 189961550 | 189960747 | 189960839 | 189961821 | 189961932 | 4 | 0 |
| SE | 50408 | ENSG00000164306 | PRIMPOL | chr4 | + | 184657084 | 184657320 | 184652022 | 184652100 | 184659339 | 184659437 | 30 | 29 |
| SE | 50411 | ENSG00000164306 | PRIMPOL | chr4 | + | 184665916 | 184666064 | 184659339 | 184659437 | 184672172 | 184672268 | 51 | 59 |
| SE | 50412 | ENSG00000164306 | PRIMPOL | chr4 | + | 184665916 | 184666064 | 184661773 | 184661903 | 184672172 | 184672268 | 68 | 92 |
| SE | 50413 | ENSG00000164306 | PRIMPOL | chr4 | + | 184678231 | 184678394 | 184672172 | 184672460 | 184682247 | 184682336 | 41 | 39 |
| SE | 50415 | ENSG00000164306 | PRIMPOL | chr4 | + | 184682247 | 184682336 | 184672172 | 184672460 | 184685408 | 184685498 | 4 | 18 |
| SE | 50418 | ENSG00000164306 | PRIMPOL | chr4 | + | 184682247 | 184682336 | 184678231 | 184678394 | 184685408 | 184685498 | 26 | 40 |
| SE | 50419 | ENSG00000164306 | PRIMPOL | chr4 | + | 184682247 | 184682336 | 184678231 | 184678394 | 184685411 | 184685498 | 24 | 23 |
| SE | 50420 | ENSG00000164306 | PRIMPOL | chr4 | + | 184685408 | 184685498 | 184672172 | 184672460 | 184685575 | 184685684 | 30 | 26 |
| SE | 50421 | ENSG00000164306 | PRIMPOL | chr4 | + | 184685408 | 184685498 | 184678276 | 184678394 | 184691498 | 184691581 | 1 | 0 |
| SE | 50423 | ENSG00000164306 | PRIMPOL | chr4 | + | 184685411 | 184685498 | 184672172 | 184672460 | 184685575 | 184685684 | 30 | 26 |
| SE | 50425 | ENSG00000164306 | PRIMPOL | chr4 | + | 184685411 | 184685498 | 184682247 | 184682336 | 184685575 | 184685684 | 38 | 33 |
| SE | 50426 | ENSG00000164306 | PRIMPOL | chr4 | + | 184685575 | 184685684 | 184685408 | 184685498 | 184691498 | 184691581 | 53 | 61 |
| SE | 50427 | ENSG00000114933 | INO80D | chr2 | - | 206007283 | 206007441 | 205993720 | 206005533 | 206009576 | 206009794 | 72 | 44 |
| SE | 50428 | ENSG00000114933 | INO80D | chr2 | - | 206062798 | 206062954 | 206056197 | 206056943 | 206063150 | 206063244 | 81 | 59 |
| SE | 50429 | ENSG00000114933 | INO80D | chr2 | - | 206062798 | 206062954 | 206056197 | 206056943 | 206085900 | 206086303 | 77 | 55 |
| SE | 50430 | ENSG00000114933 | INO80D | chr2 | - | 206062798 | 206063045 | 206056197 | 206056943 | 206063150 | 206063244 | 111 | 82 |
| SE | 50431 | ENSG00000114933 | INO80D | chr2 | - | 206062798 | 206063045 | 206056197 | 206056943 | 206085900 | 206086174 | 77 | 57 |
| SE | 50432 | ENSG00000114933 | INO80D | chr2 | - | 206063150 | 206063244 | 206056197 | 206056943 | 206085900 | 206086174 | 11 | 6 |
| SE | 50433 | ENSG00000114933 | INO80D | chr2 | - | 206063150 | 206063244 | 206062798 | 206062954 | 206085900 | 206086174 | 13 | 10 |
| SE | 50434 | ENSG00000114933 | INO80D | chr2 | - | 206063150 | 206063244 | 206062798 | 206063045 | 206085900 | 206086174 | 43 | 27 |
| SE | 50435 | ENSG00000114933 | INO80D | chr2 | - | 206063568 | 206063684 | 206063150 | 206063244 | 206085900 | 206086303 | 2 | 0 |
| SE | 50437 | ENSG00000162639 | HENMT1 | chr1 | - | 108654715 | 108654850 | 108651029 | 108651209 | 108657450 | 108657579 | 137 | 59 |
| SE | 50438 | ENSG00000162639 | HENMT1 | chr1 | - | 108655585 | 108655698 | 108654715 | 108654850 | 108657450 | 108657579 | 159 | 75 |
| SE | 50439 | ENSG00000162639 | HENMT1 | chr1 | - | 108657450 | 108657579 | 108651029 | 108651209 | 108659863 | 108659962 | 82 | 49 |
| SE | 50440 | ENSG00000162639 | HENMT1 | chr1 | - | 108659863 | 108659962 | 108657450 | 108657579 | 108660962 | 108661055 | 51 | 41 |
| SE | 50441 | ENSG00000162639 | HENMT1 | chr1 | - | 108659863 | 108659962 | 108657450 | 108657579 | 108661183 | 108661526 | 84 | 42 |
| SE | 50445 | ENSG00000121594 | CD80 | chr3 | - | 119527732 | 119527841 | 119524292 | 119525749 | 119529841 | 119529937 | 4 | 3 |
| SE | 50446 | ENSG00000121594 | CD80 | chr3 | - | 119529841 | 119529937 | 119527463 | 119527841 | 119537136 | 119537418 | 5 | 3 |
| SE | 50447 | ENSG00000120889 | TNFRSF10B | chr8 | - | 23024187 | 23024260 | 23020132 | 23022984 | 23027132 | 23027288 | 212 | 185 |
| SE | 50448 | ENSG00000120889 | TNFRSF10B | chr8 | - | 23027721 | 23027753 | 23027132 | 23027288 | 23028330 | 23028441 | 176 | 149 |
| SE | 50449 | ENSG00000120889 | TNFRSF10B | chr8 | - | 23043137 | 23043243 | 23030758 | 23030872 | 23068750 | 23069021 | 123 | 51 |
| SE | 50450 | ENSG00000171574 | ZNF584 | chr19 | + | 58409940 | 58410091 | 58408648 | 58409165 | 58415523 | 58415646 | 5 | 1 |
| SE | 50451 | ENSG00000171574 | ZNF584 | chr19 | + | 58409964 | 58410091 | 58408776 | 58409165 | 58415523 | 58415646 | 5 | 2 |
| SE | 50452 | ENSG00000171574 | ZNF584 | chr19 | + | 58410005 | 58410091 | 58408687 | 58409165 | 58415523 | 58415646 | 4 | 0 |
| SE | 50453 | ENSG00000171574 | ZNF584 | chr19 | + | 58415792 | 58415940 | 58415523 | 58415646 | 58416810 | 58417011 | 1 | 0 |

| | | | | | | | | | | | | | | |
|---|---|---|---|---|---|---|---|---|---|---|---|---|---|---|
| SE | 50458 | ENSG00000161618 | ALDH16A1 | chr19 | + | 49458485 | 49458588 | 49453224 | 49453421 | 49458959 | 49459086 | 56 | 35 | [illegible] |
| SE | 50459 | ENSG00000161618 | ALDH16A1 | chr19 | + | 49458485 | 49458588 | 49453224 | 49453421 | 49459669 | 49459730 | 14 | 4 | [illegible] |
| SE | 50460 | ENSG00000161618 | ALDH16A1 | chr19 | + | 49458959 | 49459086 | 49453224 | 49453421 | 49459669 | 49459848 | 40 | 18 | [illegible] |
| SE | 50461 | ENSG00000161618 | ALDH16A1 | chr19 | + | 49458959 | 49459086 | 49458485 | 49458588 | 49459669 | 49459848 | 86 | 49 | [illegible] |
| SE | 50465 | ENSG00000165671 | NSD1 | chr5 | + | 177135940 | 177136030 | 177135086 | 177135200 | 177191883 | 177192019 | 201 | 198 | [illegible] |
| SE | 50468 | ENSG00000165671 | NSD1 | chr5 | + | 177191883 | 177192019 | 177135086 | 177136030 | 177204119 | 177204236 | 507 | 520 | [illegible] |
| SE | 50469 | ENSG00000165671 | NSD1 | chr5 | + | 177204119 | 177204292 | 177191883 | 177192019 | 177209635 | 177209672 | 723 | 698 | [illegible] |
| SE | 50474 | ENSG00000165671 | NSD1 | chr5 | + | 177259988 | 177260168 | 177256950 | 177257151 | 177267561 | 177267718 | 143 | 99 | [illegible] |
| SE | 50475 | ENSG00000165671 | NSD1 | chr5 | + | 177273671 | 177273784 | 177269601 | 177269807 | 177280564 | 177280583 | 199 | 149 | [illegible] |
| SE | 50476 | ENSG00000165671 | NSD1 | chr5 | + | 177280564 | 177280834 | 177273671 | 177273784 | 177282464 | 177282581 | 245 | 204 | [illegible] |
| SE | 50479 | ENSG00000165671 | NSD1 | chr5 | + | 177283786 | 177283928 | 177280564 | 177280834 | 177291953 | 177292158 | 3 | 1 | [illegible] |
| SE | 50480 | ENSG00000165671 | NSD1 | chr5 | + | 177288818 | 177288925 | 177280564 | 177280834 | 177291953 | 177292158 | 110 | 97 | [illegible] |
| SE | 50481 | ENSG00000165671 | NSD1 | chr5 | + | 177288818 | 177288925 | 177283786 | 177283928 | 177291953 | 177292158 | 311 | 250 | [illegible] |
| SE | 50482 | ENSG00000165671 | NSD1 | chr5 | + | 177291953 | 177292158 | 177288818 | 177288925 | 177293831 | 177295762 | 262 | 203 | [illegible] |
| SE | 50486 | ENSG00000135069 | PSAT1 | chr9 | + | 78300601 | 78300662 | 78297124 | 78297270 | 78301953 | 78302023 | 3 | 13 | [illegible] |
| SE | 50487 | ENSG00000135069 | PSAT1 | chr9 | + | 78308413 | 78308583 | 78306313 | 78306486 | 78317675 | 78317804 | 18 | 6 | [illegible] |
| SE | 50488 | ENSG00000135069 | PSAT1 | chr9 | + | 78328050 | 78328188 | 78317675 | 78317804 | 78328980 | 78330093 | 32 | 6 | [illegible] |
| SE | 50492 | ENSG00000134326 | CMPK2 | chr2 | - | 6851449 | 6851683 | 6848315 | 6849973 | 6861183 | 6861385 | 63 | 138 | [illegible] |
| SE | 50493 | ENSG00000134326 | CMPK2 | chr2 | - | 6863463 | 6863578 | 6861183 | 6861385 | 6865021 | 6865712 | 39 | 59 | [illegible] |
| SE | 50494 | ENSG00000134326 | CMPK2 | chr2 | - | 6863463 | 6863578 | 6861183 | 6861385 | 6866425 | 6866635 | 37 | 56 | [illegible] |
| SE | 50495 | ENSG00000134326 | CMPK2 | chr2 | - | 6864331 | 6864443 | 6863463 | 6863578 | 6866425 | 6866635 | 0 | 0 | [illegible] |
| SE | 50503 | ENSG00000240401 | NA | chr2 | + | 55329421 | 55329713 | 55308831 | 55308927 | 55339502 | 55339941 | 1 | 0 | [illegible] |
| SE | 50524 | ENSG00000115685 | PPP1R7 | chr2 | + | 241158465 | 241158549 | 241157806 | 241157862 | 241159212 | 241159343 | 173 | 93 | [illegible] |
| SE | 50525 | ENSG00000115685 | PPP1R7 | chr2 | + | 241158465 | 241158549 | 241157806 | 241157862 | 241160331 | 241160433 | 1 | 1 | [illegible] |
| SE | 50527 | ENSG00000115685 | PPP1R7 | chr2 | + | 241158483 | 241158549 | 241157806 | 241157862 | 241159212 | 241159303 | 388 | 255 | [illegible] |
| SE | 50528 | ENSG00000115685 | PPP1R7 | chr2 | + | 241158483 | 241158549 | 241157806 | 241157862 | 241160331 | 241160494 | 35 | 35 | [illegible] |
| SE | 50529 | ENSG00000115685 | PPP1R7 | chr2 | + | 241159212 | 241159343 | 241157806 | 241157862 | 241160331 | 241160421 | 364 | 239 | [illegible] |
| SE | 50531 | ENSG00000115685 | PPP1R7 | chr2 | + | 241160331 | 241160494 | 241158483 | 241158549 | 241163284 | 241163401 | 367 | 252 | [illegible] |
| SE | 50532 | ENSG00000115685 | PPP1R7 | chr2 | + | 241163284 | 241163401 | 241160331 | 241160494 | 241166336 | 241166439 | 517 | 344 | [illegible] |
| SE | 50533 | ENSG00000115685 | PPP1R7 | chr2 | + | 241169780 | 241169867 | 241166336 | 241166441 | 241182646 | 241182740 | 176 | 116 | [illegible] |
| SE | 50537 | ENSG00000173436 | MICOS10 | chr1 | + | 19617257 | 19617336 | 19597000 | 19597109 | 19622099 | 19622147 | 1 | 3 | [illegible] |
| SE | 50539 | ENSG00000173436 | MICOS10 | chr1 | + | 19623473 | 19623583 | 19596976 | 19597109 | 19626386 | 19626606 | 190 | 114 | [illegible] |
| SE | 50542 | ENSG00000135617 | PRADC1 | chr2 | - | 73229460 | 73229570 | 73228794 | 73228962 | 73230112 | 73230213 | 29 | 6 | [illegible] |
| SE | 50543 | ENSG00000120370 | GORAB | chr1 | + | 170533463 | 170533651 | 170532150 | 170532284 | 170539209 | 170539567 | 4 | 16 | [illegible] |
| SE | 50544 | ENSG00000120370 | GORAB | chr1 | + | 170536309 | 170536421 | 170533463 | 170533651 | 170539209 | 170539567 | 19 | 12 | [illegible] |
| SE | 50546 | ENSG00000001461 | NIPAL3 | chr1 | + | 24433103 | 24433213 | 24419290 | 24419640 | 24440171 | 24440240 | 2 | 0 | [illegible] |
| SE | 50547 | ENSG00000001461 | NIPAL3 | chr1 | + | 24449480 | 24449626 | 24445184 | 24445244 | 24453407 | 24453504 | 37 | 33 | [illegible] |
| SE | 50548 | ENSG00000001461 | NIPAL3 | chr1 | + | 24453407 | 24453504 | 24449480 | 24449626 | 24456137 | 24456273 | 55 | 34 | [illegible] |
| SE | 50549 | ENSG00000001461 | NIPAL3 | chr1 | + | 24456137 | 24456273 | 24453407 | 24453504 | 24458887 | 24458976 | 54 | 33 | [illegible] |
| SE | 50553 | ENSG00000204131 | NHSL2 | chrX | + | 72132078 | 72132234 | 72079775 | 72080266 | 72138440 | 72140771 | 0 | 0 | [illegible] |
| SE | 50554 | ENSG00000204131 | NHSL2 | chrX | + | 72134090 | 72134218 | 72132074 | 72132234 | 72138440 | 72140771 | 13 | 11 | [illegible] |
| SE | 50555 | ENSG00000204131 | NHSL2 | chrX | + | 72134508 | 72134704 | 72134090 | 72134218 | 72138440 | 72141257 | 36 | 10 | [illegible] |
| SE | 50557 | ENSG00000204131 | NHSL2 | chrX | + | 72138440 | 72140771 | 72132074 | 72132234 | 72142231 | 72142364 | 131 | 68 | [illegible] |
| SE | 50558 | ENSG00000204131 | NHSL2 | chrX | + | 72142231 | 72142364 | 72138440 | 72140771 | 72143252 | 72143574 | 225 | 141 | [illegible] |
| SE | 50559 | ENSG00000137575 | SDCBP | chr8 | + | 58560283 | 58561795 | 58553253 | 58553303 | 58565018 | 58565084 | 11 | 5 | [illegible] |
| SE | 50563 | ENSG00000137575 | SDCBP | chr8 | + | 58575899 | 58576061 | 58572204 | 58572314 | 58578032 | 58578070 | 1132 | 805 | [illegible] |
| SE | 50564 | ENSG00000137575 | SDCBP | chr8 | + | 58575902 | 58576061 | 58572204 | 58572314 | 58578032 | 58578208 | 813 | 543 | [illegible] |
| SE | 50566 | ENSG00000139636 | LMBR1L | chr12 | - | 49097943 | 49098105 | 49097135 | 49097739 | 49100387 | 49100454 | 60 | 42 | [illegible] |
| SE | 50567 | ENSG00000139636 | LMBR1L | chr12 | - | 49101128 | 49101323 | 49100387 | 49100454 | 49101471 | 49101658 | 2 | 3 | [illegible] |
| SE | 50568 | ENSG00000139636 | LMBR1L | chr12 | - | 49101471 | 49101549 | 49100387 | 49101323 | 49102119 | 49102176 | 7 | 10 | [illegible] |
| SE | 50569 | ENSG00000139636 | LMBR1L | chr12 | - | 49101471 | 49101549 | 49101102 | 49101263 | 49102119 | 49102176 | 3 | 8 | [illegible] |
| SE | 50571 | ENSG00000139636 | LMBR1L | chr12 | - | 49104447 | 49104551 | 49103686 | 49103813 | 49104745 | 49104885 | 37 | 36 | [illegible] |
| SE | 50575 | ENSG00000139636 | LMBR1L | chr12 | - | 49104745 | 49104885 | 49104447 | 49104551 | 49105923 | 49105957 | 47 | 60 | [illegible] |
| SE | 50576 | ENSG00000139636 | LMBR1L | chr12 | - | 49105923 | 49105957 | 49103686 | 49103813 | 49106960 | 49107045 | 10 | 7 | [illegible] |
| SE | 50585 | ENSG00000139636 | LMBR1L | chr12 | - | 49109726 | 49109881 | 49106960 | 49107045 | 49110483 | 49110877 | 2 | 0 | [illegible] |
| SE | 50586 | ENSG00000154447 | SH3RF1 | chr4 | - | 169120818 | 169120989 | 169117522 | 169117782 | 169122099 | 169122199 | 7 | 22 | [illegible] |
| SE | 50595 | ENSG00000174130 | TLR6 | chr4 | - | 38843638 | 38843791 | 38829199 | 38829537 | 38856760 | 38856817 | 0 | 2 | [illegible] |
| SE | 50596 | ENSG00000085871 | MGST2 | chr4 | + | 139678542 | 139678642 | 139665767 | 139666077 | 139687659 | 139687740 | 79 | 56 | [illegible] |
| SE | 50598 | ENSG00000085871 | MGST2 | chr4 | + | 139678542 | 139678642 | 139665767 | 139666077 | 139703454 | 139703536 | 79 | 61 | [illegible] |
| SE | 50599 | ENSG00000085871 | MGST2 | chr4 | + | 139687659 | 139687740 | 139666027 | 139666077 | 139695196 | 139695267 | 2 | 0 | [illegible] |
| SE | 50604 | ENSG00000134574 | DDB2 | chr11 | + | 47232813 | 47232959 | 47216857 | 47217049 | 47234572 | 47234653 | 78 | 37 | [illegible] |
| SE | 50605 | ENSG00000134574 | DDB2 | chr11 | + | 47233226 | 47233474 | 47232813 | 47232959 | 47234572 | 47234672 | 0 | 0 | [illegible] |
| SE | 50610 | ENSG00000133275 | CSNK1G2 | chr19 | + | 1978601 | 1978750 | 1978304 | 1978511 | 1978858 | 1979093 | 414 | 180 | [illegible] |
| SE | 50611 | ENSG00000133275 | CSNK1G2 | chr19 | + | 1978858 | 1979093 | 1978304 | 1978511 | 1979162 | 1979248 | 223 | 122 | [illegible] |
| SE | 50613 | ENSG00000143033 | MTF2 | chr1 | + | 93110229 | 93110246 | 93079234 | 93079531 | 93110326 | 93110428 | 8 | 14 | [illegible] |
| SE | 50614 | ENSG00000143033 | MTF2 | chr1 | + | 93115469 | 93115618 | 93114987 | 93115088 | 93118344 | 93118440 | 218 | 200 | [illegible] |
| SE | 50616 | ENSG00000143033 | MTF2 | chr1 | + | 93127231 | 93127299 | 93120548 | 93120672 | 93133702 | 93133808 | 70 | 49 | [illegible] |
| SE | 50617 | ENSG00000143033 | MTF2 | chr1 | + | 93129277 | 93129448 | 93127231 | 93127299 | 93133702 | 93133808 | 102 | 70 | [illegible] |
| SE | 50618 | ENSG00000143033 | MTF2 | chr1 | + | 93133927 | 93133980 | 93133702 | 93133808 | 93134090 | 93134195 | 112 | 104 | [illegible] |
| SE | 50620 | ENSG00000162971 | TYW5 | chr2 | - | 199936404 | 199936492 | 199935930 | 199936047 | 199938932 | 199939070 | 29 | 15 | [illegible] |
| SE | 50621 | ENSG00000162971 | TYW5 | chr2 | - | 199940088 | 199940133 | 199938932 | 199939070 | 199948336 | 199948472 | 16 | 21 | [illegible] |
| SE | 50622 | ENSG00000162971 | TYW5 | chr2 | - | 199943764 | 199943834 | 199940088 | 199940133 | 199948336 | 199948472 | 25 | 12 | [illegible] |
| SE | 50625 | ENSG00000176463 | SLCO3A1 | chr15 | + | 91915992 | 91916458 | 91853707 | 91854088 | 92094880 | 92094979 | 120 | 80 | [illegible] |
| SE | 50626 | ENSG00000176463 | SLCO3A1 | chr15 | + | 92104278 | 92104542 | 92094880 | 92094979 | 92120464 | 92120629 | 114 | 80 | [illegible] |
| SE | 50627 | ENSG00000176463 | SLCO3A1 | chr15 | + | 92150949 | 92151014 | 92146983 | 92147159 | 92162755 | 92162998 | 58 | 31 | [illegible] |
| SE | 50628 | ENSG00000176463 | SLCO3A1 | chr15 | + | 92150954 | 92151014 | 92146983 | 92147159 | 92162755 | 92162946 | 8 | 5 | [illegible] |
| SE | 50631 | ENSG00000158869 | FCER1G | chr1 | + | 161218240 | 161218276 | 161215233 | 161215370 | 161218702 | 161218723 | 361 | 173 | [illegible] |
| SE | 50632 | ENSG00000151023 | ENKUR | chr10 | - | 24984209 | 24984376 | 24981978 | 24982770 | 24984735 | 24984905 | 8 | 40 | [illegible] |
| SE | 50633 | ENSG00000151023 | ENKUR | chr10 | - | 24984735 | 24984905 | 24981979 | 24984376 | 24990462 | 24990609 | 92 | 447 | [illegible] |
| SE | 50634 | ENSG00000151023 | ENKUR | chr10 | - | 24990462 | 24990609 | 24984401 | 24984905 | 24995645 | 24995869 | 180 | 771 | [illegible] |
| SE | 50635 | ENSG00000151023 | ENKUR | chr10 | - | 24995645 | 24995869 | 24990462 | 24990609 | 24999400 | 24999546 | 109 | 411 | [illegible] |
| SE | 50637 | ENSG00000151023 | ENKUR | chr10 | - | 24995645 | 24995869 | 24990462 | 24990609 | 25061111 | 25061369 | 96 | 366 | [illegible] |
| SE | 50640 | ENSG00000087095 | NLK | chr17 | + | 28122602 | 28122732 | 28042676 | 28043331 | 28132619 | 28132675 | 129 | 93 | [illegible] |
| SE | 50641 | ENSG00000087095 | NLK | chr17 | + | 28122602 | 28122732 | 28042676 | 28043331 | 28163542 | 28163628 | 36 | 20 | [illegible] |
| SE | 50643 | ENSG00000087095 | NLK | chr17 | + | 28194581 | 28194708 | 28192119 | 28192213 | 28196179 | 28196381 | 34 | 46 | [illegible] |
| SE | 50644 | ENSG00000182372 | CLN8 | chr8 | + | 1758248 | 1758306 | 1755787 | 1756082 | 1759296 | 1759511 | 1 | 0 | [illegible] |
| SE | 50646 | ENSG00000182372 | CLN8 | chr8 | + | 1772923 | 1772985 | 1770931 | 1771597 | 1780249 | 1780772 | 0 | 0 | [illegible] |
| SE | 50647 | ENSG00000142784 | WDTC1 | chr1 | + | 27263151 | 27263235 | 27260955 | 27261102 | 27282238 | 27282285 | 417 | 308 | [illegible] |
| SE | 50648 | ENSG00000142784 | WDTC1 | chr1 | + | 27282238 | 27282285 | 27263151 | 27263235 | 27283337 | 27283449 | 357 | 257 | [illegible] |
| SE | 50649 | ENSG00000142784 | WDTC1 | chr1 | + | 27282238 | 27282285 | 27263151 | 27263235 | 27287673 | 27287861 | 213 | 174 | [illegible] |
| SE | 50650 | ENSG00000142784 | WDTC1 | chr1 | + | 27283337 | 27283449 | 27263151 | 27263235 | 27287673 | 27287861 | 169 | 73 | [illegible] |
| SE | 50651 | ENSG00000142784 | WDTC1 | chr1 | + | 27283337 | 27283449 | 27282238 | 27282285 | 27287673 | 27287861 | 323 | 162 | [illegible] |
| SE | 50652 | ENSG00000142784 | WDTC1 | chr1 | + | 27292214 | 27292397 | 27287673 | 27287861 | 27294021 | 27294116 | 262 | 98 | [illegible] |
| SE | 50654 | ENSG00000142784 | WDTC1 | chr1 | + | 27294513 | 27294629 | 27294021 | 27294116 | 27296325 | 27296401 | 188 | 92 | [illegible] |
| SE | 50655 | ENSG00000142784 | WDTC1 | chr1 | + | 27301225 | 27301461 | 27297937 | 27298111 | 27305000 | 27305193 | 94 | 27 | [illegible] |
| SE | 50657 | ENSG00000142784 | WDTC1 | chr1 | + | 27303620 | 27305193 | 27301225 | 27301461 | 27306185 | 27308619 | 216 | 94 | [illegible] |
| SE | 50658 | ENSG00000142784 | WDTC1 | chr1 | + | 27305000 | 27305193 | 27301225 | 27301461 | 27306185 | 27308244 | 98 | 55 | [illegible] |
| SE | 50659 | ENSG00000121988 | ZRANB3 | chr2 | - | 135202831 | 135202963 | 135196968 | 135200440 | 135207433 | 135207836 | 10 | 17 | [illegible] |
| SE | 50660 | ENSG00000121988 | ZRANB3 | chr2 | - | 135224425 | 135224517 | 135219076 | 135219178 | 135227811 | 135228015 | 13 | 4 | [illegible] |
| SE | 50661 | ENSG00000121988 | ZRANB3 | chr2 | - | 135224425 | 135224517 | 135219076 | 135219178 | 135227817 | 135228015 | 4 | 1 | [illegible] |
| SE | 50662 | ENSG00000121988 | ZRANB3 | chr2 | - | 135268961 | 135269141 | 135265533 | 135265686 | 135271767 | 135271887 | 3 | 16 | [illegible] |
| SE | 50664 | ENSG00000121988 | ZRANB3 | chr2 | - | 135390801 | 135390820 | 135353320 | 135353628 | 135504328 | 135504496 | 21 | 11 | [illegible] |
| SE | 50667 | ENSG00000121988 | ZRANB3 | chr2 | - | 135504328 | 135504496 | 135390801 | 135390820 | 135531126 | 135531218 | 30 | 11 | [illegible] |
| SE | 50671 | ENSG00000142687 | KIAA0319L | chr1 | - | 35444160 | 35444303 | 35442905 | 35443028 | 35449866 | 35450005 | 79 | 36 | [illegible] |
| SE | 50673 | ENSG00000142687 | KIAA0319L | chr1 | - | 35448172 | 35448332 | 35442905 | 35443028 | 35449866 | 35450005 | 53 | 34 | [illegible] |
| SE | 50674 | ENSG00000142687 | KIAA0319L | chr1 | - | 35448172 | 35448332 | 35442905 | 35443028 | 35451627 | 35451776 | 1 | 0 | [illegible] |
| SE | 50676 | ENSG00000142687 | KIAA0319L | chr1 | - | 35448172 | 35448332 | 35444160 | 35444303 | 35451627 | 35451776 | 50 | 31 | [illegible] |
| SE | 50677 | ENSG00000142687 | KIAA0319L | chr1 | - | 35449866 | 35450005 | 35442905 | 35443028 | 35451627 | 35451776 | 0 | 0 | [illegible] |
| SE | 50678 | ENSG00000142687 | KIAA0319L | chr1 | - | 35449866 | 35450005 | 35444160 | 35444303 | 35450357 | 35450509 | 77 | 43 | [illegible] |
| SE | 50679 | ENSG00000142687 | KIAA0319L | chr1 | - | 35449866 | 35450005 | 35444160 | 35444303 | 35451627 | 35451776 | 3 | 0 | [illegible] |
| SE | 50680 | ENSG00000142687 | KIAA0319L | chr1 | - | 35449866 | 35450005 | 35448172 | 35448332 | 35451627 | 35451776 | 52 | 34 | [illegible] |
| SE | 50681 | ENSG00000142687 | KIAA0319L | chr1 | - | 35450357 | 35450509 | 35444160 | 35444303 | 35451627 | 35451776 | 36 | 22 | [illegible] |
| SE | 50682 | ENSG00000142687 | KIAA0319L | chr1 | - | 35450357 | 35450509 | 35444160 | 35444303 | 35453556 | 35453689 | 0 | 1 | [illegible] |
| SE | 50683 | ENSG00000142687 | KIAA0319L | chr1 | - | 35450357 | 35450509 | 35449866 | 35450005 | 35451627 | 35451776 | 110 | 65 | [illegible] |
| SE | 50684 | ENSG00000142687 | KIAA0319L | chr1 | - | 35451627 | 35451776 | 35444160 | 35444303 | 35453556 | 35453689 | 41 | 36 | [illegible] |
| SE | 50685 | ENSG00000142687 | KIAA0319L | chr1 | - | 35451627 | 35451776 | 35450357 | 35450509 | 35453556 | 35453689 | 77 | 58 | [illegible] |
| SE | 50687 | ENSG00000142687 | KIAA0319L | chr1 | - | 35460304 | 35460437 | 35454361 | 35454485 | 35462620 | 35462713 | 158 | 111 | [illegible] |
| SE | 50692 | ENSG00000142687 | KIAA0319L | chr1 | - | 35470862 | 35470960 | 35466607 | 35466695 | 35474804 | 35474906 | 207 | 151 | [illegible] |
| SE | 50693 | ENSG00000142687 | KIAA0319L | chr1 | - | 35474804 | 35474906 | 35470862 | 35470960 | 35478965 | 35479212 | 165 | 118 | [illegible] |
| SE | 50694 | ENSG00000142687 | KIAA0319L | chr1 | - | 35474804 | 35474906 | 35470862 | 35470960 | 35506611 | 35507135 | 86 | 56 | [illegible] |
| SE | 50695 | ENSG00000142687 | KIAA0319L | chr1 | - | 35478965 | 35479212 | 35470862 | 35470960 | 35506611 | 35506867 | 135 | 73 | [illegible] |
| SE | 50697 | ENSG00000142687 | KIAA0319L | chr1 | - | 35478965 | 35479212 | 35474804 | 35474906 | 35554349 | 35554519 | 94 | 69 | [illegible] |
| SE | 50698 | ENSG00000142687 | KIAA0319L | chr1 | - | 35506611 | 35507135 | 35474804 | 35474906 | 35554349 | 35554519 | 76 | 44 | [illegible] |
| SE | 50703 | ENSG00000113318 | MSH3 | chr5 | + | 80741463 | 80741548 | 80728850 | 80728965 | 80744505 | 80744615 | 84 | 57 | [illegible] |
| SE | 50704 | ENSG00000113318 | MSH3 | chr5 | + | 80741463 | 80741548 | 80728850 | 80728965 | 80761545 | 80761610 | 31 | 20 | [illegible] |
| SE | 50706 | ENSG00000113318 | MSH3 | chr5 | + | 80744505 | 80744615 | 80741463 | 80741548 | 80761545 | 80761610 | 93 | 59 | [illegible] |
| SE | 50708 | ENSG00000113318 | MSH3 | chr5 | + | 80761545 | 80761678 | 80744505 | 80744615 | 80775693 | 80775758 | 65 | 38 | [illegible] |
| SE | 50709 | ENSG00000113318 | MSH3 | chr5 | + | 80775693 | 80775758 | 80768834 | 80769003 | 80778719 | 80778836 | 229 | 184 | [illegible] |
| SE | 50710 | ENSG00000113318 | MSH3 | chr5 | + | 80775693 | 80775758 | 80768834 | 80769003 | 80787564 | 80787672 | 35 | 17 | [illegible] |
| SE | 50711 | ENSG00000113318 | MSH3 | chr5 | + | 80778719 | 80778836 | 80768834 | 80769003 | 80787564 | 80787672 | 41 | 29 | [illegible] |
| SE | 50712 | ENSG00000113318 | MSH3 | chr5 | + | 80778719 | 80778836 | 80775693 | 80775758 | 80787564 | 80787672 | 234 | 200 | [illegible] |
| SE | 50713 | ENSG00000113318 | MSH3 | chr5 | + | 80830636 | 80830740 | 80813624 | 80813741 | 80848447 | 80848612 | 1 | 1 | [illegible] |
| SE | 50715 | ENSG00000113318 | MSH3 | chr5 | + | 80848447 | 80848612 | 80813583 | 80813741 | 80854129 | 80854316 | 4 | 12 | [illegible] |
| SE | 50717 | ENSG00000113318 | MSH3 | chr5 | + | 80864812 | 80864942 | 80854129 | 80854316 | 80873115 | 80873287 | 80 | 52 | [illegible] |
| SE | 50721 | ENSG00000165046 | LETM2 | chr8 | + | 38394097 | 38394241 | 38392541 | 38392995 | 38400271 | 38400409 | 3 | 1 | [illegible] |
| SE | 50722 | ENSG00000165046 | LETM2 | chr8 | + | 38394182 | 38394241 | 38392541 | 38392995 | 38400271 | 38400409 | 6 | 0 | [illegible] |
| SE | 50723 | ENSG00000165046 | LETM2 | chr8 | + | 38406945 | 38407038 | 38404339 | 38404506 | 38407361 | 38407463 | 4 | 0 | [illegible] |
| SE | 50728 | ENSG00000173039 | RELA | chr11 | - | 65658717 | 65658822 | 65658286 | 65658499 | 65659665 | 65659788 | 232 | 143 | [illegible] |
| SE | 50734 | ENSG00000173039 | RELA | chr11 | - | 65661686 | 65661835 | 65660051 | 65660215 | 65662178 | 65662205 | 173 | 104 | [illegible] |
| SE | 50736 | ENSG00000173039 | RELA | chr11 | - | 65661686 | 65662088 | 65660123 | 65660215 | 65662178 | 65662205 | 211 | 117 | [illegible] |
| SE | 50737 | ENSG00000173039 | RELA | chr11 | - | 65661936 | 65662088 | 65660123 | 65660215 | 65662178 | 65662205 | 38 | 13 | [illegible] |
| SE | 50746 | ENSG00000137198 | GMPR | chr6 | + | 16250283 | 16250367 | 16246841 | 16246961 | 16254561 | 16254735 | 26 | 60 | [illegible] |
| SE | 50747 | ENSG00000137198 | GMPR | chr6 | + | 16250283 | 16250367 | 16246841 | 16246961 | 16274414 | 16274496 | 10 | 10 | [illegible] |
| SE | 50748 | ENSG00000137198 | GMPR | chr6 | + | 16254561 | 16254735 | 16246841 | 16246961 | 16274414 | 16274496 | 28 | 86 | [illegible] |
| SE | 50749 | ENSG00000137198 | GMPR | chr6 | + | 16254561 | 16254735 | 16250283 | 16250367 | 16274414 | 16274496 | 45 | 136 | [illegible] |
| SE | 50750 | ENSG00000137198 | GMPR | chr6 | + | 16285792 | 16285835 | 16278783 | 16278890 | 16290461 | 16290621 | 64 | 156 | [illegible] |
| SE | 50764 | ENSG00000112531 | QKI | chr6 | + | 163455278 | 163455421 | 163413999 | 163415335 | 163478779 | 163478802 | 414 | 342 | [illegible] |
| SE | 50766 | ENSG00000112531 | QKI | chr6 | + | 163478779 | 163478896 | 163415237 | 163415335 | 163534981 | 163534984 | 275 | 199 | [illegible] |
| SE | 50767 | ENSG00000112531 | QKI | chr6 | + | 163478779 | 163478896 | 163455215 | 163455421 | 163534981 | 163534984 | 486 | 374 | [illegible] |
| SE | 50768 | ENSG00000112531 | QKI | chr6 | + | 163478779 | 163478896 | 163455215 | 163455421 | 163561981 | 163562069 | 135 | 117 | [illegible] |
| SE | 50769 | ENSG00000112531 | QKI | chr6 | + | 163534981 | 163535125 | 163455330 | 163455421 | 163561981 | 163562069 | 351 | 288 | [illegible] |
| SE | 50770 | ENSG00000112531 | QKI | chr6 | + | 163534981 | 163535125 | 163478779 | 163478896 | 163561981 | 163562069 | 704 | 548 | [illegible] |
| SE | 50773 | ENSG00000165185 | KIAA1958 | chr9 | + | 112645649 | 112645822 | 112574056 | 112575251 | 112659262 | 112665309 | 14 | 8 | [illegible] |
| SE | 50774 | ENSG00000257698 | GIHCG | chr12 | - | 57935297 | 57935553 | 57931527 | 57931737 | 57935838 | 57936175 | 21 | 6 | [illegible] |
| SE | 50775 | ENSG00000257698 | GIHCG | chr12 | - | 57935394 | 57935553 | 57931227 | 57931737 | 57935838 | 57936155 | 21 | 6 | [illegible] |
| SE | 50776 | ENSG00000257698 | GIHCG | chr12 | - | 57935469 | 57935553 | 57930114 | 57931737 | 57935838 | 57935914 | 51 | 18 | [illegible] |
| SE | 50777 | ENSG00000257698 | GIHCG | chr12 | - | 57935469 | 57935572 | 57931452 | 57931737 | 57935838 | 57936122 | 21 | 8 | [illegible] |
| SE | 50779 | ENSG00000103351 | CLUAP1 | chr16 | + | 3504719 | 3504831 | 3500975 | 3501089 | 3506330 | 3506415 | 34 | 20 | [illegible] |
| SE | 50781 | ENSG00000103351 | CLUAP1 | chr16 | + | 3506330 | 3506415 | 3504719 | 3504831 | 3508288 | 3508464 | 35 | 35 | [illegible] |
| SE | 50784 | ENSG00000103351 | CLUAP1 | chr16 | + | 3526411 | 3526484 | 3523157 | 3523299 | 3530567 | 3530590 | 40 | 60 | [illegible] |
| SE | 50786 | ENSG00000082512 | TRAF5 | chr1 | + | 211356366 | 211356468 | 211354409 | 211354467 | 211359911 | 211360076 | 123 | 85 | [illegible] |
| SE | 50789 | ENSG00000082512 | TRAF5 | chr1 | + | 211360701 | 211360779 | 211356366 | 211356468 | 211361087 | 211361148 | 92 | 57 | [illegible] |
| SE | 50790 | ENSG00000082512 | TRAF5 | chr1 | + | 211360701 | 211360779 | 211356366 | 211356468 | 211365375 | 211365468 | 2 | 0 | [illegible] |
| SE | 50791 | ENSG00000082512 | TRAF5 | chr1 | + | 211361087 | 211361162 | 211356366 | 211356468 | 211365375 | 211365468 | 39 | 55 | [illegible] |
| SE | 50792 | ENSG00000082512 | TRAF5 | chr1 | + | 211361087 | 211361162 | 211356366 | 211360779 | 211365375 | 211365468 | 179 | 155 | [illegible] |
| SE | 50794 | ENSG00000082512 | TRAF5 | chr1 | + | 211365375 | 211365468 | 211361087 | 211361162 | 211371301 | 211371470 | 90 | 68 | [illegible] |
| SE | 50795 | ENSG00000082512 | TRAF5 | chr1 | + | 211369451 | 211369592 | 211361087 | 211361162 | 211371301 | 211371470 | 168 | 120 | [illegible] |
| SE | 50796 | ENSG00000082512 | TRAF5 | chr1 | + | 211369451 | 211369592 | 211365375 | 211365468 | 211371301 | 211371470 | 223 | 169 | [illegible] |
| SE | 50797 | ENSG00000082512 | TRAF5 | chr1 | + | 211371301 | 211371470 | 211368928 | 211369592 | 211372127 | 211372795 | 299 | 263 | [illegible] |
| SE | 50798 | ENSG00000178425 | NT5DC1 | chr6 | + | 116117860 | 116117945 | 116115690 | 116115770 | 116221053 | 116221228 | 90 | 57 | [illegible] |
| SE | 50800 | ENSG00000178425 | NT5DC1 | chr6 | + | 116238954 | 116239123 | 116238186 | 116238348 | 116243908 | 116249497 | 186 | 121 | [illegible] |
| SE | 50803 | ENSG00000211454 | AKR7L | chr1 | - | 19269271 | 19269371 | 19268571 | 19268701 | 19269582 | 19269679 | 1 | 0 | [illegible] |
| SE | 50805 | ENSG00000146540 | C7orf50 | chr7 | - | 1000469 | 1000551 | 996985 | 997804 | 1009949 | 1010143 | 68 | 47 | [illegible] |
| SE | 50808 | ENSG00000117481 | NSUN4 | chr1 | + | 46344800 | 46345144 | 46340788 | 46340919 | 46352867 | 46353028 | 29 | 4 | [illegible] |
| SE | 50810 | ENSG00000117481 | NSUN4 | chr1 | + | 46346920 | 46347075 | 46344781 | 46345144 | 46360703 | 46360828 | 14 | 8 | [illegible] |
| SE | 50812 | ENSG00000117481 | NSUN4 | chr1 | + | 46352867 | 46353028 | 46344800 | 46345144 | 46360703 | 46360828 | 18 | 19 | [illegible] |
| SE | 50813 | ENSG00000117481 | NSUN4 | chr1 | + | 46352867 | 46353028 | 46346920 | 46347075 | 46360703 | 46360828 | 37 | 28 | [illegible] |
| SE | 50823 | ENSG00000215788 | TNFRSF25 | chr1 | - | 6462862 | 6462956 | 6462628 | 6462666 | 6464374 | 6464453 | 13 | 7 | [illegible] |
| SE | 50826 | ENSG00000215788 | TNFRSF25 | chr1 | - | 6462862 | 6462970 | 6462628 | 6462666 | 6464374 | 6464453 | 13 | 8 | [illegible] |
| SE | 50827 | ENSG00000215788 | TNFRSF25 | chr1 | - | 6462862 | 6463127 | 6462628 | 6462666 | 6464374 | 6464453 | 17 | 10 | [illegible] |
| SE | 50828 | ENSG00000215788 | TNFRSF25 | chr1 | - | 6462862 | 6463147 | 6462628 | 6462666 | 6464374 | 6464453 | 16 | 7 | [illegible] |
| SE | 50830 | ENSG00000215788 | TNFRSF25 | chr1 | - | 6463071 | 6463127 | 6462628 | 6462663 | 6464374 | 6464453 | 2 | 3 | [illegible] |
| SE | 50832 | ENSG00000215788 | TNFRSF25 | chr1 | - | 6463071 | 6463127 | 6462862 | 6462970 | 6464374 | 6464453 | 8 | 3 | [illegible] |
| SE | 50833 | ENSG00000215788 | TNFRSF25 | chr1 | - | 6465087 | 6465222 | 6462862 | 6464719 | 6465439 | 6465470 | 60 | 36 | [illegible] |
| SE | 50835 | ENSG00000048052 | HDAC9 | chr7 | + | 18325511 | 18325592 | 18290440 | 18290515 | 18395114 | 18395183 | 0 | 1 | [illegible] |
| SE | 50837 | ENSG00000048052 | HDAC9 | chr7 | + | 18395114 | 18395183 | 18290091 | 18290515 | 18496261 | 18496324 | 9 | 3 | [illegible] |
| SE | 50839 | ENSG00000048052 | HDAC9 | chr7 | + | 18458841 | 18458901 | 18290449 | 18290515 | 18496261 | 18496324 | 0 | 1 | [illegible] |
| SE | 50840 | ENSG00000048052 | HDAC9 | chr7 | + | 18496261 | 18496324 | 18290421 | 18290515 | 18585280 | 18585326 | 83 | 47 | [illegible] |
| SE | 50843 | ENSG00000048052 | HDAC9 | chr7 | + | 18543168 | 18543674 | 18496261 | 18496324 | 18585280 | 18585522 | 1 | 0 | [illegible] |
| SE | 50844 | ENSG00000048052 | HDAC9 | chr7 | + | 18543168 | 18543674 | 18502607 | 18502639 | 18585280 | 18585522 | 0 | 0 | [illegible] |
| SE | 50845 | ENSG00000048052 | HDAC9 | chr7 | + | 18590335 | 18590486 | 18585280 | 18585522 | 18591515 | 18591544 | 183 | 154 | [illegible] |
| SE | 50846 | ENSG00000048052 | HDAC9 | chr7 | + | 18590344 | 18590486 | 18585280 | 18585522 | 18591515 | 18591533 | 85 | 53 | [illegible] |
| SE | 50849 | ENSG00000048052 | HDAC9 | chr7 | + | 18634626 | 18634742 | 18593907 | 18594029 | 18644670 | 18644793 | 97 | 76 | [illegible] |
| SE | 50850 | ENSG00000048052 | HDAC9 | chr7 | + | 18634626 | 18634742 | 18629349 | 18629481 | 18644670 | 18644793 | 134 | 89 | [illegible] |
| SE | 50851 | ENSG00000048052 | HDAC9 | chr7 | + | 18647784 | 18647998 | 18644670 | 18644793 | 18648465 | 18648683 | 235 | 167 | [illegible] |
| SE | 50853 | ENSG00000048052 | HDAC9 | chr7 | + | 18656906 | 18656987 | 18648465 | 18648683 | 18666212 | 18668131 | 0 | 0 | [illegible] |
| SE | 50855 | ENSG00000048052 | HDAC9 | chr7 | + | 18666212 | 18666476 | 18648465 | 18648683 | 18727579 | 18727757 | 101 | 94 | [illegible] |
| SE | 50856 | ENSG00000048052 | HDAC9 | chr7 | + | 18727579 | 18727757 | 18666212 | 18666476 | 18749004 | 18749138 | 6 | 18 | [illegible] |
| SE | 50857 | ENSG00000048052 | HDAC9 | chr7 | + | 18762156 | 18762277 | 18749004 | 18749138 | 18767105 | 18767155 | 8 | 7 | [illegible] |
| SE | 50858 | ENSG00000048052 | HDAC9 | chr7 | + | 18793344 | 18793452 | 18767105 | 18767155 | 18829160 | 18829216 | 16 | 6 | [illegible] |
| SE | 50859 | ENSG00000048052 | HDAC9 | chr7 | + | 18975805 | 18975953 | 18954145 | 18954230 | 18996022 | 18996197 | 6 | 4 | [illegible] |
| SE | 50860 | ENSG00000084652 | TXLNA | chr1 | + | 32180313 | 32180514 | 32179674 | 32179776 | 32181241 | 32181577 | 178 | 133 | [illegible] |
| SE | 50861 | ENSG00000084652 | TXLNA | chr1 | + | 32181241 | 32181577 | 32179674 | 32179776 | 32184524 | 32184616 | 170 | 107 | [illegible] |
| SE | 50863 | ENSG00000084652 | TXLNA | chr1 | + | 32187953 | 32188124 | 32184524 | 32184616 | 32190054 | 32190249 | 378 | 260 | [illegible] |
| SE | 50871 | ENSG00000115568 | ZNF142 | chr2 | - | 218658979 | 218659035 | 218650358 | 218650526 | 218659368 | 218659524 | 4 | 0 | [illegible] |
| SE | 50872 | ENSG00000115568 | ZNF142 | chr2 | - | 218658979 | 218659035 | 218658700 | 218658876 | 218659368 | 218659524 | 6 | 9 | [illegible] |
| SE | 50875 | ENSG00000115568 | ZNF142 | chr2 | - | 218658979 | 218659130 | 218658700 | 218658876 | 218659368 | 218659476 | 35 | 20 | [illegible] |
| SE | 50876 | ENSG00000181790 | ADGRB1 | chr8 | + | 142510931 | 142511073 | 142490771 | 142490815 | 142518137 | 142518241 | 0 | 3 | [illegible] |
| SE | 50888 | ENSG00000156515 | HK1 | chr10 | + | 69364782 | 69364902 | 69359896 | 69360045 | 69368535 | 69368631 | 1015 | 841 | [illegible] |

| | | | | | | | | | | | | | |
|---|---|---|---|---|---|---|---|---|---|---|---|---|---|
| SE | 50889 | ENSG00000156515 | HK1 | chr10 | + | 69368535 | 69368631 | 69364782 | 69364902 | 69369236 | 69369336 | 1062 | 900 |
| SE | 50891 | ENSG00000156515 | HK1 | chr10 | + | 69376933 | 69377089 | 69369440 | 69369624 | 69379861 | 69380095 | 475 | 300 |
| SE | 50892 | ENSG00000156515 | HK1 | chr10 | + | 69376933 | 69377089 | 69369440 | 69369624 | 69382486 | 69382791 | 253 | 146 |
| SE | 50893 | ENSG00000156515 | HK1 | chr10 | + | 69379861 | 69380095 | 69369440 | 69369624 | 69382486 | 69382791 | 142 | 96 |
| SE | 50894 | ENSG00000156515 | HK1 | chr10 | + | 69379861 | 69380095 | 69376933 | 69377089 | 69382486 | 69382791 | 364 | 249 |
| SE | 50897 | ENSG00000156515 | HK1 | chr10 | + | 69384795 | 69384915 | 69379861 | 69380095 | 69386322 | 69386418 | 361 | 291 |
| SE | 50898 | ENSG00000156515 | HK1 | chr10 | + | 69384795 | 69384915 | 69384332 | 69384481 | 69386322 | 69386418 | 651 | 542 |
| SE | 50899 | ENSG00000156515 | HK1 | chr10 | + | 69386322 | 69386418 | 69384795 | 69384915 | 69389196 | 69389296 | 745 | 578 |
| SE | 50900 | ENSG00000156515 | HK1 | chr10 | + | 69389196 | 69389296 | 69386322 | 69386418 | 69392124 | 69392308 | 474 | 336 |
| SE | 50901 | ENSG00000156515 | HK1 | chr10 | + | 69392124 | 69392308 | 69389196 | 69389296 | 69394949 | 69395105 | 575 | 367 |
| SE | 50902 | ENSG00000156515 | HK1 | chr10 | + | 69394949 | 69395105 | 69392124 | 69392308 | 69398594 | 69398828 | 436 | 281 |
| SE | 50904 | ENSG00000116641 | DOCK7 | chr1 | - | 62473981 | 62474088 | 62457537 | 62457705 | 62475207 | 62475351 | 17 | 16 |
| SE | 50905 | ENSG00000116641 | DOCK7 | chr1 | - | 62528150 | 62528305 | 62513715 | 62513898 | 62529276 | 62529446 | 16 | 24 |
| SE | 50911 | ENSG00000116641 | DOCK7 | chr1 | - | 62552731 | 62552901 | 62543655 | 62543745 | 62555824 | 62555989 | 30 | 35 |
| SE | 50912 | ENSG00000116641 | DOCK7 | chr1 | - | 62583183 | 62583254 | 62578827 | 62578966 | 62586506 | 62586624 | 21 | 20 |
| SE | 50916 | ENSG00000123064 | DDX54 | chr12 | - | 113172352 | 113172563 | 113169769 | 113169904 | 113174639 | 113174771 | 165 | 57 |
| SE | 50917 | ENSG00000123064 | DDX54 | chr12 | - | 113178976 | 113179026 | 113177051 | 113177093 | 113179142 | 113179331 | 117 | 53 |
| SE | 50922 | ENSG00000000460 | C1orf112 | chr1 | + | 169827050 | 169827199 | 169823407 | 169823472 | 169827710 | 169827840 | 12 | 7 |
| SE | 50923 | ENSG00000000460 | C1orf112 | chr1 | + | 169842416 | 169842539 | 169836926 | 169837112 | 169843673 | 169843769 | 21 | 20 |
| SE | 50924 | ENSG00000000460 | C1orf112 | chr1 | + | 169850264 | 169850345 | 169849499 | 169849604 | 169851816 | 169851936 | 25 | 28 |
| SE | 50928 | ENSG00000066056 | TIE1 | chr1 | + | 43307795 | 43307924 | 43307431 | 43307572 | 43308985 | 43309131 | 1 | 0 |
| SE | 50931 | ENSG00000204370 | SDHD | chr11 | + | 112087856 | 112087973 | 112086823 | 112086959 | 112088866 | 112089004 | 309 | 138 |
| SE | 50932 | ENSG00000255292 | NA | chr11 | + | 112170622 | 112170674 | 112170418 | 112170530 | 112179277 | 112179360 | 1 | 2 |
| SE | 50935 | ENSG00000133789 | SWAP70 | chr11 | + | 9703395 | 9703448 | 9694145 | 9694286 | 9713465 | 9713639 | 1 | 0 |
| SE | 50937 | ENSG00000133789 | SWAP70 | chr11 | + | 9713465 | 9713639 | 9694145 | 9694286 | 9728052 | 9728071 | 97 | 46 |
| SE | 50938 | ENSG00000133789 | SWAP70 | chr11 | + | 9724657 | 9724885 | 9694145 | 9694286 | 9728052 | 9728071 | 193 | 136 |
| SE | 50940 | ENSG00000133789 | SWAP70 | chr11 | + | 9728052 | 9728199 | 9694145 | 9694286 | 9729342 | 9729451 | 158 | 134 |
| SE | 50942 | ENSG00000164466 | SFXN1 | chr5 | + | 175512110 | 175512196 | 175511435 | 175511526 | 175513462 | 175513590 | 53 | 37 |
| SE | 50945 | ENSG00000205323 | SARNP | chr12 | - | 55757382 | 55757553 | 55752462 | 55752840 | 55760550 | 55760640 | 139 | 61 |
| SE | 50946 | ENSG00000205323 | SARNP | chr12 | - | 55760550 | 55760640 | 55757269 | 55757553 | 55789074 | 55789143 | 639 | 457 |
| SE | 50947 | ENSG00000205323 | SARNP | chr12 | - | 55794806 | 55794880 | 55792541 | 55794387 | 55796024 | 55796076 | 784 | 542 |
| SE | 50948 | ENSG00000205323 | SARNP | chr12 | - | 55796024 | 55796076 | 55794806 | 55794880 | 55800561 | 55800629 | 528 | 362 |
| SE | 50949 | ENSG00000205323 | SARNP | chr12 | - | 55800853 | 55800900 | 55800561 | 55800629 | 55803628 | 55803728 | 226 | 181 |
| SE | 50950 | ENSG00000205323 | SARNP | chr12 | - | 55803628 | 55803728 | 55800853 | 55800900 | 55817665 | 55817709 | 170 | 111 |
| SE | 50951 | ENSG00000075340 | ADD2 | chr2 | - | 70673240 | 70673326 | 70672877 | 70673006 | 70674677 | 70674825 | 0 | 0 |
| [illegible] | | | | | | | | | | | | | |

| | | | | | | | | | | | | | | |
|---|---|---|---|---|---|---|---|---|---|---|---|---|---|---|
| SE | 51389 | ENSG00000066427 | ATXN3 | chr14 | - | 92096673 | 92096838 | 92093745 | 92093831 | 92106528 | 92106535 | 42 | 25 | [illegible] |
| SE | 51390 | ENSG00000066427 | ATXN3 | chr14 | - | 92096673 | 92096838 | 92096092 | 92096134 | 92106528 | 92106535 | 41 | 27 | [illegible] |
| SE | 51393 | ENSG00000067606 | PRKCZ | chr1 | + | 2098346 | 2098555 | 2059540 | 2059591 | 2135261 | 2135347 | 0 | 0 | [illegible] |
| SE | 51398 | ENSG00000067606 | PRKCZ | chr1 | + | 2155994 | 2156092 | 2150789 | 2150978 | 2169517 | 2169604 | 91 | 44 | [illegible] |
| SE | 51400 | ENSG00000067606 | PRKCZ | chr1 | + | 2172300 | 2172388 | 2172054 | 2172190 | 2173896 | 2173951 | 58 | 30 | [illegible] |
| SE | 51404 | ENSG00000197302 | ZNF720 | chr16 | + | 31714366 | 31714417 | 31713305 | 31713435 | 31722625 | 31722752 | 0 | 0 | [illegible] |
| SE | 51405 | ENSG00000197302 | ZNF720 | chr16 | + | 31722625 | 31722752 | 31713228 | 31713448 | 31723257 | 31723353 | 39 | 22 | [illegible] |
| SE | 51407 | ENSG00000197302 | ZNF720 | chr16 | + | 31723257 | 31723353 | 31713228 | 31713448 | 31753765 | 31753856 | 27 | 27 | [illegible] |
| SE | 51409 | ENSG00000197302 | ZNF720 | chr16 | + | 31733510 | 31733627 | 31723257 | 31723353 | 31753765 | 31753900 | 3 | 2 | [illegible] |
| SE | 51412 | ENSG00000197302 | ZNF720 | chr16 | + | 31793334 | 31793411 | 31792629 | 31792730 | 31793701 | 31793707 | 0 | 1 | [illegible] |
| SE | 51413 | ENSG00000165417 | GTF2A1 | chr14 | - | 81185530 | 81185620 | 81175451 | 81180330 | 81192518 | 81192839 | 231 | 175 | [illegible] |
| SE | 51415 | ENSG00000143621 | ILF2 | chr1 | - | 153662704 | 153662795 | 153661787 | 153662556 | 153663018 | 153663133 | 678 | 344 | [illegible] |
| SE | 51416 | ENSG00000143621 | ILF2 | chr1 | - | 153662704 | 153662795 | 153661787 | 153662556 | 153663214 | 153663276 | 331 | 193 | [illegible] |
| SE | 51417 | ENSG00000143621 | ILF2 | chr1 | - | 153663018 | 153663133 | 153661787 | 153662556 | 153663214 | 153663276 | 196 | 88 | [illegible] |
| SE | 51419 | ENSG00000143621 | ILF2 | chr1 | - | 153663018 | 153663276 | 153662704 | 153662795 | 153664042 | 153664130 | 711 | 454 | [illegible] |
| SE | 51420 | ENSG00000143621 | ILF2 | chr1 | - | 153663214 | 153663276 | 153662704 | 153662795 | 153664042 | 153664130 | 415 | 319 | [illegible] |
| SE | 51422 | ENSG00000143621 | ILF2 | chr1 | - | 153664395 | 153664474 | 153664042 | 153664130 | 153665219 | 153665336 | 908 | 708 | [illegible] |
| SE | 51423 | ENSG00000143621 | ILF2 | chr1 | - | 153667999 | 153668077 | 153667282 | 153667657 | 153668452 | 153668557 | 755 | 538 | [illegible] |
| SE | 51425 | ENSG00000143621 | ILF2 | chr1 | - | 153670170 | 153670230 | 153668452 | 153668557 | 153670917 | 153670982 | 2 | 1 | [illegible] |
| SE | 51426 | ENSG00000107745 | MICU1 | chr10 | - | 72375782 | 72375872 | 72367326 | 72368355 | 72407928 | 72408037 | 202 | 150 | [illegible] |
| SE | 51427 | ENSG00000107745 | MICU1 | chr10 | - | 72379463 | 72379583 | 72375782 | 72375872 | 72407928 | 72408037 | 0 | 6 | [illegible] |
| SE | 51437 | ENSG00000139180 | NDUFA9 | chr12 | + | 4662532 | 4662635 | 4659035 | 4659177 | 4668456 | 4668475 | 255 | 151 | [illegible] |
| SE | 51438 | ENSG00000139180 | NDUFA9 | chr12 | + | 4667536 | 4667654 | 4662532 | 4662635 | 4668456 | 4668710 | 0 | 1 | [illegible] |
| SE | 51439 | ENSG00000137513 | NARS2 | chr11 | - | 78441090 | 78441117 | 78435967 | 78436814 | 78443660 | 78443758 | 19 | 11 | [illegible] |
| SE | 51440 | ENSG00000137513 | NARS2 | chr11 | - | 78478437 | 78478475 | 78469246 | 78469313 | 78478584 | 78478683 | 61 | 41 | [illegible] |
| SE | 51441 | ENSG00000137513 | NARS2 | chr11 | - | 78478584 | 78478683 | 78469246 | 78469313 | 78528841 | 78528936 | 9 | 0 | [illegible] |
| SE | 51443 | ENSG00000137513 | NARS2 | chr11 | - | 78559538 | 78559619 | 78528841 | 78528936 | 78566131 | 78566267 | 59 | 32 | [illegible] |
| SE | 51444 | ENSG00000265491 | RNF115 | chr1 | - | 145750406 | 145750500 | 145747994 | 145748110 | 145751437 | 145751510 | 192 | 122 | [illegible] |
| SE | 51445 | ENSG00000265491 | RNF115 | chr1 | - | 145750406 | 145750500 | 145747994 | 145748110 | 145752977 | 145753049 | 112 | 85 | [illegible] |
| SE | 51446 | ENSG00000265491 | RNF115 | chr1 | - | 145751437 | 145751510 | 145747994 | 145748110 | 145752977 | 145753049 | 54 | 32 | [illegible] |
| SE | 51453 | ENSG00000172432 | GTPBP2 | chr6 | - | 43622632 | 43622804 | 43620489 | 43622167 | 43623736 | 43623795 | 89 | 74 | [illegible] |
| SE | 51455 | ENSG00000172432 | GTPBP2 | chr6 | - | 43625362 | 43625560 | 43624887 | 43625062 | 43626225 | 43626386 | 56 | 73 | [illegible] |
| SE | 51456 | ENSG00000172432 | GTPBP2 | chr6 | - | 43625755 | 43625864 | 43624887 | 43625062 | 43626225 | 43626386 | 139 | 169 | [illegible] |
| SE | 51457 | ENSG00000172432 | GTPBP2 | chr6 | - | 43625755 | 43625864 | 43624887 | 43625560 | 43626225 | 43626386 | 366 | 422 | [illegible] |
| SE | 51458 | ENSG00000172432 | GTPBP2 | chr6 | - | 43626921 | 43626948 | 43626225 | 43626410 | 43628976 | 43629162 | 69 | 80 | [illegible] |
| SE | 51462 | ENSG00000162222 | TTC9C | chr11 | + | 62733117 | 62733199 | 62728822 | 62729086 | 62735381 | 62736243 | 19 | 7 | [illegible] |
| SE | 51463 | ENSG00000162517 | PEF1 | chr1 | - | 31632494 | 31632638 | 31629865 | 31630842 | 31635221 | 31635260 | 184 | 89 | [illegible] |
| SE | 51464 | ENSG00000162517 | PEF1 | chr1 | - | 31633158 | 31633314 | 31632494 | 31632638 | 31635221 | 31635260 | 204 | 71 | [illegible] |
| SE | 51468 | ENSG00000115602 | IL1RL1 | chr2 | + | 102340665 | 102340828 | 102340097 | 102340272 | 102342222 | 102342294 | 3 | 4 | [illegible] |
| SE | 51470 | ENSG00000133121 | STARD13 | chr13 | - | 33111777 | 33111892 | 33110685 | 33110907 | 33112720 | 33112931 | 0 | 5 | [illegible] |
| SE | 51471 | ENSG00000132604 | TERF2 | chr16 | - | 69357517 | 69357561 | 69355566 | 69357056 | 69361403 | 69361489 | 155 | 95 | [illegible] |
| SE | 51474 | ENSG00000132604 | TERF2 | chr16 | - | 69370482 | 69370629 | 69368375 | 69368595 | 69372268 | 69372282 | 120 | 49 | [illegible] |
| SE | 51475 | ENSG00000132604 | TERF2 | chr16 | - | 69372268 | 69372355 | 69370210 | 69370629 | 69384579 | 69384710 | 239 | 99 | [illegible] |
| SE | 51476 | ENSG00000132604 | TERF2 | chr16 | - | 69385390 | 69385486 | 69384579 | 69384710 | 69385592 | 69385599 | 131 | 64 | [illegible] |
| SE | 51477 | ENSG00000011009 | LYPLA2 | chr1 | + | 23793007 | 23793039 | 23792551 | 23792760 | 23793150 | 23793216 | 131 | 64 | [illegible] |
| SE | 51484 | ENSG00000160087 | UBE2J2 | chr1 | - | 1263345 | 1263386 | 1257207 | 1257310 | 1267861 | 1267992 | 122 | 82 | [illegible] |
| SE | 51487 | ENSG00000160087 | UBE2J2 | chr1 | - | 1266097 | 1266290 | 1263345 | 1263386 | 1267732 | 1267992 | 1 | 1 | [illegible] |
| SE | 51488 | ENSG00000160087 | UBE2J2 | chr1 | - | 1266097 | 1266290 | 1263345 | 1263386 | 1267861 | 1267992 | 1 | 1 | [illegible] |
| SE | 51489 | ENSG00000160087 | UBE2J2 | chr1 | - | 1267732 | 1267992 | 1263345 | 1263386 | 1273665 | 1273815 | 10 | 5 | [illegible] |
| SE | 51493 | ENSG00000160087 | UBE2J2 | chr1 | - | 1267861 | 1267992 | 1262852 | 1263386 | 1273665 | 1273833 | 73 | 48 | [illegible] |
| SE | 51497 | ENSG00000108021 | TASOR2 | chr10 | + | 5709529 | 5709663 | 5692815 | 5692918 | 5712822 | 5712918 | 17 | 28 | [illegible] |
| SE | 51501 | ENSG00000108021 | TASOR2 | chr10 | + | 5712822 | 5712918 | 5684837 | 5685175 | 5714134 | 5714176 | 49 | 33 | [illegible] |
| SE | 51502 | ENSG00000108021 | TASOR2 | chr10 | + | 5712822 | 5712918 | 5684837 | 5685175 | 5717658 | 5717750 | 11 | 11 | [illegible] |
| SE | 51504 | ENSG00000108021 | TASOR2 | chr10 | + | 5712822 | 5712918 | 5684837 | 5685175 | 5720754 | 5720774 | 6 | 2 | [illegible] |
| SE | 51505 | ENSG00000108021 | TASOR2 | chr10 | + | 5714134 | 5714207 | 5684947 | 5685175 | 5717658 | 5717750 | 34 | 22 | [illegible] |
| SE | 51506 | ENSG00000108021 | TASOR2 | chr10 | + | 5714134 | 5714207 | 5684947 | 5685175 | 5720543 | 5720668 | 4 | 6 | [illegible] |
| SE | 51512 | ENSG00000108021 | TASOR2 | chr10 | + | 5720543 | 5720668 | 5684837 | 5685175 | 5720754 | 5720774 | 121 | 125 | [illegible] |
| SE | 51513 | ENSG00000108021 | TASOR2 | chr10 | + | 5720543 | 5720668 | 5712822 | 5712918 | 5720754 | 5720774 | 129 | 125 | [illegible] |
| SE | 51514 | ENSG00000108021 | TASOR2 | chr10 | + | 5724429 | 5724533 | 5723676 | 5723777 | 5726884 | 5726957 | 253 | 253 | [illegible] |
| SE | 51515 | ENSG00000108021 | TASOR2 | chr10 | + | 5730486 | 5731203 | 5727060 | 5727123 | 5735303 | 5735546 | 161 | 145 | [illegible] |
| SE | 51516 | ENSG00000108021 | TASOR2 | chr10 | + | 5735303 | 5735546 | 5730486 | 5731203 | 5739617 | 5740497 | 161 | 161 | [illegible] |
| SE | 51521 | ENSG00000069248 | NUP133 | chr1 | - | 229449125 | 229449190 | 229444913 | 229445002 | 229450524 | 229450605 | 328 | 193 | [illegible] |
| SE | 51522 | ENSG00000069248 | NUP133 | chr1 | - | 229458160 | 229458296 | 229452524 | 229452643 | 229463542 | 229463676 | 105 | 56 | [illegible] |
| SE | 51523 | ENSG00000069248 | NUP133 | chr1 | - | 229460610 | 229460769 | 229458160 | 229458296 | 229463542 | 229463676 | 129 | 83 | [illegible] |
| SE | 51524 | ENSG00000069248 | NUP133 | chr1 | - | 229463542 | 229463676 | 229460610 | 229460769 | 229464623 | 229464875 | 181 | 161 | [illegible] |
| SE | 51526 | ENSG00000069248 | NUP133 | chr1 | - | 229484053 | 229484145 | 229477596 | 229477760 | 229486370 | 229486528 | 219 | 205 | [illegible] |
| SE | 51527 | ENSG00000069248 | NUP133 | chr1 | - | 229486370 | 229486528 | 229484053 | 229484145 | 229489954 | 229490102 | 168 | 157 | [illegible] |
| SE | 51528 | ENSG00000069248 | NUP133 | chr1 | - | 229487465 | 229487613 | 229486370 | 229486528 | 229489954 | 229490102 | 174 | 130 | [illegible] |
| SE | 51532 | ENSG00000176393 | RNPEP | chr1 | + | 201997318 | 201997554 | 201996146 | 201996263 | 201999901 | 201999945 | 181 | 99 | [illegible] |
| SE | 51533 | ENSG00000176393 | RNPEP | chr1 | + | 201997318 | 201997554 | 201996146 | 201996263 | 202001375 | 202001488 | 85 | 43 | [illegible] |
| SE | 51534 | ENSG00000176393 | RNPEP | chr1 | + | 201997318 | 201997554 | 201996146 | 201996409 | 201999901 | 201999945 | 96 | 56 | [illegible] |
| SE | 51535 | ENSG00000176393 | RNPEP | chr1 | + | 201999901 | 202000015 | 201996146 | 201996263 | 202001375 | 202001399 | 53 | 41 | [illegible] |
| SE | 51536 | ENSG00000176393 | RNPEP | chr1 | + | 201999901 | 202000015 | 201997318 | 201997554 | 202001375 | 202001399 | 146 | 97 | [illegible] |
| SE | 51539 | ENSG00000036549 | ZZZ3 | chr1 | - | 77578773 | 77578869 | 77576067 | 77576220 | 77579526 | 77579628 | 215 | 123 | [illegible] |
| SE | 51540 | ENSG00000036549 | ZZZ3 | chr1 | - | 77584766 | 77584829 | 77584516 | 77584655 | 77631849 | 77633405 | 0 | 3 | [illegible] |
| SE | 51542 | ENSG00000036549 | ZZZ3 | chr1 | - | 77631849 | 77633405 | 77584516 | 77584655 | 77639448 | 77639602 | 188 | 133 | [illegible] |
| SE | 51543 | ENSG00000036549 | ZZZ3 | chr1 | - | 77631849 | 77633405 | 77584516 | 77584655 | 77641383 | 77641446 | 54 | 46 | [illegible] |
| SE | 51544 | ENSG00000036549 | ZZZ3 | chr1 | - | 77631849 | 77633405 | 77584516 | 77584655 | 77641521 | 77641655 | 50 | 39 | [illegible] |
| SE | 51545 | ENSG00000036549 | ZZZ3 | chr1 | - | 77631849 | 77633405 | 77584766 | 77584829 | 77639448 | 77639602 | 138 | 94 | [illegible] |
| SE | 51547 | ENSG00000036549 | ZZZ3 | chr1 | - | 77639448 | 77639602 | 77584516 | 77584655 | 77641521 | 77641655 | 5 | 7 | [illegible] |
| SE | 51549 | ENSG00000036549 | ZZZ3 | chr1 | - | 77639448 | 77639602 | 77631849 | 77633405 | 77641521 | 77641655 | 142 | 100 | [illegible] |
| SE | 51550 | ENSG00000036549 | ZZZ3 | chr1 | - | 77641383 | 77641446 | 77584516 | 77584655 | 77641521 | 77641655 | 29 | 15 | [illegible] |
| SE | 51551 | ENSG00000036549 | ZZZ3 | chr1 | - | 77641383 | 77641446 | 77631849 | 77633405 | 77641521 | 77641655 | 31 | 20 | [illegible] |
| SE | 51555 | ENSG00000143801 | PSEN2 | chr1 | + | 226880468 | 226880724 | 226875364 | 226875550 | 226881887 | 226882048 | 0 | 0 | [illegible] |
| SE | 51561 | ENSG00000143801 | PSEN2 | chr1 | + | 226889757 | 226889926 | 226888828 | 226889049 | 226890034 | 226890133 | 0 | 0 | [illegible] |
| SE | 51563 | ENSG00000143801 | PSEN2 | chr1 | + | 226890655 | 226890965 | 226888828 | 226890133 | 226891277 | 226891361 | 0 | 3 | [illegible] |
| SE | 51564 | ENSG00000143801 | PSEN2 | chr1 | + | 226894006 | 226894125 | 226891742 | 226891844 | 226897719 | 226897829 | 7 | 5 | [illegible] |
| SE | 51565 | ENSG00000143801 | PSEN2 | chr1 | + | 226895423 | 226895587 | 226894006 | 226894125 | 226897719 | 226897829 | 14 | 9 | [illegible] |
| SE | 51571 | ENSG00000163564 | PYHIN1 | chr1 | + | 158936890 | 158937175 | 158931551 | 158931776 | 158938396 | 158938542 | 914 | 461 | [illegible] |
| SE | 51572 | ENSG00000163564 | PYHIN1 | chr1 | + | 158938396 | 158938542 | 158936890 | 158937175 | 158939079 | 158939120 | 589 | 293 | [illegible] |
| SE | 51573 | ENSG00000163564 | PYHIN1 | chr1 | + | 158938423 | 158938542 | 158936890 | 158937175 | 158939079 | 158939247 | 217 | 88 | [illegible] |
| SE | 51574 | ENSG00000163564 | PYHIN1 | chr1 | + | 158939079 | 158939247 | 158938396 | 158938542 | 158941976 | 158942399 | 231 | 86 | [illegible] |
| SE | 51575 | ENSG00000163564 | PYHIN1 | chr1 | + | 158943789 | 158943978 | 158941976 | 158942399 | 158944874 | 158945042 | 1928 | 1138 | [illegible] |
| SE | 51577 | ENSG00000155545 | MIER3 | chr5 | - | 56923690 | 56923833 | 56919601 | 56923585 | 56923914 | 56924042 | 66 | 76 | [illegible] |
| SE | 51578 | ENSG00000155545 | MIER3 | chr5 | - | 56936893 | 56936936 | 56935743 | 56935751 | 56937577 | 56937698 | 0 | 0 | [illegible] |
| SE | 51579 | ENSG00000155545 | MIER3 | chr5 | - | 56938295 | 56938338 | 56937577 | 56937698 | 56938882 | 56938971 | 2 | 2 | [illegible] |
| SE | 51580 | ENSG00000155545 | MIER3 | chr5 | - | 56950627 | 56950652 | 56946925 | 56947071 | 56952093 | 56952112 | 13 | 6 | [illegible] |
| SE | 51583 | ENSG00000135821 | GLUL | chr1 | - | 182385356 | 182385556 | 182378097 | 182384723 | 182385759 | 182385842 | 1101 | 637 | [illegible] |
| SE | 51584 | ENSG00000135821 | GLUL | chr1 | - | 182385759 | 182385887 | 182385356 | 182385556 | 182386255 | 182386402 | 974 | 640 | [illegible] |
| SE | 51585 | ENSG00000135821 | GLUL | chr1 | - | 182386255 | 182386402 | 182385759 | 182385887 | 182387130 | 182387292 | 1237 | 876 | [illegible] |
| SE | 51586 | ENSG00000135821 | GLUL | chr1 | - | 182386255 | 182386402 | 182385759 | 182385887 | 182388571 | 182388750 | 589 | 418 | [illegible] |
| SE | 51588 | ENSG00000135821 | GLUL | chr1 | - | 182386361 | 182386402 | 182385698 | 182385887 | 182388571 | 182388750 | 3 | 2 | [illegible] |
| SE | 51589 | ENSG00000135821 | GLUL | chr1 | - | 182387130 | 182387292 | 182385698 | 182385887 | 182388571 | 182388750 | 419 | 273 | [illegible] |
| SE | 51590 | ENSG00000135821 | GLUL | chr1 | - | 182387130 | 182387292 | 182386194 | 182386402 | 182388571 | 182388750 | 1071 | 742 | [illegible] |
| SE | 51593 | ENSG00000162694 | EXTL2 | chr1 | - | 100876793 | 100876864 | 100872386 | 100874430 | 100877475 | 100877903 | 9 | 0 | [illegible] |
| SE | 51594 | ENSG00000162694 | EXTL2 | chr1 | - | 100878393 | 100878447 | 100877518 | 100877903 | 100888752 | 100888864 | 0 | 0 | [illegible] |
| SE | 51595 | ENSG00000162694 | EXTL2 | chr1 | - | 100878423 | 100878447 | 100877475 | 100877903 | 100888752 | 100888828 | 1 | 0 | [illegible] |
| SE | 51596 | ENSG00000204217 | BMPR2 | chr2 | + | 202467518 | 202467689 | 202464808 | 202464979 | 202513718 | 202513829 | 126 | 92 | [illegible] |
| SE | 51597 | ENSG00000204217 | BMPR2 | chr2 | + | 202467518 | 202467689 | 202464808 | 202464979 | 202514887 | 202514979 | 91 | 59 | [illegible] |
| SE | 51598 | ENSG00000204217 | BMPR2 | chr2 | + | 202513718 | 202513829 | 202464808 | 202464979 | 202514887 | 202514979 | 49 | 42 | [illegible] |
| SE | 51600 | ENSG00000204217 | BMPR2 | chr2 | + | 202518821 | 202519052 | 202514887 | 202514979 | 202520086 | 202520201 | 217 | 175 | [illegible] |
| SE | 51601 | ENSG00000204217 | BMPR2 | chr2 | + | 202520086 | 202520201 | 202518821 | 202519052 | 202530793 | 202530954 | 253 | 249 | [illegible] |
| SE | 51602 | ENSG00000204217 | BMPR2 | chr2 | + | 202530793 | 202530954 | 202520086 | 202520201 | 202542310 | 202542447 | 113 | 120 | [illegible] |
| SE | 51603 | ENSG00000204217 | BMPR2 | chr2 | + | 202532584 | 202532732 | 202530793 | 202530954 | 202542310 | 202542447 | 167 | 134 | [illegible] |
| SE | 51604 | ENSG00000204217 | BMPR2 | chr2 | + | 202555251 | 202556531 | 202552715 | 202552888 | 202559695 | 202560341 | 104 | 92 | [illegible] |
| SE | 51616 | ENSG00000127922 | SEM1 | chr7 | - | 96694797 | 96694891 | 96673456 | 96673859 | 96709687 | 96709775 | 93 | 66 | [illegible] |
| SE | 51617 | ENSG00000127922 | SEM1 | chr7 | - | 96708147 | 96708214 | 96694797 | 96694891 | 96709687 | 96709873 | 0 | 0 | [illegible] |
| SE | 51618 | ENSG00000121864 | ZNF639 | chr3 | + | 179324949 | 179324993 | 179323245 | 179323291 | 179327560 | 179327631 | 0 | 0 | [illegible] |
| SE | 51620 | ENSG00000121864 | ZNF639 | chr3 | + | 179325041 | 179325322 | 179323075 | 179323291 | 179328282 | 179328351 | 2 | 1 | [illegible] |
| SE | 51621 | ENSG00000121864 | ZNF639 | chr3 | + | 179327560 | 179327631 | 179323030 | 179323291 | 179328282 | 179328351 | 80 | 64 | [illegible] |
| SE | 51623 | ENSG00000121864 | ZNF639 | chr3 | + | 179329617 | 179329728 | 179328117 | 179328351 | 179332988 | 179333123 | 309 | 270 | [illegible] |
| SE | 51625 | ENSG00000100628 | ASB2 | chr14 | - | 93953351 | 93953507 | 93950998 | 93951244 | 93954316 | 93954483 | 4 | 2 | [illegible] |
| SE | 51630 | ENSG00000181523 | SGSH | chr17 | - | 80213803 | 80213885 | 80211533 | 80212274 | 80214171 | 80214328 | 49 | 31 | [illegible] |
| SE | 51637 | ENSG00000132128 | LRRC41 | chr1 | - | 46279491 | 46279614 | 46279181 | 46279257 | 46280191 | 46280290 | 84 | 44 | [illegible] |
| SE | 51638 | ENSG00000132128 | LRRC41 | chr1 | - | 46280191 | 46280290 | 46279181 | 46279614 | 46281124 | 46281161 | 33 | 14 | [illegible] |
| SE | 51640 | ENSG00000132128 | LRRC41 | chr1 | - | 46281124 | 46281385 | 46280191 | 46280290 | 46285361 | 46286112 | 34 | 12 | [illegible] |
| SE | 51641 | ENSG00000132128 | LRRC41 | chr1 | - | 46285361 | 46286499 | 46281124 | 46281385 | 46297562 | 46297633 | 88 | 31 | [illegible] |
| SE | 51653 | ENSG00000081154 | PCNP | chr3 | + | 101590214 | 101590270 | 101585436 | 101585511 | 101592626 | 101592771 | 428 | 377 | [illegible] |
| SE | 51665 | ENSG00000136643 | RPS6KC1 | chr1 | + | 213071005 | 213071041 | 213051232 | 213051509 | 213077695 | 213077816 | 25 | 35 | [illegible] |
| SE | 51668 | ENSG00000136643 | RPS6KC1 | chr1 | + | 213104453 | 213104569 | 213077695 | 213077816 | 213117316 | 213117410 | 106 | 115 | [illegible] |
| SE | 51670 | ENSG00000136643 | RPS6KC1 | chr1 | + | 213117316 | 213117410 | 213104453 | 213104569 | 213167857 | 213167973 | 54 | 62 | [illegible] |
| SE | 51672 | ENSG00000136643 | RPS6KC1 | chr1 | + | 213129526 | 213129889 | 213104453 | 213104569 | 213167857 | 213167973 | 55 | 52 | [illegible] |
| SE | 51674 | ENSG00000136643 | RPS6KC1 | chr1 | + | 213176399 | 213176492 | 213167857 | 213167973 | 213230496 | 213230544 | 45 | 34 | [illegible] |
| SE | 51675 | ENSG00000136643 | RPS6KC1 | chr1 | + | 213176399 | 213176492 | 213167857 | 213167973 | 213240701 | 213242297 | 15 | 15 | [illegible] |
| SE | 51676 | ENSG00000136643 | RPS6KC1 | chr1 | + | 213230496 | 213230544 | 213167857 | 213167973 | 213232122 | 213232255 | 8 | 8 | [illegible] |
| SE | 51677 | ENSG00000136643 | RPS6KC1 | chr1 | + | 213232122 | 213232255 | 213167857 | 213167973 | 213240701 | 213242297 | 39 | 30 | [illegible] |
| SE | 51678 | ENSG00000136643 | RPS6KC1 | chr1 | + | 213240701 | 213242297 | 213176399 | 213176492 | 213261557 | 213261640 | 0 | 0 | [illegible] |
| SE | 51679 | ENSG00000136643 | RPS6KC1 | chr1 | + | 213240701 | 213242297 | 213232122 | 213232255 | 213261557 | 213261640 | 42 | 31 | [illegible] |
| SE | 51680 | ENSG00000136643 | RPS6KC1 | chr1 | + | 213242568 | 213242658 | 213240701 | 213242297 | 213261557 | 213261640 | 64 | 48 | [illegible] |
| SE | 51681 | ENSG00000136643 | RPS6KC1 | chr1 | + | 213262720 | 213262816 | 213261557 | 213261640 | 213272523 | 213273464 | 143 | 124 | [illegible] |
| SE | 51682 | ENSG00000198837 | DENND4B | chr1 | - | 153930691 | 153930857 | 153930538 | 153930603 | 153930946 | 153931064 | 122 | 24 | [illegible] |
| SE | 51685 | ENSG00000198837 | DENND4B | chr1 | - | 153932641 | 153932777 | 153932203 | 153932440 | 153932860 | 153932893 | 149 | 61 | [illegible] |
| SE | 51686 | ENSG00000198837 | DENND4B | chr1 | - | 153934134 | 153934302 | 153933482 | 153933871 | 153934759 | 153934964 | 61 | 27 | [illegible] |
| SE | 51689 | ENSG00000198837 | DENND4B | chr1 | - | 153940903 | 153941048 | 153940430 | 153940606 | 153941230 | 153941289 | 78 | 46 | [illegible] |
| SE | 51691 | ENSG00000198837 | DENND4B | chr1 | - | 153942186 | 153942356 | 153941868 | 153942113 | 153942555 | 153942625 | 78 | 41 | [illegible] |
| SE | 51694 | ENSG00000198837 | DENND4B | chr1 | - | 153942555 | 153942625 | 153942186 | 153942356 | 153942877 | 153943130 | 29 | 22 | [illegible] |
| SE | 51695 | ENSG00000143224 | PPOX | chr1 | + | 161166839 | 161166934 | 161166425 | 161166672 | 161170619 | 161170769 | 3 | 0 | [illegible] |
| SE | 51698 | ENSG00000143224 | PPOX | chr1 | + | 161167354 | 161167486 | 161167099 | 161167234 | 161167994 | 161168127 | 1 | 1 | [illegible] |
| SE | 51700 | ENSG00000143224 | PPOX | chr1 | + | 161167370 | 161167486 | 161167099 | 161167234 | 161167994 | 161168095 | 2 | 1 | [illegible] |
| SE | 51705 | ENSG00000143224 | PPOX | chr1 | + | 161167994 | 161168127 | 161167099 | 161167486 | 161168431 | 161168545 | 7 | 1 | [illegible] |
| SE | 51707 | ENSG00000143224 | PPOX | chr1 | + | 161168412 | 161168576 | 161167994 | 161168127 | 161168992 | 161169183 | 5 | 0 | [illegible] |
| SE | 51708 | ENSG00000143224 | PPOX | chr1 | + | 161168431 | 161168576 | 161167994 | 161168127 | 161168992 | 161169183 | 9 | 0 | [illegible] |
| SE | 51709 | ENSG00000143224 | PPOX | chr1 | + | 161168444 | 161168576 | 161167994 | 161168127 | 161168992 | 161169183 | 5 | 2 | [illegible] |
| SE | 51710 | ENSG00000143224 | PPOX | chr1 | + | 161168992 | 161169183 | 161168394 | 161168576 | 161170408 | 161170519 | 5 | 0 | [illegible] |
| SE | 51712 | ENSG00000143224 | PPOX | chr1 | + | 161169905 | 161170024 | 161169659 | 161169720 | 161170408 | 161170519 | 13 | 11 | [illegible] |
| SE | 51714 | ENSG00000143224 | PPOX | chr1 | + | 161170408 | 161170519 | 161169283 | 161170024 | 161170619 | 161170769 | 18 | 14 | [illegible] |
| SE | 51715 | ENSG00000143224 | PPOX | chr1 | + | 161170906 | 161170949 | 161170619 | 161170769 | 161176848 | 161178013 | 9 | 7 | [illegible] |
| SE | 51721 | ENSG00000166441 | RPL27A | chr11 | + | 8684685 | 8684892 | 8684005 | 8684081 | 8685677 | 8685848 | 6848 | 5122 | [illegible] |
| SE | 51722 | ENSG00000166441 | RPL27A | chr11 | + | 8684717 | 8684892 | 8683862 | 8684081 | 8685677 | 8685845 | 15783 | 10014 | [illegible] |
| SE | 51723 | ENSG00000166441 | RPL27A | chr11 | + | 8684822 | 8684892 | 8684005 | 8684081 | 8685677 | 8685848 | 4056 | 3244 | [illegible] |
| SE | 51725 | ENSG00000037637 | FBXO42 | chr1 | - | 16253095 | 16253152 | 16246839 | 16251785 | 16253634 | 16253731 | 22 | 8 | [illegible] |
| SE | 51727 | ENSG00000037637 | FBXO42 | chr1 | - | 16256605 | 16256759 | 16255710 | 16255821 | 16294782 | 16294917 | 127 | 62 | [illegible] |
| SE | 51728 | ENSG00000037637 | FBXO42 | chr1 | - | 16305802 | 16305919 | 16292313 | 16294917 | 16315168 | 16315367 | 127 | 67 | [illegible] |
| SE | 51729 | ENSG00000037637 | FBXO42 | chr1 | - | 16315168 | 16315435 | 16305802 | 16305919 | 16352254 | 16352480 | 106 | 86 | [illegible] |
| SE | 51732 | ENSG00000137941 | TTLL7 | chr1 | - | 83882668 | 83882787 | 83865027 | 83870082 | 83882962 | 83883136 | 0 | 0 | [illegible] |
| SE | 51733 | ENSG00000137941 | TTLL7 | chr1 | - | 83882962 | 83883136 | 83865023 | 83870082 | 83890320 | 83890481 | 5 | 16 | [illegible] |
| SE | 51735 | ENSG00000137941 | TTLL7 | chr1 | - | 83887176 | 83887284 | 83882962 | 83883136 | 83890320 | 83890342 | 0 | 1 | [illegible] |
| SE | 51738 | ENSG00000137941 | TTLL7 | chr1 | - | 83912443 | 83912503 | 83911164 | 83911363 | 83917603 | 83917690 | 0 | 0 | [illegible] |
| SE | 51744 | ENSG00000137941 | TTLL7 | chr1 | - | 83948627 | 83948695 | 83947123 | 83947282 | 83949864 | 83949986 | 2 | 11 | [illegible] |
| SE | 51747 | ENSG00000114841 | DNAH1 | chr3 | + | 52326734 | 52326891 | 52326139 | 52326314 | 52327881 | 52328014 | 48 | 36 | [illegible] |
| SE | 51748 | ENSG00000114841 | DNAH1 | chr3 | + | 52345494 | 52345706 | 52344489 | 52344647 | 52346471 | 52346770 | 26 | 21 | [illegible] |
| SE | 51749 | ENSG00000114841 | DNAH1 | chr3 | + | 52351961 | 52352103 | 52350507 | 52350590 | 52352551 | 52352707 | 58 | 44 | [illegible] |
| SE | 51752 | ENSG00000114841 | DNAH1 | chr3 | + | 52352551 | 52352707 | 52351961 | 52352103 | 52353102 | 52353301 | 58 | 53 | [illegible] |
| SE | 51753 | ENSG00000114841 | DNAH1 | chr3 | + | 52359245 | 52359386 | 52358557 | 52358737 | 52359915 | 52360079 | 32 | 26 | [illegible] |
| SE | 51756 | ENSG00000114841 | DNAH1 | chr3 | + | 52366732 | 52366887 | 52366456 | 52366548 | 52368740 | 52368918 | 43 | 27 | [illegible] |
| SE | 51760 | ENSG00000114841 | DNAH1 | chr3 | + | 52379904 | 52380135 | 52375954 | 52375993 | 52381639 | 52381836 | 31 | 21 | [illegible] |
| SE | 51761 | ENSG00000114841 | DNAH1 | chr3 | + | 52381639 | 52381836 | 52379904 | 52380135 | 52382319 | 52382455 | 47 | 29 | [illegible] |
| SE | 51762 | ENSG00000114841 | DNAH1 | chr3 | + | 52388166 | 52388334 | 52386661 | 52386853 | 52388417 | 52388609 | 66 | 43 | [illegible] |
| SE | 51765 | ENSG00000114841 | DNAH1 | chr3 | + | 52395307 | 52395466 | 52394914 | 52395059 | 52395546 | 52395678 | 71 | 17 | [illegible] |
| SE | 51767 | ENSG00000114841 | DNAH1 | chr3 | + | 52399544 | 52399779 | 52398849 | 52399201 | 52400324 | 52400487 | 38 | 22 | [illegible] |
| SE | 51768 | ENSG00000114841 | DNAH1 | chr3 | + | 52399620 | 52399779 | 52398849 | 52399201 | 52400324 | 52400487 | 24 | 17 | [illegible] |
| SE | 51772 | ENSG00000042429 | MED17 | chr11 | + | 93794249 | 93794439 | 93793727 | 93794035 | 93794907 | 93795060 | 2 | 1 | [illegible] |
| SE | 51773 | ENSG00000042429 | MED17 | chr11 | + | 93794907 | 93795060 | 93793173 | 93794035 | 93796409 | 93796443 | 276 | 236 | [illegible] |
| SE | 51777 | ENSG00000042429 | MED17 | chr11 | + | 93801834 | 93801972 | 93796409 | 93796540 | 93811852 | 93811888 | 11 | 23 | [illegible] |
| SE | 51778 | ENSG00000042429 | MED17 | chr11 | + | 93801834 | 93801972 | 93796936 | 93797719 | 93807517 | 93807635 | 138 | 92 | [illegible] |
| SE | 51780 | ENSG00000042429 | MED17 | chr11 | + | 93807500 | 93809876 | 93801834 | 93801972 | 93811852 | 93812347 | 69 | 41 | [illegible] |
| SE | 51781 | ENSG00000042429 | MED17 | chr11 | + | 93807517 | 93807635 | 93796409 | 93796540 | 93811852 | 93812310 | 0 | 0 | [illegible] |
| SE | 51783 | ENSG00000042429 | MED17 | chr11 | + | 93807517 | 93807635 | 93800660 | 93801972 | 93811852 | 93812310 | 73 | 45 | [illegible] |
| SE | 51784 | ENSG00000042429 | MED17 | chr11 | + | 93809716 | 93809876 | 93801834 | 93801972 | 93811852 | 93811888 | 70 | 43 | [illegible] |
| SE | 51787 | ENSG00000181135 | ZNF707 | chr8 | + | 143684623 | 143684755 | 143684472 | 143684542 | 143690056 | 143690123 | 1 | 0 | [illegible] |
| SE | 51789 | ENSG00000181135 | ZNF707 | chr8 | + | 143684755 | 143685067 | 143684486 | 143684542 | 143689203 | 143689301 | 0 | 0 | [illegible] |
| SE | 51790 | ENSG00000181135 | ZNF707 | chr8 | + | 143684755 | 143685067 | 143684486 | 143684542 | 143690056 | 143690123 | 0 | 0 | [illegible] |
| SE | 51792 | ENSG00000181135 | ZNF707 | chr8 | + | 143686916 | 143687014 | 143684507 | 143684542 | 143690056 | 143690123 | 0 | 0 | [illegible] |
| SE | 51796 | ENSG00000181135 | ZNF707 | chr8 | + | 143689203 | 143689301 | 143684478 | 143684542 | 143690056 | 143690123 | 5 | 2 | [illegible] |
| SE | 51797 | ENSG00000181135 | ZNF707 | chr8 | + | 143689203 | 143689301 | 143684623 | 143684755 | 143690056 | 143690123 | 4 | 2 | [illegible] |
| SE | 51802 | ENSG00000181135 | ZNF707 | chr8 | + | 143691072 | 143691199 | 143690056 | 143690123 | 143691599 | 143691694 | 19 | 11 | [illegible] |
| SE | 51804 | ENSG00000181135 | ZNF707 | chr8 | + | 143691072 | 143691232 | 143690056 | 143690123 | 143691599 | 143691700 | 24 | 6 | [illegible] |
| SE | 51806 | ENSG00000181135 | ZNF707 | chr8 | + | 143691596 | 143691713 | 143691072 | 143691199 | 143693670 | 143693830 | 19 | 5 | [illegible] |
| SE | 51807 | ENSG00000181135 | ZNF707 | chr8 | + | 143691596 | 143691713 | 143691072 | 143691232 | 143693670 | 143693830 | 19 | 5 | [illegible] |
| SE | 51809 | ENSG00000181135 | ZNF707 | chr8 | + | 143691599 | 143691713 | 143691072 | 143691232 | 143693670 | 143693711 | 26 | 7 | [illegible] |
| SE | 51810 | ENSG00000181135 | ZNF707 | chr8 | + | 143691609 | 143691713 | 143691072 | 143691199 | 143693670 | 143693827 | 18 | 5 | [illegible] |
| SE | 51811 | ENSG00000181135 | ZNF707 | chr8 | + | 143692000 | 143692107 | 143691599 | 143691713 | 143692244 | 143692330 | 1 | 0 | [illegible] |
| SE | 51812 | ENSG00000181135 | ZNF707 | chr8 | + | 143692000 | 143692107 | 143691599 | 143691713 | 143693670 | 143693703 | 0 | 0 | [illegible] |
| SE | 51814 | ENSG00000181135 | ZNF707 | chr8 | + | 143692244 | 143692330 | 143692000 | 143692107 | 143693670 | 143693703 | 4 | 0 | [illegible] |
| SE | 51816 | ENSG00000260314 | MRC1 | chr10 | + | 17861386 | 17861502 | 17856241 | 17856352 | 17863533 | 17863682 | 19 | 10 | [illegible] |
| SE | 51819 | ENSG00000135637 | CCDC142 | chr2 | - | 74481451 | 74481538 | 74481222 | 74481372 | 74481816 | 74482189 | 5 | 1 | [illegible] |
| SE | 51821 | ENSG00000113812 | ACTR8 | chr3 | - | 53873031 | 53873127 | 53872383 | 53872524 | 53874210 | 53874364 | 153 | 81 | [illegible] |
| SE | 51822 | ENSG00000113812 | ACTR8 | chr3 | - | 53873031 | 53873146 | 53872383 | 53872524 | 53874210 | 53874364 | 67 | 43 | [illegible] |
| SE | 51823 | ENSG00000113812 | ACTR8 | chr3 | - | 53874210 | 53874364 | 53873031 | 53873146 | 53875947 | 53876080 | 87 | 47 | [illegible] |

| | | | | | | | | | | | | | |
|---|---|---|---|---|---|---|---|---|---|---|---|---|---|
| SE | 51825 | ENSG00000113812 | ACTR8 | chr3 | - | 53877213 | 53877387 | 53875947 | 53876080 | 53877646 | 53877751 | 96 | 72 |
| SE | 51826 | ENSG00000113812 | ACTR8 | chr3 | - | 53878356 | 53878467 | 53877646 | 53877751 | 53879938 | 53880072 | 95 | 74 |
| SE | 51827 | ENSG00000113812 | ACTR8 | chr3 | - | 53879938 | 53880072 | 53878356 | 53878467 | 53881978 | 53882152 | 27 | 26 |
| SE | 51828 | ENSG00000113812 | ACTR8 | chr3 | - | 53879938 | 53880109 | 53878356 | 53878467 | 53881978 | 53882152 | 56 | 51 |
| SE | 51831 | ENSG00000169752 | NRG4 | chr15 | - | 75971165 | 75971248 | 75969128 | 75969224 | 76009199 | 76009293 | 0 | 0 |
| SE | 51835 | ENSG00000087206 | UIMC1 | chr5 | - | 176908522 | 176908694 | 176907113 | 176907177 | 176911310 | 176911389 | 137 | 133 |
| SE | 51836 | ENSG00000087206 | UIMC1 | chr5 | - | 176911310 | 176911389 | 176908522 | 176908694 | 176943334 | 176943488 | 137 | 128 |
| SE | 51838 | ENSG00000087206 | UIMC1 | chr5 | - | 176943334 | 176943488 | 176911310 | 176911389 | 176951473 | 176951577 | 446 | 345 |
| SE | 51839 | ENSG00000087206 | UIMC1 | chr5 | - | 176951473 | 176951577 | 176943334 | 176943488 | 176955958 | 176956035 | 572 | 418 |
| SE | 51844 | ENSG00000087206 | UIMC1 | chr5 | - | 176969596 | 176969706 | 176968554 | 176969291 | 176970741 | 176970866 | 73 | 54 |
| SE | 51846 | ENSG00000087206 | UIMC1 | chr5 | - | 176969600 | 176969706 | 176968554 | 176969291 | 176970741 | 176970793 | 69 | 51 |
| SE | 51847 | ENSG00000087206 | UIMC1 | chr5 | - | 176970741 | 176970866 | 176958092 | 176958154 | 176975395 | 176975480 | 93 | 46 |
| SE | 51848 | ENSG00000087206 | UIMC1 | chr5 | - | 176975395 | 176975480 | 176969940 | 176970866 | 176982468 | 176982623 | 186 | 91 |
| SE | 51849 | ENSG00000087206 | UIMC1 | chr5 | - | 176982468 | 176982623 | 176975395 | 176975480 | 177006649 | 177006715 | 188 | 95 |
| SE | 51850 | ENSG00000087206 | UIMC1 | chr5 | - | 176982468 | 176982623 | 176975395 | 176975480 | 177022463 | 177022602 | 137 | 71 |
| SE | 51852 | ENSG00000087206 | UIMC1 | chr5 | - | 177001997 | 177002111 | 176982520 | 176982623 | 177006538 | 177006570 | 0 | 0 |
| SE | 51853 | ENSG00000162496 | DHRS3 | chr1 | - | 12572727 | 12572853 | 12567909 | 12568424 | 12578717 | 12578956 | 107 | 22 |
| SE | 51854 | ENSG00000162496 | DHRS3 | chr1 | - | 12572727 | 12572853 | 12567909 | 12568424 | 12580522 | 12580666 | 63 | 9 |
| SE | 51857 | ENSG00000162496 | DHRS3 | chr1 | - | 12580522 | 12580666 | 12572727 | 12572853 | 12617153 | 12618210 | 114 | 33 |
| SE | 51869 | ENSG00000138448 | ITGAV | chr2 | + | 186630796 | 186630858 | 186625472 | 186625587 | 186633328 | 186633374 | 35 | 35 |
| SE | 51870 | ENSG00000138448 | ITGAV | chr2 | + | 186633328 | 186633374 | 186625472 | 186625587 | 186636081 | 186636207 | 8 | 6 |
| SE | 51873 | ENSG00000138448 | ITGAV | chr2 | + | 186675603 | 186675717 | 186669700 | 186669814 | 186675819 | 186675927 | 83 | 52 |
| SE | 51874 | ENSG00000138448 | ITGAV | chr2 | + | 186675819 | 186675927 | 186669700 | 186669814 | 186676812 | 186676935 | 26 | 19 |
| SE | 51875 | ENSG00000138448 | ITGAV | chr2 | + | 186676812 | 186676935 | 186675819 | 186675927 | 186677196 | 186677441 | 90 | 63 |
| SE | 51876 | ENSG00000147576 | ADHFE1 | chr8 | + | 66444366 | 66444420 | 66432504 | 66432575 | 66444593 | 66444648 | 10 | 10 |
| SE | 51878 | ENSG00000147576 | ADHFE1 | chr8 | + | 66447263 | 66447341 | 66444593 | 66444748 | 66448864 | 66448970 | 37 | 20 |
| SE | 51881 | ENSG00000285791 | NA | chr8 | + | 66474792 | 66474952 | 66473785 | 66473938 | 66496436 | 66496492 | 0 | 1 |
| SE | 51882 | ENSG00000285791 | NA | chr8 | + | 66496436 | 66496492 | 66473783 | 66473938 | 66505374 | 66505468 | 3 | 5 |
| SE | 51883 | ENSG00000184005 | ST6GALNAC3 | chr1 | + | 76313804 | 76313999 | 76074745 | 76074884 | 76412007 | 76412417 | 16 | 10 |

[illegible]

| | | | | | | | | | | | | | | | | | | | | | | | | | | | | | | | | | | | | | | | | | | | | | | |
|---|---|---|---|---|---|---|---|---|---|---|---|---|---|---|---|---|---|---|---|---|---|---|---|---|---|---|---|---|---|---|---|---|---|---|---|---|---|---|---|---|---|---|---|---|---|---|
| SE | 52219 | ENSG00000155363 | MOV10 | chr1 | + | 112685034 | 112685162 | 112674847 | 112675049 | 112686965 | 112687106 | 0 | 0 | 0 | 0 | 4 | 9 | 4 | 5 | 11 | 8 | 0 | 0 | 0 | 0 | 0 | 30 | 13 | 4 | 5 | 41 | 277 | 149 | 1 | 1 | 0 | 0 | 0 | 0 | 0.212 | 0 | 0 | 0 | 0 | 0 | 0.042 |
| SE | 52220 | ENSG00000155363 | MOV10 | chr1 | + | 112685034 | 112685162 | 112674847 | 112675049 | 112688934 | 112689138 | 0 | 0 | 0 | 0 | 2 | 12 | 13 | 17 | 19 | 11 | 0 | 1 | 0 | 0 | 0 | 39 | 23 | 2 | 27 | 21 | 277 | 149 | 1 | 1 | 0 | 0 | 0 | 0 | 0.089 | 0 | 0.023 | 0 | 0 | 0 | 0.013 |
| SE | 52223 | ENSG00000155363 | MOV10 | chr1 | + | 112686965 | 112687106 | 112685034 | 112685162 | 112688934 | 112689138 | 2 | 3 | 6 | 2 | 11 | 0 | 0 | 0 | 0 | 0 | 0 | 9 | 0 | 9 | 11 | 0 | 1 | 0 | 0 | 0 | 290 | 149 | 1 | 1 | 1 | 1 | 1 | 1 | 1 | NA | 0.822 | NA | 1 | 1 | 0.059 |
| SE | 52224 | ENSG00000155363 | MOV10 | chr1 | + | 112691664 | 112691799 | 112689839 | 112690098 | 112692760 | 112692929 | 99 | 60 | 87 | 88 | 74 | 1 | 0 | 0 | 0 | 0 | 227 | 132 | 82 | 142 | 132 | 0 | 0 | 0 | 0 | 0 | 284 | 149 | 1 | 1 | 0.981 | 1 | 1 | 1 | 1 | 1 | 1 | 1 | 1 | 1 | -0.004 |
| SE | 52226 | ENSG00000155363 | MOV10 | chr1 | + | 112695415 | 112695574 | 112694452 | 112694629 | 112696147 | 112696251 | 50 | 25 | 59 | 63 | 66 | 0 | 0 | 0 | 0 | 0 | 73 | 60 | 44 | 49 | 79 | 0 | 0 | 0 | 1 | 0 | 298 | 149 | 1 | 1 | 1 | 1 | 1 | 1 | 1 | 1 | 1 | 1 | 0.961 | 1 | 0.008 |
| SE | 52227 | ENSG00000155363 | MOV10 | chr1 | + | 112696436 | 112696534 | 112696147 | 112696251 | 112696629 | 112696846 | 61 | 32 | 73 | 57 | 56 | 0 | 0 | 1 | 0 | 0 | 93 | 42 | 42 | 99 | 54 | 0 | 0 | 0 | 0 | 0 | 247 | 149 | 1 | 1 | 1 | 1 | 0.978 | 1 | 1 | 1 | 1 | 1 | 1 | 1 | -0.004 |
| SE | 52231 | ENSG00000189171 | S100A13 | chr1 | - | 153630449 | 153630656 | 153628120 | 153628237 | 153631010 | 153631360 | 0 | 0 | 1 | 0 | 0 | 5 | 3 | 12 | 3 | 3 | 0 | 0 | 1 | 0 | 0 | 0 | 2 | 0 | 3 | 6 | 298 | 149 | 1 | 1 | 0 | 0 | 0.04 | 0 | 0 | NA | 0 | 1 | 0 | 0 | -0.242 |
| SE | 52233 | ENSG00000085185 | BCORL1 | chrX | + | 130012949 | 130016213 | 130012577 | 130012668 | 130020984 | 130021150 | 21 | 15 | 29 | 39 | 16 | 0 | 0 | 0 | 0 | 0 | 57 | 28 | 21 | 43 | 71 | 0 | 1 | 0 | 0 | 0 | 298 | 149 | 1 | 1 | 1 | 1 | 1 | 1 | 1 | 1 | 0.933 | 1 | 1 | 1 | 0.013 |
| SE | 52235 | ENSG00000085185 | BCORL1 | chrX | + | 130034454 | 130034676 | 130028634 | 130028861 | 130037366 | 130037533 | 33 | 36 | 33 | 41 | 70 | 0 | 1 | 0 | 0 | 0 | 33 | 23 | 8 | 62 | 50 | 0 | 0 | 0 | 0 | 0 | 298 | 149 | 1 | 1 | 1 | 0.947 | 1 | 1 | 1 | 1 | 1 | 1 | 1 | 1 | -0.011 |
| SE | 52238 | ENSG00000166825 | ANPEP | chr15 | - | 89790952 | 89791093 | 89790459 | 89790541 | 89792159 | 89792327 | 510 | 215 | 179 | 204 | 274 | 0 | 0 | 0 | 0 | 0 | 1592 | 794 | 469 | 589 | 679 | 0 | 0 | 1 | 0 | 0 | 290 | 149 | 1 | 1 | 1 | 1 | 1 | 1 | 1 | 1 | 1 | 0.996 | 1 | 1 | 0.001 |
| SE | 52240 | ENSG00000166825 | ANPEP | chr15 | - | 89799259 | 89799315 | 89793034 | 89793126 | 89799425 | 89799559 | 92 | 30 | 25 | 24 | 87 | 0 | 0 | 0 | 1 | 0 | 222 | 146 | 64 | 123 | 128 | 0 | 0 | 0 | 0 | 0 | 205 | 149 | 1 | 1 | 1 | 1 | 1 | 0.946 | 1 | 1 | 1 | 1 | 1 | 1 | -0.011 |
| SE | 52241 | ENSG00000166825 | ANPEP | chr15 | - | 89804945 | 89805217 | 89804490 | 89804617 | 89805320 | 89805372 | 104 | 36 | 38 | 30 | 60 | 0 | 0 | 0 | 0 | 0 | 229 | 165 | 80 | 120 | 222 | 0 | 0 | 1 | 0 | 0 | 298 | 149 | 1 | 1 | 1 | 1 | 1 | 1 | 1 | 1 | 1 | 0.976 | 1 | 1 | 0.005 |
| SE | 52242 | ENSG00000166825 | ANPEP | chr15 | - | 89805077 | 89805217 | 89804490 | 89804617 | 89805320 | 89805463 | 170 | 60 | 52 | 47 | 127 | 0 | 0 | 0 | 0 | 0 | 416 | 205 | 104 | 178 | 359 | 0 | 0 | 1 | 0 | 0 | 289 | 149 | 1 | 1 | 1 | 1 | 1 | 1 | 1 | 1 | 1 | 0.982 | 1 | 1 | 0.004 |
| SE | 52244 | ENSG00000166825 | ANPEP | chr15 | - | 89814113 | 89814294 | 89805969 | 89806945 | 89814771 | 89814852 | 2 | 2 | 1 | 0 | 1 | 1 | 0 | 1 | 0 | 0 | 15 | 1 | 6 | 0 | 0 | 0 | 4 | 3 | 0 | 0 | 298 | 149 | 1 | 1 | 0.5 | 1 | 0.333 | NA | 1 | 1 | 0.111 | 0.5 | NA | NA | 0.171 |
| SE | 52245 | ENSG00000117016 | RIMS3 | chr1 | - | 40641708 | 40641956 | 40635915 | 40636057 | 40647667 | 40647762 | 7 | 6 | 0 | 5 | 6 | 0 | 0 | 1 | 0 | 0 | 0 | 10 | 2 | 0 | 7 | 0 | 0 | 0 | 0 | 0 | 298 | 149 | 1 | 1 | 1 | 1 | 0 | 1 | 1 | NA | 1 | 1 | NA | 1 | -0.2 |
| SE | 52248 | ENSG00000092621 | PHGDH | chr1 | + | 119724318 | 119724469 | 119723375 | 119723441 | 119726850 | 119726905 | 0 | 0 | 0 | 0 | 3 | 2 | 2 | 18 | 7 | 5 | 0 | 0 | 0 | 0 | 0 | 10 | 2 | 8 | 3 | 21 | 298 | 149 | 1 | 1 | 0 | 0 | 0 | 0 | 0.231 | 0 | 0 | 0 | 0 | 0 | 0.046 |
| SE | 52253 | ENSG00000092621 | PHGDH | chr1 | + | 119729508 | 119729633 | 119727003 | 119727102 | 119734633 | 119734682 | 0 | 1 | 1 | 0 | 1 | 5 | 2 | 8 | 2 | 18 | 0 | 0 | 0 | 0 | 0 | 6 | 0 | 3 | 4 | 11 | 274 | 149 | 1 | 1 | 0 | 0.214 | 0.064 | 0 | 0.029 | 0 | NA | 0 | 0 | 0 | 0.061 |
| SE | 52259 | ENSG00000169607 | CKAP2L | chr2 | - | 112760712 | 112760764 | 112756774 | 112757155 | 112762502 | 112762569 | 0 | 1 | 4 | 0 | 0 | 0 | 0 | 0 | 0 | 0 | 0 | 0 | 0 | 0 | 0 | 0 | 0 | 1 | 0 | 0 | 201 | 149 | 1 | 1 | NA | 1 | 1 | NA | NA | NA | NA | 0 | NA | NA | 1 |
| SE | 52262 | ENSG00000132321 | IQCA1 | chr2 | - | 236487999 | 236488191 | 236486828 | 236486900 | 236493767 | 236493887 | 5 | 0 | 3 | 0 | 1 | 0 | 0 | 0 | 0 | 3 | 0 | 2 | 0 | 0 | 0 | 0 | 0 | 0 | 0 | 0 | 298 | 149 | 1 | 1 | 1 | NA | 1 | NA | 0.143 | NA | 1 | NA | NA | NA | -0.286 |
| SE | 52265 | ENSG00000267787 | NA | chr18 | + | 57731413 | 57732023 | 57641558 | 57642052 | 57734025 | 57734096 | 13 | 12 | 10 | 13 | 20 | 1 | 2 | 0 | 0 | 2 | 12 | 18 | 6 | 0 | 15 | 0 | 0 | 3 | 0 | 0 | 298 | 149 | 1 | 1 | 0.867 | 0.75 | 1 | 1 | 0.833 | 1 | 1 | 0.5 | NA | 1 | 0.015 |
| [illegible] | | | | | | | | | | | | | | | | | | | | | | | | | | | | | | | | | | | | | | | | | | | | | | |

| | | | | | | | | | | | | | | |
|---|---|---|---|---|---|---|---|---|---|---|---|---|---|---|
| SE | 52606 | ENSG00000115548 | KDM3A | chr2 | + | 86451102 | 86451213 | 86449806 | 86449962 | 86455084 | 86455143 | 175 | 108 | [illegible] |
| SE | 52608 | ENSG00000115548 | KDM3A | chr2 | + | 86474775 | 86474990 | 86470203 | 86470408 | 86477876 | 86478029 | 331 | 229 | [illegible] |
| SE | 52611 | ENSG00000164983 | TMEM65 | chr8 | - | 124330747 | 124330792 | 124323320 | 124323375 | 124371853 | 124372701 | 9 | 5 | [illegible] |
| SE | 52612 | ENSG00000164983 | TMEM65 | chr8 | - | 124330747 | 124330792 | 124327353 | 124327421 | 124371853 | 124372701 | 76 | 38 | [illegible] |
| SE | 52613 | ENSG00000197872 | CYRIA | chr2 | - | 16555068 | 16555139 | 16549458 | 16552999 | 16559459 | 16559586 | 194 | 133 | [illegible] |
| SE | 52615 | ENSG00000197872 | CYRIA | chr2 | - | 16562004 | 16562141 | 16561455 | 16561533 | 16565645 | 16565767 | 151 | 104 | [illegible] |
| SE | 52616 | ENSG00000197872 | CYRIA | chr2 | - | 16562004 | 16562141 | 16561455 | 16561533 | 16588049 | 16588129 | 149 | 102 | [illegible] |
| SE | 52617 | ENSG00000197872 | CYRIA | chr2 | - | 16563988 | 16564094 | 16561455 | 16561533 | 16565645 | 16565767 | 118 | 83 | [illegible] |
| SE | 52618 | ENSG00000197872 | CYRIA | chr2 | - | 16563988 | 16564094 | 16561455 | 16561533 | 16588049 | 16588129 | 7 | 2 | [illegible] |
| SE | 52620 | ENSG00000197872 | CYRIA | chr2 | - | 16563988 | 16564094 | 16562004 | 16562141 | 16588049 | 16588129 | 153 | 112 | [illegible] |
| SE | 52621 | ENSG00000197872 | CYRIA | chr2 | - | 16565645 | 16565767 | 16561455 | 16561533 | 16588049 | 16588129 | 101 | 68 | [illegible] |
| SE | 52622 | ENSG00000197872 | CYRIA | chr2 | - | 16565645 | 16565767 | 16562004 | 16562141 | 16588049 | 16588129 | 102 | 70 | [illegible] |
| SE | 52624 | ENSG00000197872 | CYRIA | chr2 | - | 16565645 | 16565767 | 16563988 | 16564094 | 16623863 | 16624019 | 108 | 66 | [illegible] |
| SE | 52625 | ENSG00000197872 | CYRIA | chr2 | - | 16588049 | 16588129 | 16563988 | 16564094 | 16623863 | 16624019 | 84 | 38 | [illegible] |
| SE | 52628 | ENSG00000197872 | CYRIA | chr2 | - | 16625727 | 16625769 | 16623863 | 16624019 | 16665779 | 16665834 | 2 | 0 | [illegible] |
| SE | 52629 | ENSG00000197943 | PLCG2 | chr16 | + | 81755869 | 81755966 | 81739096 | 81739385 | 81785942 | 81786182 | 9 | 1 | [illegible] |
| SE | 52630 | ENSG00000197943 | PLCG2 | chr16 | + | 81852049 | 81852126 | 81785989 | 81786182 | 81854443 | 81854587 | 0 | 1 | [illegible] |
| SE | 52631 | ENSG00000197943 | PLCG2 | chr16 | + | 81854443 | 81854587 | 81785745 | 81786182 | 81858262 | 81858356 | 265 | 86 | [illegible] |
| SE | 52640 | ENSG00000197943 | PLCG2 | chr16 | + | 81905402 | 81905507 | 81900611 | 81900780 | 81908415 | 81908591 | 79 | 46 | [illegible] |
| SE | 52641 | ENSG00000197943 | PLCG2 | chr16 | + | 81907684 | 81907774 | 81900611 | 81900780 | 81908415 | 81908591 | 181 | 109 | [illegible] |
| SE | 52642 | ENSG00000197943 | PLCG2 | chr16 | + | 81907684 | 81907774 | 81905402 | 81905507 | 81908415 | 81908591 | 379 | 234 | [illegible] |
| SE | 52643 | ENSG00000197943 | PLCG2 | chr16 | + | 81908415 | 81908591 | 81907684 | 81907774 | 81912596 | 81912716 | 322 | 191 | [illegible] |
| SE | 52644 | ENSG00000197943 | PLCG2 | chr16 | + | 81910519 | 81910720 | 81908415 | 81908591 | 81912596 | 81912716 | 145 | 46 | [illegible] |
| SE | 52650 | ENSG00000197943 | PLCG2 | chr16 | + | 81946174 | 81946263 | 81939891 | 81940059 | 81956694 | 81956730 | 351 | 212 | [illegible] |
| SE | 52651 | ENSG00000253250 | C8orf88 | chr8 | - | 90978578 | 90978652 | 90971065 | 90971141 | 90980362 | 90980461 | 2 | 3 | [illegible] |
| SE | 52652 | ENSG00000112232 | KHDRBS2 | chr6 | - | 61707329 | 61707403 | 61697194 | 61697253 | 61732681 | 61732764 | 0 | 0 | [illegible] |
| SE | 52653 | ENSG00000112232 | KHDRBS2 | chr6 | - | 62177184 | 62177312 | 62047877 | 62047994 | 62285857 | 62286225 | 15 | 9 | [illegible] |
| SE | 52654 | ENSG00000123416 | TUBA1B | chr12 | - | 49130118 | 49130274 | 49129499 | 49129722 | 49131297 | 49131397 | 2 | 0 | [illegible] |
| SE | 52655 | ENSG00000123416 | TUBA1B | chr12 | - | 49130118 | 49130344 | 49129499 | 49129722 | 49131297 | 49131395 | 2 | 0 | [illegible] |
| SE | 52656 | ENSG00000123416 | TUBA1B | chr12 | - | 49130118 | 49130402 | 49129499 | 49129722 | 49131297 | 49131395 | 2 | 0 | [illegible] |
| SE | 52657 | ENSG00000203705 | TATDN3 | chr1 | + | 212796516 | 212796590 | 212794869 | 212795127 | 212802700 | 212802763 | 8 | 6 | [illegible] |
| SE | 52658 | ENSG00000203705 | TATDN3 | chr1 | + | 212797111 | 212797196 | 212796516 | 212796590 | 212802700 | 212802751 | 49 | 36 | [illegible] |
| SE | 52660 | ENSG00000203705 | TATDN3 | chr1 | + | 212802700 | 212802763 | 212797111 | 212797196 | 212807735 | 212807848 | 7 | 6 | [illegible] |
| SE | 52664 | ENSG00000203705 | TATDN3 | chr1 | + | 212812226 | 212812328 | 212807735 | 212807848 | 212815046 | 212815597 | 2 | 1 | [illegible] |
| SE | 52665 | ENSG00000203705 | TATDN3 | chr1 | + | 212812247 | 212812328 | 212807735 | 212807848 | 212815012 | 212815415 | 56 | 26 | [illegible] |
| SE | 52669 | ENSG00000198860 | TSEN15 | chr1 | + | 184054727 | 184054863 | 184054353 | 184054435 | 184072156 | 184072194 | 122 | 50 | [illegible] |
| SE | 52670 | ENSG00000198860 | TSEN15 | chr1 | + | 184054727 | 184054863 | 184054353 | 184054435 | 184072826 | 184074212 | 87 | 42 | [illegible] |
| SE | 52671 | ENSG00000198860 | TSEN15 | chr1 | + | 184054727 | 184054863 | 184054353 | 184054435 | 184095689 | 184097485 | 63 | 38 | [illegible] |
| SE | 52677 | ENSG00000198860 | TSEN15 | chr1 | + | 184072156 | 184072298 | 184054353 | 184054435 | 184072826 | 184072986 | 50 | 32 | [illegible] |
| SE | 52681 | ENSG00000198860 | TSEN15 | chr1 | + | 184093624 | 184093724 | 184054727 | 184054863 | 184095030 | 184095092 | 2 | 2 | [illegible] |
| SE | 52682 | ENSG00000198860 | TSEN15 | chr1 | + | 184093624 | 184093724 | 184054727 | 184054863 | 184095689 | 184095845 | 2 | 2 | [illegible] |
| SE | 52683 | ENSG00000198860 | TSEN15 | chr1 | + | 184095030 | 184095092 | 184054727 | 184054863 | 184095689 | 184095845 | 0 | 0 | [illegible] |
| SE | 52685 | ENSG00000198860 | TSEN15 | chr1 | + | 184112864 | 184112960 | 184054727 | 184054863 | 184119719 | 184119835 | 3 | 2 | [illegible] |
| SE | 52687 | ENSG00000196235 | SUPT5H | chr19 | + | 39445803 | 39445965 | 39445440 | 39445637 | 39453355 | 39453442 | 83 | 54 | [illegible] |
| [illegible] | | | | | | | | | | | | | | |

| | | | | | | | | | | | | | | |
|---|---|---|---|---|---|---|---|---|---|---|---|---|---|---|
| SE | 53006 | ENSG00000158941 | CCAR2 | chr8 | + | 22617695 | 22617778 | 22617419 | 22617564 | 22618348 | 22618495 | 171 | 94 | [illegible] |
| SE | 53007 | ENSG00000158941 | CCAR2 | chr8 | + | 22617698 | 22617778 | 22617387 | 22617564 | 22618348 | 22618495 | 58 | 39 | [illegible] |
| SE | 53035 | ENSG00000197183 | NOL4L | chr20 | - | 32520810 | 32520922 | 32511346 | 32511456 | 32527757 | 32527913 | 137 | 84 | [illegible] |
| SE | 53036 | ENSG00000103657 | HERC1 | chr15 | - | 63616429 | 63616682 | 63615767 | 63615920 | 63622814 | 63622891 | 221 | 143 | [illegible] |
| SE | 53038 | ENSG00000103657 | HERC1 | chr15 | - | 63632290 | 63632605 | 63630531 | 63630635 | 63632708 | 63632811 | 0 | 0 | [illegible] |
| SE | 53040 | ENSG00000103657 | HERC1 | chr15 | - | 63633847 | 63633970 | 63630465 | 63630635 | 63634732 | 63634888 | 71 | 51 | [illegible] |
| SE | 53041 | ENSG00000103657 | HERC1 | chr15 | - | 63634732 | 63634888 | 63633847 | 63633970 | 63635960 | 63636142 | 145 | 89 | [illegible] |
| SE | 53042 | ENSG00000103657 | HERC1 | chr15 | - | 63634732 | 63634888 | 63633847 | 63633970 | 63636987 | 63637134 | 89 | 50 | [illegible] |
| SE | 53043 | ENSG00000103657 | HERC1 | chr15 | - | 63634732 | 63634888 | 63633847 | 63633970 | 63637504 | 63637550 | 89 | 52 | [illegible] |
| SE | 53044 | ENSG00000103657 | HERC1 | chr15 | - | 63635960 | 63636142 | 63633847 | 63633970 | 63636987 | 63637134 | 3 | 1 | [illegible] |
| SE | 53045 | ENSG00000103657 | HERC1 | chr15 | - | 63635960 | 63636142 | 63633847 | 63633970 | 63637504 | 63637643 | 114 | 71 | [illegible] |
| SE | 53046 | ENSG00000103657 | HERC1 | chr15 | - | 63635960 | 63636142 | 63633847 | 63633970 | 63638410 | 63638536 | 0 | 0 | [illegible] |
| SE | 53047 | ENSG00000103657 | HERC1 | chr15 | - | 63635960 | 63636142 | 63634732 | 63634888 | 63636987 | 63637134 | 62 | 40 | [illegible] |
| SE | 53049 | ENSG00000103657 | HERC1 | chr15 | - | 63636987 | 63637134 | 63633962 | 63633970 | 63637504 | 63637550 | 1 | 0 | [illegible] |
| SE | 53050 | ENSG00000103657 | HERC1 | chr15 | - | 63636987 | 63637134 | 63634732 | 63634888 | 63637504 | 63637550 | 3 | 0 | [illegible] |
| SE | 53052 | ENSG00000103657 | HERC1 | chr15 | - | 63637504 | 63637643 | 63633847 | 63633970 | 63638410 | 63638536 | 94 | 58 | [illegible] |
| SE | 53053 | ENSG00000103657 | HERC1 | chr15 | - | 63637504 | 63637643 | 63635960 | 63636142 | 63638410 | 63638536 | 207 | 125 | [illegible] |
| SE | 53054 | ENSG00000103657 | HERC1 | chr15 | - | 63638410 | 63638536 | 63637504 | 63637643 | 63638710 | 63638776 | 158 | 83 | [illegible] |
| SE | 53055 | ENSG00000103657 | HERC1 | chr15 | - | 63642956 | 63643058 | 63641469 | 63641643 | 63643403 | 63643550 | 150 | 138 | [illegible] |
| SE | 53057 | ENSG00000103657 | HERC1 | chr15 | - | 63648068 | 63648199 | 63645482 | 63645682 | 63651252 | 63651380 | 242 | 189 | [illegible] |
| SE | 53059 | ENSG00000103657 | HERC1 | chr15 | - | 63649724 | 63649925 | 63648068 | 63648199 | 63651252 | 63651380 | 352 | 275 | [illegible] |
| SE | 53060 | ENSG00000103657 | HERC1 | chr15 | - | 63662983 | 63663204 | 63661752 | 63662021 | 63664469 | 63664594 | 255 | 233 | [illegible] |
| SE | 53062 | ENSG00000103657 | HERC1 | chr15 | - | 63689588 | 63689699 | 63686358 | 63686535 | 63690540 | 63690647 | 328 | 316 | [illegible] |
| SE | 53063 | ENSG00000103657 | HERC1 | chr15 | - | 63692410 | 63692566 | 63690540 | 63690647 | 63693963 | 63694157 | 258 | 219 | [illegible] |
| SE | 53064 | ENSG00000103657 | HERC1 | chr15 | - | 63694773 | 63694894 | 63694311 | 63694549 | 63696123 | 63696339 | 152 | 104 | [illegible] |
| SE | 53070 | ENSG00000103657 | HERC1 | chr15 | - | 63718782 | 63718897 | 63718428 | 63718694 | 63723181 | 63723257 | 579 | 492 | [illegible] |
| SE | 53071 | ENSG00000103657 | HERC1 | chr15 | - | 63723181 | 63723355 | 63716301 | 63716473 | 63727646 | 63727838 | 0 | 1 | [illegible] |
| SE | 53076 | ENSG00000103657 | HERC1 | chr15 | - | 63727646 | 63727838 | 63718782 | 63718897 | 63729235 | 63729368 | 190 | 134 | [illegible] |
| SE | 53077 | ENSG00000103657 | HERC1 | chr15 | - | 63734723 | 63734849 | 63732923 | 63733145 | 63746917 | 63747083 | 234 | 188 | [illegible] |
| SE | 53079 | ENSG00000103657 | HERC1 | chr15 | - | 63747723 | 63747858 | 63734642 | 63734849 | 63749366 | 63749449 | 211 | 207 | [illegible] |
| SE | 53080 | ENSG00000103657 | HERC1 | chr15 | - | 63756436 | 63756748 | 63755228 | 63755325 | 63758174 | 63758369 | 469 | 473 | [illegible] |
| SE | 53081 | ENSG00000103657 | HERC1 | chr15 | - | 63756436 | 63756748 | 63755228 | 63755325 | 63774693 | 63775649 | 338 | 340 | [illegible] |
| SE | 53082 | ENSG00000103657 | HERC1 | chr15 | - | 63758174 | 63758369 | 63755228 | 63755325 | 63774693 | 63775649 | 1 | 3 | [illegible] |
| SE | 53083 | ENSG00000103657 | HERC1 | chr15 | - | 63758174 | 63758369 | 63756436 | 63756748 | 63764095 | 63764191 | 437 | 369 | [illegible] |
| SE | 53084 | ENSG00000103657 | HERC1 | chr15 | - | 63758174 | 63758369 | 63756436 | 63756748 | 63774693 | 63775649 | 132 | 133 | [illegible] |
| SE | 53085 | ENSG00000103657 | HERC1 | chr15 | - | 63764095 | 63764191 | 63756436 | 63756748 | 63774693 | 63775649 | 191 | 168 | [illegible] |
| SE | 53086 | ENSG00000103657 | HERC1 | chr15 | - | 63764095 | 63764191 | 63758174 | 63758369 | 63774693 | 63775649 | 494 | 403 | [illegible] |
| SE | 53093 | ENSG00000103657 | HERC1 | chr15 | - | 63780461 | 63780552 | 63779582 | 63780009 | 63833826 | 63833948 | 0 | 1 | [illegible] |
| SE | 53095 | ENSG00000006756 | ARSD | chrX | - | 2910658 | 2910793 | 2909816 | 2909979 | 2915555 | 2915692 | 147 | 147 | [illegible] |
| SE | 53097 | ENSG00000006756 | ARSD | chrX | - | 2915555 | 2915692 | 2913613 | 2914773 | 2917803 | 2918227 | 83 | 66 | [illegible] |
| SE | 53101 | ENSG00000166352 | IFTAP | chr11 | + | 36597643 | 36597730 | 36594368 | 36594592 | 36610080 | 36610239 | 0 | 0 | [illegible] |
| SE | 53102 | ENSG00000166352 | IFTAP | chr11 | + | 36597643 | 36597730 | 36594498 | 36594641 | 36610080 | 36610239 | 0 | 0 | [illegible] |
| SE | 53104 | ENSG00000166352 | IFTAP | chr11 | + | 36597643 | 36597730 | 36594545 | 36594915 | 36610080 | 36610239 | 0 | 0 | [illegible] |
| SE | 53106 | ENSG00000166352 | IFTAP | chr11 | + | 36610080 | 36610239 | 36594492 | 36594761 | 36633283 | 36633438 | 26 | 19 | [illegible] |
| SE | 53107 | ENSG00000166352 | IFTAP | chr11 | + | 36610080 | 36610239 | 36594501 | 36594641 | 36633283 | 36633438 | 30 | 21 | [illegible] |
| SE | 53108 | ENSG00000166352 | IFTAP | chr11 | + | 36610080 | 36610239 | 36594501 | 36594641 | 36648015 | 36648155 | 8 | 4 | [illegible] |
| SE | 53109 | ENSG00000166352 | IFTAP | chr11 | + | 36633283 | 36633438 | 36594498 | 36594641 | 36648015 | 36648155 | 1 | 0 | [illegible] |
| SE | 53110 | ENSG00000166352 | IFTAP | chr11 | + | 36633283 | 36633438 | 36610080 | 36610239 | 36636050 | 36636117 | 55 | 38 | [illegible] |
| SE | 53112 | ENSG00000166352 | IFTAP | chr11 | + | 36633283 | 36633438 | 36610080 | 36610239 | 36659018 | 36659268 | 26 | 19 | [illegible] |
| SE | 53113 | ENSG00000166352 | IFTAP | chr11 | + | 36633285 | 36633438 | 36610080 | 36610239 | 36636050 | 36636117 | 29 | 19 | [illegible] |
| SE | 53114 | ENSG00000166352 | IFTAP | chr11 | + | 36633285 | 36633438 | 36610080 | 36610239 | 36659018 | 36659291 | 0 | 0 | [illegible] |
| SE | 53116 | ENSG00000166352 | IFTAP | chr11 | + | 36636050 | 36636117 | 36610103 | 36610239 | 36659018 | 36659268 | 0 | 0 | [illegible] |
| SE | 53117 | ENSG00000166352 | IFTAP | chr11 | + | 36636050 | 36636117 | 36633283 | 36633438 | 36648015 | 36648155 | 35 | 27 | [illegible] |
| SE | 53118 | ENSG00000166352 | IFTAP | chr11 | + | 36636050 | 36636117 | 36633283 | 36633438 | 36659018 | 36659268 | 24 | 15 | [illegible] |
| SE | 53119 | ENSG00000166352 | IFTAP | chr11 | + | 36648015 | 36648155 | 36610080 | 36610239 | 36659018 | 36659201 | 12 | 14 | [illegible] |
| SE | 53120 | ENSG00000166352 | IFTAP | chr11 | + | 36648015 | 36648155 | 36633283 | 36633438 | 36659018 | 36659201 | 9 | 12 | [illegible] |
| SE | 53121 | ENSG00000166352 | IFTAP | chr11 | + | 36648015 | 36648155 | 36635428 | 36636117 | 36659018 | 36659201 | 19 | 24 | [illegible] |
| SE | 53123 | ENSG00000185436 | IFNLR1 | chr1 | - | 24159473 | 24159633 | 24157098 | 24157891 | 24169416 | 24169601 | 0 | 1 | [illegible] |
| SE | 53124 | ENSG00000185436 | IFNLR1 | chr1 | - | 24161541 | 24161684 | 24159473 | 24159633 | 24169416 | 24169601 | 32 | 3 | [illegible] |
| SE | 53125 | ENSG00000185436 | IFNLR1 | chr1 | - | 24180730 | 24180854 | 24169416 | 24169601 | 24187190 | 24187248 | 4 | 0 | [illegible] |
| SE | 53135 | ENSG00000071205 | ARHGAP10 | chr4 | + | 147847150 | 147847222 | 147822895 | 147822957 | 147857552 | 147857654 | 94 | 46 | [illegible] |
| SE | 53136 | ENSG00000071205 | ARHGAP10 | chr4 | + | 147857552 | 147857654 | 147847150 | 147847222 | 147864845 | 147864956 | 95 | 63 | [illegible] |
| SE | 53137 | ENSG00000071205 | ARHGAP10 | chr4 | + | 147881837 | 147881932 | 147879231 | 147879338 | 147906637 | 147906719 | 218 | 118 | [illegible] |
| SE | 53139 | ENSG00000071205 | ARHGAP10 | chr4 | + | 147909731 | 147909777 | 147881837 | 147881932 | 147913073 | 147913139 | 7 | 7 | [illegible] |
| SE | 53140 | ENSG00000071205 | ARHGAP10 | chr4 | + | 147939824 | 147939899 | 147913073 | 147913139 | 147946616 | 147946704 | 84 | 22 | [illegible] |
| SE | 53142 | ENSG00000071205 | ARHGAP10 | chr4 | + | 148064415 | 148064507 | 148046891 | 148047051 | 148071992 | 148072775 | 30 | 10 | [illegible] |
| SE | 53143 | ENSG00000124574 | ABCC10 | chr6 | + | 43427967 | 43428139 | 43427540 | 43427757 | 43430747 | 43430866 | 10 | 4 | [illegible] |
| SE | 53144 | ENSG00000124574 | ABCC10 | chr6 | + | 43427967 | 43428139 | 43427540 | 43427757 | 43432141 | 43433360 | 16 | 6 | [illegible] |
| SE | 53145 | ENSG00000124574 | ABCC10 | chr6 | + | 43430747 | 43430866 | 43427540 | 43427757 | 43432141 | 43432176 | 3 | 0 | [illegible] |
| SE | 53146 | ENSG00000124574 | ABCC10 | chr6 | + | 43430747 | 43430866 | 43427855 | 43427967 | 43432141 | 43432176 | 2 | 0 | [illegible] |
| SE | 53148 | ENSG00000124574 | ABCC10 | chr6 | + | 43432141 | 43433360 | 43427855 | 43427967 | 43434620 | 43434848 | 16 | 6 | [illegible] |
| SE | 53149 | ENSG00000124574 | ABCC10 | chr6 | + | 43433234 | 43433360 | 43427855 | 43427967 | 43434620 | 43434848 | 11 | 5 | [illegible] |
| SE | 53150 | ENSG00000124574 | ABCC10 | chr6 | + | 43435750 | 43435907 | 43434620 | 43434848 | 43436137 | 43436247 | 48 | 27 | [illegible] |
| SE | 53152 | ENSG00000124574 | ABCC10 | chr6 | + | 43442969 | 43443159 | 43441861 | 43441960 | 43443932 | 43444010 | 73 | 30 | [illegible] |
| SE | 53154 | ENSG00000124574 | ABCC10 | chr6 | + | 43445124 | 43445314 | 43444787 | 43444938 | 43445598 | 43445942 | 66 | 15 | [illegible] |
| SE | 53156 | ENSG00000124574 | ABCC10 | chr6 | + | 43446276 | 43446446 | 43445598 | 43445942 | 43447247 | 43447408 | 19 | 5 | [illegible] |
| SE | 53159 | ENSG00000105220 | GPI | chr19 | + | 34368582 | 34368702 | 34366782 | 34366851 | 34377502 | 34377578 | 301 | 139 | [illegible] |
| SE | 53161 | ENSG00000105220 | GPI | chr19 | + | 34377734 | 34377881 | 34368582 | 34368702 | 34378933 | 34379005 | 339 | 146 | [illegible] |
| SE | 53162 | ENSG00000105220 | GPI | chr19 | + | 34379517 | 34379562 | 34378933 | 34379005 | 34381465 | 34381519 | 320 | 169 | [illegible] |
| SE | 53164 | ENSG00000105220 | GPI | chr19 | + | 34396300 | 34396430 | 34393266 | 34394066 | 34396580 | 34396656 | 332 | 200 | [illegible] |
| SE | 53165 | ENSG00000105220 | GPI | chr19 | + | 34396300 | 34396430 | 34393266 | 34394066 | 34399206 | 34399335 | 178 | 101 | [illegible] |
| SE | 53166 | ENSG00000105220 | GPI | chr19 | + | 34396580 | 34396657 | 34393266 | 34394066 | 34399206 | 34399335 | 62 | 25 | [illegible] |
| SE | 53170 | ENSG00000144381 | HSPD1 | chr2 | - | 197490196 | 197490296 | 197489001 | 197489247 | 197490937 | 197491042 | 156 | 110 | [illegible] |
| SE | 53171 | ENSG00000144381 | HSPD1 | chr2 | - | 197490196 | 197490296 | 197489001 | 197489247 | 197493323 | 197493492 | 344 | 251 | [illegible] |
| SE | 53172 | ENSG00000144381 | HSPD1 | chr2 | - | 197490526 | 197490664 | 197490196 | 197490296 | 197490937 | 197491042 | 2 | 0 | [illegible] |
| SE | 53173 | ENSG00000144381 | HSPD1 | chr2 | - | 197490526 | 197490664 | 197490196 | 197490296 | 197493323 | 197493492 | 1 | 0 | [illegible] |
| SE | 53174 | ENSG00000144381 | HSPD1 | chr2 | - | 197490937 | 197491042 | 197489001 | 197489247 | 197493323 | 197493492 | 4 | 0 | [illegible] |
| SE | 53176 | ENSG00000144381 | HSPD1 | chr2 | - | 197490937 | 197491042 | 197490526 | 197490664 | 197493323 | 197493492 | 1 | 0 | [illegible] |
| SE | 53178 | ENSG00000144381 | HSPD1 | chr2 | - | 197494656 | 197494752 | 197494149 | 197494250 | 197495293 | 197495376 | 457 | 410 | [illegible] |
| SE | 53179 | ENSG00000144381 | HSPD1 | chr2 | - | 197494662 | 197494752 | 197494156 | 197494250 | 197495293 | 197495376 | 289 | 299 | [illegible] |
| SE | 53183 | ENSG00000161057 | PSMC2 | chr7 | + | 103353920 | 103353958 | 103347509 | 103347781 | 103354867 | 103354949 | 385 | 216 | [illegible] |
| SE | 53185 | ENSG00000161057 | PSMC2 | chr7 | + | 103355693 | 103355793 | 103354867 | 103354949 | 103356169 | 103356209 | 151 | 93 | [illegible] |
| SE | 53186 | ENSG00000161057 | PSMC2 | chr7 | + | 103355693 | 103355793 | 103354867 | 103354949 | 103361956 | 103362057 | 463 | 266 | [illegible] |
| SE | 53187 | ENSG00000161057 | PSMC2 | chr7 | + | 103356169 | 103356209 | 103354867 | 103354949 | 103361956 | 103362088 | 1 | 0 | [illegible] |
| SE | 53188 | ENSG00000161057 | PSMC2 | chr7 | + | 103356169 | 103356209 | 103355693 | 103355793 | 103361956 | 103362088 | 2 | 1 | [illegible] |
| SE | 53189 | ENSG00000161057 | PSMC2 | chr7 | + | 103364142 | 103364307 | 103363343 | 103363439 | 103366075 | 103366163 | 308 | 178 | [illegible] |
| SE | 53190 | ENSG00000174885 | NLRP6 | chr11 | + | 284474 | 284642 | 284229 | 284400 | 285165 | 285304 | 0 | 0 | [illegible] |
| SE | 53192 | ENSG00000149527 | PLCH2 | chr1 | + | 2487597 | 2487718 | 2487172 | 2487376 | 2489206 | 2489378 | 32 | 12 | [illegible] |
| SE | 53195 | ENSG00000131828 | PDHA1 | chrX | + | 19345735 | 19345819 | 19343929 | 19344094 | 19349311 | 19349371 | 0 | 0 | [illegible] |
| SE | 53198 | ENSG00000131828 | PDHA1 | chrX | + | 19353081 | 19353173 | 19351259 | 19351407 | 19355348 | 19355504 | 60 | 33 | [illegible] |
| SE | 53199 | ENSG00000131828 | PDHA1 | chrX | + | 19354490 | 19354583 | 19351259 | 19351407 | 19355348 | 19355445 | 109 | 67 | [illegible] |
| SE | 53201 | ENSG00000131828 | PDHA1 | chrX | + | 19357651 | 19357719 | 19355581 | 19355757 | 19358915 | 19359024 | 120 | 115 | [illegible] |
| SE | 53202 | ENSG00000259522 | NA | chr14 | - | 24188718 | 24189808 | 24188551 | 24188638 | 24190379 | 24190416 | 23 | 6 | [illegible] |
| SE | 53203 | ENSG00000083544 | TDRD3 | chr13 | + | 60439687 | 60439772 | 60397219 | 60397405 | 60460379 | 60460540 | 2 | 4 | [illegible] |
| SE | 53204 | ENSG00000083544 | TDRD3 | chr13 | + | 60444682 | 60444748 | 60439687 | 60439772 | 60460379 | 60460540 | 153 | 118 | [illegible] |
| SE | 53205 | ENSG00000083544 | TDRD3 | chr13 | + | 60444686 | 60444748 | 60439687 | 60439772 | 60460379 | 60460540 | 76 | 56 | [illegible] |
| SE | 53206 | ENSG00000083544 | TDRD3 | chr13 | + | 60460379 | 60460540 | 60444682 | 60444748 | 60467237 | 60467379 | 347 | 223 | [illegible] |
| SE | 53207 | ENSG00000083544 | TDRD3 | chr13 | + | 60467237 | 60467379 | 60460379 | 60460540 | 60483774 | 60483846 | 256 | 141 | [illegible] |
| SE | 53210 | ENSG00000083544 | TDRD3 | chr13 | + | 60494434 | 60494575 | 60485798 | 60485948 | 60509762 | 60509919 | 196 | 150 | [illegible] |
| SE | 53211 | ENSG00000083544 | TDRD3 | chr13 | + | 60528366 | 60529217 | 60510629 | 60510755 | 60535107 | 60535233 | 149 | 126 | [illegible] |
| SE | 53212 | ENSG00000162384 | CZIB | chr1 | - | 53218171 | 53218203 | 53216781 | 53216859 | 53218866 | 53218923 | 10 | 12 | [illegible] |
| SE | 53213 | ENSG00000162384 | CZIB | chr1 | - | 53218413 | 53218495 | 53218171 | 53218203 | 53218866 | 53218923 | 215 | 120 | [illegible] |
| SE | 53214 | ENSG00000162384 | CZIB | chr1 | - | 53218866 | 53218923 | 53218413 | 53218495 | 53220260 | 53220344 | 103 | 60 | [illegible] |
| SE | 53219 | ENSG00000158769 | F11R | chr1 | - | 161002539 | 161002613 | 161001284 | 161001353 | 161021009 | 161021143 | 0 | 0 | [illegible] |
| SE | 53223 | ENSG00000052723 | SIKE1 | chr1 | - | 114780109 | 114780215 | 114779097 | 114779284 | 114780436 | 114780537 | 37 | 33 | [illegible] |
| SE | 53224 | ENSG00000052723 | SIKE1 | chr1 | - | 114780109 | 114780215 | 114779097 | 114779284 | 114780448 | 114780634 | 52 | 41 | [illegible] |
| SE | 53228 | ENSG00000181852 | RNF41 | chr12 | - | 56210296 | 56210568 | 56208147 | 56208298 | 56213957 | 56214070 | 147 | 89 | [illegible] |
| SE | 53232 | ENSG00000181852 | RNF41 | chr12 | - | 56216428 | 56216613 | 56213957 | 56214070 | 56221759 | 56221860 | 123 | 65 | [illegible] |
| SE | 53233 | ENSG00000138688 | KIAA1109 | chr4 | + | 122170614 | 122170744 | 122154024 | 122154112 | 122173035 | 122173152 | 353 | 244 | [illegible] |
| SE | 53234 | ENSG00000138688 | KIAA1109 | chr4 | + | 122170614 | 122170744 | 122154024 | 122154112 | 122174573 | 122174652 | 119 | 81 | [illegible] |
| SE | 53235 | ENSG00000138688 | KIAA1109 | chr4 | + | 122173035 | 122173152 | 122154024 | 122154112 | 122174573 | 122174652 | 206 | 177 | [illegible] |
| SE | 53236 | ENSG00000138688 | KIAA1109 | chr4 | + | 122173035 | 122173152 | 122170602 | 122170744 | 122174573 | 122174652 | 439 | 342 | [illegible] |
| SE | 53237 | ENSG00000138688 | KIAA1109 | chr4 | + | 122174573 | 122174652 | 122170602 | 122170744 | 122175849 | 122175914 | 168 | 106 | [illegible] |
| SE | 53238 | ENSG00000138688 | KIAA1109 | chr4 | + | 122189999 | 122190079 | 122187894 | 122188088 | 122192228 | 122192379 | 831 | 784 | [illegible] |
| SE | 53241 | ENSG00000138688 | KIAA1109 | chr4 | + | 122192228 | 122192379 | 122189999 | 122190079 | 122196634 | 122196791 | 636 | 542 | [illegible] |
| SE | 53242 | ENSG00000138688 | KIAA1109 | chr4 | + | 122199358 | 122199450 | 122197195 | 122197274 | 122201008 | 122201126 | 271 | 214 | [illegible] |
| SE | 53244 | ENSG00000138688 | KIAA1109 | chr4 | + | 122207540 | 122207638 | 122207109 | 122207265 | 122209817 | 122209924 | 59 | 41 | [illegible] |
| SE | 53245 | ENSG00000138688 | KIAA1109 | chr4 | + | 122209158 | 122209204 | 122207109 | 122207265 | 122209817 | 122209924 | 5 | 6 | [illegible] |
| SE | 53247 | ENSG00000138688 | KIAA1109 | chr4 | + | 122209158 | 122209337 | 122207109 | 122207265 | 122209817 | 122209924 | 187 | 196 | [illegible] |
| SE | 53251 | ENSG00000138688 | KIAA1109 | chr4 | + | 122243853 | 122244027 | 122242997 | 122243068 | 122245019 | 122245140 | 507 | 521 | [illegible] |
| SE | 53252 | ENSG00000138688 | KIAA1109 | chr4 | + | 122263441 | 122263600 | 122262763 | 122262983 | 122264251 | 122264427 | 239 | 170 | [illegible] |
| SE | 53253 | ENSG00000138688 | KIAA1109 | chr4 | + | 122271017 | 122271724 | 122266782 | 122266958 | 122272159 | 122272405 | 378 | 297 | [illegible] |
| SE | 53255 | ENSG00000138688 | KIAA1109 | chr4 | + | 122289086 | 122289192 | 122286540 | 122286785 | 122301299 | 122301397 | 230 | 217 | [illegible] |
| SE | 53256 | ENSG00000138688 | KIAA1109 | chr4 | + | 122304797 | 122304983 | 122301299 | 122301397 | 122305876 | 122306045 | 186 | 144 | [illegible] |
| SE | 53257 | ENSG00000138688 | KIAA1109 | chr4 | + | 122313627 | 122313687 | 122309265 | 122309464 | 122315461 | 122315688 | 285 | 320 | [illegible] |
| SE | 53258 | ENSG00000138688 | KIAA1109 | chr4 | + | 122316347 | 122316548 | 122315461 | 122315688 | 122316731 | 122316794 | 0 | 0 | [illegible] |
| SE | 53259 | ENSG00000138688 | KIAA1109 | chr4 | + | 122316731 | 122316862 | 122315461 | 122315688 | 122318146 | 122318270 | 415 | 441 | [illegible] |
| SE | 53260 | ENSG00000138688 | KIAA1109 | chr4 | + | 122325854 | 122325917 | 122325229 | 122325320 | 122328103 | 122328357 | 83 | 77 | [illegible] |
| SE | 53261 | ENSG00000138688 | KIAA1109 | chr4 | + | 122331325 | 122331541 | 122327871 | 122328357 | 122333628 | 122333809 | 292 | 227 | [illegible] |
| SE | 53264 | ENSG00000138688 | KIAA1109 | chr4 | + | 122356620 | 122356755 | 122355795 | 122355990 | 122359556 | 122359722 | 192 | 163 | [illegible] |
| SE | 53268 | ENSG00000178537 | SLC25A20 | chr3 | - | 48879357 | 48879448 | 48862541 | 48862659 | 48883996 | 48884124 | 170 | 101 | [illegible] |
| SE | 53269 | ENSG00000178537 | SLC25A20 | chr3 | - | 48879357 | 48879448 | 48862541 | 48862659 | 48891979 | 48892072 | 46 | 40 | [illegible] |
| SE | 53270 | ENSG00000178537 | SLC25A20 | chr3 | - | 48883996 | 48884124 | 48862541 | 48862659 | 48891979 | 48892072 | 58 | 31 | [illegible] |
| SE | 53273 | ENSG00000157796 | WDR19 | chr4 | + | 39185725 | 39185817 | 39182503 | 39182563 | 39186538 | 39186604 | 9 | 7 | [illegible] |
| SE | 53274 | ENSG00000157796 | WDR19 | chr4 | + | 39186538 | 39186604 | 39185725 | 39185817 | 39189655 | 39189781 | 14 | 18 | [illegible] |
| SE | 53275 | ENSG00000157796 | WDR19 | chr4 | + | 39189655 | 39189781 | 39186538 | 39186604 | 39194543 | 39194659 | 16 | 19 | [illegible] |
| SE | 53276 | ENSG00000157796 | WDR19 | chr4 | + | 39194543 | 39194659 | 39189655 | 39189781 | 39199477 | 39199593 | 18 | 20 | [illegible] |
| SE | 53278 | ENSG00000157796 | WDR19 | chr4 | + | 39205153 | 39205266 | 39203641 | 39203722 | 39205562 | 39205572 | 17 | 31 | [illegible] |
| SE | 53279 | ENSG00000164151 | ICE1 | chr5 | + | 5454551 | 5454638 | 5447840 | 5447897 | 5457331 | 5457741 | 186 | 161 | [illegible] |
| SE | 53292 | ENSG00000256128 | LINC00944 | chr12 | - | 126747930 | 126748127 | 126745338 | 126746332 | 126760931 | 126761057 | 30 | 29 | [illegible] |
| SE | 53293 | ENSG00000256128 | LINC00944 | chr12 | - | 126747930 | 126748127 | 126745338 | 126746332 | 126770192 | 126770254 | 26 | 16 | [illegible] |
| SE | 53294 | ENSG00000256128 | LINC00944 | chr12 | - | 126747930 | 126748127 | 126745338 | 126746332 | 126772078 | 126772411 | 31 | 19 | [illegible] |
| SE | 53297 | ENSG00000256128 | LINC00944 | chr12 | - | 126760931 | 126761057 | 126747930 | 126748127 | 126770192 | 126770254 | 6 | 15 | [illegible] |
| SE | 53298 | ENSG00000256128 | LINC00944 | chr12 | - | 126760931 | 126761057 | 126747930 | 126748127 | 126772078 | 126772262 | 9 | 13 | [illegible] |
| SE | 53299 | ENSG00000256128 | LINC00944 | chr12 | - | 126763414 | 126763498 | 126760931 | 126761057 | 126770192 | 126770254 | 8 | 4 | [illegible] |
| SE | 53300 | ENSG00000256128 | LINC00944 | chr12 | - | 126763414 | 126763498 | 126760931 | 126761057 | 126772078 | 126772233 | 2 | 0 | [illegible] |
| SE | 53302 | ENSG00000256128 | LINC00944 | chr12 | - | 126770192 | 126770254 | 126747930 | 126748127 | 126772078 | 126772192 | 2 | 0 | [illegible] |
| SE | 53304 | ENSG00000256128 | LINC00944 | chr12 | - | 126770192 | 126770254 | 126763414 | 126763498 | 126772078 | 126772192 | 4 | 1 | [illegible] |
| SE | 53306 | ENSG00000162594 | IL23R | chr1 | + | 67200736 | 67200897 | 67182835 | 67182959 | 67206909 | 67207055 | 7 | 6 | [illegible] |
| SE | 53307 | ENSG00000162594 | IL23R | chr1 | + | 67200741 | 67200897 | 67182835 | 67182959 | 67206909 | 67207055 | 5 | 1 | [illegible] |
| SE | 53309 | ENSG00000162594 | IL23R | chr1 | + | 67240178 | 67240281 | 67236712 | 67236802 | 67255836 | 67255927 | 3 | 4 | [illegible] |
| SE | 53341 | ENSG00000131778 | CHD1L | chr1 | + | 147259836 | 147259918 | 147256530 | 147256562 | 147264421 | 147264532 | 91 | 45 | [illegible] |
| SE | 53342 | ENSG00000237188 | NA | chr1 | + | 147259836 | 147259918 | 147256530 | 147256562 | 147264428 | 147264584 | 26 | 17 | [illegible] |
| SE | 53343 | ENSG00000131778 | CHD1L | chr1 | + | 147264421 | 147264584 | 147259836 | 147259918 | 147265931 | 147266087 | 136 | 96 | [illegible] |
| SE | 53345 | ENSG00000237188 | NA | chr1 | + | 147268781 | 147268878 | 147267425 | 147267518 | 147272170 | 147272281 | 32 | 28 | [illegible] |
| SE | 53352 | ENSG00000237188 | NA | chr1 | + | 147284350 | 147284499 | 147280025 | 147280191 | 147285323 | 147285487 | 219 | 261 | [illegible] |
| SE | 53354 | ENSG00000130939 | UBE4B | chr1 | + | 10095460 | 10095596 | 10072027 | 10072214 | 10101107 | 10101195 | 154 | 77 | [illegible] |
| SE | 53355 | ENSG00000130939 | UBE4B | chr1 | + | 10101107 | 10101195 | 10095460 | 10095596 | 10102947 | 10103092 | 152 | 78 | [illegible] |
| SE | 53356 | ENSG00000130939 | UBE4B | chr1 | + | 10102947 | 10103092 | 10101107 | 10101195 | 10105515 | 10105744 | 172 | 64 | [illegible] |
| SE | 53359 | ENSG00000130939 | UBE4B | chr1 | + | 10117458 | 10117600 | 10105515 | 10105744 | 10119512 | 10119613 | 176 | 87 | [illegible] |
| SE | 53360 | ENSG00000130939 | UBE4B | chr1 | + | 10117458 | 10117600 | 10105515 | 10105744 | 10126793 | 10126877 | 64 | 22 | [illegible] |
| SE | 53361 | ENSG00000130939 | UBE4B | chr1 | + | 10119512 | 10119613 | 10105515 | 10105744 | 10126793 | 10126877 | 1 | 0 | [illegible] |
| SE | 53362 | ENSG00000130939 | UBE4B | chr1 | + | 10119512 | 10119613 | 10117458 | 10117600 | 10126793 | 10126877 | 69 | 37 | [illegible] |
| SE | 53365 | ENSG00000130939 | UBE4B | chr1 | + | 10132368 | 10132482 | 10130714 | 10130813 | 10134987 | 10135186 | 106 | 77 | [illegible] |
| SE | 53366 | ENSG00000130939 | UBE4B | chr1 | + | 10137066 | 10137205 | 10134987 | 10135186 | 10144939 | 10145039 | 125 | 85 | [illegible] |
| SE | 53367 | ENSG00000130939 | UBE4B | chr1 | + | 10144939 | 10145039 | 10137066 | 10137205 | 10146962 | 10147090 | 83 | 45 | [illegible] |
| SE | 53370 | ENSG00000130939 | UBE4B | chr1 | + | 10146962 | 10147090 | 10144939 | 10145039 | 10149183 | 10149282 | 93 | 47 | [illegible] |
| SE | 53372 | ENSG00000130939 | UBE4B | chr1 | + | 10151325 | 10151561 | 10149183 | 10149282 | 10161141 | 10161286 | 63 | 69 | [illegible] |
| SE | 53373 | ENSG00000130939 | UBE4B | chr1 | + | 10158355 | 10158482 | 10149183 | 10149282 | 10161141 | 10161286 | 165 | 108 | [illegible] |
| SE | 53375 | ENSG00000130939 | UBE4B | chr1 | + | 10166243 | 10166342 | 10161141 | 10161286 | 10168135 | 10168270 | 0 | 0 | [illegible] |
| SE | 53376 | ENSG00000130939 | UBE4B | chr1 | + | 10178643 | 10178818 | 10170955 | 10171329 | 10179415 | 10179562 | 193 | 136 | [illegible] |
| SE | 53378 | ENSG00000135763 | URB2 | chr1 | + | 229647509 | 229647752 | 229645858 | 229645969 | 229651234 | 229651322 | 27 | 8 | [illegible] |
| SE | 53386 | ENSG00000135837 | CEP350 | chr1 | + | 179992061 | 179992221 | 179987242 | 179987286 | 179996552 | 179996963 | 248 | 176 | [illegible] |
| SE | 53387 | ENSG00000135837 | CEP350 | chr1 | + | 179992061 | 179992221 | 179990506 | 179990621 | 179996552 | 179996963 | 357 | 273 | [illegible] |
| SE | 53388 | ENSG00000135837 | CEP350 | chr1 | + | 180003173 | 180003287 | 179996552 | 179997175 | 180006453 | 180006567 | 411 | 413 | [illegible] |
| SE | 53390 | ENSG00000135837 | CEP350 | chr1 | + | 180006453 | 180006567 | 179996552 | 179997175 | 180011928 | 180012075 | 93 | 99 | [illegible] |
| SE | 53392 | ENSG00000135837 | CEP350 | chr1 | + | 180013846 | 180014505 | 180011928 | 180012075 | 180015848 | 180015970 | 366 | 291 | [illegible] |
| SE | 53396 | ENSG00000135837 | CEP350 | chr1 | + | 180053749 | 180053934 | 180052969 | 180053166 | 180054414 | 180054502 | 295 | 283 | [illegible] |
| SE | 53397 | ENSG00000135837 | CEP350 | chr1 | + | 180053749 | 180053934 | 180052969 | 180053166 | 180062219 | 180062366 | 177 | 157 | [illegible] |
| SE | 53398 | ENSG00000135837 | CEP350 | chr1 | + | 180054414 | 180054502 | 180052969 | 180053166 | 180062219 | 180062366 | 105 | 145 | [illegible] |
| SE | 53399 | ENSG00000135837 | CEP350 | chr1 | + | 180054414 | 180054502 | 180053749 | 180053934 | 180062219 | 180062366 | 223 | 267 | [illegible] |
| SE | 53400 | ENSG00000135837 | CEP350 | chr1 | + | 180080516 | 180080661 | 180078462 | 180078674 | 180084017 | 180084178 | 252 | 159 | [illegible] |
| SE | 53402 | ENSG00000135837 | CEP350 | chr1 | + | 180087577 | 180087717 | 180084017 | 180084178 | 180090713 | 180090748 | 140 | 139 | [illegible] |
| SE | 53403 | ENSG00000135837 | CEP350 | chr1 | + | 180090713 | 180090796 | 180087577 | 180087717 | 180092613 | 180094616 | 185 | 210 | [illegible] |
| SE | 53404 | ENSG00000135837 | CEP350 | chr1 | + | 180110996 | 180111195 | 180098862 | 180098985 | 180112436 | 180113380 | 208 | 205 | [illegible] |
| SE | 53405 | ENSG00000151304 | SRFBP1 | chr5 | + | 121974195 | 121974284 | 121961974 | 121962068 | 121975314 | 121975387 | 26 | 38 | [illegible] |
| SE | 53407 | ENSG00000151304 | SRFBP1 | chr5 | + | 122020087 | 122020802 | 121994598 | 121994670 | 122022369 | 122022407 | 83 | 111 | [illegible] |
| SE | 53408 | ENSG00000151304 | SRFBP1 | chr5 | + | 122022369 | 122022407 | 121994598 | 121994670 | 122026941 | 122028633 | 6 | 1 | [illegible] |
| SE | 53409 | ENSG00000151304 | SRFBP1 | chr5 | + | 122022369 | 122022407 | 121994598 | 121994670 | 122075314 | 122075570 | 2 | 2 | [illegible] |
| SE | 53411 | ENSG00000172531 | PPP1CA | chr11 | - | 67398939 | 67399163 | 67398721 | 67398856 | 67399560 | 67399665 | 602 | 210 | [illegible] |
| SE | 53426 | ENSG00000111358 | GTF2H3 | chr12 | + | 123639263 | 123639343 | 123633738 | 123633872 | 123645454 | 123645561 | 44 | 31 | [illegible] |
| SE | 53427 | ENSG00000111358 | GTF2H3 | chr12 | + | 123639293 | 123639343 | 123633828 | 123633872 | 123645454 | 123645561 | 17 | 9 | [illegible] |
| SE | 53428 | ENSG00000111358 | GTF2H3 | chr12 | + | 123639956 | 123640001 | 123639263 | 123639343 | 123645454 | 123645561 | 0 | 0 | [illegible] |
| SE | 53429 | ENSG00000111358 | GTF2H3 | chr12 | + | 123647962 | 123648126 | 123645454 | 123645561 | 123650993 | 123651056 | 93 | 83 | [illegible] |
| SE | 53431 | ENSG00000111358 | GTF2H3 | chr12 | + | 123655770 | 123655824 | 123654923 | 123654998 | 123659515 | 123659584 | 65 | 36 | [illegible] |
| SE | 53434 | ENSG00000183918 | SH2D1A | chrX | + | 124367589 | 124367661 | 124365760 | 124365824 | 124370175 | 124370320 | 0 | 0 | [illegible] |
| SE | 53436 | ENSG00000135953 | MFSD9 | chr2 | - | 102726844 | 102726974 | 102723699 | 102723909 | 102731024 | 102731068 | 25 | 3 | [illegible] |
| SE | 53437 | ENSG00000135953 | MFSD9 | chr2 | - | 102732023 | 102732138 | 102731024 | 102731068 | 102732321 | 102732392 | 4 | 4 | [illegible] |
| SE | 53444 | ENSG00000145020 | AMT | chr3 | - | 49418970 | 49419151 | 49417817 | 49417973 | 49419259 | 49419335 | 36 | 20 | [illegible] |

| | | | | | | | | | | | | | | | | | | | | | | | | | | | | | | | | | | | | | | | | | | | | | | |
|---|---|---|---|---|---|---|---|---|---|---|---|---|---|---|---|---|---|---|---|---|---|---|---|---|---|---|---|---|---|---|---|---|---|---|---|---|---|---|---|---|---|---|---|---|---|---|
| SE | 53445 | ENSG00000145020 | AMT | chr3 | - | 49419259 | 49419335 | 49418970 | 49419151 | 49419709 | 49419788 | 2 | 1 | 1 | 6 | 1 | 0 | 0 | 0 | 0 | 0 | 0 | 4 | 4 | 6 | 12 | 0 | 0 | 4 | 5 | 0 | 225 | 149 | 1 | 1 | 1 | 1 | 1 | 1 | 1 | NA | 1 | 0.398 | 0.443 | 1 | 0.29 |
| SE | 53446 | ENSG00000145020 | AMT | chr3 | - | 49419259 | 49419393 | 49418970 | 49419151 | 49419709 | 49419788 | 4 | 1 | 2 | 9 | 5 | 0 | 0 | 0 | 0 | 0 | 0 | 11 | 7 | 11 | 19 | 0 | 0 | 4 | 5 | 0 | 283 | 149 | 1 | 1 | 1 | 1 | 1 | 1 | 1 | NA | 1 | 0.48 | 0.537 | 1 | 0.246 |
| SE | 53451 | ENSG00000145020 | AMT | chr3 | - | 49419709 | 49419788 | 49418970 | 49419405 | 49420210 | 49420342 | 13 | 7 | 6 | 17 | 9 | 0 | 0 | 0 | 0 | 0 | 31 | 24 | 48 | 31 | 40 | 0 | 0 | 0 | 0 | 2 | 228 | 149 | 1 | 1 | 1 | 1 | 1 | 1 | 1 | 1 | 1 | 1 | 1 | 0.929 | 0.014 |
| SE | 53460 | ENSG00000249915 | PDCD6 | chr5 | + | 304176 | 304221 | 272710 | 272772 | 311292 | 311402 | 15 | 0 | 4 | 7 | 0 | 3 | 0 | 0 | 0 | 0 | 5 | 14 | 3 | 12 | 3 | 0 | 2 | 1 | 0 | 0 | 194 | 149 | 1 | 1 | 0.793 | NA | 1 | 1 | NA | 1 | 0.843 | 0.697 | 1 | 1 | 0.023 |
| SE | 53461 | ENSG00000249915 | PDCD6 | chr5 | + | 304176 | 304221 | 272710 | 272772 | 314416 | 314806 | 17 | 7 | 3 | 10 | 0 | 0 | 0 | 2 | 1 | 0 | 5 | 10 | 3 | 12 | 20 | 0 | 2 | 0 | 0 | 0 | 194 | 149 | 1 | 1 | 1 | 1 | 0.535 | 0.885 | NA | 1 | 0.793 | 1 | 1 | 1 | -0.104 |
| SE | 53462 | ENSG00000249915 | PDCD6 | chr5 | + | 304176 | 306754 | 272710 | 272772 | 311292 | 311402 | 57 | 19 | 27 | 24 | 15 | 3 | 0 | 0 | 0 | 0 | 63 | 27 | 14 | 23 | 63 | 0 | 2 | 1 | 0 | 0 | 298 | 149 | 1 | 1 | 0.905 | 1 | 1 | 1 | 1 | 1 | 0.871 | 0.875 | 1 | 1 | 0.032 |
| SE | 53463 | ENSG00000249915 | PDCD6 | chr5 | + | 304176 | 306754 | 272710 | 272772 | 314416 | 314969 | 17 | 5 | 6 | 6 | 4 | 0 | 0 | 2 | 1 | 0 | 5 | 6 | 3 | 10 | 18 | 0 | 2 | 0 | 0 | 0 | 298 | 149 | 1 | 1 | 1 | 1 | 0.6 | 0.75 | 1 | 1 | 0.6 | 1 | 1 | 1 | -0.05 |
| SE | 53468 | ENSG00000249915 | PDCD6 | chr5 | + | 311292 | 311402 | 272710 | 272772 | 314416 | 314686 | 46 | 27 | 50 | 50 | 44 | 0 | 0 | 2 | 1 | 0 | 78 | 22 | 31 | 56 | 42 | 0 | 2 | 0 | 0 | 0 | 259 | 149 | 1 | 1 | 1 | 1 | 0.935 | 0.966 | 1 | 1 | 0.864 | 1 | 1 | 1 | 0.007 |
| SE | 53471 | ENSG00000249915 | PDCD6 | chr5 | + | 311292 | 311402 | 306601 | 306760 | 314416 | 314686 | 197 | 103 | 152 | 191 | 170 | 0 | 0 | 1 | 0 | 0 | 278 | 118 | 82 | 165 | 262 | 0 | 1 | 0 | 0 | 0 | 259 | 149 | 1 | 1 | 1 | 1 | 0.989 | 1 | 1 | 1 | 0.985 | 1 | 1 | 1 | 0.001 |
| SE | 53472 | ENSG00000249915 | PDCD6 | chr5 | + | 313700 | 313850 | 311292 | 311402 | 314416 | 314954 | 0 | 1 | 2 | 0 | 0 | 79 | 40 | 73 | 75 | 75 | 0 | 2 | 0 | 0 | 0 | 97 | 39 | 31 | 71 | 74 | 298 | 149 | 1 | 1 | 0 | 0.012 | 0.014 | 0 | 0 | 0 | 0.025 | 0 | 0 | 0 | 0 |
| SE | 53475 | ENSG00000136098 | NEK3 | chr13 | - | 52135728 | 52135863 | 52133688 | 52133815 | 52136799 | 52136902 | 38 | 33 | 21 | 35 | 51 | 0 | 0 | 0 | 4 | 0 | 32 | 30 | 62 | 33 | 3 | 0 | 2 | 0 | 0 | 0 | 284 | 149 | 1 | 1 | 1 | 1 | 1 | 0.821 | 1 | 1 | 0.887 | 1 | 1 | 1 | -0.013 |
| SE | 53476 | ENSG00000136098 | NEK3 | chr13 | - | 52136115 | 52136232 | 52135728 | 52135863 | 52136799 | 52136902 | 19 | 28 | 16 | 15 | 39 | 1 | 0 | 0 | 1 | 0 | 16 | 17 | 39 | 23 | 0 | 0 | 0 | 6 | 3 | 0 | 266 | 149 | 1 | 1 | 0.914 | 1 | 1 | 0.894 | 1 | 1 | 1 | 0.785 | 0.811 | NA | 0.063 |
| SE | 53477 | ENSG00000136098 | NEK3 | chr13 | - | 52136115 | 52136259 | 52133688 | 52133815 | 52136799 | 52136902 | 35 | 38 | 33 | 53 | 75 | 0 | 0 | 0 | 4 | 0 | 14 | 35 | 61 | 26 | 2 | 0 | 2 | 0 | 0 | 0 | 293 | 149 | 1 | 1 | 1 | 1 | 1 | 0.871 | 1 | 1 | 0.899 | 1 | 1 | 1 | -0.006 |
| SE | 53479 | ENSG00000136098 | NEK3 | chr13 | - | 52141019 | 52141069 | 52136799 | 52136902 | 52143914 | 52143980 | 5 | 2 | 12 | 10 | 28 | 1 | 0 | 0 | 5 | 0 | 0 | 8 | 29 | 6 | 0 | 0 | 0 | 4 | 0 | 0 | 199 | 149 | 1 | 1 | 0.789 | 1 | 1 | 0.6 | 1 | NA | 1 | 0.844 | 1 | NA | -0.07 |
| SE | 53480 | ENSG00000136098 | NEK3 | chr13 | - | 52141019 | 52141069 | 52136799 | 52136902 | 52143915 | 52143987 | 10 | 3 | 5 | 10 | 22 | 2 | 4 | 1 | 1 | 0 | 0 | 5 | 17 | 10 | 0 | 0 | 0 | 0 | 7 | 0 | 199 | 149 | 1 | 1 | 0.789 | 0.36 | 0.789 | 0.882 | 1 | NA | 1 | 1 | 0.517 | NA | -0.075 |
| SE | 53481 | ENSG00000136098 | NEK3 | chr13 | - | 52148414 | 52148469 | 52144690 | 52144891 | 52151145 | 52151232 | 31 | 23 | 14 | 34 | 68 | 0 | 1 | 0 | 0 | 0 | 49 | 15 | 19 | 20 | 0 | 0 | 0 | 0 | 0 | 0 | 204 | 149 | 1 | 1 | 1 | 0.944 | 1 | 1 | 1 | 1 | 1 | 1 | 1 | NA | -0.011 |
| SE | 53482 | ENSG00000136098 | NEK3 | chr13 | - | 52154079 | 52154173 | 52153894 | 52153992 | 52156074 | 52156248 | 25 | 14 | 24 | 24 | 31 | 0 | 0 | 0 | 0 | 0 | 8 | 46 | 27 | 24 | 0 | 0 | 0 | 1 | 0 | 0 | 243 | 149 | 1 | 1 | 1 | 1 | 1 | 1 | 1 | 1 | 1 | 0.943 | 1 | NA | 0.014 |
| SE | 53483 | ENSG00000136098 | NEK3 | chr13 | - | 52156074 | 52156248 | 52154079 | 52154173 | 52159542 | 52159550 | 23 | 11 | 31 | 30 | 29 | 1 | 0 | 0 | 0 | 0 | 16 | 50 | 56 | 18 | 0 | 0 | 0 | 0 | 0 | 0 | 298 | 149 | 1 | 1 | 0.92 | 1 | 1 | 1 | 1 | 1 | 1 | 1 | 1 | NA | -0.016 |
| SE | 53486 | ENSG00000233621 | LINC01137 | chr1 | - | 37456366 | 37456459 | 37454878 | 37455912 | 37474143 | 37474411 | 6 | 4 | 11 | 19 | 1 | 2 | 0 | 1 | 0 | 1 | 27 | 13 | 8 | 0 | 2 | 3 | 0 | 0 | 0 | 0 | 242 | 149 | 1 | 1 | 0.649 | 1 | 0.871 | 1 | 0.381 | 0.847 | 1 | 1 | NA | 1 | -0.182 |
| SE | 53487 | ENSG00000143951 | WDPCP | chr2 | - | 63259306 | 63259409 | 63174669 | 63174832 | 63313247 | 63313311 | 17 | 8 | 9 | 8 | 16 | 0 | 4 | 0 | 1 | 2 | 0 | 0 | 1 | 0 | 0 | 0 | 0 | 1 | 3 | 13 | 252 | 149 | 1 | 1 | 1 | 0.542 | 1 | 0.825 | 0.825 | NA | NA | 0.372 | 0 | 0 | 0.714 |
| SE | 53488 | ENSG00000143951 | WDPCP | chr2 | - | 63404047 | 63404657 | 63381905 | 63382094 | 63433744 | 63433930 | 72 | 35 | 49 | 23 | 66 | 0 | 0 | 1 | 0 | 1 | 12 | 33 | 21 | 27 | 22 | 0 | 0 | 2 | 0 | 0 | 298 | 149 | 1 | 1 | 1 | 1 | 0.961 | 1 | 0.971 | 1 | 1 | 0.84 | 1 | 1 | 0.018 |
| SE | 53489 | ENSG00000143951 | WDPCP | chr2 | - | 63404047 | 63404657 | 63381905 | 63382094 | 63484603 | 63484663 | 37 | 16 | 24 | 11 | 24 | 0 | 0 | 0 | 0 | 4 | 0 | 12 | 12 | 17 | 9 | 0 | 2 | 2 | 0 | 0 | 298 | 149 | 1 | 1 | 1 | 1 | 1 | 1 | 0.75 | NA | 0.75 | 0.75 | 1 | 1 | 0.075 |
| SE | 53490 | ENSG00000143951 | WDPCP | chr2 | - | 63439756 | 63439855 | 63437452 | 63437554 | 63484603 | 63484663 | 14 | 6 | 20 | 9 | 24 | 1 | 0 | 3 | 1 | 5 | 5 | 12 | 4 | 6 | 0 | 0 | 3 | 4 | 0 | 0 | 248 | 149 | 1 | 1 | 0.894 | 1 | 0.8 | 0.844 | 0.743 | 1 | 0.706 | 0.375 | 1 | NA | 0.086 |
| SE | 53492 | ENSG00000143951 | WDPCP | chr2 | - | 63486541 | 63486586 | 63484916 | 63484987 | 63487446 | 63487494 | 25 | 34 | 23 | 19 | 64 | 1 | 0 | 1 | 1 | 1 | 12 | 23 | 21 | 14 | 61 | 0 | 3 | 0 | 0 | 0 | 194 | 149 | 1 | 1 | 0.95 | 1 | 0.946 | 0.936 | 0.98 | 1 | 0.855 | 1 | 1 | 1 | -0.009 |
| SE | 53497 | ENSG00000139174 | PRICKLE1 | chr12 | - | 42483757 | 42483985 | 42472384 | 42472564 | 42484273 | 42484298 | 0 | 0 | 1 | 4 | 2 | 0 | 2 | 0 | 0 | 0 | 0 | 2 | 0 | 0 | 0 | 0 | 0 | 0 | 0 | 0 | 298 | 149 | 1 | 1 | NA | 0 | 1 | 1 | 1 | NA | 1 | NA | NA | NA | -0.25 |
| SE | 53504 | ENSG00000233008 | LINC01725 | chr1 | - | 83849906 | 83850022 | 83750197 | 83750299 | 83860818 | 83860938 | 0 | 0 | 0 | 1 | 3 | 0 | 0 | 0 | 1 | 0 | 0 | 1 | 0 | 0 | 0 | 0 | 0 | 0 | 0 | 0 | 265 | 149 | 1 | 1 | NA | NA | NA | 0.36 | 1 | NA | 1 | NA | NA | NA | -0.32 |
| SE | 53505 | ENSG00000233008 | LINC01725 | chr1 | - | 83849906 | 83850022 | 83750197 | 83750299 | 83860969 | 83861023 | 0 | 0 | 0 | 1 | 3 | 0 | 0 | 1 | 0 | 0 | 0 | 1 | 0 | 0 | 0 | 0 | 0 | 0 | 0 | 0 | 265 | 149 | 1 | 1 | NA | NA | 0 | 1 | 1 | NA | 1 | NA | NA | NA | -0.333 |
| SE | 53509 | ENSG00000145390 | USP53 | chr4 | + | 119214069 | 119214222 | 119212635 | 119212873 | 119217549 | 119217673 | 18 | 18 | 23 | 18 | 5 | 11 | 11 | 18 | 8 | 12 | 0 | 17 | 8 | 8 | 0 | 0 | 3 | 0 | 0 | 0 | 298 | 149 | 1 | 1 | 0.45 | 0.45 | 0.39 | 0.529 | 0.172 | NA | 0.739 | 1 | 1 | NA | -0.515 |
| SE | 53510 | ENSG00000145390 | USP53 | chr4 | + | 119235289 | 119235411 | 119217549 | 119217673 | 119239217 | 119239312 | 1 | 2 | 13 | 6 | 10 | 20 | 26 | 14 | 13 | 15 | 0 | 0 | 2 | 0 | 0 | 0 | 3 | 1 | 3 | 0 | 271 | 149 | 1 | 1 | 0.027 | 0.041 | 0.338 | 0.202 | 0.268 | NA | 0 | 0.524 | 0 | NA | 0.001 |
| [illegible] | | | | | | | | | | | | | | | | | | | | | | | | | | | | | | | | | | | | | | | | | | | | | | |

| | | | | | | | | | | | | | | |
|---|---|---|---|---|---|---|---|---|---|---|---|---|---|---|
| SE | 53939 | ENSG00000040275 | SPDL1 | chr5 | + | 169587260 | 169587372 | 169583737 | 169583889 | 169588393 | 169588575 | 3 | 9 | [illegible] |
| SE | 53940 | ENSG00000040275 | SPDL1 | chr5 | + | 169590752 | 169590829 | 169588534 | 169588575 | 169591047 | 169591224 | 0 | 0 | [illegible] |
| SE | 53942 | ENSG00000040275 | SPDL1 | chr5 | + | 169596560 | 169596701 | 169594570 | 169594681 | 169598475 | 169598579 | 33 | 27 | [illegible] |
| SE | 53943 | ENSG00000040275 | SPDL1 | chr5 | + | 169596584 | 169596701 | 169594570 | 169594681 | 169598475 | 169598579 | 16 | 7 | [illegible] |
| SE | 53946 | ENSG00000101977 | MCF2 | chrX | - | 139629694 | 139629844 | 139626622 | 139626756 | 139632334 | 139632454 | 1 | 1 | [illegible] |
| SE | 53947 | ENSG00000101977 | MCF2 | chrX | - | 139631394 | 139631511 | 139629694 | 139629844 | 139632334 | 139632454 | 8 | 0 | [illegible] |
| SE | 53949 | ENSG00000138101 | DTNB | chr2 | - | 25383835 | 25383889 | 25379289 | 25379323 | 25387288 | 25387378 | 3 | 8 | [illegible] |
| SE | 53950 | ENSG00000138101 | DTNB | chr2 | - | 25383835 | 25383889 | 25379289 | 25379323 | 25388201 | 25388361 | 2 | 2 | [illegible] |
| SE | 53951 | ENSG00000138101 | DTNB | chr2 | - | 25387288 | 25387378 | 25379289 | 25379323 | 25388201 | 25388361 | 6 | 6 | [illegible] |
| SE | 53952 | ENSG00000138101 | DTNB | chr2 | - | 25387288 | 25387378 | 25383835 | 25383889 | 25388201 | 25388361 | 4 | 13 | [illegible] |
| SE | 53954 | ENSG00000138101 | DTNB | chr2 | - | 25427534 | 25427631 | 25388201 | 25388361 | 25432885 | 25432999 | 23 | 10 | [illegible] |
| SE | 53955 | ENSG00000138101 | DTNB | chr2 | - | 25432885 | 25432999 | 25427534 | 25427631 | 25433909 | 25433995 | 33 | 12 | [illegible] |
| SE | 53958 | ENSG00000138101 | DTNB | chr2 | - | 25531472 | 25531597 | 25482795 | 25482873 | 25576837 | 25577004 | 51 | 15 | [illegible] |
| SE | 53961 | ENSG00000138101 | DTNB | chr2 | - | 25607235 | 25607321 | 25580720 | 25580826 | 25628170 | 25628384 | 9 | 12 | [illegible] |
| SE | 53969 | ENSG00000188641 | DPYD | chr1 | - | 97234851 | 97234994 | 97193068 | 97193248 | 97305258 | 97305378 | 234 | 186 | [illegible] |
| SE | 53970 | ENSG00000188641 | DPYD | chr1 | - | 97305258 | 97305378 | 97234851 | 97234994 | 97306176 | 97306297 | 246 | 242 | [illegible] |
| SE | 53971 | ENSG00000188641 | DPYD | chr1 | - | 97373560 | 97373644 | 97306176 | 97306297 | 97382392 | 97382461 | 298 | 293 | [illegible] |
| SE | 53972 | ENSG00000188641 | DPYD | chr1 | - | 97382392 | 97382461 | 97373560 | 97373644 | 97450058 | 97450223 | 265 | 242 | [illegible] |
| SE | 53973 | ENSG00000188641 | DPYD | chr1 | - | 97450058 | 97450223 | 97382392 | 97382461 | 97515725 | 97515941 | 486 | 391 | [illegible] |
| SE | 53975 | ENSG00000188641 | DPYD | chr1 | - | 97515725 | 97515941 | 97450058 | 97450223 | 97573759 | 97573970 | 269 | 219 | [illegible] |
| SE | 53976 | ENSG00000188641 | DPYD | chr1 | - | 97549559 | 97549744 | 97450058 | 97450223 | 97573759 | 97573970 | 269 | 245 | [illegible] |
| SE | 53977 | ENSG00000188641 | DPYD | chr1 | - | 97549559 | 97549744 | 97515725 | 97515941 | 97573759 | 97573970 | 555 | 529 | [illegible] |
| SE | 53981 | ENSG00000188641 | DPYD | chr1 | - | 97595058 | 97595166 | 97593217 | 97593387 | 97679094 | 97679182 | 833 | 926 | [illegible] |
| SE | 53982 | ENSG00000188641 | DPYD | chr1 | - | 97679094 | 97679182 | 97595058 | 97595166 | 97828113 | 97828196 | 179 | 203 | [illegible] |
| SE | 53983 | ENSG00000188641 | DPYD | chr1 | - | 97691716 | 97691798 | 97679094 | 97679182 | 97721509 | 97721671 | 473 | 535 | [illegible] |
| SE | 53984 | ENSG00000188641 | DPYD | chr1 | - | 97699350 | 97699547 | 97691206 | 97691798 | 97721509 | 97721671 | 599 | 604 | [illegible] |
| SE | 53987 | ENSG00000188641 | DPYD | chr1 | - | 97740391 | 97740479 | 97721509 | 97721671 | 97828113 | 97828196 | 1045 | 1136 | [illegible] |
| SE | 53988 | ENSG00000188641 | DPYD | chr1 | - | 97828113 | 97828196 | 97679094 | 97679182 | 97883263 | 97883374 | 354 | 377 | [illegible] |
| SE | 53989 | ENSG00000188641 | DPYD | chr1 | - | 97828113 | 97828196 | 97679094 | 97679182 | 97920883 | 97921034 | 1 | 2 | [illegible] |
| SE | 53991 | ENSG00000188641 | DPYD | chr1 | - | 97828113 | 97828196 | 97740391 | 97740479 | 97920883 | 97921034 | 348 | 339 | [illegible] |
| SE | 53992 | ENSG00000188641 | DPYD | chr1 | - | 97883263 | 97883374 | 97679094 | 97679182 | 97920883 | 97920997 | 180 | 129 | [illegible] |
| SE | 53993 | ENSG00000188641 | DPYD | chr1 | - | 97883263 | 97883374 | 97740391 | 97740479 | 97920883 | 97920997 | 192 | 136 | [illegible] |
| SE | 53994 | ENSG00000188641 | DPYD | chr1 | - | 97883263 | 97883374 | 97828113 | 97828196 | 97920883 | 97920997 | 882 | 864 | [illegible] |
| SE | 53995 | ENSG00000176927 | EFCAB5 | chr17 | + | 29950558 | 29950653 | 29943564 | 29943649 | 29968790 | 29968879 | 1 | 0 | [illegible] |
| SE | 53997 | ENSG00000176927 | EFCAB5 | chr17 | + | 30034229 | 30034385 | 29999905 | 29999976 | 30051117 | 30051217 | 2 | 5 | [illegible] |
| SE | 54006 | ENSG00000116205 | TCEANC2 | chr1 | + | 54054380 | 54054524 | 54053583 | 54053758 | 54068755 | 54068897 | 66 | 36 | [illegible] |
| SE | 54008 | ENSG00000116205 | TCEANC2 | chr1 | + | 54096284 | 54096844 | 54088596 | 54088790 | 54111051 | 54112504 | 37 | 45 | [illegible] |
| SE | 54009 | ENSG00000197989 | SNHG12 | chr1 | - | 28579532 | 28579587 | 28578537 | 28578680 | 28579911 | 28579981 | 71 | 33 | [illegible] |
| SE | 54014 | ENSG00000105609 | LILRB5 | chr19 | - | 54254017 | 54254068 | 54252870 | 54252987 | 54254364 | 54254415 | 0 | 0 | [illegible] |
| SE | 54016 | ENSG00000027001 | MIPEP | chr13 | - | 23809849 | 23809924 | 23805949 | 23806069 | 23836239 | 23836349 | 7 | 7 | [illegible] |
| SE | 54017 | ENSG00000027001 | MIPEP | chr13 | - | 23841334 | 23841488 | 23839648 | 23839726 | 23858859 | 23858912 | 20 | 17 | [illegible] |
| SE | 54019 | ENSG00000027001 | MIPEP | chr13 | - | 23858859 | 23858912 | 23839648 | 23839726 | 23862301 | 23862362 | 6 | 3 | [illegible] |
| SE | 54021 | ENSG00000027001 | MIPEP | chr13 | - | 23858859 | 23858912 | 23842269 | 23842569 | 23862301 | 23862362 | 6 | 3 | [illegible] |
| SE | 54022 | ENSG00000027001 | MIPEP | chr13 | - | 23864140 | 23864189 | 23862301 | 23862362 | 23869291 | 23869448 | 15 | 10 | [illegible] |
| SE | 54024 | ENSG00000027001 | MIPEP | chr13 | - | 23874845 | 23874909 | 23870012 | 23870195 | 23879267 | 23879354 | 35 | 50 | [illegible] |
| SE | 54030 | ENSG00000174132 | FAM174A | chr5 | + | 100562053 | 100562188 | 100535373 | 100535964 | 100586180 | 100586741 | 122 | 77 | [illegible] |
| SE | 54032 | ENSG00000180353 | HCLS1 | chr3 | - | 121632100 | 121632184 | 121631398 | 121631982 | 121632331 | 121632563 | 597 | 253 | [illegible] |
| SE | 54033 | ENSG00000180353 | HCLS1 | chr3 | - | 121632100 | 121632184 | 121631398 | 121631982 | 121633066 | 121633171 | 203 | 74 | [illegible] |
| SE | 54034 | ENSG00000180353 | HCLS1 | chr3 | - | 121632331 | 121632563 | 121631398 | 121631982 | 121633066 | 121633171 | 572 | 535 | [illegible] |
| SE | 54035 | ENSG00000180353 | HCLS1 | chr3 | - | 121632331 | 121632563 | 121632100 | 121632184 | 121633066 | 121633171 | 968 | 715 | [illegible] |
| SE | 54036 | ENSG00000180353 | HCLS1 | chr3 | - | 121635734 | 121635804 | 121634206 | 121634418 | 121636433 | 121636489 | 1693 | 1563 | [illegible] |
| SE | 54039 | ENSG00000180353 | HCLS1 | chr3 | - | 121644817 | 121644928 | 121642926 | 121642981 | 121657278 | 121657352 | 1363 | 843 | [illegible] |
| SE | 54041 | ENSG00000180353 | HCLS1 | chr3 | - | 121647318 | 121647448 | 121644229 | 121644928 | 121657278 | 121657352 | 2699 | 2015 | [illegible] |
| SE | 54042 | ENSG00000180353 | HCLS1 | chr3 | - | 121647318 | 121647448 | 121644229 | 121644928 | 121658263 | 121658347 | 1477 | 1226 | [illegible] |
| SE | 54043 | ENSG00000180353 | HCLS1 | chr3 | - | 121647318 | 121647482 | 121644817 | 121644928 | 121657278 | 121657352 | 1480 | 1229 | [illegible] |
| SE | 54044 | ENSG00000180353 | HCLS1 | chr3 | - | 121657201 | 121657352 | 121647318 | 121647448 | 121658263 | 121658347 | 613 | 362 | [illegible] |
| SE | 54045 | ENSG00000180353 | HCLS1 | chr3 | - | 121657201 | 121657352 | 121647318 | 121647448 | 121660819 | 121660878 | 4 | 2 | [illegible] |
| SE | 54046 | ENSG00000180353 | HCLS1 | chr3 | - | 121657278 | 121657352 | 121644229 | 121644928 | 121658263 | 121658347 | 598 | 340 | [illegible] |
| SE | 54047 | ENSG00000180353 | HCLS1 | chr3 | - | 121657278 | 121657352 | 121647318 | 121647448 | 121658263 | 121658347 | 1817 | 1129 | [illegible] |
| SE | 54048 | ENSG00000180353 | HCLS1 | chr3 | - | 121657278 | 121657352 | 121647318 | 121647448 | 121660819 | 121660883 | 520 | 367 | [illegible] |
| SE | 54049 | ENSG00000180353 | HCLS1 | chr3 | - | 121658263 | 121658347 | 121647318 | 121647448 | 121660819 | 121660878 | 236 | 110 | [illegible] |
| SE | 54050 | ENSG00000180353 | HCLS1 | chr3 | - | 121658263 | 121658347 | 121656890 | 121657352 | 121660819 | 121660878 | 1537 | 882 | [illegible] |
| SE | 54055 | ENSG00000158796 | DEDD | chr1 | - | 161124413 | 161124526 | 161124132 | 161124210 | 161130814 | 161130847 | 19 | 10 | [illegible] |
| SE | 54056 | ENSG00000158796 | DEDD | chr1 | - | 161130814 | 161130843 | 161124137 | 161124526 | 161132390 | 161132476 | 12 | 7 | [illegible] |
| SE | 54058 | ENSG00000158796 | DEDD | chr1 | - | 161130814 | 161130847 | 161124137 | 161124526 | 161132390 | 161132450 | 25 | 11 | [illegible] |
| SE | 54061 | ENSG00000158796 | DEDD | chr1 | - | 161131005 | 161131182 | 161130814 | 161130847 | 161132390 | 161132476 | 0 | 0 | [illegible] |
| SE | 54063 | ENSG00000235162 | C12orf75 | chr12 | + | 105348601 | 105348626 | 105330635 | 105330937 | 105365806 | 105365842 | 60 | 28 | [illegible] |
| SE | 54069 | ENSG00000107021 | TBC1D13 | chr9 | + | 128803249 | 128803448 | 128797054 | 128797214 | 128803955 | 128804119 | 66 | 31 | [illegible] |
| SE | 54070 | ENSG00000107021 | TBC1D13 | chr9 | + | 128803249 | 128803460 | 128797054 | 128797214 | 128803955 | 128804119 | 114 | 54 | [illegible] |
| SE | 54074 | ENSG00000188761 | BCL2L15 | chr1 | - | 113881772 | 113881997 | 113876815 | 113881140 | 113886536 | 113886658 | 1 | 8 | [illegible] |
| SE | 54075 | ENSG00000188761 | BCL2L15 | chr1 | - | 113881772 | 113881997 | 113876815 | 113881140 | 113887248 | 113887305 | 1 | 3 | [illegible] |
| SE | 54076 | ENSG00000188761 | BCL2L15 | chr1 | - | 113886536 | 113886658 | 113881122 | 113881140 | 113887248 | 113887375 | 0 | 2 | [illegible] |
| SE | 54077 | ENSG00000188761 | BCL2L15 | chr1 | - | 113886536 | 113886658 | 113881763 | 113881997 | 113887248 | 113887375 | 0 | 5 | [illegible] |
| SE | 54078 | ENSG00000102218 | RP2 | chrX | + | 46877504 | 46877590 | 46859987 | 46860102 | 46879685 | 46882358 | 147 | 122 | [illegible] |
| SE | 54080 | ENSG00000092847 | AGO1 | chr1 | + | 35893096 | 35893278 | 35892556 | 35892677 | 35893591 | 35893810 | 61 | 39 | [illegible] |
| SE | 54083 | ENSG00000092847 | AGO1 | chr1 | + | 35894036 | 35894171 | 35893591 | 35893810 | 35894314 | 35894402 | 93 | 58 | [illegible] |
| SE | 54085 | ENSG00000092847 | AGO1 | chr1 | + | 35906934 | 35907119 | 35902203 | 35902337 | 35913841 | 35914001 | 286 | 137 | [illegible] |
| SE | 54090 | ENSG00000115935 | WIPF1 | chr2 | - | 174581309 | 174581439 | 174571168 | 174572446 | 174585522 | 174585589 | 595 | 388 | [illegible] |
| SE | 54091 | ENSG00000115935 | WIPF1 | chr2 | - | 174581309 | 174581439 | 174575203 | 174575380 | 174585522 | 174585589 | 993 | 638 | [illegible] |
| SE | 54092 | ENSG00000115935 | WIPF1 | chr2 | - | 174585522 | 174585611 | 174581139 | 174581439 | 174597600 | 174597765 | 1111 | 676 | [illegible] |
| SE | 54093 | ENSG00000115935 | WIPF1 | chr2 | - | 174597600 | 174597858 | 174585522 | 174585611 | 174598170 | 174598206 | 631 | 364 | [illegible] |
| SE | 54094 | ENSG00000115935 | WIPF1 | chr2 | - | 174597600 | 174597858 | 174585522 | 174585611 | 174598175 | 174598211 | 632 | 366 | [illegible] |
| SE | 54100 | ENSG00000115935 | WIPF1 | chr2 | - | 174625820 | 174625960 | 174623480 | 174623508 | 174634517 | 174634605 | 36 | 24 | [illegible] |
| SE | 54103 | ENSG00000115935 | WIPF1 | chr2 | - | 174629665 | 174629868 | 174625623 | 174625960 | 174630381 | 174630429 | 28 | 30 | [illegible] |
| SE | 54109 | ENSG00000115935 | WIPF1 | chr2 | - | 174630381 | 174630696 | 174625709 | 174625960 | 174634517 | 174634536 | 0 | 0 | [illegible] |
| SE | 54111 | ENSG00000115935 | WIPF1 | chr2 | - | 174677044 | 174677124 | 174676336 | 174676666 | 174682773 | 174682916 | 0 | 0 | [illegible] |
| SE | 54120 | ENSG00000124126 | PREX1 | chr20 | - | 48632276 | 48632391 | 48630727 | 48630794 | 48632495 | 48632639 | 522 | 145 | [illegible] |
| SE | 54121 | ENSG00000124126 | PREX1 | chr20 | - | 48639765 | 48639894 | 48637710 | 48637752 | 48642167 | 48642258 | 823 | 405 | [illegible] |
| SE | 54122 | ENSG00000124126 | PREX1 | chr20 | - | 48639765 | 48639894 | 48637710 | 48637752 | 48642406 | 48642489 | 596 | 316 | [illegible] |
| SE | 54123 | ENSG00000124126 | PREX1 | chr20 | - | 48642167 | 48642258 | 48637710 | 48637752 | 48642406 | 48642489 | 375 | 165 | [illegible] |
| SE | 54126 | ENSG00000124126 | PREX1 | chr20 | - | 48649995 | 48650206 | 48649299 | 48649576 | 48650893 | 48651055 | 510 | 192 | [illegible] |
| SE | 54127 | ENSG00000124126 | PREX1 | chr20 | - | 48649995 | 48650206 | 48649299 | 48649576 | 48651395 | 48651583 | 113 | 52 | [illegible] |
| SE | 54128 | ENSG00000124126 | PREX1 | chr20 | - | 48650893 | 48651055 | 48649299 | 48649576 | 48651395 | 48651583 | 364 | 125 | [illegible] |
| SE | 54133 | ENSG00000124126 | PREX1 | chr20 | - | 48666282 | 48666355 | 48659918 | 48660061 | 48676192 | 48676268 | 844 | 367 | [illegible] |
| SE | 54134 | ENSG00000124126 | PREX1 | chr20 | - | 48679359 | 48679409 | 48676192 | 48676268 | 48679650 | 48679754 | 652 | 311 | [illegible] |
| SE | 54135 | ENSG00000124126 | PREX1 | chr20 | - | 48708259 | 48708421 | 48700752 | 48700886 | 48726289 | 48726391 | 678 | 308 | [illegible] |
| SE | 54136 | ENSG00000124126 | PREX1 | chr20 | - | 48726289 | 48726391 | 48708259 | 48708421 | 48734545 | 48734650 | 491 | 250 | [illegible] |
| SE | 54137 | ENSG00000124126 | PREX1 | chr20 | - | 48726289 | 48726391 | 48708259 | 48708421 | 48745024 | 48745147 | 150 | 87 | [illegible] |
| SE | 54138 | ENSG00000124126 | PREX1 | chr20 | - | 48734545 | 48734650 | 48708259 | 48708421 | 48745024 | 48745147 | 218 | 107 | [illegible] |
| SE | 54139 | ENSG00000124126 | PREX1 | chr20 | - | 48734545 | 48734650 | 48726289 | 48726391 | 48745024 | 48745147 | 558 | 272 | [illegible] |
| SE | 54140 | ENSG00000124126 | PREX1 | chr20 | - | 48745024 | 48745147 | 48734545 | 48734650 | 48827641 | 48827999 | 296 | 155 | [illegible] |
| SE | 54141 | ENSG00000124126 | PREX1 | chr20 | - | 48747808 | 48747880 | 48745024 | 48745147 | 48827641 | 48827999 | 171 | 85 | [illegible] |
| SE | 54142 | ENSG00000142765 | SYTL1 | chr1 | + | 27349651 | 27349765 | 27349397 | 27349498 | 27349971 | 27350132 | 117 | 31 | [illegible] |
| SE | 54144 | ENSG00000142765 | SYTL1 | chr1 | + | 27349971 | 27350132 | 27349397 | 27349498 | 27350388 | 27350485 | 45 | 7 | [illegible] |
| SE | 54145 | ENSG00000142765 | SYTL1 | chr1 | + | 27349971 | 27350132 | 27349475 | 27349765 | 27350388 | 27350485 | 96 | 21 | [illegible] |
| SE | 54146 | ENSG00000142765 | SYTL1 | chr1 | + | 27350793 | 27350952 | 27349971 | 27350485 | 27351257 | 27351336 | 197 | 60 | [illegible] |
| SE | 54151 | ENSG00000266053 | NDUFV2-AS1 | chr18 | - | 9133390 | 9133510 | 9121264 | 9121456 | 9136363 | 9136645 | 6 | 2 | [illegible] |
| SE | 54152 | ENSG00000237638 | LINC02245 | chr2 | - | 64955892 | 64955984 | 64953252 | 64954044 | 64970571 | 64970785 | 0 | 2 | [illegible] |
| SE | 54154 | ENSG00000237638 | LINC02245 | chr2 | - | 64970571 | 64970785 | 64955892 | 64955984 | 65047953 | 65048179 | 0 | 4 | [illegible] |
| SE | 54156 | ENSG00000136122 | BORA | chr13 | + | 72728925 | 72729093 | 72727748 | 72728007 | 72734959 | 72735005 | 8 | 6 | [illegible] |
| SE | 54157 | ENSG00000136122 | BORA | chr13 | + | 72728925 | 72729093 | 72727748 | 72728007 | 72737961 | 72738043 | 6 | 7 | [illegible] |
| SE | 54158 | ENSG00000136122 | BORA | chr13 | + | 72731280 | 72731387 | 72727922 | 72728007 | 72734959 | 72735005 | 6 | 5 | [illegible] |
| SE | 54160 | ENSG00000136122 | BORA | chr13 | + | 72731280 | 72731387 | 72728925 | 72729093 | 72734959 | 72735005 | 13 | 18 | [illegible] |
| SE | 54161 | ENSG00000136122 | BORA | chr13 | + | 72731280 | 72731387 | 72728925 | 72729093 | 72737961 | 72738043 | 6 | 11 | [illegible] |
| SE | 54162 | ENSG00000136122 | BORA | chr13 | + | 72734959 | 72735005 | 72727748 | 72728007 | 72737961 | 72738043 | 2 | 4 | [illegible] |
| SE | 54163 | ENSG00000136122 | BORA | chr13 | + | 72734959 | 72735005 | 72728925 | 72729093 | 72737961 | 72738043 | 4 | 4 | [illegible] |
| SE | 54164 | ENSG00000136122 | BORA | chr13 | + | 72734959 | 72735005 | 72731280 | 72731387 | 72737961 | 72738043 | 10 | 11 | [illegible] |
| SE | 54166 | ENSG00000136122 | BORA | chr13 | + | 72744980 | 72745207 | 72744504 | 72744561 | 72745943 | 72746076 | 41 | 37 | [illegible] |
| SE | 54168 | ENSG00000164300 | SERINC5 | chr5 | - | 80113571 | 80113625 | 80111650 | 80111702 | 80114513 | 80114651 | 0 | 0 | [illegible] |
| SE | 54170 | ENSG00000164300 | SERINC5 | chr5 | - | 80116137 | 80116323 | 80113571 | 80113625 | 80146089 | 80146234 | 1 | 0 | [illegible] |
| SE | 54172 | ENSG00000164300 | SERINC5 | chr5 | - | 80150881 | 80150948 | 80147244 | 80147284 | 80158835 | 80158956 | 198 | 149 | [illegible] |
| SE | 54173 | ENSG00000164300 | SERINC5 | chr5 | - | 80158835 | 80158956 | 80150881 | 80150948 | 80166382 | 80166478 | 146 | 117 | [illegible] |
| SE | 54174 | ENSG00000164300 | SERINC5 | chr5 | - | 80158835 | 80158962 | 80150881 | 80150948 | 80166382 | 80166478 | 274 | 200 | [illegible] |
| SE | 54175 | ENSG00000164300 | SERINC5 | chr5 | - | 80166382 | 80166478 | 80158835 | 80158962 | 80174953 | 80175047 | 107 | 71 | [illegible] |
| SE | 54178 | ENSG00000175376 | EIF1AD | chr11 | - | 65999349 | 65999397 | 65996544 | 65998743 | 65999566 | 65999675 | 165 | 102 | [illegible] |
| SE | 54179 | ENSG00000175376 | EIF1AD | chr11 | - | 65999566 | 65999675 | 65996544 | 65998743 | 66002039 | 66002156 | 0 | 0 | [illegible] |
| SE | 54180 | ENSG00000175376 | EIF1AD | chr11 | - | 66000052 | 66000161 | 65999566 | 65999675 | 66002039 | 66002161 | 49 | 49 | [illegible] |
| SE | 54185 | ENSG00000163781 | TOPBP1 | chr3 | - | 133612388 | 133612552 | 133611003 | 133611141 | 133616813 | 133616925 | 307 | 289 | [illegible] |
| SE | 54187 | ENSG00000163781 | TOPBP1 | chr3 | - | 133620154 | 133620347 | 133617159 | 133617326 | 133623090 | 133623193 | 97 | 85 | [illegible] |
| SE | 54188 | ENSG00000163781 | TOPBP1 | chr3 | - | 133624051 | 133624175 | 133623310 | 133623457 | 133628361 | 133628391 | 149 | 101 | [illegible] |
| SE | 54192 | ENSG00000172667 | ZMAT3 | chr3 | - | 178965444 | 178965496 | 178960120 | 178961563 | 179004060 | 179004184 | 3 | 0 | [illegible] |
| SE | 54193 | ENSG00000172667 | ZMAT3 | chr3 | - | 178982368 | 178982465 | 178965444 | 178965496 | 179004060 | 179004184 | 0 | 0 | [illegible] |
| SE | 54194 | ENSG00000172667 | ZMAT3 | chr3 | - | 179030879 | 179030999 | 179027645 | 179027812 | 179067482 | 179067809 | 116 | 82 | [illegible] |
| SE | 54195 | ENSG00000172667 | ZMAT3 | chr3 | - | 179067482 | 179067809 | 179030879 | 179030999 | 179071594 | 179071809 | 64 | 42 | [illegible] |
| SE | 54196 | ENSG00000172667 | ZMAT3 | chr3 | - | 179071326 | 179071447 | 179067482 | 179067809 | 179071594 | 179072215 | 3 | 1 | [illegible] |
| SE | 54211 | ENSG00000168395 | ING5 | chr2 | + | 241711977 | 241712071 | 241711376 | 241711488 | 241719482 | 241719612 | 59 | 51 | [illegible] |
| SE | 54213 | ENSG00000166619 | BLCAP | chr20 | - | 37521318 | 37521409 | 37518939 | 37519350 | 37527792 | 37527878 | 2 | 0 | [illegible] |
| SE | 54214 | ENSG00000166619 | BLCAP | chr20 | - | 37521318 | 37521451 | 37517417 | 37519350 | 37527792 | 37527878 | 2 | 0 | [illegible] |
| SE | 54215 | ENSG00000143643 | TTC13 | chr1 | - | 230923840 | 230923933 | 230921420 | 230921504 | 230924840 | 230924863 | 322 | 218 | [illegible] |
| SE | 54217 | ENSG00000143643 | TTC13 | chr1 | - | 230931297 | 230931472 | 230928936 | 230929093 | 230931735 | 230931877 | 288 | 205 | [illegible] |
| SE | 54218 | ENSG00000143643 | TTC13 | chr1 | - | 230931735 | 230931877 | 230928936 | 230929093 | 230933778 | 230933861 | 94 | 72 | [illegible] |
| SE | 54224 | ENSG00000143643 | TTC13 | chr1 | - | 230939385 | 230939496 | 230936157 | 230936264 | 230940439 | 230940556 | 80 | 37 | [illegible] |
| SE | 54228 | ENSG00000143643 | TTC13 | chr1 | - | 230940439 | 230940556 | 230936157 | 230936264 | 230943805 | 230943898 | 106 | 91 | [illegible] |
| SE | 54229 | ENSG00000143643 | TTC13 | chr1 | - | 230940439 | 230940556 | 230936157 | 230936264 | 230945388 | 230945454 | 13 | 5 | [illegible] |
| SE | 54231 | ENSG00000143643 | TTC13 | chr1 | - | 230943805 | 230943898 | 230936157 | 230936264 | 230945388 | 230945454 | 50 | 23 | [illegible] |
| SE | 54235 | ENSG00000143643 | TTC13 | chr1 | - | 230954332 | 230954403 | 230940439 | 230940556 | 230958223 | 230958299 | 98 | 50 | [illegible] |
| SE | 54237 | ENSG00000143643 | TTC13 | chr1 | - | 230958223 | 230958299 | 230954332 | 230954403 | 230961208 | 230961303 | 237 | 127 | [illegible] |
| SE | 54240 | ENSG00000143643 | TTC13 | chr1 | - | 230961208 | 230961303 | 230958223 | 230958299 | 230978559 | 230978796 | 127 | 60 | [illegible] |
| SE | 54241 | ENSG00000011405 | PIK3C2A | chr11 | - | 17094260 | 17094385 | 17092158 | 17092276 | 17097056 | 17097264 | 110 | 90 | [illegible] |
| SE | 54242 | ENSG00000011405 | PIK3C2A | chr11 | - | 17094260 | 17094385 | 17092158 | 17092276 | 17099859 | 17099969 | 81 | 55 | [illegible] |
| SE | 54243 | ENSG00000011405 | PIK3C2A | chr11 | - | 17097056 | 17097264 | 17092158 | 17092276 | 17099859 | 17099969 | 79 | 51 | [illegible] |
| SE | 54245 | ENSG00000011405 | PIK3C2A | chr11 | - | 17101277 | 17101434 | 17099859 | 17099969 | 17102661 | 17102831 | 95 | 79 | [illegible] |
| SE | 54247 | ENSG00000011405 | PIK3C2A | chr11 | - | 17112573 | 17112666 | 17110431 | 17110561 | 17114360 | 17114465 | 91 | 62 | [illegible] |
| SE | 54248 | ENSG00000011405 | PIK3C2A | chr11 | - | 17117490 | 17117671 | 17114360 | 17114465 | 17118644 | 17118739 | 94 | 79 | [illegible] |
| SE | 54249 | ENSG00000011405 | PIK3C2A | chr11 | - | 17119219 | 17119313 | 17118644 | 17118739 | 17119785 | 17119974 | 63 | 84 | [illegible] |
| SE | 54250 | ENSG00000011405 | PIK3C2A | chr11 | - | 17119785 | 17119974 | 17118644 | 17118739 | 17122187 | 17122333 | 54 | 63 | [illegible] |
| SE | 54251 | ENSG00000011405 | PIK3C2A | chr11 | - | 17129299 | 17129467 | 17122701 | 17122813 | 17131915 | 17132038 | 103 | 115 | [illegible] |
| SE | 54252 | ENSG00000011405 | PIK3C2A | chr11 | - | 17135110 | 17135159 | 17134818 | 17135029 | 17136481 | 17136625 | 98 | 65 | [illegible] |
| SE | 54254 | ENSG00000011405 | PIK3C2A | chr11 | - | 17148666 | 17148787 | 17147516 | 17147628 | 17150497 | 17150655 | 159 | 189 | [illegible] |
| SE | 54255 | ENSG00000011405 | PIK3C2A | chr11 | - | 17155525 | 17155629 | 17150497 | 17150655 | 17168676 | 17169741 | 128 | 128 | [illegible] |
| SE | 54258 | ENSG00000011405 | PIK3C2A | chr11 | - | 17168676 | 17169806 | 17155525 | 17155629 | 17207847 | 17207947 | 115 | 110 | [illegible] |
| SE | 54259 | ENSG00000011405 | PIK3C2A | chr11 | - | 17194388 | 17194414 | 17169351 | 17169806 | 17207847 | 17207950 | 0 | 0 | [illegible] |
| SE | 54260 | ENSG00000011405 | PIK3C2A | chr11 | - | 17204306 | 17204375 | 17169351 | 17169806 | 17207847 | 17207983 | 4 | 2 | [illegible] |
| SE | 54262 | ENSG00000090097 | PCBP4 | chr3 | - | 51959894 | 51960023 | 51959576 | 51959651 | 51960188 | 51960320 | 48 | 10 | [illegible] |
| SE | 54264 | ENSG00000090097 | PCBP4 | chr3 | - | 51961748 | 51962088 | 51961159 | 51961304 | 51967325 | 51967448 | 0 | 0 | [illegible] |
| SE | 54265 | ENSG00000090097 | PCBP4 | chr3 | - | 51961940 | 51962088 | 51961159 | 51961304 | 51962809 | 51962892 | 6 | 3 | [illegible] |
| SE | 54266 | ENSG00000090097 | PCBP4 | chr3 | - | 51961940 | 51962088 | 51961159 | 51961304 | 51967325 | 51967454 | 3 | 1 | [illegible] |
| SE | 54267 | ENSG00000090097 | PCBP4 | chr3 | - | 51962809 | 51962892 | 51961159 | 51961304 | 51967325 | 51967434 | 0 | 1 | [illegible] |
| SE | 54268 | ENSG00000090097 | PCBP4 | chr3 | - | 51962809 | 51962892 | 51961940 | 51962088 | 51967325 | 51967434 | 3 | 2 | [illegible] |
| SE | 54281 | ENSG00000143442 | POGZ | chr1 | - | 151408099 | 151408240 | 151407234 | 151407291 | 151408408 | 151408581 | 145 | 77 | [illegible] |
| SE | 54283 | ENSG00000143442 | POGZ | chr1 | - | 151408693 | 151408828 | 151408099 | 151408240 | 151411624 | 151411771 | 192 | 125 | [illegible] |
| SE | 54286 | ENSG00000143442 | POGZ | chr1 | - | 151412295 | 151412396 | 151408693 | 151408828 | 151423396 | 151423421 | 114 | 90 | [illegible] |
| SE | 54287 | ENSG00000143442 | POGZ | chr1 | - | 151423396 | 151423551 | 151412295 | 151412396 | 151440927 | 151441086 | 160 | 115 | [illegible] |
| SE | 54288 | ENSG00000143442 | POGZ | chr1 | - | 151423948 | 151424286 | 151423396 | 151423551 | 151424954 | 151425061 | 354 | 232 | [illegible] |
| SE | 54289 | ENSG00000143442 | POGZ | chr1 | - | 151423948 | 151424286 | 151423396 | 151423551 | 151427822 | 151427980 | 105 | 64 | [illegible] |
| SE | 54290 | ENSG00000143442 | POGZ | chr1 | - | 151423948 | 151424286 | 151423396 | 151423551 | 151428122 | 151428413 | 100 | 63 | [illegible] |
| SE | 54291 | ENSG00000143442 | POGZ | chr1 | - | 151423948 | 151424286 | 151423396 | 151423551 | 151429602 | 151429711 | 100 | 63 | [illegible] |
| SE | 54292 | ENSG00000143442 | POGZ | chr1 | - | 151423948 | 151424286 | 151423396 | 151423551 | 151430665 | 151430841 | 100 | 63 | [illegible] |
| SE | 54293 | ENSG00000143442 | POGZ | chr1 | - | 151424954 | 151425061 | 151423396 | 151423551 | 151427822 | 151428014 | 134 | 93 | [illegible] |
| SE | 54294 | ENSG00000143442 | POGZ | chr1 | - | 151424954 | 151425061 | 151423396 | 151423551 | 151428122 | 151428413 | 2 | 1 | [illegible] |
| SE | 54297 | ENSG00000143442 | POGZ | chr1 | - | 151424954 | 151425061 | 151423948 | 151424286 | 151428122 | 151428413 | 200 | 144 | [illegible] |
| SE | 54299 | ENSG00000143442 | POGZ | chr1 | - | 151427822 | 151428014 | 151423396 | 151423551 | 151428122 | 151428413 | 4 | 8 | [illegible] |
| SE | 54301 | ENSG00000143442 | POGZ | chr1 | - | 151427822 | 151428041 | 151423396 | 151423551 | 151428122 | 151428413 | 37 | 33 | [illegible] |
| SE | 54303 | ENSG00000143442 | POGZ | chr1 | - | 151428122 | 151428413 | 151423396 | 151423551 | 151429602 | 151429711 | 176 | 113 | [illegible] |
| SE | 54304 | ENSG00000143442 | POGZ | chr1 | - | 151428122 | 151428413 | 151423396 | 151423551 | 151430665 | 151430841 | 2 | 2 | [illegible] |
| SE | 54305 | ENSG00000143442 | POGZ | chr1 | - | 151428122 | 151428413 | 151423396 | 151423551 | 151440927 | 151441086 | 1 | 0 | [illegible] |
| SE | 54306 | ENSG00000143442 | POGZ | chr1 | - | 151428122 | 151428413 | 151423948 | 151424286 | 151429602 | 151429711 | 176 | 113 | [illegible] |
| SE | 54307 | ENSG00000143442 | POGZ | chr1 | - | 151428122 | 151428413 | 151423948 | 151424286 | 151430665 | 151430841 | 2 | 2 | [illegible] |
| SE | 54308 | ENSG00000143442 | POGZ | chr1 | - | 151428122 | 151428413 | 151424954 | 151425061 | 151430665 | 151430841 | 2 | 3 | [illegible] |
| SE | 54309 | ENSG00000143442 | POGZ | chr1 | - | 151429602 | 151429711 | 151423396 | 151423551 | 151430665 | 151430841 | 83 | 55 | [illegible] |
| SE | 54312 | ENSG00000143442 | POGZ | chr1 | - | 151430665 | 151430841 | 151423396 | 151423551 | 151440927 | 151441086 | 169 | 98 | [illegible] |
| SE | 54313 | ENSG00000143442 | POGZ | chr1 | - | 151430665 | 151430841 | 151428122 | 151428413 | 151440927 | 151441086 | 171 | 99 | [illegible] |
| SE | 54315 | ENSG00000143442 | POGZ | chr1 | - | 151450909 | 151451047 | 151442080 | 151442205 | 151459151 | 151459179 | 0 | 0 | [illegible] |
| SE | 54318 | ENSG00000117139 | KDM5B | chr1 | - | 202756357 | 202756516 | 202754996 | 202755452 | 202758390 | 202758510 | 112 | 80 | [illegible] |
| SE | 54320 | ENSG00000117139 | KDM5B | chr1 | - | 202760414 | 202760573 | 202758390 | 202758510 | 202773117 | 202773284 | 50 | 33 | [illegible] |
| SE | 54322 | ENSG00000117139 | KDM5B | chr1 | - | 202762698 | 202762808 | 202760414 | 202760573 | 202773117 | 202773284 | 49 | 30 | [illegible] |
| SE | 54323 | ENSG00000117139 | KDM5B | chr1 | - | 202764048 | 202764145 | 202760414 | 202760573 | 202773117 | 202773288 | 2 | 0 | [illegible] |
| SE | 54324 | ENSG00000117139 | KDM5B | chr1 | - | 202764048 | 202764145 | 202762698 | 202762808 | 202773117 | 202773288 | 17 | 5 | [illegible] |
| SE | 54327 | ENSG00000117139 | KDM5B | chr1 | - | 202766925 | 202767060 | 202764048 | 202764145 | 202773117 | 202773284 | 57 | 33 | [illegible] |
| SE | 54328 | ENSG00000117139 | KDM5B | chr1 | - | 202773117 | 202773284 | 202766925 | 202767060 | 202774612 | 202774735 | 43 | 18 | [illegible] |
| SE | 54329 | ENSG00000117139 | KDM5B | chr1 | - | 202773117 | 202773288 | 202764073 | 202767060 | 202774612 | 202774735 | 141 | 113 | [illegible] |
| SE | 54335 | ENSG00000176095 | IP6K1 | chr3 | - | 49738211 | 49738422 | 49732790 | 49732972 | 49747817 | 49748168 | 161 | 68 | [illegible] |
| SE | 54340 | ENSG00000178199 | ZC3H12D | chr6 | - | 149452615 | 149452722 | 149446794 | 149451479 | 149456665 | 149456900 | 23 | 11 | [illegible] |
| SE | 54341 | ENSG00000178199 | ZC3H12D | chr6 | - | 149474238 | 149474613 | 149461667 | 149461970 | 149484812 | 149485014 | 35 | 18 | [illegible] |
| SE | 54344 | ENSG00000049618 | ARID1B | chr6 | + | 157084661 | 157084905 | 156935465 | 156935576 | 157110471 | 157110561 | 113 | 57 | [illegible] |
| SE | 54347 | ENSG00000049618 | ARID1B | chr6 | + | 157133027 | 157133207 | 157110471 | 157110561 | 157167039 | 157167185 | 64 | 38 | [illegible] |
| SE | 54348 | ENSG00000049618 | ARID1B | chr6 | + | 157148623 | 157148951 | 157110471 | 157110561 | 157167039 | 157167185 | 78 | 43 | [illegible] |
| SE | 54350 | ENSG00000049618 | ARID1B | chr6 | + | 157150724 | 157150853 | 157148623 | 157148951 | 157164638 | 157164748 | 0 | 0 | [illegible] |
| SE | 54352 | ENSG00000049618 | ARID1B | chr6 | + | 157164638 | 157164748 | 157133027 | 157133207 | 157167039 | 157167185 | 0 | 2 | [illegible] |

| | | | | | | | | | | | | |
|---|---|---|---|---|---|---|---|---|---|---|---|---|
| SE | 54355 | ENSG00000049618 | ARID1B | chr6 | + | 157174007 | 157174117 | 157167039 | 157167185 | 157180968 | 157181178 | [illegible] |
| SE | 54358 | ENSG00000049618 | ARID1B | chr6 | + | 157203865 | 157203996 | 157200704 | 157201488 | 157206166 | 157207868 | [illegible] |
| SE | 54359 | ENSG00000189266 | PNRC2 | chr1 | + | 23960928 | 23961134 | 23959827 | 23959998 | 23961439 | 23961465 | [illegible] |
| SE | 54360 | ENSG00000189266 | PNRC2 | chr1 | + | 23960932 | 23961134 | 23959827 | 23959998 | 23961439 | 23961880 | [illegible] |
| SE | 54365 | ENSG00000170271 | FAXDC2 | chr5 | - | 154822471 | 154822577 | 154818493 | 154821426 | 154823386 | 154823592 | [illegible] |
| SE | 54366 | ENSG00000170271 | FAXDC2 | chr5 | - | 154823307 | 154823592 | 154822471 | 154822577 | 154830800 | 154830922 | [illegible] |
| SE | 54367 | ENSG00000170271 | FAXDC2 | chr5 | - | 154823386 | 154823592 | 154822469 | 154822577 | 154830800 | 154830922 | [illegible] |
| SE | 54369 | ENSG00000170271 | FAXDC2 | chr5 | - | 154830800 | 154830922 | 154823307 | 154823592 | 154834842 | 154835088 | [illegible] |
| SE | 54371 | ENSG00000170271 | FAXDC2 | chr5 | - | 154834624 | 154834728 | 154823307 | 154823592 | 154834842 | 154834934 | [illegible] |
| SE | 54391 | ENSG00000189229 | NA | chr3 | + | 6710452 | 6710543 | 6706635 | 6706728 | 6717568 | 6717667 | [illegible] |
| SE | 54392 | ENSG00000189229 | NA | chr3 | + | 6720086 | 6720271 | 6717568 | 6717667 | 6724449 | 6724546 | [illegible] |
| SE | 54395 | ENSG00000141127 | PRPSAP2 | chr17 | + | 18857783 | 18857853 | 18856340 | 18856429 | 18865801 | 18865952 | [illegible] |
| SE | 54396 | ENSG00000141127 | PRPSAP2 | chr17 | + | 18859339 | 18859475 | 18856341 | 18856476 | 18865468 | 18865564 | [illegible] |
| SE | 54397 | ENSG00000141127 | PRPSAP2 | chr17 | + | 18859339 | 18859475 | 18856341 | 18856476 | 18865801 | 18865900 | [illegible] |
| SE | 54398 | ENSG00000141127 | PRPSAP2 | chr17 | + | 18859339 | 18859475 | 18858103 | 18858261 | 18865468 | 18865564 | [illegible] |
| SE | 54400 | ENSG00000141127 | PRPSAP2 | chr17 | + | 18860565 | 18860691 | 18859339 | 18859475 | 18865468 | 18865564 | [illegible] |
| SE | 54401 | ENSG00000141127 | PRPSAP2 | chr17 | + | 18864867 | 18864941 | 18858130 | 18858261 | 18865468 | 18865564 | [illegible] |
| SE | 54403 | ENSG00000141127 | PRPSAP2 | chr17 | + | 18865468 | 18865564 | 18856341 | 18856476 | 18865801 | 18865900 | [illegible] |
| SE | 54411 | ENSG00000141127 | PRPSAP2 | chr17 | + | 18911102 | 18911251 | 18889821 | 18889877 | 18923913 | 18923984 | [illegible] |
| SE | 54412 | ENSG00000141127 | PRPSAP2 | chr17 | + | 18917431 | 18917536 | 18911102 | 18911251 | 18923913 | 18923984 | [illegible] |
| SE | 54413 | ENSG00000141127 | PRPSAP2 | chr17 | + | 18923913 | 18923984 | 18911102 | 18911251 | 18928810 | 18928957 | [illegible] |
| SE | 54416 | ENSG00000169554 | ZEB2 | chr2 | - | 144396411 | 144396592 | 144384080 | 144390028 | 144398300 | 144398318 | [illegible] |
| SE | 54417 | ENSG00000169554 | ZEB2 | chr2 | - | 144396411 | 144396592 | 144384080 | 144390028 | 144401198 | 144401307 | [illegible] |
| SE | 54419 | ENSG00000169554 | ZEB2 | chr2 | - | 144398300 | 144400270 | 144384080 | 144390028 | 144401198 | 144401307 | [illegible] |
| SE | 54421 | ENSG00000169554 | ZEB2 | chr2 | - | 144398300 | 144400270 | 144396411 | 144396592 | 144403915 | 144404130 | [illegible] |
| SE | 54422 | ENSG00000169554 | ZEB2 | chr2 | - | 144398300 | 144400345 | 144384080 | 144390028 | 144401198 | 144401307 | [illegible] |
| SE | 54424 | ENSG00000169554 | ZEB2 | chr2 | - | 144401198 | 144401307 | 144396411 | 144396592 | 144403915 | 144404130 | [illegible] |
| SE | 54425 | ENSG00000169554 | ZEB2 | chr2 | - | 144401198 | 144401307 | 144398300 | 144400270 | 144403915 | 144404130 | [illegible] |
| SE | 54426 | ENSG00000169554 | ZEB2 | chr2 | - | 144403915 | 144404130 | 144396411 | 144396592 | 144424795 | 144424864 | [illegible] |
| SE | 54428 | ENSG00000169554 | ZEB2 | chr2 | - | 144404835 | 144405024 | 144403915 | 144404130 | 144429768 | 144430026 | [illegible] |
| SE | 54429 | ENSG00000169554 | ZEB2 | chr2 | - | 144424795 | 144424864 | 144403915 | 144404130 | 144429768 | 144430026 | [illegible] |
| SE | 54433 | ENSG00000169554 | ZEB2 | chr2 | - | 144424795 | 144424953 | 144403915 | 144404130 | 144429768 | 144430026 | [illegible] |
| SE | 54435 | ENSG00000169554 | ZEB2 | chr2 | - | 144429768 | 144430026 | 144404835 | 144405024 | 144517277 | 144517419 | [illegible] |
| SE | 54436 | ENSG00000169554 | ZEB2 | chr2 | - | 144464165 | 144464342 | 144429981 | 144430026 | 144517277 | 144517419 | [illegible] |
| SE | 54437 | ENSG00000169554 | ZEB2 | chr2 | - | 144464165 | 144464342 | 144429981 | 144430026 | 144519938 | 144520398 | [illegible] |
| SE | 54438 | ENSG00000169554 | ZEB2 | chr2 | - | 144464165 | 144464342 | 144461832 | 144462337 | 144517277 | 144517419 | [illegible] |
| SE | 54439 | ENSG00000169554 | ZEB2 | chr2 | - | 144497707 | 144497764 | 144429768 | 144430026 | 144517277 | 144517419 | [illegible] |
| SE | 54440 | ENSG00000169554 | ZEB2 | chr2 | - | 144516457 | 144516553 | 144511388 | 144513797 | 144517277 | 144517419 | [illegible] |
| SE | 54442 | ENSG00000169554 | ZEB2 | chr2 | - | 144517277 | 144517419 | 144464165 | 144464342 | 144519938 | 144520006 | [illegible] |
| SE | 54443 | ENSG00000169554 | ZEB2 | chr2 | - | 144517277 | 144517419 | 144511384 | 144513797 | 144519938 | 144520006 | [illegible] |
| SE | 54445 | ENSG00000169554 | ZEB2 | chr2 | - | 144519938 | 144520208 | 144517277 | 144517419 | 144520351 | 144520381 | [illegible] |
| SE | 54447 | ENSG00000164305 | CASP3 | chr4 | - | 184635293 | 184635418 | 184632267 | 184632396 | 184638400 | 184638468 | [illegible] |
| SE | 54450 | ENSG00000164305 | CASP3 | chr4 | - | 184648502 | 184648631 | 184638400 | 184638468 | 184649394 | 184649447 | [illegible] |
| SE | 54458 | ENSG00000118242 | MREG | chr2 | - | 215996305 | 215996465 | 215947022 | 215947113 | 216013232 | 216013551 | [illegible] |
| SE | 54459 | ENSG00000112303 | VNN2 | chr6 | - | 132751144 | 132751278 | 132749694 | 132749865 | 132755842 | 132756035 | [illegible] |
| SE | 54460 | ENSG00000112303 | VNN2 | chr6 | - | 132751144 | 132751518 | 132749694 | 132749865 | 132755842 | 132756035 | [illegible] |
| SE | 54466 | ENSG00000112303 | VNN2 | chr6 | - | 132752672 | 132752749 | 132752460 | 132752577 | 132755842 | 132756035 | [illegible] |
| SE | 54467 | ENSG00000112303 | VNN2 | chr6 | - | 132753417 | 132753491 | 132751144 | 132751518 | 132755842 | 132756035 | [illegible] |
| SE | 54468 | ENSG00000112303 | VNN2 | chr6 | - | 132753417 | 132753491 | 132752713 | 132752749 | 132753875 | 132753949 | [illegible] |
| SE | 54471 | ENSG00000112303 | VNN2 | chr6 | - | 132753875 | 132753949 | 132752723 | 132752749 | 132755842 | 132756035 | [illegible] |
| SE | 54472 | ENSG00000112303 | VNN2 | chr6 | - | 132753875 | 132753949 | 132753417 | 132753491 | 132755842 | 132756035 | [illegible] |
| SE | 54474 | ENSG00000112303 | VNN2 | chr6 | - | 132757670 | 132757759 | 132757415 | 132757546 | 132763382 | 132763430 | [illegible] |
| SE | 54475 | ENSG00000112303 | VNN2 | chr6 | - | 132757670 | 132757902 | 132757415 | 132757546 | 132763382 | 132763459 | [illegible] |
| SE | 54477 | ENSG00000112303 | VNN2 | chr6 | - | 132760631 | 132760803 | 132757670 | 132757902 | 132763382 | 132763431 | [illegible] |
| SE | 54484 | ENSG00000272462 | NA | chr6 | + | 25993996 | 25994142 | 25992691 | 25992918 | 25997902 | 25998812 | [illegible] |
| SE | 54486 | ENSG00000164002 | EXO5 | chr1 | + | 40509450 | 40509625 | 40508740 | 40508877 | 40509732 | 40509790 | [illegible] |
| SE | 54487 | ENSG00000164002 | EXO5 | chr1 | + | 40509450 | 40509625 | 40508766 | 40508889 | 40509732 | 40509790 | [illegible] |
| SE | 54489 | ENSG00000164002 | EXO5 | chr1 | + | 40509457 | 40509625 | 40508760 | 40508908 | 40509732 | 40509790 | [illegible] |
| SE | 54492 | ENSG00000164002 | EXO5 | chr1 | + | 40509732 | 40509790 | 40508797 | 40508908 | 40514514 | 40516036 | [illegible] |
| SE | 54495 | ENSG00000151414 | NEK7 | chr1 | + | 198253039 | 198253180 | 198232552 | 198232637 | 198262574 | 198262623 | [illegible] |
| SE | 54497 | ENSG00000151414 | NEK7 | chr1 | + | 198278953 | 198279061 | 198264124 | 198264235 | 198292944 | 198293039 | [illegible] |
| SE | 54498 | ENSG00000151414 | NEK7 | chr1 | + | 198292944 | 198293039 | 198264124 | 198264235 | 198297126 | 198297240 | [illegible] |
| SE | 54499 | ENSG00000151414 | NEK7 | chr1 | + | 198292944 | 198293039 | 198264124 | 198264235 | 198319411 | 198319522 | [illegible] |
| SE | 54504 | ENSG00000092853 | CLSPN | chr1 | - | 35737338 | 35737421 | 35736913 | 35737075 | 35737991 | 35738097 | [illegible] |
| SE | 54506 | ENSG00000092853 | CLSPN | chr1 | - | 35762403 | 35762481 | 35761997 | 35762070 | 35763159 | 35763321 | [illegible] |
| SE | 54507 | ENSG00000092853 | CLSPN | chr1 | - | 35765217 | 35765326 | 35764265 | 35764714 | 35769846 | 35769928 | [illegible] |
| SE | 54508 | ENSG00000257379 | NA | chr12 | + | 53443390 | 53443773 | 53442694 | 53442733 | 53454725 | 53454869 | [illegible] |
| SE | 54509 | ENSG00000197111 | PCBP2 | chr12 | + | 53455346 | 53455370 | 53454383 | 53454869 | 53455460 | 53455477 | [illegible] |
| SE | 54510 | ENSG00000257379 | NA | chr12 | + | 53455460 | 53455493 | 53455346 | 53455370 | 53455884 | 53456001 | [illegible] |
| SE | 54511 | ENSG00000257379 | NA | chr12 | + | 53455884 | 53456001 | 53455460 | 53455493 | 53459271 | 53459403 | [illegible] |
| SE | 54513 | ENSG00000257379 | NA | chr12 | + | 53461014 | 53461143 | 53455884 | 53456001 | 53462492 | 53462567 | [illegible] |
| SE | 54520 | ENSG00000204220 | PFDN6 | chr6 | + | 33290325 | 33290450 | 33290173 | 33290244 | 33290715 | 33290862 | [illegible] |
| SE | 54521 | ENSG00000204220 | PFDN6 | chr6 | + | 33290325 | 33290450 | 33290173 | 33290244 | 33296828 | 33298401 | [illegible] |
| SE | 54525 | ENSG00000097096 | SYDE2 | chr1 | - | 85169043 | 85169225 | 85164525 | 85164757 | 85182097 | 85183200 | [illegible] |
| SE | 54526 | ENSG00000097096 | SYDE2 | chr1 | - | 85178145 | 85178272 | 85169043 | 85169225 | 85182097 | 85183200 | [illegible] |
| SE | 54531 | ENSG00000161929 | SCIMP | chr17 | - | 5223332 | 5223456 | 5221286 | 5221350 | 5234734 | 5234860 | [illegible] |
| SE | 54533 | ENSG00000161929 | SCIMP | chr17 | - | 5223353 | 5223456 | 5221286 | 5221350 | 5234734 | 5234823 | [illegible] |
| SE | 54534 | ENSG00000106028 | SSBP1 | chr7 | + | 141739123 | 141739190 | 141738357 | 141738410 | 141742168 | 141742229 | [illegible] |
| SE | 54535 | ENSG00000106028 | SSBP1 | chr7 | + | 141739123 | 141739190 | 141738357 | 141738503 | 141742168 | 141742229 | [illegible] |
| SE | 54537 | ENSG00000106028 | SSBP1 | chr7 | + | 141741748 | 141741798 | 141739123 | 141739190 | 141742168 | 141742229 | [illegible] |
| SE | 54545 | ENSG00000162441 | LZIC | chr1 | - | 9931890 | 9931972 | 9926292 | 9930457 | 9932802 | 9932898 | [illegible] |
| SE | 54547 | ENSG00000162441 | LZIC | chr1 | - | 9936518 | 9936627 | 9935491 | 9935627 | 9942623 | 9942768 | [illegible] |
| SE | 54548 | ENSG00000142089 | IFITM3 | chr11 | - | 320112 | 320248 | 319678 | 319990 | 320564 | 320841 | [illegible] |
| SE | 54552 | ENSG00000179912 | R3HDM2 | chr12 | - | 57283823 | 57284056 | 57280357 | 57280530 | 57289008 | 57289040 | [illegible] |
| SE | 54561 | ENSG00000179912 | R3HDM2 | chr12 | - | 57303175 | 57303217 | 57300094 | 57300181 | 57310263 | 57310451 | [illegible] |
| SE | 54563 | ENSG00000179912 | R3HDM2 | chr12 | - | 57310263 | 57310460 | 57303175 | 57303217 | 57395748 | 57395818 | [illegible] |
| SE | 54564 | ENSG00000179912 | R3HDM2 | chr12 | - | 57310263 | 57310463 | 57303175 | 57303217 | 57395748 | 57395818 | [illegible] |
| SE | 54569 | ENSG00000117305 | HMGCL | chr1 | - | 23804399 | 23804525 | 23801884 | 23802564 | 23808134 | 23808323 | [illegible] |
| SE | 54571 | ENSG00000117305 | HMGCL | chr1 | - | 23808134 | 23808215 | 23803587 | 23804525 | 23816674 | 23816770 | [illegible] |
| SE | 54572 | ENSG00000117305 | HMGCL | chr1 | - | 23808134 | 23808323 | 23804399 | 23804525 | 23816674 | 23816770 | [illegible] |
| SE | 54574 | ENSG00000117305 | HMGCL | chr1 | - | 23810735 | 23810799 | 23808134 | 23808323 | 23816674 | 23816770 | [illegible] |
| SE | 54575 | ENSG00000117305 | HMGCL | chr1 | - | 23814189 | 23814338 | 23804399 | 23804525 | 23816674 | 23816770 | [illegible] |
| SE | 54577 | ENSG00000117305 | HMGCL | chr1 | - | 23820509 | 23820593 | 23817475 | 23817583 | 23825355 | 23825402 | [illegible] |
| SE | 54579 | ENSG00000100092 | SH3BP1 | chr22 | + | 37642538 | 37642737 | 37641373 | 37641478 | 37642894 | 37643006 | [illegible] |
| SE | 54581 | ENSG00000100092 | SH3BP1 | chr22 | + | 37650085 | 37650249 | 37648318 | 37648435 | 37650541 | 37650725 | [illegible] |
| SE | 54582 | ENSG00000100092 | SH3BP1 | chr22 | + | 37655271 | 37655315 | 37653778 | 37653873 | 37655925 | 37656042 | [illegible] |
| SE | 54587 | ENSG00000164048 | ZNF589 | chr3 | + | 48256710 | 48257022 | 48247624 | 48247677 | 48260812 | 48260939 | [illegible] |
| SE | 54593 | ENSG00000133985 | TTC9 | chr14 | + | 70667563 | 70667746 | 70641915 | 70642535 | 70671075 | 70675366 | [illegible] |
| SE | 54594 | ENSG00000197520 | FAM177B | chr1 | + | 222742536 | 222742619 | 222737903 | 222738021 | 222746530 | 222746719 | [illegible] |
| SE | 54595 | ENSG00000197520 | FAM177B | chr1 | + | 222742536 | 222742619 | 222739683 | 222739832 | 222746530 | 222746719 | [illegible] |
| SE | 54597 | ENSG00000197520 | FAM177B | chr1 | + | 222748960 | 222749562 | 222747014 | 222747081 | 222749920 | 222750805 | [illegible] |
| SE | 54598 | ENSG00000197520 | FAM177B | chr1 | + | 222749464 | 222749562 | 222747014 | 222747081 | 222749920 | 222750196 | [illegible] |
| SE | 54600 | ENSG00000151422 | FER | chr5 | + | 108768092 | 108768238 | 108748961 | 108748993 | 108782662 | 108782755 | [illegible] |
| SE | 54602 | ENSG00000151422 | FER | chr5 | + | 108782662 | 108782755 | 108748966 | 108748993 | 108798123 | 108798389 | [illegible] |
| SE | 54606 | ENSG00000151422 | FER | chr5 | + | 108871364 | 108871502 | 108867766 | 108867950 | 108872092 | 108872212 | [illegible] |
| SE | 54607 | ENSG00000151422 | FER | chr5 | + | 108946129 | 108946222 | 108897647 | 108897848 | 108954728 | 108954932 | [illegible] |
| SE | 54608 | ENSG00000151422 | FER | chr5 | + | 108946129 | 108946222 | 108924605 | 108924789 | 108954728 | 108954932 | [illegible] |
| SE | 54609 | ENSG00000151422 | FER | chr5 | + | 108954728 | 108954932 | 108946129 | 108946222 | 108959224 | 108959347 | [illegible] |
| SE | 54612 | ENSG00000188897 | NA | chr16 | - | 11411245 | 11411327 | 11407199 | 11407337 | 11412840 | 11412953 | [illegible] |
| SE | 54613 | ENSG00000188897 | NA | chr16 | - | 11456858 | 11457039 | 11450608 | 11450820 | 11459895 | 11460054 | [illegible] |
| SE | 54617 | ENSG00000073578 | SDHA | chr5 | + | 225418 | 225562 | 224359 | 224521 | 225882 | 226047 | [illegible] |
| SE | 54618 | ENSG00000073578 | SDHA | chr5 | + | 225882 | 226047 | 225418 | 225562 | 228184 | 228333 | [illegible] |
| SE | 54621 | ENSG00000073578 | SDHA | chr5 | + | 253007 | 253080 | 251340 | 251468 | 254392 | 254506 | [illegible] |
| SE | 54622 | ENSG00000073578 | SDHA | chr5 | + | 254357 | 254506 | 251336 | 251468 | 256333 | 256682 | [illegible] |
| SE | 54623 | ENSG00000073578 | SDHA | chr5 | + | 254392 | 254506 | 251337 | 251468 | 256333 | 256648 | [illegible] |
| SE | 54625 | ENSG00000138802 | SEC24B | chr4 | + | 109462900 | 109463644 | 109433812 | 109434002 | 109473003 | 109473186 | [illegible] |
| SE | 54626 | ENSG00000138802 | SEC24B | chr4 | + | 109473003 | 109473186 | 109462900 | 109463644 | 109481676 | 109481781 | [illegible] |
| SE | 54629 | ENSG00000138802 | SEC24B | chr4 | + | 109481676 | 109481781 | 109462900 | 109463644 | 109494614 | 109494856 | [illegible] |
| SE | 54631 | ENSG00000138802 | SEC24B | chr4 | + | 109491326 | 109491407 | 109462900 | 109463644 | 109494614 | 109494856 | [illegible] |
| SE | 54632 | ENSG00000138802 | SEC24B | chr4 | + | 109491326 | 109491407 | 109481676 | 109481781 | 109494614 | 109494856 | [illegible] |
| SE | 54633 | ENSG00000138802 | SEC24B | chr4 | + | 109513746 | 109513856 | 109511956 | 109512083 | 109516527 | 109516640 | [illegible] |
| SE | 54636 | ENSG00000140992 | PDPK1 | chr16 | + | 2566355 | 2566453 | 2561479 | 2561522 | 2577424 | 2577500 | [illegible] |
| SE | 54637 | ENSG00000140992 | PDPK1 | chr16 | + | 2577424 | 2577500 | 2561479 | 2561522 | 2581294 | 2581363 | [illegible] |
| SE | 54638 | ENSG00000140992 | PDPK1 | chr16 | + | 2583411 | 2583585 | 2581606 | 2581703 | 2586675 | 2586893 | [illegible] |
| SE | 54639 | ENSG00000140992 | PDPK1 | chr16 | + | 2595792 | 2595850 | 2586675 | 2586893 | 2597122 | 2597275 | [illegible] |
| SE | 54642 | ENSG00000140992 | PDPK1 | chr16 | + | 2597122 | 2597275 | 2595792 | 2595850 | 2597650 | 2597928 | [illegible] |
| SE | 54644 | ENSG00000131791 | PRKAB2 | chr1 | - | 147166497 | 147166618 | 147162439 | 147162573 | 147166845 | 147166939 | [illegible] |
| SE | 54645 | ENSG00000131791 | PRKAB2 | chr1 | - | 147166845 | 147166939 | 147162439 | 147162573 | 147171988 | 147172167 | [illegible] |
| SE | 54646 | ENSG00000131791 | PRKAB2 | chr1 | - | 147166845 | 147166939 | 147166497 | 147166618 | 147167766 | 147167933 | [illegible] |
| SE | 54649 | ENSG00000151694 | ADAM17 | chr2 | - | 9491100 | 9491151 | 9488485 | 9490518 | 9492897 | 9492986 | [illegible] |
| SE | 54650 | ENSG00000151694 | ADAM17 | chr2 | - | 9492897 | 9492986 | 9491100 | 9491151 | 9493746 | 9493825 | [illegible] |
| SE | 54651 | ENSG00000151694 | ADAM17 | chr2 | - | 9494636 | 9494767 | 9493746 | 9493825 | 9497113 | 9497248 | [illegible] |
| SE | 54652 | ENSG00000151694 | ADAM17 | chr2 | - | 9502172 | 9502276 | 9497004 | 9497248 | 9505165 | 9505318 | [illegible] |
| SE | 54654 | ENSG00000151694 | ADAM17 | chr2 | - | 9509978 | 9510131 | 9505165 | 9505365 | 9517900 | 9517989 | [illegible] |
| SE | 54657 | ENSG00000151694 | ADAM17 | chr2 | - | 9523248 | 9523338 | 9521202 | 9521321 | 9526110 | 9526244 | [illegible] |
| SE | 54658 | ENSG00000151694 | ADAM17 | chr2 | - | 9535833 | 9535922 | 9527500 | 9527954 | 9536697 | 9536828 | [illegible] |
| SE | 54660 | ENSG00000151694 | ADAM17 | chr2 | - | 9543152 | 9543226 | 9527785 | 9527954 | 9555508 | 9555788 | [illegible] |
| SE | 54662 | ENSG00000151694 | ADAM17 | chr2 | - | 9543695 | 9543760 | 9543152 | 9543226 | 9555508 | 9555778 | [illegible] |
| SE | 54665 | ENSG00000163877 | SNIP1 | chr1 | - | 37548820 | 37548987 | 37540156 | 37540755 | 37554005 | 37554293 | [illegible] |
| SE | 54667 | ENSG00000163877 | SNIP1 | chr1 | - | 37552644 | 37552747 | 37548820 | 37548987 | 37554005 | 37554293 | [illegible] |
| SE | 54669 | ENSG00000070010 | UFD1 | chr22 | - | 19455679 | 19455768 | 19450620 | 19454830 | 19456586 | 19456634 | [illegible] |
| SE | 54670 | ENSG00000070010 | UFD1 | chr22 | - | 19458070 | 19458139 | 19456852 | 19456918 | 19465201 | 19465274 | [illegible] |
| SE | 54671 | ENSG00000070010 | UFD1 | chr22 | - | 19471686 | 19471808 | 19467402 | 19468003 | 19475067 | 19475100 | [illegible] |
| SE | 54672 | ENSG00000070010 | UFD1 | chr22 | - | 19471686 | 19471808 | 19467402 | 19468003 | 19475469 | 19475602 | [illegible] |
| SE | 54674 | ENSG00000070010 | UFD1 | chr22 | - | 19475067 | 19475100 | 19467402 | 19468003 | 19475469 | 19475602 | [illegible] |
| SE | 54675 | ENSG00000070010 | UFD1 | chr22 | - | 19475067 | 19475100 | 19467402 | 19468003 | 19479082 | 19479172 | [illegible] |
| SE | 54676 | ENSG00000070010 | UFD1 | chr22 | - | 19475067 | 19475100 | 19471686 | 19471808 | 19475469 | 19475602 | [illegible] |
| SE | 54694 | ENSG00000109667 | SLC2A9 | chr4 | - | 9826419 | 9826600 | 9779806 | 9780065 | 9834880 | 9835008 | [illegible] |
| SE | 54696 | ENSG00000109667 | SLC2A9 | chr4 | - | 9890609 | 9890711 | 9834880 | 9835008 | 9908234 | 9908345 | [illegible] |
| SE | 54697 | ENSG00000109667 | SLC2A9 | chr4 | - | 9920384 | 9920572 | 9908234 | 9908345 | 9980591 | 9980737 | [illegible] |
| SE | 54698 | ENSG00000109667 | SLC2A9 | chr4 | - | 9941912 | 9942045 | 9920384 | 9920572 | 9980591 | 9980737 | [illegible] |
| SE | 54700 | ENSG00000109667 | SLC2A9 | chr4 | - | 9985668 | 9985793 | 9980591 | 9980737 | 9996780 | 9996941 | [illegible] |
| SE | 54701 | ENSG00000109667 | SLC2A9 | chr4 | - | 9996780 | 9996941 | 9980591 | 9980737 | 10018974 | 10019073 | [illegible] |
| SE | 54703 | ENSG00000109667 | SLC2A9 | chr4 | - | 10018974 | 10019073 | 9996780 | 9996941 | 10021279 | 10021497 | [illegible] |
| SE | 54707 | ENSG00000137965 | IFI44 | chr1 | + | 78654242 | 78654279 | 78650185 | 78650652 | 78655013 | 78655209 | [illegible] |
| SE | 54709 | ENSG00000137965 | IFI44 | chr1 | + | 78660553 | 78660654 | 78659311 | 78659483 | 78663764 | 78664073 | [illegible] |
| SE | 54713 | ENSG00000137965 | IFI44 | chr1 | + | 78662703 | 78662878 | 78660553 | 78660654 | 78663764 | 78663897 | [illegible] |
| SE | 54714 | ENSG00000137965 | IFI44 | chr1 | + | 78662703 | 78662878 | 78660553 | 78660691 | 78663764 | 78663897 | [illegible] |
| SE | 54716 | ENSG00000177409 | SAMD9L | chr7 | - | 93144731 | 93144833 | 93130055 | 93135991 | 93145908 | 93146040 | [illegible] |
| SE | 54717 | ENSG00000177409 | SAMD9L | chr7 | - | 93144731 | 93144833 | 93130055 | 93135991 | 93146882 | 93147146 | [illegible] |
| SE | 54719 | ENSG00000177409 | SAMD9L | chr7 | - | 93145384 | 93145532 | 93130762 | 93135991 | 93145908 | 93146040 | [illegible] |
| SE | 54720 | ENSG00000177409 | SAMD9L | chr7 | - | 93145384 | 93145532 | 93130762 | 93135991 | 93146882 | 93147146 | [illegible] |
| SE | 54721 | ENSG00000177409 | SAMD9L | chr7 | - | 93145384 | 93145532 | 93130762 | 93135991 | 93147087 | 93147146 | [illegible] |
| SE | 54722 | ENSG00000177409 | SAMD9L | chr7 | - | 93145384 | 93145532 | 93130762 | 93135991 | 93148193 | 93148345 | [illegible] |
| SE | 54723 | ENSG00000177409 | SAMD9L | chr7 | - | 93145384 | 93145532 | 93144731 | 93144833 | 93145908 | 93146040 | [illegible] |
| SE | 54724 | ENSG00000177409 | SAMD9L | chr7 | - | 93145384 | 93145532 | 93144731 | 93144833 | 93146882 | 93147146 | [illegible] |
| SE | 54726 | ENSG00000177409 | SAMD9L | chr7 | - | 93145384 | 93146040 | 93130055 | 93135991 | 93146882 | 93147146 | [illegible] |
| SE | 54727 | ENSG00000177409 | SAMD9L | chr7 | - | 93145384 | 93146040 | 93130055 | 93135991 | 93147087 | 93147146 | [illegible] |
| SE | 54728 | ENSG00000177409 | SAMD9L | chr7 | - | 93145384 | 93146040 | 93144731 | 93144833 | 93146882 | 93147146 | [illegible] |
| SE | 54729 | ENSG00000177409 | SAMD9L | chr7 | - | 93145908 | 93146040 | 93130762 | 93135991 | 93146882 | 93147146 | [illegible] |
| SE | 54730 | ENSG00000177409 | SAMD9L | chr7 | - | 93145908 | 93146040 | 93130762 | 93135991 | 93147087 | 93147146 | [illegible] |
| SE | 54731 | ENSG00000177409 | SAMD9L | chr7 | - | 93145908 | 93146040 | 93130762 | 93135991 | 93148193 | 93148345 | [illegible] |
| SE | 54732 | ENSG00000177409 | SAMD9L | chr7 | - | 93145908 | 93146040 | 93144731 | 93144833 | 93146882 | 93147146 | [illegible] |
| SE | 54733 | ENSG00000177409 | SAMD9L | chr7 | - | 93145908 | 93146040 | 93144731 | 93144833 | 93148193 | 93148345 | [illegible] |
| SE | 54734 | ENSG00000177409 | SAMD9L | chr7 | - | 93145908 | 93146040 | 93145384 | 93145532 | 93146882 | 93147146 | [illegible] |
| SE | 54735 | ENSG00000177409 | SAMD9L | chr7 | - | 93145908 | 93146040 | 93145384 | 93145532 | 93147087 | 93147146 | [illegible] |
| SE | 54736 | ENSG00000177409 | SAMD9L | chr7 | - | 93145908 | 93146040 | 93145384 | 93145532 | 93148193 | 93148345 | [illegible] |
| SE | 54737 | ENSG00000177409 | SAMD9L | chr7 | - | 93146882 | 93147146 | 93130055 | 93135991 | 93148193 | 93148345 | [illegible] |
| SE | 54738 | ENSG00000177409 | SAMD9L | chr7 | - | 93146882 | 93147146 | 93144731 | 93144833 | 93148193 | 93148345 | [illegible] |
| SE | 54739 | ENSG00000177409 | SAMD9L | chr7 | - | 93146882 | 93147146 | 93145384 | 93145532 | 93148193 | 93148345 | [illegible] |
| SE | 54740 | ENSG00000177409 | SAMD9L | chr7 | - | 93146882 | 93147146 | 93145384 | 93146040 | 93148193 | 93148345 | [illegible] |
| SE | 54741 | ENSG00000177409 | SAMD9L | chr7 | - | 93147087 | 93147146 | 93130762 | 93135991 | 93148193 | 93148345 | [illegible] |
| SE | 54742 | ENSG00000177409 | SAMD9L | chr7 | - | 93147087 | 93147146 | 93145384 | 93145532 | 93148193 | 93148345 | [illegible] |
| SE | 54744 | ENSG00000083720 | OXCT1 | chr5 | - | 41749526 | 41749607 | 41739389 | 41739491 | 41762110 | 41762200 | [illegible] |
| SE | 54745 | ENSG00000083720 | OXCT1 | chr5 | - | 41749526 | 41749607 | 41739389 | 41739491 | 41794002 | 41794078 | [illegible] |
| SE | 54746 | ENSG00000083720 | OXCT1 | chr5 | - | 41762110 | 41762200 | 41739389 | 41739491 | 41794002 | 41794078 | [illegible] |
| SE | 54747 | ENSG00000083720 | OXCT1 | chr5 | - | 41762110 | 41762200 | 41749526 | 41749607 | 41794002 | 41794078 | [illegible] |
| SE | 54750 | ENSG00000083720 | OXCT1 | chr5 | - | 41803068 | 41803163 | 41801021 | 41801070 | 41840450 | 41840511 | [illegible] |
| SE | 54751 | ENSG00000083720 | OXCT1 | chr5 | - | 41805566 | 41805681 | 41803068 | 41803163 | 41807330 | 41807438 | [illegible] |
| SE | 54753 | ENSG00000083720 | OXCT1 | chr5 | - | 41807330 | 41807438 | 41801021 | 41801070 | 41840450 | 41840511 | [illegible] |
| SE | 54755 | ENSG00000083720 | OXCT1 | chr5 | - | 41807330 | 41807438 | 41805566 | 41805681 | 41840450 | 41840511 | [illegible] |
| SE | 54756 | ENSG00000148459 | PDSS1 | chr10 | + | 26704676 | 26704741 | 26702161 | 26702194 | 26705285 | 26705394 | [illegible] |
| SE | 54757 | ENSG00000148459 | PDSS1 | chr10 | + | 26705285 | 26705394 | 26704676 | 26704741 | 26709637 | 26709768 | [illegible] |
| SE | 54758 | ENSG00000148459 | PDSS1 | chr10 | + | 26709637 | 26709768 | 26705285 | 26705394 | 26720134 | 26720359 | [illegible] |
| SE | 54759 | ENSG00000148459 | PDSS1 | chr10 | + | 26709637 | 26709768 | 26705285 | 26705394 | 26720217 | 26720359 | [illegible] |
| SE | 54760 | ENSG00000148459 | PDSS1 | chr10 | + | 26740644 | 26740817 | 26735465 | 26735579 | 26742496 | 26742577 | [illegible] |
| SE | 54761 | ENSG00000138642 | HERC6 | chr4 | + | 88383184 | 88383380 | 88379063 | 88379120 | 88385498 | 88385575 | [illegible] |
| SE | 54762 | ENSG00000138642 | HERC6 | chr4 | + | 88383220 | 88383380 | 88378738 | 88379120 | 88385498 | 88385575 | [illegible] |
| SE | 54763 | ENSG00000138642 | HERC6 | chr4 | + | 88383231 | 88383380 | 88378910 | 88379120 | 88385498 | 88385575 | [illegible] |
| SE | 54765 | ENSG00000138642 | HERC6 | chr4 | + | 88396850 | 88396987 | 88396014 | 88396142 | 88398141 | 88398209 | [illegible] |
| SE | 54766 | ENSG00000138642 | HERC6 | chr4 | + | 88404875 | 88404997 | 88398141 | 88398209 | 88405553 | 88405613 | [illegible] |
| SE | 54767 | ENSG00000138642 | HERC6 | chr4 | + | 88413076 | 88413266 | 88408523 | 88408617 | 88423859 | 88423973 | [illegible] |
| SE | 54769 | ENSG00000138642 | HERC6 | chr4 | + | 88423859 | 88423973 | 88417424 | 88417579 | 88428579 | 88428750 | [illegible] |
| SE | 54770 | ENSG00000138642 | HERC6 | chr4 | + | 88424594 | 88424702 | 88423859 | 88423973 | 88428579 | 88428750 | [illegible] |
| SE | 54771 | ENSG00000138642 | HERC6 | chr4 | + | 88428579 | 88428750 | 88424594 | 88424702 | 88431161 | 88431305 | [illegible] |
| SE | 54772 | ENSG00000138642 | HERC6 | chr4 | + | 88431161 | 88431305 | 88428579 | 88428750 | 88435724 | 88435891 | [illegible] |
| SE | 54773 | ENSG00000138642 | HERC6 | chr4 | + | 88437710 | 88437781 | 88436904 | 88436971 | 88439873 | 88440057 | [illegible] |
| SE | 54774 | ENSG00000138642 | HERC6 | chr4 | + | 88440147 | 88440250 | 88439873 | 88440057 | 88442233 | 88443096 | [illegible] |
| SE | 54775 | ENSG00000153904 | DDAH1 | chr1 | - | 85496165 | 85496281 | 85358747 | 85358847 | 85577983 | 85578363 | [illegible] |

| | | | | | | | | | | | | | | | | | | | | | | | | | | | | | | | | | | | | | | | | | | | | | | |
|---|---|---|---|---|---|---|---|---|---|---|---|---|---|---|---|---|---|---|---|---|---|---|---|---|---|---|---|---|---|---|---|---|---|---|---|---|---|---|---|---|---|---|---|---|---|---|
| SE | 54776 | ENSG00000280670 | CCDC163 | chr1 | - | 45495417 | 45495473 | 45494391 | 45495166 | 45496555 | 45496623 | 1 | 0 | 2 | 0 | 0 | 2 | 0 | 2 | 3 | 0 | 0 | 3 | 0 | 0 | 0 | 0 | 0 | 0 | 0 | 0 | 205 | 149 | 1 | 1 | 0.267 | NA | 0.421 | 0 | NA | NA | 1 | NA | NA | NA | -0.771 |
| SE | 54777 | ENSG00000280670 | CCDC163 | chr1 | - | 45495417 | 45495545 | 45493865 | 45495166 | 45496555 | 45496623 | 5 | 1 | 3 | 5 | 1 | 2 | 0 | 2 | 3 | 0 | 0 | 3 | 6 | 0 | 6 | 0 | 0 | 0 | 0 | 0 | 277 | 149 | 1 | 1 | 0.574 | 1 | 0.447 | 0.473 | 1 | NA | 1 | 1 | NA | 1 | -0.301 |
| SE | 54778 | ENSG00000280670 | CCDC163 | chr1 | - | 45496555 | 45496623 | 45495417 | 45495545 | 45497298 | 45497383 | 2 | 1 | 2 | 5 | 0 | 0 | 0 | 0 | 0 | 1 | 0 | 3 | 4 | 0 | 6 | 0 | 0 | 0 | 0 | 0 | 217 | 149 | 1 | 1 | 1 | 1 | 1 | 1 | 0 | NA | 1 | 1 | NA | 1 | -0.2 |
| SE | 54779 | ENSG00000280670 | CCDC163 | chr1 | - | 45497298 | 45497383 | 45496210 | 45496623 | 45499344 | 45499440 | 4 | 3 | 7 | 8 | 0 | 1 | 0 | 0 | 2 | 0 | 0 | 3 | 0 | 0 | 0 | 0 | 0 | 0 | 0 | 0 | 234 | 149 | 1 | 1 | 0.718 | 1 | 1 | 0.718 | NA | NA | 1 | NA | NA | NA | -0.141 |
| SE | 54780 | ENSG00000280670 | CCDC163 | chr1 | - | 45497298 | 45497383 | 45496210 | 45496623 | 45499531 | 45499853 | 3 | 1 | 3 | 6 | 0 | 0 | 0 | 0 | 0 | 0 | 0 | 0 | 2 | 0 | 0 | 0 | 0 | 2 | 0 | 0 | 234 | 149 | 1 | 1 | 1 | 1 | 1 | 1 | NA | NA | NA | 0.389 | NA | NA | 0.611 |
| SE | 54783 | ENSG00000280670 | CCDC163 | chr1 | - | 45499344 | 45499440 | 45497298 | 45497383 | 45499531 | 45500053 | 1 | 0 | 3 | 0 | 1 | 0 | 0 | 0 | 0 | 0 | 0 | 3 | 0 | 0 | 0 | 0 | 0 | 2 | 0 | 0 | 245 | 149 | 1 | 1 | 1 | NA | 1 | NA | 1 | NA | 1 | 0 | NA | NA | 0.5 |
| SE | 54784 | ENSG00000280670 | CCDC163 | chr1 | - | 45499344 | 45499443 | 45496555 | 45496623 | 45499531 | 45499853 | 4 | 0 | 0 | 4 | 0 | 0 | 0 | 0 | 0 | 0 | 0 | 0 | 0 | 1 | 6 | 0 | 0 | 2 | 0 | 0 | 248 | 149 | 1 | 1 | 1 | NA | NA | 1 | NA | NA | NA | 0 | 1 | 1 | 0.333 |
| SE | 54785 | ENSG00000280670 | CCDC163 | chr1 | - | 45499344 | 45499443 | 45497298 | 45497383 | 45499531 | 45499853 | 4 | 2 | 4 | 4 | 1 | 0 | 0 | 0 | 0 | 0 | 0 | 3 | 0 | 1 | 6 | 0 | 0 | 2 | 0 | 0 | 248 | 149 | 1 | 1 | 1 | 1 | 1 | 1 | 1 | NA | 1 | 0 | 1 | 1 | 0.25 |
| SE | 54815 | ENSG00000204138 | PHACTR4 | chr1 | + | 28432930 | 28433046 | 28407409 | 28407463 | 28457831 | 28457859 | 1 | 0 | 2 | 1 | 0 | 5 | 2 | 12 | 7 | 14 | 0 | 0 | 0 | 0 | 6 | 4 | 6 | 4 | 4 | 13 | 265 | 149 | 1 | 1 | 0.101 | 0 | 0.086 | 0.074 | 0 | 0 | 0 | 0 | 0 | 0.206 | 0.011 |
| SE | 54820 | ENSG00000204138 | PHACTR4 | chr1 | + | 28457831 | 28457859 | 28432930 | 28433046 | 28459084 | 28459239 | 0 | 0 | 0 | 0 | 0 | 3 | 0 | 3 | 2 | 3 | 0 | 0 | 2 | 0 | 6 | 0 | 0 | 0 | 0 | 0 | 177 | 149 | 1 | 1 | 0 | NA | 0 | 0 | 0 | NA | NA | 1 | NA | 1 | -1 |
| SE | 54821 | ENSG00000204138 | PHACTR4 | chr1 | + | 28460211 | 28460292 | 28459084 | 28459258 | 28465684 | 28465849 | 73 | 52 | 47 | 67 | 34 | 0 | 3 | 0 | 2 | 0 | 73 | 50 | 64 | 91 | 144 | 0 | 0 | 0 | 0 | 0 | 230 | 149 | 1 | 1 | 1 | 0.918 | 1 | 0.956 | 1 | 1 | 1 | 1 | 1 | 1 | -0.025 |
| SE | 54823 | ENSG00000204138 | PHACTR4 | chr1 | + | 28476106 | 28476291 | 28473553 | 28474151 | 28480450 | 28480604 | 238 | 189 | 211 | 141 | 241 | 0 | 0 | 0 | 1 | 0 | 427 | 341 | 201 | 359 | 364 | 0 | 0 | 0 | 0 | 0 | 298 | 149 | 1 | 1 | 1 | 1 | 1 | 0.986 | 1 | 1 | 1 | 1 | 1 | 1 | -0.003 |
| SE | 54824 | ENSG00000204138 | PHACTR4 | chr1 | + | 28480450 | 28480604 | 28476106 | 28476291 | 28489169 | 28489225 | 164 | 152 | 174 | 110 | 280 | 0 | 0 | 0 | 0 | 0 | 433 | 256 | 173 | 321 | 239 | 0 | 0 | 0 | 1 | 0 | 298 | 149 | 1 | 1 | 1 | 1 | 1 | 1 | 1 | 1 | 1 | 1 | 0.994 | 1 | 0.001 |
| SE | 54826 | ENSG00000204138 | PHACTR4 | chr1 | + | 28490950 | 28491012 | 28489169 | 28489225 | 28496533 | 28497156 | 35 | 34 | 45 | 27 | 47 | 0 | 1 | 0 | 0 | 3 | 144 | 61 | 42 | 85 | 67 | 0 | 0 | 2 | 0 | 0 | 211 | 149 | 1 | 1 | 1 | 0.96 | 1 | 1 | 0.917 | 1 | 1 | 0.937 | 1 | 1 | -0.012 |
| SE | 54827 | ENSG00000204138 | PHACTR4 | chr1 | + | 28494229 | 28494322 | 28493014 | 28493091 | 28496533 | 28497156 | 3 | 0 | 1 | 1 | 4 | 97 | 38 | 68 | 65 | 135 | 0 | 0 | 1 | 0 | 4 | 122 | 60 | 31 | 63 | 86 | 242 | 149 | 1 | 1 | 0.019 | 0 | 0.009 | 0.009 | 0.018 | 0 | 0 | 0.019 | 0 | 0.028 | 0.002 |
| SE | 54828 | ENSG00000097021 | ACOT7 | chr1 | - | 6281101 | 6281286 | 6264268 | 6264695 | 6294863 | 6294980 | 58 | 16 | 21 | 31 | 22 | 0 | 0 | 1 | 1 | 0 | 101 | 59 | 47 | 35 | 281 | 0 | 0 | 0 | 0 | 0 | 298 | 149 | 1 | 1 | 1 | 1 | 0.913 | 0.939 | 1 | 1 | 1 | 1 | 1 | 1 | -0.03 |
| SE | 54831 | ENSG00000259030 | FPGT-TNNI3K | chr1 | + | 74204390 | 74204456 | 74201317 | 74201410 | 74236101 | 74236210 | 22 | 8 | 12 | 9 | 23 | 0 | 0 | 5 | 0 | 7 | 0 | 3 | 5 | 2 | 0 | 0 | 14 | 0 | 0 | 0 | 215 | 149 | 1 | 1 | 1 | 1 | 0.625 | 1 | 0.695 | NA | 0.129 | 1 | 1 | NA | 0.154 |
| SE | 54833 | ENSG00000259030 | FPGT-TNNI3K | chr1 | + | 74260693 | 74260754 | 74250671 | 74250769 | 74271597 | 74271708 | 0 | 3 | 0 | 0 | 0 | 2 | 3 | 1 | 3 | 2 | 0 | 7 | 0 | 0 | 0 | 0 | 1 | 0 | 0 | 0 | 210 | 149 | 1 | 1 | 0 | 0.415 | 0 | 0 | 0 | NA | 0.832 | NA | NA | NA | -0.749 |
| SE | 54843 | ENSG00000072135 | PTPN18 | chr2 | + | 130359607 | 130359646 | 130359396 | 130359492 | 130369132 | 130369201 | 31 | 11 | 24 | 35 | 25 | 0 | 0 | 0 | 2 | 1 | 31 | 42 | 23 | 48 | 84 | 0 | 0 | 0 | 8 | 0 | 188 | 149 | 1 | 1 | 1 | 1 | 1 | 0.933 | 0.952 | 1 | 1 | 1 | 0.826 | 1 | 0.012 |
| SE | 54844 | ENSG00000072135 | PTPN18 | chr2 | + | 130370704 | 130370782 | 130370309 | 130370623 | 130370874 | 130370964 | 84 | 58 | 57 | 83 | 55 | 1 | 0 | 3 | 1 | 0 | 192 | 135 | 54 | 135 | 287 | 0 | 0 | 0 | 0 | 0 | 227 | 149 | 1 | 1 | 0.982 | 1 | 0.926 | 0.982 | 1 | 1 | 1 | 1 | 1 | 1 | -0.022 |
| SE | 54845 | ENSG00000072135 | PTPN18 | chr2 | + | 130372872 | 130372947 | 130371997 | 130372483 | 130373156 | 130373220 | 336 | 162 | 166 | 352 | 115 | 3 | 5 | 1 | 2 | 0 | 1140 | 570 | 408 | 605 | 1483 | 2 | 7 | 2 | 0 | 6 | 224 | 149 | 1 | 1 | 0.987 | 0.956 | 0.991 | 0.992 | 1 | 0.997 | 0.982 | 0.993 | 1 | 0.994 | -0.008 |
| SE | 54852 | ENSG00000133056 | PIK3C2B | chr1 | - | 204433315 | 204433425 | 204432199 | 204432401 | 204434438 | 204434608 | 26 | 19 | 31 | 41 | 43 | 0 | 2 | 0 | 0 | 0 | 87 | 29 | 27 | 27 | 85 | 0 | 0 | 0 | 0 | 0 | 259 | 149 | 1 | 1 | 1 | 0.845 | 1 | 1 | 1 | 1 | 1 | 1 | 1 | 1 | -0.031 |
| SE | 54853 | ENSG00000133056 | PIK3C2B | chr1 | - | 204433315 | 204433949 | 204432199 | 204432401 | 204434438 | 204434608 | 46 | 24 | 63 | 64 | 88 | 0 | 2 | 0 | 0 | 0 | 104 | 60 | 40 | 27 | 118 | 0 | 0 | 0 | 0 | 0 | 298 | 149 | 1 | 1 | 1 | 0.857 | 1 | 1 | 1 | 1 | 1 | 1 | 1 | 1 | -0.029 |
| SE | 54854 | ENSG00000133056 | PIK3C2B | chr1 | - | 204433792 | 204433949 | 204433315 | 204433425 | 204434438 | 204434608 | 40 | 22 | 57 | 52 | 63 | 1 | 1 | 0 | 2 | 0 | 54 | 68 | 29 | 17 | 66 | 0 | 3 | 5 | 5 | 0 | 298 | 149 | 1 | 1 | 0.952 | 0.917 | 1 | 0.929 | 1 | 1 | 0.919 | 0.744 | 0.63 | 1 | 0.101 |
| SE | 54855 | ENSG00000133056 | PIK3C2B | chr1 | - | 204449184 | 204449296 | 204447435 | 204447578 | 204454668 | 204454791 | 12 | 8 | 13 | 17 | 7 | 0 | 0 | 0 | 0 | 0 | 27 | 10 | 7 | 26 | 20 | 0 | 2 | 0 | 0 | 0 | 261 | 149 | 1 | 1 | 1 | 1 | 1 | 1 | 1 | 1 | 0.741 | 1 | 1 | 1 | 0.052 |
| SE | 54856 | ENSG00000133056 | PIK3C2B | chr1 | - | 204449849 | 204450017 | 204449184 | 204449296 | 204454668 | 204454791 | 20 | 29 | 33 | 38 | 19 | 0 | 0 | 0 | 0 | 0 | 3 | 33 | 18 | 59 | 39 | 0 | 1 | 0 | 0 | 0 | 298 | 149 | 1 | 1 | 1 | 1 | 1 | 1 | 1 | 1 | 0.943 | 1 | 1 | 1 | 0.011 |
| SE | 54869 | ENSG00000141456 | PELP1 | chr17 | - | 4676035 | 4676162 | 4675273 | 4675884 | 4676356 | 4676507 | 17 | 9 | 23 | 17 | 8 | 0 | 2 | 0 | 2 | 0 | 2 | 30 | 5 | 31 | 36 | 0 | 0 | 0 | 0 | 0 | 276 | 149 | 1 | 1 | 1 | 0.708 | 1 | 0.821 | 1 | 1 | 1 | 1 | 1 | 1 | -0.094 |
| SE | 54873 | ENSG00000083123 | BCKDHB | chr6 | + | 80171281 | 80171390 | 80168874 | 80169030 | 80200933 | 80201031 | 95 | 68 | 117 | 112 | 129 | 0 | 1 | 0 | 0 | 0 | 74 | 35 | 34 | 65 | 41 | 0 | 0 | 3 | 0 | 0 | 258 | 149 | 1 | 1 | 1 | 0.975 | 1 | 1 | 1 | 1 | 1 | 0.867 | 1 | 1 | 0.022 |
| SE | 54877 | ENSG00000158825 | CDA | chr1 | + | 20604927 | 20605039 | 20589085 | 20589283 | 20613841 | 20613899 | 69 | 41 | 54 | 31 | 70 | 0 | 0 | 1 | 0 | 0 | 295 | 181 | 75 | 267 | 154 | 2 | 0 | 4 | 0 | 0 | 261 | 149 | 1 | 1 | 1 | 1 | 0.969 | 1 | 1 | 0.988 | 1 | 0.915 | 1 | 1 | 0.013 |
| SE | 54881 | ENSG00000189362 | NEMP2 | chr2 | - | 190516269 | 190516384 | 190514451 | 190514678 | 190517519 | 190517613 | 74 | 40 | 47 | 77 | 82 | 0 | 2 | 3 | 0 | 0 | 47 | 37 | 12 | 5 | 21 | 0 | 4 | 0 | 0 | 0 | 264 | 149 | 1 | 1 | 1 | 0.919 | 0.898 | 1 | 1 | 1 | 0.839 | 1 | 1 | 1 | -0.004 |
| SE | 54882 | ENSG00000189362 | NEMP2 | chr2 | - | 190516269 | 190516384 | 190514451 | 190514678 | 190518735 | 190518808 | 52 | 24 | 36 | 50 | 45 | 1 | 0 | 0 | 0 | 0 | 27 | 26 | 11 | 2 | 13 | 0 | 2 | 0 | 0 | 0 | 264 | 149 | 1 | 1 | 0.967 | 1 | 1 | 1 | 1 | 1 | 0.88 | 1 | 1 | 1 | 0.017 |
| SE | 54883 | ENSG00000189362 | NEMP2 | chr2 | - | 190517519 | 190517613 | 190514452 | 190514678 | 190518735 | 190518808 | 28 | 25 | 21 | 26 | 39 | 1 | 0 | 0 | 0 | 0 | 5 | 19 | 2 | 26 | 0 | 0 | 2 | 0 | 0 | 0 | 243 | 149 | 1 | 1 | 0.945 | 1 | 1 | 1 | 1 | 1 | 0.853 | 1 | 1 | NA | 0.026 |
| SE | 54884 | ENSG00000189362 | NEMP2 | chr2 | - | 190517519 | 190517613 | 190514452 | 190514678 | 190518951 | 190519022 | 0 | 2 | 3 | 2 | 0 | 0 | 0 | 0 | 0 | 0 | 0 | 4 | 0 | 0 | 0 | 0 | 0 | 0 | 3 | 0 | 243 | 149 | 1 | 1 | NA | 1 | 1 | 1 | NA | NA | 1 | NA | 0 | NA | 0.5 |
| SE | 54885 | ENSG00000189362 | NEMP2 | chr2 | - | 190517519 | 190517613 | 190516269 | 190516384 | 190518735 | 190518808 | 53 | 39 | 29 | 53 | 76 | 3 | 0 | 0 | 0 | 0 | 25 | 26 | 3 | 29 | 8 | 0 | 0 | 0 | 0 | 0 | 243 | 149 | 1 | 1 | 0.915 | 1 | 1 | 1 | 1 | 1 | 1 | 1 | 1 | 1 | -0.017 |
| SE | 54886 | ENSG00000189362 | NEMP2 | chr2 | - | 190518735 | 190518808 | 190514451 | 190514678 | 190518951 | 190519022 | 10 | 6 | 6 | 13 | 9 | 0 | 0 | 0 | 0 | 0 | 8 | 10 | 2 | 8 | 0 | 0 | 0 | 0 | 3 | 0 | 222 | 149 | 1 | 1 | 1 | 1 | 1 | 1 | 1 | 1 | 1 | 1 | 0.642 | NA | 0.09 |
| SE | 54887 | ENSG00000189362 | NEMP2 | chr2 | - | 190518735 | 190518808 | 190517519 | 190517613 | 190518951 | 190519022 | 41 | 35 | 25 | 42 | 56 | 0 | 0 | 0 | 2 | 0 | 13 | 23 | 4 | 44 | 3 | 0 | 0 | 0 | 0 | 0 | 222 | 149 | 1 | 1 | 1 | 1 | 1 | 0.934 | 1 | 1 | 1 | 1 | 1 | 1 | -0.013 |
| SE | 54889 | ENSG00000189362 | NEMP2 | chr2 | - | 190525262 | 190525378 | 190518951 | 190519183 | 190534558 | 190534678 | 18 | 12 | 31 | 20 | 39 | 2 | 0 | 0 | 0 | 0 | 0 | 3 | 11 | 21 | 0 | 0 | 0 | 0 | 0 | 0 | 265 | 149 | 1 | 1 | 0.835 | 1 | 1 | 1 | 1 | NA | 1 | 1 | 1 | NA | -0.033 |
| SE | 54892 | ENSG00000106261 | ZKSCAN1 | chr7 | + | 100030248 | 100030375 | 100029860 | 100029952 | 100033304 | 100034271 | 256 | 262 | 270 | 255 | 234 | 0 | 0 | 0 | 1 | 0 | 190 | 307 | 193 | 259 | 97 | 0 | 0 | 2 | 0 | 0 | 276 | 149 | 1 | 1 | 1 | 1 | 1 | 0.993 | 1 | 1 | 1 | 0.981 | 1 | 1 | 0.002 |
| SE | 54895 | ENSG00000156983 | BRPF1 | chr3 | + | 9733172 | 9733370 | 9731787 | 9732138 | 9734130 | 9734739 | 0 | 0 | 0 | 1 | 0 | 10 | 7 | 22 | 11 | 10 | 0 | 0 | 0 | 3 | 0 | 4 | 22 | 10 | 29 | 38 | 298 | 149 | 1 | 1 | 0 | 0 | 0 | 0.043 | 0 | 0 | 0 | 0 | 0.049 | 0 | -0.001 |
| SE | 54900 | ENSG00000156983 | BRPF1 | chr3 | + | 9745572 | 9745709 | 9745007 | 9745155 | 9746299 | 9746454 | 59 | 34 | 51 | 55 | 72 | 0 | 0 | 0 | 1 | 4 | 99 | 53 | 52 | 113 | 182 | 0 | 0 | 0 | 0 | 0 | 286 | 149 | 1 | 1 | 1 | 1 | 1 | 0.966 | 0.904 | 1 | 1 | 1 | 1 | 1 | -0.026 |
| SE | 54901 | ENSG00000156983 | BRPF1 | chr3 | + | 9745572 | 9745930 | 9745007 | 9745155 | 9746299 | 9746454 | 111 | 60 | 112 | 127 | 148 | 0 | 0 | 0 | 1 | 4 | 169 | 118 | 96 | 197 | 277 | 0 | 0 | 0 | 0 | 0 | 298 | 149 | 1 | 1 | 1 | 1 | 1 | 0.984 | 0.949 | 1 | 1 | 1 | 1 | 1 | -0.013 |
| SE | 54904 | ENSG00000010803 | SCMH1 | chr1 | - | 41033982 | 41034048 | 41028583 | 41028726 | 41037361 | 41037452 | 11 | 14 | 20 | 14 | 11 | 1 | 0 | 0 | 0 | 0 | 0 | 7 | 10 | 13 | 22 | 0 | 0 | 0 | 6 | 0 | 215 | 149 | 1 | 1 | 0.884 | 1 | 1 | 1 | 1 | NA | 1 | 1 | 0.6 | 1 | 0.077 |
| SE | 54906 | ENSG00000010803 | SCMH1 | chr1 | - | 41070594 | 41070721 | 41048689 | 41048890 | 41075218 | 41075451 | 51 | 31 | 53 | 73 | 96 | 0 | 0 | 0 | 0 | 0 | 46 | 88 | 41 | 77 | 96 | 0 | 0 | 0 | 3 | 0 | 276 | 149 | 1 | 1 | 1 | 1 | 1 | 1 | 1 | 1 | 1 | 1 | 0.933 | 1 | 0.013 |
| SE | 54907 | ENSG00000010803 | SCMH1 | chr1 | - | 41070594 | 41070721 | 41048689 | 41048890 | 41113282 | 41113526 | 31 | 13 | 27 | 24 | 23 | 2 | 0 | 0 | 0 | 0 | 23 | 35 | 22 | 34 | 53 | 0 | 0 | 0 | 0 | 0 | 276 | 149 | 1 | 1 | 0.893 | 1 | 1 | 1 | 1 | 1 | 1 | 1 | 1 | 1 | -0.021 |
| SE | 54908 | ENSG00000010803 | SCMH1 | chr1 | - | 41075218 | 41075451 | 41048689 | 41048890 | 41113282 | 41113526 | 12 | 10 | 11 | 9 | 19 | 2 | 0 | 0 | 0 | 0 | 0 | 11 | 12 | 12 | 6 | 0 | 0 | 0 | 0 | 0 | 298 | 149 | 1 | 1 | 0.75 | 1 | 1 | 1 | 1 | NA | 1 | 1 | 1 | 1 | -0.05 |
| SE | 54910 | ENSG00000010803 | SCMH1 | chr1 | - | 41142877 | 41143112 | 41116921 | 41117010 | 41160874 | 41160898 | 21 | 13 | 31 | 10 | 17 | 0 | 1 | 0 | 0 | 0 | 15 | 25 | 31 | 11 | 21 | 0 | 0 | 0 | 0 | 0 | 298 | 149 | 1 | 1 | 1 | 0.867 | 1 | 1 | 1 | 1 | 1 | 1 | 1 | 1 | -0.027 |
| SE | 54911 | ENSG00000010803 | SCMH1 | chr1 | - | 41151613 | 41151684 | 41142877 | 41143112 | 41152598 | 41152674 | 17 | 19 | 12 | 17 | 22 | 0 | 0 | 0 | 0 | 0 | 67 | 19 | 13 | 32 | 31 | 0 | 0 | 0 | 0 | 3 | 220 | 149 | 1 | 1 | 1 | 1 | 1 | 1 | 1 | 1 | 1 | 1 | 1 | 0.875 | 0.025 |
| SE | 54920 | ENSG00000010803 | SCMH1 | chr1 | - | 41159724 | 41159933 | 41152598 | 41152741 | 41160874 | 41160898 | 23 | 29 | 30 | 35 | 47 | 0 | 1 | 0 | 1 | 0 | 11 | 13 | 10 | 18 | 34 | 0 | 0 | 0 | 2 | 0 | 298 | 149 | 1 | 1 | 1 | 0.935 | 1 | 0.946 | 1 | 1 | 1 | 1 | 0.818 | 1 | 0.013 |
| SE | 54924 | ENSG00000124731 | TREM1 | chr6 | - | 41282394 | 41282751 | 41275620 | 41276230 | 41286606 | 41286665 | 195 | 181 | 104 | 209 | 274 | 1 | 0 | 0 | 0 | 0 | 322 | 193 | 263 | 194 | 134 | 0 | 0 | 0 | 0 | 0 | 298 | 149 | 1 | 1 | 0.99 | 1 | 1 | 1 | 1 | 1 | 1 | 1 | 1 | 1 | -0.002 |
| SE | 54925 | ENSG00000172164 | SNTB1 | chr8 | - | 120551130 | 120551251 | 120548761 | 120548958 | 120575085 | 120575225 | 1 | 0 | 2 | 0 | 0 | 76 | 76 | 107 | 109 | 84 | 0 | 6 | 0 | 0 | 0 | 191 | 121 | 93 | 187 | 141 | 270 | 149 | 1 | 1 | 0.007 | 0 | 0.01 | 0 | 0 | 0 | 0.027 | 0 | 0 | 0 | -0.002 |
| SE | 54926 | ENSG00000172164 | SNTB1 | chr8 | - | 120632443 | 120632651 | 120575085 | 120575225 | 120693691 | 120693908 | 243 | 171 | 228 | 230 | 169 | 1 | 0 | 0 | 0 | 0 | 285 | 374 | 217 | 424 | 219 | 0 | 0 | 0 | 2 | 0 | 298 | 149 | 1 | 1 | 0.992 | 1 | 1 | 1 | 1 | 1 | 1 | 1 | 0.991 | 1 | 0 |
| SE | 54928 | ENSG00000172164 | SNTB1 | chr8 | - | 120693691 | 120693908 | 120632443 | 120632651 | 120811272 | 120811563 | 170 | 111 | 195 | 159 | 132 | 0 | 0 | 0 | 0 | 0 | 168 | 253 | 131 | 243 | 125 | 0 | 0 | 3 | 0 | 0 | 298 | 149 | 1 | 1 | 1 | 1 | 1 | 1 | 1 | 1 | 1 | 0.956 | 1 | 1 | 0.009 |
| SE | 54931 | ENSG00000030419 | IKZF2 | chr2 | - | 213013790 | 213013934 | 212999690 | 213008084 | 213021992 | 213022130 | 202 | 122 | 159 | 173 | 319 | 4 | 1 | 6 | 7 | 4 | 43 | 93 | 63 | 17 | 0 | 0 | 3 | 9 | 0 | 0 | 293 | 149 | 1 | 1 | 0.963 | 0.984 | 0.931 | 0.926 | 0.976 | 1 | 0.94 | 0.781 | 1 | NA | 0.026 |
| SE | 54933 | ENSG00000030419 | IKZF2 | chr2 | - | 213021643 | 213021720 | 213013790 | 213013934 | 213021992 | 213022130 | 0 | 0 | 4 | 0 | 0 | 109 | 60 | 82 | 97 | 128 | 0 | 1 | 0 | 0 | 0 | 18 | 53 | 39 | 6 | 0 | 226 | 149 | 1 | 1 | 0 | 0 | 0.031 | 0 | 0 | 0 | 0.012 | 0 | 0 | NA | 0.003 |
| SE | 54935 | ENSG00000030419 | IKZF2 | chr2 | - | 213056565 | 213056744 | 213049712 | 213049880 | 213056832 | 213057099 | 1 | 0 | 1 | 2 | 5 | 49 | 13 | 28 | 40 | 92 | 0 | 0 | 0 | 0 | 0 | 0 | 15 | 5 | 0 | 5 | 298 | 149 | 1 | 1 | 0.01 | 0 | 0.018 | 0.024 | 0.026 | NA | 0 | 0 | NA | 0 | 0.016 |
| SE | 54936 | ENSG00000030419 | IKZF2 | chr2 | - | 213056565 | 213056744 | 213049712 | 213049880 | 213056910 | 213057099 | 0 | 0 | 3 | 1 | 3 | 27 | 13 | 34 | 25 | 50 | 0 | 0 | 0 | 0 | 0 | 13 | 7 | 16 | 5 | 0 | 298 | 149 | 1 | 1 | 0 | 0 | 0.042 | 0.02 | 0.029 | 0 | 0 | 0 | 0 | NA | 0.018 |
| SE | 54937 | ENSG00000030419 | IKZF2 | chr2 | - | 213056832 | 213057099 | 213049712 | 213049880 | 213147707 | 213147812 | 130 | 83 | 100 | 96 | 221 | 0 | 0 | 0 | 1 | 0 | 2 | 63 | 44 | 18 | 5 | 0 | 0 | 0 | 0 | 0 | 298 | 149 | 1 | 1 | 1 | 1 | 1 | 0.98 | 1 | 1 | 1 | 1 | 1 | 1 | -0.004 |
| SE | 54939 | ENSG00000030419 | IKZF2 | chr2 | - | 213056910 | 213057099 | 213049712 | 213049880 | 213147707 | 213147812 | 108 | 83 | 106 | 81 | 179 | 0 | 0 | 0 | 1 | 0 | 15 | 55 | 55 | 23 | 0 | 0 | 0 | 0 | 0 | 0 | 298 | 149 | 1 | 1 | 1 | 1 | 1 | 0.976 | 1 | 1 | 1 | 1 | 1 | NA | -0.005 |
| SE | 54940 | ENSG00000030419 | IKZF2 | chr2 | - | 213081169 | 213081238 | 213056832 | 213057099 | 213097876 | 213097960 | 3 | 0 | 1 | 0 | 1 | 1 | 0 | 1 | 1 | 4 | 0 | 0 | 0 | 0 | 0 | 0 | 0 | 1 | 0 | 0 | 218 | 149 | 1 | 1 | 0.672 | NA | 0.406 | 0 | 0.146 | NA | NA | 0 | NA | NA | 0.306 |
| SE | 54942 | ENSG00000030419 | IKZF2 | chr2 | - | 213097876 | 213097960 | 213056910 | 213057099 | 213147707 | 213147812 | 1 | 0 | 1 | 2 | 2 | 81 | 70 | 72 | 56 | 129 | 0 | 0 | 1 | 0 | 0 | 2 | 48 | 39 | 18 | 0 | 233 | 149 | 1 | 1 | 0.008 | 0 | 0.009 | 0.022 | 0.01 | 0 | 0 | 0.016 | 0 | NA | 0.006 |
| SE | 54943 | ENSG00000030419 | IKZF2 | chr2 | - | 213150143 | 213150247 | 213148595 | 213148644 | 213151412 | 213151603 | 30 | 33 | 16 | 33 | 64 | 18 | 18 | 11 | 20 | 37 | 0 | 9 | 10 | 0 | 0 | 0 | 3 | 11 | 0 | 0 | 253 | 149 | 1 | 1 | 0.495 | 0.519 | 0.461 | 0.493 | 0.505 | NA | 0.639 | 0.349 | NA | NA | 0.001 |
| SE | 54944 | ENSG00000030419 | IKZF2 | chr2 | - | 213150143 | 213150247 | 213148595 | 213148644 | 213152125 | 213152427 | 32 | 34 | 16 | 33 | 63 | 11 | 11 | 18 | 7 | 27 | 0 | 5 | 10 | 0 | 0 | 0 | 10 | 3 | 0 | 0 | 253 | 149 | 1 | 1 | 0.631 | 0.645 | 0.344 | 0.735 | 0.579 | NA | 0.227 | 0.663 | NA | NA | 0.142 |
| SE | 54948 | ENSG00000144214 | LYG1 | chr2 | - | 99292535 | 99292640 | 99291236 | 99291421 | 99295627 | 99295702 | 5 | 0 | 3 | 11 | 10 | 0 | 0 | 0 | 0 | 1 | 15 | 14 | 5 | 2 | 2 | 0 | 0 | 0 | 2 | 0 | 254 | 149 | 1 | 1 | 1 | NA | 1 | 1 | 0.854 | 1 | 1 | 1 | 0.37 | 1 | 0.09 |
| SE | 54949 | ENSG00000088448 | ANKRD10 | chr13 | - | 110883697 | 110883793 | 110878539 | 110880112 | 110906032 | 110906124 | 77 | 48 | 65 | 75 | 90 | 0 | 0 | 0 | 0 | 2 | 41 | 47 | 16 | 22 | 56 | 0 | 0 | 0 | 0 | 0 | 245 | 149 | 1 | 1 | 1 | 1 | 1 | 1 | 0.965 | 1 | 1 | 1 | 1 | 1 | -0.007 |
| SE | 54957 | ENSG00000088448 | ANKRD10 | chr13 | - | 110910617 | 110910770 | 110906032 | 110906124 | 110914724 | 110914803 | 194 | 115 | 174 | 187 | 100 | 0 | 0 | 0 | 1 | 0 | 162 | 276 | 187 | 126 | 169 | 0 | 0 | 2 | 0 | 0 | 298 | 149 | 1 | 1 | 1 | 1 | 1 | 0.989 | 1 | 1 | 1 | 0.979 | 1 | 1 | 0.002 |
| SE | 54958 | ENSG00000169991 | IFFO2 | chr1 | - | 18910341 | 18910472 | 18904279 | 18908666 | 18911383 | 18911476 | 147 | 74 | 179 | 174 | 163 | 1 | 0 | 0 | 0 | 0 | 174 | 115 | 67 | 92 | 104 | 0 | 0 | 0 | 0 | 0 | 280 | 149 | 1 | 1 | 0.987 | 1 | 1 | 1 | 1 | 1 | 1 | 1 | 1 | 1 | -0.003 |
| SE | 54959 | ENSG00000169991 | IFFO2 | chr1 | - | 18911383 | 18911476 | 18910341 | 18910472 | 18911962 | 18912083 | 109 | 67 | 138 | 131 | 97 | 1 | 2 | 0 | 0 | 0 | 192 | 103 | 50 | 91 | 110 | 1 | 2 | 0 | 0 | 0 | 242 | 149 | 1 | 1 | 0.985 | 0.954 | 1 | 1 | 1 | 0.992 | 0.969 | 1 | 1 | 1 | -0.004 |
| SE | 54961 | ENSG00000169991 | IFFO2 | chr1 | - | 18918361 | 18918502 | 18916902 | 18917042 | 18921060 | 18921121 | 196 | 124 | 165 | 182 | 121 | 0 | 2 | 3 | 0 | 0 | 281 | 132 | 128 | 132 | 294 | 0 | 0 | 0 | 0 | 0 | 290 | 149 | 1 | 1 | 1 | 0.97 | 0.966 | 1 | 1 | 1 | 1 | 1 | 1 | 1 | -0.013 |
| SE | 54964 | ENSG00000135842 | NIBAN1 | chr1 | - | 184803584 | 184803692 | 184798078 | 184798190 | 184805945 | 184805985 | 358 | 310 | 305 | 399 | 595 | 0 | 0 | 0 | 0 | 0 | 592 | 468 | 659 | 510 | 245 | 0 | 0 | 7 | 7 | 0 | 257 | 149 | 1 | 1 | 1 | 1 | 1 | 1 | 1 | 1 | 1 | 0.982 | 0.977 | 1 | 0.008 |
| SE | 54965 | ENSG00000135842 | NIBAN1 | chr1 | - | 184803584 | 184803692 | 184798078 | 184798190 | 184808073 | 184808235 | 198 | 179 | 169 | 203 | 392 | 0 | 0 | 0 | 0 | 0 | 335 | 238 | 386 | 291 | 138 | 0 | 1 | 2 | 0 | 0 | 257 | 149 | 1 | 1 | 1 | 1 | 1 | 1 | 1 | 1 | 0.993 | 0.991 | 1 | 1 | 0.003 |
| SE | 54966 | ENSG00000135842 | NIBAN1 | chr1 | - | 184805945 | 184806056 | 184798078 | 184798190 | 184808073 | 184808235 | 66 | 51 | 73 | 92 | 121 | 0 | 0 | 0 | 0 | 0 | 118 | 112 | 115 | 99 | 52 | 0 | 1 | 2 | 0 | 0 | 260 | 149 | 1 | 1 | 1 | 1 | 1 | 1 | 1 | 1 | 0.985 | 0.971 | 1 | 1 | 0.009 |
| SE | 54968 | ENSG00000135842 | NIBAN1 | chr1 | - | 184808073 | 184808235 | 184798078 | 184798190 | 184818637 | 184818825 | 151 | 91 | 81 | 115 | 179 | 0 | 0 | 0 | 0 | 0 | 125 | 120 | 200 | 95 | 106 | 0 | 0 | 3 | 0 | 0 | 298 | 149 | 1 | 1 | 1 | 1 | 1 | 1 | 1 | 1 | 1 | 0.971 | 1 | 1 | 0.006 |
| SE | 54969 | ENSG00000135842 | NIBAN1 | chr1 | - | 184808073 | 184808235 | 184803584 | 184803692 | 184823166 | 184823329 | 3 | 3 | 3 | 6 | 10 | 0 | 0 | 0 | 1 | 0 | 0 | 15 | 27 | 8 | 20 | 0 | 0 | 0 | 0 | 0 | 298 | 149 | 1 | 1 | 1 | 1 | 1 | 0.75 | 1 | NA | 1 | 1 | 1 | 1 | -0.05 |
| SE | 54971 | ENSG00000135842 | NIBAN1 | chr1 | - | 184823166 | 184823329 | 184818637 | 184818825 | 184831846 | 184831962 | 126 | 91 | 97 | 118 | 176 | 0 | 0 | 0 | 0 | 0 | 178 | 153 | 219 | 152 | 81 | 0 | 0 | 2 | 0 | 0 | 298 | 149 | 1 | 1 | 1 | 1 | 1 | 1 | 1 | 1 | 1 | 0.982 | 1 | 1 | 0.004 |
| SE | 54973 | ENSG00000135842 | NIBAN1 | chr1 | - | 184894074 | 184894206 | 184890107 | 184890222 | 184899178 | 184899309 | 283 | 283 | 238 | 284 | 294 | 0 | 0 | 0 | 0 | 0 | 722 | 391 | 848 | 506 | 232 | 0 | 0 | 0 | 0 | 2 | 281 | 149 | 1 | 1 | 1 | 1 | 1 | 1 | 1 | 1 | 1 | 1 | 1 | 0.984 | 0.003 |
| SE | 54979 | ENSG00000117859 | OSBPL9 | chr1 | + | 51614304 | 51614419 | 51577178 | 51577256 | 51616873 | 51617221 | 2 | 0 | 2 | 0 | 4 | 0 | 0 | 0 | 0 | 0 | 0 | 4 | 0 | 0 | 0 | 0 | 0 | 0 | 1 | 0 | 264 | 149 | 1 | 1 | 1 | NA | 1 | NA | 1 | NA | 1 | NA | 0 | NA | 0.5 |
| SE | 54980 | ENSG00000117859 | OSBPL9 | chr1 | + | 51651990 | 51652041 | 51616873 | 51617221 | 51669433 | 51669512 | 63 | 31 | 42 | 66 | 60 | 0 | 1 | 0 | 0 | 0 | 2 | 74 | 76 | 41 | 8 | 0 | 0 | 0 | 0 | 0 | 200 | 149 | 1 | 1 | 1 | 0.958 | 1 | 1 | 1 | 1 | 1 | 1 | 1 | 1 | -0.008 |
| SE | 54981 | ENSG00000117859 | OSBPL9 | chr1 | + | 51651990 | 51652041 | 51616873 | 51617221 | 51714002 | 51714079 | 32 | 18 | 25 | 29 | 28 | 0 | 0 | 0 | 0 | 1 | 2 | 27 | 36 | 20 | 8 | 0 | 0 | 0 | 0 | 0 | 200 | 149 | 1 | 1 | 1 | 1 | 1 | 1 | 0.954 | 1 | 1 | 1 | 1 | 1 | -0.009 |
| SE | 54982 | ENSG00000117859 | OSBPL9 | chr1 | + | 51669433 | 51669512 | 51617108 | 51617221 | 51714002 | 51714079 | 37 | 18 | 33 | 30 | 36 | 0 | 0 | 0 | 0 | 1 | 22 | 55 | 51 | 25 | 10 | 0 | 0 | 0 | 0 | 0 | 228 | 149 | 1 | 1 | 1 | 1 | 1 | 1 | 0.959 | 1 | 1 | 1 | 1 | 1 | -0.008 |
| SE | 54984 | ENSG00000117859 | OSBPL9 | chr1 | + | 51669433 | 51669814 | 51651990 | 51652041 | 51714002 | 51714079 | 11 | 4 | 8 | 16 | 6 | 11 | 7 | 13 | 11 | 21 | 0 | 14 | 24 | 10 | 0 | 0 | 0 | 19 | 8 | 0 | 298 | 149 | 1 | 1 | 0.333 | 0.222 | 0.235 | 0.421 | 0.125 | NA | 1 | 0.387 | 0.385 | NA | -0.323 |
| SE | 54985 | ENSG00000117859 | OSBPL9 | chr1 | + | 51740142 | 51740211 | 51714002 | 51714079 | 51745535 | 51745631 | 1 | 0 | 0 | 1 | 0 | 87 | 49 | 77 | 91 | 144 | 0 | 0 | 0 | 0 | 0 | 22 | 72 | 86 | 39 | 18 | 218 | 149 | 1 | 1 | 0.008 | 0 | 0 | 0.007 | 0 | 0 | 0 | 0 | 0 | 0 | 0.003 |
| SE | 54987 | ENSG00000117859 | OSBPL9 | chr1 | + | 51746709 | 51746757 | 51745089 | 51745631 | 51750144 | 51750195 | 33 | 31 | 29 | 39 | 75 | 0 | 0 | 2 | 0 | 0 | 0 | 26 | 30 | 12 | 0 | 0 | 0 | 0 | 0 | 0 | 197 | 149 | 1 | 1 | 1 | 1 | 0.916 | 1 | 1 | NA | 1 | 1 | 1 | NA | -0.017 |
| SE | 54988 | ENSG00000117859 | OSBPL9 | chr1 | + | 51748368 | 51748398 | 51746709 | 51746757 | 51750144 | 51750195 | 215 | 133 | 128 | 175 | 300 | 3 | 4 | 0 | 3 | 0 | 87 | 157 | 183 | 69 | 52 | 0 | 1 | 0 | 0 | 0 | 179 | 149 | 1 | 1 | 0.984 | 0.965 | 1 | 0.98 | 1 | 1 | 0.992 | 1 | 1 | 1 | -0.013 |
| SE | 54990 | ENSG00000117859 | OSBPL9 | chr1 | + | 51750144 | 51750195 | 51748368 | 51748398 | 51756319 | 51756358 | 144 | 95 | 80 | 105 | 176 | 0 | 1 | 1 | 0 | 0 | 72 | 98 | 124 | 36 | 45 | 0 | 0 | 0 | 0 | 0 | 200 | 149 | 1 | 1 | 1 | 0.986 | 0.983 | 1 | 1 | 1 | 1 | 1 | 1 | 1 | -0.006 |
| SE | 54991 | ENSG00000117859 | OSBPL9 | chr1 | + | 51752502 | 51752725 | 51750144 | 51750195 | 51756319 | 51756657 | 1 | 0 | 5 | 2 | 0 | 113 | 73 | 64 | 90 | 119 | 0 | 0 | 1 | 0 | 0 | 60 | 90 | 98 | 33 | 34 | 298 | 149 | 1 | 1 | 0.004 | 0 | 0.038 | 0.011 | 0 | 0 | 0 | 0.005 | 0 | 0 | 0.01 |
| SE | 54992 | ENSG00000117859 | OSBPL9 | chr1 | + | 51752502 | 51752725 | 51750144 | 51750195 | 51760689 | 51760740 | 1 | 0 | 3 | 1 | 2 | 30 | 21 | 33 | 25 | 65 | 0 | 2 | 2 | 0 | 0 | 10 | 26 | 41 | 11 | 7 | 298 | 149 | 1 | 1 | 0.016 | 0 | 0.043 | 0.02 | 0.015 | 0 | 0.037 | 0.024 | 0 | 0 | 0.007 |
| SE | 54993 | ENSG00000117859 | OSBPL9 | chr1 | + | 51756319 | 51756358 | 51750144 | 51750195 | 51759516 | 51759546 | 64 | 35 | 33 | 58 | 52 | 0 | 0 | 2 | 2 | 0 | 39 | 69 | 71 | 29 | 30 | 0 | 0 | 0 | 0 | 0 | 188 | 149 | 1 | 1 | 1 | 1 | 0.929 | 0.958 | 1 | 1 | 1 | 1 | 1 | 1 | -0.023 |
| SE | 54995 | ENSG00000117859 | OSBPL9 | chr1 | + | 51756940 | 51756995 | 51756319 | 51756358 | 51760689 | 51760771 | 0 | 0 | 0 | 2 | 0 | 48 | 41 | 36 | 47 | 86 | 0 | 0 | 0 | 0 | 0 | 21 | 26 | 38 | 7 | 14 | 204 | 149 | 1 | 1 | 0 | 0 | 0 | 0.03 | 0 | 0 | 0 | 0 | 0 | 0 | 0.006 |
| SE | 54997 | ENSG00000117859 | OSBPL9 | chr1 | + | 51761866 | 51761971 | 51759623 | 51760780 | 51765821 | 51765856 | 219 | 166 | 196 | 242 | 303 | 2 | 0 | 0 | 0 | 0 | 72 | 176 | 222 | 104 | 77 | 0 | 0 | 3 | 0 | 0 | 254 | 149 | 1 | 1 | 0.985 | 1 | 1 | 1 | 1 | 1 | 1 | 0.977 | 1 | 1 | 0.002 |
| SE | 54999 | ENSG00000117859 | OSBPL9 | chr1 | + | 51776832 | 51776918 | 51772604 | 51772723 | 51781163 | 51781335 | 120 | 96 | 122 | 129 | 204 | 1 | 0 | 0 | 1 | 0 | 25 | 75 | 71 | 52 | 4 | 0 | 0 | 0 | 0 | 0 | 235 | 149 | 1 | 1 | 0.987 | 1 | 1 | 0.988 | 1 | 1 | 1 | 1 | 1 | 1 | -0.005 |
| SE | 55000 | ENSG00000117859 | OSBPL9 | chr1 | + | 51781163 | 51781335 | 51776832 | 51776918 | 51783914 | 51784025 | 52 | 36 | 62 | 51 | 131 | 0 | 0 | 0 | 0 | 0 | 14 | 42 | 23 | 24 | 0 | 4 | 0 | 3 | 0 | 0 | 298 | 149 | 1 | 1 | 1 | 1 | 1 | 1 | 1 | 0.636 | 1 | 0.793 | 1 | NA | 0.143 |
| SE | 55001 | ENSG00000117859 | OSBPL9 | chr1 | + | 51782558 | 51782643 | 51781163 | 51781335 | 51783914 | 51784025 | 192 | 133 | 210 | 130 | 293 | 0 | 0 | 0 | 0 | 0 | 16 | 155 | 178 | 98 | 18 | 0 | 4 | 0 | 0 | 0 | 234 | 149 | 1 | 1 | 1 | 1 | 1 | 1 | 1 | 1 | 0.961 | 1 | 1 | 1 | 0.008 |
| SE | 55002 | ENSG00000117859 | OSBPL9 | chr1 | + | 51784263 | 51784327 | 51783914 | 51784025 | 51785807 | 51785886 | 47 | 25 | 53 | 48 | 48 | 0 | 0 | 0 | 0 | 0 | 0 | 17 | 49 | 16 | 0 | 0 | 0 | 1 | 0 | 0 | 213 | 149 | 1 | 1 | 1 | 1 | 1 | 1 | 1 | NA | 1 | 0.972 | 1 | NA | 0.009 |
| SE | 55003 | ENSG00000117859 | OSBPL9 | chr1 | + | 51784441 | 51784582 | 51784263 | 51784327 | 51785807 | 51785886 | 384 | 286 | 345 | 374 | 501 | 0 | 0 | 0 | 0 | 0 | 24 | 297 | 385 | 146 | 0 | 0 | 0 | 3 | 4 | 0 | 290 | 149 | 1 | 1 | 1 | 1 | 1 | 1 | 1 | 1 | 1 | 0.985 | 0.949 | NA | 0.017 |
| SE | 55004 | ENSG00000138641 | HERC3 | chr4 | + | 88595556 | 88595614 | 88592433 | 88592574 | 88605794 | 88606049 | 323 | 241 | 316 | 305 | 229 | 0 | 0 | 1 | 0 | 0 | 247 | 353 | 286 | 280 | 179 | 0 | 0 | 0 | 0 | 0 | 207 | 149 | 1 | 1 | 1 | 1 | 0.996 | 1 | 1 | 1 | 1 | 1 | 1 | 1 | -0.001 |
| SE | 55006 | ENSG00000138641 | HERC3 | chr4 | + | 88597968 | 88598087 | 88595556 | 88595614 | 88649839 | 88649895 | 4 | 2 | 2 | 2 | 2 | 0 | 1 | 2 | 2 | 0 | 4 | 0 | 1 | 0 | 0 | 0 | 0 | 9 | 0 | 0 | 268 | 149 | 1 | 1 | 1 | 0.527 | 0.357 | 0.357 | 1 | 1 | NA | 0.058 | NA | NA | 0.119 |
| SE | 55007 | ENSG00000138641 | HERC3 | chr4 | + | 88670138 | 88670252 | 88669859 | 88670023 | 88676217 | 88676241 | 187 | 160 | 167 | 131 | 181 | 0 | 2 | 0 | 0 | 0 | 91 | 170 | 151 | 118 | 33 | 0 | 3 | 0 | 0 | 0 | 263 | 149 | 1 | 1 | 1 | 0.978 | 1 | 1 | 1 | 1 | 0.97 | 1 | 1 | 1 | 0.002 |
| SE | 55009 | ENSG00000152223 | EPG5 | chr18 | - | 45855572 | 45855687 | 45847608 | 45852649 | 45857852 | 45858068 | 60 | 40 | 86 | 85 | 88 | 3 | 0 | 0 | 1 | 1 | 26 | 56 | 45 | 43 | 19 | 0 | 0 | 0 | 0 | 0 | 264 | 149 | 1 | 1 | 0.919 | 1 | 1 | 0.98 | 0.98 | 1 | 1 | 1 | 1 | 1 | -0.024 |
| SE | 55010 | ENSG00000152223 | EPG5 | chr18 | - | 45857852 | 45858068 | 45855352 | 45855687 | 45858565 | 45858651 | 76 | 59 | 66 | 87 | 75 | 0 | 0 | 0 | 0 | 0 | 1 | 41 | 41 | 37 | 19 | 0 | 3 | 0 | 0 | 0 | 298 | 149 | 1 | 1 | 1 | 1 | 1 | 1 | 1 | 1 | 0.872 | 1 | 1 | 1 | 0.026 |
| SE | 55011 | ENSG00000152223 | EPG5 | chr18 | - | 45865614 | 45865759 | 45860103 | 45860346 | 45866797 | 45867007 | 180 | 81 | 120 | 169 | 187 | 0 | 0 | 0 | 0 | 1 | 137 | 105 | 122 | 110 | 29 | 0 | 0 | 0 | 6 | 0 | 294 | 149 | 1 | 1 | 1 | 1 | 1 | 1 | 0.99 | 1 | 1 | 1 | 0.903 | 1 | 0.017 |
| SE | 55012 | ENSG00000152223 | EPG5 | chr18 | - | 45866797 | 45867007 | 45865614 | 45865759 | 45867562 | 45867748 | 218 | 107 | 142 | 186 | 200 | 3 | 1 | 0 | 0 | 0 | 140 | 126 | 135 | 137 | 6 | 0 | 0 | 0 | 0 | 0 | 298 | 149 | 1 | 1 | 0.973 | 0.982 | 1 | 1 | 1 | 1 | 1 | 1 | 1 | 1 | -0.009 |
| SE | 55013 | ENSG00000152223 | EPG5 | chr18 | - | 45867562 | 45867748 | 45866797 | 45867007 | 45870566 | 45870742 | 193 | 90 | 107 | 133 | 154 | 0 | 0 | 0 | 0 | 0 | 83 | 86 | 117 | 79 | 32 | 0 | 0 | 1 | 0 | 0 | 298 | 149 | 1 | 1 | 1 | 1 | 1 | 1 | 1 | 1 | 1 | 0.983 | 1 | 1 | 0.003 |
| SE | 55014 | ENSG00000152223 | EPG5 | chr18 | - | 45870566 | 45870742 | 45867562 | 45867748 | 45876235 | 45876342 | 125 | 58 | 86 | 105 | 102 | 0 | 0 | 0 | 0 | 0 | 37 | 68 | 87 | 45 | 60 | 0 | 0 | 1 | 0 | 0 | 298 | 149 | 1 | 1 | 1 | 1 | 1 | 1 | 1 | 1 | 1 | 0.978 | 1 | 1 | 0.004 |
| SE | 55016 | ENSG00000152223 | EPG5 | chr18 | - | 45876235 | 45876342 | 45867562 | 45867748 | 45878375 | 45878448 | 41 | 26 | 27 | 37 | 41 | 0 | 2 | 0 | 0 | 0 | 9 | 24 | 30 | 14 | 0 | 0 | 3 | 3 | 0 | 0 | 256 | 149 | 1 | 1 | 1 | 0.883 | 1 | 1 | 1 | 1 | 0.823 | 0.853 | 1 | NA | 0.058 |
| SE | 55018 | ENSG00000152223 | EPG5 | chr18 | - | 45880074 | 45880223 | 45879012 | 45879214 | 45882273 | 45882431 | 40 | 18 | 23 | 19 | 8 | 0 | 0 | 0 | 0 | 0 | 27 | 16 | 22 | 15 | 3 | 0 | 0 | 1 | 0 | 0 | 298 | 149 | 1 | 1 | 1 | 1 | 1 | 1 | 1 | 1 | 1 | 0.917 | 1 | 1 | 0.017 |
| SE | 55020 | ENSG00000152223 | EPG5 | chr18 | - | 45887750 | 45887907 | 45884616 | 45884811 | 45899403 | 45899566 | 34 | 13 | 22 | 26 | 17 | 3 | 2 | 4 | 0 | 0 | 0 | 7 | 22 | 15 | 0 | 0 | 2 | 8 | 7 | 0 | 298 | 149 | 1 | 1 | 0.85 | 0.765 | 0.733 | 1 | 1 | NA | 0.636 | 0.579 | 0.517 | NA | 0.292 |
| SE | 55023 | ENSG00000152223 | EPG5 | chr18 | - | 45900995 | 45901167 | 45899403 | 45899566 | 45903972 | 45904117 | 88 | 34 | 59 | 65 | 70 | 0 | 0 | 0 | 0 | 0 | 69 | 45 | 66 | 44 | 17 | 0 | 0 | 0 | 2 | 0 | 298 | 149 | 1 | 1 | 1 | 1 | 1 | 1 | 1 | 1 | 1 | 1 | 0.917 | 1 | 0.017 |
| SE | 55024 | ENSG00000152223 | EPG5 | chr18 | - | 45903972 | 45904117 | 45900995 | 45901167 | 45907957 | 45908081 | 103 | 40 | 40 | 81 | 103 | 0 | 0 | 0 | 0 | 0 | 44 | 59 | 60 | 53 | 12 | 0 | 0 | 0 | 3 | 0 | 294 | 149 | 1 | 1 | 1 | 1 | 1 | 1 | 1 | 1 | 1 | 1 | 0.9 | 1 | 0.02 |
| SE | 55026 | ENSG00000152223 | EPG5 | chr18 | - | 45907957 | 45908081 | 45903972 | 45904117 | 45910520 | 45910742 | 96 | 39 | 40 | 63 | 83 | 0 | 0 | 0 | 0 | 0 | 30 | 74 | 51 | 64 | 15 | 0 | 1 | 0 | 0 | 0 | 273 | 149 | 1 | 1 | 1 | 1 | 1 | 1 | 1 | 1 | 0.976 | 1 | 1 | 1 | 0.005 |
| SE | 55027 | ENSG00000152223 | EPG5 | chr18 | - | 45912289 | 45912456 | 45910520 | 45910742 | 45913705 | 45913828 | 145 | 56 | 69 | 69 | 64 | 0 | 4 | 0 | 0 | 0 | 48 | 60 | 40 | 24 | 34 | 0 | 0 | 0 | 1 | 0 | 298 | 149 | 1 | 1 | 1 | 0.875 | 1 | 1 | 1 | 1 | 1 | 1 | 0.923 | 1 | -0.01 |
| SE | 55028 | ENSG00000152223 | EPG5 | chr18 | - | 45915510 | 45915621 | 45913705 | 45913828 | 45916008 | 45916206 | 99 | 56 | 69 | 72 | 35 | 0 | 0 | 0 | 1 | 0 | 46 | 34 | 37 | 44 | 15 | 0 | 0 | 0 | 3 | 0 | 260 | 149 | 1 | 1 | 1 | 1 | 1 | 0.976 | 1 | 1 | 1 | 1 | 0.894 | 1 | 0.016 |
| SE | 55029 | ENSG00000152223 | EPG5 | chr18 | - | 45916008 | 45916206 | 45913705 | 45913828 | 45923267 | 45923387 | 0 | 1 | 1 | 1 | 0 | 0 | 1 | 0 | 0 | 2 | 0 | 1 | 1 | 3 | 0 | 0 | 2 | 2 | 0 | 0 | 298 | 149 | 1 | 1 | NA | 0.333 | 1 | 1 | 0 | NA | 0.2 | 0.2 | 1 | NA | 0.117 |
| SE | 55030 | ENSG00000152223 | EPG5 | chr18 | - | 45916008 | 45916206 | 45915510 | 45915621 | 45916437 | 45916454 | 104 | 67 | 93 | 89 | 56 | 2 | 1 | 1 | 0 | 0 | 35 | 49 | 60 | 58 | 38 | 0 | 0 | 2 | 0 | 0 | 298 | 149 | 1 | 1 | 0.963 | 0.971 | 0.979 | 1 | 1 | 1 | 1 | 0.938 | 1 | 1 | -0.005 |
| SE | 55031 | ENSG00000152223 | EPG5 | chr18 | - | 45916437 | 45916582 | 45916008 | 45916206 | 45922340 | 45922600 | 58 | 42 | 56 | 56 | 36 | 0 | 0 | 2 | 0 | 0 | 10 | 41 | 39 | 44 | 25 | 0 | 0 | 0 | 0 | 0 | 294 | 149 | 1 | 1 | 1 | 1 | 0.934 | 1 | 1 | 1 | 1 | 1 | 1 | 1 | -0.013 |
| SE | 55032 | ENSG00000152223 | EPG5 | chr18 | - | 45916437 | 45916582 | 45916008 | 45916206 | 45923267 | 45923387 | 55 | 35 | 55 | 55 | 31 | 0 | 1 | 1 | 0 | 0 | 10 | 42 | 32 | 40 | 28 | 0 | 1 | 1 | 0 | 0 | 294 | 149 | 1 | 1 | 1 | 0.947 | 0.965 | 1 | 1 | 1 | 0.955 | 0.942 | 1 | 1 | 0.003 |
| SE | 55035 | ENSG00000152223 | EPG5 | chr18 | - | 45922340 | 45922600 | 45916008 | 45916206 | 45923267 | 45923387 | 104 | 48 | 54 | 86 | 98 | 0 | 1 | 1 | 0 | 0 | 50 | 50 | 30 | 56 | 22 | 0 | 1 | 1 | 0 | 0 | 298 | 149 | 1 | 1 | 1 | 0.96 | 0.964 | 1 | 1 | 1 | 0.962 | 0.938 | 1 | 1 | 0.005 |
| SE | 55038 | ENSG00000152223 | EPG5 | chr18 | - | 45930675 | 45930830 | 45928868 | 45929009 | 45934808 | 45934966 | 124 | 76 | 103 | 129 | 160 | 0 | 0 | 0 | 0 | 0 | 147 | 105 | 131 | 95 | 36 | 0 | 0 | 0 | 4 | 0 | 298 | 149 | 1 | 1 | 1 | 1 | 1 | 1 | 1 | 1 | 1 | 1 | 0.922 | 1 | 0.016 |
| SE | 55039 | ENSG00000152223 | EPG5 | chr18 | - | 45934808 | 45934966 | 45930675 | 45930830 | 45939599 | 45939755 | 136 | 89 | 104 | 105 | 176 | 0 | 2 | 1 | 0 | 3 | 80 | 77 | 87 | 68 | 46 | 0 | 0 | 4 | 0 | 0 | 298 | 149 | 1 | 1 | 1 | 0.957 | 0.981 | 1 | 0.967 | 1 | 1 | 0.916 | 1 | 1 | -0.002 |
| SE | 55041 | ENSG00000152223 | EPG5 | chr18 | - | 45946662 | 45946768 | 45944004 | 45944119 | 45949483 | 45949591 | 42 | 11 | 28 | 21 | 32 | 0 | 1 | 0 | 0 | 0 | 7 | 7 | 8 | 24 | 13 | 0 | 0 | 1 | 0 | 0 | 255 | 149 | 1 | 1 | 1 | 0.865 | 1 | 1 | 1 | 1 | 1 | 0.824 | 1 | 1 | 0.008 |
| SE | 55042 | ENSG00000152223 | EPG5 | chr18 | - | 45948502 | 45948576 | 45944004 | 45944119 | 45949483 | 45949591 | 20 | 3 | 20 | 12 | 14 | 0 | 1 | 0 | 0 | 0 | 0 | 14 | 9 | 22 | 0 | 0 | 0 | 1 | 0 | 0 | 223 | 149 | 1 | 1 | 1 | 0.667 | 1 | 1 | 1 | NA | 1 | 0.857 | 1 | NA | -0.019 |
| SE | 55043 | ENSG00000152223 | EPG5 | chr18 | - | 45948502 | 45948576 | 45946662 | 45946768 | 45949483 | 45949591 | 47 | 11 | 56 | 29 | 35 | 0 | 0 | 0 | 0 | 0 | 48 | 34 | 37 | 47 | 10 | 0 | 0 | 1 | 1 | 0 | 223 | 149 | 1 | 1 | 1 | 1 | 1 | 1 | 1 | 1 | 1 | 0.961 | 0.969 | 1 | 0.014 |
| SE | 55044 | ENSG00000117713 | ARID1A | chr1 | + | 26760855 | 26761096 | 26732675 | 26732792 | 26761383 | 26761473 | 297 | 143 | 266 | 331 | 273 | 4 | 0 | 0 | 0 | 2 | 347 | 296 | 230 | 304 | 477 | 0 | 0 | 0 | 0 | 7 | 298 | 149 | 1 | 1 | 0.974 | 1 | 1 | 1 | 0.986 | 1 | 1 | 1 | 1 | 0.971 | -0.002 |
| SE | 55045 | ENSG00000117713 | ARID1A | chr1 | + | 26761383 | 26761473 | 26732675 | 26732792 | 26766220 | 26766366 | 2 | 0 | 0 | 0 | 1 | 0 | 0 | 0 | 0 | 0 | 0 | 0 | 0 | 0 | 7 | 0 | 0 | 1 | 0 | 0 | 239 | 149 | 1 | 1 | 1 | NA | NA | NA | 1 | NA | NA | 0 | NA | | 0.5 |
| SE | 55047 | ENSG00000117713 | ARID1A | chr1 | + | 26762972 | 26763285 | 26761383 | 26761473 | 26766220 | 26766366 | 274 | 173 | 255 | 229 | 194 | 2 | 0 | 0 | 0 | 0 | 404 | 368 | 203 | 305 | 469 | 0 | 0 | 0 | 0 | 0 | 298 | 149 | 1 | 1 | 0.986 | 1 | 1 | 1 | 1 | 1 | 1 | 1 | 1 | 1 | -0.003 |
| SE | 55049 | ENSG00000117713 | ARID1A | chr1 | + | 26771118 | 26771326 | 26766456 | 26766566 | 26772499 | 26772632 | 497 | 262 | 333 | 411 | 621 | 0 | 0 | 0 | 0 | 0 | 653 | 416 | 295 | 465 | 604 | 0 | 0 | 0 | 3 | 0 | 298 | 149 | 1 | 1 | 1 | 1 | 1 | 1 | 1 | 1 | 1 | 1 | 0.987 | 1 | 0.003 |
| SE | 55050 | ENSG00000117713 | ARID1A | chr1 | + | 26772499 | 26772632 | 26771118 | 26771326 | 26772811 | 26772987 | 592 | 329 | 444 | 491 | 679 | 0 | 0 | 0 | 0 | 0 | 743 | 475 | 363 | 534 | 704 | 0 | 0 | 0 | 4 | 0 | 282 | 149 | 1 | 1 | 1 | 1 | 1 | 1 | 1 | 1 | 1 | 1 | 0.986 | 1 | 0.003 |
| SE | 55051 | ENSG00000117713 | ARID1A | chr1 | + | 26772499 | 26772632 | 26771118 | 26771326 | 26772814 | 26772987 | 598 | 307 | 420 | 535 | 667 | 0 | 0 | 0 | 1 | 0 | 862 | 550 | 359 | 541 | 741 | 0 | 0 | 0 | 0 | 0 | 282 | 149 | 1 | 1 | 1 | 1 | 1 | 0.996 | 1 | 1 | 1 | 1 | 1 | 1 | -0.001 |
| SE | 55052 | ENSG00000117713 | ARID1A | chr1 | + | 26773579 | 26773717 | 26773345 | 26773496 | 26773801 | 26773898 | 436 | 248 | 403 | 427 | 403 | 0 | 0 | 0 | 0 | 0 | 688 | 473 | 296 | 471 | 478 | 0 | 2 | 0 | 0 | 0 | 287 | 149 | 1 | 1 | 1 | 1 | 1 | 1 | 1 | 1 | 0.992 | 1 | 1 | 1 | 0.002 |
| SE | 55053 | ENSG00000117713 | ARID1A | chr1 | + | 26774115 | 26774203 | 26773801 | 26773898 | 26774328 | 26774901 | 0 | 0 | 1 | 1 | 0 | 105 | 63 | 99 | 75 | 46 | 0 | 0 | 0 | 0 | 0 | 114 | 139 | 86 | 103 | 144 | 237 | 149 | 1 | 1 | 0 | 0 | 0.006 | 0.008 | 0 | 0 | 0 | 0 | 0 | 0 | 0.003 |
| SE | 55057 | ENSG00000117713 | ARID1A | chr1 | + | 26775816 | 26775956 | 26775576 | 26775707 | 26779022 | 26780983 | 1 | 0 | 7 | 1 | 1 | 322 | 192 | 258 | 392 | 160 | 9 | 5 | 6 | 1 | 15 | 559 | 470 | 282 | 396 | 603 | 289 | 149 | 1 | 1 | 0.002 | 0 | 0.014 | 0.001 | 0.003 | 0.008 | 0.005 | 0.011 | 0.001 | 0.013 | -0.004 |
| SE | 55072 | ENSG00000104894 | CD37 | chr19 | + | 49335885 | 49336029 | 49335713 | 49335786 | 49336908 | 49337033 | 7 | 2 | 4 | 2 | 5 | 686 | 274 | 463 | 539 | 546 | 0 | 4 | 0 | 0 | 4 | 998 | 839 | 453 | 750 | 1716 | 293 | 149 | 1 | 1 | 0.005 | 0.004 | 0.004 | 0.002 | 0.005 | 0 | 0.002 | 0 | 0 | 0.001 | 0.003 |
| SE | 55075 | ENSG00000086200 | IPO11 | chr5 | + | 62422504 | 62422568 | 62412660 | 62412929 | 62437273 | 62437417 | 0 | 0 | 2 | 0 | 0 | 29 | 15 | 14 | 18 | 20 | 0 | 0 | 0 | 0 | 0 | 0 | 16 | 10 | 0 | 0 | 213 | 149 | 1 | 1 | 0 | 0 | 0.091 | 0 | 0 | NA | 0 | 0 | NA | NA | 0.018 |
| SE | 55076 | ENSG00000086200 | IPO11 | chr5 | + | 62451729 | 62451933 | 62449730 | 62449999 | 62467130 | 62467207 | 75 | 53 | 64 | 47 | 92 | 1 | 0 | 0 | 4 | 0 | 46 | 27 | 41 | 22 | 10 | 0 | 0 | 0 | 0 | 0 | 298 | 149 | 1 | 1 | 0.974 | 1 | 1 | 0.855 | 1 | 1 | 1 | 1 | 1 | 1 | -0.034 |
| SE | 55077 | ENSG00000086200 | IPO11 | chr5 | + | 62470249 | 62470308 | 62467130 | 62467263 | 62474415 | 62474464 | 59 | 31 | 53 | 33 | 43 | 0 | 0 | 0 | 0 | 0 | 7 | 25 | 19 | 36 | 0 | 0 | 0 | 1 | 0 | 0 | 208 | 149 | 1 | 1 | 1 | 1 | 1 | 1 | 1 | 1 | 1 | 0.932 | 1 | NA | 0.017 |
| SE | 55079 | ENSG00000086200 | IPO11 | chr5 | + | 62493997 | 62494124 | 62490114 | 62490220 | 62504666 | 62504700 | 70 | 58 | 64 | 67 | 48 | 1 | 0 | 0 | 0 | 0 | 24 | 37 | 30 | 24 | 6 | 0 | 0 | 0 | 0 | 0 | 276 | 149 | 1 | 1 | 0.974 | 1 | 1 | 1 | 1 | 1 | 1 | 1 | 1 | 1 | -0.005 |
| SE | 55082 | ENSG00000086200 | IPO11 | chr5 | + | 62530708 | 62530785 | 62526141 | 62526257 | 62550366 | 62550462 | 4 | 5 | 5 | 1 | 11 | 0 | 0 | 1 | 0 | 0 | 0 | 0 | 4 | 3 | 0 | 0 | 0 | 0 | 0 | 0 | 226 | 149 | 1 | 1 | 1 | 1 | 0.767 | 1 | 1 | NA | NA | 1 | 1 | NA | -0.047 |
| SE | 55084 | ENSG00000086200 | IPO11 | chr5 | + | 62536701 | 62536781 | 62526141 | 62526257 | 62550366 | 62550462 | 1 | 0 | 1 | 2 | 0 | 0 | 0 | 1 | 0 | 0 | 0 | 0 | 0 | 1 | 0 | 0 | 0 | 0 | 0 | 0 | 229 | 149 | 1 | 1 | 1 | NA | 0.394 | 1 | NA | NA | NA | NA | 1 | NA | -0.202 |
| SE | 55085 | ENSG00000086200 | IPO11 | chr5 | + | 62536701 | 62536781 | 62530708 | 62530785 | 62550366 | 62550462 | 15 | 20 | 14 | 16 | 33 | 0 | 0 | 2 | 0 | 1 | 0 | 1 | 3 | 3 | 0 | 0 | 0 | 0 | 0 | 0 | 229 | 149 | 1 | 1 | 1 | 1 | 0.82 | 1 | 0.955 | NA | 1 | 1 | 1 | NA | -0.045 |
| SE | 55086 | ENSG00000086200 | IPO11 | chr5 | + | 62537208 | 62537289 | 62536701 | 62536781 | 62550366 | 62550462 | 83 | 88 | 85 | 70 | 98 | 1 | 0 | 1 | 2 | 0 | 14 | 51 | 34 | 36 | 22 | 0 | 0 | 0 | 0 | 0 | 230 | 149 | 1 | 1 | 0.982 | 1 | 0.982 | 0.958 | 1 | 1 | 1 | 1 | 1 | 1 | -0.016 |
| SE | 55087 | ENSG00000086200 | IPO11 | chr5 | + | 62578703 | 62578935 | 62561135 | 62561257 | 62580453 | 62581467 | 1 | 0 | 1 | 3 | 2 | 0 | 0 | 0 | 0 | 0 | 0 | 0 | 0 | 1 | 0 | 0 | 0 | 0 | 1 | 0 | 298 | 149 | 1 | 1 | 1 | NA | 1 | 1 | 1 | NA | NA | NA | 0.333 | NA | 0.667 |
| SE | 55088 | ENSG00000086200 | IPO11 | chr5 | + | 62579400 | 62579483 | 62561135 | 62561257 | 62580453 | 62580680 | 0 | 0 | 0 | 2 | 1 | 0 | 0 | 0 | 0 | 0 | 8 | 4 | 0 | 0 | 6 | 0 | 0 | 0 | 1 | 0 | 232 | 149 | 1 | 1 | NA | NA | NA | 1 | 1 | 1 | 1 | NA | 0 | 1 | 0.25 |
| SE | 55089 | ENSG00000086200 | IPO11 | chr5 | + | 62579400 | 62579483 | 62578820 | 62578935 | 62580453 | 62580680 | 4 | 4 | 4 | 10 | 7 | 1 | 0 | 0 | 2 | 2 | 15 | 6 | 0 | 0 | 6 | 0 | 0 | 0 | 1 | 0 | 232 | 149 | 1 | 1 | 0.72 | 1 | 1 | 0.763 | 0.692 | 1 | 1 | NA | 0 | 1 | 0.085 |
| SE | 55102 | ENSG00000072110 | ACTN1 | chr14 | - | 68878457 | 68878520 | 68877081 | 68877240 | 68879961 | 68880108 | 1 | 4 | 0 | 4 | 2 | 0 | 0 | 0 | 2 | 0 | 11 | 1 | 6 | 2 | 14 | 0 | 0 | 0 | 0 | 0 | 212 | 149 | 1 | 1 | 1 | 1 | NA | 0.584 | 1 | 1 | 1 | 1 | 1 | 1 | -0.104 |
| SE | 55104 | ENSG00000072110 | ACTN1 | chr14 | - | 68878457 | 68878523 | 68877081 | 68877240 | 68879961 | 68880108 | 6 | 6 | 1 | 12 | 3 | 0 | 0 | 0 | 2 | 0 | 36 | 2 | 11 | 6 | 21 | 0 | 0 | 0 | 0 | 0 | 215 | 149 | 1 | 1 | 1 | 1 | 1 | 0.806 | 1 | 1 | 1 | 1 | 1 | 1 | -0.039 |
| SE | 55105 | ENSG00000072110 | ACTN1 | chr14 | - | 68878457 | 68879069 | 68877081 | 68877240 | 68879961 | 68880108 | 107 | 138 | 87 | 119 | 184 | 0 | 0 | 0 | 2 | 0 | 282 | 164 | 85 | 210 | 1298 | 0 | 0 | 0 | 0 | 0 | 298 | 149 | 1 | 1 | 1 | 1 | 1 | 0.967 | 1 | 1 | 1 | 1 | 1 | 1 | -0.007 |
| SE | 55106 | ENSG00000072110 | ACTN1 | chr14 | - | 68878988 | 68879069 | 68877081 | 68877240 | 68879961 | 68880108 | 218 | 273 | 229 | 233 | 329 | 0 | 0 | 0 | 2 | 0 | 621 | 275 | 191 | 409 | 2571 | 0 | 0 | 0 | 0 | 0 | 230 | 149 | 1 | 1 | 1 | 1 | 1 | 0.987 | 1 | 1 | 1 | 1 | 1 | 1 | -0.003 |
| SE | 55110 | ENSG00000072110 | ACTN1 | chr14 | - | 68884774 | 68884883 | 68884167 | 68884308 | 68885424 | 68885575 | 327 | 271 | 362 | 429 | 407 | 0 | 3 | 0 | 0 | 0 | 841 | 359 | 413 | 807 | 3953 | 0 | 3 | 0 | 2 | 0 | 258 | 149 | 1 | 1 | 1 | 0.981 | 1 | 1 | 1 | 1 | 0.986 | 1 | 0.996 | 1 | 0 |
| SE | 55112 | ENSG00000072110 | ACTN1 | chr14 | - | 68900861 | 68900946 | 68893654 | 68893747 | 68902476 | 68902562 | 0 | 2 | 2 | 1 | 0 | 52 | 50 | 59 | 76 | 115 | 0 | 0 | 5 | 6 | 0 | 235 | 93 | 84 | 200 | 839 | 234 | 149 | 1 | 1 | 0 | 0.025 | 0.021 | 0.008 | 0 | 0 | 0 | 0.037 | 0.019 | 0 | 0 |
| SE | 55113 | ENSG00000072110 | ACTN1 | chr14 | - | 68902476 | 68902562 | 68893654 | 68893747 | 68904654 | 68904736 | 178 | 162 | 218 | 248 | 312 | 0 | 0 | 0 | 0 | 0 | 576 | 199 | 205 | 419 | 2209 | 0 | 0 | 3 | 0 | 0 | 235 | 149 | 1 | 1 | 1 | 1 | 1 | 1 | 1 | 1 | 1 | 0.977 | 1 | 1 | 0.005 |
| SE | 55115 | ENSG00000072110 | ACTN1 | chr14 | - | 68921005 | 68921125 | 68912155 | 68912242 | 68925557 | 68925672 | 330 | 335 | 406 | 522 | 427 | 1 | 0 | 0 | 0 | 0 | 950 | 410 | 400 | 928 | 3684 | 0 | 0 | 0 | 0 | 0 | 269 | 149 | 1 | 1 | 0.995 | 1 | 1 | 1 | 1 | 1 | 1 | 1 | 1 | 1 | -0.001 |
| SE | 55116 | ENSG00000072110 | ACTN1 | chr14 | - | 68925557 | 68925672 | 68921005 | 68921125 | 68978951 | 68979076 | 239 | 255 | 272 | 357 | 164 | 0 | 0 | 0 | 0 | 0 | 486 | 287 | 260 | 631 | 2472 | 0 | 0 | 0 | 3 | 0 | 264 | 149 | 1 | 1 | 1 | 1 | 1 | 1 | 1 | 1 | 1 | 1 | 0.992 | 1 | 0.002 |
| SE | 55118 | ENSG00000116221 | MRPL37 | chr1 | + | 54207359 | 54207504 | 54205294 | 54205410 | 54209945 | 54210131 | 1 | 0 | 0 | 3 | 0 | 188 | 106 | 179 | 183 | 485 | 0 | 0 | 0 | 0 | 0 | 381 | 239 | 183 | 204 | 452 | 294 | 149 | 1 | 1 | 0.003 | 0 | 0 | 0.008 | 0 | 0 | 0 | 0 | 0 | 0 | 0.002 |
| SE | 55125 | ENSG00000119725 | ZNF410 | chr14 | + | 73893775 | 73893932 | 73892026 | 73892208 | 73896315 | 73896506 | 172 | 113 | 158 | 161 | 191 | 1 | 0 | 0 | 3 | 0 | 304 | 141 | 149 | 214 | 231 | 0 | 0 | 0 | 0 | 3 | 298 | 149 | 1 | 1 | 0.989 | 1 | 1 | 0.964 | 1 | 1 | 1 | 1 | 1 | 0.975 | -0.004 |
| SE | 55128 | ENSG00000119725 | ZNF410 | chr14 | + | 73896347 | 73896534 | 73893796 | 73893932 | 73898070 | 73898262 | 111 | 55 | 109 | 77 | 116 | 2 | 1 | 0 | 0 | 2 | 149 | 144 | 103 | 112 | 108 | 0 | 0 | 0 | 11 | 5 | 298 | 149 | 1 | 1 | 0.965 | 0.965 | 1 | 1 | 0.967 | 1 | 1 | 1 | 0.836 | 0.915 | 0.029 |
| SE | 55130 | ENSG00000119725 | ZNF410 | chr14 | + | 73922065 | 73922206 | 73909340 | 73909430 | 73923394 | 73923522 | 52 | 34 | 44 | 45 | 96 | 0 | 0 | 0 | 0 | 0 | 57 | 27 | 31 | 21 | 34 | 0 | 1 | 0 | 0 | 0 | 290 | 149 | 1 | 1 | 1 | 1 | 1 | 1 | 1 | 1 | 0.933 | 1 | 1 | 1 | 0.013 |
| SE | 55132 | ENSG00000119725 | ZNF410 | chr14 | + | 73927106 | 73927228 | 73923394 | 73923522 | 73931502 | 73931664 | 0 | 1 | 1 | 0 | 0 | 102 | 58 | 85 | 86 | 173 | 0 | 0 | 0 | 0 | 0 | 152 | 65 | 86 | 81 | 84 | 271 | 149 | 1 | 1 | 0 | 0.009 | 0.006 | 0 | 0 | 0 | 0 | 0 | 0 | 0 | 0.003 |
| SE | 55133 | ENSG00000214530 | STARD10 | chr11 | - | 72755700 | 72755753 | 72754728 | 72755142 | 72757766 | 72757884 | 17 | 18 | 22 | 13 | 22 | 0 | 0 | 0 | 1 | 0 | 49 | 5 | 22 | 16 | 59 | 0 | 0 | 0 | 0 | 0 | 202 | 149 | 1 | 1 | 1 | 1 | 1 | 0.906 | 1 | 1 | 1 | 1 | 1 | 1 | -0.019 |
| SE | 55135 | ENSG00000214530 | STARD10 | chr11 | - | 72758529 | 72758633 | 72757766 | 72757884 | 72759233 | 72759309 | 22 | 9 | 22 | 13 | 20 | 0 | 0 | 0 | 1 | 0 | 17 | 15 | 12 | 41 | 44 | 0 | 0 | 0 | 0 | 0 | 253 | 149 | 1 | 1 | 1 | 1 | 1 | 0.884 | 1 | 1 | 1 | 1 | 1 | 1 | -0.023 |
| SE | 55138 | ENSG00000232931 | LINC00342 | chr2 | - | 95813315 | 95813957 | 95807051 | 95807302 | 95814049 | 95815156 | 1 | 3 | 1 | 6 | 1 | 18 | 15 | 8 | 7 | 23 | 0 | 0 | 5 | 1 | 0 | 0 | 6 | 2 | 3 | 0 | 298 | 149 | 1 | 1 | 0.027 | 0.091 | 0.059 | 0.3 | 0.021 | NA | 0 | 0.556 | 0.143 | NA | -0.133 |
| SE | 55140 | ENSG00000143797 | MBOAT2 | chr2 | - | 8862270 | 8862322 | 8860612 | 8860764 | 8862589 | 8862722 | 2 | 0 | 0 | 2 | 4 | 4 | 8 | 11 | 8 | 12 | 0 | 0 | 0 | 1 | 0 | 0 | 3 | 11 | 25 | 27 | 201 | 149 | 1 | 1 | 0.27 | 0 | 0 | 0.156 | 0.198 | NA | 0 | 0 | 0.029 | 0 | 0.118 |
| SE | 55141 | ENSG00000143797 | MBOAT2 | chr2 | - | 8862270 | 8862322 | 8860612 | 8860764 | 8864169 | 8864234 | 0 | 1 | 0 | 0 | 1 | 0 | 0 | 0 | 0 | 0 | 0 | 2 | 0 | 2 | 0 | 0 | 0 | 0 | 1 | 0 | 201 | 149 | 1 | 1 | NA | 1 | NA | NA | 1 | NA | 1 | NA | 0.597 | NA | 0.202 |
| SE | 55142 | ENSG00000143797 | MBOAT2 | chr2 | - | 8862589 | 8862722 | 8860612 | 8860764 | 8864169 | 8864234 | 20 | 42 | 33 | 26 | 69 | 0 | 0 | 0 | 0 | 0 | 44 | 65 | 54 | 175 | 118 | 0 | 0 | 0 | 1 | 0 | 282 | 149 | 1 | 1 | 1 | 1 | 1 | 1 | 1 | 1 | 1 | 1 | 0.989 | 1 | 0.002 |
| SE | 55143 | ENSG00000143797 | MBOAT2 | chr2 | - | 8862589 | 8862722 | 8862270 | 8862322 | 8864169 | 8864234 | 17 | 34 | 22 | 20 | 60 | 0 | 1 | 0 | 0 | 0 | 44 | 62 | 43 | 151 | 91 | 0 | 2 | 0 | 2 | 0 | 282 | 149 | 1 | 1 | 1 | 0.947 | 1 | 1 | 1 | 1 | 0.942 | 1 | 0.976 | 1 | 0.006 |
| SE | 55144 | ENSG00000143797 | MBOAT2 | chr2 | - | 8876986 | 8877213 | 8873193 | 8873300 | 8882510 | 8882565 | 28 | 34 | 14 | 27 | 46 | 0 | 0 | 0 | 1 | 0 | 19 | 36 | 53 | 146 | 69 | 0 | 0 | 0 | 0 | 0 | 298 | 149 | 1 | 1 | 1 | 1 | 1 | 0.931 | 1 | 1 | 1 | 1 | 1 | 1 | -0.014 |
| SE | 55145 | ENSG00000143797 | MBOAT2 | chr2 | - | 8877029 | 8877213 | 8873107 | 8873300 | 8882510 | 8882565 | 43 | 47 | 15 | 40 | 59 | 0 | 0 | 0 | 1 | 0 | 20 | 66 | 64 | 182 | 88 | 0 | 0 | 0 | 0 | 0 | 298 | 149 | 1 | 1 | 1 | 1 | 1 | 0.952 | 1 | 1 | 1 | 1 | 1 | 1 | -0.01 |
| SE | 55147 | ENSG00000143797 | MBOAT2 | chr2 | - | 8943186 | 8943264 | 8908620 | 8908716 | 9003539 | 9003658 | 22 | 13 | 5 | 15 | 42 | 0 | 0 | 2 | 0 | 0 | 43 | 23 | 19 | 58 | 44 | 0 | 0 | 3 | 2 | 0 | 227 | 149 | 1 | 1 | 1 | 1 | 0.621 | 1 | 1 | 1 | 1 | 0.806 | 0.95 | 1 | -0.027 |
| SE | 55149 | ENSG00000143797 | MBOAT2 | chr2 | - | 8958496 | 8958642 | 8943186 | 8943264 | 9003539 | 9003658 | 78 | 74 | 36 | 102 | 127 | 1 | 1 | 0 | 0 | 0 | 265 | 71 | 87 | 178 | 367 | 0 | 0 | 0 | 0 | 0 | 295 | 149 | 1 | 1 | 0.975 | 0.974 | 1 | 1 | 1 | 1 | 1 | 1 | 1 | 1 | -0.01 |
| SE | 55151 | ENSG00000143797 | MBOAT2 | chr2 | - | 8974358 | 8974435 | 8958496 | 8958642 | 9003539 | 9003665 | 1 | 2 | 0 | 8 | 3 | 0 | 0 | 0 | 0 | 0 | 0 | 2 | 1 | 4 | 25 | 0 | 0 | 5 | 0 | 0 | 226 | 149 | 1 | 1 | 1 | 1 | NA | 1 | 1 | NA | 1 | 0.116 | 1 | 1 | 0.221 |
| SE | 55156 | ENSG00000116586 | LAMTOR2 | chr1 | + | 156057977 | 156058067 | 156055495 | 156055675 | 156058314 | 156058506 | 132 | 136 | 150 | 126 | 43 | 3 | 0 | 0 | 0 | 0 | 472 | 227 | 122 | 227 | 493 | 0 | 0 | 3 | 0 | 0 | 239 | 149 | 1 | 1 | 0.965 | 1 | 1 | 1 | 1 | 1 | 1 | 0.962 | 1 | 1 | 0.001 |
| SE | 55157 | ENSG00000143595 | AQP10 | chr1 | + | 154321932 | 154322059 | 154321092 | 154321260 | 154322981 | 154323119 | 5 | 10 | 5 | 11 | 2 | 1 | 1 | 2 | 0 | 0 | 0 | 22 | 14 | 8 | 140 | 0 | 0 | 0 | 0 | 3 | 276 | 149 | 1 | 1 | 0.73 | 0.844 | 0.574 | 1 | 1 | NA | 1 | 1 | 1 | 0.962 | -0.161 |
| SE | 55166 | ENSG00000024526 | DEPDC1 | chr1 | - | 68482045 | 68482897 | 68481439 | 68481612 | 68483949 | 68484090 | 2 | 0 | 3 | 0 | 2 | 2 | 0 | 2 | 4 | 1 | 2 | 4 | 0 | 0 | 0 | 0 | 1 | 1 | 0 | 0 | 298 | 149 | 1 | 1 | 0.333 | NA | 0.429 | 0 | 0.5 | 1 | 0.667 | 0 | NA | NA | -0.24 |
| SE | 55168 | ENSG00000024526 | DEPDC1 | chr1 | - | [illegible] | [illegible] | [illegible] | [illegible] | [illegible] | [illegible] | [illegible] | [illegible] | [illegible] | [illegible] | [illegible] | [illegible] | [illegible] | [illegible] | [illegible] | [illegible] | [illegible] | [illegible] | [illegible] | [illegible] | [illegible] | [illegible] | [illegible] | [illegible] | [illegible] | [illegible] | [illegible] | 149 | 1 | 1 | [illegible] | [illegible] | [illegible] | [illegible] | [illegible] | [illegible] | [illegible] | [illegible] | [illegible] | [illegible] | -0.036 |
| SE | 55169 | ENSG00000054267 | ARID4B | chr1 | - | [illegible] | [illegible] | [illegible] | [illegible] | [illegible] | [illegible] | [illegible] | [illegible] | [illegible] | [illegible] | [illegible] | [illegible] | [illegible] | [illegible] | [illegible] | [illegible] | [illegible] | [illegible] | [illegible] | [illegible] | [illegible] | [illegible] | [illegible] | [illegible] | [illegible] | [illegible] | [illegible] | 149 | 1 | 1 | [illegible] | [illegible] | [illegible] | [illegible] | [illegible] | [illegible] | [illegible] | [illegible] | [illegible] | [illegible] | [illegible] |
| SE | 55170 | ENSG00000054267 | ARID4B | chr1 | - | [illegible] | [illegible] | [illegible] | [illegible] | [illegible] | [illegible] | [illegible] | [illegible] | [illegible] | [illegible] | [illegible] | [illegible] | [illegible] | [illegible] | [illegible] | [illegible] | [illegible] | [illegible] | [illegible] | [illegible] | [illegible] | [illegible] | [illegible] | [illegible] | [illegible] | [illegible] | [illegible] | 149 | 1 | 1 | [illegible] | [illegible] | [illegible] | [illegible] | [illegible] | [illegible] | [illegible] | [illegible] | [illegible] | [illegible] | [illegible] |
| SE | 55173 | ENSG00000054267 | ARID4B | chr1 | - | [illegible] | [illegible] | [illegible] | [illegible] | [illegible] | [illegible] | [illegible] | [illegible] | [illegible] | [illegible] | [illegible] | [illegible] | [illegible] | [illegible] | [illegible] | [illegible] | [illegible] | [illegible] | [illegible] | [illegible] | [illegible] | [illegible] | [illegible] | [illegible] | [illegible] | [illegible] | [illegible] | 149 | 1 | 1 | [illegible] | [illegible] | [illegible] | [illegible] | [illegible] | [illegible] | [illegible] | [illegible] | [illegible] | [illegible] | [illegible] |
| SE | 55175 | ENSG00000054267 | ARID4B | chr1 | - | [illegible] | [illegible] | [illegible] | [illegible] | [illegible] | [illegible] | [illegible] | [illegible] | [illegible] | [illegible] | [illegible] | [illegible] | [illegible] | [illegible] | [illegible] | [illegible] | [illegible] | [illegible] | [illegible] | [illegible] | [illegible] | [illegible] | [illegible] | [illegible] | [illegible] | [illegible] | [illegible] | 149 | 1 | 1 | [illegible] | [illegible] | [illegible] | [illegible] | [illegible] | [illegible] | [illegible] | [illegible] | [illegible] | [illegible] | [illegible] |
| SE | 55176 | ENSG00000054267 | ARID4B | chr1 | - | [illegible] | [illegible] | [illegible] | [illegible] | [illegible] | [illegible] | [illegible] | [illegible] | [illegible] | [illegible] | [illegible] | [illegible] | [illegible] | [illegible] | [illegible] | [illegible] | [illegible] | [illegible] | [illegible] | [illegible] | [illegible] | [illegible] | [illegible] | [illegible] | [illegible] | [illegible] | [illegible] | 149 | 1 | 1 | [illegible] | [illegible] | [illegible] | [illegible] | [illegible] | [illegible] | [illegible] | [illegible] | [illegible] | [illegible] | [illegible] |
| SE | 55177 | ENSG00000054267 | ARID4B | chr1 | - | [illegible] | [illegible] | [illegible] | [illegible] | [illegible] | [illegible] | [illegible] | [illegible] | [illegible] | [illegible] | [illegible] | [illegible] | [illegible] | [illegible] | [illegible] | [illegible] | [illegible] | [illegible] | [illegible] | [illegible] | [illegible] | [illegible] | [illegible] | [illegible] | [illegible] | [illegible] | [illegible] | 149 | 1 | 1 | [illegible] | [illegible] | [illegible] | [illegible] | [illegible] | [illegible] | [illegible] | [illegible] | [illegible] | [illegible] | [illegible] |
| SE | 55178 | ENSG00000054267 | ARID4B | chr1 | - | [illegible] | [illegible] | [illegible] | [illegible] | [illegible] | [illegible] | [illegible] | [illegible] | [illegible] | [illegible] | [illegible] | [illegible] | [illegible] | [illegible] | [illegible] | [illegible] | [illegible] | [illegible] | [illegible] | [illegible] | [illegible] | [illegible] | [illegible] | [illegible] | [illegible] | [illegible] | [illegible] | 149 | 1 | 1 | [illegible] | [illegible] | [illegible] | [illegible] | [illegible] | [illegible] | [illegible] | [illegible] | [illegible] | [illegible] | [illegible] |
| SE | 55179 | ENSG00000054267 | ARID4B | chr1 | - | [illegible] | [illegible] | [illegible] | [illegible] | [illegible] | [illegible] | [illegible] | [illegible] | [illegible] | [illegible] | [illegible] | [illegible] | [illegible] | [illegible] | [illegible] | [illegible] | [illegible] | [illegible] | [illegible] | [illegible] | [illegible] | [illegible] | [illegible] | [illegible] | [illegible] | [illegible] | [illegible] | 149 | 1 | 1 | [illegible] | [illegible] | [illegible] | [illegible] | [illegible] | [illegible] | [illegible] | [illegible] | [illegible] | [illegible] | [illegible] |
| SE | 55180 | ENSG00000054267 | ARID4B | chr1 | - | [illegible] | [illegible] | [illegible] | [illegible] | [illegible] | [illegible] | [illegible] | [illegible] | [illegible] | [illegible] | [illegible] | [illegible] | [illegible] | [illegible] | [illegible] | [illegible] | [illegible] | [illegible] | [illegible] | [illegible] | [illegible] | [illegible] | [illegible] | [illegible] | [illegible] | [illegible] | [illegible] | 149 | 1 | 1 | [illegible] | [illegible] | [illegible] | [illegible] | [illegible] | [illegible] | [illegible] | [illegible] | [illegible] | [illegible] | [illegible] |
| SE | 55182 | ENSG00000143158 | MPC2 | chr1 | - | [illegible] | [illegible] | [illegible] | [illegible] | [illegible] | [illegible] | [illegible] | [illegible] | [illegible] | [illegible] | [illegible] | [illegible] | [illegible] | [illegible] | [illegible] | [illegible] | [illegible] | [illegible] | [illegible] | [illegible] | [illegible] | [illegible] | [illegible] | [illegible] | [illegible] | [illegible] | [illegible] | 149 | 1 | 1 | [illegible] | [illegible] | [illegible] | [illegible] | [illegible] | [illegible] | [illegible] | [illegible] | [illegible] | [illegible] | [illegible] |
| SE | 55183 | ENSG00000143158 | MPC2 | chr1 | - | [illegible] | [illegible] | [illegible] | [illegible] | [illegible] | [illegible] | [illegible] | [illegible] | [illegible] | [illegible] | [illegible] | [illegible] | [illegible] | [illegible] | [illegible] | [illegible] | [illegible] | [illegible] | [illegible] | [illegible] | [illegible] | [illegible] | [illegible] | [illegible] | [illegible] | [illegible] | [illegible] | 149 | 1 | 1 | [illegible] | [illegible] | [illegible] | [illegible] | [illegible] | [illegible] | [illegible] | [illegible] | [illegible] | [illegible] | [illegible] |
| SE | 55186 | ENSG00000143158 | MPC2 | chr1 | - | [illegible] | [illegible] | [illegible] | [illegible] | [illegible] | [illegible] | [illegible] | [illegible] | [illegible] | [illegible] | [illegible] | [illegible] | [illegible] | [illegible] | [illegible] | [illegible] | [illegible] | [illegible] | [illegible] | [illegible] | [illegible] | [illegible] | [illegible] | [illegible] | [illegible] | [illegible] | [illegible] | 149 | 1 | 1 | [illegible] | [illegible] | [illegible] | [illegible] | [illegible] | [illegible] | [illegible] | [illegible] | [illegible] | [illegible] | [illegible] |
| SE | 55188 | ENSG00000162779 | AXDND1 | chr1 | + | [illegible] | [illegible] | [illegible] | [illegible] | [illegible] | [illegible] | [illegible] | [illegible] | [illegible] | [illegible] | [illegible] | [illegible] | [illegible] | [illegible] | [illegible] | [illegible] | [illegible] | [illegible] | [illegible] | [illegible] | [illegible] | [illegible] | [illegible] | [illegible] | [illegible] | [illegible] | [illegible] | 149 | 1 | 1 | [illegible] | [illegible] | [illegible] | [illegible] | [illegible] | [illegible] | [illegible] | [illegible] | [illegible] | [illegible] | [illegible] |
| SE | 55199 | ENSG00000132881 | CPLANE2 | chr1 | - | [illegible] | [illegible] | [illegible] | [illegible] | [illegible] | [illegible] | [illegible] | [illegible] | [illegible] | [illegible] | [illegible] | [illegible] | [illegible] | [illegible] | [illegible] | [illegible] | [illegible] | [illegible] | [illegible] | [illegible] | [illegible] | [illegible] | [illegible] | [illegible] | [illegible] | [illegible] | [illegible] | 149 | 1 | 1 | [illegible] | [illegible] | [illegible] | [illegible] | [illegible] | [illegible] | [illegible] | [illegible] | [illegible] | [illegible] | [illegible] |
| SE | [illegible] | [illegible] | [illegible] | [illegible] | [illegible] | [illegible] | [illegible] | [illegible] | [illegible] | [illegible] | [illegible] | [illegible] | [illegible] | [illegible] | [illegible] | [illegible] | [illegible] | [illegible] | [illegible] | [illegible] | [illegible] | [illegible] | [illegible] | [illegible] | [illegible] | [illegible] | [illegible] | [illegible] | [illegible] | [illegible] | [illegible] | [illegible] | [illegible] | [illegible] | [illegible] | [illegible] | [illegible] | [illegible] | [illegible] | [illegible] | [illegible] | [illegible] | [illegible] | [illegible] | [illegible] | [illegible] | [illegible] |

| | | | | | | | | | | | | | | |
|---|---|---|---|---|---|---|---|---|---|---|---|---|---|---|
| SE | 55233 | ENSG00000168610 | STAT3 | chr17 | - | 42348388 | 42348539 | 42346568 | 42346713 | 42388278 | 42388433 | 221 | 68 | [illegible] |
| SE | 55234 | ENSG00000168610 | STAT3 | chr17 | - | 42374058 | 42374206 | 42348388 | 42348517 | 42388278 | 42388373 | 2 | 0 | [illegible] |
| SE | 55235 | ENSG00000168610 | STAT3 | chr17 | - | 42374058 | 42374206 | 42348388 | 42348539 | 42388278 | 42388373 | 2 | 0 | [illegible] |
| SE | 55245 | ENSG00000158161 | EYA3 | chr1 | - | 28035543 | 28035680 | 28017153 | 28017239 | 28038838 | 28038905 | 62 | 63 | [illegible] |
| SE | 55248 | ENSG00000158161 | EYA3 | chr1 | - | 28057993 | 28058094 | 28048382 | 28048426 | 28088523 | 28088564 | 81 | 48 | [illegible] |
| SE | 55249 | ENSG00000135905 | DOCK10 | chr2 | - | 224768871 | 224768947 | 224765103 | 224765837 | 224770210 | 224770349 | 0 | 0 | [illegible] |
| SE | 55250 | ENSG00000135905 | DOCK10 | chr2 | - | 224774904 | 224775115 | 224773156 | 224773347 | 224778137 | 224778284 | 297 | 281 | [illegible] |
| SE | 55251 | ENSG00000135905 | DOCK10 | chr2 | - | 224778137 | 224778284 | 224773156 | 224773347 | 224787021 | 224787135 | 223 | 172 | [illegible] |
| SE | 55253 | ENSG00000135905 | DOCK10 | chr2 | - | 224792973 | 224793072 | 224789063 | 224789170 | 224793399 | 224793457 | 337 | 278 | [illegible] |
| SE | 55255 | ENSG00000135905 | DOCK10 | chr2 | - | 224801915 | 224802040 | 224800150 | 224800263 | 224804111 | 224804213 | 171 | 159 | [illegible] |
| SE | 55256 | ENSG00000135905 | DOCK10 | chr2 | - | 224804793 | 224804841 | 224804111 | 224804213 | 224805057 | 224805124 | 224 | 226 | [illegible] |
| SE | 55257 | ENSG00000135905 | DOCK10 | chr2 | - | 224814319 | 224814364 | 224807910 | 224808086 | 224816616 | 224816652 | 288 | 237 | [illegible] |
| SE | 55259 | ENSG00000135905 | DOCK10 | chr2 | - | 224837761 | 224837831 | 224834149 | 224834263 | 224839953 | 224840072 | 348 | 333 | [illegible] |
| SE | 55261 | ENSG00000135905 | DOCK10 | chr2 | - | 224852376 | 224852442 | 224849506 | 224849599 | 224852934 | 224853122 | 302 | 347 | [illegible] |
| SE | 55262 | ENSG00000135905 | DOCK10 | chr2 | - | 224852934 | 224853122 | 224852376 | 224852442 | 224854962 | 224855042 | 441 | 470 | [illegible] |
| SE | 55263 | ENSG00000135905 | DOCK10 | chr2 | - | 224856859 | 224856982 | 224854962 | 224855042 | 224862663 | 224862746 | 454 | 535 | [illegible] |
| SE | 55264 | ENSG00000135905 | DOCK10 | chr2 | - | 224862663 | 224862746 | 224856859 | 224856982 | 224864552 | 224864675 | 313 | 420 | [illegible] |
| SE | 55267 | ENSG00000135905 | DOCK10 | chr2 | - | 224886458 | 224886526 | 224886062 | 224886185 | 224896294 | 224896377 | 91 | 104 | [illegible] |
| SE | 55268 | ENSG00000135905 | DOCK10 | chr2 | - | 224886458 | 224886531 | 224886062 | 224886185 | 224896294 | 224896377 | 212 | 208 | [illegible] |
| SE | 55269 | ENSG00000135905 | DOCK10 | chr2 | - | 224916694 | 224916784 | 224896294 | 224896377 | 224931548 | 224931668 | 440 | 472 | [illegible] |
| SE | 55273 | ENSG00000260269 | NA | chr15 | - | 75597339 | 75597480 | 75527178 | 75527261 | 75598278 | 75598681 | 0 | 1 | [illegible] |
| SE | 55274 | ENSG00000260269 | NA | chr15 | - | 75598278 | 75598681 | 75527149 | 75527261 | 75601137 | 75601184 | 66 | 25 | [illegible] |
| SE | 55276 | ENSG00000242282 | NA | chr2 | - | 3533107 | 3533190 | 3531983 | 3532310 | 3534112 | 3534143 | 0 | 2 | [illegible] |
| SE | 55287 | ENSG00000178585 | CTNNBIP1 | chr1 | - | 9883704 | 9883738 | 9877904 | 9877989 | 9893008 | 9893237 | 1 | 0 | [illegible] |
| SE | 55288 | ENSG00000178585 | CTNNBIP1 | chr1 | - | 9883704 | 9883738 | 9877904 | 9877989 | 9910094 | 9910269 | 3 | 0 | [illegible] |
| SE | 55294 | ENSG00000162591 | MEGF6 | chr1 | - | 3493770 | 3493899 | 3492638 | 3492767 | 3493995 | 3494124 | 2 | 1 | [illegible] |
| SE | 55295 | ENSG00000162591 | MEGF6 | chr1 | - | 3496987 | 3497119 | 3496654 | 3496783 | 3497232 | 3497361 | 0 | 0 | [illegible] |
| SE | 55298 | ENSG00000132763 | MMACHC | chr1 | + | 45507355 | 45507550 | 45500299 | 45500413 | 45508211 | 45508364 | 7 | 10 | [illegible] |
| SE | 55301 | ENSG00000065457 | ADAT1 | chr16 | - | 75608842 | 75608988 | 75608223 | 75608323 | 75612242 | 75612861 | 56 | 48 | [illegible] |
| SE | 55302 | ENSG00000065457 | ADAT1 | chr16 | - | 75612242 | 75612861 | 75603084 | 75603171 | 75617141 | 75617272 | 69 | 65 | [illegible] |
| SE | 55303 | ENSG00000065457 | ADAT1 | chr16 | - | 75617141 | 75617272 | 75612242 | 75612861 | 75620265 | 75620334 | 65 | 66 | [illegible] |
| SE | 55306 | ENSG00000065457 | ADAT1 | chr16 | - | 75619631 | 75619743 | 75618585 | 75618640 | 75620265 | 75620334 | 0 | 0 | [illegible] |
| SE | 55308 | ENSG00000065457 | ADAT1 | chr16 | - | 75622402 | 75622464 | 75620630 | 75620820 | 75623237 | 75623275 | 38 | 15 | [illegible] |
| SE | 55313 | ENSG00000166938 | DIS3L | chr15 | + | 66314038 | 66314117 | 66308708 | 66308844 | 66315035 | 66315074 | 19 | 10 | [illegible] |
| SE | 55316 | ENSG00000166938 | DIS3L | chr15 | + | 66325830 | 66326364 | 66323492 | 66323585 | 66328969 | 66329124 | 138 | 77 | [illegible] |
| SE | 55317 | ENSG00000164111 | ANXA5 | chr4 | - | 121669601 | 121669724 | 121667945 | 121668527 | 121669953 | 121670012 | 1454 | 1291 | [illegible] |
| SE | 55318 | ENSG00000164111 | ANXA5 | chr4 | - | 121671546 | 121671642 | 121669953 | 121670012 | 121672532 | 121672558 | 1396 | 1155 | [illegible] |
| SE | 55324 | ENSG00000164111 | ANXA5 | chr4 | - | 121686287 | 121686372 | 121684676 | 121684771 | 121696580 | 121696624 | 417 | 409 | [illegible] |
| SE | 55329 | ENSG00000132635 | PCED1A | chr20 | - | 2838595 | 2838746 | 2838231 | 2838478 | 2839191 | 2839271 | 55 | 25 | [illegible] |
| SE | 55330 | ENSG00000132635 | PCED1A | chr20 | - | 2838843 | 2838929 | 2838595 | 2838746 | 2839191 | 2839271 | 59 | 25 | [illegible] |
| SE | 55331 | ENSG00000132635 | PCED1A | chr20 | - | 2838843 | 2839082 | 2838595 | 2838746 | 2839191 | 2839271 | 75 | 36 | [illegible] |
| SE | 55334 | ENSG00000204469 | PRRC2A | chr6 | + | 31622689 | 31622901 | 31620714 | 31620858 | 31623731 | 31623909 | 111 | 76 | [illegible] |
| SE | 55335 | ENSG00000204469 | PRRC2A | chr6 | + | 31622689 | 31622901 | 31620714 | 31620858 | 31624260 | 31624360 | 17 | 11 | [illegible] |
| SE | 55336 | ENSG00000204469 | PRRC2A | chr6 | + | 31622726 | 31622901 | 31620719 | 31620858 | 31623731 | 31623909 | 133 | 78 | [illegible] |
| SE | 55337 | ENSG00000204469 | PRRC2A | chr6 | + | 31622726 | 31622901 | 31620719 | 31620858 | 31624260 | 31624360 | 39 | 13 | [illegible] |
| SE | 55338 | ENSG00000204469 | PRRC2A | chr6 | + | 31623731 | 31623909 | 31620714 | 31620858 | 31624260 | 31624360 | 189 | 131 | [illegible] |
| SE | 55340 | ENSG00000204469 | PRRC2A | chr6 | + | 31624449 | 31624522 | 31624260 | 31624360 | 31625170 | 31625314 | 325 | 160 | [illegible] |
| SE | 55342 | ENSG00000204469 | PRRC2A | chr6 | + | 31626019 | 31626162 | 31625459 | 31625611 | 31626771 | 31626862 | 182 | 113 | [illegible] |
| SE | 55344 | ENSG00000204469 | PRRC2A | chr6 | + | 31629143 | 31629334 | 31627764 | 31628239 | 31629547 | 31629845 | 181 | 69 | [illegible] |
| SE | 55346 | ENSG00000204469 | PRRC2A | chr6 | + | 31634471 | 31634557 | 31634235 | 31634365 | 31634752 | 31634977 | 297 | 156 | [illegible] |
| SE | 55347 | ENSG00000204469 | PRRC2A | chr6 | + | 31634752 | 31634977 | 31634471 | 31634557 | 31635131 | 31635272 | 501 | 267 | [illegible] |
| SE | 55348 | ENSG00000204469 | PRRC2A | chr6 | + | 31635581 | 31635749 | 31635393 | 31635465 | 31635966 | 31636049 | 611 | 283 | [illegible] |
| SE | 55357 | ENSG00000131127 | ZNF141 | chr4 | + | 343781 | 343908 | 337813 | 337986 | 372663 | 373007 | 15 | 15 | [illegible] |
| SE | 55359 | ENSG00000131127 | ZNF141 | chr4 | + | 372663 | 373007 | 344334 | 344430 | 383094 | 383971 | 53 | 64 | [illegible] |
| SE | 55362 | ENSG00000149054 | ZNF215 | chr11 | + | 6943545 | 6943641 | 6941570 | 6941653 | 6955689 | 6955782 | 0 | 0 | [illegible] |
| SE | 55365 | ENSG00000159423 | ALDH4A1 | chr1 | - | 18881699 | 18881887 | 18879299 | 18879373 | 18883123 | 18883198 | 0 | 4 | [illegible] |
| SE | 55368 | ENSG00000169116 | PARM1 | chr4 | + | 75033882 | 75033961 | 75012424 | 75013150 | 75046162 | 75050113 | 1 | 5 | [illegible] |
| SE | 55369 | ENSG00000135241 | PNPLA8 | chr7 | - | 108479183 | 108479379 | 108470416 | 108472675 | 108487758 | 108487953 | 158 | 76 | [illegible] |
| SE | 55370 | ENSG00000135241 | PNPLA8 | chr7 | - | 108487758 | 108487953 | 108479183 | 108479379 | 108491409 | 108491467 | 191 | 148 | [illegible] |
| SE | 55371 | ENSG00000135241 | PNPLA8 | chr7 | - | 108487944 | 108487953 | 108479183 | 108479379 | 108491409 | 108491467 | 27 | 9 | [illegible] |
| SE | 55372 | ENSG00000135241 | PNPLA8 | chr7 | - | 108491409 | 108491467 | 108487758 | 108487953 | 108496583 | 108496755 | 149 | 130 | [illegible] |
| SE | 55373 | ENSG00000135241 | PNPLA8 | chr7 | - | 108496583 | 108496755 | 108491409 | 108491467 | 108497482 | 108497577 | 282 | 210 | [illegible] |
| SE | 55374 | ENSG00000135241 | PNPLA8 | chr7 | - | 108496583 | 108496755 | 108491409 | 108491467 | 108502490 | 108502642 | 117 | 98 | [illegible] |
| SE | 55375 | ENSG00000135241 | PNPLA8 | chr7 | - | 108497482 | 108497577 | 108491409 | 108491467 | 108502490 | 108502642 | 64 | 44 | [illegible] |
| SE | 55376 | ENSG00000135241 | PNPLA8 | chr7 | - | 108497482 | 108497577 | 108496426 | 108496755 | 108502490 | 108502642 | 226 | 155 | [illegible] |
| SE | 55377 | ENSG00000135241 | PNPLA8 | chr7 | - | 108502490 | 108502642 | 108497480 | 108497577 | 108514143 | 108514293 | 206 | 148 | [illegible] |
| SE | 55379 | ENSG00000135241 | PNPLA8 | chr7 | - | 108514435 | 108515447 | 108514143 | 108514293 | 108521475 | 108521521 | 239 | 161 | [illegible] |
| SE | 55381 | ENSG00000135241 | PNPLA8 | chr7 | - | 108514435 | 108515574 | 108514143 | 108514293 | 108521475 | 108521521 | 304 | 250 | [illegible] |
| SE | 55383 | ENSG00000135241 | PNPLA8 | chr7 | - | 108518189 | 108518301 | 108514435 | 108515447 | 108521475 | 108521521 | 4 | 0 | [illegible] |
| SE | 55384 | ENSG00000135241 | PNPLA8 | chr7 | - | 108518189 | 108518301 | 108514435 | 108515574 | 108521475 | 108521521 | 1 | 0 | [illegible] |
| SE | 55386 | ENSG00000135241 | PNPLA8 | chr7 | - | 108521475 | 108521521 | 108514435 | 108515447 | 108526028 | 108526135 | 12 | 12 | [illegible] |
| SE | 55387 | ENSG00000135241 | PNPLA8 | chr7 | - | 108521475 | 108521521 | 108514435 | 108515574 | 108526028 | 108526135 | 77 | 99 | [illegible] |
| SE | 55389 | ENSG00000130717 | UCK1 | chr9 | - | 131528943 | 131529038 | 131525928 | 131525977 | 131529127 | 131529270 | 46 | 31 | [illegible] |
| SE | 55391 | ENSG00000178607 | ERN1 | chr17 | - | 64049054 | 64049202 | 64047857 | 64047985 | 64052779 | 64052979 | 209 | 105 | [illegible] |
| SE | 55392 | ENSG00000178607 | ERN1 | chr17 | - | 64052779 | 64052979 | 64049054 | 64049202 | 64053271 | 64053371 | 152 | 65 | [illegible] |
| SE | 55393 | ENSG00000178607 | ERN1 | chr17 | - | 64053271 | 64053371 | 64052779 | 64052979 | 64054249 | 64054439 | 160 | 65 | [illegible] |
| SE | 55394 | ENSG00000178607 | ERN1 | chr17 | - | 64060468 | 64060587 | 64057801 | 64057993 | 64063985 | 64064151 | 194 | 81 | [illegible] |
| SE | 55395 | ENSG00000178607 | ERN1 | chr17 | - | 64065208 | 64065287 | 64063985 | 64064151 | 64071980 | 64072103 | 76 | 26 | [illegible] |
| SE | 55400 | ENSG00000178607 | ERN1 | chr17 | - | 64080774 | 64080808 | 64079661 | 64079734 | 64098120 | 64098241 | 136 | 49 | [illegible] |
| SE | 55401 | ENSG00000163041 | H3-3A | chr1 | + | 226064328 | 226064479 | 226062713 | 226062811 | 226065655 | 226065809 | 2791 | 1635 | [illegible] |
| SE | 55402 | ENSG00000163041 | H3-3A | chr1 | + | 226064328 | 226064479 | 226062713 | 226062811 | 226065734 | 226065809 | 742 | 373 | [illegible] |
| SE | 55403 | ENSG00000163041 | H3-3A | chr1 | + | 226064328 | 226064479 | 226062713 | 226062811 | 226071350 | 226071521 | 743 | 372 | [illegible] |
| SE | 55404 | ENSG00000163041 | H3-3A | chr1 | + | 226064328 | 226064479 | 226062749 | 226062948 | 226071350 | 226071521 | 8 | 4 | [illegible] |
| SE | 55405 | ENSG00000163041 | H3-3A | chr1 | + | 226065655 | 226065809 | 226062715 | 226062811 | 226071350 | 226071521 | 2548 | 1517 | [illegible] |
| SE | 55406 | ENSG00000163041 | H3-3A | chr1 | + | 226065655 | 226065809 | 226063976 | 226064479 | 226071350 | 226071521 | 4597 | 2780 | [illegible] |
| SE | 55408 | ENSG00000163041 | H3-3A | chr1 | + | 226065655 | 226065817 | 226064328 | 226064479 | 226071350 | 226071542 | 2053 | 1266 | [illegible] |
| SE | 55409 | ENSG00000163041 | H3-3A | chr1 | + | 226065734 | 226065809 | 226062782 | 226062811 | 226071350 | 226071846 | 1690 | 988 | [illegible] |
| SE | 55410 | ENSG00000163041 | H3-3A | chr1 | + | 226065734 | 226065809 | 226064328 | 226064479 | 226071350 | 226071846 | 1690 | 988 | [illegible] |
| SE | 55411 | ENSG00000185885 | IFITM1 | chr11 | + | 314642 | 314809 | 314039 | 314356 | 314921 | 315272 | 22 | 4 | [illegible] |
| SE | 55413 | ENSG00000158966 | CACHD1 | chr1 | + | 64582155 | 64582304 | 64550593 | 64550656 | 64602805 | 64602912 | 14 | 17 | [illegible] |
| SE | 55415 | ENSG00000158966 | CACHD1 | chr1 | + | 64632598 | 64632743 | 64629354 | 64629481 | 64634043 | 64634260 | 1 | 12 | [illegible] |
| SE | 55416 | ENSG00000158966 | CACHD1 | chr1 | + | 64647800 | 64648034 | 64641819 | 64641969 | 64652160 | 64652310 | 8 | 8 | [illegible] |
| SE | 55417 | ENSG00000070540 | WIPI1 | chr17 | - | 68428828 | 68428936 | 68427134 | 68427253 | 68429995 | 68430160 | 104 | 60 | [illegible] |
| SE | 55420 | ENSG00000070540 | WIPI1 | chr17 | - | 68429995 | 68430160 | 68428828 | 68428936 | 68433467 | 68433571 | 110 | 56 | [illegible] |
| SE | 55422 | ENSG00000070540 | WIPI1 | chr17 | - | 68452909 | 68452992 | 68450727 | 68450897 | 68457341 | 68457496 | 29 | 14 | [illegible] |
| SE | 55424 | ENSG00000116688 | MFN2 | chr1 | + | 11981809 | 11981853 | 11980298 | 11980484 | 11981973 | 11982114 | 2 | 1 | [illegible] |
| SE | 55432 | ENSG00000116688 | MFN2 | chr1 | + | 11982522 | 11982590 | 11981809 | 11982114 | 11996155 | 11996318 | 4 | 3 | [illegible] |
| SE | 55433 | ENSG00000116688 | MFN2 | chr1 | + | 11988426 | 11988516 | 11980215 | 11980484 | 11989164 | 11989343 | 2 | 0 | [illegible] |
| SE | 55434 | ENSG00000116688 | MFN2 | chr1 | + | 11988426 | 11988516 | 11981969 | 11982114 | 11989164 | 11989343 | 2 | 0 | [illegible] |
| SE | 55438 | ENSG00000116688 | MFN2 | chr1 | + | 11989164 | 11989343 | 11981809 | 11982114 | 11992554 | 11992594 | 377 | 283 | [illegible] |
| SE | 55439 | ENSG00000116688 | MFN2 | chr1 | + | 11992278 | 11992463 | 11989164 | 11989343 | 11992554 | 11992640 | 2 | 1 | [illegible] |
| SE | 55441 | ENSG00000116688 | MFN2 | chr1 | + | 11996155 | 11996318 | 11992251 | 11992690 | 11997296 | 11997421 | 141 | 140 | [illegible] |
| SE | 55444 | ENSG00000116688 | MFN2 | chr1 | + | 11998987 | 11999095 | 11998679 | 11998878 | 12001400 | 12001554 | 170 | 160 | [illegible] |
| SE | 55445 | ENSG00000116688 | MFN2 | chr1 | + | 12003757 | 12003889 | 12001981 | 12002103 | 12003991 | 12004118 | 2 | 12 | [illegible] |
| SE | 55446 | ENSG00000116688 | MFN2 | chr1 | + | 12004824 | 12004927 | 12004508 | 12004613 | 12005710 | 12005931 | 225 | 170 | [illegible] |
| SE | 55447 | ENSG00000116688 | MFN2 | chr1 | + | 12006537 | 12006693 | 12005710 | 12005931 | 12007052 | 12007249 | 145 | 105 | [illegible] |
| SE | 55457 | ENSG00000254138 | NA | chr5 | - | 31095610 | 31095641 | 31093976 | 31094059 | 31115546 | 31115581 | 0 | 2 | [illegible] |
| SE | 55462 | ENSG00000100612 | DHRS7 | chr14 | - | 60149352 | 60149568 | 60144118 | 60145013 | 60152938 | 60153178 | 810 | 493 | [illegible] |
| SE | 55464 | ENSG00000100612 | DHRS7 | chr14 | - | 60150064 | 60150117 | 60149458 | 60149568 | 60152938 | 60153178 | 375 | 198 | [illegible] |
| SE | 55465 | ENSG00000100612 | DHRS7 | chr14 | - | 60150064 | 60150187 | 60148947 | 60149568 | 60152938 | 60153174 | 1318 | 693 | [illegible] |
| SE | 55466 | ENSG00000100612 | DHRS7 | chr14 | - | 60152938 | 60153178 | 60148947 | 60149568 | 60153958 | 60154005 | 466 | 182 | [illegible] |
| SE | 55468 | ENSG00000100612 | DHRS7 | chr14 | - | 60152938 | 60153178 | 60150064 | 60150187 | 60155999 | 60156152 | 499 | 213 | [illegible] |
| SE | 55469 | ENSG00000100612 | DHRS7 | chr14 | - | 60153958 | 60154065 | 60150099 | 60150187 | 60155999 | 60156152 | 163 | 91 | [illegible] |
| SE | 55470 | ENSG00000100612 | DHRS7 | chr14 | - | 60153958 | 60154065 | 60152688 | 60153178 | 60155999 | 60156152 | 612 | 265 | [illegible] |
| SE | 55473 | ENSG00000114948 | ADAM23 | chr2 | + | 206562202 | 206562293 | 206561127 | 206561212 | 206565019 | 206565068 | 6 | 3 | [illegible] |
| SE | 55474 | ENSG00000114948 | ADAM23 | chr2 | + | 206605789 | 206605880 | 206596050 | 206596162 | 206617578 | 206617842 | 1 | 1 | [illegible] |
| SE | 55475 | ENSG00000114948 | ADAM23 | chr2 | + | 206609909 | 206610000 | 206596050 | 206596162 | 206617578 | 206621127 | 0 | 1 | [illegible] |
| SE | 55479 | ENSG00000121964 | GTDC1 | chr2 | - | 143952772 | 143952843 | 143951981 | 143952073 | 143957194 | 143957339 | 72 | 38 | [illegible] |
| SE | 55480 | ENSG00000121964 | GTDC1 | chr2 | - | 143952772 | 143952843 | 143951981 | 143952073 | 143970652 | 143970762 | 26 | 15 | [illegible] |
| SE | 55481 | ENSG00000121964 | GTDC1 | chr2 | - | 143952772 | 143952843 | 143951981 | 143952073 | 144007181 | 144007535 | 21 | 12 | [illegible] |
| SE | 55482 | ENSG00000121964 | GTDC1 | chr2 | - | 143957194 | 143957339 | 143951981 | 143952073 | 143970652 | 143970762 | 26 | 18 | [illegible] |
| SE | 55483 | ENSG00000121964 | GTDC1 | chr2 | - | 143957194 | 143957339 | 143951981 | 143952073 | 144007181 | 144007535 | 0 | 0 | [illegible] |
| SE | 55485 | ENSG00000121964 | GTDC1 | chr2 | - | 143957194 | 143957339 | 143952772 | 143952843 | 144007181 | 144007535 | 53 | 30 | [illegible] |
| SE | 55486 | ENSG00000121964 | GTDC1 | chr2 | - | 143970652 | 143970762 | 143951981 | 143952073 | 144007181 | 144007535 | 17 | 5 | [illegible] |
| SE | 55488 | ENSG00000121964 | GTDC1 | chr2 | - | 143970652 | 143970762 | 143957194 | 143957339 | 144007181 | 144007535 | 43 | 21 | [illegible] |
| SE | 55489 | ENSG00000121964 | GTDC1 | chr2 | - | 143970652 | 143970762 | 143957194 | 143957339 | 144141881 | 144142055 | 24 | 10 | [illegible] |
| SE | 55490 | ENSG00000121964 | GTDC1 | chr2 | - | 144007181 | 144007535 | 143957194 | 143957339 | 144141881 | 144142055 | 59 | 49 | [illegible] |
| SE | 55492 | ENSG00000121964 | GTDC1 | chr2 | - | 144145571 | 144145739 | 144141881 | 144142055 | 144208602 | 144208804 | 212 | 116 | [illegible] |
| SE | 55493 | ENSG00000121964 | GTDC1 | chr2 | - | 144177134 | 144177240 | 144145571 | 144145739 | 144178834 | 144178960 | 3 | 7 | [illegible] |
| SE | 55497 | ENSG00000121964 | GTDC1 | chr2 | - | 144235675 | 144235737 | 144211514 | 144211579 | 144317478 | 144317588 | 70 | 50 | [illegible] |
| SE | 55498 | ENSG00000121964 | GTDC1 | chr2 | - | 144235675 | 144235737 | 144211514 | 144211579 | 144332438 | 144332512 | 74 | 55 | [illegible] |
| SE | 55499 | ENSG00000121964 | GTDC1 | chr2 | - | 144317478 | 144317588 | 144235675 | 144235737 | 144319718 | 144319870 | 23 | 15 | [illegible] |
| SE | 55501 | ENSG00000124181 | PLCG1 | chr20 | + | 41159869 | 41159963 | 41159605 | 41159758 | 41160105 | 41160153 | 185 | 111 | [illegible] |
| SE | 55502 | ENSG00000124181 | PLCG1 | chr20 | + | 41162957 | 41162992 | 41162641 | 41162725 | 41163202 | 41163275 | 54 | 18 | [illegible] |
| SE | 55507 | ENSG00000124181 | PLCG1 | chr20 | + | 41174123 | 41174311 | 41173922 | 41174011 | 41182930 | 41182989 | 65 | 26 | [illegible] |
| SE | 55509 | ENSG00000124181 | PLCG1 | chr20 | + | 41174466 | 41174539 | 41173922 | 41174011 | 41182930 | 41182960 | 0 | 1 | [illegible] |
| SE | 55511 | ENSG00000124181 | PLCG1 | chr20 | + | 41174466 | 41174967 | 41173922 | 41174011 | 41175752 | 41175820 | 1 | 2 | [illegible] |
| SE | 55513 | ENSG00000124181 | PLCG1 | chr20 | + | 41174466 | 41174967 | 41173922 | 41174011 | 41182930 | 41182960 | 2 | 2 | [illegible] |
| SE | 55519 | ENSG00000124181 | PLCG1 | chr20 | + | 41182930 | 41182988 | 41180455 | 41180664 | 41188432 | 41188552 | 0 | 0 | [illegible] |
| SE | 55521 | ENSG00000010818 | HIVEP2 | chr6 | - | 142764798 | 142764974 | 142761463 | 142761565 | 142768381 | 142768536 | 430 | 460 | [illegible] |
| SE | 55522 | ENSG00000010818 | HIVEP2 | chr6 | - | 142768381 | 142768536 | 142764798 | 142764974 | 142776146 | 142776191 | 173 | 155 | [illegible] |
| SE | 55523 | ENSG00000010818 | HIVEP2 | chr6 | - | 142769551 | 142775125 | 142768381 | 142768536 | 142776146 | 142776191 | 368 | 330 | [illegible] |
| SE | 55524 | ENSG00000010818 | HIVEP2 | chr6 | - | 142783520 | 142783615 | 142776146 | 142776191 | 142836934 | 142837047 | 172 | 181 | [illegible] |
| SE | 55525 | ENSG00000010818 | HIVEP2 | chr6 | - | 142836934 | 142837047 | 142783520 | 142783615 | 142945098 | 142945176 | 71 | 67 | [illegible] |
| SE | 55526 | ENSG00000188986 | NELFB | chr9 | + | 137256823 | 137257054 | 137256328 | 137256428 | 137263036 | 137263222 | 106 | 42 | [illegible] |
| SE | 55527 | ENSG00000082014 | SMARCD3 | chr7 | - | 151242704 | 151242843 | 151242480 | 151242603 | 151243658 | 151243701 | 29 | 18 | [illegible] |
| SE | 55528 | ENSG00000082014 | SMARCD3 | chr7 | - | 151242720 | 151242843 | 151242136 | 151242603 | 151243658 | 151243686 | 83 | 49 | [illegible] |
| SE | 55529 | ENSG00000082014 | SMARCD3 | chr7 | - | 151243658 | 151243701 | 151242480 | 151242603 | 151275113 | 151275252 | 0 | 0 | [illegible] |
| SE | 55530 | ENSG00000082014 | SMARCD3 | chr7 | - | 151245459 | 151245671 | 151243658 | 151243701 | 151275113 | 151275251 | 20 | 11 | [illegible] |
| SE | 55531 | ENSG00000107282 | APBA1 | chr9 | - | 69452121 | 69452301 | 69449583 | 69449796 | 69456246 | 69456432 | 2 | 2 | [illegible] |
| SE | 55533 | ENSG00000107282 | APBA1 | chr9 | - | 69458155 | 69458188 | 69457052 | 69457139 | 69467822 | 69467968 | 2 | 1 | [illegible] |
| SE | 55534 | ENSG00000107282 | APBA1 | chr9 | - | 69476047 | 69476143 | 69471655 | 69471695 | 69516010 | 69517279 | 1 | 1 | [illegible] |
| SE | 55535 | ENSG00000287514 | NA | chr9 | + | 36781469 | 36781893 | 36779474 | 36779877 | 36793993 | 36796516 | 1 | 2 | [illegible] |
| SE | 55536 | ENSG00000287514 | NA | chr9 | + | 36781469 | 36781897 | 36779640 | 36779877 | 36793993 | 36796516 | 1 | 1 | [illegible] |
| SE | 55546 | ENSG00000117010 | ZNF684 | chr1 | + | 40540585 | 40540712 | 40533142 | 40533181 | 40541614 | 40541710 | 24 | 13 | [illegible] |
| SE | 55547 | ENSG00000117010 | ZNF684 | chr1 | + | 40541614 | 40541710 | 40533142 | 40533181 | 40546561 | 40546765 | 9 | 0 | [illegible] |
| SE | 55549 | ENSG00000185379 | RAD51D | chr17 | - | 35106987 | 35107061 | 35106385 | 35106481 | 35107365 | 35107447 | 4 | 5 | [illegible] |
| SE | 55551 | ENSG00000185379 | RAD51D | chr17 | - | 35106987 | 35107122 | 35106185 | 35106481 | 35118500 | 35118619 | 11 | 7 | [illegible] |
| SE | 55553 | ENSG00000185379 | RAD51D | chr17 | - | 35107365 | 35107447 | 35106127 | 35106481 | 35118500 | 35118619 | 3 | 4 | [illegible] |
| SE | 55555 | ENSG00000185379 | RAD51D | chr17 | - | 35107365 | 35107447 | 35106987 | 35107122 | 35118500 | 35118619 | 13 | 9 | [illegible] |
| SE | 55556 | ENSG00000185379 | RAD51D | chr17 | - | 35116858 | 35117037 | 35106440 | 35106481 | 35118500 | 35118619 | 0 | 0 | [illegible] |
| SE | 55557 | ENSG00000185379 | RAD51D | chr17 | - | 35116858 | 35117037 | 35106440 | 35106481 | 35119110 | 35119172 | 0 | 2 | [illegible] |
| SE | 55558 | ENSG00000185379 | RAD51D | chr17 | - | 35116858 | 35117037 | 35106987 | 35107122 | 35118500 | 35118619 | 0 | 0 | [illegible] |
| SE | 55559 | ENSG00000185379 | RAD51D | chr17 | - | 35116858 | 35117037 | 35106987 | 35107122 | 35119110 | 35119172 | 0 | 2 | [illegible] |
| SE | 55561 | ENSG00000185379 | RAD51D | chr17 | - | 35116858 | 35117037 | 35107365 | 35107447 | 35119110 | 35119172 | 3 | 2 | [illegible] |
| SE | 55562 | ENSG00000185379 | RAD51D | chr17 | - | 35118500 | 35118619 | 35106440 | 35106481 | 35119110 | 35119157 | 9 | 3 | [illegible] |
| SE | 55563 | ENSG00000185379 | RAD51D | chr17 | - | 35118500 | 35118619 | 35106987 | 35107122 | 35119110 | 35119157 | 9 | 3 | [illegible] |
| SE | 55565 | ENSG00000185379 | RAD51D | chr17 | - | 35118500 | 35118619 | 35116858 | 35117037 | 35119110 | 35119157 | 9 | 3 | [illegible] |
| SE | 55572 | ENSG00000081692 | JMJD4 | chr1 | - | 227732880 | 227733027 | 227731190 | 227732676 | 227733413 | 227733681 | 29 | 8 | [illegible] |
| SE | 55573 | ENSG00000081692 | JMJD4 | chr1 | - | 227732928 | 227733027 | 227731233 | 227732676 | 227733413 | 227733681 | 3 | 1 | [illegible] |
| SE | 55574 | ENSG00000081692 | JMJD4 | chr1 | - | 227733413 | 227733681 | 227731190 | 227732676 | 227733906 | 227734032 | 33 | 11 | [illegible] |
| SE | 55577 | ENSG00000137275 | RIPK1 | chr6 | + | 3076763 | 3076987 | 3063823 | 3064059 | 3077778 | 3077935 | 203 | 92 | [illegible] |
| SE | 55578 | ENSG00000137275 | RIPK1 | chr6 | + | 3076763 | 3076987 | 3068493 | 3068661 | 3077778 | 3077935 | 224 | 101 | [illegible] |
| SE | 55579 | ENSG00000137275 | RIPK1 | chr6 | + | 3077778 | 3077935 | 3063990 | 3064059 | 3083084 | 3083313 | 1 | 2 | [illegible] |
| SE | 55580 | ENSG00000137275 | RIPK1 | chr6 | + | 3077778 | 3077935 | 3076763 | 3076987 | 3080978 | 3081116 | 335 | 146 | [illegible] |
| SE | 55581 | ENSG00000137275 | RIPK1 | chr6 | + | 3080978 | 3081116 | 3077778 | 3077935 | 3083084 | 3083313 | 267 | 128 | [illegible] |
| SE | 55587 | ENSG00000137275 | RIPK1 | chr6 | + | 3083084 | 3083313 | 3080978 | 3081116 | 3104224 | 3104315 | 144 | 78 | [illegible] |
| SE | 55589 | ENSG00000137275 | RIPK1 | chr6 | + | 3085258 | 3085408 | 3080978 | 3081116 | 3104224 | 3104315 | 11 | 8 | [illegible] |
| SE | 55591 | ENSG00000137275 | RIPK1 | chr6 | + | 3085258 | 3085408 | 3082546 | 3083313 | 3104224 | 3104315 | 62 | 31 | [illegible] |
| SE | 55592 | ENSG00000137275 | RIPK1 | chr6 | + | 3089580 | 3089657 | 3082546 | 3083313 | 3104224 | 3104315 | 55 | 41 | [illegible] |
| SE | 55595 | ENSG00000137275 | RIPK1 | chr6 | + | 3110802 | 3110955 | 3105481 | 3106051 | 3113052 | 3115150 | 293 | 167 | [illegible] |
| SE | 55596 | ENSG00000089685 | BIRC5 | chr17 | + | 78215965 | 78216034 | 78214679 | 78214789 | 78216663 | 78216781 | 0 | 0 | [illegible] |
| SE | 55603 | ENSG00000113070 | HBEGF | chr5 | - | 140334657 | 140334748 | 140332842 | 140334280 | 140335871 | 140336027 | 59 | 47 | [illegible] |
| SE | 55605 | ENSG00000113070 | HBEGF | chr5 | - | 140345910 | 140346084 | 140342634 | 140342812 | 140346282 | 140346603 | 37 | 12 | [illegible] |
| SE | 55608 | ENSG00000144021 | CIAO1 | chr2 | + | 96268456 | 96268658 | 96267835 | 96267924 | 96271110 | 96274173 | 218 | 118 | [illegible] |
| SE | 55609 | ENSG00000144021 | CIAO1 | chr2 | + | 96268456 | 96269355 | 96267835 | 96267924 | 96271110 | 96273349 | 357 | 200 | [illegible] |
| SE | 55611 | ENSG00000213366 | GSTM2 | chr1 | + | 109668924 | 109668989 | 109668424 | 109668500 | 109669289 | 109669371 | 63 | 44 | [illegible] |
| SE | 55613 | ENSG00000149474 | KAT14 | chr20 | + | 18150820 | 18150942 | 18145098 | 18145351 | 18159083 | 18159265 | 94 | 61 | [illegible] |
| SE | 55614 | ENSG00000149474 | KAT14 | chr20 | + | 18159083 | 18159265 | 18150820 | 18150942 | 18161819 | 18162239 | 117 | 81 | [illegible] |
| SE | 55619 | ENSG00000184860 | SDR42E1 | chr16 | - | 82000300 | 82000417 | 81999698 | 82000224 | 82000790 | 82000884 | 2 | 5 | [illegible] |
| SE | 55620 | ENSG00000184860 | SDR42E1 | chr16 | - | 82007640 | 82007868 | 82000790 | 82000884 | 82011386 | 82011465 | 1 | 5 | [illegible] |
| SE | 55622 | ENSG00000184368 | MAP7D2 | chrX | - | 20050823 | 20050946 | 20044363 | 20044524 | 20052877 | 20052988 | 1 | 0 | [illegible] |
| SE | 55624 | ENSG00000115590 | IL1R2 | chr2 | + | 102008514 | 102008642 | 101991959 | 101992011 | 102009561 | 102009826 | 3 | 4 | [illegible] |
| SE | 55625 | ENSG00000115590 | IL1R2 | chr2 | + | 102009561 | 102009826 | 101991959 | 101992011 | 102015870 | 102016051 | 2 | 0 | [illegible] |
| SE | 55626 | ENSG00000115590 | IL1R2 | chr2 | + | 102015870 | 102016051 | 102009561 | 102009826 | 102019637 | 102019812 | 10 | 12 | [illegible] |
| SE | 55627 | ENSG00000115590 | IL1R2 | chr2 | + | 102022186 | 102022249 | 102019637 | 102019812 | 102024532 | 102024668 | 8 | 11 | [illegible] |
| SE | 55628 | ENSG00000115590 | IL1R2 | chr2 | + | 102022186 | 102022249 | 102019637 | 102019812 | 102026110 | 102026253 | 4 | 8 | [illegible] |
| SE | 55629 | ENSG00000115590 | IL1R2 | chr2 | + | 102024532 | 102024668 | 102019637 | 102019812 | 102026110 | 102026253 | 0 | 2 | [illegible] |
| SE | 55631 | ENSG00000115590 | IL1R2 | chr2 | + | 102026110 | 102026253 | 102022062 | 102022249 | 102028225 | 102028414 | 5 | 2 | [illegible] |
| SE | 55632 | ENSG00000117410 | ATP6V0B | chr1 | + | 43975664 | 43975848 | 43974959 | 43975107 | 43976089 | 43976173 | 12 | 0 | [illegible] |
| SE | 55633 | ENSG00000117410 | ATP6V0B | chr1 | + | 43975799 | 43975848 | 43974943 | 43975107 | 43976089 | 43976173 | 30 | 4 | [illegible] |
| SE | 55635 | ENSG00000117410 | ATP6V0B | chr1 | + | 43977025 | 43977216 | 43976772 | 43976824 | 43977980 | 43978023 | 357 | 75 | [illegible] |
| SE | 55637 | ENSG00000121481 | RNF2 | chr1 | + | 185098071 | 185098344 | 185091578 | 185091739 | 185099790 | 185099962 | 112 | 51 | [illegible] |
| SE | 55638 | ENSG00000121481 | RNF2 | chr1 | + | 185099790 | 185099962 | 185098071 | 185098344 | 185100199 | 185101057 | 192 | 125 | [illegible] |
| SE | 55639 | ENSG00000114491 | UMPS | chr3 | + | 124731505 | 124731604 | 124730444 | 124730627 | 124735092 | 124735246 | 0 | 0 | [illegible] |
| SE | 55640 | ENSG00000114491 | UMPS | chr3 | + | 124731505 | 124731604 | 124730444 | 124730627 | 124737567 | 124738239 | 0 | 0 | [illegible] |
| SE | 55643 | ENSG00000114491 | UMPS | chr3 | + | 124740023 | 124740199 | 124737567 | 124738239 | 124742151 | 124742266 | 149 | 86 | [illegible] |
| SE | 55645 | ENSG00000114491 | UMPS | chr3 | + | 124740023 | 124740199 | 124738624 | 124738735 | 124742151 | 124742266 | 62 | 41 | [illegible] |
| SE | 55648 | ENSG00000163463 | KRTCAP2 | chr1 | - | 155169786 | 155169857 | 155169407 | 155169560 | 155172564 | 155172628 | 88 | 24 | [illegible] |

| | | | | | | | | | | | | | | |
|---|---|---|---|---|---|---|---|---|---|---|---|---|---|---|
| SE | 55649 | ENSG00000163463 | KRTCAP2 | chr1 | - | 155169790 | 155172489 | 155169408 | 155169560 | 155172564 | 155172892 | 42 | 21 | [illegible] |
| SE | 55651 | ENSG00000163463 | KRTCAP2 | chr1 | - | 155172564 | 155172892 | 155169790 | 155169857 | 155173220 | 155173251 | 162 | 45 | [illegible] |
| SE | 55653 | ENSG00000163463 | KRTCAP2 | chr1 | - | 155172737 | 155172892 | 155172564 | 155172628 | 155173316 | 155173475 | 138 | 48 | [illegible] |
| SE | 55654 | ENSG00000138119 | MYOF | chr10 | - | 93313019 | 93313210 | 93310533 | 93310643 | 93316713 | 93316813 | 181 | 253 | [illegible] |
| SE | 55656 | ENSG00000138119 | MYOF | chr10 | - | 93341912 | 93341969 | 93340152 | 93340164 | 93343855 | 93343932 | 0 | 0 | [illegible] |
| SE | 55658 | ENSG00000138119 | MYOF | chr10 | - | 93351196 | 93351295 | 93349807 | 93349969 | 93351412 | 93351571 | 78 | 97 | [illegible] |
| SE | 55659 | ENSG00000138119 | MYOF | chr10 | - | 93361451 | 93361557 | 93359832 | 93359978 | 93363960 | 93364075 | 73 | 83 | [illegible] |
| SE | 55660 | ENSG00000138119 | MYOF | chr10 | - | 93377322 | 93377429 | 93374762 | 93374955 | 93379862 | 93379987 | 77 | 97 | [illegible] |
| SE | 55661 | ENSG00000138119 | MYOF | chr10 | - | 93377322 | 93377429 | 93374762 | 93374955 | 93381218 | 93381396 | 28 | 30 | [illegible] |
| SE | 55662 | ENSG00000138119 | MYOF | chr10 | - | 93379862 | 93379987 | 93374762 | 93374955 | 93381218 | 93381396 | 51 | 58 | [illegible] |
| SE | 55664 | ENSG00000138119 | MYOF | chr10 | - | 93392916 | 93392955 | 93389029 | 93389154 | 93396141 | 93396224 | 95 | 112 | [illegible] |
| SE | 55666 | ENSG00000138119 | MYOF | chr10 | - | 93409572 | 93409739 | 93408786 | 93408915 | 93431407 | 93431516 | 32 | 51 | [illegible] |
| SE | 55667 | ENSG00000138119 | MYOF | chr10 | - | 93426070 | 93426158 | 93408786 | 93408915 | 93431407 | 93431516 | 23 | 19 | [illegible] |
| SE | 55668 | ENSG00000138119 | MYOF | chr10 | - | 93426070 | 93426158 | 93409572 | 93409739 | 93431407 | 93431516 | 60 | 72 | [illegible] |
| SE | 55669 | ENSG00000138119 | MYOF | chr10 | - | 93478179 | 93478373 | 93456881 | 93456937 | 93482106 | 93482194 | 1 | 0 | [illegible] |
| SE | 55672 | ENSG00000243406 | MRPS31P5 | chr13 | - | 52186682 | 52186823 | 52180581 | 52180654 | 52189638 | 52189914 | 38 | 46 | [illegible] |
| SE | 55673 | ENSG00000243406 | MRPS31P5 | chr13 | - | 52193110 | 52193233 | 52189632 | 52189914 | 52194253 | 52194412 | 0 | 0 | [illegible] |
| SE | 55677 | ENSG00000272886 | DCP1A | chr3 | - | 53344901 | 53344936 | 53319406 | 53319473 | 53347382 | 53347524 | 2 | 0 | [illegible] |
| SE | 55678 | ENSG00000272886 | DCP1A | chr3 | - | 53344901 | 53344942 | 53319406 | 53319473 | 53347382 | 53347525 | 5 | 1 | [illegible] |
| SE | 55684 | ENSG00000215883 | CYB5RL | chr1 | - | 54186180 | 54186286 | 54184160 | 54184265 | 54187651 | 54187739 | 1 | 0 | [illegible] |
| SE | 55686 | ENSG00000256407 | NA | chr1 | - | 54187651 | 54187739 | 54184160 | 54184265 | 54195418 | 54195715 | 2 | 2 | [illegible] |
| SE | 55687 | ENSG00000215883 | CYB5RL | chr1 | - | 54190737 | 54190896 | 54184160 | 54184265 | 54195418 | 54195436 | 1 | 3 | [illegible] |
| SE | 55688 | ENSG00000215883 | CYB5RL | chr1 | - | 54190737 | 54190896 | 54187651 | 54187739 | 54195418 | 54195436 | 3 | 4 | [illegible] |
| SE | 55689 | ENSG00000215883 | CYB5RL | chr1 | - | 54190747 | 54190896 | 54187651 | 54187739 | 54195418 | 54195681 | 1 | 3 | [illegible] |
| SE | 55690 | ENSG00000135749 | PCNX2 | chr1 | - | 233016920 | 233017154 | 233014664 | 233014777 | 233054267 | 233054483 | 18 | 14 | [illegible] |
| SE | 55694 | ENSG00000135749 | PCNX2 | chr1 | - | 233112844 | 233112966 | 233065460 | 233065661 | 233135012 | 233135190 | 12 | 12 | [illegible] |
| SE | 55697 | ENSG00000135749 | PCNX2 | chr1 | - | 233161270 | 233161363 | 233139713 | 233139855 | 233177801 | 233177835 | 8 | 4 | [illegible] |
| SE | 55698 | ENSG00000135749 | PCNX2 | chr1 | - | 233200153 | 233200264 | 233198938 | 233199030 | 233208517 | 233208610 | 76 | 34 | [illegible] |
| SE | 55699 | ENSG00000135749 | PCNX2 | chr1 | - | 233208517 | 233208610 | 233200153 | 233200264 | 233217898 | 233217931 | 26 | 14 | [illegible] |
| SE | 55700 | ENSG00000135749 | PCNX2 | chr1 | - | 233208517 | 233208689 | 233200153 | 233200264 | 233217898 | 233217931 | 59 | 33 | [illegible] |
| SE | 55701 | ENSG00000135749 | PCNX2 | chr1 | - | 233208517 | 233208689 | 233200153 | 233200264 | 233227225 | 233227269 | 33 | 19 | [illegible] |
| SE | 55702 | ENSG00000135749 | PCNX2 | chr1 | - | 233217898 | 233217931 | 233208517 | 233208689 | 233218030 | 233218087 | 68 | 40 | [illegible] |
| SE | 55703 | ENSG00000135749 | PCNX2 | chr1 | - | 233217898 | 233218184 | 233208658 | 233208689 | 233227225 | 233227347 | 85 | 61 | [illegible] |
| SE | 55704 | ENSG00000135749 | PCNX2 | chr1 | - | 233218030 | 233218184 | 233208517 | 233208689 | 233227225 | 233227269 | 82 | 56 | [illegible] |
| SE | 55705 | ENSG00000135749 | PCNX2 | chr1 | - | 233218030 | 233218184 | 233208517 | 233208689 | 233236844 | 233236980 | 7 | 7 | [illegible] |
| SE | 55706 | ENSG00000135749 | PCNX2 | chr1 | - | 233227225 | 233227371 | 233208517 | 233208689 | 233236844 | 233236980 | 41 | 48 | [illegible] |
| SE | 55716 | ENSG00000188199 | NUTM2B | chr10 | + | 79709799 | 79709939 | 79708698 | 79708827 | 79710381 | 79710764 | 0 | 0 | [illegible] |
| SE | 55723 | ENSG00000225470 | JPX | chrX | + | 73944583 | 73944662 | 73944183 | 73944465 | 73994839 | 73994961 | 41 | 22 | [illegible] |
| SE | 55726 | ENSG00000225470 | JPX | chrX | + | 73946998 | 73947096 | 73944583 | 73944662 | 73998767 | 74000496 | 8 | 3 | [illegible] |
| SE | 55727 | ENSG00000225470 | JPX | chrX | + | 73946998 | 73947374 | 73944326 | 73944662 | 73962730 | 73962801 | 12 | 3 | [illegible] |
| SE | 55729 | ENSG00000225470 | JPX | chrX | + | 73946998 | 73947374 | 73944326 | 73944662 | 73994839 | 73994961 | 14 | 3 | [illegible] |
| SE | 55730 | ENSG00000225470 | JPX | chrX | + | 73946998 | 73947374 | 73944326 | 73944662 | 73998767 | 73998844 | 13 | 3 | [illegible] |
| SE | 55732 | ENSG00000225470 | JPX | chrX | + | 73946998 | 73947374 | 73944326 | 73944662 | 74004269 | 74004542 | 12 | 3 | [illegible] |
| SE | 55733 | ENSG00000225470 | JPX | chrX | + | 73962730 | 73962801 | 73944583 | 73944662 | 73994839 | 73994961 | 2 | 1 | [illegible] |
| SE | 55734 | ENSG00000225470 | JPX | chrX | + | 73962730 | 73962801 | 73944583 | 73944662 | 73998767 | 73998844 | 1 | 0 | [illegible] |
| SE | 55735 | ENSG00000225470 | JPX | chrX | + | 73962730 | 73962801 | 73946998 | 73947374 | 73994839 | 73994961 | 1 | 1 | [illegible] |
| SE | 55737 | ENSG00000225470 | JPX | chrX | + | 73962734 | 73962801 | 73944583 | 73944662 | 73994839 | 73994961 | 1 | 1 | [illegible] |
| SE | 55739 | ENSG00000225470 | JPX | chrX | + | 73962734 | 73962801 | 73944583 | 73944662 | 73998767 | 73998844 | 0 | 0 | [illegible] |
| SE | 55741 | ENSG00000225470 | JPX | chrX | + | 73962734 | 73962801 | 73946998 | 73947374 | 73994839 | 73994961 | 1 | 1 | [illegible] |
| SE | 55745 | ENSG00000225470 | JPX | chrX | + | 73984443 | 73984541 | 73944583 | 73944662 | 73994839 | 73994961 | 1 | 0 | [illegible] |
| SE | 55746 | ENSG00000225470 | JPX | chrX | + | 73984443 | 73984541 | 73944583 | 73944662 | 73998767 | 73998844 | 1 | 0 | [illegible] |
| SE | 55747 | ENSG00000225470 | JPX | chrX | + | 73994839 | 73994961 | 73944181 | 73944662 | 73998767 | 73998844 | 28 | 16 | [illegible] |
| SE | 55750 | ENSG00000225470 | JPX | chrX | + | 73994839 | 73994961 | 73944181 | 73944662 | 74004269 | 74007684 | 24 | 20 | [illegible] |
| SE | 55751 | ENSG00000225470 | JPX | chrX | + | 73994839 | 73994961 | 73944181 | 73944662 | 74062035 | 74062148 | 18 | 14 | [illegible] |
| SE | 55752 | ENSG00000225470 | JPX | chrX | + | 73994839 | 73994961 | 73944181 | 73944662 | 74069168 | 74070408 | 18 | 14 | [illegible] |
| SE | 55754 | ENSG00000225470 | JPX | chrX | + | 73994839 | 73994961 | 73946998 | 73947374 | 73998767 | 73998844 | 9 | 2 | [illegible] |
| SE | 55757 | ENSG00000225470 | JPX | chrX | + | 73994839 | 73994961 | 73962730 | 73962801 | 73998767 | 73998844 | 8 | 3 | [illegible] |
| SE | 55759 | ENSG00000225470 | JPX | chrX | + | 73994839 | 73994961 | 73984443 | 73984541 | 73998767 | 73998844 | 8 | 2 | [illegible] |
| SE | 55760 | ENSG00000225470 | JPX | chrX | + | 73994863 | 73994961 | 73944583 | 73944662 | 73998767 | 73998844 | 7 | 1 | [illegible] |
| SE | 55761 | ENSG00000225470 | JPX | chrX | + | 73994863 | 73994961 | 73962734 | 73962801 | 73998767 | 73998844 | 7 | 1 | [illegible] |
| SE | 55762 | ENSG00000225470 | JPX | chrX | + | 73997040 | 73997166 | 73994839 | 73994961 | 73998767 | 73998844 | 3 | 0 | [illegible] |
| SE | 55763 | ENSG00000225470 | JPX | chrX | + | 73997043 | 73997166 | 73994839 | 73994961 | 73998767 | 73998844 | 4 | 1 | [illegible] |
| SE | 55764 | ENSG00000225470 | JPX | chrX | + | 73997043 | 73997166 | 73994839 | 73994961 | 74004269 | 74009948 | 2 | 1 | [illegible] |
| SE | 55767 | ENSG00000225470 | JPX | chrX | + | 73998767 | 73998844 | 73944341 | 73944662 | 74004269 | 74004529 | 17 | 15 | [illegible] |
| SE | 55771 | ENSG00000225470 | JPX | chrX | + | 73998767 | 73998844 | 73994557 | 73994961 | 73998953 | 73999051 | 16 | 8 | [illegible] |
| SE | 55772 | ENSG00000225470 | JPX | chrX | + | 73998767 | 73998844 | 73994557 | 73994961 | 73999354 | 73999810 | 5 | 4 | [illegible] |
| SE | 55774 | ENSG00000225470 | JPX | chrX | + | 73998767 | 73998844 | 73994557 | 73994961 | 74004269 | 74004529 | 25 | 17 | [illegible] |
| SE | 55775 | ENSG00000225470 | JPX | chrX | + | 73998767 | 73998844 | 73997040 | 73997166 | 74004269 | 74004529 | 17 | 14 | [illegible] |
| SE | 55776 | ENSG00000225470 | JPX | chrX | + | 73998767 | 74000496 | 73994839 | 73994961 | 74004269 | 74007534 | 32 | 35 | [illegible] |
| SE | 55777 | ENSG00000225470 | JPX | chrX | + | 73998953 | 74000496 | 73994839 | 73994961 | 74004269 | 74007534 | 27 | 32 | [illegible] |
| SE | 55779 | ENSG00000225470 | JPX | chrX | + | 73999354 | 74000496 | 73944583 | 73944662 | 74004269 | 74004543 | 27 | 32 | [illegible] |
| SE | 55781 | ENSG00000225470 | JPX | chrX | + | 73999354 | 74000496 | 73994839 | 73994961 | 74004269 | 74004543 | 27 | 32 | [illegible] |
| SE | 55782 | ENSG00000225470 | JPX | chrX | + | 73999354 | 74000496 | 73998767 | 73998844 | 74004269 | 74004543 | 27 | 33 | [illegible] |
| SE | 55785 | ENSG00000012174 | MBTPS2 | chrX | + | 21868466 | 21868585 | 21853375 | 21853503 | 21869497 | 21869678 | 46 | 34 | [illegible] |
| SE | 55790 | ENSG00000187231 | SESTD1 | chr2 | - | 179117531 | 179117613 | 179115064 | 179115256 | 179121769 | 179121929 | 23 | 21 | [illegible] |
| SE | 55792 | ENSG00000187231 | SESTD1 | chr2 | - | 179149296 | 179149394 | 179146401 | 179146457 | 179151277 | 179151391 | 132 | 138 | [illegible] |
| SE | 55793 | ENSG00000187231 | SESTD1 | chr2 | - | 179172119 | 179172233 | 179151277 | 179151391 | 179176447 | 179176538 | 126 | 98 | [illegible] |
| SE | 55795 | ENSG00000186364 | NUDT17 | chr1 | + | 145846432 | 145846458 | 145846012 | 145846196 | 145846597 | 145846690 | 0 | 0 | [illegible] |
| SE | 55798 | ENSG00000144218 | AFF3 | chr2 | - | 99727076 | 99727128 | 99672537 | 99672589 | 99744103 | 99744140 | 70 | 32 | [illegible] |
| SE | 55799 | ENSG00000144218 | AFF3 | chr2 | - | 99752220 | 99752301 | 99744103 | 99744140 | 99837476 | 99837524 | 80 | 28 | [illegible] |
| SE | 55800 | ENSG00000144218 | AFF3 | chr2 | - | 99837476 | 99837524 | 99752220 | 99752301 | 100006631 | 100006775 | 36 | 14 | [illegible] |
| SE | 55801 | ENSG00000144218 | AFF3 | chr2 | - | 100011496 | 100011571 | 100008811 | 100008932 | 100063125 | 100063249 | 3 | 2 | [illegible] |
| SE | 55803 | ENSG00000144218 | AFF3 | chr2 | - | 100063125 | 100063249 | 100008886 | 100008932 | 100104401 | 100104525 | 1 | 0 | [illegible] |
| SE | 55804 | ENSG00000144218 | AFF3 | chr2 | - | 100063125 | 100063249 | 100008886 | 100008932 | 100129223 | 100129306 | 1 | 0 | [illegible] |
| SE | 55807 | ENSG00000144218 | AFF3 | chr2 | - | 100105503 | 100105583 | 100104401 | 100104518 | 100129223 | 100129306 | 10 | 4 | [illegible] |
| SE | 55810 | ENSG00000144218 | AFF3 | chr2 | - | 100112385 | 100112524 | 100105503 | 100105583 | 100142483 | 100142583 | 18 | 8 | [illegible] |
| SE | 55812 | ENSG00000144218 | AFF3 | chr2 | - | 100129223 | 100129306 | 100105503 | 100105583 | 100142483 | 100142583 | 19 | 8 | [illegible] |
| SE | 55814 | ENSG00000106615 | RHEB | chr7 | - | 151471548 | 151471605 | 151471393 | 151471441 | 151477332 | 151477415 | 516 | 435 | [illegible] |
| SE | 55816 | ENSG00000106615 | RHEB | chr7 | - | 151498088 | 151498180 | 151490942 | 151491014 | 151517890 | 151518069 | 4 | 7 | [illegible] |
| SE | 55818 | ENSG00000143641 | GALNT2 | chr1 | + | 230203136 | 230203290 | 230178217 | 230178311 | 230236013 | 230236112 | 71 | 61 | [illegible] |
| SE | 55819 | ENSG00000143641 | GALNT2 | chr1 | + | 230236352 | 230236420 | 230236013 | 230236112 | 230236659 | 230236725 | 95 | 81 | [illegible] |
| SE | 55820 | ENSG00000143641 | GALNT2 | chr1 | + | 230236659 | 230236725 | 230236352 | 230236420 | 230243305 | 230243427 | 117 | 89 | [illegible] |
| SE | 55826 | ENSG00000143641 | GALNT2 | chr1 | + | 230255217 | 230255344 | 230250456 | 230250560 | 230274444 | 230274564 | 35 | 31 | [illegible] |
| SE | 55828 | ENSG00000143641 | GALNT2 | chr1 | + | 230274444 | 230274564 | 230265240 | 230265367 | 230279302 | 230282120 | 339 | 193 | [illegible] |
| SE | 55835 | ENSG00000184178 | SCFD2 | chr4 | - | 53313635 | 53313763 | 53273825 | 53274001 | 53352597 | 53352766 | 103 | 146 | [illegible] |
| SE | 55836 | ENSG00000188186 | LAMTOR4 | chr7 | + | 100149498 | 100149565 | 100148933 | 100148975 | 100153399 | 100153517 | 2 | 1 | [illegible] |
| SE | 55837 | ENSG00000188186 | LAMTOR4 | chr7 | + | 100149498 | 100149579 | 100148906 | 100148975 | 100153399 | 100153517 | 60 | 37 | [illegible] |
| SE | 55839 | ENSG00000188186 | LAMTOR4 | chr7 | + | 100152715 | 100152757 | 100149498 | 100149565 | 100153399 | 100153517 | 1 | 0 | [illegible] |
| SE | 55840 | ENSG00000102383 | ZDHHC15 | chrX | - | 75379119 | 75379198 | 75368426 | 75372945 | 75417086 | 75417190 | 2 | 0 | [illegible] |
| SE | 55841 | ENSG00000102383 | ZDHHC15 | chrX | - | 75424651 | 75424784 | 75421863 | 75421990 | 75429947 | 75429980 | 1 | 0 | [illegible] |
| SE | 55842 | ENSG00000102383 | ZDHHC15 | chrX | - | 75429077 | 75429198 | 75424651 | 75424784 | 75429947 | 75429980 | 1 | 0 | [illegible] |
| SE | 55844 | ENSG00000186416 | NKRF | chrX | - | 119603461 | 119603559 | 119592382 | 119592513 | 119605730 | 119605883 | 0 | 0 | [illegible] |
| SE | 55856 | ENSG00000177200 | CHD9 | chr16 | + | 53222643 | 53222755 | 53209481 | 53209813 | 53226365 | 53226512 | 168 | 248 | [illegible] |
| SE | 55858 | ENSG00000177200 | CHD9 | chr16 | + | 53227547 | 53227603 | 53227395 | 53227464 | 53228982 | 53229100 | 326 | 368 | [illegible] |
| SE | 55859 | ENSG00000177200 | CHD9 | chr16 | + | 53228982 | 53229100 | 53227547 | 53227603 | 53235184 | 53235306 | 252 | 283 | [illegible] |
| SE | 55860 | ENSG00000177200 | CHD9 | chr16 | + | 53231418 | 53231505 | 53228982 | 53229100 | 53235184 | 53235306 | 30 | 21 | [illegible] |
| SE | 55861 | ENSG00000177200 | CHD9 | chr16 | + | 53231646 | 53231784 | 53231418 | 53231505 | 53235184 | 53235306 | 317 | 348 | [illegible] |
| SE | 55862 | ENSG00000177200 | CHD9 | chr16 | + | 53254437 | 53254605 | 53249870 | 53250066 | 53255599 | 53255779 | 196 | 187 | [illegible] |
| SE | 55864 | ENSG00000177200 | CHD9 | chr16 | + | 53267293 | 53267490 | 53262986 | 53263097 | 53267926 | 53268126 | 489 | 532 | [illegible] |
| SE | 55865 | ENSG00000177200 | CHD9 | chr16 | + | 53273625 | 53273785 | 53267926 | 53268126 | 53274212 | 53274302 | 245 | 249 | [illegible] |
| SE | 55866 | ENSG00000177200 | CHD9 | chr16 | + | 53296955 | 53297158 | 53292832 | 53293052 | 53303719 | 53304625 | 217 | 184 | [illegible] |
| SE | 55869 | ENSG00000177200 | CHD9 | chr16 | + | 53307680 | 53307905 | 53306236 | 53306397 | 53308685 | 53308854 | 111 | 91 | [illegible] |
| SE | 55870 | ENSG00000177200 | CHD9 | chr16 | + | 53307680 | 53307953 | 53306233 | 53306397 | 53308685 | 53308854 | 143 | 99 | [illegible] |
| SE | 55871 | ENSG00000177200 | CHD9 | chr16 | + | 53314822 | 53315044 | 53314376 | 53314516 | 53318211 | 53318340 | 143 | 136 | [illegible] |
| SE | 55872 | ENSG00000177200 | CHD9 | chr16 | + | 53319812 | 53319951 | 53318211 | 53318340 | 53321162 | 53321217 | 1 | 1 | [illegible] |
| SE | 55874 | ENSG00000177200 | CHD9 | chr16 | + | 53321162 | 53321217 | 53318211 | 53318340 | 53321525 | 53321630 | 5 | 2 | [illegible] |
| SE | 55875 | ENSG00000177200 | CHD9 | chr16 | + | 53321162 | 53321217 | 53319812 | 53319951 | 53321525 | 53321630 | 0 | 0 | [illegible] |
| SE | 55876 | ENSG00000177200 | CHD9 | chr16 | + | 53321525 | 53321630 | 53318211 | 53318340 | 53324019 | 53324141 | 129 | 127 | [illegible] |
| SE | 55877 | ENSG00000153162 | BMP6 | chr6 | + | 7845139 | 7845332 | 7726098 | 7727619 | 7861450 | 7861599 | 21 | 25 | [illegible] |
| SE | 55883 | ENSG00000100024 | UPB1 | chr22 | + | 24515200 | 24515370 | 24513323 | 24513485 | 24520386 | 24520468 | 0 | 0 | [illegible] |
| SE | 55884 | ENSG00000100024 | UPB1 | chr22 | + | 24521985 | 24522028 | 24520173 | 24520468 | 24523618 | 24523700 | 0 | 0 | [illegible] |
| SE | 55885 | ENSG00000100024 | UPB1 | chr22 | + | 24523618 | 24523773 | 24520386 | 24520468 | 24525710 | 24526668 | 5 | 0 | [illegible] |
| SE | 55886 | ENSG00000074201 | CLNS1A | chr11 | - | 77611314 | 77611374 | 77585361 | 77585459 | 77619605 | 77619695 | 0 | 0 | [illegible] |
| SE | 55887 | ENSG00000074201 | CLNS1A | chr11 | - | 77612297 | 77612370 | 77611314 | 77611374 | 77619605 | 77619695 | 9 | 3 | [illegible] |
| SE | 55888 | ENSG00000074201 | CLNS1A | chr11 | - | 77621865 | 77622044 | 77619605 | 77619695 | 77622499 | 77622531 | 0 | 0 | [illegible] |
| SE | 55891 | ENSG00000074201 | CLNS1A | chr11 | - | 77625716 | 77625818 | 77624962 | 77625070 | 77629762 | 77629899 | 1169 | 899 | [illegible] |
| SE | 55892 | ENSG00000074201 | CLNS1A | chr11 | - | 77625716 | 77625818 | 77624962 | 77625070 | 77637589 | 77637782 | 634 | 514 | [illegible] |
| SE | 55893 | ENSG00000074201 | CLNS1A | chr11 | - | 77629762 | 77629899 | 77624962 | 77625070 | 77637589 | 77637751 | 54 | 51 | [illegible] |
| SE | 55894 | ENSG00000074201 | CLNS1A | chr11 | - | 77629762 | 77629899 | 77625716 | 77625818 | 77637589 | 77637751 | 587 | 439 | [illegible] |
| SE | 55896 | ENSG00000132510 | KDM6B | chr17 | + | 7840283 | 7840370 | 7839903 | 7840024 | 7844900 | 7845020 | 0 | 0 | [illegible] |
| SE | 55897 | ENSG00000132510 | KDM6B | chr17 | + | 7845549 | 7845691 | 7845313 | 7845456 | 7845871 | 7845970 | 46 | 18 | [illegible] |
| SE | 55900 | ENSG00000136866 | ZFP37 | chr9 | - | 113049361 | 113049496 | 113038376 | 113044268 | 113049790 | 113049872 | 20 | 3 | [illegible] |
| SE | 55903 | ENSG00000251669 | FAM86EP | chr4 | - | 3947143 | 3948220 | 3941759 | 3943372 | 3953110 | 3953173 | 0 | 0 | [illegible] |
| SE | 55905 | ENSG00000251669 | FAM86EP | chr4 | - | 3948089 | 3948220 | 3942910 | 3943372 | 3953110 | 3953173 | 0 | 0 | [illegible] |
| SE | 55909 | ENSG00000251669 | FAM86EP | chr4 | - | 3954959 | 3955033 | 3953110 | 3953173 | 3955323 | 3955394 | 0 | 0 | [illegible] |
| SE | 55921 | ENSG00000122678 | POLM | chr7 | - | 44074130 | 44074530 | 44073782 | 44074025 | 44076508 | 44076629 | 13 | 6 | [illegible] |
| SE | 55922 | ENSG00000122678 | POLM | chr7 | - | 44074397 | 44074522 | 44073782 | 44074025 | 44076508 | 44076629 | 3 | 3 | [illegible] |
| SE | 55923 | ENSG00000122678 | POLM | chr7 | - | 44074397 | 44074530 | 44073782 | 44074025 | 44076508 | 44076629 | 9 | 4 | [illegible] |
| SE | 55925 | ENSG00000122678 | POLM | chr7 | - | 44076508 | 44076629 | 44073782 | 44074530 | 44078755 | 44078811 | 8 | 6 | [illegible] |
| SE | 55926 | ENSG00000122678 | POLM | chr7 | - | 44076508 | 44076629 | 44074397 | 44074522 | 44078755 | 44078811 | 1 | 3 | [illegible] |
| SE | 55929 | ENSG00000122678 | POLM | chr7 | - | 44078321 | 44078438 | 44076508 | 44076629 | 44078755 | 44078811 | 2 | 0 | [illegible] |
| SE | 55931 | ENSG00000122678 | POLM | chr7 | - | 44079860 | 44079959 | 44079570 | 44079741 | 44080732 | 44080916 | 18 | 6 | [illegible] |
| SE | 55932 | ENSG00000122678 | POLM | chr7 | - | 44080364 | 44080471 | 44079860 | 44079959 | 44080732 | 44080916 | 0 | 0 | [illegible] |
| SE | 55945 | ENSG00000187605 | TET3 | chr2 | + | 74003109 | 74003166 | 73985979 | 73986706 | 74046277 | 74048411 | 122 | 86 | [illegible] |
| SE | 55946 | ENSG00000187605 | TET3 | chr2 | + | 74003109 | 74003166 | 73985979 | 73986706 | 74073548 | 74073639 | 45 | 27 | [illegible] |
| SE | 55947 | ENSG00000187605 | TET3 | chr2 | + | 74046277 | 74048411 | 73985979 | 73986706 | 74073548 | 74073639 | 60 | 34 | [illegible] |
| SE | 55949 | ENSG00000187605 | TET3 | chr2 | + | 74087829 | 74088038 | 74080497 | 74080591 | 74089896 | 74090047 | 159 | 67 | [illegible] |
| SE | 55956 | ENSG00000136161 | RCBTB2 | chr13 | - | 48491651 | 48491715 | 48489896 | 48490251 | 48492439 | 48492573 | 0 | 2 | [illegible] |
| SE | 55957 | ENSG00000136161 | RCBTB2 | chr13 | - | 48496190 | 48496321 | 48488962 | 48490251 | 48499620 | 48499760 | 79 | 67 | [illegible] |
| SE | 55959 | ENSG00000136161 | RCBTB2 | chr13 | - | 48502723 | 48502914 | 48501741 | 48501868 | 48511769 | 48511877 | 78 | 59 | [illegible] |
| SE | 55961 | ENSG00000136161 | RCBTB2 | chr13 | - | 48511769 | 48511877 | 48502723 | 48502914 | 48512015 | 48512122 | 24 | 12 | [illegible] |
| SE | 55962 | ENSG00000136161 | RCBTB2 | chr13 | - | 48512015 | 48512174 | 48502723 | 48502914 | 48512728 | 48512895 | 74 | 34 | [illegible] |
| SE | 55967 | ENSG00000136161 | RCBTB2 | chr13 | - | 48521897 | 48521962 | 48515585 | 48515741 | 48524658 | 48524757 | 25 | 16 | [illegible] |
| SE | 55972 | ENSG00000136161 | RCBTB2 | chr13 | - | 48522307 | 48522403 | 48521897 | 48521962 | 48533027 | 48533080 | 18 | 11 | [illegible] |
| SE | 55974 | ENSG00000136161 | RCBTB2 | chr13 | - | 48524658 | 48524757 | 48521897 | 48521962 | 48533027 | 48533080 | 15 | 7 | [illegible] |
| SE | 55975 | ENSG00000136161 | RCBTB2 | chr13 | - | 48524658 | 48524757 | 48522307 | 48522403 | 48533027 | 48533080 | 16 | 21 | [illegible] |
| SE | 55977 | ENSG00000121892 | PDS5A | chr4 | - | 39837855 | 39838208 | 39822862 | 39825488 | 39841947 | 39842056 | 689 | 499 | [illegible] |
| SE | 55979 | ENSG00000121892 | PDS5A | chr4 | - | 39844655 | 39844801 | 39841947 | 39842056 | 39845817 | 39845880 | 824 | 644 | [illegible] |
| SE | 55981 | ENSG00000121892 | PDS5A | chr4 | - | 39849519 | 39849652 | 39845817 | 39845880 | 39869393 | 39869462 | 3 | 4 | [illegible] |
| SE | 55982 | ENSG00000121892 | PDS5A | chr4 | - | 39866860 | 39866997 | 39863335 | 39863459 | 39869393 | 39869462 | 454 | 372 | [illegible] |
| SE | 55983 | ENSG00000121892 | PDS5A | chr4 | - | 39866860 | 39866997 | 39863335 | 39863459 | 39872985 | 39873144 | 247 | 199 | [illegible] |
| SE | 55984 | ENSG00000121892 | PDS5A | chr4 | - | 39869393 | 39869462 | 39863335 | 39863459 | 39872985 | 39873144 | 282 | 204 | [illegible] |
| SE | 55985 | ENSG00000121892 | PDS5A | chr4 | - | 39869393 | 39869462 | 39866860 | 39866997 | 39872985 | 39873144 | 490 | 372 | [illegible] |
| SE | 55986 | ENSG00000121892 | PDS5A | chr4 | - | 39872985 | 39873144 | 39869393 | 39869462 | 39874288 | 39874408 | 669 | 525 | [illegible] |
| SE | 55987 | ENSG00000121892 | PDS5A | chr4 | - | 39872985 | 39873144 | 39869393 | 39869462 | 39876992 | 39877153 | 399 | 298 | [illegible] |
| SE | 55990 | ENSG00000121892 | PDS5A | chr4 | - | 39890248 | 39890364 | 39879727 | 39879833 | 39898388 | 39898528 | 705 | 645 | [illegible] |
| SE | 55991 | ENSG00000121892 | PDS5A | chr4 | - | 39900425 | 39900507 | 39898776 | 39898825 | 39902346 | 39902460 | 531 | 460 | [illegible] |
| SE | 55992 | ENSG00000121892 | PDS5A | chr4 | - | 39902346 | 39902460 | 39900425 | 39900507 | 39904039 | 39904191 | 572 | 425 | [illegible] |
| SE | 55993 | ENSG00000121892 | PDS5A | chr4 | - | 39904039 | 39904191 | 39902346 | 39902460 | 39913610 | 39913726 | 303 | 202 | [illegible] |
| SE | 55994 | ENSG00000121892 | PDS5A | chr4 | - | 39908394 | 39908540 | 39904039 | 39904191 | 39913610 | 39913726 | 107 | 106 | [illegible] |
| SE | 55996 | ENSG00000121892 | PDS5A | chr4 | - | 39920318 | 39920399 | 39917047 | 39917188 | 39922621 | 39922748 | 381 | 245 | [illegible] |
| SE | 56005 | ENSG00000186409 | CCDC30 | chr1 | + | 42480460 | 42480566 | 42473121 | 42473303 | 42482662 | 42482816 | 6 | 4 | [illegible] |
| SE | 56006 | ENSG00000186409 | CCDC30 | chr1 | + | 42506946 | 42507069 | 42498817 | 42498916 | 42566295 | 42566475 | 2 | 0 | [illegible] |
| SE | 56007 | ENSG00000186409 | CCDC30 | chr1 | + | 42521162 | 42521256 | 42498817 | 42498916 | 42534888 | 42534989 | 2 | 0 | [illegible] |
| SE | 56008 | ENSG00000186409 | CCDC30 | chr1 | + | 42521162 | 42521256 | 42498817 | 42498916 | 42566295 | 42566475 | 1 | 0 | [illegible] |
| SE | 56010 | ENSG00000186409 | CCDC30 | chr1 | + | 42534888 | 42534989 | 42498817 | 42498916 | 42536475 | 42536601 | 2 | 7 | [illegible] |
| SE | 56011 | ENSG00000186409 | CCDC30 | chr1 | + | 42534888 | 42534989 | 42498817 | 42498916 | 42545409 | 42545579 | 1 | 7 | [illegible] |
| SE | 56012 | ENSG00000186409 | CCDC30 | chr1 | + | 42534888 | 42534989 | 42498817 | 42498916 | 42566295 | 42566475 | 1 | 7 | [illegible] |
| SE | 56015 | ENSG00000186409 | CCDC30 | chr1 | + | 42536475 | 42536601 | 42498817 | 42498916 | 42545409 | 42545579 | 1 | 3 | [illegible] |
| SE | 56016 | ENSG00000186409 | CCDC30 | chr1 | + | 42536475 | 42536601 | 42498817 | 42498916 | 42566295 | 42566475 | 1 | 2 | [illegible] |
| SE | 56017 | ENSG00000186409 | CCDC30 | chr1 | + | 42536475 | 42536601 | 42534888 | 42534989 | 42545409 | 42545579 | 2 | 1 | [illegible] |
| SE | 56018 | ENSG00000186409 | CCDC30 | chr1 | + | 42536475 | 42536601 | 42534888 | 42534989 | 42566295 | 42566475 | 2 | 0 | [illegible] |
| SE | 56020 | ENSG00000186409 | CCDC30 | chr1 | + | 42539172 | 42539310 | 42536475 | 42536601 | 42545409 | 42545519 | 0 | 0 | [illegible] |
| SE | 56021 | ENSG00000186409 | CCDC30 | chr1 | + | 42545409 | 42545579 | 42498817 | 42498916 | 42566295 | 42566470 | 1 | 2 | [illegible] |
| SE | 56022 | ENSG00000186409 | CCDC30 | chr1 | + | 42545409 | 42545579 | 42534888 | 42534989 | 42566295 | 42566470 | 1 | 2 | [illegible] |
| SE | 56023 | ENSG00000186409 | CCDC30 | chr1 | + | 42545409 | 42545579 | 42536475 | 42536601 | 42566295 | 42566470 | 2 | 3 | [illegible] |
| SE | 56026 | ENSG00000186409 | CCDC30 | chr1 | + | 42566295 | 42566475 | 42545409 | 42545579 | 42577019 | 42577229 | 29 | 12 | [illegible] |
| SE | 56029 | ENSG00000186409 | CCDC30 | chr1 | + | 42581359 | 42581514 | 42577019 | 42577229 | 42589320 | 42589461 | 7 | 4 | [illegible] |
| SE | 56030 | ENSG00000186409 | CCDC30 | chr1 | + | 42581426 | 42581514 | 42577019 | 42577229 | 42589320 | 42590252 | 2 | 1 | [illegible] |
| SE | 56038 | ENSG00000166311 | SMPD1 | chr11 | + | 6393895 | 6394041 | 6393616 | 6393693 | 6394197 | 6394778 | 209 | 118 | [illegible] |
| SE | 56039 | ENSG00000060688 | SNRNP40 | chr1 | - | 31261528 | 31261632 | 31259567 | 31260121 | 31267870 | 31267932 | 270 | 115 | [illegible] |
| SE | 56040 | ENSG00000060688 | SNRNP40 | chr1 | - | 31267870 | 31267932 | 31261528 | 31261632 | 31269157 | 31269240 | 233 | 102 | [illegible] |
| SE | 56041 | ENSG00000126215 | XRCC3 | chr14 | - | 103712874 | 103712931 | 103711465 | 103711567 | 103715423 | 103715451 | 1 | 0 | [illegible] |
| SE | 56042 | ENSG00000126215 | XRCC3 | chr14 | - | 103712874 | 103712941 | 103711465 | 103711567 | 103715423 | 103715504 | 1 | 0 | [illegible] |
| SE | 56043 | ENSG00000126215 | XRCC3 | chr14 | - | 103712874 | 103712968 | 103711465 | 103711567 | 103715423 | 103715443 | 2 | 0 | [illegible] |
| SE | 56045 | ENSG00000152382 | TADA1 | chr1 | - | 166860185 | 166860337 | 166858118 | 166858281 | 166862202 | 166862412 | 103 | 42 | [illegible] |
| SE | 56046 | ENSG00000152382 | TADA1 | chr1 | - | 166863823 | 166863921 | 166862202 | 166862412 | 166869444 | 166869510 | 53 | 22 | [illegible] |
| SE | 56047 | ENSG00000152382 | TADA1 | chr1 | - | 166869762 | 166869854 | 166869444 | 166869510 | 166876159 | 166876264 | 37 | 15 | [illegible] |
| SE | 56048 | ENSG00000156675 | RAB11FIP1 | chr8 | - | 37870419 | 37870528 | 37858617 | 37863113 | 37874514 | 37875322 | 198 | 140 | [illegible] |
| SE | 56050 | ENSG00000156675 | RAB11FIP1 | chr8 | - | 37877108 | 37877551 | 37874514 | 37875322 | 37899070 | 37899467 | 456 | 246 | [illegible] |
| SE | 56051 | ENSG00000164402 | SEPTIN8 | chr5 | - | 132761799 | 132761896 | 132761457 | 132761626 | 132762483 | 132762645 | 28 | 26 | [illegible] |
| SE | 56053 | ENSG00000133063 | CHIT1 | chr1 | - | 203217301 | 203217372 | 203216669 | 203217133 | 203217738 | 203217865 | 0 | 0 | [illegible] |
| SE | 56054 | ENSG00000133063 | CHIT1 | chr1 | - | 203217738 | 203217865 | 203212826 | 203213045 | 203219215 | 203219329 | 0 | 0 | [illegible] |
| SE | 56055 | ENSG00000133063 | CHIT1 | chr1 | - | 203217934 | 203218091 | 203217738 | 203217865 | 203219215 | 203219329 | 0 | 0 | [illegible] |
| SE | 56056 | ENSG00000133063 | CHIT1 | chr1 | - | 203219215 | 203219329 | 203212826 | 203213045 | 203219663 | 203219849 | 1 | 0 | [illegible] |
| SE | 56057 | ENSG00000133063 | CHIT1 | chr1 | - | 203223134 | 203223259 | 203222201 | 203222325 | 203223494 | 203223660 | 0 | 0 | [illegible] |
| SE | 56058 | ENSG00000133063 | CHIT1 | chr1 | - | 203225047 | 203225104 | 203223494 | 203223660 | 203225668 | 203225870 | 2 | 0 | [illegible] |
| SE | 56059 | ENSG00000133063 | CHIT1 | chr1 | - | 203225668 | 203225870 | 203223494 | 203223660 | 203228532 | 203228562 | 0 | 0 | [illegible] |
| SE | 56061 | ENSG00000103152 | MPG | chr16 | + | 79138 | 79331 | 78256 | 78333 | 79424 | 79700 | 0 | 0 | [illegible] |

| | | | | | | | | | |
|---|---|---|---|---|---|---|---|---|---|
| SE | 56063 | ENSG00000084733 | RAB10 | chr2 | + | 26098661 | 26098722 | 26034083 | 26034735 |
| SE | 56064 | ENSG00000084733 | RAB10 | chr2 | + | 26098661 | 26098722 | 26034083 | 26034735 |
| SE | 56065 | ENSG00000084733 | RAB10 | chr2 | + | 26109767 | 26109906 | 26034083 | 26034735 |
| SE | 56066 | ENSG00000084733 | RAB10 | chr2 | + | 26109767 | 26109906 | 26098661 | 26098722 |
| SE | 56067 | ENSG00000160229 | ZNF66 | chr19 | + | 20792511 | 20792638 | 20776303 | 20776450 |
| SE | 56069 | ENSG00000160229 | ZNF66 | chr19 | + | 20793782 | 20793878 | 20776303 | 20776450 |
| SE | 56070 | ENSG00000173947 | PIFO | chr1 | + | 111347673 | 111347772 | 111346909 | 111347049 |
| SE | 56071 | ENSG00000160679 | CHTOP | chr1 | + | 153636571 | 153636653 | 153633981 | 153634343 |
| SE | 56072 | ENSG00000160679 | CHTOP | chr1 | + | 153638294 | 153638448 | 153636571 | 153636653 |
| SE | 56073 | ENSG00000160679 | CHTOP | chr1 | + | 153638294 | 153638448 | 153636571 | 153636653 |
| SE | 56074 | ENSG00000160679 | CHTOP | chr1 | + | 153642242 | 153642429 | 153638294 | 153638448 |
| SE | 56075 | ENSG00000160679 | CHTOP | chr1 | + | 153642245 | 153642429 | 153638083 | 153638448 |
| SE | 56080 | ENSG00000155016 | CYP2U1 | chr4 | + | 107944969 | 107945605 | 107931548 | 107932133 |
| SE | 56084 | ENSG00000153207 | AHCTF1 | chr1 | - | 246843794 | 246843928 | 246839097 | 246840998 |
| SE | 56086 | ENSG00000153207 | AHCTF1 | chr1 | - | 246851904 | 246852038 | 246849870 | 246851442 |
| SE | 56087 | ENSG00000153207 | AHCTF1 | chr1 | - | 246857690 | 246857814 | 246855729 | 246855827 |
| SE | 56088 | ENSG00000153207 | AHCTF1 | chr1 | - | 246863923 | 246864116 | 246861958 | 246862153 |
| SE | 56089 | ENSG00000153207 | AHCTF1 | chr1 | - | 246863923 | 246864116 | 246861958 | 246862153 |
| SE | 56091 | ENSG00000153207 | AHCTF1 | chr1 | - | 246865254 | 246865333 | 246863923 | 246864116 |
| SE | 56092 | ENSG00000153207 | AHCTF1 | chr1 | - | 246867243 | 246867351 | 246863923 | 246864116 |
| SE | 56093 | ENSG00000153207 | AHCTF1 | chr1 | - | 246876036 | 246876187 | 246867660 | 246867811 |
| SE | 56094 | ENSG00000153207 | AHCTF1 | chr1 | - | 246890955 | 246891060 | 246889965 | 246890059 |
| SE | 56095 | ENSG00000153207 | AHCTF1 | chr1 | - | 246890955 | 246891060 | 246889965 | 246890059 |
| SE | 56096 | ENSG00000153207 | AHCTF1 | chr1 | - | 246891778 | 246891919 | 246889965 | 246890059 |
| SE | 56097 | ENSG00000153207 | AHCTF1 | chr1 | - | 246891778 | 246891919 | 246890955 | 246891060 |
| SE | 56098 | ENSG00000153207 | AHCTF1 | chr1 | - | 246894658 | 246894748 | 246891778 | 246891919 |
| SE | 56099 | ENSG00000153207 | AHCTF1 | chr1 | - | 246895834 | 246895925 | 246891778 | 246891919 |
| SE | 56100 | ENSG00000153207 | AHCTF1 | chr1 | - | 246907550 | 246907758 | 246905540 | 246905657 |
| SE | 56103 | ENSG00000165689 | ENTR1 | chr9 | - | 136404054 | 136404194 | 136401921 | 136402887 |
| SE | 56104 | ENSG00000165689 | ENTR1 | chr9 | - | 136405904 | 136405978 | 136405090 | 136405202 |
| SE | 56105 | ENSG00000165689 | ENTR1 | chr9 | - | 136407144 | 136407561 | 136405904 | 136405978 |
| SE | 56109 | ENSG00000165689 | ENTR1 | chr9 | - | 136410089 | 136410239 | 136408998 | 136409067 |
| SE | 56110 | ENSG00000158109 | TPRG1L | chr1 | + | 3625423 | 3625515 | 3625014 | 3625273 |
| SE | 56112 | ENSG00000163001 | CFAP36 | chr2 | + | 55520404 | 55520479 | 55519603 | 55519916 |
| SE | 56114 | ENSG00000163001 | CFAP36 | chr2 | + | 55523720 | 55523822 | 55522101 | 55522166 |
| SE | 56115 | ENSG00000163001 | CFAP36 | chr2 | + | 55528877 | 55528992 | 55523720 | 55523822 |
| SE | 56117 | ENSG00000163001 | CFAP36 | chr2 | + | 55537482 | 55537585 | 55535711 | 55535763 |
| SE | 56121 | ENSG00000134070 | IRAK2 | chr3 | + | 10238747 | 10239039 | 10234458 | 10234659 |
| SE | 56123 | ENSG00000204406 | MBD5 | chr2 | + | 148021647 | 148021684 | 148021010 | 148021046 |
| SE | 56124 | ENSG00000204406 | MBD5 | chr2 | + | 148053932 | 148054050 | 148021026 | 148021046 |
| SE | 56129 | ENSG00000204406 | MBD5 | chr2 | + | 148342213 | 148342336 | 148233244 | 148233395 |
| SE | 56130 | ENSG00000204406 | MBD5 | chr2 | + | 148447566 | 148447740 | 148342213 | 148342336 |
| SE | 56131 | ENSG00000204406 | MBD5 | chr2 | + | 148462581 | 148462684 | 148458202 | 148458871 |
| SE | 56132 | ENSG00000204406 | MBD5 | chr2 | + | 148463738 | 148463919 | 148458202 | 148458871 |
| SE | 56133 | ENSG00000204406 | MBD5 | chr2 | + | 148489385 | 148489500 | 148485741 | 148485950 |
| SE | 56134 | ENSG00000204406 | MBD5 | chr2 | + | 148489385 | 148490594 | 148485115 | 148485950 |
| SE | 56135 | ENSG00000204406 | MBD5 | chr2 | + | 148489385 | 148490787 | 148485741 | 148485950 |
| SE | 56138 | ENSG00000204406 | MBD5 | chr2 | + | 148510059 | 148510135 | 148502396 | 148502509 |
| SE | 56142 | ENSG00000120948 | TARDBP | chr1 | + | 11020428 | 11020599 | 11018732 | 11018873 |
| SE | 56144 | ENSG00000120948 | TARDBP | chr1 | + | 11024063 | 11024335 | 11023192 | 11023329 |
| SE | 56145 | ENSG00000120948 | TARDBP | chr1 | + | 11024301 | 11024335 | 11022817 | 11023329 |
| SE | 56146 | ENSG00000120948 | TARDBP | chr1 | + | 11029686 | 11029781 | 11025109 | 11025208 |
| SE | 56156 | ENSG00000213079 | SCAF8 | chr6 | + | 154733794 | 154733930 | 154733377 | 154733503 |
| SE | 56157 | ENSG00000213079 | SCAF8 | chr6 | + | 154741955 | 154742018 | 154733846 | 154733930 |
| SE | 56158 | ENSG00000213079 | SCAF8 | chr6 | + | 154773988 | 154774072 | 154733415 | 154733930 |
| SE | 56159 | ENSG00000213079 | SCAF8 | chr6 | + | 154778000 | 154778045 | 154773988 | 154774072 |
| SE | 56160 | ENSG00000213079 | SCAF8 | chr6 | + | 154787860 | 154788022 | 154733415 | 154733930 |
| SE | 56163 | ENSG00000213079 | SCAF8 | chr6 | + | 154795008 | 154795139 | 154778000 | 154778045 |
| SE | 56164 | ENSG00000213079 | SCAF8 | chr6 | + | 154801970 | 154802147 | 154795008 | 154795139 |
| SE | 56165 | ENSG00000213079 | SCAF8 | chr6 | + | 154803543 | 154803623 | 154795008 | 154795139 |
| SE | 56166 | ENSG00000213079 | SCAF8 | chr6 | + | 154808069 | 154808201 | 154805368 | 154805486 |
| SE | 56167 | ENSG00000213079 | SCAF8 | chr6 | + | 154818478 | 154818592 | 154815715 | 154815816 |
| SE | 56169 | ENSG00000213079 | SCAF8 | chr6 | + | 154822275 | 154822409 | 154820176 | 154820333 |
| SE | 56170 | ENSG00000213079 | SCAF8 | chr6 | + | 154824233 | 154824378 | 154820176 | 154820333 |
| SE | 56172 | ENSG00000213079 | SCAF8 | chr6 | + | 154830921 | 154831140 | 154827171 | 154827240 |
| SE | 56173 | ENSG00000094975 | SUCO | chr1 | + | 172551511 | 172551626 | 172533117 | 172533497 |
| SE | 56176 | ENSG00000094975 | SUCO | chr1 | + | 172573890 | 172573998 | 172570662 | 172570730 |
| SE | 56177 | ENSG00000094975 | SUCO | chr1 | + | 172575517 | 172575623 | 172573890 | 172573998 |
| SE | 56178 | ENSG00000094975 | SUCO | chr1 | + | 172577538 | 172577559 | 172575517 | 172575623 |
| SE | 56179 | ENSG00000094975 | SUCO | chr1 | + | 172585017 | 172585086 | 172579201 | 172579267 |
| SE | 56180 | ENSG00000094975 | SUCO | chr1 | + | 172585857 | 172585948 | 172579201 | 172579267 |
| SE | 56181 | ENSG00000094975 | SUCO | chr1 | + | 172600063 | 172600168 | 172590983 | 172591071 |
| SE | 56194 | ENSG00000143106 | PSMA5 | chr1 | - | 109409927 | 109410014 | 109399041 | 109402090 |
| SE | 56200 | ENSG00000124942 | AHNAK | chr11 | - | 62535944 | 62536098 | 62535002 | 62535190 |
| SE | 56201 | ENSG00000124942 | AHNAK | chr11 | - | 62535944 | 62536098 | 62535002 | 62535190 |
| SE | 56206 | ENSG00000106462 | EZH2 | chr7 | - | 148809070 | 148809155 | 148807256 | 148807706 |
| SE | 56207 | ENSG00000106462 | EZH2 | chr7 | - | 148813958 | 148814137 | 148811624 | 148811720 |
| SE | 56208 | ENSG00000106462 | EZH2 | chr7 | - | 148813958 | 148814137 | 148811624 | 148811720 |
| SE | 56209 | ENSG00000106462 | EZH2 | chr7 | - | 148813958 | 148815546 | 148811624 | 148811720 |
| SE | 56210 | ENSG00000106462 | EZH2 | chr7 | - | 148814913 | 148815039 | 148813958 | 148814137 |
| SE | 56213 | ENSG00000106462 | EZH2 | chr7 | - | 148816683 | 148816778 | 148815505 | 148815546 |
| SE | 56214 | ENSG00000106462 | EZH2 | chr7 | - | 148816683 | 148816778 | 148815505 | 148815546 |
| SE | 56218 | ENSG00000106462 | EZH2 | chr7 | - | 148817221 | 148817412 | 148815505 | 148815546 |
| SE | 56221 | ENSG00000106462 | EZH2 | chr7 | - | 148832633 | 148832750 | 148829727 | 148829848 |
| SE | 56222 | ENSG00000106462 | EZH2 | chr7 | - | 148832633 | 148832750 | 148829727 | 148829848 |
| SE | 56229 | ENSG00000131013 | PPIL4 | chr6 | - | 149526651 | 149526776 | 149525142 | 149525209 |
| SE | 56230 | ENSG00000131013 | PPIL4 | chr6 | - | 149533457 | 149533574 | 149526651 | 149526776 |
| SE | 56231 | ENSG00000131013 | PPIL4 | chr6 | - | 149541518 | 149541586 | 149541366 | 149541431 |
| SE | 56232 | ENSG00000108523 | RNF167 | chr17 | + | 4940486 | 4940642 | 4940119 | 4940230 |
| SE | 56233 | ENSG00000108523 | RNF167 | chr17 | + | 4940486 | 4940726 | 4940007 | 4940230 |
| SE | 56234 | ENSG00000108523 | RNF167 | chr17 | + | 4940486 | 4940726 | 4940007 | 4940230 |
| SE | 56235 | ENSG00000108523 | RNF167 | chr17 | + | 4940486 | 4940726 | 4940007 | 4940230 |
| SE | 56237 | ENSG00000108523 | RNF167 | chr17 | + | 4940872 | 4940993 | 4940082 | 4940230 |
| SE | 56239 | ENSG00000108523 | RNF167 | chr17 | + | 4940872 | 4940993 | 4940486 | 4940726 |
| SE | 56240 | ENSG00000108523 | RNF167 | chr17 | + | 4940922 | 4940993 | 4940119 | 4940230 |
| SE | 56241 | ENSG00000108523 | RNF167 | chr17 | + | 4940922 | 4940993 | 4940486 | 4940726 |
| SE | 56243 | ENSG00000108523 | RNF167 | chr17 | + | 4942340 | 4942466 | 4941076 | 4941157 |
| SE | 56245 | ENSG00000108523 | RNF167 | chr17 | + | 4943178 | 4943284 | 4942850 | 4942941 |
| SE | 56249 | ENSG00000108523 | RNF167 | chr17 | + | 4944557 | 4944638 | 4943067 | 4943519 |
| SE | 56250 | ENSG00000171960 | PPIH | chr1 | + | 42659521 | 42659566 | 42659227 | 42659251 |
| SE | 56252 | ENSG00000171960 | PPIH | chr1 | + | 42660861 | 42660904 | 42659521 | 42659566 |
| SE | 56253 | ENSG00000171960 | PPIH | chr1 | + | 42663417 | 42663543 | 42660861 | 42660904 |
| SE | 56254 | ENSG00000171960 | PPIH | chr1 | + | 42664862 | 42664955 | 42660861 | 42660904 |
| SE | 56255 | ENSG00000171960 | PPIH | chr1 | + | 42668046 | 42668151 | 42667350 | 42667440 |
| SE | 56257 | ENSG00000171960 | PPIH | chr1 | + | 42675287 | 42675327 | 42667350 | 42667440 |
| SE | 56259 | ENSG00000143995 | MEIS1 | chr2 | + | 66439842 | 66439984 | 66437392 | 66437963 |
| SE | 56261 | ENSG00000143995 | MEIS1 | chr2 | + | 66440561 | 66440612 | 66439320 | 66439671 |
| SE | 56262 | ENSG00000143995 | MEIS1 | chr2 | + | 66440561 | 66440612 | 66440096 | 66440345 |
| SE | 56266 | ENSG00000143995 | MEIS1 | chr2 | + | 66512148 | 66512294 | 66496025 | 66496200 |
| SE | 56269 | ENSG00000248429 | FAM198B-AS1 | chr4 | + | 158172251 | 158172416 | 158170751 | 158171480 |
| SE | 56274 | ENSG00000248429 | FAM198B-AS1 | chr4 | + | 158179660 | 158179735 | 158171227 | 158171480 |
| SE | 56276 | ENSG00000248429 | FAM198B-AS1 | chr4 | + | 158199087 | 158199486 | 158198787 | 158198870 |
| SE | 56277 | ENSG00000165637 | VDAC2 | chr10 | + | 75211481 | 75211659 | 75211133 | 75211189 |
| SE | 56278 | ENSG00000165637 | VDAC2 | chr10 | + | 75214020 | 75214070 | 75212229 | 75212298 |
| SE | 56282 | ENSG00000182183 | SHISAL2A | chr1 | + | 52642862 | 52643002 | 52633167 | 52633675 |
| SE | 56285 | ENSG00000287684 | NA | chr1 | - | 222769749 | 222769798 | 222762118 | 222762208 |
| SE | 56286 | ENSG00000134242 | PTPN22 | chr1 | - | 113819576 | 113819654 | 113813810 | 113814969 |
| SE | 56289 | ENSG00000134242 | PTPN22 | chr1 | - | 113829591 | 113829707 | 113825141 | 113825172 |
| SE | 56290 | ENSG00000134242 | PTPN22 | chr1 | - | 113829948 | 113830029 | 113829591 | 113829707 |
| SE | 56291 | ENSG00000134242 | PTPN22 | chr1 | - | 113834909 | 113834993 | 113834308 | 113834439 |
| SE | 56292 | ENSG00000134242 | PTPN22 | chr1 | - | 113834909 | 113834993 | 113834308 | 113834439 |
| SE | 56293 | ENSG00000134242 | PTPN22 | chr1 | - | 113837589 | 113838407 | 113834308 | 113834439 |
| SE | 56294 | ENSG00000134242 | PTPN22 | chr1 | - | 113837589 | 113838407 | 113834909 | 113834993 |
| SE | 56295 | ENSG00000134242 | PTPN22 | chr1 | - | 113837589 | 113838407 | 113834909 | 113834993 |
| SE | 56296 | ENSG00000134242 | PTPN22 | chr1 | - | 113838543 | 113838620 | 113834909 | 113834993 |
| SE | 56297 | ENSG00000134242 | PTPN22 | chr1 | - | 113838543 | 113838620 | 113837589 | 113838407 |
| SE | 56299 | ENSG00000134242 | PTPN22 | chr1 | - | 113848539 | 113848626 | 113838543 | 113838620 |
| SE | 56300 | ENSG00000134242 | PTPN22 | chr1 | - | 113852026 | 113852104 | 113838543 | 113838620 |
| SE | 56301 | ENSG00000134242 | PTPN22 | chr1 | - | 113852026 | 113852104 | 113838543 | 113838620 |
| SE | 56303 | ENSG00000134242 | PTPN22 | chr1 | - | 113852026 | 113852104 | 113848539 | 113848626 |
| SE | 56304 | ENSG00000134242 | PTPN22 | chr1 | - | 113854470 | 113854537 | 113838543 | 113838620 |
| SE | 56305 | ENSG00000134242 | PTPN22 | chr1 | - | 113854470 | 113854537 | 113852026 | 113852104 |
| SE | 56307 | ENSG00000134242 | PTPN22 | chr1 | - | 113854906 | 113855049 | 113852026 | 113852104 |
| SE | 56310 | ENSG00000134242 | PTPN22 | chr1 | - | 113856547 | 113856619 | 113856381 | 113856441 |
| SE | 56312 | ENSG00000134242 | PTPN22 | chr1 | - | 113858477 | 113858573 | 113857737 | 113857776 |
| SE | 56313 | ENSG00000134242 | PTPN22 | chr1 | - | 113859001 | 113859078 | 113858477 | 113858573 |
| SE | 56315 | ENSG00000134242 | PTPN22 | chr1 | - | 113864197 | 113864382 | 113859351 | 113859460 |
| SE | 56317 | ENSG00000010072 | SPRTN | chr1 | + | 231347796 | 231347925 | 231339768 | 231339868 |
| SE | 56319 | ENSG00000134278 | SPIRE1 | chr18 | - | 12459754 | 12459928 | 12454345 | 12454483 |
| SE | 56321 | ENSG00000134278 | SPIRE1 | chr18 | - | 12506476 | 12506641 | 12496015 | 12496102 |
| SE | 56331 | ENSG00000131165 | CHMP1A | chr16 | - | 89653903 | 89653923 | 89651568 | 89651646 |
| SE | 56334 | ENSG00000104970 | KIR3DX1 | chr19 | + | 54535585 | 54535647 | 54533731 | 54534010 |
| SE | 56335 | ENSG00000104970 | KIR3DX1 | chr19 | + | 54535585 | 54535891 | 54533731 | 54534010 |
| SE | 56336 | ENSG00000104970 | KIR3DX1 | chr19 | + | 54535585 | 54535891 | 54533731 | 54534010 |
| SE | 56337 | ENSG00000104970 | KIR3DX1 | chr19 | + | 54542523 | 54542646 | 54537974 | 54538025 |
| SE | 56338 | ENSG00000117602 | RCAN3 | chr1 | + | 24514313 | 24514567 | 24502350 | 24502463 |
| SE | 56342 | ENSG00000117602 | RCAN3 | chr1 | + | 24533082 | 24533254 | 24514313 | 24514567 |
| SE | 56346 | ENSG00000142875 | PRKACB | chr1 | + | 84157334 | 84157405 | 84144321 | 84144548 |
| SE | 56350 | ENSG00000142875 | PRKACB | chr1 | + | 84175032 | 84175056 | 84144321 | 84144548 |
| SE | 56351 | ENSG00000142875 | PRKACB | chr1 | + | 84175032 | 84175056 | 84144321 | 84144548 |
| SE | 56353 | ENSG00000142875 | PRKACB | chr1 | + | 84175032 | 84175056 | 84164393 | 84164422 |
| SE | 56354 | ENSG00000142875 | PRKACB | chr1 | + | 84175032 | 84175056 | 84164393 | 84164422 |
| SE | 56355 | ENSG00000142875 | PRKACB | chr1 | + | 84175032 | 84175056 | 84164903 | 84165013 |
| SE | 56356 | ENSG00000142875 | PRKACB | chr1 | + | 84175032 | 84175056 | 84164903 | 84165013 |
| SE | 56357 | ENSG00000142875 | PRKACB | chr1 | + | 84175798 | 84175807 | 84144321 | 84144548 |
| SE | 56358 | ENSG00000142875 | PRKACB | chr1 | + | 84175798 | 84175807 | 84164381 | 84164422 |
| SE | 56359 | ENSG00000142875 | PRKACB | chr1 | + | 84175798 | 84175807 | 84164694 | 84165013 |
| SE | 56360 | ENSG00000142875 | PRKACB | chr1 | + | 84175798 | 84175807 | 84175032 | 84175056 |
| SE | 56362 | ENSG00000142875 | PRKACB | chr1 | + | 84197728 | 84197824 | 84196615 | 84196742 |
| SE | 56364 | ENSG00000142875 | PRKACB | chr1 | + | 84202682 | 84202805 | 84197728 | 84197824 |
| SE | 56365 | ENSG00000142875 | PRKACB | chr1 | + | 84202682 | 84202805 | 84197728 | 84197824 |
| SE | 56366 | ENSG00000142875 | PRKACB | chr1 | + | 84202682 | 84202805 | 84197728 | 84197824 |
| SE | 56367 | ENSG00000142875 | PRKACB | chr1 | + | 84214152 | 84214317 | 84197728 | 84197824 |
| SE | 56368 | ENSG00000142875 | PRKACB | chr1 | + | 84214152 | 84214317 | 84202682 | 84202805 |
| SE | 56369 | ENSG00000085999 | RAD54L | chr1 | + | 46248317 | 46248408 | 46247699 | 46247746 |
| SE | 56370 | ENSG00000085999 | RAD54L | chr1 | + | 46248343 | 46248408 | 46247694 | 46247746 |
| SE | 56371 | ENSG00000085999 | RAD54L | chr1 | + | 46257741 | 46257817 | 46249999 | 46250119 |
| SE | 56372 | ENSG00000085999 | RAD54L | chr1 | + | 46258685 | 46258746 | 46249999 | 46250119 |
| SE | 56375 | ENSG00000085999 | RAD54L | chr1 | + | 46271367 | 46271442 | 46270658 | 46270785 |
| SE | 56379 | ENSG00000185946 | RNPC3 | chr1 | + | 103527694 | 103527742 | 103525690 | 103526262 |
| SE | 56380 | ENSG00000185946 | RNPC3 | chr1 | + | 103543295 | 103543447 | 103541349 | 103541475 |
| SE | 56381 | ENSG00000185946 | RNPC3 | chr1 | + | 103543298 | 103543447 | 103541349 | 103541475 |
| SE | 56382 | ENSG00000185946 | RNPC3 | chr1 | + | 103544940 | 103545102 | 103543295 | 103543447 |
| SE | 56383 | ENSG00000185946 | RNPC3 | chr1 | + | 103544976 | 103545102 | 103543295 | 103543447 |
| SE | 56384 | ENSG00000031698 | SARS1 | chr1 | + | 109223977 | 109224048 | 109213917 | 109214128 |
| SE | 56385 | ENSG00000031698 | SARS1 | chr1 | + | 109230877 | 109231021 | 109229413 | 109229572 |
| SE | 56387 | ENSG00000031698 | SARS1 | chr1 | + | 109235976 | 109236106 | 109235209 | 109235431 |
| SE | 56390 | ENSG00000031698 | SARS1 | chr1 | + | 109237243 | 109237373 | 109236390 | 109236548 |
| SE | 56393 | ENSG00000151838 | CCDC175 | chr14 | - | 59525281 | 59525434 | 59521573 | 59521676 |
| SE | 56395 | ENSG00000151838 | CCDC175 | chr14 | - | 59527094 | 59527174 | 59525281 | 59525434 |
| SE | 56399 | ENSG00000151838 | CCDC175 | chr14 | - | 59540674 | 59540746 | 59538704 | 59538840 |
| SE | 56401 | ENSG00000151838 | CCDC175 | chr14 | - | 59543343 | 59543454 | 59538704 | 59538840 |
| SE | 56405 | ENSG00000151838 | CCDC175 | chr14 | - | 59545162 | 59545299 | 59538704 | 59538840 |
| SE | 56406 | ENSG00000151838 | CCDC175 | chr14 | - | 59545162 | 59545299 | 59540674 | 59540746 |
| SE | 56408 | ENSG00000151838 | CCDC175 | chr14 | - | 59551354 | 59551422 | 59540682 | 59540746 |
| SE | 56410 | ENSG00000151838 | CCDC175 | chr14 | - | 59568244 | 59568380 | 59565046 | 59565275 |
| SE | 56411 | ENSG00000151838 | CCDC175 | chr14 | - | 59572701 | 59572813 | 59565046 | 59565275 |
| SE | 56412 | ENSG00000104714 | ERICH1 | chr8 | - | 625674 | 627238 | 614745 | 615284 |
| SE | 56414 | ENSG00000104714 | ERICH1 | chr8 | - | 640582 | 640834 | 637535 | 637595 |
| SE | 56416 | ENSG00000104714 | ERICH1 | chr8 | - | 692477 | 692612 | 673288 | 674047 |
| SE | 56419 | ENSG00000104714 | ERICH1 | chr8 | - | 715860 | 716007 | 692477 | 692612 |
| SE | 56420 | ENSG00000160654 | CD3G | chr11 | + | 118349026 | 118349050 | 118344343 | 118344478 |
| SE | 56421 | ENSG00000160654 | CD3G | chr11 | + | 118349026 | 118349171 | 118344364 | 118344478 |
| SE | 56422 | ENSG00000160654 | CD3G | chr11 | + | 118349263 | 118349322 | 118349026 | 118349050 |
| SE | 56423 | ENSG00000160654 | CD3G | chr11 | + | 118349263 | 118349322 | 118349026 | 118349171 |
| SE | 56428 | ENSG00000050748 | MAPK9 | chr5 | - | 180238331 | 180238403 | 180233142 | 180236526 |
| SE | 56429 | ENSG00000050748 | MAPK9 | chr5 | - | 180238331 | 180238403 | 180233593 | 180236531 |
| SE | 56435 | ENSG00000158186 | MRAS | chr3 | + | 138400533 | 138400613 | 138398468 | 138398568 |
| SE | 56445 | ENSG00000116560 | SFPQ | chr1 | - | 35177318 | 35177399 | 35176377 | 35176474 |
| SE | 56446 | ENSG00000116560 | SFPQ | chr1 | - | 35177988 | 35178036 | 35177318 | 35177399 |
| SE | 56447 | ENSG00000116560 | SFPQ | chr1 | - | 35190693 | 35190995 | 35190497 | 35190593 |
| SE | 56454 | ENSG00000158156 | XKR8 | chr1 | + | 27965718 | 27965928 | 27963496 | 27963693 |
| SE | 56456 | ENSG00000127337 | YEATS4 | chr12 | + | 69361316 | 69361433 | 69359719 | 69360023 |
| SE | 56457 | ENSG00000127337 | YEATS4 | chr12 | + | 69362787 | 69362907 | 69359709 | 69360023 |
| SE | 56465 | ENSG00000106665 | CLIP2 | chr7 | + | 74386520 | 74386604 | 74380023 | 74380863 |
| SE | 56475 | ENSG00000162664 | ZNF326 | chr1 | + | 90020791 | 90020922 | 90018684 | 90018784 |
| SE | 56480 | ENSG00000136108 | CKAP2 | chr13 | + | 52460898 | 52460974 | 52456522 | 52456607 |
| SE | 56483 | ENSG00000114812 | VIPR1 | chr3 | + | 42512750 | 42512949 | 42502663 | 42502813 |
| SE | 56484 | ENSG00000114812 | VIPR1 | chr3 | + | 42513748 | 42513854 | 42502622 | 42502813 |

| | | | | | | | | | | | | | | | | | | |
|---|---|---|---|---|---|---|---|---|---|---|---|---|---|---|---|---|---|---|
| SE | 56490 | ENSG00000114812 | VIPR1 | chr3 | + | 42535342 | 42535384 | 42532865 | 42535104 | 42536089 | 42536394 | 11 | 26 | 21 | 26 | 21 | 0 | 0 |
| SE | 56492 | ENSG00000176853 | FAM91A1 | chr8 | + | 123778658 | 123778772 | 123777264 | 123777322 | 123779984 | 123780075 | 78 | 72 | 76 | 106 | 78 | 0 | 0 |
| SE | 56493 | ENSG00000176853 | FAM91A1 | chr8 | + | 123778658 | 123778772 | 123777264 | 123777322 | 123780479 | 123780542 | 2 | 3 | 2 | 5 | 2 | 0 | 0 |
| SE | 56494 | ENSG00000176853 | FAM91A1 | chr8 | + | 123779984 | 123780075 | 123777264 | 123777322 | 123780479 | 123780542 | 73 | 72 | 54 | 85 | 100 | 0 | 0 |
| SE | 56495 | ENSG00000176853 | FAM91A1 | chr8 | + | 123779984 | 123780075 | 123778658 | 123778772 | 123780479 | 123780542 | 166 | 161 | 133 | 212 | 206 | 0 | 0 |
| SE | 56496 | ENSG00000176853 | FAM91A1 | chr8 | + | 123779984 | 123780082 | 123777264 | 123777322 | 123780479 | 123780542 | 3 | 1 | 8 | 4 | 7 | 0 | 0 |
| SE | 56497 | ENSG00000176853 | FAM91A1 | chr8 | + | 123779984 | 123780082 | 123778658 | 123778772 | 123780479 | 123780542 | 70 | 71 | 67 | 115 | 94 | 0 | 0 |
| SE | 56498 | ENSG00000176853 | FAM91A1 | chr8 | + | 123786003 | 123786151 | 123785628 | 123785741 | 123786494 | 123786610 | 1 | 1 | 6 | 3 | 0 | 116 | 79 |
| SE | 56499 | ENSG00000176853 | FAM91A1 | chr8 | + | 123786003 | 123786151 | 123785628 | 123785741 | 123787260 | 123787355 | 0 | 0 | 0 | 0 | 0 | 1 | 1 |
| SE | 56502 | ENSG00000176853 | FAM91A1 | chr8 | + | 123787663 | 123787750 | 123787260 | 123787373 | 123789612 | 123789745 | 237 | 124 | 174 | 178 | 224 | 0 | 0 |
| SE | 56504 | ENSG00000176853 | FAM91A1 | chr8 | + | 123798089 | 123798238 | 123789612 | 123789745 | 123799519 | 123799654 | 159 | 124 | 151 | 205 | 212 | 0 | 0 |
| SE | 56505 | ENSG00000176853 | FAM91A1 | chr8 | + | 123805266 | 123805339 | 123799771 | 123799885 | 123806079 | 123806229 | 250 | 206 | 183 | 199 | 321 | 0 | 0 |
| SE | 56506 | ENSG00000176853 | FAM91A1 | chr8 | + | 123808271 | 123808376 | 123806079 | 123806229 | 123808892 | 123809016 | 240 | 177 | 157 | 210 | 287 | 1 | 0 |
| SE | 56507 | ENSG00000176853 | FAM91A1 | chr8 | + | 123812055 | 123812177 | 123810281 | 123810351 | 123812518 | 123812842 | 0 | 0 | 0 | 4 | 0 | 78 | 51 |
| SE | 56508 | ENSG00000105676 | ARMC6 | chr19 | + | 19034130 | 19034238 | 19033602 | 19033930 | 19037551 | 19037707 | 7 | 2 | 5 | 4 | 4 | 0 | 2 |
| SE | 56511 | ENSG00000105676 | ARMC6 | chr19 | + | 19034130 | 19034238 | 19033602 | 19033930 | 19042715 | 19042877 | 6 | 2 | 5 | 4 | 2 | 1 | 0 |
| SE | 56517 | ENSG00000105676 | ARMC6 | chr19 | + | 19039321 | 19039414 | 19033602 | 19033930 | 19042710 | 19042779 | 1 | 0 | 0 | 0 | 0 | 20 | 3 |
| SE | 56518 | ENSG00000105676 | ARMC6 | chr19 | + | 19039321 | 19039414 | 19033911 | 19034013 | 19042710 | 19042779 | 1 | 0 | 0 | 0 | 0 | 3 | 0 |
| SE | 56519 | ENSG00000105676 | ARMC6 | chr19 | + | 19039321 | 19039414 | 19034130 | 19034238 | 19042710 | 19042779 | 1 | 0 | 0 | 0 | 0 | 20 | 10 |
| SE | 56526 | ENSG00000143183 | TMCO1 | chr1 | - | 165743166 | 165743311 | 165724292 | 165728121 | 165752101 | 165752169 | 786 | 490 | 591 | 693 | 1761 | 0 | 0 |
| SE | 56527 | ENSG00000143183 | TMCO1 | chr1 | - | 165752101 | 165752169 | 165743166 | 165743311 | 165754227 | 165754274 | 450 | 273 | 320 | 380 | 980 | 1 | 0 |
| SE | 56532 | ENSG00000103489 | XYLT1 | chr16 | - | 17117645 | 17117979 | 17101768 | 17109017 | 17127665 | 17127861 | 78 | 34 | 74 | 62 | 41 | 2 | 0 |
| SE | 56535 | ENSG00000103489 | XYLT1 | chr16 | - | 17258987 | 17259498 | 17200481 | 17200654 | 17358011 | 17358050 | 83 | 73 | 79 | 107 | 60 | 0 | 0 |
| SE | 56539 | ENSG00000069702 | TGFBR3 | chr1 | - | 91698088 | 91698130 | 91695489 | 91695779 | 91708662 | 91708783 | 208 | 48 | 84 | 169 | 143 | 1 | 0 |
| SE | 56542 | ENSG00000069702 | TGFBR3 | chr1 | - | 91734775 | 91734959 | 91729804 | 91729973 | 91758612 | 91758750 | 696 | 226 | 283 | 553 | 388 | 2 | 0 |
| SE | 56543 | ENSG00000069702 | TGFBR3 | chr1 | - | 91758612 | 91758750 | 91734775 | 91734959 | 91786185 | 91786227 | 297 | 91 | 120 | 247 | 115 | 0 | 1 |
| SE | 56547 | ENSG00000069702 | TGFBR3 | chr1 | - | 91783205 | 91783264 | 91758612 | 91758750 | 91786185 | 91786227 | 0 | 0 | 2 | 0 | 0 | 1 | 4 |
| SE | 56548 | ENSG00000069702 | TGFBR3 | chr1 | - | 91783205 | 91783264 | 91758612 | 91758750 | 91797286 | 91797471 | 1 | 0 | 0 | 0 | 0 | 170 | 45 |
| SE | 56549 | ENSG00000069702 | TGFBR3 | chr1 | - | 91783205 | 91783264 | 91770841 | 91770970 | 91797286 | 91797471 | 1 | 0 | 0 | 0 | 0 | 1 | 0 |
| SE | 56550 | ENSG00000069702 | TGFBR3 | chr1 | - | 91786185 | 91786227 | 91734775 | 91734959 | 91797286 | 91797471 | 3 | 0 | 0 | 0 | 1 | 1 | 0 |
| SE | 56552 | ENSG00000069702 | TGFBR3 | chr1 | - | 91786185 | 91786227 | 91783205 | 91783264 | 91797286 | 91797471 | 3 | 0 | 2 | 0 | 1 | 1 | 0 |
| SE | 56554 | ENSG00000069702 | TGFBR3 | chr1 | - | 91800958 | 91801055 | 91797286 | 91797471 | 91861470 | 91861644 | 6 | 1 | 0 | 0 | 0 | 126 | 34 |
| SE | 56555 | ENSG00000069702 | TGFBR3 | chr1 | - | 91866755 | 91866842 | 91861470 | 91861644 | 91885877 | 91886122 | 0 | 0 | 0 | 0 | 0 | 69 | 3 |
| SE | 56564 | ENSG00000279457 | WASH9P | chr1 | - | 188129 | 188266 | 187375 | 187577 | 188438 | 188584 | 1 | 2 | 2 | 13 | 0 | 0 | 0 |
| SE | 56578 | ENSG00000105700 | KXD1 | chr19 | + | 18559172 | 18559285 | 18557837 | 18557914 | 18562035 | 18562157 | 0 | 0 | 0 | 0 | 0 | 41 | 13 |
| SE | 56583 | ENSG00000105700 | KXD1 | chr19 | + | 18564868 | 18565021 | 18561519 | 18562157 | 18567131 | 18567178 | 219 | 140 | 186 | 235 | 283 | 0 | 0 |
| SE | 56585 | ENSG00000105700 | KXD1 | chr19 | + | 18567131 | 18567178 | 18564868 | 18565021 | 18568401 | 18568495 | 117 | 73 | 101 | 132 | 152 | 2 | 0 |
| SE | 56586 | ENSG00000185697 | MYBL1 | chr8 | - | 66566063 | 66566243 | 66562174 | 66564825 | 66566683 | 66566788 | 248 | 108 | 72 | 168 | 359 | 64 | 24 |
| SE | 56588 | ENSG00000185697 | MYBL1 | chr8 | - | 66576006 | 66576375 | 66573363 | 66573506 | 66592439 | 66592544 | 379 | 164 | 159 | 307 | 441 | 0 | 0 |
| SE | 56589 | ENSG00000185697 | MYBL1 | chr8 | - | 66580132 | 66580366 | 66573363 | 66573506 | 66592439 | 66592544 | 317 | 174 | 168 | 287 | 486 | 0 | 0 |
| SE | 56590 | ENSG00000185697 | MYBL1 | chr8 | - | 66580132 | 66580366 | 66576006 | 66576375 | 66592439 | 66592544 | 793 | 428 | 381 | 737 | 1159 | 2 | 0 |
| SE | 56591 | ENSG00000185697 | MYBL1 | chr8 | - | 66601697 | 66601769 | 66599049 | 66599142 | 66602417 | 66602523 | 880 | 460 | 331 | 658 | 472 | 0 | 0 |
| SE | 56593 | ENSG00000183386 | FHL3 | chr1 | - | 37999256 | 37999432 | 37998717 | 37999148 | 38005356 | 38005514 | 14 | 17 | 9 | 22 | 3 | 0 | 0 |
| SE | 56595 | ENSG00000143367 | TUFT1 | chr1 | + | 151562090 | 151562165 | 151540308 | 151540426 | 151562584 | 151562686 | 3 | 0 | 1 | 0 | 0 | 0 | 0 |
| SE | 56596 | ENSG00000143367 | TUFT1 | chr1 | + | 151562584 | 151562686 | 151540304 | 151540426 | 151563903 | 151563990 | 1 | 1 | 2 | 5 | 3 | 0 | 0 |
| SE | 56599 | ENSG00000075151 | EIF4G3 | chr1 | - | 20825099 | 20825198 | 20817391 | 20817538 | 20827616 | 20827698 | 102 | 71 | 56 | 120 | 154 | 1 | 0 |
| SE | 56600 | ENSG00000075151 | EIF4G3 | chr1 | - | 20825099 | 20825198 | 20817391 | 20817538 | 20829146 | 20829272 | 52 | 29 | 33 | 63 | 85 | 1 | 0 |
| SE | 56601 | ENSG00000075151 | EIF4G3 | chr1 | - | 20827616 | 20827698 | 20817391 | 20817538 | 20829146 | 20829272 | 14 | 7 | 7 | 18 | 13 | 1 | 0 |
| SE | 56603 | ENSG00000075151 | EIF4G3 | chr1 | - | 20840855 | 20841028 | 20829146 | 20829272 | 20849414 | 20849530 | 209 | 132 | 126 | 217 | 341 | 0 | 0 |
| SE | 56605 | ENSG00000075151 | EIF4G3 | chr1 | - | 20853559 | 20853677 | 20851257 | 20851478 | 20854977 | 20855071 | 360 | 270 | 279 | 405 | 337 | 0 | 0 |
| SE | 56606 | ENSG00000075151 | EIF4G3 | chr1 | - | 20862227 | 20862332 | 20860384 | 20860517 | 20864475 | 20864712 | 189 | 156 | 100 | 218 | 165 | 2 | 0 |
| SE | 56607 | ENSG00000075151 | EIF4G3 | chr1 | - | 20886200 | 20886371 | 20879322 | 20879520 | 20893516 | 20893636 | 214 | 124 | 138 | 241 | 272 | 0 | 1 |
| SE | 56609 | ENSG00000075151 | EIF4G3 | chr1 | - | 20893516 | 20893636 | 20892618 | 20892729 | 20895367 | 20895501 | 146 | 64 | 68 | 117 | 113 | 0 | 0 |
| SE | 56610 | ENSG00000075151 | EIF4G3 | chr1 | - | 20899696 | 20899943 | 20895367 | 20895501 | 20904882 | 20904971 | 342 | 283 | 185 | 339 | 334 | 0 | 0 |
| SE | 56611 | ENSG00000075151 | EIF4G3 | chr1 | - | 20904882 | 20904971 | 20899696 | 20899943 | 20941490 | 20942330 | 240 | 188 | 132 | 233 | 222 | 1 | 0 |
| SE | 56612 | ENSG00000075151 | EIF4G3 | chr1 | - | 20941490 | 20942330 | 20904882 | 20904971 | 20950002 | 20950108 | 171 | 141 | 113 | 193 | 175 | 0 | 0 |
| SE | 56615 | ENSG00000075151 | EIF4G3 | chr1 | - | 20981047 | 20981227 | 20980333 | 20980448 | 21001198 | 21001312 | 37 | 20 | 25 | 44 | 28 | 0 | 2 |
| SE | 56616 | ENSG00000075151 | EIF4G3 | chr1 | - | 20981047 | 20981227 | 20980333 | 20980448 | 21002712 | 21002808 | 48 | 21 | 27 | 38 | 25 | 0 | 0 |
| SE | 56621 | ENSG00000075151 | EIF4G3 | chr1 | - | 20997600 | 20997633 | 20981047 | 20981227 | 21001198 | 21001307 | 0 | 0 | 0 | 0 | 0 | 7 | 5 |
| SE | 56625 | ENSG00000075151 | EIF4G3 | chr1 | - | 21001198 | 21001307 | 20980333 | 20980448 | 21002712 | 21002808 | 0 | 3 | 0 | 0 | 0 | 0 | 0 |
| SE | 56626 | ENSG00000075151 | EIF4G3 | chr1 | - | 21001198 | 21001307 | 20997600 | 20997633 | 21002712 | 21002808 | 0 | 1 | 0 | 0 | 0 | 0 | 0 |
| SE | 56627 | ENSG00000075151 | EIF4G3 | chr1 | - | 21001198 | 21001312 | 20980333 | 20980448 | 21002712 | 21002808 | 1 | 3 | 1 | 0 | 2 | 0 | 0 |
| SE | 56629 | ENSG00000075151 | EIF4G3 | chr1 | - | 21001198 | 21001324 | 20980333 | 20980448 | 21002712 | 21002808 | 0 | 5 | 4 | 0 | 0 | 0 | 0 |
| SE | 56632 | ENSG00000075151 | EIF4G3 | chr1 | - | 21002712 | 21002808 | 20981047 | 20981227 | 21050865 | 21050950 | 34 | 32 | 11 | 29 | 39 | 0 | 0 |
| SE | 56633 | ENSG00000075151 | EIF4G3 | chr1 | - | 21002712 | 21002808 | 20981047 | 20981227 | 21089137 | 21089213 | 12 | 6 | 4 | 6 | 10 | 2 | 0 |
| SE | 56634 | ENSG00000075151 | EIF4G3 | chr1 | - | 21050865 | 21050994 | 20981047 | 20981227 | 21089137 | 21089213 | 13 | 11 | 16 | 25 | 8 | 2 | 0 |
| SE | 56635 | ENSG00000075151 | EIF4G3 | chr1 | - | 21050865 | 21050994 | 21002712 | 21002808 | 21089137 | 21089213 | 35 | 38 | 23 | 45 | 40 | 0 | 0 |
| SE | 56636 | ENSG00000075151 | EIF4G3 | chr1 | - | 21064759 | 21064991 | 21050865 | 21050994 | 21089137 | 21089213 | 0 | 0 | 0 | 1 | 0 | 13 | 11 |
| SE | 56637 | ENSG00000075151 | EIF4G3 | chr1 | - | 21089137 | 21089213 | 21050865 | 21050994 | 21111322 | 21111383 | 16 | 3 | 14 | 19 | 11 | 0 | 0 |
| SE | 56644 | ENSG00000155893 | PXYLP1 | chr3 | + | 141234215 | 141234318 | 141231860 | 141231911 | 141260122 | 141260254 | 1 | 7 | 0 | 2 | 3 | 11 | 0 |
| SE | 56645 | ENSG00000155893 | PXYLP1 | chr3 | + | 141234885 | 141234956 | 141234215 | 141234318 | 141260122 | 141260254 | 6 | 1 | 0 | 5 | 2 | 1 | 6 |
| SE | 56647 | ENSG00000155893 | PXYLP1 | chr3 | + | 141291587 | 141291732 | 141287358 | 141287453 | 141292267 | 141292316 | 0 | 0 | 0 | 2 | 0 | 29 | 35 |
| SE | 56648 | ENSG00000155893 | PXYLP1 | chr3 | + | 141305755 | 141307776 | 141287313 | 141287453 | 141308203 | 141308407 | 0 | 0 | 0 | 0 | 0 | 2 | 0 |
| SE | 56650 | ENSG00000155307 | SAMSN1 | chr21 | - | 14521149 | 14521221 | 14512443 | 14512573 | 14546204 | 14546315 | 129 | 76 | 82 | 86 | 72 | 0 | 0 |
| SE | 56651 | ENSG00000155307 | SAMSN1 | chr21 | - | 14523270 | 14523394 | 14521149 | 14521221 | 14546204 | 14546313 | 7 | 5 | 0 | 3 | 0 | 306 | 196 |
| SE | 56661 | ENSG00000203697 | CAPN8 | chr1 | - | 223642682 | 223642833 | 223628661 | 223628780 | 223654329 | 223654399 | 0 | 0 | 2 | 0 | 0 | 0 | 0 |
| SE | 56663 | ENSG00000144749 | LRIG1 | chr3 | - | 66394039 | 66394203 | 66385980 | 66386301 | 66398111 | 66398183 | 101 | 69 | 121 | 85 | 178 | 0 | 0 |
| SE | 56666 | ENSG00000144749 | LRIG1 | chr3 | - | 66410128 | 66410272 | 66407347 | 66407491 | 66412870 | 66413014 | 49 | 18 | 55 | 35 | 54 | 0 | 1 |
| SE | 56668 | ENSG00000130638 | ATXN10 | chr22 | + | 45689711 | 45689903 | 45671833 | 45672179 | 45700281 | 45700322 | 24 | 12 | 30 | 31 | 9 | 0 | 0 |
| SE | 56669 | ENSG00000130638 | ATXN10 | chr22 | + | 45692995 | 45693078 | 45671797 | 45672179 | 45700281 | 45700322 | 273 | 118 | 218 | 197 | 250 | 0 | 0 |
| SE | 56670 | ENSG00000130638 | ATXN10 | chr22 | + | 45692995 | 45693078 | 45689711 | 45689903 | 45700281 | 45700322 | 430 | 177 | 346 | 308 | 364 | 2 | 0 |
| SE | 56671 | ENSG00000130638 | ATXN10 | chr22 | + | 45692995 | 45693078 | 45689711 | 45689903 | 45729424 | 45729590 | 79 | 25 | 73 | 57 | 60 | 0 | 2 |
| SE | 56672 | ENSG00000130638 | ATXN10 | chr22 | + | 45729424 | 45729590 | 45718412 | 45718493 | 45738730 | 45738839 | 296 | 154 | 235 | 299 | 464 | 0 | 0 |
| SE | 56673 | ENSG00000130638 | ATXN10 | chr22 | + | 45806958 | 45807022 | 45740368 | 45740538 | 45842990 | 45843178 | 219 | 146 | 189 | 210 | 272 | 0 | 0 |
| SE | 56674 | ENSG00000060138 | YBX3 | chr12 | - | 10703522 | 10703619 | 10701959 | 10702134 | 10704050 | 10704148 | 0 | 0 | 2 | 0 | 0 | 220 | 103 |
| SE | 56676 | ENSG00000060138 | YBX3 | chr12 | - | 10717712 | 10717777 | 10715694 | 10715783 | 10718087 | 10718114 | 0 | 0 | 2 | 0 | 0 | 180 | 78 |
| SE | 56677 | ENSG00000060138 | YBX3 | chr12 | - | 10718087 | 10718121 | 10715693 | 10715783 | 10719079 | 10719143 | 195 | 82 | 97 | 137 | 158 | 0 | 0 |
| SE | 56682 | ENSG00000008086 | CDKL5 | chrX | + | 18564476 | 18564522 | 18510819 | 18510854 | 18575353 | 18575390 | 27 | 34 | 20 | 27 | 30 | 2 | 1 |
| SE | 56684 | ENSG00000196437 | ZNF569 | chr19 | - | 37431558 | 37431647 | 37426338 | 37426378 | 37444906 | 37444964 | 0 | 1 | 0 | 1 | 0 | 38 | 21 |
| SE | 56685 | ENSG00000196437 | ZNF569 | chr19 | - | 37454833 | 37455073 | 37444906 | 37444964 | 37465312 | 37465473 | 13 | 5 | 8 | 10 | 11 | 33 | 21 |
| SE | 56686 | ENSG00000196437 | ZNF569 | chr19 | - | 37466738 | 37466804 | 37465312 | 37465473 | 37467083 | 37467290 | 2 | 5 | 6 | 4 | 1 | 13 | 7 |
| SE | 56687 | ENSG00000019582 | CD74 | chr5 | - | 150402562 | 150402625 | 150400040 | 150402250 | 150403120 | 150403312 | 4114 | 2079 | 2408 | 2717 | 4423 | 0 | 0 |
| SE | 56688 | ENSG00000019582 | CD74 | chr5 | - | 150402562 | 150402625 | 150400040 | 150402250 | 150404679 | 150404767 | 8792 | 4725 | 5522 | 6244 | 9344 | 1 | 0 |
| SE | 56691 | ENSG00000019582 | CD74 | chr5 | - | 150403120 | 150403312 | 150402562 | 150402625 | 150405084 | 150405180 | 836 | 461 | 504 | 560 | 909 | 2 | 0 |
| SE | 56692 | ENSG00000019582 | CD74 | chr5 | - | 150403120 | 150403312 | 150402562 | 150402625 | 150406258 | 150406321 | 836 | 462 | 504 | 560 | 907 | 4 | 0 |
| SE | 56693 | ENSG00000019582 | CD74 | chr5 | - | 150403120 | 150404767 | 150402562 | 150402625 | 150405084 | 150405180 | 1587 | 857 | 935 | 1150 | 1534 | 2 | 0 |
| SE | 56694 | ENSG00000019582 | CD74 | chr5 | - | 150404679 | 150404767 | 150402562 | 150402625 | 150405084 | 150405180 | 6119 | 3439 | 3950 | 4584 | 6090 | 2 | 0 |
| SE | 56695 | ENSG00000019582 | CD74 | chr5 | - | 150404679 | 150404767 | 150403120 | 150403312 | 150405084 | 150405180 | 1210 | 638 | 699 | 914 | 1007 | 0 | 0 |
| SE | 56696 | ENSG00000019582 | CD74 | chr5 | - | 150405084 | 150405180 | 150402562 | 150402625 | 150406258 | 150406321 | 3107 | 1316 | 2223 | 2800 | 1510 | 4 | 0 |
| SE | 56697 | ENSG00000019582 | CD74 | chr5 | - | 150405084 | 150405180 | 150403120 | 150403312 | 150406258 | 150406321 | 3105 | 1316 | 2218 | 2799 | 1510 | 0 | 1 |
| SE | 56698 | ENSG00000019582 | CD74 | chr5 | - | 150405084 | 150405671 | 150402562 | 150402625 | 150406258 | 150406321 | 4 | 0 | 2 | 1 | 2 | 4 | 0 |
| SE | 56699 | ENSG00000019582 | CD74 | chr5 | - | 150405084 | 150405671 | 150403120 | 150403312 | 150406258 | 150406321 | 2 | 0 | 0 | 0 | 2 | 0 | 1 |
| SE | 56700 | ENSG00000019582 | CD74 | chr5 | - | 150406258 | 150406321 | 150405084 | 150405180 | 150407151 | 150407324 | 435 | 226 | 259 | 372 | 326 | 0 | 0 |
| SE | 56702 | ENSG00000019582 | CD74 | chr5 | - | 150406880 | 150407324 | 150406038 | 150406321 | 150412624 | 150412912 | 11279 | 5618 | 8307 | 9997 | 7917 | 0 | 0 |
| SE | 56703 | ENSG00000019582 | CD74 | chr5 | - | 150407151 | 150407324 | 150406258 | 150406321 | 150412624 | 150412746 | 8220 | 4326 | 6131 | 7260 | 6491 | 0 | 0 |
| SE | 56709 | ENSG00000174827 | PDZK1 | chr1 | - | 145682499 | 145682636 | 145680911 | 145681107 | 145686476 | 145686726 | 0 | 0 | 0 | 0 | 0 | 1 | 0 |
| SE | 56710 | ENSG00000174827 | PDZK1 | chr1 | - | 145686476 | 145686726 | 145682499 | 145682636 | 145687811 | 145688023 | 7 | 0 | 2 | 9 | 3 | 2 | 0 |
| SE | 56716 | ENSG00000196782 | MAML3 | chr4 | - | 139730415 | 139730667 | 139725750 | 139725835 | 139889356 | 139890967 | 17 | 16 | 7 | 10 | 1 | 0 | 1 |
| SE | 56718 | ENSG00000134255 | CEPT1 | chr1 | + | 111161154 | 111161296 | 111159379 | 111159527 | 111174878 | 111174963 | 297 | 134 | 171 | 216 | 231 | 2 | 0 |
| SE | 56723 | ENSG00000144668 | ITGA9 | chr3 | + | 37494500 | 37494568 | 37481483 | 37481607 | 37503177 | 37503307 | 0 | 1 | 2 | 5 | 0 | 0 | 0 |
| SE | 56726 | ENSG00000184220 | CMSS1 | chr3 | + | 100141529 | 100141645 | 99817882 | 99818043 | 100146972 | 100147061 | 2 | 0 | 1 | 0 | 0 | 11 | 5 |
| SE | 56727 | ENSG00000184220 | CMSS1 | chr3 | + | 100171838 | 100171899 | 100167737 | 100167840 | 100172315 | 100172403 | 48 | 39 | 79 | 48 | 84 | 0 | 0 |
| SE | 56729 | ENSG00000184220 | CMSS1 | chr3 | + | 100176326 | 100176415 | 100172315 | 100172403 | 100178304 | 100178355 | 64 | 37 | 69 | 47 | 77 | 0 | 0 |
| SE | 56733 | ENSG00000270231 | NBPF8 | chr1 | + | 120427705 | 120427847 | 120419849 | 120420118 | 120436231 | 120436385 | 27 | 61 | 47 | 28 | 56 | 0 | 0 |
| SE | 56734 | ENSG00000270231 | NBPF8 | chr1 | + | 120440741 | 120440814 | 120439475 | 120439690 | 120441277 | 120441489 | 0 | 4 | 4 | 6 | 0 | 6 | 4 |
| SE | 56735 | ENSG00000270231 | NBPF8 | chr1 | + | 120444896 | 120444999 | 120442840 | 120443050 | 120445828 | 120446043 | 35 | 51 | 0 | 0 | 60 | 0 | 2 |
| SE | 56737 | ENSG00000270231 | NBPF8 | chr1 | + | 120452116 | 120452331 | 120451179 | 120451282 | 120453371 | 120453444 | 34 | 44 | 10 | 26 | 44 | 0 | 0 |
| SE | 56740 | ENSG00000270231 | NBPF8 | chr1 | + | 120460572 | 120460624 | 120459366 | 120459530 | 120461253 | 120461426 | 55 | 35 | 33 | 19 | 35 | 0 | 0 |
| SE | 56741 | ENSG00000117616 | RSRP1 | chr1 | - | 25243549 | 25243633 | 25242248 | 25242705 | 25245149 | 25245301 | 420 | 339 | 347 | 422 | 785 | 0 | 0 |
| SE | 56742 | ENSG00000117616 | RSRP1 | chr1 | - | 25243549 | 25244224 | 25242254 | 25242705 | 25245149 | 25245301 | 683 | 563 | 549 | 656 | 1155 | 0 | 0 |
| SE | 56747 | ENSG00000117616 | RSRP1 | chr1 | - | 25306895 | 25306949 | 25304682 | 25304821 | 25338068 | 25338213 | 9 | 2 | 5 | 2 | 1 | 2 | 2 |
| SE | 56751 | ENSG00000117616 | RSRP1 | chr1 | - | 25318062 | 25318271 | 25306895 | 25306949 | 25338068 | 25338213 | 8 | 9 | 3 | 0 | 0 | 2 | 0 |
| SE | 56762 | ENSG00000120800 | UTP20 | chr12 | + | 101285748 | 101285881 | 101285569 | 101285636 | 101286320 | 101286509 | 77 | 54 | 72 | 63 | 137 | 0 | 1 |
| SE | 56763 | ENSG00000120800 | UTP20 | chr12 | + | 101288959 | 101289041 | 101286320 | 101286509 | 101290732 | 101290888 | 19 | 11 | 24 | 19 | 39 | 0 | 0 |
| SE | 56765 | ENSG00000120800 | UTP20 | chr12 | + | 101293167 | 101293245 | 101291969 | 101292104 | 101295479 | 101295658 | 18 | 21 | 36 | 21 | 84 | 0 | 0 |
| SE | 56766 | ENSG00000120800 | UTP20 | chr12 | + | 101305914 | 101306065 | 101302447 | 101302553 | 101306698 | 101306761 | 73 | 106 | 90 | 113 | 113 | 0 | 3 |
| SE | 56767 | ENSG00000120800 | UTP20 | chr12 | + | 101346450 | 101346588 | 101345553 | 101345694 | 101352054 | 101352194 | 70 | 67 | 81 | 44 | 77 | 0 | 0 |
| SE | 56768 | ENSG00000120800 | UTP20 | chr12 | + | 101354831 | 101355118 | 101353046 | 101353129 | 101356553 | 101356693 | 98 | 101 | 87 | 108 | 145 | 0 | 0 |
| SE | 56769 | ENSG00000120800 | UTP20 | chr12 | + | 101365458 | 101365625 | 101363575 | 101363743 | 101366557 | 101366699 | 125 | 104 | 139 | 102 | 289 | 0 | 0 |
| SE | 56770 | ENSG00000120800 | UTP20 | chr12 | + | 101369720 | 101369891 | 101367859 | 101367976 | 101370431 | 101370563 | 48 | 27 | 50 | 46 | 34 | 0 | 0 |
| SE | 56771 | ENSG00000228956 | SATB1-AS1 | chr3 | + | 18465293 | 18465358 | 18464286 | 18464339 | 18465962 | 18466004 | 0 | 0 | 2 | 3 | 0 | 46 | 45 |
| SE | 56772 | ENSG00000228956 | SATB1-AS1 | chr3 | + | 18465962 | 18466039 | 18464286 | 18464339 | 18526386 | 18526539 | 120 | 92 | 187 | 156 | 397 | 0 | 0 |
| SE | 56773 | ENSG00000228956 | SATB1-AS1 | chr3 | + | 18472361 | 18472459 | 18465982 | 18466039 | 18473711 | 18473858 | 9 | 1 | 4 | 3 | 11 | 3 | 2 |
| SE | 56777 | ENSG00000228956 | SATB1-AS1 | chr3 | + | 18473711 | 18473858 | 18465962 | 18466039 | 18526386 | 18526540 | 5 | 3 | 3 | 3 | 5 | 107 | 79 |
| SE | 56778 | ENSG00000228956 | SATB1-AS1 | chr3 | + | 18473711 | 18473858 | 18472361 | 18472459 | 18526386 | 18526540 | 9 | 1 | 6 | 6 | 13 | 7 | 1 |
| SE | 56780 | ENSG00000228956 | SATB1-AS1 | chr3 | + | 18516044 | 18516178 | 18465982 | 18466039 | 18526386 | 18526540 | 0 | 0 | 0 | 0 | 1 | 107 | 79 |
| SE | 56781 | ENSG00000228956 | SATB1-AS1 | chr3 | + | 18527784 | 18528139 | 18527173 | 18527326 | 18529980 | 18530132 | 7 | 6 | 6 | 3 | 18 | 34 | 37 |
| SE | 56783 | ENSG00000228956 | SATB1-AS1 | chr3 | + | 18527950 | 18528139 | 18527173 | 18527326 | 18585386 | 18585442 | 4 | 5 | 5 | 5 | 19 | 0 | 2 |
| SE | 56784 | ENSG00000228956 | SATB1-AS1 | chr3 | + | 18528075 | 18528139 | 18527173 | 18527326 | 18529980 | 18530185 | 4 | 4 | 1 | 2 | 5 | 34 | 37 |
| SE | 56786 | ENSG00000228956 | SATB1-AS1 | chr3 | + | 18529980 | 18530185 | 18527173 | 18527326 | 18534686 | 18534763 | 34 | 37 | 69 | 54 | 171 | 3 | 5 |
| SE | 56787 | ENSG00000228956 | SATB1-AS1 | chr3 | + | 18529980 | 18530185 | 18527173 | 18527326 | 18585386 | 18585442 | 34 | 38 | 68 | 56 | 171 | 0 | 2 |
| SE | 56788 | ENSG00000228956 | SATB1-AS1 | chr3 | + | 18529980 | 18530185 | 18527173 | 18527326 | 18585619 | 18585762 | 34 | 37 | 68 | 54 | 171 | 0 | 0 |
| SE | 56789 | ENSG00000228956 | SATB1-AS1 | chr3 | + | 18529980 | 18530185 | 18527173 | 18527326 | 18606998 | 18607186 | 34 | 37 | 68 | 54 | 171 | 0 | 0 |
| SE | 56791 | ENSG00000228956 | SATB1-AS1 | chr3 | + | 18529980 | 18530326 | 18527173 | 18527326 | 18606998 | 18607186 | 34 | 37 | 68 | 54 | 171 | 0 | 0 |
| SE | 56794 | ENSG00000228956 | SATB1-AS1 | chr3 | + | 18548063 | 18548193 | 18527173 | 18527326 | 18585386 | 18585442 | 1 | 0 | 4 | 3 | 4 | 0 | 2 |
| SE | 56797 | ENSG00000228956 | SATB1-AS1 | chr3 | + | 18548063 | 18548193 | 18529983 | 18530185 | 18585386 | 18585442 | 0 | 0 | 1 | 0 | 1 | 0 | 1 |
| SE | 56814 | ENSG00000101842 | VSIG1 | chrX | + | 108061427 | 108061535 | 108058037 | 108058201 | 108066935 | 108067118 | 6 | 2 | 0 | 6 | 7 | 26 | 26 |
| SE | 56815 | ENSG00000101842 | VSIG1 | chrX | + | 108066935 | 108067134 | 108058037 | 108058201 | 108072676 | 108072832 | 37 | 29 | 51 | 41 | 78 | 0 | 0 |
| SE | 56816 | ENSG00000101842 | VSIG1 | chrX | + | 108073249 | 108073369 | 108072676 | 108072832 | 108076076 | 108076218 | 42 | 22 | 49 | 46 | 86 | 0 | 0 |
| SE | 56819 | ENSG00000119950 | MXI1 | chr10 | + | 110245742 | 110245818 | 110244827 | 110244857 | 110279179 | 110279294 | 0 | 0 | 1 | 0 | 0 | 65 | 46 |
| SE | 56821 | ENSG00000119950 | MXI1 | chr10 | + | 110279489 | 110279683 | 110279179 | 110279294 | 110279913 | 110280040 | 0 | 0 | 0 | 1 | 0 | 60 | 72 |
| SE | 56825 | ENSG00000261971 | MMP25-AS1 | chr16 | - | 3058539 | 3058702 | 3055735 | 3055945 | 3058995 | 3059085 | 5 | 6 | 8 | 4 | 2 | 0 | 0 |
| SE | 56828 | ENSG00000143398 | PIP5K1A | chr1 | + | 151224244 | 151224279 | 151198551 | 151199081 | 151227319 | 151227361 | 23 | 14 | 30 | 15 | 65 | 0 | 0 |
| SE | 56829 | ENSG00000143398 | PIP5K1A | chr1 | + | 151224247 | 151224279 | 151198550 | 151199081 | 151227319 | 151227400 | 55 | 36 | 60 | 41 | 106 | 0 | 0 |
| SE | 56831 | ENSG00000143398 | PIP5K1A | chr1 | + | 151231670 | 151231801 | 151227266 | 151227400 | 151232247 | 151232307 | 159 | 116 | 182 | 132 | 311 | 0 | 0 |
| SE | 56834 | ENSG00000143398 | PIP5K1A | chr1 | + | 151239954 | 151240039 | 151239129 | 151239178 | 151242122 | 151242269 | 103 | 56 | 100 | 113 | 97 | 0 | 0 |
| SE | 56837 | ENSG00000143398 | PIP5K1A | chr1 | + | 151246919 | 151246965 | 151242437 | 151242567 | 151247862 | 151248149 | 32 | 26 | 37 | 44 | 16 | 0 | 0 |
| SE | 56840 | ENSG00000214711 | CAPN14 | chr2 | - | 31197248 | 31197334 | 31194408 | 31194483 | 31199469 | 31199532 | 0 | 2 | 0 | 6 | 0 | 1 | 0 |
| SE | 56841 | ENSG00000214711 | CAPN14 | chr2 | - | 31201861 | 31201998 | 31200450 | 31200625 | 31202133 | 31202252 | 2 | 1 | 0 | 0 | 4 | 1 | 0 |
| SE | 56845 | ENSG00000133619 | KRBA1 | chr7 | + | 149715238 | 149715349 | 149714780 | 149715066 | 149719545 | 149719672 | 0 | 0 | 0 | 0 | 1 | 0 | 0 |
| SE | 56847 | ENSG00000133619 | KRBA1 | chr7 | + | 149729187 | 149729370 | 149728522 | 149728675 | 149730146 | 149730341 | 19 | 12 | 11 | 8 | 5 | 1 | 0 |
| SE | 56848 | ENSG00000133619 | KRBA1 | chr7 | + | 149730146 | 149730341 | 149729187 | 149729370 | 149730427 | 149730568 | 8 | 13 | 3 | 9 | 0 | 2 | 0 |
| SE | 56849 | ENSG00000173744 | AGFG1 | chr2 | + | 227491546 | 227491640 | 227472151 | 227472588 | 227519947 | 227520063 | 205 | 183 | 135 | 226 | 326 | 0 | 0 |
| SE | 56850 | ENSG00000173744 | AGFG1 | chr2 | + | 227519947 | 227520063 | 227491546 | 227491640 | 227523762 | 227523925 | 353 | 281 | 228 | 340 | 621 | 3 | 3 |
| SE | 56851 | ENSG00000173744 | AGFG1 | chr2 | + | 227523762 | 227523925 | 227519947 | 227520063 | 227524761 | 227524904 | 373 | 258 | 263 | 360 | 662 | 0 | 3 |
| SE | 56852 | ENSG00000173744 | AGFG1 | chr2 | + | 227531090 | 227531210 | 227524761 | 227524915 | 227533548 | 227533681 | 244 | 163 | 197 | 231 | 335 | 0 | 0 |
| SE | 56853 | ENSG00000173744 | AGFG1 | chr2 | + | 227532137 | 227532209 | 227531090 | 227531210 | 227533548 | 227533758 | 0 | 0 | 1 | 0 | 7 | 69 | 52 |
| SE | 56854 | ENSG00000173744 | AGFG1 | chr2 | + | 227536900 | 227536993 | 227536624 | 227536704 | 227551958 | 227552117 | 168 | 64 | 125 | 131 | 139 | 1 | 0 |
| SE | 56856 | ENSG00000117360 | PRPF3 | chr1 | + | 150324894 | 150325087 | 150321478 | 150321592 | 150325750 | 150325881 | 133 | 135 | 174 | 118 | 126 | 0 | 0 |
| SE | 56857 | ENSG00000117360 | PRPF3 | chr1 | + | 150325750 | 150325881 | 150324894 | 150325087 | 150328319 | 150328466 | 182 | 163 | 203 | 143 | 164 | 4 | 0 |
| SE | 56860 | ENSG00000117360 | PRPF3 | chr1 | + | 150332683 | 150332767 | 150327774 | 150328466 | 150332978 | 150333199 | 183 | 98 | 164 | 142 | 177 | 0 | 0 |
| SE | 56861 | ENSG00000117360 | PRPF3 | chr1 | + | 150334934 | 150335241 | 150332978 | 150333199 | 150338159 | 150338326 | 257 | 211 | 245 | 257 | 255 | 0 | 0 |
| SE | 56863 | ENSG00000117360 | PRPF3 | chr1 | + | 150338159 | 150338326 | 150334934 | 150335241 | 150343308 | 150343452 | 155 | 147 | 157 | 155 | 156 | 0 | 0 |
| SE | 56864 | ENSG00000117360 | PRPF3 | chr1 | + | 150340397 | 150340477 | 150334934 | 150335241 | 150343308 | 150343452 | 172 | 147 | 164 | 166 | 288 | 0 | 0 |
| SE | 56865 | ENSG00000117360 | PRPF3 | chr1 | + | 150340397 | 150340477 | 150338159 | 150338326 | 150343308 | 150343452 | 314 | 297 | 321 | 324 | 487 | 2 | 0 |
| SE | 56866 | ENSG00000117360 | PRPF3 | chr1 | + | 150344375 | 150344547 | 150344161 | 150344261 | 150346017 | 150346136 | 134 | 91 | 147 | 117 | 129 | 0 | 0 |
| SE | 56867 | ENSG00000117360 | PRPF3 | chr1 | + | 150344375 | 150344547 | 150344161 | 150344261 | 150346407 | 150346491 | 26 | 17 | 38 | 26 | 44 | 0 | 0 |
| SE | 56868 | ENSG00000117360 | PRPF3 | chr1 | + | 150344433 | 150344547 | 150344161 | 150344261 | 150346017 | 150346136 | 189 | 149 | 213 | 200 | 217 | 0 | 0 |
| SE | 56869 | ENSG00000117360 | PRPF3 | chr1 | + | 150344433 | 150344547 | 150344161 | 150344261 | 150346407 | 150346491 | 87 | 73 | 101 | 101 | 125 | 0 | 0 |
| SE | 56870 | ENSG00000117360 | PRPF3 | chr1 | + | 150346017 | 150346136 | 150344161 | 150344261 | 150346407 | 150346491 | 197 | 145 | 171 | 159 | 182 | 0 | 0 |
| SE | 56872 | ENSG00000117360 | PRPF3 | chr1 | + | 150346017 | 150346136 | 150344375 | 150344547 | 150349156 | 150349218 | 99 | 61 | 102 | 100 | 112 | 1 | 0 |
| SE | 56873 | ENSG00000117360 | PRPF3 | chr1 | + | 150346407 | 150346491 | 150344375 | 150344547 | 150349156 | 150349218 | 65 | 36 | 61 | 79 | 126 | 1 | 0 |
| SE | 56874 | ENSG00000117360 | PRPF3 | chr1 | + | 150346407 | 150346491 | 150346017 | 150346136 | 150349156 | 150349218 | 269 | 195 | 244 | 240 | 288 | 0 | 0 |
| SE | 56877 | ENSG00000110108 | TMEM109 | chr11 | + | 60919685 | 60919930 | 60914157 | 60914268 | 60921773 | 60923443 | 92 | 31 | 69 | 74 | 27 | 0 | 0 |
| SE | 56881 | ENSG00000125686 | MED1 | chr17 | - | 39427700 | 39427790 | 39424626 | 39424738 | 39431114 | 39431188 | 178 | 161 | 174 | 185 | 343 | 0 | 0 |
| SE | 56882 | ENSG00000125686 | MED1 | chr17 | - | 39431114 | 39431188 | 39427700 | 39427790 | 39431941 | 39432016 | 125 | 97 | 111 | 123 | 221 | 1 | 2 |
| SE | 56883 | ENSG00000125686 | MED1 | chr17 | - | 39431114 | 39431188 | 39427700 | 39427790 | 39434248 | 39434320 | 31 | 27 | 36 | 32 | 62 | 2 | 0 |
| SE | 56884 | ENSG00000125686 | MED1 | chr17 | - | 39431941 | 39432016 | 39427700 | 39427790 | 39434248 | 39434320 | 25 | 19 | 29 | 28 | 48 | 2 | 0 |
| SE | 56885 | ENSG00000125686 | MED1 | chr17 | - | 39431941 | 39432016 | 39431114 | 39431188 | 39434248 | 39434320 | 121 | 89 | 106 | 128 | 240 | 0 | 0 |
| SE | 56886 | ENSG00000125686 | MED1 | chr17 | - | 39434248 | 39434320 | 39431941 | 39432016 | 39439164 | 39439193 | 139 | 82 | 138 | 130 | 179 | 2 | 1 |
| SE | 56888 | ENSG00000125686 | MED1 | chr17 | - | 39443549 | 39443628 | 39440622 | 39440677 | 39447797 | 39447904 | 179 | 131 | 168 | 176 | 225 | 4 | 1 |
| SE | 56890 | ENSG00000125686 | MED1 | chr17 | - | 39447797 | 39447904 | 39445410 | 39445569 | 39451037 | 39451263 | 56 | 33 | 43 | 25 | 35 | 0 | 0 |
| SE | 56898 | ENSG00000085063 | CD59 | chr11 | - | 33731423 | 33731468 | 33722378 | 33722463 | 33736381 | 33736423 | 4 | 2 | 10 | 2 | 10 | 30 | 2 |
| SE | 56903 | ENSG00000179593 | ALOX15B | chr17 | + | 8046667 | 8046754 | 8045482 | 8045686 | 8046906 | 8047076 | 0 | 0 | 2 | 0 | 0 | 1 | 0 |
| SE | 56904 | ENSG00000111605 | CPSF6 | chr12 | + | 69252085 | 69252216 | 69251128 | 69251338 | 69253050 | 69253114 | 0 | 1 | 0 | 0 | 1 | 359 | 275 |
| SE | 56905 | ENSG00000111605 | CPSF6 | chr12 | + | 69253050 | 69253154 | 69251128 | 69251338 | 69256696 | 69256842 | 647 | 477 | 503 | 578 | 950 | 1 | 0 |
| SE | 56906 | ENSG00000111605 | CPSF6 | chr12 | + | 69255001 | 69256052 | 69253050 | 69253154 | 69256696 | 69256842 | 2 | 2 | 1 | 0 | 7 | 428 | 302 |
| SE | 56907 | ENSG00000111605 | CPSF6 | chr12 | + | 69257731 | 69257905 | 69256696 | 69256842 | 69258589 | 69259094 | 382 | 201 | 336 | 302 | 593 | 3 | 0 |
| SE | 56915 | ENSG00000269713 | NBPF9 | chr1 | - | 149060522 | 149061383 | 149059762 | 149059808 | 149062092 | 149062265 | 5 | 3 | 0 | 8 | 6 | 0 | 0 |
| SE | 56916 | ENSG00000269713 | NBPF9 | chr1 | - | 149065525 | 149065689 | 149064430 | 149064482 | 149069593 | 149069645 | 19 | 14 | 32 | 14 | 61 | 0 | 1 |
| SE | 56917 | ENSG00000269713 | NBPF9 | chr1 | - | 149065636 | 149065689 | 149064430 | 149064482 | 149069593 | 149069645 | 13 | 11 | 28 | 16 | 40 | 0 | 1 |
| SE | 56918 | ENSG00000269713 | NBPF9 | chr1 | - | 149070933 | 149071139 | 149069593 | 149069645 | 149071603 | 149071676 | 36 | 20 | 52 | 31 | 75 | 0 | 0 |
| SE | 56919 | ENSG00000269713 | NBPF9 | chr1 | - | 149071603 | 149071676 | 149069593 | 149069645 | 149072717 | 149072932 | 0 | 0 | 1 | 3 | 6 | 0 | 0 |
| SE | 56920 | ENSG00000269713 | NBPF9 | chr1 | - | 149073767 | 149073870 | 149072717 | 149072932 | 149075654 | 149075864 | 17 | 22 | 39 | 32 | 43 | 0 | 0 |
| SE | 56921 | ENSG00000269713 | NBPF9 | chr1 | - | 149080052 | 149080155 | 149079006 | 149079221 | 149081964 | 149082174 | 27 | 44 | 54 | 57 | 65 | 0 | 1 |
| SE | 56924 | ENSG00000269713 | NBPF9 | chr1 | - | 149090752 | 149090894 | 149082276 | 149082430 | 149098437 | 149098706 | 10 | 12 | 36 | 21 | 27 | 0 | 1 |
| SE | 56931 | ENSG00000118197 | DDX59 | chr1 | - | 200648438 | 200648567 | 200640821 | 200641256 | 200649073 | 200649149 | 93 | 30 | 59 | 82 | 99 | 0 | 0 |
| SE | 56933 | ENSG00000118197 | DDX59 | chr1 | - | 200649073 | 200649226 | 200644036 | 200644517 | 200659026 | 200659116 | 19 | 13 | 23 | 11 | 29 | 0 | 0 |
| SE | 56935 | ENSG00000118197 | DDX59 | chr1 | - | 200650424 | 200650676 | 200649073 | 200649226 | 200654805 | 200654869 | 106 | 49 | 84 | 81 | 150 | 0 | 0 |

| | | | | | | | | | | | | | | | | | | | | | | | | | | | | | | | | | | | | | | | | | | | | | | |
|---|---|---|---|---|---|---|---|---|---|---|---|---|---|---|---|---|---|---|---|---|---|---|---|---|---|---|---|---|---|---|---|---|---|---|---|---|---|---|---|---|---|---|---|---|---|---|
| SE | 56938 | ENSG00000118197 | DDX59 | chr1 | - | 200654440 | 200654550 | 200650442 | 200650676 | 200654805 | 200655104 | 2 | 1 | 2 | 2 | 0 | 3 | 0 | 0 | 0 | 4 | 0 | 6 | 0 | 12 | 0 | 0 | 0 | 2 | 0 | 0 | 259 | 149 | 1 | 1 | 0.277 | 1 | 1 | 1 | 0 | NA | 1 | 0 | 1 | NA | -0.011 |
| SE | 56939 | ENSG00000118197 | DDX59 | chr1 | - | 200654440 | 200654582 | 200649073 | 200649226 | 200654805 | 200654869 | 0 | 3 | 0 | 0 | 0 | 0 | 0 | 0 | 1 | 0 | 4 | 0 | 6 | 4 | 0 | 0 | 0 | 3 | 1 | 0 | 291 | 149 | 1 | 1 | NA | 1 | NA | 0 | NA | 1 | NA | 0.506 | 0.672 | NA | -0.226 |
| SE | 56941 | ENSG00000118197 | DDX59 | chr1 | - | 200654440 | 200654582 | 200650424 | 200650676 | 200654805 | 200654869 | 1 | 4 | 2 | 2 | 1 | 3 | 0 | 0 | 0 | 4 | 0 | 6 | 0 | 12 | 0 | 0 | 0 | 2 | 0 | 0 | 291 | 149 | 1 | 1 | 0.146 | 1 | 1 | 1 | 0.113 | NA | 1 | 0 | 1 | NA | -0.015 |
| SE | 56944 | ENSG00000118197 | DDX59 | chr1 | - | 200659026 | 200659116 | 200650424 | 200650676 | 200663918 | 200664086 | 132 | 56 | 68 | 72 | 126 | 2 | 0 | 1 | 2 | 0 | 45 | 52 | 47 | 43 | 19 | 0 | 0 | 4 | 0 | 0 | 239 | 149 | 1 | 1 | 0.976 | 1 | 0.977 | 0.957 | 1 | 1 | 1 | 0.88 | 1 | 1 | 0.006 |
| SE | 56945 | ENSG00000118197 | DDX59 | chr1 | - | 200663918 | 200664086 | 200659026 | 200659116 | 200665936 | 200666751 | 202 | 73 | 125 | 130 | 210 | 2 | 0 | 0 | 0 | 0 | 89 | 93 | 151 | 109 | 21 | 0 | 0 | 0 | 0 | 0 | 298 | 149 | 1 | 1 | 0.981 | 1 | 1 | 1 | 1 | 1 | 1 | 1 | 1 | 1 | -0.004 |
| SE | 56946 | ENSG00000188290 | HES4 | chr1 | - | 999691 | 999787 | 999525 | 999613 | 999865 | 1000097 | 4 | 1 | 2 | 1 | 0 | 1 | 0 | 0 | 0 | 0 | 0 | 0 | 0 | 1 | 0 | 0 | 0 | 0 | 0 | 0 | 245 | 149 | 1 | 1 | 0.709 | 1 | 1 | 1 | NA | NA | NA | NA | 1 | NA | -0.073 |
| SE | 56951 | ENSG00000244005 | NFS1 | chr20 | - | 35680736 | 35680871 | 35675044 | 35675202 | 35681887 | 35681981 | 76 | 59 | 44 | 71 | 50 | 0 | 0 | 0 | 1 | 0 | 96 | 90 | 29 | 44 | 56 | 0 | 0 | 0 | 0 | 0 | 284 | 149 | 1 | 1 | 1 | 1 | 1 | 0.974 | 1 | 1 | 1 | 1 | 1 | 1 | -0.005 |
| SE | 56952 | ENSG00000244005 | NFS1 | chr20 | - | 35690412 | 35690565 | 35681887 | 35681981 | 35696376 | 35696460 | 76 | 40 | 46 | 52 | 57 | 0 | 2 | 0 | 0 | 0 | 79 | 95 | 42 | 53 | 71 | 0 | 0 | 0 | 0 | 0 | 298 | 149 | 1 | 1 | 1 | 0.909 | 1 | 1 | 1 | 1 | 1 | 1 | 1 | 1 | -0.018 |
| SE | 56958 | ENSG00000120725 | SIL1 | chr5 | - | 139042619 | 139042719 | 139026800 | 139026992 | 139050937 | 139051046 | 60 | 53 | 58 | 61 | 70 | 2 | 0 | 0 | 0 | 2 | 156 | 152 | 80 | 107 | 130 | 0 | 0 | 0 | 5 | 0 | 249 | 149 | 1 | 1 | 0.947 | 1 | 1 | 1 | 0.954 | 1 | 1 | 1 | 0.928 | 1 | -0.005 |
| SE | 56959 | ENSG00000120725 | SIL1 | chr5 | - | 139127943 | 139128052 | 139127738 | 139127853 | 139196440 | 139196528 | 0 | 0 | 0 | 1 | 0 | 2 | 0 | 2 | 1 | 0 | 0 | 0 | 0 | 0 | 6 | 11 | 5 | 5 | 5 | 0 | 258 | 149 | 1 | 1 | 0 | NA | 0 | 0.366 | NA | 0 | 0 | 0 | 0 | 1 | -0.078 |
| SE | 56960 | ENSG00000120725 | SIL1 | chr5 | - | 139127943 | 139128052 | 139127738 | 139127853 | 139198268 | 139198304 | 0 | 0 | 0 | 2 | 0 | 0 | 0 | 3 | 2 | 2 | 0 | 0 | 0 | 0 | 6 | 0 | 15 | 6 | 14 | 13 | 258 | 149 | 1 | 1 | NA | NA | 0 | 0.366 | 0 | NA | 0 | 0 | 0 | 0.21 | 0.07 |
| SE | 56961 | ENSG00000120725 | SIL1 | chr5 | - | 139133333 | 139133471 | 139127738 | 139127853 | 139196440 | 139196528 | 0 | 0 | 1 | 0 | 0 | 2 | 0 | 2 | 1 | 0 | 2 | 0 | 0 | 3 | 0 | 11 | 5 | 5 | 5 | 0 | 287 | 149 | 1 | 1 | 0 | NA | 0.206 | 0 | NA | 0.086 | 0 | 0 | 0.238 | NA | -0.012 |
| SE | 56962 | ENSG00000120725 | SIL1 | chr5 | - | 139133333 | 139133471 | 139127738 | 139127853 | 139198268 | 139198306 | 0 | 0 | 1 | 0 | 0 | 2 | 0 | 3 | 2 | 2 | 2 | 0 | 0 | 0 | 0 | 0 | 15 | 6 | 14 | 13 | 287 | 149 | 1 | 1 | NA | NA | 0.148 | 0 | 0 | 1 | 0 | 0 | 0 | 0 | -0.151 |
| SE | 56963 | ENSG00000120725 | SIL1 | chr5 | - | 139137328 | 139137474 | 139127738 | 139127853 | 139196440 | 139196521 | 0 | 1 | 0 | 0 | 0 | 2 | 0 | 2 | 1 | 0 | 0 | 0 | 0 | 0 | 0 | 11 | 5 | 5 | 5 | 0 | 295 | 149 | 1 | 1 | 0 | 1 | 0 | 0 | NA | 0 | 0 | 0 | 0 | NA | 0.25 |
| SE | 56964 | ENSG00000120725 | SIL1 | chr5 | - | 139196440 | 139196528 | 139127738 | 139127853 | 139198268 | 139198297 | 2 | 0 | 2 | 1 | 0 | 2 | 0 | 3 | 2 | 2 | 11 | 5 | 5 | 5 | 0 | 0 | 15 | 6 | 14 | 13 | 237 | 149 | 1 | 1 | 1 | NA | 0.295 | 0.239 | 0 | 1 | 0.173 | 0.344 | 0.183 | 0 | 0.043 |
| SE | 56967 | ENSG00000130775 | THEMIS2 | chr1 | + | 27876587 | 27876728 | 27872542 | 27872665 | 27879643 | 27880054 | 179 | 71 | 110 | 112 | 25 | 2 | 1 | 2 | 0 | 0 | 290 | 167 | 154 | 230 | 233 | 0 | 0 | 0 | 0 | 2 | 290 | 149 | 1 | 1 | 0.979 | 0.973 | 0.966 | 1 | 1 | 1 | 1 | 1 | 1 | 0.984 | -0.013 |
| SE | 56972 | ENSG00000130775 | THEMIS2 | chr1 | + | 27881970 | 27882077 | 27879730 | 27880054 | 27882512 | 27883043 | 98 | 44 | 52 | 45 | 42 | 0 | 0 | 1 | 1 | 0 | 111 | 71 | 102 | 60 | 123 | 0 | 0 | 2 | 3 | 0 | 256 | 149 | 1 | 1 | 1 | 1 | 0.968 | 0.963 | 1 | 1 | 1 | 0.967 | 0.921 | 1 | 0.009 |
| SE | 56973 | ENSG00000130775 | THEMIS2 | chr1 | + | 27881970 | 27882125 | 27879643 | 27880054 | 27882512 | 27883043 | 114 | 51 | 57 | 49 | 45 | 0 | 0 | 1 | 1 | 0 | 80 | 100 | 86 | 62 | 153 | 0 | 0 | 2 | 3 | 0 | 298 | 149 | 1 | 1 | 1 | 1 | 0.966 | 0.961 | 1 | 1 | 1 | 0.956 | 0.912 | 1 | 0.012 |
| SE | 56978 | ENSG00000171603 | CLSTN1 | chr1 | - | 9737497 | 9737554 | 9735884 | 9736042 | 9741093 | 9741256 | 0 | 0 | 0 | 1 | 0 | 219 | 126 | 172 | 217 | 221 | 0 | 0 | 0 | 0 | 0 | 139 | 204 | 145 | 92 | 259 | 206 | 149 | 1 | 1 | 0 | 0 | 0 | 0.003 | 0 | 0 | 0 | 0 | 0 | 0 | 0.001 |
| SE | 56980 | ENSG00000171603 | CLSTN1 | chr1 | - | 9755113 | 9755309 | 9751472 | 9751681 | 9773271 | 9773394 | 463 | 276 | 450 | 432 | 638 | 0 | 0 | 2 | 3 | 0 | 501 | 691 | 311 | 378 | 689 | 0 | 0 | 0 | 0 | 0 | 298 | 149 | 1 | 1 | 1 | 1 | 0.991 | 0.986 | 1 | 1 | 1 | 1 | 1 | 1 | -0.005 |
| SE | 56981 | ENSG00000171603 | CLSTN1 | chr1 | - | 9756480 | 9756510 | 9755113 | 9755309 | 9773271 | 9773394 | 0 | 1 | 0 | 0 | 0 | 106 | 66 | 133 | 140 | 122 | 0 | 3 | 1 | 0 | 0 | 113 | 208 | 85 | 71 | 168 | 179 | 149 | 1 | 1 | 0 | 0.012 | 0 | 0 | 0 | 0 | 0.012 | 0.01 | 0 | 0 | -0.002 |
| SE | 56987 | ENSG00000131725 | WDR44 | chrX | + | 118378418 | 118378452 | 118346072 | 118346580 | 118392631 | 118393271 | 0 | 2 | 1 | 4 | 3 | 0 | 0 | 0 | 3 | 0 | 0 | 1 | 0 | 1 | 0 | 0 | 0 | 0 | 0 | 0 | 183 | 149 | 1 | 1 | NA | 1 | 1 | 0.521 | 1 | NA | 1 | NA | 1 | NA | -0.12 |
| SE | 56988 | ENSG00000131725 | WDR44 | chrX | + | 118387339 | 118387414 | 118346198 | 118346580 | 118392631 | 118393215 | 33 | 24 | 30 | 38 | 52 | 0 | 0 | 0 | 3 | 0 | 62 | 30 | 24 | 23 | 6 | 0 | 0 | 0 | 0 | 0 | 224 | 149 | 1 | 1 | 1 | 1 | 1 | 0.894 | 1 | 1 | 1 | 1 | 1 | 1 | -0.021 |
| SE | 56992 | ENSG00000131725 | WDR44 | chrX | + | 118392631 | 118393271 | 118387339 | 118387414 | 118394054 | 118394185 | 156 | 140 | 112 | 156 | 150 | 2 | 0 | 0 | 0 | 0 | 156 | 132 | 106 | 88 | 103 | 0 | 0 | 4 | 0 | 0 | 298 | 149 | 1 | 1 | 0.975 | 1 | 1 | 1 | 1 | 1 | 1 | 0.93 | 1 | 1 | 0.009 |
| SE | 56993 | ENSG00000131725 | WDR44 | chrX | + | 118404337 | 118404444 | 118398386 | 118398470 | 118406874 | 118407026 | 154 | 161 | 113 | 155 | 217 | 0 | 0 | 0 | 2 | 0 | 51 | 180 | 118 | 73 | 72 | 0 | 0 | 0 | 0 | 0 | 256 | 149 | 1 | 1 | 1 | 1 | 1 | 0.978 | 1 | 1 | 1 | 1 | 1 | 1 | -0.004 |
| SE | 56994 | ENSG00000131725 | WDR44 | chrX | + | 118436701 | 118436824 | 118432780 | 118432894 | 118441367 | 118441559 | 116 | 125 | 125 | 163 | 182 | 0 | 2 | 1 | 0 | 0 | 63 | 67 | 104 | 84 | 24 | 0 | 2 | 0 | 0 | 0 | 272 | 149 | 1 | 1 | 1 | 0.972 | 0.986 | 1 | 1 | 1 | 0.948 | 1 | 1 | 1 | 0.002 |
| SE | 56995 | ENSG00000131725 | WDR44 | chrX | + | 118441367 | 118441559 | 118436701 | 118436824 | 118442243 | 118442345 | 175 | 195 | 146 | 214 | 230 | 1 | 0 | 1 | 4 | 0 | 117 | 162 | 187 | 153 | 35 | 0 | 0 | 0 | 0 | 0 | 298 | 149 | 1 | 1 | 0.989 | 1 | 0.986 | 0.964 | 1 | 1 | 1 | 1 | 1 | 1 | -0.012 |
| SE | 56996 | ENSG00000131725 | WDR44 | chrX | + | 118444359 | 118444494 | 118443559 | 118443687 | 118448892 | 118449937 | 186 | 168 | 130 | 168 | 183 | 0 | 0 | 0 | 0 | 0 | 65 | 108 | 107 | 92 | 37 | 0 | 3 | 0 | 0 | 0 | 284 | 149 | 1 | 1 | 1 | 1 | 1 | 1 | 1 | 1 | 0.95 | 1 | 1 | 1 | 0.01 |
| SE | 56997 | ENSG00000131725 | WDR44 | chrX | + | 118447059 | 118447114 | 118444359 | 118444494 | 118448892 | 118449336 | 2 | 0 | 0 | 1 | 0 | 101 | 83 | 68 | 90 | 77 | 0 | 4 | 0 | 0 | 0 | 27 | 52 | 49 | 37 | 13 | 204 | 149 | 1 | 1 | 0.014 | 0 | 0 | 0.008 | 0 | 0 | 0.053 | 0 | 0 | 0 | -0.006 |
| SE | 57000 | ENSG00000198298 | ZNF485 | chr10 | + | 43608613 | 43608740 | 43606996 | 43607074 | 43609254 | 43609350 | 7 | 6 | 5 | 5 | 4 | 0 | 0 | 2 | 3 | 0 | 0 | 0 | 1 | 0 | 0 | 0 | 0 | 0 | 0 | 0 | 276 | 149 | 1 | 1 | 1 | 1 | 0.574 | 0.474 | 1 | NA | NA | 1 | NA | NA | -0.19 |
| SE | 57015 | ENSG00000127124 | HIVEP3 | chr1 | - | 41628748 | 41628947 | 41579736 | 41585318 | 41918412 | 41918707 | 21 | 18 | 15 | 28 | 31 | 0 | 0 | 0 | 0 | 2 | 85 | 52 | 21 | 25 | 42 | 0 | 0 | 0 | 0 | 0 | 298 | 149 | 1 | 1 | 1 | 1 | 1 | 1 | 0.886 | 1 | 1 | 1 | 1 | 1 | -0.023 |
| SE | 57016 | ENSG00000127124 | HIVEP3 | chr1 | - | 41700915 | 41700995 | 41628748 | 41628947 | 41918412 | 41918922 | 10 | 0 | 12 | 4 | 5 | 1 | 0 | 0 | 0 | 0 | 0 | 24 | 10 | 8 | 9 | 7 | 6 | 0 | 0 | 0 | 229 | 149 | 1 | 1 | 0.867 | NA | 1 | 1 | 1 | 0 | 0.722 | 1 | 1 | 1 | 0.222 |
| SE | 57017 | ENSG00000127124 | HIVEP3 | chr1 | - | 41848870 | 41849089 | 41847190 | 41848445 | 41869546 | 41869630 | 20 | 12 | 15 | 5 | 23 | 1 | 0 | 0 | 0 | 0 | 0 | 18 | 6 | 15 | 29 | 0 | 0 | 0 | 0 | 0 | 298 | 149 | 1 | 1 | 0.909 | 1 | 1 | 1 | 1 | NA | 1 | 1 | 1 | 1 | -0.018 |
| SE | 57018 | ENSG00000127124 | HIVEP3 | chr1 | - | 41848870 | 41849089 | 41847190 | 41848445 | 41918412 | 41918524 | 24 | 16 | 22 | 8 | 26 | 0 | 0 | 0 | 0 | 1 | 0 | 23 | 14 | 23 | 38 | 0 | 0 | 0 | 0 | 0 | 298 | 149 | 1 | 1 | 1 | 1 | 1 | 1 | 0.929 | NA | 1 | 1 | 1 | 1 | -0.014 |
| SE | 57019 | ENSG00000127124 | HIVEP3 | chr1 | - | 41869546 | 41869630 | 41847947 | 41848445 | 41918412 | 41918922 | 3 | 0 | 1 | 0 | 0 | 0 | 0 | 0 | 0 | 1 | 0 | 5 | 0 | 0 | 0 | 0 | 0 | 0 | 0 | 0 | 233 | 149 | 1 | 1 | 1 | NA | 1 | NA | 0 | NA | 1 | NA | NA | NA | -0.333 |
| SE | 57020 | ENSG00000127124 | HIVEP3 | chr1 | - | 41869546 | 41869630 | 41848870 | 41849089 | 41918412 | 41918922 | 4 | 0 | 3 | 0 | 0 | 6 | 4 | 9 | 3 | 3 | 0 | 5 | 0 | 0 | 0 | 0 | 5 | 8 | 8 | 9 | 233 | 149 | 1 | 1 | 0.299 | 0 | 0.176 | 0 | 0 | NA | 0.39 | 0 | 0 | 0 | -0.003 |
| SE | 57021 | ENSG00000127124 | HIVEP3 | chr1 | - | 41869546 | 41869630 | 41865547 | 41865615 | 41918412 | 41918922 | 9 | 0 | 4 | 0 | 0 | 0 | 0 | 0 | 0 | 0 | 0 | 16 | 0 | 0 | 0 | 0 | 5 | 1 | 0 | 0 | 233 | 149 | 1 | 1 | 1 | NA | 1 | NA | NA | NA | 0.672 | 0 | NA | NA | 0.664 |
| SE | 57023 | ENSG00000259673 | IQCH-AS1 | chr15 | - | 67430320 | 67430400 | 67421208 | 67421498 | 67459691 | 67459975 | 2 | 1 | 4 | 4 | 0 | 0 | 0 | 1 | 1 | 0 | 0 | 0 | 3 | 0 | 0 | 0 | 0 | 0 | 0 | 0 | 229 | 149 | 1 | 1 | 1 | 1 | 0.722 | 0.722 | NA | NA | NA | 1 | NA | NA | -0.139 |
| SE | 57025 | ENSG00000259673 | IQCH-AS1 | chr15 | - | 67459691 | 67459975 | 67430320 | 67430400 | 67518999 | 67519102 | 2 | 1 | 0 | 4 | 2 | 1 | 0 | 0 | 0 | 0 | 0 | 0 | 0 | 4 | 0 | 0 | 0 | 0 | 0 | 0 | 298 | 149 | 1 | 1 | 0.5 | 1 | NA | 1 | 1 | NA | NA | NA | 1 | NA | -0.125 |
| SE | 57027 | ENSG00000268043 | NBPF12 | chr1 | + | 146951347 | 146951489 | 146943290 | 146943562 | 146959858 | 146960006 | 2 | 6 | 6 | 2 | 2 | 0 | 0 | 0 | 0 | 0 | 4 | 2 | 5 | 9 | 3 | 0 | 0 | 2 | 0 | 0 | 291 | 149 | 1 | 1 | 1 | 1 | 1 | 1 | 1 | 1 | 1 | 0.561 | 1 | 1 | 0.088 |
| SE | 57030 | ENSG00000268043 | NBPF12 | chr1 | + | 146951813 | 146952037 | 146951347 | 146951489 | 146960108 | 146960318 | 2 | 0 | 6 | 0 | 8 | 0 | 0 | 0 | 0 | 0 | 8 | 3 | 0 | 16 | 6 | 0 | 0 | 0 | 5 | 7 | 298 | 149 | 1 | 1 | 1 | NA | 1 | NA | 1 | 1 | 1 | NA | 0.615 | 0.3 | 0.271 |
| SE | 57034 | ENSG00000268043 | NBPF12 | chr1 | + | 146963094 | 146963309 | 146960108 | 146960318 | 146968447 | 146968550 | 0 | 2 | 0 | 1 | 1 | 0 | 0 | 1 | 0 | 0 | 0 | 7 | 5 | 16 | 0 | 0 | 0 | 0 | 0 | 0 | 298 | 149 | 1 | 1 | NA | 1 | 0 | 1 | 1 | NA | 1 | 1 | 1 | NA | -0.25 |
| SE | 57035 | ENSG00000268043 | NBPF12 | chr1 | + | 146964892 | 146965104 | 146964356 | 146964429 | 146966463 | 146966673 | 99 | 37 | 117 | 61 | 95 | 0 | 0 | 0 | 0 | 0 | 95 | 68 | 64 | 143 | 43 | 0 | 0 | 0 | 1 | 0 | 298 | 149 | 1 | 1 | 1 | 1 | 1 | 1 | 1 | 1 | 1 | 1 | 0.986 | 1 | 0.003 |
| SE | 57036 | ENSG00000268043 | NBPF12 | chr1 | + | 146968447 | 146968550 | 146963094 | 146963309 | 146970646 | 146970719 | 0 | 1 | 0 | 0 | 0 | 0 | 0 | 0 | 0 | 0 | 0 | 0 | 0 | 1 | 0 | 0 | 0 | 0 | 2 | 0 | 252 | 149 | 1 | 1 | NA | 1 | NA | NA | NA | NA | NA | NA | 0.228 | NA | 0.772 |
| SE | 57037 | ENSG00000268043 | NBPF12 | chr1 | + | 146969381 | 146969596 | 146968447 | 146968550 | 146970646 | 146970719 | 65 | 43 | 72 | 67 | 108 | 1 | 0 | 0 | 0 | 0 | 37 | 81 | 96 | 130 | 80 | 0 | 0 | 0 | 1 | 0 | 298 | 149 | 1 | 1 | 0.97 | 1 | 1 | 1 | 1 | 1 | 1 | 1 | 0.985 | 1 | -0.003 |
| SE | 57038 | ENSG00000268043 | NBPF12 | chr1 | + | 146969381 | 146969596 | 146968447 | 146968550 | 146976928 | 146977001 | 32 | 23 | 33 | 31 | 46 | 1 | 0 | 0 | 2 | 0 | 12 | 45 | 40 | 66 | 38 | 0 | 0 | 4 | 0 | 0 | 298 | 149 | 1 | 1 | 0.941 | 1 | 1 | 0.886 | 1 | 1 | 1 | 0.833 | 1 | 1 | -0.001 |
| SE | 57039 | ENSG00000268043 | NBPF12 | chr1 | + | 146974738 | 146974841 | 146968447 | 146968550 | 146976928 | 146977001 | 0 | 0 | 0 | 0 | 0 | 1 | 0 | 0 | 2 | 0 | 0 | 0 | 1 | 2 | 0 | 0 | 0 | 4 | 0 | 0 | 252 | 149 | 1 | 1 | 0 | NA | NA | 0 | NA | NA | NA | 0.129 | 1 | NA | -0.565 |
| SE | 57040 | ENSG00000268043 | NBPF12 | chr1 | + | 146975676 | 146975891 | 146974738 | 146974841 | 146976928 | 146977001 | 32 | 11 | 25 | 19 | 13 | 0 | 0 | 0 | 0 | 0 | 8 | 20 | 17 | 35 | 18 | 1 | 0 | 1 | 2 | 0 | 298 | 149 | 1 | 1 | 1 | 1 | 1 | 1 | 1 | 0.8 | 1 | 0.895 | 0.897 | 1 | 0.082 |
| SE | 57046 | ENSG00000268043 | NBPF12 | chr1 | + | 146990465 | 146990517 | 146989573 | 146989746 | 146991112 | 146991285 | 42 | 76 | 59 | 44 | 96 | 1 | 0 | 0 | 0 | 0 | 30 | 72 | 96 | 70 | 63 | 0 | 0 | 2 | 4 | 0 | 201 | 149 | 1 | 1 | 0.969 | 1 | 1 | 1 | 1 | 1 | 1 | 0.973 | 0.928 | 1 | 0.014 |
| SE | 57047 | ENSG00000268043 | NBPF12 | chr1 | + | 146991112 | 146991285 | 146988840 | 146988949 | 146991994 | 146992046 | 22 | 35 | 19 | 25 | 51 | 0 | 0 | 0 | 0 | 0 | 25 | 41 | 31 | 22 | 20 | 0 | 0 | 1 | 0 | 0 | 298 | 149 | 1 | 1 | 1 | 1 | 1 | 1 | 1 | 1 | 1 | 0.939 | 1 | 1 | 0.012 |
| SE | 57048 | ENSG00000268043 | NBPF12 | chr1 | + | 146991112 | 146991285 | 146989573 | 146989746 | 146993598 | 146993707 | 1 | 0 | 0 | 0 | 2 | 0 | 0 | 0 | 0 | 0 | 0 | 0 | 2 | 4 | 0 | 0 | 0 | 0 | 3 | 0 | 298 | 149 | 1 | 1 | 1 | NA | NA | NA | 1 | NA | NA | 1 | 0.4 | NA | 0.3 |
| SE | 57049 | ENSG00000268043 | NBPF12 | chr1 | + | 146991994 | 146992046 | 146988840 | 146988949 | 146992711 | 146992884 | 0 | 1 | 0 | 0 | 1 | 0 | 0 | 0 | 0 | 1 | 0 | 1 | 2 | 0 | 2 | 12 | 0 | 0 | 1 | 0 | 201 | 149 | 1 | 1 | NA | 1 | NA | NA | 0.426 | 0 | 1 | 1 | 0 | 1 | 0.113 |
| SE | 57051 | ENSG00000003756 | RBM5 | chr3 | + | 50092978 | 50093042 | 50092042 | 50092208 | 50093719 | 50093875 | 0 | 0 | 3 | 1 | 0 | 330 | 316 | 287 | 318 | 379 | 0 | 0 | 5 | 0 | 0 | 449 | 309 | 376 | 530 | 298 | 213 | 149 | 1 | 1 | 0 | 0 | 0.007 | 0.002 | 0 | 0 | 0 | 0.009 | 0 | 0 | 0 |
| SE | 57054 | ENSG00000003756 | RBM5 | chr3 | + | 50105076 | 50105142 | 50102725 | 50103166 | 50106766 | 50106864 | 25 | 18 | 8 | 22 | 28 | 0 | 0 | 0 | 0 | 0 | 58 | 30 | 40 | 46 | 25 | 0 | 0 | 1 | 0 | 0 | 215 | 149 | 1 | 1 | 1 | 1 | 1 | 1 | 1 | 1 | 1 | 0.965 | 1 | 1 | 0.007 |
| SE | 57056 | ENSG00000003756 | RBM5 | chr3 | + | 50105548 | 50105709 | 50104576 | 50105142 | 50107481 | 50107569 | 369 | 279 | 357 | 430 | 602 | 0 | 1 | 0 | 0 | 0 | 555 | 422 | 409 | 561 | 324 | 0 | 0 | 0 | 0 | 0 | 298 | 149 | 1 | 1 | 1 | 0.993 | 1 | 1 | 1 | 1 | 1 | 1 | 1 | 1 | -0.001 |
| SE | 57057 | ENSG00000003756 | RBM5 | chr3 | + | 50105548 | 50105709 | 50104576 | 50105142 | 50109602 | 50109688 | 369 | 278 | 357 | 431 | 602 | 0 | 0 | 0 | 0 | 0 | 555 | 422 | 409 | 561 | 324 | 0 | 0 | 0 | 2 | 0 | 298 | 149 | 1 | 1 | 1 | 1 | 1 | 1 | 1 | 1 | 1 | 1 | 0.993 | 1 | 0.001 |
| SE | 57058 | ENSG00000003756 | RBM5 | chr3 | + | 50106766 | 50106864 | 50104973 | 50105142 | 50107481 | 50107569 | 163 | 112 | 113 | 129 | 261 | 0 | 1 | 0 | 0 | 0 | 230 | 88 | 110 | 125 | 25 | 0 | 0 | 0 | 0 | 0 | 247 | 149 | 1 | 1 | 1 | 0.985 | 1 | 1 | 1 | 1 | 1 | 1 | 1 | 1 | -0.003 |
| SE | 57059 | ENSG00000003756 | RBM5 | chr3 | + | 50106766 | 50106864 | 50105548 | 50105709 | 50107481 | 50107569 | 311 | 187 | 246 | 262 | 400 | 0 | 1 | 0 | 0 | 0 | 411 | 243 | 242 | 367 | 113 | 0 | 0 | 0 | 0 | 0 | 247 | 149 | 1 | 1 | 1 | 0.991 | 1 | 1 | 1 | 1 | 1 | 1 | 1 | 1 | -0.002 |
| SE | 57060 | ENSG00000003756 | RBM5 | chr3 | + | 50107017 | 50107087 | 50106766 | 50106864 | 50107481 | 50107569 | 2 | 1 | 2 | 3 | 6 | 215 | 137 | 170 | 149 | 309 | 0 | 0 | 3 | 0 | 5 | 280 | 129 | 152 | 188 | 59 | 219 | 149 | 1 | 1 | 0.006 | 0.005 | 0.008 | 0.014 | 0.013 | 0 | 0 | 0.013 | 0 | 0.055 | -0.004 |
| SE | 57061 | ENSG00000003756 | RBM5 | chr3 | + | 50107481 | 50107569 | 50104973 | 50105142 | 50108069 | 50108147 | 86 | 55 | 78 | 46 | 78 | 0 | 2 | 0 | 0 | 1 | 79 | 65 | 90 | 95 | 47 | 0 | 0 | 0 | 0 | 0 | 237 | 149 | 1 | 1 | 1 | 0.945 | 1 | 1 | 0.98 | 1 | 1 | 1 | 1 | 1 | -0.015 |
| SE | 57062 | ENSG00000003756 | RBM5 | chr3 | + | 50107481 | 50107569 | 50104973 | 50105142 | 50109602 | 50109688 | 2 | 0 | 2 | 2 | 0 | 0 | 0 | 0 | 0 | 0 | 0 | 0 | 0 | 3 | 0 | 0 | 0 | 0 | 2 | 0 | 237 | 149 | 1 | 1 | 1 | NA | 1 | 1 | NA | NA | NA | NA | 0.485 | NA | 0.515 |
| SE | 57063 | ENSG00000003756 | RBM5 | chr3 | + | 50107481 | 50107569 | 50105548 | 50105709 | 50109602 | 50109688 | 2 | 1 | 2 | 2 | 0 | 0 | 0 | 0 | 1 | 0 | 0 | 0 | 0 | 3 | 0 | 0 | 0 | 0 | 0 | 0 | 237 | 149 | 1 | 1 | 1 | 1 | 1 | 0.557 | NA | NA | NA | NA | 1 | NA | -0.111 |
| SE | 57064 | ENSG00000003756 | RBM5 | chr3 | + | 50109602 | 50109688 | 50108231 | 50108304 | 50110678 | 50110757 | 275 | 197 | 247 | 243 | 368 | 0 | 0 | 1 | 0 | 0 | 524 | 312 | 304 | 400 | 187 | 4 | 0 | 0 | 0 | 5 | 235 | 149 | 1 | 1 | 1 | 1 | 0.994 | 1 | 1 | 0.988 | 1 | 1 | 1 | 0.96 | 0.009 |
| SE | 57066 | ENSG00000003756 | RBM5 | chr3 | + | 50110678 | 50110770 | 50109602 | 50109688 | 50113382 | 50113489 | 186 | 178 | 222 | 220 | 158 | 1 | 0 | 0 | 1 | 1 | 384 | 296 | 287 | 327 | 218 | 0 | 0 | 0 | 0 | 0 | 241 | 149 | 1 | 1 | 0.991 | 1 | 1 | 0.993 | 0.99 | 1 | 1 | 1 | 1 | 1 | -0.005 |
| SE | 57067 | ENSG00000003756 | RBM5 | chr3 | + | 50113382 | 50113544 | 50110678 | 50110770 | 50115427 | 50115607 | 233 | 184 | 268 | 254 | 203 | 0 | 0 | 0 | 0 | 0 | 392 | 339 | 264 | 333 | 201 | 0 | 7 | 0 | 0 | 0 | 298 | 149 | 1 | 1 | 1 | 1 | 1 | 1 | 1 | 1 | 0.96 | 1 | 1 | 1 | 0.008 |
| SE | 57071 | ENSG00000003756 | RBM5 | chr3 | + | 50115427 | 50115607 | 50114179 | 50114251 | 50115905 | 50115943 | 762 | 624 | 678 | 828 | 828 | 0 | 0 | 0 | 0 | 0 | 1024 | 740 | 874 | 953 | 631 | 0 | 0 | 0 | 4 | 0 | 298 | 149 | 1 | 1 | 1 | 1 | 1 | 1 | 1 | 1 | 1 | 1 | 0.992 | 1 | 0.002 |
| SE | 57086 | ENSG00000272602 | ZNF595 | chr4 | + | 59429 | 59556 | 53294 | 53491 | 85730 | 87521 | 32 | 8 | 16 | 32 | 7 | 2 | 0 | 1 | 1 | 0 | 0 | 13 | 21 | 8 | 0 | 12 | 2 | 0 | 0 | 0 | 276 | 149 | 1 | 1 | 0.896 | 1 | 0.896 | 0.945 | 1 | 0 | 0.778 | 1 | 1 | NA | 0.253 |
| SE | 57087 | ENSG00000272602 | ZNF595 | chr4 | + | 60057 | 60153 | 59429 | 59556 | 85730 | 88208 | 103 | 34 | 47 | 62 | 38 | 1 | 0 | 0 | 0 | 0 | 0 | 28 | 26 | 11 | 7 | 0 | 0 | 0 | 0 | 0 | 245 | 149 | 1 | 1 | 0.984 | 1 | 1 | 1 | 1 | NA | 1 | 1 | 1 | 1 | -0.003 |
| SE | 57089 | ENSG00000176407 | KCMF1 | chr2 | + | 85027888 | 85028056 | 84971160 | 84971467 | 85035015 | 85035125 | 241 | 141 | 179 | 254 | 201 | 0 | 0 | 0 | 1 | 1 | 241 | 221 | 214 | 244 | 422 | 0 | 0 | 0 | 0 | 0 | 298 | 149 | 1 | 1 | 1 | 1 | 1 | 0.992 | 0.99 | 1 | 1 | 1 | 1 | 1 | -0.004 |
| SE | 57090 | ENSG00000176407 | KCMF1 | chr2 | + | 85035015 | 85035155 | 85027888 | 85028056 | 85043563 | 85043665 | 413 | 211 | 263 | 360 | 353 | 0 | 0 | 0 | 3 | 0 | 418 | 320 | 292 | 347 | 577 | 0 | 0 | 0 | 0 | 0 | 289 | 149 | 1 | 1 | 1 | 1 | 1 | 0.984 | 1 | 1 | 1 | 1 | 1 | 1 | -0.003 |
| SE | 57091 | ENSG00000176407 | KCMF1 | chr2 | + | 85049365 | 85049648 | 85035015 | 85035155 | 85053147 | 85059472 | 273 | 196 | 207 | 270 | 187 | 0 | 3 | 0 | 0 | 0 | 490 | 277 | 302 | 305 | 588 | 0 | 0 | 0 | 0 | 0 | 298 | 149 | 1 | 1 | 1 | 0.97 | 1 | 1 | 1 | 1 | 1 | 1 | 1 | 1 | -0.006 |
| SE | 57092 | ENSG00000124571 | XPO5 | chr6 | - | 43533906 | 43534007 | 43531478 | 43531575 | 43546570 | 43546752 | 74 | 24 | 43 | 78 | 54 | 0 | 0 | 1 | 0 | 0 | 68 | 48 | 27 | 51 | 18 | 0 | 0 | 0 | 0 | 0 | 250 | 149 | 1 | 1 | 1 | 1 | 0.962 | 1 | 1 | 1 | 1 | 1 | 1 | 1 | -0.008 |
| SE | 57093 | ENSG00000124571 | XPO5 | chr6 | - | 43547315 | 43547453 | 43546570 | 43546752 | 43547607 | 43547675 | 3 | 0 | 0 | 3 | 1 | 30 | 36 | 37 | 37 | 42 | 0 | 1 | 0 | 1 | 0 | 76 | 51 | 24 | 52 | 37 | 287 | 149 | 1 | 1 | 0.049 | 0 | 0 | 0.04 | 0.012 | 0 | 0.01 | 0 | 0.01 | 0 | 0.016 |
| SE | 57095 | ENSG00000124571 | XPO5 | chr6 | - | 43551297 | 43551453 | 43549892 | 43549934 | 43555835 | 43555964 | 104 | 72 | 72 | 82 | 123 | 0 | 2 | 0 | 0 | 0 | 161 | 110 | 75 | 108 | 82 | 0 | 0 | 0 | 0 | 0 | 298 | 149 | 1 | 1 | 1 | 0.947 | 1 | 1 | 1 | 1 | 1 | 1 | 1 | 1 | -0.011 |
| SE | 57096 | ENSG00000124571 | XPO5 | chr6 | - | 43553372 | 43553503 | 43549892 | 43549934 | 43555835 | 43555964 | 41 | 36 | 36 | 39 | 45 | 0 | 2 | 0 | 0 | 0 | 40 | 25 | 37 | 21 | 24 | 0 | 0 | 0 | 0 | 0 | 280 | 149 | 1 | 1 | 1 | 0.905 | 1 | 1 | 1 | 1 | 1 | 1 | 1 | 1 | -0.019 |
| SE | 57097 | ENSG00000124571 | XPO5 | chr6 | - | 43553372 | 43553503 | 43551297 | 43551453 | 43555835 | 43555964 | 106 | 82 | 96 | 116 | 139 | 0 | 1 | 1 | 2 | 2 | 118 | 141 | 115 | 98 | 111 | 0 | 1 | 0 | 0 | 0 | 280 | 149 | 1 | 1 | 1 | 0.978 | 0.981 | 0.969 | 0.974 | 1 | 0.987 | 1 | 1 | 1 | -0.017 |
| SE | 57098 | ENSG00000124571 | XPO5 | chr6 | - | 43555835 | 43555964 | 43551297 | 43551453 | 43560177 | 43560303 | 0 | 2 | 3 | 2 | 2 | 2 | 0 | 0 | 0 | 0 | 0 | 1 | 0 | 0 | 0 | 0 | 0 | 0 | 0 | 0 | 278 | 149 | 1 | 1 | 0 | 1 | 1 | 1 | 1 | NA | 1 | NA | NA | NA | -0.2 |
| SE | 57099 | ENSG00000124571 | XPO5 | chr6 | - | 43558500 | 43558591 | 43555835 | 43555964 | 43560177 | 43560303 | 29 | 20 | 47 | 32 | 35 | 0 | 1 | 2 | 0 | 0 | 26 | 26 | 16 | 10 | 3 | 0 | 0 | 0 | 0 | 0 | 240 | 149 | 1 | 1 | 1 | 0.925 | 0.936 | 1 | 1 | 1 | 1 | 1 | 1 | 1 | -0.028 |
| SE | 57100 | ENSG00000124571 | XPO5 | chr6 | - | 43572505 | 43572578 | 43570856 | 43570994 | 43573479 | 43573601 | 128 | 104 | 111 | 141 | 120 | 1 | 0 | 0 | 0 | 0 | 159 | 88 | 119 | 163 | 227 | 0 | 0 | 0 | 1 | 0 | 222 | 149 | 1 | 1 | 0.988 | 1 | 1 | 1 | 1 | 1 | 1 | 1 | 0.991 | 1 | -0.001 |
| SE | 57101 | ENSG00000124571 | XPO5 | chr6 | - | 43573397 | 43573601 | 43572505 | 43572578 | 43575759 | 43575985 | 17 | 11 | 15 | 11 | 6 | 0 | 0 | 0 | 1 | 0 | 8 | 32 | 10 | 22 | 40 | 0 | 0 | 0 | 0 | 0 | 298 | 149 | 1 | 1 | 1 | 1 | 1 | 0.846 | 1 | 1 | 1 | 1 | 1 | 1 | -0.031 |
| SE | 57102 | ENSG00000124571 | XPO5 | chr6 | - | 43573479 | 43573601 | 43572505 | 43572578 | 43575759 | 43576038 | 88 | 75 | 80 | 89 | 65 | 0 | 0 | 0 | 1 | 0 | 84 | 91 | 72 | 104 | 159 | 0 | 0 | 0 | 0 | 0 | 271 | 149 | 1 | 1 | 1 | 1 | 1 | 0.98 | 1 | 1 | 1 | 1 | 1 | 1 | -0.004 |
| SE | 57105 | ENSG00000162654 | GBP4 | chr1 | - | 89186999 | 89187102 | 89186332 | 89186526 | 89188581 | 89188794 | 358 | 241 | 210 | 216 | 388 | 0 | 0 | 0 | 0 | 0 | 1149 | 215 | 207 | 378 | 348 | 8 | 0 | 3 | 0 | 0 | 252 | 149 | 1 | 1 | 1 | 1 | 1 | 1 | 1 | 0.988 | 1 | 0.976 | 1 | 1 | 0.007 |
| SE | 57106 | ENSG00000162654 | GBP4 | chr1 | - | 89188581 | 89188794 | 89186999 | 89187102 | 89190037 | 89190318 | 440 | 318 | 326 | 367 | 408 | 0 | 0 | 0 | 0 | 0 | 1320 | 385 | 237 | 578 | 461 | 0 | 4 | 0 | 0 | 0 | 298 | 149 | 1 | 1 | 1 | 1 | 1 | 1 | 1 | 1 | 0.98 | 1 | 1 | 1 | 0.004 |
| SE | 57107 | ENSG00000162654 | GBP4 | chr1 | - | 89188581 | 89188794 | 89186999 | 89187102 | 89191260 | 89191506 | 182 | 138 | 123 | 132 | 198 | 0 | 0 | 0 | 1 | 0 | 623 | 130 | 106 | 234 | 200 | 0 | 0 | 0 | 0 | 0 | 298 | 149 | 1 | 1 | 1 | 1 | 1 | 0.985 | 1 | 1 | 1 | 1 | 1 | 1 | -0.003 |
| SE | 57108 | ENSG00000162654 | GBP4 | chr1 | - | 89190037 | 89190318 | 89186999 | 89187102 | 89191260 | 89191506 | 308 | 226 | 179 | 237 | 325 | 0 | 0 | 0 | 1 | 0 | 627 | 231 | 138 | 305 | 149 | 0 | 0 | 0 | 0 | 0 | 298 | 149 | 1 | 1 | 1 | 1 | 1 | 0.992 | 1 | 1 | 1 | 1 | 1 | 1 | -0.002 |
| SE | 57109 | ENSG00000162654 | GBP4 | chr1 | - | 89190037 | 89190318 | 89188581 | 89188794 | 89191260 | 89191506 | 566 | 409 | 382 | 473 | 535 | 0 | 3 | 0 | 1 | 0 | 1324 | 482 | 269 | 648 | 410 | 0 | 0 | 0 | 0 | 0 | 298 | 149 | 1 | 1 | 1 | 0.986 | 1 | 0.996 | 1 | 1 | 1 | 1 | 1 | 1 | -0.004 |
| SE | 57110 | ENSG00000162654 | GBP4 | chr1 | - | 89191260 | 89191506 | 89190037 | 89190318 | 89192903 | 89193100 | 467 | 325 | 270 | 352 | 470 | 0 | 0 | 0 | 0 | 0 | 889 | 361 | 189 | 453 | 243 | 0 | 0 | 2 | 0 | 3 | 298 | 149 | 1 | 1 | 1 | 1 | 1 | 1 | 1 | 1 | 1 | 0.979 | 1 | 0.976 | 0.009 |
| SE | 57111 | ENSG00000120437 | ACAT2 | chr6 | + | 159767004 | 159767186 | 159762918 | 159763053 | 159775169 | 159775307 | 26 | 10 | 39 | 41 | 40 | 0 | 1 | 0 | 2 | 0 | 42 | 27 | 31 | 26 | 80 | 0 | 0 | 0 | 0 | 0 | 298 | 149 | 1 | 1 | 1 | 0.833 | 1 | 0.911 | 1 | 1 | 1 | 1 | 1 | 1 | -0.051 |
| SE | 57112 | ENSG00000120437 | ACAT2 | chr6 | + | 159768510 | 159768628 | 159767004 | 159767186 | 159775169 | 159775307 | 90 | 60 | 115 | 116 | 118 | 0 | 0 | 0 | 1 | 0 | 115 | 84 | 66 | 78 | 64 | 0 | 0 | 0 | 0 | 0 | 267 | 149 | 1 | 1 | 1 | 1 | 1 | 0.985 | 1 | 1 | 1 | 1 | 1 | 1 | -0.003 |
| SE | 57114 | ENSG00000120437 | ACAT2 | chr6 | + | 159775169 | 159775313 | 159768510 | 159768628 | 159776149 | 159776272 | 95 | 60 | 136 | 140 | 189 | 0 | 0 | 0 | 0 | 0 | 123 | 81 | 78 | 106 | 48 | 0 | 3 | 0 | 0 | 0 | 293 | 149 | 1 | 1 | 1 | 1 | 1 | 1 | 1 | 1 | 0.932 | 1 | 1 | 1 | 0.014 |
| SE | 57117 | ENSG00000184305 | CCSER1 | chr4 | + | 90403923 | 90404019 | 90400035 | 90400129 | 90468233 | 90468448 | 1 | 0 | 5 | 2 | 19 | 1 | 4 | 7 | 12 | 4 | 10 | 10 | 0 | 0 | 0 | 12 | 13 | 0 | 0 | 0 | 245 | 149 | 1 | 1 | 0.378 | 0 | 0.303 | 0.092 | 0.743 | 0.336 | 0.319 | NA | NA | NA | -0.024 |
| SE | 57118 | ENSG00000184305 | CCSER1 | chr4 | + | 90628024 | 90628232 | 90468233 | 90468354 | 90723913 | 90723991 | 13 | 18 | 8 | 20 | 33 | 1 | 0 | 0 | 0 | 0 | 22 | 37 | 2 | 19 | 16 | 0 | 0 | 0 | 0 | 0 | 298 | 149 | 1 | 1 | 0.867 | 1 | 1 | 1 | 1 | 1 | 1 | 1 | 1 | 1 | -0.027 |
| SE | 57121 | ENSG00000184305 | CCSER1 | chr4 | + | 90727203 | 90727297 | 90723913 | 90723991 | 90780441 | 90780568 | 0 | 0 | 0 | 2 | 0 | 0 | 0 | 0 | 1 | 0 | 0 | 0 | 0 | 0 | 0 | 0 | 0 | 0 | 2 | 0 | 243 | 149 | 1 | 1 | NA | NA | NA | 0.551 | NA | NA | NA | NA | 0 | NA | 0.551 |
| SE | 57127 | ENSG00000172426 | RSPH9 | chr6 | + | 43666423 | 43666520 | 43656576 | 43656723 | 43670788 | 43672599 | 0 | 0 | 0 | 2 | 0 | 2 | 0 | 0 | 0 | 0 | 0 | 0 | 0 | 0 | 19 | 0 | 3 | 0 | 1 | 19 | 246 | 149 | 1 | 1 | 0 | NA | NA | 1 | NA | NA | 0 | NA | 0 | 0.377 | 0.374 |
| SE | 57130 | ENSG00000023902 | PLEKHO1 | chr1 | + | 150156065 | 150156206 | 150150606 | 150151058 | 150156910 | 150157015 | 910 | 788 | 590 | 875 | 1556 | 0 | 0 | 4 | 0 | 0 | 2560 | 1394 | 399 | 1609 | 3842 | 10 | 0 | 2 | 14 | 0 | 290 | 149 | 1 | 1 | 1 | 1 | 0.987 | 1 | 1 | 0.992 | 1 | 0.99 | 0.983 | 1 | 0.004 |
| SE | 57131 | ENSG00000023902 | PLEKHO1 | chr1 | + | 150156065 | 150156206 | 150150606 | 150151058 | 150158818 | 150159043 | 314 | 233 | 204 | 301 | 343 | 0 | 0 | 0 | 0 | 0 | 759 | 452 | 143 | 492 | 1315 | 0 | 0 | 0 | 0 | 7 | 290 | 149 | 1 | 1 | 1 | 1 | 1 | 1 | 1 | 1 | 1 | 1 | 1 | 0.99 | 0.002 |
| SE | 57132 | ENSG00000023902 | PLEKHO1 | chr1 | + | 150156910 | 150157015 | 150150606 | 150151058 | 150158818 | 150159630 | 1 | 0 | 9 | 3 | 2 | 0 | 0 | 0 | 0 | 0 | 13 | 3 | 9 | 29 | 33 | 0 | 0 | 0 | 0 | 7 | 254 | 149 | 1 | 1 | 1 | NA | 1 | 1 | 1 | 1 | 1 | 1 | 1 | 0.734 | 0.053 |
| SE | 57133 | ENSG00000023902 | PLEKHO1 | chr1 | + | 150156910 | 150157015 | 150153482 | 150156206 | 150157384 | 150157486 | 838 | 745 | 574 | 825 | 1718 | 1 | 0 | 0 | 0 | 1 | 2468 | 1301 | 371 | 1416 | 3491 | 6 | 0 | 1 | 2 | 0 | 254 | 149 | 1 | 1 | 0.998 | 1 | 1 | 1 | 0.999 | 0.996 | 1 | 0.995 | 0.998 | 1 | 0.002 |
| SE | 57134 | ENSG00000023902 | PLEKHO1 | chr1 | + | 150156910 | 150157015 | 150153482 | 150156206 | 150158818 | 150159630 | 507 | 492 | 348 | 522 | 1060 | 0 | 0 | 1 | 1 | 0 | 1621 | 838 | 247 | 1026 | 2233 | 0 | 0 | 0 | 0 | 0 | 254 | 149 | 1 | 1 | 1 | 1 | 0.995 | 0.997 | 1 | 1 | 1 | 1 | 1 | 1 | -0.002 |
| SE | 57136 | ENSG00000023902 | PLEKHO1 | chr1 | + | 150157384 | 150157486 | 150153482 | 150156206 | 150158818 | 150159043 | 172 | 94 | 112 | 154 | 141 | 0 | 0 | 1 | 1 | 0 | 405 | 172 | 76 | 150 | 560 | 0 | 0 | 0 | 0 | 0 | 251 | 149 | 1 | 1 | 1 | 1 | 0.985 | 0.989 | 1 | 1 | 1 | 1 | 1 | 1 | -0.005 |
| SE | 57138 | ENSG00000172939 | OXSR1 | chr3 | + | 38190730 | 38190839 | 38183002 | 38183115 | 38198721 | 38198863 | 224 | 175 | 142 | 212 | 267 | 0 | 0 | 0 | 0 | 0 | 223 | 255 | 202 | 234 | 18 | 8 | 0 | 0 | 0 | 0 | 258 | 149 | 1 | 1 | 1 | 1 | 1 | 1 | 1 | 0.942 | 1 | 1 | 1 | 1 | 0.012 |
| SE | 57140 | ENSG00000172939 | OXSR1 | chr3 | + | 38216095 | 38216151 | 38198721 | 38198863 | 38221577 | 38221687 | 135 | 114 | 139 | 172 | 153 | 0 | 0 | 0 | 1 | 0 | 213 | 145 | 126 | 159 | 31 | 0 | 0 | 0 | 0 | 0 | 205 | 149 | 1 | 1 | 1 | 1 | 1 | 0.992 | 1 | 1 | 1 | 1 | 1 | 1 | -0.002 |
| SE | 57142 | ENSG00000172939 | OXSR1 | chr3 | + | 38229686 | 38229735 | 38224570 | 38224704 | 38230364 | 38230430 | 230 | 203 | 197 | 269 | 398 | 1 | 4 | 1 | 4 | 2 | 424 | 278 | 287 | 383 | 82 | 0 | 1 | 0 | 6 | 0 | 198 | 149 | 1 | 1 | 0.994 | 0.974 | 0.993 | 0.981 | 0.993 | 1 | 0.995 | 1 | 0.98 | 1 | -0.008 |
| SE | 57143 | ENSG00000172939 | OXSR1 | chr3 | + | 38230364 | 38230430 | 38224570 | 38224704 | 38236838 | 38236961 | 71 | 74 | 58 | 88 | 124 | 0 | 0 | 0 | 0 | 0 | 52 | 75 | 99 | 100 | 27 | 0 | 0 | 1 | 0 | 0 | 215 | 149 | 1 | 1 | 1 | 1 | 1 | 1 | 1 | 1 | 1 | 0.986 | 1 | 1 | 0.003 |
| SE | 57144 | ENSG00000172939 | OXSR1 | chr3 | + | 38236838 | 38236961 | 38230364 | 38230430 | 38242742 | 38242778 | 238 | 207 | 211 | 274 | 410 | 0 | 0 | 0 | 0 | 0 | 178 | 245 | 249 | 256 | 65 | 0 | 0 | 2 | 0 | 0 | 272 | 149 | 1 | 1 | 1 | 1 | 1 | 1 | 1 | 1 | 1 | 0.986 | 1 | 1 | 0.003 |
| SE | 57145 | ENSG00000172939 | OXSR1 | chr3 | + | 38242742 | 38242778 | 38236660 | 38236961 | 38246074 | 38246221 | 162 | 139 | 142 | 168 | 325 | 4 | 0 | 1 | 0 | 4 | 82 | 148 | 108 | 120 | 27 | 0 | 1 | 2 | 1 | 0 | 185 | 149 | 1 | 1 | 0.97 | 1 | 0.991 | 1 | 0.985 | 1 | 0.992 | 0.978 | 0.99 | 1 | -0.003 |
| SE | 57146 | ENSG00000172939 | OXSR1 | chr3 | + | 38249965 | 38250018 | 38247667 | 38247732 | 38251402 | 38251412 | 82 | 54 | 81 | 91 | 201 | 0 | 0 | 2 | 0 | 1 | 56 | 67 | 40 | 109 | 13 | 0 | 0 | 0 | 0 | 0 | 202 | 149 | 1 | 1 | 1 | 1 | 0.968 | 1 | 0.993 | 1 | 1 | 1 | 1 | 1 | -0.008 |
| SE | 57149 | ENSG00000173276 | ZBTB21 | chr21 | - | 41992492 | 41994108 | 41986830 | 41991889 | 42002896 | 42002961 | 10 | 12 | 8 | 9 | 3 | 0 | 0 | 0 | 1 | 0 | 0 | 0 | 5 | 12 | 0 | 0 | 0 | 0 | 0 | 0 | 298 | 149 | 1 | 1 | 1 | 1 | 1 | 0.818 | 1 | NA | NA | 1 | 1 | NA | -0.036 |
| SE | 57150 | ENSG00000173276 | ZBTB21 | chr21 | - | 42002349 | 42002503 | 41988636 | 41994108 | 42002896 | 42002961 | 8 | 4 | 6 | 6 | 9 | 5 | 4 | 1 | 4 | 2 | 0 | 14 | 22 | 3 | 0 | 0 | 0 | 1 | 0 | 0 | 298 | 149 | 1 | 1 | 0.444 | 0.333 | 0.75 | 0.429 | 0.692 | NA | 1 | 0.917 | 1 | NA | -0.443 |
| SE | 57151 | ENSG00000173276 | ZBTB21 | chr21 | - | 42002349 | 42002563 | 41993947 | 41994108 | 42002896 | 42002961 | 9 | 4 | 8 | 6 | 14 | 5 | 4 | 1 | 4 | 2 | 0 | 14 | 28 | 3 | 0 | 0 | 0 | 1 | 0 | 0 | 298 | 149 | 1 | 1 | 0.474 | 0.333 | 0.8 | 0.429 | 0.778 | NA | 1 | 0.933 | 1 | NA | -0.415 |
| SE | 57152 | ENSG00000173276 | ZBTB21 | chr21 | - | 42002896 | 42002961 | 41986830 | 41994108 | 42010251 | 42010333 | 5 | 3 | 2 | 7 | 3 | 7 | 16 | 18 | 16 | 15 | 0 | 3 | 1 | 0 | 0 | 0 | 3 | 9 | 0 | 0 | 214 | 149 | 1 | 1 | 0.332 | 0.115 | 0.072 | 0.233 | 0.122 | NA | 0.41 | 0.072 | NA | NA | -0.066 |
| SE | 57153 | ENSG00000173276 | ZBTB21 | chr21 | - | 42009466 | 42009597 | 42002896 | 42002961 | 42010251 | 42010357 | 0 | 0 | 0 | 0 | 1 | 1 | 0 | 3 | 4 | 1 | 0 | 0 | 0 | 0 | 0 | 0 | 5 | 4 | 0 | 0 | 280 | 149 | 1 | 1 | 0 | NA | 0 | 0 | 0.347 | NA | 0 | 0 | NA | NA | 0.087 |
| SE | 57154 | ENSG00000159377 | PSMB4 | chr1 | + | 151399980 | 151400185 | 151399589 | 151399727 | 151400441 | 151401355 | 533 | 233 | 531 | 403 | 498 | 0 | 0 | 0 | 4 | 0 | 553 | 547 | 408 | 424 | 627 | 0 | 0 | 2 | 5 | 0 | 298 | 149 | 1 | 1 | 1 | 1 | 1 | 0.981 | 1 | 1 | 1 | 0.99 | 0.977 | 1 | 0.003 |
| SE | 57155 | ENSG00000159377 | PSMB4 | chr1 | + | 151399980 | 151400187 | 151399559 | 151399727 | 151400441 | 151400588 | 1087 | 475 | 1030 | 808 | 901 | 0 | 0 | 0 | 4 | 0 | 1222 | 1180 | 812 | 827 | 1261 | 0 | 0 | 2 | 5 | 0 | 298 | 149 | 1 | 1 | 1 | 1 | 1 | 0.99 | 1 | 1 | 1 | 0.995 | 0.988 | 1 | 0.001 |
| SE | 57157 | ENSG00000159377 | PSMB4 | chr1 | + | 151400763 | 151400845 | 151400441 | 151400588 | 151401541 | 151401630 | 225 | 82 | 184 | 204 | 148 | 0 | 0 | 0 | 0 | 0 | 254 | 207 | 105 | 167 | 240 | 0 | 4 | 0 | 0 | 0 | 231 | 149 | 1 | 1 | 1 | 1 | 1 | 1 | 1 | 1 | 0.971 | 1 | 1 | 1 | 0.006 |
| SE | 57158 | ENSG00000159377 | PSMB4 | chr1 | + | 151401238 | 151401355 | 151400441 | 151400588 | 151401541 | 151401630 | 677 | 314 | 608 | 739 | 480 | 0 | 0 | 0 | 0 | 0 | 969 | 732 | 458 | 553 | 926 | 0 | 4 | 0 | 0 | 0 | 266 | 149 | 1 | 1 | 1 | 1 | 1 | 1 | 1 | 1 | 0.99 | 1 | 1 | 1 | 0.002 |
| SE | 57159 | ENSG00000159377 | PSMB4 | chr1 | + | 151401238 | 151401355 | 151400763 | 151400845 | 151401541 | 151401630 | 1228 | 549 | 1063 | 1246 | 875 | 0 | 0 | 2 | 5 | 0 | 1552 | 1204 | 707 | 832 | 1528 | 0 | 5 | 0 | 0 | 0 | 266 | 149 | 1 | 1 | 1 | 1 | 0.997 | 0.993 | 1 | 1 | 0.993 | 1 | 1 | 1 | -0.001 |
| SE | 57160 | ENSG00000133561 | GIMAP6 | chr7 | - | 150630057 | 150630142 | 150627675 | 150628288 | 150632168 | 150632346 | 324 | 222 | 291 | 252 | 313 | 0 | 0 | 0 | 0 | 0 | 155 | 150 | 104 | 110 | 118 | 0 | 0 | 2 | 0 | 0 | 234 | 149 | 1 | 1 | 1 | 1 | 1 | 1 | 1 | 1 | 1 | 0.971 | 1 | 1 | 0.006 |
| SE | 57167 | ENSG00000011021 | CLCN6 | chr1 | + | 11843703 | 11844073 | 11841016 | 11841207 | 11845546 | 11845766 | 9 | 8 | 5 | 12 | 7 | 0 | 0 | 0 | 0 | 0 | 50 | 26 | 13 | 25 | 35 | 0 | 0 | 2 | 0 | 0 | 298 | 149 | 1 | 1 | 1 | 1 | 1 | 1 | 1 | 1 | 1 | 0.765 | 1 | 1 | 0.047 |
| SE | 57168 | ENSG00000011021 | CLCN6 | chr1 | + | 11843733 | 11844073 | 11841057 | 11841207 | 11845546 | 11845766 | 10 | 8 | 5 | 12 | 10 | 0 | 0 | 0 | 0 | 0 | 47 | 24 | 14 | 28 | 31 | 0 | 0 | 2 | 0 | 0 | 298 | 149 | 1 | 1 | 1 | 1 | 1 | 1 | 1 | 1 | 1 | 0.778 | 1 | 1 | 0.044 |
| SE | 57170 | ENSG00000129084 | PSMA1 | chr11 | - | 14507655 | 14507766 | 14504873 | 14505248 | 14510871 | 14510951 | 852 | 619 | 705 | 709 | 842 | 2 | 2 | 2 | 0 | 0 | 563 | 719 | 608 | 639 | 234 | 0 | 0 | 0 | 6 | 0 | 260 | 149 | 1 | 1 | 0.996 | 0.994 | 0.995 | 1 | 1 | 1 | 1 | 1 | 0.984 | 1 | 0 |
| SE | 57172 | ENSG00000129084 | PSMA1 | chr11 | - | 14513816 | 14514491 | 14513663 | 14513699 | 14517641 | 14517673 | 472 | 298 | 284 | 402 | 447 | 0 | 0 | 0 | 0 | 0 | 283 | 256 | 281 | 342 | 82 | 0 | 0 | 3 | 0 | 0 | 298 | 149 | 1 | 1 | 1 | 1 | 1 | 1 | 1 | 1 | 1 | 0.979 | 1 | 1 | 0.004 |
| SE | 57174 | ENSG00000129084 | PSMA1 | chr11 | - | 14610965 | 14611151 | 14518996 | 14519041 | 14643454 | 14643617 | 10 | 14 | 37 | 15 | 42 | 0 | 0 | 2 | 2 | 1 | 21 | 14 | 22 | 13 | 0 | 0 | 0 | 0 | 0 | 0 | 298 | 149 | 1 | 1 | 1 | 1 | 0.902 | 0.789 | 0.955 | 1 | 1 | 1 | 1 | NA | -0.071 |
| SE | 57175 | ENSG00000206341 | HLA-H | chr6 | + | 29888466 | 29888742 | 29887954 | 29888224 | 29890263 | 29890296 | 0 | 0 | 0 | 2 | 0 | 0 | 0 | 0 | 0 | 0 | 0 | 0 | 4 | 1 | 0 | 0 | 0 | 0 | 3 | 0 | 298 | 149 | 1 | 1 | NA | NA | NA | 1 | NA | NA | NA | 1 | 0.143 | NA | 0.428 |
| SE | 57176 | ENSG00000206341 | HLA-H | chr6 | + | 29889327 | 29889604 | 29888466 | 29888742 | 29890263 | 29890296 | 6 | 0 | 2 | 6 | 4 | 0 | 0 | 0 | 0 | 0 | 0 | 2 | 5 | 8 | 8 | 0 | 0 | 0 | 1 | 0 | 298 | 149 | 1 | 1 | 1 | NA | 1 | 1 | 1 | NA | 1 | 1 | 0.8 | 1 | 0.05 |
| SE | 57177 | ENSG00000206341 | HLA-H | chr6 | + | 29889706 | 29889823 | 29889327 | 29889604 | 29890263 | 29890296 | 943 | 112 | 553 | 216 | 705 | 4 | 0 | 2 | 1 | 4 | 575 | 655 | 48 | 519 | 1666 | 0 | 1 | 0 | 0 | 8 | 266 | 149 | 1 | 1 | 0.992 | 1 | 0.994 | 0.992 | 0.99 | 1 | 0.997 | 1 | 1 | 0.992 | -0.004 |
| SE | 57179 | ENSG00000197323 | TRIM33 | chr1 | - | 114401388 | 114401463 | 114399456 | 114399609 | 114402759 | 114402883 | 171 | 104 | 171 | 143 | 289 | 3 | 1 | 2 | 1 | 4 | 10 | 70 | 94 | 50 | 0 | 0 | 11 | 24 | 3 | 0 | 224 | 149 | 1 | 1 | 0.974 | 0.986 | 0.983 | 0.99 | 0.98 | 1 | 0.809 | 0.723 | 0.917 | NA | 0.12 |
| SE | 57180 | ENSG00000197323 | TRIM33 | chr1 | - | 114402759 | 114402883 | 114399456 | 114399609 | 114405409 | 114405759 | 182 | 179 | 170 | 242 | 285 | 0 | 0 | 0 | 0 | 1 | 58 | 248 | 232 | 135 | 32 | 0 | 0 | 0 | 0 | 0 | 273 | 149 | 1 | 1 | 1 | 1 | 1 | 1 | 0.994 | 1 | 1 | 1 | 1 | 1 | -0.001 |
| SE | 57181 | ENSG00000197323 | TRIM33 | chr1 | - | 114408676 | 114408740 | 114406940 | 114407100 | 114410183 | 114410316 | 172 | 126 | 153 | 150 | 97 | 0 | 0 | 0 | 0 | 1 | 35 | 206 | 137 | 86 | 14 | 0 | 6 | 0 | 0 | 0 | 213 | 149 | 1 | 1 | 1 | 1 | 1 | 1 | 1 | 1 | 0.96 | 1 | 1 | 1 | 0.008 |
| SE | 57182 | ENSG00000197323 | TRIM33 | chr1 | - | 114410183 | 114410316 | 114408676 | 114408740 | 114421435 | 114421636 | 155 | 102 | 146 | 120 | 68 | 0 | 1 | 1 | 0 | 0 | 23 | 175 | 86 | 38 | 4 | 0 | 0 | 0 | 0 | 0 | 282 | 149 | 1 | 1 | 1 | 0.982 | 0.987 | 1 | 1 | 1 | 1 | 1 | 1 | 1 | -0.006 |
| SE | 57183 | ENSG00000197323 | TRIM33 | chr1 | - | 114420379 | 114420451 | 114410183 | 114410316 | 114421435 | 114421636 | 16 | 8 | 17 | 21 | 11 | 104 | 79 | 104 | 92 | 44 | 8 | 61 | 32 | 13 | 0 | 10 | 125 | 42 | 23 | 0 | 221 | 149 | 1 | 1 | 0.094 | 0.064 | 0.099 | 0.133 | 0.144 | 0.35 | 0.248 | 0.339 | 0.276 | NA | -0.196 |
| SE | 57184 | ENSG00000197323 | TRIM33 | chr1 | - | 114421435 | 114421636 | 114410183 | 114410316 | 114424590 | 114424755 | 208 | 149 | 206 | 223 | 220 | 1 | 0 | 0 | 0 | 0 | 20 | 269 | 116 | 56 | 0 | 0 | 0 | 0 | 0 | 0 | 298 | 149 | 1 | 1 | 0.99 | 1 | 1 | 1 | 1 | 1 | 1 | 1 | 1 | NA | -0.002 |
| SE | 57185 | ENSG00000197323 | TRIM33 | chr1 | - | 114424590 | 114424755 | 114421435 | 114421636 | 114425448 | 114425723 | 314 | 243 | 247 | 353 | 410 | 1 | 0 | 0 | 0 | 0 | 56 | 368 | 236 | 124 | 14 | 0 | 0 | 0 | 0 | 0 | 298 | 149 | 1 | 1 | 0.994 | 1 | 1 | 1 | 1 | 1 | 1 | 1 | 1 | 1 | -0.001 |
| SE | 57186 | ENSG00000197323 | TRIM33 | chr1 | - | 114427176 | 114427294 | 114425448 | 114425723 | 114427747 | 114427894 | 176 | 158 | 171 | 209 | 380 | 0 | 0 | 0 | 0 | 0 | 19 | 149 | 173 | 76 | 0 | 0 | 1 | 0 | 0 | 0 | 267 | 149 | 1 | 1 | 1 | 1 | 1 | 1 | 1 | 1 | 0.988 | 1 | 1 | NA | 0.003 |
| SE | 57188 | ENSG00000197323 | TRIM33 | chr1 | - | 114463103 | 114463236 | 114433616 | 114433733 | 114464269 | 114464388 | 229 | 222 | 168 | 267 | 320 | 0 | 0 | 0 | 1 | 0 | 34 | 146 | 203 | 88 | 32 | 0 | 0 | 0 | 0 | 1 | 282 | 149 | 1 | 1 | 1 | 1 | 1 | 0.993 | 1 | 1 | 1 | 1 | 1 | 0.944 | 0.01 |
| SE | 57191 | ENSG00000115271 | GCA | chr2 | + | 162359493 | 162359552 | 162359043 | 162359157 | 162371305 | 162371595 | 131 | 86 | 81 | 80 | 189 | 0 | 3 | 0 | 0 | 0 | 19 | 102 | 74 | 122 | 12 | 0 | 0 | 0 | 0 | 0 | 208 | 149 | 1 | 1 | 1 | 0.954 | 1 | 1 | 1 | 1 | 1 | 1 | 1 | 1 | -0.009 |
| SE | 57192 | ENSG00000083845 | RPS5 | chr19 | + | 58387031 | 58387339 | 58386399 | 58386622 | 58388136 | 58388245 | 606 | 287 | 851 | 571 | 251 | 0 | 0 | 0 | 0 | 0 | 263 | 424 | 284 | 150 | 588 | 0 | 0 | 2 | 0 | 0 | 298 | 149 | 1 | 1 | 1 | 1 | 1 | 1 | 1 | 1 | 1 | 0.986 | 1 | 1 | 0.003 |
| SE | 57193 | ENSG00000083845 | RPS5 | chr19 | + | 58388417 | 58388480 | 58388136 | 58388245 | 58392975 | 58393185 | 0 | 0 | 1 | 1 | 7 | 2675 | 1126 | 2700 | 2467 | 2292 | 0 | 5 | 1 | 0 | 7 | 804 | 1163 | 947 | 587 | 1389 | 212 | 149 | 1 | 1 | 0 | 0 | 0 | 0 | 0.002 | 0 | 0.003 | 0.001 | 0 | 0.004 | -0.001 |
| SE | 57194 | ENSG00000083845 | RPS5 | chr19 | + | 58392975 | 58393185 | 58387288 | 58388245 | 58393358 | 58393487 | 5569 | 2506 | 5792 | 5129 | 4788 | 4 | 1 | 7 | 1 | 2 | 2269 | 2689 | 2118 | 1324 | 3500 | 5 | 0 | 2 | 0 | 20 | 298 | 149 | 1 | 1 | 0.999 | 0.999 | 0.998 | 1 | 0.999 | 0.996 | 1 | 0.998 | 1 | 0.989 | 0.002 |
| SE | 57197 | ENSG00000084112 | SSH1 | chr12 | - | 108792285 | 108792829 | 108778190 | 108789244 | 108800779 | 108800926 | 41 | 25 | 27 | 46 | 63 | 0 | 0 | 0 | 0 | 0 | 67 | 51 | 57 | 91 | 100 | 0 | 3 | 0 | 0 | 0 | 298 | 149 | 1 | 1 | 1 | 1 | 1 | 1 | 1 | 1 | 0.895 | 1 | 1 | 1 | 0.021 |
| SE | 57198 | ENSG00000084112 | SSH1 | chr12 | - | 108798999 | 108799200 | 108778190 | 108789244 | 108800779 | 108800926 | 64 | 59 | 53 | 91 | 72 | 0 | 0 | 0 | 0 | 0 | 129 | 81 | 54 | 61 | 138 | 0 | 3 | 0 | 0 | 0 | 298 | 149 | 1 | 1 | 1 | 1 | 1 | 1 | 1 | 1 | 0.931 | 1 | 1 | 1 | 0.014 |
| SE | 57201 | ENSG00000084112 | SSH1 | chr12 | - | 108809692 | 108809758 | 108807611 | 108807827 | 108811259 | 108811328 | 133 | 94 | 101 | 186 | 57 | 1 | 0 | 0 | 0 | 0 | 86 | 159 | 82 | 127 | 351 | 0 | 0 | 3 | 0 | 0 | 215 | 149 | 1 | 1 | 0.989 | 1 | 1 | 1 | 1 | 1 | 1 | 0.95 | 1 | 1 | 0.008 |
| SE | 57202 | ENSG00000084112 | SSH1 | chr12 | - | 108811259 | 108811328 | 108809692 | 108809758 | 108817037 | 108817159 | 127 | 87 | 90 | 158 | 40 | 1 | 0 | 0 | 0 | 0 | 98 | 147 | 87 | 140 | 302 | 0 | 0 | 3 | 0 | 0 | 218 | 149 | 1 | 1 | 0.989 | 1 | 1 | 1 | 1 | 1 | 1 | 1 | 1 | 1 | -0.002 |
| SE | 57205 | ENSG00000151611 | MMAA | chr4 | + | 145642362 | 145642485 | 145639079 | 145639866 | 145645985 | 145646156 | 51 | 38 | 37 | 40 | 61 | 0 | 0 | 0 | 0 | 1 | 12 | 29 | 51 | 16 | 5 | 0 | 0 | 2 | 0 | 0 | 272 | 149 | 1 | 1 | 1 | 1 | 1 | 1 | 0.971 | 1 | 1 | 1 | 1 | 1 | -0.006 |
| SE | 57213 | ENSG00000136231 | IGF2BP3 | chr7 | - | 23361533 | 23361597 | 23351304 | 23351586 | 23468481 | 23468542 | 1 | 0 | 0 | 0 | 4 | 0 | 0 | 0 | 0 | 0 | 0 | 7 | 3 | 0 | 0 | 0 | 0 | 2 | 0 | 0 | 213 | 149 | 1 | 1 | 1 | NA | NA | NA | 1 | NA | 1 | 0.512 | NA | NA | 0.244 |
| SE | 57218 | ENSG00000172915 | NBEA | chr13 | + | 35168986 | 35168995 | 35164355 | 35164509 | 35171271 | 35171452 | 0 | 0 | 2 | 1 | 0 | 2 | 2 | 0 | 0 | 0 | 0 | 0 | 0 | 0 | 0 | 2 | 0 | 2 | 3 | 0 | 158 | 149 | 1 | 1 | 0 | 0 | 1 | 1 | NA | 0 | NA | 0 | 0 | NA | 0.5 |
| SE | 57219 | ENSG00000172915 | NBEA | chr13 | + | 35183975 | 35184071 | 35182359 | 35182528 | 35195863 | 35196302 | 21 | 16 | 25 | 7 | 19 | 1 | 0 | 0 | 0 | 0 | 0 | 3 | 16 | 0 | 0 | 0 | 0 | 0 | 0 | 0 | 245 | 149 | 1 | 1 | 0.927 | 1 | 1 | 1 | 1 | NA | 1 | 1 | NA | NA | -0.015 |
| SE | 57220 | ENSG00000172915 | NBEA | chr13 | + | 35290388 | 35290450 | 35232491 | 35232619 | 35309527 | 35309592 | 13 | 23 | 19 | 12 | 13 | 0 | 1 | 0 | 0 | 0 | 23 | 9 | 5 | 3 | 0 | 0 | 0 | 0 | 0 | 0 | 211 | 149 | 1 | 1 | 1 | 0.942 | 1 | 1 | 1 | 1 | 1 | 1 | 1 | NA | -0.012 |
| SE | 57223 | ENSG00000223519 | KIF28P | chr1 | - | 246776539 | 246776684 | 246771836 | 246771991 | 246776933 | 246777040 | 3 | 5 | 4 | 25 | 5 | 0 | 0 | 0 | 0 | 0 | 15 | 15 | 5 | 20 | 54 | 0 | 2 | 0 | 0 | 0 | 294 | 149 | 1 | 1 | 1 | 1 | 1 | 1 | 1 | 1 | 0.792 | 1 | 1 | 1 | 0.042 |
| SE | 57226 | ENSG00000223519 | KIF28P | chr1 | - | 246780948 | 246781057 | 246777273 | 246777440 | 246781589 | 246781685 | 2 | 2 | 0 | 13 | 0 | 0 | 0 | 1 | 0 | 0 | 28 | 2 | 2 | 15 | 15 | 0 | 0 | 3 | 0 | 0 | 258 | 149 | 1 | 1 | 1 | 1 | 0 | 1 | NA | 1 | 1 | 0.278 | 1 | 1 | -0.106 |
| SE | 57228 | ENSG00000178691 | SUZ12 | chr17 | + | 31940285 | 31940332 | 31937006 | 31937520 | 31947616 | 31947685 | 4 | 0 | 4 | 1 | 0 | 0 | 0 | 0 | 0 | 0 | 0 | 0 | 3 | 0 | 0 | 0 | 3 | 0 | 0 | 0 | 196 | 149 | 1 | 1 | 1 | NA | 1 | 1 | NA | NA | 0 | 1 | NA | NA | 0.5 |
| SE | 57229 | ENSG00000178691 | SUZ12 | chr17 | + | 31940421 | 31940486 | 31940285 | 31940332 | 31947616 | 31947685 | 238 | 180 | 252 | 246 | 527 | 0 | 0 | 2 | 0 | 0 | 80 | 140 | 158 | 85 | 44 | 0 | 0 | 0 | 0 | 0 | 214 | 149 | 1 | 1 | 1 | 1 | 0.989 | 1 | 1 | 1 | 1 | 1 | 1 | 1 | -0.002 |
| SE | 57230 | ENSG00000178691 | SUZ12 | chr17 | + | 31947616 | 31947685 | 31937006 | 31937520 | 31966146 | 31966196 | 153 | 115 | 133 | 186 | 340 | 0 | 0 | 0 | 3 | 0 | 55 | 65 | 64 | 20 | 37 | 0 | 0 | 0 | 0 | 0 | 218 | 149 | 1 | 1 | 1 | 1 | 1 | 0.977 | 1 | 1 | 1 | 1 | 1 | 1 | -0.005 |
| SE | 57236 | ENSG00000143815 | LBR | chr1 | - | 225410290 | 225410416 | 225406561 | 225406832 | 225412453 | 225412645 | 177 | 91 | 148 | 177 | 182 | 0 | 0 | 2 | 0 | 0 | 52 | 184 | 108 | 151 | 6 | 0 | 0 | 4 | 0 | 0 | 275 | 149 | 1 | 1 | 1 | 1 | 0.976 | 1 | 1 | 1 | 1 | 0.936 | 1 | 1 | 0.008 |
| SE | 57238 | ENSG00000143815 | LBR | chr1 | - | 225411336 | 225411440 | 225406663 | 225406832 | 225412453 | 225412645 | 511 | 257 | 503 | 417 | 722 | 0 | 0 | 2 | 0 | 0 | 271 | 407 | 290 | 432 | 60 | 0 | 0 | 4 | 0 | 0 | 253 | 149 | 1 | 1 | 1 | 1 | 0.993 | 1 | 1 | 1 | 1 | 0.977 | 1 | 1 | 0.003 |
| SE | 57240 | ENSG00000143815 | LBR | chr1 | - | 225412453 | 225412645 | 225411336 | 225411440 | 225415277 | 225415332 | 1070 | 537 | 1032 | 887 | 1515 | 0 | 0 | 0 | 0 | 0 | 509 | 880 | 595 | 893 | 228 | 0 | 0 | 0 | 1 | 0 | 298 | 149 | 1 | 1 | 1 | 1 | 1 | 1 | 1 | 1 | 1 | 1 | 0.998 | 1 | 0 |
| SE | 57241 | ENSG00000143815 | LBR | chr1 | - | 225415277 | 225415332 | 225411336 | 225411440 | 225417983 | 225418180 | 66 | 31 | 57 | 62 | 142 | 0 | 0 | 0 | 1 | 0 | 60 | 92 | 45 | 56 | 18 | 0 | 0 | 2 | 0 | 0 | 204 | 149 | 1 | 1 | 1 | 1 | 1 | 0.978 | 1 | 1 | 1 | 0.943 | 1 | 1 | 0.007 |
| SE | 57242 | ENSG00000143815 | LBR | chr1 | - | 225423910 | 225424089 | 225422076 | 225422277 | 225427953 | 225428038 | 568 | 292 | 524 | 553 | 965 | 0 | 0 | 0 | 1 | 0 | 372 | 664 | 521 | 544 | 124 | 0 | 0 | 0 | 0 | 0 | 298 | 149 | 1 | 1 | 1 | 1 | 1 | 0.996 | 1 | 1 | 1 | 1 | 1 | 1 | -0.001 |
| SE | 57243 | ENSG00000162385 | MAGOH | chr1 | - | 53228871 | 53228954 | 53226899 | 53227144 | 53233541 | 53233652 | 246 | 156 | 188 | 226 | 215 | 0 | 0 | 1 | 0 | 0 | 273 | 222 | 225 | 232 | 129 | 0 | 0 | 0 | 0 | 0 | 232 | 149 | 1 | 1 | 1 | 1 | 0.992 | 1 | 1 | 1 | 1 | 1 | 1 | 1 | -0.002 |
| SE | 57250 | ENSG00000143119 | CD53 | chr1 | + | 110891391 | 110891471 | 110873147 | 110873248 | 110892344 | 110892533 | 3421 | 1895 | 2150 | 2795 | 4103 | 0 | 0 | 0 | 0 | 4 | 3709 | 3290 | 1890 | 2943 | 3845 | 0 | 0 | 0 | 0 | 0 | 229 | 149 | 1 | 1 | 1 | 1 | 1 | 1 | 0.999 | 1 | 1 | 1 | 1 | 1 | 0 |
| SE | 57251 | ENSG00000143119 | CD53 | chr1 | + | 110891391 | 110891471 | 110873147 | 110873248 | 110894326 | 110894401 | 1146 | 619 | 729 | 935 | 981 | 0 | 0 | 0 | 0 | 0 | 1160 | 1156 | 738 | 1029 | 1303 | 0 | 3 | 0 | 0 | 5 | 229 | 149 | 1 | 1 | 1 | 1 | 1 | 1 | 1 | 1 | 0.996 | 1 | 1 | 0.994 | 0.002 |
| SE | 57252 | ENSG00000143119 | CD53 | chr1 | + | 110891391 | 110891471 | 110873147 | 110873248 | 110897808 | 110897892 | 1143 | 620 | 729 | 935 | 979 | 0 | 1 | 0 | 0 | 0 | 1160 | 1149 | 727 | 1005 | 1299 | 0 | 0 | 3 | 0 | 0 | 229 | 149 | 1 | 1 | 1 | 0.998 | 1 | 1 | 1 | 1 | 1 | 0.994 | 1 | 1 | 0.001 |
| SE | 57253 | ENSG00000143119 | CD53 | chr1 | + | 110891391 | 110891471 | 110873147 | 110873248 | 110899123 | 110899514 | 1158 | 629 | 743 | 953 | 1015 | 3 | 0 | 0 | 2 | 6 | 1184 | 1162 | 781 | 1038 | 1308 | 0 | 0 | 7 | 9 | 3 | 229 | 149 | 1 | 1 | 0.996 | 1 | 1 | 0.997 | 0.991 | 1 | 1 | 0.986 | 0.987 | 0.996 | 0.003 |
| SE | 57254 | ENSG00000143119 | CD53 | chr1 | + | 110892344 | 110892533 | 110873147 | 110873248 | 110894326 | 110894401 | 1204 | 632 | 981 | 1206 | 1596 | 0 | 0 | 0 | 0 | 0 | 1199 | 1075 | 637 | 945 | 1133 | 0 | 3 | 0 | 0 | 5 | 298 | 149 | 1 | 1 | 1 | 1 | 1 | 1 | 1 | 1 | 0.994 | 1 | 1 | 0.991 | 0.003 |
| SE | 57255 | ENSG00000143119 | CD53 | chr1 | + | 110892344 | 110892533 | 110873147 | 110873248 | 110897808 | 110897892 | 1 | 0 | 1 | 0 | 4 | 0 | 1 | 0 | 0 | 0 | 0 | 1 | 1 | 0 | 0 | 0 | 0 | 3 | 0 | 0 | 298 | 149 | 1 | 1 | 1 | 0 | 1 | NA | 1 | NA | 1 | 0.143 | NA | NA | 0.178 |
| SE | 57258 | ENSG00000143119 | CD53 | chr1 | + | 110892344 | 110892533 | 110891391 | 110891471 | 110894959 | 110895055 | 2285 | 1276 | 1426 | 1867 | 3151 | 0 | 0 | 0 | 1 | 0 | 2549 | 2146 | 1163 | 1945 | 2546 | 0 | 0 | 5 | 0 | 0 | 298 | 149 | 1 | 1 | 1 | 1 | 1 | 0.999 | 1 | 1 | 1 | 0.991 | 1 | 1 | 0.002 |
| SE | 57259 | ENSG00000143119 | CD53 | chr1 | + | 110892344 | 110892533 | 110891391 | 110891471 | 110897808 | 110897892 | 2283 | 1276 | 1426 | 1866 | 3151 | 0 | 1 | 1 | 2 | 0 | 2549 | 2147 | 1164 | 1939 | 2546 | 0 | 2 | 0 | 1 | 0 | 298 | 149 | 1 | 1 | 1 | 0.998 | 0.999 | 0.998 | 1 | 1 | 0.998 | 1 | 0.999 | 1 | 0 |
| SE | 57261 | ENSG00000143119 | CD53 | chr1 | + | 110894326 | 110894401 | 110873147 | 110873248 | 110897808 | 110897892 | 3 | 0 | 5 | 0 | 1 | 0 | 1 | 0 | 0 | 0 | 1 | 6 | 11 | 0 | 5 | 0 | 0 | 3 | 0 | 0 | 224 | 149 | 1 | 1 | 1 | 0 | 1 | NA | 1 | 1 | 1 | 0.709 | NA | 1 | -0.177 |
| SE | 57263 | ENSG00000143119 | CD53 | chr1 | + | 110894326 | 110894401 | 110891391 | 110891471 | 110894959 | 110895055 | 281 | 114 | 173 | 252 | 359 | 0 | 0 | 0 | 1 | 0 | 249 | 255 | 100 | 188 | 311 | 0 | 0 | 5 | 0 | 0 | 224 | 149 | 1 | 1 | 1 | 1 | 1 | 0.994 | 1 | 1 | 1 | 0.93 | 1 | 1 | 0.013 |
| SE | 57266 | ENSG00000143119 | CD53 | chr1 | + | 110894326 | 110894401 | 110892344 | 110892533 | 110894959 | 110895055 | 1447 | 713 | 1120 | 1409 | 1890 | 3 | 0 | 1 | 1 | 0 | 1407 | 1289 | 655 | 1069 | 1401 | 0 | 0 | 0 | 6 | 0 | 224 | 149 | 1 | 1 | 0.997 | 1 | 0.999 | 0.999 | 1 | 1 | 1 | 1 | 0.992 | 1 | 0.001 |
| SE | 57267 | ENSG00000143119 | CD53 | chr1 | + | 110894326 | 110894401 | 110892344 | 110892533 | 110896652 | 110896733 | 648 | 387 | 536 | 654 | 789 | 0 | 0 | 0 | 0 | 0 | 680 | 547 | 338 | 529 | 557 | 0 | 0 | 1 | 0 | 0 | 224 | 149 | 1 | 1 | 1 | 1 | 1 | 1 | 1 | 1 | 1 | 0.996 | 1 | 1 | 0.001 |
| SE | 57268 | ENSG00000143119 | CD53 | chr1 | + | 110894326 | 110894401 | 110892344 | 110892533 | 110897808 | 110897892 | 647 | 385 | 532 | 653 | 780 | 1 | 0 | 1 | 0 | 0 | 697 | 534 | 328 | 530 | 548 | 0 | 1 | 1 | 0 | 0 | 224 | 149 | 1 | 1 | 0.998 | 1 | 0.997 | 1 | 1 | 1 | 0.997 | 0.995 | 1 | 1 | 0.001 |
| SE | 57270 | ENSG00000143119 | CD53 | chr1 | + | 110894959 | 110895055 | 110891391 | 110891471 | 110897808 | 110897892 | 17 | 8 | 10 | 6 | 12 | 0 | 1 | 1 | 2 | 0 | 15 | 20 | 13 | 9 | 46 | 0 | 2 | 0 | 1 | 0 | 245 | 149 | 1 | 1 | 1 | 0.83 | 0.859 | 0.646 | 1 | 1 | 0.859 | 1 | 0.846 | 1 | -0.074 |
| SE | 57272 | ENSG00000143119 | CD53 | chr1 | + | 110894959 | 110895055 | 110892344 | 110892533 | 110896652 | 110896733 | 851 | 372 | 592 | 709 | 557 | 0 | 0 | 0 | 0 | 0 | 739 | 735 | 322 | 456 | 677 | 0 | 0 | 1 | 0 | 0 | 245 | 149 | 1 | 1 | 1 | 1 | 1 | 1 | 1 | 1 | 1 | 0.995 | 1 | 1 | 0.001 |
| SE | 57273 | ENSG00000143119 | CD53 | chr1 | + | 110894959 | 110895055 | 110892344 | 110892533 | 110897808 | 110897892 | 17 | 8 | 11 | 6 | 12 | 1 | 0 | 1 | 0 | 0 | 15 | 20 | 8 | 9 | 46 | 0 | 1 | 1 | 0 | 0 | 245 | 149 | 1 | 1 | 0.912 | 1 | 0.87 | 1 | 1 | 1 | 0.924 | 0.83 | 1 | 1 | 0.006 |
| SE | 57278 | ENSG00000143119 | CD53 | chr1 | + | 110896652 | 110896733 | 110892344 | 110892533 | 110897808 | 110897892 | 809 | 355 | 476 | 633 | 896 | 1 | 0 | 1 | 0 | 0 | 846 | 571 | 350 | 486 | 826 | 0 | 1 | 1 | 0 | 0 | 230 | 149 | 1 | 1 | 0.998 | 1 | 0.997 | 1 | 1 | 1 | 0.997 | 0.996 | 1 | 1 | 0 |
| SE | 57291 | ENSG00000174123 | TLR10 | chr4 | - | 38775774 | 38775963 | 38772237 | 38775652 | 38782920 | 38782990 | 28 | 23 | 14 | 22 | 53 | 3 | 8 | 1 | 10 | 10 | 6 | 6 | 2 | 0 | 0 | 0 | 0 | 0 | 0 | 0 | 298 | 149 | 1 | 1 | 0.824 | 0.59 | 0.875 | 0.524 | 0.726 | 1 | 1 | 1 | NA | NA | -0.292 |
| SE | 57292 | ENSG00000174123 | TLR10 | chr4 | - | 38776109 | 38776426 | 38772238 | 38775652 | 38782920 | 38782989 | 30 | 31 | 12 | 16 | 43 | 3 | 8 | 1 | 10 | 10 | 0 | 11 | 1 | 1 | 6 | 0 | 0 | 0 | 0 | 0 | 298 | 149 | 1 | 1 | 0.833 | 0.66 | 0.857 | 0.444 | 0.683 | NA | 1 | 1 | 1 | 1 | -0.305 |
| SE | 57293 | ENSG00000174123 | TLR10 | chr4 | - | 38776109 | 38776426 | 38775774 | 38775963 | 38782920 | 38782989 | 58 | 25 | 24 | 39 | 72 | 4 | 0 | 0 | 2 | 0 | 0 | 13 | 0 | 14 | 0 | 0 | 0 | 0 | 0 | 0 | 298 | 149 | 1 | 1 | 0.879 | 1 | 1 | 0.907 | 1 | NA | 1 | NA | 1 | NA | -0.043 |
| SE | 57294 | ENSG00000174123 | TLR10 | chr4 | - | 38778970 | 38779051 | 38776109 | 38776426 | 38782920 | 38782990 | 0 | 1 | 0 | 0 | 4 | 26 | 11 | 7 | 15 | 18 | 0 | 1 | 0 | 3 | 0 | 0 | 8 | 0 | 1 | 0 | 230 | 149 | 1 | 1 | 0 | 0.056 | 0 | 0 | 0.126 | NA | 0.075 | NA | 0.66 | NA | -0.331 |
| SE | 57296 | ENSG00000131584 | ACAP3 | chr1 | - | 1296014 | 1296109 | 1295735 | 1295938 | 1296210 | 1296280 | 19 | 7 | 5 | 28 | 16 | 0 | 0 | 0 | 0 | 0 | 30 | 34 | 7 | 22 | 29 | 0 | 0 | 1 | 0 | 0 | 244 | 149 | 1 | 1 | 1 | 1 | 1 | 1 | 1 | 1 | 1 | 0.81 | 1 | 1 | 0.038 |
| SE | 57297 | ENSG00000131584 | ACAP3 | chr1 | - | 1299972 | 1300068 | 1298743 | 1299905 | 1300157 | 1300202 | 27 | 24 | 28 | 36 | 15 | 0 | 0 | 0 | 2 | 0 | 83 | 24 | 28 | 39 | 57 | 0 | 0 | 0 | 2 | 0 | 245 | 149 | 1 | 1 | 1 | 1 | 1 | 0.916 | 1 | 1 | 1 | 1 | 0.922 | 1 | -0.001 |
| SE | 57298 | ENSG00000131584 | ACAP3 | chr1 | - | 1300157 | 1300202 | 1299972 | 1300068 | 1301987 | 1302046 | 3 | 5 | 6 | 10 | 9 | 0 | 0 | 0 | 0 | 0 | 15 | 7 | 5 | 2 | 18 | 0 | 0 | 1 | 0 | 0 | 194 | 149 | 1 | 1 | 1 | 1 | 1 | 1 | 1 | 1 | 1 | 0.793 | 1 | 1 | 0.041 |
| SE | 57299 | ENSG00000131584 | ACAP3 | chr1 | - | 1300508 | 1300692 | 1300157 | 1300202 | 1301987 | 1302046 | 68 | 35 | 38 | 80 | 21 | 0 | 0 | 0 | 2 | 0 | 87 | 58 | 30 | 71 | 112 | 0 | 0 | 0 | 0 | 0 | 298 | 149 | 1 | 1 | 1 | 1 | 1 | 0.952 | 1 | 1 | 1 | 1 | 1 | 1 | -0.01 |
| SE | 57304 | ENSG00000015532 | XYLT2 | chr17 | + | 50357056 | 50357252 | 50356510 | 50356773 | 50358206 | 50358540 | 28 | 17 | 39 | 19 | 15 | 2 | 0 | 0 | 2 | 0 | 34 | 40 | 6 | 15 | 87 | 0 | 0 | 0 | 2 | 0 | 298 | 149 | 1 | 1 | 0.875 | 1 | 1 | 0.826 | 1 | 1 | 1 | 1 | 0.789 | 1 | -0.018 |
| SE | 57311 | ENSG00000226435 | ANKRD18DP | chr3 | - | 198072755 | 198072830 | 198063717 | 198063788 | 198076795 | 198077012 | 0 | 0 | 1 | 0 | 0 | 0 | 0 | 0 | 0 | 0 | 0 | 0 | 0 | 0 | 0 | 6 | 0 | 0 | 0 | 0 | 224 | 149 | 1 | 1 | NA | NA | 1 | NA | NA | 0 | NA | NA | NA | NA | 1 |
| SE | 57317 | ENSG00000181722 | ZBTB20 | chr3 | - | 114500351 | 114500391 | 114389004 | 114389105 | 114693527 | 114693575 | 74 | 80 | 95 | 72 | 153 | 0 | 0 | 1 | 0 | 1 | 12 | 57 | 59 | 59 | 38 | 0 | 0 | 1 | 0 | 0 | 189 | 149 | 1 | 1 | 1 | 1 | 0.987 | 1 | 0.992 | 1 | 1 | 0.979 | 1 | 1 | 0 |
| SE | 57318 | ENSG00000181722 | ZBTB20 | chr3 | - | 114502797 | 114502864 | 114500351 | 114500391 | 114582113 | 114582266 | 3 | 0 | 2 | 0 | 0 | 1 | 3 | 0 | 3 | 3 | 0 | 0 | 2 | 0 | 0 | 0 | 1 | 7 | 16 | 0 | 216 | 149 | 1 | 1 | 0.674 | 0 | 1 | 0 | 0 | NA | 0 | 0.165 | 0 | NA | 0.28 |
| SE | 57319 | ENSG00000181722 | ZBTB20 | chr3 | - | 114502797 | 114502864 | 114500351 | 114500391 | 114693527 | 114693575 | 2 | 0 | 0 | 0 | 1 | 78 | 78 | 95 | 73 | 144 | 0 | 0 | 4 | 1 | 0 | 16 | 57 | 56 | 48 | 36 | 216 | 149 | 1 | 1 | 0.017 | 0 | 0 | 0 | 0.005 | 0 | 0 | 0.047 | 0.014 | 0 | -0.008 |
| SE | 57320 | ENSG00000181722 | ZBTB20 | chr3 | - | 114582113 | 114582266 | 114500351 | 114500391 | 114599250 | 114599303 | 2 | 3 | 0 | 3 | 3 | 1 | 3 | 2 | 0 | 0 | 0 | 1 | 7 | 16 | 0 | 0 | 6 | 0 | 0 | 0 | 298 | 149 | 1 | 1 | 0.5 | 0.333 | 0 | 1 | 1 | NA | 0.077 | 1 | 1 | NA | -0.126 |
| SE | 57322 | ENSG00000181722 | ZBTB20 | chr3 | - | 114582113 | 114582266 | 114502797 | 114502864 | 114693527 | 114693575 | 2 | 1 | 5 | 5 | 1 | 0 | 0 | 0 | 1 | 0 | 0 | 1 | 2 | 0 | 0 | 0 | 0 | 2 | 1 | 0 | 298 | 149 | 1 | 1 | 1 | 1 | 1 | 1 | 0.333 | NA | 1 | 0.333 | 0 | NA | 0.422 |
| SE | 57324 | ENSG00000181722 | ZBTB20 | chr3 | - | 114693527 | 114693575 | 114500351 | 114500391 | 114801100 | 114801174 | 69 | 68 | 91 | 62 | 141 | 0 | 0 | 1 | 0 | 0 | 0 | 51 | 45 | 47 | 8 | 0 | 4 | 0 | 0 | 0 | 197 | 149 | 1 | 1 | 1 | 1 | 0.986 | 1 | 1 | NA | 0.906 | 1 | 1 | 1 | 0.021 |
| SE | 57328 | ENSG00000181722 | ZBTB20 | chr3 | - | 114710267 | 114710313 | 114693527 | 114693575 | 114900303 | 114900342 | 15 | 18 | 22 | 24 | 29 | 1 | 0 | 0 | 0 | 0 | 6 | 9 | 14 | 6 | 5 | 0 | 0 | 2 | 0 | 2 | 195 | 149 | 1 | 1 | 0.92 | 1 | 1 | 1 | 1 | 1 | 1 | 0.842 | 1 | 0.656 | 0.084 |
| SE | 57329 | ENSG00000181722 | ZBTB20 | chr3 | - | 114713975 | 114714015 | 114693527 | 114693575 | 114743711 | 114743779 | 2 | 2 | 0 | 1 | 0 | 1 | 0 | 0 | 0 | 0 | 0 | 0 | 4 | 0 | 0 | 10 | 0 | 0 | 0 | 0 | 189 | 149 | 1 | 1 | 0.612 | 1 | NA | 1 | NA | 0 | NA | 1 | NA | NA | 0.371 |
| SE | 57331 | ENSG00000181722 | ZBTB20 | chr3 | - | 114713975 | 114714015 | 114693527 | 114693575 | 114900303 | 114900342 | 2 | 2 | 0 | 1 | 0 | 1 | 0 | 0 | 0 | 0 | 0 | 0 | 4 | 0 | 0 | 0 | 0 | 2 | 0 | 2 | 189 | 149 | 1 | 1 | 0.612 | 1 | NA | 1 | NA | NA | NA | 0.612 | NA | 0 | 0.565 |
| SE | 57333 | ENSG00000181722 | ZBTB20 | chr3 | - | 114713975 | 114714015 | 114710267 | 114710313 | 114900303 | 114900342 | 14 | 12 | 18 | 9 | 22 | 0 | 0 | 0 | 1 | 0 | 0 | 11 | 3 | 1 | 15 | 0 | 0 | 0 | 0 | 0 | 189 | 149 | 1 | 1 | 1 | 1 | 1 | 0.876 | 1 | NA | 1 | 1 | 1 | 1 | -0.025 |
| SE | 57335 | ENSG00000181722 | ZBTB20 | chr3 | - | 114743711 | 114743779 | 114713975 | 114714015 | 114801100 | 114801177 | 0 | 4 | 0 | 1 | 0 | 22 | 39 | 25 | 19 | 33 | 0 | 0 | 0 | 0 | 0 | 9 | 10 | 13 | 12 | 0 | 217 | 149 | 1 | 1 | 0 | 0.066 | 0 | 0.035 | 0 | 0 | 0 | 0 | 0 | NA | 0.02 |
| SE | 57336 | ENSG00000181722 | ZBTB20 | chr3 | - | 114743711 | 114743779 | 114713975 | 114714015 | 114874038 | 114874108 | 0 | 2 | 0 | 0 | 0 | 0 | 0 | 0 | 0 | 0 | 0 | 0 | 0 | 0 | 0 | 0 | 0 | 1 | 0 | 0 | 217 | 149 | 1 | 1 | NA | 1 | NA | NA | NA | NA | NA | 0 | NA | NA | 1 |
| SE | 57337 | ENSG00000181722 | ZBTB20 | chr3 | - | 114801100 | 114801174 | 114693527 | 114693575 | 114900303 | 114900342 | 141 | 112 | 152 | 137 | 215 | 1 | 0 | 0 | 0 | 0 | 6 | 87 | 61 | 73 | 26 | 0 | 0 | 2 | 0 | 2 | 223 | 149 | 1 | 1 | 0.989 | 1 | 1 | 1 | 1 | 1 | 1 | 0.953 | 1 | 0.897 | 0.028 |
| SE | 57338 | ENSG00000181722 | ZBTB20 | chr3 | - | 114801100 | 114801174 | 114710267 | 114710313 | 114900303 | 114900342 | 66 | 55 | 70 | 70 | 86 | 0 | 0 | 0 | 1 | 0 | 0 | 42 | 19 | 45 | 30 | 0 | 0 | 0 | 0 | 0 | 223 | 149 | 1 | 1 | 1 | 1 | 1 | 0.979 | 1 | NA | 1 | 1 | 1 | 1 | -0.004 |
| SE | 57339 | ENSG00000181722 | ZBTB20 | chr3 | - | 114801100 | 114801174 | 114713975 | 114714015 | 114874038 | 114874108 | 11 | 19 | 15 | 3 | 19 | 0 | 0 | 0 | 0 | 0 | 9 | 0 | 7 | 0 | 0 | 0 | 0 | 1 | 0 | 0 | 223 | 149 | 1 | 1 | 1 | 1 | 1 | 1 | 1 | 1 | NA | 0.824 | NA | NA | 0.088 |
| SE | 57340 | ENSG00000181722 | ZBTB20 | chr3 | - | 114801100 | 114801174 | 114713975 | 114714015 | 114900303 | 114900342 | 83 | 92 | 94 | 86 | 116 | 0 | 0 | 1 | 0 | 0 | 9 | 48 | 29 | 50 | 23 | 0 | 0 | 0 | 0 | 0 | 223 | 149 | 1 | 1 | 1 | 1 | 0.984 | 1 | 1 | 1 | 1 | 1 | 1 | 1 | -0.003 |
| SE | 57341 | ENSG00000181722 | ZBTB20 | chr3 | - | 114874038 | 114874108 | 114713975 | 114714015 | 114900303 | 114900342 | 0 | 3 | 5 | 3 | 5 | 0 | 0 | 1 | 0 | 0 | 6 | 3 | 1 | 0 | 0 | 0 | 0 | 0 | 0 | 0 | 219 | 149 | 1 | 1 | NA | 1 | 0.773 | 1 | 1 | 1 | 1 | 1 | NA | NA | -0.057 |
| SE | 57343 | ENSG00000181722 | ZBTB20 | chr3 | - | 114900303 | 114900342 | 114801100 | 114801174 | 114974365 | 114974416 | 123 | 114 | 143 | 111 | 170 | 0 | 0 | 0 | 0 | 1 | 12 | 62 | 42 | 69 | 28 | 0 | 0 | 0 | 0 | 0 | 188 | 149 | 1 | 1 | 1 | 1 | 1 | 1 | 0.993 | 1 | 1 | 1 | 1 | 1 | -0.001 |
| SE | 57345 | ENSG00000181722 | ZBTB20 | chr3 | - | 114974365 | 114974416 | 114900303 | 114900342 | 115100174 | 115100295 | 59 | 56 | 71 | 38 | 88 | 0 | 0 | 0 | 0 | 1 | 6 | 21 | 25 | 32 | 4 | 0 | 0 | 0 | 0 | 0 | 200 | 149 | 1 | 1 | 1 | 1 | 1 | 1 | 0.985 | 1 | 1 | 1 | 1 | 1 | -0.003 |
| SE | 57346 | ENSG00000181722 | ZBTB20 | chr3 | - | 115027455 | 115027574 | 114974391 | 114974416 | 115071218 | 115071414 | 13 | 2 | 13 | 9 | 19 | 34 | 7 | 55 | 49 | 53 | 0 | 5 | 4 | 5 | 0 | 0 | 27 | 4 | 20 | 33 | 268 | 149 | 1 | 1 | 0.175 | 0.137 | 0.116 | 0.093 | 0.166 | NA | 0.093 | 0.357 | 0.122 | 0 | -0.006 |
| SE | 57347 | ENSG00000181722 | ZBTB20 | chr3 | - | 115027455 | 115027574 | 114974391 | 114974416 | 115100174 | 115100295 | 3 | 2 | 7 | 3 | 6 | 13 | 16 | 14 | 4 | 14 | 0 | 0 | 3 | 4 | 0 | 6 | 1 | 3 | 1 | 0 | 268 | 149 | 1 | 1 | 0.114 | 0.065 | 0.218 | 0.294 | 0.192 | 0 | 0 | 0.357 | 0.69 | NA | -0.085 |
| SE | 57348 | ENSG00000181722 | ZBTB20 | chr3 | - | 115027455 | 115027574 | 114974391 | 114974416 | 115144812 | 115144909 | 4 | 3 | 7 | 3 | 8 | 3 | 13 | 4 | 5 | 2 | 0 | 0 | 5 | 2 | 0 | 17 | 1 | 2 | 0 | 0 | 268 | 149 | 1 | 1 | 0.426 | 0.114 | 0.493 | 0.25 | 0.69 | 0 | 0 | 0.582 | 1 | NA | -0.001 |
| SE | 57349 | ENSG00000181722 | ZBTB20 | chr3 | - | 115027455 | 115027574 | 114974391 | 114974416 | 115147218 | 115147288 | 4 | 3 | 10 | 4 | 6 | 19 | 13 | 10 | 13 | 20 | 0 | 0 | 3 | 2 | 0 | 0 | 1 | 6 | 0 | 0 | 268 | 149 | 1 | 1 | 0.105 | 0.114 | 0.357 | 0.146 | 0.143 | NA | 0 | 0.218 | 1 | NA | -0.233 |

| | | | | | | | | | | | | | | |
|---|---|---|---|---|---|---|---|---|---|---|---|---|---|---|
| SE | 57352 | ENSG00000181722 | ZBTB20 | chr3 | - | 115071218 | 115071414 | 114974365 | 114974416 | 115147218 | 115147288 | 56 | 11 | [illegible] |
| SE | 57353 | ENSG00000181722 | ZBTB20 | chr3 | - | 115071218 | 115071414 | 115027455 | 115027574 | 115100174 | 115100295 | 17 | 0 | [illegible] |
| SE | 57354 | ENSG00000181722 | ZBTB20 | chr3 | - | 115071218 | 115071414 | 115027455 | 115027574 | 115144812 | 115144909 | 19 | 1 | [illegible] |
| SE | 57355 | ENSG00000181722 | ZBTB20 | chr3 | - | 115071218 | 115071414 | 115027455 | 115027574 | 115147218 | 115147288 | 32 | 4 | [illegible] |
| SE | 57356 | ENSG00000227057 | WDR46 | chr6 | - | 33287090 | 33287226 | 33286794 | 33286894 | 33287354 | 33287429 | 68 | 42 | [illegible] |
| SE | 57357 | ENSG00000227057 | WDR46 | chr6 | - | 33287613 | 33287718 | 33287354 | 33287505 | 33287964 | 33288002 | 70 | 50 | [illegible] |
| SE | 57359 | ENSG00000163430 | FSTL1 | chr3 | - | 120450683 | 120450746 | 120415922 | 120416027 | 120450896 | 120450960 | 3 | 4 | [illegible] |
| SE | 57360 | ENSG00000163430 | FSTL1 | chr3 | - | 120450683 | 120450797 | 120415922 | 120416027 | 120450896 | 120450991 | 2 | 1 | [illegible] |
| SE | 57362 | ENSG00000116815 | CD58 | chr1 | - | 116519230 | 116519267 | 116514533 | 116514822 | 116535964 | 116536228 | 8 | 1 | [illegible] |
| SE | 57364 | ENSG00000116815 | CD58 | chr1 | - | 116521905 | 116521983 | 116514533 | 116514822 | 116535964 | 116536228 | 140 | 120 | [illegible] |
| SE | 57365 | ENSG00000116815 | CD58 | chr1 | - | 116521905 | 116521983 | 116518699 | 116519267 | 116535964 | 116536228 | 284 | 225 | [illegible] |
| SE | 57366 | ENSG00000116815 | CD58 | chr1 | - | 116535964 | 116536228 | 116521786 | 116521983 | 116544310 | 116544604 | 627 | 445 | [illegible] |
| SE | 57382 | ENSG00000101040 | ZMYND8 | chr20 | - | 47227202 | 47227281 | 47224316 | 47224556 | 47229725 | 47229806 | 54 | 55 | [illegible] |
| SE | 57385 | ENSG00000101040 | ZMYND8 | chr20 | - | 47236325 | 47236516 | 47229725 | 47229806 | 47249286 | 47249439 | 39 | 18 | [illegible] |
| SE | 57387 | ENSG00000101040 | ZMYND8 | chr20 | - | 47238757 | 47239138 | 47236325 | 47236516 | 47249286 | 47249439 | 7 | 1 | [illegible] |
| SE | 57390 | ENSG00000101040 | ZMYND8 | chr20 | - | 47246007 | 47246517 | 47229725 | 47229806 | 47249286 | 47249439 | 94 | 74 | [illegible] |
| SE | 57392 | ENSG00000101040 | ZMYND8 | chr20 | - | 47246007 | 47246517 | 47237309 | 47239138 | 47249286 | 47249439 | 126 | 100 | [illegible] |
| SE | 57393 | ENSG00000101040 | ZMYND8 | chr20 | - | 47249286 | 47249439 | 47246007 | 47246517 | 47262287 | 47262428 | 143 | 115 | [illegible] |
| SE | 57394 | ENSG00000101040 | ZMYND8 | chr20 | - | 47262287 | 47262428 | 47249286 | 47249439 | 47276313 | 47276795 | 142 | 94 | [illegible] |
| SE | 57395 | ENSG00000101040 | ZMYND8 | chr20 | - | 47262287 | 47262428 | 47249286 | 47249439 | 47282101 | 47282217 | 50 | 45 | [illegible] |
| SE | 57396 | ENSG00000101040 | ZMYND8 | chr20 | - | 47276313 | 47276795 | 47249286 | 47249439 | 47282101 | 47282217 | 134 | 92 | [illegible] |
| SE | 57399 | ENSG00000101040 | ZMYND8 | chr20 | - | 47290186 | 47290274 | 47287228 | 47287284 | 47291795 | 47291888 | 230 | 194 | [illegible] |
| SE | 57400 | ENSG00000101040 | ZMYND8 | chr20 | - | 47290186 | 47290274 | 47287228 | 47287284 | 47294665 | 47294779 | 23 | 13 | [illegible] |
| SE | 57401 | ENSG00000101040 | ZMYND8 | chr20 | - | 47291795 | 47291888 | 47287228 | 47287284 | 47294665 | 47294779 | 113 | 120 | [illegible] |
| SE | 57403 | ENSG00000101040 | ZMYND8 | chr20 | - | 47291795 | 47291888 | 47290186 | 47290274 | 47298728 | 47298872 | 195 | 186 | [illegible] |
| [illegible] | | | | | | | | | | | | | | |

| | | | | | | | | | | | | | |
|---|---|---|---|---|---|---|---|---|---|---|---|---|---|
| SE | 57720 | ENSG00000001497 | LAS1L | chrX | - | 65531356 | 65531438 | 65529593 | 65529878 | 65532560 | 65532630 | 131 | 85 |
| SE | 57721 | ENSG00000001497 | LAS1L | chrX | - | 65532560 | 65532630 | 65531356 | 65531438 | 65533609 | 65533735 | 120 | 73 |
| SE | 57724 | ENSG00000001497 | LAS1L | chrX | - | 65534479 | 65534572 | 65533609 | 65533735 | 65534717 | 65534776 | 41 | 19 |
| SE | 57728 | ENSG00000114354 | TFG | chr3 | + | 100713642 | 100713869 | 100709294 | 100709431 | 100719974 | 100720058 | 207 | 143 |
| SE | 57729 | ENSG00000114354 | TFG | chr3 | + | 100713642 | 100713869 | 100709413 | 100709721 | 100719974 | 100720058 | 211 | 152 |
| SE | 57730 | ENSG00000114354 | TFG | chr3 | + | 100713653 | 100713869 | 100709294 | 100709431 | 100719974 | 100720058 | 185 | 140 |
| SE | 57731 | ENSG00000114354 | TFG | chr3 | + | 100713653 | 100713869 | 100709456 | 100709721 | 100719974 | 100720058 | 185 | 142 |
| SE | 57732 | ENSG00000114354 | TFG | chr3 | + | 100731741 | 100731895 | 100728711 | 100728858 | 100732507 | 100732672 | 0 | 0 |
| SE | 57733 | ENSG00000114354 | TFG | chr3 | + | 100739360 | 100739485 | 100736575 | 100736716 | 100744832 | 100744931 | 0 | 0 |
| SE | 57734 | ENSG00000114354 | TFG | chr3 | + | 100742155 | 100742200 | 100736575 | 100736716 | 100744832 | 100744931 | 1 | 0 |
| SE | 57736 | ENSG00000114354 | TFG | chr3 | + | 100747360 | 100747454 | 100744832 | 100744931 | 100748148 | 100748964 | 0 | 0 |
| SE | 57737 | ENSG00000114354 | TFG | chr3 | + | 100747413 | 100747454 | 100744832 | 100744931 | 100748148 | 100748964 | 0 | 0 |
| SE | 57739 | ENSG00000118007 | STAG1 | chr3 | - | 136357719 | 136357848 | 136348788 | 136349363 | 136359147 | 136359189 | 199 | 136 |
| SE | 57740 | ENSG00000118007 | STAG1 | chr3 | - | 136357719 | 136357848 | 136348788 | 136349363 | 136363365 | 136363467 | 76 | 66 |
| SE | 57741 | ENSG00000118007 | STAG1 | chr3 | - | 136359147 | 136359296 | 136349157 | 136349363 | 136363365 | 136363467 | 105 | 84 |
| SE | 57742 | ENSG00000118007 | STAG1 | chr3 | - | 136359147 | 136359296 | 136349157 | 136349363 | 136366942 | 136367082 | 2 | 1 |
| SE | 57743 | ENSG00000118007 | STAG1 | chr3 | - | 136359147 | 136359296 | 136357719 | 136357848 | 136363365 | 136363467 | 232 | 157 |
| SE | 57744 | ENSG00000118007 | STAG1 | chr3 | - | 136363365 | 136363467 | 136349157 | 136349363 | 136366942 | 136367082 | 69 | 45 |
| SE | 57746 | ENSG00000118007 | STAG1 | chr3 | - | 136363365 | 136363467 | 136359147 | 136359296 | 136377659 | 136377752 | 85 | 72 |
| SE | 57747 | ENSG00000118007 | STAG1 | chr3 | - | 136366942 | 136367082 | 136359147 | 136359296 | 136377659 | 136377752 | 2 | 0 |

[illegible]

| | | | | | | | | | | | | | | | | | | | | | | | | | | | | | | | | | | | | | | | | | | | | | | | | |
|---|---|---|---|---|---|---|---|---|---|---|---|---|---|---|---|---|---|---|---|---|---|---|---|---|---|---|---|---|---|---|---|---|---|---|---|---|---|---|---|---|---|---|---|---|---|---|---|---|
| SE | 58150 | ENSG00000273136 | NBPF26 | chr1 | + | 120813890 | 120813993 | 120811885 | 120812095 | 120814828 | 120815043 | 65 | 74 | 57 | 66 | 76 | 0 | 0 | 1 | 0 | 1 | 27 | 36 | 72 | 50 | 22 | 0 | 0 | 0 | 0 | 0 | 0 | 0 | 252 | 149 | 1 | 1 | 1 | 1 | 0.971 | 1 | 0.978 | 1 | 1 | 1 | 1 | 1 | -0.01 |
| SE | 58151 | ENSG00000273136 | NBPF26 | chr1 | + | 120814828 | 120815043 | 120813890 | 120813993 | 120816084 | 120816157 | 43 | 51 | 33 | 37 | 55 | 0 | 0 | 0 | 3 | 0 | 25 | 20 | 71 | 81 | 14 | 0 | 0 | 0 | 0 | 0 | 0 | 0 | 298 | 149 | 1 | 1 | 1 | 1 | 1 | 0.86 | 1 | 1 | 1 | 1 | 1 | 1 | -0.028 |
| SE | 58152 | ENSG00000273136 | NBPF26 | chr1 | + | 120816621 | 120816827 | 120816084 | 120816157 | 120818122 | 120818174 | 84 | 123 | 32 | 50 | 87 | 0 | 0 | 1 | 1 | 0 | 198 | 56 | 193 | 322 | 64 | 0 | 0 | 0 | 0 | 0 | 2 | 0 | 298 | 149 | 1 | 1 | 1 | 1 | 0.941 | 0.962 | 1 | 1 | 1 | 1 | 0.988 | 1 | -0.017 |
| SE | 58153 | ENSG00000273136 | NBPF26 | chr1 | + | 120818122 | 120818174 | 120816621 | 120816827 | 120822103 | 120822156 | 47 | 83 | 21 | 39 | 53 | 0 | 0 | 0 | 2 | 0 | 121 | 39 | 130 | 221 | 34 | 0 | 0 | 0 | 0 | 0 | 0 | 0 | 201 | 149 | 1 | 1 | 1 | 1 | 1 | 0.935 | 1 | 1 | 1 | 1 | 1 | 1 | -0.013 |
| SE | 58158 | ENSG00000273136 | NBPF26 | chr1 | + | 120837168 | 120837341 | 120833602 | 120833775 | 120838050 | 120838102 | 36 | 24 | 19 | 20 | 46 | 2 | 0 | 0 | 0 | 0 | 32 | 14 | 42 | 31 | 23 | 0 | 0 | 0 | 0 | 0 | 0 | 0 | 298 | 149 | 1 | 1 | 0.9 | 1 | 1 | 1 | 1 | 1 | 1 | 1 | 1 | 1 | -0.02 |
| SE | 58161 | ENSG00000160688 | FLAD1 | chr1 | + | 154988104 | 154988849 | 154983343 | 154984066 | 154992712 | 154992786 | 48 | 15 | 27 | 25 | 57 | 0 | 1 | 2 | 1 | 0 | 41 | 24 | 9 | 21 | 21 | 0 | 3 | 0 | 3 | 0 | 0 | 0 | 298 | 149 | 1 | 1 | 1 | 0.882 | 0.871 | 0.926 | 1 | 1 | 0.8 | 1 | 1 | 1 | -0.024 |
| SE | 58163 | ENSG00000160688 | FLAD1 | chr1 | + | 154990158 | 154990257 | 154983343 | 154984066 | 154992712 | 154992786 | 1 | 0 | 0 | 0 | 0 | 0 | 1 | 2 | 1 | 0 | 0 | 1 | 4 | 1 | 0 | 0 | 3 | 0 | 3 | 0 | 0 | 0 | 248 | 149 | 1 | 1 | 1 | 0 | 0 | 0 | NA | NA | 0.167 | 1 | 1 | NA | -0.472 |
| SE | 58164 | ENSG00000160688 | FLAD1 | chr1 | + | 154990158 | 154990257 | 154988056 | 154988849 | 154992712 | 154992786 | 7 | 3 | 3 | 5 | 3 | 0 | 0 | 2 | 0 | 0 | 0 | 4 | 6 | 3 | 0 | 0 | 0 | 0 | 0 | 0 | 0 | 0 | 248 | 149 | 1 | 1 | 1 | 1 | 0.474 | 1 | 1 | NA | 1 | 1 | 1 | NA | -0.105 |
| SE | 58166 | ENSG00000160688 | FLAD1 | chr1 | + | 154990338 | 154990528 | 154990158 | 154990257 | 154992712 | 154992786 | 41 | 23 | 29 | 29 | 17 | 3 | 1 | 0 | 0 | 0 | 17 | 36 | 13 | 17 | 63 | 0 | 1 | 0 | 1 | 0 | 1 | 0 | 298 | 149 | 1 | 1 | 0.872 | 0.92 | 1 | 1 | 1 | 1 | 0.947 | 1 | 0.895 | 1 | -0.01 |
| SE | 58170 | ENSG00000172965 | MIR4435-2HG | chr2 | - | 111208487 | 111208661 | 111196342 | 111197194 | 111233627 | 111233739 | 0 | 2 | 1 | 8 | 3 | 4 | 0 | 0 | 0 | 1 | 5 | 2 | 0 | 0 | 0 | 8 | 3 | 0 | 3 | 0 | 0 | 0 | 298 | 149 | 1 | 1 | 0 | 1 | 1 | 1 | 0.6 | 0.238 | 0.25 | NA | NA | NA | 0.476 |
| SE | 58179 | ENSG00000172965 | MIR4435-2HG | chr2 | - | 111233627 | 111233739 | 111195921 | 111197194 | 111248147 | 111248241 | 5 | 0 | 0 | 0 | 3 | 0 | 0 | 0 | 0 | 4 | 14 | 3 | 0 | 1 | 0 | 0 | 0 | 0 | 0 | 0 | 0 | 0 | 261 | 149 | 1 | 1 | 1 | NA | NA | NA | 0.3 | 1 | 1 | NA | 1 | NA | -0.35 |
| SE | 58181 | ENSG00000172965 | MIR4435-2HG | chr2 | - | 111233627 | 111233739 | 111207754 | 111208661 | 111239801 | 111239996 | 0 | 0 | 1 | 15 | 5 | 0 | 1 | 0 | 0 | 0 | 29 | 5 | 6 | 0 | 0 | 0 | 0 | 0 | 0 | 0 | 0 | 7 | 261 | 149 | 1 | 1 | NA | 0 | 1 | 1 | 1 | 1 | 1 | 1 | NA | 0 | 0 |
| SE | 58184 | ENSG00000172965 | MIR4435-2HG | chr2 | - | 111233627 | 111233739 | 111208755 | 111208936 | 111239801 | 111239996 | 1 | 0 | 0 | 11 | 3 | 0 | 2 | 0 | 0 | 0 | 24 | 5 | 12 | 0 | 0 | 0 | 1 | 0 | 1 | 0 | 0 | 0 | 261 | 149 | 1 | 1 | 1 | 0 | NA | 1 | 1 | 1 | 0.741 | 1 | NA | NA | -0.164 |
| SE | 58185 | ENSG00000172965 | MIR4435-2HG | chr2 | - | 111233627 | 111233739 | 111208755 | 111208936 | 111248147 | 111248241 | 1 | 0 | 0 | 3 | 2 | 1 | 0 | 0 | 1 | 0 | 6 | 0 | 6 | 1 | 0 | 0 | 0 | 0 | 0 | 0 | 4 | 8 | 261 | 149 | 1 | 1 | 0.363 | NA | NA | 0.631 | 1 | 1 | NA | 1 | 0.125 | 0 | 0.133 |
| SE | 58186 | ENSG00000172965 | MIR4435-2HG | chr2 | - | 111233627 | 111233739 | 111208755 | 111208936 | 111344060 | 111344237 | 3 | 0 | 1 | 11 | 5 | 1 | 0 | 0 | 3 | 1 | 12 | 3 | 8 | 12 | 0 | 0 | 0 | 2 | 0 | 2 | 1 | 0 | 261 | 149 | 1 | 1 | 0.631 | NA | 1 | 0.677 | 0.741 | 1 | 1 | 0.695 | 0.873 | NA | -0.13 |
| SE | 58187 | ENSG00000172965 | MIR4435-2HG | chr2 | - | 111233627 | 111233739 | 111208755 | 111208977 | 111239801 | 111239996 | 2 | 0 | 2 | 8 | 3 | 0 | 0 | 0 | 0 | 0 | 24 | 12 | 8 | 0 | 2 | 0 | 0 | 0 | 0 | 0 | 3 | 0 | 261 | 149 | 1 | 1 | 1 | NA | 1 | 1 | 1 | 1 | 1 | 1 | 0 | 1 | 0.2 |
| SE | 58188 | ENSG00000172965 | MIR4435-2HG | chr2 | - | 111233627 | 111233739 | 111208755 | 111208977 | 111248147 | 111248241 | 3 | 0 | 2 | 0 | 2 | 0 | 0 | 3 | 0 | 0 | 6 | 7 | 2 | 1 | 2 | 0 | 0 | 0 | 0 | 0 | 0 | 0 | 261 | 149 | 1 | 1 | 1 | NA | 0.276 | NA | 1 | 1 | 1 | 1 | 1 | 1 | -0.241 |
| SE | 58189 | ENSG00000172965 | MIR4435-2HG | chr2 | - | 111233627 | 111233739 | 111208755 | 111208977 | 111344060 | 111344237 | 5 | 0 | 3 | 8 | 5 | 0 | 0 | 0 | 3 | 2 | 12 | 10 | 4 | 12 | 2 | 0 | 0 | 1 | 0 | 1 | 5 | 0 | 261 | 149 | 1 | 1 | 1 | NA | 1 | 0.604 | 0.588 | 1 | 1 | 0.695 | 0.578 | 1 | -0.057 |
| SE | 58192 | ENSG00000172965 | MIR4435-2HG | chr2 | - | 111239801 | 111239996 | 111208755 | 111208936 | 111248147 | 111248241 | 3 | 3 | 6 | 1 | 0 | 1 | 0 | 0 | 1 | 0 | 15 | 11 | 6 | 10 | 5 | 0 | 0 | 0 | 0 | 0 | 4 | 8 | 298 | 149 | 1 | 1 | 0.6 | 1 | 1 | 0.333 | NA | 1 | 1 | 1 | 0.556 | 0.238 | -0.026 |
| SE | 58193 | ENSG00000172965 | MIR4435-2HG | chr2 | - | 111239801 | 111239996 | 111208755 | 111208936 | 111344060 | 111344237 | 2 | 5 | 1 | 10 | 10 | 1 | 0 | 0 | 3 | 1 | 14 | 1 | 3 | 7 | 0 | 0 | 0 | 2 | 0 | 2 | 1 | 0 | 298 | 149 | 1 | 1 | 0.5 | 1 | 1 | 0.625 | 0.833 | 1 | 1 | 0.429 | 0.778 | NA | -0.01 |
| SE | 58194 | ENSG00000172965 | MIR4435-2HG | chr2 | - | 111239801 | 111239996 | 111208755 | 111208977 | 111248147 | 111248241 | 3 | 1 | 6 | 1 | 0 | 0 | 0 | 3 | 0 | 0 | 15 | 10 | 6 | 13 | 5 | 0 | 0 | 0 | 0 | 0 | 0 | 0 | 298 | 149 | 1 | 1 | 1 | 1 | 0.5 | 1 | NA | 1 | 1 | 1 | 1 | 1 | -0.125 |
| SE | 58195 | ENSG00000172965 | MIR4435-2HG | chr2 | - | 111239801 | 111239996 | 111208755 | 111208977 | 111344060 | 111344237 | 2 | 3 | 1 | 10 | 10 | 0 | 0 | 0 | 3 | 2 | 14 | 0 | 3 | 10 | 0 | 0 | 0 | 1 | 0 | 1 | 5 | 0 | 298 | 149 | 1 | 1 | 1 | 1 | 1 | 0.625 | 0.714 | 1 | NA | 0.6 | 0.5 | NA | 0.168 |
| SE | 58197 | ENSG00000172965 | MIR4435-2HG | chr2 | - | 111239801 | 111239996 | 111233627 | 111233739 | 111344060 | 111344237 | 3 | 3 | 1 | 18 | 13 | 3 | 1 | 1 | 8 | 5 | 38 | 5 | 9 | 7 | 0 | 12 | 3 | 2 | 3 | 2 | 12 | 0 | 298 | 149 | 1 | 1 | 0.333 | 0.6 | 0.333 | 0.529 | 0.565 | 0.613 | 0.455 | 0.692 | 0.226 | NA | -0.025 |
| SE | 58199 | ENSG00000172965 | MIR4435-2HG | chr2 | - | 111247429 | 111247558 | 111239801 | 111239996 | 111344060 | 111344237 | 0 | 0 | 1 | 0 | 0 | 2 | 3 | 1 | 10 | 10 | 0 | 0 | 0 | 0 | 0 | 14 | 0 | 3 | 0 | 3 | 7 | 0 | 278 | 149 | 1 | 1 | 0 | 0 | 0.349 | 0 | 0 | 0 | NA | 0 | 0 | NA | 0.07 |
| SE | 58206 | ENSG00000172965 | MIR4435-2HG | chr2 | - | 111248147 | 111248241 | 111245615 | 111247558 | 111344060 | 111344237 | 25 | 18 | 9 | 27 | 25 | 0 | 0 | 1 | 0 | 0 | 31 | 37 | 28 | 15 | 57 | 0 | 0 | 0 | 0 | 0 | 0 | 0 | 243 | 149 | 1 | 1 | 1 | 1 | 0.847 | 1 | 1 | 1 | 1 | 1 | 1 | 1 | -0.031 |
| SE | 58207 | ENSG00000172965 | MIR4435-2HG | chr2 | - | 111274706 | 111274861 | 111208755 | 111208936 | 111344060 | 111344237 | 0 | 0 | 0 | 0 | 0 | 1 | 0 | 0 | 3 | 1 | 0 | 0 | 0 | 0 | 7 | 0 | 0 | 2 | 0 | 2 | 1 | 0 | 298 | 149 | 1 | 1 | 0 | NA | NA | 0 | 0 | NA | NA | 0 | 0 | 1 | -0.333 |
| SE | 58209 | ENSG00000172965 | MIR4435-2HG | chr2 | - | 111344060 | 111344237 | 111208755 | 111208936 | 111367486 | 111367575 | 16 | 5 | 10 | 19 | 24 | 0 | 0 | 1 | 0 | 0 | 5 | 4 | 11 | 9 | 10 | 0 | 0 | 0 | 0 | 0 | 0 | 0 | 298 | 149 | 1 | 1 | 1 | 1 | 0.833 | 1 | 1 | 1 | 1 | 1 | 1 | 1 | -0.033 |
| SE | 58211 | ENSG00000172965 | MIR4435-2HG | chr2 | - | 111344060 | 111344237 | 111314103 | 111314562 | 111365560 | 111365563 | 30 | 9 | 13 | 37 | 35 | 0 | 0 | 0 | 0 | 0 | 42 | 36 | 24 | 45 | 63 | 0 | 0 | 2 | 0 | 2 | 0 | 0 | 298 | 149 | 1 | 1 | 1 | 1 | 1 | 1 | 1 | 1 | 1 | 0.857 | 1 | 1 | 0.029 |
| SE | 58217 | ENSG00000172965 | MIR4435-2HG | chr2 | - | 111447211 | 111447349 | 111367486 | 111367575 | 111479906 | 111480061 | 0 | 0 | 0 | 0 | 2 | 0 | 0 | 0 | 0 | 0 | 0 | 1 | 0 | 0 | 0 | 0 | 0 | 0 | 0 | 0 | 0 | 6 | 287 | 149 | 1 | 1 | NA | NA | NA | NA | 1 | NA | 1 | NA | NA | 0 | 0.5 |
| SE | 58218 | ENSG00000172965 | MIR4435-2HG | chr2 | - | 111447211 | 111447349 | 111367486 | 111367575 | 111494884 | 111495048 | 1 | 0 | 0 | 1 | 2 | 99 | 51 | 49 | 94 | 90 | 0 | 1 | 0 | 0 | 6 | 164 | 150 | 92 | 150 | 92 | 115 | 246 | 287 | 149 | 1 | 1 | 0.005 | 0 | 0 | 0.005 | 0.011 | 0 | 0.003 | 0 | 0 | 0.013 | 0.001 |
| SE | 58219 | ENSG00000172965 | MIR4435-2HG | chr2 | - | 111447211 | 111447349 | 111429317 | 111429620 | 111479906 | 111480061 | 1 | 0 | 0 | 4 | 0 | 1 | 0 | 0 | 0 | 0 | 8 | 0 | 0 | 0 | 6 | 0 | 0 | 3 | 0 | 3 | 0 | 0 | 287 | 149 | 1 | 1 | 0.342 | NA | NA | 1 | NA | 1 | NA | 0 | NA | 1 | 0.004 |
| SE | 58222 | ENSG00000172965 | MIR4435-2HG | chr2 | - | 111469786 | 111469987 | 111367514 | 111367575 | 111479906 | 111480061 | 1 | 0 | 0 | 0 | 0 | 0 | 0 | 0 | 0 | 0 | 0 | 0 | 0 | 0 | 0 | 0 | 0 | 0 | 0 | 0 | 0 | 6 | 298 | 149 | 1 | 1 | 1 | NA | NA | NA | NA | NA | NA | NA | NA | 0 | 1 |
| SE | 58223 | ENSG00000172965 | MIR4435-2HG | chr2 | - | 111469786 | 111469987 | 111429308 | 111429620 | 111479906 | 111480061 | 1 | 5 | 2 | 1 | 0 | 1 | 0 | 0 | 0 | 0 | 0 | 6 | 8 | 0 | 2 | 0 | 0 | 3 | 0 | 3 | 0 | 0 | 298 | 149 | 1 | 1 | 0.333 | 1 | 1 | 1 | NA | NA | 1 | 0.571 | NA | 1 | -0.024 |
| SE | 58224 | ENSG00000172965 | MIR4435-2HG | chr2 | - | 111469786 | 111469987 | 111429308 | 111429620 | 111494556 | 111494685 | 1 | 5 | 2 | 1 | 0 | 0 | 0 | 0 | 1 | 0 | 0 | 6 | 8 | 0 | 2 | 0 | 0 | 0 | 0 | 0 | 0 | 9 | 298 | 149 | 1 | 1 | 1 | 1 | 1 | 0.333 | NA | NA | 1 | 1 | NA | 0.1 | 0.133 |
| SE | 58226 | ENSG00000172965 | MIR4435-2HG | chr2 | - | 111471988 | 111472097 | 111469271 | 111469987 | 111494884 | 111495034 | 0 | 0 | 0 | 0 | 0 | 3 | 0 | 4 | 4 | 0 | 0 | 0 | 6 | 1 | 0 | 8 | 0 | 1 | 0 | 1 | 1 | 4 | 258 | 149 | 1 | 1 | 0 | NA | 0 | 0 | NA | 0 | NA | 0.776 | 0.366 | 0 | -0.285 |
| SE | 58227 | ENSG00000172965 | MIR4435-2HG | chr2 | - | 111477335 | 111477481 | 111429311 | 111429620 | 111479906 | 111480061 | 0 | 2 | 0 | 1 | 0 | 1 | 0 | 0 | 0 | 0 | 0 | 0 | 4 | 6 | 5 | 0 | 0 | 3 | 0 | 3 | 0 | 0 | 295 | 149 | 1 | 1 | 0 | 1 | NA | 1 | NA | NA | NA | 0.402 | 1 | 1 | -0.134 |
| SE | 58228 | ENSG00000172965 | MIR4435-2HG | chr2 | - | 111477335 | 111477481 | 111429311 | 111429620 | 111494884 | 111494988 | 0 | 2 | 0 | 0 | 0 | 65 | 73 | 44 | 70 | 91 | 0 | 0 | 0 | 2 | 4 | 243 | 122 | 43 | 122 | 43 | 115 | 344 | 295 | 149 | 1 | 1 | 0 | 0.014 | 0 | 0 | 0 | 0 | 0 | 0 | 0.009 | 0.006 | 0 |
| SE | 58232 | ENSG00000172965 | MIR4435-2HG | chr2 | - | 111479906 | 111480061 | 111447211 | 111447349 | 111494884 | 111495029 | 11 | 0 | 4 | 1 | 4 | 1 | 0 | 0 | 1 | 0 | 0 | 4 | 7 | 3 | 0 | 0 | 0 | 0 | 0 | 0 | 0 | 6 | 298 | 149 | 1 | 1 | 0.846 | NA | 1 | 0.333 | 1 | NA | 1 | 1 | 1 | 0 | 0.045 |
| SE | 58233 | ENSG00000172965 | MIR4435-2HG | chr2 | - | 111479906 | 111480061 | 111469786 | 111469987 | 111494884 | 111495029 | 11 | 0 | 4 | 1 | 4 | 3 | 0 | 4 | 4 | 0 | 0 | 4 | 7 | 3 | 0 | 8 | 0 | 1 | 0 | 1 | 1 | 4 | 298 | 149 | 1 | 1 | 0.647 | NA | 0.333 | 0.111 | 1 | 0 | 1 | 0.778 | 0.6 | 0 | 0.047 |
| SE | 58234 | ENSG00000172965 | MIR4435-2HG | chr2 | - | 111479906 | 111480061 | 111471988 | 111472097 | 111494884 | 111495029 | 11 | 0 | 4 | 1 | 4 | 0 | 0 | 1 | 0 | 0 | 0 | 4 | 7 | 3 | 0 | 6 | 0 | 3 | 0 | 3 | 1 | 0 | 298 | 149 | 1 | 1 | 1 | NA | 0.667 | 1 | 1 | 0 | 1 | 0.538 | 0.6 | NA | 0.382 |
| SE | 58235 | ENSG00000172965 | MIR4435-2HG | chr2 | - | 111479906 | 111480061 | 111477335 | 111477481 | 111494884 | 111495029 | 11 | 0 | 4 | 2 | 4 | 0 | 0 | 0 | 0 | 0 | 0 | 4 | 11 | 7 | 5 | 0 | 0 | 0 | 0 | 0 | 0 | 4 | 298 | 149 | 1 | 1 | 1 | NA | 1 | 1 | 1 | NA | 1 | 1 | 1 | 0.385 | 0.154 |
| SE | 58236 | ENSG00000172965 | MIR4435-2HG | chr2 | - | 111494556 | 111494778 | 111429313 | 111429620 | 111494884 | 111495015 | 0 | 0 | 0 | 2 | 0 | 65 | 73 | 44 | 70 | 91 | 6 | 0 | 0 | 0 | 9 | 243 | 122 | 43 | 122 | 43 | 115 | 344 | 298 | 149 | 1 | 1 | 0 | 0 | 0 | 0.014 | 0 | 0.012 | 0 | 0 | 0 | 0.013 | -0.002 |
| SE | 58239 | ENSG00000089127 | OAS1 | chr12 | + | 112908535 | 112908824 | 112906652 | 112907219 | 112911050 | 112911235 | 380 | 318 | 234 | 404 | 369 | 2 | 0 | 0 | 0 | 0 | 1438 | 828 | 238 | 1303 | 578 | 0 | 0 | 2 | 0 | 2 | 3 | 0 | 298 | 149 | 1 | 1 | 0.99 | 1 | 1 | 1 | 1 | 1 | 1 | 0.983 | 0.995 | 1 | 0.002 |
| SE | 58242 | ENSG00000089127 | OAS1 | chr12 | + | 112908622 | 112908824 | 112906967 | 112907219 | 112911050 | 112911235 | 93 | 89 | 60 | 110 | 69 | 2 | 0 | 0 | 0 | 0 | 434 | 240 | 85 | 368 | 224 | 0 | 0 | 2 | 0 | 2 | 3 | 0 | 298 | 149 | 1 | 1 | 0.959 | 1 | 1 | 1 | 1 | 1 | 1 | 0.955 | 0.984 | 1 | 0.004 |
| SE | 58259 | ENSG00000089127 | OAS1 | chr12 | + | 112918596 | 112918617 | 112917546 | 112917700 | 112919388 | 112919903 | 7 | 3 | 1 | 5 | 1 | 0 | 0 | 0 | 0 | 140 | 0 | 7 | 0 | 2 | 0 | 326 | 0 | 0 | 0 | 0 | 0 | 0 | 170 | 149 | 1 | 1 | 1 | 1 | 1 | 1 | 0.006 | 0 | 1 | NA | 1 | NA | 0.135 |
| SE | 58261 | ENSG00000089127 | OAS1 | chr12 | + | 112919388 | 112919432 | 112917546 | 112917700 | 112931877 | 112932142 | 0 | 0 | 0 | 0 | 16 | 2 | 0 | 0 | 7 | 0 | 85 | 0 | 0 | 0 | 0 | 0 | 1 | 3 | 1 | 3 | 3 | 0 | 193 | 149 | 1 | 1 | 0 | NA | NA | 0 | 1 | 1 | 0 | 0 | 0 | NA | 0.083 |
| SE | 58264 | ENSG00000065413 | ANKRD44 | chr2 | - | 196995378 | 196995461 | 196993582 | 196993674 | 196998336 | 196998419 | 277 | 198 | 291 | 259 | 307 | 1 | 0 | 0 | 0 | 6 | 41 | 137 | 98 | 100 | 90 | 0 | 0 | 0 | 0 | 0 | 0 | 0 | 232 | 149 | 1 | 1 | 0.994 | 1 | 1 | 1 | 0.97 | 1 | 1 | 1 | 1 | 1 | -0.007 |
| SE | 58267 | ENSG00000065413 | ANKRD44 | chr2 | - | 197001752 | 197001840 | 197000418 | 197000502 | 197005693 | 197005910 | 387 | 206 | 356 | 322 | 310 | 3 | 0 | 0 | 0 | 0 | 105 | 265 | 150 | 133 | 97 | 0 | 0 | 0 | 0 | 0 | 0 | 0 | 237 | 149 | 1 | 1 | 0.988 | 1 | 1 | 1 | 1 | 1 | 1 | 1 | 1 | 1 | -0.002 |
| SE | 58268 | ENSG00000065413 | ANKRD44 | chr2 | - | 197005693 | 197005910 | 197001752 | 197001840 | 197007805 | 197007923 | 575 | 352 | 504 | 500 | 464 | 0 | 0 | 0 | 1 | 0 | 182 | 390 | 238 | 278 | 138 | 0 | 0 | 0 | 0 | 0 | 0 | 0 | 298 | 149 | 1 | 1 | 1 | 1 | 1 | 0.996 | 1 | 1 | 1 | 1 | 1 | 1 | -0.001 |
| SE | 58270 | ENSG00000065413 | ANKRD44 | chr2 | - | 197007805 | 197007923 | 197005693 | 197005910 | 197008943 | 197009031 | 713 | 537 | 674 | 615 | 694 | 0 | 0 | 0 | 0 | 0 | 331 | 589 | 483 | 414 | 148 | 0 | 0 | 0 | 0 | 0 | 1 | 0 | 267 | 149 | 1 | 1 | 1 | 1 | 1 | 1 | 1 | 1 | 1 | 1 | 0.996 | 1 | 0.001 |
| SE | 58271 | ENSG00000065413 | ANKRD44 | chr2 | - | 197008943 | 197009031 | 197007805 | 197007923 | 197013510 | 197013712 | 704 | 585 | 682 | 655 | 739 | 1 | 1 | 0 | 0 | 0 | 401 | 636 | 563 | 393 | 190 | 0 | 0 | 0 | 0 | 0 | 0 | 0 | 237 | 149 | 1 | 1 | 0.998 | 0.997 | 1 | 1 | 1 | 1 | 1 | 1 | 1 | 1 | -0.001 |
| SE | 58273 | ENSG00000065413 | ANKRD44 | chr2 | - | 197025195 | 197025267 | 197013510 | 197013712 | 197078702 | 197078814 | 519 | 398 | 488 | 466 | 1094 | 2 | 0 | 0 | 0 | 0 | 281 | 461 | 326 | 264 | 225 | 0 | 0 | 0 | 0 | 0 | 0 | 0 | 221 | 149 | 1 | 1 | 0.994 | 1 | 1 | 1 | 1 | 1 | 1 | 1 | 1 | 1 | -0.001 |
| SE | 58274 | ENSG00000065413 | ANKRD44 | chr2 | - | 197078702 | 197078814 | 197025195 | 197025267 | 197081644 | 197081725 | 661 | 506 | 675 | 622 | 947 | 0 | 0 | 0 | 2 | 0 | 297 | 510 | 411 | 314 | 249 | 0 | 0 | 0 | 0 | 0 | 0 | 0 | 261 | 149 | 1 | 1 | 1 | 1 | 1 | 0.994 | 1 | 1 | 1 | 1 | 1 | 1 | -0.001 |
| SE | 58275 | ENSG00000065413 | ANKRD44 | chr2 | - | 197078702 | 197078814 | 197025195 | 197025267 | 197083368 | 197083509 | 267 | 194 | 225 | 234 | 483 | 0 | 0 | 0 | 0 | 0 | 160 | 196 | 152 | 160 | 103 | 0 | 0 | 1 | 0 | 1 | 0 | 0 | 261 | 149 | 1 | 1 | 1 | 1 | 1 | 1 | 1 | 1 | 1 | 0.989 | 1 | 1 | 0.002 |
| SE | 58276 | ENSG00000065413 | ANKRD44 | chr2 | - | 197081644 | 197081725 | 197025195 | 197025267 | 197083368 | 197083509 | 255 | 235 | 307 | 324 | 310 | 0 | 0 | 0 | 0 | 0 | 98 | 220 | 191 | 130 | 87 | 0 | 0 | 1 | 0 | 1 | 0 | 0 | 230 | 149 | 1 | 1 | 1 | 1 | 1 | 1 | 1 | 1 | 1 | 0.992 | 1 | 1 | 0.002 |
| SE | 58277 | ENSG00000065413 | ANKRD44 | chr2 | - | 197081644 | 197081725 | 197077811 | 197078814 | 197083368 | 197083509 | 652 | 526 | 756 | 704 | 800 | 1 | 0 | 1 | 2 | 4 | 221 | 526 | 450 | 282 | 219 | 0 | 1 | 1 | 1 | 1 | 2 | 0 | 230 | 149 | 1 | 1 | 0.998 | 1 | 0.998 | 0.996 | 0.992 | 1 | 0.997 | 0.997 | 0.989 | 1 | 0 |
| SE | 58278 | ENSG00000065413 | ANKRD44 | chr2 | - | 197083368 | 197083509 | 197025195 | 197025267 | 197099815 | 197099930 | 1 | 0 | 0 | 2 | 0 | 2 | 0 | 1 | 1 | 0 | 0 | 0 | 1 | 3 | 0 | 0 | 0 | 5 | 0 | 5 | 6 | 0 | 290 | 149 | 1 | 1 | 0.204 | NA | 0 | 0.507 | NA | NA | NA | 0.093 | 0.204 | NA | 0.088 |
| SE | 58279 | ENSG00000065413 | ANKRD44 | chr2 | - | 197083368 | 197083509 | 197077811 | 197078814 | 197086679 | 197086748 | 328 | 258 | 365 | 312 | 287 | 0 | 0 | 1 | 0 | 0 | 99 | 280 | 271 | 156 | 140 | 0 | 0 | 0 | 0 | 0 | 0 | 0 | 290 | 149 | 1 | 1 | 1 | 1 | 0.995 | 1 | 1 | 1 | 1 | 1 | 1 | 1 | -0.001 |
| SE | 58280 | ENSG00000065413 | ANKRD44 | chr2 | - | 197086679 | 197086748 | 197083368 | 197083509 | 197088710 | 197088774 | 478 | 383 | 527 | 478 | 431 | 2 | 5 | 2 | 0 | 4 | 176 | 392 | 347 | 256 | 212 | 0 | 2 | 2 | 2 | 2 | 0 | 0 | 218 | 149 | 1 | 1 | 0.994 | 0.981 | 0.994 | 1 | 0.987 | 1 | 0.993 | 0.992 | 1 | 1 | -0.006 |
| SE | 58281 | ENSG00000065413 | ANKRD44 | chr2 | - | 197086679 | 197086748 | 197083368 | 197083509 | 197099815 | 197099930 | 174 | 142 | 225 | 168 | 125 | 1 | 0 | 0 | 2 | 0 | 27 | 142 | 132 | 99 | 77 | 0 | 0 | 0 | 0 | 0 | 3 | 0 | 218 | 149 | 1 | 1 | 0.992 | 1 | 1 | 0.983 | 1 | 1 | 1 | 1 | 0.958 | 1 | 0.003 |
| SE | 58282 | ENSG00000065413 | ANKRD44 | chr2 | - | 197110765 | 197110844 | 197099476 | 197099930 | 197121331 | 197121544 | 578 | 456 | 618 | 621 | 691 | 0 | 0 | 0 | 2 | 0 | 252 | 520 | 263 | 291 | 144 | 0 | 0 | 0 | 0 | 0 | 0 | 0 | 228 | 149 | 1 | 1 | 1 | 1 | 1 | 0.995 | 1 | 1 | 1 | 1 | 1 | 1 | -0.001 |
| SE | 58284 | ENSG00000065413 | ANKRD44 | chr2 | - | 197122649 | 197122792 | 197121331 | 197121544 | 197136591 | 197136662 | 191 | 110 | 207 | 191 | 109 | 0 | 0 | 0 | 0 | 1 | 81 | 161 | 109 | 131 | 87 | 0 | 0 | 0 | 0 | 0 | 0 | 0 | 292 | 149 | 1 | 1 | 1 | 1 | 1 | 1 | 0.982 | 1 | 1 | 1 | 1 | 1 | -0.004 |
| SE | 58285 | ENSG00000065413 | ANKRD44 | chr2 | - | 197125380 | 197125468 | 197121331 | 197121544 | 197125836 | 197126037 | 350 | 275 | 389 | 380 | 162 | 0 | 0 | 0 | 0 | 0 | 120 | 312 | 222 | 173 | 188 | 0 | 1 | 0 | 1 | 0 | 0 | 0 | 237 | 149 | 1 | 1 | 1 | 1 | 1 | 1 | 1 | 1 | 0.995 | 1 | 1 | 1 | 0.001 |
| SE | 58286 | ENSG00000065413 | ANKRD44 | chr2 | - | 197125380 | 197125468 | 197121331 | 197121544 | 197136591 | 197136662 | 0 | 0 | 0 | 0 | 4 | 0 | 0 | 0 | 0 | 1 | 0 | 1 | 0 | 1 | 0 | 0 | 0 | 0 | 0 | 0 | 0 | 0 | 237 | 149 | 1 | 1 | NA | NA | NA | NA | 0.715 | NA | 1 | NA | 1 | NA | -0.285 |
| SE | 58287 | ENSG00000065413 | ANKRD44 | chr2 | - | 197125380 | 197125468 | 197122649 | 197122792 | 197136591 | 197136662 | 273 | 184 | 282 | 290 | 100 | 0 | 0 | 0 | 1 | 0 | 152 | 246 | 215 | 83 | 124 | 0 | 0 | 0 | 0 | 0 | 0 | 0 | 237 | 149 | 1 | 1 | 1 | 1 | 1 | 0.995 | 1 | 1 | 1 | 1 | 1 | 1 | -0.001 |
| SE | 58288 | ENSG00000065413 | ANKRD44 | chr2 | - | 197125836 | 197126037 | 197121331 | 197121544 | 197136591 | 197136662 | 496 | 363 | 488 | 437 | 339 | 0 | 0 | 0 | 0 | 1 | 311 | 334 | 289 | 250 | 277 | 0 | 0 | 0 | 0 | 0 | 0 | 0 | 298 | 149 | 1 | 1 | 1 | 1 | 1 | 1 | 0.994 | 1 | 1 | 1 | 1 | 1 | -0.001 |
| SE | 58289 | ENSG00000065413 | ANKRD44 | chr2 | - | 197125836 | 197126037 | 197125380 | 197125468 | 197136591 | 197136662 | 919 | 683 | 956 | 913 | 529 | 0 | 0 | 0 | 0 | 4 | 475 | 702 | 576 | 451 | 491 | 0 | 0 | 0 | 0 | 0 | 0 | 0 | 298 | 149 | 1 | 1 | 1 | 1 | 1 | 1 | 0.985 | 1 | 1 | 1 | 1 | 1 | -0.003 |
| SE | 58290 | ENSG00000065413 | ANKRD44 | chr2 | - | 197136591 | 197136662 | 197125836 | 197126037 | 197147026 | 197147105 | 690 | 509 | 678 | 608 | 483 | 0 | 0 | 0 | 0 | 0 | 420 | 501 | 436 | 310 | 370 | 0 | 0 | 0 | 0 | 0 | 5 | 0 | 220 | 149 | 1 | 1 | 1 | 1 | 1 | 1 | 1 | 1 | 1 | 1 | 0.977 | 1 | 0.005 |
| SE | 58291 | ENSG00000065413 | ANKRD44 | chr2 | - | 197147026 | 197147105 | 197133970 | 197136662 | 197187022 | 197187106 | 858 | 677 | 830 | 771 | 616 | 0 | 1 | 0 | 0 | 0 | 570 | 706 | 566 | 439 | 507 | 0 | 0 | 0 | 0 | 0 | 0 | 0 | 228 | 149 | 1 | 1 | 1 | 0.998 | 1 | 1 | 1 | 1 | 1 | 1 | 1 | 1 | 0 |
| SE | 58292 | ENSG00000065413 | ANKRD44 | chr2 | - | 197187022 | 197187106 | 197147026 | 197147105 | 197310577 | 197310646 | 609 | 482 | 585 | 548 | 411 | 0 | 0 | 0 | 2 | 0 | 368 | 547 | 426 | 267 | 301 | 0 | 0 | 3 | 0 | 3 | 0 | 0 | 233 | 149 | 1 | 1 | 1 | 1 | 1 | 0.994 | 1 | 1 | 1 | 0.989 | 1 | 1 | 0.001 |
| SE | 58293 | ENSG00000196526 | AFAP1 | chr4 | - | 7772819 | 7773010 | 7768843 | 7769008 | 7774738 | 7774903 | 19 | 67 | 53 | 73 | 60 | 0 | 0 | 1 | 0 | 0 | 171 | 32 | 41 | 45 | 369 | 0 | 0 | 0 | 0 | 0 | 0 | 5 | 298 | 149 | 1 | 1 | 1 | 1 | 0.964 | 1 | 1 | 1 | 1 | 1 | 1 | 0.974 | -0.002 |
| SE | 58295 | ENSG00000196526 | AFAP1 | chr4 | - | 7786193 | 7786311 | 7778761 | 7778876 | 7793680 | 7793826 | 13 | 28 | 31 | 29 | 20 | 0 | 0 | 0 | 1 | 0 | 79 | 30 | 22 | 12 | 250 | 0 | 0 | 0 | 0 | 0 | 0 | 0 | 267 | 149 | 1 | 1 | 1 | 1 | 1 | 0.942 | 1 | 1 | 1 | 1 | 1 | 1 | -0.012 |
| SE | 58296 | ENSG00000196526 | AFAP1 | chr4 | - | 7855465 | 7855574 | 7843138 | 7843350 | 7868621 | 7868719 | 6 | 9 | 27 | 6 | 7 | 0 | 0 | 1 | 0 | 0 | 0 | 8 | 11 | 4 | 26 | 0 | 0 | 0 | 0 | 0 | 0 | 0 | 258 | 149 | 1 | 1 | 1 | 1 | 0.94 | 1 | 1 | NA | 1 | 1 | 1 | 1 | -0.012 |
| SE | 58299 | ENSG00000117262 | GPR89A | chr1 | + | 145612201 | 145612247 | 145608612 | 145608675 | 145616233 | 145616293 | 18 | 2 | 16 | 6 | 14 | 3 | 0 | 1 | 0 | 0 | 12 | 21 | 14 | 8 | 0 | 0 | 5 | 11 | 5 | 11 | 8 | 0 | 195 | 149 | 1 | 1 | 0.821 | 1 | 0.924 | 1 | 1 | 1 | 0.762 | 0.493 | 0.433 | NA | 0.277 |
| SE | 58301 | ENSG00000117262 | GPR89A | chr1 | + | 145630686 | 145630807 | 145623592 | 145623714 | 145631663 | 145631744 | 5 | 1 | 1 | 5 | 4 | 1 | 0 | 1 | 0 | 1 | 0 | 4 | 0 | 0 | 4 | 0 | 0 | 0 | 0 | 0 | 0 | 0 | 270 | 149 | 1 | 1 | 0.734 | 1 | 0.356 | 1 | 0.688 | NA | 1 | NA | NA | 1 | -0.244 |
| SE | 58302 | ENSG00000117262 | GPR89A | chr1 | + | 145645646 | 145645743 | 145643868 | 145643978 | 145646183 | 145646272 | 2 | 0 | 1 | 0 | 1 | 0 | 0 | 1 | 0 | 0 | 0 | 0 | 0 | 7 | 2 | 0 | 0 | 0 | 0 | 0 | 0 | 0 | 246 | 149 | 1 | 1 | 1 | NA | 0.377 | NA | 1 | NA | NA | NA | 1 | 1 | -0.208 |
| SE | 58305 | ENSG00000052795 | FNIP2 | chr4 | + | 158833527 | 158833628 | 158832066 | 158832138 | 158835404 | 158835476 | 66 | 64 | 43 | 60 | 50 | 0 | 0 | 0 | 0 | 2 | 46 | 30 | 74 | 74 | 36 | 0 | 1 | 0 | 1 | 0 | 0 | 0 | 250 | 149 | 1 | 1 | 1 | 1 | 1 | 1 | 0.937 | 1 | 0.947 | 1 | 1 | 1 | -0.002 |
| SE | 58308 | ENSG00000052795 | FNIP2 | chr4 | + | 158895749 | 158895865 | 158891445 | 158891646 | 158904465 | 158908050 | 91 | 73 | 80 | 57 | 85 | 2 | 0 | 0 | 0 | 0 | 31 | 44 | 53 | 54 | 0 | 0 | 0 | 1 | 0 | 1 | 0 | 0 | 265 | 149 | 1 | 1 | 0.962 | 1 | 1 | 1 | 1 | 1 | 1 | 0.968 | 1 | NA | 0 |
| SE | 58312 | ENSG00000265818 | EEF1E1-BLOC1S5 | chr6 | - | 8026366 | 8026425 | 8015725 | 8015828 | 8041138 | 8041268 | 123 | 72 | 95 | 97 | 182 | 1 | 0 | 0 | 2 | 1 | 47 | 47 | 36 | 41 | 56 | 0 | 3 | 0 | 3 | 0 | 2 | 0 | 208 | 149 | 1 | 1 | 0.989 | 1 | 1 | 0.972 | 0.992 | 1 | 0.918 | 1 | 0.936 | 1 | 0.02 |
| SE | 58313 | ENSG00000170089 | NA | chr5 | + | 177832701 | 177832792 | 177809480 | 177809521 | 177836660 | 177836800 | 0 | 3 | 0 | 0 | 0 | 0 | 0 | 0 | 2 | 5 | 0 | 0 | 0 | 0 | 0 | 0 | 3 | 10 | 3 | 10 | 0 | 0 | 240 | 149 | 1 | 1 | NA | 1 | NA | 0 | 0 | NA | 0 | 0 | NA | NA | 0.333 |
| SE | 58314 | ENSG00000170089 | NA | chr5 | + | 177836660 | 177836800 | 177809480 | 177809521 | 177876531 | 177876688 | 8 | 4 | 4 | 6 | 6 | 2 | 0 | 0 | 0 | 0 | 8 | 11 | 20 | 3 | 0 | 0 | 0 | 0 | 0 | 0 | 0 | 0 | 289 | 149 | 1 | 1 | 0.673 | 1 | 1 | 1 | 1 | 1 | 1 | 1 | 1 | NA | -0.065 |
| SE | 58315 | ENSG00000170089 | NA | chr5 | + | 177836660 | 177836800 | 177809480 | 177809521 | 177882368 | 177882383 | 0 | 3 | 0 | 3 | 5 | 2 | 0 | 0 | 0 | 4 | 0 | 3 | 10 | 0 | 0 | 0 | 0 | 0 | 0 | 0 | 0 | 0 | 289 | 149 | 1 | 1 | 0 | 1 | NA | 1 | 0.392 | NA | 1 | 1 | NA | NA | -0.402 |
| SE | 58316 | ENSG00000170089 | NA | chr5 | + | 177836660 | 177836800 | 177832701 | 177832792 | 177876531 | 177876688 | 8 | 8 | 4 | 4 | 1 | 0 | 0 | 0 | 0 | 0 | 8 | 8 | 10 | 3 | 0 | 0 | 0 | 8 | 0 | 8 | 0 | 0 | 289 | 149 | 1 | 1 | 1 | 1 | 1 | 1 | 1 | 1 | 1 | 0.392 | 1 | NA | 0.152 |
| SE | 58317 | ENSG00000170089 | NA | chr5 | + | 177876531 | 177876688 | 177809480 | 177809521 | 177882368 | 177882383 | 2 | 0 | 0 | 0 | 0 | 2 | 0 | 0 | 0 | 4 | 0 | 0 | 0 | 3 | 0 | 0 | 0 | 0 | 0 | 0 | 0 | 0 | 298 | 149 | 1 | 1 | 0.333 | NA | NA | NA | 0 | NA | NA | NA | 1 | NA | -0.834 |
| SE | 58318 | ENSG00000170089 | NA | chr5 | + | 177876531 | 177876688 | 177836660 | 177836800 | 177882368 | 177882383 | 8 | 4 | 4 | 4 | 1 | 0 | 3 | 0 | 1 | 0 | 8 | 8 | 10 | 6 | 0 | 0 | 0 | 0 | 0 | 0 | 0 | 0 | 298 | 149 | 1 | 1 | 1 | 0.4 | 1 | 0.667 | 1 | 1 | 1 | 1 | 1 | NA | -0.187 |
| SE | 58319 | ENSG00000170089 | NA | chr5 | + | 177876531 | 177876688 | 177875521 | 177875860 | 177878643 | 177878848 | 18 | 3 | 8 | 7 | 5 | 2 | 0 | 0 | 0 | 0 | 19 | 32 | 9 | 0 | 0 | 0 | 0 | 0 | 0 | 0 | 0 | 0 | 298 | 149 | 1 | 1 | 0.818 | 1 | 1 | 1 | 1 | 1 | 1 | 1 | NA | NA | -0.036 |
| SE | 58320 | ENSG00000170089 | NA | chr5 | + | 177878643 | 177878848 | 177876531 | 177876688 | 177882368 | 177882383 | 9 | 3 | 8 | 7 | 6 | 0 | 0 | 0 | 0 | 0 | 19 | 25 | 13 | 0 | 0 | 0 | 0 | 0 | 0 | 0 | 3 | 0 | 298 | 149 | 1 | 1 | 1 | 1 | 1 | 1 | 1 | 1 | 1 | 1 | 0 | NA | 0.25 |
| SE | 58323 | ENSG00000084090 | STARD7 | chr2 | - | 96193301 | 96193352 | 96193077 | 96193205 | 96194957 | 96195007 | 108 | 75 | 87 | 97 | 145 | 0 | 0 | 0 | 0 | 1 | 141 | 123 | 57 | 85 | 158 | 0 | 0 | 0 | 0 | 0 | 0 | 0 | 200 | 149 | 1 | 1 | 1 | 1 | 1 | 1 | 0.991 | 1 | 1 | 1 | 1 | 1 | -0.002 |
| SE | 58324 | ENSG00000116667 | C1orf21 | chr1 | + | 184477385 | 184477603 | 184387028 | 184387368 | 184590738 | 184590815 | 84 | 32 | 25 | 54 | 32 | 0 | 0 | 0 | 1 | 0 | 43 | 75 | 55 | 23 | 46 | 0 | 0 | 0 | 0 | 0 | 0 | 0 | 298 | 149 | 1 | 1 | 1 | 1 | 1 | 0.964 | 1 | 1 | 1 | 1 | 1 | 1 | -0.007 |
| SE | 58325 | ENSG00000116667 | C1orf21 | chr1 | + | 184507587 | 184507682 | 184477385 | 184477603 | 184590738 | 184590815 | 361 | 109 | 80 | 257 | 220 | 2 | 3 | 0 | 3 | 0 | 161 | 228 | 189 | 88 | 193 | 0 | 0 | 0 | 0 | 0 | 0 | 8 | 244 | 149 | 1 | 1 | 0.991 | 0.957 | 1 | 0.981 | 1 | 1 | 1 | 1 | 1 | 0.936 | -0.001 |
| SE | 58329 | ENSG00000116667 | C1orf21 | chr1 | + | 184590738 | 184590815 | 184507587 | 184507682 | 184598400 | 184598461 | 320 | 96 | 79 | 199 | 218 | 0 | 0 | 0 | 4 | 0 | 107 | 213 | 111 | 74 | 143 | 0 | 0 | 0 | 0 | 0 | 0 | 0 | 226 | 149 | 1 | 1 | 1 | 1 | 1 | 0.97 | 1 | 1 | 1 | 1 | 1 | 1 | -0.006 |
| SE | 58330 | ENSG00000116667 | C1orf21 | chr1 | + | 184592224 | 184592290 | 184590738 | 184590815 | 184598400 | 184598461 | 2 | 0 | 0 | 0 | 0 | 222 | 78 | 58 | 141 | 149 | 0 | 0 | 0 | 2 | 0 | 86 | 170 | 88 | 170 | 88 | 56 | 89 | 215 | 149 | 1 | 1 | 0.006 | 0 | 0 | 0 | 0 | 0 | 0 | 0 | 0.024 | 0 | -0.004 |
| SE | 58333 | ENSG00000115207 | GTF3C2 | chr2 | - | 27337480 | 27337558 | 27337243 | 27337342 | 27337925 | 27338020 | 98 | 57 | 107 | 61 | 124 | 0 | 0 | 0 | 0 | 0 | 151 | 74 | 83 | 98 | 188 | 0 | 0 | 3 | 0 | 3 | 0 | 0 | 227 | 149 | 1 | 1 | 1 | 1 | 1 | 1 | 1 | 1 | 1 | 0.948 | 1 | 1 | 0.01 |
| SE | 58334 | ENSG00000115207 | GTF3C2 | chr2 | - | 27341947 | 27342233 | 27337925 | 27338020 | 27356738 | 27357034 | 59 | 48 | 85 | 67 | 78 | 0 | 0 | 0 | 0 | 0 | 64 | 70 | 50 | 71 | 61 | 0 | 0 | 2 | 0 | 2 | 0 | 0 | 298 | 149 | 1 | 1 | 1 | 1 | 1 | 1 | 1 | 1 | 1 | 0.926 | 1 | 1 | 0.015 |
| SE | 58336 | ENSG00000115207 | GTF3C2 | chr2 | - | 27356086 | 27356186 | 27343307 | 27343578 | 27356738 | 27356767 | 0 | 0 | 0 | 1 | 0 | 23 | 20 | 33 | 25 | 14 | 0 | 0 | 0 | 0 | 0 | 35 | 16 | 13 | 16 | 13 | 22 | 20 | 249 | 149 | 1 | 1 | 0 | 0 | 0 | 0.023 | 0 | 0 | 0 | 0 | 0 | 0 | 0.005 |
| SE | 58342 | ENSG00000154174 | TOMM70 | chr3 | - | 100375017 | 100375152 | 100373537 | 100373645 | 100377704 | 100377912 | 298 | 233 | 310 | 267 | 271 | 0 | 0 | 0 | 0 | 0 | 83 | 132 | 240 | 171 | 46 | 0 | 0 | 0 | 0 | 0 | 1 | 0 | 284 | 149 | 1 | 1 | 1 | 1 | 1 | 1 | 1 | 1 | 1 | 1 | 0.989 | 1 | 0.002 |
| SE | 58343 | ENSG00000154174 | TOMM70 | chr3 | - | 100377704 | 100377912 | 100375017 | 100375152 | 100381614 | 100381763 | 225 | 181 | 262 | 219 | 323 | 0 | 0 | 1 | 0 | 0 | 68 | 88 | 189 | 103 | 36 | 0 | 0 | 0 | 0 | 0 | 0 | 0 | 298 | 149 | 1 | 1 | 1 | 1 | 0.992 | 1 | 1 | 1 | 1 | 1 | 1 | 1 | -0.002 |
| SE | 58344 | ENSG00000154174 | TOMM70 | chr3 | - | 100386217 | 100386344 | 100384478 | 100384588 | 100386804 | 100386978 | 207 | 137 | 251 | 195 | 309 | 0 | 2 | 1 | 0 | 4 | 85 | 133 | 187 | 125 | 15 | 0 | 0 | 0 | 0 | 0 | 0 | 0 | 276 | 149 | 1 | 1 | 1 | 0.974 | 0.993 | 1 | 0.977 | 1 | 1 | 1 | 1 | 1 | -0.011 |
| SE | 58345 | ENSG00000110090 | CPT1A | chr11 | - | 68759568 | 68759661 | 68754882 | 68757730 | 68760224 | 68760338 | 159 | 90 | 104 | 166 | 235 | 1 | 0 | 0 | 0 | 0 | 151 | 200 | 71 | 150 | 286 | 0 | 0 | 0 | 0 | 0 | 0 | 0 | 242 | 149 | 1 | 1 | 0.99 | 1 | 1 | 1 | 1 | 1 | 1 | 1 | 1 | 1 | -0.002 |
| SE | 58346 | ENSG00000110090 | CPT1A | chr11 | - | 68761534 | 68761687 | 68760224 | 68760338 | 68762626 | 68762761 | 260 | 142 | 107 | 236 | 186 | 1 | 0 | 0 | 3 | 0 | 258 | 282 | 112 | 160 | 456 | 0 | 0 | 6 | 0 | 6 | 1 | 0 | 298 | 149 | 1 | 1 | 0.992 | 1 | 1 | 0.975 | 1 | 1 | 1 | 0.903 | 0.988 | 1 | 0.015 |
| SE | 58347 | ENSG00000110090 | CPT1A | chr11 | - | 68761534 | 68761687 | 68760224 | 68760338 | 68773264 | 68773429 | 169 | 105 | 72 | 158 | 139 | 0 | 0 | 0 | 0 | 1 | 200 | 199 | 79 | 104 | 305 | 0 | 0 | 0 | 0 | 0 | 0 | 0 | 298 | 149 | 1 | 1 | 1 | 1 | 1 | 1 | 0.986 | 1 | 1 | 1 | 1 | 1 | -0.003 |
| SE | 58348 | ENSG00000110090 | CPT1A | chr11 | - | 68762626 | 68762761 | 68760224 | 68760338 | 68773264 | 68773429 | 140 | 89 | 63 | 140 | 113 | 0 | 0 | 0 | 0 | 1 | 319 | 124 | 74 | 134 | 335 | 0 | 0 | 0 | 0 | 0 | 0 | 0 | 284 | 149 | 1 | 1 | 1 | 1 | 1 | 1 | 0.983 | 1 | 1 | 1 | 1 | 1 | -0.003 |
| SE | 58349 | ENSG00000110090 | CPT1A | chr11 | - | 68762626 | 68762761 | 68761534 | 68761687 | 68773264 | 68773429 | 230 | 125 | 97 | 217 | 160 | 0 | 0 | 0 | 2 | 0 | 375 | 207 | 101 | 189 | 486 | 0 | 0 | 0 | 0 | 0 | 0 | 0 | 284 | 149 | 1 | 1 | 1 | 1 | 1 | 0.983 | 1 | 1 | 1 | 1 | 1 | 1 | -0.003 |
| SE | 58350 | ENSG00000110090 | CPT1A | chr11 | - | 68793314 | 68793402 | 68784814 | 68785010 | 68794803 | 68794911 | 370 | 209 | 187 | 388 | 236 | 0 | 2 | 0 | 0 | 0 | 329 | 342 | 214 | 365 | 966 | 0 | 0 | 0 | 0 | 0 | 0 | 0 | 237 | 149 | 1 | 1 | 1 | 0.985 | 1 | 1 | 1 | 1 | 1 | 1 | 1 | 1 | -0.003 |
| SE | 58351 | ENSG00000110090 | CPT1A | chr11 | - | 68794803 | 68794911 | 68793314 | 68793402 | 68796855 | 68796933 | 470 | 277 | 240 | 499 | 373 | 1 | 0 | 0 | 0 | 0 | 441 | 463 | 281 | 511 | 1342 | 0 | 0 | 0 | 0 | 0 | 0 | 0 | 257 | 149 | 1 | 1 | 0.996 | 1 | 1 | 1 | 1 | 1 | 1 | 1 | 1 | 1 | -0.001 |
| SE | 58352 | ENSG00000110090 | CPT1A | chr11 | - | 68796855 | 68796933 | 68794803 | 68794911 | 68799217 | 68799268 | 355 | 240 | 184 | 357 | 310 | 0 | 0 | 1 | 2 | 0 | 456 | 313 | 209 | 389 | 1100 | 0 | 0 | 0 | 0 | 0 | 0 | 0 | 227 | 149 | 1 | 1 | 1 | 1 | 0.992 | 0.992 | 1 | 1 | 1 | 1 | 1 | 1 | -0.003 |
| SE | 58353 | ENSG00000110090 | CPT1A | chr11 | - | 68799217 | 68799355 | 68796855 | 68796933 | 68803999 | 68804101 | 377 | 231 | 184 | 380 | 272 | 0 | 0 | 0 | 0 | 0 | 592 | 279 | 241 | 348 | 1049 | 0 | 0 | 1 | 0 | 1 | 0 | 0 | 287 | 149 | 1 | 1 | 1 | 1 | 1 | 1 | 1 | 1 | 1 | 0.992 | 1 | 1 | 0.002 |
| SE | 58354 | ENSG00000110090 | CPT1A | chr11 | - | 68807466 | 68807638 | 68803999 | 68804101 | 68815333 | 68815487 | 115 | 67 | 49 | 127 | 36 | 0 | 1 | 1 | 0 | 0 | 133 | 104 | 49 | 107 | 235 | 0 | 0 | 0 | 0 | 0 | 0 | 0 | 298 | 149 | 1 | 1 | 1 | 0.971 | 0.961 | 1 | 1 | 1 | 1 | 1 | 1 | 1 | -0.014 |
| SE | 58355 | ENSG00000110090 | CPT1A | chr11 | - | 68812436 | 68812576 | 68807466 | 68807638 | 68815333 | 68815487 | 204 | 145 | 89 | 152 | 85 | 0 | 0 | 1 | 0 | 0 | 108 | 196 | 51 | 158 | 280 | 0 | 0 | 0 | 0 | 0 | 0 | 0 | 289 | 149 | 1 | 1 | 1 | 1 | 0.979 | 1 | 1 | 1 | 1 | 1 | 1 | 1 | -0.004 |
| SE | 58357 | ENSG00000163864 | NMNAT3 | chr3 | - | 139572106 | 139572233 | 139560222 | 139561392 | 139573597 | 139573680 | 1 | 0 | 0 | 0 | 0 | 3 | 1 | 2 | 2 | 1 | 0 | 0 | 0 | 0 | 5 | 0 | 14 | 2 | 14 | 2 | 10 | 28 | 276 | 149 | 1 | 1 | 0.153 | 0 | 0 | 0 | 0 | NA | 0 | 0 | 0 | 0.088 | 0.009 |
| SE | 58358 | ENSG00000163864 | NMNAT3 | chr3 | - | 139573597 | 139573680 | 139560179 | 139561392 | 139578871 | 139579055 | 6 | 2 | 2 | 3 | 7 | 1 | 0 | 0 | 1 | 0 | 4 | 14 | 4 | 10 | 41 | 0 | 0 | 0 | 0 | 0 | 0 | 2 | 232 | 149 | 1 | 1 | 0.794 | 1 | 1 | 0.658 | 1 | 1 | 1 | 1 | 1 | 0.929 | -0.095 |
| SE | 58359 | ENSG00000163864 | NMNAT3 | chr3 | - | 139575582 | 139575994 | 139573597 | 139573680 | 139578871 | 139579055 | 0 | 0 | 0 | 0 | 0 | 5 | 1 | 0 | 2 | 6 | 0 | 0 | 2 | 0 | 0 | 4 | 12 | 4 | 12 | 4 | 9 | 39 | 298 | 149 | 1 | 1 | 0 | 0 | NA | 0 | 0 | 0 | 0 | 0.2 | 0 | 0 | -0.04 |
| SE | 58361 | ENSG00000163864 | NMNAT3 | chr3 | - | 139582926 | 139583208 | 139578871 | 139579055 | 139627615 | 139627764 | 0 | 2 | 0 | 4 | 4 | 3 | 1 | 0 | 2 | 3 | 0 | 0 | 1 | 0 | 24 | 0 | 3 | 2 | 3 | 2 | 5 | 17 | 298 | 149 | 1 | 1 | 0 | 0.5 | NA | 0.5 | 0.4 | NA | 0 | 0.2 | 0 | 0.414 | 0.196 |
| SE | 58362 | ENSG00000163864 | NMNAT3 | chr3 | - | 139627615 | 139627764 | 139578871 | 139579055 | 139637962 | 139638062 | 4 | 1 | 0 | 2 | 3 | 0 | 0 | 0 | 0 | 0 | 4 | 3 | 2 | 5 | 20 | 0 | 0 | 0 | 0 | 0 | 0 | 10 | 298 | 149 | 1 | 1 | 1 | 1 | NA | 1 | 1 | 1 | 1 | 1 | 1 | 0.5 | 0.1 |
| SE | 58363 | ENSG00000163864 | NMNAT3 | chr3 | - | 139627615 | 139627764 | 139578871 | 139579055 | 139677704 | 139677850 | 5 | 1 | 0 | 2 | 3 | 0 | 0 | 0 | 0 | 0 | 0 | 3 | 4 | 5 | 17 | 0 | 0 | 0 | 0 | 0 | 0 | 3 | 298 | 149 | 1 | 1 | 1 | 1 | NA | 1 | 1 | NA | 1 | 1 | 1 | 0.739 | 0.065 |
| SE | 58364 | ENSG00000163864 | NMNAT3 | chr3 | - | 139634610 | 139634647 | 139627615 | 139627764 | 139637962 | 139638062 | 0 | 0 | 0 | 0 | 0 | 1 | 0 | 0 | 0 | 0 | 0 | 8 | 0 | 0 | 6 | 4 | 0 | 0 | 0 | 0 | 0 | 3 | 186 | 149 | 1 | 1 | 0 | NA | NA | NA | NA | 0 | 1 | NA | NA | 0.616 | -0.539 |
| SE | 58366 | ENSG00000163864 | NMNAT3 | chr3 | - | 139637962 | 139638062 | 139627615 | 139627764 | 139677704 | 139677718 | 1 | 0 | 0 | 0 | 0 | 2 | 0 | 0 | 0 | 0 | 4 | 0 | 0 | 0 | 9 | 0 | 0 | 2 | 0 | 2 | 0 | 0 | 249 | 149 | 1 | 1 | 0.23 | NA | NA | NA | NA | 1 | NA | 0 | NA | 1 | -0.437 |
| SE | 58369 | ENSG00000104946 | TBC1D17 | chr19 | + | 49879322 | 49879504 | 49878497 | 49878572 | 49880278 | 49880402 | 14 | 7 | 14 | 7 | 7 | 41 | 10 | 22 | 32 | 31 | 0 | 7 | 6 | 16 | 29 | 0 | 18 | 12 | 18 | 12 | 12 | 57 | 298 | 149 | 1 | 1 | 0.146 | 0.259 | 0.241 | 0.099 | 0.101 | NA | 0.163 | 0.2 | 0.4 | 0.203 | -0.072 |
| SE | 58370 | ENSG00000104946 | TBC1D17 | chr19 | + | 49882040 | 49882152 | 49881267 | 49881475 | 49882241 | 49882400 | 62 | 19 | 23 | 53 | 42 | 2 | 0 | 0 | 0 | 0 | 22 | 79 | 16 | 28 | 35 | 0 | 0 | 0 | 0 | 0 | 0 | 0 | 261 | 149 | 1 | 1 | 0.947 | 1 | 1 | 1 | 1 | 1 | 1 | 1 | 1 | 1 | -0.011 |
| SE | 58373 | ENSG00000135404 | CD63 | chr12 | - | 55726120 | 55726261 | 55725811 | 55725896 | 55726699 | 55726795 | 1299 | 667 | 730 | 970 | 961 | 0 | 0 | 0 | 0 | 0 | 3387 | 1859 | 1231 | 3392 | 3460 | 0 | 0 | 3 | 0 | 3 | 6 | 0 | 290 | 149 | 1 | 1 | 1 | 1 | 1 | 1 | 1 | 1 | 1 | 0.995 | 0.997 | 1 | 0.002 |
| SE | 58374 | ENSG00000135404 | CD63 | chr12 | - | 55726699 | 55726795 | 55726120 | 55726261 | 55726889 | 55726963 | 1162 | 602 | 675 | 891 | 895 | 1 | 0 | 0 | 0 | 0 | 3175 | 1758 | 1095 | 3259 | 3071 | 0 | 0 | 1 | 0 | 1 | 10 | 0 | 245 | 149 | 1 | 1 | 0.999 | 1 | 1 | 1 | 1 | 1 | 1 | 0.999 | 0.995 | 1 | 0.001 |
| SE | 58375 | ENSG00000263513 | FAM72C | chr1 | - | 143964854 | 143964979 | 143955286 | 143956481 | 143968923 | 143969001 | 1 | 3 | 3 | 3 | 2 | 2 | 0 | 0 | 0 | 0 | 6 | 27 | 12 | 3 | 0 | 12 | 1 | 1 | 1 | 1 | 0 | 0 | 274 | 149 | 1 | 1 | 0.214 | 1 | 1 | 1 | 1 | 0.214 | 0.936 | 0.867 | 1 | NA | 0.089 |
| SE | 58378 | ENSG00000171105 | INSR | chr19 | - | 7167967 | 7168094 | 7166153 | 7166404 | 7170536 | 7170751 | 104 | 86 | 52 | 48 | 93 | 0 | 0 | 0 | 1 | 0 | 164 | 131 | 109 | 180 | 196 | 0 | 0 | 0 | 0 | 0 | 0 | 0 | 276 | 149 | 1 | 1 | 1 | 1 | 1 | 0.963 | 1 | 1 | 1 | 1 | 1 | 1 | -0.007 |
| SE | 58381 | ENSG00000169504 | CLIC4 | chr1 | + | 24797741 | 24797851 | 24745381 | 24745625 | 24814093 | 24814219 | 62 | 64 | 58 | 93 | 42 | 1 | 0 | 0 | 0 | 0 | 73 | 81 | 65 | 118 | 44 | 0 | 0 | 0 | 0 | 0 | 0 | 0 | 259 | 149 | 1 | 1 | 0.973 | 1 | 1 | 1 | 1 | 1 | 1 | 1 | 1 | 1 | -0.005 |
| SE | 58382 | ENSG00000232956 | SNHG15 | chr7 | - | 44984356 | 44984426 | 44983022 | 44984046 | 44985881 | 44986097 | 23 | 10 | 38 | 27 | 22 | 0 | 0 | 0 | 0 | 0 | 35 | 23 | 23 | 25 | 33 | 0 | 2 | 0 | 2 | 0 | 0 | 0 | 219 | 149 | 1 | 1 | 1 | 1 | 1 | 1 | 1 | 1 | 0.887 | 1 | 1 | 1 | 0.023 |
| SE | 58390 | ENSG00000284719 | NA | chr1 | + | 39800325 | 39800391 | 39799421 | 39799581 | 39804877 | 39805452 | 9 | 0 | 1 | 3 | 0 | 1 | 0 | 0 | 0 | 0 | 9 | 3 | 3 | 16 | 0 | 0 | 0 | 0 | 0 | 0 | 0 | 3 | 215 | 149 | 1 | 1 | 0.862 | NA | 1 | 1 | NA | 1 | 1 | 1 | 1 | 0 | 0.154 |
| SE | 58391 | ENSG00000060140 | STYK1 | chr12 | - | 10637070 | 10637196 | 10634566 | 10634686 | 10673515 | 10673658 | 0 | 1 | 0 | 2 | 8 | 0 | 0 | 0 | 0 | 0 | 0 | 0 | 2 | 0 | 10 | 0 | 0 | 2 | 0 | 2 | 0 | 0 | 275 | 149 | 1 | 1 | NA | 1 | NA | 1 | 1 | NA | NA | 0.351 | NA | 1 | 0.325 |
| SE | 58392 | ENSG00000060140 | STYK1 | chr12 | - | 10637070 | 10637196 | 10634566 | 10634686 | 10673965 | 10674052 | 0 | 1 | 0 | 4 | 4 | 1 | 0 | 0 | 1 | 0 | 0 | 0 | 2 | 0 | 10 | 0 | 3 | 0 | 3 | 0 | 0 | 0 | 275 | 149 | 1 | 1 | 0 | 1 | NA | 0.684 | 1 | NA | 0 | 1 | NA | 1 | 0.004 |
| SE | 58396 | ENSG00000244588 | RAD21L1 | chr20 | + | 1234084 | 1234191 | 1231525 | 1231619 | 1238043 | 1238214 | 0 | 0 | 0 | 0 | 0 | 0 | 1 | 0 | 0 | 0 | 0 | 0 | 0 | 1 | 0 | 0 | 0 | 0 | 0 | 0 | 0 | 0 | 256 | 149 | 1 | 1 | NA | 0 | NA | NA | NA | NA | NA | NA | 1 | NA | -1 |
| SE | 58401 | ENSG00000164615 | CAMLG | chr5 | + | 134741062 | 134741523 | 134738512 | 134738792 | 134750758 | 134752157 | 39 | 27 | 45 | 42 | 28 | 0 | 0 | 2 | 2 | 0 | 11 | 26 | 23 | 37 | 16 | 2 | 0 | 3 | 0 | 3 | 0 | 0 | 298 | 149 | 1 | 1 | 1 | 1 | 0.918 | 0.913 | 1 | 0.733 | 1 | 0.793 | 1 | 1 | 0.061 |
| SE | 58403 | ENSG00000164615 | CAMLG | chr5 | + | 134741065 | 134741523 | 134738494 | 134738792 | 134750758 | 134751963 | 9 | 5 | 4 | 10 | 0 | 0 | 0 | 2 | 2 | 0 | 0 | 5 | 2 | 1 | 7 | 2 | 0 | 3 | 0 | 3 | 0 | 0 | 298 | 149 | 1 | 1 | 1 | 1 | 0.5 | 0.714 | NA | 0 | 1 | 0.25 | 1 | 1 | 0.153 |
| SE | 58405 | ENSG00000164615 | CAMLG | chr5 | + | 134743986 | 134744052 | 134738582 | 134738792 | 134750758 | 134751377 | 50 | 28 | 51 | 22 | 23 | 0 | 0 | 2 | 2 | 0 | 27 | 30 | 26 | 31 | 19 | 2 | 0 | 3 | 0 | 3 | 0 | 0 | 215 | 149 | 1 | 1 | 1 | 1 | 0.946 | 0.884 | 1 | 0.903 | 1 | 0.857 | 1 | 1 | 0.014 |
| SE | 58406 | ENSG00000271425 | NBPF10 | chr1 | - | 146068771 | 146068823 | 146068002 | 146068175 | 146069542 | 146069715 | 15 | 29 | 25 | 10 | 23 | 0 | 0 | 0 | 0 | 0 | 61 | 24 | 32 | 25 | 14 | 0 | 0 | 0 | 0 | 0 | 1 | 0 | 201 | 149 | 1 | 1 | 1 | 1 | 1 | 1 | 1 | 1 | 1 | 1 | 0.949 | 1 | 0.01 |
| SE | 58407 | ENSG00000271425 | NBPF10 | chr1 | - | 146128118 | 146128282 | 146127027 | 146127079 | 146132185 | 146132237 | 20 | 51 | 33 | 41 | 39 | 0 | 1 | 0 | 0 | 0 | 54 | 48 | 58 | 26 | 61 | 0 | 0 | 0 | 0 | 0 | 0 | 0 | 298 | 149 | 1 | 1 | 1 | 0.962 | 1 | 1 | 1 | 1 | 1 | 1 | 1 | 1 | -0.008 |
| SE | 58408 | ENSG00000271425 | NBPF10 | chr1 | - | 146133522 | 146133728 | 146132185 | 146132237 | 146134192 | 146134265 | 103 | 123 | 70 | 91 | 118 | 0 | 0 | 1 | 2 | 0 | 114 | 34 | 79 | 5 | 139 | 0 | 0 | 2 | 0 | 2 | 0 | 0 | 298 | 149 | 1 | 1 | 1 | 1 | 0.972 | 0.958 | 1 | 1 | 1 | 0.952 | 1 | 1 | -0.004 |
| SE | 58409 | ENSG00000271425 | NBPF10 | chr1 | - | 146133522 | 146133728 | 146132185 | 146132237 | 146136352 | 146136455 | 38 | 53 | 30 | 39 | 49 | 0 | 0 | 0 | 0 | 0 | 55 | 34 | 44 | 6 | 61 | 0 | 0 | 2 | 0 | 2 | 0 | 0 | 298 | 149 | 1 | 1 | 1 | 1 | 1 | 1 | 1 | 1 | 1 | 0.917 | 1 | 1 | 0.017 |
| SE | 58410 | ENSG00000271425 | NBPF10 | chr1 | - | 146134192 | 146134265 | 146132185 | 146132237 | 146136352 | 146136455 | 0 | 0 | 1 | 2 | 0 | 0 | 0 | 0 | 0 | 0 | 0 | 0 | 1 | 0 | 0 | 0 | 0 | 2 | 0 | 2 | 0 | 0 | 222 | 149 | 1 | 1 | NA | NA | 1 | 1 | NA | NA | NA | 0.251 | NA | NA | 0.749 |
| SE | 58412 | ENSG00000271425 | NBPF10 | chr1 | - | 146135306 | 146135521 | 146134192 | 146134265 | 146136352 | 146136455 | 104 | 94 | 76 | 93 | 82 | 0 | 0 | 0 | 0 | 0 | 168 | 43 | 45 | 45 | 80 | 0 | 3 | 0 | 3 | 0 | 0 | 0 | 298 | 149 | 1 | 1 | 1 | 1 | 1 | 1 | 1 | 1 | 0.878 | 1 | 1 | 1 | 0.024 |
| SE | 58413 | ENSG00000271425 | NBPF10 | chr1 | - | 146136352 | 146136455 | 146134192 | 146134265 | 146144594 | 146144804 | 56 | 29 | 0 | 0 | 13 | 0 | 0 | 0 | 0 | 0 | 0 | 3 | 35 | 0 | 60 | 0 | 0 | 2 | 0 | 2 | 0 | 0 | 252 | 149 | 1 | 1 | 1 | 1 | NA | NA | 1 | NA | 1 | 0.912 | NA | 1 | 0.029 |
| SE | 58414 | ENSG00000271425 | NBPF10 | chr1 | - | 146136352 | 146136455 | 146135306 | 146135521 | 146138235 | 146138445 | 62 | 80 | 79 | 113 | 80 | 0 | 1 | 0 | 0 | 0 | 187 | 81 | 49 | 97 | 33 | 5 | 0 | 0 | 0 | 0 | 0 | 0 | 252 | 149 | 1 | 1 | 1 | 0.979 | 1 | 1 | 1 | 0.957 | 1 | 1 | 1 | 1 | 0.004 |
| SE | 58416 | ENSG00000271425 | NBPF10 | chr1 | - | 146141603 | 146141818 | 146140483 | 146140556 | 146144594 | 146144804 | 1 | 10 | 24 | 13 | 7 | 0 | 0 | 0 | 0 | 0 | 27 | 34 | 17 | 37 | 0 | 0 | 1 | 1 | 1 | 1 | 2 | 0 | 298 | 149 | 1 | 1 | 1 | 1 | 1 | 1 | 1 | 1 | 0.944 | 0.895 | 0.902 | NA | 0.065 |
| SE | 58431 | ENSG00000226674 | TEX41 | chr2 | + | 144766369 | 144766530 | 144671601 | 144671664 | 144768097 | 144768280 | 0 | 0 | 0 | 0 | 0 | 0 | 0 | 0 | 1 | 0 | 6 | 0 | 0 | 0 | 0 | 0 | 0 | 0 | 0 | 0 | 0 | 0 | 298 | 149 | 1 | 1 | NA | NA | NA | 0 | NA | 1 | NA | NA | NA | NA | -1 |
| SE | 58432 | ENSG00000226674 | TEX41 | chr2 | + | 144766369 | 144766530 | 144691038 | 144691110 | 144768097 | 144768280 | 0 | 0 | 0 | 0 | 0 | 0 | 1 | 0 | 0 | 0 | 6 | 0 | 0 | 0 | 0 | 0 | 0 | 0 | 0 | 0 | 0 | 0 | 298 | 149 | 1 | 1 | NA | 0 | NA | NA | NA | 1 | NA | NA | NA | NA | -1 |
| SE | 58570 | ENSG00000072858 | SIDT1 | chr3 | + | 113567539 | 113567710 | 113532554 | 113533243 | 113584697 | 113584769 | 2 | 5 | 1 | 7 | 0 | 0 | 0 | 0 | 1 | 0 | 0 | 1 | 0 | 0 | 5 | 0 | 0 | 0 | 0 | 0 | 0 | 0 | 298 | 149 | 1 | 1 | 1 | 1 | 1 | 0.778 | NA | NA | 1 | NA | NA | 1 | -0.055 |
| SE | 58573 | ENSG00000072858 | SIDT1 | chr3 | + | 113580607 | 113580709 | 113567539 | 113567710 | 113583408 | 113583496 | 2 | 0 | 2 | 2 | 3 | 0 | 0 | 0 | 0 | 0 | 0 | 4 | 3 | 0 | 0 | 0 | 4 | 2 | 4 | 2 | 0 | 0 | 251 | 149 | 1 | 1 | 1 | NA | 1 | 1 | 1 | NA | 0.372 | 0.471 | NA | NA | 0.579 |
| SE | 58574 | ENSG00000072858 | SIDT1 | chr3 | + | 113580607 | 113580709 | 113576921 | 113576967 | 113583408 | 113583496 | 18 | 3 | 9 | 14 | 18 | 1 | 2 | 4 | 0 | 3 | 0 | 14 | 2 | 2 | 8 | 0 | 8 | 1 | 8 | 1 | 0 | 1 | 251 | 149 | 1 | 1 | 0.914 | 0.471 | 0.572 | 1 | 0.781 | NA | 0.51 | 0.543 | 1 | 0.826 | 0.028 |
| SE | 58576 | ENSG00000072858 | SIDT1 | chr3 | + | 113583408 | 113583496 | 113567539 | 113567710 | 113584697 | 113584769 | 46 | 12 | 28 | 23 | 72 | 0 | 0 | 0 | 1 | 0 | 0 | 30 | 20 | 9 | 9 | 0 | 0 | 0 | 0 | 0 | 0 | 0 | 237 | 149 | 1 | 1 | 1 | 1 | 1 | 0.935 | 1 | NA | 1 | 1 | 1 | 1 | -0.013 |
| SE | 58580 | ENSG00000072858 | SIDT1 | chr3 | + | 113612085 | 113612194 | 113608418 | 113608536 | 113616099 | 113616176 | 10 | 12 | 28 | 16 | 15 | 0 | 0 | 0 | 1 | 1 | 0 | 11 | 12 | 3 | 21 | 0 | 1 | 0 | 1 | 0 | 0 | 0 | 258 | 149 | 1 | 1 | 1 | 1 | 1 | 0.902 | 0.897 | NA | 0.864 | 1 | 1 | 1 | -0.006 |
| SE | 58581 | ENSG00000072858 | SIDT1 | chr3 | + | 113612085 | 113612194 | 113611007 | 113611144 | 113619679 | 113619726 | 20 | 8 | 24 | 17 | 35 | 0 | 0 | 0 | 1 | 0 | 0 | 26 | 4 | 18 | 17 | 0 | 0 | 1 | 0 | 1 | 0 | 0 | 258 | 149 | 1 | 1 | 1 | 1 | 1 | 0.908 | 1 | NA | 1 | 0.698 | 1 | 1 | 0.057 |
| SE | 58582 | ENSG00000072858 | SIDT1 | chr3 | + | 113615062 | 113615077 | 113612085 | 113612194 | 113616096 | 113616176 | 1 | 0 | 0 | 0 | 0 | 2 | 1 | 0 | 0 | 3 | 0 | 0 | 0 | 0 | 0 | 0 | 3 | 0 | 3 | 0 | 0 | 0 | 164 | 149 | 1 | 1 | 0.312 | 0 | NA | NA | 0 | NA | 0 | NA | NA | NA | 0.104 |
| SE | 58583 | ENSG00000072858 | SIDT1 | chr3 | + | 113615062 | 113615077 | 113612085 | 113612194 | 113616099 | 113616176 | 1 | 0 | 0 | 0 | 0 | 19 | 14 | 43 | 36 | 18 | 0 | 1 | 0 | 0 | 0 | 14 | 19 | 10 | 19 | 10 | 3 | 22 | 164 | 149 | 1 | 1 | 0.046 | 0 | 0 | 0 | 0 | 0 | 0.046 | 0 | 0 | 0 | 0 |
| SE | 58584 | ENSG00000072858 | SIDT1 | chr3 | + | 113616096 | 113616176 | 113612085 | 113612194 | 113619679 | 113619726 | 19 | 10 | 38 | 21 | 12 | 0 | 0 | 2 | 1 | 0 | 13 | 24 | 11 | 6 | 15 | 0 | 9 | 0 | 9 | 0 | 2 | 0 | 229 | 149 | 1 | 1 | 1 | 1 | 0.925 | 0.932 | 1 | 1 | 0.634 | 1 | 0.661 | 1 | 0.112 |
| SE | 58587 | ENSG00000186854 | TRABD2A | chr2 | - | 84823952 | 84824204 | 84821649 | 84822100 | 84832054 | 84832145 | 57 | 49 | 140 | 83 | 104 | 0 | 0 | 3 | 0 | 0 | 11 | 22 | 12 | 35 | 26 | 0 | 0 | 0 | 0 | 0 | 0 | 0 | 298 | 149 | 1 | 1 | 1 | 1 | 0.959 | 1 | 1 | 1 | 1 | 1 | 1 | 1 | -0.008 |
| SE | 58588 | ENSG00000186854 | TRABD2A | chr2 | - | 84832054 | 84832145 | 84823952 | 84824204 | 84839148 | 84839323 | 21 | 30 | 93 | 55 | 57 | 0 | 0 | 0 | 0 | 3 | 11 | 2 | 6 | 15 | 3 | 0 | 0 | 0 | 0 | 0 | 0 | 0 | 240 | 149 | 1 | 1 | 1 | 1 | 1 | 1 | 0.922 | 1 | 1 | 1 | 1 | 1 | -0.016 |
| SE | 58589 | ENSG00000186854 | TRABD2A | chr2 | - | 84839148 | 84839323 | 84832054 | 84832145 | 84841860 | 84842007 | 71 | 58 | 196 | 120 | 120 | 0 | 0 | 0 | 1 | 0 | 28 | 11 | 20 | 16 | 24 | 0 | 0 | 0 | 0 | 0 | 0 | 0 | 298 | 149 | 1 | 1 | 1 | 1 | 1 | 0.984 | 1 | 1 | 1 | 1 | 1 | 1 | -0.003 |
| SE | 58590 | ENSG00000186854 | TRABD2A | chr2 | - | 84839148 | 84839323 | 84832054 | 84832145 | 84870224 | 84870785 | 8 | 14 | 53 | 25 | 37 | 2 | 0 | 3 | 1 | 1 | 0 | 2 | 2 | 0 | 10 | 0 | 0 | 0 | 0 | 0 | 0 | 0 | 298 | 149 | 1 | 1 | 0.667 | 1 | 0.898 | 0.926 | 0.949 | NA | 1 | 1 | NA | 1 | -0.112 |
| SE | 58591 | ENSG00000186854 | TRABD2A | chr2 | - | 84839148 | 84839323 | 84835918 | 84838241 | 84841860 | 84842007 | 89 | 77 | 237 | 144 | 149 | 0 | 0 | 0 | 1 | 0 | 34 | 24 | 48 | 25 | 33 | 0 | 0 | 0 | 0 | 0 | 0 | 0 | 298 | 149 | 1 | 1 | 1 | 1 | 1 | 0.986 | 1 | 1 | 1 | 1 | 1 | 1 | -0.003 |
| SE | 58594 | ENSG00000186854 | TRABD2A | chr2 | - | 84841860 | 84842007 | 84835918 | 84838241 | 84870224 | 84870785 | 71 | 75 | 177 | 121 | 110 | 0 | 1 | 3 | 0 | 1 | 1 | 28 | 28 | 32 | 26 | 0 | 0 | 0 | 0 | 0 | 0 | 0 | 296 | 149 | 1 | 1 | 1 | 0.974 | 0.967 | 1 | 0.982 | 1 | 1 | 1 | 1 | 1 | -0.015 |
| SE | 58596 | ENSG00000120694 | HSPH1 | chr13 | - | 31143791 | 31143923 | 31140183 | 31141259 | 31145562 | 31145768 | 288 | 215 | 212 | 359 | 294 | 2 | 0 | 1 | 1 | 0 | 7 | 73 | 58 | 21 | 32 | 0 | 4 | 0 | 4 | 0 | 0 | 0 | 281 | 149 | 1 | 1 | 0.987 | 1 | 0.991 | 0.995 | 1 | 1 | 0.906 | 1 | 1 | 1 | 0.013 |
| SE | 58597 | ENSG00000120694 | HSPH1 | chr13 | - | 31147958 | 31148092 | 31145562 | 31145768 | 31148373 | 31148480 | 299 | 274 | 262 | 386 | 448 | 0 | 1 | 0 | 0 | 0 | 2 | 74 | 83 | 37 | 10 | 0 | 0 | 0 | 0 | 0 | 0 | 0 | 283 | 149 | 1 | 1 | 1 | 0.993 | 1 | 1 | 1 | 1 | 1 | 1 | 1 | 1 | -0.001 |
| SE | 58598 | ENSG00000120694 | HSPH1 | chr13 | - | 31147958 | 31148092 | 31145562 | 31145768 | 31149953 | 31150182 | 155 | 141 | 149 | 220 | 226 | 3 | 0 | 1 | 0 | 2 | 1 | 54 | 62 | 21 | 10 | 0 | 0 | 0 | 0 | 0 | 0 | 0 | 283 | 149 | 1 | 1 | 0.965 | 1 | 0.987 | 1 | 0.983 | 1 | 1 | 1 | 1 | 1 | -0.013 |
| SE | 58599 | ENSG00000120694 | HSPH1 | chr13 | - | 31148373 | 31148480 | 31145562 | 31145768 | 31149953 | 31150182 | 85 | 66 | 73 | 126 | 104 | 3 | 0 | 1 | 0 | 2 | 0 | 6 | 16 | 2 | 0 | 0 | 0 | 0 | 0 | 0 | 0 | 0 | 256 | 149 | 1 | 1 | 0.943 | 1 | 0.977 | 1 | 0.968 | NA | 1 | 1 | 1 | NA | -0.022 |
| SE | 58600 | ENSG00000120694 | HSPH1 | chr13 | - | 31148373 | 31148480 | 31147958 | 31148092 | 31149953 | 31150182 | 239 | 215 | 194 | 304 | 340 | 7 | 8 | 8 | 9 | 13 | 1 | 32 | 45 | 22 | 0 | 0 | 6 | 8 | 6 | 8 | 4 | 0 | 256 | 149 | 1 | 1 | 0.952 | 0.94 | 0.934 | 0.952 | 0.938 | 1 | 0.756 | 0.766 | 0.762 | NA | 0.122 |
| SE | 58601 | ENSG00000120694 | HSPH1 | chr13 | - | 31150946 | 31151191 | 31149953 | 31150182 | 31152851 | 31152951 | 185 | 112 | 144 | 206 | 242 | 0 | 0 | 0 | 0 | 0 | 9 | 36 | 39 | 24 | 2 | 0 | 0 | 0 | 0 | 0 | 0 | 2 | 298 | 149 | 1 | 1 | 1 | 1 | 1 | 1 | 1 | 1 | 1 | 1 | 0.857 | 1 | 0.029 |
| SE | 58602 | ENSG00000120694 | HSPH1 | chr13 | - | 31151608 | 31151742 | 31150946 | 31151191 | 31152851 | 31152951 | 279 | 166 | 207 | 315 | 286 | 4 | 2 | 4 | 1 | 19 | 0 | 56 | 14 | 14 | 3 | 0 | 0 | 3 | 0 | 3 | 0 | 0 | 283 | 149 | 1 | 1 | 0.973 | 0.978 | 0.965 | 0.994 | 0.888 | NA | 1 | 0.711 | 1 | 1 | 0.032 |
| SE | 58603 | ENSG00000120694 | HSPH1 | chr13 | - | 31151608 | 31151742 | 31150946 | 31151191 | 31154632 | 31154755 | 130 | 83 | 101 | 145 | 153 | 3 | 5 | 4 | 19 | 0 | 0 | 28 | 6 | 11 | 0 | 0 | 0 | 0 | 0 | 0 | 0 | 0 | 283 | 149 | 1 | 1 | 0.958 | 0.897 | 0.93 | 0.801 | 1 | NA | 1 | 1 | 1 | NA | -0.083 |
| SE | 58605 | ENSG00000120694 | HSPH1 | chr13 | - | 31152851 | 31152951 | 31151608 | 31151742 | 31154632 | 31154755 | 260 | 141 | 157 | 288 | 242 | 0 | 1 | 2 | 0 | 0 | 12 | 44 | 12 | 16 | 6 | 0 | 0 | 0 | 0 | 0 | 0 | 0 | 249 | 149 | 1 | 1 | 1 | 0.988 | 0.979 | 1 | 1 | 1 | 1 | 1 | 1 | 1 | -0.007 |
| SE | 58606 | ENSG00000132906 | CASP9 | chr1 | - | 15493891 | 15494001 | 15491400 | 15493035 | 15495272 | 15495452 | 39 | 17 | 48 | 61 | 98 | 1 | 0 | 0 | 3 | 0 | 39 | 94 | 39 | 83 | 108 | 0 | 0 | 3 | 0 | 3 | 0 | 0 | 259 | 149 | 1 | 1 | 0.957 | 1 | 1 | 0.921 | 1 | 1 | 1 | 0.882 | 1 | 1 | -0.001 |
| SE | 58608 | ENSG00000132906 | CASP9 | chr1 | - | 15504610 | 15504758 | 15495272 | 15495452 | 15505989 | 15506079 | 31 | 24 | 21 | 32 | 42 | 0 | 0 | 0 | 0 | 0 | 39 | 52 | 24 | 22 | 52 | 0 | 0 | 1 | 0 | 1 | 0 | 0 | 297 | 149 | 1 | 1 | 1 | 1 | 1 | 1 | 1 | 1 | 1 | 0.923 | 1 | 1 | 0.015 |
| SE | 58609 | ENSG00000132906 | CASP9 | chr1 | - | 15505989 | 15506079 | 15495272 | 15495452 | 15506898 | 15507052 | 9 | 3 | 0 | 7 | 12 | 0 | 0 | 0 | 1 | 0 | 0 | 10 | 10 | 9 | 4 | 0 | 0 | 0 | 0 | 0 | 0 | 0 | 239 | 149 | 1 | 1 | 1 | 1 | NA | 0.814 | 1 | NA | 1 | 1 | 1 | 1 | -0.046 |
| SE | 58610 | ENSG00000132906 | CASP9 | chr1 | - | 15505989 | 15506079 | 15504610 | 15504758 | 15506898 | 15507052 | 18 | 15 | 11 | 26 | 32 | 1 | 0 | 0 | 0 | 0 | 18 | 41 | 28 | 15 | 23 | 0 | 0 | 0 | 0 | 0 | 0 | 0 | 239 | 149 | 1 | 1 | 0.918 | 1 | 1 | 1 | 1 | 1 | 1 | 1 | 1 | 1 | -0.016 |
| SE | 58611 | ENSG00000132906 | CASP9 | chr1 | - | 15518109 | 15518395 | 15495272 | 15495452 | 15524068 | 15524192 | 16 | 8 | 10 | 14 | 4 | 0 | 0 | 0 | 0 | 1 | 0 | 14 | 6 | 11 | 46 | 0 | 0 | 0 | 0 | 0 | 0 | 0 | 298 | 149 | 1 | 1 | 1 | 1 | 1 | 1 | 0.667 | NA | 1 | 1 | 1 | 1 | -0.067 |
| SE | 58612 | ENSG00000115594 | IL1R1 | chr2 | + | 102156145 | 102156246 | 102153962 | 102154017 | 102157718 | 102157785 | 0 | 4 | 3 | 0 | 9 | 2 | 0 | 6 | 4 | 1 | 4 | 0 | 20 | 5 | 0 | 0 | 4 | 9 | 4 | 9 | 12 | 0 | 250 | 149 | 1 | 1 | 0 | 1 | 0.23 | 0 | 0.843 | 1 | 0 | 0.57 | 0.199 | NA | -0.028 |
| SE | 58615 | ENSG00000115594 | IL1R1 | chr2 | + | 102165224 | 102165304 | 102164773 | 102165003 | 102166112 | 102166281 | 7 | 4 | 5 | 5 | 5 | 0 | 0 | 1 | 0 | 1 | 1 | 4 | 12 | 28 | 0 | 0 | 0 | 0 | 0 | 0 | 0 | 0 | 229 | 149 | 1 | 1 | 1 | 1 | 0.765 | 1 | 0.765 | 1 | 1 | 1 | 1 | NA | -0.094 |
| SE | 58616 | ENSG00000115594 | IL1R1 | chr2 | + | 102166112 | 102166281 | 102165114 | 102165304 | 102168597 | 102168663 | 16 | 6 | 10 | 12 | 17 | 0 | 0 | 1 | 0 | 4 | 1 | 4 | 18 | 78 | 0 | 0 | 0 | 4 | 0 | 4 | 0 | 0 | 298 | 149 | 1 | 1 | 1 | 1 | 0.833 | 1 | 0.68 | 1 | 1 | 0.692 | 1 | NA | -0.02 |
| SE | 58617 | ENSG00000115594 | IL1R1 | chr2 | + | 102168597 | 102168663 | 102166112 | 102166281 | 102171800 | 102171829 | 9 | 5 | 6 | 13 | 19 | 0 | 0 | 0 | 0 | 0 | 4 | 4 | 28 | 68 | 0 | 0 | 0 | 2 | 0 | 2 | 0 | 0 | 215 | 149 | 1 | 1 | 1 | 1 | 1 | 1 | 1 | 1 | 1 | 0.907 | 1 | NA | 0.023 |
| SE | 58621 | ENSG00000246526 | LINC02481 | chr4 | - | 6689308 | 6689459 | 6687447 | 6688896 | 6690298 | 6690519 | 11 | 3 | 3 | 9 | 6 | 16 | 1 | 7 | 6 | 1 | 0 | 5 | 1 | 5 | 5 | 0 | 0 | 0 | 0 | 0 | 0 | 0 | 298 | 149 | 1 | 1 | 0.256 | 0.6 | 0.176 | 0.429 | 0.75 | NA | 1 | 1 | 1 | 1 | -0.558 |
| SE | 58623 | ENSG00000100271 | TTLL1 | chr22 | - | 43046409 | 43046573 | 43039515 | 43039905 | 43059383 | 43059527 | 19 | 15 | 19 | 14 | 13 | 0 | 0 | 1 | 0 | 0 | 30 | 24 | 12 | 22 | 29 | 0 | 0 | 0 | 0 | 0 | 0 | 0 | 298 | 149 | 1 | 1 | 1 | 1 | 0.905 | 1 | 1 | 1 | 1 | 1 | 1 | 1 | -0.019 |
| SE | 58625 | ENSG00000100271 | TTLL1 | chr22 | - | 43059383 | 43059527 | 43051800 | 43051887 | 43063812 | 43063921 | 43 | 22 | 52 | 51 | 40 | 0 | 1 | 0 | 0 | 0 | 40 | 54 | 17 | 39 | 69 | 0 | 0 | 0 | 0 | 0 | 0 | 0 | 293 | 149 | 1 | 1 | 1 | 0.918 | 1 | 1 | 1 | 1 | 1 | 1 | 1 | 1 | -0.016 |
| SE | 58626 | ENSG00000100271 | TTLL1 | chr22 | - | 43063812 | 43063921 | 43051800 | 43051887 | 43068409 | 43068590 | 0 | 3 | 0 | 1 | 0 | 0 | 0 | 0 | 0 | 2 | 6 | 2 | 0 | 0 | 5 | 0 | 0 | 0 | 0 | 0 | 0 | 0 | 258 | 149 | 1 | 1 | NA | 1 | NA | 1 | 0 | 1 | 1 | NA | NA | 1 | -0.333 |
| SE | 58627 | ENSG00000100271 | TTLL1 | chr22 | - | 43064189 | 43064324 | 43063812 | 43063921 | 43068409 | 43068590 | 75 | 63 | 98 | 101 | 149 | 0 | 2 | 0 | 1 | 2 | 57 | 36 | 42 | 29 | 84 | 6 | 2 | 0 | 2 | 0 | 0 | 5 | 284 | 149 | 1 | 1 | 1 | 0.943 | 1 | 0.981 | 0.975 | 0.833 | 0.904 | 1 | 1 | 0.898 | 0.053 |
| SE | 58630 | ENSG00000134759 | ELP2 | chr18 | + | 36138269 | 36138426 | 36136306 | 36136377 | 36138794 | 36138872 | 133 | 103 | 112 | 111 | 152 | 0 | 1 | 0 | 1 | 0 | 46 | 118 | 56 | 99 | 7 | 0 | 0 | 0 | 0 | 0 | 0 | 0 | 298 | 149 | 1 | 1 | 1 | 0.981 | 1 | 0.982 | 1 | 1 | 1 | 1 | 1 | 1 | -0.007 |
| SE | 58631 | ENSG00000134759 | ELP2 | chr18 | + | 36138294 | 36138426 | 36136306 | 36136377 | 36138794 | 36138872 | 63 | 44 | 45 | 59 | 82 | 0 | 1 | 0 | 1 | 0 | 12 | 52 | 22 | 27 | 7 | 0 | 0 | 0 | 0 | 0 | 0 | 0 | 281 | 149 | 1 | 1 | 1 | 0.959 | 1 | 0.969 | 1 | 1 | 1 | 1 | 1 | 1 | -0.014 |
| SE | 58634 | ENSG00000134759 | ELP2 | chr18 | + | 36139418 | 36139613 | 36138269 | 36138426 | 36141136 | 36141201 | 4 | 6 | 14 | 5 | 14 | 0 | 0 | 1 | 6 | 2 | 0 | 0 | 4 | 3 | 0 | 0 | 0 | 0 | 0 | 0 | 0 | 0 | 298 | 149 | 1 | 1 | 1 | 1 | 0.875 | 0.294 | 0.778 | NA | NA | 1 | 1 | NA | -0.211 |
| SE | 58639 | ENSG00000134759 | ELP2 | chr18 | + | 36142825 | 36142966 | 36141136 | 36141201 | 36144938 | 36145034 | 147 | 91 | 108 | 104 | 113 | 0 | 0 | 0 | 0 | 0 | 47 | 67 | 52 | 67 | 27 | 0 | 0 | 2 | 0 | 2 | 0 | 0 | 290 | 149 | 1 | 1 | 1 | 1 | 1 | 1 | 1 | 1 | 1 | 0.93 | 1 | 1 | 0.014 |
| SE | 58640 | ENSG00000134759 | ELP2 | chr18 | + | 36142825 | 36142966 | 36141136 | 36141201 | 36154849 | 36154999 | 10 | 11 | 7 | 8 | 6 | 1 | 0 | 0 | 2 | 0 | 0 | 3 | 9 | 41 | 0 | 0 | 0 | 0 | 0 | 0 | 0 | 0 | 290 | 149 | 1 | 1 | 0.837 | 1 | 1 | 0.673 | 1 | NA | 1 | 1 | 1 | NA | -0.098 |
| SE | 58641 | ENSG00000134759 | ELP2 | chr18 | + | 36144938 | 36145034 | 36142825 | 36142966 | 36145947 | 36146048 | 179 | 106 | 125 | 134 | 163 | 0 | 0 | 0 | 0 | 0 | 78 | 81 | 39 | 71 | 38 | 0 | 0 | 0 | 0 | 0 | 1 | 0 | 245 | 149 | 1 | 1 | 1 | 1 | 1 | 1 | 1 | 1 | 1 | 1 | 0.977 | 1 | 0.005 |
| SE | 58643 | ENSG00000134759 | ELP2 | chr18 | + | 36145947 | 36146048 | 36142825 | 36142966 | 36146249 | 36146376 | 47 | 26 | 51 | 51 | 78 | 2 | 1 | 1 | 12 | 5 | 0 | 31 | 15 | 15 | 0 | 0 | 11 | 11 | 11 | 11 | 13 | 26 | 250 | 149 | 1 | 1 | 0.933 | 0.939 | 0.968 | 0.717 | 0.903 | NA | 0.627 | 0.448 | 0.407 | 0 | 0.522 |
| SE | 58644 | ENSG00000134759 | ELP2 | chr18 | + | 36145947 | 36146048 | 36142825 | 36142966 | 36154849 | 36154999 | 0 | 3 | 3 | 0 | 0 | 7 | 1 | 0 | 1 | 0 | 0 | 0 | 0 | 1 | 0 | 0 | 1 | 1 | 1 | 1 | 36 | 0 | 250 | 149 | 1 | 1 | 0 | 0.641 | 1 | 0 | NA | NA | 0 | 0 | 0.016 | NA | 0.405 |
| SE | 58648 | ENSG00000134759 | ELP2 | chr18 | + | 36158834 | 36158904 | 36156465 | 36156654 | 36159734 | 36159830 | 143 | 98 | 133 | 122 | 179 | 0 | 0 | 0 | 0 | 0 | 57 | 86 | 91 | 76 | 29 | 0 | 3 | 0 | 3 | 0 | 0 | 0 | 219 | 149 | 1 | 1 | 1 | 1 | 1 | 1 | 1 | 1 | 0.951 | 1 | 1 | 1 | 0.01 |
| SE | 58651 | ENSG00000134759 | ELP2 | chr18 | + | 36164474 | 36164667 | 36160931 | 36161004 | 36167100 | 36167222 | 116 | 96 | 129 | 101 | 179 | 0 | 0 | 0 | 1 | 0 | 45 | 77 | 54 | 56 | 14 | 0 | 4 | 0 | 4 | 0 | 0 | 0 | 298 | 149 | 1 | 1 | 1 | 1 | 1 | 0.981 | 1 | 1 | 0.906 | 1 | 1 | 1 | 0.015 |
| SE | 58652 | ENSG00000134759 | ELP2 | chr18 | + | 36167100 | 36167222 | 36160931 | 36161004 | 36170062 | 36170196 | 53 | 37 | 53 | 52 | 86 | 0 | 0 | 0 | 1 | 0 | 12 | 40 | 31 | 17 | 0 | 0 | 0 | 0 | 0 | 0 | 0 | 0 | 271 | 149 | 1 | 1 | 1 | 1 | 1 | 0.966 | 1 | 1 | 1 | 1 | 1 | NA | -0.007 |
| SE | 58654 | ENSG00000125633 | CCDC93 | chr2 | - | 117939028 | 117939111 | 117936511 | 117936739 | 117945528 | 117945582 | 34 | 36 | 32 | 22 | 66 | 1 | 0 | 0 | 0 | 0 | 22 | 44 | 48 | 27 | 0 | 0 | 0 | 0 | 0 | 0 | 0 | 0 | 232 | 149 | 1 | 1 | 0.956 | 1 | 1 | 1 | 1 | 1 | 1 | 1 | 1 | NA | -0.009 |
| SE | 58655 | ENSG00000125633 | CCDC93 | chr2 | - | 117941188 | 117941297 | 117939028 | 117939111 | 117945528 | 117945582 | 132 | 86 | 89 | 82 | 111 | 0 | 0 | 0 | 0 | 0 | 38 | 105 | 130 | 90 | 19 | 0 | 1 | 1 | 1 | 1 | 0 | 0 | 258 | 149 | 1 | 1 | 1 | 1 | 1 | 1 | 1 | 1 | 0.984 | 0.987 | 1 | 1 | 0.006 |
| SE | 58657 | ENSG00000125633 | CCDC93 | chr2 | - | 117944732 | 117944797 | 117944023 | 117944086 | 117945528 | 117945582 | 1 | 0 | 1 | 1 | 2 | 89 | 81 | 100 | 85 | 95 | 0 | 0 | 0 | 0 | 0 | 5 | 60 | 122 | 60 | 122 | 85 | 23 | 214 | 149 | 1 | 1 | 0.008 | 0 | 0.007 | 0.008 | 0.014 | 0 | 0 | 0 | 0 | 0 | 0.007 |
| SE | 58658 | ENSG00000125633 | CCDC93 | chr2 | - | 117945528 | 117945582 | 117939028 | 117939111 | 117946810 | 117946882 | 17 | 9 | 20 | 18 | 22 | 1 | 0 | 0 | 0 | 0 | 0 | 11 | 22 | 21 | 6 | 0 | 0 | 3 | 0 | 3 | 0 | 0 | 203 | 149 | 1 | 1 | 0.926 | 1 | 1 | 1 | 1 | NA | 1 | 0.843 | 1 | 1 | 0.024 |
| SE | 58659 | ENSG00000125633 | CCDC93 | chr2 | - | 117958364 | 117958478 | 117952372 | 117952435 | 117973907 | 117973994 | 55 | 46 | 61 | 77 | 74 | 0 | 0 | 0 | 0 | 0 | 35 | 32 | 79 | 68 | 7 | 0 | 2 | 0 | 2 | 0 | 0 | 0 | 263 | 149 | 1 | 1 | 1 | 1 | 1 | 1 | 1 | 1 | 0.901 | 1 | 1 | 1 | 0.02 |
| SE | 58660 | ENSG00000125633 | CCDC93 | chr2 | - | 117958364 | 117958481 | 117952372 | 117952435 | 117973907 | 117973994 | 113 | 82 | 104 | 135 | 138 | 0 | 0 | 0 | 0 | 0 | 53 | 72 | 118 | 106 | 28 | 0 | 2 | 0 | 2 | 0 | 0 | 0 | 266 | 149 | 1 | 1 | 1 | 1 | 1 | 1 | 1 | 1 | 0.953 | 1 | 1 | 1 | 0.009 |
| SE | 58662 | ENSG00000125633 | CCDC93 | chr2 | - | 117977993 | 117978027 | 117975187 | 117975280 | 117985968 | 117986069 | 37 | 32 | 41 | 35 | 48 | 0 | 0 | 1 | 0 | 2 | 30 | 48 | 71 | 59 | 22 | 0 | 0 | 2 | 0 | 2 | 1 | 0 | 183 | 149 | 1 | 1 | 1 | 1 | 0.971 | 1 | 0.951 | 1 | 1 | 0.967 | 0.98 | 1 | -0.005 |
| SE | 58663 | ENSG00000125633 | CCDC93 | chr2 | - | 117977993 | 117978030 | 117975187 | 117975280 | 117985968 | 117986069 | 159 | 89 | 126 | 121 | 186 | 0 | 0 | 1 | 0 | 2 | 52 | 179 | 209 | 168 | 47 | 0 | 0 | 2 | 0 | 2 | 1 | 0 | 186 | 149 | 1 | 1 | 1 | 1 | 0.99 | 1 | 0.987 | 1 | 1 | 0.988 | 0.993 | 1 | -0.001 |
| SE | 58664 | ENSG00000125633 | CCDC93 | chr2 | - | 117985968 | 117986069 | 117977993 | 117978030 | 117995445 | 117995502 | 232 | 147 | 183 | 171 | 273 | 0 | 1 | 2 | 1 | 2 | 156 | 280 | 277 | 270 | 58 | 0 | 0 | 2 | 0 | 2 | 0 | 0 | 250 | 149 | 1 | 1 | 1 | 0.989 | 0.982 | 0.99 | 0.988 | 1 | 1 | 0.988 | 1 | 1 | -0.008 |
| SE | 58667 | ENSG00000014919 | COX15 | chr10 | - | 99729552 | 99729734 | 99727440 | 99727563 | 99731959 | 99732100 | 148 | 94 | 145 | 137 | 111 | 0 | 0 | 0 | 0 | 0 | 140 | 132 | 70 | 167 | 115 | 0 | 1 | 0 | 1 | 0 | 0 | 0 | 298 | 149 | 1 | 1 | 1 | 1 | 1 | 1 | 1 | 1 | 0.985 | 1 | 1 | 1 | 0.003 |
| SE | 58669 | ENSG00000141458 | NPC1 | chr18 | - | 23534445 | 23534559 | 23533354 | 23533517 | 23535468 | 23535622 | 162 | 50 | 75 | 97 | 32 | 0 | 0 | 0 | 1 | 0 | 56 | 66 | 68 | 50 | 81 | 0 | 0 | 1 | 0 | 1 | 0 | 0 | 263 | 149 | 1 | 1 | 1 | 1 | 1 | 0.982 | 1 | 1 | 1 | 0.975 | 1 | 1 | 0.001 |
| SE | 58672 | ENSG00000141458 | NPC1 | chr18 | - | 23543454 | 23543569 | 23541305 | 23541433 | 23544343 | 23544526 | 63 | 26 | 42 | 41 | 13 | 0 | 0 | 0 | 0 | 0 | 4 | 64 | 83 | 31 | 34 | 0 | 0 | 1 | 0 | 1 | 0 | 0 | 264 | 149 | 1 | 1 | 1 | 1 | 1 | 1 | 1 | 1 | 1 | 0.979 | 1 | 1 | 0.004 |
| SE | 58674 | ENSG00000141458 | NPC1 | chr18 | - | 23551626 | 23551727 | 23544959 | 23545149 | 23554757 | 23554984 | 94 | 27 | 47 | 54 | 28 | 0 | 0 | 1 | 2 | 1 | 36 | 42 | 35 | 21 | 45 | 2 | 0 | 0 | 0 | 0 | 1 | 7 | 250 | 149 | 1 | 1 | 1 | 1 | 0.966 | 0.941 | 0.943 | 0.915 | 1 | 1 | 0.926 | 0.793 | 0.043 |
| SE | 58675 | ENSG00000141458 | NPC1 | chr18 | - | 23551626 | 23551727 | 23548005 | 23548108 | 23556242 | 23556613 | 35 | 16 | 21 | 30 | 12 | 0 | 1 | 0 | 0 | 0 | 0 | 25 | 11 | 29 | 20 | 0 | 6 | 0 | 6 | 0 | 0 | 0 | 250 | 149 | 1 | 1 | 1 | 0.905 | 1 | 1 | 1 | NA | 0.713 | 1 | 1 | 1 | 0.053 |
| SE | 58676 | ENSG00000141458 | NPC1 | chr18 | - | 23554757 | 23554984 | 23551626 | 23551727 | 23556242 | 23556613 | 238 | 57 | 128 | 156 | 116 | 0 | 0 | 0 | 0 | 0 | 98 | 87 | 76 | 39 | 114 | 0 | 0 | 0 | 0 | 0 | 3 | 0 | 298 | 149 | 1 | 1 | 1 | 1 | 1 | 1 | 1 | 1 | 1 | 1 | 0.867 | 1 | 0.027 |
| SE | 58677 | ENSG00000141458 | NPC1 | chr18 | - | 23557116 | 23557190 | 23556242 | 23556613 | 23561359 | 23561527 | 46 | 17 | 23 | 27 | 15 | 0 | 1 | 0 | 0 | 0 | 23 | 52 | 33 | 17 | 0 | 0 | 0 | 0 | 0 | 0 | 0 | 0 | 223 | 149 | 1 | 1 | 1 | 0.919 | 1 | 1 | 1 | 1 | 1 | 1 | 1 | NA | -0.016 |
| SE | 58679 | ENSG00000141458 | NPC1 | chr18 | - | 23561359 | 23561527 | 23560230 | 23560480 | 23568822 | 23568998 | 477 | 214 | 255 | 379 | 409 | 1 | 0 | 0 | 0 | 0 | 199 | 242 | 214 | 116 | 222 | 0 | 0 | 0 | 0 | 0 | 0 | 0 | 298 | 149 | 1 | 1 | 0.996 | 1 | 1 | 1 | 1 | 1 | 1 | 1 | 1 | 1 | -0.001 |

| | | | | | | | | | | | | | | | | | | | | | | | | | | | | | | | | | | | | | | | | | | | | | | |
|---|---|---|---|---|---|---|---|---|---|---|---|---|---|---|---|---|---|---|---|---|---|---|---|---|---|---|---|---|---|---|---|---|---|---|---|---|---|---|---|---|---|---|---|---|---|---|
| SE | 58680 | ENSG00000141458 | NPC1 | chr18 | - | 23572073 | 23572180 | 23568822 | 23568998 | 23573451 | 23573574 | 280 | 131 | 143 | 185 | 115 | 0 | 3 | 0 | 2 | 0 | 81 | 132 | 93 | 71 | 99 | 0 | 0 | 4 | 0 | 0 | 256 | 149 | 1 | 1 | 1 | 0.962 | 1 | 0.982 | 1 | 1 | 1 | 0.931 | 1 | 1 | 0.003 |
| SE | 58682 | ENSG00000227403 | LINC01806 | chr2 | + | 161245423 | 161245467 | 161244729 | 161244895 | 161246874 | 161249049 | 2 | 1 | 17 | 3 | 37 | 0 | 0 | 1 | 4 | 2 | 0 | 2 | 0 | 0 | 0 | 0 | 0 | 3 | 3 | 0 | 193 | 149 | 1 | 1 | 1 | 1 | 0.929 | 0.367 | 0.935 | NA | 1 | 0 | 0 | NA | 0.513 |
| SE | 58683 | ENSG00000173409 | ARV1 | chr1 | + | 230988319 | 230988439 | 230979093 | 230979279 | 230990109 | 230990263 | 53 | 15 | 43 | 24 | 94 | 0 | 0 | 3 | 0 | 0 | 0 | 7 | 18 | 35 | 14 | 0 | 0 | 0 | 0 | 0 | 269 | 149 | 1 | 1 | 1 | 1 | 0.888 | 1 | 1 | NA | 1 | 1 | 1 | 1 | -0.022 |
| SE | 58684 | ENSG00000173409 | ARV1 | chr1 | + | 230988319 | 230988439 | 230979093 | 230979279 | 230995759 | 230995984 | 16 | 8 | 10 | 5 | 20 | 0 | 0 | 2 | 0 | 0 | 0 | 4 | 2 | 13 | 7 | 0 | 3 | 0 | 0 | 0 | 269 | 149 | 1 | 1 | 1 | 1 | 0.735 | 1 | 1 | NA | 0.425 | 1 | 1 | 1 | 0.091 |
| SE | 58685 | ENSG00000173409 | ARV1 | chr1 | + | 230990109 | 230990263 | 230979093 | 230979279 | 230995759 | 230995984 | 29 | 8 | 19 | 23 | 61 | 0 | 0 | 2 | 0 | 0 | 0 | 3 | 5 | 5 | 0 | 0 | 3 | 0 | 0 | 0 | 298 | 149 | 1 | 1 | 1 | 1 | 0.826 | 1 | 1 | NA | 0.333 | 1 | 1 | NA | 0.188 |
| SE | 58686 | ENSG00000173409 | ARV1 | chr1 | + | 230990109 | 230990263 | 230988319 | 230988439 | 230995759 | 230995984 | 67 | 15 | 49 | 42 | 135 | 1 | 0 | 0 | 0 | 0 | 0 | 6 | 21 | 27 | 7 | 0 | 0 | 0 | 0 | 0 | 298 | 149 | 1 | 1 | 0.971 | 1 | 1 | 1 | 1 | NA | 1 | 1 | 1 | 1 | -0.006 |
| SE | 58687 | ENSG00000173409 | ARV1 | chr1 | + | 230990599 | 230990698 | 230990109 | 230990263 | 230995759 | 230995984 | 0 | 0 | 2 | 0 | 4 | 29 | 8 | 16 | 23 | 61 | 0 | 0 | 0 | 0 | 0 | 0 | 3 | 5 | 5 | 0 | 248 | 149 | 1 | 1 | 0 | 0 | 0.07 | 0 | 0.038 | NA | 0 | 0 | 0 | NA | 0.022 |
| SE | 58688 | ENSG00000173409 | ARV1 | chr1 | + | 230995759 | 230995984 | 230990109 | 230990263 | 231000137 | 231000595 | 29 | 8 | 16 | 23 | 62 | 1 | 3 | 0 | 0 | 3 | 0 | 3 | 5 | 5 | 0 | 0 | 0 | 0 | 0 | 0 | 298 | 149 | 1 | 1 | 0.935 | 0.571 | 1 | 1 | 0.912 | NA | 1 | 1 | 1 | NA | -0.116 |
| SE | 58689 | ENSG00000173409 | ARV1 | chr1 | + | 230997120 | 230997267 | 230995759 | 230995984 | 231000137 | 231000595 | 71 | 66 | 73 | 75 | 204 | 0 | 0 | 0 | 0 | 1 | 8 | 52 | 30 | 28 | 5 | 0 | 0 | 0 | 0 | 0 | 296 | 149 | 1 | 1 | 1 | 1 | 1 | 1 | 0.99 | 1 | 1 | 1 | 1 | 1 | -0.002 |
| SE | 58694 | ENSG00000198898 | CAPZA2 | chr7 | + | 116888126 | 116888190 | 116862586 | 116862650 | 116892993 | 116893045 | 350 | 252 | 226 | 370 | 482 | 0 | 0 | 0 | 0 | 1 | 15 | 169 | 118 | 55 | 61 | 0 | 0 | 0 | 0 | 0 | 213 | 149 | 1 | 1 | 1 | 1 | 1 | 1 | 0.997 | 1 | 1 | 1 | 1 | 1 | -0.001 |
| SE | 58697 | ENSG00000198898 | CAPZA2 | chr7 | + | 116912068 | 116912140 | 116909759 | 116910311 | 116916059 | 116916110 | 1009 | 957 | 546 | 1153 | 1988 | 0 | 0 | 0 | 4 | 0 | 238 | 584 | 409 | 125 | 150 | 0 | 0 | 0 | 0 | 0 | 221 | 149 | 1 | 1 | 1 | 1 | 1 | 0.995 | 1 | 1 | 1 | 1 | 1 | 1 | -0.001 |
| SE | 58700 | ENSG00000025039 | RRAGD | chr6 | - | 89379223 | 89379338 | 89377670 | 89377813 | 89387294 | 89387590 | 15 | 13 | 9 | 18 | 25 | 0 | 0 | 0 | 0 | 3 | 38 | 11 | 35 | 74 | 40 | 0 | 0 | 0 | 0 | 0 | 264 | 149 | 1 | 1 | 1 | 1 | 1 | 1 | 0.825 | 1 | 1 | 1 | 1 | 1 | -0.035 |
| SE | 58726 | ENSG00000213963 | NA | chr2 | - | 177331571 | 177331656 | 177323238 | 177323396 | 177334392 | 177334529 | 0 | 0 | 0 | 1 | 0 | 15 | 3 | 8 | 9 | 5 | 0 | 0 | 0 | 0 | 0 | 0 | 1 | 3 | 16 | 35 | 234 | 149 | 1 | 1 | 0 | 0 | 0.066 | 0 | 0 | NA | 0 | 0 | 0 | 0 | 0.013 |
| SE | 58730 | ENSG00000162415 | ZSWIM5 | chr1 | - | 45035687 | 45035823 | 45034311 | 45034469 | 45036038 | 45036299 | 7 | 8 | 13 | 4 | 6 | 0 | 0 | 1 | 0 | 0 | 0 | 0 | 0 | 0 | 12 | 0 | 0 | 0 | 0 | 0 | 285 | 149 | 1 | 1 | 1 | 1 | 0.872 | 1 | 1 | NA | NA | NA | NA | 1 | -0.026 |
| SE | 58731 | ENSG00000162415 | ZSWIM5 | chr1 | - | 45040391 | 45040538 | 45038935 | 45039073 | 45043218 | 45043395 | 12 | 2 | 8 | 6 | 8 | 0 | 0 | 0 | 1 | 0 | 0 | 0 | 9 | 4 | 0 | 0 | 0 | 3 | 0 | 11 | 296 | 149 | 1 | 1 | 1 | 1 | 1 | 0.751 | 1 | NA | NA | 0.602 | 1 | 0 | 0.416 |
| SE | 58732 | ENSG00000162415 | ZSWIM5 | chr1 | - | 45043218 | 45043395 | 45040391 | 45040538 | 45051073 | 45051253 | 5 | 2 | 5 | 4 | 1 | 0 | 0 | 0 | 0 | 0 | 0 | 0 | 5 | 2 | 4 | 0 | 0 | 4 | 0 | 7 | 298 | 149 | 1 | 1 | 1 | 1 | 1 | 1 | 1 | NA | NA | 0.385 | 1 | 0.222 | 0.464 |
| SE | 58735 | ENSG00000129968 | ABHD17A | chr19 | - | 1880895 | 1881048 | 1879920 | 1880115 | 1881234 | 1881804 | 0 | 0 | 0 | 1 | 0 | 95 | 9 | 40 | 55 | 3 | 0 | 0 | 0 | 0 | 0 | 56 | 28 | 37 | 39 | 126 | 298 | 149 | 1 | 1 | 0 | 0 | 0 | 0.009 | 0 | 0 | 0 | 0 | 0 | 0 | 0.002 |
| SE | 58736 | ENSG00000129968 | ABHD17A | chr19 | - | 1881234 | 1881804 | 1879920 | 1880115 | 1885351 | 1885433 | 130 | 10 | 43 | 59 | 7 | 1 | 1 | 0 | 1 | 0 | 56 | 31 | 39 | 48 | 134 | 0 | 0 | 0 | 0 | 0 | 298 | 149 | 1 | 1 | 0.985 | 0.833 | 1 | 0.967 | 1 | 1 | 1 | 1 | 1 | 1 | -0.043 |
| SE | 58737 | ENSG00000129968 | ABHD17A | chr19 | - | 1881261 | 1881804 | 1879920 | 1880115 | 1885351 | 1885496 | 35 | 2 | 3 | 4 | 4 | 1 | 1 | 0 | 1 | 0 | 0 | 3 | 2 | 9 | 22 | 0 | 0 | 0 | 0 | 0 | 298 | 149 | 1 | 1 | 0.946 | 0.5 | 1 | 0.667 | 1 | NA | 1 | 1 | 1 | 1 | -0.177 |
| SE | 58738 | ENSG00000129968 | ABHD17A | chr19 | - | 1884468 | 1884543 | 1881234 | 1881804 | 1885351 | 1885496 | 2 | 0 | 0 | 0 | 0 | 35 | 1 | 3 | 4 | 4 | 0 | 0 | 0 | 0 | 0 | 0 | 3 | 2 | 9 | 8 | 224 | 149 | 1 | 1 | 0.037 | 0 | 0 | 0 | 0 | NA | 0 | 0 | 0 | 0 | 0.007 |
| SE | 58741 | ENSG00000225936 | NA | chr10 | - | 117241660 | 117241783 | 117239599 | 117240181 | 117241884 | 117241923 | 25 | 23 | 1 | 54 | 18 | 2 | 0 | 1 | 0 | 0 | 24 | 5 | 22 | 16 | 125 | 0 | 0 | 0 | 0 | 3 | 272 | 149 | 1 | 1 | 0.873 | 1 | 0.354 | 1 | 1 | 1 | 1 | 1 | 1 | 0.958 | -0.146 |
| SE | 58743 | ENSG00000163710 | PCOLCE2 | chr3 | - | 142823531 | 142823615 | 142820877 | 142821045 | 142829691 | 142829731 | 0 | 0 | 0 | 1 | 0 | 0 | 0 | 0 | 0 | 0 | 7 | 5 | 4 | 44 | 19 | 0 | 0 | 0 | 2 | 8 | 233 | 149 | 1 | 1 | NA | NA | NA | 1 | NA | 1 | 1 | 1 | 0.934 | 0.603 | 0.093 |
| SE | 58745 | ENSG00000163710 | PCOLCE2 | chr3 | - | 142842923 | 142843048 | 142838769 | 142838906 | 142848216 | 142848322 | 2 | 0 | 0 | 1 | 0 | 0 | 0 | 0 | 0 | 0 | 32 | 1 | 0 | 84 | 35 | 0 | 0 | 0 | 2 | 0 | 274 | 149 | 1 | 1 | 1 | NA | NA | 1 | NA | 1 | 1 | NA | 0.958 | 1 | 0.01 |
| SE | 58751 | ENSG00000177628 | GBA | chr1 | - | 155237340 | 155237578 | 155236181 | 155236469 | 155238133 | 155238306 | 115 | 46 | 56 | 51 | 69 | 1 | 0 | 0 | 1 | 2 | 185 | 210 | 66 | 276 | 207 | 0 | 0 | 0 | 0 | 0 | 298 | 149 | 1 | 1 | 0.983 | 1 | 1 | 0.962 | 0.945 | 1 | 1 | 1 | 1 | 1 | -0.022 |
| SE | 58753 | ENSG00000177628 | GBA | chr1 | - | 155239885 | 155239962 | 155238516 | 155238650 | 155240629 | 155240717 | 1 | 0 | 0 | 1 | 0 | 0 | 0 | 0 | 0 | 0 | 4 | 5 | 0 | 3 | 0 | 0 | 3 | 0 | 0 | 0 | 226 | 149 | 1 | 1 | 1 | NA | NA | 1 | NA | 1 | 0.524 | NA | 1 | NA | 0.159 |
| SE | 58754 | ENSG00000177628 | GBA | chr1 | - | 155239885 | 155240077 | 155238516 | 155238650 | 155240629 | 155240717 | 117 | 45 | 74 | 76 | 83 | 0 | 0 | 0 | 0 | 0 | 276 | 208 | 58 | 308 | 281 | 0 | 3 | 0 | 0 | 0 | 298 | 149 | 1 | 1 | 1 | 1 | 1 | 1 | 1 | 1 | 0.972 | 1 | 1 | 1 | 0.006 |
| SE | 58755 | ENSG00000177628 | GBA | chr1 | - | 155240176 | 155240299 | 155239959 | 155240077 | 155240629 | 155240717 | 0 | 0 | 2 | 0 | 0 | 117 | 45 | 74 | 76 | 83 | 1 | 1 | 0 | 3 | 0 | 276 | 208 | 58 | 308 | 281 | 272 | 149 | 1 | 1 | 0 | 0 | 0.015 | 0 | 0 | 0.002 | 0.003 | 0 | 0.005 | 0 | 0.001 |
| SE | 58756 | ENSG00000177628 | GBA | chr1 | - | 155240629 | 155240717 | 155238516 | 155238650 | 155241085 | 155241180 | 82 | 31 | 42 | 53 | 42 | 0 | 0 | 0 | 0 | 0 | 205 | 126 | 38 | 225 | 138 | 0 | 0 | 0 | 4 | 0 | 237 | 149 | 1 | 1 | 1 | 1 | 1 | 1 | 1 | 1 | 1 | 1 | 0.973 | 1 | 0.005 |
| SE | 58758 | ENSG00000177628 | GBA | chr1 | - | 155241085 | 155241180 | 155240629 | 155240717 | 155244505 | 155244670 | 40 | 12 | 22 | 24 | 25 | 0 | 0 | 0 | 0 | 0 | 55 | 71 | 22 | 96 | 67 | 0 | 5 | 0 | 0 | 0 | 244 | 149 | 1 | 1 | 1 | 1 | 1 | 1 | 1 | 1 | 0.897 | 1 | 1 | 1 | 0.021 |
| SE | 58759 | ENSG00000177628 | GBA | chr1 | - | 155244094 | 155244230 | 155241085 | 155241180 | 155244505 | 155244606 | 5 | 0 | 0 | 0 | 3 | 7 | 0 | 4 | 0 | 0 | 6 | 0 | 0 | 0 | 7 | 0 | 1 | 2 | 7 | 0 | 285 | 149 | 1 | 1 | 0.272 | NA | 0 | NA | 1 | 1 | 0 | 0 | 0 | 1 | 0.024 |
| SE | 58760 | ENSG00000177628 | GBA | chr1 | - | 155244094 | 155244233 | 155241085 | 155241180 | 155244505 | 155244636 | 6 | 0 | 0 | 0 | 3 | 7 | 0 | 4 | 0 | 0 | 11 | 0 | 0 | 0 | 14 | 0 | 1 | 2 | 7 | 0 | 288 | 149 | 1 | 1 | 0.307 | NA | 0 | NA | 1 | 1 | 0 | 0 | 0 | 1 | 0.036 |
| SE | 58762 | ENSG00000163157 | TMOD4 | chr1 | - | 151170518 | 151170663 | 151169986 | 151170103 | 151170919 | 151171059 | 1 | 1 | 6 | 2 | 4 | 0 | 1 | 2 | 0 | 1 | 0 | 4 | 0 | 0 | 0 | 0 | 2 | 0 | 0 | 0 | 294 | 149 | 1 | 1 | 1 | 0.336 | 0.603 | 1 | 0.67 | NA | 0.503 | NA | NA | NA | 0.219 |
| SE | 58763 | ENSG00000163157 | TMOD4 | chr1 | - | 151170919 | 151171059 | 151170518 | 151170663 | 151171632 | 151171763 | 0 | 0 | 6 | 2 | 2 | 0 | 0 | 0 | 0 | 2 | 0 | 3 | 2 | 0 | 0 | 0 | 0 | 0 | 0 | 0 | 289 | 149 | 1 | 1 | NA | NA | 1 | 1 | 0.34 | NA | 1 | 1 | NA | NA | -0.22 |
| SE | 58764 | ENSG00000163157 | TMOD4 | chr1 | - | 151170919 | 151171063 | 151170518 | 151170663 | 151171632 | 151171763 | 0 | 0 | 4 | 2 | 3 | 0 | 0 | 0 | 0 | 2 | 0 | 3 | 0 | 0 | 0 | 0 | 0 | 0 | 0 | 0 | 293 | 149 | 1 | 1 | NA | NA | 1 | 1 | 0.433 | NA | 1 | NA | NA | NA | -0.189 |
| SE | 58765 | ENSG00000163157 | TMOD4 | chr1 | - | 151171432 | 151171540 | 151170919 | 151171059 | 151171632 | 151171763 | 3 | 0 | 3 | 0 | 2 | 0 | 0 | 2 | 0 | 0 | 0 | 0 | 0 | 0 | 0 | 0 | 0 | 2 | 0 | 0 | 257 | 149 | 1 | 1 | 1 | NA | 0.465 | NA | 1 | NA | NA | 0 | NA | NA | 0.822 |
| SE | 58766 | ENSG00000163157 | TMOD4 | chr1 | - | 151171432 | 151171540 | 151170919 | 151171063 | 151171632 | 151171763 | 3 | 0 | 5 | 4 | 3 | 0 | 0 | 0 | 0 | 1 | 0 | 12 | 2 | 0 | 0 | 0 | 0 | 0 | 0 | 0 | 257 | 149 | 1 | 1 | 1 | NA | 1 | 1 | 0.635 | NA | 1 | 1 | NA | NA | -0.091 |
| SE | 58767 | ENSG00000163157 | TMOD4 | chr1 | - | 151171632 | 151171763 | 151170919 | 151171063 | 151172267 | 151172315 | 2 | 4 | 6 | 6 | 3 | 0 | 0 | 0 | 0 | 2 | 0 | 2 | 5 | 2 | 17 | 0 | 0 | 0 | 0 | 0 | 280 | 149 | 1 | 1 | 1 | 1 | 1 | 1 | 0.444 | NA | 1 | 1 | 1 | 1 | -0.111 |
| SE | 58779 | ENSG00000152439 | ZNF773 | chr19 | + | 57508519 | 57508564 | 57505301 | 57505400 | 57513021 | 57513151 | 3 | 0 | 3 | 1 | 1 | 0 | 0 | 0 | 0 | 2 | 0 | 5 | 2 | 2 | 0 | 0 | 0 | 0 | 0 | 2 | 194 | 149 | 1 | 1 | 1 | NA | 1 | 1 | 0.277 | NA | 1 | 1 | 1 | 0 | 0.069 |
| SE | 58780 | ENSG00000158571 | PFKFB1 | chrX | - | 54960823 | 54960851 | 54959826 | 54959893 | 54963256 | 54963382 | 0 | 0 | 2 | 0 | 0 | 1 | 0 | 0 | 0 | 0 | 0 | 0 | 0 | 0 | 0 | 0 | 0 | 0 | 2 | 0 | 177 | 149 | 1 | 1 | 0 | NA | 1 | NA | NA | NA | NA | NA | 0 | NA | 0.5 |
| SE | 58781 | ENSG00000158571 | PFKFB1 | chrX | - | 54960823 | 54960917 | 54959826 | 54959893 | 54963256 | 54963382 | 2 | 0 | 1 | 0 | 0 | 1 | 0 | 0 | 0 | 0 | 0 | 6 | 0 | 0 | 0 | 0 | 0 | 0 | 2 | 0 | 243 | 149 | 1 | 1 | 0.551 | NA | 1 | NA | NA | NA | 1 | NA | 0 | NA | 0.276 |
| SE | 58784 | ENSG00000077254 | USP33 | chr1 | - | 77713199 | 77713281 | 77711651 | 77711855 | 77715741 | 77715868 | 180 | 125 | 116 | 126 | 158 | 0 | 1 | 0 | 0 | 0 | 80 | 123 | 76 | 109 | 68 | 0 | 0 | 0 | 0 | 0 | 231 | 149 | 1 | 1 | 1 | 0.988 | 1 | 1 | 1 | 1 | 1 | 1 | 1 | 1 | -0.002 |
| SE | 58789 | ENSG00000077254 | USP33 | chr1 | - | 77721830 | 77721901 | 77721171 | 77721205 | 77722023 | 77722196 | 140 | 72 | 118 | 123 | 204 | 0 | 0 | 0 | 0 | 1 | 16 | 70 | 54 | 50 | 29 | 0 | 0 | 0 | 0 | 0 | 220 | 149 | 1 | 1 | 1 | 1 | 1 | 1 | 0.993 | 1 | 1 | 1 | 1 | 1 | -0.001 |
| SE | 58790 | ENSG00000077254 | USP33 | chr1 | - | 77721830 | 77721925 | 77721171 | 77721205 | 77722023 | 77722196 | 189 | 109 | 160 | 156 | 252 | 0 | 0 | 0 | 0 | 1 | 30 | 101 | 77 | 68 | 60 | 0 | 0 | 0 | 0 | 0 | 244 | 149 | 1 | 1 | 1 | 1 | 1 | 1 | 0.994 | 1 | 1 | 1 | 1 | 1 | -0.001 |
| SE | 58791 | ENSG00000077254 | USP33 | chr1 | - | 77728294 | 77728712 | 77725621 | 77725762 | 77729859 | 77729938 | 565 | 471 | 514 | 573 | 1028 | 0 | 2 | 0 | 0 | 0 | 142 | 400 | 389 | 320 | 68 | 0 | 0 | 0 | 0 | 0 | 298 | 149 | 1 | 1 | 1 | 0.992 | 1 | 1 | 1 | 1 | 1 | 1 | 1 | 1 | -0.002 |
| SE | 58792 | ENSG00000077254 | USP33 | chr1 | - | 77729859 | 77729938 | 77728294 | 77728712 | 77730617 | 77730731 | 561 | 496 | 512 | 637 | 936 | 1 | 0 | 1 | 6 | 0 | 153 | 452 | 437 | 346 | 87 | 0 | 6 | 1 | 0 | 0 | 228 | 149 | 1 | 1 | 0.997 | 1 | 0.997 | 0.986 | 1 | 1 | 0.98 | 0.997 | 1 | 1 | 0.001 |
| SE | 58793 | ENSG00000077254 | USP33 | chr1 | - | 77734346 | 77734416 | 77730617 | 77730731 | 77736055 | 77736158 | 289 | 226 | 278 | 274 | 491 | 0 | 0 | 0 | 3 | 0 | 59 | 217 | 148 | 121 | 42 | 0 | 7 | 0 | 0 | 0 | 219 | 149 | 1 | 1 | 1 | 1 | 1 | 0.984 | 1 | 1 | 0.955 | 1 | 1 | 1 | 0.006 |
| SE | 58796 | ENSG00000077254 | USP33 | chr1 | - | 77736055 | 77736158 | 77734346 | 77734416 | 77739264 | 77739417 | 312 | 250 | 299 | 301 | 593 | 0 | 0 | 0 | 6 | 0 | 94 | 252 | 192 | 133 | 56 | 0 | 0 | 0 | 0 | 0 | 252 | 149 | 1 | 1 | 1 | 1 | 1 | 0.967 | 1 | 1 | 1 | 1 | 1 | 1 | -0.007 |
| SE | 58797 | ENSG00000077254 | USP33 | chr1 | - | 77736055 | 77736158 | 77734346 | 77734416 | 77740876 | 77740939 | 119 | 101 | 148 | 145 | 270 | 0 | 0 | 0 | 0 | 5 | 38 | 107 | 79 | 51 | 24 | 0 | 0 | 0 | 0 | 0 | 252 | 149 | 1 | 1 | 1 | 1 | 1 | 1 | 0.97 | 1 | 1 | 1 | 1 | 1 | -0.006 |
| SE | 58798 | ENSG00000077254 | USP33 | chr1 | - | 77739264 | 77739417 | 77734346 | 77734416 | 77740876 | 77740939 | 271 | 228 | 180 | 215 | 494 | 0 | 0 | 0 | 0 | 5 | 79 | 183 | 135 | 158 | 55 | 0 | 0 | 0 | 0 | 0 | 298 | 149 | 1 | 1 | 1 | 1 | 1 | 1 | 0.98 | 1 | 1 | 1 | 1 | 1 | -0.004 |
| SE | 58799 | ENSG00000077254 | USP33 | chr1 | - | 77739264 | 77739417 | 77736055 | 77736158 | 77740876 | 77740939 | 470 | 383 | 334 | 367 | 821 | 1 | 2 | 0 | 0 | 0 | 135 | 334 | 259 | 242 | 87 | 0 | 0 | 5 | 2 | 0 | 298 | 149 | 1 | 1 | 0.996 | 0.99 | 1 | 1 | 1 | 1 | 1 | 0.963 | 0.984 | 1 | 0.008 |
| SE | 58800 | ENSG00000077254 | USP33 | chr1 | - | 77745420 | 77745599 | 77741616 | 77741748 | 77746398 | 77746556 | 44 | 44 | 19 | 43 | 49 | 0 | 1 | 1 | 0 | 0 | 21 | 30 | 24 | 31 | 10 | 0 | 0 | 0 | 0 | 0 | 298 | 149 | 1 | 1 | 1 | 0.957 | 0.905 | 1 | 1 | 1 | 1 | 1 | 1 | 1 | -0.028 |
| SE | 58802 | ENSG00000077254 | USP33 | chr1 | - | 77746398 | 77746556 | 77741616 | 77741748 | 77759642 | 77759689 | 0 | 2 | 2 | 0 | 0 | 24 | 6 | 21 | 18 | 11 | 0 | 1 | 0 | 0 | 0 | 0 | 4 | 6 | 10 | 0 | 298 | 149 | 1 | 1 | 0 | 0.143 | 0.045 | 0 | 0 | NA | 0.111 | 0 | 0 | NA | 0.001 |
| SE | 58803 | ENSG00000077254 | USP33 | chr1 | - | 77746398 | 77746556 | 77745420 | 77745599 | 77759642 | 77759689 | 1 | 4 | 2 | 4 | 0 | 11 | 6 | 5 | 8 | 4 | 0 | 2 | 3 | 4 | 7 | 0 | 0 | 13 | 8 | 0 | 298 | 149 | 1 | 1 | 0.043 | 0.25 | 0.167 | 0.2 | 0 | NA | 1 | 0.103 | 0.2 | 1 | -0.444 |
| SE | 58805 | ENSG00000140416 | TPM1 | chr15 | + | 63044026 | 63044152 | 63043705 | 63043831 | 63056984 | 63057118 | 1 | 1 | 0 | 1 | 3 | 1 | 0 | 0 | 0 | 0 | 4 | 0 | 0 | 0 | 15 | 9 | 0 | 1 | 0 | 53 | 275 | 149 | 1 | 1 | 0.351 | 1 | NA | 1 | 1 | 0.194 | NA | 0 | NA | 0.133 | 0.729 |
| SE | 58812 | ENSG00000140416 | TPM1 | chr15 | + | 63062214 | 63062277 | 63061197 | 63061273 | 63071089 | 63071190 | 26 | 7 | 11 | 10 | 14 | 1 | 0 | 0 | 0 | 0 | 60 | 28 | 15 | 48 | 302 | 0 | 0 | 0 | 0 | 0 | 212 | 149 | 1 | 1 | 0.948 | 1 | 1 | 1 | 1 | 1 | 1 | 1 | 1 | 1 | -0.01 |
| SE | 58813 | ENSG00000140416 | TPM1 | chr15 | + | 63062575 | 63062645 | 63061197 | 63062277 | 63071089 | 63071190 | 103 | 160 | 31 | 143 | 176 | 0 | 0 | 1 | 0 | 0 | 394 | 191 | 82 | 182 | 1209 | 0 | 0 | 0 | 0 | 0 | 219 | 149 | 1 | 1 | 1 | 1 | 0.955 | 1 | 1 | 1 | 1 | 1 | 1 | 1 | -0.009 |
| SE | 58814 | ENSG00000140416 | TPM1 | chr15 | + | 63064063 | 63064142 | 63062575 | 63062645 | 63065895 | 63066091 | 8 | 2 | 1 | 0 | 3 | 0 | 0 | 0 | 0 | 0 | 0 | 15 | 0 | 2 | 13 | 0 | 0 | 1 | 0 | 0 | 228 | 149 | 1 | 1 | 1 | 1 | 1 | NA | 1 | NA | 1 | 0 | 1 | 1 | 0.25 |
| SE | 58815 | ENSG00000140416 | TPM1 | chr15 | + | 63064063 | 63064190 | 63062575 | 63062645 | 63065895 | 63066093 | 7 | 2 | 1 | 2 | 6 | 0 | 0 | 0 | 0 | 0 | 0 | 15 | 0 | 6 | 13 | 0 | 0 | 1 | 0 | 0 | 276 | 149 | 1 | 1 | 1 | 1 | 1 | 1 | 1 | NA | 1 | 0 | 1 | 1 | 0.25 |
| SE | 58816 | ENSG00000140416 | TPM1 | chr15 | + | 63064063 | 63064982 | 63062575 | 63062645 | 63065895 | 63066035 | 8 | 2 | 2 | 2 | 7 | 0 | 0 | 0 | 0 | 0 | 0 | 18 | 1 | 9 | 50 | 0 | 0 | 1 | 0 | 0 | 298 | 149 | 1 | 1 | 1 | 1 | 1 | 1 | 1 | NA | 1 | 0.333 | 1 | 1 | 0.167 |
| SE | 58819 | ENSG00000164828 | SUN1 | chr7 | + | 817279 | 817499 | 816614 | 816673 | 832504 | 832601 | 1 | 0 | 1 | 1 | 0 | 25 | 18 | 37 | 32 | 14 | 0 | 0 | 0 | 0 | 0 | 13 | 12 | 11 | 14 | 9 | 298 | 149 | 1 | 1 | 0.02 | 0 | 0.013 | 0.015 | 0 | 0 | 0 | 0 | 0 | 0 | 0.01 |
| SE | 58823 | ENSG00000164828 | SUN1 | chr7 | + | 838797 | 838986 | 832475 | 832601 | 841945 | 842019 | 158 | 96 | 158 | 148 | 276 | 0 | 0 | 0 | 0 | 0 | 131 | 93 | 102 | 104 | 190 | 0 | 0 | 3 | 0 | 0 | 298 | 149 | 1 | 1 | 1 | 1 | 1 | 1 | 1 | 1 | 1 | 0.944 | 1 | 1 | 0.011 |
| SE | 58826 | ENSG00000164828 | SUN1 | chr7 | + | 848417 | 848615 | 843340 | 843520 | 851383 | 851482 | 90 | 65 | 68 | 87 | 94 | 0 | 3 | 0 | 0 | 0 | 140 | 55 | 49 | 96 | 113 | 0 | 0 | 2 | 0 | 0 | 298 | 149 | 1 | 1 | 1 | 0.915 | 1 | 1 | 1 | 1 | 1 | 0.925 | 1 | 1 | -0.002 |
| SE | 58829 | ENSG00000164828 | SUN1 | chr7 | + | 849519 | 849603 | 843340 | 843520 | 851383 | 851482 | 0 | 0 | 1 | 0 | 0 | 0 | 3 | 0 | 0 | 0 | 0 | 0 | 0 | 0 | 0 | 0 | 0 | 2 | 0 | 0 | 233 | 149 | 1 | 1 | NA | 0 | 1 | NA | NA | NA | NA | 0 | NA | NA | 0.5 |
| SE | 58834 | ENSG00000164828 | SUN1 | chr7 | + | 849922 | 850033 | 843340 | 843520 | 851383 | 851481 | 30 | 11 | 11 | 17 | 16 | 0 | 3 | 0 | 0 | 0 | 12 | 17 | 13 | 10 | 14 | 0 | 0 | 2 | 0 | 0 | 260 | 149 | 1 | 1 | 1 | 0.678 | 1 | 1 | 1 | 1 | 1 | 0.788 | 1 | 1 | -0.022 |
| SE | 58837 | ENSG00000164828 | SUN1 | chr7 | + | 849922 | 850033 | 848417 | 848615 | 851949 | 852043 | 10 | 3 | 2 | 0 | 4 | 1 | 0 | 6 | 0 | 0 | 28 | 2 | 15 | 4 | 9 | 0 | 2 | 1 | 0 | 3 | 260 | 149 | 1 | 1 | 0.851 | 1 | 0.16 | NA | 1 | 1 | 0.364 | 0.896 | 1 | 0.632 | -0.026 |
| SE | 58843 | ENSG00000164828 | SUN1 | chr7 | + | 851383 | 851482 | 849800 | 850033 | 851949 | 852043 | 62 | 24 | 43 | 35 | 28 | 0 | 1 | 0 | 0 | 0 | 43 | 35 | 20 | 28 | 34 | 0 | 0 | 0 | 0 | 0 | 248 | 149 | 1 | 1 | 1 | 0.935 | 1 | 1 | 1 | 1 | 1 | 1 | 1 | 1 | -0.013 |
| SE | 58845 | ENSG00000164828 | SUN1 | chr7 | + | 857827 | 857957 | 856357 | 856401 | 860127 | 860382 | 86 | 72 | 112 | 99 | 85 | 2 | 0 | 1 | 0 | 0 | 205 | 91 | 55 | 89 | 103 | 4 | 0 | 0 | 2 | 0 | 279 | 149 | 1 | 1 | 0.958 | 1 | 0.984 | 1 | 1 | 0.965 | 1 | 1 | 0.96 | 1 | 0.003 |
| SE | 58846 | ENSG00000164828 | SUN1 | chr7 | + | 872469 | 872562 | 868473 | 869516 | 873214 | 874899 | 102 | 104 | 107 | 114 | 131 | 1 | 2 | 0 | 3 | 0 | 264 | 140 | 98 | 165 | 152 | 0 | 0 | 0 | 0 | 0 | 242 | 149 | 1 | 1 | 0.984 | 0.97 | 1 | 0.959 | 1 | 1 | 1 | 1 | 1 | 1 | -0.017 |
| SE | 58855 | ENSG00000182552 | RWDD4 | chr4 | - | 183655880 | 183655961 | 183650983 | 183651131 | 183658928 | 183659178 | 9 | 6 | 5 | 5 | 12 | 0 | 0 | 0 | 0 | 2 | 0 | 5 | 9 | 0 | 0 | 10 | 0 | 0 | 0 | 0 | 230 | 149 | 1 | 1 | 1 | 1 | 1 | 1 | 0.795 | 0 | 1 | 1 | NA | NA | 0.292 |
| SE | 58858 | ENSG00000196154 | S100A4 | chr1 | - | 153544653 | 153544809 | 153543612 | 153543923 | 153545752 | 153545806 | 4425 | 1820 | 2363 | 3087 | 1990 | 0 | 0 | 0 | 0 | 0 | 6126 | 4151 | 2170 | 4664 | 6590 | 0 | 0 | 0 | 0 | 4 | 298 | 149 | 1 | 1 | 1 | 1 | 1 | 1 | 1 | 1 | 1 | 1 | 1 | 0.999 | 0 |
| SE | 58859 | ENSG00000196154 | S100A4 | chr1 | - | 153545469 | 153545518 | 153544653 | 153544809 | 153545752 | 153545802 | 2 | 1 | 1 | 8 | 0 | 1388 | 574 | 779 | 1009 | 364 | 10 | 5 | 2 | 0 | 1 | 2184 | 1490 | 774 | 1670 | 2398 | 198 | 149 | 1 | 1 | 0.001 | 0.001 | 0.001 | 0.006 | 0 | 0.003 | 0.003 | 0.002 | 0 | 0 | 0 |
| SE | 58860 | ENSG00000105402 | NAPA | chr19 | - | 47489710 | 47489761 | 47487636 | 47488389 | 47490787 | 47490856 | 293 | 167 | 238 | 249 | 251 | 0 | 0 | 0 | 0 | 0 | 728 | 484 | 225 | 592 | 892 | 0 | 0 | 0 | 0 | 4 | 200 | 149 | 1 | 1 | 1 | 1 | 1 | 1 | 1 | 1 | 1 | 1 | 1 | 0.994 | 0.001 |
| SE | 58861 | ENSG00000105402 | NAPA | chr19 | - | 47492014 | 47492119 | 47490787 | 47490856 | 47492960 | 47493045 | 547 | 330 | 405 | 485 | 353 | 0 | 0 | 0 | 0 | 0 | 1290 | 898 | 418 | 975 | 1813 | 0 | 4 | 0 | 0 | 9 | 254 | 149 | 1 | 1 | 1 | 1 | 1 | 1 | 1 | 1 | 0.992 | 1 | 1 | 0.992 | 0.003 |
| SE | 58862 | ENSG00000105402 | NAPA | chr19 | - | 47493415 | 47493493 | 47492014 | 47492119 | 47495549 | 47495596 | 61 | 23 | 70 | 52 | 39 | 0 | 0 | 0 | 0 | 0 | 154 | 104 | 90 | 144 | 232 | 0 | 0 | 0 | 2 | 0 | 227 | 149 | 1 | 1 | 1 | 1 | 1 | 1 | 1 | 1 | 1 | 1 | 0.979 | 1 | 0.004 |
| SE | 58863 | ENSG00000105402 | NAPA | chr19 | - | 47493415 | 47493493 | 47492585 | 47493174 | 47495549 | 47495596 | 367 | 228 | 316 | 337 | 338 | 0 | 0 | 0 | 1 | 0 | 926 | 627 | 372 | 798 | 1141 | 0 | 0 | 0 | 4 | 0 | 227 | 149 | 1 | 1 | 1 | 1 | 1 | 0.995 | 1 | 1 | 1 | 1 | 0.992 | 1 | 0.001 |
| SE | 58864 | ENSG00000105402 | NAPA | chr19 | - | 47495549 | 47495596 | 47493070 | 47493493 | 47514842 | 47515022 | 99 | 69 | 86 | 95 | 98 | 0 | 0 | 0 | 0 | 0 | 286 | 183 | 125 | 236 | 360 | 0 | 0 | 0 | 0 | 7 | 196 | 149 | 1 | 1 | 1 | 1 | 1 | 1 | 1 | 1 | 1 | 1 | 1 | 0.975 | 0.005 |
| SE | 58865 | ENSG00000105402 | NAPA | chr19 | - | 47500632 | 47500749 | 47495549 | 47495596 | 47503422 | 47503502 | 337 | 184 | 303 | 360 | 248 | 0 | 0 | 2 | 1 | 1 | 821 | 513 | 343 | 668 | 1138 | 0 | 0 | 0 | 0 | 0 | 266 | 149 | 1 | 1 | 1 | 1 | 0.988 | 0.995 | 0.993 | 1 | 1 | 1 | 1 | 1 | -0.005 |
| SE | 58871 | ENSG00000105402 | NAPA | chr19 | - | 47503422 | 47503502 | 47500632 | 47500749 | 47514842 | 47515020 | 406 | 237 | 323 | 418 | 279 | 2 | 2 | 0 | 2 | 0 | 1077 | 622 | 348 | 726 | 1304 | 0 | 0 | 5 | 3 | 11 | 229 | 149 | 1 | 1 | 0.992 | 0.987 | 1 | 0.993 | 1 | 1 | 1 | 0.978 | 0.994 | 0.987 | 0.003 |
| SE | 58872 | ENSG00000105402 | NAPA | chr19 | - | 47505336 | 47505543 | 47503422 | 47503502 | 47514842 | 47515043 | 0 | 0 | 0 | 0 | 0 | 235 | 132 | 177 | 237 | 161 | 0 | 0 | 0 | 6 | 0 | 695 | 363 | 184 | 399 | 696 | 298 | 149 | 1 | 1 | 0 | 0 | 0 | 0 | 0 | 0 | 0 | 0 | 0.007 | 0 | -0.001 |
| SE | 58873 | ENSG00000105402 | NAPA | chr19 | - | 47506831 | 47506974 | 47503422 | 47503502 | 47514842 | 47515005 | 0 | 0 | 0 | 0 | 0 | 235 | 132 | 177 | 237 | 161 | 0 | 0 | 0 | 5 | 0 | 695 | 363 | 184 | 399 | 696 | 292 | 149 | 1 | 1 | 0 | 0 | 0 | 0 | 0 | 0 | 0 | 0 | 0.006 | 0 | -0.001 |
| SE | 58875 | ENSG00000185252 | ZNF74 | chr22 | + | 20395332 | 20395418 | 20394150 | 20394662 | 20400631 | 20400758 | 15 | 4 | 4 | 6 | 6 | 1 | 0 | 3 | 1 | 0 | 0 | 7 | 4 | 0 | 7 | 0 | 1 | 0 | 4 | 0 | 235 | 149 | 1 | 1 | 0.905 | 1 | 0.458 | 0.792 | 1 | NA | 0.816 | 1 | 0 | 1 | 0.127 |
| SE | 58879 | ENSG00000246430 | LINC00968 | chr8 | - | 56520120 | 56520567 | 56496245 | 56496333 | 56559452 | 56559671 | 2 | 0 | 3 | 2 | 0 | 2 | 1 | 0 | 1 | 3 | 0 | 3 | 1 | 0 | 0 | 0 | 13 | 0 | 2 | 0 | 298 | 149 | 1 | 1 | 0.333 | 0 | 1 | 0.5 | 0 | NA | 0.103 | 1 | 0 | NA | -0.001 |
| SE | 58880 | ENSG00000246430 | LINC00968 | chr8 | - | 56534945 | 56535009 | 56520118 | 56520567 | 56559452 | 56559497 | 1 | 0 | 0 | 1 | 0 | 0 | 0 | 1 | 2 | 0 | 0 | 0 | 0 | 0 | 0 | 0 | 0 | 1 | 0 | 0 | 213 | 149 | 1 | 1 | 1 | NA | 0 | 0.259 | NA | NA | NA | 0 | NA | NA | 0.42 |
| SE | 58884 | ENSG00000151233 | GXYLT1 | chr12 | - | 42105817 | 42106069 | 42097909 | 42098033 | 42109565 | 42109691 | 148 | 109 | 105 | 116 | 154 | 0 | 0 | 0 | 3 | 0 | 49 | 81 | 62 | 40 | 19 | 0 | 0 | 0 | 0 | 0 | 298 | 149 | 1 | 1 | 1 | 1 | 1 | 0.951 | 1 | 1 | 1 | 1 | 1 | 1 | -0.01 |
| SE | 58885 | ENSG00000151233 | GXYLT1 | chr12 | - | 42109565 | 42109691 | 42097909 | 42098033 | 42118999 | 42119171 | 27 | 29 | 16 | 30 | 56 | 0 | 0 | 0 | 1 | 0 | 0 | 2 | 15 | 7 | 0 | 0 | 0 | 0 | 0 | 0 | 275 | 149 | 1 | 1 | 1 | 1 | 1 | 0.942 | 1 | NA | 1 | 1 | 1 | NA | -0.012 |
| SE | 58886 | ENSG00000151233 | GXYLT1 | chr12 | - | 42129758 | 42129851 | 42118999 | 42119171 | 42144425 | 42144874 | 7 | 9 | 4 | 9 | 13 | 4 | 4 | 2 | 6 | 0 | 0 | 0 | 6 | 5 | 0 | 0 | 0 | 2 | 0 | 0 | 242 | 149 | 1 | 1 | 0.519 | 0.581 | 0.552 | 0.48 | 1 | NA | NA | 0.649 | 1 | NA | -0.198 |
| SE | 58888 | ENSG00000143156 | NME7 | chr1 | - | 169190642 | 169190705 | 169169446 | 169169554 | 169230717 | 169230819 | 0 | 3 | 0 | 6 | 4 | 4 | 5 | 9 | 7 | 11 | 0 | 0 | 2 | 0 | 0 | 0 | 2 | 0 | 0 | 0 | 212 | 149 | 1 | 1 | 0 | 0.297 | 0 | 0.376 | 0.204 | NA | 0 | 1 | NA | NA | -0.325 |
| SE | 58890 | ENSG00000143156 | NME7 | chr1 | - | 169287302 | 169287408 | 169237622 | 169237687 | 169298555 | 169298763 | 18 | 14 | 24 | 40 | 15 | 0 | 2 | 2 | 0 | 2 | 0 | 4 | 0 | 10 | 15 | 0 | 0 | 4 | 0 | 0 | 255 | 149 | 1 | 1 | 1 | 0.804 | 0.875 | 1 | 0.814 | NA | 1 | 0 | 1 | 1 | 0.149 |
| SE | 58891 | ENSG00000143156 | NME7 | chr1 | - | 169301969 | 169302053 | 169298604 | 169298763 | 169303144 | 169303195 | 0 | 0 | 0 | 1 | 0 | 5 | 18 | 17 | 21 | 11 | 0 | 0 | 0 | 0 | 0 | 8 | 4 | 16 | 4 | 24 | 233 | 149 | 1 | 1 | 0 | 0 | 0 | 0.03 | 0 | 0 | 0 | 0 | 0 | 0 | 0.006 |
| SE | 58893 | ENSG00000143156 | NME7 | chr1 | - | 169323116 | 169323283 | 169309969 | 169310080 | 169324392 | 169324450 | 54 | 67 | 56 | 98 | 88 | 0 | 0 | 0 | 0 | 2 | 17 | 41 | 47 | 4 | 19 | 0 | 0 | 0 | 0 | 0 | 298 | 149 | 1 | 1 | 1 | 1 | 1 | 1 | 0.957 | 1 | 1 | 1 | 1 | 1 | -0.009 |
| SE | 58894 | ENSG00000143156 | NME7 | chr1 | - | 169324392 | 169324500 | 169309969 | 169310080 | 169367707 | 169367730 | 25 | 21 | 21 | 17 | 16 | 0 | 0 | 0 | 0 | 1 | 10 | 10 | 17 | 6 | 0 | 0 | 0 | 0 | 0 | 0 | 257 | 149 | 1 | 1 | 1 | 1 | 1 | 1 | 0.903 | 1 | 1 | 1 | 1 | NA | -0.019 |
| SE | 58895 | ENSG00000143297 | FCRL5 | chr1 | - | 157520430 | 157520547 | 157519742 | 157519770 | 157521016 | 157521292 | 2 | 0 | 7 | 27 | 17 | 0 | 0 | 0 | 1 | 0 | 1 | 38 | 41 | 18 | 2 | 0 | 4 | 0 | 0 | 0 | 266 | 149 | 1 | 1 | 1 | NA | 1 | 0.938 | 1 | 1 | 0.842 | 1 | 1 | 1 | 0.016 |
| SE | 58896 | ENSG00000143297 | FCRL5 | chr1 | - | 157520430 | 157520561 | 157519742 | 157519770 | 157521016 | 157522794 | 1 | 0 | 5 | 10 | 10 | 0 | 0 | 0 | 1 | 0 | 1 | 31 | 16 | 9 | 1 | 0 | 4 | 0 | 0 | 0 | 280 | 149 | 1 | 1 | 1 | NA | 1 | 0.842 | 1 | 1 | 0.805 | 1 | 1 | 1 | 0 |
| SE | 58898 | ENSG00000143297 | FCRL5 | chr1 | - | 157534613 | 157534892 | 157527616 | 157527895 | 157539085 | 157539364 | 12 | 8 | 8 | 14 | 15 | 1 | 0 | 0 | 1 | 0 | 11 | 11 | 29 | 10 | 0 | 0 | 0 | 0 | 0 | 0 | 298 | 149 | 1 | 1 | 0.857 | 1 | 1 | 0.875 | 1 | 1 | 1 | 1 | 1 | NA | -0.054 |
| SE | 58899 | ENSG00000143297 | FCRL5 | chr1 | - | 157542858 | 157543137 | 157539085 | 157539364 | 157544261 | 157544546 | 12 | 10 | 14 | 23 | 36 | 0 | 0 | 0 | 0 | 1 | 41 | 40 | 17 | 11 | 12 | 0 | 0 | 0 | 0 | 0 | 298 | 149 | 1 | 1 | 1 | 1 | 1 | 1 | 0.947 | 1 | 1 | 1 | 1 | 1 | -0.011 |
| SE | 58901 | ENSG00000143297 | FCRL5 | chr1 | - | 157546187 | 157546280 | 157544830 | 157545082 | 157549559 | 157549580 | 3 | 6 | 1 | 5 | 1 | 0 | 0 | 0 | 0 | 0 | 19 | 9 | 13 | 0 | 10 | 0 | 0 | 1 | 0 | 0 | 242 | 149 | 1 | 1 | 1 | 1 | 1 | 1 | 1 | 1 | 1 | 0.889 | NA | 1 | 0.028 |
| SE | 58902 | ENSG00000143297 | FCRL5 | chr1 | - | 157546942 | 157547197 | 157544830 | 157545082 | 157549559 | 157549580 | 35 | 16 | 27 | 28 | 33 | 0 | 0 | 0 | 0 | 0 | 52 | 56 | 55 | 24 | 50 | 0 | 0 | 1 | 0 | 0 | 298 | 149 | 1 | 1 | 1 | 1 | 1 | 1 | 1 | 1 | 1 | 0.965 | 1 | 1 | 0.007 |
| SE | 58904 | ENSG00000109016 | DHRS7B | chr17 | + | 21172017 | 21172196 | 21126651 | 21126991 | 21183593 | 21183810 | 22 | 1 | 8 | 12 | 16 | 2 | 0 | 0 | 0 | 0 | 43 | 8 | 4 | 8 | 14 | 0 | 0 | 0 | 0 | 0 | 298 | 149 | 1 | 1 | 0.846 | 1 | 1 | 1 | 1 | 1 | 1 | 1 | 1 | 1 | -0.031 |
| SE | 58905 | ENSG00000109016 | DHRS7B | chr17 | + | 21178232 | 21178342 | 21171673 | 21172196 | 21183593 | 21183663 | 30 | 3 | 11 | 17 | 15 | 0 | 0 | 0 | 0 | 0 | 38 | 11 | 5 | 6 | 3 | 0 | 3 | 0 | 0 | 0 | 259 | 149 | 1 | 1 | 1 | 1 | 1 | 1 | 1 | 1 | 0.678 | 1 | 1 | 1 | 0.064 |
| SE | 58906 | ENSG00000109016 | DHRS7B | chr17 | + | 21184370 | 21184463 | 21183593 | 21183810 | 21188710 | 21188863 | 12 | 1 | 15 | 14 | 19 | 1 | 1 | 0 | 0 | 0 | 21 | 14 | 8 | 17 | 20 | 0 | 0 | 0 | 0 | 0 | 242 | 149 | 1 | 1 | 0.881 | 0.381 | 1 | 1 | 1 | 1 | 1 | 1 | 1 | 1 | -0.148 |
| SE | 58907 | ENSG00000109016 | DHRS7B | chr17 | + | 21188710 | 21188863 | 21184370 | 21184463 | 21190947 | 21191266 | 25 | 11 | 37 | 24 | 46 | 1 | 0 | 0 | 0 | 0 | 33 | 24 | 14 | 27 | 33 | 0 | 0 | 0 | 0 | 0 | 298 | 149 | 1 | 1 | 0.926 | 1 | 1 | 1 | 1 | 1 | 1 | 1 | 1 | 1 | -0.015 |
| SE | 58908 | ENSG00000127957 | PMS2P3 | chr7 | - | 75510930 | 75511114 | 75507746 | 75507867 | 75512308 | 75512421 | 93 | 50 | 51 | 85 | 212 | 0 | 0 | 0 | 1 | 0 | 62 | 70 | 64 | 76 | 87 | 0 | 0 | 0 | 0 | 0 | 298 | 149 | 1 | 1 | 1 | 1 | 1 | 0.977 | 1 | 1 | 1 | 1 | 1 | 1 | -0.005 |
| SE | 58911 | ENSG00000127957 | PMS2P3 | chr7 | - | 75525904 | 75525963 | 75516093 | 75516220 | 75527564 | 75527579 | 30 | 16 | 8 | 21 | 13 | 1 | 0 | 0 | 0 | 1 | 32 | 30 | 35 | 49 | 26 | 0 | 0 | 0 | 0 | 0 | 208 | 149 | 1 | 1 | 0.956 | 1 | 1 | 1 | 0.903 | 1 | 1 | 1 | 1 | 1 | -0.028 |
| SE | 58912 | ENSG00000127957 | PMS2P3 | chr7 | - | 75525904 | 75526007 | 75516093 | 75516220 | 75527564 | 75528088 | 31 | 17 | 5 | 26 | 11 | 1 | 0 | 0 | 0 | 1 | 22 | 24 | 34 | 31 | 33 | 0 | 0 | 0 | 0 | 0 | 252 | 149 | 1 | 1 | 0.948 | 1 | 1 | 1 | 0.867 | 1 | 1 | 1 | 1 | 1 | -0.037 |
| SE | 58913 | ENSG00000158428 | CATIP | chr2 | + | 218362734 | 218362902 | 218360572 | 218360659 | 218364627 | 218364752 | 2 | 1 | 1 | 2 | 0 | 1 | 0 | 0 | 0 | 0 | 0 | 2 | 0 | 1 | 11 | 0 | 0 | 0 | 3 | 0 | 298 | 149 | 1 | 1 | 0.5 | 1 | 1 | 1 | NA | NA | 1 | NA | 0.143 | 1 | 0.161 |
| SE | 58916 | ENSG00000135336 | ORC3 | chr6 | + | 87594526 | 87594613 | 87594352 | 87594407 | 87603383 | 87603528 | 8 | 17 | 12 | 5 | 21 | 1 | 0 | 0 | 1 | 5 | 0 | 6 | 12 | 4 | 0 | 0 | 4 | 0 | 0 | 0 | 236 | 149 | 1 | 1 | 0.835 | 1 | 1 | 0.759 | 0.726 | NA | 0.486 | 1 | 1 | NA | 0.035 |
| SE | 58917 | ENSG00000135336 | ORC3 | chr6 | + | 87601783 | 87601881 | 87594352 | 87594407 | 87603383 | 87603528 | 143 | 97 | 186 | 152 | 177 | 1 | 0 | 0 | 1 | 5 | 35 | 159 | 82 | 39 | 37 | 0 | 4 | 0 | 0 | 0 | 247 | 149 | 1 | 1 | 0.989 | 1 | 1 | 0.989 | 0.955 | 1 | 0.96 | 1 | 1 | 1 | -0.005 |
| SE | 58918 | ENSG00000135336 | ORC3 | chr6 | + | 87601783 | 87601881 | 87594526 | 87594613 | 87603383 | 87603528 | 82 | 62 | 113 | 80 | 102 | 1 | 4 | 1 | 0 | 0 | 10 | 107 | 51 | 21 | 19 | 0 | 0 | 0 | 0 | 0 | 247 | 149 | 1 | 1 | 0.98 | 0.903 | 0.986 | 1 | 1 | 1 | 1 | 1 | 1 | 1 | -0.026 |
| SE | 58919 | ENSG00000135336 | ORC3 | chr6 | + | 87603383 | 87603528 | 87601783 | 87601881 | 87605916 | 87606021 | 195 | 162 | 254 | 209 | 262 | 0 | 0 | 0 | 1 | 0 | 66 | 197 | 108 | 59 | 26 | 0 | 0 | 0 | 0 | 0 | 294 | 149 | 1 | 1 | 1 | 1 | 1 | 0.991 | 1 | 1 | 1 | 1 | 1 | 1 | -0.002 |
| SE | 58920 | ENSG00000135336 | ORC3 | chr6 | + | 87612088 | 87612248 | 87609095 | 87609229 | 87616313 | 87616427 | 245 | 183 | 270 | 221 | 536 | 3 | 0 | 0 | 0 | 0 | 26 | 134 | 108 | 36 | 22 | 0 | 0 | 0 | 0 | 0 | 298 | 149 | 1 | 1 | 0.976 | 1 | 1 | 1 | 1 | 1 | 1 | 1 | 1 | 1 | -0.005 |
| SE | 58922 | ENSG00000135336 | ORC3 | chr6 | + | 87634844 | 87634961 | 87621949 | 87622013 | 87636406 | 87636486 | 139 | 98 | 138 | 112 | 226 | 1 | 1 | 0 | 3 | 0 | 7 | 79 | 19 | 3 | 9 | 0 | 0 | 0 | 0 | 0 | 266 | 149 | 1 | 1 | 0.987 | 0.982 | 1 | 0.954 | 1 | 1 | 1 | 1 | 1 | 1 | -0.015 |
| SE | 58923 | ENSG00000135336 | ORC3 | chr6 | + | 87634844 | 87634961 | 87621949 | 87622013 | 87653115 | 87653249 | 78 | 42 | 84 | 57 | 83 | 2 | 1 | 5 | 0 | 0 | 7 | 34 | 15 | 0 | 9 | 0 | 0 | 2 | 0 | 0 | 266 | 149 | 1 | 1 | 0.956 | 0.959 | 0.904 | 1 | 1 | 1 | 1 | 0.808 | NA | 1 | 0.012 |
| SE | 58924 | ENSG00000135336 | ORC3 | chr6 | + | 87636406 | 87636486 | 87621949 | 87622013 | 87653115 | 87653249 | 36 | 38 | 45 | 39 | 114 | 2 | 1 | 5 | 0 | 0 | 0 | 17 | 18 | 0 | 0 | 0 | 0 | 2 | 0 | 0 | 229 | 149 | 1 | 1 | 0.921 | 0.961 | 0.854 | 1 | 1 | NA | 1 | 0.854 | NA | NA | 0.02 |
| SE | 58925 | ENSG00000135336 | ORC3 | chr6 | + | 87636406 | 87636486 | 87634844 | 87634961 | 87653115 | 87653249 | 88 | 91 | 94 | 88 | 243 | 0 | 0 | 0 | 1 | 0 | 0 | 62 | 21 | 3 | 0 | 0 | 0 | 0 | 0 | 0 | 229 | 149 | 1 | 1 | 1 | 1 | 1 | 0.983 | 1 | NA | 1 | 1 | 1 | NA | -0.003 |
| SE | 58926 | ENSG00000135336 | ORC3 | chr6 | + | 87653115 | 87653249 | 87636406 | 87636486 | 87656902 | 87656982 | 68 | 79 | 82 | 56 | 219 | 2 | 0 | 0 | 0 | 0 | 0 | 53 | 28 | 6 | 0 | 0 | 0 | 0 | 0 | 0 | 283 | 149 | 1 | 1 | 0.947 | 1 | 1 | 1 | 1 | NA | 1 | 1 | 1 | NA | -0.011 |
| SE | 58927 | ENSG00000135336 | ORC3 | chr6 | + | 87656902 | 87656982 | 87653115 | 87653249 | 87657920 | 87658018 | 38 | 47 | 45 | 42 | 132 | 0 | 4 | 0 | 0 | 2 | 0 | 31 | 12 | 8 | 0 | 0 | 0 | 1 | 0 | 0 | 229 | 149 | 1 | 1 | 1 | 0.884 | 1 | 1 | 0.977 | NA | 1 | 0.886 | 1 | NA | 0.01 |
| SE | 58928 | ENSG00000135336 | ORC3 | chr6 | + | 87656905 | 87656982 | 87653115 | 87653249 | 87657920 | 87658018 | 52 | 53 | 60 | 49 | 158 | 0 | 4 | 0 | 0 | 2 | 0 | 27 | 27 | 7 | 4 | 0 | 0 | 1 | 0 | 0 | 226 | 149 | 1 | 1 | 1 | 0.897 | 1 | 1 | 0.981 | NA | 1 | 0.947 | 1 | 1 | -0.011 |
| SE | 58930 | ENSG00000144655 | CSRNP1 | chr3 | - | 39144136 | 39144451 | 39141854 | 39144044 | 39144996 | 39145256 | 69 | 30 | 105 | 157 | 80 | 0 | 1 | 1 | 1 | 0 | 320 | 143 | 114 | 234 | 177 | 0 | 0 | 0 | 1 | 0 | 298 | 149 | 1 | 1 | 1 | 0.938 | 0.981 | 0.987 | 1 | 1 | 1 | 1 | 0.992 | 1 | -0.017 |
| SE | 58934 | ENSG00000288560 | NA | chr6 | + | 113064695 | 113064849 | 113050373 | 113050513 | 113080103 | 113080212 | 1 | 2 | 0 | 0 | 3 | 0 | 1 | 0 | 0 | 0 | 0 | 22 | 0 | 13 | 0 | 0 | 0 | 0 | 0 | 0 | 298 | 149 | 1 | 1 | 1 | 0.5 | NA | NA | 1 | NA | 1 | NA | 1 | NA | -0.167 |
| SE | 58935 | ENSG00000288560 | NA | chr6 | + | 113065242 | 113065341 | 113064695 | 113064849 | 113080103 | 113080212 | 1 | 1 | 0 | 0 | 0 | 0 | 1 | 0 | 0 | 0 | 0 | 8 | 0 | 3 | 0 | 0 | 8 | 0 | 2 | 0 | 248 | 149 | 1 | 1 | 1 | 0.375 | NA | NA | NA | NA | 0.375 | NA | 0.474 | NA | 0.263 |
| SE | 58938 | ENSG00000131778 | CHD1L | chr1 | + | 147252622 | 147252735 | 147242676 | 147242830 | 147255812 | 147255927 | 3 | 0 | 1 | 4 | 0 | 1 | 0 | 0 | 0 | 0 | 0 | 1 | 0 | 0 | 18 | 0 | 0 | 0 | 3 | 6 | 262 | 149 | 1 | 1 | 0.63 | NA | 1 | 1 | NA | NA | 1 | NA | 0 | 0.63 | 0.333 |
| SE | 58947 | ENSG00000131778 | CHD1L | chr1 | + | 147264428 | 147264584 | 147242680 | 147242830 | 147265931 | 147266087 | 63 | 66 | 48 | 71 | 78 | 0 | 0 | 0 | 0 | 1 | 78 | 78 | 58 | 101 | 228 | 0 | 0 | 0 | 0 | 0 | 298 | 149 | 1 | 1 | 1 | 1 | 1 | 1 | 0.975 | 1 | 1 | 1 | 1 | 1 | -0.005 |
| SE | 58952 | ENSG00000131778 | CHD1L | chr1 | + | 147272170 | 147272281 | 147270931 | 147271005 | 147275353 | 147275409 | 70 | 39 | 67 | 56 | 68 | 0 | 0 | 1 | 0 | 0 | 83 | 64 | 38 | 57 | 181 | 0 | 0 | 0 | 0 | 0 | 260 | 149 | 1 | 1 | 1 | 1 | 0.975 | 1 | 1 | 1 | 1 | 1 | 1 | 1 | -0.005 |
| SE | 58953 | ENSG00000131778 | CHD1L | chr1 | + | 147272170 | 147272630 | 147267425 | 147267518 | 147275353 | 147275468 | 4 | 1 | 1 | 2 | 2 | 0 | 0 | 0 | 0 | 1 | 0 | 7 | 3 | 8 | 0 | 0 | 0 | 6 | 0 | 0 | 298 | 149 | 1 | 1 | 1 | 1 | 1 | 1 | 0.5 | NA | 1 | 0.2 | 1 | NA | 0.167 |
| SE | 58957 | ENSG00000090889 | KIF4A | chrX | + | 70296997 | 70297188 | 70290690 | 70290805 | 70299112 | 70299202 | 0 | 1 | 3 | 10 | 2 | 0 | 0 | 0 | 0 | 0 | 6 | 22 | 9 | 23 | 0 | 0 | 0 | 0 | 5 | 0 | 298 | 149 | 1 | 1 | NA | 1 | 1 | 1 | 1 | 1 | 1 | 1 | 0.697 | NA | 0.076 |
| SE | 58958 | ENSG00000090889 | KIF4A | chrX | + | 70343702 | 70343761 | 70341798 | 70341931 | 70343876 | 70343982 | 6 | 2 | 1 | 3 | 7 | 0 | 0 | 0 | 2 | 0 | 5 | 4 | 5 | 29 | 0 | 0 | 0 | 4 | 0 | 0 | 208 | 149 | 1 | 1 | 1 | 1 | 1 | 0.518 | 1 | 1 | 1 | 0.472 | 1 | NA | 0.036 |
| SE | 58959 | ENSG00000090889 | KIF4A | chrX | + | 70419660 | 70419783 | 70417887 | 70418004 | 70420061 | 70420886 | 1 | 1 | 6 | 6 | 7 | 0 | 0 | 0 | 0 | 0 | 10 | 10 | 10 | 42 | 7 | 0 | 0 | 0 | 2 | 0 | 272 | 149 | 1 | 1 | 1 | 1 | 1 | 1 | 1 | 1 | 1 | 1 | 0.92 | 1 | 0.016 |
| SE | 58960 | ENSG00000197044 | ZNF441 | chr19 | + | 11778115 | 11778179 | 11777610 | 11777737 | 11778329 | 11778393 | 1 | 0 | 2 | 0 | 0 | 22 | 6 | 23 | 4 | 17 | 0 | 0 | 0 | 0 | 0 | 0 | 10 | 1 | 0 | 0 | 213 | 149 | 1 | 1 | 0.031 | 0 | 0.057 | 0 | 0 | NA | 0 | 0 | NA | NA | 0.018 |
| SE | 59021 | ENSG00000159348 | CYB5R1 | chr1 | - | 202965370 | 202965500 | 202964611 | 202964695 | 202965886 | 202965993 | 161 | 86 | 115 | 125 | 145 | 0 | 0 | 0 | 1 | 0 | 226 | 262 | 95 | 139 | 551 | 0 | 4 | 0 | 1 | 0 | 279 | 149 | 1 | 1 | 1 | 1 | 1 | 0.985 | 1 | 1 | 0.972 | 1 | 0.987 | 1 | 0.005 |
| SE | 59022 | ENSG00000159348 | CYB5R1 | chr1 | - | 202966527 | 202966600 | 202965886 | 202965993 | 202966748 | 202966898 | 98 | 27 | 36 | 48 | 66 | 2 | 0 | 2 | 0 | 0 | 54 | 82 | 45 | 49 | 265 | 0 | 0 | 0 | 0 | 0 | 222 | 149 | 1 | 1 | 0.97 | 1 | 0.924 | 1 | 1 | 1 | 1 | 1 | 1 | 1 | -0.021 |
| SE | 59023 | ENSG00000159348 | CYB5R1 | chr1 | - | 202966527 | 202966600 | 202965886 | 202966109 | 202966748 | 202966898 | 27 | 13 | 5 | 15 | 22 | 0 | 0 | 1 | 1 | 0 | 13 | 26 | 13 | 19 | 63 | 0 | 0 | 1 | 0 | 0 | 222 | 149 | 1 | 1 | 1 | 1 | 0.77 | 0.91 | 1 | 1 | 1 | 0.897 | 1 | 1 | -0.043 |
| SE | 59024 | ENSG00000159348 | CYB5R1 | chr1 | - | 202966748 | 202966898 | 202965886 | 202965993 | 202967190 | 202967225 | 44 | 16 | 24 | 28 | 39 | 0 | 0 | 0 | 0 | 0 | 21 | 66 | 23 | 27 | 135 | 0 | 0 | 2 | 0 | 0 | 298 | 149 | 1 | 1 | 1 | 1 | 1 | 1 | 1 | 1 | 1 | 0.852 | 1 | 1 | 0.03 |
| SE | 59025 | ENSG00000159348 | CYB5R1 | chr1 | - | 202966748 | 202966898 | 202965886 | 202966109 | 202967190 | 202967225 | 42 | 16 | 23 | 29 | 39 | 1 | 0 | 0 | 0 | 0 | 21 | 66 | 24 | 27 | 135 | 0 | 0 | 0 | 0 | 0 | 298 | 149 | 1 | 1 | 0.955 | 1 | 1 | 1 | 1 | 1 | 1 | 1 | 1 | 1 | -0.009 |
| SE | 59030 | ENSG00000234810 | NA | chr1 | + | 55673259 | 55673317 | 55614920 | 55615002 | 55737646 | 55738391 | 1 | 2 | 0 | 2 | 3 | 0 | 1 | 0 | 5 | 3 | 0 | 0 | 0 | 2 | 9 | 2 | 0 | 0 | 2 | 8 | 207 | 149 | 1 | 1 | 1 | 0.59 | NA | 0.224 | 0.419 | 0 | NA | NA | 0.419 | 0.447 | 0.27 |
| SE | 59034 | ENSG00000166984 | TCP10L2 | chr6 | + | 167149956 | 167150073 | 167149479 | 167149484 | 167164400 | 167164560 | 0 | 0 | 0 | 0 | 0 | 0 | 0 | 0 | 2 | 2 | 0 | 10 | 0 | 0 | 0 | 0 | 0 | 0 | 0 | 0 | 266 | 149 | 1 | 1 | NA | NA | NA | 0 | 0 | NA | 1 | NA | NA | NA | -1 |
| SE | 59036 | ENSG00000019186 | CYP24A1 | chr20 | - | 54172908 | 54173099 | 54171576 | 54171670 | 54173321 | 54173649 | 1 | 0 | 0 | 0 | 0 | 0 | 0 | 0 | 0 | 0 | 0 | 0 | 0 | 3 | 0 | 0 | 0 | 0 | 0 | 10 | 298 | 149 | 1 | 1 | 1 | NA | NA | NA | NA | NA | NA | NA | 1 | 0 | 0.5 |
| SE | 59042 | ENSG00000147813 | NAPRT | chr8 | - | 143575190 | 143575345 | 143574786 | 143575093 | 143575422 | 143575525 | 86 | 14 | 76 | 38 | 9 | 1 | 0 | 0 | 0 | 0 | 50 | 48 | 18 | 62 | 88 | 0 | 0 | 0 | 0 | 0 | 298 | 149 | 1 | 1 | 0.977 | 1 | 1 | 1 | 1 | 1 | 1 | 1 | 1 | 1 | -0.005 |
| SE | 59043 | ENSG00000147813 | NAPRT | chr8 | - | 143575229 | 143575345 | 143574784 | 143575093 | 143575422 | 143575525 | 37 | 4 | 33 | 13 | 1 | 1 | 0 | 0 | 0 | 0 | 32 | 14 | 11 | 25 | 28 | 0 | 0 | 0 | 0 | 0 | 265 | 149 | 1 | 1 | 0.954 | 1 | 1 | 1 | 1 | 1 | 1 | 1 | 1 | 1 | -0.009 |
| SE | 59044 | ENSG00000147813 | NAPRT | chr8 | - | 143575621 | 143575702 | 143575422 | 143575525 | 143576077 | 143576162 | 59 | 8 | 39 | 16 | 2 | 0 | 0 | 1 | 1 | 0 | 61 | 37 | 15 | 61 | 65 | 0 | 0 | 0 | 0 | 0 | 230 | 149 | 1 | 1 | 1 | 1 | 0.962 | 0.912 | 1 | 1 | 1 | 1 | 1 | 1 | -0.025 |
| SE | 59045 | ENSG00000147813 | NAPRT | chr8 | - | 143576645 | 143576842 | 143576431 | 143576572 | 143577268 | 143577399 | 23 | 7 | 10 | 10 | 2 | 0 | 0 | 0 | 0 | 0 | 100 | 24 | 5 | 31 | 58 | 0 | 0 | 1 | 0 | 0 | 298 | 149 | 1 | 1 | 1 | 1 | 1 | 1 | 1 | 1 | 1 | 0.714 | 1 | 1 | 0.057 |
| SE | 59046 | ENSG00000147813 | NAPRT | chr8 | - | 143577061 | 143577177 | 143576645 | 143576842 | 143577268 | 143577399 | 4 | 0 | 4 | 9 | 0 | 1 | 0 | 0 | 0 | 0 | 0 | 10 | 1 | 0 | 16 | 20 | 2 | 0 | 0 | 0 | 265 | 149 | 1 | 1 | 0.692 | NA | 1 | 1 | NA | 0 | 0.738 | 1 | NA | 1 | 0.213 |
| SE | 59061 | ENSG00000103365 | GGA2 | chr16 | - | 23474903 | 23475061 | 23469995 | 23470165 | 23478367 | 23478501 | 240 | 133 | 268 | 224 | 211 | 0 | 0 | 0 | 0 | 0 | 187 | 259 | 112 | 108 | 294 | 0 | 0 | 0 | 0 | 7 | 298 | 149 | 1 | 1 | 1 | 1 | 1 | 1 | 1 | 1 | 1 | 1 | 1 | 0.955 | 0.009 |
| SE | 59062 | ENSG00000103365 | GGA2 | chr16 | - | 23478367 | 23478501 | 23474903 | 23475061 | 23478882 | 23478911 | 199 | 123 | 163 | 170 | 149 | 1 | 0 | 3 | 1 | 0 | 210 | 232 | 85 | 93 | 329 | 0 | 0 | 0 | 0 | 0 | 283 | 149 | 1 | 1 | 0.991 | 1 | 0.966 | 0.989 | 1 | 1 | 1 | 1 | 1 | 1 | -0.011 |
| SE | 59063 | ENSG00000103365 | GGA2 | chr16 | - | 23478882 | 23478911 | 23478367 | 23478501 | 23479764 | 23479887 | 120 | 67 | 88 | 130 | 94 | 0 | 0 | 0 | 0 | 2 | 225 | 129 | 57 | 53 | 249 | 0 | 2 | 0 | 3 | 0 | 178 | 149 | 1 | 1 | 1 | 1 | 1 | 1 | 0.975 | 1 | 0.982 | 1 | 0.937 | 1 | 0.011 |
| SE | 59064 | ENSG00000103365 | GGA2 | chr16 | - | 23480644 | 23480770 | 23479764 | 23479887 | 23482922 | 23483004 | 168 | 91 | 161 | 191 | 127 | 1 | 0 | 0 | 0 | 1 | 276 | 228 | 146 | 94 | 309 | 0 | 0 | 0 | 0 | 0 | 275 | 149 | 1 | 1 | 0.989 | 1 | 1 | 1 | 0.986 | 1 | 1 | 1 | 1 | 1 | -0.005 |
| SE | 59065 | ENSG00000103365 | GGA2 | chr16 | - | 23491676 | 23491800 | 23488605 | 23488709 | 23495693 | 23495778 | 230 | 125 | 197 | 257 | 476 | 1 | 0 | 1 | 0 | 0 | 544 | 354 | 230 | 378 | 486 | 0 | 0 | 0 | 0 | 0 | 273 | 149 | 1 | 1 | 0.992 | 1 | 0.991 | 1 | 1 | 1 | 1 | 1 | 1 | 1 | -0.003 |
| SE | 59067 | ENSG00000103365 | GGA2 | chr16 | - | 23493359 | 23493458 | 23491676 | 23491800 | 23495693 | 23495772 | 166 | 73 | 129 | 138 | 307 | 0 | 0 | 1 | 0 | 0 | 338 | 253 | 153 | 235 | 275 | 0 | 0 | 0 | 0 | 0 | 248 | 149 | 1 | 1 | 1 | 1 | 0.987 | 1 | 1 | 1 | 1 | 1 | 1 | 1 | -0.003 |
| SE | 59069 | ENSG00000103365 | GGA2 | chr16 | - | 23493359 | 23494378 | 23491675 | 23491800 | 23495693 | 23495778 | 200 | 96 | 151 | 156 | 310 | 0 | 0 | 1 | 0 | 0 | 350 | 279 | 169 | 255 | 312 | 0 | 0 | 0 | 0 | 0 | 298 | 149 | 1 | 1 | 1 | 1 | 0.987 | 1 | 1 | 1 | 1 | 1 | 1 | 1 | -0.003 |
| SE | 59070 | ENSG00000103365 | GGA2 | chr16 | - | 23494298 | 23494378 | 23493359 | 23493458 | 23495693 | 23495772 | 25 | 13 | 12 | 9 | 2 | 0 | 1 | 0 | 0 | 0 | 2 | 4 | 8 | 16 | 18 | 0 | 0 | 0 | 0 | 0 | 229 | 149 | 1 | 1 | 1 | 0.894 | 1 | 1 | 1 | 1 | 1 | 1 | 1 | 1 | -0.021 |
| SE | 59071 | ENSG00000103365 | GGA2 | chr16 | - | 23494302 | 23494378 | 23491676 | 23491800 | 23495693 | 23495778 | 25 | 10 | 11 | 9 | 2 | 0 | 0 | 1 | 0 | 0 | 2 | 4 | 8 | 18 | 18 | 0 | 0 | 0 | 0 | 0 | 225 | 149 | 1 | 1 | 1 | 1 | 0.879 | 1 | 1 | 1 | 1 | 1 | 1 | 1 | -0.024 |
| SE | 59072 | ENSG00000103365 | GGA2 | chr16 | - | 23494302 | 23494378 | 23493359 | 23493458 | 23495693 | 23495778 | 137 | 44 | 116 | 134 | 126 | 0 | 1 | 0 | 0 | 0 | 125 | 142 | 80 | 113 | 170 | 0 | 0 | 0 | 0 | 0 | 225 | 149 | 1 | 1 | 1 | 0.967 | 1 | 1 | 1 | 1 | 1 | 1 | 1 | 1 | -0.007 |
| SE | 59074 | ENSG00000181409 | AATK | chr17 | - | 81126657 | 81126817 | 81126445 | 81126560 | 81127582 | 81127670 | 0 | 0 | 0 | 0 | 0 | 4 | 0 | 0 | 0 | 1 | 0 | 0 | 2 | 0 | 0 | 0 | 5 | 3 | 4 | 0 | 298 | 149 | 1 | 1 | 0 | NA | NA | NA | 0 | NA | 0 | 0.25 | 0 | NA | -0.083 |
| SE | 59076 | ENSG00000107819 | SFXN3 | chr10 | + | 101032313 | 101032482 | 101031233 | 101031534 | 101034691 | 101034855 | 68 | 42 | 31 | 54 | 45 | 2 | 0 | 0 | 0 | 0 | 157 | 66 | 32 | 35 | 154 | 0 | 0 | 0 | 0 | 0 | 298 | 149 | 1 | 1 | 0.944 | 1 | 1 | 1 | 1 | 1 | 1 | 1 | 1 | 1 | -0.011 |
| SE | 59077 | ENSG00000107819 | SFXN3 | chr10 | + | 101035496 | 101035667 | 101034691 | 101034855 | 101036002 | 101036101 | 135 | 119 | 123 | 142 | 117 | 0 | 0 | 0 | 0 | 2 | 298 | 176 | 84 | 117 | 394 | 0 | 0 | 0 | 0 | 0 | 298 | 149 | 1 | 1 | 1 | 1 | 1 | 1 | 0.967 | 1 | 1 | 1 | 1 | 1 | -0.007 |
| SE | 59078 | ENSG00000282246 | NA | chr10 | + | 13611614 | 13611683 | 13610046 | 13610185 | 13613740 | 13613881 | 162 | 135 | 152 | 137 | 259 | 0 | 3 | 3 | 0 | 0 | 8 | 124 | 156 | 114 | 6 | 0 | 0 | 1 | 0 | 0 | 218 | 149 | 1 | 1 | 1 | 0.969 | 0.972 | 1 | 1 | 1 | 1 | 0.991 | 1 | 1 | -0.01 |
| SE | 59079 | ENSG00000282246 | NA | chr10 | + | 13611614 | 13611683 | 13610046 | 13610185 | 13614014 | 13614086 | 70 | 52 | 70 | 75 | 88 | 0 | 1 | 0 | 0 | 0 | 0 | 47 | 73 | 53 | 0 | 0 | 3 | 1 | 0 | 0 | 218 | 149 | 1 | 1 | 1 | 0.973 | 1 | 1 | 1 | NA | 0.915 | 0.98 | 1 | NA | 0.03 |
| SE | 59080 | ENSG00000282246 | NA | chr10 | + | 13611614 | 13611683 | 13610046 | 13610185 | 13616397 | 13616553 | 53 | 43 | 50 | 49 | 66 | 0 | 0 | 0 | 1 | 0 | 0 | 45 | 63 | 53 | 0 | 0 | 0 | 0 | 0 | 0 | 218 | 149 | 1 | 1 | 1 | 1 | 1 | 0.971 | 1 | NA | 1 | 1 | 1 | NA | -0.006 |
| SE | 59081 | ENSG00000282246 | NA | chr10 | + | 13613740 | 13613881 | 13610046 | 13610185 | 13614014 | 13614086 | 153 | 115 | 144 | 136 | 154 | 0 | 1 | 0 | 0 | 0 | 5 | 72 | 91 | 47 | 7 | 0 | 3 | 1 | 0 | 0 | 290 | 149 | 1 | 1 | 1 | 0.983 | 1 | 1 | 1 | 1 | 0.925 | 0.979 | 1 | 1 | 0.016 |
| SE | 59083 | ENSG00000282246 | NA | chr10 | + | 13614014 | 13614086 | 13610046 | 13610185 | 13616397 | 13616553 | 110 | 94 | 114 | 128 | 98 | 0 | 0 | 0 | 1 | 0 | 10 | 71 | 70 | 53 | 0 | 0 | 0 | 0 | 0 | 0 | 221 | 149 | 1 | 1 | 1 | 1 | 1 | 0.989 | 1 | 1 | 1 | 1 | 1 | NA | -0.002 |
| SE | 59084 | ENSG00000282246 | NA | chr10 | + | 13614014 | 13614086 | 13611614 | 13611683 | 13616397 | 13616553 | 125 | 102 | 134 | 155 | 118 | 0 | 0 | 0 | 1 | 0 | 10 | 73 | 76 | 53 | 0 | 0 | 0 | 0 | 0 | 0 | 221 | 149 | 1 | 1 | 1 | 1 | 1 | 0.991 | 1 | 1 | 1 | 1 | 1 | NA | -0.002 |
| SE | 59085 | ENSG00000282246 | NA | chr10 | + | 13630259 | 13630465 | 13616530 | 13616553 | 13637996 | 13638120 | 138 | 48 | 119 | 101 | 184 | 1 | 0 | 0 | 0 | 0 | 7 | 43 | 72 | 24 | 8 | 0 | 0 | 0 | 0 | 0 | 298 | 149 | 1 | 1 | 0.986 | 1 | 1 | 1 | 1 | 1 | 1 | 1 | 1 | 1 | -0.003 |
| SE | 59087 | ENSG00000180182 | MED14 | chrX | - | 40654363 | 40654556 | 40648304 | 40651879 | 40654934 | 40655060 | 94 | 67 | 136 | 84 | 110 | 0 | 2 | 0 | 0 | 0 | 28 | 53 | 47 | 47 | 29 | 0 | 0 | 0 | 0 | 0 | 298 | 149 | 1 | 1 | 1 | 0.944 | 1 | 1 | 1 | 1 | 1 | 1 | 1 | 1 | -0.011 |
| SE | 59088 | ENSG00000180182 | MED14 | chrX | - | 40654934 | 40655060 | 40653871 | 40654556 | 40659226 | 40659334 | 90 | 70 | 111 | 64 | 81 | 0 | 0 | 0 | 0 | 0 | 18 | 78 | 50 | 51 | 35 | 0 | 0 | 1 | 0 | 0 | 275 | 149 | 1 | 1 | 1 | 1 | 1 | 1 | 1 | 1 | 1 | 0.964 | 1 | 1 | 0.007 |
| SE | 59091 | ENSG00000180182 | MED14 | chrX | - | 40679863 | 40680133 | 40675220 | 40675361 | 40726745 | 40726851 | 140 | 83 | 101 | 122 | 185 | 0 | 1 | 0 | 0 | 0 | 97 | 115 | 75 | 78 | 11 | 0 | 0 | 0 | 0 | 0 | 298 | 149 | 1 | 1 | 1 | 0.976 | 1 | 1 | 1 | 1 | 1 | 1 | 1 | 1 | -0.005 |
| SE | 59092 | ENSG00000180182 | MED14 | chrX | - | 40680757 | 40680910 | 40679863 | 40680133 | 40681851 | 40681943 | 316 | 219 | 277 | 304 | 613 | 0 | 0 | 0 | 0 | 1 | 192 | 215 | 196 | 170 | 39 | 0 | 0 | 0 | 0 | 0 | 298 | 149 | 1 | 1 | 1 | 1 | 1 | 1 | 0.997 | 1 | 1 | 1 | 1 | 1 | -0.001 |
| SE | 59093 | ENSG00000180182 | MED14 | chrX | - | 40682835 | 40682996 | 40682602 | 40682749 | 40688453 | 40688530 | 150 | 78 | 110 | 155 | 191 | 0 | 1 | 0 | 3 | 0 | 38 | 83 | 83 | 71 | 39 | 0 | 0 | 0 | 0 | 0 | 298 | 149 | 1 | 1 | 1 | 0.975 | 1 | 0.963 | 1 | 1 | 1 | 1 | 1 | 1 | -0.012 |
| SE | 59094 | ENSG00000180182 | MED14 | chrX | - | 40697023 | 40697183 | 40692707 | 40692902 | 40701164 | 40701243 | 144 | 96 | 168 | 112 | 283 | 1 | 0 | 0 | 1 | 0 | 76 | 146 | 120 | 107 | 54 | 0 | 0 | 1 | 0 | 0 | 298 | 149 | 1 | 1 | 0.986 | 1 | 1 | 0.982 | 1 | 1 | 1 | 0.984 | 1 | 1 | -0.003 |
| SE | 59095 | ENSG00000180182 | MED14 | chrX | - | 40701164 | 40701243 | 40697023 | 40697183 | 40703443 | 40703569 | 108 | 62 | 115 | 84 | 217 | 0 | 0 | 0 | 0 | 1 | 24 | 79 | 85 | 64 | 29 | 0 | 0 | 0 | 0 | 0 | 228 | 149 | 1 | 1 | 1 | 1 | 1 | 1 | 0.993 | 1 | 1 | 1 | 1 | 1 | -0.001 |
| SE | 59096 | ENSG00000180182 | MED14 | chrX | - | 40714536 | 40714710 | 40713777 | 40713907 | 40726745 | 40726851 | 174 | 103 | 116 | 149 | 251 | 0 | 1 | 0 | 0 | 0 | 120 | 155 | 159 | 102 | 58 | 0 | 0 | 0 | 0 | 0 | 298 | 149 | 1 | 1 | 1 | 0.981 | 1 | 1 | 1 | 1 | 1 | 1 | 1 | 1 | -0.004 |
| SE | 59101 | ENSG00000214941 | ZSWIM7 | chr17 | - | 15977799 | 15977913 | 15977578 | 15977700 | 15978043 | 15978163 | 8 | 16 | 18 | 9 | 15 | 0 | 0 | 0 | 0 | 0 | 23 | 10 | 4 | 18 | 36 | 0 | 2 | 0 | 0 | 0 | 263 | 149 | 1 | 1 | 1 | 1 | 1 | 1 | 1 | 1 | 0.739 | 1 | 1 | 1 | 0.052 |
| SE | 59102 | ENSG00000214941 | ZSWIM7 | chr17 | - | 15978043 | 15978163 | 15976559 | 15977092 | 15987265 | 15987368 | 27 | 11 | 4 | 5 | 15 | 0 | 3 | 0 | 0 | 0 | 42 | 11 | 2 | 15 | 26 | 0 | 0 | 0 | 0 | 0 | 269 | 149 | 1 | 1 | 1 | 0.67 | 1 | 1 | 1 | 1 | 1 | 1 | 1 | 1 | -0.066 |
| SE | 59104 | ENSG00000214941 | ZSWIM7 | chr17 | - | 15981039 | 15981144 | 15976559 | 15978163 | 15987265 | 15987368 | 102 | 66 | 80 | 72 | 156 | 0 | 1 | 0 | 2 | 2 | 22 | 117 | 41 | 54 | 193 | 0 | 0 | 0 | 0 | 0 | 254 | 149 | 1 | 1 | 1 | 0.975 | 1 | 0.955 | 0.979 | 1 | 1 | 1 | 1 | 1 | -0.018 |
| SE | 59105 | ENSG00000214941 | ZSWIM7 | chr17 | - | 15986799 | 15986886 | 15981039 | 15981144 | 15987265 | 15987368 | 0 | 6 | 0 | 0 | 1 | 47 | 28 | 38 | 50 | 88 | 0 | 0 | 0 | 0 | 7 | 10 | 62 | 22 | 32 | 78 | 236 | 149 | 1 | 1 | 0 | 0.119 | 0 | 0 | 0.007 | 0 | 0 | 0 | 0 | 0.054 | 0.014 |
| SE | 59106 | ENSG00000214941 | ZSWIM7 | chr17 | - | 15986799 | 15986892 | 15977726 | 15978163 | 15987265 | 15987368 | 0 | 0 | 0 | 0 | 2 | 0 | 1 | 0 | 2 | 2 | 0 | 3 | 0 | 3 | 5 | 0 | 0 | 0 | 0 | 0 | 242 | 149 | 1 | 1 | NA | 0 | NA | 0 | 0.381 | NA | 1 | NA | 1 | 1 | -0.873 |
| SE | 59108 | ENSG00000214941 | ZSWIM7 | chr17 | - | 15987265 | 15987368 | 15981039 | 15981144 | 15993756 | 15993778 | 56 | 32 | 48 | 53 | 94 | 0 | 0 | 0 | 0 | 0 | 31 | 64 | 25 | 38 | 105 | 0 | 0 | 2 | 0 | 0 | 252 | 149 | 1 | 1 | 1 | 1 | 1 | 1 | 1 | 1 | 1 | 0.881 | 1 | 1 | 0.024 |
| SE | 59109 | ENSG00000214941 | ZSWIM7 | chr17 | - | 15990864 | 15991104 | 15987265 | 15987368 | 15993756 | 15993778 | 0 | 0 | 2 | 0 | 4 | 36 | 21 | 34 | 38 | 56 | 0 | 0 | 0 | 0 | 0 | 28 | 21 | 12 | 26 | 93 | 298 | 149 | 1 | 1 | 0 | 0 | 0.029 | 0 | 0.034 | 0 | 0 | 0 | 0 | 0 | 0.013 |
| SE | 59111 | ENSG00000214941 | ZSWIM7 | chr17 | - | 15993756 | 15993778 | 15987265 | 15987368 | 15999518 | 15999589 | 24 | 17 | 25 | 31 | 50 | 1 | 0 | 0 | 0 | 0 | 17 | 19 | 9 | 20 | 75 | 0 | 0 | 0 | 0 | 1 | 171 | 149 | 1 | 1 | 0.954 | 1 | 1 | 1 | 1 | 1 | 1 | 1 | 1 | 0.985 | -0.006 |
| SE | 59113 | ENSG00000109674 | NEIL3 | chr4 | + | 177341475 | 177341642 | 177339782 | 177339857 | 177351379 | 177351549 | 2 | 3 | 3 | 5 | 9 | 0 | 0 | 0 | 0 | 1 | 8 | 16 | 8 | 4 | 0 | 0 | 0 | 0 | 0 | 0 | 298 | 149 | 1 | 1 | 1 | 1 | 1 | 1 | 0.818 | 1 | 1 | 1 | 1 | NA | -0.036 |
| SE | 59114 | ENSG00000159658 | EFCAB14 | chr1 | - | 46680059 | 46680138 | 46677894 | 46678636 | 46683299 | 46683425 | 0 | 1 | 0 | 0 | 0 | 191 | 107 | 151 | 120 | 269 | 0 | 0 | 0 | 0 | 0 | 151 | 157 | 108 | 94 | 80 | 228 | 149 | 1 | 1 | 0 | 0.006 | 0 | 0 | 0 | 0 | 0 | 0 | 0 | 0 | 0.001 |
| SE | 59115 | ENSG00000159658 | EFCAB14 | chr1 | - | 46685082 | 46685144 | 46684490 | 46684602 | 46686783 | 46686870 | 1 | 0 | 1 | 0 | 1 | 550 | 400 | 463 | 444 | 1048 | 6 | 2 | 2 | 2 | 0 | 558 | 483 | 388 | 567 | 272 | 211 | 149 | 1 | 1 | 0.001 | 0 | 0.002 | 0 | 0.001 | 0.008 | 0.003 | 0.004 | 0.002 | 0 | -0.003 |
| SE | 59116 | ENSG00000159658 | EFCAB14 | chr1 | - | 46686783 | 46686870 | 46684155 | 46684602 | 46689586 | 46689636 | 683 | 483 | 593 | 533 | 1308 | 1 | 0 | 0 | 0 | 0 | 726 | 640 | 494 | 669 | 399 | 0 | 0 | 0 | 0 | 0 | 236 | 149 | 1 | 1 | 0.998 | 1 | 1 | 1 | 1 | 1 | 1 | 1 | 1 | 1 | 0 |
| SE | 59118 | ENSG00000159658 | EFCAB14 | chr1 | - | 46696550 | 46696649 | 46691826 | 46691937 | 46707905 | 46708051 | 558 | 343 | 423 | 507 | 1052 | 0 | 0 | 0 | 2 | 0 | 562 | 445 | 344 | 440 | 337 | 0 | 0 | 0 | 0 | 0 | 248 | 149 | 1 | 1 | 1 | 1 | 1 | 0.993 | 1 | 1 | 1 | 1 | 1 | 1 | -0.001 |
| SE | 59120 | ENSG00000159658 | EFCAB14 | chr1 | - | 46716294 | 46716443 | 46707235 | 46708051 | 46717902 | 46718661 | 498 | 260 | 405 | 474 | 659 | 0 | 0 | 1 | 1 | 0 | 440 | 378 | 290 | 336 | 186 | 0 | 3 | 7 | 0 | 0 | 298 | 149 | 1 | 1 | 1 | 1 | 0.995 | 0.996 | 1 | 1 | 0.984 | 0.954 | 1 | 1 | 0.011 |
| SE | 59121 | ENSG00000047410 | TPR | chr1 | - | 186318732 | 186318828 | 186318446 | 186318603 | 186320311 | 186320418 | 283 | 173 | 280 | 272 | 344 | 0 | 0 | 0 | 1 | 0 | 30 | 96 | 75 | 15 | 6 | 0 | 0 | 0 | 0 | 0 | 245 | 149 | 1 | 1 | 1 | 1 | 1 | 0.994 | 1 | 1 | 1 | 1 | 1 | 1 | -0.001 |
| SE | 59122 | ENSG00000047410 | TPR | chr1 | - | 186326103 | 186326235 | 186325763 | 186325854 | 186327459 | 186327660 | 400 | 370 | 435 | 472 | 552 | 1 | 0 | 0 | 2 | 0 | 40 | 84 | 54 | 36 | 78 | 0 | 0 | 0 | 0 | 0 | 281 | 149 | 1 | 1 | 0.995 | 1 | 1 | 0.992 | 1 | 1 | 1 | 1 | 1 | 1 | -0.003 |
| SE | 59123 | ENSG00000047410 | TPR | chr1 | - | 186335067 | 186335129 | 186334324 | 186334533 | 186335337 | 186335543 | 272 | 269 | 255 | 327 | 493 | 1 | 2 | 3 | 0 | 8 | 0 | 18 | 55 | 19 | 23 | 0 | 0 | 0 | 0 | 0 | 211 | 149 | 1 | 1 | 0.995 | 0.99 | 0.984 | 1 | 0.978 | NA | 1 | 1 | 1 | 1 | -0.011 |
| SE | 59124 | ENSG00000047410 | TPR | chr1 | - | 186339641 | 186339772 | 186338032 | 186338243 | 186341027 | 186341159 | 414 | 403 | 418 | 444 | 912 | 0 | 0 | 0 | 0 | 1 | 5 | 95 | 52 | 22 | 0 | 0 | 0 | 0 | 0 | 0 | 280 | 149 | 1 | 1 | 1 | 1 | 1 | 1 | 0.998 | 1 | 1 | 1 | 1 | NA | 0 |
| SE | 59125 | ENSG00000047410 | TPR | chr1 | - | 186341251 | 186341389 | 186341027 | 186341159 | 186343325 | 186343473 | 325 | 259 | 268 | 348 | 424 | 1 | 1 | 0 | 0 | 0 | 31 | 79 | 47 | 6 | 0 | 0 | 0 | 1 | 0 | 0 | 287 | 149 | 1 | 1 | 0.994 | 0.993 | 1 | 1 | 1 | 1 | 1 | 0.961 | 1 | NA | 0.007 |
| SE | 59126 | ENSG00000047410 | TPR | chr1 | - | 186350222 | 186350388 | 186347291 | 186347458 | 186351329 | 186351470 | 649 | 745 | 615 | 661 | 1090 | 0 | 1 | 0 | 0 | 0 | 45 | 89 | 66 | 15 | 27 | 0 | 0 | 0 | 0 | 0 | 298 | 149 | 1 | 1 | 1 | 0.997 | 1 | 1 | 1 | 1 | 1 | 1 | 1 | 1 | -0.001 |
| SE | 59129 | ENSG00000047410 | TPR | chr1 | - | 186361621 | 186361709 | 186360764 | 186360905 | 186361788 | 186361869 | 414 | 358 | 395 | 448 | 431 | 0 | 0 | 0 | 4 | 0 | 18 | 76 | 59 | 30 | 10 | 0 | 3 | 0 | 0 | 0 | 237 | 149 | 1 | 1 | 1 | 1 | 1 | 0.986 | 1 | 1 | 0.941 | 1 | 1 | 1 | 0.009 |
| SE | 59130 | ENSG00000047410 | TPR | chr1 | - | 186361788 | 186361869 | 186361621 | 186361709 | 186362287 | 186362380 | 423 | 350 | 373 | 412 | 554 | 0 | 0 | 0 | 0 | 0 | 19 | 73 | 70 | 25 | 5 | 0 | 0 | 0 | 0 | 1 | 230 | 149 | 1 | 1 | 1 | 1 | 1 | 1 | 1 | 1 | 1 | 1 | 1 | 0.764 | 0.047 |
| SE | 59132 | ENSG00000228215 | LINC02770 | chr1 | + | 191964393 | 191964489 | 191823431 | 191823692 | 191965321 | 191965413 | 0 | 0 | 0 | 1 | 0 | 1 | 5 | 2 | 0 | 3 | 0 | 0 | 0 | 0 | 0 | 0 | 0 | 0 | 10 | 0 | 245 | 149 | 1 | 1 | 0 | 0 | 0 | 1 | 0 | NA | NA | NA | 0 | NA | 0.2 |
| SE | 59134 | ENSG00000168421 | RHOH | chr4 | + | 40197100 | 40197300 | 40193662 | 40193812 | 40242713 | 40242834 | 289 | 161 | 349 | 224 | 589 | 1 | 0 | 1 | 1 | 0 | 80 | 224 | 162 | 145 | 118 | 0 | 0 | 0 | 0 | 0 | 298 | 149 | 1 | 1 | 0.993 | 1 | 0.994 | 0.991 | 1 | 1 | 1 | 1 | 1 | 1 | -0.004 |
| SE | 59137 | ENSG00000168421 | RHOH | chr4 | + | 40197120 | 40197300 | 40193662 | 40193812 | 40242713 | 40242834 | 289 | 163 | 347 | 224 | 588 | 1 | 0 | 1 | 1 | 0 | 80 | 228 | 162 | 145 | 118 | 0 | 0 | 0 | 0 | 0 | 298 | 149 | 1 | 1 | 0.993 | 1 | 0.994 | 0.991 | 1 | 1 | 1 | 1 | 1 | 1 | -0.004 |
| SE | 59138 | ENSG00000168421 | RHOH | chr4 | + | 40200511 | 40200565 | 40196906 | 40197300 | 40209462 | 40209669 | 0 | 0 | 3 | 0 | 0 | 12 | 4 | 16 | 8 | 7 | 0 | 0 | 0 | 0 | 0 | 0 | 14 | 6 | 0 | 7 | 203 | 149 | 1 | 1 | 0 | 0 | 0.121 | 0 | 0 | NA | 0 | 0 | NA | 0 | 0.024 |
| SE | 59139 | ENSG00000168421 | RHOH | chr4 | + | 40200511 | 40200565 | 40196906 | 40197300 | 40242713 | 40242834 | 3 | 0 | 1 | 0 | 0 | 289 | 161 | 347 | 224 | 588 | 0 | 0 | 1 | 0 | 0 | 80 | 224 | 162 | 145 | 118 | 203 | 149 | 1 | 1 | 0.008 | 0 | 0.002 | 0 | 0 | 0 | 0 | 0.005 | 0 | 0 | 0.001 |
| SE | 59140 | ENSG00000168421 | RHOH | chr4 | + | 40205692 | 40205832 | 40197060 | 40197300 | 40209462 | 40209669 | 0 | 0 | 0 | 1 | 0 | 12 | 4 | 16 | 8 | 7 | 0 | 0 | 0 | 6 | 0 | 0 | 14 | 6 | 0 | 7 | 289 | 149 | 1 | 1 | 0 | 0 | 0 | 0.061 | 0 | NA | 0 | 0 | 1 | 0 | -0.238 |
| SE | 59141 | ENSG00000168421 | RHOH | chr4 | + | 40205692 | 40205832 | 40197060 | 40197300 | 40242713 | 40243187 | 7 | 5 | 0 | 2 | 0 | 289 | 161 | 347 | 224 | 588 | 0 | 0 | 1 | 0 | 0 | 80 | 224 | 162 | 145 | 118 | 289 | 149 | 1 | 1 | 0.012 | 0.016 | 0 | 0.005 | 0 | 0 | 0 | 0.003 | 0 | 0 | 0.006 |
| SE | 59142 | ENSG00000168421 | RHOH | chr4 | + | 40212663 | 40212791 | 40196906 | 40197300 | 40242713 | 40242834 | 5 | 2 | 1 | 0 | 2 | 289 | 161 | 347 | 224 | 588 | 0 | 0 | 0 | 1 | 0 | 80 | 224 | 162 | 145 | 118 | 277 | 149 | 1 | 1 | 0.009 | 0.007 | 0.002 | 0 | 0.002 | 0 | 0 | 0 | 0.004 | 0 | 0.003 |
| SE | 59143 | ENSG00000168421 | RHOH | chr4 | + | 40212663 | 40212791 | 40200344 | 40200565 | 40242713 | 40242834 | 2 | 0 | 0 | 0 | 0 | 5 | 2 | 3 | 0 | 0 | 0 | 0 | 0 | 1 | 0 | 0 | 4 | 1 | 0 | 0 | 277 | 149 | 1 | 1 | 0.177 | 0 | 0 | NA | NA | NA | 0 | 0 | 1 | NA | -0.274 |
| SE | 59147 | ENSG00000168421 | RHOH | chr4 | + | 40240127 | 40240345 | 40197032 | 40197300 | 40242713 | 40242801 | 0 | 1 | 0 | 0 | 0 | 289 | 161 | 347 | 224 | 588 | 0 | 0 | 0 | 0 | 0 | 80 | 224 | 162 | 145 | 118 | 298 | 149 | 1 | 1 | 0 | 0.003 | 0 | 0 | 0 | 0 | 0 | 0 | 0 | 0 | 0.001 |
| SE | 59148 | ENSG00000168421 | RHOH | chr4 | + | 40242713 | 40242834 | 40197030 | 40197300 | 40243177 | 40243193 | 565 | 290 | 688 | 474 | 997 | 0 | 0 | 1 | 0 | 0 | 129 | 480 | 307 | 270 | 245 | 0 | 7 | 0 | 0 | 0 | 270 | 149 | 1 | 1 | 1 | 1 | 0.997 | 1 | 1 | 1 | 0.974 | 1 | 1 | 1 | 0.005 |
| SE | 59154 | ENSG00000171492 | LRRC8D | chr1 | + | 89929807 | 89929878 | 89846579 | 89846658 | 89933066 | 89933160 | 6 | 1 | 2 | 10 | 1 | 0 | 1 | 1 | 5 | 0 | 13 | 0 | 0 | 0 | 0 | 0 | 2 | 1 | 7 | 0 | 220 | 149 | 1 | 1 | 1 | 0.404 | 0.575 | 0.575 | 1 | 1 | 0 | 0 | 0 | NA | 0.461 |
| SE | 59155 | ENSG00000115204 | MPV17 | chr2 | - | 27311898 | 27311951 | 27309491 | 27309981 | 27312213 | 27312246 | 85 | 52 | 66 | 74 | 94 | 1 | 3 | 0 | 0 | 0 | 190 | 128 | 52 | 82 | 266 | 0 | 0 | 0 | 0 | 0 | 202 | 149 | 1 | 1 | 0.984 | 0.927 | 1 | 1 | 1 | 1 | 1 | 1 | 1 | 1 | -0.018 |
| SE | 59156 | ENSG00000115204 | MPV17 | chr2 | - | 27311898 | 27311951 | 27309491 | 27309981 | 27312493 | 27312589 | 40 | 16 | 28 | 34 | 31 | 0 | 2 | 0 | 0 | 0 | 67 | 59 | 15 | 44 | 116 | 0 | 2 | 0 | 0 | 0 | 202 | 149 | 1 | 1 | 1 | 0.855 | 1 | 1 | 1 | 1 | 0.956 | 1 | 1 | 1 | -0.02 |
| SE | 59157 | ENSG00000115204 | MPV17 | chr2 | - | 27312213 | 27312246 | 27309491 | 27309981 | 27312493 | 27312589 | 16 | 6 | 7 | 7 | 21 | 0 | 2 | 0 | 0 | 0 | 31 | 8 | 3 | 15 | 7 | 0 | 2 | 0 | 0 | 0 | 182 | 149 | 1 | 1 | 1 | 0.711 | 1 | 1 | 1 | 1 | 0.766 | 1 | 1 | 1 | -0.011 |
| SE | 59158 | ENSG00000115204 | MPV17 | chr2 | - | 27312213 | 27312246 | 27311898 | 27311951 | 27312493 | 27312589 | 58 | 34 | 42 | 41 | 76 | 0 | 0 | 0 | 0 | 0 | 119 | 69 | 35 | 61 | 132 | 0 | 1 | 0 | 1 | 0 | 182 | 149 | 1 | 1 | 1 | 1 | 1 | 1 | 1 | 1 | 0.983 | 1 | 0.98 | 1 | 0.007 |
| SE | 59161 | ENSG00000115204 | MPV17 | chr2 | - | 27312679 | 27312772 | 27312213 | 27312246 | 27322447 | 27322522 | 0 | 1 | 1 | 3 | 2 | 0 | 0 | 0 | 0 | 0 | 0 | 0 | 4 | 6 | 12 | 0 | 0 | 0 | 0 | 11 | 242 | 149 | 1 | 1 | NA | 1 | 1 | 1 | 1 | NA | NA | 1 | 1 | 0.402 | 0.199 |
| SE | 59162 | ENSG00000115204 | MPV17 | chr2 | - | 27312993 | 27313074 | 27312679 | 27312772 | 27317077 | 27317234 | 28 | 16 | 16 | 15 | 8 | 0 | 0 | 1 | 0 | 0 | 34 | 45 | 12 | 10 | 46 | 0 | 0 | 0 | 0 | 3 | 230 | 149 | 1 | 1 | 1 | 1 | 0.912 | 1 | 1 | 1 | 1 | 1 | 1 | 0.909 | 0.001 |

| | | | | | | | | | | | | | | | | | | | | | | | | | | | | | | | | | | | | | | | | | | | | | | |
|---|---|---|---|---|---|---|---|---|---|---|---|---|---|---|---|---|---|---|---|---|---|---|---|---|---|---|---|---|---|---|---|---|---|---|---|---|---|---|---|---|---|---|---|---|---|---|
| SE | 59165 | ENSG00000115204 | MPV17 | chr2 | - | 27312993 | 27313109 | 27312679 | 27312772 | 27317077 | 27317234 | 41 | 27 | 22 | 40 | 22 | 0 | 0 | 1 | 0 | 0 | 79 | 63 | 21 | 15 | 66 | 0 | 0 | 0 | 0 | 3 | 265 | 149 | 1 | 1 | 1 | 1 | 0.925 | 1 | 1 | 1 | 1 | 1 | 1 | 0.925 | 0 |
| SE | 59168 | ENSG00000115204 | MPV17 | chr2 | - | 27316158 | 27316186 | 27312993 | 27313074 | 27317077 | 27317173 | 0 | 0 | 1 | 0 | 0 | 3 | 0 | 1 | 0 | 0 | 0 | 0 | 2 | 1 | 0 | 0 | 0 | 2 | 0 | 0 | 177 | 149 | 1 | 1 | 0 | NA | 0.457 | NA | NA | NA | NA | 0.457 | 1 | NA | -0.5 |
| SE | 59169 | ENSG00000115204 | MPV17 | chr2 | - | 27316158 | 27316186 | 27312993 | 27313109 | 27317077 | 27317173 | 0 | 0 | 0 | 0 | 0 | 2 | 0 | 0 | 10 | 5 | 0 | 0 | 5 | 1 | 0 | 21 | 6 | 4 | 0 | 6 | 177 | 149 | 1 | 1 | 0 | NA | NA | 0 | 0 | 0 | 0 | 0.513 | 1 | 0 | -0.303 |
| SE | 59173 | ENSG00000115204 | MPV17 | chr2 | - | 27322447 | 27322522 | 27312993 | 27313109 | 27323051 | 27323054 | 32 | 10 | 31 | 11 | 10 | 0 | 0 | 0 | 3 | 0 | 25 | 27 | 13 | 11 | 34 | 0 | 3 | 0 | 0 | 0 | 224 | 149 | 1 | 1 | 1 | 1 | 1 | 0.709 | 1 | 1 | 0.857 | 1 | 1 | 1 | -0.03 |
| SE | 59176 | ENSG00000080819 | CPOX | chr3 | - | 98590631 | 98590742 | 98588712 | 98588854 | 98591011 | 98591155 | 98 | 60 | 91 | 86 | 158 | 0 | 0 | 0 | 2 | 0 | 57 | 51 | 35 | 82 | 63 | 0 | 3 | 0 | 0 | 0 | 260 | 149 | 1 | 1 | 1 | 1 | 1 | 0.961 | 1 | 1 | 0.907 | 1 | 1 | 1 | 0.011 |
| SE | 59178 | ENSG00000075702 | WDR62 | chr19 | + | 36071555 | 36071716 | 36067827 | 36068010 | 36073341 | 36073525 | 2 | 0 | 2 | 6 | 4 | 0 | 0 | 0 | 0 | 0 | 0 | 0 | 2 | 6 | 0 | 0 | 0 | 1 | 0 | 0 | 298 | 149 | 1 | 1 | 1 | NA | 1 | 1 | 1 | NA | NA | 0.5 | 1 | NA | 0.25 |
| SE | 59183 | ENSG00000075702 | WDR62 | chr19 | + | 36075054 | 36075161 | 36073341 | 36073531 | 36081432 | 36081570 | 0 | 0 | 0 | 0 | 0 | 0 | 0 | 0 | 0 | 4 | 0 | 0 | 0 | 7 | 0 | 8 | 1 | 4 | 0 | 8 | 256 | 149 | 1 | 1 | NA | NA | NA | NA | 0 | 0 | 0 | 0 | 1 | 0 | -0.2 |
| SE | 59185 | ENSG00000075702 | WDR62 | chr19 | + | 36081793 | 36081935 | 36081432 | 36081570 | 36083062 | 36083241 | 0 | 0 | 0 | 0 | 0 | 1 | 1 | 2 | 1 | 5 | 0 | 0 | 1 | 0 | 0 | 8 | 9 | 11 | 2 | 8 | 291 | 149 | 1 | 1 | 0 | 0 | 0 | 0 | 0 | 0 | 0 | 0.044 | 0 | 0 | -0.009 |
| SE | 59187 | ENSG00000075702 | WDR62 | chr19 | + | 36091199 | 36091311 | 36090444 | 36090520 | 36091401 | 36091465 | 3 | 1 | 1 | 0 | 0 | 0 | 0 | 0 | 0 | 0 | 0 | 0 | 13 | 17 | 6 | 0 | 0 | 0 | 1 | 0 | 261 | 149 | 1 | 1 | 1 | 1 | 1 | NA | NA | NA | NA | 1 | 0.907 | 1 | 0.031 |
| SE | 59190 | ENSG00000075702 | WDR62 | chr19 | + | 36091401 | 36091465 | 36091199 | 36091311 | 36092688 | 36092811 | 1 | 0 | 1 | 0 | 0 | 0 | 0 | 0 | 0 | 0 | 0 | 4 | 9 | 20 | 7 | 0 | 0 | 0 | 0 | 4 | 213 | 149 | 1 | 1 | 1 | NA | 1 | NA | NA | NA | 1 | 1 | 1 | 0.55 | 0.113 |
| SE | 59191 | ENSG00000075702 | WDR62 | chr19 | + | 36100747 | 36100875 | 36099398 | 36099617 | 36101213 | 36101317 | 4 | 0 | 2 | 1 | 0 | 0 | 0 | 0 | 0 | 0 | 24 | 2 | 0 | 7 | 7 | 0 | 0 | 3 | 0 | 0 | 277 | 149 | 1 | 1 | 1 | NA | 1 | 1 | NA | 1 | 1 | 0 | 1 | 1 | 0.2 |
| SE | 59192 | ENSG00000075702 | WDR62 | chr19 | + | 36100751 | 36100875 | 36099398 | 36099617 | 36101213 | 36101317 | 3 | 0 | 1 | 1 | 0 | 0 | 0 | 0 | 0 | 0 | 24 | 0 | 0 | 5 | 7 | 0 | 0 | 3 | 0 | 0 | 273 | 149 | 1 | 1 | 1 | NA | 1 | 1 | NA | 1 | NA | 0 | 1 | 1 | 0.25 |
| SE | 59195 | ENSG00000142634 | EFHD2 | chr1 | + | 15425870 | 15426018 | 15409887 | 15410279 | 15428592 | 15429081 | 1036 | 455 | 446 | 761 | 1008 | 0 | 0 | 0 | 0 | 1 | 1584 | 810 | 660 | 955 | 1658 | 0 | 0 | 0 | 0 | 0 | 297 | 149 | 1 | 1 | 1 | 1 | 1 | 1 | 0.998 | 1 | 1 | 1 | 1 | 1 | 0 |
| SE | 59196 | ENSG00000142634 | EFHD2 | chr1 | + | 15427149 | 15427284 | 15425870 | 15426018 | 15428592 | 15430339 | 3698 | 1683 | 1691 | 2482 | 2680 | 20 | 11 | 7 | 17 | 13 | 5677 | 2437 | 1976 | 3058 | 5602 | 25 | 6 | 4 | 18 | 24 | 284 | 149 | 1 | 1 | 0.99 | 0.988 | 0.992 | 0.987 | 0.991 | 0.992 | 0.995 | 0.996 | 0.989 | 0.992 | -0.003 |
| SE | 59197 | ENSG00000142634 | EFHD2 | chr1 | + | 15427897 | 15428009 | 15427149 | 15427284 | 15428592 | 15429081 | 0 | 0 | 2 | 1 | 1 | 2377 | 1052 | 1072 | 1627 | 1788 | 10 | 0 | 1 | 0 | 0 | 3566 | 1585 | 1298 | 1946 | 3579 | 261 | 149 | 1 | 1 | 0 | 0 | 0.001 | 0 | 0 | 0.002 | 0 | 0 | 0 | 0 | 0 |
| SE | 59200 | ENSG00000113615 | SEC24A | chr5 | + | 134674614 | 134674775 | 134671808 | 134671886 | 134675044 | 134675217 | 77 | 60 | 64 | 66 | 133 | 0 | 0 | 0 | 0 | 0 | 85 | 65 | 123 | 102 | 56 | 0 | 2 | 0 | 0 | 0 | 298 | 149 | 1 | 1 | 1 | 1 | 1 | 1 | 1 | 1 | 0.942 | 1 | 1 | 1 | 0.012 |
| SE | 59201 | ENSG00000113615 | SEC24A | chr5 | + | 134674614 | 134674775 | 134671808 | 134671886 | 134679601 | 134679728 | 18 | 13 | 12 | 14 | 33 | 0 | 0 | 0 | 0 | 0 | 17 | 14 | 55 | 31 | 22 | 0 | 0 | 1 | 0 | 0 | 298 | 149 | 1 | 1 | 1 | 1 | 1 | 1 | 1 | 1 | 1 | 0.965 | 1 | 1 | 0.007 |
| SE | 59202 | ENSG00000113615 | SEC24A | chr5 | + | 134675044 | 134675217 | 134671808 | 134671886 | 134679601 | 134679728 | 1 | 0 | 0 | 2 | 0 | 0 | 0 | 0 | 0 | 0 | 0 | 2 | 0 | 6 | 0 | 0 | 0 | 1 | 0 | 0 | 298 | 149 | 1 | 1 | 1 | NA | NA | 1 | NA | NA | 1 | 0 | 1 | NA | 0.333 |
| SE | 59203 | ENSG00000113615 | SEC24A | chr5 | + | 134675044 | 134675217 | 134674614 | 134674775 | 134679601 | 134679728 | 60 | 47 | 52 | 54 | 100 | 0 | 0 | 0 | 0 | 0 | 68 | 51 | 68 | 79 | 34 | 0 | 0 | 0 | 2 | 0 | 298 | 149 | 1 | 1 | 1 | 1 | 1 | 1 | 1 | 1 | 1 | 1 | 0.952 | 1 | 0.01 |
| SE | 59205 | ENSG00000113615 | SEC24A | chr5 | + | 134679601 | 134679728 | 134676022 | 134676125 | 134682372 | 134682482 | 147 | 98 | 125 | 145 | 214 | 0 | 0 | 0 | 1 | 0 | 53 | 128 | 183 | 108 | 15 | 0 | 0 | 0 | 0 | 0 | 276 | 149 | 1 | 1 | 1 | 1 | 1 | 0.987 | 1 | 1 | 1 | 1 | 1 | 1 | -0.003 |
| SE | 59206 | ENSG00000113615 | SEC24A | chr5 | + | 134682372 | 134682482 | 134679601 | 134679728 | 134686789 | 134686902 | 159 | 93 | 117 | 128 | 217 | 0 | 1 | 0 | 1 | 0 | 83 | 114 | 176 | 106 | 11 | 0 | 0 | 0 | 0 | 0 | 259 | 149 | 1 | 1 | 1 | 0.982 | 1 | 0.987 | 1 | 1 | 1 | 1 | 1 | 1 | -0.006 |
| SE | 59207 | ENSG00000113615 | SEC24A | chr5 | + | 134693726 | 134693933 | 134692601 | 134692657 | 134697125 | 134697246 | 140 | 70 | 118 | 106 | 182 | 1 | 0 | 1 | 0 | 0 | 30 | 107 | 130 | 65 | 32 | 0 | 0 | 3 | 0 | 0 | 298 | 149 | 1 | 1 | 0.986 | 1 | 0.983 | 1 | 1 | 1 | 1 | 0.956 | 1 | 1 | 0.003 |
| SE | 59208 | ENSG00000113615 | SEC24A | chr5 | + | 134705326 | 134705437 | 134703758 | 134703932 | 134708712 | 134708888 | 52 | 30 | 38 | 55 | 70 | 0 | 0 | 0 | 0 | 0 | 16 | 28 | 81 | 38 | 4 | 0 | 0 | 0 | 1 | 0 | 260 | 149 | 1 | 1 | 1 | 1 | 1 | 1 | 1 | 1 | 1 | 1 | 0.956 | 1 | 0.009 |
| SE | 59209 | ENSG00000113615 | SEC24A | chr5 | + | 134708712 | 134708888 | 134705326 | 134705437 | 134715023 | 134715161 | 87 | 53 | 58 | 63 | 95 | 1 | 1 | 1 | 1 | 2 | 36 | 63 | 102 | 50 | 5 | 0 | 1 | 2 | 1 | 0 | 298 | 149 | 1 | 1 | 0.978 | 0.964 | 0.967 | 0.969 | 0.96 | 1 | 0.969 | 0.962 | 0.962 | 1 | -0.011 |
| SE | 59210 | ENSG00000113615 | SEC24A | chr5 | + | 134715023 | 134715161 | 134708712 | 134708888 | 134718068 | 134718173 | 111 | 87 | 99 | 102 | 136 | 1 | 0 | 1 | 0 | 0 | 38 | 86 | 136 | 39 | 6 | 0 | 2 | 2 | 0 | 0 | 287 | 149 | 1 | 1 | 0.983 | 1 | 0.981 | 1 | 1 | 1 | 0.957 | 0.972 | 1 | 1 | 0.007 |
| SE | 59212 | ENSG00000109436 | TBC1D9 | chr4 | - | 140657096 | 140657226 | 140639329 | 140639428 | 140659587 | 140659705 | 59 | 45 | 61 | 60 | 102 | 0 | 0 | 2 | 0 | 0 | 50 | 57 | 53 | 78 | 24 | 0 | 0 | 0 | 0 | 0 | 279 | 149 | 1 | 1 | 1 | 1 | 0.942 | 1 | 1 | 1 | 1 | 1 | 1 | 1 | -0.012 |
| SE | 59213 | ENSG00000109436 | TBC1D9 | chr4 | - | 140657526 | 140657812 | 140657096 | 140657226 | 140659587 | 140659705 | 178 | 126 | 138 | 175 | 164 | 0 | 0 | 0 | 0 | 0 | 160 | 140 | 81 | 128 | 92 | 0 | 0 | 0 | 2 | 0 | 298 | 149 | 1 | 1 | 1 | 1 | 1 | 1 | 1 | 1 | 1 | 1 | 0.97 | 1 | 0.006 |
| SE | 59214 | ENSG00000109436 | TBC1D9 | chr4 | - | 140670719 | 140670926 | 140669633 | 140669804 | 140676893 | 140677101 | 348 | 229 | 187 | 255 | 530 | 0 | 1 | 0 | 0 | 0 | 252 | 242 | 122 | 210 | 203 | 0 | 0 | 0 | 0 | 0 | 298 | 149 | 1 | 1 | 1 | 0.991 | 1 | 1 | 1 | 1 | 1 | 1 | 1 | 1 | -0.002 |
| SE | 59233 | ENSG00000117153 | KLHL12 | chr1 | - | 202894183 | 202894282 | 202891115 | 202892659 | 202894590 | 202894749 | 46 | 18 | 35 | 32 | 29 | 2 | 0 | 0 | 2 | 0 | 21 | 26 | 18 | 32 | 21 | 0 | 4 | 12 | 6 | 4 | 248 | 149 | 1 | 1 | 0.933 | 1 | 1 | 0.906 | 1 | 1 | 0.796 | 0.474 | 0.762 | 0.759 | 0.21 |
| SE | 59234 | ENSG00000117153 | KLHL12 | chr1 | - | 202895521 | 202895717 | 202894590 | 202894749 | 202896853 | 202896960 | 193 | 96 | 137 | 153 | 159 | 0 | 0 | 1 | 1 | 4 | 188 | 131 | 103 | 71 | 105 | 0 | 1 | 0 | 0 | 0 | 298 | 149 | 1 | 1 | 1 | 1 | 0.986 | 0.987 | 0.952 | 1 | 0.985 | 1 | 1 | 1 | -0.012 |
| SE | 59238 | ENSG00000117153 | KLHL12 | chr1 | - | 202911053 | 202911203 | 202909009 | 202909124 | 202918170 | 202918388 | 185 | 147 | 132 | 205 | 288 | 0 | 0 | 1 | 0 | 0 | 195 | 206 | 164 | 232 | 228 | 0 | 0 | 0 | 0 | 0 | 298 | 149 | 1 | 1 | 1 | 1 | 0.985 | 1 | 1 | 1 | 1 | 1 | 1 | 1 | -0.003 |
| SE | 59243 | ENSG00000011007 | ELOA | chr1 | + | 23749841 | 23749948 | 23749020 | 23749077 | 23750844 | 23750944 | 180 | 169 | 145 | 169 | 153 | 0 | 0 | 0 | 0 | 0 | 377 | 226 | 172 | 225 | 221 | 0 | 0 | 2 | 1 | 0 | 256 | 149 | 1 | 1 | 1 | 1 | 1 | 1 | 1 | 1 | 1 | 0.98 | 0.992 | 1 | 0.006 |
| SE | 59245 | ENSG00000011007 | ELOA | chr1 | + | 23756273 | 23756385 | 23755842 | 23756023 | 23759511 | 23759893 | 104 | 73 | 113 | 95 | 185 | 0 | 1 | 0 | 0 | 0 | 144 | 151 | 137 | 110 | 112 | 0 | 0 | 0 | 0 | 0 | 261 | 149 | 1 | 1 | 1 | 0.977 | 1 | 1 | 1 | 1 | 1 | 1 | 1 | 1 | -0.005 |
| SE | 59246 | ENSG00000011007 | ELOA | chr1 | + | 23756952 | 23757125 | 23755842 | 23756023 | 23759511 | 23759893 | 35 | 30 | 25 | 40 | 44 | 0 | 1 | 0 | 0 | 0 | 24 | 69 | 49 | 55 | 26 | 0 | 0 | 0 | 0 | 0 | 298 | 149 | 1 | 1 | 1 | 0.938 | 1 | 1 | 1 | 1 | 1 | 1 | 1 | 1 | -0.012 |
| SE | 59248 | ENSG00000168118 | RAB4A | chr1 | + | 229286485 | 229286566 | 229271074 | 229271370 | 229288728 | 229288843 | 99 | 70 | 71 | 79 | 112 | 1 | 0 | 0 | 0 | 0 | 88 | 99 | 76 | 65 | 206 | 0 | 0 | 0 | 2 | 0 | 230 | 149 | 1 | 1 | 0.985 | 1 | 1 | 1 | 1 | 1 | 1 | 1 | 0.955 | 1 | 0.006 |
| SE | 59252 | ENSG00000168118 | RAB4A | chr1 | + | 229295847 | 229295910 | 229288728 | 229288843 | 229297481 | 229297636 | 202 | 121 | 129 | 214 | 195 | 0 | 0 | 0 | 0 | 0 | 278 | 214 | 144 | 158 | 673 | 0 | 0 | 0 | 0 | 12 | 212 | 149 | 1 | 1 | 1 | 1 | 1 | 1 | 1 | 1 | 1 | 1 | 1 | 0.975 | 0.005 |
| SE | 59254 | ENSG00000162909 | CAPN2 | chr1 | + | 223717761 | 223717831 | 223712538 | 223712877 | 223744099 | 223744218 | 309 | 200 | 255 | 304 | 91 | 0 | 0 | 0 | 0 | 0 | 700 | 424 | 262 | 512 | 1139 | 0 | 9 | 0 | 0 | 0 | 219 | 149 | 1 | 1 | 1 | 1 | 1 | 1 | 1 | 1 | 0.97 | 1 | 1 | 1 | 0.006 |
| SE | 59255 | ENSG00000162909 | CAPN2 | chr1 | + | 223744099 | 223744218 | 223717761 | 223717831 | 223745305 | 223745439 | 386 | 227 | 307 | 411 | 211 | 0 | 0 | 0 | 0 | 0 | 773 | 543 | 361 | 656 | 1383 | 0 | 0 | 0 | 1 | 0 | 268 | 149 | 1 | 1 | 1 | 1 | 1 | 1 | 1 | 1 | 1 | 1 | 0.997 | 1 | 0.001 |
| SE | 59256 | ENSG00000162909 | CAPN2 | chr1 | + | 223745305 | 223745439 | 223744099 | 223744218 | 223746996 | 223747063 | 479 | 248 | 313 | 489 | 255 | 1 | 0 | 3 | 1 | 0 | 820 | 632 | 419 | 644 | 1274 | 0 | 0 | 0 | 1 | 11 | 283 | 149 | 1 | 1 | 0.996 | 1 | 0.982 | 0.996 | 1 | 1 | 1 | 1 | 0.997 | 0.984 | -0.001 |
| SE | 59257 | ENSG00000162909 | CAPN2 | chr1 | + | 223746996 | 223747165 | 223745305 | 223745439 | 223749038 | 223749122 | 734 | 412 | 503 | 728 | 399 | 0 | 0 | 0 | 0 | 1 | 1507 | 852 | 679 | 932 | 1758 | 0 | 0 | 0 | 2 | 0 | 298 | 149 | 1 | 1 | 1 | 1 | 1 | 1 | 0.995 | 1 | 1 | 1 | 0.996 | 1 | 0 |
| SE | 59258 | ENSG00000162909 | CAPN2 | chr1 | + | 223749038 | 223749122 | 223746996 | 223747165 | 223750889 | 223750975 | 564 | 395 | 452 | 623 | 401 | 0 | 0 | 1 | 0 | 0 | 1511 | 696 | 554 | 813 | 1519 | 10 | 0 | 0 | 7 | 0 | 233 | 149 | 1 | 1 | 1 | 1 | 0.997 | 1 | 1 | 0.99 | 1 | 1 | 0.987 | 1 | 0.004 |
| SE | 59259 | ENSG00000162909 | CAPN2 | chr1 | + | 223752795 | 223752956 | 223751996 | 223752071 | 223755479 | 223755649 | 950 | 778 | 745 | 1004 | 1348 | 0 | 0 | 0 | 1 | 0 | 3370 | 1217 | 864 | 1804 | 2683 | 0 | 0 | 1 | 5 | 0 | 298 | 149 | 1 | 1 | 1 | 1 | 1 | 0.998 | 1 | 1 | 1 | 0.998 | 0.994 | 1 | 0.001 |
| SE | 59260 | ENSG00000162909 | CAPN2 | chr1 | + | 223755479 | 223755649 | 223752795 | 223752956 | 223757368 | 223757380 | 733 | 565 | 611 | 836 | 902 | 0 | 0 | 0 | 0 | 4 | 2340 | 958 | 682 | 1441 | 2178 | 0 | 0 | 0 | 0 | 4 | 298 | 149 | 1 | 1 | 1 | 1 | 1 | 1 | 0.991 | 1 | 1 | 1 | 1 | 0.996 | -0.001 |
| SE | 59261 | ENSG00000162909 | CAPN2 | chr1 | + | 223757368 | 223757380 | 223755479 | 223755649 | 223759269 | 223759481 | 325 | 175 | 261 | 316 | 322 | 0 | 0 | 0 | 0 | 3 | 594 | 318 | 217 | 451 | 758 | 0 | 0 | 0 | 0 | 0 | 161 | 149 | 1 | 1 | 1 | 1 | 1 | 1 | 0.99 | 1 | 1 | 1 | 1 | 1 | -0.002 |
| SE | 59262 | ENSG00000162909 | CAPN2 | chr1 | + | 223766366 | 223766431 | 223764149 | 223764207 | 223769840 | 223769909 | 641 | 401 | 528 | 784 | 955 | 0 | 0 | 0 | 0 | 0 | 1093 | 629 | 441 | 731 | 1418 | 0 | 0 | 0 | 1 | 0 | 214 | 149 | 1 | 1 | 1 | 1 | 1 | 1 | 1 | 1 | 1 | 1 | 0.998 | 1 | 0 |
| SE | 59263 | ENSG00000162909 | CAPN2 | chr1 | + | 223771808 | 223771925 | 223769840 | 223770525 | 223772180 | 223772239 | 819 | 552 | 571 | 818 | 1345 | 1 | 0 | 0 | 0 | 0 | 1565 | 887 | 618 | 938 | 1556 | 0 | 0 | 0 | 0 | 0 | 266 | 149 | 1 | 1 | 0.998 | 1 | 1 | 1 | 1 | 1 | 1 | 1 | 1 | 1 | 0 |
| SE | 59273 | ENSG00000181744 | DIPK2A | chr3 | + | 143985542 | 143985846 | 143971822 | 143972989 | 144048667 | 144048719 | 20 | 17 | 16 | 15 | 10 | 0 | 3 | 0 | 0 | 0 | 5 | 22 | 3 | 3 | 7 | 0 | 0 | 0 | 0 | 0 | 298 | 149 | 1 | 1 | 1 | 0.739 | 1 | 1 | 1 | 1 | 1 | 1 | 1 | 1 | -0.052 |
| SE | 59275 | ENSG00000119004 | CYP20A1 | chr2 | + | 203245845 | 203245895 | 203238976 | 203239134 | 203246754 | 203246921 | 71 | 50 | 74 | 61 | 22 | 0 | 0 | 2 | 0 | 0 | 49 | 38 | 36 | 35 | 10 | 0 | 0 | 0 | 0 | 0 | 199 | 149 | 1 | 1 | 1 | 1 | 0.965 | 1 | 1 | 1 | 1 | 1 | 1 | 1 | -0.007 |
| SE | 59277 | ENSG00000119004 | CYP20A1 | chr2 | + | 203245845 | 203245895 | 203238976 | 203239134 | 203266513 | 203266681 | 20 | 18 | 29 | 16 | 1 | 0 | 0 | 0 | 2 | 0 | 10 | 9 | 22 | 18 | 1 | 0 | 0 | 0 | 0 | 0 | 199 | 149 | 1 | 1 | 1 | 1 | 1 | 0.857 | 1 | 1 | 1 | 1 | 1 | 1 | -0.029 |
| SE | 59281 | ENSG00000119004 | CYP20A1 | chr2 | + | 203246754 | 203246921 | 203245845 | 203245895 | 203266513 | 203266681 | 52 | 33 | 46 | 46 | 22 | 0 | 0 | 0 | 1 | 0 | 39 | 35 | 16 | 17 | 9 | 0 | 0 | 0 | 0 | 0 | 298 | 149 | 1 | 1 | 1 | 1 | 1 | 0.958 | 1 | 1 | 1 | 1 | 1 | 1 | -0.008 |
| SE | 59282 | ENSG00000119004 | CYP20A1 | chr2 | + | 203251966 | 203252109 | 203238981 | 203239134 | 203266513 | 203266681 | 50 | 60 | 78 | 53 | 118 | 0 | 0 | 0 | 2 | 0 | 60 | 37 | 49 | 44 | 29 | 0 | 0 | 0 | 0 | 0 | 292 | 149 | 1 | 1 | 1 | 1 | 1 | 0.931 | 1 | 1 | 1 | 1 | 1 | 1 | -0.014 |
| SE | 59283 | ENSG00000119004 | CYP20A1 | chr2 | + | 203251966 | 203252109 | 203245845 | 203245895 | 203266513 | 203266681 | 54 | 60 | 76 | 55 | 116 | 0 | 0 | 0 | 1 | 0 | 60 | 42 | 52 | 39 | 29 | 0 | 0 | 0 | 0 | 0 | 292 | 149 | 1 | 1 | 1 | 1 | 1 | 0.966 | 1 | 1 | 1 | 1 | 1 | 1 | -0.007 |
| SE | 59284 | ENSG00000119004 | CYP20A1 | chr2 | + | 203251966 | 203252109 | 203246754 | 203246921 | 203266513 | 203266681 | 102 | 88 | 134 | 106 | 153 | 1 | 0 | 0 | 0 | 0 | 99 | 99 | 67 | 58 | 45 | 0 | 0 | 0 | 0 | 0 | 292 | 149 | 1 | 1 | 0.981 | 1 | 1 | 1 | 1 | 1 | 1 | 1 | 1 | 1 | -0.004 |
| SE | 59285 | ENSG00000119004 | CYP20A1 | chr2 | + | 203266513 | 203266681 | 203251966 | 203252109 | 203272669 | 203272748 | 95 | 128 | 144 | 121 | 241 | 1 | 0 | 0 | 0 | 0 | 104 | 70 | 67 | 103 | 50 | 0 | 0 | 0 | 0 | 0 | 298 | 149 | 1 | 1 | 0.979 | 1 | 1 | 1 | 1 | 1 | 1 | 1 | 1 | 1 | -0.004 |
| SE | 59286 | ENSG00000119004 | CYP20A1 | chr2 | + | 203266513 | 203266681 | 203251966 | 203252109 | 203278572 | 203278688 | 67 | 73 | 90 | 65 | 151 | 0 | 0 | 0 | 0 | 1 | 97 | 51 | 72 | 79 | 35 | 0 | 0 | 0 | 0 | 0 | 298 | 149 | 1 | 1 | 1 | 1 | 1 | 1 | 0.987 | 1 | 1 | 1 | 1 | 1 | -0.003 |
| SE | 59288 | ENSG00000119004 | CYP20A1 | chr2 | + | 203272645 | 203272748 | 203266513 | 203266681 | 203280058 | 203280113 | 0 | 0 | 2 | 1 | 0 | 8 | 2 | 3 | 0 | 0 | 0 | 0 | 0 | 0 | 0 | 3 | 2 | 0 | 0 | 0 | 252 | 149 | 1 | 1 | 0 | 0 | 0.283 | 1 | NA | 0 | 0 | NA | NA | NA | 0.321 |
| SE | 59289 | ENSG00000119004 | CYP20A1 | chr2 | + | 203272669 | 203272748 | 203251966 | 203252109 | 203278572 | 203278688 | 36 | 31 | 39 | 47 | 86 | 0 | 0 | 0 | 0 | 1 | 26 | 18 | 22 | 21 | 6 | 0 | 0 | 0 | 0 | 0 | 228 | 149 | 1 | 1 | 1 | 1 | 1 | 1 | 0.983 | 1 | 1 | 1 | 1 | 1 | -0.003 |
| SE | 59293 | ENSG00000119004 | CYP20A1 | chr2 | + | 203278572 | 203278688 | 203272645 | 203272748 | 203280058 | 203280113 | 138 | 104 | 157 | 168 | 313 | 0 | 0 | 2 | 0 | 0 | 130 | 110 | 89 | 141 | 61 | 0 | 0 | 0 | 0 | 0 | 265 | 149 | 1 | 1 | 1 | 1 | 0.978 | 1 | 1 | 1 | 1 | 1 | 1 | 1 | -0.004 |
| SE | 59294 | ENSG00000119004 | CYP20A1 | chr2 | + | 203280058 | 203280113 | 203278572 | 203278688 | 203285611 | 203285679 | 124 | 58 | 112 | 130 | 240 | 0 | 0 | 0 | 0 | 1 | 98 | 93 | 56 | 110 | 69 | 0 | 0 | 0 | 0 | 0 | 204 | 149 | 1 | 1 | 1 | 1 | 1 | 1 | 0.994 | 1 | 1 | 1 | 1 | 1 | -0.001 |
| SE | 59296 | ENSG00000119004 | CYP20A1 | chr2 | + | 203289764 | 203289876 | 203285611 | 203285732 | 203292261 | 203292326 | 112 | 69 | 100 | 156 | 202 | 1 | 1 | 1 | 0 | 1 | 90 | 74 | 76 | 91 | 55 | 0 | 0 | 0 | 0 | 0 | 261 | 149 | 1 | 1 | 0.985 | 0.975 | 0.983 | 1 | 0.991 | 1 | 1 | 1 | 1 | 1 | -0.013 |
| SE | 59297 | ENSG00000119004 | CYP20A1 | chr2 | + | 203292261 | 203292326 | 203285611 | 203285732 | 203296473 | 203296563 | 50 | 31 | 29 | 51 | 66 | 0 | 1 | 0 | 0 | 0 | 26 | 28 | 37 | 11 | 6 | 0 | 0 | 0 | 0 | 0 | 214 | 149 | 1 | 1 | 1 | 0.956 | 1 | 1 | 1 | 1 | 1 | 1 | 1 | 1 | -0.009 |
| SE | 59298 | ENSG00000119004 | CYP20A1 | chr2 | + | 203296473 | 203296563 | 203292261 | 203292326 | 203296757 | 203297014 | 90 | 60 | 63 | 121 | 148 | 0 | 0 | 0 | 0 | 0 | 37 | 49 | 51 | 30 | 8 | 0 | 0 | 0 | 3 | 0 | 239 | 149 | 1 | 1 | 1 | 1 | 1 | 1 | 1 | 1 | 1 | 1 | 0.862 | 1 | 0.028 |
| SE | 59299 | ENSG00000153015 | CWC27 | chr5 | + | 64785480 | 64785579 | 64783835 | 64783979 | 64786523 | 64786627 | 283 | 264 | 211 | 240 | 272 | 0 | 0 | 1 | 0 | 0 | 44 | 153 | 148 | 93 | 20 | 0 | 0 | 0 | 0 | 0 | 248 | 149 | 1 | 1 | 1 | 1 | 0.992 | 1 | 1 | 1 | 1 | 1 | 1 | 1 | -0.002 |
| SE | 59300 | ENSG00000153015 | CWC27 | chr5 | + | 64977134 | 64977238 | 64971702 | 64971812 | 65018158 | 65018591 | 89 | 104 | 146 | 133 | 174 | 0 | 0 | 2 | 0 | 3 | 19 | 114 | 80 | 62 | 0 | 0 | 0 | 0 | 0 | 0 | 253 | 149 | 1 | 1 | 1 | 1 | 0.977 | 1 | 0.972 | 1 | 1 | 1 | 1 | NA | -0.01 |
| SE | 59308 | ENSG00000187922 | LCN10 | chr9 | - | 136740835 | 136741014 | 136739949 | 136740048 | 136741880 | 136742020 | 3 | 5 | 1 | 4 | 7 | 3 | 6 | 8 | 11 | 4 | 10 | 1 | 0 | 3 | 0 | 0 | 6 | 0 | 2 | 0 | 298 | 149 | 1 | 1 | 0.333 | 0.294 | 0.059 | 0.154 | 0.467 | 1 | 0.077 | NA | 0.429 | NA | -0.241 |
| SE | 59309 | ENSG00000187922 | LCN10 | chr9 | - | 136741251 | 136741361 | 136739949 | 136740048 | 136741880 | 136742020 | 1 | 0 | 0 | 0 | 0 | 3 | 6 | 8 | 11 | 4 | 0 | 0 | 0 | 0 | 0 | 0 | 6 | 0 | 2 | 0 | 259 | 149 | 1 | 1 | 0.161 | 0 | 0 | 0 | 0 | NA | 0 | NA | 0 | NA | 0.032 |
| SE | 59311 | ENSG00000187922 | LCN10 | chr9 | - | 136741251 | 136741361 | 136740835 | 136740943 | 136741880 | 136742020 | 1 | 0 | 1 | 0 | 0 | 4 | 6 | 0 | 5 | 1 | 0 | 0 | 0 | 0 | 0 | 32 | 2 | 0 | 5 | 0 | 259 | 149 | 1 | 1 | 0.126 | 0 | 1 | 0 | 0 | 0 | 0 | NA | 0 | NA | 0.225 |
| SE | 59313 | ENSG00000058453 | CROCC | chr1 | + | 16924324 | 16924479 | 16922662 | 16922798 | 16929845 | 16930031 | 3 | 2 | 0 | 0 | 0 | 0 | 0 | 0 | 0 | 0 | 6 | 0 | 0 | 5 | 0 | 0 | 0 | 0 | 1 | 0 | 298 | 149 | 1 | 1 | 1 | 1 | NA | NA | NA | 1 | NA | NA | 0.714 | NA | 0.143 |
| SE | 59314 | ENSG00000058453 | CROCC | chr1 | + | 16931290 | 16931397 | 16930428 | 16930594 | 16936636 | 16936873 | 35 | 22 | 18 | 16 | 13 | 1 | 0 | 0 | 0 | 0 | 32 | 5 | 18 | 10 | 18 | 0 | 0 | 0 | 0 | 0 | 256 | 149 | 1 | 1 | 0.953 | 1 | 1 | 1 | 1 | 1 | 1 | 1 | 1 | 1 | -0.009 |
| SE | 59316 | ENSG00000058453 | CROCC | chr1 | + | 16938399 | 16938483 | 16937640 | 16937737 | 16938908 | 16939142 | 23 | 9 | 21 | 9 | 22 | 1 | 0 | 0 | 0 | 0 | 18 | 6 | 8 | 14 | 14 | 0 | 0 | 0 | 0 | 0 | 233 | 149 | 1 | 1 | 0.936 | 1 | 1 | 1 | 1 | 1 | 1 | 1 | 1 | 1 | -0.013 |
| SE | 59318 | ENSG00000058453 | CROCC | chr1 | + | 16958582 | 16958750 | 16955996 | 16956156 | 16960757 | 16961078 | 39 | 14 | 26 | 18 | 7 | 0 | 1 | 0 | 0 | 0 | 18 | 10 | 13 | 21 | 51 | 0 | 0 | 0 | 0 | 0 | 298 | 149 | 1 | 1 | 1 | 0.875 | 1 | 1 | 1 | 1 | 1 | 1 | 1 | 1 | -0.025 |
| SE | 59319 | ENSG00000058453 | CROCC | chr1 | + | 16960757 | 16961130 | 16958582 | 16958750 | 16965722 | 16965892 | 3 | 1 | 6 | 1 | 7 | 0 | 0 | 1 | 0 | 0 | 0 | 1 | 4 | 0 | 16 | 0 | 0 | 0 | 0 | 0 | 298 | 149 | 1 | 1 | 1 | 1 | 0.75 | 1 | 1 | NA | 1 | 1 | NA | 1 | -0.05 |
| SE | 59320 | ENSG00000058453 | CROCC | chr1 | + | 16966407 | 16966571 | 16965998 | 16966119 | 16968202 | 16968418 | 17 | 17 | 9 | 9 | 0 | 0 | 0 | 0 | 0 | 0 | 4 | 0 | 4 | 5 | 18 | 0 | 1 | 0 | 0 | 0 | 298 | 149 | 1 | 1 | 1 | 1 | 1 | 1 | NA | 1 | 0 | 1 | 1 | 1 | 0.2 |
| SE | 59321 | ENSG00000058453 | CROCC | chr1 | + | 16971464 | 16971647 | 16970635 | 16970767 | 16972359 | 16972964 | 10 | 4 | 7 | 7 | 0 | 0 | 0 | 1 | 0 | 3 | 0 | 4 | 2 | 2 | 0 | 0 | 1 | 0 | 0 | 0 | 298 | 149 | 1 | 1 | 1 | 1 | 0.778 | 1 | 0 | NA | 0.667 | 1 | 1 | NA | -0.133 |
| SE | 59322 | ENSG00000196591 | HDAC2 | chr6 | - | 113949167 | 113949260 | 113948978 | 113949087 | 113953276 | 113953418 | 229 | 168 | 199 | 184 | 271 | 0 | 0 | 0 | 0 | 0 | 63 | 80 | 64 | 101 | 6 | 0 | 4 | 0 | 0 | 0 | 242 | 149 | 1 | 1 | 1 | 1 | 1 | 1 | 1 | 1 | 0.925 | 1 | 1 | 1 | 0.015 |
| SE | 59324 | ENSG00000196591 | HDAC2 | chr6 | - | 113956618 | 113956693 | 113955926 | 113956151 | 113958648 | 113958766 | 447 | 239 | 412 | 467 | 477 | 0 | 0 | 0 | 1 | 0 | 140 | 266 | 124 | 199 | 66 | 0 | 0 | 0 | 0 | 0 | 224 | 149 | 1 | 1 | 1 | 1 | 1 | 0.997 | 1 | 1 | 1 | 1 | 1 | 1 | -0.001 |
| SE | 59327 | ENSG00000196591 | HDAC2 | chr6 | - | 113962366 | 113962394 | 113959905 | 113960018 | 113970856 | 113971157 | 0 | 0 | 0 | 0 | 2 | 63 | 38 | 77 | 75 | 37 | 0 | 0 | 0 | 0 | 0 | 16 | 54 | 16 | 17 | 0 | 177 | 149 | 1 | 1 | 0 | 0 | 0 | 0 | 0.044 | 0 | 0 | 0 | 0 | NA | 0.009 |
| SE | 59332 | ENSG00000213064 | SFT2D2 | chr1 | + | 168229601 | 168229712 | 168225937 | 168226142 | 168231513 | 168231600 | 0 | 0 | 0 | 3 | 0 | 105 | 61 | 67 | 82 | 10 | 0 | 0 | 2 | 0 | 0 | 115 | 58 | 89 | 137 | 53 | 260 | 149 | 1 | 1 | 0 | 0 | 0 | 0.021 | 0 | 0 | 0 | 0.013 | 0 | 0 | 0.002 |
| SE | 59336 | ENSG00000213064 | SFT2D2 | chr1 | + | 168231833 | 168231919 | 168231513 | 168231600 | 168239130 | 168239160 | 69 | 46 | 49 | 63 | 18 | 0 | 0 | 0 | 0 | 0 | 39 | 32 | 64 | 71 | 46 | 0 | 0 | 0 | 2 | 0 | 235 | 149 | 1 | 1 | 1 | 1 | 1 | 1 | 1 | 1 | 1 | 1 | 0.957 | 1 | 0.009 |
| SE | 59338 | ENSG00000213064 | SFT2D2 | chr1 | + | 168235100 | 168235182 | 168231513 | 168231600 | 168236711 | 168236770 | 7 | 2 | 0 | 2 | 1 | 0 | 0 | 0 | 0 | 1 | 19 | 10 | 4 | 41 | 13 | 8 | 0 | 3 | 3 | 3 | 231 | 149 | 1 | 1 | 1 | 1 | NA | 1 | 0.392 | 0.605 | 1 | 0.462 | 0.898 | 0.737 | 0.108 |
| SE | 59339 | ENSG00000213064 | SFT2D2 | chr1 | + | 168235100 | 168235182 | 168231833 | 168231919 | 168236588 | 168236624 | 181 | 95 | 142 | 129 | 93 | 1 | 0 | 0 | 0 | 1 | 104 | 82 | 110 | 56 | 82 | 0 | 0 | 1 | 0 | 0 | 231 | 149 | 1 | 1 | 0.992 | 1 | 1 | 1 | 0.984 | 1 | 1 | 0.986 | 1 | 1 | -0.002 |
| SE | 59342 | ENSG00000213064 | SFT2D2 | chr1 | + | 168236711 | 168236770 | 168231513 | 168231600 | 168239130 | 168239160 | 30 | 16 | 27 | 41 | 27 | 0 | 0 | 0 | 0 | 0 | 38 | 15 | 29 | 33 | 3 | 0 | 0 | 0 | 2 | 0 | 208 | 149 | 1 | 1 | 1 | 1 | 1 | 1 | 1 | 1 | 1 | 1 | 0.922 | 1 | 0.016 |
| SE | 59343 | ENSG00000168216 | LMBRD1 | chr6 | - | 69705500 | 69705619 | 69701888 | 69701953 | 69713644 | 69713797 | 1 | 0 | 0 | 0 | 0 | 138 | 133 | 130 | 135 | 193 | 0 | 0 | 0 | 0 | 0 | 72 | 59 | 65 | 44 | 31 | 268 | 149 | 1 | 1 | 0.004 | 0 | 0 | 0 | 0 | 0 | 0 | 0 | 0 | 0 | 0.001 |
| SE | 59345 | ENSG00000168216 | LMBRD1 | chr6 | - | 69709156 | 69709231 | 69701888 | 69701953 | 69713644 | 69713797 | 1 | 2 | 1 | 0 | 2 | 138 | 133 | 130 | 135 | 193 | 0 | 1 | 2 | 0 | 0 | 72 | 59 | 65 | 44 | 31 | 224 | 149 | 1 | 1 | 0.005 | 0.01 | 0.005 | 0 | 0.007 | 0 | 0.011 | 0.02 | 0 | 0 | -0.001 |
| SE | 59346 | ENSG00000168216 | LMBRD1 | chr6 | - | 69710424 | 69710444 | 69701888 | 69701953 | 69713644 | 69713797 | 0 | 0 | 1 | 0 | 0 | 138 | 133 | 130 | 135 | 193 | 0 | 0 | 0 | 0 | 0 | 72 | 59 | 65 | 44 | 31 | 169 | 149 | 1 | 1 | 0 | 0 | 0.007 | 0 | 0 | 0 | 0 | 0 | 0 | 0 | 0.001 |
| SE | 59348 | ENSG00000168216 | LMBRD1 | chr6 | - | 69743783 | 69743881 | 69741788 | 69741877 | 69749340 | 69749408 | 1 | 1 | 1 | 0 | 1 | 188 | 132 | 127 | 159 | 389 | 0 | 0 | 0 | 0 | 0 | 77 | 95 | 67 | 82 | 78 | 247 | 149 | 1 | 1 | 0.003 | 0.005 | 0.005 | 0 | 0.002 | 0 | 0 | 0 | 0 | 0 | 0.003 |
| SE | 59352 | ENSG00000168216 | LMBRD1 | chr6 | - | 69748223 | 69748289 | 69741788 | 69741877 | 69749340 | 69749408 | 1 | 0 | 1 | 2 | 3 | 188 | 132 | 127 | 159 | 389 | 0 | 0 | 0 | 0 | 0 | 77 | 95 | 67 | 82 | 78 | 215 | 149 | 1 | 1 | 0.004 | 0 | 0.005 | 0.009 | 0.005 | 0 | 0 | 0 | 0 | 0 | 0.005 |
| SE | 59357 | ENSG00000168216 | LMBRD1 | chr6 | - | 69752258 | 69752356 | 69749340 | 69749408 | 69780493 | 69780538 | 496 | 328 | 358 | 344 | 611 | 0 | 1 | 0 | 0 | 0 | 266 | 286 | 186 | 200 | 215 | 0 | 0 | 0 | 0 | 0 | 247 | 149 | 1 | 1 | 1 | 0.995 | 1 | 1 | 1 | 1 | 1 | 1 | 1 | 1 | -0.001 |
| SE | 59370 | ENSG00000116127 | ALMS1 | chr2 | + | 73432128 | 73432291 | 73426452 | 73426553 | 73447959 | 73454067 | 64 | 60 | 52 | 47 | 105 | 0 | 2 | 1 | 0 | 0 | 1 | 22 | 51 | 14 | 5 | 0 | 0 | 0 | 0 | 0 | 298 | 149 | 1 | 1 | 1 | 0.938 | 0.963 | 1 | 1 | 1 | 1 | 1 | 1 | 1 | -0.02 |
| SE | 59371 | ENSG00000116127 | ALMS1 | chr2 | + | 73432197 | 73432291 | 73424429 | 73424902 | 73447959 | 73454067 | 55 | 38 | 39 | 44 | 102 | 0 | 0 | 0 | 0 | 0 | 0 | 19 | 33 | 14 | 5 | 0 | 0 | 9 | 0 | 0 | 243 | 149 | 1 | 1 | 1 | 1 | 1 | 1 | 1 | NA | 1 | 0.692 | 1 | 1 | 0.077 |
| SE | 59372 | ENSG00000116127 | ALMS1 | chr2 | + | 73432197 | 73432291 | 73424429 | 73424902 | 73489633 | 73491498 | 0 | 1 | 3 | 2 | 0 | 0 | 0 | 0 | 0 | 0 | 0 | 0 | 6 | 0 | 0 | 0 | 0 | 1 | 0 | 0 | 243 | 149 | 1 | 1 | NA | 1 | 1 | 1 | NA | NA | NA | 0.786 | NA | NA | 0.214 |
| SE | 59373 | ENSG00000116127 | ALMS1 | chr2 | + | 73432197 | 73432291 | 73426452 | 73426553 | 73447959 | 73454067 | 131 | 102 | 105 | 93 | 202 | 0 | 2 | 1 | 0 | 0 | 43 | 55 | 81 | 40 | 27 | 0 | 0 | 0 | 0 | 0 | 243 | 149 | 1 | 1 | 1 | 0.969 | 0.985 | 1 | 1 | 1 | 1 | 1 | 1 | 1 | -0.009 |
| SE | 59374 | ENSG00000116127 | ALMS1 | chr2 | + | 73447959 | 73454067 | 73432128 | 73432291 | 73455161 | 73455295 | 203 | 189 | 184 | 177 | 235 | 1 | 0 | 1 | 0 | 0 | 30 | 119 | 181 | 37 | 29 | 0 | 0 | 0 | 0 | 0 | 298 | 149 | 1 | 1 | 0.99 | 1 | 0.989 | 1 | 1 | 1 | 1 | 1 | 1 | 1 | -0.004 |
| SE | 59379 | ENSG00000116127 | ALMS1 | chr2 | + | 73550266 | 73550437 | 73534823 | 73534949 | 73557219 | 73557354 | 112 | 61 | 76 | 81 | 112 | 3 | 5 | 5 | 2 | 13 | 8 | 49 | 79 | 25 | 0 | 0 | 0 | 10 | 1 | 0 | 298 | 149 | 1 | 1 | 0.949 | 0.859 | 0.884 | 0.953 | 0.812 | 1 | 1 | 0.798 | 0.926 | NA | -0.04 |
| SE | 59380 | ENSG00000116127 | ALMS1 | chr2 | + | 73572261 | 73573424 | 73558971 | 73559142 | 73599400 | 73599521 | 105 | 75 | 83 | 113 | 153 | 0 | 2 | 1 | 0 | 0 | 1 | 33 | 55 | 16 | 0 | 0 | 1 | 0 | 0 | 0 | 298 | 149 | 1 | 1 | 1 | 0.949 | 0.976 | 1 | 1 | 1 | 0.943 | 1 | 1 | NA | -0.001 |
| SE | 59381 | ENSG00000116127 | ALMS1 | chr2 | + | 73593748 | 73593887 | 73572261 | 73573424 | 73599400 | 73599521 | 0 | 0 | 0 | 1 | 0 | 34 | 29 | 40 | 56 | 39 | 0 | 0 | 0 | 0 | 0 | 0 | 17 | 30 | 3 | 0 | 288 | 149 | 1 | 1 | 0 | 0 | 0 | 0.009 | 0 | NA | 0 | 0 | 0 | NA | 0.002 |
| SE | 59390 | ENSG00000141510 | TP53 | chr17 | - | 7674858 | 7674971 | 7674180 | 7674290 | 7675052 | 7675236 | 311 | 135 | 233 | 225 | 342 | 2 | 0 | 0 | 0 | 0 | 151 | 156 | 59 | 175 | 312 | 0 | 0 | 0 | 0 | 0 | 262 | 149 | 1 | 1 | 0.989 | 1 | 1 | 1 | 1 | 1 | 1 | 1 | 1 | 1 | -0.002 |
| SE | 59395 | ENSG00000169641 | LUZP1 | chr1 | - | 23109584 | 23109650 | 23109021 | 23109127 | 23168765 | 23168845 | 5 | 4 | 4 | 7 | 8 | 2 | 4 | 6 | 3 | 5 | 5 | 7 | 1 | 19 | 0 | 5 | 20 | 0 | 2 | 0 | 215 | 149 | 1 | 1 | 0.634 | 0.409 | 0.316 | 0.618 | 0.526 | 0.409 | 0.195 | 1 | 0.868 | NA | -0.117 |
| SE | 59398 | ENSG00000139631 | CSAD | chr12 | - | 53166204 | 53166327 | 53164621 | 53164755 | 53170071 | 53170126 | 1 | 0 | 0 | 0 | 0 | 3 | 5 | 9 | 3 | 3 | 0 | 0 | 0 | 8 | 10 | 0 | 8 | 7 | 11 | 5 | 272 | 149 | 1 | 1 | 0.154 | 0 | 0 | 0 | 0 | NA | 0 | 0 | 0.285 | 0.523 | -0.171 |
| SE | 59401 | ENSG00000139631 | CSAD | chr12 | - | 53171272 | 53171441 | 53170422 | 53170502 | 53173344 | 53173476 | 1 | 1 | 0 | 7 | 0 | 0 | 0 | 0 | 0 | 0 | 0 | 0 | 0 | 0 | 6 | 0 | 0 | 0 | 4 | 0 | 298 | 149 | 1 | 1 | 1 | 1 | NA | 1 | NA | NA | NA | NA | 0 | 1 | 0.5 |
| SE | 59403 | ENSG00000139631 | CSAD | chr12 | - | 53171325 | 53171441 | 53170422 | 53170502 | 53173344 | 53173476 | 9 | 1 | 2 | 5 | 0 | 0 | 0 | 0 | 0 | 0 | 19 | 1 | 2 | 1 | 15 | 0 | 0 | 0 | 4 | 0 | 265 | 149 | 1 | 1 | 1 | 1 | 1 | 1 | NA | 1 | 1 | 1 | 0.123 | 1 | 0.175 |
| SE | 59404 | ENSG00000139631 | CSAD | chr12 | - | 53171877 | 53171988 | 53171325 | 53171441 | 53173344 | 53173476 | 0 | 1 | 3 | 0 | 6 | 1 | 0 | 0 | 0 | 0 | 0 | 0 | 3 | 0 | 6 | 0 | 0 | 0 | 0 | 0 | 260 | 149 | 1 | 1 | 0 | 1 | 1 | NA | 1 | NA | NA | 1 | NA | 1 | -0.25 |
| SE | 59405 | ENSG00000139631 | CSAD | chr12 | - | 53171881 | 53171988 | 53170422 | 53170502 | 53172345 | 53172371 | 22 | 22 | 20 | 16 | 8 | 1 | 0 | 0 | 0 | 0 | 36 | 23 | 37 | 26 | 25 | 0 | 0 | 0 | 0 | 0 | 256 | 149 | 1 | 1 | 0.928 | 1 | 1 | 1 | 1 | 1 | 1 | 1 | 1 | 1 | -0.014 |
| SE | 59406 | ENSG00000139631 | CSAD | chr12 | - | 53172345 | 53172436 | 53170422 | 53170502 | 53173344 | 53173374 | 1 | 0 | 0 | 1 | 0 | 0 | 0 | 0 | 0 | 0 | 0 | 0 | 0 | 0 | 0 | 0 | 0 | 0 | 4 | 0 | 240 | 149 | 1 | 1 | 1 | NA | NA | 1 | NA | NA | NA | NA | 0 | NA | 1 |
| SE | 59408 | ENSG00000139631 | CSAD | chr12 | - | 53172521 | 53172648 | 53172345 | 53172436 | 53173344 | 53173374 | 52 | 58 | 64 | 44 | 39 | 0 | 0 | 0 | 2 | 0 | 114 | 30 | 67 | 81 | 99 | 0 | 0 | 1 | 0 | 0 | 276 | 149 | 1 | 1 | 1 | 1 | 1 | 0.922 | 1 | 1 | 1 | 0.973 | 1 | 1 | -0.01 |
| SE | 59411 | ENSG00000139631 | CSAD | chr12 | - | 53179776 | 53180012 | 53173727 | 53173770 | 53180566 | 53180596 | 0 | 1 | 1 | 0 | 0 | 2 | 0 | 0 | 0 | 0 | 5 | 0 | 0 | 0 | 0 | 0 | 0 | 0 | 0 | 0 | 298 | 149 | 1 | 1 | 0 | 1 | 1 | NA | NA | 1 | NA | NA | NA | NA | -0.333 |
| SE | 59413 | ENSG00000116863 | ADPRS | chr1 | + | 36091243 | 36091340 | 36088891 | 36089115 | 36091617 | 36091825 | 5 | 1 | 6 | 1 | 6 | 0 | 0 | 0 | 0 | 0 | 0 | 3 | 0 | 5 | 2 | 0 | 0 | 0 | 0 | 3 | 246 | 149 | 1 | 1 | 1 | 1 | 1 | 1 | 1 | NA | 1 | NA | 1 | 0.288 | 0.237 |
| SE | 59414 | ENSG00000115946 | PNO1 | chr2 | + | 68161214 | 68161362 | 68158379 | 68158529 | 68161682 | 68161766 | 0 | 0 | 0 | 0 | 0 | 79 | 66 | 80 | 71 | 94 | 0 | 0 | 0 | 2 | 0 | 102 | 45 | 79 | 98 | 43 | 297 | 149 | 1 | 1 | 0 | 0 | 0 | 0 | 0 | 0 | 0 | 0 | 0.01 | 0 | -0.002 |
| SE | 59415 | ENSG00000115946 | PNO1 | chr2 | + | 68173346 | 68173417 | 68162545 | 68162663 | 68174734 | 68176238 | 35 | 26 | 53 | 30 | 78 | 0 | 0 | 1 | 0 | 0 | 17 | 47 | 49 | 41 | 0 | 0 | 0 | 0 | 0 | 0 | 220 | 149 | 1 | 1 | 1 | 1 | 0.973 | 1 | 1 | 1 | 1 | 1 | 1 | NA | -0.005 |
| SE | 59417 | ENSG00000163116 | STPG2 | chr4 | - | 98134346 | 98134459 | 98128427 | 98128592 | 98143041 | 98143476 | 0 | 0 | 0 | 1 | 3 | 0 | 1 | 0 | 0 | 0 | 0 | 1 | 0 | 0 | 0 | 0 | 0 | 0 | 0 | 0 | 262 | 149 | 1 | 1 | NA | 0 | NA | 1 | 1 | NA | 1 | NA | NA | NA | -0.333 |
| SE | 59418 | ENSG00000136240 | KDELR2 | chr7 | - | 6466070 | 6466323 | 6461088 | 6463175 | 6469595 | 6469754 | 341 | 204 | 372 | 347 | 373 | 3 | 0 | 0 | 1 | 0 | 401 | 341 | 252 | 372 | 465 | 0 | 0 | 1 | 0 | 0 | 298 | 149 | 1 | 1 | 0.983 | 1 | 1 | 0.994 | 1 | 1 | 1 | 0.992 | 1 | 1 | -0.003 |
| SE | 59419 | ENSG00000136240 | KDELR2 | chr7 | - | 6469595 | 6469754 | 6466070 | 6466323 | 6474183 | 6474284 | 330 | 201 | 318 | 365 | 375 | 2 | 0 | 0 | 0 | 0 | 304 | 302 | 208 | 328 | 421 | 0 | 0 | 0 | 0 | 0 | 298 | 149 | 1 | 1 | 0.988 | 1 | 1 | 1 | 1 | 1 | 1 | 1 | 1 | 1 | -0.002 |
| SE | 59420 | ENSG00000136240 | KDELR2 | chr7 | - | 6482391 | 6482589 | 6474183 | 6474284 | 6483966 | 6484149 | 1 | 2 | 1 | 2 | 0 | 66 | 47 | 52 | 71 | 36 | 0 | 0 | 0 | 0 | 0 | 78 | 84 | 38 | 48 | 55 | 298 | 149 | 1 | 1 | 0.008 | 0.021 | 0.01 | 0.014 | 0 | 0 | 0 | 0 | 0 | 0 | 0.011 |
| SE | 59436 | ENSG00000112339 | HBS1L | chr6 | - | 135039572 | 135039767 | 135002733 | 135002842 | 135042000 | 135042126 | 376 | 260 | 388 | 349 | 481 | 1 | 0 | 1 | 0 | 0 | 65 | 220 | 296 | 134 | 30 | 0 | 0 | 0 | 0 | 0 | 298 | 149 | 1 | 1 | 0.995 | 1 | 0.995 | 1 | 1 | 1 | 1 | 1 | 1 | 1 | -0.002 |
| SE | 59437 | ENSG00000112339 | HBS1L | chr6 | - | 135039572 | 135039767 | 135002733 | 135002842 | 135050581 | 135050647 | 100 | 71 | 110 | 94 | 218 | 0 | 0 | 3 | 0 | 0 | 19 | 55 | 93 | 49 | 3 | 0 | 0 | 0 | 0 | 0 | 298 | 149 | 1 | 1 | 1 | 1 | 0.948 | 1 | 1 | 1 | 1 | 1 | 1 | 1 | -0.01 |
| SE | 59438 | ENSG00000112339 | HBS1L | chr6 | - | 135042000 | 135042126 | 135002733 | 135002842 | 135050581 | 135050644 | 130 | 113 | 136 | 128 | 167 | 0 | 0 | 3 | 0 | 0 | 6 | 88 | 129 | 51 | 18 | 0 | 0 | 0 | 0 | 0 | 275 | 149 | 1 | 1 | 1 | 1 | 0.961 | 1 | 1 | 1 | 1 | 1 | 1 | 1 | -0.008 |
| SE | 59439 | ENSG00000112339 | HBS1L | chr6 | - | 135042000 | 135042126 | 135039572 | 135039767 | 135050581 | 135050644 | 408 | 301 | 415 | 384 | 433 | 3 | 1 | 3 | 2 | 3 | 52 | 252 | 335 | 139 | 45 | 0 | 0 | 3 | 3 | 0 | 275 | 149 | 1 | 1 | 0.987 | 0.994 | 0.987 | 0.99 | 0.987 | 1 | 1 | 0.984 | 0.962 | 1 | 0 |
| SE | 59442 | ENSG00000112339 | HBS1L | chr6 | - | 135050581 | 135050647 | 135042000 | 135042126 | 135054648 | 135054800 | 172 | 139 | 195 | 190 | 199 | 0 | 2 | 1 | 0 | 2 | 6 | 103 | 173 | 54 | 23 | 0 | 0 | 0 | 0 | 0 | 215 | 149 | 1 | 1 | 1 | 0.98 | 0.993 | 1 | 0.986 | 1 | 1 | 1 | 1 | 1 | -0.008 |
| SE | 59444 | ENSG00000145246 | ATP10D | chr4 | + | 47535508 | 47535615 | 47525556 | 47525642 | 47535901 | 47536033 | 128 | 83 | 87 | 91 | 79 | 0 | 0 | 0 | 1 | 0 | 68 | 99 | 33 | 94 | 27 | 0 | 0 | 0 | 0 | 0 | 256 | 149 | 1 | 1 | 1 | 1 | 1 | 0.981 | 1 | 1 | 1 | 1 | 1 | 1 | -0.004 |
| SE | 59446 | ENSG00000145246 | ATP10D | chr4 | + | 47558922 | 47559029 | 47557663 | 47558273 | 47560948 | 47561075 | 152 | 138 | 114 | 134 | 134 | 0 | 1 | 1 | 0 | 1 | 130 | 157 | 74 | 163 | 25 | 0 | 0 | 0 | 0 | 0 | 256 | 149 | 1 | 1 | 1 | 0.988 | 0.985 | 1 | 0.987 | 1 | 1 | 1 | 1 | 1 | -0.008 |
| SE | 59448 | ENSG00000145246 | ATP10D | chr4 | + | 47582868 | 47582961 | 47582030 | 47582064 | 47587018 | 47587206 | 3 | 2 | 6 | 3 | 9 | 12 | 17 | 13 | 32 | 36 | 0 | 8 | 1 | 7 | 0 | 0 | 0 | 5 | 8 | 0 | 242 | 149 | 1 | 1 | 0.133 | 0.068 | 0.221 | 0.055 | 0.133 | NA | 1 | 0.11 | 0.35 | NA | -0.365 |
| SE | 59450 | ENSG00000115042 | FAHD2A | chr2 | + | 95405276 | 95405402 | 95402790 | 95402992 | 95405552 | 95405803 | 22 | 9 | 13 | 12 | 12 | 1 | 0 | 1 | 1 | 0 | 2 | 15 | 1 | 7 | 0 | 0 | 0 | 0 | 0 | 0 | 275 | 149 | 1 | 1 | 0.923 | 1 | 0.876 | 0.867 | 1 | 1 | 1 | 1 | 1 | NA | -0.067 |
| SE | 59451 | ENSG00000115042 | FAHD2A | chr2 | + | 95405552 | 95405803 | 95402707 | 95402872 | 95412433 | 95412542 | 4 | 0 | 1 | 3 | 1 | 1 | 0 | 0 | 0 | 0 | 0 | 3 | 0 | 3 | 9 | 0 | 0 | 0 | 0 | 0 | 298 | 149 | 1 | 1 | 0.667 | NA | 1 | 1 | 1 | NA | 1 | NA | 1 | 1 | -0.083 |
| SE | 59453 | ENSG00000115042 | FAHD2A | chr2 | + | 95410526 | 95410586 | 95402707 | 95402872 | 95412433 | 95412542 | 7 | 0 | 2 | 5 | 0 | 1 | 0 | 0 | 0 | 0 | 6 | 0 | 0 | 0 | 4 | 0 | 0 | 0 | 0 | 0 | 209 | 149 | 1 | 1 | 0.833 | NA | 1 | 1 | NA | 1 | NA | NA | NA | 1 | -0.056 |
| SE | 59457 | ENSG00000113312 | TTC1 | chr5 | + | 160010475 | 160010858 | 160009112 | 160009193 | 160035139 | 160035200 | 51 | 50 | 65 | 80 | 100 | 2 | 1 | 0 | 0 | 0 | 240 | 68 | 72 | 107 | 200 | 0 | 0 | 0 | 0 | 0 | 298 | 149 | 1 | 1 | 0.927 | 0.962 | 1 | 1 | 1 | 1 | 1 | 1 | 1 | 1 | -0.022 |
| SE | 59458 | ENSG00000113312 | TTC1 | chr5 | + | 160010499 | 160010858 | 160009124 | 160009193 | 160035139 | 160035200 | 196 | 145 | 199 | 243 | 242 | 2 | 1 | 0 | 0 | 0 | 417 | 170 | 147 | 202 | 329 | 0 | 0 | 0 | 0 | 0 | 298 | 149 | 1 | 1 | 0.98 | 0.986 | 1 | 1 | 1 | 1 | 1 | 1 | 1 | 1 | -0.007 |
| SE | 59459 | ENSG00000113312 | TTC1 | chr5 | + | 160010585 | 160010858 | 160009145 | 160009193 | 160035139 | 160035200 | 53 | 50 | 65 | 81 | 100 | 2 | 1 | 0 | 0 | 0 | 240 | 68 | 72 | 107 | 200 | 0 | 0 | 0 | 0 | 0 | 298 | 149 | 1 | 1 | 0.93 | 0.962 | 1 | 1 | 1 | 1 | 1 | 1 | 1 | 1 | -0.022 |
| SE | 59462 | ENSG00000113312 | TTC1 | chr5 | + | 160043132 | 160043169 | 160036505 | 160036803 | 160049513 | 160049662 | 83 | 68 | 88 | 99 | 123 | 0 | 2 | 0 | 0 | 0 | 103 | 63 | 73 | 114 | 104 | 0 | 0 | 0 | 0 | 0 | 186 | 149 | 1 | 1 | 1 | 0.965 | 1 | 1 | 1 | 1 | 1 | 1 | 1 | 1 | -0.007 |
| SE | 59463 | ENSG00000113312 | TTC1 | chr5 | + | 160049513 | 160049662 | 160041342 | 160043169 | 160051128 | 160051183 | 131 | 99 | 116 | 153 | 244 | 0 | 0 | 2 | 0 | 3 | 141 | 107 | 109 | 167 | 101 | 0 | 0 | 1 | 1 | 4 | 298 | 149 | 1 | 1 | 1 | 1 | 0.967 | 1 | 0.976 | 1 | 1 | 0.982 | 0.988 | 0.927 | 0.009 |
| SE | 59464 | ENSG00000113312 | TTC1 | chr5 | + | 160049513 | 160049662 | 160041342 | 160043169 | 160064931 | 160065511 | 63 | 53 | 69 | 77 | 104 | 0 | 0 | 0 | 0 | 0 | 82 | 56 | 58 | 82 | 79 | 2 | 0 | 0 | 0 | 0 | 298 | 149 | 1 | 1 | 1 | 1 | 1 | 1 | 1 | 0.953 | 1 | 1 | 1 | 1 | 0.009 |
| SE | 59467 | ENSG00000113312 | TTC1 | chr5 | + | 160064931 | 160065022 | 160051128 | 160051183 | 160065276 | 160065487 | 50 | 37 | 32 | 57 | 108 | 0 | 2 | 0 | 0 | 0 | 47 | 47 | 34 | 63 | 30 | 0 | 0 | 0 | 0 | 0 | 240 | 149 | 1 | 1 | 1 | 0.92 | 1 | 1 | 1 | 1 | 1 | 1 | 1 | 1 | -0.016 |
| SE | 59468 | ENSG00000164332 | UBLCP1 | chr5 | + | 159270359 | 159270445 | 159269907 | 159269999 | 159270527 | 159270643 | 162 | 127 | 145 | 143 | 165 | 1 | 0 | 0 | 1 | 0 | 26 | 63 | 66 | 16 | 0 | 0 | 0 | 0 | 0 | 0 | 235 | 149 | 1 | 1 | 0.99 | 1 | 1 | 0.989 | 1 | 1 | 1 | 1 | 1 | NA | -0.004 |
| SE | 59469 | ENSG00000164332 | UBLCP1 | chr5 | + | 159275147 | 159275246 | 159274541 | 159274622 | 159278237 | 159278354 | 443 | 302 | 303 | 345 | 426 | 2 | 0 | 0 | 0 | 7 | 51 | 66 | 191 | 16 | 8 | 0 | 0 | 0 | 0 | 0 | 248 | 149 | 1 | 1 | 0.993 | 1 | 1 | 1 | 0.973 | 1 | 1 | 1 | 1 | 1 | -0.007 |
| SE | 59480 | ENSG00000160310 | PRMT2 | chr21 | + | 46643534 | 46643639 | 46636895 | 46636990 | 46648457 | 46648619 | 379 | 172 | 183 | 290 | 284 | 0 | 0 | 1 | 1 | 1 | 258 | 303 | 145 | 162 | 377 | 0 | 0 | 2 | 3 | 0 | 254 | 149 | 1 | 1 | 1 | 1 | 0.991 | 0.994 | 0.994 | 1 | 1 | 0.977 | 0.969 | 1 | 0.007 |
| SE | 59481 | ENSG00000160310 | PRMT2 | chr21 | + | 46644305 | 46644488 | 46643285 | 46643639 | 46648457 | 46648538 | 1024 | 438 | 722 | 950 | 850 | 2 | 0 | 0 | 0 | 0 | 567 | 874 | 417 | 459 | 807 | 0 | 0 | 0 | 0 | 0 | 298 | 149 | 1 | 1 | 0.996 | 1 | 1 | 1 | 1 | 1 | 1 | 1 | 1 | 1 | -0.001 |
| SE | 59482 | ENSG00000160310 | PRMT2 | chr21 | + | 46649573 | 46649739 | 46648539 | 46648619 | 46658744 | 46658798 | 196 | 60 | 141 | 191 | 99 | 0 | 2 | 0 | 0 | 0 | 23 | 84 | 51 | 33 | 134 | 0 | 0 | 0 | 0 | 0 | 298 | 149 | 1 | 1 | 1 | 0.938 | 1 | 1 | 1 | 1 | 1 | 1 | 1 | 1 | -0.012 |
| SE | 59483 | ENSG00000160310 | PRMT2 | chr21 | + | 46649574 | 46649739 | 46648457 | 46648619 | 46658744 | 46658920 | 706 | 289 | 485 | 673 | 519 | 0 | 2 | 0 | 0 | 0 | 361 | 542 | 276 | 258 | 584 | 0 | 0 | 0 | 0 | 0 | 298 | 149 | 1 | 1 | 1 | 0.986 | 1 | 1 | 1 | 1 | 1 | 1 | 1 | 1 | -0.003 |
| SE | 59484 | ENSG00000160310 | PRMT2 | chr21 | + | 46658744 | 46658920 | 46649574 | 46649739 | 46660832 | 46660962 | 387 | 104 | 235 | 324 | 199 | 0 | 2 | 0 | 0 | 3 | 51 | 153 | 82 | 86 | 232 | 0 | 2 | 0 | 0 | 0 | 298 | 149 | 1 | 1 | 1 | 0.963 | 1 | 1 | 0.971 | 1 | 0.975 | 1 | 1 | 1 | -0.008 |
| SE | 59490 | ENSG00000148660 | CAMK2G | chr10 | - | 73819531 | 73819617 | 73817478 | 73817554 | 73821681 | 73821730 | 4 | 1 | 0 | 1 | 2 | 112 | 92 | 149 | 132 | 158 | 0 | 4 | 4 | 0 | 6 | 223 | 146 | 146 | 171 | 248 | 235 | 149 | 1 | 1 | 0.022 | 0.007 | 0 | 0.005 | 0.008 | 0 | 0.017 | 0.017 | 0 | 0.015 | -0.001 |
| SE | 59501 | ENSG00000148660 | CAMK2G | chr10 | - | 73828088 | 73828121 | 73825278 | 73825347 | 73837467 | 73837511 | 1 | 0 | 0 | 1 | 0 | 141 | 74 | 130 | 134 | 140 | 0 | 0 | 0 | 0 | 0 | 237 | 138 | 118 | 252 | 239 | 182 | 149 | 1 | 1 | 0.006 | 0 | 0 | 0.006 | 0 | 0 | 0 | 0 | 0 | 0 | 0.002 |
| SE | 59503 | ENSG00000148660 | CAMK2G | chr10 | - | 73847224 | 73847347 | 73842457 | 73842541 | 73848525 | 73848609 | 108 | 77 | 131 | 140 | 148 | 0 | 0 | 0 | 0 | 0 | 156 | 149 | 120 | 137 | 246 | 0 | 0 | 0 | 0 | 7 | 272 | 149 | 1 | 1 | 1 | 1 | 1 | 1 | 1 | 1 | 1 | 1 | 1 | 0.951 | 0.01 |
| SE | 59504 | ENSG00000148660 | CAMK2G | chr10 | - | 73847224 | 73847347 | 73842457 | 73842541 | 73849012 | 73849115 | 107 | 74 | 127 | 138 | 147 | 0 | 0 | 1 | 0 | 0 | 156 | 139 | 122 | 136 | 242 | 0 | 0 | 2 | 5 | 0 | 272 | 149 | 1 | 1 | 1 | 1 | 0.986 | 1 | 1 | 1 | 1 | 0.971 | 0.937 | 1 | 0.016 |
| SE | 59510 | ENSG00000148660 | CAMK2G | chr10 | - | 73852253 | 73852319 | 73849260 | 73849333 | 73853191 | 73853246 | 142 | 105 | 141 | 170 | 134 | 0 | 0 | 0 | 3 | 0 | 354 | 127 | 127 | 214 | 254 | 0 | 0 | 0 | 0 | 0 | 215 | 149 | 1 | 1 | 1 | 1 | 1 | 0.975 | 1 | 1 | 1 | 1 | 1 | 1 | -0.005 |
| SE | 59519 | ENSG00000048162 | NOP16 | chr5 | - | 176386839 | 176386909 | 176384642 | 176385327 | 176388234 | 176388343 | 24 | 29 | 25 | 51 | 27 | 0 | 1 | 0 | 2 | 0 | 76 | 50 | 17 | 25 | 41 | 12 | 10 | 0 | 2 | 6 | 219 | 149 | 1 | 1 | 1 | 0.952 | 1 | 0.946 | 1 | 0.812 | 0.773 | 1 | 0.895 | 0.823 | 0.119 |
| SE | 59524 | ENSG00000048162 | NOP16 | chr5 | - | 176388234 | 176388343 | 176386023 | 176386909 | 176388432 | 176388568 | 24 | 23 | 30 | 41 | 24 | 0 | 0 | 0 | 1 | 0 | 44 | 38 | 18 | 22 | 45 | 0 | 0 | 0 | 0 | 0 | 258 | 149 | 1 | 1 | 1 | 1 | 1 | 0.959 | 1 | 1 | 1 | 1 | 1 | 1 | -0.008 |
| SE | 59528 | ENSG00000137101 | CD72 | chr9 | - | 35618741 | 35618865 | 35618013 | 35618121 | 35646402 | 35646810 | 1 | 1 | 5 | 3 | 0 | 0 | 0 | 0 | 1 | 1 | 0 | 0 | 8 | 0 | 3 | 0 | 2 | 4 | 0 | 0 | 273 | 149 | 1 | 1 | 1 | 1 | 1 | 0.621 | 0 | NA | 0 | 0.522 | NA | 1 | 0.217 |
| SE | 59534 | ENSG00000087903 | RFX2 | chr19 | - | 6026162 | 6026237 | 6016089 | 6016271 | 6039979 | 6040241 | 1 | 3 | 1 | 3 | 10 | 0 | 0 | 0 | 0 | 0 | 0 | 0 | 15 | 1 | 21 | 0 | 0 | 0 | 0 | 2 | 224 | 149 | 1 | 1 | 1 | 1 | 1 | 1 | 1 | NA | NA | 1 | 1 | 0.875 | 0.042 |
| SE | 59542 | ENSG00000135605 | TEC | chr4 | - | 48141354 | 48141419 | 48138903 | 48139022 | 48145078 | 48145295 | 3 | 12 | 10 | 11 | 11 | 0 | 0 | 1 | 1 | 0 | 0 | 6 | 6 | 8 | 0 | 0 | 0 | 3 | 1 | 0 | 214 | 149 | 1 | 1 | 1 | 1 | 0.874 | 0.885 | 1 | NA | 1 | 0.582 | 0.848 | NA | 0.142 |
| SE | 59544 | ENSG00000135605 | TEC | chr4 | - | 48145407 | 48145579 | 48141354 | 48141419 | 48146324 | 48146399 | 25 | 12 | 12 | 17 | 22 | 0 | 1 | 0 | 0 | 0 | 23 | 13 | 37 | 14 | 0 | 0 | 0 | 0 | 0 | 0 | 298 | 149 | 1 | 1 | 1 | 0.857 | 1 | 1 | 1 | 1 | 1 | 1 | 1 | NA | -0.029 |
| SE | 59545 | ENSG00000135605 | TEC | chr4 | - | 48146324 | 48146399 | 48145407 | 48145579 | 48149556 | 48149685 | 33 | 18 | 19 | 22 | 28 | 0 | 0 | 1 | 0 | 0 | 28 | 20 | 46 | 23 | 3 | 0 | 0 | 0 | 0 | 0 | 224 | 149 | 1 | 1 | 1 | 1 | 0.927 | 1 | 1 | 1 | 1 | 1 | 1 | 1 | -0.015 |
| SE | 59547 | ENSG00000135605 | TEC | chr4 | - | 48170247 | 48170376 | 48168585 | 48168626 | 48171367 | 48171408 | 65 | 43 | 42 | 66 | 92 | 0 | 1 | 0 | 0 | 0 | 114 | 106 | 56 | 91 | 121 | 0 | 0 | 0 | 0 | 0 | 278 | 149 | 1 | 1 | 1 | 0.958 | 1 | 1 | 1 | 1 | 1 | 1 | 1 | 1 | -0.008 |
| SE | 59548 | ENSG00000135605 | TEC | chr4 | - | 48171367 | 48171408 | 48170247 | 48170376 | 48176081 | 48176186 | 1 | 0 | 2 | 4 | 0 | 1 | 0 | 0 | 0 | 1 | 7 | 3 | 1 | 4 | 0 | 0 | 0 | 0 | 0 | 0 | 190 | 149 | 1 | 1 | 0.44 | NA | 1 | 1 | 0 | 1 | 1 | 1 | 1 | NA | -0.39 |
| SE | 59549 | ENSG00000135605 | TEC | chr4 | - | 48171367 | 48171449 | 48170247 | 48170376 | 48176081 | 48176186 | 44 | 29 | 27 | 51 | 78 | 1 | 0 | 0 | 0 | 1 | 93 | 61 | 47 | 71 | 60 | 0 | 0 | 0 | 0 | 0 | 231 | 149 | 1 | 1 | 0.966 | 1 | 1 | 1 | 0.981 | 1 | 1 | 1 | 1 | 1 | -0.011 |
| SE | 59589 | ENSG00000118816 | CCNI | chr4 | - | 77055961 | 77056100 | 77055322 | 77055380 | 77066248 | 77066405 | 1714 | 1025 | 1545 | 1490 | 4298 | 0 | 0 | 1 | 2 | 0 | 1552 | 1160 | 1076 | 1172 | 1899 | 0 | 0 | 0 | 4 | 0 | 288 | 149 | 1 | 1 | 1 | 1 | 0.999 | 0.997 | 1 | 1 | 1 | 1 | 0.993 | 1 | 0.001 |
| SE | 59590 | ENSG00000118816 | CCNI | chr4 | - | 77058506 | 77058635 | 77055149 | 77055380 | 77066248 | 77066405 | 810 | 437 | 800 | 673 | 1267 | 0 | 0 | 1 | 2 | 0 | 600 | 587 | 470 | 556 | 916 | 0 | 0 | 0 | 4 | 0 | 278 | 149 | 1 | 1 | 1 | 1 | 0.998 | 0.994 | 1 | 1 | 1 | 1 | 0.987 | 1 | 0.001 |
| SE | 59592 | ENSG00000118816 | CCNI | chr4 | - | 77066248 | 77066405 | 77056248 | 77056323 | 77075471 | 77075693 | 600 | 226 | 513 | 457 | 536 | 0 | 0 | 0 | 1 | 0 | 415 | 413 | 297 | 379 | 519 | 0 | 0 | 0 | 0 | 0 | 298 | 149 | 1 | 1 | 1 | 1 | 1 | 0.996 | 1 | 1 | 1 | 1 | 1 | 1 | -0.001 |
| SE | 59593 | ENSG00000118816 | CCNI | chr4 | - | 77066248 | 77066405 | 77058506 | 77058635 | 77075471 | 77075693 | 1450 | 685 | 1333 | 1163 | 1917 | 3 | 0 | 3 | 0 | 4 | 1012 | 1023 | 808 | 982 | 1475 | 0 | 5 | 2 | 0 | 0 | 298 | 149 | 1 | 1 | 0.996 | 1 | 0.996 | 1 | 0.996 | 1 | 0.99 | 0.995 | 1 | 1 | 0.001 |
| SE | 59595 | ENSG00000171121 | KCNMB3 | chr3 | - | 179250742 | 179250934 | 179244494 | 179244693 | 179258947 | 179259208 | 4 | 5 | 3 | 2 | 8 | 0 | 0 | 0 | 0 | 0 | 0 | 18 | 0 | 14 | 2 | 0 | 0 | 0 | 2 | 0 | 298 | 149 | 1 | 1 | 1 | 1 | 1 | 1 | 1 | NA | 1 | NA | 0.778 | 1 | 0.074 |
| SE | 59598 | ENSG00000085377 | PREP | chr6 | - | 105288757 | 105288894 | 105282450 | 105282582 | 105323664 | 105323768 | 61 | 30 | 30 | 24 | 40 | 0 | 2 | 0 | 0 | 0 | 112 | 64 | 37 | 38 | 55 | 0 | 0 | 0 | 2 | 0 | 286 | 149 | 1 | 1 | 1 | 0.887 | 1 | 1 | 1 | 1 | 1 | 1 | 0.908 | 1 | -0.004 |
| SE | 59599 | ENSG00000085377 | PREP | chr6 | - | 105288757 | 105288894 | 105282450 | 105282582 | 105328828 | 105329026 | 3 | 4 | 2 | 2 | 0 | 0 | 0 | 0 | 1 | 0 | 13 | 11 | 6 | 10 | 0 | 0 | 0 | 0 | 0 | 0 | 286 | 149 | 1 | 1 | 1 | 1 | 1 | 0.51 | NA | 1 | 1 | 1 | 1 | NA | -0.123 |
| SE | 59600 | ENSG00000085377 | PREP | chr6 | - | 105323664 | 105323768 | 105282450 | 105282582 | 105328828 | 105329026 | 103 | 70 | 92 | 82 | 58 | 0 | 0 | 0 | 1 | 0 | 130 | 79 | 78 | 100 | 77 | 0 | 0 | 0 | 0 | 0 | 253 | 149 | 1 | 1 | 1 | 1 | 1 | 0.98 | 1 | 1 | 1 | 1 | 1 | 1 | -0.004 |
| SE | 59601 | ENSG00000085377 | PREP | chr6 | - | 105323664 | 105323768 | 105288757 | 105288894 | 105328828 | 105329026 | 165 | 95 | 124 | 105 | 98 | 1 | 0 | 0 | 0 | 0 | 229 | 138 | 112 | 129 | 132 | 0 | 0 | 2 | 0 | 0 | 253 | 149 | 1 | 1 | 0.99 | 1 | 1 | 1 | 1 | 1 | 1 | 0.971 | 1 | 1 | 0.004 |
| SE | 59602 | ENSG00000085377 | PREP | chr6 | - | 105323664 | 105323768 | 105322254 | 105323111 | 105328828 | 105329026 | 113 | 73 | 104 | 97 | 78 | 0 | 0 | 1 | 0 | 0 | 138 | 86 | 93 | 115 | 77 | 0 | 0 | 2 | 0 | 0 | 253 | 149 | 1 | 1 | 1 | 1 | 0.984 | 1 | 1 | 1 | 1 | 0.965 | 1 | 1 | 0.004 |
| SE | 59604 | ENSG00000085377 | PREP | chr6 | - | 105368902 | 105369024 | 105352971 | 105353077 | 105373368 | 105373578 | 471 | 258 | 355 | 414 | 522 | 0 | 0 | 0 | 0 | 0 | 407 | 447 | 226 | 358 | 394 | 0 | 0 | 2 | 0 | 0 | 271 | 149 | 1 | 1 | 1 | 1 | 1 | 1 | 1 | 1 | 1 | 0.984 | 1 | 1 | 0.003 |
| SE | 59606 | ENSG00000144026 | ZNF514 | chr2 | - | 95157350 | 95157439 | 95153132 | 95153259 | 95165030 | 95165413 | 4 | 3 | 2 | 4 | 0 | 0 | 0 | 1 | 0 | 0 | 6 | 0 | 1 | 0 | 0 | 0 | 0 | 0 | 0 | 0 | 238 | 149 | 1 | 1 | 1 | 1 | 0.556 | 1 | NA | 1 | NA | 1 | NA | NA | -0.111 |
| SE | 59608 | ENSG00000257093 | DENND11 | chr7 | - | 141666286 | 141666386 | 141665186 | 141665318 | 141674066 | 141674220 | 41 | 17 | 36 | 43 | 38 | 0 | 0 | 0 | 0 | 0 | 27 | 16 | 26 | 15 | 24 | 0 | 0 | 1 | 0 | 0 | 249 | 149 | 1 | 1 | 1 | 1 | 1 | 1 | 1 | 1 | 1 | 0.94 | 1 | 1 | 0.012 |
| SE | 59609 | ENSG00000257093 | DENND11 | chr7 | - | 141666286 | 141666425 | 141665186 | 141665318 | 141674066 | 141674220 | 147 | 63 | 131 | 146 | 149 | 0 | 0 | 0 | 0 | 0 | 75 | 66 | 91 | 57 | 54 | 0 | 0 | 1 | 0 | 0 | 288 | 149 | 1 | 1 | 1 | 1 | 1 | 1 | 1 | 1 | 1 | 0.979 | 1 | 1 | 0.004 |
| SE | 59610 | ENSG00000257093 | DENND11 | chr7 | - | 141674066 | 141674220 | 141666286 | 141666425 | 141685477 | 141685636 | 160 | 82 | 175 | 143 | 166 | 0 | 0 | 0 | 0 | 0 | 59 | 81 | 93 | 70 | 29 | 0 | 0 | 2 | 0 | 0 | 298 | 149 | 1 | 1 | 1 | 1 | 1 | 1 | 1 | 1 | 1 | 0.959 | 1 | 1 | 0.008 |
| SE | 59614 | ENSG00000165996 | HACD1 | chr10 | - | 17599289 | 17599411 | 17594204 | 17594383 | 17603559 | 17603648 | 13 | 4 | 11 | 7 | 0 | 2 | 0 | 0 | 0 | 0 | 1 | 13 | 6 | 0 | 8 | 0 | 0 | 0 | 0 | 3 | 271 | 149 | 1 | 1 | 0.781 | 1 | 1 | 1 | NA | 1 | 1 | 1 | NA | 0.595 | 0.047 |
| SE | 59620 | ENSG00000111801 | BTN3A3 | chr6 | + | 26448247 | 26448448 | 26445537 | 26445985 | 26448727 | 26448754 | 107 | 66 | 173 | 196 | 86 | 0 | 0 | 2 | 0 | 0 | 151 | 190 | 72 | 122 | 128 | 0 | 0 | 3 | 1 | 0 | 298 | 149 | 1 | 1 | 1 | 1 | 0.977 | 1 | 1 | 1 | 1 | 0.923 | 0.984 | 1 | 0.014 |
| SE | 59623 | ENSG00000111801 | BTN3A3 | chr6 | + | 26448727 | 26448754 | 26448616 | 26448637 | 26449661 | 26449688 | 513 | 298 | 384 | 475 | 446 | 0 | 0 | 0 | 0 | 0 | 561 | 367 | 212 | 240 | 472 | 0 | 0 | 0 | 5 | 0 | 176 | 149 | 1 | 1 | 1 | 1 | 1 | 1 | 1 | 1 | 1 | 1 | 0.976 | 1 | 0.005 |
| SE | 59625 | ENSG00000138279 | ANXA7 | chr10 | - | 73378910 | 73379023 | 73375100 | 73376217 | 73379878 | 73379954 | 290 | 251 | 283 | 270 | 387 | 1 | 0 | 0 | 1 | 3 | 249 | 240 | 221 | 309 | 453 | 0 | 0 | 0 | 0 | 3 | 262 | 149 | 1 | 1 | 0.994 | 1 | 1 | 0.994 | 0.987 | 1 | 1 | 1 | 1 | 0.988 | -0.003 |
| SE | 59626 | ENSG00000138279 | ANXA7 | chr10 | - | 73379878 | 73379954 | 73378910 | 73379023 | 73380030 | 73380201 | 296 | 246 | 262 | 259 | 392 | 0 | 0 | 0 | 0 | 0 | 226 | 296 | 240 | 276 | 495 | 1 | 0 | 0 | 0 | 0 | 225 | 149 | 1 | 1 | 1 | 1 | 1 | 1 | 1 | 0.993 | 1 | 1 | 1 | 1 | 0.001 |
| SE | 59627 | ENSG00000138279 | ANXA7 | chr10 | - | 73387688 | 73387783 | 73383576 | 73383690 | 73388311 | 73388414 | 456 | 389 | 466 | 487 | 704 | 2 | 0 | 3 | 0 | 2 | 1242 | 613 | 668 | 808 | 1405 | 0 | 0 | 3 | 0 | 0 | 244 | 149 | 1 | 1 | 0.993 | 1 | 0.99 | 1 | 0.995 | 1 | 1 | 0.993 | 1 | 1 | -0.003 |
| SE | 59629 | ENSG00000138279 | ANXA7 | chr10 | - | 73398180 | 73398385 | 73397163 | 73397274 | 73414011 | 73414076 | 117 | 59 | 112 | 100 | 31 | 0 | 0 | 0 | 0 | 0 | 268 | 159 | 131 | 112 | 320 | 0 | 0 | 1 | 0 | 0 | 298 | 149 | 1 | 1 | 1 | 1 | 1 | 1 | 1 | 1 | 1 | 0.985 | 1 | 1 | 0.003 |
| SE | 59630 | ENSG00000138279 | ANXA7 | chr10 | - | 73400802 | 73400857 | 73398180 | 73398385 | 73414011 | 73414058 | 39 | 14 | 36 | 31 | 17 | 3 | 0 | 1 | 0 | 0 | 50 | 60 | 41 | 53 | 92 | 18 | 0 | 4 | 2 | 0 | 204 | 149 | 1 | 1 | 0.905 | 1 | 0.963 | 1 | 1 | 0.67 | 1 | 0.882 | 0.951 | 1 | 0.073 |
| SE | 59631 | ENSG00000182827 | ACBD3 | chr1 | - | 226152334 | 226152619 | 226144678 | 226146821 | 226159183 | 226159358 | 73 | 32 | 55 | 63 | 80 | 2 | 0 | 0 | 0 | 0 | 60 | 73 | 75 | 56 | 0 | 0 | 0 | 0 | 0 | 0 | 298 | 149 | 1 | 1 | 0.948 | 1 | 1 | 1 | 1 | 1 | 1 | 1 | 1 | NA | -0.01 |
| SE | 59632 | ENSG00000182827 | ACBD3 | chr1 | - | 226154646 | 226154833 | 226152334 | 226152619 | 226159183 | 226159358 | 250 | 177 | 261 | 297 | 332 | 0 | 0 | 0 | 1 | 0 | 126 | 237 | 270 | 195 | 94 | 0 | 0 | 0 | 0 | 0 | 298 | 149 | 1 | 1 | 1 | 1 | 1 | 0.993 | 1 | 1 | 1 | 1 | 1 | 1 | -0.001 |
| SE | 59636 | ENSG00000163466 | ARPC2 | chr2 | + | 218225919 | 218225954 | 218217209 | 218217544 | 218228737 | 218228850 | 1370 | 911 | 978 | 1171 | 905 | 0 | 0 | 0 | 0 | 0 | 2241 | 1413 | 1190 | 1898 | 1797 | 0 | 1 | 2 | 3 | 0 | 184 | 149 | 1 | 1 | 1 | 1 | 1 | 1 | 1 | 1 | 0.999 | 0.998 | 0.998 | 1 | 0.001 |
| SE | 59638 | ENSG00000163466 | ARPC2 | chr2 | + | 218228737 | 218228850 | 218225918 | 218225954 | 218234351 | 218234397 | 2846 | 1892 | 1869 | 2395 | 2882 | 1 | 0 | 0 | 1 | 0 | 5030 | 2667 | 2164 | 3534 | 3908 | 0 | 0 | 0 | 0 | 0 | 262 | 149 | 1 | 1 | 0.999 | 1 | 1 | 0.999 | 1 | 1 | 1 | 1 | 1 | 1 | 0 |
| SE | 59641 | ENSG00000163466 | ARPC2 | chr2 | + | 218234848 | 218235044 | 218234351 | 218234397 | 218238663 | 218238723 | 12 | 13 | 12 | 18 | 27 | 2386 | 1668 | 1504 | 2001 | 3169 | 2 | 7 | 37 | 29 | 27 | 5150 | 2211 | 1670 | 2764 | 3498 | 298 | 149 | 1 | 1 | 0.003 | 0.004 | 0.004 | 0.004 | 0.004 | 0 | 0.002 | 0.011 | 0.005 | 0.004 | -0.001 |
| [illegible] | | | | | | | | | | | | | | | | | | | | | | | | | | | | | | | | | | | | | | | | | | | | | | |

| | | | | | | | | | | | | | | | | | | | | | | | | | | | | | | | | | | | | | | | | | | | | | | |
|---|---|---|---|---|---|---|---|---|---|---|---|---|---|---|---|---|---|---|---|---|---|---|---|---|---|---|---|---|---|---|---|---|---|---|---|---|---|---|---|---|---|---|---|---|---|---|
| SE | 59691 | ENSG00000152683 | SLC30A6 | chr2 | + | 32204589 | 32204692 | 32197706 | 32197826 | 32206885 | 32206933 | 84 | 49 | 66 | 60 | 87 | 1 | 0 | 0 | 2 | 0 | 20 | 29 | 18 | 32 | 9 | 0 | 0 | 3 | 0 | 0 | 252 | 149 | 1 | 1 | 0.98 | 1 | 1 | 0.947 | 1 | 1 | 1 | 0.78 | 1 | 1 | 0.029 |
| SE | 59692 | ENSG00000152683 | SLC30A6 | chr2 | + | 32204589 | 32204692 | 32197706 | 32197826 | 32220212 | 32221524 | 38 | 26 | 24 | 31 | 35 | 1 | 0 | 2 | 1 | 1 | 0 | 5 | 6 | 16 | 0 | 0 | 5 | 3 | 0 | 0 | 252 | 149 | 1 | 1 | 0.957 | 1 | 0.876 | 0.948 | 0.954 | NA | 0.372 | 0.542 | 1 | NA | 0.309 |
| SE | 59693 | ENSG00000152683 | SLC30A6 | chr2 | + | 32206885 | 32206933 | 32197706 | 32197826 | 32220212 | 32224112 | 3 | 1 | 0 | 4 | 0 | 1 | 0 | 2 | 1 | 1 | 0 | 3 | 5 | 0 | 0 | 0 | 5 | 3 | 0 | 0 | 197 | 149 | 1 | 1 | 0.694 | 1 | 0 | 0.752 | 0 | NA | 0.312 | 0.558 | NA | NA | 0.054 |
| SE | 59694 | ENSG00000152683 | SLC30A6 | chr2 | + | 32206885 | 32206933 | 32204589 | 32204692 | 32209492 | 32209561 | 60 | 38 | 63 | 29 | 72 | 0 | 1 | 0 | 1 | 0 | 20 | 30 | 16 | 31 | 9 | 0 | 0 | 0 | 0 | 0 | 197 | 149 | 1 | 1 | 1 | 0.966 | 1 | 0.956 | 1 | 1 | 1 | 1 | 1 | 1 | -0.016 |
| SE | 59695 | ENSG00000152683 | SLC30A6 | chr2 | + | 32206885 | 32206933 | 32204589 | 32204692 | 32220212 | 32224112 | 35 | 26 | 33 | 27 | 31 | 0 | 0 | 0 | 0 | 2 | 20 | 21 | 12 | 19 | 9 | 0 | 0 | 0 | 0 | 0 | 197 | 149 | 1 | 1 | 1 | 1 | 1 | 1 | 0.921 | 1 | 1 | 1 | 1 | 1 | -0.016 |
| SE | 59696 | ENSG00000152683 | SLC30A6 | chr2 | + | 32209492 | 32209561 | 32204589 | 32204692 | 32220212 | 32221524 | 16 | 7 | 6 | 8 | 36 | 0 | 0 | 0 | 0 | 2 | 0 | 7 | 4 | 2 | 0 | 0 | 0 | 0 | 0 | 0 | 218 | 149 | 1 | 1 | 1 | 1 | 1 | 1 | 0.925 | NA | 1 | 1 | 1 | NA | -0.015 |
| SE | 59697 | ENSG00000152683 | SLC30A6 | chr2 | + | 32209492 | 32209561 | 32206885 | 32206933 | 32220212 | 32221524 | 40 | 22 | 35 | 15 | 84 | 3 | 1 | 1 | 3 | 5 | 0 | 18 | 11 | 14 | 0 | 0 | 3 | 4 | 0 | 0 | 218 | 149 | 1 | 1 | 0.901 | 0.938 | 0.96 | 0.774 | 0.92 | NA | 0.804 | 0.653 | 1 | NA | 0.08 |
| SE | 59699 | ENSG00000143162 | CREG1 | chr1 | - | 167546100 | 167546285 | 167541012 | 167542301 | 167548001 | 167548121 | 258 | 81 | 112 | 160 | 312 | 0 | 0 | 0 | 0 | 0 | 172 | 233 | 185 | 253 | 168 | 0 | 0 | 0 | 4 | 0 | 298 | 149 | 1 | 1 | 1 | 1 | 1 | 1 | 1 | 1 | 1 | 1 | 0.969 | 1 | 0.006 |
| SE | 59704 | ENSG00000113368 | LMNB1 | chr5 | + | 126777360 | 126777867 | 126777160 | 126777252 | 126804775 | 126804914 | 54 | 26 | 66 | 59 | 37 | 0 | 0 | 2 | 0 | 0 | 181 | 73 | 63 | 133 | 37 | 0 | 1 | 0 | 0 | 0 | 298 | 149 | 1 | 1 | 1 | 1 | 0.943 | 1 | 1 | 1 | 0.973 | 1 | 1 | 1 | -0.006 |
| SE | 59706 | ENSG00000113368 | LMNB1 | chr5 | + | 126822780 | 126822885 | 126820909 | 126821135 | 126825987 | 126826107 | 349 | 306 | 399 | 388 | 469 | 0 | 1 | 1 | 3 | 0 | 1759 | 635 | 741 | 1161 | 410 | 10 | 2 | 0 | 1 | 0 | 254 | 149 | 1 | 1 | 1 | 0.994 | 0.996 | 0.987 | 1 | 0.99 | 0.995 | 1 | 0.999 | 1 | -0.001 |
| SE | 59707 | ENSG00000113368 | LMNB1 | chr5 | + | 126825987 | 126826107 | 126822780 | 126822885 | 126832693 | 126832801 | 297 | 295 | 357 | 324 | 379 | 1 | 0 | 0 | 0 | 0 | 1796 | 583 | 638 | 1070 | 448 | 0 | 1 | 0 | 0 | 0 | 269 | 149 | 1 | 1 | 0.994 | 1 | 1 | 1 | 1 | 1 | 0.997 | 1 | 1 | 1 | -0.001 |
| SE | 59716 | ENSG00000255197 | NA | chr11 | - | 47395489 | 47395577 | 47381508 | 47383832 | 47405509 | 47405547 | 38 | 32 | 25 | 28 | 19 | 0 | 0 | 3 | 0 | 0 | 27 | 21 | 18 | 78 | 56 | 0 | 0 | 0 | 0 | 0 | 237 | 149 | 1 | 1 | 1 | 1 | 0.84 | 1 | 1 | 1 | 1 | 1 | 1 | 1 | -0.032 |
| SE | 59721 | ENSG00000143633 | C1orf131 | chr1 | - | 231226972 | 231227068 | 231226385 | 231226825 | 231229149 | 231229206 | 211 | 151 | 176 | 168 | 287 | 2 | 0 | 0 | 0 | 2 | 156 | 152 | 119 | 162 | 51 | 0 | 0 | 0 | 0 | 0 | 245 | 149 | 1 | 1 | 0.985 | 1 | 1 | 1 | 0.989 | 1 | 1 | 1 | 1 | 1 | -0.005 |
| SE | 59722 | ENSG00000143633 | C1orf131 | chr1 | - | 231229149 | 231229206 | 231226972 | 231227068 | 231238856 | 231238893 | 99 | 97 | 77 | 77 | 138 | 0 | 0 | 2 | 0 | 4 | 87 | 79 | 33 | 72 | 14 | 0 | 0 | 0 | 0 | 0 | 206 | 149 | 1 | 1 | 1 | 1 | 0.965 | 1 | 0.961 | 1 | 1 | 1 | 1 | 1 | -0.015 |
| SE | 59723 | ENSG00000143633 | C1orf131 | chr1 | - | 231229149 | 231229209 | 231226972 | 231227068 | 231238856 | 231239188 | 117 | 94 | 84 | 103 | 142 | 0 | 0 | 2 | 0 | 4 | 84 | 97 | 82 | 70 | 32 | 0 | 0 | 0 | 0 | 0 | 209 | 149 | 1 | 1 | 1 | 1 | 0.968 | 1 | 0.962 | 1 | 1 | 1 | 1 | 1 | -0.014 |
| SE | 59724 | ENSG00000149231 | CCDC82 | chr11 | - | 96364979 | 96365150 | 96358992 | 96359178 | 96371012 | 96371137 | 359 | 290 | 272 | 346 | 574 | 0 | 0 | 0 | 1 | 0 | 2 | 107 | 76 | 18 | 0 | 0 | 0 | 0 | 0 | 0 | 298 | 149 | 1 | 1 | 1 | 1 | 1 | 0.994 | 1 | 1 | 1 | 1 | 1 | NA | -0.001 |
| SE | 59725 | ENSG00000149231 | CCDC82 | chr11 | - | 96365126 | 96365150 | 96358992 | 96359178 | 96371012 | 96371137 | 126 | 115 | 106 | 108 | 220 | 0 | 0 | 0 | 1 | 0 | 2 | 39 | 37 | 12 | 0 | 0 | 0 | 0 | 0 | 0 | 173 | 149 | 1 | 1 | 1 | 1 | 1 | 0.989 | 1 | 1 | 1 | 1 | 1 | NA | -0.002 |
| SE | 59726 | ENSG00000149231 | CCDC82 | chr11 | - | 96383961 | 96384761 | 96380538 | 96383473 | 96387529 | 96387613 | 414 | 414 | 314 | 444 | 603 | 0 | 0 | 0 | 0 | 7 | 36 | 122 | 81 | 38 | 42 | 0 | 0 | 0 | 0 | 0 | 298 | 149 | 1 | 1 | 1 | 1 | 1 | 1 | 0.977 | 1 | 1 | 1 | 1 | 1 | -0.005 |
| SE | 59728 | ENSG00000149231 | CCDC82 | chr11 | - | 96383961 | 96386293 | 96380547 | 96383473 | 96387529 | 96387613 | 414 | 418 | 317 | 450 | 602 | 0 | 0 | 0 | 0 | 7 | 36 | 118 | 81 | 38 | 42 | 0 | 0 | 0 | 0 | 0 | 298 | 149 | 1 | 1 | 1 | 1 | 1 | 1 | 0.977 | 1 | 1 | 1 | 1 | 1 | -0.005 |
| SE | 59731 | ENSG00000149231 | CCDC82 | chr11 | - | 96385936 | 96386020 | 96383961 | 96384761 | 96386253 | 96386293 | 1 | 1 | 2 | 0 | 3 | 98 | 74 | 70 | 99 | 90 | 0 | 0 | 5 | 0 | 0 | 6 | 32 | 7 | 14 | 0 | 233 | 149 | 1 | 1 | 0.006 | 0.009 | 0.018 | 0 | 0.021 | 0 | 0 | 0.314 | 0 | NA | -0.068 |
| SE | 59732 | ENSG00000149231 | CCDC82 | chr11 | - | 96385936 | 96386020 | 96383961 | 96384761 | 96387529 | 96387613 | 0 | 1 | 2 | 0 | 2 | 2 | 0 | 1 | 2 | 2 | 0 | 0 | 3 | 0 | 0 | 0 | 4 | 0 | 0 | 0 | 233 | 149 | 1 | 1 | 0 | 1 | 0.561 | 0 | 0.39 | NA | 0 | 1 | NA | NA | -0.11 |
| SE | 59733 | ENSG00000149231 | CCDC82 | chr11 | - | 96386253 | 96386293 | 96383961 | 96384761 | 96387529 | 96387613 | 96 | 65 | 71 | 105 | 87 | 2 | 0 | 1 | 2 | 2 | 6 | 32 | 7 | 14 | 0 | 0 | 4 | 0 | 0 | 0 | 189 | 149 | 1 | 1 | 0.974 | 1 | 0.982 | 0.976 | 0.972 | 1 | 0.863 | 1 | 1 | NA | 0.015 |
| SE | 59734 | ENSG00000149231 | CCDC82 | chr11 | - | 96386253 | 96386293 | 96383961 | 96384761 | 96389843 | 96389865 | 18 | 14 | 9 | 11 | 45 | 2 | 5 | 3 | 1 | 0 | 4 | 0 | 0 | 7 | 0 | 0 | 2 | 0 | 0 | 0 | 189 | 149 | 1 | 1 | 0.876 | 0.688 | 0.703 | 0.897 | 1 | 1 | 0 | NA | 1 | NA | 0.166 |
| SE | 59735 | ENSG00000149231 | CCDC82 | chr11 | - | 96386253 | 96386293 | 96385936 | 96386020 | 96387529 | 96387613 | 2 | 4 | 4 | 6 | 2 | 0 | 1 | 0 | 0 | 0 | 0 | 0 | 2 | 0 | 0 | 0 | 0 | 0 | 0 | 0 | 189 | 149 | 1 | 1 | 1 | 0.759 | 1 | 1 | 1 | NA | NA | 1 | NA | NA | -0.048 |
| SE | 59736 | ENSG00000149231 | CCDC82 | chr11 | - | 96387529 | 96387613 | 96383268 | 96383473 | 96389843 | 96389865 | 0 | 0 | 0 | 0 | 7 | 2 | 0 | 0 | 0 | 0 | 0 | 0 | 0 | 0 | 0 | 0 | 0 | 7 | 0 | 0 | 233 | 149 | 1 | 1 | 0 | NA | NA | NA | 1 | NA | NA | 0 | NA | NA | 0.5 |
| SE | 59737 | ENSG00000149231 | CCDC82 | chr11 | - | 96387529 | 96387613 | 96383961 | 96384761 | 96389364 | 96389523 | 1 | 0 | 1 | 2 | 2 | 1 | 0 | 0 | 0 | 0 | 0 | 4 | 0 | 0 | 0 | 0 | 0 | 0 | 0 | 0 | 233 | 149 | 1 | 1 | 0.39 | NA | 1 | 1 | 1 | NA | 1 | NA | NA | NA | -0.152 |
| SE | 59738 | ENSG00000149231 | CCDC82 | chr11 | - | 96387529 | 96387613 | 96383961 | 96384761 | 96389843 | 96389865 | 2 | 0 | 1 | 2 | 2 | 2 | 5 | 3 | 1 | 0 | 0 | 4 | 0 | 0 | 0 | 0 | 2 | 0 | 0 | 0 | 233 | 149 | 1 | 1 | 0.39 | 0 | 0.176 | 0.561 | 1 | NA | 0.561 | NA | NA | NA | -0.136 |
| SE | 59739 | ENSG00000149231 | CCDC82 | chr11 | - | 96387529 | 96387613 | 96383961 | 96386293 | 96389843 | 96389865 | 78 | 51 | 58 | 96 | 44 | 0 | 0 | 0 | 0 | 3 | 2 | 32 | 7 | 7 | 0 | 0 | 0 | 0 | 0 | 0 | 233 | 149 | 1 | 1 | 1 | 1 | 1 | 1 | 0.904 | 1 | 1 | 1 | 1 | NA | -0.019 |
| SE | 59740 | ENSG00000149231 | CCDC82 | chr11 | - | 96387529 | 96387653 | 96386253 | 96386293 | 96389843 | 96389905 | 68 | 47 | 52 | 78 | 45 | 0 | 0 | 0 | 0 | 3 | 2 | 32 | 8 | 7 | 0 | 0 | 0 | 0 | 0 | 0 | 273 | 149 | 1 | 1 | 1 | 1 | 1 | 1 | 0.891 | 1 | 1 | 1 | 1 | NA | -0.022 |
| SE | 59742 | ENSG00000149231 | CCDC82 | chr11 | - | 96389364 | 96389523 | 96387529 | 96387613 | 96389843 | 96389891 | 0 | 0 | 1 | 0 | 1 | 13 | 4 | 9 | 19 | 0 | 0 | 0 | 0 | 0 | 0 | 0 | 0 | 1 | 0 | 0 | 298 | 149 | 1 | 1 | 0 | 0 | 0.053 | 0 | 1 | NA | NA | 0 | NA | NA | 0.211 |
| SE | 59743 | ENSG00000135974 | C2orf49 | chr2 | + | 105342847 | 105343223 | 105339582 | 105339749 | 105345314 | 105349211 | 134 | 88 | 122 | 116 | 237 | 3 | 0 | 0 | 0 | 0 | 47 | 109 | 119 | 102 | 60 | 0 | 0 | 0 | 0 | 0 | 298 | 149 | 1 | 1 | 0.957 | 1 | 1 | 1 | 1 | 1 | 1 | 1 | 1 | 1 | -0.009 |
| SE | 59744 | ENSG00000176160 | HSF5 | chr17 | - | 58462781 | 58463303 | 58458767 | 58458945 | 58466884 | 58466979 | 12 | 3 | 11 | 7 | 12 | 0 | 1 | 2 | 0 | 0 | 4 | 7 | 5 | 4 | 0 | 0 | 0 | 1 | 0 | 0 | 298 | 149 | 1 | 1 | 1 | 0.6 | 0.733 | 1 | 1 | 1 | 1 | 0.714 | 1 | NA | -0.062 |
| SE | 59745 | ENSG00000067704 | IARS2 | chr1 | + | 220102362 | 220102412 | 220102128 | 220102277 | 220102494 | 220102604 | 156 | 105 | 120 | 93 | 250 | 1 | 0 | 0 | 0 | 0 | 45 | 62 | 61 | 90 | 32 | 0 | 0 | 0 | 0 | 0 | 199 | 149 | 1 | 1 | 0.992 | 1 | 1 | 1 | 1 | 1 | 1 | 1 | 1 | 1 | -0.002 |
| SE | 59746 | ENSG00000067704 | IARS2 | chr1 | + | 220103446 | 220103562 | 220102686 | 220102777 | 220105890 | 220106060 | 260 | 139 | 231 | 187 | 405 | 0 | 0 | 2 | 0 | 0 | 39 | 246 | 116 | 221 | 45 | 0 | 0 | 0 | 0 | 0 | 265 | 149 | 1 | 1 | 1 | 1 | 0.985 | 1 | 1 | 1 | 1 | 1 | 1 | 1 | -0.003 |
| SE | 59747 | ENSG00000067704 | IARS2 | chr1 | + | 220134401 | 220134510 | 220126749 | 220126843 | 220136808 | 220136911 | 157 | 79 | 147 | 129 | 189 | 0 | 0 | 0 | 0 | 0 | 41 | 136 | 79 | 133 | 3 | 0 | 3 | 0 | 0 | 0 | 258 | 149 | 1 | 1 | 1 | 1 | 1 | 1 | 1 | 1 | 0.963 | 1 | 1 | 1 | 0.007 |
| SE | 59748 | ENSG00000044459 | CNTLN | chr9 | + | 17298189 | 17298352 | 17273732 | 17273866 | 17309057 | 17309252 | 15 | 57 | 22 | 19 | 38 | 0 | 0 | 1 | 2 | 0 | 12 | 26 | 23 | 10 | 0 | 0 | 0 | 0 | 0 | 0 | 298 | 149 | 1 | 1 | 1 | 1 | 0.917 | 0.826 | 1 | 1 | 1 | 1 | 1 | NA | -0.051 |
| SE | 59749 | ENSG00000044459 | CNTLN | chr9 | + | 17309057 | 17309252 | 17298189 | 17298352 | 17330631 | 17330808 | 18 | 55 | 19 | 27 | 73 | 0 | 0 | 0 | 2 | 0 | 51 | 31 | 20 | 14 | 46 | 0 | 0 | 0 | 1 | 0 | 298 | 149 | 1 | 1 | 1 | 1 | 1 | 0.871 | 1 | 1 | 1 | 1 | 0.875 | 1 | -0.001 |
| SE | 59750 | ENSG00000044459 | CNTLN | chr9 | + | 17366616 | 17366717 | 17342324 | 17342444 | 17388161 | 17388253 | 11 | 17 | 13 | 9 | 14 | 0 | 0 | 0 | 0 | 0 | 26 | 12 | 10 | 9 | 22 | 0 | 1 | 0 | 0 | 0 | 250 | 149 | 1 | 1 | 1 | 1 | 1 | 1 | 1 | 1 | 0.877 | 1 | 1 | 1 | 0.025 |
| SE | 59751 | ENSG00000044459 | CNTLN | chr9 | + | 17444196 | 17444246 | 17415965 | 17416189 | 17457523 | 17457628 | 3 | 13 | 7 | 3 | 7 | 11 | 13 | 1 | 1 | 4 | 9 | 0 | 0 | 0 | 0 | 0 | 2 | 0 | 0 | 0 | 199 | 149 | 1 | 1 | 0.17 | 0.428 | 0.84 | 0.692 | 0.567 | 1 | 0 | NA | NA | NA | 0.039 |
| SE | 59757 | ENSG00000158711 | ELK4 | chr1 | - | 205618956 | 205619073 | 205607942 | 205616644 | 205619965 | 205620838 | 794 | 447 | 724 | 770 | 1134 | 0 | 1 | 1 | 0 | 0 | 404 | 651 | 517 | 276 | 642 | 0 | 0 | 4 | 4 | 0 | 266 | 149 | 1 | 1 | 1 | 0.996 | 0.998 | 1 | 1 | 1 | 1 | 0.986 | 0.975 | 1 | 0.007 |
| SE | 59758 | ENSG00000158711 | ELK4 | chr1 | - | 205619965 | 205620838 | 205607942 | 205616644 | 205623675 | 205623891 | 398 | 268 | 356 | 365 | 523 | 0 | 0 | 0 | 0 | 0 | 266 | 288 | 282 | 204 | 307 | 0 | 0 | 0 | 0 | 4 | 298 | 149 | 1 | 1 | 1 | 1 | 1 | 1 | 1 | 1 | 1 | 1 | 1 | 0.975 | 0.005 |
| SE | 59759 | ENSG00000158711 | ELK4 | chr1 | - | 205623675 | 205623891 | 205619267 | 205620838 | 205631631 | 205631893 | 412 | 277 | 370 | 381 | 531 | 0 | 0 | 0 | 1 | 0 | 284 | 306 | 295 | 210 | 307 | 0 | 0 | 0 | 0 | 0 | 298 | 149 | 1 | 1 | 1 | 1 | 1 | 0.995 | 1 | 1 | 1 | 1 | 1 | 1 | -0.001 |
| SE | 59761 | ENSG00000127074 | RGS13 | chr1 | + | 192638163 | 192638203 | 192637589 | 192637660 | 192644330 | 192644399 | 3 | 0 | 0 | 1 | 0 | 2 | 2 | 3 | 2 | 0 | 0 | 0 | 10 | 0 | 0 | 0 | 0 | 10 | 0 | 0 | 189 | 149 | 1 | 1 | 0.542 | 0 | 0 | 0.283 | NA | NA | NA | 0.441 | NA | NA | -0.235 |
| SE | 59763 | ENSG00000163444 | TMEM183A | chr1 | + | 203014888 | 203015048 | 203008642 | 203008810 | 203015959 | 203016094 | 395 | 203 | 336 | 366 | 695 | 0 | 1 | 1 | 0 | 4 | 609 | 428 | 305 | 465 | 715 | 0 | 0 | 0 | 6 | 0 | 298 | 149 | 1 | 1 | 1 | 0.99 | 0.994 | 1 | 0.989 | 1 | 1 | 1 | 0.975 | 1 | 0 |
| SE | 59764 | ENSG00000163444 | TMEM183A | chr1 | + | 203014888 | 203015048 | 203008642 | 203008810 | 203018480 | 203018561 | 198 | 96 | 179 | 213 | 429 | 0 | 0 | 0 | 0 | 0 | 330 | 263 | 177 | 272 | 360 | 0 | 0 | 0 | 1 | 0 | 298 | 149 | 1 | 1 | 1 | 1 | 1 | 1 | 1 | 1 | 1 | 1 | 0.993 | 1 | 0.001 |
| SE | 59765 | ENSG00000163444 | TMEM183A | chr1 | + | 203015959 | 203016140 | 203008985 | 203008810 | 203018480 | 203018561 | 101 | 47 | 92 | 87 | 92 | 0 | 0 | 0 | 0 | 0 | 63 | 89 | 72 | 115 | 153 | 0 | 0 | 0 | 1 | 0 | 298 | 149 | 1 | 1 | 1 | 1 | 1 | 1 | 1 | 1 | 1 | 1 | 0.983 | 1 | 0.003 |
| SE | 59766 | ENSG00000163444 | TMEM183A | chr1 | + | 203015959 | 203016140 | 203008985 | 203015048 | 203017755 | 203017919 | 212 | 128 | 180 | 185 | 275 | 0 | 0 | 0 | 0 | 1 | 330 | 194 | 165 | 227 | 376 | 0 | 0 | 0 | 0 | 0 | 298 | 149 | 1 | 1 | 1 | 1 | 1 | 1 | 0.993 | 1 | 1 | 1 | 1 | 1 | -0.001 |
| SE | 59767 | ENSG00000163444 | TMEM183A | chr1 | + | 203015959 | 203016140 | 203014888 | 203015048 | 203018480 | 203018561 | 298 | 153 | 248 | 239 | 354 | 0 | 0 | 0 | 0 | 0 | 342 | 254 | 200 | 307 | 508 | 0 | 0 | 0 | 5 | 0 | 298 | 149 | 1 | 1 | 1 | 1 | 1 | 1 | 1 | 1 | 1 | 1 | 0.968 | 1 | 0.006 |
| SE | 59768 | ENSG00000163444 | TMEM183A | chr1 | + | 203017755 | 203017919 | 203014888 | 203015048 | 203018480 | 203018561 | 13 | 6 | 8 | 10 | 8 | 0 | 0 | 0 | 0 | 0 | 0 | 12 | 30 | 39 | 7 | 0 | 0 | 0 | 5 | 0 | 298 | 149 | 1 | 1 | 1 | 1 | 1 | 1 | 1 | NA | 1 | 1 | 0.796 | 1 | 0.051 |
| SE | 59770 | ENSG00000204516 | MICB | chr6 | + | 31506142 | 31506430 | 31505616 | 31505871 | 31507021 | 31507300 | 136 | 80 | 104 | 107 | 111 | 1 | 0 | 0 | 0 | 0 | 274 | 229 | 113 | 156 | 231 | 0 | 1 | 0 | 0 | 0 | 298 | 149 | 1 | 1 | 0.986 | 1 | 1 | 1 | 1 | 1 | 0.991 | 1 | 1 | 1 | -0.001 |
| SE | 59771 | ENSG00000204516 | MICB | chr6 | + | 31506271 | 31506430 | 31505616 | 31505871 | 31507021 | 31507300 | 83 | 39 | 70 | 72 | 74 | 1 | 0 | 0 | 0 | 0 | 192 | 150 | 70 | 106 | 143 | 0 | 1 | 0 | 0 | 0 | 298 | 149 | 1 | 1 | 0.976 | 1 | 1 | 1 | 1 | 1 | 0.987 | 1 | 1 | 1 | -0.002 |
| SE | 59772 | ENSG00000117592 | PRDX6 | chr1 | + | 173485360 | 173485507 | 173481198 | 173481482 | 173486254 | 173486401 | 704 | 642 | 773 | 938 | 1443 | 0 | 0 | 0 | 4 | 0 | 1050 | 1197 | 621 | 953 | 4432 | 0 | 0 | 0 | 0 | 0 | 296 | 149 | 1 | 1 | 1 | 1 | 1 | 0.992 | 1 | 1 | 1 | 1 | 1 | 1 | -0.002 |
| SE | 59774 | ENSG00000155542 | SETD9 | chr5 | + | 56911168 | 56911502 | 56909623 | 56909743 | 56913010 | 56913116 | 1 | 1 | 1 | 2 | 2 | 0 | 0 | 1 | 1 | 8 | 0 | 0 | 1 | 0 | 0 | 0 | 0 | 0 | 0 | 0 | 298 | 149 | 1 | 1 | 1 | 1 | 0.333 | 0.5 | 0.111 | NA | NA | 1 | NA | NA | -0.411 |
| SE | 59775 | ENSG00000155542 | SETD9 | chr5 | + | 56911168 | 56911536 | 56909351 | 56909743 | 56913010 | 56913134 | 7 | 9 | 12 | 13 | 27 | 0 | 0 | 1 | 1 | 8 | 5 | 7 | 5 | 1 | 0 | 0 | 0 | 0 | 0 | 0 | 298 | 149 | 1 | 1 | 1 | 1 | 0.857 | 0.867 | 0.628 | 1 | 1 | 1 | 1 | NA | -0.13 |
| SE | 59776 | ENSG00000155542 | SETD9 | chr5 | + | 56914860 | 56914966 | 56913873 | 56913989 | 56916814 | 56916919 | 4 | 15 | 23 | 19 | 82 | 0 | 0 | 0 | 1 | 5 | 0 | 3 | 7 | 2 | 10 | 0 | 0 | 0 | 0 | 0 | 255 | 149 | 1 | 1 | 1 | 1 | 1 | 0.917 | 0.906 | NA | 1 | 1 | 1 | 1 | -0.035 |
| SE | 59777 | ENSG00000181817 | LSM10 | chr1 | - | 36392357 | 36392487 | 36391237 | 36391604 | 36393765 | 36394153 | 14 | 6 | 20 | 7 | 6 | 0 | 0 | 0 | 0 | 0 | 40 | 10 | 11 | 16 | 19 | 0 | 0 | 1 | 0 | 0 | 279 | 149 | 1 | 1 | 1 | 1 | 1 | 1 | 1 | 1 | 1 | 0.855 | 1 | 1 | 0.029 |
| SE | 59778 | ENSG00000181817 | LSM10 | chr1 | - | 36392357 | 36392487 | 36391237 | 36391604 | 36397766 | 36397887 | 12 | 4 | 13 | 6 | 1 | 1 | 0 | 0 | 0 | 0 | 25 | 10 | 10 | 9 | 15 | 0 | 0 | 0 | 0 | 0 | 279 | 149 | 1 | 1 | 0.865 | 1 | 1 | 1 | 1 | 1 | 1 | 1 | 1 | 1 | -0.027 |
| SE | 59779 | ENSG00000181817 | LSM10 | chr1 | - | 36393765 | 36394153 | 36391237 | 36391604 | 36397766 | 36397887 | 54 | 24 | 40 | 31 | 22 | 1 | 0 | 0 | 0 | 0 | 58 | 37 | 18 | 40 | 49 | 0 | 0 | 0 | 0 | 0 | 298 | 149 | 1 | 1 | 0.964 | 1 | 1 | 1 | 1 | 1 | 1 | 1 | 1 | 1 | -0.007 |
| SE | 59780 | ENSG00000181817 | LSM10 | chr1 | - | 36393765 | 36394153 | 36392357 | 36392487 | 36397766 | 36397887 | 56 | 26 | 48 | 33 | 28 | 0 | 0 | 0 | 1 | 0 | 73 | 37 | 18 | 47 | 53 | 0 | 0 | 0 | 0 | 0 | 298 | 149 | 1 | 1 | 1 | 1 | 1 | 0.943 | 1 | 1 | 1 | 1 | 1 | 1 | -0.011 |
| SE | 59781 | ENSG00000185163 | DDX51 | chr12 | - | 132139860 | 132139924 | 132139634 | 132139769 | 132140097 | 132140199 | 36 | 20 | 43 | 44 | 35 | 0 | 0 | 0 | 0 | 0 | 21 | 50 | 20 | 31 | 53 | 0 | 0 | 0 | 4 | 0 | 213 | 149 | 1 | 1 | 1 | 1 | 1 | 1 | 1 | 1 | 1 | 1 | 0.844 | 1 | 0.031 |
| SE | 59782 | ENSG00000185163 | DDX51 | chr12 | - | 132141102 | 132141420 | 132140830 | 132141020 | 132141497 | 132142190 | 20 | 9 | 12 | 14 | 10 | 0 | 0 | 0 | 0 | 0 | 0 | 20 | 15 | 4 | 0 | 0 | 0 | 2 | 0 | 0 | 298 | 149 | 1 | 1 | 1 | 1 | 1 | 1 | 1 | NA | 1 | 0.789 | 1 | NA | 0.07 |
| SE | 59783 | ENSG00000185163 | DDX51 | chr12 | - | 132141274 | 132141420 | 132140830 | 132141020 | 132141497 | 132141606 | 23 | 11 | 15 | 17 | 13 | 0 | 0 | 0 | 0 | 0 | 0 | 34 | 20 | 11 | 0 | 0 | 0 | 2 | 0 | 0 | 295 | 149 | 1 | 1 | 1 | 1 | 1 | 1 | 1 | NA | 1 | 0.835 | 1 | NA | 0.055 |
| SE | 59784 | ENSG00000185163 | DDX51 | chr12 | - | 132143694 | 132143909 | 132142727 | 132142878 | 132143992 | 132144319 | 2 | 3 | 2 | 1 | 3 | 0 | 0 | 0 | 0 | 0 | 0 | 0 | 0 | 2 | 3 | 5 | 0 | 0 | 4 | 0 | 298 | 149 | 1 | 1 | 1 | 1 | 1 | 1 | 1 | 0 | NA | NA | 0.2 | 1 | 0.6 |
| SE | 59786 | ENSG00000174748 | RPL15 | chr3 | + | 23918439 | 23918576 | 23917704 | 23918031 | 23919195 | 23919246 | 6643 | 3220 | 7017 | 5986 | 8174 | 1 | 4 | 0 | 2 | 0 | 4949 | 4110 | 3214 | 3508 | 7462 | 0 | 0 | 9 | 6 | 4 | 286 | 149 | 1 | 1 | 1 | 0.998 | 1 | 0.999 | 1 | 1 | 1 | 0.995 | 0.997 | 0.999 | 0.001 |
| SE | 59798 | ENSG00000166377 | ATP9B | chr18 | + | 79096475 | 79096649 | 79069391 | 79069529 | 79110354 | 79110505 | 168 | 123 | 181 | 172 | 183 | 0 | 0 | 0 | 2 | 0 | 169 | 134 | 110 | 98 | 143 | 0 | 0 | 1 | 0 | 0 | 298 | 149 | 1 | 1 | 1 | 1 | 1 | 0.977 | 1 | 1 | 1 | 0.982 | 1 | 1 | -0.001 |
| SE | 59799 | ENSG00000166377 | ATP9B | chr18 | + | 79110354 | 79110505 | 79096475 | 79096649 | 79113240 | 79113354 | 418 | 355 | 477 | 479 | 460 | 0 | 0 | 0 | 0 | 0 | 254 | 295 | 314 | 253 | 467 | 0 | 0 | 1 | 0 | 0 | 298 | 149 | 1 | 1 | 1 | 1 | 1 | 1 | 1 | 1 | 1 | 0.994 | 1 | 1 | 0.001 |
| SE | 59801 | ENSG00000166377 | ATP9B | chr18 | + | 79126266 | 79126375 | 79096475 | 79096649 | 79143801 | 79143859 | 82 | 105 | 142 | 106 | 101 | 0 | 0 | 1 | 0 | 0 | 125 | 99 | 103 | 92 | 181 | 0 | 0 | 0 | 0 | 0 | 258 | 149 | 1 | 1 | 1 | 1 | 0.988 | 1 | 1 | 1 | 1 | 1 | 1 | 1 | -0.002 |
| SE | 59805 | ENSG00000166377 | ATP9B | chr18 | + | 79194385 | 79194492 | 79193182 | 79193263 | 79206936 | 79207012 | 0 | 0 | 0 | 1 | 2 | 55 | 66 | 77 | 52 | 76 | 0 | 0 | 0 | 0 | 0 | 41 | 63 | 39 | 44 | 90 | 256 | 149 | 1 | 1 | 0 | 0 | 0 | 0.011 | 0.015 | 0 | 0 | 0 | 0 | 0 | 0.005 |
| SE | 59808 | ENSG00000166377 | ATP9B | chr18 | + | 79213961 | 79214038 | 79206935 | 79207012 | 79277053 | 79277196 | 19 | 22 | 28 | 23 | 37 | 1 | 0 | 0 | 0 | 0 | 42 | 28 | 16 | 21 | 36 | 0 | 0 | 1 | 0 | 0 | 226 | 149 | 1 | 1 | 0.926 | 1 | 1 | 1 | 1 | 1 | 1 | 0.913 | 1 | 1 | 0.003 |
| SE | 59809 | ENSG00000166377 | ATP9B | chr18 | + | 79253380 | 79253541 | 79206976 | 79207012 | 79277053 | 79277196 | 116 | 65 | 97 | 81 | 117 | 1 | 0 | 0 | 0 | 0 | 30 | 48 | 37 | 65 | 85 | 0 | 0 | 1 | 0 | 0 | 298 | 149 | 1 | 1 | 0.983 | 1 | 1 | 1 | 1 | 1 | 1 | 0.949 | 1 | 1 | 0.007 |
| SE | 59810 | ENSG00000166377 | ATP9B | chr18 | + | 79253380 | 79253541 | 79213961 | 79214038 | 79277053 | 79277196 | 147 | 94 | 117 | 88 | 168 | 0 | 0 | 0 | 2 | 0 | 53 | 42 | 53 | 88 | 103 | 0 | 0 | 0 | 0 | 0 | 298 | 149 | 1 | 1 | 1 | 1 | 1 | 0.957 | 1 | 1 | 1 | 1 | 1 | 1 | -0.009 |
| SE | 59813 | ENSG00000166377 | ATP9B | chr18 | + | 79277053 | 79277196 | 79253380 | 79253541 | 79303603 | 79303716 | 138 | 79 | 111 | 96 | 177 | 0 | 2 | 0 | 0 | 0 | 42 | 47 | 61 | 77 | 87 | 0 | 0 | 3 | 0 | 0 | 292 | 149 | 1 | 1 | 1 | 0.953 | 1 | 1 | 1 | 1 | 1 | 0.912 | 1 | 1 | 0.008 |
| SE | 59815 | ENSG00000166377 | ATP9B | chr18 | + | 79329140 | 79329302 | 79306985 | 79307234 | 79330011 | 79330104 | 79 | 70 | 82 | 131 | 82 | 0 | 5 | 0 | 0 | 0 | 52 | 65 | 104 | 62 | 74 | 0 | 0 | 0 | 4 | 0 | 298 | 149 | 1 | 1 | 1 | 0.875 | 1 | 1 | 1 | 1 | 1 | 1 | 0.886 | 1 | -0.002 |
| SE | 59816 | ENSG00000166377 | ATP9B | chr18 | + | 79337278 | 79337449 | 79336627 | 79336711 | 79342267 | 79342366 | 69 | 41 | 63 | 71 | 10 | 0 | 0 | 1 | 1 | 0 | 61 | 23 | 35 | 34 | 67 | 0 | 0 | 0 | 0 | 0 | 298 | 149 | 1 | 1 | 1 | 1 | 0.969 | 0.973 | 1 | 1 | 1 | 1 | 1 | 1 | -0.012 |
| SE | 59817 | ENSG00000166377 | ATP9B | chr18 | + | 79337316 | 79337449 | 79336625 | 79336711 | 79342267 | 79342366 | 46 | 23 | 43 | 42 | 8 | 0 | 0 | 1 | 1 | 0 | 43 | 15 | 14 | 15 | 38 | 0 | 0 | 0 | 0 | 0 | 282 | 149 | 1 | 1 | 1 | 1 | 0.958 | 0.957 | 1 | 1 | 1 | 1 | 1 | 1 | -0.017 |
| SE | 59818 | ENSG00000166377 | ATP9B | chr18 | + | 79348131 | 79348196 | 79347769 | 79347925 | 79359353 | 79359462 | 10 | 5 | 18 | 10 | 13 | 0 | 2 | 3 | 0 | 0 | 0 | 8 | 15 | 6 | 13 | 0 | 2 | 1 | 0 | 8 | 214 | 149 | 1 | 1 | 1 | 0.635 | 0.807 | 1 | 1 | NA | 0.736 | 0.913 | 1 | 0.531 | 0.093 |
| SE | 59821 | ENSG00000166377 | ATP9B | chr18 | + | 79372824 | 79372882 | 79359353 | 79359462 | 79373897 | 79374027 | 19 | 3 | 18 | 20 | 11 | 0 | 0 | 0 | 0 | 0 | 29 | 15 | 9 | 4 | 11 | 0 | 0 | 1 | 0 | 0 | 207 | 149 | 1 | 1 | 1 | 1 | 1 | 1 | 1 | 1 | 1 | 0.866 | 1 | 1 | 0.027 |
| SE | 59822 | ENSG00000166377 | ATP9B | chr18 | + | 79375393 | 79375426 | 79373897 | 79374101 | 79377246 | 79378142 | 1 | 2 | 7 | 5 | 4 | 12 | 1 | 5 | 6 | 5 | 0 | 3 | 5 | 7 | 3 | 0 | 4 | 7 | 0 | 8 | 182 | 149 | 1 | 1 | 0.064 | 0.621 | 0.534 | 0.406 | 0.396 | NA | 0.38 | 0.369 | 1 | 0.235 | -0.092 |
| SE | 59824 | ENSG00000159176 | CSRP1 | chr1 | - | 201490175 | 201490344 | 201485282 | 201488984 | 201496191 | 201496261 | 147 | 39 | 77 | 80 | 112 | 0 | 0 | 0 | 0 | 1 | 202 | 111 | 76 | 197 | 609 | 0 | 0 | 0 | 0 | 8 | 298 | 149 | 1 | 1 | 1 | 1 | 1 | 1 | 0.982 | 1 | 1 | 1 | 1 | 0.974 | 0.002 |
| SE | 59829 | ENSG00000169682 | SPNS1 | chr16 | + | 28977907 | 28978044 | 28975491 | 28975557 | 28979154 | 28979244 | 64 | 28 | 35 | 65 | 14 | 1 | 0 | 0 | 0 | 0 | 91 | 76 | 23 | 37 | 107 | 0 | 0 | 0 | 0 | 0 | 286 | 149 | 1 | 1 | 0.971 | 1 | 1 | 1 | 1 | 1 | 1 | 1 | 1 | 1 | -0.006 |
| SE | 59838 | ENSG00000120913 | PDLIM2 | chr8 | + | 22581378 | 22581530 | 22580470 | 22580697 | 22584820 | 22584890 | 110 | 50 | 97 | 87 | 82 | 0 | 0 | 0 | 0 | 2 | 82 | 120 | 44 | 55 | 125 | 0 | 0 | 0 | 0 | 0 | 298 | 149 | 1 | 1 | 1 | 1 | 1 | 1 | 0.953 | 1 | 1 | 1 | 1 | 1 | -0.009 |
| SE | 59841 | ENSG00000138709 | LARP1B | chr4 | + | 128077787 | 128077962 | 128074933 | 128074993 | 128082164 | 128082305 | 115 | 97 | 94 | 115 | 222 | 2 | 0 | 0 | 0 | 0 | 85 | 81 | 132 | 103 | 116 | 0 | 0 | 0 | 0 | 0 | 298 | 149 | 1 | 1 | 0.966 | 1 | 1 | 1 | 1 | 1 | 1 | 1 | 1 | 1 | -0.007 |
| SE | 59842 | ENSG00000138709 | LARP1B | chr4 | + | 128082164 | 128082305 | 128077787 | 128077962 | 128091000 | 128091144 | 110 | 99 | 119 | 97 | 217 | 0 | 0 | 1 | 3 | 1 | 77 | 80 | 146 | 84 | 68 | 0 | 0 | 0 | 0 | 0 | 290 | 149 | 1 | 1 | 1 | 1 | 0.984 | 0.943 | 0.991 | 1 | 1 | 1 | 1 | 1 | -0.016 |
| SE | 59845 | ENSG00000138709 | LARP1B | chr4 | + | 128091346 | 128091512 | 128091000 | 128091144 | 128107138 | 128107313 | 25 | 20 | 41 | 25 | 31 | 0 | 0 | 0 | 0 | 0 | 8 | 16 | 20 | 14 | 0 | 0 | 0 | 1 | 0 | 0 | 298 | 149 | 1 | 1 | 1 | 1 | 1 | 1 | 1 | 1 | 1 | 0.909 | 1 | NA | 0.023 |
| SE | 59846 | ENSG00000138709 | LARP1B | chr4 | + | 128098185 | 128098330 | 128091000 | 128091144 | 128107138 | 128107313 | 54 | 35 | 72 | 52 | 111 | 0 | 0 | 0 | 0 | 0 | 16 | 47 | 47 | 13 | 19 | 0 | 0 | 1 | 0 | 0 | 294 | 149 | 1 | 1 | 1 | 1 | 1 | 1 | 1 | 1 | 1 | 0.96 | 1 | 1 | 0.008 |
| SE | 59847 | ENSG00000138709 | LARP1B | chr4 | + | 128098185 | 128098330 | 128091346 | 128091512 | 128107138 | 128107313 | 84 | 54 | 106 | 87 | 193 | 0 | 1 | 0 | 0 | 0 | 16 | 77 | 72 | 17 | 24 | 0 | 0 | 1 | 0 | 0 | 294 | 149 | 1 | 1 | 1 | 0.965 | 1 | 1 | 1 | 1 | 1 | 0.973 | 1 | 1 | -0.002 |
| SE | 59848 | ENSG00000138709 | LARP1B | chr4 | + | 128114569 | 128114742 | 128107138 | 128107313 | 128121825 | 128122188 | 35 | 25 | 26 | 35 | 67 | 3 | 0 | 0 | 0 | 0 | 63 | 6 | 27 | 15 | 21 | 0 | 0 | 0 | 0 | 0 | 298 | 149 | 1 | 1 | 0.854 | 1 | 1 | 1 | 1 | 1 | 1 | 1 | 1 | 1 | -0.029 |
| SE | 59849 | ENSG00000138709 | LARP1B | chr4 | + | 128162193 | 128162317 | 128121825 | 128122188 | 128178430 | 128178642 | 4 | 2 | 4 | 5 | 13 | 2 | 0 | 0 | 1 | 3 | 8 | 4 | 1 | 0 | 0 | 0 | 0 | 0 | 0 | 0 | 273 | 149 | 1 | 1 | 0.522 | 1 | 1 | 0.732 | 0.703 | 1 | 1 | 1 | NA | NA | -0.209 |
| SE | 59850 | ENSG00000138709 | LARP1B | chr4 | + | 128176871 | 128176907 | 128162193 | 128162317 | 128178430 | 128178642 | 8 | 0 | 1 | 7 | 0 | 0 | 3 | 3 | 2 | 7 | 0 | 4 | 1 | 1 | 0 | 8 | 0 | 1 | 0 | 0 | 185 | 149 | 1 | 1 | 1 | 0 | 0.212 | 0.738 | 0 | 0 | 1 | 0.446 | 1 | NA | -0.221 |
| SE | 59851 | ENSG00000138709 | LARP1B | chr4 | + | 128178430 | 128178642 | 128122116 | 128122188 | 128179405 | 128179410 | 8 | 6 | 13 | 11 | 22 | 0 | 0 | 0 | 0 | 2 | 22 | 1 | 6 | 4 | 13 | 0 | 0 | 0 | 0 | 0 | 298 | 149 | 1 | 1 | 1 | 1 | 1 | 1 | 0.846 | 1 | 1 | 1 | 1 | 1 | -0.031 |
| SE | 59852 | ENSG00000138709 | LARP1B | chr4 | + | 128179405 | 128179512 | 128178430 | 128178642 | 128199438 | 128199599 | 7 | 8 | 17 | 18 | 19 | 0 | 0 | 0 | 1 | 0 | 22 | 4 | 7 | 7 | 19 | 0 | 0 | 0 | 0 | 0 | 256 | 149 | 1 | 1 | 1 | 1 | 1 | 0.913 | 1 | 1 | 1 | 1 | 1 | 1 | -0.017 |
| SE | 59853 | ENSG00000138709 | LARP1B | chr4 | + | 128206427 | 128206537 | 128200520 | 128200665 | 128207255 | 128207383 | 11 | 5 | 8 | 12 | 18 | 0 | 0 | 0 | 3 | 2 | 0 | 6 | 1 | 1 | 23 | 0 | 0 | 0 | 0 | 0 | 259 | 149 | 1 | 1 | 1 | 1 | 1 | 0.697 | 0.838 | NA | 1 | 1 | 1 | | -0.093 |
| SE | 59854 | ENSG00000138709 | LARP1B | chr4 | + | 128209855 | 128209993 | 128207255 | 128207383 | 128220354 | 128220440 | 5 | 6 | 8 | 9 | 21 | 0 | 0 | 0 | 0 | 0 | 0 | 6 | 0 | 0 | 24 | 0 | 0 | 1 | 0 | 0 | 287 | 149 | 1 | 1 | 1 | 1 | 1 | 1 | 1 | NA | 1 | 0 | NA | 1 | 0.333 |
| SE | 59857 | ENSG00000138709 | LARP1B | chr4 | + | 128220354 | 128220440 | 128209855 | 128209993 | 128222348 | 128222595 | 1 | 0 | 0 | 0 | 0 | 0 | 0 | 0 | 0 | 1 | 0 | 0 | 2 | 0 | 1 | 0 | 0 | 0 | 0 | 0 | 235 | 149 | 1 | 1 | 1 | NA | NA | NA | 0 | NA | NA | 1 | NA | 1 | -0.5 |
| SE | 59858 | ENSG00000117479 | SLC19A2 | chr1 | - | 169469963 | 169470186 | 169468643 | 169468836 | 169477154 | 169477757 | 10 | 6 | 23 | 9 | 11 | 0 | 0 | 0 | 0 | 0 | 8 | 8 | 4 | 0 | 5 | 0 | 0 | 2 | 0 | 0 | 298 | 149 | 1 | 1 | 1 | 1 | 1 | 1 | 1 | 1 | 1 | 0.5 | NA | 1 | 0.125 |
| SE | 59862 | ENSG00000103168 | TAF1C | chr16 | - | 84180867 | 84180995 | 84180169 | 84180344 | 84181327 | 84181663 | 0 | 0 | 0 | 0 | 0 | 1 | 0 | 0 | 0 | 0 | 0 | 1 | 0 | 0 | 0 | 0 | 0 | 0 | 0 | 0 | 277 | 149 | 1 | 1 | 0 | NA | NA | NA | NA | NA | 1 | NA | NA | NA | -1 |
| SE | 59863 | ENSG00000103168 | TAF1C | chr16 | - | 84181042 | 84181186 | 84180169 | 84180344 | 84181327 | 84181418 | 34 | 12 | 29 | 36 | 14 | 1 | 0 | 0 | 0 | 0 | 36 | 35 | 41 | 18 | 50 | 0 | 0 | 0 | 0 | 0 | 293 | 149 | 1 | 1 | 0.945 | 1 | 1 | 1 | 1 | 1 | 1 | 1 | 1 | 1 | -0.011 |
| SE | 59864 | ENSG00000103168 | TAF1C | chr16 | - | 84183071 | 84183149 | 84182201 | 84182440 | 84183243 | 84183333 | 10 | 8 | 13 | 26 | 5 | 0 | 0 | 2 | 0 | 0 | 3 | 19 | 4 | 7 | 4 | 0 | 0 | 0 | 0 | 0 | 227 | 149 | 1 | 1 | 1 | 1 | 0.81 | 1 | 1 | 1 | 1 | 1 | 1 | 1 | -0.038 |
| SE | 59865 | ENSG00000103168 | TAF1C | chr16 | - | 84183075 | 84183149 | 84182201 | 84182440 | 84183243 | 84183328 | 43 | 22 | 35 | 39 | 29 | 0 | 0 | 2 | 0 | 0 | 15 | 54 | 37 | 36 | 49 | 0 | 0 | 0 | 0 | 0 | 223 | 149 | 1 | 1 | 1 | 1 | 0.921 | 1 | 1 | 1 | 1 | 1 | 1 | 1 | -0.016 |
| SE | 59867 | ENSG00000103168 | TAF1C | chr16 | - | 84183696 | 84183778 | 84183409 | 84183507 | 84186900 | 84187020 | 14 | 8 | 12 | 24 | 1 | 1 | 0 | 0 | 0 | 0 | 9 | 17 | 10 | 16 | 4 | 0 | 0 | 0 | 0 | 0 | 231 | 149 | 1 | 1 | 0.9 | 1 | 1 | 1 | 1 | 1 | 1 | 1 | 1 | 1 | -0.02 |
| SE | 59870 | ENSG00000103168 | TAF1C | chr16 | - | 84184850 | 84185060 | 84183409 | 84183507 | 84186900 | 84186981 | 17 | 2 | 3 | 14 | 0 | 1 | 0 | 0 | 0 | 0 | 0 | 11 | 10 | 4 | 0 | 0 | 0 | 0 | 0 | 0 | 298 | 149 | 1 | 1 | 0.895 | 1 | 1 | 1 | NA | NA | 1 | 1 | 1 | NA | -0.026 |
| SE | 59873 | ENSG00000175054 | ATR | chr3 | - | 142450465 | 142450602 | 142449006 | 142449602 | 142453127 | 142453233 | 24 | 19 | 25 | 38 | 31 | 27 | 25 | 15 | 32 | 35 | 0 | 3 | 9 | 3 | 0 | 0 | 11 | 2 | 0 | 0 | 286 | 149 | 1 | 1 | 0.317 | 0.284 | 0.465 | 0.382 | 0.316 | NA | 0.124 | 0.701 | 1 | NA | -0.256 |
| SE | 59874 | ENSG00000175054 | ATR | chr3 | - | 142465096 | 142465240 | 142461939 | 142462090 | 142466323 | 142466533 | 109 | 128 | 122 | 143 | 112 | 1 | 0 | 0 | 1 | 0 | 10 | 47 | 42 | 2 | 10 | 0 | 0 | 0 | 0 | 0 | 293 | 149 | 1 | 1 | 0.982 | 1 | 1 | 0.986 | 1 | 1 | 1 | 1 | 1 | 1 | -0.006 |
| SE | 59875 | ENSG00000175054 | ATR | chr3 | - | 142466323 | 142466533 | 142465096 | 142465240 | 142467933 | 142468068 | 90 | 110 | 118 | 116 | 138 | 1 | 0 | 0 | 0 | 0 | 10 | 24 | 32 | 4 | 0 | 0 | 0 | 0 | 0 | 0 | 298 | 149 | 1 | 1 | 0.978 | 1 | 1 | 1 | 1 | 1 | 1 | 1 | 1 | NA | -0.004 |
| SE | 59876 | ENSG00000175054 | ATR | chr3 | - | 142470085 | 142470183 | 142466323 | 142469569 | 142485139 | 142485195 | 85 | 68 | 89 | 81 | 111 | 0 | 0 | 0 | 0 | 0 | 0 | 26 | 35 | 3 | 8 | 0 | 0 | 3 | 0 | 0 | 247 | 149 | 1 | 1 | 1 | 1 | 1 | 1 | 1 | NA | 1 | 0.876 | 1 | 1 | 0.031 |
| SE | 59877 | ENSG00000175054 | ATR | chr3 | - | 142496360 | 142496520 | 142491142 | 142493311 | 142497012 | 142497192 | 65 | 74 | 79 | 80 | 118 | 1 | 0 | 1 | 2 | 0 | 10 | 34 | 21 | 9 | 6 | 0 | 0 | 3 | 0 | 0 | 298 | 149 | 1 | 1 | 0.97 | 1 | 0.975 | 0.952 | 1 | 1 | 1 | 0.778 | 1 | 1 | 0.024 |
| SE | 59878 | ENSG00000175054 | ATR | chr3 | - | 142497012 | 142497192 | 142496360 | 142496520 | 142498596 | 142498774 | 101 | 91 | 90 | 99 | 126 | 0 | 0 | 1 | 2 | 0 | 5 | 47 | 18 | 18 | 6 | 0 | 0 | 0 | 0 | 0 | 298 | 149 | 1 | 1 | 1 | 1 | 0.978 | 0.961 | 1 | 1 | 1 | 1 | 1 | 1 | -0.012 |
| SE | 59879 | ENSG00000175054 | ATR | chr3 | - | 142497133 | 142497192 | 142496396 | 142496520 | 142498596 | 142498774 | 30 | 30 | 27 | 28 | 27 | 0 | 0 | 1 | 2 | 0 | 0 | 15 | 7 | 4 | 0 | 0 | 0 | 0 | 0 | 0 | 208 | 149 | 1 | 1 | 1 | 1 | 0.951 | 0.909 | 1 | NA | 1 | 1 | 1 | NA | -0.028 |
| SE | 59880 | ENSG00000175054 | ATR | chr3 | - | 142498596 | 142498774 | 142497012 | 142497192 | 142499626 | 142499718 | 159 | 134 | 117 | 141 | 153 | 0 | 0 | 0 | 3 | 0 | 8 | 72 | 29 | 33 | 7 | 0 | 0 | 0 | 0 | 0 | 298 | 149 | 1 | 1 | 1 | 1 | 1 | 0.959 | 1 | 1 | 1 | 1 | 1 | 1 | -0.008 |
| SE | 59881 | ENSG00000175054 | ATR | chr3 | - | 142503361 | 142503453 | 142499626 | 142499718 | 142505138 | 142505303 | 162 | 141 | 144 | 174 | 143 | 1 | 0 | 0 | 1 | 0 | 4 | 100 | 30 | 33 | 1 | 0 | 0 | 0 | 0 | 0 | 241 | 149 | 1 | 1 | 0.99 | 1 | 1 | 0.991 | 1 | 1 | 1 | 1 | 1 | 1 | -0.004 |
| SE | 59882 | ENSG00000175054 | ATR | chr3 | - | 142503361 | 142503453 | 142499626 | 142499718 | 142507930 | 142508109 | 75 | 56 | 74 | 86 | 58 | 2 | 3 | 0 | 0 | 0 | 4 | 51 | 14 | 27 | 1 | 0 | 0 | 0 | 0 | 0 | 241 | 149 | 1 | 1 | 0.959 | 0.92 | 1 | 1 | 1 | 1 | 1 | 1 | 1 | 1 | -0.024 |
| SE | 59883 | ENSG00000175054 | ATR | chr3 | - | 142505138 | 142505303 | 142499626 | 142499718 | 142507930 | 142508109 | 61 | 43 | 55 | 55 | 58 | 2 | 3 | 0 | 0 | 0 | 10 | 25 | 10 | 19 | 0 | 0 | 0 | 0 | 0 | 0 | 298 | 149 | 1 | 1 | 0.938 | 0.878 | 1 | 1 | 1 | 1 | 1 | 1 | 1 | NA | -0.037 |
| SE | 59884 | ENSG00000175054 | ATR | chr3 | - | 142505138 | 142505303 | 142503361 | 142503453 | 142507930 | 142508109 | 148 | 133 | 129 | 144 | 144 | 0 | 0 | 0 | 0 | 1 | 10 | 74 | 26 | 25 | 0 | 0 | 0 | 0 | 0 | 0 | 298 | 149 | 1 | 1 | 1 | 1 | 1 | 1 | 0.986 | 1 | 1 | 1 | 1 | NA | -0.003 |
| SE | 59886 | ENSG00000175054 | ATR | chr3 | - | 142515394 | 142515515 | 142513500 | 142513638 | 142519668 | 142519784 | 153 | 130 | 114 | 135 | 168 | 0 | 0 | 1 | 0 | 0 | 72 | 103 | 41 | 8 | 20 | 0 | 0 | 0 | 0 | 0 | 270 | 149 | 1 | 1 | 1 | 1 | 0.984 | 1 | 1 | 1 | 1 | 1 | 1 | 1 | -0.003 |
| SE | 59887 | ENSG00000175054 | ATR | chr3 | - | 142523992 | 142524199 | 142522727 | 142522841 | 142535079 | 142535205 | 211 | 195 | 204 | 199 | 201 | 1 | 1 | 0 | 0 | 4 | 11 | 110 | 92 | 39 | 15 | 0 | 0 | 0 | 0 | 0 | 298 | 149 | 1 | 1 | 0.991 | 0.99 | 1 | 1 | 0.962 | 1 | 1 | 1 | 1 | 1 | -0.011 |
| SE | 59888 | ENSG00000175054 | ATR | chr3 | - | 142538481 | 142538625 | 142536107 | 142536201 | 142540903 | 142541034 | 211 | 158 | 185 | 230 | 257 | 0 | 0 | 0 | 0 | 0 | 2 | 81 | 71 | 28 | 14 | 0 | 1 | 0 | 0 | 0 | 293 | 149 | 1 | 1 | 1 | 1 | 1 | 1 | 1 | 1 | 0.976 | 1 | 1 | 1 | 0.005 |
| SE | 59891 | ENSG00000175054 | ATR | chr3 | - | 142549478 | 142549673 | 142547724 | 142547910 | 142550131 | 142550302 | 228 | 195 | 146 | 239 | 175 | 0 | 0 | 2 | 0 | 0 | 0 | 88 | 67 | 17 | 0 | 0 | 0 | 1 | 0 | 0 | 298 | 149 | 1 | 1 | 1 | 1 | 0.973 | 1 | 1 | NA | 1 | 0.971 | 1 | NA | 0.004 |
| SE | 59892 | ENSG00000175054 | ATR | chr3 | - | 142553226 | 142553398 | 142550131 | 142550302 | 142553639 | 142553740 | 107 | 120 | 96 | 116 | 152 | 0 | 0 | 0 | 0 | 0 | 0 | 39 | 39 | 7 | 0 | 12 | 5 | 0 | 0 | 0 | 298 | 149 | 1 | 1 | 1 | 1 | 1 | 1 | 1 | 0 | 0.796 | 1 | 1 | NA | 0.301 |
| SE | 59893 | ENSG00000175054 | ATR | chr3 | - | 142556382 | 142556575 | 142555876 | 142556139 | 142558623 | 142558776 | 140 | 94 | 122 | 155 | 205 | 0 | 0 | 0 | 0 | 0 | 20 | 50 | 45 | 21 | 2 | 0 | 2 | 0 | 0 | 0 | 298 | 149 | 1 | 1 | 1 | 1 | 1 | 1 | 1 | 1 | 0.926 | 1 | 1 | 1 | 0.015 |
| SE | 59894 | ENSG00000175054 | ATR | chr3 | - | 142558623 | 142558776 | 142556382 | 142556575 | 142559250 | 142559441 | 133 | 66 | 119 | 123 | 169 | 0 | 0 | 0 | 0 | 0 | 0 | 49 | 46 | 25 | 1 | 0 | 4 | 0 | 0 | 0 | 298 | 149 | 1 | 1 | 1 | 1 | 1 | 1 | 1 | NA | 0.86 | 1 | 1 | 1 | 0.035 |
| SE | 59896 | ENSG00000175054 | ATR | chr3 | - | 142562231 | 142562348 | 142561242 | 142561421 | 142566120 | 142566261 | 26 | 31 | 41 | 32 | 55 | 0 | 1 | 2 | 4 | 0 | 0 | 9 | 14 | 11 | 0 | 0 | 0 | 0 | 0 | 0 | 266 | 149 | 1 | 1 | 1 | 0.946 | 0.92 | 0.818 | 1 | NA | 1 | 1 | 1 | NA | -0.063 |
| SE | 59898 | ENSG00000175054 | ATR | chr3 | - | 142566120 | 142566261 | 142562231 | 142563109 | 142568062 | 142568154 | 107 | 87 | 86 | 102 | 187 | 0 | 0 | 1 | 0 | 0 | 12 | 41 | 52 | 24 | 14 | 0 | 0 | 0 | 0 | 0 | 290 | 149 | 1 | 1 | 1 | 1 | 0.978 | 1 | 1 | 1 | 1 | 1 | 1 | 1 | -0.004 |
| SE | 59899 | ENSG00000155367 | PPM1J | chr1 | - | 112712344 | 112712457 | 112711970 | 112712055 | 112712743 | 112712803 | 4 | 2 | 6 | 7 | 0 | 0 | 0 | 0 | 1 | 3 | 11 | 0 | 0 | 0 | 0 | 0 | 0 | 0 | 2 | 0 | 262 | 149 | 1 | 1 | 1 | 1 | 1 | 0.799 | 0 | 1 | NA | NA | 0 | NA | 0.26 |
| SE | 59900 | ENSG00000155367 | PPM1J | chr1 | - | 112712344 | 112712460 | 112711970 | 112712055 | 112712743 | 112713031 | 2 | 1 | 2 | 4 | 0 | 0 | 0 | 0 | 1 | 3 | 11 | 0 | 0 | 0 | 0 | 0 | 0 | 0 | 2 | 0 | 265 | 149 | 1 | 1 | 1 | 1 | 1 | 0.692 | 0 | 1 | NA | NA | 0 | NA | 0.238 |
| SE | 59902 | ENSG00000221823 | PPP3R1 | chr2 | - | 68187254 | 68187314 | 68186467 | 68186652 | 68188513 | 68188690 | 1848 | 1455 | 1269 | 1748 | 2400 | 1 | 1 | 2 | 2 | 0 | 3374 | 1690 | 2018 | 2094 | 2281 | 0 | 0 | 0 | 0 | 10 | 209 | 149 | 1 | 1 | 0.999 | 0.999 | 0.998 | 0.998 | 1 | 1 | 1 | 1 | 1 | 0.994 | 0 |
| SE | 59903 | ENSG00000221823 | PPP3R1 | chr2 | - | 68188513 | 68188690 | 68187254 | 68187314 | 68217091 | 68217131 | 2315 | 1768 | 1648 | 2195 | 2298 | 1 | 2 | 3 | 0 | 1 | 4043 | 2209 | 2609 | 2594 | 2909 | 0 | 0 | 0 | 4 | 7 | 298 | 149 | 1 | 1 | 0.999 | 0.998 | 0.996 | 1 | 0.999 | 1 | 1 | 1 | 0.997 | 0.995 | 0 |
| SE | 59904 | ENSG00000221823 | PPP3R1 | chr2 | - | 68217091 | 68217131 | 68188513 | 68188690 | 68252124 | 68252532 | 509 | 387 | 441 | 525 | 315 | 2 | 0 | 0 | 1 | 0 | 765 | 583 | 654 | 585 | 706 | 0 | 4 | 2 | 9 | 7 | 189 | 149 | 1 | 1 | 0.995 | 1 | 1 | 0.998 | 1 | 1 | 0.991 | 0.996 | 0.981 | 0.988 | 0.007 |
| SE | 59905 | ENSG00000170606 | HSPA4 | chr5 | + | 133067416 | 133067557 | 133064979 | 133065037 | 133070373 | 133070496 | 170 | 107 | 186 | 149 | 149 | 0 | 0 | 0 | 0 | 1 | 69 | 208 | 110 | 168 | 52 | 0 | 0 | 0 | 0 | 0 | 290 | 149 | 1 | 1 | 1 | 1 | 1 | 1 | 0.987 | 1 | 1 | 1 | 1 | 1 | -0.003 |
| SE | 59906 | ENSG00000170606 | HSPA4 | chr5 | + | 133070373 | 133070496 | 133067416 | 133067557 | 133073229 | 133073329 | 343 | 237 | 269 | 281 | 260 | 0 | 0 | 0 | 0 | 0 | 180 | 288 | 215 | 240 | 64 | 0 | 2 | 5 | 0 | 0 | 272 | 149 | 1 | 1 | 1 | 1 | 1 | 1 | 1 | 1 | 0.987 | 0.959 | 1 | 1 | 0.011 |
| SE | 59907 | ENSG00000170606 | HSPA4 | chr5 | + | 133073992 | 133074126 | 133073229 | 133073329 | 133076653 | 133076898 | 597 | 498 | 479 | 578 | 496 | 1 | 0 | 0 | 1 | 0 | 282 | 439 | 496 | 451 | 77 | 0 | 0 | 0 | 0 | 0 | 283 | 149 | 1 | 1 | 0.997 | 1 | 1 | 0.997 | 1 | 1 | 1 | 1 | 1 | 1 | -0.001 |
| SE | 59910 | ENSG00000164144 | ARFIP1 | chr4 | + | 152807203 | 152807248 | 152779992 | 152780091 | 152829624 | 152829726 | 4 | 0 | 0 | 1 | 0 | 17 | 7 | 7 | 8 | 12 | 0 | 0 | 0 | 0 | 0 | 0 | 2 | 5 | 0 | 0 | 194 | 149 | 1 | 1 | 0.153 | 0 | 0 | 0.088 | 0 | NA | 0 | 0 | NA | NA | 0.048 |
| SE | 59911 | ENSG00000164144 | ARFIP1 | chr4 | + | 152807203 | 152807248 | 152780108 | 152780226 | 152809672 | 152809733 | 2 | 0 | 0 | 0 | 0 | 0 | 0 | 0 | 0 | 2 | 0 | 2 | 0 | 0 | 0 | 0 | 0 | 0 | 0 | 0 | 194 | 149 | 1 | 1 | 1 | NA | NA | NA | 0 | NA | 1 | NA | NA | NA | -0.5 |
| SE | 59912 | ENSG00000164144 | ARFIP1 | chr4 | + | 152807203 | 152807248 | 152780108 | 152780226 | 152810079 | 152810386 | 0 | 0 | 0 | 2 | 0 | 1 | 0 | 0 | 0 | 2 | 0 | 0 | 0 | 3 | 0 | 0 | 0 | 0 | 0 | 0 | 194 | 149 | 1 | 1 | 0 | NA | NA | 1 | 0 | NA | NA | NA | 1 | NA | -0.667 |
| SE | 59914 | ENSG00000164144 | ARFIP1 | chr4 | + | 152809672 | 152809733 | 152780108 | 152780226 | 152810079 | 152810386 | 1 | 1 | 0 | 1 | 2 | 1 | 0 | 0 | 0 | 2 | 0 | 0 | 1 | 0 | 0 | 0 | 0 | 0 | 0 | 0 | 210 | 149 | 1 | 1 | 0.415 | 1 | NA | 1 | 0.415 | NA | NA | 1 | NA | NA | -0.292 |
| SE | 59915 | ENSG00000164144 | ARFIP1 | chr4 | + | 152809672 | 152809733 | 152780108 | 152780226 | 152829624 | 152829726 | 0 | 0 | 1 | 0 | 0 | 58 | 37 | 35 | 30 | 22 | 0 | 0 | 0 | 0 | 0 | 3 | 20 | 5 | 25 | 2 | 210 | 149 | 1 | 1 | 0 | 0 | 0.02 | 0 | 0 | 0 | 0 | 0 | 0 | 0 | 0.004 |
| SE | 59916 | ENSG00000164144 | ARFIP1 | chr4 | + | 152809672 | 152809733 | 152807203 | 152807248 | 152810079 | 152810386 | 3 | 1 | 0 | 1 | 0 | 0 | 0 | 0 | 2 | 0 | 0 | 2 | 1 | 0 | 0 | 0 | 0 | 0 | 3 | 0 | 210 | 149 | 1 | 1 | 1 | 1 | NA | 0.262 | NA | NA | 1 | 1 | 0 | NA | 0.087 |
| SE | 59917 | ENSG00000164144 | ARFIP1 | chr4 | + | 152809672 | 152809733 | 152807203 | 152807248 | 152829624 | 152829726 | 0 | 0 | 1 | 0 | 0 | 6 | 0 | 1 | 1 | 0 | 0 | 2 | 0 | 0 | 0 | 0 | 0 | 0 | 0 | 0 | 210 | 149 | 1 | 1 | 0 | NA | 0.415 | 0 | NA | NA | 1 | NA | NA | NA | -0.862 |
| SE | 59918 | ENSG00000164144 | ARFIP1 | chr4 | + | 152863605 | 152863714 | 152829624 | 152829726 | 152872451 | 152872564 | 186 | 122 | 102 | 101 | 109 | 0 | 0 | 0 | 1 | 0 | 6 | 53 | 30 | 30 | 2 | 0 | 0 | 1 | 0 | 0 | 258 | 149 | 1 | 1 | 1 | 1 | 1 | 0.983 | 1 | 1 | 1 | 0.945 | 1 | 1 | 0.008 |
| SE | 59920 | ENSG00000164144 | ARFIP1 | chr4 | + | 152872451 | 152872564 | 152863605 | 152863714 | 152880962 | 152881111 | 223 | 143 | 116 | 156 | 220 | 0 | 0 | 0 | 1 | 0 | 3 | 55 | 53 | 37 | 3 | 0 | 0 | 0 | 0 | 0 | 262 | 149 | 1 | 1 | 1 | 1 | 1 | 0.989 | 1 | 1 | 1 | 1 | 1 | 1 | -0.002 |
| SE | 59922 | ENSG00000170043 | TRAPPC1 | chr17 | - | 7931010 | 7931149 | 7930344 | 7930734 | 7931730 | 7931829 | 460 | 253 | 300 | 396 | 901 | 0 | 0 | 0 | 0 | 0 | 912 | 523 | 326 | 408 | 2402 | 5 | 0 | 0 | 0 | 0 | 288 | 149 | 1 | 1 | 1 | 1 | 1 | 1 | 1 | 0.99 | 1 | 1 | 1 | 1 | 0.002 |
| SE | 59923 | ENSG00000170043 | TRAPPC1 | chr17 | - | 7931505 | 7931576 | 7931010 | 7931149 | 7931730 | 7931829 | 536 | 331 | 411 | 544 | 518 | 0 | 0 | 0 | 0 | 0 | 1009 | 745 | 322 | 573 | 2972 | 0 | 9 | 1 | 0 | 14 | 220 | 149 | 1 | 1 | 1 | 1 | 1 | 1 | 1 | 1 | 0.982 | 0.995 | 1 | 0.993 | 0.006 |
| SE | 59930 | ENSG00000159363 | ATP13A2 | chr1 | - | 16986462 | 16986632 | 16985957 | 16986358 | 16986804 | 16986956 | 29 | 2 | 13 | 5 | 7 | 2 | 0 | 2 | 0 | 1 | 0 | 10 | 8 | 20 | 16 | 0 | 0 | 0 | 18 | 0 | 298 | 149 | 1 | 1 | 0.879 | 1 | 0.765 | 1 | 0.778 | NA | 1 | 1 | 0.357 | 1 | 0.045 |
| SE | 59931 | ENSG00000159363 | ATP13A2 | chr1 | - | 16989886 | 16990003 | 16989690 | 16989770 | 16990126 | 16990287 | 41 | 12 | 26 | 38 | 10 | 0 | 0 | 1 | 0 | 0 | 10 | 11 | 13 | 9 | 74 | 0 | 0 | 0 | 2 | 2 | 266 | 149 | 1 | 1 | 1 | 1 | 0.936 | 1 | 1 | 1 | 1 | 1 | 0.716 | 0.954 | 0.053 |
| SE | 59933 | ENSG00000159363 | ATP13A2 | chr1 | - | 16996385 | 16996478 | 16996253 | 16996300 | 16997019 | 16997175 | 25 | 10 | 12 | 27 | 38 | 1 | 0 | 2 | 0 | 0 | 11 | 18 | 18 | 40 | 26 | 0 | 0 | 0 | 0 | 0 | 242 | 149 | 1 | 1 | 0.939 | 1 | 0.787 | 1 | 1 | 1 | 1 | 1 | 1 | 1 | -0.055 |
| SE | 59934 | ENSG00000159363 | ATP13A2 | chr1 | - | 16996385 | 16996496 | 16996253 | 16996300 | 16997019 | 16997175 | 54 | 19 | 34 | 54 | 47 | 1 | 0 | 2 | 0 | 0 | 53 | 47 | 26 | 54 | 68 | 0 | 0 | 0 | 0 | 0 | 260 | 149 | 1 | 1 | 0.969 | 1 | 0.907 | 1 | 1 | 1 | 1 | 1 | 1 | 1 | -0.025 |
| SE | 59935 | ENSG00000159363 | ATP13A2 | chr1 | - | 17000245 | 17000312 | 17000010 | 17000142 | 17000399 | 17000534 | 43 | 19 | 21 | 54 | 18 | 1 | 0 | 0 | 0 | 0 | 106 | 34 | 24 | 53 | 73 | 0 | 0 | 0 | 0 | 0 | 216 | 149 | 1 | 1 | 0.967 | 1 | 1 | 1 | 1 | 1 | 1 | 1 | 1 | 1 | -0.007 |
| SE | 59939 | ENSG00000142327 | RNPEPL1 | chr2 | + | 240573109 | 240573261 | 240572422 | 240572563 | 240573774 | 240573891 | 151 | 43 | 112 | 68 | 52 | 0 | 2 | 0 | 0 | 0 | 41 | 78 | 58 | 74 | 175 | 0 | 0 | 0 | 0 | 0 | 298 | 149 | 1 | 1 | 1 | 0.915 | 1 | 1 | 1 | 1 | 1 | 1 | 1 | 1 | -0.017 |
| SE | 59940 | ENSG00000142327 | RNPEPL1 | chr2 | + | 240573774 | 240573891 | 240573109 | 240573261 | 240574112 | 240574348 | 145 | 33 | 116 | 61 | 60 | 0 | 0 | 1 | 0 | 0 | 58 | 80 | 46 | 63 | 212 | 0 | 0 | 4 | 1 | 0 | 266 | 149 | 1 | 1 | 1 | 1 | 0.985 | 1 | 1 | 1 | 1 | 0.866 | 0.972 | 1 | 0.029 |
| SE | 59942 | ENSG00000142327 | RNPEPL1 | chr2 | + | 240576847 | 240576990 | 240574112 | 240576765 | 240577598 | 240577642 | 182 | 64 | 90 | 127 | 55 | 0 | 0 | 2 | 3 | 0 | 91 | 117 | 100 | 116 | 228 | 0 | 0 | 0 | 0 | 0 | 292 | 149 | 1 | 1 | 1 | 1 | 0.958 | 0.956 | 1 | 1 | 1 | 1 | 1 | 1 | -0.017 |
| SE | 59947 | ENSG00000117280 | RAB29 | chr1 | - | 205770732 | 205770854 | 205767985 | 205770453 | 205774832 | 205775086 | 158 | 63 | 118 | 130 | 271 | 0 | 0 | 0 | 1 | 0 | 60 | 75 | 69 | 83 | 94 | 0 | 0 | 0 | 0 | 0 | 271 | 149 | 1 | 1 | 1 | 1 | 1 | 0.986 | 1 | 1 | 1 | 1 | 1 | 1 | -0.003 |
| SE | 59948 | ENSG00000117280 | RAB29 | chr1 | - | 205771471 | 205771653 | 205770732 | 205770854 | 205772495 | 205772567 | 273 | 193 | 226 | 220 | 246 | 3 | 0 | 0 | 1 | 0 | 239 | 257 | 115 | 162 | 240 | 0 | 0 | 0 | 0 | 0 | 298 | 149 | 1 | 1 | 0.978 | 1 | 1 | 0.991 | 1 | 1 | 1 | 1 | 1 | 1 | -0.006 |
| SE | 59953 | ENSG00000117280 | RAB29 | chr1 | - | 205772495 | 205772567 | 205771061 | 205771648 | 205774832 | 205775086 | 46 | 11 | 27 | 29 | 18 | 0 | 0 | 1 | 0 | 0 | 19 | 29 | 8 | 14 | 18 | 0 | 0 | 1 | 0 | 0 | 221 | 149 | 1 | 1 | 1 | 1 | 0.948 | 1 | 1 | 1 | 1 | 0.844 | 1 | 1 | 0.021 |
| SE | 59955 | ENSG00000117280 | RAB29 | chr1 | - | 205772495 | 205772567 | 205771471 | 205771653 | 205775272 | 205775451 | 88 | 57 | 55 | 67 | 71 | 0 | 0 | 0 | 0 | 0 | 53 | 71 | 38 | 77 | 61 | 0 | 0 | 2 | 0 | 0 | 221 | 149 | 1 | 1 | 1 | 1 | 1 | 1 | 1 | 1 | 1 | 0.928 | 1 | 1 | 0.014 |
| SE | 59956 | ENSG00000117280 | RAB29 | chr1 | - | 205772495 | 205772567 | 205771471 | 205771653 | 205775373 | 205775449 | 85 | 55 | 55 | 67 | 67 | 0 | 0 | 2 | 0 | 0 | 53 | 69 | 38 | 77 | 61 | 0 | 0 | 0 | 0 | 0 | 221 | 149 | 1 | 1 | 1 | 1 | 0.949 | 1 | 1 | 1 | 1 | 1 | 1 | 1 | -0.01 |
| SE | 59962 | ENSG00000103671 | TRIP4 | chr15 | + | 64393340 | 64393476 | 64387861 | 64387964 | 64393945 | 64394115 | 0 | 0 | 2 | 0 | 0 | 80 | 43 | 36 | 35 | 45 | 0 | 0 | 0 | 0 | 0 | 65 | 83 | 52 | 84 | 15 | 285 | 149 | 1 | 1 | 0 | 0 | 0.028 | 0 | 0 | 0 | 0 | 0 | 0 | 0 | 0.006 |
| SE | 59964 | ENSG00000103671 | TRIP4 | chr15 | + | 64397605 | 64397818 | 64395397 | 64395531 | 64400742 | 64400821 | 155 | 112 | 125 | 131 | 87 | 0 | 0 | 0 | 0 | 3 | 125 | 177 | 100 | 152 | 44 | 0 | 0 | 0 | 4 | 0 | 298 | 149 | 1 | 1 | 1 | 1 | 1 | 1 | 0.935 | 1 | 1 | 1 | 0.95 | 1 | -0.003 |
| SE | 59967 | ENSG00000103671 | TRIP4 | chr15 | + | 64418540 | 64418728 | 64409612 | 64409828 | 64424030 | 64424155 | 128 | 95 | 77 | 111 | 184 | 0 | 0 | 0 | 2 | 0 | 67 | 97 | 61 | 120 | 37 | 0 | 0 | 0 | 0 | 0 | 298 | 149 | 1 | 1 | 1 | 1 | 1 | 0.965 | 1 | 1 | 1 | 1 | 1 | 1 | -0.007 |
| SE | 59970 | ENSG00000103671 | TRIP4 | chr15 | + | 64445005 | 64445108 | 64425539 | 64425631 | 64454996 | 64455236 | 221 | 118 | 136 | 160 | 175 | 0 | 2 | 0 | 0 | 0 | 29 | 199 | 57 | 119 | 43 | 0 | 0 | 0 | 0 | 0 | 252 | 149 | 1 | 1 | 1 | 0.972 | 1 | 1 | 1 | 1 | 1 | 1 | 1 | 1 | -0.006 |
| SE | 59972 | ENSG00000076770 | MBNL3 | chrX | - | 132384662 | 132384698 | 132382177 | 132382272 | 132386660 | 132386706 | 57 | 32 | 43 | 65 | 43 | 1 | 0 | 0 | 0 | 0 | 1 | 40 | 26 | 20 | 35 | 0 | 0 | 0 | 0 | 0 | 185 | 149 | 1 | 1 | 0.979 | 1 | 1 | 1 | 1 | 1 | 1 | 1 | 1 | 1 | -0.004 |
| SE | 59976 | ENSG00000188976 | NOC2L | chr1 | - | 946172 | 946286 | 945517 | 945653 | 946401 | 946545 | 227 | 118 | 224 | 244 | 141 | 0 | 0 | 0 | 0 | 0 | 578 | 211 | 240 | 335 | 544 | 0 | 0 | 0 | 0 | 10 | 263 | 149 | 1 | 1 | 1 | 1 | 1 | 1 | 1 | 1 | 1 | 1 | 1 | 0.969 | 0.006 |
| SE | 59977 | ENSG00000188976 | NOC2L | chr1 | - | 946172 | 946286 | 945517 | 945653 | 948130 | 948232 | 118 | 67 | 108 | 142 | 86 | 0 | 0 | 0 | 2 | 0 | 330 | 124 | 127 | 177 | 305 | 0 | 0 | 0 | 0 | 0 | 263 | 149 | 1 | 1 | 1 | 1 | 1 | 0.976 | 1 | 1 | 1 | 1 | 1 | 1 | -0.005 |
| SE | 59978 | ENSG00000188976 | NOC2L | chr1 | - | 946401 | 946545 | 945517 | 945653 | 948130 | 948232 | 44 | 12 | 53 | 35 | 7 | 0 | 0 | 0 | 2 | 0 | 26 | 46 | 28 | 50 | 49 | 0 | 0 | 0 | 0 | 0 | 293 | 149 | 1 | 1 | 1 | 1 | 1 | 0.899 | 1 | 1 | 1 | 1 | 1 | 1 | -0.02 |
| SE | 59979 | ENSG00000188976 | NOC2L | chr1 | - | 946401 | 946545 | 946172 | 946286 | 948130 | 948232 | 151 | 64 | 170 | 137 | 61 | 0 | 2 | 0 | 0 | 0 | 274 | 133 | 141 | 208 | 274 | 0 | 0 | 0 | 0 | 0 | 293 | 149 | 1 | 1 | 1 | 0.942 | 1 | 1 | 1 | 1 | 1 | 1 | 1 | 1 | -0.012 |
| SE | 59981 | ENSG00000230487 | PSMG3-AS1 | chr7 | + | 1574714 | 1574863 | 1570072 | 1570279 | 1575786 | 1575888 | 12 | 3 | 12 | 11 | 10 | 0 | 0 | 0 | 0 | 0 | 3 | 16 | 1 | 1 | 28 | 0 | 0 | 1 | 0 | 0 | 298 | 149 | 1 | 1 | 1 | 1 | 1 | 1 | 1 | 1 | 1 | 0.333 | 1 | 1 | 0.133 |
| SE | 59983 | ENSG00000230487 | PSMG3-AS1 | chr7 | + | 1579286 | 1579404 | 1575786 | 1575888 | 1583161 | 1583231 | 2 | 2 | 1 | 0 | 5 | 1 | 0 | 2 | 3 | 6 | 0 | 2 | 0 | 0 | 6 | 6 | 5 | 1 | 0 | 27 | 267 | 149 | 1 | 1 | 0.527 | 1 | 0.218 | 0 | 0.317 | 0 | 0.182 | 0 | NA | 0.11 | 0.339 |
| SE | 59984 | ENSG00000230487 | PSMG3-AS1 | chr7 | + | 1579286 | 1579404 | 1575786 | 1575888 | 1584383 | 1584418 | 0 | 2 | 1 | 1 | 9 | 7 | 2 | 4 | 5 | 5 | 0 | 0 | 0 | 0 | 3 | 0 | 1 | 2 | 1 | 1 | 267 | 149 | 1 | 1 | 0 | 0.358 | 0.122 | 0.1 | 0.501 | NA | 0 | 0 | 0 | 0.626 | 0.06 |
| SE | 59986 | ENSG00000230487 | PSMG3-AS1 | chr7 | + | 1583161 | 1583269 | 1579286 | 1579404 | 1584383 | 1584632 | 3 | 1 | 1 | 0 | 0 | 0 | 1 | 0 | 1 | 4 | 0 | 4 | 1 | 0 | 25 | 0 | 0 | 0 | 0 | 0 | 257 | 149 | 1 | 1 | 1 | 0.367 | 1 | 0 | 0 | NA | 1 | 1 | NA | 1 | -0.527 |
| SE | 59989 | ENSG00000123453 | SARDH | chr9 | - | 133690379 | 133690527 | 133685192 | 133685286 | 133694257 | 133694371 | 3 | 5 | 5 | 4 | 0 | 0 | 0 | 0 | 0 | 0 | 0 | 0 | 4 | 0 | 0 | 0 | 0 | 2 | 0 | 0 | 297 | 149 | 1 | 1 | 1 | 1 | 1 | 1 | NA | NA | NA | 0.501 | NA | NA | 0.499 |
| SE | 59992 | ENSG00000116574 | RHOU | chr1 | + | 228737672 | 228737731 | 228735478 | 228736004 | 228743284 | 228746664 | 54 | 23 | 32 | 25 | 28 | 7 | 5 | 1 | 8 | 2 | 57 | 53 | 53 | 40 | 23 | 5 | 18 | 0 | 2 | 9 | 208 | 149 | 1 | 1 | 0.847 | 0.767 | 0.958 | 0.691 | 0.909 | 0.891 | 0.678 | 1 | 0.935 | 0.647 | 0.004 |
| SE | 59993 | ENSG00000159346 | ADIPOR1 | chr1 | - | 202942024 | 202942218 | 202940825 | 202941701 | 202943757 | 202943863 | 128 | 60 | 62 | 174 | 132 | 0 | 0 | 0 | 0 | 0 | 200 | 207 | 110 | 195 | 1090 | 0 | 0 | 0 | 5 | 0 | 298 | 149 | 1 | 1 | 1 | 1 | 1 | 1 | 1 | 1 | 1 | 1 | 0.951 | 1 | 0.01 |
| SE | 59994 | ENSG00000159346 | ADIPOR1 | chr1 | - | 202943757 | 202943863 | 202941489 | 202941701 | 202944982 | 202945169 | 3 | 0 | 0 | 1 | 1 | 0 | 0 | 0 | 0 | 0 | 0 | 7 | 5 | 5 | 7 | 0 | 0 | 3 | 0 | 0 | 255 | 149 | 1 | 1 | 1 | NA | NA | 1 | 1 | NA | 1 | 0.493 | 1 | 1 | 0.127 |
| SE | 59995 | ENSG00000159346 | ADIPOR1 | chr1 | - | 202943757 | 202943945 | 202940831 | 202941701 | 202944982 | 202945169 | 222 | 111 | 149 | 254 | 221 | 0 | 0 | 0 | 0 | 0 | 411 | 393 | 259 | 372 | 1245 | 0 | 0 | 3 | 0 | 0 | 298 | 149 | 1 | 1 | 1 | 1 | 1 | 1 | 1 | 1 | 1 | 0.977 | 1 | 1 | 0.005 |
| SE | 59997 | ENSG00000159346 | ADIPOR1 | chr1 | - | 202944982 | 202945169 | 202943757 | 202943945 | 202946438 | 202946610 | 491 | 233 | 321 | 541 | 384 | 0 | 0 | 2 | 0 | 0 | 664 | 766 | 491 | 647 | 2656 | 0 | 0 | 0 | 4 | 0 | 298 | 149 | 1 | 1 | 1 | 1 | 0.988 | 1 | 1 | 1 | 1 | 1 | 0.988 | 1 | 0 |
| SE | 59998 | ENSG00000159346 | ADIPOR1 | chr1 | - | 202946438 | 202946610 | 202944982 | 202945169 | 202948303 | 202948420 | 532 | 254 | 333 | 597 | 616 | 0 | 2 | 0 | 0 | 0 | 525 | 720 | 548 | 686 | 3248 | 0 | 0 | 2 | 0 | 0 | 298 | 149 | 1 | 1 | 1 | 0.984 | 1 | 1 | 1 | 1 | 1 | 0.993 | 1 | 1 | -0.002 |
| SE | 60004 | ENSG00000137496 | IL18BP | chr11 | + | 71999616 | 71999662 | 71999227 | 71999370 | 71999926 | 72000012 | 2 | 2 | 0 | 0 | 0 | 9 | 7 | 3 | 5 | 11 | 0 | 0 | 2 | 0 | 0 | 0 | 7 | 4 | 0 | 15 | 195 | 149 | 1 | 1 | 0.145 | 0.179 | 0 | 0 | 0 | NA | 0 | 0.276 | NA | 0 | -0.027 |
| SE | 60006 | ENSG00000137496 | IL18BP | chr11 | + | 72001200 | 72001324 | 72000350 | 72000557 | 72001783 | 72001948 | 29 | 9 | 17 | 17 | 18 | 0 | 0 | 0 | 0 | 2 | 37 | 15 | 7 | 9 | 13 | 0 | 3 | 0 | 0 | 0 | 273 | 149 | 1 | 1 | 1 | 1 | 1 | 1 | 0.831 | 1 | 0.732 | 1 | 1 | 1 | 0.02 |
| SE | 60007 | ENSG00000137496 | IL18BP | chr11 | + | 72001404 | 72001424 | 72001200 | 72001324 | 72001783 | 72002804 | 1 | 3 | 0 | 7 | 0 | 0 | 0 | 0 | 1 | 0 | 8 | 0 | 3 | 0 | 6 | 0 | 0 | 0 | 0 | 0 | 169 | 149 | 1 | 1 | 1 | 1 | NA | 0.861 | NA | 1 | NA | 1 | NA | 1 | -0.046 |
| SE | 60008 | ENSG00000137496 | IL18BP | chr11 | + | 72001404 | 72001552 | 72001200 | 72001324 | 72001783 | 72001948 | 54 | 19 | 30 | 31 | 13 | 0 | 0 | 0 | 1 | 0 | 36 | 35 | 22 | 8 | 83 | 0 | 0 | 0 | 0 | 0 | 297 | 149 | 1 | 1 | 1 | 1 | 1 | 0.94 | 1 | 1 | 1 | 1 | 1 | 1 | -0.012 |
| SE | 60010 | ENSG00000272645 | GTF2IP20 | chr1 | - | 223986574 | 223986640 | 223984381 | 223984440 | 223988044 | 223988154 | 36 | 24 | 18 | 31 | 49 | 0 | 0 | 0 | 0 | 4 | 23 | 27 | 31 | 14 | 11 | 0 | 0 | 0 | 0 | 0 | 215 | 149 | 1 | 1 | 1 | 1 | 1 | 1 | 0.895 | 1 | 1 | 1 | 1 | 1 | -0.021 |
| SE | 60032 | ENSG00000139173 | TMEM117 | chr12 | + | 44083588 | 44083681 | 43944209 | 43944342 | 44143524 | 44143624 | 3 | 0 | 0 | 0 | 0 | 5 | 2 | 8 | 5 | 1 | 0 | 0 | 0 | 8 | 0 | 0 | 0 | 6 | 2 | 0 | 242 | 149 | 1 | 1 | 0.27 | 0 | 0 | 0 | 0 | NA | NA | 0 | 0.711 | NA | -0.301 |
| SE | 60034 | ENSG00000139173 | TMEM117 | chr12 | + | 44143524 | 44143624 | 43944209 | 43944342 | 44211289 | 44211387 | 13 | 6 | 12 | 7 | 1 | 1 | 0 | 1 | 1 | 1 | 0 | 0 | 6 | 2 | 0 | 0 | 0 | 0 | 0 | 0 | 249 | 149 | 1 | 1 | 0.886 | 1 | 0.878 | 0.807 | 0.374 | NA | NA | 1 | 1 | NA | -0.211 |
| SE | 60050 | ENSG00000096093 | EFHC1 | chr6 | + | 52423686 | 52423814 | 52420106 | 52420473 | 52423945 | 52424167 | 0 | 2 | 0 | 0 | 0 | 5 | 0 | 6 | 3 | 0 | 1 | 0 | 0 | 0 | 0 | 4 | 1 | 2 | 0 | 0 | 277 | 149 | 1 | 1 | 0 | 1 | 0 | 0 | NA | 0.119 | 0 | 0 | NA | NA | 0.21 |
| SE | 60052 | ENSG00000096093 | EFHC1 | chr6 | + | 52423686 | 52423828 | 52420344 | 52420473 | 52423945 | 52424167 | 0 | 1 | 0 | 0 | 0 | 5 | 0 | 6 | 3 | 0 | 0 | 0 | 0 | 0 | 0 | 4 | 1 | 2 | 0 | 0 | 291 | 149 | 1 | 1 | 0 | 1 | 0 | 0 | NA | 0 | 0 | 0 | NA | NA | 0.25 |
| SE | 60054 | ENSG00000096093 | EFHC1 | chr6 | + | 52429968 | 52430131 | 52423945 | 52424167 | 52438303 | 52438327 | 0 | 0 | 3 | 0 | 0 | 15 | 8 | 14 | 10 | 3 | 0 | 1 | 0 | 0 | 0 | 4 | 26 | 3 | 5 | 0 | 298 | 149 | 1 | 1 | 0 | 0 | 0.097 | 0 | 0 | 0 | 0.019 | 0 | 0 | NA | 0.015 |
| SE | 60056 | ENSG00000096093 | EFHC1 | chr6 | + | 52437561 | 52437678 | 52423945 | 52424167 | 52438303 | 52438352 | 0 | 0 | 5 | 0 | 0 | 15 | 8 | 14 | 10 | 3 | 0 | 0 | 0 | 0 | 0 | 4 | 26 | 3 | 5 | 0 | 266 | 149 | 1 | 1 | 0 | 0 | 0.167 | 0 | 0 | 0 | 0 | 0 | 0 | NA | 0.033 |
| SE | 60061 | ENSG00000096093 | EFHC1 | chr6 | + | 52464894 | 52465115 | 52452687 | 52454287 | 52469332 | 52469473 | 30 | 15 | 24 | 24 | 54 | 0 | 0 | 0 | 0 | 2 | 6 | 6 | 7 | 9 | 4 | 0 | 0 | 0 | 0 | 0 | 298 | 149 | 1 | 1 | 1 | 1 | 1 | 1 | 0.931 | 1 | 1 | 1 | 1 | 1 | -0.014 |
| SE | 60062 | ENSG00000096093 | EFHC1 | chr6 | + | 52469332 | 52469473 | 52454205 | 52454287 | 52479036 | 52479250 | 20 | 8 | 16 | 14 | 28 | 0 | 0 | 2 | 0 | 0 | 26 | 11 | 4 | 3 | 4 | 0 | 0 | 0 | 0 | 0 | 290 | 149 | 1 | 1 | 1 | 1 | 0.804 | 1 | 1 | 1 | 1 | 1 | 1 | 1 | -0.039 |
| SE | 60063 | ENSG00000096093 | EFHC1 | chr6 | + | 52481545 | 52481662 | 52479639 | 52479787 | 52488556 | 52488675 | 0 | 0 | 2 | 1 | 1 | 0 | 1 | 4 | 1 | 1 | 0 | 1 | 3 | 0 | 2 | 0 | 0 | 0 | 0 | 0 | 266 | 149 | 1 | 1 | NA | 0 | 0.219 | 0.359 | 0.359 | NA | 1 | 1 | NA | 1 | -0.766 |
| SE | 60064 | ENSG00000096093 | EFHC1 | chr6 | + | 52481545 | 52481662 | 52479639 | 52479787 | 52490139 | 52490350 | 0 | 0 | 2 | 2 | 5 | 12 | 25 | 8 | 26 | 15 | 0 | 1 | 3 | 0 | 2 | 0 | 0 | 13 | 7 | 0 | 266 | 149 | 1 | 1 | 0 | 0 | 0.123 | 0.041 | 0.157 | NA | 1 | 0.114 | 0 | 1 | -0.464 |
| SE | 60065 | ENSG00000096093 | EFHC1 | chr6 | + | 52488556 | 52488675 | 52479639 | 52479787 | 52490139 | 52490350 | 0 | 1 | 4 | 1 | 1 | 12 | 25 | 8 | 26 | 15 | 0 | 0 | 0 | 0 | 0 | 0 | 0 | 13 | 7 | 0 | 268 | 149 | 1 | 1 | 0 | 0.022 | 0.218 | 0.021 | 0.036 | NA | NA | 0 | 0 | NA | 0.059 |
| SE | 60067 | ENSG00000149532 | CPSF7 | chr11 | - | 61416363 | 61416519 | 61416104 | 61416162 | 61419948 | 61420094 | 146 | 115 | 148 | 170 | 320 | 0 | 0 | 0 | 0 | 1 | 223 | 135 | 93 | 118 | 241 | 0 | 0 | 0 | 0 | 6 | 298 | 149 | 1 | 1 | 1 | 1 | 1 | 1 | 0.994 | 1 | 1 | 1 | 1 | 0.953 | 0.008 |
| SE | 60068 | ENSG00000149532 | CPSF7 | chr11 | - | 61419921 | 61420094 | 61416104 | 61416519 | 61420469 | 61420573 | 305 | 167 | 271 | 282 | 541 | 2 | 0 | 3 | 2 | 0 | 334 | 298 | 130 | 203 | 242 | 0 | 0 | 2 | 2 | 0 | 298 | 149 | 1 | 1 | 0.987 | 1 | 0.978 | 0.986 | 1 | 1 | 1 | 0.97 | 0.981 | 1 | 0 |
| SE | 60069 | ENSG00000149532 | CPSF7 | chr11 | - | 61419948 | 61420094 | 61416104 | 61416519 | 61420469 | 61420573 | 361 | 232 | 338 | 361 | 638 | 2 | 0 | 3 | 2 | 0 | 488 | 336 | 176 | 270 | 408 | 0 | 0 | 2 | 2 | 0 | 295 | 149 | 1 | 1 | 0.989 | 1 | 0.983 | 0.989 | 1 | 1 | 1 | 0.978 | 0.986 | 1 | -0.001 |
| SE | 60086 | ENSG00000075651 | PLD1 | chr3 | - | 171686684 | 171686798 | 171677565 | 171677694 | 171687370 | 171687584 | 18 | 4 | 8 | 10 | 32 | 0 | 2 | 0 | 0 | 0 | 0 | 11 | 10 | 14 | 30 | 0 | 0 | 2 | 3 | 0 | 263 | 149 | 1 | 1 | 1 | 0.531 | 1 | 1 | 1 | NA | 1 | 0.739 | 0.726 | 1 | 0.04 |
| SE | 60087 | ENSG00000075651 | PLD1 | chr3 | - | 171687370 | 171687584 | 171686684 | 171686798 | 171692331 | 171692442 | 12 | 1 | 6 | 7 | 19 | 0 | 0 | 0 | 0 | 5 | 0 | 12 | 6 | 5 | 16 | 0 | 0 | 0 | 0 | 0 | 298 | 149 | 1 | 1 | 1 | 1 | 1 | 1 | 0.655 | NA | 1 | 1 | 1 | 1 | -0.069 |
| SE | 60089 | ENSG00000075651 | PLD1 | chr3 | - | 171699744 | 171699826 | 171692331 | 171692442 | 171708754 | 171708838 | 36 | 9 | 16 | 40 | 17 | 0 | 0 | 0 | 0 | 0 | 24 | 28 | 11 | 54 | 0 | 0 | 0 | 0 | 3 | 0 | 231 | 149 | 1 | 1 | 1 | 1 | 1 | 1 | 1 | 1 | 1 | 1 | 0.921 | NA | 0.02 |
| SE | 60091 | ENSG00000075651 | PLD1 | chr3 | - | 171734864 | 171734970 | 171726003 | 171726076 | 171737531 | 171737659 | 7 | 0 | 0 | 0 | 0 | 0 | 0 | 0 | 0 | 0 | 0 | 0 | 0 | 0 | 0 | 0 | 0 | 0 | 0 | 2 | 255 | 149 | 1 | 1 | 1 | NA | NA | NA | NA | NA | NA | NA | NA | 0 | 1 |
| SE | 60092 | ENSG00000075651 | PLD1 | chr3 | - | 171735491 | 171735637 | 171734864 | 171734970 | 171737531 | 171737659 | 88 | 63 | 53 | 54 | 68 | 1 | 0 | 0 | 0 | 0 | 62 | 62 | 60 | 143 | 20 | 0 | 0 | 0 | 0 | 0 | 295 | 149 | 1 | 1 | 0.978 | 1 | 1 | 1 | 1 | 1 | 1 | 1 | 1 | 1 | -0.004 |
| SE | 60094 | ENSG00000075651 | PLD1 | chr3 | - | 171737891 | 171738082 | 171737531 | 171737659 | 171810398 | 171810437 | 52 | 23 | 33 | 38 | 39 | 0 | 0 | 0 | 0 | 0 | 29 | 46 | 29 | 74 | 23 | 0 | 0 | 0 | 3 | 0 | 298 | 149 | 1 | 1 | 1 | 1 | 1 | 1 | 1 | 1 | 1 | 1 | 0.925 | 1 | 0.015 |
| SE | 60096 | ENSG00000075651 | PLD1 | chr3 | - | 171789840 | 171789957 | 171789645 | 171789701 | 171810398 | 171810714 | 0 | 1 | 0 | 0 | 0 | 0 | 1 | 0 | 0 | 0 | 10 | 2 | 0 | 9 | 0 | 0 | 0 | 2 | 6 | 0 | 266 | 149 | 1 | 1 | NA | 0.359 | NA | NA | NA | 1 | 1 | 0 | 0.457 | NA | -0.255 |
| SE | 60098 | ENSG00000231682 | LINC01891 | chr2 | - | 234450117 | 234450398 | 234444589 | 234444735 | 234454484 | 234454595 | 0 | 0 | 1 | 0 | 16 | 0 | 1 | 0 | 0 | 0 | 0 | 0 | 0 | 0 | 0 | 0 | 0 | 1 | 0 | 0 | 298 | 149 | 1 | 1 | NA | 0 | 1 | NA | 1 | NA | NA | 0 | NA | NA | 0.667 |
| SE | 60099 | ENSG00000160326 | SLC2A6 | chr9 | - | 133473414 | 133473600 | 133473104 | 133473250 | 133473979 | 133474088 | 6 | 5 | 5 | 11 | 4 | 0 | 4 | 0 | 0 | 0 | 0 | 4 | 3 | 10 | 7 | 0 | 3 | 0 | 0 | 0 | 298 | 149 | 1 | 1 | 1 | 0.385 | 1 | 1 | 1 | NA | 0.4 | 1 | 1 | 1 | 0.027 |
| SE | 60102 | ENSG00000139055 | ERP27 | chr12 | - | 14917177 | 14917303 | 14915488 | 14915686 | 14920931 | 14921048 | 37 | 38 | 37 | 43 | 22 | 0 | 0 | 2 | 0 | 0 | 25 | 25 | 15 | 4 | 8 | 0 | 0 | 2 | 0 | 0 | 275 | 149 | 1 | 1 | 1 | 1 | 0.909 | 1 | 1 | 1 | 1 | 0.803 | 1 | 1 | 0.021 |
| SE | 60104 | ENSG00000116288 | PARK7 | chr1 | + | 7969084 | 7969148 | 7965323 | 7965425 | 7969344 | 7969404 | 0 | 4 | 2 | 2 | 3 | 598 | 396 | 408 | 484 | 645 | 1 | 12 | 6 | 0 | 0 | 846 | 489 | 384 | 523 | 1301 | 213 | 149 | 1 | 1 | 0 | 0.007 | 0.003 | 0.003 | 0.003 | 0.001 | 0.017 | 0.011 | 0 | 0 | -0.003 |
| SE | 60105 | ENSG00000116288 | PARK7 | chr1 | + | 7969344 | 7969404 | 7965323 | 7965425 | 7970893 | 7970963 | 886 | 563 | 615 | 738 | 1097 | 0 | 5 | 3 | 3 | 0 | 1162 | 702 | 514 | 742 | 1893 | 0 | 0 | 0 | 3 | 4 | 209 | 149 | 1 | 1 | 1 | 0.988 | 0.993 | 0.994 | 1 | 1 | 1 | 1 | 0.994 | 0.997 | -0.003 |
| SE | 60107 | ENSG00000203879 | GDI1 | chrX | + | 154438215 | 154438386 | 154437165 | 154437299 | 154438522 | 154438630 | 0 | 0 | 0 | 0 | 1 | 258 | 140 | 189 | 234 | 129 | 0 | 0 | 0 | 0 | 0 | 483 | 323 | 266 | 364 | 1023 | 298 | 149 | 1 | 1 | 0 | 0 | 0 | 0 | 0.004 | 0 | 0 | 0 | 0 | 0 | 0.001 |
| SE | 60108 | ENSG00000203879 | GDI1 | chrX | + | 154438475 | 154438630 | 154437153 | 154437299 | 154438764 | 154438864 | 92 | 56 | 53 | 86 | 65 | 1 | 0 | 0 | 0 | 0 | 100 | 105 | 65 | 97 | 313 | 0 | 0 | 0 | 0 | 0 | 298 | 149 | 1 | 1 | 0.979 | 1 | 1 | 1 | 1 | 1 | 1 | 1 | 1 | 1 | -0.004 |
| SE | 60109 | ENSG00000203879 | GDI1 | chrX | + | 154438522 | 154438630 | 154436912 | 154437299 | 154438764 | 154438864 | 349 | 196 | 242 | 320 | 194 | 1 | 0 | 0 | 0 | 0 | 583 | 426 | 331 | 461 | 1336 | 0 | 0 | 0 | 0 | 0 | 257 | 149 | 1 | 1 | 0.995 | 1 | 1 | 1 | 1 | 1 | 1 | 1 | 1 | 1 | -0.001 |
| SE | 60110 | ENSG00000203879 | GDI1 | chrX | + | 154439709 | 154439797 | 154439005 | 154439140 | 154439940 | 154440189 | 0 | 0 | 0 | 0 | 0 | 161 | 72 | 102 | 163 | 99 | 0 | 0 | 0 | 2 | 0 | 165 | 147 | 108 | 204 | 544 | 237 | 149 | 1 | 1 | 0 | 0 | 0 | 0 | 0 | 0 | 0 | 0 | 0.006 | 0 | -0.001 |

| | | | | | | | | | | | | | |
|---|---|---|---|---|---|---|---|---|---|---|---|---|---|
| SE | 60111 | ENSG00000203879 | GDI1 | chrX | + | 154442110 | 154442255 | 154440523 | 154441794 | 154442373 | 154442428 | 387 | 279 |
| SE | 60112 | ENSG00000122223 | CD244 | chr1 | - | 160832518 | 160832575 | 160830159 | 160831427 | 160834050 | 160834116 | 198 | 106 |
| SE | 60113 | ENSG00000122223 | CD244 | chr1 | - | 160832518 | 160832719 | 160831270 | 160831427 | 160834050 | 160834116 | 68 | 26 |
| SE | 60114 | ENSG00000122223 | CD244 | chr1 | - | 160834050 | 160834116 | 160830159 | 160831427 | 160836194 | 160836254 | 122 | 72 |
| SE | 60115 | ENSG00000122223 | CD244 | chr1 | - | 160834050 | 160834116 | 160830159 | 160831427 | 160838450 | 160838518 | 2 | 0 |
| SE | 60116 | ENSG00000122223 | CD244 | chr1 | - | 160836194 | 160836254 | 160830159 | 160831427 | 160838450 | 160838518 | 120 | 59 |
| SE | 60118 | ENSG00000122223 | CD244 | chr1 | - | 160838938 | 160839049 | 160838450 | 160838518 | 160841209 | 160841485 | 596 | 348 |
| SE | 60119 | ENSG00000122223 | CD244 | chr1 | - | 160838938 | 160839049 | 160838450 | 160838518 | 160841583 | 160841901 | 274 | 139 |
| SE | 60120 | ENSG00000122223 | CD244 | chr1 | - | 160838938 | 160839049 | 160838450 | 160838518 | 160862616 | 160862700 | 260 | 115 |
| SE | 60121 | ENSG00000122223 | CD244 | chr1 | - | 160841209 | 160841485 | 160838450 | 160838518 | 160841583 | 160841901 | 200 | 92 |
| SE | 60123 | ENSG00000122223 | CD244 | chr1 | - | 160841209 | 160841500 | 160838450 | 160838518 | 160841583 | 160841901 | 10 | 3 |
| SE | 60125 | ENSG00000122223 | CD244 | chr1 | - | 160841583 | 160841901 | 160838450 | 160838518 | 160862616 | 160862700 | 133 | 35 |
| SE | 60127 | ENSG00000162148 | PPP1R32 | chr11 | + | 61486343 | 61486403 | 61485798 | 61485911 | 61486523 | 61486610 | 0 | 0 |
| SE | 60128 | ENSG00000162148 | PPP1R32 | chr11 | + | 61486523 | 61486610 | 61486343 | 61486403 | 61486947 | 61487061 | 0 | 4 |
| SE | 60129 | ENSG00000162148 | PPP1R32 | chr11 | + | 61489781 | 61489946 | 61487145 | 61487240 | 61490500 | 61490584 | 5 | 1 |
| SE | 60132 | ENSG00000162373 | BEND5 | chr1 | - | 48761336 | 48761470 | 48758899 | 48759284 | 48776605 | 48776891 | 3 | 7 |
| SE | 60133 | ENSG00000162373 | BEND5 | chr1 | - | 48761336 | 48761470 | 48758899 | 48759366 | 48776605 | 48776891 | 4 | 8 |
| SE | 60138 | ENSG00000171848 | RRM2 | chr2 | + | 10126874 | 10126969 | 10124716 | 10124850 | 10128847 | 10128952 | 38 | 30 |
| SE | 60141 | ENSG00000169031 | COL4A3 | chr2 | + | 227277448 | 227277553 | 227276384 | 227276477 | 227279792 | 227279890 | 4 | 2 |
| SE | 60146 | ENSG00000236810 | ELOA-AS1 | chr1 | - | 23772267 | 23772426 | 23772110 | 23772166 | 23778198 | 23778287 | 8 | 3 |
| SE | 60147 | ENSG00000236810 | ELOA-AS1 | chr1 | - | 23772271 | 23772426 | 23772110 | 23772166 | 23778198 | 23778296 | 8 | 2 |
| SE | 60149 | ENSG00000183963 | SMTN | chr22 | + | 31082909 | 31083002 | 31081317 | 31081456 | 31083178 | 31083309 | 0 | 0 |
| SE | 60150 | ENSG00000183963 | SMTN | chr22 | + | 31082915 | 31083002 | 31081317 | 31081456 | 31083178 | 31083309 | 0 | 0 |
| SE | 60152 | ENSG00000183963 | SMTN | chr22 | + | 31093783 | 31093876 | 31091674 | 31091847 | 31095302 | 31095455 | 0 | 0 |
| SE | 60153 | ENSG00000183963 | SMTN | chr22 | + | 31095851 | 31096897 | 31095533 | 31095609 | 31096997 | 31097338 | 1 | 1 |
| SE | 60154 | ENSG00000183963 | SMTN | chr22 | + | 31096732 | 31096897 | 31095533 | 31095609 | 31096997 | 31097060 | 5 | 1 |
| SE | 60155 | ENSG00000183963 | SMTN | chr22 | + | 31096997 | 31097060 | 31095533 | 31095609 | 31097268 | 31097338 | 0 | 1 |
| SE | 60159 | ENSG00000099250 | NRP1 | chr10 | - | 33202890 | 33202995 | 33197620 | 33197709 | 33207571 | 33207625 | 15 | 19 |
| SE | 60160 | ENSG00000099250 | NRP1 | chr10 | - | 33202911 | 33202995 | 33197649 | 33197709 | 33207571 | 33207716 | 4 | 9 |

| | | | | | | | | | | | | | | | | | | | | | | | | | | | | | | | | | | | | | | | | | | | | | | |
|---|---|---|---|---|---|---|---|---|---|---|---|---|---|---|---|---|---|---|---|---|---|---|---|---|---|---|---|---|---|---|---|---|---|---|---|---|---|---|---|---|---|---|---|---|---|---|
| SE | 60546 | ENSG00000116984 | MTR | chr1 | + | 236874725 | 236874846 | 236873772 | 236873840 | 236880754 | 236880836 | 194 | 189 | 241 | 207 | 373 | 0 | 0 | 0 | 0 | 0 | 143 | 177 | 230 | 144 | 70 | 0 | 0 | 0 | 2 | 0 | 270 | 149 | 1 | 1 | 1 | 1 | 1 | 1 | 1 | 1 | 1 | 1 | 0.975 | 1 | 0.005 |
| SE | 60547 | ENSG00000116984 | MTR | chr1 | + | 236880754 | 236880836 | 236874725 | 236874846 | 236885120 | 236885219 | 163 | 154 | 248 | 195 | 344 | 0 | 0 | 0 | 0 | 1 | 142 | 183 | 229 | 137 | 75 | 0 | 0 | 0 | 0 | 0 | 231 | 149 | 1 | 1 | 1 | 1 | 1 | 1 | 0.996 | 1 | 1 | 1 | 1 | 1 | -0.001 |
| SE | 60548 | ENSG00000116984 | MTR | chr1 | + | 236889180 | 236889336 | 236886291 | 236886367 | 236891132 | 236891329 | 119 | 82 | 133 | 100 | 132 | 0 | 2 | 0 | 0 | 0 | 96 | 79 | 119 | 95 | 101 | 0 | 0 | 4 | 0 | 0 | 298 | 149 | 1 | 1 | 1 | 0.953 | 1 | 1 | 1 | 1 | 1 | 0.937 | 1 | 1 | 0.003 |
| SE | 60549 | ENSG00000116984 | MTR | chr1 | + | 236894356 | 236894557 | 236891132 | 236891329 | 236895357 | 236895550 | 68 | 51 | 100 | 103 | 81 | 0 | 0 | 0 | 1 | 0 | 67 | 72 | 45 | 29 | 31 | 0 | 0 | 0 | 0 | 0 | 298 | 149 | 1 | 1 | 1 | 1 | 1 | 0.981 | 1 | 1 | 1 | 1 | 1 | 1 | -0.004 |
| SE | 60551 | ENSG00000116984 | MTR | chr1 | + | 236907044 | 236907206 | 236897557 | 236899109 | 236913297 | 236913424 | 4 | 6 | 2 | 3 | 5 | 0 | 0 | 0 | 0 | 1 | 7 | 2 | 0 | 0 | 3 | 0 | 0 | 0 | 0 | 0 | 298 | 149 | 1 | 1 | 1 | 1 | 1 | 1 | 0.714 | 1 | 1 | NA | NA | 1 | -0.057 |
| SE | 60552 | ENSG00000100410 | PHF5A | chr22 | - | 41467447 | 41467614 | 41459716 | 41460487 | 41468123 | 41468147 | 104 | 49 | 92 | 115 | 95 | 0 | 0 | 0 | 0 | 0 | 121 | 119 | 82 | 125 | 67 | 0 | 2 | 0 | 0 | 0 | 298 | 149 | 1 | 1 | 1 | 1 | 1 | 1 | 1 | 1 | 0.967 | 1 | 1 | 1 | 0.007 |
| SE | 60553 | ENSG00000100410 | PHF5A | chr22 | - | 41468019 | 41468147 | 41467447 | 41467614 | 41468601 | 41468625 | 1 | 0 | 1 | 1 | 1 | 0 | 2 | 0 | 1 | 0 | 0 | 7 | 4 | 0 | 22 | 0 | 0 | 0 | 3 | 0 | 277 | 149 | 1 | 1 | 1 | 0 | 1 | 0.35 | 1 | NA | 1 | 1 | 0 | 1 | -0.08 |
| SE | 60554 | ENSG00000100410 | PHF5A | chr22 | - | 41468123 | 41468147 | 41467447 | 41467614 | 41468601 | 41468692 | 69 | 22 | 62 | 67 | 52 | 0 | 2 | 0 | 1 | 0 | 77 | 76 | 61 | 84 | 59 | 0 | 0 | 0 | 3 | 0 | 173 | 149 | 1 | 1 | 1 | 0.905 | 1 | 0.983 | 1 | 1 | 1 | 1 | 0.96 | 1 | -0.014 |
| SE | 60555 | ENSG00000160094 | ZNF362 | chr1 | + | 33270486 | 33270612 | 33256491 | 33256654 | 33276099 | 33276163 | 7 | 5 | 5 | 1 | 11 | 0 | 0 | 0 | 2 | 0 | 26 | 23 | 4 | 4 | 20 | 0 | 0 | 0 | 0 | 0 | 275 | 149 | 1 | 1 | 1 | 1 | 1 | 0.213 | 1 | 1 | 1 | 1 | 1 | 1 | -0.157 |
| SE | 60563 | ENSG00000173406 | DAB1 | chr1 | - | 57010679 | 57010790 | 56994777 | 56998128 | 57011144 | 57011272 | 0 | 1 | 0 | 0 | 0 | 1 | 1 | 1 | 0 | 1 | 0 | 0 | 0 | 0 | 8 | 0 | 0 | 0 | 0 | 0 | 260 | 149 | 1 | 1 | 0 | 0.364 | 0 | NA | 0 | NA | NA | NA | NA | 1 | -0.909 |
| SE | 60565 | ENSG00000173406 | DAB1 | chr1 | - | 57033385 | 57033433 | 57025980 | 57026043 | 57033522 | 57033573 | 0 | 0 | 1 | 0 | 0 | 2 | 0 | 0 | 0 | 0 | 0 | 0 | 0 | 0 | 4 | 0 | 0 | 0 | 0 | 0 | 197 | 149 | 1 | 1 | 0 | NA | 1 | NA | NA | NA | NA | NA | NA | 1 | -0.5 |
| SE | 60580 | ENSG00000288637 | NA | chr4 | + | 153394261 | 153394336 | 153358473 | 153358622 | 153397218 | 153397333 | 3 | 5 | 7 | 8 | 3 | 0 | 0 | 0 | 0 | 0 | 18 | 0 | 6 | 18 | 0 | 0 | 0 | 0 | 1 | 0 | 224 | 149 | 1 | 1 | 1 | 1 | 1 | 1 | 1 | 1 | NA | 1 | 0.923 | NA | 0.026 |
| SE | 60581 | ENSG00000288637 | NA | chr4 | + | 153394261 | 153394336 | 153358473 | 153358622 | 153408970 | 153409015 | 4 | 1 | 3 | 3 | 0 | 0 | 2 | 0 | 0 | 0 | 0 | 4 | 2 | 4 | 0 | 0 | 0 | 0 | 0 | 0 | 224 | 149 | 1 | 1 | 1 | 0.25 | 1 | 1 | NA | NA | 1 | 1 | 1 | NA | -0.188 |
| SE | 60582 | ENSG00000288637 | NA | chr4 | + | 153397218 | 153397333 | 153358473 | 153358622 | 153408970 | 153409015 | 0 | 2 | 1 | 2 | 3 | 0 | 2 | 0 | 0 | 0 | 0 | 0 | 8 | 1 | 0 | 0 | 0 | 0 | 0 | 0 | 264 | 149 | 1 | 1 | NA | 0.361 | 1 | 1 | 1 | NA | NA | 1 | 1 | NA | -0.16 |
| SE | 60583 | ENSG00000121211 | MND1 | chr4 | + | 153397218 | 153397333 | 153394261 | 153394336 | 153408970 | 153409015 | 0 | 6 | 5 | 7 | 6 | 1 | 0 | 0 | 0 | 1 | 18 | 0 | 14 | 14 | 0 | 0 | 4 | 0 | 0 | 0 | 264 | 149 | 1 | 1 | 0 | 1 | 1 | 1 | 0.772 | 1 | 0 | 1 | 1 | NA | 0.004 |
| SE | 60584 | ENSG00000288637 | NA | chr4 | + | 153408970 | 153409015 | 153397218 | 153397333 | 153414750 | 153414955 | 1 | 2 | 1 | 6 | 6 | 0 | 0 | 0 | 0 | 0 | 0 | 0 | 13 | 3 | 0 | 0 | 0 | 0 | 0 | 4 | 194 | 149 | 1 | 1 | 1 | 1 | 1 | 1 | 1 | NA | NA | 1 | 1 | 0 | 0.333 |
| SE | 60585 | ENSG00000142632 | ARHGEF19 | chr1 | - | 16205537 | 16205667 | 16205350 | 16205425 | 16205930 | 16206083 | 10 | 1 | 6 | 14 | 7 | 1 | 0 | 0 | 0 | 0 | 17 | 1 | 10 | 5 | 0 | 0 | 0 | 0 | 0 | 0 | 279 | 149 | 1 | 1 | 0.842 | 1 | 1 | 1 | 1 | 1 | 1 | 1 | 1 | NA | -0.032 |
| SE | 60586 | ENSG00000070961 | ATP2B1 | chr12 | - | 89598589 | 89598743 | 89588048 | 89591295 | 89599116 | 89599299 | 0 | 0 | 0 | 0 | 1 | 264 | 145 | 162 | 201 | 500 | 0 | 0 | 0 | 0 | 0 | 8 | 33 | 67 | 25 | 0 | 298 | 149 | 1 | 1 | 0 | 0 | 0 | 0 | 0.001 | 0 | 0 | 0 | 0 | NA | 0 |
| SE | 60587 | ENSG00000070961 | ATP2B1 | chr12 | - | 89598656 | 89598743 | 89590527 | 89591295 | 89599116 | 89599299 | 0 | 0 | 0 | 0 | 1 | 264 | 145 | 162 | 201 | 500 | 0 | 0 | 3 | 0 | 0 | 8 | 33 | 67 | 25 | 0 | 236 | 149 | 1 | 1 | 0 | 0 | 0 | 0 | 0.001 | 0 | 0 | 0.027 | 0 | NA | -0.007 |
| SE | 60588 | ENSG00000070961 | ATP2B1 | chr12 | - | 89599116 | 89599299 | 89588048 | 89591295 | 89604154 | 89604346 | 264 | 145 | 162 | 202 | 500 | 0 | 0 | 0 | 0 | 0 | 8 | 33 | 67 | 25 | 0 | 0 | 0 | 1 | 0 | 0 | 298 | 149 | 1 | 1 | 1 | 1 | 1 | 1 | 1 | 1 | 1 | 0.971 | 1 | NA | 0.007 |
| SE | 60590 | ENSG00000070961 | ATP2B1 | chr12 | - | 89603711 | 89603925 | 89599116 | 89599299 | 89604154 | 89604346 | 107 | 85 | 94 | 104 | 183 | 0 | 0 | 0 | 1 | 0 | 0 | 23 | 42 | 4 | 0 | 0 | 0 | 0 | 0 | 0 | 298 | 149 | 1 | 1 | 1 | 1 | 1 | 0.981 | 1 | NA | 1 | 1 | 1 | NA | -0.004 |
| SE | 60593 | ENSG00000070961 | ATP2B1 | chr12 | - | 89619998 | 89620240 | 89616801 | 89617039 | 89621548 | 89621791 | 550 | 490 | 403 | 543 | 663 | 0 | 0 | 0 | 0 | 0 | 36 | 246 | 267 | 36 | 18 | 0 | 0 | 2 | 0 | 0 | 298 | 149 | 1 | 1 | 1 | 1 | 1 | 1 | 1 | 1 | 1 | 0.985 | 1 | 1 | 0.003 |
| SE | 60594 | ENSG00000070961 | ATP2B1 | chr12 | - | 89621548 | 89621791 | 89619998 | 89620240 | 89624182 | 89624397 | 489 | 392 | 379 | 506 | 706 | 1 | 1 | 1 | 1 | 0 | 39 | 177 | 260 | 41 | 54 | 0 | 0 | 0 | 0 | 0 | 298 | 149 | 1 | 1 | 0.996 | 0.995 | 0.995 | 0.996 | 1 | 1 | 1 | 1 | 1 | 1 | -0.004 |
| SE | 60595 | ENSG00000070961 | ATP2B1 | chr12 | - | 89624182 | 89624397 | 89621548 | 89621791 | 89626453 | 89626615 | 439 | 316 | 352 | 515 | 597 | 0 | 0 | 0 | 0 | 0 | 34 | 141 | 188 | 55 | 45 | 0 | 0 | 1 | 0 | 0 | 298 | 149 | 1 | 1 | 1 | 1 | 1 | 1 | 1 | 1 | 1 | 0.989 | 1 | 1 | 0.002 |
| SE | 60596 | ENSG00000070961 | ATP2B1 | chr12 | - | 89627677 | 89627716 | 89626453 | 89626615 | 89630504 | 89630645 | 161 | 128 | 109 | 152 | 122 | 1 | 0 | 0 | 0 | 0 | 8 | 38 | 66 | 13 | 13 | 0 | 0 | 0 | 0 | 0 | 188 | 149 | 1 | 1 | 0.992 | 1 | 1 | 1 | 1 | 1 | 1 | 1 | 1 | 1 | -0.002 |
| SE | 60597 | ENSG00000070961 | ATP2B1 | chr12 | - | 89630504 | 89630645 | 89627677 | 89627716 | 89634777 | 89634903 | 470 | 420 | 324 | 457 | 633 | 3 | 1 | 0 | 1 | 2 | 23 | 133 | 208 | 50 | 21 | 0 | 0 | 2 | 0 | 0 | 290 | 149 | 1 | 1 | 0.988 | 0.995 | 1 | 0.996 | 0.994 | 1 | 1 | 0.982 | 1 | 1 | -0.002 |
| SE | 60598 | ENSG00000070961 | ATP2B1 | chr12 | - | 89634996 | 89635251 | 89634777 | 89634903 | 89655678 | 89656067 | 407 | 319 | 325 | 349 | 615 | 4 | 0 | 0 | 0 | 0 | 23 | 126 | 150 | 38 | 39 | 0 | 0 | 0 | 0 | 0 | 298 | 149 | 1 | 1 | 0.981 | 1 | 1 | 1 | 1 | 1 | 1 | 1 | 1 | 1 | -0.004 |
| SE | 60600 | ENSG00000070961 | ATP2B1 | chr12 | - | 89655678 | 89656107 | 89642157 | 89642355 | 89708595 | 89708838 | 256 | 248 | 180 | 279 | 199 | 2 | 0 | 0 | 0 | 0 | 48 | 77 | 115 | 26 | 40 | 0 | 0 | 0 | 0 | 0 | 298 | 149 | 1 | 1 | 0.985 | 1 | 1 | 1 | 1 | 1 | 1 | 1 | 1 | 1 | -0.003 |
| SE | 60603 | ENSG00000147121 | KRBOX4 | chrX | + | 46463197 | 46463293 | 46450430 | 46450475 | 46472749 | 46473669 | 4 | 11 | 9 | 15 | 16 | 1 | 0 | 0 | 1 | 0 | 7 | 1 | 0 | 0 | 0 | 0 | 0 | 0 | 0 | 0 | 245 | 149 | 1 | 1 | 0.709 | 1 | 1 | 0.901 | 1 | 1 | 1 | NA | NA | NA | -0.078 |
| SE | 60604 | ENSG00000147121 | KRBOX4 | chrX | + | 46463197 | 46463308 | 46450430 | 46450475 | 46472749 | 46474639 | 22 | 7 | 5 | 12 | 19 | 1 | 0 | 0 | 1 | 0 | 14 | 14 | 10 | 2 | 16 | 0 | 0 | 0 | 0 | 0 | 260 | 149 | 1 | 1 | 0.927 | 1 | 1 | 0.873 | 1 | 1 | 1 | 1 | 1 | 1 | -0.04 |
| SE | 60605 | ENSG00000147121 | KRBOX4 | chrX | + | 46471141 | 46471185 | 46463197 | 46463293 | 46472749 | 46474639 | 6 | 4 | 1 | 5 | 8 | 4 | 12 | 11 | 23 | 33 | 0 | 2 | 2 | 3 | 0 | 0 | 1 | 1 | 0 | 0 | 193 | 149 | 1 | 1 | 0.537 | 0.205 | 0.066 | 0.144 | 0.158 | NA | 0.607 | 0.607 | 1 | NA | -0.516 |
| SE | 60606 | ENSG00000147121 | KRBOX4 | chrX | + | 46471141 | 46471185 | 46463197 | 46463293 | 46472752 | 46473594 | 0 | 1 | 0 | 0 | 0 | 0 | 1 | 0 | 1 | 0 | 0 | 0 | 1 | 0 | 0 | 0 | 0 | 0 | 0 | 0 | 193 | 149 | 1 | 1 | NA | 0.436 | NA | 0 | NA | NA | NA | 1 | NA | NA | -0.782 |
| SE | 60608 | ENSG00000147121 | KRBOX4 | chrX | + | 46471141 | 46471185 | 46463197 | 46463308 | 46472752 | 46473594 | 1 | 0 | 0 | 1 | 2 | 0 | 0 | 0 | 1 | 0 | 0 | 0 | 0 | 0 | 10 | 0 | 0 | 0 | 4 | 0 | 193 | 149 | 1 | 1 | 1 | NA | NA | 0.436 | 1 | NA | NA | NA | 0 | 1 | 0.312 |
| SE | 60609 | ENSG00000163399 | ATP1A1 | chr1 | + | 116390781 | 116390891 | 116390212 | 116390411 | 116392853 | 116392988 | 850 | 483 | 843 | 786 | 1487 | 0 | 0 | 2 | 0 | 0 | 898 | 772 | 519 | 596 | 1159 | 0 | 0 | 2 | 3 | 0 | 259 | 149 | 1 | 1 | 1 | 1 | 0.996 | 1 | 1 | 1 | 1 | 0.993 | 0.991 | 1 | 0.002 |
| SE | 60611 | ENSG00000163399 | ATP1A1 | chr1 | + | 116397887 | 116398038 | 116396597 | 116396734 | 116398620 | 116398789 | 749 | 450 | 858 | 770 | 985 | 0 | 0 | 0 | 0 | 0 | 575 | 741 | 510 | 540 | 947 | 0 | 1 | 0 | 0 | 0 | 298 | 149 | 1 | 1 | 1 | 1 | 1 | 1 | 1 | 1 | 0.997 | 1 | 1 | 1 | 0.001 |
| SE | 60612 | ENSG00000163399 | ATP1A1 | chr1 | + | 116403883 | 116403975 | 116401553 | 116401655 | 116404415 | 116404772 | 724 | 330 | 705 | 634 | 972 | 0 | 0 | 0 | 0 | 0 | 589 | 655 | 446 | 440 | 604 | 0 | 0 | 3 | 0 | 0 | 241 | 149 | 1 | 1 | 1 | 1 | 1 | 1 | 1 | 1 | 1 | 0.989 | 1 | 1 | 0.002 |
| SE | 60620 | ENSG00000111196 | MAGOHB | chr12 | - | 10612814 | 10612978 | 10610621 | 10610680 | 10613438 | 10613573 | 0 | 0 | 2 | 2 | 0 | 29 | 13 | 22 | 26 | 9 | 0 | 0 | 0 | 0 | 0 | 0 | 25 | 6 | 7 | 5 | 298 | 149 | 1 | 1 | 0 | 0 | 0.043 | 0.037 | 0 | NA | 0 | 0 | 0 | 0 | 0.016 |
| SE | 60621 | ENSG00000128191 | DGCR8 | chr22 | + | 20089668 | 20089811 | 20085819 | 20087321 | 20089975 | 20090258 | 79 | 51 | 48 | 68 | 111 | 0 | 0 | 0 | 0 | 0 | 73 | 46 | 23 | 76 | 87 | 0 | 3 | 0 | 0 | 0 | 292 | 149 | 1 | 1 | 1 | 1 | 1 | 1 | 1 | 1 | 0.887 | 1 | 1 | 1 | 0.023 |
| SE | 60623 | ENSG00000128191 | DGCR8 | chr22 | + | 20106176 | 20106277 | 20094711 | 20094795 | 20106591 | 20106685 | 85 | 48 | 80 | 109 | 177 | 0 | 0 | 2 | 0 | 3 | 287 | 184 | 105 | 109 | 231 | 0 | 0 | 1 | 0 | 0 | 250 | 149 | 1 | 1 | 1 | 1 | 0.96 | 1 | 0.972 | 1 | 1 | 0.984 | 1 | 1 | -0.01 |
| SE | 60624 | ENSG00000128191 | DGCR8 | chr22 | + | 20106591 | 20106685 | 20094712 | 20094795 | 20107270 | 20107398 | 0 | 0 | 2 | 1 | 13 | 0 | 0 | 0 | 0 | 4 | 0 | 0 | 1 | 0 | 5 | 0 | 0 | 0 | 0 | 0 | 243 | 149 | 1 | 1 | NA | NA | 1 | 1 | 0.666 | NA | NA | 1 | NA | 1 | -0.111 |
| SE | 60625 | ENSG00000128191 | DGCR8 | chr22 | + | 20106591 | 20106698 | 20094712 | 20094795 | 20107270 | 20107398 | 66 | 60 | 55 | 78 | 57 | 0 | 0 | 0 | 0 | 4 | 164 | 131 | 53 | 107 | 164 | 0 | 0 | 0 | 0 | 0 | 256 | 149 | 1 | 1 | 1 | 1 | 1 | 1 | 0.892 | 1 | 1 | 1 | 1 | 1 | -0.022 |
| SE | 60627 | ENSG00000159247 | TUBBP5 | chr9 | + | 138175600 | 138175901 | 138175362 | 138175471 | 138176206 | 138176508 | 0 | 0 | 3 | 0 | 0 | 0 | 0 | 0 | 0 | 0 | 0 | 2 | 11 | 0 | 0 | 0 | 0 | 1 | 0 | 0 | 298 | 149 | 1 | 1 | NA | NA | 1 | NA | NA | NA | 1 | 0.846 | NA | NA | 0.077 |
| SE | 60628 | ENSG00000135829 | DHX9 | chr1 | + | 182842544 | 182842677 | 182839346 | 182839456 | 182843293 | 182843434 | 549 | 460 | 556 | 585 | 704 | 1 | 0 | 0 | 0 | 0 | 215 | 475 | 468 | 469 | 94 | 0 | 0 | 0 | 0 | 0 | 282 | 149 | 1 | 1 | 0.997 | 1 | 1 | 1 | 1 | 1 | 1 | 1 | 1 | 1 | -0.001 |
| SE | 60629 | ENSG00000135829 | DHX9 | chr1 | + | 182842544 | 182842677 | 182839346 | 182839456 | 182852232 | 182852344 | 257 | 206 | 274 | 259 | 222 | 2 | 0 | 1 | 0 | 0 | 72 | 202 | 217 | 219 | 33 | 1 | 0 | 0 | 3 | 0 | 282 | 149 | 1 | 1 | 0.985 | 1 | 0.993 | 1 | 1 | 0.974 | 1 | 1 | 0.975 | 1 | 0.006 |
| SE | 60630 | ENSG00000135829 | DHX9 | chr1 | + | 182843293 | 182843434 | 182839346 | 182839456 | 182852232 | 182852344 | 510 | 364 | 466 | 314 | 925 | 2 | 0 | 1 | 0 | 0 | 265 | 403 | 361 | 379 | 84 | 1 | 0 | 0 | 3 | 0 | 290 | 149 | 1 | 1 | 0.992 | 1 | 0.996 | 1 | 1 | 0.993 | 1 | 1 | 0.985 | 1 | 0.002 |
| SE | 60631 | ENSG00000135829 | DHX9 | chr1 | + | 182843293 | 182843434 | 182842544 | 182842677 | 182852232 | 182852344 | 803 | 618 | 747 | 640 | 1408 | 0 | 0 | 0 | 1 | 2 | 408 | 674 | 612 | 635 | 145 | 0 | 0 | 0 | 0 | 0 | 290 | 149 | 1 | 1 | 1 | 1 | 1 | 0.997 | 0.997 | 1 | 1 | 1 | 1 | 1 | -0.001 |
| SE | 60632 | ENSG00000135829 | DHX9 | chr1 | + | 182852232 | 182852344 | 182839346 | 182839456 | 182853305 | 182853418 | 382 | 263 | 292 | 280 | 600 | 0 | 0 | 0 | 1 | 0 | 295 | 390 | 289 | 301 | 59 | 0 | 0 | 0 | 0 | 0 | 261 | 149 | 1 | 1 | 1 | 1 | 1 | 0.994 | 1 | 1 | 1 | 1 | 1 | 1 | -0.001 |
| SE | 60633 | ENSG00000135829 | DHX9 | chr1 | + | 182853305 | 182853418 | 182852232 | 182852344 | 182854029 | 182854142 | 768 | 533 | 552 | 508 | 1012 | 0 | 0 | 1 | 0 | 0 | 403 | 589 | 493 | 557 | 97 | 0 | 0 | 0 | 0 | 0 | 262 | 149 | 1 | 1 | 1 | 1 | 0.997 | 1 | 1 | 1 | 1 | 1 | 1 | 1 | -0.001 |
| SE | 60635 | ENSG00000135829 | DHX9 | chr1 | + | 182866443 | 182866585 | 182859992 | 182860184 | 182866960 | 182867043 | 836 | 417 | 671 | 771 | 1457 | 0 | 0 | 0 | 1 | 0 | 108 | 487 | 538 | 468 | 31 | 0 | 0 | 0 | 0 | 0 | 291 | 149 | 1 | 1 | 1 | 1 | 1 | 0.997 | 1 | 1 | 1 | 1 | 1 | 1 | -0.001 |
| SE | 60636 | ENSG00000135829 | DHX9 | chr1 | + | 182866960 | 182867043 | 182859992 | 182860184 | 182872336 | 182872493 | 462 | 238 | 294 | 373 | 837 | 0 | 0 | 0 | 1 | 0 | 45 | 256 | 253 | 241 | 0 | 0 | 0 | 0 | 0 | 0 | 232 | 149 | 1 | 1 | 1 | 1 | 1 | 0.996 | 1 | 1 | 1 | 1 | 1 | NA | -0.001 |
| SE | 60637 | ENSG00000135829 | DHX9 | chr1 | + | 182872336 | 182872493 | 182866960 | 182867043 | 182874853 | 182874954 | 1073 | 616 | 798 | 1045 | 2388 | 0 | 0 | 0 | 0 | 0 | 221 | 732 | 797 | 623 | 25 | 0 | 0 | 1 | 0 | 0 | 298 | 149 | 1 | 1 | 1 | 1 | 1 | 1 | 1 | 1 | 1 | 0.997 | 1 | 1 | 0.001 |
| SE | 60638 | ENSG00000135829 | DHX9 | chr1 | + | 182879249 | 182879410 | 182877417 | 182878173 | 182880496 | 182880608 | 520 | 270 | 493 | 564 | 771 | 0 | 0 | 0 | 1 | 0 | 148 | 367 | 322 | 272 | 15 | 0 | 0 | 0 | 0 | 0 | 298 | 149 | 1 | 1 | 1 | 1 | 1 | 0.996 | 1 | 1 | 1 | 1 | 1 | 1 | -0.001 |
| SE | 60639 | ENSG00000135829 | DHX9 | chr1 | + | 182880496 | 182880608 | 182879249 | 182879410 | 182881263 | 182881425 | 319 | 175 | 277 | 282 | 340 | 0 | 0 | 2 | 0 | 0 | 125 | 229 | 161 | 165 | 15 | 0 | 0 | 0 | 0 | 0 | 261 | 149 | 1 | 1 | 1 | 1 | 0.988 | 1 | 1 | 1 | 1 | 1 | 1 | 1 | -0.002 |
| SE | 60641 | ENSG00000135829 | DHX9 | chr1 | + | 182883519 | 182883635 | 182883138 | 182883368 | 182884612 | 182884813 | 865 | 569 | 743 | 859 | 1706 | 0 | 0 | 0 | 0 | 0 | 238 | 623 | 721 | 500 | 29 | 0 | 0 | 3 | 0 | 0 | 265 | 149 | 1 | 1 | 1 | 1 | 1 | 1 | 1 | 1 | 1 | 0.993 | 1 | 1 | 0.001 |
| SE | 60642 | ENSG00000135829 | DHX9 | chr1 | + | 182884612 | 182884813 | 182883519 | 182883635 | 182887082 | 182887747 | 689 | 505 | 634 | 771 | 902 | 0 | 0 | 0 | 0 | 2 | 264 | 540 | 735 | 519 | 27 | 0 | 0 | 0 | 0 | 0 | 298 | 149 | 1 | 1 | 1 | 1 | 1 | 1 | 0.996 | 1 | 1 | 1 | 1 | 1 | -0.001 |
| SE | 60643 | ENSG00000147099 | HDAC8 | chrX | - | 72330657 | 72330768 | 72329537 | 72330076 | 72351732 | 72351838 | 0 | 1 | 0 | 1 | 0 | 18 | 9 | 2 | 9 | 13 | 0 | 0 | 0 | 0 | 0 | 3 | 18 | 4 | 15 | 20 | 260 | 149 | 1 | 1 | 0 | 0.06 | 0 | 0.06 | 0 | 0 | 0 | 0 | 0 | 0 | 0.024 |
| SE | 60644 | ENSG00000147099 | HDAC8 | chrX | - | 72330690 | 72330768 | 72329556 | 72330076 | 72351732 | 72351838 | 0 | 2 | 0 | 2 | 0 | 18 | 9 | 2 | 9 | 13 | 0 | 0 | 0 | 0 | 0 | 3 | 18 | 4 | 15 | 20 | 227 | 149 | 1 | 1 | 0 | 0.127 | 0 | 0.127 | 0 | 0 | 0 | 0 | 0 | 0 | 0.051 |
| SE | 60651 | ENSG00000147099 | HDAC8 | chrX | - | 72462361 | 72462442 | 72462003 | 72462102 | 72464558 | 72464564 | 0 | 0 | 3 | 0 | 0 | 0 | 0 | 0 | 0 | 1 | 0 | 3 | 0 | 0 | 0 | 0 | 0 | 0 | 0 | 0 | 230 | 149 | 1 | 1 | NA | NA | 1 | NA | 0 | NA | 1 | NA | NA | NA | -0.5 |
| SE | 60656 | ENSG00000147099 | HDAC8 | chrX | - | 72490928 | 72491000 | 72488932 | 72489041 | 72495155 | 72495268 | 9 | 6 | 5 | 8 | 4 | 0 | 1 | 0 | 2 | 0 | 5 | 1 | 5 | 5 | 0 | 0 | 0 | 0 | 0 | 0 | 221 | 149 | 1 | 1 | 1 | 0.802 | 1 | 0.729 | 1 | 1 | 1 | 1 | 1 | NA | -0.094 |
| SE | 60657 | ENSG00000147099 | HDAC8 | chrX | - | 72490928 | 72491006 | 72484850 | 72489041 | 72495155 | 72495179 | 23 | 20 | 18 | 16 | 23 | 0 | 1 | 0 | 2 | 0 | 12 | 18 | 14 | 13 | 7 | 0 | 0 | 0 | 0 | 0 | 227 | 149 | 1 | 1 | 1 | 0.929 | 1 | 0.84 | 1 | 1 | 1 | 1 | 1 | 1 | -0.046 |
| SE | 60658 | ENSG00000147099 | HDAC8 | chrX | - | 72490928 | 72491006 | 72484850 | 72489041 | 72567888 | 72568030 | 10 | 7 | 7 | 10 | 8 | 0 | 1 | 0 | 0 | 0 | 5 | 9 | 5 | 5 | 0 | 0 | 0 | 0 | 0 | 0 | 227 | 149 | 1 | 1 | 1 | 0.821 | 1 | 1 | 1 | 1 | 1 | 1 | 1 | NA | -0.036 |
| SE | 60664 | ENSG00000147099 | HDAC8 | chrX | - | 72495155 | 72495268 | 72488932 | 72489041 | 72567888 | 72568030 | 18 | 5 | 11 | 14 | 4 | 0 | 1 | 0 | 0 | 0 | 0 | 7 | 7 | 2 | 8 | 0 | 0 | 0 | 0 | 0 | 262 | 149 | 1 | 1 | 1 | 0.74 | 1 | 1 | 1 | NA | 1 | 1 | 1 | 1 | -0.052 |
| SE | 60668 | ENSG00000147099 | HDAC8 | chrX | - | 72567888 | 72568030 | 72495155 | 72495268 | 72568753 | 72568791 | 73 | 50 | 61 | 50 | 52 | 0 | 0 | 1 | 0 | 0 | 54 | 45 | 55 | 67 | 99 | 0 | 0 | 0 | 0 | 0 | 291 | 149 | 1 | 1 | 1 | 1 | 0.969 | 1 | 1 | 1 | 1 | 1 | 1 | 1 | -0.006 |
| SE | 60672 | ENSG00000147099 | HDAC8 | chrX | - | 72572056 | 72572109 | 72567888 | 72568030 | 72572650 | 72572659 | 17 | 13 | 14 | 14 | 15 | 0 | 2 | 0 | 0 | 0 | 23 | 40 | 10 | 24 | 14 | 0 | 0 | 0 | 0 | 0 | 202 | 149 | 1 | 1 | 1 | 0.827 | 1 | 1 | 1 | 1 | 1 | 1 | 1 | 1 | -0.035 |
| SE | 60674 | ENSG00000117322 | CR2 | chr1 | + | 207473544 | 207473721 | 207472771 | 207473179 | 207473800 | 207473885 | 11 | 2 | 3 | 6 | 4 | 17 | 5 | 25 | 8 | 25 | 0 | 3 | 0 | 0 | 0 | 5 | 22 | 0 | 3 | 0 | 298 | 149 | 1 | 1 | 0.244 | 0.167 | 0.057 | 0.273 | 0.074 | 0 | 0.064 | NA | 0 | NA | 0.142 |
| SE | 60675 | ENSG00000117322 | CR2 | chr1 | + | 207476233 | 207476419 | 207474823 | 207475216 | 207477884 | 207478070 | 54 | 38 | 77 | 33 | 91 | 0 | 0 | 1 | 0 | 0 | 12 | 34 | 4 | 12 | 0 | 0 | 0 | 0 | 0 | 0 | 298 | 149 | 1 | 1 | 1 | 1 | 0.975 | 1 | 1 | 1 | 1 | 1 | 1 | NA | -0.005 |
| SE | 60677 | ENSG00000166750 | SLFN5 | chr17 | + | 35258650 | 35259702 | 35243070 | 35243143 | 35260970 | 35261096 | 387 | 414 | 733 | 504 | 483 | 1 | 0 | 0 | 1 | 0 | 95 | 256 | 249 | 121 | 31 | 0 | 0 | 0 | 0 | 0 | 298 | 149 | 1 | 1 | 0.995 | 1 | 1 | 0.996 | 1 | 1 | 1 | 1 | 1 | 1 | -0.002 |
| SE | 60678 | ENSG00000166750 | SLFN5 | chr17 | + | 35260970 | 35261096 | 35258650 | 35259702 | 35264182 | 35264903 | 496 | 608 | 978 | 651 | 748 | 0 | 0 | 0 | 2 | 0 | 111 | 401 | 281 | 172 | 27 | 3 | 0 | 0 | 0 | 0 | 275 | 149 | 1 | 1 | 1 | 1 | 1 | 0.994 | 1 | 0.952 | 1 | 1 | 1 | 1 | 0.008 |
| SE | 60690 | ENSG00000118961 | LDAH | chr2 | - | 20790254 | 20790398 | 20774809 | 20774979 | 20823036 | 20823104 | 36 | 26 | 48 | 46 | 60 | 0 | 1 | 2 | 1 | 0 | 6 | 45 | 26 | 19 | 3 | 5 | 0 | 0 | 0 | 0 | 293 | 149 | 1 | 1 | 1 | 0.93 | 0.924 | 0.959 | 1 | 0.379 | 1 | 1 | 1 | 1 | 0.087 |
| SE | 60692 | ENSG00000118961 | LDAH | chr2 | - | 20801291 | 20801465 | 20790254 | 20790398 | 20823036 | 20823089 | 22 | 8 | 10 | 17 | 9 | 0 | 0 | 5 | 2 | 0 | 0 | 15 | 8 | 4 | 10 | 0 | 3 | 0 | 5 | 0 | 298 | 149 | 1 | 1 | 1 | 1 | 0.5 | 0.81 | 1 | NA | 0.714 | 1 | 0.286 | 1 | 0.112 |
| SE | 60694 | ENSG00000118961 | LDAH | chr2 | - | 20801309 | 20801465 | 20790254 | 20790398 | 20823036 | 20823101 | 43 | 17 | 26 | 34 | 56 | 0 | 0 | 5 | 2 | 0 | 0 | 28 | 10 | 18 | 13 | 0 | 3 | 0 | 5 | 0 | 298 | 149 | 1 | 1 | 1 | 1 | 0.722 | 0.895 | 1 | NA | 0.824 | 1 | 0.643 | 1 | 0.057 |
| SE | 60695 | ENSG00000183671 | GPR1 | chr2 | - | 206203893 | 206203948 | 206176565 | 206177275 | 206213306 | 206213371 | 0 | 0 | 0 | 0 | 0 | 2 | 0 | 0 | 0 | 0 | 0 | 0 | 0 | 0 | 7 | 0 | 24 | 1 | 0 | 0 | 204 | 149 | 1 | 1 | 0 | NA | NA | NA | NA | NA | 0 | 0 | NA | 1 | -0.333 |
| SE | 60698 | ENSG00000235750 | KIAA0040 | chr1 | - | 175166561 | 175166737 | 175156985 | 175161146 | 175176399 | 175176474 | 119 | 60 | 73 | 120 | 148 | 0 | 0 | 0 | 0 | 1 | 129 | 82 | 49 | 114 | 109 | 0 | 0 | 0 | 0 | 0 | 298 | 149 | 1 | 1 | 1 | 1 | 1 | 1 | 0.987 | 1 | 1 | 1 | 1 | 1 | -0.003 |
| SE | 60699 | ENSG00000235750 | KIAA0040 | chr1 | - | 175166561 | 175166737 | 175156985 | 175161146 | 175177610 | 175177684 | 194 | 96 | 158 | 191 | 210 | 0 | 0 | 0 | 0 | 0 | 188 | 171 | 89 | 163 | 156 | 0 | 0 | 2 | 0 | 0 | 298 | 149 | 1 | 1 | 1 | 1 | 1 | 1 | 1 | 1 | 1 | 0.957 | 1 | 1 | 0.009 |
| SE | 60700 | ENSG00000235750 | KIAA0040 | chr1 | - | 175166561 | 175166737 | 175156985 | 175161146 | 175192639 | 175192754 | 132 | 69 | 78 | 137 | 152 | 0 | 0 | 0 | 0 | 0 | 141 | 80 | 51 | 114 | 129 | 0 | 0 | 1 | 0 | 0 | 298 | 149 | 1 | 1 | 1 | 1 | 1 | 1 | 1 | 1 | 1 | 0.962 | 1 | 1 | 0.008 |
| SE | 60701 | ENSG00000235750 | KIAA0040 | chr1 | - | 175176399 | 175176474 | 175161026 | 175161146 | 175177610 | 175177684 | 0 | 0 | 0 | 1 | 1 | 0 | 0 | 0 | 0 | 0 | 0 | 0 | 0 | 0 | 0 | 0 | 0 | 2 | 0 | 0 | 224 | 149 | 1 | 1 | NA | NA | NA | 1 | 1 | NA | NA | 0 | NA | NA | 1 |
| SE | 60703 | ENSG00000235750 | KIAA0040 | chr1 | - | 175177610 | 175177684 | 175156985 | 175161146 | 175192639 | 175192716 | 27 | 10 | 27 | 18 | 8 | 0 | 0 | 0 | 0 | 0 | 7 | 26 | 14 | 14 | 17 | 0 | 0 | 1 | 0 | 0 | 223 | 149 | 1 | 1 | 1 | 1 | 1 | 1 | 1 | 1 | 1 | 0.903 | 1 | 1 | 0.019 |
| SE | 60707 | ENSG00000241399 | CD302 | chr2 | - | 159781434 | 159781532 | 159780881 | 159780998 | 159783358 | 159783469 | 0 | 1 | 0 | 0 | 0 | 106 | 98 | 60 | 94 | 192 | 0 | 0 | 0 | 0 | 6 | 88 | 137 | 84 | 71 | 52 | 247 | 149 | 1 | 1 | 0 | 0.006 | 0 | 0 | 0 | 0 | 0 | 0 | 0 | 0.065 | -0.012 |
| SE | 60709 | ENSG00000106346 | USP42 | chr7 | + | 6111796 | 6111951 | 6111124 | 6111374 | 6115322 | 6115381 | 0 | 0 | 0 | 0 | 0 | 34 | 13 | 36 | 8 | 50 | 0 | 1 | 1 | 0 | 0 | 0 | 11 | 23 | 23 | 37 | 298 | 149 | 1 | 1 | 0 | 0 | 0 | 0 | 0 | NA | 0.043 | 0.021 | 0 | 0 | -0.016 |
| SE | 60710 | ENSG00000106346 | USP42 | chr7 | + | 6115322 | 6115523 | 6111124 | 6111374 | 6135840 | 6135951 | 71 | 38 | 82 | 47 | 142 | 0 | 0 | 0 | 2 | 0 | 0 | 24 | 66 | 37 | 76 | 0 | 0 | 3 | 0 | 0 | 298 | 149 | 1 | 1 | 1 | 1 | 1 | 0.922 | 1 | NA | 1 | 0.917 | 1 | 1 | 0.005 |
| SE | 60711 | ENSG00000106346 | USP42 | chr7 | + | 6116794 | 6116939 | 6115322 | 6115523 | 6128137 | 6128230 | 0 | 1 | 0 | 1 | 1 | 0 | 0 | 0 | 1 | 0 | 0 | 4 | 0 | 0 | 0 | 9 | 0 | 0 | 0 | 0 | 294 | 149 | 1 | 1 | NA | 1 | NA | 0.336 | 1 | 0 | 1 | NA | NA | NA | 0.279 |
| SE | 60714 | ENSG00000106346 | USP42 | chr7 | + | 6135840 | 6135951 | 6116479 | 6116939 | 6139091 | 6139194 | 62 | 44 | 59 | 56 | 147 | 0 | 0 | 0 | 0 | 0 | 65 | 69 | 116 | 124 | 44 | 0 | 1 | 0 | 0 | 0 | 260 | 149 | 1 | 1 | 1 | 1 | 1 | 1 | 1 | 1 | 0.975 | 1 | 1 | 1 | 0.005 |
| SE | 60719 | ENSG00000166278 | C2 | chr6 | + | 31927954 | 31928164 | 31927524 | 31927798 | 31933609 | 31933783 | 5 | 1 | 0 | 2 | 0 | 3 | 3 | 0 | 1 | 0 | 63 | 0 | 6 | 33 | 22 | 10 | 2 | 0 | 0 | 0 | 298 | 149 | 1 | 1 | 0.455 | 0.143 | NA | 0.5 | NA | 0.759 | 0 | 1 | 1 | 1 | -0.386 |
| SE | 60720 | ENSG00000166278 | C2 | chr6 | + | 31928731 | 31928917 | 31927954 | 31928164 | 31933609 | 31933783 | 10 | 3 | 0 | 7 | 0 | 2 | 0 | 0 | 0 | 0 | 53 | 12 | 9 | 45 | 47 | 0 | 0 | 0 | 4 | 1 | 298 | 149 | 1 | 1 | 0.714 | 1 | NA | 1 | NA | 1 | 1 | 1 | 0.849 | 0.959 | -0.057 |
| SE | 60722 | ENSG00000115541 | HSPE1 | chr2 | + | 197501073 | 197501238 | 197500378 | 197500439 | 197503038 | 197503128 | 340 | 214 | 290 | 387 | 275 | 0 | 0 | 0 | 0 | 0 | 516 | 212 | 336 | 345 | 305 | 0 | 0 | 1 | 0 | 0 | 298 | 149 | 1 | 1 | 1 | 1 | 1 | 1 | 1 | 1 | 1 | 0.994 | 1 | 1 | 0.001 |
| SE | 60723 | ENSG00000214194 | SMIM30 | chr7 | - | 113118067 | 113118161 | 113116717 | 113117607 | 113118439 | 113118560 | 47 | 23 | 48 | 50 | 125 | 0 | 0 | 0 | 1 | 0 | 74 | 70 | 31 | 28 | 44 | 0 | 0 | 0 | 0 | 0 | 243 | 149 | 1 | 1 | 1 | 1 | 1 | 0.968 | 1 | 1 | 1 | 1 | 1 | 1 | -0.006 |
| SE | 60724 | ENSG00000152380 | FAM151B | chr5 | + | 80495052 | 80495283 | 80487985 | 80488148 | 80499884 | 80499985 | 0 | 1 | 1 | 2 | 2 | 1 | 0 | 0 | 0 | 0 | 0 | 2 | 1 | 0 | 3 | 0 | 0 | 0 | 0 | 0 | 298 | 149 | 1 | 1 | 0 | 1 | 1 | 1 | 1 | NA | 1 | 1 | NA | 1 | -0.2 |
| SE | 60727 | ENSG00000152380 | FAM151B | chr5 | + | 80499884 | 80499985 | 80495052 | 80495283 | 80501791 | 80501917 | 4 | 1 | 2 | 0 | 1 | 5 | 1 | 1 | 0 | 0 | 0 | 4 | 2 | 1 | 6 | 0 | 8 | 6 | 0 | 10 | 250 | 149 | 1 | 1 | 0.323 | 0.373 | 0.544 | NA | 1 | NA | 0.23 | 0.166 | 1 | 0.263 | 0.145 |
| SE | 60728 | ENSG00000152380 | FAM151B | chr5 | + | 80501791 | 80501917 | 80487968 | 80488148 | 80513603 | 80513661 | 5 | 0 | 0 | 5 | 6 | 0 | 2 | 0 | 0 | 0 | 4 | 3 | 2 | 15 | 14 | 0 | 0 | 0 | 0 | 0 | 275 | 149 | 1 | 1 | 1 | 0 | NA | 1 | 1 | 1 | 1 | 1 | 1 | 1 | -0.25 |
| SE | 60729 | ENSG00000152380 | FAM151B | chr5 | + | 80505942 | 80506076 | 80501791 | 80501917 | 80510713 | 80511086 | 2 | 0 | 3 | 0 | 2 | 0 | 0 | 0 | 0 | 0 | 3 | 0 | 2 | 4 | 0 | 0 | 1 | 0 | 0 | 0 | 283 | 149 | 1 | 1 | 1 | NA | 1 | NA | 1 | 1 | 0 | 1 | 1 NA | 1 | 0.25 |
| SE | 60730 | ENSG00000152380 | FAM151B | chr5 | + | 80505942 | 80506076 | 80501791 | 80501917 | 80513603 | 80513769 | 4 | 0 | 6 | 0 | 0 | 1 | 0 | 0 | 3 | 5 | 6 | 2 | 4 | 15 | 0 | 0 | 2 | 0 | 10 | 0 | 283 | 149 | 1 | 1 | 0.678 | NA | 1 | 0 | 0 | 1 | 0.345 | 1 | 0.441 | NA | -0.277 |
| SE | 60731 | ENSG00000152380 | FAM151B | chr5 | + | 80510713 | 80511086 | 80501791 | 80501917 | 80513603 | 80513769 | 0 | 0 | 0 | 0 | 0 | 1 | 0 | 0 | 3 | 5 | 0 | 1 | 0 | 0 | 0 | 0 | 2 | 0 | 10 | 0 | 298 | 149 | 1 | 1 | 0 | NA | NA | 0 | 0 | NA | 0.2 | NA | 0 | NA | -0.1 |
| SE | 60732 | ENSG00000152380 | FAM151B | chr5 | + | 80510713 | 80511086 | 80505942 | 80506076 | 80513603 | 80513769 | 0 | 0 | 0 | 0 | 2 | 2 | 0 | 3 | 0 | 0 | 0 | 0 | 0 | 0 | 0 | 3 | 2 | 2 | 11 | 0 | 298 | 149 | 1 | 1 | 0 | NA | 0 | NA | 1 | 0 | 0 | 0 | 0 | NA | 0.333 |
| SE | 60733 | ENSG00000152380 | FAM151B | chr5 | + | 80518763 | 80518930 | 80513603 | 80513769 | 80519692 | 80519910 | 5 | 0 | 0 | 0 | 0 | 3 | 3 | 3 | 1 | 2 | 0 | 0 | 0 | 0 | 0 | 19 | 4 | 12 | 12 | 0 | 298 | 149 | 1 | 1 | 0.455 | 0 | 0 | 0 | 0 | 0 | 0 | 0 | 0 | NA | 0.091 |
| SE | 60734 | ENSG00000152380 | FAM151B | chr5 | + | 80522002 | 80522138 | 80519692 | 80519910 | 80541672 | 80542042 | 40 | 27 | 20 | 28 | 27 | 0 | 1 | 0 | 2 | 2 | 44 | 33 | 44 | 69 | 0 | 0 | 1 | 6 | 0 | 0 | 285 | 149 | 1 | 1 | 1 | 0.934 | 1 | 0.88 | 0.876 | 1 | 0.945 | 0.793 | 1 | NA | 0.004 |
| SE | 60737 | ENSG00000143303 | RRNAD1 | chr1 | + | 156732984 | 156733047 | 156731990 | 156732473 | 156733376 | 156733492 | 82 | 39 | 62 | 80 | 63 | 0 | 0 | 0 | 0 | 0 | 116 | 59 | 50 | 74 | 189 | 0 | 1 | 0 | 0 | 5 | 212 | 149 | 1 | 1 | 1 | 1 | 1 | 1 | 1 | 1 | 0.976 | 1 | 1 | 0.964 | 0.012 |
| SE | 60742 | ENSG00000160818 | GPATCH4 | chr1 | - | 156597799 | 156597922 | 156596386 | 156596478 | 156598034 | 156598121 | 2 | 0 | 0 | 0 | 0 | 14 | 12 | 11 | 15 | 19 | 0 | 0 | 0 | 0 | 0 | 20 | 10 | 19 | 14 | 0 | 272 | 149 | 1 | 1 | 0.073 | 0 | 0 | 0 | 0 | 0 | 0 | 0 | 0 | NA | 0.015 |
| SE | 60743 | ENSG00000160818 | GPATCH4 | chr1 | - | 156598034 | 156598121 | 156596293 | 156596478 | 156598226 | 156598319 | 27 | 26 | 22 | 29 | 28 | 0 | 2 | 2 | 1 | 0 | 21 | 12 | 28 | 17 | 0 | 0 | 0 | 1 | 2 | 0 | 236 | 149 | 1 | 1 | 1 | 0.891 | 0.874 | 0.948 | 1 | 1 | 1 | 0.946 | 0.843 | NA | -0.005 |
| SE | 60745 | ENSG00000160818 | GPATCH4 | chr1 | - | 156598967 | 156599104 | 156597763 | 156598319 | 156601408 | 156601462 | 18 | 10 | 10 | 24 | 12 | 0 | 0 | 0 | 1 | 0 | 15 | 14 | 8 | 34 | 32 | 0 | 1 | 0 | 0 | 0 | 286 | 149 | 1 | 1 | 1 | 1 | 1 | 0.926 | 1 | 1 | 0.879 | 1 | 1 | 1 | 0.009 |
| SE | 60746 | ENSG00000198216 | CACNA1E | chr1 | + | 181580594 | 181580776 | 181579071 | 181579224 | 181651337 | 181651441 | 0 | 0 | 0 | 0 | 2 | 0 | 0 | 0 | 1 | 0 | 0 | 0 | 4 | 2 | 0 | 0 | 0 | 0 | 0 | 0 | 298 | 149 | 1 | 1 | NA | NA | NA | 0 | 1 | NA | NA | 1 | 1 | NA | -0.5 |
| SE | 60753 | ENSG00000166025 | AMOTL1 | chr11 | + | 94795010 | 94795160 | 94768355 | 94768560 | 94799389 | 94799539 | 12 | 7 | 9 | 4 | 5 | 1 | 0 | 0 | 2 | 1 | 0 | 0 | 2 | 9 | 2 | 0 | 0 | 8 | 0 | 0 | 298 | 149 | 1 | 1 | 0.857 | 1 | 1 | 0.5 | 0.714 | NA | NA | 0.111 | 1 | 1 | 0.111 |
| SE | 60754 | ENSG00000141524 | TMC6 | chr17 | - | 78117268 | 78117347 | 78113547 | 78113624 | 78117467 | 78117644 | 346 | 143 | 361 | 339 | 196 | 0 | 1 | 0 | 0 | 0 | 269 | 367 | 184 | 172 | 471 | 0 | 0 | 0 | 0 | 0 | 228 | 149 | 1 | 1 | 1 | 0.989 | 1 | 1 | 1 | 1 | 1 | 1 | 1 | 1 | -0.002 |
| SE | 60755 | ENSG00000141524 | TMC6 | chr17 | - | 78117467 | 78117644 | 78113547 | 78113624 | 78118970 | 78119046 | 0 | 1 | 0 | 3 | 0 | 0 | 0 | 0 | 3 | 0 | 0 | 0 | 1 | 4 | 0 | 0 | 0 | 0 | 1 | 4 | 298 | 149 | 1 | 1 | NA | 1 | NA | 0.333 | NA | NA | NA | 1 | 0.667 | 0 | 0.111 |
| SE | 60756 | ENSG00000141524 | TMC6 | chr17 | - | 78117801 | 78117935 | 78117467 | 78117644 | 78118970 | 78119046 | 157 | 68 | 140 | 138 | 146 | 0 | 0 | 0 | 3 | 0 | 50 | 88 | 55 | 42 | 120 | 0 | 0 | 1 | 4 | 0 | 283 | 149 | 1 | 1 | 1 | 1 | 1 | 0.96 | 1 | 1 | 1 | 0.967 | 0.847 | 1 | 0.029 |
| SE | 60757 | ENSG00000141524 | TMC6 | chr17 | - | 78118970 | 78119046 | 78113547 | 78113624 | 78119296 | 78119392 | 101 | 48 | 78 | 103 | 118 | 3 | 0 | 0 | 0 | 0 | 12 | 68 | 25 | 52 | 118 | 0 | 0 | 0 | 0 | 0 | 225 | 149 | 1 | 1 | 0.957 | 1 | 1 | 1 | 1 | 1 | 1 | 1 | 1 | 1 | -0.009 |
| SE | 60765 | ENSG00000187741 | FANCA | chr16 | - | 89739128 | 89739289 | 89738743 | 89738974 | 89739477 | 89739553 | 15 | 8 | 5 | 10 | 15 | 0 | 2 | 0 | 0 | 0 | 31 | 7 | 6 | 9 | 35 | 0 | 0 | 0 | 0 | 0 | 298 | 149 | 1 | 1 | 1 | 0.667 | 1 | 1 | 1 | 1 | 1 | 1 | 1 | 1 | -0.067 |
| SE | 60766 | ENSG00000187741 | FANCA | chr16 | - | 89739132 | 89739289 | 89738881 | 89738974 | 89739477 | 89739553 | 26 | 8 | 12 | 10 | 16 | 0 | 2 | 0 | 0 | 0 | 24 | 15 | 7 | 10 | 35 | 0 | 0 | 0 | 0 | 0 | 298 | 149 | 1 | 1 | 1 | 0.667 | 1 | 1 | 1 | 1 | 1 | 1 | 1 | 1 | -0.067 |
| SE | 60767 | ENSG00000187741 | FANCA | chr16 | - | 89739477 | 89739553 | 89739015 | 89739289 | 89739993 | 89740078 | 29 | 10 | 10 | 21 | 17 | 0 | 1 | 0 | 0 | 0 | 56 | 9 | 10 | 9 | 41 | 0 | 0 | 0 | 0 | 0 | 225 | 149 | 1 | 1 | 1 | 0.869 | 1 | 1 | 1 | 1 | 1 | 1 | 1 | 1 | -0.026 |
| SE | 60772 | ENSG00000187741 | FANCA | chr16 | - | 89749729 | 89749902 | 89748658 | 89748767 | 89758576 | 89758705 | 27 | 16 | 12 | 16 | 4 | 0 | 0 | 0 | 1 | 0 | 39 | 21 | 18 | 22 | 54 | 0 | 0 | 0 | 0 | 0 | 298 | 149 | 1 | 1 | 1 | 1 | 1 | 0.889 | 1 | 1 | 1 | 1 | 1 | 1 | -0.022 |
| SE | 60773 | ENSG00000187741 | FANCA | chr16 | - | 89752137 | 89752222 | 89748658 | 89748767 | 89758576 | 89758705 | 12 | 12 | 3 | 5 | 7 | 0 | 0 | 0 | 1 | 0 | 33 | 3 | 3 | 4 | 13 | 0 | 0 | 0 | 0 | 0 | 234 | 149 | 1 | 1 | 1 | 1 | 1 | 0.761 | 1 | 1 | 1 | 1 | 1 | 1 | -0.048 |
| SE | 60776 | ENSG00000187741 | FANCA | chr16 | - | 89762637 | 89762793 | 89761948 | 89762022 | 89764806 | 89764947 | 5 | 3 | 5 | 2 | 4 | 0 | 0 | 1 | 2 | 0 | 0 | 6 | 2 | 5 | 9 | 0 | 0 | 0 | 0 | 3 | 298 | 149 | 1 | 1 | 1 | 1 | 0.714 | 0.333 | 1 | NA | 1 | 1 | 1 | 0.6 | -0.091 |
| SE | 60780 | ENSG00000187741 | FANCA | chr16 | - | 89795905 | 89796018 | 89792470 | 89792547 | 89799165 | 89799232 | 1 | 2 | 2 | 5 | 0 | 0 | 0 | 0 | 0 | 0 | 0 | 0 | 2 | 4 | 6 | 2 | 0 | 3 | 0 | 0 | 262 | 149 | 1 | 1 | 1 | 1 | 1 | 1 | NA | 0 | NA | 0.275 | 1 | 1 | 0.431 |
| SE | 60781 | ENSG00000187741 | FANCA | chr16 | - | 89803258 | 89803341 | 89799604 | 89799638 | 89805279 | 89805392 | 5 | 7 | 7 | 10 | 6 | 0 | 0 | 0 | 0 | 1 | 17 | 24 | 21 | 26 | 8 | 0 | 0 | 0 | 0 | 0 | 232 | 149 | 1 | 1 | 1 | 1 | 1 | 1 | 0.794 | 1 | 1 | 1 | 1 | 1 | -0.041 |
| SE | 60782 | ENSG00000187741 | FANCA | chr16 | - | 89805279 | 89805392 | 89799604 | 89799638 | 89808293 | 89808366 | 3 | 2 | 1 | 6 | 2 | 0 | 0 | 0 | 1 | 0 | 0 | 6 | 5 | 17 | 2 | 0 | 0 | 0 | 0 | 0 | 262 | 149 | 1 | 1 | 1 | 1 | 1 | 0.773 | 1 | NA | 1 | 1 | 1 | 1 | -0.045 |
| SE | 60784 | ENSG00000187741 | FANCA | chr16 | - | 89810928 | 89811057 | 89810706 | 89810802 | 89814519 | 89814613 | 3 | 1 | 0 | 5 | 1 | 0 | 0 | 0 | 1 | 0 | 4 | 11 | 7 | 37 | 1 | 0 | 0 | 0 | 0 | 0 | 278 | 149 | 1 | 1 | 1 | 1 | NA | 0.728 | 1 | 1 | 1 | 1 | 1 | 1 | -0.068 |
| SE | 60785 | ENSG00000187741 | FANCA | chr16 | - | 89810928 | 89811071 | 89810571 | 89810802 | 89814519 | 89814613 | 4 | 1 | 1 | 11 | 1 | 0 | 0 | 0 | 1 | 0 | 4 | 22 | 13 | 47 | 8 | 0 | 0 | 0 | 0 | 0 | 292 | 149 | 1 | 1 | 1 | 1 | 1 | 0.849 | 1 | 1 | 1 | 1 | 1 | 1 | -0.03 |
| SE | 60790 | ENSG00000160714 | UBE2Q1 | chr1 | - | 154550937 | 154551004 | 154548576 | 154550469 | 154551770 | 154551819 | 57 | 32 | 58 | 61 | 118 | 1 | 0 | 0 | 0 | 0 | 108 | 77 | 93 | 96 | 176 | 0 | 0 | 2 | 0 | 0 | 216 | 149 | 1 | 1 | 0.975 | 1 | 1 | 1 | 1 | 1 | 1 | 0.97 | 1 | 1 | 0.001 |
| SE | 60791 | ENSG00000160714 | UBE2Q1 | chr1 | - | 154551396 | 154551492 | 154548576 | 154550469 | 154551770 | 154551819 | 178 | 86 | 154 | 155 | 374 | 1 | 0 | 0 | 0 | 0 | 192 | 162 | 84 | 100 | 268 | 0 | 0 | 2 | 0 | 0 | 245 | 149 | 1 | 1 | 0.991 | 1 | 1 | 1 | 1 | 1 | 1 | 0.962 | 1 | 1 | 0.006 |
| SE | 60794 | ENSG00000160714 | UBE2Q1 | chr1 | - | 154551920 | 154551979 | 154551770 | 154551819 | 154552092 | 154552183 | 361 | 199 | 334 | 355 | 817 | 3 | 0 | 1 | 1 | 0 | 484 | 406 | 227 | 273 | 588 | 0 | 0 | 1 | 2 | 0 | 208 | 149 | 1 | 1 | 0.989 | 1 | 0.996 | 0.996 | 1 | 1 | 1 | 0.994 | 0.99 | 1 | -0.001 |
| SE | 60797 | ENSG00000160714 | UBE2Q1 | chr1 | - | 154553031 | 154553172 | 154552735 | 154552820 | 154554734 | 154554785 | 632 | 406 | 520 | 611 | 695 | 0 | 0 | 0 | 0 | 0 | 1170 | 713 | 512 | 821 | 1258 | 0 | 0 | 0 | 3 | 0 | 290 | 149 | 1 | 1 | 1 | 1 | 1 | 1 | 1 | 1 | 1 | 1 | 0.993 | 1 | 0.001 |
| SE | 60798 | ENSG00000160714 | UBE2Q1 | chr1 | - | 154554734 | 154554785 | 154553031 | 154553172 | 154555427 | 154555532 | 603 | 419 | 468 | 556 | 490 | 0 | 0 | 0 | 0 | 0 | 1145 | 692 | 520 | 827 | 1206 | 4 | 0 | 0 | 0 | 0 | 200 | 149 | 1 | 1 | 1 | 1 | 1 | 1 | 1 | 0.995 | 1 | 1 | 1 | 1 | 0.001 |
| SE | 60799 | ENSG00000237298 | TTN-AS1 | chr2 | + | 178523451 | 178523636 | 178523182 | 178523257 | 178535731 | 178535828 | 31 | 23 | 29 | 50 | 67 | 0 | 0 | 0 | 3 | 0 | 32 | 35 | 13 | 50 | 0 | 0 | 0 | 0 | 0 | 0 | 298 | 149 | 1 | 1 | 1 | 1 | 1 | 0.893 | 1 | 1 | 1 | 1 | 1 | NA | -0.021 |
| SE | 60800 | ENSG00000237298 | TTN-AS1 | chr2 | + | 178523451 | 178523636 | 178523182 | 178523257 | 178591138 | 178591314 | 0 | 1 | 0 | 2 | 2 | 0 | 0 | 1 | 0 | 0 | 0 | 4 | 1 | 1 | 0 | 0 | 0 | 0 | 0 | 0 | 298 | 149 | 1 | 1 | NA | 1 | 0 | 1 | 1 | NA | 1 | 1 | 1 | NA | -0.25 |
| SE | 60802 | ENSG00000237298 | TTN-AS1 | chr2 | + | 178528723 | 178528813 | 178523166 | 178523636 | 178531313 | 178531559 | 0 | 0 | 0 | 0 | 3 | 12 | 5 | 7 | 6 | 19 | 0 | 0 | 0 | 0 | 0 | 6 | 12 | 7 | 38 | 0 | 239 | 149 | 1 | 1 | 0 | 0 | 0 | 0 | 0.09 | 0 | 0 | 0 | 0 | NA | 0.018 |
| SE | 60803 | ENSG00000237298 | TTN-AS1 | chr2 | + | 178528723 | 178528813 | 178523166 | 178523636 | 178535731 | 178535828 | 0 | 0 | 0 | 2 | 0 | 31 | 23 | 29 | 49 | 65 | 0 | 0 | 0 | 3 | 0 | 32 | 35 | 13 | 50 | 0 | 239 | 149 | 1 | 1 | 0 | 0 | 0 | 0.025 | 0 | 0 | 0 | 0 | 0.036 | NA | -0.004 |
| SE | 60808 | ENSG00000237298 | TTN-AS1 | chr2 | + | 178531313 | 178531578 | 178523166 | 178523636 | 178537603 | 178537691 | 13 | 5 | 7 | 6 | 19 | 0 | 0 | 1 | 0 | 0 | 6 | 12 | 7 | 38 | 0 | 0 | 0 | 0 | 3 | 0 | 298 | 149 | 1 | 1 | 1 | 1 | 0.778 | 1 | 1 | 1 | 1 | 1 | 0.864 | NA | -0.01 |
| SE | 60810 | ENSG00000237298 | TTN-AS1 | chr2 | + | 178531313 | 178531578 | 178528723 | 178528813 | 178535731 | 178535828 | 26 | 5 | 18 | 11 | 35 | 0 | 0 | 0 | 2 | 0 | 13 | 22 | 14 | 61 | 14 | 0 | 0 | 0 | 3 | 0 | 298 | 149 | 1 | 1 | 1 | 1 | 1 | 0.733 | 1 | 1 | 1 | 1 | 0.91 | 1 | -0.035 |
| SE | 60811 | ENSG00000237298 | TTN-AS1 | chr2 | + | 178535731 | 178535828 | 178523212 | 178523636 | 178537603 | 178537691 | 91 | 57 | 75 | 108 | 186 | 0 | 0 | 1 | 0 | 0 | 52 | 99 | 32 | 156 | 47 | 0 | 0 | 0 | 3 | 0 | 246 | 149 | 1 | 1 | 1 | 1 | 0.978 | 1 | 1 | 1 | 1 | 1 | 0.969 | 1 | 0.002 |
| SE | 60814 | ENSG00000237298 | TTN-AS1 | chr2 | + | 178535731 | 178535828 | 178531313 | 178531578 | 178537603 | 178537691 | 85 | 38 | 61 | 74 | 154 | 1 | 0 | 0 | 0 | 0 | 33 | 89 | 35 | 154 | 61 | 0 | 0 | 0 | 0 | 0 | 246 | 149 | 1 | 1 | 0.981 | 1 | 1 | 1 | 1 | 1 | 1 | 1 | 1 | 1 | -0.004 |
| SE | 60815 | ENSG00000237298 | TTN-AS1 | chr2 | + | 178536333 | 178536462 | 178535731 | 178535828 | 178536878 | 178537127 | 0 | 1 | 1 | 5 | 2 | 0 | 0 | 1 | 2 | 0 | 0 | 0 | 5 | 0 | 0 | 0 | 6 | 0 | 13 | 0 | 278 | 149 | 1 | 1 | NA | 1 | 0.349 | 0.573 | 1 | NA | 0 | 1 | 0 | NA | 0.397 |
| SE | 60816 | ENSG00000237298 | TTN-AS1 | chr2 | + | 178536333 | 178536462 | 178535731 | 178535828 | 178537603 | 178537691 | 0 | 0 | 0 | 4 | 0 | 71 | 38 | 58 | 82 | 130 | 0 | 0 | 1 | 0 | 0 | 32 | 74 | 26 | 124 | 47 | 278 | 149 | 1 | 1 | 0 | 0 | 0 | 0.025 | 0 | 0 | 0 | 0.02 | 0 | 0 | 0.001 |
| SE | 60819 | ENSG00000237298 | TTN-AS1 | chr2 | + | 178537603 | 178537691 | 178523214 | 178523636 | 178538697 | 178538770 | 19 | 9 | 13 | 22 | 29 | 0 | 0 | 0 | 0 | 0 | 12 | 7 | 9 | 29 | 7 | 0 | 0 | 0 | 3 | 0 | 237 | 149 | 1 | 1 | 1 | 1 | 1 | 1 | 1 | 1 | 1 | 1 | 0.859 | 1 | 0.028 |
| SE | 60820 | ENSG00000237298 | TTN-AS1 | chr2 | + | 178537603 | 178537691 | 178523214 | 178523636 | 178540106 | 178540169 | 0 | 0 | 0 | 0 | 0 | 0 | 0 | 0 | 3 | 0 | 0 | 0 | 0 | 1 | 0 | 0 | 0 | 0 | 0 | 0 | 237 | 149 | 1 | 1 | NA | NA | NA | NA | 0 | NA | NA | NA | 1 | NA | -1 |
| SE | 60821 | ENSG00000237298 | TTN-AS1 | chr2 | + | 178537603 | 178537691 | 178523214 | 178523636 | 178597595 | 178597681 | 0 | 0 | 0 | 0 | 0 | 0 | 1 | 0 | 1 | 8 | 0 | 0 | 0 | 1 | 0 | 0 | 0 | 0 | 0 | 5 | 237 | 149 | 1 | 1 | NA | 0 | NA | 0 | 0 | NA | NA | NA | 1 | 0 | -0.5 |
| SE | 60824 | ENSG00000237298 | TTN-AS1 | chr2 | + | 178538697 | 178539890 | 178523451 | 178523636 | 178540106 | 178540169 | 2 | 3 | 2 | 1 | 0 | 0 | 0 | 0 | 3 | 0 | 0 | 1 | 0 | 4 | 0 | 0 | 0 | 0 | 0 | 0 | 298 | 149 | 1 | 1 | 1 | 1 | 1 | 1 | 0 | NA | 1 | NA | 1 | NA | -0.2 |
| SE | 60826 | ENSG00000237298 | TTN-AS1 | chr2 | + | 178540106 | 178540169 | 178523451 | 178523636 | 178542221 | 178542361 | 18 | 11 | 23 | 19 | 21 | 0 | 0 | 2 | 0 | 0 | 24 | 16 | 11 | 15 | 47 | 0 | 0 | 0 | 0 | 0 | 212 | 149 | 1 | 1 | 1 | 1 | 0.89 | 1 | 1 | 1 | 1 | 1 | 1 | 1 | -0.022 |
| SE | 60827 | ENSG00000237298 | TTN-AS1 | chr2 | + | 178540106 | 178540169 | 178539804 | 178539978 | 178542221 | 178542361 | 33 | 24 | 38 | 36 | 30 | 0 | 0 | 0 | 0 | 0 | 38 | 29 | 17 | 43 | 70 | 0 | 3 | 0 | 0 | 0 | 212 | 149 | 1 | 1 | 1 | 1 | 1 | 1 | 1 | 1 | 0.872 | 1 | 1 | 1 | 0.026 |
| SE | 60828 | ENSG00000237298 | TTN-AS1 | chr2 | + | 178542221 | 178542361 | 178523291 | 178523636 | 178571299 | 178571449 | 15 | 0 | 8 | 1 | 16 | 3 | 1 | 5 | 1 | 0 | 0 | 3 | 4 | 9 | 0 | 0 | 2 | 0 | 0 | 0 | 289 | 149 | 1 | 1 | 0.721 | 0 | 0.452 | 0.34 | 1 | NA | 0.436 | 1 | 1 | NA | -0.309 |
| SE | 60830 | ENSG00000237298 | TTN-AS1 | chr2 | + | 178542221 | 178542361 | 178523291 | 178523636 | 178583712 | 178583921 | 0 | 0 | 2 | 0 | 0 | 0 | 0 | 0 | 0 | 0 | 0 | 3 | 0 | 0 | 0 | 0 | 0 | 0 | 4 | 0 | 289 | 149 | 1 | 1 | NA | NA | 1 | NA | NA | NA | 1 | NA | 0 | NA | 0.5 |
| SE | 60831 | ENSG00000237298 | TTN-AS1 | chr2 | + | 178542221 | 178542361 | 178523291 | 178523636 | 178591138 | 178591314 | 5 | 2 | 3 | 1 | 0 | 0 | 1 | 0 | 1 | 0 | 0 | 0 | 0 | 0 | 0 | 0 | 4 | 1 | 1 | 0 | 289 | 149 | 1 | 1 | 1 | 0.508 | 1 | 0.34 | NA | NA | 0 | 0 | 0 | NA | 0.712 |
| SE | 60832 | ENSG00000237298 | TTN-AS1 | chr2 | + | 178542221 | 178542361 | 178523291 | 178523636 | 178597595 | 178597900 | 0 | 4 | 8 | 4 | 2 | 0 | 1 | 0 | 1 | 8 | 0 | 6 | 1 | 0 | 0 | 0 | 0 | 0 | 5 | 0 | 289 | 149 | 1 | 1 | NA | 0.673 | 1 | 0.673 | 0.114 | NA | 1 | 1 | NA | 0 | -0.052 |
| SE | 60837 | ENSG00000237298 | TTN-AS1 | chr2 | + | 178542221 | 178542361 | 178540106 | 178540169 | 178583523 | 178583648 | 28 | 20 | 35 | 39 | 31 | 0 | 0 | 0 | 2 | 0 | 41 | 19 | 18 | 45 | 72 | 0 | 0 | 0 | 0 | 0 | 289 | 149 | 1 | 1 | 1 | 1 | 1 | 0.91 | 1 | 1 | 1 | 1 | 1 | 1 | -0.018 |
| SE | 60838 | ENSG00000237298 | TTN-AS1 | chr2 | + | 178571299 | 178571449 | 178523223 | 178523636 | 178582571 | 178582663 | 6 | 1 | 5 | 1 | 0 | 1 | 0 | 0 | 0 | 0 | 0 | 2 | 0 | 0 | 0 | 0 | 0 | 0 | 0 | 0 | 298 | 149 | 1 | 1 | 0.75 | 1 | 1 | 1 | NA | NA | 1 | NA | NA | NA | -0.062 |
| SE | 60839 | ENSG00000237298 | TTN-AS1 | chr2 | + | 178571299 | 178571449 | 178523223 | 178523636 | 178591138 | 178591314 | 4 | 3 | 6 | 1 | 5 | 0 | 1 | 0 | 1 | 0 | 0 | 4 | 0 | 3 | 0 | 0 | 4 | 1 | 1 | 0 | 298 | 149 | 1 | 1 | 1 | 0.6 | 1 | 0.333 | 1 | NA | 0.333 | 0 | 0.6 | NA | 0.476 |
| SE | 60840 | ENSG00000237298 | TTN-AS1 | chr2 | + | 178571299 | 178571449 | 178523223 | 178523636 | 178597595 | 178597681 | 3 | 1 | 5 | 2 | 0 | 0 | 1 | 0 | 1 | 8 | 0 | 2 | 0 | 0 | 0 | 0 | 0 | 0 | 5 | 0 | 298 | 149 | 1 | 1 | 1 | 0.333 | 1 | 0.5 | 0 | NA | 1 | NA | NA | 0 | 0.067 |
| SE | 60844 | ENSG00000237298 | TTN-AS1 | chr2 | + | 178571299 | 178571449 | 178542221 | 178542361 | 178575979 | 178576557 | 16 | 1 | 6 | 2 | 16 | 0 | 0 | 0 | 0 | 0 | 0 | 3 | 4 | 9 | 0 | 0 | 0 | 0 | 3 | 0 | 298 | 149 | 1 | 1 | 1 | 1 | 1 | 1 | 1 | NA | 1 | 1 | 0.6 | NA | 0.133 |
| SE | 60846 | ENSG00000237298 | TTN-AS1 | chr2 | + | 178571299 | 178571449 | 178542221 | 178542361 | 178583523 | 178583921 | 15 | 0 | 6 | 2 | 16 | 0 | 0 | 0 | 1 | 0 | 0 | 3 | 4 | 9 | 4 | 0 | 0 | 0 | 0 | 0 | 298 | 149 | 1 | 1 | 1 | NA | 1 | 0.5 | 1 | NA | 1 | 1 | 1 | 1 | -0.125 |
| SE | 60847 | ENSG00000237298 | TTN-AS1 | chr2 | + | 178571299 | 178571449 | 178542221 | 178542361 | 178591138 | 178591314 | 16 | 2 | 7 | 2 | 21 | 5 | 2 | 1 | 1 | 0 | 0 | 5 | 4 | 12 | 0 | 0 | 0 | 0 | 0 | 0 | 298 | 149 | 1 | 1 | 0.615 | 0.333 | 0.778 | 0.5 | 1 | NA | 1 | 1 | 1 | NA | -0.355 |
| SE | 60848 | ENSG00000237298 | TTN-AS1 | chr2 | + | 178571299 | 178571449 | 178542221 | 178542361 | 178597595 | 178597681 | 15 | 0 | 6 | 3 | 16 | 0 | 4 | 6 | 4 | 3 | 0 | 3 | 4 | 9 | 0 | 0 | 6 | 1 | 0 | 0 | 298 | 149 | 1 | 1 | 1 | 0 | 0.333 | 0.273 | 0.727 | NA | 0.2 | 0.667 | 1 | NA | -0.156 |
| SE | 60849 | ENSG00000237298 | TTN-AS1 | chr2 | + | 178575112 | 178575326 | 178571299 | 178571449 | 178575979 | 178576557 | 7 | 1 | 3 | 9 | 9 | 1 | 1 | 0 | 0 | 0 | 0 | 11 | 0 | 33 | 6 | 0 | 0 | 0 | 0 | 0 | 298 | 149 | 1 | 1 | 0.778 | 0.333 | 1 | 1 | 1 | NA | 1 | NA | 1 | 1 | -0.178 |
| SE | 60850 | ENSG00000237298 | TTN-AS1 | chr2 | + | 178582571 | 178582706 | 178523451 | 178523636 | 178583712 | 178583921 | 1 | 0 | 0 | 0 | 0 | 0 | 0 | 0 | 0 | 0 | 0 | 1 | 0 | 0 | 0 | 0 | 0 | 0 | 4 | 0 | 284 | 149 | 1 | 1 | 1 | NA | NA | NA | NA | NA | 1 | NA | 0 | NA | 0.5 |
| SE | 60851 | ENSG00000237298 | TTN-AS1 | chr2 | + | 178582571 | 178582706 | 178542221 | 178542361 | 178583523 | 178583648 | 0 | 0 | 0 | 0 | 1 | 0 | 0 | 0 | 1 | 0 | 0 | 0 | 0 | 4 | 0 | 0 | 0 | 0 | 0 | 0 | 284 | 149 | 1 | 1 | NA | NA | NA | 0 | 1 | NA | NA | NA | 1 | NA | -0.5 |
| SE | 60855 | ENSG00000237298 | TTN-AS1 | chr2 | + | 178583523 | 178583648 | 178542221 | 178542361 | 178597595 | 178597681 | 0 | 0 | 0 | 1 | 0 | 0 | 4 | 6 | 4 | 3 | 0 | 0 | 0 | 0 | 0 | 0 | 6 | 1 | 0 | 0 | 274 | 149 | 1 | 1 | NA | 0 | 0 | 0.12 | 0 | NA | 0 | 0 | NA | NA | 0.03 |
| SE | 60857 | ENSG00000237298 | TTN-AS1 | chr2 | + | 178584572 | 178584993 | 178523291 | 178523636 | 178591138 | 178591314 | 2 | 0 | 0 | 0 | 1 | 0 | 1 | 0 | 1 | 0 | 0 | 0 | 0 | 0 | 11 | 0 | 4 | 1 | 1 | 0 | 298 | 149 | 1 | 1 | 1 | 0 | NA | 0 | 1 | NA | 0 | 0 | 0 | 1 | 0.25 |
| SE | 60858 | ENSG00000237298 | TTN-AS1 | chr2 | + | 178584572 | 178584993 | 178523291 | 178523636 | 178597595 | 178597681 | 0 | 0 | 1 | 0 | 0 | 0 | 1 | 0 | 1 | 8 | 0 | 0 | 0 | 0 | 0 | 0 | 0 | 0 | 0 | 5 | 298 | 149 | 1 | 1 | NA | 0 | 1 | 0 | 0 | NA | NA | NA | NA | 0 | 0.25 |
| SE | 60864 | ENSG00000237298 | TTN-AS1 | chr2 | + | 178591138 | 178591314 | 178571299 | 178571449 | 178597595 | 178597681 | 22 | 8 | 14 | 16 | 29 | 0 | 0 | 0 | 1 | 0 | 59 | 21 | 16 | 15 | 13 | 0 | 0 | 0 | 0 | 0 | 298 | 149 | 1 | 1 | 1 | 1 | 1 | 0.889 | 1 | 1 | 1 | 1 | 1 | 1 | -0.022 |
| SE | 60865 | ENSG00000237298 | TTN-AS1 | chr2 | + | 178591138 | 178591314 | 178584572 | 178584993 | 178597595 | 178597681 | 23 | 6 | 13 | 16 | 25 | 0 | 0 | 1 | 0 | 0 | 59 | 19 | 16 | 12 | 24 | 0 | 0 | 0 | 0 | 0 | 298 | 149 | 1 | 1 | 1 | 1 | 0.867 | 1 | 1 | 1 | 1 | 1 | 1 | 1 | -0.027 |
| SE | 60868 | ENSG00000237298 | TTN-AS1 | chr2 | + | 178599593 | 178599750 | 178597595 | 178597681 | 178600475 | 178600667 | 5 | 2 | 7 | 10 | 3 | 0 | 0 | 0 | 0 | 0 | 29 | 2 | 5 | 8 | 13 | 0 | 0 | 1 | 0 | 0 | 298 | 149 | 1 | 1 | 1 | 1 | 1 | 1 | 1 | 1 | 1 | 0.714 | 1 | 1 | 0.057 |
| SE | 60871 | ENSG00000237298 | TTN-AS1 | chr2 | + | 178600557 | 178600667 | 178597595 | 178597681 | 178615322 | 178615374 | 6 | 4 | 4 | 9 | 5 | 2 | 0 | 0 | 0 | 0 | 0 | 0 | 9 | 5 | 0 | 0 | 0 | 0 | 0 | 0 | 259 | 149 | 1 | 1 | 0.633 | 1 | 1 | 1 | 1 | NA | NA | 1 | 1 | NA | -0.073 |
| SE | 60873 | ENSG00000237298 | TTN-AS1 | chr2 | + | 178600557 | 178600667 | 178597595 | 178597681 | 178636573 | 178636698 | 4 | 4 | 2 | 9 | 5 | 0 | 0 | 1 | 0 | 0 | 0 | 0 | 9 | 5 | 0 | 0 | 0 | 0 | 0 | 0 | 259 | 149 | 1 | 1 | 1 | 1 | 0.535 | 1 | 1 | NA | NA | 1 | 1 | NA | -0.093 |
| SE | 60874 | ENSG00000237298 | TTN-AS1 | chr2 | + | 178600557 | 178600667 | 178597595 | 178597681 | 178713753 | 178714053 | 4 | 4 | 2 | 9 | 5 | 0 | 0 | 1 | 0 | 0 | 0 | 0 | 9 | 5 | 0 | 0 | 0 | 0 | 0 | 0 | 259 | 149 | 1 | 1 | 1 | 1 | 0.535 | 1 | 1 | NA | NA | 1 | 1 | NA | -0.093 |
| SE | 60876 | ENSG00000237298 | TTN-AS1 | chr2 | + | 178608166 | 178608266 | 178600557 | 178600667 | 178614034 | 178614060 | 1 | 1 | 0 | 1 | 0 | 0 | 0 | 0 | 2 | 0 | 0 | 0 | 0 | 3 | 0 | 0 | 0 | 0 | 0 | 0 | 249 | 149 | 1 | 1 | 1 | 1 | NA | 0.23 | NA | NA | NA | NA | 1 | NA | -0.257 |
| SE | 60878 | ENSG00000237298 | TTN-AS1 | chr2 | + | 178613211 | 178613311 | 178608201 | 178608266 | 178614034 | 178614173 | 0 | 0 | 0 | 1 | 0 | 0 | 0 | 0 | 0 | 0 | 0 | 0 | 0 | 0 | 0 | 0 | 0 | 0 | 3 | 0 | 249 | 149 | 1 | 1 | NA | NA | NA | 1 | NA | NA | NA | NA | 0 | NA | 1 |
| SE | 60892 | ENSG00000144674 | GOLGA4 | chr3 | + | 37266841 | 37266940 | 37251394 | 37251484 | 37273535 | 37273601 | 0 | 0 | 0 | 0 | 4 | 54 | 34 | 39 | 53 | 66 | 0 | 0 | 0 | 0 | 0 | 7 | 17 | 9 | 6 | 15 | 248 | 149 | 1 | 1 | 0 | 0 | 0 | 0 | 0.035 | 0 | 0 | 0 | 0 | 0 | 0.007 |
| SE | 60894 | ENSG00000144674 | GOLGA4 | chr3 | + | 37281957 | 37282272 | 37273535 | 37273601 | 37286013 | 37286061 | 249 | 224 | 214 | 230 | 243 | 0 | 2 | 0 | 0 | 0 | 80 | 120 | 78 | 45 | 68 | 0 | 0 | 0 | 0 | 0 | 298 | 149 | 1 | 1 | 1 | 0.982 | 1 | 1 | 1 | 1 | 1 | 1 | 1 | 1 | -0.004 |
| SE | 60897 | ENSG00000144674 | GOLGA4 | chr3 | + | 37287526 | 37287612 | 37286013 | 37286061 | 37289234 | 37289291 | 0 | 0 | 0 | 0 | 0 | 116 | 106 | 124 | 119 | 129 | 0 | 2 | 0 | 0 | 0 | 55 | 51 | 35 | 19 | 31 | 235 | 149 | 1 | 1 | 0 | 0 | 0 | 0 | 0 | 0 | 0.024 | 0 | 0 | 0 | -0.005 |
| SE | 60898 | ENSG00000144674 | GOLGA4 | chr3 | + | 37289234 | 37289289 | 37281957 | 37282272 | 37294978 | 37295077 | 1 | 1 | 1 | 2 | 3 | 2 | 3 | 1 | 3 | 1 | 15 | 5 | 1 | 0 | 0 | 0 | 0 | 0 | 3 | 0 | 204 | 149 | 1 | 1 | 0.268 | 0.196 | 0.422 | 0.327 | 0.687 | 1 | 1 | 1 | 0 | NA | -0.37 |
| SE | 60900 | ENSG00000144674 | GOLGA4 | chr3 | + | 37289234 | 37289291 | 37286013 | 37286061 | 37294978 | 37295077 | 210 | 188 | 183 | 203 | 204 | 0 | 0 | 1 | 0 | 1 | 68 | 91 | 61 | 31 | 34 | 0 | 0 | 0 | 0 | 0 | 206 | 149 | 1 | 1 | 1 | 1 | 0.993 | 1 | 0.993 | 1 | 1 | 1 | 1 | 1 | -0.003 |
| SE | 60901 | ENSG00000144674 | GOLGA4 | chr3 | + | 37298832 | 37299020 | 37296086 | 37296219 | 37299287 | 37299371 | 232 | 213 | 153 | 217 | 352 | 0 | 0 | 0 | 0 | 0 | 127 | 156 | 134 | 72 | 56 | 0 | 0 | 4 | 0 | 0 | 298 | 149 | 1 | 1 | 1 | 1 | 1 | 1 | 1 | 1 | 1 | 0.944 | 1 | 1 | 0.011 |
| SE | 60906 | ENSG00000144674 | GOLGA4 | chr3 | + | 37337142 | 37337163 | 37335052 | 37335166 | 37337665 | 37337734 | 135 | 132 | 136 | 121 | 144 | 6 | 0 | 1 | 1 | 0 | 0 | 44 | 27 | 13 | 5 | 0 | 0 | 5 | 0 | 0 | 170 | 149 | 1 | 1 | 0.952 | 1 | 0.992 | 0.991 | 1 | NA | 1 | 0.826 | 1 | 1 | 0.031 |
| SE | 60907 | ENSG00000144674 | GOLGA4 | chr3 | + | 37337665 | 37337734 | 37335052 | 37335166 | 37340123 | 37340199 | 29 | 22 | 18 | 20 | 39 | 0 | 0 | 1 | 0 | 0 | 0 | 7 | 9 | 2 | 0 | 0 | 0 | 0 | 0 | 0 | 218 | 149 | 1 | 1 | 1 | 1 | 0.925 | 1 | 1 | NA | 1 | 1 | 1 | NA | -0.015 |
| SE | 60908 | ENSG00000144674 | GOLGA4 | chr3 | + | 37347192 | 37347296 | 37340123 | 37340199 | 37355100 | 37355187 | 127 | 95 | 81 | 101 | 163 | 0 | 0 | 0 | 0 | 0 | 17 | 31 | 11 | 3 | 10 | 0 | 1 | 0 | 0 | 0 | 253 | 149 | 1 | 1 | 1 | 1 | 1 | 1 | 1 | 1 | 0.948 | 1 | 1 | 1 | 0.01 |
| SE | 60909 | ENSG00000144674 | GOLGA4 | chr3 | + | 37361242 | 37361305 | 37355100 | 37355187 | 37366079 | 37366751 | 18 | 20 | 16 | 23 | 33 | 87 | 80 | 57 | 64 | 102 | 8 | 13 | 9 | 1 | 0 | 18 | 29 | 33 | 9 | 0 | 212 | 149 | 1 | 1 | 0.127 | 0.149 | 0.165 | 0.202 | 0.185 | 0.238 | 0.24 | 0.161 | 0.072 | NA | -0.012 |
| SE | 60914 | ENSG00000130713 | EXOSC2 | chr9 | + | 130703052 | 130703181 | 130700841 | 130702310 | 130703693 | 130703774 | 239 | 124 | 221 | 215 | 173 | 0 | 0 | 0 | 0 | 0 | 282 | 230 | 192 | 268 | 372 | 0 | 0 | 4 | 0 | 0 | 278 | 149 | 1 | 1 | 1 | 1 | 1 | 1 | 1 | 1 | 1 | 0.963 | 1 | 1 | 0.007 |
| SE | 60916 | ENSG00000138398 | PPIG | chr2 | + | 169603641 | 169603694 | 169584341 | 169584490 | 169604025 | 169604102 | 153 | 93 | 130 | 149 | 94 | 0 | 0 | 0 | 1 | 0 | 38 | 54 | 40 | 41 | 52 | 0 | 1 | 1 | 0 | 0 | 202 | 149 | 1 | 1 | 1 | 1 | 1 | 0.991 | 1 | 1 | 0.976 | 0.967 | 1 | 1 | 0.01 |
| SE | 60918 | ENSG00000138398 | PPIG | chr2 | + | 169603641 | 169604102 | 169584352 | 169584490 | 169606038 | 169606146 | 41 | 20 | 29 | 37 | 20 | 0 | 0 | 0 | 1 | 0 | 21 | 10 | 3 | 6 | 2 | 0 | 0 | 0 | 0 | 0 | 298 | 149 | 1 | 1 | 1 | 1 | 1 | 0.949 | 1 | 1 | 1 | 1 | 1 | 1 | -0.01 |
| SE | 60919 | ENSG00000138398 | PPIG | chr2 | + | 169604025 | 169604102 | 169584341 | 169584490 | 169606038 | 169606146 | 0 | 0 | 0 | 0 | 2 | 0 | 0 | 0 | 1 | 0 | 0 | 0 | 1 | 0 | 0 | 0 | 0 | 0 | 0 | 0 | 226 | 149 | 1 | 1 | NA | NA | NA | 0 | 1 | NA | NA | 1 | NA | NA | -0.5 |
| SE | 60921 | ENSG00000138398 | PPIG | chr2 | + | 169604186 | 169604261 | 169603641 | 169604102 | 169606038 | 169606146 | 567 | 355 | 407 | 504 | 701 | 0 | 0 | 0 | 0 | 2 | 137 | 242 | 323 | 130 | 149 | 0 | 0 | 0 | 0 | 0 | 224 | 149 | 1 | 1 | 1 | 1 | 1 | 1 | 0.996 | 1 | 1 | 1 | 1 | 1 | -0.001 |
| SE | 60922 | ENSG00000138398 | PPIG | chr2 | + | 169606038 | 169606146 | 169604186 | 169604261 | 169607103 | 169607148 | 731 | 504 | 496 | 650 | 1192 | 3 | 0 | 0 | 0 | 0 | 177 | 292 | 446 | 162 | 180 | 0 | 2 | 0 | 0 | 0 | 257 | 149 | 1 | 1 | 0.993 | 1 | 1 | 1 | 1 | 1 | 0.988 | 1 | 1 | 1 | 0.001 |
| SE | 60923 | ENSG00000138398 | PPIG | chr2 | + | 169606102 | 169606146 | 169604186 | 169604261 | 169607103 | 169607148 | 67 | 45 | 45 | 75 | 146 | 3 | 0 | 0 | 0 | 0 | 30 | 15 | 35 | 24 | 38 | 0 | 2 | 0 | 0 | 0 | 193 | 149 | 1 | 1 | 0.945 | 1 | 1 | 1 | 1 | 1 | 0.853 | 1 | 1 | 1 | 0.018 |
| SE | 60924 | ENSG00000138398 | PPIG | chr2 | + | 169607103 | 169607148 | 169606038 | 169606146 | 169608670 | 169608758 | 498 | 327 | 357 | 446 | 808 | 0 | 1 | 0 | 0 | 0 | 100 | 206 | 289 | 108 | 136 | 0 | 0 | 0 | 0 | 0 | 194 | 149 | 1 | 1 | 1 | 0.996 | 1 | 1 | 1 | 1 | 1 | 1 | 1 | 1 | -0.001 |
| SE | 60925 | ENSG00000138398 | PPIG | chr2 | + | 169608670 | 169608758 | 169607103 | 169607148 | 169614463 | 169614493 | 338 | 244 | 243 | 299 | 763 | 2 | 0 | 0 | 1 | 0 | 65 | 132 | 188 | 78 | 94 | 0 | 0 | 0 | 0 | 0 | 237 | 149 | 1 | 1 | 0.991 | 1 | 1 | 0.995 | 1 | 1 | 1 | 1 | 1 | 1 | -0.003 |
| SE | 60928 | ENSG00000138398 | PPIG | chr2 | + | 169633159 | 169633247 | 169630773 | 169631933 | 169636091 | 169636162 | 150 | 154 | 153 | 138 | 244 | 0 | 0 | 0 | 2 | 0 | 25 | 54 | 70 | 36 | 10 | 0 | 0 | 2 | 0 | 0 | 237 | 149 | 1 | 1 | 1 | 1 | 1 | 0.977 | 1 | 1 | 1 | 0.957 | 1 | 1 | 0.004 |
| SE | 60929 | ENSG00000138078 | PREPL | chr2 | - | 44321826 | 44321900 | 44317606 | 44321445 | 44322730 | 44322854 | 125 | 91 | 108 | 81 | 137 | 0 | 1 | 0 | 0 | 0 | 3 | 31 | 22 | 6 | 0 | 0 | 0 | 0 | 0 | 0 | 223 | 149 | 1 | 1 | 1 | 0.984 | 1 | 1 | 1 | 1 | 1 | 1 | 1 | NA | -0.003 |
| SE | 60930 | ENSG00000138078 | PREPL | chr2 | - | 44326711 | 44326928 | 44323261 | 44323411 | 44328936 | 44329112 | 157 | 97 | 177 | 141 | 189 | 2 | 0 | 0 | 0 | 0 | 2 | 101 | 48 | 41 | 2 | 0 | 0 | 3 | 0 | 0 | 298 | 149 | 1 | 1 | 0.975 | 1 | 1 | 1 | 1 | 1 | 1 | 0.889 | 1 | 1 | 0.017 |
| SE | 60931 | ENSG00000138078 | PREPL | chr2 | - | 44328936 | 44329112 | 44326711 | 44326928 | 44332458 | 44332656 | 160 | 90 | 163 | 125 | 139 | 1 | 0 | 0 | 0 | 0 | 7 | 71 | 59 | 37 | 1 | 0 | 0 | 0 | 0 | 0 | 298 | 149 | 1 | 1 | 0.988 | 1 | 1 | 1 | 1 | 1 | 1 | 1 | 1 | 1 | -0.002 |
| SE | 60932 | ENSG00000138078 | PREPL | chr2 | - | 44332458 | 44332656 | 44328936 | 44329112 | 44338350 | 44338536 | 230 | 187 | 220 | 229 | 387 | 0 | 0 | 0 | 0 | 1 | 12 | 101 | 81 | 74 | 10 | 0 | 0 | 0 | 0 | 0 | 298 | 149 | 1 | 1 | 1 | 1 | 1 | 1 | 0.995 | 1 | 1 | 1 | 1 | 1 | -0.001 |
| SE | 60935 | ENSG00000138078 | PREPL | chr2 | - | 44346267 | 44346390 | 44344519 | 44344586 | 44359496 | 44359749 | 175 | 176 | 164 | 139 | 346 | 0 | 2 | 0 | 0 | 0 | 2 | 104 | 75 | 24 | 26 | 0 | 0 | 0 | 0 | 0 | 272 | 149 | 1 | 1 | 1 | 0.98 | 1 | 1 | 1 | 1 | 1 | 1 | 1 | 1 | -0.004 |
| SE | 60937 | ENSG00000138078 | PREPL | chr2 | - | 44359496 | 44359750 | 44346267 | 44346390 | 44361416 | 44361561 | 63 | 76 | 62 | 39 | 165 | 0 | 2 | 0 | 1 | 0 | 2 | 55 | 42 | 7 | 0 | 0 | 0 | 0 | 0 | 0 | 298 | 149 | 1 | 1 | 1 | 0.95 | 1 | 0.951 | 1 | 1 | 1 | 1 | 1 | NA | -0.02 |
| SE | 60938 | ENSG00000138078 | PREPL | chr2 | - | 44359496 | 44359750 | 44346267 | 44346390 | 44361523 | 44361805 | 63 | 76 | 63 | 38 | 164 | 6 | 8 | 6 | 3 | 3 | 2 | 55 | 45 | 7 | 0 | 0 | 0 | 6 | 5 | 0 | 298 | 149 | 1 | 1 | 0.84 | 0.826 | 0.84 | 0.864 | 0.965 | 1 | 1 | 0.789 | 0.412 | NA | 0.067 |
| SE | 60940 | ENSG00000138078 | PREPL | chr2 | - | 44360618 | 44360676 | 44359496 | 44359750 | 44361379 | 44361493 | 0 | 0 | 2 | 0 | 1 | 5 | 1 | 4 | 1 | 5 | 0 | 0 | 5 | 0 | 0 | 0 | 4 | 0 | 0 | 0 | 207 | 149 | 1 | 1 | 0 | 0 | 0.265 | 0 | 0.126 | NA | 0 | 1 | NA | NA | -0.422 |
| SE | 60943 | ENSG00000117155 | SSX2IP | chr1 | - | 84649670 | 84649948 | 84647338 | 84647607 | 84650361 | 84650527 | 1 | 0 | 0 | 0 | 0 | 99 | 96 | 24 | 109 | 119 | 0 | 0 | 0 | 0 | 0 | 9 | 53 | 56 | 17 | 36 | 298 | 149 | 1 | 1 | 0.005 | 0 | 0 | 0 | 0 | 0 | 0 | 0 | 0 | 0 | 0.001 |
| SE | 60945 | ENSG00000117155 | SSX2IP | chr1 | - | 84651882 | 84651997 | 84643705 | 84647607 | 84655831 | 84656005 | 21 | 37 | 4 | 46 | 49 | 0 | 1 | 0 | 1 | 0 | 9 | 12 | 16 | 0 | 0 | 0 | 0 | 2 | 0 | 0 | 264 | 149 | 1 | 1 | 1 | 0.954 | 1 | 0.963 | 1 | 1 | 1 | 0.819 | NA | NA | 0.044 |
| SE | 60948 | ENSG00000117155 | SSX2IP | chr1 | - | 84666121 | 84666232 | 84662197 | 84662530 | 84669680 | 84669893 | 64 | 101 | 54 | 170 | 156 | 0 | 0 | 2 | 1 | 0 | 28 | 54 | 63 | 9 | 18 | 0 | 0 | 0 | 0 | 0 | 260 | 149 | 1 | 1 | 1 | 1 | 0.939 | 0.99 | 1 | 1 | 1 | 1 | 1 | 1 | -0.014 |
| SE | 60949 | ENSG00000117155 | SSX2IP | chr1 | - | 84671176 | 84671308 | 84670645 | 84670815 | 84690153 | 84690304 | 29 | 13 | 9 | 27 | 28 | 0 | 0 | 1 | 1 | 0 | 0 | 10 | 16 | 3 | 4 | 0 | 0 | 0 | 0 | 0 | 281 | 149 | 1 | 1 | 1 | 1 | 0.827 | 0.935 | 1 | NA | 1 | 1 | 1 | 1 | -0.048 |
| SE | 60950 | ENSG00000117155 | SSX2IP | chr1 | - | 84671176 | 84671308 | 84670645 | 84670815 | 84690370 | 84690461 | 32 | 14 | 9 | 26 | 28 | 0 | 2 | 0 | 1 | 0 | 0 | 11 | 16 | 5 | 4 | 0 | 1 | 1 | 0 | 0 | 281 | 149 | 1 | 1 | 1 | 0.788 | 1 | 0.932 | 1 | NA | 0.854 | 0.895 | 1 | 1 | 0.007 |
| SE | 60951 | ENSG00000117155 | SSX2IP | chr1 | - | 84671188 | 84671308 | 84670645 | 84670815 | 84680337 | 84680387 | 0 | 0 | 0 | 0 | 0 | 1 | 1 | 0 | 4 | 0 | 0 | 0 | 0 | 0 | 5 | 0 | 0 | 0 | 0 | 0 | 269 | 149 | 1 | 1 | 0 | 0 | NA | 0 | NA | NA | NA | NA | NA | 1 | -1 |
| SE | 60952 | ENSG00000117155 | SSX2IP | chr1 | - | 84680337 | 84680387 | 84670645 | 84670815 | 84690370 | 84690461 | 1 | 1 | 0 | 5 | 0 | 0 | 2 | 0 | 1 | 0 | 0 | 0 | 0 | 0 | 0 | 0 | 1 | 1 | 0 | 0 | 199 | 149 | 1 | 1 | 1 | 0.272 | NA | 0.789 | NA | NA | 0 | 0 | NA | NA | 0.687 |
| SE | 60953 | ENSG00000117155 | SSX2IP | chr1 | - | 84680337 | 84680387 | 84671188 | 84671308 | 84690370 | 84690461 | 0 | 0 | 0 | 1 | 0 | 4 | 1 | 1 | 0 | 0 | 0 | 0 | 0 | 0 | 0 | 0 | 2 | 0 | 2 | 0 | 199 | 149 | 1 | 1 | 0 | 0 | 0 | 1 | NA | NA | 0 | NA | 0 | NA | 0.25 |
| SE | 60955 | ENSG00000255104 | NA | chr17 | + | 15700047 | 15700168 | 15699740 | 15699777 | 15701151 | 15701240 | 0 | 2 | 4 | 1 | 0 | 3 | 0 | 1 | 0 | 0 | 0 | 0 | 7 | 0 | 0 | 0 | 0 | 0 | 0 | 0 | 270 | 149 | 1 | 1 | 0 | 1 | 0.688 | 1 | NA | NA | NA | 1 | NA | NA | -0.328 |
| SE | 60956 | ENSG00000255104 | NA | chr17 | + | 15700272 | 15700366 | 15700047 | 15700168 | 15701151 | 15701240 | 6 | 3 | 7 | 7 | 3 | 0 | 0 | 1 | 0 | 0 | 0 | 3 | 2 | 0 | 4 | 0 | 0 | 0 | 0 | 0 | 243 | 149 | 1 | 1 | 1 | 1 | 0.811 | 1 | 1 | NA | 1 | 1 | NA | 1 | -0.038 |
| SE | 60960 | ENSG00000177542 | SLC25A22 | chr11 | - | 796042 | 796367 | 794986 | 795169 | 798216 | 798267 | 3 | 4 | 0 | 5 | 0 | 1 | 0 | 0 | 0 | 0 | 4 | 0 | 3 | 7 | 0 | 0 | 0 | 0 | 0 | 0 | 298 | 149 | 1 | 1 | 0.6 | 1 | NA | 1 | NA | 1 | NA | 1 | 1 | NA | -0.133 |
| SE | 60962 | ENSG00000130517 | PGPEP1 | chr19 | + | 18342858 | 18342911 | 18340597 | 18340715 | 18348947 | 18349068 | 7 | 1 | 3 | 0 | 0 | 0 | 0 | 0 | 0 | 0 | 0 | 2 | 4 | 3 | 9 | 0 | 0 | 0 | 1 | 0 | 202 | 149 | 1 | 1 | 1 | 1 | 1 | NA | NA | NA | 1 | 1 | 0.689 | 1 | 0.078 |
| SE | 60963 | ENSG00000130517 | PGPEP1 | chr19 | + | 18342858 | 18342911 | 18340597 | 18340715 | 18355894 | 18356011 | 40 | 13 | 21 | 27 | 7 | 1 | 0 | 0 | 1 | 2 | 0 | 36 | 32 | 37 | 24 | 0 | 0 | 0 | 0 | 0 | 202 | 149 | 1 | 1 | 0.967 | 1 | 1 | 0.952 | 0.721 | NA | 1 | 1 | 1 | 1 | -0.072 |
| SE | 60964 | ENSG00000130517 | PGPEP1 | chr19 | + | 18348947 | 18349068 | 18340610 | 18340715 | 18355894 | 18356011 | 2 | 0 | 3 | 0 | 0 | 1 | 0 | 0 | 1 | 2 | 0 | 2 | 2 | 3 | 6 | 0 | 0 | 0 | 0 | 0 | 270 | 149 | 1 | 1 | 0.525 | NA | 1 | 0 | 0 | NA | 1 | 1 | 1 | 1 | -0.619 |
| SE | 60965 | ENSG00000130517 | PGPEP1 | chr19 | + | 18348947 | 18349068 | 18342858 | 18342911 | 18355894 | 18356011 | 3 | 0 | 4 | 0 | 0 | 34 | 12 | 19 | 27 | 8 | 0 | 2 | 5 | 2 | 6 | 0 | 34 | 31 | 34 | 15 | 270 | 149 | 1 | 1 | 0.046 | 0 | 0.104 | 0 | 0 | NA | 0.031 | 0.082 | 0.031 | 0.181 | -0.051 |
| SE | 60969 | ENSG00000130517 | PGPEP1 | chr19 | + | 18358005 | 18358070 | 18357505 | 18357615 | 18369705 | 18369948 | 0 | 0 | 1 | 0 | 0 | 8 | 0 | 1 | 1 | 5 | 0 | 5 | 0 | 6 | 0 | 0 | 0 | 0 | 0 | 0 | 214 | 149 | 1 | 1 | 0 | NA | 0.41 | 0 | 0 | NA | 1 | NA | 1 | NA | -0.897 |
| SE | 60970 | ENSG00000151693 | ASAP2 | chr2 | + | 9297299 | 9297445 | 9279316 | 9279389 | 9320287 | 9320337 | 3 | 13 | 4 | 25 | 7 | 0 | 0 | 0 | 0 | 2 | 24 | 8 | 16 | 16 | 119 | 0 | 0 | 0 | 0 | 0 | 295 | 149 | 1 | 1 | 1 | 1 | 1 | 1 | 0.639 | 1 | 1 | 1 | 1 | 1 | -0.072 |
| SE | 60972 | ENSG00000151693 | ASAP2 | chr2 | + | 9335092 | 9335179 | 9334737 | 9334813 | 9344531 | 9344635 | 48 | 104 | 7 | 119 | 109 | 1 | 0 | 0 | 5 | 0 | 329 | 98 | 88 | 188 | 462 | 0 | 0 | 0 | 1 | 0 | 236 | 149 | 1 | 1 | 0.968 | 1 | 1 | 0.938 | 1 | 1 | 1 | 1 | 0.992 | 1 | -0.017 |

| | | | | | | | | | | | | | | |
|---|---|---|---|---|---|---|---|---|---|---|---|---|---|---|
| SE | 60974 | ENSG00000151693 | ASAP2 | chr2 | + | 9344730 | 9344800 | 9344531 | 9344635 | 9350807 | 9350895 | 39 | 89 | [illegible] |
| SE | 60975 | ENSG00000151693 | ASAP2 | chr2 | + | 9391061 | 9391196 | 9388293 | 9388546 | 9393481 | 9393647 | 16 | 26 | [illegible] |
| SE | 60976 | ENSG00000151693 | ASAP2 | chr2 | + | 9393481 | 9393647 | 9388293 | 9388546 | 9400022 | 9400072 | 12 | 11 | [illegible] |
| SE | 60977 | ENSG00000151693 | ASAP2 | chr2 | + | 9401273 | 9401396 | 9400741 | 9400830 | 9403252 | 9405677 | 9 | 13 | [illegible] |
| SE | 60978 | ENSG00000178295 | GEN1 | chr2 | + | 17768737 | 17768811 | 17766578 | 17766689 | 17772633 | 17772784 | 18 | 34 | [illegible] |
| SE | 60979 | ENSG00000178295 | GEN1 | chr2 | + | 17771195 | 17771287 | 17768737 | 17768811 | 17772633 | 17772784 | 20 | 19 | [illegible] |
| SE | 60990 | ENSG00000109381 | ELF2 | chr4 | - | 139060323 | 139060674 | 139057219 | 139059607 | 139067683 | 139067770 | 121 | 80 | [illegible] |
| SE | 60994 | ENSG00000109381 | ELF2 | chr4 | - | 139116648 | 139116807 | 139101590 | 139102353 | 139125163 | 139125329 | 5 | 1 | [illegible] |
| SE | 60995 | ENSG00000109381 | ELF2 | chr4 | - | 139117828 | 139117950 | 139116648 | 139116807 | 139125163 | 139125329 | 2 | 0 | [illegible] |
| SE | 60996 | ENSG00000109381 | ELF2 | chr4 | - | 139125163 | 139125329 | 139073453 | 139073567 | 139137629 | 139137867 | 197 | 134 | [illegible] |
| SE | 60997 | ENSG00000109381 | ELF2 | chr4 | - | 139137629 | 139137867 | 139125163 | 139125329 | 139139412 | 139139476 | 229 | 141 | [illegible] |
| SE | 60998 | ENSG00000085563 | ABCB1 | chr7 | - | 87509274 | 87509481 | 87505896 | 87506043 | 87515230 | 87515428 | 131 | 40 | [illegible] |
| SE | 60999 | ENSG00000085563 | ABCB1 | chr7 | - | 87516508 | 87516665 | 87515227 | 87515428 | 87519325 | 87519466 | 166 | 26 | [illegible] |
| SE | 61000 | ENSG00000085563 | ABCB1 | chr7 | - | 87520775 | 87520876 | 87519325 | 87519466 | 87531293 | 87531497 | 212 | 69 | [illegible] |
| SE | 61001 | ENSG00000085563 | ABCB1 | chr7 | - | 87536457 | 87536541 | 87531293 | 87531497 | 87539267 | 87539345 | 101 | 29 | [illegible] |
| SE | 61002 | ENSG00000085563 | ABCB1 | chr7 | - | 87549347 | 87549518 | 87545862 | 87546024 | 87549850 | 87550054 | 81 | 50 | [illegible] |
| SE | 61003 | ENSG00000085563 | ABCB1 | chr7 | - | 87566069 | 87566241 | 87561262 | 87561387 | 87570171 | 87570223 | 144 | 68 | [illegible] |
| SE | 61004 | ENSG00000085563 | ABCB1 | chr7 | - | 87566784 | 87566976 | 87566069 | 87566241 | 87570171 | 87570223 | 197 | 85 | [illegible] |
| SE | 61006 | ENSG00000078018 | MAP2 | chr2 | + | 209507591 | 209507641 | 209424046 | 209424276 | 209580035 | 209580100 | 2 | 0 | [illegible] |
| SE | 61017 | ENSG00000162616 | DNAJB4 | chr1 | + | 78013050 | 78013619 | 78004938 | 78005321 | 78016013 | 78016274 | 59 | 50 | [illegible] |
| SE | 61027 | ENSG00000163191 | S100A11 | chr1 | - | 152033647 | 152033800 | 152032505 | 152032823 | 152036912 | 152037004 | 1366 | 775 | [illegible] |
| SE | 61034 | ENSG00000066027 | PPP2R5A | chr1 | + | 212345802 | 212345933 | 212333496 | 212333598 | 212356625 | 212356676 | 5 | 3 | [illegible] |
| SE | 61035 | ENSG00000066027 | PPP2R5A | chr1 | + | 212345802 | 212345933 | 212342187 | 212342280 | 212347346 | 212347406 | 441 | 261 | [illegible] |
| SE | 61037 | ENSG00000066027 | PPP2R5A | chr1 | + | 212347346 | 212347406 | 212345802 | 212345933 | 212349188 | 212349242 | 94 | 54 | [illegible] |
| SE | 61039 | ENSG00000066027 | PPP2R5A | chr1 | + | 212348388 | 212348497 | 212345802 | 212345933 | 212349188 | 212349242 | 373 | 223 | [illegible] |
| SE | 61040 | ENSG00000066027 | PPP2R5A | chr1 | + | 212348388 | 212348497 | 212345802 | 212345933 | 212356625 | 212356676 | 3 | 5 | [illegible] |
| SE | 61041 | ENSG00000066027 | PPP2R5A | chr1 | + | 212348388 | 212348497 | 212347346 | 212347406 | 212349188 | 212349242 | 590 | 340 | [illegible] |
| SE | 61053 | ENSG00000039560 | RAI14 | chr5 | + | 34813573 | 34813660 | 34812179 | 34812208 | 34814582 | 34814669 | 4 | 4 | [illegible] |
| SE | 61058 | ENSG00000078808 | SDF4 | chr1 | - | 1218768 | 1218927 | 1218457 | 1218633 | 1223243 | 1223357 | 803 | 414 | [illegible] |
| SE | 61059 | ENSG00000078808 | SDF4 | chr1 | - | 1218768 | 1218943 | 1218457 | 1218633 | 1223243 | 1223357 | 518 | 260 | [illegible] |
| SE | 61060 | ENSG00000078808 | SDF4 | chr1 | - | 1218768 | 1219005 | 1218522 | 1218633 | 1223243 | 1223357 | 517 | 258 | [illegible] |
| SE | 61061 | ENSG00000078808 | SDF4 | chr1 | - | 1228467 | 1228946 | 1223831 | 1223968 | 1231891 | 1231968 | 488 | 198 | [illegible] |
| SE | 61066 | ENSG00000235194 | PPP1R3E | chr14 | - | 23300106 | 23300139 | 23299400 | 23299541 | 23300757 | 23300864 | 3 | 1 | [illegible] |
| SE | 61069 | ENSG00000173214 | MFSD4B | chr6 | + | 111263829 | 111263946 | 111262256 | 111262425 | 111265875 | 111267310 | 37 | 28 | [illegible] |
| SE | 61071 | ENSG00000173214 | MFSD4B | chr6 | + | 111273329 | 111273422 | 111265875 | 111267837 | 111276716 | 111278732 | 1 | 2 | [illegible] |
| SE | 61072 | ENSG00000173214 | MFSD4B | chr6 | + | 111273329 | 111273422 | 111265875 | 111267837 | 111277999 | 111278158 | 3 | 2 | [illegible] |
| SE | 61075 | ENSG00000173214 | MFSD4B | chr6 | + | 111277999 | 111278158 | 111273329 | 111273422 | 111295703 | 111295775 | 3 | 0 | [illegible] |
| SE | 61077 | ENSG00000143093 | STRIP1 | chr1 | + | 110039171 | 110039306 | 110038682 | 110038757 | 110040634 | 110040703 | 36 | 16 | [illegible] |
| SE | 61078 | ENSG00000143093 | STRIP1 | chr1 | + | 110039394 | 110039515 | 110039171 | 110039306 | 110040634 | 110040703 | 68 | 20 | [illegible] |
| SE | 61080 | ENSG00000143093 | STRIP1 | chr1 | + | 110040634 | 110040703 | 110039394 | 110039515 | 110041535 | 110041642 | 58 | 16 | [illegible] |
| SE | 61082 | ENSG00000143093 | STRIP1 | chr1 | + | 110049111 | 110049238 | 110047771 | 110047869 | 110049459 | 110049560 | 112 | 39 | [illegible] |
| SE | 61088 | ENSG00000198160 | MIER1 | chr1 | + | 66946149 | 66946295 | 66940027 | 66940052 | 66958058 | 66958220 | 319 | 274 | [illegible] |
| SE | 61089 | ENSG00000198160 | MIER1 | chr1 | + | 66959678 | 66959743 | 66958850 | 66958983 | 66963087 | 66963160 | 428 | 312 | [illegible] |
| SE | 61090 | ENSG00000198160 | MIER1 | chr1 | + | 66963087 | 66963160 | 66959678 | 66959743 | 66970807 | 66970959 | 410 | 321 | [illegible] |
| SE | 61091 | ENSG00000198160 | MIER1 | chr1 | + | 66972896 | 66972991 | 66971654 | 66971736 | 66976594 | 66976722 | 437 | 271 | [illegible] |
| SE | 61092 | ENSG00000198160 | MIER1 | chr1 | + | 66976594 | 66976722 | 66972896 | 66972991 | 66981778 | 66981918 | 638 | 415 | [illegible] |
| SE | 61093 | ENSG00000067225 | PKM | chr15 | - | 72200473 | 72200655 | 72199028 | 72199756 | 72202453 | 72202620 | 989 | 487 | [illegible] |
| SE | 61094 | ENSG00000067225 | PKM | chr15 | - | 72202453 | 72202620 | 72200473 | 72200655 | 72206727 | 72206880 | 1724 | 1160 | [illegible] |
| SE | 61097 | ENSG00000067225 | PKM | chr15 | - | 72217408 | 72217500 | 72210059 | 72210478 | 72218943 | 72219110 | 906 | 551 | [illegible] |
| SE | 61098 | ENSG00000067225 | PKM | chr15 | - | 72221167 | 72221284 | 72218943 | 72219110 | 72230895 | 72231113 | 2 | 0 | [illegible] |
| SE | 61099 | ENSG00000067225 | PKM | chr15 | - | 72221167 | 72221284 | 72218943 | 72219110 | 72231115 | 72231190 | 2 | 0 | [illegible] |
| SE | 61104 | ENSG00000115806 | GORASP2 | chr2 | + | 170929740 | 170929941 | 170929266 | 170929403 | 170948349 | 170948430 | 1 | 0 | [illegible] |
| SE | 61105 | ENSG00000115806 | GORASP2 | chr2 | + | 170929816 | 170929941 | 170929256 | 170929403 | 170948349 | 170948430 | 1 | 0 | [illegible] |
| SE | 61106 | ENSG00000115806 | GORASP2 | chr2 | + | 170936623 | 170936727 | 170929320 | 170929403 | 170948349 | 170948430 | 1 | 0 | [illegible] |
| SE | 61107 | ENSG00000115806 | GORASP2 | chr2 | + | 170954649 | 170954782 | 170951327 | 170951458 | 170956435 | 170956514 | 341 | 178 | [illegible] |
| SE | 61110 | ENSG00000115806 | GORASP2 | chr2 | + | 170956435 | 170956559 | 170953817 | 170954782 | 170961662 | 170961749 | 329 | 179 | [illegible] |
| SE | 61111 | ENSG00000129933 | MAU2 | chr19 | + | 19338844 | 19338939 | 19337169 | 19337265 | 19340845 | 19340873 | 75 | 52 | [illegible] |
| SE | 61113 | ENSG00000175866 | BAIAP2 | chr17 | + | 81036862 | 81036961 | 81035150 | 81035308 | 81053667 | 81053743 | 2 | 0 | [illegible] |
| SE | 61115 | ENSG00000175866 | BAIAP2 | chr17 | + | 81110913 | 81110959 | 81108474 | 81108509 | 81115769 | 81116224 | 1 | 0 | [illegible] |
| SE | 61116 | ENSG00000150782 | IL18 | chr11 | - | 112150996 | 112151072 | 112150071 | 112150206 | 112153591 | 112153603 | 0 | 0 | [illegible] |
| SE | 61120 | ENSG00000150782 | IL18 | chr11 | - | 112154974 | 112155061 | 112150071 | 112150206 | 112163905 | 112164074 | 25 | 4 | [illegible] |
| SE | 61125 | ENSG00000170522 | ELOVL6 | chr4 | - | 110093105 | 110093134 | 110059602 | 110059754 | 110105496 | 110105628 | 0 | 0 | [illegible] |
| SE | 61126 | ENSG00000170522 | ELOVL6 | chr4 | - | 110105496 | 110105628 | 110059602 | 110059754 | 110198246 | 110198367 | 51 | 39 | [illegible] |
| SE | 61127 | ENSG00000198039 | ZNF273 | chr7 | + | 64917029 | 64917118 | 64903287 | 64903419 | 64917580 | 64917707 | 4 | 10 | [illegible] |
| SE | 61128 | ENSG00000198039 | ZNF273 | chr7 | + | 64917029 | 64917118 | 64903287 | 64903419 | 64927653 | 64929124 | 0 | 5 | [illegible] |
| SE | 61129 | ENSG00000198039 | ZNF273 | chr7 | + | 64917580 | 64917707 | 64903246 | 64903419 | 64927653 | 64929124 | 6 | 3 | [illegible] |
| SE | 61130 | ENSG00000198039 | ZNF273 | chr7 | + | 64917580 | 64917707 | 64917029 | 64917118 | 64918196 | 64918292 | 10 | 13 | [illegible] |
| SE | 61131 | ENSG00000198039 | ZNF273 | chr7 | + | 64917580 | 64917707 | 64917029 | 64917118 | 64927653 | 64929124 | 10 | 10 | [illegible] |
| SE | 61132 | ENSG00000198039 | ZNF273 | chr7 | + | 64918196 | 64918292 | 64917029 | 64917118 | 64927653 | 64929124 | 18 | 13 | [illegible] |
| SE | 61133 | ENSG00000198039 | ZNF273 | chr7 | + | 64918196 | 64918292 | 64917580 | 64917707 | 64927653 | 64929124 | 24 | 17 | [illegible] |
| SE | 61145 | ENSG00000111711 | GOLT1B | chr12 | + | 21506884 | 21506976 | 21501788 | 21501948 | 21512294 | 21512376 | 9 | 11 | [illegible] |
| SE | 61146 | ENSG00000111711 | GOLT1B | chr12 | + | 21507116 | 21507151 | 21506884 | 21506976 | 21507943 | 21508109 | 2 | 0 | [illegible] |
| SE | 61148 | ENSG00000111711 | GOLT1B | chr12 | + | 21507116 | 21507151 | 21506884 | 21506976 | 21512294 | 21512376 | 2 | 0 | [illegible] |
| SE | 61150 | ENSG00000111711 | GOLT1B | chr12 | + | 21507943 | 21508109 | 21506884 | 21506976 | 21512294 | 21512376 | 6 | 2 | [illegible] |
| SE | 61152 | ENSG00000111711 | GOLT1B | chr12 | + | 21507943 | 21508109 | 21507116 | 21507151 | 21512294 | 21512376 | 1 | 0 | [illegible] |
| SE | 61155 | ENSG00000111711 | GOLT1B | chr12 | + | 21508382 | 21508561 | 21507116 | 21507151 | 21512294 | 21512376 | 116 | 70 | [illegible] |
| SE | 61156 | ENSG00000111711 | GOLT1B | chr12 | + | 21508382 | 21508561 | 21507943 | 21508109 | 21512294 | 21512376 | 105 | 65 | [illegible] |
| SE | 61162 | ENSG00000028528 | SNX1 | chr15 | + | 64118787 | 64118854 | 64118116 | 64118244 | 64123502 | 64123546 | 564 | 359 | [illegible] |
| SE | 61165 | ENSG00000028528 | SNX1 | chr15 | + | 64123502 | 64123546 | 64118787 | 64118854 | 64127173 | 64127252 | 150 | 91 | [illegible] |
| SE | 61166 | ENSG00000028528 | SNX1 | chr15 | + | 64126078 | 64126220 | 64118787 | 64118854 | 64127173 | 64127252 | 203 | 173 | [illegible] |
| SE | 61168 | ENSG00000028528 | SNX1 | chr15 | + | 64130227 | 64130321 | 64129872 | 64130029 | 64131686 | 64131892 | 348 | 225 | [illegible] |
| SE | 61169 | ENSG00000028528 | SNX1 | chr15 | + | 64131686 | 64131892 | 64130227 | 64130321 | 64134663 | 64134807 | 331 | 235 | [illegible] |
| SE | 61170 | ENSG00000028528 | SNX1 | chr15 | + | 64134663 | 64134807 | 64131686 | 64131892 | 64136329 | 64136410 | 256 | 176 | [illegible] |
| SE | 61171 | ENSG00000028528 | SNX1 | chr15 | + | 64138041 | 64138097 | 64136860 | 64136932 | 64142667 | 64143308 | 1 | 5 | [illegible] |
| SE | 61176 | ENSG00000152292 | SH2D6 | chr2 | + | 85422202 | 85422314 | 85418930 | 85419244 | 85422436 | 85422490 | 0 | 0 | [illegible] |
| SE | 61183 | ENSG00000198873 | GRK5 | chr10 | + | 119326515 | 119326611 | 119207570 | 119207969 | 119380814 | 119380927 | 104 | 68 | [illegible] |
| SE | 61185 | ENSG00000198873 | GRK5 | chr10 | + | 119396694 | 119396772 | 119380814 | 119380927 | 119423165 | 119423266 | 171 | 88 | [illegible] |
| SE | 61186 | ENSG00000198873 | GRK5 | chr10 | + | 119423165 | 119423266 | 119396694 | 119396772 | 119424992 | 119425085 | 266 | 104 | [illegible] |
| SE | 61187 | ENSG00000198873 | GRK5 | chr10 | + | 119423165 | 119423266 | 119396694 | 119396772 | 119430374 | 119430438 | 77 | 33 | [illegible] |
| SE | 61189 | ENSG00000198873 | GRK5 | chr10 | + | 119424992 | 119425085 | 119423165 | 119423266 | 119430374 | 119430438 | 300 | 139 | [illegible] |
| SE | 61190 | ENSG00000198873 | GRK5 | chr10 | + | 119430374 | 119430438 | 119396694 | 119396772 | 119431386 | 119431527 | 60 | 41 | [illegible] |
| SE | 61201 | ENSG00000131408 | NR1H2 | chr19 | + | 50377732 | 50377870 | 50377433 | 50377648 | 50378148 | 50378312 | 176 | 91 | [illegible] |
| SE | 61207 | ENSG00000162402 | USP24 | chr1 | - | 55078537 | 55078651 | 55077234 | 55077300 | 55079537 | 55079659 | 202 | 92 | [illegible] |
| SE | 61208 | ENSG00000162402 | USP24 | chr1 | - | 55078537 | 55078651 | 55077234 | 55077300 | 55081321 | 55081424 | 111 | 54 | [illegible] |
| SE | 61209 | ENSG00000162402 | USP24 | chr1 | - | 55079537 | 55079659 | 55077234 | 55077300 | 55081321 | 55081424 | 109 | 32 | [illegible] |
| SE | 61210 | ENSG00000162402 | USP24 | chr1 | - | 55079537 | 55079659 | 55078537 | 55078651 | 55081321 | 55081424 | 202 | 72 | [illegible] |
| SE | 61212 | ENSG00000162402 | USP24 | chr1 | - | 55092820 | 55092916 | 55092022 | 55092126 | 55093936 | 55094087 | 277 | 150 | [illegible] |
| SE | 61214 | ENSG00000162402 | USP24 | chr1 | - | 55100838 | 55100964 | 55099770 | 55099869 | 55101583 | 55101703 | 254 | 193 | [illegible] |
| SE | 61216 | ENSG00000162402 | USP24 | chr1 | - | 55124468 | 55124628 | 55123446 | 55123602 | 55125319 | 55125548 | 239 | 101 | [illegible] |
| SE | 61217 | ENSG00000162402 | USP24 | chr1 | - | 55134327 | 55134413 | 55134069 | 55134163 | 55137514 | 55137688 | 441 | 241 | [illegible] |
| SE | 61218 | ENSG00000162402 | USP24 | chr1 | - | 55138607 | 55138718 | 55137805 | 55137904 | 55138943 | 55139010 | 207 | 111 | [illegible] |
| SE | 61220 | ENSG00000162402 | USP24 | chr1 | - | 55144126 | 55144203 | 55142978 | 55143119 | 55145997 | 55146109 | 423 | 286 | [illegible] |
| SE | 61222 | ENSG00000162402 | USP24 | chr1 | - | 55156947 | 55157051 | 55154670 | 55154778 | 55157255 | 55157370 | 820 | 662 | [illegible] |
| SE | 61223 | ENSG00000162402 | USP24 | chr1 | - | 55176375 | 55176443 | 55172376 | 55172520 | 55177966 | 55178132 | 328 | 295 | [illegible] |
| SE | 61226 | ENSG00000167578 | RAB4B | chr19 | + | 40780018 | 40780499 | 40778273 | 40778391 | 40783777 | 40783840 | 182 | 80 | [illegible] |
| SE | 61227 | ENSG00000167578 | RAB4B | chr19 | + | 40783777 | 40783840 | 40780018 | 40780499 | 40786664 | 40786760 | 26 | 16 | [illegible] |
| SE | 61234 | ENSG00000171503 | ETFDH | chr4 | + | 158680466 | 158680607 | 158672205 | 158672490 | 158681570 | 158681614 | 16 | 9 | [illegible] |
| SE | 61236 | ENSG00000171503 | ETFDH | chr4 | + | 158680466 | 158680607 | 158680111 | 158680182 | 158681570 | 158681614 | 2 | 1 | [illegible] |
| SE | 61237 | ENSG00000171503 | ETFDH | chr4 | + | 158680466 | 158680607 | 158680111 | 158680182 | 158682194 | 158682424 | 11 | 21 | [illegible] |
| SE | 61238 | ENSG00000171503 | ETFDH | chr4 | + | 158680466 | 158681614 | 158680123 | 158680182 | 158682194 | 158682424 | 13 | 5 | [illegible] |
| SE | 61240 | ENSG00000171503 | ETFDH | chr4 | + | 158680527 | 158680607 | 158672124 | 158672490 | 158682194 | 158682424 | 8 | 14 | [illegible] |
| SE | 61241 | ENSG00000171503 | ETFDH | chr4 | + | 158680527 | 158680607 | 158680123 | 158680182 | 158682194 | 158682424 | 8 | 16 | [illegible] |
| SE | 61242 | ENSG00000171503 | ETFDH | chr4 | + | 158681570 | 158681614 | 158672306 | 158672490 | 158682194 | 158682424 | 4 | 4 | [illegible] |
| SE | 61243 | ENSG00000171503 | ETFDH | chr4 | + | 158681570 | 158681614 | 158680123 | 158680182 | 158682194 | 158682424 | 4 | 4 | [illegible] |
| SE | 61244 | ENSG00000171503 | ETFDH | chr4 | + | 158681570 | 158681614 | 158680466 | 158680607 | 158682194 | 158682424 | 4 | 5 | [illegible] |
| SE | 61246 | ENSG00000171503 | ETFDH | chr4 | + | 158685100 | 158685219 | 158684591 | 158684673 | 158690347 | 158690425 | 101 | 94 | [illegible] |
| SE | 61247 | ENSG00000171503 | ETFDH | chr4 | + | 158685151 | 158685219 | 158684591 | 158684673 | 158690347 | 158690425 | 28 | 28 | [illegible] |
| SE | 61248 | ENSG00000171503 | ETFDH | chr4 | + | 158690520 | 158690655 | 158690347 | 158690425 | 158695496 | 158695643 | 0 | 0 | [illegible] |
| SE | 61249 | ENSG00000171503 | ETFDH | chr4 | + | 158697558 | 158697670 | 158695496 | 158695643 | 158698986 | 158699130 | 25 | 40 | [illegible] |
| SE | 61255 | ENSG00000171503 | ETFDH | chr4 | + | 158700874 | 158700919 | 158698780 | 158699130 | 158703422 | 158703591 | 0 | 0 | [illegible] |
| SE | 61256 | ENSG00000171503 | ETFDH | chr4 | + | 158700878 | 158700919 | 158698986 | 158699130 | 158703422 | 158703591 | 0 | 0 | [illegible] |
| SE | 61257 | ENSG00000171503 | ETFDH | chr4 | + | 158703689 | 158706371 | 158703422 | 158703591 | 158706628 | 158706850 | 70 | 75 | [illegible] |
| SE | 61258 | ENSG00000171503 | ETFDH | chr4 | + | 158706188 | 158706371 | 158699980 | 158703591 | 158706628 | 158706850 | 133 | 138 | [illegible] |
| SE | 61259 | ENSG00000171503 | ETFDH | chr4 | + | 158706628 | 158706850 | 158703689 | 158706371 | 158708363 | 158708568 | 103 | 122 | [illegible] |
| SE | 61262 | ENSG00000197056 | ZMYM1 | chr1 | + | 35093319 | 35093441 | 35079385 | 35079442 | 35093913 | 35094083 | 1 | 9 | [illegible] |
| SE | 61263 | ENSG00000197056 | ZMYM1 | chr1 | + | 35093393 | 35093441 | 35079390 | 35079442 | 35093913 | 35094083 | 1 | 2 | [illegible] |
| SE | 61265 | ENSG00000197056 | ZMYM1 | chr1 | + | 35093913 | 35094083 | 35079384 | 35079442 | 35095818 | 35095891 | 29 | 25 | [illegible] |
| SE | 61266 | ENSG00000197056 | ZMYM1 | chr1 | + | 35093913 | 35094083 | 35079384 | 35079442 | 35097316 | 35097478 | 9 | 1 | [illegible] |
| SE | 61267 | ENSG00000197056 | ZMYM1 | chr1 | + | 35095818 | 35095891 | 35079390 | 35079442 | 35097316 | 35097478 | 19 | 25 | [illegible] |
| SE | 61269 | ENSG00000197056 | ZMYM1 | chr1 | + | 35111771 | 35111912 | 35110293 | 35110447 | 35112086 | 35112130 | 95 | 56 | [illegible] |
| SE | 61270 | ENSG00000197056 | ZMYM1 | chr1 | + | 35111775 | 35111912 | 35110293 | 35110447 | 35112086 | 35112130 | 64 | 35 | [illegible] |
| SE | 61277 | ENSG00000163882 | POLR2H | chr3 | + | 184362872 | 184363565 | 184361717 | 184362146 | 184364965 | 184365049 | 131 | 70 | [illegible] |
| SE | 61278 | ENSG00000163882 | POLR2H | chr3 | + | 184363225 | 184363565 | 184361788 | 184362146 | 184364965 | 184365049 | 131 | 70 | [illegible] |
| SE | 61279 | ENSG00000163882 | POLR2H | chr3 | + | 184364965 | 184365049 | 184362872 | 184363565 | 184365132 | 184365226 | 272 | 159 | [illegible] |
| SE | 61280 | ENSG00000163882 | POLR2H | chr3 | + | 184366716 | 184366800 | 184364954 | 184365226 | 184368176 | 184368253 | 137 | 84 | [illegible] |
| SE | 61281 | ENSG00000134058 | CDK7 | chr5 | + | 69250744 | 69250831 | 69235393 | 69235453 | 69252417 | 69252451 | 0 | 0 | [illegible] |
| SE | 61282 | ENSG00000134058 | CDK7 | chr5 | + | 69252417 | 69252451 | 69235116 | 69235453 | 69254601 | 69254667 | 53 | 15 | [illegible] |
| SE | 61285 | ENSG00000134058 | CDK7 | chr5 | + | 69254601 | 69254669 | 69252417 | 69252451 | 69258042 | 69258153 | 8 | 9 | [illegible] |
| SE | 61286 | ENSG00000134058 | CDK7 | chr5 | + | 69255459 | 69255528 | 69252417 | 69252451 | 69258042 | 69258153 | 21 | 7 | [illegible] |
| SE | 61287 | ENSG00000134058 | CDK7 | chr5 | + | 69255459 | 69255528 | 69254601 | 69254669 | 69258042 | 69258153 | 85 | 25 | [illegible] |
| SE | 61291 | ENSG00000134058 | CDK7 | chr5 | + | 69272891 | 69273041 | 69269206 | 69269293 | 69276542 | 69276569 | 107 | 67 | [illegible] |
| SE | 61294 | ENSG00000067208 | EVI5 | chr1 | - | 92563641 | 92563737 | 92508695 | 92513970 | 92605306 | 92605402 | 29 | 23 | [illegible] |
| SE | 61295 | ENSG00000067208 | EVI5 | chr1 | - | 92605306 | 92605402 | 92563641 | 92563737 | 92624175 | 92624334 | 7 | 9 | [illegible] |
| SE | 61296 | ENSG00000067208 | EVI5 | chr1 | - | 92607580 | 92607727 | 92605306 | 92605402 | 92624175 | 92624334 | 71 | 97 | [illegible] |
| SE | 61297 | ENSG00000067208 | EVI5 | chr1 | - | 92624175 | 92624334 | 92607448 | 92607727 | 92625793 | 92625934 | 94 | 121 | [illegible] |
| SE | 61298 | ENSG00000067208 | EVI5 | chr1 | - | 92624175 | 92624334 | 92607448 | 92607727 | 92636201 | 92636336 | 51 | 66 | [illegible] |
| SE | 61299 | ENSG00000067208 | EVI5 | chr1 | - | 92625793 | 92625934 | 92607448 | 92607727 | 92636201 | 92636336 | 58 | 65 | [illegible] |
| SE | 61300 | ENSG00000067208 | EVI5 | chr1 | - | 92625793 | 92625934 | 92607448 | 92607727 | 92663419 | 92663452 | 1 | 3 | [illegible] |
| SE | 61302 | ENSG00000067208 | EVI5 | chr1 | - | 92636201 | 92636336 | 92607580 | 92607727 | 92663419 | 92663452 | 11 | 13 | [illegible] |
| SE | 61303 | ENSG00000067208 | EVI5 | chr1 | - | 92636201 | 92636336 | 92625689 | 92625934 | 92662718 | 92662865 | 56 | 63 | [illegible] |
| SE | 61304 | ENSG00000067208 | EVI5 | chr1 | - | 92636201 | 92636336 | 92625689 | 92625934 | 92663419 | 92663452 | 61 | 74 | [illegible] |
| SE | 61305 | ENSG00000067208 | EVI5 | chr1 | - | 92662718 | 92662865 | 92625793 | 92625934 | 92663419 | 92663452 | 0 | 2 | [illegible] |
| SE | 61306 | ENSG00000067208 | EVI5 | chr1 | - | 92662718 | 92662865 | 92636201 | 92636336 | 92663419 | 92663452 | 5 | 4 | [illegible] |
| SE | 61309 | ENSG00000067208 | EVI5 | chr1 | - | 92663419 | 92663452 | 92662718 | 92662865 | 92665938 | 92665955 | 2 | 11 | [illegible] |
| SE | 61310 | ENSG00000067208 | EVI5 | chr1 | - | 92702140 | 92702215 | 92697859 | 92697985 | 92703394 | 92703619 | 78 | 30 | [illegible] |
| SE | 61311 | ENSG00000067208 | EVI5 | chr1 | - | 92736397 | 92736627 | 92704554 | 92704744 | 92784835 | 92785107 | 53 | 36 | [illegible] |
| SE | 61312 | ENSG00000067208 | EVI5 | chr1 | - | 92773905 | 92774027 | 92736559 | 92736627 | 92784835 | 92784966 | 1 | 1 | [illegible] |
| SE | 61314 | ENSG00000102786 | INTS6 | chr13 | - | 51379461 | 51379572 | 51378238 | 51378454 | 51382028 | 51382123 | 299 | 244 | [illegible] |
| SE | 61315 | ENSG00000102786 | INTS6 | chr13 | - | 51382028 | 51382123 | 51379461 | 51379572 | 51383328 | 51383461 | 221 | 167 | [illegible] |
| SE | 61316 | ENSG00000102786 | INTS6 | chr13 | - | 51387385 | 51387540 | 51383588 | 51383741 | 51430293 | 51430383 | 61 | 36 | [illegible] |
| SE | 61318 | ENSG00000102786 | INTS6 | chr13 | - | 51389318 | 51389444 | 51387385 | 51387540 | 51423029 | 51423090 | 57 | 40 | [illegible] |
| SE | 61319 | ENSG00000102786 | INTS6 | chr13 | - | 51389318 | 51389444 | 51387385 | 51387540 | 51430293 | 51430383 | 67 | 43 | [illegible] |
| SE | 61320 | ENSG00000102786 | INTS6 | chr13 | - | 51395299 | 51395483 | 51387385 | 51387540 | 51421111 | 51421334 | 5 | 6 | [illegible] |
| SE | 61321 | ENSG00000102786 | INTS6 | chr13 | - | 51395299 | 51395483 | 51387385 | 51387540 | 51423029 | 51423090 | 4 | 0 | [illegible] |
| SE | 61323 | ENSG00000102786 | INTS6 | chr13 | - | 51395299 | 51395483 | 51389318 | 51389444 | 51423029 | 51423090 | 155 | 87 | [illegible] |
| SE | 61325 | ENSG00000102786 | INTS6 | chr13 | - | 51421111 | 51421334 | 51387385 | 51387540 | 51423029 | 51423090 | 4 | 0 | [illegible] |
| SE | 61326 | ENSG00000102786 | INTS6 | chr13 | - | 51421111 | 51421334 | 51395299 | 51395483 | 51423029 | 51423090 | 7 | 6 | [illegible] |
| SE | 61327 | ENSG00000102786 | INTS6 | chr13 | - | 51423029 | 51423090 | 51387385 | 51387540 | 51430293 | 51430383 | 17 | 15 | [illegible] |
| SE | 61328 | ENSG00000102786 | INTS6 | chr13 | - | 51423029 | 51423090 | 51389318 | 51389444 | 51430293 | 51430383 | 16 | 15 | [illegible] |
| SE | 61332 | ENSG00000124772 | CPNE5 | chr6 | - | 36744267 | 36744325 | 36743688 | 36743762 | 36745047 | 36745150 | 14 | 3 | [illegible] |
| SE | 61333 | ENSG00000124772 | CPNE5 | chr6 | - | 36744267 | 36744526 | 36743688 | 36743762 | 36745047 | 36745150 | 7 | 1 | [illegible] |
| SE | 61346 | ENSG00000142082 | SIRT3 | chr11 | - | 233910 | 234004 | 233342 | 233534 | 236615 | 236859 | 2 | 0 | [illegible] |
| SE | 61347 | ENSG00000117528 | ABCD3 | chr1 | + | 94475142 | 94475240 | 94473765 | 94473835 | 94475613 | 94475694 | 191 | 115 | [illegible] |
| SE | 61348 | ENSG00000117528 | ABCD3 | chr1 | + | 94515145 | 94515202 | 94506537 | 94506642 | 94517051 | 94518663 | 154 | 143 | [illegible] |
| SE | 61349 | ENSG00000189319 | FAM53B | chr10 | - | 124681606 | 124682379 | 124619291 | 124623604 | 124696157 | 124696212 | 405 | 129 | [illegible] |
| SE | 61351 | ENSG00000189319 | FAM53B | chr10 | - | 124696157 | 124696212 | 124681606 | 124682379 | 124744012 | 124744269 | 98 | 40 | [illegible] |
| SE | 61352 | ENSG00000189319 | FAM53B | chr10 | - | 124706635 | 124706887 | 124681606 | 124682379 | 124744012 | 124744269 | 77 | 25 | [illegible] |
| SE | 61353 | ENSG00000189319 | FAM53B | chr10 | - | 124706635 | 124706887 | 124696157 | 124696212 | 124742427 | 124742970 | 256 | 81 | [illegible] |
| SE | 61355 | ENSG00000035115 | SH3YL1 | chr2 | - | 224863 | 224920 | 218143 | 219001 | 229965 | 230044 | 31 | 28 | [illegible] |
| SE | 61356 | ENSG00000035115 | SH3YL1 | chr2 | - | 224867 | 224920 | 218796 | 219001 | 229965 | 230044 | 5 | 6 | [illegible] |
| SE | 61357 | ENSG00000035115 | SH3YL1 | chr2 | - | 233100 | 233229 | 230377 | 231191 | 234159 | 234272 | 125 | 61 | [illegible] |
| SE | 61358 | ENSG00000035115 | SH3YL1 | chr2 | - | 242797 | 242871 | 234159 | 234272 | 243502 | 243562 | 17 | 3 | [illegible] |
| SE | 61359 | ENSG00000035115 | SH3YL1 | chr2 | - | 242797 | 242871 | 234159 | 234272 | 244626 | 244738 | 8 | 0 | [illegible] |
| SE | 61360 | ENSG00000035115 | SH3YL1 | chr2 | - | 242797 | 242871 | 234159 | 234272 | 247537 | 247602 | 10 | 0 | [illegible] |
| SE | 61361 | ENSG00000035115 | SH3YL1 | chr2 | - | 243502 | 243562 | 234159 | 234272 | 247537 | 247602 | 0 | 1 | [illegible] |
| SE | 61362 | ENSG00000035115 | SH3YL1 | chr2 | - | 243502 | 243562 | 242797 | 242871 | 247537 | 247602 | 6 | 4 | [illegible] |
| SE | 61363 | ENSG00000035115 | SH3YL1 | chr2 | - | 244626 | 244738 | 234159 | 234272 | 247537 | 247602 | 6 | 0 | [illegible] |
| SE | 61364 | ENSG00000035115 | SH3YL1 | chr2 | - | 244626 | 244738 | 242797 | 242871 | 247537 | 247602 | 5 | 0 | [illegible] |
| SE | 61365 | ENSG00000035115 | SH3YL1 | chr2 | - | 256206 | 256690 | 253004 | 253115 | 260084 | 261323 | 0 | 0 | [illegible] |
| SE | 61366 | ENSG00000035115 | SH3YL1 | chr2 | - | 260084 | 260702 | 253004 | 253115 | 262630 | 262786 | 6 | 18 | [illegible] |
| SE | 61367 | ENSG00000035115 | SH3YL1 | chr2 | - | 260084 | 261130 | 253004 | 253115 | 262630 | 262786 | 8 | 21 | [illegible] |
| SE | 61372 | ENSG00000035115 | SH3YL1 | chr2 | - | 262630 | 262786 | 260084 | 260702 | 263210 | 263368 | 1 | 1 | [illegible] |
| SE | 61378 | ENSG00000205707 | ETFRF1 | chr12 | + | 25203724 | 25203823 | 25195271 | 25195337 | 25203919 | 25204007 | 0 | 1 | [illegible] |
| SE | 61381 | ENSG00000151893 | CACUL1 | chr10 | - | 118686597 | 118686641 | 118676410 | 118686168 | 118691264 | 118691403 | 313 | 281 | [illegible] |
| SE | 61382 | ENSG00000151893 | CACUL1 | chr10 | - | 118691264 | 118691403 | 118676410 | 118686168 | 118701305 | 118701408 | 3 | 2 | [illegible] |
| SE | 61384 | ENSG00000151893 | CACUL1 | chr10 | - | 118693701 | 118693749 | 118691264 | 118691403 | 118695140 | 118695230 | 0 | 2 | [illegible] |
| SE | 61385 | ENSG00000151893 | CACUL1 | chr10 | - | 118695140 | 118695230 | 118691264 | 118691403 | 118701305 | 118701408 | 266 | 229 | [illegible] |
| SE | 61386 | ENSG00000151893 | CACUL1 | chr10 | - | 118701305 | 118701408 | 118694920 | 118695230 | 118707491 | 118707513 | 212 | 170 | [illegible] |
| SE | 61391 | ENSG00000136891 | TEX10 | chr9 | - | 100327786 | 100327962 | 100326301 | 100326479 | 100340256 | 100340369 | 49 | 12 | [illegible] |
| SE | 61394 | ENSG00000136891 | TEX10 | chr9 | - | 100340256 | 100340369 | 100327786 | 100327962 | 100346071 | 100346315 | 42 | 21 | [illegible] |
| SE | 61395 | ENSG00000136891 | TEX10 | chr9 | - | 100349183 | 100349369 | 100346693 | 100347406 | 100352771 | 100352942 | 83 | 115 | [illegible] |
| SE | 61396 | ENSG00000136891 | TEX10 | chr9 | - | 100349183 | 100349372 | 100346693 | 100347406 | 100352771 | 100352942 | 110 | 123 | [illegible] |
| SE | 61398 | ENSG00000168772 | CXXC4 | chr4 | - | 104494505 | 104494581 | 104490743 | 104492059 | 104494700 | 104494901 | 4 | 0 | [illegible] |
| SE | 61399 | ENSG00000118292 | C1orf54 | chr1 | + | 150274086 | 150274170 | 150272779 | 150272863 | 150275740 | 150275799 | 30 | 29 | [illegible] |
| SE | 61400 | ENSG00000118292 | C1orf54 | chr1 | + | 150275740 | 150275799 | 150274086 | 150274170 | 150276521 | 150276632 | 35 | 31 | [illegible] |
| SE | 61402 | ENSG00000185630 | PBX1 | chr1 | + | 164625928 | 164626096 | 164563237 | 164563311 | 164792493 | 164792738 | 3 | 0 | [illegible] |
| SE | 61415 | ENSG00000053900 | ANAPC4 | chr4 | + | 25377417 | 25377556 | 25377262 | 25377344 | 25380373 | 25380445 | 42 | 35 | [illegible] |
| SE | 61416 | ENSG00000053900 | ANAPC4 | chr4 | + | 25388837 | 25388882 | 25388716 | 25388743 | 25390135 | 25390183 | 69 | 46 | [illegible] |
| SE | 61419 | ENSG00000053900 | ANAPC4 | chr4 | + | 25414323 | 25414385 | 25413644 | 25413742 | 25414465 | 25414504 | 152 | 130 | [illegible] |

| | | | | | | | | | | | | | | | | | | | | | | | | | | | | | | | | | | | | | | | | | | | | | | |
|---|---|---|---|---|---|---|---|---|---|---|---|---|---|---|---|---|---|---|---|---|---|---|---|---|---|---|---|---|---|---|---|---|---|---|---|---|---|---|---|---|---|---|---|---|---|---|
| SE | 61420 | ENSG00000053900 | ANAPC4 | chr4 | + | 25416424 | 25416598 | 25415465 | 25415540 | 25417615 | 25417739 | 241 | 256 | 247 | 263 | 381 | 0 | 1 | 0 | 1 | 0 | 81 | 107 | 121 | 48 | 9 | 0 | 0 | 0 | 0 | 0 | 298 | 149 | 1 | 1 | 1 | 0.992 | 1 | 0.992 | 1 | 1 | 1 | 1 | 1 | 1 | -0.003 |
| SE | 61468 | ENSG00000096080 | MRPS18A | chr6 | - | 43675201 | 43675617 | 43671783 | 43671906 | 43678517 | 43678625 | 100 | 51 | 87 | 85 | 155 | 0 | 0 | 0 | 1 | 0 | 92 | 155 | 71 | 102 | 156 | 0 | 0 | 0 | 0 | 0 | 298 | 149 | 1 | 1 | 1 | 1 | 1 | 0.977 | 1 | 1 | 1 | 1 | 1 | 1 | -0.005 |
| SE | 61469 | ENSG00000096080 | MRPS18A | chr6 | - | 43675493 | 43675617 | 43671509 | 43671906 | 43678517 | 43678625 | 84 | 40 | 76 | 62 | 132 | 0 | 0 | 0 | 1 | 0 | 66 | 130 | 59 | 96 | 107 | 0 | 0 | 0 | 0 | 0 | 273 | 149 | 1 | 1 | 1 | 1 | 1 | 0.971 | 1 | 1 | 1 | 1 | 1 | 1 | -0.006 |
| SE | 61472 | ENSG00000018699 | TTC27 | chr2 | + | 32633875 | 32634005 | 32630522 | 32630700 | 32640269 | 32640368 | 138 | 95 | 121 | 123 | 115 | 0 | 1 | 3 | 0 | 0 | 127 | 121 | 100 | 76 | 145 | 0 | 0 | 0 | 0 | 0 | 279 | 149 | 1 | 1 | 1 | 0.981 | 0.956 | 1 | 1 | 1 | 1 | 1 | 1 | 1 | -0.013 |
| SE | 61473 | ENSG00000018699 | TTC27 | chr2 | + | 32640269 | 32640410 | 32633875 | 32634005 | 32650130 | 32650233 | 127 | 135 | 136 | 155 | 155 | 2 | 0 | 0 | 0 | 0 | 156 | 122 | 84 | 99 | 156 | 0 | 0 | 0 | 0 | 0 | 290 | 149 | 1 | 1 | 0.97 | 1 | 1 | 1 | 1 | 1 | 1 | 1 | 1 | 1 | -0.006 |
| SE | 61476 | ENSG00000018699 | TTC27 | chr2 | + | 32678855 | 32678922 | 32672271 | 32672384 | 32702806 | 32702920 | 119 | 102 | 117 | 140 | 130 | 1 | 0 | 1 | 0 | 4 | 43 | 83 | 93 | 30 | 71 | 0 | 0 | 0 | 0 | 0 | 216 | 149 | 1 | 1 | 0.988 | 1 | 0.988 | 1 | 0.957 | 1 | 1 | 1 | 1 | 1 | -0.013 |
| SE | 61477 | ENSG00000018699 | TTC27 | chr2 | + | 32702806 | 32702920 | 32678855 | 32678922 | 32733827 | 32733921 | 114 | 88 | 136 | 144 | 159 | 0 | 0 | 0 | 1 | 0 | 44 | 89 | 91 | 38 | 59 | 0 | 0 | 0 | 2 | 0 | 263 | 149 | 1 | 1 | 1 | 1 | 1 | 0.988 | 1 | 1 | 1 | 1 | 0.915 | 1 | 0.015 |
| SE | 61478 | ENSG00000018699 | TTC27 | chr2 | + | 32736693 | 32736816 | 32733827 | 32733923 | 32758291 | 32758519 | 78 | 55 | 108 | 96 | 109 | 0 | 0 | 2 | 0 | 4 | 31 | 50 | 53 | 46 | 30 | 0 | 2 | 0 | 0 | 0 | 272 | 149 | 1 | 1 | 1 | 1 | 0.967 | 1 | 0.937 | 1 | 0.932 | 1 | 1 | 1 | -0.006 |
| SE | 61479 | ENSG00000018699 | TTC27 | chr2 | + | 32766467 | 32766549 | 32758291 | 32758519 | 32777881 | 32777980 | 1 | 0 | 1 | 3 | 0 | 8 | 13 | 18 | 19 | 25 | 0 | 0 | 1 | 2 | 0 | 0 | 16 | 7 | 4 | 0 | 231 | 149 | 1 | 1 | 0.075 | 0 | 0.035 | 0.092 | 0 | NA | 0 | 0.084 | 0.244 | NA | -0.069 |
| SE | 61480 | ENSG00000018699 | TTC27 | chr2 | + | 32777881 | 32777980 | 32758291 | 32758519 | 32782625 | 32782678 | 19 | 20 | 23 | 35 | 36 | 0 | 0 | 2 | 1 | 2 | 0 | 18 | 12 | 12 | 4 | 0 | 0 | 0 | 0 | 0 | 248 | 149 | 1 | 1 | 1 | 1 | 0.874 | 0.955 | 0.915 | NA | 1 | | 1 | 1 | -0.051 |
| SE | 61481 | ENSG00000018699 | TTC27 | chr2 | + | 32780053 | 32780112 | 32777881 | 32777980 | 32782625 | 32782678 | 2 | 1 | 10 | 1 | 1 | 12 | 9 | 5 | 31 | 22 | 0 | 0 | 0 | 0 | 0 | 0 | 9 | 5 | 8 | 4 | 208 | 149 | 1 | 1 | 0.107 | 0.074 | 0.589 | 0.023 | 0.032 | NA | 0 | 0 | 0 | 0 | 0.165 |
| SE | 61482 | ENSG00000018699 | TTC27 | chr2 | + | 32811023 | 32811221 | 32786983 | 32787149 | 32812503 | 32812615 | 130 | 75 | 84 | 103 | 101 | 0 | 0 | 0 | 3 | 1 | 40 | 90 | 38 | 73 | 19 | 0 | 0 | 0 | 0 | 0 | 298 | 149 | 1 | 1 | 1 | 1 | 1 | 0.945 | 0.981 | 1 | 1 | 1 | 1 | 1 | -0.015 |
| SE | 61483 | ENSG00000018699 | TTC27 | chr2 | + | 32811023 | 32811221 | 32786983 | 32787149 | 32817456 | 32817557 | 48 | 38 | 40 | 44 | 47 | 2 | 2 | 8 | 1 | 6 | 0 | 51 | 12 | 23 | 11 | 0 | 0 | 0 | 0 | 0 | 298 | 149 | 1 | 1 | 0.923 | 0.905 | 0.714 | 0.957 | 0.797 | NA | 1 | | 1 | 1 | -0.141 |
| SE | 61485 | ENSG00000018699 | TTC27 | chr2 | + | 32812503 | 32812615 | 32811023 | 32811221 | 32817456 | 32817557 | 156 | 79 | 115 | 131 | 120 | 0 | 1 | 1 | 0 | 0 | 76 | 75 | 54 | 99 | 22 | 0 | 4 | 0 | 0 | 0 | 261 | 149 | 1 | 1 | 1 | 0.978 | 0.985 | 1 | 1 | 1 | 0.915 | 1 | | 1 | 0.01 |
| SE | 61486 | ENSG00000124588 | NQO2 | chr6 | + | 2989527 | 2989710 | 2987986 | 2988031 | 2993930 | 2994053 | 0 | 0 | 0 | 3 | 1 | 1 | 0 | 0 | 0 | 0 | 0 | 0 | 0 | 9 | 13 | 0 | 0 | 0 | 0 | 0 | 298 | 149 | 1 | 1 | 0 | NA | NA | 1 | 1 | NA | NA | NA | 1 | 1 | -0.333 |
| SE | 61490 | ENSG00000124588 | NQO2 | chr6 | + | 3003668 | 3003789 | 2999893 | 3000085 | 3004494 | 3004651 | 1 | 0 | 0 | 2 | 0 | 1 | 0 | 0 | 0 | 0 | 25 | 7 | 1 | 3 | 7 | 0 | 5 | 0 | 5 | 0 | 270 | 149 | 1 | 1 | 0.356 | NA | NA | 1 | NA | 1 | 0.436 | 1 | 0.249 | 1 | -0.059 |
| SE | 61492 | ENSG00000124588 | NQO2 | chr6 | + | 3003668 | 3003789 | 3002049 | 3002286 | 3006467 | 3006559 | 1 | 5 | 4 | 0 | 1 | 0 | 0 | 0 | 0 | 1 | 3 | 6 | 8 | 6 | 0 | 0 | 5 | 0 | 4 | 7 | 270 | 149 | 1 | 1 | 1 | 1 | 1 | NA | 0.356 | 1 | 0.398 | 1 | 0.453 | 0 | 0.269 |
| SE | 61498 | ENSG00000124588 | NQO2 | chr6 | + | 3010024 | 3010189 | 3006467 | 3006559 | 3012543 | 3012561 | 136 | 33 | 88 | 131 | 133 | 0 | 0 | 1 | 0 | 0 | 102 | 183 | 84 | 424 | 324 | 0 | 0 | 0 | 0 | 0 | 298 | 149 | 1 | 1 | 1 | 1 | 0.978 | 1 | 1 | 1 | 1 | 1 | 1 | 1 | -0.004 |
| SE | 61500 | ENSG00000124588 | NQO2 | chr6 | + | 3016883 | 3016985 | 3012543 | 3012674 | 3019478 | 3019755 | 54 | 15 | 32 | 50 | 39 | 2 | 0 | 0 | 2 | 0 | 59 | 89 | 32 | 244 | 134 | 0 | 0 | 0 | 0 | 0 | 251 | 149 | 1 | 1 | 0.941 | 1 | 1 | 0.937 | 1 | 1 | 1 | 1 | 1 | 1 | -0.024 |
| SE | 61501 | ENSG00000213199 | ASIC3 | chr7 | + | 151051171 | 151051319 | 151051038 | 151051095 | 151051809 | 151051901 | 0 | 0 | 3 | 0 | 1 | 0 | 0 | 0 | 0 | 0 | 0 | 1 | 0 | 0 | 0 | 0 | 1 | 0 | 0 | 0 | 297 | 149 | 1 | 1 | NA | NA | 1 | NA | 1 | NA | 0.334 | NA | NA | NA | 0.666 |
| SE | 61502 | ENSG00000213199 | ASIC3 | chr7 | + | 151052415 | 151052474 | 151052165 | 151052237 | 151052573 | 151052753 | 0 | 0 | 0 | 4 | 4 | 0 | 0 | 0 | 0 | 0 | 0 | 1 | 0 | 2 | 0 | 0 | 1 | 0 | 0 | 0 | 208 | 149 | 1 | 1 | NA | NA | NA | 1 | 1 | NA | 0.417 | NA | 1 | NA | 0.291 |
| SE | 61508 | ENSG00000083520 | DIS3 | chr13 | - | 72761362 | 72761521 | 72760528 | 72760651 | 72761645 | 72761814 | 240 | 201 | 179 | 202 | 445 | 0 | 0 | 0 | 2 | 0 | 27 | 83 | 139 | 16 | 4 | 0 | 0 | 0 | 0 | 0 | 298 | 149 | 1 | 1 | 1 | 1 | 1 | 0.981 | 1 | 1 | 1 | 1 | 1 | 1 | -0.004 |
| SE | 61510 | ENSG00000083520 | DIS3 | chr13 | - | 72775924 | 72776092 | 72775210 | 72775375 | 72777419 | 72777493 | 279 | 225 | 199 | 255 | 324 | 1 | 0 | 2 | 0 | 0 | 59 | 178 | 224 | 67 | 54 | 0 | 0 | 0 | 0 | 0 | 298 | 149 | 1 | 1 | 0.993 | 1 | 0.98 | 1 | 1 | 1 | 1 | 1 | 1 | 1 | -0.005 |
| SE | 61512 | ENSG00000083520 | DIS3 | chr13 | - | 72780845 | 72781003 | 72778186 | 72778380 | 72781288 | 72781356 | 40 | 23 | 27 | 36 | 42 | 12 | 7 | 11 | 5 | 6 | 2 | 19 | 32 | 0 | 4 | 0 | 2 | 1 | 0 | 2 | 298 | 149 | 1 | 1 | 0.625 | 0.622 | 0.551 | 0.783 | 0.778 | 1 | 0.826 | 0.941 | NA | 0.5 | -0.145 |
| SE | 61514 | ENSG00000083520 | DIS3 | chr13 | - | 72781288 | 72781356 | 72778186 | 72778380 | 72781604 | 72781925 | 19 | 8 | 14 | 14 | 6 | 1 | 0 | 5 | 1 | 0 | 0 | 3 | 4 | 0 | 2 | 0 | 0 | 0 | 0 | 0 | 217 | 149 | 1 | 1 | 0.929 | 1 | 0.658 | 0.906 | 1 | NA | 1 | 1 | NA | 1 | -0.101 |
| SE | 61524 | ENSG00000138757 | G3BP2 | chr4 | - | 75654592 | 75654745 | 75648638 | 75648741 | 75655065 | 75655246 | 5 | 0 | 2 | 3 | 7 | 450 | 348 | 421 | 563 | 623 | 0 | 1 | 0 | 0 | 0 | 341 | 397 | 777 | 502 | 120 | 298 | 149 | 1 | 1 | 0.006 | 0 | 0.002 | 0.003 | 0.006 | 0 | 0.001 | 0 | 0 | 0 | 0.003 |
| SE | 61529 | ENSG00000138757 | G3BP2 | chr4 | - | 75656923 | 75657014 | 75655767 | 75655870 | 75658842 | 75658924 | 78 | 59 | 88 | 89 | 213 | 0 | 0 | 1 | 0 | 3 | 32 | 80 | 118 | 65 | 0 | 0 | 0 | 0 | 1 | 0 | 240 | 149 | 1 | 1 | 1 | 1 | 0.982 | 1 | 0.978 | 1 | 1 | 1 | 0.976 | NA | -0.002 |
| SE | 61531 | ENSG00000138757 | G3BP2 | chr4 | - | 75657556 | 75657730 | 75655767 | 75655870 | 75658842 | 75658924 | 282 | 211 | 293 | 347 | 384 | 0 | 0 | 1 | 0 | 3 | 117 | 295 | 337 | 282 | 100 | 0 | 0 | 0 | 1 | 0 | 298 | 149 | 1 | 1 | 1 | 1 | 0.993 | 1 | 0.985 | 1 | 1 | 1 | 0.993 | 1 | -0.003 |
| SE | 61533 | ENSG00000138757 | G3BP2 | chr4 | - | 75657556 | 75657730 | 75656923 | 75657014 | 75658842 | 75658924 | 654 | 543 | 626 | 790 | 1185 | 0 | 4 | 0 | 0 | 0 | 278 | 602 | 938 | 629 | 202 | 0 | 4 | 2 | 0 | 0 | 298 | 149 | 1 | 1 | 1 | 0.985 | 1 | 1 | 1 | 1 | 0.987 | 0.996 | 1 | 1 | 0 |
| SE | 61534 | ENSG00000138757 | G3BP2 | chr4 | - | 75657800 | 75657893 | 75655767 | 75655870 | 75658842 | 75658924 | 1 | 0 | 0 | 0 | 0 | 0 | 0 | 1 | 0 | 3 | 0 | 0 | 0 | 0 | 0 | 0 | 0 | 0 | 1 | 0 | 242 | 149 | 1 | 1 | 1 | NA | 0 | NA | 0 | NA | NA | NA | 0 | NA | 0.333 |
| SE | 61535 | ENSG00000138757 | G3BP2 | chr4 | - | 75657800 | 75657893 | 75656923 | 75657014 | 75658842 | 75658924 | 1 | 0 | 0 | 0 | 0 | 0 | 4 | 0 | 0 | 0 | 0 | 0 | 0 | 0 | 0 | 0 | 4 | 2 | 0 | 0 | 242 | 149 | 1 | 1 | 1 | 0 | NA | NA | NA | NA | 0 | 0 | NA | NA | 0.5 |
| SE | 61538 | ENSG00000138757 | G3BP2 | chr4 | - | 75658842 | 75658924 | 75657556 | 75657730 | 75661930 | 75662049 | 368 | 251 | 397 | 445 | 456 | 0 | 0 | 0 | 0 | 1 | 125 | 283 | 374 | 308 | 122 | 0 | 0 | 0 | 0 | 0 | 231 | 149 | 1 | 1 | 1 | 1 | 1 | 1 | 0.997 | 1 | 1 | 1 | 1 | 1 | -0.001 |
| SE | 61541 | ENSG00000138757 | G3BP2 | chr4 | - | 75661930 | 75662049 | 75658842 | 75658924 | 75673207 | 75673298 | 331 | 192 | 310 | 371 | 254 | 0 | 0 | 0 | 0 | 0 | 53 | 207 | 273 | 224 | 81 | 0 | 0 | 1 | 0 | 3 | 268 | 149 | 1 | 1 | 1 | 1 | 1 | 1 | 1 | 1 | 1 | 0.993 | 1 | 0.938 | 0.014 |
| SE | 61543 | ENSG00000138757 | G3BP2 | chr4 | - | 75661930 | 75662073 | 75658842 | 75658924 | 75673207 | 75673414 | 212 | 137 | 218 | 249 | 211 | 0 | 0 | 0 | 0 | 0 | 39 | 167 | 205 | 173 | 63 | 0 | 0 | 1 | 0 | 3 | 292 | 149 | 1 | 1 | 1 | 1 | 1 | 1 | 1 | 1 | 1 | 0.991 | 1 | 0.915 | 0.019 |
| SE | 61544 | ENSG00000138757 | G3BP2 | chr4 | - | 75662212 | 75662520 | 75661930 | 75662049 | 75673207 | 75673364 | 2 | 1 | 0 | 0 | 0 | 126 | 59 | 99 | 129 | 53 | 0 | 0 | 1 | 0 | 0 | 14 | 40 | 68 | 55 | 18 | 298 | 149 | 1 | 1 | 0.008 | 0.008 | 0 | 0 | 0 | 0 | 0 | 0.007 | 0 | 0 | 0.002 |
| SE | 61545 | ENSG00000138757 | G3BP2 | chr4 | - | 75662212 | 75662525 | 75661930 | 75662049 | 75673018 | 75673157 | 2 | 1 | 0 | 0 | 0 | 2 | 0 | 4 | 7 | 4 | 0 | 0 | 1 | 0 | 0 | 0 | 7 | 3 | 1 | 0 | 298 | 149 | 1 | 1 | 0.333 | 1 | 0 | 0 | 0 | NA | 0 | 0.143 | 0 | NA | 0.219 |
| SE | 61565 | ENSG00000138757 | G3BP2 | chr4 | - | 75720876 | 75720969 | 75661930 | 75662049 | 75724055 | 75724176 | 0 | 0 | 0 | 0 | 2 | 0 | 0 | 4 | 0 | 1 | 0 | 0 | 0 | 0 | 0 | 0 | 0 | 1 | 0 | 0 | 242 | 149 | 1 | 1 | NA | NA | 0 | NA | 0.552 | NA | NA | 0 | NA | NA | 0.276 |
| SE | 61566 | ENSG00000138757 | G3BP2 | chr4 | - | 75720876 | 75720969 | 75661930 | 75662049 | 75724275 | 75724438 | 0 | 0 | 1 | 2 | 0 | 7 | 4 | 13 | 9 | 4 | 0 | 0 | 0 | 0 | 0 | 0 | 1 | 0 | 0 | 0 | 242 | 149 | 1 | 1 | 0 | 0 | 0.045 | 0.12 | 0 | NA | 0 | NA | NA | NA | 0.033 |
| SE | 61571 | ENSG00000138757 | G3BP2 | chr4 | - | 75724055 | 75724176 | 75661930 | 75662049 | 75724275 | 75724419 | 4 | 0 | 10 | 3 | 26 | 7 | 4 | 13 | 9 | 4 | 0 | 4 | 2 | 0 | 0 | 0 | 1 | 0 | 0 | 0 | 270 | 149 | 1 | 1 | 0.24 | 0 | 0.298 | 0.155 | 0.782 | NA | 0.688 | 1 | NA | NA | -0.549 |
| SE | 61575 | ENSG00000138757 | G3BP2 | chr4 | - | 75724055 | 75724176 | 75720876 | 75720969 | 75724275 | 75724419 | 4 | 0 | 6 | 3 | 27 | 0 | 0 | 1 | 2 | 0 | 0 | 4 | 1 | 0 | 0 | 0 | 0 | 0 | 0 | 0 | 270 | 149 | 1 | 1 | 1 | NA | 0.768 | 0.453 | 1 | NA | 1 | 1 | NA | NA | -0.195 |
| SE | 61577 | ENSG00000131876 | SNRPA1 | chr15 | - | 101287655 | 101287702 | 101285725 | 101287010 | 101291961 | 101292040 | 249 | 210 | 288 | 290 | 254 | 4 | 0 | 0 | 1 | 0 | 157 | 302 | 329 | 243 | 71 | 0 | 0 | 0 | 3 | 0 | 196 | 149 | 1 | 1 | 0.979 | 1 | 1 | 0.995 | 1 | 1 | 1 | 1 | 0.984 | 1 | -0.002 |
| SE | 61578 | ENSG00000131876 | SNRPA1 | chr15 | - | 101287655 | 101287702 | 101285725 | 101287010 | 101293024 | 101293172 | 65 | 65 | 65 | 90 | 72 | 2 | 0 | 0 | 0 | 0 | 52 | 99 | 101 | 86 | 26 | 0 | 0 | 1 | 1 | 0 | 196 | 149 | 1 | 1 | 0.961 | 1 | 1 | 1 | 1 | 1 | 1 | 0.987 | 0.985 | 1 | -0.002 |
| SE | 61581 | ENSG00000131876 | SNRPA1 | chr15 | - | 101291961 | 101292040 | 101285725 | 101287010 | 101293024 | 101293172 | 62 | 52 | 78 | 58 | 86 | 2 | 0 | 0 | 0 | 0 | 25 | 61 | 104 | 60 | 12 | 0 | 0 | 1 | 1 | 0 | 228 | 149 | 1 | 1 | 0.953 | 1 | 1 | 1 | 1 | 1 | 1 | 0.985 | 0.975 | 1 | -0.001 |
| SE | 61582 | ENSG00000131876 | SNRPA1 | chr15 | - | 101291961 | 101292040 | 101285725 | 101287010 | 101295096 | 101295204 | 4 | 0 | 0 | 1 | 0 | 1 | 0 | 3 | 5 | 0 | 0 | 0 | 0 | 0 | 0 | 6 | 0 | 30 | 6 | 0 | 228 | 149 | 1 | 1 | 0.723 | NA | 0 | 0.116 | NA | 0 | NA | 0 | 0 | NA | 0.28 |
| SE | 61583 | ENSG00000131876 | SNRPA1 | chr15 | - | 101291961 | 101292040 | 101287655 | 101287702 | 101293024 | 101293172 | 241 | 192 | 300 | 256 | 268 | 1 | 0 | 0 | 2 | 0 | 130 | 264 | 332 | 218 | 57 | 0 | 0 | 0 | 1 | 0 | 228 | 149 | 1 | 1 | 0.994 | 1 | 1 | 0.988 | 1 | | 1 | 1 | 0.993 | 1 | -0.002 |
| SE | 61585 | ENSG00000131876 | SNRPA1 | chr15 | - | 101293024 | 101293172 | 101285725 | 101287010 | 101295096 | 101295204 | 51 | 22 | 45 | 32 | 22 | 1 | 0 | 3 | 5 | 0 | 4 | 43 | 48 | 17 | 0 | 6 | 0 | 30 | 6 | 0 | 297 | 149 | 1 | 1 | 0.962 | 1 | 0.883 | 0.763 | 1 | 0.251 | 1 | 0.445 | 0.587 | NA | 0.351 |
| SE | 61586 | ENSG00000131876 | SNRPA1 | chr15 | - | 101293024 | 101293172 | 101287655 | 101287702 | 101295096 | 101295204 | 50 | 22 | 45 | 34 | 22 | 1 | 6 | 5 | 8 | 0 | 4 | 43 | 47 | 17 | 0 | 0 | 4 | 8 | 9 | 0 | 297 | 149 | 1 | 1 | 0.962 | 0.648 | 0.819 | 0.681 | 1 | 1 | 0.844 | 0.747 | 0.487 | NA | 0.052 |
| SE | 61587 | ENSG00000131876 | SNRPA1 | chr15 | - | 101293024 | 101293172 | 101291961 | 101292040 | 101295096 | 101295204 | 186 | 135 | 226 | 192 | 214 | 0 | 0 | 0 | 2 | 0 | 69 | 242 | 266 | 163 | 39 | 0 | 0 | 0 | 0 | 0 | 297 | 149 | 1 | 1 | 1 | 1 | 1 | 0.98 | 1 | 1 | 1 | 1 | 1 | 1 | -0.004 |
| SE | 61589 | ENSG00000114021 | NIT2 | chr3 | + | 100339814 | 100339935 | 100339086 | 100339205 | 100341072 | 100341143 | 78 | 44 | 61 | 65 | 58 | 1 | 0 | 2 | 1 | 0 | 42 | 12 | 28 | 65 | 19 | 0 | 0 | 0 | 0 | 0 | 270 | 149 | 1 | 1 | 0.977 | 1 | 0.944 | 0.973 | 1 | 1 | 1 | 1 | 1 | 1 | -0.021 |
| SE | 61590 | ENSG00000114021 | NIT2 | chr3 | + | 100339814 | 100339954 | 100339086 | 100339205 | 100341072 | 100341161 | 35 | 22 | 16 | 27 | 6 | 1 | 0 | 2 | 1 | 0 | 10 | 3 | 11 | 22 | 2 | 0 | 0 | 0 | 0 | 0 | 289 | 149 | 1 | 1 | 0.947 | 1 | 0.805 | 0.933 | 1 | 1 | 1 | 1 | 1 | 1 | -0.063 |
| SE | 61591 | ENSG00000114021 | NIT2 | chr3 | + | 100345584 | 100345678 | 100341072 | 100341161 | 100346180 | 100346255 | 122 | 45 | 83 | 67 | 88 | 1 | 0 | 2 | 0 | 0 | 73 | 44 | 34 | 108 | 46 | 0 | 0 | 0 | 0 | 0 | 243 | 149 | 1 | 1 | 0.987 | 1 | 0.962 | 1 | 1 | 1 | 1 | 1 | 1 | 1 | -0.01 |
| SE | 61594 | ENSG00000114021 | NIT2 | chr3 | + | 100346180 | 100346255 | 100341072 | 100341161 | 100348802 | 100348881 | 37 | 10 | 22 | 13 | 24 | 0 | 0 | 2 | 1 | 3 | 0 | 21 | 18 | 23 | 3 | 0 | 0 | 1 | 0 | 0 | 224 | 149 | 1 | 1 | 1 | 1 | 0.88 | 0.896 | 0.842 | NA | 1 | 0.923 | 1 | 1 | -0.057 |
| SE | 61595 | ENSG00000114021 | NIT2 | chr3 | + | 100346180 | 100346255 | 100345584 | 100345678 | 100348802 | 100348881 | 91 | 30 | 60 | 39 | 59 | 0 | 1 | 0 | 0 | 0 | 33 | 51 | 35 | 72 | 28 | 0 | 0 | 0 | 0 | 0 | 224 | 149 | 1 | 1 | 1 | 0.952 | 1 | 1 | 1 | 1 | 1 | 1 | 1 | 1 | -0.01 |
| SE | 61596 | ENSG00000061273 | HDAC7 | chr12 | - | 47784078 | 47784217 | 47782721 | 47783886 | 47785386 | 47785471 | 186 | 101 | 237 | 171 | 231 | 0 | 0 | 0 | 0 | 0 | 186 | 167 | 61 | 144 | 303 | 0 | 7 | 0 | 0 | 0 | 288 | 149 | 1 | 1 | 1 | 1 | 1 | 1 | 1 | 1 | 0.925 | 1 | 1 | 1 | 0.015 |
| SE | 61600 | ENSG00000061273 | HDAC7 | chr12 | - | 47789260 | 47789348 | 47788044 | 47788164 | 47789522 | 47789578 | 89 | 41 | 94 | 132 | 80 | 0 | 0 | 0 | 2 | 0 | 205 | 63 | 46 | 60 | 106 | 0 | 0 | 0 | 0 | 0 | 237 | 149 | 1 | 1 | 1 | 1 | 1 | 0.976 | 1 | 1 | 1 | 1 | 1 | 1 | -0.005 |
| SE | 61606 | ENSG00000061273 | HDAC7 | chr12 | - | 47796214 | 47796298 | 47795586 | 47795767 | 47797016 | 47797142 | 15 | 7 | 21 | 15 | 4 | 2 | 3 | 0 | 0 | 0 | 27 | 18 | 12 | 10 | 34 | 0 | 4 | 0 | 0 | 3 | 233 | 149 | 1 | 1 | 0.827 | 0.599 | 1 | 1 | 1 | 1 | 0.742 | 1 | 1 | 0.879 | -0.039 |
| SE | 61609 | ENSG00000061273 | HDAC7 | chr12 | - | 47802223 | 47802274 | 47798784 | 47798972 | 47819766 | 47819866 | 4 | 2 | 8 | 4 | 4 | 3 | 0 | 1 | 3 | 0 | 0 | 7 | 8 | 7 | 2 | 0 | 8 | 0 | 4 | 0 | 200 | 149 | 1 | 1 | 0.498 | 1 | 0.856 | 0.498 | 1 | NA | 0.395 | 1 | 0.566 | 1 | 0.03 |
| SE | 61613 | ENSG00000106804 | C5 | chr9 | - | 120957284 | 120957368 | 120953729 | 120953868 | 120960247 | 120960337 | 9 | 8 | 9 | 8 | 19 | 0 | 0 | 0 | 0 | 4 | 0 | 1 | 1 | 0 | 0 | 0 | 0 | 0 | 0 | 0 | 233 | 149 | 1 | 1 | 1 | 1 | 1 | 1 | 0.752 | NA | 1 | 1 | NA | NA | -0.05 |
| SE | 61614 | ENSG00000106804 | C5 | chr9 | - | 120982654 | 120982814 | 120981843 | 120981939 | 120989045 | 120989121 | 28 | 10 | 12 | 23 | 36 | 0 | 0 | 0 | 0 | 0 | 0 | 4 | 11 | 5 | 0 | 0 | 4 | 0 | 0 | 0 | 298 | 149 | 1 | 1 | 1 | 1 | 1 | 1 | 1 | NA | 0.333 | 1 | 1 | NA | 0.222 |
| SE | 61615 | ENSG00000106804 | C5 | chr9 | - | 121032112 | 121032195 | 121030396 | 121030487 | 121034802 | 121034826 | 23 | 17 | 16 | 12 | 40 | 0 | 0 | 0 | 1 | 1 | 8 | 5 | 9 | 12 | 8 | 0 | 0 | 0 | 0 | 0 | 232 | 149 | 1 | 1 | 1 | 1 | 1 | 0.885 | 0.963 | 1 | 1 | 1 | 1 | 1 | -0.03 |
| SE | 61616 | ENSG00000106804 | C5 | chr9 | - | 121043003 | 121043166 | 121037880 | 121037951 | 121046190 | 121046383 | 32 | 28 | 20 | 12 | 59 | 0 | 0 | 0 | 0 | 0 | 13 | 19 | 18 | 9 | 24 | 0 | 0 | 2 | 0 | 0 | 298 | 149 | 1 | 1 | 1 | 1 | 1 | 1 | 1 | 1 | 1 | 0.818 | 1 | 1 | 0.036 |
| SE | 61620 | ENSG00000234127 | TRIM26 | chr6 | - | 30198665 | 30199153 | 30190012 | 30190035 | 30213304 | 30213385 | 2 | 0 | 0 | 2 | 0 | 0 | 0 | 0 | 0 | 0 | 16 | 0 | 0 | 1 | 13 | 0 | 0 | 0 | 1 | 5 | 298 | 149 | 1 | 1 | 1 | NA | NA | 1 | NA | 1 | NA | NA | 0.333 | 0.565 | 0.367 |
| SE | 61621 | ENSG00000234127 | TRIM26 | chr6 | - | 30201034 | 30201083 | 30198665 | 30199153 | 30204655 | 30204765 | 16 | 5 | 15 | 26 | 7 | 0 | 2 | 0 | 0 | 0 | 9 | 11 | 17 | 18 | 21 | 0 | 0 | 0 | 0 | 0 | 198 | 149 | 1 | 1 | 1 | 0.653 | 1 | 1 | 1 | 1 | 1 | 1 | 1 | 1 | -0.069 |
| SE | 61623 | ENSG00000234127 | TRIM26 | chr6 | - | 30201034 | 30201138 | 30198665 | 30199153 | 30204655 | 30204765 | 198 | 103 | 133 | 134 | 130 | 0 | 2 | 0 | 0 | 0 | 94 | 118 | 115 | 135 | 161 | 0 | 0 | 0 | 0 | 0 | 253 | 149 | 1 | 1 | 1 | 0.968 | 1 | 1 | 1 | 1 | 1 | 1 | 1 | 1 | -0.006 |
| SE | 61626 | ENSG00000234127 | TRIM26 | chr6 | - | 30201034 | 30201138 | 30198737 | 30199147 | 30213304 | 30213427 | 10 | 4 | 6 | 9 | 4 | 1 | 0 | 0 | 0 | 0 | 5 | 15 | 6 | 4 | 0 | 0 | 0 | 0 | 0 | 0 | 253 | 149 | 1 | 1 | 0.855 | 1 | 1 | 1 | 1 | 1 | 1 | 1 | 1 | NA | -0.029 |
| SE | 61628 | ENSG00000234127 | TRIM26 | chr6 | - | 30204655 | 30204765 | 30198737 | 30199147 | 30213304 | 30213365 | 42 | 11 | 25 | 26 | 40 | 1 | 0 | 0 | 0 | 0 | 11 | 20 | 32 | 18 | 25 | 0 | 0 | 0 | 0 | 0 | 259 | 149 | 1 | 1 | 0.96 | 1 | 1 | 1 | 1 | 1 | 1 | 1 | 1 | 1 | -0.008 |
| SE | 61629 | ENSG00000234127 | TRIM26 | chr6 | - | 30204655 | 30204765 | 30201034 | 30201083 | 30213304 | 30213365 | 42 | 11 | 25 | 26 | 40 | 1 | 0 | 0 | 3 | 0 | 11 | 20 | 32 | 18 | 25 | 0 | 3 | 0 | 0 | 0 | 259 | 149 | 1 | 1 | 1 | 1 | 1 | 0.833 | 1 | 1 | 0.793 | 1 | 1 | 1 | 0.008 |
| SE | 61631 | ENSG00000213281 | NRAS | chr1 | - | 114709568 | 114709728 | 114708530 | 114708654 | 114713799 | 114713978 | 391 | 303 | 314 | 481 | 632 | 0 | 1 | 0 | 0 | 0 | 134 | 331 | 419 | 260 | 119 | 0 | 0 | 0 | 0 | 0 | 298 | 149 | 1 | 1 | 1 | 0.993 | 1 | 1 | 1 | 1 | 1 | 1 | 1 | 1 | -0.001 |
| SE | 61632 | ENSG00000213281 | NRAS | chr1 | - | 114709568 | 114709728 | 114708530 | 114708654 | 114716049 | 114716177 | 158 | 108 | 115 | 203 | 343 | 0 | 0 | 0 | 0 | 1 | 57 | 101 | 144 | 80 | 45 | 0 | 0 | 0 | 0 | 0 | 298 | 149 | 1 | 1 | 1 | 1 | 1 | 1 | 0.994 | 1 | 1 | 1 | 1 | 1 | -0.001 |
| SE | 61633 | ENSG00000213281 | NRAS | chr1 | - | 114713799 | 114713978 | 114708530 | 114708654 | 114716049 | 114716177 | 217 | 155 | 143 | 211 | 236 | 0 | 0 | 0 | 0 | 1 | 81 | 199 | 190 | 120 | 66 | 0 | 0 | 0 | 0 | 0 | 298 | 149 | 1 | 1 | 1 | 1 | 1 | 1 | 0.992 | 1 | 1 | 1 | 1 | 1 | -0.002 |
| SE | 61634 | ENSG00000213281 | NRAS | chr1 | - | 114713799 | 114713978 | 114709568 | 114709728 | 114716049 | 114716177 | 450 | 350 | 342 | 492 | 525 | 0 | 1 | 0 | 3 | 0 | 158 | 429 | 465 | 300 | 140 | 0 | 0 | 0 | 0 | 0 | 298 | 149 | 1 | 1 | 1 | 0.994 | 1 | 0.988 | 1 | 1 | 1 | 1 | 1 | 1 | -0.004 |
| SE | 61635 | ENSG00000213281 | NRAS | chr1 | - | 114716049 | 114716177 | 114713799 | 114713978 | 114716657 | 114716771 | 370 | 293 | 287 | 423 | 330 | 0 | 1 | 1 | 0 | 0 | 158 | 386 | 409 | 239 | 109 | 0 | 0 | 0 | 0 | 0 | 277 | 149 | 1 | 1 | 1 | 0.994 | 0.994 | 1 | 1 | 1 | 1 | 1 | 1 | 1 | -0.002 |
| SE | 61637 | ENSG00000009307 | CSDE1 | chr1 | - | 114720538 | 114720717 | 114718612 | 114718745 | 114723882 | 114724002 | 1992 | 1640 | 1978 | 2042 | 4134 | 1 | 0 | 0 | 0 | 0 | 406 | 1707 | 2556 | 1612 | 80 | 0 | 0 | 0 | 0 | 0 | 298 | 149 | 1 | 1 | 0.999 | 1 | 1 | 1 | 1 | 1 | 1 | 1 | 1 | 1 | 0 |
| SE | 61638 | ENSG00000009307 | CSDE1 | chr1 | - | 114720538 | 114720717 | 114718612 | 114718745 | 114726210 | 114726386 | 3 | 2 | 3 | 0 | 3 | 0 | 0 | 0 | 0 | 0 | 0 | 23 | 1 | 0 | 0 | 0 | 0 | 3 | 0 | 0 | 298 | 149 | 1 | 1 | 1 | 1 | 1 | NA | 1 | NA | 1 | 0.143 | NA | NA | 0.428 |
| SE | 61639 | ENSG00000009307 | CSDE1 | chr1 | - | 114723882 | 114724002 | 114718612 | 114718745 | 114726210 | 114726348 | 4 | 0 | 2 | 0 | 0 | 0 | 0 | 0 | 0 | 0 | 3 | 10 | 4 | 9 | 0 | 0 | 0 | 3 | 0 | 0 | 269 | 149 | 1 | 1 | 1 | NA | 1 | NA | NA | 1 | 1 | 0.425 | 1 | NA | 0.144 |
| SE | 61640 | ENSG00000009307 | CSDE1 | chr1 | - | 114723882 | 114724002 | 114720215 | 114720717 | 114725220 | 114725333 | 3739 | 2965 | 3662 | 3774 | 7395 | 1 | 4 | 2 | 3 | 5 | 733 | 2974 | 4621 | 2737 | 183 | 0 | 3 | 10 | 12 | 0 | 269 | 149 | 1 | 1 | 1 | 0.998 | 0.999 | 0.999 | 0.999 | 1 | 0.998 | 0.996 | 0.992 | 1 | 0.002 |
| SE | 61641 | ENSG00000009307 | CSDE1 | chr1 | - | 114723882 | 114724002 | 114720215 | 114720717 | 114726210 | 114726348 | 1722 | 1389 | 1699 | 1788 | 3381 | 2 | 0 | 0 | 0 | 0 | 387 | 1474 | 2256 | 1392 | 71 | 0 | 0 | 0 | 0 | 0 | 269 | 149 | 1 | 1 | 0.998 | 1 | 1 | 1 | 1 | 1 | 1 | 1 | 1 | 1 | 0 |
| SE | 61642 | ENSG00000009307 | CSDE1 | chr1 | - | 114725220 | 114725333 | 114720215 | 114720717 | 114726210 | 114726348 | 1805 | 1473 | 1655 | 1795 | 2741 | 2 | 0 | 0 | 0 | 0 | 316 | 1646 | 2276 | 1165 | 69 | 0 | 0 | 0 | 0 | 0 | 262 | 149 | 1 | 1 | 0.998 | 1 | 1 | 1 | 1 | 1 | 1 | 1 | 1 | 1 | 0 |
| SE | 61643 | ENSG00000009307 | CSDE1 | chr1 | - | 114725220 | 114725333 | 114723882 | 114724002 | 114726210 | 114726348 | 3815 | 3031 | 3587 | 3777 | 6765 | 3 | 0 | 2 | 0 | 5 | 669 | 3183 | 4647 | 2495 | 189 | 3 | 14 | 8 | 9 | 0 | 262 | 149 | 1 | 1 | 0.999 | 1 | 0.999 | 1 | 0.999 | 0.992 | 0.992 | 0.997 | 0.994 | 1 | 0.004 |
| SE | 61644 | ENSG00000009307 | CSDE1 | chr1 | - | 114726210 | 114726386 | 114723882 | 114724002 | 114732603 | 114732816 | 5 | 0 | 2 | 0 | 5 | 0 | 0 | 0 | 0 | 0 | 3 | 14 | 8 | 9 | 0 | 0 | 1 | 1 | 0 | 0 | 298 | 149 | 1 | 1 | 1 | NA | 1 | NA | 1 | 1 | 0.875 | 0.8 | 1 | NA | 0.081 |
| SE | 61645 | ENSG00000009307 | CSDE1 | chr1 | - | 114726210 | 114726386 | 114725220 | 114725333 | 114726982 | 114727090 | 4288 | 3280 | 3982 | 4259 | 6723 | 0 | 0 | 0 | 0 | 0 | 728 | 3502 | 4846 | 2599 | 157 | 0 | 8 | 0 | 0 | 0 | 298 | 149 | 1 | 1 | 1 | 1 | 1 | 1 | 1 | 1 | 0.995 | 1 | 1 | 1 | 0.001 |
| SE | 61646 | ENSG00000009307 | CSDE1 | chr1 | - | 114726982 | 114727090 | 114726210 | 114726386 | 114730257 | 114730422 | 3213 | 2384 | 3083 | 3221 | 3873 | 0 | 0 | 1 | 0 | 0 | 494 | 2387 | 3191 | 1763 | 109 | 0 | 0 | 0 | 0 | 0 | 257 | 149 | 1 | 1 | 1 | 1 | 0.999 | 1 | 1 | 1 | 1 | 1 | 1 | 1 | 0 |
| SE | 61647 | ENSG00000009307 | CSDE1 | chr1 | - | 114726982 | 114727090 | 114726210 | 114726386 | 114732603 | 114732816 | 1297 | 914 | 1123 | 1243 | 1844 | 2 | 0 | 0 | 0 | 0 | 196 | 1033 | 1335 | 670 | 45 | 0 | 0 | 0 | 0 | 0 | 257 | 149 | 1 | 1 | 0.997 | 1 | 1 | 1 | 1 | 1 | 1 | 1 | 1 | 1 | -0.001 |
| SE | 61648 | ENSG00000009307 | CSDE1 | chr1 | - | 114730257 | 114730422 | 114726210 | 114726386 | 114732603 | 114732816 | 0 | 0 | 1 | 0 | 12 | 2 | 0 | 0 | 0 | 0 | 0 | 0 | 4 | 0 | 0 | 0 | 0 | 0 | 0 | 0 | 298 | 149 | 1 | 1 | 0 | NA | 1 | NA | 1 | NA | NA | 1 | NA | NA | -0.333 |
| SE | 61649 | ENSG00000009307 | CSDE1 | chr1 | - | 114730257 | 114730422 | 114726982 | 114727090 | 114730507 | 114730648 | 3334 | 2518 | 3298 | 3371 | 4408 | 0 | 1 | 1 | 1 | 6 | 483 | 2214 | 3118 | 1828 | 134 | 0 | 5 | 3 | 0 | 0 | 298 | 149 | 1 | 1 | 1 | 0.999 | 0.999 | 0.999 | 0.997 | 1 | 0.996 | 0.998 | 1 | 1 | 0 |
| SE | 61652 | ENSG00000009307 | CSDE1 | chr1 | - | 114730507 | 114730648 | 114730257 | 114730422 | 114732603 | 114732816 | 2432 | 1764 | 2303 | 2407 | 4112 | 0 | 0 | 0 | 0 | 12 | 307 | 1520 | 2045 | 1220 | 101 | 0 | 0 | 4 | 0 | 0 | 290 | 149 | 1 | 1 | 1 | 1 | 1 | 1 | 0.994 | 1 | 1 | 0.996 | 1 | 1 | 0 |
| SE | 61653 | ENSG00000009307 | CSDE1 | chr1 | - | 114732603 | 114732816 | 114730507 | 114730648 | 114733731 | 114733857 | 2442 | 1790 | 2292 | 2370 | 3880 | 0 | 5 | 1 | 2 | 0 | 322 | 1763 | 2026 | 1276 | 149 | 0 | 8 | 0 | 0 | 0 | 298 | 149 | 1 | 1 | 1 | 0.994 | 0.999 | 0.998 | 1 | 1 | 0.991 | 1 | 1 | 1 | 0 |
| SE | 61654 | ENSG00000009307 | CSDE1 | chr1 | - | 114733988 | 114734117 | 114733731 | 114733857 | 114734441 | 114734523 | 3426 | 2720 | 3143 | 3158 | 6830 | 0 | 0 | 0 | 0 | 0 | 574 | 2817 | 3283 | 1953 | 248 | 0 | 5 | 0 | 0 | 0 | 278 | 149 | 1 | 1 | 1 | 1 | 1 | 1 | 1 | 1 | 0.997 | 1 | 1 | 1 | 0.001 |
| SE | 61658 | ENSG00000009307 | CSDE1 | chr1 | - | 114739691 | 114739890 | 114737962 | 114738072 | 114757924 | 114757974 | 1032 | 736 | 995 | 1017 | 1980 | 0 | 0 | 1 | 0 | 0 | 160 | 849 | 1016 | 634 | 149 | 0 | 0 | 0 | 0 | 0 | 298 | 149 | 1 | 1 | 1 | 1 | 0.998 | 1 | 1 | 1 | 1 | 1 | 1 | 1 | 0 |
| SE | 61659 | ENSG00000009307 | CSDE1 | chr1 | - | 114741526 | 114741673 | 114739691 | 114739890 | 114749820 | 114750190 | 0 | 6 | 1 | 2 | 5 | 1129 | 727 | 1073 | 1144 | 2431 | 1 | 3 | 4 | 2 | 0 | 196 | 844 | 911 | 529 | 164 | 296 | 149 | 1 | 1 | 0 | 0.004 | 0 | 0.001 | 0.001 | 0.003 | 0.002 | 0.002 | 0.002 | 0 | -0.001 |
| SE | 61660 | ENSG00000009307 | CSDE1 | chr1 | - | 114749820 | 114750207 | 114737962 | 114738072 | 114757924 | 114757983 | 104 | 63 | 109 | 98 | 124 | 0 | 0 | 1 | 0 | 0 | 16 | 77 | 86 | 90 | 17 | 0 | 0 | 0 | 0 | 0 | 298 | 149 | 1 | 1 | 1 | 1 | 0.982 | 1 | 1 | 1 | 1 | 1 | 1 | 1 | -0.004 |
| SE | 61661 | ENSG00000009307 | CSDE1 | chr1 | - | 114749820 | 114750207 | 114739691 | 114739890 | 114757924 | 114757983 | 1220 | 777 | 1156 | 1227 | 2500 | 6 | 3 | 7 | 7 | 0 | 206 | 910 | 963 | 601 | 171 | 0 | 5 | 1 | 3 | 0 | 298 | 149 | 1 | 1 | 0.99 | 0.992 | 0.988 | 0.989 | 1 | 1 | 0.989 | 0.998 | 0.99 | 1 | -0.004 |
| SE | 61666 | ENSG00000227232 | WASH7P | chr1 | - | 18267 | 18366 | 17914 | 18061 | 24737 | 24891 | 25 | 13 | 29 | 16 | 18 | 1 | 0 | 0 | 0 | 0 | 26 | 7 | 15 | 13 | 27 | 0 | 0 | 0 | 0 | 0 | 248 | 149 | 1 | 1 | 0.938 | 1 | 1 | 1 | 1 | 1 | 1 | 1 | 1 | 1 | -0.012 |
| SE | 61667 | ENSG00000186652 | PRG2 | chr11 | - | 57389009 | 57389317 | 57388576 | 57388708 | 57389886 | 57389956 | 1 | 4 | 2 | 1 | 4 | 0 | 0 | 0 | 0 | 0 | 14 | 8 | 6 | 17 | 0 | 0 | 0 | 0 | 2 | 0 | 298 | 149 | 1 | 1 | 1 | 1 | 1 | 1 | 1 | 1 | 1 | 1 | 0.81 | NA | 0.047 |
| SE | 61668 | ENSG00000186652 | PRG2 | chr11 | - | 57389042 | 57389317 | 57388576 | 57388708 | 57389886 | 57389956 | 0 | 2 | 2 | 1 | 3 | 0 | 0 | 0 | 0 | 0 | 6 | 4 | 1 | 14 | 0 | 0 | 0 | 0 | 2 | 0 | 298 | 149 | 1 | 1 | NA | 1 | 1 | 1 | 1 | 1 | | 1 | 0.778 | NA | 0.055 |
| SE | 61669 | ENSG00000233757 | NA | chr2 | + | 95211617 | 95211720 | 95208647 | 95208736 | 95212192 | 95212282 | 6 | 9 | 13 | 15 | 14 | 0 | 0 | 1 | 0 | 0 | 0 | 5 | 3 | 2 | 0 | 0 | 4 | 0 | 1 | 0 | 252 | 149 | 1 | 1 | 1 | 1 | 0.885 | 1 | 1 | NA | 0.425 | 1 | 0.542 | NA | 0.321 |
| SE | 61670 | ENSG00000233757 | NA | chr2 | + | 95212192 | 95212282 | 95208647 | 95208736 | 95214335 | 95215603 | 5 | 3 | 2 | 10 | 8 | 0 | 3 | 0 | 0 | 0 | 0 | 4 | 7 | 1 | 0 | 0 | 0 | 0 | 0 | 0 | 239 | 149 | 1 | 1 | 1 | 0.384 | 1 | 1 | 1 | NA | 1 | 1 | 1 | NA | -0.123 |
| SE | 61672 | ENSG00000150938 | CRIM1 | chr2 | + | 36396613 | 36396787 | 36355777 | 36356623 | 36441257 | 36441500 | 56 | 20 | 19 | 31 | 20 | 1 | 0 | 0 | 0 | 0 | 1 | 26 | 21 | 10 | 5 | 0 | 0 | 0 | 0 | 0 | 298 | 149 | 1 | 1 | 0.966 | 1 | 1 | 1 | 1 | 1 | 1 | 1 | 1 | 1 | -0.007 |
| SE | 61673 | ENSG00000150938 | CRIM1 | chr2 | + | 36464533 | 36464655 | 36442614 | 36442735 | 36476888 | 36477071 | 27 | 11 | 15 | 10 | 14 | 0 | 0 | 1 | 2 | 0 | 0 | 2 | 7 | 5 | 0 | 0 | 0 | 0 | 0 | 4 | 271 | 149 | 1 | 1 | 1 | 1 | 0.892 | 0.733 | 1 | NA | 1 | 1 | 1 | 0 | 0.175 |
| SE | 61675 | ENSG00000150938 | CRIM1 | chr2 | + | 36522091 | 36522313 | 36517326 | 36517542 | 36537351 | 36537546 | 56 | 17 | 23 | 27 | 20 | 1 | 1 | 3 | 1 | 2 | 43 | 28 | 19 | 28 | 0 | 0 | 3 | 0 | 0 | 0 | 298 | 149 | 1 | 1 | 0.966 | 0.895 | 0.793 | 0.931 | 0.833 | 1 | 0.824 | 1 | 1 | NA | -0.072 |
| SE | 61676 | ENSG00000150938 | CRIM1 | chr2 | + | 36522091 | 36522313 | 36517326 | 36517542 | 36544375 | 36544498 | 16 | 7 | 5 | 14 | 4 | 1 | 0 | 0 | 0 | 0 | 21 | 9 | 2 | 10 | 0 | 0 | 0 | 0 | 0 | 0 | 298 | 149 | 1 | 1 | 0.889 | 1 | 1 | 1 | 1 | 1 | 1 | 1 | 1 | NA | -0.022 |
| SE | 61677 | ENSG00000150938 | CRIM1 | chr2 | + | 36537351 | 36537546 | 36517326 | 36517542 | 36544375 | 36544498 | 37 | 13 | 23 | 20 | 18 | 1 | 0 | 0 | 0 | 0 | 10 | 21 | 14 | 22 | 0 | 0 | 0 | 0 | 0 | 0 | 298 | 149 | 1 | 1 | 0.949 | 1 | 1 | 1 | 1 | 1 | 1 | 1 | 1 | NA | -0.01 |
| SE | 61678 | ENSG00000150938 | CRIM1 | chr2 | + | 36537351 | 36537546 | 36522091 | 36522313 | 36544375 | 36544498 | 76 | 22 | 39 | 32 | 32 | 0 | 0 | 1 | 0 | 0 | 42 | 37 | 31 | 40 | 0 | 10 | 0 | 0 | 0 | 0 | 298 | 149 | 1 | 1 | 1 | 1 | 0.951 | 1 | 1 | 0.677 | 1 | 1 | 1 | NA | 0.071 |
| SE | 61681 | ENSG00000187098 | MITF | chr3 | + | 69939097 | 69939146 | 69937821 | 69938049 | 69941235 | 69941331 | 1 | 0 | 2 | 2 | 0 | 0 | 0 | 3 | 0 | 0 | 12 | 16 | 6 | 11 | 0 | 9 | 6 | 8 | 17 | 0 | 198 | 149 | 1 | 1 | 1 | NA | 0.334 | 1 | NA | 0.501 | 0.667 | 0.361 | 0.327 | NA | 0.314 |
| SE | 61683 | ENSG00000230084 | NA | chr3 | - | 42632628 | 42632695 | 42623384 | 42623812 | 42653993 | 42654156 | 3 | 0 | 1 | 1 | 0 | 0 | 0 | 0 | 1 | 0 | 0 | 2 | 4 | 4 | 0 | 0 | 0 | 1 | 0 | 9 | 216 | 149 | 1 | 1 | 1 | NA | 1 | 0.408 | NA | NA | 1 | 0.734 | 1 | 0 | 0.119 |
| SE | 61684 | ENSG00000231500 | RPS18 | chr6 | + | 33275499 | 33275657 | 33272627 | 33272726 | 33275796 | 33275883 | 8 | 0 | 4 | 8 | 14 | 3045 | 1460 | 3414 | 2807 | 2601 | 0 | 16 | 5 | 4 | 3 | 1893 | 1856 | 1882 | 1644 | 2880 | 298 | 149 | 1 | 1 | 0.001 | 0 | 0.001 | 0.001 | 0.003 | 0 | 0.004 | 0.001 | 0.001 | 0.001 | 0 |
| SE | 61685 | ENSG00000231500 | RPS18 | chr6 | + | 33275796 | 33275883 | 33272131 | 33272726 | 33275964 | 33276066 | 5333 | 2639 | 5947 | 4970 | 6699 | 3 | 2 | 1 | 0 | 0 | 3247 | 3341 | 3315 | 2917 | 5126 | 0 | 3 | 6 | 0 | 3 | 236 | 149 | 1 | 1 | 0.999 | 0.999 | 1 | 1 | 1 | 1 | 0.999 | 0.997 | 1 | 0.999 | 0.001 |
| SE | 61686 | ENSG00000231500 | RPS18 | chr6 | + | 33276175 | 33276267 | 33275865 | 33276066 | 33276390 | 33276510 | 6129 | 2952 | 7107 | 5382 | 11625 | 0 | 0 | 0 | 4 | 0 | 2329 | 2741 | 2377 | 2181 | 3792 | 0 | 0 | 0 | 0 | 0 | 241 | 149 | 1 | 1 | 1 | 1 | 1 | 0.999 | 1 | 1 | 1 | 1 | 1 | 1 | 0 |
| SE | 61697 | ENSG00000163382 | NAXE | chr1 | + | 156592100 | 156592209 | 156591761 | 156591986 | 156592364 | 156592475 | 123 | 45 | 97 | 100 | 85 | 0 | 0 | 2 | 0 | 0 | 54 | 97 | 63 | 70 | 113 | 0 | 0 | 0 | 0 | 0 | 258 | 149 | 1 | 1 | 1 | 1 | 0.966 | 1 | 1 | 1 | 1 | 1 | 1 | 1 | -0.007 |
| SE | 61699 | ENSG00000163382 | NAXE | chr1 | + | 156592556 | 156592670 | 156592100 | 156592475 | 156593407 | 156593555 | 125 | 77 | 107 | 122 | 110 | 0 | 0 | 0 | 0 | 2 | 113 | 75 | 110 | 109 | 119 | 0 | 0 | 0 | 0 | 0 | 263 | 149 | 1 | 1 | 1 | 1 | 1 | 1 | 0.969 | 1 | 1 | 1 | 1 | 1 | -0.006 |
| SE | 61700 | ENSG00000163382 | NAXE | chr1 | + | 156593407 | 156593555 | 156592556 | 156592670 | 156593881 | 156594090 | 140 | 87 | 117 | 137 | 177 | 0 | 0 | 0 | 0 | 3 | 103 | 77 | 116 | 130 | 133 | 0 | 0 | 0 | 0 | 0 | 297 | 149 | 1 | 1 | 1 | 1 | 1 | 1 | 0.967 | 1 | 1 | 1 | 1 | 1 | -0.007 |
| SE | 61701 | ENSG00000163382 | NAXE | chr1 | + | 156593463 | 156593555 | 156592556 | 156592670 | 156593881 | 156594131 | 45 | 24 | 39 | 47 | 53 | 0 | 0 | 0 | 0 | 3 | 27 | 29 | 17 | 35 | 63 | 0 | 0 | 0 | 0 | 0 | 241 | 149 | 1 | 1 | 1 | 1 | 1 | 1 | 0.916 | 1 | 1 | 1 | 1 | 1 | -0.017 |
| SE | 61702 | ENSG00000163382 | NAXE | chr1 | + | 156593881 | 156594090 | 156592556 | 156593555 | 156596706 | 156599795 | 75 | 36 | 52 | 65 | 116 | 0 | 0 | 1 | 3 | 0 | 61 | 46 | 42 | 62 | 67 | 0 | 0 | 0 | 0 | 0 | 298 | 149 | 1 | 1 | 1 | 1 | 0.963 | 0.915 | 1 | 1 | 1 | 1 | 1 | 1 | -0.024 |
| SE | 61704 | ENSG00000163382 | NAXE | chr1 | + | 156593944 | 156594090 | 156593407 | 156593555 | 156596706 | 156598309 | 0 | 0 | 0 | 0 | 1 | 0 | 0 | 1 | 3 | 0 | 0 | 0 | 0 | 3 | 0 | 0 | 0 | 0 | 0 | 0 | 295 | 149 | 1 | 1 | NA | NA | 0 | 0 | 1 | NA | NA | NA | 1 | NA | -0.667 |
| SE | 61706 | ENSG00000198836 | OPA1 | chr3 | + | 193596799 | 193596946 | 193593216 | 193593409 | 193614722 | 193615041 | 0 | 1 | 1 | 1 | 2 | 90 | 60 | 70 | 46 | 52 | 6 | 0 | 0 | 0 | 0 | 47 | 52 | 47 | 9 | 0 | 296 | 149 | 1 | 1 | 0 | 0.008 | 0.007 | 0.011 | 0.019 | 0.06 | 0 | 0 | 0 | NA | -0.006 |
| SE | 61707 | ENSG00000198836 | OPA1 | chr3 | + | 193603499 | 193603557 | 193593217 | 193593409 | 193614722 | 193615041 | 2 | 2 | 3 | 1 | 1 | 90 | 60 | 70 | 46 | 52 | 0 | 0 | 2 | 0 | 0 | 47 | 52 | 47 | 9 | 0 | 207 | 149 | 1 | 1 | 0.016 | 0.023 | 0.03 | 0.015 | 0.014 | 0 | 0 | 0.03 | 0 | NA | 0.012 |
| SE | 61708 | ENSG00000198836 | OPA1 | chr3 | + | 193617177 | 193617285 | 193615673 | 193615770 | 193617783 | 193617837 | 164 | 132 | 153 | 145 | 301 | 1 | 0 | 0 | 0 | 0 | 9 | 48 | 120 | 35 | 14 | 0 | 0 | 0 | 0 | 0 | 257 | 149 | 1 | 1 | 0.99 | 1 | 1 | 1 | 1 | 1 | 1 | 1 | 1 | 1 | -0.002 |
| SE | 61710 | ENSG00000198836 | OPA1 | chr3 | + | 193617783 | 193617837 | 193615673 | 193615770 | 193618868 | 193618936 | 16 | 6 | 13 | 14 | 51 | 8 | 1 | 7 | 7 | 13 | 0 | 0 | 17 | 5 | 6 | 0 | 2 | 4 | 0 | 0 | 203 | 149 | 1 | 1 | 0.595 | 0.815 | 0.577 | 0.595 | 0.742 | NA | 0 | 0.757 | 1 | 1 | -0.024 |
| SE | 61715 | ENSG00000198836 | OPA1 | chr3 | + | 193626091 | 193626202 | 193618702 | 193618936 | 193635417 | 193635471 | 64 | 42 | 72 | 58 | 100 | 0 | 0 | 0 | 0 | 1 | 7 | 25 | 45 | 5 | 20 | 0 | 0 | 0 | 0 | 0 | 260 | 149 | 1 | 1 | 1 | 1 | 1 | 1 | 0.983 | 1 | 1 | 1 | 1 | 1 | -0.003 |
| SE | 61716 | ENSG00000198836 | OPA1 | chr3 | + | 193628508 | 193628616 | 193618868 | 193618936 | 193631611 | 193631665 | 1 | 4 | 10 | 4 | 6 | 52 | 37 | 74 | 61 | 147 | 0 | 0 | 0 | 0 | 0 | 14 | 40 | 46 | 0 | 0 | 257 | 149 | 1 | 1 | 0.011 | 0.059 | 0.073 | 0.037 | 0.023 | 0 | 0 | 0 | NA | NA | 0.041 |
| SE | 61718 | ENSG00000198836 | OPA1 | chr3 | + | 193631611 | 193631665 | 193618702 | 193618936 | 193635417 | 193635471 | 107 | 66 | 126 | 90 | 215 | 0 | 0 | 0 | 0 | 1 | 14 | 60 | 65 | 7 | 9 | 0 | 0 | 0 | 0 | 0 | 203 | 149 | 1 | 1 | 1 | 1 | 1 | 1 | 0.994 | 1 | 1 | 1 | 1 | 1 | -0.001 |
| SE | 61719 | ENSG00000198836 | OPA1 | chr3 | + | 193631611 | 193631665 | 193626057 | 193626202 | 193635417 | 193635471 | 104 | 56 | 97 | 61 | 124 | 0 | 0 | 0 | 0 | 0 | 14 | 40 | 48 | 10 | 17 | 0 | 0 | 3 | 0 | 0 | 203 | 149 | 1 | 1 | 1 | 1 | 1 | 1 | 1 | 1 | 1 | 0.922 | 1 | 1 | 0.016 |
| SE | 61720 | ENSG00000198836 | OPA1 | chr3 | + | 193631754 | 193631769 | 193631611 | 193631665 | 193635417 | 193635522 | 2 | 0 | 2 | 1 | 0 | 125 | 83 | 151 | 108 | 207 | 0 | 0 | 3 | 0 | 0 | 22 | 72 | 78 | 7 | 17 | 164 | 149 | 1 | 1 | 0.014 | 0 | 0.012 | 0.008 | 0 | 0 | 0 | 0.034 | 0 | 0 | 0 |
| SE | 61721 | ENSG00000198836 | OPA1 | chr3 | + | 193642764 | 193642845 | 193637951 | 193638065 | 193643372 | 193643444 | 70 | 64 | 68 | 64 | 78 | 1 | 0 | 0 | 0 | 0 | 19 | 50 | 40 | 5 | 11 | 0 | 0 | 0 | 0 | 0 | 230 | 149 | 1 | 1 | 0.978 | 1 | 1 | 1 | 1 | 1 | 1 | 1 | 1 | 1 | -0.004 |
| SE | 61723 | ENSG00000198836 | OPA1 | chr3 | + | 193643372 | 193643444 | 193642974 | 193643049 | 193643527 | 193643627 | 218 | 173 | 171 | 217 | 274 | 0 | 0 | 1 | 0 | 0 | 0 | 117 | 168 | 43 | 20 | 0 | 0 | 0 | 0 | 0 | 221 | 149 | 1 | 1 | 1 | 1 | 0.991 | 1 | 1 | NA | 1 | 1 | 1 | 1 | -0.002 |
| SE | 61724 | ENSG00000198836 | OPA1 | chr3 | + | 193645552 | 193645625 | 193643951 | 193644105 | 193645727 | 193645800 | 205 | 161 | 150 | 198 | 281 | 0 | 1 | 2 | 2 | 0 | 6 | 70 | 121 | 13 | 0 | 0 | 0 | 0 | 0 | 0 | 222 | 149 | 1 | 1 | 1 | 0.991 | 0.981 | 0.985 | 1 | 1 | 1 | 1 | 1 | NA | -0.009 |
| SE | 61726 | ENSG00000198836 | OPA1 | chr3 | + | 193648069 | 193648134 | 193647064 | 193647180 | 193648794 | 193648871 | 210 | 116 | 146 | 191 | 211 | 0 | 0 | 1 | 0 | 0 | 9 | 77 | 84 | 16 | 0 | 0 | 0 | 0 | 0 | 0 | 214 | 149 | 1 | 1 | 1 | 1 | 0.99 | 1 | 1 | 1 | 1 | 1 | 1 | NA | -0.002 |
| SE | 61727 | ENSG00000198836 | OPA1 | chr3 | + | 193654861 | 193655027 | 193648765 | 193648871 | 193657079 | 193657232 | 343 | 241 | 292 | 335 | 447 | 0 | 0 | 0 | 0 | 1 | 15 | 166 | 167 | 30 | 0 | 0 | 0 | 2 | 0 | 0 | 298 | 149 | 1 | 1 | 1 | 1 | 1 | 1 | 0.996 | 1 | 1 | 0.977 | 1 | NA | 0.005 |
| SE | 61728 | ENSG00000198836 | OPA1 | chr3 | + | 193654861 | 193655052 | 193648794 | 193648871 | 193657079 | 193657232 | 202 | 143 | 170 | 205 | 276 | 0 | 0 | 0 | 0 | 1 | 15 | 97 | 84 | 14 | 0 | 0 | 0 | 2 | 0 | 0 | 298 | 149 | 1 | 1 | 1 | 1 | 1 | 1 | 0.993 | 1 | 1 | 0.955 | 1 | NA | 0.01 |
| SE | 61729 | ENSG00000198836 | OPA1 | chr3 | + | 193667370 | 193667455 | 193667169 | 193667280 | 193668314 | 193668931 | 0 | 0 | 0 | 1 | 0 | 1 | 0 | 0 | 0 | 0 | 0 | 0 | 0 | 0 | 0 | 0 | 0 | 2 | 0 | 0 | 234 | 149 | 1 | 1 | 0 | NA | NA | 1 | NA | NA | NA | 0 | NA | NA | 0.5 |
| SE | 61730 | ENSG00000198836 | OPA1 | chr3 | + | 193668314 | 193668484 | 193667169 | 193667280 | 193692062 | 193692132 | 1 | 0 | 0 | 0 | 2 | 158 | 130 | 163 | 124 | 274 | 0 | 0 | 2 | 0 | 0 | 14 | 72 | 73 | 17 | 5 | 298 | 149 | 1 | 1 | 0.003 | 0 | 0 | 0 | 0.004 | 0 | 0 | 0.014 | 0 | 0 | -0.001 |
| SE | 61731 | ENSG00000198836 | OPA1 | chr3 | + | 193689068 | 193689170 | 193667169 | 193667280 | 193692062 | 193692132 | 0 | 2 | 0 | 7 | 5 | 158 | 130 | 163 | 124 | 274 | 0 | 0 | 0 | 0 | 0 | 14 | 72 | 73 | 17 | 5 | 251 | 149 | 1 | 1 | 0 | 0.009 | 0 | 0.032 | 0.011 | 0 | 0 | 0 | 0 | 0 | 0.01 |
| SE | 61733 | ENSG00000221947 | XKR9 | chr8 | + | 70678029 | 70678139 | 70674817 | 70674899 | 70680780 | 70681330 | 0 | 0 | 0 | 0 | 0 | 2 | 0 | 3 | 0 | 7 | 0 | 0 | 0 | 2 | 0 | 0 | 0 | 0 | 0 | 0 | 259 | 149 | 1 | 1 | 0 | NA | 0 | NA | 0 | NA | NA | NA | 1 | NA | -1 |
| SE | 61734 | ENSG00000221947 | XKR9 | chr8 | + | 70680780 | 70681330 | 70674817 | 70674899 | 70706932 | 70707153 | 7 | 1 | 5 | 1 | 11 | 0 | 0 | 1 | 0 | 0 | 0 | 0 | 3 | 0 | 8 | 0 | 0 | 0 | 1 | 0 | 298 | 149 | 1 | 1 | 1 | 1 | 0.714 | 1 | 1 | NA | NA | 1 | NA | 1 | -0.057 |
| SE | 61737 | ENSG00000132485 | ZRANB2 | chr1 | - | 71072120 | 71072255 | 71070826 | 71070996 | 71072471 | 71072548 | 935 | 929 | 1048 | 1191 | 1969 | 0 | 0 | 0 | 0 | 0 | 83 | 421 | 463 | 97 | 49 | 0 | 0 | 0 | 1 | 0 | 284 | 149 | 1 | 1 | 1 | 1 | 1 | 1 | 1 | 1 | 1 | 1 | 0.981 | 1 | 0.004 |
| SE | 61738 | ENSG00000132485 | ZRANB2 | chr1 | - | 71076794 | 71076877 | 71072471 | 71072548 | 71078456 | 71078565 | 1014 | 1118 | 1147 | 1246 | 2062 | 0 | 2 | 0 | 2 | 0 | 123 | 423 | 541 | 202 | 75 | 0 | 0 | 1 | 0 | 0 | 232 | 149 | 1 | 1 | 1 | 0.997 | 1 | 0.998 | 1 | 1 | 1 | 0.997 | 1 | 1 | 0 |
| SE | 61739 | ENSG00000143727 | ACP1 | chr2 | + | 269300 | 269347 | 264899 | 265007 | 271865 | 271939 | 0 | 2 | 5 | 0 | 0 | 62 | 28 | 73 | 62 | 90 | 0 | 0 | 1 | 0 | 0 | 45 | 45 | 55 | 46 | 68 | 196 | 149 | 1 | 1 | 0 | 0.052 | 0.049 | 0 | 0 | 0 | 0 | 0.014 | 0 | 0 | 0.017 |
| SE | 61741 | ENSG00000143727 | ACP1 | chr2 | + | 272036 | 272150 | 271865 | 271939 | 276979 | 277085 | 55 | 28 | 75 | 62 | 102 | 2 | 0 | 0 | 0 | 0 | 54 | 51 | 53 | 51 | 76 | 0 | 0 | 0 | 0 | 0 | 263 | 149 | 1 | 1 | 0.94 | 1 | 1 | 1 | 1 | 1 | 1 | 1 | 1 | 1 | -0.012 |
| SE | 61742 | ENSG00000143727 | ACP1 | chr2 | + | 272191 | 272305 | 271865 | 271939 | 276979 | 277085 | 31 | 18 | 52 | 35 | 23 | 2 | 0 | 0 | 0 | 0 | 25 | 37 | 31 | 21 | 24 | 0 | 0 | 0 | 0 | 0 | 263 | 149 | 1 | 1 | 0.898 | 1 | 1 | 1 | 1 | 1 | 1 | 1 | 1 | 1 | -0.02 |
| SE | 61744 | ENSG00000143727 | ACP1 | chr2 | + | 275139 | 275201 | 272191 | 272305 | 276979 | 277085 | 232 | 164 | 269 | 248 | 347 | 0 | 0 | 1 | 0 | 0 | 205 | 202 | 130 | 180 | 260 | 0 | 0 | 6 | 2 | 0 | 211 | 149 | 1 | 1 | 1 | 1 | 0.995 | 1 | 1 | 1 | 1 | 0.939 | 0.985 | 1 | 0.014 |
| SE | 61745 | ENSG00000162437 | RAVER2 | chr1 | + | 64807205 | 64807474 | 64804990 | 64805105 | 64812737 | 64812849 | 16 | 3 | 8 | 18 | 22 | 0 | 0 | 0 | 0 | 0 | 0 | 4 | 6 | 17 | 0 | 0 | 0 | 0 | 3 | 1 | 298 | 149 | 1 | 1 | 1 | 1 | 1 | 1 | 1 | NA | 1 | 1 | 0.739 | 0 | 0.315 |
| SE | 61752 | ENSG00000153029 | MR1 | chr1 | + | 181036996 | 181037112 | 181034009 | 181034074 | 181055224 | 181055284 | 2 | 1 | 1 | 0 | 6 | 0 | 4 | 0 | 0 | 0 | 0 | 0 | 5 | 4 | 0 | 0 | 0 | 0 | 2 | 0 | 265 | 149 | 1 | 1 | 1 | 0.123 | 1 | NA | 1 | NA | NA | 0.584 | 1 | NA | -0.011 |
| SE | 61754 | ENSG00000153029 | MR1 | chr1 | + | 181049051 | 181049312 | 181036996 | 181037112 | 181053572 | 181053677 | 3 | 0 | 0 | 0 | 3 | 0 | 0 | 0 | 0 | 0 | 0 | 12 | 3 | 8 | 0 | 0 | 0 | 0 | 4 | 0 | 298 | 149 | 1 | 1 | 1 | NA | NA | NA | 1 | NA | 1 | 1 | 0.5 | NA | 0.167 |
| SE | 61755 | ENSG00000153029 | MR1 | chr1 | + | 181050010 | 181050286 | 181033886 | 181034074 | 181055224 | 181055589 | 4 | 3 | 5 | 14 | 24 | 0 | 4 | 0 | 0 | 0 | 38 | 24 | 24 | 2 | 0 | 0 | 0 | 2 | 0 | 0 | 298 | 149 | 1 | 1 | 1 | 0.273 | 1 | 1 | 1 | 1 | 1 | 0.857 | 1 | NA | -0.11 |
| SE | 61756 | ENSG00000153029 | MR1 | chr1 | + | 181050010 | 181050286 | 181049051 | 181049312 | 181053572 | 181053677 | 167 | 120 | 170 | 165 | 186 | 1 | 0 | 0 | 0 | 0 | 269 | 176 | 152 | 140 | 193 | 0 | 0 | 0 | 0 | 0 | 298 | 149 | 1 | 1 | 0.988 | 1 | 1 | 1 | 1 | 1 | 1 | 1 | 1 | 1 | -0.002 |
| SE | 61759 | ENSG00000153029 | MR1 | chr1 | + | 181053572 | 181053677 | 181036996 | 181037112 | 181055224 | 181055284 | 99 | 64 | 84 | 71 | 165 | 0 | 0 | 0 | 0 | 1 | 106 | 50 | 52 | 128 | 26 | 0 | 0 | 0 | 0 | 0 | 254 | 149 | 1 | 1 | 1 | 1 | 1 | 1 | 0.99 | 1 | 1 | 1 | 1 | 1 | -0.002 |
| SE | 61762 | ENSG00000158710 | TAGLN2 | chr1 | - | 159919660 | 159919835 | 159918750 | 159919376 | 159920329 | 159920537 | 2980 | 2084 | 1956 | 4115 | 3357 | 0 | 0 | 0 | 0 | 0 | 5771 | 3663 | 2377 | 3104 | 24965 | 2 | 3 | 1 | 8 | 0 | 298 | 149 | 1 | 1 | 1 | 1 | 1 | 1 | 1 | 0.999 | 0.998 | 0.999 | 0.995 | 1 | 0.002 |
| SE | 61763 | ENSG00000158710 | TAGLN2 | chr1 | - | 159920329 | 159920537 | 159918750 | 159919376 | 159925449 | 159925491 | 229 | 118 | 142 | 286 | 85 | 2 | 0 | 0 | 1 | 0 | 505 | 366 | 266 | 316 | 2800 | 0 | 0 | 0 | 0 | 0 | 298 | 149 | 1 | 1 | 0.983 | 1 | 1 | 0.993 | 1 | 1 | 1 | 1 | 1 | 1 | -0.005 |
| SE | 61776 | ENSG00000277425 | NA | chr20 | - | 5067208 | 5067369 | 5066081 | 5066341 | 5067966 | 5068154 | 1 | 0 | 1 | 0 | 0 | 1 | 0 | 0 | 0 | 0 | 0 | 0 | 0 | 3 | 14 | 0 | 0 | 0 | 0 | 0 | 298 | 149 | 1 | 1 | 0.333 | NA | 1 | NA | NA | NA | NA | NA | 1 | 1 | -0.334 |
| SE | 61778 | ENSG00000173041 | ZNF680 | chr7 | - | 64543706 | 64543802 | 64519877 | 64522500 | 64562924 | 64563022 | 24 | 21 | 13 | 18 | 43 | 5 | 0 | 11 | 3 | 0 | 0 | 0 | 8 | 7 | 0 | 0 | 1 | 2 | 0 | 0 | 245 | 149 | 1 | 1 | 0.745 | 1 | 0.418 | 0.785 | 1 | NA | 0 | 0.709 | 1 | NA | 0.22 |
| SE | 61779 | ENSG00000173041 | ZNF680 | chr7 | - | 64544305 | 64544432 | 64519877 | 64522500 | 64562924 | 64563041 | 8 | 2 | 2 | 3 | 0 | 5 | 0 | 11 | 3 | 0 | 23 | 7 | 2 | 0 | 0 | 0 | 1 | 2 | 0 | 0 | 276 | 149 | 1 | 1 | 0.463 | 1 | 0.089 | 0.351 | NA | 1 | 0.791 | 0.351 | NA | NA | -0.238 |
| SE | 61781 | ENSG00000173041 | ZNF680 | chr7 | - | 64560945 | 64561273 | 64544345 | 64544432 | 64562924 | 64563075 | 1 | 1 | 2 | 0 | 2 | 4 | 2 | 2 | 4 | 4 | 9 | 0 | 0 | 1 | 0 | 23 | 4 | 1 | 0 | 0 | 298 | 149 | 1 | 1 | 0.111 | 0.2 | 0.333 | 0 | 0.2 | 0.164 | 0 | 0 | 1 | NA | -0.122 |
| SE | 61783 | ENSG00000104343 | UBE2W | chr8 | - | 73795790 | 73795914 | 73794041 | 73794115 | 73805650 | 73805726 | 0 | 0 | 0 | 2 | 0 | 59 | 81 | 41 | 64 | 96 | 4 | 0 | 1 | 0 | 0 | 0 | 23 | 26 | 8 | 0 | 273 | 149 | 1 | 1 | 0 | 0 | 0 | 0.017 | 0 | 1 | 0 | 0.021 | 0 | NA | -0.252 |
| SE | 61785 | ENSG00000104343 | UBE2W | chr8 | - | 73796606 | 73796690 | 73793823 | 73794115 | 73805650 | 73805726 | 0 | 1 | 0 | 0 | 0 | 59 | 81 | 41 | 64 | 96 | 0 | 0 | 0 | 0 | 0 | 0 | 23 | 26 | 8 | 0 | 233 | 149 | 1 | 1 | 0 | 0.008 | 0 | 0 | 0 | NA | 0 | 0 | 0 | NA | 0.002 |
| SE | 61786 | ENSG00000104343 | UBE2W | chr8 | - | 73796606 | 73796690 | 73795790 | 73795914 | 73805650 | 73805726 | 0 | 0 | 0 | 0 | 2 | 0 | 0 | 0 | 0 | 0 | 0 | 0 | 0 | 0 | 0 | 4 | 0 | 1 | 0 | 0 | 233 | 149 | 1 | 1 | NA | NA | NA | NA | 1 | 0 | NA | 0 | NA | NA | 1 |
| SE | 61788 | ENSG00000104343 | UBE2W | chr8 | - | 73830380 | 73830472 | 73825146 | 73825249 | 73870268 | 73870301 | 147 | 89 | 97 | 117 | 204 | 0 | 1 | 0 | 0 | 0 | 46 | 79 | 59 | 31 | 16 | 0 | 0 | 0 | 0 | 0 | 241 | 149 | 1 | 1 | 1 | 0.982 | 1 | 1 | 1 | 1 | 1 | 1 | 1 | 1 | -0.004 |
| SE | 61790 | ENSG00000174371 | EXO1 | chr1 | + | 241849108 | 241849216 | 241848950 | 241849027 | 241850408 | 241850586 | 1 | 2 | 3 | 0 | 0 | 1 | 0 | 0 | 1 | 2 | 0 | 2 | 3 | 5 | 0 | 0 | 0 | 6 | 1 | 0 | 257 | 149 | 1 | 1 | 0.367 | 1 | 1 | 0 | 0 | NA | 1 | 0.225 | 0.744 | NA | -0.183 |
| SE | 61791 | ENSG00000174371 | EXO1 | chr1 | + | 241849111 | 241849216 | 241848730 | 241849027 | 241850408 | 241850586 | 0 | 0 | 2 | 0 | 0 | 1 | 0 | 0 | 1 | 2 | 0 | 2 | 0 | 4 | 0 | 0 | 0 | 6 | 1 | 0 | 254 | 149 | 1 | 1 | 0 | NA | 1 | 0 | 0 | NA | 1 | 0 | 0.701 | NA | -0.317 |
| SE | 61792 | ENSG00000174371 | EXO1 | chr1 | + | 241852291 | 241852411 | 241850408 | 241850586 | 241853357 | 241853481 | 6 | 0 | 22 | 4 | 4 | 0 | 0 | 0 | 1 | 2 | 0 | 22 | 20 | 10 | 0 | 11 | 0 | 3 | 2 | 0 | 269 | 149 | 1 | 1 | 1 | NA | 1 | 0.689 | 0.526 | 0 | 1 | 0.787 | 0.735 | NA | 0.173 |
| SE | 61801 | ENSG00000196975 | ANXA4 | chr2 | + | 69810593 | 69810673 | 69807981 | 69808189 | 69812652 | 69812709 | 40 | 27 | 31 | 39 | 28 | 0 | 0 | 0 | 0 | 0 | 57 | 50 | 32 | 111 | 37 | 0 | 3 | 0 | 0 | 0 | 229 | 149 | 1 | 1 | 1 | 1 | 1 | 1 | 1 | 1 | 0.916 | 1 | 1 | 1 | 0.017 |
| SE | 61802 | ENSG00000196975 | ANXA4 | chr2 | + | 69820698 | 69820821 | 69819279 | 69819338 | 69825455 | 69825774 | 410 | 264 | 247 | 248 | 436 | 0 | 0 | 0 | 0 | 0 | 558 | 413 | 215 | 663 | 504 | 10 | 0 | 0 | 0 | 0 | 272 | 149 | 1 | 1 | 1 | 1 | 1 | 1 | 1 | 0.968 | 1 | 1 | 1 | 1 | 0.006 |
| SE | 61810 | ENSG00000152760 | DYNLT5 | chr1 | + | 66755679 | 66755773 | 66754655 | 66754777 | 66770386 | 66770507 | 0 | 0 | 0 | 0 | 0 | 0 | 2 | 0 | 1 | 0 | 0 | 0 | 0 | 1 | 0 | 0 | 0 | 6 | 2 | 0 | 243 | 149 | 1 | 1 | NA | 0 | NA | 0 | NA | NA | NA | 0 | 0.235 | NA | -0.117 |
| SE | 61813 | ENSG00000164930 | FZD6 | chr8 | + | 103328267 | 103328416 | 103324480 | 103325498 | 103329654 | 103329971 | 3 | 22 | 29 | 20 | 48 | 0 | 1 | 2 | 9 | 8 | 0 | 42 | 8 | 19 | 0 | 0 | 0 | 8 | 0 | 0 | 298 | 149 | 1 | 1 | 1 | 0.917 | 0.879 | 0.526 | 0.75 | NA | 1 | 0.333 | 1 | NA | 0.037 |
| SE | 61835 | ENSG00000255418 | LINC02718 | chr11 | + | 23164990 | 23165093 | 23163493 | 23163602 | 23165189 | 23165333 | 0 | 0 | 1 | 0 | 2 | 6 | 2 | 0 | 1 | 0 | 0 | 0 | 0 | 0 | 0 | 0 | 0 | 3 | 0 | 5 | 252 | 149 | 1 | 1 | 0 | 0 | 1 | 0 | 1 | NA | 0 | NA | 0 | NA | 0.4 |
| SE | 61836 | ENSG00000255418 | LINC02718 | chr11 | + | 23189998 | 23190046 | 23183399 | 23183474 | 23192012 | 23192084 | 0 | 1 | 0 | 0 | 0 | 0 | 1 | 0 | 1 | 0 | 0 | 0 | 0 | 0 | 0 | 0 | 0 | 1 | 0 | 6 | 197 | 149 | 1 | 1 | NA | 0.431 | NA | 0 | NA | NA | NA | 0 | NA | 0 | 0.215 |
| SE | 61837 | ENSG00000170871 | KIAA0232 | chr4 | + | 6804522 | 6804606 | 6782726 | 6782841 | 6824184 | 6824684 | 0 | 0 | 0 | 2 | 0 | 7 | 1 | 4 | 3 | 3 | 0 | 0 | 2 | 3 | 0 | 0 | 5 | 6 | 5 | 0 | 233 | 149 | 1 | 1 | 0 | 0 | 0 | 0.299 | 0 | NA | 0 | 0.176 | 0.277 | NA | -0.091 |
| SE | 61838 | ENSG00000170871 | KIAA0232 | chr4 | + | 6860900 | 6864183 | 6842066 | 6842204 | 6871573 | 6871682 | 78 | 61 | 34 | 51 | 74 | 0 | 0 | 0 | 0 | 0 | 10 | 6 | 14 | 12 | 4 | 0 | 1 | 0 | 0 | 0 | 298 | 149 | 1 | 1 | 1 | 1 | 1 | 1 | 1 | 1 | 0.75 | 1 | 1 | 1 | 0.05 |
| SE | 61841 | ENSG00000162929 | KIAA1841 | chr2 | + | 61069651 | 61069698 | 61068848 | 61068985 | 61070341 | 61070500 | 0 | 4 | 2 | 4 | 0 | 9 | 11 | 5 | 7 | 23 | 0 | 0 | 0 | 0 | 0 | 6 | 5 | 7 | 5 | 0 | 196 | 149 | 1 | 1 | 0 | 0.217 | 0.233 | 0.303 | 0 | 0 | 0 | 0 | 0 | NA | 0.151 |
| SE | 61842 | ENSG00000162929 | KIAA1841 | chr2 | + | 61071605 | 61071792 | 61070325 | 61070500 | 61073457 | 61073551 | 46 | 19 | 20 | 29 | 73 | 0 | 3 | 0 | 0 | 0 | 13 | 22 | 22 | 29 | 14 | 0 | 0 | 0 | 0 | 0 | 298 | 149 | 1 | 1 | 1 | 0.76 | 1 | 1 | 1 | 1 | 1 | 1 | 1 | 1 | -0.048 |
| SE | 61845 | ENSG00000162929 | KIAA1841 | chr2 | + | 61092463 | 61092587 | 61088357 | 61088468 | 61103852 | 61103998 | 17 | 11 | 9 | 7 | 4 | 1 | 0 | 0 | 1 | 0 | 9 | 0 | 4 | 2 | 9 | 0 | 0 | 2 | 0 | 0 | 273 | 149 | 1 | 1 | 0.903 | 1 | 1 | 0.793 | 1 | 1 | NA | 0.522 | 1 | 1 | 0.059 |
| SE | 61850 | ENSG00000143393 | PI4KB | chr1 | - | 151298807 | 151299073 | 151294408 | 151294541 | 151301843 | 151301967 | 151 | 62 | 176 | 157 | 232 | 0 | 0 | 0 | 1 | 0 | 164 | 207 | 135 | 133 | 274 | 0 | 0 | 0 | 0 | 0 | 298 | 149 | 1 | 1 | 1 | 1 | 1 | 0.987 | 1 | 1 | 1 | 1 | 1 | 1 | -0.003 |
| SE | 61851 | ENSG00000143393 | PI4KB | chr1 | - | 151306135 | 151306363 | 151303540 | 151303650 | 151307573 | 151307801 | 296 | 204 | 246 | 323 | 395 | 0 | 0 | 0 | 1 | 0 | 409 | 355 | 230 | 316 | 488 | 0 | 0 | 0 | 2 | 0 | 298 | 149 | 1 | 1 | 1 | 1 | 1 | 0.994 | 1 | 1 | 1 | 1 | 0.988 | 1 | 0.001 |
| SE | 61852 | ENSG00000143393 | PI4KB | chr1 | - | 151307573 | 151307801 | 151306135 | 151306363 | 151315572 | 151315672 | 257 | 147 | 204 | 245 | 290 | 0 | 0 | 3 | 0 | 3 | 270 | 270 | 166 | 246 | 397 | 0 | 0 | 3 | 0 | 0 | 298 | 149 | 1 | 1 | 1 | 1 | 0.971 | 1 | 0.98 | 1 | 1 | 0.965 | 1 | 1 | -0.003 |
| SE | 61855 | ENSG00000143393 | PI4KB | chr1 | - | 151315572 | 151315672 | 151307573 | 151307801 | 151327270 | 151327400 | 120 | 67 | 90 | 101 | 141 | 0 | 0 | 0 | 0 | 2 | 98 | 121 | 68 | 102 | 148 | 0 | 1 | 0 | 3 | 0 | 249 | 149 | 1 | 1 | 1 | 1 | 1 | 1 | 0.977 | 1 | 0.986 | 1 | 0.953 | 1 | 0.008 |
| SE | 61856 | ENSG00000143393 | PI4KB | chr1 | - | 151315572 | 151316509 | 151307573 | 151307801 | 151324744 | 151324917 | 191 | 94 | 130 | 164 | 199 | 6 | 0 | 0 | 0 | 0 | 187 | 175 | 96 | 150 | 266 | 0 | 0 | 0 | 0 | 0 | 298 | 149 | 1 | 1 | 0.941 | 1 | 1 | 1 | 1 | 1 | 1 | 1 | 1 | 1 | -0.012 |
| SE | 61858 | ENSG00000143393 | PI4KB | chr1 | - | 151315572 | 151316509 | 151307573 | 151307801 | 151327270 | 151327416 | 174 | 88 | 136 | 127 | 169 | 0 | 0 | 0 | 0 | 2 | 124 | 160 | 91 | 147 | 236 | 0 | 1 | 0 | 3 | 0 | 298 | 149 | 1 | 1 | 1 | 1 | 1 | 1 | 0.977 | 1 | 0.988 | 1 | 0.961 | 1 | 0.006 |
| SE | 61859 | ENSG00000143393 | PI4KB | chr1 | - | 151315572 | 151316509 | 151310210 | 151310255 | 151326171 | 151326373 | 96 | 52 | 74 | 91 | 104 | 1 | 0 | 0 | 1 | 0 | 274 | 247 | 115 | 241 | 247 | 0 | 1 | 0 | 0 | 0 | 298 | 149 | 1 | 1 | 0.98 | 1 | 1 | 0.978 | 1 | 1 | 1 | 1 | 1 | 1 | -0.008 |
| SE | 61861 | ENSG00000143393 | PI4KB | chr1 | - | 151324744 | 151324917 | 151307573 | 151307801 | 151327270 | 151327377 | 12 | 1 | 3 | 0 | 2 | 0 | 0 | 0 | 0 | 2 | 0 | 0 | 2 | 4 | 5 | 0 | 1 | 0 | 3 | 0 | 298 | 149 | 1 | 1 | 1 | 1 | 1 | NA | 0.333 | NA | 0 | 1 | 0.4 | 1 | 0.233 |
| SE | 61864 | ENSG00000143393 | PI4KB | chr1 | - | 151326171 | 151326373 | 151307573 | 151307801 | 151327270 | 151327416 | 33 | 5 | 20 | 17 | 14 | 0 | 0 | 0 | 0 | 2 | 71 | 68 | 34 | 46 | 53 | 0 | 1 | 0 | 3 | 0 | 298 | 149 | 1 | 1 | 1 | 1 | 1 | 1 | 0.778 | 1 | 0.971 | 1 | 0.885 | 1 | -0.016 |
| SE | 61868 | ENSG00000076242 | MLH1 | chr3 | + | 36994829 | 36994953 | 36993594 | 36993663 | 36996618 | 36996709 | 0 | 0 | 0 | 1 | 0 | 63 | 24 | 45 | 55 | 38 | 0 | 0 | 0 | 0 | 0 | 35 | 37 | 44 | 49 | 13 | 273 | 149 | 1 | 1 | 0 | 0 | 0 | 0.01 | 0 | 0 | 0 | 0 | 0 | 0 | 0.002 |
| SE | 61873 | ENSG00000076242 | MLH1 | chr3 | + | 36997155 | 36997300 | 36996618 | 36996709 | 37000954 | 37001053 | 2 | 2 | 0 | 4 | 0 | 84 | 91 | 130 | 130 | 113 | 0 | 1 | 2 | 0 | 0 | 105 | 151 | 113 | 137 | 21 | 294 | 149 | 1 | 1 | 0.012 | 0.011 | 0 | 0.015 | 0 | 0 | 0.003 | 0.009 | 0 | 0 | 0.005 |
| SE | 61875 | ENSG00000076242 | MLH1 | chr3 | + | 37000954 | 37001053 | 36996618 | 36996709 | 37004400 | 37004474 | 170 | 164 | 227 | 208 | 227 | 0 | 0 | 0 | 1 | 0 | 249 | 217 | 215 | 251 | 40 | 8 | 0 | 0 | 0 | 0 | 248 | 149 | 1 | 1 | 1 | 1 | 1 | 0.992 | 1 | 0.949 | 1 | 1 | 1 | 1 | 0.009 |
| SE | 61876 | ENSG00000076242 | MLH1 | chr3 | + | 37004400 | 37004474 | 37000954 | 37001048 | 37006990 | 37007063 | 130 | 138 | 168 | 168 | 151 | 1 | 0 | 0 | 0 | 0 | 210 | 169 | 158 | 164 | 44 | 0 | 0 | 0 | 0 | 0 | 223 | 149 | 1 | 1 | 0.989 | 1 | 1 | 1 | 1 | 1 | 1 | 1 | 1 | 1 | -0.002 |
| SE | 61884 | ENSG00000076242 | MLH1 | chr3 | + | 37017505 | 37017599 | 37012010 | 37012099 | 37025636 | 37026007 | 0 | 1 | 1 | 3 | 8 | 6 | 2 | 8 | 1 | 0 | 7 | 0 | 7 | 5 | 0 | 0 | 4 | 4 | 6 | 0 | 243 | 149 | 1 | 1 | 0 | 0.235 | 0.071 | 0.648 | 1 | 1 | 0 | 0.518 | 0.338 | NA | -0.073 |
| SE | 61885 | ENSG00000076242 | MLH1 | chr3 | + | 37017505 | 37017599 | 37012010 | 37012099 | 37028783 | 37028847 | 0 | 1 | 1 | 3 | 8 | 0 | 0 | 3 | 2 | 1 | 7 | 0 | 7 | 5 | 0 | 0 | 0 | 2 | 0 | 0 | 243 | 149 | 1 | 1 | NA | 1 | 0.17 | 0.479 | 0.831 | 1 | NA | 0.682 | 1 | NA | -0.274 |
| SE | 61892 | ENSG00000076242 | MLH1 | chr3 | + | 37020309 | 37020463 | 37017333 | 37017599 | 37028783 | 37028932 | 120 | 118 | 141 | 132 | 180 | 0 | 0 | 2 | 0 | 0 | 219 | 84 | 112 | 156 | 42 | 0 | 0 | 0 | 0 | 0 | 298 | 149 | 1 | 1 | 1 | 1 | 0.972 | 1 | 1 | 1 | 1 | 1 | 1 | 1 | -0.006 |
| SE | 61895 | ENSG00000076242 | MLH1 | chr3 | + | 37025636 | 37026007 | 37017505 | 37017599 | 37028783 | 37028847 | 58 | 31 | 59 | 39 | 73 | 0 | 0 | 2 | 0 | 0 | 71 | 42 | 46 | 47 | 23 | 0 | 0 | 0 | 0 | 0 | 298 | 149 | 1 | 1 | 1 | 1 | 0.937 | 1 | 1 | 1 | 1 | 1 | 1 | 1 | -0.013 |
| SE | 61898 | ENSG00000076242 | MLH1 | chr3 | + | 37042267 | 37042331 | 37040185 | 37040294 | 37047518 | 37047607 | 82 | 25 | 46 | 44 | 61 | 0 | 0 | 1 | 0 | 2 | 29 | 17 | 44 | 58 | 19 | 0 | 0 | 0 | 0 | 0 | 213 | 149 | 1 | 1 | 1 | 1 | 0.97 | 1 | 0.955 | 1 | 1 | 1 | 1 | 1 | -0.015 |
| SE | 61901 | ENSG00000076242 | MLH1 | chr3 | + | 37047518 | 37047683 | 37042267 | 37042331 | 37048516 | 37048609 | 110 | 44 | 76 | 71 | 100 | 2 | 1 | 0 | 0 | 2 | 63 | 34 | 58 | 113 | 26 | 0 | 1 | 1 | 2 | 0 | 298 | 149 | 1 | 1 | 0.965 | 0.957 | 1 | 1 | 0.962 | 1 | 0.944 | 0.967 | 0.966 | 1 | 0.001 |
| SE | 61902 | ENSG00000076242 | MLH1 | chr3 | + | 37047518 | 37047683 | 37042267 | 37042331 | 37048903 | 37049017 | 61 | 24 | 43 | 36 | 46 | 0 | 0 | 0 | 0 | 4 | 26 | 18 | 32 | 53 | 19 | 0 | 0 | 0 | 0 | 0 | 298 | 149 | 1 | 1 | 1 | 1 | 1 | 1 | 0.852 | 1 | 1 | 1 | 1 | 1 | -0.03 |
| SE | 61903 | ENSG00000076242 | MLH1 | chr3 | + | 37047518 | 37047683 | 37042267 | 37042331 | 37050485 | 37050844 | 59 | 23 | 40 | 33 | 47 | 0 | 0 | 0 | 3 | 0 | 26 | 17 | 29 | 51 | 19 | 0 | 0 | 0 | 0 | 0 | 298 | 149 | 1 | 1 | 1 | 1 | 1 | 0.846 | 1 | 1 | 1 | 1 | 1 | 1 | -0.031 |
| SE | 61904 | ENSG00000076242 | MLH1 | chr3 | + | 37048516 | 37048609 | 37042267 | 37042331 | 37048903 | 37049017 | 30 | 17 | 20 | 13 | 16 | 0 | 0 | 0 | 0 | 4 | 10 | 24 | 19 | 7 | 0 | 0 | 0 | 0 | 0 | 0 | 242 | 149 | 1 | 1 | 1 | 1 | 1 | 1 | 0.711 | 1 | 1 | 1 | 1 | NA | -0.058 |
| SE | 61906 | ENSG00000076242 | MLH1 | chr3 | + | 37048903 | 37049017 | 37042267 | 37042331 | 37050485 | 37050706 | 20 | 10 | 22 | 9 | 26 | 0 | 0 | 0 | 3 | 0 | 8 | 20 | 6 | 14 | 10 | 0 | 0 | 0 | 0 | 0 | 263 | 149 | 1 | 1 | 1 | 1 | 1 | 0.63 | 1 | 1 | 1 | 1 | 1 | 1 | -0.074 |
| SE | 61907 | ENSG00000076242 | MLH1 | chr3 | + | 37048903 | 37049017 | 37047518 | 37047683 | 37050485 | 37050706 | 22 | 11 | 25 | 12 | 22 | 0 | 0 | 0 | 0 | 1 | 8 | 21 | 9 | 16 | 10 | 0 | 0 | 0 | 0 | 0 | 263 | 149 | 1 | 1 | 1 | 1 | 1 | 1 | 0.926 | 1 | 1 | 1 | 1 | 1 | -0.015 |
| SE | 61908 | ENSG00000124067 | SLC12A4 | chr16 | - | 67946437 | 67946633 | 67946170 | 67946340 | 67946936 | 67947105 | 41 | 13 | 33 | 31 | 10 | 0 | 0 | 0 | 0 | 0 | 28 | 43 | 16 | 30 | 80 | 0 | 0 | 0 | 2 | 0 | 298 | 149 | 1 | 1 | 1 | 1 | 1 | 1 | 1 | 1 | 1 | 0.8 | 1 | 1 | 0.04 |
| SE | 61909 | ENSG00000124067 | SLC12A4 | chr16 | - | 67947330 | 67947435 | 67946936 | 67947105 | 67947668 | 67947788 | 28 | 7 | 27 | 22 | 15 | 0 | 0 | 1 | 3 | 0 | 44 | 37 | 16 | 31 | 75 | 0 | 0 | 0 | 2 | 0 | 254 | 149 | 1 | 1 | 1 | 1 | 0.941 | 0.811 | 1 | 1 | 1 | 1 | 0.901 | 1 | -0.03 |
| SE | 61919 | ENSG00000080947 | CROCCP3 | chr1 | - | 16476398 | 16476504 | 16475824 | 16476062 | 16476929 | 16477074 | 7 | 10 | 5 | 12 | 0 | 0 | 1 | 1 | 1 | 0 | 0 | 8 | 8 | 3 | 21 | 0 | 0 | 0 | 2 | 0 | 255 | 149 | 1 | 1 | 1 | 0.854 | 0.745 | 0.875 | NA | NA | 1 | 1 | 0.467 | 1 | 0.002 |
| SE | 61920 | ENSG00000080947 | CROCCP3 | chr1 | - | 16484822 | 16484906 | 16484232 | 16484464 | 16485568 | 16485665 | 13 | 5 | 10 | 14 | 0 | 2 | 1 | 4 | 6 | 2 | 0 | 1 | 3 | 8 | 1 | 0 | 1 | 0 | 0 | 0 | 233 | 149 | 1 | 1 | 0.806 | 0.762 | 0.615 | 0.599 | 0 | NA | 0.39 | 1 | 1 | 1 | -0.291 |
| SE | 61922 | ENSG00000080947 | CROCCP3 | chr1 | - | 16489684 | 16489791 | 16486425 | 16486661 | 16490617 | 16490783 | 12 | 15 | 15 | 6 | 2 | 3 | 2 | 4 | 6 | 0 | 0 | 3 | 4 | 4 | 30 | 0 | 0 | 0 | 3 | 0 | 256 | 149 | 1 | 1 | 0.7 | 0.814 | 0.686 | 0.368 | 1 | NA | 1 | 1 | 0.437 | 1 | -0.146 |
| SE | 61923 | ENSG00000080947 | CROCCP3 | chr1 | - | 16491009 | 16491316 | 16490864 | 16490926 | 16499202 | 16499257 | 5 | 3 | 6 | 0 | 0 | 0 | 0 | 0 | 4 | 0 | 0 | 0 | 3 | 2 | 13 | 11 | 1 | 2 | 0 | 3 | 298 | 149 | 1 | 1 | 1 | 1 | 1 | 0 | NA | 0 | 0 | 0.429 | 1 | 0.684 | 0.327 |
| SE | 61925 | ENSG00000175449 | RFESD | chr5 | + | 95652136 | 95652331 | 95646753 | 95646817 | 95654060 | 95654236 | 4 | 2 | 9 | 1 | 5 | 0 | 0 | 1 | 0 | 0 | 0 | 19 | 0 | 2 | 0 | 0 | 0 | 0 | 0 | 0 | 298 | 149 | 1 | 1 | 1 | 1 | 0.818 | 1 | 1 | NA | 1 | NA | 1 | NA | -0.036 |
| SE | 61926 | ENSG00000175449 | RFESD | chr5 | + | 95653116 | 95653214 | 95652136 | 95652331 | 95654060 | 95654236 | 8 | 2 | 10 | 0 | 4 | 3 | 2 | 4 | 1 | 5 | 0 | 0 | 0 | 1 | 1 | 0 | 15 | 0 | 2 | 0 | 247 | 149 | 1 | 1 | 0.617 | 0.376 | 0.601 | 0 | 0.326 | NA | 0 | NA | 0.232 | 1 | -0.027 |
| SE | 61927 | ENSG00000175449 | RFESD | chr5 | + | 95654332 | 95654367 | 95654045 | 95654236 | 95656045 | 95656190 | 12 | 10 | 22 | 4 | 32 | 0 | 0 | 3 | 0 | 3 | 0 | 13 | 3 | 7 | 0 | 3 | 0 | 0 | 0 | 0 | 184 | 149 | 1 | 1 | 1 | 1 | 0.856 | 1 | 0.896 | 0 | 1 | 1 | 1 | NA | 0.2 |
| SE | 61930 | ENSG00000205643 | CDPF1 | chr22 | - | 46247105 | 46247221 | 46244050 | 46245238 | 46248171 | 46248285 | 6 | 0 | 7 | 7 | 1 | 0 | 0 | 0 | 0 | 0 | 8 | 0 | 0 | 0 | 0 | 0 | 6 | 0 | 2 | 0 | 265 | 149 | 1 | 1 | 1 | NA | 1 | 1 | 1 | 1 | 0 | NA | 0 | NA | 0.667 |
| SE | 61931 | ENSG00000205643 | CDPF1 | chr22 | - | 46247109 | 46247221 | 46244012 | 46245238 | 46248171 | 46248284 | 18 | 2 | 14 | 18 | 8 | 0 | 0 | 0 | 0 | 0 | 8 | 16 | 2 | 0 | 0 | 0 | 6 | 0 | 2 | 0 | 261 | 149 | 1 | 1 | 1 | 1 | 1 | 1 | 1 | 1 | 0.604 | 1 | 0 | NA | 0.349 |
| SE | 61932 | ENSG00000205643 | CDPF1 | chr22 | - | 46247109 | 46247221 | 46244012 | 46245238 | 46250255 | 46250311 | 14 | 2 | 10 | 12 | 7 | 3 | 0 | 0 | 0 | 1 | 0 | 16 | 2 | 6 | 0 | 7 | 0 | 0 | 0 | 0 | 261 | 149 | 1 | 1 | 0.727 | 1 | 1 | 1 | 0.8 | 0 | 1 | 1 | 1 | NA | 0.155 |
| SE | 61933 | ENSG00000205643 | CDPF1 | chr22 | - | 46247124 | 46247221 | 46244010 | 46245238 | 46248171 | 46248285 | 5 | 0 | 3 | 5 | 2 | 0 | 0 | 0 | 0 | 0 | 8 | 0 | 2 | 0 | 0 | 0 | 6 | 0 | 2 | 0 | 246 | 149 | 1 | 1 | 1 | NA | 1 | 1 | 1 | 1 | 0 | 1 | 0 | NA | 0.5 |
| SE | 61934 | ENSG00000205643 | CDPF1 | chr22 | - | 46248171 | 46248284 | 46244012 | 46245238 | 46250255 | 46250289 | 0 | 0 | 0 | 0 | 0 | 3 | 0 | 0 | 0 | 1 | 0 | 6 | 0 | 2 | 0 | 7 | 0 | 0 | 0 | 0 | 262 | 149 | 1 | 1 | 0 | NA | NA | NA | 0 | 0 | 1 | NA | 1 | NA | -0.667 |
| SE | 61935 | ENSG00000205643 | CDPF1 | chr22 | - | 46248171 | 46248284 | 46247109 | 46247221 | 46250255 | 46250289 | 5 | 0 | 4 | 6 | 1 | 1 | 0 | 0 | 0 | 0 | 8 | 0 | 0 | 0 | 0 | 0 | 0 | 0 | 6 | 0 | 262 | 149 | 1 | 1 | 0.74 | NA | 1 | 1 | 1 | 1 | NA | NA | 0 | NA | 0.435 |

| | | | | | | | | | | | | | | | | | | | | | | | | | | | | | | | | | | | | | | | | | | | | | | |
|---|---|---|---|---|---|---|---|---|---|---|---|---|---|---|---|---|---|---|---|---|---|---|---|---|---|---|---|---|---|---|---|---|---|---|---|---|---|---|---|---|---|---|---|---|---|---|
| SE | 61939 | ENSG00000132846 | ZBED3 | chr5 | - | 77078593 | 77078728 | 77072071 | 77077895 | 77087110 | 77087285 | 8 | 0 | 7 | 1 | 0 | 0 | 0 | 0 | 0 | 0 | 5 | 0 | 1 | 0 | 0 | 0 | 0 | 0 | 2 | 0 | 284 | 149 | 1 | 1 | 1 | NA | 1 | 1 | NA | 1 | NA | 1 | 0 | NA | 0.333 |
| SE | 61942 | ENSG00000136932 | TRMO | chr9 | - | 97913400 | 97913558 | 97909959 | 97910616 | 97922417 | 97922468 | 22 | 18 | 26 | 42 | 53 | 0 | 0 | 1 | 0 | 0 | 16 | 23 | 40 | 22 | 0 | 0 | 0 | 0 | 0 | 0 | 298 | 149 | 1 | 1 | 1 | 1 | 0.929 | 1 | 1 | 1 | 1 | 1 | 1 | NA | -0.014 |
| SE | 61943 | ENSG00000136932 | TRMO | chr9 | - | 97913978 | 97914026 | 97913400 | 97913558 | 97916163 | 97916338 | 2 | 1 | 2 | 2 | 0 | 55 | 43 | 73 | 65 | 72 | 0 | 0 | 0 | 0 | 0 | 12 | 27 | 30 | 15 | 6 | 197 | 149 | 1 | 1 | 0.027 | 0.017 | 0.02 | 0.023 | 0 | 0 | 0 | 0 | 0 | 0 | 0.017 |
| SE | 61945 | ENSG00000112514 | CUTA | chr6 | - | 33416919 | 33416969 | 33416441 | 33416776 | 33417096 | 33417142 | 395 | 164 | 383 | 299 | 366 | 2 | 0 | 0 | 0 | 0 | 267 | 444 | 196 | 199 | 474 | 7 | 0 | 0 | 0 | 0 | 199 | 149 | 1 | 1 | 0.993 | 1 | 1 | 1 | 1 | 0.966 | 1 | 1 | 1 | 1 | 0.005 |
| SE | 61948 | ENSG00000163904 | SENP2 | chr3 | + | 185598411 | 185598545 | 185590113 | 185590169 | 185598991 | 185599024 | 15 | 9 | 31 | 15 | 18 | 0 | 0 | 0 | 0 | 0 | 11 | 18 | 16 | 6 | 26 | 0 | 0 | 2 | 1 | 0 | 283 | 149 | 1 | 1 | 1 | 1 | 1 | 1 | 1 | 1 | 1 | 0.808 | 0.76 | 1 | 0.086 |
| SE | 61949 | ENSG00000163904 | SENP2 | chr3 | + | 185598411 | 185598545 | 185590113 | 185590169 | 185600764 | 185600855 | 15 | 8 | 31 | 14 | 15 | 1 | 0 | 0 | 0 | 0 | 11 | 14 | 14 | 1 | 26 | 0 | 0 | 0 | 0 | 0 | 283 | 149 | 1 | 1 | 0.888 | 1 | 1 | 1 | 1 | 1 | 1 | 1 | 1 | 1 | -0.022 |
| SE | 61950 | ENSG00000163904 | SENP2 | chr3 | + | 185598957 | 185599024 | 185590113 | 185590169 | 185600764 | 185600825 | 45 | 37 | 58 | 66 | 60 | 1 | 0 | 0 | 0 | 0 | 49 | 72 | 49 | 111 | 36 | 0 | 0 | 0 | 0 | 0 | 216 | 149 | 1 | 1 | 0.969 | 1 | 1 | 1 | 1 | 1 | 1 | 1 | 1 | 1 | -0.006 |
| SE | 61951 | ENSG00000163904 | SENP2 | chr3 | + | 185598957 | 185599024 | 185590113 | 185590169 | 185606329 | 185606490 | 0 | 0 | 0 | 0 | 0 | 1 | 0 | 0 | 0 | 1 | 0 | 0 | 0 | 11 | 0 | 0 | 0 | 0 | 0 | 0 | 216 | 149 | 1 | 1 | 0 | NA | NA | NA | 0 | NA | NA | NA | 1 | NA | -1 |
| SE | 61952 | ENSG00000163904 | SENP2 | chr3 | + | 185598957 | 185599024 | 185598393 | 185598545 | 185600764 | 185600825 | 70 | 65 | 99 | 111 | 111 | 0 | 0 | 2 | 0 | 0 | 74 | 120 | 82 | 122 | 40 | 0 | 0 | 0 | 0 | 0 | 216 | 149 | 1 | 1 | 1 | 1 | 0.972 | 1 | 1 | 1 | 1 | 1 | 1 | 1 | -0.006 |
| SE | 61953 | ENSG00000163904 | SENP2 | chr3 | + | 185598991 | 185599024 | 185590113 | 185590169 | 185600764 | 185600855 | 30 | 33 | 41 | 42 | 40 | 1 | 0 | 0 | 0 | 0 | 34 | 43 | 40 | 79 | 22 | 0 | 0 | 0 | 0 | 0 | 182 | 149 | 1 | 1 | 0.961 | 1 | 1 | 1 | 1 | 1 | 1 | 1 | 1 | 1 | -0.008 |
| SE | 61955 | ENSG00000163904 | SENP2 | chr3 | + | 185598991 | 185599024 | 185598411 | 185598545 | 185600764 | 185600855 | 32 | 34 | 42 | 42 | 43 | 0 | 0 | 2 | 0 | 0 | 34 | 47 | 40 | 83 | 22 | 0 | 0 | 0 | 0 | 0 | 182 | 149 | 1 | 1 | 1 | 1 | 0.945 | 1 | 1 | 1 | 1 | 1 | 1 | 1 | -0.011 |
| SE | 61956 | ENSG00000163904 | SENP2 | chr3 | + | 185600764 | 185600855 | 185590113 | 185590169 | 185606329 | 185606490 | 44 | 36 | 44 | 63 | 72 | 1 | 0 | 0 | 0 | 1 | 76 | 53 | 55 | 94 | 29 | 0 | 0 | 0 | 0 | 0 | 240 | 149 | 1 | 1 | 0.965 | 1 | 1 | 1 | 0.978 | 1 | 1 | 1 | 1 | 1 | -0.011 |
| SE | 61957 | ENSG00000163904 | SENP2 | chr3 | + | 185600764 | 185600855 | 185598957 | 185599024 | 185606329 | 185606490 | 100 | 88 | 129 | 166 | 159 | 0 | 0 | 0 | 0 | 0 | 156 | 177 | 132 | 227 | 77 | 0 | 0 | 0 | 6 | 0 | 240 | 149 | 1 | 1 | 1 | 1 | 1 | 1 | 1 | 1 | 1 | 1 | 0.959 | 1 | 0.008 |
| SE | 61958 | ENSG00000163904 | SENP2 | chr3 | + | 185609246 | 185609350 | 185606329 | 185606498 | 185611650 | 185611745 | 271 | 180 | 206 | 260 | 511 | 0 | 0 | 0 | 0 | 0 | 415 | 298 | 405 | 443 | 222 | 0 | 0 | 0 | 8 | 0 | 253 | 149 | 1 | 1 | 1 | 1 | 1 | 1 | 1 | 1 | 1 | 1 | 0.97 | 1 | 0.006 |
| SE | 61960 | ENSG00000163904 | SENP2 | chr3 | + | 185614563 | 185614740 | 185613344 | 185613408 | 185617479 | 185617509 | 226 | 184 | 231 | 196 | 336 | 0 | 1 | 0 | 0 | 1 | 304 | 293 | 388 | 388 | 154 | 0 | 0 | 0 | 4 | 0 | 298 | 149 | 1 | 1 | 1 | 0.989 | 1 | 1 | 0.994 | 1 | 1 | 1 | 0.98 | 1 | 0.001 |
| SE | 61961 | ENSG00000163904 | SENP2 | chr3 | + | 185614563 | 185614740 | 185613344 | 185613408 | 185619298 | 185619502 | 84 | 60 | 88 | 77 | 148 | 0 | 0 | 0 | 0 | 0 | 141 | 142 | 180 | 225 | 102 | 0 | 2 | 0 | 0 | 0 | 298 | 149 | 1 | 1 | 1 | 1 | 1 | 1 | 1 | 1 | 0.973 | 1 | 1 | 1 | 0.005 |
| SE | 61962 | ENSG00000163904 | SENP2 | chr3 | + | 185617479 | 185617611 | 185613344 | 185613408 | 185619298 | 185619502 | 225 | 194 | 197 | 178 | 257 | 0 | 0 | 0 | 0 | 0 | 180 | 203 | 305 | 237 | 112 | 0 | 2 | 0 | 0 | 0 | 281 | 149 | 1 | 1 | 1 | 1 | 1 | 1 | 1 | 1 | 0.982 | 1 | 1 | 1 | 0.004 |
| SE | 61963 | ENSG00000163904 | SENP2 | chr3 | + | 185617479 | 185617611 | 185613344 | 185613408 | 185621825 | 185621905 | 1 | 7 | 0 | 0 | 2 | 0 | 0 | 0 | 0 | 0 | 0 | 9 | 0 | 4 | 0 | 0 | 0 | 0 | 1 | 0 | 281 | 149 | 1 | 1 | 1 | 1 | NA | NA | 1 | NA | 1 | NA | 0.68 | NA | 0.16 |
| SE | 61965 | ENSG00000163904 | SENP2 | chr3 | + | 185617479 | 185617611 | 185614502 | 185614740 | 185623997 | 185624082 | 130 | 120 | 134 | 114 | 177 | 0 | 0 | 0 | 2 | 0 | 157 | 145 | 190 | 161 | 51 | 0 | 0 | 0 | 0 | 0 | 281 | 149 | 1 | 1 | 1 | 1 | 1 | 0.968 | 1 | 1 | 1 | 1 | 1 | 1 | -0.006 |
| SE | 61966 | ENSG00000163904 | SENP2 | chr3 | + | 185619298 | 185619502 | 185613344 | 185613408 | 185621825 | 185621905 | 62 | 43 | 69 | 77 | 109 | 0 | 0 | 0 | 0 | 0 | 36 | 90 | 63 | 78 | 35 | 0 | 0 | 0 | 1 | 0 | 298 | 149 | 1 | 1 | 1 | 1 | 1 | 1 | 1 | 1 | 1 | 1 | 0.975 | 1 | 0.005 |
| SE | 61967 | ENSG00000163904 | SENP2 | chr3 | + | 185619298 | 185619502 | 185614563 | 185614740 | 185623997 | 185624082 | 2 | 2 | 5 | 2 | 3 | 0 | 0 | 0 | 2 | 0 | 0 | 9 | 8 | 3 | 13 | 0 | 0 | 0 | 0 | 0 | 298 | 149 | 1 | 1 | 1 | 1 | 1 | 0.333 | 1 | NA | 1 | 1 | 1 | 1 | -0.133 |
| SE | 61968 | ENSG00000163904 | SENP2 | chr3 | + | 185619298 | 185619502 | 185617479 | 185617611 | 185621825 | 185621905 | 304 | 240 | 278 | 263 | 383 | 1 | 6 | 0 | 0 | 1 | 225 | 310 | 391 | 316 | 172 | 0 | 9 | 0 | 0 | 0 | 298 | 149 | 1 | 1 | 0.993 | 0.952 | 1 | 1 | 0.995 | 1 | 0.945 | 1 | 1 | 1 | -0.001 |
| SE | 61972 | ENSG00000163904 | SENP2 | chr3 | + | 185623997 | 185624082 | 185617479 | 185617611 | 185626297 | 185626307 | 29 | 11 | 17 | 23 | 57 | 0 | 0 | 0 | 0 | 0 | 35 | 64 | 25 | 49 | 12 | 0 | 0 | 3 | 0 | 0 | 234 | 149 | 1 | 1 | 1 | 1 | 1 | 1 | 1 | 1 | 1 | 1 | 0.912 | 1 | 0.018 |
| SE | 61974 | ENSG00000205423 | CNEP1R1 | chr16 | + | 50029724 | 50029798 | 50026159 | 50026467 | 50033396 | 50033506 | 50 | 32 | 40 | 37 | 45 | 2 | 0 | 0 | 0 | 0 | 33 | 28 | 40 | 34 | 22 | 0 | 0 | 0 | 0 | 0 | 223 | 149 | 1 | 1 | 0.944 | 1 | 1 | 1 | 1 | 1 | 1 | 1 | 1 | 1 | -0.011 |
| SE | 61978 | ENSG00000044115 | CTNNA1 | chr5 | + | 138782289 | 138782403 | 138781922 | 138782029 | 138783176 | 138783372 | 2 | 0 | 0 | 4 | 0 | 591 | 437 | 280 | 388 | 213 | 0 | 4 | 3 | 0 | 5 | 2343 | 763 | 489 | 1201 | 2423 | 263 | 149 | 1 | 1 | 0.002 | 0 | 0 | 0.006 | 0 | 0 | 0.003 | 0.003 | 0 | 0.001 | 0 |
| SE | 61980 | ENSG00000044115 | CTNNA1 | chr5 | + | 138783176 | 138783372 | 138781922 | 138782029 | 138824529 | 138824588 | 586 | 435 | 278 | 388 | 213 | 0 | 0 | 0 | 0 | 0 | 2343 | 763 | 489 | 1201 | 2423 | 0 | 3 | 3 | 0 | 0 | 298 | 149 | 1 | 1 | 1 | 1 | 1 | 1 | 1 | 1 | 0.992 | 0.988 | 1 | 1 | 0.004 |
| SE | 61981 | ENSG00000044115 | CTNNA1 | chr5 | + | 138810037 | 138810204 | 138781922 | 138782029 | 138824529 | 138824588 | 7 | 0 | 1 | 2 | 7 | 0 | 0 | 0 | 0 | 0 | 10 | 0 | 0 | 4 | 0 | 0 | 3 | 3 | 0 | 0 | 298 | 149 | 1 | 1 | 1 | NA | 1 | 1 | 1 | 1 | 0 | 0 | 1 | NA | 0.5 |
| SE | 61982 | ENSG00000044115 | CTNNA1 | chr5 | + | 138810037 | 138810204 | 138783176 | 138783372 | 138812182 | 138812302 | 348 | 196 | 172 | 251 | 415 | 1 | 0 | 3 | 0 | 0 | 573 | 268 | 153 | 463 | 627 | 0 | 0 | 0 | 0 | 0 | 298 | 149 | 1 | 1 | 0.994 | 1 | 0.966 | 1 | 1 | 1 | 1 | 1 | 1 | 1 | -0.008 |
| SE | 61983 | ENSG00000044115 | CTNNA1 | chr5 | + | 138810037 | 138810204 | 138783176 | 138783372 | 138824529 | 138824588 | 130 | 87 | 71 | 86 | 184 | 1 | 0 | 0 | 0 | 0 | 222 | 93 | 47 | 213 | 235 | 0 | 0 | 0 | 0 | 0 | 298 | 149 | 1 | 1 | 0.985 | 1 | 1 | 1 | 1 | 1 | 1 | 1 | 1 | 1 | -0.003 |
| SE | 61984 | ENSG00000044115 | CTNNA1 | chr5 | + | 138812182 | 138812302 | 138783176 | 138783372 | 138824529 | 138824588 | 335 | 261 | 203 | 267 | 378 | 1 | 0 | 0 | 0 | 0 | 935 | 412 | 186 | 367 | 868 | 0 | 0 | 0 | 0 | 0 | 269 | 149 | 1 | 1 | 0.995 | 1 | 1 | 1 | 1 | 1 | 1 | 1 | 1 | 1 | -0.001 |
| SE | 61985 | ENSG00000044115 | CTNNA1 | chr5 | + | 138812182 | 138812302 | 138810037 | 138810204 | 138824529 | 138824588 | 552 | 370 | 300 | 432 | 609 | 0 | 0 | 0 | 0 | 2 | 1286 | 587 | 292 | 617 | 1254 | 0 | 0 | 0 | 0 | 0 | 269 | 149 | 1 | 1 | 1 | 1 | 1 | 1 | 0.994 | 1 | 1 | 1 | 1 | 1 | -0.001 |
| SE | 61986 | ENSG00000044115 | CTNNA1 | chr5 | + | 138824529 | 138824799 | 138812182 | 138812302 | 138827514 | 138827718 | 609 | 428 | 353 | 471 | 640 | 0 | 0 | 0 | 0 | 0 | 1300 | 719 | 366 | 647 | 1648 | 0 | 0 | 0 | 3 | 0 | 298 | 149 | 1 | 1 | 1 | 1 | 1 | 1 | 1 | 1 | 1 | 1 | 0.991 | 1 | 0.002 |
| SE | 61995 | ENSG00000044115 | CTNNA1 | chr5 | + | 138881041 | 138881137 | 138827514 | 138827718 | 138886211 | 138886292 | 0 | 0 | 0 | 0 | 0 | 166 | 98 | 91 | 141 | 116 | 0 | 5 | 0 | 0 | 0 | 319 | 202 | 123 | 216 | 514 | 245 | 149 | 1 | 1 | 0 | 0 | 0 | 0 | 0 | 0 | 0.015 | 0 | 0 | 0 | -0.003 |
| SE | 61999 | ENSG00000044115 | CTNNA1 | chr5 | + | 138891182 | 138891211 | 138887489 | 138887642 | 138904348 | 138904441 | 0 | 0 | 0 | 0 | 4 | 141 | 91 | 72 | 95 | 135 | 0 | 0 | 0 | 0 | 0 | 301 | 115 | 72 | 165 | 293 | 178 | 149 | 1 | 1 | 0 | 0 | 0 | 0 | 0.024 | 0 | 0 | 0 | 0 | 0 | 0.005 |
| SE | 62000 | ENSG00000044115 | CTNNA1 | chr5 | + | 138904348 | 138904441 | 138887489 | 138887642 | 138917741 | 138917779 | 263 | 176 | 142 | 191 | 244 | 0 | 0 | 0 | 0 | 0 | 574 | 232 | 134 | 296 | 513 | 0 | 0 | 1 | 3 | 0 | 242 | 149 | 1 | 1 | 1 | 1 | 1 | 1 | 1 | 1 | 1 | 0.988 | 0.984 | 1 | 0.006 |
| SE | 62002 | ENSG00000044115 | CTNNA1 | chr5 | + | 138929245 | 138929356 | 138925255 | 138925407 | 138930472 | 138930509 | 319 | 224 | 153 | 278 | 308 | 0 | 0 | 0 | 2 | 0 | 614 | 357 | 175 | 384 | 743 | 0 | 0 | 0 | 0 | 0 | 260 | 149 | 1 | 1 | 1 | 1 | 1 | 0.988 | 1 | 1 | 1 | 1 | 1 | 1 | -0.002 |
| SE | 62004 | ENSG00000044115 | CTNNA1 | chr5 | + | 138932887 | 138932958 | 138932577 | 138932712 | 138933801 | 138934831 | 2 | 1 | 0 | 0 | 0 | 127 | 57 | 55 | 88 | 96 | 0 | 0 | 0 | 0 | 0 | 335 | 91 | 39 | 143 | 239 | 220 | 149 | 1 | 1 | 0.011 | 0.012 | 0 | 0 | 0 | 0 | 0 | 0 | 0 | 0 | 0.005 |
| SE | 62009 | ENSG00000182885 | ADGRG3 | chr16 | + | 57676199 | 57676338 | 57673320 | 57673468 | 57678169 | 57678282 | 11 | 0 | 12 | 14 | 10 | 0 | 0 | 0 | 3 | 0 | 42 | 18 | 165 | 290 | 87 | 0 | 0 | 8 | 5 | 3 | 288 | 149 | 1 | 1 | 1 | NA | 1 | 0.707 | 1 | 1 | 1 | 0.914 | 0.968 | 0.938 | -0.037 |
| SE | 62010 | ENSG00000182885 | ADGRG3 | chr16 | + | 57679176 | 57679311 | 57678169 | 57678316 | 57679815 | 57679855 | 3 | 0 | 6 | 4 | 8 | 0 | 0 | 0 | 0 | 0 | 6 | 23 | 82 | 138 | 65 | 0 | 0 | 0 | 0 | 4 | 284 | 149 | 1 | 1 | 1 | NA | 1 | 1 | 1 | 1 | 1 | 1 | 1 | 0.895 | 0.021 |
| SE | 62011 | ENSG00000182885 | ADGRG3 | chr16 | + | 57680504 | 57680617 | 57680264 | 57680365 | 57684389 | 57684483 | 10 | 0 | 1 | 10 | 0 | 0 | 0 | 0 | 0 | 0 | 0 | 0 | 26 | 82 | 24 | 0 | 0 | 1 | 0 | 0 | 262 | 149 | 1 | 1 | 1 | NA | 1 | 1 | NA | NA | NA | 0.937 | 1 | 1 | 0.021 |
| SE | 62012 | ENSG00000182885 | ADGRG3 | chr16 | + | 57683931 | 57684212 | 57680504 | 57680617 | 57684389 | 57684483 | 7 | 0 | 9 | 6 | 1 | 0 | 0 | 0 | 0 | 0 | 0 | 28 | 30 | 96 | 53 | 0 | 0 | 9 | 2 | 0 | 298 | 149 | 1 | 1 | 1 | NA | 1 | 1 | 1 | NA | 1 | 0.625 | 0.96 | 1 | 0.104 |
| SE | 62013 | ENSG00000182885 | ADGRG3 | chr16 | + | 57684389 | 57684483 | 57680504 | 57680617 | 57685642 | 57685926 | 1 | 0 | 5 | 5 | 7 | 0 | 0 | 0 | 0 | 0 | 2 | 7 | 24 | 24 | 14 | 0 | 2 | 0 | 0 | 0 | 243 | 149 | 1 | 1 | 1 | NA | 1 | 1 | 1 | 1 | 0.682 | 1 | 1 | 1 | 0.064 |
| SE | 62014 | ENSG00000182885 | ADGRG3 | chr16 | + | 57685642 | 57685926 | 57680504 | 57680617 | 57688351 | 57688695 | 2 | 0 | 4 | 8 | 10 | 0 | 0 | 0 | 1 | 0 | 2 | 8 | 48 | 57 | 27 | 0 | 0 | 0 | 0 | 0 | 298 | 149 | 1 | 1 | 1 | NA | 1 | 0.8 | 1 | 1 | 1 | 1 | 1 | 1 | -0.05 |
| SE | 62017 | ENSG00000136930 | PSMB7 | chr9 | - | 124384076 | 124384208 | 124383595 | 124383950 | 124384597 | 124384656 | 0 | 0 | 0 | 3 | 0 | 0 | 0 | 0 | 1 | 0 | 0 | 2 | 2 | 7 | 0 | 0 | 0 | 0 | 0 | 0 | 281 | 149 | 1 | 1 | NA | NA | NA | 0.614 | NA | NA | 1 | 1 | 1 | NA | -0.386 |
| SE | 62018 | ENSG00000136930 | PSMB7 | chr9 | - | 124384597 | 124384656 | 124356763 | 124356915 | 124405316 | 124405432 | 391 | 177 | 300 | 306 | 589 | 0 | 0 | 0 | 3 | 0 | 631 | 495 | 347 | 535 | 746 | 0 | 0 | 1 | 0 | 0 | 208 | 149 | 1 | 1 | 1 | 1 | 1 | 0.986 | 1 | 1 | 1 | 0.996 | 1 | 1 | -0.002 |
| SE | 62025 | ENSG00000135766 | EGLN1 | chr1 | - | 231373979 | 231374099 | 231370561 | 231370698 | 231420997 | 231421110 | 279 | 161 | 255 | 327 | 492 | 0 | 2 | 3 | 1 | 0 | 330 | 221 | 204 | 189 | 128 | 7 | 0 | 1 | 0 | 0 | 269 | 149 | 1 | 1 | 1 | 0.978 | 0.979 | 0.995 | 1 | 0.963 | 1 | 1 | 1 | 1 | -0.002 |
| SE | 62028 | ENSG00000124313 | IQSEC2 | chrX | - | 53235782 | 53235832 | 53232874 | 53235184 | 53236321 | 53236495 | 2 | 0 | 0 | 1 | 0 | 6 | 2 | 2 | 4 | 5 | 4 | 0 | 0 | 0 | 0 | 9 | 4 | 2 | 0 | 26 | 199 | 149 | 1 | 1 | 0.2 | 0 | 0 | 0.158 | 0 | 0.25 | 0 | 0 | NA | 0 | 0.009 |
| SE | 62030 | ENSG00000124313 | IQSEC2 | chrX | - | 53246968 | 53247135 | 53243053 | 53243471 | 53248113 | 53248156 | 26 | 24 | 14 | 38 | 24 | 0 | 0 | 0 | 1 | 0 | 91 | 9 | 17 | 24 | 41 | 0 | 0 | 0 | 0 | 0 | 298 | 149 | 1 | 1 | 1 | 1 | 1 | 0.95 | 1 | 1 | 1 | 1 | 1 | 1 | -0.01 |
| SE | 62031 | ENSG00000124313 | IQSEC2 | chrX | - | 53279563 | 53279659 | 53255799 | 53256061 | 53281495 | 53281535 | 1 | 0 | 0 | 0 | 4 | 0 | 0 | 0 | 2 | 0 | 9 | 0 | 0 | 0 | 8 | 0 | 0 | 0 | 0 | 0 | 245 | 149 | 1 | 1 | 1 | NA | NA | 0 | 1 | 1 | NA | NA | NA | 1 | -0.333 |
| SE | 62032 | ENSG00000124313 | IQSEC2 | chrX | - | 53279563 | 53279659 | 53255799 | 53256061 | 53291894 | 53291916 | 0 | 0 | 0 | 0 | 2 | 3 | 2 | 2 | 3 | 3 | 1 | 0 | 0 | 0 | 0 | 0 | 9 | 5 | 0 | 11 | 245 | 149 | 1 | 1 | 0 | 0 | 0 | 0 | 0.288 | 1 | 0 | 0 | NA | 0 | -0.192 |
| SE | 62034 | ENSG00000124313 | IQSEC2 | chrX | - | 53279563 | 53279659 | 53266327 | 53267064 | 53281495 | 53281535 | 3 | 0 | 0 | 0 | 2 | 0 | 2 | 0 | 1 | 0 | 8 | 5 | 0 | 1 | 10 | 0 | 0 | 1 | 0 | 0 | 245 | 149 | 1 | 1 | 1 | 0 | NA | 0 | 1 | 1 | 1 | 0 | 1 | 1 | -0.3 |
| SE | 62035 | ENSG00000124313 | IQSEC2 | chrX | - | 53279563 | 53279776 | 53266641 | 53267064 | 53281495 | 53281581 | 0 | 0 | 0 | 0 | 2 | 0 | 2 | 0 | 1 | 0 | 0 | 5 | 0 | 1 | 1 | 0 | 0 | 1 | 0 | 0 | 298 | 149 | 1 | 1 | NA | 0 | NA | 0 | 1 | NA | 1 | 0 | 1 | 1 | -0.417 |
| SE | 62037 | ENSG00000218510 | LINC00339 | chr1 | + | 22027413 | 22027534 | 22025492 | 22025945 | 22030279 | 22030374 | 3 | 0 | 0 | 1 | 0 | 0 | 0 | 2 | 4 | 0 | 0 | 0 | 0 | 0 | 0 | 0 | 2 | 0 | 0 | 0 | 270 | 149 | 1 | 1 | 1 | NA | 0 | 0.121 | NA | NA | 0 | NA | NA | NA | 0.374 |
| SE | 62038 | ENSG00000218510 | LINC00339 | chr1 | + | 22027413 | 22027534 | 22025510 | 22025639 | 22030279 | 22030374 | 4 | 0 | 0 | 5 | 0 | 0 | 1 | 2 | 0 | 0 | 9 | 0 | 0 | 0 | 0 | 0 | 0 | 0 | 0 | 0 | 270 | 149 | 1 | 1 | 1 | 0 | 0 | 1 | NA | 1 | NA | NA | NA | NA | -0.5 |
| SE | 62039 | ENSG00000218510 | LINC00339 | chr1 | + | 22027413 | 22027534 | 22025521 | 22025643 | 22030279 | 22030374 | 5 | 0 | 0 | 2 | 0 | 2 | 0 | 2 | 13 | 6 | 0 | 1 | 0 | 0 | 0 | 0 | 4 | 2 | 6 | 9 | 270 | 149 | 1 | 1 | 0.58 | NA | 0 | 0.078 | 0 | NA | 0.121 | 0 | 0 | 0 | 0.134 |
| SE | 62040 | ENSG00000218510 | LINC00339 | chr1 | + | 22027413 | 22028379 | 22025552 | 22025639 | 22030279 | 22030617 | 4 | 0 | 0 | 4 | 0 | 0 | 1 | 2 | 0 | 0 | 9 | 0 | 2 | 0 | 0 | 0 | 0 | 0 | 0 | 0 | 298 | 149 | 1 | 1 | 1 | 0 | 0 | 1 | NA | 1 | NA | 1 | NA | NA | -0.5 |
| SE | 62042 | ENSG00000218510 | LINC00339 | chr1 | + | 22028527 | 22028660 | 22025531 | 22025643 | 22030279 | 22031223 | 0 | 1 | 0 | 0 | 0 | 2 | 0 | 2 | 13 | 6 | 0 | 0 | 0 | 0 | 0 | 0 | 4 | 2 | 6 | 9 | 282 | 149 | 1 | 1 | 0 | 1 | 0 | 0 | 0 | NA | 0 | 0 | 0 | 0 | 0.2 |
| SE | 62043 | ENSG00000144040 | SFXN5 | chr2 | - | 72956979 | 72957103 | 72944739 | 72945099 | 72961130 | 72961248 | 2 | 0 | 4 | 5 | 1 | 13 | 1 | 14 | 21 | 0 | 0 | 0 | 0 | 0 | 0 | 0 | 14 | 18 | 21 | 38 | 273 | 149 | 1 | 1 | 0.077 | 0 | 0.135 | 0.115 | 1 | NA | 0 | 0 | 0 | 0 | 0.265 |
| SE | 62044 | ENSG00000144040 | SFXN5 | chr2 | - | 72956979 | 72957103 | 72944739 | 72945099 | 72968447 | 72968533 | 1 | 0 | 3 | 4 | 1 | 2 | 0 | 1 | 1 | 2 | 0 | 0 | 0 | 0 | 4 | 0 | 0 | 2 | 3 | 14 | 273 | 149 | 1 | 1 | 0.214 | NA | 0.621 | 0.686 | 0.214 | NA | NA | 0 | 0 | 0.135 | 0.389 |
| SE | 62045 | ENSG00000144040 | SFXN5 | chr2 | - | 72961130 | 72961248 | 72942035 | 72945099 | 72968447 | 72968533 | 15 | 2 | 18 | 22 | 0 | 2 | 0 | 1 | 1 | 2 | 0 | 14 | 17 | 21 | 38 | 0 | 0 | 2 | 3 | 14 | 267 | 149 | 1 | 1 | 0.807 | 1 | 0.909 | 0.925 | 0 | NA | 1 | 0.826 | 0.796 | 0.602 | -0.078 |
| SE | 62046 | ENSG00000144040 | SFXN5 | chr2 | - | 72961130 | 72961248 | 72942035 | 72945099 | 72988257 | 72988348 | 9 | 1 | 11 | 16 | 1 | 1 | 0 | 0 | 0 | 0 | 0 | 10 | 14 | 20 | 38 | 0 | 0 | 0 | 0 | 0 | 267 | 149 | 1 | 1 | 0.834 | 1 | 1 | 1 | 1 | NA | 1 | 1 | 1 | 1 | -0.033 |
| SE | 62047 | ENSG00000144040 | SFXN5 | chr2 | - | 72961130 | 72961248 | 72956979 | 72957103 | 72968447 | 72968533 | 3 | 1 | 7 | 4 | 0 | 0 | 0 | 0 | 0 | 0 | 0 | 0 | 0 | 0 | 0 | 0 | 0 | 0 | 0 | 4 | 267 | 149 | 1 | 1 | 1 | 1 | 1 | 1 | NA | NA | NA | NA | NA | 0 | 1 |
| SE | 62048 | ENSG00000144040 | SFXN5 | chr2 | - | 72967236 | 72967323 | 72945021 | 72945099 | 72968447 | 72968533 | 0 | 1 | 3 | 6 | 4 | 2 | 0 | 1 | 1 | 2 | 0 | 6 | 0 | 6 | 6 | 0 | 0 | 2 | 3 | 14 | 236 | 149 | 1 | 1 | 0 | 1 | 0.654 | 0.791 | 0.558 | NA | 1 | 0 | 0.558 | 0.213 | 0.158 |
| SE | 62049 | ENSG00000144040 | SFXN5 | chr2 | - | 72967236 | 72967323 | 72961201 | 72961248 | 72968447 | 72968533 | 2 | 1 | 3 | 6 | 0 | 7 | 1 | 7 | 7 | 0 | 0 | 6 | 4 | 5 | 6 | 0 | 4 | 3 | 1 | 0 | 236 | 149 | 1 | 1 | 0.153 | 0.387 | 0.213 | 0.351 | NA | NA | 0.486 | 0.457 | 0.759 | 1 | -0.399 |
| SE | 62050 | ENSG00000144040 | SFXN5 | chr2 | - | 72968447 | 72968533 | 72942035 | 72945099 | 72988257 | 72988348 | 0 | 0 | 1 | 0 | 0 | 1 | 0 | 0 | 0 | 0 | 11 | 0 | 2 | 4 | 14 | 0 | 0 | 0 | 0 | 0 | 235 | 149 | 1 | 1 | 0 | NA | 1 | NA | NA | 1 | NA | 1 | 1 | 1 | -0.5 |
| SE | 62051 | ENSG00000144040 | SFXN5 | chr2 | - | 72968447 | 72968533 | 72961130 | 72961248 | 72988257 | 72988348 | 6 | 0 | 5 | 4 | 0 | 0 | 0 | 0 | 0 | 1 | 11 | 4 | 3 | 2 | 0 | 0 | 0 | 0 | 0 | 0 | 235 | 149 | 1 | 1 | 1 | NA | 1 | 1 | 0 | 1 | 1 | 1 | 1 | NA | -0.25 |
| SE | 62053 | ENSG00000144040 | SFXN5 | chr2 | - | 73022521 | 73022576 | 73019446 | 73020264 | 73023187 | 73023214 | 35 | 25 | 21 | 31 | 16 | 2 | 0 | 0 | 0 | 0 | 36 | 62 | 25 | 64 | 76 | 0 | 0 | 0 | 0 | 0 | 204 | 149 | 1 | 1 | 0.927 | 1 | 1 | 1 | 1 | 1 | 1 | 1 | 1 | 1 | -0.015 |
| SE | 62056 | ENSG00000144040 | SFXN5 | chr2 | - | 73058527 | 73058596 | 73040853 | 73040931 | 73070459 | 73070567 | 16 | 5 | 9 | 5 | 7 | 0 | 2 | 0 | 0 | 0 | 16 | 42 | 15 | 12 | 63 | 0 | 0 | 0 | 0 | 0 | 218 | 149 | 1 | 1 | 1 | 0.631 | 1 | 1 | 1 | 1 | 1 | 1 | 1 | 1 | -0.074 |
| SE | 62058 | ENSG00000169507 | SLC38A11 | chr2 | - | 164907954 | 164908090 | 164898607 | 164898730 | 164908639 | 164908771 | 2 | 0 | 0 | 0 | 0 | 1 | 0 | 0 | 0 | 2 | 0 | 0 | 0 | 0 | 0 | 0 | 0 | 6 | 0 | 0 | 285 | 149 | 1 | 1 | 0.511 | NA | NA | NA | 0 | NA | 0 | NA | NA | NA | 0.256 |
| SE | 62060 | ENSG00000169507 | SLC38A11 | chr2 | - | 164944568 | 164944634 | 164939449 | 164939556 | 164945592 | 164945727 | 5 | 1 | 0 | 0 | 2 | 1 | 0 | 0 | 0 | 0 | 0 | 4 | 0 | 0 | 0 | 0 | 0 | 0 | 0 | 0 | 215 | 149 | 1 | 1 | 0.776 | 1 | NA | NA | 1 | NA | 1 | NA | NA | NA | -0.075 |
| SE | 62073 | ENSG00000196550 | FAM72A | chr1 | - | 206201873 | 206201889 | 206199806 | 206199884 | 206201993 | 206202446 | 0 | 3 | 1 | 0 | 3 | 0 | 0 | 2 | 0 | 0 | 0 | 0 | 0 | 4 | 0 | 0 | 0 | 0 | 1 | 2 | 0 | 165 | 149 | 1 | 1 | NA | 1 | 0.311 | NA | 1 | NA | NA | 0 | 0.644 | NA | 0.448 |
| SE | 62076 | ENSG00000263528 | IKBKE | chr1 | + | 206478159 | 206478339 | 206477748 | 206477859 | 206478942 | 206479133 | 33 | 23 | 29 | 32 | 27 | 0 | 0 | 0 | 1 | 0 | 40 | 34 | 13 | 51 | 58 | 0 | 0 | 0 | 0 | 0 | 298 | 149 | 1 | 1 | 1 | 1 | 1 | 0.941 | 1 | 1 | 1 | 1 | 1 | 1 | -0.012 |
| SE | 62081 | ENSG00000263528 | IKBKE | chr1 | + | 206493265 | 206493378 | 206491647 | 206491749 | 206496111 | 206496274 | 0 | 0 | 1 | 0 | 0 | 0 | 0 | 0 | 3 | 2 | 6 | 0 | 1 | 0 | 0 | 16 | 5 | 0 | 1 | 6 | 262 | 149 | 1 | 1 | NA | NA | 1 | 0 | 0 | 0.176 | 0 | 1 | 0 | 0 | 0.098 |
| SE | 62086 | ENSG00000263528 | IKBKE | chr1 | + | 206493919 | 206493991 | 206493265 | 206493378 | 206496111 | 206496274 | 89 | 39 | 85 | 80 | 85 | 0 | 0 | 1 | 0 | 0 | 218 | 69 | 47 | 105 | 79 | 0 | 0 | 0 | 0 | 0 | 221 | 149 | 1 | 1 | 1 | 1 | 0.983 | 1 | 1 | 1 | 1 | 1 | 1 | 1 | -0.003 |
| SE | 62087 | ENSG00000143486 | EIF2D | chr1 | - | 206593618 | 206593793 | 206580934 | 206581162 | 206595717 | 206595838 | 68 | 50 | 88 | 50 | 137 | 0 | 0 | 2 | 0 | 0 | 38 | 49 | 35 | 34 | 58 | 0 | 0 | 0 | 0 | 0 | 298 | 149 | 1 | 1 | 1 | 1 | 0.957 | 1 | 1 | 1 | 1 | 1 | 1 | 1 | -0.009 |
| SE | 62089 | ENSG00000143486 | EIF2D | chr1 | - | 206599462 | 206599612 | [illegible] | 206599092 | 206600262 | 206600308 | 70 | 55 | 65 | 71 | 75 | 1 | 0 | 1 | 2 | 0 | 45 | 38 | 41 | 33 | 135 | 22 | 3 | 0 | 11 | 0 | 298 | 149 | 1 | 1 | 0.972 | 1 | 0.97 | 0.947 | 1 | 0.506 | 0.864 | 1 | 0.6 | 1 | 0.184 |
| SE | 62091 | ENSG00000143486 | EIF2D | chr1 | - | 206599732 | 206599836 | 206599462 | 206599612 | 206600262 | 206600308 | 195 | 117 | 160 | 149 | 207 | 1 | 1 | 0 | 0 | 1 | 145 | 182 | 123 | 116 | 189 | 0 | 0 | 0 | 0 | 0 | 253 | 149 | 1 | 1 | 0.991 | 0.986 | 1 | 1 | 0.992 | 1 | 1 | 1 | 1 | 1 | -0.006 |
| SE | 62092 | ENSG00000143486 | EIF2D | chr1 | - | 206602335 | 206602453 | 206600262 | 206600308 | 206602950 | 206602958 | 278 | 153 | 186 | 204 | 256 | 0 | 1 | 0 | 1 | 0 | 292 | 231 | 149 | 204 | 256 | 0 | 0 | 0 | 0 | 0 | 267 | 149 | 1 | 1 | 1 | 0.988 | 1 | 0.991 | 1 | 1 | 1 | 1 | 1 | 1 | -0.004 |
| SE | 62094 | ENSG00000143486 | EIF2D | chr1 | - | 206604608 | 206604729 | 206603082 | 206603204 | 206605399 | 206605507 | 1 | 0 | 3 | 4 | 0 | 58 | 29 | 66 | 46 | 86 | 4 | 0 | 0 | 0 | 0 | 35 | 76 | 18 | 28 | 76 | 270 | 149 | 1 | 1 | 0.009 | 0 | 0.024 | 0.046 | 0 | 0.059 | 0 | 0 | 0 | 0 | 0.004 |
| SE | 62097 | ENSG00000125498 | KIR2DL1 | chr19 | + | 54780088 | 54780166 | 54778611 | 54778662 | 54782921 | 54783023 | 0 | 0 | 0 | 1 | 0 | 10 | 3 | 4 | 1 | 14 | 0 | 0 | 0 | 2 | 0 | 0 | 0 | 10 | 12 | 0 | 227 | 149 | 1 | 1 | 0 | 0 | 0 | 0.396 | 0 | NA | NA | 0 | 0.099 | NA | 0.03 |
| SE | 62098 | ENSG00000092529 | CAPN3 | chr15 | + | 42405925 | 42405943 | 42403740 | 42403777 | 42408210 | 42408324 | 0 | 0 | 0 | 1 | 2 | 12 | 17 | 3 | 7 | 9 | 0 | 0 | 3 | 1 | 0 | 12 | 20 | 10 | 44 | 19 | 167 | 149 | 1 | 1 | 0 | 0 | 0 | 0.113 | 0.165 | 0 | 0 | 0.211 | 0.02 | 0 | 0.009 |
| SE | 62099 | ENSG00000092529 | CAPN3 | chr15 | + | 42405925 | 42405943 | 42403740 | 42403777 | 42409302 | 42409380 | 0 | 0 | 0 | 0 | 2 | 8 | 0 | 7 | 7 | 2 | 0 | 0 | 3 | 0 | 0 | 6 | 11 | 0 | 9 | 0 | 167 | 149 | 1 | 1 | 0 | NA | 0 | 0 | 0.472 | 0 | 0 | 1 | 0 | NA | -0.132 |
| SE | 62118 | ENSG00000148488 | ST8SIA6 | chr10 | - | 17323064 | 17323157 | 17315420 | 17321346 | 17327013 | 17327109 | 59 | 30 | 24 | 40 | 20 | 0 | 4 | 0 | 0 | 0 | 31 | 10 | 26 | 23 | 12 | 0 | 0 | 1 | 0 | 0 | 242 | 149 | 1 | 1 | 1 | 0.822 | 1 | 1 | 1 | 1 | 1 | 0.941 | 1 | 1 | -0.024 |
| SE | 62119 | ENSG00000148488 | ST8SIA6 | chr10 | - | 17323064 | 17323157 | 17315420 | 17321346 | 17331407 | 17331552 | 37 | 13 | 16 | 26 | 7 | 1 | 0 | 0 | 0 | 0 | 31 | 4 | 18 | 19 | 6 | 0 | 0 | 0 | 0 | 0 | 242 | 149 | 1 | 1 | 0.958 | 1 | 1 | 1 | 1 | 1 | 1 | 1 | 1 | 1 | -0.008 |
| SE | 62120 | ENSG00000148488 | ST8SIA6 | chr10 | - | 17327013 | 17327126 | 17315420 | 17321346 | 17331407 | 17331552 | 8 | 6 | 2 | 17 | 1 | 1 | 0 | 0 | 0 | 0 | 22 | 4 | 8 | 2 | 4 | 0 | 0 | 0 | 0 | 0 | 262 | 149 | 1 | 1 | 0.82 | 1 | 1 | 1 | 1 | 1 | 1 | 1 | 1 | 1 | -0.036 |
| SE | 62122 | ENSG00000148488 | ST8SIA6 | chr10 | - | 17379356 | 17379425 | 17359513 | 17359600 | 17390530 | 17390620 | 6 | 1 | 0 | 8 | 0 | 2 | 7 | 1 | 9 | 0 | 0 | 0 | 0 | 3 | 6 | 0 | 2 | 2 | 4 | 0 | 218 | 149 | 1 | 1 | 0.672 | 0.089 | 0 | 0.378 | NA | NA | 0 | 0 | 0.339 | 1 | -0.05 |
| SE | 62129 | ENSG00000182179 | UBA7 | chr3 | - | 49807735 | 49807927 | 49806072 | 49806165 | 49808019 | 49808112 | 503 | 317 | 529 | 528 | 489 | 0 | 0 | 1 | 0 | 0 | 627 | 556 | 253 | 507 | 824 | 0 | 0 | 0 | 0 | 0 | 298 | 149 | 1 | 1 | 1 | 1 | 0.996 | 1 | 1 | 1 | 1 | 1 | 1 | 1 | -0.001 |
| SE | 62130 | ENSG00000182179 | UBA7 | chr3 | - | 49808385 | 49808468 | 49808019 | 49808112 | 49809541 | 49809725 | 185 | 85 | 180 | 157 | 187 | 0 | 0 | 1 | 0 | 0 | 182 | 135 | 84 | 165 | 281 | 0 | 0 | 0 | 0 | 0 | 232 | 149 | 1 | 1 | 1 | 1 | 0.991 | 1 | 1 | 1 | 1 | 1 | 1 | 1 | -0.002 |
| SE | 62131 | ENSG00000182179 | UBA7 | chr3 | - | 49808975 | 49809159 | 49808385 | 49808468 | 49809541 | 49809725 | 87 | 41 | 93 | 88 | 56 | 0 | 0 | 2 | 0 | 0 | 170 | 135 | 52 | 133 | 143 | 0 | 0 | 0 | 0 | 6 | 298 | 149 | 1 | 1 | 1 | 1 | 0.959 | 1 | 1 | 1 | 1 | 1 | 1 | 0.923 | 0.007 |
| SE | 62137 | ENSG00000182179 | UBA7 | chr3 | - | 49810516 | 49810672 | 49810262 | 49810428 | 49810751 | 49810832 | 158 | 110 | 97 | 167 | 117 | 0 | 0 | 2 | 1 | 0 | 347 | 178 | 80 | 132 | 418 | 0 | 3 | 0 | 0 | 0 | 298 | 149 | 1 | 1 | 1 | 1 | 0.96 | 0.988 | 1 | 1 | 0.967 | 1 | 1 | 1 | -0.004 |
| SE | 62144 | ENSG00000162650 | ATXN7L2 | chr1 | + | 109486056 | 109486122 | 109483904 | 109484080 | 109486505 | 109486610 | 5 | 3 | 6 | 3 | 3 | 0 | 0 | 0 | 0 | 0 | 10 | 0 | 5 | 0 | 0 | 13 | 0 | 0 | 0 | 0 | 215 | 149 | 1 | 1 | 1 | 1 | 1 | 1 | 1 | 0.348 | NA | 1 | NA | NA | 0.326 |
| SE | 62145 | ENSG00000162650 | ATXN7L2 | chr1 | + | 109486056 | 109486304 | 109483953 | 109484080 | 109486505 | 109486610 | 3 | 0 | 2 | 1 | 0 | 0 | 0 | 0 | 0 | 0 | 0 | 2 | 3 | 0 | 0 | 13 | 0 | 0 | 0 | 0 | 298 | 149 | 1 | 1 | 1 | NA | 1 | 1 | NA | 0 | 1 | 1 | NA | NA | 0.333 |
| SE | 62146 | ENSG00000162650 | ATXN7L2 | chr1 | + | 109488382 | 109488465 | 109487517 | 109487804 | 109488846 | 109489100 | 16 | 1 | 18 | 6 | 16 | 0 | 0 | 0 | 0 | 0 | 0 | 11 | 4 | 7 | 2 | 0 | 0 | 2 | 2 | 0 | 232 | 149 | 1 | 1 | 1 | 1 | 1 | 1 | 1 | NA | 1 | 0.562 | 0.692 | 1 | 0.186 |
| SE | 62147 | ENSG00000162650 | ATXN7L2 | chr1 | + | 109489282 | 109489706 | 109488846 | 109489100 | 109489929 | 109490128 | 0 | 0 | 0 | 1 | 0 | 11 | 3 | 9 | 11 | 7 | 0 | 0 | 0 | 0 | 0 | 17 | 39 | 7 | 4 | 3 | 298 | 149 | 1 | 1 | 0 | 0 | 0 | 0.043 | 0 | 0 | 0 | 0 | 0 | 0 | 0.009 |
| SE | 62154 | ENSG00000117625 | RCOR3 | chr1 | + | 211274209 | 211274262 | 211271231 | 211271309 | 211276256 | 211276335 | 100 | 98 | 93 | 115 | 117 | 0 | 0 | 2 | 0 | 0 | 102 | 75 | 105 | 116 | 90 | 0 | 0 | 0 | 0 | 0 | 202 | 149 | 1 | 1 | 1 | 1 | 0.972 | 1 | 1 | 1 | 1 | 1 | 1 | 1 | -0.006 |
| SE | 62155 | ENSG00000117625 | RCOR3 | chr1 | + | 211278116 | 211278241 | 211276256 | 211276418 | 211289177 | 211289396 | 202 | 190 | 178 | 217 | 210 | 0 | 0 | 0 | 0 | 2 | 175 | 197 | 196 | 212 | 174 | 0 | 0 | 0 | 0 | 0 | 274 | 149 | 1 | 1 | 1 | 1 | 1 | 1 | 0.983 | 1 | 1 | 1 | 1 | 1 | -0.003 |
| SE | 62156 | ENSG00000117625 | RCOR3 | chr1 | + | 211279237 | 211279316 | 211278116 | 211278241 | 211289177 | 211289396 | 119 | 127 | 128 | 97 | 203 | 0 | 1 | 0 | 0 | 0 | 82 | 104 | 111 | 132 | 53 | 0 | 0 | 0 | 0 | 0 | 228 | 149 | 1 | 1 | 1 | 0.988 | 1 | 1 | 1 | 1 | 1 | 1 | 1 | 1 | -0.002 |
| SE | 62160 | ENSG00000117625 | RCOR3 | chr1 | + | 211295675 | 211295753 | 211289177 | 211289396 | 211313423 | 211313474 | 51 | 65 | 58 | 54 | 107 | 0 | 2 | 0 | 0 | 0 | 54 | 65 | 54 | 70 | 52 | 0 | 0 | 0 | 0 | 0 | 227 | 149 | 1 | 1 | 1 | 0.955 | 1 | 1 | 1 | 1 | 1 | 1 | 1 | 1 | -0.009 |
| SE | 62161 | ENSG00000117625 | RCOR3 | chr1 | + | 211295675 | 211295753 | 211291523 | 211291599 | 211304082 | 211304140 | 89 | 58 | 57 | 72 | 230 | 0 | 0 | 0 | 3 | 0 | 115 | 87 | 68 | 98 | 48 | 0 | 0 | 0 | 0 | 0 | 227 | 149 | 1 | 1 | 1 | 1 | 1 | 0.94 | 1 | 1 | 1 | 1 | 1 | 1 | -0.012 |
| SE | 62162 | ENSG00000117625 | RCOR3 | chr1 | + | 211304082 | 211304140 | 211289177 | 211289396 | 211313423 | 211314591 | 18 | 11 | 21 | 16 | 39 | 0 | 2 | 0 | 0 | 0 | 8 | 24 | 8 | 14 | 25 | 0 | 0 | 0 | 0 | 0 | 207 | 149 | 1 | 1 | 1 | 0.798 | 1 | 1 | 1 | 1 | 1 | 1 | 1 | 1 | -0.04 |
| SE | 62163 | ENSG00000117625 | RCOR3 | chr1 | + | 211304082 | 211304140 | 211295675 | 211295753 | 211312354 | 211312487 | 106 | 77 | 71 | 77 | 139 | 0 | 0 | 0 | 1 | 0 | 49 | 61 | 61 | 63 | 76 | 0 | 0 | 0 | 0 | 0 | 207 | 149 | 1 | 1 | 1 | 1 | 1 | 0.982 | 1 | 1 | 1 | 1 | 1 | 1 | -0.004 |
| SE | 62166 | ENSG00000117625 | RCOR3 | chr1 | + | 211312354 | 211312487 | 211295675 | 211295753 | 211312719 | 211312819 | 44 | 29 | 28 | 42 | 72 | 0 | 1 | 1 | 3 | 0 | 0 | 34 | 28 | 53 | 21 | 0 | 0 | 1 | 4 | 5 | 282 | 149 | 1 | 1 | 1 | 0.939 | 0.937 | 0.881 | 1 | NA | 1 | 0.937 | 0.875 | 0.689 | 0.076 |
| SE | 62168 | ENSG00000117625 | RCOR3 | chr1 | + | 211312719 | 211312835 | 211295675 | 211295753 | 211313423 | 211313672 | 0 | 1 | 1 | 3 | 0 | 2 | 5 | 1 | 4 | 3 | 0 | 0 | 1 | 4 | 5 | 8 | 0 | 0 | 0 | 7 | 265 | 149 | 1 | 1 | 0 | 0.101 | 0.36 | 0.297 | 0 | 0 | NA | 1 | 1 | 0.287 | -0.42 |
| SE | 62176 | ENSG00000204677 | FAM153CP | chr5 | + | 178016585 | 178016616 | 178008959 | 178009381 | 178020259 | 178020453 | 5 | 13 | 3 | 11 | 4 | 0 | 1 | 2 | 2 | 2 | 0 | 0 | 31 | 5 | 0 | 0 | 0 | 0 | 0 | 0 | 180 | 149 | 1 | 1 | 1 | 0.915 | 0.554 | 0.82 | 0.623 | NA | NA | 1 | 1 | NA | -0.218 |
| SE | 62177 | ENSG00000204677 | FAM153CP | chr5 | + | 178020259 | 178020453 | 178009076 | 178009381 | 178035095 | 178035154 | 0 | 2 | 4 | 10 | 4 | 0 | 4 | 1 | 2 | 0 | 1 | 0 | 0 | 0 | 0 | 0 | 0 | 0 | 0 | 0 | 298 | 149 | 1 | 1 | NA | 0.2 | 0.667 | 0.714 | 1 | 1 | NA | NA | NA | NA | -0.355 |
| SE | 62178 | ENSG00000204677 | FAM153CP | chr5 | + | 178023232 | 178023332 | 178020259 | 178020453 | 178035095 | 178035154 | 5 | 5 | 9 | 8 | 2 | 0 | 1 | 2 | 8 | 2 | 1 | 0 | 3 | 0 | 0 | 1 | 0 | 0 | 0 | 0 | 249 | 149 | 1 | 1 | 1 | 0.749 | 0.729 | 0.374 | 0.374 | 0.374 | NA | 1 | NA | NA | -0.042 |
| SE | 62181 | ENSG00000204677 | FAM153CP | chr5 | + | 178052802 | 178052989 | 178048664 | 178052687 | 178053839 | 178053973 | 3 | 2 | 0 | 1 | 1 | 0 | 1 | 2 | 6 | 2 | 0 | 0 | 0 | 0 | 0 | 0 | 10 | 4 | 0 | 3 | 298 | 149 | 1 | 1 | 1 | 0.5 | 0 | 0.077 | 0.2 | NA | 0 | 0 | NA | 0 | 0.355 |
| SE | 62182 | ENSG00000204677 | FAM153CP | chr5 | + | 178052908 | 178052989 | 178052531 | 178052687 | 178053839 | 178053973 | 1 | 0 | 0 | 0 | 1 | 0 | 1 | 2 | 6 | 2 | 0 | 0 | 0 | 0 | 0 | 0 | 10 | 4 | 0 | 3 | 230 | 149 | 1 | 1 | 1 | 0 | 0 | 0 | 0.245 | NA | 0 | 0 | NA | 0 | 0.249 |
| SE | 62188 | ENSG00000113569 | NUP155 | chr5 | - | 37292878 | 37292985 | 37288136 | 37292038 | 37294328 | 37294465 | 77 | 39 | 70 | 36 | 62 | 0 | 0 | 0 | 2 | 0 | 6 | 54 | 54 | 35 | 6 | 0 | 0 | 0 | 0 | 0 | 256 | 149 | 1 | 1 | 1 | 1 | 1 | 0.913 | 1 | 1 | 1 | 1 | 1 | 1 | -0.017 |
| SE | 62189 | ENSG00000113569 | NUP155 | chr5 | - | 37294328 | 37294465 | 37292878 | 37292985 | 37298867 | 37298978 | 211 | 70 | 153 | 136 | 116 | 0 | 0 | 1 | 0 | 2 | 21 | 90 | 108 | 80 | 1 | 0 | 0 | 2 | 1 | 0 | 286 | 149 | 1 | 1 | 1 | 1 | 0.988 | 1 | 0.968 | 1 | 1 | 0.966 | 0.977 | 1 | 0.003 |
| SE | 62192 | ENSG00000113569 | NUP155 | chr5 | - | 37309403 | 37309572 | 37309128 | 37309267 | 37310551 | 37310668 | 0 | 0 | 0 | 0 | 2 | 65 | 33 | 81 | 79 | 125 | 0 | 0 | 0 | 0 | 0 | 18 | 79 | 65 | 38 | 18 | 298 | 149 | 1 | 1 | 0 | 0 | 0 | 0 | 0.008 | 0 | 0 | 0 | 0 | 0 | 0.002 |
| SE | 62193 | ENSG00000113569 | NUP155 | chr5 | - | 37310551 | 37310743 | 37309128 | 37309267 | 37314197 | 37314328 | 170 | 114 | 205 | 200 | 278 | 0 | 0 | 0 | 1 | 0 | 82 | 168 | 188 | 163 | 29 | 0 | 0 | 0 | 0 | 0 | 298 | 149 | 1 | 1 | 1 | 1 | 1 | 0.99 | 1 | 1 | 1 | 1 | 1 | 1 | -0.002 |
| SE | 62195 | ENSG00000113569 | NUP155 | chr5 | - | 37330037 | 37330132 | 37329189 | 37329278 | 37331684 | 37331795 | 76 | 52 | 87 | 87 | 231 | 0 | 0 | 0 | 0 | 0 | 28 | 64 | 66 | 45 | 7 | 0 | 0 | 1 | 2 | 0 | 244 | 149 | 1 | 1 | 1 | 1 | 1 | 1 | 1 | 1 | 1 | 0.976 | 0.932 | 1 | 0.018 |
| SE | 62196 | ENSG00000113569 | NUP155 | chr5 | - | 37341089 | 37341242 | 37337817 | 37337918 | 37342548 | 37342646 | 141 | 80 | 139 | 102 | 132 | 0 | 0 | 0 | 1 | 1 | 124 | 106 | 105 | 80 | 13 | 0 | 0 | 0 | 1 | 0 | 298 | 149 | 1 | 1 | 1 | 1 | 1 | 0.981 | 0.985 | 1 | 1 | 1 | 0.976 | 1 | -0.002 |
| SE | 62198 | ENSG00000113569 | NUP155 | chr5 | - | 37348504 | 37348596 | 37342396 | 37342646 | 37349171 | 37349245 | 88 | 65 | 76 | 62 | 172 | 0 | 1 | 0 | 0 | 0 | 120 | 74 | 58 | 35 | 12 | 0 | 0 | 0 | 0 | 0 | 241 | 149 | 1 | 1 | 1 | 0.976 | 1 | 1 | 1 | 1 | 1 | 1 | 1 | 1 | -0.005 |
| SE | 62200 | ENSG00000187800 | PEAR1 | chr1 | + | 156902187 | 156902285 | 156893719 | 156893837 | 156903917 | 156904027 | 0 | 2 | 0 | 0 | 6 | 0 | 0 | 0 | 0 | 0 | 18 | 14 | 16 | 2 | 43 | 0 | 0 | 0 | 0 | 4 | 247 | 149 | 1 | 1 | NA | 1 | NA | NA | 1 | 1 | 1 | 1 | 1 | 0.866 | 0.027 |
| SE | 62201 | ENSG00000187800 | PEAR1 | chr1 | + | 156904943 | 156904998 | 156904747 | 156904852 | 156905323 | 156905424 | 0 | 3 | 0 | 6 | 1 | 2 | 1 | 0 | 2 | 0 | 7 | 1 | 3 | 7 | 32 | 11 | 1 | 2 | 10 | 10 | 204 | 149 | 1 | 1 | 0 | 0.687 | NA | 0.687 | 1 | 0.317 | 0.422 | 0.523 | 0.338 | 0.7 | 0.134 |
| SE | 62204 | ENSG00000119778 | ATAD2B | chr2 | - | 23757417 | 23758101 | 23754646 | 23754774 | 23762208 | 23762346 | 129 | 108 | 95 | 122 | 126 | 6 | 1 | 1 | 1 | 5 | 0 | 70 | 138 | 29 | 0 | 0 | 0 | 0 | 0 | 0 | 298 | 149 | 1 | 1 | 0.915 | 0.982 | 0.979 | 0.984 | 0.926 | NA | 1 | 1 | 1 | NA | -0.043 |
| SE | 62205 | ENSG00000119778 | ATAD2B | chr2 | - | 23765505 | 23765628 | 23762208 | 23762346 | 23782868 | 23783028 | 234 | 218 | 181 | 240 | 367 | 0 | 0 | 1 | 0 | 0 | 46 | 163 | 197 | 66 | 32 | 0 | 0 | 1 | 0 | 0 | 272 | 149 | 1 | 1 | 1 | 1 | 0.99 | 1 | 1 | 1 | 1 | 0.991 | 1 | 1 | 0 |
| SE | 62206 | ENSG00000119778 | ATAD2B | chr2 | - | 23788511 | 23788647 | 23786026 | 23786223 | 23798137 | 23798323 | 119 | 112 | 105 | 110 | 168 | 1 | 1 | 0 | 0 | 0 | 20 | 71 | 85 | 27 | 17 | 0 | 3 | 1 | 0 | 0 | 285 | 149 | 1 | 1 | 0.984 | 0.983 | 1 | 1 | 1 | 1 | 0.925 | 0.978 | 1 | 1 | 0.013 |
| SE | 62207 | ENSG00000119778 | ATAD2B | chr2 | - | 23798137 | 23798323 | 23788511 | 23788647 | 23810315 | 23810502 | 131 | 96 | 123 | 112 | 194 | 0 | 0 | 0 | 0 | 0 | 22 | 63 | 55 | 42 | 27 | 0 | 0 | 2 | 0 | 0 | 298 | 149 | 1 | 1 | 1 | 1 | 1 | 1 | 1 | 1 | 1 | 0.932 | 1 | 1 | 0.014 |
| SE | 62208 | ENSG00000119778 | ATAD2B | chr2 | - | 23799824 | 23799993 | 23798137 | 23798323 | 23806170 | 23806228 | 5 | 0 | 1 | 5 | 0 | 1 | 3 | 1 | 0 | 5 | 0 | 0 | 0 | 0 | 0 | 2 | 2 | 3 | 0 | 7 | 298 | 149 | 1 | 1 | 0.714 | 0 | 0.333 | 1 | 0 | 0 | 0 | 0 | NA | 0 | 0.409 |
| SE | 62210 | ENSG00000119778 | ATAD2B | chr2 | - | 23806170 | 23806228 | 23798137 | 23798323 | 23808988 | 23809176 | 0 | 2 | 0 | 0 | 5 | 0 | 0 | 1 | 0 | 0 | 0 | 2 | 3 | 0 | 7 | 0 | 0 | 0 | 0 | 0 | 207 | 149 | 1 | 1 | NA | 1 | 0 | NA | 1 | NA | 1 | 1 | NA | 1 | -0.333 |
| SE | 62212 | ENSG00000119778 | ATAD2B | chr2 | - | 23808988 | 23809176 | 23806170 | 23806228 | 23810315 | 23810502 | 6 | 5 | 0 | 3 | 5 | 0 | 0 | 2 | 0 | 0 | 0 | 2 | 6 | 0 | 7 | 2 | 0 | 2 | 0 | 0 | 298 | 149 | 1 | 1 | 1 | 1 | 0 | 1 | 1 | 0 | 1 | 0.6 | NA | 1 | 0.15 |
| SE | 62213 | ENSG00000119778 | ATAD2B | chr2 | - | 23823257 | 23823569 | 23819746 | 23819882 | 23828848 | 23828939 | 161 | 176 | 147 | 187 | 251 | 0 | 2 | 2 | 1 | 2 | 72 | 246 | 169 | 93 | 89 | 0 | 0 | 0 | 0 | 0 | 298 | 149 | 1 | 1 | 1 | 0.978 | 0.974 | 0.989 | 0.984 | 1 | 1 | 1 | 1 | 1 | -0.015 |
| SE | 62215 | ENSG00000119778 | ATAD2B | chr2 | - | 23833918 | 23834078 | 23828848 | 23828939 | 23857414 | 23857503 | 210 | 235 | 207 | 249 | 200 | 0 | 1 | 0 | 0 | 0 | 159 | 146 | 167 | 64 | 98 | 0 | 0 | 3 | 0 | 0 | 298 | 149 | 1 | 1 | 1 | 0.992 | 1 | 1 | 1 | 1 | 1 | 0.965 | 1 | 1 | 0.005 |
| SE | 62217 | ENSG00000119778 | ATAD2B | chr2 | - | 23857414 | 23857503 | 23833918 | 23834078 | 23864808 | 23864924 | 30 | 31 | 45 | 45 | 50 | 0 | 1 | 0 | 0 | 0 | 22 | 38 | 31 | 2 | 0 | 0 | 0 | 0 | 0 | 0 | 238 | 149 | 1 | 1 | 1 | 0.951 | 1 | 1 | 1 | 1 | 1 | 1 | 1 | NA | -0.01 |
| SE | 62218 | ENSG00000119778 | ATAD2B | chr2 | - | 23857414 | 23857503 | 23856008 | 23856442 | 23863380 | 23863555 | 82 | 83 | 43 | 64 | 49 | 0 | 0 | 1 | 0 | 0 | 61 | 47 | 94 | 47 | 45 | 0 | 0 | 0 | 0 | 0 | 238 | 149 | 1 | 1 | 1 | 1 | 0.964 | 1 | 1 | 1 | 1 | 1 | 1 | 1 | -0.007 |
| SE | 62219 | ENSG00000119778 | ATAD2B | chr2 | - | 23863380 | 23863555 | 23833918 | 23834078 | 23864808 | 23864924 | 101 | 116 | 114 | 131 | 82 | 0 | 1 | 0 | 0 | 0 | 55 | 50 | 41 | 21 | 24 | 0 | 0 | 0 | 0 | 0 | 298 | 149 | 1 | 1 | 1 | 0.983 | 1 | 1 | 1 | 1 | 1 | 1 | 1 | 1 | -0.003 |
| SE | 62221 | ENSG00000119778 | ATAD2B | chr2 | - | 23863380 | 23863555 | 23857414 | 23857503 | 23867834 | 23867887 | 179 | 188 | 128 | 167 | 87 | 0 | 0 | 0 | 0 | 0 | 161 | 117 | 184 | 86 | 52 | 0 | 1 | 0 | 0 | 0 | 298 | 149 | 1 | 1 | 1 | 1 | 1 | 1 | 1 | 1 | 0.983 | 1 | 1 | 1 | 0.003 |
| SE | 62222 | ENSG00000119778 | ATAD2B | chr2 | - | 23864808 | 23864924 | 23857414 | 23857503 | 23867834 | 23867887 | 81 | 70 | 57 | 82 | 72 | 0 | 0 | 0 | 0 | 0 | 28 | 47 | 41 | 20 | 12 | 0 | 1 | 0 | 0 | 0 | 265 | 149 | 1 | 1 | 1 | 1 | 1 | 1 | 1 | 1 | 0.964 | 1 | 1 | 1 | 0.007 |
| SE | 62223 | ENSG00000119778 | ATAD2B | chr2 | - | 23864808 | 23864924 | 23863380 | 23863555 | 23867834 | 23867887 | 175 | 181 | 166 | 207 | 152 | 2 | 0 | 1 | 0 | 0 | 73 | 93 | 82 | 41 | 36 | 0 | 0 | 2 | 0 | 0 | 265 | 149 | 1 | 1 | 0.98 | 1 | 0.989 | 1 | 1 | 1 | 1 | 0.958 | 1 | 1 | 0.002 |
| SE | 62224 | ENSG00000119778 | ATAD2B | chr2 | - | 23880638 | 23880755 | 23875828 | 23875904 | 23884764 | 23884873 | 440 | 505 | 363 | 467 | 462 | 0 | 0 | 0 | 0 | 0 | 618 | 588 | 615 | 417 | 215 | 0 | 0 | 3 | 3 | 0 | 266 | 149 | 1 | 1 | 1 | 1 | 1 | 1 | 1 | 1 | 1 | 0.991 | 0.987 | 1 | 0.004 |
| SE | 62226 | ENSG00000119778 | ATAD2B | chr2 | - | 23884764 | 23884873 | 23880638 | 23880755 | 23885726 | 23885829 | 509 | 612 | 412 | 545 | 589 | 0 | 0 | 0 | 0 | 0 | 801 | 715 | 702 | 556 | 342 | 0 | 0 | 0 | 0 | 8 | 258 | 149 | 1 | 1 | 1 | 1 | 1 | 1 | 1 | 1 | 1 | 1 | 1 | 0.961 | 0.008 |
| SE | 62227 | ENSG00000119778 | ATAD2B | chr2 | - | 23885726 | 23885829 | 23880638 | 23880755 | 23895818 | 23895970 | 0 | 0 | 1 | 0 | 0 | 0 | 0 | 0 | 0 | 0 | 0 | 0 | 0 | 0 | 4 | 0 | 5 | 0 | 0 | 0 | 252 | 149 | 1 | 1 | NA | NA | 1 | NA | NA | NA | 0 | NA | NA | 1 | 0.5 |
| SE | 62228 | ENSG00000119778 | ATAD2B | chr2 | - | 23887831 | 23887985 | 23885726 | 23885829 | 23888349 | 23888399 | 381 | 429 | 280 | 363 | 449 | 0 | 2 | 0 | 2 | 0 | 529 | 551 | 505 | 381 | 177 | 0 | 2 | 0 | 2 | 0 | 298 | 149 | 1 | 1 | 1 | 0.991 | 1 | 0.989 | 1 | 1 | 0.993 | 1 | 0.99 | 1 | -0.001 |
| SE | 62229 | ENSG00000119778 | ATAD2B | chr2 | - | 23888349 | 23888399 | 23885726 | 23885829 | 23895818 | 23895970 | 11 | 9 | 19 | 15 | 7 | 1 | 0 | 1 | 0 | 0 | 4 | 8 | 16 | 7 | 4 | 0 | 0 | 0 | 0 | 0 | 199 | 149 | 1 | 1 | 0.892 | 1 | 0.934 | 1 | 1 | 1 | 1 | 1 | 1 | 1 | -0.035 |
| SE | 62230 | ENSG00000119778 | ATAD2B | chr2 | - | 23895818 | 23895970 | 23888349 | 23888399 | 23926554 | 23927065 | 78 | 65 | 69 | 68 | 54 | 0 | 0 | 0 | 0 | 0 | 54 | 80 | 92 | 68 | 17 | 0 | 1 | 0 | 0 | 0 | 298 | 149 | 1 | 1 | 1 | 1 | 1 | 1 | 1 | 1 | 1 | 0.979 | 1 | 1 | 0.004 |
| SE | 62234 | ENSG00000197043 | ANXA6 | chr5 | - | 151108454 | 151108550 | 151105244 | 151105303 | 151109752 | 151109846 | 1342 | 735 | 1191 | 1170 | 1145 | 0 | 0 | 1 | 2 | 0 | 1224 | 1842 | 719 | 1037 | 1309 | 0 | 3 | 0 | 0 | 0 | 245 | 149 | 1 | 1 | 1 | 1 | 0.999 | 0.997 | 1 | 1 | 0.997 | 1 | 1 | 1 | 0 |
| SE | 62236 | ENSG00000197043 | ANXA6 | chr5 | - | 151109752 | 151109846 | 151108454 | 151108550 | 151110626 | 151110644 | 1284 | 687 | 1175 | 1305 | 1309 | 1 | 0 | 0 | 0 | 0 | 1375 | 1952 | 840 | 1172 | 1534 | 0 | 0 | 0 | 0 | 0 | 243 | 149 | 1 | 1 | 0.999 | 1 | 1 | 1 | 1 | 1 | 1 | 1 | 1 | 1 | 0 |
| SE | 62238 | ENSG00000197043 | ANXA6 | chr5 | - | 151119299 | 151119390 | 151117757 | 151117837 | 151122146 | 151122260 | 1725 | 1089 | 1451 | 1791 | 1622 | 7 | 2 | 4 | 7 | 2 | 2523 | 2855 | 1343 | 2037 | 2492 | 2 | 12 | 3 | 1 | 0 | 240 | 149 | 1 | 1 | 0.994 | 0.997 | 0.996 | 0.994 | 0.998 | 0.999 | 0.993 | 0.996 | 0.999 | 1 | -0.002 |
| SE | 62239 | ENSG00000197043 | ANXA6 | chr5 | - | 151126401 | 151126480 | 151124285 | 151124367 | 151128180 | 151128239 | 1112 | 638 | 937 | 943 | 973 | 2 | 0 | 0 | 3 | 0 | 681 | 1201 | 499 | 899 | 1035 | 0 | 6 | 0 | 2 | 7 | 228 | 149 | 1 | 1 | 0.997 | 1 | 1 | 0.995 | 1 | 1 | 0.992 | 1 | 0.997 | 0.99 | 0.003 |
| SE | 62240 | ENSG00000197043 | ANXA6 | chr5 | - | 151133093 | 151133187 | 151132475 | 151132571 | 151134426 | 151134483 | 1649 | 960 | 1283 | 1540 | 1334 | 0 | 0 | 1 | 0 | 0 | 1750 | 2359 | 1045 | 1636 | 2231 | 0 | 13 | 1 | 0 | 4 | 243 | 149 | 1 | 1 | 1 | 1 | 0.999 | 1 | 1 | 1 | 0.991 | 0.998 | 1 | 0.997 | 0.003 |
| SE | 62241 | ENSG00000197043 | ANXA6 | chr5 | - | 151137230 | 151137321 | 151136255 | 151136335 | 151138677 | 151138791 | 3006 | 1957 | 2308 | 2740 | 2983 | 0 | 0 | 0 | 0 | 0 | 3804 | 4051 | 1958 | 2931 | 4205 | 0 | 0 | 1 | 0 | 0 | 240 | 149 | 1 | 1 | 1 | 1 | 1 | 1 | 1 | 1 | 1 | 0.999 | 1 | 1 | 0 |
| SE | 62250 | ENSG00000196220 | SRGAP3 | chr3 | - | 9049260 | 9049469 | 9047390 | 9047475 | 9053026 | 9053224 | 0 | 0 | 0 | 0 | 0 | 4 | 1 | 1 | 1 | 3 | 0 | 0 | 7 | 0 | 0 | 0 | 1 | 0 | 0 | 0 | 298 | 149 | 1 | 1 | 0 | 0 | 0 | 0 | 0 | NA | NA | 1 | NA | NA | -1 |
| SE | 62260 | ENSG00000091831 | ESR1 | chr6 | + | 152094384 | 152094568 | 152060990 | 152061124 | 152098731 | 152098920 | 1 | 0 | 0 | 1 | 0 | 0 | 0 | 1 | 1 | 0 | 12 | 0 | 2 | 3 | 0 | 0 | 1 | 0 | 0 | 0 | 298 | 149 | 1 | 1 | 1 | NA | 0 | 0.333 | NA | 1 | NA | 1 | 1 | NA | -0.556 |
| SE | 62267 | ENSG00000254126 | CD8B2 | chr2 | + | 106496172 | 106496262 | 106490873 | 106491233 | 106502473 | 106502563 | 0 | 0 | 0 | 0 | 0 | 0 | 0 | 1 | 0 | 0 | 0 | 0 | 1 | 0 | 0 | 0 | 0 | 0 | 0 | 0 | 239 | 149 | 1 | 1 | NA | NA | 0 | NA | NA | NA | NA | 1 | NA | NA | -1 |
| SE | 62268 | ENSG00000172888 | ZNF621 | chr3 | + | 40529318 | 40529445 | 40525778 | 40525864 | 40530208 | 40530316 | 26 | 11 | 44 | 37 | 32 | 1 | 2 | 6 | 1 | 0 | 0 | 26 | 22 | 6 | 18 | 0 | 0 | 7 | 2 | 0 | 276 | 149 | 1 | 1 | 0.933 | 0.748 | 0.798 | 0.952 | 1 | NA | 1 | 0.629 | 0.618 | 1 | 0.074 |
| SE | 62269 | ENSG00000172888 | ZNF621 | chr3 | + | 40529494 | 40529575 | 40525778 | 40525864 | 40530208 | 40530316 | 4 | 4 | 6 | 2 | 0 | 1 | 2 | 6 | 1 | 0 | 12 | 7 | 6 | 14 | 8 | 0 | 0 | 7 | 2 | 0 | 230 | 149 | 1 | 1 | 0.722 | 0.564 | 0.393 | 0.564 | NA | 1 | 1 | 0.357 | 0.819 | 1 | -0.274 |
| SE | 62276 | ENSG00000261236 | BOP1 | chr8 | - | 144263691 | 144263761 | 144263477 | 144263610 | 144263830 | 144263911 | 15 | 6 | 6 | 6 | 4 | 0 | 0 | 0 | 0 | 0 | 0 | 11 | 6 | 8 | 5 | 0 | 0 | 0 | 2 | 0 | 219 | 149 | 1 | 1 | 1 | 1 | 1 | 1 | 1 | NA | 1 | 1 | 0.731 | 1 | 0.067 |
| SE | 62278 | ENSG00000261236 | BOP1 | chr8 | - | 144276223 | 144276304 | 144264916 | 144265071 | 144289094 | 144289304 | 118 | 72 | 90 | 117 | 63 | 0 | 0 | 0 | 0 | 0 | 120 | 113 | 36 | 115 | 198 | 0 | 4 | 0 | 0 | 0 | 230 | 149 | 1 | 1 | 1 | 1 | 1 | 1 | 1 | 1 | 0.948 | 1 | 1 | 1 | 0.01 |
| SE | 62283 | ENSG00000112031 | MTRF1L | chr6 | - | 152994512 | 152994676 | 152992856 | 152992974 | 152998549 | 152998629 | 46 | 54 | 46 | 71 | 70 | 0 | 0 | 1 | 0 | 0 | 62 | 91 | 88 | 112 | 56 | 0 | 0 | 0 | 0 | 0 | 298 | 149 | 1 | 1 | 1 | 1 | 0.958 | 1 | 1 | 1 | 1 | 1 | 1 | 1 | -0.008 |
| SE | 62284 | ENSG00000112031 | MTRF1L | chr6 | - | 152995135 | 152995294 | 152992856 | 152992974 | 152998549 | 152998629 | 51 | 42 | 46 | 47 | 55 | 0 | 0 | 1 | 0 | 0 | 43 | 101 | 74 | 91 | 12 | 0 | 0 | 0 | 0 | 0 | 298 | 149 | 1 | 1 | 1 | 1 | 0.958 | 1 | 1 | 1 | 1 | 1 | 1 | 1 | -0.008 |
| SE | 62286 | ENSG00000112031 | MTRF1L | chr6 | - | 152995135 | 152995319 | 152992856 | 152992974 | 152998549 | 152998629 | 47 | 36 | 32 | 49 | 49 | 0 | 0 | 1 | 0 | 0 | 50 | 91 | 30 | 58 | 28 | 0 | 0 | 0 | 0 | 0 | 298 | 149 | 1 | 1 | 1 | 1 | 0.941 | 1 | 1 | 1 | 1 | 1 | 1 | 1 | -0.012 |
| SE | 62290 | ENSG00000039319 | ZFYVE16 | chr5 | + | 80411645 | 80411773 | 80408023 | 80408153 | 80434108 | 80434217 | 0 | 0 | 0 | 2 | 0 | 9 | 9 | 5 | 19 | 13 | 0 | 0 | 0 | 0 | 0 | 0 | 3 | 0 | 0 | 0 | 277 | 149 | 1 | 1 | 0 | 0 | 0 | 0.054 | 0 | NA | 0 | NA | NA | NA | 0.011 |
| SE | 62291 | ENSG00000039319 | ZFYVE16 | chr5 | + | 80427491 | 80427545 | 80408063 | 80408153 | 80434108 | 80434217 | 27 | 11 | 4 | 14 | 4 | 9 | 9 | 5 | 19 | 13 | 0 | 16 | 3 | 8 | 0 | 0 | 3 | 0 | 0 | 0 | 203 | 149 | 1 | 1 | 0.688 | 0.473 | 0.37 | 0.351 | 0.184 | NA | 0.797 | 1 | 1 | NA | -0.519 |
| SE | 62292 | ENSG00000039319 | ZFYVE16 | chr5 | + | 80434108 | 80434217 | 80427491 | 80427545 | 80436755 | 80439007 | 86 | 70 | 23 | 54 | 29 | 0 | 1 | 0 | 0 | 0 | 0 | 29 | 15 | 12 | 0 | 0 | 0 | 0 | 0 | 0 | 258 | 149 | 1 | 1 | 1 | 0.976 | 1 | 1 | 1 | NA | 1 | 1 | 1 | NA | -0.005 |
| SE | 62293 | ENSG00000039319 | ZFYVE16 | chr5 | + | 80435761 | 80435854 | 80434108 | 80434217 | 80436755 | 80437159 | 0 | 0 | 0 | 0 | 0 | 77 | 61 | 21 | 48 | 34 | 0 | 2 | 0 | 0 | 0 | 9 | 27 | 14 | 6 | 0 | 242 | 149 | 1 | 1 | 0 | 0 | 0 | 0 | 0 | 0 | 0.044 | 0 | 0 | NA | -0.011 |
| SE | 62294 | ENSG00000039319 | ZFYVE16 | chr5 | + | 80439935 | 80440032 | 80436755 | 80439007 | 80443122 | 80443284 | 93 | 70 | 72 | 100 | 131 | 0 | 1 | 0 | 0 | 0 | 0 | 32 | 36 | 14 | 0 | 0 | 0 | 0 | 0 | 0 | 246 | 149 | 1 | 1 | 1 | 0.977 | 1 | 1 | 1 | NA | 1 | 1 | 1 | NA | -0.005 |
| SE | 62295 | ENSG00000039319 | ZFYVE16 | chr5 | + | 80443793 | 80443920 | 80443151 | 80443284 | 80445262 | 80445405 | 6 | 4 | 6 | 2 | 3 | 40 | 47 | 22 | 35 | 46 | 0 | 8 | 1 | 0 | 0 | 0 | 17 | 17 | 7 | 0 | 276 | 149 | 1 | 1 | 0.075 | 0.044 | 0.128 | 0.03 | 0.034 | NA | 0.203 | 0.031 | 0 | NA | -0.016 |
| SE | 62296 | ENSG00000039319 | ZFYVE16 | chr5 | + | 80448025 | 80448404 | 80445262 | 80445405 | 80449590 | 80449713 | 106 | 86 | 69 | 107 | 175 | 12 | 7 | 14 | 33 | 19 | 0 | 29 | 21 | 6 | 7 | 0 | 9 | 21 | 0 | 0 | 298 | 149 | 1 | 1 | 0.815 | 0.86 | 0.711 | 0.618 | 0.822 | NA | 0.617 | 0.333 | 1 | 1 | 0.028 |
| SE | 62297 | ENSG00000039319 | ZFYVE16 | chr5 | + | 80448282 | 80448404 | 80445262 | 80445405 | 80449590 | 80449713 | 61 | 34 | 36 | 53 | 115 | 12 | 7 | 14 | 33 | 19 | 0 | 10 | 9 | 2 | 7 | 0 | 9 | 21 | 0 | 0 | 271 | 149 | 1 | 1 | 0.736 | 0.728 | 0.586 | 0.469 | 0.769 | NA | 0.379 | 0.191 | 1 | 1 | 0.015 |
| SE | 62298 | ENSG00000039319 | ZFYVE16 | chr5 | + | 80456460 | 80456565 | 80455691 | 80455774 | 80456944 | 80457092 | 147 | 118 | 55 | 120 | 204 | 0 | 0 | 0 | 1 | 0 | 30 | 17 | 26 | 23 | 0 | 0 | 0 | 0 | 0 | 0 | 254 | 149 | 1 | 1 | 1 | 1 | 1 | 0.986 | 1 | 1 | 1 | 1 | 1 | NA | -0.003 |
| SE | 62299 | ENSG00000039319 | ZFYVE16 | chr5 | + | 80472760 | 80472923 | 80459413 | 80459494 | 80473753 | 80473859 | 163 | 188 | 93 | 136 | 333 | 0 | 1 | 1 | 7 | 8 | 0 | 38 | 41 | 4 | 19 | 0 | 0 | 0 | 0 | 0 | 298 | 149 | 1 | 1 | 1 | 0.989 | 0.979 | 0.907 | 0.954 | NA | 1 | 1 | 1 | 1 | -0.034 |
| SE | 62300 | ENSG00000039319 | ZFYVE16 | chr5 | + | 80474662 | 80474830 | 80473753 | 80473859 | 80477218 | 80477311 | 144 | 130 | 91 | 128 | 273 | 0 | 0 | 0 | 0 | 1 | 0 | 33 | 20 | 0 | 22 | 0 | 0 | 0 | 0 | 0 | 298 | 149 | 1 | 1 | 1 | 1 | 1 | 1 | 0.993 | NA | 1 | 1 | NA | 1 | -0.001 |
| SE | 62307 | ENSG00000134686 | PHC2 | chr1 | - | 33332274 | 33332404 | 33331624 | 33331699 | 33334184 | 33334290 | 17 | 6 | 6 | 17 | 10 | 0 | 0 | 0 | 0 | 0 | 25 | 54 | 68 | 39 | 31 | 2 | 0 | 0 | 0 | 0 | 279 | 149 | 1 | 1 | 1 | 1 | 1 | 1 | 1 | 0.87 | 1 | 1 | 1 | 1 | 0.026 |
| SE | 62308 | ENSG00000134686 | PHC2 | chr1 | - | 33333843 | 33333932 | 33332274 | 33332404 | 33334184 | 33334292 | 2 | 0 | 0 | 0 | 0 | 13 | 5 | 5 | 15 | 8 | 2 | 0 | 0 | 0 | 0 | 31 | 43 | 30 | 9 | 29 | 238 | 149 | 1 | 1 | 0.088 | 0 | 0 | 0 | 0 | 0.039 | 0 | 0 | 0 | 0 | 0.01 |
| SE | 62309 | ENSG00000134686 | PHC2 | chr1 | - | 33334089 | 33334292 | 33332274 | 33332404 | 33354400 | 33354566 | 79 | 56 | 85 | 75 | 126 | 0 | 0 | 0 | 0 | 0 | 58 | 297 | 332 | 160 | 223 | 0 | 7 | 0 | 0 | 0 | 298 | 149 | 1 | 1 | 1 | 1 | 1 | 1 | 1 | 1 | 0.955 | 1 | 1 | 1 | 0.009 |
| SE | 62310 | ENSG00000134686 | PHC2 | chr1 | - | 33368535 | 33368622 | 33367115 | 33367428 | 33370420 | 33370585 | 0 | 0 | 1 | 0 | 0 | 0 | 0 | 0 | 0 | 0 | 9 | 0 | 0 | 0 | 0 | 5 | 0 | 0 | 1 | 0 | 236 | 149 | 1 | 1 | NA | NA | 1 | NA | NA | 0.532 | NA | NA | 0 | NA | 0.734 |
| SE | 62311 | ENSG00000223466 | LINC01825 | chr2 | + | 195538040 | 195538132 | 195532944 | 195533243 | 195538311 | 195540287 | 11 | 0 | 8 | 5 | 7 | 1 | 0 | 0 | 0 | 2 | 0 | 6 | 0 | 0 | 0 | 0 | 0 | 0 | 3 | 0 | 241 | 149 | 1 | 1 | 0.872 | NA | 1 | 1 | 0.684 | NA | 1 | NA | 0 | NA | 0.389 |
| SE | 62315 | ENSG00000185811 | IKZF1 | chr7 | + | 50368005 | 50368065 | 50327637 | 50327757 | 50369521 | 50369601 | 65 | 53 | 91 | 93 | 132 | 1 | 0 | 0 | 0 | 0 | 124 | 97 | 83 | 147 | 136 | 0 | 0 | 0 | 0 | 0 | 209 | 149 | 1 | 1 | 0.979 | 1 | 1 | 1 | 1 | 1 | 1 | 1 | 1 | 1 | -0.004 |
| SE | 62322 | ENSG00000185811 | IKZF1 | chr7 | + | 50376532 | 50376793 | 50327378 | 50327757 | 50399917 | 50405098 | 355 | 211 | 274 | 289 | 161 | 0 | 0 | 1 | 0 | 0 | 152 | 226 | 137 | 176 | 202 | 0 | 3 | 0 | 0 | 0 | 298 | 149 | 1 | 1 | 1 | 1 | 0.993 | 1 | 1 | 1 | 0.974 | 1 | 1 | 1 | 0.004 |
| SE | 62325 | ENSG00000185811 | IKZF1 | chr7 | + | 50382539 | 50382707 | 50327637 | 50327757 | 50399917 | 50405098 | 136 | 72 | 140 | 96 | 93 | 0 | 0 | 1 | 0 | 0 | 56 | 77 | 28 | 51 | 43 | 0 | 3 | 0 | 0 | 0 | 298 | 149 | 1 | 1 | 1 | 1 | 0.986 | 1 | 1 | 1 | 0.928 | 1 | 1 | 1 | 0.012 |
| SE | 62326 | ENSG00000185811 | IKZF1 | chr7 | + | 50382539 | 50382707 | 50376532 | 50376793 | 50387344 | 50387470 | 544 | 361 | 477 | 426 | 478 | 0 | 0 | 1 | 0 | 1 | 332 | 427 | 222 | 237 | 232 | 0 | 0 | 0 | 6 | 0 | 298 | 149 | 1 | 1 | 1 | 1 | 0.996 | 1 | 0.996 | 1 | 1 | 1 | 0.952 | 1 | 0.008 |
| SE | 62327 | ENSG00000185811 | IKZF1 | chr7 | + | 50382539 | 50382707 | 50376532 | 50376793 | 50391728 | 50391863 | 311 | 211 | 226 | 212 | 317 | 0 | 0 | 0 | 0 | 2 | 224 | 203 | 142 | 135 | 97 | 0 | 0 | 0 | 0 | 0 | 298 | 149 | 1 | 1 | 1 | 1 | 1 | 1 | 0.988 | 1 | 1 | 1 | 1 | 1 | -0.002 |
| SE | 62331 | ENSG00000185811 | IKZF1 | chr7 | + | 50387344 | 50387470 | 50327378 | 50327757 | 50391728 | 50391863 | 635 | 419 | 604 | 548 | 990 | 5 | 0 | 1 | 3 | 0 | 529 | 551 | 377 | 329 | 407 | 0 | 4 | 5 | 3 | 0 | 275 | 149 | 1 | 1 | 0.986 | 1 | 0.997 | 0.99 | 1 | 1 | 0.987 | 0.976 | 0.983 | 1 | 0.005 |
| SE | 62332 | ENSG00000185811 | IKZF1 | chr7 | + | 50387344 | 50387470 | 50327378 | 50327757 | 50399917 | 50400627 | 4 | 8 | 2 | 7 | 5 | 0 | 0 | 1 | 0 | 0 | 9 | 6 | 3 | 4 | 0 | 0 | 3 | 0 | 0 | 0 | 275 | 149 | 1 | 1 | 1 | 1 | 0.52 | 1 | 1 | 1 | 0.52 | 1 | 1 | NA | 0.024 |
| SE | 62333 | ENSG00000185811 | IKZF1 | chr7 | + | 50387344 | 50387470 | 50376532 | 50376793 | 50391728 | 50391863 | 633 | 418 | 603 | 544 | 986 | 0 | 0 | 0 | 0 | 2 | 529 | 545 | 377 | 332 | 407 | 0 | 0 | 0 | 0 | 0 | 275 | 149 | 1 | 1 | 1 | 1 | 1 | 1 | 0.996 | 1 | 1 | 1 | 1 | 1 | -0.001 |
| SE | 62338 | ENSG00000185811 | IKZF1 | chr7 | + | 50391728 | 50391863 | 50327378 | 50327757 | 50399917 | 50399966 | 822 | 440 | 673 | 633 | 1091 | 0 | 0 | 1 | 0 | 0 | 505 | 687 | 377 | 373 | 401 | 0 | 3 | 0 | 0 | 0 | 284 | 149 | 1 | 1 | 1 | 1 | 0.997 | 1 | 1 | 1 | 0.992 | 1 | 1 | 1 | 0.001 |
| SE | 62341 | ENSG00000185811 | IKZF1 | chr7 | + | 50391728 | 50391863 | 50387344 | 50387470 | 50399917 | 50399966 | 1551 | 927 | 1372 | 1254 | 2133 | 5 | 7 | 0 | 4 | 3 | 1052 | 1273 | 784 | 724 | 828 | 9 | 0 | 3 | 1 | 0 | 284 | 149 | 1 | 1 | 0.994 | 0.986 | 1 | 0.994 | 0.997 | 0.984 | 1 | 0.993 | 0.997 | 1 | -0.001 |
| SE | 62342 | ENSG00000078246 | TULP3 | chr12 | + | 2920762 | 2920922 | 2909528 | 2909580 | 2922261 | 2922402 | 18 | 18 | 21 | 29 | 41 | 5 | 2 | 3 | 3 | 4 | 0 | 16 | 18 | 31 | 11 | 0 | 0 | 2 | 0 | 5 | 298 | 149 | 1 | 1 | 0.643 | 0.818 | 0.778 | 0.829 | 0.837 | NA | 1 | 0.818 | 1 | 0.524 | -0.055 |
| SE | 62343 | ENSG00000078246 | TULP3 | chr12 | + | 2930247 | 2930345 | 2922261 | 2922402 | 2931036 | 2931240 | 32 | 30 | 37 | 18 | 28 | 0 | 0 | 1 | 0 | 0 | 0 | 23 | 19 | 28 | 27 | 0 | 0 | 0 | 0 | 0 | 247 | 149 | 1 | 1 | 1 | 1 | 0.957 | 1 | 1 | NA | 1 | 1 | 1 | 1 | -0.009 |
| SE | 62344 | ENSG00000078246 | TULP3 | chr12 | + | 2930905 | 2930939 | 2930247 | 2930345 | 2931036 | 2931240 | 0 | 1 | 1 | 2 | 0 | 22 | 18 | 21 | 14 | 11 | 0 | 4 | 1 | 3 | 0 | 0 | 12 | 9 | 18 | 13 | 183 | 149 | 1 | 1 | 0 | 0.043 | 0.037 | 0.104 | 0 | NA | 0.213 | 0.083 | 0.119 | 0 | -0.067 |
| SE | 62346 | ENSG00000078246 | TULP3 | chr12 | + | 2937630 | 2937729 | 2934446 | 2934561 | 2938113 | 2938285 | 9 | 7 | 14 | 7 | 14 | 0 | 0 | 1 | 0 | 0 | 0 | 7 | 8 | 7 | 0 | 0 | 3 | 0 | 0 | 0 | 248 | 149 | 1 | 1 | 1 | 1 | 0.894 | 1 | 1 | NA | 0.584 | 1 | 1 | NA | 0.117 |
| SE | 62347 | ENSG00000078246 | TULP3 | chr12 | + | 2938113 | 2938285 | 2937630 | 2937729 | 2939310 | 2939345 | 20 | 24 | 21 | 24 | 47 | 0 | 0 | 0 | 0 | 0 | 14 | 8 | 36 | 27 | 5 | 0 | 1 | 0 | 0 | 0 | 298 | 149 | 1 | 1 | 1 | 1 | 1 | 1 | 1 | 1 | 0.8 | 1 | 1 | 1 | 0.04 |
| SE | 62348 | ENSG00000078246 | TULP3 | chr12 | + | 2940021 | 2940081 | 2939310 | 2939424 | 2940194 | 2940338 | 0 | 0 | 0 | 1 | 0 | 2 | 0 | 0 | 3 | 3 | 0 | 0 | 0 | 0 | 7 | 7 | 9 | 4 | 0 | 0 | 209 | 149 | 1 | 1 | 0 | NA | NA | 0.192 | 0 | 0 | 0 | 0 | NA | 1 | -0.186 |
| SE | 62357 | ENSG00000197956 | S100A6 | chr1 | - | 153535201 | 153535360 | 153534598 | 153534830 | 153535948 | 153535991 | 2613 | 1019 | 1591 | 1954 | 1182 | 0 | 0 | 0 | 0 | 0 | 3952 | 2813 | 1532 | 3674 | 4901 | 0 | 0 | 0 | 2 | 0 | 298 | 149 | 1 | 1 | 1 | 1 | 1 | 1 | 1 | 1 | 1 | 1 | 0.999 | 1 | 0 |
| SE | 62358 | ENSG00000197956 | S100A6 | chr1 | - | 153535201 | 153535375 | 153534598 | 153534830 | 153535948 | 153536215 | 1985 | 838 | 1201 | 1470 | 1008 | 0 | 0 | 0 | 0 | 0 | 3292 | 1934 | 1169 | 2888 | 3616 | 0 | 0 | 0 | 2 | 0 | 298 | 149 | 1 | 1 | 1 | 1 | 1 | 1 | 1 | 1 | 1 | 1 | 0.999 | 1 | 0 |
| SE | 62359 | ENSG00000197956 | S100A6 | chr1 | - | 153535788 | 153535933 | 153535201 | 153535360 | 153536107 | 153536244 | 3 | 2 | 4 | 2 | 1 | 1 | 0 | 0 | 1 | 0 | 0 | 0 | 0 | 4 | 0 | 0 | 7 | 0 | 0 | 16 | 294 | 149 | 1 | 1 | 0.603 | 1 | 1 | 0.503 | 1 | NA | 0 | NA | 1 | 0 | 0.488 |
| SE | 62361 | ENSG00000214193 | SH3D21 | chr1 | + | 36307516 | 36307607 | 36307109 | 36307285 | 36307769 | 36307825 | 0 | 0 | 0 | 0 | 4 | 0 | 0 | 0 | 2 | 0 | 0 | 2 | 2 | 0 | 0 | 0 | 0 | 0 | 0 | 0 | 240 | 149 | 1 | 1 | NA | NA | NA | 0 | 1 | NA | 1 | 1 | NA | NA | -0.5 |
| SE | 62362 | ENSG00000214193 | SH3D21 | chr1 | + | 36319070 | 36319164 | 36309547 | 36309590 | 36319674 | 36320798 | 0 | 0 | 0 | 3 | 1 | 0 | 0 | 0 | 0 | 0 | 0 | 5 | 0 | 2 | 18 | 0 | 0 | 4 | 0 | 0 | 243 | 149 | 1 | 1 | NA | NA | NA | 1 | 1 | NA | 1 | 0 | 1 | 1 | 0.25 |
| SE | 62363 | ENSG00000214193 | SH3D21 | chr1 | + | 36319259 | 36319312 | 36319070 | 36319164 | 36319674 | 36320751 | 0 | 1 | 0 | 0 | 0 | 0 | 0 | 0 | 0 | 0 | 0 | 0 | 0 | 6 | 15 | 0 | 1 | 0 | 0 | 9 | 202 | 149 | 1 | 1 | NA | 1 | NA | NA | NA | NA | 0 | NA | 1 | 0.551 | 0.483 |
| SE | 62364 | ENSG00000214193 | SH3D21 | chr1 | + | 36319441 | 36319536 | 36319259 | 36319312 | 36319674 | 36320798 | 3 | 3 | 4 | 1 | 0 | 0 | 0 | 0 | 0 | 0 | 0 | 0 | 0 | 0 | 0 | 0 | 4 | 0 | 0 | 0 | 244 | 149 | 1 | 1 | 1 | 1 | 1 | 1 | NA | NA | 0 | NA | NA | NA | 1 |
| SE | 62365 | ENSG00000281404 | LINC01176 | chr7 | + | 30402911 | 30403022 | 30390884 | 30391035 | 30403601 | 30404055 | 0 | 2 | 0 | 1 | 0 | 0 | 0 | 0 | 0 | 1 | 0 | 5 | 6 | 4 | 0 | 0 | 0 | 0 | 0 | 0 | 260 | 149 | 1 | 1 | NA | 1 | NA | 1 | 0 | NA | 1 | 1 | 1 | NA | -0.333 |
| SE | 62367 | ENSG00000168826 | ZBTB49 | chr4 | + | 4306137 | 4306184 | 4301988 | 4303091 | 4320639 | 4321783 | 1 | 4 | 1 | 5 | 4 | 1 | 0 | 0 | 0 | 2 | 0 | 0 | 0 | 0 | 0 | 9 | 0 | 0 | 9 | 0 | 196 | 149 | 1 | 1 | 0.432 | 1 | 1 | 1 | 0.603 | 0 | NA | NA | 0 | NA | 0.807 |
| SE | 62368 | ENSG00000168826 | ZBTB49 | chr4 | + | 4313040 | 4313114 | 4302310 | 4303091 | 4315635 | 4315718 | 31 | 15 | 14 | 21 | 16 | 0 | 0 | 0 | 0 | 1 | 29 | 9 | 22 | 26 | 4 | 0 | 0 | 0 | 4 | 0 | 223 | 149 | 1 | 1 | 1 | 1 | 1 | 1 | 0.914 | 1 | 1 | 1 | 0.813 | 1 | 0.02 |
| SE | 62370 | ENSG00000168826 | ZBTB49 | chr4 | + | 4313040 | 4313114 | 4306137 | 4306184 | 4320639 | 4320697 | 18 | 0 | 6 | 7 | 6 | 0 | 0 | 0 | 2 | 0 | 0 | 2 | 4 | 0 | 7 | 0 | 0 | 0 | 0 | 0 | 223 | 149 | 1 | 1 | 1 | NA | 1 | 0.7 | 1 | NA | 1 | 1 | NA | 1 | -0.075 |
| SE | 62371 | ENSG00000168826 | ZBTB49 | chr4 | + | 4315635 | 4315718 | 4301988 | 4303091 | 4320639 | 4321398 | 0 | 0 | 0 | 0 | 1 | 1 | 0 | 0 | 0 | 2 | 0 | 0 | 0 | 4 | 0 | 9 | 0 | 0 | 9 | 0 | 232 | 149 | 1 | 1 | 0 | NA | NA | NA | 0.243 | 0 | NA | NA | 0.222 | NA | 0.01 |
| SE | 62375 | ENSG00000116885 | OSCP1 | chr1 | - | 36420475 | 36420615 | 36418990 | 36419054 | 36422149 | 36422181 | 1 | 0 | 1 | 2 | 8 | 0 | 0 | 1 | 1 | 0 | 4 | 6 | 4 | 0 | 4 | 0 | 0 | 0 | 0 | 0 | 289 | 149 | 1 | 1 | 1 | NA | 0.34 | 0.508 | 1 | 1 | 1 | 1 | NA | 1 | -0.288 |
| SE | 62390 | ENSG00000051596 | THOC3 | chr5 | - | 175967110 | 175967267 | 175961272 | 175961434 | 176007001 | 176007141 | 1 | 2 | 2 | 0 | 2 | 0 | 0 | 0 | 0 | 1 | 5 | 0 | 2 | 0 | 0 | 0 | 0 | 0 | 2 | 0 | 298 | 149 | 1 | 1 | 1 | 1 | 1 | NA | 0.5 | 1 | NA | 1 | 0 | NA | 0.208 |
| SE | 62391 | ENSG00000051596 | THOC3 | chr5 | - | 175967110 | 175967267 | 175961272 | 175961434 | 176034565 | 176034612 | 11 | 3 | 13 | 10 | 2 | 0 | 0 | 0 | 1 | 0 | 15 | 1 | 2 | 0 | 8 | 0 | 0 | 0 | 0 | 8 | 298 | 149 | 1 | 1 | 1 | 1 | 1 | 0.833 | 1 | 1 | 1 | 1 | NA | 0.333 | 0.133 |
| SE | 62393 | ENSG00000051596 | THOC3 | chr5 | - | 176007001 | 176007141 | 175961419 | 175961904 | 176034565 | 176034606 | 0 | 0 | 1 | 0 | 1 | 0 | 0 | 0 | 1 | 0 | 0 | 0 | 0 | 2 | 0 | 0 | 0 | 0 | 0 | 8 | 289 | 149 | 1 | 1 | NA | NA | 1 | 0 | 1 | NA | NA | NA | 1 | 0 | 0.167 |
| SE | 62394 | ENSG00000051596 | THOC3 | chr5 | - | 176007001 | 176007141 | 175967110 | 175967267 | 176034565 | 176034606 | 1 | 1 | 3 | 0 | 2 | 11 | 2 | 13 | 10 | 2 | 0 | 0 | 2 | 0 | 0 | 10 | 1 | 2 | 0 | 8 | 289 | 149 | 1 | 1 | 0.045 | 0.205 | 0.106 | 0 | 0.34 | 0 | 0 | 0.34 | NA | 0 | 0.054 |
| SE | 62395 | ENSG00000051596 | THOC3 | chr5 | - | 176011008 | 176011099 | 176007001 | 176007141 | 176034565 | 176034606 | 0 | 0 | 0 | 1 | 1 | 0 | 0 | 1 | 0 | 0 | 0 | 0 | 0 | 0 | 3 | 0 | 0 | 0 | 0 | 0 | 240 | 149 | 1 | 1 | NA | NA | 0 | 1 | 1 | NA | NA | NA | NA | 1 | -0.333 |
| SE | 62396 | ENSG00000122912 | SLC25A16 | chr10 | - | 68487143 | 68487212 | 68477997 | 68483588 | 68493131 | 68493198 | 32 | 35 | 40 | 24 | 61 | 0 | 0 | 1 | 0 | 0 | 20 | 34 | 76 | 40 | 11 | 0 | 0 | 0 | 0 | 0 | 218 | 149 | 1 | 1 | 1 | 1 | 0.965 | 1 | 1 | 1 | 1 | 1 | 1 | 1 | -0.007 |
| SE | 62397 | ENSG00000122912 | SLC25A16 | chr10 | - | 68488466 | 68488629 | 68487143 | 68487212 | 68493131 | 68493198 | 132 | 102 | 116 | 84 | 124 | 1 | 0 | 0 | 2 | 1 | 65 | 106 | 123 | 142 | 62 | 0 | 0 | 4 | 0 | 0 | 298 | 149 | 1 | 1 | 0.985 | 1 | 1 | 0.955 | 0.984 | 1 | 1 | 0.939 | 1 | 1 | -0.003 |
| SE | 62398 | ENSG00000122912 | SLC25A16 | chr10 | - | 68493448 | 68493570 | 68493131 | 68493198 | 68496433 | 68496512 | 41 | 38 | 30 | 37 | 64 | 1 | 0 | 0 | 0 | 0 | 25 | 22 | 16 | 27 | 4 | 0 | 0 | 0 | 2 | 0 | 271 | 149 | 1 | 1 | 0.958 | 1 | 1 | 1 | 1 | 1 | 1 | 1 | 0.881 | 1 | 0.015 |
| SE | 62400 | ENSG00000122912 | SLC25A16 | chr10 | - | 68496433 | 68496512 | 68493173 | 68493198 | 68503631 | 68503695 | 4 | 4 | 0 | 2 | 0 | 0 | 0 | 2 | 1 | 5 | 0 | 0 | 0 | 2 | 0 | 0 | 0 | 0 | 0 | 0 | 228 | 149 | 1 | 1 | 1 | 1 | 0 | 0.567 | 0 | NA | NA | NA | 1 | NA | -0.487 |
| SE | 62402 | ENSG00000122912 | SLC25A16 | chr10 | - | 68506584 | 68506718 | 68503631 | 68503695 | 68516750 | 68516843 | 45 | 51 | 55 | 62 | 113 | 0 | 0 | 4 | 0 | 0 | 18 | 21 | 48 | 55 | 37 | 0 | 2 | 0 | 0 | 0 | 283 | 149 | 1 | 1 | 1 | 1 | 0.879 | 1 | 1 | 1 | 0.847 | 1 | 1 | 1 | 0.006 |

| | | | | | | | | | | | | | | |
|---|---|---|---|---|---|---|---|---|---|---|---|---|---|---|
| SE | 62403 | ENSG00000122912 | SLC25A16 | chr10 | - | 68516750 | 68516843 | 68506584 | 68506718 | 68527245 | 68527523 | 16 | 12 | [illegible] |
| SE | 62404 | ENSG00000122912 | SLC25A16 | chr10 | - | 68517083 | 68517185 | 68516750 | 68516843 | 68527245 | 68527438 | 5 | 6 | [illegible] |
| SE | 62405 | ENSG00000122912 | SLC25A16 | chr10 | - | 68517083 | 68517265 | 68516750 | 68516843 | 68527245 | 68527416 | 9 | 6 | [illegible] |
| SE | 62407 | ENSG00000033867 | SLC4A7 | chr3 | - | 27389930 | 27390104 | 27385891 | 27386023 | 27391739 | 27391808 | 260 | 193 | [illegible] |
| SE | 62409 | ENSG00000033867 | SLC4A7 | chr3 | - | 27421621 | 27421779 | 27420699 | 27420787 | 27422780 | 27422840 | 74 | 35 | [illegible] |
| SE | 62415 | ENSG00000033867 | SLC4A7 | chr3 | - | 27436387 | 27436548 | 27433915 | 27434104 | 27437387 | 27437526 | 450 | 261 | [illegible] |
| SE | 62417 | ENSG00000113761 | ZNF346 | chr5 | + | 177041125 | 177041229 | 177022695 | 177022913 | 177062057 | 177062151 | 13 | 2 | [illegible] |
| SE | 62422 | ENSG00000113761 | ZNF346 | chr5 | + | 177044388 | 177044533 | 177041777 | 177041870 | 177050702 | 177050936 | 35 | 33 | [illegible] |
| SE | 62423 | ENSG00000113761 | ZNF346 | chr5 | + | 177044388 | 177044533 | 177041777 | 177041870 | 177050750 | 177050936 | 67 | 60 | [illegible] |
| SE | 62425 | ENSG00000113761 | ZNF346 | chr5 | + | 177044388 | 177044533 | 177041777 | 177041870 | 177064511 | 177064601 | 30 | 35 | [illegible] |
| SE | 62426 | ENSG00000113761 | ZNF346 | chr5 | + | 177044391 | 177044533 | 177041125 | 177041229 | 177062057 | 177062151 | 3 | 6 | [illegible] |
| SE | 62427 | ENSG00000113761 | ZNF346 | chr5 | + | 177044391 | 177044533 | 177041777 | 177041870 | 177050750 | 177050936 | 39 | 28 | [illegible] |
| SE | 62429 | ENSG00000113761 | ZNF346 | chr5 | + | 177044391 | 177044533 | 177041777 | 177041870 | 177064511 | 177064966 | 2 | 3 | [illegible] |
| SE | 62430 | ENSG00000113761 | ZNF346 | chr5 | + | 177050702 | 177050936 | 177022695 | 177022913 | 177062057 | 177062151 | 10 | 6 | [illegible] |
| SE | 62432 | ENSG00000113761 | ZNF346 | chr5 | + | 177050702 | 177050936 | 177041777 | 177041870 | 177064511 | 177065241 | 0 | 1 | [illegible] |
| SE | 62434 | ENSG00000113761 | ZNF346 | chr5 | + | 177050702 | 177050936 | 177044388 | 177044533 | 177064511 | 177065241 | 5 | 1 | [illegible] |
| SE | 62436 | ENSG00000113761 | ZNF346 | chr5 | + | 177050750 | 177050936 | 177041777 | 177041870 | 177064511 | 177064601 | 1 | 1 | [illegible] |
| SE | 62438 | ENSG00000113761 | ZNF346 | chr5 | + | 177050750 | 177050936 | 177044388 | 177044533 | 177064511 | 177064601 | 37 | 28 | [illegible] |
| SE | 62439 | ENSG00000113761 | ZNF346 | chr5 | + | 177062057 | 177062151 | 177041777 | 177041870 | 177064511 | 177064601 | 24 | 22 | [illegible] |
| SE | 62440 | ENSG00000113761 | ZNF346 | chr5 | + | 177062057 | 177062151 | 177041777 | 177041870 | 177079381 | 177080554 | 3 | 7 | [illegible] |
| SE | 62441 | ENSG00000113761 | ZNF346 | chr5 | + | 177062057 | 177062151 | 177044388 | 177044533 | 177064511 | 177064601 | 22 | 18 | [illegible] |
| SE | 62443 | ENSG00000113761 | ZNF346 | chr5 | + | 177064511 | 177064601 | 177041777 | 177041870 | 177079381 | 177080554 | 1 | 1 | [illegible] |
| SE | 62444 | ENSG00000113761 | ZNF346 | chr5 | + | 177064511 | 177064601 | 177062057 | 177062151 | 177079381 | 177080554 | 18 | 13 | [illegible] |
| SE | 62445 | ENSG00000286905 | NA | chr2 | - | 3536197 | 3536313 | 3533107 | 3534255 | 3536723 | 3536911 | 1 | 4 | [illegible] |
| SE | 62446 | ENSG00000120265 | PCMT1 | chr6 | + | 149765690 | 149765753 | 149749815 | 149749956 | 149771161 | 149771266 | 2 | 0 | [illegible] |
| SE | 62447 | ENSG00000120265 | PCMT1 | chr6 | + | 149771161 | 149771266 | 149749442 | 149749956 | 149773137 | 149773169 | 316 | 210 | [illegible] |
| SE | 62448 | ENSG00000120265 | PCMT1 | chr6 | + | 149771161 | 149771266 | 149749442 | 149749956 | 149789953 | 149790056 | 69 | 52 | [illegible] |
| SE | 62449 | ENSG00000120265 | PCMT1 | chr6 | + | 149772013 | 149772247 | 149771161 | 149771266 | 149773137 | 149773150 | 24 | 11 | [illegible] |
| SE | 62454 | ENSG00000120265 | PCMT1 | chr6 | + | 149773137 | 149773169 | 149749694 | 149749956 | 149789953 | 149790056 | 37 | 34 | [illegible] |
| SE | 62455 | ENSG00000120265 | PCMT1 | chr6 | + | 149773137 | 149773169 | 149771161 | 149771266 | 149789953 | 149790056 | 286 | 191 | [illegible] |
| SE | 62456 | ENSG00000120265 | PCMT1 | chr6 | + | 149789953 | 149790058 | 149773137 | 149773169 | 149793548 | 149793632 | 494 | 383 | [illegible] |
| SE | 62457 | ENSG00000120265 | PCMT1 | chr6 | + | 149796414 | 149796500 | 149793548 | 149793669 | 149802199 | 149802369 | 292 | 244 | [illegible] |
| SE | 62461 | ENSG00000146856 | AGBL3 | chr7 | + | 134989249 | 134989310 | 134987874 | 134987996 | 134993492 | 134993678 | 1 | 3 | [illegible] |
| SE | 62462 | ENSG00000146856 | AGBL3 | chr7 | + | 134989249 | 134989310 | 134987874 | 134987996 | 135010041 | 135012511 | 0 | 1 | [illegible] |
| SE | 62463 | ENSG00000146856 | AGBL3 | chr7 | + | 134993492 | 134993678 | 134987874 | 134987996 | 135010041 | 135012511 | 0 | 0 | [illegible] |
| SE | 62464 | ENSG00000146856 | AGBL3 | chr7 | + | 134993492 | 134993678 | 134989249 | 134989310 | 135010041 | 135012511 | 1 | 2 | [illegible] |
| SE | 62465 | ENSG00000146856 | AGBL3 | chr7 | + | 134993492 | 134993678 | 134989249 | 134989310 | 135017051 | 135017159 | 1 | 2 | [illegible] |
| SE | 62467 | ENSG00000130508 | PXDN | chr2 | - | 1638845 | 1638978 | 1635407 | 1635521 | 1639301 | 1639422 | 0 | 0 | [illegible] |
| SE | 62471 | ENSG00000130508 | PXDN | chr2 | - | 1687631 | 1687703 | 1684079 | 1684151 | 1691927 | 1691999 | 0 | 0 | [illegible] |
| SE | 62472 | ENSG00000116752 | BCAS2 | chr1 | - | 114570699 | 114570750 | 114569991 | 114570072 | 114575589 | 114575751 | 239 | 218 | [illegible] |
| SE | 62473 | ENSG00000116752 | BCAS2 | chr1 | - | 114575589 | 114575751 | 114570699 | 114570750 | 114576687 | 114576758 | 335 | 281 | [illegible] |
| SE | 62474 | ENSG00000116752 | BCAS2 | chr1 | - | 114576687 | 114576758 | 114575589 | 114575751 | 114581298 | 114581391 | 176 | 115 | [illegible] |
| SE | 62501 | ENSG00000183943 | PRKX | chrX | - | 3626418 | 3626514 | 3621258 | 3621316 | 3641851 | 3641971 | 206 | 108 | [illegible] |
| SE | 62503 | ENSG00000183943 | PRKX | chrX | - | 3655148 | 3655412 | 3626418 | 3626514 | 3670178 | 3670215 | 3 | 6 | [illegible] |
| SE | 62509 | ENSG00000049167 | ERCC8 | chr5 | - | 60896458 | 60896515 | 60890888 | 60891086 | 60898275 | 60898400 | 0 | 1 | [illegible] |
| SE | 62513 | ENSG00000049167 | ERCC8 | chr5 | - | 60904787 | 60904873 | 60903647 | 60903716 | 60918264 | 60918388 | 11 | 8 | [illegible] |
| SE | 62514 | ENSG00000049167 | ERCC8 | chr5 | - | 60904791 | 60904873 | 60903359 | 60903716 | 60918264 | 60918388 | 39 | 24 | [illegible] |
| SE | 62515 | ENSG00000049167 | ERCC8 | chr5 | - | 60909647 | 60909957 | 60904791 | 60904873 | 60918264 | 60918388 | 1 | 0 | [illegible] |
| SE | 62516 | ENSG00000049167 | ERCC8 | chr5 | - | 60915185 | 60915286 | 60904791 | 60904873 | 60918264 | 60918388 | 0 | 0 | [illegible] |
| SE | 62518 | ENSG00000049167 | ERCC8 | chr5 | - | 60924290 | 60924356 | 60922053 | 60922155 | 60928863 | 60928959 | 0 | 2 | [illegible] |
| SE | 62519 | ENSG00000049167 | ERCC8 | chr5 | - | 60924290 | 60924367 | 60922053 | 60922155 | 60928863 | 60928959 | 4 | 0 | [illegible] |
| SE | 62523 | ENSG00000049167 | ERCC8 | chr5 | - | 60928863 | 60928959 | 60922053 | 60922155 | 60944931 | 60945008 | 31 | 18 | [illegible] |
| SE | 62524 | ENSG00000049167 | ERCC8 | chr5 | - | 60932106 | 60932148 | 60928863 | 60928959 | 60944931 | 60945064 | 0 | 0 | [illegible] |
| SE | 62529 | ENSG00000004866 | ST7 | chr7 | + | 117098612 | 117098815 | 117014935 | 117015014 | 117099761 | 117099844 | 2 | 2 | [illegible] |
| SE | 62532 | ENSG00000004866 | ST7 | chr7 | + | 117119560 | 117119720 | 117099761 | 117099844 | 117129792 | 117129809 | 97 | 30 | [illegible] |
| SE | 62534 | ENSG00000004866 | ST7 | chr7 | + | 117136080 | 117136235 | 117131884 | 117131960 | 117138434 | 117138532 | 64 | 39 | [illegible] |
| SE | 62538 | ENSG00000004866 | ST7 | chr7 | + | 117190833 | 117190936 | 117189320 | 117189393 | 117209786 | 117209922 | 45 | 43 | [illegible] |
| SE | 62547 | ENSG00000144028 | SNRNP200 | chr2 | - | 96276462 | 96276724 | 96275256 | 96275349 | 96276903 | 96276985 | 0 | 0 | [illegible] |
| SE | 62548 | ENSG00000144028 | SNRNP200 | chr2 | - | 96276903 | 96276985 | 96275256 | 96275349 | 96277080 | 96277241 | 337 | 164 | [illegible] |
| SE | 62549 | ENSG00000144028 | SNRNP200 | chr2 | - | 96279450 | 96279559 | 96278808 | 96278998 | 96281813 | 96281922 | 294 | 107 | [illegible] |
| SE | 62550 | ENSG00000144028 | SNRNP200 | chr2 | - | 96281813 | 96281922 | 96279450 | 96279559 | 96283200 | 96283352 | 234 | 72 | [illegible] |
| SE | 62551 | ENSG00000144028 | SNRNP200 | chr2 | - | 96287862 | 96287969 | 96287438 | 96287557 | 96288662 | 96288746 | 375 | 218 | [illegible] |
| SE | 62553 | ENSG00000144028 | SNRNP200 | chr2 | - | 96305392 | 96305544 | 96304704 | 96304868 | 96307309 | 96307406 | 182 | 99 | [illegible] |
| SE | 62556 | ENSG00000092330 | TINF2 | chr14 | - | 24241203 | 24241311 | 24241019 | 24241116 | 24241674 | 24241776 | 332 | 160 | [illegible] |
| SE | 62558 | ENSG00000092330 | TINF2 | chr14 | - | 24241674 | 24241776 | 24241019 | 24241116 | 24242140 | 24242605 | 9 | 4 | [illegible] |
| SE | 62561 | ENSG00000092330 | TINF2 | chr14 | - | 24241889 | 24241994 | 24241674 | 24241776 | 24242376 | 24242585 | 166 | 70 | [illegible] |
| SE | 62562 | ENSG00000107614 | TRDMT1 | chr10 | - | 17157440 | 17157784 | 17154676 | 17154734 | 17159145 | 17159229 | 79 | 32 | [illegible] |
| SE | 62563 | ENSG00000107614 | TRDMT1 | chr10 | - | 17160304 | 17160374 | 17159145 | 17159229 | 17161482 | 17161548 | 118 | 63 | [illegible] |
| SE | 62564 | ENSG00000107614 | TRDMT1 | chr10 | - | 17160304 | 17160374 | 17159145 | 17159229 | 17162165 | 17162237 | 35 | 19 | [illegible] |
| SE | 62565 | ENSG00000107614 | TRDMT1 | chr10 | - | 17161482 | 17161548 | 17159145 | 17159229 | 17162165 | 17162237 | 43 | 24 | [illegible] |
| SE | 62567 | ENSG00000107614 | TRDMT1 | chr10 | - | 17161482 | 17161548 | 17160304 | 17160374 | 17168840 | 17168917 | 54 | 31 | [illegible] |
| SE | 62568 | ENSG00000107614 | TRDMT1 | chr10 | - | 17161482 | 17161548 | 17160304 | 17160374 | 17174550 | 17174660 | 53 | 31 | [illegible] |
| SE | 62569 | ENSG00000107614 | TRDMT1 | chr10 | - | 17162165 | 17162237 | 17160304 | 17160374 | 17168840 | 17168917 | 16 | 13 | [illegible] |
| SE | 62573 | ENSG00000107614 | TRDMT1 | chr10 | - | 17168840 | 17168917 | 17160304 | 17160374 | 17174550 | 17174660 | 14 | 4 | [illegible] |
| SE | 62576 | ENSG00000107614 | TRDMT1 | chr10 | - | 17169309 | 17169539 | 17168840 | 17168917 | 17174550 | 17174660 | 22 | 8 | [illegible] |
| SE | 62577 | ENSG00000255028 | NA | chr11 | - | 109415923 | 109416171 | 109413253 | 109413556 | 109512578 | 109512725 | 0 | 0 | [illegible] |
| SE | 62582 | ENSG00000183150 | GPR19 | chr12 | - | 12688512 | 12688932 | 12684397 | 12684507 | 12695458 | 12695511 | 0 | 0 | [illegible] |
| SE | 62585 | ENSG00000173597 | SULT1B1 | chr4 | - | 69733412 | 69733507 | 69730500 | 69730681 | 69734137 | 69734264 | 251 | 188 | [illegible] |
| SE | 62586 | ENSG00000173597 | SULT1B1 | chr4 | - | 69749720 | 69749818 | 69734137 | 69734264 | 69754669 | 69754798 | 277 | 198 | [illegible] |
| SE | 62587 | ENSG00000173597 | SULT1B1 | chr4 | - | 69749720 | 69749818 | 69734137 | 69734264 | 69760458 | 69760598 | 108 | 87 | [illegible] |
| SE | 62588 | ENSG00000173597 | SULT1B1 | chr4 | - | 69754669 | 69754798 | 69734137 | 69734264 | 69755069 | 69755201 | 268 | 241 | [illegible] |
| SE | 62589 | ENSG00000173597 | SULT1B1 | chr4 | - | 69754669 | 69754798 | 69734137 | 69734264 | 69760458 | 69760571 | 4 | 2 | [illegible] |
| SE | 62590 | ENSG00000173597 | SULT1B1 | chr4 | - | 69754669 | 69754798 | 69749720 | 69749818 | 69760458 | 69760571 | 172 | 110 | [illegible] |
| SE | 62591 | ENSG00000173597 | SULT1B1 | chr4 | - | 69755069 | 69755201 | 69754669 | 69754798 | 69760207 | 69760250 | 279 | 266 | [illegible] |
| SE | 62593 | ENSG00000173597 | SULT1B1 | chr4 | - | 69755069 | 69755261 | 69734137 | 69734264 | 69760458 | 69760620 | 145 | 91 | [illegible] |
| SE | 62594 | ENSG00000173597 | SULT1B1 | chr4 | - | 69755069 | 69755261 | 69754669 | 69754798 | 69760207 | 69760250 | 272 | 252 | [illegible] |
| SE | 62596 | ENSG00000173597 | SULT1B1 | chr4 | - | 69760207 | 69760250 | 69754669 | 69754798 | 69760458 | 69760598 | 10 | 6 | [illegible] |
| SE | 62597 | ENSG00000173597 | SULT1B1 | chr4 | - | 69760207 | 69760250 | 69755069 | 69755201 | 69760458 | 69760598 | 30 | 36 | [illegible] |
| SE | 62599 | ENSG00000143622 | RIT1 | chr1 | - | 155904310 | 155904502 | 155897807 | 155900618 | 155904730 | 155904804 | 358 | 316 | [illegible] |
| SE | 62602 | ENSG00000143622 | RIT1 | chr1 | - | 155904602 | 155904610 | 155904310 | 155904502 | 155904730 | 155904804 | 0 | 0 | [illegible] |
| SE | 62604 | ENSG00000143622 | RIT1 | chr1 | - | 155904730 | 155904804 | 155904310 | 155904502 | 155910449 | 155910506 | 178 | 133 | [illegible] |
| SE | 62605 | ENSG00000143622 | RIT1 | chr1 | - | 155910655 | 155910804 | 155910449 | 155910506 | 155910877 | 155910915 | 83 | 75 | [illegible] |
| SE | 62609 | ENSG00000015676 | NUDCD3 | chr7 | - | 44413190 | 44413283 | 44404439 | 44404583 | 44425899 | 44426279 | 7 | 2 | [illegible] |
| SE | 62611 | ENSG00000015676 | NUDCD3 | chr7 | - | 44413190 | 44413283 | 44404439 | 44404583 | 44481291 | 44481462 | 7 | 2 | [illegible] |
| SE | 62612 | ENSG00000015676 | NUDCD3 | chr7 | - | 44425899 | 44426279 | 44413190 | 44413283 | 44427570 | 44427703 | 0 | 0 | [illegible] |
| SE | 62614 | ENSG00000015676 | NUDCD3 | chr7 | - | 44427570 | 44427703 | 44404439 | 44404583 | 44481291 | 44481462 | 77 | 41 | [illegible] |
| SE | 62616 | ENSG00000015676 | NUDCD3 | chr7 | - | 44427570 | 44427703 | 44413190 | 44413283 | 44481291 | 44481462 | 7 | 8 | [illegible] |
| SE | 62617 | ENSG00000015676 | NUDCD3 | chr7 | - | 44485530 | 44485622 | 44484967 | 44485284 | 44489822 | 44490265 | 12 | 4 | [illegible] |
| SE | 62620 | ENSG00000033178 | UBA6 | chr4 | - | 67626359 | 67626477 | 67624993 | 67625187 | 67629070 | 67629142 | 149 | 89 | [illegible] |
| SE | 62621 | ENSG00000033178 | UBA6 | chr4 | - | 67631856 | 67631908 | 67631707 | 67631771 | 67633344 | 67633473 | 124 | 78 | [illegible] |
| SE | 62624 | ENSG00000033178 | UBA6 | chr4 | - | 67663138 | 67663215 | 67661662 | 67662255 | 67663884 | 67663947 | 136 | 70 | [illegible] |
| SE | 62625 | ENSG00000033178 | UBA6 | chr4 | - | 67663884 | 67663947 | 67663138 | 67663215 | 67665188 | 67665292 | 171 | 95 | [illegible] |
| SE | 62626 | ENSG00000033178 | UBA6 | chr4 | - | 67673696 | 67673777 | 67670469 | 67670592 | 67677610 | 67677722 | 288 | 318 | [illegible] |
| SE | 62627 | ENSG00000033178 | UBA6 | chr4 | - | 67677610 | 67677722 | 67673696 | 67673777 | 67678438 | 67678533 | 264 | 237 | [illegible] |
| SE | 62628 | ENSG00000033178 | UBA6 | chr4 | - | 67696644 | 67696707 | 67682118 | 67682213 | 67701048 | 67701142 | 85 | 57 | [illegible] |
| SE | 62637 | ENSG00000128815 | WDFY4 | chr10 | + | 48709715 | 48709966 | 48684875 | 48685001 | 48720010 | 48720125 | 89 | 70 | [illegible] |
| SE | 62638 | ENSG00000128815 | WDFY4 | chr10 | + | 48727469 | 48727659 | 48725880 | 48726070 | 48729431 | 48729589 | 64 | 43 | [illegible] |
| SE | 62639 | ENSG00000128815 | WDFY4 | chr10 | + | 48760346 | 48760440 | 48742967 | 48743548 | 48774457 | 48774672 | 73 | 42 | [illegible] |
| SE | 62642 | ENSG00000128815 | WDFY4 | chr10 | + | 48790726 | 48790917 | 48789873 | 48789985 | 48796297 | 48796450 | 126 | 73 | [illegible] |
| SE | 62644 | ENSG00000128815 | WDFY4 | chr10 | + | 48811538 | 48811708 | 48807858 | 48807958 | 48813956 | 48814082 | 39 | 25 | [illegible] |
| SE | 62648 | ENSG00000128815 | WDFY4 | chr10 | + | 48867264 | 48867342 | 48832572 | 48832709 | 48873490 | 48873697 | 236 | 181 | [illegible] |
| SE | 62649 | ENSG00000128815 | WDFY4 | chr10 | + | 48873490 | 48873697 | 48867264 | 48867342 | 48875088 | 48875140 | 146 | 94 | [illegible] |
| SE | 62650 | ENSG00000128815 | WDFY4 | chr10 | + | 48875088 | 48875140 | 48873490 | 48873697 | 48877032 | 48877199 | 99 | 52 | [illegible] |
| SE | 62651 | ENSG00000128815 | WDFY4 | chr10 | + | 48957128 | 48957282 | 48946859 | 48946969 | 48959721 | 48959813 | 75 | 37 | [illegible] |
| SE | 62652 | ENSG00000128815 | WDFY4 | chr10 | + | 48959721 | 48959813 | 48946859 | 48946969 | 48963841 | 48964054 | 36 | 22 | [illegible] |
| SE | 62653 | ENSG00000128815 | WDFY4 | chr10 | + | 48981366 | 48981478 | 48978308 | 48978393 | 48982508 | 48982956 | 113 | 81 | [illegible] |
| SE | 62654 | ENSG00000183762 | KREMEN1 | chr22 | + | 29121356 | 29121481 | 29098861 | 29098953 | 29125262 | 29125416 | 0 | 0 | [illegible] |
| SE | 62658 | ENSG00000161640 | SIGLEC11 | chr19 | - | 49959772 | 49959820 | 49959359 | 49959623 | 49960136 | 49960421 | 2 | 0 | [illegible] |
| SE | 62660 | ENSG00000229809 | ZNF688 | chr16 | - | 30571009 | 30571123 | 30569671 | 30570436 | 30571433 | 30571652 | 52 | 27 | [illegible] |
| SE | 62661 | ENSG00000229809 | ZNF688 | chr16 | - | 30571009 | 30571123 | 30570733 | 30570940 | 30571433 | 30571652 | 26 | 6 | [illegible] |
| SE | 62662 | ENSG00000187726 | DNAJB13 | chr11 | + | 73964877 | 73965035 | 73959493 | 73959655 | 73966137 | 73966251 | 0 | 0 | [illegible] |
| SE | 62666 | ENSG00000115525 | ST3GAL5 | chr2 | - | 85846376 | 85846424 | 85839188 | 85840392 | 85847860 | 85848204 | 2 | 0 | [illegible] |
| SE | 62667 | ENSG00000115525 | ST3GAL5 | chr2 | - | 85846376 | 85846563 | 85839148 | 85840392 | 85847860 | 85848084 | 40 | 50 | [illegible] |
| SE | 62668 | ENSG00000115525 | ST3GAL5 | chr2 | - | 85847381 | 85847562 | 85846376 | 85846424 | 85847860 | 85847984 | 19 | 8 | [illegible] |
| SE | 62670 | ENSG00000115525 | ST3GAL5 | chr2 | - | 85847860 | 85848084 | 85846376 | 85846563 | 85861180 | 85861292 | 37 | 48 | [illegible] |
| SE | 62671 | ENSG00000115525 | ST3GAL5 | chr2 | - | 85847860 | 85848204 | 85846376 | 85846424 | 85861180 | 85861292 | 103 | 107 | [illegible] |
| SE | 62672 | ENSG00000115525 | ST3GAL5 | chr2 | - | 85847860 | 85848204 | 85846376 | 85846563 | 85861180 | 85861292 | 134 | 149 | [illegible] |
| SE | 62673 | ENSG00000115525 | ST3GAL5 | chr2 | - | 85847860 | 85848381 | 85846376 | 85846424 | 85861180 | 85861292 | 19 | 9 | [illegible] |
| SE | 62674 | ENSG00000115525 | ST3GAL5 | chr2 | - | 85847860 | 85848381 | 85846376 | 85846563 | 85861180 | 85861292 | 50 | 51 | [illegible] |
| SE | 62678 | ENSG00000115525 | ST3GAL5 | chr2 | - | 85851263 | 85851703 | 85847860 | 85848204 | 85863361 | 85863369 | 5 | 15 | [illegible] |
| SE | 62680 | ENSG00000115525 | ST3GAL5 | chr2 | - | 85851263 | 85851703 | 85847860 | 85850834 | 85861180 | 85861292 | 3 | 2 | [illegible] |
| SE | 62690 | ENSG00000115525 | ST3GAL5 | chr2 | - | 85861180 | 85861292 | 85847860 | 85848204 | 85863361 | 85863369 | 148 | 153 | [illegible] |
| SE | 62691 | ENSG00000115525 | ST3GAL5 | chr2 | - | 85861180 | 85861292 | 85851263 | 85851703 | 85863361 | 85863369 | 51 | 47 | [illegible] |
| SE | 62693 | ENSG00000115525 | ST3GAL5 | chr2 | - | 85867844 | 85868058 | 85863361 | 85863485 | 85870071 | 85870258 | 0 | 0 | [illegible] |
| SE | 62696 | ENSG00000115525 | ST3GAL5 | chr2 | - | 85875414 | 85875545 | 85863361 | 85863485 | 85888823 | 85888910 | 0 | 0 | [illegible] |
| SE | 62697 | ENSG00000236008 | LINC01814 | chr2 | - | 8542784 | 8542974 | 8541076 | 8542225 | 8545147 | 8545236 | 0 | 0 | [illegible] |
| SE | 62700 | ENSG00000236008 | LINC01814 | chr2 | - | 8570447 | 8570581 | 8545147 | 8545236 | 8576910 | 8577141 | 0 | 0 | [illegible] |
| SE | 62705 | ENSG00000049247 | UTS2 | chr1 | - | 7847803 | 7847882 | 7846691 | 7846766 | 7849639 | 7849683 | 2 | 29 | [illegible] |
| SE | 62706 | ENSG00000049247 | UTS2 | chr1 | - | 7847803 | 7847882 | 7846691 | 7846766 | 7850811 | 7850922 | 0 | 6 | [illegible] |
| SE | 62707 | ENSG00000049247 | UTS2 | chr1 | - | 7849639 | 7849683 | 7846691 | 7846766 | 7850811 | 7850922 | 7 | 21 | [illegible] |
| SE | 62708 | ENSG00000049247 | UTS2 | chr1 | - | 7849639 | 7849683 | 7847611 | 7847882 | 7850811 | 7850922 | 8 | 44 | [illegible] |
| SE | 62709 | ENSG00000049247 | UTS2 | chr1 | - | 7852900 | 7852923 | 7850811 | 7850922 | 7853306 | 7853431 | 0 | 4 | [illegible] |
| SE | 62710 | ENSG00000049247 | UTS2 | chr1 | - | 7852900 | 7853118 | 7850811 | 7850922 | 7853306 | 7853512 | 1 | 9 | [illegible] |
| SE | 62713 | ENSG00000151779 | NBAS | chr2 | - | 15238467 | 15238686 | 15234544 | 15234747 | 15275483 | 15275818 | 53 | 49 | [illegible] |
| SE | 62714 | ENSG00000151779 | NBAS | chr2 | - | 15276850 | 15277101 | 15275483 | 15275818 | 15287072 | 15287183 | 172 | 129 | [illegible] |
| SE | 62716 | ENSG00000151779 | NBAS | chr2 | - | 15353552 | 15353710 | 15351991 | 15352081 | 15356302 | 15356416 | 104 | 47 | [illegible] |
| SE | 62717 | ENSG00000151779 | NBAS | chr2 | - | 15366579 | 15366693 | 15356302 | 15356416 | 15374607 | 15374720 | 152 | 120 | [illegible] |
| SE | 62718 | ENSG00000151779 | NBAS | chr2 | - | 15383214 | 15383317 | 15379601 | 15379831 | 15394226 | 15394349 | 93 | 51 | [illegible] |
| SE | 62719 | ENSG00000151779 | NBAS | chr2 | - | 15415545 | 15415719 | 15402167 | 15402301 | 15417526 | 15417581 | 251 | 186 | [illegible] |
| SE | 62721 | ENSG00000151779 | NBAS | chr2 | - | 15478225 | 15478289 | 15475686 | 15475880 | 15488893 | 15489022 | 191 | 159 | [illegible] |
| SE | 62722 | ENSG00000151779 | NBAS | chr2 | - | 15483261 | 15483368 | 15478244 | 15478289 | 15488893 | 15489022 | 1 | 0 | [illegible] |
| SE | 62724 | ENSG00000151779 | NBAS | chr2 | - | 15504144 | 15504213 | 15488893 | 15489022 | 15534542 | 15534641 | 74 | 60 | [illegible] |
| SE | 62725 | ENSG00000151779 | NBAS | chr2 | - | 15511211 | 15511350 | 15504144 | 15504213 | 15534542 | 15534641 | 303 | 245 | [illegible] |
| SE | 62728 | ENSG00000151779 | NBAS | chr2 | - | 15551492 | 15551536 | 15534542 | 15534641 | 15553425 | 15553473 | 20 | 8 | [illegible] |
| SE | 62734 | ENSG00000101333 | PLCB4 | chr20 | + | 9373046 | 9373104 | 9372302 | 9372403 | 9380053 | 9380162 | 0 | 0 | [illegible] |
| SE | 62738 | ENSG00000145416 | MARCHF1 | chr4 | - | 163555848 | 163555973 | 163545595 | 163545743 | 163585748 | 163585929 | 2 | 0 | [illegible] |
| SE | 62748 | ENSG00000162961 | DPY30 | chr2 | - | 31869847 | 31869952 | 31867808 | 31868492 | 31891991 | 31892134 | 101 | 74 | [illegible] |
| SE | 62749 | ENSG00000162961 | DPY30 | chr2 | - | 31869847 | 31869952 | 31867808 | 31868492 | 31917925 | 31918037 | 95 | 63 | [illegible] |
| SE | 62752 | ENSG00000162961 | DPY30 | chr2 | - | 31891991 | 31892134 | 31867808 | 31868492 | 31917925 | 31918037 | 106 | 93 | [illegible] |
| SE | 62755 | ENSG00000162961 | DPY30 | chr2 | - | 31891991 | 31892134 | 31883385 | 31883462 | 31932066 | 31932135 | 171 | 116 | [illegible] |
| SE | 62756 | ENSG00000162961 | DPY30 | chr2 | - | 31932066 | 31932135 | 31891991 | 31892134 | 31943301 | 31943383 | 30 | 11 | [illegible] |
| SE | 62759 | ENSG00000162961 | DPY30 | chr2 | - | 32029593 | 32029736 | 32023904 | 32024256 | 32039278 | 32039326 | 254 | 227 | [illegible] |
| SE | 62760 | ENSG00000088543 | C3orf18 | chr3 | - | 50560916 | 50561064 | 50558024 | 50559737 | 50561721 | 50561747 | 38 | 19 | [illegible] |
| SE | 62763 | ENSG00000088543 | C3orf18 | chr3 | - | 50565465 | 50565861 | 50561721 | 50561747 | 50566005 | 50566094 | 14 | 0 | [illegible] |
| SE | 62764 | ENSG00000088543 | C3orf18 | chr3 | - | 50565465 | 50565861 | 50561721 | 50561747 | 50567462 | 50567665 | 14 | 6 | [illegible] |
| SE | 62767 | ENSG00000088543 | C3orf18 | chr3 | - | 50566005 | 50566094 | 50561721 | 50561747 | 50567462 | 50567625 | 0 | 2 | [illegible] |
| SE | 62768 | ENSG00000088543 | C3orf18 | chr3 | - | 50566005 | 50566094 | 50565465 | 50565861 | 50567462 | 50567625 | 0 | 0 | [illegible] |
| SE | 62769 | ENSG00000134243 | SORT1 | chr1 | - | 109314260 | 109314384 | 109309567 | 109314057 | 109314671 | 109314778 | 86 | 104 | [illegible] |
| SE | 62770 | ENSG00000134243 | SORT1 | chr1 | - | 109316849 | 109316958 | 109314671 | 109314778 | 109317852 | 109317969 | 321 | 284 | [illegible] |
| SE | 62771 | ENSG00000134243 | SORT1 | chr1 | - | 109317852 | 109317969 | 109316849 | 109316958 | 109322931 | 109323121 | 267 | 261 | [illegible] |
| SE | 62772 | ENSG00000134243 | SORT1 | chr1 | - | 109342013 | 109342158 | 109340723 | 109340879 | 109345750 | 109345878 | 328 | 369 | [illegible] |
| SE | 62773 | ENSG00000134243 | SORT1 | chr1 | - | 109397206 | 109397340 | 109369529 | 109369589 | 109397586 | 109397861 | 0 | 0 | [illegible] |
| SE | 62774 | ENSG00000135338 | LCA5 | chr6 | - | 79513211 | 79513741 | 79493612 | 79493750 | 79518704 | 79519085 | 3 | 2 | [illegible] |
| SE | 62779 | ENSG00000163785 | RYK | chr3 | - | 134175929 | 134176039 | 134159236 | 134159373 | 134183001 | 134183071 | 27 | 14 | [illegible] |
| SE | 62781 | ENSG00000163785 | RYK | chr3 | - | 134191848 | 134191974 | 134188836 | 134188923 | 134195081 | 134195182 | 173 | 121 | [illegible] |
| SE | 62782 | ENSG00000163785 | RYK | chr3 | - | 134191848 | 134191983 | 134188836 | 134188923 | 134195081 | 134195182 | 177 | 139 | [illegible] |
| SE | 62783 | ENSG00000163785 | RYK | chr3 | - | 134202729 | 134202874 | 134195081 | 134195182 | 134207471 | 134207525 | 137 | 109 | [illegible] |
| SE | 62785 | ENSG00000138646 | HERC5 | chr4 | + | 88463531 | 88463623 | 88460094 | 88460171 | 88467058 | 88467204 | 5 | 0 | [illegible] |
| SE | 62787 | ENSG00000138646 | HERC5 | chr4 | + | 88468345 | 88468422 | 88467058 | 88467204 | 88469156 | 88469260 | 154 | 195 | [illegible] |
| SE | 62788 | ENSG00000138646 | HERC5 | chr4 | + | 88470613 | 88470673 | 88469156 | 88469260 | 88472408 | 88472502 | 71 | 76 | [illegible] |
| SE | 62789 | ENSG00000138646 | HERC5 | chr4 | + | 88472408 | 88472502 | 88470613 | 88470673 | 88475840 | 88476030 | 93 | 81 | [illegible] |
| SE | 62790 | ENSG00000138646 | HERC5 | chr4 | + | 88486114 | 88486228 | 88479352 | 88479507 | 88487068 | 88487179 | 110 | 144 | [illegible] |
| SE | 62792 | ENSG00000138646 | HERC5 | chr4 | + | 88487068 | 88487179 | 88479352 | 88479507 | 88489165 | 88489336 | 52 | 56 | [illegible] |
| SE | 62793 | ENSG00000138646 | HERC5 | chr4 | + | 88487068 | 88487179 | 88479352 | 88479507 | 88493011 | 88493155 | 0 | 1 | [illegible] |
| SE | 62794 | ENSG00000138646 | HERC5 | chr4 | + | 88487068 | 88487179 | 88486114 | 88486228 | 88489165 | 88489336 | 132 | 152 | [illegible] |
| SE | 62796 | ENSG00000138646 | HERC5 | chr4 | + | 88489165 | 88489336 | 88487068 | 88487179 | 88493011 | 88493155 | 111 | 136 | [illegible] |
| SE | 62797 | ENSG00000138646 | HERC5 | chr4 | + | 88493011 | 88493155 | 88489165 | 88489336 | 88494164 | 88494331 | 113 | 114 | [illegible] |
| SE | 62798 | ENSG00000138646 | HERC5 | chr4 | + | 88499925 | 88499992 | 88494164 | 88494331 | 88500914 | 88500985 | 135 | 122 | [illegible] |
| SE | 62799 | ENSG00000138646 | HERC5 | chr4 | + | 88500914 | 88500985 | 88494164 | 88494331 | 88504231 | 88504415 | 20 | 39 | [illegible] |
| SE | 62801 | ENSG00000169246 | NPIPB3 | chr16 | - | 21419489 | 21419766 | 21414956 | 21415085 | 21424978 | 21425136 | 0 | 0 | [illegible] |
| SE | 62802 | ENSG00000169246 | NPIPB3 | chr16 | - | 21419634 | 21419766 | 21414956 | 21415085 | 21424978 | 21425136 | 0 | 0 | [illegible] |
| SE | 62804 | ENSG00000169246 | NPIPB3 | chr16 | - | 21425289 | 21425378 | 21424978 | 21425136 | 21431504 | 21431553 | 0 | 0 | [illegible] |
| SE | 62805 | ENSG00000101882 | NKAP | chrX | - | 119934493 | 119934557 | 119932106 | 119932216 | 119936296 | 119936428 | 31 | 40 | [illegible] |
| SE | 62826 | ENSG00000196387 | ZNF140 | chr12 | + | 133080821 | 133081072 | 133080473 | 133080551 | 133081272 | 133081329 | 3 | 1 | [illegible] |
| SE | 62827 | ENSG00000196387 | ZNF140 | chr12 | + | 133081272 | 133081329 | 133080821 | 133081072 | 133083102 | 133083229 | 4 | 2 | [illegible] |
| SE | 62829 | ENSG00000196387 | ZNF140 | chr12 | + | 133093414 | 133093505 | 133083462 | 133083561 | 133100986 | 133101053 | 0 | 2 | [illegible] |
| SE | 62831 | ENSG00000196387 | ZNF140 | chr12 | + | 133100986 | 133101053 | 133083465 | 133083561 | 133105509 | 133107428 | 19 | 16 | [illegible] |
| SE | 62832 | ENSG00000196387 | ZNF140 | chr12 | + | 133100986 | 133101053 | 133093414 | 133093505 | 133105509 | 133107428 | 6 | 8 | [illegible] |
| SE | 62836 | ENSG00000197451 | HNRNPAB | chr5 | + | 178209329 | 178209432 | 178207093 | 178207225 | 178210131 | 178210272 | 407 | 315 | [illegible] |
| SE | 62837 | ENSG00000197451 | HNRNPAB | chr5 | + | 178209329 | 178209432 | 178207093 | 178207225 | 178210552 | 178211155 | 407 | 315 | [illegible] |
| SE | 62838 | ENSG00000197451 | HNRNPAB | chr5 | + | 178209329 | 178209441 | 178207093 | 178207225 | 178210552 | 178211119 | 409 | 317 | [illegible] |
| SE | 62839 | ENSG00000197451 | HNRNPAB | chr5 | + | 178209329 | 178209447 | 178207093 | 178207225 | 178210131 | 178210272 | 426 | 326 | [illegible] |
| SE | 62840 | ENSG00000197451 | HNRNPAB | chr5 | + | 178209329 | 178209447 | 178207093 | 178207225 | 178210552 | 178211142 | 422 | 323 | [illegible] |
| SE | 62844 | ENSG00000237190 | CDKN2AIPNL | chr5 | - | 134409902 | 134410002 | 134402064 | 134402926 | 134411615 | 134411881 | 39 | 17 | [illegible] |
| SE | 62858 | ENSG00000112335 | SNX3 | chr6 | - | 108214497 | 108214622 | 108211221 | 108212254 | 108222233 | 108222320 | 899 | 727 | [illegible] |
| SE | 62862 | ENSG00000112335 | SNX3 | chr6 | - | 108222949 | 108223045 | 108214497 | 108214622 | 108260759 | 108261040 | 758 | 567 | [illegible] |
| SE | 62863 | ENSG00000112335 | SNX3 | chr6 | - | 108222949 | 108223045 | 108214497 | 108214622 | 108260825 | 108261244 | 408 | 319 | [illegible] |
| SE | 62865 | ENSG00000140612 | SEC11A | chr15 | - | 84670724 | 84670782 | 84669537 | 84670069 | 84680712 | 84680832 | 734 | 418 | [illegible] |
| SE | 62867 | ENSG00000140612 | SEC11A | chr15 | - | 84679931 | 84679969 | 84670724 | 84670782 | 84680712 | 84680832 | 1 | 0 | [illegible] |

| | | | | | | | | | | | | | | |
|---|---|---|---|---|---|---|---|---|---|---|---|---|---|---|
| SE | 62868 | ENSG00000140612 | SEC11A | chr15 | - | 84680712 | 84680832 | 84669543 | 84670069 | 84691534 | 84691644 | 3 | 0 | [illegible] |
| SE | 62870 | ENSG00000140612 | SEC11A | chr15 | - | 84680712 | 84680832 | 84670724 | 84670782 | 84716024 | 84716089 | 285 | 159 | [illegible] |
| SE | 62871 | ENSG00000140612 | SEC11A | chr15 | - | 84687624 | 84687774 | 84670724 | 84670782 | 84716024 | 84716063 | 27 | 11 | [illegible] |
| SE | 62873 | ENSG00000140612 | SEC11A | chr15 | - | 84687624 | 84687774 | 84680712 | 84680832 | 84716024 | 84716063 | 678 | 403 | [illegible] |
| SE | 62874 | ENSG00000140612 | SEC11A | chr15 | - | 84691534 | 84691644 | 84680712 | 84680832 | 84716024 | 84716063 | 168 | 59 | [illegible] |
| SE | 62875 | ENSG00000140612 | SEC11A | chr15 | - | 84691534 | 84691644 | 84687624 | 84687774 | 84716024 | 84716063 | 651 | 358 | [illegible] |
| SE | 62876 | ENSG00000140612 | SEC11A | chr15 | - | 84691969 | 84692120 | 84691534 | 84691644 | 84716024 | 84716067 | 0 | 0 | [illegible] |
| SE | 62879 | ENSG00000114120 | SLC25A36 | chr3 | + | 140959462 | 140959540 | 140956524 | 140956691 | 140963126 | 140963227 | 77 | 39 | [illegible] |
| SE | 62880 | ENSG00000114120 | SLC25A36 | chr3 | + | 140966347 | 140966434 | 140963126 | 140963227 | 140966933 | 140967036 | 7 | 2 | [illegible] |
| SE | 62881 | ENSG00000114120 | SLC25A36 | chr3 | + | 140966347 | 140966434 | 140963126 | 140963227 | 140966999 | 140967036 | 7 | 2 | [illegible] |
| SE | 62882 | ENSG00000114120 | SLC25A36 | chr3 | + | 140966347 | 140966434 | 140963126 | 140963227 | 140970926 | 140970993 | 15 | 6 | [illegible] |
| SE | 62883 | ENSG00000114120 | SLC25A36 | chr3 | + | 140966933 | 140967036 | 140963126 | 140963227 | 140970926 | 140970993 | 14 | 10 | [illegible] |
| SE | 62884 | ENSG00000114120 | SLC25A36 | chr3 | + | 140966933 | 140967036 | 140966347 | 140966434 | 140970926 | 140970993 | 11 | 8 | [illegible] |
| SE | 62885 | ENSG00000114120 | SLC25A36 | chr3 | + | 140966999 | 140967036 | 140963126 | 140963227 | 140970926 | 140970993 | 9 | 4 | [illegible] |
| SE | 62886 | ENSG00000114120 | SLC25A36 | chr3 | + | 140966999 | 140967036 | 140966347 | 140966434 | 140970926 | 140970993 | 9 | 3 | [illegible] |
| SE | 62887 | ENSG00000114120 | SLC25A36 | chr3 | + | 140968682 | 140968730 | 140966933 | 140967036 | 140970926 | 140970993 | 0 | 0 | [illegible] |
| SE | 62888 | ENSG00000012963 | UBR7 | chr14 | + | 93209823 | 93209957 | 93207240 | 93207441 | 93210647 | 93210708 | 42 | 38 | [illegible] |
| SE | 62890 | ENSG00000012963 | UBR7 | chr14 | + | 93215175 | 93215281 | 93214928 | 93214982 | 93219211 | 93219361 | 29 | 49 | [illegible] |
| SE | 62892 | ENSG00000012963 | UBR7 | chr14 | + | 93219211 | 93219361 | 93215175 | 93215281 | 93222312 | 93222374 | 0 | 0 | [illegible] |
| SE | 62894 | ENSG00000012963 | UBR7 | chr14 | + | 93220248 | 93220411 | 93219211 | 93219361 | 93222312 | 93222374 | 160 | 108 | [illegible] |
| SE | 62895 | ENSG00000012963 | UBR7 | chr14 | + | 93222312 | 93222374 | 93220248 | 93220411 | 93226942 | 93227024 | 68 | 59 | [illegible] |
| SE | 62897 | ENSG00000119771 | KLHL29 | chr2 | + | 23703185 | 23703379 | 23696332 | 23696513 | 23703718 | 23703863 | 0 | 4 | [illegible] |
| SE | 62920 | ENSG00000185420 | SMYD3 | chr1 | - | 246186742 | 246186942 | 245929869 | 245929937 | 246193674 | 246193727 | 1 | 0 | [illegible] |
| SE | 62922 | ENSG00000185420 | SMYD3 | chr1 | - | 246335366 | 246335474 | 246330479 | 246330537 | 246355030 | 246355094 | 57 | 31 | [illegible] |
| SE | 62923 | ENSG00000185420 | SMYD3 | chr1 | - | 246355030 | 246355094 | 246335366 | 246335474 | 246507053 | 246507144 | 26 | 5 | [illegible] |
| SE | 62925 | ENSG00000163235 | TGFA | chr2 | - | 70461628 | 70461701 | 70456365 | 70456488 | 70465615 | 70465736 | 0 | 0 | [illegible] |
| SE | 62926 | ENSG00000142892 | PIGK | chr1 | - | 77161294 | 77161405 | 77153535 | 77154621 | 77161593 | 77161711 | 243 | 164 | [illegible] |
| SE | 62929 | ENSG00000142892 | PIGK | chr1 | - | 77169259 | 77169395 | 77166718 | 77166830 | 77210435 | 77210489 | 267 | 195 | [illegible] |
| SE | 62930 | ENSG00000142892 | PIGK | chr1 | - | 77169259 | 77169395 | 77166718 | 77166830 | 77219309 | 77219429 | 264 | 196 | [illegible] |
| SE | 62932 | ENSG00000142892 | PIGK | chr1 | - | 77206639 | 77206731 | 77169259 | 77169395 | 77210435 | 77210489 | 189 | 169 | [illegible] |
| SE | 62933 | ENSG00000142892 | PIGK | chr1 | - | 77210435 | 77210489 | 77166718 | 77166830 | 77219309 | 77219426 | 20 | 3 | [illegible] |
| SE | 62934 | ENSG00000142892 | PIGK | chr1 | - | 77210435 | 77210489 | 77169259 | 77169395 | 77219309 | 77219426 | 20 | 1 | [illegible] |
| SE | 62935 | ENSG00000177683 | THAP5 | chr7 | - | 108565829 | 108566022 | 108562131 | 108565105 | 108569098 | 108569453 | 32 | 20 | [illegible] |
| SE | 62936 | ENSG00000177683 | THAP5 | chr7 | - | 108565829 | 108566022 | 108562131 | 108565105 | 108569489 | 108569706 | 56 | 40 | [illegible] |
| SE | 62937 | ENSG00000177683 | THAP5 | chr7 | - | 108565951 | 108566022 | 108564569 | 108565105 | 108569098 | 108569256 | 4 | 8 | [illegible] |
| SE | 62938 | ENSG00000177683 | THAP5 | chr7 | - | 108565951 | 108566022 | 108564569 | 108565105 | 108569489 | 108569750 | 18 | 14 | [illegible] |
| SE | 62940 | ENSG00000186235 | LINC02610 | chr2 | - | 238229315 | 238229668 | 238227683 | 238227798 | 238231333 | 238231370 | 0 | 1 | [illegible] |
| SE | 62947 | ENSG00000130595 | TNNT3 | chr11 | + | 1923047 | 1923061 | 1922778 | 1922891 | 1923554 | 1923572 | 1 | 2 | [illegible] |
| SE | 62951 | ENSG00000130595 | TNNT3 | chr11 | + | 1925098 | 1925116 | 1923554 | 1923572 | 1925255 | 1925276 | 1 | 6 | [illegible] |
| SE | 62955 | ENSG00000130595 | TNNT3 | chr11 | + | 1926469 | 1926481 | 1925255 | 1925276 | 1929809 | 1929828 | 2 | 11 | [illegible] |
| SE | 62957 | ENSG00000130595 | TNNT3 | chr11 | + | 1926694 | 1926709 | 1926469 | 1926481 | 1929809 | 1929828 | 0 | 0 | [illegible] |
| SE | 62964 | ENSG00000117222 | RBBP5 | chr1 | - | 205105027 | 205105168 | 205103856 | 205104019 | 205114788 | 205114961 | 113 | 57 | [illegible] |
| SE | 62965 | ENSG00000117222 | RBBP5 | chr1 | - | 205114788 | 205114961 | 205105027 | 205105168 | 205121854 | 205121966 | 46 | 22 | [illegible] |
| SE | 62966 | ENSG00000117222 | RBBP5 | chr1 | - | 205115857 | 205115883 | 205114788 | 205114961 | 205121854 | 205121978 | 48 | 8 | [illegible] |
| SE | 62971 | ENSG00000228590 | MIR4432HG | chr2 | - | 60339315 | 60339481 | 60336445 | 60338720 | 60339601 | 60339708 | 0 | 0 | [illegible] |
| SE | 62972 | ENSG00000228590 | MIR4432HG | chr2 | - | 60339601 | 60339708 | 60336445 | 60338720 | 60343087 | 60343168 | 2 | 1 | [illegible] |
| SE | 62973 | ENSG00000228590 | MIR4432HG | chr2 | - | 60342816 | 60342996 | 60336445 | 60338720 | 60343087 | 60343168 | 1 | 0 | [illegible] |
| SE | 62974 | ENSG00000228590 | MIR4432HG | chr2 | - | 60342816 | 60342996 | 60340165 | 60340238 | 60343087 | 60343168 | 1 | 0 | [illegible] |
| SE | 62976 | ENSG00000228590 | MIR4432HG | chr2 | - | 60351808 | 60352308 | 60343087 | 60343168 | 60369908 | 60370037 | 0 | 0 | [illegible] |
| SE | 62977 | ENSG00000228590 | MIR4432HG | chr2 | - | 60351808 | 60352308 | 60343087 | 60343168 | 60402087 | 60402210 | 1 | 0 | [illegible] |
| SE | 62978 | ENSG00000228590 | MIR4432HG | chr2 | - | 60359269 | 60359848 | 60343087 | 60343168 | 60369908 | 60370037 | 0 | 1 | [illegible] |
| SE | 62980 | ENSG00000228590 | MIR4432HG | chr2 | - | 60369908 | 60370037 | 60343087 | 60343168 | 60402087 | 60402210 | 1 | 4 | [illegible] |
| SE | 62981 | ENSG00000228590 | MIR4432HG | chr2 | - | 60369908 | 60370037 | 60351808 | 60352308 | 60402087 | 60402210 | 1 | 4 | [illegible] |
| SE | 62982 | ENSG00000228590 | MIR4432HG | chr2 | - | 60369908 | 60370037 | 60359093 | 60359848 | 60402087 | 60402210 | 1 | 5 | [illegible] |
| SE | 62983 | ENSG00000228590 | MIR4432HG | chr2 | - | 60378501 | 60378620 | 60369908 | 60370037 | 60402087 | 60402210 | 0 | 0 | [illegible] |
| SE | 62985 | ENSG00000066697 | MSANTD3 | chr9 | + | 100444127 | 100445237 | 100442148 | 100442356 | 100450556 | 100451229 | 0 | 0 | [illegible] |
| SE | 62990 | ENSG00000137497 | NUMA1 | chr11 | - | 72012894 | 72016260 | 72010785 | 72010854 | 72016407 | 72016530 | 236 | 124 | [illegible] |
| SE | 62991 | ENSG00000137497 | NUMA1 | chr11 | - | 72016407 | 72016530 | 72010785 | 72010854 | 72017686 | 72017827 | 322 | 231 | [illegible] |
| SE | 62992 | ENSG00000137497 | NUMA1 | chr11 | - | 72021203 | 72021291 | 72019493 | 72019617 | 72022338 | 72022419 | 457 | 210 | [illegible] |
| SE | 62993 | ENSG00000137497 | NUMA1 | chr11 | - | 72024273 | 72024353 | 72023064 | 72023147 | 72029204 | 72029290 | 516 | 276 | [illegible] |
| SE | 62997 | ENSG00000137497 | NUMA1 | chr11 | - | 72050661 | 72050833 | 72035901 | 72035975 | 72069841 | 72069911 | 0 | 0 | [illegible] |
| SE | 62998 | ENSG00000137497 | NUMA1 | chr11 | - | 72050661 | 72050833 | 72049418 | 72049539 | 72069841 | 72069911 | 0 | 0 | [illegible] |
| SE | 62999 | ENSG00000137497 | NUMA1 | chr11 | - | 72062588 | 72062686 | 72035901 | 72035975 | 72069841 | 72069911 | 0 | 0 | [illegible] |
| SE | 63001 | ENSG00000137497 | NUMA1 | chr11 | - | 72068104 | 72068231 | 72035901 | 72035975 | 72068495 | 72068617 | 6 | 1 | [illegible] |
| SE | 63004 | ENSG00000137497 | NUMA1 | chr11 | - | 72068495 | 72068617 | 72035901 | 72035975 | 72069841 | 72069911 | 1 | 0 | [illegible] |
| SE | 63006 | ENSG00000137497 | NUMA1 | chr11 | - | 72069841 | 72069911 | 72035901 | 72035975 | 72080457 | 72080494 | 41 | 18 | [illegible] |
| SE | 63028 | ENSG00000156113 | KCNMA1 | chr10 | - | 77108480 | 77108572 | 77090399 | 77090510 | 77110172 | 77110343 | 2 | 0 | [illegible] |
| SE | 63035 | ENSG00000163026 | WDCP | chr2 | - | 24032828 | 24032946 | 24029346 | 24031162 | 24037676 | 24039512 | 56 | 29 | [illegible] |
| SE | 63036 | ENSG00000285533 | RELA-DT | chr11 | + | 65663426 | 65663683 | 65663005 | 65663148 | 65665420 | 65665543 | 1 | 0 | [illegible] |
| SE | 63037 | ENSG00000285533 | RELA-DT | chr11 | + | 65663426 | 65665543 | 65663006 | 65663148 | 65671422 | 65671704 | 0 | 0 | [illegible] |
| SE | 63038 | ENSG00000285533 | RELA-DT | chr11 | + | 65665420 | 65665543 | 65663005 | 65663148 | 65671422 | 65671716 | 1 | 2 | [illegible] |
| SE | 63039 | ENSG00000285533 | RELA-DT | chr11 | + | 65665420 | 65665543 | 65663426 | 65663683 | 65666130 | 65666217 | 7 | 5 | [illegible] |
| SE | 63042 | ENSG00000267316 | NA | chr18 | - | 61573893 | 61573950 | 61571341 | 61571544 | 61577027 | 61577151 | 0 | 0 | [illegible] |
| SE | 63043 | ENSG00000267316 | NA | chr18 | - | 61577027 | 61577211 | 61571347 | 61571544 | 61579406 | 61579456 | 6 | 13 | [illegible] |
| SE | 63045 | ENSG00000116741 | RGS2 | chr1 | + | 192809457 | 192809535 | 192809142 | 192809181 | 192810165 | 192810267 | 0 | 0 | [illegible] |
| SE | 63046 | ENSG00000116741 | RGS2 | chr1 | + | 192810165 | 192810267 | 192809038 | 192809181 | 192810369 | 192810431 | 756 | 662 | [illegible] |
| SE | 63050 | ENSG00000116741 | RGS2 | chr1 | + | 192810369 | 192810431 | 192810165 | 192810267 | 192810980 | 192811032 | 1063 | 952 | [illegible] |
| SE | 63051 | ENSG00000116741 | RGS2 | chr1 | + | 192810980 | 192811147 | 192810369 | 192810431 | 192811401 | 192812275 | 2720 | 2559 | [illegible] |
| SE | 63054 | ENSG00000006652 | IFRD1 | chr7 | + | 112456913 | 112457038 | 112456001 | 112456086 | 112458860 | 112458912 | 171 | 154 | [illegible] |
| SE | 63055 | ENSG00000006652 | IFRD1 | chr7 | + | 112456913 | 112457038 | 112456001 | 112456086 | 112461865 | 112461916 | 64 | 54 | [illegible] |
| SE | 63057 | ENSG00000006652 | IFRD1 | chr7 | + | 112458860 | 112459018 | 112456913 | 112457038 | 112461865 | 112461916 | 142 | 128 | [illegible] |
| SE | 63061 | ENSG00000006652 | IFRD1 | chr7 | + | 112465638 | 112465838 | 112462269 | 112462378 | 112467980 | 112468115 | 0 | 3 | [illegible] |
| SE | 63071 | ENSG00000132953 | XPO4 | chr13 | - | 20786964 | 20787057 | 20777328 | 20783919 | 20790461 | 20790580 | 46 | 29 | [illegible] |
| SE | 63072 | ENSG00000132953 | XPO4 | chr13 | - | 20787480 | 20787598 | 20786964 | 20787057 | 20790461 | 20790580 | 37 | 41 | [illegible] |
| SE | 63073 | ENSG00000132953 | XPO4 | chr13 | - | 20788485 | 20788616 | 20787480 | 20787598 | 20790461 | 20790580 | 107 | 82 | [illegible] |
| SE | 63074 | ENSG00000132953 | XPO4 | chr13 | - | 20790461 | 20790580 | 20777328 | 20783919 | 20796075 | 20796256 | 82 | 64 | [illegible] |
| SE | 63075 | ENSG00000132953 | XPO4 | chr13 | - | 20790461 | 20790580 | 20788485 | 20788616 | 20796075 | 20796256 | 130 | 100 | [illegible] |
| SE | 63077 | ENSG00000132953 | XPO4 | chr13 | - | 20800155 | 20800325 | 20796763 | 20797057 | 20807456 | 20807634 | 1 | 5 | [illegible] |
| SE | 63078 | ENSG00000132953 | XPO4 | chr13 | - | 20800830 | 20800990 | 20800155 | 20800325 | 20807456 | 20807634 | 128 | 88 | [illegible] |
| SE | 63079 | ENSG00000132953 | XPO4 | chr13 | - | 20807456 | 20807634 | 20796763 | 20797057 | 20808435 | 20808581 | 56 | 44 | [illegible] |
| SE | 63080 | ENSG00000132953 | XPO4 | chr13 | - | 20807456 | 20807634 | 20800830 | 20800990 | 20808435 | 20808581 | 116 | 77 | [illegible] |
| SE | 63081 | ENSG00000132953 | XPO4 | chr13 | - | 20843769 | 20843886 | 20842894 | 20843048 | 20855626 | 20855765 | 88 | 82 | [illegible] |
| SE | 63084 | ENSG00000132953 | XPO4 | chr13 | - | 20868595 | 20868701 | 20862968 | 20863123 | 20902669 | 20902810 | 32 | 21 | [illegible] |
| SE | 63093 | ENSG00000131037 | EPS8L1 | chr19 | + | 55083766 | 55084642 | 55083615 | 55083644 | 55085840 | 55085892 | 0 | 0 | [illegible] |
| SE | 63094 | ENSG00000093217 | XYLB | chr3 | + | 38360338 | 38360408 | 38348549 | 38348632 | 38365198 | 38365285 | 6 | 0 | [illegible] |
| SE | 63095 | ENSG00000093217 | XYLB | chr3 | + | 38362936 | 38363017 | 38360338 | 38360408 | 38365198 | 38365285 | 2 | 3 | [illegible] |
| SE | 63098 | ENSG00000093217 | XYLB | chr3 | + | 38379245 | 38379342 | 38376917 | 38376991 | 38395504 | 38395563 | 3 | 5 | [illegible] |
| SE | 63101 | ENSG00000138050 | THUMPD2 | chr2 | - | 39768900 | 39769093 | 39768423 | 39768501 | 39769709 | 39770119 | 7 | 9 | [illegible] |
| SE | 63102 | ENSG00000138050 | THUMPD2 | chr2 | - | 39768900 | 39769093 | 39768423 | 39768501 | 39771504 | 39771640 | 3 | 2 | [illegible] |
| SE | 63103 | ENSG00000138050 | THUMPD2 | chr2 | - | 39769709 | 39770119 | 39768423 | 39768501 | 39771504 | 39771640 | 30 | 39 | [illegible] |
| SE | 63105 | ENSG00000138050 | THUMPD2 | chr2 | - | 39769709 | 39770119 | 39768902 | 39769191 | 39771504 | 39771640 | 28 | 34 | [illegible] |
| SE | 63106 | ENSG00000138050 | THUMPD2 | chr2 | - | 39769757 | 39770119 | 39768423 | 39768501 | 39771514 | 39771640 | 8 | 11 | [illegible] |
| SE | 63107 | ENSG00000138050 | THUMPD2 | chr2 | - | 39771504 | 39771640 | 39769709 | 39770119 | 39779113 | 39779152 | 26 | 29 | [illegible] |
| SE | 63108 | ENSG00000138050 | THUMPD2 | chr2 | - | 39771514 | 39771640 | 39769757 | 39770119 | 39779113 | 39779221 | 8 | 13 | [illegible] |
| SE | 63110 | ENSG00000115761 | NOL10 | chr2 | - | 10607184 | 10607311 | 10603077 | 10603157 | 10644319 | 10644372 | 126 | 96 | [illegible] |
| SE | 63111 | ENSG00000115761 | NOL10 | chr2 | - | 10644319 | 10644372 | 10607184 | 10607311 | 10659171 | 10659250 | 24 | 22 | [illegible] |
| SE | 63116 | ENSG00000115761 | NOL10 | chr2 | - | 10659171 | 10659250 | 10657741 | 10657891 | 10662958 | 10663044 | 114 | 111 | [illegible] |
| SE | 63117 | ENSG00000115761 | NOL10 | chr2 | - | 10662958 | 10663044 | 10659171 | 10659250 | 10667217 | 10667278 | 148 | 141 | [illegible] |
| SE | 63120 | ENSG00000115761 | NOL10 | chr2 | - | 10681970 | 10682069 | 10673519 | 10673557 | 10689795 | 10689975 | 1 | 0 | [illegible] |
| SE | 63121 | ENSG00000115761 | NOL10 | chr2 | - | 10681970 | 10682069 | 10675793 | 10675871 | 10684566 | 10684612 | 115 | 65 | [illegible] |
| SE | 63122 | ENSG00000115761 | NOL10 | chr2 | - | 10684566 | 10684612 | 10681970 | 10682069 | 10689795 | 10689972 | 72 | 37 | [illegible] |
| SE | 63125 | ENSG00000164134 | NAA15 | chr4 | + | 139334173 | 139334258 | 139301504 | 139301831 | 139336847 | 139336952 | 143 | 127 | [illegible] |
| SE | 63126 | ENSG00000164134 | NAA15 | chr4 | + | 139342825 | 139342960 | 139340911 | 139341069 | 139344185 | 139344339 | 289 | 228 | [illegible] |
| SE | 63127 | ENSG00000164134 | NAA15 | chr4 | + | 139344185 | 139344339 | 139340911 | 139341069 | 139349461 | 139349581 | 140 | 150 | [illegible] |
| SE | 63128 | ENSG00000164134 | NAA15 | chr4 | + | 139354025 | 139354098 | 139351504 | 139351611 | 139357385 | 139357555 | 168 | 86 | [illegible] |
| SE | 63130 | ENSG00000164134 | NAA15 | chr4 | + | 139357385 | 139357555 | 139354025 | 139354098 | 139359742 | 139359857 | 294 | 193 | [illegible] |
| SE | 63131 | ENSG00000164134 | NAA15 | chr4 | + | 139370210 | 139370404 | 139361723 | 139361937 | 139376364 | 139376473 | 180 | 163 | [illegible] |
| SE | 63132 | ENSG00000164134 | NAA15 | chr4 | + | 139376364 | 139376473 | 139370210 | 139370404 | 139378755 | 139378854 | 89 | 58 | [illegible] |
| SE | 63133 | ENSG00000164134 | NAA15 | chr4 | + | 139378755 | 139378854 | 139376364 | 139376473 | 139384831 | 139384978 | 191 | 144 | [illegible] |
| SE | 63134 | ENSG00000164134 | NAA15 | chr4 | + | 139384831 | 139384978 | 139378755 | 139378854 | 139386132 | 139386230 | 351 | 272 | [illegible] |
| SE | 63135 | ENSG00000164134 | NAA15 | chr4 | + | 139384834 | 139384978 | 139378755 | 139378854 | 139386132 | 139386230 | 229 | 167 | [illegible] |
| SE | 63136 | ENSG00000164134 | NAA15 | chr4 | + | 139386132 | 139386230 | 139384831 | 139384978 | 139387883 | 139391368 | 290 | 219 | [illegible] |
| SE | 63137 | ENSG00000164134 | NAA15 | chr4 | + | 139393590 | 139393722 | 139386167 | 139386230 | 139404397 | 139404490 | 2 | 1 | [illegible] |
| SE | 63139 | ENSG00000224875 | EML4-AS1 | chr2 | - | 42149934 | 42150032 | 42147650 | 42147840 | 42169969 | 42170231 | 4 | 0 | [illegible] |
| SE | 63140 | ENSG00000228262 | LINC01320 | chr2 | + | 33886371 | 33886474 | 33855687 | 33855753 | 33926839 | 33926968 | 0 | 2 | [illegible] |
| SE | 63156 | ENSG00000251474 | RPL32P3 | chr3 | - | 129397039 | 129397140 | 129393606 | 129393795 | 129399380 | 129399495 | 1 | 0 | [illegible] |
| SE | 63160 | ENSG00000168061 | SAC3D1 | chr11 | + | 65043024 | 65043164 | 65041244 | 65041866 | 65044224 | 65044673 | 2 | 0 | [illegible] |
| SE | 63166 | ENSG00000175662 | TOM1L2 | chr17 | - | 17848822 | 17848859 | 17843510 | 17847783 | 17862730 | 17862848 | 1 | 4 | [illegible] |
| SE | 63167 | ENSG00000175662 | TOM1L2 | chr17 | - | 17850892 | 17850952 | 17848822 | 17848859 | 17861475 | 17861551 | 35 | 33 | [illegible] |
| SE | 63168 | ENSG00000175662 | TOM1L2 | chr17 | - | 17850892 | 17850952 | 17848822 | 17848859 | 17866295 | 17866419 | 20 | 3 | [illegible] |
| SE | 63169 | ENSG00000175662 | TOM1L2 | chr17 | - | 17857768 | 17857855 | 17850892 | 17850952 | 17861475 | 17861551 | 1 | 0 | [illegible] |
| SE | 63170 | ENSG00000175662 | TOM1L2 | chr17 | - | 17861475 | 17861551 | 17848822 | 17848859 | 17862730 | 17862848 | 14 | 8 | [illegible] |
| SE | 63172 | ENSG00000175662 | TOM1L2 | chr17 | - | 17861475 | 17861551 | 17850892 | 17850952 | 17866295 | 17866419 | 5 | 19 | [illegible] |
| SE | 63173 | ENSG00000175662 | TOM1L2 | chr17 | - | 17862730 | 17862848 | 17848822 | 17848859 | 17866295 | 17866419 | 16 | 6 | [illegible] |
| SE | 63174 | ENSG00000175662 | TOM1L2 | chr17 | - | 17862730 | 17862848 | 17861475 | 17861551 | 17866295 | 17866419 | 47 | 24 | [illegible] |
| SE | 63175 | ENSG00000175662 | TOM1L2 | chr17 | - | 17866295 | 17866419 | 17862730 | 17862848 | 17866875 | 17866924 | 55 | 31 | [illegible] |
| SE | 63176 | ENSG00000175662 | TOM1L2 | chr17 | - | 17866295 | 17866419 | 17862730 | 17862848 | 17869339 | 17869473 | 21 | 9 | [illegible] |
| SE | 63177 | ENSG00000175662 | TOM1L2 | chr17 | - | 17866875 | 17866924 | 17862730 | 17862848 | 17869339 | 17869473 | 4 | 5 | [illegible] |
| SE | 63178 | ENSG00000175662 | TOM1L2 | chr17 | - | 17866875 | 17866924 | 17866295 | 17866419 | 17869339 | 17869473 | 35 | 20 | [illegible] |
| SE | 63180 | ENSG00000175662 | TOM1L2 | chr17 | - | 17882704 | 17882863 | 17879626 | 17879743 | 17884633 | 17884768 | 48 | 43 | [illegible] |
| SE | 63184 | ENSG00000175662 | TOM1L2 | chr17 | - | 17884633 | 17884768 | 17879626 | 17879743 | 17898595 | 17898674 | 4 | 0 | [illegible] |
| SE | 63191 | ENSG00000175662 | TOM1L2 | chr17 | - | 17907446 | 17907531 | 17898595 | 17898674 | 17972261 | 17972397 | 77 | 49 | [illegible] |
| SE | 63201 | ENSG00000118960 | HS1BP3 | chr2 | - | 20623895 | 20624031 | 20612296 | 20619245 | 20624731 | 20624773 | 102 | 31 | [illegible] |
| SE | 63204 | ENSG00000118960 | HS1BP3 | chr2 | - | 20640972 | 20641180 | 20634715 | 20638652 | 20651031 | 20651082 | 108 | 45 | [illegible] |
| SE | 63205 | ENSG00000118960 | HS1BP3 | chr2 | - | 20645339 | 20645505 | 20638435 | 20638652 | 20651031 | 20651078 | 17 | 6 | [illegible] |
| SE | 63210 | ENSG00000204564 | C6orf136 | chr6 | + | 30647605 | 30647846 | 30647185 | 30647303 | 30649557 | 30649913 | 2 | 5 | [illegible] |
| SE | 63222 | ENSG00000068784 | SRBD1 | chr2 | - | 45392944 | 45393129 | 45388679 | 45389599 | 45413113 | 45413293 | 201 | 205 | [illegible] |
| SE | 63223 | ENSG00000068784 | SRBD1 | chr2 | - | 45413113 | 45413293 | 45392944 | 45393129 | 45418364 | 45418541 | 194 | 157 | [illegible] |
| SE | 63224 | ENSG00000068784 | SRBD1 | chr2 | - | 45419787 | 45419894 | 45418364 | 45418541 | 45488239 | 45488331 | 45 | 37 | [illegible] |
| SE | 63225 | ENSG00000068784 | SRBD1 | chr2 | - | 45476992 | 45477075 | 45419787 | 45419894 | 45488239 | 45488331 | 93 | 39 | [illegible] |
| SE | 63226 | ENSG00000068784 | SRBD1 | chr2 | - | 45476992 | 45477075 | 45419787 | 45419894 | 45546731 | 45546839 | 39 | 27 | [illegible] |
| SE | 63227 | ENSG00000068784 | SRBD1 | chr2 | - | 45488239 | 45488331 | 45419787 | 45419894 | 45546731 | 45546839 | 27 | 30 | [illegible] |
| SE | 63228 | ENSG00000068784 | SRBD1 | chr2 | - | 45488239 | 45488331 | 45476992 | 45477075 | 45546731 | 45546839 | 78 | 33 | [illegible] |
| SE | 63231 | ENSG00000068784 | SRBD1 | chr2 | - | 45553622 | 45553730 | 45547521 | 45547612 | 45567935 | 45568021 | 2 | 5 | [illegible] |
| SE | 63235 | ENSG00000068784 | SRBD1 | chr2 | - | 45585607 | 45585774 | 45581692 | 45581810 | 45599448 | 45599835 | 81 | 71 | [illegible] |
| SE | 63240 | ENSG00000106105 | GARS1 | chr7 | + | 30606325 | 30606397 | 30603495 | 30603572 | 30609584 | 30609730 | 0 | 1 | [illegible] |
| SE | 63242 | ENSG00000106105 | GARS1 | chr7 | + | 30609584 | 30609730 | 30602341 | 30603572 | 30615895 | 30616058 | 121 | 98 | [illegible] |
| SE | 63243 | ENSG00000106105 | GARS1 | chr7 | + | 30612095 | 30612245 | 30603495 | 30609730 | 30615895 | 30616058 | 617 | 437 | [illegible] |
| SE | 63244 | ENSG00000106105 | GARS1 | chr7 | + | 30613444 | 30613597 | 30612095 | 30612245 | 30615895 | 30616058 | 2 | 1 | [illegible] |
| SE | 63245 | ENSG00000106105 | GARS1 | chr7 | + | 30626233 | 30626319 | 30621392 | 30622462 | 30628559 | 30628669 | 170 | 141 | [illegible] |
| SE | 63246 | ENSG00000106105 | GARS1 | chr7 | + | 30627056 | 30627190 | 30626233 | 30626319 | 30628559 | 30628669 | 2 | 4 | [illegible] |
| SE | 63252 | ENSG00000105185 | PDCD5 | chr19 | + | 32586857 | 32586929 | 32585815 | 32585907 | 32587252 | 32587414 | 46 | 33 | [illegible] |
| SE | 63263 | ENSG00000013810 | TACC3 | chr4 | + | 1727707 | 1728787 | 1723727 | 1723870 | 1730886 | 1730962 | 249 | 137 | [illegible] |
| SE | 63264 | ENSG00000106153 | CHCHD2 | chr7 | - | 56104225 | 56104475 | 56102635 | 56103011 | 56106363 | 56106431 | 392 | 119 | [illegible] |
| SE | 63268 | ENSG00000263482 | ANTXRLP1 | chr10 | - | 46263509 | 46263608 | 46254020 | 46254096 | 46269381 | 46269461 | 33 | 67 | [illegible] |
| SE | 63269 | ENSG00000263482 | ANTXRLP1 | chr10 | - | 46268625 | 46268686 | 46263509 | 46263608 | 46268776 | 46268857 | 10 | 0 | [illegible] |
| SE | 63273 | ENSG00000095002 | MSH2 | chr2 | + | 47429741 | 47429941 | 47416295 | 47416429 | 47445547 | 47445657 | 149 | 94 | [illegible] |
| SE | 63274 | ENSG00000095002 | MSH2 | chr2 | + | 47466657 | 47466808 | 47463030 | 47463154 | 47470964 | 47471062 | 152 | 109 | [illegible] |
| SE | 63275 | ENSG00000095002 | MSH2 | chr2 | + | 47470964 | 47471062 | 47466657 | 47466808 | 47475024 | 47475270 | 121 | 98 | [illegible] |
| SE | 63276 | ENSG00000095002 | MSH2 | chr2 | + | 47470987 | 47471062 | 47466657 | 47466808 | 47475024 | 47475270 | 41 | 35 | [illegible] |
| SE | 63277 | ENSG00000095002 | MSH2 | chr2 | + | 47476366 | 47476571 | 47475024 | 47475270 | 47478271 | 47478519 | 89 | 65 | [illegible] |
| SE | 63279 | ENSG00000177084 | POLE | chr12 | - | 132632314 | 132632437 | 132626310 | 132626317 | 132632663 | 132632795 | 25 | 8 | [illegible] |
| SE | 63280 | ENSG00000177084 | POLE | chr12 | - | 132632314 | 132632488 | 132626266 | 132626317 | 132632663 | 132632795 | 26 | 10 | [illegible] |
| SE | 63281 | ENSG00000177084 | POLE | chr12 | - | 132632314 | 132632508 | 132626116 | 132626317 | 132632663 | 132632795 | 43 | 24 | [illegible] |
| SE | 63282 | ENSG00000177084 | POLE | chr12 | - | 132643406 | 132643560 | 132643223 | 132643330 | 132643836 | 132643977 | 15 | 13 | [illegible] |
| SE | 63283 | ENSG00000177084 | POLE | chr12 | - | 132648928 | 132649072 | 132643836 | 132643977 | 132649305 | 132649515 | 15 | 16 | [illegible] |
| SE | 63286 | ENSG00000177084 | POLE | chr12 | - | 132675397 | 132675517 | 132673574 | 132673707 | 132675734 | 132675820 | 44 | 22 | [illegible] |
| SE | 63287 | ENSG00000177084 | POLE | chr12 | - | 132678451 | 132678569 | 132677577 | 132677719 | 132679496 | 132679651 | 0 | 1 | [illegible] |
| SE | 63289 | ENSG00000177084 | POLE | chr12 | - | 132680606 | 132680687 | 132680177 | 132680222 | 132681137 | 132681279 | 36 | 23 | [illegible] |
| SE | 63290 | ENSG00000171132 | PRKCE | chr2 | + | 45677831 | 45677916 | 45652400 | 45652448 | 45842999 | 45843063 | 0 | 0 | [illegible] |
| SE | 63293 | ENSG00000171132 | PRKCE | chr2 | + | 45840379 | 45840551 | 45701382 | 45701524 | 45842999 | 45843063 | 4 | 0 | [illegible] |
| SE | 63294 | ENSG00000171132 | PRKCE | chr2 | + | 46004541 | 46004638 | 46001403 | 46001546 | 46007461 | 46007661 | 55 | 41 | [illegible] |
| SE | 63295 | ENSG00000171132 | PRKCE | chr2 | + | 46086207 | 46086362 | 46010343 | 46010517 | 46145092 | 46145231 | 28 | 16 | [illegible] |
| SE | 63300 | ENSG00000138018 | SELENOI | chr2 | + | 26367145 | 26367220 | 26364831 | 26364940 | 26373366 | 26373506 | 48 | 19 | [illegible] |
| SE | 63303 | ENSG00000138018 | SELENOI | chr2 | + | 26375039 | 26375148 | 26373366 | 26373629 | 26389004 | 26389067 | 11 | 1 | [illegible] |
| SE | 63304 | ENSG00000138018 | SELENOI | chr2 | + | 26383298 | 26383347 | 26373366 | 26373629 | 26384958 | 26385139 | 15 | 1 | [illegible] |
| SE | 63306 | ENSG00000138018 | SELENOI | chr2 | + | 26383298 | 26383347 | 26375039 | 26375148 | 26389004 | 26389067 | 6 | 0 | [illegible] |
| SE | 63307 | ENSG00000138018 | SELENOI | chr2 | + | 26384958 | 26385139 | 26373366 | 26373629 | 26389004 | 26389067 | 3 | 0 | [illegible] |
| SE | 63308 | ENSG00000138018 | SELENOI | chr2 | + | 26384958 | 26385139 | 26383298 | 26383347 | 26386353 | 26386536 | 19 | 12 | [illegible] |
| SE | 63309 | ENSG00000138018 | SELENOI | chr2 | + | 26384958 | 26385139 | 26383298 | 26383347 | 26389004 | 26389067 | 19 | 4 | [illegible] |
| SE | 63312 | ENSG00000115825 | PRKD3 | chr2 | - | 37254203 | 37254289 | 37250501 | 37253350 | 37256661 | 37256929 | 133 | 94 | [illegible] |
| SE | 63313 | ENSG00000115825 | PRKD3 | chr2 | - | 37256661 | 37256929 | 37254203 | 37254289 | 37259582 | 37259681 | 211 | 138 | [illegible] |
| SE | 63314 | ENSG00000115825 | PRKD3 | chr2 | - | 37259582 | 37259681 | 37256661 | 37256929 | 37260222 | 37260384 | 190 | 119 | [illegible] |
| SE | 63315 | ENSG00000115825 | PRKD3 | chr2 | - | 37259582 | 37259681 | 37256661 | 37256929 | 37267429 | 37267536 | 92 | 56 | [illegible] |
| SE | 63316 | ENSG00000115825 | PRKD3 | chr2 | - | 37260222 | 37260384 | 37256661 | 37256929 | 37267429 | 37267536 | 85 | 37 | [illegible] |
| SE | 63320 | ENSG00000115825 | PRKD3 | chr2 | - | 37269614 | 37269687 | 37260222 | 37260384 | 37272379 | 37272423 | 43 | 36 | [illegible] |
| SE | 63324 | ENSG00000115825 | PRKD3 | chr2 | - | 37279745 | 37279929 | 37277865 | 37277989 | 37282541 | 37282619 | 345 | 184 | [illegible] |
| SE | 63326 | ENSG00000115825 | PRKD3 | chr2 | - | 37289355 | 37289513 | 37286176 | 37286369 | 37293132 | 37293271 | 90 | 60 | [illegible] |
| SE | 63327 | ENSG00000115825 | PRKD3 | chr2 | - | 37290867 | 37290999 | 37286176 | 37286369 | 37293132 | 37293271 | 118 | 70 | [illegible] |
| SE | 63328 | ENSG00000115825 | PRKD3 | chr2 | - | 37290867 | 37290999 | 37289355 | 37289513 | 37293132 | 37293271 | 275 | 228 | [illegible] |
| SE | 63329 | ENSG00000115825 | PRKD3 | chr2 | - | 37293132 | 37293271 | 37290867 | 37290999 | 37316236 | 37317023 | 185 | 137 | [illegible] |
| SE | 63330 | ENSG00000115825 | PRKD3 | chr2 | - | 37316236 | 37317179 | 37293132 | 37293271 | 37324680 | 37324833 | 99 | 65 | [illegible] |
| SE | 63332 | ENSG00000138028 | CGREF1 | chr2 | - | 27110672 | 27110968 | 27104286 | 27104377 | 27118845 | 27119030 | 1 | 0 | [illegible] |
| SE | 63335 | ENSG00000151092 | NGLY1 | chr3 | - | 25722656 | 25722899 | 25720013 | 25720191 | 25729132 | 25729318 | 2 | 5 | [illegible] |
| SE | 63336 | ENSG00000151092 | NGLY1 | chr3 | - | 25729132 | 25729318 | 25720013 | 25720191 | 25733871 | 25733982 | 60 | 37 | [illegible] |
| SE | 63339 | ENSG00000151092 | NGLY1 | chr3 | - | 25736003 | 25736149 | 25733871 | 25733982 | 25737333 | 25737455 | 271 | 211 | [illegible] |
| SE | 63344 | ENSG00000151092 | NGLY1 | chr3 | - | 25736308 | 25736400 | 25733871 | 25733982 | 25739576 | 25739799 | 4 | 8 | [illegible] |
| SE | 63345 | ENSG00000151092 | NGLY1 | chr3 | - | 25736308 | 25736400 | 25733871 | 25733982 | 25751097 | 25751263 | 5 | 8 | [illegible] |
| SE | 63347 | ENSG00000151092 | NGLY1 | chr3 | - | 25736308 | 25736400 | 25736003 | 25736149 | 25739576 | 25739799 | 2 | 7 | [illegible] |

| | | | | | | | | | | | | | | | | | | | | | | | | | | | | | | | | | | | | | | | | | | | | | | |
|---|---|---|---|---|---|---|---|---|---|---|---|---|---|---|---|---|---|---|---|---|---|---|---|---|---|---|---|---|---|---|---|---|---|---|---|---|---|---|---|---|---|---|---|---|---|---|
| SE | 63348 | ENSG00000151092 | NGLY1 | chr3 | - | 25736308 | 25736400 | 25736003 | 25736149 | 25751097 | 25751263 | 3 | 7 | 10 | 9 | 10 | 0 | 1 | 2 | 1 | 0 | 0 | 10 | 4 | 0 | 0 | 0 | 0 | 3 | 0 | 0 | 241 | 149 | 1 | 1 | 1 | 0.812 | 0.756 | 0.848 | 1 | NA | 1 | 0.452 | NA | NA | 0.157 |
| SE | 63351 | ENSG00000151092 | NGLY1 | chr3 | - | 25737333 | 25737455 | 25733871 | 25733982 | 25740402 | 25740450 | 0 | 2 | 3 | 1 | 0 | 0 | 0 | 1 | 0 | 3 | 0 | 0 | 6 | 0 | 0 | 0 | 3 | 5 | 6 | 0 | 271 | 149 | 1 | 1 | NA | 1 | 0.623 | 1 | 0 | NA | 0 | 0.398 | 0 | NA | 0.523 |
| SE | 63352 | ENSG00000151092 | NGLY1 | chr3 | - | 25737333 | 25737455 | 25733871 | 25733982 | 25751097 | 25751263 | 2 | 2 | 2 | 1 | 0 | 7 | 5 | 9 | 7 | 10 | 0 | 0 | 4 | 0 | 0 | 0 | 0 | 2 | 10 | 0 | 271 | 149 | 1 | 1 | 0.136 | 0.18 | 0.109 | 0.073 | 0 | NA | NA | 0.524 | 0 | NA | -0.162 |
| SE | 63354 | ENSG00000151092 | NGLY1 | chr3 | - | 25737333 | 25737455 | 25736003 | 25736149 | 25740402 | 25740450 | 119 | 101 | 96 | 133 | 115 | 0 | 0 | 0 | 0 | 2 | 42 | 107 | 54 | 98 | 12 | 0 | 2 | 0 | 0 | 0 | 271 | 149 | 1 | 1 | 1 | 1 | 1 | 1 | 0.969 | 1 | 0.967 | 1 | 1 | 1 | 0 |
| SE | 63355 | ENSG00000151092 | NGLY1 | chr3 | - | 25737333 | 25737455 | 25736003 | 25736149 | 25751097 | 25751263 | 121 | 101 | 95 | 133 | 115 | 0 | 1 | 2 | 1 | 0 | 42 | 107 | 52 | 98 | 12 | 0 | 0 | 3 | 0 | 0 | 271 | 149 | 1 | 1 | 1 | 0.982 | 0.963 | 0.987 | 1 | 1 | 1 | 0.905 | 1 | 1 | 0.005 |
| SE | 63357 | ENSG00000151092 | NGLY1 | chr3 | - | 25737333 | 25737455 | 25736308 | 25736400 | 25751097 | 25751263 | 2 | 2 | 5 | 10 | 4 | 1 | 0 | 0 | 0 | 0 | 6 | 0 | 8 | 4 | 14 | 0 | 0 | 0 | 0 | 0 | 271 | 149 | 1 | 1 | 0.524 | 1 | 1 | 1 | 1 | 1 | NA | 1 | 1 | 1 | -0.095 |
| SE | 63358 | ENSG00000151092 | NGLY1 | chr3 | - | 25739576 | 25739799 | 25733871 | 25733982 | 25740402 | 25740450 | 4 | 3 | 4 | 4 | 4 | 0 | 0 | 1 | 0 | 3 | 0 | 0 | 3 | 6 | 0 | 0 | 3 | 5 | 6 | 0 | 298 | 149 | 1 | 1 | 1 | 1 | 0.667 | 1 | 0.4 | NA | 0 | 0.231 | 0.333 | NA | 0.625 |
| SE | 63359 | ENSG00000151092 | NGLY1 | chr3 | - | 25739576 | 25739799 | 25733871 | 25733982 | 25751097 | 25751263 | 65 | 53 | 50 | 61 | 80 | 7 | 5 | 9 | 7 | 10 | 18 | 35 | 22 | 60 | 0 | 0 | 0 | 2 | 10 | 0 | 298 | 149 | 1 | 1 | 0.823 | 0.841 | 0.735 | 0.813 | 0.8 | 1 | 1 | 0.846 | 0.75 | NA | -0.097 |
| SE | 63360 | ENSG00000151092 | NGLY1 | chr3 | - | 25739576 | 25739799 | 25736003 | 25736149 | 25740402 | 25740450 | 6 | 6 | 4 | 2 | 4 | 0 | 0 | 0 | 0 | 2 | 0 | 10 | 5 | 0 | 0 | 0 | 2 | 0 | 0 | 0 | 298 | 149 | 1 | 1 | 1 | 1 | 1 | 1 | 0.5 | NA | 0.714 | 1 | NA | NA | 0.043 |
| SE | 63361 | ENSG00000151092 | NGLY1 | chr3 | - | 25739576 | 25739799 | 25736003 | 25736149 | 25751097 | 25751263 | 67 | 56 | 50 | 59 | 80 | 0 | 1 | 2 | 1 | 0 | 18 | 45 | 24 | 54 | 0 | 0 | 0 | 3 | 0 | 0 | 298 | 149 | 1 | 1 | 1 | 0.966 | 0.926 | 0.967 | 1 | 1 | 1 | 0.8 | 1 | NA | 0.022 |
| SE | 63362 | ENSG00000151092 | NGLY1 | chr3 | - | 25739576 | 25739799 | 25736308 | 25736400 | 25751097 | 25751263 | 64 | 53 | 50 | 59 | 80 | 1 | 0 | 0 | 0 | 0 | 18 | 35 | 22 | 54 | 0 | 0 | 0 | 0 | 0 | 0 | 298 | 149 | 1 | 1 | 0.97 | 1 | 1 | 1 | 1 | 1 | 1 | 1 | 1 | NA | -0.006 |
| SE | 63363 | ENSG00000151092 | NGLY1 | chr3 | - | 25739576 | 25739799 | 25736308 | 25737455 | 25740402 | 25740450 | 92 | 64 | 72 | 88 | 130 | 0 | 0 | 1 | 0 | 0 | 21 | 34 | 64 | 41 | 3 | 0 | 0 | 2 | 0 | 0 | 298 | 149 | 1 | 1 | 1 | 1 | 0.973 | 1 | 1 | 1 | 1 | 0.941 | 1 | 1 | 0.006 |
| SE | 63364 | ENSG00000151092 | NGLY1 | chr3 | - | 25739576 | 25739799 | 25736308 | 25737455 | 25751097 | 25751263 | 153 | 114 | 118 | 145 | 206 | 2 | 0 | 0 | 0 | 0 | 39 | 69 | 83 | 95 | 3 | 0 | 0 | 0 | 0 | 0 | 298 | 149 | 1 | 1 | 0.975 | 1 | 1 | 1 | 1 | 1 | 1 | 1 | 1 | 1 | -0.005 |
| SE | 63365 | ENSG00000151092 | NGLY1 | chr3 | - | 25740402 | 25740450 | 25733871 | 25733982 | 25751097 | 25751263 | 2 | 1 | 0 | 2 | 6 | 7 | 5 | 9 | 7 | 10 | 0 | 11 | 6 | 6 | 0 | 0 | 0 | 2 | 10 | 0 | 197 | 149 | 1 | 1 | 0.178 | 0.131 | 0 | 0.178 | 0.312 | NA | 1 | 0.694 | 0.312 | NA | -0.509 |
| SE | 63366 | ENSG00000151092 | NGLY1 | chr3 | - | 25740402 | 25740450 | 25736003 | 25736149 | 25751097 | 25751263 | 2 | 1 | 0 | 2 | 5 | 0 | 1 | 2 | 1 | 0 | 0 | 10 | 1 | 0 | 0 | 0 | 0 | 3 | 0 | 0 | 197 | 149 | 1 | 1 | 1 | 0.431 | 0 | 0.602 | 1 | NA | 1 | 0.201 | NA | NA | 0.006 |
| SE | 63367 | ENSG00000151092 | NGLY1 | chr3 | - | 25740402 | 25740450 | 25737333 | 25737455 | 25751097 | 25751263 | 2 | 1 | 0 | 2 | 3 | 2 | 0 | 0 | 0 | 0 | 0 | 8 | 3 | 0 | 0 | 0 | 0 | 0 | 0 | 0 | 197 | 149 | 1 | 1 | 0.431 | 1 | NA | 1 | 1 | NA | 1 | 1 | NA | NA | -0.142 |
| SE | 63368 | ENSG00000151092 | NGLY1 | chr3 | - | 25740402 | 25740450 | 25739576 | 25739799 | 25751097 | 25751263 | 5 | 4 | 2 | 4 | 7 | 64 | 53 | 50 | 59 | 80 | 0 | 8 | 4 | 0 | 0 | 18 | 35 | 22 | 54 | 0 | 197 | 149 | 1 | 1 | 0.056 | 0.054 | 0.029 | 0.049 | 0.062 | 0 | 0.147 | 0.121 | 0 | NA | -0.017 |
| SE | 63369 | ENSG00000151092 | NGLY1 | chr3 | - | 25761677 | 25761708 | 25751097 | 25751263 | 25764065 | 25764311 | 0 | 1 | 1 | 2 | 2 | 128 | 114 | 91 | 147 | 108 | 0 | 5 | 0 | 0 | 0 | 86 | 73 | 51 | 48 | 49 | 180 | 149 | 1 | 1 | 0 | 0.007 | 0.009 | 0.011 | 0.015 | 0 | 0.054 | 0 | 0 | 0 | -0.002 |
| SE | 63375 | ENSG00000237803 | LINC00211 | chr2 | - | 37828932 | 37829038 | 37826246 | 37827878 | 37829247 | 37829432 | 0 | 0 | 2 | 0 | 1 | 0 | 0 | 0 | 0 | 0 | 16 | 5 | 10 | 2 | 12 | 0 | 0 | 2 | 0 | 0 | 255 | 149 | 1 | 1 | NA | NA | 1 | NA | 1 | 1 | 1 | 0.745 | 1 | 1 | 0.051 |
| SE | 63376 | ENSG00000237803 | LINC00211 | chr2 | - | 37828932 | 37829038 | 37826246 | 37827878 | 37865113 | 37865278 | 0 | 0 | 2 | 0 | 1 | 0 | 1 | 0 | 1 | 0 | 16 | 5 | 10 | 2 | 12 | 0 | 6 | 0 | 1 | 0 | 255 | 149 | 1 | 1 | NA | 0 | 1 | 0 | 1 | 1 | 0.327 | 1 | 0.539 | 1 | -0.273 |
| SE | 63377 | ENSG00000237803 | LINC00211 | chr2 | - | 37865113 | 37865278 | 37827009 | 37827878 | 37873485 | 37873542 | 2 | 7 | 0 | 3 | 0 | 0 | 0 | 0 | 0 | 0 | 0 | 14 | 0 | 8 | 47 | 0 | 7 | 0 | 0 | 0 | 298 | 149 | 1 | 1 | 1 | 1 | NA | 1 | NA | NA | 0.5 | NA | 1 | 1 | 0.167 |
| SE | 63378 | ENSG00000237803 | LINC00211 | chr2 | - | 37871807 | 37871888 | 37868028 | 37868272 | 37873485 | 37873542 | 0 | 0 | 0 | 0 | 0 | 0 | 0 | 0 | 1 | 0 | 0 | 0 | 0 | 0 | 2 | 0 | 3 | 0 | 2 | 10 | 230 | 149 | 1 | 1 | NA | NA | NA | 0 | NA | NA | 0 | NA | 0 | 0.115 | -0.038 |
| SE | 63379 | ENSG00000237803 | LINC00211 | chr2 | - | 37873485 | 37873542 | 37865113 | 37865278 | 37875742 | 37875883 | 5 | 14 | 0 | 4 | 0 | 0 | 0 | 0 | 1 | 0 | 0 | 11 | 0 | 17 | 73 | 0 | 0 | 4 | 0 | 0 | 206 | 149 | 1 | 1 | 1 | 1 | NA | 0.743 | NA | NA | 1 | 0 | 1 | 1 | 0.164 |
| SE | 63381 | ENSG00000126870 | DYNC2I1 | chr7 | + | 158884563 | 158884619 | 158879683 | 158879989 | 158887020 | 158887075 | 34 | 24 | 18 | 11 | 36 | 2 | 0 | 0 | 2 | 0 | 17 | 53 | 17 | 9 | 0 | 0 | 0 | 5 | 8 | 0 | 205 | 149 | 1 | 1 | 0.925 | 1 | 1 | 0.8 | 1 | 1 | 1 | 0.712 | 0.45 | NA | 0.154 |
| SE | 63382 | ENSG00000126870 | DYNC2I1 | chr7 | + | 158905988 | 158906091 | 158902375 | 158902595 | 158911549 | 158911679 | 56 | 37 | 45 | 36 | 78 | 5 | 4 | 1 | 0 | 3 | 14 | 36 | 62 | 39 | 0 | 0 | 5 | 4 | 3 | 0 | 252 | 149 | 1 | 1 | 0.869 | 0.845 | 0.964 | 1 | 0.939 | 1 | 0.81 | 0.902 | 0.885 | NA | 0.024 |
| SE | 63383 | ENSG00000126870 | DYNC2I1 | chr7 | + | 158922376 | 158922549 | 158918739 | 158918869 | 158923570 | 158923733 | 39 | 25 | 39 | 34 | 42 | 0 | 0 | 0 | 0 | 0 | 30 | 40 | 21 | 25 | 33 | 0 | 1 | 0 | 0 | 0 | 298 | 149 | 1 | 1 | 1 | 1 | 1 | 1 | 1 | 1 | 0.952 | 1 | 1 | 1 | 0.01 |
| SE | 63384 | ENSG00000126870 | DYNC2I1 | chr7 | + | 158934417 | 158934549 | 158934128 | 158934228 | 158941924 | 158942148 | 16 | 14 | 47 | 18 | 49 | 2 | 0 | 0 | 0 | 2 | 22 | 12 | 9 | 0 | 16 | 0 | 0 | 7 | 0 | 0 | 281 | 149 | 1 | 1 | 0.809 | 1 | 1 | 1 | 0.929 | 1 | 1 | 0.405 | NA | 1 | 0.096 |
| SE | 63386 | ENSG00000118263 | KLF7 | chr2 | - | 207123773 | 207124404 | 207088457 | 207088581 | 207165466 | 207165948 | 73 | 75 | 118 | 109 | 179 | 0 | 0 | 0 | 0 | 0 | 119 | 103 | 212 | 154 | 74 | 5 | 0 | 0 | 0 | 0 | 298 | 149 | 1 | 1 | 1 | 1 | 1 | 1 | 1 | 0.922 | 1 | 1 | 1 | 1 | 0.016 |
| SE | 63387 | ENSG00000118263 | KLF7 | chr2 | - | 207123773 | 207124404 | 207088457 | 207088581 | 207167139 | 207167246 | 43 | 50 | 68 | 63 | 109 | 0 | 0 | 0 | 4 | 0 | 81 | 56 | 138 | 98 | 47 | 0 | 0 | 0 | 0 | 0 | 298 | 149 | 1 | 1 | 1 | 1 | 1 | 0.887 | 1 | 1 | 1 | 1 | 1 | 1 | -0.023 |

[illegible]

| | | | | | | | | | | | | | | | | | | | | | | | | | | | | | | | | | | | | | | | | | | | | | | |
|---|---|---|---|---|---|---|---|---|---|---|---|---|---|---|---|---|---|---|---|---|---|---|---|---|---|---|---|---|---|---|---|---|---|---|---|---|---|---|---|---|---|---|---|---|---|---|
| SE | 63720 | ENSG00000166986 | MARS1 | chr12 | + | 57498156 | 57498273 | 57490537 | 57490644 | 57498419 | 57498623 | 213 | 159 | 212 | 270 | 251 | 0 | 0 | 1 | 2 | 0 | 367 | 231 | 212 | 256 | 370 | 0 | 0 | 0 | 0 | 9 | 266 | 149 | 1 | 1 | 1 | 1 | 0.992 | 0.987 | 1 | 1 | 1 | 1 | 1 | 0.958 | 0.004 |
| SE | 63721 | ENSG00000166986 | MARS1 | chr12 | + | 57500320 | 57500522 | 57498419 | 57498623 | 57504224 | 57504299 | 236 | 156 | 321 | 332 | 231 | 1 | 1 | 1 | 0 | 0 | 328 | 321 | 218 | 258 | 464 | 0 | 1 | 0 | 1 | 0 | 298 | 149 | 1 | 1 | 0.992 | 0.987 | 0.994 | 1 | 1 | 1 | 0.994 | 1 | 0.992 | 1 | -0.003 |
| SE | 63722 | ENSG00000166986 | MARS1 | chr12 | + | 57504224 | 57504299 | 57500320 | 57500522 | 57511697 | 57511868 | 184 | 93 | 189 | 209 | 142 | 0 | 0 | 0 | 0 | 0 | 198 | 206 | 125 | 196 | 348 | 0 | 1 | 0 | 0 | 0 | 224 | 149 | 1 | 1 | 1 | 1 | 1 | 1 | 1 | 1 | 0.993 | 1 | 1 | 1 | 0.001 |
| SE | 63724 | ENSG00000166986 | MARS1 | chr12 | + | 57512302 | 57512353 | 57512007 | 57512103 | 57512750 | 57512964 | 26 | 14 | 22 | 49 | 12 | 2 | 0 | 0 | 2 | 0 | 32 | 26 | 29 | 48 | 95 | 0 | 0 | 4 | 9 | 0 | 200 | 149 | 1 | 1 | 0.906 | 1 | 1 | 0.948 | 1 | 1 | 1 | 0.844 | 0.799 | 1 | 0.042 |
| SE | 63726 | ENSG00000115840 | SLC25A12 | chr2 | - | 171787570 | 171787661 | 171783404 | 171785475 | 171787788 | 171787947 | 66 | 33 | 42 | 38 | 47 | 0 | 0 | 0 | 3 | 0 | 31 | 36 | 22 | 8 | 37 | 0 | 0 | 0 | 0 | 0 | 240 | 149 | 1 | 1 | 1 | 1 | 1 | 0.887 | 1 | 1 | 1 | 1 | 1 | 1 | -0.023 |
| SE | 63728 | ENSG00000115840 | SLC25A12 | chr2 | - | 171815120 | 171815202 | 171810223 | 171810276 | 171826797 | 171826882 | 27 | 13 | 21 | 19 | 12 | 0 | 0 | 0 | 0 | 0 | 20 | 17 | 12 | 3 | 4 | 0 | 0 | 2 | 0 | 0 | 231 | 149 | 1 | 1 | 1 | 1 | 1 | 1 | 1 | 1 | 1 | 0.795 | 1 | 1 | 0.041 |
| SE | 63729 | ENSG00000115840 | SLC25A12 | chr2 | - | 171826797 | 171826882 | 171815108 | 171815202 | 171833962 | 171834056 | 77 | 37 | 68 | 48 | 29 | 0 | 0 | 0 | 1 | 0 | 25 | 90 | 29 | 23 | 30 | 0 | 0 | 0 | 0 | 0 | 234 | 149 | 1 | 1 | 1 | 1 | 1 | 0.968 | 1 | 1 | 1 | 1 | 1 | 1 | -0.006 |
| SE | 63731 | ENSG00000115840 | SLC25A12 | chr2 | - | 171848172 | 171848276 | 171844368 | 171844508 | 171855833 | 171855949 | 2 | 0 | 0 | 0 | 2 | 116 | 39 | 55 | 59 | 113 | 0 | 0 | 0 | 0 | 0 | 26 | 64 | 47 | 46 | 21 | 253 | 149 | 1 | 1 | 0.01 | 0 | 0 | 0 | 0.01 | 0 | 0 | 0 | 0 | 0 | 0.004 |
| SE | 63733 | ENSG00000115840 | SLC25A12 | chr2 | - | 171855833 | 171855949 | 171844368 | 171844508 | 171868680 | 171868823 | 185 | 54 | 101 | 94 | 178 | 0 | 0 | 0 | 3 | 0 | 31 | 89 | 79 | 92 | 23 | 0 | 0 | 0 | 0 | 0 | 265 | 149 | 1 | 1 | 1 | 1 | 1 | 0.946 | 1 | 1 | 1 | 1 | 1 | 1 | -0.011 |
| SE | 63737 | ENSG00000152061 | RABGAP1L | chr1 | + | 174241482 | 174241657 | 174231144 | 174231355 | 174250474 | 174250632 | 990 | 728 | 542 | 838 | 781 | 0 | 0 | 0 | 0 | 1 | 186 | 1109 | 666 | 315 | 114 | 0 | 0 | 0 | 0 | 0 | 298 | 149 | 1 | 1 | 1 | 1 | 1 | 1 | 0.997 | 1 | 1 | 1 | 1 | 1 | -0.001 |
| SE | 63738 | ENSG00000152061 | RABGAP1L | chr1 | + | 174252443 | 174252590 | 174250474 | 174250632 | 174272413 | 174272480 | 241 | 183 | 149 | 223 | 319 | 1 | 0 | 0 | 0 | 0 | 53 | 252 | 236 | 93 | 26 | 0 | 0 | 0 | 0 | 0 | 296 | 149 | 1 | 1 | 0.992 | 1 | 1 | 1 | 1 | 1 | 1 | 1 | 1 | 1 | -0.002 |
| SE | 63739 | ENSG00000152061 | RABGAP1L | chr1 | + | 174252479 | 174252590 | 174250474 | 174250632 | 174272413 | 174272480 | 576 | 472 | 365 | 504 | 625 | 1 | 0 | 0 | 0 | 0 | 116 | 574 | 524 | 211 | 45 | 0 | 0 | 0 | 0 | 0 | 260 | 149 | 1 | 1 | 0.997 | 1 | 1 | 1 | 1 | 1 | 1 | 1 | 1 | 1 | -0.001 |
| SE | 63742 | ENSG00000152061 | RABGAP1L | chr1 | + | 174370978 | 174371072 | 174304985 | 174305127 | 174393994 | 174394145 | 225 | 128 | 127 | 202 | 162 | 3 | 0 | 0 | 0 | 0 | 35 | 214 | 151 | 84 | 28 | 0 | 0 | 0 | 0 | 0 | 243 | 149 | 1 | 1 | 0.979 | 1 | 1 | 1 | 1 | 1 | 1 | 1 | 1 | 1 | -0.004 |
| SE | 63743 | ENSG00000152061 | RABGAP1L | chr1 | + | 174393994 | 174394145 | 174304985 | 174305127 | 174544843 | 174545911 | 3 | 2 | 0 | 4 | 1 | 0 | 0 | 0 | 0 | 0 | 0 | 0 | 0 | 0 | 0 | 0 | 2 | 0 | 0 | 0 | 298 | 149 | 1 | 1 | 1 | 1 | NA | 1 | 1 | NA | 0 | NA | NA | NA | 1 |
| SE | 63747 | ENSG00000152061 | RABGAP1L | chr1 | + | 174811831 | 174811960 | 174799931 | 174800500 | 174957456 | 174957529 | 328 | 159 | 121 | 368 | 384 | 1 | 0 | 0 | 2 | 7 | 961 | 287 | 132 | 595 | 1253 | 8 | 0 | 0 | 0 | 7 | 278 | 149 | 1 | 1 | 0.994 | 1 | 1 | 0.99 | 0.967 | 0.985 | 1 | 1 | 1 | 0.99 | -0.005 |
| SE | 63750 | ENSG00000152061 | RABGAP1L | chr1 | + | 174877391 | 174877605 | 174875545 | 174875713 | 174957456 | 174957549 | 39 | 41 | 23 | 54 | 23 | 0 | 0 | 2 | 0 | 0 | 151 | 37 | 17 | 97 | 73 | 0 | 0 | 0 | 0 | 0 | 298 | 149 | 1 | 1 | 1 | 1 | 0.852 | 1 | 1 | 1 | 1 | 1 | 1 | 1 | -0.03 |
| SE | 63752 | ENSG00000152061 | RABGAP1L | chr1 | + | 174976077 | 174976182 | 174969276 | 174969387 | 174978806 | 174978890 | 341 | 279 | 181 | 301 | 334 | 0 | 0 | 0 | 1 | 0 | 121 | 134 | 117 | 187 | 130 | 0 | 0 | 1 | 0 | 0 | 254 | 149 | 1 | 1 | 1 | 1 | 1 | 0.994 | 1 | 1 | 1 | 0.986 | 1 | 1 | 0.002 |
| SE | 63756 | ENSG00000141429 | GALNT1 | chr18 | + | 35683390 | 35683598 | 35663627 | 35663802 | 35687015 | 35687186 | 263 | 194 | 210 | 245 | 445 | 0 | 0 | 0 | 0 | 0 | 42 | 192 | 198 | 68 | 28 | 0 | 0 | 2 | 0 | 0 | 298 | 149 | 1 | 1 | 1 | 1 | 1 | 1 | 1 | 1 | 1 | 0.98 | 1 | 1 | 0.004 |
| SE | 63757 | ENSG00000141429 | GALNT1 | chr18 | + | 35689172 | 35689290 | 35687015 | 35687186 | 35691011 | 35691192 | 720 | 547 | 586 | 668 | 568 | 0 | 0 | 0 | 0 | 0 | 92 | 467 | 505 | 135 | 46 | 0 | 0 | 1 | 0 | 0 | 267 | 149 | 1 | 1 | 1 | 1 | 1 | 1 | 1 | 1 | 1 | 0.996 | 1 | 1 | 0.001 |
| SE | 63758 | ENSG00000141429 | GALNT1 | chr18 | + | 35702896 | 35702995 | 35692180 | 35692320 | 35703508 | 35703643 | 648 | 547 | 500 | 483 | 1139 | 0 | 0 | 0 | 0 | 0 | 43 | 333 | 342 | 113 | 19 | 0 | 0 | 2 | 0 | 0 | 248 | 149 | 1 | 1 | 1 | 1 | 1 | 1 | 1 | 1 | 1 | 0.99 | 1 | 1 | 0.002 |

[illegible]

| | | | | | | | | | | | | | | | | | | | | | | | | | | | | | | | | | | | | | | | | | | | | | | |
|---|---|---|---|---|---|---|---|---|---|---|---|---|---|---|---|---|---|---|---|---|---|---|---|---|---|---|---|---|---|---|---|---|---|---|---|---|---|---|---|---|---|---|---|---|---|---|
| SE | 64075 | ENSG00000110066 | KMT5B | chr11 | - | 68174087 | 68174210 | 68173803 | 68173913 | 68175017 | 68175183 | 6 | 0 | 2 | 7 | 2 | 291 | 229 | 249 | 228 | 337 | 0 | 0 | 3 | 0 | 0 | 33 | 135 | 159 | 113 | 30 | 272 | 149 | 1 | 1 | 0.011 | 0 | 0.004 | 0.017 | 0.003 | 0 | 0 | 0.01 | 0 | 0 | 0.005 |
| SE | 64077 | ENSG00000110066 | KMT5B | chr11 | - | 68179469 | 68179537 | 68175017 | 68175183 | 68185780 | 68185928 | 21 | 16 | 17 | 9 | 17 | 12 | 8 | 10 | 12 | 14 | 13 | 22 | 15 | 6 | 0 | 0 | 13 | 21 | 20 | 0 | 217 | 149 | 1 | 1 | 0.546 | 0.579 | 0.539 | 0.34 | 0.455 | 1 | 0.537 | 0.329 | 0.171 | NA | -0.017 |
| SE | 64079 | ENSG00000110066 | KMT5B | chr11 | - | 68180131 | 68180200 | 68179469 | 68179537 | 68185780 | 68185928 | 79 | 68 | 108 | 69 | 87 | 0 | 0 | 2 | 0 | 0 | 23 | 62 | 42 | 59 | 55 | 0 | 3 | 0 | 0 | 0 | 218 | 149 | 1 | 1 | 1 | 1 | 0.974 | 1 | 1 | 1 | 0.934 | 1 | 1 | 1 | 0.008 |
| SE | 64080 | ENSG00000110066 | KMT5B | chr11 | - | 68189916 | 68190152 | 68185780 | 68185928 | 68213137 | 68213286 | 170 | 150 | 181 | 204 | 246 | 0 | 1 | 0 | 2 | 4 | 70 | 137 | 92 | 99 | 8 | 0 | 1 | 0 | 0 | 0 | 298 | 149 | 1 | 1 | 1 | 0.987 | 1 | 0.981 | 0.969 | 1 | 0.986 | 1 | 1 | 1 | -0.01 |
| SE | 64082 | ENSG00000283050 | GTF2IP12 | chr4 | - | 119410516 | 119410658 | 119410231 | 119410289 | 119412351 | 119412542 | 0 | 0 | 0 | 0 | 4 | 0 | 1 | 0 | 0 | 3 | 7 | 0 | 0 | 0 | 6 | 0 | 0 | 3 | 0 | 0 | 291 | 149 | 1 | 1 | NA | 0 | NA | NA | 0.406 | 1 | NA | 0 | NA | 1 | -0.464 |
| SE | 64083 | ENSG00000283050 | GTF2IP12 | chr4 | - | 119410516 | 119410712 | 119410231 | 119410289 | 119412351 | 119412542 | 0 | 0 | 0 | 0 | 0 | 0 | 1 | 0 | 0 | 3 | 7 | 0 | 0 | 0 | 6 | 0 | 0 | 3 | 0 | 0 | 298 | 149 | 1 | 1 | NA | 0 | NA | NA | 0 | 1 | NA | 0 | NA | 1 | -0.667 |
| SE | 64084 | ENSG00000283050 | GTF2IP12 | chr4 | - | 119413288 | 119413407 | 119412351 | 119412542 | 119419041 | 119419106 | 0 | 1 | 0 | 0 | 10 | 0 | 2 | 4 | 4 | 12 | 0 | 0 | 0 | 0 | 3 | 12 | 1 | 0 | 4 | 0 | 268 | 149 | 1 | 1 | NA | 0.218 | 0 | 0 | 0.317 | 0 | 0 | NA | 0 | 1 | -0.116 |
| SE | 64085 | ENSG00000283050 | GTF2IP12 | chr4 | - | 119420493 | 119420590 | 119419958 | 119420155 | 119422450 | 119422574 | 2 | 3 | 3 | 2 | 1 | 0 | 0 | 3 | 0 | 1 | 5 | 0 | 0 | 0 | 0 | 0 | 0 | 1 | 0 | 0 | 246 | 149 | 1 | 1 | 1 | 1 | 0.377 | 1 | 0.377 | 1 | NA | 0 | NA | NA | 0.251 |
| SE | 64086 | ENSG00000283050 | GTF2IP12 | chr4 | - | 119429419 | 119429609 | 119422450 | 119422574 | 119431836 | 119431901 | 4 | 4 | 3 | 3 | 3 | 2 | 3 | 0 | 3 | 0 | 13 | 1 | 0 | 6 | 0 | 0 | 0 | 1 | 0 | 0 | 298 | 149 | 1 | 1 | 0.5 | 0.4 | 1 | 0.333 | 1 | 1 | 1 | 0 | 1 | NA | -0.103 |
| SE | 64087 | ENSG00000283050 | GTF2IP12 | chr4 | - | 119440559 | 119441054 | 119434739 | 119434975 | 119442157 | 119442217 | 28 | 17 | 19 | 20 | 10 | 0 | 0 | 0 | 0 | 0 | 41 | 39 | 4 | 7 | 50 | 4 | 0 | 0 | 0 | 0 | 298 | 149 | 1 | 1 | 1 | 1 | 1 | 1 | 1 | 0.837 | 1 | 1 | 1 | 1 | 0.033 |
| SE | 64089 | ENSG00000283050 | GTF2IP12 | chr4 | - | 119444873 | 119445027 | 119444367 | 119444433 | 119445311 | 119445527 | 0 | 0 | 1 | 1 | 0 | 6 | 3 | 6 | 1 | 0 | 0 | 7 | 0 | 0 | 0 | 18 | 4 | 0 | 2 | 11 | 298 | 149 | 1 | 1 | 0 | 0 | 0.077 | 0.333 | NA | 0 | 0.467 | NA | 0 | 0 | -0.014 |
| SE | 64090 | ENSG00000283050 | GTF2IP12 | chr4 | - | 119450040 | 119450125 | 119448827 | 119448959 | 119454245 | 119454630 | 1 | 1 | 0 | 0 | 0 | 2 | 3 | 4 | 5 | 0 | 13 | 1 | 0 | 2 | 0 | 6 | 2 | 0 | 0 | 7 | 234 | 149 | 1 | 1 | 0.241 | 0.175 | 0 | 0 | NA | 0.58 | 0.241 | NA | 1 | 0 | -0.351 |
| SE | 64091 | ENSG00000283050 | GTF2IP12 | chr4 | - | 119450040 | 119450150 | 119448827 | 119448959 | 119454245 | 119454638 | 1 | 1 | 2 | 1 | 0 | 2 | 3 | 4 | 5 | 0 | 25 | 1 | 0 | 0 | 0 | 6 | 2 | 0 | 0 | 7 | 259 | 149 | 1 | 1 | 0.223 | 0.161 | 0.223 | 0.103 | NA | 0.706 | 0.223 | NA | NA | 0 | -0.132 |
| SE | 64092 | ENSG00000267796 | LIN37 | chr19 | + | 35752803 | 35752833 | 35752433 | 35752484 | 35752913 | 35752999 | 38 | 22 | 42 | 44 | 32 | 0 | 0 | 0 | 1 | 4 | 78 | 69 | 39 | 44 | 23 | 0 | 0 | 1 | 0 | 0 | 179 | 149 | 1 | 1 | 1 | 1 | 1 | 0.973 | 0.869 | 1 | 1 | 0.97 | 1 | 1 | -0.026 |
| SE | 64093 | ENSG00000115919 | KYNU | chr2 | + | 142879462 | 142879666 | 142877683 | 142877736 | 142885348 | 142885536 | 1 | 0 | 0 | 0 | 0 | 92 | 74 | 47 | 91 | 51 | 0 | 0 | 3 | 0 | 0 | 10 | 47 | 58 | 52 | 13 | 298 | 149 | 1 | 1 | 0.005 | 0 | 0 | 0 | 0 | 0 | 0 | 0.025 | 0 | 0 | -0.004 |
| SE | 64095 | ENSG00000115919 | KYNU | chr2 | + | 142881344 | 142881430 | 142877663 | 142877736 | 142918608 | 142918729 | 1 | 3 | 0 | 2 | 0 | 1 | 0 | 0 | 0 | 0 | 0 | 1 | 0 | 0 | 0 | 0 | 0 | 0 | 0 | 0 | 235 | 149 | 1 | 1 | 0.388 | 1 | NA | 1 | NA | NA | 1 | NA | NA | NA | -0.204 |
| SE | 64096 | ENSG00000115919 | KYNU | chr2 | + | 142885348 | 142885536 | 142877656 | 142877736 | 142918608 | 142918729 | 288 | 283 | 143 | 267 | 171 | 1 | 0 | 0 | 0 | 0 | 51 | 157 | 175 | 104 | 49 | 0 | 0 | 0 | 0 | 0 | 298 | 149 | 1 | 1 | 0.993 | 1 | 1 | 1 | 1 | 1 | 1 | 1 | 1 | 1 | -0.001 |
| SE | 64097 | ENSG00000115919 | KYNU | chr2 | + | 142885348 | 142885536 | 142881344 | 142881430 | 142918608 | 142918729 | 203 | 232 | 99 | 187 | 136 | 0 | 1 | 0 | 0 | 0 | 64 | 130 | 121 | 52 | 36 | 0 | 0 | 0 | 0 | 0 | 298 | 149 | 1 | 1 | 1 | 0.991 | 1 | 1 | 1 | 1 | 1 | 1 | 1 | 1 | -0.002 |
| SE | 64098 | ENSG00000115919 | KYNU | chr2 | + | 142918608 | 142918729 | 142885348 | 142885536 | 142927658 | 142927741 | 383 | 396 | 191 | 343 | 304 | 0 | 3 | 2 | 0 | 0 | 95 | 212 | 215 | 122 | 45 | 0 | 0 | 0 | 0 | 0 | 270 | 149 | 1 | 1 | 1 | 0.986 | 0.981 | 1 | 1 | 1 | 1 | 1 | 1 | 1 | -0.007 |
| SE | 64099 | ENSG00000115919 | KYNU | chr2 | + | 142954021 | 142954067 | 142927658 | 142927741 | 142954809 | 142954871 | 0 | 3 | 0 | 0 | 4 | 170 | 204 | 79 | 162 | 189 | 0 | 0 | 0 | 0 | 0 | 58 | 74 | 80 | 47 | 16 | 195 | 149 | 1 | 1 | 0 | 0.011 | 0 | 0 | 0.016 | 0 | 0 | 0 | 0 | 0 | 0.005 |
| SE | 64100 | ENSG00000115919 | KYNU | chr2 | + | 142960623 | 142960770 | 142957640 | 142957715 | 142985083 | 142985182 | 169 | 176 | 97 | 174 | 208 | 0 | 2 | 1 | 1 | 0 | 42 | 99 | 107 | 69 | 8 | 0 | 3 | 1 | 0 | 0 | 296 | 149 | 1 | 1 | 1 | 0.978 | 0.98 | 0.989 | 1 | 1 | 0.943 | 0.982 | 1 | 1 | 0.004 |
| SE | 64101 | ENSG00000115919 | KYNU | chr2 | + | 143029626 | 143029679 | 142985947 | 142986021 | 143033235 | 143033321 | 72 | 60 | 50 | 64 | 82 | 3 | 0 | 2 | 0 | 9 | 0 | 24 | 29 | 33 | 0 | 0 | 0 | 0 | 0 | 0 | 202 | 149 | 1 | 1 | 0.947 | 1 | 0.949 | 1 | 0.87 | NA | 1 | 1 | 1 | NA | -0.047 |
| SE | 64102 | ENSG00000115919 | KYNU | chr2 | + | 143033235 | 143033321 | 143029626 | 143029679 | 143040427 | 143040658 | 92 | 75 | 49 | 66 | 109 | 0 | 2 | 0 | 0 | 0 | 0 | 34 | 33 | 26 | 0 | 0 | 0 | 0 | 0 | 0 | 235 | 149 | 1 | 1 | 1 | 0.96 | 1 | 1 | 1 | NA | 1 | 1 | 1 | NA | -0.008 |
| SE | 64103 | ENSG00000283602 | NA | chr5 | - | 140591063 | 140591150 | 140590874 | 140590991 | 140591669 | 140591752 | 1 | 0 | 0 | 0 | 0 | 6 | 0 | 0 | 1 | 5 | 0 | 0 | 0 | 0 | 0 | 0 | 0 | 0 | 0 | 30 | 236 | 149 | 1 | 1 | 0.095 | NA | NA | 0 | 0 | NA | NA | NA | NA | 0 | 0.032 |
| SE | 64105 | ENSG00000283602 | NA | chr5 | - | 140591073 | 140591150 | 140590874 | 140590991 | 140591669 | 140591752 | 2 | 0 | 2 | 0 | 1 | 6 | 0 | 0 | 1 | 5 | 0 | 0 | 2 | 2 | 0 | 0 | 0 | 0 | 0 | 30 | 226 | 149 | 1 | 1 | 0.18 | NA | 1 | 0 | 0.116 | NA | NA | 1 | 1 | 0 | -0.343 |
| SE | 64106 | ENSG00000283602 | NA | chr5 | - | 140591073 | 140591150 | 140590874 | 140590991 | 140592023 | 140592101 | 0 | 0 | 2 | 0 | 0 | 17 | 9 | 6 | 4 | 4 | 0 | 0 | 2 | 0 | 5 | 0 | 20 | 9 | 25 | 37 | 226 | 149 | 1 | 1 | 0 | 0 | 0.18 | 0 | 0 | NA | 0 | 0.128 | 0 | 0.082 | -0.017 |
| SE | 64108 | ENSG00000283602 | NA | chr5 | - | 140591669 | 140591752 | 140591063 | 140591150 | 140592023 | 140592101 | 1 | 2 | 0 | 2 | 1 | 0 | 0 | 0 | 0 | 0 | 8 | 0 | 1 | 2 | 0 | 0 | 0 | 0 | 0 | 5 | 232 | 149 | 1 | 1 | 1 | 1 | NA | 1 | 1 | 1 | NA | 1 | 1 | 0 | 0.25 |
| SE | 64110 | ENSG00000050555 | LAMC3 | chr9 | + | 131052490 | 131052683 | 131049019 | 131049130 | 131052849 | 131052965 | 1 | 0 | 0 | 0 | 0 | 0 | 0 | 0 | 0 | 0 | 0 | 7 | 1 | 0 | 0 | 0 | 0 | 0 | 5 | 0 | 298 | 149 | 1 | 1 | 1 | NA | NA | NA | NA | NA | 1 | 1 | 0 | NA | 0.333 |
| SE | 64115 | ENSG00000215717 | TMEM167B | chr1 | + | 109091659 | 109091748 | 109090779 | 109090882 | 109092889 | 109093021 | 1 | 0 | 0 | 0 | 0 | 183 | 118 | 166 | 197 | 144 | 0 | 0 | 0 | 2 | 0 | 199 | 232 | 286 | 202 | 50 | 238 | 149 | 1 | 1 | 0.003 | 0 | 0 | 0 | 0 | 0 | 0 | 0 | 0.006 | 0 | -0.001 |
| SE | 64119 | ENSG00000135205 | CCDC146 | chr7 | + | 77241690 | 77241900 | 77236946 | 77237029 | 77254505 | 77254563 | 31 | 22 | 27 | 47 | 15 | 1 | 2 | 5 | 0 | 0 | 16 | 9 | 8 | 0 | 6 | 0 | 1 | 2 | 0 | 0 | 298 | 149 | 1 | 1 | 0.939 | 0.846 | 0.73 | 1 | 1 | 1 | 0.818 | 0.667 | NA | 1 | 0.032 |
| SE | 64127 | ENSG00000163126 | ANKRD23 | chr2 | - | 96838498 | 96839644 | 96833310 | 96833540 | 96839726 | 96839825 | 7 | 2 | 4 | 3 | 1 | 0 | 0 | 0 | 1 | 1 | 0 | 3 | 2 | 0 | 0 | 0 | 0 | 0 | 0 | 0 | 298 | 149 | 1 | 1 | 1 | 1 | 1 | 0.6 | 0.333 | NA | 1 | 1 | NA | NA | -0.213 |
| SE | 64128 | ENSG00000163126 | ANKRD23 | chr2 | - | 96839726 | 96839828 | 96833310 | 96833540 | 96839996 | 96840095 | 0 | 0 | 0 | 1 | 1 | 2 | 1 | 2 | 1 | 0 | 0 | 0 | 0 | 0 | 0 | 0 | 0 | 2 | 0 | 0 | 251 | 149 | 1 | 1 | 0 | 0 | 0 | 0.372 | 1 | NA | NA | 0 | NA | NA | 0.274 |
| SE | 64132 | ENSG00000168906 | MAT2A | chr2 | + | 85541828 | 85541972 | 85541632 | 85541745 | 85542154 | 85542373 | 853 | 599 | 793 | 861 | 1192 | 0 | 0 | 0 | 2 | 0 | 669 | 542 | 324 | 583 | 507 | 0 | 0 | 0 | 5 | 0 | 293 | 149 | 1 | 1 | 1 | 1 | 1 | 0.995 | 1 | 1 | 1 | 1 | 0.983 | 1 | 0.002 |
| SE | 64134 | ENSG00000117643 | MAN1C1 | chr1 | + | 25686439 | 25686536 | 25616790 | 25618337 | 25746667 | 25746783 | 58 | 32 | 88 | 50 | 88 | 0 | 2 | 0 | 0 | 0 | 44 | 24 | 8 | 29 | 33 | 0 | 0 | 0 | 0 | 0 | 246 | 149 | 1 | 1 | 1 | 0.906 | 1 | 1 | 1 | 1 | 1 | 1 | 1 | 1 | -0.019 |
| SE | 64136 | ENSG00000117643 | MAN1C1 | chr1 | + | 25749254 | 25749335 | 25686439 | 25686536 | 25758591 | 25758709 | 0 | 1 | 6 | 2 | 3 | 0 | 0 | 0 | 0 | 0 | 5 | 0 | 0 | 3 | 0 | 0 | 0 | 5 | 0 | 0 | 230 | 149 | 1 | 1 | NA | 1 | 1 | 1 | 1 | 1 | NA | 0 | 1 | NA | 0.333 |
| SE | 64138 | ENSG00000117643 | MAN1C1 | chr1 | + | 25763873 | 25763967 | 25758591 | 25758709 | 25771656 | 25771772 | 121 | 72 | 174 | 116 | 105 | 0 | 0 | 0 | 2 | 1 | 70 | 43 | 45 | 46 | 59 | 0 | 0 | 1 | 0 | 4 | 243 | 149 | 1 | 1 | 1 | 1 | 1 | 0.973 | 0.985 | 1 | 1 | 0.965 | 1 | 0.9 | 0.019 |
| SE | 64139 | ENSG00000117643 | MAN1C1 | chr1 | + | 25775931 | 25776029 | 25771656 | 25771772 | 25778104 | 25778324 | 0 | 4 | 2 | 0 | 0 | 103 | 55 | 136 | 96 | 59 | 0 | 0 | 0 | 0 | 0 | 54 | 22 | 24 | 36 | 45 | 247 | 149 | 1 | 1 | 0 | 0.042 | 0.009 | 0 | 0 | 0 | 0 | 0 | 0 | 0 | 0.01 |
| SE | 64140 | ENSG00000117643 | MAN1C1 | chr1 | + | 25778104 | 25778324 | 25771656 | 25771772 | 25780939 | 25781103 | 110 | 64 | 171 | 117 | 71 | 0 | 0 | 1 | 0 | 0 | 54 | 22 | 27 | 36 | 61 | 0 | 0 | 0 | 0 | 0 | 298 | 149 | 1 | 1 | 1 | 1 | 0.988 | 1 | 1 | 1 | 1 | 1 | 1 | 1 | -0.002 |
| SE | 64141 | ENSG00000117643 | MAN1C1 | chr1 | + | 25778104 | 25778324 | 25771656 | 25771772 | 25782584 | 25782700 | 103 | 58 | 137 | 96 | 61 | 0 | 0 | 0 | 1 | 0 | 54 | 22 | 24 | 36 | 48 | 0 | 0 | 0 | 0 | 0 | 298 | 149 | 1 | 1 | 1 | 1 | 1 | 0.98 | 1 | 1 | 1 | 1 | 1 | 1 | -0.004 |
| SE | 64142 | ENSG00000117643 | MAN1C1 | chr1 | + | 25780939 | 25781112 | 25771656 | 25771772 | 25782584 | 25782700 | 14 | 16 | 28 | 16 | 29 | 0 | 0 | 0 | 1 | 0 | 0 | 0 | 3 | 0 | 4 | 0 | 0 | 0 | 0 | 0 | 298 | 149 | 1 | 1 | 1 | 1 | 1 | 0.889 | 1 | NA | NA | 1 | NA | 1 | -0.022 |
| SE | 64143 | ENSG00000117643 | MAN1C1 | chr1 | + | 25780939 | 25781112 | 25778104 | 25778324 | 25782584 | 25782700 | 21 | 25 | 62 | 37 | 41 | 0 | 3 | 1 | 0 | 2 | 0 | 0 | 6 | 0 | 20 | 0 | 0 | 0 | 0 | 3 | 298 | 149 | 1 | 1 | 1 | 0.806 | 0.969 | 1 | 0.911 | NA | NA | 1 | NA | 0.769 | 0.053 |
| SE | 64147 | ENSG00000032444 | PNPLA6 | chr19 | + | 7536190 | 7536273 | 7535716 | 7536020 | 7536448 | 7536546 | 92 | 60 | 78 | 76 | 84 | 0 | 0 | 2 | 0 | 0 | 198 | 147 | 63 | 68 | 327 | 0 | 0 | 0 | 0 | 0 | 232 | 149 | 1 | 1 | 1 | 1 | 0.962 | 1 | 1 | 1 | 1 | 1 | 1 | 1 | -0.008 |
| SE | 64148 | ENSG00000032444 | PNPLA6 | chr19 | + | 7541983 | 7542067 | 7541447 | 7541684 | 7542560 | 7542670 | 19 | 13 | 18 | 23 | 2 | 0 | 0 | 0 | 0 | 0 | 39 | 12 | 6 | 21 | 19 | 0 | 0 | 0 | 2 | 0 | 233 | 149 | 1 | 1 | 1 | 1 | 1 | 1 | 1 | 1 | 1 | 1 | 0.87 | 1 | 0.026 |
| SE | 64150 | ENSG00000032444 | PNPLA6 | chr19 | + | 7553871 | 7554015 | 7551361 | 7551437 | 7554208 | 7554272 | 40 | 39 | 23 | 43 | 16 | 0 | 0 | 0 | 0 | 0 | 80 | 53 | 25 | 19 | 91 | 0 | 0 | 1 | 0 | 0 | 293 | 149 | 1 | 1 | 1 | 1 | 1 | 1 | 1 | 1 | 1 | 0.927 | 1 | 1 | 0.015 |
| SE | 64151 | ENSG00000032444 | PNPLA6 | chr19 | + | 7553874 | 7554015 | 7551361 | 7551437 | 7554208 | 7554243 | 54 | 44 | 28 | 39 | 15 | 0 | 0 | 0 | 0 | 0 | 82 | 57 | 24 | 31 | 92 | 0 | 0 | 1 | 0 | 0 | 290 | 149 | 1 | 1 | 1 | 1 | 1 | 1 | 1 | 1 | 1 | 0.925 | 1 | 1 | 0.015 |
| SE | 64155 | ENSG00000032444 | PNPLA6 | chr19 | + | 7561013 | 7561110 | 7560590 | 7560764 | 7561207 | 7561317 | 277 | 217 | 257 | 243 | 188 | 3 | 0 | 1 | 0 | 0 | 493 | 390 | 234 | 316 | 487 | 0 | 6 | 1 | 3 | 0 | 246 | 149 | 1 | 1 | 0.982 | 1 | 0.994 | 1 | 1 | 1 | 0.975 | 0.993 | 0.985 | 1 | 0.005 |
| SE | 64156 | ENSG00000032444 | PNPLA6 | chr19 | + | 7561013 | 7561110 | 7560590 | 7560764 | 7561487 | 7561761 | 145 | 117 | 126 | 112 | 88 | 0 | 0 | 0 | 0 | 0 | 170 | 167 | 96 | 106 | 234 | 0 | 7 | 0 | 0 | 0 | 246 | 149 | 1 | 1 | 1 | 1 | 1 | 1 | 1 | 1 | 0.935 | 1 | 1 | 1 | 0.013 |
| SE | 64157 | ENSG00000032444 | PNPLA6 | chr19 | + | 7561207 | 7561317 | 7560590 | 7560764 | 7561487 | 7561761 | 141 | 93 | 111 | 113 | 99 | 0 | 0 | 0 | 0 | 0 | 200 | 163 | 101 | 129 | 390 | 0 | 7 | 0 | 0 | 0 | 259 | 149 | 1 | 1 | 1 | 1 | 1 | 1 | 1 | 1 | 0.931 | 1 | 1 | 1 | 0.014 |
| SE | 64158 | ENSG00000032444 | PNPLA6 | chr19 | + | 7561207 | 7561317 | 7561013 | 7561110 | 7561487 | 7561761 | 270 | 193 | 242 | 245 | 199 | 0 | 0 | 1 | 0 | 0 | 523 | 380 | 236 | 336 | 643 | 0 | 0 | 0 | 0 | 0 | 259 | 149 | 1 | 1 | 1 | 1 | 0.993 | 1 | 1 | 1 | 1 | 1 | 1 | 1 | -0.001 |
| SE | 64159 | ENSG00000143387 | CTSK | chr1 | - | 150799167 | 150799273 | 150794879 | 150796898 | 150799543 | 150799705 | 48 | 49 | 29 | 43 | 67 | 0 | 0 | 0 | 0 | 1 | 14 | 12 | 63 | 29 | 4 | 0 | 1 | 2 | 2 | 0 | 255 | 149 | 1 | 1 | 1 | 1 | 1 | 1 | 0.975 | 1 | 0.875 | 0.948 | 0.894 | 1 | 0.052 |
| SE | 64160 | ENSG00000143387 | CTSK | chr1 | - | 150799167 | 150799285 | 150796207 | 150796898 | 150799543 | 150799709 | 22 | 22 | 13 | 15 | 31 | 0 | 0 | 0 | 0 | 1 | 10 | 13 | 33 | 11 | 0 | 0 | 1 | 2 | 2 | 0 | 267 | 149 | 1 | 1 | 1 | 1 | 1 | 1 | 0.945 | 1 | 0.879 | 0.902 | 0.754 | NA | 0.105 |
| SE | 64162 | ENSG00000143387 | CTSK | chr1 | - | 150806101 | 150806224 | 150805860 | 150806016 | 150806685 | 150806806 | 32 | 31 | 38 | 35 | 52 | 0 | 1 | 0 | 0 | 0 | 30 | 58 | 42 | 64 | 28 | 0 | 0 | 0 | 0 | 0 | 272 | 149 | 1 | 1 | 1 | 0.944 | 1 | 1 | 1 | 1 | 1 | 1 | 1 | 1 | -0.011 |
| SE | 64164 | ENSG00000143387 | CTSK | chr1 | - | 150807248 | 150807341 | 150806685 | 150806806 | 150808213 | 150808260 | 0 | 0 | 1 | 0 | 0 | 17 | 5 | 15 | 17 | 7 | 0 | 0 | 1 | 0 | 0 | 15 | 11 | 11 | 14 | 0 | 242 | 149 | 1 | 1 | 0 | 0 | 0.039 | 0 | 0 | 0 | 0 | 0.053 | 0 | NA | -0.005 |
| SE | 64165 | ENSG00000138035 | PNPT1 | chr2 | - | 55640626 | 55640705 | 55637551 | 55637599 | 55643157 | 55643213 | 58 | 53 | 51 | 113 | 68 | 0 | 0 | 0 | 0 | 0 | 13 | 60 | 50 | 62 | 0 | 0 | 4 | 0 | 0 | 0 | 228 | 149 | 1 | 1 | 1 | 1 | 1 | 1 | 1 | 1 | 0.907 | 1 | 1 | NA | 0.023 |
| SE | 64166 | ENSG00000138035 | PNPT1 | chr2 | - | 55644636 | 55644720 | 55643318 | 55643425 | 55645348 | 55645432 | 136 | 143 | 159 | 197 | 177 | 1 | 0 | 0 | 0 | 0 | 65 | 159 | 122 | 176 | 0 | 0 | 0 | 0 | 0 | 0 | 233 | 149 | 1 | 1 | 0.989 | 1 | 1 | 1 | 1 | 1 | 1 | 1 | 1 | NA | -0.002 |
| SE | 64167 | ENSG00000138035 | PNPT1 | chr2 | - | 55646414 | 55646486 | 55646258 | 55646322 | 55647346 | 55647382 | 103 | 92 | 98 | 106 | 133 | 0 | 0 | 0 | 0 | 0 | 38 | 111 | 70 | 141 | 7 | 0 | 0 | 0 | 4 | 0 | 221 | 149 | 1 | 1 | 1 | 1 | 1 | 1 | 1 | 1 | 1 | 1 | 0.96 | 1 | 0.008 |
| SE | 64169 | ENSG00000138035 | PNPT1 | chr2 | - | 55660156 | 55660193 | 55656304 | 55656371 | 55661955 | 55662026 | 89 | 83 | 116 | 117 | 232 | 1 | 0 | 0 | 0 | 0 | 51 | 103 | 45 | 104 | 10 | 0 | 0 | 0 | 0 | 0 | 186 | 149 | 1 | 1 | 0.986 | 1 | 1 | 1 | 1 | 1 | 1 | 1 | 1 | 1 | -0.003 |
| SE | 64170 | ENSG00000138035 | PNPT1 | chr2 | - | 55667861 | 55667958 | 55666990 | 55667093 | 55671318 | 55671368 | 233 | 282 | 222 | 270 | 301 | 0 | 2 | 1 | 0 | 0 | 114 | 167 | 126 | 273 | 15 | 0 | 0 | 0 | 0 | 0 | 246 | 149 | 1 | 1 | 1 | 0.988 | 0.993 | 1 | 1 | 1 | 1 | 1 | 1 | 1 | -0.004 |
| SE | 64171 | ENSG00000138035 | PNPT1 | chr2 | - | 55672892 | 55673079 | 55671994 | 55672046 | 55679681 | 55679795 | 256 | 267 | 162 | 285 | 313 | 0 | 5 | 0 | 0 | 0 | 166 | 236 | 181 | 312 | 19 | 0 | 0 | 1 | 0 | 0 | 298 | 149 | 1 | 1 | 1 | 0.964 | 1 | 1 | 1 | 1 | 1 | 0.989 | 1 | 1 | -0.005 |
| SE | 64172 | ENSG00000138035 | PNPT1 | chr2 | - | 55680711 | 55680759 | 55679681 | 55679795 | 55680854 | 55680918 | 57 | 49 | 52 | 49 | 60 | 0 | 0 | 0 | 1 | 0 | 31 | 35 | 45 | 69 | 8 | 0 | 0 | 0 | 0 | 0 | 197 | 149 | 1 | 1 | 1 | 1 | 1 | 0.974 | 1 | 1 | 1 | 1 | 1 | 1 | -0.005 |
| SE | 64173 | ENSG00000138035 | PNPT1 | chr2 | - | 55686369 | 55686444 | 55684942 | 55685048 | 55693662 | 55693823 | 13 | 14 | 9 | 15 | 12 | 3 | 0 | 0 | 0 | 0 | 0 | 1 | 9 | 2 | 0 | 0 | 0 | 0 | 3 | 0 | 224 | 149 | 1 | 1 | 0.742 | 1 | 1 | 1 | 1 | NA | 1 | 1 | 0.307 | NA | 0.179 |
| SE | 64174 | ENSG00000138035 | PNPT1 | chr2 | - | 55687644 | 55687705 | 55686369 | 55686444 | 55693662 | 55693823 | 21 | 9 | 19 | 15 | 7 | 0 | 2 | 0 | 0 | 0 | 6 | 7 | 8 | 18 | 0 | 0 | 0 | 0 | 0 | 0 | 210 | 149 | 1 | 1 | 1 | 0.761 | 1 | 1 | 1 | 1 | 1 | 1 | 1 | NA | -0.048 |
| SE | 64175 | ENSG00000048991 | R3HDM1 | chr2 | + | 135565736 | 135565925 | 135531454 | 135531633 | 135602499 | 135602670 | 0 | 0 | 0 | 0 | 0 | 19 | 9 | 28 | 17 | 6 | 0 | 0 | 1 | 0 | 0 | 5 | 23 | 6 | 4 | 18 | 298 | 149 | 1 | 1 | 0 | 0 | 0 | 0 | 0 | 0 | 0 | 0.077 | 0 | 0 | -0.015 |
| SE | 64182 | ENSG00000048991 | R3HDM1 | chr2 | + | 135622653 | 135622732 | 135621493 | 135621608 | 135631717 | 135631777 | 99 | 81 | 92 | 117 | 234 | 0 | 0 | 3 | 0 | 3 | 91 | 119 | 132 | 98 | 65 | 0 | 0 | 0 | 0 | 2 | 228 | 149 | 1 | 1 | 1 | 1 | 0.952 | 1 | 0.981 | 1 | 1 | 1 | 1 | 0.955 | -0.004 |
| SE | 64183 | ENSG00000048991 | R3HDM1 | chr2 | + | 135638617 | 135638655 | 135636087 | 135636183 | 135638738 | 135638789 | 94 | 70 | 96 | 84 | 179 | 0 | 0 | 2 | 0 | 1 | 88 | 109 | 69 | 81 | 91 | 0 | 0 | 0 | 0 | 0 | 187 | 149 | 1 | 1 | 1 | 1 | 0.975 | 1 | 0.993 | 1 | 1 | 1 | 1 | 1 | -0.006 |
| SE | 64187 | ENSG00000048991 | R3HDM1 | chr2 | + | 135661269 | 135661393 | 135651729 | 135652032 | 135675331 | 135675486 | 23 | 24 | 29 | 31 | 17 | 0 | 0 | 0 | 0 | 0 | 38 | 58 | 54 | 53 | 36 | 0 | 2 | 0 | 0 | 0 | 273 | 149 | 1 | 1 | 1 | 1 | 1 | 1 | 1 | 1 | 0.941 | 1 | 1 | 1 | 0.012 |
| SE | 64188 | ENSG00000048991 | R3HDM1 | chr2 | + | 135680172 | 135680324 | 135675331 | 135675486 | 135709432 | 135709536 | 201 | 126 | 188 | 212 | 315 | 0 | 0 | 0 | 0 | 0 | 122 | 220 | 160 | 140 | 153 | 0 | 0 | 0 | 6 | 0 | 298 | 149 | 1 | 1 | 1 | 1 | 1 | 1 | 1 | 1 | 1 | 1 | 0.921 | 1 | 0.016 |
| SE | 64189 | ENSG00000048991 | R3HDM1 | chr2 | + | 135715549 | 135715694 | 135710058 | 135710231 | 135721923 | 135722006 | 56 | 58 | 52 | 55 | 62 | 2 | 0 | 0 | 1 | 1 | 51 | 33 | 51 | 30 | 33 | 0 | 0 | 0 | 0 | 0 | 294 | 149 | 1 | 1 | 0.934 | 1 | 1 | 0.965 | 0.969 | 1 | 1 | 1 | 1 | 1 | -0.026 |
| SE | 64190 | ENSG00000048991 | R3HDM1 | chr2 | + | 135721923 | 135722006 | 135715549 | 135715694 | 135722468 | 135722537 | 44 | 27 | 34 | 35 | 32 | 0 | 1 | 0 | 0 | 0 | 33 | 33 | 33 | 23 | 19 | 0 | 0 | 0 | 0 | 0 | 232 | 149 | 1 | 1 | 1 | 0.945 | 1 | 1 | 1 | 1 | 1 | 1 | 1 | 1 | -0.011 |
| SE | 64193 | ENSG00000171824 | EXOSC10 | chr1 | - | 11077364 | 11077443 | 11076841 | 11076948 | 11077600 | 11077651 | 83 | 56 | 85 | 86 | 139 | 0 | 0 | 1 | 0 | 0 | 47 | 67 | 61 | 55 | 54 | 0 | 0 | 0 | 3 | 0 | 228 | 149 | 1 | 1 | 1 | 1 | 0.982 | 1 | 1 | 1 | 1 | 1 | 0.923 | 1 | 0.012 |
| SE | 64194 | ENSG00000171824 | EXOSC10 | chr1 | - | 11079710 | 11079822 | 11076841 | 11077651 | 11080498 | 11080514 | 90 | 50 | 77 | 86 | 97 | 0 | 0 | 0 | 0 | 0 | 60 | 69 | 60 | 61 | 57 | 0 | 0 | 1 | 0 | 0 | 261 | 149 | 1 | 1 | 1 | 1 | 1 | 1 | 1 | 1 | 1 | 0.972 | 1 | 1 | 0.006 |
| SE | 64195 | ENSG00000171824 | EXOSC10 | chr1 | - | 11080498 | 11080514 | 11079710 | 11079822 | 11080763 | 11080912 | 10 | 8 | 10 | 10 | 15 | 0 | 2 | 1 | 0 | 0 | 0 | 5 | 4 | 14 | 5 | 0 | 0 | 0 | 0 | 0 | 165 | 149 | 1 | 1 | 1 | 0.783 | 0.9 | 1 | 1 | NA | 1 | 1 | 1 | 1 | -0.063 |
| SE | 64196 | ENSG00000171824 | EXOSC10 | chr1 | - | 11080498 | 11080549 | 11079710 | 11079822 | 11080763 | 11080912 | 57 | 34 | 52 | 55 | 65 | 0 | 2 | 1 | 0 | 0 | 12 | 51 | 22 | 48 | 27 | 0 | 0 | 0 | 0 | 0 | 200 | 149 | 1 | 1 | 1 | 0.927 | 0.975 | 1 | 1 | 1 | 1 | 1 | 1 | 1 | -0.02 |
| SE | 64199 | ENSG00000171824 | EXOSC10 | chr1 | - | 11095757 | 11095881 | 11091492 | 11091597 | 11098019 | 11098156 | 224 | 153 | 166 | 262 | 305 | 0 | 0 | 0 | 1 | 0 | 287 | 184 | 169 | 188 | 187 | 0 | 0 | 0 | 2 | 0 | 273 | 149 | 1 | 1 | 1 | 1 | 1 | 0.993 | 1 | 1 | 1 | 1 | 0.981 | 1 | 0.002 |
| SE | 64200 | ENSG00000163806 | SPDYA | chr2 | + | 28788657 | 28788809 | 28783906 | 28783978 | 28793862 | 28793997 | 1286 | 941 | 991 | 1128 | 1366 | 1 | 0 | 0 | 0 | 0 | 205 | 528 | 809 | 252 | 79 | 0 | 0 | 0 | 0 | 0 | 298 | 149 | 1 | 1 | 0.998 | 1 | 1 | 1 | 1 | 1 | 1 | 1 | 1 | 1 | 0 |
| SE | 64201 | ENSG00000163806 | SPDYA | chr2 | + | 28793862 | 28793997 | 28783906 | 28783978 | 28799198 | 28799299 | 745 | 595 | 559 | 747 | 1659 | 1 | 0 | 5 | 6 | 1 | 100 | 265 | 423 | 161 | 42 | 0 | 0 | 4 | 0 | 0 | 284 | 149 | 1 | 1 | 0.997 | 1 | 0.983 | 0.985 | 0.999 | 1 | 1 | 0.982 | 1 | 1 | -0.004 |
| SE | 64204 | ENSG00000138032 | PPM1B | chr2 | + | 44201185 | 44202045 | 44168874 | 44169274 | 44209209 | 44209327 | 362 | 256 | 336 | 376 | 395 | 0 | 0 | 0 | 0 | 0 | 165 | 283 | 471 | 218 | 76 | 0 | 0 | 5 | 0 | 0 | 298 | 149 | 1 | 1 | 1 | 1 | 1 | 1 | 1 | 1 | 1 | 0.979 | 1 | 1 | 0.004 |
| SE | 64206 | ENSG00000285542 | NA | chr2 | + | 44217966 | 44218078 | 44209209 | 44209327 | 44218479 | 44218537 | 471 | 431 | 433 | 509 | 626 | 0 | 0 | 0 | 0 | 0 | 230 | 398 | 878 | 598 | 68 | 0 | 0 | 1 | 0 | 0 | 261 | 149 | 1 | 1 | 1 | 1 | 1 | 1 | 1 | 1 | 1 | 0.998 | 1 | 1 | 0 |
| SE | 64210 | ENSG00000117697 | NSL1 | chr1 | - | 212739533 | 212739601 | 212726152 | 212727179 | 212782371 | 212782426 | 36 | 49 | 46 | 39 | 113 | 2 | 0 | 0 | 0 | 0 | 11 | 47 | 23 | 15 | 0 | 0 | 0 | 0 | 0 | 0 | 217 | 149 | 1 | 1 | 0.925 | 1 | 1 | 1 | 1 | 1 | 1 | 1 | 1 | NA | -0.015 |
| SE | 64213 | ENSG00000117697 | NSL1 | chr1 | - | 212766188 | 212766273 | 212738561 | 212738686 | 212782371 | 212782426 | 12 | 17 | 12 | 12 | 17 | 4 | 2 | 5 | 4 | 0 | 12 | 10 | 1 | 3 | 0 | 0 | 0 | 0 | 0 | 0 | 234 | 149 | 1 | 1 | 0.656 | 0.844 | 0.604 | 0.656 | 1 | 1 | 1 | 1 | 1 | NA | -0.248 |
| SE | 64216 | ENSG00000117697 | NSL1 | chr1 | - | 212787558 | 212787637 | 212784140 | 212784493 | 212791529 | 212791581 | 153 | 97 | 126 | 119 | 149 | 0 | 0 | 1 | 0 | 0 | 52 | 115 | 51 | 98 | 45 | 0 | 2 | 0 | 0 | 0 | 228 | 149 | 1 | 1 | 1 | 1 | 0.988 | 1 | 1 | 1 | 0.974 | 1 | 1 | 1 | 0.003 |
| SE | 64219 | ENSG00000102524 | TNFSF13B | chr13 | + | 108303252 | 108303365 | 108286802 | 108286859 | 108303453 | 108303604 | 298 | 314 | 194 | 293 | 380 | 1 | 0 | 1 | 0 | 0 | 327 | 318 | 137 | 247 | 60 | 0 | 0 | 0 | 0 | 0 | 262 | 149 | 1 | 1 | 0.994 | 1 | 0.991 | 1 | 1 | 1 | 1 | 1 | 1 | 1 | -0.003 |
| SE | 64220 | ENSG00000064225 | ST3GAL6 | chr3 | + | 98756368 | 98756540 | 98732235 | 98732532 | 98768429 | 98768529 | 0 | 5 | 0 | 1 | 0 | 0 | 0 | 0 | 1 | 0 | 8 | 0 | 1 | 10 | 3 | 19 | 3 | 0 | 2 | 3 | 298 | 149 | 1 | 1 | NA | 1 | NA | 0.333 | NA | 0.174 | 0 | 1 | 0.714 | 0.333 | 0.222 |
| SE | 64227 | ENSG00000064225 | ST3GAL6 | chr3 | + | 98773919 | 98773983 | 98772812 | 98772916 | 98783562 | 98783739 | 13 | 4 | 6 | 16 | 26 | 0 | 0 | 1 | 0 | 0 | 24 | 11 | 8 | 11 | 4 | 0 | 0 | 0 | 0 | 0 | 213 | 149 | 1 | 1 | 1 | 1 | 0.808 | 1 | 1 | 1 | 1 | 1 | 1 | 1 | -0.038 |
| SE | 64229 | ENSG00000064225 | ST3GAL6 | chr3 | + | 98773919 | 98773983 | 98772812 | 98772916 | 98791840 | 98791993 | 15 | 6 | 6 | 15 | 19 | 2 | 0 | 0 | 0 | 0 | 24 | 3 | 8 | 11 | 4 | 0 | 0 | 0 | 0 | 0 | 213 | 149 | 1 | 1 | 0.84 | 1 | 1 | 1 | 1 | 1 | 1 | 1 | 1 | 1 | -0.032 |
| SE | 64230 | ENSG00000064225 | ST3GAL6 | chr3 | + | 98773919 | 98773983 | 98772812 | 98772916 | 98793674 | 98793765 | 13 | 5 | 6 | 20 | 17 | 0 | 2 | 0 | 0 | 0 | 24 | 3 | 8 | 11 | 4 | 0 | 0 | 0 | 0 | 0 | 213 | 149 | 1 | 1 | 1 | 0.636 | 1 | 1 | 1 | 1 | 1 | 1 | 1 | 1 | -0.073 |
| SE | 64231 | ENSG00000064225 | ST3GAL6 | chr3 | + | 98784944 | 98785040 | 98772812 | 98772916 | 98791840 | 98791993 | 1 | 0 | 3 | 2 | 1 | 2 | 0 | 0 | 0 | 0 | 4 | 0 | 1 | 2 | 13 | 0 | 0 | 0 | 0 | 0 | 245 | 149 | 1 | 1 | 0.233 | NA | 1 | 1 | 1 | 1 | NA | 1 | 1 | 1 | -0.192 |
| SE | 64233 | ENSG00000064225 | ST3GAL6 | chr3 | + | 98784944 | 98785040 | 98773919 | 98773983 | 98791840 | 98791993 | 15 | 10 | 14 | 13 | 27 | 2 | 2 | 0 | 1 | 2 | 4 | 6 | 13 | 0 | 15 | 0 | 0 | 0 | 0 | 0 | 245 | 149 | 1 | 1 | 0.82 | 0.753 | 1 | 0.888 | 0.891 | 1 | 1 | 1 | NA | 1 | -0.13 |
| SE | 64234 | ENSG00000064225 | ST3GAL6 | chr3 | + | 98791840 | 98791993 | 98772812 | 98772916 | 98793674 | 98793944 | 11 | 20 | 13 | 10 | 42 | 0 | 2 | 0 | 0 | 0 | 19 | 5 | 6 | 15 | 6 | 0 | 0 | 0 | 0 | 0 | 298 | 149 | 1 | 1 | 1 | 0.833 | 1 | 1 | 1 | 1 | 1 | 1 | 1 | 1 | -0.033 |
| SE | 64237 | ENSG00000088280 | ASAP3 | chr1 | - | 23439160 | 23439230 | 23438746 | 23438834 | 23441101 | 23441211 | 4 | 1 | 0 | 1 | 0 | 0 | 0 | 0 | 0 | 0 | 0 | 0 | 0 | 0 | 0 | 0 | 2 | 0 | 0 | 0 | 219 | 149 | 1 | 1 | 1 | 1 | NA | 1 | NA | NA | 0 | NA | NA | NA | 1 |
| SE | 64239 | ENSG00000175104 | TRAF6 | chr11 | - | 36494975 | 36495047 | 36492550 | 36492628 | 36498489 | 36498640 | 49 | 20 | 33 | 37 | 49 | 2 | 0 | 0 | 0 | 0 | 25 | 34 | 48 | 103 | 24 | 0 | 0 | 0 | 0 | 0 | 221 | 149 | 1 | 1 | 0.943 | 1 | 1 | 1 | 1 | 1 | 1 | 1 | 1 | 1 | -0.011 |
| SE | 64241 | ENSG00000175104 | TRAF6 | chr11 | - | 36498489 | 36498640 | 36497107 | 36497266 | 36501219 | 36501537 | 71 | 40 | 44 | 83 | 74 | 0 | 0 | 0 | 0 | 0 | 35 | 58 | 67 | 72 | 12 | 0 | 0 | 0 | 2 | 0 | 298 | 149 | 1 | 1 | 1 | 1 | 1 | 1 | 1 | 1 | 1 | 1 | 0.947 | 1 | 0.011 |
| SE | 64242 | ENSG00000175104 | TRAF6 | chr11 | - | 36501219 | 36501537 | 36498489 | 36498640 | 36510047 | 36510272 | 64 | 28 | 35 | 60 | 56 | 2 | 0 | 0 | 0 | 0 | 38 | 52 | 56 | 35 | 8 | 0 | 0 | 0 | 0 | 0 | 298 | 149 | 1 | 1 | 0.941 | 1 | 1 | 1 | 1 | 1 | 1 | 1 | 1 | 1 | -0.012 |
| SE | 64243 | ENSG00000175104 | TRAF6 | chr11 | - | 36501718 | 36501811 | 36501219 | 36501537 | 36510047 | 36510256 | 0 | 0 | 3 | 0 | 0 | 31 | 15 | 21 | 31 | 35 | 0 | 0 | 0 | 0 | 0 | 24 | 38 | 31 | 20 | 8 | 242 | 149 | 1 | 1 | 0 | 0 | 0.081 | 0 | 0 | 0 | 0 | 0 | 0 | 0 | 0.016 |
| SE | 64245 | ENSG00000185499 | MUC1 | chr1 | - | 155187454 | 155187544 | 155187224 | 155187374 | 155187721 | 155187767 | 0 | 0 | 1 | 0 | 1 | 0 | 0 | 0 | 2 | 0 | 0 | 7 | 6 | 3 | 10 | 0 | 0 | 0 | 0 | 0 | 239 | 149 | 1 | 1 | NA | NA | 1 | 0 | 1 | NA | 1 | 1 | 1 | 1 | -0.333 |
| SE | 64247 | ENSG00000185499 | MUC1 | chr1 | - | 155187454 | 155187576 | 155186707 | 155187374 | 155187721 | 155187831 | 2 | 1 | 2 | 0 | 1 | 0 | 0 | 0 | 2 | 0 | 0 | 14 | 11 | 4 | 20 | 0 | 0 | 0 | 0 | 0 | 271 | 149 | 1 | 1 | 1 | 1 | 1 | 0 | 1 | NA | 1 | 1 | 1 | 1 | -0.2 |
| SE | 64251 | ENSG00000185499 | MUC1 | chr1 | - | 155187721 | 155187858 | 155187454 | 155187576 | 155188162 | 155188212 | 1 | 1 | 0 | 0 | 0 | 0 | 0 | 1 | 0 | 0 | 0 | 7 | 0 | 1 | 10 | 0 | 0 | 0 | 0 | 0 | 286 | 149 | 1 | 1 | 1 | 1 | 0 | NA | NA | NA | 1 | NA | 1 | 1 | -0.333 |
| SE | 64254 | ENSG00000185499 | MUC1 | chr1 | - | 155188007 | 155188063 | 155187454 | 155187576 | 155188162 | 155188212 | 1 | 1 | 0 | 0 | 2 | 0 | 0 | 1 | 0 | 0 | 6 | 6 | 6 | 7 | 0 | 0 | 0 | 0 | 0 | 0 | 205 | 149 | 1 | 1 | 1 | 1 | 0 | NA | 1 | 1 | 1 | 1 | 1 | NA | -0.25 |
| SE | 64261 | ENSG00000213699 | SLC35F6 | chr2 | + | 26774250 | 26774323 | 26764283 | 26764426 | 26775043 | 26775215 | 48 | 14 | 50 | 37 | 7 | 2 | 1 | 0 | 0 | 0 | 0 | 53 | 32 | 26 | 109 | 0 | 0 | 0 | 0 | 8 | 222 | 149 | 1 | 1 | 0.942 | 0.904 | 1 | 1 | 1 | NA | 1 | 1 | 1 | 0.901 | -0.006 |
| SE | 64262 | ENSG00000213699 | SLC35F6 | chr2 | + | 26774250 | 26774323 | 26764283 | 26764426 | 26775463 | 26775676 | 29 | 12 | 27 | 20 | 4 | 0 | 0 | 0 | 2 | 0 | 0 | 19 | 12 | 11 | 23 | 0 | 1 | 0 | 0 | 0 | 222 | 149 | 1 | 1 | 1 | 1 | 1 | 0.87 | 1 | NA | 0.927 | 1 | 1 | 1 | -0.008 |
| SE | 64263 | ENSG00000213699 | SLC35F6 | chr2 | + | 26775043 | 26775215 | 26764283 | 26764426 | 26775463 | 26775676 | 25 | 9 | 12 | 20 | 23 | 0 | 0 | 0 | 2 | 0 | 8 | 28 | 12 | 6 | 63 | 0 | 1 | 0 | 0 | 0 | 298 | 149 | 1 | 1 | 1 | 1 | 1 | 0.833 | 1 | 1 | 0.933 | 1 | 1 | 1 | -0.02 |
| SE | 64265 | ENSG00000213699 | SLC35F6 | chr2 | + | 26775463 | 26775676 | 26764289 | 26764426 | 26778041 | 26778219 | 0 | 1 | 0 | 5 | 0 | 1 | 0 | 1 | 0 | 0 | 0 | 1 | 3 | 3 | 0 | 0 | 0 | 0 | 0 | 0 | 298 | 149 | 1 | 1 | 0 | 1 | 0 | 1 | NA | NA | 1 | 1 | 1 | NA | -0.5 |
| SE | 64267 | ENSG00000168887 | C2orf68 | chr2 | - | 85609434 | 85609586 | 85605253 | 85609067 | 85611667 | 85611786 | 114 | 68 | 113 | 111 | 92 | 1 | 1 | 3 | 0 | 0 | 99 | 98 | 69 | 86 | 234 | 0 | 0 | 0 | 4 | 24 | 298 | 149 | 1 | 1 | 0.983 | 0.971 | 0.95 | 1 | 1 | 1 | 1 | 1 | 0.915 | 0.83 | 0.032 |
| SE | 64269 | ENSG00000145919 | BOD1 | chr5 | - | 173609237 | 173609434 | 173607144 | 173608292 | 173616199 | 173616358 | 38 | 19 | 46 | 51 | 50 | 0 | 2 | 0 | 0 | 1 | 33 | 21 | 14 | 16 | 33 | 0 | 2 | 0 | 1 | 0 | 298 | 149 | 1 | 1 | 1 | 0.826 | 1 | 1 | 0.962 | 1 | 0.84 | 1 | 0.889 | 1 | 0.012 |
| SE | 64270 | ENSG00000145919 | BOD1 | chr5 | - | 173613130 | 173613255 | 173607520 | 173608292 | 173616199 | 173616588 | 16 | 9 | 27 | 35 | 16 | 0 | 2 | 0 | 0 | 1 | 67 | 17 | 6 | 32 | 30 | 0 | 2 | 0 | 1 | 0 | 274 | 149 | 1 | 1 | 1 | 0.71 | 1 | 1 | 0.897 | 1 | 0.822 | 1 | 0.946 | 1 | -0.032 |
| SE | 64271 | ENSG00000145919 | BOD1 | chr5 | - | 173613130 | 173613255 | 173609237 | 173609434 | 173616199 | 173616588 | 43 | 21 | 61 | 46 | 51 | 1 | 0 | 0 | 1 | 1 | 28 | 15 | 13 | 9 | 66 | 0 | 3 | 0 | 0 | 2 | 274 | 149 | 1 | 1 | 0.959 | 1 | 1 | 0.962 | 0.965 | 1 | 0.731 | 1 | 1 | 0.947 | 0.042 |
| SE | 64272 | ENSG00000068878 | PSME4 | chr2 | - | 53869375 | 53869538 | 53866746 | 53866880 | 53874338 | 53874494 | 78 | 59 | 50 | 86 | 141 | 2 | 1 | 1 | 1 | 3 | 26 | 63 | 52 | 34 | 0 | 0 | 0 | 0 | 3 | 0 | 298 | 149 | 1 | 1 | 0.951 | 0.967 | 0.962 | 0.977 | 0.959 | 1 | 1 | 1 | 0.85 | NA | 0.001 |
| SE | 64274 | ENSG00000068878 | PSME4 | chr2 | - | 53875626 | 53875755 | 53874338 | 53874494 | 53885689 | 53885775 | 281 | 123 | 197 | 191 | 273 | 0 | 0 | 1 | 0 | 0 | 40 | 156 | 201 | 105 | 33 | 0 | 0 | 0 | 0 | 0 | 278 | 149 | 1 | 1 | 1 | 1 | 0.991 | 1 | 1 | 1 | 1 | 1 | 1 | 1 | -0.002 |
| SE | 64275 | ENSG00000068878 | PSME4 | chr2 | - | 53887258 | 53887467 | 53885689 | 53885775 | 53887857 | 53887989 | 443 | 228 | 328 | 337 | 549 | 0 | 0 | 1 | 0 | 0 | 32 | 235 | 339 | 175 | 42 | 0 | 0 | 0 | 0 | 0 | 298 | 149 | 1 | 1 | 1 | 1 | 0.994 | 1 | 1 | 1 | 1 | 1 | 1 | 1 | -0.001 |
| SE | 64276 | ENSG00000068878 | PSME4 | chr2 | - | 53887857 | 53887989 | 53887258 | 53887467 | 53888720 | 53888812 | 314 | 186 | 223 | 306 | 461 | 0 | 0 | 0 | 0 | 0 | 59 | 200 | 333 | 147 | 45 | 0 | 0 | 0 | 3 | 0 | 281 | 149 | 1 | 1 | 1 | 1 | 1 | 1 | 1 | 1 | 1 | 1 | 0.963 | 1 | 0.007 |
| SE | 64278 | ENSG00000068878 | PSME4 | chr2 | - | 53892807 | 53892960 | 53890103 | 53890208 | 53895006 | 53895076 | 47 | 25 | 53 | 51 | 115 | 1 | 0 | 0 | 0 | 0 | 6 | 49 | 61 | 59 | 16 | 0 | 0 | 0 | 0 | 0 | 298 | 149 | 1 | 1 | 0.959 | 1 | 1 | 1 | 1 | 1 | 1 | 1 | 1 | 1 | -0.008 |
| SE | 64280 | ENSG00000068878 | PSME4 | chr2 | - | 53893673 | 53893799 | 53892807 | 53892960 | 53895582 | 53895736 | 116 | 104 | 107 | 118 | 145 | 0 | 2 | 0 | 0 | 0 | 22 | 149 | 114 | 94 | 20 | 0 | 1 | 2 | 0 | 0 | 275 | 149 | 1 | 1 | 1 | 0.966 | 1 | 1 | 1 | 1 | 0.988 | 0.969 | 1 | 1 | 0.002 |
| SE | 64281 | ENSG00000068878 | PSME4 | chr2 | - | 53895006 | 53895076 | 53892807 | 53892960 | 53895582 | 53895736 | 44 | 40 | 49 | 45 | 49 | 0 | 2 | 0 | 0 | 0 | 6 | 40 | 27 | 40 | 2 | 0 | 1 | 2 | 0 | 0 | 219 | 149 | 1 | 1 | 1 | 0.932 | 1 | 1 | 1 | 1 | 0.965 | 0.902 | 1 | 1 | 0.013 |
| SE | 64282 | ENSG00000068878 | PSME4 | chr2 | - | 53895006 | 53895076 | 53893673 | 53893799 | 53895582 | 53895736 | 141 | 98 | 111 | 160 | 180 | 1 | 2 | 0 | 0 | 2 | 30 | 145 | 102 | 87 | 9 | 0 | 0 | 0 | 0 | 0 | 219 | 149 | 1 | 1 | 0.99 | 0.971 | 1 | 1 | 0.984 | 1 | 1 | 1 | 1 | 1 | -0.011 |
| SE | 64283 | ENSG00000068878 | PSME4 | chr2 | - | 53897869 | 53897999 | 53896803 | 53896885 | 53898300 | 53898354 | 353 | 278 | 246 | 369 | 547 | 0 | 0 | 0 | 0 | 1 | 212 | 329 | 358 | 332 | 42 | 0 | 0 | 0 | 0 | 0 | 279 | 149 | 1 | 1 | 1 | 1 | 1 | 1 | 0.997 | 1 | 1 | 1 | 1 | 1 | -0.001 |
| SE | 64286 | ENSG00000068878 | PSME4 | chr2 | - | 53910074 | 53910130 | 53908783 | 53908840 | 53919150 | 53919246 | 116 | 76 | 122 | 153 | 202 | 0 | 0 | 0 | 0 | 0 | 34 | 126 | 176 | 105 | 36 | 0 | 0 | 0 | 1 | 0 | 205 | 149 | 1 | 1 | 1 | 1 | 1 | 1 | 1 | 1 | 1 | 1 | 0.987 | 1 | 0.003 |
| SE | 64287 | ENSG00000068878 | PSME4 | chr2 | - | 53925538 | 53925689 | 53923320 | 53923419 | 53925958 | 53926023 | 402 | 246 | 278 | 363 | 470 | 2 | 0 | 0 | 0 | 0 | 118 | 398 | 387 | 213 | 43 | 0 | 0 | 0 | 0 | 0 | 298 | 149 | 1 | 1 | 0.99 | 1 | 1 | 1 | 1 | 1 | 1 | 1 | 1 | 1 | -0.002 |
| SE | 64288 | ENSG00000068878 | PSME4 | chr2 | - | 53925958 | 53926023 | 53925538 | 53925689 | 53927393 | 53927483 | 253 | 163 | 184 | 245 | 268 | 0 | 0 | 0 | 0 | 0 | 97 | 250 | 251 | 157 | 29 | 0 | 0 | 0 | 0 | 2 | 214 | 149 | 1 | 1 | 1 | 1 | 1 | 1 | 1 | 1 | 1 | 1 | 1 | 0.91 | 0.018 |
| SE | 64289 | ENSG00000068878 | PSME4 | chr2 | - | 53936763 | 53936827 | 53936086 | 53936161 | 53937390 | 53937540 | 106 | 63 | 71 | 120 | 140 | 0 | 2 | 0 | 1 | 0 | 52 | 104 | 145 | 99 | 21 | 0 | 0 | 0 | 3 | 0 | 213 | 149 | 1 | 1 | 1 | 0.957 | 1 | 0.988 | 1 | 1 | 1 | 1 | 0.958 | 1 | -0.003 |
| SE | 64290 | ENSG00000068878 | PSME4 | chr2 | - | 53942158 | 53942224 | 53939955 | 53940000 | 53948420 | 53948537 | 0 | 0 | 1 | 4 | 0 | 98 | 68 | 68 | 95 | 187 | 0 | 0 | 6 | 0 | 0 | 40 | 89 | 117 | 71 | 3 | 215 | 149 | 1 | 1 | 0 | 0 | 0.01 | 0.028 | 0 | 0 | 0 | 0.034 | 0 | 0 | 0.001 |
| SE | 64293 | ENSG00000177000 | MTHFR | chr1 | - | 11794728 | 11794863 | 11794357 | 11794538 | 11795097 | 11795348 | 59 | 40 | 41 | 58 | 53 | 0 | 0 | 0 | 1 | 0 | 231 | 86 | 30 | 77 | 135 | 0 | 0 | 0 | 0 | 0 | 284 | 149 | 1 | 1 | 1 | 1 | 1 | 0.968 | 1 | 1 | 1 | 1 | 1 | 1 | -0.006 |
| SE | 64296 | ENSG00000177000 | MTHFR | chr1 | - | 11800211 | 11800322 | 11796205 | 11796399 | 11801160 | 11801399 | 44 | 7 | 27 | 43 | 34 | 0 | 0 | 0 | 0 | 0 | 55 | 21 | 30 | 56 | 60 | 0 | 0 | 1 | 0 | 0 | 260 | 149 | 1 | 1 | 1 | 1 | 1 | 1 | 1 | 1 | 1 | 0.945 | 1 | 1 | 0.011 |
| SE | 64300 | ENSG00000131873 | CHSY1 | chr15 | - | 101235081 | 101235577 | 101175726 | 101178980 | 101251136 | 101252048 | 84 | 42 | 53 | 77 | 57 | 0 | 0 | 0 | 0 | 0 | 117 | 100 | 90 | 186 | 76 | 0 | 0 | 3 | 0 | 0 | 298 | 149 | 1 | 1 | 1 | 1 | 1 | 1 | 1 | 1 | 1 | 0.938 | 1 | 1 | 0.012 |
| SE | 64303 | ENSG00000101336 | HCK | chr20 | + | 32069687 | 32069751 | 32052241 | 32052486 | 32071661 | 32071782 | 4 | 0 | 0 | 0 | 0 | 350 | 245 | 162 | 236 | 204 | 0 | 0 | 0 | 0 | 0 | 787 | 301 | 170 | 429 | 880 | 213 | 149 | 1 | 1 | 0.008 | 0 | 0 | 0 | 0 | 0 | 0 | 0 | 0 | 0 | 0.002 |
| SE | 64304 | ENSG00000101336 | HCK | chr20 | + | 32073318 | 32073361 | 32071661 | 32071782 | 32073715 | 32073818 | 484 | 350 | 253 | 368 | 316 | 0 | 0 | 0 | 0 | 0 | 1112 | 526 | 234 | 573 | 1367 | 0 | 0 | 1 | 0 | 0 | 192 | 149 | 1 | 1 | 1 | 1 | 1 | 1 | 1 | 1 | 1 | 0.995 | 1 | 1 | 0.001 |
| SE | 64305 | ENSG00000101336 | HCK | chr20 | + | 32073318 | 32073361 | 32071661 | 32071782 | 32073718 | 32073818 | 285 | 235 | 136 | 209 | 158 | 0 | 0 | 0 | 0 | 0 | 695 | 304 | 171 | 365 | 896 | 0 | 0 | 1 | 0 | 1 | 192 | 149 | 1 | 1 | 1 | 1 | 1 | 1 | 1 | 1 | 1 | 0.993 | 1 | 0.999 | 0.002 |
| SE | 64310 | ENSG00000101336 | HCK | chr20 | + | 32099003 | 32099135 | 32093862 | 32094016 | 32101316 | 32101519 | 953 | 710 | 452 | 804 | 434 | 0 | 0 | 0 | 0 | 0 | 2684 | 1115 | 519 | 1464 | 2717 | 0 | 0 | 0 | 3 | 0 | 281 | 149 | 1 | 1 | 1 | 1 | 1 | 1 | 1 | 1 | 1 | 1 | 0.996 | 1 | 0.001 |
| SE | 64311 | ENSG00000231259 | ANAPC1P2 | chr2 | - | 87034151 | 87034243 | 87032068 | 87032203 | 87037293 | 87037528 | 73 | 123 | 69 | 175 | 74 | 0 | 0 | 0 | 1 | 0 | 85 | 73 | 0 | 217 | 36 | 0 | 6 | 0 | 0 | 0 | 241 | 149 | 1 | 1 | 1 | 1 | 1 | 0.991 | 1 | 1 | 0.883 | NA | 1 | 1 | 0.027 |
| SE | 64312 | ENSG00000231259 | ANAPC1P2 | chr2 | - | 87043712 | 87043984 | 87037293 | 87037528 | 87049588 | 87049803 | 6 | 3 | 3 | 9 | 2 | 0 | 0 | 0 | 0 | 0 | 0 | 6 | 2 | 11 | 0 | 0 | 0 | 1 | 0 | 0 | 298 | 149 | 1 | 1 | 1 | 1 | 1 | 1 | 1 | NA | 1 | 0.5 | 1 | NA | 0.167 |
| SE | 64313 | ENSG00000231259 | ANAPC1P2 | chr2 | - | 87047187 | 87047306 | 87043712 | 87043984 | 87049588 | 87049803 | 141 | 61 | 106 | 107 | 119 | 2 | 0 | 0 | 2 | 0 | 79 | 85 | 62 | 99 | 118 | 0 | 4 | 0 | 0 | 0 | 268 | 149 | 1 | 1 | 0.975 | 1 | 1 | 0.967 | 1 | 1 | 0.922 | 1 | 1 | 1 | 0.004 |
| SE | 64314 | ENSG00000231259 | ANAPC1P2 | chr2 | - | 87047187 | 87047306 | 87043712 | 87043984 | 87052813 | 87052992 | 61 | 25 | 50 | 50 | 53 | 0 | 1 | 0 | 0 | 0 | 44 | 24 | 38 | 39 | 38 | 0 | 2 | 0 | 0 | 0 | 268 | 149 | 1 | 1 | 1 | 0.933 | 1 | 1 | 1 | 1 | 0.87 | 1 | 1 | 1 | 0.013 |
| SE | 64315 | ENSG00000231259 | ANAPC1P2 | chr2 | - | 87049588 | 87049803 | 87043712 | 87043984 | 87052813 | 87052992 | 162 | 96 | 169 | 160 | 106 | 0 | 1 | 0 | 0 | 0 | 144 | 267 | 224 | 141 | 233 | 0 | 2 | 0 | 0 | 0 | 298 | 149 | 1 | 1 | 1 | 0.98 | 1 | 1 | 1 | 1 | 0.985 | 1 | 1 | 1 | -0.001 |
| SE | 64318 | ENSG00000166949 | SMAD3 | chr15 | + | 67170553 | 67170604 | 67166530 | 67166853 | 67181240 | 67181420 | 61 | 41 | 50 | 61 | 31 | 0 | 0 | 0 | 0 | 0 | 141 | 120 | 41 | 99 | 139 | 0 | 0 | 0 | 0 | 2 | 200 | 149 | 1 | 1 | 1 | 1 | 1 | 1 | 1 | 1 | 1 | 1 | 1 | 0.981 | 0.004 |
| SE | 64319 | ENSG00000166949 | SMAD3 | chr15 | + | 67181240 | 67181453 | 67170553 | 67170604 | 67187364 | 67187447 | 53 | 19 | 32 | 47 | 26 | 1 | 0 | 0 | 2 | 0 | 81 | 70 | 32 | 67 | 75 | 0 | 0 | 0 | 0 | 0 | 298 | 149 | 1 | 1 | 0.964 | 1 | 1 | 0.922 | 1 | 1 | 1 | 1 | 1 | 1 | -0.023 |
| SE | 64320 | ENSG00000166949 | SMAD3 | chr15 | + | 67184726 | 67184864 | 67180879 | 67181453 | 67187364 | 67187509 | 112 | 46 | 72 | 98 | 29 | 0 | 0 | 0 | 1 | 0 | 196 | 211 | 117 | 107 | 170 | 0 | 0 | 1 | 0 | 4 | 287 | 149 | 1 | 1 | 1 | 1 | 1 | 0.981 | 1 | 1 | 1 | 0.984 | 1 | 0.957 | 0.008 |
| SE | 64321 | ENSG00000166949 | SMAD3 | chr15 | + | 67187019 | 67187130 | 67184774 | 67184864 | 67187364 | 67187509 | 0 | 0 | 0 | 1 | 0 | 68 | 29 | 36 | 55 | 14 | 0 | 0 | 0 | 0 | 0 | 121 | 136 | 90 | 63 | 100 | 260 | 149 | 1 | 1 | 0 | 0 | 0 | 0.01 | 0 | 0 | 0 | 0 | 0 | 0 | 0.002 |
| SE | 64325 | ENSG00000138363 | ATIC | chr2 | + | 215325986 | 215326138 | 215319664 | 215319731 | 215326821 | 215326978 | 145 | 92 | 208 | 164 | 202 | 0 | 0 | 0 | 0 | 2 | 148 | 215 | 77 | 111 | 148 | 0 | 0 | 0 | 0 | 0 | 298 | 149 | 1 | 1 | 1 | 1 | 1 | 1 | 0.981 | 1 | 1 | 1 | 1 | 1 | -0.004 |
| SE | 64326 | ENSG00000138363 | ATIC | chr2 | + | 215326821 | 215326978 | 215325986 | 215326138 | 215332381 | 215332507 | 264 | 186 | 359 | 283 | 374 | 0 | 0 | 0 | 0 | 0 | 234 | 364 | 137 | 204 | 234 | 0 | 0 | 0 | 5 | 0 | 298 | 149 | 1 | 1 | 1 | 1 | 1 | 1 | 1 | 1 | 1 | 1 | 0.953 | 1 | 0.009 |
| SE | 64327 | ENSG00000138363 | ATIC | chr2 | + | 215338778 | 215338907 | 215336021 | 215336124 | 215344778 | 215344871 | 258 | 174 | 313 | 318 | 595 | 0 | 0 | 0 | 0 | 3 | 303 | 300 | 189 | 174 | 314 | 0 | 0 | 0 | 0 | 0 | 278 | 149 | 1 | 1 | 1 | 1 | 1 | 1 | 0.991 | 1 | 1 | 1 | 1 | 1 | -0.002 |
| SE | 64350 | ENSG00000152672 | CLEC4F | chr2 | - | 70812446 | 70812598 | 70809738 | 70809857 | 70815993 | 70817112 | 0 | 47 | 1 | 39 | 11 | 0 | 6 | 0 | 5 | 0 | 7 | 0 | 0 | 0 | 0 | 0 | 0 | 0 | 0 | 0 | 298 | 149 | 1 | 1 | NA | 0.797 | 1 | 0.796 | 1 | 1 | NA | NA | NA | NA | -0.102 |
| SE | 64351 | ENSG00000152672 | CLEC4F | chr2 | - | 70819354 | 70819444 | 70815993 | 70817112 | 70819774 | 70819891 | 0 | 20 | 0 | 17 | 2 | 0 | 0 | 0 | 5 | 0 | 0 | 0 | 1 | 8 | 0 | 0 | 0 | 0 | 0 | 0 | 239 | 149 | 1 | 1 | NA | 1 | NA | 0.679 | 1 | NA | NA | 1 | 1 | NA | -0.107 |
| SE | 64353 | ENSG00000163219 | ARHGAP25 | chr2 | + | 68757637 | 68757730 | 68710504 | 68710698 | 68775220 | 68775417 | 4 | 4 | 1 | 4 | 4 | 2 | 0 | 0 | 0 | 0 | 8 | 3 | 0 | 0 | 11 | 0 | 8 | 0 | 0 | 0 | 242 | 149 | 1 | 1 | 0.552 | 1 | 1 | 1 | 1 | 1 | 0.188 | NA | NA | 1 | 0.181 |
| SE | 64354 | ENSG00000163219 | ARHGAP25 | chr2 | + | 68757637 | 68757730 | 68734951 | 68735260 | 68774845 | 68775138 | 2 | 0 | 1 | 0 | 9 | 0 | 0 | 0 | 0 | 0 | 14 | 0 | 0 | 3 | 0 | 0 | 2 | 5 | 0 | 0 | 242 | 149 | 1 | 1 | 1 | NA | 1 | NA | 1 | 1 | 0 | 0 | 1 | NA | 0.5 |
| SE | 64356 | ENSG00000163219 | ARHGAP25 | chr2 | + | 68775220 | 68775420 | 68734810 | 68735260 | 68782232 | 68782286 | 516 | 339 | 484 | 401 | 457 | 1 | 0 | 0 | 3 | 0 | 396 | 430 | 352 | 229 | 328 | 0 | 5 | 0 | 1 | 0 | 298 | 149 | 1 | 1 | 0.996 | 1 | 1 | 0.985 | 1 | 1 | 0.977 | 1 | 0.991 | 1 | 0.003 |
| SE | 64357 | ENSG00000163219 | ARHGAP25 | chr2 | + | 68775220 | 68775420 | 68734810 | 68735260 | 68807272 | 68807480 | 155 | 157 | 227 | 173 | 234 | 3 | 0 | 1 | 0 | 0 | 195 | 124 | 135 | 78 | 141 | 0 | 0 | 0 | 0 | 0 | 298 | 149 | 1 | 1 | 0.963 | 1 | 0.991 | 1 | 1 | 1 | 1 | 1 | 1 | 1 | -0.009 |
| SE | 64358 | ENSG00000163219 | ARHGAP25 | chr2 | + | 68782232 | 68782320 | 68734810 | 68735260 | 68807272 | 68807480 | 13 | 4 | 12 | 6 | 16 | 3 | 0 | 1 | 0 | 0 | 0 | 26 | 28 | 23 | 0 | 0 | 0 | 0 | 0 | 0 | 237 | 149 | 1 | 1 | 0.731 | 1 | 0.883 | 1 | 1 | NA | 1 | 1 | 1 | NA | -0.077 |
| SE | 64359 | ENSG00000163219 | ARHGAP25 | chr2 | + | 68782232 | 68782320 | 68734810 | 68735260 | 68807275 | 68807480 | 2 | 1 | 3 | 3 | 0 | 0 | 0 | 2 | 0 | 0 | 0 | 8 | 3 | 1 | 0 | 0 | 0 | 0 | 0 | 0 | 237 | 149 | 1 | 1 | 1 | 1 | 0.485 | 1 | NA | NA | 1 | 1 | 1 | NA | -0.129 |
| SE | 64360 | ENSG00000163219 | ARHGAP25 | chr2 | + | 68782232 | 68782320 | 68774800 | 68775420 | 68787839 | 68787922 | 598 | 287 | 434 | 367 | 424 | 0 | 1 | 0 | 0 | 0 | 267 | 458 | 293 | 158 | 270 | 0 | 0 | 0 | 0 | 0 | 237 | 149 | 1 | 1 | 1 | 0.994 | 1 | 1 | 1 | 1 | 1 | 1 | 1 | 1 | -0.001 |
| SE | 64361 | ENSG00000163219 | ARHGAP25 | chr2 | + | 68782232 | 68782320 | 68774800 | 68775420 | 68807272 | 68807480 | 196 | 111 | 144 | 110 | 135 | 0 | 0 | 0 | 4 | 0 | 90 | 203 | 158 | 106 | 107 | 0 | 0 | 6 | 0 | 0 | 237 | 149 | 1 | 1 | 1 | 1 | 1 | 0.945 | 1 | 1 | 1 | 0.943 | 1 | 1 | 0 |
| SE | 64365 | ENSG00000163219 | ARHGAP25 | chr2 | + | 68813286 | 68813419 | 68807230 | 68807480 | 68816153 | 68816311 | 133 | 33 | 105 | 101 | 118 | 0 | 0 | 0 | 0 | 1 | 20 | 114 | 85 | 55 | 57 | 0 | 0 | 0 | 0 | 0 | 282 | 149 | 1 | 1 | 1 | 1 | 1 | 1 | 0.984 | 1 | 1 | 1 | 1 | 1 | -0.003 |
| SE | 64368 | ENSG00000163219 | ARHGAP25 | chr2 | + | 68819122 | 68819319 | 68817872 | 68817994 | 68822339 | 68822872 | 864 | 338 | 628 | 563 | 689 | 1 | 0 | 0 | 0 | 0 | 288 | 568 | 359 | 391 | 377 | 0 | 0 | 0 | 0 | 0 | 298 | 149 | 1 | 1 | 0.998 | 1 | 1 | 1 | 1 | 1 | 1 | 1 | 1 | 1 | 0 |
| SE | 64369 | ENSG00000163219 | ARHGAP25 | chr2 | + | 68822339 | 68822872 | 68819122 | 68819319 | 68825986 | 68826302 | 531 | 219 | 396 | 387 | 419 | 1 | 0 | 0 | 4 | 0 | 211 | 330 | 239 | 222 | 266 | 0 | 0 | 0 | 0 | 0 | 298 | 149 | 1 | 1 | 0.996 | 1 | 1 | 0.98 | 1 | 1 | 1 | 1 | 1 | 1 | -0.005 |
| SE | 64372 | ENSG00000264538 | SUZ12P1 | chr17 | + | 30734896 | 30734943 | 30731885 | 30732123 | 30743300 | 30743369 | 10 | 0 | 4 | 2 | 2 | 2 | 4 | 0 | 4 | 2 | 0 | 8 | 2 | 1 | 0 | 0 | 0 | 4 | 0 | 0 | 196 | 149 | 1 | 1 | 0.792 | 0 | 1 | 0.275 | 0.432 | NA | 1 | 0.275 | 1 | NA | -0.259 |
| SE | 64376 | ENSG00000264538 | SUZ12P1 | chr17 | + | 30743300 | 30743369 | 30731705 | 30732123 | 30766502 | 30766588 | 1 | 0 | 0 | 1 | 0 | 0 | 0 | 0 | 0 | 0 | 0 | 0 | 0 | 0 | 0 | 0 | 0 | 1 | 0 | 0 | 218 | 149 | 1 | 1 | 1 | NA | NA | 1 | NA | NA | NA | 0 | NA | NA | 1 |
| SE | 64377 | ENSG00000264538 | SUZ12P1 | chr17 | + | 30743300 | 30743369 | 30735032 | 30735097 | 30758171 | 30758335 | 7 | 15 | 24 | 16 | 46 | 0 | 0 | 3 | 0 | 0 | 19 | 27 | 13 | 28 | 30 | 0 | 0 | 0 | 0 | 0 | 218 | 149 | 1 | 1 | 1 | 1 | 0.845 | 1 | 1 | 1 | 1 | 1 | 1 | 1 | -0.031 |
| SE | 64379 | ENSG00000264538 | SUZ12P1 | chr17 | + | 30743300 | 30743369 | 30735032 | 30735097 | 30766502 | 30766588 | 7 | 15 | 18 | 16 | 33 | 0 | 0 | 0 | 1 | 1 | 19 | 27 | 13 | 27 | 30 | 0 | 0 | 0 | 0 | 0 | 218 | 149 | 1 | 1 | 1 | 1 | 1 | 0.916 | 0.958 | 1 | 1 | 1 | 1 | 1 | -0.025 |
| SE | 64380 | ENSG00000264538 | SUZ12P1 | chr17 | + | 30759490 | 30759539 | 30731885 | 30732123 | 30766502 | 30766588 | 21 | 13 | 22 | 23 | 32 | 0 | 0 | 0 | 0 | 0 | 36 | 21 | 9 | 22 | 41 | 0 | 0 | 1 | 0 | 0 | 198 | 149 | 1 | 1 | 1 | 1 | 1 | 1 | 1 | 1 | 1 | 0.871 | 1 | 1 | 0.026 |
| SE | 64381 | ENSG00000264538 | SUZ12P1 | chr17 | + | 30759490 | 30759539 | 30735032 | 30735097 | 30766502 | 30766588 | 32 | 13 | 34 | 35 | 69 | 0 | 0 | 0 | 1 | 1 | 49 | 27 | 13 | 30 | 62 | 0 | 0 | 0 | 0 | 0 | 198 | 149 | 1 | 1 | 1 | 1 | 1 | 0.963 | 0.981 | 1 | 1 | 1 | 1 | 1 | -0.011 |
| SE | 64384 | ENSG00000264538 | SUZ12P1 | chr17 | + | 30769000 | 30769064 | 30768833 | 30768867 | 30769871 | 30770034 | 29 | 41 | 33 | 46 | 47 | 2 | 0 | 0 | 0 | 0 | 62 | 30 | 21 | 15 | 77 | 0 | 0 | 0 | 0 | 0 | 213 | 149 | 1 | 1 | 0.91 | 1 | 1 | 1 | 1 | 1 | 1 | 1 | 1 | 1 | -0.018 |
| SE | 64386 | ENSG00000256206 | NA | chr11 | - | 14514389 | 14514491 | 14513569 | 14513699 | 14517641 | 14517745 | 198 | 99 | 124 | 184 | 201 | 0 | 0 | 0 | 0 | 0 | 144 | 111 | 108 | 131 | 31 | 0 | 0 | 3 | 0 | 0 | 251 | 149 | 1 | 1 | 1 | 1 | 1 | 1 | 1 | 1 | 1 | 0.955 | 1 | 1 | 0.009 |
| SE | 64387 | ENSG00000101577 | LPIN2 | chr18 | - | 2922046 | 2922199 | 2921532 | 2921647 | 2923774 | 2923861 | 383 | 196 | 342 | 331 | 398 | 2 | 0 | 0 | 0 | 0 | 143 | 209 | 214 | 185 | 158 | 0 | 0 | 0 | 0 | 0 | 298 | 149 | 1 | 1 | 0.99 | 1 | 1 | 1 | 1 | 1 | 1 | 1 | 1 | 1 | -0.002 |
| SE | 64389 | ENSG00000101577 | LPIN2 | chr18 | - | 2925223 | 2925368 | 2924397 | 2924546 | 2926722 | 2926805 | 268 | 170 | 212 | 231 | 250 | 0 | 3 | 3 | 0 | 2 | 153 | 125 | 191 | 130 | 147 | 0 | 0 | 3 | 0 | 0 | 294 | 149 | 1 | 1 | 1 | 0.966 | 0.973 | 1 | 0.984 | 1 | 1 | 0.97 | 1 | 1 | -0.009 |
| SE | 64390 | ENSG00000101577 | LPIN2 | chr18 | - | 2925223 | 2925368 | 2924397 | 2924546 | 2927721 | 2927811 | 140 | 69 | 103 | 111 | 53 | 0 | 0 | 0 | 0 | 0 | 45 | 70 | 86 | 73 | 64 | 0 | 0 | 0 | 2 | 0 | 294 | 149 | 1 | 1 | 1 | 1 | 1 | 1 | 1 | 1 | 1 | 1 | 0.949 | 1 | 0.01 |
| SE | 64391 | ENSG00000101577 | LPIN2 | chr18 | - | 2925223 | 2925368 | 2924397 | 2924546 | 2929064 | 2929158 | 139 | 71 | 105 | 114 | 62 | 1 | 0 | 2 | 0 | 0 | 45 | 70 | 86 | 85 | 66 | 0 | 0 | 0 | 1 | 2 | 294 | 149 | 1 | 1 | 0.986 | 1 | 0.964 | 1 | 1 | 1 | 1 | 1 | 0.977 | 0.944 | 0.006 |
| SE | 64392 | ENSG00000101577 | LPIN2 | chr18 | - | 2926722 | 2926805 | 2924397 | 2924546 | 2927721 | 2927811 | 101 | 113 | 87 | 97 | 128 | 0 | 0 | 0 | 0 | 0 | 104 | 67 | 103 | 102 | 101 | 0 | 0 | 0 | 2 | 0 | 232 | 149 | 1 | 1 | 1 | 1 | 1 | 1 | 1 | 1 | 1 | 1 | 0.97 | 1 | 0.006 |
| SE | 64393 | ENSG00000101577 | LPIN2 | chr18 | - | 2926722 | 2926805 | 2924397 | 2924546 | 2929064 | 2929158 | 5 | 3 | 5 | 0 | 1 | 1 | 0 | 2 | 0 | 0 | 0 | 11 | 5 | 8 | 0 | 0 | 0 | 0 | 1 | 2 | 232 | 149 | 1 | 1 | 0.763 | 1 | 0.616 | NA | 1 | NA | 1 | 1 | 0.837 | 0 | 0.136 |
| SE | 64394 | ENSG00000101577 | LPIN2 | chr18 | - | 2926722 | 2926805 | 2925223 | 2925368 | 2927721 | 2927811 | 229 | 209 | 193 | 216 | 319 | 1 | 0 | 0 | 0 | 0 | 202 | 122 | 204 | 159 | 184 | 0 | 0 | 0 | 0 | 0 | 232 | 149 | 1 | 1 | 0.993 | 1 | 1 | 1 | 1 | 1 | 1 | 1 | 1 | 1 | -0.001 |
| SE | 64396 | ENSG00000101577 | LPIN2 | chr18 | - | 2926722 | 2926805 | 2925223 | 2925368 | 2931255 | 2931443 | 106 | 69 | 84 | 96 | 151 | 0 | 0 | 0 | 1 | 0 | 77 | 49 | 74 | 41 | 46 | 0 | 2 | 0 | 0 | 0 | 232 | 149 | 1 | 1 | 1 | 1 | 1 | 0.984 | 1 | 1 | 0.94 | 1 | 1 | 1 | 0.009 |
| SE | 64397 | ENSG00000101577 | LPIN2 | chr18 | - | 2927721 | 2927811 | 2924397 | 2924546 | 2929064 | 2929158 | 6 | 2 | 8 | 8 | 14 | 1 | 0 | 2 | 0 | 0 | 0 | 3 | 10 | 9 | 0 | 0 | 0 | 0 | 1 | 2 | 239 | 149 | 1 | 1 | 0.789 | 1 | 0.714 | 1 | 1 | NA | 1 | 1 | 0.849 | 0 | 0.188 |
| SE | 64398 | ENSG00000101577 | LPIN2 | chr18 | - | 2927721 | 2927811 | 2925223 | 2925368 | 2929064 | 2929158 | 6 | 2 | 8 | 8 | 14 | 0 | 2 | 2 | 3 | 9 | 0 | 3 | 10 | 7 | 0 | 0 | 0 | 0 | 12 | 2 | 239 | 149 | 1 | 1 | 1 | 0.384 | 0.714 | 0.624 | 0.492 | NA | 1 | 1 | 0.267 | 0 | 0.076 |
| SE | 64401 | ENSG00000101577 | LPIN2 | chr18 | - | 2927721 | 2927811 | 2926722 | 2926805 | 2931255 | 2931443 | 36 | 42 | 44 | 41 | 77 | 0 | 0 | 0 | 2 | 0 | 46 | 13 | 44 | 21 | 54 | 0 | 0 | 0 | 0 | 0 | 239 | 149 | 1 | 1 | 1 | 1 | 1 | 0.927 | 1 | 1 | 1 | 1 | 1 | 1 | -0.015 |
| SE | 64403 | ENSG00000101577 | LPIN2 | chr18 | - | 2929064 | 2929158 | 2925223 | 2925368 | 2931255 | 2931443 | 147 | 125 | 186 | 189 | 124 | 0 | 0 | 0 | 1 | 0 | 124 | 140 | 99 | 134 | 83 | 0 | 2 | 0 | 0 | 0 | 243 | 149 | 1 | 1 | 1 | 1 | 1 | 0.991 | 1 | 1 | 0.977 | 1 | 1 | 1 | 0.003 |
| SE | 64404 | ENSG00000101577 | LPIN2 | chr18 | - | 2929064 | 2929158 | 2926722 | 2926805 | 2931255 | 2931443 | 152 | 124 | 186 | 186 | 115 | 0 | 0 | 0 | 2 | 0 | 124 | 151 | 102 | 130 | 81 | 0 | 0 | 0 | 0 | 0 | 243 | 149 | 1 | 1 | 1 | 1 | 1 | 0.983 | 1 | 1 | 1 | 1 | 1 | 1 | -0.003 |
| SE | 64405 | ENSG00000101577 | LPIN2 | chr18 | - | 2929064 | 2929158 | 2927721 | 2927811 | 2931255 | 2931443 | 157 | 127 | 193 | 201 | 135 | 0 | 0 | 0 | 0 | 0 | 124 | 150 | 119 | 138 | 81 | 0 | 2 | 0 | 4 | 0 | 243 | 149 | 1 | 1 | 1 | 1 | 1 | 1 | 1 | 1 | 0.979 | 1 | 0.955 | 1 | 0.013 |
| SE | 64406 | ENSG00000101577 | LPIN2 | chr18 | - | 2931255 | 2931443 | 2929064 | 2929158 | 2934350 | 2934450 | 486 | 350 | 451 | 508 | 457 | 0 | 0 | 0 | 2 | 0 | 337 | 375 | 280 | 371 | 272 | 0 | 0 | 0 | 0 | 0 | 298 | 149 | 1 | 1 | 1 | 1 | 1 | 0.992 | 1 | 1 | 1 | 1 | 1 | 1 | -0.002 |
| SE | 64407 | ENSG00000101577 | LPIN2 | chr18 | - | 2934350 | 2934450 | 2931255 | 2931443 | 2937691 | 2938037 | 368 | 265 | 337 | 393 | 346 | 0 | 0 | 0 | 0 | 0 | 243 | 261 | 242 | 283 | 200 | 0 | 0 | 0 | 2 | 0 | 249 | 149 | 1 | 1 | 1 | 1 | 1 | 1 | 1 | 1 | 1 | 1 | 0.988 | 1 | 0.002 |
| SE | 64408 | ENSG00000101577 | LPIN2 | chr18 | - | 2937691 | 2938037 | 2934350 | 2934450 | 2939479 | 2939603 | 377 | 250 | 381 | 369 | 430 | 0 | 0 | 0 | 1 | 0 | 178 | 221 | 259 | 221 | 188 | 0 | 0 | 0 | 3 | 0 | 298 | 149 | 1 | 1 | 1 | 1 | 1 | 0.995 | 1 | 1 | 1 | 1 | 0.974 | 1 | 0.004 |
| SE | 64409 | ENSG00000101577 | LPIN2 | chr18 | - | 2939479 | 2939603 | 2937691 | 2938037 | 2940604 | 2940712 | 368 | 238 | 300 | 286 | 596 | 0 | 0 | 0 | 0 | 4 | 228 | 185 | 234 | 165 | 183 | 0 | 0 | 0 | 0 | 0 | 273 | 149 | 1 | 1 | 1 | 1 | 1 | 1 | 0.988 | 1 | 1 | 1 | 1 | 1 | -0.002 |
| SE | 64410 | ENSG00000101577 | LPIN2 | chr18 | - | 2951054 | 2951356 | 2940604 | 2940712 | 2954503 | 2954599 | 483 | 351 | 355 | 440 | 795 | 0 | 0 | 0 | 1 | 1 | 457 | 284 | 370 | 368 | 356 | 0 | 0 | 0 | 0 | 0 | 298 | 149 | 1 | 1 | 1 | 1 | 1 | 0.995 | 0.997 | 1 | 1 | 1 | 1 | 1 | -0.002 |
| SE | 64412 | ENSG00000145241 | CENPC | chr4 | - | 67474887 | 67474978 | 67468761 | 67472675 | 67489966 | 67490121 | 66 | 38 | 62 | 47 | 82 | 0 | 1 | 0 | 1 | 4 | 0 | 20 | 16 | 2 | 0 | 0 | 0 | 0 | 0 | 0 | 240 | 149 | 1 | 1 | 1 | 0.959 | 1 | 0.967 | 0.927 | NA | 1 | 1 | 1 | NA | -0.029 |
| SE | 64413 | ENSG00000145241 | CENPC | chr4 | - | 67492179 | 67492275 | 67489966 | 67490121 | 67492868 | 67492997 | 136 | 97 | 112 | 133 | 129 | 0 | 2 | 0 | 0 | 0 | 9 | 27 | 58 | 6 | 0 | 0 | 0 | 0 | 0 | 0 | 245 | 149 | 1 | 1 | 1 | 0.967 | 1 | 1 | 1 | 1 | 1 | 1 | 1 | NA | -0.007 |
| SE | 64414 | ENSG00000145241 | CENPC | chr4 | - | 67495158 | 67495212 | 67493404 | 67493988 | 67505204 | 67505284 | 195 | 183 | 183 | 283 | 368 | 0 | 0 | 0 | 0 | 0 | 69 | 123 | 108 | 27 | 32 | 0 | 4 | 1 | 0 | 0 | 203 | 149 | 1 | 1 | 1 | 1 | 1 | 1 | 1 | 1 | 0.958 | 0.988 | 1 | 1 | 0.011 |
| SE | 64415 | ENSG00000145241 | CENPC | chr4 | - | 67505204 | 67505284 | 67495158 | 67495212 | 67506787 | 67506934 | 231 | 221 | 241 | 328 | 479 | 0 | 0 | 0 | 0 | 0 | 73 | 137 | 152 | 48 | 30 | 0 | 3 | 0 | 0 | 0 | 229 | 149 | 1 | 1 | 1 | 1 | 1 | 1 | 1 | 1 | 0.967 | 1 | 1 | 1 | 0.007 |
| SE | 64416 | ENSG00000145241 | CENPC | chr4 | - | 67508813 | 67509105 | 67506787 | 67506934 | 67512401 | 67512569 | 347 | 277 | 325 | 349 | 751 | 0 | 0 | 0 | 0 | 6 | 21 | 145 | 166 | 26 | 15 | 0 | 0 | 0 | 0 | 0 | 298 | 149 | 1 | 1 | 1 | 1 | 1 | 1 | 0.984 | 1 | 1 | 1 | 1 | 1 | -0.003 |
| SE | 64417 | ENSG00000145241 | CENPC | chr4 | - | 67510672 | 67511025 | 67508813 | 67509105 | 67512401 | 67512569 | 4 | 0 | 0 | 3 | 0 | 145 | 91 | 120 | 144 | 291 | 0 | 2 | 2 | 0 | 0 | 0 | 50 | 67 | 4 | 4 | 298 | 149 | 1 | 1 | 0.014 | 0 | 0 | 0.01 | 0 | NA | 0.02 | 0.015 | 0 | 0 | -0.004 |
| SE | 64419 | ENSG00000145241 | CENPC | chr4 | - | 67540979 | 67541050 | 67539839 | 67539934 | 67544148 | 67544195 | 232 | 195 | 225 | 364 | 284 | 0 | 0 | 0 | 0 | 0 | 83 | 285 | 204 | 111 | 62 | 0 | 0 | 0 | 0 | 0 | 220 | 149 | 1 | 1 | 1 | 1 | 0.993 | 1 | 1 | 1 | 1 | 1 | 1 | 1 | -0.001 |
| SE | 64422 | ENSG00000143919 | CAMKMT | chr2 | + | 44704282 | 44704343 | 44549554 | 44549728 | 44707398 | 44707462 | 3 | 2 | 2 | 3 | 0 | 0 | 0 | 0 | 0 | 0 | 0 | 12 | 4 | 6 | 0 | 0 | 0 | 3 | 0 | 0 | 210 | 149 | 1 | 1 | 1 | 1 | 1 | 1 | NA | NA | 1 | 0.486 | 1 | NA | 0.171 |
| SE | 64425 | ENSG00000143919 | CAMKMT | chr2 | + | 44715286 | 44715353 | 44704282 | 44704343 | 44743621 | 44743696 | 12 | 4 | 5 | 10 | 15 | 0 | 5 | 1 | 0 | 5 | 2 | 4 | 8 | 0 | 0 | 0 | 4 | 0 | 0 | 3 | 216 | 149 | 1 | 1 | 1 | 0.356 | 0.775 | 1 | 0.674 | 1 | 0.408 | 1 | NA | 0 | 0.159 |
| SE | 64426 | ENSG00000143919 | CAMKMT | chr2 | + | 44723728 | 44723816 | 44715286 | 44715353 | 44743621 | 44743657 | 2 | 1 | 0 | 0 | 0 | 23 | 9 | 16 | 14 | 53 | 0 | 0 | 0 | 0 | 0 | 17 | 20 | 19 | 14 | 5 | 237 | 149 | 1 | 1 | 0.052 | 0.065 | 0 | 0 | 0 | 0 | 0 | 0 | 0 | 0 | 0.023 |
| SE | 64427 | ENSG00000143919 | CAMKMT | chr2 | + | 44743621 | 44743696 | 44704282 | 44704343 | 44754054 | 44754118 | 4 | 16 | 9 | 9 | 18 | 0 | 0 | 0 | 0 | 0 | 10 | 27 | 6 | 7 | 7 | 2 | 0 | 0 | 0 | 0 | 224 | 149 | 1 | 1 | 1 | 1 | 1 | 1 | 1 | 0.769 | 1 | 1 | 1 | 1 | 0.046 |
| SE | 64431 | ENSG00000162813 | BPNT1 | chr1 | - | 220067029 | 220067125 | 220062798 | 220062954 | 220067301 | 220067393 | 0 | 0 | 0 | 0 | 0 | 96 | 46 | 80 | 71 | 93 | 0 | 2 | 0 | 0 | 0 | 62 | 74 | 37 | 56 | 98 | 245 | 149 | 1 | 1 | 0 | 0 | 0 | 0 | 0 | 0 | 0.016 | 0 | 0 | 0 | -0.003 |
| SE | 64435 | ENSG00000254415 | SIGLEC14 | chr19 | - | 51645476 | 51645530 | 51643778 | 51644036 | 51645781 | 51646060 | 34 | 0 | 0 | 123 | 14 | 0 | 0 | 0 | 0 | 0 | 72 | 57 | 40 | 93 | 84 | 0 | 0 | 0 | 1 | 0 | 203 | 149 | 1 | 1 | 1 | NA | NA | 1 | 1 | 1 | 1 | 1 | 0.986 | 1 | 0.003 |
| SE | 64437 | ENSG00000189403 | HMGB1 | chr13 | - | 30463206 | 30463352 | 30462537 | 30462712 | 30463530 | 30463694 | 2254 | 1635 | 2480 | 2979 | 3379 | 2 | 1 | 1 | 5 | 0 | 1754 | 2457 | 2268 | 1648 | 848 | 0 | 4 | 0 | 0 | 0 | 295 | 149 | 1 | 1 | 0.998 | 0.999 | 0.999 | 0.997 | 1 | 1 | 0.997 | 1 | 1 | 1 | -0.001 |
| SE | 64438 | ENSG00000160352 | ZNF714 | chr19 | + | 21083977 | 21084069 | 21082173 | 21082348 | 21098184 | 21098311 | 11 | 1 | 18 | 5 | 4 | 0 | 0 | 1 | 0 | 0 | 0 | 12 | 15 | 13 | 0 | 13 | 0 | 3 | 0 | 0 | 241 | 149 | 1 | 1 | 1 | 1 | 0.918 | 1 | 1 | 0 | 1 | 0.756 | 1 | NA | 0.295 |
| SE | 64440 | ENSG00000160352 | ZNF714 | chr19 | + | 21098811 | 21098910 | 21098184 | 21098311 | 21116806 | 21118885 | 21 | 10 | 17 | 8 | 23 | 0 | 0 | 4 | 0 | 6 | 0 | 6 | 1 | 1 | 0 | 0 | 0 | 13 | 10 | 0 | 248 | 149 | 1 | 1 | 1 | 1 | 0.719 | 1 | 0.697 | NA | 1 | 0.044 | 0.057 | NA | 0.516 |
| SE | 64441 | ENSG00000160352 | ZNF714 | chr19 | + | 21098811 | 21098910 | 21098184 | 21098311 | 21116828 | 21118477 | 11 | 1 | 6 | 5 | 13 | 0 | 0 | 3 | 0 | 6 | 0 | 6 | 7 | 4 | 0 | 0 | 0 | 2 | 0 | 0 | 248 | 149 | 1 | 1 | 1 | 1 | 0.546 | 1 | 1 | NA | 1 | 0.678 | 1 | NA | 0.017 |
| SE | 64442 | ENSG00000160352 | ZNF714 | chr19 | + | 21098814 | 21098910 | 21098184 | 21098311 | 21116806 | 21117084 | 21 | 12 | 16 | 10 | 15 | 0 | 0 | 4 | 0 | 6 | 0 | 11 | 7 | 5 | 0 | 0 | 0 | 13 | 10 | 0 | 245 | 149 | 1 | 1 | 1 | 1 | 0.709 | 1 | 0.603 | NA | 1 | 0.247 | 0.233 | NA | 0.369 |
| SE | 64443 | ENSG00000160352 | ZNF714 | chr19 | + | 21109622 | 21109644 | 21098814 | 21098910 | 21116806 | 21118885 | 1 | 0 | 0 | 0 | 8 | 16 | 9 | 14 | 5 | 22 | 0 | 0 | 0 | 0 | 0 | 0 | 7 | 4 | 0 | 0 | 171 | 149 | 1 | 1 | 0.052 | 0 | 0 | 0 | 0.241 | NA | 0 | 0 | NA | NA | 0.059 |
| SE | 64446 | ENSG00000145569 | OTULINL | chr5 | + | 14608747 | 14609017 | 14607329 | 14607458 | 14610140 | 14616180 | 480 | 394 | 349 | 455 | 1178 | 3 | 0 | 0 | 0 | 0 | 426 | 470 | 480 | 725 | 107 | 0 | 0 | 0 | 0 | 0 | 298 | 149 | 1 | 1 | 0.988 | 1 | 1 | 1 | 1 | 1 | 1 | 1 | 1 | 1 | -0.002 |
| SE | 64453 | ENSG00000065150 | IPO5 | chr13 | + | 97981241 | 97981400 | 97976692 | 97976786 | 97982502 | 97982569 | 2 | 0 | 0 | 0 | 0 | 98 | 74 | 132 | 102 | 58 | 0 | 0 | 1 | 0 | 0 | 43 | 73 | 63 | 42 | 17 | 298 | 149 | 1 | 1 | 0.01 | 0 | 0 | 0 | 0 | 0 | 0 | 0.008 | 0 | 0 | 0 |
| SE | 64458 | ENSG00000065150 | IPO5 | chr13 | + | 98007361 | 98007587 | 98006129 | 98006348 | 98008058 | 98008142 | 0 | 1 | 0 | 6 | 1 | 108 | 112 | 112 | 105 | 145 | 0 | 0 | 0 | 0 | 0 | 34 | 67 | 109 | 57 | 13 | 298 | 149 | 1 | 1 | 0 | 0.004 | 0 | 0.028 | 0.003 | 0 | 0 | 0 | 0 | 0 | 0.007 |
| SE | 64459 | ENSG00000065150 | IPO5 | chr13 | + | 98016728 | 98016851 | 98015725 | 98015781 | 98018484 | 98018704 | 427 | 308 | 444 | 404 | 969 | 0 | 0 | 0 | 0 | 4 | 124 | 295 | 257 | 210 | 52 | 0 | 0 | 0 | 0 | 0 | 272 | 149 | 1 | 1 | 1 | 1 | 1 | 1 | 0.993 | 1 | 1 | 1 | 1 | 1 | -0.001 |
| SE | 64462 | ENSG00000135046 | ANXA1 | chr9 | + | 73163475 | 73163532 | 73162781 | 73162861 | 73165115 | 73165209 | 3160 | 1870 | 1953 | 2729 | 4488 | 3 | 2 | 3 | 0 | 1 | 575 | 1731 | 1391 | 1046 | 227 | 0 | 0 | 0 | 0 | 0 | 206 | 149 | 1 | 1 | 0.999 | 0.999 | 0.998 | 1 | 1 | 1 | 1 | 1 | 1 | 1 | -0.001 |
| SE | 64463 | ENSG00000135046 | ANXA1 | chr9 | + | 73166096 | 73166192 | 73165115 | 73165209 | 73167496 | 73167555 | 2100 | 1195 | 1393 | 1807 | 3681 | 0 | 0 | 0 | 0 | 3 | 155 | 981 | 792 | 522 | 73 | 0 | 0 | 0 | 0 | 0 | 245 | 149 | 1 | 1 | 1 | 1 | 1 | 1 | 0.999 | 1 | 1 | 1 | 1 | 1 | 0 |
| SE | 64464 | ENSG00000233369 | GTF2IP4 | chr7 | + | 73183114 | 73183298 | 73182076 | 73182148 | 73184660 | 73184719 | 72 | 24 | 46 | 64 | 54 | 0 | 0 | 2 | 0 | 0 | 66 | 48 | 24 | 82 | 19 | 0 | 0 | 0 | 0 | 0 | 298 | 149 | 1 | 1 | 1 | 1 | 0.92 | 1 | 1 | 1 | 1 | 1 | 1 | 1 | -0.016 |
| SE | 64466 | ENSG00000233369 | GTF2IP4 | chr7 | + | 73202111 | 73202140 | 73200448 | 73200632 | 73203422 | 73203464 | 364 | 244 | 261 | 330 | 519 | 0 | 0 | 0 | 1 | 0 | 427 | 375 | 296 | 423 | 198 | 0 | 0 | 0 | 0 | 0 | 178 | 149 | 1 | 1 | 1 | 1 | 1 | 0.996 | 1 | 1 | 1 | 1 | 1 | 1 | -0.001 |
| SE | 64470 | ENSG00000256223 | ZNF10 | chr12 | + | 133139198 | 133139249 | 133130836 | 133131014 | 133143519 | 133143622 | 0 | 0 | 0 | 0 | 0 | 0 | 1 | 0 | 0 | 0 | 0 | 0 | 0 | 0 | 14 | 0 | 0 | 0 | 0 | 0 | 200 | 149 | 1 | 1 | NA | 0 | NA | NA | NA | NA | NA | NA | NA | 1 | -1 |
| SE | 64473 | ENSG00000256223 | ZNF10 | chr12 | + | 133143519 | 133143622 | 133139198 | 133139249 | 133144433 | 133144525 | 0 | 1 | 1 | 0 | 1 | 0 | 1 | 0 | 1 | 0 | 0 | 0 | 0 | 0 | 14 | 0 | 0 | 0 | 0 | 0 | 252 | 149 | 1 | 1 | NA | 0.372 | 1 | 0 | 1 | NA | NA | NA | NA | 1 | -0.407 |
| SE | 64475 | ENSG00000256223 | ZNF10 | chr12 | + | 133151027 | 133151154 | 133143868 | 133144525 | 133155502 | 133155689 | 6 | 8 | 6 | 3 | 9 | 0 | 2 | 0 | 0 | 0 | 0 | 2 | 3 | 0 | 0 | 0 | 0 | 0 | 0 | 0 | 276 | 149 | 1 | 1 | 1 | 0.683 | 1 | 1 | 1 | NA | 1 | 1 | NA | NA | -0.063 |

| | | | | | | | | | | | | |
|---|---|---|---|---|---|---|---|---|---|---|---|---|
| SE | 64476 | ENSG00000256223 | ZNF10 | chr12 | + | 133151808 | 133151904 | 133150986 | 133151154 | 133155502 | 133155689 | 18 | 13 |
| SE | 64478 | ENSG00000146263 | MMS22L | chr6 | - | 97181903 | 97182054 | 97179407 | 97179559 | 97186496 | 97186690 | 11 | 19 |
| SE | 64479 | ENSG00000146263 | MMS22L | chr6 | - | 97233860 | 97233980 | 97230268 | 97231652 | 97246627 | 97246690 | 59 | 34 |
| SE | 64480 | ENSG00000146263 | MMS22L | chr6 | - | 97263334 | 97263448 | 97253464 | 97254733 | 97267871 | 97268002 | 32 | 24 |
| SE | 64481 | ENSG00000146263 | MMS22L | chr6 | - | 97270106 | 97270283 | 97269901 | 97269992 | 97272703 | 97272881 | 4 | 3 |
| SE | 64482 | ENSG00000146263 | MMS22L | chr6 | - | 97278848 | 97278898 | 97272974 | 97273062 | 97281236 | 97281304 | 46 | 48 |
| SE | 64484 | ENSG00000118412 | CASP8AP2 | chr6 | + | 89846561 | 89846644 | 89829893 | 89829934 | 89853165 | 89853234 | 143 | 135 |
| SE | 64485 | ENSG00000118412 | CASP8AP2 | chr6 | + | 89846561 | 89846644 | 89829899 | 89830067 | 89853165 | 89853234 | 168 | 162 |
| SE | 64486 | ENSG00000118412 | CASP8AP2 | chr6 | + | 89846561 | 89846644 | 89829938 | 89830106 | 89853165 | 89853234 | 156 | 141 |
| SE | 64487 | ENSG00000118412 | CASP8AP2 | chr6 | + | 89854816 | 89854917 | 89853165 | 89853234 | 89855469 | 89855544 | 399 | 412 |
| SE | 64489 | ENSG00000118412 | CASP8AP2 | chr6 | + | 89857083 | 89857199 | 89855469 | 89855544 | 89862125 | 89862793 | 237 | 192 |
| SE | 64490 | ENSG00000172116 | CD8B | chr2 | - | 86822320 | 86822374 | 86815346 | 86815718 | 86844921 | 86844958 | 5 | 5 |
| SE | 64491 | ENSG00000172116 | CD8B | chr2 | - | 86844921 | 86844958 | 86815449 | 86815718 | 86852996 | 86853086 | 3 | 0 |
| SE | 64492 | ENSG00000172116 | CD8B | chr2 | - | 86844921 | 86844958 | 86838166 | 86842319 | 86846683 | 86846773 | 247 | 195 |
| SE | 64493 | ENSG00000172116 | CD8B | chr2 | - | 86844921 | 86844958 | 86838166 | 86842319 | 86852996 | 86853086 | 59 | 53 |
| SE | 64494 | ENSG00000172116 | CD8B | chr2 | - | 86846683 | 86846773 | 86838166 | 86842319 | 86852996 | 86853086 | 66 | 41 |
| SE | 64496 | ENSG00000172116 | CD8B | chr2 | - | 86852996 | 86853086 | 86815338 | 86815718 | 86858056 | 86858416 | 122 | 99 |
| SE | 64497 | ENSG00000172116 | CD8B | chr2 | - | 86852996 | 86853086 | 86846683 | 86846773 | 86858056 | 86858416 | 318 | 232 |
| SE | 64499 | ENSG00000263002 | ZNF234 | chr19 | + | 44149770 | 44149976 | 44148770 | 44148897 | 44150412 | 44150505 | 0 | 0 |
| SE | 64500 | ENSG00000263002 | ZNF234 | chr19 | + | 44150412 | 44150505 | 44148770 | 44148897 | 44156251 | 44158119 | 48 | 32 |
| SE | 64501 | ENSG00000105750 | ZNF85 | chr19 | + | 20934023 | 20934150 | 20923221 | 20923403 | 20934948 | 20935047 | 8 | 8 |
| SE | 64502 | ENSG00000105750 | ZNF85 | chr19 | + | 20934023 | 20934150 | 20923221 | 20923403 | 20948743 | 20950302 | 3 | 0 |
| SE | 64504 | ENSG00000105750 | ZNF85 | chr19 | + | 20934948 | 20935047 | 20934023 | 20934150 | 20946299 | 20946389 | 4 | 6 |
| SE | 64505 | ENSG00000105750 | ZNF85 | chr19 | + | 20934948 | 20935047 | 20934023 | 20934150 | 20948743 | 20949093 | 6 | 7 |
| SE | 64506 | ENSG00000105750 | ZNF85 | chr19 | + | 20934965 | 20935047 | 20934023 | 20934150 | 20948743 | 20948915 | 1 | 0 |
| SE | 64507 | ENSG00000105750 | ZNF85 | chr19 | + | 20934969 | 20935047 | 20934023 | 20934150 | 20948743 | 20948893 | 1 | 0 |
| SE | 64508 | ENSG00000105750 | ZNF85 | chr19 | + | 20944251 | 20944296 | 20934969 | 20935047 | 20946299 | 20946340 | 2 | 3 |
| SE | 64510 | ENSG00000105750 | ZNF85 | chr19 | + | 20946299 | 20946389 | 20934023 | 20934150 | 20948743 | 20948791 | 5 | 0 |
| SE | 64511 | ENSG00000105750 | ZNF85 | chr19 | + | 20946299 | 20946389 | 20934948 | 20935047 | 20948743 | 20948791 | 4 | 1 |
| SE | 64515 | ENSG00000166801 | FAM111A | chr11 | + | 59143496 | 59143695 | 59143082 | 59143304 | 59144808 | 59144896 | 58 | 40 |
| SE | 64516 | ENSG00000166801 | FAM111A | chr11 | + | 59143496 | 59143695 | 59143082 | 59143304 | 59145826 | 59145856 | 76 | 69 |
| SE | 64518 | ENSG00000166801 | FAM111A | chr11 | + | 59143496 | 59145856 | 59143082 | 59143304 | 59148835 | 59148953 | 106 | 133 |
| SE | 64519 | ENSG00000166801 | FAM111A | chr11 | + | 59143496 | 59145856 | 59143082 | 59143304 | 59151749 | 59155039 | 54 | 38 |
| SE | 64520 | ENSG00000166801 | FAM111A | chr11 | + | 59144537 | 59144896 | 59143496 | 59143695 | 59145826 | 59145856 | 17 | 20 |
| SE | 64521 | ENSG00000166801 | FAM111A | chr11 | + | 59144808 | 59144896 | 59143082 | 59143304 | 59145826 | 59145856 | 8 | 15 |
| SE | 64523 | ENSG00000166801 | FAM111A | chr11 | + | 59145826 | 59145856 | 59143082 | 59143304 | 59148796 | 59148953 | 58 | 65 |
| SE | 64524 | ENSG00000166801 | FAM111A | chr11 | + | 59145826 | 59145856 | 59143082 | 59143304 | 59148835 | 59148953 | 27 | 43 |
| SE | 64525 | ENSG00000166801 | FAM111A | chr11 | + | 59145826 | 59145856 | 59143082 | 59143304 | 59151749 | 59153504 | 4 | 5 |
| SE | 64526 | ENSG00000166801 | FAM111A | chr11 | + | 59145826 | 59145856 | 59143496 | 59143695 | 59148703 | 59148953 | 4 | 9 |
| SE | 64527 | ENSG00000166801 | FAM111A | chr11 | + | 59145826 | 59145856 | 59143496 | 59143695 | 59148835 | 59148953 | 22 | 30 |
| SE | 64528 | ENSG00000166801 | FAM111A | chr11 | + | 59145826 | 59145856 | 59144537 | 59144896 | 59148835 | 59148953 | 14 | 26 |
| SE | 64529 | ENSG00000166801 | FAM111A | chr11 | + | 59148703 | 59148953 | 59145826 | 59145856 | 59151749 | 59153504 | 197 | 256 |
| SE | 64531 | ENSG00000166801 | FAM111A | chr11 | + | 59148796 | 59148953 | 59144861 | 59145856 | 59151749 | 59153504 | 260 | 344 |
| SE | 64533 | ENSG00000166801 | FAM111A | chr11 | + | 59148835 | 59148953 | 59143496 | 59145856 | 59151749 | 59153504 | 273 | 368 |
| SE | 64535 | ENSG00000066455 | GOLGA5 | chr14 | + | 92810253 | 92810377 | 92809299 | 92809519 | 92811550 | 92811754 | 159 | 95 |
| SE | 64536 | ENSG00000066455 | GOLGA5 | chr14 | + | 92816250 | 92816421 | 92811550 | 92811754 | 92819707 | 92819836 | 379 | 322 |
| SE | 64537 | ENSG00000066455 | GOLGA5 | chr14 | + | 92819707 | 92819836 | 92816250 | 92816421 | 92824545 | 92824644 | 226 | 202 |
| SE | 64538 | ENSG00000066455 | GOLGA5 | chr14 | + | 92833121 | 92833347 | 92824545 | 92824644 | 92835558 | 92835664 | 154 | 113 |
| SE | 64539 | ENSG00000066455 | GOLGA5 | chr14 | + | 92835558 | 92835664 | 92833121 | 92833347 | 92837385 | 92837449 | 168 | 96 |
| SE | 64540 | ENSG00000066455 | GOLGA5 | chr14 | + | 92837385 | 92837449 | 92835558 | 92835664 | 92839365 | 92839947 | 131 | 86 |
| SE | 64541 | ENSG00000147533 | GOLGA7 | chr8 | + | 41490832 | 41490965 | 41490481 | 41490617 | 41497508 | 41497661 | 153 | 82 |
| SE | 64542 | ENSG00000147533 | GOLGA7 | chr8 | + | 41490869 | 41490965 | 41490561 | 41490617 | 41497508 | 41497661 | 124 | 61 |
| SE | 64543 | ENSG00000198677 | TTC37 | chr5 | - | 95484723 | 95484845 | 95482472 | 95482631 | 95490907 | 95491036 | 216 | 169 |
| SE | 64544 | ENSG00000198677 | TTC37 | chr5 | - | 95498369 | 95498591 | 95497415 | 95497488 | 95502879 | 95503003 | 244 | 188 |
| SE | 64545 | ENSG00000198677 | TTC37 | chr5 | - | 95503813 | 95503938 | 95502879 | 95503003 | 95506933 | 95506992 | 298 | 233 |
| SE | 64547 | ENSG00000198677 | TTC37 | chr5 | - | 95525586 | 95525673 | 95525397 | 95525501 | 95528012 | 95528152 | 186 | 113 |
| SE | 64548 | ENSG00000198677 | TTC37 | chr5 | - | 95528012 | 95528152 | 95525586 | 95525673 | 95529000 | 95529102 | 213 | 146 |
| SE | 64549 | ENSG00000198677 | TTC37 | chr5 | - | 95543183 | 95543327 | 95541814 | 95541906 | 95547060 | 95547151 | 396 | 344 |
| SE | 64550 | ENSG00000198677 | TTC37 | chr5 | - | 95548559 | 95548715 | 95546973 | 95547151 | 95550580 | 95550650 | 270 | 193 |
| SE | 64551 | ENSG00000198677 | TTC37 | chr5 | - | 95553006 | 95553228 | 95550580 | 95550650 | 95554935 | 95554977 | 1 | 0 |
| SE | 64552 | ENSG00000144036 | EXOC6B | chr2 | - | 72182893 | 72182905 | 72179308 | 72179461 | 72184074 | 72184187 | 1 | 3 |
| SE | 64553 | ENSG00000144036 | EXOC6B | chr2 | - | 72480615 | 72480750 | 72461475 | 72465339 | 72492317 | 72492429 | 99 | 102 |
| SE | 64554 | ENSG00000144036 | EXOC6B | chr2 | - | 72498453 | 72498551 | 72496453 | 72496559 | 72499900 | 72499972 | 17 | 13 |
| SE | 64555 | ENSG00000144036 | EXOC6B | chr2 | - | 72513131 | 72513252 | 72499900 | 72499972 | 72514633 | 72514680 | 86 | 63 |
| SE | 64556 | ENSG00000144036 | EXOC6B | chr2 | - | 72514633 | 72514680 | 72513131 | 72513252 | 72515042 | 72515126 | 83 | 75 |
| SE | 64557 | ENSG00000144036 | EXOC6B | chr2 | - | 72559452 | 72559521 | 72515042 | 72515126 | 72575491 | 72575668 | 66 | 82 |
| SE | 64558 | ENSG00000144036 | EXOC6B | chr2 | - | 72718102 | 72718307 | 72575491 | 72575668 | 72731006 | 72731052 | 142 | 128 |
| SE | 64559 | ENSG00000144036 | EXOC6B | chr2 | - | 72731006 | 72731052 | 72718102 | 72718307 | 72741303 | 72741469 | 29 | 14 |
| SE | 64561 | ENSG00000144036 | EXOC6B | chr2 | - | 72733070 | 72733118 | 72731154 | 72731245 | 72741303 | 72741469 | 119 | 112 |
| SE | 64562 | ENSG00000144036 | EXOC6B | chr2 | - | 72741303 | 72741469 | 72718102 | 72718307 | 72825797 | 72826019 | 28 | 20 |
| SE | 64564 | ENSG00000213185 | FAM24B | chr10 | - | 122850423 | 122850520 | 122849095 | 122849439 | 122855644 | 122855786 | 12 | 10 |
| SE | 64567 | ENSG00000145214 | DGKQ | chr4 | - | 962771 | 962920 | 962434 | 962613 | 963138 | 963290 | 40 | 27 |
| SE | 64568 | ENSG00000145214 | DGKQ | chr4 | - | 964072 | 964093 | 963141 | 963290 | 965175 | 965291 | 2 | 1 |
| SE | 64570 | ENSG00000145214 | DGKQ | chr4 | - | 965927 | 966078 | 965490 | 965529 | 966465 | 966527 | 31 | 9 |
| SE | 64571 | ENSG00000145214 | DGKQ | chr4 | - | 966063 | 966301 | 965490 | 965529 | 966465 | 966527 | 1 | 3 |
| SE | 64575 | ENSG00000135624 | CCT7 | chr2 | + | 73237551 | 73237687 | 73234335 | 73234384 | 73239642 | 73239796 | 2 | 1 |
| SE | 64577 | ENSG00000135624 | CCT7 | chr2 | + | 73242972 | 73243129 | 73234335 | 73234384 | 73244543 | 73244715 | 2 | 1 |
| SE | 64578 | ENSG00000135624 | CCT7 | chr2 | + | 73242972 | 73243129 | 73234335 | 73234384 | 73247761 | 73247926 | 0 | 0 |
| SE | 64579 | ENSG00000135624 | CCT7 | chr2 | + | 73242972 | 73243129 | 73240436 | 73240543 | 73247761 | 73247926 | 1 | 0 |
| SE | 64580 | ENSG00000135624 | CCT7 | chr2 | + | 73243003 | 73243129 | 73234235 | 73234384 | 73244543 | 73244630 | 2 | 1 |
| SE | 64581 | ENSG00000135624 | CCT7 | chr2 | + | 73243003 | 73243129 | 73234235 | 73234384 | 73247761 | 73247926 | 0 | 0 |
| SE | 64582 | ENSG00000135624 | CCT7 | chr2 | + | 73243003 | 73243129 | 73240436 | 73240543 | 73247761 | 73247926 | 105 | 53 |
| SE | 64583 | ENSG00000135624 | CCT7 | chr2 | + | 73243996 | 73244049 | 73242972 | 73243129 | 73244543 | 73244630 | 251 | 151 |
| SE | 64584 | ENSG00000135624 | CCT7 | chr2 | + | 73244543 | 73244715 | 73234235 | 73234384 | 73247761 | 73247789 | 418 | 211 |
| SE | 64586 | ENSG00000135624 | CCT7 | chr2 | + | 73247761 | 73247926 | 73244543 | 73244715 | 73248990 | 73249061 | 905 | 563 |
| SE | 64587 | ENSG00000135624 | CCT7 | chr2 | + | 73248990 | 73249179 | 73247761 | 73247926 | 73249818 | 73249844 | 840 | 614 |
| SE | 64588 | ENSG00000135624 | CCT7 | chr2 | + | 73248990 | 73249179 | 73247761 | 73247926 | 73250305 | 73250438 | 488 | 352 |
| SE | 64589 | ENSG00000135624 | CCT7 | chr2 | + | 73249818 | 73249916 | 73247761 | 73247926 | 73250305 | 73250438 | 325 | 231 |
| SE | 64590 | ENSG00000135624 | CCT7 | chr2 | + | 73249818 | 73249916 | 73248990 | 73249179 | 73250305 | 73250438 | 673 | 482 |
| SE | 64598 | ENSG00000100644 | HIF1A | chr14 | + | 61727452 | 61727655 | 61726705 | 61726818 | 61732417 | 61732524 | 678 | 572 |
| SE | 64599 | ENSG00000100644 | HIF1A | chr14 | + | 61738086 | 61738373 | 61736888 | 61737109 | 61740504 | 61740627 | 756 | 465 |
| SE | 64601 | ENSG00000100644 | HIF1A | chr14 | + | 61738086 | 61738373 | 61736888 | 61737109 | 61740754 | 61741188 | 394 | 273 |
| SE | 64602 | ENSG00000100644 | HIF1A | chr14 | + | 61738086 | 61738373 | 61736888 | 61737109 | 61744704 | 61744813 | 394 | 274 |
| SE | 64603 | ENSG00000100644 | HIF1A | chr14 | + | 61740504 | 61740627 | 61736888 | 61737109 | 61740754 | 61741188 | 214 | 138 |
| SE | 64604 | ENSG00000100644 | HIF1A | chr14 | + | 61740504 | 61740627 | 61738086 | 61738373 | 61740754 | 61741188 | 558 | 325 |
| SE | 64605 | ENSG00000100644 | HIF1A | chr14 | + | 61740542 | 61740627 | 61736889 | 61737109 | 61740754 | 61741188 | 180 | 110 |
| SE | 64607 | ENSG00000100644 | HIF1A | chr14 | + | 61740754 | 61741188 | 61736888 | 61737109 | 61744704 | 61744813 | 437 | 339 |
| SE | 64608 | ENSG00000100644 | HIF1A | chr14 | + | 61740754 | 61741188 | 61738086 | 61738373 | 61744704 | 61744813 | 437 | 340 |
| SE | 64611 | ENSG00000170802 | FOXN2 | chr2 | + | 48346200 | 48346751 | 48314636 | 48314814 | 48373291 | 48373328 | 49 | 35 |
| SE | 64612 | ENSG00000170802 | FOXN2 | chr2 | + | 48359046 | 48359147 | 48346200 | 48346751 | 48362642 | 48362707 | 279 | 183 |
| SE | 64613 | ENSG00000170802 | FOXN2 | chr2 | + | 48359046 | 48359147 | 48346200 | 48346751 | 48373291 | 48373328 | 94 | 66 |
| SE | 64614 | ENSG00000170802 | FOXN2 | chr2 | + | 48362642 | 48362707 | 48346200 | 48346751 | 48373291 | 48373328 | 36 | 33 |
| SE | 64615 | ENSG00000170802 | FOXN2 | chr2 | + | 48362642 | 48362707 | 48359046 | 48359147 | 48373291 | 48373328 | 198 | 141 |
| SE | 64617 | ENSG00000148737 | TCF7L2 | chr10 | + | 112951482 | 112951607 | 112951206 | 112951273 | 113040024 | 113040126 | 95 | 130 |
| SE | 64620 | ENSG00000148737 | TCF7L2 | chr10 | + | 113089396 | 113089540 | 113040024 | 113040126 | 113141183 | 113141316 | 1 | 1 |
| SE | 64622 | ENSG00000148737 | TCF7L2 | chr10 | + | 113158666 | 113158717 | 113158020 | 113158069 | 113159919 | 113159992 | 3 | 5 |
| SE | 64632 | ENSG00000148737 | TCF7L2 | chr10 | + | 113160618 | 113160691 | 113159919 | 113159992 | 113165554 | 113165972 | 5 | 5 |
| SE | 64634 | ENSG00000148737 | TCF7L2 | chr10 | + | 113161578 | 113161603 | 113160618 | 113160691 | 113165554 | 113165612 | 0 | 2 |
| SE | 64638 | ENSG00000230773 | NA | chr2 | - | 47996715 | 47996790 | 47943530 | 47943612 | 48003299 | 48003425 | 1 | 0 |
| SE | 64641 | ENSG00000230773 | NA | chr2 | - | 48243468 | 48243619 | 48164887 | 48164934 | 48313689 | 48313764 | 1 | 3 |
| SE | 64645 | ENSG00000198755 | RPL10A | chr6 | + | 35468798 | 35468873 | 35468400 | 35468439 | 35468946 | 35469027 | 1810 | 648 |
| SE | 64646 | ENSG00000198755 | RPL10A | chr6 | + | 35468946 | 35469529 | 35468407 | 35468439 | 35470178 | 35470351 | 4680 | 2278 |
| SE | 64667 | ENSG00000167220 | HDHD2 | chr18 | - | 47130243 | 47130328 | 47115130 | 47115348 | 47134495 | 47134677 | 206 | 232 |
| SE | 64668 | ENSG00000167220 | HDHD2 | chr18 | - | 47130243 | 47130328 | 47115130 | 47115348 | 47150377 | 47150459 | 55 | 67 |
| SE | 64669 | ENSG00000167220 | HDHD2 | chr18 | - | 47134495 | 47134677 | 47130243 | 47130328 | 47136301 | 47136449 | 153 | 165 |
| SE | 64670 | ENSG00000167220 | HDHD2 | chr18 | - | 47134495 | 47134677 | 47130243 | 47130328 | 47136338 | 47136449 | 155 | 168 |
| SE | 64671 | ENSG00000167220 | HDHD2 | chr18 | - | 47134495 | 47134704 | 47130241 | 47130328 | 47136301 | 47136449 | 167 | 181 |
| SE | 64672 | ENSG00000167220 | HDHD2 | chr18 | - | 47134495 | 47134704 | 47130241 | 47130328 | 47136338 | 47136449 | 191 | 199 |
| SE | 64673 | ENSG00000167220 | HDHD2 | chr18 | - | 47136301 | 47136449 | 47130243 | 47130328 | 47150377 | 47150500 | 15 | 8 |
| SE | 64674 | ENSG00000167220 | HDHD2 | chr18 | - | 47136338 | 47136449 | 47130241 | 47130328 | 47150377 | 47150498 | 13 | 7 |
| SE | 64675 | ENSG00000167220 | HDHD2 | chr18 | - | 47137234 | 47137296 | 47136301 | 47136449 | 47150377 | 47150476 | 2 | 1 |
| SE | 64677 | ENSG00000167220 | HDHD2 | chr18 | - | 47149048 | 47149108 | 47136338 | 47136449 | 47150377 | 47150479 | 0 | 0 |
| SE | 64682 | ENSG00000136715 | SAP130 | chr2 | - | 127996349 | 127996491 | 127993186 | 127993308 | 127999740 | 127999845 | 45 | 35 |
| SE | 64683 | ENSG00000136715 | SAP130 | chr2 | - | 127996349 | 127996491 | 127993186 | 127993308 | 128000055 | 128000146 | 22 | 12 |
| SE | 64684 | ENSG00000136715 | SAP130 | chr2 | - | 127999740 | 127999845 | 127993186 | 127993308 | 128000055 | 128000146 | 32 | 30 |
| SE | 64687 | ENSG00000093144 | ECHDC1 | chr6 | - | 127314815 | 127314896 | 127288711 | 127290277 | 127327001 | 127327144 | 177 | 114 |
| SE | 64691 | ENSG00000093144 | ECHDC1 | chr6 | - | 127315933 | 127316081 | 127314815 | 127314896 | 127316449 | 127316502 | 1 | 0 |
| SE | 64693 | ENSG00000093144 | ECHDC1 | chr6 | - | 127316449 | 127316502 | 127315933 | 127315974 | 127327001 | 127327137 | 118 | 64 |
| SE | 64694 | ENSG00000093144 | ECHDC1 | chr6 | - | 127326495 | 127326574 | 127316449 | 127316502 | 127327001 | 127327144 | 0 | 0 |
| SE | 64695 | ENSG00000093144 | ECHDC1 | chr6 | - | 127327001 | 127327144 | 127316255 | 127316502 | 127343335 | 127343549 | 209 | 140 |
| SE | 64703 | ENSG00000251201 | TMED7-TICAM2 | chr5 | - | 115616317 | 115616445 | 115578650 | 115581315 | 115620434 | 115620680 | 142 | 128 |
| SE | 64704 | ENSG00000164985 | PSIP1 | chr9 | - | 15468629 | 15468843 | 15466747 | 15466859 | 15468956 | 15469058 | 528 | 544 |
| SE | 64705 | ENSG00000164985 | PSIP1 | chr9 | - | 15472631 | 15472750 | 15469937 | 15469993 | 15474008 | 15474237 | 165 | 168 |
| SE | 64707 | ENSG00000164985 | PSIP1 | chr9 | - | 15489269 | 15489440 | 15486826 | 15486931 | 15489985 | 15490124 | 3 | 3 |
| SE | 64708 | ENSG00000164985 | PSIP1 | chr9 | - | 15489985 | 15490124 | 15486826 | 15486931 | 15506560 | 15506637 | 590 | 505 |
| SE | 64709 | ENSG00000164985 | PSIP1 | chr9 | - | 15510116 | 15510326 | 15506560 | 15506637 | 15510816 | 15510991 | 187 | 150 |
| SE | 64710 | ENSG00000164985 | PSIP1 | chr9 | - | 15510116 | 15510329 | 15504958 | 15506637 | 15510816 | 15510946 | 186 | 146 |
| SE | 64713 | ENSG00000167895 | TMC8 | chr17 | + | 78132787 | 78132870 | 78132358 | 78132508 | 78133405 | 78133542 | 106 | 65 |
| SE | 64719 | ENSG00000167895 | TMC8 | chr17 | + | 78137716 | 78137814 | 78137234 | 78137323 | 78138004 | 78138071 | 197 | 125 |
| SE | 64720 | ENSG00000167895 | TMC8 | chr17 | + | 78138348 | 78138479 | 78138004 | 78138188 | 78138573 | 78138732 | 293 | 126 |
| SE | 64733 | ENSG00000042493 | CAPG | chr2 | - | 85395537 | 85395626 | 85394752 | 85394958 | 85398019 | 85398131 | 783 | 344 |
| SE | 64734 | ENSG00000042493 | CAPG | chr2 | - | 85395979 | 85396010 | 85395574 | 85395626 | 85398019 | 85398152 | 0 | 1 |
| SE | 64736 | ENSG00000042493 | CAPG | chr2 | - | 85401528 | 85401683 | 85401164 | 85401329 | 85401784 | 85401957 | 577 | 252 |
| SE | 64742 | ENSG00000108387 | SEPTIN4 | chr17 | - | 58526681 | 58526978 | 58526219 | 58526313 | 58531479 | 58531551 | 2 | 1 |
| SE | 64743 | ENSG00000108387 | SEPTIN4 | chr17 | - | 58526681 | 58526978 | 58526219 | 58526313 | 58531960 | 58532046 | 4 | 2 |
| SE | 64744 | ENSG00000108387 | SEPTIN4 | chr17 | - | 58526681 | 58526978 | 58526219 | 58526313 | 58539111 | 58539208 | 2 | 1 |
| SE | 64750 | ENSG00000108387 | SEPTIN4 | chr17 | - | 58531479 | 58531551 | 58526681 | 58526978 | 58531960 | 58532066 | 0 | 0 |
| SE | 64754 | ENSG00000157470 | FAM81A | chr15 | + | 59516644 | 59516840 | 59514288 | 59514424 | 59521253 | 59523555 | 2 | 11 |
| SE | 64755 | ENSG00000144677 | CTDSPL | chr3 | + | 37911672 | 37911827 | 37862177 | 37862278 | 37947056 | 37947171 | 0 | 0 |
| SE | 64756 | ENSG00000144677 | CTDSPL | chr3 | + | 37947056 | 37947211 | 37861879 | 37862278 | 37964570 | 37964672 | 3 | 8 |
| SE | 64758 | ENSG00000144677 | CTDSPL | chr3 | + | 37971406 | 37971499 | 37967825 | 37967882 | 37975708 | 37975786 | 38 | 25 |
| SE | 64761 | ENSG00000050628 | PTGER3 | chr1 | - | 70974296 | 70974388 | 70953762 | 70953789 | 71012304 | 71012484 | 0 | 0 |
| SE | 64771 | ENSG00000111780 | NA | chr12 | + | 120457075 | 120457179 | 120446656 | 120446829 | 120459906 | 120460006 | 63 | 27 |
| SE | 64772 | ENSG00000163006 | CCDC138 | chr2 | + | 108788031 | 108788089 | 108786756 | 108786915 | 108788851 | 108788966 | 6 | 12 |
| SE | 64773 | ENSG00000163006 | CCDC138 | chr2 | + | 108788031 | 108788089 | 108786772 | 108786804 | 108788851 | 108788966 | 3 | 4 |
| SE | 64774 | ENSG00000163006 | CCDC138 | chr2 | + | 108791554 | 108791802 | 108788851 | 108788966 | 108794539 | 108794721 | 11 | 13 |
| SE | 64775 | ENSG00000163006 | CCDC138 | chr2 | + | 108791674 | 108791802 | 108788851 | 108788966 | 108794539 | 108794721 | 17 | 25 |
| SE | 64776 | ENSG00000163006 | CCDC138 | chr2 | + | 108791679 | 108791802 | 108788851 | 108788966 | 108794539 | 108794712 | 10 | 13 |
| SE | 64777 | ENSG00000163006 | CCDC138 | chr2 | + | 108791711 | 108791802 | 108788851 | 108788966 | 108794539 | 108794670 | 6 | 10 |
| SE | 64778 | ENSG00000163006 | CCDC138 | chr2 | + | 108798427 | 108798586 | 108794539 | 108794721 | 108804891 | 108805008 | 13 | 36 |
| SE | 64779 | ENSG00000163006 | CCDC138 | chr2 | + | 108839184 | 108839301 | 108815940 | 108816105 | 108846737 | 108846780 | 26 | 17 |
| SE | 64780 | ENSG00000163006 | CCDC138 | chr2 | + | 108846737 | 108846930 | 108815940 | 108816105 | 108873450 | 108873589 | 3 | 3 |
| SE | 64781 | ENSG00000163006 | CCDC138 | chr2 | + | 108856793 | 108856970 | 108846737 | 108846930 | 108873450 | 108873589 | 22 | 9 |
| SE | 64782 | ENSG00000163006 | CCDC138 | chr2 | + | 108873450 | 108873589 | 108856793 | 108856970 | 108876087 | 108876578 | 16 | 10 |
| SE | 64795 | ENSG00000228486 | C2orf92 | chr2 | + | 97671190 | 97671306 | 97669775 | 97669834 | 97674455 | 97674557 | 1 | 0 |
| SE | 64796 | ENSG00000228486 | C2orf92 | chr2 | + | 97671193 | 97671306 | 97669792 | 97669834 | 97674455 | 97674557 | 2 | 0 |
| SE | 64798 | ENSG00000228486 | C2orf92 | chr2 | + | 97671219 | 97671306 | 97669792 | 97669834 | 97674455 | 97674557 | 1 | 0 |
| SE | 64799 | ENSG00000228486 | C2orf92 | chr2 | + | 97671330 | 97671481 | 97669816 | 97669834 | 97674455 | 97674557 | 0 | 0 |
| SE | 64800 | ENSG00000228486 | C2orf92 | chr2 | + | 97671330 | 97671499 | 97669745 | 97669834 | 97674455 | 97674557 | 1 | 2 |
| SE | 64801 | ENSG00000228486 | C2orf92 | chr2 | + | 97671330 | 97671499 | 97669745 | 97669834 | 97675844 | 97675928 | 0 | 0 |
| SE | 64802 | ENSG00000228486 | C2orf92 | chr2 | + | 97671330 | 97671574 | 97669792 | 97669834 | 97674455 | 97674557 | 0 | 0 |
| SE | 64803 | ENSG00000228486 | C2orf92 | chr2 | + | 97674455 | 97674557 | 97669742 | 97669834 | 97675844 | 97675928 | 1 | 0 |
| SE | 64807 | ENSG00000228486 | C2orf92 | chr2 | + | 97674455 | 97674561 | 97666956 | 97667054 | 97675844 | 97675928 | 0 | 0 |
| SE | 64813 | ENSG00000228486 | C2orf92 | chr2 | + | 97688894 | 97688993 | 97675844 | 97675928 | 97699025 | 97699136 | 4 | 2 |
| SE | 64815 | ENSG00000228486 | C2orf92 | chr2 | + | 97690255 | 97690327 | 97675820 | 97675928 | 97699025 | 97699114 | 3 | 6 |
| SE | 64816 | ENSG00000228486 | C2orf92 | chr2 | + | 97690255 | 97690327 | 97688894 | 97688993 | 97699025 | 97699114 | 2 | 2 |
| SE | 64817 | ENSG00000228486 | C2orf92 | chr2 | + | 97694446 | 97694599 | 97675844 | 97675928 | 97699025 | 97699048 | 2 | 0 |
| SE | 64820 | ENSG00000228486 | C2orf92 | chr2 | + | 97699025 | 97699136 | 97690255 | 97690327 | 97702668 | 97702725 | 3 | 7 |
| SE | 64821 | ENSG00000228486 | C2orf92 | chr2 | + | 97701153 | 97701304 | 97699025 | 97699136 | 97702668 | 97702704 | 4 | 3 |
| SE | 64823 | ENSG00000198324 | PHETA1 | chr12 | - | 111365563 | 111365651 | 111360650 | 111363463 | 111366112 | 111366257 | 2 | 1 |
| SE | 64828 | ENSG00000104907 | TRMT1 | chr19 | - | 13112895 | 13113011 | 13110157 | 13110306 | 13115278 | 13115402 | 19 | 19 |
| SE | 64829 | ENSG00000104907 | TRMT1 | chr19 | - | 13112908 | 13113011 | 13110157 | 13110306 | 13115278 | 13115466 | 17 | 11 |
| SE | 64830 | ENSG00000104907 | TRMT1 | chr19 | - | 13115625 | 13115768 | 13115278 | 13115402 | 13115996 | 13116027 | 76 | 70 |
| SE | 64831 | ENSG00000104907 | TRMT1 | chr19 | - | 13115625 | 13115768 | 13115278 | 13115466 | 13115996 | 13116027 | 161 | 105 |
| SE | 64834 | ENSG00000104907 | TRMT1 | chr19 | - | 13115996 | 13116052 | 13115625 | 13115768 | 13116145 | 13116340 | 89 | 96 |
| SE | 64840 | ENSG00000137185 | ZSCAN9 | chr6 | + | 28230335 | 28230488 | 28227689 | 28227837 | 28232561 | 28233466 | 1 | 0 |
| SE | 64841 | ENSG00000137185 | ZSCAN9 | chr6 | + | 28230401 | 28230488 | 28227689 | 28227837 | 28232561 | 28233015 | 1 | 0 |
| SE | 64842 | ENSG00000158435 | CNOT11 | chr2 | + | 101257790 | 101257955 | 101252885 | 101253478 | 101262538 | 101262691 | 201 | 82 |
| SE | 64843 | ENSG00000158435 | CNOT11 | chr2 | + | 101262538 | 101262691 | 101257790 | 101257955 | 101264844 | 101265047 | 297 | 142 |
| SE | 64845 | ENSG00000138032 | PPM1B | chr2 | + | 44218479 | 44218537 | 44209209 | 44209327 | 44230412 | 44231472 | 26 | 25 |
| SE | 64847 | ENSG00000163069 | SGCB | chr4 | - | 52027967 | 52028099 | 52020705 | 52024160 | 52028729 | 52028921 | 36 | 20 |
| SE | 64848 | ENSG00000163069 | SGCB | chr4 | - | 52031851 | 52031961 | 52029677 | 52029863 | 52033430 | 52033640 | 3 | 2 |
| SE | 64849 | ENSG00000163069 | SGCB | chr4 | - | 52031864 | 52031961 | 52029677 | 52029863 | 52033430 | 52033640 | 1 | 1 |
| SE | 64850 | ENSG00000163069 | SGCB | chr4 | - | 52033430 | 52033640 | 52029677 | 52029863 | 52038226 | 52038245 | 24 | 12 |
| SE | 64851 | ENSG00000206530 | CFAP44 | chr3 | - | 113293951 | 113294038 | 113290852 | 113291748 | 113294686 | 113294821 | 5 | 11 |
| SE | 64852 | ENSG00000206530 | CFAP44 | chr3 | - | 113296724 | 113296885 | 113294686 | 113294821 | 113303915 | 113304117 | 9 | 14 |
| SE | 64853 | ENSG00000206530 | CFAP44 | chr3 | - | 113296724 | 113296902 | 113294686 | 113294821 | 113303915 | 113304067 | 6 | 2 |
| SE | 64854 | ENSG00000206530 | CFAP44 | chr3 | - | 113327615 | 113327819 | 113326444 | 113326640 | 113330167 | 113330668 | 28 | 51 |
| SE | 64855 | ENSG00000206530 | CFAP44 | chr3 | - | 113333405 | 113333583 | 113330167 | 113330668 | 113341743 | 113341918 | 7 | 9 |
| SE | 64856 | ENSG00000206530 | CFAP44 | chr3 | - | 113333405 | 113333583 | 113330167 | 113330668 | 113358744 | 113358875 | 3 | 7 |
| SE | 64858 | ENSG00000206530 | CFAP44 | chr3 | - | 113341743 | 113341918 | 113333405 | 113333583 | 113358744 | 113358875 | 4 | 4 |
| SE | 64859 | ENSG00000206530 | CFAP44 | chr3 | - | 113344515 | 113344712 | 113341743 | 113341918 | 113358744 | 113358875 | 13 | 23 |
| SE | 64861 | ENSG00000206530 | CFAP44 | chr3 | - | 113373410 | 113373556 | 113366038 | 113366309 | 113379305 | 113379551 | 0 | 0 |
| SE | 64862 | ENSG00000206530 | CFAP44 | chr3 | - | 113379305 | 113379551 | 113363476 | 113363532 | 113380898 | 113381060 | 36 | 33 |
| SE | 64867 | ENSG00000170310 | STX8 | chr17 | - | 9378551 | 9378653 | 9250470 | 9250645 | 9491828 | 9491921 | 163 | 163 |
| SE | 64870 | ENSG00000170310 | STX8 | chr17 | - | 9568370 | 9568470 | 9557433 | 9557528 | 9575791 | 9575811 | 311 | 191 |
| SE | 64872 | ENSG00000115109 | EPB41L5 | chr2 | + | 120093248 | 120093276 | 120091554 | 120091661 | 120100243 | 120100286 | 14 | 6 |
| SE | 64873 | ENSG00000115109 | EPB41L5 | chr2 | + | 120143002 | 120143131 | 120131617 | 120131715 | 120146224 | 120146289 | 30 | 15 |
| SE | 64874 | ENSG00000115109 | EPB41L5 | chr2 | + | 120164835 | 120164910 | 120160566 | 120160974 | 120167465 | 120167507 | 14 | 11 |

| | | | | | | | | | | | | | |
|---|---|---|---|---|---|---|---|---|---|---|---|---|---|
| SE | 64875 | ENSG00000115109 | EPB41L5 | chr2 | + | 120167465 | 120167507 | 120164835 | 120164910 | 120174840 | 120175861 | 4 | 4 |
| SE | 64876 | ENSG00000115109 | EPB41L5 | chr2 | + | 120167876 | 120168007 | 120167465 | 120167507 | 120174840 | 120179119 | 7 | 4 |
| SE | 64877 | ENSG00000115109 | EPB41L5 | chr2 | + | 120167879 | 120168007 | 120167465 | 120167507 | 120174840 | 120179118 | 3 | 1 |
| SE | 64879 | ENSG00000162722 | TRIM58 | chr1 | + | 247864704 | 247864935 | 247860616 | 247860712 | 247867844 | 247867867 | 81 | 114 |
| SE | 64880 | ENSG00000119777 | TMEM214 | chr2 | + | 27034066 | 27034266 | 27032909 | 27033166 | 27035134 | 27035285 | 142 | 55 |
| SE | 64881 | ENSG00000119777 | TMEM214 | chr2 | + | 27035593 | 27035728 | 27035134 | 27035285 | 27035969 | 27036052 | 181 | 54 |
| SE | 64885 | ENSG00000119777 | TMEM214 | chr2 | + | 27040029 | 27040198 | 27039046 | 27039164 | 27040344 | 27040492 | 26 | 4 |
| SE | 64887 | ENSG00000105997 | HOXA3 | chr7 | - | 27126885 | 27127070 | 27122558 | 27122642 | 27140082 | 27140186 | 3 | 1 |
| SE | 64888 | ENSG00000254999 | BRK1 | chr3 | + | 10125625 | 10125708 | 10115674 | 10115819 | 10126268 | 10127190 | 852 | 653 |
| SE | 64889 | ENSG00000123684 | LPGAT1 | chr1 | - | 211783228 | 211783502 | 211778917 | 211779044 | 211787631 | 211787727 | 873 | 700 |
| SE | 64890 | ENSG00000123684 | LPGAT1 | chr1 | - | 211783228 | 211783502 | 211778917 | 211779044 | 211793071 | 211793190 | 134 | 79 |
| SE | 64891 | ENSG00000123684 | LPGAT1 | chr1 | - | 211787631 | 211787727 | 211778917 | 211779044 | 211793071 | 211793190 | 530 | 463 |
| SE | 64896 | ENSG00000135945 | REV1 | chr2 | - | 99403694 | 99403815 | 99402888 | 99403106 | 99404443 | 99404677 | 80 | 80 |
| SE | 64897 | ENSG00000135945 | REV1 | chr2 | - | 99406324 | 99406431 | 99405927 | 99406106 | 99408028 | 99408131 | 39 | 22 |
| SE | 64898 | ENSG00000135945 | REV1 | chr2 | - | 99406324 | 99406490 | 99405909 | 99406106 | 99408028 | 99408131 | 79 | 39 |
| SE | 64899 | ENSG00000135945 | REV1 | chr2 | - | 99410694 | 99410867 | 99408028 | 99408131 | 99412730 | 99412951 | 165 | 89 |
| SE | 64900 | ENSG00000135945 | REV1 | chr2 | - | 99434331 | 99434448 | 99429839 | 99429945 | 99438600 | 99438727 | 17 | 11 |
| SE | 64901 | ENSG00000135945 | REV1 | chr2 | - | 99435833 | 99435941 | 99434331 | 99434448 | 99438600 | 99438727 | 130 | 98 |
| SE | 64902 | ENSG00000135945 | REV1 | chr2 | - | 99436646 | 99436791 | 99435836 | 99435941 | 99438600 | 99438777 | 2 | 1 |
| SE | 64903 | ENSG00000135945 | REV1 | chr2 | - | 99442316 | 99442469 | 99438573 | 99439310 | 99449335 | 99449504 | 124 | 88 |
| SE | 64904 | ENSG00000135945 | REV1 | chr2 | - | 99442316 | 99442469 | 99438573 | 99439310 | 99462495 | 99462622 | 50 | 34 |
| SE | 64905 | ENSG00000135945 | REV1 | chr2 | - | 99449335 | 99449504 | 99438573 | 99439310 | 99462495 | 99462622 | 177 | 140 |
| SE | 64906 | ENSG00000135945 | REV1 | chr2 | - | 99449335 | 99449504 | 99442316 | 99442469 | 99462495 | 99462622 | 251 | 193 |
| SE | 64908 | ENSG00000135945 | REV1 | chr2 | - | 99462495 | 99462622 | 99451344 | 99451501 | 99464921 | 99464985 | 92 | 58 |
| SE | 64909 | ENSG00000246985 | SOCS2-AS1 | chr12 | - | 93565612 | 93565744 | 93545070 | 93545358 | 93566743 | 93566970 | 1 | 3 |
| SE | 64911 | ENSG00000246985 | SOCS2-AS1 | chr12 | - | 93565612 | 93565744 | 93551432 | 93551512 | 93566743 | 93566970 | 1 | 3 |
| SE | 64912 | ENSG00000246985 | SOCS2-AS1 | chr12 | - | 93567063 | 93567159 | 93566743 | 93566970 | 93570817 | 93570854 | 6 | 0 |
| SE | 64913 | ENSG00000246985 | SOCS2-AS1 | chr12 | - | 93567063 | 93567159 | 93566743 | 93566970 | 93571566 | 93571768 | 4 | 0 |
| SE | 64914 | ENSG00000048342 | CC2D2A | chr4 | + | 15473221 | 15473255 | 15469870 | 15470057 | 15475914 | 15475971 | 0 | 0 |
| SE | 64924 | ENSG00000204642 | HLA-F | chr6 | + | 29725030 | 29725306 | 29724156 | 29724448 | 29725446 | 29725563 | 1950 | 1038 |
| SE | 64925 | ENSG00000204642 | HLA-F | chr6 | + | 29725446 | 29725563 | 29724172 | 29724448 | 29738121 | 29738188 | 1 | 5 |
| SE | 64931 | ENSG00000198400 | NTRK1 | chr1 | + | 156868103 | 156868249 | 156866909 | 156866978 | 156868504 | 156868647 | 2 | 5 |
| SE | 64944 | ENSG00000163644 | PPM1K | chr4 | - | 88265000 | 88265135 | 88257619 | 88262726 | 88268740 | 88268906 | 195 | 100 |
| SE | 64945 | ENSG00000163644 | PPM1K | chr4 | - | 88265000 | 88265135 | 88257619 | 88262726 | 88277142 | 88277243 | 192 | 100 |
| SE | 64946 | ENSG00000163644 | PPM1K | chr4 | - | 88268189 | 88268334 | 88261870 | 88262726 | 88268740 | 88268906 | 87 | 55 |
| SE | 64948 | ENSG00000163644 | PPM1K | chr4 | - | 88268740 | 88268906 | 88257619 | 88262726 | 88277142 | 88277243 | 91 | 50 |
| SE | 64949 | ENSG00000163644 | PPM1K | chr4 | - | 88268740 | 88268906 | 88265000 | 88265135 | 88277142 | 88277243 | 94 | 50 |
| SE | 64950 | ENSG00000163644 | PPM1K | chr4 | - | 88270999 | 88271189 | 88268761 | 88268906 | 88277142 | 88277243 | 0 | 1 |
| SE | 64951 | ENSG00000163644 | PPM1K | chr4 | - | 88277142 | 88277243 | 88265000 | 88265135 | 88284405 | 88284631 | 2 | 6 |
| SE | 64954 | ENSG00000121579 | NAA50 | chr3 | - | 113723421 | 113723541 | 113722905 | 113722972 | 113723958 | 113724092 | 318 | 215 |
| SE | 64956 | ENSG00000053371 | AKR7A2 | chr1 | - | 19307001 | 19307101 | 19306017 | 19306147 | 19308454 | 19308505 | 28 | 14 |
| SE | 64958 | ENSG00000053371 | AKR7A2 | chr1 | - | 19307313 | 19307410 | 19307001 | 19307101 | 19308454 | 19308642 | 9 | 9 |
| SE | 64959 | ENSG00000053371 | AKR7A2 | chr1 | - | 19308157 | 19308262 | 19306017 | 19306147 | 19308454 | 19308505 | 28 | 16 |
| SE | 64960 | ENSG00000053371 | AKR7A2 | chr1 | - | 19308157 | 19308262 | 19307001 | 19307101 | 19308454 | 19308505 | 31 | 19 |
| SE | 64966 | ENSG00000182973 | CNOT10 | chr3 | + | 32704810 | 32704972 | 32703790 | 32703962 | 32708669 | 32708775 | 242 | 149 |
| SE | 64968 | ENSG00000182973 | CNOT10 | chr3 | + | 32717153 | 32717237 | 32716224 | 32716311 | 32720113 | 32720231 | 137 | 92 |
| SE | 64970 | ENSG00000182973 | CNOT10 | chr3 | + | 32734799 | 32734976 | 32733422 | 32733544 | 32737409 | 32737490 | 245 | 172 |
| SE | 64975 | ENSG00000182973 | CNOT10 | chr3 | + | 32762732 | 32762863 | 32759457 | 32759571 | 32764454 | 32764490 | 94 | 75 |
| SE | 64976 | ENSG00000182973 | CNOT10 | chr3 | + | 32764454 | 32764490 | 32762732 | 32762863 | 32764681 | 32764809 | 49 | 38 |
| SE | 64977 | ENSG00000182973 | CNOT10 | chr3 | + | 32764681 | 32764809 | 32762732 | 32762863 | 32773456 | 32773616 | 4 | 2 |
| SE | 64978 | ENSG00000182973 | CNOT10 | chr3 | + | 32765049 | 32765177 | 32764681 | 32764809 | 32773456 | 32773616 | 0 | 0 |
| SE | 64979 | ENSG00000182973 | CNOT10 | chr3 | + | 32769886 | 32769962 | 32764681 | 32764809 | 32773456 | 32773714 | 75 | 47 |
| SE | 64980 | ENSG00000120675 | DNAJC15 | chr13 | + | 43065685 | 43065737 | 43023202 | 43023734 | 43078611 | 43078688 | 25 | 17 |
| SE | 64981 | ENSG00000120675 | DNAJC15 | chr13 | + | 43068929 | 43069003 | 43065685 | 43065737 | 43078611 | 43078688 | 200 | 116 |
| SE | 64985 | ENSG00000159840 | ZYX | chr7 | + | 143382592 | 143382685 | 143382247 | 143382447 | 143382800 | 143382925 | 732 | 333 |
| SE | 64987 | ENSG00000159840 | ZYX | chr7 | + | 143384089 | 143384245 | 143382999 | 143383322 | 143388218 | 143388339 | 0 | 0 |
| SE | 64990 | ENSG00000159840 | ZYX | chr7 | + | 143388218 | 143388339 | 143382800 | 143383322 | 143388488 | 143388500 | 472 | 224 |
| SE | 65006 | ENSG00000167107 | ACSF2 | chr17 | + | 50461632 | 50461686 | 50461241 | 50461287 | 50462183 | 50462302 | 8 | 2 |
| SE | 65007 | ENSG00000167107 | ACSF2 | chr17 | + | 50461632 | 50461686 | 50461241 | 50461370 | 50462183 | 50462302 | 28 | 5 |
| SE | 65011 | ENSG00000167107 | ACSF2 | chr17 | + | 50472427 | 50472579 | 50470930 | 50471135 | 50473664 | 50473790 | 45 | 10 |
| SE | 65016 | ENSG00000157184 | CPT2 | chr1 | + | 53202322 | 53202429 | 53200718 | 53200799 | 53210014 | 53211250 | 93 | 51 |
| SE | 65020 | ENSG00000157184 | CPT2 | chr1 | + | 53212780 | 53212816 | 53210014 | 53211250 | 53213263 | 53213846 | 0 | 0 |
| SE | 65021 | ENSG00000157184 | CPT2 | chr1 | + | 53212780 | 53212816 | 53210014 | 53211319 | 53213263 | 53213846 | 1 | 0 |
| SE | 65027 | ENSG00000091409 | ITGA6 | chr2 | + | 172469124 | 172469380 | 172465538 | 172465663 | 172474054 | 172474265 | 3 | 1 |
| SE | 65028 | ENSG00000091409 | ITGA6 | chr2 | + | 172469124 | 172469380 | 172467480 | 172467560 | 172470973 | 172471105 | 203 | 157 |
| SE | 65029 | ENSG00000091409 | ITGA6 | chr2 | + | 172469124 | 172469380 | 172467480 | 172467560 | 172474054 | 172474265 | 91 | 60 |
| SE | 65031 | ENSG00000091409 | ITGA6 | chr2 | + | 172470973 | 172471105 | 172467480 | 172467560 | 172474054 | 172474112 | 179 | 180 |
| SE | 65033 | ENSG00000091409 | ITGA6 | chr2 | + | 172470973 | 172471105 | 172468997 | 172469380 | 172474054 | 172474112 | 293 | 277 |
| SE | 65035 | ENSG00000091409 | ITGA6 | chr2 | + | 172472773 | 172472890 | 172470973 | 172471105 | 172474054 | 172474265 | 0 | 0 |
| SE | 65036 | ENSG00000091409 | ITGA6 | chr2 | + | 172479989 | 172480051 | 172479640 | 172479739 | 172484781 | 172484942 | 190 | 197 |
| SE | 65037 | ENSG00000091409 | ITGA6 | chr2 | + | 172487263 | 172487453 | 172487022 | 172487138 | 172487546 | 172487630 | 410 | 408 |
| SE | 65041 | ENSG00000068697 | LAPTM4A | chr2 | - | 20034316 | 20034415 | 20032649 | 20033279 | 20034966 | 20035062 | 988 | 568 |
| SE | 65044 | ENSG00000111275 | ALDH2 | chr12 | + | 111783157 | 111783298 | 111781917 | 111782022 | 111785266 | 111785346 | 165 | 109 |
| SE | 65046 | ENSG00000111275 | ALDH2 | chr12 | + | 111790433 | 111790562 | 111789822 | 111789934 | 111791305 | 111791419 | 321 | 130 |
| SE | 65047 | ENSG00000111275 | ALDH2 | chr12 | + | 111790433 | 111790562 | 111789822 | 111789934 | 111792060 | 111792163 | 62 | 15 |
| SE | 65048 | ENSG00000111275 | ALDH2 | chr12 | + | 111791305 | 111791419 | 111789822 | 111789934 | 111792060 | 111792163 | 195 | 124 |
| SE | 65049 | ENSG00000111275 | ALDH2 | chr12 | + | 111791305 | 111791419 | 111790433 | 111790562 | 111792060 | 111792163 | 454 | 240 |
| SE | 65050 | ENSG00000111275 | ALDH2 | chr12 | + | 111792060 | 111792163 | 111791305 | 111791419 | 111792597 | 111792782 | 463 | 277 |
| SE | 65051 | ENSG00000111275 | ALDH2 | chr12 | + | 111792597 | 111792782 | 111791305 | 111791419 | 111798077 | 111798242 | 62 | 62 |
| SE | 65052 | ENSG00000111275 | ALDH2 | chr12 | + | 111799905 | 111800063 | 111798077 | 111798242 | 111803858 | 111803973 | 350 | 282 |
| SE | 65053 | ENSG00000111275 | ALDH2 | chr12 | + | 111803858 | 111803973 | 111799905 | 111800063 | 111809542 | 111809630 | 334 | 304 |
| SE | 65054 | ENSG00000115355 | CCDC88A | chr2 | - | 55299827 | 55299919 | 55296257 | 55296523 | 55301205 | 55301277 | 62 | 72 |
| SE | 65055 | ENSG00000115355 | CCDC88A | chr2 | - | 55299827 | 55299919 | 55296257 | 55296523 | 55308808 | 55309023 | 1 | 0 |
| SE | 65056 | ENSG00000115355 | CCDC88A | chr2 | - | 55299838 | 55299919 | 55296257 | 55296523 | 55300956 | 55300996 | 44 | 49 |
| SE | 65057 | ENSG00000115355 | CCDC88A | chr2 | - | 55299838 | 55299919 | 55296257 | 55296523 | 55301205 | 55301277 | 147 | 152 |
| SE | 65058 | ENSG00000115355 | CCDC88A | chr2 | - | 55299838 | 55299919 | 55296257 | 55296523 | 55308808 | 55309023 | 45 | 49 |
| SE | 65060 | ENSG00000115355 | CCDC88A | chr2 | - | 55300956 | 55300996 | 55299838 | 55299919 | 55301205 | 55301272 | 1 | 0 |
| SE | 65062 | ENSG00000115355 | CCDC88A | chr2 | - | 55301205 | 55301277 | 55299827 | 55299919 | 55308808 | 55309023 | 52 | 43 |
| SE | 65063 | ENSG00000115355 | CCDC88A | chr2 | - | 55301871 | 55302072 | 55299838 | 55301277 | 55303068 | 55303152 | 129 | 156 |
| SE | 65064 | ENSG00000115355 | CCDC88A | chr2 | - | 55301871 | 55302072 | 55299838 | 55301277 | 55308808 | 55309023 | 123 | 137 |
| SE | 65065 | ENSG00000115355 | CCDC88A | chr2 | - | 55303068 | 55303152 | 55299838 | 55301277 | 55308808 | 55309023 | 30 | 51 |
| SE | 65067 | ENSG00000115355 | CCDC88A | chr2 | - | 55308808 | 55309023 | 55302662 | 55303152 | 55309161 | 55309207 | 200 | 320 |
| SE | 65068 | ENSG00000115355 | CCDC88A | chr2 | - | 55309161 | 55309254 | 55303068 | 55303152 | 55315927 | 55315952 | 3 | 13 |
| SE | 65071 | ENSG00000115355 | CCDC88A | chr2 | - | 55317563 | 55317841 | 55315927 | 55316114 | 55318842 | 55318904 | 176 | 150 |
| SE | 65072 | ENSG00000115355 | CCDC88A | chr2 | - | 55322527 | 55322692 | 55318842 | 55319004 | 55328293 | 55328425 | 238 | 293 |
| SE | 65073 | ENSG00000115355 | CCDC88A | chr2 | - | 55334093 | 55335164 | 55332130 | 55332693 | 55336680 | 55336818 | 159 | 312 |
| SE | 65074 | ENSG00000115355 | CCDC88A | chr2 | - | 55334093 | 55335164 | 55332130 | 55332693 | 55339463 | 55339648 | 53 | 116 |
| SE | 65075 | ENSG00000115355 | CCDC88A | chr2 | - | 55336680 | 55336818 | 55332130 | 55332693 | 55339463 | 55339648 | 134 | 192 |
| SE | 65076 | ENSG00000115355 | CCDC88A | chr2 | - | 55336680 | 55336818 | 55334093 | 55335164 | 55339463 | 55339648 | 240 | 388 |
| SE | 65077 | ENSG00000115355 | CCDC88A | chr2 | - | 55339463 | 55339648 | 55336680 | 55336818 | 55343647 | 55343792 | 218 | 290 |
| SE | 65079 | ENSG00000115355 | CCDC88A | chr2 | - | 55349517 | 55349599 | 55346174 | 55346333 | 55362307 | 55362448 | 75 | 132 |
| SE | 65081 | ENSG00000115355 | CCDC88A | chr2 | - | 55355578 | 55355751 | 55346174 | 55346333 | 55362307 | 55362448 | 83 | 79 |
| SE | 65082 | ENSG00000115355 | CCDC88A | chr2 | - | 55355578 | 55355751 | 55349517 | 55349599 | 55362307 | 55362448 | 192 | 184 |
| SE | 65083 | ENSG00000115355 | CCDC88A | chr2 | - | 55355578 | 55355751 | 55349517 | 55349599 | 55372451 | 55372510 | 109 | 105 |
| SE | 65084 | ENSG00000115355 | CCDC88A | chr2 | - | 55362307 | 55362448 | 55355578 | 55355751 | 55363949 | 55364019 | 181 | 238 |
| SE | 65086 | ENSG00000115355 | CCDC88A | chr2 | - | 55363949 | 55364033 | 55355578 | 55355751 | 55372451 | 55372510 | 60 | 82 |
| SE | 65088 | ENSG00000115355 | CCDC88A | chr2 | - | 55372451 | 55372510 | 55363949 | 55364033 | 55374813 | 55374883 | 121 | 185 |
| SE | 65089 | ENSG00000115355 | CCDC88A | chr2 | - | 55374813 | 55374883 | 55363949 | 55364033 | 55388777 | 55388844 | 57 | 73 |
| SE | 65090 | ENSG00000265808 | SEC22B | chr1 | - | 120168839 | 120168949 | 120163209 | 120163370 | 120176306 | 120176520 | 560 | 479 |
| SE | 65096 | ENSG00000178038 | ALS2CL | chr3 | - | 46683129 | 46683326 | 46682028 | 46682094 | 46683781 | 46683848 | 4 | 17 |
| SE | 65097 | ENSG00000178038 | ALS2CL | chr3 | - | 46683133 | 46683326 | 46682028 | 46682094 | 46683781 | 46683848 | 2 | 7 |
| SE | 65098 | ENSG00000203756 | TMEM244 | chr6 | - | 129833459 | 129833585 | 129831243 | 129831386 | 129843529 | 129843603 | 2 | 2 |
| SE | 65103 | ENSG00000186073 | CDIN1 | chr15 | + | 36644277 | 36644323 | 36579625 | 36579961 | 36645222 | 36645287 | 4 | 4 |
| SE | 65106 | ENSG00000186073 | CDIN1 | chr15 | + | 36647568 | 36647658 | 36645222 | 36645287 | 36654097 | 36654158 | 0 | 0 |
| SE | 65107 | ENSG00000186073 | CDIN1 | chr15 | + | 36658260 | 36658294 | 36657832 | 36657905 | 36691684 | 36691764 | 0 | 0 |
| SE | 65108 | ENSG00000186073 | CDIN1 | chr15 | + | 36668110 | 36668219 | 36657832 | 36657905 | 36691684 | 36691764 | 1 | 0 |
| SE | 65111 | ENSG00000186073 | CDIN1 | chr15 | + | 36790207 | 36790358 | 36709855 | 36709961 | 36808323 | 36808457 | 0 | 1 |
| SE | 65112 | ENSG00000282572 | NA | chr7 | - | 26288 | 26402 | 20833 | 21029 | 31059 | 31194 | 3 | 7 |
| SE | 65117 | ENSG00000173960 | UBXN2A | chr2 | + | 23968918 | 23969035 | 23958300 | 23958355 | 23971275 | 23971414 | 4 | 0 |
| SE | 65118 | ENSG00000173960 | UBXN2A | chr2 | + | 23984672 | 23984831 | 23982895 | 23983033 | 23999671 | 24000203 | 85 | 96 |
| SE | 65122 | ENSG00000104728 | ARHGEF10 | chr8 | + | 1880047 | 1880164 | 1876570 | 1876734 | 1882634 | 1882749 | 1 | 5 |
| SE | 65123 | ENSG00000104728 | ARHGEF10 | chr8 | + | 1926376 | 1926463 | 1925282 | 1925404 | 1928426 | 1928650 | 5 | 1 |
| SE | 65126 | ENSG00000163576 | EFHB | chr3 | - | 19920504 | 19920567 | 19919832 | 19919976 | 19936140 | 19936334 | 0 | 0 |
| SE | 65128 | ENSG00000138801 | PAPSS1 | chr4 | - | 107631630 | 107631860 | 107613665 | 107614387 | 107644801 | 107645070 | 127 | 55 |
| SE | 65129 | ENSG00000138801 | PAPSS1 | chr4 | - | 107656895 | 107657007 | 107654694 | 107654900 | 107659958 | 107660072 | 226 | 107 |
| SE | 65130 | ENSG00000138801 | PAPSS1 | chr4 | - | 107687038 | 107687177 | 107682014 | 107682133 | 107693770 | 107694006 | 325 | 180 |
| SE | 65131 | ENSG00000138801 | PAPSS1 | chr4 | - | 107693770 | 107694006 | 107687038 | 107687177 | 107701170 | 107701285 | 296 | 144 |
| SE | 65133 | ENSG00000138801 | PAPSS1 | chr4 | - | 107718341 | 107718396 | 107701170 | 107701285 | 107720119 | 107720226 | 2 | 0 |
| SE | 65137 | ENSG00000152689 | RASGRP3 | chr2 | + | 33493053 | 33493172 | 33447819 | 33447943 | 33511709 | 33511842 | 0 | 0 |
| SE | 65139 | ENSG00000152689 | RASGRP3 | chr2 | + | 33511709 | 33511842 | 33476596 | 33476707 | 33515009 | 33515206 | 23 | 30 |
| SE | 65140 | ENSG00000152689 | RASGRP3 | chr2 | + | 33511811 | 33511842 | 33476648 | 33476707 | 33515009 | 33515206 | 7 | 10 |
| SE | 65141 | ENSG00000152689 | RASGRP3 | chr2 | + | 33534322 | 33534400 | 33527136 | 33527412 | 33539093 | 33539210 | 24 | 25 |
| SE | 65142 | ENSG00000152689 | RASGRP3 | chr2 | + | 33543511 | 33543627 | 33539650 | 33540108 | 33549603 | 33549751 | 11 | 18 |
| SE | 65143 | ENSG00000152689 | RASGRP3 | chr2 | + | 33555530 | 33555567 | 33549603 | 33549751 | 33558210 | 33558336 | 28 | 25 |
| SE | 65144 | ENSG00000152689 | RASGRP3 | chr2 | + | 33558671 | 33559030 | 33558210 | 33558336 | 33562728 | 33564605 | 33 | 26 |
| SE | 65145 | ENSG00000002822 | MAD1L1 | chr7 | - | 2149151 | 2149238 | 2069193 | 2069338 | 2213211 | 2213234 | 100 | 81 |
| SE | 65146 | ENSG00000002822 | MAD1L1 | chr7 | - | 2201850 | 2202098 | 2200264 | 2200557 | 2213211 | 2213273 | 0 | 0 |
| SE | 65155 | ENSG00000196912 | ANKRD36B | chr2 | - | 97507994 | 97508065 | 97507563 | 97507657 | 97509580 | 97509733 | 1 | 4 |
| SE | 65156 | ENSG00000196912 | ANKRD36B | chr2 | - | 97523325 | 97523467 | 97515731 | 97515945 | 97532310 | 97532384 | 32 | 14 |
| SE | 65157 | ENSG00000196912 | ANKRD36B | chr2 | - | 97551288 | 97551361 | 97549585 | 97549614 | 97551451 | 97551480 | 277 | 208 |
| SE | 65158 | ENSG00000196912 | ANKRD36B | chr2 | - | 97558965 | 97558994 | 97558798 | 97558871 | 97560831 | 97560860 | 55 | 11 |
| SE | 65166 | ENSG00000136842 | TMOD1 | chr9 | + | 97546184 | 97546341 | 97524140 | 97524308 | 97553280 | 97553400 | 0 | 0 |
| SE | 65171 | ENSG00000268362 | NA | chr19 | + | 24036252 | 24036382 | 24033432 | 24033621 | 24036745 | 24037405 | 6 | 7 |
| SE | 65172 | ENSG00000268362 | NA | chr19 | + | 24036252 | 24036382 | 24033432 | 24033621 | 24042442 | 24042722 | 2 | 0 |
| SE | 65179 | ENSG00000268362 | NA | chr19 | + | 24048936 | 24049014 | 24046183 | 24046279 | 24049409 | 24049548 | 0 | 3 |
| SE | 65182 | ENSG00000268362 | NA | chr19 | + | 24049661 | 24049839 | 24049404 | 24049548 | 24057930 | 24058106 | 0 | 1 |
| SE | 65184 | ENSG00000268362 | NA | chr19 | + | 24050184 | 24050302 | 24049404 | 24049548 | 24057930 | 24058106 | 0 | 1 |
| SE | 65186 | ENSG00000268362 | NA | chr19 | + | 24057930 | 24058106 | 24049404 | 24049548 | 24065462 | 24065648 | 0 | 2 |
| SE | 65189 | ENSG00000268362 | NA | chr19 | + | 24057930 | 24058230 | 24049404 | 24049548 | 24065462 | 24066726 | 0 | 1 |
| SE | 65190 | ENSG00000268362 | NA | chr19 | + | 24057930 | 24058300 | 24049404 | 24049548 | 24065462 | 24065830 | 0 | 1 |
| SE | 65198 | ENSG00000010244 | ZNF207 | chr17 | + | 32360891 | 32360967 | 32360597 | 32360765 | 32361467 | 32361515 | 667 | 393 |
| SE | 65199 | ENSG00000010244 | ZNF207 | chr17 | + | 32360891 | 32360967 | 32360597 | 32360765 | 32362913 | 32362984 | 249 | 146 |
| SE | 65200 | ENSG00000010244 | ZNF207 | chr17 | + | 32361467 | 32361515 | 32360597 | 32360765 | 32362913 | 32362984 | 56 | 73 |
| SE | 65202 | ENSG00000010244 | ZNF207 | chr17 | + | 32365329 | 32365487 | 32362913 | 32362984 | 32367771 | 32368014 | 761 | 628 |
| SE | 65204 | ENSG00000230606 | NA | chr2 | - | 97422438 | 97422656 | 97416220 | 97416405 | 97422954 | 97423084 | 2 | 0 |
| SE | 65210 | ENSG00000230606 | NA | chr2 | - | 97423801 | 97423918 | 97423176 | 97423272 | 97425877 | 97426058 | 1 | 1 |
| SE | 65211 | ENSG00000230606 | NA | chr2 | - | 97424294 | 97424405 | 97423801 | 97423918 | 97433299 | 97433527 | 0 | 0 |
| SE | 65214 | ENSG00000143870 | PDIA6 | chr2 | - | 10790718 | 10790833 | 10789748 | 10789889 | 10791794 | 10791925 | 1237 | 637 |
| SE | 65215 | ENSG00000143870 | PDIA6 | chr2 | - | 10793095 | 10793202 | 10791794 | 10791925 | 10797080 | 10797207 | 838 | 461 |
| SE | 65216 | ENSG00000143870 | PDIA6 | chr2 | - | 10797080 | 10797207 | 10793095 | 10793202 | 10802498 | 10802640 | 521 | 310 |
| SE | 65217 | ENSG00000143870 | PDIA6 | chr2 | - | 10797699 | 10797757 | 10797080 | 10797207 | 10802498 | 10802640 | 733 | 451 |
| SE | 65218 | ENSG00000143870 | PDIA6 | chr2 | - | 10802498 | 10802640 | 10797699 | 10797757 | 10812677 | 10812785 | 496 | 314 |
| SE | 65219 | ENSG00000143870 | PDIA6 | chr2 | - | 10810242 | 10810420 | 10802498 | 10802640 | 10812677 | 10812754 | 0 | 0 |
| SE | 65228 | ENSG00000169379 | ARL13B | chr3 | + | 93988671 | 93988711 | 93980141 | 93980482 | 93995873 | 93995944 | 0 | 0 |
| SE | 65230 | ENSG00000169379 | ARL13B | chr3 | + | 93988858 | 93989116 | 93988671 | 93988711 | 93995873 | 93995944 | 0 | 0 |
| SE | 65234 | ENSG00000169379 | ARL13B | chr3 | + | 93995873 | 93995944 | 93980141 | 93980482 | 94035330 | 94035436 | 2 | 0 |
| SE | 65235 | ENSG00000169379 | ARL13B | chr3 | + | 93995873 | 93995944 | 93980141 | 93980482 | 94036551 | 94036754 | 2 | 0 |
| SE | 65238 | ENSG00000169379 | ARL13B | chr3 | + | 93996547 | 93996684 | 93995873 | 93995944 | 93997079 | 93997136 | 10 | 1 |
| SE | 65239 | ENSG00000169379 | ARL13B | chr3 | + | 93996547 | 93996684 | 93995873 | 93995944 | 94003658 | 94003908 | 1 | 1 |
| SE | 65241 | ENSG00000169379 | ARL13B | chr3 | + | 93996562 | 93996684 | 93995873 | 93995944 | 94035330 | 94035436 | 5 | 0 |
| SE | 65242 | ENSG00000169379 | ARL13B | chr3 | + | 93997079 | 93997136 | 93995873 | 93995944 | 93997246 | 93997301 | 1 | 3 |
| SE | 65244 | ENSG00000169379 | ARL13B | chr3 | + | 93997079 | 93997136 | 93995873 | 93995944 | 94035330 | 94035436 | 0 | 0 |
| SE | 65245 | ENSG00000169379 | ARL13B | chr3 | + | 93997079 | 93997136 | 93996547 | 93996684 | 94003658 | 94003908 | 18 | 4 |
| SE | 65246 | ENSG00000169379 | ARL13B | chr3 | + | 93997079 | 93997136 | 93996547 | 93996684 | 94035330 | 94035436 | 4 | 1 |
| SE | 65250 | ENSG00000169379 | ARL13B | chr3 | + | 93997246 | 93997301 | 93997079 | 93997136 | 94035330 | 94035436 | 3 | 1 |
| SE | 65251 | ENSG00000169379 | ARL13B | chr3 | + | 94003658 | 94003908 | 93980138 | 93980482 | 94035330 | 94035436 | 60 | 29 |
| SE | 65253 | ENSG00000169379 | ARL13B | chr3 | + | 94003658 | 94003908 | 93995873 | 93995944 | 94035330 | 94035436 | 66 | 32 |
| SE | 65255 | ENSG00000169379 | ARL13B | chr3 | + | 94003658 | 94003908 | 93996547 | 93996684 | 94035330 | 94035436 | 61 | 30 |
| SE | 65256 | ENSG00000169379 | ARL13B | chr3 | + | 94003658 | 94003908 | 93997079 | 93997136 | 94035330 | 94035436 | 75 | 32 |
| SE | 65257 | ENSG00000169379 | ARL13B | chr3 | + | 94003658 | 94003908 | 93997246 | 93997301 | 94035330 | 94035436 | 69 | 33 |
| SE | 65258 | ENSG00000169379 | ARL13B | chr3 | + | 94035330 | 94035436 | 93980141 | 93980482 | 94036551 | 94036754 | 21 | 11 |
| SE | 65260 | ENSG00000169379 | ARL13B | chr3 | + | 94035330 | 94035436 | 94003658 | 94003908 | 94036551 | 94036754 | 81 | 40 |
| SE | 65261 | ENSG00000169379 | ARL13B | chr3 | + | 94037931 | 94038018 | 94036551 | 94036754 | 94039879 | 94039988 | 0 | 0 |
| SE | 65262 | ENSG00000169379 | ARL13B | chr3 | + | 94050823 | 94050892 | 94049405 | 94049522 | 94053186 | 94053300 | 51 | 18 |
| SE | 65276 | ENSG00000171219 | CDC42BPG | chr11 | - | 64827685 | 64827783 | 64827526 | 64827611 | 64829470 | 64830070 | 8 | 0 |
| SE | 65280 | ENSG00000152291 | TGOLN2 | chr2 | - | 85324914 | 85324998 | 85318026 | 85322741 | 85326507 | 85326804 | 500 | 308 |
| SE | 65283 | ENSG00000115295 | CLIP4 | chr2 | + | 29121373 | 29121521 | 29114877 | 29115149 | 29131257 | 29131397 | 51 | 34 |
| SE | 65284 | ENSG00000115295 | CLIP4 | chr2 | + | 29121373 | 29121521 | 29115414 | 29115665 | 29131257 | 29131397 | 62 | 38 |
| SE | 65286 | ENSG00000115295 | CLIP4 | chr2 | + | 29145231 | 29145367 | 29143708 | 29143945 | 29152684 | 29152828 | 192 | 92 |
| SE | 65287 | ENSG00000115295 | CLIP4 | chr2 | + | 29156353 | 29156443 | 29152684 | 29152828 | 29157203 | 29157347 | 416 | 305 |
| SE | 65289 | ENSG00000115295 | CLIP4 | chr2 | + | 29163830 | 29163954 | 29160332 | 29160467 | 29167475 | 29167540 | 151 | 95 |
| SE | 65290 | ENSG00000115295 | CLIP4 | chr2 | + | 29174372 | 29174445 | 29167475 | 29167540 | 29181571 | 29181697 | 179 | 88 |
| SE | 65295 | ENSG00000149499 | EML3 | chr11 | - | 62604112 | 62604201 | 62603943 | 62604041 | 62605112 | 62605180 | 103 | 44 |
| SE | 65296 | ENSG00000149499 | EML3 | chr11 | - | 62605641 | 62605773 | 62605112 | 62605180 | 62605854 | 62605980 | 72 | 16 |
| SE | 65301 | ENSG00000169621 | APLF | chr2 | + | 68490189 | 68490261 | 68467584 | 68467827 | 68502730 | 68502903 | 46 | 28 |
| SE | 65302 | ENSG00000169621 | APLF | chr2 | + | 68502730 | 68502903 | 68490189 | 68490261 | 68513547 | 68513680 | 66 | 46 |
| SE | 65307 | ENSG00000169621 | APLF | chr2 | + | 68535282 | 68535383 | 68526060 | 68526242 | 68537871 | 68538227 | 0 | 0 |
| SE | 65310 | ENSG00000285756 | NA | chrX | - | 3818441 | 3818500 | 3817527 | 3817778 | 3818656 | 3818703 | 24 | 26 |
| SE | 65311 | ENSG00000285756 | NA | chrX | - | 3818441 | 3818500 | 3817527 | 3817778 | 3822373 | 3822580 | 40 | 28 |
| SE | 65313 | ENSG00000285756 | NA | chrX | - | 3818656 | 3818703 | 3817527 | 3817778 | 3822373 | 3822580 | 2 | 2 |
| SE | 65317 | ENSG00000285756 | NA | chrX | - | 3818656 | 3818703 | 3818441 | 3818500 | 3825087 | 3825231 | 7 | 7 |
| SE | 65322 | ENSG00000285756 | NA | chrX | - | 3822373 | 3822580 | 3817527 | 3817778 | 3828505 | 3828597 | 0 | 3 |
| SE | 65323 | ENSG00000285756 | NA | chrX | - | 3822373 | 3822580 | 3818441 | 3818500 | 3825087 | 3825231 | 63 | 31 |
| SE | 65326 | ENSG00000285756 | NA | chrX | - | 3822373 | 3822580 | 3818656 | 3818703 | 3826600 | 3826631 | 5 | 3 |
| SE | 65327 | ENSG00000285756 | NA | chrX | - | 3822373 | 3822580 | 3818656 | 3818703 | 3828505 | 3828597 | 4 | 6 |
| SE | 65328 | ENSG00000285756 | NA | chrX | - | 3825087 | 3825231 | 3818656 | 3818703 | 3826600 | 3826631 | 1 | 0 |
| SE | 65329 | ENSG00000285756 | NA | chrX | - | 3825087 | 3825231 | 3818656 | 3818703 | 3828505 | 3828979 | 0 | 0 |
| SE | 65333 | ENSG00000285756 | NA | chrX | - | 3826600 | 3826729 | 3818656 | 3818703 | 3828505 | 3828979 | 3 | 0 |
| SE | 65334 | ENSG00000285756 | NA | chrX | - | 3826600 | 3826729 | 3822373 | 3822580 | 3828505 | 3828979 | 4 | 0 |

| | | | | | | | | | | | | | | | | | | | | | | | | | | | | | | | | | | | | | | | | | | | | | | |
|---|---|---|---|---|---|---|---|---|---|---|---|---|---|---|---|---|---|---|---|---|---|---|---|---|---|---|---|---|---|---|---|---|---|---|---|---|---|---|---|---|---|---|---|---|---|---|
| SE | 65335 | ENSG00000285756 | NA | chrX | - | 3826600 | 3826729 | 3825087 | 3825231 | 3828505 | 3828979 | 4 | 0 | 19 | 6 | 5 | 0 | 0 | 0 | 0 | 1 | 12 | 0 | 3 | 3 | 12 | 0 | 0 | 0 | 0 | 2 | 278 | 149 | 1 | 1 | 1 | NA | 1 | 1 | 0.728 | 1 | NA | 1 | 1 | 0.763 | -0.009 |
| SE | 65336 | ENSG00000285756 | NA | chrX | - | 3828505 | 3829392 | 3825087 | 3825231 | 3854691 | 3854927 | 0 | 17 | 0 | 0 | 1 | 0 | 3 | 0 | 0 | 0 | 0 | 0 | 9 | 0 | 97 | 0 | 0 | 0 | 0 | 0 | 298 | 149 | 1 | 1 | NA | 0.739 | NA | NA | 1 | NA | NA | 1 | NA | 1 | -0.131 |
| SE | 65338 | ENSG00000285756 | NA | chrX | - | 3854691 | 3854927 | 3828505 | 3829392 | 3860789 | 3860996 | 0 | 19 | 1 | 0 | 2 | 0 | 0 | 1 | 0 | 1 | 0 | 0 | 9 | 0 | 102 | 0 | 0 | 0 | 0 | 0 | 298 | 149 | 1 | 1 | NA | 1 | 0.333 | NA | 0.5 | NA | NA | 1 | NA | 1 | -0.389 |
| SE | 65339 | ENSG00000285756 | NA | chrX | - | 3860789 | 3860996 | 3853009 | 3854927 | 3866921 | 3867541 | 0 | 2 | 1 | 0 | 0 | 0 | 1 | 1 | 0 | 0 | 0 | 0 | 0 | 0 | 6 | 0 | 0 | 1 | 0 | 0 | 298 | 149 | 1 | 1 | NA | 0.5 | 0.333 | NA | NA | NA | NA | 0 | NA | 1 | -0.084 |
| SE | 65340 | ENSG00000285756 | NA | chrX | - | 3860789 | 3860996 | 3853009 | 3854927 | 3866949 | 3867445 | 0 | 3 | 1 | 0 | 0 | 0 | 2 | 2 | 0 | 0 | 0 | 0 | 0 | 0 | 6 | 0 | 0 | 0 | 0 | 9 | 298 | 149 | 1 | 1 | NA | 0.429 | 0.2 | NA | NA | NA | NA | NA | NA | 0.25 | 0.065 |
| SE | 65341 | ENSG00000285756 | NA | chrX | - | 3863501 | 3863645 | 3860798 | 3860996 | 3866949 | 3867806 | 0 | 9 | 0 | 0 | 0 | 0 | 1 | 0 | 0 | 0 | 0 | 0 | 0 | 0 | 2 | 0 | 0 | 0 | 0 | 0 | 293 | 149 | 1 | 1 | NA | 0.821 | NA | NA | NA | NA | NA | NA | NA | 1 | -0.179 |
| SE | 65342 | ENSG00000285756 | NA | chrX | - | 3866921 | 3867806 | 3854691 | 3854927 | 3893115 | 3893351 | 4 | 34 | 10 | 10 | 5 | 0 | 0 | 0 | 0 | 0 | 17 | 0 | 3 | 10 | 28 | 0 | 0 | 0 | 0 | 4 | 298 | 149 | 1 | 1 | 1 | 1 | 1 | 1 | 1 | 1 | NA | 1 | 1 | 0.778 | 0.055 |
| SE | 65343 | ENSG00000285756 | NA | chrX | - | 3866949 | 3867806 | 3863501 | 3863645 | 3893115 | 3893351 | 4 | 42 | 9 | 10 | 5 | 2 | 1 | 0 | 1 | 0 | 17 | 0 | 2 | 10 | 30 | 0 | 0 | 0 | 1 | 0 | 298 | 149 | 1 | 1 | 0.5 | 0.955 | 1 | 0.833 | 1 | 1 | NA | 1 | 0.833 | 1 | -0.101 |
| SE | 65344 | ENSG00000285756 | NA | chrX | - | 3893115 | 3893351 | 3866921 | 3867806 | 3899215 | 3899422 | 13 | 37 | 14 | 12 | 7 | 0 | 34 | 0 | 0 | 0 | 17 | 0 | 2 | 17 | 31 | 0 | 0 | 6 | 0 | 41 | 298 | 149 | 1 | 1 | 1 | 0.352 | 1 | 1 | 1 | 1 | NA | 0.143 | 1 | 0.274 | 0.266 |
| SE | 65345 | ENSG00000285756 | NA | chrX | - | 3893115 | 3893351 | 3866921 | 3867806 | 3901927 | 3902071 | 4 | 33 | 10 | 11 | 5 | 0 | 9 | 0 | 0 | 0 | 32 | 0 | 3 | 13 | 58 | 0 | 0 | 0 | 0 | 1 | 298 | 149 | 1 | 1 | 1 | 0.647 | 1 | 1 | 1 | 1 | NA | 1 | 1 | 0.967 | -0.062 |
| SE | 65346 | ENSG00000285756 | NA | chrX | - | 3899215 | 3899422 | 3866921 | 3867806 | 3901927 | 3902071 | 0 | 38 | 1 | 0 | 0 | 0 | 9 | 0 | 0 | 0 | 0 | 0 | 7 | 0 | 51 | 0 | 0 | 0 | 0 | 1 | 298 | 149 | 1 | 1 | NA | 0.679 | 1 | NA | NA | NA | NA | 1 | NA | 0.962 | -0.141 |
| SE | 65347 | ENSG00000285756 | NA | chrX | - | 3899215 | 3899422 | 3891437 | 3893351 | 3901927 | 3902071 | 9 | 8 | 6 | 2 | 2 | 0 | 0 | 1 | 1 | 0 | 0 | 0 | 1 | 7 | 13 | 15 | 0 | 1 | 3 | 30 | 298 | 149 | 1 | 1 | 1 | 1 | 0.75 | 0.5 | 1 | 0 | NA | 0.333 | 0.538 | 0.178 | 0.588 |
| SE | 65350 | ENSG00000105948 | TTC26 | chr7 | + | 139168343 | 139168397 | 139166839 | 139166895 | 139169288 | 139169352 | 0 | 0 | 1 | 1 | 1 | 0 | 0 | 1 | 0 | 0 | 0 | 0 | 2 | 0 | 0 | 0 | 0 | 0 | 0 | 0 | 203 | 149 | 1 | 1 | NA | NA | 0.423 | 1 | 1 | NA | NA | 1 | NA | NA | -0.192 |
| SE | 65355 | ENSG00000153214 | TMEM87B | chr2 | + | 112066935 | 112067067 | 112059976 | 112060037 | 112074911 | 112074962 | 206 | 135 | 149 | 179 | 315 | 0 | 0 | 0 | 0 | 0 | 116 | 171 | 79 | 95 | 75 | 0 | 0 | 0 | 0 | 4 | 281 | 149 | 1 | 1 | 1 | 1 | 1 | 1 | 1 | 1 | 1 | 1 | 1 | 0.909 | 0.018 |
| SE | 65356 | ENSG00000153214 | TMEM87B | chr2 | + | 112066935 | 112067067 | 112059976 | 112060037 | 112081056 | 112081118 | 2 | 1 | 5 | 6 | 1 | 0 | 0 | 0 | 1 | 0 | 0 | 8 | 4 | 1 | 0 | 0 | 0 | 0 | 0 | 0 | 281 | 149 | 1 | 1 | 1 | 1 | 1 | 0.761 | 1 | NA | 1 | 1 | 1 | NA | -0.048 |
| SE | 65357 | ENSG00000153214 | TMEM87B | chr2 | + | 112074911 | 112074962 | 112059976 | 112060037 | 112081056 | 112081118 | 1 | 0 | 0 | 0 | 0 | 0 | 0 | 0 | 1 | 0 | 0 | 0 | 1 | 0 | 4 | 0 | 0 | 0 | 0 | 0 | 200 | 149 | 1 | 1 | 1 | NA | NA | 0 | NA | NA | NA | 1 | NA | 1 | -0.5 |
| SE | 65359 | ENSG00000153214 | TMEM87B | chr2 | + | 112074911 | 112074962 | 112066935 | 112067067 | 112081056 | 112081118 | 50 | 32 | 23 | 60 | 69 | 2 | 0 | 1 | 3 | 0 | 11 | 30 | 30 | 45 | 0 | 0 | 0 | 2 | 1 | 0 | 200 | 149 | 1 | 1 | 0.949 | 1 | 0.945 | 0.937 | 1 | 1 | 1 | 0.918 | 0.971 | NA | -0.006 |
| SE | 65360 | ENSG00000153214 | TMEM87B | chr2 | + | 112077191 | 112077282 | 112066935 | 112067067 | 112081056 | 112081118 | 83 | 83 | 60 | 98 | 154 | 2 | 0 | 1 | 3 | 0 | 96 | 37 | 27 | 41 | 51 | 0 | 0 | 2 | 1 | 0 | 240 | 149 | 1 | 1 | 0.963 | 1 | 0.974 | 0.953 | 1 | 1 | 1 | 0.893 | 0.962 | 1 | 0.007 |
| SE | 65361 | ENSG00000153214 | TMEM87B | chr2 | + | 112077191 | 112077282 | 112066935 | 112067067 | 112086004 | 112086104 | 1 | 3 | 1 | 3 | 2 | 0 | 0 | 0 | 0 | 0 | 7 | 2 | 1 | 4 | 0 | 0 | 0 | 1 | 0 | 0 | 240 | 149 | 1 | 1 | 1 | 1 | 1 | 1 | 1 | 1 | 1 | 0.383 | 1 | NA | 0.154 |
| SE | 65362 | ENSG00000153214 | TMEM87B | chr2 | + | 112077191 | 112077282 | 112074911 | 112074962 | 112081056 | 112081118 | 278 | 203 | 224 | 262 | 463 | 2 | 2 | 0 | 0 | 0 | 190 | 210 | 84 | 104 | 149 | 0 | 1 | 1 | 0 | 0 | 240 | 149 | 1 | 1 | 0.989 | 0.984 | 1 | 1 | 1 | 1 | 0.992 | 0.981 | 1 | 1 | 0 |
| SE | 65363 | ENSG00000153214 | TMEM87B | chr2 | + | 112081056 | 112081118 | 112066935 | 112067067 | 112086004 | 112086104 | 4 | 6 | 4 | 11 | 8 | 0 | 0 | 0 | 0 | 0 | 0 | 4 | 6 | 11 | 27 | 0 | 0 | 1 | 0 | 0 | 211 | 149 | 1 | 1 | 1 | 1 | 1 | 1 | 1 | NA | 1 | 0.809 | 1 | 1 | 0.048 |
| SE | 65364 | ENSG00000153214 | TMEM87B | chr2 | + | 112081056 | 112081118 | 112077191 | 112077282 | 112086004 | 112086104 | 44 | 36 | 56 | 64 | 82 | 0 | 2 | 0 | 0 | 0 | 41 | 48 | 19 | 32 | 72 | 0 | 0 | 0 | 0 | 0 | 211 | 149 | 1 | 1 | 1 | 0.927 | 1 | 1 | 1 | 1 | 1 | 1 | 1 | 1 | -0.015 |
| SE | 65366 | ENSG00000153214 | TMEM87B | chr2 | + | 112097232 | 112097291 | 112097043 | 112097152 | 112098594 | 112098698 | 207 | 163 | 156 | 198 | 172 | 0 | 0 | 0 | 0 | 0 | 110 | 202 | 127 | 126 | 47 | 0 | 0 | 0 | 0 | 5 | 208 | 149 | 1 | 1 | 1 | 1 | 1 | 1 | 1 | 1 | 1 | 1 | 1 | 0.871 | 0.026 |
| SE | 65367 | ENSG00000153214 | TMEM87B | chr2 | + | 112107787 | 112107840 | 112106001 | 112106075 | 112116083 | 112119273 | 34 | 14 | 29 | 30 | 90 | 0 | 0 | 1 | 0 | 0 | 13 | 18 | 14 | 15 | 0 | 0 | 0 | 0 | 0 | 0 | 202 | 149 | 1 | 1 | 1 | 1 | 0.955 | 1 | 1 | 1 | 1 | 1 | 1 | NA | -0.009 |
| SE | 65368 | ENSG00000153214 | TMEM87B | chr2 | + | 112112898 | 112112929 | 112107787 | 112107840 | 112116083 | 112119273 | 141 | 152 | 148 | 138 | 318 | 0 | 0 | 0 | 0 | 1 | 36 | 98 | 106 | 116 | 68 | 3 | 0 | 0 | 0 | 0 | 180 | 149 | 1 | 1 | 1 | 1 | 1 | 1 | 0.996 | 0.909 | 1 | 1 | 1 | 1 | 0.017 |
| SE | 65369 | ENSG00000087008 | ACOX3 | chr4 | - | 8370907 | 8370994 | 8366281 | 8367080 | 8373560 | 8373628 | 20 | 5 | 20 | 6 | 11 | 0 | 0 | 0 | 0 | 0 | 7 | 9 | 7 | 11 | 45 | 0 | 3 | 0 | 0 | 0 | 236 | 149 | 1 | 1 | 1 | 1 | 1 | 1 | 1 | 1 | 0.654 | 1 | 1 | 1 | 0.069 |
| SE | 65370 | ENSG00000087008 | ACOX3 | chr4 | - | 8370907 | 8370994 | 8366281 | 8367080 | 8374977 | 8375152 | 12 | 3 | 13 | 7 | 10 | 0 | 0 | 0 | 0 | 0 | 7 | 4 | 0 | 8 | 15 | 0 | 0 | 0 | 0 | 8 | 236 | 149 | 1 | 1 | 1 | 1 | 1 | 1 | 1 | 1 | 1 | NA | 1 | 0.542 | 0.115 |
| SE | 65371 | ENSG00000087008 | ACOX3 | chr4 | - | 8370907 | 8370994 | 8366281 | 8367080 | 8381491 | 8381607 | 9 | 1 | 15 | 3 | 5 | 0 | 1 | 3 | 0 | 0 | 7 | 4 | 0 | 0 | 10 | 0 | 0 | 1 | 0 | 0 | 236 | 149 | 1 | 1 | 1 | 0.387 | 0.759 | 1 | 1 | 1 | 1 | 0 | NA | 1 | 0.079 |
| SE | 65375 | ENSG00000087008 | ACOX3 | chr4 | - | 8374977 | 8375152 | 8370907 | 8370994 | 8381491 | 8381607 | 13 | 14 | 8 | 13 | 19 | 0 | 0 | 2 | 0 | 0 | 9 | 3 | 0 | 16 | 41 | 0 | 0 | 0 | 0 | 0 | 298 | 149 | 1 | 1 | 1 | 1 | 0.667 | 1 | 1 | 1 | 1 | NA | 1 | 1 | -0.067 |
| SE | 65376 | ENSG00000087008 | ACOX3 | chr4 | - | 8392332 | 8392453 | 8389611 | 8389734 | 8394619 | 8394742 | 30 | 11 | 12 | 20 | 14 | 1 | 2 | 0 | 0 | 0 | 0 | 31 | 12 | 18 | 31 | 0 | 0 | 0 | 0 | 0 | 270 | 149 | 1 | 1 | 0.943 | 0.752 | 1 | 1 | 1 | NA | 1 | 1 | 1 | 1 | -0.061 |
| SE | 65377 | ENSG00000087008 | ACOX3 | chr4 | - | 8399555 | 8399652 | 8396936 | 8397119 | 8405954 | 8406043 | 22 | 9 | 30 | 21 | 17 | 0 | 0 | 0 | 0 | 0 | 0 | 24 | 18 | 22 | 40 | 0 | 6 | 0 | 0 | 0 | 246 | 149 | 1 | 1 | 1 | 1 | 1 | 1 | 1 | NA | 0.708 | 1 | 1 | 1 | 0.073 |
| SE | 65378 | ENSG00000087008 | ACOX3 | chr4 | - | 8410211 | 8410355 | 8405954 | 8406043 | 8414291 | 8414381 | 24 | 9 | 21 | 25 | 27 | 0 | 0 | 1 | 0 | 0 | 29 | 64 | 15 | 25 | 29 | 0 | 0 | 0 | 0 | 0 | 293 | 149 | 1 | 1 | 1 | 1 | 0.914 | 1 | 1 | 1 | 1 | 1 | 1 | 1 | -0.017 |
| SE | 65380 | ENSG00000087008 | ACOX3 | chr4 | - | 8416377 | 8416535 | 8415765 | 8415999 | 8440647 | 8440723 | 18 | 1 | 18 | 6 | 12 | 1 | 0 | 0 | 0 | 0 | 0 | 34 | 6 | 13 | 20 | 0 | 0 | 0 | 0 | 0 | 298 | 149 | 1 | 1 | 0.9 | 1 | 1 | 1 | 1 | NA | 1 | 1 | 1 | 1 | -0.02 |
| SE | 65388 | ENSG00000140265 | ZSCAN29 | chr15 | - | 43369595 | 43370025 | 43368922 | 43369127 | 43370557 | 43371043 | 88 | 38 | 68 | 53 | 18 | 1 | 0 | 0 | 0 | 0 | 87 | 54 | 19 | 42 | 18 | 0 | 0 | 0 | 0 | 0 | 298 | 149 | 1 | 1 | 0.978 | 1 | 1 | 1 | 1 | 1 | 1 | 1 | 1 | 1 | -0.004 |
| SE | 65389 | ENSG00000143942 | CHAC2 | chr2 | + | 53771906 | 53771942 | 53767803 | 53768021 | 53774141 | 53775196 | 5 | 2 | 2 | 5 | 16 | 0 | 6 | 0 | 0 | 0 | 0 | 12 | 0 | 0 | 0 | 0 | 0 | 0 | 0 | 0 | 185 | 149 | 1 | 1 | 1 | 0.212 | 1 | 1 | 1 | NA | 1 | NA | NA | NA | -0.158 |
| SE | 65390 | ENSG00000164172 | MOCS2 | chr5 | - | 53101358 | 53101509 | 53100410 | 53100534 | 53102096 | 53102224 | 92 | 80 | 135 | 127 | 250 | 0 | 0 | 3 | 2 | 1 | 61 | 107 | 60 | 95 | 38 | 0 | 0 | 0 | 0 | 0 | 298 | 149 | 1 | 1 | 1 | 1 | 0.957 | 0.969 | 0.992 | 1 | 1 | 1 | 1 | 1 | -0.016 |
| SE | 65394 | ENSG00000164172 | MOCS2 | chr5 | - | 53107076 | 53107221 | 53102096 | 53102224 | 53108521 | 53108643 | 168 | 76 | 148 | 147 | 207 | 0 | 0 | 3 | 2 | 0 | 56 | 114 | 67 | 91 | 22 | 0 | 0 | 0 | 0 | 0 | 294 | 149 | 1 | 1 | 1 | 1 | 1 | 0.974 | 1 | 1 | 1 | 1 | 1 | 1 | -0.005 |
| SE | 65395 | ENSG00000129083 | COPB1 | chr11 | - | 14453224 | 14453305 | 14451834 | 14451936 | 14457783 | 14457883 | 3 | 0 | 1 | 0 | 0 | 0 | 0 | 0 | 0 | 1 | 0 | 0 | 1 | 7 | 0 | 0 | 0 | 0 | 1 | 0 | 230 | 149 | 1 | 1 | 1 | NA | 1 | NA | 0 | NA | NA | 1 | 0.819 | NA | -0.243 |
| SE | 65396 | ENSG00000129083 | COPB1 | chr11 | - | 14457783 | 14457883 | 14451834 | 14451936 | 14458531 | 14458687 | 50 | 51 | 52 | 47 | 113 | 0 | 0 | 1 | 0 | 0 | 11 | 13 | 25 | 4 | 0 | 0 | 0 | 0 | 0 | 0 | 249 | 149 | 1 | 1 | 1 | 1 | 0.969 | 1 | 1 | 1 | 1 | 1 | 1 | NA | -0.006 |
| SE | 65397 | ENSG00000129083 | COPB1 | chr11 | - | 14468680 | 14468860 | 14466281 | 14466426 | 14469335 | 14469563 | 543 | 357 | 410 | 453 | 846 | 0 | 0 | 0 | 0 | 6 | 26 | 231 | 277 | 62 | 47 | 0 | 1 | 0 | 0 | 0 | 298 | 149 | 1 | 1 | 1 | 1 | 1 | 1 | 0.986 | 1 | 0.991 | 1 | 1 | 1 | -0.001 |
| SE | 65398 | ENSG00000129083 | COPB1 | chr11 | - | 14469335 | 14469563 | 14468680 | 14468860 | 14474494 | 14474615 | 618 | 432 | 438 | 463 | 944 | 0 | 0 | 0 | 0 | 0 | 30 | 323 | 370 | 73 | 44 | 0 | 0 | 1 | 0 | 0 | 298 | 149 | 1 | 1 | 1 | 1 | 1 | 1 | 1 | 1 | 1 | 0.995 | 1 | 1 | 0.001 |
| SE | 65399 | ENSG00000129083 | COPB1 | chr11 | - | 14475784 | 14475945 | 14474494 | 14474615 | 14476918 | 14477015 | 402 | 288 | 362 | 366 | 889 | 0 | 0 | 1 | 0 | 3 | 26 | 235 | 246 | 53 | 28 | 0 | 0 | 0 | 0 | 0 | 298 | 149 | 1 | 1 | 1 | 1 | 0.995 | 1 | 0.993 | 1 | 1 | 1 | 1 | 1 | -0.002 |
| SE | 65400 | ENSG00000129083 | COPB1 | chr11 | - | 14479568 | 14479714 | 14476916 | 14477015 | 14480758 | 14480905 | 614 | 424 | 522 | 541 | 909 | 0 | 0 | 0 | 1 | 0 | 40 | 309 | 357 | 74 | 21 | 0 | 0 | 1 | 0 | 0 | 295 | 149 | 1 | 1 | 1 | 1 | 1 | 0.996 | 1 | 1 | 1 | 0.994 | 1 | 1 | 0 |
| SE | 65401 | ENSG00000129083 | COPB1 | chr11 | - | 14494531 | 14494727 | 14494316 | 14494439 | 14498837 | 14498985 | 0 | 0 | 0 | 2 | 1 | 105 | 45 | 70 | 72 | 97 | 0 | 4 | 1 | 1 | 0 | 17 | 40 | 40 | 20 | 2 | 298 | 149 | 1 | 1 | 0 | 0 | 0 | 0.014 | 0.005 | 0 | 0.048 | 0.012 | 0.024 | 0 | -0.013 |
| SE | 65403 | ENSG00000245937 | LINC01184 | chr5 | - | 127956803 | 127956921 | 127939151 | 127941220 | 127966162 | 127966509 | 6 | 2 | 3 | 3 | 0 | 0 | 3 | 0 | 0 | 0 | 0 | 0 | 0 | 7 | 0 | 0 | 0 | 0 | 0 | 0 | 267 | 149 | 1 | 1 | 1 | 0.271 | 1 | 1 | NA | NA | NA | NA | 1 | NA | -0.182 |
| SE | 65405 | ENSG00000245937 | LINC01184 | chr5 | - | 128061143 | 128061244 | 127965887 | 127966928 | 128081972 | 128082161 | 8 | 2 | 14 | 9 | 26 | 0 | 0 | 0 | 0 | 1 | 0 | 24 | 1 | 5 | 13 | 0 | 0 | 0 | 0 | 0 | 250 | 149 | 1 | 1 | 1 | 1 | 1 | 1 | 0.939 | NA | 1 | 1 | 1 | 1 | -0.012 |
| SE | 65406 | ENSG00000245937 | LINC01184 | chr5 | - | 128061143 | 128061244 | 127965887 | 127966928 | 128082734 | 128083100 | 12 | 4 | 18 | 18 | 48 | 0 | 0 | 1 | 0 | 7 | 0 | 9 | 0 | 8 | 13 | 0 | 0 | 0 | 0 | 0 | 250 | 149 | 1 | 1 | 1 | 1 | 0.915 | 1 | 0.803 | NA | 1 | NA | 1 | 1 | -0.056 |
| SE | 65407 | ENSG00000245937 | LINC01184 | chr5 | - | 128061143 | 128061244 | 128021527 | 128024298 | 128081972 | 128082161 | 14 | 1 | 14 | 6 | 12 | 0 | 1 | 0 | 0 | 0 | 0 | 19 | 4 | 0 | 0 | 0 | 0 | 0 | 0 | 0 | 250 | 149 | 1 | 1 | 1 | 0.373 | 1 | 1 | 1 | NA | 1 | 1 | NA | NA | -0.125 |
| SE | 65408 | ENSG00000245937 | LINC01184 | chr5 | - | 128062369 | 128062500 | 128061143 | 128061244 | 128082734 | 128083047 | 0 | 0 | 0 | 0 | 0 | 9 | 4 | 8 | 14 | 31 | 0 | 0 | 0 | 3 | 0 | 0 | 2 | 1 | 3 | 0 | 280 | 149 | 1 | 1 | 0 | 0 | 0 | 0 | 0 | NA | 0 | 0 | 0.347 | NA | -0.116 |
| SE | 65409 | ENSG00000245937 | LINC01184 | chr5 | - | 128062369 | 128062500 | 128061143 | 128061244 | 128082969 | 128083090 | 0 | 0 | 0 | 0 | 0 | 3 | 0 | 2 | 2 | 0 | 0 | 0 | 0 | 3 | 0 | 0 | 0 | 0 | 0 | 0 | 280 | 149 | 1 | 1 | 0 | NA | 0 | 0 | NA | NA | NA | NA | 1 | NA | -1 |
| SE | 65413 | ENSG00000245937 | LINC01184 | chr5 | - | 128064736 | 128064850 | 128061143 | 128061244 | 128076946 | 128077093 | 0 | 0 | 2 | 0 | 0 | 0 | 0 | 4 | 1 | 2 | 0 | 0 | 0 | 0 | 10 | 0 | 8 | 0 | 1 | 0 | 263 | 149 | 1 | 1 | NA | NA | 0.221 | 0 | 0 | NA | 0 | NA | 0 | 1 | -0.26 |
| SE | 65414 | ENSG00000245937 | LINC01184 | chr5 | - | 128064736 | 128064850 | 128061143 | 128061244 | 128081972 | 128082194 | 0 | 0 | 2 | 0 | 0 | 7 | 1 | 5 | 5 | 7 | 0 | 0 | 0 | 1 | 5 | 0 | 17 | 1 | 0 | 0 | 263 | 149 | 1 | 1 | 0 | 0 | 0.185 | 0 | 0 | NA | 0 | 0 | 1 | 1 | -0.463 |
| SE | 65415 | ENSG00000245937 | LINC01184 | chr5 | - | 128064736 | 128064850 | 128061143 | 128061244 | 128082734 | 128083070 | 0 | 1 | 2 | 0 | 2 | 9 | 4 | 8 | 14 | 31 | 0 | 0 | 0 | 0 | 5 | 0 | 2 | 1 | 3 | 0 | 263 | 149 | 1 | 1 | 0 | 0.124 | 0.124 | 0 | 0.035 | NA | 0 | 0 | 0 | 1 | -0.193 |
| SE | 65418 | ENSG00000245937 | LINC01184 | chr5 | - | 128076946 | 128077093 | 127965884 | 127966928 | 128081972 | 128082194 | 1 | 0 | 0 | 0 | 1 | 0 | 0 | 0 | 0 | 1 | 0 | 0 | 1 | 0 | 0 | 0 | 0 | 0 | 0 | 0 | 296 | 149 | 1 | 1 | 1 | NA | NA | NA | 0.335 | NA | NA | 1 | NA | NA | -0.333 |
| SE | 65420 | ENSG00000245937 | LINC01184 | chr5 | - | 128076946 | 128077093 | 128061143 | 128061244 | 128081972 | 128082194 | 1 | 0 | 4 | 1 | 3 | 7 | 1 | 5 | 5 | 7 | 0 | 8 | 1 | 1 | 0 | 0 | 17 | 1 | 0 | 0 | 296 | 149 | 1 | 1 | 0.067 | 0 | 0.287 | 0.091 | 0.177 | NA | 0.192 | 0.335 | 1 | NA | -0.385 |
| SE | 65421 | ENSG00000245937 | LINC01184 | chr5 | - | 128076946 | 128077093 | 128061143 | 128061244 | 128082734 | 128082881 | 4 | 0 | 10 | 4 | 2 | 9 | 4 | 8 | 14 | 31 | 0 | 8 | 0 | 1 | 0 | 0 | 2 | 1 | 3 | 0 | 296 | 149 | 1 | 1 | 0.183 | 0 | 0.386 | 0.126 | 0.031 | NA | 0.668 | 0 | 0.144 | NA | -0.125 |
| SE | 65422 | ENSG00000245937 | LINC01184 | chr5 | - | 128076946 | 128077093 | 128061143 | 128061244 | 128082969 | 128083070 | 0 | 0 | 4 | 1 | 2 | 3 | 0 | 2 | 2 | 0 | 0 | 8 | 0 | 1 | 0 | 0 | 0 | 0 | 0 | 0 | 296 | 149 | 1 | 1 | 0 | NA | 0.502 | 0.201 | 1 | NA | 1 | NA | 1 | NA | -0.574 |
| SE | 65425 | ENSG00000245937 | LINC01184 | chr5 | - | 128076946 | 128077093 | 128064736 | 128064850 | 128081972 | 128082194 | 1 | 0 | 0 | 0 | 1 | 0 | 0 | 0 | 0 | 0 | 0 | 0 | 1 | 3 | 5 | 0 | 0 | 0 | 1 | 0 | 296 | 149 | 1 | 1 | 1 | NA | NA | NA | 1 | NA | NA | 1 | 0.602 | 1 | 0.133 |
| SE | 65426 | ENSG00000245937 | LINC01184 | chr5 | - | 128076946 | 128077093 | 128064736 | 128064850 | 128082734 | 128082881 | 4 | 0 | 6 | 3 | 0 | 0 | 1 | 1 | 0 | 2 | 0 | 0 | 0 | 3 | 5 | 0 | 0 | 0 | 0 | 0 | 296 | 149 | 1 | 1 | 1 | 0 | 0.751 | 1 | 0 | NA | NA | | 1 | 1 | -0.45 |
| SE | 65428 | ENSG00000245937 | LINC01184 | chr5 | - | 128081972 | 128082191 | 128061143 | 128061244 | 128082734 | 128082974 | 7 | 1 | 5 | 5 | 7 | 9 | 4 | 8 | 14 | 31 | 0 | 17 | 1 | 0 | 0 | 0 | 2 | 1 | 3 | 0 | 298 | 149 | 1 | 1 | 0.28 | 0.111 | 0.238 | 0.152 | 0.101 | NA | 0.81 | 0.333 | 0 | NA | -0.205 |
| SE | 65430 | ENSG00000245937 | LINC01184 | chr5 | - | 128081972 | 128082194 | 128061143 | 128061244 | 128082734 | 128082960 | 7 | 1 | 5 | 5 | 7 | 9 | 4 | 8 | 14 | 31 | 0 | 17 | 1 | 0 | 0 | 0 | 2 | 1 | 3 | 0 | 298 | 149 | 1 | 1 | 0.28 | 0.111 | 0.238 | 0.152 | 0.101 | NA | 0.81 | 0.333 | 0 | NA | -0.205 |
| SE | 65431 | ENSG00000245937 | LINC01184 | chr5 | - | 128081972 | 128082194 | 128064736 | 128064850 | 128082734 | 128082960 | 0 | 0 | 0 | 0 | 0 | 0 | 1 | 1 | 0 | 2 | 0 | 0 | 0 | 1 | 0 | 0 | 0 | 0 | 0 | 0 | 298 | 149 | 1 | 1 | NA | 0 | 0 | NA | 0 | NA | NA | NA | 1 | NA | -1 |
| SE | 65432 | ENSG00000245937 | LINC01184 | chr5 | - | 128081972 | 128082194 | 128076946 | 128077093 | 128082734 | 128082960 | 1 | 0 | 0 | 0 | 1 | 4 | 0 | 6 | 3 | 0 | 0 | 0 | 0 | 1 | 0 | 0 | 0 | 0 | 0 | 0 | 298 | 149 | 1 | 1 | 0.111 | NA | 0 | 0 | 1 | NA | NA | 1 | NA | NA | -0.722 |
| SE | 65433 | ENSG00000164919 | COX6C | chr8 | - | 99887489 | 99887618 | 99877864 | 99878265 | 99891907 | 99892047 | 1570 | 708 | 1040 | 1073 | 1217 | 0 | 0 | 0 | 0 | 0 | 1733 | 993 | 880 | 1248 | 1218 | 0 | 4 | 0 | 0 | 0 | 278 | 149 | 1 | 1 | 1 | 1 | 1 | 1 | 1 | 1 | 0.993 | 1 | 1 | 1 | 0.001 |
| SE | 65434 | ENSG00000164919 | COX6C | chr8 | - | 99891907 | 99892047 | 99887489 | 99887618 | 99893638 | 99893681 | 945 | 577 | 762 | 810 | 873 | 0 | 0 | 0 | 0 | 1 | 1401 | 773 | 690 | 1064 | 1051 | 0 | 0 | 0 | 0 | 0 | 289 | 149 | 1 | 1 | 1 | 1 | 1 | 1 | 0.998 | 1 | 1 | 1 | 1 | 1 | 0 |
| SE | 65435 | ENSG00000164919 | COX6C | chr8 | - | 99891907 | 99892052 | 99887060 | 99887618 | 99893638 | 99893680 | 1756 | 967 | 1307 | 1317 | 1377 | 0 | 0 | 0 | 0 | 1 | 2250 | 1302 | 1189 | 1639 | 1683 | 0 | 0 | 0 | 0 | 0 | 294 | 149 | 1 | 1 | 1 | 1 | 1 | 1 | 0.999 | 1 | 1 | 1 | 1 | 1 | 0 |
| SE | 65438 | ENSG00000166164 | BRD7 | chr16 | - | 50320662 | 50320774 | 50320247 | 50320391 | 50321981 | 50322038 | 260 | 185 | 224 | 241 | 324 | 0 | 0 | 0 | 1 | 0 | 125 | 239 | 383 | 308 | 125 | 0 | 0 | 0 | 0 | 0 | 261 | 149 | 1 | 1 | 1 | 1 | 1 | 0.993 | 1 | 1 | 1 | 1 | 1 | 1 | -0.001 |
| SE | 65439 | ENSG00000166164 | BRD7 | chr16 | - | 50320662 | 50320777 | 50320247 | 50320391 | 50321981 | 50322038 | 189 | 143 | 175 | 163 | 263 | 0 | 0 | 0 | 1 | 0 | 100 | 145 | 230 | 213 | 72 | 0 | 0 | 0 | 0 | 0 | 264 | 149 | 1 | 1 | 1 | 1 | 1 | 0.989 | 1 | 1 | 1 | 1 | 1 | 1 | -0.002 |
| SE | 65440 | ENSG00000166164 | BRD7 | chr16 | - | 50321981 | 50322038 | 50320662 | 50320774 | 50325747 | 50325883 | 76 | 41 | 55 | 68 | 107 | 0 | 0 | 0 | 0 | 0 | 53 | 112 | 212 | 147 | 32 | 0 | 0 | 1 | 0 | 0 | 206 | 149 | 1 | 1 | 1 | 1 | 1 | 1 | 1 | 1 | 1 | 0.994 | 1 | 1 | 0.001 |
| SE | 65442 | ENSG00000166164 | BRD7 | chr16 | - | 50325747 | 50325883 | 50321981 | 50322038 | 50328668 | 50328744 | 54 | 22 | 38 | 51 | 55 | 0 | 0 | 1 | 1 | 0 | 43 | 71 | 158 | 92 | 14 | 0 | 0 | 0 | 0 | 0 | 285 | 149 | 1 | 1 | 1 | 1 | 0.952 | 0.964 | 1 | 1 | 1 | 1 | 1 | 1 | -0.017 |
| SE | 65445 | ENSG00000166164 | BRD7 | chr16 | - | 50339975 | 50340086 | 50334710 | 50334895 | 50350022 | 50350167 | 301 | 311 | 222 | 328 | 381 | 0 | 1 | 0 | 0 | 0 | 683 | 341 | 651 | 563 | 216 | 0 | 0 | 0 | 0 | 0 | 260 | 149 | 1 | 1 | 1 | 0.994 | 1 | 1 | 1 | 1 | 1 | 1 | 1 | 1 | -0.001 |
| SE | 65446 | ENSG00000166164 | BRD7 | chr16 | - | 50350022 | 50350167 | 50339975 | 50340086 | 50354424 | 50354482 | 389 | 355 | 287 | 352 | 490 | 0 | 0 | 0 | 2 | 2 | 704 | 392 | 748 | 631 | 260 | 0 | 0 | 0 | 0 | 0 | 294 | 149 | 1 | 1 | 1 | 1 | 1 | 1 | 0.992 | 1 | 1 | 1 | 1 | 1 | -0.002 |
| SE | 65448 | ENSG00000131508 | UBE2D2 | chr5 | + | 139623367 | 139623461 | 139614860 | 139614966 | 139626755 | 139626801 | 945 | 439 | 786 | 851 | 990 | 1 | 0 | 0 | 2 | 5 | 724 | 671 | 469 | 509 | 801 | 0 | 0 | 1 | 0 | 0 | 243 | 149 | 1 | 1 | 0.998 | 1 | 1 | 0.996 | 0.992 | 1 | 1 | 0.997 | 1 | 1 | -0.002 |
| SE | 65455 | ENSG00000204344 | STK19 | chr6 | + | 31979413 | 31980548 | 31978902 | 31978998 | 31980653 | 31980801 | 25 | 4 | 17 | 25 | 18 | 0 | 1 | 0 | 0 | 0 | 40 | 34 | 28 | 21 | 13 | 0 | 0 | 0 | 0 | 0 | 298 | 149 | 1 | 1 | 1 | 0.667 | 1 | 1 | 1 | 1 | 1 | 1 | 1 | 1 | -0.067 |
| SE | 65456 | ENSG00000204344 | STK19 | chr6 | + | 31980450 | 31980548 | 31978902 | 31978998 | 31980653 | 31980724 | 10 | 4 | 10 | 8 | 9 | 0 | 1 | 0 | 0 | 0 | 33 | 0 | 0 | 9 | 4 | 0 | 0 | 0 | 0 | 0 | 247 | 149 | 1 | 1 | 1 | 0.707 | 1 | 1 | 1 | 1 | NA | NA | 1 | 1 | -0.059 |
| SE | 65457 | ENSG00000147234 | FRMPD3 | chrX | + | 107526581 | 107526736 | 107449651 | 107450005 | 107530408 | 107530511 | 7 | 1 | 2 | 3 | 0 | 0 | 0 | 0 | 0 | 0 | 0 | 7 | 1 | 0 | 0 | 0 | 0 | 0 | 2 | 0 | 298 | 149 | 1 | 1 | 1 | 1 | 1 | 1 | NA | NA | 1 | 1 | 0 | NA | 0.333 |
| SE | 65458 | ENSG00000147234 | FRMPD3 | chrX | + | 107533504 | 107533550 | 107530408 | 107530511 | 107545736 | 107545841 | 10 | 0 | 1 | 6 | 0 | 5 | 0 | 2 | 2 | 0 | 9 | 17 | 4 | 4 | 9 | 0 | 4 | 0 | 2 | 11 | 195 | 149 | 1 | 1 | 0.604 | NA | 0.276 | 0.696 | NA | 1 | 0.765 | 1 | 0.604 | 0.385 | -0.225 |
| SE | 65459 | ENSG00000147234 | FRMPD3 | chrX | + | 107545736 | 107545841 | 107530408 | 107530511 | 107550048 | 107550156 | 10 | 0 | 2 | 5 | 0 | 0 | 0 | 0 | 0 | 0 | 0 | 12 | 0 | 2 | 22 | 0 | 0 | 0 | 3 | 0 | 254 | 149 | 1 | 1 | 1 | NA | 1 | 1 | NA | NA | 1 | NA | 0.281 | 1 | 0.24 |
| SE | 65460 | ENSG00000147234 | FRMPD3 | chrX | + | 107550048 | 107550156 | 107544582 | 107545841 | 107552794 | 107552926 | 26 | 6 | 3 | 21 | 8 | 1 | 0 | 0 | 1 | 0 | 0 | 40 | 4 | 4 | 27 | 0 | 0 | 0 | 0 | 0 | 257 | 149 | 1 | 1 | 0.938 | 1 | 1 | 1 | 0.823 | NA | 1 | 1 | 1 | 1 | -0.048 |
| SE | 65461 | ENSG00000147234 | FRMPD3 | chrX | + | 107576314 | 107576459 | 107564886 | 107565066 | 107597320 | 107598142 | 11 | 3 | 3 | 12 | 0 | 0 | 0 | 0 | 0 | 0 | 8 | 10 | 1 | 1 | 17 | 0 | 0 | 2 | 0 | 0 | 294 | 149 | 1 | 1 | 1 | 1 | 1 | 1 | NA | 1 | 1 | 0.202 | 1 | 1 | 0.16 |
| SE | 65462 | ENSG00000287625 | NA | chr2 | + | 84941213 | 84941299 | 84926018 | 84927396 | 84962829 | 84963145 | 11 | 6 | 3 | 0 | 9 | 0 | 0 | 1 | 0 | 0 | 33 | 4 | 0 | 4 | 28 | 0 | 0 | 0 | 0 | 0 | 235 | 149 | 1 | 1 | 1 | 1 | 0.655 | NA | 1 | 1 | 1 | NA | 1 | 1 | -0.086 |
| SE | 65463 | ENSG00000118508 | RAB32 | chr6 | + | 146549463 | 146549741 | 146543832 | 146544121 | 146554455 | 146554953 | 337 | 248 | 155 | 365 | 385 | 0 | 2 | 0 | 0 | 0 | 1431 | 527 | 326 | 911 | 2737 | 0 | 0 | 0 | 11 | 0 | 298 | 149 | 1 | 1 | 1 | 0.984 | 1 | 1 | 1 | 1 | 1 | 1 | 0.976 | 1 | 0.002 |
| SE | 65464 | ENSG00000150873 | C2orf50 | chr2 | + | 11140509 | 11140549 | 11133127 | 11133591 | 11143919 | 11144142 | 0 | 0 | 0 | 0 | 0 | 0 | 1 | 0 | 0 | 0 | 0 | 0 | 0 | 1 | 0 | 0 | 0 | 0 | 0 | 0 | 189 | 149 | 1 | 1 | NA | 0 | NA | NA | NA | NA | NA | NA | 1 | NA | -1 |
| SE | 65466 | ENSG00000130348 | QRSL1 | chr6 | + | 106640348 | 106640508 | 106629577 | 106629705 | 106640822 | 106640862 | 138 | 112 | 110 | 118 | 180 | 0 | 0 | 0 | 2 | 0 | 109 | 122 | 149 | 183 | 74 | 0 | 0 | 0 | 0 | 0 | 298 | 149 | 1 | 1 | 1 | 1 | 1 | 0.967 | 1 | 1 | 1 | 1 | 1 | 1 | -0.007 |
| SE | 65467 | ENSG00000130348 | QRSL1 | chr6 | + | 106642993 | 106643090 | 106640822 | 106640921 | 106649024 | 106649201 | 195 | 161 | 183 | 179 | 260 | 0 | 0 | 0 | 0 | 0 | 187 | 163 | 162 | 205 | 90 | 0 | 0 | 4 | 0 | 0 | 246 | 149 | 1 | 1 | 1 | 1 | 1 | 1 | 1 | 1 | 1 | 0.961 | 1 | 1 | 0.008 |
| SE | 65470 | ENSG00000264324 | NA | chr2 | - | 74342457 | 74342537 | 74338854 | 74338903 | 74361003 | 74361156 | 5 | 7 | 4 | 5 | 0 | 0 | 0 | 2 | 0 | 0 | 0 | 2 | 6 | 0 | 10 | 0 | 0 | 0 | 0 | 0 | 229 | 149 | 1 | 1 | 1 | 1 | 0.565 | 1 | NA | NA | 1 | 1 | NA | 1 | -0.109 |
| SE | 65471 | ENSG00000264324 | NA | chr2 | - | 74342457 | 74342537 | 74338854 | 74338903 | 74361412 | 74361636 | 5 | 7 | 4 | 6 | 0 | 0 | 0 | 0 | 1 | 0 | 0 | 2 | 4 | 0 | 10 | 0 | 0 | 0 | 0 | 0 | 229 | 149 | 1 | 1 | 1 | 1 | 1 | 0.796 | NA | NA | 1 | 1 | NA | 1 | -0.051 |
| SE | 65472 | ENSG00000264324 | NA | chr2 | - | 74361003 | 74361156 | 74338854 | 74338903 | 74361412 | 74361636 | 0 | 0 | 3 | 1 | 0 | 0 | 0 | 0 | 1 | 0 | 0 | 0 | 1 | 0 | 0 | 0 | 0 | 0 | 0 | 0 | 298 | 149 | 1 | 1 | NA | NA | 1 | 0.333 | NA | NA | NA | 1 | NA | NA | -0.334 |
| SE | 65473 | ENSG00000264324 | NA | chr2 | - | 74361003 | 74361156 | 74342457 | 74342537 | 74361412 | 74361636 | 0 | 0 | 1 | 1 | 0 | 0 | 0 | 0 | 1 | 0 | 0 | 0 | 3 | 0 | 0 | 0 | 0 | 0 | 0 | 0 | 298 | 149 | 1 | 1 | NA | NA | 1 | 0.333 | NA | NA | NA | 1 | NA | NA | -0.334 |
| SE | 65474 | ENSG00000264324 | NA | chr2 | - | 74362051 | 74362141 | 74361412 | 74361636 | 74362649 | 74362729 | 247 | 138 | 205 | 249 | 230 | 0 | 0 | 0 | 1 | 0 | 340 | 305 | 162 | 248 | 451 | 0 | 0 | 0 | 0 | 0 | 239 | 149 | 1 | 1 | 1 | 1 | 1 | 0.994 | 1 | 1 | 1 | 1 | 1 | 1 | -0.001 |
| SE | 65476 | ENSG00000246228 | CASC8 | chr8 | - | 127323014 | 127323154 | 127289816 | 127290060 | 127420828 | 127421374 | 22 | 0 | 5 | 1 | 6 | 17 | 0 | 4 | 1 | 2 | 0 | 0 | 0 | 11 | 0 | 0 | 0 | 0 | 0 | 0 | 289 | 149 | 1 | 1 | 0.4 | NA | 0.392 | 0.34 | 0.607 | NA | NA | NA | 1 | NA | -0.565 |
| SE | 65477 | ENSG00000163541 | SUCLG1 | chr2 | - | 84431507 | 84431659 | 84425413 | 84425603 | 84433351 | 84433435 | 399 | 212 | 282 | 227 | 332 | 0 | 0 | 0 | 1 | 0 | 441 | 321 | 186 | 340 | 336 | 0 | 0 | 1 | 4 | 0 | 298 | 149 | 1 | 1 | 1 | 1 | 1 | 0.991 | 1 | 1 | 1 | 0.989 | 0.977 | 1 | 0.005 |
| SE | 65486 | ENSG00000125875 | TBC1D20 | chr20 | - | 440247 | 440317 | 439107 | 439295 | 441587 | 441689 | 89 | 52 | 87 | 119 | 107 | 0 | 0 | 0 | 0 | 0 | 96 | 142 | 44 | 78 | 214 | 0 | 3 | 0 | 0 | 0 | 219 | 149 | 1 | 1 | 1 | 1 | 1 | 1 | 1 | 1 | 0.97 | 1 | 1 | 1 | 0.006 |
| SE | 65487 | ENSG00000125875 | TBC1D20 | chr20 | - | 440247 | 440389 | 439107 | 439295 | 441587 | 441689 | 240 | 132 | 208 | 306 | 272 | 0 | 0 | 0 | 0 | 0 | 249 | 282 | 112 | 211 | 453 | 0 | 3 | 0 | 0 | 0 | 291 | 149 | 1 | 1 | 1 | 1 | 1 | 1 | 1 | 1 | 0.98 | 1 | 1 | 1 | 0.004 |
| SE | 65488 | ENSG00000125875 | TBC1D20 | chr20 | - | 440964 | 441060 | 440247 | 440389 | 441587 | 441689 | 0 | 0 | 1 | 0 | 2 | 111 | 66 | 96 | 149 | 136 | 0 | 0 | 0 | 0 | 0 | 125 | 108 | 55 | 113 | 161 | 245 | 149 | 1 | 1 | 0 | 0 | 0.006 | 0 | 0.009 | 0 | 0 | 0 | 0 | 0 | 0.003 |
| SE | 65490 | ENSG00000125875 | TBC1D20 | chr20 | - | 445049 | 445130 | 439107 | 442043 | 447888 | 448074 | 113 | 62 | 98 | 130 | 125 | 0 | 3 | 0 | 0 | 0 | 96 | 99 | 71 | 106 | 285 | 0 | 0 | 0 | 0 | 0 | 230 | 149 | 1 | 1 | 1 | 0.93 | 1 | 1 | 1 | 1 | 1 | 1 | 1 | 1 | -0.014 |
| SE | 65495 | ENSG00000121989 | ACVR2A | chr2 | + | 147915190 | 147915334 | 147899743 | 147899898 | 147917282 | 147917426 | 56 | 48 | 55 | 66 | 65 | 1 | 1 | 0 | 0 | 0 | 12 | 22 | 48 | 10 | 22 | 0 | 0 | 0 | 0 | 0 | 293 | 149 | 1 | 1 | 0.966 | 0.961 | 1 | 1 | 1 | 1 | 1 | 1 | 1 | 1 | -0.015 |
| SE | 65496 | ENSG00000121989 | ACVR2A | chr2 | + | 147918446 | 147918592 | 147917282 | 147917426 | 147920229 | 147920344 | 72 | 72 | 87 | 65 | 76 | 0 | 1 | 0 | 0 | 0 | 90 | 36 | 46 | 7 | 0 | 0 | 0 | 0 | 0 | 0 | 295 | 149 | 1 | 1 | 1 | 0.973 | 1 | 1 | 1 | 1 | 1 | 1 | 1 | NA | -0.005 |
| SE | 65497 | ENSG00000121989 | ACVR2A | chr2 | + | 147918446 | 147918592 | 147917282 | 147917426 | 147926030 | 147926161 | 44 | 40 | 50 | 40 | 38 | 3 | 0 | 0 | 2 | 0 | 49 | 29 | 26 | 4 | 0 | 0 | 0 | 0 | 0 | 0 | 295 | 149 | 1 | 1 | 0.881 | 1 | 1 | 0.91 | 1 | 1 | 1 | 1 | 1 | NA | -0.042 |
| SE | 65498 | ENSG00000121989 | ACVR2A | chr2 | + | 147920229 | 147920344 | 147917282 | 147917426 | 147926030 | 147926161 | 5 | 1 | 2 | 3 | 3 | 3 | 0 | 0 | 2 | 0 | 0 | 0 | 3 | 5 | 0 | 0 | 0 | 0 | 0 | 0 | 264 | 149 | 1 | 1 | 0.485 | 1 | 1 | 0.458 | 1 | NA | NA | 1 | 1 | NA | -0.211 |
| SE | 65499 | ENSG00000121989 | ACVR2A | chr2 | + | 147920229 | 147920344 | 147918446 | 147918592 | 147926030 | 147926161 | 36 | 30 | 30 | 25 | 36 | 1 | 0 | 0 | 0 | 0 | 43 | 7 | 21 | 8 | 0 | 10 | 0 | 0 | 0 | 0 | 264 | 149 | 1 | 1 | 0.953 | 1 | 1 | 1 | 1 | 0.708 | 1 | 1 | 1 | NA | 0.064 |
| SE | 65500 | ENSG00000121989 | ACVR2A | chr2 | + | 147922972 | 147923111 | 147920229 | 147920344 | 147926030 | 147926161 | 26 | 15 | 26 | 24 | 34 | 5 | 0 | 3 | 3 | 4 | 24 | 0 | 18 | 1 | 0 | 0 | 0 | 3 | 6 | 0 | 288 | 149 | 1 | 1 | 0.729 | 1 | 0.818 | 0.805 | 0.815 | 1 | NA | 0.756 | 0.079 | NA | 0.222 |
| SE | 65501 | ENSG00000140386 | SCAPER | chr15 | - | 76376161 | 76376311 | 76353948 | 76354140 | 76381377 | 76381615 | 14 | 11 | 21 | 11 | 40 | 0 | 0 | 1 | 0 | 0 | 0 | 8 | 1 | 2 | 0 | 0 | 0 | 2 | 0 | 0 | 298 | 149 | 1 | 1 | 1 | 1 | 0.913 | 1 | 1 | NA | 1 | 0.2 | 1 | NA | 0.249 |
| SE | 65502 | ENSG00000140386 | SCAPER | chr15 | - | 76381377 | 76381615 | 76376161 | 76376311 | 76404523 | 76404679 | 38 | 33 | 51 | 49 | 60 | 0 | 0 | 0 | 0 | 1 | 11 | 12 | 7 | 14 | 0 | 0 | 0 | 0 | 0 | 0 | 298 | 149 | 1 | 1 | 1 | 1 | 1 | 1 | 0.968 | 1 | 1 | 1 | 1 | NA | -0.006 |
| SE | 65503 | ENSG00000140386 | SCAPER | chr15 | - | 76404523 | 76404679 | 76379708 | 76381615 | 76434077 | 76434310 | 68 | 45 | 72 | 89 | 73 | 0 | 0 | 1 | 0 | 0 | 15 | 16 | 34 | 26 | 0 | 0 | 0 | 0 | 0 | 0 | 298 | 149 | 1 | 1 | 1 | 1 | 0.973 | 1 | 1 | 1 | 1 | 1 | 1 | NA | -0.005 |
| SE | 65504 | ENSG00000140386 | SCAPER | chr15 | - | 76434077 | 76434310 | 76404523 | 76404679 | 76471211 | 76471335 | 68 | 38 | 70 | 88 | 60 | 0 | 2 | 0 | 0 | 0 | 4 | 12 | 42 | 14 | 0 | 0 | 0 | 0 | 0 | 0 | 298 | 149 | 1 | 1 | 1 | 0.905 | 1 | 1 | 1 | 1 | 1 | 1 | 1 | NA | -0.019 |
| SE | 65505 | ENSG00000140386 | SCAPER | chr15 | - | 76567269 | 76567405 | 76504858 | 76504974 | 76574157 | 76574284 | 2 | 2 | 4 | 0 | 6 | 34 | 19 | 45 | 45 | 36 | 0 | 0 | 0 | 6 | 0 | 0 | 19 | 13 | 10 | 0 | 285 | 149 | 1 | 1 | 0.03 | 0.052 | 0.044 | 0 | 0.08 | NA | 0 | 0 | 0.239 | NA | -0.038 |
| SE | 65506 | ENSG00000140386 | SCAPER | chr15 | - | 76574157 | 76574284 | 76504858 | 76504974 | 76621763 | 76621829 | 104 | 68 | 92 | 107 | 124 | 0 | 0 | 0 | 2 | 0 | 6 | 38 | 32 | 12 | 14 | 0 | 0 | 1 | 0 | 0 | 276 | 149 | 1 | 1 | 1 | 1 | 1 | 0.967 | 1 | 1 | 1 | 0.945 | 1 | 1 | 0.004 |
| SE | 65507 | ENSG00000140386 | SCAPER | chr15 | - | 76621763 | 76621829 | 76574157 | 76574284 | 76665652 | 76665658 | 113 | 77 | 69 | 90 | 145 | 0 | 0 | 0 | 0 | 0 | 12 | 29 | 28 | 2 | 14 | 0 | 0 | 0 | 2 | 0 | 215 | 149 | 1 | 1 | 1 | 1 | 1 | 1 | 1 | 1 | 1 | 1 | 0.409 | 1 | 0.118 |
| SE | 65508 | ENSG00000140386 | SCAPER | chr15 | - | 76665652 | 76665789 | 76621763 | 76621829 | 76701757 | 76701865 | 104 | 111 | 92 | 119 | 159 | 0 | 0 | 2 | 0 | 0 | 0 | 26 | 37 | 6 | 7 | 0 | 0 | 0 | 0 | 0 | 286 | 149 | 1 | 1 | 1 | 1 | 0.96 | 1 | 1 | NA | 1 | 1 | 1 | 1 | -0.008 |
| SE | 65509 | ENSG00000140386 | SCAPER | chr15 | - | 76705902 | 76705984 | 76702849 | 76703002 | 76728594 | 76728737 | 67 | 67 | 81 | 47 | 77 | 0 | 0 | 0 | 0 | 0 | 5 | 33 | 45 | 4 | 1 | 0 | 0 | 2 | 0 | 0 | 231 | 149 | 1 | 1 | 1 | 1 | 1 | 1 | 1 | 1 | 1 | 0.936 | 1 | 1 | 0.013 |
| SE | 65512 | ENSG00000140386 | SCAPER | chr15 | - | 76774367 | 76774409 | 76771741 | 76771954 | 76774854 | 76775064 | 3 | 0 | 0 | 0 | 0 | 50 | 41 | 55 | 50 | 71 | 0 | 0 | 0 | 0 | 0 | 12 | 8 | 24 | 12 | 10 | 191 | 149 | 1 | 1 | 0.045 | 0 | 0 | 0 | 0 | 0 | 0 | 0 | 0 | 0 | 0.009 |
| SE | 65513 | ENSG00000140386 | SCAPER | chr15 | - | 76793232 | 76793391 | 76774854 | 76775117 | 76795279 | 76795440 | 0 | 2 | 5 | 3 | 0 | 31 | 42 | 48 | 41 | 33 | 10 | 0 | 0 | 0 | 0 | 0 | 20 | 26 | 14 | 0 | 298 | 149 | 1 | 1 | 0 | 0.023 | 0.05 | 0.035 | 0 | 1 | 0 | 0 | 0 | NA | -0.228 |
| SE | 65514 | ENSG00000140386 | SCAPER | chr15 | - | 76795240 | 76795440 | 76774854 | 76775117 | 76800247 | 76800267 | 109 | 125 | 118 | 119 | 148 | 2 | 0 | 0 | 4 | 0 | 18 | 86 | 108 | 18 | 19 | 0 | 0 | 3 | 0 | 0 | 298 | 149 | 1 | 1 | 0.965 | 1 | 1 | 0.937 | 1 | 1 | 1 | 0.947 | 1 | 1 | -0.009 |
| SE | 65515 | ENSG00000140386 | SCAPER | chr15 | - | 76795279 | 76795440 | 76774854 | 76775117 | 76800247 | 76800364 | 140 | 166 | 166 | 156 | 181 | 2 | 0 | 0 | 4 | 0 | 18 | 104 | 134 | 29 | 15 | 0 | 0 | 3 | 0 | 0 | 298 | 149 | 1 | 1 | 0.972 | 1 | 1 | 0.951 | 1 | 1 | 1 | 0.957 | 1 | 1 | -0.007 |
| SE | 65518 | ENSG00000140386 | SCAPER | chr15 | - | 76857808 | 76857879 | 76841733 | 76841931 | 76862415 | 76862533 | 129 | 128 | 129 | 122 | 167 | 0 | 1 | 0 | 1 | 0 | 26 | 49 | 45 | 48 | 49 | 0 | 0 | 0 | 0 | 0 | 220 | 149 | 1 | 1 | 1 | 0.989 | 1 | 0.988 | 1 | 1 | 1 | 1 | 1 | 1 | -0.005 |
| SE | 65520 | ENSG00000140386 | SCAPER | chr15 | - | 76883811 | 76883876 | 76862415 | 76862533 | 76904613 | 76904791 | 44 | 37 | 29 | 26 | 39 | 0 | 0 | 0 | 1 | 0 | 6 | 10 | 34 | 15 | 10 | 0 | 0 | 1 | 0 | 0 | 214 | 149 | 1 | 1 | 1 | 1 | 1 | 1 | 1 | 1 | 1 | 0.959 | 1 | 1 | 0.008 |
| SE | 65522 | ENSG00000140386 | SCAPER | chr15 | - | 76904613 | 76904791 | 76883811 | 76883876 | 76905298 | 76905337 | 1 | 0 | 2 | 0 | 0 | 39 | 32 | 30 | 48 | 22 | 0 | 0 | 1 | 0 | 0 | 4 | 18 | 20 | 12 | 28 | 298 | 149 | 1 | 1 | 0.013 | 0 | 0.032 | 0 | 0 | 0 | 0 | 0.024 | 0 | 0 | 0.004 |
| SE | 65526 | ENSG00000119820 | YIPF4 | chr2 | + | 32292176 | 32292348 | 32290482 | 32290636 | 32298233 | 32298311 | 182 | 153 | 228 | 187 | 320 | 0 | 0 | 3 | 0 | 0 | 81 | 115 | 76 | 60 | 25 | 0 | 0 | 0 | 0 | 0 | 298 | 149 | 1 | 1 | 1 | 1 | 0.974 | 1 | 1 | 1 | 1 | 1 | 1 | 1 | -0.005 |
| SE | 65528 | ENSG00000119820 | YIPF4 | chr2 | + | 32298233 | 32298311 | 32292176 | 32292348 | 32301442 | 32301495 | 33 | 29 | 38 | 30 | 70 | 1 | 0 | 1 | 1 | 0 | 26 | 31 | 7 | 9 | 0 | 0 | 3 | 3 | 0 | 0 | 227 | 149 | 1 | 1 | 0.956 | 1 | 0.961 | 0.952 | 1 | 1 | 0.872 | 0.605 | 1 | NA | 0.105 |
| SE | 65529 | ENSG00000119820 | YIPF4 | chr2 | + | 32301381 | 32301495 | 32298233 | 32298311 | 32305488 | 32305866 | 173 | 138 | 179 | 166 | 312 | 0 | 0 | 0 | 0 | 1 | 48 | 91 | 67 | 64 | 17 | 0 | 0 | 0 | 0 | 0 | 263 | 149 | 1 | 1 | 1 | 1 | 1 | 1 | 0.994 | 1 | 1 | 1 | 1 | 1 | -0.001 |
| SE | 65530 | ENSG00000284292 | NA | chr7 | + | 99333324 | 99333417 | 99325878 | 99326004 | 99338180 | 99338285 | 156 | 152 | 112 | 165 | 196 | 0 | 0 | 1 | 0 | 0 | 317 | 190 | 175 | 235 | 288 | 0 | 0 | 0 | 0 | 0 | 242 | 149 | 1 | 1 | 1 | 1 | 0.986 | 1 | 1 | 1 | 1 | 1 | 1 | 1 | -0.003 |
| SE | 65533 | ENSG00000284292 | NA | chr7 | + | 99339857 | 99340009 | 99338180 | 99338285 | 99344292 | 99344515 | 0 | 0 | 0 | 0 | 0 | 93 | 69 | 75 | 110 | 122 | 0 | 2 | 0 | 2 | 0 | 92 | 104 | 66 | 85 | 159 | 298 | 149 | 1 | 1 | 0 | 0 | 0 | 0 | 0 | 0 | 0.01 | 0 | 0.012 | 0 | -0.004 |
| SE | 65536 | ENSG00000284292 | NA | chr7 | + | 99344292 | 99344515 | 99338180 | 99338285 | 99353908 | 99354121 | 97 | 71 | 78 | 113 | 123 | 0 | 0 | 2 | 0 | 0 | 108 | 109 | 70 | 97 | 165 | 0 | 0 | 5 | 0 | 0 | 298 | 149 | 1 | 1 | 1 | 1 | 0.951 | 1 | 1 | 1 | 1 | 0.875 | 1 | 1 | 0.015 |
| SE | 65540 | ENSG00000284292 | NA | chr7 | + | 99358339 | 99358415 | 99353908 | 99354121 | 99359544 | 99359738 | 164 | 52 | 128 | 90 | 73 | 1 | 0 | 0 | 0 | 0 | 157 | 171 | 128 | 121 | 216 | 0 | 0 | 0 | 0 | 0 | 225 | 149 | 1 | 1 | 0.991 | 1 | 1 | 1 | 1 | 1 | 1 | 1 | 1 | 1 | -0.002 |
| SE | 65543 | ENSG00000155097 | ATP6V1C1 | chr8 | + | 103040797 | 103040968 | 103021082 | 103021225 | 103042339 | 103042407 | 122 | 84 | 99 | 143 | 137 | 3 | 0 | 0 | 1 | 0 | 42 | 84 | 94 | 78 | 0 | 12 | 0 | 0 | 0 | 0 | 298 | 149 | 1 | 1 | 0.953 | 1 | 1 | 0.986 | 1 | 0.636 | 1 | 1 | 1 | NA | 0.079 |
| SE | 65544 | ENSG00000155097 | ATP6V1C1 | chr8 | + | 103040797 | 103040968 | 103021082 | 103021225 | 103048869 | 103048955 | 44 | 53 | 43 | 67 | 56 | 1 | 0 | 1 | 1 | 0 | 37 | 34 | 40 | 44 | 0 | 0 | 0 | 0 | 0 | 0 | 298 | 149 | 1 | 1 | 0.957 | 1 | 0.956 | 0.971 | 1 | 1 | 1 | 1 | 1 | NA | -0.023 |
| SE | 65545 | ENSG00000155097 | ATP6V1C1 | chr8 | + | 103042339 | 103042407 | 103021087 | 103021225 | 103048869 | 103048955 | 33 | 17 | 23 | 35 | 41 | 1 | 0 | 1 | 1 | 0 | 38 | 2 | 13 | 14 | 0 | 0 | 0 | 0 | 0 | 0 | 217 | 149 | 1 | 1 | 0.958 | 1 | 0.94 | 0.96 | 1 | 1 | 1 | 1 | 1 | NA | -0.028 |
| SE | 65546 | ENSG00000155097 | ATP6V1C1 | chr8 | + | 103042339 | 103042407 | 103040797 | 103040968 | 103048869 | 103048955 | 96 | 43 | 71 | 98 | 117 | 1 | 0 | 0 | 0 | 0 | 31 | 36 | 59 | 36 | 0 | 0 | 0 | 0 | 0 | 0 | 217 | 149 | 1 | 1 | 0.985 | 1 | 1 | 1 | 1 | 1 | 1 | 1 | 1 | NA | -0.003 |
| SE | 65547 | ENSG00000155097 | ATP6V1C1 | chr8 | + | 103064713 | 103064811 | 103063134 | 103063228 | 103066320 | 103066447 | 150 | 95 | 114 | 120 | 300 | 0 | 2 | 0 | 1 | 0 | 7 | 104 | 73 | 76 | 0 | 0 | 0 | 0 | 0 | 0 | 247 | 149 | 1 | 1 | 1 | 0.966 | 1 | 0.986 | 1 | 1 | 1 | 1 | 1 | NA | -0.01 |
| SE | 65548 | ENSG00000155097 | ATP6V1C1 | chr8 | + | 103066320 | 103066447 | 103064559 | 103064811 | 103068651 | 103068825 | 175 | 135 | 138 | 168 | 380 | 0 | 0 | 0 | 0 | 1 | 5 | 103 | 101 | 73 | 4 | 0 | 0 | 0 | 0 | 0 | 276 | 149 | 1 | 1 | 1 | 1 | 1 | 1 | 0.995 | 1 | 1 | 1 | 1 | 1 | -0.001 |
| SE | 65554 | ENSG00000153201 | RANBP2 | chr2 | + | 108735531 | 108735762 | 108731321 | 108731474 | 108736103 | 108736249 | 288 | 245 | 258 | 266 | 383 | 0 | 0 | 1 | 0 | 0 | 90 | 195 | 220 | 163 | 126 | 0 | 0 | 0 | 0 | 0 | 298 | 149 | 1 | 1 | 1 | 1 | 0.992 | 1 | 1 | 1 | 1 | 1 | 1 | 1 | -0.002 |
| SE | 65557 | ENSG00000153201 | RANBP2 | chr2 | + | 108762100 | 108762195 | 108758412 | 108758548 | 108771700 | 108771871 | 180 | 126 | 170 | 183 | 319 | 0 | 0 | 0 | 0 | 0 | 0 | 57 | 36 | 20 | 0 | 0 | 0 | 1 | 2 | 0 | 244 | 149 | 1 | 1 | 1 | 1 | 1 | 1 | 1 | NA | 1 | 0.956 | 0.859 | NA | 0.062 |
| SE | 65558 | ENSG00000153201 | RANBP2 | chr2 | + | 108763236 | 108768388 | 108758412 | 108758548 | 108771700 | 108771871 | 138 | 125 | 115 | 136 | 324 | 0 | 0 | 0 | 0 | 0 | 6 | 14 | 30 | 15 | 8 | 0 | 0 | 1 | 2 | 0 | 298 | 149 | 1 | 1 | 1 | 1 | 1 | 1 | 1 | 1 | 1 | 0.938 | 0.789 | 1 | 0.055 |
| SE | 65559 | ENSG00000153201 | RANBP2 | chr2 | + | 108763236 | 108768388 | 108762100 | 108762195 | 108771700 | 108771871 | 217 | 172 | 208 | 192 | 457 | 0 | 0 | 1 | 0 | 0 | 1 | 38 | 34 | 7 | 2 | 0 | 0 | 0 | 0 | 0 | 298 | 149 | 1 | 1 | 1 | 1 | 0.99 | 1 | 1 | 1 | 1 | 1 | 1 | 1 | -0.002 |
| SE | 65560 | ENSG00000153201 | RANBP2 | chr2 | + | 108775731 | 108775936 | 108772867 | 108773046 | 108777129 | 108777231 | 261 | 198 | 189 | 211 | 300 | 0 | 0 | 0 | 5 | 16 | 0 | 62 | 50 | 5 | 2 | 0 | 0 | 0 | 0 | 0 | 298 | 149 | 1 | 1 | 1 | 1 | 1 | 0.955 | 0.904 | NA | 1 | 1 | 1 | 1 | -0.028 |
| SE | 65561 | ENSG00000153201 | RANBP2 | chr2 | + | 108777129 | 108777231 | 108775731 | 108775936 | 108781268 | 108781429 | 371 | 275 | 287 | 328 | 407 | 1 | 1 | 0 | 1 | 2 | 0 | 89 | 69 | 10 | 2 | 0 | 3 | 2 | 0 | 0 | 251 | 149 | 1 | 1 | 0.995 | 0.994 | 1 | 0.995 | 0.992 | NA | 0.946 | 0.953 | 1 | 1 | 0.02 |
| SE | 65566 | ENSG00000147526 | TACC1 | chr8 | + | 38788703 | 38788819 | 38757280 | 38757409 | 38819521 | 38819631 | 103 | 72 | 75 | 134 | 70 | 0 | 0 | 0 | 1 | 0 | 76 | 131 | 103 | 114 | 57 | 0 | 0 | 0 | 0 | 0 | 265 | 149 | 1 | 1 | 1 | 1 | 1 | 0.987 | 1 | 1 | 1 | 1 | 1 | 1 | -0.003 |
| SE | 65568 | ENSG00000147526 | TACC1 | chr8 | + | 38788703 | 38788819 | 38787235 | 38787743 | 38819521 | 38819631 | 214 | 102 | 161 | 241 | 108 | 0 | 0 | 0 | 0 | 0 | 102 | 205 | 172 | 214 | 52 | 0 | 1 | 0 | 0 | 0 | 265 | 149 | 1 | 1 | 1 | 1 | 1 | 1 | 1 | 1 | 0.991 | 1 | 1 | 1 | 0.002 |
| SE | 65571 | ENSG00000147526 | TACC1 | chr8 | + | 38819521 | 38820635 | 38787235 | 38787743 | 38825307 | 38825368 | 227 | 205 | 157 | 267 | 251 | 2 | 2 | 0 | 2 | 0 | 220 | 268 | 277 | 249 | 206 | 0 | 0 | 0 | 0 | 3 | 298 | 149 | 1 | 1 | 0.983 | 0.981 | 1 | 0.985 | 1 | 1 | 1 | 1 | 1 | 0.972 | -0.005 |
| SE | 65572 | ENSG00000147526 | TACC1 | chr8 | + | 38819521 | 38820635 | 38788703 | 38788819 | 38825307 | 38825368 | 371 | 290 | 256 | 423 | 331 | 0 | 0 | 0 | 0 | 0 | 302 | 429 | 421 | 408 | 243 | 0 | 0 | 3 | 0 | 0 | 298 | 149 | 1 | 1 | 1 | 1 | 1 | 1 | 1 | 1 | 1 | 0.986 | 1 | 1 | 0.003 |
| SE | 65573 | ENSG00000147526 | TACC1 | chr8 | + | 38819521 | 38820635 | 38805451 | 38805644 | 38825307 | 38825368 | 238 | 219 | 166 | 283 | 257 | 0 | 2 | 0 | 0 | 0 | 220 | 267 | 280 | 253 | 209 | 0 | 0 | 0 | 0 | 8 | 298 | 149 | 1 | 1 | 1 | 0.982 | 1 | 1 | 1 | 1 | 1 | 1 | 1 | 0.929 | 0.011 |
| SE | 65575 | ENSG00000147526 | TACC1 | chr8 | + | 38823989 | 38824025 | 38819920 | 38820635 | 38825307 | 38825368 | 0 | 2 | 0 | 0 | 0 | 227 | 205 | 157 | 267 | 251 | 0 | 0 | 0 | 1 | 0 | 220 | 267 | 277 | 249 | 206 | 185 | 149 | 1 | 1 | 0 | 0.008 | 0 | 0 | 0 | 0 | 0 | 0 | 0.003 | 0 | 0.001 |
| SE | 65577 | ENSG00000147526 | TACC1 | chr8 | + | 38825307 | 38825368 | 38819521 | 38820635 | 38831124 | 38831177 | 102 | 96 | 57 | 112 | 89 | 0 | 0 | 0 | 0 | 0 | 110 | 93 | 121 | 95 | 64 | 0 | 0 | 3 | 0 | 0 | 210 | 149 | 1 | 1 | 1 | 1 | 1 | 1 | 1 | 1 | 1 | 0.966 | 1 | 1 | 0.007 |
| SE | 65580 | ENSG00000147526 | TACC1 | chr8 | + | 38827167 | 38827288 | 38825307 | 38825368 | 38831124 | 38831177 | 328 | 290 | 270 | 358 | 361 | 0 | 0 | 0 | 0 | 1 | 330 | 373 | 416 | 434 | 309 | 0 | 4 | 0 | 0 | 0 | 270 | 149 | 1 | 1 | 1 | 1 | 1 | 1 | 0.995 | 1 | 0.981 | 1 | 1 | 1 | 0.003 |
| SE | 65581 | ENSG00000147526 | TACC1 | chr8 | + | 38827167 | 38827375 | 38820546 | 38820635 | 38831124 | 38831177 | 126 | 143 | 107 | 167 | 151 | 0 | 0 | 0 | 0 | 0 | 149 | 146 | 184 | 150 | 84 | 0 | 0 | 3 | 0 | 0 | 298 | 149 | 1 | 1 | 1 | 1 | 1 | 1 | 1 | 1 | 1 | 0.968 | 1 | 1 | 0.006 |
| SE | 65582 | ENSG00000147526 | TACC1 | chr8 | + | 38827167 | 38827375 | 38825307 | 38825368 | 38831124 | 38831177 | 344 | 337 | 299 | 396 | 404 | 0 | 0 | 0 | 0 | 1 | 355 | 400 | 424 | 426 | 281 | 0 | 4 | 0 | 0 | 0 | 298 | 149 | 1 | 1 | 1 | 1 | 1 | 1 | 0.995 | 1 | 0.98 | 1 | 1 | 1 | 0.003 |
| SE | 65583 | ENSG00000147526 | TACC1 | chr8 | + | 38827167 | 38827375 | 38825307 | 38825368 | 38836161 | 38836287 | 218 | 194 | 193 | 229 | 253 | 0 | 0 | 1 | 0 | 0 | 206 | 254 | 240 | 276 | 197 | 0 | 0 | 0 | 0 | 0 | 298 | 149 | 1 | 1 | 1 | 1 | 0.99 | 1 | 1 | 1 | 1 | 1 | 1 | 1 | -0.002 |
| SE | 65584 | ENSG00000147526 | TACC1 | chr8 | + | 38831124 | 38831177 | 38825307 | 38825368 | 38836161 | 38836287 | 95 | 79 | 86 | 103 | 102 | 0 | 0 | 1 | 0 | 0 | 63 | 102 | 121 | 134 | 54 | 0 | 0 | 0 | 0 | 0 | 202 | 149 | 1 | 1 | 1 | 1 | 0.984 | 1 | 1 | 1 | 1 | 1 | 1 | 1 | -0.003 |
| SE | 65585 | ENSG00000147526 | TACC1 | chr8 | + | 38831124 | 38831177 | 38827167 | 38827375 | 38836161 | 38836287 | 221 | 222 | 192 | 270 | 251 | 0 | 0 | 1 | 0 | 0 | 208 | 244 | 305 | 284 | 138 | 0 | 0 | 0 | 0 | 0 | 202 | 149 | 1 | 1 | 1 | 1 | 0.993 | 1 | 1 | 1 | 1 | 1 | 1 | 1 | -0.001 |
| SE | 65586 | ENSG00000147526 | TACC1 | chr8 | + | 38836161 | 38836287 | 38831124 | 38831177 | 38838469 | 38838507 | 259 | 234 | 245 | 289 | 267 | 0 | 0 | 1 | 0 | 3 | 242 | 285 | 419 | 392 | 199 | 0 | 5 | 0 | 0 | 0 | 275 | 149 | 1 | 1 | 1 | 1 | 0.993 | 1 | 0.98 | 1 | 0.969 | 1 | 1 | 1 | 0.001 |
| SE | 65587 | ENSG00000147526 | TACC1 | chr8 | + | 38836161 | 38836287 | 38831124 | 38831177 | 38840223 | 38840267 | 155 | 123 | 143 | 179 | 146 | 0 | 0 | 0 | 0 | 0 | 150 | 177 | 228 | 226 | 111 | 0 | 0 | 0 | 1 | 0 | 275 | 149 | 1 | 1 | 1 | 1 | 1 | 1 | 1 | 1 | 1 | 1 | 0.992 | 1 | 0.002 |
| SE | 65588 | ENSG00000147526 | TACC1 | chr8 | + | 38838469 | 38838546 | 38831124 | 38831177 | 38840223 | 38840267 | 178 | 121 | 129 | 149 | 240 | 0 | 0 | 0 | 0 | 0 | 141 | 107 | 159 | 113 | 91 | 0 | 0 | 0 | 1 | 0 | 226 | 149 | 1 | 1 | 1 | 1 | 1 | 1 | 1 | 1 | 1 | 1 | 0.987 | 1 | 0.003 |
| SE | 65589 | ENSG00000147526 | TACC1 | chr8 | + | 38838469 | 38838546 | 38836161 | 38836287 | 38840223 | 38840267 | 280 | 230 | 230 | 258 | 355 | 0 | 0 | 0 | 0 | 0 | 234 | 200 | 348 | 273 | 175 | 0 | 0 | 3 | 0 | 0 | 226 | 149 | 1 | 1 | 1 | 1 | 1 | 1 | 1 | 1 | 1 | 0.987 | 1 | 1 | 0.003 |
| SE | 65590 | ENSG00000147526 | TACC1 | chr8 | + | 38840223 | 38840267 | 38838469 | 38838546 | 38842286 | 38842447 | 257 | 209 | 189 | 212 | 381 | 2 | 0 | 0 | 0 | 0 | 159 | 130 | 243 | 170 | 137 | 0 | 0 | 0 | 0 | 0 | 193 | 149 | 1 | 1 | 0.99 | 1 | 1 | 1 | 1 | 1 | 1 | 1 | 1 | 1 | -0.002 |
| SE | 65593 | ENSG00000198574 | SH2D1B | chr1 | - | 162398922 | 162399087 | 162395267 | 162397315 | 162411882 | 162412136 | 424 | 145 | 139 | 169 | 182 | 3 | 0 | 1 | 1 | 0 | 21 | 61 | 62 | 29 | 14 | 0 | 0 | 0 | 0 | 0 | 298 | 149 | 1 | 1 | 0.986 | 1 | 0.986 | 0.988 | 1 | 1 | 1 | 1 | 1 | 1 | -0.008 |
| SE | 65594 | ENSG00000198574 | SH2D1B | chr1 | - | 162402738 | 162402802 | 162395267 | 162397315 | 162411882 | 162412136 | 89 | 29 | 30 | 31 | 19 | 3 | 0 | 0 | 1 | 0 | 13 | 14 | 17 | 7 | 3 | 0 | 0 | 0 | 0 | 0 | 213 | 149 | 1 | 1 | 0.954 | 1 | 0.955 | 0.956 | 1 | 1 | 1 | 1 | 1 | 1 | -0.027 |
| SE | 65595 | ENSG00000198574 | SH2D1B | chr1 | - | 162402738 | 162402802 | 162398922 | 162399087 | 162411882 | 162412136 | 283 | 102 | 94 | 143 | 76 | 0 | 0 | 0 | 1 | 0 | 25 | 54 | 43 | 15 | 15 | 0 | 0 | 0 | 0 | 0 | 213 | 149 | 1 | 1 | 1 | 1 | 1 | 0.99 | 1 | 1 | 1 | 1 | 1 | 1 | -0.002 |
| SE | 65596 | ENSG00000112078 | KCTD20 | chr6 | + | 36443417 | 36443530 | 36443016 | 36443111 | 36470051 | 36470257 | 0 | 0 | 0 | 0 | 0 | 23 | 17 | 12 | 29 | 11 | 0 | 0 | 3 | 0 | 0 | 10 | 13 | 16 | 43 | 46 | 262 | 149 | 1 | 1 | 0 | 0 | 0 | 0 | 0 | 0 | 0 | 0.096 | 0 | 0 | -0.019 |
| SE | 65599 | ENSG00000112078 | KCTD20 | chr6 | + | 36452563 | 36452680 | 36451533 | 36451632 | 36470051 | 36470257 | 0 | 0 | 1 | 0 | 0 | 3 | 0 | 1 | 0 | 0 | 0 | 0 | 0 | 0 | 0 | 0 | 7 | 1 | 5 | 6 | 266 | 149 | 1 | 1 | 0 | NA | 0.359 | NA | NA | NA | 0 | 0 | 0 | 0 | 0.179 |
| SE | 65601 | ENSG00000112078 | KCTD20 | chr6 | + | 36470051 | 36470257 | 36442766 | 36443111 | 36474788 | 36474889 | 158 | 122 | 80 | 185 | 227 | 0 | 0 | 1 | 1 | 0 | 176 | 130 | 95 | 169 | 133 | 0 | 0 | 0 | 0 | 0 | 298 | 149 | 1 | 1 | 1 | 1 | 0.976 | 1 | 1 | 1 | 1 | 1 | 1 | 1 | -0.005 |
| SE | 65603 | ENSG00000112078 | KCTD20 | chr6 | + | 36470051 | 36470257 | 36442766 | 36443111 | 36481561 | 36481759 | 23 | 17 | 13 | 31 | 14 | 0 | 0 | 0 | 1 | 0 | 14 | 17 | 32 | 48 | 46 | 0 | 0 | 0 | 0 | 0 | 298 | 149 | 1 | 1 | 1 | 1 | 1 | 0.939 | 1 | 1 | 1 | 1 | 1 | 1 | -0.012 |
| SE | 65604 | ENSG00000112078 | KCTD20 | chr6 | + | 36470051 | 36470257 | 36451533 | 36451632 | 36479120 | 36479223 | 17 | 13 | 12 | 7 | 19 | 0 | 0 | 0 | 0 | 1 | 18 | 14 | 17 | 14 | 26 | 0 | 0 | 0 | 0 | 0 | 298 | 149 | 1 | 1 | 1 | 1 | 1 | 1 | 0.905 | 1 | 1 | 1 | 1 | 1 | -0.019 |
| SE | 65605 | ENSG00000112078 | KCTD20 | chr6 | + | 36474788 | 36475062 | 36442999 | 36443111 | 36479120 | 36479223 | 142 | 111 | 74 | 123 | 198 | 3 | 1 | 1 | 0 | 0 | 118 | 102 | 78 | 108 | 140 | 0 | 0 | 0 | 0 | 0 | 298 | 149 | 1 | 1 | 0.959 | 0.982 | 0.974 | 1 | 1 | 1 | 1 | 1 | 1 | 1 | -0.017 |
| SE | 65606 | ENSG00000112078 | KCTD20 | chr6 | + | 36474788 | 36475062 | 36442999 | 36443111 | 36481561 | 36481759 | 6 | 4 | 4 | 5 | 17 | 0 | 0 | 0 | 1 | 0 | 0 | 3 | 8 | 7 | 8 | 0 | 0 | 0 | 0 | 0 | 298 | 149 | 1 | 1 | 1 | 1 | 1 | 0.714 | 1 | NA | 1 | 1 | 1 | 1 | -0.057 |
| SE | 65610 | ENSG00000112078 | KCTD20 | chr6 | + | 36474788 | 36475062 | 36470051 | 36470257 | 36484713 | 36484824 | 138 | 105 | 71 | 156 | 216 | 0 | 2 | 1 | 2 | 0 | 173 | 117 | 79 | 126 | 87 | 0 | 0 | 1 | 0 | 0 | 298 | 149 | 1 | 1 | 1 | 0.963 | 0.973 | 0.975 | 1 | 1 | 1 | 0.975 | 1 | 1 | -0.013 |
| SE | 65611 | ENSG00000112078 | KCTD20 | chr6 | + | 36479120 | 36479223 | 36442766 | 36443111 | 36481561 | 36481759 | 7 | 4 | 0 | 8 | 7 | 0 | 0 | 0 | 1 | 0 | 31 | 17 | 12 | 26 | 61 | 0 | 0 | 0 | 0 | 0 | 252 | 149 | 1 | 1 | 1 | 1 | NA | 0.825 | 1 | 1 | 1 | 1 | 1 | 1 | -0.044 |
| SE | 65619 | ENSG00000112078 | KCTD20 | chr6 | + | 36479590 | 36479711 | 36470051 | 36470257 | 36484713 | 36484824 | 2 | 3 | 2 | 2 | 7 | 0 | 2 | 1 | 2 | 0 | 4 | 3 | 3 | 15 | 0 | 0 | 0 | 1 | 0 | 0 | 270 | 149 | 1 | 1 | 1 | 0.453 | 0.525 | 0.356 | 1 | 1 | 1 | 0.623 | 1 | NA | -0.239 |
| SE | 65621 | ENSG00000112078 | KCTD20 | chr6 | + | 36479590 | 36479711 | 36474788 | 36475062 | 36484713 | 36484824 | 2 | 1 | 3 | 1 | 1 | 3 | 0 | 3 | 0 | 0 | 7 | 0 | 0 | 0 | 7 | 0 | 0 | 3 | 0 | 0 | 270 | 149 | 1 | 1 | 0.269 | 1 | 0.356 | 1 | 1 | 0.356 | NA | NA | NA | NA | 0.369 |
| SE | 65623 | ENSG00000112078 | KCTD20 | chr6 | + | 36479590 | 36479711 | 36479120 | 36479223 | 36484713 | 36484824 | 161 | 166 | 124 | 189 | 348 | 0 | 0 | 0 | 0 | 0 | 277 | 125 | 131 | 182 | 135 | 3 | 0 | 3 | 0 | 0 | 270 | 149 | 1 | 1 | 1 | 1 | 1 | 1 | 1 | 0.981 | 1 | 0.96 | 1 | 1 | 0.012 |
| SE | 65624 | ENSG00000112078 | KCTD20 | chr6 | + | 36481561 | 36481759 | 36470051 | 36470257 | 36484713 | 36484824 | 243 | 194 | 217 | 282 | 319 | 0 | 2 | 1 | 2 | 0 | 474 | 290 | 331 | 346 | 331 | 0 | 0 | 1 | 0 | 0 | 298 | 149 | 1 | 1 | 1 | 0.98 | 0.991 | 0.986 | 1 | 1 | 1 | 0.994 | 1 | 1 | -0.007 |
| SE | 65625 | ENSG00000112078 | KCTD20 | chr6 | + | 36481561 | 36481759 | 36474788 | 36475062 | 36484713 | 36484824 | 249 | 198 | 219 | 285 | 333 | 3 | 0 | 3 | 0 | 0 | 470 | 289 | 323 | 348 | 339 | 7 | 0 | 0 | 0 | 0 | 298 | 149 | 1 | 1 | 0.976 | 1 | 0.973 | 1 | 1 | 0.971 | 1 | 1 | 1 | 1 | -0.004 |
| SE | 65626 | ENSG00000112078 | KCTD20 | chr6 | + | 36481561 | 36481759 | 36479120 | 36479223 | 36484713 | 36484824 | 252 | 198 | 216 | 291 | 323 | 0 | 0 | 0 | 0 | 0 | 508 | 303 | 327 | 368 | 405 | 3 | 0 | 3 | 0 | 0 | 298 | 149 | 1 | 1 | 1 | 1 | 1 | 1 | 1 | 0.988 | 1 | 0.982 | 1 | 1 | 0.006 |
| SE | 65627 | ENSG00000112078 | KCTD20 | chr6 | + | 36481561 | 36481759 | 36479590 | 36479711 | 36484713 | 36484824 | 461 | 406 | 359 | 532 | 685 | 0 | 0 | 1 | 0 | 0 | 895 | 518 | 550 | 678 | 537 | 2 | 0 | 0 | 0 | 0 | 298 | 149 | 1 | 1 | 1 | 1 | 0.994 | 1 | 1 | 0.996 | 1 | 1 | 1 | 1 | 0 |
| SE | 65629 | ENSG00000142856 | ITGB3BP | chr1 | - | 63446805 | 63446856 | 63442509 | 63443195 | 63453917 | 63453974 | 17 | 15 | 6 | 15 | 31 | 0 | 0 | 0 | 0 | 3 | 29 | 7 | 6 | 0 | 27 | 0 | 0 | 0 | 0 | 0 | 200 | 149 | 1 | 1 | 1 | 1 | 1 | 1 | 0.885 | 1 | 1 | 1 | NA | 1 | -0.023 |
| SE | 65631 | ENSG00000142856 | ITGB3BP | chr1 | - | 63454379 | 63454473 | 63453614 | 63453974 | 63454889 | 63454949 | 118 | 102 | 48 | 100 | 233 | 0 | 0 | 2 | 1 | 2 | 64 | 51 | 66 | 54 | 123 | 0 | 0 | 0 | 0 | 2 | 243 | 149 | 1 | 1 | 1 | 1 | 0.936 | 0.984 | 0.986 | 1 | 1 | 1 | 1 | 0.974 | -0.014 |
| SE | 65632 | ENSG00000142856 | ITGB3BP | chr1 | - | 63454889 | 63454968 | 63454379 | 63454473 | 63508527 | 63508570 | 27 | 20 | 13 | 16 | 74 | 0 | 0 | 0 | 0 | 0 | 6 | 8 | 14 | 8 | 25 | 0 | 0 | 0 | 2 | 0 | 228 | 149 | 1 | 1 | 1 | 1 | 1 | 1 | 1 | 1 | 1 | 1 | 0.723 | 1 | 0.055 |
| SE | 65635 | ENSG00000142856 | ITGB3BP | chr1 | - | 63478763 | 63478833 | 63457683 | 63458373 | 63490082 | 63490214 | 69 | 40 | 14 | 55 | 81 | 0 | 0 | 0 | 1 | 0 | 56 | 19 | 12 | 32 | 28 | 0 | 0 | 0 | 0 | 0 | 219 | 149 | 1 | 1 | 1 | 1 | 1 | 0.974 | 1 | 1 | 1 | 1 | 1 | 1 | -0.005 |
| SE | 65636 | ENSG00000142856 | ITGB3BP | chr1 | - | 63478763 | 63478833 | 63457683 | 63458373 | 63508527 | 63508570 | 17 | 6 | 2 | 10 | 11 | 0 | 0 | 0 | 2 | 0 | 0 | 0 | 4 | 0 | 1 | 0 | 0 | 0 | 0 | 0 | 219 | 149 | 1 | 1 | 1 | 1 | 1 | 0.773 | 1 | NA | NA | 1 | NA | 1 | -0.045 |
| SE | 65638 | ENSG00000142856 | ITGB3BP | chr1 | - | 63490082 | 63490218 | 63457683 | 63458373 | 63508527 | 63508570 | 83 | 51 | 45 | 82 | 103 | 0 | 0 | 0 | 2 | 0 | 24 | 19 | 23 | 19 | 63 | 0 | 0 | 0 | 0 | 0 | 285 | 149 | 1 | 1 | 1 | 1 | 1 | 0.955 | 1 | 1 | 1 | 1 | 1 | 1 | -0.009 |
| SE | 65640 | ENSG00000142856 | ITGB3BP | chr1 | - | 63508527 | 63508570 | 63490082 | 63490218 | 63523128 | 63523182 | 79 | 49 | 37 | 78 | 95 | 0 | 0 | 0 | 1 | 0 | 22 | 19 | 21 | 17 | 35 | 0 | 0 | 0 | 0 | 0 | 192 | 149 | 1 | 1 | 1 | 1 | 1 | 0.984 | 1 | 1 | 1 | 1 | 1 | 1 | -0.003 |
| SE | 65642 | ENSG00000142856 | ITGB3BP | chr1 | - | 63510064 | 63510181 | 63508527 | 63508570 | 63523128 | 63523194 | 2 | 0 | 4 | 1 | 5 | 59 | 26 | 17 | 57 | 45 | 2 | 0 | 0 | 0 | 0 | 18 | 12 | 28 | 11 | 0 | 266 | 149 | 1 | 1 | 0.019 | 0 | 0.116 | 0.01 | 0.059 | 0.059 | 0 | 0 | 0 | NA | 0.026 |
| SE | 65643 | ENSG00000142856 | ITGB3BP | chr1 | - | 63527878 | 63527970 | 63508527 | 63508570 | 63529130 | 63529186 | 2 | 0 | 2 | 1 | 1 | 2 | 7 | 4 | 3 | 18 | 0 | 0 | 0 | 0 | 0 | 0 | 3 | 0 | 4 | 30 | 241 | 149 | 1 | 1 | 0.382 | 0 | 0.236 | 0.171 | 0.033 | NA | 0 | NA | 0 | 0 | 0.164 |
| SE | 65644 | ENSG00000142856 | ITGB3BP | chr1 | - | 63529130 | 63529186 | 63508527 | 63508570 | 63593524 | 63593721 | 2 | 7 | 4 | 3 | 17 | 0 | 0 | 0 | 0 | 0 | 0 | 3 | 0 | 4 | 34 | 0 | 0 | 0 | 2 | 0 | 205 | 149 | 1 | 1 | 1 | 1 | 1 | 1 | 1 | NA | 1 | NA | 0.592 | 1 | 0.136 |
| SE | 65645 | ENSG00000143924 | EML4 | chr2 | + | 42207928 | 42208082 | 42169390 | 42169636 | 42245504 | 42245688 | 0 | 0 | 0 | 1 | 1 | 96 | 49 | 74 | 91 | 57 | 0 | 1 | 1 | 0 | 0 | 10 | 56 | 71 | 51 | 29 | 298 | 149 | 1 | 1 | 0 | 0 | 0 | 0.005 | 0.009 | 0 | 0.009 | 0.007 | 0 | 0 | 0 |
| SE | 65646 | ENSG00000143924 | EML4 | chr2 | + | 42256500 | 42256630 | 42245504 | 42245687 | 42261120 | 42261294 | 330 | 316 | 444 | 384 | 955 | 0 | 0 | 0 | 0 | 0 | 158 | 274 | 368 | 274 | 50 | 0 | 0 | 1 | 0 | 0 | 279 | 149 | 1 | 1 | 1 | 1 | 1 | 1 | 1 | 1 | 1 | 0.995 | 1 | 1 | 0.001 |
| SE | 65647 | ENSG00000143924 | EML4 | chr2 | + | 42261120 | 42261294 | 42256500 | 42256630 | 42263177 | 42263306 | 329 | 303 | 435 | 390 | 1089 | 0 | 2 | 1 | 0 | 0 | 157 | 275 | 406 | 251 | 48 | 0 | 0 | 0 | 0 | 0 | 298 | 149 | 1 | 1 | 1 | 0.987 | 0.995 | 1 | 1 | 1 | 1 | 1 | 1 | 1 | -0.004 |
| SE | 65649 | ENSG00000143924 | EML4 | chr2 | + | 42282822 | 42282972 | 42280849 | 42280973 | 42284633 | 42284703 | 489 | 432 | 510 | 481 | 581 | 0 | 0 | 3 | 0 | 0 | 65 | 292 | 332 | 267 | 200 | 0 | 0 | 0 | 0 | 5 | 298 | 149 | 1 | 1 | 1 | 1 | 0.988 | 1 | 1 | 1 | 1 | 1 | 1 | 0.952 | 0.007 |
| SE | 65651 | ENSG00000143924 | EML4 | chr2 | + | 42303314 | 42303446 | 42303103 | 42303229 | 42304483 | 42304551 | 483 | 438 | 590 | 541 | 664 | 0 | 0 | 1 | 0 | 0 | 271 | 364 | 597 | 290 | 66 | 0 | 0 | 0 | 0 | 0 | 281 | 149 | 1 | 1 | 1 | 1 | 0.997 | 1 | 1 | 1 | 1 | 1 | 1 | 1 | -0.001 |
| SE | 65653 | ENSG00000143924 | EML4 | chr2 | + | 42315961 | 42316050 | 42304483 | 42304551 | 42317426 | 42317524 | 156 | 127 | 195 | 216 | 273 | 0 | 0 | 1 | 0 | 0 | 57 | 114 | 167 | 98 | 37 | 0 | 0 | 2 | 0 | 0 | 238 | 149 | 1 | 1 | 1 | 1 | 1 | 1 | 1 | 1 | 1 | 0.981 | 1 | 1 | 0.004 |
| SE | 65656 | ENSG00000143924 | EML4 | chr2 | + | 42317426 | 42317524 | 42315961 | 42316050 | 42325466 | 42325554 | 174 | 122 | 216 | 218 | 238 | 1 | 1 | 0 | 0 | 0 | 52 | 140 | 181 | 139 | 33 | 0 | 0 | 0 | 0 | 0 | 247 | 149 | 1 | 1 | 0.991 | 0.987 | 1 | 1 | 1 | 1 | 1 | 1 | 1 | 1 | -0.004 |
| SE | 65657 | ENSG00000143924 | EML4 | chr2 | + | 42328885 | 42329016 | 42326153 | 42326252 | 42329733 | 42330744 | 441 | 302 | 548 | 480 | 1000 | 0 | 0 | 1 | 0 | 0 | 113 | 251 | 326 | 196 | 49 | 0 | 0 | 0 | 0 | 0 | 280 | 149 | 1 | 1 | 1 | 1 | 0.997 | 1 | 1 | 1 | 1 | 1 | 1 | 1 | -0.001 |
| SE | 65660 | ENSG00000111729 | CLEC4A | chr12 | + | 8125560 | 8125677 | 8123616 | 8123960 | 8136787 | 8136903 | 194 | 160 | 150 | 135 | 275 | 1 | 0 | 0 | 0 | 0 | 280 | 117 | 70 | 211 | 77 | 0 | 0 | 0 | 0 | 0 | 266 | 149 | 1 | 1 | 0.991 | 1 | 1 | 1 | 1 | 1 | 1 | 1 | 1 | 1 | -0.002 |
| SE | 65662 | ENSG00000111729 | CLEC4A | chr12 | + | 8129263 | 8129362 | 8123878 | 8123960 | 8136787 | 8136903 | 30 | 18 | 12 | 8 | 2 | 1 | 0 | 0 | 0 | 0 | 0 | 0 | 3 | 5 | 0 | 0 | 0 | 0 | 0 | 0 | 248 | 149 | 1 | 1 | 0.947 | 1 | 1 | 1 | 1 | NA | NA | 1 | 1 | NA | -0.011 |
| SE | 65665 | ENSG00000111729 | CLEC4A | chr12 | + | 8135584 | 8135736 | 8123852 | 8123960 | 8136787 | 8136903 | 149 | 80 | 62 | 81 | 164 | 1 | 0 | 0 | 0 | 0 | 124 | 72 | 36 | 72 | 63 | 0 | 0 | 0 | 0 | 0 | 298 | 149 | 1 | 1 | 0.987 | 1 | 1 | 1 | 1 | 1 | 1 | 1 | 1 | 1 | -0.003 |
| SE | 65667 | ENSG00000111729 | CLEC4A | chr12 | + | 8135584 | 8135736 | 8129263 | 8129362 | 8136787 | 8136903 | 422 | 273 | 214 | 225 | 272 | 0 | 0 | 1 | 0 | 0 | 461 | 191 | 129 | 382 | 155 | 0 | 0 | 0 | 0 | 0 | 298 | 149 | 1 | 1 | 1 | 1 | 0.991 | 1 | 1 | 1 | 1 | 1 | 1 | 1 | -0.002 |
| SE | 65668 | ENSG00000111729 | CLEC4A | chr12 | + | 8136787 | 8136903 | 8135584 | 8135736 | 8138139 | 8138206 | 273 | 141 | 147 | 156 | 298 | 0 | 0 | 1 | 0 | 0 | 244 | 142 | 80 | 200 | 110 | 0 | 0 | 0 | 0 | 0 | 265 | 149 | 1 | 1 | 1 | 1 | 0.988 | 1 | 1 | 1 | 1 | 1 | 1 | 1 | -0.002 |
| SE | 65670 | ENSG00000138336 | TET1 | chr10 | + | 68651845 | 68651936 | 68644697 | 68647005 | 68652500 | 68652594 | 4 | 9 | 12 | 15 | 35 | 0 | 0 | 3 | 3 | 1 | 14 | 8 | 4 | 1 | 0 | 0 | 10 | 0 | 0 | 0 | 240 | 149 | 1 | 1 | 1 | 1 | 0.713 | 0.756 | 0.956 | 1 | 0.332 | 1 | 1 | NA | 0.052 |
| SE | 65671 | ENSG00000138336 | TET1 | chr10 | + | 68652500 | 68652594 | 68644697 | 68647005 | 68667044 | 68667256 | 3 | 3 | 8 | 8 | 11 | 0 | 0 | 1 | 1 | 0 | 0 | 11 | 2 | 0 | 0 | 0 | 0 | 0 | 0 | 0 | 243 | 149 | 1 | 1 | 1 | 1 | 0.831 | 1 | 1 | NA | 1 | 1 | NA | NA | -0.034 |
| SE | 65676 | ENSG00000151502 | VPS26B | chr11 | + | 134234896 | 134235053 | 134224670 | 134225345 | 134243118 | 134243294 | 104 | 68 | 120 | 155 | 261 | 3 | 1 | 0 | 0 | 0 | 84 | 143 | 57 | 77 | 218 | 0 | 0 | 0 | 0 | 0 | 298 | 149 | 1 | 1 | 0.945 | 0.971 | 1 | 1 | 1 | 1 | 1 | 1 | 1 | 1 | -0.017 |
| SE | 65677 | ENSG00000151502 | VPS26B | chr11 | + | 134236341 | 134236746 | 134234961 | 134235053 | 134239990 | 134240155 | 2 | 2 | 3 | 5 | 6 | 213 | 126 | 171 | 214 | 286 | 0 | 4 | 0 | 4 | 0 | 315 | 250 | 122 | 179 | 456 | 298 | 149 | 1 | 1 | 0.005 | 0.008 | 0.009 | 0.012 | 0.01 | 0 | 0.008 | 0 | 0.011 | 0 | 0.005 |
| SE | 65678 | ENSG00000151502 | VPS26B | chr11 | + | 134239990 | 134240155 | 134225193 | 134225345 | 134243118 | 134243196 | 319 | 208 | 186 | 285 | 413 | 3 | 1 | 0 | 0 | 0 | 554 | 347 | 215 | 342 | 774 | 0 | 0 | 0 | 0 | 0 | 298 | 149 | 1 | 1 | 0.982 | 0.99 | 1 | 1 | 1 | 1 | 1 | 1 | 1 | 1 | -0.006 |
| SE | 65679 | ENSG00000151502 | VPS26B | chr11 | + | 134239990 | 134240155 | 134234896 | 134235053 | 134243118 | 134243196 | 531 | 333 | 357 | 499 | 698 | 1 | 2 | 4 | 2 | 1 | 855 | 597 | 336 | 521 | 1230 | 0 | 0 | 0 | 0 | 10 | 298 | 149 | 1 | 1 | 0.996 | 0.988 | 0.978 | 0.992 | 0.997 | 1 | 1 | 1 | 1 | 0.984 | -0.007 |
| SE | 65681 | ENSG00000179104 | TMTC2 | chr12 | + | 82985924 | 82986046 | 82930430 | 82930545 | 83050903 | 83051018 | 0 | 1 | 1 | 1 | 0 | 0 | 0 | 0 | 0 | 0 | 0 | 0 | 0 | 3 | 0 | 1 | 0 | 0 | 0 | 0 | 271 | 149 | 1 | 1 | NA | 1 | 1 | 1 | NA | 0 | NA | NA | 1 | NA | 0.5 |
| SE | 65682 | ENSG00000179104 | TMTC2 | chr12 | + | 83030797 | 83030879 | 82985924 | 82986046 | 83050903 | 83051018 | 9 | 13 | 13 | 5 | 6 | 0 | 1 | 1 | 1 | 0 | 0 | 2 | 13 | 23 | 0 | 0 | 0 | 0 | 0 | 0 | 231 | 149 | 1 | 1 | 1 | 0.893 | 0.893 | 0.763 | 1 | NA | 1 | 1 | 1 | NA | -0.09 |
| SE | 65684 | ENSG00000186141 | POLR3C | chr1 | - | 145836493 | 145836574 | 145833489 | 145833582 | 145836814 | 145836854 | 93 | 84 | 106 | 113 | 159 | 2 | 0 | 1 | 0 | 0 | 132 | 168 | 89 | 88 | 108 | 0 | 0 | 0 | 0 | 0 | 230 | 149 | 1 | 1 | 0.968 | 1 | 0.986 | 1 | 1 | 1 | 1 | 1 | 1 | 1 | -0.009 |
| SE | 65687 | ENSG00000186141 | POLR3C | chr1 | - | 145838055 | 145838206 | 145833489 | 145833582 | 145839889 | 145839991 | 107 | 79 | 83 | 107 | 149 | 0 | 0 | 1 | 1 | 1 | 231 | 106 | 135 | 101 | 125 | 6 | 0 | 0 | 0 | 0 | 298 | 149 | 1 | 1 | 1 | 1 | 1 | 0.982 | 0.987 | 0.951 | 1 | 1 | 1 | 1 | 0.004 |
| SE | 65689 | ENSG00000186141 | POLR3C | chr1 | - | 145838055 | 145838206 | 145836493 | 145836574 | 145842338 | 145842493 | 7 | 0 | 6 | 6 | 4 | 0 | 0 | 0 | 1 | 0 | 5 | 7 | 5 | 2 | 9 | 0 | 0 | 0 | 0 | 0 | 298 | 149 | 1 | 1 | 1 | NA | 1 | 0.75 | 1 | 1 | 1 | 1 | 1 | 1 | -0.062 |
| SE | 65690 | ENSG00000186141 | POLR3C | chr1 | - | 145839889 | 145839991 | 145836493 | 145836574 | 145842338 | 145844402 | 2 | 0 | 2 | 4 | 2 | 0 | 0 | 0 | 1 | 0 | 0 | 5 | 5 | 0 | 14 | 0 | 0 | 0 | 0 | 0 | 251 | 149 | 1 | 1 | 1 | NA | 1 | 0.704 | 1 | NA | 1 | 1 | NA | 1 | -0.074 |

| | | | | | | | | | | | | | | |
|---|---|---|---|---|---|---|---|---|---|---|---|---|---|---|
| SE | 65691 | ENSG00000186141 | POLR3C | chr1 | + | 145839889 | 145839991 | 145838055 | 145838206 | 145842338 | 145844402 | 89 | 77 | [illegible] |
| SE | 65692 | ENSG00000186141 | POLR3C | chr1 | + | 145840115 | 145840165 | 145839889 | 145839991 | 145842338 | 145844402 | 17 | 5 | [illegible] |
| SE | 65693 | ENSG00000186141 | POLR3C | chr1 | + | 145840921 | 145841071 | 145840115 | 145840165 | 145842338 | 145844402 | 202 | 106 | [illegible] |
| SE | 65694 | ENSG00000100243 | CYB5R3 | chr22 | - | 42627604 | 42627688 | 42627303 | 42627389 | 42628151 | 42628281 | 273 | 101 | [illegible] |
| SE | 65695 | ENSG00000100243 | CYB5R3 | chr22 | - | 42628151 | 42628281 | 42627604 | 42627688 | 42630881 | 42630988 | 265 | 112 | [illegible] |
| SE | 65696 | ENSG00000100243 | CYB5R3 | chr22 | - | 42646640 | 42646701 | 42636714 | 42636846 | 42646885 | 42646960 | 0 | 0 | [illegible] |
| SE | 65699 | ENSG00000102900 | NUP93 | chr16 | + | 56748233 | 56748426 | 56730128 | 56730211 | 56758537 | 56758655 | 70 | 31 | [illegible] |
| SE | 65700 | ENSG00000102900 | NUP93 | chr16 | + | 56758537 | 56758655 | 56748233 | 56748426 | 56798475 | 56798538 | 80 | 68 | [illegible] |
| SE | 65701 | ENSG00000102900 | NUP93 | chr16 | + | 56798475 | 56798538 | 56758537 | 56758655 | 56805503 | 56805552 | 66 | 66 | [illegible] |
| SE | 65702 | ENSG00000102900 | NUP93 | chr16 | + | 56805503 | 56805632 | 56798475 | 56798538 | 56818663 | 56818709 | 94 | 103 | [illegible] |
| SE | 65703 | ENSG00000102900 | NUP93 | chr16 | + | 56837607 | 56837726 | 56836600 | 56836717 | 56838951 | 56839069 | 148 | 96 | [illegible] |
| SE | 65707 | ENSG00000102900 | NUP93 | chr16 | + | 56842577 | 56842695 | 56841704 | 56841833 | 56844498 | 56844769 | 0 | 0 | [illegible] |
| SE | 65708 | ENSG00000138434 | ITPRID2 | chr2 | + | 181896029 | 181896079 | 181892614 | 181892660 | 181896907 | 181896964 | 168 | 151 | [illegible] |
| SE | 65709 | ENSG00000138434 | ITPRID2 | chr2 | + | 181900695 | 181900904 | 181899013 | 181899112 | 181901765 | 181901974 | 545 | 491 | [illegible] |
| SE | 65711 | ENSG00000138434 | ITPRID2 | chr2 | + | 181918322 | 181918470 | 181916721 | 181917005 | 181918597 | 181918675 | 5 | 8 | [illegible] |
| SE | 65712 | ENSG00000138434 | ITPRID2 | chr2 | + | 181918754 | 181918882 | 181915215 | 181918675 | 181919295 | 181919375 | 52 | 66 | [illegible] |
| SE | 65717 | ENSG00000198554 | WDHD1 | chr14 | - | 55007279 | 55007375 | 55002092 | 55002185 | 55008315 | 55008366 | 6 | 2 | [illegible] |
| SE | 65718 | ENSG00000198554 | WDHD1 | chr14 | - | 55010308 | 55010460 | 55008607 | 55008719 | 55013484 | 55013596 | 24 | 16 | [illegible] |
| SE | 65719 | ENSG00000198554 | WDHD1 | chr14 | - | 55013484 | 55013596 | 55010308 | 55010460 | 55026710 | 55026803 | 18 | 13 | [illegible] |
| SE | 65720 | ENSG00000198554 | WDHD1 | chr14 | - | 55026710 | 55026803 | 55013484 | 55013596 | 55027027 | 55027099 | 7 | 2 | [illegible] |
| SE | 65721 | ENSG00000144909 | OSBPL11 | chr3 | - | 125547405 | 125547592 | 125538450 | 125538633 | 125560378 | 125560519 | 129 | 115 | [illegible] |
| SE | 65723 | ENSG00000144909 | OSBPL11 | chr3 | - | 125560378 | 125560519 | 125552180 | 125552679 | 125563697 | 125563843 | 218 | 177 | [illegible] |
| SE | 65724 | ENSG00000144909 | OSBPL11 | chr3 | - | 125563697 | 125563843 | 125560378 | 125560519 | 125567393 | 125567595 | 355 | 253 | [illegible] |
| SE | 65725 | ENSG00000144909 | OSBPL11 | chr3 | - | 125576188 | 125576365 | 125567393 | 125567595 | 125578959 | 125579039 | 278 | 217 | [illegible] |
| SE | 65726 | ENSG00000148303 | RPL7A | chr9 | + | 133349386 | 133349700 | 133348936 | 133349042 | 133349911 | 133350052 | 10179 | 6031 | [illegible] |
| SE | 65727 | ENSG00000148303 | RPL7A | chr9 | + | 133349550 | 133349700 | 133348921 | 133349042 | 133349911 | 133350052 | 22414 | 12812 | [illegible] |
| SE | 65728 | ENSG00000148303 | RPL7A | chr9 | + | 133350239 | 133350319 | 133349911 | 133350052 | 133350596 | 133350695 | 11790 | 6557 | [illegible] |
| SE | 65730 | ENSG00000138073 | PREB | chr2 | - | 27131673 | 27131831 | 27130790 | 27131508 | 27132009 | 27132082 | 19 | 11 | [illegible] |
| SE | 65732 | ENSG00000138073 | PREB | chr2 | - | 27132842 | 27132923 | 27132510 | 27132727 | 27133116 | 27133337 | 278 | 131 | [illegible] |
| [illegible] | | | | | | | | | | | | | | |

| | | | | | | | | | | | | | |
|---|---|---|---|---|---|---|---|---|---|---|---|---|---|
| SE | 66098 | ENSG00000000938 | FGR | chr1 | - | 27621558 | 27621657 | 27616856 | 27617006 | 27625088 | 27625151 | 2 | 6 |
| SE | 66099 | ENSG00000000938 | FGR | chr1 | - | 27621558 | 27621657 | 27616856 | 27617006 | 27635064 | 27635185 | 2 | 5 |
| SE | 66100 | ENSG00000000938 | FGR | chr1 | - | 27621558 | 27621657 | 27617192 | 27617296 | 27623041 | 27623144 | 1882 | 1081 |
| SE | 66101 | ENSG00000000938 | FGR | chr1 | - | 27621558 | 27621657 | 27617192 | 27617296 | 27623690 | 27623929 | 1097 | 665 |
| SE | 66102 | ENSG00000000938 | FGR | chr1 | - | 27621558 | 27621657 | 27617192 | 27617296 | 27625088 | 27625151 | 1090 | 666 |
| SE | 66103 | ENSG00000000938 | FGR | chr1 | - | 27621558 | 27621657 | 27617192 | 27617296 | 27635064 | 27635185 | 1089 | 665 |
| SE | 66104 | ENSG00000000938 | FGR | chr1 | - | 27623041 | 27623144 | 27616856 | 27617006 | 27623690 | 27623929 | 359 | 176 |
| SE | 66105 | ENSG00000000938 | FGR | chr1 | - | 27623041 | 27623144 | 27616856 | 27617006 | 27625088 | 27625151 | 0 | 0 |
| SE | 66106 | ENSG00000000938 | FGR | chr1 | - | 27623041 | 27623144 | 27617192 | 27617296 | 27623690 | 27623929 | 360 | 176 |
| SE | 66107 | ENSG00000000938 | FGR | chr1 | - | 27623041 | 27623144 | 27617192 | 27617296 | 27625088 | 27625151 | 1 | 0 |
| SE | 66108 | ENSG00000000938 | FGR | chr1 | - | 27623041 | 27623144 | 27621558 | 27621657 | 27623690 | 27623929 | 1112 | 573 |
| SE | 66113 | ENSG00000000938 | FGR | chr1 | - | 27623690 | 27623929 | 27621558 | 27621657 | 27626045 | 27626135 | 11 | 4 |
| SE | 66114 | ENSG00000000938 | FGR | chr1 | - | 27623690 | 27623929 | 27623041 | 27623144 | 27625088 | 27625151 | 871 | 402 |
| SE | 66115 | ENSG00000000938 | FGR | chr1 | - | 27625088 | 27625151 | 27616856 | 27617006 | 27635064 | 27635134 | 19 | 5 |
| SE | 66116 | ENSG00000000938 | FGR | chr1 | - | 27625088 | 27625151 | 27617192 | 27617296 | 27635064 | 27635134 | 20 | 6 |
| SE | 66117 | ENSG00000000938 | FGR | chr1 | - | 27625088 | 27625151 | 27621558 | 27621657 | 27626045 | 27626135 | 22 | 10 |
| SE | 66118 | ENSG00000000938 | FGR | chr1 | - | 27625088 | 27625151 | 27621558 | 27621657 | 27635064 | 27635134 | 20 | 5 |
| SE | 66119 | ENSG00000000938 | FGR | chr1 | - | 27625088 | 27625151 | 27623383 | 27623929 | 27626045 | 27626135 | 206 | 98 |
| SE | 66127 | ENSG00000234323 | LINC01505 | chr9 | + | 106564924 | 106565022 | 106558471 | 106558583 | 106588376 | 106588464 | 0 | 0 |
| SE | 66134 | ENSG00000234323 | LINC01505 | chr9 | + | 106588827 | 106588940 | 106588376 | 106588464 | 106603755 | 106604801 | 0 | 0 |
| SE | 66135 | ENSG00000234323 | LINC01505 | chr9 | + | 106588827 | 106589013 | 106588376 | 106588464 | 106603755 | 106604513 | 0 | 0 |
| SE | 66142 | ENSG00000115641 | FHL2 | chr2 | - | 105389875 | 105390227 | 105386457 | 105386540 | 105396646 | 105396785 | 0 | 0 |
| SE | 66144 | ENSG00000115641 | FHL2 | chr2 | - | 105396646 | 105396697 | 105386360 | 105386540 | 105398841 | 105399051 | 0 | 0 |
| SE | 66145 | ENSG00000115641 | FHL2 | chr2 | - | 105396646 | 105396697 | 105386360 | 105386540 | 105398969 | 105399061 | 0 | 0 |
| SE | 66148 | ENSG00000088179 | PTPN4 | chr2 | + | 119809836 | 119809991 | 119759921 | 119760384 | 119862535 | 119862622 | 121 | 68 |
| SE | 66149 | ENSG00000088179 | PTPN4 | chr2 | + | 119877322 | 119877365 | 119862535 | 119862643 | 119877463 | 119877542 | 107 | 77 |
| SE | 66153 | ENSG00000088179 | PTPN4 | chr2 | + | 119885794 | 119885882 | 119882502 | 119882623 | 119915178 | 119915242 | 97 | 62 |
| SE | 66154 | ENSG00000088179 | PTPN4 | chr2 | + | 119885794 | 119885882 | 119882502 | 119882623 | 119932423 | 119932549 | 95 | 62 |
| SE | 66155 | ENSG00000088179 | PTPN4 | chr2 | + | 119900717 | 119900806 | 119882502 | 119882623 | 119915178 | 119915242 | 83 | 96 |

[illegible]

| | | | | | | | | | | | | | |
|---|---|---|---|---|---|---|---|---|---|---|---|---|---|
| SE | 66514 | ENSG00000081320 | STK17B | chr2 | - | 196139619 | 196139799 | 196133582 | 196137729 | 196141248 | 196141297 | 4158 | 3890 [illegible] |
| SE | 66515 | ENSG00000081320 | STK17B | chr2 | - | 196141248 | 196141297 | 196139619 | 196139799 | 196143559 | 196143686 | 3081 | 3039 [illegible] |
| SE | 66516 | ENSG00000081320 | STK17B | chr2 | - | 196145910 | 196146055 | 196143559 | 196143686 | 196156438 | 196156651 | 4209 | 3531 [illegible] |
| SE | 66517 | ENSG00000081320 | STK17B | chr2 | - | 196156438 | 196156651 | 196145910 | 196146055 | 196163261 | 196163427 | 3558 | 2777 [illegible] |
| SE | 66518 | ENSG00000081320 | STK17B | chr2 | - | 196163261 | 196163427 | 196156438 | 196156651 | 196171332 | 196171578 | 2131 | 1705 [illegible] |
| SE | 66519 | ENSG00000081320 | STK17B | chr2 | - | 196165582 | 196165734 | 196163363 | 196163427 | 196166093 | 196166277 | 17 | 14 [illegible] |
| SE | 66521 | ENSG00000081320 | STK17B | chr2 | - | 196174248 | 196174407 | 196163261 | 196163427 | 196176225 | 196176327 | 3 | 3 [illegible] |
| SE | 66537 | ENSG00000223508 | RPL23AP53 | chr8 | - | 219289 | 219417 | 213185 | 213710 | 230803 | 230907 | 7 | 1 [illegible] |
| SE | 66538 | ENSG00000223508 | RPL23AP53 | chr8 | - | 219289 | 219417 | 213185 | 213710 | 232128 | 232211 | 10 | 1 [illegible] |
| SE | 66539 | ENSG00000223508 | RPL23AP53 | chr8 | - | 230803 | 230907 | 213185 | 213710 | 232128 | 232231 | 0 | 0 [illegible] |
| SE | 66541 | ENSG00000144792 | ZNF660 | chr3 | + | 44587129 | 44587250 | 44586104 | 44586213 | 44594013 | 44594295 | 0 | 1 [illegible] |
| SE | 66543 | ENSG00000170917 | NUDT6 | chr4 | - | 122897623 | 122897678 | 122892574 | 122893225 | 122912567 | 122912623 | 15 | 8 [illegible] |
| SE | 66547 | ENSG00000168438 | CDC40 | chr6 | + | 110228831 | 110228976 | 110226166 | 110226243 | 110229953 | 110230333 | 307 | 276 [illegible] |
| SE | 66548 | ENSG00000133872 | SARAF | chr8 | - | 30065987 | 30066067 | 30063794 | 30063913 | 30066776 | 30066918 | 674 | 422 [illegible] |
| SE | 66549 | ENSG00000133872 | SARAF | chr8 | - | 30065987 | 30066139 | 30063002 | 30063913 | 30066776 | 30066918 | 2950 | 1725 [illegible] |
| SE | 66550 | ENSG00000133872 | SARAF | chr8 | - | 30066027 | 30066139 | 30063125 | 30063913 | 30066776 | 30066918 | 1187 | 693 [illegible] |
| SE | 66551 | ENSG00000133872 | SARAF | chr8 | - | 30066776 | 30066918 | 30065711 | 30066139 | 30069641 | 30069949 | 2867 | 1748 [illegible] |
| SE | 66559 | ENSG00000269293 | ZSCAN16-AS1 | chr6 | - | 28125622 | 28125819 | 28121794 | 28123211 | 28136744 | 28137293 | 3 | 0 [illegible] |
| SE | 66560 | ENSG00000136371 | MTHFS | chr15 | - | 79839480 | 79839709 | 79833584 | 79833653 | 79841209 | 79841336 | 0 | 0 [illegible] |
| SE | 66561 | ENSG00000136371 | MTHFS | chr15 | - | 79841209 | 79841336 | 79833584 | 79833653 | 79845157 | 79845327 | 0 | 0 [illegible] |
| SE | 66562 | ENSG00000136371 | MTHFS | chr15 | - | 79889092 | 79889354 | 79843546 | 79845442 | 79896871 | 79897014 | 94 | 48 [illegible] |
| SE | 66581 | ENSG00000226259 | GTF2H2B | chr5 | + | 70417821 | 70417919 | 70415351 | 70415389 | 70420248 | 70420337 | 2 | 0 [illegible] |
| SE | 66583 | ENSG00000226259 | GTF2H2B | chr5 | + | 70421734 | 70421771 | 70421279 | 70421357 | 70422112 | 70422199 | 0 | 0 [illegible] |
| SE | 66584 | ENSG00000226259 | GTF2H2B | chr5 | + | 70421736 | 70421771 | 70421279 | 70421357 | 70422112 | 70422199 | 33 | 0 [illegible] |
| SE | 66585 | ENSG00000226259 | GTF2H2B | chr5 | + | 70422872 | 70422923 | 70422112 | 70422199 | 70423298 | 70423353 | 31 | 0 [illegible] |
| SE | 66586 | ENSG00000226259 | GTF2H2B | chr5 | + | 70427584 | 70427690 | 70423298 | 70423353 | 70433869 | 70433960 | 70 | 0 [illegible] |
| SE | 66587 | ENSG00000226259 | GTF2H2B | chr5 | + | 70437530 | 70437594 | 70434914 | 70435022 | 70440755 | 70440859 | 43 | 12 [illegible] |
| SE | 66588 | ENSG00000180488 | MIGA1 | chr1 | + | 77803269 | 77803406 | 77801330 | 77801508 | 77806974 | 77807101 | 210 | 158 [illegible] |
| SE | 66589 | ENSG00000180488 | MIGA1 | chr1 | + | 77803269 | 77803406 | 77801330 | 77801508 | 77813733 | 77813867 | 93 | 58 [illegible] |
| SE | 66590 | ENSG00000180488 | MIGA1 | chr1 | + | 77806974 | 77807101 | 77801330 | 77801508 | 77813733 | 77813867 | 42 | 42 [illegible] |
| SE | 66591 | ENSG00000180488 | MIGA1 | chr1 | + | 77806974 | 77807101 | 77803269 | 77803406 | 77813733 | 77813867 | 160 | 141 [illegible] |
| SE | 66592 | ENSG00000180488 | MIGA1 | chr1 | + | 77813733 | 77813867 | 77806974 | 77807101 | 77815107 | 77815231 | 96 | 80 [illegible] |
| SE | 66593 | ENSG00000180488 | MIGA1 | chr1 | + | 77843306 | 77843407 | 77815107 | 77815231 | 77858937 | 77859056 | 77 | 58 [illegible] |
| SE | 66594 | ENSG00000180488 | MIGA1 | chr1 | + | 77858937 | 77859056 | 77843306 | 77843407 | 77861223 | 77861322 | 55 | 47 [illegible] |
| SE | 66596 | ENSG00000180488 | MIGA1 | chr1 | + | 77859313 | 77859386 | 77858937 | 77859056 | 77861223 | 77861322 | 23 | 26 [illegible] |
| SE | 66599 | ENSG00000180488 | MIGA1 | chr1 | + | 77861223 | 77861322 | 77843306 | 77843407 | 77863893 | 77864028 | 46 | 32 [illegible] |
| SE | 66600 | ENSG00000180488 | MIGA1 | chr1 | + | 77861223 | 77861322 | 77843306 | 77843407 | 77866337 | 77866391 | 0 | 0 [illegible] |
| SE | 66601 | ENSG00000180488 | MIGA1 | chr1 | + | 77861223 | 77861322 | 77859313 | 77859386 | 77863893 | 77864028 | 55 | 38 [illegible] |
| SE | 66602 | ENSG00000180488 | MIGA1 | chr1 | + | 77863893 | 77864028 | 77843306 | 77843407 | 77866337 | 77866391 | 26 | 15 [illegible] |
| SE | 66603 | ENSG00000180488 | MIGA1 | chr1 | + | 77863893 | 77864028 | 77861223 | 77861322 | 77866337 | 77866391 | 88 | 50 [illegible] |
| SE | 66604 | ENSG00000180488 | MIGA1 | chr1 | + | 77866337 | 77866391 | 77863893 | 77864028 | 77873000 | 77873120 | 17 | 8 [illegible] |
| SE | 66605 | ENSG00000180488 | MIGA1 | chr1 | + | 77873000 | 77873120 | 77866337 | 77866391 | 77874845 | 77878376 | 48 | 28 [illegible] |
| SE | 66606 | ENSG00000180488 | MIGA1 | chr1 | + | 77873003 | 77873120 | 77866337 | 77866391 | 77874845 | 77876163 | 51 | 33 [illegible] |
| SE | 66608 | ENSG00000115687 | PASK | chr2 | - | 241123948 | 241124133 | 241122731 | 241122899 | 241126195 | 241127451 | 13 | 10 [illegible] |
| SE | 66612 | ENSG00000115687 | PASK | chr2 | - | 241142836 | 241143074 | 241138653 | 241138794 | 241149413 | 241149435 | 0 | 0 [illegible] |
| SE | 66618 | ENSG00000170445 | HARS1 | chr5 | - | 140677907 | 140678015 | 140677654 | 140677753 | 140683099 | 140683219 | 67 | 27 [illegible] |
| SE | 66620 | ENSG00000170445 | HARS1 | chr5 | - | 140683099 | 140683219 | 140679077 | 140679883 | 140690854 | 140690944 | 184 | 84 [illegible] |
| SE | 66621 | ENSG00000170445 | HARS1 | chr5 | - | 140683545 | 140683664 | 140683099 | 140683219 | 140690854 | 140690944 | 0 | 0 [illegible] |
| SE | 66622 | ENSG00000170445 | HARS1 | chr5 | - | 140683545 | 140683667 | 140683099 | 140683219 | 140690854 | 140690944 | 0 | 0 [illegible] |
| SE | 66626 | ENSG00000206418 | RAB12 | chr18 | + | 8635532 | 8635622 | 8633188 | 8633327 | 8636252 | 8636357 | 223 | 215 [illegible] |
| SE | 66630 | ENSG00000237037 | NDUFA6-DT | chr22 | + | 42122792 | 42122888 | 42091351 | 42091600 | 42124046 | 42124130 | 0 | 0 [illegible] |
| SE | 66640 | ENSG00000064703 | DDX20 | chr1 | + | 111760986 | 111761125 | 111760705 | 111760848 | 111761225 | 111761284 | 69 | 47 [illegible] |
| SE | 66641 | ENSG00000064703 | DDX20 | chr1 | + | 111762254 | 111762337 | 111760973 | 111761284 | 111762676 | 111762782 | 79 | 77 [illegible] |
| SE | 66647 | ENSG00000084623 | EIF3I | chr1 | + | 32226104 | 32226320 | 32224409 | 32224475 | 32226402 | 32226530 | 277 | 137 [illegible] |
| SE | 66648 | ENSG00000084623 | EIF3I | chr1 | + | 32226148 | 32226320 | 32224409 | 32224475 | 32226402 | 32226530 | 278 | 137 [illegible] |
| SE | 66649 | ENSG00000084623 | EIF3I | chr1 | + | 32226170 | 32226283 | 32224409 | 32224475 | 32226402 | 32226530 | 151 | 78 [illegible] |
| SE | 66652 | ENSG00000084623 | EIF3I | chr1 | + | 32226170 | 32226320 | 32224033 | 32224475 | 32226402 | 32226530 | 445 | 233 [illegible] |
| SE | 66663 | ENSG00000005483 | KMT2E | chr7 | + | 105038384 | 105038525 | 105038125 | 105038199 | 105039152 | 105039269 | 6 | 4 [illegible] |
| SE | 66665 | ENSG00000005483 | KMT2E | chr7 | + | 105039152 | 105039269 | 105038125 | 105038199 | 105040838 | 105040856 | 0 | 2 [illegible] |
| SE | 66666 | ENSG00000005483 | KMT2E | chr7 | + | 105039152 | 105039269 | 105038384 | 105038525 | 105040838 | 105040856 | 0 | 1 [illegible] |
| SE | 66667 | ENSG00000005483 | KMT2E | chr7 | + | 105062163 | 105062278 | 105040838 | 105041023 | 105063350 | 105063377 | 382 | 235 [illegible] |
| SE | 66668 | ENSG00000005483 | KMT2E | chr7 | + | 105063350 | 105063580 | 105040838 | 105041023 | 105066726 | 105066807 | 359 | 284 [illegible] |
| SE | 66669 | ENSG00000005483 | KMT2E | chr7 | + | 105063350 | 105063580 | 105040838 | 105041023 | 105073618 | 105073677 | 11 | 2 [illegible] |
| SE | 66671 | ENSG00000005483 | KMT2E | chr7 | + | 105066726 | 105066807 | 105040838 | 105041023 | 105073618 | 105073677 | 83 | 83 [illegible] |
| SE | 66672 | ENSG00000005483 | KMT2E | chr7 | + | 105066726 | 105066807 | 105063350 | 105063580 | 105068474 | 105068545 | 125 | 101 [illegible] |
| SE | 66677 | ENSG00000005483 | KMT2E | chr7 | + | 105081687 | 105081797 | 105077302 | 105077433 | 105090008 | 105090106 | 262 | 182 [illegible] |
| SE | 66678 | ENSG00000005483 | KMT2E | chr7 | + | 105090008 | 105090273 | 105081687 | 105081797 | 105091215 | 105091314 | 573 | 465 [illegible] |
| SE | 66680 | ENSG00000005483 | KMT2E | chr7 | + | 105101885 | 105102194 | 105091215 | 105091314 | 105105438 | 105105693 | 275 | 238 [illegible] |
| SE | 66682 | ENSG00000005483 | KMT2E | chr7 | + | 105106521 | 105106772 | 105105858 | 105106003 | 105107165 | 105107222 | 777 | 766 [illegible] |
| SE | 66683 | ENSG00000005483 | KMT2E | chr7 | + | 105107165 | 105107222 | 105106521 | 105106772 | 105107361 | 105107925 | 465 | 430 [illegible] |
| SE | 66685 | ENSG00000005483 | KMT2E | chr7 | + | 105110279 | 105110368 | 105108941 | 105109228 | 105110770 | 105110867 | 310 | 227 [illegible] |
| SE | 66686 | ENSG00000005483 | KMT2E | chr7 | + | 105110476 | 105110602 | 105110279 | 105110368 | 105110770 | 105110868 | 419 | 372 [illegible] |
| SE | 66687 | ENSG00000137992 | DBT | chr1 | - | 100206444 | 100206636 | 100206229 | 100206301 | 100210693 | 100210771 | 41 | 45 [illegible] |
| SE | 66688 | ENSG00000137992 | DBT | chr1 | - | 100210693 | 100210771 | 100206444 | 100206636 | 100214816 | 100214983 | 34 | 26 [illegible] |
| SE | 66689 | ENSG00000137992 | DBT | chr1 | - | 100213498 | 100213624 | 100210693 | 100210771 | 100214816 | 100214983 | 0 | 0 [illegible] |
| SE | 66690 | ENSG00000137992 | DBT | chr1 | - | 100214816 | 100214983 | 100210693 | 100210771 | 100218625 | 100218747 | 25 | 13 [illegible] |
| SE | 66691 | ENSG00000137992 | DBT | chr1 | - | 100215982 | 100216199 | 100214816 | 100214983 | 100218625 | 100218747 | 81 | 54 [illegible] |
| SE | 66693 | ENSG00000137992 | DBT | chr1 | - | 100248898 | 100249085 | 100240760 | 100240884 | 100249769 | 100249833 | 3 | 1 [illegible] |
| SE | 66717 | ENSG00000186522 | SEPTIN10 | chr2 | - | 109553086 | 109553219 | 109545801 | 109546237 | 109564365 | 109564534 | 14 | 8 [illegible] |
| SE | 66718 | ENSG00000186522 | SEPTIN10 | chr2 | - | 109567814 | 109567976 | 109565762 | 109565859 | 109574580 | 109574767 | 32 | 18 [illegible] |
| SE | 66719 | ENSG00000186522 | SEPTIN10 | chr2 | - | 109574580 | 109574767 | 109567814 | 109567976 | 109585125 | 109585321 | 19 | 9 [illegible] |
| SE | 66729 | ENSG00000155729 | KCTD18 | chr2 | - | 200504747 | 200504959 | 200498890 | 200499084 | 200506856 | 200507091 | 156 | 101 [illegible] |
| SE | 66730 | ENSG00000155729 | KCTD18 | chr2 | - | 200504747 | 200504959 | 200498890 | 200499084 | 200510010 | 200510068 | 113 | 72 [illegible] |
| SE | 66731 | ENSG00000155729 | KCTD18 | chr2 | - | 200504769 | 200504959 | 200498890 | 200499084 | 200506856 | 200507091 | 54 | 44 [illegible] |
| SE | 66732 | ENSG00000155729 | KCTD18 | chr2 | - | 200504769 | 200504959 | 200498890 | 200499084 | 200510010 | 200510068 | 11 | 15 [illegible] |
| SE | 66733 | ENSG00000155729 | KCTD18 | chr2 | - | 200506856 | 200507091 | 200498890 | 200499084 | 200510010 | 200510068 | 6 | 2 [illegible] |
| SE | 66734 | ENSG00000155729 | KCTD18 | chr2 | - | 200506856 | 200507091 | 200504747 | 200504959 | 200510010 | 200510068 | 47 | 31 [illegible] |
| SE | 66735 | ENSG00000167747 | C19orf48 | chr19 | - | 50799271 | 50799357 | 50798703 | 50798958 | 50802216 | 50802249 | 3 | 0 [illegible] |
| SE | 66736 | ENSG00000167747 | C19orf48 | chr19 | - | 50799293 | 50799357 | 50797703 | 50798958 | 50802216 | 50802257 | 9 | 4 [illegible] |
| SE | 66745 | ENSG00000138468 | SENP7 | chr3 | - | 101458954 | 101459052 | 101417592 | 101417790 | 101493872 | 101493968 | 231 | 249 [illegible] |
| SE | 66748 | ENSG00000129292 | PHF20L1 | chr8 | + | 132794409 | 132794581 | 132777791 | 132777911 | 132794732 | 132794817 | 317 | 338 [illegible] |
| SE | 66749 | ENSG00000129292 | PHF20L1 | chr8 | + | 132794409 | 132794581 | 132777791 | 132777911 | 132803818 | 132804032 | 168 | 169 [illegible] |
| SE | 66750 | ENSG00000129292 | PHF20L1 | chr8 | + | 132794732 | 132794817 | 132777788 | 132777911 | 132803818 | 132804032 | 1 | 0 [illegible] |
| SE | 66751 | ENSG00000129292 | PHF20L1 | chr8 | + | 132794732 | 132794817 | 132794409 | 132794581 | 132798771 | 132798816 | 278 | 272 [illegible] |
| SE | 66754 | ENSG00000129292 | PHF20L1 | chr8 | + | 132798771 | 132798860 | 132794732 | 132794817 | 132799082 | 132799172 | 162 | 163 [illegible] |
| SE | 66755 | ENSG00000129292 | PHF20L1 | chr8 | + | 132798771 | 132798860 | 132794732 | 132794817 | 132799094 | 132799172 | 163 | 169 [illegible] |
| SE | 66761 | ENSG00000129292 | PHF20L1 | chr8 | + | 132807722 | 132807783 | 132804614 | 132804740 | 132811045 | 132811128 | 2 | 0 [illegible] |
| SE | 66765 | ENSG00000129292 | PHF20L1 | chr8 | + | 132844155 | 132844318 | 132841552 | 132842875 | 132845780 | 132845923 | 252 | 234 [illegible] |
| SE | 66775 | ENSG00000131653 | TRAF7 | chr16 | + | 2172190 | 2172374 | 2171571 | 2171605 | 2172464 | 2172589 | 125 | 44 [illegible] |
| SE | 66777 | ENSG00000002330 | BAD | chr11 | - | 64271612 | 64271803 | 64269829 | 64270337 | 64284181 | 64284376 | 36 | 26 [illegible] |
| SE | 66778 | ENSG00000002330 | BAD | chr11 | - | 64276903 | 64277043 | 64271612 | 64271803 | 64284181 | 64284376 | 0 | 0 [illegible] |
| SE | 66786 | ENSG00000169432 | SCN9A | chr2 | - | 166243489 | 166243529 | 166242574 | 166242656 | 166251764 | 166251885 | 0 | 1 [illegible] |
| SE | 66789 | ENSG00000169432 | SCN9A | chr2 | - | 166293230 | 166293372 | 166288436 | 166288643 | 166294598 | 166294662 | 10 | 10 [illegible] |
| SE | 66790 | ENSG00000169432 | SCN9A | chr2 | - | 166294598 | 166294662 | 166288436 | 166288643 | 166303089 | 166303302 | 1 | 0 [illegible] |
| SE | 66792 | ENSG00000169432 | SCN9A | chr2 | - | 166304237 | 166304329 | 166302062 | 166303302 | 166305791 | 166305920 | 8 | 5 [illegible] |
| SE | 66796 | ENSG00000135929 | CYP27A1 | chr2 | + | 218809576 | 218809767 | 218781748 | 218782437 | 218812221 | 218812421 | 36 | 17 [illegible] |
| SE | 66797 | ENSG00000205485 | NA | chr7 | + | 76615493 | 76615663 | 76551463 | 76551522 | 76621115 | 76621388 | 10 | 0 [illegible] |
| SE | 66800 | ENSG00000137075 | RNF38 | chr9 | - | 36344831 | 36344953 | 36342324 | 36342424 | 36351114 | 36351199 | 221 | 96 [illegible] |
| SE | 66801 | ENSG00000137075 | RNF38 | chr9 | - | 36352741 | 36352848 | 36351114 | 36351199 | 36353169 | 36353331 | 97 | 74 [illegible] |
| SE | 66802 | ENSG00000137075 | RNF38 | chr9 | - | 36356302 | 36356473 | 36353169 | 36353331 | 36357774 | 36357942 | 127 | 85 [illegible] |
| SE | 66803 | ENSG00000137075 | RNF38 | chr9 | - | 36357774 | 36357942 | 36353169 | 36353331 | 36369718 | 36369932 | 78 | 72 [illegible] |
| SE | 66804 | ENSG00000137075 | RNF38 | chr9 | - | 36369718 | 36369932 | 36357774 | 36357942 | 36375933 | 36376127 | 142 | 117 [illegible] |
| SE | 66805 | ENSG00000137075 | RNF38 | chr9 | - | 36372495 | 36372593 | 36369718 | 36369932 | 36375933 | 36376127 | 0 | 2 [illegible] |
| SE | 66806 | ENSG00000137075 | RNF38 | chr9 | - | 36375933 | 36376127 | 36369718 | 36369932 | 36390466 | 36390616 | 118 | 82 [illegible] |
| SE | 66807 | ENSG00000137075 | RNF38 | chr9 | - | 36375933 | 36376127 | 36369718 | 36369932 | 36400553 | 36400872 | 75 | 51 [illegible] |
| SE | 66808 | ENSG00000137075 | RNF38 | chr9 | - | 36390466 | 36390616 | 36369718 | 36369932 | 36400553 | 36400923 | 1 | 0 [illegible] |
| SE | 66809 | ENSG00000137075 | RNF38 | chr9 | - | 36390466 | 36390616 | 36375933 | 36376127 | 36400096 | 36400269 | 54 | 34 [illegible] |
| SE | 66811 | ENSG00000137075 | RNF38 | chr9 | - | 36390466 | 36390616 | 36375933 | 36376127 | 36400935 | 36401198 | 55 | 35 [illegible] |
| SE | 66814 | ENSG00000137075 | RNF38 | chr9 | - | 36424612 | 36424683 | 36390466 | 36390616 | 36487307 | 36487548 | 15 | 11 [illegible] |
| SE | 66817 | ENSG00000031003 | FAM13B | chr5 | - | 137948954 | 137949184 | 137946227 | 137946311 | 137952627 | 137952709 | 336 | 344 [illegible] |
| SE | 66818 | ENSG00000031003 | FAM13B | chr5 | - | 137954165 | 137954376 | 137953335 | 137953465 | 137959615 | 137959763 | 417 | 375 [illegible] |
| SE | 66820 | ENSG00000031003 | FAM13B | chr5 | - | 137959615 | 137959763 | 137954165 | 137954376 | 137960165 | 137960214 | 386 | 322 [illegible] |
| SE | 66822 | ENSG00000031003 | FAM13B | chr5 | - | 137960165 | 137960214 | 137959615 | 137959763 | 137962404 | 137962469 | 243 | 213 [illegible] |
| SE | 66823 | ENSG00000031003 | FAM13B | chr5 | - | 137962404 | 137962469 | 137960165 | 137960214 | 137985256 | 137985389 | 396 | 355 [illegible] |
| SE | 66824 | ENSG00000031003 | FAM13B | chr5 | - | 137962404 | 137962469 | 137960165 | 137960214 | 138006989 | 138007147 | 95 | 89 [illegible] |
| SE | 66825 | ENSG00000031003 | FAM13B | chr5 | - | 137985256 | 137985389 | 137960165 | 137960214 | 138006989 | 138007147 | 1 | 0 [illegible] |
| SE | 66826 | ENSG00000031003 | FAM13B | chr5 | - | 137985256 | 137985389 | 137962404 | 137962469 | 137987460 | 137987616 | 822 | 656 [illegible] |
| SE | 66827 | ENSG00000031003 | FAM13B | chr5 | - | 137985256 | 137985389 | 137962404 | 137962469 | 138006989 | 138007147 | 291 | 264 [illegible] |
| SE | 66829 | ENSG00000031003 | FAM13B | chr5 | - | 137988273 | 137988315 | 137985256 | 137985389 | 138006989 | 138007147 | 75 | 59 [illegible] |
| SE | 66830 | ENSG00000031003 | FAM13B | chr5 | - | 138011007 | 138011149 | 138006989 | 138007147 | 138011767 | 138011945 | 501 | 435 [illegible] |
| SE | 66831 | ENSG00000031003 | FAM13B | chr5 | - | 138011767 | 138011945 | 138011007 | 138011149 | 138018301 | 138018514 | 309 | 256 [illegible] |
| SE | 66832 | ENSG00000031003 | FAM13B | chr5 | - | 138018301 | 138018514 | 138011767 | 138011945 | 138021030 | 138021197 | 130 | 100 [illegible] |
| SE | 66847 | ENSG00000198598 | MMP17 | chr12 | + | 131844535 | 131844633 | 131843996 | 131844081 | 131845117 | 131845200 | 1 | 0 [illegible] |
| SE | 66849 | ENSG00000156869 | FRRS1 | chr1 | - | 99711354 | 99711532 | 99710918 | 99710949 | 99712104 | 99712163 | 0 | 1 [illegible] |
| SE | 66851 | ENSG00000156869 | FRRS1 | chr1 | - | 99717409 | 99717525 | 99715585 | 99715672 | 99719533 | 99719647 | 12 | 10 [illegible] |
| SE | 66852 | ENSG00000156869 | FRRS1 | chr1 | - | 99740792 | 99740940 | 99738085 | 99738268 | 99742178 | 99742273 | 30 | 6 [illegible] |
| SE | 66853 | ENSG00000211689 | TRGC1 | chr7 | - | 38262170 | 38262218 | 38257878 | 38260230 | 38265348 | 38265678 | 187 | 82 [illegible] |
| SE | 66854 | ENSG00000082781 | ITGB5 | chr3 | - | 124796387 | 124796817 | 124773689 | 124773912 | 124809021 | 124809156 | 83 | 79 [illegible] |
| SE | 66859 | ENSG00000082781 | ITGB5 | chr3 | - | 124819738 | 124819834 | 124817620 | 124817710 | 124821312 | 124821458 | 63 | 117 [illegible] |
| SE | 66861 | ENSG00000082781 | ITGB5 | chr3 | - | 124857231 | 124857313 | 124848308 | 124848558 | 124859241 | 124859446 | 0 | 0 [illegible] |
| SE | 66890 | ENSG00000140470 | ADAMTS17 | chr15 | - | 99976044 | 99976222 | 99971436 | 99974562 | 99993047 | 99993200 | 0 | 1 [illegible] |
| SE | 66891 | ENSG00000140470 | ADAMTS17 | chr15 | - | 100053896 | 100054054 | 100051571 | 100051731 | 100063678 | 100063759 | 2 | 0 [illegible] |
| SE | 66892 | ENSG00000140470 | ADAMTS17 | chr15 | - | 100063678 | 100063759 | 100053896 | 100054054 | 100096355 | 100096476 | 0 | 0 [illegible] |
| SE | 66893 | ENSG00000140470 | ADAMTS17 | chr15 | - | 100262351 | 100262435 | 100261478 | 100261636 | 100281228 | 100281401 | 13 | 4 [illegible] |
| SE | 66896 | ENSG00000213145 | CRIP1 | chr14 | + | 105487212 | 105487299 | 105486866 | 105486972 | 105488165 | 105488260 | 181 | 77 [illegible] |
| SE | 66897 | ENSG00000196504 | PRPF40A | chr2 | - | 152658947 | 152659009 | 152657888 | 152658020 | 152659092 | 152659189 | 294 | 273 [illegible] |
| SE | 66898 | ENSG00000196504 | PRPF40A | chr2 | - | 152669136 | 152669270 | 152664137 | 152664252 | 152670230 | 152670381 | 559 | 515 [illegible] |
| SE | 66899 | ENSG00000196504 | PRPF40A | chr2 | - | 152676404 | 152676779 | 152673597 | 152673706 | 152677450 | 152677475 | 526 | 516 [illegible] |
| SE | 66902 | ENSG00000196504 | PRPF40A | chr2 | - | 152693157 | 152693476 | 152693037 | 152693068 | 152694541 | 152694622 | 3 | 3 [illegible] |
| SE | 66904 | ENSG00000285417 | NA | chr1 | + | 206041565 | 206041690 | 206035252 | 206035299 | 206051852 | 206052001 | 3 | 2 [illegible] |
| SE | 66905 | ENSG00000285417 | NA | chr1 | + | 206060400 | 206060449 | 206052844 | 206053265 | 206061344 | 206061397 | 0 | 2 [illegible] |
| SE | 66907 | ENSG00000285417 | NA | chr1 | + | 206061344 | 206061397 | 206060400 | 206060449 | 206061661 | 206061820 | 1 | 4 [illegible] |
| SE | 66910 | ENSG00000152520 | PAN3 | chr13 | + | 28174271 | 28174393 | 28138505 | 28139087 | 28176492 | 28176559 | 160 | 138 [illegible] |
| SE | 66911 | ENSG00000152520 | PAN3 | chr13 | + | 28177864 | 28177935 | 28176492 | 28176559 | 28197184 | 28197346 | 135 | 111 [illegible] |
| SE | 66915 | ENSG00000152520 | PAN3 | chr13 | + | 28220230 | 28220378 | 28177864 | 28177935 | 28260446 | 28260551 | 98 | 47 [illegible] |
| SE | 66916 | ENSG00000152520 | PAN3 | chr13 | + | 28220230 | 28220378 | 28197184 | 28197346 | 28256291 | 28256539 | 165 | 119 [illegible] |
| SE | 66917 | ENSG00000152520 | PAN3 | chr13 | + | 28256291 | 28256539 | 28177864 | 28177935 | 28260446 | 28260551 | 292 | 138 [illegible] |
| SE | 66918 | ENSG00000152520 | PAN3 | chr13 | + | 28256291 | 28256539 | 28220230 | 28220378 | 28260446 | 28260551 | 392 | 189 [illegible] |
| SE | 66919 | ENSG00000152520 | PAN3 | chr13 | + | 28260446 | 28260551 | 28256291 | 28256539 | 28261400 | 28261458 | 456 | 212 [illegible] |
| SE | 66920 | ENSG00000152520 | PAN3 | chr13 | + | 28267094 | 28267211 | 28266714 | 28266876 | 28267299 | 28267401 | 275 | 121 [illegible] |
| SE | 66921 | ENSG00000152520 | PAN3 | chr13 | + | 28267299 | 28267401 | 28266714 | 28266876 | 28270700 | 28270866 | 65 | 38 [illegible] |
| SE | 66923 | ENSG00000152520 | PAN3 | chr13 | + | 28280411 | 28280541 | 28277236 | 28277376 | 28281314 | 28281379 | 415 | 319 [illegible] |
| SE | 66925 | ENSG00000152520 | PAN3 | chr13 | + | 28281314 | 28281379 | 28280411 | 28280541 | 28292381 | 28295335 | 181 | 143 [illegible] |
| SE | 66926 | ENSG00000152520 | PAN3 | chr13 | + | 28287983 | 28288122 | 28280411 | 28280541 | 28292381 | 28295335 | 72 | 78 [illegible] |
| SE | 66931 | ENSG00000167658 | EEF2 | chr19 | - | 3977214 | 3977347 | 3976055 | 3976747 | 3977427 | 3977610 | 3685 | 1538 [illegible] |
| SE | 66932 | ENSG00000167658 | EEF2 | chr19 | - | 3977818 | 3978172 | 3977427 | 3977610 | 3979328 | 3979436 | 5110 | 2286 [illegible] |
| SE | 66933 | ENSG00000167658 | EEF2 | chr19 | - | 3979328 | 3979436 | 3977427 | 3977610 | 3980513 | 3980709 | 1 | 2 [illegible] |
| SE | 66935 | ENSG00000167658 | EEF2 | chr19 | - | 3981946 | 3982052 | 3981338 | 3981452 | 3982245 | 3982313 | 5877 | 3806 [illegible] |
| SE | 66942 | ENSG00000172493 | AFF1 | chr4 | + | 87046165 | 87046286 | 86948497 | 86948571 | 87046694 | 87046890 | 247 | 166 [illegible] |
| SE | 66943 | ENSG00000172493 | AFF1 | chr4 | + | 87046165 | 87046286 | 87006752 | 87007424 | 87046694 | 87046890 | 203 | 144 [illegible] |
| SE | 66944 | ENSG00000172493 | AFF1 | chr4 | + | 87046694 | 87047594 | 87006752 | 87007424 | 87084119 | 87084164 | 83 | 69 [illegible] |
| SE | 66946 | ENSG00000172493 | AFF1 | chr4 | + | 87089983 | 87090070 | 87084119 | 87084164 | 87091792 | 87091824 | 161 | 160 [illegible] |
| SE | 66947 | ENSG00000172493 | AFF1 | chr4 | + | 87094914 | 87094969 | 87091792 | 87091829 | 87105627 | 87105682 | 108 | 123 [illegible] |
| SE | 66948 | ENSG00000172493 | AFF1 | chr4 | + | 87105627 | 87105682 | 87094914 | 87094969 | 87108158 | 87108300 | 26 | 32 [illegible] |
| SE | 66949 | ENSG00000172493 | AFF1 | chr4 | + | 87105807 | 87105845 | 87105627 | 87105682 | 87108158 | 87108300 | 97 | 116 [illegible] |
| SE | 66955 | ENSG00000172493 | AFF1 | chr4 | + | 87134470 | 87134694 | 87132270 | 87132408 | 87135579 | 87141035 | 217 | 206 [illegible] |
| SE | 66957 | ENSG00000172493 | AFF1 | chr4 | + | 87134473 | 87134694 | 87132270 | 87132408 | 87135579 | 87141038 | 89 | 98 [illegible] |
| SE | 66958 | ENSG00000135960 | EDAR | chr2 | - | 108930963 | 108931032 | 108930119 | 108930242 | 108988959 | 108989220 | 8 | 5 [illegible] |
| SE | 66961 | ENSG00000286239 | NA | chr2 | - | 223880123 | 223880232 | 223878528 | 223878730 | 223881941 | 223882072 | 279 | 187 [illegible] |
| SE | 66962 | ENSG00000286239 | NA | chr2 | - | 223880123 | 223880232 | 223878528 | 223878730 | 223884647 | 223884749 | 38 | 31 [illegible] |
| SE | 66963 | ENSG00000286239 | NA | chr2 | - | 223881941 | 223882072 | 223878528 | 223878730 | 223884647 | 223884749 | 182 | 132 [illegible] |
| SE | 66965 | ENSG00000286239 | NA | chr2 | - | 223884647 | 223884749 | 223881941 | 223882072 | 223894233 | 223894339 | 299 | 220 [illegible] |
| SE | 66966 | ENSG00000286239 | NA | chr2 | - | 223884647 | 223884749 | 223881941 | 223882072 | 223898957 | 223899051 | 116 | 70 [illegible] |
| SE | 66969 | ENSG00000286239 | NA | chr2 | - | 223912252 | 223912326 | 223905946 | 223906001 | 223917942 | 223918010 | 104 | 50 [illegible] |
| SE | 66970 | ENSG00000132570 | PCBD2 | chr5 | + | 134910334 | 134910466 | 134905119 | 134905223 | 134959039 | 134959095 | 27 | 23 [illegible] |
| SE | 66971 | ENSG00000132570 | PCBD2 | chr5 | + | 134969938 | 134970092 | 134910334 | 134910466 | 134996272 | 134996512 | 3 | 0 [illegible] |
| SE | 66972 | ENSG00000132570 | PCBD2 | chr5 | + | 134975137 | 134975324 | 134969938 | 134970092 | 134996272 | 134996512 | 17 | 0 [illegible] |
| SE | 66974 | ENSG00000162645 | GBP2 | chr1 | - | 89114015 | 89114296 | 89112471 | 89112684 | 89116991 | 89117234 | 1050 | 841 [illegible] |
| SE | 66975 | ENSG00000162645 | GBP2 | chr1 | - | 89116991 | 89117234 | 89114015 | 89114296 | 89117576 | 89117773 | 864 | 540 [illegible] |
| SE | 66976 | ENSG00000167393 | PPP2R3B | chrX | - | 340764 | 340940 | 338612 | 338896 | 341306 | 341396 | 33 | 28 [illegible] |
| SE | 66977 | ENSG00000167393 | PPP2R3B | chrX | - | 362254 | 362332 | 361404 | 361590 | 386367 | 386955 | 0 | 2 [illegible] |
| SE | 66979 | ENSG00000137161 | CNPY3 | chr6 | + | 42935573 | 42935670 | 42929479 | 42929721 | 42938089 | 42938207 | 6 | 0 [illegible] |
| SE | 66980 | ENSG00000137161 | CNPY3 | chr6 | + | 42937716 | 42937839 | 42935573 | 42935670 | 42938089 | 42938170 | 988 | 651 [illegible] |
| SE | 66981 | ENSG00000137161 | CNPY3 | chr6 | + | 42938089 | 42938207 | 42937716 | 42937839 | 42938567 | 42939294 | 1232 | 961 [illegible] |
| SE | 66983 | ENSG00000135930 | EIF4E2 | chr2 | + | 232556415 | 232556530 | 232550682 | 232550744 | 232564246 | 232564351 | 49 | 30 [illegible] |
| SE | 66986 | ENSG00000135930 | EIF4E2 | chr2 | + | 232564246 | 232564351 | 232556415 | 232556530 | 232566828 | 232566981 | 378 | 285 [illegible] |
| SE | 66987 | ENSG00000135930 | EIF4E2 | chr2 | + | 232564246 | 232564351 | 232557883 | 232558018 | 232566828 | 232566981 | 667 | 515 [illegible] |
| SE | 66989 | ENSG00000135930 | EIF4E2 | chr2 | + | 232574262 | 232574341 | 232574073 | 232574136 | 232580903 | 232581332 | 8 | 3 [illegible] |
| SE | 67000 | ENSG00000121542 | SEC22A | chr3 | + | 123209198 | 123209399 | 123201926 | 123201986 | 123223558 | 123223680 | 66 | 82 [illegible] |
| SE | 67001 | ENSG00000121542 | SEC22A | chr3 | + | 123223643 | 123223722 | 123209198 | 123209399 | 123225102 | 123225297 | 48 | 32 [illegible] |
| SE | 67004 | ENSG00000121542 | SEC22A | chr3 | + | 123245898 | 123246014 | 123225102 | 123225297 | 123259523 | 123259589 | 47 | 17 [illegible] |
| SE | 67005 | ENSG00000121542 | SEC22A | chr3 | + | 123245898 | 123246014 | 123225102 | 123225297 | 123271521 | 123271664 | 23 | 8 [illegible] |
| SE | 67006 | ENSG00000121542 | SEC22A | chr3 | + | 123259523 | 123259589 | 123209198 | 123209399 | 123271521 | 123271664 | 4 | 10 [illegible] |
| SE | 67007 | ENSG00000121542 | SEC22A | chr3 | + | 123259523 | 123259589 | 123225102 | 123225297 | 123271521 | 123271664 | 5 | 11 [illegible] |
| SE | 67008 | ENSG00000121542 | SEC22A | chr3 | + | 123259523 | 123259589 | 123245898 | 123246014 | 123271521 | 123271664 | 28 | 17 [illegible] |
| SE | 67016 | ENSG00000112658 | SRF | chr6 | + | 43175705 | 43175967 | 43173846 | 43174113 | 43178293 | 43178485 | 44 | 25 [illegible] |
| SE | 67018 | ENSG00000112658 | SRF | chr6 | + | 43176547 | 43176667 | 43175705 | 43175967 | 43178293 | 43178485 | 88 | 38 [illegible] |
| SE | 67019 | ENSG00000112658 | SRF | chr6 | + | 43178293 | 43178485 | 43173846 | 43174113 | 43179094 | 43181506 | 0 | 2 [illegible] |
| SE | 67020 | ENSG00000112658 | SRF | chr6 | + | 43178805 | 43178882 | 43178293 | 43178485 | 43179094 | 43181506 | 33 | 17 [illegible] |

| | | | | | | | | | | |
|---|---|---|---|---|---|---|---|---|---|---|
| SE | 67023 | ENSG00000225434 | LINC01504 | chr9 | + | 72324988 | 72325054 | 72314985 | 72315090 | 72335391 |
| SE | 67024 | ENSG00000225434 | LINC01504 | chr9 | + | 72324988 | 72325054 | 72314985 | 72315090 | 72341591 |
| SE | 67025 | ENSG00000225434 | LINC01504 | chr9 | + | 72328663 | 72328721 | 72314985 | 72315090 | 72335391 |
| SE | 67026 | ENSG00000225434 | LINC01504 | chr9 | + | 72328663 | 72328721 | 72314985 | 72315090 | 72336765 |
| SE | 67027 | ENSG00000225434 | LINC01504 | chr9 | + | 72328663 | 72328721 | 72314985 | 72315090 | 72341591 |
| SE | 67028 | ENSG00000225434 | LINC01504 | chr9 | + | 72328663 | 72328721 | 72324988 | 72325054 | 72335391 |
| SE | 67029 | ENSG00000225434 | LINC01504 | chr9 | + | 72328663 | 72328721 | 72324988 | 72325054 | 72341591 |
| SE | 67030 | ENSG00000225434 | LINC01504 | chr9 | + | 72335391 | 72335497 | 72314985 | 72315090 | 72336765 |
| SE | 67031 | ENSG00000225434 | LINC01504 | chr9 | + | 72335391 | 72335497 | 72314985 | 72315090 | 72341591 |
| SE | 67032 | ENSG00000225434 | LINC01504 | chr9 | + | 72335391 | 72335497 | 72324988 | 72325054 | 72341591 |
| SE | 67033 | ENSG00000225434 | LINC01504 | chr9 | + | 72335391 | 72335497 | 72328663 | 72328721 | 72336765 |
| SE | 67034 | ENSG00000225434 | LINC01504 | chr9 | + | 72335391 | 72335497 | 72328663 | 72328721 | 72341591 |
| SE | 67036 | ENSG00000225434 | LINC01504 | chr9 | + | 72336765 | 72336864 | 72328663 | 72328721 | 72341591 |
| SE | 67037 | ENSG00000225434 | LINC01504 | chr9 | + | 72336765 | 72336864 | 72335391 | 72335497 | 72341591 |
| SE | 67039 | ENSG00000169087 | HSPBAP1 | chr3 | - | 122752590 | 122752674 | 122741002 | 122741113 | 122755259 |
| SE | 67040 | ENSG00000169087 | HSPBAP1 | chr3 | - | 122755259 | 122755431 | 122741002 | 122741113 | 122759223 |
| SE | 67041 | ENSG00000169087 | HSPBAP1 | chr3 | - | 122758771 | 122758853 | 122755323 | 122755431 | 122759223 |
| SE | 67042 | ENSG00000169087 | HSPBAP1 | chr3 | - | 122768700 | 122768882 | 122759219 | 122759360 | 122777720 |
| SE | 67043 | ENSG00000169087 | HSPBAP1 | chr3 | - | 122768700 | 122768882 | 122759219 | 122759360 | 122793616 |
| SE | 67044 | ENSG00000169087 | HSPBAP1 | chr3 | - | 122777720 | 122777906 | 122759219 | 122759360 | 122793616 |
| SE | 67045 | ENSG00000169087 | HSPBAP1 | chr3 | - | 122777720 | 122777906 | 122768700 | 122768882 | 122793616 |
| SE | 67046 | ENSG00000169087 | HSPBAP1 | chr3 | - | 122786408 | 122786490 | 122777720 | 122777906 | 122793616 |
| SE | 67047 | ENSG00000242258 | LINC00996 | chr7 | + | 150435029 | 150435157 | 150433729 | 150433829 | 150442548 |
| SE | 67048 | ENSG00000242258 | LINC00996 | chr7 | + | 150446699 | 150446889 | 150442548 | 150443160 | 150447075 |
| SE | 67049 | ENSG00000170906 | NDUFA3 | chr19 | + | 54103113 | 54103188 | 54102727 | 54102888 | 54105933 |
| SE | 67051 | ENSG00000170906 | NDUFA3 | chr19 | + | 54106352 | 54106446 | 54105933 | 54106011 | 54106763 |
| SE | 67058 | ENSG00000186567 | CEACAM19 | chr19 | + | 44671745 | 44671986 | 44666957 | 44667321 | 44672595 |
| SE | 67059 | ENSG00000186567 | CEACAM19 | chr19 | + | 44673711 | 44673845 | 44672477 | 44672964 | 44676270 |
| SE | 67060 | ENSG00000186567 | CEACAM19 | chr19 | + | 44678852 | 44678936 | 44676270 | 44676421 | 44680287 |
| SE | 67062 | ENSG00000127720 | METTL25 | chr12 | + | 82438717 | 82438791 | 82434694 | 82434724 | 82456726 |
| SE | 67063 | ENSG00000127720 | METTL25 | chr12 | + | 82456726 | 82456820 | 82438514 | 82438791 | 82476643 |
| SE | 67065 | ENSG00000215022 | NA | chr6 | - | 13281428 | 13281553 | 13279294 | 13281156 | 13294795 |
| SE | 67066 | ENSG00000188177 | ZC3H6 | chr2 | + | 112324963 | 112325197 | 112324151 | 112324663 | 112331004 |
| SE | 67068 | ENSG00000225205 | NA | chr2 | - | 172512333 | 172512518 | 172503438 | 172504142 | 172556088 |
| SE | 67069 | ENSG00000115233 | PSMD14 | chr2 | + | 161367783 | 161367903 | 161367477 | 161367549 | 161370106 |
| SE | 67070 | ENSG00000168899 | VAMP5 | chr2 | + | 85591724 | 85591862 | 85584430 | 85584493 | 85592947 |
| SE | 67074 | ENSG00000204681 | GABBR1 | chr6 | - | 29609228 | 29609379 | 29608600 | 29608733 | 29610923 |
| SE | 67076 | ENSG00000204681 | GABBR1 | chr6 | - | 29629086 | 29629107 | 29627485 | 29627646 | 29631395 |
| SE | 67077 | ENSG00000204681 | GABBR1 | chr6 | - | 29629086 | 29629247 | 29627485 | 29627646 | 29631395 |
| SE | 67079 | ENSG00000204681 | GABBR1 | chr6 | - | 29630457 | 29630643 | 29629086 | 29629247 | 29631395 |
| SE | 67081 | ENSG00000204681 | GABBR1 | chr6 | - | 29632300 | 29632385 | 29631395 | 29631599 | 29632849 |
| SE | 67084 | ENSG00000162910 | MRPL55 | chr1 | - | 228108234 | 228108318 | 228107666 | 228107869 | 228108954 |
| SE | 67085 | ENSG00000162910 | MRPL55 | chr1 | - | 228108234 | 228108321 | 228107667 | 228107869 | 228108954 |
| SE | 67086 | ENSG00000162910 | MRPL55 | chr1 | - | 228108436 | 228108474 | 228108234 | 228108318 | 228108954 |
| SE | 67087 | ENSG00000162910 | MRPL55 | chr1 | - | 228108436 | 228108474 | 228108234 | 228108318 | 228109148 |
| SE | 67089 | ENSG00000162910 | MRPL55 | chr1 | - | 228108436 | 228108474 | 228108234 | 228108321 | 228108954 |
| SE | 67090 | ENSG00000162910 | MRPL55 | chr1 | - | 228108436 | 228108474 | 228108234 | 228108321 | 228109148 |
| SE | 67094 | ENSG00000162910 | MRPL55 | chr1 | - | 228108436 | 228109021 | 228108246 | 228108321 | 228109148 |
| SE | 67096 | ENSG00000162910 | MRPL55 | chr1 | - | 228108954 | 228109021 | 228108234 | 228108318 | 228109148 |
| SE | 67098 | ENSG00000162910 | MRPL55 | chr1 | - | 228108954 | 228109021 | 228108436 | 228108474 | 228109148 |
| SE | 67100 | ENSG00000198369 | SPRED2 | chr2 | - | 65316733 | 65316883 | 65310850 | 65314169 | 65331986 |
| SE | 67103 | ENSG00000120594 | PLXDC2 | chr10 | + | 20001774 | 20001986 | 19816238 | 19817191 | 20046868 |
| SE | 67104 | ENSG00000120594 | PLXDC2 | chr10 | + | 20046868 | 20047015 | 20001774 | 20001986 | 20068169 |
| SE | 67105 | ENSG00000120594 | PLXDC2 | chr10 | + | 20068169 | 20068239 | 20046868 | 20047015 | 20143294 |
| SE | 67106 | ENSG00000120594 | PLXDC2 | chr10 | + | 20143294 | 20143417 | 20068169 | 20068239 | 20147783 |
| SE | 67107 | ENSG00000120594 | PLXDC2 | chr10 | + | 20211668 | 20211729 | 20177327 | 20177409 | 20217425 |
| SE | 67108 | ENSG00000120594 | PLXDC2 | chr10 | + | 20219063 | 20219102 | 20217425 | 20217576 | 20245344 |
| SE | 67110 | ENSG00000146414 | SHPRH | chr6 | - | 145888019 | 145888100 | 145864244 | 145864491 | 145910447 |
| SE | 67111 | ENSG00000146414 | SHPRH | chr6 | - | 145893214 | 145893393 | 145888019 | 145888100 | 145894149 |
| SE | 67113 | ENSG00000146414 | SHPRH | chr6 | - | 145894884 | 145894977 | 145894149 | 145894236 | 145910447 |
| SE | 67114 | ENSG00000146414 | SHPRH | chr6 | - | 145922285 | 145922348 | 145921166 | 145921392 | 145922662 |
| SE | 67122 | ENSG00000273143 | DUSP5-DT | chr10 | - | 110435123 | 110435207 | 110428839 | 110429010 | 110442210 |
| SE | 67123 | ENSG00000123136 | DDX39A | chr19 | - | 14409535 | 14409645 | 14409302 | 14409447 | 14409741 |
| SE | 67126 | ENSG00000123136 | DDX39A | chr19 | - | 14412550 | 14413224 | 14411217 | 14411598 | 14419269 |
| SE | 67130 | ENSG00000170854 | RIOX2 | chr3 | - | 97949843 | 97950012 | 97947360 | 97947449 | 97950785 |
| SE | 67131 | ENSG00000170854 | RIOX2 | chr3 | - | 97949843 | 97950015 | 97947360 | 97947449 | 97950785 |
| SE | 67132 | ENSG00000170854 | RIOX2 | chr3 | - | 97950785 | 97950888 | 97947360 | 97947449 | 97954391 |
| SE | 67133 | ENSG00000170854 | RIOX2 | chr3 | - | 97952151 | 97952252 | 97950785 | 97950888 | 97954391 |
| SE | 67134 | ENSG00000170854 | RIOX2 | chr3 | - | 97959050 | 97959179 | 97954006 | 97954495 | 97961588 |
| SE | 67135 | ENSG00000115866 | DARS1 | chr2 | - | 135912485 | 135912566 | 135911381 | 135911493 | 135914468 |
| SE | 67137 | ENSG00000115866 | DARS1 | chr2 | - | 135922783 | 135922918 | 135920452 | 135920600 | 135924386 |
| SE | 67138 | ENSG00000115866 | DARS1 | chr2 | - | 135924386 | 135924498 | 135920452 | 135920600 | 135932782 |
| SE | 67139 | ENSG00000115866 | DARS1 | chr2 | - | 135932782 | 135932842 | 135924386 | 135924498 | 135933909 |
| SE | 67140 | ENSG00000115866 | DARS1 | chr2 | - | 135979273 | 135979366 | 135961395 | 135961498 | 135985402 |
| SE | 67141 | ENSG00000115866 | DARS1 | chr2 | - | 135983396 | 135983454 | 135979273 | 135979366 | 135985402 |
| SE | 67142 | ENSG00000118257 | NRP2 | chr2 | + | 205763673 | 205763936 | 205752834 | 205752975 | 205765473 |
| SE | 67147 | ENSG00000089060 | SLC8B1 | chr12 | - | 113334442 | 113334835 | 113332722 | 113332960 | 113334992 |
| SE | 67148 | ENSG00000089060 | SLC8B1 | chr12 | - | 113334442 | 113334838 | 113332722 | 113332960 | 113334992 |
| SE | 67149 | ENSG00000152117 | SMPD4BP | chr2 | + | 131497209 | 131497293 | 131492812 | 131493140 | 131498866 |
| SE | 67150 | ENSG00000152117 | SMPD4BP | chr2 | + | 131497209 | 131497293 | 131492812 | 131493140 | 131500203 |
| SE | 67151 | ENSG00000152117 | SMPD4BP | chr2 | + | 131498866 | 131498953 | 131492812 | 131493140 | 131500203 |
| SE | 67152 | ENSG00000152117 | SMPD4BP | chr2 | + | 131498866 | 131498953 | 131497209 | 131497293 | 131500203 |
| SE | 67153 | ENSG00000152117 | SMPD4BP | chr2 | + | 131501357 | 131501508 | 131501180 | 131501233 | 131506311 |
| SE | 67154 | ENSG00000152117 | SMPD4BP | chr2 | + | 131506311 | 131506442 | 131501357 | 131501508 | 131508536 |
| SE | 67155 | ENSG00000152117 | SMPD4BP | chr2 | + | 131515601 | 131515747 | 131508536 | 131508608 | 131516313 |
| SE | 67157 | ENSG00000152117 | SMPD4BP | chr2 | + | 131519813 | 131519942 | 131519534 | 131519666 | 131520096 |
| SE | 67159 | ENSG00000153250 | RBMS1 | chr2 | - | 160275629 | 160275714 | 160272150 | 160274764 | 160277302 |
| SE | 67165 | ENSG00000153250 | RBMS1 | chr2 | - | 160303329 | 160303487 | 160300650 | 160300730 | 160313155 |
| SE | 67166 | ENSG00000153250 | RBMS1 | chr2 | - | 160313155 | 160313247 | 160303329 | 160303487 | 160318168 |
| SE | 67167 | ENSG00000153250 | RBMS1 | chr2 | - | 160318168 | 160318227 | 160313155 | 160313247 | 160367215 |
| SE | 67168 | ENSG00000153250 | RBMS1 | chr2 | - | 160367215 | 160367391 | 160318168 | 160318227 | 160493288 |
| SE | 67170 | ENSG00000153250 | RBMS1 | chr2 | - | 160432367 | 160432477 | 160367215 | 160367391 | 160493288 |
| SE | 67181 | ENSG00000135002 | RFK | chr9 | - | 76388553 | 76388656 | 76385525 | 76387529 | 76392417 |
| SE | 67182 | ENSG00000135002 | RFK | chr9 | - | 76392417 | 76392569 | 76388553 | 76388656 | 76394089 |
| SE | 67183 | ENSG00000197226 | TBC1D9B | chr5 | - | 179865837 | 179865888 | 179865253 | 179865360 | 179867777 |
| SE | 67185 | ENSG00000197226 | TBC1D9B | chr5 | - | 179869768 | 179869834 | 179867303 | 179867849 | 179870254 |
| SE | 67186 | ENSG00000197226 | TBC1D9B | chr5 | - | 179870254 | 179870495 | 179869768 | 179869834 | 179871461 |
| SE | 67187 | ENSG00000197226 | TBC1D9B | chr5 | - | 179875919 | 179876037 | 179874901 | 179875187 | 179878308 |
| SE | 67188 | ENSG00000197226 | TBC1D9B | chr5 | - | 179878308 | 179878523 | 179874901 | 179875187 | 179888102 |
| SE | 67189 | ENSG00000197226 | TBC1D9B | chr5 | - | 179879046 | 179879197 | 179878308 | 179878523 | 179888102 |
| SE | 67201 | ENSG00000124151 | NCOA3 | chr20 | + | 47583182 | 47583261 | 47501886 | 47502019 | 47622228 |
| SE | 67203 | ENSG00000124151 | NCOA3 | chr20 | + | 47627001 | 47627176 | 47625380 | 47625481 | 47627560 |
| SE | 67204 | ENSG00000124151 | NCOA3 | chr20 | + | 47633495 | 47633636 | 47627921 | 47628023 | 47634017 |
| SE | 67206 | ENSG00000124151 | NCOA3 | chr20 | + | 47634017 | 47634195 | 47633495 | 47633636 | 47635321 |
| SE | 67207 | ENSG00000124151 | NCOA3 | chr20 | + | 47634047 | 47634195 | 47633495 | 47633636 | 47635321 |
| SE | 67209 | ENSG00000124151 | NCOA3 | chr20 | + | 47639007 | 47639202 | 47637647 | 47637783 | 47639576 |
| SE | 67211 | ENSG00000124151 | NCOA3 | chr20 | + | 47639576 | 47639822 | 47639007 | 47639202 | 47639924 |
| SE | 67212 | ENSG00000124151 | NCOA3 | chr20 | + | 47639621 | 47639822 | 47639007 | 47639202 | 47639924 |
| SE | 67213 | ENSG00000124151 | NCOA3 | chr20 | + | 47639924 | 47640051 | 47639007 | 47639202 | 47642212 |
| SE | 67217 | ENSG00000144118 | RALB | chr2 | + | 120280834 | 120280907 | 120278617 | 120278778 | 120293140 |
| SE | 67219 | ENSG00000144118 | RALB | chr2 | + | 120285873 | 120286082 | 120280834 | 120280907 | 120289579 |
| SE | 67220 | ENSG00000144118 | RALB | chr2 | + | 120289579 | 120289757 | 120278614 | 120278778 | 120293140 |
| SE | 67221 | ENSG00000144118 | RALB | chr2 | + | 120289579 | 120289757 | 120280834 | 120280907 | 120293140 |
| SE | 67232 | ENSG00000003509 | NDUFAF7 | chr2 | + | 37242634 | 37242693 | 37241577 | 37241637 | 37243862 |
| SE | 67233 | ENSG00000003509 | NDUFAF7 | chr2 | + | 37242634 | 37242693 | 37241577 | 37241791 | 37243862 |
| SE | 67238 | ENSG00000123338 | NCKAP1L | chr12 | + | 54508388 | 54508531 | 54507852 | 54507909 | 54509668 |
| SE | 67240 | ENSG00000123338 | NCKAP1L | chr12 | + | 54516238 | 54516295 | 54511948 | 54512105 | 54516895 |
| SE | 67241 | ENSG00000123338 | NCKAP1L | chr12 | + | 54517532 | 54517642 | 54516895 | 54516992 | 54517805 |
| SE | 67243 | ENSG00000123338 | NCKAP1L | chr12 | + | 54518650 | 54518732 | 54517805 | 54517938 | 54518913 |
| SE | 67244 | ENSG00000123338 | NCKAP1L | chr12 | + | 54518913 | 54518972 | 54517805 | 54517938 | 54520693 |
| SE | 67245 | ENSG00000123338 | NCKAP1L | chr12 | + | 54518913 | 54518972 | 54518595 | 54518732 | 54519186 |
| SE | 67246 | ENSG00000123338 | NCKAP1L | chr12 | + | 54519186 | 54519332 | 54518913 | 54518972 | 54520693 |
| SE | 67247 | ENSG00000123338 | NCKAP1L | chr12 | + | 54523393 | 54523539 | 54521118 | 54521238 | 54526527 |
| SE | 67248 | ENSG00000123338 | NCKAP1L | chr12 | + | 54523824 | 54523956 | 54523393 | 54523539 | 54526527 |
| SE | 67249 | ENSG00000123338 | NCKAP1L | chr12 | + | 54526527 | 54526746 | 54523824 | 54523956 | 54528246 |
| SE | 67252 | ENSG00000123338 | NCKAP1L | chr12 | + | 54531742 | 54531825 | 54531490 | 54531584 | 54532169 |
| SE | 67253 | ENSG00000101695 | RNF125 | chr18 | + | 32045641 | 32045732 | 32041905 | 32042273 | 32065901 |
| SE | 67257 | ENSG00000162636 | FAM102B | chr1 | + | 108612177 | 108612294 | 108606222 | 108606252 | 108616381 |
| SE | 67258 | ENSG00000162636 | FAM102B | chr1 | + | 108628676 | 108628804 | 108624664 | 108628584 | 108629587 |
| SE | 67266 | ENSG00000152056 | AP1S3 | chr2 | - | 223776048 | 223776158 | 223765267 | 223765350 | 223777690 |
| SE | 67269 | ENSG00000082269 | FAM135A | chr6 | + | 70415290 | 70415376 | 70413420 | 70413702 | 70426438 |
| SE | 67270 | ENSG00000082269 | FAM135A | chr6 | + | 70418374 | 70418492 | 70415290 | 70415376 | 70426438 |
| SE | 67272 | ENSG00000082269 | FAM135A | chr6 | + | 70426438 | 70426532 | 70415290 | 70415376 | 70428303 |
| SE | 67273 | ENSG00000082269 | FAM135A | chr6 | + | 70426569 | 70426709 | 70415290 | 70415376 | 70428303 |
| SE | 67274 | ENSG00000082269 | FAM135A | chr6 | + | 70426569 | 70426764 | 70415290 | 70415376 | 70428303 |
| SE | 67277 | ENSG00000082269 | FAM135A | chr6 | + | 70491033 | 70491083 | 70482000 | 70482154 | 70502635 |
| SE | 67278 | ENSG00000082269 | FAM135A | chr6 | + | 70523966 | 70524121 | 70522512 | 70522586 | 70524342 |
| SE | 67279 | ENSG00000082269 | FAM135A | chr6 | + | 70523966 | 70524121 | 70522512 | 70522586 | 70524930 |
| SE | 67280 | ENSG00000082269 | FAM135A | chr6 | + | 70524930 | 70525191 | 70523966 | 70524121 | 70525863 |
| SE | 67281 | ENSG00000082269 | FAM135A | chr6 | + | 70528291 | 70528452 | 70524342 | 70526698 | 70533159 |
| SE | 67282 | ENSG00000082269 | FAM135A | chr6 | + | 70533159 | 70533251 | 70528291 | 70528452 | 70533756 |
| SE | 67285 | ENSG00000116199 | FAM20B | chr1 | + | 179043714 | 179044224 | 179025893 | 179026098 | 179050278 |
| SE | 67287 | ENSG00000174483 | BBS1 | chr11 | + | 66511204 | 66511239 | 66510633 | 66511089 | 66514405 |
| SE | 67289 | ENSG00000256349 | NA | chr11 | + | 66529818 | 66529952 | 66526648 | 66526807 | 66530893 |
| SE | 67293 | ENSG00000164117 | FBXO8 | chr4 | - | 174259698 | 174259825 | 174241099 | 174241218 | 174262763 |
| SE | 67299 | ENSG00000204764 | RANBP17 | chr5 | + | 170916464 | 170916584 | 170914166 | 170914240 | 170918712 |
| SE | 67302 | ENSG00000204764 | RANBP17 | chr5 | + | 171028871 | 171028989 | 170968241 | 170968377 | 171170129 |
| SE | 67308 | ENSG00000205702 | CYP2D7 | chr22 | - | 42141533 | 42141675 | 42141151 | 42141396 | 42141867 |
| SE | 67310 | ENSG00000157570 | TSPAN18 | chr11 | + | 44860327 | 44860469 | 44764425 | 44764512 | 44906406 |
| SE | 67312 | ENSG00000157570 | TSPAN18 | chr11 | + | 44906406 | 44906479 | 44860327 | 44860469 | 44917971 |
| SE | 67323 | ENSG00000163728 | TTC14 | chr3 | + | 180603123 | 180603323 | 180602890 | 180603015 | 180604851 |
| SE | 67324 | ENSG00000163728 | TTC14 | chr3 | + | 180603123 | 180603323 | 180602890 | 180603015 | 180605765 |
| SE | 67325 | ENSG00000163728 | TTC14 | chr3 | + | 180604224 | 180604309 | 180602890 | 180603015 | 180604851 |
| SE | 67326 | ENSG00000163728 | TTC14 | chr3 | + | 180604224 | 180604309 | 180602890 | 180603015 | 180605765 |
| SE | 67327 | ENSG00000163728 | TTC14 | chr3 | + | 180604224 | 180604309 | 180603123 | 180603323 | 180604851 |
| SE | 67331 | ENSG00000163728 | TTC14 | chr3 | + | 180604851 | 180605007 | 180602890 | 180603015 | 180605765 |
| SE | 67335 | ENSG00000163728 | TTC14 | chr3 | + | 180605765 | 180605837 | 180602890 | 180603015 | 180606252 |
| SE | 67336 | ENSG00000163728 | TTC14 | chr3 | + | 180608700 | 180608810 | 180607647 | 180607765 | 180609629 |
| SE | 67337 | ENSG00000163728 | TTC14 | chr3 | + | 180608700 | 180608810 | 180608162 | 180608271 | 180609629 |
| SE | 67338 | ENSG00000106636 | YKT6 | chr7 | + | 44204567 | 44204650 | 44200958 | 44201239 | 44206384 |
| SE | 67339 | ENSG00000106636 | YKT6 | chr7 | + | 44204574 | 44204650 | 44200982 | 44201239 | 44206384 |
| SE | 67368 | ENSG00000159479 | MED8 | chr1 | - | 43386588 | 43386670 | 43385672 | 43386226 | 43386857 |
| SE | 67370 | ENSG00000159479 | MED8 | chr1 | - | 43387502 | 43387647 | 43386857 | 43386998 | 43389758 |
| SE | 67373 | ENSG00000187688 | TRPV2 | chr17 | + | 16423468 | 16423767 | 16422598 | 16422889 | 16427448 |
| SE | 67374 | ENSG00000187688 | TRPV2 | chr17 | + | 16426098 | 16426269 | 16423468 | 16423767 | 16426721 |
| SE | 67377 | ENSG00000187688 | TRPV2 | chr17 | + | 16431783 | 16431850 | 16428816 | 16428982 | 16434889 |
| SE | 67378 | ENSG00000187688 | TRPV2 | chr17 | + | 16433573 | 16433698 | 16428816 | 16428982 | 16434889 |
| SE | 67379 | ENSG00000187688 | TRPV2 | chr17 | + | 16433573 | 16433698 | 16428816 | 16428982 | 16436788 |
| SE | 67380 | ENSG00000187688 | TRPV2 | chr17 | + | 16433573 | 16433698 | 16431965 | 16432300 | 16434889 |
| SE | 67381 | ENSG00000187688 | TRPV2 | chr17 | + | 16434858 | 16434969 | 16433573 | 16433698 | 16436788 |
| SE | 67382 | ENSG00000187688 | TRPV2 | chr17 | + | 16434889 | 16434969 | 16428816 | 16428982 | 16436788 |
| SE | 67385 | ENSG00000250131 | NA | chr4 | + | 177593471 | 177593546 | 177444978 | 177445443 | 177596460 |
| SE | 67398 | ENSG00000287151 | C2orf27A | chr2 | + | 131688165 | 131688303 | 131686329 | 131686441 | 131722416 |
| SE | 67400 | ENSG00000287151 | C2orf27A | chr2 | + | 131722416 | 131722648 | 131688165 | 131688303 | 131733832 |
| SE | 67401 | ENSG00000287151 | C2orf27A | chr2 | + | 131733832 | 131733952 | 131688165 | 131688303 | 131750632 |
| SE | 67404 | ENSG00000214960 | CRPPA | chr7 | - | 16216065 | 16216197 | 16087524 | 16091799 | 16258389 |
| SE | 67405 | ENSG00000214960 | CRPPA | chr7 | - | 16301420 | 16301466 | 16278128 | 16278226 | 16308522 |
| SE | 67406 | ENSG00000214960 | CRPPA | chr7 | - | 16301420 | 16301466 | 16278128 | 16278226 | 16376091 |
| SE | 67407 | ENSG00000214960 | CRPPA | chr7 | - | 16308522 | 16308627 | 16278128 | 16278226 | 16376091 |
| SE | 67408 | ENSG00000214960 | CRPPA | chr7 | - | 16308522 | 16308627 | 16301420 | 16301466 | 16376091 |
| SE | 67415 | ENSG00000224043 | CCNT2-AS1 | chr2 | - | 134877519 | 134877643 | 134868943 | 134869041 | 134918562 |
| SE | 67417 | ENSG00000226696 | LENG8-AS1 | chr19 | - | 54447194 | 54447260 | 54445920 | 54446031 | 54448164 |
| SE | 67418 | ENSG00000204160 | ZDHHC18 | chr1 | + | 26850300 | 26850438 | 26848607 | 26848757 | 26850557 |
| SE | 67419 | ENSG00000204160 | ZDHHC18 | chr1 | + | 26856184 | 26856262 | 26853725 | 26853825 | 26856847 |
| SE | 67429 | ENSG00000184371 | CSF1 | chr1 | + | 109914258 | 109914381 | 109910812 | 109911062 | 109915633 |
| SE | 67432 | ENSG00000232125 | DYTN | chr2 | - | 206698755 | 206698971 | 206694815 | 206694877 | 206699726 |
| SE | 67442 | ENSG00000152359 | POC5 | chr5 | - | 75677773 | 75677950 | 75674123 | 75674578 | 75689011 |
| SE | 67443 | ENSG00000152359 | POC5 | chr5 | - | 75685206 | 75685484 | 75674430 | 75674578 | 75689011 |
| SE | 67444 | ENSG00000152359 | POC5 | chr5 | - | 75685206 | 75685484 | 75677773 | 75677950 | 75689011 |
| SE | 67447 | ENSG00000152359 | POC5 | chr5 | - | 75707736 | 75707875 | 75705238 | 75705787 | 75712606 |
| SE | 67450 | ENSG00000152359 | POC5 | chr5 | - | 75712606 | 75712665 | 75705238 | 75705787 | 75712853 |
| SE | 67451 | ENSG00000152359 | POC5 | chr5 | - | 75712606 | 75712665 | 75707483 | 75707875 | 75712853 |
| SE | 67453 | ENSG00000152359 | POC5 | chr5 | - | 75712853 | 75712951 | 75707736 | 75707875 | 75717305 |
| SE | 67454 | ENSG00000005381 | MPO | chr17 | - | 58272747 | 58272918 | 58271654 | 58271892 | 58273413 |
| SE | 67456 | ENSG00000162923 | WDR26 | chr1 | - | 224389832 | 224389860 | 224385665 | 224386502 | 224393827 |
| SE | 67459 | ENSG00000162923 | WDR26 | chr1 | - | 224393827 | 224394013 | 224385665 | 224386502 | 224398096 |
| SE | 67461 | ENSG00000162923 | WDR26 | chr1 | - | 224398096 | 224398226 | 224389183 | 224389860 | 224398888 |
| SE | 67463 | ENSG00000162923 | WDR26 | chr1 | - | 224398514 | 224398593 | 224385145 | 224389860 | 224398888 |
| SE | 67465 | ENSG00000162923 | WDR26 | chr1 | - | 224398888 | 224399034 | 224398514 | 224398593 | 224400949 |
| SE | 67467 | ENSG00000162923 | WDR26 | chr1 | - | 224404429 | 224404570 | 224400949 | 224401069 | 224411426 |
| SE | 67468 | ENSG00000162923 | WDR26 | chr1 | - | 224411426 | 224411565 | 224404429 | 224404570 | 224418259 |
| SE | 67469 | ENSG00000162923 | WDR26 | chr1 | - | 224411426 | 224411565 | 224404429 | 224404570 | 224419517 |
| SE | 67471 | ENSG00000162923 | WDR26 | chr1 | - | 224418259 | 224418398 | 224404429 | 224404570 | 224419517 |
| SE | 67473 | ENSG00000162923 | WDR26 | chr1 | - | 224418259 | 224418416 | 224404429 | 224404570 | 224419517 |
| SE | 67475 | ENSG00000162923 | WDR26 | chr1 | - | 224424517 | 224424654 | 224419517 | 224419615 | 224431476 |
| SE | 67476 | ENSG00000162923 | WDR26 | chr1 | - | 224424517 | 224424654 | 224419517 | 224419615 | 224431524 |
| SE | 67482 | ENSG00000255529 | POLR2M | chr15 | + | 57707466 | 57707566 | 57706713 | 57706955 | 57708713 |
| SE | 67484 | ENSG00000120159 | CAAP1 | chr9 | - | 26866633 | 26866780 | 26861065 | 26861139 | 26869925 |
| SE | 67487 | ENSG00000120159 | CAAP1 | chr9 | - | 26887312 | 26887513 | 26886103 | 26886188 | 26892258 |

| | | | | | | | | | | | | | | | | | | | | | | | | | | | | | | | | | | | | | | | | | | | | | |
|---|---|---|---|---|---|---|---|---|---|---|---|---|---|---|---|---|---|---|---|---|---|---|---|---|---|---|---|---|---|---|---|---|---|---|---|---|---|---|---|---|---|---|---|---|---|---|
| SE | 67488 | ENSG00000120159 | CAAP1 | chr9 | - | 26887312 | 26887513 | 26886103 | 26886188 | 26892412 | 26892604 | 70 | 81 | 85 | 87 | 83 | 0 | 0 | 0 | 0 | 3 | 111 | 45 | 73 | 60 | 15 | 0 | 0 | 0 | 0 | 0 | 298 | 149 | 1 | 1 | 1 | 1 | 1 | 1 | 0.933 | 1 | 1 | 1 | 1 | 1 | -0.013 |
| SE | 67492 | ENSG00000151553 | FAM160B1 | chr10 | + | 114836123 | 114836246 | 114833232 | 114833402 | 114842932 | 114843226 | 81 | 78 | 56 | 79 | 117 | 0 | 0 | 0 | 0 | 0 | 91 | 89 | 139 | 95 | 109 | 0 | 0 | 1 | 0 | 0 | 272 | 149 | 1 | 1 | 1 | 1 | 1 | 1 | 1 | 1 | 1 | 0.987 | 1 | 1 | 0.003 |
| SE | 67493 | ENSG00000151553 | FAM160B1 | chr10 | + | 114846558 | 114846728 | 114846174 | 114846367 | 114847089 | 114847233 | 186 | 113 | 133 | 161 | 352 | 0 | 0 | 1 | 0 | 0 | 57 | 100 | 175 | 97 | 63 | 0 | 0 | 0 | 0 | 0 | 298 | 149 | 1 | 1 | 1 | 1 | 0.985 | 1 | 1 | 1 | 1 | 1 | 1 | 1 | -0.003 |
| SE | 67494 | ENSG00000151553 | FAM160B1 | chr10 | + | 114848646 | 114848737 | 114847089 | 114847233 | 114855196 | 114855340 | 135 | 114 | 95 | 156 | 204 | 0 | 0 | 0 | 3 | 0 | 37 | 114 | 146 | 75 | 29 | 0 | 0 | 1 | 0 | 0 | 240 | 149 | 1 | 1 | 1 | 1 | 1 | 0.97 | 1 | 1 | 1 | 0.989 | 1 | 1 | -0.004 |
| SE | 67496 | ENSG00000068654 | POLR1A | chr2 | - | 86041888 | 86042103 | 86040391 | 86040559 | 86042973 | 86043195 | 50 | 56 | 54 | 55 | 80 | 0 | 0 | 0 | 0 | 0 | 93 | 58 | 44 | 47 | 32 | 0 | 0 | 2 | 0 | 0 | 298 | 149 | 1 | 1 | 1 | 1 | 1 | 1 | 1 | 1 | 1 | 0.917 | 1 | 1 | 0.017 |
| SE | 67498 | ENSG00000068654 | POLR1A | chr2 | - | 86045277 | 86045360 | 86042973 | 86043195 | 86045616 | 86045769 | 12 | 3 | 9 | 4 | 8 | 2 | 0 | 2 | 0 | 0 | 27 | 12 | 10 | 9 | 0 | 0 | 0 | 4 | 0 | 0 | 232 | 149 | 1 | 1 | 0.794 | 1 | 0.743 | 1 | 1 | 1 | 1 | 0.616 | 1 | NA | 0.003 |
| SE | 67502 | ENSG00000068654 | POLR1A | chr2 | - | 86081600 | 86081706 | 86080815 | 86080978 | 86083081 | 86083168 | 76 | 36 | 57 | 63 | 57 | 0 | 1 | 0 | 0 | 0 | 74 | 50 | 15 | 24 | 62 | 0 | 0 | 0 | 1 | 0 | 255 | 149 | 1 | 1 | 1 | 0.955 | 1 | 1 | 1 | 1 | 1 | 1 | 0.933 | 1 | 0.004 |
| SE | 67504 | ENSG00000068654 | POLR1A | chr2 | - | 86098610 | 86098760 | 86089633 | 86089929 | 86105699 | 86105873 | 31 | 24 | 54 | 52 | 40 | 0 | 0 | 2 | 0 | 0 | 54 | 41 | 11 | 31 | 18 | 0 | 0 | 0 | 0 | 0 | 298 | 149 | 1 | 1 | 1 | 1 | 0.931 | 1 | 1 | 1 | 1 | 1 | 1 | 1 | -0.014 |
| SE | 67506 | ENSG00000165714 | BORCS5 | chr12 | + | 12361205 | 12361349 | 12357077 | 12357509 | 12435627 | 12435785 | 69 | 15 | 45 | 47 | 23 | 2 | 0 | 1 | 0 | 0 | 49 | 47 | 26 | 46 | 55 | 0 | 0 | 0 | 0 | 0 | 293 | 149 | 1 | 1 | 0.946 | 1 | 0.958 | 1 | 1 | 1 | 1 | 1 | 1 | 1 | -0.019 |
| SE | 67507 | ENSG00000153574 | RPIA | chr2 | + | 88698483 | 88698544 | 88691672 | 88691983 | 88729277 | 88729337 | 1 | 1 | 1 | 2 | 0 | 0 | 0 | 1 | 0 | 0 | 0 | 6 | 6 | 4 | 1 | 0 | 0 | 0 | 0 | 0 | 210 | 149 | 1 | 1 | 1 | 1 | 0.415 | 1 | NA | NA | 1 | 1 | 1 | 1 | -0.146 |
| SE | 67508 | ENSG00000153574 | RPIA | chr2 | + | 88700008 | 88700064 | 88698483 | 88698544 | 88729277 | 88729337 | 51 | 33 | 109 | 65 | 98 | 0 | 0 | 0 | 1 | 0 | 105 | 71 | 59 | 53 | 58 | 0 | 0 | 0 | 0 | 0 | 205 | 149 | 1 | 1 | 1 | 1 | 1 | 0.979 | 1 | 1 | 1 | 1 | 1 | 1 | -0.004 |
| SE | 67509 | ENSG00000153574 | RPIA | chr2 | + | 88729277 | 88729337 | 88700008 | 88700064 | 88734551 | 88734616 | 59 | 44 | 111 | 83 | 95 | 0 | 0 | 0 | 6 | 0 | 93 | 63 | 57 | 60 | 65 | 0 | 5 | 1 | 0 | 0 | 209 | 149 | 1 | 1 | 1 | 1 | 1 | 0.908 | 1 | 1 | 0.9 | 0.976 | 1 | 1 | 0.006 |
| SE | 67510 | ENSG00000153574 | RPIA | chr2 | + | 88736534 | 88736676 | 88735668 | 88735737 | 88749980 | 88750929 | 65 | 63 | 104 | 104 | 90 | 0 | 0 | 1 | 0 | 0 | 96 | 83 | 68 | 115 | 58 | 0 | 0 | 0 | 0 | 0 | 291 | 149 | 1 | 1 | 1 | 1 | 0.982 | 1 | 1 | 1 | 1 | 1 | 1 | 1 | -0.004 |
| SE | 67511 | ENSG00000153574 | RPIA | chr2 | + | 88737976 | 88738076 | 88736534 | 88736676 | 88749980 | 88750929 | 114 | 61 | 111 | 89 | 153 | 1 | 0 | 2 | 2 | 0 | 134 | 62 | 89 | 170 | 117 | 0 | 0 | 4 | 0 | 0 | 249 | 149 | 1 | 1 | 0.986 | 1 | 0.971 | 0.964 | 1 | 1 | 1 | 0.93 | 1 | 1 | -0.002 |
| SE | 67512 | ENSG00000148426 | PROSER2 | chr10 | + | 11851996 | 11852215 | 11823338 | 11823470 | 11866530 | 11866647 | 1 | 3 | 10 | 5 | 2 | 0 | 0 | 0 | 0 | 0 | 6 | 7 | 12 | 8 | 10 | 0 | 0 | 0 | 0 | 1 | 298 | 149 | 1 | 1 | 1 | 1 | 1 | 1 | 1 | 1 | 1 | 1 | 1 | 0.833 | 0.033 |
| SE | 67514 | ENSG00000115159 | GPD2 | chr2 | + | 156476097 | 156476207 | 156436378 | 156436513 | 156496043 | 156496215 | 21 | 18 | 9 | 29 | 18 | 1 | 0 | 0 | 0 | 0 | 7 | 31 | 11 | 16 | 0 | 0 | 0 | 0 | 0 | 0 | 259 | 149 | 1 | 1 | 0.924 | 1 | 1 | 1 | 1 | 1 | 1 | 1 | 1 | NA | -0.015 |
| SE | 67516 | ENSG00000115159 | GPD2 | chr2 | + | 156496043 | 156496215 | 156476097 | 156476207 | 156510795 | 156510834 | 74 | 53 | 40 | 66 | 47 | 0 | 0 | 0 | 1 | 0 | 33 | 84 | 58 | 58 | 29 | 0 | 0 | 0 | 0 | 0 | 298 | 149 | 1 | 1 | 1 | 1 | 1 | 0.971 | 1 | 1 | 1 | 1 | 1 | 1 | -0.006 |
| SE | 67517 | ENSG00000115159 | GPD2 | chr2 | + | 156570086 | 156570218 | 156569362 | 156569538 | 156571133 | 156571292 | 139 | 90 | 68 | 162 | 131 | 0 | 0 | 0 | 0 | 2 | 51 | 144 | 98 | 129 | 18 | 0 | 0 | 0 | 0 | 0 | 281 | 149 | 1 | 1 | 1 | 1 | 1 | 1 | 0.972 | 1 | 1 | 1 | 1 | 1 | -0.006 |
| SE | 67518 | ENSG00000115159 | GPD2 | chr2 | + | 156571133 | 156571292 | 156570086 | 156570218 | 156578888 | 156579001 | 172 | 97 | 76 | 176 | 168 | 0 | 0 | 0 | 0 | 0 | 54 | 146 | 132 | 114 | 0 | 0 | 0 | 0 | 2 | 0 | 298 | 149 | 1 | 1 | 1 | 1 | 1 | 1 | 1 | 1 | 1 | 1 | 0.966 | NA | 0.008 |
| SE | 67519 | ENSG00000115159 | GPD2 | chr2 | + | 156579085 | 156579164 | 156578888 | 156579001 | 156579689 | 156579788 | 135 | 94 | 52 | 152 | 330 | 0 | 0 | 0 | 1 | 2 | 45 | 58 | 107 | 100 | 41 | 0 | 0 | 0 | 0 | 0 | 228 | 149 | 1 | 1 | 1 | 1 | 1 | 0.99 | 0.991 | 1 | 1 | 1 | 1 | 1 | -0.004 |
| [illegible] | [illegible] | [illegible] | [illegible] | [illegible] | [illegible] | [illegible] | [illegible] | [illegible] | [illegible] | [illegible] | [illegible] | [illegible] | [illegible] | [illegible] | [illegible] | [illegible] | [illegible] | [illegible] | [illegible] | [illegible] | [illegible] | [illegible] | [illegible] | [illegible] | [illegible] | [illegible] | [illegible] | [illegible] | [illegible] | [illegible] | [illegible] | [illegible] | [illegible] | [illegible] | [illegible] | [illegible] | [illegible] | [illegible] | [illegible] | [illegible] | [illegible] | [illegible] | [illegible] | [illegible] | [illegible] | [illegible] |

| | | | | | | | | | | | | | | |
|---|---|---|---|---|---|---|---|---|---|---|---|---|---|---|
| SE | 67861 | ENSG00000018280 | SLC11A1 | chr2 | + | 218385146 | 218385266 | 218384166 | 218384365 | 218385411 | 218385485 | 151 | 75 | [illegible] |
| SE | 67862 | ENSG00000018280 | SLC11A1 | chr2 | + | 218385146 | 218385266 | 218384166 | 218384365 | 218386634 | 218386741 | 195 | 98 | [illegible] |
| SE | 67863 | ENSG00000018280 | SLC11A1 | chr2 | + | 218385146 | 218385485 | 218383686 | 218384365 | 218386634 | 218387230 | 135 | 69 | [illegible] |
| SE | 67867 | ENSG00000133627 | ACTR3B | chr7 | + | 152783186 | 152783242 | 152759748 | 152759926 | 152800530 | 152800655 | 10 | 7 | [illegible] |
| SE | 67868 | ENSG00000133627 | ACTR3B | chr7 | + | 152800530 | 152800655 | 152783186 | 152783242 | 152801620 | 152801731 | 37 | 35 | [illegible] |
| SE | 67870 | ENSG00000133627 | ACTR3B | chr7 | + | 152816480 | 152816588 | 152814549 | 152814645 | 152820298 | 152820442 | 94 | 77 | [illegible] |
| SE | 67873 | ENSG00000133627 | ACTR3B | chr7 | + | 152853493 | 152853577 | 152825029 | 152825122 | 152854457 | 152854949 | 3 | 4 | [illegible] |
| SE | 67875 | ENSG00000164164 | OTUD4 | chr4 | - | 145141378 | 145141639 | 145139950 | 145139991 | 145143364 | 145143445 | 97 | 71 | [illegible] |
| SE | 67876 | ENSG00000164164 | OTUD4 | chr4 | - | 145142195 | 145142334 | 145141378 | 145141639 | 145143364 | 145143445 | 149 | 144 | [illegible] |
| SE | 67878 | ENSG00000164164 | OTUD4 | chr4 | - | 145146266 | 145146429 | 145144310 | 145144434 | 145150512 | 145150699 | 334 | 282 | [illegible] |
| SE | 67879 | ENSG00000164164 | OTUD4 | chr4 | - | 145152540 | 145152635 | 145150801 | 145150905 | 145155410 | 145155475 | 262 | 237 | [illegible] |
| SE | 67881 | ENSG00000164164 | OTUD4 | chr4 | - | 145155935 | 145155996 | 145155568 | 145155686 | 145159502 | 145159635 | 124 | 107 | [illegible] |
| SE | 67882 | ENSG00000164164 | OTUD4 | chr4 | - | 145159502 | 145159635 | 145155935 | 145155996 | 145162639 | 145162721 | 261 | 210 | [illegible] |
| SE | 67883 | ENSG00000164164 | OTUD4 | chr4 | - | 145162639 | 145162721 | 145159136 | 145159635 | 145164153 | 145164226 | 221 | 173 | [illegible] |
| SE | 67886 | ENSG00000115145 | STAM2 | chr2 | - | 152133401 | 152133484 | 152133172 | 152133260 | 152135508 | 152135603 | 305 | 283 | [illegible] |
| SE | 67887 | ENSG00000115145 | STAM2 | chr2 | - | 152150144 | 152150229 | 152148224 | 152148300 | 152175602 | 152175729 | 54 | 56 | [illegible] |
| SE | 67889 | ENSG00000250075 | NA | chr4 | - | 67419893 | 67420049 | 67418541 | 67418991 | 67420897 | 67421054 | 0 | 0 | [illegible] |
| SE | 67891 | ENSG00000039650 | PNKP | chr19 | - | 49861607 | 49861695 | 49861252 | 49861510 | 49861771 | 49861776 | 20 | 11 | [illegible] |
| SE | 67892 | ENSG00000039650 | PNKP | chr19 | - | 49861771 | 49861881 | 49861607 | 49861695 | 49862043 | 49862105 | 62 | 16 | [illegible] |
| SE | 67893 | ENSG00000039650 | PNKP | chr19 | - | 49862043 | 49862105 | 49861607 | 49861695 | 49862184 | 49862209 | 25 | 10 | [illegible] |
| SE | 67900 | ENSG00000039650 | PNKP | chr19 | - | 49865126 | 49865426 | 49864178 | 49864236 | 49866398 | 49866445 | 22 | 7 | [illegible] |
| SE | 67901 | ENSG00000039650 | PNKP | chr19 | - | 49867468 | 49867532 | 49867195 | 49867217 | 49867678 | 49867701 | 2 | 1 | [illegible] |
| SE | 67903 | ENSG00000079156 | OSBPL6 | chr2 | + | 178331551 | 178331605 | 178328255 | 178328378 | 178332640 | 178332701 | 1 | 1 | [illegible] |
| SE | 67908 | ENSG00000168301 | KCTD6 | chr3 | + | 58494152 | 58494408 | 58492118 | 58492169 | 58498712 | 58498782 | 1 | 0 | [illegible] |
| SE | 67909 | ENSG00000168301 | KCTD6 | chr3 | + | 58500310 | 58500403 | 58498712 | 58498782 | 58500945 | 58501167 | 0 | 0 | [illegible] |
| SE | 67916 | ENSG00000112130 | RNF8 | chr6 | + | 37361337 | 37361558 | 37360445 | 37360574 | 37368377 | 37369218 | 0 | 0 | [illegible] |
| SE | 67920 | ENSG00000112130 | RNF8 | chr6 | + | 37376925 | 37377033 | 37371511 | 37371574 | 37381149 | 37381354 | 31 | 25 | [illegible] |
| SE | 67922 | ENSG00000112130 | RNF8 | chr6 | + | 37381149 | 37381354 | 37371511 | 37371574 | 37390741 | 37391168 | 24 | 21 | [illegible] |
| SE | 67924 | ENSG00000091490 | SEL1L3 | chr4 | - | 25757533 | 25757606 | 25747432 | 25748564 | 25758940 | 25759068 | 36 | 19 | [illegible] |
| SE | 67931 | ENSG00000091490 | SEL1L3 | chr4 | - | 25784227 | 25784290 | 25782241 | 25782418 | 25788223 | 25788364 | 167 | 133 | [illegible] |
| SE | 67932 | ENSG00000091490 | SEL1L3 | chr4 | - | 25788223 | 25788364 | 25784227 | 25784290 | 25790454 | 25790574 | 239 | 154 | [illegible] |
| SE | 67933 | ENSG00000091490 | SEL1L3 | chr4 | - | 25802282 | 25802462 | 25790454 | 25790574 | 25804540 | 25804752 | 239 | 127 | [illegible] |
| SE | 67934 | ENSG00000091490 | SEL1L3 | chr4 | - | 25821995 | 25822128 | 25819807 | 25819940 | 25830097 | 25830156 | 445 | 350 | [illegible] |
| SE | 67938 | ENSG00000091490 | SEL1L3 | chr4 | - | 25832994 | 25833110 | 25830097 | 25830156 | 25833447 | 25833569 | 434 | 339 | [illegible] |
| SE | 67939 | ENSG00000091490 | SEL1L3 | chr4 | - | 25832994 | 25833110 | 25830097 | 25830156 | 25835196 | 25835323 | 273 | 208 | [illegible] |
| SE | 67941 | ENSG00000091490 | SEL1L3 | chr4 | - | 25833447 | 25833569 | 25830097 | 25830156 | 25835196 | 25835323 | 109 | 85 | [illegible] |
| SE | 67943 | ENSG00000091490 | SEL1L3 | chr4 | - | 25833447 | 25833569 | 25832994 | 25833110 | 25835196 | 25835323 | 266 | 213 | [illegible] |
| SE | 67945 | ENSG00000091490 | SEL1L3 | chr4 | - | 25835196 | 25835323 | 25830097 | 25830156 | 25847293 | 25847864 | 96 | 74 | [illegible] |
| SE | 67955 | ENSG00000123374 | CDK2 | chr12 | + | 55968777 | 55968948 | 55968048 | 55968169 | 55971043 | 55971099 | 17 | 20 | [illegible] |
| SE | 67959 | ENSG00000188282 | RUFY4 | chr2 | + | 218058594 | 218058681 | 218034959 | 218035394 | 218068594 | 218069558 | 3 | 0 | [illegible] |
| SE | 67961 | ENSG00000188282 | RUFY4 | chr2 | + | 218073815 | 218073885 | 218073242 | 218073386 | 218075092 | 218075740 | 8 | 2 | [illegible] |
| SE | 67962 | ENSG00000188282 | RUFY4 | chr2 | + | 218073815 | 218073945 | 218073242 | 218073386 | 218075092 | 218075740 | 3 | 1 | [illegible] |
| SE | 67964 | ENSG00000141522 | ARHGDIA | chr17 | - | 81869172 | 81869236 | 81867720 | 81869075 | 81869541 | 81869625 | 348 | 164 | [illegible] |
| SE | 67966 | ENSG00000141522 | ARHGDIA | chr17 | - | 81869541 | 81869625 | 81867720 | 81869075 | 81869740 | 81869957 | 371 | 132 | [illegible] |
| SE | 67967 | ENSG00000166762 | CATSPER2 | chr15 | - | 43631429 | 43631538 | 43630570 | 43630732 | 43632198 | 43632363 | 0 | 0 | [illegible] |
| SE | 67968 | ENSG00000166762 | CATSPER2 | chr15 | - | 43632716 | 43632928 | 43632198 | 43632363 | 43635359 | 43635416 | 0 | 3 | [illegible] |
| SE | 67969 | ENSG00000166762 | CATSPER2 | chr15 | - | 43632716 | 43632934 | 43632198 | 43632363 | 43635359 | 43635416 | 0 | 3 | [illegible] |
| SE | 67971 | ENSG00000166762 | CATSPER2 | chr15 | - | 43635726 | 43635826 | 43635359 | 43635416 | 43636040 | 43636219 | 0 | 0 | [illegible] |
| SE | 67974 | ENSG00000166762 | CATSPER2 | chr15 | - | 43647916 | 43648063 | 43647293 | 43647467 | 43664417 | 43664543 | 0 | 0 | [illegible] |
| SE | 67976 | ENSG00000166762 | CATSPER2 | chr15 | - | 43649726 | 43649879 | 43647916 | 43648063 | 43664417 | 43664543 | 0 | 2 | [illegible] |
| SE | 67977 | ENSG00000166762 | CATSPER2 | chr15 | - | 43649726 | 43649879 | 43647916 | 43648066 | 43668027 | 43668118 | 0 | 3 | [illegible] |
| SE | 67978 | ENSG00000166762 | CATSPER2 | chr15 | - | 43664417 | 43664543 | 43649726 | 43649879 | 43668027 | 43668118 | 0 | 2 | [illegible] |
| SE | 67983 | ENSG00000197959 | DNM3 | chr1 | + | 172131174 | 172131288 | 172093695 | 172093725 | 172308727 | 172308839 | 8 | 30 | [illegible] |
| SE | 67985 | ENSG00000197959 | DNM3 | chr1 | + | 172308727 | 172308839 | 172131174 | 172131288 | 172379017 | 172379182 | 36 | 71 | [illegible] |
| SE | 67988 | ENSG00000157077 | ZFYVE9 | chr1 | + | 52266654 | 52266831 | 52263772 | 52263872 | 52268462 | 52268632 | 2 | 8 | [illegible] |
| SE | 67989 | ENSG00000157077 | ZFYVE9 | chr1 | + | 52332767 | 52332918 | 52303820 | 52303925 | 52334687 | 52334768 | 22 | 19 | [illegible] |
| SE | 67990 | ENSG00000157077 | ZFYVE9 | chr1 | + | 52334687 | 52334768 | 52332767 | 52332918 | 52337771 | 52337934 | 11 | 9 | [illegible] |
| SE | 67991 | ENSG00000147124 | ZNF41 | chrX | - | 47467409 | 47467760 | 47456271 | 47456398 | 47482458 | 47482585 | 4 | 2 | [illegible] |
| SE | 67995 | ENSG00000235703 | LINC00894 | chrX | + | 149941056 | 149941239 | 149940658 | 149940886 | 149944017 | 149944100 | 3 | 2 | [illegible] |
| SE | 67996 | ENSG00000235703 | LINC00894 | chrX | + | 149941789 | 149941847 | 149940658 | 149940886 | 149944017 | 149944100 | 0 | 1 | [illegible] |
| SE | 67998 | ENSG00000235703 | LINC00894 | chrX | + | 149942183 | 149942329 | 149941789 | 149941847 | 149943650 | 149943762 | 0 | 0 | [illegible] |
| SE | 67999 | ENSG00000235703 | LINC00894 | chrX | + | 149942183 | 149942329 | 149941789 | 149941847 | 149944017 | 149944100 | 0 | 0 | [illegible] |
| SE | 68000 | ENSG00000235703 | LINC00894 | chrX | + | 149943650 | 149943762 | 149941789 | 149941847 | 149944017 | 149944100 | 4 | 0 | [illegible] |
| SE | 68002 | ENSG00000235703 | LINC00894 | chrX | + | 149944290 | 149944570 | 149944017 | 149944100 | 149945347 | 149948175 | 2 | 7 | [illegible] |
| SE | 68004 | ENSG00000235703 | LINC00894 | chrX | + | 149948077 | 149948175 | 149945857 | 149947347 | 149948305 | 149948394 | 6 | 4 | [illegible] |
| SE | 68005 | ENSG00000235703 | LINC00894 | chrX | + | 149962582 | 149962716 | 149960764 | 149960948 | 150015950 | 150016140 | 11 | 8 | [illegible] |
| SE | 68009 | ENSG00000214517 | PPME1 | chr11 | + | 74222311 | 74222369 | 74204352 | 74204445 | 74225204 | 74225256 | 45 | 13 | [illegible] |
| SE | 68011 | ENSG00000137414 | FAM8A1 | chr6 | + | 17604905 | 17605029 | 17602589 | 17602710 | 17605873 | 17606013 | 188 | 92 | [illegible] |
| SE | 68014 | ENSG00000187257 | RSBN1L | chr7 | + | 77768660 | 77768803 | 77765490 | 77765632 | 77773146 | 77773314 | 454 | 299 | [illegible] |
| SE | 68015 | ENSG00000172890 | NADSYN1 | chr11 | + | 71458427 | 71458544 | 71455109 | 71455170 | 71460721 | 71460947 | 74 | 50 | [illegible] |
| SE | 68016 | ENSG00000172890 | NADSYN1 | chr11 | + | 71458427 | 71458544 | 71455109 | 71455170 | 71463431 | 71463485 | 168 | 103 | [illegible] |
| SE | 68018 | ENSG00000172890 | NADSYN1 | chr11 | + | 71460040 | 71460213 | 71458427 | 71458544 | 71460721 | 71460947 | 2 | 2 | [illegible] |
| SE | 68019 | ENSG00000172890 | NADSYN1 | chr11 | + | 71460040 | 71460213 | 71458427 | 71458544 | 71463431 | 71463485 | 3 | 0 | [illegible] |
| SE | 68027 | ENSG00000172890 | NADSYN1 | chr11 | + | 71497482 | 71497611 | 71491833 | 71491903 | 71498351 | 71498528 | 144 | 64 | [illegible] |
| SE | 68028 | ENSG00000172890 | NADSYN1 | chr11 | + | 71501301 | 71501426 | 71498351 | 71498528 | 71505567 | 71505872 | 87 | 36 | [illegible] |
| SE | 68029 | ENSG00000172890 | NADSYN1 | chr11 | + | 71501301 | 71501426 | 71498351 | 71498528 | 71521089 | 71521204 | 86 | 36 | [illegible] |
| SE | 68030 | ENSG00000172890 | NADSYN1 | chr11 | + | 71503089 | 71503303 | 71501301 | 71501426 | 71505567 | 71505872 | 1 | 0 | [illegible] |
| SE | 68034 | ENSG00000181544 | FANCB | chrX | - | 14844855 | 14845286 | 14844502 | 14844740 | 14850504 | 14850674 | 32 | 17 | [illegible] |
| SE | 68036 | ENSG00000181544 | FANCB | chrX | - | 14857861 | 14857954 | 14853038 | 14853167 | 14859181 | 14859247 | 54 | 22 | [illegible] |
| SE | 68037 | ENSG00000181544 | FANCB | chrX | - | 14862251 | 14862332 | 14859181 | 14859334 | 14864559 | 14865580 | 2 | 0 | [illegible] |
| SE | 68038 | ENSG00000181544 | FANCB | chrX | - | 14868922 | 14869043 | 14864559 | 14865580 | 14872985 | 14873052 | 17 | 28 | [illegible] |
| SE | 68039 | ENSG00000064989 | CALCRL | chr2 | - | 187387328 | 187387465 | 187385544 | 187385631 | 187387665 | 187387756 | 19 | 4 | [illegible] |
| SE | 68040 | ENSG00000064989 | CALCRL | chr2 | - | 187387328 | 187387493 | 187385544 | 187385631 | 187387665 | 187387756 | 23 | 6 | [illegible] |
| SE | 68041 | ENSG00000064989 | CALCRL | chr2 | - | 187431296 | 187431364 | 187387665 | 187387756 | 187448038 | 187448146 | 5 | 5 | [illegible] |
| SE | 68042 | ENSG00000064989 | CALCRL | chr2 | - | 187436597 | 187436656 | 187431296 | 187431364 | 187448038 | 187448252 | 0 | 5 | [illegible] |
| SE | 68045 | ENSG00000213341 | CHUK | chr10 | - | 100199970 | 100200020 | 100194424 | 100194521 | 100200670 | 100200780 | 98 | 63 | [illegible] |
| SE | 68046 | ENSG00000213341 | CHUK | chr10 | - | 100202087 | 100202149 | 100200670 | 100200780 | 100204505 | 100204657 | 82 | 91 | [illegible] |
| SE | 68047 | ENSG00000213341 | CHUK | chr10 | - | 100207229 | 100207332 | 100205075 | 100205199 | 100209594 | 100209789 | 245 | 159 | [illegible] |
| SE | 68048 | ENSG00000213341 | CHUK | chr10 | - | 100209594 | 100209789 | 100207229 | 100207332 | 100217994 | 100218130 | 349 | 258 | [illegible] |
| SE | 68050 | ENSG00000213341 | CHUK | chr10 | - | 100222111 | 100222181 | 100220587 | 100220676 | 100222865 | 100222980 | 98 | 78 | [illegible] |
| SE | 68051 | ENSG00000162999 | DUSP19 | chr2 | + | 183083507 | 183083554 | 183078558 | 183079159 | 183087039 | 183087192 | 13 | 11 | [illegible] |
| SE | 68053 | ENSG00000162999 | DUSP19 | chr2 | + | 183087157 | 183087192 | 183083507 | 183083554 | 183095430 | 183095903 | 2 | 6 | [illegible] |
| SE | 68054 | ENSG00000230438 | SERPINB9P1 | chr6 | - | 2857910 | 2857991 | 2854657 | 2855690 | 2876292 | 2876425 | 3 | 7 | [illegible] |
| SE | 68055 | ENSG00000230438 | SERPINB9P1 | chr6 | - | 2857910 | 2857995 | 2854201 | 2855690 | 2872816 | 2873086 | 3 | 7 | [illegible] |
| SE | 68056 | ENSG00000230438 | SERPINB9P1 | chr6 | - | 2857910 | 2857995 | 2854201 | 2855690 | 2876292 | 2876425 | 3 | 7 | [illegible] |
| SE | 68057 | ENSG00000230438 | SERPINB9P1 | chr6 | - | 2872816 | 2873086 | 2854574 | 2855690 | 2876292 | 2876425 | 0 | 0 | [illegible] |
| SE | 68059 | ENSG00000230438 | SERPINB9P1 | chr6 | - | 2876292 | 2876425 | 2854244 | 2855690 | 2881284 | 2881286 | 0 | 0 | [illegible] |
| SE | 68062 | ENSG00000143933 | CALM2 | chr2 | - | 47170733 | 47170764 | 47162518 | 47162662 | 47176440 | 47176500 | 1663 | 1323 | [illegible] |
| SE | 68063 | ENSG00000143933 | CALM2 | chr2 | - | 47172413 | 47172459 | 47170733 | 47170764 | 47176235 | 47176453 | 4 | 8 | [illegible] |
| SE | 68064 | ENSG00000143933 | CALM2 | chr2 | - | 47172413 | 47172459 | 47170733 | 47170764 | 47176276 | 47176470 | 4 | 8 | [illegible] |
| SE | 68066 | ENSG00000110880 | CORO1C | chr12 | - | 108648962 | 108649020 | 108648604 | 108648850 | 108652271 | 108652345 | 583 | 528 | [illegible] |
| SE | 68067 | ENSG00000110880 | CORO1C | chr12 | - | 108652271 | 108652417 | 108648962 | 108649020 | 108654305 | 108654410 | 640 | 620 | [illegible] |
| SE | 68069 | ENSG00000110880 | CORO1C | chr12 | - | 108657303 | 108657423 | 108654305 | 108654410 | 108658737 | 108658822 | 368 | 327 | [illegible] |
| SE | 68071 | ENSG00000110880 | CORO1C | chr12 | - | 108662028 | 108662158 | 108654305 | 108654410 | 108678271 | 108678394 | 143 | 106 | [illegible] |
| SE | 68072 | ENSG00000110880 | CORO1C | chr12 | - | 108678271 | 108678394 | 108662022 | 108662158 | 108701123 | 108701323 | 233 | 148 | [illegible] |
| SE | 68073 | ENSG00000110880 | CORO1C | chr12 | - | 108678271 | 108678394 | 108662022 | 108662158 | 108731428 | 108731518 | 102 | 73 | [illegible] |
| SE | 68074 | ENSG00000110880 | CORO1C | chr12 | - | 108701123 | 108701323 | 108662022 | 108662158 | 108731428 | 108731518 | 22 | 25 | [illegible] |
| SE | 68075 | ENSG00000110880 | CORO1C | chr12 | - | 108701123 | 108701323 | 108678271 | 108678394 | 108731428 | 108731518 | 153 | 100 | [illegible] |
| SE | 68079 | ENSG00000163577 | EIF5A2 | chr3 | - | 170907641 | 170907799 | 170906988 | 170907093 | 170908542 | 170908604 | 40 | 19 | [illegible] |
| SE | 68080 | ENSG00000163577 | EIF5A2 | chr3 | - | 170907641 | 170907841 | 170906988 | 170907093 | 170908542 | 170908598 | 47 | 24 | [illegible] |
| SE | 68086 | ENSG00000135838 | NPL | chr1 | + | 182794355 | 182794439 | 182792231 | 182792286 | 182812155 | 182812190 | 45 | 52 | [illegible] |
| SE | 68088 | ENSG00000135838 | NPL | chr1 | + | 182803697 | 182803771 | 182792230 | 182792286 | 182812155 | 182812190 | 15 | 1 | [illegible] |
| SE | 68091 | ENSG00000135838 | NPL | chr1 | + | 182806144 | 182806232 | 182794349 | 182794439 | 182812155 | 182812190 | 28 | 21 | [illegible] |
| SE | 68092 | ENSG00000135838 | NPL | chr1 | + | 182806144 | 182806232 | 182803697 | 182803771 | 182809842 | 182809970 | 124 | 71 | [illegible] |
| SE | 68096 | ENSG00000135838 | NPL | chr1 | + | 182809842 | 182809970 | 182803697 | 182803771 | 182812155 | 182812213 | 3 | 0 | [illegible] |
| SE | 68097 | ENSG00000135838 | NPL | chr1 | + | 182809842 | 182809970 | 182806144 | 182806232 | 182812155 | 182812213 | 7 | 0 | [illegible] |
| SE | 68104 | ENSG00000135838 | NPL | chr1 | + | 182822114 | 182822199 | 182818540 | 182818689 | 182829602 | 182830384 | 5 | 0 | [illegible] |
| SE | 68106 | ENSG00000135838 | NPL | chr1 | + | 182825780 | 182825820 | 182818540 | 182818689 | 182829602 | 182829679 | 0 | 1 | [illegible] |
| SE | 68107 | ENSG00000135838 | NPL | chr1 | + | 182825780 | 182825820 | 182822114 | 182822199 | 182829602 | 182829679 | 8 | 4 | [illegible] |
| SE | 68109 | ENSG00000142621 | FHAD1 | chr1 | + | 15289398 | 15289666 | 15272322 | 15272529 | 15296683 | 15296762 | 2 | 1 | [illegible] |
| SE | 68115 | ENSG00000142621 | FHAD1 | chr1 | + | 15380700 | 15380796 | 15375602 | 15375730 | 15381230 | 15381451 | 0 | 0 | [illegible] |
| SE | 68125 | ENSG00000082153 | BZW1 | chr2 | + | 200815666 | 200815761 | 200815340 | 200815517 | 200816324 | 200816390 | 1034 | 503 | [illegible] |
| SE | 68126 | ENSG00000082153 | BZW1 | chr2 | + | 200817105 | 200817241 | 200816324 | 200816390 | 200817973 | 200818083 | 687 | 369 | [illegible] |
| SE | 68127 | ENSG00000082153 | BZW1 | chr2 | + | 200821182 | 200821305 | 200819981 | 200820120 | 200822146 | 200822243 | 846 | 383 | [illegible] |
| SE | 68130 | ENSG00000086288 | NME8 | chr7 | + | 37850377 | 37850435 | 37850259 | 37850299 | 37850628 | 37850735 | 8 | 3 | [illegible] |
| SE | 68131 | ENSG00000086288 | NME8 | chr7 | + | 37850628 | 37850735 | 37850259 | 37850299 | 37857273 | 37857345 | 0 | 2 | [illegible] |
| SE | 68132 | ENSG00000086288 | NME8 | chr7 | + | 37857273 | 37857345 | 37850628 | 37850735 | 37862027 | 37862144 | 11 | 8 | [illegible] |
| SE | 68133 | ENSG00000086288 | NME8 | chr7 | + | 37857273 | 37857345 | 37850628 | 37850735 | 37863395 | 37863462 | 4 | 4 | [illegible] |
| SE | 68134 | ENSG00000086288 | NME8 | chr7 | + | 37857273 | 37857345 | 37850628 | 37850735 | 37884302 | 37884447 | 3 | 2 | [illegible] |
| SE | 68135 | ENSG00000086288 | NME8 | chr7 | + | 37862027 | 37862144 | 37850628 | 37850735 | 37863395 | 37863462 | 5 | 7 | [illegible] |
| SE | 68140 | ENSG00000086288 | NME8 | chr7 | + | 37864347 | 37864421 | 37863395 | 37863462 | 37867701 | 37867898 | 15 | 14 | [illegible] |
| SE | 68141 | ENSG00000086288 | NME8 | chr7 | + | 37864347 | 37864421 | 37863395 | 37863462 | 37876831 | 37877007 | 10 | 7 | [illegible] |
| SE | 68142 | ENSG00000086288 | NME8 | chr7 | + | 37864347 | 37864421 | 37863395 | 37863462 | 37884302 | 37884447 | 10 | 7 | [illegible] |
| SE | 68143 | ENSG00000086288 | NME8 | chr7 | + | 37865524 | 37865617 | 37864347 | 37864421 | 37867701 | 37867898 | 4 | 3 | [illegible] |
| SE | 68144 | ENSG00000086288 | NME8 | chr7 | + | 37867701 | 37867898 | 37863395 | 37863462 | 37876831 | 37877007 | 4 | 6 | [illegible] |
| SE | 68145 | ENSG00000086288 | NME8 | chr7 | + | 37867701 | 37867898 | 37863395 | 37863462 | 37884302 | 37884447 | 5 | 1 | [illegible] |
| SE | 68146 | ENSG00000086288 | NME8 | chr7 | + | 37867701 | 37867898 | 37864347 | 37864421 | 37876831 | 37877007 | 7 | 12 | [illegible] |
| SE | 68147 | ENSG00000086288 | NME8 | chr7 | + | 37867701 | 37867898 | 37864347 | 37864421 | 37884302 | 37884447 | 8 | 7 | [illegible] |
| SE | 68148 | ENSG00000086288 | NME8 | chr7 | + | 37876831 | 37877007 | 37857273 | 37857345 | 37884302 | 37884447 | 5 | 5 | [illegible] |
| SE | 68149 | ENSG00000086288 | NME8 | chr7 | + | 37876831 | 37877007 | 37863395 | 37863462 | 37884302 | 37884447 | 6 | 5 | [illegible] |
| SE | 68150 | ENSG00000086288 | NME8 | chr7 | + | 37876831 | 37877007 | 37864347 | 37864421 | 37884302 | 37884447 | 5 | 5 | [illegible] |
| SE | 68151 | ENSG00000086288 | NME8 | chr7 | + | 37876831 | 37877007 | 37867701 | 37867898 | 37884302 | 37884447 | 7 | 10 | [illegible] |
| SE | 68152 | ENSG00000086288 | NME8 | chr7 | + | 37894465 | 37894610 | 37888276 | 37888428 | 37896869 | 37897092 | 34 | 13 | [illegible] |
| SE | 68153 | ENSG00000168758 | SEMA4C | chr2 | - | 96864237 | 96864382 | 96863925 | 96864148 | 96864704 | 96864880 | 61 | 35 | [illegible] |
| SE | 68154 | ENSG00000168758 | SEMA4C | chr2 | - | 96866282 | 96866431 | 96865866 | 96865929 | 96867777 | 96867923 | 23 | 16 | [illegible] |
| SE | 68155 | ENSG00000168758 | SEMA4C | chr2 | - | 96866282 | 96866431 | 96865866 | 96865929 | 96870377 | 96870757 | 18 | 10 | [illegible] |
| SE | 68156 | ENSG00000168758 | SEMA4C | chr2 | - | 96867777 | 96867923 | 96865866 | 96865929 | 96870377 | 96870757 | 1 | 2 | [illegible] |
| SE | 68157 | ENSG00000168758 | SEMA4C | chr2 | - | 96867777 | 96867923 | 96866282 | 96866431 | 96870377 | 96870757 | 7 | 8 | [illegible] |
| SE | 68158 | ENSG00000168758 | SEMA4C | chr2 | - | 96867777 | 96867926 | 96865866 | 96865929 | 96870377 | 96870757 | 1 | 0 | [illegible] |
| SE | 68159 | ENSG00000168758 | SEMA4C | chr2 | - | 96867777 | 96867926 | 96866282 | 96866431 | 96870377 | 96870757 | 7 | 6 | [illegible] |
| SE | 68161 | ENSG00000047230 | CTPS2 | chrX | - | 16617149 | 16617246 | 16609540 | 16609685 | 16620276 | 16620332 | 38 | 31 | [illegible] |
| SE | 68162 | ENSG00000047230 | CTPS2 | chrX | - | 16639146 | 16639243 | 16620276 | 16620332 | 16667513 | 16667557 | 38 | 28 | [illegible] |
| SE | 68163 | ENSG00000047230 | CTPS2 | chrX | - | 16670579 | 16670674 | 16667661 | 16667724 | 16678361 | 16678450 | 32 | 32 | [illegible] |
| SE | 68164 | ENSG00000047230 | CTPS2 | chrX | - | 16678361 | 16678450 | 16670579 | 16670674 | 16683093 | 16683226 | 52 | 57 | [illegible] |
| SE | 68166 | ENSG00000047230 | CTPS2 | chrX | - | 16693370 | 16693487 | 16693140 | 16693224 | 16698235 | 16698336 | 36 | 26 | [illegible] |
| SE | 68167 | ENSG00000047230 | CTPS2 | chrX | - | 16702736 | 16702941 | 16698922 | 16699093 | 16712721 | 16712936 | 26 | 34 | [illegible] |
| SE | 68172 | ENSG00000169629 | RGPD8 | chr2 | - | 112412430 | 112412626 | 112406296 | 112406384 | 112417683 | 112417914 | 2 | 0 | [illegible] |
| SE | 68178 | ENSG00000020922 | MRE11 | chr11 | - | 94456275 | 94456338 | 94447218 | 94447438 | 94459407 | 94459581 | 162 | 176 | [illegible] |
| SE | 68179 | ENSG00000020922 | MRE11 | chr11 | - | 94464112 | 94464239 | 94460935 | 94461036 | 94467812 | 94467893 | 238 | 244 | [illegible] |
| SE | 68180 | ENSG00000020922 | MRE11 | chr11 | - | 94464112 | 94464239 | 94460935 | 94461036 | 94470470 | 94470642 | 92 | 121 | [illegible] |
| SE | 68182 | ENSG00000020922 | MRE11 | chr11 | - | 94467812 | 94467893 | 94460935 | 94461036 | 94470470 | 94470642 | 75 | 60 | [illegible] |
| SE | 68183 | ENSG00000020922 | MRE11 | chr11 | - | 94467812 | 94467893 | 94460935 | 94461036 | 94471573 | 94471759 | 0 | 5 | [illegible] |
| SE | 68184 | ENSG00000020922 | MRE11 | chr11 | - | 94467812 | 94467893 | 94464112 | 94464239 | 94470470 | 94470642 | 221 | 179 | [illegible] |
| SE | 68185 | ENSG00000020922 | MRE11 | chr11 | - | 94470470 | 94470642 | 94460935 | 94461036 | 94471573 | 94471759 | 110 | 84 | [illegible] |
| SE | 68186 | ENSG00000020922 | MRE11 | chr11 | - | 94470470 | 94470642 | 94467812 | 94467893 | 94471573 | 94471759 | 213 | 172 | [illegible] |
| SE | 68187 | ENSG00000020922 | MRE11 | chr11 | - | 94471573 | 94471759 | 94460935 | 94461036 | 94476288 | 94476403 | 110 | 82 | [illegible] |
| SE | 68188 | ENSG00000020922 | MRE11 | chr11 | - | 94471573 | 94471759 | 94460935 | 94461036 | 94478734 | 94478876 | 1 | 0 | [illegible] |
| SE | 68189 | ENSG00000020922 | MRE11 | chr11 | - | 94476288 | 94476403 | 94460935 | 94461036 | 94478734 | 94478876 | 108 | 88 | [illegible] |
| SE | 68191 | ENSG00000020922 | MRE11 | chr11 | - | 94476288 | 94476403 | 94471573 | 94471759 | 94485923 | 94486084 | 87 | 65 | [illegible] |
| SE | 68193 | ENSG00000020922 | MRE11 | chr11 | - | 94492641 | 94492906 | 94490832 | 94490965 | 94493790 | 94493843 | 43 | 22 | [illegible] |
| SE | 68194 | ENSG00000020922 | MRE11 | chr11 | - | 94492781 | 94492897 | 94490832 | 94490965 | 94493683 | 94493812 | 110 | 92 | [illegible] |
| SE | 68195 | ENSG00000020922 | MRE11 | chr11 | - | 94492781 | 94492897 | 94490832 | 94490965 | 94493790 | 94493885 | 111 | 94 | [illegible] |
| SE | 68196 | ENSG00000020922 | MRE11 | chr11 | - | 94492781 | 94492906 | 94490832 | 94490965 | 94493683 | 94493813 | 186 | 135 | [illegible] |
| SE | 68197 | ENSG00000020922 | MRE11 | chr11 | - | 94492781 | 94492906 | 94490832 | 94490965 | 94493790 | 94493834 | 173 | 128 | [illegible] |
| SE | 68206 | ENSG00000103707 | MTFMT | chr15 | - | 65021513 | 65021616 | 65020196 | 65020272 | 65023671 | 65023794 | 69 | 52 | [illegible] |
| SE | 68207 | ENSG00000103707 | MTFMT | chr15 | - | 65021513 | 65021616 | 65020196 | 65020272 | 65026830 | 65026931 | 46 | 29 | [illegible] |
| SE | 68210 | ENSG00000107643 | MAPK8 | chr10 | + | 48308094 | 48308288 | 48306654 | 48306821 | 48325896 | 48325965 | 4 | 6 | [illegible] |
| SE | 68211 | ENSG00000107643 | MAPK8 | chr10 | + | 48325896 | 48325965 | 48306654 | 48306821 | 48401611 | 48401782 | 10 | 9 | [illegible] |
| SE | 68212 | ENSG00000107643 | MAPK8 | chr10 | + | 48325896 | 48325965 | 48306654 | 48306821 | 48404851 | 48404981 | 2 | 0 | [illegible] |
| SE | 68213 | ENSG00000107643 | MAPK8 | chr10 | + | 48325896 | 48325965 | 48308094 | 48308288 | 48401611 | 48401782 | 10 | 12 | [illegible] |
| SE | 68215 | ENSG00000107643 | MAPK8 | chr10 | + | 48401611 | 48401782 | 48306638 | 48306821 | 48404851 | 48404981 | 112 | 64 | [illegible] |
| SE | 68216 | ENSG00000107643 | MAPK8 | chr10 | + | 48401611 | 48401782 | 48325896 | 48325965 | 48404851 | 48404981 | 113 | 70 | [illegible] |
| SE | 68218 | ENSG00000107643 | MAPK8 | chr10 | + | 48420154 | 48420320 | 48410029 | 48410168 | 48425178 | 48425219 | 83 | 45 | [illegible] |
| SE | 68219 | ENSG00000107643 | MAPK8 | chr10 | + | 48426379 | 48426504 | 48425887 | 48426070 | 48427079 | 48427143 | 140 | 115 | [illegible] |
| SE | 68220 | ENSG00000107643 | MAPK8 | chr10 | + | 48431192 | 48431270 | 48425188 | 48427143 | 48434878 | 48434898 | 83 | 66 | [illegible] |
| SE | 68221 | ENSG00000107643 | MAPK8 | chr10 | + | 48431192 | 48431270 | 48425188 | 48427143 | 48434883 | 48439357 | 77 | 61 | [illegible] |
| SE | 68222 | ENSG00000090273 | NUDC | chr1 | + | 26911127 | 26911235 | 26902281 | 26902366 | 26924088 | 26924166 | 0 | 1 | [illegible] |
| SE | 68227 | ENSG00000114054 | PCCB | chr3 | + | 136260262 | 136260355 | 136256554 | 136256623 | 136260478 | 136260535 | 1 | 0 | [illegible] |
| SE | 68228 | ENSG00000114054 | PCCB | chr3 | + | 136260478 | 136260535 | 136256554 | 136256623 | 136261951 | 136262010 | 142 | 58 | [illegible] |
| SE | 68230 | ENSG00000114054 | PCCB | chr3 | + | 136297951 | 136298072 | 136293755 | 136293864 | 136301029 | 136301111 | 97 | 35 | [illegible] |
| SE | 68231 | ENSG00000114054 | PCCB | chr3 | + | 136326325 | 136326385 | 136316940 | 136317064 | 136326802 | 136326910 | 1 | 0 | [illegible] |
| SE | 68232 | ENSG00000114054 | PCCB | chr3 | + | 136327154 | 136327255 | 136326802 | 136326910 | 136327633 | 136327732 | 50 | 34 | [illegible] |
| SE | 68233 | ENSG00000091972 | CD200 | chr3 | + | 112335942 | 112336017 | 112333155 | 112333224 | 112340901 | 112340983 | 1 | 0 | [illegible] |
| SE | 68235 | ENSG00000091972 | CD200 | chr3 | + | 112340901 | 112340983 | 112333067 | 112333224 | 112344961 | 112345075 | 5 | 2 | [illegible] |
| SE | 68241 | ENSG00000189223 | PAX8-AS1 | chr2 | + | 113236512 | 113236613 | 113235539 | 113235570 | 113238089 | 113238531 | 0 | 0 | [illegible] |
| SE | 68242 | ENSG00000189223 | PAX8-AS1 | chr2 | + | 113236512 | 113236613 | 113235539 | 113235570 | 113255389 | 113255549 | 0 | 0 | [illegible] |
| SE | 68245 | ENSG00000189223 | PAX8-AS1 | chr2 | + | 113238089 | 113238212 | 113235505 | 113235570 | 113255389 | 113255549 | 0 | 1 | [illegible] |
| SE | 68249 | ENSG00000189223 | PAX8-AS1 | chr2 | + | 113250766 | 113250906 | 113235544 | 113235570 | 113255389 | 113255534 | 0 | 0 | [illegible] |
| SE | 68251 | ENSG00000189223 | PAX8-AS1 | chr2 | + | 113250766 | 113250906 | 113238089 | 113238212 | 113255389 | 113255534 | 1 | 0 | [illegible] |
| SE | 68252 | ENSG00000189223 | PAX8-AS1 | chr2 | + | 113250766 | 113250922 | 113235424 | 113235570 | 113255389 | 113255549 | 0 | 0 | [illegible] |
| SE | 68254 | ENSG00000189223 | PAX8-AS1 | chr2 | + | 113255389 | 113255549 | 113235518 | 113235570 | 113263277 | 113263454 | 2 | 0 | [illegible] |
| SE | 68258 | ENSG00000189223 | PAX8-AS1 | chr2 | + | 113255389 | 113255549 | 113238089 | 113238212 | 113263277 | 113263454 | 1 | 1 | [illegible] |
| SE | 68259 | ENSG00000189223 | PAX8-AS1 | chr2 | + | 113259263 | 113259584 | 113235517 | 113235570 | 113263277 | 113263334 | 3 | 3 | [illegible] |
| SE | 68260 | ENSG00000189223 | PAX8-AS1 | chr2 | + | 113259263 | 113259584 | 113238089 | 113238212 | 113263277 | 113263334 | 3 | 3 | [illegible] |
| SE | 68261 | ENSG00000189223 | PAX8-AS1 | chr2 | + | 113259263 | 113259584 | 113255389 | 113255549 | 113263277 | 113263334 | 6 | 12 | [illegible] |
| SE | 68262 | ENSG00000189223 | PAX8-AS1 | chr2 | + | 113270200 | 113270307 | 113265475 | 113265580 | 113271835 | 113271971 | 2 | 3 | [illegible] |
| SE | 68263 | ENSG00000189223 | PAX8-AS1 | chr2 | + | 113271835 | 113271971 | 113269823 | 113270307 | 113273056 | 113273164 | 5 | 7 | [illegible] |
| SE | 68265 | ENSG00000135916 | ITM2C | chr2 | + | 230873416 | 230873557 | 230864941 | 230865145 | 230875619 | 230875808 | 55 | 45 | [illegible] |
| SE | 68266 | ENSG00000135916 | ITM2C | chr2 | + | 230876856 | 230876967 | 230875619 | 230875808 | 230877399 | 230877550 | 115 | 89 | [illegible] |

| | | | | | | | | | | | | | | |
|---|---|---|---|---|---|---|---|---|---|---|---|---|---|---|
| SE | 68267 | ENSG00000169855 | ROBO1 | chr3 | - | 78600112 | 78600309 | 78597238 | 78598927 | 78606732 | 78607041 | 1 | 1 | [illegible] |
| SE | 68268 | ENSG00000169855 | ROBO1 | chr3 | - | 78647628 | 78647655 | 78646147 | 78646190 | 78651731 | 78651929 | 0 | 0 | [illegible] |
| SE | 68271 | ENSG00000115085 | ZAP70 | chr2 | + | 97724015 | 97724438 | 97713915 | 97713994 | 97725091 | 97725252 | 134 | 50 | [illegible] |
| SE | 68272 | ENSG00000204387 | SNHG32 | chr6 | + | 31835410 | 31835451 | 31834607 | 31835200 | 31837235 | 31837435 | 676 | 477 | [illegible] |
| SE | 68273 | ENSG00000204387 | SNHG32 | chr6 | + | 31835410 | 31835455 | 31834921 | 31835200 | 31837235 | 31837435 | 143 | 106 | [illegible] |
| SE | 68277 | ENSG00000122085 | MTERF4 | chr2 | - | 241089979 | 241090044 | 241087093 | 241089365 | 241090302 | 241090449 | 15 | 12 | [illegible] |
| SE | 68278 | ENSG00000122085 | MTERF4 | chr2 | - | 241089979 | 241090044 | 241087093 | 241089365 | 241094276 | 241094429 | 58 | 32 | [illegible] |
| SE | 68280 | ENSG00000122085 | MTERF4 | chr2 | - | 241090302 | 241090449 | 241089979 | 241090044 | 241099395 | 241099894 | 0 | 1 | [illegible] |
| SE | 68281 | ENSG00000122085 | MTERF4 | chr2 | - | 241091894 | 241092114 | 241090277 | 241090449 | 241094276 | 241094432 | 1 | 0 | [illegible] |
| SE | 68283 | ENSG00000122085 | MTERF4 | chr2 | - | 241094276 | 241094429 | 241089979 | 241090044 | 241102252 | 241102332 | 43 | 23 | [illegible] |
| SE | 68287 | ENSG00000122085 | MTERF4 | chr2 | - | 241095129 | 241096438 | 241094276 | 241094432 | 241099395 | 241099894 | 0 | 5 | [illegible] |
| SE | 68288 | ENSG00000122085 | MTERF4 | chr2 | - | 241095129 | 241096438 | 241094276 | 241094432 | 241102252 | 241102295 | 0 | 3 | [illegible] |
| SE | 68291 | ENSG00000122085 | MTERF4 | chr2 | - | 241096392 | 241096438 | 241094276 | 241094432 | 241099395 | 241099426 | 0 | 2 | [illegible] |
| SE | 68292 | ENSG00000122085 | MTERF4 | chr2 | - | 241097242 | 241097427 | 241089979 | 241090044 | 241099395 | 241099426 | 36 | 36 | [illegible] |
| SE | 68293 | ENSG00000122085 | MTERF4 | chr2 | - | 241097242 | 241097427 | 241089979 | 241090044 | 241102252 | 241102299 | 20 | 7 | [illegible] |
| SE | 68294 | ENSG00000122085 | MTERF4 | chr2 | - | 241097242 | 241097427 | 241094276 | 241094429 | 241102252 | 241102299 | 20 | 7 | [illegible] |
| SE | 68298 | ENSG00000122085 | MTERF4 | chr2 | - | 241097242 | 241097427 | 241095129 | 241096438 | 241102252 | 241102299 | 153 | 112 | [illegible] |
| SE | 68299 | ENSG00000122085 | MTERF4 | chr2 | - | 241099395 | 241099894 | 241089979 | 241090044 | 241102252 | 241102287 | 16 | 7 | [illegible] |
| SE | 68300 | ENSG00000122085 | MTERF4 | chr2 | - | 241099395 | 241099894 | 241094405 | 241094432 | 241102252 | 241102287 | 16 | 7 | [illegible] |
| SE | 68301 | ENSG00000122085 | MTERF4 | chr2 | - | 241099395 | 241099894 | 241095129 | 241096438 | 241102252 | 241102287 | 16 | 9 | [illegible] |
| SE | 68304 | ENSG00000122085 | MTERF4 | chr2 | - | 241101163 | 241101221 | 241099398 | 241099894 | 241102252 | 241102293 | 0 | 3 | [illegible] |
| SE | 68306 | ENSG00000074695 | LMAN1 | chr18 | - | 59347512 | 59347571 | 59345918 | 59346051 | 59349112 | 59349236 | 215 | 171 | [illegible] |
| SE | 68308 | ENSG00000115290 | GRB14 | chr2 | - | 164525003 | 164525078 | 164521979 | 164522117 | 164527013 | 164527135 | 8 | 2 | [illegible] |
| SE | 68309 | ENSG00000115290 | GRB14 | chr2 | - | 164539765 | 164539897 | 164527013 | 164527135 | 164547659 | 164547816 | 0 | 0 | [illegible] |
| SE | 68310 | ENSG00000168016 | TRANK1 | chr3 | - | 36834386 | 36834475 | 36830872 | 36833919 | 36834761 | 36834907 | 0 | 0 | [illegible] |
| SE | 68311 | ENSG00000168016 | TRANK1 | chr3 | - | 36834761 | 36834907 | 36830872 | 36833919 | 36838612 | 36838716 | 102 | 59 | [illegible] |
| SE | 68313 | ENSG00000168016 | TRANK1 | chr3 | - | 36838612 | 36838716 | 36834761 | 36834907 | 36842621 | 36842710 | 193 | 118 | [illegible] |
| SE | 68317 | ENSG00000168016 | TRANK1 | chr3 | - | 36852145 | 36852345 | 36851718 | 36851856 | 36855172 | 36856645 | 295 | 169 | [illegible] |
| SE | 68319 | ENSG00000168016 | TRANK1 | chr3 | - | 36860905 | 36861160 | 36858717 | 36858894 | 36864318 | 36864480 | 266 | 104 | [illegible] |
| SE | 68320 | ENSG00000168016 | TRANK1 | chr3 | - | 36864318 | 36864480 | 36860905 | 36861160 | 36874125 | 36874296 | 343 | 160 | [illegible] |
| SE | 68322 | ENSG00000168016 | TRANK1 | chr3 | - | 36892201 | 36892340 | 36889828 | 36889960 | 36892900 | 36892984 | 493 | 303 | [illegible] |
| SE | 68323 | ENSG00000168016 | TRANK1 | chr3 | - | 36892201 | 36892340 | 36889828 | 36889960 | 36899108 | 36899259 | 251 | 144 | [illegible] |
| SE | 68324 | ENSG00000168016 | TRANK1 | chr3 | - | 36892900 | 36892984 | 36889828 | 36889960 | 36899108 | 36899259 | 7 | 0 | [illegible] |
| SE | 68325 | ENSG00000168016 | TRANK1 | chr3 | - | 36892900 | 36892984 | 36892201 | 36892340 | 36895639 | 36895758 | 380 | 219 | [illegible] |
| SE | 68329 | ENSG00000168016 | TRANK1 | chr3 | - | 36899108 | 36899259 | 36892900 | 36892984 | 36903148 | 36903275 | 140 | 63 | [illegible] |
| SE | 68330 | ENSG00000168016 | TRANK1 | chr3 | - | 36903148 | 36903275 | 36899106 | 36899259 | 36908322 | 36908454 | 244 | 89 | [illegible] |
| SE | 68332 | ENSG00000168016 | TRANK1 | chr3 | - | 36929314 | 36929415 | 36908322 | 36908454 | 36944786 | 36944861 | 2 | 0 | [illegible] |
| SE | 68337 | ENSG00000114942 | EEF1B2 | chr2 | + | 206160556 | 206160710 | 206159933 | 206160059 | 206161345 | 206161454 | 3105 | 1668 | [illegible] |
| SE | 68338 | ENSG00000114942 | EEF1B2 | chr2 | + | 206160587 | 206160710 | 206159894 | 206160059 | 206161345 | 206161472 | 4852 | 2468 | [illegible] |
| SE | 68339 | ENSG00000114942 | EEF1B2 | chr2 | + | 206160886 | 206161132 | 206160587 | 206160710 | 206161345 | 206161472 | 3 | 10 | [illegible] |
| SE | 68340 | ENSG00000114942 | EEF1B2 | chr2 | + | 206161345 | 206161472 | 206160556 | 206160710 | 206162037 | 206162077 | 5006 | 2319 | [illegible] |
| SE | 68341 | ENSG00000173726 | TOMM20 | chr1 | - | 235113767 | 235113910 | 235109340 | 235112108 | 235119817 | 235119899 | 412 | 222 | [illegible] |
| SE | 68342 | ENSG00000173726 | TOMM20 | chr1 | - | 235119338 | 235119406 | 235113767 | 235113910 | 235119817 | 235119899 | 1 | 1 | [illegible] |
| SE | 68343 | ENSG00000173726 | TOMM20 | chr1 | - | 235122325 | 235122372 | 235119817 | 235119899 | 235128594 | 235128837 | 179 | 95 | [illegible] |
| SE | 68348 | ENSG00000125630 | POLR1B | chr2 | + | 112548145 | 112548213 | 112547420 | 112547567 | 112549266 | 112549360 | 0 | 0 | [illegible] |
| SE | 68351 | ENSG00000125630 | POLR1B | chr2 | + | 112554523 | 112554578 | 112552644 | 112552816 | 112559292 | 112559395 | 0 | 0 | [illegible] |
| SE | 68353 | ENSG00000125630 | POLR1B | chr2 | + | 112557909 | 112558081 | 112554523 | 112554578 | 112559292 | 112559395 | 44 | 21 | [illegible] |
| SE | 68354 | ENSG00000125630 | POLR1B | chr2 | + | 112559292 | 112559574 | 112552644 | 112552816 | 112564365 | 112564499 | 17 | 14 | [illegible] |
| SE | 68355 | ENSG00000125630 | POLR1B | chr2 | + | 112564365 | 112564499 | 112559292 | 112559574 | 112567966 | 112568137 | 53 | 23 | [illegible] |
| SE | 68358 | ENSG00000114988 | LMAN2L | chr2 | - | 96713106 | 96713139 | 96711863 | 96712025 | 96733518 | 96733601 | 0 | 0 | [illegible] |
| SE | 68359 | ENSG00000114988 | LMAN2L | chr2 | - | 96713106 | 96713139 | 96711863 | 96712025 | 96734408 | 96734526 | 0 | 0 | [illegible] |
| SE | 68362 | ENSG00000114988 | LMAN2L | chr2 | - | 96737136 | 96737217 | 96734408 | 96734526 | 96737948 | 96738067 | 0 | 0 | [illegible] |
| SE | 68363 | ENSG00000114988 | LMAN2L | chr2 | - | 96737948 | 96738067 | 96734408 | 96734526 | 96739853 | 96740041 | 42 | 15 | [illegible] |
| SE | 68364 | ENSG00000185621 | LMLN | chr3 | + | 198003014 | 198003125 | 197999265 | 197999342 | 198019252 | 198019385 | 5 | 6 | [illegible] |
| SE | 68365 | ENSG00000185621 | LMLN | chr3 | + | 198019252 | 198019385 | 198003014 | 198003125 | 198021445 | 198021605 | 12 | 10 | [illegible] |
| SE | 68366 | ENSG00000185621 | LMLN | chr3 | + | 198035832 | 198036043 | 198024657 | 198024788 | 198038566 | 198038667 | 4 | 2 | [illegible] |
| SE | 68370 | ENSG00000244026 | FAM86DP | chr3 | - | 75427426 | 75427692 | 75426449 | 75426599 | 75427774 | 75427908 | 6 | 0 | [illegible] |
| SE | 68371 | ENSG00000244026 | FAM86DP | chr3 | - | 75427426 | 75427692 | 75426449 | 75426599 | 75430831 | 75430914 | 9 | 0 | [illegible] |
| SE | 68372 | ENSG00000244026 | FAM86DP | chr3 | - | 75427774 | 75427908 | 75426449 | 75426599 | 75429135 | 75429237 | 0 | 0 | [illegible] |
| SE | 68373 | ENSG00000244026 | FAM86DP | chr3 | - | 75427774 | 75427908 | 75426449 | 75426599 | 75430831 | 75430914 | 0 | 0 | [illegible] |
| SE | 68374 | ENSG00000244026 | FAM86DP | chr3 | - | 75427774 | 75427908 | 75427426 | 75427692 | 75430831 | 75430914 | 1 | 0 | [illegible] |
| SE | 68375 | ENSG00000244026 | FAM86DP | chr3 | - | 75429135 | 75429237 | 75426449 | 75426599 | 75430831 | 75430914 | 2 | 0 | [illegible] |
| SE | 68377 | ENSG00000244026 | FAM86DP | chr3 | - | 75430831 | 75430914 | 75426449 | 75426599 | 75432807 | 75432870 | 4 | 6 | [illegible] |
| SE | 68380 | ENSG00000136710 | CCDC115 | chr2 | - | 130341666 | 130341749 | 130340907 | 130341039 | 130341840 | 130341944 | 80 | 38 | [illegible] |
| SE | 68381 | ENSG00000157985 | AGAP1 | chr2 | + | 235717556 | 235717644 | 235709176 | 235709237 | 235740962 | 235741048 | 35 | 11 | [illegible] |
| SE | 68382 | ENSG00000157985 | AGAP1 | chr2 | + | 235718380 | 235718392 | 235717556 | 235717644 | 235740962 | 235741048 | 0 | 0 | [illegible] |
| SE | 68384 | ENSG00000157985 | AGAP1 | chr2 | + | 235908737 | 235908906 | 235883344 | 235883449 | 235930764 | 235930923 | 59 | 20 | [illegible] |
| SE | 68387 | ENSG00000006451 | RALA | chr7 | + | 39686630 | 39686781 | 39623571 | 39623825 | 39696684 | 39696859 | 35 | 34 | [illegible] |
| SE | 68388 | ENSG00000006451 | RALA | chr7 | + | 39690381 | 39690590 | 39686630 | 39686781 | 39696684 | 39696859 | 462 | 307 | [illegible] |
| SE | 68392 | ENSG00000115365 | LANCL1 | chr2 | - | 210436215 | 210436392 | 210435386 | 210435459 | 210437689 | 210437872 | 131 | 93 | [illegible] |
| SE | 68393 | ENSG00000115365 | LANCL1 | chr2 | - | 210436215 | 210436392 | 210435386 | 210435459 | 210440597 | 210440745 | 84 | 60 | [illegible] |
| SE | 68394 | ENSG00000115365 | LANCL1 | chr2 | - | 210437689 | 210437872 | 210435386 | 210435459 | 210440597 | 210440744 | 33 | 25 | [illegible] |
| SE | 68396 | ENSG00000115365 | LANCL1 | chr2 | - | 210441307 | 210441443 | 210440597 | 210440744 | 210455106 | 210455314 | 122 | 120 | [illegible] |
| SE | 68397 | ENSG00000115365 | LANCL1 | chr2 | - | 210455106 | 210455314 | 210441307 | 210441443 | 210471958 | 210472076 | 108 | 77 | [illegible] |
| SE | 68399 | ENSG00000115365 | LANCL1 | chr2 | - | 210471506 | 210471645 | 210455258 | 210455314 | 210471958 | 210472076 | 0 | 0 | [illegible] |
| SE | 68400 | ENSG00000115365 | LANCL1 | chr2 | - | 210476315 | 210476412 | 210471958 | 210472076 | 210476619 | 210476705 | 71 | 25 | [illegible] |
| SE | 68404 | ENSG00000131437 | KIF3A | chr5 | - | 132706450 | 132706459 | 132703462 | 132703619 | 132710958 | 132711057 | 1 | 0 | [illegible] |
| SE | 68405 | ENSG00000131437 | KIF3A | chr5 | - | 132706450 | 132706459 | 132703462 | 132703619 | 132715756 | 132715931 | 0 | 0 | [illegible] |
| SE | 68407 | ENSG00000131437 | KIF3A | chr5 | - | 132710958 | 132711057 | 132703462 | 132703619 | 132715756 | 132715931 | 33 | 24 | [illegible] |
| SE | 68408 | ENSG00000131437 | KIF3A | chr5 | - | 132710958 | 132711057 | 132706450 | 132706459 | 132715756 | 132715931 | 21 | 21 | [illegible] |
| SE | 68413 | ENSG00000020129 | NCDN | chr1 | + | 35563201 | 35563426 | 35562391 | 35562633 | 35563766 | 35563909 | 35 | 16 | [illegible] |
| SE | 68415 | ENSG00000101104 | PABPC1L | chr20 | + | 44916755 | 44916871 | 44912659 | 44912853 | 44921593 | 44921731 | 5 | 1 | [illegible] |
| SE | 68416 | ENSG00000101104 | PABPC1L | chr20 | + | 44918905 | 44919045 | 44916755 | 44916871 | 44919182 | 44919277 | 48 | 35 | [illegible] |
| SE | 68418 | ENSG00000101104 | PABPC1L | chr20 | + | 44921593 | 44921731 | 44919182 | 44919277 | 44924160 | 44924256 | 95 | 73 | [illegible] |
| SE | 68419 | ENSG00000101104 | PABPC1L | chr20 | + | 44924160 | 44924256 | 44921593 | 44921731 | 44930517 | 44930726 | 76 | 53 | [illegible] |
| SE | 68421 | ENSG00000101104 | PABPC1L | chr20 | + | 44931832 | 44932026 | 44930568 | 44930726 | 44933056 | 44933185 | 1 | 1 | [illegible] |
| SE | 68422 | ENSG00000101104 | PABPC1L | chr20 | + | 44931832 | 44932026 | 44930568 | 44930726 | 44933071 | 44933185 | 2 | 1 | [illegible] |
| SE | 68425 | ENSG00000101104 | PABPC1L | chr20 | + | 44932341 | 44932432 | 44931832 | 44932026 | 44933056 | 44933185 | 0 | 0 | [illegible] |
| SE | 68426 | ENSG00000101104 | PABPC1L | chr20 | + | 44932341 | 44932432 | 44931832 | 44932026 | 44933071 | 44933185 | 0 | 0 | [illegible] |
| SE | 68430 | ENSG00000101104 | PABPC1L | chr20 | + | 44936893 | 44936976 | 44936636 | 44936730 | 44938060 | 44938191 | 0 | 0 | [illegible] |
| SE | 68431 | ENSG00000101104 | PABPC1L | chr20 | + | 44938060 | 44938191 | 44936636 | 44936730 | 44958336 | 44958807 | 8 | 6 | [illegible] |
| SE | 68432 | ENSG00000101104 | PABPC1L | chr20 | + | 44938673 | 44938748 | 44938044 | 44938191 | 44939125 | 44939184 | 6 | 2 | [illegible] |
| SE | 68433 | ENSG00000101104 | PABPC1L | chr20 | + | 44938673 | 44938748 | 44938044 | 44938191 | 44958336 | 44958952 | 5 | 0 | [illegible] |
| SE | 68434 | ENSG00000101104 | PABPC1L | chr20 | + | 44952642 | 44952704 | 44938060 | 44938191 | 44958336 | 44959035 | 1 | 2 | [illegible] |
| SE | 68435 | ENSG00000198589 | LRBA | chr4 | - | 150266398 | 150266441 | 150264696 | 150265812 | 150277852 | 150278004 | 0 | 0 | [illegible] |
| SE | 68437 | ENSG00000198589 | LRBA | chr4 | - | 150282449 | 150282646 | 150277852 | 150278004 | 150285932 | 150286034 | 169 | 155 | [illegible] |
| SE | 68438 | ENSG00000198589 | LRBA | chr4 | - | 150302624 | 150302792 | 150285932 | 150286034 | 150310228 | 150310384 | 170 | 141 | [illegible] |
| SE | 68439 | ENSG00000198589 | LRBA | chr4 | - | 150302627 | 150302792 | 150285932 | 150286034 | 150310228 | 150310384 | 70 | 78 | [illegible] |
| SE | 68440 | ENSG00000198589 | LRBA | chr4 | - | 150349991 | 150350090 | 150325808 | 150325898 | 150415437 | 150415590 | 113 | 80 | [illegible] |
| SE | 68441 | ENSG00000198589 | LRBA | chr4 | - | 150349991 | 150350159 | 150325808 | 150325898 | 150415437 | 150415590 | 232 | 172 | [illegible] |
| SE | 68442 | ENSG00000198589 | LRBA | chr4 | - | 150353730 | 150353924 | 150349991 | 150350159 | 150415437 | 150415590 | 0 | 0 | [illegible] |
| SE | 68443 | ENSG00000198589 | LRBA | chr4 | - | 150415437 | 150415590 | 150349991 | 150350159 | 151014426 | 151014861 | 86 | 66 | [illegible] |
| SE | 68444 | ENSG00000198589 | LRBA | chr4 | - | 150588047 | 150588184 | 150490917 | 150491035 | 150590712 | 150590859 | 469 | 310 | [illegible] |
| SE | 68445 | ENSG00000198589 | LRBA | chr4 | - | 150590712 | 150590859 | 150588047 | 150588184 | 150599006 | 150599131 | 540 | 333 | [illegible] |
| SE | 68447 | ENSG00000198589 | LRBA | chr4 | - | 150735257 | 150735366 | 150683550 | 150683717 | 150761782 | 150761847 | 443 | 311 | [illegible] |
| SE | 68449 | ENSG00000198589 | LRBA | chr4 | - | 150817123 | 150817257 | 150808319 | 150808398 | 150828179 | 150828621 | 376 | 319 | [illegible] |
| SE | 68450 | ENSG00000198589 | LRBA | chr4 | - | 150849421 | 150849575 | 150848817 | 150848998 | 150850723 | 150850902 | 217 | 176 | [illegible] |
| SE | 68452 | ENSG00000198589 | LRBA | chr4 | - | 150868181 | 150868305 | 150851884 | 150852943 | 150870524 | 150870606 | 197 | 233 | [illegible] |
| SE | 68453 | ENSG00000198589 | LRBA | chr4 | - | 150872662 | 150872755 | 150871344 | 150871453 | 150893051 | 150893149 | 341 | 360 | [illegible] |
| SE | 68454 | ENSG00000198589 | LRBA | chr4 | - | 150872662 | 150872755 | 150871344 | 150871453 | 150900048 | 150900217 | 150 | 142 | [illegible] |
| SE | 68455 | ENSG00000198589 | LRBA | chr4 | - | 150893051 | 150893149 | 150872662 | 150872755 | 150896393 | 150896456 | 283 | 289 | [illegible] |
| SE | 68456 | ENSG00000198589 | LRBA | chr4 | - | 150896393 | 150896456 | 150872662 | 150872755 | 150900048 | 150900217 | 2 | 3 | [illegible] |
| SE | 68458 | ENSG00000198589 | LRBA | chr4 | - | 150906296 | 150906405 | 150905837 | 150905990 | 150908333 | 150908467 | 181 | 166 | [illegible] |
| SE | 68459 | ENSG00000198589 | LRBA | chr4 | - | 150916616 | 150916738 | 150916400 | 150916527 | 150921197 | 150921293 | 131 | 85 | [illegible] |
| SE | 68460 | ENSG00000198589 | LRBA | chr4 | - | 150921197 | 150921293 | 150916616 | 150916738 | 150928515 | 150928616 | 103 | 73 | [illegible] |
| SE | 68462 | ENSG00000143799 | PARP1 | chr1 | - | 226371015 | 226371106 | 226370433 | 226370517 | 226374225 | 226376067 | 0 | 0 | [illegible] |
| SE | 68464 | ENSG00000143799 | PARP1 | chr1 | - | 226383035 | 226383183 | 226379921 | 226381208 | 226385503 | 226385680 | 528 | 314 | [illegible] |
| SE | 68465 | ENSG00000143799 | PARP1 | chr1 | - | 226390409 | 226390624 | 226388655 | 226388755 | 226392198 | 226392314 | 336 | 189 | [illegible] |
| SE | 68466 | ENSG00000143799 | PARP1 | chr1 | - | 226392198 | 226392314 | 226388655 | 226388755 | 226402213 | 226402379 | 200 | 121 | [illegible] |
| SE | 68469 | ENSG00000119508 | NR4A3 | chr9 | + | 99825658 | 99825832 | 99821854 | 99822407 | 99828040 | 99828993 | 3 | 2 | [illegible] |
| SE | 68470 | ENSG00000119508 | NR4A3 | chr9 | + | 99826729 | 99826801 | 99825658 | 99825832 | 99828040 | 99828993 | 0 | 1 | [illegible] |
| SE | 68474 | ENSG00000127837 | AAMP | chr2 | - | 218265578 | 218265682 | 218265370 | 218265461 | 218265830 | 218265946 | 149 | 62 | [illegible] |
| SE | 68475 | ENSG00000127837 | AAMP | chr2 | - | 218265830 | 218265946 | 218265578 | 218265682 | 218266063 | 218266147 | 165 | 90 | [illegible] |
| SE | 68476 | ENSG00000127837 | AAMP | chr2 | - | 218266846 | 218266986 | 218265578 | 218266587 | 218267493 | 218267613 | 145 | 69 | [illegible] |
| SE | 68477 | ENSG00000242372 | EIF6 | chr20 | - | 35279565 | 35279747 | 35278906 | 35279206 | 35279941 | 35280118 | 212 | 72 | [illegible] |
| SE | 68479 | ENSG00000242372 | EIF6 | chr20 | - | 35284175 | 35284261 | 35279941 | 35280118 | 35284380 | 35284492 | 87 | 30 | [illegible] |
| SE | 68484 | ENSG00000177879 | AP3S1 | chr5 | + | 115866669 | 115866761 | 115841934 | 115842106 | 115870016 | 115870128 | 679 | 505 | [illegible] |
| SE | 68485 | ENSG00000177879 | AP3S1 | chr5 | + | 115866669 | 115866761 | 115841934 | 115842106 | 115895086 | 115895158 | 52 | 25 | [illegible] |
| SE | 68486 | ENSG00000177879 | AP3S1 | chr5 | + | 115870016 | 115870128 | 115841934 | 115842106 | 115895086 | 115895158 | 732 | 555 | [illegible] |
| SE | 68487 | ENSG00000177879 | AP3S1 | chr5 | + | 115870016 | 115870128 | 115866660 | 115866761 | 115895086 | 115895158 | 1361 | 1029 | [illegible] |
| SE | 68489 | ENSG00000177879 | AP3S1 | chr5 | + | 115895086 | 115895158 | 115870016 | 115870128 | 115913361 | 115913900 | 216 | 188 | [illegible] |
| SE | 68490 | ENSG00000177879 | AP3S1 | chr5 | + | 115898648 | 115898789 | 115895086 | 115895158 | 115902884 | 115904088 | 0 | 1 | [illegible] |
| SE | 68491 | ENSG00000177879 | AP3S1 | chr5 | + | 115898648 | 115898789 | 115895086 | 115895158 | 115913361 | 115914081 | 2 | 0 | [illegible] |
| SE | 68492 | ENSG00000177879 | AP3S1 | chr5 | + | 115902884 | 115902992 | 115870016 | 115870128 | 115913361 | 115914011 | 211 | 123 | [illegible] |
| SE | 68495 | ENSG00000177879 | AP3S1 | chr5 | + | 115912088 | 115912136 | 115902884 | 115902992 | 115913361 | 115913457 | 7 | 1 | [illegible] |
| SE | 68498 | ENSG00000087087 | SRRT | chr7 | + | 100881658 | 100881805 | 100881284 | 100881413 | 100882052 | 100882241 | 147 | 59 | [illegible] |
| SE | 68502 | ENSG00000087087 | SRRT | chr7 | + | 100888041 | 100888143 | 100887702 | 100887859 | 100888473 | 100888660 | 155 | 79 | [illegible] |
| SE | 68508 | ENSG00000125462 | MIR9-1HG | chr1 | - | 156407841 | 156407975 | 156404252 | 156404601 | 156414653 | 156414753 | 2 | 0 | [illegible] |
| SE | 68509 | ENSG00000125462 | MIR9-1HG | chr1 | - | 156407841 | 156407975 | 156404252 | 156404601 | 156420170 | 156420289 | 2 | 0 | [illegible] |
| SE | 68515 | ENSG00000125462 | MIR9-1HG | chr1 | - | 156414653 | 156414753 | 156404255 | 156404601 | 156420170 | 156420336 | 2 | 0 | [illegible] |
| SE | 68534 | ENSG00000131236 | CAP1 | chr1 | + | 40041428 | 40041493 | 40040881 | 40040965 | 40059336 | 40059458 | 12 | 16 | [illegible] |
| SE | 68535 | ENSG00000131236 | CAP1 | chr1 | + | 40057445 | 40057495 | 40040779 | 40040801 | 40059336 | 40059458 | 3 | 0 | [illegible] |
| SE | 68536 | ENSG00000131236 | CAP1 | chr1 | + | 40059333 | 40059458 | 40040322 | 40040467 | 40060066 | 40060130 | 1568 | 1047 | [illegible] |
| SE | 68537 | ENSG00000131236 | CAP1 | chr1 | + | 40059336 | 40059458 | 40040322 | 40040467 | 40060066 | 40060089 | 1556 | 1037 | [illegible] |
| SE | 68538 | ENSG00000131236 | CAP1 | chr1 | + | 40064226 | 40064370 | 40061734 | 40061812 | 40066214 | 40066218 | 1380 | 826 | [illegible] |
| SE | 68539 | ENSG00000131236 | CAP1 | chr1 | + | 40064473 | 40064559 | 40064226 | 40064370 | 40066214 | 40066218 | 2251 | 1419 | [illegible] |
| SE | 68540 | ENSG00000131236 | CAP1 | chr1 | + | 40066214 | 40066320 | 40064473 | 40064559 | 40067539 | 40067556 | 1634 | 1000 | [illegible] |
| SE | 68541 | ENSG00000131236 | CAP1 | chr1 | + | 40069689 | 40069874 | 40067539 | 40067717 | 40070158 | 40070282 | 2571 | 1547 | [illegible] |
| SE | 68546 | ENSG00000141013 | GAS8 | chr16 | + | 90035632 | 90035687 | 90032717 | 90032924 | 90036380 | 90036586 | 8 | 4 | [illegible] |
| SE | 68547 | ENSG00000086717 | PPEF1 | chrX | + | 18697844 | 18697931 | 18690912 | 18691094 | 18700332 | 18700435 | 1 | 0 | [illegible] |
| SE | 68551 | ENSG00000086717 | PPEF1 | chrX | + | 18825750 | 18825835 | 18823922 | 18824086 | 18827275 | 18827547 | 0 | 0 | [illegible] |
| SE | 68552 | ENSG00000224660 | SH3BP5-AS1 | chr3 | + | 15258108 | 15258183 | 15256478 | 15257606 | 15258545 | 15260582 | 7 | 4 | [illegible] |
| SE | 68553 | ENSG00000224660 | SH3BP5-AS1 | chr3 | + | 15258108 | 15258183 | 15256478 | 15257606 | 15258633 | 15260582 | 1 | 0 | [illegible] |
| SE | 68554 | ENSG00000224660 | SH3BP5-AS1 | chr3 | + | 15258545 | 15260582 | 15257225 | 15257606 | 15264295 | 15264515 | 22 | 25 | [illegible] |
| SE | 68555 | ENSG00000224660 | SH3BP5-AS1 | chr3 | + | 15258633 | 15260582 | 15254183 | 15257606 | 15264295 | 15264493 | 21 | 23 | [illegible] |
| SE | 68558 | ENSG00000138442 | WDR12 | chr2 | - | 202894580 | 202894626 | 202892616 | 202892702 | 202896064 | 202896219 | 71 | 45 | [illegible] |
| SE | 68560 | ENSG00000138442 | WDR12 | chr2 | - | 202899530 | 202899637 | 202897299 | 202897415 | 202901024 | 202901119 | 177 | 106 | [illegible] |
| SE | 68561 | ENSG00000065802 | ASB1 | chr2 | + | 238433312 | 238433447 | 238426829 | 238427119 | 238433548 | 238433695 | 2 | 0 | [illegible] |
| SE | 68562 | ENSG00000065802 | ASB1 | chr2 | + | 238433312 | 238433447 | 238426829 | 238427119 | 238433553 | 238433695 | 0 | 0 | [illegible] |
| SE | 68566 | ENSG00000140368 | PSTPIP1 | chr15 | + | 76999460 | 76999614 | 76995411 | 76995609 | 77018147 | 77018248 | 2 | 0 | [illegible] |
| SE | 68567 | ENSG00000140368 | PSTPIP1 | chr15 | + | 77007846 | 77008065 | 76995121 | 76995609 | 77018147 | 77018248 | 1 | 2 | [illegible] |
| SE | 68568 | ENSG00000140368 | PSTPIP1 | chr15 | + | 77018147 | 77018248 | 76995125 | 76995609 | 77018456 | 77018531 | 381 | 175 | [illegible] |
| SE | 68569 | ENSG00000140368 | PSTPIP1 | chr15 | + | 77023569 | 77023667 | 77018456 | 77018531 | 77024012 | 77024427 | 3 | 0 | [illegible] |
| SE | 68573 | ENSG00000140368 | PSTPIP1 | chr15 | + | 77031179 | 77031278 | 77029528 | 77029574 | 77035507 | 77035563 | 5 | 1 | [illegible] |
| SE | 68574 | ENSG00000140368 | PSTPIP1 | chr15 | + | 77032297 | 77032394 | 77031179 | 77031278 | 77032861 | 77032952 | 156 | 61 | [illegible] |
| SE | 68575 | ENSG00000140368 | PSTPIP1 | chr15 | + | 77032297 | 77032428 | 77031179 | 77031278 | 77032861 | 77032952 | 76 | 17 | [illegible] |
| SE | 68578 | ENSG00000140368 | PSTPIP1 | chr15 | + | 77032861 | 77032952 | 77032297 | 77032394 | 77035507 | 77035563 | 166 | 78 | [illegible] |
| SE | 68580 | ENSG00000140368 | PSTPIP1 | chr15 | + | 77035801 | 77035935 | 77035507 | 77035563 | 77037044 | 77037216 | 281 | 138 | [illegible] |
| SE | 68581 | ENSG00000140368 | PSTPIP1 | chr15 | + | 77035810 | 77035935 | 77035507 | 77035563 | 77037044 | 77037331 | 161 | 85 | [illegible] |
| SE | 68586 | ENSG00000169710 | FASN | chr17 | - | 82087324 | 82087504 | 82087049 | 82087253 | 82087684 | 82087861 | 66 | 43 | [illegible] |
| SE | 68587 | ENSG00000169710 | FASN | chr17 | - | 82087953 | 82088034 | 82087684 | 82087861 | 82088115 | 82088307 | 34 | 12 | [illegible] |
| SE | 68588 | ENSG00000239779 | WBP1 | chr2 | + | 74458831 | 74458919 | 74458438 | 74458671 | 74459437 | 74459562 | 2 | 0 | [illegible] |
| SE | 68590 | ENSG00000239779 | WBP1 | chr2 | + | 74458831 | 74459117 | 74458471 | 74458671 | 74459477 | 74459562 | 2 | 0 | [illegible] |
| SE | 68592 | ENSG00000239779 | WBP1 | chr2 | + | 74458996 | 74459098 | 74458399 | 74458671 | 74459437 | 74459745 | 2 | 0 | [illegible] |
| SE | 68595 | ENSG00000239779 | WBP1 | chr2 | + | 74459437 | 74459562 | 74458831 | 74458919 | 74459642 | 74459745 | 9 | 13 | [illegible] |
| SE | 68596 | ENSG00000239779 | WBP1 | chr2 | + | 74459437 | 74459745 | 74458638 | 74458671 | 74459872 | 74460049 | 36 | 21 | [illegible] |
| SE | 68598 | ENSG00000239779 | WBP1 | chr2 | + | 74459477 | 74459562 | 74458831 | 74459117 | 74459642 | 74459745 | 5 | 4 | [illegible] |
| SE | 68600 | ENSG00000239779 | WBP1 | chr2 | + | 74459642 | 74459745 | 74458437 | 74458671 | 74459872 | 74459905 | 51 | 22 | [illegible] |
| SE | 68601 | ENSG00000239779 | WBP1 | chr2 | + | 74459651 | 74459745 | 74458468 | 74458671 | 74459872 | 74460049 | 34 | 20 | [illegible] |
| SE | 68610 | ENSG00000085978 | ATG16L1 | chr2 | + | 233264891 | 233265143 | 233256101 | 233256195 | 233270001 | 233270067 | 71 | 47 | [illegible] |
| SE | 68614 | ENSG00000085978 | ATG16L1 | chr2 | + | 233277567 | 233277673 | 233274303 | 233274778 | 233281104 | 233281114 | 70 | 41 | [illegible] |
| SE | 68616 | ENSG00000085978 | ATG16L1 | chr2 | + | 233293255 | 233293357 | 233292386 | 233292434 | 233294256 | 233294519 | 46 | 36 | [illegible] |
| SE | 68617 | ENSG00000163902 | RPN1 | chr3 | - | 128625873 | 128626012 | 128625533 | 128625653 | 128626732 | 128626832 | 955 | 475 | [illegible] |
| SE | 68619 | ENSG00000163902 | RPN1 | chr3 | - | 128631947 | 128632157 | 128629950 | 128630143 | 128637798 | 128638105 | 866 | 540 | [illegible] |
| SE | 68620 | ENSG00000163902 | RPN1 | chr3 | - | 128637798 | 128638105 | 128631947 | 128632157 | 128644918 | 128644983 | 915 | 493 | [illegible] |
| SE | 68622 | ENSG00000163902 | RPN1 | chr3 | - | 128644534 | 128644650 | 128641045 | 128641134 | 128644918 | 128644983 | 0 | 0 | [illegible] |
| SE | 68624 | ENSG00000223960 | CHROMR | chr2 | + | 178426365 | 178426505 | 178413944 | 178414113 | 178430677 | 178430889 | 56 | 32 | [illegible] |
| SE | 68633 | ENSG00000197714 | ZNF460 | chr19 | + | 57284550 | 57284677 | 57280050 | 57280124 | 57290698 | 57291045 | 76 | 41 | [illegible] |
| SE | 68639 | ENSG00000166295 | ANAPC16 | chr10 | + | 72220053 | 72220379 | 72216028 | 72216072 | 72233000 | 72235858 | 15 | 1 | [illegible] |
| SE | 68641 | ENSG00000166295 | ANAPC16 | chr10 | + | 72223887 | 72224056 | 72215999 | 72216072 | 72233000 | 72235858 | 71 | 60 | [illegible] |
| SE | 68644 | ENSG00000166295 | ANAPC16 | chr10 | + | 72223887 | 72224056 | 72220053 | 72220379 | 72233000 | 72235858 | 33 | 41 | [illegible] |
| SE | 68645 | ENSG00000166295 | ANAPC16 | chr10 | + | 72230365 | 72230440 | 72216028 | 72216072 | 72233000 | 72233615 | 127 | 56 | [illegible] |
| SE | 68646 | ENSG00000166295 | ANAPC16 | chr10 | + | 72230365 | 72230440 | 72220053 | 72220379 | 72233000 | 72233615 | 138 | 59 | [illegible] |
| SE | 68647 | ENSG00000166295 | ANAPC16 | chr10 | + | 72230365 | 72230440 | 72223132 | 72224056 | 72233000 | 72233615 | 639 | 354 | [illegible] |
| SE | 68650 | ENSG00000285733 | NA | chr6 | - | 169458220 | 169458464 | 169457533 | 169457634 | 169582835 | 169582934 | 0 | 0 | [illegible] |
| SE | 68652 | ENSG00000184465 | WDR27 | chr6 | - | 169613558 | 169613656 | 169602218 | 169602321 | 169632946 | 169633068 | 10 | 3 | [illegible] |
| SE | 68653 | ENSG00000285733 | NA | chr6 | - | 169672254 | 169672396 | 169634427 | 169634525 | 169701550 | 169702047 | 0 | 1 | [illegible] |
| SE | 68654 | ENSG00000285733 | NA | chr6 | - | 169688816 | 169689012 | 169672254 | 169672396 | 169701550 | 169702047 | 37 | 45 | [illegible] |
| SE | 68696 | ENSG00000137309 | HMGA1 | chr6 | + | 34240736 | 34240882 | 34237203 | 34237317 | 34242711 | 34242795 | 77 | 27 | [illegible] |
| SE | 68697 | ENSG00000137309 | HMGA1 | chr6 | + | 34240736 | 34240915 | 34236872 | 34237317 | 34242711 | 34242795 | 104 | 41 | [illegible] |
| SE | 68698 | ENSG00000196295 | GARS1-DT | chr7 | - | 30521763 | 30521852 | 30517205 | 30517498 | 30524311 | 30524380 | 0 | 0 | [illegible] |
| SE | 68699 | ENSG00000196295 | GARS1-DT | chr7 | - | 30551317 | 30551395 | 30550567 | 30550781 | 30552099 | 30552177 | 4 | 9 | [illegible] |
| SE | 68701 | ENSG00000196295 | GARS1-DT | chr7 | - | 30551317 | 30551395 | 30550567 | 30550781 | 30568832 | 30569821 | 1 | 10 | [illegible] |
| SE | 68702 | ENSG00000196295 | GARS1-DT | chr7 | - | 30551317 | 30551479 | 30548358 | 30550781 | 30552099 | 30552179 | 4 | 10 | [illegible] |
| SE | 68705 | ENSG00000196295 | GARS1-DT | chr7 | - | 30552099 | 30552179 | 30550320 | 30550781 | 30563730 | 30563998 | 21 | 20 | [illegible] |
| SE | 68706 | ENSG00000196295 | GARS1-DT | chr7 | - | 30552099 | 30552179 | 30551317 | 30551395 | 30561465 | 30561487 | 21 | 22 | [illegible] |
| SE | 68710 | ENSG00000196295 | GARS1-DT | chr7 | - | 30563730 | 30563998 | 30550635 | 30550781 | 30568832 | 30568989 | 77 | 63 | [illegible] |
| SE | 68712 | ENSG00000196295 | GARS1-DT | chr7 | - | 30576563 | 30576710 | 30568832 | 30568989 | 30579005 | 30579128 | 5 | 2 | [illegible] |
| SE | 68713 | ENSG00000196295 | GARS1-DT | chr7 | - | 30576563 | 30576710 | 30568832 | 30568989 | 30579230 | 30579314 | 8 | 0 | [illegible] |

| | | | | | | | | | | | | | | | | | | | | | | | | | | | | | | | | | | | | | | | | | | | | | | |
|---|---|---|---|---|---|---|---|---|---|---|---|---|---|---|---|---|---|---|---|---|---|---|---|---|---|---|---|---|---|---|---|---|---|---|---|---|---|---|---|---|---|---|---|---|---|---|
| SE | 68714 | ENSG00000196295 | GARS1-DT | chr7 | - | 30576563 | 30576710 | 30568832 | 30568989 | 30585329 | 30585459 | 2 | 1 | 0 | 0 | 1 | 3 | 2 | 1 | 8 | 0 | 0 | 5 | 3 | 0 | 0 | 8 | 6 | 1 | 0 | 0 | 296 | 149 | 1 | 1 | 0.251 | 0.201 | 0 | 0 | 1 | 0 | 0.296 | 0.602 | NA | NA | -0.009 |
| SE | 68715 | ENSG00000196295 | GARS1-DT | chr7 | - | 30576563 | 30576710 | 30574441 | 30574881 | 30577648 | 30577759 | 170 | 125 | 134 | 154 | 144 | 0 | 0 | 0 | 0 | 3 | 241 | 161 | 122 | 147 | 263 | 0 | 0 | 0 | 0 | 0 | 296 | 149 | 1 | 1 | 1 | 1 | 1 | 1 | 0.96 | 1 | 1 | 1 | 1 | 1 | -0.008 |
| SE | 68716 | ENSG00000196295 | GARS1-DT | chr7 | - | 30576563 | 30576710 | 30574441 | 30574881 | 30579230 | 30579314 | 108 | 72 | 73 | 85 | 71 | 0 | 0 | 0 | 0 | 0 | 141 | 121 | 55 | 93 | 132 | 0 | 0 | 0 | 2 | 0 | 296 | 149 | 1 | 1 | 1 | 1 | 1 | 1 | 1 | 1 | 1 | 1 | 0.959 | 1 | 0.008 |
| SE | 68717 | ENSG00000196295 | GARS1-DT | chr7 | - | 30576563 | 30576710 | 30574441 | 30574881 | 30585329 | 30585459 | 102 | 73 | 73 | 85 | 71 | 0 | 0 | 2 | 0 | 0 | 141 | 122 | 55 | 89 | 132 | 0 | 0 | 0 | 0 | 0 | 296 | 149 | 1 | 1 | 1 | 1 | 0.948 | 1 | 1 | 1 | 1 | 1 | 1 | 1 | -0.01 |
| SE | 68718 | ENSG00000196295 | GARS1-DT | chr7 | - | 30577648 | 30578091 | 30574441 | 30574881 | 30579230 | 30579314 | 41 | 29 | 46 | 73 | 67 | 0 | 0 | 0 | 0 | 0 | 71 | 92 | 27 | 24 | 68 | 0 | 0 | 0 | 2 | 0 | 298 | 149 | 1 | 1 | 1 | 1 | 1 | 1 | 1 | 1 | 1 | 1 | 0.857 | 1 | 0.029 |
| SE | 68720 | ENSG00000196295 | GARS1-DT | chr7 | - | 30579005 | 30579128 | 30566421 | 30568989 | 30579230 | 30579314 | 10 | 14 | 4 | 14 | 21 | 0 | 0 | 0 | 2 | 0 | 17 | 20 | 21 | 26 | 79 | 0 | 0 | 0 | 0 | 0 | 272 | 149 | 1 | 1 | 1 | 1 | 1 | 0.793 | 1 | 1 | 1 | 1 | 1 | 1 | -0.041 |
| SE | 68721 | ENSG00000196295 | GARS1-DT | chr7 | - | 30579005 | 30579128 | 30566421 | 30568989 | 30585329 | 30585456 | 0 | 0 | 0 | 1 | 0 | 3 | 2 | 1 | 8 | 0 | 0 | 0 | 0 | 2 | 0 | 8 | 6 | 1 | 0 | 0 | 272 | 149 | 1 | 1 | 0 | 0 | 0 | 0.064 | NA | 0 | 0 | 0 | 1 | NA | -0.234 |
| SE | 68723 | ENSG00000196295 | GARS1-DT | chr7 | - | 30579005 | 30579128 | 30576563 | 30576710 | 30585329 | 30585456 | 3 | 2 | 0 | 0 | 0 | 0 | 1 | 0 | 0 | 0 | 0 | 6 | 0 | 10 | 0 | 0 | 5 | 0 | 0 | 0 | 272 | 149 | 1 | 1 | 1 | 0.523 | NA | NA | NA | NA | 0.397 | NA | 1 | NA | 0.063 |
| SE | 68725 | ENSG00000196295 | GARS1-DT | chr7 | - | 30579230 | 30579314 | 30566421 | 30568989 | 30584422 | 30584453 | 3 | 2 | 0 | 3 | 4 | 1 | 0 | 0 | 0 | 0 | 8 | 10 | 8 | 5 | 4 | 0 | 0 | 1 | 0 | 0 | 233 | 149 | 1 | 1 | 0.657 | 1 | NA | 1 | 1 | 1 | 0.865 | 1 | 1 | 1 | -0.059 |
| SE | 68729 | ENSG00000196295 | GARS1-DT | chr7 | - | 30579230 | 30579314 | 30574422 | 30574881 | 30585329 | 30585456 | 24 | 6 | 18 | 27 | 17 | 0 | 0 | 2 | 0 | 0 | 20 | 30 | 18 | 44 | 46 | 0 | 0 | 0 | 0 | 0 | 233 | 149 | 1 | 1 | 1 | 1 | 0.852 | 1 | 1 | 1 | 1 | 1 | 1 | 1 | -0.03 |
| SE | 68731 | ENSG00000196295 | GARS1-DT | chr7 | - | 30579230 | 30579314 | 30579005 | 30579128 | 30585329 | 30585456 | 40 | 25 | 23 | 46 | 43 | 0 | 0 | 0 | 0 | 0 | 45 | 59 | 45 | 82 | 110 | 0 | 0 | 0 | 2 | 0 | 233 | 149 | 1 | 1 | 1 | 1 | 1 | 1 | 1 | 1 | 1 | 1 | 0.963 | 1 | 0.007 |
| SE | 68732 | ENSG00000196295 | GARS1-DT | chr7 | - | 30584422 | 30584453 | 30568676 | 30568989 | 30584710 | 30584808 | 3 | 0 | 0 | 1 | 0 | 0 | 0 | 0 | 2 | 0 | 0 | 0 | 0 | 0 | 0 | 16 | 14 | 0 | 4 | 0 | 180 | 149 | 1 | 1 | 1 | NA | NA | 0.293 | NA | 0 | 0 | NA | 0 | NA | 0.646 |
| SE | 68733 | ENSG00000196295 | GARS1-DT | chr7 | - | 30584422 | 30584453 | 30568676 | 30568989 | 30585329 | 30585479 | 1 | 0 | 0 | 0 | 0 | 3 | 2 | 1 | 8 | 0 | 0 | 0 | 0 | 0 | 0 | 8 | 6 | 1 | 0 | 0 | 180 | 149 | 1 | 1 | 0.216 | 0 | 0 | 0 | NA | 0 | 0 | 0 | NA | NA | 0.054 |
| SE | 68739 | ENSG00000196295 | GARS1-DT | chr7 | - | 30585329 | 30585479 | 30566403 | 30568989 | 30594710 | 30594740 | 63 | 55 | 52 | 95 | 59 | 0 | 0 | 0 | 0 | 3 | 83 | 110 | 62 | 86 | 113 | 0 | 0 | 0 | 0 | 0 | 298 | 149 | 1 | 1 | 1 | 1 | 1 | 1 | 0.908 | 1 | 1 | 1 | 1 | 1 | -0.018 |
| SE | 68748 | ENSG00000196141 | SPATS2L | chr2 | + | 200372108 | 200372251 | 200329430 | 200329480 | 200389222 | 200389283 | 0 | 3 | 1 | 1 | 0 | 5 | 13 | 8 | 19 | 8 | 0 | 0 | 0 | 5 | 0 | 67 | 19 | 23 | 40 | 10 | 292 | 149 | 1 | 1 | 0 | 0.105 | 0.06 | 0.026 | 0 | 0 | 0 | 0 | 0.06 | 0 | 0.026 |
| SE | 68750 | ENSG00000196141 | SPATS2L | chr2 | + | 200378140 | 200378371 | 200329430 | 200329480 | 200389222 | 200389283 | 0 | 0 | 0 | 2 | 0 | 5 | 13 | 8 | 19 | 8 | 0 | 3 | 1 | 2 | 0 | 67 | 19 | 23 | 40 | 10 | 298 | 149 | 1 | 1 | 0 | 0 | 0 | 0.05 | 0 | 0 | 0.073 | 0.021 | 0.024 | 0 | -0.014 |
| SE | 68752 | ENSG00000196141 | SPATS2L | chr2 | + | 200383866 | 200384017 | 200378140 | 200378371 | 200389222 | 200389283 | 5 | 12 | 6 | 8 | 4 | 0 | 0 | 0 | 2 | 0 | 12 | 24 | 11 | 26 | 0 | 0 | 3 | 1 | 1 | 0 | 298 | 149 | 1 | 1 | 1 | 1 | 1 | 0.667 | 1 | 1 | 0.8 | 0.846 | 0.929 | NA | 0.04 |
| SE | 68754 | ENSG00000196141 | SPATS2L | chr2 | + | 200416378 | 200416428 | 200412310 | 200412419 | 200419249 | 200419333 | 38 | 54 | 36 | 52 | 29 | 0 | 0 | 0 | 0 | 0 | 228 | 93 | 53 | 195 | 44 | 0 | 0 | 1 | 0 | 0 | 199 | 149 | 1 | 1 | 1 | 1 | 1 | 1 | 1 | 1 | 0.986 | 1 | 1 | 1 | 0.003 |
| SE | 68757 | ENSG00000284691 | NA | chr7 | + | 150407583 | 150408488 | 150405145 | 150405317 | 150410159 | 150410249 | 4 | 0 | 1 | 2 | 0 | 0 | 0 | 0 | 0 | 0 | 0 | 0 | 1 | 0 | 0 | 0 | 0 | 2 | 0 | 0 | 298 | 149 | 1 | 1 | 1 | NA | 1 | 1 | NA | NA | 0 | 1 | NA | NA | 0.5 |
| SE | 68758 | ENSG00000163872 | YEATS2 | chr3 | + | 183715143 | 183715262 | 183697796 | 183697993 | 183717650 | 183717748 | 61 | 43 | 45 | 71 | 67 | 1 | 0 | 0 | 0 | 0 | 57 | 28 | 34 | 72 | 22 | 0 | 0 | 0 | 0 | 0 | 268 | 149 | 1 | 1 | 0.971 | 1 | 1 | 1 | 1 | 1 | 1 | 1 | 1 | 1 | -0.006 |
| SE | 68759 | ENSG00000163872 | YEATS2 | chr3 | + | 183721890 | 183722136 | 183718499 | 183718592 | 183724418 | 183724531 | 244 | 179 | 194 | 198 | 297 | 0 | 0 | 0 | 0 | 2 | 227 | 168 | 121 | 239 | 158 | 0 | 0 | 0 | 0 | 0 | 298 | 149 | 1 | 1 | 1 | 1 | 1 | 1 | 0.987 | 1 | 1 | 1 | 1 | 1 | -0.003 |
| SE | 68764 | ENSG00000163872 | YEATS2 | chr3 | + | 183756527 | 183756689 | 183754125 | 183754365 | 183758861 | 183758965 | 129 | 83 | 105 | 134 | 131 | 0 | 0 | 0 | 0 | 0 | 100 | 76 | 64 | 87 | 172 | 0 | 0 | 0 | 2 | 0 | 298 | 149 | 1 | 1 | 1 | 1 | 1 | 1 | 1 | 1 | 1 | 1 | 0.956 | 1 | 0.009 |
| SE | 68765 | ENSG00000163872 | YEATS2 | chr3 | + | 183758861 | 183758965 | 183756527 | 183756689 | 183761506 | 183761614 | 104 | 67 | 82 | 111 | 126 | 0 | 0 | 0 | 0 | 0 | 61 | 64 | 53 | 76 | 134 | 0 | 0 | 2 | 0 | 0 | 253 | 149 | 1 | 1 | 1 | 1 | 1 | 1 | 1 | 1 | 1 | 0.94 | 1 | 1 | 0.012 |
| SE | 68766 | ENSG00000163872 | YEATS2 | chr3 | + | 183761506 | 183761614 | 183758861 | 183758965 | 183762096 | 183762279 | 95 | 58 | 72 | 103 | 97 | 0 | 0 | 0 | 1 | 0 | 76 | 79 | 52 | 94 | 76 | 0 | 0 | 0 | 0 | 0 | 257 | 149 | 1 | 1 | 1 | 1 | 1 | 0.984 | 1 | 1 | 1 | 1 | 1 | 1 | -0.003 |
| SE | 68773 | ENSG00000100523 | DDHD1 | chr14 | - | 53091784 | 53091932 | 53073740 | 53073847 | 53103682 | 53103856 | 129 | 139 | 114 | 116 | 144 | 0 | 2 | 0 | 1 | 0 | 103 | 89 | 135 | 68 | 46 | 0 | 0 | 1 | 0 | 0 | 297 | 149 | 1 | 1 | 1 | 0.972 | 1 | 0.983 | 1 | 1 | 1 | 0.985 | 1 | 1 | -0.006 |
| SE | 68774 | ENSG00000100523 | DDHD1 | chr14 | - | 53092472 | 53092675 | 53091784 | 53091932 | 53093315 | 53093444 | 0 | 0 | 0 | 0 | 0 | 161 | 160 | 107 | 147 | 197 | 0 | 2 | 0 | 0 | 0 | 59 | 87 | 96 | 41 | 45 | 298 | 149 | 1 | 1 | 0 | 0 | 0 | 0 | 0 | 0 | 0.011 | 0 | 0 | 0 | -0.002 |
| SE | 68776 | ENSG00000100523 | DDHD1 | chr14 | - | 53096116 | 53096218 | 53093315 | 53093444 | 53103030 | 53103051 | 28 | 30 | 23 | 15 | 33 | 1 | 2 | 11 | 6 | 10 | 0 | 12 | 7 | 1 | 0 | 0 | 0 | 5 | 0 | 0 | 251 | 149 | 1 | 1 | 0.943 | 0.899 | 0.554 | 0.597 | 0.662 | NA | 1 | 0.454 | 1 | NA | -0.087 |
| SE | 68779 | ENSG00000100523 | DDHD1 | chr14 | - | 53103030 | 53103051 | 53096116 | 53096218 | 53103682 | 53103856 | 2 | 3 | 4 | 0 | 2 | 5 | 3 | 14 | 5 | 14 | 0 | 0 | 1 | 0 | 0 | 0 | 0 | 2 | 5 | 0 | 170 | 149 | 1 | 1 | 0.26 | 0.467 | 0.2 | 0 | 0.111 | NA | NA | 0.305 | 0 | NA | 0.055 |
| SE | 68783 | ENSG00000137947 | GTF2B | chr1 | - | 88857205 | 88857487 | 88852632 | 88853346 | 88859881 | 88860011 | 351 | 228 | 278 | 323 | 564 | 1 | 0 | 0 | 0 | 0 | 294 | 469 | 476 | 338 | 162 | 0 | 0 | 0 | 0 | 0 | 298 | 149 | 1 | 1 | 0.994 | 1 | 1 | 1 | 1 | 1 | 1 | 1 | 1 | 1 | -0.001 |

[illegible]

| | | | | | | | | | | | | | | | | | |
|---|---|---|---|---|---|---|---|---|---|---|---|---|---|---|---|---|---|---|---|---|---|---|---|
| SE | 69108 | ENSG00000105374 | NKG7 | chr19 | - | 51372160 | 51372202 | 51371619 | 51371835 | 51372378 | 51372410 | 18 | 2 | 2 | 3 | 6 | 5 | 0 | 2 | 0 | 6 | 0 | 16 |
| SE | 69111 | ENSG00000148341 | SH3GLB2 | chr9 | - | 129009105 | 129009346 | 129007035 | 129008791 | 129009770 | 129009871 | 83 | 44 | 92 | 75 | 80 | 0 | 0 | 0 | 0 | 0 | 80 | 84 |
| SE | 69113 | ENSG00000148341 | SH3GLB2 | chr9 | - | 129010669 | 129010693 | 129010119 | 129010209 | 129012235 | 129012298 | 145 | 71 | 135 | 132 | 208 | 3 | 0 | 0 | 0 | 0 | 215 | 172 |
| SE | 69114 | ENSG00000148341 | SH3GLB2 | chr9 | - | 129010669 | 129010693 | 129010119 | 129010209 | 129014410 | 129014503 | 12 | 8 | 24 | 19 | 18 | 0 | 0 | 1 | 0 | 0 | 16 | 5 |
| SE | 69115 | ENSG00000148341 | SH3GLB2 | chr9 | - | 129012235 | 129012298 | 129010119 | 129010209 | 129014410 | 129014503 | 26 | 25 | 50 | 27 | 18 | 0 | 0 | 1 | 0 | 0 | 26 | 44 |
| SE | 69117 | ENSG00000148341 | SH3GLB2 | chr9 | - | 129013007 | 129013019 | 129012235 | 129012298 | 129014410 | 129014503 | 0 | 0 | 2 | 0 | 0 | 162 | 109 | 173 | 175 | 195 | 0 | 0 |
| SE | 69118 | ENSG00000148341 | SH3GLB2 | chr9 | - | 129014410 | 129014503 | 129012229 | 129012298 | 129014770 | 129014904 | 234 | 149 | 243 | 254 | 301 | 0 | 0 | 0 | 0 | 3 | 331 | 257 |
| SE | 69119 | ENSG00000148341 | SH3GLB2 | chr9 | - | 129015739 | 129015860 | 129014770 | 129014904 | 129021090 | 129021211 | 1 | 0 | 0 | 2 | 4 | 92 | 46 | 93 | 82 | 73 | 0 | 0 |
| SE | 69121 | ENSG00000148341 | SH3GLB2 | chr9 | - | 129021090 | 129021219 | 129014770 | 129014904 | 129022281 | 129022423 | 150 | 71 | 127 | 135 | 113 | 0 | 0 | 0 | 0 | 0 | 150 | 114 |
| SE | 69122 | ENSG00000148341 | SH3GLB2 | chr9 | - | 129021090 | 129021219 | 129014770 | 129014904 | 129028091 | 129028185 | 85 | 43 | 89 | 82 | 72 | 0 | 0 | 0 | 0 | 0 | 96 | 83 |
| SE | 69123 | ENSG00000148341 | SH3GLB2 | chr9 | - | 129022281 | 129022423 | 129014770 | 129014904 | 129028091 | 129028185 | 19 | 15 | 16 | 19 | 21 | 0 | 0 | 0 | 0 | 0 | 8 | 20 |
| SE | 69124 | ENSG00000148341 | SH3GLB2 | chr9 | - | 129022281 | 129022423 | 129021090 | 129021219 | 129028091 | 129028185 | 84 | 43 | 54 | 74 | 59 | 0 | 0 | 0 | 2 | 0 | 62 | 48 |
| SE | 69127 | ENSG00000175505 | CLCF1 | chr11 | - | 67367459 | 67367626 | 67364167 | 67365630 | 67373523 | 67373598 | 4 | 3 | 5 | 3 | 0 | 0 | 0 | 1 | 0 | 0 | 16 | 3 |
| SE | 69128 | ENSG00000155252 | PI4K2A | chr10 | + | 97650940 | 97651141 | 97640685 | 97641177 | 97656284 | 97656416 | 61 | 7 | 46 | 85 | 49 | 0 | 0 | 0 | 2 | 0 | 122 | 62 |
| SE | 69129 | ENSG00000155252 | PI4K2A | chr10 | + | 97656284 | 97656416 | 97650940 | 97651141 | 97656820 | 97656974 | 96 | 11 | 62 | 123 | 94 | 0 | 0 | 1 | 0 | 0 | 165 | 92 |
| SE | 69130 | ENSG00000155252 | PI4K2A | chr10 | + | 97656284 | 97656416 | 97650940 | 97651141 | 97662906 | 97662968 | 40 | 5 | 35 | 53 | 41 | 0 | 0 | 0 | 0 | 0 | 79 | 48 |
| SE | 69131 | ENSG00000155252 | PI4K2A | chr10 | + | 97656820 | 97656974 | 97650940 | 97651141 | 97662906 | 97662968 | 44 | 7 | 33 | 36 | 36 | 0 | 0 | 0 | 0 | 0 | 26 | 44 |
| SE | 69132 | ENSG00000155252 | PI4K2A | chr10 | + | 97656820 | 97656974 | 97656284 | 97656416 | 97662906 | 97662968 | 100 | 13 | 59 | 106 | 90 | 0 | 0 | 0 | 0 | 0 | 112 | 88 |
| SE | 69133 | ENSG00000155252 | PI4K2A | chr10 | + | 97662906 | 97662968 | 97656820 | 97656974 | 97666437 | 97666571 | 15 | 0 | 11 | 9 | 10 | 0 | 0 | 0 | 0 | 0 | 7 | 5 |
| SE | 69135 | ENSG00000155252 | PI4K2A | chr10 | + | 97666437 | 97666571 | 97664884 | 97664984 | 97667060 | 97667120 | 104 | 50 | 97 | 100 | 107 | 0 | 0 | 0 | 0 | 0 | 143 | 126 |
| SE | 69136 | ENSG00000155252 | PI4K2A | chr10 | + | 97667060 | 97667120 | 97666437 | 97666571 | 97673580 | 97676434 | 88 | 38 | 78 | 79 | 84 | 0 | 0 | 0 | 0 | 0 | 108 | 96 |
| SE | 69148 | ENSG00000169499 | PLEKHA2 | chr8 | + | 38917906 | 38918070 | 38901797 | 38901892 | 38935993 | 38936050 | 187 | 95 | 181 | 203 | 183 | 1 | 0 | 2 | 3 | 1 | 235 | 223 |
| SE | 69149 | ENSG00000169499 | PLEKHA2 | chr8 | + | 38935993 | 38936050 | 38917906 | 38918070 | 38943788 | 38943837 | 231 | 154 | 235 | 256 | 241 | 0 | 0 | 0 | 0 | 0 | 256 | 295 |
| SE | 69153 | ENSG00000169499 | PLEKHA2 | chr8 | + | 38946123 | 38946221 | 38943788 | 38943837 | 38950849 | 38950990 | 421 | 254 | 375 | 413 | 639 | 1 | 0 | 0 | 0 | 0 | 478 | 550 |
| SE | 69155 | ENSG00000169499 | PLEKHA2 | chr8 | + | 38957322 | 38957386 | 38953296 | 38953367 | 38968591 | 38968669 | 536 | 266 | 370 | 422 | 535 | 6 | 0 | 0 | 1 | 2 | 719 | 652 |
| SE | 69156 | ENSG00000169499 | PLEKHA2 | chr8 | + | 38968591 | 38968669 | 38957322 | 38957386 | 38969420 | 38970227 | 498 | 231 | 329 | 371 | 512 | 2 | 0 | 2 | 0 | 2 | 606 | 587 |
| SE | 69157 | ENSG00000172757 | CFL1 | chr11 | - | 65855653 | 65855730 | 65854672 | 65855448 | 65855934 | 65856242 | 2643 | 1263 | 1957 | 2348 | 2502 | 0 | 0 | 1 | 1 | 0 | 3490 | 3114 |
| SE | 69158 | ENSG00000172757 | CFL1 | chr11 | - | 65858096 | 65858342 | 65855973 | 65856242 | 65859230 | 65859282 | 714 | 265 | 533 | 621 | 340 | 0 | 0 | 1 | 0 | 0 | 955 | 1114 |
| SE | 69159 | ENSG00000172757 | CFL1 | chr11 | - | 65858096 | 65858630 | 65855934 | 65856242 | 65859230 | 65859415 | 714 | 265 | 533 | 621 | 340 | 0 | 0 | 1 | 0 | 0 | 955 | 1114 |
| SE | 69162 | ENSG00000132530 | XAF1 | chr17 | + | 6758861 | 6758959 | 6758088 | 6758224 | 6759255 | 6759718 | 0 | 0 | 0 | 0 | 0 | 13 | 10 | 5 | 5 | 9 | 0 | 0 |
| SE | 69163 | ENSG00000132530 | XAF1 | chr17 | + | 6758861 | 6758959 | 6758088 | 6758224 | 6759661 | 6759718 | 2 | 0 | 0 | 1 | 0 | 75 | 76 | 38 | 88 | 83 | 0 | 0 |
| SE | 69165 | ENSG00000132530 | XAF1 | chr17 | + | 6759255 | 6759718 | 6758088 | 6758224 | 6760405 | 6760601 | 39 | 45 | 17 | 36 | 39 | 2 | 1 | 1 | 0 | 7 | 26 | 32 |
| SE | 69170 | ENSG00000132530 | XAF1 | chr17 | + | 6759840 | 6759881 | 6758088 | 6758224 | 6760405 | 6760601 | 0 | 0 | 0 | 0 | 0 | 2 | 1 | 1 | 0 | 7 | 0 | 0 |
| SE | 69171 | ENSG00000132530 | XAF1 | chr17 | + | 6759840 | 6759881 | 6759661 | 6759718 | 6760405 | 6760601 | 0 | 0 | 0 | 1 | 0 | 67 | 63 | 26 | 75 | 55 | 0 | 0 |
| SE | 69173 | ENSG00000132530 | XAF1 | chr17 | + | 6760405 | 6760601 | 6759255 | 6759718 | 6762154 | 6762206 | 127 | 120 | 55 | 166 | 146 | 1 | 0 | 0 | 0 | 0 | 78 | 132 |
| SE | 69183 | ENSG00000113712 | CSNK1A1 | chr5 | - | 149507026 | 149507133 | 149504845 | 149505595 | 149509878 | 149509953 | 1153 | 994 | 1032 | 1192 | 1900 | 0 | 0 | 0 | 0 | 1 | 838 | 807 |
| SE | 69184 | ENSG00000113712 | CSNK1A1 | chr5 | - | 149509878 | 149509953 | 149507026 | 149507133 | 149511793 | 149511872 | 954 | 722 | 735 | 893 | 1463 | 0 | 0 | 0 | 0 | 4 | 661 | 685 |
| SE | 69185 | ENSG00000113712 | CSNK1A1 | chr5 | - | 149511065 | 149511158 | 149509878 | 149509953 | 149511793 | 149511872 | 0 | 0 | 0 | 0 | 6 | 407 | 269 | 298 | 360 | 504 | 0 | 0 |
| SE | 69188 | ENSG00000113712 | CSNK1A1 | chr5 | - | 149515668 | 149515726 | 149513069 | 149513209 | 149520289 | 149520388 | 0 | 1 | 3 | 3 | 1 | 895 | 673 | 749 | 833 | 1165 | 0 | 6 |
| SE | 69190 | ENSG00000113712 | CSNK1A1 | chr5 | - | 149517793 | 149517877 | 149515668 | 149515726 | 149520289 | 149520388 | 28 | 18 | 26 | 21 | 39 | 0 | 1 | 1 | 2 | 0 | 12 | 49 |
| SE | 69191 | ENSG00000113712 | CSNK1A1 | chr5 | - | 149520289 | 149520388 | 149513069 | 149513209 | 149525044 | 149525171 | 1216 | 929 | 984 | 1156 | 1797 | 1 | 6 | 2 | 0 | 0 | 1107 | 1003 |
| SE | 69193 | ENSG00000113712 | CSNK1A1 | chr5 | - | 149525044 | 149525171 | 149520289 | 149520388 | 149550074 | 149550181 | 1243 | 886 | 1004 | 1114 | 1503 | 2 | 0 | 4 | 0 | 2 | 1254 | 1132 |
| SE | 69194 | ENSG00000113712 | CSNK1A1 | chr5 | - | 149529603 | 149529719 | 149525044 | 149525171 | 149531393 | 149531511 | 0 | 0 | 0 | 2 | 0 | 0 | 0 | 1 | 1 | 0 | 0 | 0 |
| SE | 69195 | ENSG00000113712 | CSNK1A1 | chr5 | - | 149529603 | 149529719 | 149525044 | 149525171 | 149550074 | 149550181 | 0 | 0 | 0 | 1 | 0 | 565 | 344 | 488 | 480 | 498 | 0 | 0 |
| SE | 69197 | ENSG00000113712 | CSNK1A1 | chr5 | - | 149531393 | 149531511 | 149525044 | 149525171 | 149550074 | 149550181 | 0 | 0 | 1 | 2 | 0 | 565 | 344 | 488 | 480 | 498 | 0 | 0 |
| SE | 69199 | ENSG00000113712 | CSNK1A1 | chr5 | - | 149550074 | 149550181 | 149525044 | 149525171 | 149550841 | 149551168 | 764 | 469 | 648 | 680 | 671 | 4 | 0 | 0 | 0 | 0 | 738 | 712 |
| SE | 69201 | ENSG00000115694 | STK25 | chr2 | - | 241497299 | 241497378 | 241496397 | 241496534 | 241497615 | 241497687 | 0 | 0 | 0 | 1 | 0 | 19 | 17 | 28 | 22 | 8 | 0 | 0 |
| SE | 69203 | ENSG00000115694 | STK25 | chr2 | - | 241500739 | 241500796 | 241499406 | 241500281 | 241501477 | 241501698 | 107 | 49 | 91 | 100 | 86 | 0 | 0 | 0 | 0 | 0 | 121 | 108 |
| SE | 69205 | ENSG00000115694 | STK25 | chr2 | - | 241500739 | 241500847 | 241500259 | 241500281 | 241501477 | 241501799 | 47 | 20 | 40 | 48 | 31 | 0 | 0 | 0 | 0 | 0 | 67 | 34 |
| SE | 69206 | ENSG00000115694 | STK25 | chr2 | - | 241500739 | 241501077 | 241500172 | 241500281 | 241501477 | 241501526 | 51 | 20 | 46 | 50 | 31 | 0 | 0 | 0 | 0 | 0 | 76 | 34 |
| SE | 69214 | ENSG00000115694 | STK25 | chr2 | - | 241508005 | 241508135 | 241501356 | 241501708 | 241508442 | 241508556 | 13 | 11 | 13 | 20 | 2 | 0 | 0 | 1 | 1 | 0 | 19 | 12 |
| SE | 69216 | ENSG00000119396 | RAB14 | chr9 | - | 121186952 | 121187019 | 121183310 | 121183398 | 121201638 | 121201869 | 645 | 545 | 526 | 732 | 736 | 1 | 0 | 0 | 0 | 0 | 1140 | 895 |
| SE | 69225 | ENSG00000115484 | CCT4 | chr2 | - | 61872081 | 61872316 | 61869439 | 61869553 | 61873001 | 61873112 | 115 | 56 | 124 | 98 | 204 | 1 | 0 | 3 | 0 | 1 | 58 | 71 |
| [illegible] | | | | | | | | | | | | | | | | | | | | | | | |

| | | | | | | | | | | | | | | | | | | | | | | | | | | | | | | | | | | | | | | | | | | | | | | |
|---|---|---|---|---|---|---|---|---|---|---|---|---|---|---|---|---|---|---|---|---|---|---|---|---|---|---|---|---|---|---|---|---|---|---|---|---|---|---|---|---|---|---|---|---|---|---|
| SE | 69568 | ENSG00000177479 | ARIH2 | chr3 | + | 48980352 | 48980496 | 48979481 | 48979633 | 48981659 | 48981728 | 421 | 207 | 269 | 444 | 426 | 0 | 0 | 1 | 0 | 0 | 470 | 374 | 276 | 375 | 670 | 0 | 0 | 0 | 0 | 0 | 293 | 149 | 1 | 1 | 1 | 1 | 0.993 | 1 | 1 | 1 | 1 | 1 | 1 | 1 | -0.001 |
| SE | 69569 | ENSG00000177479 | ARIH2 | chr3 | + | 48981659 | 48981728 | 48979481 | 48979633 | 48982895 | 48982979 | 98 | 48 | 59 | 90 | 46 | 0 | 0 | 1 | 0 | 0 | 84 | 109 | 62 | 76 | 144 | 0 | 0 | 0 | 0 | 0 | 218 | 149 | 1 | 1 | 1 | 0.97 | 1 | 1 | 1 | 1 | 1 | 1 | 1 | 1 | -0.006 |
| SE | 69570 | ENSG00000177479 | ARIH2 | chr3 | + | 48982895 | 48982979 | 48981659 | 48981728 | 48983198 | 48983620 | 203 | 125 | 134 | 215 | 159 | 0 | 1 | 0 | 0 | 0 | 221 | 208 | 144 | 187 | 303 | 0 | 0 | 0 | 0 | 0 | 233 | 149 | 1 | 1 | 1 | 0.988 | 1 | 1 | 1 | 1 | 1 | 1 | 1 | 1 | -0.002 |
| SE | 69571 | ENSG00000134262 | AP4B1 | chr1 | - | 113895756 | 113896038 | 113894193 | 113895492 | 113896257 | 113896465 | 79 | 29 | 63 | 78 | 58 | 0 | 0 | 1 | 0 | 2 | 21 | 68 | 32 | 49 | 27 | 0 | 0 | 0 | 0 | 0 | 298 | 149 | 1 | 1 | 1 | 1 | 0.969 | 1 | 0.935 | 1 | 1 | 1 | 1 | 1 | -0.019 |
| SE | 69572 | ENSG00000134262 | AP4B1 | chr1 | - | 113895756 | 113896038 | 113894193 | 113895492 | 113898717 | 113898801 | 43 | 20 | 40 | 53 | 32 | 0 | 0 | 0 | 0 | 0 | 14 | 36 | 33 | 27 | 12 | 0 | 0 | 2 | 0 | 0 | 298 | 149 | 1 | 1 | 1 | 1 | 1 | 1 | 1 | 1 | 1 | 0.892 | 1 | 1 | 0.022 |
| SE | 69575 | ENSG00000134262 | AP4B1 | chr1 | - | 113898717 | 113898801 | 113897839 | 113897943 | 113899903 | 113900245 | 119 | 66 | 118 | 139 | 97 | 2 | 0 | 0 | 0 | 0 | 134 | 84 | 73 | 96 | 70 | 0 | 0 | 0 | 1 | 0 | 233 | 149 | 1 | 1 | 0.974 | 1 | 1 | 1 | 1 | 1 | 1 | 1 | 0.984 | 1 | -0.002 |
| SE | 69576 | ENSG00000134262 | AP4B1 | chr1 | - | 113899903 | 113900400 | 113898717 | 113898801 | 113901235 | 113901383 | 144 | 73 | 119 | 103 | 134 | 0 | 0 | 0 | 0 | 0 | 118 | 124 | 111 | 153 | 97 | 0 | 0 | 0 | 3 | 0 | 298 | 149 | 1 | 1 | 1 | 1 | 1 | 1 | 1 | 1 | 1 | 1 | 0.962 | 1 | 0.008 |
| SE | 69577 | ENSG00000134262 | AP4B1 | chr1 | - | 113900269 | 113900400 | 113898717 | 113898801 | 113901235 | 113901383 | 69 | 34 | 51 | 34 | 86 | 0 | 0 | 0 | 0 | 0 | 45 | 71 | 49 | 93 | 40 | 0 | 0 | 0 | 3 | 0 | 280 | 149 | 1 | 1 | 1 | 1 | 1 | 1 | 1 | 1 | 1 | 1 | 0.943 | 1 | 0.011 |
| SE | 69578 | ENSG00000134262 | AP4B1 | chr1 | - | 113901235 | 113901383 | 113899903 | 113900400 | 113901754 | 113901885 | 165 | 129 | 131 | 111 | 173 | 0 | 0 | 0 | 3 | 0 | 130 | 181 | 175 | 214 | 116 | 0 | 0 | 4 | 1 | 0 | 297 | 149 | 1 | 1 | 1 | 1 | 1 | 0.949 | 1 | 1 | 1 | 0.956 | 0.991 | 1 | 0 |
| SE | 69580 | ENSG00000134262 | AP4B1 | chr1 | - | 113901754 | 113901885 | 113899941 | 113900400 | 113904604 | 113904793 | 4 | 0 | 0 | 3 | 0 | 0 | 0 | 0 | 4 | 0 | 0 | 0 | 5 | 1 | 0 | 0 | 0 | 0 | 0 | 0 | 280 | 149 | 1 | 1 | 1 | NA | NA | 0.285 | NA | NA | NA | 1 | 1 | NA | -0.358 |
| SE | 69583 | ENSG00000134262 | AP4B1 | chr1 | - | 113902637 | 113902862 | 113899903 | 113900400 | 113904604 | 113904793 | 12 | 2 | 11 | 19 | 7 | 0 | 0 | 0 | 4 | 0 | 0 | 5 | 10 | 3 | 26 | 0 | 0 | 0 | 0 | 0 | 298 | 149 | 1 | 1 | 1 | 1 | 1 | 0.704 | 1 | NA | | 1 | 1 | 1 | -0.059 |
| SE | 69584 | ENSG00000134262 | AP4B1 | chr1 | - | 113902637 | 113902862 | 113901754 | 113901885 | 113904604 | 113904793 | 38 | 10 | 29 | 46 | 22 | 4 | 0 | 0 | 0 | 0 | 32 | 49 | 31 | 16 | 78 | 0 | 0 | 1 | 0 | 0 | 298 | 149 | 1 | 1 | 0.826 | 1 | 1 | 1 | 1 | 1 | 1 | 0.939 | 1 | 1 | -0.023 |
| SE | 69587 | ENSG00000157379 | DHRS1 | chr14 | - | 24296737 | 24296881 | 24296508 | 24296588 | 24298956 | 24299130 | 75 | 58 | 53 | 58 | 53 | 0 | 1 | 2 | 0 | 0 | 107 | 37 | 71 | 65 | 87 | 0 | 0 | 1 | 0 | 0 | 293 | 149 | 1 | 1 | 1 | 0.967 | 0.931 | 1 | 1 | 1 | 1 | 0.973 | 1 | 1 | -0.015 |
| SE | 69588 | ENSG00000135390 | ATP5MC2 | chr12 | - | 53672575 | 53672645 | 53669870 | 53670114 | 53676052 | 53676082 | 7 | 2 | 7 | 4 | 0 | 1 | 0 | 0 | 0 | 0 | 2 | 9 | 8 | 7 | 0 | 0 | 0 | 0 | 0 | 0 | 219 | 149 | 1 | 1 | 0.826 | 1 | 1 | 1 | NA | 1 | 1 | 1 | 1 | NA | -0.043 |
| SE | 69589 | ENSG00000135390 | ATP5MC2 | chr12 | - | 53673695 | 53673812 | 53672575 | 53672645 | 53676175 | 53677408 | 1 | 0 | 0 | 0 | 0 | 0 | 0 | 2 | 0 | 0 | 0 | 0 | 0 | 5 | 0 | 0 | 0 | 0 | 0 | 0 | 266 | 149 | 1 | 1 | 1 | NA | 0 | NA | NA | NA | NA | NA | 1 | NA | -0.5 |
| SE | 69592 | ENSG00000138036 | DYNC2LI1 | chr2 | + | 43787180 | 43787250 | 43783519 | 43783554 | 43789632 | 43789721 | 7 | 22 | 13 | 18 | 23 | 0 | 0 | 1 | 0 | 0 | 0 | 0 | 5 | 10 | 0 | 0 | 0 | 0 | 0 | 0 | 219 | 149 | 1 | 1 | 1 | 1 | 0.898 | 1 | 1 | NA | NA | 1 | 1 | NA | -0.02 |
| SE | 69593 | ENSG00000138036 | DYNC2LI1 | chr2 | + | 43795886 | 43795958 | 43794456 | 43794643 | 43796717 | 43796795 | 8 | 9 | 7 | 8 | 24 | 0 | 0 | 0 | 1 | 0 | 0 | 0 | 2 | 6 | 0 | 0 | 0 | 0 | 0 | 0 | 221 | 149 | 1 | 1 | 1 | 1 | 1 | 1 | 0.942 | NA | NA | 1 | 1 | NA | -0.012 |
| SE | 69595 | ENSG00000138036 | DYNC2LI1 | chr2 | + | 43795889 | 43795958 | 43794456 | 43794643 | 43796717 | 43796795 | 9 | 7 | 8 | 8 | 29 | 0 | 0 | 0 | 0 | 1 | 0 | 2 | 9 | 4 | 0 | 0 | 0 | 0 | 0 | 0 | 218 | 149 | 1 | 1 | 1 | 1 | 1 | 1 | 0.952 | NA | 1 | 1 | 1 | NA | -0.01 |
| SE | 69596 | ENSG00000138036 | DYNC2LI1 | chr2 | + | 43795889 | 43795958 | 43794456 | 43794643 | 43800840 | 43800917 | 2 | 0 | 3 | 2 | 9 | 0 | 0 | 0 | 1 | 0 | 0 | 2 | 5 | 0 | 0 | 0 | 0 | 0 | 0 | 0 | 218 | 149 | 1 | 1 | 1 | NA | 1 | 0.578 | 1 | NA | 1 | 1 | NA | NA | -0.106 |
| SE | 69597 | ENSG00000138036 | DYNC2LI1 | chr2 | + | 43796717 | 43796795 | 43794456 | 43794643 | 43800840 | 43800917 | 11 | 8 | 1 | 8 | 22 | 0 | 0 | 0 | 1 | 0 | 0 | 4 | 0 | 5 | 0 | 0 | 0 | 0 | 0 | 0 | 227 | 149 | 1 | 1 | 1 | 1 | 1 | 0.84 | 1 | NA | 1 | NA | 1 | NA | -0.032 |
| SE | 69598 | ENSG00000138036 | DYNC2LI1 | chr2 | + | 43796717 | 43796795 | 43795886 | 43795958 | 43800840 | 43800917 | 20 | 17 | 9 | 15 | 56 | 0 | 0 | 1 | 0 | 0 | 0 | 4 | 4 | 15 | 0 | 0 | 0 | 0 | 0 | 0 | 227 | 149 | 1 | 1 | 1 | 1 | 0.855 | 1 | 1 | NA | 1 | 1 | 1 | NA | -0.029 |
| SE | 69599 | ENSG00000184922 | FMNL1 | chr17 | + | 45230603 | 45230687 | 45221884 | 45222253 | 45232366 | 45232480 | 251 | 110 | 217 | 161 | 104 | 0 | 0 | 0 | 0 | 0 | 311 | 228 | 167 | 327 | 481 | 0 | 1 | 0 | 0 | 0 | 233 | 149 | 1 | 1 | 1 | 1 | 1 | 1 | 1 | 1 | 0.993 | 1 | 1 | 1 | 0.001 |
| SE | 69600 | ENSG00000184922 | FMNL1 | chr17 | + | 45233647 | 45233731 | 45233223 | 45233297 | 45234071 | 45234200 | 829 | 322 | 721 | 749 | 521 | 0 | 0 | 0 | 0 | 0 | 1035 | 789 | 582 | 777 | 1359 | 0 | 0 | 0 | 2 | 0 | 233 | 149 | 1 | 1 | 1 | 1 | 1 | 1 | 1 | 1 | 1 | 1 | 0.996 | 1 | 0.001 |
| SE | 69603 | ENSG00000184922 | FMNL1 | chr17 | + | 45243117 | 45243320 | 45242340 | 45242465 | 45243790 | 45244025 | 1035 | 430 | 683 | 756 | 629 | 0 | 0 | 0 | 0 | 0 | 1235 | 903 | 728 | 1072 | 1449 | 0 | 0 | 0 | 2 | 0 | 298 | 149 | 1 | 1 | 1 | 1 | 1 | 1 | 1 | 1 | 1 | 1 | 0.996 | 1 | 0.001 |
| SE | 69608 | ENSG00000163412 | EIF4E3 | chr3 | - | 71693874 | 71693941 | 71690009 | 71690165 | 71696459 | 71696520 | 415 | 417 | 380 | 482 | 1001 | 0 | 0 | 0 | 0 | 0 | 1055 | 578 | 690 | 919 | 496 | 0 | 1 | 5 | 12 | 0 | 216 | 149 | 1 | 1 | 1 | 1 | 1 | 1 | 1 | 1 | 0.997 | 0.99 | 0.981 | 1 | 0.006 |
| SE | 69610 | ENSG00000163412 | EIF4E3 | chr3 | - | 71696459 | 71696520 | 71690009 | 71690165 | 71710411 | 71710484 | 0 | 0 | 1 | 1 | 0 | 1 | 2 | 0 | 0 | 0 | 0 | 1 | 6 | 5 | 0 | 0 | 0 | 1 | 0 | 6 | 210 | 149 | 1 | 1 | 0 | 0 | 1 | 1 | NA | NA | 1 | 0.81 | 1 | 0 | -0.203 |
| SE | 69611 | ENSG00000163412 | EIF4E3 | chr3 | - | 71696459 | 71696520 | 71690009 | 71690165 | 71725191 | 71725402 | 0 | 0 | 0 | 0 | 0 | 1 | 1 | 0 | 0 | 0 | 0 | 1 | 5 | 5 | 0 | 0 | 0 | 3 | 0 | 0 | 210 | 149 | 1 | 1 | 0 | 0 | NA | NA | NA | | 1 | 0.542 | 1 | NA | -0.847 |
| SE | 69612 | ENSG00000163412 | EIF4E3 | chr3 | - | 71696459 | 71696520 | 71693874 | 71693941 | 71699613 | 71699708 | 310 | 324 | 260 | 379 | 672 | 5 | 0 | 2 | 2 | 0 | 704 | 428 | 496 | 659 | 332 | 0 | 0 | 0 | 0 | 6 | 210 | 149 | 1 | 1 | 0.978 | 1 | 0.989 | 0.993 | 1 | 1 | 1 | 1 | 1 | 0.975 | -0.003 |
| SE | 69616 | ENSG00000163412 | EIF4E3 | chr3 | - | 71699613 | 71699708 | 71696459 | 71696520 | 71725191 | 71725402 | 139 | 124 | 84 | 145 | 250 | 0 | 0 | 0 | 0 | 0 | 265 | 149 | 140 | 173 | 152 | 0 | 0 | 0 | 5 | 0 | 244 | 149 | 1 | 1 | 1 | 1 | 1 | 1 | 1 | 1 | 1 | 1 | 0.955 | 1 | 0.009 |
| SE | 69617 | ENSG00000163412 | EIF4E3 | chr3 | - | 71710411 | 71710484 | 71696459 | 71696520 | 71725191 | 71725402 | 3 | 5 | 4 | 5 | 3 | 0 | 0 | 0 | 0 | 0 | 0 | 3 | 1 | 0 | 0 | 0 | 0 | 0 | 5 | 0 | 222 | 149 | 1 | 1 | 1 | 1 | 1 | 1 | 1 | NA | 1 | 1 | 0 | NA | 0.333 |
| SE | 69618 | ENSG00000163412 | EIF4E3 | chr3 | - | 71728475 | 71728623 | 71710411 | 71710484 | 71754643 | 71754773 | 3 | 6 | 0 | 1 | 1 | 1 | 0 | 0 | 0 | 0 | 0 | 1 | 13 | 4 | 4 | 0 | 0 | 0 | 0 | 0 | 297 | 149 | 1 | 1 | 0.601 | 1 | NA | 1 | 1 | NA | 1 | 1 | 1 | 1 | -0.1 |
| SE | 69620 | ENSG00000156711 | MAPK13 | chr6 | + | 36131270 | 36131400 | 36130483 | 36130701 | 36135752 | 36135832 | 9 | 5 | 19 | 19 | 1 | 0 | 0 | 0 | 0 | 0 | 8 | 13 | 11 | 9 | 6 | 0 | 0 | 2 | 0 | 0 | 279 | 149 | 1 | 1 | 1 | 1 | 1 | 1 | 1 | 1 | 1 | 0.746 | 1 | 1 | 0.051 |
| SE | 69621 | ENSG00000156711 | MAPK13 | chr6 | + | 36132620 | 36132679 | 36131270 | 36131400 | 36135752 | 36135832 | 82 | 24 | 80 | 72 | 33 | 2 | 0 | 0 | 1 | 0 | 27 | 71 | 46 | 66 | 101 | 0 | 0 | 2 | 2 | 0 | 208 | 149 | 1 | 1 | 0.967 | 1 | 1 | 0.981 | 1 | 1 | 1 | 0.943 | 0.959 | 1 | 0.009 |
| SE | 69624 | ENSG00000183617 | MRPL54 | chr19 | + | 3765165 | 3765331 | 3762681 | 3762818 | 3767260 | 3767565 | 125 | 58 | 104 | 126 | 87 | 1 | 1 | 0 | 0 | 0 | 115 | 130 | 113 | 110 | 151 | 0 | 0 | 0 | 0 | 0 | 298 | 149 | 1 | 1 | 0.984 | 0.967 | 1 | 1 | 1 | 1 | 1 | 1 | 1 | 1 | -0.01 |
| SE | 69627 | ENSG00000197622 | CDC42SE1 | chr1 | - | 151055015 | 151055126 | 151054230 | 151054321 | 151059478 | 151059615 | 786 | 320 | 660 | 621 | 1267 | 2 | 1 | 0 | 0 | 2 | 371 | 545 | 352 | 427 | 495 | 0 | 0 | 0 | 0 | 0 | 260 | 149 | 1 | 1 | 0.996 | 0.995 | 1 | 1 | 0.997 | 1 | 1 | 1 | 1 | 1 | -0.002 |
| SE | 69628 | ENSG00000197622 | CDC42SE1 | chr1 | - | 151055015 | 151055993 | 151054230 | 151054321 | 151059478 | 151059574 | 1434 | 662 | 1271 | 1199 | 1525 | 2 | 1 | 0 | 0 | 2 | 910 | 1234 | 912 | 847 | 1278 | 0 | 0 | 0 | 0 | 0 | 298 | 149 | 1 | 1 | 0.997 | 0.997 | 1 | 1 | 0.997 | 1 | 1 | 1 | 1 | 1 | -0.002 |
| SE | 69629 | ENSG00000197622 | CDC42SE1 | chr1 | - | 151055676 | 151055993 | 151054230 | 151054321 | 151059478 | 151059622 | 640 | 338 | 587 | 566 | 215 | 2 | 1 | 0 | 0 | 2 | 551 | 695 | 563 | 408 | 751 | 0 | 0 | 0 | 0 | 0 | 298 | 149 | 1 | 1 | 0.994 | 0.994 | 1 | 1 | 0.982 | 1 | 1 | 1 | 1 | 1 | -0.006 |
| SE | 69630 | ENSG00000197622 | CDC42SE1 | chr1 | - | 151055676 | 151055993 | 151055015 | 151055126 | 151056655 | 151056786 | 1354 | 702 | 1046 | 1037 | 1220 | 9 | 0 | 5 | 0 | 0 | 953 | 1073 | 842 | 773 | 989 | 0 | 5 | 6 | 3 | 0 | 298 | 149 | 1 | 1 | 0.987 | 1 | 0.991 | 1 | 1 | 1 | 0.991 | 0.986 | 0.992 | 1 | 0.002 |
| SE | 69639 | ENSG00000120451 | SNX19 | chr11 | - | 130903254 | 130903384 | 130880621 | 130880806 | 130905952 | 130905999 | 146 | 96 | 126 | 121 | 205 | 0 | 0 | 0 | 0 | 0 | 163 | 146 | 90 | 106 | 175 | 0 | 0 | 1 | 0 | 0 | 279 | 149 | 1 | 1 | 1 | 1 | 1 | 1 | 1 | 1 | 1 | 0.98 | 1 | 1 | 0.004 |
| SE | 69643 | ENSG00000258181 | NA | chr12 | + | 47249325 | 47249428 | 47249135 | 47249231 | 47257461 | 47257539 | 6 | 6 | 11 | 3 | 17 | 5 | 2 | 2 | 5 | 13 | 0 | 0 | 2 | 0 | 0 | 9 | 0 | 1 | 1 | 0 | 252 | 149 | 1 | 1 | 0.415 | 0.639 | 0.765 | 0.262 | 0.436 | 0 | NA | 0.542 | 0 | NA | 0.323 |
| SE | 69650 | ENSG00000143390 | RFX5 | chr1 | - | 151343680 | 151343882 | 151343341 | 151343442 | 151344196 | 151344278 | 168 | 102 | 159 | 181 | 157 | 0 | 0 | 0 | 0 | 0 | 195 | 183 | 123 | 223 | 171 | 0 | 0 | 2 | 5 | 0 | 298 | 149 | 1 | 1 | 1 | 1 | 1 | 1 | 1 | 1 | 1 | 0.969 | 0.957 | 1 | 0.015 |
| SE | 69651 | ENSG00000143390 | RFX5 | chr1 | - | 151344727 | 151344847 | 151344332 | 151344536 | 151345105 | 151345177 | 191 | 82 | 160 | 133 | 202 | 0 | 0 | 0 | 0 | 0 | 185 | 200 | 75 | 132 | 264 | 0 | 0 | 0 | 4 | 0 | 269 | 149 | 1 | 1 | 1 | 1 | 1 | 1 | 1 | 1 | 1 | 1 | 0.948 | 1 | 0.01 |
| SE | 69655 | ENSG00000143390 | RFX5 | chr1 | - | 151345927 | 151345961 | 151344988 | 151345188 | 151346214 | 151346333 | 28 | 11 | 22 | 25 | 31 | 0 | 0 | 0 | 0 | 0 | 4 | 18 | 15 | 15 | 25 | 0 | 0 | 0 | 2 | 0 | 183 | 149 | 1 | 1 | 1 | 1 | 1 | 1 | 1 | 1 | 1 | 1 | 0.859 | 1 | 0.028 |
| SE | 69662 | ENSG00000184708 | EIF4ENIF1 | chr22 | - | 31448152 | 31448232 | 31447425 | 31447568 | 31449347 | 31449383 | 48 | 26 | 42 | 53 | 44 | 0 | 0 | 1 | 0 | 0 | 34 | 62 | 28 | 48 | 81 | 0 | 0 | 0 | 0 | 0 | 229 | 149 | 1 | 1 | 1 | 1 | 0.965 | 1 | 1 | 1 | 1 | 1 | 1 | 1 | -0.007 |
| SE | 69663 | ENSG00000184708 | EIF4ENIF1 | chr22 | - | 31450288 | 31450360 | 31449347 | 31449408 | 31454143 | 31454376 | 35 | 33 | 32 | 25 | 92 | 0 | 0 | 0 | 0 | 0 | 26 | 25 | 12 | 26 | 53 | 0 | 0 | 0 | 4 | 0 | 221 | 149 | 1 | 1 | 1 | 1 | 1 | 1 | 1 | 1 | 1 | 1 | 0.814 | 1 | 0.037 |
| SE | 69665 | ENSG00000184708 | EIF4ENIF1 | chr22 | - | 31455851 | 31455987 | 31455135 | 31455315 | 31462931 | 31463133 | 100 | 54 | 80 | 109 | 159 | 0 | 0 | 0 | 1 | 0 | 137 | 102 | 66 | 64 | 134 | 0 | 0 | 0 | 0 | 0 | 285 | 149 | 1 | 1 | 1 | 1 | 1 | 0.983 | 1 | 1 | 1 | 1 | 1 | 1 | -0.003 |
| SE | 69667 | ENSG00000184708 | EIF4ENIF1 | chr22 | - | 31462931 | 31463133 | 31458474 | 31458650 | 31463680 | 31463967 | 103 | 86 | 91 | 66 | 86 | 0 | 0 | 1 | 0 | 0 | 112 | 80 | 52 | 86 | 124 | 0 | 0 | 0 | 0 | 0 | 298 | 149 | 1 | 1 | 1 | 1 | 0.978 | 1 | 1 | 1 | 1 | 1 | 1 | 1 | -0.004 |
| SE | 69668 | ENSG00000184708 | EIF4ENIF1 | chr22 | - | 31462931 | 31463133 | 31458474 | 31458650 | 31468174 | 31468302 | 29 | 23 | 26 | 23 | 30 | 0 | 1 | 0 | 0 | 0 | 42 | 11 | 12 | 28 | 46 | 0 | 0 | 0 | 0 | 0 | 298 | 149 | 1 | 1 | 1 | 0.92 | 1 | 1 | 1 | 1 | 1 | 1 | 1 | 1 | -0.016 |
| SE | 69669 | ENSG00000184708 | EIF4ENIF1 | chr22 | - | 31463680 | 31463967 | 31458474 | 31458650 | 31468174 | 31468302 | 55 | 42 | 53 | 70 | 58 | 0 | 1 | 0 | 0 | 0 | 38 | 80 | 50 | 56 | 64 | 0 | 0 | 0 | 0 | 0 | 298 | 149 | 1 | 1 | 1 | 0.955 | 1 | 1 | 1 | 1 | 1 | 1 | 1 | 1 | -0.009 |
| SE | 69671 | ENSG00000184708 | EIF4ENIF1 | chr22 | - | 31463680 | 31463967 | 31462931 | 31463133 | 31471843 | 31471917 | 74 | 63 | 65 | 47 | 60 | 0 | 0 | 0 | 0 | 0 | 73 | 69 | 41 | 68 | 84 | 0 | 0 | 0 | 3 | 0 | 298 | 149 | 1 | 1 | 1 | 1 | 1 | 1 | 1 | 1 | 1 | 1 | 0.919 | 1 | 0.016 |
| SE | 69672 | ENSG00000184708 | EIF4ENIF1 | chr22 | - | 31468174 | 31468302 | 31462931 | 31463133 | 31471843 | 31471917 | 39 | 42 | 51 | 52 | 66 | 0 | 0 | 0 | 0 | 0 | 71 | 63 | 34 | 75 | 72 | 0 | 0 | 0 | 3 | 0 | 277 | 149 | 1 | 1 | 1 | 1 | 1 | 1 | 1 | 1 | 1 | 1 | 0.931 | 1 | 0.014 |
| SE | 69673 | ENSG00000184708 | EIF4ENIF1 | chr22 | - | 31468174 | 31468302 | 31463680 | 31463967 | 31471843 | 31471917 | 93 | 84 | 103 | 118 | 124 | 0 | 0 | 0 | 2 | 4 | 106 | 143 | 83 | 126 | 130 | 0 | 0 | 0 | 5 | 0 | 277 | 149 | 1 | 1 | 1 | 1 | 1 | 0.969 | 0.943 | 1 | 1 | 1 | 0.931 | 1 | -0.004 |
| SE | 69676 | ENSG00000004897 | CDC27 | chr17 | - | 47122443 | 47122600 | 47117702 | 47121017 | 47129392 | 47129521 | 72 | 65 | 67 | 84 | 116 | 0 | 0 | 0 | 0 | 1 | 0 | 29 | 29 | 9 | 0 | 0 | 0 | 0 | 0 | 0 | 298 | 149 | 1 | 1 | 1 | 1 | 1 | 1 | 0.935 | NA | 1 | 1 | 1 | NA | -0.013 |
| SE | 69677 | ENSG00000004897 | CDC27 | chr17 | - | 47123885 | 47123960 | 47122443 | 47122600 | 47129392 | 47129521 | 75 | 50 | 64 | 84 | 60 | 4 | 4 | 4 | 4 | 2 | 0 | 49 | 61 | 30 | 0 | 0 | 3 | 2 | 0 | 0 | 224 | 149 | 1 | 1 | 0.926 | 0.893 | 0.914 | 0.933 | 0.952 | NA | 0.916 | 0.953 | 1 | NA | -0.033 |
| SE | 69678 | ENSG00000004897 | CDC27 | chr17 | - | 47132256 | 47132374 | 47129392 | 47129521 | 47137151 | 47137244 | 220 | 129 | 154 | 257 | 385 | 0 | 0 | 0 | 2 | 4 | 57 | 147 | 113 | 57 | 7 | 0 | 2 | 0 | 0 | 0 | 267 | 149 | 1 | 1 | 1 | 1 | 1 | 0.986 | 0.982 | 1 | 0.976 | 1 | 1 | 1 | -0.002 |
| SE | 69679 | ENSG00000004897 | CDC27 | chr17 | - | 47132256 | 47132374 | 47129392 | 47129521 | 47138738 | 47138891 | 86 | 60 | 78 | 107 | 209 | 0 | 0 | 1 | 0 | 0 | 19 | 56 | 46 | 26 | 0 | 0 | 0 | 0 | 0 | 0 | 267 | 149 | 1 | 1 | 1 | 1 | 0.978 | 1 | 1 | 1 | 1 | 1 | 1 | NA | -0.004 |
| SE | 69680 | ENSG00000004897 | CDC27 | chr17 | - | 47137151 | 47137360 | 47129392 | 47129521 | 47138738 | 47138891 | 137 | 125 | 119 | 137 | 196 | 0 | 0 | 1 | 0 | 0 | 53 | 122 | 140 | 64 | 17 | 0 | 0 | 0 | 0 | 0 | 298 | 149 | 1 | 1 | 1 | 1 | 0.983 | 1 | 1 | 1 | 1 | 1 | 1 | 1 | -0.003 |
| SE | 69681 | ENSG00000004897 | CDC27 | chr17 | - | 47137151 | 47137360 | 47132256 | 47132374 | 47138738 | 47138891 | 272 | 194 | 196 | 285 | 371 | 0 | 0 | 0 | 0 | 2 | 91 | 222 | 208 | 95 | 24 | 0 | 0 | 1 | 0 | 0 | 298 | 149 | 1 | 1 | 1 | 1 | 1 | 1 | 0.989 | 1 | 1 | 0.99 | 1 | 1 | 0 |
| SE | 69682 | ENSG00000004897 | CDC27 | chr17 | - | 47139770 | 47139973 | 47138738 | 47138891 | 47141852 | 47142022 | 4 | 0 | 0 | 1 | 2 | 126 | 115 | 117 | 101 | 186 | 0 | 0 | 0 | 0 | 0 | 38 | 53 | 108 | 56 | 21 | 298 | 149 | 1 | 1 | 0.016 | 0 | 0 | 0.005 | 0.005 | 0 | 0 | 0 | 0 | 0 | 0.005 |
| SE | 69686 | ENSG00000004897 | CDC27 | chr17 | - | 47151805 | 47151918 | 47143882 | 47143982 | 47154671 | 47154786 | 294 | 241 | 238 | 253 | 885 | 0 | 1 | 0 | 0 | 0 | 75 | 223 | 250 | 86 | 49 | 0 | 0 | 0 | 0 | 0 | 262 | 149 | 1 | 1 | 1 | 0.993 | 1 | 1 | 1 | 1 | 1 | 1 | 1 | 1 | -0.001 |
| SE | 69687 | ENSG00000004897 | CDC27 | chr17 | - | 47151805 | 47151936 | 47143882 | 47143982 | 47154671 | 47154786 | 123 | 103 | 135 | 116 | 372 | 0 | 1 | 0 | 0 | 0 | 33 | 85 | 119 | 34 | 11 | 0 | 0 | 0 | 0 | 0 | 280 | 149 | 1 | 1 | 1 | 0.982 | 1 | 1 | 1 | 1 | 1 | 1 | 1 | 1 | -0.004 |
| SE | 69688 | ENSG00000004897 | CDC27 | chr17 | - | 47156912 | 47156983 | 47154697 | 47154786 | 47157229 | 47157384 | 197 | 161 | 103 | 184 | 375 | 1 | 1 | 0 | 2 | 2 | 75 | 201 | 170 | 77 | 2 | 0 | 0 | 1 | 0 | 0 | 220 | 149 | 1 | 1 | 0.993 | 0.991 | 1 | 0.984 | 0.992 | 1 | 1 | 0.991 | 1 | 1 | -0.006 |
| SE | 69689 | ENSG00000004897 | CDC27 | chr17 | - | 47156912 | 47157124 | 47154671 | 47154786 | 47157229 | 47157267 | 369 | 304 | 249 | 356 | 977 | 1 | 1 | 0 | 2 | 2 | 208 | 377 | 318 | 132 | 36 | 0 | 0 | 1 | 0 | 0 | 298 | 149 | 1 | 1 | 0.995 | 0.993 | 1 | 0.989 | 0.996 | 1 | 1 | 0.994 | 1 | 1 | -0.004 |
| SE | 69691 | ENSG00000004897 | CDC27 | chr17 | - | 47158205 | 47158303 | 47154671 | 47154786 | 47181561 | 47181637 | 0 | 3 | 0 | 1 | 1 | 0 | 0 | 0 | 0 | 0 | 0 | 4 | 0 | 0 | 0 | 0 | 0 | 2 | 0 | 0 | 247 | 149 | 1 | 1 | NA | 1 | NA | 1 | 1 | NA | 1 | 0 | NA | NA | 0.5 |
| SE | 69692 | ENSG00000004897 | CDC27 | chr17 | - | 47158205 | 47158303 | 47157229 | 47157384 | 47169916 | 47169977 | 337 | 281 | 250 | 363 | 556 | 0 | 0 | 1 | 0 | 3 | 211 | 350 | 259 | 139 | 39 | 0 | 1 | 1 | 0 | 0 | 247 | 149 | 1 | 1 | 1 | 1 | 0.993 | 1 | 0.991 | 1 | 0.995 | 0.994 | 1 | 1 | -0.001 |
| SE | 69693 | ENSG00000004897 | CDC27 | chr17 | - | 47169916 | 47170042 | 47158205 | 47158303 | 47171916 | 47172047 | 290 | 218 | 247 | 286 | 378 | 0 | 0 | 0 | 2 | 0 | 141 | 250 | 155 | 109 | 32 | 0 | 0 | 0 | 0 | 0 | 275 | 149 | 1 | 1 | 1 | 1 | 1 | 0.987 | 1 | 1 | 1 | 1 | 1 | 1 | -0.003 |
| SE | 69697 | ENSG00000198901 | PRC1 | chr15 | - | 90969078 | 90969120 | 90966039 | 90967202 | 90969446 | 90969623 | 0 | 1 | 0 | 3 | 2 | 0 | 0 | 0 | 1 | 0 | 0 | 0 | 5 | 1 | 0 | 0 | 8 | 5 | 9 | 0 | 191 | 149 | 1 | 1 | NA | 1 | NA | 0.701 | 1 | NA | 0 | 0.438 | 0.08 | NA | 0.728 |
| SE | 69698 | ENSG00000198901 | PRC1 | chr15 | - | 90969078 | 90969120 | 90966039 | 90967202 | 90969523 | 90969623 | 0 | 1 | 0 | 3 | 0 | 0 | 1 | 1 | 0 | 0 | 0 | 0 | 10 | 0 | 0 | 0 | 0 | 4 | 0 | 0 | 191 | 149 | 1 | 1 | NA | 0.438 | 0 | 1 | NA | NA | NA | 0.661 | NA | NA | -0.182 |
| SE | 69699 | ENSG00000198901 | PRC1 | chr15 | - | 90969078 | 90969120 | 90966464 | 90966653 | 90969446 | 90969623 | 0 | 0 | 0 | 0 | 0 | 0 | 0 | 2 | 0 | 0 | 0 | 0 | 0 | 1 | 0 | 0 | 0 | 1 | 0 | 0 | 191 | 149 | 1 | 1 | NA | NA | 0 | NA | NA | NA | NA | 0 | 1 | NA | -0.5 |
| SE | 69703 | ENSG00000198901 | PRC1 | chr15 | - | 90984017 | 90984140 | 90981498 | 90981981 | 90994406 | 90994537 | 6 | 6 | 5 | 7 | 12 | 0 | 0 | 0 | 0 | 0 | 16 | 20 | 46 | 99 | 11 | 0 | 0 | 0 | 3 | 0 | 272 | 149 | 1 | 1 | 1 | 1 | 1 | 1 | 1 | 1 | 1 | 1 | 0.948 | 1 | 0.01 |
| SE | 69704 | ENSG00000198901 | PRC1 | chr15 | - | 90984692 | 90984825 | 90981747 | 90981981 | 90994406 | 90994500 | 6 | 1 | 3 | 8 | 7 | 0 | 0 | 0 | 0 | 0 | 41 | 9 | 43 | 30 | 8 | 0 | 0 | 0 | 3 | 0 | 282 | 149 | 1 | 1 | 1 | 1 | 1 | 1 | 1 | 1 | 1 | 1 | 0.841 | 1 | 0.032 |
| SE | 69714 | ENSG00000177674 | AGTRAP | chr1 | + | 11747439 | 11747545 | 11736083 | 11736235 | 11750076 | 11750125 | 2 | 1 | 0 | 0 | 0 | 1 | 0 | 0 | 0 | 0 | 0 | 0 | 0 | 5 | 0 | 0 | 0 | 0 | 0 | 0 | 255 | 149 | 1 | 1 | 0.539 | 1 | NA | NA | NA | NA | NA | NA | 1 | NA | -0.23 |
| SE | 69717 | ENSG00000177674 | AGTRAP | chr1 | + | 11748414 | 11748610 | 11747439 | 11747545 | 11750076 | 11750125 | 110 | 35 | 55 | 79 | 17 | 2 | 1 | 0 | 0 | 0 | 106 | 93 | 30 | 125 | 176 | 0 | 0 | 0 | 5 | 0 | 298 | 149 | 1 | 1 | 0.965 | 0.946 | 1 | 1 | 1 | 1 | 1 | 1 | 0.926 | 1 | -0.003 |
| SE | 69718 | ENSG00000170312 | CDK1 | chr10 | + | 60788059 | 60788230 | 60785663 | 60785787 | 60791889 | 60791969 | 18 | 5 | 53 | 36 | 50 | 0 | 0 | 0 | 0 | 6 | 0 | 38 | 55 | 16 | 16 | 0 | 0 | 2 | 0 | 0 | 298 | 149 | 1 | 1 | 1 | 1 | 1 | 1 | 0.806 | NA | 1 | 0.932 | 1 | 1 | -0.022 |
| SE | 69720 | ENSG00000173559 | NABP1 | chr2 | + | 191708811 | 191708928 | 191685598 | 191693416 | 191709733 | 191709783 | 15 | 15 | 13 | 4 | 7 | 0 | 1 | 0 | 0 | 0 | 0 | 0 | 20 | 6 | 0 | 0 | 0 | 0 | 0 | 0 | 266 | 149 | 1 | 1 | 1 | 0.894 | 1 | 1 | 1 | NA | NA | 1 | 1 | NA | -0.021 |
| SE | 69721 | ENSG00000173559 | NABP1 | chr2 | + | 191708811 | 191708928 | 191685598 | 191693416 | 191735915 | 191736150 | 10 | 13 | 10 | 4 | 7 | 2 | 0 | 0 | 0 | 0 | 0 | 0 | 13 | 5 | 0 | 0 | 0 | 0 | 0 | 0 | 266 | 149 | 1 | 1 | 0.737 | 1 | 1 | 1 | 1 | NA | NA | 1 | 1 | NA | -0.053 |
| SE | 69723 | ENSG00000173559 | NABP1 | chr2 | + | 191709733 | 191709783 | 191708811 | 191708928 | 191735915 | 191736150 | 5 | 1 | 2 | 1 | 0 | 0 | 1 | 0 | 1 | 0 | 0 | 0 | 4 | 1 | 0 | 0 | 0 | 0 | 0 | 0 | 199 | 149 | 1 | 1 | 1 | 0.428 | 1 | 0.428 | NA | NA | NA | 1 | 1 | NA | -0.286 |
| SE | 69724 | ENSG00000078596 | ITM2A | chrX | - | 79363422 | 79363554 | 79362941 | 79363139 | 79366856 | 79366927 | 136 | 66 | 86 | 127 | 78 | 2 | 0 | 0 | 0 | 0 | 27 | 110 | 83 | 55 | 75 | 0 | 0 | 0 | 0 | 0 | 281 | 149 | 1 | 1 | 0.973 | 1 | 1 | 1 | 1 | 1 | 1 | 1 | 1 | 1 | -0.005 |
| SE | 69727 | ENSG00000130347 | RTN4IP1 | chr6 | - | 106587678 | 106587862 | 106583327 | 106583420 | 106592163 | 106592300 | 14 | 13 | 25 | 15 | 11 | 0 | 0 | 0 | 0 | 0 | 27 | 17 | 7 | 7 | 0 | 0 | 0 | 0 | 5 | 0 | 298 | 149 | 1 | 1 | 1 | 1 | 1 | 1 | 1 | 1 | 1 | 1 | 0.412 | NA | 0.147 |
| SE | 69728 | ENSG00000130347 | RTN4IP1 | chr6 | - | 106621424 | 106621493 | 106619201 | 106619326 | 106622817 | 106622969 | 21 | 4 | 16 | 14 | 17 | 0 | 2 | 0 | 0 | 3 | 10 | 7 | 4 | 5 | 0 | 0 | 0 | 2 | 0 | 0 | 218 | 149 | 1 | 1 | 1 | 0.578 | 1 | 1 | 0.795 | 1 | 1 | 0.578 | 1 | NA | -0.02 |
| SE | 69736 | ENSG00000082126 | MPP4 | chr2 | - | 201687290 | 201687371 | 201685918 | 201686050 | 201690201 | 201690279 | 0 | 0 | 0 | 1 | 0 | 0 | 0 | 0 | 0 | 0 | 0 | 0 | 0 | 0 | 0 | 0 | 3 | 0 | 0 | 0 | 230 | 149 | 1 | 1 | NA | NA | NA | 1 | NA | NA | 0 | NA | NA | NA | 1 |
| SE | 69737 | ENSG00000149782 | PLCB3 | chr11 | + | 64255233 | 64255313 | 64254897 | 64255038 | 64255395 | 64255449 | 17 | 26 | 26 | 25 | 21 | 0 | 1 | 0 | 0 | 0 | 49 | 28 | 22 | 55 | 106 | 0 | 0 | 0 | 0 | 0 | 229 | 149 | 1 | 1 | 1 | 0.944 | 1 | 1 | 1 | 1 | 1 | 1 | 1 | 1 | -0.011 |
| SE | 69738 | ENSG00000149782 | PLCB3 | chr11 | + | 64258472 | 64258713 | 64256617 | 64256764 | 64258884 | 64258969 | 48 | 21 | 15 | 40 | 14 | 0 | 0 | 0 | 0 | 0 | 49 | 13 | 11 | 44 | 72 | 0 | 0 | 0 | 0 | 4 | 298 | 149 | 1 | 1 | 1 | 1 | 1 | 1 | 1 | 1 | 1 | 1 | 1 | 0.9 | 0.02 |
| SE | 69740 | ENSG00000149782 | PLCB3 | chr11 | + | 64264020 | 64264112 | 64263690 | 64263795 | 64264950 | 64265006 | 19 | 9 | 18 | 19 | 12 | 0 | 0 | 0 | 0 | 2 | 29 | 20 | 6 | 24 | 60 | 0 | 0 | 0 | 3 | 0 | 241 | 149 | 1 | 1 | 1 | 1 | 1 | 1 | 0.788 | 1 | 1 | 1 | 0.832 | 1 | -0.009 |
| SE | 69741 | ENSG00000149782 | PLCB3 | chr11 | + | 64266494 | 64266552 | 64266314 | 64266404 | 64267184 | 64267271 | 28 | 27 | 23 | 51 | 34 | 0 | 1 | 2 | 0 | 0 | 21 | 36 | 23 | 52 | 62 | 0 | 2 | 0 | 0 | 0 | 207 | 149 | 1 | 1 | 1 | 0.951 | 0.892 | 1 | 1 | 1 | 0.928 | 1 | 1 | 1 | -0.017 |
| SE | 69746 | ENSG00000144228 | SPOPL | chr2 | + | 138550902 | 138551054 | 138550156 | 138550294 | 138559021 | 138559199 | 4 | 1 | 1 | 0 | 0 | 0 | 0 | 0 | 0 | 0 | 0 | 4 | 6 | 4 | 4 | 0 | 4 | 5 | 0 | 0 | 298 | 149 | 1 | 1 | 1 | 1 | 1 | NA | NA | NA | 0.333 | 0.375 | 1 | 1 | 0.323 |
| SE | 69747 | ENSG00000144228 | SPOPL | chr2 | + | 138550902 | 138551053 | 138550482 | 138550604 | 138552553 | 138552681 | 296 | 246 | 186 | 246 | 274 | 0 | 0 | 0 | 1 | 0 | 39 | 171 | 206 | 70 | 35 | 0 | 0 | 0 | 0 | 0 | 298 | 149 | 1 | 1 | 1 | 1 | 1 | 0.992 | 1 | 1 | 1 | 1 | 1 | 1 | -0.002 |
| SE | 69749 | ENSG00000144228 | SPOPL | chr2 | + | 138550973 | 138551054 | 138550482 | 138550604 | 138552553 | 138552582 | 87 | 62 | 68 | 68 | 95 | 0 | 0 | 0 | 1 | 0 | 0 | 72 | 93 | 19 | 27 | 0 | 0 | 0 | 0 | 0 | 230 | 149 | 1 | 1 | 1 | 1 | 1 | 0.978 | 1 | NA | 1 | 1 | 1 | 1 | -0.004 |
| SE | 69751 | ENSG00000144228 | SPOPL | chr2 | + | 138552253 | 138552681 | 138550902 | 138551054 | 138554438 | 138554535 | 215 | 147 | 117 | 179 | 186 | 1 | 0 | 0 | 0 | 0 | 9 | 111 | 156 | 58 | 15 | 0 | 0 | 0 | 0 | 0 | 277 | 149 | 1 | 1 | 0.991 | 1 | 1 | 1 | 1 | | 1 | 1 | 1 | 1 | -0.002 |
| SE | 69752 | ENSG00000144228 | SPOPL | chr2 | + | 138552553 | 138552681 | 138550902 | 138551054 | 138559021 | 138559199 | 298 | 220 | 169 | 268 | 313 | 2 | 1 | 1 | 3 | 1 | 15 | 181 | 222 | 78 | 36 | 0 | 0 | 0 | 0 | 0 | 277 | 149 | 1 | 1 | 0.988 | 0.992 | 0.989 | 0.98 | 0.994 | 1 | 1 | 1 | 1 | 1 | -0.011 |
| SE | 69753 | ENSG00000144228 | SPOPL | chr2 | + | 138554438 | 138554535 | 138550902 | 138551054 | 138559021 | 138559199 | 21 | 17 | 14 | 19 | 11 | 2 | 1 | 1 | 3 | 1 | 0 | 9 | 22 | 20 | 0 | 0 | 0 | 0 | 0 | 0 | 246 | 149 | 1 | 1 | 0.864 | 0.911 | 0.895 | 0.793 | 0.869 | NA | 1 | 1 | 1 | NA | -0.134 |
| SE | 69755 | ENSG00000144228 | SPOPL | chr2 | + | 138559021 | 138559199 | 138552553 | 138552681 | 138559281 | 138559337 | 326 | 186 | 148 | 263 | 398 | 0 | 0 | 0 | 0 | 0 | 23 | 225 | 238 | 50 | 27 | 0 | 2 | 0 | 0 | 0 | 298 | 149 | 1 | 1 | 1 | 1 | 1 | 1 | 1 | 1 | 0.983 | 1 | 1 | 1 | 0.003 |
| SE | 69756 | ENSG00000144228 | SPOPL | chr2 | + | 138560804 | 138560927 | 138559281 | 138559337 | 138564707 | 138564850 | 360 | 208 | 187 | 295 | 450 | 0 | 0 | 0 | 1 | 0 | 16 | 192 | 256 | 33 | 10 | 0 | 0 | 0 | 0 | 0 | 272 | 149 | 1 | 1 | 1 | 1 | 1 | 0.994 | 1 | 1 | 1 | 1 | 1 | 1 | -0.001 |
| SE | 69757 | ENSG00000196247 | ZNF107 | chr7 | + | 64669036 | 64669178 | 64666098 | 64666285 | 64679176 | 64679336 | 1 | 0 | 6 | 6 | 0 | 5 | 3 | 0 | 12 | 5 | 0 | 0 | 4 | 0 | 14 | 0 | 22 | 19 | 30 | 6 | 291 | 149 | 1 | 1 | 0.093 | 0 | 1 | 0.204 | 0 | NA | 0 | 0.097 | 0 | 0.544 | 0.099 |
| SE | 69758 | ENSG00000196247 | ZNF107 | chr7 | + | 64669036 | 64669178 | 64666098 | 64666285 | 64691247 | 64691374 | 0 | 0 | 2 | 1 | 0 | 5 | 3 | 3 | 1 | 2 | 0 | 0 | 2 | 0 | 7 | 8 | 0 | 4 | 5 | 0 | 291 | 149 | 1 | 1 | 0 | 0 | 0.254 | 0.339 | 0 | 0 | NA | 0.204 | 0 | 1 | -0.182 |
| SE | 69760 | ENSG00000196247 | ZNF107 | chr7 | + | 64679176 | 64679336 | 64669036 | 64669178 | 64691247 | 64691374 | 7 | 0 | 6 | 14 | 3 | 0 | 0 | 1 | 0 | 0 | 0 | 2 | 2 | 6 | 13 | 0 | 0 | 0 | 0 | 0 | 298 | 149 | 1 | 1 | 1 | NA | 0.75 | 1 | 1 | NA | 1 | 1 | 1 | 1 | -0.062 |
| SE | 69761 | ENSG00000196247 | ZNF107 | chr7 | + | 64683945 | 64684205 | 64679176 | 64679336 | 64684334 | 64684519 | 1 | 2 | 2 | 12 | 0 | 0 | 0 | 1 | 6 | 0 | 3 | 5 | 8 | 5 | 0 | 0 | 6 | 4 | 0 | 0 | 298 | 149 | 1 | 1 | 1 | 1 | 0.5 | 0.5 | NA | 1 | 0.294 | 0.5 | 1 | NA | 0.051 |
| SE | 69762 | ENSG00000196247 | ZNF107 | chr7 | + | 64684086 | 64684205 | 64679176 | 64679336 | 64684334 | 64684390 | 2 | 2 | 1 | 9 | 0 | 0 | 0 | 1 | 6 | 0 | 6 | 11 | 8 | 7 | 0 | 0 | 6 | 4 | 0 | 0 | 268 | 149 | 1 | 1 | 1 | 1 | 0.357 | 0.455 | NA | 1 | 0.505 | 0.527 | 1 | NA | -0.055 |
| SE | 69763 | ENSG00000196247 | ZNF107 | chr7 | + | 64684086 | 64684205 | 64679176 | 64679336 | 64691247 | 64691374 | 2 | 0 | 0 | 0 | 0 | 6 | 0 | 1 | 9 | 3 | 0 | 7 | 0 | 0 | 0 | 0 | 2 | 0 | 6 | 6 | 268 | 149 | 1 | 1 | 0.156 | NA | 0 | 0 | 0 | NA | 0.661 | NA | 0 | 0 | -0.181 |
| SE | 69764 | ENSG00000196247 | ZNF107 | chr7 | + | 64690398 | 64690525 | 64679176 | 64679336 | 64691247 | 64691374 | 17 | 7 | 14 | 15 | 12 | 6 | 0 | 1 | 9 | 3 | 0 | 1 | 3 | 7 | 0 | 0 | 2 | 0 | 6 | 6 | 276 | 149 | 1 | 1 | 0.605 | 1 | 0.883 | 0.474 | 0.683 | NA | 0.213 | 1 | 0.386 | 0 | 0.329 |
| SE | 69765 | ENSG00000196247 | ZNF107 | chr7 | + | 64691247 | 64691374 | 64666103 | 64666285 | 64706323 | 64708656 | 8 | 10 | 4 | 6 | 13 | 1 | 2 | 1 | 0 | 0 | 8 | 4 | 4 | 5 | 0 | 0 | 0 | 0 | 0 | 0 | 276 | 149 | 1 | 1 | 0.812 | 0.73 | 0.683 | 1 | 1 | 1 | 1 | 1 | 1 | NA | -0.155 |
| SE | 69766 | ENSG00000196247 | ZNF107 | chr7 | + | 64691864 | 64691960 | 64691144 | 64691374 | 64706323 | 64706399 | 42 | 44 | 35 | 40 | 60 | 3 | 8 | 1 | 5 | 13 | 8 | 6 | 9 | 12 | 0 | 0 | 6 | 0 | 0 | 0 | 245 | 149 | 1 | 1 | 0.895 | 0.77 | 0.955 | 0.83 | 0.737 | 1 | 0.378 | 1 | 1 | NA | -0.007 |
| SE | 69771 | ENSG00000164051 | CCDC51 | chr3 | - | 48433706 | 48433871 | 48432163 | 48433166 | 48434816 | 48435015 | 25 | 14 | 19 | 25 | 29 | 0 | 0 | 0 | 0 | 0 | 20 | 25 | 25 | 8 | 59 | 0 | 0 | 0 | 1 | 0 | 298 | 149 | 1 | 1 | 1 | 1 | 1 | 1 | 1 | 1 | 1 | 1 | 0.8 | 1 | 0.04 |
| SE | 69772 | ENSG00000164051 | CCDC51 | chr3 | - | 48433706 | 48433976 | 48432194 | 48433166 | 48434816 | 48435015 | 13 | 7 | 9 | 10 | 11 | 0 | 0 | 0 | 0 | 0 | 13 | 15 | 8 | 4 | 13 | 0 | 0 | 0 | 1 | 0 | 298 | 149 | 1 | 1 | 1 | 1 | 1 | 1 | 1 | 1 | 1 | 1 | 0.667 | 1 | 0.067 |
| SE | 69773 | ENSG00000164687 | FABP5 | chr8 | + | 81283365 | 81283538 | 81280535 | 81280674 | 81283872 | 81283974 | 36 | 21 | 48 | 25 | 26 | 0 | 0 | 0 | 0 | 0 | 61 | 35 | 56 | 66 | 64 | 0 | 0 | 0 | 0 | 3 | 298 | 149 | 1 | 1 | 1 | 1 | 1 | 1 | 1 | 1 | 1 | 1 | 1 | 0.914 | 0.017 |
| SE | 69775 | ENSG00000182004 | SNRPE | chr1 | + | 203862181 | 203862222 | 203861602 | 203861743 | 203863662 | 203863725 | 115 | 64 | 93 | 92 | 72 | 2 | 0 | 0 | 0 | 0 | 94 | 97 | 102 | 95 | 80 | 0 | 0 | 0 | 0 | 0 | 190 | 149 | 1 | 1 | 0.978 | 1 | 1 | 1 | 1 | 1 | 1 | 1 | 1 | 1 | -0.004 |
| SE | 69776 | ENSG00000182004 | SNRPE | chr1 | + | 203862181 | 203862222 | 203861849 | 203861955 | 203863662 | 203863725 | 115 | 63 | 92 | 88 | 62 | 2 | 0 | 0 | 0 | 1 | 94 | 90 | 95 | 90 | 80 | 0 | 0 | 0 | 0 | 0 | 190 | 149 | 1 | 1 | 0.978 | 1 | 1 | 1 | 0.98 | 1 | 1 | 1 | 1 | 1 | -0.008 |
| SE | 69777 | ENSG00000182004 | SNRPE | chr1 | + | 203863662 | 203863725 | 203861602 | 203862222 | 203865040 | 203865119 | 376 | 192 | 368 | 349 | 277 | 0 | 0 | 3 | 2 | 0 | 200 | 282 | 274 | 261 | 181 | 0 | 0 | 2 | 0 | 0 | 212 | 149 | 1 | 1 | 1 | 1 | 0.989 | 0.992 | 1 | 1 | 1 | 0.99 | 1 | 1 | -0.002 |
| SE | 69783 | ENSG00000065675 | PRKCQ | chr10 | - | 6464312 | 6464404 | 6462302 | 6462365 | 6478991 | 6479165 | 283 | 129 | 232 | 232 | 503 | 0 | 0 | 0 | 0 | 0 | 178 | 230 | 218 | 128 | 110 | 0 | 0 | 0 | 3 | 0 | 241 | 149 | 1 | 1 | 1 | 1 | 1 | 1 | 1 | 1 | 1 | 1 | 0.963 | 1 | 0.007 |
| SE | 69784 | ENSG00000065675 | PRKCQ | chr10 | - | 6483439 | 6483600 | 6478991 | 6479165 | 6485151 | 6485269 | 277 | 125 | 269 | 231 | 345 | 1 | 0 | 0 | 0 | 0 | 147 | 213 | 236 | 114 | 136 | 0 | 0 | 0 | 0 | 0 | 298 | 149 | 1 | 1 | 0.993 | 1 | 1 | 1 | 1 | 1 | 1 | 1 | 1 | 1 | -0.001 |
| SE | 69786 | ENSG00000065675 | PRKCQ | chr10 | - | 6485151 | 6485269 | 6478991 | 6479165 | 6491682 | 6491812 | 2 | 1 | 0 | 3 | 0 | 0 | 0 | 0 | 0 | 0 | 5 | 9 | 4 | 8 | 14 | 0 | 0 | 4 | 4 | 0 | 267 | 149 | 1 | 1 | 1 | 1 | NA | 1 | NA | 1 | 1 | 0.358 | 0.527 | 1 | 0.223 |
| SE | 69793 | ENSG00000184007 | PTP4A2 | chr1 | - | 31911695 | 31911826 | 31910037 | 31910112 | 31912980 | 31913023 | 1493 | 1099 | 1078 | 1475 | 1898 | 0 | 0 | 2 | 0 | 0 | 3678 | 1602 | 1862 | 2344 | 1835 | 0 | 3 | 0 | 4 | 0 | 280 | 149 | 1 | 1 | 1 | 1 | 0.997 | | 1 | 1 | 0.996 | 1 | 1 | 1 | 0 |
| SE | 69795 | ENSG00000184007 | PTP4A2 | chr1 | - | 31911695 | 31911826 | 31910037 | 31910112 | 31918969 | 31919368 | 1659 | 1208 | 1148 | 1631 | 2103 | 6 | 2 | 0 | 7 | 1 | 4109 | 1799 | 2004 | 2537 | 2108 | 0 | 1 | 9 | 0 | 0 | 280 | 149 | 1 | 1 | 0.993 | 0.997 | 1 | 0.992 | 0.999 | 1 | 0.999 | 0.992 | 1 | 1 | -0.002 |
| SE | 69796 | ENSG00000184007 | PTP4A2 | chr1 | - | 31911695 | 31911826 | 31910037 | 31910112 | 31937986 | 31938368 | 1490 | 1095 | 1077 | 1474 | 1878 | 0 | 0 | 0 | 2 | 0 | 3658 | 1588 | 1851 | 2349 | 1833 | 0 | 0 | 0 | 0 | 0 | 280 | 149 | 1 | 1 | 1 | 1 | 1 | 0.997 | 1 | 1 | 1 | 1 | 1 | 1 | -0.001 |
| SE | 69798 | ENSG00000184007 | PTP4A2 | chr1 | - | 31912980 | 31913023 | 31911695 | 31911826 | 31915894 | 31915987 | 11 | 6 | 5 | 12 | 30 | 2401 | 1653 | 1590 | 2080 | 3069 | 38 | 19 | 16 | 16 | 27 | 5056 | 2461 | 2423 | 3309 | 2550 | 192 | 149 | 1 | 1 | 0.004 | 0.003 | 0.002 | 0.004 | 0.008 | 0.006 | 0.006 | 0.005 | 0.004 | 0.008 | -0.002 |
| SE | 69800 | ENSG00000184007 | PTP4A2 | chr1 | - | 31915894 | 31915987 | 31910037 | 31910112 | 31918969 | 31919109 | 1285 | 994 | 952 | 1211 | 1429 | 6 | 2 | 0 | 7 | 1 | 3170 | 1462 | 1496 | 2052 | 1318 | 0 | 1 | 9 | 0 | 0 | 242 | 149 | 1 | 1 | 0.992 | 0.997 | 1 | 0.991 | 0.999 | 1 | 0.999 | 0.99 | 1 | 1 | -0.002 |
| SE | 69801 | ENSG00000184007 | PTP4A2 | chr1 | - | 31915894 | 31915987 | 31910037 | 31910112 | 31937986 | 31938368 | 37 | 38 | 28 | 41 | 54 | 0 | 0 | 0 | 2 | 0 | 234 | 89 | 108 | 159 | 11 | 0 | 0 | 0 | 0 | 0 | 242 | 149 | 1 | 1 | 1 | 1 | 1 | 0.927 | 1 | 1 | 1 | 1 | 1 | 1 | -0.015 |
| SE | 69803 | ENSG00000184007 | PTP4A2 | chr1 | - | 31915894 | 31915987 | 31911695 | 31911826 | 31937986 | 31938368 | 1522 | 1120 | 1014 | 1343 | 1783 | 3 | 2 | 4 | 7 | 0 | 3407 | 1501 | 1581 | 2121 | 1768 | 14 | 3 | 2 | 14 | 16 | 242 | 149 | 1 | 1 | 0.997 | 0.997 | 0.994 | 0.992 | 1 | 0.993 | 0.997 | 0.998 | 0.989 | 0.986 | 0.003 |
| SE | 69804 | ENSG00000184007 | PTP4A2 | chr1 | - | 31918969 | 31919658 | 31910037 | 31910112 | 31937986 | 31938368 | 369 | 203 | 248 | 332 | 60 | 0 | 0 | 0 | 2 | 0 | 427 | 327 | 269 | 370 | 215 | 0 | 1 | 0 | 0 | 0 | 298 | 149 | 1 | 1 | 1 | 1 | 1 | 0.988 | 1 | 1 | 1 | 1 | 1 | 1 | -0.002 |
| SE | 69805 | ENSG00000184007 | PTP4A2 | chr1 | - | 31918969 | 31919658 | 31915894 | 31915987 | 31937986 | 31938368 | 2502 | 1707 | 1762 | 2243 | 2755 | 0 | 0 | 0 | 0 | 0 | 5045 | 2694 | 2492 | 3457 | 2294 | 0 | 1 | 0 | 3 | 0 | 298 | 149 | 1 | 1 | 1 | 1 | 1 | 1 | 1 | 1 | 0.999 | 1 | 0.998 | 1 | 0.001 |
| SE | 69806 | ENSG00000184007 | PTP4A2 | chr1 | - | 31924017 | 31924165 | 31915894 | 31915987 | 31932671 | 31932700 | 1 | 0 | 0 | 0 | 0 | 0 | 0 | 0 | 2 | 0 | 0 | 3 | 0 | 0 | 0 | 0 | 0 | 1 | 0 | 0 | 297 | 149 | 1 | 1 | 1 | NA | NA | 0 | NA | NA | 1 | 0 | NA | NA | 0 |
| SE | 69807 | ENSG00000184007 | PTP4A2 | chr1 | - | 31930038 | 31930334 | 31918969 | 31919658 | 31930932 | 31931142 | 2 | 2 | 2 | 3 | 0 | 7 | 7 | 4 | 12 | 6 | 0 | 2 | 0 | 0 | 0 | 23 | 14 | 2 | 8 | 0 | 298 | 149 | 1 | 1 | 0.125 | 0.125 | 0.2 | 0.111 | 0 | 0 | 0.067 | 0 | 0 | NA | 0.095 |
| SE | 69808 | ENSG00000184007 | PTP4A2 | chr1 | - | 31930932 | 31931142 | 31918999 | 31919658 | 31935588 | 31935659 | 7 | 8 | 4 | 12 | 6 | 0 | 0 | 0 | 1 | 0 | 23 | 14 | 2 | 8 | 0 | 0 | 0 | 0 | 0 | 0 | 298 | 149 | 1 | 1 | 1 | 1 | 1 | 0.857 | 1 | 1 | 1 | 1 | 1 | NA | -0.029 |
| SE | 69809 | ENSG00000184007 | PTP4A2 | chr1 | - | 31930932 | 31931142 | 31918999 | 31919658 | 31937754 | 31937775 | 7 | 7 | 4 | 12 | 6 | 5 | 0 | 0 | 2 | 1 | 23 | 14 | 2 | 8 | 0 | 0 | 0 | 0 | 3 | 0 | 298 | 149 | 1 | 1 | 0.412 | 1 | 1 | 0.75 | 0.75 | 1 | 1 | 1 | 0.571 | NA | -0.11 |
| SE | 69814 | ENSG00000184007 | PTP4A2 | chr1 | - | 31935588 | 31935659 | 31919568 | 31919658 | 31937777 | 31937818 | 0 | 0 | 0 | 1 | 0 | 13 | 3 | 1 | 12 | 16 | 0 | 0 | 0 | 0 | 0 | 0 | 9 | 5 | 10 | 5 | 220 | 149 | 1 | 1 | 0 | 0 | 0 | 0.053 | 0 | NA | 0 | 0 | 0 | 0 | 0.011 |
| SE | 69816 | ENSG00000184007 | PTP4A2 | chr1 | - | 31935588 | 31935659 | 31930932 | 31931142 | 31937777 | 31937818 | 0 | 1 | 0 | 0 | 0 | 0 | 1 | 0 | 3 | 0 | 0 | 0 | 0 | 0 | 0 | 0 | 2 | 0 | 0 | 0 | 220 | 149 | 1 | 1 | NA | 0.404 | NA | 0 | NA | NA | 0 | NA | NA | NA | 0.202 |
| SE | 69817 | ENSG00000115163 | CENPA | chr2 | + | 26792501 | 26792833 | 26792130 | 26792240 | 26793144 | 26793326 | 0 | 1 | 7 | 6 | 0 | 0 | 0 | 0 | 0 | 0 | 12 | 4 | 10 | 8 | 4 | 0 | 0 | 3 | 1 | 0 | 298 | 149 | 1 | 1 | NA | 1 | 1 | 1 | NA | 1 | 1 | 0.625 | 0.8 | 1 | 0.115 |
| SE | 69818 | ENSG00000115163 | CENPA | chr2 | + | 26792755 | 26792833 | 26792130 | 26792240 | 26793144 | 26793326 | 1 | 2 | 14 | 10 | 0 | 0 | 0 | 0 | 0 | 0 | 29 | 18 | 25 | 29 | 6 | 0 | 0 | 3 | 1 | 0 | 227 | 149 | 1 | 1 | 1 | 1 | 1 | 1 | NA | 1 | 1 | 0.845 | 0.95 | 1 | 0.041 |
| SE | 69820 | ENSG00000163803 | PLB1 | chr2 | + | 28582404 | 28582505 | 28579626 | 28579707 | 28585760 | 28585842 | 0 | 0 | 0 | 0 | 3 | 0 | 0 | 0 | 0 | 1 | 0 | 0 | 1 | 0 | 0 | 0 | 0 | 0 | 0 | 0 | 250 | 149 | 1 | 1 | NA | NA | NA | NA | 0.641 | NA | NA | 1 | NA | NA | -0.359 |
| SE | 69821 | ENSG00000163803 | PLB1 | chr2 | + | 28601251 | 28601332 | 28600808 | 28600860 | 28601898 | 28601964 | 0 | 0 | 0 | 1 | 0 | 0 | 1 | 0 | 0 | 0 | 0 | 3 | 2 | 3 | 0 | 0 | 0 | 0 | 0 | 0 | 230 | 149 | 1 | 1 | NA | 0 | NA | 1 | NA | NA | 1 | 1 | 1 | NA | -0.5 |
| SE | 69822 | ENSG00000163803 | PLB1 | chr2 | + | 28603965 | 28604047 | 28602820 | 28602921 | 28604654 | 28604759 | 10 | 3 | 0 | 0 | 3 | 7 | 0 | 0 | 1 | 0 | 33 | 19 | 9 | 6 | 11 | 6 | 1 | 0 | 2 | 25 | 231 | 149 | 1 | 1 | 0.48 | 1 | NA | 0 | 1 | 0.78 | 0.925 | 1 | 0.659 | 0.221 | -0.097 |
| SE | 69823 | ENSG00000163803 | PLB1 | chr2 | + | 28603965 | 28604047 | 28602820 | 28602921 | 28605852 | 28605948 | 6 | 1 | 0 | 0 | 3 | 0 | 0 | 0 | 0 | 0 | 20 | 4 | 5 | 6 | 10 | 0 | 0 | 0 | 1 | 0 | 231 | 149 | 1 | 1 | 1 | 1 | NA | NA | 1 | 1 | 1 | 1 | 0.795 | 1 | 0.041 |
| SE | 69825 | ENSG00000163803 | PLB1 | chr2 | + | 28604654 | 28604759 | 28602820 | 28602921 | 28605852 | 28605948 | 10 | 4 | 6 | 4 | 3 | 0 | 0 | 0 | 0 | 0 | 13 | 2 | 4 | 8 | 67 | 0 | 0 | 0 | 1 | 0 | 254 | 149 | 1 | 1 | 1 | 1 | 1 | 1 | 1 | 1 | 1 | 1 | 0.824 | 1 | 0.035 |
| SE | 69828 | ENSG00000163803 | PLB1 | chr2 | + | 28604654 | 28604759 | 28603965 | 28604047 | 28606495 | 28606567 | 5 | 2 | 4 | 2 | 0 | 0 | 1 | 0 | 0 | 0 | 13 | 10 | 4 | 0 | 17 | 0 | 0 | 3 | 2 | 0 | 254 | 149 | 1 | 1 | 1 | 0.54 | 1 | 1 | NA | 1 | 1 | 0.439 | 0 | 1 | 0.197 |
| SE | 69833 | ENSG00000163803 | PLB1 | chr2 | + | 28620878 | 28620978 | 28620599 | 28620643 | 28625056 | 28625108 | 29 | 21 | 14 | 34 | 36 | 0 | 0 | 0 | 0 | 0 | 27 | 46 | 10 | 84 | 71 | 0 | 1 | 0 | 0 | 0 | 249 | 149 | 1 | 1 | 1 | 1 | 1 | 1 | 1 | 1 | 0.965 | 1 | 1 | 1 | 0.007 |
| SE | 69837 | ENSG00000144635 | DYNC1LI1 | chr3 | - | 32528445 | 32528601 | 32525973 | 32526908 | 32529539 | 32529660 | 282 | 173 | 242 | 309 | 482 | 0 | 0 | 0 | 0 | 1 | 186 | 164 | 171 | 226 | 111 | 0 | 0 | 0 | 0 | 0 | 298 | 149 | 1 | 1 | 1 | 1 | 1 | 1 | 0.996 | 1 | 1 | 1 | 1 | 1 | -0.001 |
| SE | 69838 | ENSG00000144635 | DYNC1LI1 | chr3 | - | 32529539 | 32529660 | 32528445 | 32528601 | 32530283 | 32530328 | 254 | 180 | 232 | 274 | 476 | 0 | 0 | 0 | 0 | 0 | 219 | 186 | 170 | 202 | 79 | 0 | 0 | 2 | 0 | 0 | 270 | 149 | 1 | 1 | 1 | 1 | 1 | 1 | 1 | 1 | 1 | 0.979 | 1 | 1 | 0.004 |
| SE | 69839 | ENSG00000144635 | DYNC1LI1 | chr3 | - | 32530283 | 32530328 | 32529539 | 32529660 | 32530460 | 32530520 | 162 | 114 | 142 | 161 | 335 | 0 | 0 | 0 | 0 | 2 | 140 | 112 | 88 | 117 | 47 | 0 | 0 | 0 | 0 | 0 | 194 | 149 | 1 | 1 | 1 | 1 | 1 | 1 | 0.992 | 1 | 1 | 1 | 1 | 1 | -0.002 |
| SE | 69840 | ENSG00000144635 | DYNC1LI1 | chr3 | - | 32530460 | 32530520 | 32530283 | 32530328 | 32532985 | 32533097 | 167 | 141 | 144 | 165 | 410 | 0 | 0 | 1 | 0 | 0 | 160 | 105 | 96 | 105 | 33 | 0 | 0 | 0 | 0 | 2 | 209 | 149 | 1 | 1 | 1 | 1 | 0.99 | 1 | 1 | 1 | 1 | 1 | 1 | 0.922 | 0.014 |
| SE | 69841 | ENSG00000144635 | DYNC1LI1 | chr3 | - | 32532985 | 32533097 | 32530460 | 32530520 | 32534510 | 32534646 | 250 | 199 | 209 | 256 | 657 | 0 | 1 | 0 | 1 | 2 | 328 | 151 | 136 | 149 | 60 | 0 | 0 | 0 | 0 | 0 | 261 | 149 | 1 | 1 | 1 | 0.991 | 1 | 0.993 | 0.995 | 1 | 1 | 1 | 1 | 1 | -0.004 |
| SE | 69842 | ENSG00000144635 | DYNC1LI1 | chr3 | - | 32541036 | 32541206 | 32537010 | 32537104 | 32544875 | 32545106 | 449 | 336 | 324 | 372 | 741 | 0 | 1 | 0 | 0 | 3 | 263 | 288 | 276 | 253 | 268 | 0 | 0 | 1 | 0 | 0 | 298 | 149 | 1 | 1 | 1 | 0.994 | 1 | 1 | 0.992 | 1 | 1 | 0.993 | 1 | 1 | -0.001 |
| SE | 69843 | ENSG00000144635 | DYNC1LI1 | chr3 | - | 32544875 | 32545106 | 32541036 | 32541206 | 32570345 | 32570419 | 191 | 152 | 148 | 163 | 348 | 0 | 0 | 0 | 1 | 0 | 104 | 125 | 111 | 122 | 129 | 0 | 0 | 0 | 0 | 0 | 298 | 149 | 1 | 1 | 1 | 1 | 1 | 0.988 | 1 | 1 | 1 | 1 | 1 | 1 | -0.002 |
| SE | 69844 | ENSG00000144635 | DYNC1LI1 | chr3 | - | 32545848 | 32545965 | 32544875 | 32545106 | 32570345 | 32570419 | 150 | 90 | 122 | 138 | 98 | 1 | 0 | 0 | 0 | 0 | 267 | 160 | 157 | 215 | 107 | 0 | 0 | 0 | 0 | 0 | 266 | 149 | 1 | 1 | 0.988 | 1 | 1 | 1 | 1 | 1 | 1 | 1 | 1 | 1 | -0.002 |
| SE | 69846 | ENSG00000107581 | EIF3A | chr10 | - | 119038237 | 119038439 | 119037118 | 119037309 | 119041993 | 119042772 | 468 | 400 | 309 | 416 | 400 | 0 | 1 | 1 | 0 | 0 | 95 | 325 | 586 | 299 | 76 | 0 | 0 | 0 | 0 | 0 | 298 | 149 | 1 | 1 | 1 | 0.995 | 0.994 | 1 | 1 | 1 | 1 | 1 | 1 | 1 | -0.002 |
| SE | 69848 | ENSG00000107581 | EIF3A | chr10 | - | 119056739 | 119056853 | 119051198 | 119051321 | 119056935 | 119057040 | 1233 | 1126 | 1051 | 1153 | 1921 | 0 | 1 | 0 | 0 | 0 | 287 | 1007 | 1347 | 947 | 31 | 0 | 0 | 0 | 0 | 0 | 263 | 149 | 1 | 1 | 1 | 0.998 | 1 | 1 | 1 | 1 | 1 | 1 | 1 | 1 | 0 |
| SE | 69849 | ENSG00000107581 | EIF3A | chr10 | - | 119059601 | 119059718 | 119059211 | 119059397 | 119060745 | 119060844 | 487 | 327 | 498 | 547 | 585 | 0 | 0 | 0 | 1 | 0 | 169 | 378 | 412 | 193 | 24 | 5 | 0 | 0 | 0 | 0 | 266 | 149 | 1 | 1 | 1 | 1 | 1 | 0.997 | 1 | 0.95 | 1 | 1 | 1 | 1 | 0.009 |
| SE | 69850 | ENSG00000107581 | EIF3A | chr10 | - | 119060745 | 119060844 | 119059601 | 119059718 | 119065398 | 119065570 | 218 | 140 | 213 | 212 | 278 | 0 | 0 | 0 | 0 | 0 | 124 | 181 | 227 | 96 | 15 | 0 | 3 | 0 | 0 | 0 | 248 | 149 | 1 | 1 | 1 | 1 | 1 | 1 | 1 | 1 | 0.973 | 1 | 1 | 1 | 0.005 |
| SE | 69852 | ENSG00000107581 | EIF3A | chr10 | - | 119061223 | 119061328 | 119060745 | 119060844 | 119069445 | 119069654 | 191 | 115 | 287 | 286 | 496 | 0 | 0 | 0 | 2 | 0 | 58 | 245 | 265 | 110 | 27 | 0 | 0 | 0 | 0 | 0 | 254 | 149 | 1 | 1 | 1 | 1 | 1 | 0.988 | 1 | 1 | 1 | 1 | 1 | 1 | -0.002 |
| SE | 69853 | ENSG00000107581 | EIF3A | chr10 | - | 119065398 | 119065570 | 119060745 | 119060844 | 119069445 | 119069654 | 767 | 593 | 621 | 649 | 1237 | 0 | 0 | 0 | 2 | 0 | 240 | 551 | 466 | 435 | 81 | 0 | 0 | 0 | 0 | 0 | 298 | 149 | 1 | 1 | 1 | 1 | 1 | 0.994 | 1 | 1 | 1 | 1 | 1 | 1 | -0.001 |
| SE | 69855 | ENSG00000107581 | EIF3A | chr10 | - | 119069445 | 119069654 | 119065398 | 119065570 | 119070885 | 119071085 | 1092 | 815 | 971 | 938 | 1676 | 0 | 0 | 0 | 0 | 0 | 261 | 789 | 660 | 562 | 119 | 0 | 1 | 1 | 4 | 0 | 298 | 149 | 1 | 1 | 1 | 1 | 1 | 1 | 1 | 1 | 0.997 | 0.997 | 0.986 | 1 | 0.004 |
| SE | 69856 | ENSG00000107581 | EIF3A | chr10 | - | 119072889 | 119073053 | 119070885 | 119071085 | 119073440 | 119073577 | 1557 | 1046 | 1173 | 1213 | 2059 | 0 | 1 | 0 | 2 | 0 | 264 | 839 | 925 | 579 | 194 | 0 | 0 | 0 | 0 | 0 | 298 | 149 | 1 | 1 | 1 | 0.998 | 1 | 0.997 | 1 | 1 | 1 | 1 | 1 | 1 | -0.001 |
| SE | 69857 | ENSG00000117475 | BLZF1 | chr1 | + | 169378329 | 169378529 | 169376539 | 169376979 | 169380480 | 169380562 | 127 | 112 | 121 | 146 | 204 | 4 | 1 | 1 | 2 | 12 | 0 | 148 | 50 | 15 | 0 | 0 | 1 | 0 | 3 | 0 | 298 | 149 | 1 | 1 | 0.941 | 0.982 | 0.984 | 0.973 | 0.895 | NA | 0.987 | 1 | 0.714 | NA | 0.055 |
| SE | 69858 | ENSG00000117475 | BLZF1 | chr1 | + | 169382061 | 169382281 | 169380480 | 169380609 | 169386996 | 169387209 | 129 | 139 | 106 | 158 | 236 | 0 | 0 | 2 | 0 | 7 | 0 | 115 | 73 | 45 | 0 | 0 | 0 | 0 | 0 | 0 | 298 | 149 | 1 | 1 | 1 | 1 | 0.964 | 1 | 0.944 | NA | 1 | 1 | 1 | NA | -0.018 |
| SE | 69859 | ENSG00000160766 | GBAP1 | chr1 | - | 155217518 | 155217593 | 155217239 | 155217373 | 155217963 | 155218049 | 6 | 0 | 0 | 0 | 6 | 3 | 0 | 4 | 2 | 3 | 0 | 0 | 0 | 0 | 0 | 14 | 0 | 0 | 3 | 0 | 224 | 149 | 1 | 1 | 0.571 | NA | 0 | 0 | 0.571 | 0 | NA | NA | 0 | NA | 0.285 |
| SE | 69860 | ENSG00000160766 | GBAP1 | chr1 | - | 155217704 | 155217846 | 155217239 | 155217373 | 155217963 | 155218150 | 0 | 0 | 0 | 0 | 4 | 3 | 0 | 4 | 2 | 3 | 0 | 0 | 0 | 0 | 0 | 14 | 0 | 0 | 3 | 0 | 291 | 149 | 1 | 1 | 0 | NA | 0 | 0 | 0.406 | 0 | NA | NA | 0 | NA | 0.102 |
| SE | 69861 | ENSG00000160766 | GBAP1 | chr1 | - | 155218387 | 155218479 | 155217239 | 155218049 | 155218847 | 155218943 | 6 | 0 | 2 | 0 | 3 | 1 | 0 | 2 | 0 | 0 | 0 | 0 | 0 | 9 | 2 | 0 | 0 | 0 | 0 | 0 | 241 | 149 | 1 | 1 | 0.788 | NA | 0.382 | NA | 1 | NA | NA | NA | 1 | 1 | -0.277 |
| SE | 69862 | ENSG00000160766 | GBAP1 | chr1 | - | 155224876 | 155225087 | 155217963 | 155218049 | 155227381 | 155227422 | 0 | 2 | 0 | 2 | 1 | 0 | 0 | 0 | 0 | 0 | 0 | 2 | 0 | 0 | 0 | 7 | 0 | 0 | 0 | 0 | 298 | 149 | 1 | 1 | NA | 1 | NA | 1 | 1 | 0 | 1 | NA | NA | NA | 0.5 |
| SE | 69867 | ENSG00000105136 | ZNF419 | chr19 | + | 57490146 | 57490185 | 57487717 | 57487983 | 57491470 | 57491597 | 0 | 0 | 2 | 1 | 0 | 3 | 0 | 0 | 2 | 3 | 0 | 0 | 0 | 0 | 0 | 0 | 2 | 1 | 0 | 0 | 188 | 149 | 1 | 1 | 0 | NA | 1 | 0.284 | 0 | NA | 0 | 0 | NA | NA | 0.321 |
| SE | 69868 | ENSG00000105136 | ZNF419 | chr19 | + | 57492112 | 57492211 | 57487964 | 57491597 | 57492855 | 57493783 | 5 | 0 | 10 | 8 | 9 | 4 | 1 | 6 | 7 | 1 | 0 | 3 | 2 | 0 | 0 | 0 | 1 | 0 | 0 | 0 | 248 | 149 | 1 | 1 | 0.429 | 0 | 0.5 | 0.407 | 0.844 | NA | 0.643 | 1 | NA | NA | -0.385 |
| SE | 69870 | ENSG00000105063 | PPP6R1 | chr19 | - | 55232074 | 55232211 | 55231801 | 55231982 | 55236642 | 55236821 | 973 | 596 | 718 | 876 | 852 | 0 | 0 | 1 | 0 | 0 | 1487 | 1399 | 716 | 1079 | 3284 | 9 | 6 | 4 | 0 | 0 | 286 | 149 | 1 | 1 | 1 | 1 | 0.997 | 1 | 1 | 0.989 | 0.992 | 0.989 | 1 | 1 | 0.005 |
| SE | 69871 | ENSG00000105063 | PPP6R1 | chr19 | - | 55241476 | 55241639 | 55241238 | 55241391 | 55242165 | 55242279 | 199 | 50 | 142 | 153 | 84 | 0 | 2 | 0 | 0 | 0 | 60 | 153 | 52 | 74 | 250 | 0 | 0 | 0 | 0 | 0 | 298 | 149 | 1 | 1 | 1 | 0.926 | 1 | 1 | 1 | 1 | 1 | 1 | 1 | 1 | -0.015 |
| SE | 69872 | ENSG00000176659 | C20orf197 | chr20 | + | 60058906 | 60058996 | 60055924 | 60056008 | 60069832 | 60070043 | 0 | 1 | 0 | 1 | 0 | 2 | 4 | 6 | 0 | 0 | 0 | 0 | 0 | 0 | 0 | 0 | 3 | 0 | 0 | 0 | 229 | 149 | 1 | 1 | 0 | 0.14 | 0 | 0.098 | NA | NA | 0 | NA | NA | NA | 0.06 |
| SE | 69874 | ENSG00000109171 | SLAIN2 | chr4 | + | 48377895 | 48378060 | 48369848 | 48369997 | 48382567 | 48382927 | 325 | 270 | 217 | 316 | 566 | 0 | 0 | 1 | 0 | 0 | 495 | 272 | 230 | 345 | 440 | 0 | 0 | 0 | 0 | 3 | 298 | 149 | 1 | 1 | 1 | 1 | 0.991 | 1 | 1 | 1 | 1 | 1 | 1 | 0.987 | 0.001 |
| SE | 69875 | ENSG00000109171 | SLAIN2 | chr4 | + | 48379689 | 48379848 | 48377895 | 48378060 | 48382567 | 48382927 | 914 | 578 | 620 | 878 | 1327 | 0 | 1 | 0 | 0 | 0 | 1113 | 830 | 623 | 717 | 1012 | 0 | 0 | 1 | 0 | 0 | 298 | 149 | 1 | 1 | 1 | 0.997 | 1 | 1 | 1 | 1 | 1 | 0.997 | 1 | 1 | 0 |
| SE | 69876 | ENSG00000109171 | SLAIN2 | chr4 | + | 48382567 | 48382639 | 48379826 | 48379848 | 48382727 | 48382927 | 374 | 233 | 246 | 362 | 522 | 0 | 1 | 0 | 1 | 0 | 361 | 336 | 241 | 242 | 371 | 7 | 0 | 3 | 0 | 0 | 221 | 149 | 1 | 1 | 1 | 0.994 | 1 | 0.996 | 1 | 0.972 | 1 | 0.982 | 1 | 1 | 0.007 |
| SE | 69877 | ENSG00000109171 | SLAIN2 | chr4 | + | 48382567 | 48382927 | 48379689 | 48379848 | 48383646 | 48383784 | 612 | 408 | 412 | 576 | 1060 | 0 | 0 | 0 | 0 | 4 | 590 | 525 | 424 | 477 | 630 | 0 | 0 | 0 | 0 | 0 | 298 | 149 | 1 | 1 | 1 | 1 | 1 | 1 | 0.993 | 1 | 1 | 1 | 1 | 1 | -0.001 |
| SE | 69878 | ENSG00000109171 | SLAIN2 | chr4 | + | 48382727 | 48382927 | 48379826 | 48379848 | 48383646 | 48383784 | 109 | 78 | 81 | 86 | 204 | 0 | 0 | 0 | 0 | 4 | 81 | 61 | 93 | 104 | 100 | 0 | 0 | 0 | 0 | 0 | 298 | 149 | 1 | 1 | 1 | 1 | 1 | 1 | 0.962 | 1 | 1 | 1 | 1 | 1 | -0.008 |
| SE | 69879 | ENSG00000109171 | SLAIN2 | chr4 | + | 48383646 | 48383784 | 48379689 | 48379848 | 48420124 | 48420443 | 251 | 175 | 205 | 209 | 527 | 0 | 0 | 0 | 0 | 0 | 102 | 176 | 192 | 164 | 53 | 0 | 0 | 2 | 0 | 0 | 287 | 149 | 1 | 1 | 1 | 1 | 1 | 1 | 1 | 1 | 1 | 0.98 | 1 | 1 | 0.004 |
| SE | 69880 | ENSG00000109171 | SLAIN2 | chr4 | + | 48387365 | 48387466 | 48383646 | 48383784 | 48394575 | 48394630 | 0 | 1 | 1 | 0 | 2 | 76 | 64 | 74 | 97 | 237 | 0 | 0 | 0 | 0 | 0 | 53 | 71 | 86 | 99 | 78 | 250 | 149 | 1 | 1 | 0 | 0.009 | 0.008 | 0 | 0.005 | 0 | 0 | 0 | 0 | 0 | 0.004 |
| SE | 69882 | ENSG00000109171 | SLAIN2 | chr4 | + | 48420124 | 48420443 | 48383646 | 48383784 | 48422010 | 48425015 | 379 | 281 | 322 | 347 | 670 | 0 | 0 | 0 | 2 | 4 | 149 | 271 | 292 | 234 | 108 | 0 | 0 | 0 | 7 | 0 | 298 | 149 | 1 | 1 | 1 | 1 | 1 | 0.989 | 0.988 | 1 | 1 | 1 | 0.944 | 1 | 0.007 |
| SE | 69886 | ENSG00000283761 | NA | chr1 | + | 100017681 | 100017815 | 100015301 | 100015420 | 100043072 | 100043229 | 60 | 54 | 38 | 49 | 59 | 1 | 2 | 1 | 2 | 0 | 0 | 24 | 25 | 10 | 7 | 0 | 3 | 0 | 0 | 0 | 283 | 149 | 1 | 1 | 0.969 | 0.934 | 0.952 | 0.928 | 1 | NA | 0.808 | 1 | 1 | 1 | 0.005 |
| SE | 69887 | ENSG00000283761 | NA | chr1 | + | 100017681 | 100017815 | 100015301 | 100015420 | 100049908 | 100050004 | 64 | 57 | 41 | 48 | 58 | 2 | 0 | 1 | 1 | 0 | 0 | 24 | 24 | 12 | 7 | 0 | 0 | 0 | 0 | 0 | 283 | 149 | 1 | 1 | 0.944 | 1 | 0.956 | 0.962 | 1 | NA | 1 | 1 | 1 | 1 | -0.028 |
| SE | 69889 | ENSG00000283761 | NA | chr1 | + | 100043072 | 100043229 | 100015301 | 100015420 | 100049908 | 100050004 | 14 | 8 | 10 | 13 | 5 | 2 | 0 | 1 | 1 | 0 | 13 | 11 | 8 | 0 | 0 | 0 | 0 | 0 | 0 | 0 | 298 | 149 | 1 | 1 | 0.778 | 1 | 0.833 | 0.867 | 1 | 1 | 1 | 1 | NA | NA | -0.104 |
| SE | 69890 | ENSG00000283761 | NA | chr1 | + | 100043072 | 100043229 | 100017681 | 100017815 | 100049908 | 100050004 | 13 | 6 | 9 | 12 | 6 | 4 | 3 | 3 | 0 | 0 | 13 | 8 | 9 | 0 | 0 | 0 | 0 | 0 | 2 | 0 | 298 | 149 | 1 | 1 | 0.619 | 0.5 | 0.6 | 1 | 1 | 1 | 1 | 1 | 0 | NA | -0.006 |
| SE | 69891 | ENSG00000283761 | NA | chr1 | + | 100049908 | 100050004 | 100043072 | 100043229 | 100058665 | 100058728 | 53 | 39 | 40 | 55 | 46 | 0 | 0 | 0 | 2 | 0 | 90 | 55 | 46 | 41 | 21 | 0 | 0 | 0 | 0 | 0 | 245 | 149 | 1 | 1 | 1 | 1 | 1 | 0.944 | 1 | 1 | 1 | 1 | 1 | 1 | -0.011 |
| SE | 69892 | ENSG00000109618 | SEPSECS | chr4 | - | 25127263 | 25127357 | 25125693 | 25125784 | 25144773 | 25144865 | 32 | 43 | 46 | 58 | 97 | 0 | 0 | 1 | 0 | 0 | 0 | 9 | 38 | 11 | 1 | 0 | 0 | 0 | 0 | 0 | 243 | 149 | 1 | 1 | 1 | 1 | 0.966 | 1 | 1 | NA | 1 | 1 | 1 | 1 | -0.007 |
| SE | 69894 | ENSG00000109618 | SEPSECS | chr4 | - | 25159509 | 25159764 | 25158952 | 25159107 | 25160255 | 25160546 | 0 | 0 | 0 | 1 | 1 | 10 | 5 | 8 | 0 | 17 | 0 | 0 | 0 | 2 | 0 | 0 | 7 | 2 | 2 | 3 | 298 | 149 | 1 | 1 | 0 | 0 | 0 | 1 | 0.029 | NA | 0 | 0 | 0.333 | 0 | 0.123 |
| SE | 69895 | ENSG00000109618 | SEPSECS | chr4 | - | 25159627 | 25159764 | 25158952 | 25159107 | 25160255 | 25160394 | 0 | 0 | 0 | 1 | 0 | 10 | 5 | 8 | 0 | 17 | 0 | 0 | 0 | 0 | 0 | 0 | 7 | 2 | 2 | 3 | 286 | 149 | 1 | 1 | 0 | 0 | 0 | 1 | 0 | NA | 0 | 0 | 0 | 0 | 0.2 |
| SE | 69901 | ENSG00000105851 | PIK3CG | chr7 | + | 106867549 | 106869556 | 106865478 | 106865776 | 106872536 | 106872602 | 200 | 104 | 148 | 174 | 229 | 0 | 0 | 2 | 0 | 0 | 104 | 145 | 158 | 126 | 55 | 0 | 0 | 0 | 0 | 0 | 298 | 149 | 1 | 1 | 1 | 1 | 0.974 | 1 | 1 | 1 | 1 | 1 | 1 | 1 | -0.005 |
| SE | 69903 | ENSG00000105851 | PIK3CG | chr7 | + | 106872712 | 106872938 | 106872536 | 106872602 | 106874699 | 106874803 | 371 | 194 | 300 | 313 | 516 | 0 | 0 | 0 | 2 | 1 | 181 | 378 | 365 | 331 | 168 | 6 | 0 | 0 | 0 | 0 | 298 | 149 | 1 | 1 | 1 | 1 | 1 | 0.987 | 0.996 | 0.938 | 1 | 1 | 1 | 1 | 0.009 |
| SE | 69904 | ENSG00000105851 | PIK3CG | chr7 | + | 106879518 | 106879665 | 106874699 | 106874803 | 106882116 | 106882207 | 614 | 317 | 411 | 513 | 563 | 3 | 1 | 0 | 1 | 2 | 364 | 412 | 380 | 517 | 218 | 0 | 8 | 0 | 2 | 0 | 296 | 149 | 1 | 1 | 0.99 | 0.994 | 1 | 0.996 | 0.993 | 1 | 0.963 | 1 | 0.992 | 1 | 0.004 |
| SE | 69905 | ENSG00000105851 | PIK3CG | chr7 | + | 106883032 | 106883163 | 106882116 | 106882207 | 106884154 | 106884266 | 757 | 460 | 521 | 661 | 896 | 1 | 0 | 1 | 0 | 0 | 364 | 555 | 419 | 694 | 187 | 0 | 0 | 0 | 8 | 0 | 280 | 149 | 1 | 1 | 0.998 | 1 | 0.996 | 1 | 1 | 1 | 1 | 1 | 0.979 | 1 | 0.003 |
| SE | 69914 | ENSG00000064995 | TAF11 | chr6 | - | 34882931 | 34883080 | 34880288 | 34880376 | 34887786 | 34888027 | 124 | 94 | 99 | 161 | 123 | 0 | 0 | 1 | 0 | 0 | 186 | 175 | 165 | 118 | 153 | 0 | 3 | 0 | 0 | 0 | 298 | 149 | 1 | 1 | 1 | 1 | 1 | 1 | 1 | 1 | 0.967 | 1 | 1 | 1 | 0.007 |
| SE | 69915 | ENSG00000238005 | NA | chr1 | - | 234961416 | 234961566 | 234959648 | 234959916 | 234968387 | 234968490 | 2 | 0 | 0 | 0 | 0 | 0 | 0 | 0 | 2 | 0 | 4 | 3 | 10 | 6 | 0 | 0 | 3 | 0 | 6 | 2 | 298 | 149 | 1 | 1 | 1 | NA | NA | 0 | NA | 1 | 0.333 | 1 | 0.333 | 0 | -0.033 |
| SE | 69924 | ENSG00000288710 | NA | chr12 | - | 49022279 | 49022353 | 49022042 | 49022151 | 49022589 | 49022875 | 227 | 136 | 222 | 255 | 199 | 0 | 0 | 0 | 0 | 0 | 622 | 308 | 350 | 359 | 540 | 0 | 0 | 0 | 0 | 10 | 223 | 149 | 1 | 1 | 1 | 1 | 1 | | 1 | 1 | 1 | 1 | 1 | 0.973 | 0.005 |
| SE | 69925 | ENSG00000169016 | E2F6 | chr2 | - | 11453581 | 11453798 | 11451349 | 11451806 | 11455341 | 11455474 | 52 | 47 | 21 | 35 | 47 | 0 | 0 | 1 | 0 | 0 | 30 | 26 | 19 | 24 | 36 | 0 | 0 | 0 | 0 | 0 | 298 | 149 | 1 | 1 | 1 | 1 | 0.913 | 1 | 1 | 1 | 1 | 1 | 1 | 1 | -0.017 |
| SE | 69932 | ENSG00000134987 | WDR36 | chr5 | + | 111104176 | 111104352 | 111103785 | 111103918 | 111104696 | 111104817 | 184 | 132 | 154 | 180 | 198 | 1 | 0 | 0 | 5 | 0 | 18 | 79 | 93 | 64 | 8 | 0 | 0 | 0 | 0 | 0 | 298 | 149 | 1 | 1 | 0.989 | 1 | 1 | 0.947 | 1 | 1 | 1 | 1 | 1 | 1 | -0.013 |
| SE | 69933 | ENSG00000134987 | WDR36 | chr5 | + | 111104696 | 111104817 | 111104176 | 111104352 | 111105294 | 111105360 | 223 | 155 | 195 | 208 | 192 | 4 | 1 | 0 | 1 | 0 | 4 | 97 | 95 | 58 | 0 | 0 | 0 | 0 | 0 | 0 | 270 | 149 | 1 | 1 | 0.969 | 0.988 | 1 | 0.991 | 1 | 1 | 1 | 1 | 1 | NA | -0.01 |
| SE | 69934 | ENSG00000134987 | WDR36 | chr5 | + | 111120995 | 111121141 | 111120495 | 111120593 | 111123804 | 111123924 | 224 | 146 | 185 | 225 | 499 | 0 | 2 | 0 | 1 | 0 | 0 | 107 | 76 | 24 | 2 | 0 | 0 | 0 | 0 | 0 | 295 | 149 | 1 | 1 | 1 | 0.974 | 1 | 0.991 | 1 | NA | 1 | 1 | 1 | 1 | -0.007 |
| SE | 69935 | ENSG00000134987 | WDR36 | chr5 | + | 111124107 | 111124189 | 111123804 | 111123924 | 111125607 | 111125795 | 90 | 63 | 60 | 76 | 90 | 2 | 2 | 0 | 3 | 0 | 0 | 20 | 16 | 9 | 7 | 0 | 7 | 0 | 0 | 0 | 231 | 149 | 1 | 1 | 0.967 | 0.953 | 1 | 1 | 0.951 | NA | 0.648 | 1 | 1 | 1 | 0.062 |
| SE | 69936 | ENSG00000123427 | EEF1AKMT3 | chr12 | + | 57774628 | 57774767 | 57773016 | 57773128 | 57780254 | 57780430 | 2 | 1 | 3 | 1 | 0 | 9 | 4 | 8 | 1 | 3 | 0 | 0 | 0 | 0 | 0 | 14 | 0 | 2 | 0 | 0 | 288 | 149 | 1 | 1 | 0.103 | 0.115 | 0.162 | 0.341 | 0 | 0 | NA | 0 | NA | NA | 0.144 |
| SE | 69937 | ENSG00000120254 | MTHFD1L | chr6 | + | 150887841 | 150887981 | 150885633 | 150885734 | 150905649 | 150905761 | 11 | 11 | 15 | 18 | 13 | 0 | 0 | 0 | 0 | 0 | 7 | 4 | 11 | 59 | 20 | 0 | 2 | 0 | 0 | 0 | 289 | 149 | 1 | 1 | 1 | 1 | 1 | 1 | 1 | 1 | 0.508 | 1 | 1 | 1 | 0.098 |
| SE | 69938 | ENSG00000120254 | MTHFD1L | chr6 | + | 150887844 | 150887981 | 150885633 | 150885734 | 150905649 | 150905761 | 8 | 7 | 9 | 18 | 10 | 0 | 0 | 0 | 0 | 0 | 18 | 5 | 12 | 41 | 20 | 0 | 2 | 0 | 0 | 0 | 286 | 149 | 1 | 1 | 1 | 1 | 1 | 1 | 1 | 1 | 0.566 | 1 | 1 | 1 | 0.087 |
| SE | 69939 | ENSG00000120254 | MTHFD1L | chr6 | + | 150905649 | 150905761 | 150887841 | 150887981 | 150918576 | 150918668 | 7 | 9 | 14 | 19 | 8 | 0 | 0 | 0 | 1 | 0 | 16 | 1 | 11 | 39 | 26 | 0 | 0 | 0 | 0 | 0 | 261 | 149 | 1 | 1 | 1 | 1 | 1 | 0.916 | 1 | 1 | 1 | 1 | 1 | 1 | -0.017 |
| SE | 69940 | ENSG00000120254 | MTHFD1L | chr6 | + | 150936803 | 150936940 | 150926121 | 150926295 | 150944485 | 150944593 | 4 | 4 | 11 | 9 | 11 | 0 | 0 | 3 | 1 | 0 | 33 | 4 | 4 | 38 | 17 | 0 | 0 | 0 | 0 | 0 | 286 | 149 | 1 | 1 | 1 | 1 | 0.656 | 0.824 | 1 | 1 | 1 | 1 | 1 | 1 | -0.104 |
| SE | 69943 | ENSG00000108883 | EFTUD2 | chr17 | - | 44852408 | 44852562 | 44851709 | 44851817 | 44853295 | 44853390 | 240 | 115 | 176 | 200 | 142 | 0 | 0 | 0 | 0 | 0 | 201 | 210 | 116 | 180 | 390 | 0 | 0 | 2 | 5 | 0 | 298 | 149 | 1 | 1 | 1 | 1 | 1 | 1 | 1 | 1 | 1 | 0.967 | 0.947 | 1 | 0.017 |
| SE | 69944 | ENSG00000108883 | EFTUD2 | chr17 | - | 44852408 | 44852562 | 44851709 | 44851817 | 44853516 | 44853635 | 105 | 52 | 79 | 93 | 61 | 0 | 0 | 0 | 0 | 0 | 80 | 86 | 62 | 88 | 164 | 0 | 0 | 0 | 1 | 0 | 298 | 149 | 1 | 1 | 1 | 1 | 1 | 1 | 1 | 1 | 1 | 1 | 0.978 | 1 | 0.004 |
| SE | 69945 | ENSG00000108883 | EFTUD2 | chr17 | - | 44853295 | 44853390 | 44851709 | 44851817 | 44853516 | 44853635 | 100 | 55 | 79 | 81 | 80 | 0 | 0 | 0 | 0 | 0 | 138 | 67 | 34 | 77 | 179 | 0 | 0 | 0 | 1 | 0 | 244 | 149 | 1 | 1 | 1 | 1 | 1 | 1 | 1 | 1 | 1 | 1 | 0.979 | 1 | 0.004 |
| SE | 69947 | ENSG00000108883 | EFTUD2 | chr17 | - | 44857074 | 44857157 | 44854917 | 44855004 | 44859079 | 44859181 | 226 | 121 | 130 | 211 | 335 | 1 | 0 | 2 | 1 | 0 | 312 | 223 | 155 | 229 | 486 | 0 | 0 | 0 | 0 | 0 | 232 | 149 | 1 | 1 | 0.993 | 1 | 0.977 | 0.993 | 1 | 1 | 1 | 1 | 1 | 1 | -0.007 |
| SE | 69948 | ENSG00000108883 | EFTUD2 | chr17 | - | 44859079 | 44859181 | 44857074 | 44857157 | 44859904 | 44860045 | 285 | 165 | 207 | 294 | 522 | 0 | 0 | 0 | 1 | 0 | 451 | 316 | 206 | 312 | 780 | 0 | 0 | 0 | 0 | 0 | 251 | 149 | 1 | 1 | 1 | 1 | 1 | 0.994 | 1 | 1 | 1 | 1 | 1 | 1 | -0.001 |
| SE | 69949 | ENSG00000108883 | EFTUD2 | chr17 | - | 44862712 | 44862906 | 44860431 | 44860543 | 44863654 | 44863782 | 301 | 175 | 225 | 314 | 231 | 1 | 0 | 0 | 0 | 0 | 356 | 364 | 125 | 262 | 475 | 0 | 0 | 0 | 0 | 0 | 298 | 149 | 1 | 1 | 0.993 | 1 | 1 | 1 | 1 | 1 | 1 | 1 | 1 | 1 | -0.001 |
| SE | 69950 | ENSG00000108883 | EFTUD2 | chr17 | - | 44867806 | 44867897 | 44864929 | 44865065 | 44872445 | 44872526 | 138 | 102 | 100 | 120 | 162 | 0 | 1 | 0 | 0 | 0 | 156 | 144 | 49 | 110 | 223 | 0 | 0 | 0 | 1 | 3 | 240 | 149 | 1 | 1 | 1 | 0.984 | 1 | 1 | 1 | 1 | 1 | 1 | 0.986 | 0.979 | 0.004 |
| SE | 69951 | ENSG00000108883 | EFTUD2 | chr17 | - | 44868286 | 44868350 | 44867806 | 44867897 | 44872445 | 44872570 | 152 | 91 | 127 | 128 | 127 | 0 | 0 | 1 | 1 | 0 | 142 | 122 | 90 | 105 | 249 | 0 | 0 | 0 | 0 | 7 | 213 | 149 | 1 | 1 | 1 | 1 | 0.989 | 0.989 | 1 | 1 | 1 | 1 | 1 | 0.961 | 0.003 |
| SE | 69952 | ENSG00000108883 | EFTUD2 | chr17 | - | 44868286 | 44868350 | 44867806 | 44867897 | 44875933 | 44876100 | 63 | 44 | 66 | 59 | 65 | 0 | 0 | 5 | 0 | 0 | 82 | 70 | 43 | 62 | 82 | 3 | 0 | 0 | 0 | 0 | 213 | 149 | 1 | 1 | 1 | 1 | 0.902 | 1 | 1 | 0.95 | 1 | 1 | 1 | 1 | -0.01 |
| SE | 69953 | ENSG00000108883 | EFTUD2 | chr17 | - | 44868381 | 44868479 | 44867806 | 44867897 | 44872445 | 44872526 | 0 | 0 | 0 | 2 | 0 | 0 | 0 | 1 | 1 | 0 | 0 | 2 | 4 | 6 | 4 | 0 | 0 | 0 | 0 | 7 | 247 | 149 | 1 | 1 | NA | NA | 0 | 0.547 | NA | NA | 1 | 1 | 1 | 0.256 | -0.54 |
| SE | 69954 | ENSG00000108883 | EFTUD2 | chr17 | - | 44872445 | 44872570 | 44867806 | 44867897 | 44875933 | 44876100 | 185 | 85 | 106 | 133 | 140 | 0 | 0 | 5 | 0 | 0 | 166 | 150 | 58 | 92 | 179 | 3 | 0 | 0 | 0 | 0 | 274 | 149 | 1 | 1 | 1 | 1 | 0.92 | 1 | 1 | 0.968 | 1 | 1 | 1 | 1 | -0.01 |
| SE | 69957 | ENSG00000108883 | EFTUD2 | chr17 | - | 44885255 | 44885334 | 44883555 | 44883724 | 44894416 | 44894497 | 179 | 108 | 110 | 156 | 308 | 0 | 0 | 2 | 0 | 0 | 279 | 182 | 121 | 153 | 325 | 0 | 0 | 0 | 0 | 0 | 228 | 149 | 1 | 1 | 1 | 1 | 1 | 0.981 | 1 | 1 | 1 | 1 | 1 | 1 | -0.004 |
| SE | 69958 | ENSG00000108883 | EFTUD2 | chr17 | - | 44886584 | 44886750 | 44885255 | 44885334 | 44894416 | 44894525 | 357 | 175 | 246 | 296 | 464 | 0 | 0 | 2 | 0 | 8 | 592 | 319 | 246 | 359 | 653 | 0 | 0 | 0 | 0 | 0 | 298 | 149 | 1 | 1 | 1 | 1 | 0.984 | 1 | 0.967 | 1 | 1 | 1 | 1 | 1 | -0.01 |
| SE | 69959 | ENSG00000108883 | EFTUD2 | chr17 | - | 44894244 | 44894525 | 44886584 | 44886750 | 44899368 | 44899403 | 29 | 6 | 13 | 17 | 4 | 0 | 0 | 0 | 0 | 0 | 34 | 20 | 9 | 12 | 13 | 0 | 0 | 0 | 3 | 0 | 298 | 149 | 1 | 1 | 1 | 1 | 1 | 1 | 1 | 1 | 1 | 1 | 0.667 | 1 | 0.067 |

| | | | | | | | | | | | | | |
|---|---|---|---|---|---|---|---|---|---|---|---|---|---|
| SE | 69960 | ENSG00000108883 | EFTUD2 | chr17 | - | 44894416 | 44894525 | 44886584 | 44886750 | 44899368 | 44899431 | 195 | 104 |
| SE | 69961 | ENSG00000108883 | EFTUD2 | chr17 | - | 44895825 | 44895989 | 44894416 | 44894525 | 44898974 | 44899049 | 1 | 3 |
| SE | 69963 | ENSG00000108883 | EFTUD2 | chr17 | - | 44898974 | 44899041 | 44894416 | 44894525 | 44899368 | 44899403 | 0 | 0 |
| SE | 69965 | ENSG00000106069 | CHN2 | chr7 | + | 29292924 | 29292963 | 29194924 | 29194990 | 29354624 | 29354663 | 0 | 0 |
| SE | 69969 | ENSG00000106069 | CHN2 | chr7 | + | 29400240 | 29400351 | 29398372 | 29398486 | 29400542 | 29400696 | 0 | 0 |
| SE | 69970 | ENSG00000106069 | CHN2 | chr7 | + | 29495951 | 29496036 | 29479711 | 29480356 | 29504743 | 29504821 | 6 | 4 |
| SE | 69975 | ENSG00000135775 | COG2 | chr1 | + | 230661324 | 230661557 | 230660757 | 230660823 | 230663140 | 230663221 | 0 | 0 |
| SE | 69977 | ENSG00000135775 | COG2 | chr1 | + | 230669355 | 230669535 | 230668675 | 230668784 | 230671515 | 230671640 | 371 | 251 |
| SE | 69979 | ENSG00000135775 | COG2 | chr1 | + | 230683573 | 230683635 | 230678912 | 230679052 | 230685084 | 230685236 | 72 | 31 |
| SE | 69982 | ENSG00000135775 | COG2 | chr1 | + | 230688419 | 230688562 | 230688070 | 230688143 | 230690013 | 230690153 | 150 | 104 |
| SE | 69983 | ENSG00000135775 | COG2 | chr1 | + | 230688422 | 230688562 | 230688070 | 230688143 | 230690013 | 230690153 | 116 | 88 |
| SE | 69985 | ENSG00000165695 | AK8 | chr9 | - | 132792633 | 132792775 | 132727453 | 132727534 | 132814637 | 132814727 | 0 | 2 |
| SE | 70048 | ENSG00000111644 | ACRBP | chr12 | - | 6640059 | 6640227 | 6638953 | 6639037 | 6640342 | 6640522 | 21 | 9 |
| SE | 70049 | ENSG00000111644 | ACRBP | chr12 | - | 6643538 | 6643671 | 6640342 | 6640522 | 6644136 | 6644506 | 21 | 9 |
| SE | 70050 | ENSG00000111644 | ACRBP | chr12 | - | 6645219 | 6645337 | 6644054 | 6644605 | 6646482 | 6646577 | 11 | 11 |
| SE | 70055 | ENSG00000151665 | PIGF | chr2 | - | 46591802 | 46591925 | 46588074 | 46588179 | 46592474 | 46592583 | 15 | 7 |
| SE | 70056 | ENSG00000151665 | PIGF | chr2 | - | 46592474 | 46592583 | 46580936 | 46581591 | 46612227 | 46612274 | 99 | 65 |
| SE | 70057 | ENSG00000151665 | PIGF | chr2 | - | 46613693 | 46613785 | 46612227 | 46612344 | 46614936 | 46615185 | 95 | 55 |
| SE | 70107 | ENSG00000121691 | CAT | chr11 | + | 34457775 | 34457847 | 34456664 | 34456817 | 34461250 | 34461389 | 2 | 0 |
| SE | 70109 | ENSG00000121691 | CAT | chr11 | + | 34461250 | 34461389 | 34456664 | 34456817 | 34464104 | 34464235 | 504 | 412 |
| SE | 70110 | ENSG00000121691 | CAT | chr11 | + | 34461250 | 34461389 | 34457775 | 34457847 | 34464104 | 34464235 | 128 | 126 |
| SE | 70118 | ENSG00000130561 | SAG | chr2 | + | 233346402 | 233346412 | 233342270 | 233342326 | 233346806 | 233347054 | 0 | 0 |
| SE | 70121 | ENSG00000123485 | HJURP | chr2 | - | 233847396 | 233847461 | 233845727 | 233845820 | 233849762 | 233849859 | 4 | 0 |
| SE | 70122 | ENSG00000123485 | HJURP | chr2 | - | 233847396 | 233847461 | 233845727 | 233845820 | 233852564 | 233852620 | 1 | 0 |
| SE | 70123 | ENSG00000123485 | HJURP | chr2 | - | 233849762 | 233849859 | 233845727 | 233845820 | 233852564 | 233852620 | 0 | 2 |
| SE | 70125 | ENSG00000105953 | OGDH | chr7 | + | 44622805 | 44622875 | 44606594 | 44606653 | 44624316 | 44624565 | 2 | 0 |
| SE | 70126 | ENSG00000105953 | OGDH | chr7 | + | 44624316 | 44624565 | 44606571 | 44606653 | 44645326 | 44645518 | 517 | 383 |
| SE | 70129 | ENSG00000105953 | OGDH | chr7 | + | 44645326 | 44645518 | 44624316 | 44624565 | 44673786 | 44673941 | 344 | 246 |
| SE | 70130 | ENSG00000105953 | OGDH | chr7 | + | 44647443 | 44647534 | 44645326 | 44645518 | 44666735 | 44666851 | 43 | 33 |

| | | | | | | | | | | | | | | |
|---|---|---|---|---|---|---|---|---|---|---|---|---|---|---|
| SE | 70495 | ENSG00000170266 | GLB1 | chr3 | - | 33072543 | 33072713 | 33068818 | 33068970 | 33096718 | 33096835 | 68 | 50 | [illegible] |
| SE | 70500 | ENSG00000134086 | VHL | chr3 | + | 10146513 | 10146636 | 10142545 | 10143021 | 10149786 | 10150145 | 122 | 58 | [illegible] |
| SE | 70501 | ENSG00000198162 | MAN1A2 | chr1 | + | 117402185 | 117402441 | 117367448 | 117368485 | 117405548 | 117405645 | 432 | 347 | [illegible] |
| SE | 70502 | ENSG00000198162 | MAN1A2 | chr1 | + | 117405548 | 117405645 | 117367448 | 117368485 | 117414712 | 117414831 | 917 | 467 | [illegible] |
| SE | 70503 | ENSG00000198162 | MAN1A2 | chr1 | + | 117405548 | 117405645 | 117367448 | 117368485 | 117420568 | 117420649 | 3 | 0 | [illegible] |
| SE | 70504 | ENSG00000198162 | MAN1A2 | chr1 | + | 117414712 | 117414831 | 117367942 | 117368485 | 117420568 | 117420630 | 900 | 598 | [illegible] |
| SE | 70505 | ENSG00000198162 | MAN1A2 | chr1 | + | 117414712 | 117414831 | 117405548 | 117405645 | 117420568 | 117420630 | 1881 | 1143 | [illegible] |
| SE | 70506 | ENSG00000198162 | MAN1A2 | chr1 | + | 117420568 | 117420649 | 117414712 | 117414831 | 117442230 | 117442325 | 1288 | 844 | [illegible] |
| SE | 70507 | ENSG00000198162 | MAN1A2 | chr1 | + | 117499381 | 117499554 | 117496762 | 117496982 | 117502854 | 117502970 | 176 | 131 | [illegible] |
| SE | 70510 | ENSG00000021300 | PLEKHB1 | chr11 | + | 73649011 | 73649087 | 73647548 | 73647738 | 73650552 | 73650705 | 27 | 19 | [illegible] |
| SE | 70515 | ENSG00000021300 | PLEKHB1 | chr11 | + | 73655802 | 73655907 | 73652974 | 73653014 | 73660752 | 73660756 | 0 | 1 | [illegible] |
| SE | 70523 | ENSG00000167850 | CD300C | chr17 | - | 74542860 | 74542987 | 74541072 | 74541736 | 74545721 | 74546115 | 21 | 0 | [illegible] |
| SE | 70525 | ENSG00000073849 | ST6GAL1 | chr3 | + | 186942347 | 186942427 | 186930501 | 186930834 | 186948854 | 186948938 | 1 | 0 | [illegible] |
| SE | 70527 | ENSG00000073849 | ST6GAL1 | chr3 | + | 186942347 | 186942427 | 186931351 | 186931394 | 186963784 | 186963887 | 3 | 0 | [illegible] |
| SE | 70529 | ENSG00000073849 | ST6GAL1 | chr3 | + | 186948854 | 186948938 | 186942347 | 186942427 | 186963784 | 186963926 | 8 | 2 | [illegible] |
| SE | 70530 | ENSG00000073849 | ST6GAL1 | chr3 | + | 186963784 | 186963926 | 186930324 | 186930834 | 187038741 | 187038873 | 198 | 72 | [illegible] |
| SE | 70540 | ENSG00000073849 | ST6GAL1 | chr3 | + | 187038741 | 187038873 | 186963784 | 186963926 | 187042653 | 187042831 | 405 | 171 | [illegible] |
| SE | 70543 | ENSG00000073849 | ST6GAL1 | chr3 | + | 187038741 | 187038873 | 187021855 | 187022004 | 187042653 | 187042831 | 271 | 124 | [illegible] |
| SE | 70545 | ENSG00000073849 | ST6GAL1 | chr3 | + | 187038741 | 187038873 | 187025478 | 187025517 | 187042653 | 187042831 | 275 | 145 | [illegible] |
| SE | 70549 | ENSG00000073849 | ST6GAL1 | chr3 | + | 187042653 | 187043310 | 187038741 | 187038873 | 187051248 | 187051346 | 589 | 336 | [illegible] |
| SE | 70550 | ENSG00000073849 | ST6GAL1 | chr3 | + | 187042653 | 187043310 | 187038741 | 187038873 | 187072848 | 187072947 | 268 | 139 | [illegible] |
| SE | 70551 | ENSG00000073849 | ST6GAL1 | chr3 | + | 187051248 | 187051346 | 187038741 | 187038873 | 187072848 | 187072901 | 299 | 172 | [illegible] |
| SE | 70552 | ENSG00000073849 | ST6GAL1 | chr3 | + | 187051248 | 187051346 | 187042653 | 187043310 | 187072848 | 187072901 | 610 | 337 | [illegible] |
| SE | 70554 | ENSG00000073849 | ST6GAL1 | chr3 | + | 187072848 | 187072947 | 187051248 | 187051346 | 187074158 | 187074204 | 578 | 345 | [illegible] |
| SE | 70555 | ENSG00000073849 | ST6GAL1 | chr3 | + | 187073404 | 187074333 | 187071973 | 187072947 | 187075561 | 187078551 | 275 | 130 | [illegible] |
| SE | 70556 | ENSG00000073849 | ST6GAL1 | chr3 | + | 187074158 | 187074333 | 187072848 | 187072947 | 187075561 | 187075622 | 719 | 380 | [illegible] |
| SE | 70559 | ENSG00000022840 | RNF10 | chr12 | + | 120552498 | 120552698 | 120546404 | 120546601 | 120557281 | 120557466 | 197 | 70 | [illegible] |
| SE | 70560 | ENSG00000022840 | RNF10 | chr12 | + | 120554717 | 120554808 | 120546404 | 120546601 | 120557281 | 120557466 | 116 | 68 | [illegible] |
| SE | 70563 | ENSG00000022840 | RNF10 | chr12 | + | 120562944 | 120563070 | 120560725 | 120560886 | 120563331 | 120563456 | 164 | 83 | [illegible] |
| SE | 70564 | ENSG00000022840 | RNF10 | chr12 | + | 120562944 | 120563070 | 120560725 | 120560886 | 120563346 | 120563623 | 404 | 174 | [illegible] |
| SE | 70565 | ENSG00000022840 | RNF10 | chr12 | + | 120563331 | 120563623 | 120562944 | 120563070 | 120563809 | 120563943 | 370 | 176 | [illegible] |
| SE | 70566 | ENSG00000022840 | RNF10 | chr12 | + | 120563346 | 120563623 | 120562944 | 120563070 | 120563809 | 120563943 | 612 | 267 | [illegible] |
| SE | 70567 | ENSG00000022840 | RNF10 | chr12 | + | 120563440 | 120563623 | 120562944 | 120563070 | 120563809 | 120563943 | 339 | 157 | [illegible] |
| SE | 70568 | ENSG00000022840 | RNF10 | chr12 | + | 120563809 | 120563943 | 120562944 | 120563070 | 120565427 | 120565529 | 2 | 0 | [illegible] |
| SE | 70570 | ENSG00000022840 | RNF10 | chr12 | + | 120565427 | 120565529 | 120563809 | 120563943 | 120575630 | 120575688 | 2 | 0 | [illegible] |
| SE | 70572 | ENSG00000022840 | RNF10 | chr12 | + | 120566824 | 120566980 | 120565427 | 120565529 | 120575630 | 120575688 | 240 | 112 | [illegible] |
| SE | 70573 | ENSG00000022840 | RNF10 | chr12 | + | 120571190 | 120571291 | 120565483 | 120565529 | 120575630 | 120575688 | 143 | 80 | [illegible] |
| SE | 70584 | ENSG00000188092 | GPR89B | chr1 | + | 147968331 | 147968428 | 147966553 | 147966663 | 147968874 | 147968963 | 27 | 6 | [illegible] |
| SE | 70585 | ENSG00000188092 | GPR89B | chr1 | + | 147968874 | 147968963 | 147968331 | 147968428 | 147969866 | 147969959 | 22 | 7 | [illegible] |
| SE | 70591 | ENSG00000164548 | TRA2A | chr7 | - | 23506137 | 23506266 | 23505745 | 23505813 | 23507419 | 23507535 | 492 | 387 | [illegible] |
| SE | 70592 | ENSG00000164548 | TRA2A | chr7 | - | 23507419 | 23507535 | 23505745 | 23505813 | 23512893 | 23513082 | 394 | 329 | [illegible] |
| SE | 70594 | ENSG00000164548 | TRA2A | chr7 | - | 23516362 | 23516528 | 23512893 | 23513082 | 23521706 | 23521840 | 895 | 673 | [illegible] |
| SE | 70595 | ENSG00000164548 | TRA2A | chr7 | - | 23516362 | 23516528 | 23512893 | 23513082 | 23531788 | 23531980 | 267 | 201 | [illegible] |
| SE | 70596 | ENSG00000164548 | TRA2A | chr7 | - | 23521706 | 23521840 | 23512893 | 23513082 | 23531788 | 23531981 | 153 | 86 | [illegible] |
| SE | 70597 | ENSG00000164548 | TRA2A | chr7 | - | 23521706 | 23521840 | 23516362 | 23516528 | 23531788 | 23531981 | 766 | 541 | [illegible] |
| SE | 70598 | ENSG00000164548 | TRA2A | chr7 | - | 23521706 | 23522432 | 23512893 | 23513082 | 23531788 | 23531981 | 112 | 103 | [illegible] |
| SE | 70599 | ENSG00000164548 | TRA2A | chr7 | - | 23521706 | 23522432 | 23516362 | 23516528 | 23531788 | 23531981 | 733 | 558 | [illegible] |
| SE | 70601 | ENSG00000164548 | TRA2A | chr7 | - | 23522131 | 23522432 | 23521706 | 23521840 | 23531180 | 23531270 | 29 | 16 | [illegible] |
| SE | 70606 | ENSG00000225889 | NA | chr2 | + | 64188457 | 64188562 | 64143283 | 64143346 | 64201747 | 64201894 | 0 | 0 | [illegible] |
| SE | 70608 | ENSG00000225889 | NA | chr2 | + | 64206971 | 64207045 | 64201747 | 64201894 | 64226446 | 64226492 | 0 | 1 | [illegible] |
| SE | 70609 | ENSG00000225889 | NA | chr2 | + | 64206974 | 64207045 | 64201747 | 64201894 | 64226446 | 64226492 | 1 | 1 | [illegible] |
| SE | 70610 | ENSG00000225889 | NA | chr2 | + | 64214128 | 64214254 | 64206971 | 64207045 | 64214466 | 64214534 | 0 | 0 | [illegible] |
| SE | 70611 | ENSG00000225889 | NA | chr2 | + | 64214466 | 64214534 | 64206971 | 64207045 | 64226446 | 64226492 | 1 | 0 | [illegible] |
| SE | 70612 | ENSG00000225889 | NA | chr2 | + | 64226446 | 64226492 | 64206971 | 64207045 | 64229496 | 64229640 | 0 | 1 | [illegible] |
| SE | 70620 | ENSG00000225889 | NA | chr2 | + | 64250888 | 64250987 | 64229844 | 64229958 | 64251118 | 64251983 | 2 | 0 | [illegible] |
| SE | 70625 | ENSG00000160307 | S100B | chr21 | - | 46600281 | 46600375 | 46598961 | 46599503 | 46602277 | 46602416 | 4 | 0 | [illegible] |
| SE | 70628 | ENSG00000076201 | PTPN23 | chr3 | + | 47406324 | 47406405 | 47405914 | 47406046 | 47406480 | 47406612 | 56 | 13 | [illegible] |
| SE | 70629 | ENSG00000076201 | PTPN23 | chr3 | + | 47406702 | 47406750 | 47406480 | 47406612 | 47407129 | 47407186 | 60 | 30 | [illegible] |
| SE | 70630 | ENSG00000188647 | PTAR1 | chr9 | - | 69718649 | 69718684 | 69709524 | 69718568 | 69723325 | 69723571 | 45 | 34 | [illegible] |
| SE | 70631 | ENSG00000188647 | PTAR1 | chr9 | - | 69718649 | 69718684 | 69709524 | 69718568 | 69732138 | 69732352 | 11 | 3 | [illegible] |
| SE | 70632 | ENSG00000188647 | PTAR1 | chr9 | - | 69718649 | 69718684 | 69718020 | 69718565 | 69723325 | 69723571 | 42 | 30 | [illegible] |
| SE | 70633 | ENSG00000188647 | PTAR1 | chr9 | - | 69718649 | 69718684 | 69718020 | 69718565 | 69732138 | 69732352 | 7 | 5 | [illegible] |
| SE | 70637 | ENSG00000188647 | PTAR1 | chr9 | - | 69741791 | 69741858 | 69734149 | 69734254 | 69759852 | 69760011 | 34 | 32 | [illegible] |
| SE | 70644 | ENSG00000168214 | RBPJ | chr4 | + | 26362445 | 26362628 | 26321364 | 26321871 | 26386352 | 26386391 | 144 | 136 | [illegible] |
| SE | 70645 | ENSG00000168214 | RBPJ | chr4 | + | 26362445 | 26362628 | 26322869 | 26322950 | 26386352 | 26386391 | 144 | 136 | [illegible] |
| SE | 70648 | ENSG00000168214 | RBPJ | chr4 | + | 26366603 | 26366726 | 26362520 | 26362628 | 26386352 | 26386391 | 0 | 0 | [illegible] |
| SE | 70649 | ENSG00000168214 | RBPJ | chr4 | + | 26383884 | 26383960 | 26362445 | 26362628 | 26386352 | 26386391 | 0 | 1 | [illegible] |
| SE | 70655 | ENSG00000168214 | RBPJ | chr4 | + | 26412929 | 26412987 | 26406174 | 26406270 | 26414048 | 26414155 | 0 | 1 | [illegible] |
| SE | 70656 | ENSG00000168214 | RBPJ | chr4 | + | 26412929 | 26412987 | 26406174 | 26406270 | 26415474 | 26415640 | 0 | 1 | [illegible] |
| SE | 70658 | ENSG00000168214 | RBPJ | chr4 | + | 26414048 | 26414155 | 26412929 | 26412987 | 26415474 | 26415640 | 4 | 9 | [illegible] |
| SE | 70659 | ENSG00000168214 | RBPJ | chr4 | + | 26420550 | 26420725 | 26415474 | 26415640 | 26424341 | 26424383 | 630 | 428 | [illegible] |
| SE | 70662 | ENSG00000168214 | RBPJ | chr4 | + | 26428719 | 26428860 | 26424630 | 26424743 | 26430418 | 26430522 | 223 | 176 | [illegible] |
| SE | 70664 | ENSG00000168214 | RBPJ | chr4 | + | 26429897 | 26430053 | 26428719 | 26428860 | 26430691 | 26431156 | 154 | 127 | [illegible] |
| SE | 70665 | ENSG00000168214 | RBPJ | chr4 | + | 26430418 | 26430522 | 26428719 | 26428860 | 26430691 | 26430703 | 133 | 125 | [illegible] |
| SE | 70667 | ENSG00000115762 | PLEKHB2 | chr2 | + | 131120933 | 131120978 | 131104846 | 131105398 | 131125752 | 131125905 | 97 | 52 | [illegible] |
| SE | 70668 | ENSG00000115762 | PLEKHB2 | chr2 | + | 131130720 | 131130760 | 131126683 | 131126786 | 131132901 | 131132991 | 555 | 374 | [illegible] |
| SE | 70679 | ENSG00000095059 | DHPS | chr19 | - | 12677107 | 12677211 | 12676016 | 12676078 | 12677290 | 12677396 | 142 | 63 | [illegible] |
| SE | 70682 | ENSG00000095059 | DHPS | chr19 | - | 12677290 | 12677396 | 12676016 | 12676142 | 12679456 | 12679520 | 154 | 95 | [illegible] |
| SE | 70683 | ENSG00000095059 | DHPS | chr19 | - | 12679456 | 12679543 | 12677290 | 12677396 | 12679622 | 12679719 | 372 | 204 | [illegible] |
| SE | 70684 | ENSG00000095059 | DHPS | chr19 | - | 12679800 | 12679922 | 12679622 | 12679719 | 12680149 | 12680325 | 179 | 112 | [illegible] |
| SE | 70685 | ENSG00000095059 | DHPS | chr19 | - | 12679800 | 12679922 | 12679622 | 12679719 | 12680160 | 12680220 | 268 | 153 | [illegible] |
| SE | 70687 | ENSG00000095059 | DHPS | chr19 | - | 12679800 | 12680030 | 12679456 | 12679719 | 12680160 | 12680325 | 174 | 109 | [illegible] |
| SE | 70689 | ENSG00000114268 | PFKFB4 | chr3 | - | 48535511 | 48535658 | 48525564 | 48525669 | 48536255 | 48536463 | 72 | 47 | [illegible] |
| SE | 70690 | ENSG00000114268 | PFKFB4 | chr3 | - | 48535532 | 48535658 | 48525564 | 48525669 | 48536255 | 48536463 | 30 | 14 | [illegible] |
| SE | 70691 | ENSG00000114268 | PFKFB4 | chr3 | - | 48536255 | 48536463 | 48525564 | 48525669 | 48538497 | 48538619 | 37 | 24 | [illegible] |
| SE | 70693 | ENSG00000114268 | PFKFB4 | chr3 | - | 48539253 | 48539310 | 48538497 | 48538619 | 48539696 | 48539771 | 38 | 37 | [illegible] |
| SE | 70701 | ENSG00000197275 | RAD54B | chr8 | - | 94378567 | 94378634 | 94378179 | 94378380 | 94380144 | 94380406 | 10 | 9 | [illegible] |
| SE | 70702 | ENSG00000197275 | RAD54B | chr8 | - | 94404076 | 94404239 | 94400237 | 94400463 | 94407438 | 94407720 | 20 | 23 | [illegible] |
| SE | 70703 | ENSG00000197275 | RAD54B | chr8 | - | 94432610 | 94432656 | 94411120 | 94411315 | 94458267 | 94458436 | 1 | 0 | [illegible] |
| SE | 70706 | ENSG00000197275 | RAD54B | chr8 | - | 94458267 | 94458436 | 94431159 | 94432656 | 94467404 | 94467555 | 17 | 2 | [illegible] |
| SE | 70707 | ENSG00000153896 | ZNF599 | chr19 | - | 34772335 | 34772473 | 34769428 | 34769555 | 34772823 | 34773191 | 2 | 1 | [illegible] |
| SE | 70708 | ENSG00000112110 | MRPL18 | chr6 | + | 159790667 | 159790787 | 159789885 | 159790060 | 159790939 | 159791126 | 2 | 1 | [illegible] |
| SE | 70709 | ENSG00000112110 | MRPL18 | chr6 | + | 159790939 | 159791126 | 159789885 | 159790060 | 159797286 | 159797404 | 187 | 104 | [illegible] |
| SE | 70712 | ENSG00000117362 | APH1A | chr1 | - | 150267715 | 150267789 | 150267355 | 150267478 | 150267956 | 150267991 | 179 | 83 | [illegible] |
| SE | 70714 | ENSG00000117362 | APH1A | chr1 | - | 150267956 | 150267991 | 150267355 | 150267478 | 150268697 | 150269015 | 2 | 0 | [illegible] |
| SE | 70717 | ENSG00000183283 | DAZAP2 | chr12 | + | 51240342 | 51240461 | 51238723 | 51238920 | 51240870 | 51240929 | 196 | 96 | [illegible] |
| SE | 70721 | ENSG00000183283 | DAZAP2 | chr12 | + | 51240870 | 51241116 | 51240342 | 51240461 | 51246060 | 51246717 | 172 | 83 | [illegible] |
| SE | 70738 | ENSG00000102125 | TAZ | chrX | + | 154414100 | 154414190 | 154413481 | 154413567 | 154418545 | 154418602 | 6 | 2 | [illegible] |
| SE | 70740 | ENSG00000102125 | TAZ | chrX | + | 154418545 | 154418602 | 154413551 | 154413567 | 154419542 | 154419623 | 0 | 0 | [illegible] |
| SE | 70741 | ENSG00000102125 | TAZ | chrX | + | 154418545 | 154418602 | 154414100 | 154414190 | 154419542 | 154419623 | 0 | 0 | [illegible] |
| SE | 70742 | ENSG00000102125 | TAZ | chrX | + | 154419542 | 154419623 | 154412919 | 154413567 | 154420031 | 154420094 | 20 | 10 | [illegible] |
| SE | 70744 | ENSG00000102125 | TAZ | chrX | + | 154419542 | 154419746 | 154413481 | 154413567 | 154420031 | 154420094 | 22 | 12 | [illegible] |
| SE | 70746 | ENSG00000102125 | TAZ | chrX | + | 154420211 | 154420264 | 154419542 | 154420094 | 154420657 | 154420735 | 89 | 49 | [illegible] |
| SE | 70747 | ENSG00000166681 | BEX3 | chrX | + | 103377027 | 103377106 | 103376394 | 103376613 | 103377462 | 103378077 | 16 | 7 | [illegible] |
| SE | 70748 | ENSG00000166681 | BEX3 | chrX | + | 103377027 | 103377106 | 103376394 | 103376613 | 103377509 | 103378164 | 27 | 23 | [illegible] |
| SE | 70749 | ENSG00000040933 | INPP4A | chr2 | + | 98519945 | 98520154 | 98518963 | 98519025 | 98533376 | 98533456 | 60 | 50 | [illegible] |
| SE | 70751 | ENSG00000040933 | INPP4A | chr2 | + | 98539527 | 98539675 | 98538890 | 98538981 | 98543876 | 98544007 | 525 | 267 | [illegible] |
| SE | 70752 | ENSG00000040933 | INPP4A | chr2 | + | 98543876 | 98544007 | 98539527 | 98539675 | 98545968 | 98546073 | 516 | 250 | [illegible] |
| SE | 70754 | ENSG00000040933 | INPP4A | chr2 | + | 98545968 | 98546073 | 98539527 | 98539675 | 98552785 | 98552969 | 3 | 2 | [illegible] |
| SE | 70755 | ENSG00000040933 | INPP4A | chr2 | + | 98545968 | 98546073 | 98543876 | 98544007 | 98546585 | 98546694 | 563 | 286 | [illegible] |
| SE | 70759 | ENSG00000040933 | INPP4A | chr2 | + | 98548954 | 98548969 | 98546585 | 98546694 | 98552785 | 98552969 | 1 | 1 | [illegible] |
| SE | 70761 | ENSG00000040933 | INPP4A | chr2 | + | 98565639 | 98565766 | 98564639 | 98564763 | 98566028 | 98566169 | 118 | 49 | [illegible] |
| SE | 70762 | ENSG00000144199 | FAHD2B | chr2 | - | 97085698 | 97086198 | 97084168 | 97084277 | 97090108 | 97090325 | 27 | 18 | [illegible] |
| SE | 70763 | ENSG00000144199 | FAHD2B | chr2 | - | 97090108 | 97090325 | 97084168 | 97084277 | 97091461 | 97091712 | 4 | 11 | [illegible] |
| SE | 70774 | ENSG00000128656 | CHN1 | chr2 | - | 174875787 | 174875865 | 174846879 | 174846957 | 174877839 | 174877941 | 0 | 0 | [illegible] |
| SE | 70776 | ENSG00000128656 | CHN1 | chr2 | - | 174877839 | 174878128 | 174846879 | 174846957 | 174915057 | 174915171 | 4 | 6 | [illegible] |
| SE | 70779 | ENSG00000128656 | CHN1 | chr2 | - | 174917324 | 174917431 | 174915057 | 174915171 | 174918533 | 174918565 | 0 | 1 | [illegible] |
| SE | 70783 | ENSG00000128656 | CHN1 | chr2 | - | 174945169 | 174945261 | 174944887 | 174944943 | 174952163 | 174952202 | 1 | 0 | [illegible] |
| SE | 70788 | ENSG00000064201 | TSPAN32 | chr11 | + | 2304106 | 2304204 | 2302707 | 2302958 | 2308735 | 2308810 | 116 | 27 | [illegible] |
| SE | 70790 | ENSG00000064201 | TSPAN32 | chr11 | + | 2316228 | 2316312 | 2314484 | 2314571 | 2316575 | 2316667 | 45 | 7 | [illegible] |
| SE | 70794 | ENSG00000064201 | TSPAN32 | chr11 | + | 2317345 | 2317525 | 2316228 | 2316667 | 2317862 | 2318142 | 16 | 0 | [illegible] |
| SE | 70795 | ENSG00000064201 | TSPAN32 | chr11 | + | 2317426 | 2317525 | 2316635 | 2316667 | 2317862 | 2318142 | 10 | 0 | [illegible] |
| SE | 70796 | ENSG00000213799 | ZNF845 | chr19 | + | 53340823 | 53340925 | 53333748 | 53333792 | 53341234 | 53341322 | 0 | 0 | [illegible] |
| SE | 70797 | ENSG00000213799 | ZNF845 | chr19 | + | 53345505 | 53345632 | 53341234 | 53341322 | 53350817 | 53354869 | 49 | 47 | [illegible] |
| SE | 70798 | ENSG00000122026 | RPL21 | chr13 | + | 27253764 | 27253843 | 27251361 | 27251585 | 27254219 | 27254281 | 13127 | 6957 | [illegible] |
| SE | 70799 | ENSG00000122026 | RPL21 | chr13 | + | 27253764 | 27253843 | 27251361 | 27251585 | 27255241 | 27255354 | 49 | 47 | [illegible] |
| SE | 70800 | ENSG00000122026 | RPL21 | chr13 | + | 27253764 | 27253843 | 27251361 | 27251585 | 27256183 | 27256334 | 4 | 5 | [illegible] |
| SE | 70801 | ENSG00000122026 | RPL21 | chr13 | + | 27253764 | 27253843 | 27251582 | 27251850 | 27254219 | 27254281 | 11728 | 6201 | [illegible] |
| SE | 70802 | ENSG00000122026 | RPL21 | chr13 | + | 27253764 | 27253843 | 27251596 | 27251699 | 27254219 | 27254281 | 11726 | 6209 | [illegible] |
| SE | 70803 | ENSG00000122026 | RPL21 | chr13 | + | 27254219 | 27254281 | 27251557 | 27251585 | 27255241 | 27255349 | 2751 | 1671 | [illegible] |
| SE | 70806 | ENSG00000122026 | RPL21 | chr13 | + | 27254219 | 27254281 | 27253704 | 27253843 | 27256183 | 27256307 | 1230 | 642 | [illegible] |
| SE | 70808 | ENSG00000122026 | RPL21 | chr13 | + | 27255241 | 27255349 | 27254219 | 27254281 | 27256183 | 27256334 | 13470 | 7250 | [illegible] |
| SE | 70809 | ENSG00000122026 | RPL21 | chr13 | + | 27255241 | 27255354 | 27251557 | 27251585 | 27256183 | 27256307 | 18098 | 10168 | [illegible] |
| SE | 70810 | ENSG00000122026 | RPL21 | chr13 | + | 27255241 | 27255354 | 27253764 | 27253843 | 27256183 | 27256307 | 18149 | 10216 | [illegible] |
| SE | 70811 | ENSG00000122026 | RPL21 | chr13 | + | 27255241 | 27255354 | 27254219 | 27254281 | 27256183 | 27256307 | 32885 | 18254 | [illegible] |
| SE | 70820 | ENSG00000115839 | RAB3GAP1 | chr2 | + | 135090997 | 135091130 | 135058010 | 135058086 | 135093614 | 135093693 | 98 | 77 | [illegible] |
| SE | 70821 | ENSG00000115839 | RAB3GAP1 | chr2 | + | 135093614 | 135093693 | 135058010 | 135058086 | 135113150 | 135113270 | 23 | 17 | [illegible] |
| SE | 70822 | ENSG00000115839 | RAB3GAP1 | chr2 | + | 135124164 | 135124246 | 135120818 | 135120918 | 135126180 | 135126249 | 96 | 88 | [illegible] |
| SE | 70823 | ENSG00000115839 | RAB3GAP1 | chr2 | + | 135130551 | 135130721 | 135129994 | 135130087 | 135132894 | 135132984 | 224 | 171 | [illegible] |
| SE | 70825 | ENSG00000115839 | RAB3GAP1 | chr2 | + | 135162554 | 135162651 | 135153648 | 135153876 | 135162747 | 135162851 | 191 | 163 | [illegible] |
| SE | 70826 | ENSG00000115839 | RAB3GAP1 | chr2 | + | 135164593 | 135164696 | 135162985 | 135163101 | 135168544 | 135168749 | 168 | 143 | [illegible] |
| SE | 70828 | ENSG00000115839 | RAB3GAP1 | chr2 | + | 135169733 | 135169848 | 135168544 | 135168749 | 135175326 | 135175588 | 0 | 2 | [illegible] |
| SE | 70829 | ENSG00000105325 | FZR1 | chr19 | + | 3533293 | 3533398 | 3532416 | 3532650 | 3534420 | 3534513 | 79 | 32 | [illegible] |
| SE | 70830 | ENSG00000115392 | FANCL | chr2 | - | 58161521 | 58161638 | 58160107 | 58160179 | 58163433 | 58163517 | 2 | 10 | [illegible] |
| SE | 70831 | ENSG00000115392 | FANCL | chr2 | - | 58162865 | 58162947 | 58161521 | 58161638 | 58163028 | 58163074 | 16 | 15 | [illegible] |
| SE | 70832 | ENSG00000115392 | FANCL | chr2 | - | 58162865 | 58162947 | 58161521 | 58161638 | 58163433 | 58163517 | 10 | 11 | [illegible] |
| SE | 70833 | ENSG00000115392 | FANCL | chr2 | - | 58163028 | 58163074 | 58161521 | 58161638 | 58163433 | 58163517 | 3 | 8 | [illegible] |
| SE | 70834 | ENSG00000115392 | FANCL | chr2 | - | 58163028 | 58163074 | 58162865 | 58162947 | 58163433 | 58163517 | 9 | 12 | [illegible] |
| SE | 70836 | ENSG00000115392 | FANCL | chr2 | - | 58194228 | 58194273 | 58165723 | 58165874 | 58198578 | 58198662 | 1 | 0 | [illegible] |
| SE | 70837 | ENSG00000115392 | FANCL | chr2 | - | 58194228 | 58194273 | 58165723 | 58165874 | 58198593 | 58198662 | 2 | 0 | [illegible] |
| SE | 70839 | ENSG00000115392 | FANCL | chr2 | - | 58221941 | 58222042 | 58204129 | 58204226 | 58241217 | 58241372 | 26 | 18 | [illegible] |
| SE | 70846 | ENSG00000243970 | PPIEL | chr1 | - | 39524835 | 39525057 | 39523806 | 39523896 | 39529054 | 39529164 | 0 | 1 | [illegible] |
| SE | 70847 | ENSG00000243970 | PPIEL | chr1 | - | 39524835 | 39525151 | 39523806 | 39523896 | 39529054 | 39529164 | 2 | 1 | [illegible] |
| SE | 70848 | ENSG00000243970 | PPIEL | chr1 | - | 39525698 | 39525848 | 39522313 | 39522505 | 39529015 | 39529164 | 0 | 0 | [illegible] |
| SE | 70849 | ENSG00000243970 | PPIEL | chr1 | - | 39525698 | 39525848 | 39522313 | 39522505 | 39529054 | 39529164 | 0 | 2 | [illegible] |
| SE | 70852 | ENSG00000243970 | PPIEL | chr1 | - | 39537190 | 39537343 | 39529054 | 39529164 | 39541656 | 39541841 | 1 | 0 | [illegible] |
| SE | 70856 | ENSG00000243970 | PPIEL | chr1 | - | 39548682 | 39548746 | 39547578 | 39547634 | 39548861 | 39548951 | 3 | 4 | [illegible] |
| SE | 70862 | ENSG00000120833 | SOCS2 | chr12 | + | 93572684 | 93573036 | 93569813 | 93570198 | 93574721 | 93575010 | 37 | 14 | [illegible] |
| SE | 70863 | ENSG00000174915 | PTDSS2 | chr11 | + | 451351 | 451493 | 450450 | 450637 | 460186 | 460288 | 0 | 0 | [illegible] |
| SE | 70864 | ENSG00000174915 | PTDSS2 | chr11 | + | 451384 | 451493 | 450295 | 450637 | 460186 | 460288 | 0 | 0 | [illegible] |
| SE | 70865 | ENSG00000174915 | PTDSS2 | chr11 | + | 473894 | 473977 | 458673 | 460288 | 479084 | 479090 | 15 | 8 | [illegible] |
| SE | 70866 | ENSG00000255857 | PXN-AS1 | chr12 | + | 120210929 | 120211106 | 120209838 | 120209934 | 120212374 | 120213138 | 1 | 2 | [illegible] |
| SE | 70874 | ENSG00000119912 | IDE | chr10 | - | 92463730 | 92464003 | 92461190 | 92461252 | 92468878 | 92468990 | 48 | 46 | [illegible] |
| SE | 70877 | ENSG00000119912 | IDE | chr10 | - | 92465675 | 92465843 | 92463730 | 92464003 | 92468878 | 92468990 | 74 | 52 | [illegible] |
| SE | 70878 | ENSG00000119912 | IDE | chr10 | - | 92468878 | 92468990 | 92465675 | 92465843 | 92470253 | 92470345 | 67 | 51 | [illegible] |
| SE | 70880 | ENSG00000119912 | IDE | chr10 | - | 92478644 | 92478789 | 92475883 | 92475994 | 92479276 | 92479421 | 3 | 0 | [illegible] |
| SE | 70881 | ENSG00000119912 | IDE | chr10 | - | 92478644 | 92478789 | 92475883 | 92475994 | 92480792 | 92480916 | 2 | 0 | [illegible] |
| SE | 70883 | ENSG00000119912 | IDE | chr10 | - | 92479276 | 92479421 | 92475883 | 92475994 | 92480792 | 92480916 | 22 | 27 | [illegible] |
| SE | 70887 | ENSG00000119912 | IDE | chr10 | - | 92479276 | 92479421 | 92478644 | 92478789 | 92483254 | 92483337 | 40 | 26 | [illegible] |
| SE | 70890 | ENSG00000119912 | IDE | chr10 | - | 92480792 | 92480916 | 92478644 | 92478789 | 92483254 | 92483337 | 8 | 7 | [illegible] |
| SE | 70891 | ENSG00000119912 | IDE | chr10 | - | 92480792 | 92480916 | 92479276 | 92479421 | 92483254 | 92483337 | 8 | 9 | [illegible] |
| SE | 70893 | ENSG00000119912 | IDE | chr10 | - | 92483254 | 92483337 | 92479276 | 92479421 | 92487195 | 92487318 | 71 | 43 | [illegible] |
| SE | 70894 | ENSG00000119912 | IDE | chr10 | - | 92487195 | 92487318 | 92482948 | 92483337 | 92490492 | 92490595 | 168 | 129 | [illegible] |
| SE | 70895 | ENSG00000119912 | IDE | chr10 | - | 92504793 | 92504897 | 92490492 | 92490595 | 92506441 | 92506522 | 165 | 127 | [illegible] |
| SE | 70896 | ENSG00000119912 | IDE | chr10 | - | 92507574 | 92507666 | 92506441 | 92506522 | 92508112 | 92508205 | 155 | 99 | [illegible] |
| SE | 70897 | ENSG00000119912 | IDE | chr10 | - | 92508112 | 92508205 | 92507574 | 92507666 | 92508727 | 92508890 | 205 | 133 | [illegible] |
| SE | 70898 | ENSG00000119912 | IDE | chr10 | - | 92508727 | 92508890 | 92506441 | 92508205 | 92510049 | 92510162 | 186 | 118 | [illegible] |
| SE | 70900 | ENSG00000119912 | IDE | chr10 | - | 92554880 | 92554917 | 92537365 | 92537550 | 92555371 | 92555424 | 0 | 0 | [illegible] |
| SE | 70901 | ENSG00000119912 | IDE | chr10 | - | 92554880 | 92554917 | 92537365 | 92537550 | 92559033 | 92559215 | 6 | 1 | [illegible] |
| SE | 70903 | ENSG00000119912 | IDE | chr10 | - | 92559033 | 92559215 | 92537365 | 92537550 | 92573921 | 92574056 | 10 | 10 | [illegible] |
| SE | 70954 | ENSG00000131781 | FMO5 | chr1 | - | 147208851 | 147209051 | 147201151 | 147201504 | 147212392 | 147212535 | 43 | 16 | [illegible] |
| SE | 70956 | ENSG00000111665 | CDCA3 | chr12 | - | 6849322 | 6849413 | 6849153 | 6849198 | 6849564 | 6849858 | 4 | 1 | [illegible] |
| SE | 70957 | ENSG00000111665 | CDCA3 | chr12 | - | 6849322 | 6849429 | 6848815 | 6849198 | 6849564 | 6849783 | 3 | 1 | [illegible] |
| SE | 70958 | ENSG00000111665 | CDCA3 | chr12 | - | 6849564 | 6849858 | 6844792 | 6846903 | 6850466 | 6850490 | 0 | 0 | [illegible] |
| SE | 70962 | ENSG00000178719 | GRINA | chr8 | + | 143992244 | 143992373 | 143991877 | 143992078 | 143992464 | 143992608 | 227 | 100 | [illegible] |
| SE | 70963 | ENSG00000181798 | LINC00471 | chr2 | - | 231513526 | 231513632 | 231508421 | 231509343 | 231514096 | 231514352 | 2 | 2 | [illegible] |
| SE | 70965 | ENSG00000225830 | ERCC6 | chr10 | - | 49475326 | 49475496 | 49474026 | 49474242 | 49476214 | 49476258 | 0 | 1 | [illegible] |
| SE | 70966 | ENSG00000225830 | ERCC6 | chr10 | - | 49476214 | 49476310 | 49472375 | 49474242 | 49478353 | 49478470 | 45 | 30 | [illegible] |
| SE | 70967 | ENSG00000225830 | ERCC6 | chr10 | - | 49488157 | 49488379 | 49483345 | 49483516 | 49493116 | 49493252 | 3 | 0 | [illegible] |
| SE | 70968 | ENSG00000225830 | ERCC6 | chr10 | - | 49488157 | 49488379 | 49483345 | 49483516 | 49500537 | 49506480 | 0 | 0 | [illegible] |
| SE | 70969 | ENSG00000225830 | ERCC6 | chr10 | - | 49493116 | 49493252 | 49483345 | 49483516 | 49500537 | 49500696 | 34 | 23 | [illegible] |
| SE | 70970 | ENSG00000225830 | ERCC6 | chr10 | - | 49493116 | 49493252 | 49483345 | 49483516 | 49505883 | 49506012 | 25 | 22 | [illegible] |
| SE | 70971 | ENSG00000225830 | ERCC6 | chr10 | - | 49493116 | 49493252 | 49488157 | 49488379 | 49500537 | 49500696 | 13 | 4 | [illegible] |
| SE | 70972 | ENSG00000225830 | ERCC6 | chr10 | - | 49500537 | 49500696 | 49483345 | 49483516 | 49505883 | 49505999 | 56 | 31 | [illegible] |
| SE | 70974 | ENSG00000225830 | ERCC6 | chr10 | - | 49505883 | 49506012 | 49483345 | 49483516 | 49528416 | 49528525 | 4 | 1 | [illegible] |
| SE | 70975 | ENSG00000225830 | ERCC6 | chr10 | - | 49505883 | 49506012 | 49500537 | 49500696 | 49524032 | 49524777 | 70 | 41 | [illegible] |
| SE | 70976 | ENSG00000225830 | ERCC6 | chr10 | - | 49524032 | 49524777 | 49504991 | 49506012 | 49528416 | 49528525 | 135 | 89 | [illegible] |
| SE | 70980 | ENSG00000127081 | ZNF484 | chr9 | - | 92875014 | 92875059 | 92856191 | 92856318 | 92877819 | 92878038 | 8 | 7 | [illegible] |
| SE | 70981 | ENSG00000127081 | ZNF484 | chr9 | - | 92875014 | 92875059 | 92856191 | 92856318 | 92877889 | 92878038 | 30 | 9 | [illegible] |
| SE | 70982 | ENSG00000275778 | NA | chr12 | - | 10882216 | 10882698 | 10848371 | 10848407 | 10973654 | 10973721 | 3 | 0 | [illegible] |
| SE | 70983 | ENSG00000275778 | NA | chr12 | - | 10884153 | 10884275 | 10883060 | 10883096 | 10973654 | 10973721 | 13 | 1 | [illegible] |
| SE | 70986 | ENSG00000275778 | NA | chr12 | - | 11034781 | 11034885 | 10973654 | 10973721 | 11171421 | 11171598 | 1 | 4 | [illegible] |
| SE | 71010 | ENSG00000173867 | NA | chr15 | - | 88465586 | 88465773 | 88459500 | 88459863 | 88511431 | 88511521 | 0 | 4 | [illegible] |
| SE | 71012 | ENSG00000173867 | NA | chr15 | - | 88544610 | 88544694 | 88530622 | 88531715 | 88546539 | 88546585 | 0 | 0 | [illegible] |
| SE | 71013 | ENSG00000248636 | NA | chr12 | - | 119402314 | 119402430 | 119387986 | 119388328 | 119553547 | 119553645 | 0 | 0 | [illegible] |
| SE | 71016 | ENSG00000115657 | ABCB6 | chr2 | - | 219210933 | 219211108 | 219210380 | 219210823 | 219212386 | 219212491 | 9 | 6 | [illegible] |
| SE | 71036 | ENSG00000054356 | PTPRN | chr2 | - | 219291469 | 219291523 | 219290825 | 219290890 | 219294974 | 219295028 | 0 | 0 | [illegible] |
| SE | 71039 | ENSG00000272419 | LINC01145 | chr1 | - | 145169526 | 145169615 | 145169035 | 145169086 | 145170941 | 145171185 | 8 | 4 | [illegible] |
| SE | 71040 | ENSG00000272419 | LINC01145 | chr1 | - | 145169526 | 145169615 | 145169035 | 145169086 | 145214676 | 145214731 | 1 | 1 | [illegible] |

| | | | | | | | | | | | | | | | | | | | | | | | | | | | | | | | | | | | | | | | | | | | | | | | | |
|---|---|---|---|---|---|---|---|---|---|---|---|---|---|---|---|---|---|---|---|---|---|---|---|---|---|---|---|---|---|---|---|---|---|---|---|---|---|---|---|---|---|---|---|---|---|---|---|---|
| SE | 71041 | ENSG00000272419 | LINC01145 | chr1 | - | 145169526 | 145169615 | 145169035 | 145169086 | 145215638 | 145215814 | 2 | 1 | 8 | 7 | 9 | 21 | 8 | 22 | 10 | 37 | 0 | 0 | 0 | 0 | 1 | 8 | 0 | 0 | 14 | 2 | 8 | 0 | 238 | 149 | 1 | 1 | 0.056 | 0.073 | 0.185 | 0.305 | 0.132 NA | | 0 | 0.238 | 0.385 NA | | -0.057 |
| SE | 71042 | ENSG00000272419 | LINC01145 | chr1 | - | 145170941 | 145171185 | 145164098 | 145164638 | 145201141 | 145201301 | 1 | 0 | 3 | 2 | 0 | 0 | 0 | 0 | 0 | 2 | 7 | 0 | 7 | 0 | 0 | 0 | 0 | 0 | 0 | 1 | 0 | 0 | 298 | 149 | 1 | 1 | 1 NA | | 1 | 1 | 0 | 1 NA | | 0 NA | NA | | 0.25 |
| SE | 71043 | ENSG00000272419 | LINC01145 | chr1 | - | 145170941 | 145171185 | 145164098 | 145164638 | 145214676 | 145214731 | 6 | 0 | 5 | 8 | 0 | 5 | 0 | 0 | 0 | 0 | 13 | 0 | 13 | 0 | 0 | 0 | 0 | 1 | 0 | 1 | 0 | 5 | 298 | 149 | 1 | 1 | 0.375 NA | | 1 | 1 NA | | 0.867 NA | | 0 NA | NA | 0 | 0.503 |
| SE | 71044 | ENSG00000272419 | LINC01145 | chr1 | - | 145170941 | 145171185 | 145164098 | 145164638 | 145215638 | 145215814 | 10 | 3 | 9 | 6 | 25 | 0 | 0 | 1 | 0 | 0 | 7 | 5 | 7 | 5 | 2 | 4 | 0 | 0 | 0 | 0 | 0 | 0 | 298 | 149 | 1 | 1 | 1 | 1 | 0.818 | 1 | 1 | 1 | 1 | 1 | 1 NA | | -0.036 |
| SE | 71045 | ENSG00000272419 | LINC01145 | chr1 | - | 145170941 | 145171185 | 145168404 | 145169086 | 145214676 | 145214731 | 37 | 4 | 14 | 21 | 34 | 4 | 0 | 0 | 4 | 0 | 6 | 8 | 6 | 8 | 2 | 15 | 0 | 5 | 2 | 1 | 0 | 0 | 298 | 149 | 1 | 1 | 0.822 | 1 | 1 | 0.724 | 1 | 0.375 | 0.667 | 0.5 | 1 NA | | 0.274 |
| SE | 71046 | ENSG00000272419 | LINC01145 | chr1 | - | 145170941 | 145171185 | 145168404 | 145169086 | 145215638 | 145215814 | 41 | 7 | 18 | 19 | 59 | 21 | 8 | 22 | 10 | 37 | 0 | 13 | 0 | 13 | 4 | 19 | 0 | 0 | 14 | 2 | 8 | 0 | 298 | 149 | 1 | 1 | 0.494 | 0.304 | 0.29 | 0.487 | 0.444 NA | | 0.317 | 0.5 | 0.543 NA | | -0.05 |
| SE | 71047 | ENSG00000272419 | LINC01145 | chr1 | - | 145170941 | 145171185 | 145169526 | 145169615 | 145214676 | 145214731 | 12 | 3 | 8 | 16 | 0 | 0 | 0 | 0 | 3 | 0 | 6 | 0 | 6 | 0 | 0 | 11 | 11 | 0 | 0 | 0 | 0 | 0 | 298 | 149 | 1 | 1 | 1 | 1 | 1 | 0.727 NA | | 1 NA | NA | | 1 | 1 | -0.068 |
| SE | 71048 | ENSG00000272419 | LINC01145 | chr1 | - | 145170941 | 145171185 | 145169526 | 145169615 | 145215638 | 145215814 | 16 | 6 | 12 | 14 | 25 | 1 | 0 | 0 | 0 | 1 | 0 | 5 | 0 | 5 | 2 | 15 | 11 | 0 | 0 | 0 | 0 | 0 | 298 | 149 | 1 | 1 | 0.889 | 1 | 1 | 1 | 0.926 NA | | 1 | 1 | 1 | 1 | -0.037 |
| SE | 71049 | ENSG00000272419 | LINC01145 | chr1 | - | 145201141 | 145201301 | 145164098 | 145164638 | 145215638 | 145215872 | 0 | 2 | 7 | 3 | 9 | 0 | 0 | 1 | 0 | 0 | 0 | 0 | 0 | 0 | 13 | 4 | 0 | 0 | 0 | 0 | 0 | 0 | 298 | 149 | 1 | 1 NA | | 1 | 0.778 | 1 | 1 NA | NA | | 1 | 1 NA | 1 | -0.055 |
| SE | 71050 | ENSG00000272419 | LINC01145 | chr1 | - | 145201141 | 145201301 | 145170941 | 145171185 | 145215638 | 145215872 | 0 | 2 | 8 | 3 | 7 | 9 | 3 | 7 | 4 | 25 | 0 | 0 | 0 | 0 | 12 | 4 | 0 | 0 | 5 | 2 | 4 | 0 | 298 | 149 | 1 | 1 | 0 | 0.25 | 0.364 | 0.273 | 0.123 NA | | 0 | 0.75 | 0.333 NA | | -0.159 |
| SE | 71051 | ENSG00000272419 | LINC01145 | chr1 | - | 145214676 | 145214731 | 145164098 | 145164638 | 145215638 | 145215726 | 22 | 6 | 14 | 4 | 6 | 0 | 0 | 1 | 0 | 0 | 6 | 6 | 6 | 6 | 10 | 2 | 5 | 0 | 0 | 0 | 0 | 0 | 204 | 149 | 1 | 1 | 1 | 1 | 0.911 | 1 | 1 | 1 | 1 | 1 | 1 | 1 | -0.018 |
| SE | 71052 | ENSG00000272419 | LINC01145 | chr1 | - | 145214676 | 145214731 | 145168420 | 145169086 | 145215638 | 145215726 | 21 | 6 | 14 | 8 | 6 | 21 | 8 | 22 | 10 | 37 | 10 | 8 | 10 | 8 | 10 | 2 | 0 | 0 | 14 | 2 | 8 | 0 | 204 | 149 | 1 | 1 | 0.422 | 0.354 | 0.317 | 0.369 | 0.106 | 1 | 0.294 | 0.785 | 0.154 NA | | -0.245 |
| SE | 71053 | ENSG00000272419 | LINC01145 | chr1 | - | 145214676 | 145214731 | 145169526 | 145169615 | 145215638 | 145215726 | 17 | 6 | 14 | 7 | 6 | 1 | 0 | 0 | 0 | 1 | 5 | 6 | 5 | 6 | 9 | 2 | 0 | 0 | 0 | 0 | 0 | 0 | 204 | 149 | 1 | 1 | 0.925 | 1 | 1 | 1 | 0.814 | 1 | 1 | 1 | 1 NA | | -0.052 |
| SE | 71054 | ENSG00000272419 | LINC01145 | chr1 | - | 145214676 | 145214731 | 145170941 | 145171185 | 145215638 | 145215726 | 22 | 6 | 17 | 10 | 6 | 9 | 3 | 7 | 4 | 25 | 11 | 6 | 11 | 6 | 9 | 2 | 0 | 0 | 5 | 2 | 4 | 0 | 204 | 149 | 1 | 1 | 0.641 | 0.594 | 0.639 | 0.646 | 0.149 | 1 | 0.467 | 0.767 | 0.268 NA | | -0.092 |
| SE | 71055 | ENSG00000272419 | LINC01145 | chr1 | - | 145214676 | 145214731 | 145201187 | 145201301 | 145215638 | 145215726 | 20 | 7 | 15 | 5 | 6 | 0 | 2 | 7 | 3 | 7 | 5 | 10 | 5 | 10 | 9 | 3 | 0 | 0 | 0 | 12 | 4 | 0 | 204 | 149 | 1 | 1 | 1 | 0.719 | 0.61 | 0.549 | 0.385 | 1 | 1 | 0.354 | 0.354 NA | | -0.024 |
| SE | 71056 | ENSG00000169914 | OTUD3 | chr1 | + | 19890384 | 19890533 | 19882394 | 19882734 | 19897539 | 19897662 | 9 | 5 | 8 | 11 | 9 | 1 | 0 | 0 | 0 | 0 | 0 | 17 | 0 | 17 | 17 | 5 | 4 | 0 | 0 | 0 | 0 | 0 | 298 | 149 | 1 | 1 | 0.818 | 1 | 1 | 1 | 1 NA | | 1 | 1 | 1 | 1 | -0.036 |
| SE | 71060 | ENSG00000115486 | GGCX | chr2 | - | 85554142 | 85554306 | 85553231 | 85553497 | 85555483 | 85555590 | 34 | 10 | 16 | 19 | 25 | 0 | 1 | 1 | 1 | 0 | 10 | 34 | 10 | 34 | 6 | 29 | 8 | 0 | 0 | 0 | 0 | 0 | 298 | 149 | 1 | 1 | 1 | 0.833 | 0.889 | 0.905 | 1 | 1 | 1 | 1 | 1 | 1 | -0.075 |
| SE | 71061 | ENSG00000115486 | GGCX | chr2 | - | 85555483 | 85555590 | 85553231 | 85553497 | 85558916 | 85559075 | 0 | 0 | 1 | 1 | 0 | 0 | 0 | 2 | 0 | 0 | 0 | 2 | 0 | 2 | 0 | 0 | 0 | 0 | 0 | 0 | 0 | 0 | 256 | 149 | 1 | 1 NA | NA | | 0.225 | 1 NA | NA | | 1 NA | NA | NA | | -0.387 |
| SE | 71068 | ENSG00000134490 | TMEM241 | chr18 | - | 23368714 | 23368750 | 23365633 | 23365668 | 23370215 | 23370261 | 12 | 5 | 7 | 7 | 13 | 0 | 0 | 0 | 0 | 2 | 14 | 22 | 14 | 22 | 4 | 6 | 4 | 0 | 0 | 0 | 0 | 0 | 185 | 149 | 1 | 1 | 1 | 1 | 1 | 1 | 0.84 | 1 | 1 | 1 | 1 | 1 | -0.032 |
| SE | 71070 | ENSG00000134490 | TMEM241 | chr18 | - | 23370215 | 23370261 | 23356593 | 23356663 | 23373711 | 23373763 | 1 | 0 | 0 | 0 | 4 | 0 | 1 | 0 | 0 | 0 | 5 | 5 | 5 | 5 | 0 | 0 | 0 | 0 | 0 | 0 | 0 | 0 | 195 | 149 | 1 | 1 | 1 | 0 NA | NA | | 1 | 1 | 1 NA | NA | NA | | -0.333 |
| [illegible] | | | | | | | | | | | | | | | | | | | | | | | | | | | | | | | | | | | | | | | | | | | | | | | | |

| | | | | | | | | | | | | |
|---|---|---|---|---|---|---|---|---|---|---|---|---|
| SE | 71494 | ENSG00000211460 | TSN | chr2 | + | 121761408 | 121761524 | 121758709 | 121758806 | 121763004 | 121763057 | [illegible] |
| SE | 71498 | ENSG00000170006 | TMEM154 | chr4 | - | 152644414 | 152644442 | 152643087 | 152643173 | 152652537 | 152652576 | [illegible] |
| SE | 71501 | ENSG00000102221 | JADE3 | chrX | + | 47033620 | 47033788 | 47027891 | 47028103 | 47038948 | 47039065 | [illegible] |
| SE | 71506 | ENSG00000170325 | PRDM10 | chr11 | - | 129944770 | 129945012 | 129942425 | 129942629 | 129947144 | 129947292 | [illegible] |
| SE | 71516 | ENSG00000144504 | ANKMY1 | chr2 | - | 240507559 | 240507691 | 240500451 | 240500565 | 240509347 | 240509455 | [illegible] |
| SE | 71518 | ENSG00000144504 | ANKMY1 | chr2 | - | 240507559 | 240507691 | 240500451 | 240500565 | 240512801 | 240512942 | [illegible] |
| SE | 71520 | ENSG00000144504 | ANKMY1 | chr2 | - | 240509347 | 240509455 | 240500451 | 240500565 | 240512801 | 240512942 | [illegible] |
| SE | 71521 | ENSG00000144504 | ANKMY1 | chr2 | - | 240509347 | 240509455 | 240507559 | 240507691 | 240511860 | 240512001 | [illegible] |
| SE | 71522 | ENSG00000144504 | ANKMY1 | chr2 | - | 240509347 | 240509455 | 240507559 | 240507691 | 240512801 | 240512942 | [illegible] |
| SE | 71523 | ENSG00000144504 | ANKMY1 | chr2 | - | 240511860 | 240512001 | 240500451 | 240500565 | 240512801 | 240512942 | [illegible] |
| SE | 71524 | ENSG00000144504 | ANKMY1 | chr2 | - | 240511860 | 240512001 | 240507559 | 240507691 | 240512801 | 240512942 | [illegible] |
| SE | 71525 | ENSG00000144504 | ANKMY1 | chr2 | - | 240511860 | 240512001 | 240509347 | 240509455 | 240512801 | 240512942 | [illegible] |
| SE | 71527 | ENSG00000144504 | ANKMY1 | chr2 | - | 240526228 | 240526445 | 240525684 | 240525849 | 240529036 | 240529509 | [illegible] |
| SE | 71529 | ENSG00000144504 | ANKMY1 | chr2 | - | 240529036 | 240529509 | 240525684 | 240525849 | 240560730 | 240560776 | [illegible] |
| SE | 71530 | ENSG00000144504 | ANKMY1 | chr2 | - | 240529036 | 240529509 | 240526228 | 240526445 | 240552913 | 240553057 | [illegible] |
| SE | 71532 | ENSG00000144504 | ANKMY1 | chr2 | - | 240557189 | 240557352 | 240554865 | 240555055 | 240558303 | 240558619 | [illegible] |
| SE | 71533 | ENSG00000144504 | ANKMY1 | chr2 | - | 240557189 | 240557352 | 240554865 | 240555055 | 240560730 | 240561110 | [illegible] |
| SE | 71534 | ENSG00000144504 | ANKMY1 | chr2 | - | 240558303 | 240558619 | 240554978 | 240555055 | 240560730 | 240560826 | [illegible] |
| SE | 71539 | ENSG00000145216 | FIP1L1 | chr4 | + | 53379072 | 53379117 | 53378217 | 53378528 | 53379224 | 53379264 | [illegible] |
| SE | 71540 | ENSG00000145216 | FIP1L1 | chr4 | + | 53382277 | 53382335 | 53377802 | 53379264 | 53383772 | 53383876 | [illegible] |
| SE | 71547 | ENSG00000129691 | ASH2L | chr8 | + | 38107020 | 38107166 | 38106377 | 38106444 | 38110378 | 38110467 | [illegible] |
| SE | 71548 | ENSG00000129691 | ASH2L | chr8 | + | 38114191 | 38114287 | 38110736 | 38110833 | 38114904 | 38115000 | [illegible] |
| SE | 71549 | ENSG00000129691 | ASH2L | chr8 | + | 38114904 | 38115000 | 38110736 | 38110833 | 38116649 | 38116725 | [illegible] |
| SE | 71550 | ENSG00000129691 | ASH2L | chr8 | + | 38116649 | 38116725 | 38114757 | 38115000 | 38117451 | 38117812 | [illegible] |
| SE | 71554 | ENSG00000129691 | ASH2L | chr8 | + | 38119269 | 38119363 | 38116649 | 38116725 | 38120931 | 38121149 | [illegible] |
| SE | 71557 | ENSG00000198799 | LRIG2 | chr1 | + | 113095873 | 113096017 | 113094611 | 113094755 | 113096221 | 113096365 | [illegible] |
| SE | 71561 | ENSG00000198799 | LRIG2 | chr1 | + | 113119232 | 113119523 | 113116286 | 113116436 | 113123874 | 113124997 | [illegible] |
| SE | 71578 | ENSG00000164114 | MAP9 | chr4 | - | 155355070 | 155355160 | 155353178 | 155353340 | 155357448 | 155357519 | [illegible] |
| SE | 71579 | ENSG00000164114 | MAP9 | chr4 | - | 155355715 | 155355884 | 155355070 | 155355160 | 155357448 | 155357519 | [illegible] |
| SE | 71580 | ENSG00000164114 | MAP9 | chr4 | - | 155374936 | 155375021 | 155372995 | 155373456 | 155375775 | 155375870 | [illegible] |
| SE | 71581 | ENSG00000164114 | MAP9 | chr4 | - | 155375775 | 155375870 | 155374936 | 155375021 | 155376770 | 155376813 | [illegible] |
| SE | 71583 | ENSG00000164114 | MAP9 | chr4 | - | 155375775 | 155375914 | 155374936 | 155375021 | 155376770 | 155376813 | [illegible] |
| SE | 71585 | ENSG00000115504 | EHBP1 | chr2 | + | 62858405 | 62858510 | 62831018 | 62831158 | 62859168 | 62859291 | [illegible] |
| SE | 71586 | ENSG00000115504 | EHBP1 | chr2 | + | 62948259 | 62948316 | 62943801 | 62943850 | 62955516 | 62956117 | [illegible] |
| SE | 71587 | ENSG00000115504 | EHBP1 | chr2 | + | 62948259 | 62949162 | 62943801 | 62943850 | 62955516 | 62955660 | [illegible] |
| SE | 71588 | ENSG00000115504 | EHBP1 | chr2 | + | 62949093 | 62949162 | 62948291 | 62948316 | 62955516 | 62955660 | [illegible] |
| SE | 71592 | ENSG00000115504 | EHBP1 | chr2 | + | 63038742 | 63038816 | 63037534 | 63037634 | 63045065 | 63045180 | [illegible] |
| SE | 71593 | ENSG00000100448 | CTSG | chr14 | - | 24575264 | 24575412 | 24574674 | 24574810 | 24576168 | 24576250 | [illegible] |
| SE | 71594 | ENSG00000104998 | IL27RA | chr19 | + | 14039507 | 14039665 | 14032385 | 14032503 | 14039752 | 14039910 | [illegible] |
| SE | 71595 | ENSG00000104998 | IL27RA | chr19 | + | 14039507 | 14039665 | 14032385 | 14032503 | 14042452 | 14042612 | [illegible] |
| SE | 71596 | ENSG00000104998 | IL27RA | chr19 | + | 14039752 | 14039910 | 14032385 | 14032503 | 14042452 | 14042612 | [illegible] |
| SE | 71597 | ENSG00000104998 | IL27RA | chr19 | + | 14039752 | 14039910 | 14039507 | 14039665 | 14042452 | 14042612 | [illegible] |
| SE | 71598 | ENSG00000104998 | IL27RA | chr19 | + | 14050757 | 14050883 | 14049155 | 14049314 | 14051606 | 14051700 | [illegible] |
| SE | 71601 | ENSG00000169247 | SH3TC2 | chr5 | - | 149006133 | 149006251 | 149004621 | 149004902 | 149006880 | 149006912 | [illegible] |
| SE | 71602 | ENSG00000169247 | SH3TC2 | chr5 | - | 149026059 | 149026200 | 149011793 | 149012734 | 149026571 | 149026752 | [illegible] |
| SE | 71607 | ENSG00000169247 | SH3TC2 | chr5 | - | 149036343 | 149036675 | 149031585 | 149031687 | 149038294 | 149038490 | [illegible] |
| SE | 71610 | ENSG00000169247 | SH3TC2 | chr5 | - | 149045893 | 149045961 | 149044532 | 149044638 | 149047861 | 149047989 | [illegible] |
| SE | 71612 | ENSG00000184182 | UBE2F | chr2 | + | 237973648 | 237973747 | 237973091 | 237973225 | 237994743 | 237994809 | [illegible] |
| SE | 71613 | ENSG00000184182 | UBE2F | chr2 | + | 237986161 | 237986292 | 237973091 | 237973225 | 237987962 | 237987992 | [illegible] |
| SE | 71615 | ENSG00000184182 | UBE2F | chr2 | + | 237987962 | 237987992 | 237973648 | 237973747 | 237994743 | 237994809 | [illegible] |
| SE | 71616 | ENSG00000184182 | UBE2F | chr2 | + | 237990423 | 237990552 | 237987962 | 237987992 | 237994743 | 237994809 | [illegible] |
| SE | 71619 | ENSG00000184182 | UBE2F | chr2 | + | 238003367 | 238003527 | 237994743 | 237994809 | 238016565 | 238016630 | [illegible] |
| SE | 71620 | ENSG00000184182 | UBE2F | chr2 | + | 238003367 | 238003527 | 237994743 | 237994809 | 238025341 | 238025412 | [illegible] |
| SE | 71626 | ENSG00000184182 | UBE2F | chr2 | + | 238016565 | 238016633 | 238003367 | 238003527 | 238025341 | 238025412 | [illegible] |
| SE | 71627 | ENSG00000184182 | UBE2F | chr2 | + | 238025341 | 238025412 | 237987962 | 237987992 | 238035877 | 238035940 | [illegible] |
| SE | 71629 | ENSG00000184182 | UBE2F | chr2 | + | 238025341 | 238025412 | 237994743 | 237994809 | 238035877 | 238035940 | [illegible] |
| SE | 71630 | ENSG00000184182 | UBE2F | chr2 | + | 238025341 | 238025412 | 238016532 | 238016633 | 238030555 | 238030595 | [illegible] |
| SE | 71632 | ENSG00000184182 | UBE2F | chr2 | + | 238030555 | 238030613 | 238025260 | 238025412 | 238032221 | 238032254 | [illegible] |
| SE | 71636 | ENSG00000184182 | UBE2F | chr2 | + | 238035877 | 238035940 | 238025341 | 238025412 | 238041287 | 238041293 | [illegible] |
| SE | 71637 | ENSG00000113916 | BCL6 | chr3 | - | 187724940 | 187725078 | 187721376 | 187722601 | 187725498 | 187725629 | [illegible] |
| SE | 71639 | ENSG00000113916 | BCL6 | chr3 | - | 187726730 | 187726898 | 187725498 | 187725629 | 187729049 | 187730021 | [illegible] |
| SE | 71640 | ENSG00000113916 | BCL6 | chr3 | - | 187728359 | 187728544 | 187725498 | 187725629 | 187729049 | 187730021 | [illegible] |
| SE | 71642 | ENSG00000113916 | BCL6 | chr3 | - | 187731708 | 187731868 | 187729049 | 187730021 | 187733532 | 187733703 | [illegible] |
| SE | 71643 | ENSG00000113916 | BCL6 | chr3 | - | 187731708 | 187731930 | 187729049 | 187730021 | 187733532 | 187733703 | [illegible] |
| SE | 71645 | ENSG00000163832 | ELP6 | chr3 | - | 47498285 | 47498432 | 47495639 | 47496197 | 47501649 | 47501851 | [illegible] |
| SE | 71647 | ENSG00000163832 | ELP6 | chr3 | - | 47501649 | 47501851 | 47495639 | 47496197 | 47504329 | 47504448 | [illegible] |
| SE | 71652 | ENSG00000163832 | ELP6 | chr3 | - | 47511147 | 47511226 | 47504329 | 47504448 | 47513536 | 47513709 | [illegible] |
| SE | 71654 | ENSG00000077380 | DYNC1I2 | chr2 | + | 171690146 | 171690263 | 171687408 | 171687627 | 171707286 | 171707377 | [illegible] |
| SE | 71655 | ENSG00000077380 | DYNC1I2 | chr2 | + | 171690146 | 171690263 | 171687671 | 171687873 | 171707286 | 171707377 | [illegible] |
| SE | 71660 | ENSG00000077380 | DYNC1I2 | chr2 | + | 171712766 | 171712826 | 171706889 | 171707377 | 171715327 | 171715350 | [illegible] |
| SE | 71661 | ENSG00000077380 | DYNC1I2 | chr2 | + | 171725617 | 171725713 | 171715327 | 171715443 | 171725918 | 171725946 | [illegible] |
| SE | 71663 | ENSG00000077380 | DYNC1I2 | chr2 | + | 171726790 | 171726916 | 171726193 | 171726293 | 171727820 | 171727967 | [illegible] |
| SE | 71664 | ENSG00000140398 | NEIL1 | chr15 | + | 75347202 | 75347473 | 75346954 | 75347110 | 75348883 | 75349098 | [illegible] |
| SE | 71665 | ENSG00000140398 | NEIL1 | chr15 | + | 75347246 | 75347473 | 75347052 | 75347110 | 75348883 | 75349170 | [illegible] |
| SE | 71666 | ENSG00000140398 | NEIL1 | chr15 | + | 75347834 | 75347999 | 75347434 | 75347473 | 75348883 | 75349339 | [illegible] |
| SE | 71667 | ENSG00000140398 | NEIL1 | chr15 | + | 75347958 | 75347999 | 75347068 | 75347473 | 75348883 | 75349024 | [illegible] |
| SE | 71668 | ENSG00000140398 | NEIL1 | chr15 | + | 75347958 | 75348026 | 75347202 | 75347473 | 75348883 | 75348971 | [illegible] |
| SE | 71672 | ENSG00000140398 | NEIL1 | chr15 | + | 75351300 | 75351376 | 75348883 | 75349339 | 75352110 | 75352230 | [illegible] |
| SE | 71674 | ENSG00000140398 | NEIL1 | chr15 | + | 75353738 | 75353866 | 75352601 | 75352701 | 75354249 | 75354277 | [illegible] |
| SE | 71677 | ENSG00000153823 | PID1 | chr2 | - | 229262682 | 229262893 | 229155817 | 229155964 | 229271013 | 229271265 | [illegible] |
| SE | 71678 | ENSG00000135976 | ANKRD36 | chr2 | + | 97118343 | 97118517 | 97118063 | 97118178 | 97127066 | 97127134 | [illegible] |
| SE | 71679 | ENSG00000135976 | ANKRD36 | chr2 | + | 97118343 | 97118517 | 97118063 | 97118178 | 97142639 | 97142660 | [illegible] |
| SE | 71680 | ENSG00000135976 | ANKRD36 | chr2 | + | 97122886 | 97122993 | 97118343 | 97118517 | 97124459 | 97124597 | [illegible] |
| SE | 71682 | ENSG00000135976 | ANKRD36 | chr2 | + | 97122886 | 97122993 | 97118343 | 97118517 | 97179737 | 97179766 | [illegible] |
| SE | 71684 | ENSG00000135976 | ANKRD36 | chr2 | + | 97124459 | 97124597 | 97118343 | 97118517 | 97179737 | 97179766 | [illegible] |
| SE | 71686 | ENSG00000135976 | ANKRD36 | chr2 | + | 97124459 | 97124597 | 97122886 | 97122993 | 97179737 | 97179766 | [illegible] |
| SE | 71687 | ENSG00000135976 | ANKRD36 | chr2 | + | 97127066 | 97127134 | 97118063 | 97118178 | 97142639 | 97142660 | [illegible] |
| SE | 71689 | ENSG00000135976 | ANKRD36 | chr2 | + | 97127066 | 97127134 | 97118343 | 97118517 | 97179737 | 97179766 | [illegible] |
| SE | 71690 | ENSG00000135976 | ANKRD36 | chr2 | + | 97127066 | 97127134 | 97122886 | 97122993 | 97179737 | 97179766 | [illegible] |
| SE | 71691 | ENSG00000135976 | ANKRD36 | chr2 | + | 97127066 | 97127134 | 97124459 | 97124597 | 97179737 | 97179766 | [illegible] |
| SE | 71693 | ENSG00000135976 | ANKRD36 | chr2 | + | 97144639 | 97144712 | 97144517 | 97144546 | 97146485 | 97146516 | [illegible] |
| SE | 71695 | ENSG00000135976 | ANKRD36 | chr2 | + | 97146485 | 97146516 | 97144639 | 97144712 | 97151878 | 97151939 | [illegible] |
| SE | 71698 | ENSG00000135976 | ANKRD36 | chr2 | + | 97149294 | 97149361 | 97146485 | 97146516 | 97151878 | 97151939 | [illegible] |
| SE | 71699 | ENSG00000135976 | ANKRD36 | chr2 | + | 97151878 | 97151939 | 97149294 | 97149361 | 97152503 | 97152534 | [illegible] |
| SE | 71701 | ENSG00000135976 | ANKRD36 | chr2 | + | 97154674 | 97154741 | 97152503 | 97152534 | 97158106 | 97158167 | [illegible] |
| SE | 71702 | ENSG00000135976 | ANKRD36 | chr2 | + | 97158587 | 97158655 | 97158106 | 97158167 | 97162098 | 97162138 | [illegible] |
| SE | 71705 | ENSG00000135976 | ANKRD36 | chr2 | + | 97181597 | 97181626 | 97179737 | 97179766 | 97181720 | 97181793 | [illegible] |
| SE | 71706 | ENSG00000135976 | ANKRD36 | chr2 | + | 97183581 | 97183654 | 97183458 | 97183487 | 97185315 | 97185344 | [illegible] |
| SE | 71709 | ENSG00000135976 | ANKRD36 | chr2 | + | 97198585 | 97198658 | 97198462 | 97198491 | 97200333 | 97200362 | [illegible] |
| SE | 71710 | ENSG00000135976 | ANKRD36 | chr2 | + | 97200333 | 97200362 | 97198462 | 97198491 | 97200452 | 97200525 | [illegible] |
| SE | 71713 | ENSG00000135976 | ANKRD36 | chr2 | + | 97204190 | 97204263 | 97204067 | 97204096 | 97209680 | 97209709 | [illegible] |
| SE | 71714 | ENSG00000135976 | ANKRD36 | chr2 | + | 97205939 | 97205968 | 97204190 | 97204263 | 97207810 | 97207839 | [illegible] |
| SE | 71715 | ENSG00000135976 | ANKRD36 | chr2 | + | 97206062 | 97206135 | 97205939 | 97205968 | 97207810 | 97207839 | [illegible] |
| SE | 71716 | ENSG00000135976 | ANKRD36 | chr2 | + | 97207810 | 97207839 | 97204190 | 97204263 | 97209680 | 97209709 | [illegible] |
| SE | 71718 | ENSG00000135976 | ANKRD36 | chr2 | + | 97207810 | 97207839 | 97206062 | 97206135 | 97209680 | 97209709 | [illegible] |
| SE | 71719 | ENSG00000135976 | ANKRD36 | chr2 | + | 97207933 | 97208006 | 97202201 | 97202230 | 97209680 | 97209709 | [illegible] |
| SE | 71723 | ENSG00000135976 | ANKRD36 | chr2 | + | 97209680 | 97209709 | 97204190 | 97204263 | 97211545 | 97211574 | [illegible] |
| SE | 71724 | ENSG00000135976 | ANKRD36 | chr2 | + | 97209680 | 97209709 | 97207810 | 97207839 | 97209799 | 97209872 | [illegible] |
| SE | 71725 | ENSG00000135976 | ANKRD36 | chr2 | + | 97209799 | 97209872 | 97209680 | 97209709 | 97211545 | 97211574 | [illegible] |
| SE | 71726 | ENSG00000135976 | ANKRD36 | chr2 | + | 97219173 | 97219246 | 97219049 | 97219078 | 97224808 | 97224879 | [illegible] |
| SE | 71731 | ENSG00000179051 | RCC2 | chr1 | - | 17412121 | 17412194 | 17406759 | 17409194 | 17413072 | 17413178 | [illegible] |
| SE | 71732 | ENSG00000179051 | RCC2 | chr1 | - | 17413072 | 17413178 | 17406759 | 17409194 | 17413536 | 17413717 | [illegible] |
| SE | 71733 | ENSG00000179051 | RCC2 | chr1 | - | 17413072 | 17413178 | 17406759 | 17409194 | 17416479 | 17416646 | [illegible] |
| SE | 71734 | ENSG00000179051 | RCC2 | chr1 | - | 17413536 | 17413717 | 17406759 | 17409194 | 17416479 | 17416646 | [illegible] |
| SE | 71736 | ENSG00000179051 | RCC2 | chr1 | - | 17420713 | 17420828 | 17416479 | 17416646 | 17422202 | 17422291 | [illegible] |
| SE | 71737 | ENSG00000179051 | RCC2 | chr1 | - | 17422202 | 17422291 | 17416479 | 17416646 | 17425540 | 17425684 | [illegible] |
| SE | 71738 | ENSG00000179051 | RCC2 | chr1 | - | 17422202 | 17422291 | 17420713 | 17420828 | 17422704 | 17422836 | [illegible] |
| SE | 71739 | ENSG00000179051 | RCC2 | chr1 | - | 17422704 | 17422836 | 17422202 | 17422291 | 17425540 | 17425684 | [illegible] |
| SE | 71740 | ENSG00000179051 | RCC2 | chr1 | - | 17425540 | 17425684 | 17416479 | 17416646 | 17429105 | 17429199 | [illegible] |
| SE | 71742 | ENSG00000179051 | RCC2 | chr1 | - | 17429105 | 17429199 | 17422704 | 17422836 | 17438229 | 17438522 | [illegible] |
| SE | 71744 | ENSG00000071909 | MYO3B | chr2 | + | 170200149 | 170200284 | 170199284 | 170199391 | 170214378 | 170214483 | [illegible] |
| SE | 71746 | ENSG00000071909 | MYO3B | chr2 | + | 170383072 | 170383189 | 170382015 | 170382112 | 170383709 | 170383814 | [illegible] |
| SE | 71747 | ENSG00000071909 | MYO3B | chr2 | + | 170392380 | 170392495 | 170391519 | 170391618 | 170400187 | 170400314 | [illegible] |
| SE | 71748 | ENSG00000071909 | MYO3B | chr2 | + | 170466505 | 170466711 | 170463367 | 170463445 | 170498591 | 170498703 | [illegible] |
| SE | 71750 | ENSG00000071909 | MYO3B | chr2 | + | 170514920 | 170515022 | 170501784 | 170501865 | 170519437 | 170519540 | [illegible] |
| SE | 71753 | ENSG00000128309 | MPST | chr22 | + | 37023751 | 37023927 | 37019695 | 37019774 | 37024191 | 37024810 | [illegible] |
| SE | 71754 | ENSG00000128309 | MPST | chr22 | + | 37023751 | 37023927 | 37019767 | 37019872 | 37024191 | 37024810 | [illegible] |
| SE | 71755 | ENSG00000128309 | MPST | chr22 | + | 37025001 | 37025127 | 37024191 | 37024810 | 37029215 | 37029796 | [illegible] |
| SE | 71757 | ENSG00000021574 | SPAST | chr2 | + | 32087491 | 32087578 | 32063555 | 32064246 | 32089521 | 32089605 | [illegible] |
| SE | 71759 | ENSG00000021574 | SPAST | chr2 | + | 32087494 | 32087578 | 32063781 | 32064246 | 32089521 | 32089605 | [illegible] |
| SE | 71762 | ENSG00000021574 | SPAST | chr2 | + | 32089521 | 32089605 | 32087491 | 32087578 | 32098795 | 32098891 | [illegible] |
| SE | 71764 | ENSG00000021574 | SPAST | chr2 | + | 32098800 | 32098891 | 32089521 | 32089605 | 32114637 | 32114825 | [illegible] |
| SE | 71767 | ENSG00000021574 | SPAST | chr2 | + | 32128407 | 32128479 | 32116118 | 32116212 | 32136562 | 32136638 | [illegible] |
| SE | 71769 | ENSG00000021574 | SPAST | chr2 | + | 32136876 | 32136968 | 32136562 | 32136638 | 32137108 | 32137188 | [illegible] |
| SE | 71770 | ENSG00000021574 | SPAST | chr2 | + | 32141903 | 32141946 | 32137108 | 32137188 | 32143335 | 32143349 | [illegible] |
| SE | 71771 | ENSG00000021574 | SPAST | chr2 | + | 32143335 | 32143415 | 32141903 | 32141946 | 32144936 | 32145007 | [illegible] |
| SE | 71785 | ENSG00000196586 | MYO6 | chr6 | + | 75858901 | 75858993 | 75857096 | 75857254 | 75861022 | 75861095 | [illegible] |
| SE | 71788 | ENSG00000196586 | MYO6 | chr6 | + | 75873206 | 75873300 | 75870646 | 75870685 | 75879819 | 75879950 | [illegible] |
| SE | 71790 | ENSG00000196586 | MYO6 | chr6 | + | 75895230 | 75895260 | 75892529 | 75892690 | 75907604 | 75907708 | [illegible] |
| SE | 71793 | ENSG00000260238 | PMF1-BGLAP | chr1 | + | 156236287 | 156236483 | 156233627 | 156233728 | 156242552 | 156242591 | [illegible] |
| SE | 71794 | ENSG00000260238 | PMF1-BGLAP | chr1 | + | 156236348 | 156236483 | 156233627 | 156233728 | 156242552 | 156242591 | [illegible] |
| SE | 71796 | ENSG00000258539 | NA | chr10 | - | 124765391 | 124765608 | 124760385 | 124760503 | 124774674 | 124774782 | [illegible] |
| SE | 71797 | ENSG00000258539 | NA | chr10 | - | 124774674 | 124774782 | 124765391 | 124765608 | 124789042 | 124789157 | [illegible] |
| SE | 71798 | ENSG00000258539 | NA | chr10 | - | 124789042 | 124789157 | 124774674 | 124774782 | 124790272 | 124790338 | [illegible] |
| SE | 71799 | ENSG00000258539 | NA | chr10 | - | 124789042 | 124789157 | 124774674 | 124774782 | 124791723 | 124791727 | [illegible] |
| SE | 71800 | ENSG00000258539 | NA | chr10 | - | 124790272 | 124790338 | 124774674 | 124774782 | 124791723 | 124791727 | [illegible] |
| SE | 71801 | ENSG00000258539 | NA | chr10 | - | 124790272 | 124790338 | 124789042 | 124789157 | 124791723 | 124791727 | [illegible] |
| SE | 71802 | ENSG00000124491 | F13A1 | chr6 | - | 6151812 | 6151949 | 6144083 | 6145772 | 6167457 | 6167618 | [illegible] |
| SE | 71803 | ENSG00000124491 | F13A1 | chr6 | - | 6151812 | 6151949 | 6144083 | 6145772 | 6174579 | 6174867 | [illegible] |
| SE | 71804 | ENSG00000124491 | F13A1 | chr6 | - | 6167457 | 6167618 | 6144083 | 6145772 | 6174579 | 6174867 | [illegible] |
| SE | 71805 | ENSG00000124491 | F13A1 | chr6 | - | 6167457 | 6167618 | 6151812 | 6151949 | 6174579 | 6174867 | [illegible] |
| SE | 71806 | ENSG00000124491 | F13A1 | chr6 | - | 6174579 | 6174867 | 6167457 | 6167618 | 6181987 | 6182141 | [illegible] |
| SE | 71808 | ENSG00000124491 | F13A1 | chr6 | - | 6195796 | 6195885 | 6174579 | 6174867 | 6197222 | 6197326 | [illegible] |
| SE | 71809 | ENSG00000124491 | F13A1 | chr6 | - | 6248311 | 6248419 | 6197222 | 6197326 | 6266557 | 6266809 | [illegible] |
| SE | 71810 | ENSG00000124491 | F13A1 | chr6 | - | 6250810 | 6250929 | 6248311 | 6248419 | 6266557 | 6266809 | [illegible] |
| SE | 71811 | ENSG00000124491 | F13A1 | chr6 | - | 6266557 | 6266809 | 6250810 | 6250929 | 6305350 | 6305539 | [illegible] |
| SE | 71812 | ENSG00000124491 | F13A1 | chr6 | - | 6305350 | 6305539 | 6266557 | 6266809 | 6318534 | 6318682 | [illegible] |
| SE | 71813 | ENSG00000119041 | GTF3C3 | chr2 | - | 196766564 | 196766717 | 196763034 | 196764685 | 196769914 | 196770039 | [illegible] |
| SE | 71815 | ENSG00000119041 | GTF3C3 | chr2 | - | 196773668 | 196773726 | 196773006 | 196773153 | 196776009 | 196776111 | [illegible] |
| SE | 71816 | ENSG00000119041 | GTF3C3 | chr2 | - | 196773912 | 196774140 | 196772915 | 196773153 | 196775115 | 196775251 | [illegible] |
| SE | 71817 | ENSG00000119041 | GTF3C3 | chr2 | - | 196775115 | 196775251 | 196772915 | 196773153 | 196776009 | 196776111 | [illegible] |
| SE | 71821 | ENSG00000115607 | IL18RAP | chr2 | + | 102419560 | 102419797 | 102418688 | 102418940 | 102424230 | 102424414 | [illegible] |
| SE | 71822 | ENSG00000115607 | IL18RAP | chr2 | + | 102423177 | 102423347 | 102418688 | 102418940 | 102423810 | 102423874 | [illegible] |
| SE | 71823 | ENSG00000115607 | IL18RAP | chr2 | + | 102423177 | 102423347 | 102418688 | 102418940 | 102424230 | 102424414 | [illegible] |
| SE | 71824 | ENSG00000115607 | IL18RAP | chr2 | + | 102423177 | 102423347 | 102419444 | 102419797 | 102424230 | 102424414 | [illegible] |
| SE | 71825 | ENSG00000115607 | IL18RAP | chr2 | + | 102423810 | 102424135 | 102418688 | 102418940 | 102424230 | 102424414 | [illegible] |
| SE | 71826 | ENSG00000115607 | IL18RAP | chr2 | + | 102423810 | 102424135 | 102423177 | 102423347 | 102424230 | 102424414 | [illegible] |
| SE | 71828 | ENSG00000115607 | IL18RAP | chr2 | + | 102447069 | 102447207 | 102445188 | 102445340 | 102450847 | 102451021 | [illegible] |
| SE | 71832 | ENSG00000183486 | MX2 | chr21 | + | 41372933 | 41372999 | 41361998 | 41362055 | 41376835 | 41377155 | [illegible] |
| SE | 71833 | ENSG00000183486 | MX2 | chr21 | + | 41372933 | 41372999 | 41363268 | 41363389 | 41376835 | 41377155 | [illegible] |
| SE | 71835 | ENSG00000183486 | MX2 | chr21 | + | 41372933 | 41372999 | 41370212 | 41370645 | 41376835 | 41377155 | [illegible] |
| SE | 71836 | ENSG00000183486 | MX2 | chr21 | + | 41373897 | 41374127 | 41363268 | 41363416 | 41376835 | 41376949 | [illegible] |
| SE | 71837 | ENSG00000183486 | MX2 | chr21 | + | 41373897 | 41374127 | 41370212 | 41370645 | 41376835 | 41376949 | [illegible] |
| SE | 71838 | ENSG00000183486 | MX2 | chr21 | + | 41373897 | 41374127 | 41372933 | 41372999 | 41376835 | 41376949 | [illegible] |
| SE | 71840 | ENSG00000183486 | MX2 | chr21 | + | 41382409 | 41382564 | 41377788 | 41377981 | 41395586 | 41395785 | [illegible] |
| SE | 71842 | ENSG00000183486 | MX2 | chr21 | + | 41395586 | 41395785 | 41382409 | 41382564 | 41397612 | 41397691 | [illegible] |
| SE | 71844 | ENSG00000183486 | MX2 | chr21 | + | 41399195 | 41399337 | 41398896 | 41399019 | 41406743 | 41406998 | [illegible] |
| SE | 71846 | ENSG00000183486 | MX2 | chr21 | + | 41403266 | 41403343 | 41398896 | 41399019 | 41406743 | 41406998 | [illegible] |
| SE | 71848 | ENSG00000241962 | NA | chr2 | + | 99206633 | 99206757 | 99195572 | 99195756 | 99215997 | 99216114 | [illegible] |
| SE | 71850 | ENSG00000185414 | MRPL30 | chr2 | + | 99186176 | 99186254 | 99181151 | 99181249 | 99188176 | 99188257 | [illegible] |
| SE | 71853 | ENSG00000185414 | MRPL30 | chr2 | + | 99188176 | 99188257 | 99181151 | 99181249 | 99194750 | 99194897 | [illegible] |
| SE | 71854 | ENSG00000105639 | JAK3 | chr19 | - | 17835923 | 17836051 | 17835082 | 17835215 | 17837128 | 17837213 | [illegible] |
| SE | 71855 | ENSG00000105639 | JAK3 | chr19 | - | 17837128 | 17837213 | 17835082 | 17835215 | 17837931 | 17838063 | [illegible] |
| SE | 71858 | ENSG00000064666 | CNN2 | chr19 | + | 1026976 | 1027074 | 1026585 | 1026724 | 1031070 | 1031192 | [illegible] |
| SE | 71859 | ENSG00000064666 | CNN2 | chr19 | + | 1032391 | 1032458 | 1030942 | 1031192 | 1032558 | 1032663 | [illegible] |
| SE | 71860 | ENSG00000064666 | CNN2 | chr19 | + | 1032558 | 1032663 | 1032391 | 1032458 | 1036129 | 1036246 | [illegible] |
| SE | 71861 | ENSG00000064666 | CNN2 | chr19 | + | 1032558 | 1032669 | 1032391 | 1032458 | 1036129 | 1036246 | [illegible] |
| SE | 71862 | ENSG00000064666 | CNN2 | chr19 | + | 1032558 | 1032696 | 1032391 | 1032458 | 1036129 | 1036246 | [illegible] |
| SE | 71865 | ENSG00000064666 | CNN2 | chr19 | + | 1036129 | 1036246 | 1032558 | 1032669 | 1036415 | 1036472 | [illegible] |
| SE | 71867 | ENSG00000064666 | CNN2 | chr19 | + | 1036415 | 1036562 | 1032558 | 1032696 | 1037624 | 1037733 | [illegible] |
| SE | 71877 | ENSG00000115514 | TXNDC9 | chr2 | - | 99322560 | 99322659 | 99322062 | 99322209 | 99327534 | 99327653 | [illegible] |
| SE | 71878 | ENSG00000115514 | TXNDC9 | chr2 | - | 99327534 | 99327653 | 99321533 | 99322209 | 99332983 | 99333242 | [illegible] |
| SE | 71879 | ENSG00000115514 | TXNDC9 | chr2 | - | 99327534 | 99327653 | 99321533 | 99322209 | 99333021 | 99333242 | [illegible] |
| SE | 71881 | ENSG00000115112 | TFCP2L1 | chr2 | - | 121231825 | 121231968 | 121225561 | 121225613 | 121234090 | 121234194 | [illegible] |
| SE | 71882 | ENSG00000115112 | TFCP2L1 | chr2 | - | 121248163 | 121248270 | 121246817 | 121246970 | 121248981 | 121249087 | [illegible] |
| SE | 71884 | ENSG00000074054 | CLASP1 | chr2 | - | 121387762 | 121387906 | 121387121 | 121387228 | 121397139 | 121397283 | [illegible] |
| SE | 71885 | ENSG00000074054 | CLASP1 | chr2 | - | 121401867 | 121401870 | 121401508 | 121401672 | 121404370 | 121404434 | [illegible] |
| SE | 71891 | ENSG00000074054 | CLASP1 | chr2 | - | 121430072 | 121430177 | 121425138 | 121425306 | 121447336 | 121447507 | [illegible] |
| SE | 71892 | ENSG00000074054 | CLASP1 | chr2 | - | 121444938 | 121444962 | 121430072 | 121430177 | 121447336 | 121447507 | [illegible] |
| SE | 71894 | ENSG00000074054 | CLASP1 | chr2 | - | 121459979 | 121460125 | 121458839 | 121458975 | 121461100 | 121461193 | [illegible] |
| SE | 71896 | ENSG00000074054 | CLASP1 | chr2 | - | 121525844 | 121525920 | 121503166 | 121503234 | 121605700 | 121606180 | [illegible] |
| SE | 71897 | ENSG00000074054 | CLASP1 | chr2 | - | 121525844 | 121525920 | 121515664 | 121515762 | 121528676 | 121528780 | [illegible] |
| SE | 71898 | ENSG00000074054 | CLASP1 | chr2 | - | 121527798 | 121527890 | 121525844 | 121525920 | 121528676 | 121528780 | [illegible] |
| SE | 71899 | ENSG00000074054 | CLASP1 | chr2 | - | 121527798 | 121527890 | 121525844 | 121525920 | 121605700 | 121606180 | [illegible] |
| SE | 71900 | ENSG00000074054 | CLASP1 | chr2 | - | 121528676 | 121528780 | 121525844 | 121525920 | 121605700 | 121606180 | [illegible] |
| SE | 71901 | ENSG00000074054 | CLASP1 | chr2 | - | 121528676 | 121528780 | 121527798 | 121527890 | 121530246 | 121530325 | [illegible] |
| SE | 71902 | ENSG00000074054 | CLASP1 | chr2 | - | 121528676 | 121528780 | 121527798 | 121527890 | 121605700 | 121606180 | [illegible] |
| SE | 71904 | ENSG00000074054 | CLASP1 | chr2 | - | 121530246 | 121530325 | 121527798 | 121527890 | 121605700 | 121606180 | [illegible] |
| SE | 71906 | ENSG00000074054 | CLASP1 | chr2 | - | 121605700 | 121606180 | 121530246 | 121530325 | 121649371 | 121649438 | [illegible] |

| | | | | | | | | | | | | | | |
|---|---|---|---|---|---|---|---|---|---|---|---|---|---|---|
| SE | 71910 | ENSG00000143751 | SDE2 | chr1 | - | 225987895 | 225988388 | 225982701 | 225985523 | 225991242 | 225991363 | 217 | 187 | [illegible] |
| SE | 71912 | ENSG00000143324 | XPR1 | chr1 | + | 180834873 | 180835045 | 180811406 | 180811488 | 180863707 | 180863874 | 9 | 7 | [illegible] |
| SE | 71915 | ENSG00000143324 | XPR1 | chr1 | + | 180873802 | 180873942 | 180863707 | 180863874 | 180880075 | 180880297 | 93 | 67 | [illegible] |
| SE | 71916 | ENSG00000157637 | SLC38A10 | chr17 | - | 81250069 | 81250093 | 81246884 | 81247058 | 81251492 | 81251612 | 3 | 0 | [illegible] |
| SE | 71919 | ENSG00000157637 | SLC38A10 | chr17 | - | 81250069 | 81250093 | 81246884 | 81247061 | 81252194 | 81252273 | 3 | 0 | [illegible] |
| SE | 71920 | ENSG00000157637 | SLC38A10 | chr17 | - | 81251492 | 81251612 | 81246884 | 81247058 | 81252194 | 81252272 | 112 | 30 | [illegible] |
| SE | 71922 | ENSG00000157637 | SLC38A10 | chr17 | - | 81251492 | 81251612 | 81250069 | 81250093 | 81252194 | 81252272 | 119 | 36 | [illegible] |
| SE | 71923 | ENSG00000157637 | SLC38A10 | chr17 | - | 81252194 | 81252683 | 81246884 | 81247061 | 81253072 | 81253240 | 231 | 98 | [illegible] |
| SE | 71924 | ENSG00000157637 | SLC38A10 | chr17 | - | 81253072 | 81253240 | 81246884 | 81247061 | 81260237 | 81260394 | 277 | 174 | [illegible] |
| SE | 71925 | ENSG00000157637 | SLC38A10 | chr17 | - | 81260237 | 81260394 | 81253072 | 81253240 | 81275968 | 81276151 | 277 | 174 | [illegible] |
| SE | 71928 | ENSG00000157637 | SLC38A10 | chr17 | - | 81284849 | 81284895 | 81283408 | 81283502 | 81289690 | 81289808 | 29 | 7 | [illegible] |
| SE | 71929 | ENSG00000115520 | COQ10B | chr2 | + | 197459931 | 197460081 | 197453422 | 197453664 | 197462538 | 197462731 | 165 | 109 | [illegible] |
| SE | 71931 | ENSG00000115520 | COQ10B | chr2 | + | 197470069 | 197470171 | 197462538 | 197462731 | 197473756 | 197474168 | 245 | 189 | [illegible] |
| SE | 71934 | ENSG00000115540 | MOB4 | chr2 | + | 197523623 | 197523686 | 197516065 | 197516146 | 197535529 | 197535630 | 122 | 62 | [illegible] |
| SE | 71935 | ENSG00000115540 | MOB4 | chr2 | + | 197523623 | 197523686 | 197516065 | 197516146 | 197540110 | 197540153 | 13 | 1 | [illegible] |
| SE | 71936 | ENSG00000115540 | MOB4 | chr2 | + | 197535529 | 197535605 | 197516083 | 197516146 | 197540110 | 197540153 | 2 | 0 | [illegible] |
| SE | 71937 | ENSG00000115540 | MOB4 | chr2 | + | 197535529 | 197535630 | 197516046 | 197516146 | 197540110 | 197540153 | 116 | 45 | [illegible] |
| SE | 71940 | ENSG00000144362 | PHOSPHO2 | chr2 | + | 169697892 | 169697982 | 169697360 | 169697531 | 169700945 | 169701138 | 0 | 0 | [illegible] |
| SE | 71941 | ENSG00000138385 | SSB | chr2 | + | 169800951 | 169801026 | 169798870 | 169798976 | 169805473 | 169805577 | 302 | 202 | [illegible] |
| SE | 71942 | ENSG00000138385 | SSB | chr2 | + | 169805473 | 169805577 | 169800951 | 169801026 | 169805664 | 169805839 | 565 | 390 | [illegible] |
| SE | 71943 | ENSG00000138385 | SSB | chr2 | + | 169805997 | 169806113 | 169805664 | 169805839 | 169806784 | 169806890 | 1 | 8 | [illegible] |
| SE | 71945 | ENSG00000197930 | ERO1A | chr14 | - | 52646374 | 52646461 | 52646153 | 52646287 | 52652238 | 52652308 | 595 | 531 | [illegible] |
| SE | 71946 | ENSG00000197930 | ERO1A | chr14 | - | 52652238 | 52652308 | 52646374 | 52646461 | 52653068 | 52653315 | 523 | 480 | [illegible] |
| SE | 71947 | ENSG00000197930 | ERO1A | chr14 | - | 52653068 | 52653315 | 52652238 | 52652308 | 52657916 | 52658009 | 1027 | 924 | [illegible] |
| SE | 71949 | ENSG00000197930 | ERO1A | chr14 | - | 52661292 | 52661304 | 52658123 | 52658150 | 52671629 | 52671703 | 308 | 264 | [illegible] |
| SE | 71950 | ENSG00000197930 | ERO1A | chr14 | - | 52663800 | 52663847 | 52658123 | 52658150 | 52671629 | 52671703 | 0 | 3 | [illegible] |
| SE | 71951 | ENSG00000197930 | ERO1A | chr14 | - | 52663800 | 52663847 | 52661292 | 52661304 | 52671629 | 52671703 | 81 | 82 | [illegible] |
| SE | 71952 | ENSG00000197930 | ERO1A | chr14 | - | 52666374 | 52666495 | 52663780 | 52663847 | 52671629 | 52671703 | 656 | 476 | [illegible] |
| SE | 71954 | ENSG00000144560 | VGLL4 | chr3 | - | 11671261 | 11671394 | 11601832 | 11602022 | 11702970 | 11703047 | 2 | 0 | [illegible] |
| SE | 71960 | ENSG00000160710 | ADAR | chr1 | - | 154590183 | 154590409 | 154589756 | 154589928 | 154596804 | 154596938 | 678 | 399 | [illegible] |
| SE | 71961 | ENSG00000160710 | ADAR | chr1 | - | 154590261 | 154590409 | 154589756 | 154589928 | 154596804 | 154596995 | 433 | 239 | [illegible] |
| SE | 71962 | ENSG00000160710 | ADAR | chr1 | - | 154590341 | 154590409 | 154589756 | 154589928 | 154596804 | 154596995 | 153 | 88 | [illegible] |
| SE | 71963 | ENSG00000160710 | ADAR | chr1 | - | 154597827 | 154597976 | 154597122 | 154597267 | 154598401 | 154598572 | 726 | 536 | [illegible] |
| SE | 71964 | ENSG00000160710 | ADAR | chr1 | - | 154602384 | 154602626 | 154600176 | 154602248 | 154607991 | 154608181 | 63 | 46 | [illegible] |
| SE | 71971 | ENSG00000160710 | ADAR | chr1 | - | 154625062 | 154625153 | 154601040 | 154602626 | 154627953 | 154628013 | 0 | 0 | [illegible] |
| SE | 71988 | ENSG00000197635 | DPP4 | chr2 | - | 162008561 | 162008661 | 162004823 | 162005809 | 162009240 | 162009295 | 87 | 88 | [illegible] |
| SE | 71989 | ENSG00000197635 | DPP4 | chr2 | - | 162009240 | 162009295 | 162004823 | 162005809 | 162011792 | 162011987 | 22 | 18 | [illegible] |
| SE | 71990 | ENSG00000197635 | DPP4 | chr2 | - | 162019222 | 162019276 | 162018728 | 162018850 | 162020228 | 162020296 | 69 | 76 | [illegible] |
| SE | 71991 | ENSG00000197635 | DPP4 | chr2 | - | 162022754 | 162022799 | 162020580 | 162020688 | 162024803 | 162024939 | 153 | 160 | [illegible] |
| SE | 71992 | ENSG00000197635 | DPP4 | chr2 | - | 162032034 | 162032265 | 162024803 | 162024939 | 162033540 | 162033653 | 0 | 2 | [illegible] |
| SE | 71993 | ENSG00000197635 | DPP4 | chr2 | - | 162035163 | 162035324 | 162033540 | 162033653 | 162038301 | 162038422 | 165 | 149 | [illegible] |
| SE | 71996 | ENSG00000197635 | DPP4 | chr2 | - | 162046701 | 162046805 | 162045531 | 162045612 | 162046914 | 162047006 | 0 | 1 | [illegible] |
| SE | 71999 | ENSG00000139629 | GALNT6 | chr12 | - | 51355805 | 51355958 | 51351246 | 51354492 | 51357348 | 51357450 | 33 | 21 | [illegible] |
| SE | 72006 | ENSG00000139629 | GALNT6 | chr12 | - | 51383460 | 51383582 | 51382312 | 51382509 | 51391600 | 51391675 | 2 | 5 | [illegible] |
| SE | 72009 | ENSG00000139629 | GALNT6 | chr12 | - | 51390849 | 51390937 | 51383460 | 51383582 | 51391081 | 51391195 | 4 | 0 | [illegible] |
| SE | 72010 | ENSG00000139629 | GALNT6 | chr12 | - | 51390849 | 51390937 | 51383460 | 51383582 | 51391286 | 51391390 | 2 | 0 | [illegible] |
| SE | 72011 | ENSG00000139629 | GALNT6 | chr12 | - | 51390849 | 51390937 | 51383460 | 51383582 | 51391354 | 51391434 | 2 | 0 | [illegible] |
| SE | 72012 | ENSG00000139629 | GALNT6 | chr12 | - | 51390849 | 51390937 | 51383460 | 51383582 | 51391600 | 51391629 | 4 | 0 | [illegible] |
| SE | 72014 | ENSG00000139629 | GALNT6 | chr12 | - | 51391081 | 51391195 | 51390849 | 51390937 | 51391286 | 51391591 | 2 | 0 | [illegible] |
| SE | 72016 | ENSG00000139629 | GALNT6 | chr12 | - | 51391081 | 51391195 | 51390849 | 51390937 | 51391600 | 51391652 | 2 | 0 | [illegible] |
| SE | 72017 | ENSG00000079785 | DDX1 | chr2 | + | 15602547 | 15602631 | 15599668 | 15599716 | 15603191 | 15603275 | 166 | 85 | [illegible] |
| SE | 72018 | ENSG00000079785 | DDX1 | chr2 | + | 15613223 | 15613284 | 15607174 | 15607313 | 15617243 | 15617342 | 219 | 152 | [illegible] |
| SE | 72019 | ENSG00000079785 | DDX1 | chr2 | + | 15617221 | 15617342 | 15613223 | 15613284 | 15618180 | 15618270 | 60 | 48 | [illegible] |
| SE | 72020 | ENSG00000079785 | DDX1 | chr2 | + | 15617243 | 15617342 | 15613223 | 15613284 | 15618180 | 15618270 | 213 | 156 | [illegible] |
| SE | 72022 | ENSG00000079785 | DDX1 | chr2 | + | 15628444 | 15628517 | 15627053 | 15627145 | 15629601 | 15629697 | 38 | 24 | [illegible] |
| SE | 72023 | ENSG00000079785 | DDX1 | chr2 | + | 15628444 | 15628839 | 15627053 | 15627145 | 15629601 | 15629697 | 82 | 60 | [illegible] |
| SE | 72024 | ENSG00000079785 | DDX1 | chr2 | + | 15628637 | 15628839 | 15628444 | 15628517 | 15629601 | 15629697 | 60 | 45 | [illegible] |
| SE | 72038 | ENSG00000115053 | NCL | chr2 | - | 231455400 | 231455624 | 231453530 | 231455267 | 231456009 | 231456136 | 1268 | 894 | [illegible] |
| SE | 72044 | ENSG00000115053 | NCL | chr2 | - | 231456009 | 231456764 | 231455400 | 231455624 | 231457000 | 231457124 | 3247 | 2494 | [illegible] |
| SE | 72045 | ENSG00000115053 | NCL | chr2 | - | 231456630 | 231456764 | 231454750 | 231455624 | 231457000 | 231457124 | 2517 | 1967 | [illegible] |
| SE | 72047 | ENSG00000115053 | NCL | chr2 | - | 231457642 | 231457800 | 231457000 | 231457124 | 231458265 | 231458344 | 2183 | 1786 | [illegible] |
| SE | 72048 | ENSG00000115053 | NCL | chr2 | - | 231457642 | 231457800 | 231457000 | 231457124 | 231459000 | 231459125 | 666 | 459 | [illegible] |
| SE | 72049 | ENSG00000115053 | NCL | chr2 | - | 231458265 | 231458344 | 231457000 | 231457124 | 231459000 | 231459125 | 1 | 1 | [illegible] |
| SE | 72050 | ENSG00000115053 | NCL | chr2 | - | 231458265 | 231458344 | 231457642 | 231457800 | 231459000 | 231459125 | 1050 | 910 | [illegible] |
| SE | 72051 | ENSG00000115053 | NCL | chr2 | - | 231458265 | 231458389 | 231457000 | 231457124 | 231459000 | 231459125 | 1720 | 1500 | [illegible] |
| SE | 72052 | ENSG00000115053 | NCL | chr2 | - | 231458265 | 231458389 | 231457000 | 231457800 | 231459000 | 231459125 | 3229 | 2817 | [illegible] |
| SE | 72053 | ENSG00000115053 | NCL | chr2 | - | 231458265 | 231458620 | 231457000 | 231457124 | 231459000 | 231459125 | 0 | 0 | [illegible] |
| SE | 72054 | ENSG00000115053 | NCL | chr2 | - | 231458265 | 231458620 | 231457642 | 231457800 | 231459000 | 231459125 | 1417 | 1254 | [illegible] |
| SE | 72055 | ENSG00000115053 | NCL | chr2 | - | 231458265 | 231459125 | 231456630 | 231457800 | 231460151 | 231460293 | 2222 | 1859 | [illegible] |
| SE | 72056 | ENSG00000115053 | NCL | chr2 | - | 231459000 | 231459125 | 231457000 | 231457800 | 231460151 | 231460293 | 792 | 599 | [illegible] |
| SE | 72057 | ENSG00000272980 | NA | chr6 | + | 167003192 | 167004401 | 167000059 | 167000113 | 167010812 | 167010893 | 31 | 21 | [illegible] |
| SE | 72058 | ENSG00000130414 | NDUFA10 | chr2 | - | 239960954 | 239961186 | 239957372 | 239959037 | 239990073 | 239990182 | 91 | 35 | [illegible] |
| SE | 72059 | ENSG00000130414 | NDUFA10 | chr2 | - | 239961022 | 239961186 | 239957357 | 239959037 | 239990073 | 239990182 | 91 | 35 | [illegible] |
| SE | 72060 | ENSG00000130414 | NDUFA10 | chr2 | - | 239964182 | 239964261 | 239960805 | 239961186 | 239990073 | 239990182 | 0 | 0 | [illegible] |
| SE | 72065 | ENSG00000130414 | NDUFA10 | chr2 | - | 239990073 | 239990182 | 239957370 | 239961186 | 240005209 | 240005286 | 220 | 135 | [illegible] |
| SE | 72066 | ENSG00000130414 | NDUFA10 | chr2 | - | 239990073 | 239990182 | 239957370 | 239961186 | 240011616 | 240011660 | 54 | 20 | [illegible] |
| SE | 72068 | ENSG00000130414 | NDUFA10 | chr2 | - | 240005209 | 240005295 | 239987929 | 239990182 | 240007315 | 240007370 | 248 | 193 | [illegible] |
| SE | 72069 | ENSG00000130414 | NDUFA10 | chr2 | - | 240007315 | 240007370 | 239990073 | 239990182 | 240011616 | 240011660 | 54 | 34 | [illegible] |
| SE | 72071 | ENSG00000130414 | NDUFA10 | chr2 | - | 240011616 | 240014860 | 239989251 | 239990182 | 240018552 | 240018639 | 386 | 275 | [illegible] |
| SE | 72072 | ENSG00000130414 | NDUFA10 | chr2 | - | 240011616 | 240014860 | 240007315 | 240007370 | 240018552 | 240018639 | 484 | 367 | [illegible] |
| SE | 72074 | ENSG00000130414 | NDUFA10 | chr2 | - | 240017814 | 240017905 | 240011616 | 240014860 | 240018552 | 240018639 | 3 | 1 | [illegible] |
| SE | 72075 | ENSG00000130414 | NDUFA10 | chr2 | - | 240018552 | 240018639 | 240011616 | 240014860 | 240021196 | 240021412 | 582 | 409 | [illegible] |
| SE | 72077 | ENSG00000130414 | NDUFA10 | chr2 | - | 240021770 | 240021860 | 240021196 | 240021412 | 240022171 | 240022340 | 0 | 0 | [illegible] |
| SE | 72083 | ENSG00000180228 | PRKRA | chr2 | - | 178443266 | 178443384 | 178440257 | 178441704 | 178444421 | 178444500 | 136 | 88 | [illegible] |
| SE | 72084 | ENSG00000180228 | PRKRA | chr2 | - | 178443266 | 178443403 | 178441609 | 178441704 | 178444421 | 178444500 | 55 | 49 | [illegible] |
| SE | 72085 | ENSG00000180228 | PRKRA | chr2 | - | 178443289 | 178443384 | 178441609 | 178441704 | 178444421 | 178444500 | 46 | 24 | [illegible] |
| SE | 72088 | ENSG00000180228 | PRKRA | chr2 | - | 178445425 | 178445535 | 178444421 | 178444500 | 178447455 | 178447586 | 6 | 1 | [illegible] |
| SE | 72092 | ENSG00000180228 | PRKRA | chr2 | - | 178450241 | 178450411 | 178447319 | 178447586 | 178450965 | 178451021 | 111 | 60 | [illegible] |
| SE | 72093 | ENSG00000180228 | PRKRA | chr2 | - | 178450365 | 178450411 | 178447455 | 178447586 | 178450965 | 178451146 | 2 | 5 | [illegible] |
| SE | 72095 | ENSG00000122484 | RPAP2 | chr1 | + | 92300193 | 92300239 | 92299058 | 92299146 | 92301475 | 92301590 | 41 | 30 | [illegible] |
| SE | 72097 | ENSG00000122484 | RPAP2 | chr1 | + | 92307187 | 92307276 | 92304283 | 92304349 | 92320598 | 92320634 | 173 | 142 | [illegible] |
| SE | 72098 | ENSG00000122484 | RPAP2 | chr1 | + | 92320598 | 92320634 | 92307187 | 92307276 | 92333390 | 92333473 | 11 | 10 | [illegible] |
| SE | 72099 | ENSG00000122484 | RPAP2 | chr1 | + | 92323444 | 92324375 | 92320598 | 92320634 | 92333390 | 92333473 | 190 | 166 | [illegible] |
| SE | 72101 | ENSG00000102034 | ELF4 | chrX | - | 130069299 | 130069677 | 130063954 | 130067525 | 130071039 | 130071232 | 362 | 187 | [illegible] |
| SE | 72102 | ENSG00000102034 | ELF4 | chrX | - | 130071039 | 130071232 | 130069299 | 130069677 | 130071335 | 130071419 | 629 | 404 | [illegible] |
| SE | 72103 | ENSG00000102034 | ELF4 | chrX | - | 130071039 | 130071232 | 130069299 | 130069677 | 130072225 | 130072417 | 279 | 169 | [illegible] |
| SE | 72104 | ENSG00000102034 | ELF4 | chrX | - | 130071335 | 130071419 | 130069299 | 130069677 | 130072225 | 130072417 | 271 | 179 | [illegible] |
| SE | 72105 | ENSG00000102034 | ELF4 | chrX | - | 130071335 | 130071419 | 130071039 | 130071232 | 130072225 | 130072417 | 620 | 414 | [illegible] |
| SE | 72109 | ENSG00000251569 | NA | chr2 | + | 169487068 | 169487134 | 169482250 | 169482333 | 169487805 | 169487855 | 4 | 0 | [illegible] |
| SE | 72110 | ENSG00000251569 | NA | chr2 | + | 169493740 | 169493836 | 169492873 | 169493009 | 169497626 | 169497689 | 2 | 1 | [illegible] |
| SE | 72112 | ENSG00000251569 | NA | chr2 | + | 169503094 | 169503178 | 169499485 | 169499620 | 169504302 | 169504326 | 5 | 2 | [illegible] |
| SE | 72117 | ENSG00000138399 | FASTKD1 | chr2 | - | 169560386 | 169560785 | 169557186 | 169557297 | 169563224 | 169563350 | 82 | 57 | [illegible] |
| SE | 72118 | ENSG00000138399 | FASTKD1 | chr2 | - | 169569183 | 169569252 | 169563224 | 169563350 | 169571003 | 169571239 | 25 | 17 | [illegible] |
| SE | 72119 | ENSG00000138399 | FASTKD1 | chr2 | - | 169569183 | 169569252 | 169563224 | 169563350 | 169571652 | 169572171 | 53 | 27 | [illegible] |
| SE | 72121 | ENSG00000139531 | SUOX | chr12 | + | 55997614 | 55997723 | 55997258 | 55997339 | 56002211 | 56002271 | 5 | 2 | [illegible] |
| SE | 72122 | ENSG00000139531 | SUOX | chr12 | + | 55999332 | 55999458 | 55997295 | 55997339 | 56002211 | 56002271 | 8 | 4 | [illegible] |
| SE | 72123 | ENSG00000139531 | SUOX | chr12 | + | 55999332 | 55999458 | 55997614 | 55997723 | 56002211 | 56002271 | 16 | 7 | [illegible] |
| SE | 72124 | ENSG00000139531 | SUOX | chr12 | + | 55999332 | 55999590 | 55997614 | 55997723 | 56001885 | 56001948 | 8 | 3 | [illegible] |
| SE | 72125 | ENSG00000139531 | SUOX | chr12 | + | 55999332 | 55999590 | 55997614 | 55997723 | 56002211 | 56002271 | 8 | 3 | [illegible] |
| SE | 72134 | ENSG00000172936 | MYD88 | chr3 | + | 38139863 | 38139998 | 38139151 | 38139272 | 38140387 | 38140549 | 86 | 31 | [illegible] |
| SE | 72139 | ENSG00000172936 | MYD88 | chr3 | + | 38140756 | 38140848 | 38140387 | 38140568 | 38141131 | 38141265 | 268 | 154 | [illegible] |
| SE | 72141 | ENSG00000115904 | SOS1 | chr2 | - | 38995122 | 38995387 | 38989269 | 38989314 | 38996921 | 38997038 | 128 | 90 | [illegible] |
| SE | 72142 | ENSG00000115904 | SOS1 | chr2 | - | 39007030 | 39007193 | 39006411 | 39006529 | 39010583 | 39010703 | 648 | 541 | [illegible] |
| SE | 72144 | ENSG00000115904 | SOS1 | chr2 | - | 39012125 | 39012348 | 39010583 | 39010703 | 39013459 | 39013563 | 332 | 247 | [illegible] |
| SE | 72145 | ENSG00000115904 | SOS1 | chr2 | - | 39013459 | 39013563 | 39012125 | 39012348 | 39013866 | 39013989 | 267 | 233 | [illegible] |
| SE | 72147 | ENSG00000115904 | SOS1 | chr2 | - | 39024009 | 39024137 | 39022569 | 39023225 | 39035211 | 39035310 | 413 | 305 | [illegible] |
| SE | 72148 | ENSG00000115904 | SOS1 | chr2 | - | 39035389 | 39035500 | 39035211 | 39035310 | 39051143 | 39051287 | 177 | 141 | [illegible] |
| SE | 72149 | ENSG00000115904 | SOS1 | chr2 | - | 39051143 | 39051287 | 39035389 | 39035500 | 39054613 | 39054823 | 399 | 318 | [illegible] |
| SE | 72151 | ENSG00000115904 | SOS1 | chr2 | - | 39067627 | 39067753 | 39058672 | 39058804 | 39120335 | 39120450 | 159 | 108 | [illegible] |
| SE | 72153 | ENSG00000112799 | LY86 | chr6 | + | 6626292 | 6626421 | 6624925 | 6625012 | 6649624 | 6649677 | 507 | 415 | [illegible] |
| SE | 72154 | ENSG00000112799 | LY86 | chr6 | + | 6626292 | 6626421 | 6624925 | 6625012 | 6654543 | 6654982 | 158 | 117 | [illegible] |
| SE | 72155 | ENSG00000112799 | LY86 | chr6 | + | 6649624 | 6649677 | 6624925 | 6625012 | 6654543 | 6654982 | 110 | 92 | [illegible] |
| SE | 72162 | ENSG00000134317 | GRHL1 | chr2 | + | 9998964 | 9999029 | 9996315 | 9996401 | 10000592 | 10000731 | 14 | 48 | [illegible] |
| SE | 72163 | ENSG00000172059 | KLF11 | chr2 | + | 10047649 | 10048595 | 10046149 | 10046419 | 10052226 | 10054836 | 206 | 156 | [illegible] |
| SE | 72168 | ENSG00000059588 | TARBP1 | chr1 | - | 234393361 | 234393476 | 234392415 | 234392552 | 234393645 | 234393837 | 11 | 22 | [illegible] |
| SE | 72170 | ENSG00000059588 | TARBP1 | chr1 | - | 234398381 | 234398553 | 234393645 | 234393837 | 234400852 | 234400948 | 35 | 37 | [illegible] |
| SE | 72171 | ENSG00000059588 | TARBP1 | chr1 | - | 234398381 | 234398553 | 234393645 | 234393837 | 234401180 | 234401216 | 34 | 29 | [illegible] |
| SE | 72172 | ENSG00000059588 | TARBP1 | chr1 | - | 234400852 | 234400948 | 234393645 | 234393837 | 234401180 | 234401262 | 7 | 5 | [illegible] |
| SE | 72173 | ENSG00000059588 | TARBP1 | chr1 | - | 234400852 | 234400948 | 234398381 | 234398553 | 234401180 | 234401262 | 15 | 15 | [illegible] |
| SE | 72174 | ENSG00000059588 | TARBP1 | chr1 | - | 234405311 | 234405480 | 234401180 | 234401262 | 234405902 | 234405993 | 0 | 0 | [illegible] |
| SE | 72176 | ENSG00000059588 | TARBP1 | chr1 | - | 234429135 | 234429324 | 234427575 | 234427766 | 234429415 | 234429677 | 132 | 120 | [illegible] |
| SE | 72178 | ENSG00000059588 | TARBP1 | chr1 | - | 234433409 | 234433571 | 234430086 | 234430301 | 234437274 | 234437372 | 48 | 55 | [illegible] |
| SE | 72180 | ENSG00000059588 | TARBP1 | chr1 | - | 234446802 | 234446975 | 234437274 | 234437372 | 234450427 | 234450566 | 47 | 17 | [illegible] |
| SE | 72181 | ENSG00000059588 | TARBP1 | chr1 | - | 234448479 | 234448579 | 234446802 | 234446975 | 234450427 | 234450566 | 36 | 37 | [illegible] |
| SE | 72184 | ENSG00000164902 | PHAX | chr5 | + | 126608363 | 126608484 | 126603560 | 126604183 | 126617249 | 126617333 | 285 | 201 | [illegible] |
| SE | 72195 | ENSG00000145425 | RPS3A | chr4 | + | 151100974 | 151101012 | 151099670 | 151099714 | 151102870 | 151103079 | 0 | 0 | [illegible] |
| SE | 72200 | ENSG00000145425 | RPS3A | chr4 | + | 151100974 | 151101162 | 151100426 | 151100588 | 151102870 | 151102932 | 17915 | 10459 | [illegible] |
| SE | 72201 | ENSG00000145425 | RPS3A | chr4 | + | 151104176 | 151104286 | 151102870 | 151103079 | 151104471 | 151104490 | 32311 | 18214 | [illegible] |
| SE | 72202 | ENSG00000145425 | RPS3A | chr4 | + | 151104184 | 151104286 | 151102870 | 151103079 | 151104471 | 151104637 | 17779 | 9743 | [illegible] |
| SE | 72204 | ENSG00000284825 | NA | chr6 | - | 167116742 | 167117087 | 167114956 | 167116161 | 167119185 | 167119342 | 1 | 0 | [illegible] |
| SE | 72206 | ENSG00000223546 | LINC00630 | chrX | + | 102770351 | 102770420 | 102769157 | 102769341 | 102816991 | 102817082 | 4 | 4 | [illegible] |
| SE | 72207 | ENSG00000223546 | LINC00630 | chrX | + | 102770351 | 102770420 | 102769157 | 102769341 | 102825992 | 102826169 | 3 | 1 | [illegible] |
| SE | 72208 | ENSG00000223546 | LINC00630 | chrX | + | 102770351 | 102770420 | 102769157 | 102769341 | 102826968 | 102827144 | 2 | 1 | [illegible] |
| SE | 72210 | ENSG00000223546 | LINC00630 | chrX | + | 102816991 | 102817082 | 102769186 | 102769341 | 102826968 | 102827136 | 0 | 2 | [illegible] |
| SE | 72211 | ENSG00000223546 | LINC00630 | chrX | + | 102816991 | 102817082 | 102770351 | 102770420 | 102825992 | 102826169 | 8 | 8 | [illegible] |
| SE | 72213 | ENSG00000223546 | LINC00630 | chrX | + | 102825992 | 102826169 | 102769180 | 102769341 | 102826968 | 102827144 | 10 | 9 | [illegible] |
| SE | 72219 | ENSG00000223546 | LINC00630 | chrX | + | 102845856 | 102846020 | 102839800 | 102839863 | 102865041 | 102865152 | 11 | 12 | [illegible] |
| SE | 72220 | ENSG00000223546 | LINC00630 | chrX | + | 102845856 | 102846020 | 102839907 | 102839972 | 102865041 | 102865152 | 12 | 14 | [illegible] |
| SE | 72221 | ENSG00000223546 | LINC00630 | chrX | + | 102865041 | 102865152 | 102845856 | 102846020 | 102884142 | 102884166 | 8 | 8 | [illegible] |
| SE | 72222 | ENSG00000223546 | LINC00630 | chrX | + | 102877482 | 102877611 | 102865041 | 102865152 | 102884142 | 102885406 | 1 | 0 | [illegible] |
| SE | 72223 | ENSG00000223546 | LINC00630 | chrX | + | 102878012 | 102878119 | 102877482 | 102877611 | 102884142 | 102885406 | 1 | 1 | [illegible] |
| SE | 72226 | ENSG00000164291 | ARSK | chr5 | + | 95586561 | 95586733 | 95582915 | 95583198 | 95591400 | 95591625 | 19 | 34 | [illegible] |
| SE | 72228 | ENSG00000154473 | BUB3 | chr10 | + | 123157728 | 123157880 | 123155657 | 123155727 | 123160406 | 123160565 | 648 | 483 | [illegible] |
| SE | 72230 | ENSG00000072195 | SPEG | chr2 | + | 219444024 | 219444059 | 219443107 | 219443150 | 219444652 | 219444742 | 3 | 0 | [illegible] |
| SE | 72231 | ENSG00000072195 | SPEG | chr2 | + | 219444029 | 219444059 | 219443107 | 219443150 | 219444652 | 219444742 | 0 | 0 | [illegible] |
| SE | 72233 | ENSG00000072195 | SPEG | chr2 | + | 219451624 | 219451807 | 219450669 | 219451279 | 219461881 | 219462057 | 8 | 2 | [illegible] |
| SE | 72235 | ENSG00000149016 | TUT1 | chr11 | - | 62581105 | 62581206 | 62578560 | 62579030 | 62581385 | 62581701 | 31 | 16 | [illegible] |
| SE | 72236 | ENSG00000149016 | TUT1 | chr11 | - | 62581385 | 62581701 | 62578560 | 62579030 | 62589030 | 62589221 | 35 | 23 | [illegible] |
| SE | 72238 | ENSG00000173465 | ZNRD2 | chr11 | + | 65571390 | 65571779 | 65570885 | 65570970 | 65572442 | 65572892 | 66 | 26 | [illegible] |
| SE | 72240 | ENSG00000152348 | ATG10 | chr5 | + | 81976192 | 81976327 | 81972050 | 81972306 | 81987558 | 81987678 | 8 | 2 | [illegible] |
| SE | 72243 | ENSG00000152348 | ATG10 | chr5 | + | 82178489 | 82178587 | 82164398 | 82164537 | 82252561 | 82252659 | 30 | 18 | [illegible] |
| SE | 72246 | ENSG00000152348 | ATG10 | chr5 | + | 82253313 | 82253429 | 82252561 | 82252659 | 82254067 | 82254792 | 16 | 10 | [illegible] |
| SE | 72247 | ENSG00000152348 | ATG10 | chr5 | + | 82253313 | 82253429 | 82252561 | 82252659 | 82276144 | 82276857 | 4 | 4 | [illegible] |
| SE | 72254 | ENSG00000162885 | B3GALNT2 | chr1 | - | 235458602 | 235458786 | 235455558 | 235455684 | 235470849 | 235470960 | 33 | 39 | [illegible] |
| SE | 72255 | ENSG00000162885 | B3GALNT2 | chr1 | - | 235458602 | 235458786 | 235455558 | 235455684 | 235480053 | 235480149 | 32 | 39 | [illegible] |
| SE | 72256 | ENSG00000162885 | B3GALNT2 | chr1 | - | 235465635 | 235465714 | 235455558 | 235455684 | 235470849 | 235470930 | 24 | 24 | [illegible] |
| SE | 72257 | ENSG00000162885 | B3GALNT2 | chr1 | - | 235465635 | 235465714 | 235458602 | 235458786 | 235470849 | 235470930 | 33 | 37 | [illegible] |
| SE | 72258 | ENSG00000162885 | B3GALNT2 | chr1 | - | 235470849 | 235470960 | 235455558 | 235455684 | 235480053 | 235480149 | 21 | 39 | [illegible] |
| SE | 72259 | ENSG00000162885 | B3GALNT2 | chr1 | - | 235470849 | 235470960 | 235458602 | 235458786 | 235480053 | 235480149 | 22 | 38 | [illegible] |
| SE | 72260 | ENSG00000162885 | B3GALNT2 | chr1 | - | 235489167 | 235489268 | 235484321 | 235484515 | 235494680 | 235494828 | 19 | 32 | [illegible] |
| SE | 72261 | ENSG00000162885 | B3GALNT2 | chr1 | - | 235496185 | 235496308 | 235494680 | 235494828 | 235504140 | 235504422 | 3 | 0 | [illegible] |
| SE | 72262 | ENSG00000186635 | ARAP1 | chr11 | - | 72686041 | 72686191 | 72685068 | 72685681 | 72687438 | 72687502 | 667 | 270 | [illegible] |
| SE | 72264 | ENSG00000186635 | ARAP1 | chr11 | - | 72686094 | 72686191 | 72685643 | 72685681 | 72687438 | 72687502 | 416 | 169 | [illegible] |
| SE | 72265 | ENSG00000186635 | ARAP1 | chr11 | - | 72686094 | 72686191 | 72685643 | 72685681 | 72687687 | 72687738 | 1 | 0 | [illegible] |
| SE | 72269 | ENSG00000186635 | ARAP1 | chr11 | - | 72693324 | 72693470 | 72688454 | 72688537 | 72693691 | 72693805 | 531 | 244 | [illegible] |
| SE | 72270 | ENSG00000186635 | ARAP1 | chr11 | - | 72693691 | 72693805 | 72693138 | 72693470 | 72694979 | 72695040 | 395 | 194 | [illegible] |
| SE | 72271 | ENSG00000186635 | ARAP1 | chr11 | - | 72695717 | 72695865 | 72695541 | 72695628 | 72696548 | 72696654 | 653 | 521 | [illegible] |
| SE | 72272 | ENSG00000186635 | ARAP1 | chr11 | - | 72697910 | 72698106 | 72697597 | 72697649 | 72699004 | 72699107 | 1377 | 715 | [illegible] |
| SE | 72273 | ENSG00000186635 | ARAP1 | chr11 | - | 72699004 | 72699107 | 72697910 | 72698106 | 72699416 | 72699552 | 762 | 398 | [illegible] |
| SE | 72274 | ENSG00000186635 | ARAP1 | chr11 | - | 72699416 | 72699552 | 72699004 | 72699107 | 72701648 | 72701783 | 362 | 188 | [illegible] |
| SE | 72277 | ENSG00000186635 | ARAP1 | chr11 | - | 72712195 | 72712339 | 72711429 | 72711499 | 72712437 | 72712568 | 548 | 282 | [illegible] |
| SE | 72278 | ENSG00000186635 | ARAP1 | chr11 | - | 72713175 | 72713243 | 72712437 | 72712568 | 72714151 | 72714321 | 196 | 101 | [illegible] |
| SE | 72284 | ENSG00000131368 | MRPS25 | chr3 | - | 15053379 | 15053467 | 15049211 | 15049935 | 15059368 | 15059475 | 77 | 68 | [illegible] |
| SE | 72285 | ENSG00000131368 | MRPS25 | chr3 | - | 15059368 | 15059475 | 15053290 | 15053467 | 15065060 | 15065260 | 157 | 145 | [illegible] |
| SE | 72286 | ENSG00000006062 | MAP3K14 | chr17 | - | 45267091 | 45267198 | 45266536 | 45266681 | 45267405 | 45267759 | 28 | 25 | [illegible] |
| SE | 72287 | ENSG00000006062 | MAP3K14 | chr17 | - | 45274122 | 45274254 | 45273502 | 45273607 | 45274463 | 45274593 | 33 | 11 | [illegible] |
| SE | 72289 | ENSG00000006062 | MAP3K14 | chr17 | - | 45287153 | 45287364 | 45286430 | 45287045 | 45289235 | 45289305 | 169 | 86 | [illegible] |
| SE | 72290 | ENSG00000006062 | MAP3K14 | chr17 | - | 45289235 | 45289305 | 45287153 | 45287364 | 45290489 | 45290765 | 113 | 86 | [illegible] |
| SE | 72292 | ENSG00000156521 | TYSND1 | chr10 | - | 70142667 | 70142853 | 70137980 | 70140141 | 70143841 | 70143972 | 43 | 16 | [illegible] |
| SE | 72293 | ENSG00000156521 | TYSND1 | chr10 | - | 70143841 | 70143972 | 70137980 | 70140141 | 70145420 | 70146700 | 8 | 4 | [illegible] |
| SE | 72294 | ENSG00000156521 | TYSND1 | chr10 | - | 70143841 | 70143972 | 70137980 | 70140141 | 70146532 | 70146676 | 1 | 0 | [illegible] |
| SE | 72295 | ENSG00000059377 | TBXAS1 | chr7 | + | 139780739 | 139780824 | 139778247 | 139778471 | 139782665 | 139782729 | 1 | 0 | [illegible] |
| SE | 72296 | ENSG00000059377 | TBXAS1 | chr7 | + | 139780739 | 139780824 | 139778247 | 139778471 | 139787337 | 139787426 | 2 | 0 | [illegible] |
| SE | 72297 | ENSG00000059377 | TBXAS1 | chr7 | + | 139782487 | 139782564 | 139780739 | 139780824 | 139782665 | 139782729 | 2 | 0 | [illegible] |
| SE | 72298 | ENSG00000059377 | TBXAS1 | chr7 | + | 139782665 | 139782729 | 139777050 | 139777241 | 139787337 | 139787426 | 0 | 0 | [illegible] |
| SE | 72299 | ENSG00000059377 | TBXAS1 | chr7 | + | 139782665 | 139782729 | 139778247 | 139778471 | 139787337 | 139787426 | 0 | 0 | [illegible] |
| SE | 72300 | ENSG00000059377 | TBXAS1 | chr7 | + | 139782665 | 139782729 | 139780739 | 139780824 | 139787337 | 139787426 | 1 | 0 | [illegible] |
| SE | 72302 | ENSG00000059377 | TBXAS1 | chr7 | + | 139906100 | 139906198 | 139875584 | 139875637 | 139911224 | 139911321 | 3 | 1 | [illegible] |
| SE | 72304 | ENSG00000059377 | TBXAS1 | chr7 | + | 139936190 | 139936307 | 139911224 | 139911321 | 139953367 | 139953456 | 505 | 268 | [illegible] |
| SE | 72305 | ENSG00000059377 | TBXAS1 | chr7 | + | 139952550 | 139952688 | 139936190 | 139936307 | 139953367 | 139953456 | 1 | 0 | [illegible] |
| SE | 72308 | ENSG00000059377 | TBXAS1 | chr7 | + | 139957633 | 139957764 | 139936190 | 139936307 | 139961918 | 139962233 | 328 | 213 | [illegible] |
| SE | 72310 | ENSG00000126254 | RBM42 | chr19 | + | 35631139 | 35631224 | 35629519 | 35629673 | 35631330 | 35631405 | 91 | 33 | [illegible] |
| SE | 72314 | ENSG00000126254 | RBM42 | chr19 | + | 35632935 | 35633001 | 35631330 | 35631405 | 35633100 | 35633252 | 16 | 10 | [illegible] |
| SE | 72317 | ENSG00000197299 | BLM | chr15 | + | 90747388 | 90747490 | 90717345 | 90717440 | 90749366 | 90750067 | 54 | 26 | [illegible] |
| SE | 72318 | ENSG00000197299 | BLM | chr15 | + | 90759980 | 90760279 | 90754810 | 90754938 | 90760593 | 90761255 | 7 | 6 | [illegible] |
| SE | 72319 | ENSG00000197299 | BLM | chr15 | + | 90760146 | 90760279 | 90754810 | 90754938 | 90760593 | 90761255 | 15 | 12 | [illegible] |
| SE | 72320 | ENSG00000197299 | BLM | chr15 | + | 90765295 | 90765414 | 90762965 | 90763157 | 90766909 | 90767023 | 39 | 15 | [illegible] |
| SE | 72321 | ENSG00000197299 | BLM | chr15 | + | 90769437 | 90769586 | 90769132 | 90769231 | 90782821 | 90782928 | 50 | 37 | [illegible] |
| SE | 72322 | ENSG00000197299 | BLM | chr15 | + | 90769437 | 90769586 | 90769132 | 90769231 | 90784920 | 90785081 | 27 | 21 | [illegible] |

| | | | | | | | | | | | | | | | | | | | | | | | | | | | | | | | | | | | | | | | | | | | | | | |
|---|---|---|---|---|---|---|---|---|---|---|---|---|---|---|---|---|---|---|---|---|---|---|---|---|---|---|---|---|---|---|---|---|---|---|---|---|---|---|---|---|---|---|---|---|---|---|
| SE | 72323 | ENSG00000197299 | BLM | chr15 | + | 90782821 | 90782928 | 90769132 | 90769231 | 90784920 | 90785081 | 13 | 7 | 9 | 13 | 10 | 0 | 0 | 0 | 0 | 0 | 7 | 14 | 26 | 16 | 3 | 0 | 0 | 2 | 0 | 0 | 256 | 149 | 1 | 1 | 1 | 1 | 1 | 1 | 1 | 1 | 1 | 0.883 | 1 | 1 | 0.023 |
| SE | 72325 | ENSG00000197299 | BLM | chr15 | + | 90803520 | 90803720 | 90798189 | 90798337 | 90809136 | 90809259 | 36 | 26 | 49 | 22 | 82 | 1 | 0 | 0 | 0 | 0 | 0 | 38 | 29 | 4 | 0 | 0 | 4 | 0 | 0 | 0 | 298 | 149 | 1 | 1 | 0.947 | 1 | 1 | 1 | 1 | NA | 0.826 | 1 | 1 | NA | 0.047 |
| SE | 72326 | ENSG00000197299 | BLM | chr15 | + | 90804166 | 90804359 | 90803520 | 90803720 | 90809136 | 90809259 | 52 | 25 | 38 | 56 | 62 | 0 | 0 | 0 | 1 | 0 | 0 | 54 | 50 | 23 | 0 | 0 | 6 | 0 | 0 | 0 | 298 | 149 | 1 | 1 | 1 | 1 | 1 | 0.966 | 1 | NA | 0.818 | 1 | 1 | NA | 0.054 |
| SE | 72327 | ENSG00000197299 | BLM | chr15 | + | 90811677 | 90811800 | 90811204 | 90811406 | 90815101 | 90815342 | 0 | 0 | 0 | 0 | 0 | 31 | 20 | 58 | 38 | 90 | 0 | 0 | 2 | 0 | 0 | 32 | 39 | 18 | 44 | 20 | 272 | 149 | 1 | 1 | 0 | 0 | 0 | 0 | 0 | 0 | 0 | 0.057 | 0 | 0 | -0.011 |
| SE | 72328 | ENSG00000146143 | PRIM2 | chr6 | + | 57318436 | 57318599 | 57317623 | 57317701 | 57320456 | 57320560 | 110 | 104 | 168 | 148 | 158 | 0 | 0 | 0 | 0 | 0 | 93 | 119 | 122 | 69 | 99 | 0 | 0 | 0 | 2 | 0 | 298 | 149 | 1 | 1 | 1 | 1 | 1 | 1 | 1 | 1 | 1 | 1 | 0.945 | 1 | 0.011 |
| SE | 72329 | ENSG00000146143 | PRIM2 | chr6 | + | 57324200 | 57324280 | 57320456 | 57320560 | 57325924 | 57326045 | 134 | 129 | 130 | 90 | 220 | 0 | 1 | 0 | 0 | 0 | 82 | 101 | 98 | 82 | 83 | 0 | 0 | 0 | 0 | 0 | 229 | 149 | 1 | 1 | 1 | 0.988 | 1 | 1 | 1 | 1 | 1 | 1 | 1 | 1 | -0.002 |
| SE | 72339 | ENSG00000138433 | CIR1 | chr2 | - | 174351618 | 174351725 | 174350659 | 174350745 | 174378922 | 174379040 | 279 | 215 | 215 | 214 | 553 | 0 | 0 | 0 | 1 | 0 | 38 | 142 | 287 | 43 | 9 | 0 | 0 | 0 | 0 | 0 | 256 | 149 | 1 | 1 | 1 | 1 | 1 | 0.992 | 1 | 1 | 1 | 1 | 1 | 1 | -0.002 |
| SE | 72340 | ENSG00000138433 | CIR1 | chr2 | - | 174378922 | 174379040 | 174351618 | 174351725 | 174380201 | 174380250 | 420 | 311 | 320 | 332 | 744 | 4 | 0 | 0 | 0 | 0 | 71 | 240 | 413 | 64 | 56 | 0 | 0 | 0 | 0 | 0 | 267 | 149 | 1 | 1 | 0.983 | 1 | 1 | 1 | 1 | 1 | 1 | 1 | 1 | 1 | -0.003 |
| SE | 72341 | ENSG00000125503 | PPP1R12C | chr19 | - | 55092441 | 55092544 | 55092221 | 55092326 | 55092621 | 55092653 | 133 | 69 | 98 | 89 | 69 | 1 | 1 | 0 | 0 | 0 | 150 | 123 | 86 | 113 | 268 | 0 | 0 | 0 | 0 | 0 | 252 | 149 | 1 | 1 | 0.987 | 0.976 | 1 | 1 | 1 | 1 | 1 | 1 | 1 | 1 | -0.007 |
| SE | 72342 | ENSG00000125503 | PPP1R12C | chr19 | - | 55092621 | 55092662 | 55092441 | 55092544 | 55092782 | 55092865 | 89 | 41 | 71 | 65 | 35 | 0 | 0 | 0 | 5 | 0 | 70 | 58 | 46 | 56 | 150 | 0 | 0 | 0 | 0 | 0 | 190 | 149 | 1 | 1 | 1 | 1 | 1 | 0.911 | 1 | 1 | 1 | 1 | 1 | 1 | -0.018 |
| SE | 72345 | ENSG00000125503 | PPP1R12C | chr19 | - | 55103408 | 55103568 | 55096261 | 55096335 | 55117222 | 55117413 | 0 | 0 | 0 | 2 | 0 | 0 | 0 | 0 | 0 | 0 | 0 | 2 | 1 | 0 | 6 | 0 | 4 | 0 | 0 | 0 | 298 | 149 | 1 | 1 | NA | NA | NA | 1 | NA | NA | 0.2 | 1 | NA | 1 | 0.267 |
| SE | 72346 | ENSG00000125503 | PPP1R12C | chr19 | - | 55112466 | 55112585 | 55103408 | 55103568 | 55112664 | 55112795 | 72 | 26 | 67 | 59 | 24 | 0 | 0 | 0 | 0 | 0 | 57 | 99 | 37 | 60 | 126 | 0 | 2 | 0 | 0 | 0 | 268 | 149 | 1 | 1 | 1 | 1 | 1 | 1 | 1 | 1 | 0.965 | 1 | 1 | 1 | 0.007 |
| SE | 72347 | ENSG00000125503 | PPP1R12C | chr19 | - | 55112664 | 55112795 | 55103408 | 55103568 | 55117222 | 55117413 | 6 | 6 | 8 | 8 | 2 | 0 | 0 | 0 | 0 | 0 | 10 | 10 | 4 | 4 | 34 | 0 | 0 | 0 | 0 | 2 | 280 | 149 | 1 | 1 | 1 | 1 | 1 | 1 | 1 | 1 | 1 | 1 | 1 | 0.9 | 0.02 |
| SE | 72350 | ENSG00000148835 | TAF5 | chr10 | + | 103385325 | 103385490 | 103383237 | 103383367 | 103387174 | 103387352 | 30 | 23 | 24 | 57 | 31 | 12 | 1 | 3 | 0 | 2 | 29 | 60 | 37 | 14 | 0 | 0 | 1 | 2 | 0 | 0 | 298 | 149 | 1 | 1 | 0.556 | 0.92 | 0.8 | 1 | 0.886 | 1 | 0.968 | 0.902 | 1 | NA | -0.135 |
| SE | 72354 | ENSG00000163214 | DHX57 | chr2 | - | 38802714 | 38802915 | 38797728 | 38798442 | 38806558 | 38806693 | 127 | 93 | 110 | 81 | 179 | 0 | 0 | 2 | 0 | 0 | 39 | 77 | 78 | 68 | 0 | 0 | 6 | 1 | 0 | 0 | 298 | 149 | 1 | 1 | 1 | 1 | 0.965 | 1 | 1 | 1 | 0.865 | 0.975 | 1 | NA | 0.033 |
| SE | 72355 | ENSG00000163214 | DHX57 | chr2 | - | 38806558 | 38806693 | 38802714 | 38802915 | 38813820 | 38813895 | 127 | 113 | 113 | 83 | 221 | 0 | 0 | 0 | 1 | 0 | 13 | 99 | 103 | 107 | 0 | 0 | 0 | 0 | 0 | 0 | 284 | 149 | 1 | 1 | 1 | 1 | 1 | 0.978 | 1 | 1 | 1 | 1 | 1 | NA | -0.004 |
| SE | 72356 | ENSG00000163214 | DHX57 | chr2 | - | 38818872 | 38818960 | 38815520 | 38815655 | 38819048 | 38819144 | 24 | 5 | 17 | 17 | 21 | 1 | 0 | 0 | 0 | 0 | 10 | 26 | 8 | 31 | 0 | 0 | 0 | 0 | 0 | 0 | 237 | 149 | 1 | 1 | 0.938 | 1 | 1 | 1 | 1 | 1 | 1 | 1 | 1 | NA | -0.012 |
| SE | 72357 | ENSG00000163214 | DHX57 | chr2 | - | 38818876 | 38818960 | 38815520 | 38815655 | 38819048 | 38819108 | 46 | 23 | 43 | 33 | 41 | 1 | 0 | 0 | 0 | 0 | 10 | 53 | 23 | 64 | 0 | 0 | 0 | 0 | 0 | 0 | 233 | 149 | 1 | 1 | 0.967 | 1 | 1 | 1 | 1 | 1 | 1 | 1 | 1 | NA | -0.007 |
| SE | 72359 | ENSG00000163214 | DHX57 | chr2 | - | 38854053 | 38854178 | 38848268 | 38848402 | 38855056 | 38855252 | 139 | 54 | 55 | 77 | 62 | 0 | 2 | 0 | 0 | 0 | 28 | 67 | 55 | 46 | 8 | 0 | 0 | 0 | 0 | 0 | 274 | 149 | 1 | 1 | 1 | 0.936 | 1 | 1 | 1 | 1 | 1 | 1 | 1 | 1 | -0.013 |
| SE | 72360 | ENSG00000163214 | DHX57 | chr2 | - | 38856339 | 38856461 | 38853044 | 38855252 | 38858660 | 38858836 | 142 | 85 | 122 | 93 | 409 | 0 | 0 | 0 | 0 | 3 | 13 | 91 | 48 | 82 | 0 | 0 | 0 | 0 | 0 | 0 | 271 | 149 | 1 | 1 | 1 | 1 | 1 | 1 | 0.987 | 1 | 1 | 1 | 1 | NA | -0.003 |
| SE | 72361 | ENSG00000163214 | DHX57 | chr2 | - | 38862144 | 38862333 | 38860998 | 38861837 | 38863360 | 38863519 | 86 | 52 | 62 | 82 | 100 | 0 | 0 | 0 | 1 | 0 | 57 | 86 | 76 | 86 | 45 | 0 | 0 | 0 | 5 | 0 | 298 | 149 | 1 | 1 | 1 | 1 | 1 | 0.976 | 1 | 1 | 1 | 1 | 0.896 | 1 | 0.016 |
| SE | 72364 | ENSG00000112877 | CEP72 | chr5 | + | 633768 | 633947 | 624470 | 624579 | 635371 | 635584 | 37 | 17 | 43 | 34 | 25 | 0 | 0 | 0 | 0 | 0 | 49 | 13 | 22 | 24 | 51 | 0 | 0 | 1 | 0 | 0 | 298 | 149 | 1 | 1 | 1 | 1 | 1 | 1 | 1 | 1 | 1 | 0.917 | 1 | 1 | 0.017 |
| SE | 72365 | ENSG00000112877 | CEP72 | chr5 | + | 665940 | 666099 | 665179 | 665325 | 666863 | 667168 | 4 | 0 | 0 | 3 | 0 | 1 | 0 | 0 | 2 | 0 | 0 | 0 | 2 | 15 | 0 | 0 | 0 | 0 | 0 | 0 | 298 | 149 | 1 | 1 | 0.667 | NA | NA | 0.429 | NA | NA | NA | 1 | 1 | NA | -0.452 |
| SE | 72367 | ENSG00000169045 | HNRNPH1 | chr5 | - | 179617050 | 179617110 | 179616868 | 179616958 | 179617513 | 179617649 | 2996 | 3270 | 2868 | 3034 | 3237 | 23 | 18 | 16 | 4 | 9 | 1069 | 2010 | 2272 | 1774 | 402 | 1 | 5 | 10 | 2 | 0 | 209 | 149 | 1 | 1 | 0.989 | 0.992 | 0.992 | 0.998 | 0.996 | 0.999 | 0.997 | 0.994 | 0.998 | 1 | -0.004 |
| SE | 72369 | ENSG00000169045 | HNRNPH1 | chr5 | - | 179617988 | 179618060 | 179617816 | 179617911 | 179618144 | 179618301 | 196 | 233 | 234 | 269 | 126 | 1 | 0 | 0 | 0 | 0 | 89 | 187 | 299 | 123 | 44 | 0 | 0 | 0 | 0 | 0 | 221 | 149 | 1 | 1 | 0.992 | 1 | 1 | 1 | 1 | 1 | 1 | 1 | 1 | 1 | -0.002 |
| SE | 72370 | ENSG00000169045 | HNRNPH1 | chr5 | - | 179617988 | 179618060 | 179617816 | 179617911 | 179620891 | 179620959 | 1 | 2 | 0 | 4 | 3 | 33 | 19 | 24 | 23 | 25 | 0 | 2 | 0 | 0 | 0 | 4 | 17 | 27 | 12 | 0 | 221 | 149 | 1 | 1 | 0.02 | 0.066 | 0 | 0.105 | 0.075 | 0 | 0.073 | 0 | 0 | NA | 0.035 |
| SE | 72372 | ENSG00000169045 | HNRNPH1 | chr5 | - | 179618144 | 179618323 | 179617988 | 179618060 | 179620891 | 179621035 | 752 | 861 | 894 | 1013 | 708 | 1 | 0 | 0 | 1 | 0 | 356 | 770 | 1114 | 707 | 102 | 0 | 0 | 0 | 0 | 0 | 298 | 149 | 1 | 1 | 0.997 | 1 | 1 | 0.998 | 1 | 1 | 1 | 1 | 1 | 1 | -0.001 |
| SE | 72375 | ENSG00000169045 | HNRNPH1 | chr5 | - | 179619268 | 179619407 | 179618144 | 179618323 | 179621241 | 179621397 | 390 | 349 | 370 | 357 | 559 | 0 | 1 | 0 | 0 | 0 | 160 | 322 | 259 | 294 | 70 | 0 | 0 | 0 | 0 | 0 | 288 | 149 | 1 | 1 | 1 | 0.994 | 1 | 1 | 1 | 1 | 1 | 1 | 1 | 1 | -0.001 |
| SE | 72376 | ENSG00000169045 | HNRNPH1 | chr5 | - | 179620891 | 179621035 | 179618144 | 179618323 | 179621241 | 179621335 | 1318 | 1272 | 1443 | 1562 | 2835 | 0 | 1 | 0 | 0 | 0 | 634 | 1220 | 1186 | 1131 | 262 | 0 | 0 | 0 | 0 | 0 | 293 | 149 | 1 | 1 | 1 | 0.998 | 1 | 1 | 1 | 1 | 1 | 1 | 1 | 1 | 0 |
| SE | 72377 | ENSG00000169045 | HNRNPH1 | chr5 | - | 179620891 | 179621035 | 179619100 | 179619407 | 179621241 | 179621335 | 2213 | 2145 | 2345 | 2550 | 5070 | 0 | 0 | 1 | 3 | 0 | 996 | 2055 | 1745 | 1856 | 436 | 0 | 4 | 3 | 0 | 0 | 293 | 149 | 1 | 1 | 1 | 1 | 0.999 | 0.998 | 1 | 1 | 0.996 | 0.997 | 1 | 1 | 0.001 |
| SE | 72378 | ENSG00000169045 | HNRNPH1 | chr5 | - | 179621241 | 179621397 | 179620674 | 179621035 | 179623036 | 179623164 | 2076 | 1686 | 2009 | 2211 | 3162 | 0 | 0 | 1 | 1 | 0 | 725 | 1615 | 1476 | 1342 | 443 | 0 | 0 | 0 | 0 | 0 | 298 | 149 | 1 | 1 | 1 | 1 | 0.999 | 0.999 | 1 | 1 | 1 | 1 | 1 | 1 | 0 |
| SE | 72379 | ENSG00000169045 | HNRNPH1 | chr5 | - | 179621370 | 179621397 | 179620813 | 179621035 | 179623036 | 179623164 | 125 | 86 | 115 | 138 | 48 | 0 | 0 | 1 | 1 | 0 | 15 | 64 | 89 | 71 | 40 | 0 | 0 | 0 | 0 | 0 | 176 | 149 | 1 | 1 | 1 | 1 | 0.99 | 0.992 | 1 | 1 | 1 | 1 | 1 | 1 | -0.004 |
| SE | 72380 | ENSG00000169045 | HNRNPH1 | chr5 | - | 179621842 | 179621976 | 179621241 | 179621397 | 179623036 | 179623164 | 5 | 3 | 4 | 6 | 9 | 918 | 584 | 773 | 873 | 612 | 0 | 3 | 10 | 7 | 0 | 198 | 606 | 597 | 462 | 199 | 283 | 149 | 1 | 1 | 0.003 | 0.003 | 0.003 | 0.004 | 0.008 | 0 | 0.003 | 0.009 | 0.008 | 0 | 0 |
| SE | 72381 | ENSG00000169045 | HNRNPH1 | chr5 | - | 179623036 | 179623164 | 179620891 | 179621397 | 179623594 | 179623668 | 972 | 623 | 847 | 972 | 644 | 0 | 2 | 0 | 0 | 0 | 258 | 754 | 673 | 518 | 198 | 0 | 4 | 4 | 2 | 0 | 277 | 149 | 1 | 1 | 1 | 0.994 | 1 | 1 | 1 | 1 | 0.99 | 0.989 | 0.993 | 1 | 0.004 |
| SE | 72382 | ENSG00000169045 | HNRNPH1 | chr5 | - | 179623036 | 179623164 | 179620891 | 179621397 | 179634064 | 179634157 | 806 | 507 | 673 | 752 | 572 | 0 | 1 | 4 | 0 | 0 | 183 | 547 | 520 | 393 | 163 | 0 | 0 | 0 | 0 | 0 | 277 | 149 | 1 | 1 | 1 | 0.996 | 0.989 | 1 | 1 | 1 | 1 | 1 | 1 | 1 | -0.003 |
| SE | 72383 | ENSG00000169045 | HNRNPH1 | chr5 | - | 179623036 | 179623187 | 179621241 | 179621397 | 179623594 | 179623710 | 828 | 523 | 681 | 766 | 590 | 0 | 2 | 0 | 0 | 0 | 190 | 567 | 536 | 421 | 169 | 0 | 4 | 4 | 2 | 0 | 298 | 149 | 1 | 1 | 1 | 0.992 | 1 | 1 | 1 | 1 | 0.986 | 0.985 | 0.991 | 1 | 0.006 |
| SE | 72387 | ENSG00000143434 | SEMA6C | chr1 | - | 151136447 | 151136579 | 151136010 | 151136163 | 151136856 | 151137074 | 5 | 8 | 11 | 5 | 1 | 1 | 0 | 0 | 0 | 0 | 0 | 0 | 0 | 0 | 10 | 0 | 0 | 0 | 0 | 0 | 281 | 149 | 1 | 1 | 0.726 | 1 | 1 | 1 | 1 | NA | NA | NA | NA | 1 | -0.055 |
| SE | 72389 | ENSG00000143434 | SEMA6C | chr1 | - | 151138315 | 151138406 | 151137985 | 151138105 | 151138629 | 151138731 | 2 | 7 | 0 | 6 | 7 | 0 | 0 | 0 | 1 | 0 | 0 | 0 | 1 | 2 | 0 | 0 | 0 | 0 | 0 | 0 | 240 | 149 | 1 | 1 | 1 | 1 | NA | 0.788 | 1 | NA | NA | 1 | 1 | NA | -0.053 |
| SE | 72390 | ENSG00000143434 | SEMA6C | chr1 | - | 151138629 | 151138731 | 151138008 | 151138105 | 151139424 | 151139461 | 2 | 7 | 1 | 6 | 0 | 0 | 0 | 0 | 2 | 0 | 0 | 0 | 1 | 2 | 8 | 0 | 0 | 0 | 0 | 0 | 251 | 149 | 1 | 1 | 1 | 1 | 1 | 0.64 | NA | NA | NA | 1 | 1 | 1 | -0.09 |
| SE | 72391 | ENSG00000143434 | SEMA6C | chr1 | - | 151142503 | 151142675 | 151139975 | 151140090 | 151144384 | 151144406 | 3 | 1 | 1 | 5 | 0 | 0 | 0 | 0 | 0 | 0 | 0 | 0 | 6 | 4 | 0 | 0 | 0 | 3 | 0 | 0 | 298 | 149 | 1 | 1 | 1 | 1 | 1 | 1 | NA | NA | NA | 0.5 | 1 | NA | 0.25 |
| SE | 72392 | ENSG00000138835 | RGS3 | chr9 | + | 113514457 | 113514654 | 113508540 | 113508580 | 113517540 | 113517624 | 1 | 0 | 0 | 0 | 0 | 1 | 0 | 0 | 0 | 0 | 0 | 0 | 0 | 3 | 0 | 0 | 0 | 0 | 0 | 0 | 298 | 149 | 1 | 1 | 0.333 | NA | NA | NA | NA | NA | NA | NA | 1 | NA | -0.667 |
| SE | 72393 | ENSG00000138835 | RGS3 | chr9 | + | 113522929 | 113523041 | 113517367 | 113517624 | 113529220 | 113529264 | 168 | 42 | 65 | 73 | 52 | 0 | 0 | 0 | 1 | 0 | 4 | 87 | 29 | 30 | 66 | 0 | 0 | 0 | 0 | 0 | 261 | 149 | 1 | 1 | 1 | 1 | 1 | 0.977 | 1 | 1 | 1 | 1 | 1 | 1 | -0.005 |
| SE | 72394 | ENSG00000138835 | RGS3 | chr9 | + | 113529220 | 113529264 | 113522929 | 113523041 | 113541319 | 113542089 | 27 | 10 | 13 | 15 | 6 | 1 | 0 | 0 | 0 | 0 | 0 | 16 | 6 | 1 | 14 | 0 | 0 | 0 | 0 | 0 | 193 | 149 | 1 | 1 | 0.954 | 1 | 1 | 1 | 1 | NA | 1 | 1 | 1 | 1 | -0.009 |
| SE | 72395 | ENSG00000138835 | RGS3 | chr9 | + | 113536795 | 113536918 | 113529220 | 113529264 | 113541319 | 113542089 | 165 | 28 | 53 | 72 | 57 | 0 | 0 | 0 | 1 | 0 | 8 | 63 | 16 | 30 | 76 | 0 | 0 | 0 | 0 | 0 | 272 | 149 | 1 | 1 | 1 | 1 | 1 | 0.975 | 1 | 1 | 1 | 1 | 1 | 1 | -0.005 |
| SE | 72396 | ENSG00000138835 | RGS3 | chr9 | + | 113536795 | 113536918 | 113529220 | 113529264 | 113582071 | 113582242 | 112 | 17 | 36 | 43 | 30 | 0 | 2 | 0 | 0 | 0 | 8 | 56 | 15 | 30 | 45 | 0 | 0 | 0 | 0 | 0 | 272 | 149 | 1 | 1 | 1 | 0.823 | 1 | 1 | 1 | 1 | 1 | 1 | 1 | 1 | -0.035 |
| SE | 72399 | ENSG00000138835 | RGS3 | chr9 | + | 113581558 | 113581658 | 113580927 | 113581005 | 113582071 | 113582242 | 4 | 0 | 1 | 1 | 0 | 14 | 9 | 1 | 3 | 0 | 0 | 2 | 0 | 0 | 0 | 0 | 4 | 0 | 0 | 0 | 249 | 149 | 1 | 1 | 0.146 | 0 | 0.374 | 0.166 | NA | NA | 0.23 | NA | NA | NA | -0.058 |
| SE | 72406 | ENSG00000138835 | RGS3 | chr9 | + | 113594918 | 113594980 | 113594429 | 113594531 | 113596767 | 113597723 | 58 | 20 | 21 | 48 | 50 | 0 | 0 | 0 | 1 | 0 | 19 | 50 | 15 | 25 | 41 | 0 | 0 | 0 | 0 | 0 | 211 | 149 | 1 | 1 | 1 | 1 | 1 | 0.971 | 1 | 1 | 1 | 1 | 1 | 1 | -0.006 |
| SE | 72407 | ENSG00000138835 | RGS3 | chr9 | + | 113595598 | 113595765 | 113594918 | 113594980 | 113596767 | 113597723 | 216 | 82 | 116 | 195 | 214 | 4 | 0 | 1 | 2 | 0 | 139 | 225 | 84 | 89 | 191 | 0 | 6 | 0 | 0 | 0 | 298 | 149 | 1 | 1 | 0.964 | 1 | 0.983 | 0.98 | 1 | 1 | 0.949 | 1 | 1 | 1 | -0.004 |
| SE | 72411 | ENSG00000122042 | UBL3 | chr13 | - | 29772111 | 29772198 | 29767617 | 29767695 | 29777154 | 29777263 | 491 | 359 | 403 | 508 | 832 | 0 | 0 | 0 | 2 | 0 | 153 | 298 | 382 | 254 | 62 | 0 | 0 | 0 | 0 | 0 | 236 | 149 | 1 | 1 | 1 | 1 | 1 | 0.994 | 1 | 1 | 1 | 1 | 1 | 1 | -0.001 |
| SE | 72412 | ENSG00000122042 | UBL3 | chr13 | - | 29777154 | 29777263 | 29772111 | 29772198 | 29849511 | 29850617 | 378 | 304 | 327 | 450 | 599 | 5 | 0 | 2 | 0 | 0 | 92 | 298 | 267 | 242 | 58 | 0 | 0 | 0 | 0 | 0 | 258 | 149 | 1 | 1 | 0.978 | 1 | 0.99 | 1 | 1 | 1 | 1 | 1 | 1 | 1 | -0.006 |
| SE | 72414 | ENSG00000114978 | MOB1A | chr2 | - | 74165217 | 74165351 | 74159068 | 74159254 | 74167013 | 74167107 | 2190 | 1677 | 1642 | 2153 | 3485 | 1 | 1 | 0 | 3 | 6 | 859 | 1632 | 1328 | 1213 | 777 | 0 | 4 | 1 | 0 | 0 | 283 | 149 | 1 | 1 | 0.999 | 0.999 | 1 | 0.997 | 0.997 | 1 | 0.995 | 0.999 | 1 | 1 | 0 |
| SE | 72415 | ENSG00000114978 | MOB1A | chr2 | - | 74165217 | 74165351 | 74159068 | 74159254 | 74172585 | 74172749 | 1438 | 1143 | 1067 | 1482 | 2540 | 0 | 0 | 0 | 0 | 0 | 499 | 1082 | 880 | 787 | 532 | 0 | 0 | 2 | 0 | 0 | 283 | 149 | 1 | 1 | 1 | 1 | 1 | 1 | 1 | 1 | 1 | 0.996 | 1 | 1 | 0.001 |
| SE | 72416 | ENSG00000114978 | MOB1A | chr2 | - | 74167013 | 74167107 | 74159068 | 74159254 | 74172585 | 74172749 | 401 | 256 | 280 | 367 | 572 | 0 | 0 | 0 | 0 | 0 | 159 | 318 | 231 | 202 | 138 | 0 | 0 | 2 | 0 | 0 | 243 | 149 | 1 | 1 | 1 | 1 | 1 | 1 | 1 | 1 | 1 | 0.986 | 1 | 1 | 0.003 |
| SE | 72417 | ENSG00000114978 | MOB1A | chr2 | - | 74167013 | 74167107 | 74164021 | 74165351 | 74172585 | 74172749 | 1160 | 790 | 857 | 1047 | 1515 | 4 | 0 | 0 | 5 | 2 | 519 | 869 | 683 | 634 | 387 | 0 | 0 | 1 | 3 | 5 | 243 | 149 | 1 | 1 | 0.994 | 1 | 1 | 0.992 | 0.998 | 1 | 1 | 0.998 | 0.992 | 0.979 | 0.003 |
| SE | 72421 | ENSG00000251442 | LINC01094 | chr4 | + | 78649600 | 78649709 | 78645869 | 78646263 | 78680760 | 78681664 | 8 | 7 | 2 | 2 | 19 | 0 | 1 | 0 | 0 | 1 | 14 | 0 | 0 | 0 | 0 | 0 | 0 | 0 | 0 | 0 | 258 | 149 | 1 | 1 | 1 | 0.802 | 1 | 1 | 0.916 | 1 | NA | NA | NA | NA | -0.056 |
| SE | 72422 | ENSG00000251442 | LINC01094 | chr4 | + | 78662917 | 78662972 | 78649600 | 78649709 | 78680760 | 78681704 | 0 | 0 | 0 | 0 | 4 | 5 | 4 | 2 | 0 | 9 | 0 | 0 | 0 | 0 | 0 | 14 | 0 | 0 | 0 | 0 | 204 | 149 | 1 | 1 | 0 | 0 | 0 | NA | 0.245 | 0 | NA | NA | NA | NA | 0.061 |
| SE | 72425 | ENSG00000251442 | LINC01094 | chr4 | + | 78662917 | 78663011 | 78649600 | 78649709 | 78680760 | 78681664 | 0 | 0 | 0 | 2 | 4 | 5 | 4 | 2 | 0 | 9 | 0 | 0 | 0 | 0 | 0 | 14 | 0 | 0 | 0 | 0 | 243 | 149 | 1 | 1 | 0 | 0 | 0 | 1 | 0.214 | 0 | NA | NA | NA | NA | 0.243 |
| SE | 72426 | ENSG00000251442 | LINC01094 | chr4 | + | 78673342 | 78673414 | 78649600 | 78649709 | 78680760 | 78682548 | 0 | 2 | 0 | 0 | 0 | 5 | 4 | 2 | 0 | 9 | 0 | 0 | 0 | 0 | 0 | 14 | 0 | 0 | 0 | 0 | 221 | 149 | 1 | 1 | 0 | 0.252 | 0 | NA | 0 | 0 | NA | NA | NA | NA | 0.063 |
| SE | 72430 | ENSG00000047932 | GOPC | chr6 | - | 117570859 | 117570955 | 117566853 | 117567034 | 117573466 | 117573632 | 97 | 57 | 94 | 77 | 91 | 2 | 1 | 4 | 3 | 8 | 1 | 38 | 51 | 6 | 0 | 10 | 4 | 8 | 0 | 0 | 245 | 149 | 1 | 1 | 0.967 | 0.972 | 0.935 | 0.94 | 0.874 | 0.057 | 0.852 | 0.795 | 1 | NA | 0.262 |
| SE | 72431 | ENSG00000047932 | GOPC | chr6 | - | 117570859 | 117570955 | 117569571 | 117569736 | 117573466 | 117573632 | 260 | 142 | 248 | 215 | 224 | 2 | 2 | 0 | 1 | 0 | 26 | 119 | 105 | 32 | 0 | 0 | 2 | 3 | 0 | 0 | 245 | 149 | 1 | 1 | 0.988 | 0.977 | 1 | 0.992 | 1 | 1 | 0.973 | 0.955 | 1 | NA | 0.009 |
| SE | 72433 | ENSG00000074582 | BCS1L | chr2 | + | 218660036 | 218660166 | 218659713 | 218659743 | 218660938 | 218661307 | 0 | 0 | 0 | 3 | 1 | 1 | 3 | 4 | 1 | 7 | 0 | 0 | 0 | 0 | 0 | 0 | 3 | 1 | 0 | 2 | 279 | 149 | 1 | 1 | 0 | 0 | 0 | 0.616 | 0.071 | NA | 0 | 0 | NA | 0 | 0.137 |
| SE | 72434 | ENSG00000074582 | BCS1L | chr2 | + | 218660036 | 218660218 | 218659708 | 218659743 | 218660306 | 218660482 | 0 | 1 | 1 | 0 | 0 | 0 | 0 | 0 | 0 | 3 | 0 | 0 | 0 | 0 | 0 | 0 | 0 | 0 | 2 | 0 | 298 | 149 | 1 | 1 | NA | 1 | 1 | NA | 0 | NA | NA | NA | 0 | NA | 0.667 |
| SE | 72435 | ENSG00000074582 | BCS1L | chr2 | + | 218660036 | 218660218 | 218659708 | 218659743 | 218660938 | 218661307 | 0 | 1 | 1 | 0 | 0 | 1 | 3 | 4 | 1 | 7 | 0 | 0 | 0 | 0 | 0 | 0 | 3 | 1 | 0 | 2 | 298 | 149 | 1 | 1 | 0 | 0.143 | 0.111 | 0 | 0 | NA | 0 | 0 | NA | 0 | 0.051 |
| SE | 72436 | ENSG00000074582 | BCS1L | chr2 | + | 218660036 | 218660245 | 218659682 | 218659743 | 218660938 | 218661307 | 0 | 1 | 1 | 0 | 0 | 1 | 3 | 4 | 1 | 7 | 0 | 0 | 0 | 0 | 0 | 0 | 3 | 1 | 0 | 2 | 298 | 149 | 1 | 1 | 0 | 0.143 | 0.111 | 0 | 0 | NA | 0 | 0 | NA | 0 | 0.051 |
| SE | 72437 | ENSG00000074582 | BCS1L | chr2 | + | 218660306 | 218660421 | 218658763 | 218659173 | 218660938 | 218661217 | 0 | 0 | 1 | 1 | 9 | 0 | 3 | 1 | 1 | 0 | 0 | 0 | 1 | 2 | 0 | 0 | 0 | 0 | 0 | 7 | 264 | 149 | 1 | 1 | NA | 0 | 0.361 | 0.361 | 1 | NA | NA | 1 | 1 | 0 | -0.236 |
| SE | 72438 | ENSG00000074582 | BCS1L | chr2 | + | 218660306 | 218660421 | 218659759 | 218659800 | 218660938 | 218661217 | 0 | 0 | 1 | 1 | 9 | 1 | 0 | 0 | 0 | 0 | 0 | 0 | 1 | 2 | 0 | 0 | 0 | 0 | 0 | 4 | 264 | 149 | 1 | 1 | 0 | NA | 1 | 1 | 1 | NA | NA | 1 | 1 | 0 | 0.083 |
| SE | 72439 | ENSG00000074582 | BCS1L | chr2 | + | 218660306 | 218660482 | 218659716 | 218659743 | 218660938 | 218661307 | 0 | 0 | 0 | 0 | 2 | 1 | 3 | 4 | 1 | 7 | 0 | 0 | 0 | 0 | 0 | 0 | 3 | 1 | 0 | 2 | 298 | 149 | 1 | 1 | 0 | 0 | 0 | 0 | 0.125 | NA | 0 | 0 | NA | 0 | 0.025 |
| SE | 72441 | ENSG00000074582 | BCS1L | chr2 | + | 218660938 | 218661307 | 218659682 | 218659743 | 218661405 | 218661416 | 15 | 11 | 17 | 11 | 17 | 0 | 3 | 0 | 1 | 0 | 32 | 17 | 11 | 8 | 13 | 0 | 0 | 0 | 0 | 0 | 298 | 149 | 1 | 1 | 1 | 0.647 | 1 | 0.846 | 1 | 1 | 1 | 1 | 1 | 1 | -0.101 |
| SE | 72442 | ENSG00000074582 | BCS1L | chr2 | + | 218660938 | 218661307 | 218659771 | 218660421 | 218661405 | 218661416 | 15 | 8 | 15 | 11 | 20 | 0 | 0 | 0 | 0 | 1 | 32 | 14 | 12 | 10 | 21 | 0 | 0 | 0 | 0 | 0 | 298 | 149 | 1 | 1 | 1 | 1 | 1 | 1 | 0.909 | 1 | 1 | 1 | 1 | 1 | -0.018 |
| SE | 72443 | ENSG00000074582 | BCS1L | chr2 | + | 218661405 | 218661545 | 218659655 | 218659743 | 218661758 | 218661916 | 12 | 13 | 16 | 23 | 16 | 0 | 0 | 1 | 0 | 0 | 0 | 9 | 12 | 4 | 21 | 0 | 0 | 0 | 0 | 0 | 289 | 149 | 1 | 1 | 1 | 1 | 0.892 | 1 | 1 | NA | 1 | 1 | 1 | 1 | -0.022 |
| SE | 72444 | ENSG00000074582 | BCS1L | chr2 | + | 218661405 | 218661545 | 218659771 | 218660421 | 218661758 | 218661916 | 12 | 13 | 16 | 22 | 17 | 0 | 0 | 0 | 0 | 0 | 0 | 9 | 12 | 4 | 21 | 0 | 0 | 0 | 4 | 0 | 289 | 149 | 1 | 1 | 1 | 1 | 1 | 1 | 1 | NA | 1 | 1 | 0.34 | 1 | 0.165 |
| SE | 72448 | ENSG00000163481 | RNF25 | chr2 | - | 218665154 | 218665247 | 218664738 | 218664873 | 218667911 | 218667981 | 31 | 11 | 28 | 14 | 52 | 0 | 1 | 0 | 0 | 0 | 21 | 14 | 19 | 14 | 4 | 0 | 0 | 0 | 0 | 0 | 242 | 149 | 1 | 1 | 1 | 0.871 | 1 | 1 | 1 | 1 | 1 | 1 | 1 | 1 | -0.026 |
| SE | 72449 | ENSG00000163481 | RNF25 | chr2 | - | 218665915 | 218666059 | 218664738 | 218664873 | 218667911 | 218667981 | 4 | 3 | 0 | 0 | 4 | 0 | 1 | 0 | 0 | 0 | 5 | 13 | 0 | 3 | 0 | 0 | 0 | 0 | 0 | 0 | 293 | 149 | 1 | 1 | 1 | 0.604 | NA | NA | 1 | 1 | 1 | NA | 1 | NA | -0.132 |
| SE | 72450 | ENSG00000163481 | RNF25 | chr2 | - | 218665915 | 218666059 | 218665153 | 218665247 | 218666158 | 218666230 | 58 | 35 | 53 | 59 | 182 | 0 | 0 | 1 | 0 | 2 | 83 | 51 | 58 | 36 | 38 | 0 | 0 | 0 | 2 | 0 | 293 | 149 | 1 | 1 | 1 | 1 | 0.964 | 1 | 0.979 | 1 | 1 | 1 | 0.902 | 1 | 0.008 |
| SE | 72452 | ENSG00000163481 | RNF25 | chr2 | - | 218665915 | 218666084 | 218665154 | 218665247 | 218666158 | 218666230 | 34 | 18 | 35 | 24 | 114 | 0 | 0 | 1 | 0 | 2 | 45 | 31 | 27 | 14 | 38 | 0 | 0 | 0 | 2 | 0 | 298 | 149 | 1 | 1 | 1 | 1 | 0.946 | 1 | 0.966 | 1 | 1 | 1 | 0.778 | 1 | 0.027 |
| SE | 72454 | ENSG00000163481 | RNF25 | chr2 | - | 218666158 | 218666230 | 218665915 | 218666059 | 218667911 | 218667981 | 43 | 23 | 41 | 60 | 118 | 0 | 0 | 0 | 0 | 1 | 73 | 29 | 43 | 31 | 15 | 0 | 0 | 0 | 0 | 0 | 221 | 149 | 1 | 1 | 1 | 1 | 1 | 1 | 0.988 | 1 | 1 | 1 | 1 | 1 | -0.002 |
| SE | 72456 | ENSG00000163482 | STK36 | chr2 | + | 218688390 | 218688493 | 218685084 | 218685228 | 218688696 | 218688876 | 0 | 0 | 0 | 1 | 0 | 8 | 6 | 5 | 12 | 0 | 0 | 0 | 0 | 0 | 0 | 12 | 12 | 3 | 13 | 12 | 252 | 149 | 1 | 1 | 0 | 0 | 0 | 0.047 | NA | 0 | 0 | 0 | 0 | 0 | 0.012 |
| SE | 72457 | ENSG00000163482 | STK36 | chr2 | + | 218693239 | 218693344 | 218692426 | 218692710 | 218693722 | 218693821 | 0 | 2 | 5 | 3 | 1 | 0 | 0 | 0 | 0 | 0 | 1 | 1 | 2 | 1 | 0 | 5 | 0 | 0 | 0 | 0 | 254 | 149 | 1 | 1 | NA | 1 | 1 | 1 | 1 | 0.105 | 1 | 1 | 1 | NA | 0.224 |
| SE | 72458 | ENSG00000163482 | STK36 | chr2 | + | 218694524 | 218694635 | 218694263 | 218694327 | 218696526 | 218696601 | 3 | 2 | 10 | 2 | 0 | 0 | 0 | 0 | 1 | 0 | 0 | 3 | 1 | 0 | 22 | 0 | 0 | 0 | 0 | 0 | 260 | 149 | 1 | 1 | 1 | 1 | 1 | 0.534 | NA | NA | 1 | 1 | NA | 1 | -0.117 |
| SE | 72459 | ENSG00000163482 | STK36 | chr2 | + | 218697853 | 218698001 | 218697462 | 218697610 | 218698601 | 218698701 | 7 | 7 | 5 | 3 | 6 | 1 | 0 | 0 | 0 | 0 | 1 | 0 | 13 | 12 | 0 | 3 | 9 | 2 | 2 | 0 | 297 | 149 | 1 | 1 | 0.778 | 1 | 1 | 1 | 1 | 0.143 | 0 | 0.765 | 0.751 | NA | 0.541 |
| SE | 72460 | ENSG00000163482 | STK36 | chr2 | + | 218697902 | 218698001 | 218697462 | 218697610 | 218698601 | 218699348 | 3 | 4 | 5 | 0 | 3 | 1 | 0 | 0 | 0 | 0 | 1 | 0 | 7 | 10 | 0 | 3 | 9 | 2 | 2 | 0 | 248 | 149 | 1 | 1 | 0.643 | 1 | 1 | NA | 1 | 0.167 | 0 | 0.678 | 0.75 | NA | 0.512 |
| SE | 72464 | ENSG00000023516 | AKAP11 | chr13 | + | 42285985 | 42286035 | 42272151 | 42272228 | 42286299 | 42286399 | 42 | 23 | 24 | 19 | 33 | 4 | 0 | 0 | 4 | 3 | 0 | 15 | 35 | 17 | 0 | 0 | 0 | 0 | 0 | 0 | 199 | 149 | 1 | 1 | 0.887 | 1 | 1 | 0.781 | 0.892 | NA | 1 | 1 | 1 | NA | -0.088 |
| SE | 72465 | ENSG00000023516 | AKAP11 | chr13 | + | 42297047 | 42297182 | 42295694 | 42295742 | 42298532 | 42298797 | 158 | 126 | 151 | 128 | 392 | 7 | 1 | 6 | 10 | 16 | 0 | 76 | 54 | 4 | 13 | 0 | 6 | 8 | 0 | 0 | 284 | 149 | 1 | 1 | 0.922 | 0.985 | 0.93 | 0.87 | 0.928 | NA | 0.869 | 0.78 | 1 | 1 | 0.015 |
| SE | 72467 | ENSG00000023516 | AKAP11 | chr13 | + | 42313893 | 42313940 | 42308453 | 42308609 | 42319087 | 42323261 | 37 | 36 | 64 | 60 | 91 | 0 | 2 | 0 | 0 | 0 | 0 | 24 | 27 | 7 | 0 | 0 | 0 | 0 | 0 | 0 | 196 | 149 | 1 | 1 | 1 | 0.932 | 1 | 1 | 1 | NA | 1 | 1 | 1 | NA | -0.014 |
| SE | 72470 | ENSG00000118495 | PLAGL1 | chr6 | - | 143947984 | 143948460 | 143940299 | 143942663 | 143960472 | 143960556 | 5 | 6 | 3 | 6 | 24 | 0 | 0 | 0 | 1 | 0 | 20 | 1 | 8 | 4 | 0 | 0 | 0 | 0 | 0 | 0 | 298 | 149 | 1 | 1 | 1 | 1 | 1 | 0.75 | 1 | 1 | 1 | 1 | 1 | NA | -0.05 |
| SE | 72471 | ENSG00000118495 | PLAGL1 | chr6 | - | 143947984 | 143948460 | 143940299 | 143942663 | 143964786 | 143964818 | 5 | 6 | 3 | 6 | 24 | 3 | 0 | 0 | 4 | 0 | 20 | 0 | 4 | 5 | 0 | 0 | 0 | 3 | 2 | 0 | 298 | 149 | 1 | 1 | 0.455 | 1 | 1 | 0.429 | 1 | 1 | NA | 0.4 | 0.556 | NA | 0.125 |
| SE | 72473 | ENSG00000118495 | PLAGL1 | chr6 | - | 143960468 | 143960542 | 143947984 | 143948460 | 143964786 | 143964818 | 17 | 4 | 6 | 13 | 7 | 0 | 0 | 0 | 0 | 0 | 7 | 23 | 11 | 17 | 0 | 0 | 0 | 0 | 1 | 0 | 223 | 149 | 1 | 1 | 1 | 1 | 1 | 1 | 1 | 1 | 1 | 1 | 0.919 | NA | 0.02 |
| SE | 72476 | ENSG00000118495 | PLAGL1 | chr6 | - | 143960468 | 143960556 | 143948259 | 143948460 | 143964786 | 143964818 | 3 | 12 | 5 | 5 | 14 | 0 | 0 | 0 | 0 | 0 | 4 | 1 | 3 | 7 | 6 | 0 | 0 | 0 | 1 | 0 | 237 | 149 | 1 | 1 | 1 | 1 | 1 | 1 | 1 | 1 | 1 | 1 | 0.815 | 1 | 0.037 |
| SE | 72477 | ENSG00000118495 | PLAGL1 | chr6 | - | 143960468 | 143960580 | 143947984 | 143948460 | 143964786 | 143964818 | 2 | 1 | 2 | 2 | 2 | 0 | 0 | 0 | 0 | 0 | 4 | 0 | 1 | 7 | 2 | 0 | 0 | 0 | 1 | 0 | 261 | 149 | 1 | 1 | 1 | 1 | 1 | 1 | 1 | 1 | NA | 1 | 0.8 | 1 | 0.05 |
| SE | 72478 | ENSG00000118495 | PLAGL1 | chr6 | - | 143960472 | 143960556 | 143942296 | 143942663 | 143964786 | 143964818 | 0 | 9 | 3 | 1 | 10 | 3 | 0 | 0 | 4 | 0 | 0 | 1 | 0 | 5 | 4 | 0 | 0 | 3 | 2 | 0 | 233 | 149 | 1 | 1 | 0 | 1 | 1 | 0.138 | 1 | NA | 1 | 0 | 0.615 | 1 | -0.026 |
| SE | 72479 | ENSG00000118495 | PLAGL1 | chr6 | - | 143960472 | 143960556 | 143947984 | 143948460 | 143964786 | 143964818 | 0 | 9 | 3 | 1 | 10 | 0 | 0 | 0 | 0 | 0 | 0 | 1 | 4 | 5 | 4 | 0 | 0 | 0 | 1 | 0 | 233 | 149 | 1 | 1 | NA | 1 | 1 | 1 | 1 | NA | 1 | 1 | 0.762 | 1 | 0.059 |
| SE | 72486 | ENSG00000118495 | PLAGL1 | chr6 | - | 143965107 | 143965313 | 143964786 | 143964818 | 143966157 | 143966198 | 0 | 1 | 0 | 0 | 0 | 16 | 18 | 5 | 8 | 22 | 0 | 0 | 0 | 0 | 0 | 27 | 43 | 19 | 37 | 4 | 298 | 149 | 1 | 1 | 0 | 0.027 | 0 | 0 | 0 | 0 | 0 | 0 | 0 | 0 | 0.005 |
| SE | 72489 | ENSG00000118495 | PLAGL1 | chr6 | - | 143965849 | 143966198 | 143964786 | 143964818 | 143968906 | 143968978 | 2 | 1 | 0 | 0 | 2 | 7 | 0 | 0 | 7 | 11 | 9 | 6 | 4 | 4 | 4 | 0 | 4 | 4 | 8 | 0 | 298 | 149 | 1 | 1 | 0.125 | 1 | NA | 0 | 0.083 | 1 | 0.429 | 0.333 | 0.2 | 1 | -0.29 |
| SE | 72492 | ENSG00000118495 | PLAGL1 | chr6 | - | 143966157 | 143966212 | 143964786 | 143964818 | 143968906 | 143968978 | 12 | 11 | 1 | 5 | 8 | 7 | 0 | 0 | 7 | 11 | 10 | 29 | 9 | 24 | 0 | 0 | 4 | 4 | 8 | 0 | 204 | 149 | 1 | 1 | 0.556 | 1 | 1 | 0.343 | 0.347 | 1 | 0.841 | 0.622 | 0.687 | NA | -0.138 |
| SE | 72495 | ENSG00000118495 | PLAGL1 | chr6 | - | 143985134 | 143985174 | 143968906 | 143968978 | 144008089 | 144008259 | 2 | 1 | 6 | 0 | 2 | 0 | 0 | 0 | 0 | 0 | 0 | 5 | 5 | 3 | 0 | 0 | 0 | 0 | 1 | 0 | 189 | 149 | 1 | 1 | 1 | 1 | 1 | NA | 1 | NA | 1 | 1 | 0.703 | NA | 0.099 |
| SE | 72497 | ENSG00000118495 | PLAGL1 | chr6 | - | 143985134 | 143985174 | 143968906 | 143968978 | 144064450 | 144064598 | 2 | 1 | 5 | 0 | 6 | 2 | 0 | 1 | 2 | 2 | 0 | 4 | 5 | 3 | 0 | 18 | 3 | 1 | 0 | 0 | 189 | 149 | 1 | 1 | 0.441 | 1 | 0.798 | 0 | 0.703 | 0 | 0.512 | 0.798 | 1 | NA | 0.011 |
| SE | 72498 | ENSG00000118495 | PLAGL1 | chr6 | - | 143985134 | 143985174 | 143968906 | 143968978 | 144064467 | 144064598 | 3 | 1 | 5 | 0 | 2 | 8 | 6 | 2 | 6 | 0 | 0 | 9 | 6 | 5 | 8 | 0 | 2 | 7 | 7 | 11 | 189 | 149 | 1 | 1 | 0.228 | 0.116 | 0.663 | 0 | 1 | NA | 0.78 | 0.403 | 0.36 | 0.364 | -0.075 |
| SE | 72499 | ENSG00000118495 | PLAGL1 | chr6 | - | 143985134 | 143985196 | 143968608 | 143968978 | 144064467 | 144064540 | 2 | 1 | 5 | 0 | 2 | 8 | 6 | 2 | 6 | 0 | 0 | 4 | 5 | 3 | 0 | 0 | 2 | 7 | 7 | 11 | 211 | 149 | 1 | 1 | 0.15 | 0.105 | 0.638 | 0 | 1 | NA | 0.585 | 0.335 | 0.232 | 0 | 0.091 |
| SE | 72501 | ENSG00000177842 | ZNF620 | chr3 | + | 40507842 | 40507941 | 40506303 | 40506376 | 40515859 | 40517529 | 0 | 0 | 0 | 0 | 4 | 5 | 2 | 0 | 2 | 4 | 0 | 0 | 0 | 0 | 0 | 11 | 0 | 0 | 1 | 0 | 248 | 149 | 1 | 1 | 0 | 0 | NA | 0 | 0.375 | 0 | NA | NA | 0 | NA | 0.094 |
| SE | 72503 | ENSG00000177842 | ZNF620 | chr3 | + | 40508655 | 40508774 | 40506303 | 40506376 | 40515859 | 40518736 | 0 | 0 | 1 | 0 | 0 | 5 | 2 | 0 | 2 | 4 | 0 | 2 | 1 | 0 | 0 | 11 | 0 | 0 | 1 | 0 | 268 | 149 | 1 | 1 | 0 | 0 | 1 | 0 | 0 | 0 | 1 | 1 | 0 | NA | -0.3 |
| SE | 72504 | ENSG00000177842 | ZNF620 | chr3 | + | 40511469 | 40511596 | 40506303 | 40506376 | 40512401 | 40512472 | 10 | 2 | 11 | 6 | 7 | 0 | 0 | 0 | 2 | 3 | 11 | 3 | 0 | 0 | 0 | 0 | 0 | 0 | 0 | 0 | 276 | 149 | 1 | 1 | 1 | 1 | 1 | 0.618 | 0.557 | 1 | 1 | NA | NA | NA | -0.165 |
| SE | 72505 | ENSG00000177842 | ZNF620 | chr3 | + | 40511469 | 40511596 | 40506303 | 40506376 | 40515859 | 40515944 | 5 | 0 | 1 | 1 | 2 | 5 | 2 | 0 | 2 | 4 | 0 | 5 | 9 | 1 | 0 | 11 | 0 | 0 | 1 | 0 | 276 | 149 | 1 | 1 | 0.351 | 0 | 1 | 0.213 | 0.213 | 0 | 1 | 1 | 0.351 | NA | -0.232 |
| SE | 72506 | ENSG00000177842 | ZNF620 | chr3 | + | 40511469 | 40511596 | 40507842 | 40507941 | 40515859 | 40515944 | 2 | 0 | 0 | 0 | 0 | 0 | 0 | 0 | 0 | 2 | 0 | 5 | 9 | 1 | 0 | 0 | 0 | 0 | 0 | 0 | 276 | 149 | 1 | 1 | 1 | NA | NA | NA | 0 | NA | 1 | 1 | 1 | NA | -0.5 |
| SE | 72507 | ENSG00000177842 | ZNF620 | chr3 | + | 40511469 | 40511596 | 40508655 | 40508774 | 40515859 | 40515944 | 1 | 0 | 0 | 0 | 0 | 0 | 0 | 0 | 0 | 0 | 0 | 5 | 9 | 1 | 0 | 0 | 2 | 1 | 0 | 0 | 276 | 149 | 1 | 1 | 1 | NA | NA | NA | NA | NA | 0.574 | 0.829 | 1 | NA | 0.199 |
| SE | 72508 | ENSG00000177842 | ZNF620 | chr3 | + | 40512401 | 40512515 | 40506303 | 40506376 | 40515859 | 40515944 | 4 | 6 | 2 | 6 | 3 | 5 | 2 | 0 | 2 | 4 | 0 | 0 | 3 | 4 | 0 | 11 | 0 | 0 | 1 | 0 | 263 | 149 | 1 | 1 | 0.312 | 0.63 | 1 | 0.63 | 0.298 | 0 | NA | 1 | 0.694 | NA | 0.009 |
| SE | 72509 | ENSG00000177842 | ZNF620 | chr3 | + | 40512401 | 40512515 | 40511469 | 40511596 | 40515859 | 40515944 | 10 | 8 | 12 | 7 | 5 | 1 | 0 | 0 | 0 | 0 | 11 | 3 | 3 | 9 | 0 | 0 | 5 | 9 | 1 | 0 | 263 | 149 | 1 | 1 | 0.85 | 1 | 1 | 1 | 1 | 1 | 0.254 | 0.159 | 0.836 | NA | 0.408 |
| SE | 72510 | ENSG00000163013 | FBXO41 | chr2 | - | 73263677 | 73263830 | 73263212 | 73263308 | 73263937 | 73264053 | 22 | 12 | 23 | 7 | 5 | 0 | 0 | 0 | 2 | 0 | 44 | 10 | 12 | 14 | 36 | 0 | 0 | 1 | 0 | 0 | 298 | 149 | 1 | 1 | 1 | 1 | 1 | 0.636 | 1 | 1 | 1 | 0.857 | 1 | 1 | -0.044 |
| SE | 72511 | ENSG00000163013 | FBXO41 | chr2 | - | 73265892 | 73265966 | 73265281 | 73265640 | 73266456 | 73266682 | 5 | 0 | 3 | 0 | 0 | 1 | 0 | 1 | 0 | 0 | 16 | 5 | 0 | 7 | 10 | 0 | 4 | 0 | 0 | 0 | 223 | 149 | 1 | 1 | 0.77 | NA | 0.667 | NA | NA | 1 | 0.455 | NA | 1 | 1 | -0.145 |
| SE | 72515 | ENSG00000072415 | MPP5 | chr14 | + | 67302409 | 67302571 | 67301971 | 67302118 | 67303521 | 67303599 | 54 | 68 | 69 | 65 | 107 | 0 | 0 | 0 | 0 | 3 | 30 | 28 | 59 | 31 | 56 | 0 | 0 | 0 | 0 | 0 | 298 | 149 | 1 | 1 | 1 | 1 | 1 | 1 | 0.947 | 1 | 1 | 1 | 1 | 1 | -0.011 |
| SE | 72516 | ENSG00000072415 | MPP5 | chr14 | + | 67317407 | 67317479 | 67315293 | 67316903 | 67320229 | 67320397 | 46 | 41 | 45 | 48 | 45 | 0 | 0 | 0 | 0 | 0 | 2 | 25 | 51 | 5 | 0 | 0 | 0 | 3 | 0 | 0 | 221 | 149 | 1 | 1 | 1 | 1 | 1 | 1 | 1 | 1 | 1 | 0.92 | 1 | NA | 0.02 |
| SE | 72533 | ENSG00000135426 | TESPA1 | chr12 | - | 54962430 | 54963242 | 54961167 | 54961267 | 54963741 | 54963950 | 263 | 220 | 422 | 354 | 605 | 0 | 0 | 0 | 0 | 0 | 60 | 151 | 182 | 138 | 287 | 4 | 0 | 1 | 0 | 0 | 298 | 149 | 1 | 1 | 1 | 1 | 1 | 1 | 1 | 0.882 | 1 | 0.989 | 1 | 1 | 0.026 |
| SE | 72536 | ENSG00000135426 | TESPA1 | chr12 | - | 54981773 | 54981945 | 54974399 | 54974607 | 54984584 | 54984731 | 10 | 15 | 24 | 20 | 35 | 78 | 53 | 66 | 125 | 72 | 0 | 9 | 7 | 6 | 0 | 0 | 47 | 40 | 32 | 170 | 298 | 149 | 1 | 1 | 0.06 | 0.124 | 0.154 | 0.074 | 0.196 | NA | 0.087 | 0.08 | 0.086 | 0 | 0.058 |
| SE | 72538 | ENSG00000135426 | TESPA1 | chr12 | - | 54983993 | 54984166 | 54981773 | 54981945 | 54984584 | 54984731 | 9 | 14 | 47 | 15 | 102 | 0 | 0 | 2 | 2 | 0 | 0 | 16 | 0 | 9 | 6 | 0 | 4 | 5 | 0 | 0 | 298 | 149 | 1 | 1 | 1 | 1 | 0.922 | 0.789 | 1 | NA | 0.667 | 0 | 1 | 1 | 0.275 |
| SE | 72540 | ENSG00000034152 | MAP2K3 | chr17 | + | 21295674 | 21295769 | 21284710 | 21284969 | 21296085 | 21296143 | 0 | 1 | 0 | 0 | 0 | 0 | 0 | 1 | 1 | 5 | 0 | 0 | 3 | 6 | 6 | 5 | 3 | 0 | 0 | 5 | 244 | 149 | 1 | 1 | NA | 1 | 0 | 0 | 0 | 0 | 0 | 1 | 1 | 0.423 | -0.235 |
| SE | 72541 | ENSG00000034152 | MAP2K3 | chr17 | + | 21295674 | 21295769 | 21284710 | 21284969 | 21298412 | 21298479 | 2 | 0 | 1 | 0 | 0 | 56 | 18 | 35 | 25 | 14 | 0 | 0 | 0 | 0 | 3 | 38 | 50 | 35 | 26 | 108 | 244 | 149 | 1 | 1 | 0.021 | 0 | 0.017 | 0 | 0 | 0 | 0 | 0 | 0 | 0.017 | 0.004 |
| SE | 72542 | ENSG00000034152 | MAP2K3 | chr17 | + | 21295674 | 21295769 | 21287990 | 21288091 | 21296085 | 21296143 | 0 | 1 | 0 | 0 | 0 | 2 | 0 | 0 | 0 | 0 | 0 | 0 | 3 | 6 | 5 | 0 | 0 | 0 | 0 | 0 | 244 | 149 | 1 | 1 | 0 | 1 | NA | NA | NA | NA | NA | 1 | 1 | 1 | -0.5 |
| SE | 72543 | ENSG00000034152 | MAP2K3 | chr17 | + | 21295674 | 21295769 | 21287990 | 21288091 | 21298412 | 21298479 | 2 | 0 | 1 | 0 | 0 | 0 | 0 | 0 | 2 | 0 | 0 | 0 | 0 | 0 | 2 | 7 | 1 | 2 | 0 | 0 | 244 | 149 | 1 | 1 | 1 | NA | 1 | 0 | NA | 0 | 0 | 0 | NA | 1 | 0.417 |
| SE | 72546 | ENSG00000034152 | MAP2K3 | chr17 | + | 21296085 | 21296210 | 21295674 | 21295769 | 21298412 | 21298444 | 4 | 3 | 2 | 5 | 7 | 2 | 0 | 1 | 0 | 0 | 6 | 2 | 3 | 11 | 16 | 0 | 0 | 0 | 0 | 2 | 274 | 149 | 1 | 1 | 0.521 | 1 | 0.521 | 1 | 1 | 1 | 1 | 1 | 1 | 0.813 | -0.154 |
| SE | 72548 | ENSG00000034152 | MAP2K3 | chr17 | + | 21305050 | 21305128 | 21304425 | 21304553 | 21312141 | 21312281 | 289 | 141 | 173 | 263 | 213 | 0 | 0 | 0 | 0 | 0 | 300 | 387 | 185 | 331 | 802 | 0 | 0 | 0 | 2 | 0 | 227 | 149 | 1 | 1 | 1 | 1 | 1 | 1 | 1 | 1 | 1 | 1 | 0.991 | 1 | 0.002 |
| SE | 72549 | ENSG00000034152 | MAP2K3 | chr17 | + | 21312141 | 21312281 | 21305050 | 21305128 | 21313491 | 21313537 | 307 | 143 | 168 | 278 | 186 | 1 | 0 | 3 | 4 | 0 | 256 | 310 | 219 | 355 | 785 | 0 | 1 | 0 | 3 | 2 | 289 | 149 | 1 | 1 | 0.994 | 1 | 0.967 | 0.973 | 1 | 1 | 0.994 | 1 | 0.984 | 0.995 | -0.008 |
| SE | 72550 | ENSG00000068724 | TTC7A | chr2 | + | 46917183 | 46917277 | 46916156 | 46916577 | 46950362 | 46950526 | 0 | 1 | 2 | 4 | 24 | 0 | 0 | 3 | 0 | 1 | 8 | 4 | 10 | 0 | 0 | 0 | 0 | 0 | 0 | 0 | 243 | 149 | 1 | 1 | NA | 1 | 0.29 | 1 | 0.936 | 1 | 1 | 1 | NA | NA | -0.194 |
| SE | 72551 | ENSG00000068724 | TTC7A | chr2 | + | 46951601 | 46951733 | 46950420 | 46950526 | 46956838 | 46957007 | 0 | 0 | 0 | 0 | 0 | 161 | 92 | 117 | 151 | 208 | 0 | 8 | 0 | 0 | 0 | 325 | 151 | 102 | 132 | 386 | 281 | 149 | 1 | 1 | 0 | 0 | 0 | 0 | 0 | 0 | 0.027 | 0 | 0 | 0 | -0.005 |
| SE | 72554 | ENSG00000068724 | TTC7A | chr2 | + | 46978791 | 46978907 | 46956838 | 46957007 | 46993449 | 46993528 | 123 | 76 | 81 | 104 | 86 | 2 | 0 | 0 | 0 | 2 | 258 | 148 | 102 | 121 | 170 | 0 | 0 | 0 | 0 | 0 | 265 | 149 | 1 | 1 | 0.972 | 1 | 1 | 1 | 0.96 | 1 | 1 | 1 | 1 | 1 | -0.014 |
| SE | 72556 | ENSG00000068724 | TTC7A | chr2 | + | 47005921 | 47006059 | 46995135 | 46995199 | 47011330 | 47011435 | 152 | 44 | 95 | 120 | 39 | 0 | 0 | 0 | 0 | 2 | 245 | 145 | 76 | 89 | 198 | 0 | 0 | 0 | 0 | 0 | 287 | 149 | 1 | 1 | 1 | 1 | 1 | 1 | 0.91 | 1 | 1 | 1 | 1 | 1 | -0.018 |
| SE | 72560 | ENSG00000068724 | TTC7A | chr2 | + | 47021861 | 47021979 | 47011330 | 47011435 | 47029223 | 47029384 | 45 | 11 | 20 | 36 | 15 | 0 | 0 | 0 | 3 | 0 | 47 | 14 | 25 | 16 | 37 | 0 | 0 | 0 | 0 | 0 | 267 | 149 | 1 | 1 | 1 | 1 | 1 | 0.87 | 1 | 1 | 1 | 1 | 1 | 1 | -0.026 |
| SE | 72562 | ENSG00000068724 | TTC7A | chr2 | + | 47023407 | 47023465 | 47011330 | 47011435 | 47029223 | 47029384 | 0 | 2 | 0 | 4 | 5 | 0 | 0 | 0 | 3 | 0 | 0 | 1 | 0 | 0 | 0 | 0 | 0 | 0 | 0 | 0 | 207 | 149 | 1 | 1 | NA | 1 | NA | 0.49 | 1 | NA | 1 | NA | NA | NA | -0.17 |
| SE | 72565 | ENSG00000068724 | TTC7A | chr2 | + | 47024286 | 47024359 | 47011330 | 47011435 | 47029223 | 47029384 | 41 | 11 | 22 | 18 | 15 | 0 | 0 | 0 | 3 | 0 | 8 | 26 | 12 | 9 | 39 | 0 | 0 | 0 | 0 | 0 | 222 | 149 | 1 | 1 | 1 | 1 | 1 | 0.801 | 1 | 1 | 1 | 1 | 1 | 1 | -0.04 |
| SE | 72567 | ENSG00000068724 | TTC7A | chr2 | + | 47024286 | 47024359 | 47023407 | 47023465 | 47029223 | 47029384 | 95 | 32 | 51 | 63 | 57 | 0 | 0 | 0 | 1 | 0 | 42 | 59 | 39 | 28 | 93 | 0 | 0 | 0 | 0 | 0 | 222 | 149 | 1 | 1 | 1 | 1 | 1 | 0.977 | 1 | 1 | 1 | 1 | 1 | 1 | -0.005 |
| SE | 72568 | ENSG00000068724 | TTC7A | chr2 | + | 47046314 | 47046431 | 47029223 | 47029384 | 47049948 | 47050046 | 43 | 17 | 41 | 56 | 21 | 1 | 2 | 0 | 2 | 0 | 143 | 43 | 16 | 46 | 74 | 0 | 0 | 2 | 0 | 0 | 266 | 149 | 1 | 1 | 0.96 | 0.826 | 1 | 0.94 | 1 | 1 | 1 | 0.818 | 1 | 1 | -0.018 |
| SE | 72569 | ENSG00000068724 | TTC7A | chr2 | + | 47047265 | 47047337 | 47046314 | 47046431 | 47049948 | 47050046 | 1 | 0 | 0 | 0 | 0 | 38 | 12 | 28 | 48 | 19 | 0 | 1 | 0 | 0 | 0 | 110 | 27 | 14 | 38 | 52 | 221 | 149 | 1 | 1 | 0.017 | 0 | 0 | 0 | 0 | 0 | 0.024 | 0 | 0 | 0 | -0.001 |
| SE | 72570 | ENSG00000068724 | TTC7A | chr2 | + | 47051745 | 47051880 | 47049948 | 47050046 | 47060768 | 47060956 | 68 | 32 | 61 | 69 | 32 | 1 | 0 | 0 | 3 | 0 | 99 | 81 | 58 | 34 | 81 | 4 | 3 | 0 | 0 | 12 | 284 | 149 | 1 | 1 | 0.973 | 1 | 1 | 0.923 | 1 | 0.928 | 0.934 | 1 | 1 | 0.78 | 0.051 |
| SE | 72572 | ENSG00000197223 | C1D | chr2 | - | 68046343 | 68046410 | 68045987 | 68046043 | 68047172 | 68047315 | 484 | 238 | 351 | 407 | 485 | 2 | 0 | 1 | 0 | 0 | 247 | 291 | 343 | 270 | 120 | 0 | 0 | 0 | 4 | 0 | 216 | 149 | 1 | 1 | 0.994 | 1 | 0.996 | 1 | 1 | 1 | 1 | 1 | 0.979 | 1 | 0.002 |
| SE | 72574 | ENSG00000197223 | C1D | chr2 | - | 68053060 | 68053137 | 68047172 | 68047315 | 68062957 | 68063009 | 1 | 0 | 0 | 1 | 0 | 1 | 0 | 0 | 0 | 0 | 0 | 0 | 0 | 0 | 0 | 0 | 0 | 1 | 0 | 0 | 226 | 149 | 1 | 1 | 0.397 | NA | NA | 1 | NA | NA | NA | 0 | NA | NA | 0.699 |
| SE | 72577 | ENSG00000157680 | DGKI | chr7 | - | 137412169 | 137412207 | 137407874 | 137407995 | 137444076 | 137444102 | 1 | 0 | 0 | 0 | 0 | 0 | 0 | 0 | 0 | 0 | 0 | 0 | 0 | 0 | 0 | 8 | 0 | 0 | 0 | 0 | 187 | 149 | 1 | 1 | 1 | NA | NA | NA | NA | 0 | NA | NA | NA | NA | 1 |
| SE | 72582 | ENSG00000189350 | TOGARAM2 | chr2 | + | 29017153 | 29017304 | 29014394 | 29014561 | 29017791 | 29017956 | 27 | 6 | 2 | 17 | 1 | 0 | 0 | 0 | 1 | 0 | 12 | 27 | 5 | 11 | 8 | 0 | 0 | 0 | 0 | 0 | 298 | 149 | 1 | 1 | 1 | 1 | 1 | 0.895 | 1 | 1 | 1 | 1 | 1 | 1 | -0.021 |
| SE | 72583 | ENSG00000189350 | TOGARAM2 | chr2 | + | 29017153 | 29017304 | 29014394 | 29014561 | 29022157 | 29022308 | 6 | 2 | 0 | 10 | 1 | 1 | 0 | 0 | 2 | 0 | 0 | 0 | 3 | 6 | 3 | 0 | 0 | 0 | 0 | 0 | 298 | 149 | 1 | 1 | 0.75 | 1 | NA | 0.714 | 1 | NA | NA | 1 | 1 | 1 | -0.134 |
| SE | 72584 | ENSG00000189350 | TOGARAM2 | chr2 | + | 29017153 | 29017304 | 29014394 | 29014561 | 29024138 | 29024374 | 6 | 2 | 2 | 8 | 1 | 2 | 0 | 0 | 2 | 0 | 0 | 0 | 3 | 6 | 3 | 0 | 0 | 0 | 2 | 0 | 298 | 149 | 1 | 1 | 0.6 | 1 | 1 | 0.667 | 1 | NA | NA | 1 | 0.6 | 1 | -0.013 |
| SE | 72585 | ENSG00000189350 | TOGARAM2 | chr2 | + | 29017791 | 29017956 | 29014394 | 29014561 | 29022157 | 29022308 | 16 | 1 | 2 | 9 | 0 | 1 | 0 | 0 | 2 | 0 | 0 | 16 | 6 | 5 | 0 | 0 | 0 | 0 | 0 | 0 | 298 | 149 | 1 | 1 | 0.889 | 1 | 1 | 0.692 | NA | NA | 1 | 1 | 1 | NA | -0.105 |
| SE | 72586 | ENSG00000189350 | TOGARAM2 | chr2 | + | 29017791 | 29017956 | 29014394 | 29014561 | 29023085 | 29023191 | 2 | 0 | 1 | 1 | 0 | 0 | 2 | 0 | 0 | 0 | 0 | 0 | 3 | 0 | 0 | 0 | 0 | 0 | 0 | 0 | 298 | 149 | 1 | 1 | 1 | 0 | 1 | 1 | NA | NA | NA | 1 | NA | NA | -0.25 |
| SE | 72587 | ENSG00000189350 | TOGARAM2 | chr2 | + | 29017791 | 29017956 | 29017153 | 29017304 | 29022157 | 29022308 | 37 | 5 | 4 | 17 | 0 | 0 | 0 | 0 | 2 | 0 | 12 | 43 | 8 | 10 | 5 | 0 | 0 | 0 | 0 | 0 | 298 | 149 | 1 | 1 | 1 | 1 | 1 | 0.81 | NA | 1 | 1 | 1 | 1 | 1 | -0.047 |
| SE | 72588 | ENSG00000189350 | TOGARAM2 | chr2 | + | 29022157 | 29022308 | 29014466 | 29014561 | 29023085 | 29023191 | 15 | 4 | 6 | 5 | 0 | 0 | 2 | 0 | 0 | 0 | 0 | 13 | 0 | 0 | 0 | 0 | 0 | 0 | 0 | 0 | 298 | 149 | 1 | 1 | 1 | 0.5 | 1 | 1 | NA | NA | 1 | NA | NA | NA | -0.125 |
| SE | 72589 | ENSG00000189350 | TOGARAM2 | chr2 | + | 29022157 | 29022308 | 29017791 | 29017956 | 29023085 | 29023191 | 30 | 5 | 8 | 11 | 0 | 2 | 0 | 1 | 0 | 0 | 0 | 29 | 6 | 5 | 0 | 0 | 0 | 3 | 0 | 0 | 298 | 149 | 1 | 1 | 0.882 | 1 | 0.8 | 1 | NA | NA | 1 | 0.5 | 1 | NA | 0.087 |
| SE | 72590 | ENSG00000189350 | TOGARAM2 | chr2 | + | 29023085 | 29023191 | 29014394 | 29014561 | 29024138 | 29024374 | 13 | 6 | 4 | 6 | 3 | 2 | 0 | 0 | 2 | 0 | 0 | 11 | 1 | 3 | 0 | 0 | 0 | 0 | 2 | 0 | 255 | 149 | 1 | 1 | 0.792 | 1 | 1 | 0.637 | 1 | NA | 1 | 1 | 0.467 | NA | 0.063 |
| SE | 72591 | ENSG00000189350 | TOGARAM2 | chr2 | + | 29033468 | 29033563 | 29032770 | 29033051 | 29035463 | 29035656 | 40 | 22 | 15 | 28 | 22 | 0 | 0 | 0 | 0 | 0 | 36 | 45 | 20 | 25 | 10 | 0 | 1 | 0 | 0 | 0 | 244 | 149 | 1 | 1 | 1 | 1 | 1 | 1 | 1 | 1 | 0.965 | 1 | 1 | 1 | 0.007 |
| SE | 72603 | ENSG00000115977 | AAK1 | chr2 | - | 69478950 | 69479061 | 69476879 | 69476990 | 69480859 | 69480961 | 2 | 3 | 5 | 5 | 0 | 0 | 1 | 0 | 0 | 0 | 3 | 0 | 6 | 0 | 0 | 0 | 0 | 0 | 0 | 6 | 260 | 149 | 1 | 1 | 1 | 0.632 | 1 | 1 | NA | 1 | NA | 1 | NA | 0 | 0.241 |
| SE | 72611 | ENSG00000115977 | AAK1 | chr2 | - | 69509230 | 69509460 | 69507420 | 69507578 | 69514470 | 69514749 | 78 | 48 | 84 | 108 | 58 | 0 | 0 | 2 | 0 | 0 | 64 | 65 | 79 | 49 | 21 | 0 | 0 | 0 | 0 | 0 | 298 | 149 | 1 | 1 | 1 | 1 | 0.955 | 1 | 1 | 1 | 1 | 1 | 1 | 1 | -0.009 |
| SE | 72612 | ENSG00000115977 | AAK1 | chr2 | - | 69525032 | 69525112 | 69520833 | 69520988 | 69527215 | 69527319 | 200 | 156 | 273 | 191 | 413 | 0 | 1 | 2 | 1 | 3 | 234 | 185 | 242 | 185 | 101 | 0 | 0 | 0 | 0 | 0 | 229 | 149 | 1 | 1 | 1 | 0.99 | 0.989 | 0.992 | 0.989 | 1 | 1 | 1 | 1 | 1 | -0.008 |
| SE | 72613 | ENSG00000115977 | AAK1 | chr2 | - | 69527215 | 69527319 | 69525032 | 69525112 | 69530007 | 69530140 | 617 | 511 | 681 | 625 | 1068 | 0 | 0 | 0 | 0 | 0 | 587 | 529 | 671 | 490 | 279 | 0 | 3 | 0 | 3 | 0 | 253 | 149 | 1 | 1 | 1 | 1 | 1 | 1 | 1 | 1 | 0.99 | 1 | 0.99 | 1 | 0.004 |
| SE | 72614 | ENSG00000115977 | AAK1 | chr2 | - | 69532040 | 69532162 | 69530624 | 69530706 | 69542522 | 69542665 | 555 | 531 | 669 | 559 | 1012 | 1 | 0 | 0 | 0 | 0 | 537 | 478 | 656 | 495 | 287 | 0 | 0 | 7 | 3 | 0 | 271 | 149 | 1 | 1 | 0.997 | 1 | 1 | 1 | 1 | 1 | 1 | 0.981 | 0.989 | 1 | 0.005 |
| SE | 72615 | ENSG00000115977 | AAK1 | chr2 | - | 69542522 | 69542665 | 69531716 | 69532162 | 69544435 | 69544509 | 450 | 389 | 496 | 375 | 637 | 0 | 1 | 1 | 0 | 2 | 400 | 357 | 508 | 400 | 247 | 0 | 0 | 0 | 0 | 0 | 292 | 149 | 1 | 1 | 1 | 0.995 | 0.996 | 1 | 0.994 | 1 | 1 | 1 | 1 | 1 | -0.003 |
| SE | 72616 | ENSG00000115977 | AAK1 | chr2 | - | 69556859 | 69556978 | 69544238 | 69544544 | 69642877 | 69643274 | 324 | 284 | 355 | 342 | 275 | 2 | 1 | 1 | 2 | 0 | 311 | 255 | 340 | 248 | 266 | 0 | 3 | 6 | 1 | 0 | 268 | 149 | 1 | 1 | 0.989 | 0.994 | 0.995 | 0.99 | 1 | 1 | 0.979 | 0.969 | 0.993 | 1 | 0.005 |
| SE | 72617 | ENSG00000147669 | POLR2K | chr8 | + | 100151346 | 100151416 | 100150622 | 100150709 | 100151823 | 100151916 | 319 | 215 | 261 | 248 | 717 | 0 | 0 | 2 | 0 | 2 | 231 | 238 | 204 | 245 | 200 | 0 | 0 | 2 | 2 | 0 | 219 | 149 | 1 | 1 | 1 | 1 | 0.989 | 1 | 0.996 | 1 | 1 | 0.986 | 0.988 | 1 | 0.002 |
| SE | 72618 | ENSG00000147669 | POLR2K | chr8 | + | 100151823 | 100151916 | 100151346 | 100151416 | 100153293 | 100154003 | 524 | 336 | 454 | 441 | 1481 | 0 | 0 | 0 | 0 | 2 | 362 | 384 | 333 | 424 | 287 | 0 | 0 | 2 | 0 | 0 | 242 | 149 | 1 | 1 | 1 | 1 | 1 | 1 | 0.998 | 1 | 1 | 0.99 | 1 | 1 | 0.002 |
| SE | 72619 | ENSG00000104267 | CA2 | chr8 | + | 85473388 | 85473468 | 85465271 | 85465469 | 85473692 | 85473811 | 0 | 1 | 2 | 0 | 0 | 51 | 325 | 59 | 126 | 88 | 0 | 3 | 0 | 0 | 0 | 327 | 287 | 122 | 221 | 375 | 229 | 149 | 1 | 1 | 0 | 0.002 | 0.022 | 0 | 0 | 0 | 0.007 | 0 | 0 | 0 | 0.003 |
| SE | 72620 | ENSG00000104267 | CA2 | chr8 | + | 85473692 | 85473811 | 85465271 | 85465469 | 85474323 | 85474416 | 96 | 590 | 111 | 204 | 209 | 0 | 2 | 0 | 0 | 0 | 636 | 531 | 219 | 383 | 623 | 0 | 0 | 0 | 0 | 0 | 268 | 149 | 1 | 1 | 1 | 0.994 | 1 | 1 | 1 | 1 | 1 | 1 | 1 | 1 | -0.001 |
| SE | 72621 | ENSG00000104267 | CA2 | chr8 | + | 85474323 | 85474416 | 85465271 | 85465469 | 85475797 | 85475860 | 66 | 273 | 40 | 113 | 104 | 0 | 1 | 0 | 0 | 0 | 429 | 266 | 102 | 264 | 351 | 0 | 0 | 0 | 0 | 0 | 242 | 149 | 1 | 1 | 1 | 0.994 | 1 | 1 | 1 | 1 | 1 | 1 | 1 | 1 | -0.001 |
| SE | 72624 | ENSG00000169679 | BUB1 | chr2 | - | 110657035 | 110657117 | 110655738 | 110655916 | 110657545 | 110657621 | 6 | 4 | 16 | 4 | 16 | 0 | 1 | 0 | 0 | 0 | 1 | 13 | 13 | 24 | 0 | 0 | 0 | 0 | 0 | 0 | 231 | 149 | 1 | 1 | 1 | 0.721 | 1 | 1 | 1 | 1 | 1 | 1 | 1 | NA | -0.056 |
| SE | 72625 | ENSG00000169679 | BUB1 | chr2 | - | 110672660 | 110672857 | 110670524 | 110670568 | 110674085 | 110674224 | 25 | 7 | 41 | 48 | 35 | 1 | 0 | 0 | 0 | 0 | 74 | 31 | 47 | 56 | 33 | 0 | 0 | 0 | 0 | 0 | 298 | 149 | 1 | 1 | 0.926 | 1 | 1 | 1 | 1 | 1 | 1 | 1 | 1 | 1 | -0.015 |
| SE | 72641 | ENSG00000153094 | BCL2L11 | chr2 | + | 111130131 | 111130247 | 111123745 | 111123869 | 111150043 | 111150147 | 0 | 0 | 0 | 0 | 0 | 14 | 3 | 17 | 9 | 7 | 0 | 0 | 0 | 1 | 0 | 9 | 4 | 16 | 13 | 8 | 265 | 149 | 1 | 1 | 0 | 0 | 0 | 0 | 0 | 0 | 0 | 0 | 0.041 | 0 | -0.008 |
| SE | 72642 | ENSG00000153094 | BCL2L11 | chr2 | + | 111130131 | 111130247 | 111124049 | 111124139 | 111150043 | 111150147 | 0 | 0 | 0 | 0 | 0 | 76 | 22 | 54 | 41 | 70 | 0 | 0 | 0 | 2 | 0 | 107 | 110 | 176 | 87 | 112 | 265 | 149 | 1 | 1 | 0 | 0 | 0 | 0 | 0 | 0 | 0 | 0 | 0.013 | 0 | -0.003 |
| SE | 72643 | ENSG00000153094 | BCL2L11 | chr2 | + | 111142312 | 111142391 | 111124049 | 111124139 | 111144487 | 111144520 | 0 | 0 | 0 | 0 | 0 | 0 | 0 | 3 | 0 | 0 | 0 | 0 | 1 | 0 | 0 | 0 | 1 | 3 | 0 | 0 | 228 | 149 | 1 | 1 | NA | NA | 0 | NA | NA | NA | 0 | 0.179 | NA | NA | -0.089 |
| SE | 72644 | ENSG00000153094 | BCL2L11 | chr2 | + | 111142312 | 111142391 | 111124049 | 111124139 | 111150043 | 111150147 | 0 | 1 | 0 | 0 | 0 | 76 | 22 | 54 | 41 | 70 | 0 | 8 | 0 | 0 | 0 | 107 | 110 | 176 | 87 | 112 | 228 | 149 | 1 | 1 | 0 | 0.029 | 0 | 0 | 0 | 0 | 0.045 | 0 | 0 | 0 | -0.003 |
| SE | 72647 | ENSG00000153094 | BCL2L11 | chr2 | + | 111144487 | 111144520 | 111142312 | 111142391 | 111150043 | 111150147 | 0 | 0 | 0 | 0 | 0 | 0 | 1 | 0 | 0 | 0 | 10 | 0 | 0 | 0 | 0 | 0 | 8 | 1 | 0 | 0 | 182 | 149 | 1 | 1 | NA | 0 | NA | NA | NA | 1 | 0 | 0 | NA | NA | -0.333 |
| SE | 72649 | ENSG00000153094 | BCL2L11 | chr2 | + | 111150043 | 111150147 | 111123732 | 111123869 | 111164132 | 111164231 | 37 | 20 | 37 | 31 | 34 | 0 | 0 | 0 | 2 | 0 | 35 | 62 | 100 | 53 | 85 | 0 | 4 | 0 | 0 | 0 | 253 | 149 | 1 | 1 | 1 | 1 | 1 | 0.901 | 1 | 1 | 0.901 | 1 | 1 | 1 | 0 |
| SE | 72652 | ENSG00000153094 | BCL2L11 | chr2 | + | 111150043 | 111150147 | 111144487 | 111144520 | 111164132 | 111164231 | 28 | 17 | 26 | 22 | 28 | 0 | 0 | 0 | 0 | 0 | 26 | 60 | 90 | 40 | 77 | 0 | 0 | 0 | 3 | 0 | 253 | 149 | 1 | 1 | 1 | 1 | 1 | 1 | 1 | 1 | 1 | 1 | 0.887 | 1 | 0.023 |
| SE | 72661 | ENSG00000129295 | LRRC6 | chr8 | - | 132578407 | 132578516 | 132572198 | 132572480 | 132583693 | 132583775 | 0 | 2 | 3 | 1 | 1 | 2 | 1 | 7 | 11 | 22 | 0 | 8 | 0 | 12 | 0 | 5 | 2 | 2 | 2 | 0 | 258 | 149 | 1 | 1 | 0 | 0.536 | 0.198 | 0.05 | 0.026 | 0 | 0.698 | 0 | 0.776 | NA | -0.206 |
| SE | 72662 | ENSG00000129295 | LRRC6 | chr8 | - | 132615037 | 132615097 | 132611293 | 132611363 | 132622610 | 132622688 | 6 | 3 | 12 | 9 | 16 | 0 | 0 | 0 | 2 | 1 | 0 | 3 | 0 | 0 | 0 | 0 | 0 | 0 | 0 | 0 | 209 | 149 | 1 | 1 | 1 | 1 | 1 | 0.762 | 0.919 | NA | 1 | NA | NA | NA | -0.064 |
| SE | 72663 | ENSG00000129295 | LRRC6 | chr8 | - | 132622610 | 132622688 | 132615037 | 132615097 | 132625271 | 132625387 | 5 | 2 | 12 | 13 | 19 | 0 | 0 | 0 | 1 | 0 | 0 | 6 | 0 | 0 | 0 | 0 | 0 | 0 | 0 | 0 | 227 | 149 | 1 | 1 | 1 | 1 | 1 | 0.895 | 1 | NA | 1 | NA | NA | NA | -0.021 |
| SE | 72664 | ENSG00000129295 | LRRC6 | chr8 | - | 132632739 | 132632963 | 132625271 | 132625454 | 132637934 | 132638107 | 7 | 4 | 7 | 15 | 16 | 1 | 3 | 5 | 8 | 11 | 0 | 10 | 4 | 6 | 0 | 0 | 8 | 0 | 6 | 4 | 298 | 149 | 1 | 1 | 0.778 | 0.4 | 0.412 | 0.484 | 0.421 | NA | 0.385 | 1 | 0.333 | 0 | 0.069 |
| SE | 72666 | ENSG00000129295 | LRRC6 | chr8 | - | 132643623 | 132643686 | 132637984 | 132638107 | 132656829 | 132656907 | 0 | 1 | 2 | 0 | 0 | 4 | 9 | 11 | 12 | 29 | 0 | 0 | 0 | 0 | 0 | 0 | 15 | 6 | 5 | 7 | 212 | 149 | 1 | 1 | 0 | 0.072 | 0.113 | 0 | 0 | NA | 0 | 0 | 0 | 0 | 0.037 |
| SE | 72667 | ENSG00000129295 | LRRC6 | chr8 | - | 132656829 | 132656907 | 132637934 | 132638107 | 132661459 | 132661627 | 5 | 9 | 14 | 13 | 34 | 0 | 0 | 0 | 0 | 0 | 0 | 16 | 6 | 9 | 7 | 0 | 3 | 0 | 0 | 0 | 227 | 149 | 1 | 1 | 1 | 1 | 1 | 1 | 1 | NA | 0.778 | 1 | 1 | 1 | 0.055 |
| SE | 72669 | ENSG00000113387 | SUB1 | chr5 | + | 32536637 | 32536695 | 32531848 | 32531916 | 32571090 | 32571174 | 0 | 0 | 1 | 0 | 3 | 0 | 7 | 2 | 6 | 5 | 0 | 0 | 1 | 0 | 0 | 0 | 24 | 1 | 2 | 0 | 207 | 149 | 1 | 1 | NA | 0 | 0.265 | 0 | 0.302 | NA | 0 | 0.419 | 0 | NA | 0.002 |
| SE | 72670 | ENSG00000113387 | SUB1 | chr5 | + | 32536637 | 32536695 | 32531848 | 32531916 | 32588511 | 32588584 | 5 | 1 | 1 | 3 | 3 | 4 | 5 | 3 | 7 | 12 | 0 | 1 | 3 | 0 | 7 | 0 | 4 | 4 | 7 | 0 | 207 | 149 | 1 | 1 | 0.474 | 0.126 | 0.194 | 0.236 | 0.153 | NA | 0.153 | 0.351 | 0 | 1 | -0.139 |
| SE | 72671 | ENSG00000113387 | SUB1 | chr5 | + | 32571090 | 32571174 | 32531817 | 32531916 | 32588511 | 32588584 | 7 | 11 | 12 | 12 | 22 | 4 | 5 | 3 | 7 | 12 | 0 | 32 | 3 | 4 | 0 | 0 | 4 | 4 | 7 | 0 | 233 | 149 | 1 | 1 | 0.528 | 0.585 | 0.719 | 0.523 | 0.54 | NA | 0.836 | 0.324 | 0.268 | NA | 0.103 |
| SE | 72672 | ENSG00000113387 | SUB1 | chr5 | + | 32571090 | 32571174 | 32536637 | 32536695 | 32588511 | 32588584 | 7 | 4 | 10 | 6 | 20 | 5 | 1 | 0 | 3 | 3 | 0 | 8 | 4 | 2 | 0 | 0 | 2 | 3 | 0 | 7 | 233 | 149 | 1 | 1 | 0.472 | 0.719 | 1 | 0.561 | 0.81 | NA | 0.719 | 0.46 | 1 | 0 | 0.168 |
| SE | 72673 | ENSG00000113387 | SUB1 | chr5 | + | 32586368 | 32586501 | 32585570 | 32585625 | 32588511 | 32588584 | 15 | 4 | 10 | 4 | 2 | 463 | 201 | 331 | 295 | 246 | 5 | 5 | 8 | 12 | 0 | 134 | 444 | 376 | 250 | 166 | 282 | 149 | 1 | 1 | 0.017 | 0.01 | 0.016 | 0.007 | 0.004 | 0.019 | 0.006 | 0.011 | 0.025 | 0 | -0.001 |
| SE | 72677 | ENSG00000113387 | SUB1 | chr5 | + | 32588511 | 32588584 | 32571090 | 32571174 | 32598960 | 32599069 | 4 | 9 | 7 | 3 | 11 | 0 | 0 | 0 | 0 | 0 | 0 | 26 | 0 | 2 | 0 | 0 | 0 | 2 | 0 | 0 | 222 | 149 | 1 | 1 | 1 | 1 | 1 | 1 | 1 | NA | 1 | 0 | 1 | NA | 0.333 |
| SE | 72678 | ENSG00000113387 | SUB1 | chr5 | + | 32588511 | 32588584 | 32585556 | 32585625 | 32591562 | 32591685 | 1019 | 471 | 730 | 734 | 611 | 1 | 1 | 0 | 0 | 0 | 618 | 898 | 905 | 654 | 384 | 0 | 2 | 2 | 0 | 0 | 222 | 149 | 1 | 1 | 0.999 | 0.997 | 1 | 1 | 1 | 1 | 0.997 | 0.997 | 1 | 1 | 0 |
| SE | 72679 | ENSG00000113387 | SUB1 | chr5 | + | 32591562 | 32591685 | 32571090 | 32571174 | 32598960 | 32599069 | 849 | 527 | 860 | 808 | 1007 | 0 | 0 | 0 | 0 | 0 | 651 | 1081 | 1029 | 676 | 435 | 0 | 0 | 2 | 0 | 0 | 272 | 149 | 1 | 1 | 1 | 1 | 1 | 1 | 1 | 1 | 1 | 0.996 | 1 | 1 | 0.001 |
| SE | 72680 | ENSG00000113387 | SUB1 | chr5 | + | 32591562 | 32591685 | 32588511 | 32588584 | 32598960 | 32599069 | 1889 | 1009 | 1610 | 1554 | 1646 | 0 | 0 | 3 | 0 | 0 | 1271 | 1995 | 1953 | 1350 | 823 | 0 | 0 | 5 | 0 | 0 | 272 | 149 | 1 | 1 | 1 | 1 | 0.997 | 1 | 1 | 1 | 1 | 0.995 | 1 | 1 | 0 |
| SE | 72681 | ENSG00000113387 | SUB1 | chr5 | + | 32594570 | 32594872 | 32591562 | 32591685 | 32598960 | 32599069 | 2 | 0 | 2 | 3 | 0 | 1121 | 652 | 1049 | 977 | 1144 | 4 | 2 | 12 | 6 | 0 | 815 | 1247 | 1258 | 855 | 524 | 298 | 149 | 1 | 1 | 0.001 | 0 | 0.001 | 0.002 | 0 | 0.002 | 0.001 | 0.005 | 0.003 | 0 | -0.001 |
| SE | 72685 | ENSG00000114784 | EIF1B | chr3 | + | 40310892 | 40311056 | 40309706 | 40309972 | 40311469 | 40311571 | 1146 | 929 | 975 | 1217 | 1076 | 4 | 3 | 1 | 3 | 4 | 2727 | 1950 | 1770 | 2745 | 3615 | 8 | 0 | 8 | 13 | 0 | 298 | 149 | 1 | 1 | 0.993 | 0.994 | 0.998 | 0.995 | 0.993 | 0.994 | 1 | 0.991 | 0.991 | 1 | -0.001 |
| SE | 72686 | ENSG00000104047 | DTWD1 | chr15 | + | 49623670 | 49623798 | 49621068 | 49621122 | 49625112 | 49625431 | 0 | 0 | 0 | 0 | 5 | 7 | 1 | 8 | 7 | 6 | 0 | 0 | 0 | 0 | 0 | 0 | 2 | 2 | 1 | 0 | 277 | 149 | 1 | 1 | 0 | 0 | 0 | 0 | 0.31 | NA | 0 | 0 | 0 | NA | 0.062 |
| SE | 72687 | ENSG00000104047 | DTWD1 | chr15 | + | 49625112 | 49625431 | 49621036 | 49621122 | 49634535 | 49634595 | 8 | 2 | 9 | 7 | 6 | 0 | 0 | 0 | 0 | 0 | 0 | 2 | 2 | 1 | 0 | 0 | 2 | 0 | 0 | 0 | 298 | 149 | 1 | 1 | 1 | 1 | 1 | 1 | 1 | NA | 0.333 | 1 | 1 | NA | 0.222 |
| SE | 72688 | ENSG00000104047 | DTWD1 | chr15 | + | 49626743 | 49626893 | 49625112 | 49625431 | 49629586 | 49629967 | 4 | 2 | 3 | 7 | 7 | 0 | 0 | 0 | 0 | 0 | 14 | 1 | 2 | 0 | 0 | 0 | 0 | 3 | 0 | 0 | 298 | 149 | 1 | 1 | 1 | 1 | 1 | 1 | 1 | 1 | 1 | 0.25 | NA | NA | 0.25 |
| SE | 72689 | ENSG00000104047 | DTWD1 | chr15 | + | 49626743 | 49626893 | 49625112 | 49625431 | 49630823 | 49631060 | 4 | 2 | 3 | 6 | 7 | 2 | 1 | 0 | 0 | 0 | 14 | 1 | 5 | 0 | 0 | 0 | 5 | 0 | 0 | 0 | 298 | 149 | 1 | 1 | 0.5 | 0.5 | 1 | 1 | 1 | 1 | 0.091 | 1 | NA | NA | 0.103 |
| SE | 72692 | ENSG00000104047 | DTWD1 | chr15 | + | 49626743 | 49626893 | 49625112 | 49625431 | 49632170 | 49632302 | 4 | 2 | 3 | 6 | 7 | 0 | 5 | 4 | 0 | 1 | 28 | 1 | 2 | 0 | 0 | 0 | 0 | 0 | 0 | 0 | 298 | 149 | 1 | 1 | 1 | 0.167 | 0.273 | 1 | 0.778 | 1 | 1 | 1 | NA | NA | -0.356 |
| SE | 72695 | ENSG00000104047 | DTWD1 | chr15 | + | 49631000 | 49631058 | 49626743 | 49626893 | 49632158 | 49632302 | 6 | 11 | 17 | 12 | 25 | 1 | 0 | 0 | 0 | 0 | 28 | 8 | 1 | 10 | 2 | 0 | 0 | 0 | 0 | 0 | 207 | 149 | 1 | 1 | 0.812 | 1 | 1 | 1 | 1 | 1 | 1 | 1 | 1 | 1 | -0.038 |
| SE | 72696 | ENSG00000104047 | DTWD1 | chr15 | + | 49632158 | 49632302 | 49625112 | 49625431 | 49634535 | 49634730 | 112 | 115 | 141 | 147 | 243 | 1 | 1 | 1 | 0 | 0 | 31 | 66 | 52 | 54 | 42 | 0 | 0 | 0 | 0 | 0 | 293 | 149 | 1 | 1 | 0.983 | 0.983 | 0.986 | 1 | 1 | 1 | 1 | 1 | 1 | 1 | -0.01 |
| SE | 72697 | ENSG00000104047 | DTWD1 | chr15 | + | 49632170 | 49632302 | 49625112 | 49625431 | 49634535 | 49634595 | 59 | 57 | 93 | 76 | 141 | 1 | 1 | 1 | 0 | 0 | 23 | 28 | 35 | 35 | 16 | 0 | 0 | 0 | 0 | 0 | 281 | 149 | 1 | 1 | 0.969 | 0.968 | 0.98 | 1 | 1 | 1 | 1 | 1 | 1 | 1 | -0.017 |
| SE | 72701 | ENSG00000172765 | TMCC1 | chr3 | - | 129827802 | 129828077 | 129670329 | 129671264 | 129832773 | 129832826 | 51 | 32 | 55 | 60 | 39 | 0 | 0 | 2 | 0 | 0 | 46 | 54 | 28 | 35 | 8 | 0 | 4 | 0 | 0 | 0 | 298 | 149 | 1 | 1 | 1 | 1 | 0.932 | 1 | 1 | 1 | 0.871 | 1 | 1 | 1 | 0.012 |
| SE | 72702 | ENSG00000172765 | TMCC1 | chr3 | - | 129827802 | 129828508 | 129670329 | 129671264 | 129832773 | 129832826 | 92 | 73 | 109 | 112 | 100 | 0 | 0 | 2 | 0 | 0 | 79 | 93 | 75 | 82 | 26 | 0 | 4 | 0 | 0 | 0 | 298 | 149 | 1 | 1 | 1 | 1 | 0.965 | 1 | 1 | 1 | 0.921 | 1 | 1 | 1 | 0.009 |
| SE | 72703 | ENSG00000172765 | TMCC1 | chr3 | - | 129832773 | 129832826 | 129827755 | 129828077 | 129838897 | 129839060 | 6 | 3 | 6 | 5 | 17 | 0 | 0 | 0 | 1 | 0 | 4 | 1 | 3 | 0 | 0 | 0 | 0 | 0 | 0 | 0 | 202 | 149 | 1 | 1 | 1 | 1 | 1 | 0.787 | 1 | 1 | 1 | 1 | NA | NA | -0.043 |
| SE | 72705 | ENSG00000172765 | TMCC1 | chr3 | - | 129838897 | 129839060 | 129832773 | 129832826 | 129857249 | 129857405 | 7 | 8 | 8 | 15 | 23 | 1 | 0 | 0 | 0 | 0 | 8 | 11 | 13 | 0 | 9 | 0 | 0 | 0 | 0 | 1 | 298 | 149 | 1 | 1 | 0.778 | 1 | 1 | 1 | 1 | 1 | 1 | 1 | NA | 0.818 | 0.001 |
| SE | 72706 | ENSG00000172765 | TMCC1 | chr3 | - | 129838897 | 129839060 | 129832773 | 129832826 | 129860939 | 129861012 | 7 | 8 | 8 | 15 | 26 | 1 | 0 | 0 | 0 | 1 | 8 | 11 | 13 | 0 | 9 | 0 | 0 | 9 | 0 | 0 | 298 | 149 | 1 | 1 | 0.778 | 1 | 1 | 1 | 0.929 | 1 | 1 | 0.419 | NA | 1 | 0.087 |
| SE | 72708 | ENSG00000172765 | TMCC1 | chr3 | - | 129857249 | 129857405 | 129832773 | 129832826 | 129860939 | 129861012 | 3 | 4 | 1 | 6 | 9 | 1 | 0 | 0 | 0 | 1 | 0 | 3 | 7 | 14 | 1 | 0 | 0 | 9 | 0 | 0 | 298 | 149 | 1 | 1 | 0.6 | 1 | 1 | 1 | 0.818 | NA | 1 | 0.28 | 1 | 1 | 0.064 |
| SE | 72709 | ENSG00000172765 | TMCC1 | chr3 | - | 129857249 | 129857405 | 129832773 | 129832826 | 129867009 | 129867098 | 1 | 0 | 0 | 0 | 0 | 0 | 0 | 1 | 2 | 0 | 0 | 3 | 0 | 0 | 1 | 0 | 0 | 0 | 8 | 0 | 298 | 149 | 1 | 1 | 1 | NA | 0 | 0 | NA | NA | 1 | NA | 0 | 1 | -0.333 |
| SE | 72711 | ENSG00000172765 | TMCC1 | chr3 | - | 129857249 | 129857405 | 129838897 | 129839060 | 129860939 | 129861012 | 2 | 5 | 1 | 6 | 9 | 0 | 1 | 0 | 0 | 3 | 0 | 3 | 7 | 14 | 0 | 0 | 0 | 0 | 0 | 0 | 298 | 149 | 1 | 1 | 1 | 0.714 | 1 | 1 | 0.6 | NA | 1 | 1 | 1 | NA | -0.137 |
| SE | 72712 | ENSG00000172765 | TMCC1 | chr3 | - | 129857249 | 129857405 | 129838897 | 129839060 | 129880308 | 129880559 | 8 | 8 | 5 | 14 | 9 | 1 | 0 | 1 | 2 | 5 | 0 | 2 | 3 | 6 | 0 | 0 | 2 | 3 | 1 | 0 | 298 | 149 | 1 | 1 | 0.8 | 1 | 0.714 | 0.778 | 0.474 | NA | 0.333 | 0.333 | 0.75 | NA | 0.281 |
| SE | 72713 | ENSG00000172765 | TMCC1 | chr3 | - | 129857249 | 129857405 | 129848342 | 129848412 | 129860939 | 129861012 | 20 | 12 | 6 | 22 | 26 | 14 | 20 | 13 | 17 | 34 | 7 | 6 | 26 | 30 | 0 | 36 | 13 | 65 | 34 | 0 | 298 | 149 | 1 | 1 | 0.417 | 0.231 | 0.188 | 0.393 | 0.277 | 0.089 | 0.188 | 0.167 | 0.306 | NA | 0.114 |
| SE | 72714 | ENSG00000172765 | TMCC1 | chr3 | - | 129857249 | 129857405 | 129848342 | 129848412 | 129867009 | 129867098 | 18 | 8 | 5 | 16 | 17 | 0 | 0 | 2 | 0 | 3 | 7 | 6 | 19 | 16 | 0 | 0 | 0 | 0 | 0 | 0 | 298 | 149 | 1 | 1 | 1 | 1 | 0.556 | 1 | 0.739 | 1 | 1 | 1 | 1 | NA | -0.141 |
| SE | 72716 | ENSG00000172765 | TMCC1 | chr3 | - | 129860939 | 129861012 | 129832773 | 129832826 | 129867009 | 129867100 | 0 | 0 | 0 | 0 | 1 | 0 | 0 | 1 | 2 | 0 | 0 | 0 | 4 | 7 | 0 | 0 | 0 | 0 | 8 | 0 | 222 | 149 | 1 | 1 | NA | NA | 0 | 0 | 1 | NA | NA | 1 | 0.37 | NA | -0.352 |
| SE | 72718 | ENSG00000172765 | TMCC1 | chr3 | - | 129860939 | 129861012 | 129838897 | 129839060 | 129880308 | 129880559 | 6 | 10 | 7 | 6 | 8 | 1 | 0 | 1 | 2 | 5 | 0 | 6 | 19 | 2 | 0 | 0 | 2 | 3 | 1 | 0 | 222 | 149 | 1 | 1 | 0.801 | 1 | 0.825 | 0.668 | 0.518 | NA | 0.668 | 0.81 | 0.573 | NA | 0.079 |
| SE | 72719 | ENSG00000172765 | TMCC1 | chr3 | - | 129860939 | 129861012 | 129848345 | 129848412 | 129867009 | 129867100 | 6 | 4 | 8 | 6 | 0 | 0 | 0 | 2 | 0 | 3 | 25 | 3 | 30 | 24 | 0 | 0 | 0 | 0 | 0 | 0 | 222 | 149 | 1 | 1 | 1 | 1 | 0.729 | 1 | 0 | 1 | 1 | 1 | 1 | NA | -0.254 |
| SE | 72721 | ENSG00000172765 | TMCC1 | chr3 | - | 129860939 | 129861012 | 129857249 | 129857405 | 129867009 | 129867100 | 2 | 4 | 0 | 2 | 3 | 0 | 0 | 0 | 0 | 0 | 0 | 3 | 6 | 18 | 0 | 0 | 3 | 0 | 0 | 0 | 222 | 149 | 1 | 1 | 1 | 1 | NA | 1 | 1 | NA | 0.402 | 1 | 1 | NA | 0.199 |
| SE | 72722 | ENSG00000172765 | TMCC1 | chr3 | - | 129860939 | 129861012 | 129857249 | 129857405 | 129880308 | 129880559 | 8 | 13 | 8 | 12 | 12 | 8 | 7 | 5 | 14 | 9 | 0 | 9 | 26 | 16 | 0 | 0 | 2 | 3 | 6 | 0 | 222 | 149 | 1 | 1 | 0.402 | 0.555 | 0.518 | 0.365 | 0.472 | NA | 0.751 | 0.853 | 0.642 | NA | -0.286 |
| SE | 72725 | ENSG00000172765 | TMCC1 | chr3 | - | 129867009 | 129867098 | 129857249 | 129857405 | 129880308 | 129880559 | 6 | 2 | 2 | 1 | 4 | 8 | 7 | 5 | 14 | 9 | 0 | 3 | 5 | 0 | 0 | 0 | 2 | 3 | 6 | 0 | 238 | 149 | 1 | 1 | 0.32 | 0.152 | 0.2 | 0.043 | 0.218 | NA | 0.484 | 0.511 | 0 | NA | -0.145 |
| SE | 72727 | ENSG00000172765 | TMCC1 | chr3 | - | 129880308 | 129880559 | 129848345 | 129848412 | 129893181 | 129893321 | 50 | 26 | 49 | 44 | 106 | 0 | 0 | 0 | 0 | 1 | 38 | 51 | 119 | 45 | 13 | 0 | 0 | 0 | 0 | 0 | 298 | 149 | 1 | 1 | 1 | 1 | 1 | 1 | 0.981 | 1 | 1 | 1 | 1 | 1 | -0.004 |
| SE | 72729 | ENSG00000172765 | TMCC1 | chr3 | - | 129888981 | 129889104 | 129880308 | 129880559 | 129893493 | 129893606 | 2 | 0 | 0 | 0 | 0 | 19 | 8 | 10 | 18 | 23 | 0 | 0 | 0 | 0 | 0 | 26 | 19 | 23 | 3 | 0 | 272 | 149 | 1 | 1 | 0.055 | 0 | 0 | 0 | 0 | 0 | 0 | 0 | 0 | NA | 0.011 |

| | | | | | | | | | | | | | | |
|---|---|---|---|---|---|---|---|---|---|---|---|---|---|---|
| SE | 72732 | ENSG00000112394 | SLC16A10 | chr6 | + | 111177211 | 111177665 | 111172694 | 111172839 | 111193292 | 111193324 | 5 | 10 | [illegible] |
| SE | 72733 | ENSG00000112394 | SLC16A10 | chr6 | + | 111177211 | 111177665 | 111172694 | 111172839 | 111206591 | 111206735 | 12 | 15 | [illegible] |
| SE | 72736 | ENSG00000146083 | RNF44 | chr5 | - | 176530847 | 176531021 | 176530581 | 176530743 | 176531462 | 176531630 | 26 | 15 | [illegible] |
| SE | 72742 | ENSG00000106565 | TMEM176B | chr7 | - | 150793087 | 150793315 | 150792055 | 150792175 | 150793543 | 150793600 | 1173 | 411 | [illegible] |
| SE | 72746 | ENSG00000240498 | CDKN2B-AS1 | chr9 | + | 22029432 | 22029594 | 21994790 | 21995161 | 22046750 | 22046900 | 1 | 0 | [illegible] |
| SE | 72752 | ENSG00000240498 | CDKN2B-AS1 | chr9 | + | 22056251 | 22056387 | 22049105 | 22049228 | 22092307 | 22092469 | 1 | 6 | [illegible] |
| SE | 72753 | ENSG00000240498 | CDKN2B-AS1 | chr9 | + | 22056251 | 22056387 | 22049105 | 22049228 | 22097257 | 22097364 | 1 | 7 | [illegible] |
| SE | 72757 | ENSG00000240498 | CDKN2B-AS1 | chr9 | + | 22061952 | 22062026 | 22056251 | 22056387 | 22063943 | 22064018 | 1 | 1 | [illegible] |
| SE | 72761 | ENSG00000240498 | CDKN2B-AS1 | chr9 | + | 22063943 | 22064018 | 22056251 | 22056387 | 22096371 | 22096514 | 1 | 0 | [illegible] |
| SE | 72766 | ENSG00000240498 | CDKN2B-AS1 | chr9 | + | 22092307 | 22092469 | 22049105 | 22049228 | 22096371 | 22096514 | 0 | 0 | [illegible] |
| SE | 72767 | ENSG00000240498 | CDKN2B-AS1 | chr9 | + | 22092307 | 22092469 | 22056251 | 22056387 | 22096371 | 22096514 | 0 | 0 | [illegible] |
| SE | 72768 | ENSG00000240498 | CDKN2B-AS1 | chr9 | + | 22096371 | 22096514 | 22049105 | 22049228 | 22097257 | 22097364 | 1 | 1 | [illegible] |
| SE | 72769 | ENSG00000240498 | CDKN2B-AS1 | chr9 | + | 22096371 | 22096514 | 22056326 | 22056387 | 22097257 | 22097364 | 2 | 1 | [illegible] |
| SE | 72778 | ENSG00000113407 | TARS1 | chr5 | + | 33445323 | 33445404 | 33440804 | 33441143 | 33448540 | 33448730 | 142 | 67 | [illegible] |
| SE | 72779 | ENSG00000113407 | TARS1 | chr5 | + | 33446606 | 33446764 | 33445323 | 33445404 | 33448540 | 33448731 | 4 | 0 | [illegible] |
| SE | 72780 | ENSG00000113407 | TARS1 | chr5 | + | 33446643 | 33446764 | 33445323 | 33445404 | 33448540 | 33448731 | 4 | 0 | [illegible] |
| SE | 72781 | ENSG00000113407 | TARS1 | chr5 | + | 33448540 | 33448731 | 33445323 | 33445404 | 33453288 | 33453346 | 192 | 131 | [illegible] |
| SE | 72782 | ENSG00000113407 | TARS1 | chr5 | + | 33448540 | 33448731 | 33445323 | 33445404 | 33453293 | 33453412 | 85 | 52 | [illegible] |
| SE | 72784 | ENSG00000113407 | TARS1 | chr5 | + | 33453288 | 33453412 | 33448540 | 33448731 | 33454944 | 33455066 | 226 | 158 | [illegible] |
| SE | 72785 | ENSG00000113407 | TARS1 | chr5 | + | 33453293 | 33453412 | 33448540 | 33448731 | 33454944 | 33455012 | 112 | 79 | [illegible] |
| [illegible] | | | | | | | | | | | | | | |

| | | | | | | | | | | | | |
|---|---|---|---|---|---|---|---|---|---|---|---|---|
| SE | 73152 | ENSG00000144233 | AMMECR1L | chr2 | - | 127871248 | 127871359 | 127866899 | 127866996 | 127873827 | 127874272 | 6 |
| SE | 73153 | ENSG00000144233 | AMMECR1L | chr2 | - | 127871248 | 127871359 | 127869453 | 127869544 | 127873827 | 127874272 | 13 |
| SE | 73154 | ENSG00000144233 | AMMECR1L | chr2 | - | 127871248 | 127871359 | 127869453 | 127869544 | 127884202 | 127884312 | 5 |
| SE | 73157 | ENSG00000144233 | AMMECR1L | chr2 | - | 127873827 | 127874272 | 127869453 | 127869544 | 127884202 | 127884312 | 17 |
| SE | 73158 | ENSG00000144233 | AMMECR1L | chr2 | - | 127873827 | 127874272 | 127871248 | 127871359 | 127884202 | 127884312 | 33 |
| SE | 73165 | ENSG00000145391 | SETD7 | chr4 | - | 139533164 | 139533366 | 139529030 | 139529220 | 139546919 | 139547049 | 97 |
| SE | 73167 | ENSG00000145391 | SETD7 | chr4 | - | 139548018 | 139548119 | 139546919 | 139547049 | 139556097 | 139556208 | 0 |
| SE | 73178 | ENSG00000183091 | NEB | chr2 | - | 151494160 | 151494253 | 151492386 | 151492494 | 151505477 | 151505570 | 3 |
| SE | 73184 | ENSG00000183091 | NEB | chr2 | - | 151503348 | 151503441 | 151492386 | 151492494 | 151505477 | 151505570 | 2 |
| SE | 73185 | ENSG00000183091 | NEB | chr2 | - | 151503348 | 151503441 | 151494160 | 151494253 | 151505477 | 151505570 | 0 |
| SE | 73188 | ENSG00000183091 | NEB | chr2 | - | 151514817 | 151514928 | 151514317 | 151514428 | 151516458 | 151516563 | 1 |
| SE | 73191 | ENSG00000088325 | TPX2 | chr20 | + | 31742540 | 31742647 | 31739270 | 31739621 | 31757406 | 31757582 | 10 |
| SE | 73200 | ENSG00000115307 | AUP1 | chr2 | - | 74526940 | 74527363 | 74526745 | 74526836 | 74527735 | 74528085 | 166 |
| SE | 73201 | ENSG00000115307 | AUP1 | chr2 | - | 74527470 | 74527590 | 74526940 | 74527363 | 74527735 | 74527838 | 323 |
| SE | 73202 | ENSG00000115307 | AUP1 | chr2 | - | 74528247 | 74528321 | 74527939 | 74528006 | 74528416 | 74528489 | 407 |
| SE | 73203 | ENSG00000115307 | AUP1 | chr2 | - | 74528416 | 74528489 | 74527939 | 74528006 | 74528750 | 74528935 | 63 |
| SE | 73204 | ENSG00000115307 | AUP1 | chr2 | - | 74528416 | 74528489 | 74527939 | 74528006 | 74529131 | 74529282 | 2 |
| SE | 73205 | ENSG00000115307 | AUP1 | chr2 | - | 74528750 | 74528935 | 74527939 | 74528006 | 74529131 | 74529282 | 26 |
| SE | 73207 | ENSG00000115307 | AUP1 | chr2 | - | 74528750 | 74529282 | 74528416 | 74528489 | 74529361 | 74529499 | 242 |
| SE | 73208 | ENSG00000115307 | AUP1 | chr2 | - | 74529131 | 74529282 | 74528416 | 74528489 | 74529361 | 74529499 | 67 |
| SE | 73212 | ENSG00000143499 | SMYD2 | chr1 | + | 214314761 | 214314872 | 214305186 | 214305250 | 214318078 | 214318139 | 63 |
| SE | 73213 | ENSG00000143499 | SMYD2 | chr1 | + | 214324640 | 214324708 | 214318858 | 214318983 | 214327621 | 214327724 | 147 |
| SE | 73214 | ENSG00000143499 | SMYD2 | chr1 | + | 214330167 | 214330278 | 214327493 | 214327724 | 214334199 | 214334308 | 66 |
| SE | 73215 | ENSG00000143499 | SMYD2 | chr1 | + | 214330949 | 214334308 | 214330167 | 214330278 | 214336703 | 214337130 | 107 |
| SE | 73216 | ENSG00000143499 | SMYD2 | chr1 | + | 214334199 | 214334308 | 214330167 | 214330278 | 214336703 | 214337131 | 55 |
| SE | 73229 | ENSG00000170190 | SLC16A5 | chr17 | + | 75093588 | 75093835 | 75088937 | 75089304 | 75098037 | 75098181 | 10 |
| SE | 73249 | ENSG00000114956 | DGUOK | chr2 | + | 73938909 | 73939022 | 73932597 | 73932683 | 73958145 | 73958245 | 3 |
| SE | 73254 | ENSG00000114956 | DGUOK | chr2 | + | 73946718 | 73946906 | 73938909 | 73939022 | 73957124 | 73957240 | 56 |
| SE | 73255 | ENSG00000114956 | DGUOK | chr2 | + | 73946718 | 73946906 | 73938909 | 73939022 | 73958145 | 73958245 | 128 |
| SE | 73257 | ENSG00000114956 | DGUOK | chr2 | + | 73950584 | 73950732 | 73946718 | 73946906 | 73957124 | 73957240 | 98 |
| SE | 73261 | ENSG00000114956 | DGUOK | chr2 | + | 73957124 | 73957240 | 73938909 | 73939022 | 73958145 | 73958245 | 81 |
| SE | 73269 | ENSG00000168280 | KIF5C | chr2 | + | 148981354 | 148981561 | 148978921 | 148978990 | 148983619 | 148983766 | 8 |
| SE | 73270 | ENSG00000168280 | KIF5C | chr2 | + | 148994420 | 148994538 | 148984722 | 148991198 | 148997263 | 148997340 | 32 |
| SE | 73273 | ENSG00000168280 | KIF5C | chr2 | + | 149008471 | 149008661 | 149007962 | 149008067 | 149010134 | 149010351 | 0 |
| SE | 73274 | ENSG00000168280 | KIF5C | chr2 | + | 149020175 | 149020382 | 149011569 | 149011683 | 149023077 | 149023510 | 0 |
| SE | 73279 | ENSG00000153037 | SRP19 | chr5 | + | 112862507 | 112862583 | 112861187 | 112861417 | 112864456 | 112864528 | 212 |
| SE | 73280 | ENSG00000153037 | SRP19 | chr5 | + | 112864433 | 112864528 | 112862507 | 112862583 | 112891602 | 112898371 | 0 |
| SE | 73282 | ENSG00000153037 | SRP19 | chr5 | + | 112864620 | 112864732 | 112862507 | 112862583 | 112891602 | 112891725 | 1 |
| SE | 73283 | ENSG00000153037 | SRP19 | chr5 | + | 112864620 | 112864732 | 112864456 | 112864528 | 112867371 | 112867935 | 416 |
| SE | 73285 | ENSG00000153037 | SRP19 | chr5 | + | 112864620 | 112864732 | 112864456 | 112864528 | 112891602 | 112891725 | 281 |
| SE | 73286 | ENSG00000153037 | SRP19 | chr5 | + | 112881956 | 112882049 | 112864456 | 112864528 | 112891602 | 112891637 | 0 |
| SE | 73287 | ENSG00000153037 | SRP19 | chr5 | + | 112881956 | 112882049 | 112864620 | 112864732 | 112891602 | 112891637 | 0 |
| SE | 73288 | ENSG00000011566 | MAP4K3 | chr2 | - | 39254449 | 39254520 | 39251829 | 39251885 | 39258347 | 39258440 | 29 |
| SE | 73289 | ENSG00000011566 | MAP4K3 | chr2 | - | 39272481 | 39272542 | 39271910 | 39272400 | 39278406 | 39278419 | 16 |
| SE | 73290 | ENSG00000011566 | MAP4K3 | chr2 | - | 39299742 | 39299801 | 39293229 | 39293268 | 39309460 | 39309519 | 31 |
| SE | 73292 | ENSG00000011566 | MAP4K3 | chr2 | - | 39303073 | 39303117 | 39299742 | 39299801 | 39309460 | 39309519 | 6 |
| SE | 73293 | ENSG00000011566 | MAP4K3 | chr2 | - | 39307942 | 39308005 | 39299742 | 39299801 | 39309460 | 39309519 | 22 |
| SE | 73294 | ENSG00000011566 | MAP4K3 | chr2 | - | 39307942 | 39308005 | 39303073 | 39303117 | 39309460 | 39309519 | 5 |
| SE | 73295 | ENSG00000011566 | MAP4K3 | chr2 | - | 39378065 | 39378123 | 39356164 | 39356339 | 39436891 | 39437065 | 32 |
| SE | 73299 | ENSG00000136699 | SMPD4 | chr2 | - | 130157250 | 130157396 | 130156584 | 130156675 | 130164373 | 130164445 | 62 |
| SE | 73300 | ENSG00000136699 | SMPD4 | chr2 | - | 130158183 | 130158265 | 130157250 | 130157396 | 130161185 | 130161272 | 0 |
| SE | 73302 | ENSG00000136699 | SMPD4 | chr2 | - | 130159512 | 130159634 | 130157250 | 130157396 | 130161185 | 130161272 | 2 |
| SE | 73305 | ENSG00000136699 | SMPD4 | chr2 | - | 130161185 | 130161272 | 130158183 | 130158265 | 130164373 | 130164445 | 9 |
| SE | 73306 | ENSG00000136699 | SMPD4 | chr2 | - | 130167457 | 130167590 | 130164373 | 130164445 | 130172348 | 130172455 | 110 |
| SE | 73307 | ENSG00000136699 | SMPD4 | chr2 | - | 130167457 | 130167590 | 130164373 | 130164445 | 130174913 | 130175000 | 55 |
| SE | 73314 | ENSG00000136699 | SMPD4 | chr2 | - | 130173513 | 130173656 | 130172786 | 130172895 | 130176553 | 130176637 | 0 |
| SE | 73319 | ENSG00000115970 | THADA | chr2 | - | 43232712 | 43232882 | 43230850 | 43231343 | 43279764 | 43279896 | 40 |
| SE | 73320 | ENSG00000115970 | THADA | chr2 | - | 43279764 | 43279896 | 43232712 | 43232882 | 43286907 | 43287061 | 32 |
| SE | 73321 | ENSG00000115970 | THADA | chr2 | - | 43286907 | 43287061 | 43279764 | 43279896 | 43291695 | 43291768 | 84 |
| SE | 73322 | ENSG00000115970 | THADA | chr2 | - | 43291695 | 43292222 | 43286984 | 43287061 | 43292833 | 43293066 | 87 |
| SE | 73325 | ENSG00000115970 | THADA | chr2 | - | 43370119 | 43370197 | 43344121 | 43344237 | 43371964 | 43372122 | 0 |
| SE | 73326 | ENSG00000115970 | THADA | chr2 | - | 43370119 | 43370197 | 43344121 | 43344237 | 43397970 | 43398058 | 0 |
| SE | 73327 | ENSG00000115970 | THADA | chr2 | - | 43371964 | 43372122 | 43370119 | 43370197 | 43397970 | 43398058 | 0 |
| SE | 73328 | ENSG00000115970 | THADA | chr2 | - | 43397970 | 43398139 | 43343000 | 43344237 | 43428099 | 43428231 | 101 |
| SE | 73332 | ENSG00000115970 | THADA | chr2 | - | 43485233 | 43485325 | 43430212 | 43430302 | 43498832 | 43498955 | 71 |
| SE | 73333 | ENSG00000115970 | THADA | chr2 | - | 43527878 | 43527988 | 43508647 | 43508780 | 43541158 | 43541316 | 128 |
| SE | 73335 | ENSG00000115970 | THADA | chr2 | - | 43549209 | 43549368 | 43541158 | 43541316 | 43551788 | 43551922 | 157 |
| SE | 73340 | ENSG00000115970 | THADA | chr2 | - | 43571706 | 43571862 | 43570387 | 43570510 | 43572813 | 43572965 | 264 |
| SE | 73344 | ENSG00000115970 | THADA | chr2 | - | 43581740 | 43581928 | 43578512 | 43578607 | 43586400 | 43586449 | 116 |
| SE | 73345 | ENSG00000115970 | THADA | chr2 | - | 43592843 | 43592890 | 43592316 | 43592416 | 43595646 | 43595974 | 0 |
| SE | 73347 | ENSG00000115604 | IL18R1 | chr2 | + | 102367824 | 102368068 | 102362632 | 102362718 | 102371952 | 102372118 | 72 |
| SE | 73348 | ENSG00000115604 | IL18R1 | chr2 | + | 102367824 | 102368068 | 102362632 | 102362718 | 102375906 | 102376063 | 14 |
| SE | 73349 | ENSG00000115604 | IL18R1 | chr2 | + | 102371952 | 102372118 | 102362632 | 102362718 | 102375906 | 102376063 | 84 |
| SE | 73350 | ENSG00000115604 | IL18R1 | chr2 | + | 102371952 | 102372118 | 102367824 | 102368068 | 102375906 | 102376063 | 136 |
| SE | 73352 | ENSG00000115604 | IL18R1 | chr2 | + | 102373936 | 102373993 | 102371952 | 102372118 | 102396530 | 102398777 | 25 |
| SE | 73353 | ENSG00000115604 | IL18R1 | chr2 | + | 102375906 | 102376063 | 102362632 | 102362718 | 102381619 | 102381682 | 74 |
| SE | 73354 | ENSG00000115604 | IL18R1 | chr2 | + | 102375906 | 102376063 | 102373936 | 102373993 | 102381619 | 102381682 | 104 |
| SE | 73355 | ENSG00000115604 | IL18R1 | chr2 | + | 102381619 | 102381682 | 102375906 | 102376063 | 102384877 | 102384998 | 79 |
| SE | 73356 | ENSG00000115604 | IL18R1 | chr2 | + | 102381619 | 102381682 | 102375906 | 102376063 | 102386860 | 102387000 | 21 |
| SE | 73357 | ENSG00000115604 | IL18R1 | chr2 | + | 102384877 | 102384998 | 102375906 | 102376063 | 102386860 | 102387000 | 40 |
| SE | 73358 | ENSG00000115604 | IL18R1 | chr2 | + | 102384877 | 102384998 | 102381619 | 102381682 | 102386860 | 102387000 | 97 |
| SE | 73359 | ENSG00000115604 | IL18R1 | chr2 | + | 102386860 | 102387000 | 102384877 | 102384998 | 102396530 | 102396886 | 36 |
| SE | 73360 | ENSG00000115604 | IL18R1 | chr2 | + | 102390055 | 102390217 | 102386860 | 102387000 | 102394468 | 102394627 | 105 |
| SE | 73361 | ENSG00000115604 | IL18R1 | chr2 | + | 102390055 | 102390217 | 102386860 | 102387000 | 102396530 | 102396886 | 37 |
| SE | 73362 | ENSG00000115604 | IL18R1 | chr2 | + | 102394468 | 102394627 | 102386860 | 102387000 | 102396530 | 102396886 | 27 |
| SE | 73363 | ENSG00000115604 | IL18R1 | chr2 | + | 102394468 | 102394627 | 102390055 | 102390217 | 102396530 | 102396886 | 100 |
| SE | 73365 | ENSG00000145736 | GTF2H2 | chr5 | - | 71048005 | 71048113 | 71045443 | 71045507 | 71048761 | 71048849 | 70 |
| SE | 73367 | ENSG00000145736 | GTF2H2 | chr5 | - | 71059689 | 71059744 | 71055029 | 71055457 | 71060843 | 71060930 | 39 |
| SE | 73368 | ENSG00000145736 | GTF2H2 | chr5 | - | 71060119 | 71060170 | 71059689 | 71059744 | 71060843 | 71060930 | 14 |
| SE | 73369 | ENSG00000145736 | GTF2H2 | chr5 | - | 71061271 | 71061308 | 71060506 | 71060930 | 71061687 | 71061765 | 8 |
| SE | 73377 | ENSG00000092208 | GEMIN2 | chr14 | + | 39114828 | 39114913 | 39114284 | 39114475 | 39117998 | 39118088 | 55 |
| SE | 73379 | ENSG00000092208 | GEMIN2 | chr14 | + | 39124991 | 39125036 | 39122429 | 39122543 | 39128279 | 39128341 | 28 |
| SE | 73380 | ENSG00000092208 | GEMIN2 | chr14 | + | 39124991 | 39125036 | 39122429 | 39122543 | 39131957 | 39132068 | 6 |
| SE | 73381 | ENSG00000092208 | GEMIN2 | chr14 | + | 39128279 | 39128348 | 39122429 | 39122543 | 39131957 | 39132023 | 17 |
| SE | 73382 | ENSG00000092208 | GEMIN2 | chr14 | + | 39128279 | 39128348 | 39124991 | 39125036 | 39131957 | 39132023 | 31 |
| SE | 73383 | ENSG00000092208 | GEMIN2 | chr14 | + | 39133660 | 39133719 | 39131772 | 39132068 | 39135930 | 39136155 | 14 |
| SE | 73384 | ENSG00000092208 | GEMIN2 | chr14 | + | 39133660 | 39133719 | 39131772 | 39132068 | 39136439 | 39136541 | 14 |
| SE | 73385 | ENSG00000092208 | GEMIN2 | chr14 | + | 39135930 | 39136155 | 39131957 | 39132068 | 39136439 | 39136624 | 1 |
| SE | 73389 | ENSG00000240344 | PPIL3 | chr2 | - | 200877496 | 200877604 | 200876941 | 200877037 | 200881420 | 200881488 | 4 |
| SE | 73390 | ENSG00000240344 | PPIL3 | chr2 | - | 200882341 | 200882382 | 200881420 | 200881488 | 200885261 | 200885367 | 16 |
| SE | 73391 | ENSG00000240344 | PPIL3 | chr2 | - | 200882341 | 200882382 | 200881420 | 200881488 | 200885697 | 200885772 | 16 |
| SE | 73392 | ENSG00000240344 | PPIL3 | chr2 | - | 200882341 | 200882435 | 200881420 | 200881488 | 200885261 | 200885367 | 40 |
| SE | 73394 | ENSG00000240344 | PPIL3 | chr2 | - | 200885261 | 200885343 | 200881420 | 200881488 | 200885697 | 200885772 | 6 |
| SE | 73395 | ENSG00000240344 | PPIL3 | chr2 | - | 200885261 | 200885367 | 200881420 | 200881488 | 200885697 | 200885772 | 19 |
| SE | 73396 | ENSG00000240344 | PPIL3 | chr2 | - | 200885261 | 200885367 | 200882341 | 200882382 | 200885697 | 200885772 | 16 |
| SE | 73398 | ENSG00000240344 | PPIL3 | chr2 | - | 200886913 | 200886982 | 200885697 | 200885772 | 200887612 | 200887685 | 1 |
| SE | 73400 | ENSG00000240344 | PPIL3 | chr2 | - | 200887612 | 200887685 | 200885318 | 200885772 | 200889243 | 200889266 | 164 |
| SE | 73401 | ENSG00000156510 | HKDC1 | chr10 | + | 69257035 | 69257131 | 69250532 | 69250652 | 69257326 | 69257426 | 9 |
| SE | 73402 | ENSG00000156510 | HKDC1 | chr10 | + | 69258775 | 69258959 | 69257326 | 69257426 | 69261138 | 69261294 | 16 |
| SE | 73403 | ENSG00000196290 | NIF3L1 | chr2 | + | 200890522 | 200890585 | 200889368 | 200889391 | 200891735 | 200891789 | 0 |
| SE | 73405 | ENSG00000196290 | NIF3L1 | chr2 | + | 200890522 | 200890585 | 200889368 | 200889391 | 200891968 | 200892379 | 0 |
| SE | 73408 | ENSG00000196290 | NIF3L1 | chr2 | + | 200890522 | 200890585 | 200889368 | 200889652 | 200891968 | 200892379 | 0 |
| SE | 73409 | ENSG00000196290 | NIF3L1 | chr2 | + | 200891735 | 200891789 | 200889368 | 200889652 | 200891917 | 200892379 | 1 |
| SE | 73411 | ENSG00000196290 | NIF3L1 | chr2 | + | 200897075 | 200897214 | 200895263 | 200895390 | 200899384 | 200899468 | 117 |
| SE | 73413 | ENSG00000115942 | ORC2 | chr2 | - | 200925835 | 200925932 | 200920992 | 200921139 | 200926767 | 200926900 | 61 |
| SE | 73414 | ENSG00000115942 | ORC2 | chr2 | - | 200925835 | 200925932 | 200920992 | 200921139 | 200931338 | 200931448 | 36 |
| SE | 73415 | ENSG00000115942 | ORC2 | chr2 | - | 200926767 | 200926900 | 200920992 | 200921139 | 200931338 | 200931448 | 14 |
| SE | 73417 | ENSG00000115942 | ORC2 | chr2 | - | 200933875 | 200933974 | 200931338 | 200931448 | 200935698 | 200935892 | 85 |
| SE | 73418 | ENSG00000115942 | ORC2 | chr2 | - | 200935698 | 200935892 | 200933875 | 200933974 | 200937905 | 200937966 | 156 |
| SE | 73419 | ENSG00000115942 | ORC2 | chr2 | - | 200937905 | 200937966 | 200933875 | 200933974 | 200941247 | 200941279 | 59 |
| SE | 73420 | ENSG00000115942 | ORC2 | chr2 | - | 200942684 | 200942777 | 200941247 | 200941279 | 200949553 | 200949643 | 182 |
| SE | 73421 | ENSG00000115942 | ORC2 | chr2 | - | 200949553 | 200949643 | 200942683 | 200942777 | 200957400 | 200957544 | 119 |
| SE | 73422 | ENSG00000115942 | ORC2 | chr2 | - | 200958029 | 200958133 | 200957400 | 200957544 | 200959391 | 200959440 | 175 |
| SE | 73423 | ENSG00000177463 | NR2C2 | chr3 | + | 14959156 | 14959196 | 14947704 | 14947906 | 14959535 | 14959641 | 8 |
| SE | 73424 | ENSG00000177463 | NR2C2 | chr3 | + | 14959535 | 14959641 | 14947704 | 14947906 | 14962408 | 14962492 | 9 |
| SE | 73432 | ENSG00000177463 | NR2C2 | chr3 | + | 15034669 | 15034809 | 15032378 | 15032500 | 15037999 | 15038137 | 96 |
| SE | 73433 | ENSG00000177463 | NR2C2 | chr3 | + | 15037999 | 15039227 | 15034669 | 15034809 | 15042833 | 15043349 | 98 |
| SE | 73438 | ENSG00000198721 | ECI2 | chr6 | - | 4119185 | 4119275 | 4117307 | 4117451 | 4126134 | 4126237 | 10 |
| SE | 73439 | ENSG00000198721 | ECI2 | chr6 | - | 4125249 | 4125370 | 4117307 | 4117451 | 4126134 | 4126237 | 25 |
| SE | 73444 | ENSG00000198721 | ECI2 | chr6 | - | 4126134 | 4126237 | 4125249 | 4125370 | 4127761 | 4127831 | 140 |
| SE | 73446 | ENSG00000198721 | ECI2 | chr6 | - | 4130371 | 4130560 | 4127761 | 4127831 | 4130766 | 4130865 | 241 |
| SE | 73447 | ENSG00000198721 | ECI2 | chr6 | - | 4130371 | 4130642 | 4127761 | 4127831 | 4130766 | 4130865 | 112 |
| SE | 73454 | ENSG00000143374 | TARS2 | chr1 | + | 150496809 | 150496908 | 150496481 | 150496628 | 150497529 | 150497587 | 36 |
| SE | 73456 | ENSG00000143374 | TARS2 | chr1 | + | 150498896 | 150499034 | 150498501 | 150498664 | 150499215 | 150499293 | 31 |
| SE | 73459 | ENSG00000003402 | CFLAR | chr2 | + | 201129728 | 201130146 | 201116163 | 201116481 | 201135971 | 201136107 | 99 |
| SE | 73460 | ENSG00000003402 | CFLAR | chr2 | + | 201129728 | 201130146 | 201118490 | 201118901 | 201133028 | 201133102 | 790 |
| SE | 73461 | ENSG00000003402 | CFLAR | chr2 | + | 201133028 | 201133134 | 201116153 | 201116481 | 201135971 | 201136107 | 490 |
| SE | 73462 | ENSG00000003402 | CFLAR | chr2 | + | 201133028 | 201133134 | 201129502 | 201130146 | 201135971 | 201136107 | 1112 |
| SE | 73464 | ENSG00000003402 | CFLAR | chr2 | + | 201135971 | 201136107 | 201132871 | 201133134 | 201145377 | 201145432 | 645 |
| SE | 73465 | ENSG00000003402 | CFLAR | chr2 | + | 201140356 | 201140439 | 201132871 | 201133134 | 201145377 | 201145432 | 192 |
| SE | 73466 | ENSG00000003402 | CFLAR | chr2 | + | 201140356 | 201140439 | 201135971 | 201136107 | 201145377 | 201145432 | 802 |
| SE | 73467 | ENSG00000003402 | CFLAR | chr2 | + | 201140356 | 201140439 | 201135971 | 201136107 | 201149002 | 201149052 | 356 |
| SE | 73468 | ENSG00000003402 | CFLAR | chr2 | + | 201142644 | 201142762 | 201140356 | 201140439 | 201145377 | 201145432 | 1 |
| SE | 73469 | ENSG00000003402 | CFLAR | chr2 | + | 201145377 | 201145432 | 201135971 | 201136107 | 201149002 | 201149052 | 127 |
| SE | 73475 | ENSG00000119013 | NDUFB3 | chr2 | + | 201072338 | 201072361 | 201072007 | 201072059 | 201078880 | 201079022 | 1 |
| SE | 73477 | ENSG00000119013 | NDUFB3 | chr2 | + | 201078880 | 201079022 | 201072000 | 201072059 | 201085458 | 201085615 | 323 |
| SE | 73478 | ENSG00000215559 | ANKRD20A11P | chr21 | - | 13915186 | 13915409 | 13911717 | 13911763 | 13916746 | 13916958 | 21 |
| SE | 73479 | ENSG00000215559 | ANKRD20A11P | chr21 | - | 13915186 | 13915409 | 13911717 | 13911763 | 13917554 | 13917836 | 34 |
| SE | 73482 | ENSG00000215559 | ANKRD20A11P | chr21 | - | 13927334 | 13927447 | 13925163 | 13925293 | 13932227 | 13932374 | 2 |
| SE | 73483 | ENSG00000215559 | ANKRD20A11P | chr21 | - | 13943743 | 13943862 | 13940814 | 13940978 | 13945038 | 13945109 | 2 |
| SE | 73484 | ENSG00000215559 | ANKRD20A11P | chr21 | - | 13954065 | 13954150 | 13951231 | 13951260 | 13967380 | 13967436 | 10 |
| SE | 73486 | ENSG00000215559 | ANKRD20A11P | chr21 | - | 13954065 | 13954150 | 13951231 | 13951260 | 13979701 | 13980437 | 9 |
| SE | 73487 | ENSG00000215559 | ANKRD20A11P | chr21 | - | 13958115 | 13958185 | 13954065 | 13954150 | 13967380 | 13967436 | 2 |
| SE | 73488 | ENSG00000215559 | ANKRD20A11P | chr21 | - | 13958115 | 13958185 | 13954065 | 13954150 | 13969972 | 13970110 | 5 |
| SE | 73490 | ENSG00000215559 | ANKRD20A11P | chr21 | - | 13962739 | 13962871 | 13951231 | 13951260 | 13967380 | 13967436 | 14 |
| SE | 73491 | ENSG00000215559 | ANKRD20A11P | chr21 | - | 13962739 | 13962871 | 13958115 | 13958185 | 13967380 | 13967436 | 21 |
| SE | 73493 | ENSG00000215559 | ANKRD20A11P | chr21 | - | 13967380 | 13967436 | 13954065 | 13954150 | 13969972 | 13970110 | 5 |
| SE | 73494 | ENSG00000215559 | ANKRD20A11P | chr21 | - | 13967380 | 13967436 | 13958115 | 13958185 | 13969972 | 13970110 | 5 |
| SE | 73495 | ENSG00000215559 | ANKRD20A11P | chr21 | - | 13969972 | 13970110 | 13954065 | 13954150 | 13979701 | 13979911 | 3 |
| SE | 73496 | ENSG00000215559 | ANKRD20A11P | chr21 | - | 13971166 | 13971273 | 13969972 | 13970110 | 13979701 | 13979911 | 1 |
| SE | 73501 | ENSG00000138380 | CARF | chr2 | + | 202913033 | 202913102 | 202912272 | 202912324 | 202917876 | 202918043 | 2 |
| SE | 73502 | ENSG00000138380 | CARF | chr2 | + | 202913033 | 202913114 | 202912217 | 202912324 | 202917876 | 202918043 | 1 |
| SE | 73503 | ENSG00000138380 | CARF | chr2 | + | 202924296 | 202924415 | 202917876 | 202918043 | 202934501 | 202934599 | 7 |
| SE | 73505 | ENSG00000138380 | CARF | chr2 | + | 202924296 | 202924415 | 202917876 | 202918043 | 202942739 | 202942867 | 7 |
| SE | 73506 | ENSG00000138380 | CARF | chr2 | + | 202924296 | 202924415 | 202917876 | 202918043 | 202952558 | 202952679 | 8 |
| SE | 73508 | ENSG00000138380 | CARF | chr2 | + | 202924356 | 202924415 | 202917876 | 202918043 | 202942739 | 202942967 | 4 |
| SE | 73509 | ENSG00000138380 | CARF | chr2 | + | 202934501 | 202934599 | 202917876 | 202918043 | 202938273 | 202938395 | 5 |
| SE | 73511 | ENSG00000138380 | CARF | chr2 | + | 202934501 | 202934599 | 202924296 | 202924415 | 202941859 | 202941966 | 0 |
| SE | 73512 | ENSG00000138380 | CARF | chr2 | + | 202938273 | 202938395 | 202917876 | 202918043 | 202941859 | 202941980 | 3 |
| SE | 73513 | ENSG00000138380 | CARF | chr2 | + | 202938273 | 202938395 | 202934501 | 202934599 | 202941859 | 202941980 | 1 |
| SE | 73519 | ENSG00000138380 | CARF | chr2 | + | 202942739 | 202942967 | 202924356 | 202924415 | 202952558 | 202952620 | 47 |
| SE | 73520 | ENSG00000138380 | CARF | chr2 | + | 202942739 | 202942967 | 202924356 | 202924415 | 202955673 | 202955758 | 0 |
| SE | 73521 | ENSG00000138380 | CARF | chr2 | + | 202942739 | 202942967 | 202941859 | 202941980 | 202952558 | 202952620 | 132 |
| SE | 73522 | ENSG00000138380 | CARF | chr2 | + | 202942762 | 202942967 | 202941859 | 202941980 | 202952558 | 202952679 | 47 |
| SE | 73523 | ENSG00000138380 | CARF | chr2 | + | 202955673 | 202955758 | 202942739 | 202942967 | 202961236 | 202961426 | 26 |
| SE | 73526 | ENSG00000138380 | CARF | chr2 | + | 202961236 | 202961426 | 202955673 | 202955758 | 202969918 | 202970062 | 42 |
| SE | 73527 | ENSG00000138380 | CARF | chr2 | + | 202966977 | 202967098 | 202955673 | 202955758 | 202969918 | 202970062 | 51 |
| SE | 73531 | ENSG00000138380 | CARF | chr2 | + | 202974333 | 202974496 | 202971504 | 202971738 | 202981554 | 202981685 | 68 |
| SE | 73532 | ENSG00000138380 | CARF | chr2 | + | 202977268 | 202977332 | 202971504 | 202971738 | 202981554 | 202981685 | 28 |
| SE | 73534 | ENSG00000138380 | CARF | chr2 | + | 202981554 | 202981685 | 202977268 | 202977332 | 202982071 | 202982441 | 112 |
| SE | 73535 | ENSG00000138380 | CARF | chr2 | + | 202982071 | 202982441 | 202981554 | 202981685 | 202983505 | 202983642 | 45 |
| SE | 73538 | ENSG00000146576 | C7orf26 | chr7 | + | 6599828 | 6600356 | 6594411 | 6594600 | 6602100 | 6602140 | 214 |
| SE | 73539 | ENSG00000146576 | C7orf26 | chr7 | + | 6602663 | 6602752 | 6602100 | 6602140 | 6607536 | 6607597 | 0 |
| SE | 73540 | ENSG00000115232 | ITGA4 | chr2 | + | 181458195 | 181458317 | 181457206 | 181457851 | 181474959 | 181475066 | 539 |
| SE | 73544 | ENSG00000115232 | ITGA4 | chr2 | + | 181475158 | 181475288 | 181474959 | 181475066 | 181480136 | 181480266 | 393 |
| SE | 73548 | ENSG00000115232 | ITGA4 | chr2 | + | 181480136 | 181480266 | 181474959 | 181475066 | 181481597 | 181481683 | 1172 |
| SE | 73551 | ENSG00000115232 | ITGA4 | chr2 | + | 181494721 | 181494812 | 181493033 | 181493419 | 181495370 | 181495416 | 1347 |
| SE | 73553 | ENSG00000115232 | ITGA4 | chr2 | + | 181498622 | 181498777 | 181495370 | 181495416 | 181509657 | 181509805 | 1179 |
| SE | 73555 | ENSG00000115232 | ITGA4 | chr2 | + | 181534271 | 181534370 | 181531656 | 181531776 | 181534815 | 181534935 | 735 |
| SE | 73556 | ENSG00000151164 | RAD9B | chr12 | + | 110511427 | 110511583 | 110506578 | 110506693 | 110512778 | 110512878 | 0 |
| SE | 73562 | ENSG00000230006 | ANKRD36BP2 | chr2 | + | 88797281 | 88797423 | 88792420 | 88792494 | 88801098 | 88804880 | 0 |
| SE | 73564 | ENSG00000115998 | C2orf42 | chr2 | - | 70165527 | 70165635 | 70160624 | 70160787 | 70169556 | 70169661 | 57 |
| SE | 73565 | ENSG00000115998 | C2orf42 | chr2 | - | 70175672 | 70175777 | 70169556 | 70169661 | 70179531 | 70179642 | 60 |
| SE | 73567 | ENSG00000115998 | C2orf42 | chr2 | - | 70182666 | 70182935 | 70181162 | 70181997 | 70190690 | 70190907 | 106 |
| SE | 73570 | ENSG00000115998 | C2orf42 | chr2 | - | 70190542 | 70190715 | 70182666 | 70182935 | 70190972 | 70191002 | 3 |
| SE | 73579 | ENSG00000129472 | RAB2B | chr14 | - | 21463655 | 21463767 | 21462349 | 21462418 | 21468356 | 21468449 | 130 |
| SE | 73582 | ENSG00000147443 | DOK2 | chr8 | - | 21910672 | 21910857 | 21908872 | 21909931 | 21911900 | 21911988 | 451 |
| SE | 73583 | ENSG00000147443 | DOK2 | chr8 | - | 21910672 | 21910857 | 21908872 | 21909931 | 21912228 | 21912510 | 307 |
| SE | 73584 | ENSG00000147443 | DOK2 | chr8 | - | 21910672 | 21910857 | 21909828 | 21909909 | 21911900 | 21911988 | 193 |
| SE | 73585 | ENSG00000147443 | DOK2 | chr8 | - | 21911900 | 21911988 | 21908872 | 21909931 | 21912228 | 21912510 | 51 |
| SE | 73591 | ENSG00000124356 | STAMBP | chr2 | + | 73829404 | 73829510 | 73828960 | 73828996 | 73844812 | 73844888 | 0 |
| SE | 73593 | ENSG00000124356 | STAMBP | chr2 | + | 73829404 | 73829510 | 73828966 | 73829070 | 73844812 | 73844888 | 0 |
| SE | 73596 | ENSG00000124356 | STAMBP | chr2 | + | 73830844 | 73831059 | 73828960 | 73829510 | 73844812 | 73844888 | 88 |
| SE | 73597 | ENSG00000124356 | STAMBP | chr2 | + | 73830844 | 73831059 | 73828982 | 73829070 | 73844812 | 73844888 | 79 |
| SE | 73599 | ENSG00000124356 | STAMBP | chr2 | + | 73847386 | 73847572 | 73845166 | 73845262 | 73849362 | 73849487 | 137 |
| SE | 73600 | ENSG00000124356 | STAMBP | chr2 | + | 73847386 | 73847753 | 73844812 | 73845262 | 73849362 | 73849487 | 219 |
| SE | 73601 | ENSG00000124356 | STAMBP | chr2 | + | 73847389 | 73847753 | 73845166 | 73845262 | 73849362 | 73849487 | 82 |
| SE | 73604 | ENSG00000124356 | STAMBP | chr2 | + | 73859253 | 73859366 | 73850375 | 73850513 | 73860051 | 73860151 | 162 |
| SE | 73607 | ENSG00000124356 | STAMBP | chr2 | + | 73862202 | 73862304 | 73860051 | 73860151 | 73870142 | 73870352 | 37 |

| | | | | | | | | | | | | | | | | | | | | | | | |
|---|---|---|---|---|---|---|---|---|---|---|---|---|---|---|---|---|---|---|---|---|---|---|---|
| SE | 73609 | ENSG00000124356 | STAMBP | chr2 | + | 73862202 | 73862945 | 73860051 | 73860151 | 73870142 | 73870352 | 49 | 42 | 34 | 66 | 154 | 1 | 0 | 1 | 0 | 0 | 60 | 34 | [illegible] |
| SE | 73610 | ENSG00000124356 | STAMBP | chr2 | + | 73862202 | 73867174 | 73860051 | 73860151 | 73870142 | 73870352 | 49 | 42 | 34 | 66 | 154 | 1 | 0 | 1 | 0 | 0 | 60 | 34 | [illegible] |
| SE | 73611 | ENSG00000124356 | STAMBP | chr2 | + | 73870142 | 73870352 | 73860051 | 73860151 | 73873351 | 73873601 | 4 | 0 | 1 | 5 | 0 | 0 | 0 | 0 | 0 | 0 | 7 | 2 | [illegible] |
| SE | 73612 | ENSG00000124356 | STAMBP | chr2 | + | 73870142 | 73870352 | 73862202 | 73862304 | 73873351 | 73873601 | 3 | 0 | 0 | 5 | 3 | 0 | 0 | 0 | 0 | 0 | 7 | 2 | [illegible] |
| SE | 73613 | ENSG00000124356 | STAMBP | chr2 | + | 73871792 | 73872417 | 73870142 | 73870352 | 73873351 | 73873601 | 0 | 0 | 0 | 2 | 3 | 3 | 0 | 0 | 5 | 0 | 0 | 0 | [illegible] |
| SE | 73614 | ENSG00000124356 | STAMBP | chr2 | + | 73871954 | 73872417 | 73869706 | 73870352 | 73873351 | 73873497 | 0 | 0 | 1 | 3 | 3 | 3 | 0 | 0 | 5 | 0 | 0 | 0 | [illegible] |
| SE | 73615 | ENSG00000196453 | ZNF777 | chr7 | - | 149454110 | 149454237 | 149450998 | 149451112 | 149455176 | 149456037 | 116 | 65 | 95 | 86 | 140 | 0 | 0 | 0 | 2 | 0 | 97 | 195 | [illegible] |
| SE | 73619 | ENSG00000119707 | RBM25 | chr14 | + | 73076318 | 73076368 | 73071626 | 73071747 | 73077368 | 73077536 | 132 | 105 | 158 | 121 | 126 | 1 | 0 | 0 | 0 | 0 | 63 | 88 | [illegible] |
| SE | 73620 | ENSG00000119707 | RBM25 | chr14 | + | 73077368 | 73077536 | 73076318 | 73076368 | 73083493 | 73083551 | 522 | 453 | 580 | 595 | 907 | 0 | 0 | 1 | 0 | 0 | 203 | 403 | [illegible] |
| SE | 73621 | ENSG00000119707 | RBM25 | chr14 | + | 73083493 | 73083551 | 73077368 | 73077536 | 73088000 | 73088161 | 540 | 467 | 569 | 617 | 1108 | 1 | 1 | 3 | 1 | 1 | 210 | 387 | [illegible] |
| SE | 73623 | ENSG00000119707 | RBM25 | chr14 | + | 73096914 | 73097100 | 73088000 | 73088161 | 73099379 | 73099433 | 1099 | 1069 | 1197 | 1349 | 2484 | 2 | 0 | 2 | 0 | 0 | 618 | 723 | [illegible] |
| SE | 73624 | ENSG00000119707 | RBM25 | chr14 | + | 73100300 | 73100362 | 73099666 | 73099750 | 73100713 | 73101179 | 11 | 26 | 9 | 16 | 10 | 0 | 1 | 0 | 3 | 9 | 0 | 12 | [illegible] |
| SE | 73628 | ENSG00000119707 | RBM25 | chr14 | + | 73114285 | 73114333 | 73112151 | 73112250 | 73119712 | 73120129 | 328 | 280 | 288 | 361 | 535 | 1 | 1 | 0 | 0 | 0 | 54 | 120 | [illegible] |
| SE | 73630 | ENSG00000158417 | EIF5B | chr2 | + | 99389717 | 99389849 | 99382779 | 99382921 | 99390218 | 99390401 | 198 | 165 | 218 | 226 | 222 | 1 | 0 | 1 | 0 | 0 | 0 | 53 | [illegible] |
| SE | 73631 | ENSG00000158417 | EIF5B | chr2 | + | 99389717 | 99389849 | 99382779 | 99382921 | 99390543 | 99390705 | 116 | 89 | 131 | 113 | 112 | 0 | 0 | 1 | 1 | 0 | 0 | 31 | [illegible] |
| SE | 73632 | ENSG00000158417 | EIF5B | chr2 | + | 99390218 | 99390401 | 99382779 | 99382921 | 99390543 | 99390690 | 124 | 74 | 78 | 114 | 89 | 0 | 0 | 1 | 1 | 0 | 0 | 55 | [illegible] |
| SE | 73633 | ENSG00000158417 | EIF5B | chr2 | + | 99390218 | 99390401 | 99389606 | 99389849 | 99390543 | 99390690 | 205 | 152 | 164 | 228 | 199 | 0 | 0 | 0 | 0 | 0 | 0 | 80 | [illegible] |
| SE | 73634 | ENSG00000154328 | NEIL2 | chr8 | + | 11770173 | 11770335 | 11769698 | 11769791 | 11771445 | 11771585 | 14 | 5 | 15 | 7 | 16 | 1 | 0 | 1 | 0 | 0 | 14 | 5 | [illegible] |
| SE | 73635 | ENSG00000154328 | NEIL2 | chr8 | + | 11770173 | 11770335 | 11769698 | 11769791 | 11779597 | 11779950 | 4 | 0 | 1 | 0 | 1 | 0 | 0 | 0 | 0 | 0 | 0 | 3 | [illegible] |
| SE | 73636 | ENSG00000154328 | NEIL2 | chr8 | + | 11771445 | 11771585 | 11769638 | 11769791 | 11779597 | 11779950 | 5 | 3 | 16 | 2 | 8 | 0 | 0 | 0 | 0 | 0 | 30 | 21 | [illegible] |
| SE | 73637 | ENSG00000154328 | NEIL2 | chr8 | + | 11771445 | 11771585 | 11769709 | 11770335 | 11779597 | 11779950 | 14 | 8 | 29 | 9 | 24 | 1 | 0 | 0 | 0 | 1 | 44 | 23 | [illegible] |
| SE | 73638 | ENSG00000154328 | NEIL2 | chr8 | + | 11771445 | 11771590 | 11770173 | 11770335 | 11779597 | 11779950 | 14 | 5 | 14 | 10 | 16 | 1 | 0 | 0 | 0 | 1 | 14 | 2 | [illegible] |
| SE | 73641 | ENSG00000115526 | CHST10 | chr2 | - | 100395508 | 100395614 | 100391859 | 100393782 | 100397907 | 100398142 | 21 | 12 | 8 | 34 | 21 | 4 | 1 | 0 | 0 | 0 | 0 | 54 | [illegible] |
| SE | 73642 | ENSG00000115526 | CHST10 | chr2 | - | 100402563 | 100402655 | 100397907 | 100398142 | 100406575 | 100406707 | 16 | 14 | 23 | 19 | 28 | 0 | 0 | 2 | 0 | 0 | 1 | 15 | [illegible] |
| SE | 73643 | ENSG00000115526 | CHST10 | chr2 | - | 100406575 | 100406707 | 100402563 | 100402655 | 100415040 | 100415111 | 5 | 12 | 11 | 16 | 21 | 0 | 0 | 2 | 0 | 0 | 1 | 7 | [illegible] |
| SE | 73645 | ENSG00000115526 | CHST10 | chr2 | - | 100408542 | 100408660 | 100406575 | 100406707 | 100415040 | 100415111 | 0 | 0 | 3 | 1 | 0 | 4 | 3 | 8 | 7 | 11 | 0 | 0 | [illegible] |
| SE | 73652 | ENSG00000178075 | GRAMD1C | chr3 | + | 113901030 | 113901146 | 113882751 | 113882832 | 113904139 | 113904272 | 1 | 6 | 1 | 17 | 16 | 2 | 0 | 2 | 0 | 0 | 0 | 30 | [illegible] |
| SE | 73653 | ENSG00000178075 | GRAMD1C | chr3 | + | 113901030 | 113901146 | 113897469 | 113897813 | 113904139 | 113904272 | 1 | 4 | 0 | 10 | 7 | 0 | 0 | 0 | 1 | 0 | 0 | 13 | [illegible] |
| SE | 73656 | ENSG00000178075 | GRAMD1C | chr3 | + | 113904139 | 113904272 | 113901030 | 113901146 | 113908957 | 113909120 | 1 | 11 | 0 | 11 | 5 | 0 | 0 | 0 | 0 | 2 | 0 | 19 | [illegible] |
| SE | 73658 | ENSG00000178075 | GRAMD1C | chr3 | + | 113936265 | 113936442 | 113934431 | 113934535 | 113938085 | 113938143 | 7 | 8 | 7 | 20 | 23 | 0 | 1 | 1 | 0 | 0 | 0 | 5 | [illegible] |
| SE | 73659 | ENSG00000081019 | RSBN1 | chr1 | - | 113777209 | 113777352 | 113768221 | 113768389 | 113777670 | 113777808 | 340 | 192 | 328 | 291 | 612 | 1 | 0 | 0 | 0 | 0 | 101 | 226 | [illegible] |
| SE | 73662 | ENSG00000151687 | ANKAR | chr2 | + | 189676455 | 189677091 | 189674669 | 189674830 | 189689526 | 189689964 | 4 | 1 | 6 | 1 | 8 | 18 | 13 | 7 | 10 | 10 | 0 | 1 | [illegible] |
| SE | 73663 | ENSG00000151687 | ANKAR | chr2 | + | 189692254 | 189692418 | 189689526 | 189689964 | 189693073 | 189693177 | 4 | 3 | 2 | 3 | 0 | 1 | 3 | 0 | 1 | 0 | 2 | 14 | [illegible] |
| SE | 73664 | ENSG00000151687 | ANKAR | chr2 | + | 189692254 | 189692418 | 189689526 | 189689964 | 189696149 | 189696369 | 7 | 3 | 3 | 3 | 0 | 22 | 17 | 26 | 16 | 73 | 2 | 14 | [illegible] |
| SE | 73665 | ENSG00000151687 | ANKAR | chr2 | + | 189693073 | 189693177 | 189689526 | 189689964 | 189694980 | 189695161 | 4 | 4 | 0 | 2 | 0 | 1 | 5 | 2 | 3 | 0 | 0 | 0 | [illegible] |
| SE | 73666 | ENSG00000151687 | ANKAR | chr2 | + | 189694980 | 189695161 | 189689526 | 189689964 | 189696149 | 189696369 | 9 | 9 | 9 | 3 | 1 | 22 | 17 | 26 | 16 | 73 | 0 | 2 | [illegible] |
| SE | 73667 | ENSG00000151687 | ANKAR | chr2 | + | 189705022 | 189705224 | 189689526 | 189689964 | 189706937 | 189707146 | 4 | 1 | 3 | 8 | 8 | 6 | 1 | 0 | 5 | 3 | 0 | 6 | [illegible] |
| SE | 73668 | ENSG00000151687 | ANKAR | chr2 | + | 189705022 | 189705224 | 189696149 | 189696369 | 189706937 | 189707146 | 2 | 2 | 11 | 6 | 9 | 1 | 1 | 3 | 1 | 1 | 0 | 6 | [illegible] |
| SE | 73669 | ENSG00000151687 | ANKAR | chr2 | + | 189711048 | 189711153 | 189706937 | 189707146 | 189719571 | 189719813 | 8 | 4 | 10 | 10 | 0 | 10 | 6 | 8 | 8 | 11 | 0 | 3 | [illegible] |
| SE | 73670 | ENSG00000151687 | ANKAR | chr2 | + | 189720618 | 189720787 | 189719571 | 189719813 | 189727855 | 189728097 | 33 | 32 | 36 | 20 | 64 | 0 | 1 | 0 | 1 | 0 | 24 | 30 | [illegible] |
| SE | 73671 | ENSG00000151687 | ANKAR | chr2 | + | 189727822 | 189728097 | 189720618 | 189720787 | 189728266 | 189728420 | 16 | 6 | 15 | 5 | 19 | 1 | 0 | 0 | 0 | 0 | 0 | 9 | [illegible] |
| SE | 73672 | ENSG00000151687 | ANKAR | chr2 | + | 189727855 | 189728097 | 189720618 | 189720787 | 189728266 | 189728420 | 31 | 16 | 29 | 9 | 50 | 1 | 0 | 0 | 0 | 0 | 12 | 15 | [illegible] |
| SE | 73673 | ENSG00000151687 | ANKAR | chr2 | + | 189728266 | 189728420 | 189720618 | 189720787 | 189728659 | 189728821 | 3 | 4 | 6 | 4 | 4 | 0 | 1 | 0 | 2 | 0 | 0 | 3 | [illegible] |
| SE | 73674 | ENSG00000151687 | ANKAR | chr2 | + | 189728659 | 189728821 | 189720618 | 189720787 | 189741341 | 189741451 | 5 | 6 | 7 | 9 | 19 | 0 | 0 | 0 | 1 | 0 | 18 | 9 | [illegible] |
| SE | 73675 | ENSG00000151687 | ANKAR | chr2 | + | 189728659 | 189728821 | 189728266 | 189728420 | 189733106 | 189733229 | 3 | 4 | 6 | 6 | 4 | 0 | 0 | 4 | 0 | 0 | 0 | 3 | [illegible] |
| SE | 73677 | ENSG00000151687 | ANKAR | chr2 | + | 189728659 | 189728821 | 189728266 | 189728420 | 189738564 | 189738682 | 6 | 6 | 6 | 6 | 8 | 1 | 0 | 1 | 0 | 0 | 0 | 6 | [illegible] |
| SE | 73678 | ENSG00000151687 | ANKAR | chr2 | + | 189730494 | 189730601 | 189728659 | 189728821 | 189733106 | 189733229 | 19 | 14 | 8 | 28 | 26 | 1 | 0 | 0 | 2 | 0 | 42 | 18 | [illegible] |
| SE | 73681 | ENSG00000151687 | ANKAR | chr2 | + | 189733106 | 189733229 | 189728266 | 189728420 | 189738564 | 189738682 | 2 | 6 | 6 | 5 | 6 | 1 | 0 | 1 | 0 | 0 | 0 | 9 | [illegible] |
| SE | 73683 | ENSG00000151687 | ANKAR | chr2 | + | 189733106 | 189733229 | 189728659 | 189728821 | 189738564 | 189738682 | 3 | 6 | 2 | 7 | 6 | 4 | 2 | 0 | 2 | 4 | 0 | 9 | [illegible] |
| SE | 73685 | ENSG00000151687 | ANKAR | chr2 | + | 189733106 | 189733229 | 189730494 | 189730601 | 189741341 | 189741451 | 10 | 8 | 11 | 20 | 22 | 0 | 0 | 0 | 0 | 0 | 42 | 12 | [illegible] |
| SE | 73686 | ENSG00000151687 | ANKAR | chr2 | + | 189737682 | 189737841 | 189728266 | 189728420 | 189738564 | 189738682 | 37 | 49 | 30 | 30 | 70 | 1 | 0 | 1 | 0 | 0 | 65 | 29 | [illegible] |
| SE | 73693 | ENSG00000151687 | ANKAR | chr2 | + | 189738564 | 189738682 | 189737682 | 189737841 | 189741341 | 189741451 | 54 | 60 | 39 | 34 | 98 | 0 | 1 | 0 | 1 | 1 | 84 | 39 | [illegible] |
| SE | 73694 | ENSG00000151687 | ANKAR | chr2 | + | 189743274 | 189743474 | 189741341 | 189741451 | 189744741 | 189744788 | 0 | 0 | 2 | 2 | 0 | 21 | 19 | 17 | 24 | 30 | 0 | 0 | [illegible] |
| SE | 73696 | ENSG00000157349 | DDX19B | chr16 | + | 70317495 | 70317588 | 70315968 | 70316104 | 70319600 | 70319683 | 83 | 59 | 86 | 58 | 85 | 1 | 0 | 0 | 0 | 0 | 187 | 117 | [illegible] |
| SE | 73697 | ENSG00000157349 | DDX19B | chr16 | + | 70319600 | 70319709 | 70315968 | 70316104 | 70324584 | 70324687 | 1 | 0 | 1 | 0 | 1 | 12 | 2 | 2 | 15 | 6 | 0 | 0 | [illegible] |
| SE | 73698 | ENSG00000157349 | DDX19B | chr16 | + | 70319600 | 70319709 | 70317495 | 70317588 | 70324584 | 70324687 | 3 | 0 | 1 | 2 | 1 | 128 | 98 | 129 | 95 | 210 | 0 | 4 | [illegible] |
| SE | 73699 | ENSG00000114770 | ABCC5 | chr3 | - | 183927329 | 183927443 | 183925554 | 183925719 | 183928746 | 183928825 | 97 | 44 | 50 | 108 | 141 | 0 | 0 | 0 | 1 | 0 | 116 | 92 | [illegible] |
| SE | 73700 | ENSG00000114770 | ABCC5 | chr3 | - | 183927329 | 183927443 | 183925554 | 183925719 | 183937900 | 183938060 | 47 | 19 | 36 | 45 | 85 | 0 | 0 | 1 | 0 | 0 | 60 | 35 | [illegible] |
| SE | 73701 | ENSG00000114770 | ABCC5 | chr3 | - | 183928746 | 183928825 | 183925554 | 183925719 | 183937900 | 183938060 | 48 | 16 | 28 | 32 | 52 | 0 | 0 | 1 | 0 | 0 | 84 | 59 | [illegible] |
| SE | 73702 | ENSG00000114770 | ABCC5 | chr3 | - | 183928746 | 183928825 | 183927329 | 183927443 | 183937900 | 183938060 | 98 | 41 | 42 | 89 | 106 | 0 | 0 | 0 | 0 | 0 | 140 | 116 | [illegible] |
| SE | 73704 | ENSG00000114770 | ABCC5 | chr3 | - | 183942726 | 183942916 | 183937900 | 183938060 | 183945849 | 183945939 | 57 | 22 | 45 | 64 | 57 | 0 | 0 | 0 | 0 | 0 | 62 | 77 | [illegible] |
| SE | 73711 | ENSG00000114770 | ABCC5 | chr3 | - | 184009888 | 184009977 | 183989225 | 183989383 | 184014263 | 184014447 | 1 | 0 | 1 | 0 | 4 | 71 | 57 | 55 | 64 | 99 | 0 | 0 | [illegible] |
| SE | 73712 | ENSG00000144224 | UBXN4 | chr2 | + | 135747657 | 135747765 | 135741733 | 135742011 | 135748266 | 135748369 | 0 | 0 | 2 | 0 | 0 | 59 | 14 | 33 | 36 | 39 | 0 | 0 | [illegible] |
| SE | 73714 | ENSG00000144224 | UBXN4 | chr2 | + | 135755516 | 135755691 | 135748266 | 135748369 | 135761817 | 135761881 | 274 | 217 | 231 | 342 | 483 | 0 | 0 | 0 | 0 | 0 | 174 | 173 | [illegible] |
| SE | 73716 | ENSG00000144224 | UBXN4 | chr2 | + | 135772419 | 135772547 | 135770570 | 135770735 | 135776248 | 135776351 | 495 | 380 | 448 | 481 | 660 | 0 | 0 | 0 | 1 | 0 | 89 | 288 | [illegible] |
| SE | 73717 | ENSG00000144224 | UBXN4 | chr2 | + | 135772419 | 135772547 | 135770570 | 135770735 | 135778947 | 135779079 | 228 | 216 | 204 | 240 | 301 | 0 | 0 | 0 | 1 | 0 | 68 | 209 | [illegible] |
| SE | 73718 | ENSG00000144224 | UBXN4 | chr2 | + | 135776248 | 135776351 | 135770570 | 135770735 | 135778947 | 135779079 | 212 | 123 | 200 | 191 | 237 | 0 | 0 | 0 | 1 | 0 | 51 | 114 | [illegible] |
| SE | 73720 | ENSG00000162419 | GMEB1 | chr1 | + | 28690073 | 28690186 | 28683563 | 28683740 | 28691584 | 28691709 | 54 | 50 | 76 | 62 | 139 | 0 | 0 | 0 | 1 | 0 | 98 | 51 | [illegible] |
| SE | 73722 | ENSG00000162419 | GMEB1 | chr1 | + | 28692941 | 28693045 | 28691584 | 28691709 | 28696926 | 28697084 | 119 | 77 | 117 | 117 | 219 | 1 | 0 | 0 | 0 | 0 | 157 | 138 | [illegible] |
| SE | 73723 | ENSG00000162419 | GMEB1 | chr1 | + | 28696926 | 28697084 | 28692941 | 28693045 | 28702437 | 28702569 | 139 | 60 | 90 | 116 | 166 | 2 | 0 | 1 | 1 | 1 | 146 | 132 | [illegible] |
| SE | 73724 | ENSG00000162419 | GMEB1 | chr1 | + | 28702437 | 28702569 | 28696926 | 28697084 | 28704191 | 28704329 | 164 | 86 | 99 | 138 | 160 | 0 | 0 | 0 | 1 | 0 | 180 | 149 | [illegible] |
| SE | 73725 | ENSG00000162419 | GMEB1 | chr1 | + | 28704191 | 28704329 | 28702437 | 28702569 | 28710519 | 28710642 | 161 | 111 | 118 | 169 | 222 | 1 | 1 | 0 | 1 | 0 | 247 | 229 | [illegible] |
| SE | 73730 | ENSG00000162733 | DDR2 | chr1 | + | 162755155 | 162755303 | 162754623 | 162754855 | 162755663 | 162755769 | 1 | 0 | 1 | 5 | 2 | 0 | 0 | 0 | 0 | 0 | 0 | 1 | [illegible] |
| SE | 73732 | ENSG00000166508 | MCM7 | chr7 | - | 100095386 | 100095470 | 100094172 | 100094341 | 100095773 | 100095973 | 111 | 39 | 127 | 122 | 63 | 0 | 0 | 0 | 0 | 0 | 186 | 123 | [illegible] |
| SE | 73733 | ENSG00000166508 | MCM7 | chr7 | - | 100097300 | 100097384 | 100095773 | 100096167 | 100097613 | 100097745 | 146 | 52 | 191 | 222 | 160 | 0 | 0 | 0 | 0 | 0 | 279 | 186 | [illegible] |
| SE | 73734 | ENSG00000166508 | MCM7 | chr7 | - | 100097833 | 100097948 | 100097613 | 100097745 | 100098140 | 100098290 | 250 | 184 | 361 | 312 | 259 | 0 | 0 | 0 | 0 | 0 | 696 | 412 | [illegible] |
| SE | 73735 | ENSG00000166508 | MCM7 | chr7 | - | 100099588 | 100099753 | 100099022 | 100099403 | 100100013 | 100100093 | 202 | 91 | 238 | 233 | 241 | 0 | 2 | 0 | 0 | 0 | 305 | 202 | [illegible] |
| SE | 73739 | ENSG00000119772 | DNMT3A | chr2 | - | 25239129 | 25239215 | 25236929 | 25237005 | 25240301 | 25240450 | 75 | 68 | 100 | 80 | 89 | 0 | 0 | 0 | 0 | 0 | 81 | 30 | [illegible] |
| SE | 73741 | ENSG00000119772 | DNMT3A | chr2 | - | 25240301 | 25240450 | 25239129 | 25239215 | 25240639 | 25240730 | 62 | 59 | 100 | 67 | 57 | 0 | 2 | 0 | 0 | 0 | 84 | 53 | [illegible] |
| SE | 73742 | ENSG00000119772 | DNMT3A | chr2 | - | 25241561 | 25241707 | 25240639 | 25240730 | 25243897 | 25243970 | 36 | 49 | 89 | 43 | 22 | 0 | 0 | 0 | 0 | 0 | 61 | 73 | [illegible] |
| SE | 73743 | ENSG00000119772 | DNMT3A | chr2 | - | 25241561 | 25241777 | 25240639 | 25240730 | 25243897 | 25243982 | 13 | 31 | 42 | 15 | 10 | 0 | 0 | 0 | 0 | 0 | 27 | 46 | [illegible] |
| SE | 73744 | ENSG00000119772 | DNMT3A | chr2 | - | 25247590 | 25247749 | 25247050 | 25247158 | 25248036 | 25248252 | 38 | 22 | 43 | 43 | 22 | 0 | 0 | 0 | 0 | 0 | 57 | 58 | [illegible] |
| SE | 73745 | ENSG00000119772 | DNMT3A | chr2 | - | 25249656 | 25249724 | 25248036 | 25248252 | 25252193 | 25252311 | 2 | 0 | 0 | 2 | 0 | 0 | 0 | 0 | 2 | 1 | 0 | 2 | [illegible] |
| SE | 73747 | ENSG00000119772 | DNMT3A | chr2 | - | 25249656 | 25249724 | 25248036 | 25248252 | 25282440 | 25282711 | 0 | 0 | 0 | 2 | 0 | 0 | 0 | 0 | 0 | 0 | 0 | 2 | [illegible] |
| SE | 73749 | ENSG00000119772 | DNMT3A | chr2 | - | 25275499 | 25275543 | 25248036 | 25248252 | 25282440 | 25282711 | 7 | 12 | 8 | 12 | 2 | 0 | 0 | 0 | 0 | 0 | 18 | 2 | [illegible] |
| SE | 73750 | ENSG00000119772 | DNMT3A | chr2 | - | 25282440 | 25282711 | 25275499 | 25275543 | 25300138 | 25300243 | 54 | 44 | 72 | 77 | 29 | 0 | 0 | 0 | 0 | 0 | 64 | 46 | [illegible] |
| SE | 73752 | ENSG00000081052 | COL4A4 | chr2 | - | 227077893 | 227078077 | 227062529 | 227062598 | 227080442 | 227080549 | 6 | 0 | 10 | 2 | 6 | 2 | 0 | 0 | 0 | 0 | 0 | 0 | [illegible] |
| SE | 73764 | ENSG00000146285 | SCML4 | chr6 | - | 107746688 | 107746889 | 107744948 | 107745143 | 107749683 | 107749813 | 29 | 18 | 40 | 35 | 21 | 1 | 0 | 0 | 3 | 0 | 17 | 23 | [illegible] |
| SE | 73765 | ENSG00000146285 | SCML4 | chr6 | - | 107749683 | 107749813 | 107744948 | 107746889 | 107772171 | 107772386 | 57 | 41 | 84 | 50 | 43 | 0 | 0 | 0 | 0 | 0 | 33 | 32 | [illegible] |
| SE | 73767 | ENSG00000146285 | SCML4 | chr6 | - | 107755570 | 107755664 | 107749683 | 107749813 | 107824125 | 107824294 | 2 | 4 | 2 | 3 | 3 | 14 | 0 | 17 | 10 | 3 | 0 | 0 | [illegible] |
| SE | 73807 | ENSG00000043462 | LCP2 | chr5 | - | 170250729 | 170250885 | 170246232 | 170248819 | 170252433 | 170252511 | 1664 | 2181 | 2078 | 2285 | 2308 | 0 | 0 | 0 | 0 | 0 | 4327 | 2080 | [illegible] |
| SE | 73808 | ENSG00000043462 | LCP2 | chr5 | - | 170252433 | 170252511 | 170250729 | 170250885 | 170253118 | 170253213 | 906 | 821 | 832 | 1017 | 1036 | 0 | 0 | 0 | 1 | 0 | 1893 | 968 | [illegible] |
| SE | 73809 | ENSG00000043462 | LCP2 | chr5 | - | 170252433 | 170252511 | 170250729 | 170250885 | 170256525 | 170256575 | 577 | 590 | 494 | 644 | 625 | 0 | 3 | 0 | 0 | 0 | 1228 | 609 | [illegible] |
| SE | 73810 | ENSG00000043462 | LCP2 | chr5 | - | 170253118 | 170253213 | 170250729 | 170250885 | 170256525 | 170256575 | 322 | 155 | 227 | 290 | 444 | 0 | 3 | 0 | 0 | 0 | 742 | 308 | [illegible] |
| SE | 73811 | ENSG00000043462 | LCP2 | chr5 | - | 170253118 | 170253213 | 170252433 | 170252511 | 170256525 | 170256575 | 654 | 386 | 560 | 662 | 853 | 3 | 0 | 1 | 0 | 0 | 1407 | 670 | [illegible] |
| SE | 73812 | ENSG00000043462 | LCP2 | chr5 | - | 170258036 | 170258166 | 170256525 | 170256575 | 170258865 | 170258878 | 877 | 510 | 638 | 822 | 1249 | 1 | 0 | 4 | 3 | 3 | 2088 | 923 | [illegible] |
| SE | 73813 | ENSG00000043462 | LCP2 | chr5 | - | 170258865 | 170258878 | 170258036 | 170258166 | 170261106 | 170261137 | 492 | 318 | 370 | 444 | 793 | 0 | 0 | 0 | 0 | 0 | 1205 | 566 | [illegible] |
| SE | 73814 | ENSG00000043462 | LCP2 | chr5 | - | 170261106 | 170261137 | 170258865 | 170258878 | 170262634 | 170262742 | 554 | 387 | 446 | 562 | 873 | 0 | 0 | 0 | 0 | 0 | 1374 | 665 | [illegible] |
| SE | 73816 | ENSG00000043462 | LCP2 | chr5 | - | 170268384 | 170268482 | 170267008 | 170267075 | 170270718 | 170270917 | 532 | 358 | 508 | 522 | 1522 | 2 | 0 | 0 | 0 | 0 | 970 | 673 | [illegible] |
| SE | 73817 | ENSG00000043462 | LCP2 | chr5 | - | 170268384 | 170268482 | 170267008 | 170267075 | 170274300 | 170274338 | 57 | 36 | 30 | 48 | 44 | 0 | 0 | 0 | 0 | 2 | 119 | 69 | [illegible] |
| SE | 73818 | ENSG00000043462 | LCP2 | chr5 | - | 170270718 | 170270917 | 170267008 | 170267075 | 170274300 | 170274338 | 515 | 373 | 483 | 479 | 1426 | 0 | 0 | 0 | 0 | 2 | 791 | 579 | [illegible] |
| SE | 73820 | ENSG00000043462 | LCP2 | chr5 | - | 170270718 | 170270917 | 170268384 | 170268482 | 170275319 | 170275351 | 483 | 327 | 485 | 488 | 1489 | 0 | 0 | 0 | 0 | 0 | 918 | 631 | [illegible] |
| SE | 73821 | ENSG00000043462 | LCP2 | chr5 | - | 170274300 | 170274338 | 170268384 | 170268482 | 170275319 | 170275351 | 79 | 75 | 86 | 91 | 180 | 0 | 0 | 0 | 0 | 0 | 191 | 134 | [illegible] |
| SE | 73822 | ENSG00000043462 | LCP2 | chr5 | - | 170274300 | 170274338 | 170268384 | 170268482 | 170275794 | 170275860 | 2 | 2 | 2 | 5 | 0 | 1 | 0 | 0 | 0 | 0 | 21 | 3 | [illegible] |
| SE | 73823 | ENSG00000043462 | LCP2 | chr5 | - | 170274300 | 170274338 | 170270729 | 170270917 | 170275319 | 170275351 | 579 | 437 | 550 | 550 | 1586 | 0 | 0 | 1 | 1 | 0 | 908 | 681 | [illegible] |
| SE | 73830 | ENSG00000110080 | ST3GAL4 | chr11 | + | 126408306 | 126408496 | 126408098 | 126408194 | 126409267 | 126409411 | 110 | 45 | 57 | 93 | 85 | 0 | 0 | 0 | 0 | 0 | 105 | 85 | [illegible] |
| SE | 73833 | ENSG00000167792 | NDUFV1 | chr11 | + | 67607298 | 67607616 | 67606953 | 67607076 | 67608395 | 67608478 | 1 | 0 | 0 | 0 | 1 | 65 | 35 | 66 | 54 | 30 | 0 | 0 | [illegible] |
| SE | 73834 | ENSG00000167792 | NDUFV1 | chr11 | + | 67607365 | 67607496 | 67606932 | 67607076 | 67608395 | 67608518 | 3 | 0 | 3 | 0 | 0 | 65 | 35 | 66 | 54 | 30 | 0 | 0 | [illegible] |
| SE | 73835 | ENSG00000167792 | NDUFV1 | chr11 | + | 67607365 | 67607496 | 67606932 | 67607076 | 67608399 | 67608478 | 0 | 0 | 3 | 0 | 0 | 10 | 3 | 0 | 1 | 0 | 0 | 0 | [illegible] |
| SE | 73836 | ENSG00000167792 | NDUFV1 | chr11 | + | 67607365 | 67607616 | 67606943 | 67607076 | 67608395 | 67608478 | 0 | 0 | 3 | 0 | 1 | 65 | 35 | 66 | 54 | 30 | 0 | 0 | [illegible] |
| SE | 73841 | ENSG00000167792 | NDUFV1 | chr11 | + | 67610380 | 67610570 | 67609451 | 67609635 | 67610994 | 67611207 | 105 | 44 | 103 | 59 | 67 | 0 | 2 | 0 | 0 | 0 | 65 | 101 | [illegible] |
| SE | 73846 | ENSG00000108465 | CDK5RAP3 | chr17 | + | 47973930 | 47974031 | 47973518 | 47973693 | 47974399 | 47974447 | 166 | 141 | 161 | 129 | 113 | 2 | 0 | 0 | 0 | 0 | 366 | 228 | [illegible] |
| SE | 73847 | ENSG00000108465 | CDK5RAP3 | chr17 | + | 47978834 | 47978917 | 47977831 | 47977910 | 47980592 | 47980798 | 63 | 47 | 62 | 87 | 38 | 0 | 0 | 0 | 0 | 0 | 70 | 62 | [illegible] |
| SE | 73852 | ENSG00000132196 | HSD17B7 | chr1 | + | 162795620 | 162795663 | 162792564 | 162792862 | 162796584 | 162796677 | 1 | 0 | 0 | 2 | 0 | 13 | 2 | 10 | 8 | 16 | 0 | 0 | [illegible] |
| SE | 73854 | ENSG00000132196 | HSD17B7 | chr1 | + | 162802831 | 162802949 | 162797827 | 162799937 | 162803430 | 162803535 | 1 | 1 | 7 | 2 | 0 | 26 | 22 | 19 | 13 | 27 | 0 | 0 | [illegible] |
| SE | 73855 | ENSG00000132196 | HSD17B7 | chr1 | + | 162803430 | 162803535 | 162799503 | 162799937 | 162804266 | 162804312 | 44 | 40 | 33 | 28 | 47 | 0 | 0 | 0 | 1 | 0 | 13 | 32 | [illegible] |
| SE | 73857 | ENSG00000063587 | ZNF275 | chrX | + | 153336633 | 153336710 | 153334152 | 153334285 | 153345519 | 153345621 | 10 | 5 | 10 | 8 | 0 | 1 | 0 | 0 | 0 | 0 | 12 | 4 | [illegible] |
| SE | 73860 | ENSG00000102547 | CAB39L | chr13 | - | 49401132 | 49401270 | 49382506 | 49382941 | 49433317 | 49433393 | 0 | 1 | 0 | 2 | 4 | 27 | 17 | 16 | 20 | 31 | 0 | 0 | [illegible] |
| SE | 73861 | ENSG00000102547 | CAB39L | chr13 | - | 49412911 | 49413040 | 49382799 | 49382941 | 49433317 | 49433393 | 0 | 1 | 0 | 0 | 0 | 27 | 17 | 16 | 20 | 31 | 0 | 0 | [illegible] |
| SE | 73862 | ENSG00000164609 | SLU7 | chr5 | - | 160417111 | 160417241 | 160415124 | 160415310 | 160418595 | 160419081 | 8 | 5 | 0 | 4 | 29 | 10 | 8 | 27 | 14 | 40 | 0 | 5 | [illegible] |
| SE | 73866 | ENSG00000237499 | WAKMAR2 | chr6 | - | 137860981 | 137861011 | 137857726 | 137857966 | 137862815 | 137862925 | 3 | 0 | 2 | 1 | 0 | 1 | 0 | 0 | 0 | 0 | 0 | 4 | [illegible] |
| SE | 73868 | ENSG00000237499 | WAKMAR2 | chr6 | - | 137862815 | 137862925 | 137857726 | 137857966 | 137865158 | 137865372 | 1 | 0 | 2 | 0 | 0 | 0 | 0 | 2 | 0 | 0 | 0 | 5 | [illegible] |
| SE | 73870 | ENSG00000237499 | WAKMAR2 | chr6 | - | 137862815 | 137862925 | 137860981 | 137861011 | 137865158 | 137865372 | 2 | 1 | 3 | 1 | 0 | 0 | 4 | 0 | 0 | 4 | 0 | 10 | [illegible] |
| SE | 73871 | ENSG00000237499 | WAKMAR2 | chr6 | - | 137865158 | 137865372 | 137862815 | 137862925 | 137866582 | 137866884 | 0 | 1 | 2 | 1 | 0 | 0 | 0 | 1 | 0 | 0 | 0 | 8 | [illegible] |
| SE | 73872 | ENSG00000237499 | WAKMAR2 | chr6 | - | 137866582 | 137866884 | 137862815 | 137862925 | 137867641 | 137867729 | 3 | 0 | 1 | 0 | 3 | 0 | 0 | 0 | 0 | 0 | 2 | 0 | [illegible] |
| SE | 73873 | ENSG00000243244 | STON1 | chr2 | + | 48530712 | 48530829 | 48530185 | 48530216 | 48580586 | 48582563 | 0 | 2 | 0 | 5 | 0 | 5 | 4 | 4 | 2 | 1 | 0 | 0 | [illegible] |
| SE | 73878 | ENSG00000123600 | METTL8 | chr2 | - | 171325840 | 171325906 | 171315745 | 171324362 | 171337452 | 171337502 | 22 | 14 | 12 | 26 | 24 | 2 | 0 | 0 | 1 | 0 | 0 | 4 | [illegible] |
| SE | 73879 | ENSG00000123600 | METTL8 | chr2 | - | 171326041 | 171326148 | 171325840 | 171325906 | 171330558 | 171330616 | 85 | 73 | 90 | 97 | 114 | 2 | 0 | 0 | 4 | 0 | 162 | 89 | [illegible] |
| SE | 73880 | ENSG00000123600 | METTL8 | chr2 | - | 171326041 | 171326148 | 171325840 | 171325906 | 171337452 | 171337502 | 28 | 29 | 24 | 28 | 76 | 1 | 1 | 0 | 0 | 0 | 6 | 34 | [illegible] |
| SE | 73881 | ENSG00000123600 | METTL8 | chr2 | - | 171326041 | 171326148 | 171325840 | 171325906 | 171339183 | 171339256 | 28 | 26 | 23 | 30 | 72 | 0 | 3 | 1 | 0 | 0 | 18 | 26 | [illegible] |
| SE | 73886 | ENSG00000123600 | METTL8 | chr2 | - | 171330558 | 171330698 | 171325840 | 171325906 | 171337452 | 171337502 | 2 | 0 | 0 | 1 | 0 | 1 | 1 | 0 | 0 | 0 | 0 | 2 | [illegible] |
| SE | 73887 | ENSG00000123600 | METTL8 | chr2 | - | 171330558 | 171330698 | 171326041 | 171326148 | 171331764 | 171331867 | 55 | 51 | 72 | 83 | 43 | 0 | 0 | 0 | 1 | 0 | 134 | 79 | [illegible] |
| SE | 73888 | ENSG00000123600 | METTL8 | chr2 | - | 171330558 | 171330698 | 171326041 | 171326148 | 171331803 | 171331867 | 85 | 82 | 101 | 97 | 59 | 4 | 0 | 0 | 0 | 2 | 175 | 88 | [illegible] |
| SE | 73891 | ENSG00000123600 | METTL8 | chr2 | - | 171331764 | 171331867 | 171330558 | 171330616 | 171337452 | 171337502 | 12 | 11 | 10 | 14 | 4 | 0 | 1 | 0 | 0 | 0 | 19 | 25 | [illegible] |
| SE | 73892 | ENSG00000123600 | METTL8 | chr2 | - | 171331764 | 171331867 | 171330558 | 171330698 | 171337452 | 171337502 | 11 | 12 | 15 | 21 | 4 | 0 | 0 | 0 | 0 | 0 | 19 | 31 | [illegible] |
| SE | 73894 | ENSG00000123600 | METTL8 | chr2 | - | 171331803 | 171331867 | 171330558 | 171330616 | 171337452 | 171337502 | 18 | 13 | 21 | 17 | 4 | 0 | 1 | 0 | 0 | 0 | 49 | 28 | [illegible] |
| SE | 73895 | ENSG00000123600 | METTL8 | chr2 | - | 171331803 | 171331867 | 171330558 | 171330698 | 171337452 | 171337502 | 39 | 39 | 44 | 35 | 20 | 0 | 0 | 0 | 0 | 0 | 60 | 43 | [illegible] |
| SE | 73896 | ENSG00000123600 | METTL8 | chr2 | - | 171331803 | 171331867 | 171330558 | 171330698 | 171360421 | 171360513 | 10 | 2 | 10 | 12 | 6 | 0 | 2 | 0 | 0 | 0 | 11 | 9 | [illegible] |
| SE | 73899 | ENSG00000123600 | METTL8 | chr2 | - | 171337452 | 171337502 | 171330558 | 171330616 | 171339183 | 171339213 | 13 | 6 | 7 | 10 | 4 | 3 | 0 | 0 | 0 | 0 | 18 | 22 | [illegible] |
| SE | 73902 | ENSG00000123600 | METTL8 | chr2 | - | 171339183 | 171339554 | 171330624 | 171330698 | 171360421 | 171360513 | 46 | 27 | 28 | 11 | 34 | 0 | 2 | 0 | 0 | 0 | 23 | 13 | [illegible] |
| SE | 73903 | ENSG00000123600 | METTL8 | chr2 | - | 171339183 | 171339554 | 171331807 | 171331867 | 171360421 | 171360513 | 48 | 33 | 28 | 12 | 34 | 1 | 0 | 0 | 1 | 0 | 23 | 13 | [illegible] |
| SE | 73905 | ENSG00000123600 | METTL8 | chr2 | - | 171360421 | 171360513 | 171339183 | 171339554 | 171414602 | 171414726 | 29 | 15 | 15 | 4 | 23 | 0 | 0 | 0 | 0 | 0 | 16 | 10 | [illegible] |
| SE | 73906 | ENSG00000123600 | METTL8 | chr2 | - | 171392042 | 171392197 | 171339183 | 171339554 | 171414602 | 171414726 | 3 | 2 | 3 | 8 | 4 | 0 | 0 | 0 | 0 | 0 | 0 | 0 | [illegible] |
| SE | 73907 | ENSG00000123600 | METTL8 | chr2 | - | 171392042 | 171392197 | 171360421 | 171360513 | 171414602 | 171414726 | 29 | 25 | 22 | 21 | 26 | 1 | 0 | 1 | 1 | 0 | 7 | 9 | [illegible] |
| SE | 73908 | ENSG00000123600 | METTL8 | chr2 | - | 171392042 | 171392197 | 171360421 | 171360513 | 171433882 | 171433990 | 32 | 24 | 21 | 23 | 25 | 1 | 0 | 0 | 0 | 0 | 7 | 9 | [illegible] |
| SE | 73910 | ENSG00000123600 | METTL8 | chr2 | - | 171414602 | 171414726 | 171360421 | 171360513 | 171433882 | 171434087 | 1 | 1 | 1 | 2 | 0 | 1 | 0 | 0 | 0 | 0 | 12 | 2 | [illegible] |
| SE | 73912 | ENSG00000123600 | METTL8 | chr2 | - | 171414602 | 171414726 | 171392042 | 171392197 | 171433882 | 171434087 | 3 | 3 | 3 | 5 | 4 | 6 | 1 | 2 | 6 | 3 | 6 | 2 | [illegible] |
| SE | 73913 | ENSG00000123600 | METTL8 | chr2 | - | 171414602 | 171414726 | 171392042 | 171392197 | 171434673 | 171434780 | 3 | 2 | 3 | 4 | 4 | 2 | 1 | 2 | 0 | 5 | 0 | 1 | [illegible] |
| SE | 73914 | ENSG00000123600 | METTL8 | chr2 | - | 171433025 | 171433219 | 171392042 | 171392197 | 171433882 | 171433990 | 0 | 0 | 0 | 0 | 0 | 6 | 1 | 2 | 6 | 3 | 0 | 2 | [illegible] |
| SE | 73919 | ENSG00000161692 | DBF4B | chr17 | + | 44741335 | 44741452 | 44738378 | 44738424 | 44747082 | 44747191 | 5 | 6 | 3 | 3 | 12 | 0 | 0 | 1 | 0 | 0 | 3 | 6 | [illegible] |
| SE | 73920 | ENSG00000161692 | DBF4B | chr17 | + | 44748340 | 44748465 | 44747390 | 44747515 | 44750594 | 44750677 | 2 | 2 | 0 | 7 | 0 | 3 | 0 | 1 | 1 | 0 | 0 | 9 | [illegible] |
| SE | 73925 | ENSG00000155506 | LARP1 | chr5 | + | 154795174 | 154795319 | 154794045 | 154794262 | 154799590 | 154799759 | 467 | 277 | 517 | 519 | 533 | 0 | 0 | 0 | 0 | 3 | 592 | 582 | [illegible] |
| SE | 73926 | ENSG00000155506 | LARP1 | chr5 | + | 154802006 | 154802399 | 154799872 | 154800042 | 154803289 | 154803413 | 585 | 397 | 588 | 549 | 948 | 2 | 0 | 0 | 1 | 0 | 962 | 847 | [illegible] |
| SE | 73928 | ENSG00000155506 | LARP1 | chr5 | + | 154805880 | 154806032 | 154803539 | 154803745 | 154808458 | 154808603 | 239 | 124 | 266 | 285 | 192 | 0 | 1 | 0 | 0 | 0 | 322 | 319 | [illegible] |
| SE | 73929 | ENSG00000155506 | LARP1 | chr5 | + | 154811246 | 154811356 | 154808458 | 154808603 | 154811512 | 154811640 | 367 | 284 | 460 | 421 | 499 | 0 | 0 | 0 | 0 | 0 | 652 | 427 | [illegible] |
| SE | 73931 | ENSG00000169967 | MAP3K2 | chr2 | - | 127318168 | 127318317 | 127317628 | 127317760 | 127322045 | 127322252 | 403 | 320 | 283 | 396 | 330 | 0 | 0 | 0 | 1 | 0 | 50 | 324 | [illegible] |
| SE | 73933 | ENSG00000169967 | MAP3K2 | chr2 | - | 127330391 | 127330505 | 127329920 | 127330008 | 127335869 | 127335969 | 354 | 367 | 332 | 344 | 770 | 0 | 0 | 0 | 2 | 0 | 145 | 221 | [illegible] |
| SE | 73934 | ENSG00000120896 | SORBS3 | chr8 | + | 22565825 | 22565872 | 22565265 | 22565354 | 22566344 | 22566484 | 20 | 14 | 9 | 10 | 12 | 0 | 0 | 2 | 0 | 0 | 7 | 8 | [illegible] |
| SE | 73939 | ENSG00000120896 | SORBS3 | chr8 | + | 22572339 | 22572446 | 22571691 | 22571821 | 22574666 | 22574724 | 32 | 23 | 33 | 22 | 4 | 1 | 0 | 0 | 0 | 0 | 2 | 6 | [illegible] |
| SE | 73940 | ENSG00000144134 | RABL2A | chr2 | + | 113627855 | 113628010 | 113627310 | 113627400 | 113628549 | 113628713 | 4 | 0 | 0 | 0 | 0 | 0 | 0 | 2 | 0 | 0 | 0 | 4 | [illegible] |
| SE | 73943 | ENSG00000144134 | RABL2A | chr2 | + | 113635050 | 113635130 | 113634152 | 113634232 | 113641355 | 113641450 | 15 | 1 | 3 | 1 | 9 | 0 | 0 | 0 | 1 | 0 | 5 | 17 | [illegible] |
| SE | 73944 | ENSG00000144134 | RABL2A | chr2 | + | 113640893 | 113641005 | 113634152 | 113634232 | 113641352 | 113641450 | 4 | 0 | 3 | 2 | 1 | 5 | 3 | 6 | 16 | 6 | 0 | 6 | [illegible] |
| SE | 73947 | ENSG00000144134 | RABL2A | chr2 | + | 113640893 | 113641005 | 113635050 | 113635130 | 113641355 | 113641450 | 12 | 3 | 3 | 7 | 12 | 0 | 0 | 0 | 0 | 2 | 16 | 21 | [illegible] |
| SE | 73948 | ENSG00000144134 | RABL2A | chr2 | + | 113640893 | 113641005 | 113637366 | 113637486 | 113641352 | 113641450 | 4 | 0 | 1 | 0 | 1 | 0 | 0 | 0 | 2 | 0 | 0 | 0 | [illegible] |
| SE | 73949 | ENSG00000144134 | RABL2A | chr2 | + | 113641352 | 113641450 | 113634152 | 113634232 | 113641780 | 113641864 | 8 | 7 | 14 | 22 | 20 | 0 | 0 | 2 | 0 | 0 | 2 | 2 | [illegible] |
| SE | 73950 | ENSG00000144134 | RABL2A | chr2 | + | 113641355 | 113641450 | 113634152 | 113634232 | 113641780 | 113641864 | 4 | 3 | 8 | 7 | 12 | 0 | 0 | 2 | 0 | 0 | 2 | 0 | [illegible] |
| SE | 73956 | ENSG00000166436 | TRIM66 | chr11 | - | 8618721 | 8618968 | 8617763 | 8618003 | 8619382 | 8619535 | 15 | 8 | 16 | 17 | 11 | 0 | 0 | 0 | 0 | 0 | 5 | 30 | [illegible] |
| SE | 73957 | ENSG00000166436 | TRIM66 | chr11 | - | 8618749 | 8618968 | 8612036 | 8618003 | 8619382 | 8619535 | 23 | 12 | 26 | 26 | 17 | 0 | 0 | 0 | 0 | 0 | 51 | 42 | [illegible] |
| SE | 73959 | ENSG00000166436 | TRIM66 | chr11 | - | 8620049 | 8620124 | 8619382 | 8619535 | 8621644 | 8621819 | 8 | 0 | 6 | 4 | 0 | 2 | 3 | 0 | 0 | 0 | 0 | 0 | [illegible] |
| SE | 73961 | ENSG00000166436 | TRIM66 | chr11 | - | 8620445 | 8620572 | 8619382 | 8619535 | 8621644 | 8621819 | 2 | 1 | 0 | 0 | 2 | 2 | 3 | 0 | 0 | 0 | 0 | 3 | [illegible] |
| SE | 73966 | ENSG00000166436 | TRIM66 | chr11 | - | 8621031 | 8621321 | 8620445 | 8620572 | 8621644 | 8621819 | 21 | 27 | 31 | 56 | 43 | 0 | 0 | 0 | 0 | 0 | 28 | 9 | [illegible] |
| SE | 73967 | ENSG00000166436 | TRIM66 | chr11 | - | 8643008 | 8643126 | 8640226 | 8641152 | 8646446 | 8646561 | 2 | 3 | 4 | 8 | 6 | 0 | 1 | 0 | 0 | 0 | 0 | 13 | [illegible] |
| SE | 73971 | ENSG00000166436 | TRIM66 | chr11 | - | 8654433 | 8654578 | 8651799 | 8651903 | 8663128 | 8663248 | 12 | 0 | 5 | 6 | 6 | 0 | 0 | 0 | 1 | 0 | 0 | 0 | [illegible] |
| SE | 73972 | ENSG00000166436 | TRIM66 | chr11 | - | 8663128 | 8663248 | 8651799 | 8651903 | 8671785 | 8671866 | 0 | 0 | 0 | 1 | 0 | 0 | 0 | 0 | 0 | 0 | 0 | 0 | [illegible] |
| SE | 73973 | ENSG00000166436 | TRIM66 | chr11 | - | 8674805 | 8674883 | 8672247 | 8672385 | 8679619 | 8679814 | 12 | 10 | 15 | 22 | 34 | 3 | 0 | 0 | 0 | 0 | 23 | 26 | [illegible] |
| SE | 73980 | ENSG00000259692 | NA | chr15 | - | 81915346 | 81915457 | 81818324 | 81818379 | 81925672 | 81925743 | 0 | 0 | 0 | 0 | 1 | 0 | 1 | 0 | 0 | 0 | 14 | 0 | [illegible] |
| SE | 73988 | ENSG00000072163 | LIMS2 | chr2 | - | 127640269 | 127640318 | 127640069 | 127640145 | 127640895 | 127640988 | 1 | 2 | 7 | 22 | 6 | 1 | 0 | 0 | 6 | 1 | 0 | 6 | [illegible] |
| SE | 73998 | ENSG00000096433 | ITPR3 | chr6 | + | 33677503 | 33677629 | 33677014 | 33677089 | 33678420 | 33678543 | 168 | 58 | 163 | 127 | 138 | 1 | 0 | 0 | 0 | 0 | 94 | 189 | [illegible] |
| SE | 73999 | ENSG00000096433 | ITPR3 | chr6 | + | 33686408 | 33686519 | 33686052 | 33686253 | 33687008 | 33687104 | 96 | 31 | 100 | 72 | 31 | 0 | 0 | 0 | 1 | 0 | 52 | 74 | [illegible] |
| SE | 74002 | ENSG00000228242 | XPC-AS1 | chr3 | + | 14147954 | 14148701 | 14145369 | 14145683 | 14149841 | 14149997 | 0 | 3 | 0 | 1 | 0 | 0 | 0 | 0 | 0 | 0 | 0 | 0 | [illegible] |
| SE | 74003 | ENSG00000228242 | XPC-AS1 | chr3 | + | 14149506 | 14149602 | 14148596 | 14148701 | 14149841 | 14149997 | 0 | 1 | 1 | 0 | 3 | 0 | 2 | 0 | 0 | 0 | 0 | 0 | [illegible] |
| SE | 74011 | ENSG00000206559 | ZCWPW2 | chr3 | + | 28478813 | 28478931 | 28435109 | 28435269 | 28492126 | 28492173 | 4 | 0 | 4 | 1 | 5 | 0 | 1 | 0 | 0 | 1 | 0 | 0 | [illegible] |
| SE | 74033 | ENSG00000284644 | NA | chr7 | - | 144252295 | 144252492 | 144251407 | 144251610 | 144254433 | 144254696 | 0 | 0 | 1 | 0 | 4 | 0 | 0 | 0 | 0 | 1 | 0 | 0 | [illegible] |
| SE | 74035 | ENSG00000284644 | NA | chr7 | - | 144311590 | 144311704 | 144251324 | 144251610 | 144343186 | 144343298 | 0 | 2 | 0 | 0 | 0 | 0 | 0 | 0 | 0 | 0 | 0 | 0 | [illegible] |
| SE | 74036 | ENSG00000244479 | OR2A1-AS1 | chr7 | - | 144311883 | 144311936 | 144311590 | 144311704 | 144343186 | 144343298 | 0 | 1 | 0 | 0 | 0 | 0 | 0 | 0 | 0 | 0 | 0 | 0 | [illegible] |
| SE | 74037 | ENSG00000284644 | NA | chr7 | - | 144343186 | 144343298 | 144251386 | 144251610 | 144355337 | 144355529 | 0 | 0 | 0 | 0 | 0 | 0 | 0 | 0 | 5 | 0 | 0 | 3 | [illegible] |
| SE | 74042 | ENSG00000162928 | PEX13 | chr2 | + | 61045725 | 61045851 | 61031418 | 61032113 | 61048471 | 61051990 | 183 | 132 | 127 | 142 | 265 | 2 | 0 | 0 | 0 | 0 | 65 | 150 | [illegible] |
| SE | 74043 | ENSG00000187951 | NA | chr15 | + | 30748575 | 30748670 | 30716219 | 30716342 | 30750401 | 30750556 | 4 | 1 | 6 | 2 | 4 | 1 | 0 | 0 | 0 | 0 | 12 | 7 | [illegible] |
| SE | 74044 | ENSG00000187951 | NA | chr15 | + | 30750401 | 30750556 | 30716219 | 30716342 | 30767857 | 30768126 | 10 | 2 | 6 | 1 | 2 | 0 | 0 | 1 | 0 | 0 | 15 | 1 | [illegible] |
| SE | 74046 | ENSG00000111011 | RSRC2 | chr12 | - | 122511108 | 122511188 | 122508217 | 122508447 | 122515104 | 122515227 | 240 | 198 | 178 | 323 | 205 | 0 | 0 | 0 | 1 | 0 | 38 | 196 | [illegible] |
| SE | 74047 | ENSG00000111011 | RSRC2 | chr12 | - | 122511108 | 122511188 | 122508217 | 122508447 | 122517226 | 122517430 | 143 | 117 | 110 | 199 | 130 | 0 | 0 | 1 | 0 | 0 | 23 | 121 | [illegible] |
| SE | 74048 | ENSG00000111011 | RSRC2 | chr12 | - | 122511108 | 122511188 | 122508217 | 122508450 | 122515104 | 122515227 | 209 | 196 | 170 | 174 | 201 | 0 | 0 | 0 | 1 | 0 | 55 | 134 | [illegible] |
| SE | 74049 | ENSG00000111011 | RSRC2 | chr12 | - | 122511108 | 122511188 | 122508217 | 122508450 | 122517226 | 122517430 | 122 | 127 | 101 | 96 | 123 | 0 | 0 | 2 | 0 | 0 | 38 | 87 | [illegible] |
| SE | 74053 | ENSG00000111011 | RSRC2 | chr12 | - | 122514711 | 122514728 | 122513766 | 122513874 | 122515104 | 122515227 | 0 | 2 | 3 | 2 | 0 | 2 | 2 | 1 | 13 | 10 | 0 | 2 | [illegible] |
| SE | 74054 | ENSG00000111011 | RSRC2 | chr12 | - | 122515104 | 122515227 | 122508217 | 122508447 | 122517226 | 122517430 | 17 | 33 | 32 | 45 | 27 | 0 | 0 | 1 | 0 | 0 | 11 | 44 | [illegible] |
| SE | 74055 | ENSG00000111011 | RSRC2 | chr12 | - | 122515104 | 122515227 | 122508217 | 122508450 | 122517226 | 122517430 | 17 | 33 | 32 | 45 | 27 | 0 | 0 | 2 | 0 | 0 | 11 | 44 | [illegible] |
| SE | 74056 | ENSG00000111011 | RSRC2 | chr12 | - | 122515104 | 122515227 | 122511108 | 122511188 | 122517226 | 122517430 | 180 | 160 | 155 | 210 | 156 | 1 | 0 | 2 | 0 | 0 | 39 | 138 | [illegible] |
| SE | 74059 | ENSG00000111011 | RSRC2 | chr12 | - | 122520503 | 122520581 | 122518838 | 122519051 | 122521384 | 122521428 | 35 | 32 | 38 | 55 | 64 | 0 | 0 | 0 | 1 | 2 | 18 | 42 | [illegible] |
| SE | 74060 | ENSG00000111011 | RSRC2 | chr12 | - | 122521384 | 122521428 | 122518838 | 122519029 | 122526847 | 122526960 | 33 | 20 | 17 | 22 | 36 | 0 | 2 | 0 | 0 | 0 | 0 | 16 | [illegible] |
| SE | 74061 | ENSG00000111011 | RSRC2 | chr12 | - | 122522142 | 122522299 | 122518838 | 122521428 | 122526847 | 122526936 | 127 | 86 | 84 | 126 | 126 | 4 | 0 | 0 | 1 | 0 | 50 | 161 | [illegible] |
| SE | 74062 | ENSG00000111011 | RSRC2 | chr12 | - | 122522142 | 122522329 | 122521384 | 122521428 | 122526847 | 122526949 | 105 | 79 | 63 | 110 | 118 | 4 | 0 | 0 | 1 | 0 | 50 | 135 | [illegible] |
| SE | 74064 | ENSG00000111011 | RSRC2 | chr12 | - | 122526169 | 122526293 | 122522798 | 122522906 | 122526847 | 122526940 | 0 | 2 | 1 | 1 | 0 | 9 | 0 | 0 | 15 | 2 | 0 | 6 | [illegible] |
| SE | 74066 | ENSG00000205268 | PDE7A | chr8 | - | 65723540 | 65723621 | 65714333 | 65719495 | 65724254 | 65724351 | 764 | 647 | 1085 | 879 | 1302 | 0 | 0 | 1 | 0 | 0 | 492 | 575 | [illegible] |
| SE | 74067 | ENSG00000205268 | PDE7A | chr8 | - | 65724254 | 65724351 | 65721797 | 65723621 | 65724776 | 65724921 | 982 | 926 | 1383 | 1292 | 1821 | 0 | 0 | 1 | 0 | 0 | 619 | 822 | [illegible] |
| SE | 74069 | ENSG00000205268 | PDE7A | chr8 | - | 65731560 | 65731631 | 65727169 | 65727301 | 65734793 | 65734894 | 1 | 2 | 4 | 1 | 0 | 121 | 77 | 203 | 163 | 511 | 0 | 5 | [illegible] |
| SE | 74071 | ENSG00000205268 | PDE7A | chr8 | - | 65739501 | 65739597 | 65734793 | 65734894 | 65747651 | 65747803 | 205 | 107 | 276 | 220 | 332 | 1 | 2 | 0 | 0 | 3 | 45 | 124 | [illegible] |
| SE | 74072 | ENSG00000205268 | PDE7A | chr8 | - | 65745406 | 65745470 | 65739501 | 65739597 | 65747651 | 65747803 | 297 | 178 | 407 | 323 | 755 | 0 | 0 | 0 | 0 | 0 | 118 | 183 | [illegible] |
| SE | 74074 | ENSG00000205268 | PDE7A | chr8 | - | 65788881 | 65788981 | 65782782 | 65782843 | 65838473 | 65838748 | 12 | 11 | 10 | 13 | 1 | 20 | 13 | 46 | 7 | 42 | 0 | 2 | [illegible] |
| SE | 74076 | ENSG00000205268 | PDE7A | chr8 | - | 65841370 | 65841467 | 65782782 | 65782843 | 65842081 | 65842322 | 50 | 13 | 34 | 33 | 8 | 1 | 0 | 0 | 0 | 0 | 31 | 36 | [illegible] |

| | | | | | | | | | | | | | | |
|---|---|---|---|---|---|---|---|---|---|---|---|---|---|---|
| SE | 74077 | ENSG00000085721 | RRN3 | chr16 | - | 15071120 | 15071251 | 15070069 | 15070254 | 15074722 | 15074861 | 112 | 42 | [illegible] |
| SE | 74078 | ENSG00000085721 | RRN3 | chr16 | - | 15072949 | 15073080 | 15071120 | 15071251 | 15074722 | 15074861 | 167 | 70 | [illegible] |
| SE | 74079 | ENSG00000085721 | RRN3 | chr16 | - | 15076557 | 15076650 | 15074676 | 15074861 | 15079997 | 15080096 | 131 | 87 | [illegible] |
| SE | 74081 | ENSG00000085721 | RRN3 | chr16 | - | 15084641 | 15084705 | 15079997 | 15080096 | 15085638 | 15085698 | 54 | 22 | [illegible] |
| SE | 74104 | ENSG00000255587 | RAB44 | chr6 | + | 36718027 | 36718118 | 36717272 | 36717419 | 36718492 | 36718588 | 0 | 8 | [illegible] |
| SE | 74105 | ENSG00000255587 | RAB44 | chr6 | + | 36725861 | 36725943 | 36721150 | 36722733 | 36727576 | 36727691 | 29 | 6 | [illegible] |
| SE | 74106 | ENSG00000255587 | RAB44 | chr6 | + | 36725861 | 36725943 | 36721150 | 36722733 | 36728699 | 36728801 | 13 | 1 | [illegible] |
| SE | 74109 | ENSG00000093167 | LRRFIP2 | chr3 | - | 37055085 | 37055165 | 37054410 | 37054515 | 37058789 | 37058910 | 197 | 189 | [illegible] |
| SE | 74111 | ENSG00000093167 | LRRFIP2 | chr3 | - | 37066223 | 37066325 | 37065809 | 37065942 | 37072789 | 37072882 | 0 | 0 | [illegible] |
| SE | 74114 | ENSG00000093167 | LRRFIP2 | chr3 | - | 37072789 | 37072882 | 37066223 | 37066325 | 37075023 | 37075116 | 0 | 0 | [illegible] |
| SE | 74116 | ENSG00000093167 | LRRFIP2 | chr3 | - | 37108636 | 37108684 | 37096615 | 37096660 | 37110990 | 37111065 | 0 | 0 | [illegible] |
| SE | 74117 | ENSG00000093167 | LRRFIP2 | chr3 | - | 37108636 | 37108684 | 37096615 | 37096660 | 37112914 | 37112980 | 0 | 0 | [illegible] |
| SE | 74120 | ENSG00000093167 | LRRFIP2 | chr3 | - | 37110990 | 37111065 | 37096615 | 37096660 | 37112914 | 37112980 | 0 | 1 | [illegible] |
| SE | 74123 | ENSG00000093167 | LRRFIP2 | chr3 | - | 37110990 | 37111065 | 37108636 | 37108684 | 37121491 | 37121536 | 0 | 0 | [illegible] |
| SE | 74125 | ENSG00000093167 | LRRFIP2 | chr3 | - | 37112914 | 37112980 | 37096615 | 37096660 | 37129062 | 37129149 | 0 | 0 | [illegible] |
| SE | 74126 | ENSG00000093167 | LRRFIP2 | chr3 | - | 37112914 | 37112980 | 37108636 | 37108684 | 37121491 | 37121536 | 0 | 0 | [illegible] |
| SE | 74127 | ENSG00000093167 | LRRFIP2 | chr3 | - | 37112914 | 37112980 | 37108636 | 37108684 | 37129062 | 37129149 | 0 | 0 | [illegible] |
| SE | 74128 | ENSG00000093167 | LRRFIP2 | chr3 | - | 37112914 | 37112980 | 37110990 | 37111065 | 37121491 | 37121536 | 0 | 2 | [illegible] |
| SE | 74129 | ENSG00000093167 | LRRFIP2 | chr3 | - | 37115053 | 37115095 | 37112914 | 37112980 | 37121491 | 37121536 | 0 | 0 | [illegible] |
| SE | 74131 | ENSG00000093167 | LRRFIP2 | chr3 | - | 37127629 | 37127680 | 37121634 | 37121691 | 37129062 | 37129149 | 1 | 0 | [illegible] |
| SE | 74132 | ENSG00000093167 | LRRFIP2 | chr3 | - | 37129062 | 37129149 | 37096615 | 37096660 | 37148893 | 37149038 | 225 | 229 | [illegible] |
| SE | 74134 | ENSG00000111679 | PTPN6 | chr12 | + | 6951458 | 6951520 | 6951261 | 6951291 | 6951608 | 6951731 | 63 | 28 | [illegible] |
| SE | 74136 | ENSG00000111679 | PTPN6 | chr12 | + | 6951504 | 6951731 | 6946709 | 6946739 | 6951982 | 6952177 | 215 | 102 | [illegible] |
| SE | 74137 | ENSG00000111679 | PTPN6 | chr12 | + | 6951608 | 6951731 | 6946573 | 6946739 | 6951911 | 6952177 | 8 | 0 | [illegible] |
| SE | 74138 | ENSG00000111679 | PTPN6 | chr12 | + | 6951608 | 6951731 | 6946573 | 6946739 | 6951982 | 6952177 | 212 | 99 | [illegible] |
| SE | 74141 | ENSG00000111679 | PTPN6 | chr12 | + | 6951982 | 6952177 | 6951440 | 6951731 | 6954804 | 6954853 | 1001 | 523 | [illegible] |
| SE | 74144 | ENSG00000111679 | PTPN6 | chr12 | + | 6954804 | 6954994 | 6951608 | 6952177 | 6955371 | 6955394 | 753 | 403 | [illegible] |
| SE | 74148 | ENSG00000111679 | PTPN6 | chr12 | + | 6960343 | 6960435 | 6960087 | 6960239 | 6960805 | 6960945 | 1450 | 968 | [illegible] |
| SE | 74151 | ENSG00000168785 | TSPAN5 | chr4 | - | 98478684 | 98478746 | 98476412 | 98476460 | 98482004 | 98482175 | 10 | 5 | [illegible] |
| SE | 74152 | ENSG00000168785 | TSPAN5 | chr4 | - | 98478684 | 98478810 | 98476412 | 98476460 | 98482004 | 98482175 | 57 | 21 | [illegible] |
| SE | 74153 | ENSG00000168785 | TSPAN5 | chr4 | - | 98482004 | 98482175 | 98478684 | 98478746 | 98507677 | 98507728 | 34 | 14 | [illegible] |
| SE | 74157 | ENSG00000091317 | CMTM6 | chr3 | - | 32487937 | 32488036 | 32481311 | 32484097 | 32491709 | 32491886 | 818 | 623 | [illegible] |
| SE | 74158 | ENSG00000126264 | HCST | chr19 | + | 35903771 | 35903903 | 35903350 | 35903416 | 35904119 | 35904377 | 620 | 151 | [illegible] |
| SE | 74164 | ENSG00000285906 | NA | chr3 | - | 11207241 | 11207398 | 11194992 | 11195204 | 11225699 | 11225877 | 0 | 0 | [illegible] |
| SE | 74165 | ENSG00000113658 | SMAD5 | chr5 | + | 136147831 | 136147906 | 136132844 | 136132962 | 136153591 | 136153763 | 31 | 15 | [illegible] |
| SE | 74167 | ENSG00000113658 | SMAD5 | chr5 | + | 136152674 | 136152758 | 136147831 | 136147906 | 136153591 | 136153729 | 0 | 5 | [illegible] |
| SE | 74176 | ENSG00000116698 | SMG7 | chr1 | + | 183537144 | 183537215 | 183533675 | 183533832 | 183538379 | 183538440 | 206 | 140 | [illegible] |
| SE | 74180 | ENSG00000116698 | SMG7 | chr1 | + | 183545965 | 183546337 | 183544929 | 183545312 | 183549207 | 183549288 | 180 | 117 | [illegible] |
| SE | 74181 | ENSG00000116698 | SMG7 | chr1 | + | 183545965 | 183546337 | 183544929 | 183545312 | 183549763 | 183549923 | 96 | 62 | [illegible] |
| SE | 74183 | ENSG00000116698 | SMG7 | chr1 | + | 183549207 | 183549288 | 183544929 | 183545312 | 183549763 | 183549923 | 70 | 53 | [illegible] |
| SE | 74185 | ENSG00000116698 | SMG7 | chr1 | + | 183549207 | 183549288 | 183545965 | 183546337 | 183550750 | 183550921 | 47 | 29 | [illegible] |
| SE | 74186 | ENSG00000116698 | SMG7 | chr1 | + | 183549763 | 183549923 | 183545965 | 183546337 | 183550750 | 183550921 | 103 | 90 | [illegible] |
| SE | 74187 | ENSG00000116698 | SMG7 | chr1 | + | 183549763 | 183549923 | 183549207 | 183549288 | 183550750 | 183550921 | 220 | 174 | [illegible] |
| SE | 74188 | ENSG00000116698 | SMG7 | chr1 | + | 183551044 | 183551190 | 183550750 | 183550921 | 183551817 | 183552263 | 331 | 222 | [illegible] |
| SE | 74197 | ENSG00000235257 | ITGA9-AS1 | chr3 | - | 37790842 | 37790911 | 37753928 | 37754052 | 37821013 | 37821100 | 0 | 0 | [illegible] |
| SE | 74200 | ENSG00000235257 | ITGA9-AS1 | chr3 | - | 37806078 | 37806185 | 37790842 | 37790911 | 37808703 | 37808880 | 0 | 4 | [illegible] |
| SE | 74203 | ENSG00000235257 | ITGA9-AS1 | chr3 | - | 37808703 | 37808880 | 37753928 | 37754052 | 37821013 | 37821095 | 3 | 1 | [illegible] |
| SE | 74205 | ENSG00000235257 | ITGA9-AS1 | chr3 | - | 37808703 | 37808880 | 37790750 | 37790911 | 37818809 | 37818930 | 1 | 4 | [illegible] |
| SE | 74206 | ENSG00000235257 | ITGA9-AS1 | chr3 | - | 37808703 | 37808880 | 37790750 | 37790911 | 37821013 | 37821095 | 4 | 1 | [illegible] |
| SE | 74207 | ENSG00000235257 | ITGA9-AS1 | chr3 | - | 37808703 | 37808880 | 37790750 | 37790911 | 37834678 | 37834699 | 1 | 2 | [illegible] |
| SE | 74208 | ENSG00000235257 | ITGA9-AS1 | chr3 | - | 37808703 | 37808880 | 37794885 | 37796375 | 37821013 | 37821095 | 3 | 1 | [illegible] |
| SE | 74209 | ENSG00000235257 | ITGA9-AS1 | chr3 | - | 37808703 | 37808880 | 37806077 | 37806228 | 37834678 | 37834699 | 2 | 11 | [illegible] |
| SE | 74211 | ENSG00000235257 | ITGA9-AS1 | chr3 | - | 37818809 | 37818930 | 37790842 | 37790911 | 37821013 | 37821100 | 5 | 3 | [illegible] |
| SE | 74214 | ENSG00000235257 | ITGA9-AS1 | chr3 | - | 37821013 | 37821100 | 37790750 | 37790911 | 37834678 | 37834744 | 4 | 3 | [illegible] |
| SE | 74215 | ENSG00000235257 | ITGA9-AS1 | chr3 | - | 37821013 | 37821100 | 37805828 | 37808880 | 37834678 | 37834744 | 6 | 4 | [illegible] |
| SE | 74216 | ENSG00000235257 | ITGA9-AS1 | chr3 | - | 37821013 | 37821100 | 37805828 | 37808880 | 37861700 | 37861764 | 0 | 0 | [illegible] |
| SE | 74217 | ENSG00000235257 | ITGA9-AS1 | chr3 | - | 37834678 | 37834767 | 37808495 | 37808880 | 37861700 | 37861720 | 1 | 2 | [illegible] |
| SE | 74218 | ENSG00000235257 | ITGA9-AS1 | chr3 | - | 37834678 | 37834767 | 37821013 | 37821100 | 37851423 | 37851450 | 5 | 4 | [illegible] |
| SE | 74221 | ENSG00000235257 | ITGA9-AS1 | chr3 | - | 37851423 | 37851492 | 37834678 | 37834767 | 37861700 | 37861732 | 1 | 0 | [illegible] |
| SE | 74222 | ENSG00000067248 | DHX29 | chr5 | - | 55259847 | 55259944 | 55256054 | 55256540 | 55261367 | 55261499 | 81 | 41 | [illegible] |
| SE | 74223 | ENSG00000067248 | DHX29 | chr5 | - | 55261367 | 55261499 | 55259847 | 55259944 | 55267137 | 55267231 | 39 | 27 | [illegible] |
| SE | 74224 | ENSG00000067248 | DHX29 | chr5 | - | 55262629 | 55262932 | 55261367 | 55261499 | 55267137 | 55267231 | 52 | 38 | [illegible] |
| SE | 74225 | ENSG00000067248 | DHX29 | chr5 | - | 55285695 | 55285861 | 55285292 | 55285416 | 55289269 | 55289428 | 132 | 104 | [illegible] |
| SE | 74227 | ENSG00000137478 | FCHSD2 | chr11 | - | 72840876 | 72840959 | 72836744 | 72838874 | 72841453 | 72841583 | 149 | 88 | [illegible] |
| SE | 74228 | ENSG00000137478 | FCHSD2 | chr11 | - | 72840876 | 72840959 | 72836744 | 72838874 | 72842620 | 72842841 | 36 | 19 | [illegible] |
| SE | 74230 | ENSG00000137478 | FCHSD2 | chr11 | - | 72841453 | 72841583 | 72836744 | 72838874 | 72842620 | 72842841 | 116 | 72 | [illegible] |
| SE | 74231 | ENSG00000137478 | FCHSD2 | chr11 | - | 72841453 | 72841583 | 72836744 | 72838874 | 72843150 | 72843328 | 4 | 0 | [illegible] |
| SE | 74233 | ENSG00000137478 | FCHSD2 | chr11 | - | 72841453 | 72841583 | 72840876 | 72840959 | 72843150 | 72843328 | 107 | 67 | [illegible] |
| SE | 74236 | ENSG00000137478 | FCHSD2 | chr11 | - | 72842620 | 72842841 | 72840876 | 72840959 | 72843150 | 72843328 | 183 | 171 | [illegible] |
| SE | 74238 | ENSG00000137478 | FCHSD2 | chr11 | - | 72843150 | 72843328 | 72840876 | 72840959 | 72849754 | 72849889 | 0 | 0 | [illegible] |
| SE | 74239 | ENSG00000137478 | FCHSD2 | chr11 | - | 72843150 | 72843328 | 72841453 | 72841583 | 72849754 | 72849889 | 4 | 0 | [illegible] |
| SE | 74240 | ENSG00000137478 | FCHSD2 | chr11 | - | 72843448 | 72843532 | 72842953 | 72843328 | 72849754 | 72849879 | 259 | 220 | [illegible] |
| SE | 74241 | ENSG00000137478 | FCHSD2 | chr11 | - | 72849754 | 72849879 | 72843448 | 72843532 | 72867864 | 72868026 | 141 | 126 | [illegible] |
| SE | 74242 | ENSG00000137478 | FCHSD2 | chr11 | - | 72849754 | 72849889 | 72843448 | 72843532 | 72867864 | 72868026 | 237 | 203 | [illegible] |
| SE | 74243 | ENSG00000137478 | FCHSD2 | chr11 | - | 72867864 | 72868026 | 72849754 | 72849889 | 72887469 | 72887574 | 215 | 153 | [illegible] |
| SE | 74244 | ENSG00000137478 | FCHSD2 | chr11 | - | 72887469 | 72887574 | 72867864 | 72868026 | 72889828 | 72889945 | 260 | 160 | [illegible] |
| SE | 74245 | ENSG00000137478 | FCHSD2 | chr11 | - | 72889828 | 72889945 | 72887469 | 72887574 | 72902542 | 72902638 | 332 | 185 | [illegible] |
| SE | 74248 | ENSG00000137478 | FCHSD2 | chr11 | - | 73000989 | 73001134 | 72988963 | 72989097 | 73015808 | 73015885 | 483 | 379 | [illegible] |
| SE | 74249 | ENSG00000137478 | FCHSD2 | chr11 | - | 73000989 | 73001134 | 72988963 | 72989097 | 73083694 | 73083740 | 250 | 212 | [illegible] |
| SE | 74250 | ENSG00000137478 | FCHSD2 | chr11 | - | 73015808 | 73015885 | 72988963 | 72989097 | 73083694 | 73083740 | 91 | 84 | [illegible] |
| SE | 74254 | ENSG00000137478 | FCHSD2 | chr11 | - | 73140030 | 73140128 | 73083694 | 73083740 | 73141856 | 73141913 | 193 | 151 | [illegible] |
| SE | 74256 | ENSG00000137959 | IFI44L | chr1 | + | 78620877 | 78620971 | 78620448 | 78620571 | 78628950 | 78628999 | 0 | 0 | [illegible] |
| SE | 74259 | ENSG00000137959 | IFI44L | chr1 | + | 78628950 | 78628999 | 78620381 | 78620571 | 78635336 | 78635489 | 0 | 1 | [illegible] |
| SE | 74261 | ENSG00000137959 | IFI44L | chr1 | + | 78628950 | 78628999 | 78627905 | 78628393 | 78629719 | 78629915 | 209 | 660 | [illegible] |
| SE | 74262 | ENSG00000137959 | IFI44L | chr1 | + | 78628950 | 78628999 | 78627905 | 78628393 | 78635336 | 78635489 | 77 | 271 | [illegible] |
| SE | 74266 | ENSG00000137959 | IFI44L | chr1 | + | 78629719 | 78629915 | 78625669 | 78628999 | 78635341 | 78635489 | 144 | 408 | [illegible] |
| SE | 74267 | ENSG00000137959 | IFI44L | chr1 | + | 78629719 | 78629915 | 78627905 | 78628393 | 78635336 | 78635489 | 193 | 436 | [illegible] |
| SE | 74268 | ENSG00000137959 | IFI44L | chr1 | + | 78635336 | 78635489 | 78628950 | 78628999 | 78637031 | 78637203 | 49 | 129 | [illegible] |
| SE | 74271 | ENSG00000167515 | TRAPPC2L | chr16 | + | 88859344 | 88859443 | 88858618 | 88858789 | 88859572 | 88859750 | 0 | 0 | [illegible] |
| SE | 74272 | ENSG00000167515 | TRAPPC2L | chr16 | + | 88859344 | 88859750 | 88858618 | 88858789 | 88859892 | 88859972 | 58 | 21 | [illegible] |
| SE | 74273 | ENSG00000167515 | TRAPPC2L | chr16 | + | 88859572 | 88859750 | 88858641 | 88858789 | 88859892 | 88859972 | 57 | 21 | [illegible] |
| SE | 74274 | ENSG00000154359 | LONRF1 | chr8 | - | 12725726 | 12725879 | 12721905 | 12723254 | 12728900 | 12729063 | 59 | 46 | [illegible] |
| SE | 74275 | ENSG00000154359 | LONRF1 | chr8 | - | 12731735 | 12731857 | 12729173 | 12729332 | 12735285 | 12735400 | 32 | 41 | [illegible] |
| SE | 74277 | ENSG00000125863 | MKKS | chr20 | - | 10408627 | 10408803 | 10407615 | 10407726 | 10420527 | 10420758 | 52 | 19 | [illegible] |
| SE | 74278 | ENSG00000125863 | MKKS | chr20 | - | 10412529 | 10412700 | 10408627 | 10408803 | 10413795 | 10413931 | 25 | 18 | [illegible] |
| SE | 74281 | ENSG00000125863 | MKKS | chr20 | - | 10428729 | 10428816 | 10420527 | 10420758 | 10434107 | 10434214 | 0 | 1 | [illegible] |
| SE | 74283 | ENSG00000182247 | UBE2E2 | chr3 | + | 23499607 | 23499740 | 23217261 | 23217312 | 23532553 | 23532615 | 104 | 82 | [illegible] |
| SE | 74300 | ENSG00000101187 | SLCO4A1 | chr20 | + | 62668011 | 62668184 | 62667744 | 62667910 | 62668476 | 62668541 | 0 | 0 | [illegible] |
| SE | 74303 | ENSG00000101187 | SLCO4A1 | chr20 | + | 62668476 | 62668541 | 62668011 | 62668184 | 62668929 | 62669078 | 0 | 0 | [illegible] |
| SE | 74304 | ENSG00000196074 | SYCP2 | chr20 | - | 59866292 | 59866392 | 59865806 | 59865865 | 59866494 | 59866589 | 25 | 8 | [illegible] |
| SE | 74305 | ENSG00000196074 | SYCP2 | chr20 | - | 59868412 | 59868568 | 59867710 | 59867847 | 59868834 | 59868925 | 19 | 7 | [illegible] |
| SE | 74307 | ENSG00000196074 | SYCP2 | chr20 | - | 59881436 | 59881492 | 59880965 | 59881023 | 59881944 | 59882002 | 11 | 9 | [illegible] |
| SE | 74308 | ENSG00000196074 | SYCP2 | chr20 | - | 59885102 | 59885144 | 59882094 | 59882165 | 59885927 | 59885964 | 0 | 0 | [illegible] |
| SE | 74309 | ENSG00000196074 | SYCP2 | chr20 | - | 59895436 | 59895597 | 59893523 | 59893593 | 59896428 | 59896528 | 29 | 25 | [illegible] |
| SE | 74312 | ENSG00000114166 | KAT2B | chr3 | + | 20072332 | 20072459 | 20040445 | 20040780 | 20095262 | 20095408 | 260 | 194 | [illegible] |
| SE | 74313 | ENSG00000114166 | KAT2B | chr3 | + | 20095262 | 20095408 | 20072332 | 20072459 | 20099861 | 20099954 | 569 | 519 | [illegible] |
| SE | 74315 | ENSG00000114166 | KAT2B | chr3 | + | 20101286 | 20101468 | 20099861 | 20099954 | 20111595 | 20111787 | 475 | 375 | [illegible] |
| SE | 74316 | ENSG00000114166 | KAT2B | chr3 | + | 20119597 | 20119723 | 20114881 | 20114988 | 20122667 | 20122795 | 473 | 268 | [illegible] |
| SE | 74318 | ENSG00000114166 | KAT2B | chr3 | + | 20136941 | 20137052 | 20127422 | 20127549 | 20140220 | 20140364 | 651 | 593 | [illegible] |
| SE | 74319 | ENSG00000114166 | KAT2B | chr3 | + | 20146315 | 20146430 | 20140220 | 20140364 | 20147962 | 20147999 | 647 | 517 | [illegible] |
| SE | 74320 | ENSG00000114166 | KAT2B | chr3 | + | 20147962 | 20147999 | 20146315 | 20146430 | 20148242 | 20148306 | 377 | 305 | [illegible] |
| SE | 74321 | ENSG00000114166 | KAT2B | chr3 | + | 20148402 | 20148487 | 20148242 | 20148306 | 20152331 | 20154404 | 190 | 132 | [illegible] |
| SE | 74323 | ENSG00000140025 | EFCAB11 | chr14 | - | 89924512 | 89924677 | 89797126 | 89797324 | 89931540 | 89931631 | 7 | 2 | [illegible] |
| SE | 74324 | ENSG00000151789 | ZNF385D | chr3 | - | 21441685 | 21441742 | 21436819 | 21437203 | 21510860 | 21510986 | 0 | 1 | [illegible] |
| SE | 74329 | ENSG00000151789 | ZNF385D | chr3 | - | 21563261 | 21563312 | 21437076 | 21437203 | 21564573 | 21564684 | 0 | 2 | [illegible] |
| SE | 74332 | ENSG00000151789 | ZNF385D | chr3 | - | 21564573 | 21564684 | 21510860 | 21511023 | 21750894 | 21751114 | 4 | 14 | [illegible] |
| SE | 74333 | ENSG00000151789 | ZNF385D | chr3 | - | 21664885 | 21665028 | 21564573 | 21564684 | 21750894 | 21751114 | 3 | 5 | [illegible] |
| SE | 74335 | ENSG00000187118 | CMC1 | chr3 | + | 28253730 | 28253765 | 28241623 | 28241812 | 28316332 | 28316423 | 0 | 0 | [illegible] |
| SE | 74336 | ENSG00000187118 | CMC1 | chr3 | + | 28263290 | 28263380 | 28241583 | 28241812 | 28294433 | 28294526 | 3 | 7 | [illegible] |
| SE | 74338 | ENSG00000187118 | CMC1 | chr3 | + | 28263290 | 28263380 | 28253730 | 28253765 | 28316332 | 28316423 | 13 | 25 | [illegible] |
| SE | 74339 | ENSG00000187118 | CMC1 | chr3 | + | 28292735 | 28292964 | 28263290 | 28263380 | 28294433 | 28294526 | 811 | 200 | [illegible] |
| SE | 74341 | ENSG00000187118 | CMC1 | chr3 | + | 28292735 | 28292964 | 28291800 | 28291954 | 28294433 | 28294526 | 1569 | 352 | [illegible] |
| SE | 74343 | ENSG00000187118 | CMC1 | chr3 | + | 28294433 | 28294526 | 28241623 | 28241812 | 28316332 | 28316407 | 3 | 5 | [illegible] |
| SE | 74349 | ENSG00000187118 | CMC1 | chr3 | + | 28298218 | 28298350 | 28294433 | 28294526 | 28316332 | 28316423 | 74 | 9 | [illegible] |
| SE | 74350 | ENSG00000187118 | CMC1 | chr3 | + | 28298218 | 28298350 | 28297100 | 28297161 | 28316332 | 28316423 | 91 | 8 | [illegible] |
| SE | 74353 | ENSG00000175984 | DENND2C | chr1 | - | 114595354 | 114595650 | 114594472 | 114594578 | 114595831 | 114595873 | 0 | 0 | [illegible] |
| SE | 74355 | ENSG00000175984 | DENND2C | chr1 | - | 114645447 | 114645559 | 114625178 | 114626188 | 114654504 | 114654761 | 2 | 0 | [illegible] |
| SE | 74356 | ENSG00000175984 | DENND2C | chr1 | - | 114654504 | 114654761 | 114625178 | 114626188 | 114669982 | 114670049 | 1 | 0 | [illegible] |
| SE | 74359 | ENSG00000112787 | FBRSL1 | chr12 | + | 132508152 | 132508350 | 132489550 | 132490861 | 132525733 | 132525823 | 62 | 29 | [illegible] |
| SE | 74360 | ENSG00000112787 | FBRSL1 | chr12 | + | 132525733 | 132525823 | 132508152 | 132508350 | 132548002 | 132548032 | 29 | 17 | [illegible] |
| SE | 74361 | ENSG00000112787 | FBRSL1 | chr12 | + | 132527952 | 132527988 | 132525733 | 132525823 | 132548002 | 132548032 | 41 | 10 | [illegible] |
| SE | 74362 | ENSG00000112787 | FBRSL1 | chr12 | + | 132571067 | 132571136 | 132570334 | 132570540 | 132571306 | 132571524 | 2 | 3 | [illegible] |
| SE | 74366 | ENSG00000112787 | FBRSL1 | chr12 | + | 132581438 | 132581516 | 132576798 | 132576931 | 132582061 | 132582266 | 30 | 3 | [illegible] |
| SE | 74367 | ENSG00000112787 | FBRSL1 | chr12 | + | 132581740 | 132581824 | 132581438 | 132581516 | 132582061 | 132582266 | 16 | 0 | [illegible] |
| SE | 74372 | ENSG00000177889 | UBE2N | chr12 | - | 93410733 | 93410874 | 93405683 | 93410079 | 93411052 | 93411299 | 752 | 457 | [illegible] |
| SE | 74373 | ENSG00000177889 | UBE2N | chr12 | - | 93410733 | 93410874 | 93405683 | 93410079 | 93411125 | 93411299 | 219 | 117 | [illegible] |
| SE | 74374 | ENSG00000177889 | UBE2N | chr12 | - | 93424269 | 93424390 | 93411052 | 93411299 | 93441854 | 93441947 | 0 | 0 | [illegible] |
| SE | 74375 | ENSG00000215375 | MYL5 | chr4 | + | 676072 | 676209 | 673579 | 675854 | 678657 | 678765 | 3 | 1 | [illegible] |
| SE | 74380 | ENSG00000148291 | SURF2 | chr9 | + | 133360264 | 133360434 | 133357710 | 133357814 | 133361055 | 133361158 | 3 | 2 | [illegible] |
| SE | 74381 | ENSG00000188846 | RPL14 | chr3 | + | 40461406 | 40461506 | 40458641 | 40458736 | 40461607 | 40461661 | 4153 | 2098 | [illegible] |
| SE | 74382 | ENSG00000188846 | RPL14 | chr3 | + | 40461406 | 40461661 | 40458641 | 40458736 | 40461938 | 40462132 | 2590 | 1320 | [illegible] |
| SE | 74384 | ENSG00000131378 | RFTN1 | chr3 | - | 16323375 | 16323457 | 16315844 | 16317232 | 16357931 | 16358047 | 193 | 121 | [illegible] |
| SE | 74387 | ENSG00000131378 | RFTN1 | chr3 | - | 16409374 | 16409483 | 16377717 | 16378102 | 16493724 | 16493877 | 215 | 168 | [illegible] |
| SE | 74389 | ENSG00000131378 | RFTN1 | chr3 | - | 16433850 | 16434037 | 16409374 | 16409483 | 16493724 | 16493877 | 459 | 252 | [illegible] |
| SE | 74390 | ENSG00000131378 | RFTN1 | chr3 | - | 16493724 | 16493877 | 16433850 | 16434037 | 16513441 | 16513522 | 330 | 190 | [illegible] |
| SE | 74391 | ENSG00000131378 | RFTN1 | chr3 | - | 16493724 | 16493880 | 16433850 | 16434037 | 16513441 | 16513699 | 270 | 144 | [illegible] |
| SE | 74393 | ENSG00000154781 | CCDC174 | chr3 | + | 14661529 | 14661707 | 14658870 | 14658929 | 14665027 | 14665123 | 120 | 99 | [illegible] |
| SE | 74397 | ENSG00000117691 | NENF | chr1 | + | 212442545 | 212442625 | 212432972 | 212433120 | 212444338 | 212444442 | 6 | 7 | [illegible] |
| SE | 74398 | ENSG00000117691 | NENF | chr1 | + | 212442564 | 212442625 | 212432919 | 212433120 | 212444338 | 212444442 | 8 | 8 | [illegible] |
| SE | 74399 | ENSG00000117691 | NENF | chr1 | + | 212444338 | 212444442 | 212442545 | 212442625 | 212445829 | 212446372 | 42 | 30 | [illegible] |
| SE | 74400 | ENSG00000168237 | GLYCTK | chr3 | + | 52290303 | 52290719 | 52287819 | 52287876 | 52290959 | 52291111 | 8 | 0 | [illegible] |
| SE | 74403 | ENSG00000159618 | ADGRG5 | chr16 | + | 57562383 | 57562459 | 57562055 | 57562157 | 57563090 | 57563247 | 93 | 36 | [illegible] |
| SE | 74404 | ENSG00000159618 | ADGRG5 | chr16 | + | 57570417 | 57570535 | 57567855 | 57568124 | 57574814 | 57575092 | 50 | 9 | [illegible] |
| SE | 74418 | ENSG00000163328 | GPR155 | chr2 | - | 174446610 | 174446747 | 174445080 | 174445176 | 174453736 | 174453841 | 32 | 37 | [illegible] |
| SE | 74420 | ENSG00000163328 | GPR155 | chr2 | - | 174453736 | 174453841 | 174446610 | 174446747 | 174461401 | 174461492 | 14 | 24 | [illegible] |
| SE | 74421 | ENSG00000163328 | GPR155 | chr2 | - | 174459877 | 174460088 | 174446610 | 174446747 | 174461401 | 174461492 | 42 | 43 | [illegible] |
| SE | 74422 | ENSG00000163328 | GPR155 | chr2 | - | 174459877 | 174460088 | 174453736 | 174453841 | 174461401 | 174461492 | 77 | 65 | [illegible] |
| SE | 74429 | ENSG00000163328 | GPR155 | chr2 | - | 174483009 | 174483158 | 174482872 | 174482918 | 174486872 | 174487000 | 8 | 9 | [illegible] |
| SE | 74432 | ENSG00000172940 | SLC22A13 | chr3 | + | 38276911 | 38277127 | 38276013 | 38276096 | 38277371 | 38278757 | 0 | 0 | [illegible] |
| SE | 74437 | ENSG00000072401 | UBE2D1 | chr10 | + | 58363608 | 58363686 | 58361494 | 58361526 | 58364770 | 58364876 | 246 | 93 | [illegible] |
| SE | 74438 | ENSG00000072401 | UBE2D1 | chr10 | + | 58364770 | 58364876 | 58363608 | 58363686 | 58367922 | 58368016 | 427 | 233 | [illegible] |
| SE | 74439 | ENSG00000138376 | BARD1 | chr2 | - | 214730234 | 214730280 | 214728537 | 214729008 | 214730410 | 214730508 | 0 | 4 | [illegible] |
| SE | 74448 | ENSG00000116209 | TMEM59 | chr1 | - | 54036609 | 54036718 | 54026680 | 54032305 | 54040755 | 54040837 | 563 | 231 | [illegible] |
| SE | 74449 | ENSG00000116209 | TMEM59 | chr1 | - | 54045691 | 54045786 | 54043372 | 54043525 | 54047266 | 54047372 | 873 | 460 | [illegible] |
| SE | 74450 | ENSG00000116209 | TMEM59 | chr1 | - | 54045691 | 54045786 | 54043372 | 54043525 | 54052999 | 54053204 | 298 | 148 | [illegible] |
| SE | 74453 | ENSG00000116209 | TMEM59 | chr1 | - | 54047266 | 54047372 | 54043372 | 54043525 | 54053355 | 54053504 | 9 | 8 | [illegible] |
| SE | 74454 | ENSG00000116209 | TMEM59 | chr1 | - | 54047266 | 54047372 | 54045485 | 54045786 | 54052999 | 54053192 | 693 | 351 | [illegible] |
| SE | 74455 | ENSG00000116209 | TMEM59 | chr1 | - | 54047266 | 54047372 | 54045485 | 54045786 | 54053355 | 54053504 | 524 | 282 | [illegible] |
| SE | 74456 | ENSG00000116209 | TMEM59 | chr1 | - | 54048649 | 54048682 | 54047266 | 54047372 | 54052999 | 54053405 | 1 | 1 | [illegible] |
| SE | 74457 | ENSG00000116209 | TMEM59 | chr1 | - | 54048649 | 54048682 | 54047266 | 54047372 | 54053355 | 54053412 | 0 | 0 | [illegible] |
| SE | 74458 | ENSG00000116539 | ASH1L | chr1 | - | 155341935 | 155342102 | 155339327 | 155339368 | 155343313 | 155343486 | 250 | 185 | [illegible] |
| SE | 74459 | ENSG00000116539 | ASH1L | chr1 | - | 155345962 | 155346207 | 155344182 | 155344273 | 155346382 | 155346469 | 5 | 5 | [illegible] |
| SE | 74460 | ENSG00000116539 | ASH1L | chr1 | - | 155347655 | 155347901 | 155346382 | 155346469 | 155352705 | 155352858 | 53 | 52 | [illegible] |
| SE | 74461 | ENSG00000116539 | ASH1L | chr1 | - | 155347655 | 155347904 | 155346382 | 155346469 | 155352705 | 155352858 | 54 | 53 | [illegible] |
| SE | 74463 | ENSG00000116539 | ASH1L | chr1 | - | 155349326 | 155349459 | 155347655 | 155347904 | 155349541 | 155349596 | 190 | 112 | [illegible] |
| SE | 74464 | ENSG00000116539 | ASH1L | chr1 | - | 155349326 | 155349459 | 155347655 | 155347904 | 155352705 | 155352858 | 121 | 63 | [illegible] |
| SE | 74465 | ENSG00000116539 | ASH1L | chr1 | - | 155349541 | 155349596 | 155347655 | 155347904 | 155352705 | 155352858 | 120 | 97 | [illegible] |
| SE | 74466 | ENSG00000116539 | ASH1L | chr1 | - | 155349541 | 155349596 | 155349326 | 155349459 | 155352705 | 155352858 | 189 | 147 | [illegible] |
| SE | 74467 | ENSG00000116539 | ASH1L | chr1 | - | 155357315 | 155357410 | 155354472 | 155354630 | 155360300 | 155360409 | 81 | 68 | [illegible] |
| SE | 74469 | ENSG00000116539 | ASH1L | chr1 | - | 155378502 | 155378548 | 155378280 | 155378389 | 155380042 | 155380116 | 244 | 182 | [illegible] |
| SE | 74473 | ENSG00000116539 | ASH1L | chr1 | - | 155440489 | 155440593 | 155438326 | 155439068 | 155477885 | 155482449 | 10 | 15 | [illegible] |
| SE | 74477 | ENSG00000116539 | ASH1L | chr1 | - | 155459796 | 155459898 | 155440489 | 155440593 | 155477885 | 155482449 | 91 | 80 | [illegible] |
| SE | 74479 | ENSG00000116539 | ASH1L | chr1 | - | 155477885 | 155482449 | 155459796 | 155459898 | 155521099 | 155521618 | 281 | 271 | [illegible] |
| SE | 74480 | ENSG00000116539 | ASH1L | chr1 | - | 155477885 | 155482449 | 155459796 | 155459898 | 155562152 | 155562291 | 82 | 75 | [illegible] |
| SE | 74482 | ENSG00000116539 | ASH1L | chr1 | - | 155521099 | 155521618 | 155459796 | 155459898 | 155562152 | 155562291 | 27 | 25 | [illegible] |
| SE | 74483 | ENSG00000116539 | ASH1L | chr1 | - | 155521099 | 155521618 | 155477885 | 155482449 | 155562152 | 155562291 | 226 | 221 | [illegible] |
| SE | 74484 | ENSG00000116539 | ASH1L | chr1 | - | 155521099 | 155521618 | 155477885 | 155482449 | 155562778 | 155562841 | 222 | 206 | [illegible] |
| SE | 74485 | ENSG00000116539 | ASH1L | chr1 | - | 155562152 | 155562291 | 155521099 | 155521618 | 155563072 | 155563115 | 22 | 21 | [illegible] |
| SE | 74486 | ENSG00000116539 | ASH1L | chr1 | - | 155562152 | 155562295 | 155477885 | 155482449 | 155562778 | 155562807 | 1 | 0 | [illegible] |
| SE | 74487 | ENSG00000116539 | ASH1L | chr1 | - | 155562152 | 155562295 | 155521099 | 155521618 | 155562778 | 155562807 | 25 | 23 | [illegible] |
| SE | 74488 | ENSG00000116539 | ASH1L | chr1 | - | 155562152 | 155562295 | 155521099 | 155521618 | 155563072 | 155563092 | 24 | 23 | [illegible] |
| SE | 74492 | ENSG00000123219 | CENPK | chr5 | - | 65542801 | 65542848 | 65529116 | 65529199 | 65551563 | 65551636 | 117 | 110 | [illegible] |
| SE | 74493 | ENSG00000123219 | CENPK | chr5 | - | 65551105 | 65551242 | 65542801 | 65542848 | 65551563 | 65551636 | 3 | 0 | [illegible] |
| SE | 74494 | ENSG00000123219 | CENPK | chr5 | - | 65551563 | 65551636 | 65542801 | 65542848 | 65552492 | 65552549 | 158 | 145 | [illegible] |
| SE | 74495 | ENSG00000123219 | CENPK | chr5 | - | 65554796 | 65554898 | 65552492 | 65552549 | 65561462 | 65561564 | 82 | 60 | [illegible] |
| SE | 74496 | ENSG00000123219 | CENPK | chr5 | - | 65554796 | 65554946 | 65552492 | 65552549 | 65561462 | 65561535 | 123 | 93 | [illegible] |
| SE | 74497 | ENSG00000123219 | CENPK | chr5 | - | 65562802 | 65563008 | 65561115 | 65561564 | 65563097 | 65563123 | 1 | 0 | [illegible] |
| SE | 74505 | ENSG00000135525 | MAP7 | chr6 | - | 136383670 | 136383781 | 136377754 | 136377868 | 136388392 | 136388510 | 0 | 7 | [illegible] |
| SE | 74507 | ENSG00000170364 | SETMAR | chr3 | + | 4312897 | 4313229 | 4303331 | 4303526 | 4313646 | 4313761 | 0 | 4 | [illegible] |
| SE | 74508 | ENSG00000170364 | SETMAR | chr3 | + | 4312897 | 4313229 | 4303331 | 4303526 | 4313666 | 4313761 | 0 | 4 | [illegible] |
| SE | 74509 | ENSG00000170364 | SETMAR | chr3 | + | 4312897 | 4313761 | 4303303 | 4303526 | 4316211 | 4317567 | 8 | 10 | [illegible] |
| SE | 74510 | ENSG00000170364 | SETMAR | chr3 | + | 4313646 | 4313761 | 4303336 | 4303526 | 4316211 | 4316536 | 4 | 7 | [illegible] |
| SE | 74511 | ENSG00000170364 | SETMAR | chr3 | + | 4313646 | 4313761 | 4303607 | 4303868 | 4316211 | 4316536 | 5 | 5 | [illegible] |
| SE | 74512 | ENSG00000170364 | SETMAR | chr3 | + | 4313666 | 4313761 | 4303331 | 4303526 | 4316211 | 4317265 | 6 | 4 | [illegible] |
| SE | 74513 | ENSG00000166192 | SENP8 | chr15 | + | 72139311 | 72139435 | 72118324 | 72118464 | 72139576 | 72139735 | 0 | 0 | [illegible] |
| SE | 74517 | ENSG00000064933 | PMS1 | chr2 | + | 189785345 | 189785471 | 189784488 | 189784770 | 189791789 | 189791941 | 0 | 3 | [illegible] |
| SE | 74518 | ENSG00000064933 | PMS1 | chr2 | + | 189785345 | 189785471 | 189784512 | 189784564 | 189791789 | 189791941 | 0 | 3 | [illegible] |
| SE | 74520 | ENSG00000064933 | PMS1 | chr2 | + | 189795768 | 189795951 | 189791789 | 189791941 | 189818016 | 189818180 | 123 | 95 | [illegible] |
| SE | 74521 | ENSG00000064933 | PMS1 | chr2 | + | 189805651 | 189805754 | 189791789 | 189791941 | 189818016 | 189818180 | 63 | 40 | [illegible] |

| | | | | | | | | | | | | | | |
|---|---|---|---|---|---|---|---|---|---|---|---|---|---|---|
| SE | 74524 | ENSG00000064933 | PMS1 | chr2 | + | 189852654 | 189852777 | 189843963 | 189844080 | 189853938 | 189854082 | 156 | 110 | [illegible] |
| SE | 74525 | ENSG00000064933 | PMS1 | chr2 | + | 189855828 | 189855916 | 189854238 | 189855128 | 189863742 | 189864114 | 0 | 2 | [illegible] |
| SE | 74526 | ENSG00000064933 | PMS1 | chr2 | + | 189863742 | 189863974 | 189854238 | 189855128 | 189867798 | 189867929 | 113 | 90 | [illegible] |
| SE | 74527 | ENSG00000064933 | PMS1 | chr2 | + | 189863742 | 189863974 | 189854238 | 189855128 | 189873495 | 189873528 | 112 | 86 | [illegible] |
| SE | 74528 | ENSG00000064933 | PMS1 | chr2 | + | 189863742 | 189864228 | 189854238 | 189855128 | 189867798 | 189867929 | 154 | 106 | [illegible] |
| SE | 74529 | ENSG00000064933 | PMS1 | chr2 | + | 189863742 | 189864228 | 189854238 | 189855128 | 189873495 | 189873656 | 115 | 87 | [illegible] |
| SE | 74530 | ENSG00000064933 | PMS1 | chr2 | + | 189867798 | 189867929 | 189854238 | 189855128 | 189873495 | 189873656 | 61 | 47 | [illegible] |
| SE | 74532 | ENSG00000064933 | PMS1 | chr2 | + | 189867798 | 189867929 | 189863742 | 189864228 | 189873495 | 189873656 | 102 | 67 | [illegible] |
| SE | 74534 | ENSG00000183960 | KCNH8 | chr3 | + | 19390480 | 19390638 | 19347720 | 19347965 | 19395103 | 19395311 | 12 | 4 | [illegible] |
| SE | 74536 | ENSG00000183960 | KCNH8 | chr3 | + | 19395103 | 19395311 | 19347724 | 19347965 | 19438163 | 19438361 | 5 | 3 | [illegible] |
| SE | 74537 | ENSG00000183960 | KCNH8 | chr3 | + | 19451154 | 19451404 | 19450105 | 19450305 | 19456767 | 19456982 | 8 | 8 | [illegible] |
| SE | 74538 | ENSG00000090975 | PITPNM2 | chr12 | - | 122987510 | 122987659 | 122987280 | 122987430 | 122987784 | 122987901 | 33 | 18 | [illegible] |
| SE | 74540 | ENSG00000090975 | PITPNM2 | chr12 | - | 122991754 | 122991898 | 122990544 | 122990709 | 122992498 | 122992669 | 1 | 0 | [illegible] |
| SE | 74541 | ENSG00000090975 | PITPNM2 | chr12 | - | 122992498 | 122992669 | 122990544 | 122990709 | 122994800 | 122994979 | 24 | 16 | [illegible] |
| SE | 74542 | ENSG00000090975 | PITPNM2 | chr12 | - | 123009849 | 123010077 | 123001053 | 123001158 | 123034512 | 123034685 | 0 | 0 | [illegible] |
| SE | 74543 | ENSG00000090975 | PITPNM2 | chr12 | - | 123012612 | 123012734 | 123009849 | 123010077 | 123034512 | 123034685 | 115 | 81 | [illegible] |
| SE | 74545 | ENSG00000090975 | PITPNM2 | chr12 | - | 123034512 | 123034685 | 123013827 | 123014042 | 123110384 | 123110488 | 50 | 52 | [illegible] |
| SE | 74546 | ENSG00000090975 | PITPNM2 | chr12 | - | 123110384 | 123110488 | 123034512 | 123034685 | 123150752 | 123151090 | 13 | 16 | [illegible] |
| SE | 74548 | ENSG00000165792 | METTL17 | chr14 | + | 20993117 | 20993191 | 20992540 | 20992622 | 20994542 | 20994613 | 38 | 27 | [illegible] |
| SE | 74549 | ENSG00000165792 | METTL17 | chr14 | + | 20993117 | 20994063 | 20992540 | 20992622 | 20994542 | 20994613 | 102 | 72 | [illegible] |
| SE | 74554 | ENSG00000114353 | GNAI2 | chr3 | + | 50253023 | 50253184 | 50251408 | 50252538 | 50256191 | 50256320 | 1951 | 858 | [illegible] |
| SE | 74555 | ENSG00000114353 | GNAI2 | chr3 | + | 50256936 | 50257090 | 50256722 | 50256852 | 50257499 | 50257714 | 4782 | 2582 | [illegible] |
| SE | 74561 | ENSG00000089818 | NECAP1 | chr12 | + | 8092675 | 8092784 | 8090194 | 8091850 | 8092871 | 8093043 | 170 | 77 | [illegible] |
| SE | 74565 | ENSG00000163512 | AZI2 | chr3 | - | 28341511 | 28341596 | 28340401 | 28340622 | 28348160 | 28348477 | 2 | 1 | [illegible] |
| SE | 74566 | ENSG00000163512 | AZI2 | chr3 | - | 28341511 | 28341596 | 28340401 | 28340622 | 28348600 | 28348790 | 3 | 1 | [illegible] |
| SE | 74567 | ENSG00000163512 | AZI2 | chr3 | - | 28341511 | 28341647 | 28340401 | 28340622 | 28348160 | 28348477 | 2 | 1 | [illegible] |
| SE | 74568 | ENSG00000163512 | AZI2 | chr3 | - | 28341511 | 28341647 | 28340401 | 28340622 | 28348600 | 28348819 | 2 | 1 | [illegible] |
| SE | 74578 | ENSG00000128284 | APOL3 | chr22 | - | 36149258 | 36149371 | 36149045 | 36149185 | 36160668 | 36160929 | 34 | 19 | [illegible] |
| SE | 74579 | ENSG00000128284 | APOL3 | chr22 | - | 36149770 | 36149964 | 36149074 | 36149142 | 36156843 | 36157006 | 4 | 0 | [illegible] |
| [illegible] | | | | | | | | | | | | | | |

| | | | | | | | | | | | | | | |
|---|---|---|---|---|---|---|---|---|---|---|---|---|---|---|
| SE | 75033 | ENSG00000130023 | ERMARD | chr6 | + | 169753286 | 169753372 | 169751621 | 169751663 | 169753863 | 169754032 | 5 | 1 | [illegible] |
| SE | 75034 | ENSG00000130023 | ERMARD | chr6 | + | 169753316 | 169753372 | 169751624 | 169751663 | 169753863 | 169754032 | 1 | 1 | [illegible] |
| SE | 75038 | ENSG00000130023 | ERMARD | chr6 | + | 169755282 | 169755422 | 169753863 | 169754032 | 169756337 | 169756415 | 46 | 21 | [illegible] |
| SE | 75039 | ENSG00000130023 | ERMARD | chr6 | + | 169755287 | 169755422 | 169753863 | 169754032 | 169756337 | 169756439 | 19 | 6 | [illegible] |
| SE | 75042 | ENSG00000130023 | ERMARD | chr6 | + | 169769539 | 169769713 | 169767014 | 169768171 | 169773318 | 169773402 | 121 | 90 | [illegible] |
| SE | 75045 | ENSG00000130023 | ERMARD | chr6 | + | 169776333 | 169776411 | 169775939 | 169776065 | 169779181 | 169779295 | 0 | 6 | [illegible] |
| SE | 75047 | ENSG00000130023 | ERMARD | chr6 | + | 169779181 | 169779295 | 169775939 | 169776065 | 169781329 | 169781516 | 7 | 9 | [illegible] |
| SE | 75049 | ENSG00000036054 | TBC1D23 | chr3 | + | 100283606 | 100283811 | 100281717 | 100281847 | 100295086 | 100295211 | 98 | 57 | [illegible] |
| SE | 75050 | ENSG00000036054 | TBC1D23 | chr3 | + | 100290577 | 100290701 | 100281741 | 100281847 | 100295086 | 100295211 | 77 | 59 | [illegible] |
| SE | 75051 | ENSG00000036054 | TBC1D23 | chr3 | + | 100290577 | 100290701 | 100283606 | 100283811 | 100295086 | 100295211 | 130 | 106 | [illegible] |
| SE | 75052 | ENSG00000036054 | TBC1D23 | chr3 | + | 100302066 | 100302237 | 100299238 | 100299331 | 100304845 | 100304888 | 157 | 89 | [illegible] |
| SE | 75058 | ENSG00000198223 | CSF2RA | chrX | + | 1282677 | 1282779 | 1274754 | 1274818 | 1285777 | 1285920 | 88 | 76 | [illegible] |
| SE | 75061 | ENSG00000198223 | CSF2RA | chrX | + | 1288518 | 1288642 | 1285777 | 1285920 | 1288758 | 1288888 | 275 | 275 | [illegible] |
| SE | 75062 | ENSG00000198223 | CSF2RA | chrX | + | 1288758 | 1288888 | 1285777 | 1285920 | 1290336 | 1290509 | 123 | 116 | [illegible] |
| SE | 75063 | ENSG00000198223 | CSF2RA | chrX | + | 1294327 | 1294461 | 1290336 | 1290509 | 1295426 | 1295456 | 157 | 135 | [illegible] |
| SE | 75065 | ENSG00000198223 | CSF2RA | chrX | + | 1300490 | 1300626 | 1295426 | 1295456 | 1303922 | 1304019 | 99 | 70 | [illegible] |
| SE | 75077 | ENSG00000188554 | NBR1 | chr17 | + | 43177935 | 43177998 | 43175790 | 43175901 | 43179393 | 43179412 | 448 | 380 | [illegible] |
| SE | 75078 | ENSG00000188554 | NBR1 | chr17 | + | 43180794 | 43180817 | 43179393 | 43179412 | 43186249 | 43186444 | 437 | 325 | [illegible] |
| SE | 75079 | ENSG00000188554 | NBR1 | chr17 | + | 43186249 | 43186444 | 43180794 | 43180817 | 43189041 | 43189119 | 713 | 549 | [illegible] |
| SE | 75082 | ENSG00000188554 | NBR1 | chr17 | + | 43193347 | 43193638 | 43193093 | 43193253 | 43194349 | 43194499 | 565 | 388 | [illegible] |
| SE | 75084 | ENSG00000188554 | NBR1 | chr17 | + | 43196480 | 43196591 | 43194963 | 43195039 | 43196941 | 43197106 | 402 | 286 | [illegible] |
| SE | 75085 | ENSG00000188554 | NBR1 | chr17 | + | 43196480 | 43196591 | 43194963 | 43195039 | 43200166 | 43200608 | 214 | 165 | [illegible] |
| SE | 75086 | ENSG00000188554 | NBR1 | chr17 | + | 43196941 | 43197106 | 43194963 | 43195039 | 43200166 | 43200608 | 141 | 129 | [illegible] |
| SE | 75088 | ENSG00000188554 | NBR1 | chr17 | + | 43200166 | 43200608 | 43196941 | 43197106 | 43201685 | 43201780 | 328 | 237 | [illegible] |
| SE | 75089 | ENSG00000188554 | NBR1 | chr17 | + | 43201685 | 43201780 | 43200166 | 43200608 | 43202654 | 43202712 | 258 | 152 | [illegible] |
| SE | 75090 | ENSG00000188554 | NBR1 | chr17 | + | 43201685 | 43201780 | 43200166 | 43200608 | 43203680 | 43203786 | 130 | 86 | [illegible] |
| SE | 75091 | ENSG00000188554 | NBR1 | chr17 | + | 43202654 | 43202712 | 43200166 | 43200608 | 43203680 | 43203786 | 52 | 31 | [illegible] |
| SE | 75099 | ENSG00000198948 | MFAP3L | chr4 | - | 170006602 | 170006736 | 170004990 | 170006010 | 170026859 | 170027210 | 13 | 20 | [illegible] |
| SE | 75101 | ENSG00000114423 | CBLB | chr3 | - | 105671023 | 105671046 | 105670232 | 105670352 | 105672336 | 105672462 | 2 | 3 | [illegible] |
| SE | 75103 | ENSG00000114423 | CBLB | chr3 | - | 105672336 | 105672462 | 105670232 | 105670352 | 105678430 | 105678571 | 0 | 0 | [illegible] |
| SE | 75104 | ENSG00000114423 | CBLB | chr3 | - | 105672336 | 105672462 | 105671023 | 105671046 | 105678430 | 105678571 | 14 | 4 | [illegible] |
| SE | 75107 | ENSG00000114423 | CBLB | chr3 | - | 105685319 | 105685466 | 105680991 | 105681818 | 105693493 | 105693588 | 302 | 154 | [illegible] |
| SE | 75108 | ENSG00000114423 | CBLB | chr3 | - | 105702093 | 105702459 | 105693493 | 105693588 | 105703987 | 105704173 | 176 | 127 | [illegible] |
| SE | 75109 | ENSG00000114423 | CBLB | chr3 | - | 105703987 | 105704173 | 105702093 | 105702459 | 105720046 | 105720250 | 222 | 190 | [illegible] |
| SE | 75110 | ENSG00000114423 | CBLB | chr3 | - | 105720046 | 105720250 | 105703987 | 105704173 | 105734008 | 105734140 | 186 | 133 | [illegible] |
| SE | 75111 | ENSG00000114423 | CBLB | chr3 | - | 105724025 | 105724189 | 105720046 | 105720250 | 105734008 | 105734140 | 0 | 0 | [illegible] |
| SE | 75112 | ENSG00000114423 | CBLB | chr3 | - | 105740493 | 105740631 | 105737170 | 105737258 | 105745916 | 105746038 | 429 | 284 | [illegible] |
| SE | 75115 | ENSG00000114423 | CBLB | chr3 | - | 105776395 | 105776542 | 105720046 | 105720250 | 105853413 | 105853646 | 175 | 117 | [illegible] |
| SE | 75116 | ENSG00000114423 | CBLB | chr3 | - | 105776395 | 105776542 | 105751461 | 105751618 | 105853413 | 105853646 | 297 | 190 | [illegible] |
| SE | 75118 | ENSG00000114423 | CBLB | chr3 | - | 105853413 | 105853664 | 105776395 | 105776542 | 105867409 | 105867591 | 412 | 272 | [illegible] |
| SE | 75123 | ENSG00000112343 | TRIM38 | chr6 | + | 25966334 | 25966933 | 25963048 | 25963282 | 25969324 | 25969420 | 503 | 295 | [illegible] |
| SE | 75126 | ENSG00000248124 | RRN3P1 | chr16 | - | 21800706 | 21800837 | 21799649 | 21799834 | 21802635 | 21802728 | 136 | 88 | [illegible] |
| SE | 75127 | ENSG00000248124 | RRN3P1 | chr16 | - | 21802635 | 21802728 | 21800706 | 21800837 | 21806130 | 21806229 | 97 | 90 | [illegible] |
| SE | 75128 | ENSG00000248124 | RRN3P1 | chr16 | - | 21809645 | 21809715 | 21806130 | 21806229 | 21811777 | 21811837 | 9 | 14 | [illegible] |
| SE | 75130 | ENSG00000248124 | RRN3P1 | chr16 | - | 21811777 | 21811837 | 21806130 | 21806229 | 21812285 | 21812415 | 23 | 10 | [illegible] |
| SE | 75131 | ENSG00000248124 | RRN3P1 | chr16 | - | 21819301 | 21819481 | 21818655 | 21818761 | 21820294 | 21820410 | 9 | 5 | [illegible] |
| SE | 75132 | ENSG00000124226 | RNF114 | chr20 | + | 49941106 | 49941220 | 49936398 | 49936552 | 49941560 | 49941711 | 0 | 1 | [illegible] |
| SE | 75133 | ENSG00000124226 | RNF114 | chr20 | + | 49941560 | 49941711 | 49936396 | 49936552 | 49945381 | 49945488 | 137 | 58 | [illegible] |
| SE | 75135 | ENSG00000124226 | RNF114 | chr20 | + | 49949247 | 49949355 | 49946135 | 49946250 | 49952075 | 49952152 | 321 | 198 | [illegible] |
| SE | 75138 | ENSG00000156256 | USP16 | chr21 | + | 29026566 | 29026661 | 29024655 | 29024777 | 29027872 | 29027974 | 0 | 0 | [illegible] |
| SE | 75139 | ENSG00000156256 | USP16 | chr21 | + | 29030594 | 29030773 | 29027872 | 29027974 | 29034836 | 29034919 | 503 | 457 | [illegible] |
| SE | 75140 | ENSG00000156256 | USP16 | chr21 | + | 29036270 | 29036374 | 29034836 | 29034940 | 29037278 | 29037463 | 373 | 357 | [illegible] |
| SE | 75141 | ENSG00000156256 | USP16 | chr21 | + | 29037275 | 29037463 | 29036270 | 29036374 | 29038334 | 29038430 | 356 | 308 | [illegible] |
| SE | 75142 | ENSG00000156256 | USP16 | chr21 | + | 29037278 | 29037463 | 29036270 | 29036374 | 29038334 | 29038430 | 252 | 241 | [illegible] |
| SE | 75143 | ENSG00000156256 | USP16 | chr21 | + | 29039025 | 29039156 | 29038334 | 29038430 | 29039480 | 29039568 | 278 | 189 | [illegible] |
| SE | 75144 | ENSG00000156256 | USP16 | chr21 | + | 29039480 | 29039568 | 29039025 | 29039156 | 29040608 | 29040687 | 165 | 118 | [illegible] |
| SE | 75145 | ENSG00000156256 | USP16 | chr21 | + | 29040608 | 29040687 | 29039480 | 29039568 | 29042012 | 29042104 | 163 | 142 | [illegible] |
| SE | 75146 | ENSG00000156256 | USP16 | chr21 | + | 29048760 | 29048855 | 29046666 | 29047321 | 29053801 | 29053958 | 69 | 55 | [illegible] |
| SE | 75147 | ENSG00000156256 | USP16 | chr21 | + | 29050091 | 29050178 | 29048760 | 29048855 | 29053801 | 29053958 | 249 | 182 | [illegible] |
| SE | 75148 | ENSG00000156256 | USP16 | chr21 | + | 29053801 | 29053958 | 29050091 | 29050178 | 29054065 | 29054488 | 333 | 238 | [illegible] |
| SE | 75149 | ENSG00000232810 | TNF | chr6 | + | 31576533 | 31576579 | 31575564 | 31575927 | 31576766 | 31576814 | 27 | 10 | [illegible] |
| SE | 75152 | ENSG00000227507 | LTB | chr6 | - | 31581813 | 31581859 | 31581558 | 31581630 | 31582495 | 31582522 | 280 | 172 | [illegible] |
| SE | 75155 | ENSG00000059758 | CDK17 | chr12 | - | 96286663 | 96286761 | 96286042 | 96286148 | 96289166 | 96289287 | 338 | 196 | [illegible] |
| SE | 75157 | ENSG00000059758 | CDK17 | chr12 | - | 96297626 | 96297721 | 96294998 | 96295122 | 96300303 | 96300360 | 14 | 7 | [illegible] |
| SE | 75160 | ENSG00000059758 | CDK17 | chr12 | - | 96323947 | 96324112 | 96313320 | 96313454 | 96334718 | 96334865 | 373 | 254 | [illegible] |
| SE | 75162 | ENSG00000137962 | ARHGAP29 | chr1 | - | 94184150 | 94184288 | 94179724 | 94179957 | 94184871 | 94185060 | 11 | 22 | [illegible] |
| SE | 75163 | ENSG00000204438 | GPANK1 | chr6 | - | 31664840 | 31665003 | 31663852 | 31664577 | 31665860 | 31666283 | 35 | 31 | [illegible] |
| SE | 75164 | ENSG00000204438 | GPANK1 | chr6 | - | 31664840 | 31665003 | 31663852 | 31664580 | 31665860 | 31666283 | 49 | 21 | [illegible] |
| SE | 75167 | ENSG00000072121 | ZFYVE26 | chr14 | - | 67753706 | 67753766 | 67752343 | 67752526 | 67754070 | 67754212 | 59 | 42 | [illegible] |
| SE | 75171 | ENSG00000072121 | ZFYVE26 | chr14 | - | 67755050 | 67755250 | 67754070 | 67754212 | 67761365 | 67761491 | 32 | 38 | [illegible] |
| SE | 75172 | ENSG00000072121 | ZFYVE26 | chr14 | - | 67755947 | 67756145 | 67755050 | 67755250 | 67761365 | 67761491 | 128 | 74 | [illegible] |
| SE | 75173 | ENSG00000072121 | ZFYVE26 | chr14 | - | 67761365 | 67762412 | 67755050 | 67755250 | 67762671 | 67762819 | 68 | 39 | [illegible] |
| SE | 75174 | ENSG00000072121 | ZFYVE26 | chr14 | - | 67772046 | 67772210 | 67769593 | 67769730 | 67775015 | 67775114 | 55 | 33 | [illegible] |
| SE | 75176 | ENSG00000072121 | ZFYVE26 | chr14 | - | 67802078 | 67802282 | 67798013 | 67798622 | 67804100 | 67804264 | 118 | 89 | [illegible] |
| SE | 75177 | ENSG00000072121 | ZFYVE26 | chr14 | - | 67802078 | 67802282 | 67798013 | 67798622 | 67805453 | 67805618 | 45 | 27 | [illegible] |
| SE | 75179 | ENSG00000072121 | ZFYVE26 | chr14 | - | 67804100 | 67804264 | 67798013 | 67798622 | 67805453 | 67805618 | 1 | 1 | [illegible] |
| SE | 75180 | ENSG00000072121 | ZFYVE26 | chr14 | - | 67804100 | 67804264 | 67802078 | 67802282 | 67805453 | 67805618 | 73 | 63 | [illegible] |
| SE | 75181 | ENSG00000072121 | ZFYVE26 | chr14 | - | 67805216 | 67805305 | 67798013 | 67798622 | 67805453 | 67805618 | 11 | 8 | [illegible] |
| SE | 75182 | ENSG00000072121 | ZFYVE26 | chr14 | - | 67805216 | 67805305 | 67804100 | 67804264 | 67805453 | 67805618 | 44 | 27 | [illegible] |
| SE | 75185 | ENSG00000072121 | ZFYVE26 | chr14 | - | 67806544 | 67806675 | 67805453 | 67805618 | 67807397 | 67807920 | 68 | 47 | [illegible] |
| SE | 75193 | ENSG00000147419 | CCDC25 | chr8 | - | 27764285 | 27764533 | 27762418 | 27762458 | 27765203 | 27765251 | 2 | 0 | [illegible] |
| SE | 75196 | ENSG00000204410 | MSH5 | chr6 | + | 31740453 | 31740613 | 31740004 | 31740062 | 31741162 | 31741286 | 11 | 15 | [illegible] |
| SE | 75197 | ENSG00000204410 | MSH5 | chr6 | + | 31742876 | 31742957 | 31741162 | 31741286 | 31743107 | 31743149 | 30 | 21 | [illegible] |
| SE | 75199 | ENSG00000204410 | MSH5 | chr6 | + | 31743903 | 31744025 | 31743107 | 31743170 | 31744545 | 31744581 | 15 | 12 | [illegible] |
| SE | 75200 | ENSG00000204410 | MSH5 | chr6 | + | 31743903 | 31744299 | 31743107 | 31743170 | 31744545 | 31744581 | 30 | 30 | [illegible] |
| SE | 75204 | ENSG00000204410 | MSH5 | chr6 | + | 31747386 | 31747432 | 31744545 | 31744581 | 31753300 | 31753439 | 10 | 10 | [illegible] |
| SE | 75205 | ENSG00000204410 | MSH5 | chr6 | + | 31758532 | 31758620 | 31758164 | 31758293 | 31758765 | 31758875 | 8 | 4 | [illegible] |
| SE | 75209 | ENSG00000204410 | MSH5 | chr6 | + | 31761749 | 31761780 | 31761471 | 31761615 | 31761865 | 31761955 | 1 | 0 | [illegible] |
| SE | 75210 | ENSG00000204410 | MSH5 | chr6 | + | 31761865 | 31761955 | 31761749 | 31761780 | 31762111 | 31762185 | 4 | 8 | [illegible] |
| SE | 75211 | ENSG00000255152 | MSH5-SAPCD1 | chr6 | + | 31762789 | 31763168 | 31762570 | 31763414 | 31763555 | 31763555 | 9 | 1 | [illegible] |
| SE | 75212 | ENSG00000100503 | NIN | chr14 | - | 50729522 | 50729723 | 50725826 | 50726066 | 50735515 | 50735617 | 376 | 412 | [illegible] |
| SE | 75213 | ENSG00000100503 | NIN | chr14 | - | 50729522 | 50729723 | 50727235 | 50727751 | 50735515 | 50735617 | 268 | 284 | [illegible] |
| SE | 75216 | ENSG00000100503 | NIN | chr14 | - | 50744242 | 50744365 | 50743415 | 50743529 | 50747991 | 50748105 | 876 | 782 | [illegible] |
| SE | 75220 | ENSG00000100503 | NIN | chr14 | - | 50759856 | 50760359 | 50754741 | 50754867 | 50763825 | 50763964 | 75 | 86 | [illegible] |
| SE | 75222 | ENSG00000100503 | NIN | chr14 | - | 50761789 | 50761911 | 50747991 | 50748105 | 50763825 | 50763964 | 523 | 578 | [illegible] |
| SE | 75223 | ENSG00000100503 | NIN | chr14 | - | 50761789 | 50761911 | 50754741 | 50754867 | 50763825 | 50763964 | 534 | 584 | [illegible] |
| SE | 75224 | ENSG00000100503 | NIN | chr14 | - | 50761789 | 50761911 | 50759856 | 50760359 | 50763825 | 50763964 | 664 | 711 | [illegible] |
| SE | 75225 | ENSG00000100503 | NIN | chr14 | - | 50761789 | 50761972 | 50747991 | 50748105 | 50763825 | 50763964 | 8 | 7 | [illegible] |
| SE | 75226 | ENSG00000100503 | NIN | chr14 | - | 50761789 | 50761972 | 50754741 | 50754867 | 50763825 | 50763964 | 19 | 15 | [illegible] |
| SE | 75227 | ENSG00000100503 | NIN | chr14 | - | 50761789 | 50761972 | 50759856 | 50760359 | 50763825 | 50763964 | 137 | 117 | [illegible] |
| SE | 75229 | ENSG00000100503 | NIN | chr14 | - | 50766779 | 50766890 | 50763825 | 50763964 | 50770387 | 50770562 | 125 | 149 | [illegible] |
| SE | 75231 | ENSG00000100503 | NIN | chr14 | - | 50772948 | 50773095 | 50772300 | 50772468 | 50776948 | 50777139 | 713 | 595 | [illegible] |
| SE | 75233 | ENSG00000100503 | NIN | chr14 | - | 50792711 | 50792881 | 50778764 | 50778804 | 50806736 | 50806818 | 538 | 431 | [illegible] |
| SE | 75235 | ENSG00000100503 | NIN | chr14 | - | 50806207 | 50806276 | 50792711 | 50792881 | 50806736 | 50806818 | 2 | 3 | [illegible] |
| SE | 75238 | ENSG00000065883 | CDK13 | chr7 | + | 39987598 | 39988258 | 39950120 | 39951852 | 39997493 | 39997664 | 299 | 234 | [illegible] |
| SE | 75239 | ENSG00000065883 | CDK13 | chr7 | + | 39997493 | 39997664 | 39987598 | 39988258 | 39999360 | 39999500 | 611 | 477 | [illegible] |
| SE | 75240 | ENSG00000065883 | CDK13 | chr7 | + | 39997493 | 39997664 | 39987598 | 39988258 | 40001860 | 40002031 | 282 | 223 | [illegible] |
| SE | 75242 | ENSG00000065883 | CDK13 | chr7 | + | 39999360 | 39999500 | 39987598 | 39988258 | 40001860 | 40002031 | 212 | 169 | [illegible] |
| SE | 75243 | ENSG00000065883 | CDK13 | chr7 | + | 39999360 | 39999500 | 39997493 | 39997664 | 40001860 | 40002031 | 541 | 424 | [illegible] |
| SE | 75245 | ENSG00000065883 | CDK13 | chr7 | + | 40045835 | 40046025 | 40001860 | 40002031 | 40047820 | 40047877 | 885 | 575 | [illegible] |
| SE | 75248 | ENSG00000204396 | VWA7 | chr6 | - | 31766244 | 31766384 | 31765882 | 31766057 | 31766462 | 31766764 | 2 | 0 | [illegible] |
| SE | 75249 | ENSG00000175575 | PAAF1 | chr11 | + | 73878778 | 73878819 | 73876979 | 73877068 | 73887353 | 73887457 | 23 | 23 | [illegible] |
| SE | 75250 | ENSG00000175575 | PAAF1 | chr11 | + | 73878782 | 73878819 | 73876996 | 73877068 | 73887353 | 73887457 | 13 | 14 | [illegible] |
| SE | 75257 | ENSG00000175575 | PAAF1 | chr11 | + | 73909398 | 73909593 | 73900269 | 73900420 | 73914412 | 73914447 | 75 | 68 | [illegible] |
| SE | 75259 | ENSG00000175575 | PAAF1 | chr11 | + | 73918949 | 73919032 | 73916544 | 73916660 | 73924614 | 73924697 | 22 | 21 | [illegible] |
| SE | 75261 | ENSG00000128272 | ATF4 | chr22 | + | 39521257 | 39521671 | 39520575 | 39520751 | 39521772 | 39522668 | 520 | 360 | [illegible] |
| SE | 75262 | ENSG00000128272 | ATF4 | chr22 | + | 39521353 | 39521671 | 39520177 | 39520751 | 39521772 | 39522602 | 635 | 395 | [illegible] |
| SE | 75263 | ENSG00000128272 | ATF4 | chr22 | + | 39521371 | 39521671 | 39520575 | 39520751 | 39521772 | 39522602 | 522 | 360 | [illegible] |
| SE | 75264 | ENSG00000128272 | ATF4 | chr22 | + | 39521374 | 39521671 | 39520603 | 39520751 | 39521772 | 39522602 | 520 | 360 | [illegible] |
| SE | 75265 | ENSG00000128272 | ATF4 | chr22 | + | 39521431 | 39521671 | 39520578 | 39520751 | 39521772 | 39522602 | 520 | 360 | [illegible] |
| SE | 75267 | ENSG00000100504 | PYGL | chr14 | - | 50912155 | 50912303 | 50911977 | 50912036 | 50913028 | 50913130 | 256 | 155 | [illegible] |
| SE | 75269 | ENSG00000100504 | PYGL | chr14 | - | 50915335 | 50915499 | 50914700 | 50914815 | 50915824 | 50915971 | 227 | 182 | [illegible] |
| SE | 75270 | ENSG00000100504 | PYGL | chr14 | - | 50916641 | 50916734 | 50915824 | 50915971 | 50916961 | 50917105 | 327 | 227 | [illegible] |
| SE | 75271 | ENSG00000100504 | PYGL | chr14 | - | 50920540 | 50920623 | 50916961 | 50917105 | 50920955 | 50921067 | 235 | 137 | [illegible] |
| SE | 75272 | ENSG00000100504 | PYGL | chr14 | - | 50923968 | 50924100 | 50920955 | 50921067 | 50931672 | 50931776 | 431 | 276 | [illegible] |
| SE | 75274 | ENSG00000024048 | UBR2 | chr6 | + | 42573733 | 42573993 | 42564028 | 42564397 | 42592150 | 42592229 | 351 | 272 | [illegible] |
| SE | 75278 | ENSG00000024048 | UBR2 | chr6 | + | 42594190 | 42594304 | 42592150 | 42592229 | 42603587 | 42603718 | 520 | 460 | [illegible] |
| SE | 75279 | ENSG00000024048 | UBR2 | chr6 | + | 42603587 | 42603718 | 42594190 | 42594304 | 42605720 | 42605859 | 387 | 313 | [illegible] |
| SE | 75280 | ENSG00000024048 | UBR2 | chr6 | + | 42605720 | 42605859 | 42603587 | 42603718 | 42606588 | 42606651 | 300 | 223 | [illegible] |
| SE | 75282 | ENSG00000024048 | UBR2 | chr6 | + | 42615070 | 42615178 | 42612170 | 42612291 | 42616001 | 42616090 | 317 | 262 | [illegible] |
| SE | 75283 | ENSG00000024048 | UBR2 | chr6 | + | 42616001 | 42616090 | 42615070 | 42615178 | 42617408 | 42617507 | 229 | 201 | [illegible] |
| SE | 75285 | ENSG00000024048 | UBR2 | chr6 | + | 42640208 | 42640270 | 42637010 | 42637194 | 42641581 | 42641692 | 284 | 271 | [illegible] |
| SE | 75286 | ENSG00000024048 | UBR2 | chr6 | + | 42644472 | 42644536 | 42644213 | 42644336 | 42645465 | 42645590 | 324 | 190 | [illegible] |
| SE | 75287 | ENSG00000024048 | UBR2 | chr6 | + | 42650283 | 42650386 | 42648117 | 42648170 | 42652022 | 42652071 | 375 | 326 | [illegible] |
| SE | 75288 | ENSG00000024048 | UBR2 | chr6 | + | 42655620 | 42655723 | 42652490 | 42652645 | 42658023 | 42658133 | 422 | 382 | [illegible] |
| SE | 75290 | ENSG00000024048 | UBR2 | chr6 | + | 42666166 | 42666245 | 42663257 | 42663419 | 42670091 | 42670240 | 103 | 78 | [illegible] |
| SE | 75292 | ENSG00000024048 | UBR2 | chr6 | + | 42679723 | 42679832 | 42678538 | 42678669 | 42683054 | 42683111 | 287 | 199 | [illegible] |
| SE | 75293 | ENSG00000024048 | UBR2 | chr6 | + | 42683054 | 42683111 | 42679723 | 42679832 | 42684793 | 42684871 | 226 | 192 | [illegible] |
| SE | 75295 | ENSG00000184916 | JAG2 | chr14 | - | 105144929 | 105145061 | 105143481 | 105143638 | 105145730 | 105145973 | 1 | 1 | [illegible] |
| SE | 75296 | ENSG00000184916 | JAG2 | chr14 | - | 105150990 | 105151104 | 105150864 | 105150911 | 105151282 | 105151396 | 4 | 0 | [illegible] |
| SE | 75311 | ENSG00000204256 | BRD2 | chr6 | + | 32971594 | 32972927 | 32968593 | 32969056 | 32974461 | 32974765 | 90 | 55 | [illegible] |
| SE | 75313 | ENSG00000204256 | BRD2 | chr6 | + | 32974461 | 32974765 | 32971594 | 32972927 | 32975020 | 32975112 | 322 | 240 | [illegible] |
| SE | 75315 | ENSG00000204256 | BRD2 | chr6 | + | 32976561 | 32976936 | 32975383 | 32976464 | 32977441 | 32977570 | 961 | 593 | [illegible] |
| SE | 75316 | ENSG00000204256 | BRD2 | chr6 | + | 32977259 | 32977570 | 32976561 | 32976936 | 32977756 | 32978005 | 1207 | 793 | [illegible] |
| SE | 75317 | ENSG00000204256 | BRD2 | chr6 | + | 32977259 | 32977570 | 32976561 | 32976936 | 32979827 | 32980132 | 1 | 1 | [illegible] |
| SE | 75318 | ENSG00000204256 | BRD2 | chr6 | + | 32977416 | 32977570 | 32976568 | 32976936 | 32977756 | 32977901 | 1207 | 802 | [illegible] |
| SE | 75319 | ENSG00000204256 | BRD2 | chr6 | + | 32977441 | 32977570 | 32976561 | 32976936 | 32977756 | 32978005 | 1934 | 1249 | [illegible] |
| SE | 75320 | ENSG00000204256 | BRD2 | chr6 | + | 32977441 | 32977570 | 32976561 | 32976936 | 32979827 | 32980132 | 663 | 419 | [illegible] |
| SE | 75322 | ENSG00000204256 | BRD2 | chr6 | + | 32978125 | 32978388 | 32977756 | 32978005 | 32980341 | 32980464 | 324 | 221 | [illegible] |
| SE | 75325 | ENSG00000204256 | BRD2 | chr6 | + | 32979827 | 32980132 | 32978125 | 32978388 | 32980341 | 32980464 | 544 | 413 | [illegible] |
| SE | 75326 | ENSG00000204256 | BRD2 | chr6 | + | 32979874 | 32980132 | 32978125 | 32978388 | 32980341 | 32980464 | 222 | 161 | [illegible] |
| SE | 75327 | ENSG00000196262 | PPIA | chr7 | + | 44797519 | 44797621 | 44796680 | 44796793 | 44799246 | 44799277 | 8 | 7 | [illegible] |
| SE | 75331 | ENSG00000196262 | PPIA | chr7 | + | 44798747 | 44798904 | 44797599 | 44798006 | 44799246 | 44799277 | 77 | 45 | [illegible] |
| SE | 75333 | ENSG00000196262 | PPIA | chr7 | + | 44799391 | 44799480 | 44796978 | 44799277 | 44799701 | 44799874 | 9943 | 5686 | [illegible] |
| SE | 75337 | ENSG00000196262 | PPIA | chr7 | + | 44799701 | 44799874 | 44799391 | 44799480 | 44800423 | 44800543 | 5300 | 2994 | [illegible] |
| SE | 75338 | ENSG00000196262 | PPIA | chr7 | + | 44799701 | 44799874 | 44799391 | 44799480 | 44801286 | 44801630 | 9448 | 5299 | [illegible] |
| SE | 75339 | ENSG00000196262 | PPIA | chr7 | + | 44800423 | 44800543 | 44799391 | 44799480 | 44801286 | 44801624 | 1 | 0 | [illegible] |
| SE | 75340 | ENSG00000196262 | PPIA | chr7 | + | 44800423 | 44800543 | 44799701 | 44799874 | 44801286 | 44801624 | 1 | 0 | [illegible] |
| SE | 75343 | ENSG00000180992 | MRPL14 | chr6 | - | 44116540 | 44116629 | 44113450 | 44114209 | 44127343 | 44127452 | 56 | 19 | [illegible] |
| SE | 75347 | ENSG00000171453 | POLR1C | chr6 | + | 43520274 | 43520427 | 43519705 | 43520185 | 43520624 | 43520703 | 60 | 37 | [illegible] |
| SE | 75348 | ENSG00000171453 | POLR1C | chr6 | + | 43520624 | 43520774 | 43519705 | 43520427 | 43520931 | 43521048 | 44 | 13 | [illegible] |
| SE | 75349 | ENSG00000171453 | POLR1C | chr6 | + | 43521181 | 43521317 | 43520931 | 43521048 | 43529248 | 43529404 | 8 | 6 | [illegible] |
| SE | 75350 | ENSG00000171453 | POLR1C | chr6 | + | 43529248 | 43529359 | 43520931 | 43521048 | 43550967 | 43551011 | 1 | 0 | [illegible] |
| SE | 75352 | ENSG00000066044 | ELAVL1 | chr19 | - | 7967564 | 7967790 | 7958572 | 7963807 | 7973558 | 7973625 | 201 | 86 | [illegible] |
| SE | 75355 | ENSG00000066044 | ELAVL1 | chr19 | - | 7973558 | 7973625 | 7967564 | 7967790 | 7973724 | 7973878 | 0 | 0 | [illegible] |
| SE | 75356 | ENSG00000066044 | ELAVL1 | chr19 | - | 7973724 | 7973878 | 7967564 | 7967790 | 7981082 | 7981112 | 557 | 332 | [illegible] |
| SE | 75357 | ENSG00000186063 | AIDA | chr1 | - | 222670132 | 222670250 | 222668012 | 222669989 | 222673312 | 222673435 | 664 | 566 | [illegible] |
| SE | 75358 | ENSG00000186063 | AIDA | chr1 | - | 222686929 | 222687036 | 222676095 | 222676218 | 222687594 | 222687658 | 470 | 386 | [illegible] |
| SE | 75359 | ENSG00000186063 | AIDA | chr1 | - | 222687594 | 222687658 | 222686929 | 222687036 | 222693788 | 222693843 | 399 | 338 | [illegible] |
| SE | 75360 | ENSG00000186063 | AIDA | chr1 | - | 222693788 | 222693843 | 222687594 | 222687658 | 222694209 | 222694263 | 470 | 448 | [illegible] |
| SE | 75364 | ENSG00000132434 | LANCL2 | chr7 | + | 55398422 | 55398630 | 55391792 | 55391910 | 55399956 | 55400104 | 34 | 38 | [illegible] |
| SE | 75365 | ENSG00000132434 | LANCL2 | chr7 | + | 55411906 | 55412089 | 55401173 | 55401320 | 55425253 | 55425430 | 40 | 23 | [illegible] |
| SE | 75367 | ENSG00000139746 | RBM26 | chr13 | - | 79358273 | 79358433 | 79355219 | 79355384 | 79359574 | 79359686 | 346 | 269 | [illegible] |
| SE | 75368 | ENSG00000139746 | RBM26 | chr13 | - | 79359574 | 79359686 | 79355219 | 79355384 | 79365577 | 79365718 | 112 | 106 | [illegible] |
| SE | 75369 | ENSG00000139746 | RBM26 | chr13 | - | 79370944 | 79371162 | 79368729 | 79368990 | 79371841 | 79371930 | 399 | 394 | [illegible] |
| SE | 75373 | ENSG00000113552 | GNPDA1 | chr5 | - | 142007798 | 142007900 | 142006035 | 142006326 | 142011911 | 142012041 | 70 | 46 | [illegible] |
| SE | 75374 | ENSG00000113552 | GNPDA1 | chr5 | - | 142012287 | 142012339 | 142011911 | 142012041 | 142012668 | 142012996 | 5 | 2 | [illegible] |
| SE | 75375 | ENSG00000113552 | GNPDA1 | chr5 | - | 142012287 | 142012339 | 142011911 | 142012041 | 142012994 | 142013030 | 7 | 2 | [illegible] |
| SE | 75376 | ENSG00000113552 | GNPDA1 | chr5 | - | 142012287 | 142012377 | 142011911 | 142012041 | 142012668 | 142012973 | 9 | 3 | [illegible] |
| SE | 75378 | ENSG00000156508 | EEF1A1 | chr6 | - | 73518029 | 73518264 | 73515749 | 73517934 | 73518353 | 73518610 | 58486 | 31985 | [illegible] |
| SE | 75379 | ENSG00000156508 | EEF1A1 | chr6 | - | 73518353 | 73518610 | 73489307 | 73492804 | 73518697 | 73518848 | 46816 | 27095 | [illegible] |
| SE | 75380 | ENSG00000156508 | EEF1A1 | chr6 | - | 73518353 | 73518610 | 73517514 | 73518264 | 73518697 | 73518848 | 79601 | 44086 | [illegible] |
| SE | 75381 | ENSG00000156508 | EEF1A1 | chr6 | - | 73519336 | 73519507 | 73518931 | 73519228 | 73519882 | 73520056 | 48693 | 31235 | [illegible] |
| SE | 75382 | ENSG00000156508 | EEF1A1 | chr6 | - | 73519336 | 73519516 | 73518697 | 73519228 | 73519882 | 73520056 | 105666 | 71355 | [illegible] |
| SE | 75388 | ENSG00000247134 | NA | chr8 | + | 32996200 | 32996316 | 32927920 | 32928093 | 33041634 | 33041743 | 2 | 0 | [illegible] |
| SE | 75391 | ENSG00000247134 | NA | chr8 | + | 32996200 | 32996355 | 32927976 | 32928093 | 33041634 | 33041771 | 1 | 0 | [illegible] |
| SE | 75393 | ENSG00000247134 | NA | chr8 | + | 33041634 | 33041743 | 32996177 | 32996316 | 33044366 | 33044855 | 5 | 0 | [illegible] |
| SE | 75395 | ENSG00000247134 | NA | chr8 | + | 33041634 | 33041771 | 32996177 | 32996316 | 33044366 | 33044614 | 27 | 10 | [illegible] |
| SE | 75398 | ENSG00000169826 | CSGALNACT2 | chr10 | + | 43158714 | 43158931 | 43154896 | 43155810 | 43160493 | 43160595 | 374 | 269 | [illegible] |
| SE | 75399 | ENSG00000169826 | CSGALNACT2 | chr10 | + | 43158714 | 43158931 | 43154896 | 43155810 | 43163865 | 43164044 | 183 | 166 | [illegible] |
| SE | 75400 | ENSG00000169826 | CSGALNACT2 | chr10 | + | 43158714 | 43158931 | 43154896 | 43155810 | 43167003 | 43167098 | 182 | 167 | [illegible] |
| SE | 75401 | ENSG00000169826 | CSGALNACT2 | chr10 | + | 43158714 | 43158931 | 43154896 | 43155810 | 43175950 | 43176032 | 182 | 167 | [illegible] |
| SE | 75402 | ENSG00000169826 | CSGALNACT2 | chr10 | + | 43160493 | 43160595 | 43154896 | 43155810 | 43163865 | 43164044 | 107 | 67 | [illegible] |
| SE | 75403 | ENSG00000169826 | CSGALNACT2 | chr10 | + | 43160493 | 43160595 | 43154896 | 43155810 | 43167003 | 43167098 | 4 | 1 | [illegible] |
| SE | 75404 | ENSG00000169826 | CSGALNACT2 | chr10 | + | 43160493 | 43160595 | 43154896 | 43155810 | 43175950 | 43176032 | 3 | 0 | [illegible] |
| SE | 75407 | ENSG00000169826 | CSGALNACT2 | chr10 | + | 43160493 | 43160595 | 43158714 | 43158931 | 43175950 | 43176032 | 140 | 69 | [illegible] |
| SE | 75408 | ENSG00000169826 | CSGALNACT2 | chr10 | + | 43163865 | 43164044 | 43154896 | 43155810 | 43167003 | 43167098 | 155 | 109 | [illegible] |
| SE | 75409 | ENSG00000169826 | CSGALNACT2 | chr10 | + | 43163865 | 43164044 | 43154896 | 43155810 | 43175950 | 43176032 | 7 | 5 | [illegible] |
| SE | 75411 | ENSG00000169826 | CSGALNACT2 | chr10 | + | 43163865 | 43164044 | 43158714 | 43158931 | 43175950 | 43176032 | 8 | 5 | [illegible] |
| SE | 75414 | ENSG00000169826 | CSGALNACT2 | chr10 | + | 43167003 | 43167098 | 43154896 | 43155810 | 43175950 | 43176032 | 96 | 81 | [illegible] |
| SE | 75415 | ENSG00000169826 | CSGALNACT2 | chr10 | + | 43167003 | 43167098 | 43158714 | 43158931 | 43175950 | 43176032 | 96 | 82 | [illegible] |
| SE | 75418 | ENSG00000244482 | LILRA6 | chr19 | - | 54238485 | 54238613 | 54236591 | 54237193 | 54238821 | 54239089 | 34 | 20 | [illegible] |
| SE | 75423 | ENSG00000244482 | LILRA6 | chr19 | - | 54240830 | 54241127 | 54240273 | 54240576 | 54241661 | 54241878 | 82 | 45 | [illegible] |
| SE | 75424 | ENSG00000244482 | LILRA6 | chr19 | - | 54241575 | 54241878 | 54240830 | 54241127 | 54242025 | 54242310 | 53 | 28 | [illegible] |
| SE | 75425 | ENSG00000244482 | LILRA6 | chr19 | - | 54241661 | 54241878 | 54240830 | 54241127 | 54242025 | 54242310 | 9 | 20 | [illegible] |
| SE | 75426 | ENSG00000244482 | LILRA6 | chr19 | - | 54242518 | 54242554 | 54242025 | 54242310 | 54242717 | 54242751 | 6 | 0 | [illegible] |
| SE | 75427 | ENSG00000135049 | AGTPBP1 | chr9 | - | 85575314 | 85575475 | 85546538 | 85547286 | 85578919 | 85579025 | 402 | 314 | [illegible] |
| SE | 75429 | ENSG00000135049 | AGTPBP1 | chr9 | - | 85589527 | 85589681 | 85586830 | 85586960 | 85592559 | 85592704 | 330 | 198 | [illegible] |
| SE | 75430 | ENSG00000135049 | AGTPBP1 | chr9 | - | 85596361 | 85596449 | 85592559 | 85592704 | 85618982 | 85619131 | 602 | 326 | [illegible] |

| | | | | | | | | | | | | | | | | | | | | | | | | | | | | | | | | | | | | | | | | | | | | | |
|---|---|---|---|---|---|---|---|---|---|---|---|---|---|---|---|---|---|---|---|---|---|---|---|---|---|---|---|---|---|---|---|---|---|---|---|---|---|---|---|---|---|---|---|---|---|
| SE | 75431 | ENSG00000135049 | AGTPBP1 | chr9 | - | 85621201 | 85621285 | 85619214 | 85619301 | 85632661 | 85633374 | 703 | 381 | 464 | 477 | 1061 | 1 | 0 | 0 | 0 | 0 | 421 | 417 | 565 | 276 | 159 | 0 | 0 | 2 | 0 | 0 | 233 | 149 | 1 | 1 | 0.998 | 1 | 1 | 1 | 1 | 1 | 1 | 0.994 | 1 | 1 | 0.001 |
| SE | 75432 | ENSG00000135049 | AGTPBP1 | chr9 | - | 85632661 | 85633374 | 85621201 | 85621285 | 85642826 | 85642943 | 892 | 552 | 635 | 661 | 1282 | 0 | 1 | 0 | 1 | 3 | 341 | 680 | 823 | 406 | 135 | 0 | 0 | 5 | 0 | 0 | 298 | 149 | 1 | 1 | 1 | 0.996 | 1 | 0.997 | 0.995 | 1 | 1 | 0.988 | 1 | 1 | 0 |
| SE | 75433 | ENSG00000135049 | AGTPBP1 | chr9 | - | 85632661 | 85633374 | 85621201 | 85621285 | 85655142 | 85655200 | 413 | 220 | 287 | 278 | 572 | 0 | 0 | 0 | 0 | 0 | 259 | 277 | 342 | 166 | 114 | 0 | 0 | 3 | 0 | 0 | 298 | 149 | 1 | 1 | 1 | 1 | 1 | 1 | 1 | 1 | 1 | 0.983 | 1 | 1 | 0.003 |
| SE | 75434 | ENSG00000135049 | AGTPBP1 | chr9 | - | 85632661 | 85633374 | 85621201 | 85621285 | 85669484 | 85669578 | 416 | 221 | 288 | 278 | 572 | 1 | 0 | 0 | 0 | 0 | 259 | 277 | 342 | 166 | 110 | 0 | 2 | 0 | 0 | 0 | 298 | 149 | 1 | 1 | 0.995 | 1 | 1 | 1 | 1 | 1 | 0.986 | 1 | 1 | 1 | 0.002 |
| SE | 75435 | ENSG00000135049 | AGTPBP1 | chr9 | - | 85642826 | 85642943 | 85596361 | 85596449 | 85655142 | 85655200 | 0 | 7 | 2 | 7 | 2 | 0 | 1 | 0 | 0 | 0 | 0 | 4 | 0 | 0 | 0 | 0 | 0 | 0 | 0 | 0 | 266 | 149 | 1 | 1 | NA | 0.797 | 1 | 1 | 1 | NA | 1 | NA | NA | NA | -0.051 |
| SE | 75436 | ENSG00000135049 | AGTPBP1 | chr9 | - | 85642826 | 85642943 | 85621201 | 85621285 | 85655142 | 85655200 | 0 | 4 | 0 | 7 | 5 | 0 | 0 | 0 | 0 | 0 | 0 | 4 | 5 | 0 | 0 | 0 | 0 | 3 | 0 | 0 | 266 | 149 | 1 | 1 | NA | 1 | NA | 1 | 1 | NA | 1 | 0.483 | NA | NA | 0.258 |
| SE | 75438 | ENSG00000135049 | AGTPBP1 | chr9 | - | 85646320 | 85646418 | 85642826 | 85642943 | 85655142 | 85655200 | 642 | 400 | 457 | 501 | 1196 | 0 | 3 | 0 | 6 | 2 | 158 | 471 | 664 | 336 | 60 | 0 | 4 | 0 | 0 | 0 | 247 | 149 | 1 | 1 | 1 | 0.988 | 1 | 0.981 | 0.997 | 1 | 0.986 | 1 | 1 | 1 | -0.004 |
| SE | 75441 | ENSG00000135049 | AGTPBP1 | chr9 | - | 85655142 | 85655200 | 85632661 | 85633374 | 85669484 | 85669578 | 0 | 0 | 0 | 0 | 0 | 3 | 1 | 1 | 0 | 0 | 0 | 0 | 0 | 0 | 2 | 0 | 0 | 0 | 0 | 0 | 207 | 149 | 1 | 1 | 0 | 0 | 0 | NA | NA | NA | NA | NA | NA | 1 | -1 |
| SE | 75443 | ENSG00000135049 | AGTPBP1 | chr9 | - | 85669484 | 85669578 | 85632661 | 85633374 | 85677435 | 85677582 | 2 | 3 | 1 | 3 | 0 | 1 | 0 | 0 | 0 | 0 | 0 | 6 | 0 | 0 | 2 | 0 | 0 | 0 | 0 | 0 | 243 | 149 | 1 | 1 | 0.551 | 1 | 1 | 1 | NA | NA | 1 | NA | NA | 1 | -0.112 |
| SE | 75444 | ENSG00000135049 | AGTPBP1 | chr9 | - | 85672549 | 85672681 | 85669484 | 85669578 | 85677435 | 85677582 | 868 | 598 | 629 | 755 | 1029 | 2 | 4 | 0 | 4 | 2 | 344 | 688 | 733 | 438 | 170 | 0 | 6 | 3 | 0 | 2 | 281 | 149 | 1 | 1 | 0.996 | 0.988 | 1 | 0.99 | 0.996 | 1 | 0.984 | 0.992 | 1 | 0.978 | 0.003 |
| SE | 75445 | ENSG00000135049 | AGTPBP1 | chr9 | - | 85677435 | 85677582 | 85669484 | 85669578 | 85692688 | 85692780 | 2 | 5 | 2 | 4 | 2 | 0 | 0 | 0 | 0 | 1 | 0 | 6 | 3 | 0 | 2 | 0 | 0 | 0 | 0 | 0 | 296 | 149 | 1 | 1 | 1 | 1 | 1 | 1 | 0.502 | NA | 1 | 1 | NA | 1 | -0.1 |
| SE | 75447 | ENSG00000135049 | AGTPBP1 | chr9 | - | 85678334 | 85678398 | 85677435 | 85677582 | 85692688 | 85692780 | 232 | 152 | 163 | 183 | 291 | 0 | 1 | 2 | 0 | 0 | 115 | 129 | 194 | 108 | 80 | 0 | 0 | 0 | 0 | 0 | 213 | 149 | 1 | 1 | 1 | 0.991 | 0.983 | 1 | 1 | 1 | 1 | 1 | 1 | 1 | -0.005 |
| SE | 75448 | ENSG00000135049 | AGTPBP1 | chr9 | - | 85681267 | 85681335 | 85677435 | 85677582 | 85692688 | 85692780 | 126 | 92 | 95 | 124 | 130 | 0 | 1 | 2 | 0 | 0 | 108 | 137 | 173 | 91 | 21 | 0 | 0 | 0 | 0 | 0 | 217 | 149 | 1 | 1 | 1 | 0.984 | 0.97 | 1 | 1 | 1 | 1 | 1 | 1 | 1 | -0.009 |
| SE | 75449 | ENSG00000135049 | AGTPBP1 | chr9 | - | 85681267 | 85681335 | 85678334 | 85678398 | 85692688 | 85692780 | 476 | 332 | 339 | 418 | 472 | 1 | 0 | 2 | 0 | 0 | 223 | 412 | 410 | 259 | 150 | 0 | 0 | 0 | 0 | 0 | 217 | 149 | 1 | 1 | 0.997 | 1 | 0.991 | 1 | 1 | 1 | 1 | 1 | 1 | 1 | -0.002 |
| SE | 75450 | ENSG00000135049 | AGTPBP1 | chr9 | - | 85692688 | 85692813 | 85681267 | 85681335 | 85741774 | 85742029 | 106 | 101 | 84 | 120 | 127 | 0 | 0 | 0 | 0 | 0 | 47 | 98 | 108 | 54 | 22 | 0 | 0 | 0 | 4 | 0 | 274 | 149 | 1 | 1 | 1 | 1 | 1 | 1 | 1 | 1 | 1 | 1 | 0.88 | 1 | 0.024 |
| SE | 75451 | ENSG00000135049 | AGTPBP1 | chr9 | - | 85712501 | 85712566 | 85692688 | 85692813 | 85741774 | 85741818 | 240 | 175 | 146 | 230 | 192 | 0 | 0 | 0 | 0 | 0 | 123 | 199 | 291 | 146 | 75 | 0 | 3 | 0 | 0 | 0 | 214 | 149 | 1 | 1 | 1 | 1 | 1 | 1 | 1 | 1 | 0.979 | 1 | 1 | 1 | 0.004 |
| SE | 75452 | ENSG00000074842 | MYDGF | chr19 | - | 4660668 | 4660750 | 4659687 | 4660003 | 4664875 | 4664906 | 214 | 85 | 181 | 150 | 75 | 1 | 0 | 0 | 0 | 0 | 185 | 286 | 231 | 167 | 291 | 0 | 0 | 2 | 0 | 0 | 231 | 149 | 1 | 1 | 0.993 | 1 | 1 | 1 | 1 | 1 | 1 | 0.987 | 1 | 1 | 0.001 |
| SE | 75453 | ENSG00000074842 | MYDGF | chr19 | - | 4664875 | 4664937 | 4660579 | 4660750 | 4668594 | 4668645 | 171 | 75 | 162 | 117 | 60 | 0 | 0 | 0 | 2 | 0 | 172 | 244 | 193 | 136 | 267 | 0 | 0 | 0 | 0 | 0 | 211 | 149 | 1 | 1 | 1 | 1 | 1 | 0.976 | 1 | 1 | 1 | 1 | 1 | 1 | -0.005 |
| SE | 75454 | ENSG00000197081 | IGF2R | chr6 | + | 160027184 | 160027314 | 160024571 | 160024704 | 160029549 | 160029655 | 727 | 261 | 402 | 525 | 522 | 0 | 0 | 0 | 0 | 0 | 369 | 542 | 395 | 498 | 637 | 0 | 0 | 1 | 0 | 0 | 279 | 149 | 1 | 1 | 1 | 1 | 1 | 1 | 1 | 1 | 1 | 0.995 | 1 | 1 | 0.001 |
| SE | 75457 | ENSG00000197081 | IGF2R | chr6 | + | 160045744 | 160045882 | 160044513 | 160044657 | 160046497 | 160046645 | 1518 | 743 | 838 | 1127 | 740 | 2 | 0 | 1 | 0 | 0 | 1801 | 1376 | 1480 | 1371 | 1635 | 0 | 0 | 2 | 1 | 6 | 287 | 149 | 1 | 1 | 0.997 | 1 | 0.998 | 1 | 1 | 1 | 1 | 0.997 | 0.999 | 0.993 | 0.001 |
| SE | 75458 | ENSG00000197081 | IGF2R | chr6 | + | 160046891 | 160047057 | 160046497 | 160046645 | 160047158 | 160047336 | 0 | 0 | 0 | 0 | 0 | 335 | 140 | 188 | 202 | 293 | 10 | 0 | 0 | 0 | 0 | 310 | 214 | 301 | 249 | 280 | 298 | 149 | 1 | 1 | 0 | 0 | 0 | 0 | 0 | 0.016 | 0 | 0 | 0 | 0 | -0.003 |
| SE | 75460 | ENSG00000197081 | IGF2R | chr6 | + | 160056423 | 160056525 | 160050472 | 160050652 | 160058905 | 160059098 | 316 | 104 | 164 | 169 | 194 | 0 | 0 | 0 | 0 | 0 | 191 | 197 | 214 | 234 | 274 | 0 | 3 | 0 | 5 | 0 | 251 | 149 | 1 | 1 | 1 | 1 | 1 | 1 | 1 | 1 | 0.975 | 1 | 0.965 | 1 | 0.012 |
| SE | 75461 | ENSG00000197081 | IGF2R | chr6 | + | 160058022 | 160058124 | 160056423 | 160056525 | 160058905 | 160059098 | 632 | 293 | 335 | 370 | 388 | 0 | 0 | 0 | 0 | 0 | 517 | 408 | 524 | 489 | 598 | 0 | 0 | 0 | 3 | 0 | 251 | 149 | 1 | 1 | 1 | 1 | 1 | 1 | 1 | 1 | 1 | 1 | 0.99 | 1 | 0.002 |
| SE | 75462 | ENSG00000197081 | IGF2R | chr6 | + | 160061502 | 160061646 | 160060546 | 160060717 | 160061752 | 160061928 | 634 | 291 | 361 | 476 | 383 | 0 | 1 | 0 | 0 | 0 | 387 | 574 | 424 | 416 | 500 | 0 | 0 | 0 | 0 | 0 | 293 | 149 | 1 | 1 | 1 | 0.993 | 1 | 1 | 1 | 1 | 1 | 1 | 1 | 1 | -0.001 |
| SE | 75463 | ENSG00000197081 | IGF2R | chr6 | + | 160061502 | 160061646 | 160060546 | 160060717 | 160062531 | 160062619 | 322 | 128 | 147 | 204 | 175 | 0 | 0 | 0 | 0 | 0 | 217 | 209 | 164 | 190 | 275 | 0 | 0 | 0 | 1 | 0 | 293 | 149 | 1 | 1 | 1 | 1 | 1 | 1 | 1 | 1 | 1 | 1 | 0.99 | 1 | 0.002 |
| SE | 75464 | ENSG00000197081 | IGF2R | chr6 | + | 160061752 | 160061928 | 160060546 | 160060717 | 160062531 | 160062619 | 509 | 210 | 296 | 396 | 308 | 0 | 0 | 0 | 0 | 0 | 422 | 441 | 367 | 299 | 490 | 0 | 0 | 0 | 1 | 0 | 298 | 149 | 1 | 1 | 1 | 1 | 1 | 1 | 1 | 1 | 1 | 1 | 0.993 | 1 | 0.001 |
| SE | 75465 | ENSG00000197081 | IGF2R | chr6 | + | 160061752 | 160061928 | 160061502 | 160061646 | 160062531 | 160062619 | 821 | 372 | 510 | 668 | 516 | 0 | 0 | 0 | 0 | 0 | 592 | 806 | 629 | 528 | 715 | 0 | 0 | 2 | 3 | 0 | 298 | 149 | 1 | 1 | 1 | 1 | 1 | 1 | 1 | 1 | 1 | 0.994 | 0.989 | 1 | 0.003 |
| SE | 75466 | ENSG00000197081 | IGF2R | chr6 | + | 160062531 | 160062619 | 160061752 | 160061928 | 160063414 | 160063630 | 943 | 372 | 480 | 686 | 534 | 0 | 0 | 0 | 0 | 0 | 801 | 770 | 698 | 519 | 902 | 0 | 2 | 0 | 5 | 0 | 237 | 149 | 1 | 1 | 1 | 1 | 1 | 1 | 1 | 1 | 0.996 | 1 | 0.985 | 1 | 0.004 |
| SE | 75467 | ENSG00000197081 | IGF2R | chr6 | + | 160062531 | 160062619 | 160061752 | 160061928 | 160064400 | 160064531 | 353 | 135 | 207 | 282 | 193 | 0 | 0 | 0 | 0 | 0 | 213 | 276 | 227 | 193 | 306 | 0 | 0 | 0 | 2 | 0 | 237 | 149 | 1 | 1 | 1 | 1 | 1 | 1 | 1 | 1 | 1 | 1 | 0.984 | 1 | 0.003 |

[illegible]

| | | | | | | | | | | | | | | |
|---|---|---|---|---|---|---|---|---|---|---|---|---|---|---|
| SE | 75807 | ENSG00000105671 | DDX49 | chr19 | + | 18920579 | 18920703 | 18919704 | 18919856 | 18921662 | 18921748 | 28 | 12 | [illegible] |
| SE | 75808 | ENSG00000105671 | DDX49 | chr19 | + | 18920650 | 18920703 | 18919727 | 18919856 | 18921662 | 18921748 | 10 | 4 | [illegible] |
| SE | 75811 | ENSG00000105671 | DDX49 | chr19 | + | 18921662 | 18921748 | 18920579 | 18920703 | 18921842 | 18921964 | 42 | 20 | [illegible] |
| SE | 75814 | ENSG00000153147 | SMARCA5 | chr4 | + | 143526280 | 143526460 | 143525450 | 143525551 | 143527867 | 143528023 | 1177 | 1124 | [illegible] |
| SE | 75816 | ENSG00000153147 | SMARCA5 | chr4 | + | 143528582 | 143528714 | 143526280 | 143526460 | 143530457 | 143530526 | 204 | 171 | [illegible] |
| SE | 75817 | ENSG00000153147 | SMARCA5 | chr4 | + | 143530457 | 143530526 | 143526280 | 143526460 | 143534854 | 143534964 | 152 | 120 | [illegible] |
| SE | 75820 | ENSG00000153147 | SMARCA5 | chr4 | + | 143546775 | 143546908 | 143545469 | 143545583 | 143547384 | 143547503 | 216 | 138 | [illegible] |
| SE | 75821 | ENSG00000153147 | SMARCA5 | chr4 | + | 143547927 | 143548140 | 143547384 | 143547503 | 143549996 | 143550104 | 1199 | 1157 | [illegible] |
| SE | 75822 | ENSG00000112139 | MDGA1 | chr6 | - | 37641736 | 37642049 | 37638536 | 37638667 | 37643808 | 37643943 | 1 | 1 | [illegible] |
| SE | 75823 | ENSG00000112139 | MDGA1 | chr6 | - | 37645932 | 37645956 | 37644496 | 37644649 | 37646197 | 37646375 | 0 | 10 | [illegible] |
| SE | 75824 | ENSG00000124523 | SIRT5 | chr6 | + | 13578771 | 13578833 | 13574235 | 13574306 | 13579450 | 13579609 | 0 | 3 | [illegible] |
| SE | 75825 | ENSG00000124523 | SIRT5 | chr6 | + | 13578771 | 13578833 | 13574600 | 13574744 | 13579450 | 13579609 | 0 | 3 | [illegible] |
| SE | 75829 | ENSG00000124523 | SIRT5 | chr6 | + | 13584986 | 13585127 | 13584075 | 13584225 | 13591668 | 13591894 | 0 | 0 | [illegible] |
| SE | 75830 | ENSG00000124523 | SIRT5 | chr6 | + | 13588330 | 13588464 | 13584075 | 13584225 | 13591668 | 13591894 | 47 | 44 | [illegible] |
| SE | 75834 | ENSG00000124523 | SIRT5 | chr6 | + | 13591668 | 13591894 | 13588330 | 13588464 | 13595476 | 13595564 | 39 | 24 | [illegible] |
| SE | 75835 | ENSG00000124523 | SIRT5 | chr6 | + | 13594429 | 13594845 | 13591668 | 13591894 | 13595476 | 13595564 | 0 | 0 | [illegible] |
| SE | 75837 | ENSG00000124523 | SIRT5 | chr6 | + | 13595476 | 13595564 | 13591668 | 13591894 | 13596962 | 13597016 | 31 | 15 | [illegible] |
| SE | 75839 | ENSG00000124523 | SIRT5 | chr6 | + | 13596962 | 13597016 | 13595476 | 13595564 | 13599031 | 13599155 | 32 | 11 | [illegible] |
| SE | 75841 | ENSG00000124523 | SIRT5 | chr6 | + | 13597727 | 13597864 | 13596962 | 13597016 | 13600833 | 13600949 | 0 | 0 | [illegible] |
| SE | 75842 | ENSG00000124523 | SIRT5 | chr6 | + | 13597730 | 13597864 | 13596962 | 13597016 | 13599031 | 13599155 | 0 | 0 | [illegible] |
| SE | 75843 | ENSG00000124523 | SIRT5 | chr6 | + | 13597730 | 13597864 | 13596962 | 13597016 | 13600833 | 13603253 | 0 | 0 | [illegible] |
| SE | 75845 | ENSG00000124523 | SIRT5 | chr6 | + | 13599031 | 13599155 | 13596962 | 13597016 | 13611789 | 13612287 | 18 | 7 | [illegible] |
| SE | 75846 | ENSG00000124523 | SIRT5 | chr6 | + | 13599031 | 13599155 | 13597727 | 13597864 | 13600833 | 13600949 | 18 | 4 | [illegible] |
| SE | 75847 | ENSG00000124523 | SIRT5 | chr6 | + | 13599031 | 13600949 | 13596962 | 13597016 | 13611789 | 13615158 | 39 | 12 | [illegible] |
| SE | 75849 | ENSG00000124523 | SIRT5 | chr6 | + | 13600833 | 13600949 | 13596962 | 13597016 | 13611789 | 13612287 | 22 | 8 | [illegible] |
| SE | 75854 | ENSG00000132436 | FIGNL1 | chr7 | - | 50447967 | 50448055 | 50446980 | 50447297 | 50448180 | 50448289 | 3 | 2 | [illegible] |
| SE | 75856 | ENSG00000132436 | FIGNL1 | chr7 | - | 50449117 | 50449723 | 50448180 | 50448289 | 50450300 | 50450350 | 2 | 1 | [illegible] |
| SE | 75857 | ENSG00000132436 | FIGNL1 | chr7 | - | 50449499 | 50449723 | 50448180 | 50448289 | 50450300 | 50450336 | 1 | 0 | [illegible] |
| SE | 75858 | ENSG00000132436 | FIGNL1 | chr7 | - | 50449544 | 50449723 | 50448180 | 50448289 | 50450300 | 50450336 | 4 | 0 | [illegible] |
| SE | 75859 | ENSG00000132436 | FIGNL1 | chr7 | - | 50449555 | 50449723 | 50448180 | 50448289 | 50450300 | 50450324 | 0 | 0 | [illegible] |
| SE | 75860 | ENSG00000132436 | FIGNL1 | chr7 | - | 50449562 | 50449723 | 50448180 | 50448289 | 50450300 | 50450350 | 2 | 2 | [illegible] |
| SE | 75861 | ENSG00000250264 | NA | chr6 | - | 32816860 | 32816952 | 32814319 | 32814601 | 32829399 | 32829536 | 1 | 0 | [illegible] |
| SE | 75862 | ENSG00000250264 | NA | chr6 | - | 32830617 | 32830806 | 32830266 | 32830440 | 32832332 | 32832461 | 235 | 147 | [illegible] |
| SE | 75863 | ENSG00000250264 | NA | chr6 | - | 32835153 | 32835359 | 32832626 | 32832824 | 32835642 | 32835773 | 263 | 98 | [illegible] |
| SE | 75864 | ENSG00000250264 | NA | chr6 | - | 32837536 | 32837651 | 32835642 | 32835773 | 32837740 | 32838237 | 201 | 87 | [illegible] |
| SE | 75865 | ENSG00000253704 | NA | chr8 | + | 94568048 | 94568535 | 94553730 | 94553907 | 94570109 | 94570648 | 0 | 1 | [illegible] |
| SE | 75866 | ENSG00000253704 | NA | chr8 | + | 94568142 | 94568535 | 94553769 | 94553907 | 94570109 | 94570531 | 0 | 1 | [illegible] |
| SE | 75867 | ENSG00000253704 | NA | chr8 | + | 94568142 | 94568621 | 94553712 | 94553907 | 94570109 | 94570648 | 0 | 0 | [illegible] |
| SE | 75868 | ENSG00000204264 | PSMB8 | chr6 | - | 32841530 | 32841735 | 32840716 | 32841047 | 32842133 | 32842263 | 552 | 307 | [illegible] |
| SE | 75869 | ENSG00000240065 | PSMB9 | chr6 | + | 32856137 | 32856205 | 32854191 | 32854289 | 32857262 | 32857394 | 202 | 80 | [illegible] |
| SE | 75874 | ENSG00000164430 | CGAS | chr6 | - | 73440208 | 73440445 | 73428708 | 73428811 | 73445527 | 73445747 | 197 | 138 | [illegible] |
| SE | 75875 | ENSG00000164430 | CGAS | chr6 | - | 73440208 | 73440445 | 73428708 | 73428811 | 73451524 | 73452181 | 68 | 38 | [illegible] |
| SE | 75876 | ENSG00000164430 | CGAS | chr6 | - | 73445527 | 73445747 | 73428708 | 73428811 | 73451524 | 73452181 | 76 | 57 | [illegible] |
| SE | 75879 | ENSG00000138071 | ACTR2 | chr2 | + | 65242048 | 65242063 | 65239851 | 65239962 | 65246523 | 65246739 | 0 | 0 | [illegible] |
| SE | 75880 | ENSG00000138071 | ACTR2 | chr2 | + | 65246184 | 65246307 | 65239851 | 65239962 | 65246523 | 65246718 | 4 | 10 | [illegible] |
| SE | 75881 | ENSG00000138071 | ACTR2 | chr2 | + | 65246523 | 65246739 | 65239851 | 65239962 | 65251026 | 65251099 | 3755 | 2850 | [illegible] |
| SE | 75882 | ENSG00000138071 | ACTR2 | chr2 | + | 65251026 | 65251099 | 65246523 | 65246739 | 65253727 | 65253864 | 2832 | 2133 | [illegible] |
| SE | 75883 | ENSG00000138071 | ACTR2 | chr2 | + | 65255544 | 65255694 | 65253727 | 65253864 | 65261246 | 65261392 | 4466 | 3043 | [illegible] |
| SE | 75884 | ENSG00000138071 | ACTR2 | chr2 | + | 65261246 | 65261392 | 65255544 | 65255694 | 65265042 | 65265175 | 3161 | 2059 | [illegible] |
| SE | 75885 | ENSG00000138071 | ACTR2 | chr2 | + | 65261246 | 65261392 | 65255544 | 65255694 | 65268563 | 65268755 | 1524 | 1002 | [illegible] |
| SE | 75886 | ENSG00000138071 | ACTR2 | chr2 | + | 65265042 | 65265175 | 65255544 | 65255694 | 65268563 | 65268755 | 1473 | 1093 | [illegible] |
| SE | 75887 | ENSG00000138071 | ACTR2 | chr2 | + | 65265042 | 65265175 | 65261246 | 65261392 | 65268563 | 65268755 | 3107 | 2150 | [illegible] |
| SE | 75888 | ENSG00000010704 | HFE | chr6 | + | 26090840 | 26091104 | 26087280 | 26087516 | 26091313 | 26091589 | 33 | 19 | [illegible] |
| SE | 75891 | ENSG00000010704 | HFE | chr6 | + | 26090909 | 26091104 | 26087280 | 26087516 | 26091313 | 26091589 | 25 | 13 | [illegible] |
| SE | 75894 | ENSG00000010704 | HFE | chr6 | + | 26091313 | 26091589 | 26090840 | 26091104 | 26092684 | 26092960 | 37 | 16 | [illegible] |
| SE | 75896 | ENSG00000010704 | HFE | chr6 | + | 26092608 | 26092641 | 26091313 | 26091589 | 26092726 | 26092960 | 0 | 0 | [illegible] |
| SE | 75898 | ENSG00000010704 | HFE | chr6 | + | 26093118 | 26093232 | 26092608 | 26093026 | 26094403 | 26094553 | 0 | 1 | [illegible] |
| SE | 75899 | ENSG00000010704 | HFE | chr6 | + | 26094185 | 26094218 | 26093118 | 26093232 | 26094403 | 26095020 | 5 | 2 | [illegible] |
| SE | 75900 | ENSG00000147650 | LRP12 | chr8 | - | 104531906 | 104531963 | 104508938 | 104509074 | 104588818 | 104589258 | 2 | 7 | [illegible] |
| SE | 75902 | ENSG00000188229 | TUBB4B | chr9 | + | 137241720 | 137241829 | 137241286 | 137241417 | 137241910 | 137242021 | 123 | 49 | [illegible] |
| SE | 75903 | ENSG00000188229 | TUBB4B | chr9 | + | 137241910 | 137242021 | 137241720 | 137241829 | 137242495 | 137243707 | 214 | 77 | [illegible] |
| SE | 75908 | ENSG00000134324 | LPIN1 | chr2 | + | 11764203 | 11764289 | 11763022 | 11763286 | 11765532 | 11765625 | 0 | 0 | [illegible] |
| SE | 75909 | ENSG00000134324 | LPIN1 | chr2 | + | 11767762 | 11767858 | 11765532 | 11765733 | 11771371 | 11771679 | 133 | 91 | [illegible] |
| SE | 75911 | ENSG00000134324 | LPIN1 | chr2 | + | 11782200 | 11782507 | 11779518 | 11779645 | 11783828 | 11783922 | 174 | 118 | [illegible] |
| SE | 75912 | ENSG00000134324 | LPIN1 | chr2 | + | 11784885 | 11785076 | 11783828 | 11783922 | 11787073 | 11787167 | 92 | 77 | [illegible] |
| SE | 75913 | ENSG00000134324 | LPIN1 | chr2 | + | 11787073 | 11787167 | 11783828 | 11783922 | 11788386 | 11788456 | 23 | 15 | [illegible] |
| SE | 75915 | ENSG00000134324 | LPIN1 | chr2 | + | 11795407 | 11795487 | 11791913 | 11792006 | 11802906 | 11803033 | 83 | 61 | [illegible] |
| SE | 75916 | ENSG00000134324 | LPIN1 | chr2 | + | 11802906 | 11803033 | 11795407 | 11795487 | 11804422 | 11804519 | 147 | 104 | [illegible] |
| SE | 75917 | ENSG00000134324 | LPIN1 | chr2 | + | 11805069 | 11805156 | 11804422 | 11804571 | 11815087 | 11815240 | 110 | 81 | [illegible] |
| SE | 75918 | ENSG00000134324 | LPIN1 | chr2 | + | 11819483 | 11819598 | 11815087 | 11815240 | 11820410 | 11820514 | 155 | 76 | [illegible] |
| SE | 75919 | ENSG00000135334 | AKIRIN2 | chr6 | - | 87677817 | 87677967 | 87675859 | 87675931 | 87681619 | 87681763 | 642 | 475 | [illegible] |
| SE | 75922 | ENSG00000137261 | KIAA0319 | chr6 | - | 24554540 | 24554631 | 24551433 | 24551525 | 24556606 | 24556729 | 0 | 0 | [illegible] |
| SE | 75925 | ENSG00000137261 | KIAA0319 | chr6 | - | 24579857 | 24579950 | 24578109 | 24578242 | 24580925 | 24581013 | 1 | 0 | [illegible] |
| SE | 75927 | ENSG00000173588 | CEP83 | chr12 | - | 94309917 | 94310107 | 94307697 | 94308917 | 94312913 | 94312929 | 46 | 32 | [illegible] |
| SE | 75928 | ENSG00000173588 | CEP83 | chr12 | - | 94328204 | 94328308 | 94312913 | 94313017 | 94331699 | 94331829 | 1 | 0 | [illegible] |
| SE | 75929 | ENSG00000173588 | CEP83 | chr12 | - | 94401204 | 94401281 | 94400903 | 94400981 | 94402250 | 94402336 | 0 | 3 | [illegible] |
| SE | 75931 | ENSG00000173588 | CEP83 | chr12 | - | 94434002 | 94434053 | 94412483 | 94412591 | 94435274 | 94435327 | 0 | 0 | [illegible] |
| SE | 75932 | ENSG00000173588 | CEP83 | chr12 | - | 94435274 | 94435327 | 94412317 | 94412591 | 94459556 | 94459867 | 8 | 7 | [illegible] |
| SE | 75933 | ENSG00000160185 | UBASH3A | chr21 | + | 42406307 | 42406361 | 42403861 | 42404058 | 42409421 | 42409608 | 42 | 21 | [illegible] |
| SE | 75938 | ENSG00000160185 | UBASH3A | chr21 | + | 42443153 | 42443418 | 42442451 | 42442596 | 42444533 | 42444643 | 17 | 10 | [illegible] |
| SE | 75939 | ENSG00000160185 | UBASH3A | chr21 | + | 42443311 | 42443418 | 42442451 | 42442596 | 42444533 | 42444643 | 28 | 24 | [illegible] |
| SE | 75942 | ENSG00000196730 | DAPK1 | chr9 | + | 87639649 | 87639698 | 87639353 | 87639483 | 87639788 | 87639815 | 131 | 144 | [illegible] |
| SE | 75945 | ENSG00000196730 | DAPK1 | chr9 | + | 87686550 | 87686739 | 87681403 | 87681626 | 87697006 | 87697204 | 122 | 64 | [illegible] |
| SE | 75947 | ENSG00000147383 | NSDHL | chrX | + | 152831633 | 152831700 | 152830966 | 152831117 | 152846281 | 152846432 | 0 | 0 | [illegible] |
| SE | 75948 | ENSG00000147383 | NSDHL | chrX | + | 152858769 | 152858916 | 152850264 | 152850423 | 152862595 | 152862724 | 75 | 44 | [illegible] |
| SE | 75955 | ENSG00000234745 | HLA-B | chr6 | - | 31354632 | 31354665 | 31354478 | 31354526 | 31355106 | 31355223 | 11756 | 5119 | [illegible] |
| SE | 75956 | ENSG00000234745 | HLA-B | chr6 | - | 31354632 | 31354665 | 31354478 | 31354526 | 31355316 | 31355404 | 1640 | 617 | [illegible] |
| SE | 75957 | ENSG00000234745 | HLA-B | chr6 | - | 31355106 | 31355223 | 31354478 | 31354526 | 31355316 | 31355592 | 11174 | 5086 | [illegible] |
| SE | 75959 | ENSG00000234745 | HLA-B | chr6 | - | 31356166 | 31356442 | 31355316 | 31355592 | 31356687 | 31356957 | 5660 | 1889 | [illegible] |
| SE | 75960 | ENSG00000234745 | HLA-B | chr6 | - | 31356166 | 31356957 | 31355390 | 31355592 | 31357085 | 31357179 | 4067 | 1109 | [illegible] |
| SE | 75961 | ENSG00000234745 | HLA-B | chr6 | - | 31356687 | 31356957 | 31355316 | 31355592 | 31357085 | 31357179 | 177 | 102 | [illegible] |
| SE | 75962 | ENSG00000118873 | RAB3GAP2 | chr1 | - | 220157801 | 220157876 | 220157269 | 220157488 | 220159385 | 220159421 | 94 | 73 | [illegible] |
| SE | 75963 | ENSG00000118873 | RAB3GAP2 | chr1 | - | 220159385 | 220159421 | 220157801 | 220157876 | 220162197 | 220162268 | 101 | 101 | [illegible] |
| SE | 75965 | ENSG00000118873 | RAB3GAP2 | chr1 | - | 220190376 | 220190520 | 220190063 | 220190146 | 220191067 | 220191284 | 110 | 76 | [illegible] |
| SE | 75966 | ENSG00000118873 | RAB3GAP2 | chr1 | - | 220191067 | 220191284 | 220190063 | 220190146 | 220193239 | 220193379 | 86 | 45 | [illegible] |
| SE | 75968 | ENSG00000118873 | RAB3GAP2 | chr1 | - | 220210387 | 220210489 | 220205906 | 220206006 | 220210800 | 220210876 | 154 | 140 | [illegible] |
| SE | 75969 | ENSG00000118873 | RAB3GAP2 | chr1 | - | 220210800 | 220210876 | 220210387 | 220210489 | 220210954 | 220211002 | 120 | 102 | [illegible] |
| SE | 75970 | ENSG00000118873 | RAB3GAP2 | chr1 | - | 220210954 | 220211002 | 220210800 | 220210876 | 220212886 | 220212968 | 120 | 88 | [illegible] |
| SE | 75971 | ENSG00000118873 | RAB3GAP2 | chr1 | - | 220219434 | 220219629 | 220213855 | 220213979 | 220232798 | 220232863 | 4 | 0 | [illegible] |
| SE | 75972 | ENSG00000251364 | NA | chr11 | - | 7427872 | 7428044 | 7427265 | 7427654 | 7440126 | 7440260 | 2 | 4 | [illegible] |
| SE | 75973 | ENSG00000251364 | NA | chr11 | - | 7434982 | 7435022 | 7428008 | 7428044 | 7440126 | 7441719 | 0 | 0 | [illegible] |
| SE | 75974 | ENSG00000251364 | NA | chr11 | - | 7440126 | 7440260 | 7427872 | 7428044 | 7450372 | 7450442 | 4 | 4 | [illegible] |
| SE | 75975 | ENSG00000251364 | NA | chr11 | - | 7440126 | 7441719 | 7427872 | 7428044 | 7450372 | 7450442 | 8 | 7 | [illegible] |
| SE | 75976 | ENSG00000251364 | NA | chr11 | - | 7451812 | 7451963 | 7450372 | 7450442 | 7457795 | 7457959 | 1 | 0 | [illegible] |
| SE | 75978 | ENSG00000251364 | NA | chr11 | - | 7457795 | 7457943 | 7450372 | 7450442 | 7461238 | 7461336 | 6 | 2 | [illegible] |
| SE | 75979 | ENSG00000251364 | NA | chr11 | - | 7457795 | 7457959 | 7450372 | 7450442 | 7461238 | 7461336 | 6 | 2 | [illegible] |
| SE | 75980 | ENSG00000251364 | NA | chr11 | - | 7461238 | 7461336 | 7450372 | 7450442 | 7469640 | 7469766 | 10 | 16 | [illegible] |
| SE | 75982 | ENSG00000251364 | NA | chr11 | - | 7504765 | 7504872 | 7469640 | 7469766 | 7512122 | 7513644 | 5 | 6 | [illegible] |
| SE | 75984 | ENSG00000100316 | RPL3 | chr22 | - | 39317460 | 39317629 | 39316705 | 39316841 | 39318399 | 39318592 | 11896 | 5477 | [illegible] |
| SE | 75985 | ENSG00000100316 | RPL3 | chr22 | - | 39317460 | 39317629 | 39316705 | 39317021 | 39318399 | 39318592 | 5747 | 2589 | [illegible] |
| SE | 75996 | ENSG00000213676 | ATF6B | chr6 | - | 32117586 | 32117694 | 32117322 | 32117404 | 32117858 | 32118038 | 204 | 99 | [illegible] |
| SE | 75997 | ENSG00000213676 | ATF6B | chr6 | - | 32117586 | 32117694 | 32117322 | 32117404 | 32118774 | 32118866 | 152 | 65 | [illegible] |
| SE | 75998 | ENSG00000213676 | ATF6B | chr6 | - | 32117858 | 32118038 | 32117322 | 32117404 | 32118774 | 32118866 | 186 | 114 | [illegible] |
| SE | 76004 | ENSG00000204305 | AGER | chr6 | - | 32181605 | 32181632 | 32181350 | 32181477 | 32182246 | 32182388 | 28 | 19 | [illegible] |
| SE | 76006 | ENSG00000204305 | AGER | chr6 | - | 32182246 | 32182388 | 32181523 | 32181632 | 32182840 | 32183023 | 25 | 16 | [illegible] |
| SE | 76007 | ENSG00000204305 | AGER | chr6 | - | 32182538 | 32182698 | 32181605 | 32181632 | 32182840 | 32183023 | 6 | 3 | [illegible] |
| SE | 76008 | ENSG00000204305 | AGER | chr6 | - | 32182567 | 32182698 | 32181523 | 32181632 | 32182840 | 32183023 | 6 | 6 | [illegible] |
| SE | 76009 | ENSG00000204305 | AGER | chr6 | - | 32182567 | 32182698 | 32182246 | 32182388 | 32182840 | 32183023 | 28 | 23 | [illegible] |
| SE | 76012 | ENSG00000204304 | PBX2 | chr6 | - | 32186603 | 32186690 | 32184732 | 32186474 | 32187241 | 32187395 | 130 | 71 | [illegible] |
| SE | 76014 | ENSG00000204304 | PBX2 | chr6 | - | 32187646 | 32187782 | 32186972 | 32187395 | 32187965 | 32188156 | 776 | 422 | [illegible] |
| SE | 76015 | ENSG00000204304 | PBX2 | chr6 | - | 32188722 | 32188796 | 32188256 | 32188504 | 32189694 | 32189942 | 312 | 147 | [illegible] |
| SE | 76019 | ENSG00000204301 | NOTCH4 | chr6 | - | 32196323 | 32196421 | 32194842 | 32196150 | 32196924 | 32197072 | 0 | 0 | [illegible] |
| SE | 76030 | ENSG00000196230 | TUBB | chr6 | + | 30722917 | 30723028 | 30721617 | 30722645 | 30723339 | 30725409 | 1322 | 767 | [illegible] |
| SE | 76031 | ENSG00000137312 | FLOT1 | chr6 | - | 30730021 | 30730186 | 30727708 | 30728145 | 30730427 | 30730565 | 458 | 188 | [illegible] |
| SE | 76032 | ENSG00000137312 | FLOT1 | chr6 | - | 30730918 | 30731100 | 30730681 | 30730727 | 30740157 | 30740310 | 501 | 275 | [illegible] |
| SE | 76037 | ENSG00000137312 | FLOT1 | chr6 | - | 30741613 | 30741704 | 30740678 | 30740798 | 30742146 | 30742203 | 2 | 0 | [illegible] |
| SE | 76041 | ENSG00000137312 | FLOT1 | chr6 | - | 30741661 | 30741704 | 30741189 | 30741333 | 30742146 | 30742203 | 0 | 2 | [illegible] |
| SE | 76043 | ENSG00000137312 | FLOT1 | chr6 | - | 30741791 | 30741867 | 30741189 | 30741704 | 30742146 | 30742203 | 88 | 29 | [illegible] |
| SE | 76044 | ENSG00000128590 | DNAJB9 | chr7 | + | 108571716 | 108571943 | 108569873 | 108570103 | 108572898 | 108574850 | 181 | 93 | [illegible] |
| SE | 76049 | ENSG00000140391 | TSPAN3 | chr15 | - | 77056063 | 77056255 | 77055788 | 77055863 | 77070891 | 77071109 | 77 | 53 | [illegible] |
| SE | 76050 | ENSG00000166896 | ATP23 | chr12 | + | 57945627 | 57945673 | 57941566 | 57941888 | 57946994 | 57947076 | 18 | 9 | [illegible] |
| SE | 76053 | ENSG00000180822 | PSMG4 | chr6 | + | 3263683 | 3263759 | 3258946 | 3259196 | 3267590 | 3268049 | 22 | 5 | [illegible] |
| SE | 76057 | ENSG00000170275 | CRTAP | chr3 | + | 33129938 | 33130067 | 33124407 | 33124579 | 33132554 | 33132600 | 913 | 514 | [illegible] |
| SE | 76058 | ENSG00000170275 | CRTAP | chr3 | + | 33129938 | 33130067 | 33124407 | 33124579 | 33134181 | 33134265 | 344 | 174 | [illegible] |
| SE | 76059 | ENSG00000170275 | CRTAP | chr3 | + | 33129938 | 33130067 | 33124407 | 33124579 | 33142394 | 33147773 | 344 | 174 | [illegible] |
| SE | 76060 | ENSG00000170275 | CRTAP | chr3 | + | 33132554 | 33132700 | 33124407 | 33124579 | 33134181 | 33134265 | 347 | 172 | [illegible] |
| SE | 76062 | ENSG00000170275 | CRTAP | chr3 | + | 33132554 | 33132700 | 33129938 | 33130067 | 33134181 | 33134265 | 918 | 518 | [illegible] |
| SE | 76063 | ENSG00000170275 | CRTAP | chr3 | + | 33132554 | 33132700 | 33129938 | 33130067 | 33142394 | 33143158 | 573 | 348 | [illegible] |
| SE | 76064 | ENSG00000170275 | CRTAP | chr3 | + | 33134181 | 33134265 | 33124407 | 33124579 | 33142394 | 33143158 | 246 | 102 | [illegible] |
| SE | 76065 | ENSG00000170275 | CRTAP | chr3 | + | 33134181 | 33134265 | 33129938 | 33130067 | 33142394 | 33143158 | 246 | 102 | [illegible] |
| SE | 76071 | ENSG00000137692 | DCUN1D5 | chr11 | - | 103089226 | 103089318 | 103083255 | 103083326 | 103091683 | 103091861 | 32 | 20 | [illegible] |
| SE | 76073 | ENSG00000170515 | PA2G4 | chr12 | + | 56106989 | 56107095 | 56106161 | 56106716 | 56107186 | 56107219 | 725 | 477 | [illegible] |
| SE | 76074 | ENSG00000170515 | PA2G4 | chr12 | + | 56109229 | 56109293 | 56107520 | 56107613 | 56109856 | 56109901 | 830 | 625 | [illegible] |
| SE | 76075 | ENSG00000170515 | PA2G4 | chr12 | + | 56109856 | 56109935 | 56109229 | 56109293 | 56110398 | 56110477 | 757 | 538 | [illegible] |
| SE | 76077 | ENSG00000186115 | CYP4F2 | chr19 | - | 15886241 | 15886308 | 15885923 | 15886053 | 15889422 | 15889693 | 2 | 0 | [illegible] |
| SE | 76081 | ENSG00000112697 | TMEM30A | chr6 | - | 75260121 | 75260402 | 75259346 | 75259490 | 75260823 | 75260911 | 0 | 0 | [illegible] |
| SE | 76082 | ENSG00000112697 | TMEM30A | chr6 | - | 75265230 | 75265338 | 75260823 | 75260911 | 75267640 | 75267748 | 666 | 425 | [illegible] |
| SE | 76090 | ENSG00000262943 | ALOX12P2 | chr17 | + | 6894317 | 6894513 | 6892002 | 6892075 | 6894883 | 6895006 | 4 | 0 | [illegible] |
| SE | 76091 | ENSG00000262943 | ALOX12P2 | chr17 | + | 6894317 | 6894519 | 6892002 | 6892075 | 6894883 | 6895006 | 4 | 0 | [illegible] |
| SE | 76097 | ENSG00000136044 | APPL2 | chr12 | - | 105195584 | 105195627 | 105195444 | 105195501 | 105197764 | 105197953 | 93 | 46 | [illegible] |
| SE | 76099 | ENSG00000136044 | APPL2 | chr12 | - | 105199372 | 105199524 | 105197764 | 105197953 | 105203702 | 105203785 | 65 | 33 | [illegible] |
| SE | 76100 | ENSG00000136044 | APPL2 | chr12 | - | 105199372 | 105199531 | 105197763 | 105197880 | 105203702 | 105203785 | 70 | 55 | [illegible] |
| SE | 76101 | ENSG00000136044 | APPL2 | chr12 | - | 105199372 | 105199531 | 105197764 | 105197953 | 105203702 | 105203785 | 134 | 80 | [illegible] |
| SE | 76102 | ENSG00000136044 | APPL2 | chr12 | - | 105203702 | 105203785 | 105195584 | 105195627 | 105207060 | 105207207 | 32 | 10 | [illegible] |
| SE | 76104 | ENSG00000136044 | APPL2 | chr12 | - | 105203702 | 105203785 | 105199331 | 105199531 | 105207970 | 105208001 | 43 | 37 | [illegible] |
| SE | 76105 | ENSG00000136044 | APPL2 | chr12 | - | 105207060 | 105207207 | 105199331 | 105199531 | 105207970 | 105207989 | 29 | 14 | [illegible] |
| SE | 76110 | ENSG00000100220 | RTCB | chr22 | - | 32396073 | 32396249 | 32395025 | 32395214 | 32399602 | 32399759 | 179 | 113 | [illegible] |
| SE | 76112 | ENSG00000100220 | RTCB | chr22 | - | 32399602 | 32399759 | 32396073 | 32396249 | 32401746 | 32401903 | 315 | 232 | [illegible] |
| SE | 76114 | ENSG00000100220 | RTCB | chr22 | - | 32408174 | 32408242 | 32406661 | 32406761 | 32408754 | 32408833 | 185 | 148 | [illegible] |
| SE | 76117 | ENSG00000162337 | LRP5 | chr11 | + | 68357649 | 68357847 | 68347846 | 68348243 | 68363746 | 68363943 | 4 | 1 | [illegible] |
| SE | 76119 | ENSG00000162337 | LRP5 | chr11 | + | 68404222 | 68404416 | 68403482 | 68403699 | 68406523 | 68406813 | 0 | 0 | [illegible] |
| SE | 76127 | ENSG00000135486 | HNRNPA1 | chr12 | + | 54281794 | 54281941 | 54281352 | 54281502 | 54282089 | 54282138 | 4684 | 2989 | [illegible] |
| SE | 76128 | ENSG00000135486 | HNRNPA1 | chr12 | + | 54282572 | 54282662 | 54282393 | 54282486 | 54282799 | 54282874 | 702 | 645 | [illegible] |
| SE | 76129 | ENSG00000135486 | HNRNPA1 | chr12 | + | 54282572 | 54282665 | 54282393 | 54282486 | 54282799 | 54282874 | 3504 | 3355 | [illegible] |
| SE | 76131 | ENSG00000135486 | HNRNPA1 | chr12 | + | 54282572 | 54282665 | 54282393 | 54282486 | 54283811 | 54283967 | 704 | 647 | [illegible] |
| SE | 76133 | ENSG00000135486 | HNRNPA1 | chr12 | + | 54282799 | 54282874 | 54282393 | 54282486 | 54283811 | 54283878 | 1549 | 1334 | [illegible] |
| SE | 76135 | ENSG00000135486 | HNRNPA1 | chr12 | + | 54282799 | 54282874 | 54282393 | 54282665 | 54283811 | 54283878 | 4198 | 3892 | [illegible] |
| SE | 76136 | ENSG00000135486 | HNRNPA1 | chr12 | + | 54283062 | 54283234 | 54282799 | 54282874 | 54283811 | 54283967 | 44 | 20 | [illegible] |
| SE | 76137 | ENSG00000135486 | HNRNPA1 | chr12 | + | 54283078 | 54283234 | 54282393 | 54282486 | 54283811 | 54283967 | 44 | 20 | [illegible] |
| SE | 76159 | ENSG00000135486 | HNRNPA1 | chr12 | + | 54285973 | 54286112 | 54284257 | 54284317 | 54286661 | 54286732 | 0 | 0 | [illegible] |
| SE | 76161 | ENSG00000135486 | HNRNPA1 | chr12 | + | 54286661 | 54286732 | 54284257 | 54284317 | 54286952 | 54287087 | 25 | 22 | [illegible] |
| SE | 76162 | ENSG00000135486 | HNRNPA1 | chr12 | + | 54286679 | 54286732 | 54284257 | 54284317 | 54286952 | 54287088 | 6 | 0 | [illegible] |
| SE | 76166 | ENSG00000164691 | TAGAP | chr6 | - | 159038113 | 159038228 | 159034467 | 159037124 | 159039113 | 159039309 | 1394 | 618 | [illegible] |
| SE | 76168 | ENSG00000164691 | TAGAP | chr6 | - | 159042077 | 159042244 | 159041353 | 159041515 | 159043977 | 159044031 | 520 | 435 | [illegible] |
| SE | 76169 | ENSG00000164691 | TAGAP | chr6 | - | 159043588 | 159043655 | 159041353 | 159041515 | 159043977 | 159044031 | 398 | 196 | [illegible] |
| SE | 76171 | ENSG00000132423 | COQ3 | chr6 | - | 99383007 | 99383130 | 99380188 | 99380341 | 99383697 | 99383824 | 1 | 1 | [illegible] |
| SE | 76172 | ENSG00000132424 | PNISR | chr6 | - | 99400543 | 99401630 | 99397628 | 99400311 | 99402539 | 99402710 | 241 | 273 | [illegible] |
| SE | 76173 | ENSG00000132424 | PNISR | chr6 | - | 99402539 | 99402710 | 99398988 | 99400311 | 99404602 | 99404702 | 15 | 16 | [illegible] |
| SE | 76175 | ENSG00000132424 | PNISR | chr6 | - | 99406030 | 99406168 | 99403828 | 99404702 | 99408080 | 99408271 | 909 | 1062 | [illegible] |
| SE | 76176 | ENSG00000132424 | PNISR | chr6 | - | 99409172 | 99409286 | 99408080 | 99408271 | 99410740 | 99410964 | 227 | 278 | [illegible] |
| SE | 76177 | ENSG00000132424 | PNISR | chr6 | - | 99409172 | 99409344 | 99408080 | 99408271 | 99410740 | 99410964 | 481 | 538 | [illegible] |
| SE | 76178 | ENSG00000132424 | PNISR | chr6 | - | 99409172 | 99409344 | 99408080 | 99408271 | 99412550 | 99412734 | 241 | 290 | [illegible] |
| SE | 76179 | ENSG00000132424 | PNISR | chr6 | - | 99410740 | 99410964 | 99408080 | 99408271 | 99412550 | 99412734 | 306 | 237 | [illegible] |
| SE | 76180 | ENSG00000132424 | PNISR | chr6 | - | 99410740 | 99410964 | 99409172 | 99409344 | 99412550 | 99412734 | 549 | 485 | [illegible] |
| SE | 76181 | ENSG00000132424 | PNISR | chr6 | - | 99414571 | 99414686 | 99412550 | 99412739 | 99416348 | 99416428 | 522 | 529 | [illegible] |
| SE | 76182 | ENSG00000132424 | PNISR | chr6 | - | 99414571 | 99414686 | 99412550 | 99412739 | 99425214 | 99425277 | 458 | 463 | [illegible] |
| SE | 76183 | ENSG00000132424 | PNISR | chr6 | - | 99414571 | 99414690 | 99412550 | 99412739 | 99416348 | 99416428 | 830 | 856 | [illegible] |
| SE | 76184 | ENSG00000132424 | PNISR | chr6 | - | 99414571 | 99414690 | 99412550 | 99412739 | 99425214 | 99425329 | 525 | 513 | [illegible] |
| SE | 76185 | ENSG00000132424 | PNISR | chr6 | - | 99414571 | 99416428 | 99411790 | 99412739 | 99425214 | 99425258 | 560 | 571 | [illegible] |
| SE | 76186 | ENSG00000132424 | PNISR | chr6 | - | 99415568 | 99415747 | 99414571 | 99414690 | 99416348 | 99416428 | 0 | 2 | [illegible] |
| SE | 76187 | ENSG00000132424 | PNISR | chr6 | - | 99415635 | 99415747 | 99414571 | 99414686 | 99416348 | 99416428 | 0 | 0 | [illegible] |
| SE | 76188 | ENSG00000132424 | PNISR | chr6 | - | 99415635 | 99415747 | 99414571 | 99414690 | 99416348 | 99416428 | 2 | 0 | [illegible] |
| SE | 76189 | ENSG00000132424 | PNISR | chr6 | - | 99416348 | 99416428 | 99412550 | 99412739 | 99425214 | 99425270 | 61 | 68 | [illegible] |
| SE | 76198 | ENSG00000112237 | CCNC | chr6 | - | 99549507 | 99549575 | 99546394 | 99546474 | 99550992 | 99551028 | 93 | 74 | [illegible] |
| SE | 76199 | ENSG00000112237 | CCNC | chr6 | - | 99550217 | 99550309 | 99546394 | 99546474 | 99550992 | 99551028 | 79 | 49 | [illegible] |
| SE | 76201 | ENSG00000112237 | CCNC | chr6 | - | 99561596 | 99561681 | 99561366 | 99561436 | 99562841 | 99562948 | 112 | 43 | [illegible] |
| SE | 76202 | ENSG00000112237 | CCNC | chr6 | - | 99562841 | 99562948 | 99561596 | 99561681 | 99568495 | 99568628 | 69 | 29 | [illegible] |
| SE | 76203 | ENSG00000112237 | CCNC | chr6 | - | 99566897 | 99567038 | 99562841 | 99562948 | 99568495 | 99568623 | 0 | 0 | [illegible] |
| SE | 76205 | ENSG00000146070 | PLA2G7 | chr6 | - | 46705152 | 46705301 | 46704200 | 46704696 | 46707990 | 46708161 | 307 | 219 | [illegible] |
| SE | 76208 | ENSG00000146070 | PLA2G7 | chr6 | - | 46714459 | 46714553 | 46712268 | 46712337 | 46716974 | 46717096 | 56 | 35 | [illegible] |
| SE | 76209 | ENSG00000146070 | PLA2G7 | chr6 | - | 46716383 | 46716528 | 46712268 | 46712337 | 46716974 | 46717096 | 53 | 38 | [illegible] |
| SE | 76210 | ENSG00000146070 | PLA2G7 | chr6 | - | 46716383 | 46716528 | 46714459 | 46714553 | 46716974 | 46717096 | 117 | 63 | [illegible] |
| SE | 76211 | ENSG00000146070 | PLA2G7 | chr6 | - | 46716974 | 46717096 | 46712268 | 46712337 | 46735179 | 46735375 | 2 | 0 | [illegible] |
| SE | 76212 | ENSG00000146070 | PLA2G7 | chr6 | - | 46722782 | 46722925 | 46716974 | 46717096 | 46735179 | 46735375 | 26 | 11 | [illegible] |
| SE | 76213 | ENSG00000146070 | PLA2G7 | chr6 | - | 46722782 | 46722925 | 46716974 | 46717096 | 46735549 | 46735693 | 25 | 11 | [illegible] |
| SE | 76214 | ENSG00000083097 | DOP1A | chr6 | + | 83096730 | 83096823 | 83067665 | 83067779 | 83096924 | 83097115 | 33 | 32 | [illegible] |
| SE | 76215 | ENSG00000083097 | DOP1A | chr6 | + | 83113322 | 83113421 | 83110124 | 83110314 | 83118887 | 83118987 | 29 | 36 | [illegible] |
| SE | 76216 | ENSG00000083097 | DOP1A | chr6 | + | 83121929 | 83122050 | 83120682 | 83120791 | 83122862 | 83122982 | 27 | 26 | [illegible] |
| SE | 76217 | ENSG00000083097 | DOP1A | chr6 | + | 83122862 | 83122982 | 83121929 | 83122050 | 83124704 | 83124819 | 48 | 41 | [illegible] |

| | | | | | | | | | | | | | |
|---|---|---|---|---|---|---|---|---|---|---|---|---|---|
| SE | 76219 | ENSG00000083097 | DOP1A | chr6 | + | 83130122 | 83130397 | 83128886 | 83129508 | 83132175 | 83132328 | 80 | 84 |
| SE | 76220 | ENSG00000083097 | DOP1A | chr6 | + | 83130122 | 83130397 | 83128886 | 83129508 | 83134186 | 83134287 | 35 | 27 |
| SE | 76221 | ENSG00000083097 | DOP1A | chr6 | + | 83132175 | 83132328 | 83128886 | 83129508 | 83134186 | 83134287 | 49 | 72 |
| SE | 76222 | ENSG00000083097 | DOP1A | chr6 | + | 83132175 | 83132328 | 83130122 | 83130397 | 83134186 | 83134287 | 94 | 128 |
| SE | 76223 | ENSG00000083097 | DOP1A | chr6 | + | 83135618 | 83135878 | 83134186 | 83134287 | 83137172 | 83139162 | 89 | 99 |
| SE | 76224 | ENSG00000083097 | DOP1A | chr6 | + | 83139999 | 83140111 | 83137172 | 83139162 | 83140220 | 83140403 | 89 | 96 |
| SE | 76225 | ENSG00000083097 | DOP1A | chr6 | + | 83147235 | 83147291 | 83145523 | 83145658 | 83148758 | 83148863 | 49 | 35 |
| SE | 76226 | ENSG00000083097 | DOP1A | chr6 | + | 83147235 | 83147291 | 83145523 | 83145658 | 83151592 | 83151659 | 16 | 8 |
| SE | 76227 | ENSG00000083097 | DOP1A | chr6 | + | 83148758 | 83148863 | 83145523 | 83145658 | 83151592 | 83151659 | 30 | 29 |
| SE | 76228 | ENSG00000083097 | DOP1A | chr6 | + | 83148758 | 83148863 | 83147235 | 83147291 | 83151592 | 83151659 | 66 | 57 |
| SE | 76229 | ENSG00000083097 | DOP1A | chr6 | + | 83152287 | 83152367 | 83151882 | 83152027 | 83153510 | 83153620 | 68 | 51 |
| SE | 76230 | ENSG00000083097 | DOP1A | chr6 | + | 83153893 | 83154043 | 83153510 | 83153620 | 83154179 | 83154241 | 71 | 45 |
| SE | 76231 | ENSG00000083097 | DOP1A | chr6 | + | 83155950 | 83156103 | 83154179 | 83154241 | 83157181 | 83157318 | 74 | 40 |
| SE | 76232 | ENSG00000083097 | DOP1A | chr6 | + | 83157181 | 83157318 | 83154179 | 83154241 | 83158566 | 83158622 | 59 | 36 |
| SE | 76233 | ENSG00000083097 | DOP1A | chr6 | + | 83159795 | 83159960 | 83158566 | 83158622 | 83162789 | 83162919 | 46 | 42 |
| SE | 76234 | ENSG00000083097 | DOP1A | chr6 | + | 83164659 | 83164719 | 83162789 | 83162919 | 83167861 | 83168331 | 13 | 11 |
| SE | 76236 | ENSG00000166278 | C2 | chr6 | + | 31934165 | 31934299 | 31933866 | 31933965 | 31935922 | 31935990 | 8 | 15 |
| SE | 76239 | ENSG00000204351 | SKIV2L | chr6 | + | 31960009 | 31960119 | 31959296 | 31959400 | 31960219 | 31960337 | 92 | 64 |
| SE | 76241 | ENSG00000204351 | SKIV2L | chr6 | + | 31960431 | 31960546 | 31960219 | 31960337 | 31960665 | 31960740 | 51 | 46 |
| SE | 76247 | ENSG00000204348 | DXO | chr6 | - | 31970592 | 31970825 | 31970342 | 31970378 | 31970911 | 31970923 | 6 | 0 |
| SE | 76248 | ENSG00000204348 | DXO | chr6 | - | 31971319 | 31971681 | 31970911 | 31971147 | 31971928 | 31972138 | 34 | 14 |
| SE | 76249 | ENSG00000204348 | DXO | chr6 | - | 31971319 | 31971725 | 31970897 | 31971147 | 31971928 | 31972009 | 20 | 3 |
| SE | 76252 | ENSG00000211450 | SELENOH | chr11 | + | 57741808 | 57741954 | 57741490 | 57741717 | 57742862 | 57743038 | 119 | 49 |
| SE | 76264 | ENSG00000154839 | SKA1 | chr18 | + | 50385215 | 50385353 | 50382128 | 50382226 | 50391123 | 50391293 | 8 | 5 |
| SE | 76265 | ENSG00000065427 | KARS1 | chr16 | - | 75634172 | 75634292 | 75631692 | 75631855 | 75635679 | 75635805 | 641 | 376 |
| SE | 76270 | ENSG00000065427 | KARS1 | chr16 | - | 75641563 | 75641723 | 75640183 | 75640349 | 75644282 | 75644462 | 322 | 191 |
| SE | 76276 | ENSG00000224848 | NA | chr9 | + | 96720965 | 96721130 | 96687049 | 96687324 | 96773459 | 96773977 | 1 | 5 |
| SE | 76281 | ENSG00000124508 | BTN2A2 | chr6 | + | 26383791 | 26383915 | 26383095 | 26383299 | 26388012 | 26388249 | 13 | 2 |
| SE | 76283 | ENSG00000124508 | BTN2A2 | chr6 | + | 26383791 | 26383915 | 26383101 | 26383181 | 26388012 | 26388249 | 14 | 7 |
| SE | 76285 | ENSG00000124508 | BTN2A2 | chr6 | + | 26383791 | 26383915 | 26383112 | 26383244 | 26385014 | 26385362 | 35 | 19 |
| SE | 76286 | ENSG00000124508 | BTN2A2 | chr6 | + | 26383791 | 26383915 | 26383112 | 26383244 | 26388012 | 26388249 | 10 | 3 |
| SE | 76287 | ENSG00000124508 | BTN2A2 | chr6 | + | 26383791 | 26383915 | 26383112 | 26383244 | 26390004 | 26390211 | 10 | 3 |
| SE | 76289 | ENSG00000124508 | BTN2A2 | chr6 | + | 26385014 | 26385362 | 26383107 | 26383181 | 26390004 | 26390211 | 2 | 0 |
| SE | 76290 | ENSG00000124508 | BTN2A2 | chr6 | + | 26385014 | 26385362 | 26383112 | 26383244 | 26388012 | 26388289 | 23 | 20 |
| SE | 76291 | ENSG00000124508 | BTN2A2 | chr6 | + | 26385014 | 26385362 | 26383112 | 26383244 | 26390004 | 26390211 | 0 | 0 |
| SE | 76292 | ENSG00000124508 | BTN2A2 | chr6 | + | 26385014 | 26385362 | 26383791 | 26383915 | 26388012 | 26388289 | 50 | 37 |
| SE | 76293 | ENSG00000124508 | BTN2A2 | chr6 | + | 26385014 | 26385362 | 26383791 | 26383915 | 26390004 | 26390211 | 27 | 17 |
| SE | 76295 | ENSG00000124508 | BTN2A2 | chr6 | + | 26388012 | 26388294 | 26383100 | 26383181 | 26390004 | 26390137 | 59 | 42 |
| SE | 76296 | ENSG00000124508 | BTN2A2 | chr6 | + | 26388012 | 26388294 | 26383432 | 26383915 | 26390004 | 26390137 | 54 | 43 |
| SE | 76298 | ENSG00000269313 | MAGIX | chrX | + | 49164862 | 49165090 | 49164706 | 49164789 | 49165189 | 49165361 | 0 | 2 |
| SE | 76299 | ENSG00000269313 | MAGIX | chrX | + | 49164907 | 49165090 | 49163931 | 49164789 | 49165189 | 49165361 | 2 | 2 |
| SE | 76301 | ENSG00000124549 | BTN2A3P | chr6 | + | 26422088 | 26422197 | 26421390 | 26421624 | 26422932 | 26423759 | 1 | 0 |
| SE | 76302 | ENSG00000124549 | BTN2A3P | chr6 | + | 26423398 | 26423759 | 26422932 | 26423277 | 26426241 | 26426523 | 1 | 7 |
| SE | 76303 | ENSG00000124549 | BTN2A3P | chr6 | + | 26427870 | 26428107 | 26426241 | 26426523 | 26428642 | 26428785 | 11 | 0 |
| SE | 76309 | ENSG00000167996 | FTH1 | chr11 | - | 61964986 | 61965105 | 61964565 | 61964891 | 61965368 | 61965515 | 2020 | 1894 |
| SE | 76310 | ENSG00000167996 | FTH1 | chr11 | - | 61964986 | 61965112 | 61964284 | 61964891 | 61965368 | 61965515 | 10765 | 8668 |
| SE | 76312 | ENSG00000167996 | FTH1 | chr11 | - | 61965368 | 61965515 | 61964746 | 61965112 | 61967311 | 61967453 | 14173 | 10304 |
| SE | 76313 | ENSG00000167996 | FTH1 | chr11 | - | 61965368 | 61965577 | 61959717 | 61959927 | 61967311 | 61967371 | 15 | 25 |
| SE | 76314 | ENSG00000167996 | FTH1 | chr11 | - | 61965368 | 61965577 | 61964986 | 61965112 | 61967311 | 61967371 | 8758 | 6791 |
| SE | 76315 | ENSG00000085491 | SLC25A24 | chr1 | - | 108139057 | 108139208 | 108134042 | 108136837 | 108143542 | 108143710 | 190 | 124 |
| SE | 76316 | ENSG00000085491 | SLC25A24 | chr1 | - | 108143542 | 108143710 | 108139057 | 108139208 | 108148278 | 108148386 | 210 | 166 |
| SE | 76317 | ENSG00000085491 | SLC25A24 | chr1 | - | 108156700 | 108156885 | 108154982 | 108155084 | 108157461 | 108157620 | 5 | 12 |
| SE | 76322 | ENSG00000073050 | XRCC1 | chr19 | - | 43546594 | 43546727 | 43546051 | 43546106 | 43546883 | 43546977 | 52 | 39 |
| SE | 76323 | ENSG00000073050 | XRCC1 | chr19 | - | 43551570 | 43551687 | 43546883 | 43546977 | 43552016 | 43552275 | 48 | 33 |
| SE | 76325 | ENSG00000073050 | XRCC1 | chr19 | - | 43574909 | 43575002 | 43560909 | 43561020 | 43575407 | 43575506 | 26 | 18 |
| SE | 76328 | ENSG00000136861 | CDK5RAP2 | chr9 | - | 120404035 | 120404113 | 120402805 | 120403071 | 120408346 | 120408468 | 31 | 13 |
| SE | 76330 | ENSG00000136861 | CDK5RAP2 | chr9 | - | 120411357 | 120411474 | 120409126 | 120409316 | 120415039 | 120415159 | 321 | 279 |
| SE | 76331 | ENSG00000136861 | CDK5RAP2 | chr9 | - | 120419787 | 120419960 | 120415039 | 120415159 | 120437294 | 120437527 | 76 | 52 |
| SE | 76332 | ENSG00000136861 | CDK5RAP2 | chr9 | - | 120419787 | 120419960 | 120415039 | 120415159 | 120439398 | 120439972 | 76 | 52 |
| SE | 76334 | ENSG00000136861 | CDK5RAP2 | chr9 | - | 120422692 | 120422741 | 120419787 | 120419960 | 120439398 | 120439972 | 38 | 31 |
| SE | 76335 | ENSG00000136861 | CDK5RAP2 | chr9 | - | 120437294 | 120437527 | 120415039 | 120415159 | 120439398 | 120439972 | 48 | 21 |
| SE | 76336 | ENSG00000136861 | CDK5RAP2 | chr9 | - | 120437294 | 120437527 | 120419787 | 120419960 | 120439398 | 120439972 | 48 | 21 |
| SE | 76339 | ENSG00000136861 | CDK5RAP2 | chr9 | - | 120447894 | 120448126 | 120443619 | 120443742 | 120453455 | 120453873 | 91 | 91 |
| SE | 76341 | ENSG00000136861 | CDK5RAP2 | chr9 | - | 120453455 | 120453873 | 120447894 | 120448126 | 120467859 | 120467997 | 38 | 50 |
| SE | 76342 | ENSG00000136861 | CDK5RAP2 | chr9 | - | 120458449 | 120458622 | 120453455 | 120453873 | 120467859 | 120467997 | 84 | 46 |
| SE | 76344 | ENSG00000136861 | CDK5RAP2 | chr9 | - | 120460571 | 120460747 | 120458449 | 120458622 | 120467859 | 120467997 | 44 | 27 |
| SE | 76345 | ENSG00000136861 | CDK5RAP2 | chr9 | - | 120467859 | 120467997 | 120453455 | 120453873 | 120477349 | 120477354 | 0 | 0 |
| SE | 76346 | ENSG00000136861 | CDK5RAP2 | chr9 | - | 120467859 | 120467997 | 120458449 | 120458622 | 120470110 | 120470217 | 132 | 128 |
| SE | 76347 | ENSG00000136861 | CDK5RAP2 | chr9 | - | 120467859 | 120467997 | 120460571 | 120460667 | 120477349 | 120477354 | 117 | 117 |
| SE | 76350 | ENSG00000136861 | CDK5RAP2 | chr9 | - | 120471747 | 120471878 | 120467859 | 120467997 | 120477349 | 120477354 | 109 | 73 |
| SE | 76353 | ENSG00000136861 | CDK5RAP2 | chr9 | - | 120536371 | 120536526 | 120529977 | 120530140 | 120539040 | 120539120 | 199 | 145 |
| SE | 76354 | ENSG00000136861 | CDK5RAP2 | chr9 | - | 120571973 | 120572041 | 120568320 | 120568388 | 120579919 | 120580061 | 87 | 54 |
| SE | 76355 | ENSG00000136861 | CDK5RAP2 | chr9 | - | 120571976 | 120572041 | 120568320 | 120568388 | 120579919 | 120580005 | 32 | 22 |
| SE | 76356 | ENSG00000227540 | DNAJC9-AS1 | chr10 | + | 73253190 | 73253254 | 73252790 | 73253095 | 73253892 | 73254344 | 0 | 0 |
| SE | 76357 | ENSG00000225292 | NA | chr10 | + | 112890965 | 112891072 | 112888734 | 112888807 | 112905976 | 112906111 | 3 | 0 |
| SE | 76361 | ENSG00000177150 | FAM210A | chr18 | - | 13687742 | 13687882 | 13681604 | 13682105 | 13726328 | 13726663 | 5 | 6 |
| SE | 76365 | ENSG00000164674 | SYTL3 | chr6 | + | 158665394 | 158665613 | 158662749 | 158662812 | 158682924 | 158682989 | 92 | 60 |
| SE | 76366 | ENSG00000164674 | SYTL3 | chr6 | + | 158682924 | 158682989 | 158665394 | 158665613 | 158707229 | 158707281 | 134 | 61 |
| SE | 76367 | ENSG00000164674 | SYTL3 | chr6 | + | 158707229 | 158707281 | 158665394 | 158665613 | 158708321 | 158708391 | 67 | 17 |
| SE | 76368 | ENSG00000164674 | SYTL3 | chr6 | + | 158713799 | 158713878 | 158682924 | 158682989 | 158725502 | 158725637 | 2 | 1 |
| SE | 76374 | ENSG00000164674 | SYTL3 | chr6 | + | 158751927 | 158752030 | 158725502 | 158725637 | 158760639 | 158760745 | 2 | 0 |
| SE | 76375 | ENSG00000164674 | SYTL3 | chr6 | + | 158757210 | 158757381 | 158751927 | 158752030 | 158760639 | 158760745 | 107 | 36 |
| SE | 76376 | ENSG00000164674 | SYTL3 | chr6 | + | 158762075 | 158762178 | 158760639 | 158760745 | 158763303 | 158763509 | 355 | 153 |
| SE | 76377 | ENSG00000164674 | SYTL3 | chr6 | + | 158763303 | 158763509 | 158762075 | 158762178 | 158764494 | 158764869 | 357 | 136 |
| SE | 76381 | ENSG00000288546 | NA | chrX | - | 22620923 | 22621057 | 22389314 | 22389720 | 22743228 | 22743338 | 1 | 0 |
| SE | 76382 | ENSG00000204520 | MICA | chr6 | + | 31402156 | 31402356 | 31400748 | 31400763 | 31403444 | 31403673 | 0 | 0 |
| SE | 76383 | ENSG00000204520 | MICA | chr6 | + | 31403444 | 31403702 | 31400748 | 31400763 | 31410542 | 31410715 | 2 | 0 |
| SE | 76384 | ENSG00000204520 | MICA | chr6 | + | 31410542 | 31410797 | 31399783 | 31400783 | 31411071 | 31411359 | 60 | 35 |
| SE | 76394 | ENSG00000164116 | GUCY1A1 | chr4 | + | 155668089 | 155668183 | 155667345 | 155667419 | 155696755 | 155697122 | 0 | 1 |
| SE | 76395 | ENSG00000164116 | GUCY1A1 | chr4 | + | 155696755 | 155696849 | 155667345 | 155667419 | 155696934 | 155697122 | 1 | 1 |
| SE | 76396 | ENSG00000164116 | GUCY1A1 | chr4 | + | 155703931 | 155703993 | 155696755 | 155697122 | 155708235 | 155708294 | 17 | 100 |
| SE | 76397 | ENSG00000164116 | GUCY1A1 | chr4 | + | 155708235 | 155708294 | 155703931 | 155703993 | 155710541 | 155710611 | 15 | 76 |
| SE | 76398 | ENSG00000164116 | GUCY1A1 | chr4 | + | 155710541 | 155711251 | 155708235 | 155708294 | 155713097 | 155713583 | 53 | 137 |
| SE | 76399 | ENSG00000164116 | GUCY1A1 | chr4 | + | 155710681 | 155711251 | 155708235 | 155708294 | 155713097 | 155713583 | 46 | 85 |
| SE | 76408 | ENSG00000081377 | CDC14B | chr9 | - | 96541825 | 96541892 | 96538974 | 96539140 | 96562692 | 96562785 | 29 | 32 |
| SE | 76410 | ENSG00000081377 | CDC14B | chr9 | - | 96565392 | 96565455 | 96564776 | 96564852 | 96619218 | 96619843 | 10 | 26 |
| SE | 76411 | ENSG00000081377 | CDC14B | chr9 | - | 96565392 | 96565483 | 96564776 | 96564852 | 96619218 | 96619297 | 25 | 36 |
| SE | 76414 | ENSG00000168538 | TRAPPC11 | chr4 | + | 183675163 | 183675237 | 183674712 | 183674812 | 183677457 | 183677475 | 353 | 290 |
| SE | 76415 | ENSG00000168538 | TRAPPC11 | chr4 | + | 183680119 | 183680267 | 183679352 | 183679486 | 183683974 | 183684054 | 101 | 82 |
| SE | 76416 | ENSG00000168538 | TRAPPC11 | chr4 | + | 183680207 | 183680267 | 183679352 | 183679486 | 183683974 | 183684006 | 2 | 2 |
| SE | 76419 | ENSG00000168538 | TRAPPC11 | chr4 | + | 183685270 | 183685403 | 183685083 | 183685145 | 183691315 | 183691471 | 165 | 155 |
| SE | 76421 | ENSG00000168538 | TRAPPC11 | chr4 | + | 183691315 | 183691471 | 183686617 | 183686748 | 183692959 | 183693147 | 248 | 195 |
| SE | 76422 | ENSG00000168538 | TRAPPC11 | chr4 | + | 183697502 | 183697568 | 183694603 | 183694723 | 183697678 | 183697835 | 278 | 215 |
| SE | 76424 | ENSG00000168538 | TRAPPC11 | chr4 | + | 183706806 | 183706940 | 183704978 | 183705070 | 183708406 | 183708416 | 106 | 39 |
| SE | 76425 | ENSG00000112739 | PRPF4B | chr6 | + | 4047174 | 4047248 | 4043812 | 4044023 | 4049015 | 4049106 | 253 | 177 |
| SE | 76427 | ENSG00000112739 | PRPF4B | chr6 | + | 4060412 | 4060644 | 4058727 | 4058814 | 4061014 | 4061147 | 287 | 260 |
| SE | 76428 | ENSG00000112739 | PRPF4B | chr6 | + | 4060412 | 4060686 | 4058727 | 4058814 | 4061014 | 4061147 | 315 | 271 |
| SE | 76429 | ENSG00000112739 | PRPF4B | chr6 | + | 4061014 | 4061147 | 4060412 | 4060686 | 4061654 | 4061700 | 58 | 27 |
| SE | 76430 | ENSG00000112739 | PRPF4B | chr6 | + | 4061654 | 4061700 | 4060412 | 4061147 | 4062212 | 4062299 | 30 | 12 |
| SE | 76433 | ENSG00000198492 | YTHDF2 | chr1 | + | 28737657 | 28737682 | 28736939 | 28737147 | 28738258 | 28738329 | 121 | 72 |
| SE | 76434 | ENSG00000198492 | YTHDF2 | chr1 | + | 28740302 | 28740398 | 28738258 | 28738338 | 28742402 | 28742883 | 0 | 1 |
| SE | 76447 | ENSG00000168685 | IL7R | chr5 | + | 35856031 | 35856169 | 35852694 | 35852938 | 35860851 | 35860948 | 0 | 1 |
| SE | 76448 | ENSG00000168685 | IL7R | chr5 | + | 35856976 | 35857059 | 35856031 | 35856169 | 35860851 | 35860926 | 2376 | 1229 |
| SE | 76450 | ENSG00000168685 | IL7R | chr5 | + | 35871055 | 35871213 | 35867305 | 35867463 | 35875511 | 35875587 | 3154 | 2396 |
| SE | 76451 | ENSG00000168685 | IL7R | chr5 | + | 35873479 | 35873648 | 35871055 | 35871213 | 35874448 | 35874542 | 3843 | 2507 |
| SE | 76457 | ENSG00000155906 | RMND1 | chr6 | - | 151417080 | 151417138 | 151405719 | 151405836 | 151417278 | 151417399 | 4 | 0 |
| SE | 76458 | ENSG00000155906 | RMND1 | chr6 | - | 151418532 | 151418579 | 151416590 | 151417399 | 151421244 | 151421321 | 2 | 2 |
| SE | 76459 | ENSG00000155906 | RMND1 | chr6 | - | 151422540 | 151422605 | 151421244 | 151421321 | 151423524 | 151423631 | 92 | 56 |
| SE | 76460 | ENSG00000155906 | RMND1 | chr6 | - | 151427481 | 151427582 | 151421244 | 151423631 | 151430137 | 151430177 | 45 | 40 |
| SE | 76461 | ENSG00000155906 | RMND1 | chr6 | - | 151436419 | 151436554 | 151433154 | 151433230 | 151445307 | 151445825 | 28 | 14 |
| SE | 76462 | ENSG00000155906 | RMND1 | chr6 | - | 151436445 | 151436554 | 151433154 | 151433230 | 151445307 | 151445825 | 41 | 35 |
| SE | 76463 | ENSG00000155906 | RMND1 | chr6 | - | 151436445 | 151436554 | 151433154 | 151433230 | 151452015 | 151452158 | 14 | 11 |
| SE | 76464 | ENSG00000155906 | RMND1 | chr6 | - | 151445307 | 151445825 | 151433154 | 151433230 | 151452015 | 151452018 | 15 | 4 |
| SE | 76467 | ENSG00000111817 | DSE | chr6 | + | 116254839 | 116255002 | 116254206 | 116254383 | 116257215 | 116257337 | 0 | 3 |
| SE | 76468 | ENSG00000111817 | DSE | chr6 | + | 116257215 | 116257337 | 116254172 | 116254295 | 116258445 | 116259240 | 2 | 0 |
| SE | 76469 | ENSG00000111817 | DSE | chr6 | + | 116257215 | 116257337 | 116254206 | 116254383 | 116258445 | 116259240 | 2 | 0 |
| SE | 76472 | ENSG00000111817 | DSE | chr6 | + | 116370869 | 116371121 | 116369846 | 116369927 | 116399197 | 116399666 | 9 | 13 |
| SE | 76473 | ENSG00000111817 | DSE | chr6 | + | 116430953 | 116431193 | 116426573 | 116426827 | 116435586 | 116438243 | 68 | 67 |
| SE | 76475 | ENSG00000120262 | CCDC170 | chr6 | + | 151536317 | 151536446 | 151494016 | 151494185 | 151538044 | 151538301 | 6 | 6 |
| SE | 76476 | ENSG00000120262 | CCDC170 | chr6 | + | 151573173 | 151573491 | 151548303 | 151548489 | 151585888 | 151586089 | 10 | 6 |
| SE | 76477 | ENSG00000120262 | CCDC170 | chr6 | + | 151585888 | 151586089 | 151573173 | 151573491 | 151593106 | 151593280 | 12 | 9 |
| SE | 76479 | ENSG00000144115 | THNSL2 | chr2 | + | 88185327 | 88185479 | 88182947 | 88183073 | 88185897 | 88186626 | 4 | 0 |
| SE | 76482 | ENSG00000135919 | SERPINE2 | chr2 | - | 223982680 | 223982781 | 223980310 | 223980397 | 223984751 | 223984950 | 24 | 23 |
| SE | 76483 | ENSG00000135919 | SERPINE2 | chr2 | - | 223991802 | 223992000 | 223984751 | 223984950 | 223998114 | 223998342 | 41 | 42 |
| SE | 76485 | ENSG00000143314 | MRPL24 | chr1 | - | 156737645 | 156737776 | 156737302 | 156737534 | 156738030 | 156738134 | 125 | 50 |
| SE | 76486 | ENSG00000143314 | MRPL24 | chr1 | - | 156738030 | 156738134 | 156737302 | 156737534 | 156738342 | 156738438 | 38 | 29 |
| SE | 76494 | ENSG00000153406 | NMRAL1 | chr16 | - | 4463419 | 4463512 | 4461691 | 4461959 | 4463641 | 4463850 | 0 | 0 |
| SE | 76495 | ENSG00000153406 | NMRAL1 | chr16 | - | 4463419 | 4463512 | 4461691 | 4461959 | 4463659 | 4463850 | 0 | 0 |
| SE | 76496 | ENSG00000153406 | NMRAL1 | chr16 | - | 4463641 | 4463850 | 4461693 | 4461959 | 4466152 | 4466336 | 123 | 53 |
| SE | 76497 | ENSG00000153406 | NMRAL1 | chr16 | - | 4463659 | 4463850 | 4461690 | 4461959 | 4466152 | 4466402 | 307 | 123 |
| SE | 76498 | ENSG00000153406 | NMRAL1 | chr16 | - | 4463677 | 4463850 | 4461698 | 4461959 | 4466152 | 4466174 | 121 | 53 |
| SE | 76499 | ENSG00000153406 | NMRAL1 | chr16 | - | 4466526 | 4466685 | 4466152 | 4466402 | 4469226 | 4469465 | 2 | 4 |
| SE | 76501 | ENSG00000153406 | NMRAL1 | chr16 | - | 4469226 | 4469465 | 4466152 | 4466402 | 4474092 | 4474166 | 194 | 101 |
| SE | 76504 | ENSG00000153406 | NMRAL1 | chr16 | - | 4471313 | 4471449 | 4469629 | 4469698 | 4474092 | 4474166 | 1 | 0 |
| SE | 76507 | ENSG00000034693 | PEX3 | chr6 | + | 143459084 | 143459216 | 143450804 | 143451115 | 143462915 | 143462997 | 133 | 108 |
| SE | 76509 | ENSG00000034693 | PEX3 | chr6 | + | 143479075 | 143479198 | 143474785 | 143474856 | 143489142 | 143490616 | 24 | 18 |
| SE | 76510 | ENSG00000034693 | PEX3 | chr6 | + | 143485151 | 143485248 | 143479075 | 143479198 | 143489142 | 143490616 | 54 | 46 |
| SE | 76513 | ENSG00000165672 | PRDX3 | chr10 | - | 119174450 | 119174592 | 119173736 | 119173872 | 119177020 | 119177153 | 143 | 85 |
| SE | 76514 | ENSG00000165672 | PRDX3 | chr10 | - | 119177020 | 119177153 | 119173736 | 119173872 | 119178754 | 119178812 | 44 | 20 |
| SE | 76520 | ENSG00000118507 | AKAP7 | chr6 | + | 131165080 | 131165217 | 131160058 | 131160198 | 131169112 | 131169273 | 102 | 89 |
| SE | 76521 | ENSG00000118507 | AKAP7 | chr6 | + | 131169112 | 131169273 | 131165080 | 131165217 | 131199460 | 131199573 | 99 | 69 |
| SE | 76522 | ENSG00000118507 | AKAP7 | chr6 | + | 131199460 | 131199573 | 131169112 | 131169273 | 131219660 | 131219808 | 73 | 66 |
| SE | 76523 | ENSG00000118507 | AKAP7 | chr6 | + | 131199460 | 131199573 | 131169112 | 131169273 | 131281529 | 131283532 | 26 | 23 |
| SE | 76524 | ENSG00000118507 | AKAP7 | chr6 | + | 131219660 | 131219808 | 131199460 | 131199573 | 131281529 | 131283532 | 87 | 99 |
| SE | 76527 | ENSG00000204209 | DAXX | chr6 | - | 33320735 | 33321567 | 33320379 | 33320591 | 33321718 | 33321974 | 306 | 151 |
| SE | 76532 | ENSG00000134697 | GNL2 | chr1 | - | 37574664 | 37574823 | 37574342 | 37574456 | 37575594 | 37575699 | 107 | 56 |
| SE | 76534 | ENSG00000134697 | GNL2 | chr1 | - | 37587310 | 37587495 | 37583866 | 37583933 | 37590705 | 37590845 | 244 | 141 |
| SE | 76535 | ENSG00000134697 | GNL2 | chr1 | - | 37593761 | 37593846 | 37592711 | 37592806 | 37595758 | 37595937 | 78 | 49 |
| SE | 76536 | ENSG00000197279 | ZNF165 | chr6 | + | 28086171 | 28086310 | 28085480 | 28085891 | 28088562 | 28089563 | 3 | 2 |
| SE | 76537 | ENSG00000135298 | ADGRB3 | chr6 | + | 68637390 | 68637562 | 68635281 | 68636000 | 68638660 | 68639432 | 5 | 10 |
| SE | 76540 | ENSG00000111670 | GNPTAB | chr12 | - | 101749100 | 101749191 | 101745498 | 101747241 | 101753371 | 101753539 | 776 | 391 |
| SE | 76542 | ENSG00000111670 | GNPTAB | chr12 | - | 101760029 | 101760143 | 101757571 | 101757657 | 101761126 | 101761346 | 621 | 270 |
| SE | 76544 | ENSG00000111670 | GNPTAB | chr12 | - | 101768036 | 101768160 | 101766090 | 101766294 | 101770020 | 101770191 | 663 | 290 |
| SE | 76545 | ENSG00000111670 | GNPTAB | chr12 | - | 101780556 | 101780621 | 101778289 | 101780286 | 101786011 | 101786216 | 435 | 164 |
| SE | 76546 | ENSG00000111670 | GNPTAB | chr12 | - | 101786011 | 101786217 | 101780556 | 101780621 | 101788547 | 101788589 | 552 | 178 |
| SE | 76547 | ENSG00000111670 | GNPTAB | chr12 | - | 101810526 | 101810658 | 101796676 | 101796762 | 101830558 | 101830663 | 0 | 0 |
| SE | 76550 | ENSG00000170919 | TPT1-AS1 | chr13 | + | 45344304 | 45344446 | 45341418 | 45341630 | 45360140 | 45360486 | 0 | 2 |
| SE | 76551 | ENSG00000170919 | TPT1-AS1 | chr13 | + | 45344304 | 45344446 | 45341418 | 45341630 | 45369958 | 45370182 | 0 | 2 |
| SE | 76553 | ENSG00000170919 | TPT1-AS1 | chr13 | + | 45344304 | 45344446 | 45341512 | 45341614 | 45369958 | 45370182 | 0 | 0 |
| SE | 76555 | ENSG00000170919 | TPT1-AS1 | chr13 | + | 45344328 | 45344446 | 45341428 | 45341614 | 45369958 | 45370182 | 6 | 1 |
| SE | 76556 | ENSG00000170919 | TPT1-AS1 | chr13 | + | 45344328 | 45344446 | 45341428 | 45341614 | 45370079 | 45370182 | 7 | 2 |
| SE | 76557 | ENSG00000170919 | TPT1-AS1 | chr13 | + | 45344328 | 45344446 | 45341440 | 45341630 | 45360140 | 45360239 | 0 | 0 |
| SE | 76558 | ENSG00000170919 | TPT1-AS1 | chr13 | + | 45344328 | 45344446 | 45341440 | 45341630 | 45369958 | 45370182 | 0 | 0 |
| SE | 76560 | ENSG00000170919 | TPT1-AS1 | chr13 | + | 45344378 | 45344446 | 45341512 | 45341614 | 45370079 | 45370182 | 3 | 1 |
| SE | 76561 | ENSG00000170919 | TPT1-AS1 | chr13 | + | 45344753 | 45344909 | 45344304 | 45344446 | 45351896 | 45352014 | 0 | 0 |
| SE | 76562 | ENSG00000170919 | TPT1-AS1 | chr13 | + | 45344753 | 45344909 | 45344304 | 45344446 | 45369958 | 45371812 | 0 | 0 |
| SE | 76563 | ENSG00000170919 | TPT1-AS1 | chr13 | + | 45344753 | 45344909 | 45344304 | 45344446 | 45370079 | 45370182 | 1 | 0 |
| SE | 76567 | ENSG00000170919 | TPT1-AS1 | chr13 | + | 45351896 | 45352014 | 45344304 | 45344446 | 45369958 | 45370182 | 0 | 0 |
| SE | 76577 | ENSG00000170919 | TPT1-AS1 | chr13 | + | 45370079 | 45370182 | 45344753 | 45344909 | 45377060 | 45377215 | 6 | 6 |
| SE | 76578 | ENSG00000170919 | TPT1-AS1 | chr13 | + | 45378529 | 45378671 | 45374933 | 45377215 | 45379194 | 45379274 | 26 | 18 |
| SE | 76597 | ENSG00000170919 | TPT1-AS1 | chr13 | + | 45383090 | 45383235 | 45378529 | 45380001 | 45389734 | 45389820 | 1 | 0 |
| SE | 76598 | ENSG00000170919 | TPT1-AS1 | chr13 | + | 45383090 | 45383235 | 45379194 | 45379335 | 45389734 | 45389820 | 1 | 0 |
| SE | 76599 | ENSG00000170919 | TPT1-AS1 | chr13 | + | 45383159 | 45383235 | 45379194 | 45379335 | 45389734 | 45389820 | 1 | 0 |
| SE | 76600 | ENSG00000170919 | TPT1-AS1 | chr13 | + | 45383159 | 45383235 | 45379925 | 45380001 | 45389734 | 45389820 | 1 | 0 |
| SE | 76604 | ENSG00000170919 | TPT1-AS1 | chr13 | + | 45390713 | 45390858 | 45389734 | 45389820 | 45391031 | 45391548 | 6 | 9 |
| SE | 76612 | ENSG00000112695 | COX7A2 | chr6 | - | 75240300 | 75240385 | 75237674 | 75237988 | 75241122 | 75241265 | 448 | 240 |
| SE | 76620 | ENSG00000231389 | HLA-DPA1 | chr6 | - | 33068637 | 33068804 | 33064568 | 33065347 | 33069018 | 33069300 | 2240 | 1060 |
| SE | 76621 | ENSG00000231389 | HLA-DPA1 | chr6 | - | 33069018 | 33069300 | 33068637 | 33068804 | 33069640 | 33069886 | 1911 | 949 |
| SE | 76622 | ENSG00000231389 | HLA-DPA1 | chr6 | - | 33069640 | 33069886 | 33069018 | 33069300 | 33073470 | 33073669 | 2561 | 1137 |
| SE | 76631 | ENSG00000107882 | SUFU | chr10 | + | 102599432 | 102599544 | 102597139 | 102597293 | 102615267 | 102615402 | 57 | 35 |
| SE | 76632 | ENSG00000107882 | SUFU | chr10 | + | 102617289 | 102617428 | 102615267 | 102615402 | 102627174 | 102627243 | 41 | 29 |
| SE | 76636 | ENSG00000135503 | ACVR1B | chr12 | + | 51980968 | 51981199 | 51976326 | 51976575 | 51991862 | 51991993 | 31 | 19 |
| SE | 76638 | ENSG00000135503 | ACVR1B | chr12 | + | 51983998 | 51984166 | 51976326 | 51976575 | 51986817 | 51986942 | 1 | 0 |
| SE | 76639 | ENSG00000135503 | ACVR1B | chr12 | + | 51983998 | 51984166 | 51976326 | 51976575 | 51991862 | 51991993 | 0 | 2 |
| SE | 76641 | ENSG00000135503 | ACVR1B | chr12 | + | 51983998 | 51984166 | 51980968 | 51981199 | 51986817 | 51986942 | 35 | 23 |
| SE | 76643 | ENSG00000135503 | ACVR1B | chr12 | + | 51985191 | 51985348 | 51980968 | 51981199 | 51986817 | 51986942 | 19 | 14 |
| SE | 76644 | ENSG00000135503 | ACVR1B | chr12 | + | 51985191 | 51985348 | 51980968 | 51981199 | 51991862 | 51991992 | 3 | 1 |
| SE | 76645 | ENSG00000135503 | ACVR1B | chr12 | + | 51985191 | 51985348 | 51983998 | 51984166 | 51986817 | 51986942 | 35 | 24 |
| SE | 76646 | ENSG00000135503 | ACVR1B | chr12 | + | 51985191 | 51985348 | 51983998 | 51984166 | 51991862 | 51991992 | 19 | 11 |
| SE | 76647 | ENSG00000135503 | ACVR1B | chr12 | + | 51986817 | 51986942 | 51976326 | 51976575 | 51991862 | 51991993 | 26 | 6 |
| SE | 76649 | ENSG00000135503 | ACVR1B | chr12 | + | 51986817 | 51986942 | 51983998 | 51984166 | 51991862 | 51991993 | 27 | 6 |
| SE | 76651 | ENSG00000161800 | RACGAP1 | chr12 | - | 49990683 | 49990792 | 49989161 | 49990343 | 49991997 | 49992133 | 33 | 11 |
| SE | 76655 | ENSG00000161800 | RACGAP1 | chr12 | - | 50016630 | 50016715 | 50006433 | 50006636 | 50025397 | 50025448 | 21 | 15 |
| SE | 76658 | ENSG00000161800 | RACGAP1 | chr12 | - | 50016630 | 50016719 | 50006433 | 50006636 | 50025397 | 50025461 | 27 | 17 |
| SE | 76659 | ENSG00000161800 | RACGAP1 | chr12 | - | 50016630 | 50016719 | 50006433 | 50006636 | 50031696 | 50031867 | 19 | 17 |
| SE | 76660 | ENSG00000161800 | RACGAP1 | chr12 | - | 50016634 | 50016715 | 50006433 | 50006636 | 50031696 | 50031867 | 0 | 0 |
| SE | 76662 | ENSG00000161800 | RACGAP1 | chr12 | - | 50018515 | 50018589 | 50016630 | 50016715 | 50025397 | 50025451 | 0 | 0 |
| SE | 76664 | ENSG00000161800 | RACGAP1 | chr12 | - | 50018515 | 50018589 | 50016630 | 50016719 | 50025397 | 50025451 | 0 | 3 |
| SE | 76670 | ENSG00000161800 | RACGAP1 | chr12 | - | 50021061 | 50021177 | 50018515 | 50018589 | 50025397 | 50025438 | 0 | 0 |

| | | | | | | | | | | | | | | | | | | | | | | | | | | | | | | | | | | | | | | | | | | | | | | |
|---|---|---|---|---|---|---|---|---|---|---|---|---|---|---|---|---|---|---|---|---|---|---|---|---|---|---|---|---|---|---|---|---|---|---|---|---|---|---|---|---|---|---|---|---|---|---|
| SE | 76673 | ENSG00000188807 | TMEM201 | chr1 | + | 9609839 | 9609911 | 9607556 | 9607789 | 9610505 | 9610805 | 3 | 0 | 2 | 3 | 0 | 0 | 0 | 0 | 0 | 0 | 0 | 3 | 2 | 4 | 17 | 0 | 0 | 0 | 0 | 13 | 221 | 149 | 1 | 1 | 1 | NA | 1 | 1 | NA | NA | 1 | 1 | 1 | 0.469 | 0.133 |
| SE | 76674 | ENSG00000188807 | TMEM201 | chr1 | + | 9611014 | 9611073 | 9610505 | 9610805 | 9611752 | 9611890 | 0 | 0 | 0 | 0 | 0 | 5 | 5 | 0 | 4 | 0 | 0 | 0 | 1 | 0 | 0 | 3 | 0 | 0 | 3 | 8 | 208 | 149 | 1 | 1 | 0 | 0 | NA | 0 | NA | 0 | NA | 1 | 0 | 0 | -0.25 |
| SE | 76676 | ENSG00000204227 | RING1 | chr6 | + | 33211157 | 33211547 | 33209914 | 33210130 | 33211728 | 33212002 | 88 | 41 | 78 | 89 | 60 | 1 | 0 | 0 | 0 | 0 | 109 | 95 | 45 | 48 | 124 | 0 | 0 | 0 | 0 | 0 | 298 | 149 | 1 | 1 | 0.978 | 1 | 1 | 1 | 1 | 1 | 1 | 1 | 1 | 1 | -0.004 |
| SE | 76677 | ENSG00000141642 | ELAC1 | chr18 | + | 50984095 | 50984563 | 50974396 | 50974561 | 50986618 | 50988121 | 39 | 21 | 45 | 34 | 43 | 1 | 0 | 0 | 0 | 0 | 24 | 22 | 11 | 16 | 20 | 0 | 1 | 0 | 0 | 0 | 298 | 149 | 1 | 1 | 0.951 | 1 | 1 | 1 | 1 | 1 | 0.917 | 1 | 1 | 1 | 0.007 |
| SE | 76678 | ENSG00000141642 | ELAC1 | chr18 | + | 50984486 | 50984563 | 50974396 | 50974561 | 50986618 | 50986946 | 20 | 12 | 19 | 14 | 31 | 1 | 0 | 0 | 0 | 0 | 11 | 7 | 9 | 11 | 0 | 0 | 1 | 0 | 0 | 0 | 226 | 149 | 1 | 1 | 0.93 | 1 | 1 | 1 | 1 | 1 | 0.822 | 1 | 1 | NA | 0.03 |
| SE | 76680 | ENSG00000223501 | VPS52 | chr6 | - | 33264373 | 33264900 | 33264078 | 33264103 | 33266556 | 33266712 | 98 | 45 | 87 | 86 | 76 | 0 | 0 | 1 | 0 | 0 | 111 | 96 | 79 | 98 | 133 | 0 | 0 | 0 | 0 | 0 | 298 | 149 | 1 | 1 | 1 | 1 | 0.978 | 1 | 1 | 1 | 1 | 1 | 1 | 1 | -0.004 |
| SE | 76681 | ENSG00000223501 | VPS52 | chr6 | - | 33266556 | 33266712 | 33264007 | 33264103 | 33267187 | 33267265 | 61 | 51 | 65 | 69 | 85 | 0 | 0 | 0 | 0 | 0 | 63 | 76 | 29 | 32 | 116 | 0 | 0 | 0 | 2 | 0 | 298 | 149 | 1 | 1 | 1 | 1 | 1 | 1 | 1 | 1 | 1 | 1 | 0.889 | 1 | 0.022 |
| SE | 76682 | ENSG00000223501 | VPS52 | chr6 | - | 33267187 | 33267321 | 33266556 | 33266712 | 33267681 | 33267739 | 145 | 121 | 138 | 165 | 256 | 0 | 0 | 0 | 0 | 0 | 163 | 151 | 84 | 93 | 285 | 0 | 0 | 2 | 0 | 0 | 283 | 149 | 1 | 1 | 1 | 1 | 1 | 1 | 1 | 1 | 1 | 0.957 | 1 | 1 | 0.009 |
| SE | 76684 | ENSG00000223501 | VPS52 | chr6 | - | 33268498 | 33268649 | 33268107 | 33268208 | 33269013 | 33269189 | 71 | 45 | 98 | 94 | 62 | 0 | 0 | 0 | 1 | 0 | 101 | 90 | 90 | 136 | 252 | 0 | 0 | 0 | 0 | 0 | 298 | 149 | 1 | 1 | 1 | 1 | 1 | 0.979 | 1 | 1 | 1 | 1 | 1 | 1 | -0.004 |
| SE | 76685 | ENSG00000223501 | VPS52 | chr6 | - | 33269013 | 33269189 | 33268498 | 33268649 | 33269489 | 33269557 | 195 | 122 | 203 | 198 | 130 | 0 | 0 | 0 | 0 | 0 | 301 | 223 | 205 | 199 | 456 | 0 | 0 | 0 | 1 | 0 | 298 | 149 | 1 | 1 | 1 | 1 | 1 | 1 | 1 | 1 | 1 | 1 | 0.99 | 1 | 0.002 |
| SE | 76686 | ENSG00000223501 | VPS52 | chr6 | - | 33269013 | 33269218 | 33268498 | 33268649 | 33269489 | 33269557 | 41 | 21 | 58 | 59 | 31 | 0 | 0 | 0 | 0 | 0 | 50 | 42 | 50 | 61 | 149 | 0 | 0 | 0 | 1 | 0 | 298 | 149 | 1 | 1 | 1 | 1 | 1 | 1 | 1 | 1 | 1 | 1 | 0.968 | 1 | 0.006 |
| SE | 76689 | ENSG00000237441 | RGL2 | chr6 | - | 33294687 | 33294762 | 33293794 | 33294066 | 33294976 | 33295045 | 81 | 53 | 61 | 93 | 44 | 1 | 0 | 0 | 0 | 0 | 197 | 97 | 66 | 82 | 114 | 5 | 0 | 0 | 0 | 0 | 224 | 149 | 1 | 1 | 0.982 | 1 | 1 | 1 | 1 | 0.963 | 1 | 1 | 1 | 1 | 0.004 |
| SE | 76691 | ENSG00000237441 | RGL2 | chr6 | - | 33297213 | 33297343 | 33297031 | 33297115 | 33297930 | 33297993 | 1 | 0 | 0 | 0 | 0 | 2 | 0 | 0 | 0 | 0 | 0 | 0 | 0 | 0 | 0 | 0 | 0 | 0 | 2 | 0 | 279 | 149 | 1 | 1 | 0.211 | NA | NA | NA | NA | NA | NA | NA | 0 | NA | 0.211 |
| SE | 76692 | ENSG00000237441 | RGL2 | chr6 | - | 33297213 | 33297343 | 33297031 | 33297115 | 33298454 | 33298810 | 2 | 0 | 0 | 0 | 0 | 52 | 14 | 41 | 45 | 62 | 0 | 2 | 0 | 0 | 0 | 97 | 37 | 20 | 58 | 79 | 279 | 149 | 1 | 1 | 0.02 | 0 | 0 | 0 | 0 | 0 | 0.028 | 0 | 0 | 0 | -0.002 |
| SE | 76698 | ENSG00000231925 | TAPBP | chr6 | - | 33308104 | 33308158 | 33304988 | 33305387 | 33313216 | 33313477 | 0 | 0 | 0 | 0 | 0 | 226 | 76 | 168 | 173 | 55 | 0 | 2 | 0 | 0 | 0 | 92 | 226 | 95 | 117 | 338 | 203 | 149 | 1 | 1 | 0 | 0 | 0 | 0 | 0 | 0 | 0.006 | 0 | 0 | 0 | -0.001 |
| SE | 76700 | ENSG00000231925 | TAPBP | chr6 | - | 33313693 | 33313864 | 33313216 | 33313477 | 33314004 | 33314041 | 372 | 156 | 256 | 341 | 104 | 0 | 0 | 0 | 3 | 0 | 235 | 347 | 162 | 284 | 352 | 0 | 0 | 2 | 6 | 0 | 298 | 149 | 1 | 1 | 1 | 1 | 1 | 0.983 | 1 | 1 | 1 | 0.976 | 0.959 | 1 | 0.01 |
| SE | 76702 | ENSG00000125656 | CLPP | chr19 | + | 6366257 | 6366363 | 6364451 | 6364639 | 6368537 | 6368545 | 72 | 21 | 84 | 48 | 34 | 1 | 0 | 0 | 1 | 0 | 72 | 40 | 40 | 71 | 137 | 0 | 0 | 4 | 0 | 0 | 255 | 149 | 1 | 1 | 0.977 | 1 | 1 | 0.966 | 1 | 1 | 1 | 0.854 | 1 | 1 | 0.018 |
| SE | 76703 | ENSG00000044090 | CUL7 | chr6 | - | 43046510 | 43046601 | 43046235 | 43046407 | 43046879 | 43047107 | 25 | 8 | 5 | 10 | 14 | 2 | 3 | 6 | 10 | 7 | 0 | 13 | 14 | 23 | 25 | 0 | 2 | 3 | 0 | 2 | 240 | 149 | 1 | 1 | 0.886 | 0.623 | 0.341 | 0.383 | 0.554 | NA | 0.801 | 0.743 | 1 | 0.886 | -0.3 |
| SE | 76704 | ENSG00000044090 | CUL7 | chr6 | - | 43047946 | 43047989 | 43046510 | 43047107 | 43048147 | 43048160 | 5 | 1 | 0 | 5 | 7 | 6 | 7 | 3 | 3 | 4 | 0 | 10 | 5 | 2 | 0 | 0 | 0 | 4 | 3 | 5 | 192 | 149 | 1 | 1 | 0.393 | 0.1 | 0 | 0.564 | 0.576 | NA | 1 | 0.492 | 0.341 | 0 | -0.132 |
| SE | 76705 | ENSG00000044090 | CUL7 | chr6 | - | 43048147 | 43048253 | 43046879 | 43047107 | 43048331 | 43048343 | 9 | 9 | 8 | 4 | 4 | 0 | 0 | 0 | 1 | 0 | 0 | 1 | 8 | 17 | 29 | 0 | 0 | 0 | 0 | 0 | 255 | 149 | 1 | 1 | 1 | 1 | 1 | 0.7 | 1 | NA | 1 | 1 | 1 | 1 | -0.06 |
| SE | 76706 | ENSG00000044090 | CUL7 | chr6 | - | 43050259 | 43050398 | 43049230 | 43050159 | 43050967 | 43051468 | 41 | 25 | 22 | 41 | 23 | 0 | 0 | 2 | 0 | 0 | 24 | 34 | 23 | 30 | 75 | 0 | 0 | 0 | 0 | 0 | 288 | 149 | 1 | 1 | 1 | 1 | 0.851 | 1 | 1 | 1 | 1 | 1 | 1 | 1 | -0.03 |
| SE | 76707 | ENSG00000142751 | GPN2 | chr1 | - | 26885972 | 26886133 | 26884159 | 26884290 | 26886476 | 26886525 | 22 | 9 | 21 | 12 | 7 | 0 | 0 | 1 | 0 | 0 | 32 | 34 | 25 | 12 | 21 | 0 | 0 | 0 | 0 | 0 | 298 | 149 | 1 | 1 | 1 | 1 | 0.913 | 1 | 1 | 1 | 1 | 1 | 1 | 1 | -0.017 |
| SE | 76708 | ENSG00000142751 | GPN2 | chr1 | - | 26885972 | 26886133 | 26884159 | 26884290 | 26888968 | 26889125 | 46 | 27 | 33 | 39 | 19 | 0 | 0 | 2 | 1 | 0 | 72 | 81 | 49 | 32 | 70 | 0 | 0 | 2 | 0 | 0 | 298 | 149 | 1 | 1 | 1 | 1 | 0.892 | 0.951 | 1 | 1 | 1 | 0.925 | 1 | 1 | -0.016 |
| SE | 76709 | ENSG00000142751 | GPN2 | chr1 | - | 26886476 | 26886525 | 26884159 | 26884290 | 26888968 | 26889125 | 0 | 0 | 1 | 0 | 0 | 0 | 0 | 2 | 1 | 0 | 0 | 0 | 0 | 0 | 0 | 0 | 0 | 2 | 0 | 0 | 198 | 149 | 1 | 1 | NA | NA | 0.273 | 0 | NA | NA | NA | 0 | NA | NA | 0.137 |
| SE | 76710 | ENSG00000142751 | GPN2 | chr1 | - | 26886476 | 26886525 | 26885972 | 26886133 | 26888968 | 26889125 | 0 | 2 | 0 | 0 | 0 | 24 | 20 | 12 | 27 | 12 | 0 | 0 | 0 | 0 | 0 | 40 | 47 | 24 | 20 | 49 | 198 | 149 | 1 | 1 | 0 | 0.07 | 0 | 0 | 0 | 0 | 0 | 0 | 0 | 0 | 0.014 |
| SE | 76718 | ENSG00000112511 | PHF1 | chr6 | + | 33412507 | 33412589 | 33412247 | 33412422 | 33413195 | 33413294 | 15 | 7 | 21 | 17 | 0 | 0 | 0 | 0 | 0 | 0 | 21 | 11 | 21 | 15 | 7 | 0 | 0 | 0 | 0 | 6 | 231 | 149 | 1 | 1 | 1 | 1 | 1 | 1 | NA | 1 | 1 | 1 | 1 | 0.429 | 0.114 |
| SE | 76726 | ENSG00000112511 | PHF1 | chr6 | + | 33415234 | 33415329 | 33414724 | 33414829 | 33415589 | 33415670 | 4 | 6 | 14 | 4 | 4 | 0 | 0 | 0 | 0 | 0 | 10 | 2 | 5 | 11 | 0 | 0 | 0 | 0 | 10 | 0 | 244 | 149 | 1 | 1 | 1 | 1 | 1 | 1 | 1 | 1 | 1 | 1 | 0.402 | NA | 0.149 |
| SE | 76728 | ENSG00000112511 | PHF1 | chr6 | + | 33415234 | 33415333 | 33415064 | 33415144 | 33415592 | 33415670 | 25 | 20 | 36 | 31 | 65 | 0 | 0 | 1 | 0 | 0 | 34 | 38 | 27 | 22 | 9 | 0 | 0 | 0 | 0 | 0 | 248 | 149 | 1 | 1 | 1 | 1 | 0.956 | 1 | 1 | 1 | 1 | 1 | 1 | 1 | -0.009 |
| SE | 76729 | ENSG00000112511 | PHF1 | chr6 | + | 33415260 | 33415329 | 33414724 | 33414829 | 33415589 | 33415670 | 3 | 6 | 16 | 4 | 0 | 0 | 0 | 0 | 0 | 0 | 10 | 2 | 5 | 7 | 0 | 0 | 0 | 0 | 10 | 0 | 218 | 149 | 1 | 1 | 1 | 1 | 1 | 1 | NA | 1 | 1 | 1 | 0.324 | NA | 0.169 |
| SE | 76730 | ENSG00000112511 | PHF1 | chr6 | + | 33415260 | 33415329 | 33414724 | 33414829 | 33415592 | 33415670 | 1 | 0 | 3 | 4 | 0 | 0 | 0 | 0 | 0 | 0 | 0 | 3 | 4 | 1 | 0 | 0 | 0 | 2 | 0 | 0 | 218 | 149 | 1 | 1 | 1 | NA | 1 | 1 | NA | NA | 1 | 0.578 | 1 | NA | 0.141 |
| SE | 76731 | ENSG00000197283 | SYNGAP1 | chr6 | + | 33432160 | 33432252 | 33428229 | 33428313 | 33432684 | 33432806 | 4 | 7 | 10 | 8 | 10 | 0 | 1 | 0 | 0 | 0 | 0 | 6 | 1 | 3 | 4 | 0 | 0 | 0 | 0 | 0 | 241 | 149 | 1 | 1 | 1 | 0.812 | 1 | 1 | 1 | NA | 1 | 1 | 1 | 1 | -0.038 |
| SE | 76732 | ENSG00000197283 | SYNGAP1 | chr6 | + | 33438418 | 33438563 | 33437667 | 33438291 | 33438774 | 33438919 | 132 | 64 | 166 | 127 | 267 | 0 | 0 | 0 | 0 | 0 | 105 | 92 | 103 | 33 | 181 | 0 | 0 | 0 | 1 | 0 | 294 | 149 | 1 | 1 | 1 | 1 | 1 | 1 | 1 | 1 | 1 | 1 | 0.944 | 1 | 0.011 |
| SE | 76734 | ENSG00000197283 | SYNGAP1 | chr6 | + | 33448788 | 33448868 | 33447842 | 33447933 | 33451759 | 33452395 | 22 | 12 | 38 | 16 | 31 | 4 | 1 | 13 | 6 | 7 | 0 | 64 | 18 | 24 | 14 | 0 | 4 | 2 | 1 | 5 | 229 | 149 | 1 | 1 | 0.782 | 0.886 | 0.655 | 0.634 | 0.742 | NA | 0.912 | 0.854 | 0.94 | 0.646 | -0.098 |
| SE | 76735 | ENSG00000167930 | FAM234A | chr16 | + | 246956 | 247069 | 234833 | 234857 | 248243 | 248397 | 0 | 0 | 0 | 0 | 0 | 2 | 2 | 0 | 2 | 0 | 0 | 0 | 0 | 0 | 3 | 0 | 0 | 0 | 3 | 0 | 262 | 149 | 1 | 1 | 0 | 0 | NA | 0 | NA | NA | NA | NA | 0 | 1 | -0.5 |
| SE | 76740 | ENSG00000167930 | FAM234A | chr16 | + | 253832 | 253929 | 249548 | 249654 | 254380 | 254681 | 0 | 2 | 0 | 0 | 0 | 54 | 23 | 13 | 15 | 5 | 0 | 0 | 0 | 0 | 0 | 16 | 12 | 5 | 22 | 38 | 246 | 149 | 1 | 1 | 0 | 0.05 | 0 | 0 | 0 | 0 | 0 | 0 | 0 | 0 | 0.01 |
| SE | 76746 | ENSG00000145996 | CDKAL1 | chr6 | + | 20546345 | 20546523 | 20535350 | 20535394 | 20649292 | 20649377 | 124 | 144 | 149 | 119 | 120 | 0 | 0 | 0 | 0 | 0 | 111 | 140 | 102 | 58 | 38 | 0 | 2 | 0 | 0 | 0 | 298 | 149 | 1 | 1 | 1 | 1 | 1 | 1 | 1 | 1 | 0.972 | 1 | 1 | 1 | 0.006 |
| SE | 76747 | ENSG00000145996 | CDKAL1 | chr6 | + | 20548592 | 20548705 | 20546340 | 20546523 | 20649292 | 20649377 | 297 | 286 | 278 | 332 | 450 | 0 | 2 | 0 | 4 | 0 | 444 | 305 | 267 | 271 | 127 | 0 | 6 | 4 | 0 | 0 | 262 | 149 | 1 | 1 | 1 | 0.988 | 1 | 0.979 | 1 | 1 | 0.967 | 0.974 | 1 | 1 | 0.005 |
| SE | 76748 | ENSG00000145996 | CDKAL1 | chr6 | + | 20781144 | 20781265 | 20758594 | 20758643 | 20846074 | 20846178 | 393 | 251 | 249 | 376 | 345 | 1 | 0 | 2 | 0 | 0 | 268 | 331 | 205 | 275 | 228 | 0 | 0 | 0 | 3 | 0 | 270 | 149 | 1 | 1 | 0.995 | 1 | 0.986 | 1 | 1 | 1 | 1 | 1 | 0.981 | 1 | 0 |
| SE | 76749 | ENSG00000145996 | CDKAL1 | chr6 | + | 20846074 | 20846178 | 20781144 | 20781265 | 20955418 | 20955585 | 309 | 221 | 206 | 301 | 294 | 0 | 1 | 1 | 1 | 3 | 232 | 266 | 180 | 258 | 288 | 0 | 1 | 2 | 0 | 0 | 253 | 149 | 1 | 1 | 1 | 0.992 | 0.992 | 0.994 | 0.983 | 1 | 0.994 | 0.981 | 1 | 1 | -0.003 |
| SE | 76754 | ENSG00000168036 | CTNNB1 | chr3 | + | 41224189 | 41224321 | 41224020 | 41224081 | 41224525 | 41224711 | 0 | 0 | 0 | 0 | 0 | 166 | 80 | 109 | 122 | 119 | 0 | 0 | 4 | 0 | 0 | 110 | 106 | 120 | 142 | 83 | 281 | 149 | 1 | 1 | 0 | 0 | 0 | 0 | 0 | 0 | 0 | 0.017 | 0 | 0 | -0.003 |
| SE | 76756 | ENSG00000168036 | CTNNB1 | chr3 | + | 41224953 | 41225207 | 41223991 | 41224753 | 41225333 | 41225572 | 543 | 326 | 415 | 516 | 510 | 0 | 0 | 0 | 0 | 0 | 361 | 295 | 448 | 359 | 300 | 0 | 0 | 0 | 2 | 0 | 298 | 149 | 1 | 1 | 1 | 1 | 1 | 1 | 1 | 1 | 1 | 1 | 0.989 | 1 | 0.002 |
| SE | 76762 | ENSG00000168036 | CTNNB1 | chr3 | + | 41239133 | 41239354 | 41238015 | 41238076 | 41239659 | 41239767 | 247 | 87 | 219 | 219 | 235 | 1 | 0 | 0 | 0 | 0 | 152 | 161 | 149 | 134 | 53 | 0 | 0 | 0 | 0 | 0 | 298 | 149 | 1 | 1 | 0.992 | 1 | 1 | 1 | 1 | 1 | 1 | 1 | 1 | 1 | -0.002 |
| SE | 76763 | ENSG00000099995 | SF3A1 | chr22 | - | 30340694 | 30340812 | 30340195 | 30340381 | 30341691 | 30341850 | 491 | 277 | 436 | 495 | 870 | 1 | 0 | 0 | 0 | 0 | 555 | 534 | 452 | 385 | 722 | 0 | 0 | 0 | 0 | 0 | 267 | 149 | 1 | 1 | 0.996 | 1 | 1 | 1 | 1 | 1 | 1 | 1 | 1 | 1 | -0.001 |
| SE | 76764 | ENSG00000099995 | SF3A1 | chr22 | - | 30340694 | 30340812 | 30340195 | 30340381 | 30342199 | 30342350 | 275 | 157 | 232 | 292 | 336 | 0 | 0 | 0 | 0 | 0 | 371 | 297 | 273 | 238 | 450 | 9 | 0 | 0 | 0 | 0 | 267 | 149 | 1 | 1 | 1 | 1 | 1 | 1 | 1 | 0.958 | 1 | 1 | 1 | 1 | 0.008 |
| SE | 76765 | ENSG00000099995 | SF3A1 | chr22 | - | 30341691 | 30341885 | 30340195 | 30340381 | 30342199 | 30342350 | 881 | 593 | 744 | 894 | 1539 | 0 | 0 | 0 | 0 | 0 | 1836 | 1134 | 1036 | 1036 | 1737 | 9 | 0 | 0 | 0 | 0 | 298 | 149 | 1 | 1 | 1 | 1 | 1 | 1 | 1 | 0.99 | 1 | 1 | 1 | 1 | 0.002 |
| SE | 76766 | ENSG00000099995 | SF3A1 | chr22 | - | 30341691 | 30341885 | 30340694 | 30340812 | 30342199 | 30342350 | 1096 | 713 | 948 | 1097 | 2073 | 0 | 0 | 0 | 0 | 0 | 2020 | 1382 | 1215 | 1186 | 2009 | 0 | 0 | 0 | 3 | 0 | 298 | 149 | 1 | 1 | 1 | 1 | 1 | 1 | 1 | 1 | 1 | 1 | 0.995 | 1 | 0.001 |
| SE | 76767 | ENSG00000099995 | SF3A1 | chr22 | - | 30344932 | 30345190 | 30342804 | 30342879 | 30346311 | 30346519 | 535 | 309 | 523 | 572 | 565 | 0 | 0 | 1 | 0 | 0 | 805 | 709 | 599 | 616 | 1363 | 0 | 0 | 0 | 0 | 0 | 298 | 149 | 1 | 1 | 1 | 1 | 0.996 | 1 | 1 | 1 | 1 | 1 | 1 | 1 | -0.001 |
| SE | 76778 | ENSG00000135317 | SNX14 | chr6 | - | 85530191 | 85530275 | 85528261 | 85528362 | 85533598 | 85533680 | 440 | 261 | 317 | 367 | 396 | 0 | 0 | 1 | 3 | 2 | 149 | 219 | 248 | 235 | 68 | 0 | 0 | 1 | 0 | 0 | 233 | 149 | 1 | 1 | 1 | 1 | 0.995 | 0.987 | 0.992 | 1 | 1 | 0.994 | 1 | 1 | -0.004 |
| SE | 76779 | ENSG00000135317 | SNX14 | chr6 | - | 85530191 | 85530275 | 85528261 | 85528362 | 85536791 | 85536924 | 197 | 148 | 138 | 140 | 189 | 0 | 2 | 0 | 0 | 0 | 107 | 77 | 119 | 127 | 41 | 0 | 0 | 0 | 0 | 0 | 233 | 149 | 1 | 1 | 1 | 0.979 | 1 | 1 | 1 | 1 | 1 | 1 | 1 | 1 | -0.004 |
| SE | 76780 | ENSG00000135317 | SNX14 | chr6 | - | 85533598 | 85533800 | 85528261 | 85528362 | 85536791 | 85536924 | 225 | 164 | 128 | 182 | 334 | 0 | 2 | 0 | 0 | 0 | 25 | 113 | 104 | 104 | 46 | 0 | 0 | 0 | 0 | 0 | 298 | 149 | 1 | 1 | 1 | 0.976 | 1 | 1 | 1 | 1 | 1 | 1 | 1 | 1 | -0.005 |
| SE | 76785 | ENSG00000135317 | SNX14 | chr6 | - | 85541984 | 85542043 | 85533598 | 85533800 | 85543181 | 85543306 | 186 | 162 | 103 | 147 | 280 | 2 | 0 | 3 | 3 | 1 | 56 | 136 | 85 | 84 | 30 | 0 | 0 | 2 | 1 | 0 | 208 | 149 | 1 | 1 | 0.985 | 1 | 0.961 | 0.972 | 0.995 | 1 | 1 | 0.968 | 0.984 | 1 | -0.008 |
| SE | 76786 | ENSG00000135317 | SNX14 | chr6 | - | 85541984 | 85542043 | 85536791 | 85536924 | 85543181 | 85543306 | 588 | 473 | 298 | 434 | 841 | 2 | 0 | 0 | 0 | 0 | 154 | 451 | 320 | 320 | 81 | 0 | 0 | 0 | 0 | 0 | 208 | 149 | 1 | 1 | 0.995 | 1 | 1 | 1 | 1 | 1 | 1 | 1 | 1 | 1 | -0.001 |
| SE | 76789 | ENSG00000135317 | SNX14 | chr6 | - | 85543604 | 85543760 | 85543181 | 85543306 | 85547111 | 85547226 | 492 | 374 | 274 | 374 | 651 | 1 | 0 | 0 | 0 | 0 | 190 | 327 | 232 | 277 | 88 | 0 | 0 | 0 | 0 | 0 | 298 | 149 | 1 | 1 | 0.996 | 1 | 1 | 1 | 1 | 1 | 1 | 1 | 1 | 1 | -0.001 |
| SE | 76790 | ENSG00000135317 | SNX14 | chr6 | - | 85543604 | 85545989 | 85543181 | 85543306 | 85547111 | 85547226 | 348 | 288 | 195 | 250 | 457 | 1 | 0 | 0 | 0 | 0 | 149 | 264 | 171 | 224 | 62 | 0 | 0 | 0 | 0 | 0 | 298 | 149 | 1 | 1 | 0.994 | 1 | 1 | 1 | 1 | 1 | 1 | 1 | 1 | 1 | -0.001 |
| SE | 76791 | ENSG00000135317 | SNX14 | chr6 | - | 85547111 | 85547226 | 85543181 | 85543760 | 85547505 | 85547550 | 77 | 47 | 26 | 61 | 72 | 0 | 0 | 0 | 0 | 1 | 28 | 40 | 35 | 21 | 33 | 0 | 0 | 2 | 0 | 0 | 264 | 149 | 1 | 1 | 1 | 1 | 1 | 1 | 0.976 | 1 | 1 | 0.908 | 1 | 1 | 0.014 |
| SE | 76793 | ENSG00000135317 | SNX14 | chr6 | - | 85547505 | 85547550 | 85547111 | 85547226 | 85548300 | 85548376 | 49 | 22 | 24 | 45 | 39 | 0 | 0 | 0 | 2 | 0 | 20 | 24 | 11 | 21 | 24 | 0 | 0 | 0 | 0 | 0 | 194 | 149 | 1 | 1 | 1 | 1 | 1 | 0.945 | 1 | 1 | 1 | 1 | 1 | 1 | -0.011 |
| SE | 76794 | ENSG00000135317 | SNX14 | chr6 | - | 85548300 | 85548376 | 85547401 | 85547550 | 85557975 | 85558060 | 76 | 48 | 37 | 52 | 74 | 0 | 0 | 0 | 2 | 0 | 13 | 32 | 18 | 25 | 16 | 0 | 0 | 0 | 0 | 0 | 225 | 149 | 1 | 1 | 1 | 1 | 1 | 0.945 | 1 | 1 | 1 | 1 | 1 | 1 | -0.011 |
| SE | 76798 | ENSG00000135317 | SNX14 | chr6 | - | 85557975 | 85558060 | 85549722 | 85549879 | 85572136 | 85572215 | 127 | 59 | 89 | 76 | 107 | 0 | 0 | 4 | 0 | 0 | 13 | 65 | 33 | 23 | 20 | 0 | 0 | 0 | 0 | 0 | 234 | 149 | 1 | 1 | 1 | 1 | 0.934 | 1 | 1 | 1 | 1 | 1 | 1 | 1 | -0.013 |
| SE | 76799 | ENSG00000135317 | SNX14 | chr6 | - | 85560773 | 85561340 | 85557975 | 85558060 | 85565331 | 85565419 | 0 | 0 | 0 | 3 | 0 | 147 | 102 | 81 | 108 | 222 | 0 | 9 | 0 | 0 | 0 | 116 | 97 | 52 | 102 | 39 | 298 | 149 | 1 | 1 | 0 | 0 | 0 | 0.014 | 0 | 0 | 0.044 | 0 | 0 | 0 | -0.006 |
| SE | 76800 | ENSG00000135317 | SNX14 | chr6 | - | 85561214 | 85561340 | 85557975 | 85558060 | 85565331 | 85565419 | 0 | 0 | 0 | 3 | 0 | 147 | 102 | 81 | 108 | 222 | 0 | 0 | 0 | 0 | 0 | 116 | 97 | 52 | 102 | 39 | 275 | 149 | 1 | 1 | 0 | 0 | 0 | 0.015 | 0 | 0 | 0 | 0 | 0 | 0 | 0.003 |
| SE | 76801 | ENSG00000135317 | SNX14 | chr6 | - | 85564727 | 85564985 | 85557975 | 85558060 | 85565331 | 85565419 | 4 | 0 | 0 | 0 | 0 | 147 | 102 | 81 | 108 | 222 | 0 | 1 | 0 | 0 | 0 | 116 | 97 | 52 | 102 | 39 | 298 | 149 | 1 | 1 | 0.013 | 0 | 0 | 0 | 0 | 0 | 0.005 | 0 | 0 | 0 | 0.002 |
| SE | 76804 | ENSG00000135317 | SNX14 | chr6 | - | 85564729 | 85564985 | 85557975 | 85558060 | 85565331 | 85565419 | 4 | 0 | 0 | 0 | 0 | 147 | 102 | 81 | 108 | 222 | 0 | 1 | 0 | 0 | 0 | 116 | 97 | 52 | 102 | 39 | 298 | 149 | 1 | 1 | 0.013 | 0 | 0 | 0 | 0 | 0 | 0.005 | 0 | 0 | 0 | 0.002 |
| SE | 76806 | ENSG00000135317 | SNX14 | chr6 | - | 85565331 | 85565419 | 85549722 | 85549879 | 85572136 | 85572215 | 4 | 3 | 1 | 0 | 6 | 0 | 0 | 4 | 0 | 0 | 0 | 4 | 4 | 10 | 0 | 0 | 0 | 0 | 0 | 0 | 237 | 149 | 1 | 1 | 1 | 1 | 0.136 | NA | 1 | NA | 1 | 1 | 1 | NA | -0.216 |
| SE | 76807 | ENSG00000135317 | SNX14 | chr6 | - | 85565331 | 85565419 | 85557975 | 85558060 | 85567533 | 85567577 | 179 | 123 | 113 | 134 | 306 | 2 | 0 | 0 | 1 | 1 | 187 | 112 | 61 | 118 | 63 | 0 | 0 | 0 | 0 | 0 | 237 | 149 | 1 | 1 | 0.983 | 1 | 1 | 0.988 | 0.995 | 1 | 1 | 1 | 1 | 1 | -0.007 |
| SE | 76814 | ENSG00000135317 | SNX14 | chr6 | - | 85574257 | 85574378 | 85572297 | 85572371 | 85593578 | 85593718 | 63 | 25 | 35 | 65 | 59 | 0 | 0 | 0 | 0 | 0 | 18 | 50 | 12 | 50 | 30 | 0 | 0 | 1 | 0 | 0 | 270 | 149 | 1 | 1 | 1 | 1 | 1 | 1 | 1 | 1 | 1 | 0.869 | 1 | 1 | 0.026 |
| SE | 76816 | ENSG00000135317 | SNX14 | chr6 | - | 85574257 | 85574378 | 85572297 | 85572374 | 85593578 | 85593718 | 179 | 122 | 103 | 200 | 130 | 0 | 0 | 0 | 3 | 0 | 110 | 122 | 80 | 102 | 61 | 0 | 0 | 0 | 0 | 0 | 270 | 149 | 1 | 1 | 1 | 1 | 1 | 0.974 | 1 | 1 | 1 | 1 | 1 | 1 | -0.005 |
| SE | 76817 | ENSG00000135317 | SNX14 | chr6 | - | 85581195 | 85581258 | 85574257 | 85574378 | 85588862 | 85588950 | 0 | 2 | 0 | 0 | 0 | 3 | 10 | 9 | 9 | 1 | 0 | 0 | 0 | 0 | 0 | 5 | 9 | 24 | 1 | 6 | 212 | 149 | 1 | 1 | 0 | 0.123 | 0 | 0 | 0 | 0 | 0 | 0 | 0 | 0 | 0.025 |
| SE | 76824 | ENSG00000172508 | CARNS1 | chr11 | + | 67420938 | 67421219 | 67420608 | 67420840 | 67423374 | 67424684 | 14 | 2 | 3 | 5 | 0 | 0 | 0 | 0 | 0 | 1 | 0 | 0 | 1 | 0 | 0 | 0 | 0 | 0 | 0 | 0 | 298 | 149 | 1 | 1 | 1 | 1 | 1 | 1 | 0 | NA | NA | 1 | NA | NA | -0.2 |
| SE | 76825 | ENSG00000255339 | NA | chr10 | - | 100509315 | 100509511 | 100509006 | 100509102 | 100526398 | 100526554 | 31 | 22 | 46 | 28 | 17 | 0 | 0 | 1 | 0 | 0 | 40 | 39 | 31 | 24 | 71 | 0 | 0 | 2 | 0 | 0 | 298 | 149 | 1 | 1 | 1 | 1 | 0.958 | 1 | 1 | 1 | 1 | 0.886 | 1 | 1 | 0.014 |
| SE | 76826 | ENSG00000255339 | NA | chr10 | - | 100516095 | 100516219 | 100509006 | 100509102 | 100526974 | 100527074 | 3 | 1 | 3 | 3 | 6 | 0 | 0 | 2 | 0 | 0 | 0 | 6 | 1 | 7 | 0 | 0 | 0 | 1 | 1 | 3 | 273 | 149 | 1 | 1 | 1 | 1 | 0.45 | 1 | 1 | NA | 1 | 0.353 | 0.793 | 0 | 0.354 |
| SE | 76827 | ENSG00000255339 | NA | chr10 | - | 100516873 | 100516960 | 100516095 | 100516219 | 100526974 | 100527074 | 18 | 22 | 18 | 24 | 9 | 0 | 0 | 0 | 0 | 0 | 5 | 9 | 22 | 28 | 29 | 0 | 0 | 0 | 4 | 0 | 236 | 149 | 1 | 1 | 1 | 1 | 1 | 1 | 1 | 1 | 1 | 1 | 0.815 | 1 | 0.037 |
| SE | 76829 | ENSG00000255339 | NA | chr10 | - | 100526398 | 100526554 | 100509006 | 100509102 | 100526974 | 100527074 | 184 | 118 | 241 | 207 | 144 | 0 | 0 | 2 | 0 | 0 | 177 | 227 | 150 | 122 | 360 | 0 | 0 | 1 | 1 | 3 | 298 | 149 | 1 | 1 | 1 | 1 | 0.984 | 1 | 1 | 1 | 1 | 0.987 | 0.984 | 0.984 | 0.006 |
| SE | 76831 | ENSG00000146842 | TMEM209 | chr7 | - | 130175511 | 130175609 | 130173824 | 130173939 | 130181622 | 130181719 | 36 | 21 | 29 | 35 | 80 | 0 | 2 | 0 | 0 | 0 | 33 | 35 | 26 | 27 | 8 | 0 | 0 | 0 | 0 | 0 | 247 | 149 | 1 | 1 | 1 | 0.864 | 1 | 1 | 1 | 1 | 1 | 1 | 1 | 1 | -0.027 |
| SE | 76833 | ENSG00000146842 | TMEM209 | chr7 | - | 130178401 | 130178527 | 130175511 | 130175609 | 130184183 | 130184255 | 76 | 72 | 65 | 65 | 181 | 0 | 0 | 0 | 0 | 3 | 43 | 61 | 38 | 66 | 38 | 0 | 3 | 0 | 0 | 0 | 275 | 149 | 1 | 1 | 1 | 1 | 1 | 1 | 0.97 | 1 | 0.917 | 1 | 1 | 1 | 0.011 |
| SE | 76834 | ENSG00000146842 | TMEM209 | chr7 | - | 130181622 | 130181719 | 130175511 | 130175609 | 130184183 | 130184255 | 78 | 93 | 89 | 81 | 130 | 0 | 0 | 0 | 0 | 3 | 71 | 98 | 73 | 50 | 7 | 0 | 3 | 0 | 0 | 0 | 246 | 149 | 1 | 1 | 1 | 1 | 1 | 1 | 0.963 | 1 | 0.952 | 1 | 1 | 1 | 0.002 |
| SE | 76836 | ENSG00000146842 | TMEM209 | chr7 | - | 130202531 | 130202663 | 130201849 | 130202091 | 130203787 | 130203846 | 185 | 104 | 174 | 140 | 391 | 0 | 0 | 0 | 0 | 0 | 31 | 179 | 99 | 63 | 28 | 0 | 4 | 0 | 0 | 0 | 281 | 149 | 1 | 1 | 1 | 1 | 1 | 1 | 1 | 1 | 0.96 | 1 | 1 | 1 | 0.008 |
| SE | 76838 | ENSG00000124788 | ATXN1 | chr6 | - | 16432945 | 16433096 | 16432563 | 16432733 | 16485971 | 16486109 | 1 | 1 | 3 | 3 | 9 | 0 | 0 | 0 | 0 | 0 | 0 | 24 | 28 | 12 | 21 | 0 | 0 | 3 | 0 | 0 | 298 | 149 | 1 | 1 | 1 | 1 | 1 | 1 | 1 | NA | 1 | 0.824 | 1 | 1 | 0.044 |
| SE | 76839 | ENSG00000124788 | ATXN1 | chr6 | - | 16522626 | 16522688 | 16485111 | 16486109 | 16585779 | 16585907 | 118 | 74 | 82 | 90 | 146 | 1 | 1 | 0 | 0 | 0 | 121 | 162 | 133 | 179 | 114 | 0 | 0 | 4 | 0 | 0 | 211 | 149 | 1 | 1 | 0.988 | 0.981 | 1 | 1 | 1 | 1 | 1 | 0.959 | 1 | 1 | 0.002 |
| SE | 76840 | ENSG00000124788 | ATXN1 | chr6 | - | 16585779 | 16585907 | 16522626 | 16522688 | 16657775 | 16657901 | 138 | 103 | 101 | 141 | 179 | 0 | 0 | 0 | 0 | 4 | 136 | 204 | 186 | 204 | 179 | 0 | 0 | 0 | 0 | 0 | 277 | 149 | 1 | 1 | 1 | 1 | 1 | 1 | 0.96 | 1 | 1 | 1 | 1 | 1 | -0.008 |
| SE | 76846 | ENSG00000124788 | ATXN1 | chr6 | - | 16754042 | 16754091 | 16753232 | 16753347 | 16754639 | 16754746 | 1 | 0 | 2 | 4 | 3 | 1 | 0 | 0 | 0 | 0 | 5 | 2 | 6 | 3 | 0 | 0 | 0 | 0 | 0 | 0 | 198 | 149 | 1 | 1 | 0.429 | NA | 1 | 1 | 1 | 1 | 1 | 1 | 1 | NA | -0.143 |
| SE | 76847 | ENSG00000124788 | ATXN1 | chr6 | - | 16754042 | 16754091 | 16753232 | 16753347 | 16760056 | 16760102 | 2 | 0 | 2 | 4 | 3 | 0 | 2 | 0 | 0 | 0 | 5 | 2 | 6 | 3 | 0 | 0 | 1 | 0 | 0 | 0 | 198 | 149 | 1 | 1 | 1 | 0 | 1 | 1 | 1 | 1 | 0.601 | 1 | 1 | NA | -0.1 |
| SE | 76849 | ENSG00000124788 | ATXN1 | chr6 | - | 16754639 | 16754746 | 16753232 | 16753347 | 16761297 | 16761370 | 1 | 0 | 0 | 0 | 0 | 27 | 1 | 10 | 14 | 13 | 0 | 0 | 0 | 0 | 0 | 3 | 17 | 10 | 12 | 6 | 256 | 149 | 1 | 1 | 0.021 | 0 | 0 | 0 | 0 | 0 | 0 | 0 | 0 | 0 | 0.004 |
| SE | 76851 | ENSG00000288721 | NA | chr6 | - | 41909268 | 41909522 | 41907015 | 41907287 | 41916784 | 41916939 | 25 | 18 | 31 | 12 | 18 | 3 | 1 | 5 | 1 | 2 | 0 | 5 | 27 | 21 | 20 | 0 | 0 | 6 | 0 | 0 | 298 | 149 | 1 | 1 | 0.806 | 0.9 | 0.756 | 0.857 | 0.818 | NA | 1 | 0.692 | 1 | 1 | -0.096 |
| SE | 76852 | ENSG00000124641 | MED20 | chr6 | - | 41909344 | 41909522 | 41906803 | 41907287 | 41916784 | 41916939 | 16 | 8 | 17 | 10 | 15 | 3 | 1 | 5 | 1 | 2 | 0 | 2 | 15 | 21 | 11 | 0 | 0 | 6 | 0 | 0 | 298 | 149 | 1 | 1 | 0.727 | 0.8 | 0.63 | 0.833 | 0.789 | NA | 1 | 0.556 | 1 | 1 | -0.133 |
| SE | 76855 | ENSG00000124641 | MED20 | chr6 | - | 41916784 | 41916939 | 41908951 | 41909522 | 41921004 | 41921093 | 34 | 21 | 46 | 24 | 46 | 1 | 0 | 0 | 0 | 0 | 8 | 15 | 29 | 54 | 14 | 0 | 0 | 0 | 0 | 0 | 298 | 149 | 1 | 1 | 0.944 | 1 | 1 | 1 | 1 | 1 | 1 | 1 | 1 | 1 | -0.011 |
| SE | 76856 | ENSG00000124641 | MED20 | chr6 | - | 41916784 | 41916939 | 41912984 | 41913100 | 41921004 | 41921093 | 18 | 13 | 29 | 16 | 31 | 0 | 0 | 2 | 0 | 0 | 8 | 13 | 14 | 33 | 3 | 0 | 0 | 0 | 0 | 0 | 298 | 149 | 1 | 1 | 1 | 1 | 0.879 | 1 | 1 | 1 | 1 | 1 | 1 | 1 | -0.024 |
| SE | 76858 | ENSG00000168661 | ZNF30 | chr19 | + | 34942593 | 34942673 | 34933627 | 34933723 | 34943222 | 34945168 | 9 | 6 | 7 | 7 | 10 | 5 | 0 | 3 | 6 | 2 | 0 | 3 | 1 | 4 | 0 | 0 | 0 | 0 | 4 | 0 | 229 | 149 | 1 | 1 | 0.539 | 1 | 0.603 | 0.432 | 0.765 | NA | 1 | 1 | 0.394 | NA | -0.13 |
| SE | 76861 | ENSG00000112576 | CCND3 | chr6 | - | 41937234 | 41937394 | 41935434 | 41936107 | 41940369 | 41940585 | 229 | 109 | 113 | 173 | 108 | 0 | 0 | 0 | 0 | 0 | 271 | 277 | 125 | 205 | 591 | 0 | 0 | 1 | 0 | 0 | 298 | 149 | 1 | 1 | 1 | 1 | 1 | 1 | 1 | 1 | 1 | 0.984 | 1 | 1 | 0.003 |
| SE | 76865 | ENSG00000112576 | CCND3 | chr6 | - | 41940951 | 41941161 | 41940493 | 41940585 | 42048500 | 42048688 | 0 | 0 | 0 | 4 | 0 | 55 | 24 | 52 | 61 | 14 | 0 | 1 | 3 | 3 | 23 | 0 | 45 | 13 | 5 | 5 | 298 | 149 | 1 | 1 | 0 | 0 | 0 | 0.032 | 0 | NA | 0.011 | 0.103 | 0.231 | 0.697 | -0.254 |
| SE | 76866 | ENSG00000112576 | CCND3 | chr6 | - | 42030008 | 42030190 | 41937174 | 41937394 | 42048500 | 42048686 | 3 | 0 | 4 | 5 | 1 | 47 | 6 | 31 | 25 | 12 | 0 | 15 | 13 | 0 | 0 | 0 | 46 | 31 | 11 | 15 | 298 | 149 | 1 | 1 | 0.031 | 0 | 0.061 | 0.091 | 0.04 | NA | 0.14 | 0.173 | 0 | 0 | -0.034 |
| SE | 76867 | ENSG00000112576 | CCND3 | chr6 | - | 42030008 | 42030190 | 41940409 | 41940585 | 42048500 | 42048686 | 6 | 0 | 6 | 7 | 0 | 55 | 24 | 52 | 61 | 14 | 0 | 11 | 10 | 0 | 0 | 0 | 45 | 13 | 5 | 5 | 298 | 149 | 1 | 1 | 0.052 | 0 | 0.055 | 0.054 | 0 | NA | 0.109 | 0.278 | 0 | 0 | -0.065 |
| SE | 76868 | ENSG00000112576 | CCND3 | chr6 | - | 42030008 | 42030190 | 42010837 | 42011067 | 42048500 | 42048686 | 3 | 0 | 2 | 6 | 0 | 0 | 0 | 0 | 1 | 0 | 0 | 10 | 9 | 0 | 0 | 0 | 0 | 6 | 0 | 0 | 298 | 149 | 1 | 1 | 1 | NA | 1 | 1 | 0 | NA | 1 | 0.429 | NA | NA | 0.035 |
| SE | 76873 | ENSG00000001167 | NFYA | chr6 | + | 41084045 | 41084192 | 41079028 | 41079164 | 41089578 | 41089710 | 245 | 214 | 314 | 319 | 238 | 1 | 0 | 0 | 0 | 0 | 234 | 197 | 193 | 242 | 71 | 0 | 0 | 0 | 0 | 0 | 296 | 149 | 1 | 1 | 0.992 | 1 | 1 | 1 | 1 | 1 | 1 | 1 | 1 | 1 | -0.002 |
| SE | 76874 | ENSG00000001167 | NFYA | chr6 | + | 41089578 | 41089710 | 41084045 | 41084192 | 41090203 | 41090309 | 195 | 161 | 261 | 271 | 171 | 0 | 0 | 4 | 0 | 2 | 169 | 161 | 150 | 183 | 44 | 0 | 0 | 2 | 0 | 0 | 281 | 149 | 1 | 1 | 1 | 1 | 0.972 | 1 | 0.978 | 1 | 1 | 0.975 | 1 | 1 | -0.005 |
| SE | 76875 | ENSG00000001167 | NFYA | chr6 | + | 41091527 | 41091694 | 41090203 | 41090309 | 41092911 | 41093085 | 211 | 138 | 169 | 200 | 202 | 0 | 0 | 0 | 1 | 0 | 138 | 111 | 149 | 200 | 67 | 0 | 0 | 0 | 0 | 0 | 298 | 149 | 1 | 1 | 1 | 1 | 1 | 0.99 | 1 | 1 | 1 | 1 | 1 | 1 | -0.002 |
| SE | 76876 | ENSG00000001167 | NFYA | chr6 | + | 41092911 | 41093085 | 41091527 | 41091694 | 41094395 | 41094497 | 225 | 148 | 176 | 211 | 239 | 0 | 2 | 0 | 0 | 0 | 107 | 116 | 135 | 211 | 103 | 0 | 0 | 0 | 0 | 0 | 298 | 149 | 1 | 1 | 1 | 0.974 | 1 | 1 | 1 | 1 | 1 | 1 | 1 | 1 | -0.005 |
| SE | 76881 | ENSG00000235109 | ZSCAN31 | chr6 | - | 28329302 | 28329714 | 28327382 | 28327533 | 28335829 | 28335936 | 0 | 0 | 3 | 2 | 3 | 1 | 0 | 1 | 0 | 0 | 2 | 3 | 0 | 0 | 0 | 0 | 3 | 0 | 0 | 0 | 298 | 149 | 1 | 1 | 0 | NA | 0.6 | 1 | 1 | 1 | 0.333 | NA | NA | NA | -0.016 |
| SE | 76882 | ENSG00000235109 | ZSCAN31 | chr6 | - | 28329302 | 28329714 | 28327382 | 28327533 | 28336081 | 28336198 | 0 | 0 | 3 | 2 | 3 | 0 | 0 | 0 | 1 | 0 | 2 | 3 | 0 | 0 | 0 | 0 | 0 | 1 | 0 | 0 | 298 | 149 | 1 | 1 | NA | NA | 1 | 0.5 | 1 | 1 | 1 | 0 | NA | NA | 0.167 |
| SE | 76883 | ENSG00000235109 | ZSCAN31 | chr6 | - | 28329302 | 28329778 | 28327382 | 28327533 | 28335829 | 28336134 | 0 | 0 | 3 | 2 | 3 | 1 | 0 | 1 | 0 | 0 | 2 | 3 | 0 | 0 | 0 | 0 | 3 | 0 | 0 | 0 | 298 | 149 | 1 | 1 | 0 | NA | 0.6 | 1 | 1 | 1 | 0.333 | NA | NA | NA | -0.016 |
| SE | 76884 | ENSG00000235109 | ZSCAN31 | chr6 | - | 28329302 | 28329778 | 28327382 | 28327533 | 28336081 | 28336148 | 1 | 1 | 3 | 2 | 3 | 0 | 0 | 0 | 1 | 0 | 2 | 3 | 0 | 0 | 0 | 0 | 0 | 1 | 0 | 0 | 298 | 149 | 1 | 1 | 1 | 1 | 1 | 0.5 | 1 | 1 | 1 | 0 | NA | NA | 0.233 |
| SE | 76895 | ENSG00000118515 | SGK1 | chr6 | - | 134172661 | 134172774 | 134172192 | 134172316 | 134173022 | 134173154 | 560 | 379 | 299 | 643 | 1108 | 0 | 2 | 3 | 0 | 0 | 537 | 254 | 210 | 606 | 159 | 0 | 0 | 0 | 0 | 0 | 262 | 149 | 1 | 1 | 1 | 0.991 | 0.983 | 1 | 1 | 1 | 1 | 1 | 1 | 1 | -0.005 |
| SE | 76896 | ENSG00000118515 | SGK1 | chr6 | - | 134173022 | 134173154 | 134172661 | 134172774 | 134173273 | 134173357 | 571 | 417 | 313 | 591 | 1068 | 0 | 0 | 0 | 3 | 1 | 534 | 273 | 214 | 604 | 170 | 0 | 0 | 0 | 0 | 0 | 281 | 149 | 1 | 1 | 0.997 | 1 | 1 | 0.991 | 0.998 | 1 | 1 | 1 | 1 | 1 | -0.003 |
| SE | 76901 | ENSG00000204590 | GNL1 | chr6 | - | 30553083 | 30553179 | 30547351 | 30547530 | 30554574 | 30554646 | 0 | 1 | 0 | 0 | 0 | 1 | 0 | 0 | 0 | 0 | 0 | 1 | 3 | 6 | 0 | 0 | 0 | 0 | 2 | 0 | 245 | 149 | 1 | 1 | NA | 1 | NA | NA | NA | NA | 1 | 1 | 0.646 | NA | 0.118 |
| SE | 76905 | ENSG00000151914 | DST | chr6 | - | 56464684 | 56464756 | 56463564 | 56463764 | 56466077 | 56466195 | 0 | 1 | 0 | 0 | 0 | 6 | 4 | 3 | 2 | 3 | 0 | 0 | 0 | 0 | 0 | 0 | 1 | 3 | 0 | 0 | 221 | 149 | 1 | 1 | 0 | 0.144 | 0 | 0 | 0 | NA | 0 | 0 | NA | NA | 0.029 |
| SE | 76906 | ENSG00000151914 | DST | chr6 | - | 56468981 | 56468999 | 56466077 | 56466195 | 56469882 | 56469957 | 10 | 9 | 5 | 5 | 8 | 3 | 0 | 0 | 0 | 4 | 0 | 0 | 1 | 0 | 0 | 0 | 2 | 3 | 0 | 0 | 167 | 149 | 1 | 1 | 0.748 | 1 | 1 | 1 | 0.641 | NA | 0 | 0.229 | NA | NA | 0.763 |
| SE | 76907 | ENSG00000151914 | DST | chr6 | - | 56482682 | 56482877 | 56482049 | 56482178 | 56485311 | 56485471 | 14 | 4 | 5 | 6 | 10 | 0 | 0 | 0 | 0 | 0 | 0 | 0 | 0 | 0 | 0 | 0 | 1 | 0 | 0 | 0 | 298 | 149 | 1 | 1 | 1 | 1 | 1 | 1 | 1 | NA | NA | 0 | NA | NA | 1 |
| SE | 76909 | ENSG00000151914 | DST | chr6 | - | 56598475 | 56598709 | 56597739 | 56598006 | 56600068 | 56600221 | 21 | 31 | 14 | 22 | 25 | 0 | 1 | 0 | 0 | 3 | 0 | 2 | 0 | 0 | 0 | 0 | 0 | 0 | 0 | 0 | 298 | 149 | 1 | 1 | 1 | 0.939 | 1 | 1 | 0.806 | NA | 1 | NA | NA | NA | -0.051 |
| SE | 76910 | ENSG00000151914 | DST | chr6 | - | 56601442 | 56601676 | 56600068 | 56600221 | 56602881 | 56603031 | 0 | 0 | 2 | 3 | 0 | 0 | 1 | 0 | 0 | 0 | 0 | 2 | 1 | 0 | 0 | 0 | 0 | 0 | 0 | 0 | 298 | 149 | 1 | 1 | NA | 0 | 1 | 1 | NA | NA | 1 | 1 | NA | NA | -0.333 |
| SE | 76912 | ENSG00000151914 | DST | chr6 | - | 56703647 | 56703746 | 56701887 | 56701965 | 56704279 | 56704369 | 0 | 1 | 0 | 0 | 0 | 21 | 28 | 14 | 16 | 2 | 0 | 0 | 0 | 0 | 0 | 0 | 3 | 0 | 1 | 0 | 248 | 149 | 1 | 1 | 0 | 0.021 | 0 | 0 | 0 | NA | 0 | NA | 0 | NA | 0.004 |
| SE | 76917 | ENSG00000167483 | NIBAN3 | chr19 | + | 17537375 | 17537543 | 17533478 | 17537701 | 17539149 | 17539225 | 38 | 13 | 38 | 59 | 50 | 0 | 0 | 1 | 3 | 0 | 58 | 81 | 3 | 20 | 98 | 0 | 0 | 0 | 0 | 0 | 298 | 149 | 1 | 1 | 1 | 1 | 0.95 | 0.908 | 1 | 1 | 1 | 1 | 1 | 1 | -0.028 |
| SE | 76920 | ENSG00000167483 | NIBAN3 | chr19 | + | 17543523 | 17543631 | 17543316 | 17543433 | 17551785 | 17551960 | 18 | 7 | 36 | 19 | 13 | 1 | 0 | 0 | 0 | 0 | 49 | 27 | 5 | 10 | 99 | 0 | 0 | 0 | 0 | 3 | 257 | 149 | 1 | 1 | 0.913 | 1 | 1 | 1 | 1 | 1 | 1 | 1 | 1 | 0.95 | -0.007 |
| SE | 76922 | ENSG00000167483 | NIBAN3 | chr19 | + | 17546685 | 17546797 | 17543316 | 17543433 | 17551785 | 17551960 | 4 | 4 | 1 | 3 | 8 | 1 | 0 | 0 | 0 | 0 | 6 | 13 | 0 | 12 | 50 | 0 | 0 | 0 | 0 | 3 | 261 | 149 | 1 | 1 | 0.695 | 1 | 1 | 1 | 1 | 1 | 1 | NA | 1 | 0.905 | -0.037 |
| SE | 76925 | ENSG00000167483 | NIBAN3 | chr19 | + | 17549443 | 17549527 | 17543316 | 17543433 | 17551785 | 17552024 | 33 | 29 | 36 | 55 | 26 | 1 | 0 | 0 | 0 | 0 | 82 | 69 | 8 | 56 | 167 | 0 | 0 | 0 | 0 | 3 | 233 | 149 | 1 | 1 | 0.955 | 1 | 1 | 1 | 1 | 1 | 1 | 1 | 1 | 0.973 | -0.004 |
| SE | 76931 | ENSG00000131018 | SYNE1 | chr6 | - | 152141202 | 152141329 | 152139949 | 152140161 | 152143622 | 152143765 | 249 | 106 | 198 | 247 | 162 | 0 | 2 | 0 | 0 | 0 | 7 | 163 | 165 | 32 | 11 | 0 | 0 | 1 | 0 | 0 | 276 | 149 | 1 | 1 | 1 | 0.966 | 1 | 1 | 1 | 1 | 1 | 0.989 | 1 | 1 | -0.005 |
| SE | 76934 | ENSG00000131018 | SYNE1 | chr6 | - | 152151958 | 152152141 | 152151552 | 152151690 | 152154891 | 152154934 | 352 | 239 | 331 | 325 | 352 | 0 | 0 | 0 | 0 | 0 | 50 | 243 | 285 | 126 | 53 | 0 | 0 | 1 | 0 | 0 | 298 | 149 | 1 | 1 | 1 | 1 | 1 | 1 | 1 | 1 | 1 | 0.993 | 1 | 1 | 0.001 |
| SE | 76935 | ENSG00000131018 | SYNE1 | chr6 | - | 152164162 | 152164325 | 152155909 | 152156097 | 152176393 | 152176560 | 414 | 235 | 320 | 381 | 359 | 0 | 0 | 0 | 0 | 0 | 59 | 245 | 288 | 105 | 48 | 0 | 0 | 2 | 0 | 0 | 298 | 149 | 1 | 1 | 1 | 1 | 1 | 1 | 1 | 1 | 1 | 0.986 | 1 | 1 | 0.003 |
| SE | 76937 | ENSG00000131018 | SYNE1 | chr6 | - | 152180135 | 152180294 | 152176393 | 152176560 | 152185273 | 152185411 | 390 | 267 | 317 | 378 | 436 | 0 | 2 | 0 | 0 | 2 | 109 | 273 | 257 | 135 | 34 | 0 | 0 | 0 | 0 | 0 | 298 | 149 | 1 | 1 | 1 | 0.985 | 1 | 1 | 0.991 | 1 | 1 | 1 | 1 | 1 | -0.005 |
| SE | 76938 | ENSG00000131018 | SYNE1 | chr6 | - | 152180135 | 152180294 | 152176393 | 152176560 | 152189251 | 152189407 | 424 | 311 | 306 | 425 | 440 | 2 | 0 | 0 | 2 | 0 | 108 | 343 | 301 | 153 | 48 | 0 | 0 | 0 | 0 | 0 | 298 | 149 | 1 | 1 | 0.991 | 1 | 1 | 0.991 | 1 | 1 | 1 | 1 | 1 | 1 | -0.004 |
| SE | 76939 | ENSG00000131018 | SYNE1 | chr6 | - | 152211493 | 152211588 | 152207971 | 152208206 | 152213611 | 152213759 | 498 | 296 | 344 | 351 | 395 | 0 | 0 | 0 | 0 | 0 | 147 | 384 | 351 | 146 | 44 | 0 | 0 | 4 | 0 | 0 | 244 | 149 | 1 | 1 | 1 | 1 | 1 | 1 | 1 | 1 | 1 | 0.982 | 1 | 1 | 0.004 |
| SE | 76940 | ENSG00000131018 | SYNE1 | chr6 | - | 152214905 | 152215060 | 152213611 | 152213759 | 152218256 | 152218385 | 506 | 270 | 311 | 463 | 425 | 5 | 3 | 2 | 2 | 0 | 54 | 245 | 271 | 127 | 14 | 0 | 2 | 1 | 0 | 0 | 298 | 149 | 1 | 1 | 0.981 | 0.978 | 0.987 | 0.991 | 1 | 1 | 0.984 | 0.993 | 1 | 1 | -0.008 |
| SE | 76941 | ENSG00000131018 | SYNE1 | chr6 | - | 152225720 | 152225876 | 152224493 | 152224664 | 152230546 | 152230702 | 328 | 202 | 289 | 326 | 473 | 0 | 0 | 1 | 1 | 6 | 18 | 192 | 201 | 103 | 0 | 0 | 3 | 4 | 3 | 0 | 298 | 149 | 1 | 1 | 1 | 1 | 0.993 | 0.994 | 0.975 | 1 | 0.97 | 0.962 | 0.945 | NA | 0.023 |
| SE | 76942 | ENSG00000131018 | SYNE1 | chr6 | - | 152236106 | 152236303 | 152234667 | 152234800 | 152236816 | 152236948 | 405 | 180 | 285 | 316 | 351 | 1 | 0 | 0 | 0 | 0 | 74 | 198 | 266 | 96 | 13 | 0 | 0 | 0 | 0 | 0 | 298 | 149 | 1 | 1 | 0.995 | 1 | 1 | 1 | 1 | 1 | 1 | 1 | 1 | 1 | -0.001 |
| SE | 76944 | ENSG00000131018 | SYNE1 | chr6 | - | 152244536 | 152244656 | 152242239 | 152242440 | 152249160 | 152249262 | 402 | 286 | 306 | 355 | 411 | 0 | 1 | 0 | 4 | 0 | 113 | 364 | 413 | 146 | 33 | 0 | 0 | 3 | 1 | 0 | 269 | 149 | 1 | 1 | 1 | 0.994 | 1 | 0.98 | 1 | 1 | 1 | 0.987 | 0.988 | 1 | 0 |
| SE | 76945 | ENSG00000131018 | SYNE1 | chr6 | - | 152249160 | 152249262 | 152244536 | 152244656 | 152254879 | 152255089 | 528 | 333 | 393 | 450 | 367 | 0 | 0 | 1 | 3 | 0 | 144 | 471 | 550 | 218 | 63 | 0 | 0 | 2 | 0 | 0 | 251 | 149 | 1 | 1 | 1 | 1 | 0.996 | 0.989 | 1 | 1 | 1 | 0.994 | 1 | 1 | -0.002 |
| SE | 76946 | ENSG00000131018 | SYNE1 | chr6 | - | 152262031 | 152262188 | 152256633 | 152256765 | 152278088 | 152278280 | 257 | 137 | 170 | 225 | 227 | 0 | 0 | 0 | 0 | 0 | 37 | 204 | 179 | 95 | 34 | 0 | 0 | 1 | 0 | 0 | 298 | 149 | 1 | 1 | 1 | 1 | 1 | 1 | 1 | 1 | 1 | 0.989 | 1 | 1 | 0.002 |
| SE | 76947 | ENSG00000131018 | SYNE1 | chr6 | - | 152281806 | 152281980 | 152277501 | 152278280 | 152283977 | 152284172 | 306 | 181 | 207 | 256 | 120 | 0 | 1 | 0 | 0 | 0 | 81 | 247 | 235 | 112 | 66 | 0 | 0 | 0 | 0 | 0 | 298 | 149 | 1 | 1 | 1 | 0.989 | 1 | 1 | 1 | 1 | 1 | 1 | 1 | 1 | -0.002 |
| SE | 76948 | ENSG00000131018 | SYNE1 | chr6 | - | 152283977 | 152284172 | 152281806 | 152281980 | 152293587 | 152293749 | 199 | 116 | 138 | 195 | 114 | 1 | 0 | 1 | 0 | 0 | 77 | 189 | 136 | 72 | 32 | 0 | 1 | 0 | 0 | 0 | 298 | 149 | 1 | 1 | 0.99 | 1 | 0.986 | 1 | 1 | 1 | 0.99 | 1 | 1 | 1 | -0.003 |
| SE | 76949 | ENSG00000131018 | SYNE1 | chr6 | - | 152283977 | 152284184 | 152281806 | 152281980 | 152293587 | 152293749 | 161 | 95 | 113 | 133 | 78 | 1 | 0 | 1 | 0 | 0 | 49 | 142 | 116 | 48 | 19 | 0 | 1 | 0 | 0 | 0 | 298 | 149 | 1 | 1 | 0.988 | 1 | 0.983 | 1 | 1 | 1 | 0.986 | 1 | 1 | 1 | -0.003 |
| SE | 76952 | ENSG00000131018 | SYNE1 | chr6 | - | 152326295 | 152326633 | 152325957 | 152326102 | 152329729 | 152331890 | 197 | 117 | 114 | 188 | 83 | 0 | 0 | 0 | 0 | 0 | 33 | 141 | 167 | 35 | 49 | 0 | 0 | 1 | 0 | 0 | 298 | 149 | 1 | 1 | 1 | 1 | 1 | 1 | 1 | 1 | 1 | 0.988 | 1 | 1 | 0.002 |
| SE | 76954 | ENSG00000131018 | SYNE1 | chr6 | - | 152352026 | 152352353 | 152350617 | 152350770 | 152353262 | 152353433 | 224 | 129 | 123 | 230 | 126 | 1 | 0 | 2 | 0 | 1 | 8 | 137 | 248 | 15 | 3 | 0 | 0 | 1 | 0 | 0 | 298 | 149 | 1 | 1 | 0.991 | 1 | 0.969 | 1 | 0.984 | 1 | 1 | 0.992 | 1 | 1 | -0.01 |
| SE | 76955 | ENSG00000131018 | SYNE1 | chr6 | - | 152364846 | 152365019 | 152362169 | 152362323 | 152367217 | 152367382 | 160 | 67 | 80 | 121 | 134 | 0 | 0 | 0 | 0 | 0 | 8 | 111 | 154 | 10 | 28 | 0 | 3 | 0 | 0 | 0 | 298 | 149 | 1 | 1 | 1 | 1 | 1 | 1 | 1 | 1 | 0.949 | 1 | 1 | 1 | 0.01 |
| SE | 76956 | ENSG00000131018 | SYNE1 | chr6 | - | 152368971 | 152369127 | 152366991 | 152367382 | 152369470 | 152369614 | 20 | 6 | 19 | 14 | 1 | 89 | 54 | 36 | 78 | 44 | 0 | 7 | 9 | 0 | 0 | 0 | 48 | 38 | 3 | 0 | 298 | 149 | 1 | 1 | 0.101 | 0.053 | 0.209 | 0.082 | 0.011 | NA | 0.068 | 0.106 | 0 | NA | 0.033 |
| SE | 76957 | ENSG00000131018 | SYNE1 | chr6 | - | 152385673 | 152385838 | 152381005 | 152381362 | 152387071 | 152387381 | 355 | 181 | 198 | 254 | 199 | 1 | 0 | 0 | 0 | 0 | 44 | 235 | 325 | 37 | 42 | 0 | 0 | 0 | 0 | 0 | 298 | 149 | 1 | 1 | 0.994 | 1 | 1 | 1 | 1 | 1 | 1 | 1 | 1 | 1 | -0.001 |
| SE | 76958 | ENSG00000131018 | SYNE1 | chr6 | - | 152390279 | 152390452 | 152387071 | 152387381 | 152391276 | 152391568 | 244 | 91 | 107 | 188 | 49 | 1 | 1 | 0 | 0 | 0 | 69 | 199 | 267 | 28 | 38 | 0 | 0 | 3 | 0 | 0 | 298 | 149 | 1 | 1 | 0.992 | 0.978 | 1 | 1 | 1 | 1 | 1 | 0.978 | 1 | 1 | -0.002 |
| SE | 76960 | ENSG00000131018 | SYNE1 | chr6 | - | 152413351 | 152413531 | 152409558 | 152409709 | 152416386 | 152417015 | 333 | 186 | 130 | 207 | 134 | 1 | 0 | 0 | 1 | 0 | 40 | 152 | 182 | 50 | 6 | 0 | 0 | 0 | 0 | 0 | 298 | 149 | 1 | 1 | 0.994 | 1 | 1 | 0.99 | 1 | 1 | 1 | 1 | 1 | 1 | -0.003 |
| SE | 76961 | ENSG00000131018 | SYNE1 | chr6 | - | 152419568 | 152419722 | 152416386 | 152417015 | 152425380 | 152425547 | 261 | 132 | 127 | 208 | 127 | 0 | 0 | 0 | 1 | 0 | 51 | 178 | 189 | 46 | 7 | 0 | 0 | 0 | 0 | 0 | 298 | 149 | 1 | 1 | 1 | 1 | 1 | 0.99 | 1 | 1 | 1 | 1 | 1 | 1 | -0.002 |
| SE | 76962 | ENSG00000131018 | SYNE1 | chr6 | - | 152430111 | 152430210 | 152428204 | 152428392 | 152430481 | 152430709 | 367 | 194 | 155 | 329 | 112 | 0 | 0 | 0 | 0 | 0 | 2 | 276 | 268 | 64 | 30 | 0 | 0 | 2 | 0 | 0 | 248 | 149 | 1 | 1 | 1 | 1 | 1 | 1 | 1 | 1 | 1 | 0.988 | 1 | 1 | 0.002 |
| SE | 76963 | ENSG00000131018 | SYNE1 | chr6 | - | 152433794 | 152433945 | 152430481 | 152430709 | 152435940 | 152436101 | 377 | 177 | 169 | 247 | 73 | 0 | 0 | 2 | 0 | 0 | 10 | 147 | 241 | 43 | 0 | 0 | 0 | 0 | 0 | 0 | 298 | 149 | 1 | 1 | 1 | 1 | 0.977 | 1 | 1 | 1 | 1 | 1 | 1 | NA | -0.005 |
| SE | 76964 | ENSG00000131018 | SYNE1 | chr6 | - | 152435940 | 152436101 | 152430481 | 152430709 | 152441129 | 152441270 | 210 | 102 | 97 | 159 | 93 | 0 | 0 | 0 | 0 | 0 | 0 | 106 | 139 | 28 | 10 | 0 | 0 | 2 | 0 | 0 | 298 | 149 | 1 | 1 | 1 | 1 | 1 | 1 | 1 | NA | 1 | 0.972 | 1 | 1 | 0.007 |
| SE | 76965 | ENSG00000131018 | SYNE1 | chr6 | - | 152447457 | 152447622 | 152444410 | 152444578 | 152449532 | 152449641 | 291 | 157 | 179 | 218 | 98 | 0 | 0 | 0 | 0 | 0 | 21 | 247 | 297 | 72 | 4 | 0 | 0 | 3 | 0 | 0 | 298 | 149 | 1 | 1 | 1 | 1 | 1 | 1 | 1 | 1 | 1 | 0.98 | 1 | 1 | 0.004 |
| SE | 76966 | ENSG00000131018 | SYNE1 | chr6 | - | 152450624 | 152450833 | 152449532 | 152449641 | 152451046 | 152451205 | 247 | 106 | 135 | 129 | 101 | 1 | 0 | 0 | 0 | 0 | 17 | 133 | 200 | 28 | 24 | 0 | 0 | 0 | 0 | 0 | 298 | 149 | 1 | 1 | 0.992 | 1 | 1 | 1 | 1 | 1 | 1 | 1 | 1 | 1 | -0.002 |
| SE | 76968 | ENSG00000131018 | SYNE1 | chr6 | - | 152461596 | 152461740 | 152458756 | 152458930 | 152462737 | 152462890 | 381 | 198 | 177 | 285 | 164 | 0 | 1 | 0 | 0 | 2 | 33 | 306 | 256 | 88 | 12 | 0 | 0 | 0 | 0 | 0 | 293 | 149 | 1 | 1 | 1 | 0.99 | 1 | 1 | 0.977 | 1 | 1 | 1 | 1 | 1 | -0.007 |
| SE | 76970 | ENSG00000131018 | SYNE1 | chr6 | - | 152484834 | 152484972 | 152483084 | 152483249 | 152488395 | 152488503 | 357 | 146 | 157 | 250 | 157 | 0 | 1 | 0 | 0 | 0 | 70 | 242 | 339 | 107 | 31 | 0 | 0 | 0 | 0 | 0 | 287 | 149 | 1 | 1 | 1 | 0.987 | 1 | 1 | 1 | 1 | 1 | 1 | 1 | 1 | -0.003 |
| SE | 76971 | ENSG00000131018 | SYNE1 | chr6 | - | 152498741 | 152498792 | 152488395 | 152488503 | 152502632 | 152502742 | 3 | 0 | 0 | 0 | 0 | 174 | 70 | 77 | 129 | 88 | 0 | 0 | 2 | 4 | 0 | 19 | 118 | 195 | 65 | 28 | 200 | 149 | 1 | 1 | 0.013 | 0 | 0 | 0 | 0 | 0 | 0 | 0.008 | 0.044 | 0 | -0.008 |
| SE | 76972 | ENSG00000131018 | SYNE1 | chr6 | - | 152511010 | 152511103 | 152510192 | 152510371 | 152520458 | 152520542 | 545 | 238 | 290 | 358 | 313 | 0 | 0 | 2 | 4 | 0 | 105 | 503 | 378 | 151 | 103 | 0 | 0 | 0 | 0 | 0 | 242 | 149 | 1 | 1 | 1 | 1 | 0.989 | 0.982 | 1 | 1 | 1 | 1 | 1 | 1 | -0.006 |
| SE | 76973 | ENSG00000131018 | SYNE1 | chr6 | - | 152511570 | 152511591 | 152511010 | 152511103 | 152520458 | 152520542 | 0 | 0 | 0 | 0 | 0 | 306 | 121 | 138 | 186 | 172 | 0 | 6 | 0 | 2 | 0 | 31 | 254 | 183 | 90 | 43 | 170 | 149 | 1 | 1 | 0 | 0 | 0 | 0 | 0 | 0 | 0.02 | 0 | 0.019 | 0 | -0.008 |
| SE | 76974 | ENSG00000131018 | SYNE1 | chr6 | - | 152542159 | 152542240 | 152539959 | 152540021 | 152628264 | 152628554 | 0 | 0 | 0 | 1 | 0 | 70 | 29 | 59 | 70 | 57 | 0 | 5 | 0 | 0 | 0 | 32 | 77 | 53 | 21 | 26 | 230 | 149 | 1 | 1 | 0 | 0 | 0 | 0.009 | 0 | 0 | 0.04 | 0 | 0 | 0 | -0.006 |
| SE | 76976 | ENSG00000172469 | MANEA | chr6 | + | 95589999 | 95590173 | 95586401 | 95586983 | 95596736 | 95596846 | 3 | 1 | 1 | 2 | 4 | 54 | 59 | 55 | 21 | 117 | 0 | 0 | 0 | 0 | 0 | 0 | 6 | 15 | 0 | 0 | 298 | 149 | 1 | 1 | 0.027 | 0.008 | 0.009 | 0.045 | 0.017 | NA | 0 | 0 | NA | NA | 0.021 |
| SE | 76977 | ENSG00000172469 | MANEA | chr6 | + | 95592563 | 95592646 | 95586401 | 95586983 | 95596736 | 95596846 | 0 | 0 | 0 | 9 | 5 | 54 | 59 | 55 | 21 | 117 | 0 | 0 | 0 | 0 | 0 | 0 | 6 | 15 | 0 | 0 | 232 | 149 | 1 | 1 | 0 | 0 | 0 | 0.216 | 0.027 | NA | 0 | 0 | NA | NA | 0.049 |
| SE | 76979 | ENSG00000172469 | MANEA | chr6 | + | 95596736 | 95596846 | 95586401 | 95586983 | 95604826 | 95604903 | 82 | 73 | 86 | 33 | 159 | 0 | 3 | 2 | 0 | 1 | 0 | 15 | 22 | 0 | 0 | 0 | 0 | 0 | 0 | 0 | 259 | 149 | 1 | 1 | 1 | 0.933 | 0.961 | 1 | 0.989 | NA | 1 | 1 | NA | NA | -0.023 |
| SE | 76980 | ENSG00000172469 | MANEA | chr6 | + | 95604826 | 95604903 | 95596736 | 95596846 | 95605747 | 95606023 | 86 | 50 | 87 | 52 | 171 | 0 | 0 | 0 | 1 | 0 | 0 | 16 | 20 | 0 | 0 | 0 | 0 | 0 | 0 | 0 | 226 | 149 | 1 | 1 | 1 | 1 | 1 | 0.972 | 1 | NA | 1 | 1 | NA | NA | -0.006 |
| SE | 76982 | ENSG00000170485 | NPAS2 | chr2 | + | 100949366 | 100949480 | 100948234 | 100948355 | 100964057 | 100964176 | 6 | 2 | 17 | 6 | 24 | 0 | 0 | 0 | 0 | 3 | 0 | 0 | 0 | 0 | 2 | 0 | 0 | 0 | 0 | 0 | 263 | 149 | 1 | 1 | 1 | 1 | 1 | 1 | 0.819 | NA | NA | NA | NA | 1 | -0.036 |
| SE | 76984 | ENSG00000135318 | NT5E | chr6 | + | 85490507 | 85490657 | 85489493 | 85489599 | 85491976 | 85492177 | 33 | 14 | 44 | 38 | 83 | 2 | 0 | 1 | 0 | 0 | 11 | 11 | 18 | 9 | 6 | 0 | 0 | 0 | 0 | 0 | 298 | 149 | 1 | 1 | 0.892 | 1 | 0.957 | 1 | 1 | 1 | 1 | 1 | 1 | 1 | -0.03 |
| SE | 76985 | ENSG00000135318 | NT5E | chr6 | + | 85491976 | 85492177 | 85489493 | 85489599 | 85493840 | 85495784 | 30 | 13 | 41 | 30 | 83 | 1 | 0 | 1 | 1 | 2 | 3 | 0 | 0 | 0 | 2 | 0 | 0 | 0 | 0 | 0 | 298 | 149 | 1 | 1 | 0.938 | 1 | 0.953 | 0.938 | 0.954 | 1 | NA | NA | NA | 1 | -0.043 |
| SE | 76986 | ENSG00000173511 | VEGFB | chr11 | + | 64236253 | 64236327 | 64235812 | 64236009 | 64237186 | 64237222 | 44 | 24 | 56 | 36 | 28 | 2 | 0 | 0 | 0 | 0 | 25 | 15 | 22 | 1 | 79 | 0 | 0 | 0 | 0 | 0 | 223 | 149 | 1 | 1 | 0.936 | 1 | 1 | 1 | 1 | 1 | 1 | 1 | 1 | 1 | -0.013 |
| SE | 76987 | ENSG00000173511 | VEGFB | chr11 | + | 64237186 | 64237222 | 64236253 | 64236327 | 64237419 | 64237655 | 39 | 18 | 36 | 23 | 17 | 0 | 0 | 0 | 1 | 0 | 25 | 11 | 13 | 3 | 63 | 0 | 0 | 1 | 0 | 0 | 185 | 149 | 1 | 1 | 1 | 1 | 1 | 0.949 | 1 | 1 | 1 | 0.913 | 1 | 1 | 0.007 |
| SE | 76988 | ENSG00000271793 | NA | chr6 | - | 85547115 | 85547226 | 85543604 | 85543760 | 85547505 | 85547550 | 3 | 5 | 5 | 7 | 7 | 0 | 0 | 0 | 0 | 1 | 5 | 6 | 6 | 3 | 10 | 0 | 0 | 2 | 0 | 0 | 260 | 149 | 1 | 1 | 1 | 1 | 1 | 1 | 0.8 | 1 | 1 | 0.632 | 1 | 1 | 0.034 |
| SE | 76989 | ENSG00000271793 | NA | chr6 | - | 85612831 | 85612920 | 85574257 | 85574378 | 85614980 | 85615234 | 16 | 3 | 1 | 6 | 0 | 0 | 2 | 0 | 0 | 0 | 0 | 2 | 4 | 0 | 0 | 0 | 0 | 0 | 0 | 0 | 238 | 149 | 1 | 1 | 1 | 0.484 | 1 | 1 | NA | NA | 1 | 1 | NA | NA | -0.129 |
| SE | 76990 | ENSG00000271793 | NA | chr6 | - | 85612831 | 85612923 | 85574257 | 85574378 | 85614980 | 85615347 | 80 | 34 | 55 | 42 | 63 | 0 | 2 | 0 | 0 | 0 | 6 | 33 | 41 | 20 | 0 | 0 | 0 | 0 | 0 | 0 | 241 | 149 | 1 | 1 | 1 | 0.913 | 1 | 1 | 1 | 1 | 1 | 1 | 1 | NA | -0.017 |
| SE | 76991 | ENSG00000137404 | NRM | chr6 | - | 30689275 | 30689452 | 30688046 | 30688942 | 30690024 | 30690243 | 18 | 5 | 4 | 17 | 2 | 0 | 0 | 0 | 0 | 0 | 0 | 20 | 7 | 9 | 26 | 0 | 0 | 0 | 0 | 6 | 298 | 149 | 1 | 1 | 1 | 1 | 1 | 1 | 1 | NA | 1 | 1 | 1 | 0.684 | 0.079 |
| SE | 76993 | ENSG00000137404 | NRM | chr6 | - | 30689275 | 30689452 | 30688046 | 30688942 | 30690841 | 30690967 | 18 | 5 | 4 | 17 | 2 | 2 | 0 | 0 | 0 | 0 | 0 | 20 | 7 | 9 | 29 | 0 | 6 | 0 | 0 | 7 | 298 | 149 | 1 | 1 | 0.818 | 1 | 1 | 1 | 1 | NA | 0.625 | 1 | 1 | 0.674 | 0.139 |
| SE | 76994 | ENSG00000137404 | NRM | chr6 | - | 30690024 | 30690243 | 30688046 | 30688942 | 30690841 | 30690986 | 6 | 2 | 7 | 3 | 0 | 2 | 0 | 0 | 0 | 0 | 12 | 6 | 7 | 1 | 29 | 0 | 6 | 0 | 0 | 7 | 298 | 149 | 1 | 1 | 0.6 | 1 | 1 | 1 | NA | 1 | 0.333 | 1 | 1 | 0.674 | 0.099 |
| SE | 76995 | ENSG00000137404 | NRM | chr6 | - | 30690024 | 30690243 | 30689275 | 30689452 | 30690841 | 30690986 | 6 | 2 | 7 | 3 | 0 | 0 | 0 | 0 | 0 | 0 | 12 | 6 | 7 | 1 | 23 | 0 | 0 | 0 | 0 | 3 | 298 | 149 | 1 | 1 | 1 | 1 | 1 | 1 | NA | 1 | 1 | 1 | 1 | 0.793 | 0.041 |
| SE | 76996 | ENSG00000137404 | NRM | chr6 | - | 30690046 | 30690243 | 30688046 | 30688942 | 30690841 | 30690976 | 8 | 2 | 9 | 5 | 0 | 2 | 0 | 0 | 0 | 0 | 16 | 7 | 12 | 4 | 23 | 0 | 6 | 0 | 0 | 7 | 298 | 149 | 1 | 1 | 0.667 | 1 | 1 | 1 | NA | 1 | 0.368 | 1 | 1 | 0.622 | 0.119 |
| SE | 76997 | ENSG00000137404 | NRM | chr6 | - | 30690046 | 30690243 | 30689275 | 30689452 | 30690841 | 30690976 | 18 | 2 | 16 | 4 | 4 | 0 | 0 | 0 | 0 | 0 | 12 | 9 | 15 | 6 | 49 | 0 | 0 | 0 | 0 | 3 | 298 | 149 | 1 | 1 | 1 | 1 | 1 | 1 | 1 | 1 | 1 | 1 | 1 | 0.891 | 0.022 |
| SE | 77001 | ENSG00000149657 | LSM14B | chr20 | + | 62127584 | 62127701 | 62124616 | 62124780 | 62129784 | 62129952 | 4 | 0 | 0 | 1 | 0 | 6 | 3 | 0 | 2 | 2 | 30 | 3 | 3 | 0 | 0 | 0 | 0 | 0 | 5 | 3 | 266 | 149 | 1 | 1 | 0.272 | 0 | NA | 0.219 | 0 | 1 | 1 | 1 | 0 | 0 | -0.477 |
| SE | 77011 | ENSG00000158691 | ZSCAN12 | chr6 | - | 28390595 | 28391742 | 28378954 | 28382609 | 28392901 | 28393046 | 36 | 15 | 20 | 23 | 30 | 0 | 1 | 0 | 1 | 0 | 0 | 13 | 21 | 4 | 3 | 0 | 0 | 4 | 0 | 0 | 298 | 149 | 1 | 1 | 1 | 0.882 | 1 | 0.92 | 1 | NA | 1 | 0.724 | 1 | 1 | 0.029 |
| SE | 77012 | ENSG00000158691 | ZSCAN12 | chr6 | - | 28392901 | 28393046 | 28378954 | 28382609 | 28399656 | 28399731 | 1 | 1 | 1 | 2 | 0 | 1 | 0 | 1 | 0 | 4 | 0 | 0 | 4 | 6 | 0 | 0 | 0 | 0 | 0 | 0 | 294 | 149 | 1 | 1 | 0.336 | 1 | 0.336 | 1 | 0 | NA | NA | 1 | 1 | NA | -0.466 |
| SE | 77013 | ENSG00000158691 | ZSCAN12 | chr6 | - | 28398003 | 28398473 | 28392901 | 28393046 | 28399656 | 28399731 | 25 | 8 | 30 | 15 | 14 | 1 | 0 | 1 | 1 | 0 | 19 | 9 | 26 | 13 | 0 | 0 | 0 | 0 | 6 | 0 | 298 | 149 | 1 | 1 | 0.926 | 1 | 0.938 | 0.882 | 1 | 1 | 1 | 1 | 0.52 | NA | 0.069 |
| SE | 77015 | ENSG00000107341 | UBE2R2 | chr9 | + | 33900173 | 33900271 | 33886880 | 33886967 | 33911963 | 33912098 | 683 | 497 | 529 | 643 | 711 | 0 | 0 | 0 | 0 | 6 | 1308 | 632 | 733 | 840 | 1280 | 0 | 0 | 0 | 0 | 0 | 247 | 149 | 1 | 1 | 1 | 1 | 1 | 1 | 0.986 | 1 | 1 | 1 | 1 | 1 | -0.003 |
| SE | 77017 | ENSG00000146433 | TMEM181 | chr6 | + | 158580939 | 158580995 | 158573419 | 158573523 | 158583953 | 158584044 | 105 | 54 | 77 | 89 | 103 | 0 | 0 | 0 | 0 | 0 | 26 | 52 | 69 | 28 | 53 | 0 | 0 | 2 | 0 | 0 | 205 | 149 | 1 | 1 | 1 | 1 | 1 | 1 | 1 | 1 | 1 | 0.962 | 1 | 1 | 0.008 |
| SE | 77018 | ENSG00000146433 | TMEM181 | chr6 | + | 158583953 | 158584044 | 158580939 | 158580995 | 158585303 | 158585425 | 315 | 127 | 166 | 207 | 362 | 4 | 0 | 3 | 5 | 2 | 102 | 176 | 162 | 103 | 255 | 0 | 1 | 11 | 0 | 0 | 240 | 149 | 1 | 1 | 0.98 | 1 | 0.972 | 0.963 | 0.991 | 1 | 0.991 | 0.901 | 1 | 1 | 0.003 |
| SE | 77019 | ENSG00000146433 | TMEM181 | chr6 | + | 158589671 | 158589782 | 158585303 | 158585425 | 158605266 | 158605347 | 231 | 102 | 158 | 187 | 357 | 0 | 0 | 0 | 1 | 0 | 76 | 115 | 128 | 86 | 181 | 0 | 0 | 0 | 0 | 0 | 260 | 149 | 1 | 1 | 1 | 1 | 1 | 0.991 | 1 | 1 | 1 | 1 | 1 | 1 | -0.002 |
| SE | 77020 | ENSG00000146433 | TMEM181 | chr6 | + | 158605266 | 158605347 | 158589671 | 158589782 | 158607243 | 158607343 | 185 | 73 | 139 | 170 | 240 | 0 | 0 | 4 | 0 | 0 | 31 | 76 | 106 | 37 | 95 | 0 | 2 | 0 | 0 | 0 | 230 | 149 | 1 | 1 | 1 | 1 | 0.957 | 1 | 1 | 1 | 0.961 | 1 | 1 | 1 | -0.001 |
| SE | 77021 | ENSG00000146433 | TMEM181 | chr6 | + | 158605266 | 158605347 | 158589671 | 158589782 | 158608332 | 158608463 | 81 | 27 | 53 | 62 | 103 | 2 | 0 | 0 | 0 | 0 | 13 | 31 | 51 | 17 | 36 | 0 | 3 | 0 | 1 | 0 | 230 | 149 | 1 | 1 | 0.963 | 1 | 1 | 1 | 1 | 1 | 0.87 | 1 | 0.917 | 1 | 0.035 |
| SE | 77022 | ENSG00000146433 | TMEM181 | chr6 | + | 158607243 | 158607343 | 158589671 | 158589782 | 158608332 | 158608463 | 22 | 12 | 28 | 23 | 24 | 2 | 0 | 0 | 0 | 0 | 23 | 36 | 33 | 10 | 33 | 0 | 3 | 0 | 1 | 0 | 249 | 149 | 1 | 1 | 0.868 | 1 | 1 | 1 | 1 | 1 | 0.878 | 1 | 0.857 | 1 | 0.027 |
| SE | 77024 | ENSG00000146433 | TMEM181 | chr6 | + | 158623549 | 158623607 | 158608658 | 158608750 | 158625103 | 158625206 | 62 | 20 | 46 | 72 | 51 | 2 | 0 | 0 | 1 | 2 | 10 | 39 | 29 | 6 | 46 | 0 | 0 | 0 | 0 | 0 | 207 | 149 | 1 | 1 | 0.957 | 1 | 1 | 0.981 | 0.948 | 1 | 1 | 1 | 1 | 1 | -0.023 |
| SE | 77030 | ENSG00000164663 | USP49 | chr6 | - | 41803805 | 41804010 | 41799829 | 41799938 | 41805627 | 41807011 | 7 | 14 | 10 | 8 | 2 | 0 | 0 | 0 | 0 | 0 | 0 | 0 | 17 | 29 | 0 | 0 | 0 | 5 | 0 | 0 | 298 | 149 | 1 | 1 | 1 | 1 | 1 | 1 | 1 | NA | NA | 0.63 | 1 | NA | 0.185 |
| SE | 77034 | ENSG00000014123 | UFL1 | chr6 | + | 96524381 | 96524410 | 96523145 | 96523291 | 96525296 | 96525394 | 254 | 227 | 243 | 220 | 356 | 0 | 0 | 0 | 0 | 1 | 45 | 138 | 87 | 49 | 32 | 0 | 0 | 0 | 0 | 0 | 178 | 149 | 1 | 1 | 1 | 1 | 1 | 1 | 0.997 | 1 | 1 | 1 | 1 | 1 | -0.001 |
| SE | 77036 | ENSG00000014123 | UFL1 | chr6 | + | 96536243 | 96536390 | 96534262 | 96534321 | 96537373 | 96537549 | 186 | 155 | 188 | 157 | 225 | 0 | 1 | 1 | 2 | 0 | 8 | 73 | 45 | 19 | 13 | 0 | 0 | 0 | 2 | 0 | 296 | 149 | 1 | 1 | 1 | 0.987 | 0.99 | 0.975 | 1 | 1 | 1 | 1 | 0.827 | 1 | 0.025 |
| SE | 77037 | ENSG00000014123 | UFL1 | chr6 | + | 96540534 | 96540655 | 96538630 | 96538810 | 96542893 | 96543016 | 120 | 129 | 90 | 110 | 195 | 0 | 0 | 0 | 0 | 0 | 23 | 54 | 45 | 3 | 16 | 0 | 0 | 1 | 0 | 0 | 270 | 149 | 1 | 1 | 1 | 1 | 1 | 1 | 1 | 1 | 1 | 0.961 | 1 | 1 | 0.008 |
| SE | 77039 | ENSG00000160785 | SLC25A44 | chr1 | + | 156207861 | 156208013 | 156199834 | 156200472 | 156210239 | 156212795 | 26 | 24 | 29 | 23 | 25 | 2 | 1 | 0 | 1 | 0 | 29 | 39 | 30 | 23 | 58 | 0 | 4 | 0 | 0 | 0 | 298 | 149 | 1 | 1 | 0.867 | 0.923 | 1 | 0.92 | 1 | 1 | 0.83 | 1 | 1 | 1 | -0.024 |
| SE | 77041 | ENSG00000160785 | SLC25A44 | chr1 | + | 156207885 | 156208013 | 156199834 | 156200472 | 156210239 | 156212795 | 30 | 30 | 35 | 37 | 26 | 2 | 1 | 0 | 1 | 0 | 29 | 53 | 42 | 33 | 63 | 0 | 4 | 0 | 0 | 0 | 277 | 149 | 1 | 1 | 0.89 | 0.942 | 1 | 0.952 | 1 | 1 | 0.877 | 1 | 1 | 1 | -0.019 |
| SE | 77044 | ENSG00000165568 | AKR1E2 | chr10 | + | 4833349 | 4833466 | 4830674 | 4830842 | 4835674 | 4835809 | 9 | 12 | 23 | 1 | 30 | 1 | 0 | 1 | 0 | 4 | 0 | 17 | 27 | 1 | 66 | 0 | 2 | 1 | 0 | 8 | 266 | 149 | 1 | 1 | 0.834 | 1 | 0.928 | 1 | 0.808 | NA | 0.826 | 0.938 | 1 | 0.822 | 0.018 |
| SE | 77046 | ENSG00000165568 | AKR1E2 | chr10 | + | 4839728 | 4839826 | 4837458 | 4837581 | 4841784 | 4841857 | 6 | 4 | 12 | 7 | 25 | 0 | 1 | 0 | 0 | 0 | 0 | 6 | 11 | 1 | 19 | 0 | 0 | 0 | 0 | 0 | 247 | 149 | 1 | 1 | 1 | 0.707 | 1 | 1 | 1 | NA | 1 | 1 | 1 | 1 | -0.059 |
| SE | 77048 | ENSG00000165568 | AKR1E2 | chr10 | + | 4841784 | 4841840 | 4839783 | 4839826 | 4842420 | 4842504 | 0 | 0 | 0 | 1 | 8 | 2 | 0 | 2 | 4 | 7 | 0 | 2 | 3 | 0 | 0 | 13 | 3 | 4 | 1 | 29 | 205 | 149 | 1 | 1 | 0 | NA | 0 | 0.154 | 0.454 | 0 | 0.326 | 0.353 | 0 | 0 | 0.016 |
| SE | 77051 | ENSG00000125779 | PANK2 | chr20 | + | 3901547 | 3901617 | 3891243 | 3891411 | 3907925 | 3908135 | 0 | 0 | 0 | 0 | 0 | 0 | 0 | 0 | 0 | 1 | 0 | 0 | 0 | 2 | 0 | 5 | 0 | 4 | 0 | 0 | 219 | 149 | 1 | 1 | NA | NA | NA | NA | 0 | 0 | NA | 0 | 1 | NA | -0.333 |
| SE | 77052 | ENSG00000125779 | PANK2 | chr20 | + | 3901551 | 3901617 | 3889669 | 3889728 | 3907925 | 3908278 | 0 | 0 | 0 | 0 | 0 | 40 | 23 | 29 | 24 | 8 | 0 | 0 | 0 | 2 | 0 | 11 | 42 | 21 | 25 | 57 | 215 | 149 | 1 | 1 | 0 | 0 | 0 | 0 | 0 | 0 | 0 | 0 | 0.053 | 0 | -0.011 |
| SE | 77053 | ENSG00000125779 | PANK2 | chr20 | + | 3910576 | 3910830 | 3907925 | 3908278 | 3912457 | 3912634 | 359 | 259 | 280 | 320 | 470 | 0 | 0 | 2 | 5 | 0 | 788 | 417 | 224 | 463 | 442 | 6 | 6 | 0 | 0 | 2 | 298 | 149 | 1 | 1 | 1 | 1 | 0.986 | 0.97 | 1 | 0.985 | 0.972 | 1 | 1 | 0.991 | 0.002 |
| SE | 77054 | ENSG00000125779 | PANK2 | chr20 | + | 3910576 | 3910830 | 3907925 | 3908278 | 3916926 | 3917050 | 111 | 60 | 64 | 99 | 118 | 0 | 0 | 2 | 0 | 0 | 133 | 103 | 45 | 135 | 145 | 0 | 0 | 0 | 0 | 4 | 298 | 149 | 1 | 1 | 1 | 1 | 0.941 | 1 | 1 | 1 | 1 | 1 | 1 | 0.948 | -0.001 |
| SE | 77055 | ENSG00000125779 | PANK2 | chr20 | + | 3912457 | 3912634 | 3907925 | 3908278 | 3916926 | 3917050 | 268 | 204 | 277 | 247 | 435 | 0 | 0 | 2 | 0 | 0 | 595 | 300 | 211 | 307 | 428 | 0 | 0 | 0 | 0 | 4 | 298 | 149 | 1 | 1 | 1 | 1 | 0.986 | 1 | 1 | 1 | 1 | 1 | 1 | 0.982 | 0.001 |
| SE | 77057 | ENSG00000125779 | PANK2 | chr20 | + | 3912457 | 3912634 | 3910576 | 3910830 | 3923243 | 3923631 | 252 | 204 | 218 | 226 | 357 | 0 | 0 | 0 | 0 | 1 | 682 | 315 | 183 | 336 | 314 | 0 | 0 | 0 | 0 | 0 | 298 | 149 | 1 | 1 | 1 | 1 | 1 | 1 | 0.994 | 1 | 1 | 1 | 1 | 1 | -0.001 |
| SE | 77058 | ENSG00000125779 | PANK2 | chr20 | + | 3916926 | 3917050 | 3910576 | 3910830 | 3923243 | 3923631 | 1 | 5 | 2 | 3 | 13 | 0 | 0 | 0 | 0 | 1 | 20 | 6 | 6 | 15 | 20 | 0 | 0 | 0 | 0 | 0 | 273 | 149 | 1 | 1 | 1 | 1 | 1 | 1 | 0.876 | 1 | 1 | 1 | 1 | 1 | -0.025 |
| SE | 77059 | ENSG00000125779 | PANK2 | chr20 | + | 3916926 | 3917050 | 3912457 | 3912634 | 3918670 | 3918789 | 391 | 285 | 383 | 343 | 545 | 0 | 0 | 0 | 0 | 0 | 749 | 429 | 309 | 397 | 587 | 0 | 0 | 0 | 1 | 0 | 273 | 149 | 1 | 1 | 1 | 1 | 1 | 1 | 1 | 1 | 1 | 1 | 0.995 | 1 | 0.001 |
| SE | 77063 | ENSG00000164880 | INTS1 | chr7 | - | 1472242 | 1472386 | 1471570 | 1471641 | 1473071 | 1473150 | 33 | 14 | 26 | 36 | 9 | 0 | 0 | 0 | 2 | 0 | 15 | 27 | 28 | 39 | 23 | 0 | 0 | 0 | 0 | 0 | 293 | 149 | 1 | 1 | 1 | 1 | 1 | 0.902 | 1 | 1 | 1 | 1 | 1 | 1 | -0.02 |
| SE | 77064 | ENSG00000164880 | INTS1 | chr7 | - | 1472272 | 1472386 | 1471570 | 1471641 | 1473071 | 1473125 | 75 | 30 | 70 | 81 | 23 | 0 | 0 | 0 | 2 | 0 | 46 | 45 | 65 | 54 | 35 | 0 | 0 | 0 | 0 | 0 | 263 | 149 | 1 | 1 | 1 | 1 | 1 | 0.958 | 1 | 1 | 1 | 1 | 1 | 1 | -0.008 |
| SE | 77066 | ENSG00000164880 | INTS1 | chr7 | - | 1480834 | 1480933 | 1479429 | 1479684 | 1481341 | 1481488 | 17 | 5 | 16 | 16 | 15 | 0 | 0 | 0 | 0 | 0 | 0 | 33 | 14 | 18 | 23 | 0 | 0 | 0 | 0 | 4 | 248 | 149 | 1 | 1 | 1 | 1 | 1 | 1 | 1 | NA | 1 | 1 | 1 | 0.776 | 0.056 |
| SE | 77067 | ENSG00000104312 | RIPK2 | chr8 | + | 89759321 | 89759446 | 89757816 | 89758233 | 89762828 | 89762858 | 0 | 0 | 0 | 0 | 0 | 10 | 6 | 8 | 8 | 0 | 0 | 8 | 0 | 0 | 0 | 0 | 9 | 1 | 6 | 0 | 274 | 149 | 1 | 1 | 0 | 0 | 0 | 0 | NA | NA | 0.326 | 0 | 0 | NA | -0.109 |
| SE | 77069 | ENSG00000104312 | RIPK2 | chr8 | + | 89769771 | 89769929 | 89765340 | 89765496 | 89771740 | 89771790 | 320 | 237 | 191 | 303 | 393 | 0 | 0 | 1 | 0 | 0 | 219 | 202 | 262 | 303 | 63 | 0 | 0 | 0 | 0 | 0 | 298 | 149 | 1 | 1 | 1 | 1 | 0.99 | 1 | 1 | 1 | 1 | 1 | 1 | 1 | -0.002 |
| SE | 77071 | ENSG00000086232 | EIF2AK1 | chr7 | - | 6028614 | 6028697 | 6026727 | 6026961 | 6028917 | 6029032 | 312 | 270 | 303 | 392 | 798 | 0 | 0 | 0 | 0 | 0 | 689 | 574 | 344 | 405 | 2468 | 0 | 6 | 0 | 5 | 0 | 232 | 149 | 1 | 1 | 1 | 1 | 1 | 1 | 1 | 1 | 0.984 | 1 | 0.981 | 1 | 0.007 |
| SE | 77072 | ENSG00000086232 | EIF2AK1 | chr7 | - | 6037423 | 6037524 | 6028917 | 6029032 | 6040891 | 6041219 | 270 | 242 | 253 | 451 | 876 | 0 | 0 | 0 | 0 | 1 | 568 | 496 | 297 | 403 | 2180 | 0 | 0 | 0 | 0 | 0 | 250 | 149 | 1 | 1 | 1 | 1 | 1 | 1 | 0.998 | 1 | 1 | 1 | 1 | 1 | 0 |
| SE | 77073 | ENSG00000086232 | EIF2AK1 | chr7 | - | 6038559 | 6038671 | 6037423 | 6037524 | 6040891 | 6041219 | 613 | 519 | 519 | 930 | 1171 | 2 | 0 | 0 | 2 | 0 | 1549 | 1029 | 637 | 1037 | 4878 | 0 | 0 | 0 | 7 | 0 | 261 | 149 | 1 | 1 | 0.994 | 1 | 1 | 0.996 | 1 | 1 | 1 | 1 | 0.988 | 1 | 0 |
| SE | 77074 | ENSG00000086232 | EIF2AK1 | chr7 | - | 6042932 | 6042993 | 6040891 | 6041219 | 6044561 | 6044661 | 489 | 408 | 326 | 511 | 1335 | 0 | 0 | 0 | 0 | 0 | 1132 | 728 | 477 | 654 | 3609 | 0 | 0 | 1 | 1 | 0 | 210 | 149 | 1 | 1 | 1 | 1 | 1 | 1 | 1 | 1 | 1 | 0.997 | 0.998 | 1 | 0.001 |
| SE | 77075 | ENSG00000086232 | EIF2AK1 | chr7 | - | 6044561 | 6044661 | 6042932 | 6042993 | 6046070 | 6046151 | 688 | 560 | 482 | 724 | 1959 | 0 | 0 | 2 | 0 | 0 | 1412 | 937 | 598 | 891 | 4686 | 0 | 2 | 0 | 0 | 3 | 249 | 149 | 1 | 1 | 1 | 1 | 0.993 | 1 | 1 | 1 | 0.996 | 1 | 1 | 0.999 | 0 |
| SE | 77077 | ENSG00000086232 | EIF2AK1 | chr7 | - | 6046070 | 6046151 | 6044561 | 6044661 | 6046991 | 6047091 | 569 | 494 | 410 | 661 | 1560 | 1 | 0 | 1 | 2 | 1 | 1091 | 769 | 429 | 808 | 3869 | 0 | 2 | 0 | 4 | 6 | 230 | 149 | 1 | 1 | 0.997 | 1 | 0.996 | 0.995 | 0.999 | 1 | 0.996 | 1 | 0.992 | 0.998 | 0 |
| SE | 77078 | ENSG00000086232 | EIF2AK1 | chr7 | - | 6048806 | 6048844 | 6046722 | 6047091 | 6049911 | 6050045 | 399 | 314 | 302 | 478 | 629 | 0 | 0 | 0 | 0 | 0 | 643 | 588 | 351 | 538 | 2485 | 0 | 2 | 0 | 0 | 6 | 187 | 149 | 1 | 1 | 1 | 1 | 1 | 1 | 1 | 1 | 0.996 | 1 | 1 | 1 | 0.001 |
| SE | 77079 | ENSG00000086232 | EIF2AK1 | chr7 | - | 6048806 | 6048844 | 6046722 | 6047091 | 6054545 | 6054704 | 101 | 120 | 90 | 135 | 270 | 0 | 0 | 0 | 0 | 0 | 193 | 182 | 121 | 222 | 883 | 0 | 0 | 0 | 0 | 6 | 187 | 149 | 1 | 1 | 1 | 1 | 1 | 1 | 1 | 1 | 1 | 1 | 1 | 0.992 | 0.002 |
| SE | 77080 | ENSG00000086232 | EIF2AK1 | chr7 | - | 6049911 | 6050045 | 6046722 | 6047091 | 6054545 | 6054704 | 50 | 44 | 47 | 76 | 47 | 0 | 0 | 0 | 0 | 0 | 177 | 75 | 65 | 73 | 388 | 0 | 0 | 0 | 0 | 6 | 283 | 149 | 1 | 1 | 1 | 1 | 1 | 1 | 1 | 1 | 1 | 1 | 1 | 0.971 | 0.006 |
| SE | 77086 | ENSG00000197461 | PDGFA | chr7 | - | 518938 | 519068 | 517393 | 517490 | 519959 | 520296 | 2 | 6 | 1 | 0 | 4 | 0 | 0 | 0 | 0 | 0 | 10 | 3 | 3 | 16 | 35 | 0 | 3 | 0 | 0 | 0 | 279 | 149 | 1 | 1 | 1 | 1 | 1 | NA | 1 | 1 | 0.348 | 1 | 1 | 1 | 0.13 |

| | | | | | | | | | | | | | | | | | | | | | | | | | | | | | | | | | | | | | | | | | | | | | | |
|---|---|---|---|---|---|---|---|---|---|---|---|---|---|---|---|---|---|---|---|---|---|---|---|---|---|---|---|---|---|---|---|---|---|---|---|---|---|---|---|---|---|---|---|---|---|---|
| SE | 77088 | ENSG00000178381 | ZFAND2A | chr7 | - | 1157655 | 1157750 | 1155452 | 1155584 | 1158157 | 1158257 | 186 | 79 | 115 | 148 | 116 | 2 | 3 | 0 | 2 | 0 | 438 | 212 | 166 | 232 | 388 | 0 | 2 | 0 | 4 | 3 | 244 | 149 | 1 | 1 | 0.983 | 0.941 | 1 | 0.978 | 1 | 1 | 0.985 | 1 | 0.973 | 0.987 | -0.009 |
| SE | 77089 | ENSG00000178381 | ZFAND2A | chr7 | - | 1157655 | 1157750 | 1155452 | 1155584 | 1159963 | 1160153 | 42 | 21 | 22 | 32 | 16 | 0 | 0 | 0 | 0 | 0 | 113 | 51 | 44 | 48 | 88 | 0 | 0 | 0 | 2 | 0 | 244 | 149 | 1 | 1 | 1 | 1 | 1 | 1 | 1 | 1 | 1 | 1 | 0.936 | 1 | 0.013 |
| SE | 77090 | ENSG00000178381 | ZFAND2A | chr7 | - | 1158157 | 1158257 | 1155452 | 1155584 | 1159963 | 1160153 | 59 | 31 | 34 | 68 | 44 | 0 | 0 | 0 | 0 | 0 | 159 | 84 | 53 | 121 | 161 | 0 | 0 | 0 | 2 | 0 | 249 | 149 | 1 | 1 | 1 | 1 | 1 | 1 | 1 | 1 | 1 | 1 | 0.973 | 1 | 0.005 |
| SE | 77091 | ENSG00000178381 | ZFAND2A | chr7 | - | 1158157 | 1158257 | 1157022 | 1157750 | 1159963 | 1160153 | 212 | 90 | 130 | 183 | 151 | 1 | 0 | 2 | 0 | 0 | 503 | 245 | 181 | 307 | 453 | 0 | 0 | 0 | 0 | 0 | 249 | 149 | 1 | 1 | 0.992 | 1 | 0.975 | 1 | 1 | 1 | 1 | 1 | 1 | 1 | -0.007 |
| SE | 77098 | ENSG00000164587 | RPS14 | chr5 | - | 150446801 | 150446963 | 150445608 | 150445685 | 150447584 | 150447735 | 18214 | 8093 | 20630 | 15002 | 46632 | 3 | 0 | 0 | 0 | 0 | 7665 | 8513 | 6288 | 5119 | 12387 | 0 | 0 | 0 | 0 | 4 | 298 | 149 | 1 | 1 | 1 | 1 | 1 | 1 | 1 | 1 | 1 | 1 | 1 | 0.999 | 0 |
| SE | 77099 | ENSG00000164587 | RPS14 | chr5 | - | 150447584 | 150447735 | 150446801 | 150446963 | 150449702 | 150449739 | 15726 | 7971 | 16798 | 12800 | 31618 | 2 | 0 | 0 | 1 | 6 | 7658 | 7406 | 5884 | 4627 | 11478 | 0 | 4 | 0 | 0 | 5 | 298 | 149 | 1 | 1 | 1 | 1 | 1 | 1 | 1 | 1 | 0.999 | 1 | 1 | 0.999 | 0 |
| SE | 77100 | ENSG00000122512 | PMS2 | chr7 | - | 5978595 | 5978696 | 5977587 | 5977757 | 5982823 | 5982991 | 50 | 23 | 57 | 68 | 49 | 2 | 0 | 0 | 0 | 0 | 10 | 63 | 76 | 46 | 51 | 0 | 0 | 0 | 0 | 0 | 250 | 149 | 1 | 1 | 0.937 | 1 | 1 | 1 | 1 | 1 | 1 | 1 | 1 | 1 | -0.013 |
| SE | 77103 | ENSG00000122512 | PMS2 | chr7 | - | 5991972 | 5992057 | 5986758 | 5987620 | 5995533 | 5995633 | 25 | 7 | 22 | 17 | 41 | 1 | 0 | 0 | 0 | 0 | 15 | 8 | 15 | 3 | 0 | 0 | 0 | 0 | 0 | 0 | 234 | 149 | 1 | 1 | 0.941 | 1 | 1 | 1 | 1 | 1 | 1 | 1 | 1 | NA | -0.012 |
| SE | 77104 | ENSG00000122512 | PMS2 | chr7 | - | 5995533 | 5995633 | 5991972 | 5992057 | 5997325 | 5997423 | 72 | 30 | 56 | 55 | 100 | 2 | 0 | 0 | 0 | 0 | 76 | 26 | 24 | 37 | 32 | 0 | 3 | 1 | 0 | 0 | 249 | 149 | 1 | 1 | 0.956 | 1 | 1 | 1 | 1 | 1 | 0.838 | 0.935 | 1 | 1 | 0.037 |
| SE | 77111 | ENSG00000164654 | MIOS | chr7 | + | 7567606 | 7567688 | 7566883 | 7567072 | 7568025 | 7568123 | 10 | 6 | 6 | 8 | 24 | 4 | 0 | 2 | 5 | 5 | 4 | 18 | 10 | 11 | 0 | 0 | 7 | 0 | 0 | 0 | 231 | 149 | 1 | 1 | 0.617 | 1 | 0.659 | 0.508 | 0.756 | 1 | 0.624 | 1 | 1 | NA | -0.198 |
| SE | 77112 | ENSG00000164654 | MIOS | chr7 | + | 7567606 | 7567688 | 7566883 | 7567072 | 7572435 | 7572667 | 2 | 0 | 1 | 1 | 1 | 4 | 4 | 0 | 1 | 0 | 0 | 0 | 3 | 6 | 0 | 6 | 2 | 1 | 0 | 0 | 231 | 149 | 1 | 1 | 0.244 | 0 | 1 | 0.392 | 1 | 0 | 0 | 0.659 | 1 | NA | 0.112 |
| SE | 77113 | ENSG00000164654 | MIOS | chr7 | + | 7567606 | 7567688 | 7567052 | 7567484 | 7568025 | 7568123 | 12 | 7 | 6 | 8 | 27 | 5 | 0 | 0 | 8 | 4 | 4 | 16 | 10 | 10 | 0 | 0 | 0 | 0 | 6 | 0 | 231 | 149 | 1 | 1 | 0.608 | 1 | 1 | 0.392 | 0.813 | 1 | 1 | 1 | 0.518 | NA | -0.117 |
| SE | 77114 | ENSG00000164654 | MIOS | chr7 | + | 7567606 | 7567688 | 7567052 | 7567484 | 7572435 | 7572667 | 4 | 0 | 0 | 1 | 1 | 0 | 2 | 2 | 0 | 0 | 0 | 1 | 3 | 6 | 0 | 0 | 0 | 1 | 3 | 0 | 231 | 149 | 1 | 1 | 1 | 0 | 0 | 1 | 1 | NA | 1 | 0.659 | 0.563 | NA | -0.141 |
| SE | 77116 | ENSG00000164654 | MIOS | chr7 | + | 7568025 | 7568123 | 7567052 | 7567484 | 7572435 | 7572667 | 23 | 12 | 13 | 27 | 38 | 0 | 2 | 2 | 0 | 0 | 0 | 18 | 4 | 14 | 5 | 0 | 0 | 1 | 3 | 0 | 247 | 149 | 1 | 1 | 1 | 0.784 | 0.797 | 1 | 1 | NA | 1 | 0.707 | 0.738 | 1 | 0.055 |
| SE | 77121 | ENSG00000164654 | MIOS | chr7 | + | 7585635 | 7585805 | 7568025 | 7568123 | 7588497 | 7588563 | 80 | 75 | 75 | 76 | 100 | 2 | 0 | 0 | 1 | 0 | 27 | 51 | 39 | 32 | 5 | 0 | 0 | 0 | 0 | 0 | 298 | 149 | 1 | 1 | 0.952 | 1 | 1 | 0.974 | 1 | 1 | 1 | 1 | 1 | 1 | -0.015 |
| SE | 77123 | ENSG00000164654 | MIOS | chr7 | + | 7585635 | 7585805 | 7582953 | 7583372 | 7588497 | 7588563 | 157 | 114 | 139 | 145 | 164 | 0 | 0 | 0 | 0 | 3 | 65 | 108 | 54 | 69 | 5 | 0 | 5 | 0 | 0 | 0 | 298 | 149 | 1 | 1 | 1 | 1 | 1 | 1 | 0.965 | 1 | 0.915 | 1 | 1 | 1 | 0.01 |
| SE | 77124 | ENSG00000164654 | MIOS | chr7 | + | 7589404 | 7589563 | 7588497 | 7588563 | 7594979 | 7595132 | 137 | 94 | 126 | 126 | 207 | 1 | 1 | 0 | 2 | 0 | 52 | 104 | 56 | 63 | 8 | 0 | 0 | 0 | 0 | 0 | 298 | 149 | 1 | 1 | 0.986 | 0.979 | 1 | 0.969 | 1 | 1 | 1 | 1 | 1 | 1 | -0.013 |
| SE | 77126 | ENSG00000164654 | MIOS | chr7 | + | 7594979 | 7595132 | 7588497 | 7588563 | 7596256 | 7596461 | 54 | 30 | 37 | 56 | 64 | 2 | 0 | 6 | 1 | 5 | 9 | 45 | 11 | 17 | 0 | 0 | 0 | 5 | 0 | 0 | 298 | 149 | 1 | 1 | 0.931 | 1 | 0.755 | 0.966 | 0.865 | 1 | 1 | 0.524 | 1 | NA | 0.022 |
| SE | 77127 | ENSG00000164654 | MIOS | chr7 | + | 7594979 | 7595132 | 7589404 | 7589563 | 7596256 | 7596461 | 92 | 47 | 75 | 85 | 149 | 0 | 0 | 1 | 0 | 0 | 9 | 75 | 27 | 36 | 3 | 0 | 0 | 0 | 0 | 0 | 298 | 149 | 1 | 1 | 1 | 1 | 0.974 | 1 | 1 | 1 | 1 | 1 | 1 | 1 | -0.005 |
| SE | 77128 | ENSG00000164654 | MIOS | chr7 | + | 7596256 | 7596461 | 7594979 | 7595132 | 7605941 | 7606071 | 117 | 70 | 109 | 136 | 153 | 1 | 0 | 0 | 0 | 0 | 9 | 94 | 43 | 44 | 2 | 0 | 0 | 0 | 0 | 0 | 298 | 149 | 1 | 1 | 0.983 | 1 | 1 | 1 | 1 | 1 | 1 | 1 | 1 | 1 | -0.003 |
| SE | 77133 | ENSG00000240859 | NA | chr7 | + | 152440 | 152547 | 149548 | 150132 | 153408 | 154904 | 0 | 0 | 0 | 0 | 0 | 2 | 0 | 2 | 7 | 0 | 5 | 4 | 0 | 1 | 0 | 11 | 4 | 0 | 7 | 0 | 256 | 149 | 1 | 1 | 0 | NA | 0 | 0 | NA | 0.209 | 0.368 | NA | 0.077 | NA | -0.218 |
| SE | 77134 | ENSG00000065833 | ME1 | chr6 | - | 83223759 | 83223933 | 83216497 | 83216596 | 83227334 | 83227477 | 16 | 10 | 10 | 14 | 14 | 0 | 0 | 0 | 0 | 3 | 0 | 11 | 6 | 0 | 64 | 0 | 2 | 0 | 3 | 0 | 298 | 149 | 1 | 1 | 1 | 1 | 1 | 1 | 0.7 | NA | 0.733 | 1 | 0 | 1 | 0.257 |
| SE | 77135 | ENSG00000065833 | ME1 | chr6 | - | 83407767 | 83407901 | 83398366 | 83398516 | 83430876 | 83431051 | 24 | 17 | 22 | 6 | 19 | 0 | 0 | 1 | 0 | 0 | 1 | 1 | 5 | 3 | 30 | 0 | 0 | 2 | 0 | 0 | 283 | 149 | 1 | 1 | 1 | 1 | 0.921 | 1 | 1 | 1 | 1 | 0.568 | 1 | 1 | 0.071 |
| SE | 77136 | ENSG00000172014 | ANKRD20A4P | chr9 | + | 64370716 | 64370942 | 64369410 | 64369877 | 64373186 | 64373301 | 1 | 0 | 1 | 2 | 0 | 1 | 0 | 1 | 0 | 0 | 0 | 0 | 0 | 0 | 0 | 0 | 2 | 0 | 0 | 0 | 298 | 149 | 1 | 1 | 0.333 | NA | 0.333 | 1 | NA | NA | 0 | NA | NA | NA | 0.555 |
| SE | 77138 | ENSG00000256625 | NA | chr12 | + | 27139965 | 27140097 | 27134972 | 27135086 | 27144991 | 27145084 | 2 | 2 | 9 | 2 | 4 | 2 | 0 | 4 | 0 | 1 | 0 | 2 | 0 | 6 | 7 | 11 | 3 | 0 | 0 | 0 | 281 | 149 | 1 | 1 | 0.347 | 1 | 0.544 | 1 | 0.68 | 0 | 0.261 | NA | 1 | 1 | 0.149 |
| SE | 77145 | ENSG00000106009 | BRAT1 | chr7 | - | 2547323 | 2547478 | 2544908 | 2545056 | 2554304 | 2554447 | 66 | 24 | 44 | 34 | 28 | 0 | 0 | 0 | 0 | 0 | 23 | 30 | 40 | 7 | 76 | 0 | 0 | 1 | 0 | 0 | 298 | 149 | 1 | 1 | 1 | 1 | 1 | 1 | 1 | 1 | 1 | 0.952 | 1 | 1 | 0.01 |
| SE | 77146 | ENSG00000172339 | ALG14 | chr1 | - | 95041492 | 95041612 | 95027128 | 95027260 | 95064865 | 95065017 | 4 | 0 | 1 | 0 | 0 | 9 | 3 | 8 | 9 | 6 | 0 | 0 | 2 | 0 | 7 | 19 | 13 | 5 | 0 | 4 | 269 | 149 | 1 | 1 | 0.198 | 0 | 0.065 | 0 | 0 | 0 | 0 | 0.181 | NA | 0.492 | -0.116 |
| SE | 77147 | ENSG00000231160 | KLF3-AS1 | chr4 | - | 38620071 | 38620162 | 38614624 | 38614708 | 38624421 | 38624579 | 0 | 0 | 0 | 3 | 1 | 10 | 13 | 18 | 4 | 21 | 0 | 0 | 0 | 0 | 0 | 15 | 10 | 2 | 3 | 0 | 240 | 149 | 1 | 1 | 0 | 0 | 0 | 0.318 | 0.029 | 0 | 0 | 0 | 0 | NA | 0.069 |
| SE | 77148 | ENSG00000231160 | KLF3-AS1 | chr4 | - | 38624421 | 38624579 | 38614624 | 38614708 | 38628038 | 38628102 | 10 | 13 | 17 | 4 | 21 | 0 | 0 | 1 | 0 | 0 | 15 | 10 | 2 | 3 | 0 | 0 | 0 | 0 | 0 | 0 | 298 | 149 | 1 | 1 | 1 | 1 | 0.895 | 1 | 1 | 1 | 1 | 1 | 1 | NA | -0.021 |
| SE | 77150 | ENSG00000231160 | KLF3-AS1 | chr4 | - | 38625601 | 38625762 | 38624421 | 38624579 | 38628038 | 38628102 | 6 | 0 | 5 | 2 | 4 | 0 | 0 | 1 | 0 | 0 | 0 | 5 | 1 | 2 | 0 | 0 | 0 | 0 | 0 | 0 | 298 | 149 | 1 | 1 | 1 | NA | 0.714 | 1 | 1 | NA | 1 | 1 | 1 | NA | -0.072 |
| SE | 77151 | ENSG00000231160 | KLF3-AS1 | chr4 | - | 38630499 | 38630581 | 38626407 | 38628102 | 38635955 | 38636047 | 9 | 3 | 15 | 9 | 14 | 0 | 0 | 0 | 0 | 0 | 0 | 6 | 0 | 0 | 0 | 0 | 0 | 1 | 0 | 0 | 231 | 149 | 1 | 1 | 1 | 1 | 1 | 1 | 1 | NA | 1 | 0 | NA | NA | 0.5 |
| SE | 77155 | ENSG00000136295 | TTYH3 | chr7 | + | 2658285 | 2658459 | 2656397 | 2656534 | 2658939 | 2659015 | 85 | 32 | 28 | 76 | 25 | 0 | 2 | 0 | 0 | 0 | 206 | 50 | 33 | 69 | 395 | 0 | 0 | 0 | 0 | 0 | 298 | 149 | 1 | 1 | 1 | 0.889 | 1 | 1 | 1 | 1 | 1 | 1 | 1 | 1 | -0.022 |
| SE | 77156 | ENSG00000136295 | TTYH3 | chr7 | + | 2658939 | 2659015 | 2658285 | 2658459 | 2661667 | 2664800 | 46 | 16 | 17 | 53 | 26 | 0 | 0 | 0 | 0 | 0 | 125 | 24 | 23 | 39 | 231 | 0 | 0 | 3 | 0 | 0 | 225 | 149 | 1 | 1 | 1 | 1 | 1 | 1 | 1 | 1 | 1 | 0.835 | 1 | 1 | 0.033 |
| SE | 77216 | ENSG00000137831 | UACA | chr15 | - | 70677107 | 70677140 | 70676306 | 70676591 | 70678098 | 70678206 | 9 | 7 | 2 | 8 | 7 | 0 | 1 | 0 | 0 | 4 | 0 | 4 | 3 | 14 | 0 | 0 | 0 | 0 | 0 | 10 | 182 | 149 | 1 | 1 | 1 | 0.851 | 1 | 1 | 0.589 | NA | 1 | 1 | 1 | 0 | 0.138 |
| SE | 77217 | ENSG00000137831 | UACA | chr15 | - | 70687539 | 70687646 | 70684264 | 70684446 | 70687749 | 70687820 | 15 | 29 | 14 | 21 | 16 | 0 | 2 | 0 | 0 | 3 | 21 | 21 | 9 | 0 | 15 | 0 | 0 | 0 | 0 | 0 | 256 | 149 | 1 | 1 | 1 | 0.894 | 1 | 1 | 0.756 | 1 | 1 | 1 | NA | 1 | -0.07 |
| SE | 77219 | ENSG00000229043 | ZFAND2A-DT | chr7 | + | 1162961 | 1163217 | 1160431 | 1160623 | 1163451 | 1163495 | 0 | 0 | 1 | 1 | 0 | 0 | 0 | 0 | 0 | 2 | 0 | 0 | 0 | 0 | 0 | 0 | 0 | 0 | 2 | 0 | 298 | 149 | 1 | 1 | NA | NA | 1 | 1 | 0 | NA | NA | NA | 0 | NA | 0.667 |
| SE | 77224 | ENSG00000064419 | TNPO3 | chr7 | - | 128978982 | 128979123 | 128975818 | 128975935 | 128979970 | 128980031 | 193 | 101 | 197 | 195 | 194 | 0 | 2 | 0 | 0 | 0 | 184 | 121 | 118 | 100 | 122 | 0 | 0 | 2 | 0 | 0 | 290 | 149 | 1 | 1 | 1 | 0.963 | 1 | 1 | 1 | 1 | 1 | 0.968 | 1 | 1 | -0.001 |
| SE | 77225 | ENSG00000064419 | TNPO3 | chr7 | - | 128979970 | 128980031 | 128975818 | 128975935 | 128982247 | 128982324 | 27 | 23 | 32 | 41 | 42 | 0 | 0 | 1 | 0 | 0 | 53 | 17 | 21 | 23 | 28 | 0 | 0 | 1 | 0 | 0 | 210 | 149 | 1 | 1 | 1 | 1 | 0.958 | 1 | 1 | 1 | 1 | 0.937 | 1 | 1 | 0.004 |
| SE | 77227 | ENSG00000064419 | TNPO3 | chr7 | - | 128986728 | 128986920 | 128984167 | 128984259 | 128989960 | 128990100 | 240 | 127 | 219 | 200 | 277 | 5 | 0 | 1 | 0 | 0 | 140 | 140 | 123 | 210 | 115 | 0 | 0 | 0 | 4 | 0 | 298 | 149 | 1 | 1 | 0.96 | 1 | 0.991 | 1 | 1 | 1 | 1 | 1 | 0.963 | 1 | -0.002 |
| SE | 77228 | ENSG00000064419 | TNPO3 | chr7 | - | 128986728 | 128986920 | 128984167 | 128984259 | 128997388 | 128997535 | 94 | 50 | 74 | 78 | 86 | 0 | 0 | 0 | 3 | 0 | 48 | 31 | 46 | 76 | 43 | 0 | 0 | 0 | 0 | 0 | 298 | 149 | 1 | 1 | 1 | 1 | 1 | 0.929 | 1 | 1 | 1 | 1 | 1 | 1 | -0.014 |
| SE | 77229 | ENSG00000064419 | TNPO3 | chr7 | - | 128991998 | 128992090 | 128989960 | 128990100 | 128993806 | 128993914 | 149 | 79 | 102 | 114 | 197 | 1 | 0 | 0 | 0 | 0 | 93 | 133 | 73 | 134 | 77 | 0 | 0 | 0 | 0 | 0 | 241 | 149 | 1 | 1 | 0.989 | 1 | 1 | 1 | 1 | 1 | 1 | 1 | 1 | 1 | -0.002 |
| SE | 77230 | ENSG00000064419 | TNPO3 | chr7 | - | 128993806 | 128993914 | 128991998 | 128992090 | 128997388 | 128997535 | 146 | 92 | 113 | 137 | 191 | 0 | 0 | 0 | 0 | 0 | 117 | 135 | 85 | 129 | 80 | 0 | 4 | 0 | 1 | 0 | 257 | 149 | 1 | 1 | 1 | 1 | 1 | 1 | 1 | 1 | 0.951 | 1 | 0.987 | 1 | 0.012 |
| SE | 77231 | ENSG00000064419 | TNPO3 | chr7 | - | 128997388 | 128997535 | 128986728 | 128986920 | 129000428 | 129000567 | 181 | 118 | 149 | 163 | 194 | 0 | 0 | 0 | 0 | 0 | 246 | 206 | 177 | 222 | 87 | 0 | 0 | 0 | 3 | 0 | 296 | 149 | 1 | 1 | 1 | 1 | 1 | 1 | 1 | 1 | 1 | 1 | 0.974 | 1 | 0.005 |
| SE | 77232 | ENSG00000064419 | TNPO3 | chr7 | - | 129005015 | 129005159 | 129001058 | 129001234 | 129014978 | 129015135 | 221 | 174 | 194 | 205 | 211 | 0 | 2 | 0 | 0 | 5 | 153 | 175 | 188 | 194 | 96 | 1 | 3 | 2 | 0 | 0 | 293 | 149 | 1 | 1 | 1 | 0.978 | 1 | 1 | 0.955 | 0.987 | 0.967 | 0.98 | 1 | 1 | 0 |
| SE | 77236 | ENSG00000003147 | ICA1 | chr7 | - | 8141968 | 8142055 | 8141764 | 8141817 | 8143874 | 8143972 | 0 | 0 | 0 | 2 | 0 | 11 | 6 | 8 | 9 | 14 | 0 | 0 | 0 | 0 | 0 | 44 | 8 | 8 | 33 | 2 | 236 | 149 | 1 | 1 | 0 | 0 | 0 | 0.123 | 0 | 0 | 0 | 0 | 0 | 0 | 0.025 |
| SE | 77237 | ENSG00000003147 | ICA1 | chr7 | - | 8223487 | 8223559 | 8221352 | 8221398 | 8227637 | 8227851 | 0 | 2 | 0 | 0 | 2 | 2 | 0 | 0 | 0 | 0 | 0 | 0 | 0 | 1 | 0 | 4 | 0 | 0 | 0 | 11 | 221 | 149 | 1 | 1 | 0 | 1 | NA | NA | 1 | 0 | NA | NA | 1 | 0 | 0.333 |
| SE | 77238 | ENSG00000003147 | ICA1 | chr7 | - | 8223487 | 8223559 | 8221352 | 8221398 | 8228600 | 8228673 | 0 | 2 | 0 | 0 | 0 | 15 | 12 | 8 | 26 | 68 | 0 | 0 | 2 | 4 | 0 | 131 | 40 | 18 | 129 | 126 | 221 | 149 | 1 | 1 | 0 | 0.101 | 0 | 0 | 0 | 0 | 0 | 0.07 | 0.02 | 0 | 0.002 |
| SE | 77240 | ENSG00000003147 | ICA1 | chr7 | - | 8227637 | 8227851 | 8223487 | 8223559 | 8228600 | 8228673 | 1 | 0 | 0 | 0 | 2 | 0 | 0 | 0 | 0 | 0 | 0 | 0 | 1 | 1 | 2 | 0 | 0 | 2 | 3 | 0 | 298 | 149 | 1 | 1 | 1 | NA | NA | NA | 1 | NA | NA | 0.2 | 0.143 | 1 | 0.552 |
| SE | 77241 | ENSG00000003147 | ICA1 | chr7 | - | 8232589 | 8232733 | 8228600 | 8228673 | 8235909 | 8236005 | 6 | 5 | 2 | 15 | 33 | 0 | 1 | 0 | 0 | 0 | 74 | 34 | 16 | 65 | 124 | 0 | 0 | 0 | 0 | 0 | 293 | 149 | 1 | 1 | 1 | 0.718 | 1 | 1 | 1 | 1 | 1 | 1 | 1 | 1 | -0.056 |
| SE | 77242 | ENSG00000003147 | ICA1 | chr7 | - | 8232589 | 8232752 | 8228600 | 8228673 | 8235909 | 8236005 | 7 | 5 | 2 | 17 | 33 | 0 | 1 | 0 | 0 | 0 | 74 | 34 | 17 | 68 | 141 | 0 | 0 | 0 | 0 | 0 | 298 | 149 | 1 | 1 | 1 | 0.714 | 1 | 1 | 1 | 1 | 1 | 1 | 1 | 1 | -0.057 |
| SE | 77243 | ENSG00000003147 | ICA1 | chr7 | - | 8232589 | 8232755 | 8228600 | 8228673 | 8235909 | 8235926 | 10 | 5 | 4 | 17 | 38 | 0 | 1 | 0 | 0 | 0 | 78 | 36 | 18 | 109 | 134 | 0 | 0 | 0 | 0 | 0 | 298 | 149 | 1 | 1 | 1 | 0.714 | 1 | 1 | 1 | 1 | 1 | 1 | 1 | 1 | -0.057 |
| SE | 77250 | ENSG00000283486 | FAM95C | chr9 | - | 38541455 | 38541562 | 38540568 | 38540849 | 38542311 | 38542589 | 4 | 0 | 0 | 1 | 0 | 0 | 0 | 0 | 2 | 0 | 0 | 0 | 0 | 9 | 0 | 0 | 0 | 0 | 0 | 0 | 256 | 149 | 1 | 1 | 1 | NA | NA | 0.225 | NA | NA | NA | NA | 1 | NA | -0.387 |
| SE | 77251 | ENSG00000106399 | RPA3 | chr7 | - | 7640319 | 7640581 | 7639069 | 7639144 | 7640722 | 7640972 | 57 | 17 | 53 | 53 | 53 | 0 | 0 | 0 | 0 | 0 | 45 | 33 | 15 | 86 | 43 | 0 | 0 | 0 | 1 | 0 | 298 | 149 | 1 | 1 | 1 | 1 | 1 | 1 | 1 | 1 | 1 | 1 | 0.977 | 1 | 0.005 |
| SE | 77252 | ENSG00000141664 | ZCCHC2 | chr18 | + | 62539680 | 62539792 | 62523024 | 62524363 | 62542497 | 62542574 | 146 | 133 | 151 | 192 | 184 | 0 | 0 | 0 | 0 | 0 | 420 | 250 | 311 | 394 | 164 | 2 | 0 | 0 | 0 | 0 | 261 | 149 | 1 | 1 | 1 | 1 | 1 | 1 | 1 | 0.992 | 1 | 1 | 1 | 1 | 0.002 |
| SE | 77258 | ENSG00000141664 | ZCCHC2 | chr18 | + | 62570102 | 62570231 | 62565001 | 62565096 | 62574056 | 62575550 | 156 | 113 | 146 | 187 | 317 | 0 | 0 | 0 | 0 | 2 | 250 | 270 | 224 | 359 | 102 | 0 | 0 | 0 | 0 | 0 | 278 | 149 | 1 | 1 | 1 | 1 | 1 | 1 | 0.988 | 1 | 1 | 1 | 1 | 1 | -0.002 |
| SE | 77259 | ENSG00000141664 | ZCCHC2 | chr18 | + | 62576511 | 62576652 | 62574056 | 62575550 | 62584445 | 62584595 | 47 | 40 | 23 | 53 | 36 | 0 | 0 | 0 | 2 | 0 | 45 | 97 | 79 | 91 | 40 | 0 | 0 | 0 | 0 | 0 | 290 | 149 | 1 | 1 | 1 | 1 | 1 | 0.932 | 1 | 1 | 1 | 1 | 1 | 1 | -0.014 |
| SE | 77261 | ENSG00000168944 | CEP120 | chr5 | - | 123372649 | 123372751 | 123364495 | 123364594 | 123377373 | 123377535 | 217 | 184 | 200 | 194 | 341 | 2 | 0 | 0 | 5 | 0 | 22 | 111 | 129 | 31 | 0 | 0 | 0 | 0 | 0 | 0 | 251 | 149 | 1 | 1 | 0.985 | 1 | 1 | 0.958 | 1 | 1 | 1 | 1 | 1 | NA | -0.011 |
| SE | 77262 | ENSG00000168944 | CEP120 | chr5 | - | 123372649 | 123372772 | 123364495 | 123364594 | 123377373 | 123377535 | 398 | 308 | 393 | 347 | 622 | 2 | 0 | 0 | 5 | 0 | 22 | 186 | 244 | 50 | 25 | 0 | 0 | 0 | 0 | 0 | 272 | 149 | 1 | 1 | 0.991 | 1 | 1 | 0.974 | 1 | 1 | 1 | 1 | 1 | 1 | -0.007 |
| SE | 77263 | ENSG00000168944 | CEP120 | chr5 | - | 123382110 | 123382200 | 123378335 | 123378428 | 123382736 | 123382889 | 169 | 92 | 117 | 126 | 232 | 0 | 1 | 0 | 1 | 0 | 10 | 97 | 115 | 37 | 8 | 0 | 0 | 0 | 0 | 0 | 239 | 149 | 1 | 1 | 1 | 0.983 | 1 | 0.987 | 1 | 1 | 1 | 1 | 1 | 1 | -0.006 |
| SE | 77264 | ENSG00000168944 | CEP120 | chr5 | - | 123384950 | 123385133 | 123382985 | 123383082 | 123386517 | 123386667 | 167 | 81 | 137 | 151 | 142 | 4 | 0 | 2 | 0 | 3 | 0 | 76 | 83 | 7 | 18 | 0 | 2 | 5 | 0 | 0 | 298 | 149 | 1 | 1 | 0.954 | 1 | 0.972 | 1 | 0.959 | NA | 0.95 | 0.892 | 1 | 1 | 0.016 |
| SE | 77265 | ENSG00000168944 | CEP120 | chr5 | - | 123386517 | 123386667 | 123382985 | 123383082 | 123388303 | 123388606 | 7 | 1 | 2 | 3 | 12 | 0 | 0 | 0 | 0 | 0 | 0 | 3 | 5 | 0 | 0 | 0 | 0 | 2 | 0 | 0 | 298 | 149 | 1 | 1 | 1 | 1 | 1 | 1 | 1 | NA | 1 | 0.556 | NA | NA | 0.222 |
| SE | 77266 | ENSG00000168944 | CEP120 | chr5 | - | 123391109 | 123391337 | 123389923 | 123390140 | 123393299 | 123393497 | 305 | 213 | 215 | 233 | 525 | 0 | 0 | 0 | 0 | 1 | 10 | 126 | 216 | 53 | 55 | 0 | 0 | 0 | 0 | 0 | 298 | 149 | 1 | 1 | 1 | 1 | 1 | 1 | 0.996 | 1 | 1 | 1 | 1 | 1 | -0.001 |
| SE | 77267 | ENSG00000168944 | CEP120 | chr5 | - | 123393299 | 123393497 | 123389923 | 123391337 | 123399135 | 123399284 | 287 | 180 | 208 | 228 | 497 | 1 | 0 | 0 | 2 | 0 | 17 | 153 | 191 | 49 | 27 | 0 | 0 | 0 | 0 | 0 | 298 | 149 | 1 | 1 | 0.993 | 1 | 1 | 0.983 | 1 | 1 | 1 | 1 | 1 | 1 | -0.005 |
| SE | 77268 | ENSG00000168944 | CEP120 | chr5 | - | 123403216 | 123403344 | 123399135 | 123399284 | 123403693 | 123403740 | 0 | 0 | 0 | 0 | 0 | 0 | 0 | 2 | 1 | 3 | 0 | 5 | 1 | 0 | 0 | 0 | 0 | 1 | 4 | 0 | 277 | 149 | 1 | 1 | NA | NA | 0 | 0 | 0 | NA | 1 | 0.35 | 0 | NA | -0.45 |
| SE | 77269 | ENSG00000168944 | CEP120 | chr5 | - | 123403255 | 123403344 | 123399135 | 123399284 | 123412398 | 123412540 | 0 | 0 | 0 | 2 | 0 | 94 | 47 | 89 | 82 | 122 | 0 | 0 | 0 | 3 | 0 | 6 | 51 | 37 | 10 | 21 | 238 | 149 | 1 | 1 | 0 | 0 | 0 | 0.015 | 0 | 0 | 0 | 0 | 0.158 | 0 | -0.029 |
| SE | 77270 | ENSG00000168944 | CEP120 | chr5 | - | 123403693 | 123403740 | 123399135 | 123399284 | 123412398 | 123412540 | 0 | 0 | 1 | 0 | 2 | 94 | 47 | 89 | 82 | 122 | 0 | 0 | 0 | 0 | 0 | 6 | 51 | 37 | 10 | 21 | 196 | 149 | 1 | 1 | 0 | 0 | 0.008 | 0 | 0.012 | 0 | 0 | 0 | 0 | 0 | 0.004 |
| SE | 77276 | ENSG00000005108 | THSD7A | chr7 | - | 11379069 | 11379280 | 11376569 | 11376657 | 11379629 | 11379712 | 0 | 2 | 1 | 0 | 0 | 0 | 0 | 0 | 1 | 3 | 0 | 2 | 0 | 0 | 0 | 0 | 0 | 0 | 0 | 0 | 298 | 149 | 1 | 1 | NA | 1 | 1 | 0 | 0 | NA | 1 | NA | NA | NA | -0.5 |
| SE | 77279 | ENSG00000113643 | RARS1 | chr5 | + | 168497227 | 168497348 | 168495314 | 168495436 | 168500590 | 168500720 | 270 | 178 | 192 | 213 | 220 | 0 | 0 | 0 | 1 | 4 | 136 | 204 | 137 | 113 | 94 | 0 | 0 | 0 | 0 | 0 | 270 | 149 | 1 | 1 | 1 | 1 | 1 | 0.992 | 0.968 | 1 | 1 | 1 | 1 | 1 | -0.008 |
| SE | 77280 | ENSG00000113643 | RARS1 | chr5 | + | 168497946 | 168498046 | 168497217 | 168497348 | 168500590 | 168500720 | 0 | 1 | 0 | 0 | 0 | 127 | 85 | 106 | 111 | 71 | 0 | 0 | 0 | 4 | 0 | 62 | 96 | 79 | 44 | 6 | 249 | 149 | 1 | 1 | 0 | 0.007 | 0 | 0 | 0 | 0 | 0 | 0 | 0.052 | 0 | -0.009 |
| SE | 77281 | ENSG00000113643 | RARS1 | chr5 | + | 168500590 | 168500720 | 168497011 | 168497348 | 168506020 | 168506030 | 123 | 81 | 102 | 106 | 70 | 0 | 0 | 2 | 0 | 0 | 62 | 93 | 78 | 40 | 6 | 0 | 0 | 0 | 0 | 0 | 279 | 149 | 1 | 1 | 1 | 1 | 0.965 | 1 | 1 | 1 | 1 | 1 | 1 | 1 | -0.007 |
| SE | 77282 | ENSG00000113643 | RARS1 | chr5 | + | 168502000 | 168502105 | 168500590 | 168500720 | 168506020 | 168506199 | 142 | 121 | 128 | 131 | 96 | 0 | 0 | 1 | 0 | 0 | 77 | 99 | 126 | 83 | 36 | 0 | 0 | 0 | 0 | 0 | 254 | 149 | 1 | 1 | 1 | 1 | 0.987 | 1 | 1 | 1 | 1 | 1 | 1 | 1 | -0.003 |
| SE | 77283 | ENSG00000113643 | RARS1 | chr5 | + | 168502002 | 168502105 | 168500590 | 168500720 | 168506020 | 168506030 | 52 | 43 | 53 | 57 | 32 | 0 | 0 | 1 | 0 | 0 | 4 | 46 | 60 | 33 | 0 | 0 | 0 | 0 | 0 | 0 | 252 | 149 | 1 | 1 | 1 | 1 | 0.969 | 1 | 1 | 1 | 1 | 1 | 1 | NA | -0.006 |
| SE | 77285 | ENSG00000113643 | RARS1 | chr5 | + | 168517814 | 168518062 | 168516777 | 168516950 | 168519080 | 168519298 | 154 | 92 | 106 | 136 | 202 | 0 | 0 | 1 | 0 | 0 | 21 | 53 | 55 | 26 | 16 | 0 | 0 | 0 | 0 | 0 | 298 | 149 | 1 | 1 | 1 | 1 | 0.981 | 1 | 1 | 1 | 1 | 1 | 1 | 1 | -0.004 |
| SE | 77286 | ENSG00000117000 | RLF | chr1 | + | 40189054 | 40189209 | 40161386 | 40161636 | 40190771 | 40190853 | 96 | 86 | 80 | 140 | 126 | 1 | 0 | 0 | 0 | 0 | 65 | 126 | 171 | 89 | 83 | 0 | 4 | 0 | 0 | 0 | 298 | 149 | 1 | 1 | 0.98 | 1 | 1 | 1 | 1 | 1 | 0.94 | 1 | 1 | 1 | 0.008 |
| SE | 77287 | ENSG00000117000 | RLF | chr1 | + | 40195631 | 40195764 | 40190771 | 40190853 | 40202411 | 40202614 | 180 | 135 | 157 | 224 | 339 | 0 | 0 | 0 | 1 | 0 | 187 | 222 | 302 | 151 | 181 | 0 | 0 | 0 | 0 | 0 | 282 | 149 | 1 | 1 | 1 | 1 | 1 | 0.992 | 1 | 1 | 1 | 1 | 1 | 1 | -0.002 |
| SE | 77288 | ENSG00000117000 | RLF | chr1 | + | 40222573 | 40222710 | 40202411 | 40202614 | 40231516 | 40231658 | 37 | 19 | 37 | 68 | 64 | 0 | 0 | 0 | 0 | 0 | 0 | 56 | 77 | 10 | 7 | 0 | 0 | 2 | 0 | 0 | 286 | 149 | 1 | 1 | 1 | 1 | 1 | 1 | 1 | NA | 1 | 0.953 | 1 | 1 | 0.012 |
| SE | 77289 | ENSG00000117000 | RLF | chr1 | + | 40231516 | 40231658 | 40222573 | 40222710 | 40235791 | 40240921 | 50 | 41 | 45 | 55 | 63 | 1 | 0 | 1 | 1 | 1 | 0 | 77 | 81 | 27 | 0 | 0 | 11 | 3 | 0 | 0 | 291 | 149 | 1 | 1 | 0.962 | 1 | 0.958 | 0.966 | 0.97 | NA | 0.782 | 0.933 | 1 | NA | 0.066 |
| SE | 77290 | ENSG00000106392 | C1GALT1 | chr7 | + | 7209426 | 7209566 | 7182624 | 7182820 | 7210445 | 7210567 | 2 | 0 | 0 | 0 | 0 | 0 | 0 | 0 | 2 | 0 | 0 | 3 | 0 | 0 | 0 | 0 | 0 | 0 | 0 | 0 | 289 | 149 | 1 | 1 | 1 | NA | NA | 0 | NA | NA | 1 | NA | NA | NA | -0.5 |
| SE | 77291 | ENSG00000106392 | C1GALT1 | chr7 | + | 7209426 | 7209566 | 7182624 | 7182820 | 7234302 | 7235297 | 0 | 0 | 0 | 0 | 0 | 35 | 22 | 28 | 25 | 29 | 0 | 0 | 1 | 0 | 0 | 18 | 16 | 11 | 11 | 0 | 289 | 149 | 1 | 1 | 0 | 0 | 0 | 0 | 0 | 0 | 0 | 0.045 | 0 | NA | -0.011 |
| SE | 77292 | ENSG00000099968 | BCL2L13 | chr22 | + | 17655661 | 17655832 | 17638101 | 17638700 | 17683213 | 17683321 | 143 | 116 | 164 | 147 | 181 | 0 | 0 | 0 | 0 | 2 | 231 | 205 | 95 | 231 | 322 | 0 | 0 | 1 | 0 | 0 | 298 | 149 | 1 | 1 | 1 | 1 | 1 | 1 | 0.978 | 1 | 1 | 0.979 | 1 | 1 | 0 |
| SE | 77294 | ENSG00000099968 | BCL2L13 | chr22 | + | 17683213 | 17683321 | 17655661 | 17655832 | 17688985 | 17689142 | 281 | 188 | 286 | 256 | 366 | 0 | 0 | 6 | 0 | 0 | 609 | 404 | 197 | 474 | 581 | 0 | 0 | 0 | 7 | 0 | 257 | 149 | 1 | 1 | 1 | 1 | 0.965 | 1 | 1 | 1 | 1 | 1 | 0.975 | 1 | -0.002 |
| SE | 77296 | ENSG00000099968 | BCL2L13 | chr22 | + | 17683213 | 17683321 | 17655661 | 17655832 | 17726676 | 17730855 | 100 | 79 | 123 | 117 | 124 | 0 | 0 | 1 | 0 | 0 | 195 | 141 | 65 | 171 | 255 | 0 | 0 | 0 | 0 | 0 | 257 | 149 | 1 | 1 | 1 | 1 | 0.986 | 1 | 1 | 1 | 1 | 1 | 1 | 1 | -0.003 |
| SE | 77298 | ENSG00000099968 | BCL2L13 | chr22 | + | 17688985 | 17689142 | 17655661 | 17655832 | 17726676 | 17730622 | 10 | 6 | 13 | 3 | 4 | 0 | 0 | 1 | 0 | 0 | 6 | 14 | 0 | 14 | 0 | 0 | 0 | 0 | 0 | 0 | 298 | 149 | 1 | 1 | 1 | 1 | 0.867 | 1 | 1 | 1 | 1 | NA | 1 | NA | -0.027 |
| SE | 77303 | ENSG00000099968 | BCL2L13 | chr22 | + | 17696140 | 17696210 | 17683213 | 17683321 | 17726676 | 17730855 | 0 | 1 | 7 | 5 | 10 | 0 | 0 | 0 | 0 | 0 | 0 | 10 | 7 | 1 | 8 | 14 | 0 | 0 | 3 | 0 | 219 | 149 | 1 | 1 | NA | 1 | 1 | 1 | 1 | 0 | 1 | 1 | 0.185 | 1 | 0.363 |
| SE | 77306 | ENSG00000099968 | BCL2L13 | chr22 | + | 17702242 | 17702386 | 17655661 | 17655832 | 17726676 | 17728699 | 77 | 64 | 85 | 94 | 89 | 0 | 0 | 1 | 0 | 0 | 94 | 130 | 47 | 101 | 186 | 0 | 0 | 0 | 0 | 0 | 293 | 149 | 1 | 1 | 1 | 1 | 0.977 | 1 | 1 | 1 | 1 | 1 | 1 | 1 | -0.005 |
| SE | 77320 | ENSG00000242802 | AP5Z1 | chr7 | + | 4781567 | 4781754 | 4781174 | 4781312 | 4783315 | 4783460 | 60 | 25 | 68 | 68 | 28 | 0 | 2 | 0 | 0 | 0 | 80 | 63 | 41 | 76 | 101 | 0 | 0 | 0 | 0 | 0 | 298 | 149 | 1 | 1 | 1 | 0.862 | 1 | 1 | 1 | 1 | 1 | 1 | 1 | 1 | -0.028 |
| SE | 77321 | ENSG00000242802 | AP5Z1 | chr7 | + | 4790213 | 4790591 | 4789831 | 4789929 | 4790672 | 4790887 | 18 | 5 | 26 | 20 | 6 | 0 | 0 | 0 | 0 | 0 | 38 | 30 | 13 | 28 | 2 | 0 | 0 | 0 | 2 | 0 | 298 | 149 | 1 | 1 | 1 | 1 | 1 | 1 | 1 | 1 | 1 | 1 | 0.875 | 1 | 0.025 |
| SE | 77322 | ENSG00000242802 | AP5Z1 | chr7 | + | 4790388 | 4790591 | 4789831 | 4789929 | 4790672 | 4790887 | 17 | 5 | 25 | 20 | 6 | 0 | 0 | 0 | 0 | 0 | 38 | 30 | 13 | 28 | 2 | 0 | 0 | 0 | 2 | 0 | 298 | 149 | 1 | 1 | 1 | 1 | 1 | 1 | 1 | 1 | 1 | 1 | 0.875 | 1 | 0.025 |
| SE | 77323 | ENSG00000242802 | AP5Z1 | chr7 | + | 4790458 | 4790591 | 4789831 | 4789929 | 4790672 | 4790886 | 18 | 5 | 35 | 23 | 7 | 0 | 0 | 0 | 0 | 0 | 44 | 34 | 18 | 42 | 4 | 0 | 0 | 0 | 2 | 0 | 282 | 149 | 1 | 1 | 1 | 1 | 1 | 1 | 1 | 1 | 1 | 1 | 0.917 | 1 | 0.017 |
| SE | 77333 | ENSG00000214022 | REPIN1 | chr7 | + | 150370759 | 150370884 | 150369670 | 150369868 | 150371230 | 150371632 | 7 | 9 | 2 | 11 | 1 | 7 | 2 | 6 | 11 | 8 | 0 | 1 | 0 | 0 | 4 | 0 | 7 | 9 | 4 | 7 | 274 | 149 | 1 | 1 | 0.352 | 0.71 | 0.153 | 0.352 | 0.064 | NA | 0.072 | 0 | 0 | 0.237 | 0.249 |
| SE | 77334 | ENSG00000189043 | NDUFA4 | chr7 | - | 10938089 | 10938147 | 10931942 | 10933692 | 10940015 | 10940153 | 328 | 138 | 327 | 272 | 563 | 1 | 0 | 0 | 0 | 0 | 221 | 234 | 159 | 200 | 340 | 0 | 0 | 0 | 0 | 0 | 207 | 149 | 1 | 1 | 0.996 | 1 | 1 | 1 | 1 | 1 | 1 | 1 | 1 | 1 | -0.001 |
| SE | 77346 | ENSG00000167380 | ZNF226 | chr19 | + | 44166704 | 44166882 | 44165587 | 44165807 | 44170034 | 44170095 | 2 | 0 | 1 | 0 | 0 | 33 | 34 | 41 | 30 | 23 | 0 | 5 | 0 | 0 | 0 | 20 | 15 | 17 | 28 | 18 | 298 | 149 | 1 | 1 | 0.029 | 0 | 0.012 | 0 | 0 | 0 | 0.143 | 0 | 0 | 0 | -0.02 |
| SE | 77347 | ENSG00000167380 | ZNF226 | chr19 | + | 44167818 | 44167860 | 44166704 | 44166882 | 44170034 | 44170095 | 1 | 0 | 0 | 0 | 0 | 0 | 0 | 1 | 0 | 0 | 0 | 0 | 0 | 0 | 0 | 0 | 5 | 1 | 0 | 0 | 191 | 149 | 1 | 1 | 1 | NA | 0 | NA | NA | NA | 0 | 0 | NA | NA | 0.5 |
| SE | 77353 | ENSG00000171316 | CHD7 | chr8 | + | 60780999 | 60781050 | 60741364 | 60743097 | 60794985 | 60795127 | 7 | 5 | 8 | 5 | 0 | 0 | 2 | 0 | 0 | 2 | 0 | 16 | 5 | 4 | 1 | 0 | 0 | 0 | 0 | 0 | 200 | 149 | 1 | 1 | 1 | 0.651 | 1 | 1 | 0 | NA | 1 | 1 | 1 | 1 | -0.27 |
| SE | 77354 | ENSG00000171316 | CHD7 | chr8 | + | 60780999 | 60781430 | 60741258 | 60743097 | 60794985 | 60795127 | 24 | 22 | 30 | 35 | 6 | 0 | 2 | 0 | 0 | 2 | 51 | 40 | 54 | 48 | 1 | 0 | 0 | 0 | 0 | 0 | 298 | 149 | 1 | 1 | 1 | 0.846 | 1 | 1 | 0.6 | 1 | 1 | 1 | 1 | 1 | -0.111 |
| SE | 77355 | ENSG00000171316 | CHD7 | chr8 | + | 60780999 | 60781430 | 60741258 | 60743097 | 60800387 | 60800525 | 13 | 11 | 13 | 18 | 0 | 0 | 1 | 1 | 0 | 2 | 0 | 21 | 19 | 14 | 1 | 0 | 0 | 0 | 0 | 0 | 298 | 149 | 1 | 1 | 1 | 0.846 | 0.867 | 1 | NA | NA | 1 | 1 | 1 | 1 | -0.072 |
| SE | 77356 | ENSG00000171316 | CHD7 | chr8 | + | 60794985 | 60795127 | 60741258 | 60743097 | 60800387 | 60800525 | 11 | 6 | 13 | 10 | 6 | 0 | 1 | 1 | 0 | 0 | 10 | 0 | 13 | 19 | 0 | 0 | 0 | 0 | 0 | 0 | 291 | 149 | 1 | 1 | 1 | 0.754 | 0.869 | 1 | 1 | 1 | NA | 1 | 1 | NA | -0.075 |
| SE | 77357 | ENSG00000171316 | CHD7 | chr8 | + | 60794985 | 60795127 | 60780999 | 60781430 | 60800387 | 60800525 | 22 | 15 | 31 | 27 | 10 | 0 | 0 | 1 | 0 | 0 | 61 | 19 | 49 | 53 | 0 | 0 | 0 | 1 | 0 | 0 | 291 | 149 | 1 | 1 | 1 | 1 | 0.941 | 1 | 1 | 1 | 1 | 0.962 | 1 | NA | -0.002 |
| SE | 77358 | ENSG00000171316 | CHD7 | chr8 | + | 60822502 | 60822746 | 60822023 | 60822145 | 60823839 | 60824016 | 56 | 48 | 93 | 71 | 124 | 0 | 0 | 0 | 1 | 0 | 47 | 71 | 89 | 85 | 3 | 0 | 0 | 0 | 0 | 0 | 298 | 149 | 1 | 1 | 1 | 1 | 1 | 0.973 | 1 | 1 | 1 | 1 | 1 | 1 | -0.005 |
| SE | 77359 | ENSG00000171316 | CHD7 | chr8 | + | 60837667 | 60837835 | 60836816 | 60837012 | 60838075 | 60838255 | 92 | 66 | 149 | 137 | 105 | 0 | 0 | 0 | 0 | 0 | 48 | 62 | 117 | 127 | 8 | 0 | 2 | 0 | 0 | 0 | 298 | 149 | 1 | 1 | 1 | 1 | 1 | 1 | 1 | 1 | 0.939 | 1 | 1 | 1 | 0.012 |
| SE | 77360 | ENSG00000171316 | CHD7 | chr8 | + | 60844863 | 60845063 | 60841846 | 60842052 | 60845249 | 60845409 | 41 | 51 | 78 | 48 | 99 | 0 | 0 | 0 | 0 | 0 | 56 | 44 | 133 | 58 | 0 | 0 | 0 | 5 | 0 | 0 | 298 | 149 | 1 | 1 | 1 | 1 | 1 | 1 | 1 | 1 | 1 | 0.93 | 1 | NA | 0.017 |
| SE | 77361 | ENSG00000171316 | CHD7 | chr8 | + | 60856444 | 60856888 | 60855974 | 60856202 | 60860903 | 60861125 | 56 | 42 | 87 | 101 | 125 | 0 | 0 | 0 | 0 | 3 | 67 | 73 | 109 | 137 | 22 | 0 | 0 | 0 | 0 | 0 | 298 | 149 | 1 | 1 | 1 | 1 | 1 | 1 | 0.954 | 1 | 1 | 1 | 1 | 1 | -0.009 |
| SE | 77366 | ENSG00000124422 | USP22 | chr17 | - | 21028541 | 21028674 | 21021112 | 21021226 | 21042664 | 21043011 | 199 | 130 | 207 | 246 | 143 | 3 | 0 | 0 | 0 | 2 | 421 | 221 | 185 | 175 | 568 | 0 | 0 | 2 | 0 | 4 | 282 | 149 | 1 | 1 | 0.972 | 1 | 1 | 1 | 0.974 | 1 | 1 | 0.98 | 1 | 0.987 | -0.004 |
| SE | 77367 | ENSG00000124422 | USP22 | chr17 | - | 21028541 | 21028674 | 21021112 | 21021226 | 21043320 | 21043375 | 145 | 98 | 165 | 192 | 124 | 1 | 0 | 0 | 0 | 0 | 343 | 193 | 154 | 160 | 424 | 0 | 0 | 0 | 0 | 0 | 282 | 149 | 1 | 1 | 0.987 | 1 | 1 | 1 | 1 | 1 | 1 | 1 | 1 | 1 | -0.003 |
| SE | 77372 | ENSG00000086827 | ZW10 | chr11 | - | 113739212 | 113739382 | 113738263 | 113738394 | 113741693 | 113741765 | 110 | 49 | 92 | 97 | 151 | 2 | 0 | 0 | 0 | 0 | 35 | 61 | 31 | 54 | 24 | 0 | 0 | 0 | 0 | 0 | 298 | 149 | 1 | 1 | 0.965 | 1 | 1 | 1 | 1 | 1 | 1 | 1 | 1 | 1 | -0.007 |
| SE | 77373 | ENSG00000086827 | ZW10 | chr11 | - | 113743801 | 113744040 | 113741693 | 113741765 | 113747530 | 113747713 | 69 | 37 | 51 | 65 | 100 | 5 | 0 | 0 | 1 | 1 | 33 | 32 | 16 | 32 | 0 | 0 | 0 | 0 | 3 | 0 | 298 | 149 | 1 | 1 | 0.873 | 1 | 1 | 0.97 | 0.98 | 1 | 1 | 1 | 0.842 | NA | 0.004 |
| SE | 77374 | ENSG00000086827 | ZW10 | chr11 | - | 113748256 | 113748420 | 113747530 | 113747713 | 113757661 | 113757853 | 101 | 61 | 73 | 120 | 179 | 0 | 1 | 0 | 4 | 0 | 33 | 44 | 40 | 54 | 0 | 0 | 0 | 0 | 0 | 0 | 298 | 149 | 1 | 1 | 1 | 0.968 | 1 | 0.938 | 1 | 1 | 1 | 1 | 1 | NA | -0.019 |
| SE | 77375 | ENSG00000086827 | ZW10 | chr11 | - | 113757661 | 113757853 | 113748256 | 113748420 | 113758553 | 113758706 | 80 | 56 | 57 | 112 | 131 | 1 | 0 | 0 | 0 | 0 | 14 | 19 | 29 | 36 | 0 | 0 | 0 | 0 | 0 | 0 | 298 | 149 | 1 | 1 | 0.976 | 1 | 1 | 1 | 1 | 1 | 1 | 1 | 1 | NA | -0.005 |
| SE | 77376 | ENSG00000086827 | ZW10 | chr11 | - | 113760208 | 113760368 | 113758553 | 113758706 | 113760512 | 113760590 | 98 | 58 | 66 | 52 | 120 | 1 | 0 | 0 | 0 | 0 | 46 | 45 | 38 | 42 | 26 | 0 | 0 | 0 | 0 | 0 | 298 | 149 | 1 | 1 | 0.98 | 1 | 1 | 1 | 1 | 1 | 1 | 1 | 1 | 1 | -0.004 |
| SE | 77378 | ENSG00000139154 | AEBP2 | chr12 | + | 19493799 | 19493986 | 19473247 | 19473355 | 19512397 | 19512465 | 314 | 242 | 274 | 330 | 472 | 0 | 0 | 1 | 0 | 0 | 142 | 207 | 213 | 113 | 117 | 0 | 0 | 0 | 0 | 0 | 298 | 149 | 1 | 1 | 1 | 1 | 0.993 | 1 | 1 | 1 | 1 | 1 | 1 | 1 | -0.001 |
| SE | 77379 | ENSG00000139154 | AEBP2 | chr12 | + | 19500096 | 19500221 | 19493799 | 19493986 | 19512397 | 19512465 | 379 | 266 | 337 | 386 | 432 | 2 | 0 | 1 | 0 | 1 | 161 | 215 | 290 | 134 | 107 | 0 | 0 | 0 | 0 | 0 | 274 | 149 | 1 | 1 | 0.99 | 1 | 0.995 | 1 | 0.996 | 1 | 1 | 1 | 1 | 1 | -0.004 |
| SE | 77380 | ENSG00000139154 | AEBP2 | chr12 | + | 19512397 | 19512465 | 19500092 | 19500221 | 19518086 | 19518117 | 67 | 71 | 66 | 81 | 69 | 0 | 0 | 0 | 1 | 0 | 16 | 28 | 64 | 18 | 9 | 0 | 0 | 0 | 0 | 0 | 217 | 149 | 1 | 1 | 1 | 1 | 1 | 0.982 | 1 | 1 | 1 | 1 | 1 | 1 | -0.004 |
| SE | 77388 | ENSG00000089820 | ARHGAP4 | chrX | - | 153912699 | 153912802 | 153911128 | 153911189 | 153913023 | 153913051 | 462 | 205 | 402 | 370 | 312 | 1 | 0 | 7 | 0 | 3 | 497 | 470 | 301 | 440 | 818 | 0 | 1 | 3 | 6 | 7 | 252 | 149 | 1 | 1 | 0.996 | 1 | 0.971 | 1 | 0.984 | 1 | 0.996 | 0.983 | 0.977 | 0.986 | 0.002 |
| SE | 77389 | ENSG00000089820 | ARHGAP4 | chrX | - | 153913217 | 153913302 | 153913023 | 153913051 | 153913408 | 153913600 | 295 | 144 | 284 | 264 | 149 | 1 | 0 | 0 | 0 | 0 | 251 | 231 | 189 | 271 | 452 | 0 | 0 | 0 | 0 | 0 | 234 | 149 | 1 | 1 | 0.995 | 1 | 1 | 1 | 1 | 1 | 1 | 1 | 1 | 1 | -0.001 |
| SE | 77390 | ENSG00000089820 | ARHGAP4 | chrX | - | 153913777 | 153913879 | 153913209 | 153913600 | 153918831 | 153919053 | 366 | 172 | 360 | 329 | 163 | 0 | 0 | 3 | 0 | 0 | 388 | 354 | 196 | 385 | 776 | 0 | 0 | 0 | 1 | 0 | 251 | 149 | 1 | 1 | 1 | 1 | 0.986 | 1 | 1 | 1 | 1 | 1 | 0.996 | 1 | -0.002 |
| SE | 77391 | ENSG00000089820 | ARHGAP4 | chrX | - | 153913777 | 153913879 | 153913209 | 153913600 | 153920625 | 153920631 | 111 | 77 | 113 | 104 | 104 | 0 | 0 | 3 | 0 | 0 | 83 | 77 | 49 | 122 | 149 | 0 | 0 | 0 | 0 | 0 | 251 | 149 | 1 | 1 | 1 | 1 | 0.957 | 1 | 1 | 1 | 1 | 1 | 1 | 1 | -0.009 |
| SE | 77392 | ENSG00000089820 | ARHGAP4 | chrX | - | 153913777 | 153913879 | 153913408 | 153913516 | 153918831 | 153919053 | 209 | 85 | 205 | 179 | 51 | 0 | 1 | 0 | 0 | 3 | 258 | 253 | 133 | 214 | 512 | 0 | 0 | 0 | 3 | 0 | 251 | 149 | 1 | 1 | 1 | 0.981 | 1 | 1 | 0.91 | 1 | 1 | 1 | 0.977 | 1 | -0.017 |
| SE | 77399 | ENSG00000089820 | ARHGAP4 | chrX | - | 153921096 | 153921159 | 153920411 | 153920808 | 153921364 | 153921400 | 78 | 21 | 80 | 68 | 7 | 0 | 1 | 1 | 0 | 0 | 82 | 59 | 54 | 116 | 121 | 0 | 0 | 0 | 0 | 0 | 212 | 149 | 1 | 1 | 1 | 0.937 | 0.983 | 1 | 1 | 1 | 1 | 1 | 1 | 1 | -0.016 |
| SE | 77400 | ENSG00000089820 | ARHGAP4 | chrX | - | 153921096 | 153921223 | 153920738 | 153920808 | 153921364 | 153921527 | 25 | 14 | 32 | 31 | 3 | 0 | 1 | 1 | 0 | 0 | 32 | 17 | 16 | 35 | 30 | 0 | 0 | 0 | 0 | 0 | 276 | 149 | 1 | 1 | 1 | 0.883 | 0.945 | 1 | 1 | 1 | 1 | 1 | 1 | 1 | -0.034 |
| SE | 77401 | ENSG00000089820 | ARHGAP4 | chrX | - | 153933942 | 153934001 | 153932538 | 153932584 | 153934376 | 153934475 | 160 | 98 | 147 | 123 | 255 | 0 | 0 | 0 | 1 | 0 | 186 | 176 | 102 | 181 | 327 | 0 | 0 | 0 | 0 | 0 | 208 | 149 | 1 | 1 | 1 | 1 | 1 | 0.989 | 1 | 1 | 1 | 1 | 1 | 1 | -0.002 |
| SE | 77402 | ENSG00000089820 | ARHGAP4 | chrX | - | 153933942 | 153934001 | 153932538 | 153932584 | 153934883 | 153934999 | 72 | 43 | 70 | 52 | 108 | 1 | 0 | 0 | 0 | 0 | 71 | 67 | 19 | 69 | 123 | 0 | 0 | 0 | 0 | 0 | 208 | 149 | 1 | 1 | 0.981 | 1 | 1 | 1 | 1 | 1 | 1 | 1 | 1 | 1 | -0.004 |
| SE | 77403 | ENSG00000089820 | ARHGAP4 | chrX | - | 153934376 | 153934475 | 153932538 | 153932584 | 153934883 | 153934999 | 11 | 5 | 11 | 12 | 1 | 1 | 0 | 0 | 0 | 0 | 9 | 6 | 11 | 11 | 25 | 0 | 0 | 0 | 0 | 0 | 248 | 149 | 1 | 1 | 0.869 | 1 | 1 | 1 | 1 | 1 | 1 | 1 | 1 | 1 | -0.026 |
| SE | 77404 | ENSG00000089820 | ARHGAP4 | chrX | - | 153934376 | 153934475 | 153933942 | 153934001 | 153934883 | 153934999 | 106 | 64 | 95 | 83 | 155 | 0 | 0 | 2 | 0 | 0 | 131 | 116 | 102 | 135 | 245 | 0 | 0 | 0 | 0 | 0 | 248 | 149 | 1 | 1 | 1 | 1 | 0.966 | 1 | 1 | 1 | 1 | 1 | 1 | 1 | -0.007 |
| SE | 77405 | ENSG00000105963 | ADAP1 | chr7 | - | 900532 | 900613 | 900101 | 900164 | 904125 | 904272 | 34 | 20 | 17 | 27 | 14 | 2 | 0 | 0 | 1 | 1 | 73 | 31 | 13 | 40 | 32 | 0 | 1 | 3 | 5 | 12 | 230 | 149 | 1 | 1 | 0.917 | 1 | 1 | 0.946 | 0.901 | 1 | 0.953 | 0.737 | 0.838 | 0.633 | 0.121 |
| SE | 77408 | ENSG00000164077 | MON1A | chr3 | - | 49910118 | 49910884 | 49909252 | 49909400 | 49913219 | 49913359 | 12 | 2 | 3 | 2 | 4 | 0 | 0 | 0 | 2 | 2 | 0 | 15 | 0 | 0 | 5 | 0 | 0 | 0 | 0 | 0 | 298 | 149 | 1 | 1 | 1 | 1 | 1 | 0.333 | 0.5 | NA | 1 | NA | NA | 1 | -0.233 |
| SE | 77409 | ENSG00000164077 | MON1A | chr3 | - | 49911525 | 49912011 | 49910118 | 49910884 | 49913219 | 49913359 | 34 | 16 | 17 | 25 | 31 | 1 | 0 | 0 | 1 | 4 | 0 | 19 | 1 | 15 | 5 | 0 | 1 | 0 | 0 | 0 | 298 | 149 | 1 | 1 | 0.944 | 1 | 1 | 0.926 | 0.795 | NA | 0.905 | 1 | 1 | 1 | -0.043 |
| SE | 77421 | ENSG00000047648 | ARHGAP6 | chrX | - | 11143979 | 11144248 | 11142232 | 11142313 | 11156528 | 11156626 | 23 | 63 | 7 | 73 | 65 | 1 | 0 | 0 | 0 | 0 | 45 | 39 | 39 | 47 | 262 | 0 | 0 | 0 | 0 | 0 | 298 | 149 | 1 | 1 | 0.92 | 1 | 1 | 1 | 1 | 1 | 1 | 1 | 1 | 1 | -0.016 |
| SE | 77423 | ENSG00000047648 | ARHGAP6 | chrX | - | 11148629 | 11148719 | 11142232 | 11144245 | 11156528 | 11156626 | 0 | 0 | 0 | 1 | 7 | 0 | 0 | 0 | 0 | 1 | 0 | 0 | 1 | 0 | 8 | 0 | 0 | 0 | 0 | 0 | 239 | 149 | 1 | 1 | NA | NA | NA | 1 | 0.814 | NA | NA | 1 | NA | 1 | -0.093 |
| SE | 77425 | ENSG00000047648 | ARHGAP6 | chrX | - | 11179301 | 11179452 | 11178099 | 11178248 | 11182062 | 11182074 | 49 | 72 | 21 | 151 | 91 | 0 | 0 | 0 | 0 | 0 | 168 | 104 | 67 | 84 | 448 | 0 | 0 | 1 | 0 | 0 | 298 | 149 | 1 | 1 | 1 | 1 | 1 | 1 | 1 | 1 | 1 | 0.971 | 1 | 1 | 0.006 |
| SE | 77427 | ENSG00000071189 | SNX13 | chr7 | - | 17796826 | 17796939 | 17790760 | 17794292 | 17798689 | 17798758 | 112 | 92 | 115 | 115 | 141 | 0 | 3 | 0 | 2 | 5 | 0 | 51 | 62 | 2 | 0 | 0 | 0 | 0 | 0 | 0 | 262 | 149 | 1 | 1 | 1 | 0.946 | 1 | 0.97 | 0.941 | NA | 1 | 1 | 1 | NA | -0.029 |
| SE | 77429 | ENSG00000071189 | SNX13 | chr7 | - | 17821508 | 17821648 | 17816181 | 17816289 | 17826021 | 17826091 | 160 | 191 | 163 | 168 | 227 | 0 | 1 | 0 | 0 | 0 | 9 | 69 | 79 | 18 | 9 | 0 | 0 | 0 | 0 | 0 | 289 | 149 | 1 | 1 | 1 | 0.99 | 1 | 1 | 1 | 1 | 1 | 1 | 1 | 1 | -0.002 |
| SE | 77430 | ENSG00000071189 | SNX13 | chr7 | - | 17821508 | 17821681 | 17816181 | 17816289 | 17826021 | 17826091 | 117 | 144 | 129 | 124 | 180 | 0 | 1 | 0 | 0 | 0 | 9 | 48 | 56 | 8 | 0 | 0 | 0 | 0 | 0 | 0 | 298 | 149 | 1 | 1 | 1 | 0.986 | 1 | 1 | 1 | 1 | 1 | 1 | 1 | NA | -0.003 |
| SE | 77431 | ENSG00000071189 | SNX13 | chr7 | - | 17830009 | 17830047 | 17826021 | 17826091 | 17834051 | 17834184 | 71 | 72 | 56 | 59 | 76 | 0 | 0 | 0 | 6 | 11 | 0 | 23 | 35 | 6 | 9 | 0 | 4 | 0 | 0 | 0 | 187 | 149 | 1 | 1 | 1 | 1 | 1 | 0.887 | 0.846 | NA | 0.821 | 1 | 1 | 1 | -0.009 |
| SE | 77432 | ENSG00000071189 | SNX13 | chr7 | - | 17834760 | 17834865 | 17834051 | 17834184 | 17839806 | 17840000 | 140 | 70 | 87 | 92 | 139 | 0 | 1 | 0 | 0 | 1 | 0 | 46 | 50 | 13 | 13 | 0 | 0 | 0 | 0 | 0 | 254 | 149 | 1 | 1 | 1 | 0.976 | 1 | 1 | 0.988 | NA | 1 | 1 | 1 | 1 | -0.007 |
| SE | 77433 | ENSG00000071189 | SNX13 | chr7 | - | 17845594 | 17845694 | 17839806 | 17840000 | 17868406 | 17868490 | 159 | 110 | 122 | 119 | 259 | 3 | 1 | 0 | 0 | 0 | 10 | 76 | 49 | 20 | 2 | 0 | 0 | 0 | 0 | 0 | 249 | 149 | 1 | 1 | 0.969 | 0.985 | 1 | 1 | 1 | 1 | 1 | 1 | 1 | 1 | -0.009 |
| SE | 77434 | ENSG00000071189 | SNX13 | chr7 | - | 17850346 | 17850435 | 17845594 | 17845694 | 17850825 | 17850964 | 275 | 229 | 186 | 209 | 439 | 3 | 1 | 0 | 1 | 0 | 32 | 121 | 127 | 29 | 44 | 0 | 0 | 0 | 0 | 0 | 238 | 149 | 1 | 1 | 0.983 | 0.993 | 1 | 0.992 | 1 | 1 | 1 | 1 | 1 | 1 | -0.006 |
| SE | 77435 | ENSG00000071189 | SNX13 | chr7 | - | 17850825 | 17850964 | 17845594 | 17845694 | 17868406 | 17868490 | 90 | 78 | 137 | 176 | 132 | 0 | 0 | 0 | 1 | 0 | 15 | 78 | 50 | 67 | 9 | 0 | 0 | 0 | 0 | 0 | 288 | 149 | 1 | 1 | 1 | 1 | 1 | 0.989 | 1 | 1 | 1 | 1 | 1 | 1 | -0.002 |
| SE | 77436 | ENSG00000071189 | SNX13 | chr7 | - | 17875479 | 17875581 | 17873527 | 17873616 | 17875668 | 17875790 | 69 | 49 | 168 | 257 | 220 | 0 | 0 | 2 | 0 | 1 | 125 | 99 | 102 | 98 | 155 | 0 | 0 | 0 | 0 | 0 | 251 | 149 | 1 | 1 | 1 | 1 | 0.98 | 1 | 0.992 | 1 | 1 | 1 | 1 | 1 | -0.006 |
| SE | 77439 | ENSG00000071189 | SNX13 | chr7 | - | 17897333 | 17897446 | 17893331 | 17893434 | 17940283 | 17940451 | 161 | 124 | 96 | 161 | 196 | 0 | 0 | 0 | 0 | 0 | 55 | 96 | 63 | 26 | 38 | 0 | 0 | 4 | 0 | 0 | 262 | 149 | 1 | 1 | 1 | 1 | 1 | 1 | 1 | 1 | 1 | 0.9 | 1 | 1 | 0.02 |
| SE | 77440 | ENSG00000071189 | SNX13 | chr7 | - | 17897843 | 17897884 | 17897333 | 17897446 | 17907248 | 17907340 | 3 | 1 | 1 | 0 | 0 | 1 | 0 | 1 | 1 | 0 | 10 | 3 | 3 | 0 | 0 | 0 | 0 | 0 | 0 | 0 | 190 | 149 | 1 | 1 | 0.702 | 1 | 0.44 | 0 | NA | 1 | 1 | 1 | NA | NA | -0.465 |
| SE | 77441 | ENSG00000071189 | SNX13 | chr7 | - | 17897843 | 17897884 | 17897333 | 17897446 | 17940283 | 17940431 | 1 | 1 | 1 | 0 | 1 | 51 | 31 | 29 | 62 | 45 | 0 | 0 | 2 | 0 | 0 | 6 | 36 | 13 | 1 | 0 | 190 | 149 | 1 | 1 | 0.015 | 0.025 | 0.026 | 0 | 0.017 | 0 | 0 | 0.108 | 0 | NA | -0.01 |
| SE | 77443 | ENSG00000071189 | SNX13 | chr7 | - | 17907248 | 17907340 | 17897333 | 17897446 | 17940283 | 17940488 | 1 | 0 | 1 | 2 | 0 | 51 | 31 | 29 | 62 | 45 | 0 | 2 | 0 | 0 | 0 | 6 | 36 | 13 | 1 | 0 | 241 | 149 | 1 | 1 | 0.012 | 0 | 0.021 | 0.02 | 0 | 0 | 0.033 | 0 | 0 | NA | 0.002 |
| SE | 77444 | ENSG00000071189 | SNX13 | chr7 | - | 17907248 | 17907340 | 17897843 | 17897884 | 17940283 | 17940488 | 2 | 0 | 0 | 1 | 0 | 0 | 0 | 0 | 0 | 1 | 10 | 2 | 2 | 0 | 0 | 0 | 0 | 1 | 0 | 0 | 241 | 149 | 1 | 1 | 1 | NA | NA | 1 | 0 | 1 | 1 | 0.553 | NA | NA | -0.184 |
| SE | 77447 | ENSG00000122550 | KLHL7 | chr7 | + | 23117833 | 23117964 | 23106297 | 23106794 | 23123776 | 23123879 | 4 | 2 | 13 | 0 | 11 | 3 | 0 | 0 | 0 | 0 | 7 | 9 | 0 | 0 | 26 | 0 | 0 | 0 | 0 | 0 | 280 | 149 | 1 | 1 | 0.415 | 1 | 1 | NA | 1 | 1 | 1 | NA | NA | 1 | -0.146 |
| SE | 77450 | ENSG00000122550 | KLHL7 | chr7 | + | 23143850 | 23144025 | 23140768 | 23140944 | 23152066 | 23152209 | 75 | 33 | 46 | 84 | 88 | 1 | 4 | 0 | 0 | 0 | 9 | 62 | 19 | 11 | 6 | 0 | 0 | 0 | 0 | 0 | 298 | 149 | 1 | 1 | 0.974 | 0.805 | | 1 | 1 | 1 | 1 | 1 | 1 | 1 | -0.044 |
| SE | 77451 | ENSG00000122550 | KLHL7 | chr7 | + | 23172947 | 23173045 | 23167835 | 23168037 | 23174014 | 23175302 | 59 | 48 | 38 | 46 | 40 | 0 | 0 | 0 | 0 | 0 | 0 | 22 | 7 | 33 | 3 | 0 | 1 | 0 | 0 | 0 | 247 | 149 | 1 | 1 | 1 | 1 | 1 | 1 | 1 | NA | 0.93 | 1 | 1 | 1 | 0.017 |
| SE | 77458 | ENSG00000106100 | NOD1 | chr7 | - | 30435997 | 30436081 | 30429373 | 30429457 | 30437572 | 30437656 | 19 | 15 | 21 | 12 | 26 | 0 | 0 | 0 | 2 | 0 | 37 | 21 | 14 | 30 | 20 | 0 | 0 | 0 | 0 | 0 | 233 | 149 | 1 | 1 | 1 | 1 | 1 | 0.793 | 1 | 1 | 1 | 1 | 1 | 1 | -0.041 |
| SE | 77459 | ENSG00000106100 | NOD1 | chr7 | - | 30435997 | 30436081 | 30429373 | 30429457 | 30446140 | 30446224 | 3 | 2 | 4 | 3 | 10 | 13 | 0 | 1 | 2 | 1 | 0 | 2 | 2 | 12 | 2 | 0 | 0 | 0 | 0 | 0 | 233 | 149 | 1 | 1 | 0.129 | 1 | 0.719 | 0.49 | 0.865 | NA | 1 | 1 | 1 | 1 | -0.359 |
| SE | 77463 | ENSG00000106100 | NOD1 | chr7 | - | 30459151 | 30459240 | 30456720 | 30457042 | 30459900 | 30460041 | 78 | 81 | 68 | 82 | 127 | 0 | 0 | 3 | 0 | 0 | 194 | 143 | 67 | 93 | 137 | 0 | 0 | 0 | 0 | 0 | 238 | 149 | 1 | 1 | 1 | 1 | 0.934 | 1 | 1 | 1 | 1 | 1 | 1 | 1 | -0.013 |
| SE | 77464 | ENSG00000106100 | NOD1 | chr7 | - | 30463608 | 30463698 | 30459900 | 30460041 | 30475847 | 30475988 | 6 | 0 | 6 | 3 | 8 | 2 | 0 | 0 | 0 | 2 | 7 | 6 | 4 | 7 | 6 | 8 | 2 | 0 | 0 | 0 | 239 | 149 | 1 | 1 | 0.652 | NA | 1 | 1 | 0.714 | 0.353 | 0.652 | 1 | 1 | 1 | 0.041 |
| SE | 77466 | ENSG00000106100 | NOD1 | chr7 | - | 30463985 | 30464061 | 30459900 | 30460041 | 30478605 | 30478636 | 0 | 0 | 0 | 0 | 0 | 9 | 7 | 5 | 6 | 8 | 0 | 0 | 0 | 2 | 0 | 23 | 26 | 3 | 5 | 0 | 225 | 149 | 1 | 1 | 0 | 0 | 0 | 0 | 0 | 0 | 0 | 0 | 0.209 | NA | -0.052 |
| SE | 77467 | ENSG00000106100 | NOD1 | chr7 | - | 30475847 | 30475988 | 30459900 | 30460041 | 30478605 | 30478728 | 4 | 0 | 0 | 0 | 2 | 9 | 7 | 5 | 6 | 8 | 8 | 2 | 0 | 0 | 0 | 23 | 26 | 3 | 5 | 0 | 290 | 149 | 1 | 1 | 0.186 | 0 | 0 | 0 | 0.114 | 0.152 | 0.038 | 0 | 0 | NA | 0.012 |
| SE | 77468 | ENSG00000106100 | NOD1 | chr7 | - | 30475847 | 30475988 | 30463608 | 30463698 | 30478605 | 30478728 | 3 | 0 | 1 | 0 | 0 | 2 | 0 | 2 | 3 | 4 | 0 | 0 | 0 | 0 | 0 | 16 | 1 | 3 | 5 | 6 | 290 | 149 | 1 | 1 | 0.435 | NA | 0.204 | 0 | 0 | 0 | 0 | 0 | 0 | 0 | 0.16 |
| SE | 77471 | ENSG00000008405 | CRY1 | chr12 | - | 107049770 | 107049852 | 107022083 | 107022192 | 107092803 | 107093549 | 3 | 0 | 0 | 0 | 0 | 40 | 16 | 32 | 22 | 19 | 0 | 0 | 0 | 0 | 0 | 0 | 8 | 19 | 30 | 16 | 231 | 149 | 1 | 1 | 0.046 | 0 | 0 | 0 | 0 | NA | 0 | 0 | 0 | 0 | 0.009 |
| SE | 77472 | ENSG00000229358 | DPY19L1P1 | chr7 | - | 32590878 | 32591004 | 32585976 | 32586136 | 32593183 | 32593306 | 1 | 0 | 0 | 1 | 0 | 3 | 1 | 0 | 0 | 0 | 0 | 0 | 0 | 17 | 0 | 0 | 0 | 0 | 1 | 0 | 275 | 149 | 1 | 1 | 0.153 | 0 | NA | 1 | NA | NA | NA | NA | 0.902 | NA | -0.518 |
| SE | 77473 | ENSG00000229358 | DPY19L1P1 | chr7 | - | 32590878 | 32591004 | 32585976 | 32586136 | 32620403 | 32620484 | 1 | 0 | 0 | 0 | 0 | 0 | 0 | 0 | 0 | 0 | 0 | 0 | 0 | 9 | 0 | 0 | 0 | 0 | 8 | 0 | 275 | 149 | 1 | 1 | 1 | NA | NA | NA | NA | NA | NA | NA | 0.379 | NA | 0.621 |
| SE | 77474 | ENSG00000229358 | DPY19L1P1 | chr7 | - | 32593183 | 32593306 | 32585976 | 32586136 | 32620403 | 32620484 | 3 | 1 | 1 | 0 | 0 | 0 | 0 | 0 | 0 | 0 | 0 | 0 | 0 | 4 | 0 | 0 | 0 | 0 | 8 | 0 | 272 | 149 | 1 | 1 | 1 | 1 | 1 | NA | NA | NA | NA | NA | 0.215 | NA | 0.785 |
| SE | 77475 | ENSG00000229358 | DPY19L1P1 | chr7 | - | 32593183 | 32593306 | 32590878 | 32591004 | 32620403 | 32620484 | 1 | 0 | 1 | 0 | 0 | 1 | 0 | 0 | 0 | 0 | 0 | 0 | 0 | 14 | 0 | 0 | 0 | 0 | 0 | 0 | 272 | 149 | 1 | 1 | 0.354 | NA | 1 | NA | NA | NA | NA | NA | 1 | NA | -0.323 |
| SE | 77476 | ENSG00000229358 | DPY19L1P1 | chr7 | - | 32623360 | 32623439 | 32621012 | 32621072 | 32632542 | 32632620 | 0 | 0 | 3 | 2 | 5 | 0 | 1 | 0 | 1 | 0 | 0 | 3 | 0 | 32 | 14 | 0 | 1 | 0 | 7 | 0 | 228 | 149 | 1 | 1 | NA | 0 | 1 | 1 | 0.766 | NA | 0.662 | NA | 0.749 | 1 | -0.112 |
| SE | 77477 | ENSG00000229358 | DPY19L1P1 | chr7 | - | 32639273 | 32639365 | 32635251 | 32635351 | 32655986 | 32656044 | 8 | 6 | 10 | 21 | 14 | 0 | 0 | 0 | 0 | 2 | 11 | 34 | 15 | 105 | 97 | 0 | 0 | 2 | 0 | 0 | 241 | 149 | 1 | 1 | 1 | 1 | 1 | 1 | 0.812 | 1 | 1 | 0.823 | 1 | 1 | -0.002 |
| SE | 77478 | ENSG00000229358 | DPY19L1P1 | chr7 | - | 32716717 | 32716809 | 32678992 | 32679130 | 32719135 | 32719193 | 2 | 0 | 1 | 1 | 3 | 0 | 0 | 0 | 0 | 0 | 0 | 7 | 1 | 0 | 4 | 0 | 0 | 0 | 1 | 0 | 241 | 149 | 1 | 1 | 1 | NA | 1 | 1 | 1 | NA | 1 | 1 | 0 | 1 | 0.25 |
| SE | 77479 | ENSG00000227240 | NA | chr1 | + | 193886320 | 193886417 | 193673209 | 193673256 | 193899286 | 193899367 | 2 | 5 | 0 | 1 | 1 | 0 | 0 | 0 | 2 | 0 | 0 | 3 | 2 | 0 | 0 | 0 | 0 | 0 | 0 | 0 | 246 | 149 | 1 | 1 | 1 | 1 | NA | 0.232 | 1 | NA | 1 | 1 | NA | NA | -0.192 |
| SE | 77481 | ENSG00000227240 | NA | chr1 | + | 194149702 | 194149856 | 194126139 | 194126185 | 194197522 | 194197587 | 1 | 3 | 3 | 0 | 0 | 0 | 0 | 0 | 1 | 0 | 0 | 0 | 0 | 5 | 0 | 0 | 0 | 0 | 0 | 0 | 298 | 149 | 1 | 1 | 1 | 1 | 1 | 0 | NA | NA | NA | NA | 1 | NA | -0.25 |
| SE | 77482 | ENSG00000127540 | UQCR11 | chr19 | - | 1602293 | 1602426 | 1599411 | 1599560 | 1605359 | 1605443 | 0 | 0 | 0 | 3 | 0 | 100 | 31 | 80 | 67 | 53 | 0 | 1 | 0 | 0 | 0 | 31 | 79 | 19 | 59 | 133 | 282 | 149 | 1 | 1 | 0 | 0 | 0 | 0.023 | 0 | 0 | 0.007 | 0 | 0 | 0 | 0.003 |
| SE | 77489 | ENSG00000164620 | RELL2 | chr5 | + | 141639463 | 141639649 | 141638794 | 141638860 | 141639919 | 141640295 | 25 | 11 | 26 | 34 | 22 | 0 | 0 | 0 | 0 | 0 | 33 | 33 | 29 | 26 | 62 | 0 | 3 | 0 | 0 | 0 | 298 | 149 | 1 | 1 | 1 | 1 | 1 | 1 | 1 | 1 | 0.846 | 1 | 1 | 1 | 0.031 |
| SE | 77490 | ENSG00000164620 | RELL2 | chr5 | + | 141639919 | 141640295 | 141638794 | 141638860 | 141640411 | 141640445 | 25 | 0 | 14 | 28 | 15 | 0 | 0 | 0 | 0 | 0 | 5 | 55 | 25 | 10 | 36 | 0 | 0 | 0 | 0 | 6 | 298 | 149 | 1 | 1 | 1 | NA | 1 | 1 | 1 | 1 | 1 | 1 | 1 | 0.75 | 0.05 |
| SE | 77494 | ENSG00000160799 | CCDC12 | chr3 | - | 46923328 | 46923363 | 46922235 | 46922312 | 46923606 | 46923668 | 253 | 142 | 226 | 183 | 219 | 0 | 0 | 1 | 0 | 0 | 97 | 155 | 151 | 200 | 316 | 0 | 0 | 0 | 0 | 0 | 184 | 149 | 1 | 1 | 1 | 1 | 0.995 | 1 | 1 | 1 | 1 | 1 | 1 | 1 | -0.001 |
| SE | 77496 | ENSG00000160799 | CCDC12 | chr3 | - | 46940997 | 46941065 | 46924919 | 46925535 | 46976636 | 46976745 | 86 | 63 | 105 | 82 | 156 | 0 | 0 | 1 | 3 | 0 | 214 | 112 | 104 | 207 | 220 | 0 | 0 | 0 | 0 | 0 | 217 | 149 | 1 | 1 | 1 | 1 | 0.986 | 0.949 | 1 | 1 | 1 | 1 | 1 | 1 | -0.013 |
| SE | 77499 | ENSG00000106415 | GLCCI1 | chr7 | + | 8022482 | 8022569 | 8003896 | 8004059 | 8055435 | 8055549 | 71 | 43 | 99 | 65 | 90 | 0 | 0 | 0 | 2 | 0 | 71 | 91 | 65 | 57 | 40 | 0 | 0 | 0 | 0 | 0 | 236 | 149 | 1 | 1 | 1 | 1 | 1 | 0.954 | 1 | 1 | 1 | 1 | 1 | 1 | -0.009 |
| SE | 77503 | ENSG00000106415 | GLCCI1 | chr7 | + | 8055432 | 8055549 | 8022482 | 8022569 | 8060095 | 8060109 | 140 | 97 | 211 | 127 | 198 | 0 | 0 | 0 | 0 | 0 | 69 | 181 | 111 | 80 | 73 | 0 | 0 | 0 | 0 | 3 | 266 | 149 | 1 | 1 | 1 | 1 | 1 | 1 | 1 | 1 | 1 | 1 | 1 | 0.932 | 0.014 |
| SE | 77505 | ENSG00000106415 | GLCCI1 | chr7 | + | 8055435 | 8055549 | 8022482 | 8022569 | 8060095 | 8060232 | 149 | 100 | 218 | 142 | 198 | 0 | 0 | 0 | 0 | 0 | 73 | 184 | 113 | 103 | 102 | 0 | 0 | 0 | 0 | 3 | 263 | 149 | 1 | 1 | 1 | 1 | 1 | 1 | 1 | 1 | 1 | 1 | 1 | 0.951 | 0.01 |
| SE | 77506 | ENSG00000106415 | GLCCI1 | chr7 | + | 8060095 | 8060248 | 8022482 | 8022569 | 8084896 | 8085017 | 0 | 0 | 0 | 0 | 3 | 1 | 0 | 0 | 0 | 0 | 0 | 3 | 0 | 0 | 3 | 0 | 0 | 0 | 0 | 0 | 298 | 149 | 1 | 1 | 0 | NA | NA | NA | 1 | NA | 1 | NA | NA | 1 | -0.5 |
| SE | 77507 | ENSG00000106415 | GLCCI1 | chr7 | + | 8070920 | 8071131 | 8060095 | 8060248 | 8084896 | 8085017 | 133 | 85 | 163 | 111 | 234 | 0 | 0 | 0 | 0 | 3 | 80 | 139 | 88 | 84 | 37 | 0 | 3 | 0 | 0 | 0 | 298 | 149 | 1 | 1 | 1 | 1 | 1 | 1 | 0.975 | 1 | 0.959 | 1 | 1 | 1 | 0.003 |
| SE | 77508 | ENSG00000106415 | GLCCI1 | chr7 | + | 8070920 | 8071131 | 8060095 | 8060248 | 8086192 | 8086939 | 105 | 59 | 125 | 81 | 163 | 0 | 1 | 0 | 0 | 0 | 78 | 122 | 50 | 70 | 19 | 0 | 0 | 0 | 2 | 0 | 298 | 149 | 1 | 1 | 1 | 0.967 | 1 | 1 | 1 | 1 | 1 | 1 | 0.946 | 1 | 0.004 |
| SE | 77509 | ENSG00000106415 | GLCCI1 | chr7 | + | 8084896 | 8085017 | 8060095 | 8060248 | 8086192 | 8089080 | 52 | 44 | 62 | 51 | 109 | 0 | 1 | 0 | 0 | 0 | 1 | 66 | 34 | 31 | 26 | 0 | 0 | 0 | 2 | 0 | 270 | 149 | 1 | 1 | 1 | 0.96 | 1 | 1 | 1 | 1 | 1 | 1 | 0.895 | 1 | 0.013 |
| SE | 77511 | ENSG00000136243 | NUP42 | chr7 | + | 23185069 | 23185298 | 23182023 | 23182206 | 23187051 | 23187146 | 109 | 83 | 100 | 115 | 70 | 0 | 0 | 1 | 1 | 0 | 51 | 88 | 52 | 57 | 55 | 0 | 0 | 0 | 4 | 0 | 298 | 149 | 1 | 1 | 1 | 1 | 0.98 | 0.983 | 1 | 1 | 1 | 1 | 0.877 | 1 | 0.017 |
| SE | 77513 | ENSG00000136243 | NUP42 | chr7 | + | 23196679 | 23196766 | 23187051 | 23187146 | 23199457 | 23199542 | 18 | 13 | 10 | 14 | 38 | 4 | 0 | 0 | 0 | 1 | 0 | 3 | 4 | 8 | 0 | 0 | 0 | 0 | 0 | 0 | 236 | 149 | 1 | 1 | 0.74 | 1 | 1 | 1 | 0.96 | NA | 1 | 1 | 1 | NA | -0.06 |
| SE | 77519 | ENSG00000109339 | MAPK10 | chr4 | - | 86067772 | 86067955 | 86064265 | 86064390 | 86159297 | 86159467 | 0 | 0 | 1 | 0 | 0 | 0 | 0 | 0 | 0 | 1 | 6 | 3 | 0 | 2 | 0 | 0 | 0 | 0 | 0 | 0 | 298 | 149 | 1 | 1 | NA | NA | 1 | NA | 0 | 1 | 1 | NA | 1 | NA | -0.5 |
| SE | 77526 | ENSG00000109339 | MAPK10 | chr4 | - | 86103185 | 86103244 | 86101893 | 86102032 | 86107222 | 86107243 | 4 | 1 | 0 | 4 | 0 | 0 | 0 | 0 | 2 | 0 | 18 | 7 | 0 | 0 | 8 | 7 | 7 | 0 | 0 | 5 | 208 | 149 | 1 | 1 | 1 | 1 | NA | 0.589 | NA | 0.648 | 0.417 | NA | NA | 0.534 | 0.33 |
| SE | 77531 | ENSG00000109339 | MAPK10 | chr4 | - | 86152462 | 86152629 | 86107222 | 86107352 | 86159297 | 86159467 | 1 | 0 | 0 | 1 | 0 | 0 | 0 | 0 | 0 | 0 | 10 | 6 | 0 | 0 | 19 | 7 | 2 | 0 | 0 | 1 | 298 | 149 | 1 | 1 | 1 | NA | NA | 1 | NA | 0.417 | 0.6 | NA | NA | 0.905 | 0.359 |
| SE | 77533 | ENSG00000109339 | MAPK10 | chr4 | - | 86152462 | 86152629 | 86151990 | 86152196 | 86159297 | 86159467 | 3 | 0 | 0 | 2 | 0 | 0 | 0 | 0 | 0 | 0 | 10 | 7 | 0 | 0 | 19 | 0 | 0 | 0 | 0 | 5 | 298 | 149 | 1 | 1 | 1 | NA | NA | 1 | NA | 1 | 1 | NA | NA | 0.655 | 0.115 |
| SE | 77535 | ENSG00000109339 | MAPK10 | chr4 | - | 86159297 | 86159467 | 86107222 | 86107352 | 86194335 | 86194407 | 3 | 0 | 0 | 0 | 0 | 0 | 0 | 0 | 0 | 0 | 30 | 5 | 0 | 0 | 9 | 0 | 0 | 0 | 0 | 5 | 298 | 149 | 1 | 1 | 1 | NA | NA | NA | NA | 1 | 1 | NA | NA | 0.474 | 0.175 |
| SE | 77539 | ENSG00000109339 | MAPK10 | chr4 | - | 86194308 | 86194407 | 86107222 | 86107352 | 86325901 | 86326557 | 2 | 0 | 0 | 0 | 0 | 0 | 0 | 0 | 0 | 0 | 15 | 4 | 0 | 0 | 0 | 0 | 1 | 0 | 0 | 0 | 248 | 149 | 1 | 1 | 1 | NA | NA | NA | NA | 1 | 0.706 | NA | NA | NA | 0.147 |
| SE | 77541 | ENSG00000109339 | MAPK10 | chr4 | - | 86194308 | 86194407 | 86159049 | 86159467 | 86325901 | 86326557 | 3 | 1 | 0 | 1 | 0 | 0 | 0 | 0 | 0 | 0 | 23 | 8 | 0 | 0 | 27 | 0 | 3 | 0 | 0 | 0 | 248 | 149 | 1 | 1 | 1 | 1 | NA | 1 | NA | 1 | 0.616 | NA | NA | 1 | 0.128 |
| SE | 77552 | ENSG00000109339 | MAPK10 | chr4 | - | 86321843 | 86322029 | 86194308 | 86194407 | 86325901 | 86326557 | 2 | 0 | 2 | 1 | 3 | 4 | 0 | 0 | 0 | 0 | 0 | 0 | 0 | 2 | 4 | 23 | 4 | 0 | 0 | 4 | 298 | 149 | 1 | 1 | 0.2 | NA | 1 | | 1 | 0 | 0 | NA | 1 | 0.333 | 0.467 |
| SE | 77592 | ENSG00000127947 | PTPN12 | chr7 | + | 77538764 | 77538845 | 77537553 | 77537645 | 77571077 | 77571186 | 8 | 2 | 1 | 2 | 2 | 336 | 272 | 173 | 314 | 248 | 0 | 0 | 8 | 0 | 0 | 188 | 284 | 320 | 206 | 134 | 230 | 149 | 1 | 1 | 0.015 | 0.005 | 0.004 | 0.004 | 0.005 | 0 | 0 | 0.016 | 0 | 0 | 0.003 |
| SE | 77593 | ENSG00000127947 | PTPN12 | chr7 | + | 77538764 | 77538845 | 77538028 | 77538106 | 77571077 | 77571186 | 4 | 2 | 1 | 2 | 1 | 6 | 0 | 0 | 3 | 0 | 0 | 0 | 4 | 0 | 0 | 0 | 0 | 0 | 0 | 0 | 230 | 149 | 1 | 1 | 0.302 | 1 | 1 | 0.302 | 1 | NA | NA | 1 | NA | NA | -0.279 |
| SE | 77596 | ENSG00000127947 | PTPN12 | chr7 | + | 77571077 | 77571186 | 77537294 | 77537645 | 77585542 | 77585581 | 245 | 214 | 130 | 222 | 192 | 1 | 0 | 0 | 0 | 0 | 146 | 232 | 249 | 153 | 112 | 0 | 0 | 0 | 0 | 0 | 258 | 149 | 1 | 1 | 0.993 | 1 | 1 | 1 | 1 | 1 | 1 | 1 | 1 | 1 | -0.001 |
| SE | 77597 | ENSG00000127947 | PTPN12 | chr7 | + | 77571077 | 77571186 | 77537294 | 77537645 | 77592184 | 77592256 | 243 | 212 | 127 | 220 | 192 | 0 | 2 | 0 | 0 | 0 | 149 | 228 | 248 | 153 | 112 | 0 | 0 | 0 | 0 | 0 | 258 | 149 | 1 | 1 | 1 | 0.984 | 1 | 1 | 1 | 1 | 1 | 1 | 1 | 1 | -0.003 |
| SE | 77598 | ENSG00000127947 | PTPN12 | chr7 | + | 77581426 | 77581503 | 77537294 | 77537645 | 77585542 | 77585581 | 3 | 5 | 3 | 8 | 2 | 1 | 0 | 0 | 0 | 0 | 9 | 0 | 11 | 2 | 18 | 0 | 0 | 0 | 0 | 0 | 226 | 149 | 1 | 1 | 0.664 | 1 | 1 | 1 | 1 | 1 | NA | 1 | 1 | 1 | -0.067 |
| SE | 77602 | ENSG00000127947 | PTPN12 | chr7 | + | 77583554 | 77583650 | 77571152 | 77571186 | 77592184 | 77592256 | 1 | 4 | 0 | 1 | 1 | 0 | 0 | 0 | 0 | 0 | 0 | 0 | 7 | 0 | 9 | 3 | 0 | 0 | 0 | 0 | 245 | 149 | 1 | 1 | 1 | 1 | NA | 1 | 1 | 0 | NA | 1 | NA | 1 | 0.333 |
| SE | 77604 | ENSG00000127947 | PTPN12 | chr7 | + | 77585542 | 77585581 | 77537294 | 77537645 | 77592184 | 77592256 | 24 | 26 | 37 | 35 | 55 | 0 | 2 | 0 | 0 | 0 | 40 | 24 | 19 | 24 | 42 | 0 | 0 | 0 | 0 | 0 | 188 | 149 | 1 | 1 | 1 | 0.912 | 1 | 1 | 1 | 1 | 1 | 1 | 1 | 1 | -0.018 |
| SE | 77605 | ENSG00000127947 | PTPN12 | chr7 | + | 77585542 | 77585581 | 77537294 | 77537645 | 77597841 | 77597901 | 0 | 0 | 0 | 0 | 3 | 0 | 0 | 0 | 0 | 0 | 0 | 0 | 1 | 0 | 0 | 0 | 0 | 2 | 4 | 0 | 188 | 149 | 1 | 1 | NA | NA | NA | NA | 1 | NA | NA | 0.284 | 0 | NA | 0.858 |
| SE | 77606 | ENSG00000127947 | PTPN12 | chr7 | + | 77585542 | 77585581 | 77571077 | 77571186 | 77592184 | 77592256 | 25 | 28 | 40 | 37 | 55 | 0 | 0 | 0 | 0 | 0 | 40 | 28 | 20 | 24 | 42 | 3 | 0 | 0 | 0 | 0 | 188 | 149 | 1 | 1 | 1 | 1 | 1 | 1 | 1 | 0.914 | 1 | 1 | 1 | 1 | 0.017 |
| SE | 77607 | ENSG00000127947 | PTPN12 | chr7 | + | 77585542 | 77585581 | 77583554 | 77583650 | 77592184 | 77592256 | 421 | 340 | 216 | 435 | 418 | 0 | 2 | 0 | 1 | 0 | 244 | 342 | 310 | 278 | 324 | 0 | 0 | 0 | 0 | 0 | 188 | 149 | 1 | 1 | 1 | 0.993 | 1 | 0.997 | 1 | 1 | 1 | 1 | 1 | 1 | -0.002 |
| SE | 77608 | ENSG00000127947 | PTPN12 | chr7 | + | 77585542 | 77585581 | 77583554 | 77583650 | 77597841 | 77597901 | 233 | 200 | 111 | 222 | 177 | 1 | 0 | 0 | 0 | 0 | 114 | 198 | 181 | 148 | 183 | 0 | 0 | 0 | 1 | 0 | 188 | 149 | 1 | 1 | 0.995 | 1 | 1 | 1 | 1 | 1 | 1 | 1 | 0.992 | 1 | 0.001 |
| SE | 77609 | ENSG00000127947 | PTPN12 | chr7 | + | 77592184 | 77592256 | 77537294 | 77537645 | 77597841 | 77597901 | 213 | 231 | 158 | 336 | 366 | 0 | 0 | 0 | 0 | 0 | 317 | 209 | 228 | 184 | 241 | 0 | 0 | 2 | 4 | 0 | 221 | 149 | 1 | 1 | 1 | 1 | 1 | 1 | 1 | 1 | 1 | 0.987 | 0.969 | 1 | 0.009 |
| SE | 77610 | ENSG00000127947 | PTPN12 | chr7 | + | 77592184 | 77592256 | 77537294 | 77537645 | 77607234 | 77607301 | 0 | 2 | 0 | 0 | 0 | 0 | 0 | 0 | 0 | 0 | 0 | 0 | 1 | 0 | 0 | 0 | 0 | 2 | 0 | 0 | 221 | 149 | 1 | 1 | NA | 1 | NA | NA | | NA | NA | 0.252 | NA | NA | 0.748 |
| SE | 77611 | ENSG00000127947 | PTPN12 | chr7 | + | 77592184 | 77592256 | 77583554 | 77583650 | 77597841 | 77597901 | 213 | 231 | 158 | 337 | 366 | 1 | 0 | 0 | 0 | 0 | 317 | 209 | 228 | 184 | 241 | 0 | 0 | 0 | 1 | 0 | 221 | 149 | 1 | 1 | 0.993 | 1 | 1 | 1 | 1 | 1 | 1 | 1 | 0.992 | 1 | 0 |
| SE | 77612 | ENSG00000127947 | PTPN12 | chr7 | + | 77592184 | 77592256 | 77585542 | 77585581 | 77597841 | 77597901 | 472 | 421 | 307 | 614 | 618 | 0 | 0 | 0 | 3 | 3 | 479 | 396 | 436 | 358 | 472 | 0 | 1 | 2 | 0 | 0 | 221 | 149 | 1 | 1 | 1 | 1 | 1 | 0.993 | 0.993 | 1 | 0.996 | 0.993 | 1 | 1 | -0.001 |
| SE | 77613 | ENSG00000127947 | PTPN12 | chr7 | + | 77597841 | 77597901 | 77537294 | 77537645 | 77607234 | 77607301 | 4 | 0 | 0 | 1 | 2 | 0 | 0 | 0 | 0 | 0 | 0 | 3 | 4 | 0 | 0 | 0 | 0 | 2 | 0 | 0 | 209 | 149 | 1 | 1 | 1 | NA | NA | 1 | 1 | NA | 1 | 0.588 | NA | NA | 0.206 |
| SE | 77614 | ENSG00000127947 | PTPN12 | chr7 | + | 77597841 | 77597901 | 77592184 | 77592256 | 77600663 | 77600728 | 521 | 486 | 366 | 707 | 877 | 0 | 0 | 0 | 0 | 0 | 689 | 462 | 517 | 453 | 544 | 0 | 0 | 0 | 1 | 0 | 209 | 149 | 1 | 1 | 1 | 1 | 1 | 1 | 1 | 1 | 1 | 1 | 0.997 | 1 | 0.001 |
| SE | 77615 | ENSG00000127947 | PTPN12 | chr7 | + | 77597841 | 77597901 | 77592184 | 77592256 | 77607234 | 77607301 | 173 | 133 | 98 | 173 | 163 | 0 | 0 | 0 | 0 | 0 | 110 | 106 | 151 | 121 | 151 | 0 | 0 | 1 | 0 | 0 | 209 | 149 | 1 | 1 | 1 | 1 | 1 | 1 | 1 | 1 | 1 | 0.991 | 1 | 1 | 0.002 |
| SE | 77616 | ENSG00000127947 | PTPN12 | chr7 | + | 77600663 | 77600806 | 77592184 | 77592256 | 77607234 | 77607301 | 502 | 393 | 260 | 477 | 846 | 0 | 0 | 0 | 0 | 0 | 276 | 321 | 374 | 203 | 182 | 0 | 0 | 1 | 0 | 0 | 292 | 149 | 1 | 1 | 1 | 1 | 1 | 1 | 1 | 1 | 1 | 0.995 | 1 | 1 | 0.001 |
| SE | 77617 | ENSG00000127947 | PTPN12 | chr7 | + | 77600663 | 77600806 | 77597841 | 77597901 | 77604878 | 77604935 | 353 | 352 | 275 | 544 | 721 | 0 | 0 | 0 | 0 | 0 | 572 | 360 | 365 | 337 | 400 | 0 | 0 | 2 | 0 | 0 | 292 | 149 | 1 | 1 | 1 | 1 | 1 | 1 | 1 | 1 | 1 | 0.989 | 1 | 1 | 0.002 |
| SE | 77618 | ENSG00000127947 | PTPN12 | chr7 | + | 77600663 | 77600806 | 77597841 | 77597901 | 77607234 | 77607301 | 856 | 747 | 529 | 1018 | 1574 | 5 | 1 | 1 | 5 | 11 | 846 | 677 | 738 | 539 | 582 | 0 | 3 | 5 | 0 | 7 | 292 | 149 | 1 | 1 | 0.989 | 0.997 | 0.996 | 0.99 | 0.986 | 1 | 0.991 | 0.987 | 1 | 0.977 | 0.001 |
| SE | 77619 | ENSG00000127947 | PTPN12 | chr7 | + | 77600663 | 77600806 | 77597841 | 77597901 | 77610764 | 77610802 | 352 | 349 | 270 | 542 | 720 | 0 | 0 | 0 | 0 | 3 | 592 | 356 | 376 | 332 | 400 | 0 | 0 | 0 | 2 | 0 | 292 | 149 | 1 | 1 | 1 | 1 | 1 | 1 | 0.992 | 1 | 1 | 1 | 0.988 | 1 | 0.001 |
| SE | 77620 | ENSG00000127947 | PTPN12 | chr7 | + | 77604878 | 77604935 | 77597841 | 77597901 | 77607234 | 77607301 | 2 | 1 | 1 | 0 | 6 | 5 | 1 | 1 | 5 | 11 | 0 | 0 | 3 | 0 | 0 | 0 | 3 | 5 | 0 | 7 | 206 | 149 | 1 | 1 | 0.224 | 0.42 | 0.42 | 0 | 0.283 | NA | 0 | 0.303 | NA | 0 | 0.168 |
| SE | 77621 | ENSG00000127947 | PTPN12 | chr7 | + | 77604878 | 77604935 | 77600663 | 77600806 | 77607234 | 77607301 | 6 | 4 | 9 | 5 | 10 | 507 | 398 | 262 | 479 | 857 | 2 | 4 | 6 | 6 | 0 | 276 | 321 | 376 | 208 | 182 | 206 | 149 | 1 | 1 | 0.008 | 0.007 | 0.024 | 0.007 | 0.008 | 0.005 | 0.009 | 0.011 | 0.02 | 0 | 0.002 |

| | | | | | | | | | | | | | | |
|---|---|---|---|---|---|---|---|---|---|---|---|---|---|---|
| SE | 77622 | ENSG00000127947 | PTPN12 | chr7 | + | 77607234 | 77607301 | 77597841 | 77597901 | 77610764 | 77610802 | 175 | 181 | [illegible] |
| SE | 77624 | ENSG00000127947 | PTPN12 | chr7 | + | 77618479 | 77618565 | 77610947 | 77611046 | 77626704 | 77627675 | 774 | 669 | [illegible] |
| SE | 77626 | ENSG00000127947 | PTPN12 | chr7 | + | 77635781 | 77635849 | 77632347 | 77632425 | 77637017 | 77637048 | 1018 | 808 | [illegible] |
| SE | 77627 | ENSG00000127947 | PTPN12 | chr7 | + | 77637017 | 77637048 | 77635781 | 77635849 | 77638623 | 77638708 | 978 | 737 | [illegible] |
| SE | 77628 | ENSG00000127947 | PTPN12 | chr7 | + | 77638623 | 77638731 | 77636764 | 77637048 | 77639218 | 77639280 | 1442 | 1072 | [illegible] |
| SE | 77634 | ENSG00000105877 | DNAH11 | chr7 | + | 21564185 | 21564397 | 21561070 | 21561170 | 21570068 | 21570299 | 2 | 0 | [illegible] |
| SE | 77638 | ENSG00000104863 | LIN7B | chr19 | + | 49115259 | 49115331 | 49114465 | 49114967 | 49116262 | 49116472 | 1 | 0 | [illegible] |
| SE | 77639 | ENSG00000104863 | LIN7B | chr19 | + | 49115824 | 49116024 | 49115259 | 49115331 | 49116262 | 49116472 | 0 | 0 | [illegible] |
| SE | 77645 | ENSG00000214021 | TTLL3 | chr3 | + | 9816073 | 9816202 | 9813247 | 9813345 | 9826996 | 9827072 | 22 | 15 | [illegible] |
| SE | 77647 | ENSG00000214021 | TTLL3 | chr3 | + | 9817644 | 9817759 | 9813247 | 9813345 | 9818821 | 9818871 | 21 | 17 | [illegible] |
| SE | 77649 | ENSG00000214021 | TTLL3 | chr3 | + | 9817644 | 9817759 | 9813247 | 9813345 | 9826996 | 9827072 | 19 | 15 | [illegible] |
| SE | 77655 | ENSG00000214021 | TTLL3 | chr3 | + | 9817689 | 9817759 | 9813247 | 9813345 | 9818821 | 9818921 | 2 | 2 | [illegible] |
| SE | 77662 | ENSG00000214021 | TTLL3 | chr3 | + | 9820545 | 9820741 | 9813247 | 9813345 | 9826996 | 9827072 | 8 | 1 | [illegible] |
| SE | 77663 | ENSG00000214021 | TTLL3 | chr3 | + | 9820545 | 9820741 | 9816073 | 9816202 | 9826996 | 9827072 | 5 | 1 | [illegible] |
| SE | 77664 | ENSG00000214021 | TTLL3 | chr3 | + | 9820545 | 9820741 | 9817644 | 9817759 | 9826996 | 9827072 | 7 | 3 | [illegible] |
| SE | 77665 | ENSG00000214021 | TTLL3 | chr3 | + | 9825799 | 9825948 | 9813247 | 9813345 | 9826996 | 9827072 | 47 | 29 | [illegible] |
| SE | 77667 | ENSG00000214021 | TTLL3 | chr3 | + | 9827214 | 9827240 | 9826996 | 9827072 | 9828959 | 9829395 | 3 | 1 | [illegible] |
| SE | 77670 | ENSG00000214021 | TTLL3 | chr3 | + | 9834426 | 9834503 | 9833103 | 9833245 | 9834680 | 9834907 | 4 | 3 | [illegible] |
| SE | 77675 | ENSG00000117009 | KMO | chr1 | + | 241536462 | 241536564 | 241532337 | 241532498 | 241548828 | 241548898 | 1 | 2 | [illegible] |
| SE | 77676 | ENSG00000117009 | KMO | chr1 | + | 241536471 | 241536564 | 241532368 | 241532498 | 241548828 | 241548898 | 0 | 0 | [illegible] |
| SE | 77677 | ENSG00000117009 | KMO | chr1 | + | 241564986 | 241565058 | 241562052 | 241562332 | 241566490 | 241566575 | 22 | 23 | [illegible] |
| SE | 77690 | ENSG00000011426 | ANLN | chr7 | + | 36417079 | 36417190 | 36415757 | 36415884 | 36419243 | 36419479 | 3 | 8 | [illegible] |
| SE | 77691 | ENSG00000011426 | ANLN | chr7 | + | 36422632 | 36422809 | 36421856 | 36421992 | 36423816 | 36423943 | 6 | 6 | [illegible] |
| SE | 77692 | ENSG00000011426 | ANLN | chr7 | + | 36424544 | 36424593 | 36423816 | 36423943 | 36424685 | 36424742 | 0 | 3 | [illegible] |
| SE | 77693 | ENSG00000011426 | ANLN | chr7 | + | 36428306 | 36428360 | 36426915 | 36427028 | 36439203 | 36439290 | 0 | 0 | [illegible] |
| SE | 77694 | ENSG00000011426 | ANLN | chr7 | + | 36443754 | 36443862 | 36439203 | 36439290 | 36449664 | 36449678 | 6 | 5 | [illegible] |
| SE | 77711 | ENSG00000136205 | TNS3 | chr7 | - | 47283696 | 47283865 | 47280285 | 47280354 | 47291954 | 47292032 | 43 | 16 | [illegible] |
| SE | 77712 | ENSG00000136205 | TNS3 | chr7 | - | 47292827 | 47292905 | 47291954 | 47292032 | 47293732 | 47293828 | 92 | 54 | [illegible] |
| SE | 77715 | ENSG00000136205 | TNS3 | chr7 | - | 47346186 | 47346356 | 47344923 | 47345038 | 47368364 | 47369621 | 38 | 18 | [illegible] |
| SE | 77716 | ENSG00000136205 | TNS3 | chr7 | - | 47368364 | 47369621 | 47346186 | 47346356 | 47396799 | 47396904 | 80 | 50 | [illegible] |
| SE | 77719 | ENSG00000136205 | TNS3 | chr7 | - | 47437262 | 47437313 | 47435281 | 47435404 | 47439486 | 47439658 | 37 | 27 | [illegible] |
| SE | 77720 | ENSG00000136205 | TNS3 | chr7 | - | 47481102 | 47481141 | 47442002 | 47442055 | 47506906 | 47506944 | 63 | 26 | [illegible] |
| SE | 77721 | ENSG00000136205 | TNS3 | chr7 | - | 47529035 | 47529147 | 47506906 | 47506944 | 47582050 | 47582111 | 55 | 20 | [illegible] |
| SE | 77722 | ENSG00000136205 | TNS3 | chr7 | - | 47557036 | 47557150 | 47529035 | 47529147 | 47582050 | 47582111 | 1 | 2 | [illegible] |
| SE | 77725 | ENSG00000288700 | NA | chr3 | + | 134755658 | 134755704 | 134615183 | 134615438 | 134779737 | 134779899 | 0 | 0 | [illegible] |
| SE | 77726 | ENSG00000288700 | NA | chr3 | + | 134755658 | 134755704 | 134615183 | 134615438 | 134925815 | 134925880 | 0 | 0 | [illegible] |
| SE | 77727 | ENSG00000288700 | NA | chr3 | + | 134779737 | 134779881 | 134615364 | 134615438 | 134925815 | 134925880 | 0 | 6 | [illegible] |
| SE | 77735 | ENSG00000250565 | ATP6V1E2 | chr2 | - | 46537375 | 46537576 | 46536610 | 46536703 | 46541389 | 46541589 | 0 | 0 | [illegible] |
| SE | 77736 | ENSG00000164168 | TMEM184C | chr4 | + | 147631392 | 147631505 | 147629598 | 147629692 | 147633764 | 147633936 | 26 | 15 | [illegible] |
| SE | 77737 | ENSG00000164168 | TMEM184C | chr4 | + | 147632902 | 147633002 | 147631392 | 147631505 | 147633764 | 147633936 | 76 | 55 | [illegible] |
| SE | 77738 | ENSG00000164168 | TMEM184C | chr4 | + | 147632954 | 147633002 | 147631392 | 147631505 | 147633764 | 147633936 | 20 | 19 | [illegible] |
| SE | 77775 | ENSG00000152457 | DCLRE1C | chr10 | - | 14922980 | 14923069 | 14919737 | 14919832 | 14926842 | 14926897 | 93 | 99 | [illegible] |
| SE | 77777 | ENSG00000152457 | DCLRE1C | chr10 | - | 14926842 | 14926897 | 14922980 | 14923069 | 14932853 | 14932955 | 41 | 51 | [illegible] |
| SE | 77779 | ENSG00000152457 | DCLRE1C | chr10 | - | 14934379 | 14934520 | 14932853 | 14932955 | 14934702 | 14934775 | 133 | 68 | [illegible] |
| SE | 77781 | ENSG00000152457 | DCLRE1C | chr10 | - | 14936537 | 14936593 | 14935462 | 14935564 | 14945104 | 14945189 | 61 | 34 | [illegible] |
| SE | 77782 | ENSG00000152457 | DCLRE1C | chr10 | - | 14939809 | 14939869 | 14935462 | 14935564 | 14945104 | 14945189 | 39 | 26 | [illegible] |
| SE | 77783 | ENSG00000152457 | DCLRE1C | chr10 | - | 14939809 | 14939869 | 14936537 | 14936593 | 14945104 | 14945189 | 110 | 84 | [illegible] |
| SE | 77792 | ENSG00000134250 | NOTCH2 | chr1 | - | 119921712 | 119921809 | 119920228 | 119920397 | 119922635 | 119922778 | 181 | 142 | [illegible] |
| SE | 77793 | ENSG00000134250 | NOTCH2 | chr1 | - | 119922235 | 119922446 | 119921712 | 119921809 | 119922635 | 119922778 | 438 | 296 | [illegible] |
| SE | 77794 | ENSG00000134250 | NOTCH2 | chr1 | - | 119922235 | 119922446 | 119921712 | 119921809 | 119923636 | 119923984 | 221 | 153 | [illegible] |
| SE | 77795 | ENSG00000134250 | NOTCH2 | chr1 | - | 119922635 | 119922778 | 119921712 | 119921809 | 119923636 | 119923984 | 283 | 171 | [illegible] |
| SE | 77796 | ENSG00000134250 | NOTCH2 | chr1 | - | 119922635 | 119922778 | 119922235 | 119922446 | 119923636 | 119923984 | 499 | 314 | [illegible] |
| SE | 77797 | ENSG00000134250 | NOTCH2 | chr1 | - | 119923636 | 119923984 | 119922235 | 119922446 | 119925304 | 119925810 | 131 | 70 | [illegible] |
| SE | 77798 | ENSG00000134250 | NOTCH2 | chr1 | - | 119925304 | 119925810 | 119923636 | 119923984 | 119926498 | 119926611 | 355 | 206 | [illegible] |
| SE | 77799 | ENSG00000134250 | NOTCH2 | chr1 | - | 119941525 | 119941754 | 119940554 | 119940756 | 119948413 | 119948566 | 446 | 354 | [illegible] |
| SE | 77800 | ENSG00000134250 | NOTCH2 | chr1 | - | 119941525 | 119941754 | 119940554 | 119940756 | 119950723 | 119950837 | 328 | 288 | [illegible] |
| SE | 77801 | ENSG00000134250 | NOTCH2 | chr1 | - | 119948413 | 119948566 | 119940554 | 119940756 | 119950723 | 119950837 | 0 | 0 | [illegible] |
| SE | 77802 | ENSG00000134250 | NOTCH2 | chr1 | - | 119948413 | 119948566 | 119941525 | 119941754 | 119949006 | 119949126 | 333 | 219 | [illegible] |
| SE | 77805 | ENSG00000134250 | NOTCH2 | chr1 | - | 119949006 | 119949126 | 119948413 | 119948566 | 119950723 | 119950837 | 381 | 253 | [illegible] |
| SE | 77807 | ENSG00000134250 | NOTCH2 | chr1 | - | 119965452 | 119965566 | 119963573 | 119963807 | 119966375 | 119966489 | 878 | 763 | [illegible] |
| SE | 77808 | ENSG00000134250 | NOTCH2 | chr1 | - | 119966375 | 119966489 | 119965452 | 119965566 | 119967432 | 119967621 | 979 | 841 | [illegible] |
| SE | 77809 | ENSG00000134250 | NOTCH2 | chr1 | - | 119967432 | 119967621 | 119966375 | 119966489 | 119969510 | 119969744 | 496 | 388 | [illegible] |
| SE | 77811 | ENSG00000134250 | NOTCH2 | chr1 | - | 119969510 | 119969744 | 119967432 | 119967621 | 119986959 | 119987082 | 182 | 126 | [illegible] |
| SE | 77813 | ENSG00000134250 | NOTCH2 | chr1 | - | 119986959 | 119987082 | 119969510 | 119969744 | 119996996 | 119997332 | 531 | 328 | [illegible] |
| SE | 77814 | ENSG00000134250 | NOTCH2 | chr1 | - | 119996996 | 119997332 | 119986959 | 119987082 | 120005328 | 120005588 | 538 | 308 | [illegible] |
| SE | 77821 | ENSG00000134250 | NOTCH2 | chr1 | - | 120097179 | 120097279 | 120051483 | 120051779 | 120100653 | 120100779 | 10 | 6 | [illegible] |
| SE | 77827 | ENSG00000169490 | TM2D2 | chr8 | - | 38993544 | 38993660 | 38988807 | 38991545 | 38995317 | 38995405 | 144 | 103 | [illegible] |
| SE | 77828 | ENSG00000188191 | PRKAR1B | chr7 | - | 584507 | 584568 | 579255 | 579377 | 596145 | 596304 | 25 | 31 | [illegible] |
| SE | 77831 | ENSG00000235618 | FAM21EP | chr10 | - | 50056783 | 50056911 | 50054637 | 50054724 | 50059100 | 50059273 | 0 | 0 | [illegible] |
| SE | 77833 | ENSG00000256043 | CTSO | chr4 | - | 155926783 | 155926881 | 155924120 | 155926070 | 155928335 | 155928428 | 0 | 0 | [illegible] |
| SE | 77836 | ENSG00000256043 | CTSO | chr4 | - | 155928338 | 155928428 | 155924117 | 155926070 | 155929541 | 155929705 | 94 | 34 | [illegible] |
| SE | 77837 | ENSG00000256043 | CTSO | chr4 | - | 155928610 | 155928689 | 155928335 | 155928428 | 155929541 | 155929705 | 1 | 0 | [illegible] |
| SE | 77838 | ENSG00000256043 | CTSO | chr4 | - | 155937361 | 155937483 | 155929541 | 155929705 | 155939370 | 155939538 | 695 | 578 | [illegible] |
| SE | 77840 | ENSG00000088727 | KIF9 | chr3 | - | 47243050 | 47243245 | 47240800 | 47241015 | 47244790 | 47244924 | 1 | 1 | [illegible] |
| SE | 77843 | ENSG00000088727 | KIF9 | chr3 | - | 47271236 | 47271461 | 47267179 | 47267263 | 47275324 | 47275490 | 8 | 2 | [illegible] |
| SE | 77844 | ENSG00000088727 | KIF9 | chr3 | - | 47273551 | 47273658 | 47267179 | 47267263 | 47275324 | 47275490 | 5 | 5 | [illegible] |
| SE | 77845 | ENSG00000088727 | KIF9 | chr3 | - | 47273551 | 47273658 | 47271236 | 47271461 | 47275324 | 47275490 | 6 | 3 | [illegible] |
| SE | 77846 | ENSG00000088727 | KIF9 | chr3 | - | 47277281 | 47277379 | 47275227 | 47275490 | 47282494 | 47282537 | 4 | 5 | [illegible] |
| SE | 77847 | ENSG00000088727 | KIF9 | chr3 | - | 47280951 | 47281195 | 47277281 | 47277379 | 47282139 | 47282198 | 0 | 0 | [illegible] |
| SE | 77848 | ENSG00000088727 | KIF9 | chr3 | - | 47282139 | 47282385 | 47277281 | 47277379 | 47282494 | 47282577 | 0 | 0 | [illegible] |
| SE | 77849 | ENSG00000078053 | AMPH | chr7 | - | 38417824 | 38417950 | 38394004 | 38394214 | 38422420 | 38422477 | 0 | 0 | [illegible] |
| SE | 77851 | ENSG00000078053 | AMPH | chr7 | - | 38422420 | 38422477 | 38394004 | 38394214 | 38426953 | 38426986 | 0 | 0 | [illegible] |
| SE | 77852 | ENSG00000078053 | AMPH | chr7 | - | 38429055 | 38429577 | 38426953 | 38426986 | 38429841 | 38429865 | 0 | 0 | [illegible] |
| SE | 77867 | ENSG00000198815 | FOXJ3 | chr1 | - | 42188736 | 42188928 | 42181916 | 42182024 | 42191302 | 42191719 | 303 | 241 | [illegible] |
| SE | 77868 | ENSG00000198815 | FOXJ3 | chr1 | - | 42189302 | 42189404 | 42188736 | 42188928 | 42191302 | 42191719 | 418 | 323 | [illegible] |
| SE | 77869 | ENSG00000198815 | FOXJ3 | chr1 | - | 42194889 | 42195064 | 42191302 | 42191719 | 42199101 | 42199230 | 499 | 401 | [illegible] |
| SE | 77870 | ENSG00000198815 | FOXJ3 | chr1 | - | 42199101 | 42199230 | 42194889 | 42195064 | 42205761 | 42205863 | 334 | 265 | [illegible] |
| SE | 77871 | ENSG00000198815 | FOXJ3 | chr1 | - | 42199101 | 42199230 | 42194889 | 42195064 | 42227882 | 42227966 | 246 | 197 | [illegible] |
| SE | 77872 | ENSG00000198815 | FOXJ3 | chr1 | - | 42205761 | 42205863 | 42194889 | 42195064 | 42227882 | 42227966 | 77 | 51 | [illegible] |
| SE | 77874 | ENSG00000198815 | FOXJ3 | chr1 | - | 42205761 | 42205863 | 42199101 | 42199230 | 42265114 | 42265189 | 103 | 72 | [illegible] |
| SE | 77875 | ENSG00000198815 | FOXJ3 | chr1 | - | 42227882 | 42227966 | 42199101 | 42199230 | 42265114 | 42265189 | 212 | 139 | [illegible] |
| SE | 77876 | ENSG00000198815 | FOXJ3 | chr1 | - | 42227882 | 42227966 | 42205761 | 42205863 | 42265114 | 42265189 | 283 | 187 | [illegible] |
| SE | 77877 | ENSG00000198815 | FOXJ3 | chr1 | - | 42265114 | 42265189 | 42227882 | 42227966 | 42278347 | 42278672 | 336 | 243 | [illegible] |
| SE | 77884 | ENSG00000106609 | TMEM248 | chr7 | + | 66944975 | 66945261 | 66941847 | 66942024 | 66948543 | 66948694 | 385 | 174 | [illegible] |
| SE | 77885 | ENSG00000106609 | TMEM248 | chr7 | + | 66948543 | 66948694 | 66944975 | 66945261 | 66950951 | 66951135 | 413 | 187 | [illegible] |
| SE | 77886 | ENSG00000106609 | TMEM248 | chr7 | + | 66950951 | 66951135 | 66948543 | 66948694 | 66953225 | 66953369 | 513 | 282 | [illegible] |
| SE | 77887 | ENSG00000106609 | TMEM248 | chr7 | + | 66950951 | 66951135 | 66948543 | 66948694 | 66955501 | 66958551 | 241 | 124 | [illegible] |
| SE | 77888 | ENSG00000106609 | TMEM248 | chr7 | + | 66953225 | 66953369 | 66948543 | 66948694 | 66955501 | 66956153 | 172 | 75 | [illegible] |
| SE | 77893 | ENSG00000160188 | RSPH1 | chr21 | - | 42492965 | 42493079 | 42492757 | 42492863 | 42496132 | 42496224 | 0 | 4 | [illegible] |
| SE | 77903 | ENSG00000106524 | ANKMY2 | chr7 | - | 16603548 | 16603660 | 16602379 | 16602509 | 16604720 | 16604849 | 4 | 0 | [illegible] |
| SE | 77904 | ENSG00000106524 | ANKMY2 | chr7 | - | 16609629 | 16609765 | 16604720 | 16604849 | 16610548 | 16610763 | 72 | 40 | [illegible] |
| SE | 77905 | ENSG00000106524 | ANKMY2 | chr7 | - | 16615743 | 16615904 | 16610548 | 16610763 | 16624982 | 16625081 | 73 | 61 | [illegible] |
| SE | 77906 | ENSG00000106524 | ANKMY2 | chr7 | - | 16624982 | 16625081 | 16615743 | 16615904 | 16627039 | 16627178 | 76 | 77 | [illegible] |
| SE | 77907 | ENSG00000106524 | ANKMY2 | chr7 | - | 16644625 | 16644812 | 16636390 | 16636455 | 16645506 | 16645662 | 1 | 1 | [illegible] |
| SE | 77908 | ENSG00000106524 | ANKMY2 | chr7 | - | 16644684 | 16644812 | 16636390 | 16636455 | 16645506 | 16645816 | 4 | 5 | [illegible] |
| SE | 77910 | ENSG00000234585 | CCT6P3 | chr7 | + | 65072144 | 65072247 | 65070908 | 65071042 | 65073440 | 65073514 | 0 | 0 | [illegible] |
| SE | 77911 | ENSG00000146731 | CCT6A | chr7 | + | 56054368 | 56054503 | 56052421 | 56052485 | 56055623 | 56055797 | 387 | 231 | [illegible] |
| SE | 77912 | ENSG00000146731 | CCT6A | chr7 | + | 56056310 | 56056414 | 56055623 | 56055797 | 56057992 | 56058103 | 727 | 529 | [illegible] |
| SE | 77913 | ENSG00000146731 | CCT6A | chr7 | + | 56058361 | 56058521 | 56057992 | 56058103 | 56058619 | 56058702 | 331 | 264 | [illegible] |
| SE | 77914 | ENSG00000146731 | CCT6A | chr7 | + | 56058619 | 56058702 | 56057992 | 56058103 | 56059543 | 56059640 | 38 | 17 | [illegible] |
| SE | 77915 | ENSG00000146731 | CCT6A | chr7 | + | 56059543 | 56059640 | 56058619 | 56058702 | 56060268 | 56060416 | 500 | 302 | [illegible] |
| SE | 77917 | ENSG00000146731 | CCT6A | chr7 | + | 56062682 | 56062755 | 56060806 | 56060940 | 56063012 | 56063178 | 148 | 110 | [illegible] |
| SE | 77918 | ENSG00000168056 | LTBP3 | chr11 | - | 65540012 | 65540153 | 65539719 | 65539881 | 65540244 | 65540382 | 55 | 6 | [illegible] |
| SE | 77919 | ENSG00000168056 | LTBP3 | chr11 | - | 65551547 | 65551564 | 65551401 | 65551474 | 65551971 | 65552157 | 12 | 10 | [illegible] |
| SE | 77922 | ENSG00000225940 | C5orf67 | chr5 | - | 56513761 | 56513925 | 56511566 | 56511736 | 56533081 | 56533155 | 3 | 0 | [illegible] |
| SE | 77923 | ENSG00000095015 | MAP3K1 | chr5 | + | 56864733 | 56864934 | 56859714 | 56859915 | 56865339 | 56865456 | 929 | 656 | [illegible] |
| SE | 77924 | ENSG00000095015 | MAP3K1 | chr5 | + | 56865828 | 56865977 | 56865339 | 56865456 | 56871909 | 56872031 | 599 | 414 | [illegible] |
| SE | 77926 | ENSG00000095015 | MAP3K1 | chr5 | + | 56875031 | 56875310 | 56872824 | 56873005 | 56880710 | 56880802 | 421 | 288 | [illegible] |
| SE | 77927 | ENSG00000095015 | MAP3K1 | chr5 | + | 56878979 | 56879101 | 56872824 | 56873005 | 56880710 | 56880802 | 124 | 73 | [illegible] |
| SE | 77928 | ENSG00000095015 | MAP3K1 | chr5 | + | 56878979 | 56879101 | 56875031 | 56875310 | 56880710 | 56880802 | 206 | 112 | [illegible] |
| SE | 77929 | ENSG00000095015 | MAP3K1 | chr5 | + | 56880710 | 56880802 | 56878979 | 56879101 | 56881082 | 56881272 | 315 | 171 | [illegible] |
| SE | 77930 | ENSG00000095015 | MAP3K1 | chr5 | + | 56881569 | 56882866 | 56881082 | 56881272 | 56883526 | 56883679 | 1142 | 942 | [illegible] |
| SE | 77933 | ENSG00000095015 | MAP3K1 | chr5 | + | 56887377 | 56888357 | 56885952 | 56886063 | 56893530 | 56896152 | 805 | 611 | [illegible] |
| SE | 77935 | ENSG00000095015 | MAP3K1 | chr5 | + | 56888225 | 56888357 | 56887377 | 56887520 | 56893530 | 56896152 | 528 | 359 | [illegible] |
| SE | 77936 | ENSG00000111832 | RWDD1 | chr6 | + | 116572421 | 116572478 | 116571562 | 116571655 | 116572857 | 116572985 | 0 | 0 | [illegible] |
| SE | 77937 | ENSG00000111832 | RWDD1 | chr6 | + | 116572421 | 116572478 | 116571562 | 116571655 | 116574057 | 116574171 | 1 | 0 | [illegible] |
| SE | 77941 | ENSG00000111832 | RWDD1 | chr6 | + | 116572857 | 116572985 | 116572421 | 116572478 | 116574057 | 116574171 | 6 | 8 | [illegible] |
| SE | 77942 | ENSG00000111832 | RWDD1 | chr6 | + | 116572857 | 116572985 | 116572421 | 116572478 | 116580294 | 116580360 | 1 | 0 | [illegible] |
| SE | 77946 | ENSG00000111832 | RWDD1 | chr6 | + | 116590887 | 116590950 | 116590271 | 116590404 | 116592979 | 116593588 | 186 | 127 | [illegible] |
| SE | 77947 | ENSG00000221829 | FANCG | chr9 | - | 35074370 | 35074494 | 35073834 | 35074216 | 35074926 | 35075082 | 21 | 5 | [illegible] |
| SE | 77948 | ENSG00000221829 | FANCG | chr9 | - | 35074370 | 35074494 | 35073834 | 35074216 | 35075278 | 35075325 | 8 | 0 | [illegible] |
| SE | 77951 | ENSG00000221829 | FANCG | chr9 | - | 35075961 | 35076028 | 35075464 | 35075754 | 35076431 | 35076583 | 1 | 0 | [illegible] |
| SE | 77952 | ENSG00000221829 | FANCG | chr9 | - | 35078140 | 35078343 | 35077263 | 35077399 | 35078604 | 35078736 | 3 | 1 | [illegible] |
| SE | 77953 | ENSG00000120314 | WDR55 | chr5 | + | 140668414 | 140668502 | 140668233 | 140668334 | 140668611 | 140668791 | 166 | 80 | [illegible] |
| SE | 77956 | ENSG00000281103 | TRG-AS1 | chr7 | + | 38350345 | 38350450 | 38341614 | 38341799 | 38375556 | 38378627 | 12 | 15 | [illegible] |
| SE | 77958 | ENSG00000173200 | PARP15 | chr3 | + | 122594746 | 122594870 | 122577649 | 122577853 | 122610493 | 122610730 | 4 | 0 | [illegible] |
| SE | 77960 | ENSG00000173200 | PARP15 | chr3 | + | 122605935 | 122606055 | 122594746 | 122594870 | 122610493 | 122610730 | 186 | 164 | [illegible] |
| SE | 77961 | ENSG00000173200 | PARP15 | chr3 | + | 122613040 | 122613268 | 122610493 | 122610730 | 122615285 | 122615320 | 92 | 36 | [illegible] |
| SE | 77962 | ENSG00000173200 | PARP15 | chr3 | + | 122613040 | 122613268 | 122610493 | 122610730 | 122615778 | 122615857 | 78 | 30 | [illegible] |
| SE | 77963 | ENSG00000173200 | PARP15 | chr3 | + | 122613040 | 122613268 | 122610493 | 122610730 | 122617085 | 122617164 | 72 | 26 | [illegible] |
| SE | 77964 | ENSG00000173200 | PARP15 | chr3 | + | 122615285 | 122615320 | 122610493 | 122610730 | 122615778 | 122615857 | 13 | 5 | [illegible] |
| SE | 77965 | ENSG00000173200 | PARP15 | chr3 | + | 122615285 | 122615320 | 122610493 | 122610730 | 122617085 | 122617164 | 8 | 1 | [illegible] |
| SE | 77967 | ENSG00000173200 | PARP15 | chr3 | + | 122615285 | 122615320 | 122613040 | 122613268 | 122617085 | 122617164 | 8 | 3 | [illegible] |
| SE | 77968 | ENSG00000173200 | PARP15 | chr3 | + | 122615778 | 122615857 | 122610493 | 122610730 | 122617085 | 122617164 | 3 | 13 | [illegible] |
| SE | 77969 | ENSG00000173200 | PARP15 | chr3 | + | 122615778 | 122615857 | 122613040 | 122613268 | 122617014 | 122617164 | 130 | 76 | [illegible] |
| SE | 77970 | ENSG00000173200 | PARP15 | chr3 | + | 122615778 | 122615857 | 122613040 | 122613268 | 122617085 | 122617164 | 5 | 6 | [illegible] |
| SE | 77971 | ENSG00000173200 | PARP15 | chr3 | + | 122615778 | 122615857 | 122615285 | 122615320 | 122617085 | 122617164 | 7 | 7 | [illegible] |
| SE | 77973 | ENSG00000173200 | PARP15 | chr3 | + | 122632085 | 122632219 | 122621443 | 122621611 | 122635019 | 122635194 | 158 | 110 | [illegible] |
| SE | 77984 | ENSG00000106635 | BCL7B | chr7 | - | 73552166 | 73552242 | 73543547 | 73543644 | 73557486 | 73557609 | 96 | 73 | [illegible] |
| SE | 77985 | ENSG00000106635 | BCL7B | chr7 | - | 73553547 | 73553669 | 73552166 | 73552242 | 73557040 | 73557251 | 4 | 0 | [illegible] |
| SE | 77989 | ENSG00000182022 | CHST15 | chr10 | - | 124045666 | 124046724 | 124044579 | 124044919 | 124092323 | 124092414 | 216 | 189 | [illegible] |
| SE | 77990 | ENSG00000146729 | NIPSNAP2 | chr7 | + | 55978349 | 55978395 | 55978125 | 55978265 | 55981472 | 55981498 | 162 | 101 | [illegible] |
| SE | 77991 | ENSG00000146729 | NIPSNAP2 | chr7 | + | 55978349 | 55978395 | 55978125 | 55978265 | 55982209 | 55982280 | 9 | 0 | [illegible] |
| SE | 77993 | ENSG00000146729 | NIPSNAP2 | chr7 | + | 55979762 | 55979886 | 55978349 | 55978395 | 55981472 | 55981503 | 1 | 0 | [illegible] |
| SE | 77994 | ENSG00000146729 | NIPSNAP2 | chr7 | + | 55981472 | 55981567 | 55978125 | 55978265 | 55982209 | 55982280 | 78 | 56 | [illegible] |
| SE | 77995 | ENSG00000146729 | NIPSNAP2 | chr7 | + | 55981472 | 55981567 | 55978349 | 55978395 | 55982209 | 55982280 | 231 | 154 | [illegible] |
| SE | 77997 | ENSG00000146729 | NIPSNAP2 | chr7 | + | 55984846 | 55984878 | 55983727 | 55983868 | 55997365 | 55997449 | 77 | 46 | [illegible] |
| SE | 77999 | ENSG00000146729 | NIPSNAP2 | chr7 | + | 55994893 | 55994988 | 55984846 | 55984878 | 55997365 | 55997449 | 153 | 81 | [illegible] |
| SE | 78000 | ENSG00000146729 | NIPSNAP2 | chr7 | + | 55997365 | 55997449 | 55983727 | 55983868 | 55999007 | 55999304 | 79 | 69 | [illegible] |
| SE | 78001 | ENSG00000146733 | PSPH | chr7 | - | 56015022 | 56015171 | 56011063 | 56011869 | 56017233 | 56017379 | 11 | 9 | [illegible] |
| SE | 78004 | ENSG00000146733 | PSPH | chr7 | - | 56019599 | 56019734 | 56017233 | 56017379 | 56021072 | 56021231 | 14 | 15 | [illegible] |
| SE | 78005 | ENSG00000146733 | PSPH | chr7 | - | 56019599 | 56019808 | 56017276 | 56017379 | 56021072 | 56021231 | 5 | 6 | [illegible] |
| SE | 78006 | ENSG00000146733 | PSPH | chr7 | - | 56033380 | 56033494 | 56031928 | 56032054 | 56033960 | 56034103 | 0 | 0 | [illegible] |
| SE | 78007 | ENSG00000146733 | PSPH | chr7 | - | 56043147 | 56043261 | 56033960 | 56034154 | 56051137 | 56051604 | 0 | 1 | [illegible] |
| SE | 78011 | ENSG00000173890 | GPR160 | chr3 | + | 170079773 | 170079897 | 170038914 | 170039043 | 170083904 | 170084071 | 189 | 73 | [illegible] |
| SE | 78016 | ENSG00000125977 | EIF2S2 | chr20 | - | 34098497 | 34098633 | 34097415 | 34097516 | 34103461 | 34103565 | 876 | 718 | [illegible] |
| SE | 78017 | ENSG00000125977 | EIF2S2 | chr20 | - | 34105367 | 34105545 | 34103461 | 34103565 | 34112095 | 34112243 | 310 | 271 | [illegible] |
| SE | 78022 | ENSG00000136250 | AOAH | chr7 | - | 36530417 | 36530514 | 36522038 | 36522115 | 36532146 | 36532206 | 471 | 319 | [illegible] |
| SE | 78023 | ENSG00000136250 | AOAH | chr7 | - | 36532146 | 36532206 | 36530417 | 36530514 | 36532285 | 36532344 | 428 | 275 | [illegible] |
| SE | 78024 | ENSG00000136250 | AOAH | chr7 | - | 36532285 | 36532344 | 36532146 | 36532206 | 36540318 | 36540491 | 453 | 289 | [illegible] |
| SE | 78025 | ENSG00000136250 | AOAH | chr7 | - | 36540318 | 36540491 | 36532285 | 36532344 | 36549438 | 36549475 | 325 | 200 | [illegible] |
| SE | 78028 | ENSG00000136250 | AOAH | chr7 | - | 36576573 | 36576656 | 36549438 | 36549475 | 36594338 | 36594430 | 557 | 477 | [illegible] |
| SE | 78029 | ENSG00000136250 | AOAH | chr7 | - | 36632035 | 36632106 | 36623189 | 36623250 | 36637850 | 36637910 | 828 | 611 | [illegible] |
| SE | 78030 | ENSG00000136250 | AOAH | chr7 | - | 36632035 | 36632106 | 36623189 | 36623250 | 36673942 | 36674009 | 342 | 260 | [illegible] |
| SE | 78031 | ENSG00000136250 | AOAH | chr7 | - | 36637850 | 36637910 | 36623189 | 36623250 | 36673942 | 36674009 | 14 | 7 | [illegible] |
| SE | 78032 | ENSG00000136250 | AOAH | chr7 | - | 36637850 | 36637910 | 36632035 | 36632106 | 36659165 | 36659265 | 912 | 686 | [illegible] |
| SE | 78033 | ENSG00000136250 | AOAH | chr7 | - | 36637850 | 36637910 | 36632035 | 36632106 | 36673942 | 36674009 | 354 | 262 | [illegible] |
| SE | 78034 | ENSG00000136250 | AOAH | chr7 | - | 36659165 | 36659265 | 36632035 | 36632106 | 36673942 | 36674009 | 400 | 268 | [illegible] |
| SE | 78036 | ENSG00000136250 | AOAH | chr7 | - | 36659165 | 36659265 | 36637850 | 36637910 | 36724021 | 36724449 | 308 | 256 | [illegible] |
| SE | 78037 | ENSG00000136250 | AOAH | chr7 | - | 36673942 | 36674009 | 36637850 | 36637910 | 36724021 | 36724500 | 35 | 26 | [illegible] |
| SE | 78038 | ENSG00000136250 | AOAH | chr7 | - | 36673942 | 36674009 | 36659165 | 36659265 | 36724021 | 36724500 | 401 | 295 | [illegible] |
| SE | 78041 | ENSG00000151116 | UEVLD | chr11 | - | 18534329 | 18534453 | 18529608 | 18532487 | 18536405 | 18536469 | 58 | 45 | [illegible] |
| SE | 78042 | ENSG00000151116 | UEVLD | chr11 | - | 18534329 | 18534453 | 18529608 | 18532487 | 18544622 | 18544796 | 34 | 29 | [illegible] |
| SE | 78043 | ENSG00000151116 | UEVLD | chr11 | - | 18536405 | 18536469 | 18531200 | 18532487 | 18544622 | 18544796 | 16 | 7 | [illegible] |
| SE | 78044 | ENSG00000151116 | UEVLD | chr11 | - | 18536405 | 18536469 | 18534329 | 18534453 | 18544622 | 18544796 | 35 | 22 | [illegible] |
| SE | 78045 | ENSG00000151116 | UEVLD | chr11 | - | 18546879 | 18547050 | 18544622 | 18544796 | 18558227 | 18558330 | 55 | 45 | [illegible] |
| SE | 78046 | ENSG00000151116 | UEVLD | chr11 | - | 18564891 | 18565010 | 18558227 | 18558330 | 18570213 | 18570377 | 45 | 36 | [illegible] |
| SE | 78047 | ENSG00000151116 | UEVLD | chr11 | - | 18566346 | 18566482 | 18564891 | 18565010 | 18570213 | 18570377 | 87 | 107 | [illegible] |
| SE | 78048 | ENSG00000151116 | UEVLD | chr11 | - | 18566346 | 18566482 | 18564891 | 18565010 | 18575346 | 18575412 | 44 | 77 | [illegible] |
| SE | 78049 | ENSG00000151116 | UEVLD | chr11 | - | 18570213 | 18570377 | 18564891 | 18565010 | 18575346 | 18575412 | 24 | 28 | [illegible] |
| SE | 78052 | ENSG00000151116 | UEVLD | chr11 | - | 18570620 | 18570738 | 18570344 | 18570377 | 18575346 | 18575412 | 1 | 4 | [illegible] |
| SE | 78053 | ENSG00000151116 | UEVLD | chr11 | - | 18570620 | 18570738 | 18570344 | 18570377 | 18578723 | 18578808 | 1 | 3 | [illegible] |
| SE | 78054 | ENSG00000151116 | UEVLD | chr11 | - | 18575346 | 18575412 | 18566346 | 18566482 | 18578723 | 18578808 | 28 | 12 | [illegible] |
| SE | 78055 | ENSG00000151116 | UEVLD | chr11 | - | 18575346 | 18575412 | 18569897 | 18570377 | 18578723 | 18578808 | 48 | 36 | [illegible] |
| SE | 78056 | ENSG00000151116 | UEVLD | chr11 | - | 18575346 | 18575412 | 18570620 | 18570738 | 18578723 | 18578808 | 24 | 9 | [illegible] |
| SE | 78061 | ENSG00000127952 | STYXL1 | chr7 | - | 76013741 | 76013887 | 76005258 | 76005404 | 76028641 | 76028671 | 64 | 18 | [illegible] |
| SE | 78062 | ENSG00000127952 | STYXL1 | chr7 | - | 76021850 | 76021992 | 76005258 | 76005404 | 76028641 | 76028671 | 64 | 34 | [illegible] |
| SE | 78063 | ENSG00000127952 | STYXL1 | chr7 | - | 76021850 | 76021992 | 76013741 | 76013887 | 76028641 | 76028671 | 114 | 63 | [illegible] |
| SE | 78075 | ENSG00000196923 | PDLIM7 | chr5 | - | 177489770 | 177489832 | 177489392 | 177489627 | 177490869 | 177490906 | 17 | 8 | [illegible] |
| SE | 78076 | ENSG00000196923 | PDLIM7 | chr5 | - | 177489770 | 177490031 | 177489392 | 177489627 | 177490869 | 177490906 | 7 | 2 | [illegible] |
| SE | 78082 | ENSG00000196923 | PDLIM7 | chr5 | - | 177491806 | 177491976 | 177491071 | 177491420 | 177492404 | 177492435 | 2 | 0 | [illegible] |
| SE | 78083 | ENSG00000196923 | PDLIM7 | chr5 | - | 177491806 | 177491995 | 177491403 | 177491420 | 177492404 | 177492435 | 0 | 0 | [illegible] |
| SE | 78084 | ENSG00000196923 | PDLIM7 | chr5 | - | 177492404 | 177492435 | 177491009 | 177491146 | 177492525 | 177492572 | 7 | 1 | [illegible] |
| SE | 78085 | ENSG00000005020 | SKAP2 | chr7 | - | 26684735 | 26684848 | 26670090 | 26670192 | 26690284 | 26690362 | 620 | 732 | [illegible] |
| SE | 78087 | ENSG00000164543 | STK17A | chr7 | + | 43619596 | 43619723 | 43608255 | 43608400 | 43623571 | 43623620 | 783 | 435 | [illegible] |
| SE | 78088 | ENSG00000164543 | STK17A | chr7 | + | 43623571 | 43623620 | 43619596 | 43619723 | 43623708 | 43623888 | 586 | 367 | [illegible] |
| SE | 78089 | ENSG00000164543 | STK17A | chr7 | + | 43626477 | 43626586 | 43624517 | 43625035 | 43650557 | 43650713 | 8 | 12 | [illegible] |
| SE | 78099 | ENSG00000141258 | SGSM2 | chr17 | + | 2343544 | 2343620 | 2337500 | 2337745 | 2361636 | 2361799 | 59 | 43 | [illegible] |
| SE | 78102 | ENSG00000141258 | SGSM2 | chr17 | + | 2371261 | 2371415 | 2365214 | 2365341 | 2372189 | 2372254 | 81 | 48 | [illegible] |

| | | | | | | | | | | | | | | | | | | | | | | | | | | | | | | | | | | | | | | | | | | | | | | |
|---|---|---|---|---|---|---|---|---|---|---|---|---|---|---|---|---|---|---|---|---|---|---|---|---|---|---|---|---|---|---|---|---|---|---|---|---|---|---|---|---|---|---|---|---|---|---|
| SE | 78104 | ENSG00000141258 | SGSM2 | chr17 | + | 2379035 | 2379203 | 2377856 | 2377953 | 2379431 | 2381050 | 131 | 68 | 132 | 101 | 93 | 0 | 0 | 0 | 0 | 0 | 147 | 127 | 79 | 117 | 168 | 0 | 0 | 0 | 4 | 0 | 298 | 149 | 1 | 1 | 1 | 1 | 1 | 1 | 1 | 1 | 1 | 1 | 0.936 | 1 | 0.013 |
| SE | 78105 | ENSG00000141258 | SGSM2 | chr17 | + | 2379431 | 2379493 | 2379167 | 2379203 | 2380264 | 2380810 | 19 | 16 | 16 | 14 | 24 | 0 | 0 | 1 | 0 | 0 | 19 | 10 | 14 | 26 | 29 | 0 | 0 | 0 | 0 | 0 | 211 | 149 | 1 | 1 | 1 | 1 | 0.919 | 1 | 1 | 1 | 1 | 1 | 1 | 1 | -0.016 |
| SE | 78116 | ENSG00000254319 | NA | chr8 | - | 2705214 | 2705286 | 2673501 | 2674845 | 2728329 | 2728386 | 1 | 0 | 0 | 1 | 1 | 0 | 0 | 0 | 0 | 0 | 0 | 0 | 0 | 0 | 0 | 7 | 0 | 0 | 1 | 0 | 221 | 149 | 1 | 1 | 1 | NA | NA | 1 | 1 | 0 | NA | NA | 0 | NA | 1 |
| SE | 78121 | ENSG00000106638 | TBL2 | chr7 | - | 73576540 | 73576701 | 73574382 | 73574513 | 73577972 | 73578267 | 2 | 1 | 3 | 5 | 0 | 2 | 0 | 0 | 0 | 0 | 0 | 0 | 2 | 0 | 0 | 0 | 0 | 0 | 0 | 0 | 298 | 149 | 1 | 1 | 0.333 | 1 | 1 | 1 | NA | NA | NA | 1 | NA | NA | -0.167 |
| SE | 78123 | ENSG00000106638 | TBL2 | chr7 | - | 73576540 | 73576701 | 73574382 | 73574513 | 73578419 | 73578619 | 2 | 2 | 4 | 4 | 0 | 9 | 8 | 2 | 13 | 0 | 0 | 0 | 2 | 0 | 0 | 22 | 0 | 2 | 3 | 13 | 298 | 149 | 1 | 1 | 0.1 | 0.111 | 0.5 | 0.133 | NA | 0 | NA | 0.333 | 0 | 0 | 0.128 |
| SE | 78126 | ENSG00000126524 | SBDS | chr7 | - | 66991136 | 66991301 | 66987679 | 66988499 | 66993216 | 66993417 | 269 | 248 | 395 | 442 | 782 | 1 | 0 | 2 | 3 | 1 | 276 | 535 | 564 | 360 | 93 | 0 | 2 | 2 | 4 | 0 | 298 | 149 | 1 | 1 | 0.993 | 1 | 0.99 | 0.987 | 0.997 | 1 | 0.993 | 0.993 | 0.978 | 1 | 0.001 |
| SE | 78127 | ENSG00000126524 | SBDS | chr7 | - | 66993216 | 66993417 | 66991060 | 66991301 | 66994211 | 66994341 | 469 | 362 | 580 | 655 | 950 | 0 | 0 | 2 | 2 | 0 | 575 | 803 | 693 | 671 | 384 | 0 | 0 | 0 | 0 | 0 | 298 | 149 | 1 | 1 | 1 | 1 | 0.993 | 0.994 | 1 | 1 | 1 | 1 | 1 | 1 | -0.003 |
| SE | 78128 | ENSG00000126524 | SBDS | chr7 | - | 66994211 | 66994341 | 66991136 | 66991301 | 66995289 | 66995586 | 120 | 57 | 90 | 129 | 149 | 0 | 0 | 1 | 0 | 0 | 48 | 131 | 165 | 68 | 34 | 0 | 0 | 0 | 0 | 0 | 279 | 149 | 1 | 1 | 1 | 1 | 0.98 | 1 | 1 | 1 | 1 | 1 | 1 | 1 | -0.004 |
| SE | 78130 | ENSG00000268750 | NA | chr19 | + | 57820493 | 57820549 | 57819718 | 57819945 | 57856103 | 57856233 | 18 | 9 | 12 | 12 | 0 | 1 | 0 | 0 | 1 | 0 | 8 | 13 | 19 | 2 | 28 | 8 | 0 | 0 | 1 | 0 | 205 | 149 | 1 | 1 | 0.929 | 1 | 1 | 0.897 | NA | 0.421 | 1 | 1 | 0.592 | 1 | 0.154 |
| SE | 78131 | ENSG00000268750 | NA | chr19 | + | 57820493 | 57820549 | 57819718 | 57819945 | 57856106 | 57856233 | 18 | 13 | 15 | 11 | 0 | 2 | 0 | 1 | 2 | 0 | 27 | 11 | 20 | 2 | 28 | 0 | 0 | 0 | 0 | 12 | 205 | 149 | 1 | 1 | 0.867 | 1 | 0.916 | 0.8 | NA | 1 | 1 | 1 | 1 | 0.629 | -0.03 |
| SE | 78132 | ENSG00000268750 | NA | chr19 | + | 57839022 | 57839149 | 57819718 | 57819945 | 57856103 | 57856233 | 10 | 1 | 7 | 8 | 1 | 1 | 0 | 0 | 1 | 0 | 0 | 2 | 5 | 1 | 0 | 8 | 0 | 0 | 1 | 0 | 276 | 149 | 1 | 1 | 0.844 | 1 | 1 | 0.812 | 1 | 0 | 1 | 1 | 0.351 | NA | 0.343 |
| SE | 78133 | ENSG00000268750 | NA | chr19 | + | 57839022 | 57839149 | 57819718 | 57819945 | 57856106 | 57856233 | 10 | 1 | 7 | 8 | 1 | 2 | 0 | 1 | 2 | 0 | 0 | 0 | 7 | 1 | 0 | 0 | 0 | 0 | 0 | 12 | 276 | 149 | 1 | 1 | 0.73 | 1 | 0.791 | 0.683 | 1 | NA | NA | 1 | 1 | 0 | 0.174 |
| SE | 78134 | ENSG00000268750 | NA | chr19 | + | 57839022 | 57839149 | 57820493 | 57820549 | 57856103 | 57856233 | 12 | 2 | 2 | 10 | 0 | 0 | 0 | 0 | 1 | 0 | 0 | 2 | 8 | 11 | 5 | 0 | 2 | 0 | 0 | 0 | 276 | 149 | 1 | 1 | 1 | 1 | 1 | 0.844 | NA | NA | 0.351 | 1 | 1 | 1 | 0.123 |
| SE | 78135 | ENSG00000268750 | NA | chr19 | + | 57839022 | 57839149 | 57820493 | 57820549 | 57856106 | 57856233 | 11 | 2 | 2 | 10 | 0 | 0 | 4 | 3 | 0 | 0 | 0 | 0 | 10 | 11 | 5 | 19 | 0 | 1 | 0 | 0 | 276 | 149 | 1 | 1 | 1 | 0.213 | 0.265 | 1 | NA | 0 | NA | 0.844 | 1 | 1 | -0.091 |
| SE | 78138 | ENSG00000214765 | SEPTIN7P2 | chr7 | - | 45726183 | 45726236 | 45724049 | 45724480 | 45728177 | 45728313 | 5 | 7 | 4 | 11 | 15 | 0 | 4 | 3 | 1 | 0 | 9 | 3 | 0 | 0 | 0 | 0 | 0 | 0 | 0 | 0 | 202 | 149 | 1 | 1 | 1 | 0.563 | 0.496 | 0.89 | 1 | 1 | 1 | NA | NA | NA | -0.21 |
| SE | 78140 | ENSG00000214765 | SEPTIN7P2 | chr7 | - | 45736786 | 45736910 | 45727826 | 45728313 | 45744996 | 45745048 | 117 | 109 | 123 | 123 | 150 | 0 | 0 | 0 | 0 | 0 | 173 | 89 | 208 | 126 | 47 | 0 | 0 | 1 | 0 | 0 | 273 | 149 | 1 | 1 | 1 | 1 | 1 | 1 | 1 | 1 | 1 | 0.991 | 1 | 1 | 0.002 |
| SE | 78142 | ENSG00000214765 | SEPTIN7P2 | chr7 | - | 45747214 | 45747311 | 45746826 | 45746948 | 45748625 | 45748664 | 27 | 37 | 38 | 26 | 21 | 0 | 0 | 0 | 0 | 0 | 60 | 32 | 39 | 54 | 15 | 0 | 0 | 1 | 0 | 0 | 246 | 149 | 1 | 1 | 1 | 1 | 1 | 1 | 1 | 1 | 1 | 0.959 | 1 | 1 | 0.008 |
| SE | 78143 | ENSG00000214765 | SEPTIN7P2 | chr7 | - | 45751528 | 45751646 | 45748625 | 45748714 | 45756908 | 45756992 | 74 | 63 | 92 | 95 | 73 | 1 | 0 | 0 | 1 | 0 | 84 | 51 | 102 | 86 | 14 | 2 | 0 | 0 | 0 | 0 | 267 | 149 | 1 | 1 | 0.976 | 1 | 1 | 0.981 | 1 | 0.959 | 1 | 1 | 1 | 1 | 0 |
| SE | 78145 | ENSG00000214765 | SEPTIN7P2 | chr7 | - | 45765705 | 45765812 | 45759073 | 45759174 | 45768659 | 45768985 | 0 | 0 | 4 | 0 | 0 | 7 | 2 | 20 | 8 | 1 | 0 | 0 | 0 | 0 | 0 | 1 | 9 | 10 | 9 | 0 | 256 | 149 | 1 | 1 | 0 | 0 | 0.104 | 0 | 0 | 0 | 0 | 0 | 0 | NA | 0.021 |
| SE | 78151 | ENSG00000181704 | YIPF6 | chrX | + | 68510678 | 68510776 | 68499050 | 68499123 | 68511848 | 68511977 | 2 | 0 | 1 | 0 | 0 | 9 | 2 | 12 | 9 | 7 | 0 | 0 | 0 | 0 | 0 | 4 | 6 | 0 | 2 | 4 | 247 | 149 | 1 | 1 | 0.118 | 0 | 0.048 | 0 | 0 | 0 | 0 | NA | 0 | 0 | 0.033 |
| SE | 78153 | ENSG00000181704 | YIPF6 | chrX | + | 68511848 | 68511977 | 68499050 | 68499123 | 68513326 | 68513405 | 33 | 19 | 50 | 46 | 32 | 1 | 0 | 0 | 0 | 0 | 36 | 30 | 25 | 10 | 14 | 0 | 0 | 0 | 0 | 0 | 278 | 149 | 1 | 1 | 0.946 | 1 | 1 | 1 | 1 | 1 | 1 | 1 | 1 | 1 | -0.011 |
| SE | 78154 | ENSG00000181704 | YIPF6 | chrX | + | 68518769 | 68518812 | 68513326 | 68513405 | 68521371 | 68521497 | 55 | 20 | 53 | 50 | 51 | 1 | 0 | 0 | 0 | 0 | 73 | 28 | 32 | 2 | 11 | 0 | 0 | 1 | 0 | 0 | 192 | 149 | 1 | 1 | 0.977 | 1 | 1 | 1 | 1 | 1 | 1 | 0.961 | 1 | 1 | 0.003 |
| SE | 78157 | ENSG00000239713 | APOBEC3G | chr22 | + | 39080932 | 39081227 | 39078931 | 39079085 | 39086278 | 39086567 | 459 | 187 | 209 | 293 | 336 | 0 | 0 | 0 | 0 | 0 | 333 | 291 | 134 | 178 | 364 | 0 | 0 | 1 | 0 | 0 | 298 | 149 | 1 | 1 | 1 | 1 | 1 | 1 | 1 | 1 | 1 | 0.985 | 1 | 1 | 0.003 |
| SE | 78158 | ENSG00000239713 | APOBEC3G | chr22 | + | 39081470 | 39081585 | 39080932 | 39081227 | 39086278 | 39086567 | 315 | 142 | 172 | 263 | 193 | 1 | 0 | 0 | 0 | 0 | 150 | 212 | 104 | 153 | 228 | 0 | 0 | 0 | 0 | 0 | 264 | 149 | 1 | 1 | 0.994 | 1 | 1 | 1 | 1 | 1 | 1 | 1 | 1 | 1 | -0.001 |
| SE | 78159 | ENSG00000239713 | APOBEC3G | chr22 | + | 39083730 | 39083884 | 39081470 | 39081585 | 39086278 | 39086567 | 543 | 216 | 270 | 459 | 456 | 3 | 0 | 1 | 4 | 2 | 312 | 479 | 135 | 288 | 291 | 0 | 2 | 1 | 0 | 0 | 298 | 149 | 1 | 1 | 0.989 | 1 | 0.993 | 0.983 | 0.991 | 1 | 0.992 | 0.985 | 1 | 1 | -0.004 |
| SE | 78160 | ENSG00000223705 | NSUN5P1 | chr7 | + | 75410605 | 75410726 | 75410361 | 75410460 | 75414884 | 75415023 | 5 | 0 | 4 | 8 | 0 | 2 | 0 | 0 | 0 | 0 | 0 | 0 | 0 | 1 | 8 | 0 | 6 | 6 | 5 | 0 | 270 | 149 | 1 | 1 | 0.58 | NA | 1 | 1 | NA | NA | 0 | 0 | 0.099 | 1 | 0.585 |
| SE | 78163 | ENSG00000223705 | NSUN5P1 | chr7 | + | 75412788 | 75412932 | 75411654 | 75411759 | 75414884 | 75415023 | 53 | 23 | 47 | 38 | 72 | 0 | 0 | 0 | 0 | 0 | 20 | 48 | 16 | 19 | 38 | 0 | 0 | 2 | 0 | 0 | 293 | 149 | 1 | 1 | 1 | 1 | 1 | 1 | 1 | 1 | 1 | 0.803 | 1 | 1 | 0.039 |
| SE | 78173 | ENSG00000227486 | NA | chrX | + | 55985998 | 55986224 | 55908231 | 55908800 | 56002824 | 56002983 | 1 | 1 | 0 | 6 | 0 | 0 | 0 | 1 | 0 | 0 | 0 | 1 | 0 | 0 | 0 | 0 | 0 | 0 | 0 | 0 | 298 | 149 | 1 | 1 | 1 | 1 | 0 | 1 | NA | NA | 1 | NA | NA | NA | -0.25 |
| SE | 78174 | ENSG00000227486 | NA | chrX | + | 55985998 | 55986224 | 55908231 | 55908800 | 56014844 | 56021946 | 1 | 0 | 0 | 5 | 5 | 8 | 3 | 8 | 8 | 23 | 0 | 1 | 0 | 0 | 0 | 0 | 13 | 13 | 3 | 0 | 298 | 149 | 1 | 1 | 0.059 | 0 | 0 | 0.238 | 0.098 | NA | 0.037 | 0 | 0 | NA | 0.067 |
| SE | 78175 | ENSG00000227486 | NA | chrX | + | 55985998 | 55986224 | 55908282 | 55908517 | 56002824 | 56002983 | 0 | 1 | 0 | 4 | 0 | 0 | 0 | 0 | 0 | 0 | 0 | 0 | 0 | 0 | 0 | 0 | 0 | 1 | 0 | 0 | 298 | 149 | 1 | 1 | NA | 1 | NA | 1 | NA | NA | NA | 0 | NA | NA | 1 |
| SE | 78176 | ENSG00000227486 | NA | chrX | + | 55985998 | 55986224 | 55908282 | 55908517 | 56014844 | 56021946 | 0 | 0 | 0 | 3 | 5 | 0 | 0 | 0 | 0 | 0 | 0 | 0 | 0 | 0 | 0 | 0 | 0 | 1 | 0 | 0 | 298 | 149 | 1 | 1 | NA | NA | NA | 1 | 1 | NA | NA | 0 | NA | NA | 1 |
| SE | 78177 | ENSG00000227486 | NA | chrX | + | 56002824 | 56002983 | 55908152 | 55908800 | 56014844 | 56014961 | 0 | 0 | 1 | 0 | 0 | 8 | 3 | 8 | 8 | 23 | 0 | 0 | 1 | 4 | 0 | 0 | 13 | 13 | 3 | 0 | 298 | 149 | 1 | 1 | 0 | 0 | 0.059 | 0 | 0 | NA | 0 | 0.037 | 0.4 | NA | -0.134 |
| SE | 78179 | ENSG00000227486 | NA | chrX | + | 56002824 | 56002983 | 55985998 | 55986224 | 56014844 | 56014961 | 0 | 1 | 0 | 3 | 0 | 0 | 0 | 0 | 2 | 5 | 0 | 0 | 1 | 4 | 0 | 0 | 0 | 0 | 0 | 0 | 298 | 149 | 1 | 1 | NA | 1 | NA | 0.429 | 0 | NA | NA | 1 | 1 | NA | -0.524 |
| SE | 78180 | ENSG00000227486 | NA | chrX | + | 56014844 | 56014961 | 55908242 | 55908800 | 56204972 | 56205253 | 3 | 1 | 6 | 8 | 21 | 0 | 0 | 0 | 1 | 0 | 0 | 13 | 18 | 2 | 0 | 0 | 0 | 0 | 0 | 0 | 266 | 149 | 1 | 1 | 1 | 1 | 1 | 0.818 | 1 | NA | 1 | 1 | 1 | NA | -0.036 |
| SE | 78182 | ENSG00000196715 | VKORC1L1 | chr7 | + | 65948670 | 65948780 | 65873073 | 65873565 | 65954073 | 65959558 | 38 | 26 | 43 | 44 | 63 | 0 | 0 | 1 | 1 | 0 | 14 | 42 | 18 | 32 | 97 | 0 | 0 | 0 | 0 | 0 | 259 | 149 | 1 | 1 | 1 | 1 | 0.961 | 0.962 | 1 | 1 | 1 | 1 | 1 | 1 | -0.015 |
| SE | 78183 | ENSG00000136273 | HUS1 | chr7 | - | 47965299 | 47965438 | 47964319 | 47964632 | 47967805 | 47967925 | 64 | 78 | 143 | 103 | 129 | 0 | 0 | 0 | 0 | 0 | 48 | 60 | 67 | 78 | 52 | 0 | 0 | 0 | 2 | 0 | 288 | 149 | 1 | 1 | 1 | 1 | 1 | 1 | 1 | 1 | 1 | 1 | 0.953 | 1 | 0.009 |
| SE | 78184 | ENSG00000136273 | HUS1 | chr7 | - | 47967805 | 47967925 | 47963287 | 47965438 | 47969218 | 47969318 | 170 | 127 | 218 | 197 | 307 | 0 | 0 | 0 | 1 | 0 | 99 | 127 | 131 | 131 | 109 | 0 | 0 | 0 | 0 | 0 | 269 | 149 | 1 | 1 | 1 | 1 | 1 | 0.991 | 1 | 1 | 1 | 1 | 1 | 1 | -0.002 |
| SE | 78185 | ENSG00000136273 | HUS1 | chr7 | - | 47969218 | 47969318 | 47963287 | 47965438 | 47975612 | 47975687 | 70 | 48 | 65 | 75 | 128 | 0 | 1 | 0 | 0 | 0 | 52 | 48 | 38 | 57 | 63 | 0 | 0 | 0 | 0 | 0 | 249 | 149 | 1 | 1 | 1 | 0.966 | 1 | 1 | 1 | 1 | 1 | 1 | 1 | 1 | -0.007 |
| SE | 78186 | ENSG00000136273 | HUS1 | chr7 | - | 47971259 | 47971516 | 47968818 | 47969318 | 47975612 | 47975687 | 0 | 0 | 0 | 0 | 0 | 108 | 91 | 121 | 126 | 211 | 0 | 3 | 1 | 2 | 0 | 78 | 63 | 65 | 89 | 70 | 298 | 149 | 1 | 1 | 0 | 0 | 0 | 0 | 0 | 0 | 0.023 | 0.008 | 0.011 | 0 | -0.008 |
| SE | 78189 | ENSG00000124486 | USP9X | chrX | + | 41123470 | 41123724 | 41085444 | 41086109 | 41128999 | 41129145 | 357 | 283 | 363 | 367 | 258 | 0 | 0 | 2 | 1 | 0 | 259 | 407 | 320 | 208 | 86 | 0 | 0 | 0 | 3 | 0 | 298 | 149 | 1 | 1 | 1 | 1 | 0.989 | 0.995 | 1 | 1 | 1 | 1 | 0.972 | 1 | 0.002 |
| SE | 78190 | ENSG00000124486 | USP9X | chrX | + | 41128999 | 41129145 | 41123470 | 41123724 | 41131456 | 41131536 | 405 | 332 | 404 | 450 | 247 | 0 | 0 | 1 | 0 | 0 | 310 | 438 | 384 | 255 | 117 | 0 | 0 | 0 | 1 | 0 | 295 | 149 | 1 | 1 | 1 | 1 | 0.995 | 1 | 1 | 1 | 1 | 1 | 0.992 | 1 | 0.001 |
| SE | 78193 | ENSG00000124486 | USP9X | chrX | + | 41140965 | 41141217 | 41136803 | 41137022 | 41141292 | 41141431 | 386 | 343 | 303 | 343 | 416 | 0 | 0 | 0 | 1 | 0 | 85 | 338 | 309 | 228 | 20 | 3 | 0 | 0 | 0 | 0 | 298 | 149 | 1 | 1 | 1 | 1 | 1 | 0.994 | 1 | 0.934 | 1 | 1 | 1 | 1 | 0.012 |
| SE | 78194 | ENSG00000124486 | USP9X | chrX | + | 41140965 | 41141217 | 41140655 | 41140771 | 41141292 | 41141431 | 612 | 486 | 421 | 532 | 595 | 0 | 2 | 0 | 2 | 0 | 126 | 423 | 430 | 289 | 34 | 0 | 0 | 0 | 0 | 0 | 298 | 149 | 1 | 1 | 1 | 0.992 | 1 | 0.993 | 1 | 1 | 1 | 1 | 1 | 1 | -0.003 |
| SE | 78195 | ENSG00000124486 | USP9X | chrX | + | 41143290 | 41143443 | 41141292 | 41141431 | 41144521 | 41144626 | 671 | 468 | 471 | 545 | 762 | 7 | 0 | 0 | 0 | 0 | 168 | 606 | 576 | 264 | 93 | 0 | 0 | 0 | 0 | 0 | 298 | 149 | 1 | 1 | 0.98 | 1 | 1 | 1 | 1 | 1 | 1 | 1 | 1 | 1 | -0.004 |
| SE | 78197 | ENSG00000124486 | USP9X | chrX | + | 41152947 | 41153081 | 41148368 | 41148575 | 41162789 | 41162877 | 398 | 278 | 295 | 373 | 458 | 0 | 0 | 0 | 0 | 0 | 114 | 235 | 315 | 196 | 7 | 0 | 0 | 0 | 2 | 0 | 283 | 149 | 1 | 1 | 1 | 1 | 1 | 1 | 1 | 1 | 1 | 1 | 0.981 | 1 | 0.004 |
| SE | 78201 | ENSG00000124486 | USP9X | chrX | + | 41186516 | 41186642 | 41184396 | 41184675 | 41187991 | 41188117 | 174 | 134 | 181 | 147 | 220 | 0 | 0 | 0 | 0 | 0 | 8 | 106 | 162 | 52 | 4 | 0 | 0 | 0 | 1 | 0 | 275 | 149 | 1 | 1 | 1 | 1 | 1 | 1 | 1 | 1 | 1 | 1 | 0.966 | 1 | 0.007 |
| SE | 78202 | ENSG00000124486 | USP9X | chrX | + | 41187991 | 41188117 | 41186516 | 41186642 | 41189308 | 41189475 | 330 | 204 | 231 | 260 | 358 | 0 | 0 | 0 | 0 | 0 | 45 | 207 | 276 | 81 | 4 | 0 | 1 | 0 | 4 | 0 | 275 | 149 | 1 | 1 | 1 | 1 | 1 | 1 | 1 | 1 | 0.991 | 1 | 0.916 | 1 | 0.019 |
| SE | 78203 | ENSG00000124486 | USP9X | chrX | + | 41189308 | 41189475 | 41187991 | 41188117 | 41196250 | 41196359 | 772 | 483 | 554 | 583 | 676 | 0 | 0 | 0 | 0 | 0 | 152 | 486 | 581 | 191 | 17 | 0 | 2 | 0 | 0 | 0 | 298 | 149 | 1 | 1 | 1 | 1 | 1 | 1 | 1 | 1 | 0.992 | 1 | 1 | 1 | 0.002 |
| SE | 78204 | ENSG00000124486 | USP9X | chrX | + | 41198527 | 41198750 | 41197363 | 41197510 | 41201059 | 41201280 | 589 | 393 | 481 | 466 | 803 | 0 | 0 | 0 | 0 | 1 | 60 | 322 | 329 | 89 | 18 | 0 | 0 | 0 | 0 | 0 | 298 | 149 | 1 | 1 | 1 | 1 | 1 | 1 | 0.998 | 1 | 1 | 1 | 1 | 1 | 0 |
| SE | 78205 | ENSG00000124486 | USP9X | chrX | + | 41201059 | 41201280 | 41198527 | 41198750 | 41205302 | 41205493 | 518 | 315 | 461 | 427 | 688 | 4 | 0 | 3 | 2 | 0 | 34 | 250 | 250 | 76 | 14 | 0 | 0 | 2 | 0 | 0 | 298 | 149 | 1 | 1 | 0.985 | 1 | 0.987 | 0.991 | 1 | 1 | 1 | 0.984 | 1 | 1 | -0.004 |
| SE | 78206 | ENSG00000124486 | USP9X | chrX | + | 41210508 | 41210682 | 41205302 | 41205493 | 41214567 | 41214709 | 331 | 185 | 240 | 229 | 441 | 0 | 0 | 0 | 2 | 0 | 0 | 134 | 164 | 42 | 2 | 0 | 0 | 0 | 0 | 0 | 298 | 149 | 1 | 1 | 1 | 1 | 1 | 0.983 | 1 | NA | 1 | 1 | 1 | 1 | -0.003 |
| SE | 78207 | ENSG00000124486 | USP9X | chrX | + | 41214567 | 41214709 | 41210508 | 41210682 | 41215898 | 41216652 | 475 | 345 | 375 | 337 | 571 | 1 | 1 | 0 | 3 | 0 | 8 | 194 | 337 | 65 | 1 | 0 | 0 | 2 | 0 | 0 | 291 | 149 | 1 | 1 | 0.996 | 0.994 | 1 | 0.983 | 1 | 1 | 1 | 0.989 | 1 | 1 | -0.003 |
| SE | 78208 | ENSG00000124486 | USP9X | chrX | + | 41219101 | 41219231 | 41218371 | 41218597 | 41223216 | 41223402 | 98 | 88 | 95 | 110 | 239 | 2 | 1 | 3 | 9 | 2 | 6 | 49 | 69 | 42 | 0 | 0 | 18 | 6 | 0 | 0 | 279 | 149 | 1 | 1 | 0.963 | 0.979 | 0.944 | 0.867 | 0.985 | 1 | 0.592 | 0.86 | 1 | NA | 0.085 |
| SE | 78209 | ENSG00000124486 | USP9X | chrX | + | 41223216 | 41223402 | 41219101 | 41219231 | 41224741 | 41224962 | 197 | 156 | 187 | 212 | 300 | 0 | 0 | 0 | 0 | 7 | 0 | 99 | 110 | 51 | 0 | 0 | 0 | 0 | 0 | 0 | 298 | 149 | 1 | 1 | 1 | 1 | 1 | 1 | 0.955 | NA | 1 | 1 | 1 | NA | -0.009 |
| SE | 78210 | ENSG00000196937 | FAM3C | chr7 | - | 121351142 | 121351269 | 121348877 | 121350550 | 121360042 | 121360127 | 181 | 149 | 126 | 180 | 249 | 0 | 0 | 0 | 2 | 0 | 97 | 219 | 154 | 82 | 24 | 0 | 0 | 0 | 0 | 0 | 276 | 149 | 1 | 1 | 1 | 1 | 1 | 0.98 | 1 | 1 | 1 | 1 | 1 | 1 | -0.004 |
| SE | 78211 | ENSG00000196937 | FAM3C | chr7 | - | 121362896 | 121362947 | 121360042 | 121360127 | 121364129 | 121364188 | 149 | 112 | 140 | 162 | 282 | 1 | 0 | 0 | 0 | 1 | 49 | 176 | 134 | 35 | 31 | 0 | 1 | 0 | 2 | 0 | 200 | 149 | 1 | 1 | 0.991 | 1 | 1 | 1 | 0.995 | 1 | 0.992 | 1 | 0.929 | 1 | 0.013 |
| SE | 78212 | ENSG00000196937 | FAM3C | chr7 | - | 121380208 | 121380264 | 121378909 | 121379014 | 121382956 | 121383010 | 1 | 0 | 0 | 1 | 0 | 76 | 58 | 60 | 83 | 90 | 0 | 4 | 2 | 0 | 0 | 16 | 106 | 82 | 34 | 11 | 205 | 149 | 1 | 1 | 0.009 | 0 | 0 | 0.009 | 0 | 0 | 0.027 | 0.017 | 0 | 0 | -0.005 |
| SE | 78216 | ENSG00000183696 | UPP1 | chr7 | + | 48101823 | 48101870 | 48094762 | 48094827 | 48103781 | 48103991 | 5 | 3 | 4 | 5 | 0 | 1 | 0 | 0 | 0 | 0 | 7 | 25 | 9 | 23 | 9 | 0 | 0 | 0 | 0 | 6 | 196 | 149 | 1 | 1 | 0.792 | 1 | 1 | 1 | NA | 1 | 1 | 1 | 1 | 0.533 | 0.041 |
| SE | 78234 | ENSG00000106605 | BLVRA | chr7 | + | 43760016 | 43760110 | 43758686 | 43758734 | 43771137 | 43771170 | 1 | 1 | 0 | 1 | 0 | 6 | 5 | 5 | 4 | 0 | 8 | 0 | 0 | 0 | 8 | 21 | 13 | 7 | 21 | 11 | 243 | 149 | 1 | 1 | 0.093 | 0.109 | 0 | 0.133 | NA | 0.189 | 0 | 0 | 0 | 0.308 | -0.016 |
| SE | 78235 | ENSG00000106605 | BLVRA | chr7 | + | 43787903 | 43788025 | 43771134 | 43771170 | 43791248 | 43791368 | 130 | 145 | 159 | 148 | 52 | 0 | 0 | 0 | 0 | 0 | 499 | 236 | 89 | 348 | 437 | 0 | 0 | 1 | 0 | 0 | 271 | 149 | 1 | 1 | 1 | 1 | 1 | 1 | 1 | 1 | 1 | 0.98 | 1 | 1 | 0.004 |
| SE | 78236 | ENSG00000106605 | BLVRA | chr7 | + | 43803675 | 43803847 | 43800464 | 43800572 | 43806976 | 43807342 | 241 | 285 | 191 | 210 | 306 | 0 | 0 | 0 | 1 | 0 | 1901 | 437 | 219 | 1211 | 913 | 0 | 0 | 0 | 3 | 0 | 298 | 149 | 1 | 1 | 1 | 1 | 1 | 0.991 | 1 | 1 | 1 | 1 | 0.995 | 1 | -0.001 |
| SE | 78239 | ENSG00000107438 | PDLIM1 | chr10 | - | 95247214 | 95247366 | 95238567 | 95238685 | 95263863 | 95264063 | 309 | 353 | 192 | 554 | 338 | 0 | 1 | 0 | 7 | 0 | 1617 | 786 | 296 | 483 | 6368 | 0 | 4 | 2 | 2 | 4 | 298 | 149 | 1 | 1 | 1 | 0.994 | 1 | 0.975 | 1 | 1 | 0.99 | 0.987 | 0.992 | 0.999 | 0 |
| SE | 78244 | ENSG00000154978 | VOPP1 | chr7 | - | 55561853 | 55562051 | 55521071 | 55521130 | 55572270 | 55572454 | 0 | 0 | 0 | 0 | 0 | 46 | 10 | 20 | 35 | 1 | 0 | 4 | 0 | 0 | 0 | 7 | 21 | 23 | 16 | 14 | 298 | 149 | 1 | 1 | 0 | 0 | 0 | 0 | 0 | 0 | 0.087 | 0 | 0 | 0 | -0.017 |
| SE | 78247 | ENSG00000158560 | DYNC1I1 | chr7 | + | 95813195 | 95813337 | 95810391 | 95810506 | 95869882 | 95869998 | 1 | 1 | 0 | 0 | 0 | 0 | 0 | 0 | 0 | 0 | 6 | 0 | 1 | 0 | 6 | 0 | 0 | 2 | 0 | 0 | 291 | 149 | 1 | 1 | 1 | 1 | NA | NA | NA | 1 | NA | 0.204 | NA | 1 | 0.265 |
| SE | 78248 | ENSG00000158560 | DYNC1I1 | chr7 | + | 95813246 | 95813337 | 95810391 | 95810506 | 95869882 | 95869998 | 2 | 1 | 2 | 8 | 1 | 0 | 0 | 0 | 0 | 0 | 22 | 1 | 3 | 6 | 36 | 0 | 0 | 2 | 0 | 0 | 240 | 149 | 1 | 1 | 1 | 1 | 1 | 1 | 1 | 1 | 1 | 0.482 | 1 | 1 | 0.104 |
| SE | 78249 | ENSG00000158560 | DYNC1I1 | chr7 | + | 95828056 | 95828116 | 95813195 | 95813337 | 95869882 | 95869998 | 2 | 4 | 1 | 3 | 0 | 2 | 1 | 0 | 0 | 0 | 0 | 8 | 0 | 6 | 45 | 22 | 0 | 1 | 1 | 13 | 209 | 149 | 1 | 1 | 0.416 | 0.74 | 1 | 1 | NA | 0 | 1 | 0 | 0.811 | 0.712 | 0.284 |
| SE | 78251 | ENSG00000158560 | DYNC1I1 | chr7 | + | 96028174 | 96028321 | 95995947 | 95996073 | 96032666 | 96032780 | 0 | 1 | 1 | 1 | 0 | 0 | 0 | 0 | 0 | 0 | 0 | 0 | 0 | 9 | 0 | 0 | 0 | 1 | 0 | 0 | 296 | 149 | 1 | 1 | NA | 1 | 1 | 1 | NA | NA | NA | 0 | 1 | NA | 0.5 |
| SE | 78252 | ENSG00000272391 | POM121C | chr7 | - | 75424048 | 75424225 | 75421508 | 75423203 | 75424525 | 75424628 | 238 | 96 | 178 | 197 | 450 | 1 | 0 | 0 | 0 | 0 | 404 | 242 | 141 | 192 | 242 | 0 | 0 | 0 | 0 | 0 | 298 | 149 | 1 | 1 | 0.992 | 1 | 1 | 1 | 1 | 1 | 1 | 1 | 1 | 1 | -0.002 |
| SE | 78255 | ENSG00000173715 | C11orf80 | chr11 | + | 66748378 | 66748517 | 66744735 | 66744919 | 66756352 | 66756505 | 18 | 4 | 7 | 9 | 18 | 2 | 0 | 0 | 3 | 3 | 0 | 2 | 1 | 22 | 2 | 0 | 1 | 0 | 0 | 0 | 288 | 149 | 1 | 1 | 0.823 | 1 | 1 | 0.608 | 0.756 | NA | 0.509 | 1 | 1 | 1 | -0.04 |
| SE | 78256 | ENSG00000173715 | C11orf80 | chr11 | + | 66748378 | 66748517 | 66744735 | 66744919 | 66759042 | 66759096 | 6 | 1 | 1 | 14 | 5 | 9 | 1 | 3 | 0 | 0 | 0 | 0 | 0 | 5 | 2 | 0 | 2 | 1 | 0 | 0 | 288 | 149 | 1 | 1 | 0.256 | 0.341 | 0.147 | 1 | 1 | NA | 0 | 0 | 1 | 1 | 0.049 |
| SE | 78257 | ENSG00000173715 | C11orf80 | chr11 | + | 66756352 | 66756505 | 66744735 | 66744919 | 66759042 | 66759084 | 13 | 6 | 2 | 15 | 8 | 9 | 1 | 3 | 0 | 0 | 0 | 7 | 7 | 10 | 10 | 0 | 2 | 1 | 0 | 0 | 298 | 149 | 1 | 1 | 0.419 | 0.75 | 0.25 | 1 | 1 | NA | 0.636 | 0.778 | 1 | 1 | -0.17 |
| SE | 78258 | ENSG00000173715 | C11orf80 | chr11 | + | 66756352 | 66756505 | 66748378 | 66748517 | 66759042 | 66759084 | 23 | 9 | 9 | 14 | 20 | 0 | 1 | 1 | 7 | 2 | 0 | 8 | 8 | 32 | 10 | 0 | 0 | 0 | 0 | 0 | 298 | 149 | 1 | 1 | 1 | 0.818 | 0.818 | 0.5 | 0.833 | NA | 1 | 1 | 1 | 1 | -0.206 |
| SE | 78259 | ENSG00000173715 | C11orf80 | chr11 | + | 66759042 | 66759096 | 66744768 | 66744919 | 66788159 | 66788269 | 8 | 2 | 6 | 12 | 5 | 0 | 0 | 0 | 0 | 1 | 0 | 2 | 5 | 4 | 6 | 0 | 0 | 0 | 0 | 0 | 203 | 149 | 1 | 1 | 1 | 1 | 1 | 1 | 0.786 | NA | 1 | 1 | 1 | 1 | -0.043 |
| SE | 78261 | ENSG00000173715 | C11orf80 | chr11 | + | 66788159 | 66788269 | 66759042 | 66759096 | 66796280 | 66796364 | 8 | 5 | 9 | 17 | 10 | 0 | 0 | 0 | 0 | 8 | 0 | 2 | 13 | 6 | 24 | 0 | 0 | 0 | 0 | 0 | 259 | 149 | 1 | 1 | 1 | 1 | 1 | 1 | 0.418 | NA | 1 | 1 | 1 | 1 | -0.116 |
| SE | 78262 | ENSG00000173715 | C11orf80 | chr11 | + | 66788159 | 66788269 | 66759042 | 66759096 | 66800640 | 66800696 | 2 | 5 | 4 | 10 | 6 | 3 | 0 | 0 | 0 | 0 | 0 | 2 | 7 | 2 | 6 | 0 | 0 | 0 | 0 | 0 | 259 | 149 | 1 | 1 | 0.277 | 1 | 1 | 1 | 1 | NA | 1 | 1 | 1 | 1 | -0.145 |
| SE | 78263 | ENSG00000173715 | C11orf80 | chr11 | + | 66788159 | 66788273 | 66759042 | 66759096 | 66796280 | 66796364 | 2 | 5 | 5 | 11 | 5 | 0 | 0 | 0 | 0 | 8 | 0 | 2 | 7 | 2 | 6 | 0 | 0 | 0 | 0 | 0 | 263 | 149 | 1 | 1 | 1 | 1 | 1 | 1 | 0.261 | NA | 1 | 1 | 1 | 1 | -0.148 |
| SE | 78264 | ENSG00000173715 | C11orf80 | chr11 | + | 66796280 | 66796364 | 66759042 | 66759096 | 66800640 | 66800686 | 6 | 4 | 7 | 15 | 22 | 3 | 0 | 0 | 0 | 0 | 2 | 9 | 4 | 2 | 0 | 0 | 0 | 0 | 0 | 0 | 233 | 149 | 1 | 1 | 0.561 | 1 | 1 | 1 | 1 | 1 | 1 | 1 | 1 | NA | -0.088 |
| SE | 78265 | ENSG00000173715 | C11orf80 | chr11 | + | 66796280 | 66796364 | 66788159 | 66788269 | 66800640 | 66800686 | 12 | 4 | 12 | 22 | 24 | 0 | 0 | 0 | 0 | 1 | 2 | 9 | 8 | 11 | 18 | 0 | 0 | 0 | 0 | 0 | 233 | 149 | 1 | 1 | 1 | 1 | 1 | 1 | 0.939 | 1 | 1 | 1 | 1 | 1 | -0.012 |
| SE | 78268 | ENSG00000173715 | C11orf80 | chr11 | + | 66821642 | 66821764 | 66815833 | 66816199 | 66822563 | 66822660 | 29 | 21 | 68 | 46 | 97 | 0 | 0 | 0 | 0 | 0 | 54 | 49 | 48 | 84 | 65 | 0 | 2 | 0 | 0 | 0 | 271 | 149 | 1 | 1 | 1 | 1 | 1 | 1 | 1 | 1 | 0.931 | 1 | 1 | 1 | 0.014 |
| SE | 78271 | ENSG00000173715 | C11orf80 | chr11 | + | 66822563 | 66822674 | 66821642 | 66821764 | 66828265 | 66828364 | 32 | 25 | 77 | 59 | 102 | 1 | 0 | 1 | 2 | 0 | 49 | 86 | 64 | 84 | 64 | 0 | 1 | 1 | 6 | 4 | 260 | 149 | 1 | 1 | 0.948 | 1 | 0.978 | 0.944 | 1 | 1 | 0.98 | 0.973 | 0.889 | 0.902 | 0.025 |
| SE | 78272 | ENSG00000173715 | C11orf80 | chr11 | + | 66839131 | 66839207 | 66838369 | 66838445 | 66842848 | 66843069 | 0 | 0 | 0 | 2 | 0 | 10 | 15 | 18 | 10 | 5 | 0 | 0 | 1 | 0 | 0 | 13 | 28 | 13 | 12 | 27 | 225 | 149 | 1 | 1 | 0 | 0 | 0 | 0.117 | 0 | 0 | 0 | 0.048 | 0 | 0 | 0.014 |
| SE | 78273 | ENSG00000105829 | BET1 | chr7 | - | 93972574 | 93972663 | 93962761 | 93965687 | 93975903 | 93976100 | 4 | 3 | 10 | 6 | 14 | 0 | 4 | 0 | 2 | 1 | 17 | 2 | 4 | 4 | 0 | 8 | 7 | 4 | 0 | 0 | 238 | 149 | 1 | 1 | 1 | 0.32 | 1 | 0.653 | 0.898 | 0.571 | 0.152 | 0.385 | 1 | NA | 0.247 |
| SE | 78276 | ENSG00000164715 | LMTK2 | chr7 | + | 98137314 | 98137442 | 98106861 | 98107280 | 98141396 | 98141541 | 67 | 44 | 47 | 69 | 78 | 0 | 0 | 1 | 0 | 0 | 79 | 65 | 64 | 96 | 28 | 0 | 0 | 0 | 0 | 0 | 277 | 149 | 1 | 1 | 1 | 1 | 0.962 | 1 | 1 | 1 | 1 | 1 | 1 | 1 | -0.008 |
| SE | 78278 | ENSG00000164715 | LMTK2 | chr7 | + | 98185050 | 98185135 | 98171540 | 98171674 | 98186876 | 98186998 | 31 | 16 | 68 | 38 | 59 | 0 | 0 | 0 | 0 | 0 | 76 | 27 | 39 | 78 | 43 | 0 | 0 | 2 | 0 | 0 | 234 | 149 | 1 | 1 | 1 | 1 | 1 | 1 | 1 | 1 | 1 | 0.925 | 1 | 1 | 0.015 |
| SE | 78279 | ENSG00000164715 | LMTK2 | chr7 | + | 98190727 | 98190877 | 98186876 | 98186998 | 98191613 | 98194572 | 76 | 30 | 71 | 66 | 101 | 0 | 0 | 0 | 0 | 0 | 60 | 62 | 54 | 54 | 100 | 0 | 0 | 0 | 0 | 0 | 298 | 149 | 1 | 1 | 1 | 1 | 0.973 | 1 | 1 | 1 | 1 | 1 | 1 | 1 | -0.005 |
| SE | 78280 | ENSG00000164715 | LMTK2 | chr7 | + | 98191613 | 98194572 | 98190727 | 98190877 | 98203573 | 98203706 | 75 | 43 | 79 | 78 | 118 | 0 | 1 | 4 | 0 | 0 | 79 | 60 | 66 | 106 | 96 | 0 | 0 | 0 | 0 | 0 | 298 | 149 | 1 | 1 | 1 | 0.956 | 0.908 | 1 | 1 | 1 | 1 | 1 | 1 | 1 | -0.027 |
| SE | 78281 | ENSG00000164715 | LMTK2 | chr7 | + | 98203943 | 98204186 | 98203573 | 98203706 | 98205463 | 98209638 | 46 | 32 | 50 | 51 | 51 | 0 | 3 | 0 | 0 | 0 | 61 | 25 | 81 | 82 | 35 | 0 | 0 | 0 | 0 | 0 | 298 | 149 | 1 | 1 | 1 | 0.842 | 1 | 1 | 1 | 1 | 1 | 1 | 1 | 1 | -0.032 |
| SE | 78282 | ENSG00000062725 | APPBP2 | chr17 | - | 60451879 | 60452045 | 60443157 | 60447834 | 60454301 | 60454492 | 214 | 137 | 164 | 207 | 300 | 2 | 0 | 0 | 0 | 1 | 34 | 198 | 152 | 77 | 27 | 0 | 1 | 0 | 1 | 0 | 298 | 149 | 1 | 1 | 0.982 | 1 | 1 | 1 | 0.993 | 1 | 0.99 | 1 | 0.975 | 1 | 0.002 |
| SE | 78283 | ENSG00000062725 | APPBP2 | chr17 | - | 60460662 | 60460787 | 60456295 | 60456381 | 60461809 | 60461915 | 195 | 120 | 112 | 173 | 332 | 6 | 1 | 0 | 0 | 1 | 33 | 44 | 92 | 48 | 29 | 0 | 0 | 0 | 2 | 0 | 274 | 149 | 1 | 1 | 0.946 | 0.985 | 1 | 1 | 0.994 | 1 | 1 | 1 | 0.929 | 1 | -0.001 |
| SE | 78284 | ENSG00000062725 | APPBP2 | chr17 | - | 60461809 | 60461915 | 60456295 | 60456381 | 60461993 | 60462061 | 54 | 37 | 19 | 43 | 82 | 4 | 1 | 0 | 5 | 1 | 7 | 9 | 22 | 10 | 0 | 0 | 6 | 1 | 0 | 0 | 255 | 149 | 1 | 1 | 0.887 | 0.956 | 1 | 0.834 | 0.98 | 1 | 0.467 | 0.928 | 1 | NA | 0.083 |
| SE | 78285 | ENSG00000062725 | APPBP2 | chr17 | - | 60466290 | 60466459 | 60464020 | 60464110 | 60479147 | 60479271 | 271 | 166 | 206 | 235 | 245 | 0 | 0 | 1 | 0 | 0 | 6 | 143 | 162 | 107 | 11 | 0 | 0 | 0 | 0 | 0 | 298 | 149 | 1 | 1 | 1 | 1 | 0.99 | 1 | 1 | 1 | 1 | 1 | 1 | 1 | -0.002 |
| SE | 78286 | ENSG00000062725 | APPBP2 | chr17 | - | 60468438 | 60468510 | 60466350 | 60466459 | 60477468 | 60477506 | 3 | 2 | 1 | 6 | 2 | 1 | 1 | 0 | 1 | 0 | 0 | 3 | 1 | 1 | 0 | 0 | 0 | 0 | 0 | 0 | 221 | 149 | 1 | 1 | 0.669 | 0.574 | 1 | 0.802 | 1 | NA | 1 | 1 | 1 | NA | -0.191 |
| SE | 78287 | ENSG00000062725 | APPBP2 | chr17 | - | 60468438 | 60468510 | 60466350 | 60466459 | 60479147 | 60479271 | 4 | 2 | 1 | 8 | 4 | 77 | 43 | 59 | 57 | 39 | 0 | 4 | 1 | 1 | 0 | 0 | 27 | 59 | 28 | 3 | 221 | 149 | 1 | 1 | 0.034 | 0.03 | 0.011 | 0.086 | 0.065 | NA | 0.091 | 0.011 | 0.024 | 0 | 0.014 |
| SE | 78288 | ENSG00000062725 | APPBP2 | chr17 | - | 60477468 | 60477506 | 60466350 | 60466459 | 60479147 | 60479271 | 1 | 1 | 0 | 1 | 0 | 77 | 43 | 59 | 57 | 39 | 0 | 0 | 0 | 0 | 0 | 0 | 27 | 59 | 28 | 3 | 187 | 149 | 1 | 1 | 0.01 | 0.018 | 0 | 0.014 | 0 | NA | 0 | 0 | 0 | 0 | 0.008 |
| SE | 78290 | ENSG00000062725 | APPBP2 | chr17 | - | 60479147 | 60479271 | 60466290 | 60466459 | 60494465 | 60494617 | 134 | 65 | 105 | 94 | 54 | 0 | 0 | 0 | 1 | 0 | 0 | 42 | 73 | 42 | 3 | 0 | 0 | 0 | 0 | 0 | 273 | 149 | 1 | 1 | 1 | 1 | 1 | 0.981 | 1 | NA | 1 | 1 | 1 | 1 | -0.004 |
| SE | 78291 | ENSG00000062725 | APPBP2 | chr17 | - | 60496412 | 60496441 | 60494465 | 60494617 | 60500398 | 60500487 | 4 | 1 | 3 | 5 | 7 | 38 | 15 | 16 | 29 | 23 | 0 | 0 | 0 | 7 | 0 | 0 | 12 | 18 | 12 | 12 | 178 | 149 | 1 | 1 | 0.081 | 0.053 | 0.136 | 0.126 | 0.203 | NA | 0 | 0 | 0.328 | 0 | 0.038 |
| SE | 78295 | ENSG00000134882 | UBAC2 | chr13 | + | 99314096 | 99314220 | 99238426 | 99238554 | 99340319 | 99340565 | 0 | 0 | 0 | 1 | 1 | 0 | 0 | 0 | 0 | 0 | 0 | 0 | 1 | 0 | 0 | 0 | 0 | 0 | 4 | 0 | 273 | 149 | 1 | 1 | NA | NA | NA | 1 | 1 | NA | NA | 1 | 0 | NA | 0.5 |
| SE | 78296 | ENSG00000134882 | UBAC2 | chr13 | + | 99314096 | 99314220 | 99244514 | 99244624 | 99318021 | 99318069 | 418 | 264 | 341 | 395 | 529 | 1 | 1 | 1 | 6 | 4 | 335 | 322 | 205 | 314 | 553 | 0 | 0 | 3 | 0 | 0 | 273 | 149 | 1 | 1 | 0.996 | 0.993 | 0.995 | 0.973 | 0.986 | 1 | 1 | 0.974 | 1 | 1 | -0.006 |
| SE | 78297 | ENSG00000134882 | UBAC2 | chr13 | + | 99314096 | 99314220 | 99244514 | 99244624 | 99340319 | 99340565 | 226 | 132 | 183 | 206 | 257 | 0 | 0 | 0 | 0 | 0 | 153 | 167 | 89 | 144 | 229 | 0 | 0 | 2 | 3 | 0 | 273 | 149 | 1 | 1 | 1 | 1 | 1 | 1 | 1 | 1 | 1 | 0.96 | 0.963 | 1 | 0.015 |
| SE | 78298 | ENSG00000134882 | UBAC2 | chr13 | + | 99318021 | 99318069 | 99244514 | 99244624 | 99340319 | 99340565 | 94 | 69 | 82 | 114 | 120 | 0 | 0 | 0 | 0 | 0 | 155 | 94 | 56 | 69 | 214 | 0 | 0 | 2 | 3 | 0 | 197 | 149 | 1 | 1 | 1 | 1 | 1 | 1 | 1 | 1 | 1 | 0.955 | 0.946 | 1 | 0.02 |
| SE | 78299 | ENSG00000134882 | UBAC2 | chr13 | + | 99318021 | 99318069 | 99314096 | 99314220 | 99340319 | 99340565 | 271 | 195 | 231 | 296 | 367 | 0 | 0 | 0 | 0 | 1 | 332 | 229 | 152 | 241 | 542 | 0 | 0 | 1 | 0 | 0 | 197 | 149 | 1 | 1 | 1 | 1 | 1 | 1 | 0.996 | 1 | 1 | 0.991 | 1 | 1 | 0.001 |
| SE | 78300 | ENSG00000134882 | UBAC2 | chr13 | + | 99340319 | 99340565 | 99318021 | 99318069 | 99367786 | 99367906 | 347 | 235 | 267 | 359 | 388 | 0 | 0 | 2 | 1 | 0 | 369 | 258 | 181 | 235 | 630 | 0 | 2 | 2 | 0 | 0 | 298 | 149 | 1 | 1 | 1 | 1 | 0.985 | 0.994 | 1 | 1 | 0.985 | 0.978 | 1 | 1 | 0.003 |
| SE | 78301 | ENSG00000134882 | UBAC2 | chr13 | + | 99340319 | 99340565 | 99318021 | 99318069 | 99385227 | 99385315 | 259 | 183 | 205 | 274 | 344 | 1 | 0 | 0 | 0 | 0 | 302 | 218 | 143 | 201 | 475 | 0 | 0 | 0 | 0 | 0 | 298 | 149 | 1 | 1 | 0.992 | 1 | 1 | 1 | 1 | 1 | 1 | 1 | 1 | 1 | -0.002 |
| SE | 78302 | ENSG00000134882 | UBAC2 | chr13 | + | 99367786 | 99367906 | 99318021 | 99318069 | 99385227 | 99385315 | 104 | 45 | 67 | 74 | 51 | 1 | 0 | 0 | 0 | 0 | 58 | 77 | 24 | 24 | 120 | 0 | 0 | 0 | 0 | 0 | 269 | 149 | 1 | 1 | 0.983 | 1 | 1 | 1 | 1 | 1 | 1 | 1 | 1 | 1 | -0.003 |
| SE | 78307 | ENSG00000122547 | EEPD1 | chr7 | + | 36296992 | 36297187 | 36287588 | 36287777 | 36299006 | 36299192 | 64 | 29 | 40 | 39 | 48 | 0 | 1 | 0 | 0 | 0 | 54 | 5 | 32 | 62 | 46 | 0 | 0 | 0 | 0 | 0 | 298 | 149 | 1 | 1 | 1 | 0.935 | 1 | 1 | 1 | 1 | 1 | 1 | 1 | 1 | -0.013 |
| SE | 78308 | ENSG00000186187 | ZNRF1 | chr16 | + | 75084428 | 75084637 | 74999344 | 75000095 | 75093571 | 75093667 | 0 | 0 | 0 | 3 | 0 | 2 | 0 | 0 | 3 | 2 | 0 | 0 | 0 | 0 | 0 | 0 | 0 | 2 | 1 | 6 | 298 | 149 | 1 | 1 | 0 | NA | NA | 0.333 | 0 | NA | NA | 0 | 0 | 0 | 0.111 |
| SE | 78309 | ENSG00000186187 | ZNRF1 | chr16 | + | 75093571 | 75093667 | 74999023 | 75000095 | 75104783 | 75104889 | 7 | 2 | 7 | 6 | 4 | 2 | 0 | 0 | 0 | 0 | 9 | 0 | 2 | 11 | 26 | 0 | 0 | 0 | 0 | 0 | 245 | 149 | 1 | 1 | 0.68 | 1 | 1 | 1 | 1 | 1 | NA | 1 | 1 | 1 | -0.064 |
| SE | 78317 | ENSG00000228409 | CCT6P1 | chr7 | + | 65752163 | 65752202 | 65751141 | 65751278 | 65754526 | 65754661 | 0 | 0 | 0 | 0 | 0 | 0 | 1 | 1 | 1 | 0 | 0 | 1 | 0 | 0 | 0 | 0 | 0 | 0 | 0 | 0 | 188 | 149 | 1 | 1 | NA | 0 | 0 | 0 | NA | NA | 1 | NA | NA | NA | -1 |
| SE | 78318 | ENSG00000228409 | CCT6P1 | chr7 | + | 65752163 | 65752310 | 65751141 | 65751278 | 65754526 | 65754661 | 4 | 1 | 12 | 6 | 3 | 0 | 1 | 1 | 1 | 0 | 0 | 11 | 13 | 8 | 4 | 0 | 0 | 0 | 0 | 0 | 296 | 149 | 1 | 1 | 1 | 0.335 | 0.858 | 0.751 | 1 | NA | 1 | 1 | 1 | 1 | -0.211 |
| SE | 78319 | ENSG00000228409 | CCT6P1 | chr7 | + | 65752163 | 65752310 | 65751141 | 65751278 | 65755779 | 65755952 | 2 | 3 | 8 | 1 | 0 | 4 | 2 | 1 | 9 | 8 | 0 | 1 | 2 | 0 | 0 | 0 | 0 | 1 | 0 | 2 | 296 | 149 | 1 | 1 | 0.201 | 0.43 | 0.801 | 0.053 | 0 | NA | 1 | 0.502 | NA | 0 | -0.204 |
| SE | 78320 | ENSG00000228409 | CCT6P1 | chr7 | + | 65754526 | 65754661 | 65751141 | 65751278 | 65755779 | 65755952 | 7 | 7 | 11 | 10 | 12 | 4 | 2 | 1 | 9 | 8 | 0 | 13 | 6 | 8 | 0 | 0 | 0 | 1 | 0 | 2 | 284 | 149 | 1 | 1 | 0.479 | 0.647 | 0.852 | 0.368 | 0.44 | NA | 1 | 0.759 | 1 | 0 | -0.133 |
| SE | 78321 | ENSG00000228409 | CCT6P1 | chr7 | + | 65754526 | 65754661 | 65752163 | 65752310 | 65755779 | 65755952 | 11 | 6 | 18 | 15 | 15 | 2 | 2 | 4 | 0 | 0 | 0 | 23 | 19 | 16 | 4 | 0 | 0 | 2 | 0 | 0 | 284 | 149 | 1 | 1 | 0.743 | 0.611 | 0.702 | 1 | 1 | NA | 1 | 0.833 | 1 | 1 | -0.147 |
| SE | 78324 | ENSG00000228409 | CCT6P1 | chr7 | + | 65758235 | 65758317 | 65756468 | 65756574 | 65759174 | 65759271 | 5 | 1 | 6 | 5 | 13 | 0 | 1 | 0 | 0 | 0 | 0 | 7 | 7 | 0 | 3 | 0 | 0 | 0 | 0 | 0 | 231 | 149 | 1 | 1 | 1 | 0.392 | 1 | 1 | 1 | NA | 1 | 1 | NA | 1 | -0.122 |
| SE | 78331 | ENSG00000178741 | COX5A | chr15 | - | 74923647 | 74923770 | 74919790 | 74920442 | 74926765 | 74926887 | 127 | 88 | 150 | 145 | 147 | 1 | 0 | 0 | 0 | 0 | 258 | 148 | 113 | 209 | 334 | 0 | 2 | 0 | 0 | 0 | 272 | 149 | 1 | 1 | 0.986 | 1 | 1 | 1 | 1 | 1 | 0.976 | 1 | 1 | 1 | 0.002 |
| SE | 78334 | ENSG00000178741 | COX5A | chr15 | - | 74926765 | 74926887 | 74923593 | 74923770 | 74929115 | 74929232 | 225 | 132 | 247 | 230 | 206 | 0 | 0 | 0 | 1 | 0 | 397 | 264 | 158 | 339 | 483 | 0 | 0 | 0 | 0 | 0 | 271 | 149 | 1 | 1 | 1 | 1 | 1 | 0.992 | 1 | 1 | 1 | 1 | 1 | 1 | -0.002 |
| SE | 78335 | ENSG00000178741 | COX5A | chr15 | - | 74929115 | 74929232 | 74926765 | 74926887 | 74937914 | 74938066 | 160 | 87 | 187 | 147 | 129 | 0 | 0 | 0 | 2 | 0 | 319 | 216 | 99 | 214 | 302 | 0 | 0 | 0 | 4 | 0 | 266 | 149 | 1 | 1 | 1 | 1 | 1 | 0.976 | 1 | 1 | 1 | 1 | 0.968 | 1 | 0.002 |
| SE | 78360 | ENSG00000072609 | CHFR | chr12 | - | 132872284 | 132872394 | 132861466 | 132861634 | 132887195 | 132887340 | 7 | 7 | 10 | 8 | 13 | 0 | 0 | 1 | 0 | 0 | 30 | 18 | 19 | 23 | 24 | 0 | 2 | 0 | 0 | 0 | 259 | 149 | 1 | 1 | 1 | 1 | 0.852 | 1 | 1 | 1 | 0.838 | 1 | 1 | 1 | 0.003 |
| SE | 78362 | ENSG00000072609 | CHFR | chr12 | - | 132887195 | 132887340 | 132877554 | 132877654 | 132887546 | 132887564 | 109 | 65 | 80 | 90 | 49 | 0 | 0 | 0 | 0 | 1 | 92 | 123 | 63 | 77 | 191 | 0 | 0 | 0 | 0 | 0 | 294 | 149 | 1 | 1 | 1 | 1 | 1 | 1 | 0.961 | 1 | 1 | 1 | 1 | 1 | -0.008 |
| SE | 78363 | ENSG00000072609 | CHFR | chr12 | - | 132898999 | 132899213 | 132877554 | 132877654 | 132908775 | 132909147 | 0 | 0 | 2 | 0 | 0 | 0 | 0 | 2 | 0 | 0 | 0 | 0 | 0 | 0 | 0 | 0 | 2 | 0 | 0 | 0 | 298 | 149 | 1 | 1 | NA | NA | 0.333 | NA | NA | NA | 0 | NA | NA | NA | 0.333 |
| SE | 78364 | ENSG00000132432 | SEC61G | chr7 | - | 54755778 | 54755881 | 54752249 | 54752420 | 54757494 | 54757594 | 578 | 364 | 499 | 487 | 236 | 0 | 0 | 0 | 0 | 0 | 402 | 556 | 548 | 499 | 229 | 0 | 0 | 0 | 6 | 0 | 252 | 149 | 1 | 1 | 1 | 1 | 1 | 1 | 1 | 1 | 1 | 1 | 0.98 | 1 | 0.004 |
| SE | 78366 | ENSG00000132432 | SEC61G | chr7 | - | 54757494 | 54757594 | 54754791 | 54755881 | 54759161 | 54759198 | 190 | 117 | 147 | 130 | 128 | 2 | 0 | 0 | 0 | 0 | 117 | 207 | 180 | 172 | 32 | 0 | 0 | 0 | 0 | 0 | 249 | 149 | 1 | 1 | 0.983 | 1 | 1 | 1 | 1 | 1 | 1 | 1 | 1 | 1 | -0.003 |
| SE | 78367 | ENSG00000132432 | SEC61G | chr7 | - | 54759157 | 54759334 | 54757494 | 54757594 | 54759790 | 54759974 | 177 | 110 | 166 | 155 | 45 | 0 | 0 | 0 | 0 | 0 | 140 | 166 | 165 | 121 | 75 | 0 | 0 | 2 | 0 | 0 | 298 | 149 | 1 | 1 | 1 | 1 | 1 | 1 | 1 | 1 | 1 | 0.976 | 1 | 1 | 0.005 |
| SE | 78368 | ENSG00000176102 | CSTF3 | chr11 | - | 33090531 | 33090727 | 33086987 | 33087141 | 33092270 | 33092340 | 53 | 41 | 39 | 63 | 159 | 1 | 0 | 1 | 0 | 0 | 0 | 32 | 24 | 18 | 5 | 0 | 0 | 0 | 0 | 0 | 298 | 149 | 1 | 1 | 0.964 | 1 | 0.951 | 1 | 1 | NA | 1 | 1 | 1 | 1 | -0.017 |
| SE | 78369 | ENSG00000176102 | CSTF3 | chr11 | - | 33092270 | 33092340 | 33090531 | 33090727 | 33096305 | 33096408 | 36 | 27 | 32 | 42 | 91 | 0 | 0 | 1 | 0 | 0 | 9 | 31 | 13 | 15 | 0 | 0 | 0 | 0 | 0 | 0 | 219 | 149 | 1 | 1 | 1 | 1 | 0.956 | 1 | 1 | 1 | 1 | 1 | 1 | NA | -0.009 |
| SE | 78370 | ENSG00000176102 | CSTF3 | chr11 | - | 33099607 | 33099717 | 33099033 | 33099150 | 33102176 | 33102339 | 34 | 36 | 41 | 35 | 46 | 1 | 0 | 2 | 0 | 2 | 9 | 15 | 10 | 22 | 0 | 0 | 6 | 7 | 0 | 0 | 259 | 149 | 1 | 1 | 0.951 | 1 | 0.922 | 1 | 0.93 | 1 | 0.59 | 0.451 | 1 | NA | 0.2 |
| SE | 78371 | ENSG00000176102 | CSTF3 | chr11 | - | 33144869 | 33144973 | 33141884 | 33141986 | 33161298 | 33161495 | 0 | 0 | 2 | 0 | 0 | 34 | 26 | 37 | 39 | 51 | 0 | 0 | 0 | 0 | 0 | 13 | 11 | 20 | 17 | 0 | 253 | 149 | 1 | 1 | 0 | 0 | 0.031 | 0 | 0 | 0 | 0 | 0 | 0 | NA | 0.006 |
| SE | 78380 | ENSG00000198900 | TOP1 | chr20 | + | 41061393 | 41061490 | 41029430 | 41029455 | 41076170 | 41076294 | 76 | 64 | 80 | 83 | 59 | 1 | 2 | 0 | 0 | 0 | 51 | 110 | 184 | 99 | 15 | 0 | 0 | 0 | 0 | 0 | 246 | 149 | 1 | 1 | 0.979 | 0.951 | 1 | 1 | 1 | 1 | 1 | 1 | 1 | 1 | -0.014 |
| SE | 78382 | ENSG00000198900 | TOP1 | chr20 | + | 41077581 | 41077637 | 41076170 | 41076294 | 41080084 | 41080180 | 73 | 79 | 61 | 85 | 102 | 0 | 0 | 0 | 1 | 0 | 209 | 98 | 247 | 168 | 58 | 0 | 0 | 0 | 0 | 0 | 205 | 149 | 1 | 1 | 1 | 1 | 1 | 0.984 | 1 | 1 | 1 | 1 | 1 | 1 | -0.003 |
| SE | 78383 | ENSG00000198900 | TOP1 | chr20 | + | 41080084 | 41080180 | 41076182 | 41077637 | 41081164 | 41081240 | 90 | 93 | 84 | 118 | 119 | 0 | 0 | 0 | 0 | 0 | 270 | 124 | 332 | 195 | 94 | 0 | 0 | 0 | 2 | 0 | 245 | 149 | 1 | 1 | 1 | 1 | 1 | 1 | 1 | 1 | 1 | 1 | 0.983 | 1 | 0.003 |
| SE | 78386 | ENSG00000198900 | TOP1 | chr20 | + | 41115370 | 41115439 | 41113969 | 41114155 | 41116277 | 41116392 | 181 | 115 | 163 | 162 | 162 | 0 | 0 | 4 | 1 | 0 | 71 | 161 | 177 | 71 | 20 | 0 | 0 | 0 | 0 | 0 | 218 | 149 | 1 | 1 | 1 | 1 | 0.965 | 0.991 | 1 | 1 | 1 | 1 | 1 | 1 | -0.009 |
| SE | 78388 | ENSG00000172115 | CYCS | chr7 | - | 25123950 | 25124127 | 25118655 | 25123849 | 25125199 | 25125260 | 501 | 322 | 495 | 528 | 748 | 4 | 3 | 0 | 0 | 0 | 324 | 572 | 553 | 565 | 205 | 0 | 2 | 3 | 7 | 0 | 298 | 149 | 1 | 1 | 0.984 | 0.982 | 1 | 1 | 1 | 1 | 0.993 | 0.989 | 0.976 | 1 | 0.002 |
| SE | 78395 | ENSG00000169919 | GUSB | chr7 | - | 65979398 | 65979541 | 65976014 | 65976202 | 65979726 | 65979911 | 161 | 86 | 156 | 168 | 60 | 3 | 0 | 1 | 0 | 0 | 212 | 182 | 168 | 252 | 318 | 0 | 2 | 2 | 0 | 0 | 292 | 149 | 1 | 1 | 0.965 | 1 | 0.988 | 1 | 1 | 1 | 0.979 | 1 | 1 | 1 | -0.005 |
| SE | 78402 | ENSG00000127980 | PEX1 | chr7 | - | 92503040 | 92503195 | 92501506 | 92501673 | 92504731 | 92504902 | 26 | 9 | 28 | 20 | 20 | 0 | 0 | 0 | 0 | 0 | 0 | 12 | 13 | 6 | 0 | 0 | 4 | 0 | 0 | 0 | 298 | 149 | 1 | 1 | 1 | 1 | 1 | 1 | 1 | NA | 0.6 | 1 | 1 | NA | 0.133 |
| SE | 78403 | ENSG00000127980 | PEX1 | chr7 | - | 92504731 | 92504902 | 92503040 | 92503195 | 92506247 | 92506344 | 35 | 30 | 46 | 43 | 66 | 1 | 2 | 4 | 0 | 0 | 18 | 31 | 14 | 2 | 0 | 0 | 1 | 0 | 0 | 0 | 298 | 149 | 1 | 1 | 0.946 | 0.882 | 0.852 | 1 | 1 | 1 | 0.939 | 1 | 1 | NA | -0.049 |
| SE | 78404 | ENSG00000127980 | PEX1 | chr7 | - | 92509328 | 92509411 | 92506993 | 92507126 | 92510943 | 92511047 | 35 | 13 | 27 | 41 | 71 | 13 | 15 | 21 | 28 | 16 | 10 | 24 | 6 | 11 | 0 | 0 | 5 | 4 | 0 | 0 | 232 | 149 | 1 | 1 | 0.634 | 0.358 | 0.452 | 0.485 | 0.74 | 1 | 0.755 | 0.491 | 1 | NA | -0.278 |
| SE | 78405 | ENSG00000127980 | PEX1 | chr7 | - | 92513847 | 92513967 | 92511579 | 92511703 | 92517275 | 92517415 | 96 | 88 | 105 | 113 | 306 | 0 | 0 | 0 | 2 | 0 | 6 | 68 | 47 | 35 | 0 | 0 | 0 | 0 | 0 | 0 | 269 | 149 | 1 | 1 | 1 | 1 | 1 | 0.969 | 1 | 1 | 1 | 1 | 1 | NA | -0.006 |
| SE | 78406 | ENSG00000127980 | PEX1 | chr7 | - | 92518140 | 92518255 | 92517275 | 92518042 | 92518994 | 92519078 | 197 | 152 | 224 | 181 | 381 | 0 | 0 | 0 | 0 | 2 | 30 | 119 | 133 | 84 | 61 | 0 | 0 | 0 | 0 | 0 | 264 | 149 | 1 | 1 | 1 | 1 | 1 | 1 | 0.991 | 1 | 1 | 1 | 1 | 1 | -0.002 |
| SE | 78407 | ENSG00000127980 | PEX1 | chr7 | - | 92518959 | 92519078 | 92518140 | 92518255 | 92522101 | 92522245 | 48 | 32 | 40 | 44 | 97 | 1 | 0 | 0 | 0 | 0 | 0 | 26 | 21 | 21 | 8 | 0 | 0 | 5 | 0 | 0 | 268 | 149 | 1 | 1 | 0.964 | 1 | 1 | 1 | 1 | NA | 1 | 0.7 | 1 | 1 | 0.068 |
| SE | 78409 | ENSG00000161405 | IKZF3 | chr17 | - | 39777650 | 39777767 | 39757717 | 39766493 | 39788257 | 39788374 | 559 | 245 | 423 | 485 | 525 | 0 | 0 | 2 | 0 | 0 | 188 | 527 | 302 | 324 | 290 | 0 | 0 | 2 | 0 | 0 | 266 | 149 | 1 | 1 | 1 | 1 | 0.992 | 1 | 1 | 1 | 1 | 0.988 | 1 | 1 | 0.001 |
| SE | 78419 | ENSG00000161405 | IKZF3 | chr17 | - | 39788257 | 39788374 | 39777650 | 39777767 | 39829386 | 39829488 | 149 | 53 | 121 | 122 | 162 | 0 | 0 | 0 | 0 | 0 | 41 | 117 | 67 | 80 | 38 | 0 | 0 | 3 | 0 | 0 | 266 | 149 | 1 | 1 | 1 | 1 | 1 | 1 | 1 | 1 | 1 | 0.926 | 1 | 1 | 0.015 |
| SE | 78423 | ENSG00000161405 | IKZF3 | chr17 | - | 39791415 | 39791583 | 39777650 | 39777767 | 39829386 | 39829488 | 2 | 2 | 1 | 5 | 4 | 0 | 0 | 0 | 0 | 0 | 0 | 7 | 7 | 15 | 0 | 0 | 0 | 3 | 0 | 0 | 298 | 149 | 1 | 1 | 1 | 1 | 1 | 1 | 1 | NA | 1 | 0.538 | 1 | NA | 0.154 |
| SE | 78427 | ENSG00000161405 | IKZF3 | chr17 | - | 39792672 | 39792933 | 39777650 | 39777767 | 39829386 | 39829488 | 111 | 60 | 69 | 75 | 59 | 0 | 0 | 0 | 0 | 0 | 51 | 107 | 65 | 86 | 56 | 0 | 0 | 3 | 0 | 0 | 298 | 149 | 1 | 1 | 1 | 1 | 1 | 1 | 1 | 1 | 1 | 0.915 | 1 | 1 | 0.017 |
| SE | 78431 | ENSG00000161405 | IKZF3 | chr17 | - | 39832097 | 39832151 | 39792672 | 39792933 | 39864119 | 39864126 | 36 | 9 | 31 | 32 | 2 | 0 | 0 | 0 | 0 | 0 | 21 | 23 | 15 | 22 | 16 | 0 | 0 | 0 | 0 | 0 | 203 | 149 | 1 | 1 | 1 | 1 | 1 | 0.959 | 1 | 1 | 1 | 1 | 1 | 1 | -0.008 |
| SE | 78433 | ENSG00000255036 | SUGT1P4-STRA6LP-CCDC180 | chr9 | + | 97290853 | 97290902 | 97290583 | 97290665 | 97307735 | 97307806 | 7 | 4 | 8 | 4 | 8 | 0 | 0 | 0 | 1 | 0 | 0 | 8 | 4 | 3 | 0 | 0 | 0 | 2 | 2 | 0 | 198 | 149 | 1 | 1 | 1 | 1 | 1 | 0.751 | 1 | NA | 1 | 0.601 | 0.53 | NA | 0.24 |
| SE | 78434 | ENSG00000255036 | SUGT1P4-STRA6LP-CCDC180 | chr9 | + | 97291370 | 97291487 | 97290853 | 97290902 | 97307735 | 97307806 | 3 | 3 | 5 | 1 | 5 | 1 | 0 | 2 | 1 | 0 | 0 | 4 | 3 | 0 | 5 | 0 | 4 | 0 | 0 | 0 | 266 | 149 | 1 | 1 | 0.627 | 1 | 0.583 | 0.359 | 1 | NA | 0.359 | 1 | NA | 1 | -0.073 |
| SE | 78436 | ENSG00000255036 | SUGT1P4-STRA6LP-CCDC180 | chr9 | + | 97294844 | 97294991 | 97293946 | 97294110 | 97307735 | 97307806 | 8 | 5 | 13 | 10 | 5 | 0 | 0 | 0 | 1 | 0 | 0 | 16 | 2 | 20 | 5 | 0 | 0 | 0 | 0 | 0 | 296 | 149 | 1 | 1 | 1 | 1 | 1 | 0.834 | 1 | NA | 1 | 1 | 1 | 1 | -0.033 |
| SE | 78438 | ENSG00000255036 | SUGT1P4-STRA6LP-CCDC180 | chr9 | + | 97317064 | 97317228 | 97314850 | 97314946 | 97318462 | 97318582 | 17 | 23 | 10 | 6 | 25 | 1 | 0 | 0 | 0 | 0 | 0 | 33 | 11 | 15 | 6 | 0 | 0 | 0 | 0 | 0 | 298 | 149 | 1 | 1 | 0.895 | 1 | 1 | 1 | 1 | NA | 1 | 1 | 1 | 1 | -0.021 |
| SE | 78439 | ENSG00000255036 | SUGT1P4-STRA6LP-CCDC180 | chr9 | + | 97318462 | 97318582 | 97317064 | 97317228 | 97320125 | 97320205 | 8 | 6 | 10 | 10 | 5 | 0 | 0 | 0 | 0 | 0 | 0 | 19 | 7 | 11 | 10 | 0 | 0 | 0 | 2 | 0 | 269 | 149 | 1 | 1 | 1 | 1 | 1 | 1 | 1 | NA | 1 | 1 | 0.753 | 1 | 0.062 |
| SE | 78441 | ENSG00000255036 | SUGT1P4-STRA6LP-CCDC180 | chr9 | + | 97343339 | 97343563 | 97330321 | 97330767 | 97349110 | 97349291 | 8 | 11 | 11 | 11 | 15 | 0 | 0 | 0 | 1 | 0 | 0 | 0 | 9 | 8 | 10 | 0 | 0 | 0 | 0 | 0 | 298 | 149 | 1 | 1 | 1 | 1 | 1 | 0.846 | 1 | NA | NA | 1 | 1 | 1 | -0.031 |
| SE | 78442 | ENSG00000255036 | SUGT1P4-STRA6LP-CCDC180 | chr9 | + | 97343339 | 97343563 | 97330321 | 97330767 | 97354568 | 97354713 | 9 | 14 | 13 | 13 | 26 | 0 | 0 | 0 | 0 | 3 | 8 | 0 | 9 | 14 | 13 | 0 | 1 | 0 | 0 | 0 | 298 | 149 | 1 | 1 | 1 | 1 | 1 | 1 | 0.812 | 1 | 0 | 1 | 1 | 1 | 0.162 |
| SE | 78443 | ENSG00000255036 | SUGT1P4-STRA6LP-CCDC180 | chr9 | + | 97347313 | 97347489 | 97343339 | 97343563 | 97349110 | 97349291 | 17 | 6 | 16 | 5 | 9 | 1 | 0 | 0 | 2 | 0 | 28 | 4 | 8 | 9 | 9 | 0 | 0 | 0 | 0 | 0 | 298 | 149 | 1 | 1 | 0.895 | 1 | 1 | 0.556 | 1 | 1 | 1 | 1 | 1 | 1 | -0.11 |
| SE | 78444 | ENSG00000255036 | SUGT1P4-STRA6LP-CCDC180 | chr9 | + | 97347313 | 97347666 | 97343339 | 97343563 | 97349110 | 97349291 | 15 | 7 | 14 | 3 | 7 | 1 | 0 | 0 | 2 | 0 | 19 | 7 | 13 | 15 | 19 | 0 | 0 | 0 | 0 | 0 | 298 | 149 | 1 | 1 | 0.882 | 1 | 1 | 0.429 | 1 | 1 | 1 | 1 | 1 | 1 | -0.138 |
| SE | 78445 | ENSG00000255036 | SUGT1P4-STRA6LP-CCDC180 | chr9 | + | 97347313 | 97347666 | 97343339 | 97343563 | 97350408 | 97350555 | 13 | 6 | 9 | 3 | 7 | 0 | 0 | 0 | 1 | 0 | 14 | 3 | 8 | 10 | 0 | 0 | 5 | 3 | 0 | 0 | 298 | 149 | 1 | 1 | 1 | 1 | 1 | 0.6 | 1 | 1 | 0.231 | 0.571 | 1 | NA | 0.219 |
| SE | 78446 | ENSG00000255036 | SUGT1P4-STRA6LP-CCDC180 | chr9 | + | 97349090 | 97349291 | 97347313 | 97347666 | 97350408 | 97350555 | 2 | 3 | 2 | 3 | 1 | 0 | 0 | 0 | 0 | 0 | 0 | 0 | 0 | 0 | 7 | 0 | 0 | 0 | 1 | 0 | 298 | 149 | 1 | 1 | 1 | 1 | 1 | 1 | 1 | NA | NA | NA | 0 | 1 | 0.5 |
| SE | 78447 | ENSG00000255036 | SUGT1P4-STRA6LP-CCDC180 | chr9 | + | 97349110 | 97349291 | 97343339 | 97343563 | 97350408 | 97350555 | 3 | 2 | 2 | 4 | 1 | 0 | 0 | 0 | 1 | 0 | 0 | 0 | 0 | 0 | 7 | 0 | 5 | 3 | 0 | 0 | 298 | 149 | 1 | 1 | 1 | 1 | 1 | 0.667 | 1 | NA | 0 | 0 | NA | 1 | 0.6 |
| SE | 78448 | ENSG00000255036 | SUGT1P4-STRA6LP-CCDC180 | chr9 | + | 97349110 | 97349291 | 97347313 | 97347666 | 97350408 | 97350555 | 4 | 3 | 7 | 2 | 1 | 0 | 0 | 0 | 0 | 0 | 5 | 4 | 5 | 6 | 26 | 0 | 0 | 0 | 1 | 0 | 298 | 149 | 1 | 1 | 1 | 1 | 1 | 1 | 1 | 1 | 1 | 1 | 0.75 | 1 | 0.05 |
| SE | 78450 | ENSG00000255036 | SUGT1P4-STRA6LP-CCDC180 | chr9 | + | 97359981 | 97360101 | 97357626 | 97357725 | 97361725 | 97361898 | 13 | 12 | 16 | 11 | 10 | 0 | 0 | 0 | 1 | 0 | 7 | 1 | 7 | 9 | 12 | 0 | 0 | 0 | 0 | 0 | 269 | 149 | 1 | 1 | 1 | 1 | 1 | 0.859 | 1 | 1 | 1 | 1 | 1 | 1 | -0.028 |
| SE | 78451 | ENSG00000255036 | SUGT1P4-STRA6LP-CCDC180 | chr9 | + | 97359985 | 97360101 | 97357626 | 97357725 | 97361725 | 97361898 | 8 | 12 | 9 | 10 | 10 | 0 | 0 | 0 | 1 | 0 | 0 | 1 | 3 | 11 | 27 | 0 | 0 | 0 | 0 | 0 | 265 | 149 | 1 | 1 | 1 | 1 | 1 | 0.849 | 1 | NA | 1 | 1 | 1 | 1 | -0.03 |
| SE | 78452 | ENSG00000255036 | SUGT1P4-STRA6LP-CCDC180 | chr9 | + | 97362195 | 97362251 | 97361725 | 97361898 | 97362362 | 97362441 | 2 | 10 | 4 | 17 | 4 | 0 | 0 | 0 | 0 | 0 | 7 | 1 | 6 | 10 | 0 | 0 | 1 | 0 | 0 | 0 | 205 | 149 | 1 | 1 | 1 | 1 | 1 | 1 | 1 | 1 | 0.421 | 1 | 1 | NA | 0.145 |
| SE | 78455 | ENSG00000255036 | SUGT1P4-STRA6LP-CCDC180 | chr9 | + | 97365672 | 97365739 | 97364017 | 97364128 | 97369921 | 97370082 | 1 | 1 | 4 | 1 | 1 | 0 | 0 | 0 | 0 | 0 | 0 | 0 | 0 | 0 | 0 | 0 | 2 | 0 | 2 | 0 | 216 | 149 | 1 | 1 | 1 | 1 | 1 | 1 | 1 | NA | 0 | NA | 0 | NA | 1 |
| SE | 78456 | ENSG00000255036 | SUGT1P4-STRA6LP-CCDC180 | chr9 | + | 97366558 | 97366700 | 97365672 | 97365739 | 97369921 | 97370082 | 14 | 18 | 22 | 27 | 13 | 0 | 0 | 1 | 0 | 0 | 20 | 13 | 14 | 14 | 9 | 0 | 0 | 0 | 0 | 0 | 291 | 149 | 1 | 1 | 1 | 1 | 0.918 | 1 | 1 | 1 | 1 | 1 | 1 | 1 | -0.016 |
| SE | 78457 | ENSG00000255036 | SUGT1P4-STRA6LP-CCDC180 | chr9 | + | 97371594 | 97371706 | 97370640 | 97370778 | 97374542 | 97374648 | 3 | 4 | 7 | 3 | 1 | 0 | 0 | 0 | 0 | 0 | 0 | 2 | 0 | 2 | 22 | 0 | 0 | 0 | 8 | 0 | 261 | 149 | 1 | 1 | 1 | 1 | 1 | 1 | 1 | NA | 1 | NA | 0.125 | 1 | 0.292 |
| SE | 78459 | ENSG00000006715 | VPS41 | chr7 | - | 38728340 | 38728434 | 38726908 | 38726988 | 38728541 | 38728586 | 0 | 7 | 7 | 1 | 0 | 122 | 60 | 75 | 65 | 87 | 0 | 0 | 0 | 0 | 0 | 9 | 61 | 40 | 39 | 0 | 243 | 149 | 1 | 1 | 0 | 0.067 | 0.054 | 0.009 | 0 | 0 | 0 | 0 | 0 | NA | 0.026 |
| SE | 78461 | ENSG00000006715 | VPS41 | chr7 | - | 38817816 | 38817882 | 38796744 | 38796864 | 38830253 | 38830328 | 54 | 29 | 45 | 43 | 43 | 0 | 0 | 0 | 5 | 0 | 0 | 44 | 43 | 49 | 3 | 0 | 0 | 0 | 0 | 0 | 215 | 149 | 1 | 1 | 1 | 1 | 1 | 0.856 | 1 | NA | 1 | 1 | 1 | 1 | -0.029 |
| SE | 78462 | ENSG00000006715 | VPS41 | chr7 | - | 38821202 | 38821265 | 38817802 | 38817882 | 38830253 | 38830328 | 343 | 265 | 246 | 288 | 251 | 0 | 0 | 0 | 0 | 0 | 111 | 280 | 244 | 293 | 96 | 0 | 0 | 1 | 0 | 0 | 212 | 149 | 1 | 1 | 1 | 1 | 1 | 1 | 1 | 1 | 1 | 0.994 | 1 | 1 | 0.001 |
| SE | 78467 | ENSG00000177156 | TALDO1 | chr11 | + | 755878 | 756002 | 747427 | 747578 | 759008 | 759057 | 144 | 68 | 84 | 126 | 16 | 0 | 2 | 2 | 0 | 0 | 309 | 174 | 127 | 210 | 477 | 4 | 3 | 2 | 0 | 0 | 273 | 149 | 1 | 1 | 1 | 0.949 | 0.958 | 1 | 1 | 0.977 | 0.969 | 0.972 | 1 | 1 | -0.002 |
| SE | 78468 | ENSG00000177156 | TALDO1 | chr11 | + | 755878 | 756002 | 752263 | 752473 | 758949 | 759057 | 241 | 120 | 166 | 192 | 88 | 0 | 0 | 0 | 0 | 0 | 664 | 324 | 226 | 506 | 905 | 0 | 1 | 0 | 0 | 0 | 273 | 149 | 1 | 1 | 1 | 1 | 1 | 1 | 1 | 1 | 0.994 | 1 | 1 | 1 | 0.001 |
| SE | 78469 | ENSG00000177156 | TALDO1 | chr11 | + | 763746 | 763940 | 763343 | 763519 | 764287 | 764433 | 351 | 167 | 218 | 318 | 392 | 0 | 0 | 0 | 0 | 0 | 763 | 597 | 335 | 725 | 1502 | 0 | 2 | 0 | 0 | 0 | 298 | 149 | 1 | 1 | 1 | 1 | 1 | 1 | 1 | 1 | 0.993 | 1 | 1 | 1 | 0.001 |
| SE | 78470 | ENSG00000177156 | TALDO1 | chr11 | + | 763746 | 763944 | 763343 | 763519 | 764287 | 764433 | 1045 | 516 | 708 | 929 | 853 | 0 | 0 | 0 | 0 | 0 | 2946 | 1756 | 1088 | 2489 | 4949 | 0 | 2 | 0 | 0 | 0 | 298 | 149 | 1 | 1 | 1 | 1 | 1 | 1 | 1 | 1 | 0.998 | 1 | 1 | 1 | 0 |
| SE | 78477 | ENSG00000164659 | ELAPOR2 | chr7 | - | 86911621 | 86911695 | 86909811 | 86910001 | 86912071 | 86912202 | 1 | 0 | 0 | 0 | 0 | 30 | 21 | 34 | 11 | 7 | 0 | 0 | 0 | 0 | 0 | 0 | 16 | 9 | 8 | 1 | 223 | 149 | 1 | 1 | 0.022 | 0 | 0 | 0 | 0 | NA | 0 | 0 | 0 | 0 | 0.004 |
| SE | 78478 | ENSG00000164659 | ELAPOR2 | chr7 | - | 86912071 | 86912171 | 86909811 | 86910001 | 86912940 | 86913065 | 25 | 18 | 30 | 11 | 2 | 0 | 0 | 0 | 0 | 0 | 0 | 16 | 9 | 8 | 1 | 0 | 0 | 0 | 1 | 0 | 249 | 149 | 1 | 1 | 1 | 1 | 1 | 1 | 1 | NA | 1 | 1 | 0.827 | 1 | 0.043 |
| SE | 78479 | ENSG00000164659 | ELAPOR2 | chr7 | - | 86912071 | 86912245 | 86909811 | 86910001 | 86912940 | 86913204 | 60 | 43 | 70 | 30 | 17 | 0 | 0 | 0 | 0 | 0 | 9 | 33 | 19 | 16 | 2 | 0 | 0 | 0 | 1 | 0 | 298 | 149 | 1 | 1 | 1 | 1 | 1 | 1 | 1 | 1 | 1 | 1 | 0.889 | 1 | 0.022 |
| SE | 78480 | ENSG00000164659 | ELAPOR2 | chr7 | - | 86914722 | 86914860 | 86912940 | 86913204 | 86918441 | 86918544 | 31 | 10 | 39 | 18 | 39 | 0 | 0 | 0 | 1 | 0 | 9 | 9 | 5 | 20 | 7 | 0 | 0 | 0 | 0 | 0 | 287 | 149 | 1 | 1 | 1 | 1 | 1 | 0.903 | 1 | 1 | 1 | 1 | 1 | 1 | -0.019 |
| SE | 78481 | ENSG00000164659 | ELAPOR2 | chr7 | - | 86940009 | 86940115 | 86938807 | 86938960 | 86942017 | 86942104 | 65 | 11 | 66 | 19 | 33 | 0 | 0 | 0 | 0 | 0 | 12 | 24 | 8 | 12 | 25 | 0 | 1 | 0 | 0 | 0 | 255 | 149 | 1 | 1 | 1 | 1 | 1 | 1 | 1 | 1 | 0.933 | 1 | 1 | 1 | 0.013 |
| SE | 78482 | ENSG00000164659 | ELAPOR2 | chr7 | - | 86941351 | 86941385 | 86940009 | 86940115 | 86942017 | 86942104 | 2 | 0 | 2 | 0 | 0 | 40 | 9 | 39 | 8 | 23 | 0 | 0 | 4 | 0 | 0 | 0 | 10 | 4 | 4 | 25 | 183 | 149 | 1 | 1 | 0.039 | 0 | 0.04 | 0 | 0 | NA | 0 | 0.449 | 0 | 0 | -0.096 |
| SE | 78483 | ENSG00000164659 | ELAPOR2 | chr7 | - | 86947726 | 86947922 | 86944898 | 86945046 | 86964903 | 86965024 | 29 | 19 | 34 | 18 | 25 | 1 | 1 | 1 | 0 | 0 | 3 | 18 | 6 | 2 | 11 | 0 | 4 | 0 | 0 | 0 | 298 | 149 | 1 | 1 | 0.935 | 0.905 | 0.944 | 1 | 1 | 1 | 0.692 | 1 | 1 | 1 | 0.018 |
| SE | 78491 | ENSG00000125434 | SLC25A35 | chr17 | - | 8292522 | 8292588 | 8291332 | 8291485 | 8294432 | 8294807 | 10 | 5 | 8 | 7 | 5 | 0 | 0 | 0 | 0 | 0 | 0 | 31 | 6 | 5 | 2 | 0 | 0 | 1 | 0 | 0 | 215 | 149 | 1 | 1 | 1 | 1 | 1 | 1 | 1 | NA | 1 | 0.806 | 1 | 1 | 0.048 |
| SE | 78492 | ENSG00000108825 | PTGES3L-AARSD1 | chr17 | - | 42979549 | 42979773 | 42979377 | 42979444 | 42980045 | 42980086 | 0 | 1 | 2 | 4 | 1 | 0 | 0 | 1 | 0 | 0 | 5 | 3 | 2 | 0 | 101 | 0 | 0 | 0 | 0 | 0 | 298 | 149 | 1 | 1 | NA | 1 | 0.5 | 1 | 1 | 1 | 1 | 1 | NA | 1 | -0.125 |
| SE | 78493 | ENSG00000106336 | FBXO24 | chr7 | + | 100589976 | 100590075 | 100584689 | 100584785 | 100590173 | 100590315 | 1 | 1 | 1 | 0 | 0 | 0 | 0 | 0 | 0 | 0 | 0 | 0 | 1 | 0 | 0 | 0 | 0 | 2 | 0 | 0 | 248 | 149 | 1 | 1 | 1 | 1 | 1 | NA | NA | NA | NA | 0.231 | NA | NA | 0.769 |
| SE | 78495 | ENSG00000106336 | FBXO24 | chr7 | + | 100590173 | 100590315 | 100589976 | 100590075 | 100591666 | 100591902 | 4 | 1 | 2 | 1 | 0 | 2 | 0 | 0 | 0 | 0 | 0 | 0 | 1 | 0 | 0 | 0 | 0 | 0 | 0 | 0 | 291 | 149 | 1 | 1 | 0.506 | 1 | 1 | 1 | NA | NA | NA | 1 | NA | NA | -0.123 |
| SE | 78496 | ENSG00000106336 | FBXO24 | chr7 | + | 100590173 | 100590357 | 100589976 | 100590075 | 100591666 | 100591846 | 4 | 1 | 4 | 3 | 0 | 2 | 0 | 0 | 0 | 0 | 0 | 0 | 1 | 0 | 0 | 0 | 0 | 0 | 0 | 0 | 298 | 149 | 1 | 1 | 0.5 | 1 | 1 | 1 | NA | NA | NA | 1 | NA | NA | -0.125 |
| SE | 78497 | ENSG00000106336 | FBXO24 | chr7 | + | 100595101 | 100595223 | 100594382 | 100594541 | 100595574 | 100595706 | 1 | 0 | 1 | 0 | 0 | 0 | 0 | 0 | 2 | 0 | 0 | 0 | 0 | 2 | 10 | 0 | 0 | 0 | 0 | 0 | 271 | 149 | 1 | 1 | 1 | NA | 1 | NA | 0 | NA | NA | NA | 1 | 1 | -0.333 |
| SE | 78502 | ENSG00000112592 | TBP | chr6 | + | 170556878 | 170557083 | 170554378 | 170554463 | 170561790 | 170562188 | 34 | 22 | 21 | 28 | 55 | 0 | 0 | 0 | 0 | 0 | 26 | 16 | 18 | 28 | 25 | 0 | 0 | 3 | 0 | 0 | 298 | 149 | 1 | 1 | 1 | 1 | 1 | 1 | 1 | 1 | 1 | 0.75 | 1 | 1 | 0.05 |
| SE | 78503 | ENSG00000112592 | TBP | chr6 | + | 170556881 | 170557083 | 170554301 | 170554463 | 170561790 | 170561910 | 35 | 26 | 28 | 32 | 60 | 0 | 0 | 0 | 0 | 0 | 20 | 16 | 19 | 28 | 21 | 0 | 0 | 3 | 0 | 0 | 298 | 149 | 1 | 1 | 1 | 1 | 1 | 1 | 1 | 1 | 1 | 0.76 | 1 | 1 | 0.048 |
| SE | 78505 | ENSG00000112592 | TBP | chr6 | + | 170571409 | 170571504 | 170569611 | 170569740 | 170572185 | 170572832 | 45 | 18 | 27 | 18 | 43 | 2 | 0 | 0 | 0 | 0 | 7 | 14 | 4 | 5 | 18 | 0 | 0 | 0 | 0 | 0 | 244 | 149 | 1 | 1 | 0.932 | 1 | 1 | 1 | 1 | 1 | 1 | 1 | 1 | 1 | -0.014 |
| SE | 78508 | ENSG00000126353 | CCR7 | chr17 | - | 40558892 | 40558942 | 40553768 | 40555818 | 40565399 | 40565472 | 128 | 68 | 200 | 96 | 110 | 4 | 0 | 0 | 0 | 0 | 0 | 37 | 42 | 12 | 16 | 0 | 0 | 0 | 0 | 0 | 199 | 149 | 1 | 1 | 0.96 | 1 | 1 | 1 | 1 | NA | 1 | 1 | 1 | 1 | -0.008 |
| SE | 78511 | ENSG00000115423 | DNAH6 | chr2 | + | 84547491 | 84547612 | 84547267 | 84547402 | 84548287 | 84548417 | 1 | 0 | 0 | 2 | 0 | 0 | 0 | 0 | 0 | 0 | 0 | 0 | 0 | 0 | 0 | 0 | 3 | 0 | 0 | 0 | 270 | 149 | 1 | 1 | 1 | NA | NA | 1 | NA | NA | 0 | NA | NA | NA | 1 |
| SE | 78512 | ENSG00000115423 | DNAH6 | chr2 | + | 84577256 | 84577408 | 84573466 | 84573587 | 84579526 | 84579679 | 0 | 4 | 0 | 1 | 10 | 1 | 0 | 0 | 0 | 1 | 0 | 0 | 1 | 0 | 0 | 0 | 0 | 0 | 0 | 0 | 298 | 149 | 1 | 1 | 0 | 1 | NA | 1 | 0.833 | NA | NA | 1 | NA | NA | -0.292 |
| SE | 78515 | ENSG00000115423 | DNAH6 | chr2 | + | 84659025 | 84659169 | 84658291 | 84658474 | 84669288 | 84669510 | 0 | 2 | 5 | 0 | 4 | 0 | 0 | 0 | 0 | 2 | 0 | 0 | 2 | 12 | 0 | 0 | 0 | 0 | 0 | 0 | 293 | 149 | 1 | 1 | NA | 1 | 1 | NA | 0.504 | NA | NA | 1 | 1 | NA | -0.165 |
| SE | 78516 | ENSG00000115423 | DNAH6 | chr2 | + | 84685325 | 84685472 | 84681356 | 84681528 | 84686483 | 84686557 | 0 | 1 | 2 | 0 | 2 | 1 | 0 | 0 | 0 | 0 | 6 | 0 | 0 | 0 | 0 | 0 | 0 | 0 | 0 | 0 | 296 | 149 | 1 | 1 | 0 | 1 | 1 | NA | 1 | 1 | NA | NA | NA | NA | -0.25 |
| SE | 78517 | ENSG00000115423 | DNAH6 | chr2 | + | 84727668 | 84727902 | 84722624 | 84722804 | 84733443 | 84733579 | 32 | 19 | 29 | 16 | 48 | 0 | 2 | 0 | 0 | 0 | 0 | 0 | 2 | 0 | 0 | 0 | 0 | 0 | 0 | 0 | 298 | 149 | 1 | 1 | 1 | 0.826 | 1 | 1 | 1 | NA | NA | 1 | NA | NA | -0.035 |
| SE | 78519 | ENSG00000019991 | HGF | chr7 | - | 81743352 | 81743471 | 81729604 | 81729779 | 81752119 | 81752247 | 10 | 10 | 12 | 13 | 7 | 0 | 0 | 2 | 0 | 0 | 21 | 18 | 5 | 6 | 0 | 0 | 0 | 0 | 0 | 0 | 268 | 149 | 1 | 1 | 1 | 1 | 0.769 | 1 | 1 | 1 | 1 | 1 | 1 | NA | -0.046 |
| SE | 78520 | ENSG00000019991 | HGF | chr7 | - | 81744999 | 81745120 | 81743352 | 81743471 | 81752119 | 81752247 | 54 | 12 | 25 | 32 | 51 | 0 | 0 | 0 | 1 | 0 | 20 | 53 | 23 | 32 | 10 | 0 | 0 | 0 | 0 | 0 | 270 | 149 | 1 | 1 | 1 | 1 | 1 | 0.946 | 1 | 1 | 1 | 1 | 1 | 1 | -0.011 |
| SE | 78521 | ENSG00000019991 | HGF | chr7 | - | 81752119 | 81752247 | 81744999 | 81745120 | 81757188 | 81757303 | 42 | 11 | 16 | 19 | 44 | 0 | 1 | 0 | 0 | 1 | 7 | 42 | 22 | 23 | 15 | 0 | 0 | 0 | 0 | 0 | 277 | 149 | 1 | 1 | 1 | 0.855 | 1 | 1 | 0.959 | 1 | 1 | 1 | 1 | 1 | -0.037 |
| SE | 78522 | ENSG00000019991 | HGF | chr7 | - | 81752119 | 81752262 | 81744999 | 81745120 | 81757188 | 81757303 | 49 | 11 | 12 | 30 | 45 | 0 | 1 | 0 | 0 | 1 | 13 | 45 | 25 | 35 | 14 | 0 | 0 | 0 | 0 | 0 | 292 | 149 | 1 | 1 | 1 | 0.849 | 1 | 1 | 0.958 | 1 | 1 | 1 | 1 | 1 | -0.039 |
| SE | 78523 | ENSG00000180233 | ZNRF2 | chr7 | + | 30323641 | 30323737 | 30284596 | 30285826 | 30355727 | 30355833 | 124 | 115 | 105 | 144 | 158 | 0 | 1 | 0 | 0 | 0 | 133 | 113 | 125 | 101 | 63 | 0 | 0 | 0 | 0 | 0 | 245 | 149 | 1 | 1 | 1 | 0.986 | 1 | 1 | 1 | 1 | 1 | 1 | 1 | 1 | -0.003 |
| SE | 78524 | ENSG00000180233 | ZNRF2 | chr7 | + | 30355727 | 30355833 | 30323641 | 30323737 | 30362376 | 30362456 | 274 | 208 | 204 | 309 | 389 | 0 | 0 | 0 | 0 | 0 | 254 | 219 | 268 | 226 | 107 | 0 | 0 | 0 | 1 | 4 | 255 | 149 | 1 | 1 | 1 | 1 | 1 | 1 | 1 | 1 | 1 | 1 | 0.992 | 0.94 | 0.014 |
| SE | 78525 | ENSG00000180233 | ZNRF2 | chr7 | + | 30355727 | 30355833 | 30323641 | 30323737 | 30366034 | 30367689 | 65 | 67 | 64 | 97 | 72 | 0 | 0 | 1 | 0 | 0 | 70 | 67 | 73 | 51 | 36 | 0 | 0 | 0 | 0 | 0 | 255 | 149 | 1 | 1 | 1 | 1 | 0.974 | 1 | 1 | 1 | 1 | 1 | 1 | 1 | -0.005 |
| SE | 78526 | ENSG00000180233 | ZNRF2 | chr7 | + | 30362376 | 30362456 | 30323641 | 30323737 | 30366034 | 30367689 | 97 | 80 | 61 | 87 | 145 | 0 | 0 | 1 | 0 | 0 | 71 | 80 | 73 | 109 | 28 | 0 | 0 | 0 | 0 | 0 | 229 | 149 | 1 | 1 | 1 | 1 | 0.975 | 1 | 1 | 1 | 1 | 1 | 1 | 1 | -0.005 |
| SE | 78529 | ENSG00000164054 | SHISA5 | chr3 | - | 48469727 | 48469843 | 48467875 | 48469573 | 48479176 | 48479257 | 300 | 138 | 178 | 246 | 413 | 0 | 0 | 0 | 0 | 0 | 360 | 356 | 188 | 445 | 452 | 0 | 0 | 2 | 0 | 2 | 265 | 149 | 1 | 1 | 1 | 1 | 1 | 1 | 1 | 1 | 1 | 0.981 | 1 | 1 | 0.004 |
| SE | 78531 | ENSG00000164054 | SHISA5 | chr3 | - | 48479176 | 48479257 | 48469727 | 48469843 | 48501136 | 48501293 | 72 | 31 | 21 | 36 | 66 | 0 | 0 | 0 | 0 | 0 | 114 | 86 | 59 | 108 | 135 | 0 | 0 | 0 | 2 | 0 | 230 | 149 | 1 | 1 | 1 | 1 | 1 | 1 | 1 | 1 | 1 | 1 | 0.972 | 1 | 0.006 |
| SE | 78532 | ENSG00000164054 | SHISA5 | chr3 | - | 48502320 | 48502452 | 48501136 | 48501293 | 48504018 | 48504154 | 0 | 0 | 0 | 0 | 0 | 15 | 2 | 2 | 5 | 8 | 0 | 0 | 0 | 0 | 4 | 24 | 16 | 15 | 11 | 17 | 281 | 149 | 1 | 1 | 0 | 0 | 0 | 0 | 0 | 0 | 0 | 0 | 0 | 0.111 | -0.022 |
| SE | 78533 | ENSG00000164054 | SHISA5 | chr3 | - | 48502320 | 48502543 | 48501178 | 48501293 | 48503094 | 48503193 | 0 | 0 | 0 | 0 | 0 | 4 | 0 | 2 | 3 | 0 | 0 | 0 | 0 | 0 | 4 | 0 | 12 | 2 | 5 | 9 | 298 | 149 | 1 | 1 | 0 | NA | 0 | 0 | NA | NA | 0 | 0 | 0 | 0.182 | -0.045 |
| SE | 78535 | ENSG00000164054 | SHISA5 | chr3 | - | 48503136 | 48503193 | 48501136 | 48501293 | 48504422 | 48504776 | 5 | 0 | 2 | 3 | 0 | 5 | 1 | 2 | 0 | 0 | 0 | 6 | 1 | 6 | 9 | 0 | 0 | 0 | 4 | 16 | 248 | 149 | 1 | 1 | 0.375 | 0 | 0.375 | 1 | NA | NA | 1 | 1 | 0.474 | 0.253 | -0.244 |
| SE | 78537 | ENSG00000188786 | MTF1 | chr1 | - | 37817418 | 37817482 | 37809573 | 37815566 | 37822120 | 37822716 | 67 | 49 | 89 | 62 | 122 | 0 | 0 | 0 | 1 | 0 | 110 | 62 | 73 | 157 | 93 | 0 | 0 | 0 | 6 | 0 | 213 | 149 | 1 | 1 | 1 | 1 | 1 | 0.977 | 1 | 1 | 1 | 1 | 0.948 | 1 | 0.006 |
| SE | 78538 | ENSG00000188786 | MTF1 | chr1 | - | 37822120 | 37822716 | 37817418 | 37817482 | 37823709 | 37823812 | 141 | 114 | 140 | 143 | 391 | 0 | 0 | 0 | 0 | 1 | 234 | 113 | 140 | 273 | 94 | 0 | 0 | 0 | 0 | 0 | 298 | 149 | 1 | 1 | 1 | 1 | 1 | 1 | 0.995 | 1 | 1 | 1 | 1 | 1 | -0.001 |
| SE | 78539 | ENSG00000188786 | MTF1 | chr1 | - | 37835078 | 37835215 | 37832244 | 37832322 | 37838624 | 37838756 | 136 | 124 | 128 | 148 | 283 | 1 | 0 | 0 | 0 | 0 | 291 | 183 | 188 | 321 | 100 | 0 | 0 | 0 | 0 | 0 | 286 | 149 | 1 | 1 | 0.986 | 1 | 1 | 1 | 1 | 1 | 1 | 1 | 1 | 1 | -0.003 |
| SE | 78540 | ENSG00000188786 | MTF1 | chr1 | - | 37835670 | 37835744 | 37835078 | 37835215 | 37838624 | 37838756 | 151 | 104 | 130 | 115 | 155 | 0 | 0 | 0 | 0 | 0 | 219 | 151 | 161 | 310 | 89 | 0 | 0 | 2 | 0 | 0 | 223 | 149 | 1 | 1 | 1 | 1 | 1 | 1 | 1 | 1 | 1 | 0.982 | 1 | 1 | 0.004 |
| SE | 78541 | ENSG00000188786 | MTF1 | chr1 | - | 37835670 | 37835744 | 37835078 | 37835215 | 37839919 | 37840158 | 78 | 60 | 71 | 67 | 88 | 0 | 1 | 0 | 0 | 0 | 124 | 88 | 105 | 214 | 60 | 0 | 0 | 0 | 0 | 0 | 223 | 149 | 1 | 1 | 1 | 0.976 | 1 | 1 | 1 | 1 | 1 | 1 | 1 | 1 | -0.005 |

| | | | | | | | | | | | | | | |
|---|---|---|---|---|---|---|---|---|---|---|---|---|---|---|
| SE | 78542 | ENSG00000188786 | MTF1 | chr1 | - | 37838624 | 37838756 | 37835078 | 37835215 | 37839919 | 37840158 | 150 | 140 | [illegible] |
| SE | 78543 | ENSG00000188786 | MTF1 | chr1 | - | 37838624 | 37838756 | 37835670 | 37835744 | 37839919 | 37840158 | 224 | 184 | [illegible] |
| SE | 78545 | ENSG00000241685 | ARPC1A | chr7 | + | 99363542 | 99363633 | 99359533 | 99359738 | 99365890 | 99366214 | 123 | 55 | [illegible] |
| SE | 78546 | ENSG00000234545 | FAM133B | chr7 | - | 92566009 | 92566061 | 92561136 | 92562368 | 92569822 | 92569915 | 26 | 48 | [illegible] |
| SE | 78547 | ENSG00000234545 | FAM133B | chr7 | - | 92566013 | 92566061 | 92560757 | 92562368 | 92569822 | 92569915 | 48 | 90 | [illegible] |
| SE | 78548 | ENSG00000234545 | FAM133B | chr7 | - | 92585297 | 92585396 | 92581505 | 92581603 | 92590267 | 92590393 | 0 | 5 | [illegible] |
| SE | 78549 | ENSG00000169902 | TPST1 | chr7 | + | 66205793 | 66205921 | 66205316 | 66205522 | 66240324 | 66240550 | 0 | 0 | [illegible] |
| SE | 78550 | ENSG00000169902 | TPST1 | chr7 | + | 66209852 | 66209904 | 66205384 | 66205522 | 66240324 | 66241270 | 0 | 0 | [illegible] |
| SE | 78551 | ENSG00000169902 | TPST1 | chr7 | + | 66240324 | 66241270 | 66205333 | 66205522 | 66286510 | 66286709 | 5 | 7 | [illegible] |
| SE | 78553 | ENSG00000169902 | TPST1 | chr7 | + | 66286510 | 66286709 | 66240324 | 66241270 | 66356824 | 66356871 | 5 | 7 | [illegible] |
| SE | 78554 | ENSG00000169902 | TPST1 | chr7 | + | 66352504 | 66352555 | 66240324 | 66241270 | 66356824 | 66356871 | 3 | 3 | [illegible] |
| SE | 78555 | ENSG00000169902 | TPST1 | chr7 | + | 66352504 | 66352555 | 66286510 | 66286709 | 66356824 | 66356871 | 8 | 8 | [illegible] |
| SE | 78560 | ENSG00000088038 | CNOT3 | chr19 | + | 54144005 | 54144134 | 54143659 | 54143749 | 54144236 | 54144332 | 295 | 210 | [illegible] |
| SE | 78561 | ENSG00000088038 | CNOT3 | chr19 | + | 54145541 | 54145817 | 54144236 | 54144332 | 54145909 | 54146043 | 97 | 61 | [illegible] |
| SE | 78562 | ENSG00000088038 | CNOT3 | chr19 | + | 54145594 | 54145817 | 54144236 | 54144332 | 54145909 | 54146043 | 120 | 62 | [illegible] |
| SE | 78563 | ENSG00000088038 | CNOT3 | chr19 | + | 54145597 | 54145817 | 54144236 | 54144332 | 54145909 | 54146043 | 201 | 132 | [illegible] |
| SE | 78572 | ENSG00000091129 | NRCAM | chr7 | - | 108225644 | 108225701 | 108223724 | 108223836 | 108226207 | 108226378 | 0 | 0 | [illegible] |
| SE | 78573 | ENSG00000091129 | NRCAM | chr7 | - | 108232325 | 108232522 | 108231030 | 108231153 | 108234582 | 108234688 | 4 | 10 | [illegible] |
| SE | 78574 | ENSG00000091129 | NRCAM | chr7 | - | 108237751 | 108237769 | 108234582 | 108234688 | 108239958 | 108240049 | 0 | 1 | [illegible] |
| SE | 78575 | ENSG00000091129 | NRCAM | chr7 | - | 108239958 | 108240170 | 108234582 | 108234688 | 108312664 | 108312731 | 18 | 18 | [illegible] |
| SE | 78576 | ENSG00000088930 | XRN2 | chr20 | + | 21326278 | 21326406 | 21303330 | 21303473 | 21326489 | 21326601 | 748 | 584 | [illegible] |
| SE | 78579 | ENSG00000088930 | XRN2 | chr20 | + | 21328558 | 21328670 | 21326489 | 21326601 | 21330615 | 21330705 | 584 | 426 | [illegible] |
| SE | 78581 | ENSG00000088930 | XRN2 | chr20 | + | 21330480 | 21330539 | 21328558 | 21328670 | 21330615 | 21330705 | 598 | 368 | [illegible] |
| SE | 78582 | ENSG00000088930 | XRN2 | chr20 | + | 21331560 | 21331633 | 21330615 | 21330705 | 21331767 | 21331818 | 347 | 224 | [illegible] |
| SE | 78583 | ENSG00000088930 | XRN2 | chr20 | + | 21333543 | 21333618 | 21332282 | 21332440 | 21333703 | 21333837 | 778 | 708 | [illegible] |
| SE | 78584 | ENSG00000088930 | XRN2 | chr20 | + | 21333543 | 21333618 | 21332282 | 21332440 | 21333936 | 21333994 | 192 | 184 | [illegible] |
| SE | 78585 | ENSG00000088930 | XRN2 | chr20 | + | 21333703 | 21333837 | 21332282 | 21332440 | 21333936 | 21333994 | 881 | 713 | [illegible] |
| SE | 78586 | ENSG00000088930 | XRN2 | chr20 | + | 21333703 | 21333837 | 21333543 | 21333618 | 21333936 | 21333994 | 1469 | 1223 | [illegible] |
| SE | 78587 | ENSG00000088930 | XRN2 | chr20 | + | 21333703 | 21333837 | 21333543 | 21333618 | 21339043 | 21339088 | 558 | 492 | [illegible] |
| SE | 78588 | ENSG00000088930 | XRN2 | chr20 | + | 21348145 | 21348253 | 21346414 | 21346550 | 21349388 | 21349461 | 161 | 96 | [illegible] |
| SE | 78589 | ENSG00000088930 | XRN2 | chr20 | + | 21348340 | 21348430 | 21348145 | 21348253 | 21349388 | 21349461 | 341 | 210 | [illegible] |
| SE | 78590 | ENSG00000088930 | XRN2 | chr20 | + | 21349388 | 21349461 | 21346414 | 21346550 | 21354788 | 21354872 | 143 | 73 | [illegible] |
| SE | 78591 | ENSG00000088930 | XRN2 | chr20 | + | 21354788 | 21354872 | 21349388 | 21349461 | 21356079 | 21356177 | 248 | 154 | [illegible] |
| SE | 78592 | ENSG00000088930 | XRN2 | chr20 | + | 21356585 | 21356665 | 21356079 | 21356177 | 21357735 | 21357792 | 425 | 271 | [illegible] |
| SE | 78593 | ENSG00000088930 | XRN2 | chr20 | + | 21356585 | 21356665 | 21356079 | 21356177 | 21365420 | 21365489 | 52 | 40 | [illegible] |
| SE | 78594 | ENSG00000088930 | XRN2 | chr20 | + | 21357735 | 21357792 | 21356079 | 21356177 | 21365420 | 21365489 | 170 | 125 | [illegible] |
| SE | 78596 | ENSG00000088930 | XRN2 | chr20 | + | 21386867 | 21387006 | 21381993 | 21382057 | 21389272 | 21389825 | 302 | 144 | [illegible] |
| SE | 78599 | ENSG00000083168 | KAT6A | chr8 | - | 41949221 | 41949363 | 41946512 | 41947912 | 41955295 | 41955411 | 511 | 297 | [illegible] |
| SE | 78600 | ENSG00000083168 | KAT6A | chr8 | - | 41949221 | 41949363 | 41946512 | 41947912 | 41974703 | 41974734 | 277 | 176 | [illegible] |
| SE | 78601 | ENSG00000083168 | KAT6A | chr8 | - | 41955295 | 41955411 | 41946512 | 41947912 | 41974703 | 41974822 | 160 | 76 | [illegible] |
| SE | 78602 | ENSG00000083168 | KAT6A | chr8 | - | 41955295 | 41955411 | 41949221 | 41949363 | 41974703 | 41974822 | 389 | 196 | [illegible] |
| SE | 78603 | ENSG00000083168 | KAT6A | chr8 | - | 41957048 | 41957180 | 41955295 | 41955411 | 41974703 | 41974734 | 0 | 0 | [illegible] |
| SE | 78604 | ENSG00000083168 | KAT6A | chr8 | - | 41980845 | 41980927 | 41978620 | 41978777 | 41981838 | 41981954 | 404 | 243 | [illegible] |
| SE | 78605 | ENSG00000083168 | KAT6A | chr8 | - | 41987454 | 41987563 | 41981838 | 41981954 | 42048377 | 42049302 | 400 | 272 | [illegible] |
| SE | 78606 | ENSG00000083168 | KAT6A | chr8 | - | 42048377 | 42049302 | 41987454 | 41987563 | 42051900 | 42051987 | 329 | 230 | [illegible] |
| SE | 78611 | ENSG00000042317 | SPATA7 | chr14 | + | 88393392 | 88393488 | 88391380 | 88391455 | 88396155 | 88396184 | 0 | 0 | [illegible] |
| SE | 78613 | ENSG00000042317 | SPATA7 | chr14 | + | 88414673 | 88414732 | 88396155 | 88396203 | 88415204 | 88415266 | 1 | 0 | [illegible] |
| SE | 78614 | ENSG00000042317 | SPATA7 | chr14 | + | 88414673 | 88414732 | 88396155 | 88396203 | 88416710 | 88416761 | 1 | 0 | [illegible] |
| SE | 78615 | ENSG00000042317 | SPATA7 | chr14 | + | 88415204 | 88415266 | 88396155 | 88396203 | 88416710 | 88416844 | 11 | 10 | [illegible] |
| SE | 78616 | ENSG00000042317 | SPATA7 | chr14 | + | 88415204 | 88415266 | 88414673 | 88414732 | 88416710 | 88416844 | 6 | 5 | [illegible] |
| SE | 78617 | ENSG00000042317 | SPATA7 | chr14 | + | 88426231 | 88426704 | 88415925 | 88416844 | 88427629 | 88427696 | 38 | 37 | [illegible] |
| SE | 78627 | ENSG00000196357 | ZNF565 | chr19 | - | 36194232 | 36194328 | 36182275 | 36183733 | 36195029 | 36195156 | 37 | 12 | [illegible] |
| SE | 78629 | ENSG00000196357 | ZNF565 | chr19 | - | 36201976 | 36202050 | 36194826 | 36195156 | 36214887 | 36215081 | 10 | 2 | [illegible] |
| SE | 78630 | ENSG00000168014 | C2CD3 | chr11 | - | 74016598 | 74016801 | 74013443 | 74013525 | 74028286 | 74028398 | 0 | 0 | [illegible] |
| SE | 78637 | ENSG00000168014 | C2CD3 | chr11 | - | 74085521 | 74085886 | 74084880 | 74084970 | 74090812 | 74090936 | 34 | 20 | [illegible] |
| SE | 78638 | ENSG00000168014 | C2CD3 | chr11 | - | 74085617 | 74085886 | 74084880 | 74084970 | 74090812 | 74090936 | 92 | 91 | [illegible] |
| SE | 78639 | ENSG00000168014 | C2CD3 | chr11 | - | 74090812 | 74090936 | 74085521 | 74085886 | 74092415 | 74092588 | 58 | 36 | [illegible] |
| SE | 78640 | ENSG00000168014 | C2CD3 | chr11 | - | 74093815 | 74093999 | 74092415 | 74092588 | 74095227 | 74095408 | 96 | 63 | [illegible] |
| SE | 78641 | ENSG00000168014 | C2CD3 | chr11 | - | 74100524 | 74100676 | 74098008 | 74098255 | 74103130 | 74103625 | 82 | 41 | [illegible] |
| SE | 78642 | ENSG00000168014 | C2CD3 | chr11 | - | 74103130 | 74103625 | 74100524 | 74100676 | 74106370 | 74106489 | 80 | 48 | [illegible] |
| SE | 78643 | ENSG00000168014 | C2CD3 | chr11 | - | 74109033 | 74109152 | 74106370 | 74106493 | 74113779 | 74113892 | 49 | 51 | [illegible] |
| SE | 78645 | ENSG00000168014 | C2CD3 | chr11 | - | 74114383 | 74114593 | 74109033 | 74109152 | 74118227 | 74118382 | 31 | 18 | [illegible] |
| SE | 78649 | ENSG00000168014 | C2CD3 | chr11 | - | 74139604 | 74139828 | 74137057 | 74138967 | 74161398 | 74161556 | 108 | 75 | [illegible] |
| SE | 78653 | ENSG00000274272 | NA | chr7 | - | 100575870 | 100576008 | 100573296 | 100575782 | 100576237 | 100576323 | 285 | 159 | [illegible] |
| SE | 78654 | ENSG00000077454 | LRCH4 | chr7 | - | 100577085 | 100577154 | 100576901 | 100577005 | 100577272 | 100577389 | 101 | 52 | [illegible] |
| SE | 78656 | ENSG00000274272 | NA | chr7 | - | 100577085 | 100577154 | 100576693 | 100577005 | 100577663 | 100577740 | 37 | 20 | [illegible] |
| SE | 78657 | ENSG00000274272 | NA | chr7 | - | 100577272 | 100577389 | 100576693 | 100577005 | 100577496 | 100577558 | 166 | 133 | [illegible] |
| SE | 78658 | ENSG00000274272 | NA | chr7 | - | 100577272 | 100577389 | 100576693 | 100577005 | 100577663 | 100577740 | 0 | 7 | [illegible] |
| SE | 78661 | ENSG00000274272 | NA | chr7 | - | 100577496 | 100577558 | 100576693 | 100577005 | 100577663 | 100577740 | 88 | 46 | [illegible] |
| SE | 78663 | ENSG00000274272 | NA | chr7 | - | 100577496 | 100577558 | 100577272 | 100577389 | 100577663 | 100577740 | 247 | 176 | [illegible] |
| SE | 78664 | ENSG00000274272 | NA | chr7 | - | 100578158 | 100578258 | 100577821 | 100577912 | 100578398 | 100578511 | 313 | 262 | [illegible] |
| SE | 78665 | ENSG00000274272 | NA | chr7 | - | 100578398 | 100578511 | 100577821 | 100577912 | 100578649 | 100578700 | 72 | 16 | [illegible] |
| SE | 78668 | ENSG00000135218 | CD36 | chr7 | + | 80656539 | 80656700 | 80646445 | 80646860 | 80661062 | 80661210 | 713 | 471 | [illegible] |
| SE | 78669 | ENSG00000135218 | CD36 | chr7 | + | 80662989 | 80663169 | 80661062 | 80661210 | 80664405 | 80664497 | 1279 | 1001 | [illegible] |
| SE | 78673 | ENSG00000135218 | CD36 | chr7 | + | 80671921 | 80672040 | 80669952 | 80671164 | 80672769 | 80672843 | 1396 | 1082 | [illegible] |
| SE | 78674 | ENSG00000135218 | CD36 | chr7 | + | 80672769 | 80672843 | 80671921 | 80672040 | 80673354 | 80673409 | 1066 | 809 | [illegible] |
| SE | 78675 | ENSG00000135218 | CD36 | chr7 | + | 80673354 | 80673409 | 80672769 | 80672843 | 80673982 | 80674119 | 1067 | 828 | [illegible] |
| SE | 78678 | ENSG00000248919 | ATP5MF-PTCD1 | chr7 | - | 99447951 | 99448188 | 99435030 | 99435268 | 99460085 | 99460193 | 0 | 0 | [illegible] |
| SE | 78681 | ENSG00000248919 | ATP5MF-PTCD1 | chr7 | - | 99460085 | 99460193 | 99434789 | 99435268 | 99466128 | 99466162 | 2 | 0 | [illegible] |
| SE | 78683 | ENSG00000087258 | GNAO1 | chr16 | + | 56235012 | 56235384 | 56192492 | 56192616 | 56275930 | 56276072 | 0 | 0 | [illegible] |
| SE | 78684 | ENSG00000087258 | GNAO1 | chr16 | + | 56255520 | 56255591 | 56192573 | 56192616 | 56275930 | 56276072 | 2 | 0 | [illegible] |
| SE | 78691 | ENSG00000083642 | PDS5B | chr13 | + | 32658238 | 32658325 | 32651803 | 32652007 | 32658433 | 32658531 | 299 | 245 | [illegible] |
| SE | 78692 | ENSG00000083642 | PDS5B | chr13 | + | 32667763 | 32667844 | 32659153 | 32659280 | 32673215 | 32673356 | 231 | 227 | [illegible] |
| SE | 78693 | ENSG00000083642 | PDS5B | chr13 | + | 32673215 | 32673356 | 32667763 | 32667844 | 32675843 | 32675959 | 357 | 310 | [illegible] |
| SE | 78694 | ENSG00000083642 | PDS5B | chr13 | + | 32687133 | 32687285 | 32683877 | 32684023 | 32688455 | 32688569 | 332 | 255 | [illegible] |
| SE | 78695 | ENSG00000083642 | PDS5B | chr13 | + | 32687133 | 32687285 | 32683877 | 32684023 | 32694222 | 32694304 | 143 | 114 | [illegible] |
| SE | 78696 | ENSG00000083642 | PDS5B | chr13 | + | 32688455 | 32688569 | 32683877 | 32684023 | 32694222 | 32694304 | 99 | 71 | [illegible] |
| SE | 78697 | ENSG00000083642 | PDS5B | chr13 | + | 32688455 | 32688569 | 32687133 | 32687285 | 32694222 | 32694304 | 288 | 212 | [illegible] |
| SE | 78698 | ENSG00000083642 | PDS5B | chr13 | + | 32699729 | 32699869 | 32696853 | 32696902 | 32701322 | 32701438 | 169 | 152 | [illegible] |
| SE | 78699 | ENSG00000083642 | PDS5B | chr13 | + | 32742590 | 32742727 | 32741079 | 32741148 | 32745976 | 32746100 | 292 | 213 | [illegible] |
| SE | 78700 | ENSG00000083642 | PDS5B | chr13 | + | 32742590 | 32742727 | 32741079 | 32741148 | 32753331 | 32753536 | 158 | 119 | [illegible] |
| SE | 78701 | ENSG00000083642 | PDS5B | chr13 | + | 32745976 | 32746100 | 32741079 | 32741148 | 32753331 | 32753536 | 125 | 95 | [illegible] |
| SE | 78702 | ENSG00000083642 | PDS5B | chr13 | + | 32745976 | 32746100 | 32742590 | 32742727 | 32753331 | 32753536 | 258 | 186 | [illegible] |
| SE | 78703 | ENSG00000083642 | PDS5B | chr13 | + | 32755841 | 32755956 | 32753331 | 32753536 | 32758086 | 32758219 | 195 | 188 | [illegible] |
| SE | 78704 | ENSG00000083642 | PDS5B | chr13 | + | 32770653 | 32770761 | 32770120 | 32770560 | 32773188 | 32773324 | 120 | 134 | [illegible] |
| SE | 78711 | ENSG00000075975 | MKRN2 | chr3 | + | 12568874 | 12568997 | 12557089 | 12557176 | 12570070 | 12570252 | 8 | 3 | [illegible] |
| SE | 78712 | ENSG00000075975 | MKRN2 | chr3 | + | 12568874 | 12569003 | 12557056 | 12557176 | 12570070 | 12570252 | 39 | 17 | [illegible] |
| SE | 78713 | ENSG00000075975 | MKRN2 | chr3 | + | 12570324 | 12570583 | 12570070 | 12570252 | 12572068 | 12572373 | 0 | 0 | [illegible] |
| SE | 78715 | ENSG00000171777 | RASGRP4 | chr19 | - | 38414847 | 38415123 | 38413393 | 38413474 | 38417051 | 38417147 | 93 | 23 | [illegible] |
| SE | 78717 | ENSG00000171777 | RASGRP4 | chr19 | - | 38421968 | 38422153 | 38421093 | 38421200 | 38426068 | 38426162 | 73 | 32 | [illegible] |
| SE | 78718 | ENSG00000166886 | NAB2 | chr12 | + | 57092447 | 57092581 | 57091124 | 57091998 | 57092916 | 57092968 | 56 | 34 | [illegible] |
| SE | 78725 | ENSG00000147145 | LPAR4 | chrX | + | 78750729 | 78750792 | 78750173 | 78750223 | 78751209 | 78751294 | 0 | 0 | [illegible] |
| SE | 78726 | ENSG00000225762 | LINC01389 | chr1 | - | 47386247 | 47386356 | 47380927 | 47381143 | 47408319 | 47408477 | 5 | 1 | [illegible] |
| SE | 78730 | ENSG00000127995 | CASD1 | chr7 | + | 94518202 | 94518323 | 94517559 | 94517656 | 94527161 | 94527206 | 86 | 53 | [illegible] |
| SE | 78731 | ENSG00000127995 | CASD1 | chr7 | + | 94533204 | 94533249 | 94528187 | 94528250 | 94533678 | 94533758 | 96 | 63 | [illegible] |
| SE | 78732 | ENSG00000127995 | CASD1 | chr7 | + | 94535308 | 94535523 | 94533678 | 94533802 | 94537471 | 94537894 | 185 | 109 | [illegible] |
| SE | 78733 | ENSG00000127995 | CASD1 | chr7 | + | 94535308 | 94535523 | 94533678 | 94533802 | 94538966 | 94539056 | 97 | 72 | [illegible] |
| SE | 78734 | ENSG00000127995 | CASD1 | chr7 | + | 94537471 | 94537894 | 94533678 | 94533802 | 94538966 | 94539056 | 48 | 50 | [illegible] |
| SE | 78735 | ENSG00000127995 | CASD1 | chr7 | + | 94537471 | 94537894 | 94535308 | 94535523 | 94538966 | 94539056 | 136 | 85 | [illegible] |
| SE | 78736 | ENSG00000127995 | CASD1 | chr7 | + | 94545544 | 94545701 | 94544410 | 94544530 | 94547095 | 94547175 | 166 | 87 | [illegible] |
| SE | 78737 | ENSG00000275111 | ZNF2 | chr2 | + | 95176187 | 95176259 | 95165788 | 95165860 | 95177482 | 95177609 | 29 | 8 | [illegible] |
| SE | 78738 | ENSG00000275111 | ZNF2 | chr2 | + | 95180158 | 95180272 | 95177482 | 95177609 | 95181102 | 95184317 | 47 | 18 | [illegible] |
| SE | 78741 | ENSG00000179406 | LINC00174 | chr7 | - | 66395878 | 66395956 | 66388324 | 66390250 | 66399691 | 66399795 | 5 | 1 | [illegible] |
| SE | 78743 | ENSG00000179406 | LINC00174 | chr7 | - | 66400321 | 66400437 | 66399691 | 66399795 | 66400922 | 66401027 | 1 | 3 | [illegible] |
| SE | 78744 | ENSG00000179406 | LINC00174 | chr7 | - | 66400922 | 66401027 | 66395878 | 66395956 | 66402456 | 66402545 | 2 | 0 | [illegible] |
| SE | 78745 | ENSG00000179406 | LINC00174 | chr7 | - | 66400922 | 66401027 | 66395878 | 66395956 | 66407289 | 66407364 | 2 | 0 | [illegible] |
| SE | 78746 | ENSG00000179406 | LINC00174 | chr7 | - | 66401194 | 66401394 | 66400922 | 66401027 | 66402456 | 66402545 | 10 | 7 | [illegible] |
| SE | 78747 | ENSG00000179406 | LINC00174 | chr7 | - | 66401194 | 66401394 | 66400922 | 66401027 | 66407289 | 66407364 | 2 | 5 | [illegible] |
| SE | 78748 | ENSG00000179406 | LINC00174 | chr7 | - | 66402456 | 66402545 | 66395878 | 66395956 | 66407289 | 66407364 | 9 | 3 | [illegible] |
| SE | 78750 | ENSG00000179406 | LINC00174 | chr7 | - | 66402456 | 66402545 | 66401194 | 66401394 | 66407289 | 66407364 | 17 | 6 | [illegible] |
| SE | 78751 | ENSG00000179406 | LINC00174 | chr7 | - | 66407289 | 66407364 | 66395878 | 66395956 | 66409059 | 66409244 | 7 | 2 | [illegible] |
| SE | 78752 | ENSG00000179406 | LINC00174 | chr7 | - | 66427948 | 66428040 | 66411614 | 66411702 | 66429678 | 66429757 | 11 | 9 | [illegible] |
| SE | 78753 | ENSG00000179406 | LINC00174 | chr7 | - | 66427948 | 66428040 | 66411614 | 66411702 | 66439134 | 66439304 | 8 | 3 | [illegible] |
| SE | 78754 | ENSG00000179406 | LINC00174 | chr7 | - | 66427948 | 66428040 | 66411614 | 66411702 | 66450070 | 66450236 | 5 | 3 | [illegible] |
| SE | 78755 | ENSG00000179406 | LINC00174 | chr7 | - | 66439134 | 66439304 | 66411614 | 66411702 | 66450070 | 66450236 | 15 | 5 | [illegible] |
| SE | 78758 | ENSG00000179406 | LINC00174 | chr7 | - | 66455658 | 66455853 | 66450070 | 66450236 | 66467681 | 66467719 | 7 | 2 | [illegible] |
| SE | 78760 | ENSG00000179406 | LINC00174 | chr7 | - | 66455658 | 66455853 | 66450070 | 66450236 | 66479008 | 66479096 | 11 | 2 | [illegible] |
| SE | 78762 | ENSG00000179406 | LINC00174 | chr7 | - | 66467681 | 66467719 | 66450070 | 66450236 | 66479008 | 66479096 | 1 | 0 | [illegible] |
| SE | 78763 | ENSG00000179406 | LINC00174 | chr7 | - | 66467681 | 66467719 | 66455658 | 66455853 | 66476258 | 66476327 | 2 | 0 | [illegible] |
| SE | 78764 | ENSG00000179406 | LINC00174 | chr7 | - | 66467681 | 66467719 | 66455658 | 66455853 | 66479008 | 66479096 | 0 | 0 | [illegible] |
| SE | 78769 | ENSG00000179406 | LINC00174 | chr7 | - | 66485094 | 66485153 | 66481540 | 66481599 | 66487948 | 66487990 | 26 | 20 | [illegible] |
| SE | 78771 | ENSG00000179406 | LINC00174 | chr7 | - | 66487948 | 66488038 | 66481540 | 66481599 | 66489937 | 66490059 | 10 | 3 | [illegible] |
| SE | 78774 | ENSG00000179406 | LINC00174 | chr7 | - | 66492273 | 66492339 | 66491349 | 66491471 | 66493215 | 66493517 | 2 | 1 | [illegible] |
| SE | 78775 | ENSG00000173114 | LRRN3 | chr7 | + | 111097026 | 111097133 | 111091005 | 111091504 | 111099880 | 111099962 | 26 | 14 | [illegible] |
| SE | 78776 | ENSG00000173114 | LRRN3 | chr7 | + | 111097026 | 111097133 | 111091005 | 111091504 | 111122414 | 111123593 | 11 | 7 | [illegible] |
| SE | 78777 | ENSG00000173114 | LRRN3 | chr7 | + | 111099880 | 111099962 | 111091126 | 111091504 | 111122414 | 111123593 | 48 | 31 | [illegible] |
| SE | 78778 | ENSG00000173114 | LRRN3 | chr7 | + | 111099880 | 111099962 | 111097026 | 111097133 | 111122414 | 111123593 | 35 | 22 | [illegible] |
| SE | 78781 | ENSG00000131023 | LATS1 | chr6 | - | 149683078 | 149684592 | 149679874 | 149680457 | 149701778 | 149701963 | 133 | 127 | [illegible] |
| SE | 78782 | ENSG00000131023 | LATS1 | chr6 | - | 149695073 | 149695221 | 149679874 | 149680457 | 149701778 | 149701963 | 105 | 79 | [illegible] |
| SE | 78783 | ENSG00000131023 | LATS1 | chr6 | - | 149695073 | 149695221 | 149679874 | 149680457 | 149717848 | 149718034 | 1 | 0 | [illegible] |
| SE | 78784 | ENSG00000131023 | LATS1 | chr6 | - | 149695073 | 149695221 | 149682918 | 149684592 | 149701778 | 149701963 | 199 | 170 | [illegible] |
| SE | 78788 | ENSG00000106245 | BUD31 | chr7 | + | 99410033 | 99410169 | 99408640 | 99409004 | 99411063 | 99411186 | 212 | 140 | [illegible] |
| SE | 78792 | ENSG00000106245 | BUD31 | chr7 | + | 99411063 | 99411186 | 99410033 | 99410169 | 99416137 | 99416260 | 555 | 406 | [illegible] |
| SE | 78795 | ENSG00000146802 | TMEM168 | chr7 | - | 112772780 | 112772818 | 112767174 | 112767744 | 112775175 | 112775318 | 8 | 12 | [illegible] |
| SE | 78796 | ENSG00000146802 | TMEM168 | chr7 | - | 112772780 | 112773055 | 112762376 | 112767744 | 112775175 | 112775318 | 85 | 86 | [illegible] |
| SE | 78797 | ENSG00000146802 | TMEM168 | chr7 | - | 112783697 | 112783783 | 112775175 | 112775318 | 112790159 | 112790390 | 27 | 13 | [illegible] |
| SE | 78798 | ENSG00000146802 | TMEM168 | chr7 | - | 112783697 | 112784953 | 112775175 | 112775318 | 112790159 | 112790393 | 113 | 50 | [illegible] |
| SE | 78799 | ENSG00000146802 | TMEM168 | chr7 | - | 112788453 | 112788544 | 112783697 | 112784953 | 112790159 | 112790393 | 3 | 2 | [illegible] |
| SE | 78802 | ENSG00000135269 | TES | chr7 | + | 116250160 | 116250496 | 116249019 | 116249272 | 116257293 | 116258783 | 840 | 568 | [illegible] |
| SE | 78803 | ENSG00000135269 | TES | chr7 | + | 116251759 | 116251975 | 116234533 | 116234619 | 116257293 | 116257668 | 0 | 2 | [illegible] |
| SE | 78806 | ENSG00000135269 | TES | chr7 | + | 116252317 | 116252476 | 116251506 | 116251975 | 116257293 | 116257576 | 929 | 399 | [illegible] |
| SE | 78811 | ENSG00000224897 | POT1-AS1 | chr7 | + | 125080711 | 125080813 | 125020332 | 125020413 | 125099403 | 125099513 | 0 | 2 | [illegible] |
| SE | 78812 | ENSG00000224897 | POT1-AS1 | chr7 | + | 125080711 | 125080813 | 125020332 | 125020413 | 125116873 | 125117028 | 0 | 3 | [illegible] |
| SE | 78813 | ENSG00000224897 | POT1-AS1 | chr7 | + | 125080711 | 125080813 | 125020332 | 125020413 | 125125824 | 125125977 | 0 | 3 | [illegible] |
| SE | 78816 | ENSG00000224897 | POT1-AS1 | chr7 | + | 125089689 | 125089776 | 125020332 | 125020413 | 125099403 | 125099513 | 2 | 1 | [illegible] |
| SE | 78817 | ENSG00000224897 | POT1-AS1 | chr7 | + | 125089689 | 125089776 | 125020332 | 125020413 | 125125824 | 125125977 | 0 | 1 | [illegible] |
| SE | 78819 | ENSG00000224897 | POT1-AS1 | chr7 | + | 125089689 | 125089776 | 125080711 | 125080813 | 125125824 | 125125977 | 0 | 1 | [illegible] |
| SE | 78820 | ENSG00000224897 | POT1-AS1 | chr7 | + | 125099403 | 125099513 | 125020332 | 125020413 | 125116873 | 125117028 | 4 | 1 | [illegible] |
| SE | 78822 | ENSG00000224897 | POT1-AS1 | chr7 | + | 125116873 | 125117028 | 125020332 | 125020413 | 125125824 | 125125907 | 0 | 0 | [illegible] |
| SE | 78823 | ENSG00000224897 | POT1-AS1 | chr7 | + | 125116873 | 125117028 | 125080711 | 125080813 | 125125824 | 125125907 | 0 | 1 | [illegible] |
| SE | 78834 | ENSG00000148834 | GSTO1 | chr10 | + | 104259575 | 104259798 | 104254985 | 104255271 | 104266083 | 104266190 | 232 | 112 | [illegible] |
| SE | 78838 | ENSG00000105854 | PON2 | chr7 | - | 95415147 | 95415272 | 95412323 | 95412477 | 95416171 | 95416297 | 2 | 1 | [illegible] |
| SE | 78841 | ENSG00000105854 | PON2 | chr7 | - | 95434258 | 95434333 | 95424514 | 95424585 | 95434877 | 95434951 | 0 | 2 | [illegible] |
| SE | 78842 | ENSG00000158604 | TMED4 | chr7 | - | 44581092 | 44581239 | 44577893 | 44579628 | 44581448 | 44581574 | 325 | 176 | [illegible] |
| SE | 78849 | ENSG00000105971 | CAV2 | chr7 | + | 116500259 | 116500447 | 116499737 | 116499931 | 116505970 | 116506093 | 3 | 1 | [illegible] |
| SE | 78859 | ENSG00000158321 | AUTS2 | chr7 | + | 69825779 | 69825875 | 69824570 | 69824687 | 69899285 | 69899498 | 17 | 14 | [illegible] |
| SE | 78863 | ENSG00000158321 | AUTS2 | chr7 | + | 70771548 | 70771644 | 70766113 | 70766334 | 70777102 | 70777135 | 6 | 0 | [illegible] |
| SE | 78866 | ENSG00000158321 | AUTS2 | chr7 | + | 70781614 | 70781756 | 70777102 | 70777174 | 70784941 | 70785019 | 122 | 30 | [illegible] |
| SE | 78867 | ENSG00000158321 | AUTS2 | chr7 | + | 70787085 | 70787137 | 70785954 | 70786038 | 70789747 | 70790109 | 2 | 2 | [illegible] |
| SE | 78868 | ENSG00000158321 | AUTS2 | chr7 | + | 70787208 | 70787431 | 70785954 | 70786038 | 70789747 | 70789926 | 84 | 27 | [illegible] |
| SE | 78869 | ENSG00000196792 | STRN3 | chr14 | - | 30905417 | 30905558 | 30902535 | 30902643 | 30906876 | 30907044 | 204 | 159 | [illegible] |
| SE | 78870 | ENSG00000196792 | STRN3 | chr14 | - | 30906876 | 30907044 | 30902535 | 30902643 | 30911040 | 30911162 | 34 | 35 | [illegible] |
| SE | 78871 | ENSG00000196792 | STRN3 | chr14 | - | 30913523 | 30913657 | 30912006 | 30912182 | 30918965 | 30919106 | 36 | 28 | [illegible] |
| SE | 78872 | ENSG00000196792 | STRN3 | chr14 | - | 30913523 | 30913657 | 30912006 | 30912182 | 30929200 | 30929311 | 25 | 22 | [illegible] |
| SE | 78874 | ENSG00000196792 | STRN3 | chr14 | - | 30918965 | 30919106 | 30912006 | 30912182 | 30929200 | 30929311 | 2 | 2 | [illegible] |
| SE | 78875 | ENSG00000196792 | STRN3 | chr14 | - | 30918965 | 30919106 | 30912006 | 30912182 | 30935162 | 30935304 | 2 | 3 | [illegible] |
| SE | 78876 | ENSG00000196792 | STRN3 | chr14 | - | 30918965 | 30919106 | 30913523 | 30913657 | 30929200 | 30929311 | 16 | 8 | [illegible] |
| SE | 78882 | ENSG00000196792 | STRN3 | chr14 | - | 30947089 | 30947263 | 30936494 | 30936624 | 30955619 | 30955693 | 75 | 79 | [illegible] |
| SE | 78883 | ENSG00000196792 | STRN3 | chr14 | - | 30950862 | 30950944 | 30947089 | 30947263 | 30955619 | 30955693 | 169 | 151 | [illegible] |
| SE | 78885 | ENSG00000198874 | TYW1 | chr7 | + | 67009582 | 67009684 | 66998816 | 66998954 | 67014366 | 67014561 | 93 | 89 | [illegible] |
| SE | 78887 | ENSG00000198874 | TYW1 | chr7 | + | 67024899 | 67025022 | 67009582 | 67009684 | 67049948 | 67050066 | 50 | 43 | [illegible] |
| SE | 78888 | ENSG00000198874 | TYW1 | chr7 | + | 67049948 | 67050066 | 67009582 | 67009684 | 67055834 | 67055887 | 40 | 29 | [illegible] |
| SE | 78890 | ENSG00000198874 | TYW1 | chr7 | + | 67083429 | 67083539 | 67067137 | 67067403 | 67098540 | 67098718 | 92 | 73 | [illegible] |
| SE | 78895 | ENSG00000173598 | NUDT4 | chr12 | + | 93380987 | 93381031 | 93378549 | 93378693 | 93394608 | 93394719 | 4 | 3 | [illegible] |
| SE | 78898 | ENSG00000173598 | NUDT4 | chr12 | + | 93395488 | 93395533 | 93394608 | 93394719 | 93398770 | 93398855 | 61 | 29 | [illegible] |
| SE | 78900 | ENSG00000184307 | ZDHHC23 | chr3 | + | 113948819 | 113948963 | 113948685 | 113948734 | 113953699 | 113954410 | 8 | 10 | [illegible] |
| SE | 78901 | ENSG00000184307 | ZDHHC23 | chr3 | + | 113956338 | 113956506 | 113953699 | 113954410 | 113958362 | 113958538 | 8 | 5 | [illegible] |
| SE | 78903 | ENSG00000184307 | ZDHHC23 | chr3 | + | 113959520 | 113959600 | 113958362 | 113958538 | 113961051 | 113962978 | 0 | 0 | [illegible] |
| SE | 78912 | ENSG00000198839 | ZNF277 | chr7 | + | 112286872 | 112287074 | 112206694 | 112206807 | 112295868 | 112295957 | 126 | 85 | [illegible] |
| SE | 78913 | ENSG00000198839 | ZNF277 | chr7 | + | 112286876 | 112287074 | 112206694 | 112206807 | 112295868 | 112295957 | 101 | 82 | [illegible] |
| SE | 78915 | ENSG00000198839 | ZNF277 | chr7 | + | 112296228 | 112296311 | 112286872 | 112287074 | 112318181 | 112318273 | 160 | 192 | [illegible] |
| SE | 78916 | ENSG00000198839 | ZNF277 | chr7 | + | 112318181 | 112318273 | 112286872 | 112287074 | 112327716 | 112327827 | 209 | 147 | [illegible] |
| SE | 78918 | ENSG00000198839 | ZNF277 | chr7 | + | 112336103 | 112336171 | 112330083 | 112330216 | 112337729 | 112337820 | 246 | 154 | [illegible] |
| SE | 78920 | ENSG00000197363 | ZNF517 | chr8 | + | 144803640 | 144803767 | 144802869 | 144802947 | 144804124 | 144804238 | 9 | 4 | [illegible] |
| SE | 78923 | ENSG00000184903 | IMMP2L | chr7 | - | 111521312 | 111521449 | 111487237 | 111487341 | 111532517 | 111532585 | 40 | 39 | [illegible] |
| SE | 78924 | ENSG00000184903 | IMMP2L | chr7 | - | 111521312 | 111521449 | 111487237 | 111487341 | 111561850 | 111561901 | 74 | 44 | [illegible] |
| SE | 78925 | ENSG00000184903 | IMMP2L | chr7 | - | 111532517 | 111532585 | 111487239 | 111487341 | 111561850 | 111561923 | 0 | 0 | [illegible] |
| SE | 78934 | ENSG00000004059 | ARF5 | chr7 | + | 127589082 | 127589163 | 127588385 | 127588565 | 127589484 | 127589574 | 400 | 172 | [illegible] |
| SE | 78935 | ENSG00000004059 | ARF5 | chr7 | + | 127589484 | 127589594 | 127589082 | 127589163 | 127590065 | 127590137 | 561 | 246 | [illegible] |
| SE | 78976 | ENSG00000283149 | NA | chr3 | + | 186582715 | 186582773 | 186581994 | 186582077 | 186583864 | 186583976 | 155 | 113 | [illegible] |
| SE | 78982 | ENSG00000183735 | TBK1 | chr12 | + | 64457907 | 64458018 | 64455839 | 64455957 | 64460188 | 64460329 | 1 | 3 | [illegible] |
| SE | 78983 | ENSG00000183735 | TBK1 | chr12 | + | 64460188 | 64460329 | 64455815 | 64455957 | 64464333 | 64464399 | 275 | 219 | [illegible] |
| SE | 78984 | ENSG00000183735 | TBK1 | chr12 | + | 64460246 | 64460329 | 64455839 | 64455957 | 64464333 | 64464463 | 71 | 40 | [illegible] |
| SE | 78985 | ENSG00000183735 | TBK1 | chr12 | + | 64462472 | 64462521 | 64460188 | 64460329 | 64464333 | 64464463 | 0 | 0 | [illegible] |
| SE | 78986 | ENSG00000183735 | TBK1 | chr12 | + | 64464572 | 64464657 | 64464333 | 64464463 | 64466900 | 64467082 | 0 | 0 | [illegible] |
| SE | 78987 | ENSG00000183735 | TBK1 | chr12 | + | 64466900 | 64467082 | 64464333 | 64464463 | 64474229 | 64474390 | 309 | 209 | [illegible] |
| SE | 78988 | ENSG00000183735 | TBK1 | chr12 | + | 64468572 | 64468672 | 64466900 | 64467082 | 64474229 | 64474390 | 2 | 0 | [illegible] |
| SE | 78991 | ENSG00000183735 | TBK1 | chr12 | + | 64471743 | 64471995 | 64466900 | 64467082 | 64474229 | 64474390 | 0 | 0 | [illegible] |
| SE | 78992 | ENSG00000183735 | TBK1 | chr12 | + | 64474229 | 64474390 | 64466900 | 64467082 | 64480011 | 64480122 | 220 | 172 | [illegible] |

| | | | | | | |
|---|---|---|---|---|---|---|
| SE | 78993 | ENSG00000183735 | TBK1 | chr12 | + | [illegible] |
| SE | 78994 | ENSG00000183735 | TBK1 | chr12 | + | [illegible] |
| SE | 78995 | ENSG00000183735 | TBK1 | chr12 | + | [illegible] |
| SE | 78996 | ENSG00000183735 | TBK1 | chr12 | + | [illegible] |
| SE | 78999 | ENSG00000183735 | TBK1 | chr12 | + | [illegible] |
| SE | 79000 | ENSG00000183735 | TBK1 | chr12 | + | [illegible] |
| SE | 79001 | ENSG00000183735 | TBK1 | chr12 | + | [illegible] |
| SE | 79005 | ENSG00000183735 | TBK1 | chr12 | + | [illegible] |
| SE | 79015 | ENSG00000273297 | NA | chr7 | - | [illegible] |
| SE | 79018 | ENSG00000160218 | TRAPPC10 | chr21 | + | [illegible] |
| SE | 79019 | ENSG00000160218 | TRAPPC10 | chr21 | + | [illegible] |
| SE | 79020 | ENSG00000160218 | TRAPPC10 | chr21 | + | [illegible] |
| SE | 79023 | ENSG00000160218 | TRAPPC10 | chr21 | + | [illegible] |
| SE | 79024 | ENSG00000160218 | TRAPPC10 | chr21 | + | [illegible] |
| SE | 79026 | ENSG00000160218 | TRAPPC10 | chr21 | + | [illegible] |
| SE | 79027 | ENSG00000160218 | TRAPPC10 | chr21 | + | [illegible] |
| SE | 79028 | ENSG00000160218 | TRAPPC10 | chr21 | + | [illegible] |
| SE | 79029 | ENSG00000160218 | TRAPPC10 | chr21 | + | [illegible] |
| SE | 79031 | ENSG00000160218 | TRAPPC10 | chr21 | + | [illegible] |
| SE | 79033 | ENSG00000160218 | TRAPPC10 | chr21 | + | [illegible] |
| SE | 79035 | ENSG00000160218 | TRAPPC10 | chr21 | + | [illegible] |
| SE | 79049 | ENSG00000136045 | PWP1 | chr12 | + | [illegible] |
| SE | 79050 | ENSG00000136045 | PWP1 | chr12 | + | [illegible] |
| SE | 79051 | ENSG00000136045 | PWP1 | chr12 | + | [illegible] |
| SE | 79052 | ENSG00000136045 | PWP1 | chr12 | + | [illegible] |
| SE | 79062 | ENSG00000128536 | CDHR3 | chr7 | + | [illegible] |
| SE | 79063 | ENSG00000158636 | EMSY | chr11 | + | [illegible] |
| SE | 79066 | ENSG00000158636 | EMSY | chr11 | + | [illegible] |
| SE | 79067 | ENSG00000158636 | EMSY | chr11 | + | [illegible] |
| SE | 79068 | ENSG00000158636 | EMSY | chr11 | + | [illegible] |
| SE | 79069 | ENSG00000158636 | EMSY | chr11 | + | [illegible] |
| SE | 79071 | ENSG00000158636 | EMSY | chr11 | + | [illegible] |
| SE | 79073 | ENSG00000158636 | EMSY | chr11 | + | [illegible] |
| SE | 79074 | ENSG00000158636 | EMSY | chr11 | + | [illegible] |
| SE | 79075 | ENSG00000158636 | EMSY | chr11 | + | [illegible] |
| SE | 79084 | ENSG00000106086 | PLEKHA8 | chr7 | + | [illegible] |
| SE | 79085 | ENSG00000106086 | PLEKHA8 | chr7 | + | [illegible] |
| SE | 79086 | ENSG00000106086 | PLEKHA8 | chr7 | + | [illegible] |
| SE | 79087 | ENSG00000106086 | PLEKHA8 | chr7 | + | [illegible] |
| SE | 79089 | ENSG00000106086 | PLEKHA8 | chr7 | + | [illegible] |
| SE | 79090 | ENSG00000106086 | PLEKHA8 | chr7 | + | [illegible] |
| SE | 79091 | ENSG00000269955 | FMC1-LUC7L2 | chr7 | + | [illegible] |
| SE | 79092 | ENSG00000269955 | FMC1-LUC7L2 | chr7 | + | [illegible] |
| SE | 79093 | ENSG00000269955 | FMC1-LUC7L2 | chr7 | + | [illegible] |
| SE | 79094 | ENSG00000269955 | FMC1-LUC7L2 | chr7 | + | [illegible] |
| SE | 79095 | ENSG00000269955 | FMC1-LUC7L2 | chr7 | + | [illegible] |
| SE | 79096 | ENSG00000269955 | FMC1-LUC7L2 | chr7 | + | [illegible] |
| SE | 79123 | ENSG00000160219 | GAB3 | chrX | - | [illegible] |
| SE | 79124 | ENSG00000160219 | GAB3 | chrX | - | [illegible] |
| SE | 79125 | ENSG00000160219 | GAB3 | chrX | - | [illegible] |
| SE | 79130 | ENSG00000026950 | BTN3A1 | chr6 | + | [illegible] |
| SE | 79136 | ENSG00000169660 | HEXD | chr17 | + | [illegible] |
| SE | 79138 | ENSG00000169660 | HEXD | chr17 | + | [illegible] |
| SE | 79139 | ENSG00000169660 | HEXD | chr17 | + | [illegible] |
| SE | 79141 | ENSG00000169660 | HEXD | chr17 | + | [illegible] |
| SE | 79142 | ENSG00000169660 | HEXD | chr17 | + | [illegible] |
| SE | 79146 | ENSG00000104299 | INTS9 | chr8 | - | [illegible] |
| SE | 79147 | ENSG00000104299 | INTS9 | chr8 | - | [illegible] |
| SE | 79161 | ENSG00000145191 | EIF2B5 | chr3 | + | [illegible] |
| SE | 79162 | ENSG00000145191 | EIF2B5 | chr3 | + | [illegible] |
| SE | 79163 | ENSG00000145191 | EIF2B5 | chr3 | + | [illegible] |
| SE | 79164 | ENSG00000145191 | EIF2B5 | chr3 | + | [illegible] |
| SE | 79171 | ENSG00000134291 | TMEM106C | chr12 | + | [illegible] |
| SE | 79173 | ENSG00000170632 | ARMC10 | chr7 | + | [illegible] |
| SE | 79176 | ENSG00000170632 | ARMC10 | chr7 | + | [illegible] |
| SE | 79177 | ENSG00000170632 | ARMC10 | chr7 | + | [illegible] |
| SE | 79184 | ENSG00000170632 | ARMC10 | chr7 | + | [illegible] |
| SE | 79186 | ENSG00000170632 | ARMC10 | chr7 | + | [illegible] |
| SE | 79187 | ENSG00000160949 | TONSL | chr8 | - | [illegible] |
| SE | 79188 | ENSG00000160949 | TONSL | chr8 | - | [illegible] |
| SE | 79189 | ENSG00000160949 | TONSL | chr8 | - | [illegible] |
| SE | 79192 | ENSG00000236753 | MKLN1-AS | chr7 | - | [illegible] |
| SE | 79193 | ENSG00000236753 | MKLN1-AS | chr7 | - | [illegible] |
| SE | 79195 | ENSG00000054277 | OPN3 | chr1 | - | [illegible] |
| SE | 79197 | ENSG00000054277 | OPN3 | chr1 | - | [illegible] |
| SE | 79198 | ENSG00000054277 | OPN3 | chr1 | - | [illegible] |
| SE | 79199 | ENSG00000242588 | NA | chr7 | + | [illegible] |
| SE | 79202 | ENSG00000242588 | NA | chr7 | + | [illegible] |
| SE | 79204 | ENSG00000242588 | NA | chr7 | + | [illegible] |
| SE | 79205 | ENSG00000242588 | NA | chr7 | + | [illegible] |
| SE | 79207 | ENSG00000242588 | NA | chr7 | + | [illegible] |
| SE | 79209 | ENSG00000242588 | NA | chr7 | + | [illegible] |
| SE | 79210 | ENSG00000242588 | NA | chr7 | + | [illegible] |
| SE | 79217 | ENSG00000242588 | NA | chr7 | + | [illegible] |
| SE | 79218 | ENSG00000242588 | NA | chr7 | + | [illegible] |
| SE | 79223 | ENSG00000132823 | OSER1 | chr20 | - | [illegible] |
| SE | 79224 | ENSG00000132823 | OSER1 | chr20 | - | [illegible] |
| SE | 79225 | ENSG00000132823 | OSER1 | chr20 | - | [illegible] |
| SE | 79226 | ENSG00000132823 | OSER1 | chr20 | - | [illegible] |
| SE | 79228 | ENSG00000116679 | IVNS1ABP | chr1 | - | [illegible] |
| SE | 79231 | ENSG00000116679 | IVNS1ABP | chr1 | - | [illegible] |
| SE | 79233 | ENSG00000116679 | IVNS1ABP | chr1 | - | [illegible] |
| SE | 79237 | ENSG00000196511 | TPK1 | chr7 | - | [illegible] |
| SE | 79238 | ENSG00000196511 | TPK1 | chr7 | - | [illegible] |
| SE | 79241 | ENSG00000196511 | TPK1 | chr7 | - | [illegible] |
| SE | 79250 | ENSG00000133606 | MKRN1 | chr7 | - | [illegible] |
| SE | 79251 | ENSG00000133606 | MKRN1 | chr7 | - | [illegible] |
| SE | 79252 | ENSG00000133606 | MKRN1 | chr7 | - | [illegible] |
| SE | 79253 | ENSG00000133606 | MKRN1 | chr7 | - | [illegible] |
| SE | 79255 | ENSG00000133606 | MKRN1 | chr7 | - | [illegible] |
| SE | 79256 | ENSG00000133606 | MKRN1 | chr7 | - | [illegible] |
| SE | 79258 | ENSG00000133606 | MKRN1 | chr7 | - | [illegible] |
| SE | 79260 | ENSG00000133606 | MKRN1 | chr7 | - | [illegible] |
| SE | 79262 | ENSG00000133606 | MKRN1 | chr7 | - | [illegible] |
| SE | 79264 | ENSG00000133606 | MKRN1 | chr7 | - | [illegible] |
| SE | 79268 | ENSG00000133574 | GIMAP4 | chr7 | + | [illegible] |
| SE | 79269 | ENSG00000133574 | GIMAP4 | chr7 | + | [illegible] |
| SE | 79272 | ENSG00000128607 | KLHDC10 | chr7 | + | [illegible] |
| SE | 79274 | ENSG00000128607 | KLHDC10 | chr7 | + | [illegible] |
| SE | 79278 | ENSG00000154803 | FLCN | chr17 | - | [illegible] |
| SE | 79281 | ENSG00000154803 | FLCN | chr17 | - | [illegible] |
| SE | 79282 | ENSG00000128578 | STRIP2 | chr7 | + | [illegible] |
| SE | 79283 | ENSG00000128578 | STRIP2 | chr7 | + | [illegible] |
| SE | 79285 | ENSG00000102390 | PBDC1 | chrX | + | [illegible] |
| SE | 79286 | ENSG00000102390 | PBDC1 | chrX | + | [illegible] |
| SE | 79292 | ENSG00000001631 | KRIT1 | chr7 | - | [illegible] |
| SE | 79293 | ENSG00000001631 | KRIT1 | chr7 | - | [illegible] |
| SE | 79294 | ENSG00000001631 | KRIT1 | chr7 | - | [illegible] |
| SE | 79295 | ENSG00000001631 | KRIT1 | chr7 | - | [illegible] |
| SE | 79296 | ENSG00000001631 | KRIT1 | chr7 | - | [illegible] |
| SE | 79297 | ENSG00000001631 | KRIT1 | chr7 | - | [illegible] |
| SE | 79299 | ENSG00000001631 | KRIT1 | chr7 | - | [illegible] |
| SE | 79300 | ENSG00000001631 | KRIT1 | chr7 | - | [illegible] |
| SE | 79301 | ENSG00000001631 | KRIT1 | chr7 | - | [illegible] |
| SE | 79302 | ENSG00000001631 | KRIT1 | chr7 | - | [illegible] |
| SE | 79303 | ENSG00000001631 | KRIT1 | chr7 | - | [illegible] |
| SE | 79305 | ENSG00000001631 | KRIT1 | chr7 | - | [illegible] |
| SE | 79306 | ENSG00000001631 | KRIT1 | chr7 | - | [illegible] |
| SE | 79307 | ENSG00000001631 | KRIT1 | chr7 | - | [illegible] |
| SE | 79308 | ENSG00000001631 | KRIT1 | chr7 | - | [illegible] |
| SE | 79309 | ENSG00000001631 | KRIT1 | chr7 | - | [illegible] |
| SE | 79310 | ENSG00000001631 | KRIT1 | chr7 | - | [illegible] |
| SE | 79312 | ENSG00000001631 | KRIT1 | chr7 | - | [illegible] |
| SE | 79313 | ENSG00000001631 | KRIT1 | chr7 | - | [illegible] |
| SE | 79318 | ENSG00000162600 | OMA1 | chr1 | - | [illegible] |
| SE | 79319 | ENSG00000162600 | OMA1 | chr1 | - | [illegible] |
| SE | 79320 | ENSG00000162600 | OMA1 | chr1 | - | [illegible] |
| SE | 79323 | ENSG00000162600 | OMA1 | chr1 | - | [illegible] |
| SE | 79324 | ENSG00000162600 | OMA1 | chr1 | - | [illegible] |
| SE | 79325 | ENSG00000162600 | OMA1 | chr1 | - | [illegible] |
| SE | 79327 | ENSG00000162600 | OMA1 | chr1 | - | [illegible] |
| SE | 79328 | ENSG00000162600 | OMA1 | chr1 | - | [illegible] |
| SE | 79329 | ENSG00000162600 | OMA1 | chr1 | - | [illegible] |
| SE | 79331 | ENSG00000162600 | OMA1 | chr1 | - | [illegible] |
| SE | 79333 | ENSG00000162600 | OMA1 | chr1 | - | [illegible] |
| SE | 79334 | ENSG00000162600 | OMA1 | chr1 | - | [illegible] |
| SE | 79335 | ENSG00000180758 | GPR157 | chr1 | - | [illegible] |
| SE | 79336 | ENSG00000231721 | LINC-PINT | chr7 | - | [illegible] |
| SE | 79341 | ENSG00000231721 | LINC-PINT | chr7 | - | [illegible] |
| SE | 79344 | ENSG00000231721 | LINC-PINT | chr7 | - | [illegible] |
| SE | 79345 | ENSG00000231721 | LINC-PINT | chr7 | - | [illegible] |
| SE | 79346 | ENSG00000231721 | LINC-PINT | chr7 | - | [illegible] |
| SE | 79350 | ENSG00000231721 | LINC-PINT | chr7 | - | [illegible] |
| SE | 79353 | ENSG00000231721 | LINC-PINT | chr7 | - | [illegible] |
| SE | 79357 | ENSG00000231721 | LINC-PINT | chr7 | - | [illegible] |
| SE | 79358 | ENSG00000280828 | NA | chr7 | + | [illegible] |
| SE | 79365 | ENSG00000105887 | MTPN | chr7 | - | [illegible] |
| SE | 79366 | ENSG00000105887 | MTPN | chr7 | - | [illegible] |
| SE | 79368 | ENSG00000101574 | METTL4 | chr18 | - | [illegible] |
| SE | 79369 | ENSG00000101574 | METTL4 | chr18 | - | [illegible] |
| SE | 79370 | ENSG00000101574 | METTL4 | chr18 | - | [illegible] |
| SE | 79371 | ENSG00000101574 | METTL4 | chr18 | - | [illegible] |
| SE | 79372 | ENSG00000101574 | METTL4 | chr18 | - | [illegible] |
| SE | 79373 | ENSG00000101574 | METTL4 | chr18 | - | [illegible] |
| SE | 79374 | ENSG00000122482 | ZNF644 | chr1 | - | [illegible] |
| SE | 79376 | ENSG00000122482 | ZNF644 | chr1 | - | [illegible] |
| SE | 79377 | ENSG00000122482 | ZNF644 | chr1 | - | [illegible] |
| SE | 79378 | ENSG00000122482 | ZNF644 | chr1 | - | [illegible] |
| SE | 79379 | ENSG00000122482 | ZNF644 | chr1 | - | [illegible] |
| SE | 79381 | ENSG00000054523 | KIF1B | chr1 | + | [illegible] |
| SE | 79383 | ENSG00000054523 | KIF1B | chr1 | + | [illegible] |
| SE | 79385 | ENSG00000054523 | KIF1B | chr1 | + | [illegible] |
| SE | 79387 | ENSG00000054523 | KIF1B | chr1 | + | [illegible] |
| SE | 79393 | ENSG00000054523 | KIF1B | chr1 | + | [illegible] |
| SE | 79394 | ENSG00000054523 | KIF1B | chr1 | + | [illegible] |
| SE | 79399 | ENSG00000221866 | PLXNA4 | chr7 | - | [illegible] |
| SE | 79400 | ENSG00000221866 | PLXNA4 | chr7 | - | [illegible] |
| SE | 79401 | ENSG00000065970 | FOXJ2 | chr12 | + | [illegible] |
| SE | 79405 | ENSG00000133624 | ZNF767P | chr7 | - | [illegible] |
| SE | 79406 | ENSG00000133624 | ZNF767P | chr7 | - | [illegible] |
| SE | 79407 | ENSG00000133624 | ZNF767P | chr7 | - | [illegible] |
| SE | 79410 | ENSG00000139637 | MYG1 | chr12 | + | [illegible] |
| SE | 79411 | ENSG00000139637 | MYG1 | chr12 | + | [illegible] |
| SE | 79413 | ENSG00000125388 | GRK4 | chr4 | + | [illegible] |
| SE | 79414 | ENSG00000125388 | GRK4 | chr4 | + | [illegible] |
| SE | 79415 | ENSG00000125388 | GRK4 | chr4 | + | [illegible] |
| SE | 79416 | ENSG00000125388 | GRK4 | chr4 | + | [illegible] |
| SE | 79417 | ENSG00000125388 | GRK4 | chr4 | + | [illegible] |
| SE | 79426 | ENSG00000170260 | ZNF212 | chr7 | + | [illegible] |
| SE | 79427 | ENSG00000170260 | ZNF212 | chr7 | + | [illegible] |
| SE | 79429 | ENSG00000163513 | TGFBR2 | chr3 | + | [illegible] |
| SE | 79430 | ENSG00000163513 | TGFBR2 | chr3 | + | [illegible] |
| SE | 79431 | ENSG00000163513 | TGFBR2 | chr3 | + | [illegible] |
| SE | 79441 | ENSG00000147894 | C9orf72 | chr9 | - | [illegible] |
| SE | 79442 | ENSG00000147894 | C9orf72 | chr9 | - | [illegible] |
| SE | 79443 | ENSG00000147894 | C9orf72 | chr9 | - | [illegible] |
| SE | 79444 | ENSG00000147894 | C9orf72 | chr9 | - | [illegible] |
| SE | 79446 | ENSG00000147894 | C9orf72 | chr9 | - | [illegible] |
| SE | 79447 | ENSG00000147894 | C9orf72 | chr9 | - | [illegible] |
| SE | 79448 | ENSG00000147894 | C9orf72 | chr9 | - | [illegible] |
| SE | 79449 | ENSG00000147894 | C9orf72 | chr9 | - | [illegible] |
| SE | 79451 | ENSG00000121680 | PEX16 | chr11 | - | [illegible] |
| SE | 79452 | ENSG00000121680 | PEX16 | chr11 | - | [illegible] |
| SE | 79454 | ENSG00000121680 | PEX16 | chr11 | - | [illegible] |
| SE | 79457 | ENSG00000121680 | PEX16 | chr11 | - | [illegible] |
| SE | 79461 | ENSG00000166135 | HIF1AN | chr10 | + | [illegible] |
| SE | 79462 | ENSG00000166135 | HIF1AN | chr10 | + | [illegible] |
| SE | 79463 | ENSG00000166135 | HIF1AN | chr10 | + | [illegible] |
| SE | 79464 | ENSG00000166135 | HIF1AN | chr10 | + | [illegible] |

| | | | | | | | | | | | | | | | | | | | | | | | | | | | | | | | | | | | | | | | | | | | | | | |
|---|---|---|---|---|---|---|---|---|---|---|---|---|---|---|---|---|---|---|---|---|---|---|---|---|---|---|---|---|---|---|---|---|---|---|---|---|---|---|---|---|---|---|---|---|---|---|
| SE | 79465 | ENSG00000130021 | PUDP | chrX | - | 7077219 | 7077449 | 7048919 | 7050472 | 7105619 | 7105838 | 103 | 101 | 91 | 109 | 105 | 0 | 0 | 1 | 0 | 0 | 249 | 95 | 42 | 125 | 70 | 0 | 0 | 0 | 0 | 0 | 298 | 149 | 1 | 1 | 1 | 1 | 0.978 | 1 | 1 | 1 | 1 | 1 | 1 | 1 | -0.004 |
| SE | 79466 | ENSG00000130021 | PUDP | chrX | - | 7116965 | 7117034 | 7105619 | 7105838 | 7148052 | 7148158 | 1 | 1 | 1 | 0 | 3 | 51 | 50 | 54 | 49 | 34 | 0 | 0 | 0 | 0 | 0 | 102 | 60 | 17 | 46 | 31 | 218 | 149 | 1 | 1 | 0.013 | 0.013 | 0.012 | 0 | 0.057 | 0 | 0 | 0 | 0 | 0 | 0.019 |
| SE | 79470 | ENSG00000253882 | NA | chr7 | - | 143833246 | 143833883 | 143811968 | 143813779 | 143836639 | 143836708 | 2 | 0 | 0 | 0 | 0 | 1 | 1 | 0 | 0 | 0 | 0 | 0 | 5 | 0 | 13 | 0 | 0 | 0 | 0 | 0 | 298 | 149 | 1 | 1 | 0.5 | 0 | NA | NA | NA | NA | NA | 1 | NA | 1 | -0.75 |
| SE | 79476 | ENSG00000145725 | PPIP5K2 | chr5 | + | 103129305 | 103129703 | 103120199 | 103120488 | 103133452 | 103133513 | 296 | 208 | 242 | 288 | 293 | 0 | 0 | 1 | 0 | 0 | 10 | 130 | 90 | 60 | 33 | 0 | 0 | 0 | 0 | 0 | 298 | 149 | 1 | 1 | 1 | 1 | 0.992 | 1 | 1 | 1 | 1 | 1 | 1 | 1 | -0.002 |
| SE | 79477 | ENSG00000145725 | PPIP5K2 | chr5 | + | 103129305 | 103129703 | 103120199 | 103120488 | 103136731 | 103136822 | 172 | 129 | 131 | 149 | 157 | 0 | 0 | 1 | 0 | 0 | 6 | 72 | 71 | 32 | 19 | 0 | 0 | 0 | 0 | 0 | 298 | 149 | 1 | 1 | 1 | 1 | 0.985 | 1 | 1 | 1 | 1 | 1 | 1 | 1 | -0.003 |
| SE | 79478 | ENSG00000145725 | PPIP5K2 | chr5 | + | 103133452 | 103133648 | 103120199 | 103120488 | 103136731 | 103136822 | 161 | 81 | 120 | 155 | 215 | 0 | 0 | 1 | 0 | 0 | 10 | 71 | 42 | 7 | 13 | 0 | 0 | 0 | 0 | 0 | 298 | 149 | 1 | 1 | 1 | 1 | 0.984 | 1 | 1 | 1 | 1 | 1 | 1 | 1 | -0.003 |
| SE | 79479 | ENSG00000145725 | PPIP5K2 | chr5 | + | 103133452 | 103133648 | 103129305 | 103129703 | 103136731 | 103136822 | 285 | 160 | 230 | 295 | 351 | 0 | 0 | 0 | 1 | 0 | 14 | 129 | 61 | 35 | 27 | 0 | 0 | 0 | 0 | 0 | 298 | 149 | 1 | 1 | 1 | 1 | 1 | 0.993 | 1 | 1 | 1 | 1 | 1 | 1 | -0.001 |
| SE | 79480 | ENSG00000145725 | PPIP5K2 | chr5 | + | 103135992 | 103136114 | 103120261 | 103120488 | 103136731 | 103136822 | 0 | 0 | 0 | 0 | 5 | 0 | 0 | 1 | 0 | 0 | 0 | 8 | 1 | 0 | 0 | 0 | 0 | 0 | 0 | 0 | 271 | 149 | 1 | 1 | NA | NA | 0 | NA | 1 | NA | 1 | 1 | NA | NA | -0.5 |
| SE | 79482 | ENSG00000145725 | PPIP5K2 | chr5 | + | 103159145 | 103159328 | 103158451 | 103158573 | 103167178 | 103167320 | 195 | 127 | 156 | 185 | 358 | 0 | 0 | 0 | 1 | 0 | 18 | 51 | 57 | 4 | 8 | 0 | 0 | 0 | 0 | 0 | 298 | 149 | 1 | 1 | 1 | 1 | 1 | 0.989 | 1 | 1 | 1 | 1 | 1 | 1 | -0.002 |
| SE | 79483 | ENSG00000145725 | PPIP5K2 | chr5 | + | 103167178 | 103167320 | 103159145 | 103159328 | 103173154 | 103173282 | 82 | 61 | 73 | 85 | 186 | 0 | 0 | 1 | 0 | 0 | 11 | 21 | 26 | 0 | 5 | 0 | 0 | 0 | 0 | 0 | 291 | 149 | 1 | 1 | 1 | 1 | 0.974 | 1 | 1 | 1 | 1 | 1 | NA | 1 | -0.005 |
| SE | 79484 | ENSG00000145725 | PPIP5K2 | chr5 | + | 103168071 | 103168295 | 103167178 | 103167320 | 103173154 | 103173282 | 173 | 155 | 132 | 209 | 360 | 1 | 1 | 1 | 0 | 2 | 11 | 68 | 53 | 18 | 0 | 0 | 0 | 0 | 0 | 0 | 298 | 149 | 1 | 1 | 0.989 | 0.987 | 0.985 | 1 | 0.989 | 1 | 1 | 1 | 1 | NA | -0.01 |
| SE | 79485 | ENSG00000145725 | PPIP5K2 | chr5 | + | 103176865 | 103176910 | 103173857 | 103173972 | 103177666 | 103177774 | 22 | 9 | 18 | 15 | 28 | 102 | 95 | 60 | 96 | 145 | 11 | 13 | 17 | 0 | 0 | 0 | 33 | 32 | 0 | 0 | 194 | 149 | 1 | 1 | 0.142 | 0.068 | 0.187 | 0.107 | 0.129 | 1 | 0.232 | 0.29 | NA | NA | -0.381 |
| SE | 79487 | ENSG00000145725 | PPIP5K2 | chr5 | + | 103186319 | 103186439 | 103184671 | 103184744 | 103190841 | 103190982 | 18 | 24 | 26 | 33 | 40 | 75 | 52 | 49 | 71 | 167 | 0 | 0 | 8 | 0 | 0 | 0 | 31 | 15 | 5 | 0 | 269 | 149 | 1 | 1 | 0.117 | 0.204 | 0.227 | 0.205 | 0.117 | NA | 0 | 0.228 | 0 | NA | 0.098 |
| SE | 79488 | ENSG00000145725 | PPIP5K2 | chr5 | + | 103187313 | 103187376 | 103186319 | 103186439 | 103190841 | 103190982 | 13 | 6 | 6 | 15 | 17 | 9 | 15 | 14 | 20 | 15 | 0 | 2 | 0 | 0 | 0 | 0 | 0 | 0 | 0 | 0 | 212 | 149 | 1 | 1 | 0.504 | 0.219 | 0.231 | 0.345 | 0.443 | NA | 1 | NA | NA | NA | -0.652 |
| SE | 79490 | ENSG00000145725 | PPIP5K2 | chr5 | + | 103194899 | 103195025 | 103159145 | 103159328 | 103201521 | 103201533 | 32 | 53 | 32 | 43 | 79 | 0 | 2 | 0 | 0 | 0 | 0 | 14 | 17 | 2 | 0 | 0 | 0 | 0 | 0 | 0 | 275 | 149 | 1 | 1 | 1 | 0.935 | 1 | 1 | 1 | NA | 1 | 1 | 1 | NA | -0.013 |
| SE | 79491 | ENSG00000222011 | FAM185A | chr7 | + | 102751691 | 102751801 | 102749009 | 102749417 | 102757853 | 102757876 | 23 | 11 | 31 | 17 | 9 | 0 | 0 | 2 | 2 | 0 | 0 | 11 | 8 | 2 | 3 | 0 | 0 | 4 | 0 | 0 | 259 | 149 | 1 | 1 | 1 | 1 | 0.899 | 0.83 | 1 | NA | 1 | 0.535 | 1 | 1 | 0.062 |
| SE | 79493 | ENSG00000222011 | FAM185A | chr7 | + | 102787334 | 102787469 | 102777252 | 102777348 | 102808289 | 102808840 | 20 | 12 | 16 | 19 | 20 | 0 | 1 | 0 | 0 | 1 | 12 | 20 | 11 | 13 | 11 | 0 | 0 | 0 | 0 | 0 | 284 | 149 | 1 | 1 | 1 | 0.863 | 1 | 1 | 0.913 | 1 | 1 | 1 | 1 | 1 | -0.045 |
| SE | 79500 | ENSG00000066422 | ZBTB11 | chr3 | - | 101664963 | 101665808 | 101664537 | 101664714 | 101671129 | 101671361 | 265 | 191 | 227 | 234 | 257 | 1 | 0 | 0 | 0 | 0 | 38 | 113 | 157 | 42 | 10 | 0 | 0 | 0 | 0 | 0 | 298 | 149 | 1 | 1 | 0.993 | 1 | 1 | 1 | 1 | 1 | 1 | 1 | 1 | 1 | -0.001 |
| SE | 79501 | ENSG00000066422 | ZBTB11 | chr3 | - | 101671129 | 101671361 | 101664963 | 101665808 | 101671977 | 101672213 | 191 | 144 | 152 | 150 | 210 | 0 | 0 | 2 | 2 | 0 | 28 | 98 | 66 | 54 | 51 | 0 | 0 | 2 | 0 | 0 | 298 | 149 | 1 | 1 | 1 | 1 | 0.974 | 0.974 | 1 | 1 | 1 | 0.943 | 1 | 1 | 0.001 |
| SE | 79502 | ENSG00000066422 | ZBTB11 | chr3 | - | 101671977 | 101672213 | 101664963 | 101665808 | 101676604 | 101677132 | 68 | 64 | 55 | 46 | 39 | 1 | 0 | 0 | 0 | 1 | 14 | 41 | 25 | 26 | 12 | 0 | 0 | 0 | 0 | 0 | 298 | 149 | 1 | 1 | 0.971 | 1 | 1 | 1 | 0.951 | 1 | 1 | 1 | 1 | 1 | -0.016 |
| SE | 79505 | ENSG00000146918 | NCAPG2 | chr7 | - | 158645518 | 158645619 | 158644288 | 158644388 | 158646459 | 158646563 | 23 | 75 | 71 | 54 | 98 | 0 | 0 | 0 | 2 | 0 | 58 | 62 | 61 | 106 | 34 | 0 | 0 | 0 | 3 | 0 | 250 | 149 | 1 | 1 | 1 | 1 | 1 | 0.941 | 1 | 1 | 1 | 1 | 0.955 | 1 | -0.003 |
| SE | 79507 | ENSG00000146918 | NCAPG2 | chr7 | - | 158652292 | 158652480 | 158646459 | 158646563 | 158654594 | 158654694 | 13 | 25 | 12 | 18 | 4 | 2 | 0 | 0 | 0 | 0 | 0 | 7 | 24 | 16 | 0 | 0 | 0 | 1 | 1 | 0 | 298 | 149 | 1 | 1 | 0.765 | 1 | 1 | 1 | 1 | NA | 1 | 0.923 | 0.889 | NA | 0.016 |
| SE | 79508 | ENSG00000146918 | NCAPG2 | chr7 | - | 158655338 | 158655455 | 158655117 | 158655258 | 158656259 | 158656433 | 26 | 12 | 28 | 22 | 18 | 1 | 0 | 0 | 0 | 0 | 9 | 8 | 16 | 39 | 5 | 0 | 0 | 0 | 0 | 0 | 266 | 149 | 1 | 1 | 0.936 | 1 | 1 | 1 | 1 | 1 | 1 | 1 | 1 | 1 | -0.013 |
| SE | 79514 | ENSG00000146918 | NCAPG2 | chr7 | - | 158703649 | 158703702 | 158701821 | 158701938 | 158704723 | 158704804 | 1 | 0 | 2 | 0 | 3 | 2 | 1 | 12 | 3 | 5 | 0 | 2 | 19 | 2 | 0 | 0 | 1 | 0 | 4 | 1 | 202 | 149 | 1 | 1 | 0.269 | 0 | 0.109 | 0 | 0.307 | NA | 0.596 | 1 | 0.269 | 0 | -0.329 |
| SE | 79515 | ENSG00000179218 | CALR | chr19 | + | 12939427 | 12939631 | 12938577 | 12939235 | 12940052 | 12940147 | 2531 | 1410 | 2344 | 2332 | 2931 | 1 | 2 | 5 | 0 | 2 | 4051 | 2675 | 1958 | 2675 | 4088 | 5 | 4 | 8 | 6 | 0 | 298 | 149 | 1 | 1 | 0.999 | 0.997 | 0.996 | 1 | 0.999 | 0.998 | 0.997 | 0.992 | 0.996 | 1 | 0.002 |
| SE | 79517 | ENSG00000185055 | EFCAB10 | chr7 | - | 105567466 | 105567490 | 105565232 | 105565615 | 105569202 | 105569290 | 0 | 0 | 0 | 0 | 0 | 2 | 0 | 0 | 0 | 0 | 0 | 5 | 1 | 0 | 0 | 0 | 0 | 0 | 0 | 0 | 173 | 149 | 1 | 1 | 0 | NA | NA | NA | NA | NA | 1 | 1 | NA | NA | -1 |
| SE | 79520 | ENSG00000105778 | AVL9 | chr7 | + | 32553726 | 32553767 | 32552228 | 32552295 | 32554557 | 32554596 | 145 | 106 | 116 | 120 | 170 | 0 | 0 | 0 | 0 | 0 | 32 | 98 | 98 | 134 | 73 | 0 | 3 | 0 | 0 | 0 | 190 | 149 | 1 | 1 | 1 | 1 | 1 | 1 | 1 | 1 | 0.962 | 1 | 1 | 1 | 0.008 |
| SE | 79522 | ENSG00000105778 | AVL9 | chr7 | + | 32570019 | 32570154 | 32558928 | 32559464 | 32573198 | 32573418 | 155 | 131 | 181 | 187 | 284 | 3 | 0 | 0 | 0 | 0 | 72 | 216 | 115 | 193 | 37 | 0 | 0 | 0 | 2 | 0 | 284 | 149 | 1 | 1 | 0.964 | 1 | 1 | 1 | 1 | 1 | 1 | 1 | 0.981 | 1 | -0.003 |
| SE | 79523 | ENSG00000105778 | AVL9 | chr7 | + | 32575954 | 32576072 | 32573198 | 32573418 | 32580218 | 32580272 | 154 | 101 | 172 | 143 | 181 | 1 | 0 | 0 | 0 | 0 | 112 | 140 | 136 | 133 | 26 | 0 | 0 | 0 | 0 | 0 | 267 | 149 | 1 | 1 | 0.988 | 1 | 1 | 1 | 1 | 1 | 1 | 1 | 1 | 1 | -0.002 |
| [illegible] | | | | | | | | | | | | | | | | | | | | | | | | | | | | | | | | | | | | | | | | | | | | | | |

| | | | | | | | | | | | | | | |
|---|---|---|---|---|---|---|---|---|---|---|---|---|---|---|
| SE | 79902 | ENSG00000068885 | IFT80 | chr3 | - | 160343650 | 160343815 | 160307662 | 160307781 | 160356012 | 160356150 | 3 | 2 | [illegible] |
| SE | 79903 | ENSG00000068885 | IFT80 | chr3 | - | 160343650 | 160343815 | 160319759 | 160319939 | 160356012 | 160356150 | 7 | 7 | [illegible] |
| SE | 79904 | ENSG00000068885 | IFT80 | chr3 | - | 160366042 | 160366152 | 160357484 | 160357578 | 160375811 | 160375880 | 56 | 82 | [illegible] |
| SE | 79905 | ENSG00000068885 | IFT80 | chr3 | - | 160366092 | 160366152 | 160357488 | 160357578 | 160375811 | 160375880 | 30 | 48 | [illegible] |
| SE | 79906 | ENSG00000068885 | IFT80 | chr3 | - | 160377832 | 160377890 | 160377429 | 160377540 | 160381502 | 160381572 | 4 | 0 | [illegible] |
| SE | 79908 | ENSG00000068885 | IFT80 | chr3 | - | 160383436 | 160383504 | 160381502 | 160381718 | 160384405 | 160384646 | 15 | 16 | [illegible] |
| SE | 79909 | ENSG00000068885 | IFT80 | chr3 | - | 160391545 | 160391643 | 160383436 | 160384646 | 160399145 | 160399209 | 0 | 0 | [illegible] |
| SE | 79911 | ENSG00000150768 | DLAT | chr11 | + | 112028791 | 112028945 | 112028514 | 112028639 | 112037272 | 112037460 | 58 | 23 | [illegible] |
| SE | 79913 | ENSG00000150768 | DLAT | chr11 | + | 112033403 | 112033530 | 112028791 | 112028945 | 112039243 | 112039397 | 25 | 17 | [illegible] |
| SE | 79915 | ENSG00000150768 | DLAT | chr11 | + | 112037272 | 112037460 | 112028791 | 112028945 | 112039243 | 112039397 | 44 | 23 | [illegible] |
| SE | 79917 | ENSG00000150768 | DLAT | chr11 | + | 112043465 | 112043533 | 112039243 | 112039397 | 112045137 | 112045230 | 102 | 76 | [illegible] |
| SE | 79918 | ENSG00000150768 | DLAT | chr11 | + | 112043471 | 112043533 | 112039243 | 112039397 | 112045137 | 112045230 | 31 | 35 | [illegible] |
| SE | 79920 | ENSG00000150768 | DLAT | chr11 | + | 112051233 | 112051349 | 112045862 | 112045970 | 112059902 | 112060065 | 160 | 105 | [illegible] |
| SE | 79922 | ENSG00000172331 | BPGM | chr7 | + | 134647134 | 134647259 | 134646810 | 134646937 | 134661446 | 134662108 | 0 | 0 | [illegible] |
| SE | 79924 | ENSG00000172331 | BPGM | chr7 | + | 134661446 | 134662108 | 134646852 | 134646937 | 134678852 | 134679808 | 106 | 42 | [illegible] |
| SE | 79930 | ENSG00000006530 | AGK | chr7 | + | 141593145 | 141593185 | 141555438 | 141555567 | 141596561 | 141596641 | 217 | 118 | [illegible] |
| SE | 79932 | ENSG00000006530 | AGK | chr7 | + | 141594674 | 141594928 | 141593145 | 141593185 | 141596561 | 141596641 | 2 | 0 | [illegible] |
| SE | 79933 | ENSG00000006530 | AGK | chr7 | + | 141600491 | 141600557 | 141596561 | 141596641 | 141601204 | 141601280 | 3 | 0 | [illegible] |
| SE | 79938 | ENSG00000006530 | AGK | chr7 | + | 141633900 | 141633980 | 141621731 | 141621801 | 141636959 | 141636981 | 151 | 73 | [illegible] |
| SE | 79939 | ENSG00000006530 | AGK | chr7 | + | 141641247 | 141641398 | 141636959 | 141637017 | 141641810 | 141641908 | 290 | 146 | [illegible] |
| SE | 79940 | ENSG00000006530 | AGK | chr7 | + | 141651524 | 141651609 | 141649130 | 141649333 | 141652786 | 141652888 | 127 | 98 | [illegible] |
| SE | 79941 | ENSG00000158169 | FANCC | chr9 | - | 95107065 | 95107269 | 95099053 | 95101850 | 95111462 | 95111637 | 10 | 6 | [illegible] |
| SE | 79949 | ENSG00000186480 | INSIG1 | chr7 | + | 155302746 | 155302846 | 155302250 | 155302417 | 155308240 | 155308757 | 235 | 104 | [illegible] |
| SE | 79951 | ENSG00000042980 | ADAM28 | chr8 | + | 24299973 | 24300077 | 24294068 | 24294195 | 24309893 | 24309970 | 69 | 57 | [illegible] |
| SE | 79952 | ENSG00000042980 | ADAM28 | chr8 | + | 24308640 | 24308768 | 24299973 | 24300077 | 24309893 | 24309970 | 4 | 1 | [illegible] |
| SE | 79954 | ENSG00000042980 | ADAM28 | chr8 | + | 24311360 | 24311437 | 24310162 | 24310241 | 24313387 | 24313580 | 59 | 41 | [illegible] |
| SE | 79955 | ENSG00000042980 | ADAM28 | chr8 | + | 24311360 | 24311437 | 24310162 | 24310241 | 24320235 | 24320307 | 45 | 26 | [illegible] |
| SE | 79956 | ENSG00000042980 | ADAM28 | chr8 | + | 24313387 | 24313580 | 24310162 | 24310241 | 24320235 | 24320307 | 32 | 42 | [illegible] |
| SE | 79959 | ENSG00000042980 | ADAM28 | chr8 | + | 24339465 | 24339568 | 24335445 | 24335641 | 24341597 | 24341757 | 16 | 22 | [illegible] |
| SE | 79960 | ENSG00000042980 | ADAM28 | chr8 | + | 24343100 | 24343181 | 24340868 | 24341757 | 24343505 | 24343584 | 39 | 44 | [illegible] |
| SE | 79961 | ENSG00000042980 | ADAM28 | chr8 | + | 24349863 | 24349972 | 24343505 | 24343584 | 24351231 | 24351310 | 63 | 63 | [illegible] |
| SE | 79962 | ENSG00000042980 | ADAM28 | chr8 | + | 24353769 | 24353832 | 24351986 | 24352052 | 24354383 | 24354472 | 21 | 46 | [illegible] |
| SE | 79963 | ENSG00000288558 | DUS4L-BCAP29 | chr7 | + | 107564587 | 107564676 | 107563970 | 107564209 | 107567049 | 107567186 | 3 | 1 | [illegible] |
| SE | 79964 | ENSG00000288558 | DUS4L-BCAP29 | chr7 | + | 107564587 | 107564676 | 107563970 | 107564247 | 107567049 | 107567186 | 2 | 1 | [illegible] |
| SE | 79965 | ENSG00000288558 | DUS4L-BCAP29 | chr7 | + | 107571144 | 107571266 | 107567049 | 107567186 | 107575187 | 107575310 | 17 | 6 | [illegible] |
| SE | 79966 | ENSG00000288558 | DUS4L-BCAP29 | chr7 | + | 107573703 | 107573821 | 107571144 | 107571266 | 107575187 | 107575310 | 20 | 11 | [illegible] |
| SE | 79967 | ENSG00000288558 | DUS4L-BCAP29 | chr7 | + | 107573703 | 107573883 | 107571144 | 107571266 | 107575187 | 107575310 | 9 | 7 | [illegible] |
| SE | 79969 | ENSG00000197124 | ZNF682 | chr19 | - | 20015174 | 20015266 | 20007155 | 20007275 | 20015773 | 20015842 | 0 | 0 | [illegible] |
| SE | 79970 | ENSG00000197124 | ZNF682 | chr19 | - | 20015773 | 20015837 | 20007101 | 20007275 | 20023003 | 20023099 | 0 | 0 | [illegible] |
| SE | 79971 | ENSG00000124193 | SRSF6 | chr20 | + | 43458360 | 43458509 | 43457892 | 43458140 | 43459152 | 43459420 | 352 | 226 | [illegible] |
| SE | 79972 | ENSG00000124193 | SRSF6 | chr20 | + | 43458360 | 43458509 | 43457892 | 43458140 | 43459770 | 43459895 | 674 | 438 | [illegible] |
| SE | 79973 | ENSG00000124193 | SRSF6 | chr20 | + | 43459152 | 43459420 | 43457927 | 43458140 | 43459770 | 43459895 | 168 | 259 | [illegible] |
| SE | 79975 | ENSG00000124193 | SRSF6 | chr20 | + | 43460702 | 43460852 | 43460514 | 43460598 | 43463343 | 43463605 | 677 | 654 | [illegible] |
| SE | 79978 | ENSG00000118058 | KMT2A | chr11 | + | 118471661 | 118474315 | 118468774 | 118468844 | 118476804 | 118476982 | 237 | 226 | [illegible] |
| SE | 79979 | ENSG00000118058 | KMT2A | chr11 | + | 118474178 | 118474315 | 118468774 | 118468844 | 118476804 | 118476982 | 137 | 147 | [illegible] |
| SE | 79984 | ENSG00000118058 | KMT2A | chr11 | + | 118496260 | 118496367 | 118495699 | 118495893 | 118497935 | 118498073 | 393 | 247 | [illegible] |
| SE | 79986 | ENSG00000118058 | KMT2A | chr11 | + | 118507507 | 118507609 | 118506263 | 118506646 | 118509947 | 118510110 | 0 | 0 | [illegible] |
| SE | 79987 | ENSG00000118058 | KMT2A | chr11 | + | 118507528 | 118507609 | 118502397 | 118506646 | 118509135 | 118509200 | 218 | 113 | [illegible] |
| SE | 79988 | ENSG00000118058 | KMT2A | chr11 | + | 118507528 | 118507609 | 118502397 | 118506646 | 118509947 | 118510118 | 92 | 64 | [illegible] |
| SE | 79989 | ENSG00000118058 | KMT2A | chr11 | + | 118509135 | 118509200 | 118502397 | 118506646 | 118509947 | 118510110 | 66 | 20 | [illegible] |
| SE | 79994 | ENSG00000165874 | SHLD2P1 | chr10 | - | 46625806 | 46626013 | 46621605 | 46621834 | 46630952 | 46631087 | 5 | 0 | [illegible] |
| SE | 79995 | ENSG00000204934 | ATP6V0E2-AS1 | chr7 | - | 149870241 | 149870351 | 149867772 | 149869858 | 149871641 | 149871796 | 10 | 5 | [illegible] |
| SE | 79998 | ENSG00000084070 | SMAP2 | chr1 | + | 40362299 | 40362436 | 40344849 | 40344910 | 40413015 | 40413102 | 59 | 43 | [illegible] |
| SE | 79999 | ENSG00000084070 | SMAP2 | chr1 | + | 40406735 | 40406869 | 40344849 | 40344910 | 40408652 | 40408738 | 876 | 668 | [illegible] |
| SE | 80000 | ENSG00000084070 | SMAP2 | chr1 | + | 40406735 | 40406869 | 40362299 | 40362436 | 40408652 | 40408738 | 943 | 715 | [illegible] |
| SE | 80001 | ENSG00000084070 | SMAP2 | chr1 | + | 40406735 | 40406869 | 40373726 | 40374223 | 40408652 | 40408738 | 1777 | 1022 | [illegible] |
| SE | 80002 | ENSG00000084070 | SMAP2 | chr1 | + | 40414158 | 40414240 | 40413015 | 40413102 | 40415271 | 40415381 | 653 | 386 | [illegible] |
| SE | 80003 | ENSG00000084070 | SMAP2 | chr1 | + | 40415271 | 40415381 | 40413015 | 40413102 | 40416175 | 40416341 | 459 | 251 | [illegible] |
| SE | 80004 | ENSG00000084070 | SMAP2 | chr1 | + | 40416175 | 40416341 | 40415271 | 40415381 | 40416779 | 40417096 | 1307 | 661 | [illegible] |
| SE | 80010 | ENSG00000183742 | MACC1 | chr7 | - | 20158203 | 20160245 | 20154192 | 20154381 | 20161747 | 20161870 | 9 | 10 | [illegible] |
| SE | 80011 | ENSG00000183742 | MACC1 | chr7 | - | 20164255 | 20164399 | 20161747 | 20161870 | 20170713 | 20170778 | 2 | 0 | [illegible] |
| SE | 80012 | ENSG00000183742 | MACC1 | chr7 | - | 20170713 | 20170778 | 20161747 | 20161870 | 20184988 | 20185078 | 2 | 1 | [illegible] |
| SE | 80013 | ENSG00000183742 | MACC1 | chr7 | - | 20170713 | 20170778 | 20161747 | 20161870 | 20217298 | 20217384 | 2 | 2 | [illegible] |
| SE | 80014 | ENSG00000183742 | MACC1 | chr7 | - | 20170713 | 20170781 | 20164368 | 20164399 | 20198524 | 20198670 | 2 | 0 | [illegible] |
| SE | 80015 | ENSG00000183742 | MACC1 | chr7 | - | 20184988 | 20185078 | 20161747 | 20161870 | 20217298 | 20217404 | 2 | 2 | [illegible] |
| SE | 80016 | ENSG00000183742 | MACC1 | chr7 | - | 20184988 | 20185078 | 20170713 | 20170778 | 20217298 | 20217404 | 1 | 2 | [illegible] |
| SE | 80018 | ENSG00000183742 | MACC1 | chr7 | - | 20184988 | 20185078 | 20170713 | 20170781 | 20217298 | 20217404 | 1 | 2 | [illegible] |
| SE | 80020 | ENSG00000183742 | MACC1 | chr7 | - | 20198524 | 20198670 | 20184988 | 20185078 | 20217298 | 20217357 | 0 | 1 | [illegible] |
| SE | 80021 | ENSG00000091732 | ZC3HC1 | chr7 | - | 130023510 | 130023723 | 130022318 | 130022525 | 130024262 | 130024377 | 25 | 12 | [illegible] |
| SE | 80022 | ENSG00000091732 | ZC3HC1 | chr7 | - | 130024262 | 130024377 | 130023510 | 130023723 | 130026157 | 130026312 | 20 | 11 | [illegible] |
| SE | 80023 | ENSG00000091732 | ZC3HC1 | chr7 | - | 130024262 | 130024506 | 130023510 | 130023723 | 130026157 | 130026312 | 37 | 22 | [illegible] |
| SE | 80026 | ENSG00000091732 | ZC3HC1 | chr7 | - | 130048973 | 130049144 | 130039463 | 130039547 | 130051220 | 130051369 | 8 | 3 | [illegible] |
| SE | 80027 | ENSG00000091732 | ZC3HC1 | chr7 | - | 130050442 | 130050488 | 130049032 | 130049144 | 130051220 | 130051374 | 0 | 0 | [illegible] |
| SE | 80029 | ENSG00000074800 | ENO1 | chr1 | - | 8870451 | 8870510 | 8867987 | 8868057 | 8871890 | 8871944 | 1435 | 799 | [illegible] |
| SE | 80030 | ENSG00000074800 | ENO1 | chr1 | - | 8870451 | 8870510 | 8867987 | 8868057 | 8878579 | 8878646 | 478 | 289 | [illegible] |
| SE | 80033 | ENSG00000074800 | ENO1 | chr1 | - | 8871890 | 8871986 | 8867987 | 8868057 | 8878579 | 8878646 | 2 | 0 | [illegible] |
| SE | 80034 | ENSG00000074800 | ENO1 | chr1 | - | 8871890 | 8871986 | 8870451 | 8870510 | 8874823 | 8874914 | 1470 | 758 | [illegible] |
| SE | 80035 | ENSG00000074800 | ENO1 | chr1 | - | 8871890 | 8871986 | 8870451 | 8870510 | 8878579 | 8878646 | 799 | 429 | [illegible] |
| SE | 80037 | ENSG00000074800 | ENO1 | chr1 | - | 8874823 | 8874914 | 8870451 | 8870510 | 8878579 | 8879092 | 3 | 1 | [illegible] |
| SE | 80038 | ENSG00000074800 | ENO1 | chr1 | - | 8874823 | 8874914 | 8871890 | 8871986 | 8878579 | 8879092 | 522 | 238 | [illegible] |
| SE | 80039 | ENSG00000074800 | ENO1 | chr1 | - | 8874823 | 8874917 | 8870451 | 8870510 | 8878579 | 8878686 | 556 | 234 | [illegible] |
| SE | 80040 | ENSG00000074800 | ENO1 | chr1 | - | 8874823 | 8874917 | 8870946 | 8871986 | 8878579 | 8878686 | 1226 | 559 | [illegible] |
| SE | 80041 | ENSG00000074800 | ENO1 | chr1 | - | 8875788 | 8876031 | 8874823 | 8874917 | 8878579 | 8878646 | 0 | 0 | [illegible] |
| SE | 80045 | ENSG00000142657 | PGD | chr1 | + | 10400392 | 10400572 | 10399497 | 10399704 | 10404160 | 10404279 | 399 | 232 | [illegible] |
| SE | 80046 | ENSG00000142657 | PGD | chr1 | + | 10403070 | 10403136 | 10399497 | 10399704 | 10404160 | 10404276 | 618 | 353 | [illegible] |
| SE | 80048 | ENSG00000142657 | PGD | chr1 | + | 10404160 | 10404276 | 10403070 | 10403136 | 10408070 | 10408140 | 803 | 563 | [illegible] |
| SE | 80049 | ENSG00000142657 | PGD | chr1 | + | 10404160 | 10404279 | 10403070 | 10403136 | 10408070 | 10408140 | 1699 | 1013 | [illegible] |
| SE | 80050 | ENSG00000142657 | PGD | chr1 | + | 10411417 | 10411552 | 10408070 | 10408140 | 10413061 | 10413251 | 1082 | 589 | [illegible] |
| SE | 80051 | ENSG00000142657 | PGD | chr1 | + | 10417375 | 10417509 | 10416986 | 10417117 | 10418825 | 10418925 | 823 | 506 | [illegible] |
| SE | 80052 | ENSG00000142657 | PGD | chr1 | + | 10419416 | 10419539 | 10418825 | 10418925 | 10419629 | 10419830 | 999 | 430 | [illegible] |
| SE | 80055 | ENSG00000140287 | HDC | chr15 | - | 50257424 | 50257547 | 50254529 | 50254664 | 50258403 | 50258517 | 17 | 5 | [illegible] |
| SE | 80057 | ENSG00000198198 | SZT2 | chr1 | + | 43403600 | 43403774 | 43403176 | 43403302 | 43404379 | 43404550 | 123 | 43 | [illegible] |
| SE | 80058 | ENSG00000198198 | SZT2 | chr1 | + | 43403600 | 43403774 | 43403176 | 43403302 | 43415081 | 43415213 | 55 | 15 | [illegible] |
| SE | 80059 | ENSG00000198198 | SZT2 | chr1 | + | 43403606 | 43403774 | 43403176 | 43403302 | 43404379 | 43404550 | 70 | 28 | [illegible] |
| SE | 80060 | ENSG00000198198 | SZT2 | chr1 | + | 43404379 | 43404550 | 43403176 | 43403302 | 43415081 | 43415213 | 16 | 13 | [illegible] |
| SE | 80061 | ENSG00000198198 | SZT2 | chr1 | + | 43404379 | 43404550 | 43403600 | 43403774 | 43406282 | 43406894 | 70 | 28 | [illegible] |
| SE | 80064 | ENSG00000198198 | SZT2 | chr1 | + | 43415081 | 43415213 | 43404379 | 43404550 | 43419733 | 43419944 | 16 | 9 | [illegible] |
| SE | 80065 | ENSG00000198198 | SZT2 | chr1 | + | 43415081 | 43415213 | 43404379 | 43404550 | 43420152 | 43420323 | 16 | 9 | [illegible] |
| SE | 80068 | ENSG00000198198 | SZT2 | chr1 | + | 43416534 | 43416641 | 43404379 | 43404550 | 43420152 | 43420323 | 0 | 0 | [illegible] |
| SE | 80069 | ENSG00000198198 | SZT2 | chr1 | + | 43416534 | 43416641 | 43415081 | 43415213 | 43419733 | 43419944 | 24 | 19 | [illegible] |
| SE | 80070 | ENSG00000198198 | SZT2 | chr1 | + | 43416534 | 43416641 | 43415081 | 43415213 | 43420152 | 43420323 | 1 | 0 | [illegible] |
| SE | 80071 | ENSG00000198198 | SZT2 | chr1 | + | 43419733 | 43419944 | 43404379 | 43404550 | 43420152 | 43420323 | 26 | 22 | [illegible] |
| SE | 80072 | ENSG00000198198 | SZT2 | chr1 | + | 43419733 | 43419944 | 43415081 | 43415213 | 43420152 | 43420323 | 26 | 22 | [illegible] |
| SE | 80075 | ENSG00000198198 | SZT2 | chr1 | + | 43419733 | 43419961 | 43415081 | 43415213 | 43420152 | 43420323 | 3 | 0 | [illegible] |
| SE | 80077 | ENSG00000198198 | SZT2 | chr1 | + | 43420748 | 43420983 | 43420152 | 43420323 | 43421173 | 43421303 | 65 | 60 | [illegible] |
| SE | 80083 | ENSG00000198198 | SZT2 | chr1 | + | 43426714 | 43426809 | 43426367 | 43426538 | 43427280 | 43427445 | 11 | 5 | [illegible] |
| SE | 80087 | ENSG00000198198 | SZT2 | chr1 | + | 43437170 | 43437323 | 43435199 | 43435329 | 43437405 | 43437508 | 40 | 31 | [illegible] |
| SE | 80088 | ENSG00000198198 | SZT2 | chr1 | + | 43439357 | 43439442 | 43438928 | 43439093 | 43439604 | 43439638 | 97 | 83 | [illegible] |
| SE | 80089 | ENSG00000198198 | SZT2 | chr1 | + | 43439604 | 43439769 | 43439357 | 43439442 | 43439880 | 43440048 | 180 | 159 | [illegible] |
| SE | 80092 | ENSG00000167123 | CERCAM | chr9 | + | 128421203 | 128421511 | 128420480 | 128421074 | 128422867 | 128422978 | 3 | 0 | [illegible] |
| SE | 80094 | ENSG00000167123 | CERCAM | chr9 | + | 128434101 | 128434229 | 128431169 | 128431303 | 128434409 | 128434607 | 47 | 5 | [illegible] |
| SE | 80100 | ENSG00000158525 | CPA5 | chr7 | + | 130367905 | 130367990 | 130367371 | 130367571 | 130368409 | 130368597 | 1 | 2 | [illegible] |
| SE | 80103 | ENSG00000186265 | BTLA | chr3 | - | 112469757 | 112469804 | 112463965 | 112466383 | 112479454 | 112479769 | 56 | 46 | [illegible] |
| SE | 80104 | ENSG00000186265 | BTLA | chr3 | - | 112471211 | 112471355 | 112469757 | 112469804 | 112479454 | 112479769 | 137 | 94 | [illegible] |
| SE | 80111 | ENSG00000170265 | ZNF282 | chr7 | + | 149212357 | 149212471 | 149210584 | 149210704 | 149213700 | 149213814 | 63 | 40 | [illegible] |
| SE | 80112 | ENSG00000075624 | ACTB | chr7 | - | 5529018 | 5529059 | 5528280 | 5528719 | 5529160 | 5529400 | 0 | 0 | [illegible] |
| SE | 80114 | ENSG00000075624 | ACTB | chr7 | - | 5529534 | 5529663 | 5528280 | 5529400 | 5530523 | 5530581 | 15888 | 7152 | [illegible] |
| SE | 80115 | ENSG00000075624 | ACTB | chr7 | - | 5529534 | 5529684 | 5529160 | 5529400 | 5530523 | 5530627 | 12726 | 5971 | [illegible] |
| SE | 80116 | ENSG00000075624 | ACTB | chr7 | - | 5529534 | 5529733 | 5529160 | 5529400 | 5530523 | 5530601 | 12726 | 5970 | [illegible] |
| SE | 80119 | ENSG00000197024 | ZNF398 | chr7 | + | 149166816 | 149166930 | 149153944 | 149154340 | 149176467 | 149176581 | 36 | 29 | [illegible] |
| SE | 80122 | ENSG00000122515 | ZMIZ2 | chr7 | + | 44749814 | 44749980 | 44748580 | 44748631 | 44756187 | 44756299 | 2 | 2 | [illegible] |
| SE | 80123 | ENSG00000122515 | ZMIZ2 | chr7 | + | 44749814 | 44749980 | 44748861 | 44748991 | 44756187 | 44756299 | 0 | 2 | [illegible] |
| SE | 80124 | ENSG00000122515 | ZMIZ2 | chr7 | + | 44750971 | 44751091 | 44748580 | 44748631 | 44756187 | 44756299 | 0 | 0 | [illegible] |
| SE | 80125 | ENSG00000122515 | ZMIZ2 | chr7 | + | 44750971 | 44751091 | 44749814 | 44749980 | 44756187 | 44756299 | 0 | 0 | [illegible] |
| SE | 80129 | ENSG00000122515 | ZMIZ2 | chr7 | + | 44760150 | 44760228 | 44759280 | 44759460 | 44760424 | 44760593 | 7 | 5 | [illegible] |
| SE | 80132 | ENSG00000012124 | CD22 | chr19 | + | 35331905 | 35332074 | 35329186 | 35329230 | 35332552 | 35332924 | 7 | 4 | [illegible] |
| SE | 80134 | ENSG00000012124 | CD22 | chr19 | + | 35331926 | 35332074 | 35329175 | 35329230 | 35332552 | 35332924 | 8 | 4 | [illegible] |
| SE | 80137 | ENSG00000012124 | CD22 | chr19 | + | 35332018 | 35332074 | 35329165 | 35329230 | 35332577 | 35332924 | 7 | 1 | [illegible] |
| SE | 80138 | ENSG00000012124 | CD22 | chr19 | + | 35332018 | 35332074 | 35329165 | 35329230 | 35332593 | 35332924 | 4 | 0 | [illegible] |
| SE | 80139 | ENSG00000012124 | CD22 | chr19 | + | 35332018 | 35332074 | 35329165 | 35329230 | 35332652 | 35332924 | 11 | 0 | [illegible] |
| SE | 80155 | ENSG00000012124 | CD22 | chr19 | + | 35336035 | 35336341 | 35329186 | 35329230 | 35338167 | 35338431 | 0 | 0 | [illegible] |
| SE | 80157 | ENSG00000012124 | CD22 | chr19 | + | 35336035 | 35336341 | 35332546 | 35332924 | 35337754 | 35337977 | 102 | 63 | [illegible] |
| SE | 80160 | ENSG00000012124 | CD22 | chr19 | + | 35336148 | 35336341 | 35329186 | 35329230 | 35338167 | 35338431 | 1 | 0 | [illegible] |
| SE | 80161 | ENSG00000012124 | CD22 | chr19 | + | 35336148 | 35336341 | 35332552 | 35332924 | 35337754 | 35337807 | 42 | 24 | [illegible] |
| SE | 80162 | ENSG00000012124 | CD22 | chr19 | + | 35336250 | 35336341 | 35332545 | 35332924 | 35337754 | 35338021 | 23 | 19 | [illegible] |
| SE | 80163 | ENSG00000012124 | CD22 | chr19 | + | 35337754 | 35338021 | 35329187 | 35329230 | 35338167 | 35338170 | 87 | 40 | [illegible] |
| SE | 80166 | ENSG00000012124 | CD22 | chr19 | + | 35337754 | 35338021 | 35336035 | 35336341 | 35338167 | 35338170 | 128 | 63 | [illegible] |
| SE | 80171 | ENSG00000012124 | CD22 | chr19 | + | 35341342 | 35341606 | 35340880 | 35341138 | 35341701 | 35341724 | 168 | 83 | [illegible] |
| SE | 80172 | ENSG00000012124 | CD22 | chr19 | + | 35341525 | 35341606 | 35340880 | 35341138 | 35341701 | 35341965 | 51 | 20 | [illegible] |
| SE | 80174 | ENSG00000179454 | KLHL28 | chr14 | - | 44931332 | 44931541 | 44924323 | 44929191 | 44934114 | 44934558 | 73 | 53 | [illegible] |
| SE | 80175 | ENSG00000179454 | KLHL28 | chr14 | - | 44934114 | 44934558 | 44931332 | 44931541 | 44945029 | 44945928 | 66 | 54 | [illegible] |
| SE | 80176 | ENSG00000179454 | KLHL28 | chr14 | - | 44945029 | 44945928 | 44934114 | 44934558 | 44961845 | 44961947 | 120 | 50 | [illegible] |
| SE | 80177 | ENSG00000179454 | KLHL28 | chr14 | - | 44953622 | 44953709 | 44945232 | 44945928 | 44961596 | 44961967 | 0 | 0 | [illegible] |
| SE | 80178 | ENSG00000179454 | KLHL28 | chr14 | - | 44953622 | 44953709 | 44945232 | 44945928 | 44961845 | 44961909 | 0 | 0 | [illegible] |
| SE | 80180 | ENSG00000147065 | MSN | chrX | + | 65588885 | 65589068 | 65588380 | 65588612 | 65593252 | 65593425 | 27 | 7 | [illegible] |
| SE | 80181 | ENSG00000147065 | MSN | chrX | + | 65588885 | 65589068 | 65588380 | 65588612 | 65600661 | 65600750 | 24 | 4 | [illegible] |
| SE | 80184 | ENSG00000147065 | MSN | chrX | + | 65589645 | 65589769 | 65588885 | 65589068 | 65593252 | 65593425 | 1 | 1 | [illegible] |
| SE | 80185 | ENSG00000147065 | MSN | chrX | + | 65589645 | 65589769 | 65588885 | 65589068 | 65624643 | 65625865 | 1 | 0 | [illegible] |
| SE | 80186 | ENSG00000147065 | MSN | chrX | + | 65600661 | 65600750 | 65588397 | 65588612 | 65624643 | 65625865 | 0 | 0 | [illegible] |
| SE | 80187 | ENSG00000147065 | MSN | chrX | + | 65600661 | 65600750 | 65588397 | 65588612 | 65716817 | 65716901 | 0 | 0 | [illegible] |
| SE | 80188 | ENSG00000147065 | MSN | chrX | + | 65600661 | 65600750 | 65588885 | 65589068 | 65624643 | 65625865 | 0 | 0 | [illegible] |
| SE | 80190 | ENSG00000147065 | MSN | chrX | + | 65716817 | 65716901 | 65588376 | 65588612 | 65727813 | 65727909 | 1048 | 451 | [illegible] |
| SE | 80191 | ENSG00000147065 | MSN | chrX | + | 65716817 | 65716901 | 65667653 | 65667853 | 65727813 | 65727909 | 2422 | 1032 | [illegible] |
| SE | 80193 | ENSG00000147065 | MSN | chrX | + | 65727813 | 65727909 | 65716817 | 65716901 | 65731106 | 65731190 | 646 | 293 | [illegible] |
| SE | 80194 | ENSG00000147065 | MSN | chrX | + | 65729437 | 65729712 | 65716817 | 65716901 | 65731106 | 65731190 | 1584 | 851 | [illegible] |
| SE | 80195 | ENSG00000147065 | MSN | chrX | + | 65729437 | 65729712 | 65727813 | 65727909 | 65731106 | 65731190 | 3132 | 1555 | [illegible] |
| SE | 80196 | ENSG00000147065 | MSN | chrX | + | 65731837 | 65731984 | 65731106 | 65731190 | 65733183 | 65733280 | 4758 | 2855 | [illegible] |
| SE | 80197 | ENSG00000147065 | MSN | chrX | + | 65733183 | 65733280 | 65731837 | 65731984 | 65735266 | 65735430 | 5912 | 3907 | [illegible] |
| SE | 80198 | ENSG00000147065 | MSN | chrX | + | 65736794 | 65736925 | 65735266 | 65735430 | 65737177 | 65737338 | 8274 | 5891 | [illegible] |
| SE | 80199 | ENSG00000147065 | MSN | chrX | + | 65737177 | 65737338 | 65735266 | 65735430 | 65738524 | 65738617 | 5723 | 3807 | [illegible] |
| SE | 80200 | ENSG00000147065 | MSN | chrX | + | 65738524 | 65738617 | 65737177 | 65737338 | 65738969 | 65739194 | 7978 | 5030 | [illegible] |
| SE | 80206 | ENSG00000153814 | JAZF1 | chr7 | - | 27840697 | 27840867 | 27830572 | 27832976 | 27895219 | 27895416 | 220 | 76 | [illegible] |
| SE | 80207 | ENSG00000153814 | JAZF1 | chr7 | - | 27895219 | 27895416 | 27840697 | 27840867 | 27991908 | 27991981 | 193 | 88 | [illegible] |
| SE | 80208 | ENSG00000153814 | JAZF1 | chr7 | - | 27913391 | 27913453 | 27895218 | 27895416 | 27991908 | 27991981 | 0 | 1 | [illegible] |
| SE | 80210 | ENSG00000153814 | JAZF1 | chr7 | - | 28101618 | 28101804 | 27991908 | 27991981 | 28180462 | 28180611 | 0 | 2 | [illegible] |
| SE | 80211 | ENSG00000158623 | COPG2 | chr7 | - | 130550523 | 130550649 | 130549313 | 130549376 | 130551240 | 130551344 | 101 | 60 | [illegible] |
| SE | 80212 | ENSG00000158623 | COPG2 | chr7 | - | 130554480 | 130554724 | 130552354 | 130552430 | 130555036 | 130555132 | 69 | 37 | [illegible] |
| SE | 80213 | ENSG00000010318 | PHF7 | chr3 | + | 52412810 | 52412920 | 52410659 | 52411247 | 52413995 | 52414048 | 31 | 20 | [illegible] |
| SE | 80214 | ENSG00000010318 | PHF7 | chr3 | + | 52413995 | 52414048 | 52412810 | 52412920 | 52414495 | 52414508 | 30 | 22 | [illegible] |
| SE | 80218 | ENSG00000010318 | PHF7 | chr3 | + | 52420902 | 52421062 | 52419832 | 52419934 | 52423090 | 52423283 | 0 | 1 | [illegible] |
| SE | 80219 | ENSG00000010318 | PHF7 | chr3 | + | 52421647 | 52421754 | 52419832 | 52419934 | 52423090 | 52423406 | 2 | 1 | [illegible] |
| SE | 80220 | ENSG00000010318 | PHF7 | chr3 | + | 52421647 | 52421754 | 52420902 | 52421062 | 52423090 | 52423406 | 3 | 2 | [illegible] |
| SE | 80221 | ENSG00000010318 | PHF7 | chr3 | + | 52421647 | 52422881 | 52419919 | 52419934 | 52423090 | 52423283 | 0 | 2 | [illegible] |
| SE | 80222 | ENSG00000010318 | PHF7 | chr3 | + | 52421647 | 52422881 | 52420902 | 52421062 | 52423090 | 52423283 | 1 | 3 | [illegible] |
| SE | 80228 | ENSG00000198961 | PJA2 | chr5 | - | 109344189 | 109344311 | 109334712 | 109337356 | 109344704 | 109344819 | 602 | 365 | [illegible] |
| SE | 80229 | ENSG00000198961 | PJA2 | chr5 | - | 109355914 | 109356026 | 109344704 | 109344819 | 109362839 | 109363022 | 458 | 335 | [illegible] |
| SE | 80230 | ENSG00000198961 | PJA2 | chr5 | - | 109381502 | 109381703 | 109378203 | 109379254 | 109383402 | 109383433 | 882 | 649 | [illegible] |
| SE | 80231 | ENSG00000198961 | PJA2 | chr5 | - | 109390637 | 109390758 | 109383395 | 109383520 | 109409116 | 109409314 | 4 | 2 | [illegible] |
| SE | 80235 | ENSG00000113303 | BTNL8 | chr5 | + | 180947511 | 180947625 | 180911338 | 180911614 | 180948354 | 180948375 | 1 | 0 | [illegible] |
| SE | 80236 | ENSG00000113303 | BTNL8 | chr5 | + | 180947511 | 180947755 | 180911338 | 180911614 | 180948354 | 180948375 | 0 | 0 | [illegible] |
| SE | 80237 | ENSG00000113303 | BTNL8 | chr5 | + | 180947624 | 180947755 | 180911488 | 180911614 | 180948354 | 180948375 | 0 | 0 | [illegible] |
| SE | 80238 | ENSG00000113303 | BTNL8 | chr5 | + | 180948354 | 180948375 | 180947511 | 180947625 | 180948919 | 180948946 | 1 | 0 | [illegible] |
| SE | 80240 | ENSG00000077454 | LRCH4 | chr7 | - | 100577085 | 100577389 | 100576693 | 100577005 | 100577496 | 100577558 | 206 | 156 | [illegible] |
| SE | 80241 | ENSG00000077454 | LRCH4 | chr7 | - | 100577085 | 100577389 | 100576693 | 100577005 | 100577663 | 100577740 | 38 | 27 | [illegible] |
| SE | 80243 | ENSG00000169410 | PTPN9 | chr15 | - | 75479847 | 75479914 | 75473688 | 75473767 | 75490207 | 75490301 | 174 | 148 | [illegible] |
| SE | 80244 | ENSG00000169410 | PTPN9 | chr15 | - | 75490207 | 75490301 | 75479847 | 75479914 | 75505674 | 75505816 | 218 | 174 | [illegible] |
| SE | 80245 | ENSG00000169410 | PTPN9 | chr15 | - | 75523120 | 75523245 | 75517258 | 75517364 | 75527117 | 75527261 | 81 | 55 | [illegible] |
| SE | 80250 | ENSG00000164535 | DAGLB | chr7 | - | 6416626 | 6416754 | 6412965 | 6413034 | 6416840 | 6416921 | 23 | 10 | [illegible] |
| SE | 80251 | ENSG00000164535 | DAGLB | chr7 | - | 6421726 | 6421804 | 6416840 | 6416921 | 6424751 | 6424835 | 32 | 25 | [illegible] |
| SE | 80252 | ENSG00000164535 | DAGLB | chr7 | - | 6421726 | 6421804 | 6416840 | 6416921 | 6425987 | 6426114 | 3 | 4 | [illegible] |
| SE | 80257 | ENSG00000164535 | DAGLB | chr7 | - | 6425987 | 6426114 | 6424751 | 6424835 | 6430479 | 6430607 | 54 | 58 | [illegible] |
| SE | 80258 | ENSG00000164535 | DAGLB | chr7 | - | 6430479 | 6430607 | 6425987 | 6426114 | 6432836 | 6432959 | 72 | 70 | [illegible] |
| SE | 80260 | ENSG00000164535 | DAGLB | chr7 | - | 6430479 | 6430607 | 6425987 | 6426114 | 6445952 | 6446104 | 29 | 34 | [illegible] |
| SE | 80262 | ENSG00000164535 | DAGLB | chr7 | - | 6432836 | 6432959 | 6425987 | 6426114 | 6445952 | 6446104 | 1 | 0 | [illegible] |
| SE | 80263 | ENSG00000164535 | DAGLB | chr7 | - | 6432836 | 6432959 | 6430479 | 6430607 | 6434761 | 6435020 | 100 | 82 | [illegible] |
| SE | 80264 | ENSG00000164535 | DAGLB | chr7 | - | 6432836 | 6432959 | 6430479 | 6430607 | 6436361 | 6436533 | 44 | 34 | [illegible] |
| SE | 80265 | ENSG00000164535 | DAGLB | chr7 | - | 6432836 | 6432959 | 6430479 | 6430607 | 6445952 | 6446104 | 44 | 34 | [illegible] |
| SE | 80266 | ENSG00000164535 | DAGLB | chr7 | - | 6434761 | 6435020 | 6430479 | 6430607 | 6436361 | 6436533 | 34 | 25 | [illegible] |
| SE | 80267 | ENSG00000164535 | DAGLB | chr7 | - | 6434761 | 6435020 | 6430479 | 6430607 | 6445952 | 6446104 | 2 | 6 | [illegible] |
| SE | 80270 | ENSG00000164535 | DAGLB | chr7 | - | 6436361 | 6436533 | 6426085 | 6426114 | 6445952 | 6446104 | 10 | 5 | [illegible] |
| SE | 80271 | ENSG00000164535 | DAGLB | chr7 | - | 6436361 | 6436533 | 6430479 | 6430607 | 6445952 | 6446104 | 10 | 5 | [illegible] |
| SE | 80272 | ENSG00000164535 | DAGLB | chr7 | - | 6436361 | 6436533 | 6432836 | 6432959 | 6445952 | 6446104 | 10 | 5 | [illegible] |
| [illegible] | [illegible] | [illegible] | [illegible] | [illegible] | [illegible] | [illegible] | [illegible] | [illegible] | [illegible] | [illegible] | [illegible] | [illegible] | [illegible] | [illegible] |

| | | | | | | | | | | | | | | |
|---|---|---|---|---|---|---|---|---|---|---|---|---|---|---|
| SE | 80302 | ENSG00000168137 | SETD5 | chr3 | + | 9454004 | 9454101 | 9453738 | 9453868 | 9464409 | 9464672 | 19 | 28 | [illegible] |
| SE | 80305 | ENSG00000168137 | SETD5 | chr3 | + | 9470458 | 9470929 | 9464409 | 9464672 | 9473235 | 9473372 | 140 | 121 | [illegible] |
| SE | 80307 | ENSG00000197958 | RPL12 | chr9 | - | 127448336 | 127448423 | 127447679 | 127447989 | 127449280 | 127449362 | 6916 | 4639 | [illegible] |
| SE | 80308 | ENSG00000197958 | RPL12 | chr9 | - | 127449609 | 127449708 | 127449280 | 127449362 | 127450730 | 127450804 | 5833 | 3487 | [illegible] |
| SE | 80309 | ENSG00000197958 | RPL12 | chr9 | - | 127450730 | 127450804 | 127449609 | 127449708 | 127451280 | 127451373 | 3687 | 1936 | [illegible] |
| SE | 80310 | ENSG00000241973 | PI4KA | chr22 | - | 20711340 | 20711461 | 20710698 | 20710858 | 20712485 | 20712611 | 89 | 20 | [illegible] |
| SE | 80311 | ENSG00000241973 | PI4KA | chr22 | - | 20726487 | 20726541 | 20721297 | 20721418 | 20727229 | 20727397 | 161 | 83 | [illegible] |
| SE | 80312 | ENSG00000241973 | PI4KA | chr22 | - | 20726497 | 20726541 | 20721297 | 20721418 | 20727229 | 20727357 | 63 | 34 | [illegible] |
| SE | 80314 | ENSG00000241973 | PI4KA | chr22 | - | 20732970 | 20733098 | 20729891 | 20730011 | 20733735 | 20733843 | 217 | 137 | [illegible] |
| SE | 80316 | ENSG00000241973 | PI4KA | chr22 | - | 20749904 | 20749994 | 20747483 | 20747702 | 20751292 | 20751376 | 106 | 74 | [illegible] |
| SE | 80317 | ENSG00000241973 | PI4KA | chr22 | - | 20764816 | 20764950 | 20761303 | 20761386 | 20765099 | 20765236 | 134 | 84 | [illegible] |
| SE | 80318 | ENSG00000241973 | PI4KA | chr22 | - | 20793192 | 20793243 | 20765584 | 20765693 | 20798583 | 20798687 | 50 | 33 | [illegible] |
| SE | 80320 | ENSG00000241973 | PI4KA | chr22 | - | 20798583 | 20798687 | 20796145 | 20796314 | 20799092 | 20799276 | 210 | 132 | [illegible] |
| SE | 80321 | ENSG00000241973 | PI4KA | chr22 | - | 20799092 | 20799276 | 20798477 | 20798687 | 20799670 | 20799766 | 366 | 244 | [illegible] |
| SE | 80322 | ENSG00000241973 | PI4KA | chr22 | - | 20799670 | 20799766 | 20798939 | 20799276 | 20801972 | 20802018 | 361 | 255 | [illegible] |
| SE | 80324 | ENSG00000241973 | PI4KA | chr22 | - | 20807361 | 20807458 | 20804973 | 20805165 | 20810966 | 20811032 | 817 | 524 | [illegible] |
| SE | 80326 | ENSG00000241973 | PI4KA | chr22 | - | 20810966 | 20811032 | 20807361 | 20807458 | 20818482 | 20818549 | 79 | 45 | [illegible] |
| SE | 80327 | ENSG00000241973 | PI4KA | chr22 | - | 20813357 | 20813506 | 20807361 | 20807458 | 20818482 | 20818549 | 106 | 93 | [illegible] |
| SE | 80329 | ENSG00000241973 | PI4KA | chr22 | - | 20818482 | 20818549 | 20813357 | 20813506 | 20819640 | 20819900 | 136 | 124 | [illegible] |
| SE | 80330 | ENSG00000241973 | PI4KA | chr22 | - | 20838614 | 20838731 | 20834561 | 20834655 | 20858569 | 20858811 | 48 | 28 | [illegible] |
| SE | 80336 | ENSG00000163492 | CCDC141 | chr2 | - | 178845625 | 178845742 | 178835755 | 178837744 | 178850048 | 178850161 | 41 | 100 | [illegible] |
| SE | 80337 | ENSG00000163492 | CCDC141 | chr2 | - | 178877963 | 178878143 | 178872132 | 178872312 | 178884900 | 178884951 | 73 | 55 | [illegible] |
| SE | 80338 | ENSG00000163492 | CCDC141 | chr2 | - | 178886751 | 178886871 | 178884900 | 178885092 | 178888526 | 178888668 | 23 | 26 | [illegible] |
| SE | 80340 | ENSG00000163492 | CCDC141 | chr2 | - | 178975056 | 178975165 | 178961229 | 178961483 | 178978483 | 178978675 | 30 | 27 | [illegible] |
| SE | 80346 | ENSG00000114698 | PLSCR4 | chr3 | - | 146250556 | 146250637 | 146222064 | 146222092 | 146250959 | 146251125 | 0 | 0 | [illegible] |
| SE | 80349 | ENSG00000102531 | FNDC3A | chr13 | + | 49075288 | 49075364 | 49006151 | 49006289 | 49114654 | 49114731 | 115 | 103 | [illegible] |
| SE | 80350 | ENSG00000102531 | FNDC3A | chr13 | + | 49114654 | 49114731 | 49075288 | 49075364 | 49131136 | 49131374 | 195 | 159 | [illegible] |
| SE | 80351 | ENSG00000102531 | FNDC3A | chr13 | + | 49172042 | 49172096 | 49168612 | 49168751 | 49174434 | 49174559 | 116 | 103 | [illegible] |
| SE | 80353 | ENSG00000102531 | FNDC3A | chr13 | + | 49174434 | 49174559 | 49168612 | 49168751 | 49175366 | 49175541 | 148 | 87 | [illegible] |
| SE | 80355 | ENSG00000102531 | FNDC3A | chr13 | + | 49178567 | 49178654 | 49175366 | 49175541 | 49185963 | 49186102 | 217 | 227 | [illegible] |
| SE | 80356 | ENSG00000102531 | FNDC3A | chr13 | + | 49185963 | 49186102 | 49178567 | 49178654 | 49187121 | 49187190 | 262 | 277 | [illegible] |
| SE | 80357 | ENSG00000102531 | FNDC3A | chr13 | + | 49191014 | 49191109 | 49188514 | 49188633 | 49191208 | 49191384 | 58 | 45 | [illegible] |
| SE | 80359 | ENSG00000102531 | FNDC3A | chr13 | + | 49191208 | 49191384 | 49188514 | 49188633 | 49196876 | 49196990 | 125 | 96 | [illegible] |
| SE | 80362 | ENSG00000102531 | FNDC3A | chr13 | + | 49201799 | 49201966 | 49198361 | 49198574 | 49203156 | 49203205 | 218 | 176 | [illegible] |
| SE | 80364 | ENSG00000134308 | YWHAQ | chr2 | - | 9588164 | 9588328 | 9587413 | 9587509 | 9591391 | 9591515 | 1440 | 847 | [illegible] |
| SE | 80365 | ENSG00000204371 | EHMT2 | chr6 | - | 31880672 | 31880848 | 31879758 | 31880264 | 31881013 | 31881092 | 181 | 106 | [illegible] |
| SE | 80366 | ENSG00000204371 | EHMT2 | chr6 | - | 31884890 | 31885016 | 31884644 | 31884799 | 31886580 | 31886682 | 22 | 5 | [illegible] |
| SE | 80367 | ENSG00000204371 | EHMT2 | chr6 | - | 31884911 | 31885016 | 31884644 | 31884799 | 31886580 | 31886682 | 46 | 19 | [illegible] |
| SE | 80368 | ENSG00000204371 | EHMT2 | chr6 | - | 31884911 | 31885055 | 31884644 | 31884799 | 31886580 | 31886682 | 25 | 8 | [illegible] |
| SE | 80370 | ENSG00000204371 | EHMT2 | chr6 | - | 31892825 | 31892910 | 31892693 | 31892734 | 31893361 | 31893662 | 79 | 51 | [illegible] |
| SE | 80384 | ENSG00000142303 | ADAMTS10 | chr19 | - | 8589451 | 8589574 | 8589241 | 8589365 | 8589888 | 8589991 | 10 | 4 | [illegible] |
| SE | 80385 | ENSG00000142303 | ADAMTS10 | chr19 | - | 8589451 | 8589585 | 8589241 | 8589365 | 8589888 | 8589991 | 20 | 7 | [illegible] |
| SE | 80386 | ENSG00000142303 | ADAMTS10 | chr19 | - | 8596072 | 8596219 | 8595761 | 8595903 | 8596306 | 8596412 | 18 | 13 | [illegible] |
| SE | 80387 | ENSG00000142303 | ADAMTS10 | chr19 | - | 8605622 | 8605809 | 8605011 | 8605358 | 8608133 | 8608248 | 23 | 12 | [illegible] |
| SE | 80394 | ENSG00000197694 | SPTAN1 | chr9 | + | 128568771 | 128568897 | 128566737 | 128566977 | 128574674 | 128574735 | 573 | 340 | [illegible] |
| SE | 80395 | ENSG00000197694 | SPTAN1 | chr9 | + | 128572755 | 128572824 | 128568771 | 128568897 | 128574674 | 128574815 | 2 | 0 | [illegible] |
| SE | 80396 | ENSG00000197694 | SPTAN1 | chr9 | + | 128577351 | 128577506 | 128577128 | 128577273 | 128578109 | 128578245 | 908 | 576 | [illegible] |
| SE | 80397 | ENSG00000197694 | SPTAN1 | chr9 | + | 128587605 | 128587698 | 128585747 | 128585965 | 128588808 | 128588943 | 538 | 424 | [illegible] |
| SE | 80398 | ENSG00000197694 | SPTAN1 | chr9 | + | 128588808 | 128588943 | 128587605 | 128587698 | 128591476 | 128591625 | 949 | 694 | [illegible] |
| SE | 80400 | ENSG00000197694 | SPTAN1 | chr9 | + | 128601344 | 128601400 | 128600079 | 128600115 | 128603542 | 128603590 | 0 | 0 | [illegible] |
| SE | 80401 | ENSG00000197694 | SPTAN1 | chr9 | + | 128603542 | 128603590 | 128598962 | 128600115 | 128604325 | 128604335 | 258 | 163 | [illegible] |
| SE | 80402 | ENSG00000197694 | SPTAN1 | chr9 | + | 128604325 | 128604417 | 128598962 | 128600115 | 128605033 | 128605178 | 119 | 93 | [illegible] |
| SE | 80403 | ENSG00000197694 | SPTAN1 | chr9 | + | 128607603 | 128607703 | 128605295 | 128605477 | 128607851 | 128607989 | 479 | 331 | [illegible] |
| SE | 80404 | ENSG00000197694 | SPTAN1 | chr9 | + | 128607851 | 128608049 | 128607603 | 128607703 | 128608129 | 128608276 | 473 | 267 | [illegible] |
| SE | 80406 | ENSG00000197694 | SPTAN1 | chr9 | + | 128611713 | 128611845 | 128609121 | 128609284 | 128612108 | 128612246 | 804 | 452 | [illegible] |
| SE | 80407 | ENSG00000197694 | SPTAN1 | chr9 | + | 128612108 | 128612246 | 128611601 | 128611845 | 128613380 | 128613485 | 717 | 405 | [illegible] |
| SE | 80408 | ENSG00000197694 | SPTAN1 | chr9 | + | 128617986 | 128618108 | 128617639 | 128617760 | 128618870 | 128619003 | 505 | 318 | [illegible] |
| SE | 80410 | ENSG00000197694 | SPTAN1 | chr9 | + | 128625102 | 128625179 | 128624327 | 128624487 | 128625768 | 128625978 | 464 | 252 | [illegible] |
| SE | 80411 | ENSG00000197694 | SPTAN1 | chr9 | + | 128626999 | 128627175 | 128626510 | 128626687 | 128627385 | 128628025 | 0 | 0 | [illegible] |
| SE | 80414 | ENSG00000197694 | SPTAN1 | chr9 | + | 128630320 | 128630375 | 128627924 | 128627942 | 128632126 | 128632323 | 216 | 134 | [illegible] |
| SE | 80416 | ENSG00000117906 | RCN2 | chr15 | + | 76938725 | 76938815 | 76935525 | 76935722 | 76943757 | 76943871 | 1 | 0 | [illegible] |
| SE | 80418 | ENSG00000117906 | RCN2 | chr15 | + | 76943757 | 76943871 | 76935525 | 76935722 | 76947420 | 76947489 | 305 | 243 | [illegible] |
| SE | 80419 | ENSG00000126746 | ZNF384 | chr12 | - | 6669030 | 6669189 | 6666476 | 6668115 | 6670759 | 6670838 | 179 | 109 | [illegible] |
| SE | 80425 | ENSG00000126746 | ZNF384 | chr12 | - | 6688166 | 6688226 | 6679219 | 6679525 | 6689097 | 6689292 | 71 | 41 | [illegible] |
| SE | 80426 | ENSG00000126746 | ZNF384 | chr12 | - | 6688166 | 6688226 | 6679219 | 6679525 | 6689367 | 6689454 | 57 | 37 | [illegible] |
| SE | 80427 | ENSG00000126746 | ZNF384 | chr12 | - | 6688166 | 6688226 | 6679454 | 6679521 | 6689097 | 6689292 | 2 | 3 | [illegible] |
| SE | 80432 | ENSG00000006534 | ALDH3B1 | chr11 | + | 68019714 | 68019796 | 68019065 | 68019255 | 68021484 | 68021522 | 102 | 36 | [illegible] |
| SE | 80435 | ENSG00000133216 | EPHB2 | chr1 | + | 22907952 | 22908168 | 22906709 | 22906957 | 22909021 | 22909171 | 9 | 0 | [illegible] |
| SE | 80436 | ENSG00000085788 | DDHD2 | chr8 | + | 38234393 | 38234510 | 38232986 | 38233214 | 38237537 | 38237622 | 68 | 49 | [illegible] |
| SE | 80437 | ENSG00000085788 | DDHD2 | chr8 | + | 38234393 | 38234584 | 38232986 | 38233214 | 38237537 | 38237585 | 179 | 162 | [illegible] |
| SE | 80438 | ENSG00000085788 | DDHD2 | chr8 | + | 38234399 | 38234584 | 38232986 | 38233214 | 38237537 | 38237616 | 104 | 84 | [illegible] |
| SE | 80441 | ENSG00000085788 | DDHD2 | chr8 | + | 38245741 | 38245950 | 38241964 | 38242385 | 38246232 | 38246300 | 163 | 84 | [illegible] |
| SE | 80442 | ENSG00000085788 | DDHD2 | chr8 | + | 38245741 | 38245950 | 38241964 | 38242385 | 38247712 | 38247835 | 109 | 52 | [illegible] |
| SE | 80443 | ENSG00000085788 | DDHD2 | chr8 | + | 38246232 | 38246300 | 38238088 | 38238209 | 38247712 | 38247835 | 58 | 44 | [illegible] |
| SE | 80444 | ENSG00000085788 | DDHD2 | chr8 | + | 38246232 | 38246300 | 38241964 | 38242385 | 38247712 | 38247835 | 58 | 44 | [illegible] |
| SE | 80448 | ENSG00000085788 | DDHD2 | chr8 | + | 38247712 | 38247835 | 38246183 | 38246300 | 38249707 | 38249747 | 150 | 137 | [illegible] |
| SE | 80451 | ENSG00000085788 | DDHD2 | chr8 | + | 38249707 | 38249803 | 38246803 | 38247835 | 38251911 | 38252028 | 118 | 99 | [illegible] |
| SE | 80457 | ENSG00000099901 | RANBP1 | chr22 | + | 20122579 | 20122697 | 20122263 | 20122421 | 20125307 | 20125436 | 0 | 0 | [illegible] |
| SE | 80462 | ENSG00000154511 | DIPK1A | chr1 | - | 92847182 | 92847359 | 92832736 | 92833034 | 92850847 | 92850955 | 64 | 29 | [illegible] |
| SE | 80465 | ENSG00000154511 | DIPK1A | chr1 | - | 92876295 | 92876430 | 92850847 | 92850955 | 92961375 | 92961462 | 72 | 31 | [illegible] |
| SE | 80467 | ENSG00000179818 | PCBP1-AS1 | chr2 | - | 69964230 | 69964427 | 69962262 | 69963500 | 69988741 | 69988838 | 0 | 0 | [illegible] |
| SE | 80471 | ENSG00000179818 | PCBP1-AS1 | chr2 | - | 69964368 | 69964427 | 69960115 | 69963500 | 69988741 | 69988838 | 0 | 0 | [illegible] |
| SE | 80478 | ENSG00000179818 | PCBP1-AS1 | chr2 | - | 69988445 | 69988505 | 69960107 | 69963500 | 69988741 | 69988838 | 0 | 0 | [illegible] |
| SE | 80479 | ENSG00000179818 | PCBP1-AS1 | chr2 | - | 69988445 | 69988505 | 69960107 | 69963500 | 69996685 | 69996888 | 0 | 0 | [illegible] |
| SE | 80484 | ENSG00000179818 | PCBP1-AS1 | chr2 | - | 69988741 | 69988838 | 69960103 | 69963500 | 69996685 | 69996777 | 2 | 0 | [illegible] |
| SE | 80489 | ENSG00000179818 | PCBP1-AS1 | chr2 | - | 69996685 | 69996777 | 69988741 | 69988838 | 69996857 | 69996888 | 2 | 0 | [illegible] |
| SE | 80496 | ENSG00000179818 | PCBP1-AS1 | chr2 | - | 69997912 | 69998134 | 69996751 | 69996888 | 70051202 | 70051305 | 0 | 0 | [illegible] |
| SE | 80503 | ENSG00000179818 | PCBP1-AS1 | chr2 | - | 70015447 | 70015517 | 69996685 | 69996888 | 70015837 | 70016150 | 1 | 0 | [illegible] |
| SE | 80504 | ENSG00000179818 | PCBP1-AS1 | chr2 | - | 70015447 | 70015580 | 69996685 | 69996888 | 70018014 | 70018097 | 1 | 1 | [illegible] |
| SE | 80509 | ENSG00000179818 | PCBP1-AS1 | chr2 | - | 70018014 | 70018131 | 69996685 | 69996888 | 70032132 | 70032255 | 3 | 5 | [illegible] |
| SE | 80511 | ENSG00000179818 | PCBP1-AS1 | chr2 | - | 70018014 | 70018131 | 69996685 | 69996888 | 70085546 | 70085570 | 2 | 4 | [illegible] |
| SE | 80512 | ENSG00000179818 | PCBP1-AS1 | chr2 | - | 70018014 | 70018131 | 70013248 | 70015580 | 70051202 | 70051304 | 1 | 6 | [illegible] |
| SE | 80514 | ENSG00000179818 | PCBP1-AS1 | chr2 | - | 70032132 | 70032255 | 70018014 | 70018131 | 70051202 | 70051305 | 19 | 17 | [illegible] |
| SE | 80515 | ENSG00000179818 | PCBP1-AS1 | chr2 | - | 70032132 | 70032255 | 70029997 | 70031931 | 70051202 | 70051305 | 36 | 30 | [illegible] |
| SE | 80517 | ENSG00000179818 | PCBP1-AS1 | chr2 | - | 70051202 | 70051305 | 69996685 | 69996888 | 70059642 | 70059680 | 17 | 10 | [illegible] |
| SE | 80518 | ENSG00000179818 | PCBP1-AS1 | chr2 | - | 70051202 | 70051305 | 69996685 | 69996888 | 70085546 | 70085624 | 15 | 6 | [illegible] |
| SE | 80522 | ENSG00000179818 | PCBP1-AS1 | chr2 | - | 70051202 | 70051305 | 70032132 | 70032255 | 70083537 | 70083616 | 15 | 11 | [illegible] |
| SE | 80523 | ENSG00000179818 | PCBP1-AS1 | chr2 | - | 70051202 | 70051305 | 70048686 | 70048957 | 70083537 | 70083616 | 3 | 1 | [illegible] |
| SE | 80527 | ENSG00000179818 | PCBP1-AS1 | chr2 | - | 70053730 | 70053792 | 70051044 | 70051305 | 70055655 | 70055749 | 6 | 1 | [illegible] |
| SE | 80528 | ENSG00000179818 | PCBP1-AS1 | chr2 | - | 70053730 | 70053792 | 70051044 | 70051305 | 70055713 | 70055823 | 12 | 1 | [illegible] |
| SE | 80530 | ENSG00000179818 | PCBP1-AS1 | chr2 | - | 70053730 | 70053792 | 70051044 | 70051305 | 70083506 | 70083687 | 6 | 2 | [illegible] |
| SE | 80531 | ENSG00000179818 | PCBP1-AS1 | chr2 | - | 70053730 | 70053792 | 70051044 | 70051305 | 70083537 | 70083687 | 6 | 1 | [illegible] |
| SE | 80533 | ENSG00000179818 | PCBP1-AS1 | chr2 | - | 70053730 | 70053812 | 70051202 | 70051305 | 70055713 | 70055910 | 6 | 1 | [illegible] |
| SE | 80534 | ENSG00000179818 | PCBP1-AS1 | chr2 | - | 70053730 | 70053815 | 70051202 | 70051305 | 70055655 | 70055910 | 6 | 1 | [illegible] |
| SE | 80535 | ENSG00000179818 | PCBP1-AS1 | chr2 | - | 70053730 | 70053815 | 70051202 | 70051305 | 70055713 | 70055910 | 6 | 1 | [illegible] |
| SE | 80536 | ENSG00000179818 | PCBP1-AS1 | chr2 | - | 70053730 | 70053815 | 70051202 | 70051305 | 70083537 | 70083687 | 6 | 1 | [illegible] |
| SE | 80537 | ENSG00000179818 | PCBP1-AS1 | chr2 | - | 70053730 | 70053994 | 70051202 | 70051305 | 70055655 | 70055910 | 7 | 2 | [illegible] |
| SE | 80538 | ENSG00000179818 | PCBP1-AS1 | chr2 | - | 70055655 | 70055749 | 70053730 | 70053792 | 70083506 | 70083687 | 0 | 0 | [illegible] |
| SE | 80540 | ENSG00000179818 | PCBP1-AS1 | chr2 | - | 70055655 | 70055910 | 70048708 | 70051305 | 70077300 | 70077381 | 0 | 0 | [illegible] |
| SE | 80541 | ENSG00000179818 | PCBP1-AS1 | chr2 | - | 70055655 | 70055910 | 70048708 | 70051305 | 70083537 | 70083687 | 0 | 0 | [illegible] |
| SE | 80543 | ENSG00000179818 | PCBP1-AS1 | chr2 | - | 70055655 | 70055910 | 70053730 | 70053792 | 70059642 | 70059679 | 15 | 2 | [illegible] |
| SE | 80545 | ENSG00000179818 | PCBP1-AS1 | chr2 | - | 70055655 | 70055910 | 70053730 | 70053792 | 70085546 | 70085624 | 0 | 0 | [illegible] |
| SE | 80550 | ENSG00000179818 | PCBP1-AS1 | chr2 | - | 70055713 | 70055910 | 70051016 | 70051305 | 70077300 | 70077381 | 4 | 4 | [illegible] |
| SE | 80551 | ENSG00000179818 | PCBP1-AS1 | chr2 | - | 70055713 | 70055910 | 70051016 | 70051305 | 70083506 | 70083687 | 4 | 4 | [illegible] |
| SE | 80552 | ENSG00000179818 | PCBP1-AS1 | chr2 | - | 70055713 | 70055910 | 70051016 | 70051305 | 70083537 | 70083687 | 4 | 4 | [illegible] |
| SE | 80554 | ENSG00000179818 | PCBP1-AS1 | chr2 | - | 70055713 | 70055910 | 70053730 | 70053792 | 70059642 | 70059680 | 21 | 2 | [illegible] |
| SE | 80555 | ENSG00000179818 | PCBP1-AS1 | chr2 | - | 70055713 | 70055910 | 70053730 | 70053792 | 70083506 | 70083687 | 6 | 0 | [illegible] |
| SE | 80556 | ENSG00000179818 | PCBP1-AS1 | chr2 | - | 70055713 | 70055910 | 70053730 | 70053792 | 70083537 | 70083687 | 6 | 0 | [illegible] |
| SE | 80557 | ENSG00000179818 | PCBP1-AS1 | chr2 | - | 70055713 | 70055910 | 70053730 | 70053792 | 70085546 | 70085575 | 6 | 0 | [illegible] |
| SE | 80559 | ENSG00000179818 | PCBP1-AS1 | chr2 | - | 70059642 | 70059680 | 69996685 | 69996888 | 70085546 | 70085574 | 0 | 0 | [illegible] |
| SE | 80560 | ENSG00000179818 | PCBP1-AS1 | chr2 | - | 70059642 | 70059680 | 70048706 | 70051305 | 70077300 | 70077381 | 2 | 5 | [illegible] |
| SE | 80561 | ENSG00000179818 | PCBP1-AS1 | chr2 | - | 70059642 | 70059680 | 70048706 | 70051305 | 70083506 | 70083687 | 5 | 5 | [illegible] |
| SE | 80565 | ENSG00000179818 | PCBP1-AS1 | chr2 | - | 70059642 | 70059680 | 70053730 | 70053792 | 70083506 | 70083687 | 1 | 0 | [illegible] |
| SE | 80566 | ENSG00000179818 | PCBP1-AS1 | chr2 | - | 70059642 | 70059680 | 70053730 | 70053792 | 70083537 | 70083687 | 2 | 0 | [illegible] |
| SE | 80567 | ENSG00000179818 | PCBP1-AS1 | chr2 | - | 70059642 | 70059680 | 70053730 | 70053792 | 70085546 | 70085574 | 1 | 0 | [illegible] |
| SE | 80568 | ENSG00000179818 | PCBP1-AS1 | chr2 | - | 70059642 | 70059680 | 70055616 | 70055910 | 70077300 | 70077381 | 0 | 0 | [illegible] |
| SE | 80569 | ENSG00000179818 | PCBP1-AS1 | chr2 | - | 70059642 | 70059680 | 70055616 | 70055910 | 70083506 | 70083687 | 1 | 0 | [illegible] |
| SE | 80570 | ENSG00000179818 | PCBP1-AS1 | chr2 | - | 70059642 | 70059680 | 70055616 | 70055910 | 70083537 | 70083687 | 11 | 0 | [illegible] |
| SE | 80571 | ENSG00000179818 | PCBP1-AS1 | chr2 | - | 70059642 | 70059680 | 70055616 | 70055910 | 70085546 | 70085574 | 3 | 2 | [illegible] |
| SE | 80573 | ENSG00000179818 | PCBP1-AS1 | chr2 | - | 70077300 | 70077381 | 70051202 | 70051305 | 70083506 | 70083687 | 0 | 2 | [illegible] |
| SE | 80574 | ENSG00000179818 | PCBP1-AS1 | chr2 | - | 70077300 | 70077381 | 70051202 | 70051305 | 70083537 | 70083687 | 0 | 1 | [illegible] |
| SE | 80575 | ENSG00000179818 | PCBP1-AS1 | chr2 | - | 70077300 | 70077381 | 70051202 | 70051305 | 70085546 | 70085595 | 0 | 1 | [illegible] |
| SE | 80576 | ENSG00000179818 | PCBP1-AS1 | chr2 | - | 70077300 | 70077381 | 70055616 | 70055910 | 70083506 | 70083687 | 0 | 1 | [illegible] |
| SE | 80578 | ENSG00000179818 | PCBP1-AS1 | chr2 | - | 70077300 | 70077381 | 70055616 | 70055910 | 70085546 | 70085595 | 0 | 0 | [illegible] |
| SE | 80579 | ENSG00000179818 | PCBP1-AS1 | chr2 | - | 70077300 | 70077381 | 70059642 | 70059680 | 70083506 | 70083687 | 0 | 1 | [illegible] |
| SE | 80580 | ENSG00000179818 | PCBP1-AS1 | chr2 | - | 70077300 | 70077381 | 70059642 | 70059680 | 70083537 | 70083687 | 0 | 0 | [illegible] |
| SE | 80581 | ENSG00000179818 | PCBP1-AS1 | chr2 | - | 70077300 | 70077381 | 70059642 | 70059680 | 70085546 | 70085595 | 0 | 0 | [illegible] |
| SE | 80582 | ENSG00000179818 | PCBP1-AS1 | chr2 | - | 70081369 | 70081490 | 70059642 | 70059680 | 70083506 | 70083687 | 0 | 0 | [illegible] |
| SE | 80583 | ENSG00000179818 | PCBP1-AS1 | chr2 | - | 70081369 | 70081490 | 70059642 | 70059680 | 70083537 | 70083687 | 1 | 0 | [illegible] |
| SE | 80584 | ENSG00000179818 | PCBP1-AS1 | chr2 | - | 70081369 | 70081490 | 70059642 | 70059680 | 70085546 | 70085624 | 0 | 0 | [illegible] |
| SE | 80586 | ENSG00000179818 | PCBP1-AS1 | chr2 | - | 70083506 | 70083687 | 70053730 | 70053792 | 70085546 | 70085554 | 33 | 19 | [illegible] |
| SE | 80587 | ENSG00000179818 | PCBP1-AS1 | chr2 | - | 70083506 | 70083687 | 70055713 | 70055910 | 70085546 | 70085554 | 33 | 18 | [illegible] |
| SE | 80589 | ENSG00000179818 | PCBP1-AS1 | chr2 | - | 70083506 | 70083687 | 70077300 | 70077381 | 70085546 | 70085554 | 33 | 19 | [illegible] |
| SE | 80590 | ENSG00000179818 | PCBP1-AS1 | chr2 | - | 70083506 | 70083687 | 70081369 | 70081490 | 70085546 | 70085554 | 33 | 18 | [illegible] |
| SE | 80592 | ENSG00000179818 | PCBP1-AS1 | chr2 | - | 70083537 | 70083687 | 70053730 | 70053792 | 70085546 | 70085573 | 33 | 18 | [illegible] |
| SE | 80593 | ENSG00000179818 | PCBP1-AS1 | chr2 | - | 70083537 | 70083687 | 70055616 | 70055910 | 70085546 | 70085573 | 33 | 18 | [illegible] |
| SE | 80595 | ENSG00000179818 | PCBP1-AS1 | chr2 | - | 70083537 | 70083687 | 70077300 | 70077381 | 70085546 | 70085573 | 33 | 18 | [illegible] |
| SE | 80596 | ENSG00000179818 | PCBP1-AS1 | chr2 | - | 70083537 | 70083687 | 70081369 | 70081490 | 70085546 | 70085573 | 34 | 18 | [illegible] |
| SE | 80597 | ENSG00000179818 | PCBP1-AS1 | chr2 | - | 70085815 | 70085903 | 70085546 | 70085624 | 70086054 | 70086273 | 6 | 5 | [illegible] |
| SE | 80598 | ENSG00000179818 | PCBP1-AS1 | chr2 | - | 70085815 | 70085903 | 70085546 | 70085624 | 70086123 | 70086273 | 6 | 5 | [illegible] |
| SE | 80599 | ENSG00000179818 | PCBP1-AS1 | chr2 | - | 70085815 | 70085903 | 70085546 | 70085624 | 70086251 | 70086388 | 6 | 5 | [illegible] |
| SE | 80600 | ENSG00000179818 | PCBP1-AS1 | chr2 | - | 70085815 | 70085903 | 70085546 | 70085624 | 70086430 | 70086485 | 6 | 5 | [illegible] |
| SE | 80601 | ENSG00000179818 | PCBP1-AS1 | chr2 | - | 70085815 | 70085930 | 70085546 | 70085624 | 70086977 | 70086996 | 11 | 10 | [illegible] |
| SE | 80602 | ENSG00000179818 | PCBP1-AS1 | chr2 | - | 70085815 | 70085930 | 70085546 | 70085624 | 70087280 | 70087435 | 11 | 11 | [illegible] |
| SE | 80614 | ENSG00000185164 | NOMO2 | chr16 | - | 18538858 | 18538964 | 18538525 | 18538676 | 18542171 | 18542261 | 78 | 34 | [illegible] |
| SE | 80615 | ENSG00000185164 | NOMO2 | chr16 | - | 18549520 | 18549627 | 18547227 | 18547300 | 18551438 | 18551539 | 22 | 12 | [illegible] |
| SE | 80616 | ENSG00000185164 | NOMO2 | chr16 | - | 18551438 | 18551539 | 18549520 | 18549627 | 18554806 | 18554852 | 23 | 6 | [illegible] |
| SE | 80620 | ENSG00000175600 | SUGCT | chr7 | + | 40274512 | 40274656 | 40237634 | 40237726 | 40316759 | 40316855 | 4 | 2 | [illegible] |
| SE | 80625 | ENSG00000258227 | CLEC5A | chr7 | - | 141945340 | 141945400 | 141931719 | 141931826 | 141946806 | 141946936 | 0 | 0 | [illegible] |
| SE | 80626 | ENSG00000258227 | CLEC5A | chr7 | - | 141945340 | 141945400 | 141935813 | 141935950 | 141946213 | 141946312 | 20 | 13 | [illegible] |
| SE | 80627 | ENSG00000258227 | CLEC5A | chr7 | - | 141945340 | 141945400 | 141935813 | 141935950 | 141946806 | 141946936 | 1 | 3 | [illegible] |
| SE | 80628 | ENSG00000258227 | CLEC5A | chr7 | - | 141946213 | 141946312 | 141935813 | 141935950 | 141946806 | 141946933 | 6 | 4 | [illegible] |
| SE | 80629 | ENSG00000258227 | CLEC5A | chr7 | - | 141946213 | 141946312 | 141945340 | 141945400 | 141946806 | 141946933 | 23 | 13 | [illegible] |
| SE | 80632 | ENSG00000175324 | LSM1 | chr8 | - | 38171964 | 38172033 | 38163334 | 38163840 | 38176274 | 38176488 | 59 | 38 | [illegible] |
| SE | 80633 | ENSG00000180357 | ZNF609 | chr15 | + | 64622826 | 64623052 | 64499291 | 64500166 | 64670345 | 64670433 | 323 | 200 | [illegible] |
| SE | 80634 | ENSG00000180357 | ZNF609 | chr15 | + | 64670345 | 64670433 | 64622826 | 64623052 | 64673915 | 64674771 | 212 | 118 | [illegible] |
| SE | 80635 | ENSG00000180357 | ZNF609 | chr15 | + | 64673915 | 64676256 | 64670345 | 64670433 | 64678115 | 64678482 | 243 | 178 | [illegible] |
| SE | 80636 | ENSG00000180357 | ZNF609 | chr15 | + | 64673915 | 64676256 | 64670345 | 64670433 | 64680184 | 64680360 | 94 | 43 | [illegible] |
| SE | 80637 | ENSG00000180357 | ZNF609 | chr15 | + | 64678115 | 64678482 | 64670345 | 64670433 | 64680184 | 64680360 | 100 | 53 | [illegible] |
| SE | 80638 | ENSG00000180357 | ZNF609 | chr15 | + | 64678115 | 64678482 | 64673915 | 64676256 | 64680184 | 64680360 | 248 | 185 | [illegible] |
| SE | 80639 | ENSG00000180357 | ZNF609 | chr15 | + | 64681308 | 64681387 | 64680645 | 64680862 | 64681691 | 64686068 | 60 | 40 | [illegible] |
| SE | 80640 | ENSG00000214439 | FAM185BP | chr7 | - | 77098909 | 77098951 | 77094001 | 77094097 | 77109937 | 77110076 | 6 | 7 | [illegible] |
| SE | 80646 | ENSG00000168090 | COPS6 | chr7 | + | 100090901 | 100090949 | 100090361 | 100090654 | 100091037 | 100091152 | 180 | 67 | [illegible] |
| SE | 80648 | ENSG00000166997 | CNPY4 | chr7 | + | 100122480 | 100122577 | 100122051 | 100122385 | 100124513 | 100124631 | 28 | 18 | [illegible] |
| SE | 80649 | ENSG00000166997 | CNPY4 | chr7 | + | 100122783 | 100122906 | 100122051 | 100122385 | 100124513 | 100124631 | 12 | 9 | [illegible] |
| SE | 80660 | ENSG00000149131 | SERPING1 | chr11 | + | 57606710 | 57606799 | 57606407 | 57606547 | 57611716 | 57611936 | 0 | 0 | [illegible] |
| SE | 80661 | ENSG00000149131 | SERPING1 | chr11 | + | 57614327 | 57614596 | 57611716 | 57611936 | 57618680 | 57619159 | 14 | 13 | [illegible] |
| SE | 80662 | ENSG00000063169 | BICRA | chr19 | + | 47670442 | 47670544 | 47608941 | 47609168 | 47673569 | 47673615 | 12 | 3 | [illegible] |
| SE | 80663 | ENSG00000063169 | BICRA | chr19 | + | 47681975 | 47682152 | 47678961 | 47681276 | 47694114 | 47694726 | 2 | 0 | [illegible] |
| SE | 80664 | ENSG00000213420 | GPC2 | chr7 | - | 100175571 | 100175894 | 100173457 | 100174765 | 100176206 | 100176365 | 0 | 1 | [illegible] |
| SE | 80665 | ENSG00000242294 | STAG3L5P | chr7 | + | 100345864 | 100345968 | 100338772 | 100338889 | 100349716 | 100350158 | 173 | 73 | [illegible] |
| SE | 80666 | ENSG00000086619 | ERO1B | chr1 | - | 236243421 | 236243495 | 236236277 | 236236398 | 236249884 | 236249967 | 104 | 77 | [illegible] |
| SE | 80668 | ENSG00000086619 | ERO1B | chr1 | - | 236269874 | 236269994 | 236253421 | 236253505 | 236281681 | 236281958 | 20 | 18 | [illegible] |
| SE | 80669 | ENSG00000121716 | PILRB | chr7 | + | 100353372 | 100353483 | 100353209 | 100353270 | 100353894 | 100354012 | 22 | 7 | [illegible] |
| SE | 80673 | ENSG00000121716 | PILRB | chr7 | + | 100355142 | 100355240 | 100353372 | 100354012 | 100356393 | 100356883 | 74 | 33 | [illegible] |
| SE | 80674 | ENSG00000121716 | PILRB | chr7 | + | 100356393 | 100356883 | 100355142 | 100355240 | 100358219 | 100358259 | 57 | 26 | [illegible] |
| SE | 80675 | ENSG00000121716 | PILRB | chr7 | + | 100356749 | 100356938 | 100355142 | 100355240 | 100358219 | 100358366 | 107 | 60 | [illegible] |
| SE | 80676 | ENSG00000121716 | PILRB | chr7 | + | 100358219 | 100358366 | 100356393 | 100356883 | 100358932 | 100359008 | 63 | 25 | [illegible] |
| SE | 80680 | ENSG00000085514 | PILRA | chr7 | + | 100397878 | 100397912 | 100389887 | 100390106 | 100399290 | 100399302 | 285 | 182 | [illegible] |
| SE | 80681 | ENSG00000085514 | PILRA | chr7 | + | 100399580 | 100399612 | 100399290 | 100399340 | 100399784 | 100400028 | 153 | 88 | [illegible] |
| SE | 80683 | ENSG00000175471 | MCTP1 | chr5 | - | 94708511 | 94708609 | 94703740 | 94705203 | 94710817 | 94710927 | 22 | 28 | [illegible] |
| SE | 80684 | ENSG00000175471 | MCTP1 | chr5 | - | 94710817 | 94710927 | 94708511 | 94708609 | 94714776 | 94714886 | 56 | 82 | [illegible] |
| SE | 80685 | ENSG00000175471 | MCTP1 | chr5 | - | 94710817 | 94710927 | 94708511 | 94708609 | 94779109 | 94779163 | 28 | 51 | [illegible] |
| SE | 80686 | ENSG00000175471 | MCTP1 | chr5 | - | 94714776 | 94714886 | 94708511 | 94708609 | 94779109 | 94779163 | 48 | 38 | [illegible] |
| SE | 80688 | ENSG00000175471 | MCTP1 | chr5 | - | 94779109 | 94779163 | 94714776 | 94714886 | 94799012 | 94799132 | 74 | 59 | [illegible] |
| SE | 80689 | ENSG00000175471 | MCTP1 | chr5 | - | 94779109 | 94779163 | 94714776 | 94714886 | 94940083 | 94940195 | 12 | 14 | [illegible] |
| SE | 80694 | ENSG00000175471 | MCTP1 | chr5 | - | 94870416 | 94870491 | 94867221 | 94868452 | 94870871 | 94870973 | 128 | 163 | [illegible] |
| SE | 80696 | ENSG00000175471 | MCTP1 | chr5 | - | 94894648 | 94894835 | 94888878 | 94888972 | 94909250 | 94909381 | 208 | 242 | [illegible] |
| SE | 80697 | ENSG00000175471 | MCTP1 | chr5 | - | 94912805 | 94912976 | 94909250 | 94909381 | 94923961 | 94924021 | 202 | 198 | [illegible] |
| SE | 80703 | ENSG00000175471 | MCTP1 | chr5 | - | 94931952 | 94931991 | 94799012 | 94799132 | 94940083 | 94940195 | 17 | 36 | [illegible] |
| SE | 80706 | ENSG00000175471 | MCTP1 | chr5 | - | 94953218 | 94953361 | 94942347 | 94942427 | 95017366 | 95017484 | 230 | 340 | [illegible] |
| SE | 80707 | ENSG00000175471 | MCTP1 | chr5 | - | 94976744 | 94976809 | 94953218 | 94953361 | 95017366 | 95017487 | 0 | 0 | [illegible] |
| SE | 80710 | ENSG00000147535 | PLPP5 | chr8 | - | 38267266 | 38267391 | 38266140 | 38266311 | 38267896 | 38267960 | 58 | 35 | [illegible] |
| SE | 80719 | ENSG00000160813 | PPP1R35 | chr7 | - | 100435630 | 100435767 | 100435281 | 100435533 | 100435847 | 100436064 | 84 | 30 | [illegible] |
| SE | 80720 | ENSG00000075223 | SEMA3C | chr7 | - | 80758330 | 80758488 | 80751268 | 80751336 | 80761615 | 80761657 | 32 | 13 | [illegible] |
| SE | 80721 | ENSG00000075223 | SEMA3C | chr7 | - | 80847259 | 80847345 | 80828584 | 80828745 | 80916678 | 80916819 | 2 | 0 | [illegible] |
| SE | 80722 | ENSG00000075223 | SEMA3C | chr7 | - | 80905828 | 80905916 | 80828584 | 80828745 | 80916678 | 80916819 | 4 | 2 | [illegible] |
| SE | 80724 | ENSG00000125505 | MBOAT7 | chr19 | - | 54188432 | 54188511 | 54188216 | 54188346 | 54189337 | 54189360 | 53 | 45 | [illegible] |
| SE | 80728 | ENSG00000163636 | PSMD6 | chr3 | - | 64022738 | 64022801 | 64022317 | 64022523 | 64023050 | 64023478 | 0 | 5 | [illegible] |

| | | | | | | | | | | | | | | | | | | | | | | | |
|---|---|---|---|---|---|---|---|---|---|---|---|---|---|---|---|---|---|---|---|---|---|---|---|
| SE | 80730 | ENSG00000285953 | NA | chr7 | - | 92244901 | 92245134 | 92242033 | 92242137 | 92245789 | 92245924 | 3 | 11 | 7 | 3 | 11 | 0 | 0 | 1 | 0 | 0 | 0 | 1 |
| SE | 80733 | ENSG00000151208 | DLG5 | chr10 | - | 77794848 | 77794958 | 77793543 | 77794117 | 77796060 | 77796188 | 25 | 23 | 11 | 30 | 42 | 0 | 0 | 0 | 0 | 0 | 0 | 0 |
| SE | 80734 | ENSG00000151208 | DLG5 | chr10 | - | 77821081 | 77822101 | 77819894 | 77820018 | 77824383 | 77824476 | 55 | 3 | 20 | 31 | 21 | 0 | 0 | 1 | 0 | 0 | 9 | 8 |
| SE | 80735 | ENSG00000151208 | DLG5 | chr10 | - | 77829354 | 77829530 | 77828881 | 77828985 | 77830216 | 77830344 | 53 | 17 | 31 | 48 | 15 | 0 | 0 | 0 | 0 | 2 | 0 | 28 |
| SE | 80737 | ENSG00000086300 | SNX10 | chr7 | + | 26346419 | 26346466 | 26291861 | 26292086 | 26360974 | 26361061 | 138 | 92 | 76 | 82 | 110 | 1 | 0 | 0 | 0 | 0 | 15 | 47 |
| SE | 80741 | ENSG00000086300 | SNX10 | chr7 | + | 26357006 | 26357127 | 26354056 | 26354184 | 26360974 | 26361031 | 22 | 21 | 14 | 19 | 30 | 4 | 14 | 1 | 13 | 14 | 28 | 27 |
| SE | 80742 | ENSG00000086300 | SNX10 | chr7 | + | 26360974 | 26361061 | 26346419 | 26346466 | 26364534 | 26364635 | 249 | 179 | 170 | 185 | 279 | 0 | 0 | 0 | 0 | 0 | 44 | 120 |
| SE | 80743 | ENSG00000004864 | SLC25A13 | chr7 | - | 96131742 | 96131881 | 96121838 | 96121997 | 96146555 | 96146696 | 23 | 21 | 124 | 19 | 15 | 0 | 0 | 2 | 0 | 3 | 41 | 11 |
| SE | 80747 | ENSG00000004864 | SLC25A13 | chr7 | - | 96191108 | 96191247 | 96189293 | 96189378 | 96193036 | 96193183 | 15 | 15 | 86 | 10 | 40 | 0 | 0 | 0 | 0 | 0 | 20 | 13 |
| SE | 80748 | ENSG00000004864 | SLC25A13 | chr7 | - | 96208837 | 96208977 | 96193036 | 96193183 | 96277195 | 96277338 | 81 | 61 | 229 | 60 | 89 | 0 | 0 | 2 | 0 | 0 | 75 | 85 |
| SE | 80750 | ENSG00000004864 | SLC25A13 | chr7 | - | 96234801 | 96234917 | 96208837 | 96208977 | 96296897 | 96296951 | 13 | 17 | 36 | 24 | 24 | 4 | 1 | 2 | 0 | 4 | 21 | 13 |
| SE | 80752 | ENSG00000004864 | SLC25A13 | chr7 | - | 96277195 | 96277338 | 96234801 | 96234917 | 96296897 | 96296951 | 33 | 48 | 135 | 53 | 33 | 0 | 0 | 0 | 0 | 0 | 37 | 31 |
| SE | 80753 | ENSG00000136280 | CCM2 | chr7 | + | 45029443 | 45029509 | 45000188 | 45000363 | 45038252 | 45038426 | 0 | 0 | 0 | 0 | 1 | 3 | 2 | 2 | 2 | 2 | 0 | 0 |
| SE | 80754 | ENSG00000136280 | CCM2 | chr7 | + | 45029443 | 45029509 | 45027700 | 45027797 | 45038252 | 45038426 | 1 | 0 | 0 | 0 | 1 | 79 | 66 | 91 | 97 | 133 | 0 | 0 |
| SE | 80755 | ENSG00000136280 | CCM2 | chr7 | + | 45038252 | 45038426 | 44999474 | 44999857 | 45063917 | 45064001 | 99 | 50 | 94 | 86 | 135 | 1 | 0 | 0 | 0 | 0 | 33 | 80 |
| SE | 80757 | ENSG00000136280 | CCM2 | chr7 | + | 45038252 | 45038426 | 45027014 | 45027266 | 45063917 | 45064001 | 100 | 50 | 94 | 87 | 134 | 2 | 0 | 2 | 1 | 0 | 33 | 81 |
| SE | 80761 | ENSG00000136280 | CCM2 | chr7 | + | 45038252 | 45038426 | 45027025 | 45027797 | 45073459 | 45073519 | 79 | 67 | 91 | 98 | 133 | 0 | 0 | 0 | 0 | 0 | 67 | 72 |
| SE | 80764 | ENSG00000136280 | CCM2 | chr7 | + | 45063917 | 45064001 | 45038252 | 45038426 | 45064462 | 45064494 | 156 | 63 | 157 | 127 | 191 | 1 | 0 | 0 | 3 | 0 | 95 | 133 |
| SE | 80767 | ENSG00000136280 | CCM2 | chr7 | + | 45064462 | 45064646 | 45063917 | 45064001 | 45069825 | 45069961 | 125 | 56 | 130 | 99 | 167 | 0 | 0 | 0 | 0 | 0 | 102 | 141 |
| SE | 80769 | ENSG00000136280 | CCM2 | chr7 | + | 45068442 | 45068579 | 45027025 | 45027797 | 45072725 | 45072783 | 0 | 0 | 0 | 0 | 3 | 0 | 2 | 12 | 2 | 1 | 20 | 0 |
| SE | 80770 | ENSG00000136280 | CCM2 | chr7 | + | 45068442 | 45068579 | 45038252 | 45038426 | 45072725 | 45072783 | 0 | 0 | 3 | 2 | 3 | 1 | 0 | 0 | 4 | 1 | 10 | 0 |
| SE | 80777 | ENSG00000136280 | CCM2 | chr7 | + | 45070093 | 45070168 | 45069825 | 45069961 | 45070388 | 45071872 | 0 | 0 | 1 | 0 | 0 | 4 | 1 | 0 | 5 | 5 | 0 | 0 |
| SE | 80778 | ENSG00000136280 | CCM2 | chr7 | + | 45070093 | 45070168 | 45069825 | 45069961 | 45072725 | 45072783 | 0 | 0 | 1 | 0 | 0 | 86 | 20 | 44 | 50 | 33 | 0 | 2 |
| SE | 80779 | ENSG00000136280 | CCM2 | chr7 | + | 45070388 | 45071872 | 45070093 | 45070168 | 45072725 | 45072783 | 3 | 0 | 3 | 5 | 3 | 0 | 0 | 0 | 0 | 0 | 14 | 5 |
| SE | 80781 | ENSG00000136280 | CCM2 | chr7 | + | 45072725 | 45072783 | 45027025 | 45027797 | 45073459 | 45073519 | 62 | 34 | 78 | 65 | 35 | 0 | 0 | 0 | 0 | 0 | 37 | 74 |
| SE | 80782 | ENSG00000136280 | CCM2 | chr7 | + | 45072725 | 45072783 | 45038252 | 45038426 | 45073459 | 45073519 | 63 | 32 | 67 | 67 | 34 | 0 | 1 | 0 | 1 | 0 | 32 | 70 |
| SE | 80792 | ENSG00000123191 | ATP7B | chr13 | - | 51949661 | 51949796 | 51944108 | 51944291 | 51950006 | 51950161 | 1 | 0 | 5 | 1 | 0 | 0 | 0 | 0 | 0 | 3 | 0 | 0 |
| SE | 80795 | ENSG00000123191 | ATP7B | chr13 | - | 51958310 | 51958400 | 51957515 | 51957607 | 51960147 | 51960322 | 0 | 0 | 0 | 1 | 0 | 0 | 1 | 1 | 0 | 0 | 0 | 0 |
| SE | 80796 | ENSG00000123191 | ATP7B | chr13 | - | 51958310 | 51958544 | 51957515 | 51957607 | 51960147 | 51960322 | 0 | 0 | 0 | 2 | 0 | 0 | 1 | 1 | 0 | 0 | 0 | 0 |
| SE | 80800 | ENSG00000123191 | ATP7B | chr13 | - | 51972594 | 51972832 | 51970491 | 51970749 | 51973934 | 51974175 | 1 | 0 | 0 | 0 | 0 | 2 | 3 | 0 | 2 | 4 | 0 | 0 |
| SE | 80803 | ENSG00000101082 | SLA2 | chr20 | - | 36641244 | 36641378 | 36634489 | 36634589 | 36645836 | 36645881 | 190 | 105 | 144 | 235 | 171 | 0 | 0 | 0 | 1 | 0 | 193 | 267 |
| SE | 80805 | ENSG00000151948 | GLT1D1 | chr12 | + | 128875913 | 128876062 | 128853426 | 128853649 | 128899235 | 128899287 | 12 | 4 | 4 | 7 | 1 | 0 | 0 | 0 | 0 | 0 | 8 | 18 |
| SE | 80806 | ENSG00000151948 | GLT1D1 | chr12 | + | 128888638 | 128888744 | 128853426 | 128853649 | 128899235 | 128899287 | 46 | 61 | 37 | 53 | 72 | 0 | 0 | 0 | 0 | 0 | 144 | 116 |
| SE | 80807 | ENSG00000151948 | GLT1D1 | chr12 | + | 128888638 | 128888744 | 128853426 | 128853649 | 128945325 | 128945369 | 2 | 3 | 1 | 2 | 1 | 0 | 0 | 1 | 0 | 0 | 0 | 2 |
| SE | 80809 | ENSG00000151948 | GLT1D1 | chr12 | + | 128899235 | 128899287 | 128853477 | 128853649 | 128945325 | 128945369 | 0 | 2 | 0 | 0 | 1 | 0 | 0 | 1 | 0 | 0 | 18 | 9 |
| SE | 80810 | ENSG00000151948 | GLT1D1 | chr12 | + | 128899235 | 128899287 | 128888638 | 128888744 | 128912423 | 128912460 | 32 | 48 | 31 | 57 | 55 | 1 | 1 | 1 | 0 | 0 | 166 | 105 |
| SE | 80814 | ENSG00000151948 | GLT1D1 | chr12 | + | 128899235 | 128899287 | 128888638 | 128888744 | 128945325 | 128945369 | 12 | 17 | 9 | 19 | 18 | 0 | 0 | 0 | 1 | 0 | 109 | 52 |
| SE | 80818 | ENSG00000151948 | GLT1D1 | chr12 | + | 128912423 | 128912460 | 128888638 | 128888744 | 128926358 | 128926454 | 0 | 0 | 0 | 0 | 0 | 1 | 0 | 0 | 0 | 4 | 0 | 2 |
| SE | 80822 | ENSG00000151948 | GLT1D1 | chr12 | + | 128914932 | 128914989 | 128888638 | 128888744 | 128927104 | 128927154 | 1 | 0 | 3 | 2 | 1 | 5 | 0 | 0 | 0 | 0 | 8 | 0 |
| SE | 80823 | ENSG00000151948 | GLT1D1 | chr12 | + | 128914932 | 128914989 | 128888638 | 128888744 | 128945325 | 128945369 | 0 | 2 | 0 | 1 | 0 | 0 | 0 | 0 | 1 | 0 | 7 | 0 |
| SE | 80827 | ENSG00000151948 | GLT1D1 | chr12 | + | 128914932 | 128914989 | 128912423 | 128912460 | 128926358 | 128926454 | 40 | 56 | 23 | 58 | 53 | 0 | 0 | 0 | 0 | 0 | 173 | 105 |
| SE | 80829 | ENSG00000151948 | GLT1D1 | chr12 | + | 128926358 | 128926454 | 128888638 | 128888744 | 128945325 | 128945369 | 0 | 0 | 0 | 0 | 4 | 0 | 0 | 0 | 1 | 0 | 7 | 3 |
| SE | 80830 | ENSG00000151948 | GLT1D1 | chr12 | + | 128926358 | 128926454 | 128899235 | 128899287 | 128927104 | 128927154 | 76 | 81 | 45 | 56 | 101 | 0 | 0 | 0 | 0 | 0 | 310 | 139 |
| SE | 80837 | ENSG00000151948 | GLT1D1 | chr12 | + | 128927104 | 128927154 | 128926358 | 128926454 | 128945325 | 128945369 | 106 | 99 | 53 | 80 | 94 | 0 | 0 | 0 | 0 | 0 | 406 | 169 |
| SE | 80839 | ENSG00000181016 | LSMEM1 | chr7 | + | 112486351 | 112486439 | 112484811 | 112484943 | 112486922 | 112487051 | 1 | 0 | 0 | 0 | 0 | 3 | 5 | 5 | 8 | 3 | 0 | 0 |
| SE | 80840 | ENSG00000181652 | ATG9B | chr7 | - | 151015359 | 151015713 | 151014945 | 151015123 | 151015881 | 151016023 | 0 | 0 | 0 | 2 | 2 | 0 | 0 | 0 | 2 | 0 | 0 | 0 |
| SE | 80843 | ENSG00000164896 | FASTK | chr7 | - | 151076912 | 151077236 | 151076625 | 151076832 | 151077309 | 151077401 | 30 | 18 | 27 | 46 | 29 | 0 | 0 | 1 | 0 | 0 | 33 | 74 |
| SE | 80845 | ENSG00000164896 | FASTK | chr7 | - | 151078841 | 151079021 | 151078561 | 151078701 | 151079499 | 151079922 | 54 | 18 | 52 | 30 | 16 | 1 | 0 | 0 | 1 | 0 | 11 | 20 |
| SE | 80846 | ENSG00000164896 | FASTK | chr7 | - | 151078922 | 151079021 | 151078561 | 151078701 | 151079499 | 151079922 | 13 | 14 | 24 | 15 | 8 | 1 | 0 | 0 | 1 | 0 | 0 | 9 |
| SE | 80849 | ENSG00000133612 | AGAP3 | chr7 | + | 151117370 | 151117456 | 151117094 | 151117182 | 151117635 | 151117682 | 51 | 39 | 38 | 43 | 30 | 0 | 0 | 0 | 0 | 0 | 95 | 23 |
| SE | 80851 | ENSG00000133612 | AGAP3 | chr7 | + | 151119986 | 151120145 | 151118293 | 151118632 | 151123793 | 151123876 | 60 | 42 | 34 | 37 | 23 | 0 | 0 | 1 | 1 | 0 | 83 | 15 |
| SE | 80852 | ENSG00000133612 | AGAP3 | chr7 | + | 151120519 | 151120567 | 151119986 | 151120145 | 151122724 | 151122793 | 0 | 0 | 0 | 2 | 0 | 1 | 2 | 4 | 1 | 0 | 0 | 0 |
| SE | 80853 | ENSG00000133612 | AGAP3 | chr7 | + | 151120519 | 151120567 | 151119986 | 151120145 | 151123793 | 151123886 | 1 | 0 | 0 | 1 | 0 | 16 | 7 | 6 | 14 | 3 | 0 | 0 |
| SE | 80856 | ENSG00000133612 | AGAP3 | chr7 | + | 151138142 | 151138313 | 151134399 | 151134568 | 151139978 | 151140116 | 37 | 13 | 11 | 15 | 7 | 0 | 0 | 0 | 1 | 0 | 60 | 31 |
| SE | 80857 | ENSG00000133612 | AGAP3 | chr7 | + | 151139978 | 151140116 | 151134424 | 151134568 | 151141897 | 151142052 | 22 | 7 | 1 | 9 | 5 | 0 | 0 | 4 | 0 | 2 | 0 | 4 |
| SE | 80858 | ENSG00000133612 | AGAP3 | chr7 | + | 151142411 | 151142634 | 151142162 | 151142253 | 151143340 | 151143596 | 58 | 49 | 46 | 42 | 15 | 1 | 0 | 0 | 0 | 0 | 101 | 9 |
| SE | 80860 | ENSG00000122557 | HERPUD2 | chr7 | - | 35635134 | 35635458 | 35634311 | 35634429 | 35638349 | 35638472 | 620 | 504 | 505 | 635 | 831 | 2 | 3 | 0 | 6 | 4 | 877 | 655 |
| SE | 80861 | ENSG00000122557 | HERPUD2 | chr7 | - | 35635134 | 35635458 | 35634311 | 35634429 | 35667433 | 35667588 | 352 | 341 | 264 | 364 | 247 | 0 | 0 | 1 | 0 | 0 | 575 | 351 |
| SE | 80862 | ENSG00000122557 | HERPUD2 | chr7 | - | 35638349 | 35638472 | 35634311 | 35634429 | 35667433 | 35667588 | 151 | 126 | 145 | 175 | 292 | 0 | 0 | 1 | 0 | 0 | 241 | 170 |
| SE | 80863 | ENSG00000122557 | HERPUD2 | chr7 | - | 35638349 | 35638472 | 35635134 | 35635458 | 35667433 | 35667588 | 412 | 286 | 387 | 440 | 871 | 0 | 0 | 2 | 0 | 1 | 538 | 474 |
| SE | 80864 | ENSG00000122557 | HERPUD2 | chr7 | - | 35667433 | 35667588 | 35638349 | 35638472 | 35694183 | 35694601 | 214 | 163 | 175 | 229 | 413 | 0 | 4 | 1 | 0 | 0 | 270 | 240 |
| SE | 80865 | ENSG00000122557 | HERPUD2 | chr7 | - | 35670214 | 35670328 | 35667433 | 35667588 | 35673200 | 35673278 | 69 | 62 | 344 | 385 | 359 | 2 | 2 | 1 | 0 | 1 | 468 | 324 |
| SE | 80869 | ENSG00000122557 | HERPUD2 | chr7 | - | 35694183 | 35694627 | 35673200 | 35673278 | 35694800 | 35695135 | 45 | 6 | 239 | 253 | 248 | 0 | 0 | 0 | 0 | 0 | 399 | 263 |
| SE | 80870 | ENSG00000006576 | PHTF2 | chr7 | + | 77840220 | 77840300 | 77798791 | 77798971 | 77893607 | 77893664 | 90 | 55 | 61 | 80 | 127 | 0 | 0 | 0 | 0 | 0 | 46 | 49 |
| SE | 80871 | ENSG00000006576 | PHTF2 | chr7 | + | 77854732 | 77854834 | 77840179 | 77840300 | 77893607 | 77893664 | 0 | 2 | 0 | 2 | 2 | 74 | 52 | 48 | 59 | 120 | 0 | 0 |
| SE | 80873 | ENSG00000006576 | PHTF2 | chr7 | + | 77901761 | 77901920 | 77900710 | 77900780 | 77908792 | 77908958 | 145 | 146 | 135 | 215 | 501 | 0 | 0 | 0 | 4 | 0 | 57 | 198 |
| SE | 80874 | ENSG00000006576 | PHTF2 | chr7 | + | 77907853 | 77907983 | 77901761 | 77901920 | 77908792 | 77908958 | 1 | 0 | 2 | 1 | 6 | 81 | 71 | 65 | 109 | 244 | 0 | 0 |
| SE | 80877 | ENSG00000006576 | PHTF2 | chr7 | + | 77940527 | 77940659 | 77940029 | 77940302 | 77942699 | 77942786 | 99 | 83 | 75 | 81 | 128 | 14 | 4 | 15 | 9 | 10 | 0 | 62 |
| SE | 80878 | ENSG00000006576 | PHTF2 | chr7 | + | 77942699 | 77942786 | 77940198 | 77940302 | 77949677 | 77949833 | 30 | 18 | 30 | 21 | 41 | 0 | 0 | 0 | 0 | 0 | 0 | 22 |
| SE | 80879 | ENSG00000006576 | PHTF2 | chr7 | + | 77942699 | 77942786 | 77940198 | 77940302 | 77951616 | 77951712 | 10 | 3 | 14 | 6 | 6 | 1 | 0 | 0 | 0 | 0 | 0 | 0 |
| SE | 80880 | ENSG00000006576 | PHTF2 | chr7 | + | 77949677 | 77949833 | 77940029 | 77940302 | 77951616 | 77951712 | 63 | 54 | 38 | 65 | 95 | 1 | 0 | 0 | 0 | 0 | 3 | 45 |
| SE | 80885 | ENSG00000126883 | NUP214 | chr9 | + | 131144279 | 131144754 | 131140548 | 131140710 | 131146128 | 131146304 | 100 | 49 | 65 | 75 | 153 | 0 | 0 | 0 | 1 | 0 | 138 | 48 |
| SE | 80886 | ENSG00000126883 | NUP214 | chr9 | + | 131144567 | 131144754 | 131140548 | 131140710 | 131146128 | 131146193 | 38 | 18 | 21 | 14 | 39 | 0 | 0 | 0 | 1 | 0 | 82 | 13 |
| SE | 80887 | ENSG00000126883 | NUP214 | chr9 | + | 131144576 | 131144754 | 131140548 | 131140710 | 131146128 | 131146304 | 36 | 13 | 16 | 12 | 24 | 0 | 0 | 0 | 1 | 0 | 82 | 13 |
| SE | 80888 | ENSG00000126883 | NUP214 | chr9 | + | 131146128 | 131146304 | 131144279 | 131144754 | 131147489 | 131147584 | 78 | 32 | 47 | 51 | 49 | 0 | 0 | 0 | 0 | 0 | 137 | 53 |
| [illegible] | | | | | | | | | | | | | | | | | | | | | | | |

| | | | | | | | | | | | | | | | | | | | | | | | | | | | | | | | | | | | | | | | | | | | | | | |
|---|---|---|---|---|---|---|---|---|---|---|---|---|---|---|---|---|---|---|---|---|---|---|---|---|---|---|---|---|---|---|---|---|---|---|---|---|---|---|---|---|---|---|---|---|---|---|
| SE | 81268 | ENSG00000123505 | AMD1 | chr6 | + | 110888856 | 110888983 | 110874784 | 110875215 | 110890253 | 110890356 | 340 | 276 | 313 | 558 | 360 | 0 | 3 | 0 | 0 | 0 | 116 | 275 | 207 | 167 | 93 | 0 | 0 | 0 | 0 | 0 | 276 | 149 | 1 | 1 | 1 | 0.98 | 1 | 1 | 1 | 1 | 1 | 1 | 1 | 1 | -0.004 |
| SE | 81269 | ENSG00000123505 | AMD1 | chr6 | + | 110888856 | 110888983 | 110874784 | 110875215 | 110892160 | 110892203 | 1 | 7 | 2 | 8 | 4 | 1 | 0 | 0 | 0 | 0 | 0 | 15 | 1 | 1 | 0 | 0 | 0 | 0 | 3 | 0 | 276 | 149 | 1 | 1 | 0.351 | 1 | 1 | 1 | 1 | NA | 1 | 1 | 0.153 | NA | 0.153 |
| SE | 81270 | ENSG00000123505 | AMD1 | chr6 | + | 110888856 | 110888983 | 110887504 | 110887591 | 110890253 | 110890356 | 626 | 492 | 592 | 1008 | 787 | 1 | 0 | 0 | 1 | 4 | 209 | 503 | 375 | 298 | 185 | 0 | 0 | 0 | 0 | 0 | 276 | 149 | 1 | 1 | 0.997 | 1 | 1 | 0.998 | 0.991 | 1 | 1 | 1 | 1 | 1 | -0.003 |
| SE | 81271 | ENSG00000123505 | AMD1 | chr6 | + | 110888856 | 110888983 | 110887504 | 110887591 | 110892160 | 110892203 | 276 | 212 | 268 | 427 | 414 | 0 | 0 | 0 | 1 | 0 | 87 | 236 | 161 | 114 | 92 | 0 | 0 | 0 | 0 | 0 | 276 | 149 | 1 | 1 | 1 | 1 | 1 | 0.996 | 1 | 1 | 1 | 1 | 1 | 1 | -0.001 |
| SE | 81272 | ENSG00000123505 | AMD1 | chr6 | + | 110890253 | 110890356 | 110874783 | 110875215 | 110892160 | 110892203 | 96 | 64 | 98 | 115 | 181 | 1 | 0 | 0 | 0 | 0 | 16 | 58 | 66 | 31 | 13 | 0 | 0 | 0 | 3 | 0 | 252 | 149 | 1 | 1 | 0.983 | 1 | 1 | 1 | 1 | 1 | 1 | 1 | 0.859 | 1 | 0.025 |
| SE | 81273 | ENSG00000123505 | AMD1 | chr6 | + | 110890253 | 110890356 | 110874783 | 110875215 | 110892298 | 110892443 | 0 | 4 | 0 | 0 | 0 | 0 | 0 | 0 | 3 | 1 | 0 | 0 | 0 | 0 | 0 | 0 | 0 | 0 | 2 | 0 | 252 | 149 | 1 | 1 | NA | 1 | NA | 0 | 0 | NA | NA | NA | 0 | NA | 0.333 |
| SE | 81274 | ENSG00000123505 | AMD1 | chr6 | + | 110890253 | 110890356 | 110887504 | 110887591 | 110892160 | 110892203 | 97 | 61 | 98 | 116 | 185 | 0 | 0 | 0 | 1 | 0 | 16 | 58 | 66 | 31 | 13 | 0 | 0 | 0 | 0 | 0 | 252 | 149 | 1 | 1 | 1 | | 1 | 0.986 | 1 | 1 | 1 | 1 | 1 | 1 | -0.003 |
| SE | 81275 | ENSG00000123505 | AMD1 | chr6 | + | 110890253 | 110890356 | 110888856 | 110888983 | 110892160 | 110892203 | 446 | 341 | 424 | 695 | 554 | 0 | 1 | 0 | 1 | 0 | 123 | 325 | 280 | 211 | 106 | 0 | 0 | 0 | 0 | 0 | 252 | 149 | 1 | 1 | 1 | 0.995 | 1 | 0.998 | 1 | 1 | 1 | 1 | 1 | 1 | -0.001 |
| SE | 81276 | ENSG00000123505 | AMD1 | chr6 | + | 110892160 | 110892203 | 110874769 | 110875215 | 110892298 | 110892443 | 88 | 76 | 82 | 137 | 128 | 0 | 0 | 0 | 3 | 1 | 41 | 73 | 66 | 31 | 7 | 0 | 0 | 0 | 2 | 0 | 192 | 149 | 1 | 1 | 1 | 1 | 1 | 0.973 | 0.99 | 1 | 1 | 1 | 0.923 | 1 | 0.008 |
| SE | 81277 | ENSG00000123505 | AMD1 | chr6 | + | 110892160 | 110892203 | 110888285 | 110890356 | 110892298 | 110892443 | 390 | 300 | 366 | 577 | 511 | 0 | 1 | 0 | 0 | 0 | 108 | 281 | 247 | 148 | 64 | 0 | 0 | 0 | 0 | 0 | 192 | 149 | 1 | 1 | 1 | 0.996 | 1 | 1 | 1 | 1 | 1 | 1 | 1 | 1 | -0.001 |
| SE | 81278 | ENSG00000155096 | AZIN1 | chr8 | - | 102829820 | 102829933 | 102829271 | 102829486 | 102833055 | 102833218 | 473 | 275 | 335 | 429 | 621 | 0 | 0 | 1 | 3 | 2 | 258 | 267 | 415 | 379 | 121 | 0 | 0 | 2 | 4 | 0 | 262 | 149 | 1 | 1 | 1 | 1 | 0.995 | 0.988 | 0.994 | 1 | 1 | 0.992 | 0.982 | 1 | 0.001 |
| SE | 81279 | ENSG00000155096 | AZIN1 | chr8 | - | 102829820 | 102829936 | 102829271 | 102829486 | 102833055 | 102833218 | 857 | 498 | 626 | 804 | 1221 | 0 | 0 | 1 | 3 | 2 | 491 | 545 | 741 | 705 | 251 | 0 | 0 | 2 | 4 | 0 | 265 | 149 | 1 | 1 | 1 | 1 | 0.997 | 0.993 | 0.997 | 1 | 1 | 0.995 | 0.99 | 1 | 0 |
| SE | 81280 | ENSG00000155096 | AZIN1 | chr8 | - | 102829820 | 102829936 | 102829271 | 102829486 | 102834188 | 102834263 | 494 | 277 | 338 | 442 | 634 | 0 | 0 | 0 | 0 | 0 | 272 | 275 | 414 | 386 | 116 | 0 | 0 | 5 | 0 | 0 | 265 | 149 | 1 | 1 | 1 | 1 | 1 | 1 | 1 | 1 | 1 | 0.979 | 1 | 1 | 0.004 |
| SE | 81282 | ENSG00000155096 | AZIN1 | chr8 | - | 102833055 | 102833218 | 102829271 | 102829486 | 102834188 | 102834263 | 255 | 142 | 215 | 286 | 398 | 0 | 0 | 0 | 0 | 0 | 113 | 180 | 207 | 211 | 128 | 0 | 0 | 5 | 0 | 0 | 298 | 149 | 1 | 1 | 1 | 1 | 1 | 1 | 1 | 1 | 1 | 0.954 | 1 | 1 | 0.009 |
| SE | 81284 | ENSG00000155096 | AZIN1 | chr8 | - | 102834188 | 102834263 | 102833055 | 102833218 | 102834665 | 102834747 | 591 | 375 | 523 | 619 | 792 | 3 | 0 | 3 | 3 | 1 | 276 | 464 | 558 | 504 | 280 | 0 | 0 | 3 | 3 | 0 | 224 | 149 | 1 | 1 | 0.992 | 1 | 0.991 | 0.993 | 0.998 | 1 | 1 | 0.992 | 0.991 | 1 | -0.002 |
| SE | 81285 | ENSG00000155096 | AZIN1 | chr8 | - | 102834665 | 102834747 | 102834188 | 102834263 | 102836255 | 102836390 | 566 | 376 | 504 | 581 | 732 | 0 | 0 | 1 | 6 | 2 | 234 | 423 | 570 | 481 | 275 | 0 | 1 | 3 | 4 | 0 | 231 | 149 | 1 | 1 | 1 | 1 | 0.997 | 0.984 | 0.996 | 1 | 0.996 | 0.992 | 0.987 | 1 | 0 |
| SE | 81286 | ENSG00000155096 | AZIN1 | chr8 | - | 102836255 | 102836390 | 102834665 | 102834747 | 102838743 | 102838916 | 871 | 669 | 753 | 917 | 1259 | 0 | 0 | 0 | 0 | 0 | 493 | 823 | 1029 | 859 | 476 | 0 | 0 | 2 | 0 | 0 | 284 | 149 | 1 | 1 | 1 | 1 | 1 | 1 | 1 | 1 | 1 | 0.996 | 1 | 1 | 0.001 |
| SE | 81287 | ENSG00000155096 | AZIN1 | chr8 | - | 102836255 | 102836541 | 102834665 | 102834747 | 102838743 | 102838916 | 436 | 294 | 376 | 453 | 509 | 0 | 0 | 0 | 0 | 0 | 193 | 331 | 457 | 379 | 190 | 0 | 0 | 2 | 0 | 0 | 298 | 149 | 1 | 1 | 1 | 1 | 1 | 1 | 1 | 1 | 1 | 0.991 | 1 | 1 | 0.002 |
| SE | 81288 | ENSG00000155096 | AZIN1 | chr8 | - | 102839649 | 102839823 | 102838743 | 102838916 | 102843550 | 102843729 | 870 | 634 | 758 | 861 | 1130 | 0 | 0 | 0 | 3 | 0 | 502 | 845 | 866 | 863 | 525 | 0 | 0 | 0 | 0 | 0 | 298 | 149 | 1 | 1 | 1 | 1 | 1 | 0.993 | 1 | 1 | 1 | 1 | 1 | 1 | -0.001 |
| SE | 81294 | ENSG00000155096 | AZIN1 | chr8 | - | 102858012 | 102858150 | 102843550 | 102843747 | 102863806 | 102864035 | 392 | 217 | 400 | 410 | 401 | 0 | 0 | 0 | 1 | 0 | 295 | 376 | 349 | 468 | 316 | 0 | 0 | 5 | 0 | 0 | 287 | 149 | 1 | 1 | 1 | 1 | 1 | 0.995 | 1 | 1 | 1 | 0.973 | 1 | 1 | 0.004 |
| SE | 81295 | ENSG00000155096 | AZIN1 | chr8 | - | 102859069 | 102859188 | 102858012 | 102858150 | 102863806 | 102864128 | 0 | 0 | 0 | 0 | 0 | 199 | 113 | 231 | 232 | 237 | 0 | 1 | 0 | 0 | 0 | 204 | 198 | 222 | 293 | 200 | 268 | 149 | 1 | 1 | 0 | 0 | 0 | 0 | 0 | 0 | 0.003 | 0 | 0 | 0 | -0.001 |
| SE | 81306 | ENSG00000141480 | ARRB2 | chr17 | + | 4720392 | 4720472 | 4720215 | 4720299 | 4720549 | 4720640 | 399 | 208 | 209 | 285 | 357 | 0 | 0 | 0 | 0 | 0 | 761 | 356 | 321 | 476 | 763 | 0 | 0 | 0 | 4 | 0 | 229 | 149 | 1 | 1 | 1 | 1 | 1 | 1 | 1 | 1 | 1 | 1 | 0.987 | 1 | 0.003 |
| SE | 81308 | ENSG00000141480 | ARRB2 | chr17 | + | 4720549 | 4720640 | 4720215 | 4720299 | 4720945 | 4721499 | 254 | 173 | 147 | 223 | 187 | 0 | 2 | 0 | 3 | 0 | 548 | 282 | 191 | 306 | 505 | 0 | 0 | 0 | 1 | 0 | 240 | 149 | 1 | 1 | 1 | 0.982 | 1 | 0.979 | 1 | 1 | 1 | 1 | 0.995 | 1 | -0.007 |
| SE | 81309 | ENSG00000141480 | ARRB2 | chr17 | + | 4720585 | 4720640 | 4720215 | 4720299 | 4720945 | 4721061 | 231 | 159 | 134 | 188 | 176 | 0 | 2 | 0 | 3 | 0 | 515 | 248 | 162 | 268 | 441 | 0 | 0 | 0 | 1 | 0 | 204 | 149 | 1 | 1 | 1 | 0.983 | 1 | 0.979 | 1 | 1 | 1 | 1 | 0.995 | 1 | -0.007 |
| SE | 81314 | ENSG00000106608 | URGCP | chr7 | - | 43887789 | 43887811 | 43887414 | 43887485 | 43926131 | 43926411 | 10 | 13 | 4 | 6 | 5 | 0 | 0 | 0 | 0 | 3 | 0 | 4 | 5 | 3 | 1 | 0 | 0 | 0 | 0 | 0 | 171 | 149 | 1 | 1 | 1 | 1 | 1 | 1 | 0.592 | NA | 1 | 1 | 1 | 1 | -0.082 |
| SE | 81316 | ENSG00000106608 | URGCP | chr7 | - | 43887789 | 43887816 | 43887414 | 43887485 | 43926131 | 43926411 | 12 | 14 | 5 | 6 | 7 | 0 | 0 | 0 | 0 | 3 | 0 | 4 | 6 | 3 | 1 | 0 | 0 | 0 | 0 | 0 | 176 | 149 | 1 | 1 | 1 | 1 | 1 | 1 | 0.664 | NA | 1 | 1 | 1 | 1 | -0.067 |
| SE | 81317 | ENSG00000106608 | URGCP | chr7 | - | 43910730 | 43910852 | 43887789 | 43887816 | 43926131 | 43926411 | 0 | 2 | 0 | 0 | 0 | 2 | 1 | 1 | 2 | 2 | 0 | 0 | 2 | 0 | 0 | 0 | 0 | 0 | 0 | 0 | 271 | 149 | 1 | 1 | 0 | 0.524 | 0 | 0 | 0 | NA | NA | 1 | NA | NA | -0.895 |
| SE | 81318 | ENSG00000135164 | DMTF1 | chr7 | + | 87154227 | 87154278 | 87152452 | 87152555 | 87154389 | 87154528 | 0 | 0 | 0 | 5 | 0 | 11 | 0 | 1 | 2 | 5 | 0 | 0 | 0 | 0 | 0 | 0 | 0 | 0 | 1 | 0 | 200 | 149 | 1 | 1 | 0 | NA | 0 | 0.651 | 0 | NA | NA | NA | 0 | NA | 0.163 |
| SE | 81321 | ENSG00000135164 | DMTF1 | chr7 | + | 87154389 | 87154528 | 87152435 | 87152555 | 87163238 | 87163332 | 10 | 0 | 1 | 1 | 5 | 3 | 1 | 0 | 2 | 0 | 0 | 0 | 0 | 1 | 0 | 0 | 0 | 0 | 1 | 0 | 288 | 149 | 1 | 1 | 0.633 | 0 | 1 | 0.206 | 1 | NA | NA | NA | 0.341 | NA | 0.227 |
| SE | 81322 | ENSG00000135164 | DMTF1 | chr7 | + | 87154389 | 87154528 | 87152435 | 87152555 | 87163494 | 87163617 | 18 | 5 | 6 | 5 | 7 | 14 | 10 | 18 | 20 | 13 | 0 | 0 | 0 | 1 | 0 | 0 | 23 | 12 | 0 | 22 | 288 | 149 | 1 | 1 | 0.399 | 0.206 | 0.147 | 0.115 | 0.218 | NA | 0 | 0 | 1 | 0 | -0.033 |
| SE | 81324 | ENSG00000135164 | DMTF1 | chr7 | + | 87154389 | 87154528 | 87154227 | 87154278 | 87155454 | 87155497 | 3 | 3 | 0 | 8 | 6 | 0 | 0 | 0 | 1 | 0 | 3 | 0 | 0 | 1 | 0 | 0 | 0 | 0 | 0 | 0 | 288 | 149 | 1 | 1 | 1 | 1 | NA | 0.805 | 1 | 1 | NA | NA | 1 | NA | -0.049 |
| SE | 81326 | ENSG00000135164 | DMTF1 | chr7 | + | 87155454 | 87155497 | 87154389 | 87154528 | 87163494 | 87163617 | 2 | 2 | 0 | 2 | 4 | 8 | 5 | 5 | 4 | 2 | 3 | 0 | 0 | 0 | 0 | 0 | 0 | 0 | 0 | 0 | 192 | 149 | 1 | 1 | 0.162 | 0.237 | 0 | 0.28 | 0.608 | 1 | NA | NA | NA | NA | -0.743 |
| SE | 81327 | ENSG00000135164 | DMTF1 | chr7 | + | 87163238 | 87163332 | 87152425 | 87152555 | 87163494 | 87163539 | 8 | 1 | 1 | 6 | 6 | 14 | 10 | 18 | 20 | 13 | 0 | 2 | 2 | 6 | 3 | 0 | 23 | 12 | 0 | 22 | 243 | 149 | 1 | 1 | 0.259 | 0.058 | 0.033 | 0.155 | 0.221 | NA | 0.051 | 0.093 | 1 | 0.077 | -0.16 |
| SE | 81328 | ENSG00000135164 | DMTF1 | chr7 | + | 87163238 | 87163332 | 87154389 | 87154528 | 87163494 | 87163539 | 5 | 0 | 1 | 4 | 6 | 8 | 5 | 5 | 4 | 2 | 0 | 2 | 2 | 5 | 3 | 0 | 0 | 0 | 0 | 0 | 243 | 149 | 1 | 1 | 0.277 | 0 | 0.109 | 0.38 | 0.648 | NA | 1 | 1 | 1 | 1 | -0.717 |
| SE | 81331 | ENSG00000135164 | DMTF1 | chr7 | + | 87164253 | 87164331 | 87162881 | 87163617 | 87164933 | 87165023 | 0 | 3 | 0 | 4 | 2 | 89 | 66 | 66 | 105 | 125 | 0 | 0 | 0 | 0 | 0 | 22 | 35 | 37 | 8 | 29 | 227 | 149 | 1 | 1 | 0 | 0.029 | 0 | 0.024 | 0.01 | 0 | 0 | 0 | 0 | 0 | 0.013 |
| SE | 81333 | ENSG00000135164 | DMTF1 | chr7 | + | 87164933 | 87165050 | 87163219 | 87163617 | 87170994 | 87171089 | 73 | 59 | 54 | 90 | 106 | 0 | 0 | 0 | 0 | 0 | 19 | 29 | 32 | 7 | 29 | 0 | 4 | 0 | 0 | 0 | 266 | 149 | 1 | 1 | 1 | 1 | 1 | 1 | 1 | 1 | 0.802 | 1 | 1 | 1 | 0.04 |
| SE | 81334 | ENSG00000135164 | DMTF1 | chr7 | + | 87166482 | 87166605 | 87163494 | 87163617 | 87170994 | 87171017 | 167 | 154 | 147 | 226 | 227 | 0 | 0 | 0 | 0 | 0 | 19 | 153 | 165 | 64 | 32 | 0 | 4 | 0 | 0 | 0 | 272 | 149 | 1 | 1 | 1 | 1 | 1 | 1 | 1 | 1 | 0.954 | 1 | 1 | 1 | 0.009 |
| SE | 81336 | ENSG00000135164 | DMTF1 | chr7 | + | 87179544 | 87179702 | 87174592 | 87174669 | 87181308 | 87181341 | 434 | 409 | 385 | 470 | 485 | 3 | 1 | 3 | 2 | 1 | 80 | 301 | 357 | 70 | 68 | 0 | 0 | 5 | 0 | 0 | 298 | 149 | 1 | 1 | 0.986 | 0.995 | 0.985 | 0.992 | 0.996 | 1 | 1 | 0.973 | 1 | 1 | -0.004 |
| SE | 81337 | ENSG00000135164 | DMTF1 | chr7 | + | 87181308 | 87181341 | 87179544 | 87179702 | 87182055 | 87182337 | 149 | 137 | 140 | 150 | 174 | 1 | 2 | 0 | 2 | 0 | 25 | 70 | 125 | 22 | 12 | 0 | 0 | 0 | 0 | 0 | 182 | 149 | 1 | 1 | 0.992 | 0.982 | 1 | 0.984 | 1 | 1 | 1 | 1 | 1 | 1 | -0.008 |
| SE | 81338 | ENSG00000135164 | DMTF1 | chr7 | + | 87184396 | 87184625 | 87182016 | 87182337 | 87185828 | 87185933 | 209 | 192 | 233 | 228 | 233 | 2 | 0 | 1 | 0 | 2 | 19 | 133 | 117 | 53 | 10 | 0 | 1 | 1 | 0 | 0 | 298 | 149 | 1 | 1 | 0.981 | 1 | 0.991 | 1 | 0.983 | 1 | 0.985 | 0.983 | 1 | 1 | -0.003 |
| SE | 81339 | ENSG00000135164 | DMTF1 | chr7 | + | 87184819 | 87184892 | 87184396 | 87184625 | 87185828 | 87185980 | 8 | 0 | 0 | 1 | 7 | 84 | 79 | 98 | 68 | 139 | 0 | 0 | 1 | 0 | 0 | 12 | 54 | 45 | 6 | 0 | 222 | 149 | 1 | 1 | 0.06 | 0 | 0 | 0.01 | 0.033 | 0 | 0 | 0.015 | 0 | NA | 0.017 |
| SE | 81344 | ENSG00000135164 | DMTF1 | chr7 | + | 87194683 | 87194828 | 87193724 | 87194102 | 87195030 | 87195166 | 232 | 241 | 215 | 282 | 406 | 12 | 27 | 32 | 32 | 40 | 0 | 180 | 171 | 45 | 4 | 0 | 9 | 18 | 9 | 10 | 294 | 149 | 1 | 1 | 0.907 | 0.819 | 0.773 | 0.817 | 0.837 | NA | 0.91 | 0.828 | 0.717 | 0.169 | 0.175 |
| SE | 81347 | ENSG00000173852 | DPY19L1 | chr7 | - | 34969432 | 34969532 | 34966893 | 34966971 | 34973513 | 34973605 | 54 | 27 | 33 | 32 | 41 | 0 | 0 | 0 | 0 | 0 | 17 | 31 | 26 | 24 | 21 | 0 | 0 | 0 | 2 | 0 | 249 | 149 | 1 | 1 | 1 | 1 | 1 | 1 | 1 | 1 | 1 | 1 | 0.878 | 1 | 0.024 |
| SE | 81365 | ENSG00000146757 | ZNF92 | chr7 | + | 65387901 | 65388028 | 65373798 | 65374000 | 65388805 | 65388901 | 61 | 26 | 58 | 58 | 75 | 5 | 0 | 1 | 0 | 0 | 4 | 25 | 1 | 0 | 0 | 0 | 0 | 0 | 0 | 0 | 276 | 149 | 1 | 1 | 0.868 | 1 | 0.969 | 1 | 1 | 1 | 1 | 1 | NA | NA | -0.033 |
| SE | 81366 | ENSG00000146757 | ZNF92 | chr7 | + | 65387901 | 65388028 | 65373798 | 65374000 | 65398340 | 65401135 | 24 | 10 | 24 | 25 | 32 | 7 | 3 | 8 | 8 | 0 | 0 | 5 | 0 | 0 | 2 | 10 | 1 | 0 | 0 | 0 | 276 | 149 | 1 | 1 | 0.649 | 0.643 | 0.618 | 0.628 | 1 | 0 | 0.73 | NA | NA | 1 | 0.131 |
| SE | 81367 | ENSG00000146757 | ZNF92 | chr7 | + | 65388805 | 65388901 | 65373880 | 65374000 | 65398340 | 65401081 | 24 | 12 | 16 | 25 | 24 | 7 | 3 | 8 | 8 | 0 | 2 | 9 | 0 | 0 | 0 | 10 | 1 | 0 | 0 | 0 | 245 | 149 | 1 | 1 | 0.676 | 0.709 | 0.549 | 0.655 | 1 | 0.108 | 0.846 | NA | NA | NA | 0.241 |
| SE | 81368 | ENSG00000146757 | ZNF92 | chr7 | + | 65388805 | 65388901 | 65387901 | 65388028 | 65398340 | 65401081 | 58 | 29 | 57 | 58 | 73 | 3 | 0 | 8 | 0 | 6 | 6 | 29 | 1 | 0 | 0 | 0 | 0 | 0 | 0 | 2 | 245 | 149 | 1 | 1 | 0.922 | 1 | 0.812 | 1 | 0.881 | 1 | 1 | 1 | NA | 0 | 0.173 |
| SE | 81369 | ENSG00000258733 | LINC02328 | chr14 | + | 86000860 | 86000953 | 85934709 | 85934861 | 86028368 | 86028458 | 2 | 2 | 2 | 0 | 0 | 2 | 1 | 1 | 3 | 0 | 9 | 3 | 3 | 0 | 0 | 0 | 0 | 0 | 6 | 0 | 242 | 149 | 1 | 1 | 0.381 | 0.552 | 0.552 | 0 | NA | 1 | 1 | 1 | 0 | NA | -0.379 |
| SE | 81370 | ENSG00000258733 | LINC02328 | chr14 | + | 86000860 | 86000953 | 85934709 | 85934861 | 86031135 | 86031230 | 2 | 1 | 1 | 0 | 0 | 0 | 0 | 1 | 2 | 0 | 9 | 3 | 3 | 0 | 0 | 0 | 0 | 0 | 0 | 0 | 242 | 149 | 1 | 1 | 1 | 1 | 0.381 | 0 | NA | 1 | 1 | 1 | NA | NA | -0.405 |
| SE | 81371 | ENSG00000258733 | LINC02328 | chr14 | + | 86000860 | 86000953 | 85934709 | 85934861 | 86070862 | 86070967 | 4 | 1 | 2 | 1 | 0 | 16 | 6 | 7 | 6 | 13 | 9 | 6 | 3 | 0 | 0 | 0 | 9 | 3 | 0 | 0 | 242 | 149 | 1 | 1 | 0.133 | 0.093 | 0.15 | 0.093 | 0 | 1 | 0.291 | 0.381 | NA | NA | -0.464 |
| SE | 81374 | ENSG00000258733 | LINC02328 | chr14 | + | 86000860 | 86000953 | 85983945 | 85984022 | 86070862 | 86070967 | 2 | 0 | 1 | 2 | 0 | 2 | 0 | 3 | 2 | 2 | 0 | 3 | 0 | 0 | 0 | 0 | 0 | 0 | 0 | 10 | 242 | 149 | 1 | 1 | 0.381 | NA | 0.17 | 0.381 | 0 | NA | 1 | NA | NA | 0 | -0.267 |
| SE | 81375 | ENSG00000258733 | LINC02328 | chr14 | + | 86028368 | 86028458 | 85934676 | 85934861 | 86031135 | 86031230 | 39 | 9 | 18 | 22 | 29 | 0 | 0 | 1 | 2 | 0 | 0 | 10 | 14 | 9 | 0 | 0 | 0 | 0 | 0 | 0 | 239 | 149 | 1 | 1 | 1 | 1 | 0.918 | 0.873 | 1 | NA | 1 | 1 | 1 | NA | -0.042 |
| SE | 81376 | ENSG00000258733 | LINC02328 | chr14 | + | 86028368 | 86028458 | 85934676 | 85934861 | 86070862 | 86070967 | 2 | 1 | 1 | 3 | 0 | 16 | 6 | 7 | 6 | 13 | 0 | 0 | 0 | 6 | 0 | 0 | 9 | 3 | 0 | 0 | 239 | 149 | 1 | 1 | 0.072 | 0.094 | 0.082 | 0.238 | 0 | NA | 0 | 0 | 1 | NA | -0.236 |
| SE | 81377 | ENSG00000258733 | LINC02328 | chr14 | + | 86028368 | 86028458 | 85983945 | 85984022 | 86031135 | 86031230 | 39 | 8 | 18 | 19 | 32 | 3 | 1 | 2 | 0 | 0 | 0 | 10 | 14 | 3 | 0 | 0 | 0 | 0 | 0 | 0 | 239 | 149 | 1 | 1 | 0.89 | 0.833 | 0.849 | 1 | 1 | NA | 1 | 1 | 1 | NA | -0.086 |
| SE | 81379 | ENSG00000258733 | LINC02328 | chr14 | + | 86028368 | 86028458 | 86000860 | 86000953 | 86031135 | 86031230 | 38 | 9 | 18 | 19 | 29 | 0 | 0 | 2 | 0 | 0 | 0 | 10 | 14 | 3 | 0 | 0 | 0 | 0 | 0 | 0 | 239 | 149 | 1 | 1 | 1 | 1 | 0.849 | 1 | 1 | NA | 1 | 1 | 1 | NA | -0.03 |
| SE | 81380 | ENSG00000258733 | LINC02328 | chr14 | + | 86028368 | 86028458 | 86000860 | 86000953 | 86070862 | 86070967 | 0 | 1 | 1 | 0 | 0 | 2 | 0 | 1 | 1 | 0 | 0 | 0 | 0 | 0 | 0 | 0 | 3 | 0 | 0 | 0 | 239 | 149 | 1 | 1 | 0 | 1 | 0.384 | 0 | NA | NA | 0 | NA | NA | NA | 0.346 |
| SE | 81381 | ENSG00000258733 | LINC02328 | chr14 | + | 86031135 | 86031230 | 85934625 | 85934861 | 86070862 | 86070967 | 1 | 2 | 3 | 1 | 4 | 16 | 6 | 7 | 6 | 13 | 0 | 0 | 0 | 0 | 0 | 0 | 9 | 3 | 0 | 0 | 244 | 149 | 1 | 1 | 0.037 | 0.169 | 0.207 | 0.092 | 0.158 | NA | 0 | 0 | NA | NA | 0.133 |
| SE | 81382 | ENSG00000258733 | LINC02328 | chr14 | + | 86031135 | 86031230 | 85983945 | 85984022 | 86035713 | 86035815 | 13 | 4 | 11 | 4 | 39 | 0 | 0 | 0 | 3 | 0 | 0 | 0 | 5 | 2 | 5 | 0 | 0 | 0 | 0 | 0 | 244 | 149 | 1 | 1 | 1 | 1 | 1 | 0.449 | 1 | NA | NA | 1 | 1 | 1 | -0.11 |
| SE | 81383 | ENSG00000258733 | LINC02328 | chr14 | + | 86031135 | 86031230 | 85983945 | 85984022 | 86070862 | 86070967 | 2 | 2 | 4 | 0 | 4 | 2 | 0 | 3 | 2 | 2 | 0 | 0 | 0 | 0 | 0 | 0 | 0 | 0 | 0 | 10 | 244 | 149 | 1 | 1 | 0.379 | 1 | 0.449 | 0 | 0.55 | NA | NA | NA | NA | 0 | 0.476 |
| SE | 81384 | ENSG00000258733 | LINC02328 | chr14 | + | 86031135 | 86031230 | 86000860 | 86000953 | 86070862 | 86070967 | 1 | 2 | 4 | 0 | 4 | 2 | 0 | 1 | 1 | 0 | 0 | 0 | 0 | 0 | 0 | 0 | 3 | 0 | 0 | 0 | 244 | 149 | 1 | 1 | 0.234 | 1 | 0.71 | 0 | 1 | NA | 0 | NA | NA | NA | 0.589 |
| SE | 81385 | ENSG00000258733 | LINC02328 | chr14 | + | 86031135 | 86031230 | 86028368 | 86028458 | 86070862 | 86070967 | 23 | 5 | 12 | 13 | 29 | 1 | 0 | 0 | 0 | 0 | 0 | 1 | 8 | 1 | 0 | 0 | 0 | 0 | 0 | 0 | 244 | 149 | 1 | 1 | 0.934 | 1 | 1 | 1 | 1 | NA | 1 | 1 | 1 | NA | -0.013 |
| SE | 81386 | ENSG00000258733 | LINC02328 | chr14 | + | 86035713 | 86035815 | 85983945 | 85984022 | 86070862 | 86070967 | 0 | 0 | 0 | 0 | 0 | 2 | 0 | 3 | 2 | 2 | 0 | 0 | 0 | 0 | 5 | 0 | 0 | 0 | 0 | 10 | 251 | 149 | 1 | 1 | 0 | NA | 0 | 0 | 0 | NA | NA | NA | NA | 0.229 | -0.229 |
| SE | 81387 | ENSG00000258733 | LINC02328 | chr14 | + | 86035713 | 86035815 | 86031135 | 86031230 | 86041725 | 86041833 | 49 | 17 | 30 | 32 | 83 | 0 | 0 | 0 | 0 | 2 | 6 | 12 | 11 | 4 | 19 | 0 | 0 | 0 | 0 | 0 | 251 | 149 | 1 | 1 | 1 | 1 | 1 | 1 | 0.961 | 1 | 1 | 1 | 1 | 1 | -0.008 |
| SE | 81388 | ENSG00000258733 | LINC02328 | chr14 | + | 86035713 | 86035815 | 86031135 | 86031230 | 86070862 | 86070967 | 11 | 4 | 9 | 7 | 21 | 5 | 2 | 3 | 0 | 6 | 0 | 0 | 7 | 0 | 5 | 0 | 0 | 1 | 0 | 0 | 251 | 149 | 1 | 1 | 0.566 | 0.543 | 0.64 | 1 | 0.675 | NA | NA | 0.806 | NA | 1 | -0.218 |
| SE | 81389 | ENSG00000258733 | LINC02328 | chr14 | + | 86041725 | 86041833 | 86031135 | 86031230 | 86070862 | 86070967 | 2 | 4 | 1 | 3 | 15 | 5 | 2 | 3 | 0 | 6 | 2 | 0 | 0 | 0 | 0 | 0 | 0 | 1 | 0 | 0 | 257 | 149 | 1 | 1 | 0.188 | 0.537 | 0.162 | 1 | 0.592 | 1 | NA | 0 | NA | NA | -0.004 |
| SE | 81391 | ENSG00000258733 | LINC02328 | chr14 | + | 86070862 | 86070967 | 85934665 | 85934861 | 86127949 | 86128116 | 26 | 11 | 14 | 20 | 39 | 1 | 1 | 0 | 3 | 1 | 2 | 18 | 9 | 0 | 0 | 0 | 0 | 0 | 0 | 0 | 254 | 149 | 1 | 1 | 0.938 | 0.866 | 1 | 0.796 | 0.958 | 1 | 1 | 1 | NA | NA | -0.088 |
| SE | 81392 | ENSG00000258733 | LINC02328 | chr14 | + | 86128664 | 86128773 | 86127949 | 86128116 | 86129495 | 86129778 | 8 | 14 | 6 | 6 | 20 | 21 | 9 | 5 | 8 | 33 | 0 | 8 | 3 | 7 | 0 | 2 | 6 | 2 | 0 | 0 | 258 | 149 | 1 | 1 | 0.18 | 0.473 | 0.409 | 0.302 | 0.259 | 0 | 0.435 | 0.464 | 1 | NA | -0.15 |
| SE | 81393 | ENSG00000258733 | LINC02328 | chr14 | + | 86128714 | 86128773 | 86127949 | 86128116 | 86129495 | 86129778 | 2 | 5 | 0 | 2 | 13 | 21 | 9 | 5 | 8 | 33 | 0 | 0 | 1 | 2 | 0 | 2 | 6 | 2 | 0 | 0 | 208 | 149 | 1 | 1 | 0.064 | 0.285 | 0 | 0.152 | 0.22 | 0 | 0 | 0.264 | 1 | NA | -0.172 |
| SE | 81394 | ENSG00000106404 | CLDN15 | chr7 | - | 101233872 | 101234116 | 101232832 | 101232914 | 101234277 | 101234442 | 0 | 0 | 0 | 0 | 0 | 4 | 3 | 11 | 2 | 7 | 0 | 2 | 0 | 0 | 0 | 0 | 7 | 2 | 0 | 0 | 298 | 149 | 1 | 1 | 0 | 0 | 0 | 0 | 0 | NA | 0.125 | 0 | NA | NA | -0.062 |
| SE | 81397 | ENSG00000106404 | CLDN15 | chr7 | - | 101236746 | 101236886 | 101233768 | 101234442 | 101237364 | 101237750 | 4 | 0 | 3 | 2 | 0 | 8 | 3 | 9 | 3 | 3 | 0 | 3 | 0 | 0 | 18 | 0 | 11 | 2 | 3 | 0 | 289 | 149 | 1 | 1 | 0.205 | 0 | 0.147 | 0.256 | 0 | NA | 0.123 | 0 | 0 | 1 | -0.159 |
| SE | 81398 | ENSG00000158683 | PKD1L1 | chr7 | - | 47873898 | 47874010 | 47866418 | 47866614 | 47876096 | 47876217 | 0 | 1 | 1 | 0 | 0 | 0 | 0 | 0 | 0 | 0 | 0 | 0 | 0 | 0 | 0 | 0 | 0 | 1 | 0 | 0 | 261 | 149 | 1 | 1 | NA | 1 | 1 | NA | NA | NA | NA | 0 | NA | NA | 1 |
| SE | 81399 | ENSG00000158683 | PKD1L1 | chr7 | - | 47876096 | 47876217 | 47866418 | 47866614 | 47877488 | 47877631 | 0 | 2 | 0 | 1 | 0 | 0 | 2 | 0 | 0 | 0 | 0 | 0 | 1 | 0 | 0 | 0 | 0 | 0 | 0 | 0 | 270 | 149 | 1 | 1 | NA | 0.356 | NA | 1 | NA | NA | NA | 1 | NA | NA | -0.322 |
| SE | 81403 | ENSG00000129103 | SUMF2 | chr7 | + | 56068481 | 56068638 | 56064001 | 56064378 | 56072996 | 56073111 | 207 | 102 | 193 | 150 | 277 | 0 | 0 | 2 | 0 | 0 | 143 | 216 | 120 | 127 | 242 | 0 | 0 | 0 | 0 | 0 | 298 | 149 | 1 | 1 | 1 | 1 | 0.98 | 1 | 1 | 1 | 1 | 1 | 1 | 1 | -0.004 |
| SE | 81404 | ENSG00000129103 | SUMF2 | chr7 | + | 56068481 | 56068647 | 56064254 | 56064378 | 56072996 | 56073111 | 36 | 16 | 31 | 32 | 18 | 0 | 0 | 2 | 0 | 0 | 34 | 55 | 27 | 33 | 21 | 0 | 0 | 0 | 0 | 0 | 298 | 149 | 1 | 1 | 1 | 1 | 0.886 | 1 | 1 | 1 | 1 | 1 | 1 | 1 | -0.023 |
| SE | 81405 | ENSG00000129103 | SUMF2 | chr7 | + | 56072996 | 56073111 | 56068481 | 56068638 | 56074173 | 56074218 | 317 | 167 | 288 | 249 | 493 | 2 | 0 | 2 | 0 | 4 | 250 | 327 | 187 | 159 | 454 | 0 | 3 | 1 | 0 | 0 | 264 | 149 | 1 | 1 | 0.989 | 1 | 0.988 | 1 | 0.986 | 1 | 0.984 | 0.991 | 1 | 1 | -0.002 |
| SE | 81406 | ENSG00000129103 | SUMF2 | chr7 | + | 56072996 | 56073111 | 56068481 | 56068638 | 56076833 | 56076889 | 115 | 53 | 104 | 73 | 159 | 0 | 2 | 1 | 0 | 2 | 65 | 121 | 73 | 72 | 176 | 0 | 0 | 0 | 0 | 0 | 264 | 149 | 1 | 1 | 1 | 0.937 | 0.983 | 1 | 0.978 | 1 | 1 | 1 | 1 | 1 | -0.02 |
| SE | 81408 | ENSG00000129103 | SUMF2 | chr7 | + | 56074173 | 56074218 | 56068481 | 56068638 | 56076833 | 56076889 | 2 | 0 | 2 | 1 | 1 | 0 | 2 | 1 | 0 | 2 | 0 | 5 | 5 | 4 | 0 | 0 | 0 | 0 | 0 | 0 | 194 | 149 | 1 | 1 | 1 | 0 | 0.606 | 1 | 0.277 | NA | 1 | 1 | 1 | NA | -0.423 |
| SE | 81414 | ENSG00000129103 | SUMF2 | chr7 | + | 56074585 | 56074736 | 56072996 | 56073111 | 56078101 | 56078186 | 2 | 11 | 9 | 9 | 13 | 1 | 0 | 0 | 2 | 0 | 0 | 14 | 4 | 1 | 20 | 0 | 3 | 1 | 1 | 0 | 298 | 149 | 1 | 1 | 0.5 | 1 | 1 | 0.692 | 1 | NA | 0.7 | 0.667 | 0.333 | 1 | 0.163 |
| SE | 81416 | ENSG00000129103 | SUMF2 | chr7 | + | 56074585 | 56074736 | 56074113 | 56074218 | 56078101 | 56078186 | 135 | 70 | 144 | 138 | 243 | 1 | 0 | 0 | 1 | 0 | 115 | 191 | 68 | 81 | 196 | 0 | 0 | 0 | 0 | 0 | 298 | 149 | 1 | 1 | 0.985 | 1 | 1 | 0.986 | 1 | 1 | 1 | 1 | 1 | 1 | -0.006 |
| SE | 81418 | ENSG00000129103 | SUMF2 | chr7 | + | 56076833 | 56076889 | 56074173 | 56074218 | 56078101 | 56078186 | 45 | 22 | 63 | 42 | 47 | 1 | 0 | 0 | 1 | 0 | 45 | 75 | 28 | 35 | 32 | 0 | 0 | 0 | 0 | 0 | 205 | 149 | 1 | 1 | 0.97 | 1 | 1 | 0.968 | 1 | 1 | 1 | 1 | 1 | 1 | -0.012 |
| SE | 81419 | ENSG00000129103 | SUMF2 | chr7 | + | 56076833 | 56076889 | 56074585 | 56074736 | 56078101 | 56078186 | 141 | 80 | 169 | 142 | 245 | 0 | 0 | 2 | 0 | 0 | 103 | 187 | 56 | 76 | 161 | 0 | 0 | 0 | 0 | 0 | 205 | 149 | 1 | 1 | 1 | 1 | 0.984 | 1 | 1 | 1 | 1 | 1 | 1 | 1 | -0.003 |
| SE | 81421 | ENSG00000129103 | SUMF2 | chr7 | + | 56078363 | 56078508 | 56078101 | 56078186 | 56079527 | 56079567 | 289 | 126 | 298 | 259 | 169 | 0 | 0 | 1 | 0 | 0 | 179 | 273 | 160 | 140 | 279 | 0 | 1 | 0 | 0 | 0 | 294 | 149 | 1 | 1 | 1 | 1 | 0.993 | 1 | 1 | 1 | 0.993 | 1 | 1 | 1 | 0 |
| SE | 81426 | ENSG00000011275 | RNF216 | chr7 | - | 5641153 | 5641376 | 5624055 | 5624125 | 5652412 | 5652510 | 162 | 121 | 241 | 233 | 317 | 0 | 2 | 0 | 0 | 0 | 226 | 227 | 164 | 138 | 316 | 0 | 0 | 0 | 0 | 0 | 298 | 149 | 1 | 1 | 1 | 0.968 | 1 | 1 | 1 | 1 | 1 | 1 | 1 | 1 | -0.006 |
| SE | 81427 | ENSG00000011275 | RNF216 | chr7 | - | 5652412 | 5652510 | 5641153 | 5641376 | 5711760 | 5711839 | 186 | 154 | 229 | 223 | 251 | 0 | 0 | 0 | 0 | 0 | 251 | 195 | 220 | 199 | 301 | 0 | 0 | 4 | 0 | 0 | 247 | 149 | 1 | 1 | 1 | 1 | 1 | 1 | 1 | 1 | 1 | 0.971 | 1 | 1 | 0.006 |
| SE | 81428 | ENSG00000011275 | RNF216 | chr7 | - | 5711760 | 5711839 | 5652412 | 5652510 | 5712714 | 5712863 | 163 | 123 | 175 | 175 | 152 | 0 | 1 | 0 | 0 | 0 | 189 | 119 | 181 | 138 | 270 | 0 | 0 | 0 | 0 | 0 | 228 | 149 | 1 | 1 | 1 | 0.988 | 1 | 1 | 1 | 1 | 1 | 1 | 1 | 1 | -0.002 |
| SE | 81430 | ENSG00000011275 | RNF216 | chr7 | - | 5725323 | 5725438 | 5721032 | 5721172 | 5729431 | 5729596 | 256 | 191 | 283 | 302 | 241 | 0 | 0 | 2 | 0 | 2 | 412 | 270 | 298 | 327 | 525 | 0 | 0 | 0 | 0 | 0 | 264 | 149 | 1 | 1 | 1 | 1 | 0.988 | 1 | 0.986 | 1 | 1 | 1 | 1 | 1 | -0.005 |
| SE | 81431 | ENSG00000011275 | RNF216 | chr7 | - | 5729431 | 5729596 | 5725323 | 5725438 | 5739275 | 5739352 | 103 | 94 | 129 | 148 | 140 | 0 | 0 | 0 | 0 | 0 | 197 | 153 | 165 | 199 | 331 | 0 | 0 | 0 | 5 | 0 | 298 | 149 | 1 | 1 | 1 | 1 | 1 | 1 | 1 | 1 | 1 | 1 | 0.952 | 1 | 0.01 |
| SE | 81433 | ENSG00000011275 | RNF216 | chr7 | - | 5732969 | 5733162 | 5730714 | 5730817 | 5739275 | 5739492 | 1 | 0 | 0 | 1 | 0 | 227 | 159 | 198 | 222 | 389 | 0 | 0 | 0 | 0 | 0 | 209 | 151 | 127 | 161 | 275 | 298 | 149 | 1 | 1 | 0.002 | 0 | 0 | 0.002 | 0 | 0 | 0 | 0 | 0 | 0 | 0.001 |
| SE | 81439 | ENSG00000011275 | RNF216 | chr7 | - | 5775004 | 5775179 | 5766894 | 5766939 | 5781540 | 5781620 | 6 | 0 | 0 | 0 | 0 | 0 | 1 | 1 | 0 | 0 | 0 | 0 | 0 | 0 | 0 | 6 | 6 | 1 | 2 | 5 | 298 | 149 | 1 | 1 | 1 | 0 | 0 | NA | NA | 0 | 0 | 0 | 0 | 0 | 0.333 |
| SE | 81440 | ENSG00000011275 | RNF216 | chr7 | - | 5777486 | 5777628 | 5766894 | 5766939 | 5781540 | 5781620 | 0 | 0 | 0 | 1 | 2 | 0 | 1 | 1 | 0 | 0 | 0 | 0 | 0 | 0 | 0 | 6 | 6 | 1 | 2 | 5 | 291 | 149 | 1 | 1 | NA | 0 | 0 | 1 | 1 | 0 | 0 | 0 | 0 | 0 | 0.5 |
| SE | 81450 | ENSG00000106443 | PHF14 | chr7 | + | 10982371 | 10983159 | 10974834 | 10974945 | 11013746 | 11013906 | 45 | 40 | 36 | 58 | 60 | 1 | 0 | 0 | 0 | 0 | 44 | 58 | 31 | 37 | 34 | 0 | 0 | 0 | 0 | 0 | 298 | 149 | 1 | 1 | 0.957 | 1 | 1 | 1 | 1 | 1 | 1 | 1 | 1 | 1 | -0.009 |
| SE | 81451 | ENSG00000106443 | PHF14 | chr7 | + | 10990702 | 10990847 | 10974834 | 10974945 | 11013746 | 11013906 | 104 | 93 | 105 | 109 | 233 | 1 | 0 | 0 | 0 | 0 | 67 | 64 | 47 | 47 | 16 | 0 | 0 | 0 | 0 | 0 | 294 | 149 | 1 | 1 | 0.981 | 1 | 1 | 1 | 1 | 1 | 1 | 1 | 1 | 1 | -0.004 |
| SE | 81452 | ENSG00000106443 | PHF14 | chr7 | + | 10990702 | 10990847 | 10982371 | 10983159 | 11013746 | 11013906 | 218 | 216 | 218 | 206 | 400 | 1 | 0 | 0 | 0 | 0 | 123 | 141 | 122 | 88 | 64 | 0 | 0 | 0 | 0 | 0 | 294 | 149 | 1 | 1 | 0.991 | 1 | 1 | 1 | 1 | 1 | 1 | 1 | 1 | 1 | -0.002 |
| SE | 81454 | ENSG00000106443 | PHF14 | chr7 | + | 11013746 | 11013906 | 10990702 | 10990847 | 11028680 | 11028760 | 93 | 87 | 96 | 96 | 229 | 2 | 0 | 0 | 1 | 0 | 65 | 52 | 41 | 35 | 11 | 0 | 0 | 0 | 0 | 0 | 298 | 149 | 1 | 1 | 0.959 | 1 | 1 | 0.98 | 1 | 1 | 1 | 1 | 1 | 1 | -0.012 |
| SE | 81455 | ENSG00000106443 | PHF14 | chr7 | + | 11022867 | 11022979 | 10990702 | 10990847 | 11028680 | 11028760 | 69 | 48 | 57 | 67 | 107 | 2 | 0 | 0 | 1 | 0 | 17 | 11 | 34 | 21 | 15 | 0 | 0 | 0 | 0 | 0 | 261 | 149 | 1 | 1 | 0.952 | 1 | 1 | 0.975 | 1 | 1 | 1 | 1 | 1 | 1 | -0.015 |
| SE | 81456 | ENSG00000106443 | PHF14 | chr7 | + | 11022867 | 11022979 | 11013746 | 11013906 | 11028680 | 11028760 | 146 | 114 | 128 | 126 | 271 | 0 | 1 | 1 | 1 | 0 | 39 | 40 | 65 | 43 | 22 | 0 | 0 | 0 | 0 | 0 | 261 | 149 | 1 | 1 | 1 | 0.985 | 0.986 | 0.986 | 1 | 1 | 1 | 1 | 1 | 1 | -0.009 |
| SE | 81458 | ENSG00000106443 | PHF14 | chr7 | + | 11036417 | 11036688 | 11035639 | 11035786 | 11036984 | 11037091 | 237 | 207 | 168 | 190 | 268 | 0 | 0 | 0 | 1 | 0 | 29 | 138 | 89 | 47 | 9 | 0 | 0 | 0 | 0 | 0 | 298 | 149 | 1 | 1 | 1 | 1 | 1 | 0.99 | 1 | 1 | 1 | 1 | 1 | 1 | -0.002 |
| SE | 81459 | ENSG00000106443 | PHF14 | chr7 | + | 11038759 | 11038855 | 11036984 | 11037091 | 11040671 | 11040775 | 142 | 132 | 110 | 111 | 149 | 2 | 1 | 1 | 2 | 3 | 0 | 77 | 60 | 23 | 0 | 0 | 11 | 4 | 0 | 0 | 245 | 149 | 1 | 1 | 0.977 | 0.988 | 0.985 | 0.971 | 0.968 | NA | 0.81 | 0.901 | 1 | NA | 0.074 |
| SE | 81461 | ENSG00000106443 | PHF14 | chr7 | + | 11110524 | 11110551 | 11061963 | 11062085 | 11111349 | 11111467 | 1 | 0 | 0 | 0 | 0 | 48 | 44 | 54 | 51 | 54 | 0 | 0 | 0 | 0 | 0 | 19 | 35 | 3 | 33 | 38 | 176 | 149 | 1 | 1 | 0.017 | 0 | 0 | 0 | 0 | 0 | 0 | 0 | 0 | 0 | 0.003 |
| SE | 81467 | ENSG00000106477 | CEP41 | chr7 | - | 130426628 | 130426759 | 130416918 | 130416966 | 130427954 | 130428018 | 0 | 1 | 0 | 2 | 0 | 12 | 5 | 9 | 15 | 18 | 0 | 0 | 1 | 0 | 0 | 18 | 3 | 0 | 11 | 2 | 280 | 149 | 1 | 1 | 0 | 0.096 | 0 | 0.066 | 0 | 0 | 0 | 1 | 0 | 0 | -0.168 |
| SE | 81468 | ENSG00000106477 | CEP41 | chr7 | - | 130426628 | 130426759 | 130421852 | 130422007 | 130427954 | 130428018 | 0 | 0 | 0 | 2 | 0 | 1 | 3 | 3 | 3 | 0 | 0 | 0 | 2 | 0 | 0 | 0 | 4 | 0 | 3 | 26 | 280 | 149 | 1 | 1 | 0 | 0 | 0 | 0.262 | NA | NA | 0 | 1 | 0 | 0 | -0.184 |
| SE | 81469 | ENSG00000106477 | CEP41 | chr7 | - | 130426645 | 130426759 | 130416918 | 130416966 | 130427954 | 130428018 | 0 | 0 | 0 | 2 | 0 | 12 | 5 | 9 | 15 | 18 | 0 | 0 | 1 | 0 | 0 | 18 | 3 | 0 | 11 | 2 | 263 | 149 | 1 | 1 | 0 | 0 | 0 | 0.07 | 0 | 0 | 0 | 1 | 0 | 0 | -0.186 |
| SE | 81470 | ENSG00000106477 | CEP41 | chr7 | - | 130426645 | 130426759 | 130421852 | 130422007 | 130427954 | 130428018 | 0 | 0 | 0 | 2 | 0 | 1 | 3 | 3 | 3 | 0 | 0 | 0 | 1 | 0 | 0 | 0 | 4 | 0 | 3 | 26 | 263 | 149 | 1 | 1 | 0 | 0 | 0 | 0.274 | NA | NA | 0 | 1 | 0 | 0 | -0.181 |
| SE | 81474 | ENSG00000091140 | DLD | chr7 | + | 107901737 | 107901817 | 107893199 | 107893278 | 107903477 | 107903547 | 132 | 87 | 153 | 153 | 87 | 0 | 0 | 1 | 0 | 0 | 2 | 93 | 65 | 38 | 30 | 0 | 0 | 1 | 0 | 0 | 229 | 149 | 1 | 1 | 1 | 1 | 0.99 | 1 | 1 | 1 | 1 | 0.977 | 1 | 1 | 0.003 |
| SE | 81476 | ENSG00000091140 | DLD | chr7 | + | 107904662 | 107904746 | 107903477 | 107903547 | 107904862 | 107905058 | 2 | 1 | 1 | 1 | 3 | 0 | 0 | 0 | 0 | 0 | 0 | 7 | 0 | 5 | 0 | 0 | 0 | 2 | 0 | 0 | 233 | 149 | 1 | 1 | 1 | 1 | 1 | 1 | 1 | NA | 1 | 0 | 1 | NA | 0.333 |
| SE | 81477 | ENSG00000091140 | DLD | chr7 | + | 107904662 | 107904746 | 107903477 | 107903547 | 107904957 | 107905058 | 2 | 1 | 2 | 1 | 1 | 94 | 46 | 87 | 97 | 71 | 0 | 0 | 0 | 5 | 0 | 2 | 58 | 74 | 24 | 3 | 233 | 149 | 1 | 1 | 0.013 | 0.014 | 0.014 | 0.007 | 0.009 | 0 | 0 | 0 | 0.118 | 0 | -0.012 |
| SE | 81478 | ENSG00000091140 | DLD | chr7 | + | 107904957 | 107905058 | 107903477 | 107903547 | 107906266 | 107906368 | 66 | 31 | 63 | 88 | 40 | 0 | 1 | 0 | 0 | 0 | 2 | 44 | 59 | 20 | 3 | 0 | 0 | 0 | 0 | 0 | 250 | 149 | 1 | 1 | 1 | 0.949 | 1 | 1 | 1 | 1 | 1 | 1 | 1 | 1 | -0.01 |
| SE | 81483 | ENSG00000188517 | COL25A1 | chr4 | - | 108819811 | 108819898 | 108817396 | 108817435 | 108824173 | 108824227 | 0 | 0 | 0 | 0 | 4 | 0 | 0 | 0 | 0 | 0 | 0 | 0 | 0 | 0 | 0 | 0 | 0 | 0 | 1 | 0 | 236 | 149 | 1 | 1 | NA | NA | NA | NA | 1 | NA | NA | NA | 0 | NA | 1 |
| SE | 81488 | ENSG00000188517 | COL25A1 | chr4 | - | 108860926 | 108860971 | 108859651 | 108859733 | 108863318 | 108863387 | 2 | 0 | 0 | 0 | 0 | 0 | 0 | 0 | 1 | 0 | 0 | 0 | 2 | 0 | 0 | 0 | 0 | 0 | 0 | 0 | 194 | 149 | 1 | 1 | 1 | NA | NA | 0 | NA | NA | NA | 1 | NA | NA | -0.5 |
| SE | 81494 | ENSG00000188517 | COL25A1 | chr4 | - | 109010357 | 109010375 | 108974532 | 108974559 | 109048167 | 109048175 | 0 | 1 | 0 | 0 | 0 | 0 | 0 | 0 | 1 | 0 | 5 | 0 | 0 | 0 | 7 | 0 | 0 | 0 | 0 | 0 | 167 | 149 | 1 | 1 | NA | 1 | NA | 0 | NA | 1 | NA | NA | NA | 1 | -0.5 |
| SE | 81495 | ENSG00000136193 | SCRN1 | chr7 | - | 29926451 | 29926632 | 29920103 | 29924115 | 29936555 | 29936721 | 127 | 94 | 107 | 139 | 97 | 2 | 0 | 0 | 1 | 0 | 103 | 60 | 35 | 45 | 19 | 0 | 0 | 1 | 1 | 0 | 298 | 149 | 1 | 1 | 0.969 | | 1 | 0.986 | 1 | 1 | 1 | 0.946 | 0.957 | 1 | 0.01 |
| SE | 81496 | ENSG00000136193 | SCRN1 | chr7 | - | 29940681 | 29940876 | 29936555 | 29936721 | 29943976 | 29944179 | 87 | 78 | 77 | 104 | 107 | 0 | 1 | 0 | 1 | 0 | 35 | 43 | 18 | 46 | 34 | 0 | 4 | 0 | 0 | 0 | 298 | 149 | 1 | 1 | 1 | 0.975 | 1 | 0.981 | 1 | 1 | 0.843 | 1 | 1 | 1 | 0.023 |
| SE | 81497 | ENSG00000136193 | SCRN1 | chr7 | - | 29955178 | 29955360 | 29943976 | 29944179 | 29968908 | 29969013 | 109 | 100 | 64 | 151 | 149 | 1 | 0 | 0 | 0 | 0 | 210 | 64 | 41 | 145 | 115 | 0 | 0 | 0 | 0 | 0 | 298 | 149 | 1 | 1 | 0.982 | 1 | 1 | 1 | 1 | 1 | 1 | 1 | 1 | 1 | -0.004 |
| SE | 81498 | ENSG00000136193 | SCRN1 | chr7 | - | 29968908 | 29969068 | 29955178 | 29955360 | 29989641 | 29989730 | 66 | 52 | 34 | 86 | 74 | 0 | 1 | 0 | 4 | 0 | 92 | 32 | 15 | 58 | 63 | 3 | 1 | 1 | 1 | 0 | 298 | 149 | 1 | 1 | 1 | 0.963 | 1 | 0.915 | 1 | 0.939 | 0.941 | 0.882 | 0.967 | 1 | 0.03 |
| SE | 81500 | ENSG00000106080 | FKBP14 | chr7 | - | 30020212 | 30020304 | 30018995 | 30019123 | 30022664 | 30022816 | 2 | 5 | 10 | 8 | 9 | 1 | 1 | 3 | 6 | 5 | 0 | 11 | 0 | 0 | 0 | 0 | 6 | 8 | 0 | 0 | 241 | 149 | 1 | 1 | 0.553 | 0.756 | 0.673 | 0.452 | 0.527 | NA | 0 | 0.459 | NA | NA | 0.363 |
| SE | 81503 | ENSG00000092820 | EZR | chr6 | - | 158776407 | 158776504 | 158771243 | 158771407 | 158783519 | 158783666 | 1881 | 1548 | 1632 | 1913 | 2244 | 0 | 0 | 0 | 0 | 0 | 4315 | 2584 | 3368 | 2477 | 3661 | 0 | 0 | 1 | 0 | 9 | 246 | 149 | 1 | 1 | 1 | 1 | 1 | 1 | 1 | 1 | 1 | 1 | 1 | 0.996 | 0.001 |
| SE | 81504 | ENSG00000092820 | EZR | chr6 | - | 158783519 | 158783666 | 158776407 | 158776504 | 158784643 | 158784727 | 1426 | 1093 | 1344 | 1418 | 1999 | 0 | 0 | 0 | 0 | 0 | 3256 | 2126 | 2459 | 1936 | 2570 | 0 | 1 | 1 | 1 | 0 | 296 | 149 | 1 | 1 | 1 | 1 | 1 | 1 | 1 | 1 | 0.999 | 0.999 | 0.999 | 1 | 0.001 |
| SE | 81505 | ENSG00000092820 | EZR | chr6 | - | 158784643 | 158784727 | 158783519 | 158783666 | 158785308 | 158785583 | 716 | 489 | 673 | 724 | 774 | 0 | 0 | 1 | 0 | 0 | 1293 | 1084 | 1036 | 824 | 1146 | 0 | 0 | 0 | 0 | 0 | 233 | 149 | 1 | 1 | 1 | 1 | 0.998 | 1 | 1 | 1 | 1 | 1 | 1 | 1 | 0 |
| SE | 81506 | ENSG00000092820 | EZR | chr6 | - | 158785308 | 158785583 | 158784643 | 158784727 | 158789287 | 158789371 | 465 | 302 | 378 | 460 | 469 | 0 | 0 | 0 | 0 | 0 | 874 | 711 | 632 | 530 | 687 | 0 | 0 | 0 | 0 | 6 | 298 | 149 | 1 | 1 | 1 | 1 | 1 | 1 | 1 | 1 | 1 | 1 | 1 | 0.983 | 0.003 |
| SE | 81508 | ENSG00000092820 | EZR | chr6 | - | 158788313 | 158788479 | 158787107 | 158787203 | 158789287 | 158789371 | 0 | 0 | 1 | 1 | 0 | 545 | 387 | 505 | 511 | 956 | 0 | 0 | 2 | 0 | 0 | 696 | 684 | 706 | 451 | 625 | 298 | 149 | 1 | 1 | 0 | 0 | 0.001 | 0.001 | 0 | 0 | 0 | 0.001 | 0 | 0 | 0 |
| SE | 81514 | ENSG00000131171 | SH3BGRL | chrX | + | 81276983 | 81277169 | 81202101 | 81202245 | 81278330 | 81278411 | 1739 | 1444 | 1190 | 1908 | 2178 | 0 | 1 | 0 | 0 | 0 | 1537 | 1386 | 849 | 738 | 1975 | 0 | 0 | 0 | 0 | 0 | 298 | 149 | 1 | 1 | 1 | 0.999 | 1 | 1 | 1 | 1 | 1 | 1 | 1 | 1 | 0 |
| SE | 81515 | ENSG00000071462 | BUD23 | chr7 | + | 73683766 | 73683804 | 73683570 | 73683673 | 73686635 | 73686731 | 31 | 9 | 15 | 13 | 7 | 2 | 0 | 0 | 0 | 0 | 29 | 24 | 34 | 19 | 55 | 0 | 0 | 0 | 0 | 0 | 187 | 149 | 1 | 1 | 0.925 | 1 | 1 | 1 | 1 | 1 | 1 | 1 | 1 | 1 | -0.015 |
| SE | 81516 | ENSG00000071462 | BUD23 | chr7 | + | 73683766 | 73683901 | 73683600 | 73683673 | 73686635 | 73686731 | 0 | 2 | 3 | 3 | 0 | 2 | 0 | 0 | 0 | 0 | 0 | 4 | 4 | 5 | 0 | 0 | 0 | 0 | 0 | 0 | 284 | 149 | 1 | 1 | 0 | 1 | 1 | 1 | NA | NA | 1 | 1 | 1 | NA | -0.25 |
| SE | 81517 | ENSG00000071462 | BUD23 | chr7 | + | 73683766 | 73683923 | 73683568 | 73683673 | 73686635 | 73686731 | 0 | 2 | 3 | 0 | 0 | 2 | 0 | 0 | 0 | 0 | 0 | 4 | 2 | 2 | 0 | 0 | 0 | 0 | 0 | 0 | 298 | 149 | 1 | 1 | 0 | 1 | 1 | NA | NA | NA | 1 | 1 | 1 | NA | -0.333 |
| SE | 81518 | ENSG00000071462 | BUD23 | chr7 | + | 73683766 | 73684000 | 73683625 | 73683673 | 73686635 | 73686731 | 0 | 2 | 3 | 5 | 7 | 2 | 0 | 0 | 0 | 0 | 0 | 4 | 7 | 2 | 0 | 0 | 0 | 0 | 0 | 0 | 298 | 149 | 1 | 1 | 0 | 1 | 1 | 1 | 1 | NA | 1 | 1 | 1 | NA | -0.2 |
| SE | 81523 | ENSG00000071462 | BUD23 | chr7 | + | 73693328 | 73693414 | 73692595 | 73692646 | 73693623 | 73693669 | 193 | 74 | 192 | 178 | 110 | 0 | 0 | 0 | 1 | 0 | 236 | 247 | 190 | 140 | 215 | 5 | 0 | 0 | 0 | 0 | 235 | 149 | 1 | 1 | 1 | 1 | 1 | 0.991 | 1 | 0.968 | 1 | 1 | 1 | 1 | 0.005 |
| SE | 81526 | ENSG00000071462 | BUD23 | chr7 | + | 73693623 | 73693669 | 73692595 | 73693414 | 73693991 | 73694004 | 165 | 71 | 168 | 169 | 99 | 0 | 0 | 0 | 0 | 0 | 252 | 210 | 166 | 123 | 183 | 0 | 0 | 2 | 0 | 0 | 195 | 149 | 1 | 1 | 1 | 1 | 1 | 1 | 1 | 1 | 1 | 0.984 | 1 | 1 | 0.003 |
| SE | 81530 | ENSG00000071462 | BUD23 | chr7 | + | 73697465 | 73697516 | 73693991 | 73694050 | 73697604 | 73697694 | 3 | 1 | 1 | 2 | 2 | 105 | 44 | 96 | 108 | 78 | 4 | 0 | 0 | 0 | 2 | 117 | 112 | 103 | 62 | 103 | 200 | 149 | 1 | 1 | 0.021 | 0.017 | 0.008 | 0.014 | 0.019 | 0.025 | 0 | 0 | 0 | 0.014 | 0.008 |
| SE | 81531 | ENSG00000106089 | STX1A | chr7 | - | 73716580 | 73716734 | 73709044 | 73709122 | 73719601 | 73719626 | 0 | 0 | 0 | 0 | 0 | 3 | 6 | 0 | 4 | 9 | 0 | 3 | 0 | 0 | 0 | 22 | 3 | 7 | 7 | 28 | 298 | 149 | 1 | 1 | 0 | 0 | NA | 0 | 0 | 0 | 0.333 | 0 | 0 | 0 | -0.067 |
| SE | 81533 | ENSG00000106344 | RBM28 | chr7 | - | 128317658 | 128317733 | 128314763 | 128315020 | 128317956 | 128318106 | 99 | 65 | 78 | 97 | 96 | 3 | 1 | 0 | 0 | 0 | 139 | 101 | 92 | 53 | 40 | 0 | 0 | 2 | 2 | 0 | 224 | 149 | 1 | 1 | 0.956 | 0.977 | | 1 | 1 | 1 | 1 | 0.968 | 0.946 | 1 | 0.004 |
| SE | 81536 | ENSG00000106344 | RBM28 | chr7 | - | 128337130 | 128337202 | 128335846 | 128336042 | 128338249 | 128338342 | 119 | 105 | 98 | 133 | 165 | 0 | 0 | 0 | 0 | 0 | 200 | 90 | 107 | 186 | 35 | 0 | 0 | 0 | 1 | 0 | 221 | 149 | 1 | 1 | 1 | 1 | 1 | 1 | 1 | 1 | 1 | 1 | 0.992 | 1 | 0.002 |
| SE | 81537 | ENSG00000106344 | RBM28 | chr7 | - | 128337130 | 128337202 | 128335846 | 128336042 | 128338725 | 128338801 | 32 | 28 | 29 | 43 | 63 | 0 | 0 | 0 | 0 | 0 | 54 | 13 | 17 | 53 | 4 | 0 | 0 | 0 | 4 | 0 | 221 | 149 | 1 | 1 | 1 | 1 | 1 | 1 | 1 | 1 | 1 | 1 | 0.899 | 1 | 0.02 |
| SE | 81538 | ENSG00000106344 | RBM28 | chr7 | - | 128338249 | 128338342 | 128335846 | 128336042 | 128338725 | 128338801 | 66 | 40 | 37 | 63 | 75 | 0 | 0 | 0 | 0 | 0 | 31 | 27 | 45 | 67 | 24 | 0 | 0 | 0 | 4 | 0 | 242 | 149 | 1 | 1 | 1 | 1 | 1 | 1 | 1 | 1 | 1 | 1 | 0.912 | 1 | 0.018 |
| SE | 81540 | ENSG00000106344 | RBM28 | chr7 | - | 128339226 | 128339321 | 128338725 | 128338801 | 128339632 | 128339791 | 135 | 91 | 97 | 131 | 181 | 0 | 0 | 0 | 0 | 0 | 63 | 66 | 91 | 142 | 37 | 0 | 0 | 1 | 0 | 0 | 244 | 149 | 1 | 1 | 1 | 1 | 1 | 1 | 1 | 1 | 1 | 0.982 | 1 | 1 | 0.004 |
| SE | 81541 | ENSG00000106344 | RBM28 | chr7 | - | 128341117 | 128341219 | 128339632 | 128339791 | 128343675 | 128343908 | 0 | 0 | 0 | 1 | 0 | 35 | 13 | 29 | 26 | 13 | 0 | 0 | 0 | 0 | 0 | 57 | 74 | 19 | 27 | 0 | 251 | 149 | 1 | 1 | 0 | 0 | 0 | 0.022 | 0 | 0 | 0 | 0 | 0 | NA | 0.004 |
| SE | 81543 | ENSG00000106348 | IMPDH1 | chr7 | - | 128394888 | 128395033 | 128394455 | 128394599 | 128395130 | 128395274 | 449 | 229 | 310 | 345 | 247 | 0 | 0 | 2 | 0 | 0 | 836 | 408 | 338 | 489 | 1027 | 0 | 0 | 0 | 0 | 0 | 294 | 149 | 1 | 1 | 1 | 1 | 0.987 | 1 | 1 | 1 | 1 | 1 | 1 | 1 | -0.003 |
| SE | 81544 | ENSG00000106348 | IMPDH1 | chr7 | - | 128396599 | 128396695 | 128395130 | 128395274 | 128396931 | 128397022 | 378 | 194 | 287 | 295 | 245 | 0 | 0 | 1 | 2 | 0 | 802 | 363 | 274 | 415 | 939 | 0 | 0 | 0 | 0 | 0 | 245 | 149 | 1 | 1 | 1 | 1 | 0.994 | 0.989 | 1 | 1 | 1 | 1 | 1 | 1 | -0.003 |
| SE | 81545 | ENSG00000106348 | IMPDH1 | chr7 | - | 128400816 | 128400891 | 128400332 | 128400539 | 128401014 | 128401116 | 99 | 45 | 56 | 68 | 60 | 0 | 4 | 0 | 0 | 0 | 177 | 109 | 80 | 97 | 208 | 0 | 0 | 0 | 0 | 0 | 224 | 149 | 1 | 1 | 1 | 0.882 | 1 | 1 | 1 | 1 | 1 | 1 | 1 | 1 | -0.024 |
| SE | 81547 | ENSG00000106348 | IMPDH1 | chr7 | - | 128405766 | 128405865 | 128403705 | 128403754 | 128409288 | 128409352 | 54 | 10 | 32 | 18 | 7 | 0 | 0 | 1 | 0 | 0 | 46 | 39 | 10 | 19 | 116 | 0 | 0 | 0 | 0 | 0 | 248 | 149 | 1 | 1 | 1 | 1 | 0.951 | 1 | 1 | 1 | 1 | 1 | 1 | 1 | -0.01 |
| SE | 81550 | ENSG00000106348 | IMPDH1 | chr7 | - | 128409288 | 128409352 | 128405766 | 128405865 | 128409440 | 128409484 | 4 | 1 | 4 | 1 | 0 | 0 | 0 | 0 | 0 | 0 | 6 | 5 | 0 | 2 | 33 | 0 | 1 | 0 | 0 | 0 | 213 | 149 | 1 | 1 | 1 | 1 | 1 | 1 | NA | 1 | 0.778 | NA | 1 | 1 | 0.055 |
| SE | 81554 | ENSG00000106348 | IMPDH1 | chr7 | - | 128409440 | 128409484 | 128405766 | 128405865 | 128409755 | 128409901 | 2 | 0 | 0 | 0 | 0 | 15 | 13 | 9 | 13 | 1 | 0 | 1 | 0 | 0 | 0 | 6 | 6 | 4 | 2 | 32 | 193 | 149 | 1 | 1 | 0.093 | 0 | 0 | 0 | 0 | 0 | 0.114 | 0 | 0 | 0 | -0.004 |
| SE | 81556 | ENSG00000167703 | SLC43A2 | chr17 | - | 1583203 | 1583336 | 1578249 | 1578323 | 1585912 | 1586051 | 94 | 77 | 69 | 89 | 67 | 0 | 0 | 1 | 0 | 0 | 149 | 108 | 86 | 122 | 245 | 0 | 0 | 0 | 0 | 0 | 282 | 149 | 1 | 1 | 1 | 1 | 0.973 | 1 | 1 | 1 | 1 | 1 | 1 | 1 | -0.005 |
| SE | 81563 | ENSG00000167703 | SLC43A2 | chr17 | - | 1616716 | 1616769 | 1614978 | 1615034 | 1627714 | 1627911 | 7 | 0 | 3 | 1 | 0 | 1 | 0 | 0 | 0 | 0 | 18 | 4 | 0 | 1 | 18 | 2 | 0 | 2 | 0 | 27 | 202 | 149 | 1 | 1 | 0.838 | NA | 1 | 1 | NA | 0.869 | 1 | 0 | 1 | 0.33 | 0.306 |
| SE | 81564 | ENSG00000167703 | SLC43A2 | chr17 | - | 1616716 | 1616769 | 1616561 | 1616630 | 1627714 | 1627911 | 7 | 0 | 3 | 1 | 0 | 1 | 0 | 0 | 0 | 0 | 0 | 4 | 1 | 1 | 18 | 0 | 0 | 0 | 1 | 0 | 202 | 149 | 1 | 1 | 0.838 | NA | 1 | 1 | NA | NA | 1 | 1 | 0.425 | 1 | 0.09 |
| SE | 81573 | ENSG00000132256 | TRIM5 | chr11 | - | 5683985 | 5684139 | 5679961 | 5680238 | 5684867 | 5685043 | 2 | 0 | 0 | 1 | 0 | 43 | 52 | 41 | 55 | 61 | 0 | 0 | 0 | 0 | 0 | 29 | 33 | 18 | 59 | 43 | 298 | 149 | 1 | 1 | 0.023 | 0 | 0 | 0.009 | 0 | 0 | 0 | 0 | 0 | 0 | 0.006 |
| SE | 81576 | ENSG00000165055 | METTL2B | chr7 | + | 128488100 | 128488161 | 128480646 | 128480696 | 128498035 | 128498142 | 47 | 39 | 46 | 56 | 60 | 0 | 0 | 0 | 3 | 0 | 126 | 27 | 25 | 43 | 77 | 0 | 0 | 0 | 0 | 0 | 210 | 149 | 1 | 1 | 1 | 1 | 1 | 0.93 | 1 | 1 | 1 | 1 | 1 | 1 | -0.014 |
| SE | 81577 | ENSG00000165055 | METTL2B | chr7 | + | 128493803 | 128493943 | 128480646 | 128480696 | 128498035 | 128498142 | 52 | 29 | 50 | 30 | 45 | 0 | 0 | 0 | 3 | 0 | 22 | 33 | 11 | 21 | 28 | 0 | 0 | 0 | 0 | 0 | 289 | 149 | 1 | 1 | 1 | 1 | 1 | 0.838 | 1 | 1 | 1 | 1 | 1 | 1 | -0.032 |
| SE | 81578 | ENSG00000165055 | METTL2B | chr7 | + | 128493803 | 128493943 | 128488100 | 128488161 | 128498035 | 128498142 | 104 | 45 | 100 | 52 | 116 | 0 | 0 | 0 | 0 | 0 | 34 | 72 | 12 | 41 | 88 | 0 | 0 | 0 | 1 | 0 | 289 | 149 | 1 | 1 | 1 | 1 | 1 | 1 | 1 | 1 | 1 | 1 | 0.955 | 1 | 0.009 |
| SE | 81580 | ENSG00000165055 | METTL2B | chr7 | + | 128500902 | 128500968 | 128498035 | 128498142 | 128501761 | 128501832 | 51 | 19 | 41 | 27 | 34 | 1 | 0 | 0 | 0 | 0 | 39 | 13 | 8 | 22 | 13 | 0 | 0 | 0 | 0 | 0 | 215 | 149 | 1 | 1 | 0.972 | 1 | 1 | 1 | 1 | 1 | 1 | 1 | 1 | 1 | -0.006 |
| SE | 81581 | ENSG00000253729 | PRKDC | chr8 | - | 47776843 | 47776983 | 47773110 | 47774377 | 47777685 | 47777874 | 221 | 159 | 210 | 256 | 476 | 0 | 0 | 1 | 0 | 0 | 93 | 178 | 265 | 200 | 22 | 0 | 0 | 0 | 0 | 0 | 289 | 149 | 1 | 1 | 1 | 1 | 0.991 | 1 | 1 | 1 | 1 | 1 | 1 | 1 | -0.002 |
| SE | 81582 | ENSG00000253729 | PRKDC | chr8 | - | 47777685 | 47777874 | 47776843 | 47776983 | 47778458 | 47778537 | 208 | 153 | 213 | 221 | 295 | 0 | 0 | 0 | 0 | 0 | 72 | 162 | 269 | 140 | 39 | 0 | 0 | 0 | 4 | 0 | 298 | 149 | 1 | 1 | 1 | 1 | 1 | 1 | 1 | 1 | 1 | 1 | 0.946 | 1 | 0.011 |
| SE | 81585 | ENSG00000253729 | PRKDC | chr8 | - | 47783741 | 47783809 | 47782377 | 47782598 | 47785112 | 47785317 | 215 | 149 | 191 | 253 | 306 | 0 | 0 | 0 | 1 | 0 | 53 | 177 | 162 | 165 | 26 | 0 | 2 | 0 | 1 | 0 | 217 | 149 | 1 | 1 | 1 | 1 | 1 | 0.994 | 1 | 1 | 0.984 | 1 | 0.991 | 1 | 0.004 |
| SE | 81586 | ENSG00000253729 | PRKDC | chr8 | - | 47799209 | 47799390 | 47798236 | 47798397 | 47800792 | 47800986 | 122 | 100 | 113 | 149 | 158 | 0 | 0 | 0 | 0 | 0 | 54 | 55 | 96 | 90 | 16 | 0 | 0 | 2 | 0 | 0 | 298 | 149 | 1 | 1 | 1 | 1 | 1 | 1 | 1 | 1 | 1 | 0.96 | 1 | 1 | 0.008 |
| SE | 81587 | ENSG00000253729 | PRKDC | chr8 | - | 47817449 | 47817561 | 47807136 | 47807326 | 47819401 | 47819510 | 443 | 358 | 355 | 445 | 410 | 0 | 0 | 0 | 0 | 0 | 210 | 318 | 418 | 504 | 30 | 0 | 0 | 0 | 3 | 0 | 261 | 149 | 1 | 1 | 1 | 1 | 1 | 1 | 1 | 1 | 1 | 1 | 0.99 | 1 | 0.002 |
| SE | 81588 | ENSG00000253729 | PRKDC | chr8 | - | 47820718 | 47820943 | 47819401 | 47819510 | 47821603 | 47821792 | 235 | 218 | 233 | 264 | 308 | 0 | 0 | 0 | 0 | 0 | 86 | 173 | 249 | 266 | 43 | 0 | 0 | 0 | 3 | 0 | 298 | 149 | 1 | 1 | 1 | 1 | 1 | 1 | 1 | 1 | 1 | 1 | 0.978 | 1 | 0.004 |
| SE | 81589 | ENSG00000253729 | PRKDC | chr8 | - | 47826655 | 47826861 | 47823857 | 47823996 | 47828167 | 47828347 | 158 | 107 | 133 | 171 | 156 | 0 | 2 | 0 | 0 | 4 | 46 | 131 | 155 | 99 | 12 | 0 | 3 | 0 | 1 | 0 | 298 | 149 | 1 | 1 | 1 | 0.964 | 1 | 1 | 0.951 | 1 | 0.956 | 1 | 0.98 | 1 | -0.004 |
| SE | 81590 | ENSG00000253729 | PRKDC | chr8 | - | 47836337 | 47836527 | 47834195 | 47834396 | 47837211 | 47837419 | 218 | 172 | 193 | 191 | 234 | 0 | 0 | 0 | 1 | 0 | 81 | 160 | 218 | 227 | 82 | 0 | 2 | 0 | 0 | 0 | 298 | 149 | 1 | 1 | 1 | 1 | 1 | 0.99 | 1 | 1 | 0.976 | 1 | 1 | 1 | 0.003 |
| SE | 81591 | ENSG00000253729 | PRKDC | chr8 | - | 47837211 | 47837419 | 47834195 | 47834396 | 47839147 | 47839246 | 240 | 171 | 196 | 171 | 517 | 0 | 0 | 1 | 0 | 0 | 87 | 218 | 196 | 184 | 21 | 0 | 0 | 0 | 0 | 0 | 298 | 149 | 1 | 1 | 1 | 1 | 0.99 | 1 | 1 | 1 | 1 | 1 | 1 | 1 | -0.002 |
| SE | 81592 | ENSG00000253729 | PRKDC | chr8 | - | 47839147 | 47839246 | 47837211 | 47837419 | 47840015 | 47840189 | 506 | 372 | 416 | 416 | 899 | 2 | 1 | 0 | 1 | 1 | 162 | 421 | 420 | 377 | 45 | 0 | 0 | 0 | 0 | 0 | 248 | 149 | 1 | 1 | 0.993 | 0.996 | 1 | 0.996 | 0.998 | 1 | 1 | 1 | 1 | 1 | -0.003 |
| SE | 81593 | ENSG00000253729 | PRKDC | chr8 | - | 47854082 | 47854214 | 47852672 | 47852784 | 47857155 | 47857299 | 98 | 56 | 80 | 95 | 121 | 0 | 0 | 0 | 0 | 0 | 61 | 136 | 127 | 131 | 39 | 0 | 1 | 0 | 4 | 0 | 281 | 149 | 1 | 1 | 1 | 1 | 1 | 1 | 1 | 1 | 0.986 | 1 | 0.946 | 1 | 0.014 |
| SE | 81595 | ENSG00000253729 | PRKDC | chr8 | - | 47864555 | 47864763 | 47863398 | 47863577 | 47877723 | 47877851 | 289 | 172 | 302 | 309 | 309 | 0 | 0 | 0 | 0 | 0 | 164 | 253 | 303 | 298 | 20 | 0 | 0 | 1 | 0 | 0 | 298 | 149 | 1 | 1 | 1 | 1 | 1 | 1 | 1 | 1 | 1 | 0.993 | 1 | 1 | 0.001 |
| SE | 81596 | ENSG00000253729 | PRKDC | chr8 | - | 47885943 | 47886147 | 47881911 | 47882097 | 47887546 | 47887705 | 253 | 169 | 223 | 262 | 248 | 2 | 0 | 3 | 0 | 0 | 111 | 182 | 226 | 214 | 18 | 0 | 0 | 0 | 1 | 0 | 298 | 149 | 1 | 1 | 0.984 | 1 | 0.974 | 1 | 1 | 1 | 1 | 1 | 0.991 | 1 | -0.007 |
| SE | 81597 | ENSG00000253729 | PRKDC | chr8 | - | 47885943 | 47886147 | 47881911 | 47882097 | 47888517 | 47888650 | 170 | 108 | 135 | 160 | 139 | 0 | 0 | 0 | 0 | 0 | 56 | 92 | 134 | 144 | 6 | 0 | 0 | 0 | 1 | 0 | 298 | 149 | 1 | 1 | 1 | 1 | 1 | 1 | 1 | 1 | 1 | 1 | 0.986 | 1 | 0.003 |
| SE | 81598 | ENSG00000253729 | PRKDC | chr8 | - | 47887546 | 47887705 | 47881911 | 47882097 | 47888517 | 47888650 | 78 | 31 | 58 | 67 | 64 | 0 | 0 | 0 | 0 | 0 | 9 | 29 | 61 | 45 | 7 | 0 | 0 | 0 | 1 | 0 | 298 | 149 | 1 | 1 | 1 | 1 | 1 | 1 | 1 | 1 | 1 | 1 | 0.957 | 1 | 0.009 |
| SE | 81601 | ENSG00000253729 | PRKDC | chr8 | - | 47897160 | 47897294 | 47893138 | 47893387 | 47898469 | 47898569 | 162 | 97 | 124 | 179 | 171 | 0 | 1 | 0 | 0 | 0 | 38 | 109 | 85 | 64 | 21 | 0 | 0 | 0 | 0 | 0 | 283 | 149 | 1 | 1 | 1 | 0.981 | 1 | 1 | 1 | 1 | 1 | 1 | 1 | 1 | -0.004 |
| SE | 81602 | ENSG00000253729 | PRKDC | chr8 | - | 47897160 | 47897294 | 47893138 | 47893387 | 47900372 | 47900467 | 19 | 8 | 32 | 33 | 48 | 0 | 0 | 0 | 1 | 0 | 0 | 17 | 18 | 16 | 0 | 0 | 0 | 4 | 0 | 0 | 283 | 149 | 1 | 1 | 1 | 1 | 1 | 1 | 0.962 | NA | 1 | 0.703 | 1 | NA | 0.091 |
| SE | 81603 | ENSG00000253729 | PRKDC | chr8 | - | 47898469 | 47898569 | 47893138 | 47893387 | 47900372 | 47900467 | 99 | 60 | 76 | 83 | 97 | 0 | 0 | 0 | 1 | 1 | 17 | 47 | 30 | 43 | 4 | 0 | 0 | 4 | 0 | 0 | 249 | 149 | 1 | 1 | 1 | 1 | 1 | 1 | 0.983 | 1 | 1 | 0.818 | 1 | 1 | 0.033 |
| SE | 81604 | ENSG00000253729 | PRKDC | chr8 | - | 47898469 | 47898569 | 47897160 | 47897294 | 47900372 | 47900467 | 239 | 152 | 176 | 230 | 217 | 0 | 0 | 0 | 0 | 0 | 55 | 147 | 103 | 96 | 25 | 0 | 0 | 3 | 0 | 0 | 249 | 149 | 1 | 1 | 1 | 1 | 1 | 1 | 1 | 1 | 1 | 0.954 | 1 | 1 | 0.009 |
| SE | 81606 | ENSG00000253729 | PRKDC | chr8 | - | 47927770 | 47927890 | 47926832 | 47927353 | 47929091 | 47929169 | 316 | 263 | 288 | 322 | 672 | 0 | 0 | 0 | 0 | 0 | 225 | 347 | 324 | 327 | 61 | 0 | 2 | 0 | 0 | 0 | 269 | 149 | 1 | 1 | 1 | 1 | 1 | 1 | 1 | 1 | 0.99 | 1 | 1 | 1 | 0.002 |
| SE | 81608 | ENSG00000253729 | PRKDC | chr8 | - | 47935008 | 47935058 | 47933964 | 47934090 | 47936352 | 47936517 | 52 | 32 | 42 | 41 | 47 | 0 | 0 | 0 | 0 | 0 | 5 | 33 | 61 | 46 | 21 | 0 | 0 | 2 | 0 | 0 | 199 | 149 | 1 | 1 | 1 | 1 | 1 | 1 | 1 | 1 | 1 | 0.958 | 1 | 1 | 0.008 |
| SE | 81611 | ENSG00000253729 | PRKDC | chr8 | - | 47955873 | 47955948 | 47954337 | 47954446 | 47957170 | 47957263 | 315 | 284 | 247 | 311 | 487 | 0 | 0 | 1 | 0 | 0 | 158 | 228 | 235 | 267 | 109 | 0 | 0 | 0 | 0 | 0 | 224 | 149 | 1 | 1 | 1 | 1 | 0.994 | 1 | 1 | 1 | 1 | 1 | 1 | 1 | -0.001 |
| SE | 81613 | ENSG00000161036 | LRWD1 | chr7 | + | 102472453 | 102472609 | 102472217 | 102472309 | 102472691 | 102472804 | 27 | 10 | 14 | 18 | 0 | 0 | 1 | 0 | 0 | 0 | 21 | 20 | 30 | 22 | 40 | 6 | 0 | 0 | 0 | 0 | 298 | 149 | 1 | 1 | 1 | 0.833 | 1 | 1 | NA | 0.636 | 1 | 1 | 1 | 1 | 0.031 |
| SE | 81614 | ENSG00000068976 | PYGM | chr11 | - | 64747223 | 64747358 | 64746920 | 64746987 | 64750375 | 64750583 | 20 | 8 | 19 | 16 | 18 | 0 | 1 | 0 | 0 | 0 | 17 | 31 | 14 | 2 | 33 | 0 | 0 | 0 | 0 | 0 | 284 | 149 | 1 | 1 | 1 | 0.808 | 1 | 1 | 1 | 1 | 1 | 1 | 1 | 1 | -0.038 |
| SE | 81615 | ENSG00000068976 | PYGM | chr11 | - | 64747223 | 64747358 | 64746920 | 64746987 | 64751324 | 64751466 | 16 | 7 | 14 | 9 | 12 | 0 | 0 | 0 | 1 | 0 | 17 | 30 | 14 | 3 | 21 | 0 | 0 | 0 | 0 | 0 | 284 | 149 | 1 | 1 | 1 | 1 | 1 | 0.825 | 1 | 1 | 1 | 1 | 1 | 1 | -0.035 |
| SE | 81616 | ENSG00000068976 | PYGM | chr11 | - | 64750375 | 64750583 | 64746920 | 64746987 | 64751324 | 64751466 | 7 | 6 | 8 | 5 | 7 | 0 | 0 | 0 | 1 | 0 | 6 | 5 | 10 | 3 | 3 | 0 | 0 | 0 | 0 | 0 | 298 | 149 | 1 | 1 | 1 | 1 | 1 | 0.714 | 1 | 1 | 1 | 1 | 1 | 1 | -0.057 |
| SE | 81619 | ENSG00000068976 | PYGM | chr11 | - | 64758245 | 64758349 | 64757778 | 64757910 | 64758436 | 64758515 | 6 | 0 | 6 | 10 | 1 | 0 | 0 | 0 | 1 | 0 | 0 | 1 | 5 | 0 | 0 | 4 | 0 | 0 | 0 | 0 | 253 | 149 | 1 | 1 | 1 | NA | 1 | 0.855 | 1 | 0 | 1 | 1 | NA | NA | 0.297 |
| SE | 81621 | ENSG00000135093 | USP30 | chr12 | + | 109054679 | 109054792 | 109052590 | 109052761 | 109056681 | 109056791 | 0 | 2 | 2 | 0 | 10 | 6 | 4 | 8 | 6 | 0 | 8 | 0 | 2 | 0 | 0 | 7 | 0 | 4 | 7 | 0 | 262 | 149 | 1 | 1 | 0 | 0.221 | 0.124 | 0 | 1 | 0.394 | NA | 0.221 | 0 | NA | 0.064 |
| SE | 81622 | ENSG00000135093 | USP30 | chr12 | + | 109056681 | 109056791 | 109052575 | 109052761 | 109057925 | 109058096 | 15 | 19 | 19 | 9 | 6 | 0 | 0 | 0 | 1 | 0 | 31 | 7 | 12 | 10 | 0 | 0 | 0 | 0 | 0 | 0 | 259 | 149 | 1 | 1 | 1 | 1 | 1 | 0.838 | 1 | 1 | 1 | 1 | 1 | NA | -0.032 |
| SE | 81626 | ENSG00000135093 | USP30 | chr12 | + | 109071611 | 109071710 | 109067523 | 109067627 | 109072304 | 109072347 | 32 | 18 | 24 | 23 | 102 | 0 | 0 | 0 | 1 | 0 | 87 | 25 | 27 | 51 | 36 | 0 | 0 | 0 | 0 | 0 | 248 | 149 | 1 | 1 | 1 | 1 | 1 | 0.933 | 1 | 1 | 1 | 1 | 1 | 1 | -0.013 |
| SE | 81627 | ENSG00000135093 | USP30 | chr12 | + | 109072304 | 109072350 | 109067523 | 109067627 | 109073437 | 109073532 | 3 | 6 | 5 | 5 | 6 | 0 | 0 | 1 | 0 | 0 | 43 | 3 | 3 | 7 | 0 | 0 | 0 | 6 | 0 | 0 | 195 | 149 | 1 | 1 | 1 | 1 | 0.793 | 1 | 1 | 1 | 1 | 0.276 | 1 | NA | 0.14 |
| SE | 81628 | ENSG00000135093 | USP30 | chr12 | + | 109081333 | 109081393 | 109073437 | 109073532 | 109081932 | 109082019 | 47 | 23 | 49 | 44 | 69 | 0 | 1 | 0 | 0 | 0 | 102 | 27 | 95 | 63 | 102 | 0 | 0 | 0 | 0 | 0 | 209 | 149 | 1 | 1 | 1 | 0.943 | 1 | 1 | 1 | 1 | 1 | 1 | 1 | 1 | -0.011 |
| SE | 81629 | ENSG00000135093 | USP30 | chr12 | + | 109082662 | 109082743 | 109081932 | 109082019 | 109082842 | 109082917 | 60 | 37 | 68 | 59 | 134 | 0 | 0 | 0 | 0 | 0 | 96 | 45 | 142 | 75 | 105 | 0 | 0 | 0 | 1 | 0 | 230 | 149 | 1 | 1 | 1 | 1 | 1 | 1 | 1 | 1 | 1 | 1 | 0.98 | 1 | 0.004 |
| SE | 81631 | ENSG00000130856 | ZNF236 | chr18 | + | 76871700 | 76871825 | 76868684 | 76868863 | 76875491 | 76875664 | 152 | 97 | 128 | 143 | 196 | 0 | 0 | 0 | 0 | 0 | 240 | 172 | 131 | 101 | 177 | 0 | 0 | 2 | 4 | 0 | 274 | 149 | 1 | 1 | 1 | 1 | 1 | 1 | 1 | 1 | 1 | 0.973 | 0.932 | 1 | 0.019 |
| SE | 81632 | ENSG00000130856 | ZNF236 | chr18 | + | 76881283 | 76881512 | 76880112 | 76880316 | 76895012 | 76895285 | 45 | 45 | 43 | 56 | 121 | 0 | 0 | 2 | 0 | 0 | 29 | 45 | 35 | 55 | 10 | 0 | 0 | 0 | 0 | 0 | 298 | 149 | 1 | 1 | 1 | 1 | 0.915 | 1 | 1 | 1 | 1 | 1 | 1 | 1 | -0.017 |
| SE | 81634 | ENSG00000130856 | ZNF236 | chr18 | + | 76927926 | 76928106 | 76927276 | 76927517 | 76937155 | 76937343 | 18 | 18 | 23 | 30 | 13 | 0 | 0 | 0 | 0 | 2 | 12 | 19 | 27 | 14 | 0 | 0 | 0 | 0 | 0 | 0 | 298 | 149 | 1 | 1 | 1 | 1 | 1 | 1 | 0.765 | 1 | 1 | 1 | 1 | NA | -0.047 |
| SE | 81636 | ENSG00000130856 | ZNF236 | chr18 | + | 76935979 | 76936061 | 76927926 | 76928106 | 76937155 | 76937343 | 5 | 2 | 1 | 1 | 4 | 10 | 11 | 15 | 9 | 4 | 0 | 0 | 0 | 0 | 0 | 12 | 9 | 15 | 9 | 0 | 231 | 149 | 1 | 1 | 0.244 | 0.105 | 0.041 | 0.067 | 0.392 | 0 | 0 | 0 | 0 | NA | 0.17 |
| SE | 81637 | ENSG00000130856 | ZNF236 | chr18 | + | 76947520 | 76947652 | 76937155 | 76937343 | 76959686 | 76959816 | 8 | 5 | 11 | 4 | 0 | 0 | 0 | 2 | 0 | 0 | 0 | 10 | 4 | 9 | 0 | 0 | 0 | 0 | 2 | 0 | 281 | 149 | 1 | 1 | 1 | 1 | 0.745 | 1 | NA | NA | 1 | 1 | 0.705 | NA | 0.035 |
| SE | 81640 | ENSG00000133195 | SLC39A11 | chr17 | - | 72817869 | 72817935 | 72736649 | 72736719 | 72849633 | 72849776 | 0 | 0 | 0 | 1 | 0 | 27 | 11 | 31 | 16 | 9 | 0 | 0 | 0 | 0 | 0 | 50 | 44 | 14 | 63 | 26 | 215 | 149 | 1 | 1 | 0 | 0 | 0 | 0.042 | 0 | 0 | 0 | 0 | 0 | 0 | 0.008 |
| SE | 81644 | ENSG00000133195 | SLC39A11 | chr17 | - | 73091823 | 73092116 | 73088656 | 73088775 | 73092494 | 73092687 | 1 | 2 | 0 | 0 | 0 | 0 | 0 | 0 | 2 | 0 | 6 | 0 | 0 | 0 | 0 | 0 | 9 | 2 | 2 | 0 | 298 | 149 | 1 | 1 | 1 | 1 | NA | 0 | NA | 1 | 0 | 0 | 0 | NA | 0.417 |
| SE | 81647 | ENSG00000130948 | HSD17B3 | chr9 | - | 96240757 | 96240907 | 96235305 | 96235570 | 96244290 | 96244394 | 11 | 3 | 0 | 2 | 9 | 0 | 2 | 0 | 0 | 0 | 16 | 8 | 0 | 4 | 0 | 0 | 0 | 0 | 0 | 0 | 298 | 149 | 1 | 1 | 1 | 0.429 | NA | 1 | 1 | 1 | 1 | NA | 1 | NA | -0.143 |
| SE | 81650 | ENSG00000130948 | HSD17B3 | chr9 | - | 96246555 | 96246590 | 96245344 | 96245426 | 96251417 | 96251485 | 0 | 1 | 0 | 0 | 0 | 0 | 0 | 0 | 0 | 0 | 11 | 2 | 0 | 0 | 0 | 0 | 0 | 0 | 1 | 0 | 184 | 149 | 1 | 1 | NA | 1 | NA | NA | NA | 1 | 1 | NA | 0 | NA | 0.333 |
| SE | 81651 | ENSG00000130948 | HSD17B3 | chr9 | - | 96249750 | 96249786 | 96245344 | 96245426 | 96251417 | 96251485 | 0 | 0 | 0 | 0 | 3 | 0 | 0 | 0 | 0 | 0 | 0 | 0 | 0 | 0 | 0 | 0 | 0 | 0 | 1 | 0 | 185 | 149 | 1 | 1 | NA | NA | NA | NA | 1 | NA | NA | NA | 0 | NA | 1 |
| SE | 81652 | ENSG00000130948 | HSD17B3 | chr9 | - | 96249750 | 96249786 | 96246555 | 96246590 | 96251417 | 96251485 | 3 | 3 | 1 | 0 | 3 | 0 | 0 | 0 | 0 | 0 | 0 | 3 | 0 | 0 | 0 | 0 | 2 | 0 | 1 | 0 | 185 | 149 | 1 | 1 | 1 | 1 | 1 | NA | 1 | NA | 0.547 | NA | NA | NA | 0.453 |
| SE | 81654 | ENSG00000130948 | HSD17B3 | chr9 | - | 96254867 | 96254943 | 96252802 | 96252910 | 96298415 | 96298462 | 7 | 9 | 5 | 7 | 4 | 11 | 5 | 0 | 8 | 20 | 17 | 11 | 0 | 5 | 0 | 35 | 9 | 0 | 0 | 8 | 225 | 149 | 1 | 1 | 0.296 | 0.544 | 1 | 0.367 | 0.117 | 0.243 | 0.447 | NA | 1 | 0 | 0.042 |
| SE | 81655 | ENSG00000181090 | EHMT1 | chr9 | + | 137637501 | 137637624 | 137619006 | 137619049 | 137710966 | 137711030 | 0 | 0 | 0 | 0 | 0 | 11 | 4 | 3 | 8 | 9 | 0 | 0 | 0 | 3 | 0 | 0 | 0 | 1 | 2 | 22 | 272 | 149 | 1 | 1 | 0 | 0 | 0 | 0 | 0 | NA | NA | 0 | 0.451 | 0 | -0.15 |
| SE | 81656 | ENSG00000181090 | EHMT1 | chr9 | + | 137650859 | 137651037 | 137619002 | 137619049 | 137710966 | 137711030 | 4 | 0 | 6 | 1 | 0 | 11 | 4 | 3 | 8 | 9 | 0 | 0 | 0 | 7 | 5 | 0 | 0 | 1 | 2 | 22 | 298 | 149 | 1 | 1 | 0.154 | 0 | 0.5 | 0.059 | 0 | NA | NA | 0 | 0.636 | 0.102 | -0.103 |
| SE | 81657 | ENSG00000181090 | EHMT1 | chr9 | + | 137650859 | 137651037 | 137619002 | 137619049 | 137716625 | 137717045 | 0 | 0 | 0 | 0 | 0 | 1 | 0 | 0 | 0 | 0 | 0 | 4 | 0 | 0 | 5 | 11 | 6 | 2 | 0 | 17 | 298 | 149 | 1 | 1 | 0 | NA | NA | NA | NA | 0 | 0.25 | 0 | NA | 0.128 | -0.095 |
| SE | 81658 | ENSG00000181090 | EHMT1 | chr9 | + | 137650859 | 137651037 | 137646376 | 137646438 | 137710966 | 137711030 | 5 | 0 | 8 | 2 | 0 | 0 | 1 | 0 | 3 | 0 | 0 | 0 | 0 | 14 | 0 | 0 | 0 | 0 | 6 | 2 | 298 | 149 | 1 | 1 | 1 | 0 | 1 | 0.25 | NA | NA | NA | NA | 0.538 | 0 | 0.293 |
| SE | 81659 | ENSG00000181090 | EHMT1 | chr9 | + | 137650859 | 137651037 | 137646376 | 137646438 | 137716625 | 137717045 | 1 | 0 | 2 | 1 | 0 | 0 | 0 | 0 | 2 | 0 | 0 | 4 | 0 | 7 | 0 | 0 | 0 | 3 | 5 | 0 | 298 | 149 | 1 | 1 | 1 | NA | 1 | 0.2 | NA | NA | 1 | 0 | 0.412 | NA | 0.263 |
| SE | 81661 | ENSG00000181090 | EHMT1 | chr9 | + | 137710966 | 137711030 | 137650859 | 137651037 | 137716625 | 137716953 | 19 | 6 | 19 | 19 | 6 | 0 | 0 | 0 | 0 | 0 | 0 | 1 | 15 | 30 | 43 | 0 | 4 | 0 | 0 | 0 | 213 | 149 | 1 | 1 | 1 | 1 | 1 | 1 | 1 | NA | 0.149 | 1 | 1 | 1 | 0.213 |
| SE | 81662 | ENSG00000181090 | EHMT1 | chr9 | + | 137716625 | 137717182 | 137710966 | 137711030 | 137728348 | 137728400 | 100 | 42 | 84 | 99 | 52 | 0 | 1 | 0 | 0 | 0 | 49 | 57 | 78 | 95 | 123 | 0 | 0 | 0 | 0 | 0 | 298 | 149 | 1 | 1 | 1 | 0.955 | 1 | 1 | 1 | 1 | 1 | 1 | 1 | 1 | -0.009 |
| SE | 81663 | ENSG00000181090 | EHMT1 | chr9 | + | 137728348 | 137728529 | 137716625 | 137717182 | 137743370 | 137743528 | 216 | 118 | 171 | 191 | 158 | 0 | 0 | 0 | 0 | 2 | 191 | 182 | 165 | 181 | 265 | 0 | 0 | 0 | 0 | 0 | 298 | 149 | 1 | 1 | 1 | 1 | 1 | 1 | 0.975 | 1 | 1 | 1 | 1 | 1 | -0.005 |
| SE | 81664 | ENSG00000181090 | EHMT1 | chr9 | + | 137728348 | 137728529 | 137716625 | 137717182 | 137743391 | 137743528 | 160 | 76 | 166 | 141 | 150 | 0 | 0 | 0 | 0 | 0 | 198 | 159 | 174 | 184 | 245 | 0 | 2 | 0 | 2 | 0 | 298 | 149 | 1 | 1 | 1 | 1 | 1 | 1 | 1 | 1 | 0.975 | 1 | 0.979 | 1 | 0.009 |
| SE | 81665 | ENSG00000181090 | EHMT1 | chr9 | + | 137728348 | 137728529 | 137716625 | 137717182 | 137754170 | 137754291 | 70 | 32 | 67 | 62 | 35 | 0 | 1 | 0 | 0 | 0 | 43 | 56 | 62 | 72 | 60 | 0 | 0 | 0 | 0 | 0 | 298 | 149 | 1 | 1 | 1 | 0.941 | 1 | 1 | 1 | 1 | 1 | 1 | 1 | 1 | -0.012 |
| SE | 81666 | ENSG00000181090 | EHMT1 | chr9 | + | 137728348 | 137728593 | 137716625 | 137717182 | 137743370 | 137743528 | 68 | 32 | 66 | 62 | 35 | 0 | 0 | 0 | 0 | 2 | 43 | 56 | 62 | 69 | 56 | 0 | 0 | 0 | 0 | 0 | 298 | 149 | 1 | 1 | 1 | 1 | 1 | 1 | 0.897 | 1 | 1 | 1 | 1 | 1 | -0.021 |
| SE | 81668 | ENSG00000181090 | EHMT1 | chr9 | + | 137757879 | 137758011 | 137754170 | 137754291 | 137762674 | 137762769 | 123 | 66 | 107 | 102 | 116 | 0 | 0 | 0 | 1 | 0 | 84 | 129 | 68 | 143 | 144 | 0 | 0 | 0 | 3 | 0 | 281 | 149 | 1 | 1 | 1 | 1 | 1 | 0.982 | 1 | 1 | 1 | 1 | 0.962 | 1 | 0.004 |
| SE | 81670 | ENSG00000181090 | EHMT1 | chr9 | + | 137790847 | 137790970 | 137782290 | 137782397 | 137798812 | 137798914 | 163 | 97 | 192 | 179 | 109 | 0 | 0 | 2 | 0 | 0 | 126 | 153 | 89 | 158 | 279 | 0 | 0 | 1 | 6 | 0 | 272 | 149 | 1 | 1 | 1 | 1 | 0.981 | 1 | 1 | 1 | 1 | 0.98 | 0.935 | 1 | 0.013 |
| SE | 81673 | ENSG00000181090 | EHMT1 | chr9 | + | 137811460 | 137811615 | 137800879 | 137800984 | 137813005 | 137813173 | 189 | 96 | 181 | 177 | 103 | 1 | 0 | 0 | 0 | 0 | 191 | 169 | 63 | 147 | 184 | 0 | 0 | 0 | 0 | 0 | 298 | 149 | 1 | 1 | 0.99 | 1 | 1 | 1 | 1 | 1 | 1 | 1 | 1 | 1 | -0.002 |
| SE | 81675 | ENSG00000160856 | FCRL3 | chr1 | - | 157695328 | 157695607 | 157690254 | 157690533 | 157696039 | 157696327 | 143 | 55 | 125 | 90 | 80 | 0 | 0 | 0 | 1 | 0 | 32 | 186 | 36 | 33 | 72 | 0 | 0 | 0 | 0 | 0 | 298 | 149 | 1 | 1 | 1 | 1 | 1 | 0.978 | 1 | 1 | 1 | 1 | 1 | 1 | -0.004 |

| | | | | | | | | | | | | | | | | | | | | | | | | | | | | | | | | | | | | | | | | | | | | | | | | |
|---|---|---|---|---|---|---|---|---|---|---|---|---|---|---|---|---|---|---|---|---|---|---|---|---|---|---|---|---|---|---|---|---|---|---|---|---|---|---|---|---|---|---|---|---|---|---|---|---|
| SE | 81676 | ENSG00000160856 | FCRL3 | chr1 | - | 157695328 | 157695625 | 157690254 | 157690533 | 157696039 | 157696327 | 68 | 28 | 59 | 56 | 39 | 0 | 0 | 0 | 1 | 0 | 25 | 107 | 20 | 14 | 36 | 0 | 0 | 0 | 0 | 0 | 0 | 0 | 298 | 149 | 1 | 1 | 1 | 1 | 1 | 0.966 | 1 | 1 | 1 | 1 | 1 | 1 | -0.007 |
| SE | 81679 | ENSG00000160856 | FCRL3 | chr1 | - | 157697139 | 157697424 | 157696039 | 157696327 | 157698383 | 157698629 | 135 | 43 | 65 | 73 | 75 | 4 | 0 | 0 | 0 | 0 | 33 | 83 | 17 | 35 | 40 | 0 | 0 | 0 | 0 | 0 | 0 | 0 | 298 | 149 | 1 | 1 | 0.944 | 1 | 1 | 1 | 1 | 1 | 1 | 1 | 1 | 1 | -0.011 |
| SE | 81680 | ENSG00000160856 | FCRL3 | chr1 | - | 157697139 | 157697919 | 157696039 | 157696327 | 157698383 | 157698629 | 378 | 123 | 181 | 213 | 253 | 4 | 0 | 0 | 0 | 0 | 105 | 347 | 83 | 153 | 236 | 0 | 0 | 0 | 0 | 0 | 0 | 0 | 298 | 149 | 1 | 1 | 0.979 | 1 | 1 | 1 | 1 | 1 | 1 | 1 | 1 | 1 | -0.004 |
| SE | 81681 | ENSG00000160856 | FCRL3 | chr1 | - | 157697658 | 157697919 | 157696039 | 157696327 | 157698383 | 157698629 | 253 | 82 | 118 | 142 | 189 | 4 | 0 | 0 | 0 | 0 | 76 | 273 | 80 | 119 | 196 | 0 | 0 | 0 | 0 | 0 | 0 | 0 | 298 | 149 | 1 | 1 | 0.969 | 1 | 1 | 1 | 1 | 1 | 1 | 1 | 1 | 1 | -0.006 |
| SE | 81682 | ENSG00000160856 | FCRL3 | chr1 | - | 157697658 | 157697919 | 157697139 | 157697424 | 157698383 | 157698629 | 305 | 96 | 145 | 171 | 234 | 1 | 0 | 0 | 0 | 0 | 72 | 305 | 75 | 125 | 226 | 0 | 0 | 0 | 0 | 0 | 0 | 0 | 298 | 149 | 1 | 1 | 0.993 | 1 | 1 | 1 | 1 | 1 | 1 | 1 | 1 | 1 | -0.001 |
| SE | 81683 | ENSG00000160856 | FCRL3 | chr1 | - | 157698383 | 157698629 | 157696039 | 157696327 | 157699691 | 157699712 | 100 | 29 | 42 | 40 | 67 | 0 | 0 | 0 | 1 | 0 | 39 | 105 | 27 | 20 | 56 | 0 | 0 | 0 | 0 | 0 | 0 | 0 | 298 | 149 | 1 | 1 | 1 | 1 | 1 | 0.952 | 1 | 1 | 1 | 1 | 1 | 1 | -0.01 |
| SE | 81686 | ENSG00000171681 | ATF7IP | chr12 | + | 14420301 | 14420446 | 14408489 | 14408544 | 14423908 | 14424183 | 1 | 1 | 3 | 0 | 0 | 0 | 1 | 0 | 2 | 0 | 0 | 0 | 0 | 0 | 9 | 0 | 0 | 0 | 0 | 0 | 0 | 0 | 294 | 149 | 1 | 1 | 1 | 0.336 | 1 | 0 | NA | NA | NA | NA | NA | 1 | -0.416 |
| SE | 81690 | ENSG00000171681 | ATF7IP | chr12 | + | 14423908 | 14425473 | 14385063 | 14385400 | 14434336 | 14434423 | 503 | 442 | 365 | 497 | 712 | 0 | 0 | 0 | 0 | 0 | 112 | 574 | 552 | 282 | 93 | 0 | 0 | 0 | 0 | 0 | 2 | 0 | 298 | 149 | 1 | 1 | 1 | 1 | 1 | 1 | 1 | 1 | 1 | 1 | 0.986 | 1 | 0.003 |
| SE | 81692 | ENSG00000171681 | ATF7IP | chr12 | + | 14434336 | 14434423 | 14423760 | 14425473 | 14436105 | 14436251 | 1502 | 1420 | 1160 | 1555 | 2159 | 2 | 3 | 1 | 2 | 0 | 442 | 1710 | 1766 | 988 | 281 | 0 | 0 | 0 | 0 | 0 | 0 | 0 | 236 | 149 | 1 | 1 | 0.998 | 0.997 | 0.999 | 0.998 | 1 | 1 | 1 | 1 | 1 | 1 | -0.002 |
| SE | 81693 | ENSG00000171681 | ATF7IP | chr12 | + | 14434339 | 14434423 | 14423908 | 14425473 | 14436105 | 14436251 | 1244 | 1156 | 963 | 1271 | 1711 | 2 | 3 | 1 | 2 | 0 | 363 | 1387 | 1430 | 803 | 217 | 0 | 0 | 0 | 0 | 0 | 0 | 0 | 233 | 149 | 1 | 1 | 0.997 | 0.996 | 0.998 | 0.998 | 1 | 1 | 1 | 1 | 1 | 1 | -0.002 |
| SE | 81694 | ENSG00000171681 | ATF7IP | chr12 | + | 14438129 | 14438267 | 14436105 | 14436251 | 14446987 | 14447030 | 1088 | 978 | 869 | 1174 | 1834 | 0 | 1 | 0 | 0 | 2 | 271 | 1027 | 1114 | 620 | 102 | 0 | 0 | 0 | 0 | 0 | 0 | 0 | 287 | 149 | 1 | 1 | 1 | 0.998 | 1 | 1 | 0.998 | 1 | 1 | 1 | 1 | 1 | -0.001 |
| SE | 81695 | ENSG00000171681 | ATF7IP | chr12 | + | 14438129 | 14438267 | 14436105 | 14436251 | 14456560 | 14456634 | 666 | 653 | 531 | 744 | 1176 | 0 | 0 | 1 | 0 | 0 | 155 | 743 | 789 | 482 | 68 | 0 | 0 | 0 | 0 | 0 | 0 | 0 | 287 | 149 | 1 | 1 | 1 | 1 | 0.996 | 1 | 1 | 1 | 1 | 1 | 1 | 1 | -0.001 |
| SE | 81696 | ENSG00000171681 | ATF7IP | chr12 | + | 14446987 | 14447053 | 14436105 | 14436251 | 14456560 | 14456634 | 114 | 91 | 75 | 103 | 194 | 0 | 0 | 1 | 0 | 0 | 19 | 83 | 96 | 32 | 9 | 0 | 0 | 0 | 0 | 0 | 0 | 0 | 215 | 149 | 1 | 1 | 1 | 1 | 0.981 | 1 | 1 | 1 | 1 | 1 | 1 | 1 | -0.004 |
| SE | 81698 | ENSG00000171681 | ATF7IP | chr12 | + | 14456560 | 14456634 | 14446987 | 14447053 | 14460494 | 14460567 | 199 | 150 | 153 | 188 | 328 | 1 | 0 | 0 | 0 | 0 | 58 | 151 | 156 | 77 | 19 | 0 | 0 | 0 | 0 | 0 | 0 | 0 | 223 | 149 | 1 | 1 | 0.993 | 1 | 1 | 1 | 1 | 1 | 1 | 1 | 1 | 1 | -0.001 |
| SE | 81700 | ENSG00000171681 | ATF7IP | chr12 | + | 14466525 | 14466590 | 14460494 | 14461133 | 14475889 | 14475968 | 242 | 161 | 152 | 250 | 302 | 0 | 2 | 0 | 0 | 0 | 2 | 217 | 176 | 81 | 6 | 0 | 0 | 0 | 0 | 0 | 0 | 0 | 214 | 149 | 1 | 1 | 1 | 0.982 | 1 | 1 | 1 | 1 | 1 | 1 | 1 | 1 | -0.004 |
| SE | 81710 | ENSG00000113013 | HSPA9 | chr5 | - | 138556451 | 138556592 | 138553755 | 138556114 | 138556773 | 138556821 | 816 | 840 | 791 | 992 | 1272 | 0 | 0 | 0 | 1 | 0 | 1228 | 941 | 1521 | 1511 | 174 | 0 | 0 | 0 | 0 | 0 | 1 | 0 | 290 | 149 | 1 | 1 | 1 | 1 | 1 | 0.998 | 1 | 1 | 1 | 1 | 0.999 | 1 | 0 |
| SE | 81711 | ENSG00000113013 | HSPA9 | chr5 | - | 138557868 | 138557986 | 138556773 | 138557496 | 138558552 | 138558631 | 1195 | 977 | 1124 | 1144 | 1583 | 0 | 0 | 0 | 0 | 0 | 1196 | 960 | 1615 | 1201 | 172 | 0 | 0 | 0 | 0 | 0 | 9 | 0 | 267 | 149 | 1 | 1 | 1 | 1 | 1 | 1 | 1 | 1 | 1 | 1 | 0.987 | 1 | 0.003 |
| SE | 81712 | ENSG00000113013 | HSPA9 | chr5 | - | 138558552 | 138558657 | 138557868 | 138557986 | 138559863 | 138560091 | 748 | 609 | 732 | 737 | 990 | 0 | 0 | 0 | 0 | 0 | 706 | 619 | 1018 | 778 | 103 | 0 | 1 | 0 | 1 | 0 | 0 | 0 | 254 | 149 | 1 | 1 | 1 | 1 | 1 | 1 | 1 | 1 | 0.997 | 1 | 1 | 1 | 0.001 |
| SE | 81713 | ENSG00000113013 | HSPA9 | chr5 | - | 138559863 | 138560091 | 138558552 | 138558657 | 138561579 | 138561789 | 632 | 459 | 570 | 662 | 675 | 4 | 0 | 3 | 4 | 5 | 575 | 473 | 782 | 618 | 58 | 0 | 0 | 0 | 0 | 2 | 5 | 0 | 298 | 149 | 1 | 1 | 0.988 | 1 | 0.99 | 0.988 | 0.985 | 1 | 1 | 0.995 | 0.984 | 1 | -0.006 |
| SE | 81716 | ENSG00000113013 | HSPA9 | chr5 | - | 138567648 | 138567722 | 138567454 | 138567561 | 138568924 | 138569049 | 937 | 600 | 740 | 930 | 1121 | 1 | 4 | 0 | 3 | 3 | 781 | 759 | 1009 | 860 | 226 | 0 | 0 | 0 | 0 | 4 | 1 | 0 | 223 | 149 | 1 | 1 | 0.998 | 0.99 | 1 | 0.995 | 0.996 | 1 | 1 | 0.994 | 0.998 | 1 | -0.003 |

[illegible]

| | | | | | | | | | | | | | | | | | | | | | | | |
|---|---|---|---|---|---|---|---|---|---|---|---|---|---|---|---|---|---|---|---|---|---|---|---|
| SE | 82075 | ENSG00000198286 | CARD11 | chr7 | - | 2910049 | 2910165 | 2906141 | 2906842 | 2912171 | 2912296 | 212 | 78 | 201 | 199 | 141 | 0 | 1 | 0 | 0 | 0 | 110 | 140 |
| SE | 82078 | ENSG00000198286 | CARD11 | chr7 | - | 2937906 | 2938032 | 2937036 | 2937234 | 2938678 | 2938831 | 408 | 190 | 314 | 367 | 361 | 0 | 1 | 0 | 0 | 0 | 370 | 389 |
| SE | 82079 | ENSG00000198286 | CARD11 | chr7 | - | 2958499 | 2958631 | 2947574 | 2947787 | 2980519 | 2980580 | 208 | 110 | 199 | 234 | 141 | 0 | 1 | 0 | 0 | 0 | 199 | 278 |
| SE | 82082 | ENSG00000230524 | COL6A4P1 | chr3 | - | 15173455 | 15174061 | 15169939 | 15170577 | 15174762 | 15175103 | 0 | 0 | 0 | 0 | 2 | 0 | 0 | 0 | 0 | 0 | 0 | 0 |
| SE | 82086 | ENSG00000184216 | IRAK1 | chrX | - | 154014041 | 154014278 | 154013042 | 154013343 | 154016031 | 154016073 | 82 | 41 | 65 | 42 | 76 | 1 | 0 | 0 | 0 | 0 | 71 | 112 |
| SE | 82087 | ENSG00000184216 | IRAK1 | chrX | - | 154014041 | 154014278 | 154013042 | 154013433 | 154016031 | 154016073 | 126 | 64 | 99 | 84 | 109 | 1 | 0 | 0 | 2 | 0 | 99 | 158 |
| SE | 82091 | ENSG00000211751 | TRBC1 | chr7 | + | 142792521 | 142792539 | 142791693 | 142792080 | 142792691 | 142792798 | 1210 | 447 | 1621 | 1040 | 1538 | 2 | 0 | 8 | 2 | 0 | 132 | 826 |
| SE | 82092 | ENSG00000211751 | TRBC1 | chr7 | + | 142792521 | 142792539 | 142791693 | 142792080 | 142793120 | 142793368 | 135 | 16 | 145 | 84 | 50 | 0 | 0 | 1 | 0 | 0 | 39 | 69 |
| SE | 82093 | ENSG00000211751 | TRBC1 | chr7 | + | 142792691 | 142792798 | 142791693 | 142792080 | 142793120 | 142793368 | 1138 | 451 | 1420 | 971 | 1306 | 0 | 0 | 1 | 0 | 0 | 179 | 788 |
| SE | 82095 | ENSG00000128891 | CCDC32 | chr15 | - | 40539238 | 40539355 | 40528682 | 40528790 | 40557215 | 40557372 | 0 | 4 | 6 | 2 | 1 | 1 | 1 | 3 | 0 | 2 | 0 | 6 |
| SE | 82096 | ENSG00000128891 | CCDC32 | chr15 | - | 40539238 | 40539355 | 40531340 | 40532302 | 40553594 | 40554127 | 0 | 0 | 0 | 0 | 0 | 1 | 0 | 0 | 0 | 0 | 0 | 9 |
| SE | 82098 | ENSG00000128891 | CCDC32 | chr15 | - | 40539238 | 40539355 | 40534899 | 40535010 | 40557215 | 40557372 | 0 | 4 | 7 | 2 | 1 | 0 | 0 | 0 | 0 | 2 | 0 | 6 |
| SE | 82101 | ENSG00000128891 | CCDC32 | chr15 | - | 40553436 | 40554127 | 40534342 | 40539355 | 40562771 | 40563027 | 0 | 0 | 3 | 0 | 0 | 0 | 0 | 0 | 0 | 0 | 0 | 4 |
| SE | 82104 | ENSG00000128891 | CCDC32 | chr15 | - | 40553594 | 40554127 | 40539238 | 40539355 | 40562771 | 40563027 | 0 | 0 | 2 | 0 | 0 | 0 | 0 | 0 | 0 | 0 | 0 | 8 |
| SE | 82105 | ENSG00000128891 | CCDC32 | chr15 | - | 40557215 | 40557372 | 40539228 | 40539355 | 40562771 | 40563022 | 101 | 52 | 88 | 75 | 122 | 0 | 0 | 0 | 0 | 0 | 107 | 127 |
| SE | 82113 | ENSG00000136048 | DRAM1 | chr12 | + | 101914173 | 101914232 | 101901290 | 101901433 | 101921215 | 101921542 | 1 | 0 | 1 | 1 | 0 | 0 | 0 | 0 | 2 | 0 | 11 | 6 |
| SE | 82114 | ENSG00000136048 | DRAM1 | chr12 | + | 101920108 | 101920201 | 101901290 | 101901433 | 101921215 | 101921542 | 52 | 59 | 30 | 31 | 39 | 0 | 0 | 0 | 2 | 0 | 70 | 122 |
| SE | 82117 | ENSG00000164675 | IQUB | chr7 | - | 123479794 | 123479970 | 123469213 | 123469384 | 123496695 | 123496906 | 0 | 0 | 0 | 2 | 0 | 0 | 0 | 0 | 0 | 0 | 0 | 0 |
| SE | 82118 | ENSG00000164675 | IQUB | chr7 | - | 123496695 | 123496906 | 123469213 | 123469384 | 123502596 | 123502752 | 0 | 0 | 0 | 2 | 0 | 0 | 0 | 0 | 0 | 3 | 0 | 8 |
| SE | 82119 | ENSG00000164675 | IQUB | chr7 | - | 123502596 | 123502752 | 123496695 | 123496906 | 123511943 | 123512344 | 0 | 0 | 0 | 2 | 2 | 0 | 0 | 1 | 3 | 4 | 0 | 2 |
| SE | 82128 | ENSG00000188002 | NA | chr5 | - | 1626785 | 1626926 | 1602690 | 1602735 | 1632643 | 1632751 | 6 | 0 | 10 | 9 | 13 | 2 | 0 | 1 | 6 | 8 | 5 | 9 |
| SE | 82129 | ENSG00000188002 | NA | chr5 | - | 1628754 | 1629630 | 1626785 | 1626926 | 1632643 | 1632751 | 11 | 6 | 17 | 22 | 16 | 0 | 0 | 0 | 0 | 2 | 3 | 12 |
| SE | 82130 | ENSG00000188002 | NA | chr5 | - | 1630408 | 1630566 | 1628754 | 1629630 | 1632643 | 1632751 | 3 | 3 | 4 | 0 | 2 | 3 | 2 | 2 | 3 | 11 | 0 | 18 |
| SE | 82131 | ENSG00000188002 | NA | chr5 | - | 1630408 | 1630566 | 1628754 | 1629630 | 1632844 | 1632906 | 1 | 5 | 4 | 0 | 2 | 3 | 0 | 5 | 0 | 3 | 0 | 3 |
| SE | 82133 | ENSG00000188002 | NA | chr5 | - | 1632643 | 1632751 | 1628754 | 1629630 | 1632844 | 1632906 | 2 | 1 | 1 | 3 | 9 | 3 | 0 | 5 | 0 | 3 | 7 | 12 |
| SE | 82134 | ENSG00000188002 | NA | chr5 | - | 1632643 | 1632751 | 1630408 | 1630566 | 1632844 | 1632906 | 3 | 0 | 3 | 0 | 0 | 0 | 4 | 3 | 0 | 0 | 7 | 13 |
| SE | 82136 | ENSG00000132640 | BTBD3 | chr20 | + | 11919085 | 11919176 | 11917777 | 11918601 | 11919717 | 11919836 | 10 | 7 | 7 | 14 | 26 | 2 | 1 | 2 | 2 | 7 | 18 | 8 |
| SE | 82138 | ENSG00000172354 | GNB2 | chr7 | + | 100676534 | 100676573 | 100676137 | 100676322 | 100676692 | 100676799 | 100 | 36 | 70 | 117 | 47 | 0 | 0 | 0 | 0 | 0 | 95 | 133 |
| SE | 82140 | ENSG00000172354 | GNB2 | chr7 | + | 100677351 | 100677415 | 100676692 | 100676799 | 100677497 | 100677660 | 230 | 88 | 173 | 200 | 145 | 0 | 0 | 2 | 0 | 0 | 341 | 188 |
| SE | 82141 | ENSG00000065000 | AP3D1 | chr19 | - | 2108686 | 2108766 | 2100987 | 2102268 | 2109085 | 2109125 | 257 | 172 | 256 | 287 | 158 | 1 | 0 | 0 | 0 | 0 | 411 | 362 |
| SE | 82142 | ENSG00000065000 | AP3D1 | chr19 | - | 2108686 | 2108876 | 2102236 | 2102268 | 2109085 | 2109207 | 130 | 86 | 110 | 165 | 74 | 1 | 0 | 0 | 0 | 0 | 200 | 205 |
| SE | 82143 | ENSG00000065000 | AP3D1 | chr19 | - | 2112859 | 2112931 | 2111678 | 2111828 | 2113335 | 2113413 | 79 | 62 | 74 | 93 | 99 | 0 | 0 | 0 | 0 | 0 | 100 | 135 |
| SE | 82144 | ENSG00000065000 | AP3D1 | chr19 | - | 2112859 | 2112967 | 2111678 | 2111828 | 2113335 | 2113413 | 82 | 49 | 81 | 84 | 86 | 0 | 0 | 0 | 0 | 0 | 103 | 124 |
| SE | 82148 | ENSG00000065000 | AP3D1 | chr19 | - | 2115218 | 2115418 | 2114747 | 2114821 | 2115537 | 2115613 | 236 | 158 | 219 | 226 | 205 | 0 | 0 | 0 | 0 | 0 | 346 | 244 |
| SE | 82151 | ENSG00000065000 | AP3D1 | chr19 | - | 2133396 | 2133484 | 2132470 | 2132578 | 2137010 | 2137091 | 2 | 0 | 0 | 0 | 2 | 100 | 47 | 66 | 71 | 140 | 0 | 0 |
| SE | 82153 | ENSG00000146828 | SLC12A9 | chr7 | + | 100854155 | 100854378 | 100852712 | 100852835 | 100858834 | 100858942 | 3 | 1 | 2 | 3 | 0 | 0 | 0 | 0 | 0 | 0 | 0 | 0 |
| SE | 82155 | ENSG00000146828 | SLC12A9 | chr7 | + | 100856867 | 100857176 | 100852732 | 100852835 | 100858834 | 100858942 | 18 | 11 | 14 | 30 | 4 | 0 | 0 | 0 | 0 | 0 | 48 | 23 |
| SE | 82156 | ENSG00000146828 | SLC12A9 | chr7 | + | 100856867 | 100857176 | 100854155 | 100854378 | 100858834 | 100858942 | 18 | 12 | 16 | 30 | 8 | 1 | 0 | 0 | 2 | 0 | 52 | 25 |
| SE | 82157 | ENSG00000146828 | SLC12A9 | chr7 | + | 100856867 | 100857176 | 100855705 | 100855837 | 100858834 | 100858942 | 26 | 12 | 16 | 38 | 9 | 1 | 0 | 0 | 1 | 0 | 48 | 23 |
| SE | 82159 | ENSG00000146828 | SLC12A9 | chr7 | + | 100861137 | 100861262 | 100860149 | 100860232 | 100861391 | 100861584 | 52 | 19 | 40 | 35 | 38 | 0 | 0 | 1 | 0 | 0 | 37 | 57 |
| SE | 82161 | ENSG00000168546 | GFRA2 | chr8 | - | 21782584 | 21782899 | 21774971 | 21775055 | 21788119 | 21788159 | 1 | 1 | 1 | 4 | 3 | 0 | 1 | 0 | 0 | 0 | 0 | 0 |
| SE | 82162 | ENSG00000168546 | GFRA2 | chr8 | - | 21782584 | 21783142 | 21774971 | 21775055 | 21788119 | 21788819 | 0 | 1 | 0 | 4 | 0 | 0 | 1 | 0 | 0 | 0 | 0 | 0 |
| SE | 82165 | ENSG00000087077 | TRIP6 | chr7 | + | 100871060 | 100871177 | 100870573 | 100870743 | 100871542 | 100871721 | 1 | 0 | 0 | 0 | 0 | 18 | 7 | 14 | 28 | 18 | 0 | 0 |
| SE | 82169 | ENSG00000146872 | TLK2 | chr17 | + | 62507380 | 62507449 | 62481120 | 62481206 | 62508407 | 62508511 | 0 | 0 | 0 | 0 | 0 | 1 | 3 | 2 | 10 | 5 | 0 | 2 |
| SE | 82174 | ENSG00000146872 | TLK2 | chr17 | + | 62536169 | 62536337 | 62523133 | 62523177 | 62552301 | 62552339 | 153 | 111 | 189 | 168 | 232 | 0 | 0 | 0 | 0 | 0 | 246 | 186 |
| SE | 82178 | ENSG00000146872 | TLK2 | chr17 | + | 62553662 | 62553755 | 62552301 | 62552397 | 62560015 | 62560034 | 143 | 105 | 137 | 124 | 128 | 1 | 0 | 2 | 2 | 0 | 239 | 235 |
| SE | 82182 | ENSG00000146872 | TLK2 | chr17 | + | 62578476 | 62578574 | 62576708 | 62576775 | 62586134 | 62586226 | 110 | 96 | 102 | 117 | 92 | 0 | 0 | 0 | 0 | 0 | 167 | 126 |
| SE | 82183 | ENSG00000146872 | TLK2 | chr17 | + | 62578476 | 62580192 | 62576708 | 62576775 | 62586134 | 62586226 | 218 | 185 | 195 | 239 | 251 | 0 | 0 | 0 | 0 | 0 | 284 | 218 |
| SE | 82184 | ENSG00000146872 | TLK2 | chr17 | + | 62580110 | 62580192 | 62578476 | 62578574 | 62586134 | 62586226 | 269 | 192 | 221 | 269 | 319 | 0 | 0 | 0 | 0 | 0 | 426 | 276 |
| SE | 82185 | ENSG00000146872 | TLK2 | chr17 | + | 62586134 | 62586226 | 62578476 | 62580192 | 62596584 | 62596674 | 237 | 164 | 190 | 235 | 338 | 0 | 0 | 0 | 1 | 0 | 288 | 186 |
| SE | 82186 | ENSG00000146872 | TLK2 | chr17 | + | 62596584 | 62596674 | 62585595 | 62586226 | 62600650 | 62600820 | 271 | 168 | 189 | 225 | 333 | 0 | 0 | 2 | 0 | 0 | 246 | 137 |
| SE | 82188 | ENSG00000146872 | TLK2 | chr17 | + | 62602041 | 62602180 | 62600650 | 62600820 | 62606129 | 62606188 | 89 | 65 | 108 | 109 | 115 | 0 | 0 | 0 | 0 | 3 | 22 | 81 |
| SE | 82189 | ENSG00000146872 | TLK2 | chr17 | + | 62608040 | 62608148 | 62606095 | 62606241 | 62612391 | 62612573 | 106 | 47 | 87 | 89 | 142 | 0 | 0 | 0 | 2 | 0 | 34 | 85 |

| | | | | | | | | | | | | | |
|---|---|---|---|---|---|---|---|---|---|---|---|---|---|
| SE | 82502 | ENSG00000185201 | IFITM2 | chr11 | + | 308754 | 308889 | 308508 | 308538 | 309017 | 309392 | 0 | 0 |
| SE | 82504 | ENSG00000137193 | PIM1 | chr6 | + | 37172995 | 37173172 | 37171124 | 37171491 | 37173933 | 37174247 | 798 | 350 |
| SE | 82506 | ENSG00000198315 | ZKSCAN8 | chr6 | + | 28144330 | 28144424 | 28141937 | 28142029 | 28149482 | 28149624 | 1 | 2 |
| SE | 82507 | ENSG00000198315 | ZKSCAN8 | chr6 | + | 28148315 | 28148824 | 28141882 | 28142029 | 28149482 | 28149624 | 71 | 65 |
| SE | 82509 | ENSG00000198315 | ZKSCAN8 | chr6 | + | 28148315 | 28148838 | 28141909 | 28142029 | 28149482 | 28149624 | 45 | 42 |
| SE | 82510 | ENSG00000198315 | ZKSCAN8 | chr6 | + | 28148315 | 28148838 | 28144330 | 28144424 | 28149482 | 28149624 | 11 | 5 |
| SE | 82511 | ENSG00000198315 | ZKSCAN8 | chr6 | + | 28152260 | 28152384 | 28151844 | 28151936 | 28153055 | 28153079 | 210 | 194 |
| SE | 82515 | ENSG00000061938 | TNK2 | chr3 | - | 195870113 | 195870205 | 195867360 | 195868709 | 195872275 | 195872470 | 96 | 33 |
| SE | 82516 | ENSG00000061938 | TNK2 | chr3 | - | 195870113 | 195870378 | 195868294 | 195868709 | 195872275 | 195872470 | 21 | 14 |
| SE | 82518 | ENSG00000061938 | TNK2 | chr3 | - | 195878445 | 195878592 | 195878252 | 195878347 | 195879048 | 195879175 | 251 | 129 |
| SE | 82524 | ENSG00000061938 | TNK2 | chr3 | - | 195891911 | 195892007 | 195888425 | 195888606 | 195892410 | 195892784 | 1 | 1 |
| SE | 82526 | ENSG00000061938 | TNK2 | chr3 | - | 195894315 | 195894363 | 195891911 | 195892007 | 195908484 | 195909802 | 0 | 0 |
| SE | 82528 | ENSG00000164815 | ORC5 | chr7 | - | 104136780 | 104136893 | 104126340 | 104126893 | 104161071 | 104161182 | 62 | 26 |
| SE | 82529 | ENSG00000164815 | ORC5 | chr7 | - | 104161071 | 104161182 | 104136780 | 104136893 | 104165234 | 104165282 | 94 | 43 |
| SE | 82530 | ENSG00000164815 | ORC5 | chr7 | - | 104165234 | 104165282 | 104136780 | 104136893 | 104188250 | 104188381 | 0 | 0 |
| SE | 82531 | ENSG00000164815 | ORC5 | chr7 | - | 104165234 | 104165282 | 104161071 | 104161182 | 104166771 | 104166884 | 77 | 43 |
| SE | 82533 | ENSG00000164815 | ORC5 | chr7 | - | 104168472 | 104168525 | 104165234 | 104165282 | 104188250 | 104188381 | 2 | 0 |
| SE | 82535 | ENSG00000164815 | ORC5 | chr7 | - | 104184122 | 104184171 | 104183942 | 104184033 | 104188250 | 104188381 | 84 | 67 |
| SE | 82536 | ENSG00000164815 | ORC5 | chr7 | - | 104195142 | 104195254 | 104188250 | 104188381 | 104197724 | 104197767 | 118 | 84 |
| SE | 82537 | ENSG00000164815 | ORC5 | chr7 | - | 104197724 | 104197799 | 104195142 | 104195254 | 104200757 | 104200958 | 104 | 57 |
| SE | 82538 | ENSG00000164815 | ORC5 | chr7 | - | 104202376 | 104202544 | 104200820 | 104200958 | 104204141 | 104204234 | 1 | 0 |
| SE | 82540 | ENSG00000167476 | JSRP1 | chr19 | - | 2254186 | 2254301 | 2253619 | 2253793 | 2254444 | 2254482 | 0 | 0 |
| SE | 82550 | ENSG00000139737 | SLAIN1 | chr13 | + | 77746513 | 77746855 | 77744282 | 77744432 | 77753202 | 77753241 | 186 | 105 |
| SE | 82552 | ENSG00000106263 | EIF3B | chr7 | + | 2362644 | 2362764 | 2360709 | 2360902 | 2363631 | 2363760 | 299 | 164 |
| SE | 82553 | ENSG00000106263 | EIF3B | chr7 | + | 2363069 | 2363127 | 2362644 | 2362764 | 2363631 | 2363760 | 686 | 396 |
| SE | 82555 | ENSG00000106263 | EIF3B | chr7 | + | 2371776 | 2371849 | 2369471 | 2369682 | 2372672 | 2372795 | 284 | 157 |
| SE | 82556 | ENSG00000235944 | ZNF815P | chr7 | + | 5831180 | 5831283 | 5823159 | 5823526 | 5833526 | 5833568 | 2 | 2 |
| SE | 82557 | ENSG00000235944 | ZNF815P | chr7 | + | 5831180 | 5831283 | 5823159 | 5823526 | 5839948 | 5840075 | 0 | 0 |
| SE | 82558 | ENSG00000235944 | ZNF815P | chr7 | + | 5833526 | 5833568 | 5823215 | 5823526 | 5839948 | 5840075 | 4 | 9 |
| SE | 82559 | ENSG00000235944 | ZNF815P | chr7 | + | 5833526 | 5833568 | 5823215 | 5823526 | 5840689 | 5840856 | 1 | 1 |
| SE | 82560 | ENSG00000235944 | ZNF815P | chr7 | + | 5833526 | 5833568 | 5831180 | 5831283 | 5839948 | 5840075 | 4 | 5 |
| SE | 82561 | ENSG00000235944 | ZNF815P | chr7 | + | 5839069 | 5839204 | 5833526 | 5833568 | 5839948 | 5840042 | 1 | 0 |
| SE | 82566 | ENSG00000104915 | STX10 | chr19 | - | 13145287 | 13145395 | 13144763 | 13144870 | 13149028 | 13149091 | 195 | 109 |
| SE | 82567 | ENSG00000127824 | TUBA4A | chr2 | - | 219251564 | 219251713 | 219249709 | 219251323 | 219252007 | 219252230 | 425 | 259 |
| SE | 82568 | ENSG00000127824 | TUBA4A | chr2 | - | 219251564 | 219251713 | 219249709 | 219251323 | 219253855 | 219253901 | 405 | 233 |
| SE | 82569 | ENSG00000127824 | TUBA4A | chr2 | - | 219251564 | 219251713 | 219249709 | 219251323 | 219277813 | 219277902 | 401 | 233 |
| SE | 82570 | ENSG00000127824 | TUBA4A | chr2 | - | 219252007 | 219252230 | 219249709 | 219251323 | 219253855 | 219253916 | 55 | 45 |
| SE | 82571 | ENSG00000127824 | TUBA4A | chr2 | - | 219252007 | 219252230 | 219249709 | 219251323 | 219277813 | 219277902 | 1 | 0 |
| SE | 82573 | ENSG00000127824 | TUBA4A | chr2 | - | 219252007 | 219252230 | 219251193 | 219251713 | 219254094 | 219254608 | 24 | 32 |
| SE | 82584 | ENSG00000106628 | POLD2 | chr7 | - | 44117942 | 44118064 | 44117516 | 44117742 | 44121833 | 44121905 | 59 | 32 |
| SE | 82587 | ENSG00000013374 | NUB1 | chr7 | + | 151351423 | 151351482 | 151349072 | 151349240 | 151352811 | 151352882 | 685 | 573 |
| SE | 82589 | ENSG00000013374 | NUB1 | chr7 | + | 151355767 | 151355950 | 151352811 | 151352882 | 151356127 | 151356222 | 672 | 558 |
| SE | 82591 | ENSG00000013374 | NUB1 | chr7 | + | 151355767 | 151355950 | 151352811 | 151352882 | 151366938 | 151367125 | 427 | 348 |
| SE | 82593 | ENSG00000013374 | NUB1 | chr7 | + | 151356127 | 151356222 | 151355767 | 151355950 | 151360140 | 151360209 | 454 | 372 |
| SE | 82594 | ENSG00000013374 | NUB1 | chr7 | + | 151360140 | 151360247 | 151352811 | 151352882 | 151366938 | 151367125 | 319 | 283 |
| SE | 82595 | ENSG00000013374 | NUB1 | chr7 | + | 151360140 | 151360247 | 151355767 | 151355950 | 151366938 | 151367125 | 315 | 278 |
| SE | 82596 | ENSG00000013374 | NUB1 | chr7 | + | 151367860 | 151367968 | 151366938 | 151367125 | 151374096 | 151374201 | 150 | 110 |
| SE | 82602 | ENSG00000197448 | GSTK1 | chr7 | + | 143264085 | 143264167 | 143263428 | 143263585 | 143264547 | 143264676 | 726 | 347 |
| SE | 82603 | ENSG00000197448 | GSTK1 | chr7 | + | 143264547 | 143264676 | 143264085 | 143264167 | 143264991 | 143265092 | 1264 | 644 |
| SE | 82605 | ENSG00000197448 | GSTK1 | chr7 | + | 143268090 | 143268184 | 143267616 | 143267733 | 143268787 | 143268943 | 661 | 394 |
| SE | 82606 | ENSG00000145375 | SPATA5 | chr4 | + | 122952198 | 122952270 | 122947232 | 122947488 | 122979231 | 122979386 | 0 | 0 |
| SE | 82607 | ENSG00000145375 | SPATA5 | chr4 | + | 122979231 | 122979386 | 122947232 | 122947488 | 123028185 | 123028395 | 78 | 84 |
| SE | 82608 | ENSG00000145375 | SPATA5 | chr4 | + | 123028185 | 123028395 | 122979231 | 122979386 | 123056386 | 123056440 | 99 | 113 |
| SE | 82610 | ENSG00000145375 | SPATA5 | chr4 | + | 123057208 | 123057288 | 123028185 | 123028395 | 123090578 | 123090705 | 19 | 28 |
| SE | 82611 | ENSG00000145375 | SPATA5 | chr4 | + | 123090578 | 123090705 | 123057208 | 123057288 | 123256015 | 123256180 | 55 | 50 |
| SE | 82620 | ENSG00000106144 | CASP2 | chr7 | + | 143304673 | 143304783 | 143300203 | 143303933 | 143304939 | 143305071 | 133 | 76 |
| SE | 82621 | ENSG00000171863 | RPS7 | chr2 | + | 3575816 | 3575888 | 3575324 | 3575684 | 3576486 | 3576630 | 7921 | 4249 |
| SE | 82633 | ENSG00000229153 | EPHA1-AS1 | chr7 | + | 143504773 | 143504830 | 143415067 | 143415199 | 143518281 | 143518517 | 2 | 0 |
| SE | 82634 | ENSG00000229153 | EPHA1-AS1 | chr7 | + | 143504773 | 143504830 | 143415067 | 143415199 | 143518930 | 143523449 | 0 | 0 |
| SE | 82635 | ENSG00000229153 | EPHA1-AS1 | chr7 | + | 143518281 | 143518517 | 143415067 | 143415199 | 143518930 | 143523449 | 18 | 6 |
| SE | 82636 | ENSG00000229153 | EPHA1-AS1 | chr7 | + | 143518281 | 143518517 | 143504773 | 143504830 | 143518930 | 143523449 | 16 | 4 |
| SE | 82637 | ENSG00000103145 | HCFC1R1 | chr16 | - | 3023232 | 3023361 | 3022624 | 3022998 | 3023846 | 3023961 | 22 | 2 |
| SE | 82638 | ENSG00000103145 | HCFC1R1 | chr16 | - | 3023232 | 3023412 | 3022840 | 3022998 | 3023846 | 3024273 | 21 | 2 |
| SE | 82639 | ENSG00000103145 | HCFC1R1 | chr16 | - | 3023232 | 3023530 | 3022625 | 3022998 | 3023846 | 3023968 | 36 | 9 |
| SE | 82644 | ENSG00000131373 | HACL1 | chr3 | - | 15579909 | 15580045 | 15574976 | 15575082 | 15582876 | 15582989 | 110 | 45 |
| SE | 82645 | ENSG00000131373 | HACL1 | chr3 | - | 15582876 | 15582989 | 15579899 | 15580045 | 15591599 | 15591680 | 45 | 15 |
| SE | 82647 | ENSG00000131373 | HACL1 | chr3 | - | 15585247 | 15585342 | 15582876 | 15582989 | 15589539 | 15589612 | 33 | 12 |
| SE | 82648 | ENSG00000131373 | HACL1 | chr3 | - | 15586524 | 15586602 | 15582876 | 15582989 | 15589539 | 15589612 | 24 | 8 |
| SE | 82649 | ENSG00000131373 | HACL1 | chr3 | - | 15586524 | 15586602 | 15582876 | 15582989 | 15591599 | 15591680 | 1 | 2 |
| SE | 82651 | ENSG00000131373 | HACL1 | chr3 | - | 15589539 | 15589612 | 15582876 | 15582989 | 15591599 | 15591680 | 16 | 7 |
| SE | 82652 | ENSG00000131373 | HACL1 | chr3 | - | 15589539 | 15589612 | 15586524 | 15586602 | 15591599 | 15591680 | 60 | 25 |
| SE | 82653 | ENSG00000131373 | HACL1 | chr3 | - | 15591599 | 15591680 | 15589539 | 15589612 | 15596383 | 15596424 | 52 | 22 |
| SE | 82659 | ENSG00000104447 | TRPS1 | chr8 | - | 115619131 | 115620060 | 115603872 | 115605002 | 115668544 | 115668702 | 246 | 208 |
| SE | 82660 | ENSG00000104447 | TRPS1 | chr8 | - | 115623600 | 115623758 | 115619131 | 115620060 | 115637233 | 115637330 | 8 | 8 |
| SE | 82662 | ENSG00000104447 | TRPS1 | chr8 | - | 115637233 | 115637330 | 115619912 | 115620060 | 115667873 | 115668294 | 0 | 0 |
| SE | 82664 | ENSG00000104447 | TRPS1 | chr8 | - | 115637233 | 115637330 | 115623600 | 115623758 | 115667873 | 115668294 | 1 | 0 |
| SE | 82667 | ENSG00000141380 | SS18 | chr18 | - | 26038554 | 26038659 | 26035830 | 26035923 | 26039288 | 26039456 | 134 | 72 |
| SE | 82669 | ENSG00000141380 | SS18 | chr18 | - | 26068493 | 26068557 | 26064831 | 26064871 | 26078075 | 26078160 | 5 | 5 |
| SE | 82670 | ENSG00000141380 | SS18 | chr18 | - | 26078075 | 26078160 | 26057588 | 26057742 | 26087500 | 26087577 | 258 | 210 |
| SE | 82671 | ENSG00000141380 | SS18 | chr18 | - | 26079010 | 26079125 | 26078075 | 26078160 | 26087500 | 26087577 | 1 | 1 |
| SE | 82672 | ENSG00000141380 | SS18 | chr18 | - | 26079010 | 26079146 | 26078075 | 26078160 | 26082362 | 26082528 | 1 | 1 |
| SE | 82673 | ENSG00000141380 | SS18 | chr18 | - | 26079010 | 26079146 | 26078075 | 26078160 | 26084051 | 26084175 | 1 | 1 |
| SE | 82674 | ENSG00000141380 | SS18 | chr18 | - | 26080304 | 26080423 | 26078075 | 26078160 | 26082362 | 26082528 | 2 | 0 |
| SE | 82680 | ENSG00000141380 | SS18 | chr18 | - | 26082362 | 26082528 | 26080304 | 26080423 | 26084051 | 26084175 | 1 | 0 |
| SE | 82681 | ENSG00000141380 | SS18 | chr18 | - | 26082362 | 26082528 | 26080304 | 26080423 | 26087500 | 26087577 | 4 | 0 |
| SE | 82683 | ENSG00000141380 | SS18 | chr18 | - | 26084051 | 26084175 | 26080370 | 26080423 | 26087500 | 26087577 | 1 | 1 |
| SE | 82684 | ENSG00000141380 | SS18 | chr18 | - | 26084051 | 26084175 | 26082362 | 26082528 | 26087500 | 26087577 | 1 | 1 |
| SE | 82693 | ENSG00000097033 | SH3GLB1 | chr1 | + | 86719506 | 86719635 | 86715723 | 86715865 | 86722539 | 86722673 | 399 | 316 |
| SE | 82694 | ENSG00000097033 | SH3GLB1 | chr1 | + | 86728402 | 86728441 | 86724312 | 86724405 | 86730087 | 86730111 | 0 | 0 |
| SE | 82695 | ENSG00000097033 | SH3GLB1 | chr1 | + | 86728402 | 86728489 | 86724312 | 86724405 | 86730087 | 86730111 | 0 | 5 |
| SE | 82696 | ENSG00000097033 | SH3GLB1 | chr1 | + | 86728402 | 86728489 | 86724312 | 86724405 | 86734601 | 86734691 | 3 | 1 |
| SE | 82698 | ENSG00000097033 | SH3GLB1 | chr1 | + | 86730087 | 86730111 | 86728402 | 86728489 | 86734601 | 86734691 | 0 | 0 |
| SE | 82701 | ENSG00000127914 | AKAP9 | chr7 | + | 91973710 | 91973968 | 91940839 | 91941147 | 91980288 | 91980333 | 287 | 252 |
| SE | 82706 | ENSG00000127914 | AKAP9 | chr7 | + | 91992157 | 91992211 | 91980872 | 91980958 | 91992884 | 91993055 | 49 | 40 |
| SE | 82707 | ENSG00000127914 | AKAP9 | chr7 | + | 92000847 | 92003235 | 91995602 | 91995800 | 92012428 | 92012642 | 218 | 230 |
| SE | 82708 | ENSG00000127914 | AKAP9 | chr7 | + | 92012428 | 92012642 | 92000847 | 92003235 | 92016128 | 92016267 | 164 | 153 |
| SE | 82709 | ENSG00000127914 | AKAP9 | chr7 | + | 92014248 | 92014328 | 92012428 | 92012642 | 92016128 | 92016267 | 362 | 394 |
| SE | 82710 | ENSG00000127914 | AKAP9 | chr7 | + | 92016128 | 92016267 | 92014248 | 92014328 | 92017016 | 92017102 | 654 | 701 |
| SE | 82713 | ENSG00000127914 | AKAP9 | chr7 | + | 92042667 | 92042771 | 92040673 | 92040898 | 92045007 | 92045213 | 110 | 137 |
| SE | 82714 | ENSG00000127914 | AKAP9 | chr7 | + | 92043281 | 92043377 | 92042667 | 92042771 | 92045007 | 92045213 | 2 | 1 |
| SE | 82715 | ENSG00000127914 | AKAP9 | chr7 | + | 92061259 | 92061422 | 92052725 | 92052958 | 92062273 | 92062486 | 341 | 354 |
| SE | 82717 | ENSG00000127914 | AKAP9 | chr7 | + | 92065230 | 92065463 | 92062273 | 92062486 | 92066426 | 92066546 | 124 | 165 |
| SE | 82720 | ENSG00000127914 | AKAP9 | chr7 | + | 92070904 | 92071009 | 92070029 | 92070206 | 92076854 | 92077007 | 42 | 40 |
| SE | 82721 | ENSG00000127914 | AKAP9 | chr7 | + | 92076854 | 92077007 | 92070904 | 92071009 | 92077695 | 92077875 | 104 | 141 |
| SE | 82722 | ENSG00000127914 | AKAP9 | chr7 | + | 92079078 | 92080152 | 92077695 | 92077875 | 92082521 | 92082662 | 156 | 187 |
| SE | 82723 | ENSG00000127914 | AKAP9 | chr7 | + | 92084818 | 92084940 | 92084639 | 92084703 | 92085494 | 92085686 | 144 | 135 |
| SE | 82724 | ENSG00000127914 | AKAP9 | chr7 | + | 92086227 | 92086416 | 92085494 | 92085686 | 92089384 | 92089529 | 124 | 105 |
| SE | 82726 | ENSG00000127914 | AKAP9 | chr7 | + | 92092673 | 92093316 | 92089384 | 92089529 | 92095022 | 92095173 | 137 | 173 |
| SE | 82727 | ENSG00000127914 | AKAP9 | chr7 | + | 92093096 | 92093316 | 92089384 | 92089529 | 92095022 | 92095173 | 195 | 228 |
| SE | 82730 | ENSG00000127914 | AKAP9 | chr7 | + | 92102593 | 92102826 | 92099686 | 92101056 | 92108493 | 92108633 | 40 | 48 |
| SE | 82734 | ENSG00000127914 | AKAP9 | chr7 | + | 92107292 | 92107422 | 92100855 | 92101056 | 92108493 | 92108633 | 40 | 51 |
| SE | 82736 | ENSG00000127914 | AKAP9 | chr7 | + | 92107292 | 92107422 | 92105677 | 92105763 | 92108493 | 92108633 | 91 | 114 |
| SE | 82738 | ENSG00000127914 | AKAP9 | chr7 | + | 92108493 | 92108633 | 92105677 | 92105763 | 92110121 | 92110571 | 25 | 33 |
| SE | 82739 | ENSG00000165632 | TAF3 | chr10 | + | 8013730 | 8013837 | 8009077 | 8009330 | 8014636 | 8016631 | 34 | 18 |
| SE | 82740 | ENSG00000140396 | NCOA2 | chr8 | - | 70126812 | 70127047 | 70124687 | 70124865 | 70128432 | 70128510 | 277 | 176 |
| SE | 82743 | ENSG00000140396 | NCOA2 | chr8 | - | 70144641 | 70144848 | 70141183 | 70141399 | 70157124 | 70157146 | 82 | 94 |
| SE | 82745 | ENSG00000140396 | NCOA2 | chr8 | - | 70148272 | 70148483 | 70141178 | 70141399 | 70157124 | 70157146 | 28 | 10 |
| SE | 82747 | ENSG00000140396 | NCOA2 | chr8 | - | 70148272 | 70148483 | 70144641 | 70144848 | 70157124 | 70157146 | 55 | 53 |
| SE | 82748 | ENSG00000140396 | NCOA2 | chr8 | - | 70159504 | 70159652 | 70155970 | 70157240 | 70162710 | 70162854 | 289 | 217 |
| SE | 82750 | ENSG00000140396 | NCOA2 | chr8 | - | 70213902 | 70214075 | 70174755 | 70174859 | 70216659 | 70216764 | 367 | 367 |
| SE | 82751 | ENSG00000140396 | NCOA2 | chr8 | - | 70213902 | 70214075 | 70174755 | 70174859 | 70296743 | 70296800 | 313 | 308 |
| SE | 82753 | ENSG00000140396 | NCOA2 | chr8 | - | 70216659 | 70216764 | 70213902 | 70214075 | 70296743 | 70296800 | 72 | 63 |
| SE | 82754 | ENSG00000166455 | C16orf46 | chr16 | - | 81063745 | 81063993 | 81060979 | 81062138 | 81077135 | 81077213 | 2 | 0 |
| SE | 82758 | ENSG00000237357 | NA | chr9 | - | 42568784 | 42568914 | 42566992 | 42568059 | 42569028 | 42569310 | 0 | 1 |
| SE | 82759 | ENSG00000272968 | RBAK-RBAKDN | chr7 | + | 5048032 | 5048091 | 4984909 | 4985075 | 5057294 | 5057421 | 5 | 10 |
| SE | 82762 | ENSG00000196873 | CBWD3 | chr9 | + | 68232436 | 68232495 | 68232002 | 68232198 | 68234999 | 68235128 | 21 | 19 |
| SE | 82763 | ENSG00000196873 | CBWD3 | chr9 | + | 68243182 | 68243280 | 68234999 | 68235128 | 68245212 | 68245299 | 0 | 0 |
| SE | 82764 | ENSG00000196873 | CBWD3 | chr9 | + | 68243182 | 68243280 | 68241912 | 68242185 | 68245212 | 68245299 | 0 | 0 |
| SE | 82765 | ENSG00000196873 | CBWD3 | chr9 | + | 68256920 | 68256980 | 68248805 | 68248897 | 68258531 | 68258569 | 37 | 24 |
| SE | 82768 | ENSG00000196873 | CBWD3 | chr9 | + | 68260856 | 68260996 | 68259346 | 68259434 | 68264467 | 68264514 | 3 | 2 |
| SE | 82769 | ENSG00000196873 | CBWD3 | chr9 | + | 68260856 | 68261001 | 68258531 | 68258569 | 68264467 | 68264514 | 0 | 0 |
| SE | 82770 | ENSG00000196873 | CBWD3 | chr9 | + | 68266219 | 68266306 | 68264467 | 68264514 | 68268933 | 68268978 | 20 | 16 |
| SE | 82771 | ENSG00000196873 | CBWD3 | chr9 | + | 68268933 | 68268978 | 68266219 | 68266306 | 68285988 | 68286040 | 3 | 3 |
| SE | 82772 | ENSG00000196873 | CBWD3 | chr9 | + | 68269599 | 68269722 | 68268933 | 68268978 | 68274881 | 68274938 | 2 | 1 |
| SE | 82773 | ENSG00000196873 | CBWD3 | chr9 | + | 68269599 | 68269722 | 68268933 | 68268978 | 68285988 | 68286040 | 2 | 1 |
| SE | 82774 | ENSG00000196873 | CBWD3 | chr9 | + | 68270301 | 68270518 | 68268933 | 68268978 | 68274881 | 68274938 | 0 | 0 |
| SE | 82775 | ENSG00000196873 | CBWD3 | chr9 | + | 68274881 | 68274938 | 68268933 | 68268978 | 68285988 | 68286040 | 5 | 1 |
| SE | 82776 | ENSG00000196873 | CBWD3 | chr9 | + | 68274881 | 68274938 | 68269599 | 68269722 | 68285988 | 68286040 | 2 | 0 |
| SE | 82778 | ENSG00000122591 | FAM126A | chr7 | - | 22946951 | 22947247 | 22941268 | 22946163 | 22960255 | 22960415 | 0 | 0 |
| SE | 82779 | ENSG00000122591 | FAM126A | chr7 | - | 22947032 | 22947247 | 22945982 | 22946163 | 22960255 | 22960415 | 0 | 0 |
| SE | 82780 | ENSG00000122591 | FAM126A | chr7 | - | 22960255 | 22960415 | 22934210 | 22946163 | 22964414 | 22964531 | 82 | 106 |
| SE | 82781 | ENSG00000122591 | FAM126A | chr7 | - | 22960255 | 22960415 | 22934210 | 22946163 | 22976209 | 22976305 | 81 | 102 |
| SE | 82784 | ENSG00000122591 | FAM126A | chr7 | - | 22961234 | 22961322 | 22960255 | 22960415 | 22964414 | 22964531 | 375 | 320 |
| SE | 82785 | ENSG00000122591 | FAM126A | chr7 | - | 22961234 | 22961322 | 22960255 | 22960415 | 22976209 | 22976305 | 176 | 157 |
| SE | 82786 | ENSG00000122591 | FAM126A | chr7 | - | 22964414 | 22964531 | 22934210 | 22946163 | 22976209 | 22976305 | 125 | 75 |
| SE | 82787 | ENSG00000122591 | FAM126A | chr7 | - | 22964414 | 22964531 | 22934210 | 22946163 | 23013924 | 23014084 | 0 | 1 |
| SE | 82788 | ENSG00000122591 | FAM126A | chr7 | - | 22964414 | 22964531 | 22960255 | 22960415 | 22976209 | 22976305 | 126 | 78 |
| SE | 82789 | ENSG00000122591 | FAM126A | chr7 | - | 22964414 | 22964531 | 22961234 | 22961322 | 22976209 | 22976305 | 324 | 237 |
| SE | 82790 | ENSG00000122591 | FAM126A | chr7 | - | 22976209 | 22976305 | 22964414 | 22964531 | 22976692 | 22976808 | 234 | 137 |
| SE | 82792 | ENSG00000122591 | FAM126A | chr7 | - | 22978268 | 22978404 | 22977340 | 22977421 | 22983943 | 22984045 | 124 | 86 |
| SE | 82794 | ENSG00000122591 | FAM126A | chr7 | - | 22983943 | 22984045 | 22934210 | 22946163 | 23013924 | 23014130 | 0 | 0 |
| SE | 82795 | ENSG00000122591 | FAM126A | chr7 | - | 22983943 | 22984045 | 22978163 | 22978448 | 22991060 | 22991139 | 214 | 147 |
| SE | 82797 | ENSG00000122591 | FAM126A | chr7 | - | 22991060 | 22991139 | 22980296 | 22984045 | 23013924 | 23014074 | 186 | 154 |
| SE | 82801 | ENSG00000106268 | NUDT1 | chr7 | + | 2242924 | 2243048 | 2242225 | 2242256 | 2244562 | 2244726 | 0 | 0 |
| SE | 82802 | ENSG00000106268 | NUDT1 | chr7 | + | 2244562 | 2244726 | 2242904 | 2242975 | 2249856 | 2250002 | 40 | 11 |
| SE | 82803 | ENSG00000106268 | NUDT1 | chr7 | + | 2249856 | 2250002 | 2244475 | 2244726 | 2250828 | 2251145 | 86 | 26 |
| SE | 82805 | ENSG00000136271 | DDX56 | chr7 | - | 44571491 | 44571736 | 44570014 | 44570128 | 44572346 | 44572437 | 76 | 41 |
| SE | 82806 | ENSG00000136271 | DDX56 | chr7 | - | 44571491 | 44571736 | 44570014 | 44570128 | 44572562 | 44572744 | 0 | 0 |
| SE | 82807 | ENSG00000136271 | DDX56 | chr7 | - | 44572346 | 44572437 | 44569983 | 44570128 | 44572562 | 44572744 | 3 | 1 |
| SE | 82808 | ENSG00000136271 | DDX56 | chr7 | - | 44572346 | 44572437 | 44571491 | 44571736 | 44572562 | 44572744 | 52 | 22 |
| SE | 82811 | ENSG00000106682 | EIF4H | chr7 | + | 74178655 | 74178826 | 74174357 | 74174442 | 74187610 | 74187798 | 0 | 0 |
| SE | 82812 | ENSG00000106682 | EIF4H | chr7 | + | 74179337 | 74179418 | 74174357 | 74174442 | 74187610 | 74187798 | 0 | 0 |
| SE | 82813 | ENSG00000106682 | EIF4H | chr7 | + | 74187610 | 74187798 | 74174230 | 74174442 | 74189672 | 74189737 | 956 | 659 |
| SE | 82814 | ENSG00000106682 | EIF4H | chr7 | + | 74189672 | 74189737 | 74187610 | 74187798 | 74189821 | 74189918 | 1345 | 1048 |
| SE | 82816 | ENSG00000106682 | EIF4H | chr7 | + | 74194740 | 74194878 | 74189821 | 74189918 | 74195168 | 74195494 | 1407 | 976 |
| SE | 82817 | ENSG00000106682 | EIF4H | chr7 | + | 74194740 | 74195059 | 74189821 | 74189918 | 74195168 | 74197077 | 918 | 660 |
| SE | 82818 | ENSG00000155849 | ELMO1 | chr7 | - | 36870392 | 36870475 | 36861658 | 36861736 | 36894853 | 36895017 | 39 | 21 |
| SE | 82820 | ENSG00000155849 | ELMO1 | chr7 | - | 36878009 | 36878117 | 36870392 | 36870475 | 36894853 | 36895017 | 138 | 83 |
| SE | 82821 | ENSG00000155849 | ELMO1 | chr7 | - | 36887559 | 36887672 | 36870392 | 36870475 | 36894853 | 36895017 | 89 | 55 |
| SE | 82824 | ENSG00000155849 | ELMO1 | chr7 | - | 37133129 | 37133234 | 37096618 | 37096727 | 37211385 | 37211517 | 290 | 237 |
| SE | 82826 | ENSG00000155849 | ELMO1 | chr7 | - | 37224878 | 37225030 | 37222614 | 37222693 | 37233094 | 37233194 | 276 | 246 |
| SE | 82827 | ENSG00000155849 | ELMO1 | chr7 | - | 37244355 | 37244391 | 37233094 | 37233194 | 37259180 | 37259350 | 193 | 169 |
| SE | 82828 | ENSG00000155849 | ELMO1 | chr7 | - | 37271831 | 37271882 | 37259180 | 37259350 | 37314849 | 37314922 | 264 | 240 |
| SE | 82830 | ENSG00000155849 | ELMO1 | chr7 | - | 37393821 | 37393917 | 37342612 | 37342763 | 37448859 | 37448950 | 0 | 0 |
| SE | 82831 | ENSG00000196329 | GIMAP5 | chr7 | + | 150747713 | 150747863 | 150740878 | 150740927 | 150749731 | 150750033 | 1 | 2 |
| SE | 82832 | ENSG00000162129 | CLPB | chr11 | - | 72294324 | 72294444 | 72294021 | 72294126 | 72294619 | 72294693 | 75 | 62 |
| SE | 82835 | ENSG00000162129 | CLPB | chr11 | - | 72308526 | 72308604 | 72307198 | 72307254 | 72317105 | 72317220 | 69 | 35 |
| SE | 82843 | ENSG00000181513 | ACBD4 | chr17 | + | 45133030 | 45133201 | 45132604 | 45132663 | 45135340 | 45135592 | 0 | 0 |
| SE | 82844 | ENSG00000181513 | ACBD4 | chr17 | + | 45136107 | 45136232 | 45135646 | 45135953 | 45136499 | 45136620 | 18 | 16 |
| SE | 82848 | ENSG00000181513 | ACBD4 | chr17 | + | 45139020 | 45139160 | 45137912 | 45137988 | 45143442 | 45143850 | 25 | 13 |
| SE | 82851 | ENSG00000146776 | ATXN7L1 | chr7 | - | 105610528 | 105610603 | 105607770 | 105607893 | 105613861 | 105614816 | 31 | 31 |
| SE | 82852 | ENSG00000146776 | ATXN7L1 | chr7 | - | 105613861 | 105614816 | 105610528 | 105610603 | 105624074 | 105624267 | 33 | 30 |
| SE | 82855 | ENSG00000146776 | ATXN7L1 | chr7 | - | 105624074 | 105624267 | 105620199 | 105620321 | 105638352 | 105638609 | 76 | 35 |
| SE | 82859 | ENSG00000146776 | ATXN7L1 | chr7 | - | 105665065 | 105665288 | 105642837 | 105643121 | 105788603 | 105788708 | 83 | 55 |
| SE | 82861 | ENSG00000146776 | ATXN7L1 | chr7 | - | 105788603 | 105788708 | 105665065 | 105665288 | 105876377 | 105876477 | 53 | 27 |
| SE | 82862 | ENSG00000146776 | ATXN7L1 | chr7 | - | 105788603 | 105788708 | 105760372 | 105761463 | 105875811 | 105875880 | 48 | 25 |
| SE | 82863 | ENSG00000146776 | ATXN7L1 | chr7 | - | 105875811 | 105875880 | 105665065 | 105665288 | 105876377 | 105876574 | 51 | 25 |
| SE | 82865 | ENSG00000161048 | NAPEPLD | chr7 | - | 103115059 | 103115174 | 103099775 | 103103554 | 103119576 | 103119766 | 85 | 76 |
| SE | 82867 | ENSG00000161048 | NAPEPLD | chr7 | - | 103119576 | 103120223 | 103099775 | 103103554 | 103128482 | 103128792 | 22 | 15 |
| SE | 82868 | ENSG00000161048 | NAPEPLD | chr7 | - | 103119576 | 103120223 | 103115059 | 103115174 | 103128482 | 103128792 | 64 | 38 |
| SE | 82869 | ENSG00000267673 | FDX2 | chr19 | - | 10315385 | 10315553 | 10310852 | 10310940 | 10315704 | 10315795 | 13 | 5 |
| SE | 82870 | ENSG00000237471 | NA | chr7 | + | 45977518 | 45977678 | 45940448 | 45940880 | 45978349 | 45978468 | 27 | 9 |
| SE | 82875 | ENSG00000170852 | KBTBD2 | chr7 | - | 32874991 | 32875157 | 32868171 | 32870880 | 32879434 | 32879672 | 473 | 371 |
| SE | 82876 | ENSG00000170852 | KBTBD2 | chr7 | - | 32879434 | 32879942 | 32874991 | 32875157 | 32891535 | 32891760 | 270 | 207 |
| SE | 82877 | ENSG00000205763 | RP9P | chr7 | - | 32921338 | 32921430 | 32918310 | 32918372 | 32930077 | 32930108 | 7 | 6 |
| SE | 82881 | ENSG00000122643 | NT5C3A | chr7 | - | 33017438 | 33017601 | 33014129 | 33014831 | 33024038 | 33024108 | 0 | 0 |
| SE | 82883 | ENSG00000122643 | NT5C3A | chr7 | - | 33019634 | 33019724 | 33017203 | 33017601 | 33026816 | 33026915 | 89 | 117 |
| SE | 82884 | ENSG00000122643 | NT5C3A | chr7 | - | 33021271 | 33021357 | 33019634 | 33019724 | 33022052 | 33022099 | 496 | 521 |
| SE | 82885 | ENSG00000122643 | NT5C3A | chr7 | - | 33022052 | 33022099 | 33019634 | 33019724 | 33026816 | 33026915 | 1 | 0 |
| SE | 82886 | ENSG00000122643 | NT5C3A | chr7 | - | 33024038 | 33024108 | 33017203 | 33017601 | 33026816 | 33026915 | 210 | 239 |
| SE | 82887 | ENSG00000122643 | NT5C3A | chr7 | - | 33024038 | 33024108 | 33021824 | 33022099 | 33026816 | 33026915 | 471 | 539 |
| SE | 82888 | ENSG00000122643 | NT5C3A | chr7 | - | 33026816 | 33026915 | 33023718 | 33024108 | 33035933 | 33035983 | 395 | 489 |
| SE | 82893 | ENSG00000197008 | ZNF138 | chr7 | + | 64814058 | 64814147 | 64794427 | 64794571 | 64814917 | 64815044 | 2 | 0 |
| SE | 82894 | ENSG00000197008 | ZNF138 | chr7 | + | 64814917 | 64815044 | 64794387 | 64794571 | 64831450 | 64832242 | 7 | 3 |
| SE | 82895 | ENSG00000197008 | ZNF138 | chr7 | + | 64815571 | 64815653 | 64814917 | 64815044 | 64831450 | 64833327 | 7 | 1 |
| SE | 82897 | ENSG00000197008 | ZNF138 | chr7 | + | 64815575 | 64815653 | 64814917 | 64815044 | 64831450 | 64833680 | 4 | 1 |
| SE | 82898 | ENSG00000197008 | ZNF138 | chr7 | + | 64830939 | 64831076 | 64814917 | 64815044 | 64831450 | 64831803 | 5 | 7 |
| SE | 82899 | ENSG00000197008 | ZNF138 | chr7 | + | 64830939 | 64831076 | 64815575 | 64815653 | 64831450 | 64831803 | 5 | 7 |
| SE | 82901 | ENSG00000285106 | NA | chr7 | - | 130872500 | 130872758 | 130868317 | 130868472 | 130884316 | 130884396 | 3 | 0 |

| | | | | | | | | | | | | | | | | | | | | | | | | | | | | | | | | | | | | | | | | | | | | | | |
|---|---|---|---|---|---|---|---|---|---|---|---|---|---|---|---|---|---|---|---|---|---|---|---|---|---|---|---|---|---|---|---|---|---|---|---|---|---|---|---|---|---|---|---|---|---|---|
| SE | 82902 | ENSG00000285106 | NA | chr7 | - | 130912109 | 130912152 | 130881057 | 130884396 | 130945719 | 130945835 | 3 | 2 | [illegible] | [illegible] | [illegible] | [illegible] | [illegible] | [illegible] | [illegible] | [illegible] | 0 | 0 | 4 | 0 | 0 | 0 | 0 | 0 | 0 | 0 | 192 | 149 | 1 | 1 | 0.28 | 0.437 | 1 | NA | NA | NA | NA | 1 | NA | NA | -0.428 |
| SE | 82903 | ENSG00000285106 | NA | chr7 | - | 130945719 | 130945835 | 130912109 | 130912152 | 130984052 | 130984109 | 42 | 37 | [illegible] | [illegible] | [illegible] | [illegible] | [illegible] | [illegible] | [illegible] | [illegible] | 13 | 17 | 39 | 31 | 3 | 0 | 0 | 0 | 0 | 0 | 265 | 149 | 1 | 1 | 1 | 1 | 0.962 | 1 | 1 | 1 | 1 | 1 | 1 | 1 | -0.008 |
| SE | 82905 | ENSG00000234456 | MAGI2-AS3 | chr7 | + | 79453586 | 79453655 | 79453157 | 79453191 | 79458846 | 79458999 | 0 | 2 | [illegible] | [illegible] | [illegible] | [illegible] | [illegible] | [illegible] | [illegible] | [illegible] | 0 | 1 | 3 | 0 | 0 | 0 | 0 | 0 | 0 | 0 | 218 | 149 | 1 | 1 | NA | 1 | 0.578 | 1 | NA | NA | 1 | 1 | NA | NA | -0.141 |
| SE | 82906 | ENSG00000234456 | MAGI2-AS3 | chr7 | + | 79454865 | 79455007 | 79452876 | 79453191 | 79458846 | 79458999 | 2 | 57 | 8 | 27 | 8 | 0 | 0 | 1 | 0 | 0 | 0 | 1 | 29 | 3 | 12 | 0 | 0 | 0 | 0 | 0 | 291 | 149 | 1 | 1 | 1 | 1 | 0.804 | 1 | 1 | NA | 1 | 1 | 1 | 1 | -0.039 |
| SE | 82907 | ENSG00000234456 | MAGI2-AS3 | chr7 | + | 79454865 | 79455007 | 79453571 | 79453655 | 79458846 | 79458999 | 2 | 57 | 8 | 27 | 8 | 0 | 2 | 2 | 4 | 0 | 0 | 1 | 29 | 3 | 12 | 0 | 1 | 3 | 0 | 0 | 291 | 149 | 1 | 1 | 1 | 0.936 | 0.672 | 0.776 | 1 | NA | 0.339 | 0.832 | 1 | 1 | 0.084 |
| SE | 82908 | ENSG00000234456 | MAGI2-AS3 | chr7 | + | 79454865 | 79455007 | 79453734 | 79453789 | 79458846 | 79458999 | 2 | 57 | 8 | 27 | 8 | 0 | 2 | 0 | 0 | 0 | 0 | 1 | 29 | 3 | 12 | 0 | 0 | 0 | 0 | 0 | 291 | 149 | 1 | 1 | 1 | 0.936 | 1 | 1 | 1 | NA | 1 | 1 | 1 | 1 | -0.013 |
| SE | 82909 | ENSG00000234456 | MAGI2-AS3 | chr7 | + | 79454865 | 79455007 | 79453975 | 79454244 | 79458846 | 79458999 | 2 | 64 | 8 | 28 | 8 | 0 | 1 | 0 | 2 | 0 | 0 | 1 | 30 | 3 | 14 | 0 | 0 | 0 | 0 | 0 | 291 | 149 | 1 | 1 | 1 | 0.97 | 1 | 0.878 | 1 | NA | 1 | 1 | 1 | 1 | -0.03 |
| SE | 82910 | ENSG00000234456 | MAGI2-AS3 | chr7 | + | 79454885 | 79455007 | 79453559 | 79453655 | 79458846 | 79458999 | 2 | 55 | 8 | 27 | 8 | 0 | 2 | 2 | 4 | 0 | 0 | 1 | 29 | 3 | 12 | 0 | 1 | 3 | 0 | 0 | 271 | 149 | 1 | 1 | 1 | 0.938 | 0.687 | 0.788 | 1 | NA | 0.355 | 0.842 | 1 | 1 | 0.083 |
| SE | 82911 | ENSG00000234456 | MAGI2-AS3 | chr7 | + | 79454885 | 79455007 | 79453752 | 79453789 | 79458846 | 79458999 | 2 | 55 | 10 | 28 | 8 | 0 | 2 | 0 | 0 | 0 | 0 | 1 | 29 | 3 | 12 | 0 | 0 | 0 | 0 | 0 | 271 | 149 | 1 | 1 | 1 | 0.938 | 1 | 1 | 1 | NA | 1 | 1 | 1 | 1 | -0.012 |
| SE | 82912 | ENSG00000234456 | MAGI2-AS3 | chr7 | + | 79454885 | 79455007 | 79453850 | 79454244 | 79458846 | 79458999 | 2 | 61 | 8 | 31 | 8 | 0 | 1 | 0 | 2 | 0 | 0 | 2 | 33 | 3 | 21 | 0 | 0 | 0 | 0 | 0 | 271 | 149 | 1 | 1 | 1 | 0.971 | 1 | 0.895 | 1 | NA | 1 | 1 | 1 | 1 | -0.027 |
| SE | 82913 | ENSG00000234456 | MAGI2-AS3 | chr7 | + | 79454885 | 79458730 | 79453999 | 79454244 | 79458846 | 79458999 | 0 | 8 | 0 | 4 | 0 | 0 | 1 | 0 | 2 | 0 | 0 | 1 | 2 | 0 | 9 | 0 | 0 | 0 | 0 | 0 | 298 | 149 | 1 | 1 | NA | 0.8 | NA | 0.5 | NA | NA | 1 | 1 | NA | 1 | -0.35 |
| SE | 82914 | ENSG00000234456 | MAGI2-AS3 | chr7 | + | 79455889 | 79456180 | 79454865 | 79455007 | 79458846 | 79458999 | 1 | 12 | 0 | 6 | 4 | 2 | 58 | 10 | 28 | 8 | 0 | 0 | 3 | 2 | 0 | 0 | 1 | 31 | 3 | 12 | 298 | 149 | 1 | 1 | 0.2 | 0.094 | 0 | 0.097 | 0.2 | NA | 0 | 0.046 | 0.25 | 0 | 0.044 |
| SE | 82916 | ENSG00000234456 | MAGI2-AS3 | chr7 | + | 79456089 | 79456180 | 79454828 | 79455007 | 79458846 | 79458999 | 1 | 10 | 0 | 2 | 0 | 2 | 58 | 10 | 28 | 8 | 0 | 0 | 0 | 0 | 0 | 0 | 1 | 31 | 3 | 12 | 240 | 149 | 1 | 1 | 0.237 | 0.097 | 0 | 0.042 | 0 | NA | 0 | 0 | 0 | 0 | 0.075 |
| SE | 82917 | ENSG00000234456 | MAGI2-AS3 | chr7 | + | 79458658 | 79458730 | 79454085 | 79455374 | 79458846 | 79458999 | 0 | 3 | 0 | 0 | 0 | 0 | 1 | 0 | 0 | 0 | 0 | 0 | 0 | 0 | 0 | 0 | 0 | 2 | 0 | 0 | 221 | 149 | 1 | 1 | NA | 0.669 | NA | NA | NA | NA | NA | 0 | NA | NA | 0.669 |
| SE | 82918 | ENSG00000234456 | MAGI2-AS3 | chr7 | + | 79458658 | 79458730 | 79454885 | 79455007 | 79458846 | 79458999 | 0 | 4 | 0 | 2 | 0 | 2 | 58 | 10 | 28 | 8 | 0 | 0 | 0 | 0 | 0 | 0 | 1 | 31 | 3 | 12 | 221 | 149 | 1 | 1 | 0 | 0.044 | 0 | 0.046 | 0 | NA | 0 | 0 | 0 | 0 | 0.018 |
| SE | 82919 | ENSG00000234456 | MAGI2-AS3 | chr7 | + | 79458658 | 79458730 | 79455911 | 79456180 | 79458846 | 79458999 | 0 | 5 | 0 | 0 | 0 | 1 | 11 | 0 | 3 | 2 | 0 | 0 | 0 | 0 | 0 | 0 | 0 | 2 | 1 | 0 | 221 | 149 | 1 | 1 | 0 | 0.235 | NA | 0 | 0 | NA | NA | 0 | 0 | NA | 0.059 |
| SE | 82920 | ENSG00000234456 | MAGI2-AS3 | chr7 | + | 79459406 | 79459453 | 79458846 | 79458999 | 79459535 | 79459625 | 0 | 1 | 0 | 0 | 0 | 4 | 61 | 7 | 23 | 3 | 0 | 0 | 0 | 0 | 0 | 0 | 2 | 44 | 3 | 0 | 196 | 149 | 1 | 1 | 0 | 0.012 | 0 | 0 | 0 | NA | 0 | 0 | 0 | NA | 0.002 |
| SE | 82923 | ENSG00000234456 | MAGI2-AS3 | chr7 | + | 79470593 | 79470679 | 79458846 | 79458999 | 79470782 | 79470991 | 2 | 19 | 4 | 10 | 4 | 2 | 11 | 1 | 3 | 0 | 20 | 0 | 12 | 0 | 0 | 0 | 0 | 3 | 1 | 6 | 235 | 149 | 1 | 1 | 0.388 | 0.523 | 0.717 | 0.679 | 1 | 1 | NA | 0.717 | 0 | 0 | 0.232 |
| SE | 82927 | ENSG00000122786 | CALD1 | chr7 | + | 134935687 | 134935765 | 134932987 | 134933390 | 134941091 | 134941237 | 0 | 0 | 0 | 0 | 0 | 5 | 2 | 2 | 13 | 15 | 0 | 0 | 0 | 3 | 0 | 0 | 3 | 14 | 3 | 15 | 227 | 149 | 1 | 1 | 0 | 0 | 0 | 0 | 0 | NA | 0 | 0 | 0.396 | 0 | -0.099 |
| SE | 82930 | ENSG00000148606 | POLR3A | chr10 | - | 77982652 | 77982817 | 77982153 | 77982318 | 77983919 | 77983978 | 60 | 52 | 92 | 111 | 150 | 1 | 0 | 0 | 0 | 0 | 90 | 70 | 56 | 99 | 53 | 0 | 0 | 3 | 1 | 0 | 298 | 149 | 1 | 1 | 0.968 | 1 | 1 | 1 | 1 | 1 | 1 | 0.903 | 0.98 | 1 | 0.017 |
| SE | 82931 | ENSG00000148606 | POLR3A | chr10 | - | 77982652 | 77982817 | 77982153 | 77982318 | 77984204 | 77984298 | 20 | 15 | 38 | 49 | 42 | 0 | 0 | 0 | 0 | 0 | 22 | 27 | 18 | 37 | 31 | 0 | 0 | 2 | 0 | 0 | 298 | 149 | 1 | 1 | 1 | 1 | 1 | 1 | 1 | 1 | 1 | 0.818 | 1 | 1 | 0.036 |
| SE | 82932 | ENSG00000148606 | POLR3A | chr10 | - | 77983919 | 77984012 | 77982153 | 77982318 | 77984204 | 77984298 | 19 | 12 | 31 | 28 | 23 | 0 | 0 | 0 | 0 | 0 | 17 | 8 | 13 | 29 | 17 | 0 | 0 | 2 | 0 | 0 | 242 | 149 | 1 | 1 | 1 | 1 | 1 | 1 | 1 | 1 | 1 | 0.8 | 1 | 1 | 0.04 |
| SE | 82933 | ENSG00000148606 | POLR3A | chr10 | - | 77983919 | 77984012 | 77982153 | 77982318 | 77985169 | 77985340 | 1 | 0 | 0 | 0 | 0 | 0 | 0 | 0 | 0 | 0 | 0 | 0 | 3 | 1 | 0 | 0 | 1 | 0 | 0 | 0 | 242 | 149 | 1 | 1 | 1 | NA | NA | NA | NA | NA | 0 | 1 | 1 | NA | 0.333 |
| SE | 82934 | ENSG00000148606 | POLR3A | chr10 | - | 77983919 | 77984012 | 77982652 | 77982817 | 77984204 | 77984298 | 58 | 49 | 83 | 92 | 122 | 0 | 0 | 0 | 2 | 0 | 85 | 51 | 47 | 86 | 39 | 0 | 0 | 0 | 0 | 0 | 242 | 149 | 1 | 1 | 1 | 1 | 1 | 0.966 | 1 | 1 | 1 | 1 | 1 | 1 | -0.007 |
| SE | 82935 | ENSG00000148606 | POLR3A | chr10 | - | 77984204 | 77984298 | 77982153 | 77982318 | 77985169 | 77985340 | 20 | 15 | 20 | 23 | 27 | 0 | 0 | 0 | 0 | 0 | 25 | 31 | 19 | 25 | 32 | 0 | 1 | 0 | 0 | 0 | 243 | 149 | 1 | 1 | 1 | 1 | 1 | 1 | 1 | 1 | 0.95 | 1 | 1 | 1 | 0.01 |
| SE | 82938 | ENSG00000148606 | POLR3A | chr10 | - | 77999980 | 78000118 | 77993196 | 77993367 | 78000975 | 78001094 | 84 | 36 | 49 | 40 | 48 | 0 | 3 | 0 | 0 | 0 | 103 | 71 | 27 | 81 | 30 | 0 | 0 | 0 | 0 | 0 | 287 | 149 | 1 | 1 | 1 | 0.862 | 1 | 1 | 1 | 1 | 1 | 1 | 1 | 1 | -0.028 |
| SE | 82948 | ENSG00000075142 | SRI | chr7 | - | 88209982 | 88210130 | 88208859 | 88209452 | 88210881 | 88210925 | 501 | 371 | 382 | 434 | 705 | 0 | 0 | 0 | 0 | 0 | 527 | 422 | 248 | 390 | 466 | 0 | 0 | 1 | 0 | 0 | 297 | 149 | 1 | 1 | 1 | 1 | 1 | 1 | 1 | 1 | 1 | 0.992 | 1 | 1 | 0.002 |
| SE | 82949 | ENSG00000075142 | SRI | chr7 | - | 88210881 | 88210925 | 88208859 | 88209452 | 88217121 | 88217191 | 26 | 12 | 29 | 26 | 18 | 0 | 2 | 0 | 0 | 0 | 26 | 21 | 27 | 27 | 12 | 0 | 0 | 3 | 0 | 0 | 193 | 149 | 1 | 1 | 1 | 0.822 | 1 | 1 | 1 | 1 | 1 | 0.874 | 1 | 1 | -0.01 |
| SE | 82952 | ENSG00000127954 | STEAP4 | chr7 | - | 88282640 | 88283168 | 88280914 | 88281079 | 88283813 | 88284271 | 91 | 43 | 58 | 46 | 124 | 0 | 0 | 0 | 0 | 0 | 57 | 36 | 61 | 96 | 0 | 18 | 0 | 0 | 2 | 0 | 298 | 149 | 1 | 1 | 1 | 1 | 1 | 1 | 1 | 0.613 | 1 | 1 | 0.96 | NA | 0.107 |
| SE | 82955 | ENSG00000008277 | ADAM22 | chr7 | + | 88128601 | 88128676 | 88125588 | 88125659 | 88130387 | 88130439 | 22 | 11 | 16 | 25 | 29 | 0 | 0 | 0 | 1 | 0 | 20 | 18 | 19 | 19 | 12 | 0 | 0 | 0 | 0 | 0 | 224 | 149 | 1 | 1 | 1 | 1 | 1 | 0.943 | 1 | 1 | 1 | 1 | 1 | 1 | -0.011 |
| SE | 82956 | ENSG00000008277 | ADAM22 | chr7 | + | 88131268 | 88131435 | 88130387 | 88130459 | 88132866 | 88132951 | 38 | 19 | 30 | 39 | 29 | 2 | 0 | 0 | 0 | 0 | 0 | 11 | 30 | 19 | 0 | 10 | 0 | 0 | 0 | 0 | 298 | 149 | 1 | 1 | 0.905 | 1 | 1 | 1 | 1 | 0 | 1 | 1 | 1 | NA | 0.231 |
| SE | 82957 | ENSG00000008277 | ADAM22 | chr7 | + | 88131268 | 88131435 | 88130387 | 88130459 | 88135979 | 88136031 | 27 | 10 | 20 | 26 | 23 | 0 | 0 | 0 | 0 | 0 | 0 | 9 | 16 | 16 | 0 | 0 | 6 | 0 | 0 | 0 | 298 | 149 | 1 | 1 | 1 | 1 | 1 | 1 | 1 | NA | 0.429 | 1 | 1 | NA | 0.19 |
| SE | 82958 | ENSG00000008277 | ADAM22 | chr7 | + | 88132866 | 88132951 | 88130387 | 88130459 | 88135979 | 88136031 | 2 | 0 | 0 | 0 | 0 | 0 | 0 | 0 | 0 | 0 | 10 | 0 | 0 | 0 | 0 | 0 | 6 | 0 | 0 | 0 | 234 | 149 | 1 | 1 | 1 | NA | NA | NA | NA | 1 | 0 | NA | NA | NA | 0.5 |
| SE | 82959 | ENSG00000008277 | ADAM22 | chr7 | + | 88132866 | 88132951 | 88131268 | 88131435 | 88134328 | 88134419 | 19 | 10 | 13 | 16 | 9 | 0 | 0 | 0 | 1 | 0 | 0 | 2 | 17 | 6 | 0 | 0 | 0 | 0 | 0 | 0 | 234 | 149 | 1 | 1 | 1 | 1 | 1 | 0.911 | 1 | NA | 1 | 1 | 1 | NA | -0.018 |
| SE | 82960 | ENSG00000008277 | ADAM22 | chr7 | + | 88132866 | 88132951 | 88131268 | 88131435 | 88135979 | 88136031 | 11 | 7 | 11 | 12 | 6 | 1 | 0 | 1 | 0 | 0 | 0 | 2 | 12 | 3 | 0 | 0 | 0 | 0 | 0 | 0 | 234 | 149 | 1 | 1 | 0.875 | 1 | 0.875 | 1 | 1 | NA | 1 | 1 | 1 | NA | -0.05 |
| SE | 82961 | ENSG00000008277 | ADAM22 | chr7 | + | 88132866 | 88132982 | 88131268 | 88131435 | 88134328 | 88134419 | 11 | 6 | 11 | 12 | 6 | 0 | 0 | 0 | 1 | 0 | 0 | 2 | 12 | 3 | 0 | 0 | 0 | 0 | 0 | 0 | 265 | 149 | 1 | 1 | 1 | 1 | 1 | 0.871 | 1 | NA | 1 | 1 | 1 | NA | -0.026 |
| SE | 82962 | ENSG00000008277 | ADAM22 | chr7 | + | 88134328 | 88134419 | 88131268 | 88131435 | 88135979 | 88136031 | 5 | 1 | 1 | 1 | 3 | 1 | 0 | 1 | 0 | 0 | 0 | 0 | 2 | 0 | 0 | 0 | 0 | 0 | 0 | 0 | 240 | 149 | 1 | 1 | 0.756 | 1 | 0.383 | 1 | 1 | NA | NA | 1 | NA | NA | -0.172 |
| SE | 82963 | ENSG00000008277 | ADAM22 | chr7 | + | 88134328 | 88134419 | 88132866 | 88132951 | 88135979 | 88136031 | 14 | 5 | 3 | 4 | 6 | 0 | 1 | 0 | 0 | 0 | 0 | 0 | 7 | 3 | 0 | 0 | 0 | 0 | 0 | 0 | 240 | 149 | 1 | 1 | 1 | 0.756 | 1 | 1 | 1 | NA | NA | 1 | 1 | NA | -0.049 |
| SE | 82964 | ENSG00000008277 | ADAM22 | chr7 | + | 88148976 | 88149057 | 88145414 | 88145507 | 88150980 | 88151031 | 17 | 17 | 22 | 18 | 11 | 0 | 0 | 0 | 1 | 0 | 0 | 16 | 12 | 5 | 0 | 0 | 0 | 0 | 0 | 0 | 230 | 149 | 1 | 1 | 1 | 1 | 1 | 0.921 | 1 | NA | 1 | 1 | 1 | NA | -0.016 |
| SE | 82965 | ENSG00000008277 | ADAM22 | chr7 | + | 88153220 | 88153326 | 88151256 | 88151320 | 88155886 | 88156006 | 12 | 15 | 31 | 13 | 10 | 1 | 0 | 0 | 0 | 0 | 0 | 0 | 3 | 9 | 9 | 0 | 0 | 0 | 0 | 0 | 255 | 149 | 1 | 1 | 0.875 | 1 | 1 | 1 | 1 | NA | NA | 1 | 1 | 1 | -0.025 |
| SE | 82967 | ENSG00000008277 | ADAM22 | chr7 | + | 88178934 | 88179129 | 88171543 | 88171561 | 88181504 | 88181605 | 2 | 0 | 0 | 0 | 0 | 12 | 6 | 5 | 10 | 7 | 0 | 0 | 0 | 0 | 0 | 0 | 9 | 0 | 8 | 5 | 298 | 149 | 1 | 1 | 0.077 | 0 | 0 | 0 | 0 | NA | 0 | NA | 0 | 0 | 0.015 |
| SE | 82969 | ENSG00000008277 | ADAM22 | chr7 | + | 88186614 | 88186701 | 88181957 | 88182024 | 88193115 | 88193239 | 0 | 0 | 0 | 1 | 0 | 11 | 11 | 7 | 9 | 11 | 0 | 0 | 0 | 0 | 0 | 0 | 11 | 4 | 11 | 0 | 236 | 149 | 1 | 1 | 0 | 0 | 0 | 0.066 | 0 | NA | 0 | 0 | 0 | NA | 0.013 |
| SE | 82970 | ENSG00000008277 | ADAM22 | chr7 | + | 88193115 | 88193239 | 88181957 | 88182024 | 88196470 | 88196582 | 15 | 11 | 9 | 20 | 16 | 0 | 0 | 1 | 0 | 0 | 0 | 11 | 10 | 21 | 0 | 0 | 0 | 0 | 0 | 0 | 273 | 149 | 1 | 1 | 1 | 1 | 0.831 | 1 | 1 | NA | 1 | 1 | 1 | NA | -0.034 |
| SE | 82972 | ENSG00000186088 | GSAP | chr7 | - | 77311840 | 77311940 | 77310750 | 77311449 | 77313487 | 77313549 | 121 | 60 | 61 | 72 | 132 | 0 | 0 | 0 | 0 | 1 | 30 | 47 | 11 | 7 | 16 | 0 | 0 | 0 | 0 | 0 | 249 | 149 | 1 | 1 | 1 | 1 | 1 | 1 | 0.987 | 1 | 1 | 1 | 1 | 1 | -0.003 |
| SE | 82973 | ENSG00000186088 | GSAP | chr7 | - | 77312100 | 77312202 | 77310750 | 77311449 | 77313487 | 77313549 | 77 | 43 | 55 | 31 | 36 | 0 | 0 | 0 | 0 | 1 | 10 | 24 | 23 | 1 | 9 | 0 | 0 | 0 | 0 | 0 | 251 | 149 | 1 | 1 | 1 | 1 | 1 | 1 | 0.955 | 1 | 1 | 1 | 1 | 1 | -0.009 |
| SE | 82975 | ENSG00000186088 | GSAP | chr7 | - | 77321332 | 77321403 | 77320724 | 77320819 | 77323646 | 77323742 | 226 | 143 | 91 | 138 | 178 | 0 | 0 | 6 | 0 | 0 | 11 | 98 | 52 | 50 | 8 | 0 | 0 | 0 | 0 | 0 | 220 | 149 | 1 | 1 | 1 | 1 | 0.911 | 1 | 1 | 1 | 1 | 1 | 1 | 1 | -0.018 |
| SE | 82976 | ENSG00000186088 | GSAP | chr7 | - | 77323646 | 77323742 | 77321332 | 77321403 | 77326211 | 77326246 | 152 | 100 | 80 | 115 | 192 | 5 | 0 | 0 | 1 | 1 | 2 | 70 | 40 | 34 | 6 | 0 | 0 | 0 | 0 | 0 | 245 | 149 | 1 | 1 | 0.949 | 1 | 1 | 0.986 | 0.992 | 1 | 1 | 1 | 1 | 1 | -0.015 |
| SE | 82977 | ENSG00000186088 | GSAP | chr7 | - | 77328605 | 77328637 | 77326211 | 77326273 | 77329332 | 77329391 | 133 | 134 | 101 | 114 | 148 | 0 | 0 | 0 | 0 | 3 | 16 | 125 | 47 | 31 | 2 | 0 | 0 | 0 | 0 | 0 | 181 | 149 | 1 | 1 | 1 | 1 | 1 | 1 | 0.976 | 1 | 1 | 1 | 1 | 1 | -0.005 |
| SE | 82983 | ENSG00000186088 | GSAP | chr7 | - | 77355554 | 77355647 | 77355212 | 77355430 | 77360823 | 77360901 | 362 | 235 | 244 | 224 | 481 | 0 | 0 | 0 | 1 | 0 | 96 | 185 | 92 | 83 | 90 | 0 | 0 | 0 | 0 | 0 | 242 | 149 | 1 | 1 | 1 | 1 | 1 | 0.993 | 1 | 1 | 1 | 1 | 1 | 1 | -0.001 |
| SE | 82984 | ENSG00000186088 | GSAP | chr7 | - | 77362582 | 77362660 | 77360823 | 77360901 | 77374069 | 77374155 | 573 | 397 | 404 | 438 | 793 | 0 | 0 | 0 | 1 | 0 | 202 | 327 | 182 | 178 | 159 | 6 | 0 | 0 | 0 | 0 | 227 | 149 | 1 | 1 | 1 | 1 | 1 | 1 | 1 | 0.957 | 1 | 1 | 1 | 1 | 0.009 |
| SE | 82985 | ENSG00000186088 | GSAP | chr7 | - | 77374069 | 77374155 | 77362582 | 77362660 | 77376847 | 77376907 | 176 | 117 | 112 | 108 | 217 | 2 | 0 | 1 | 1 | 0 | 29 | 97 | 50 | 56 | 31 | 0 | 0 | 0 | 0 | 0 | 235 | 149 | 1 | 1 | 0.982 | 1 | 0.986 | 0.986 | 1 | 1 | 1 | 1 | 1 | 1 | -0.009 |
| SE | 82986 | ENSG00000186088 | GSAP | chr7 | - | 77375057 | 77375101 | 77374069 | 77374155 | 77376847 | 77376907 | 639 | 433 | 370 | 496 | 776 | 0 | 0 | 0 | 1 | 0 | 277 | 391 | 202 | 258 | 234 | 0 | 0 | 0 | 0 | 2 | 193 | 149 | 1 | 1 | 1 | 1 | 1 | 0.997 | 1 | 1 | 1 | 1 | 1 | 0.989 | 0.002 |
| SE | 82988 | ENSG00000186088 | GSAP | chr7 | - | 77396981 | 77397035 | 77387359 | 77387448 | 77406028 | 77406105 | 46 | 31 | 23 | 42 | 82 | 0 | 0 | 0 | 1 | 0 | 10 | 13 | 13 | 34 | 63 | 0 | 0 | 0 | 0 | 0 | 203 | 149 | 1 | 1 | 1 | 1 | 1 | 0.969 | 1 | 1 | 1 | 1 | 1 | 1 | -0.006 |
| SE | 82989 | ENSG00000186088 | GSAP | chr7 | - | 77397345 | 77397415 | 77396981 | 77397035 | 77406028 | 77406105 | 78 | 35 | 39 | 60 | 148 | 0 | 0 | 0 | 0 | 0 | 30 | 21 | 18 | 45 | 44 | 2 | 0 | 0 | 0 | 0 | 219 | 149 | 1 | 1 | 1 | 1 | 1 | 1 | 1 | 0.911 | 1 | 1 | 1 | 1 | 0.018 |
| SE | 82990 | ENSG00000186088 | GSAP | chr7 | - | 77404558 | 77404615 | 77397345 | 77397415 | 77406028 | 77406105 | 114 | 53 | 54 | 53 | 211 | 2 | 0 | 0 | 4 | 1 | 66 | 24 | 7 | 43 | 42 | 0 | 0 | 0 | 3 | 0 | 206 | 149 | 1 | 1 | 0.976 | 1 | 1 | 0.906 | 0.993 | 1 | 1 | 1 | 0.912 | 1 | -0.007 |
| SE | 82991 | ENSG00000186088 | GSAP | chr7 | - | 77406028 | 77406105 | 77397345 | 77397415 | 77416212 | 77416349 | 3 | 0 | 1 | 4 | 1 | 0 | 2 | 0 | 0 | 0 | 0 | 0 | 0 | 3 | 0 | 0 | 0 | 0 | 0 | 0 | 226 | 149 | 1 | 1 | 1 | 0 | 1 | 1 | 1 | NA | NA | NA | 1 | NA | -0.2 |
| SE | 82993 | ENSG00000265681 | RPL17 | chr18 | - | 49491404 | 49491445 | 49490792 | 49490927 | 49491625 | 49491814 | 782 | 434 | 944 | 833 | 990 | 4 | 0 | 3 | 3 | 0 | 455 | 463 | 511 | 402 | 397 | 0 | 0 | 1 | 0 | 0 | 190 | 149 | 1 | 1 | 0.994 | 1 | 0.996 | 0.995 | 1 | 1 | 1 | 0.998 | 1 | 1 | -0.003 |
| SE | 82998 | ENSG00000265681 | RPL17 | chr18 | - | 49491745 | 49491833 | 49491531 | 49491584 | 49492457 | 49492482 | 11 | 0 | 4 | 9 | 6 | 474 | 254 | 551 | 423 | 418 | 0 | 3 | 1 | 4 | 8 | 126 | 296 | 260 | 215 | 224 | 237 | 149 | 1 | 1 | 0.014 | 0 | 0.005 | 0.013 | 0.009 | 0 | 0.006 | 0.002 | 0.012 | 0.022 | 0 |
| SE | 83000 | ENSG00000122783 | CYREN | chr7 | - | 135167731 | 135167807 | 135165781 | 135166871 | 135168785 | 135169060 | 37 | 25 | 27 | 41 | 21 | 2 | 0 | 0 | 0 | 0 | 37 | 33 | 22 | 18 | 50 | 0 | 0 | 0 | 0 | 0 | 225 | 149 | 1 | 1 | 0.925 | 1 | 1 | 1 | 1 | 1 | 1 | 1 | 1 | | -0.015 |
| SE | 83002 | ENSG00000106460 | TMEM106B | chr7 | + | 12214200 | 12214394 | 12211299 | 12211425 | 12214808 | 12214885 | 0 | 0 | 1 | 1 | 0 | 21 | 13 | 22 | 34 | 8 | 0 | 0 | 0 | 0 | 0 | 0 | 12 | 6 | 0 | 4 | 298 | 149 | 1 | 1 | 0 | 0 | 0.022 | 0.014 | 0 | NA | 0 | 0 | NA | 0 | 0.007 |
| SE | 83005 | ENSG00000106460 | TMEM106B | chr7 | + | 12218457 | 12218521 | 12214808 | 12215027 | 12229678 | 12229819 | 89 | 47 | 60 | 55 | 161 | 1 | 0 | 0 | 0 | 0 | 0 | 28 | 8 | 0 | 13 | 0 | 0 | 0 | 0 | 0 | 213 | 149 | 1 | 1 | 0.984 | 1 | 1 | 1 | 1 | NA | 1 | 1 | NA | 1 | -0.003 |
| SE | 83006 | ENSG00000106460 | TMEM106B | chr7 | + | 12218457 | 12218521 | 12214808 | 12215027 | 12230388 | 12230438 | 75 | 40 | 54 | 55 | 150 | 0 | 0 | 0 | 0 | 0 | 0 | 26 | 11 | 0 | 4 | 0 | 9 | 0 | 0 | 0 | 213 | 149 | 1 | 1 | 1 | 1 | 1 | 1 | 1 | NA | 0.669 | 1 | NA | 1 | 0.11 |
| SE | 83012 | ENSG00000106460 | TMEM106B | chr7 | + | 12231061 | 12231115 | 12230357 | 12230438 | 12231836 | 12232400 | 241 | 140 | 246 | 261 | 552 | 3 | 0 | 3 | 3 | 2 | 42 | 60 | 36 | 14 | 0 | 0 | 0 | 0 | 0 | 0 | 203 | 149 | 1 | 1 | 0.983 | 1 | 0.984 | 0.985 | 0.995 | 1 | 1 | 1 | 1 | NA | -0.011 |
| SE | 83014 | ENSG00000152484 | USP12 | chr13 | - | 27075190 | 27075388 | 27066155 | 27069384 | 27089882 | 27089966 | 185 | 197 | 141 | 188 | 360 | 0 | 0 | 1 | 0 | 0 | 82 | 141 | 157 | 69 | 32 | 0 | 0 | 0 | 0 | 0 | 298 | 149 | 1 | 1 | 1 | 1 | 0.986 | 1 | 1 | 1 | 1 | 1 | 1 | 1 | -0.003 |
| SE | 83016 | ENSG00000184083 | FAM120C | chrX | - | 54085714 | 54085916 | 54081321 | 54081460 | 54091311 | 54091426 | 7 | 2 | 1 | 4 | 6 | 0 | 0 | 3 | 7 | 0 | 0 | 0 | 2 | 3 | 0 | 3 | 0 | 2 | 6 | 0 | 298 | 149 | 1 | 1 | 1 | 1 | 0.143 | 0.222 | 1 | 0 | NA | 0.333 | 0.2 | NA | 0.495 |
| SE | 83017 | ENSG00000184083 | FAM120C | chrX | - | 54087754 | 54087964 | 54085714 | 54085916 | 54091311 | 54091426 | 23 | 15 | 30 | 24 | 13 | 0 | 1 | 0 | 0 | 0 | 0 | 3 | 15 | 14 | 0 | 0 | 0 | 0 | 0 | 0 | 298 | 149 | 1 | 1 | 1 | 0.882 | 1 | 1 | 1 | NA | 1 | 1 | 1 | NA | -0.024 |
| SE | 83019 | ENSG00000170356 | OR2A20P | chr7 | - | 144253532 | 144253588 | 144250044 | 144251610 | 144254433 | 144254696 | 23 | 2 | 9 | 0 | 6 | 0 | 0 | 0 | 0 | 0 | 6 | 11 | 8 | 29 | 0 | 0 | 1 | 0 | 0 | 0 | 205 | 149 | 1 | 1 | 1 | 1 | 1 | NA | 1 | 1 | 0.889 | 1 | 1 | NA | 0.028 |
| SE | 83020 | ENSG00000170356 | OR2A20P | chr7 | - | 144253532 | 144253588 | 144251661 | 144252492 | 144254433 | 144254696 | 21 | 7 | 32 | 3 | 24 | 0 | 0 | 1 | 0 | 0 | 25 | 21 | 15 | 40 | 9 | 0 | 0 | 0 | 0 | 0 | 205 | 149 | 1 | 1 | 1 | 1 | 0.959 | 1 | 1 | 1 | 1 | 1 | 1 | 1 | -0.008 |
| SE | 83021 | ENSG00000170356 | OR2A20P | chr7 | - | 144255508 | 144255732 | 144254433 | 144254696 | 144256180 | 144256244 | 3 | 7 | 6 | 3 | 7 | 0 | 0 | 0 | 0 | 0 | 0 | 10 | 9 | 4 | 0 | 0 | 5 | 0 | 0 | 0 | 298 | 149 | 1 | 1 | 1 | 1 | 1 | 1 | 1 | NA | 0.5 | 1 | 1 | NA | 0.167 |
| SE | 83022 | ENSG00000164292 | RHOBTB3 | chr5 | + | 95736888 | 95737075 | 95731858 | 95732084 | 95748332 | 95748441 | 81 | 26 | 29 | 65 | 64 | 1 | 1 | 0 | 0 | 0 | 24 | 24 | 19 | 11 | 17 | 0 | 0 | 0 | 0 | 0 | 298 | 149 | 1 | 1 | 0.976 | 0.929 | 1 | 1 | 1 | 1 | 1 | 1 | 1 | 1 | -0.019 |
| SE | 83024 | ENSG00000164292 | RHOBTB3 | chr5 | + | 95755395 | 95755761 | 95752238 | 95752350 | 95763507 | 95763542 | 210 | 86 | 72 | 214 | 204 | 1 | 0 | 5 | 0 | 0 | 477 | 294 | 110 | 237 | 868 | 0 | 0 | 0 | 0 | 0 | 298 | 149 | 1 | 1 | 0.991 | 1 | 0.878 | 1 | 1 | 1 | 1 | 1 | 1 | 1 | -0.026 |
| SE | 83028 | ENSG00000244198 | ARHGEF35-AS1 | chr7 | + | 144239039 | 144239095 | 144207692 | 144207804 | 144239274 | 144239388 | 1 | 0 | 1 | 0 | 0 | 1 | 0 | 0 | 0 | 0 | 0 | 0 | 0 | 8 | 0 | 0 | 0 | 0 | 3 | 0 | 205 | 149 | 1 | 1 | 1 | NA | 1 | NA | NA | NA | NA | NA | 0.66 | NA | 0.34 |
| SE | 83029 | ENSG00000244198 | ARHGEF35-AS1 | chr7 | + | 144239042 | 144239095 | 144207692 | 144207804 | 144239274 | 144239388 | 1 | 0 | 1 | 0 | 0 | 0 | 0 | 0 | 0 | 0 | 0 | 0 | 0 | 9 | 0 | 0 | 0 | 0 | 3 | 0 | 202 | 149 | 1 | 1 | 1 | NA | 1 | NA | NA | NA | NA | NA | 0.689 | NA | 0.311 |
| SE | 83032 | ENSG00000144840 | RABL3 | chr3 | - | 120690448 | 120690487 | 120684937 | 120689888 | 120694152 | 120694224 | 96 | 68 | 49 | 47 | 100 | 0 | 1 | 0 | 0 | 0 | 40 | 30 | 27 | 52 | 0 | 0 | 0 | 2 | 0 | 0 | 188 | 149 | 1 | 1 | 1 | 0.982 | 1 | 1 | 1 | 1 | 1 | 0.915 | 1 | NA | 0.018 |
| SE | 83033 | ENSG00000144840 | RABL3 | chr3 | - | 120690448 | 120690487 | 120684937 | 120689888 | 120698422 | 120698573 | 17 | 5 | 9 | 3 | 38 | 0 | 1 | 0 | 0 | 0 | 13 | 14 | 6 | 6 | 0 | 0 | 0 | 0 | 0 | 0 | 188 | 149 | 1 | 1 | 1 | 0.798 | 1 | 1 | 1 | 1 | 1 | 1 | 1 | NA | -0.04 |
| SE | 83034 | ENSG00000144840 | RABL3 | chr3 | - | 120694152 | 120694224 | 120684937 | 120689888 | 120698422 | 120698520 | 55 | 56 | 51 | 40 | 43 | 0 | 1 | 0 | 0 | 0 | 0 | 43 | 34 | 34 | 6 | 0 | 0 | 0 | 0 | 0 | 221 | 149 | 1 | 1 | 1 | 0.974 | 1 | 1 | 1 | NA | 1 | 1 | 1 | 1 | -0.005 |
| SE | 83039 | ENSG00000144840 | RABL3 | chr3 | - | 120738526 | 120738667 | 120730695 | 120730787 | 120742461 | 120742533 | 5 | 6 | 6 | 1 | 17 | 23 | 27 | 17 | 26 | 17 | 0 | 6 | 43 | 13 | 0 | 8 | 6 | 14 | 39 | 0 | 290 | 149 | 1 | 1 | 0.1 | 0.102 | 0.154 | 0.019 | 0.339 | 0 | 0.339 | 0.612 | 0.146 | NA | -0.131 |
| SE | 83040 | ENSG00000144840 | RABL3 | chr3 | - | 120738549 | 120738667 | 120730695 | 120730787 | 120742461 | 120742516 | 1 | 0 | 2 | 0 | 0 | 23 | 27 | 17 | 26 | 17 | 0 | 0 | 4 | 0 | 0 | 8 | 6 | 14 | 39 | 0 | 267 | 149 | 1 | 1 | 0.024 | 0 | 0.062 | 0 | 0 | 0 | 0 | 0.138 | 0 | NA | -0.017 |
| SE | 83041 | ENSG00000204959 | ARHGEF34P | chr7 | - | 144273959 | 144274121 | 144272444 | 144272515 | 144274745 | 144274903 | 26 | 8 | 18 | 3 | 7 | 1 | 3 | 0 | 0 | 0 | 0 | 8 | 4 | 56 | 0 | 0 | 0 | 0 | 12 | 0 | 298 | 149 | 1 | 1 | 0.929 | 0.571 | 1 | 1 | 1 | NA | 1 | 1 | 0.7 | NA | 0 |
| SE | 83043 | ENSG00000196305 | IARS1 | chr9 | - | 92229000 | 92229126 | 92223345 | 92223489 | 92240642 | 92240691 | 101 | 65 | 105 | 86 | 190 | 0 | 0 | 0 | 0 | 0 | 71 | 33 | 39 | 34 | 0 | 0 | 0 | 1 | 0 | 0 | 275 | 149 | 1 | 1 | 1 | 1 | 1 | 1 | 1 | 1 | 1 | 0.955 | 1 | NA | 0.011 |
| SE | 83044 | ENSG00000196305 | IARS1 | chr9 | - | 92229000 | 92229126 | 92223345 | 92223489 | 92240855 | 92240961 | 167 | 126 | 192 | 163 | 371 | 0 | 0 | 0 | 0 | 0 | 141 | 41 | 71 | 56 | 0 | 0 | 0 | 2 | 0 | 0 | 275 | 149 | 1 | 1 | 1 | 1 | 1 | 1 | 1 | 1 | 1 | 0.951 | 1 | NA | 0.012 |
| SE | 83045 | ENSG00000196305 | IARS1 | chr9 | - | 92240642 | 92240691 | 92223345 | 92223489 | 92240855 | 92240961 | 0 | 3 | 1 | 3 | 7 | 0 | 0 | 0 | 0 | 0 | 0 | 2 | 2 | 4 | 0 | 0 | 0 | 2 | 0 | 0 | 198 | 149 | 1 | 1 | NA | 1 | 1 | 1 | 1 | NA | 1 | 0.429 | 1 | NA | 0.19 |
| SE | 83046 | ENSG00000196305 | IARS1 | chr9 | - | 92240642 | 92240691 | 92229000 | 92229126 | 92240855 | 92240961 | 4 | 8 | 5 | 3 | 7 | 70 | 66 | 91 | 77 | 181 | 0 | 10 | 5 | 4 | 0 | 70 | 16 | 36 | 22 | 0 | 198 | 149 | 1 | 1 | 0.041 | 0.084 | 0.04 | 0.028 | 0.028 | 0 | 0.32 | 0.095 | 0.12 | NA | -0.09 |
| SE | 83047 | ENSG00000196305 | IARS1 | chr9 | - | 92240855 | 92240961 | 92228686 | 92229126 | 92242153 | 92242330 | 108 | 86 | 126 | 119 | 239 | 0 | 0 | 0 | 1 | 0 | 103 | 32 | 54 | 42 | 0 | 0 | 0 | 0 | 0 | 0 | 255 | 149 | 1 | 1 | 1 | 1 | 1 | 0.986 | 1 | 1 | 1 | 1 | 1 | NA | -0.003 |
| SE | 83049 | ENSG00000196305 | IARS1 | chr9 | - | 92243215 | 92243311 | 92242153 | 92242330 | 92244958 | 92245071 | 197 | 156 | 169 | 155 | 388 | 0 | 0 | 0 | 0 | 0 | 38 | 112 | 119 | 81 | 8 | 0 | 3 | 0 | 0 | 0 | 245 | 149 | 1 | 1 | 1 | 1 | 1 | 1 | 1 | 1 | 0.958 | 1 | 1 | 1 | 0.008 |
| SE | 83051 | ENSG00000196305 | IARS1 | chr9 | - | 92247376 | 92247551 | 92244958 | 92245071 | 92249857 | 92249941 | 171 | 158 | 238 | 193 | 299 | 0 | 0 | 0 | 0 | 0 | 75 | 151 | 150 | 136 | 8 | 0 | 0 | 0 | 0 | 2 | 298 | 149 | 1 | 1 | 1 | 1 | 1 | 1 | 1 | 1 | 1 | 1 | 1 | 0.667 | 0.067 |
| SE | 83052 | ENSG00000196305 | IARS1 | chr9 | - | 92253361 | 92253446 | 92251807 | 92251885 | 92255534 | 92255604 | 40 | 26 | 60 | 33 | 107 | 2 | 1 | 1 | 0 | 2 | 9 | 28 | 13 | 10 | 0 | 0 | 2 | 1 | 0 | 0 | 234 | 149 | 1 | 1 | 0.927 | 0.943 | 0.974 | 1 | 0.971 | 1 | 0.899 | 0.892 | 1 | NA | 0.015 |
| SE | 83053 | ENSG00000196305 | IARS1 | chr9 | - | 92253361 | 92253446 | 92251807 | 92251885 | 92256679 | 92256800 | 40 | 26 | 60 | 33 | 102 | 3 | 0 | 0 | 1 | 0 | 9 | 28 | 13 | 10 | 0 | 0 | 0 | 1 | 0 | 0 | 234 | 149 | 1 | 1 | 0.895 | 1 | 1 | 0.955 | 1 | 1 | 1 | 0.892 | 1 | NA | -0.003 |
| SE | 83054 | ENSG00000196305 | IARS1 | chr9 | - | 92253361 | 92253453 | 92251807 | 92251885 | 92255534 | 92255604 | 48 | 32 | 69 | 35 | 107 | 2 | 1 | 1 | 0 | 2 | 9 | 29 | 15 | 10 | 0 | 0 | 2 | 1 | 0 | 0 | 241 | 149 | 1 | 1 | 0.937 | 0.952 | 0.977 | 1 | 0.971 | 1 | 0.9 | 0.903 | 1 | NA | 0.017 |
| SE | 83055 | ENSG00000196305 | IARS1 | chr9 | - | 92253361 | 92253453 | 92251807 | 92251885 | 92256679 | 92256800 | 92 | 54 | 115 | 78 | 179 | 3 | 0 | 0 | 1 | 0 | 12 | 43 | 27 | 40 | 9 | 0 | 0 | 1 | 0 | 0 | 241 | 149 | 1 | 1 | 0.95 | 1 | 1 | 0.98 | 1 | 1 | 1 | 0.943 | 1 | 1 | -0.003 |
| SE | 83056 | ENSG00000196305 | IARS1 | chr9 | - | 92255534 | 92255604 | 92251807 | 92251885 | 92256679 | 92256800 | 15 | 3 | 3 | 5 | 12 | 3 | 0 | 0 | 1 | 0 | 0 | 8 | 5 | 0 | 0 | 0 | 0 | 1 | 0 | 0 | 219 | 149 | 1 | 1 | 0.773 | 1 | 1 | 0.773 | 1 | NA | 1 | 0.773 | NA | NA | 0.023 |
| SE | 83061 | ENSG00000196305 | IARS1 | chr9 | - | 92267238 | 92267409 | 92265479 | 92265553 | 92268173 | 92268300 | 1 | 0 | 0 | 0 | 0 | 85 | 51 | 108 | 61 | 87 | 0 | 0 | 0 | 0 | 5 | 41 | 53 | 87 | 41 | 16 | 298 | 149 | 1 | 1 | 0.006 | 0 | 0 | 0 | 0 | 0 | 0 | 0 | 0 | 0.135 | -0.026 |
| SE | 83067 | ENSG00000196305 | IARS1 | chr9 | - | 92290835 | 92290922 | 92289300 | 92289426 | 92293610 | 92293693 | 1 | 0 | 0 | 1 | 0 | 42 | 30 | 41 | 38 | 15 | 0 | 0 | 0 | 0 | 0 | 18 | 24 | 31 | 7 | 0 | 236 | 149 | 1 | 1 | 0.015 | 0 | 0 | 0.016 | 0 | 0 | 0 | 0 | 0 | NA | 0.006 |
| SE | 83074 | ENSG00000184863 | RBM33 | chr7 | + | 155665174 | 155665253 | 155644450 | 155644919 | 155672866 | 155672915 | 311 | 247 | 285 | 299 | 370 | 1 | 1 | 0 | 2 | 0 | 417 | 322 | 400 | 359 | 357 | 0 | 1 | 0 | 0 | 0 | 228 | 149 | 1 | 1 | 0.995 | 0.994 | 1 | 0.99 | 1 | 1 | 0.995 | 1 | 1 | 1 | -0.003 |
| SE | 83075 | ENSG00000184863 | RBM33 | chr7 | + | 155665174 | 155665253 | 155644450 | 155644919 | 155678607 | 155678684 | 22 | 11 | 17 | 27 | 3 | 0 | 0 | 1 | 0 | 0 | 11 | 27 | 17 | 25 | 12 | 0 | 0 | 0 | 0 | 0 | 228 | 149 | 1 | 1 | 1 | 1 | 0.917 | 1 | 1 | 1 | 1 | 1 | 1 | 1 | -0.017 |
| SE | 83076 | ENSG00000184863 | RBM33 | chr7 | + | 155672866 | 155672915 | 155644450 | 155644919 | 155678607 | 155678684 | 244 | 173 | 163 | 185 | 550 | 0 | 0 | 1 | 0 | 0 | 452 | 190 | 284 | 272 | 354 | 0 | 0 | 0 | 0 | 0 | 198 | 149 | 1 | 1 | 1 | 1 | 0.992 | 1 | 1 | 1 | 1 | 1 | 1 | 1 | -0.002 |
| SE | 83077 | ENSG00000184863 | RBM33 | chr7 | + | 155672866 | 155672915 | 155665174 | 155665253 | 155678607 | 155678684 | 532 | 415 | 434 | 460 | 927 | 0 | 2 | 0 | 0 | 1 | 872 | 499 | 678 | 599 | 694 | 0 | 0 | 1 | 1 | 0 | 198 | 149 | 1 | 1 | 1 | 0.994 | 1 | 1 | 0.999 | 1 | 1 | 0.998 | 0.998 | 1 | -0.001 |
| SE | 83078 | ENSG00000184863 | RBM33 | chr7 | + | 155680589 | 155680908 | 155678607 | 155678684 | 155700772 | 155700944 | 1491 | 1248 | 1115 | 1435 | 3346 | 0 | 3 | 0 | 0 | 0 | 2629 | 1606 | 1811 | 1896 | 2141 | 0 | 0 | 0 | 0 | 0 | 298 | 149 | 1 | 1 | 1 | 0.995 | 1 | 1 | 1 | 1 | 1 | 1 | 1 | 1 | -0.001 |
| SE | 83080 | ENSG00000184863 | RBM33 | chr7 | + | 155700772 | 155700944 | 155680589 | 155680908 | 155706859 | 155707068 | 636 | 531 | 480 | 670 | 925 | 0 | 2 | 0 | 0 | 0 | 907 | 807 | 850 | 846 | 710 | 0 | 0 | 0 | 0 | 0 | 298 | 149 | 1 | 1 | 1 | 0.993 | 1 | 1 | 1 | 1 | 1 | 1 | 1 | 1 | -0.001 |
| SE | 83081 | ENSG00000184863 | RBM33 | chr7 | + | 155706859 | 155707068 | 155700772 | 155700944 | 155711172 | 155711455 | 349 | 286 | 317 | 363 | 501 | 1 | 3 | 3 | 2 | 0 | 577 | 442 | 469 | 425 | 455 | 0 | 3 | 2 | 0 | 0 | 298 | 149 | 1 | 1 | 0.994 | 0.979 | 0.981 | 0.989 | 1 | 1 | 0.987 | 0.992 | 1 | 1 | -0.007 |
| SE | 83083 | ENSG00000184863 | RBM33 | chr7 | + | 155707293 | 155707380 | 155706859 | 155707068 | 155711172 | 155711455 | 0 | 3 | 2 | 0 | 0 | 26 | 9 | 66 | 42 | 50 | 0 | 0 | 0 | 0 | 0 | 28 | 23 | 10 | 19 | 4 | 236 | 149 | 1 | 1 | 0 | 0.174 | 0.019 | 0 | 0 | 0 | 0 | 0 | 0 | 0 | 0.039 |
| SE | 83084 | ENSG00000184863 | RBM33 | chr7 | + | 155707293 | 155707380 | 155706859 | 155707068 | 155711202 | 155711455 | 0 | 4 | 2 | 0 | 0 | 46 | 26 | 38 | 53 | 24 | 0 | 0 | 0 | 2 | 0 | 36 | 78 | 45 | 24 | 58 | 236 | 149 | 1 | 1 | 0 | 0.089 | 0.032 | 0 | 0 | 0 | 0 | 0 | 0.05 | 0 | 0.014 |
| SE | 83085 | ENSG00000184863 | RBM33 | chr7 | + | 155711172 | 155711455 | 155700772 | 155700944 | 155718384 | 155718443 | 30 | 37 | 28 | 29 | 37 | 0 | 0 | 0 | 0 | 0 | 78 | 17 | 57 | 71 | 27 | 0 | 0 | 0 | 4 | 0 | 298 | 149 | 1 | 1 | 1 | 1 | 1 | 1 | 1 | 1 | 1 | 1 | 0.899 | 1 | 0.02 |
| SE | 83086 | ENSG00000184863 | RBM33 | chr7 | + | 155711202 | 155711455 | 155700772 | 155700944 | 155718384 | 155718435 | 32 | 34 | 25 | 28 | 42 | 0 | 0 | 0 | 0 | 0 | 83 | 14 | 59 | 74 | 36 | 0 | 0 | 0 | 4 | 0 | 298 | 149 | 1 | 1 | 1 | 1 | 1 | 1 | 1 | 1 | 1 | 1 | 0.902 | 1 | 0.02 |
| SE | 83087 | ENSG00000184863 | RBM33 | chr7 | + | 155739714 | 155740026 | 155738059 | 155738403 | 155741818 | 155742106 | 73 | 50 | 81 | 86 | 25 | 0 | 0 | 0 | 0 | 0 | 101 | 85 | 85 | 95 | 142 | 0 | 0 | 2 | 0 | 0 | 298 | 149 | 1 | 1 | 1 | 1 | 1 | 1 | 1 | 1 | 1 | 0.955 | 1 | 1 | 0.009 |
| SE | 83090 | ENSG00000126653 | NSRP1 | chr17 | + | 30117301 | 30117332 | 30116826 | 30116863 | 30118079 | 30118173 | 2 | 0 | 0 | 1 | 4 | 10 | 9 | 14 | 5 | 3 | 5 | 0 | 2 | 0 | 0 | 0 | 8 | 4 | 2 | 0 | 180 | 149 | 1 | 1 | 0.142 | 0 | 0 | 0.142 | 0.525 | 1 | 0 | 0.293 | 0 | NA | -0.161 |
| SE | 83093 | ENSG00000126653 | NSRP1 | chr17 | + | 30118079 | 30118173 | 30116780 | 30116863 | 30172541 | 30172598 | 38 | 44 | 53 | 34 | 21 | 0 | 0 | 0 | 1 | 0 | 28 | 18 | 12 | 9 | 11 | 0 | 0 | 0 | 0 | 0 | 243 | 149 | 1 | 1 | 1 | 1 | 1 | 0.954 | 1 | 1 | 1 | 1 | 1 | 1 | -0.009 |
| SE | 83097 | ENSG00000126653 | NSRP1 | chr17 | + | 30156539 | 30156662 | 30118079 | 30118173 | 30163066 | 30163162 | 0 | 5 | 1 | 0 | 0 | 3 | 0 | 2 | 1 | 2 | 0 | 0 | 4 | 0 | 0 | 0 | 3 | 9 | 0 | 0 | 272 | 149 | 1 | 1 | 0 | 1 | 0.215 | 0 | 0 | NA | 0 | 0.196 | NA | NA | 0.145 |
| SE | 83098 | ENSG00000126653 | NSRP1 | chr17 | + | 30156539 | 30156662 | 30118079 | 30118173 | 30172541 | 30172598 | 2 | 2 | 2 | 0 | 0 | 38 | 44 | 54 | 35 | 25 | 0 | 0 | 0 | 0 | 0 | 28 | 18 | 13 | 10 | 11 | 272 | 149 | 1 | 1 | 0.028 | 0.024 | 0.02 | 0 | 0 | 0 | 0 | 0 | 0 | 0 | 0.014 |
| SE | 83099 | ENSG00000126653 | NSRP1 | chr17 | + | 30163066 | 30163162 | 30116818 | 30116863 | 30172541 | 30172598 | 7 | 1 | 2 | 4 | 3 | 0 | 0 | 0 | 1 | 0 | 3 | 7 | 8 | 0 | 0 | 0 | 0 | 0 | 0 | 0 | 245 | 149 | 1 | 1 | 1 | 1 | 1 | 0.709 | 1 | 1 | 1 | 1 | NA | NA | -0.058 |
| SE | 83101 | ENSG00000126653 | NSRP1 | chr17 | + | 30163066 | 30163162 | 30156539 | 30156662 | 30172541 | 30172598 | 7 | 6 | 3 | 4 | 4 | 2 | 2 | 2 | 0 | 0 | 3 | 7 | 13 | 0 | 0 | 0 | 0 | 0 | 0 | 0 | 245 | 149 | 1 | 1 | 0.68 | 0.646 | 0.477 | 1 | 1 | 1 | 1 | 1 | NA | NA | -0.239 |
| SE | 83103 | ENSG00000126653 | NSRP1 | chr17 | + | 30168102 | 30168214 | 30118079 | 30118173 | 30172541 | 30172592 | 0 | 2 | 1 | 0 | 0 | 38 | 44 | 54 | 35 | 25 | 0 | 0 | 0 | 0 | 0 | 28 | 18 | 13 | 10 | 11 | 261 | 149 | 1 | 1 | 0 | 0.025 | 0.01 | 0 | 0 | 0 | 0 | 0 | 0 | 0 | 0.007 |
| SE | 83104 | ENSG00000126653 | NSRP1 | chr17 | + | 30168813 | 30168925 | 30118079 | 30118173 | 30172541 | 30172592 | 0 | 0 | 0 | 1 | 4 | 38 | 44 | 54 | 35 | 25 | 0 | 0 | 0 | 0 | 0 | 28 | 18 | 13 | 10 | 11 | 261 | 149 | 1 | 1 | 0 | 0 | 0 | 0.016 | 0.084 | 0 | 0 | 0 | 0 | 0 | 0.02 |
| SE | 83110 | ENSG00000146909 | NOM1 | chr7 | + | 156969086 | 156969196 | 156963904 | 156964026 | 156969528 | 156973176 | 42 | 23 | 37 | 31 | 44 | 1 | 0 | 0 | 0 | 0 | 38 | 52 | 16 | 34 | 19 | 0 | 0 | 0 | 0 | 0 | 259 | 149 | 1 | 1 | 0.96 | 1 | 1 | 1 | 1 | 1 | 1 | 1 | 1 | 1 | -0.008 |
| SE | 83115 | ENSG00000234741 | GAS5 | chr1 | - | 173864483 | 173864506 | 173863901 | 173864304 | 173865228 | 173865282 | 87 | 53 | 120 | 61 | 130 | 4 | 0 | 0 | 2 | 0 | 25 | 44 | 30 | 20 | 24 | 0 | 0 | 0 | 0 | 0 | 172 | 149 | 1 | 1 | 0.95 | 1 | 1 | 0.964 | 1 | 1 | 1 | 1 | 1 | 1 | -0.017 |
| SE | 83116 | ENSG00000234741 | GAS5 | chr1 | - | 173864483 | 173864704 | 173864256 | 173864304 | 173865228 | 173865282 | 331 | 207 | 418 | 315 | 418 | 4 | 0 | 0 | 2 | 0 | 125 | 123 | 201 | 98 | 112 | 0 | 0 | 0 | 0 | 0 | 298 | 149 | 1 | 1 | 0.976 | 1 | 1 | 0.987 | 1 | 1 | 1 | 1 | 1 | 1 | -0.007 |
| SE | 83118 | ENSG00000234741 | GAS5 | chr1 | - | 173864674 | 173864704 | 173863900 | 173864304 | 173865228 | 173865282 | 198 | 105 | 265 | 216 | 259 | 4 | 0 | 0 | 2 | 0 | 94 | 63 | 137 | 76 | 79 | 0 | 0 | 0 | 0 | 0 | 179 | 149 | 1 | 1 | 0.976 | 1 | 1 | 0.989 | 1 | 1 | 1 | 1 | 1 | 1 | -0.007 |
| SE | 83119 | ENSG00000234741 | GAS5 | chr1 | - | 173864674 | 173864704 | 173863900 | 173864304 | 173865470 | 173865547 | 3 | 1 | 11 | 6 | 8 | 1 | 0 | 1 | 0 | 5 | 0 | 0 | 7 | 7 | 0 | 0 | 0 | 0 | 0 | 0 | 179 | 149 | 1 | 1 | 0.714 | 1 | 0.902 | 1 | 0.571 | NA | NA | 1 | 1 | NA | -0.163 |
| SE | 83120 | ENSG00000234741 | GAS5 | chr1 | - | 173864674 | 173864704 | 173863900 | 173864304 | 173865509 | 173865547 | 2 | 1 | 8 | 6 | 8 | 1 | 0 | 9 | 6 | 3 | 0 | 0 | 4 | 7 | 0 | 0 | 0 | 0 | 0 | 8 | 179 | 149 | 1 | 1 | 0.625 | 1 | 0.425 | 0.454 | 0.689 | NA | NA | 1 | 1 | 0 | -0.028 |
| SE | 83121 | ENSG00000234741 | GAS5 | chr1 | - | 173864674 | 173864704 | 173864483 | 173864506 | 173865228 | 173865282 | 756 | 389 | 904 | 661 | 1213 | 0 | 0 | 0 | 0 | 0 | 255 | 279 | 388 | 229 | 315 | 0 | 0 | 0 | 1 | 0 | 179 | 149 | 1 | 1 | 1 | 1 | 1 | 1 | 1 | 1 | 1 | 1 | 0.995 | 1 | 0.001 |
| SE | 83122 | ENSG00000234741 | GAS5 | chr1 | - | 173865228 | 173865282 | 173863900 | 173864304 | 173865470 | 173865547 | 147 | 94 | 243 | 147 | 263 | 1 | 0 | 1 | 0 | 5 | 60 | 22 | 96 | 37 | 58 | 0 | 0 | 0 | 0 | 0 | 203 | 149 | 1 | 1 | 0.991 | 1 | 0.994 | 1 | 0.975 | 1 | 1 | 1 | 1 | 1 | -0.008 |
| SE | 83125 | ENSG00000234741 | GAS5 | chr1 | - | 173865228 | 173865282 | 173864256 | 173864704 | 173865509 | 173865547 | 695 | 293 | 596 | 454 | 702 | 2 | 1 | 1 | 0 | 0 | 131 | 316 | 219 | 220 | 217 | 0 | 0 | 0 | 0 | 0 | 203 | 149 | 1 | 1 | 0.996 | 0.995 | 0.998 | 1 | 1 | 1 | 1 | 1 | 1 | 1 | -0.002 |
| SE | 83126 | ENSG00000234741 | GAS5 | chr1 | - | 173865856 | 173866206 | 173865352 | 173865547 | 173866990 | 173867043 | 417 | 189 | 481 | 325 | 414 | 0 | 0 | 0 | 3 | 0 | 138 | 138 | 243 | 100 | 150 | 0 | 0 | 0 | 0 | 0 | 298 | 149 | 1 | 1 | 1 | 1 | 1 | 0.982 | 1 | 1 | 1 | 1 | 1 | 1 | -0.004 |
| SE | 83127 | ENSG00000234741 | GAS5 | chr1 | - | 173866176 | 173866206 | 173865856 | 173865894 | 173866527 | 173866567 | 960 | 509 | 1116 | 778 | 1105 | 0 | 0 | 1 | 1 | 3 | 293 | 419 | 468 | 346 | 352 | 2 | 0 | 1 | 0 | 0 | 179 | 149 | 1 | 1 | 1 | 1 | 0.999 | 0.998 | 0.997 | 0.992 | 1 | 0.997 | 1 | 1 | 0.001 |
| SE | 83128 | ENSG00000234741 | GAS5 | chr1 | - | 173866527 | 173866796 | 173865856 | 173866206 | 173866990 | 173867043 | 203 | 104 | 209 | 160 | 186 | 1 | 0 | 0 | 0 | 1 | 35 | 121 | 142 | 93 | 52 | 0 | 0 | 1 | 0 | 0 | 298 | 149 | 1 | 1 | 0.99 | 1 | 1 | 1 | 0.989 | 1 | 1 | 1 | 1 | 1 | -0.004 |
| SE | 83138 | ENSG00000240225 | ZNF542P | chr19 | + | 56369278 | 56369337 | 56368865 | 56368936 | 56373156 | 56373283 | 2 | 6 | 4 | 4 | 2 | 13 | 14 | 10 | 31 | 10 | 0 | 14 | 2 | 2 | 24 | 75 | 15 | 0 | 4 | 113 | 208 | 149 | 1 | 1 | 0.099 | 0.235 | 0.223 | 0.085 | 0.125 | 0 | 0.401 | 1 | 0.264 | 0.132 | -0.206 |
| SE | 83144 | ENSG00000188878 | FBF1 | chr17 | - | 75914746 | 75914932 | 75914121 | 75914298 | 75915016 | 75915139 | 9 | 13 | 7 | 12 | 4 | 0 | 0 | 0 | 3 | 0 | 0 | 31 | 7 | 0 | 9 | 0 | 0 | 0 | 0 | 0 | 298 | 149 | 1 | 1 | 1 | 1 | 1 | 0.667 | 1 | NA | 1 | 1 | NA | 1 | -0.067 |
| SE | 83147 | ENSG00000188878 | FBF1 | chr17 | - | 75939817 | 75939882 | 75938172 | 75938232 | 75940847 | 75941038 | 7 | 0 | 4 | 1 | 0 | 6 | 2 | 2 | 4 | 2 | 8 | 4 | 3 | 0 | 0 | 6 | 2 | 5 | 2 | 0 | 214 | 149 | 1 | 1 | 0.448 | 0 | 0.582 | 0.148 | 0 | 0.481 | 0.582 | 0.295 | 0 | NA | -0.104 |
| SE | 83151 | ENSG00000055130 | CUL1 | chr7 | + | 148759303 | 148759354 | 148756967 | 148757150 | 148759547 | 148759638 | 216 | 154 | 188 | 222 | 233 | 0 | 2 | 1 | 0 | 0 | 277 | 233 | 106 | 300 | 18 | 5 | 3 | 0 | 2 | 0 | 200 | 149 | 1 | 1 | 1 | 0.983 | 0.993 | 1 | 1 | 0.976 | 0.983 | 1 | 0.991 | 1 | 0.005 |
| SE | 83153 | ENSG00000055130 | CUL1 | chr7 | + | 148759547 | 148759638 | 148759303 | 148759354 | 148760332 | 148760484 | 322 | 249 | 293 | 316 | 345 | 0 | 2 | 0 | 0 | 0 | 401 | 267 | 180 | 458 | 20 | 0 | 0 | 0 | 0 | 0 | 240 | 149 | 1 | 1 | 1 | 0.987 | 1 | 1 | 1 | 1 | 1 | 1 | 1 | 1 | -0.003 |
| SE | 83161 | ENSG00000055130 | CUL1 | chr7 | + | 148792725 | 148792818 | 148790309 | 148790441 | 148797811 | 148797859 | 183 | 144 | 179 | 164 | 262 | 0 | 0 | 0 | 0 | 0 | 124 | 166 | 108 | 244 | 17 | 0 | 0 | 0 | 3 | 0 | 242 | 149 | 1 | 1 | 1 | 1 | 1 | 1 | 1 | 1 | 1 | 1 | 0.98 | 1 | 0.004 |
| SE | 83169 | ENSG00000160408 | ST6GALNAC6 | chr9 | - | 127897825 | 127898010 | 127896241 | 127896332 | 127905206 | 127905408 | 0 | 0 | 5 | 1 | 0 | 90 | 28 | 62 | 62 | 39 | 0 | 0 | 1 | 0 | 0 | 32 | 35 | 58 | 39 | 31 | 298 | 149 | 1 | 1 | 0 | 0 | 0.039 | 0.008 | 0 | 0 | 0 | 0.009 | 0 | 0 | 0.008 |
| SE | 83170 | ENSG00000160408 | ST6GALNAC6 | chr9 | - | 127897955 | 127898010 | 127896241 | 127896332 | 127899502 | 127899574 | 0 | 0 | 18 | 1 | 3 | 0 | 0 | 0 | 2 | 0 | 0 | 4 | 1 | 0 | 0 | 0 | 0 | 0 | 0 | 0 | 204 | 149 | 1 | 1 | NA | NA | 1 | 0.268 | 1 | NA | 1 | 1 | NA | NA | -0.244 |
| SE | 83171 | ENSG00000197965 | MPZL1 | chr1 | + | 167765582 | 167765749 | 167721981 | 167722242 | 167787819 | 167790819 | 26 | 15 | 26 | 20 | 12 | 0 | 0 | 0 | 0 | 1 | 23 | 12 | 17 | 23 | 22 | 0 | 0 | 0 | 0 | 0 | 298 | 149 | 1 | 1 | 1 | 1 | 1 | 1 | 0.857 | 1 | 1 | 1 | 1 | 1 | -0.029 |
| SE | 83173 | ENSG00000197965 | MPZL1 | chr1 | + | 167773235 | 167773368 | 167765582 | 167765749 | 167776063 | 167776166 | 6 | 3 | 8 | 8 | 24 | 0 | 0 | 0 | 0 | 0 | 16 | 4 | 2 | 1 | 0 | 0 | 0 | 0 | 3 | 0 | 282 | 149 | 1 | 1 | 1 | 1 | 1 | 1 | | 1 | 1 | 1 | 0.15 | NA | 0.213 |
| SE | 83177 | ENSG00000205482 | SPDYE18 | chr7 | - | 77055235 | 77055321 | 77052959 | 77053203 | 77056549 | 77056608 | 3 | 3 | 0 | 1 | 1 | 4 | 0 | 0 | 0 | 0 | 0 | 12 | 0 | 0 | 0 | 0 | 0 | 0 | 0 | 0 | 235 | 149 | 1 | 1 | 0.322 | 1 | NA | 1 | 1 | NA | 1 | NA | NA | NA | -0.169 |
| SE | 83178 | ENSG00000205482 | SPDYE18 | chr7 | - | 77057636 | 77057867 | 77056549 | 77056608 | 77059179 | 77059398 | 1 | 2 | 4 | 1 | 1 | 3 | 0 | 0 | 0 | 0 | 0 | 15 | 0 | 0 | 0 | 0 | 0 | 0 | 0 | 7 | 298 | 149 | 1 | 1 | 0.143 | 1 | 1 | 1 | 1 | NA | 1 | NA | NA | 0 | 0.329 |
| SE | 83185 | ENSG00000175567 | UCP2 | chr11 | - | 73975490 | 73975671 | 73974671 | 73975121 | 73976640 | 73976742 | 1189 | 497 | 735 | 711 | 777 | 1 | 1 | 0 | 0 | 0 | 1031 | 1303 | 456 | 579 | 3069 | 0 | 0 | 3 | 2 | 0 | 298 | 149 | 1 | 1 | 0.998 | 0.996 | 1 | 1 | 1 | 1 | 1 | 0.987 | 0.993 | 1 | 0.003 |
| SE | 83186 | ENSG00000175567 | UCP2 | chr11 | - | 73976822 | 73977017 | 73976640 | 73976742 | 73977885 | 73978096 | 566 | 209 | 342 | 349 | 315 | 2 | 0 | 0 | 0 | 0 | 365 | 525 | 196 | 238 | 986 | 0 | 0 | 0 | 0 | 0 | 298 | 149 | 1 | 1 | 0.993 | 1 | 1 | 1 | 1 | 1 | 1 | 1 | 1 | 1 | -0.001 |
| SE | 83187 | ENSG00000175567 | UCP2 | chr11 | - | 73977885 | 73978096 | 73976640 | 73976742 | 73978252 | 73978477 | 401 | 149 | 299 | 245 | 231 | 1 | 0 | 0 | 0 | 0 | 203 | 299 | 140 | 148 | 562 | 0 | 0 | 0 | 0 | 0 | 298 | 149 | 1 | 1 | 0.995 | 1 | 1 | 1 | 1 | 1 | 1 | 1 | 1 | 1 | -0.001 |
| SE | 83189 | ENSG00000175567 | UCP2 | chr11 | - | 73979935 | 73980052 | 73978695 | 73978834 | 73981475 | 73981632 | 1 | 0 | 0 | 0 | 2 | 2 | 2 | 3 | 2 | 0 | 10 | 0 | 0 | 4 | 8 | 0 | 10 | 0 | 0 | 16 | 266 | 149 | 1 | 1 | 0.219 | 0 | 0 | 0 | 1 | 1 | 0 | NA | 1 | 0.219 | -0.311 |
| SE | 83190 | ENSG00000175567 | UCP2 | chr11 | - | 73980014 | 73980052 | 73978252 | 73978477 | 73981475 | 73981632 | 0 | 2 | 0 | 0 | 0 | 659 | 282 | 488 | 463 | 794 | 0 | 2 | 0 | 2 | 0 | 259 | 674 | 216 | 283 | 930 | 187 | 149 | 1 | 1 | 0 | 0.006 | 0 | 0 | 0 | 0 | 0.002 | 0 | 0.006 | 0 | 0 |
| SE | 83197 | ENSG00000183773 | AIFM3 | chr22 | + | 20977699 | 20977776 | 20977031 | 20977095 | 20977887 | 20978005 | 11 | 7 | 4 | 2 | 7 | 2 | 0 | 0 | 1 | 0 | 0 | 8 | 5 | 9 | 51 | 0 | 0 | 0 | 0 | 0 | 226 | 149 | 1 | 1 | 0.784 | 1 | 1 | 0.569 | 1 | NA | 1 | 1 | 1 | 1 | -0.129 |
| SE | 83198 | ENSG00000183773 | AIFM3 | chr22 | + | 20977705 | 20977776 | 20977031 | 20977095 | 20977887 | 20978005 | 5 | 4 | 3 | 2 | 1 | 2 | 0 | 0 | 1 | 0 | 0 | 3 | 0 | 3 | 29 | 0 | 0 | 0 | 0 | 0 | 220 | 149 | 1 | 1 | 0.629 | 1 | 1 | 0.575 | 1 | NA | 1 | NA | 1 | 1 | -0.159 |
| SE | 83199 | ENSG00000183773 | AIFM3 | chr22 | + | 20979270 | 20979369 | 20977887 | 20978005 | 20979626 | 20979702 | 16 | 3 | 2 | 5 | 2 | 0 | 0 | 0 | 0 | 0 | 0 | 2 | 1 | 4 | 26 | 0 | 0 | 3 | 0 | 0 | 248 | 149 | 1 | 1 | 1 | 1 | 1 | 1 | 1 | NA | 1 | 0.167 | 1 | 1 | 0.208 |
| SE | 83202 | ENSG00000174384 | PMS2P6 | chr7 | + | 73100617 | 73100706 | 73097383 | 73097567 | 73105292 | 73105389 | 0 | 0 | 0 | 0 | 0 | 0 | 0 | 0 | 4 | 0 | 0 | 0 | 0 | 2 | 0 | 0 | 0 | 0 | 0 | 0 | 238 | 149 | 1 | 1 | NA | NA | NA | NA | 0 | NA | NA | NA | 1 | NA | -1 |
| SE | 83204 | ENSG00000150637 | CD226 | chr18 | - | 69842283 | 69842395 | 69832535 | 69832739 | 69895700 | 69895809 | 11 | 6 | 4 | 3 | 0 | 0 | 0 | 0 | 1 | 0 | 0 | 3 | 11 | 0 | 2 | 0 | 0 | 0 | 0 | 3 | 261 | 149 | 1 | 1 | 1 | 1 | 1 | 0.631 | NA | NA | 1 | 1 | NA | 0.276 | 0.149 |
| SE | 83207 | ENSG00000150637 | CD226 | chr18 | - | 69867356 | 69867411 | 69853273 | 69864439 | 69873143 | 69873246 | 352 | 375 | 243 | 667 | 486 | 0 | 0 | 0 | 0 | 0 | 304 | 247 | 244 | 281 | 274 | 0 | 0 | 2 | 0 | 0 | 204 | 149 | 1 | 1 | 1 | 1 | 1 | 1 | 1 | 1 | 1 | 0.989 | 1 | 1 | 0.002 |
| SE | 83208 | ENSG00000150637 | CD226 | chr18 | - | 69867356 | 69867411 | 69853273 | 69864439 | 69895700 | 69896045 | 73 | 69 | 59 | 116 | 109 | 0 | 0 | 0 | 4 | 0 | 60 | 59 | 33 | 25 | 42 | 0 | 0 | 0 | 0 | 0 | 204 | 149 | 1 | 1 | 1 | 1 | 1 | 0.955 | 1 | 1 | 1 | 1 | 1 | 1 | -0.009 |
| SE | 83210 | ENSG00000150637 | CD226 | chr18 | - | 69873143 | 69873246 | 69853273 | 69864439 | 69895700 | 69896045 | 247 | 202 | 143 | 383 | 417 | 0 | 0 | 0 | 4 | 0 | 192 | 147 | 138 | 129 | 294 | 0 | 0 | 0 | 0 | 0 | 252 | 149 | 1 | 1 | 1 | 1 | 1 | 0.983 | 1 | 1 | 1 | 1 | 1 | 1 | -0.003 |
| SE | 83212 | ENSG00000150637 | CD226 | chr18 | - | 69895700 | 69896045 | 69873143 | 69873246 | 69946733 | 69947069 | 521 | 476 | 345 | 914 | 802 | 1 | 0 | 0 | 0 | 2 | 490 | 382 | 357 | 411 | 734 | 0 | 0 | 0 | 0 | 0 | 298 | 149 | 1 | 1 | 0.996 | 1 | 1 | 1 | 0.995 | 1 | 1 | 1 | 1 | 1 | -0.002 |
| SE | 83213 | ENSG00000150637 | CD226 | chr18 | - | 69895700 | 69896045 | 69873143 | 69873246 | 69961591 | 69961803 | 353 | 315 | 223 | 642 | 583 | 0 | 2 | 0 | 1 | 0 | 340 | 241 | 225 | 256 | 408 | 0 | 0 | 0 | 0 | 0 | 298 | 149 | 1 | 1 | 1 | 0.987 | 1 | 0.997 | 1 | 1 | 1 | 1 | 1 | 1 | -0.003 |
| SE | 83214 | ENSG00000150637 | CD226 | chr18 | - | 69944483 | 69944568 | 69895700 | 69896045 | 69946733 | 69946770 | 1 | 2 | 0 | 0 | 1 | 182 | 183 | 124 | 307 | 233 | 0 | 2 | 0 | 0 | 4 | 150 | 168 | 149 | 182 | 342 | 234 | 149 | 1 | 1 | 0.003 | 0.007 | 0 | 0 | 0.003 | 0 | 0.008 | 0 | 0 | 0.007 | 0 |
| SE | 83215 | ENSG00000150637 | CD226 | chr18 | - | 69946733 | 69947069 | 69895700 | 69896045 | 69947360 | 69947433 | 483 | 530 | 301 | 753 | 753 | 1 | 0 | 1 | 2 | 3 | 522 | 397 | 418 | 501 | 1174 | 0 | 2 | 2 | 0 | 0 | 298 | 149 | 1 | 1 | 0.996 | 1 | 0.993 | 0.995 | 0.992 | 1 | 0.99 | 0.991 | 1 | 1 | -0.001 |
| SE | 83216 | ENSG00000150637 | CD226 | chr18 | - | 69947360 | 69947433 | 69895700 | 69896045 | 69956613 | 69956670 | 0 | 0 | 0 | 2 | 0 | 1 | 0 | 0 | 0 | 0 | 0 | 2 | 0 | 0 | 0 | 0 | 0 | 0 | 0 | 0 | 222 | 149 | 1 | 1 | 0 | NA | NA | 1 | NA | NA | 1 | NA | NA | NA | -0.5 |
| SE | 83217 | ENSG00000150637 | CD226 | chr18 | - | 69947360 | 69947433 | 69895700 | 69896045 | 69956754 | 69956996 | 5 | 5 | 7 | 6 | 6 | 3 | 0 | 4 | 4 | 2 | 2 | 2 | 2 | 0 | 0 | 0 | 0 | 0 | 3 | 0 | 222 | 149 | 1 | 1 | 0.528 | 1 | 0.54 | 0.502 | 0.668 | 1 | 1 | 1 | 0 | NA | -0.102 |
| SE | 83219 | ENSG00000150637 | CD226 | chr18 | - | 69947360 | 69947532 | 69895700 | 69896045 | 69956613 | 69956670 | 1 | 0 | 0 | 2 | 0 | 1 | 0 | 0 | 0 | 0 | 0 | 2 | 0 | 0 | 0 | 0 | 0 | 0 | 0 | 0 | 298 | 149 | 1 | 1 | 0.333 | NA | NA | 1 | NA | NA | 1 | NA | NA | NA | -0.334 |
| SE | 83220 | ENSG00000150637 | CD226 | chr18 | - | 69947360 | 69947532 | 69895700 | 69896045 | 69956754 | 69957176 | 3 | 4 | 10 | 3 | 1 | 3 | 0 | 4 | 4 | 2 | 0 | 3 | 0 | 0 | 0 | 0 | 0 | 0 | 3 | 0 | 298 | 149 | 1 | 1 | 0.333 | 1 | 0.556 | 0.273 | 0.2 | NA | 1 | NA | 0 | NA | -0.028 |
| SE | 83222 | ENSG00000205583 | STAG3L1 | chr7 | + | 75361971 | 75362088 | 75361373 | 75361459 | 75365254 | 75365353 | 7 | 1 | 5 | 0 | 0 | 0 | 0 | 0 | 0 | 0 | 1 | 0 | 0 | 0 | 2 | 0 | 1 | 0 | 0 | 0 | 266 | 149 | 1 | 1 | 1 | 1 | 1 | NA | NA | 1 | 0 | NA | NA | 1 | 0.333 |
| SE | 83223 | ENSG00000205583 | STAG3L1 | chr7 | + | 75364632 | 75364756 | 75361971 | 75362088 | 75365254 | 75365353 | 3 | 0 | 0 | 0 | 1 | 2 | 1 | 1 | 0 | 0 | 0 | 0 | 0 | 0 | 0 | 1 | 0 | 0 | 0 | 0 | 273 | 149 | 1 | 1 | 0.45 | 0 | 0 | NA | 1 | 0 | NA | NA | | NA | 0.362 |
| SE | 83224 | ENSG00000205583 | STAG3L1 | chr7 | + | 75365563 | 75365643 | 75365254 | 75365353 | 75366323 | 75366438 | 0 | 2 | 1 | 0 | 3 | 0 | 0 | 0 | 0 | 0 | 0 | 2 | 1 | 0 | 0 | 0 | 1 | 0 | 0 | 0 | 229 | 149 | 1 | 1 | NA | 1 | 1 | NA | 1 | NA | 0.565 | 1 | NA | NA | 0.218 |
| SE | 83225 | ENSG00000205583 | STAG3L1 | chr7 | + | 75365951 | 75366059 | 75365563 | 75365643 | 75366323 | 75366438 | 1 | 0 | 1 | 0 | 1 | 0 | 0 | 0 | 0 | 1 | 0 | 0 | 0 | 0 | 0 | 0 | 2 | 1 | 0 | 0 | 257 | 149 | 1 | 1 | 1 | NA | 1 | NA | 0.367 | NA | 0 | 0 | NA | NA | 0.789 |
| SE | 83227 | ENSG00000205903 | ZNF316 | chr7 | + | 6639041 | 6639141 | 6637845 | 6638009 | 6641824 | 6641962 | 18 | 18 | 23 | 14 | 12 | 0 | 1 | 0 | 1 | 0 | 49 | 37 | 28 | 31 | 69 | 0 | 0 | 0 | 0 | 0 | 249 | 149 | 1 | 1 | 1 | 0.915 | 1 | 0.893 | 1 | 1 | 1 | 1 | 1 | 1 | -0.038 |
| SE | 83229 | ENSG00000205903 | ZNF316 | chr7 | + | 6643821 | 6643948 | 6642963 | 6643073 | 6644479 | 6644565 | 219 | 140 | 200 | 264 | 180 | 0 | 0 | 0 | 0 | 0 | 591 | 252 | 227 | 437 | 589 | 0 | 3 | 0 | 0 | 0 | 276 | 149 | 1 | 1 | 1 | 1 | 1 | 1 | 1 | 1 | 0.978 | 1 | 1 | 1 | 0.004 |
| SE | 83230 | ENSG00000164631 | ZNF12 | chr7 | - | 6705158 | 6705223 | 6697684 | 6697811 | 6706431 | 6706536 | 41 | 22 | 23 | 24 | 8 | 1 | 0 | 0 | 0 | 0 | 5 | 5 | 9 | 16 | 0 | 0 | 0 | 0 | 0 | 0 | 214 | 149 | 1 | 1 | 0.966 | 1 | 1 | 1 | 1 | 1 | 1 | 1 | 1 | NA | -0.007 |
| SE | 83231 | ENSG00000106077 | ABHD11 | chr7 | - | 73736928 | 73737110 | 73736093 | 73736691 | 73737220 | 73737360 | 22 | 13 | 15 | 11 | 5 | 0 | 0 | 0 | 1 | 0 | 0 | 38 | 14 | 17 | 34 | 0 | 0 | 0 | 0 | 23 | 298 | 149 | 1 | 1 | 1 | 1 | 1 | 0.846 | 1 | NA | 1 | 1 | 1 | 0.425 | 0.113 |
| SE | 83232 | ENSG00000106077 | ABHD11 | chr7 | - | 73736928 | 73737110 | 73736093 | 73736691 | 73737561 | 73737735 | 14 | 8 | 9 | 9 | 5 | 0 | 0 | 2 | 9 | 0 | 0 | 37 | 10 | 9 | 16 | 0 | 9 | 0 | 3 | 1 | 298 | 149 | 1 | 1 | 1 | 1 | 0.692 | 0.333 | 1 | NA | 0.673 | 1 | 0.6 | 0.889 | 0.015 |
| SE | 83233 | ENSG00000106077 | ABHD11 | chr7 | - | 73736928 | 73737220 | 73736535 | 73736691 | 73737561 | 73737735 | 14 | 8 | 9 | 10 | 5 | 0 | 0 | 2 | 9 | 0 | 0 | 35 | 10 | 9 | 16 | 0 | 9 | 0 | 3 | 1 | 298 | 149 | 1 | 1 | 1 | 1 | 0.692 | 0.357 | 1 | NA | 0.66 | 1 | 0.6 | 0.889 | 0.023 |
| SE | 83234 | ENSG00000106077 | ABHD11 | chr7 | - | 73737220 | 73737360 | 73736563 | 73736691 | 73737561 | 73737735 | 0 | 0 | 0 | 1 | 0 | 0 | 0 | 2 | 9 | 0 | 0 | 0 | 0 | 0 | 23 | 0 | 9 | 0 | 3 | 1 | 289 | 149 | 1 | 1 | NA | NA | 0 | 0.054 | NA | NA | 0 | NA | 0 | 0.922 | -0.28 |
| SE | 83235 | ENSG00000106077 | ABHD11 | chr7 | - | 73737220 | 73737360 | 73736928 | 73737110 | 73737561 | 73737735 | 8 | 5 | 6 | 2 | 0 | 0 | 0 | 0 | 0 | 0 | 0 | 5 | 4 | 8 | 18 | 0 | 4 | 0 | 0 | 0 | 289 | 149 | 1 | 1 | 1 | 1 | 1 | 1 | NA | NA | 0.392 | 1 | 1 | 1 | 0.152 |
| SE | 83236 | ENSG00000106077 | ABHD11 | chr7 | - | 73737220 | 73737391 | 73736093 | 73736691 | 73737561 | 73737735 | 4 | 0 | 2 | 1 | 7 | 0 | 0 | 2 | 9 | 0 | 0 | 5 | 3 | 0 | 34 | 0 | 9 | 0 | 3 | 1 | 298 | 149 | 1 | 1 | 1 | NA | 0.333 | 0.053 | 1 | NA | 0.217 | 1 | 0 | 0.944 | 0.056 |
| SE | 83237 | ENSG00000106077 | ABHD11 | chr7 | - | 73737220 | 73737391 | 73736928 | 73737110 | 73737561 | 73737735 | 12 | 5 | 8 | 2 | 7 | 0 | 0 | 0 | 0 | 0 | 0 | 10 | 7 | 8 | 29 | 0 | 4 | 0 | 0 | 0 | 298 | 149 | 1 | 1 | 1 | 1 | 1 | 1 | 1 | NA | 0.556 | 1 | 1 | 1 | 0.111 |
| SE | 83240 | ENSG00000137501 | SYTL2 | chr11 | - | 85704857 | 85705028 | 85700514 | 85700593 | 85707428 | 85707531 | 192 | 91 | 128 | 194 | 267 | 2 | 1 | 1 | 2 | 2 | 10 | 24 | 55 | 22 | 3 | 0 | 0 | 0 | 0 | 0 | 298 | 149 | 1 | 1 | 0.98 | 0.978 | 0.985 | 0.98 | 0.985 | 1 | 1 | 1 | 1 | 1 | -0.018 |
| SE | 83241 | ENSG00000137501 | SYTL2 | chr11 | - | 85708037 | 85708150 | 85707428 | 85707531 | 85709330 | 85709500 | 0 | 3 | 0 | 6 | 1 | 60 | 33 | 45 | 60 | 66 | 0 | 0 | 0 | 0 | 0 | 0 | 5 | 19 | 5 | 3 | 262 | 149 | 1 | 1 | 0 | 0.049 | 0 | 0.054 | 0.009 | NA | 0 | 0 | 0 | 0 | 0.022 |
| SE | 83242 | ENSG00000137501 | SYTL2 | chr11 | - | 85711112 | 85711232 | 85709330 | 85709500 | 85714412 | 85714507 | 122 | 89 | 127 | 149 | 116 | 0 | 0 | 1 | 2 | 0 | 12 | 16 | 26 | 10 | 0 | 0 | 0 | 0 | 0 | 0 | 269 | 149 | 1 | 1 | 1 | 1 | 0.986 | 0.976 | 1 | 1 | 1 | 1 | 1 | NA | -0.008 |
| SE | 83243 | ENSG00000137501 | SYTL2 | chr11 | - | 85711112 | 85711232 | 85709330 | 85709500 | 85718789 | 85719168 | 70 | 52 | 62 | 88 | 46 | 2 | 0 | 0 | 1 | 0 | 12 | 3 | 18 | 5 | 0 | 0 | 0 | 0 | 0 | 0 | 269 | 149 | 1 | 1 | 0.951 | 1 | 1 | 0.98 | 1 | 1 | 1 | 1 | 1 | NA | -0.014 |
| SE | 83244 | ENSG00000137501 | SYTL2 | chr11 | - | 85714412 | 85714507 | 85709330 | 85709500 | 85718789 | 85718843 | 41 | 35 | 32 | 54 | 129 | 2 | 0 | 0 | 1 | 0 | 0 | 17 | 20 | 15 | 0 | 0 | 0 | 0 | 0 | 0 | 244 | 149 | 1 | 1 | 0.926 | 1 | 1 | 0.971 | 1 | NA | 1 | 1 | 1 | NA | -0.021 |
| SE | 83245 | ENSG00000137501 | SYTL2 | chr11 | - | 85714412 | 85714507 | 85711112 | 85711232 | 85718789 | 85718843 | 95 | 71 | 94 | 116 | 200 | 6 | 1 | 0 | 6 | 2 | 0 | 28 | 31 | 20 | 0 | 0 | 0 | 3 | 0 | 0 | 244 | 149 | 1 | 1 | 0.906 | 0.977 | 1 | 0.922 | 0.984 | NA | 1 | 0.863 | 1 | NA | 0.003 |
| SE | 83252 | ENSG00000137501 | SYTL2 | chr11 | - | 85717482 | 85717530 | 85714412 | 85714507 | 85718789 | 85718843 | 4 | 0 | 0 | 7 | 5 | 59 | 44 | 42 | 72 | 158 | 0 | 2 | 1 | 0 | 0 | 0 | 24 | 25 | 15 | 0 | 197 | 149 | 1 | 1 | 0.049 | 0 | 0 | 0.068 | 0.023 | NA | 0.059 | 0.029 | 0 | NA | -0.001 |
| SE | 83254 | ENSG00000137501 | SYTL2 | chr11 | - | 85717482 | 85717726 | 85714412 | 85714507 | 85718789 | 85718843 | 0 | 0 | 0 | 2 | 1 | 59 | 44 | 42 | 72 | 158 | 0 | 2 | 0 | 0 | 0 | 0 | 24 | 25 | 15 | 0 | 298 | 149 | 1 | 1 | 0 | 0 | 0 | 0.014 | 0.003 | NA | 0.04 | 0 | 0 | NA | -0.01 |
| SE | 83257 | ENSG00000137501 | SYTL2 | chr11 | - | 85745636 | 85745772 | 85737574 | 85737656 | 85748271 | 85748423 | 109 | 68 | 92 | 162 | 198 | 1 | 0 | 0 | 1 | 0 | 8 | 37 | 44 | 40 | 9 | 0 | 0 | 1 | 0 | 0 | 285 | 149 | 1 | 1 | 0.983 | 1 | 1 | 0.988 | 1 | 1 | 1 | 0.958 | 1 | 1 | 0.003 |
| SE | 83258 | ENSG00000137501 | SYTL2 | chr11 | - | 85757624 | 85758114 | 85748271 | 85748423 | 85810953 | 85811129 | 219 | 158 | 150 | 249 | 229 | 0 | 0 | 0 | 4 | 0 | 27 | 75 | 110 | 40 | 12 | 0 | 0 | 0 | 0 | 0 | 298 | 149 | 1 | 1 | 1 | 1 | 1 | 0.969 | 1 | 1 | 1 | 1 | 1 | 1 | -0.006 |
| SE | 83262 | ENSG00000169871 | TRIM56 | chr7 | + | 101087005 | 101087168 | 101085480 | 101085571 | 101087311 | 101088128 | 95 | 50 | 86 | 85 | 31 | 0 | 0 | 0 | 1 | 0 | 17 | 56 | 30 | 31 | 33 | 0 | 0 | 0 | 0 | 0 | 298 | 149 | 1 | 1 | 1 | 1 | 1 | 0.977 | 1 | 1 | 1 | 1 | 1 | 1 | -0.005 |
| SE | 83263 | ENSG00000146373 | RNF217 | chr6 | + | 124971500 | 124971598 | 124963307 | 124963426 | 124996574 | 124996655 | 0 | 0 | 0 | 4 | 0 | 0 | 0 | 3 | 0 | 0 | 0 | 0 | 0 | 4 | 6 | 0 | 6 | 0 | 0 | 0 | 247 | 149 | 1 | 1 | NA | NA | 0 | 1 | NA | NA | 0 | NA | 1 | 1 | -0.167 |
| SE | 83265 | ENSG00000146373 | RNF217 | chr6 | + | 124996574 | 124996655 | 124963307 | 124963426 | 125045210 | 125045444 | 1 | 1 | 2 | 1 | 2 | 2 | 1 | 4 | 1 | 7 | 4 | 0 | 1 | 0 | 0 | 6 | 10 | 7 | 9 | 0 | 230 | 149 | 1 | 1 | 0.245 | 0.393 | 0.245 | 0.393 | 0.156 | 0.302 | 0 | 0.085 | 0 | NA | 0.19 |
| SE | 83266 | ENSG00000146373 | RNF217 | chr6 | + | 124996574 | 124996655 | 124971500 | 124971598 | 125045210 | 125045444 | 1 | 1 | 0 | 5 | 2 | 2 | 1 | 2 | 2 | 2 | 4 | 0 | 1 | 4 | 0 | 0 | 0 | 0 | 0 | 11 | 230 | 149 | 1 | 1 | 0.245 | 0.393 | 0 | 0.618 | 0.393 | 1 | NA | 1 | 1 | 0 | -0.42 |
| SE | 83267 | ENSG00000146373 | RNF217 | chr6 | + | 125006075 | 125006177 | 124996574 | 124996655 | 125009173 | 125009282 | 1 | 0 | 1 | 0 | 0 | 0 | 0 | 0 | 0 | 0 | 0 | 0 | 0 | 0 | 0 | 0 | 6 | 0 | 0 | 0 | 251 | 149 | 1 | 1 | 1 | NA | 1 | NA | NA | NA | 0 | NA | NA | NA | 1 |
| SE | 83268 | ENSG00000146373 | RNF217 | chr6 | + | 125009173 | 125009282 | 124996574 | 124996655 | 125045210 | 125045444 | 0 | 0 | 0 | 0 | 0 | 1 | 1 | 2 | 5 | 2 | 0 | 6 | 0 | 4 | 0 | 4 | 0 | 1 | 4 | 0 | 258 | 149 | 1 | 1 | 0 | 0 | 0 | 0 | 0 | 0 | 1 | 0 | 0.366 | NA | -0.342 |
| SE | 83269 | ENSG00000146373 | RNF217 | chr6 | + | 125045210 | 125045444 | 124971500 | 124971598 | 125057941 | 125058106 | 10 | 14 | 5 | 10 | 15 | 0 | 0 | 0 | 1 | 0 | 14 | 2 | 3 | 19 | 34 | 0 | 0 | 0 | 0 | 0 | 298 | 149 | 1 | 1 | 1 | 1 | 1 | 0.833 | 1 | 1 | 1 | 1 | 1 | 1 | -0.033 |
| SE | 83273 | ENSG00000233429 | HOTAIRM1 | chr7 | + | 27099097 | 27099366 | 27096123 | 27096388 | 27099778 | 27099966 | 1 | 2 | 0 | 0 | 0 | 29 | 40 | 35 | 37 | 68 | 0 | 0 | 4 | 0 | 0 | 19 | 42 | 34 | 9 | 36 | 298 | 149 | 1 | 1 | 0.017 | 0.024 | 0 | 0 | 0 | 0 | 0 | 0.056 | 0 | 0 | -0.003 |
| SE | 83275 | ENSG00000240291 | NA | chr10 | - | 18539193 | 18539299 | 18513114 | 18513660 | 18542068 | 18542177 | 7 | 6 | 9 | 5 | 0 | 7 | 7 | 2 | 1 | 6 | 0 | 0 | 8 | 0 | 0 | 0 | 0 | 1 | 0 | 0 | 255 | 149 | 1 | 1 | 0.369 | 0.334 | 0.724 | 0.745 | 0 | NA | NA | 0.824 | NA | NA | -0.39 |
| SE | 83282 | ENSG00000087237 | CETP | chr16 | + | 56973330 | 56973510 | 56971991 | 56972083 | 56975100 | 56975151 | 1 | 0 | 3 | 1 | 5 | 0 | 0 | 0 | 3 | 0 | 7 | 3 | 0 | 16 | 69 | 0 | 0 | 0 | 0 | 10 | 298 | 149 | 1 | 1 | 1 | NA | 1 | 0.143 | 1 | 1 | 1 | NA | 1 | 0.775 | -0.158 |
| SE | 83284 | ENSG00000003436 | TFPI | chr2 | - | 187484810 | 187484987 | 187483929 | 187484216 | 187488336 | 187488375 | 43 | 117 | 19 | 107 | 107 | 0 | 0 | 0 | 1 | 0 | 72 | 146 | 52 | 52 | 28 | 0 | 0 | 0 | 0 | 0 | 298 | 149 | 1 | 1 | 1 | 1 | 1 | 0.982 | 1 | 1 | 1 | 1 | 1 | 1 | -0.004 |
| SE | 83286 | ENSG00000003436 | TFPI | chr2 | - | 187513615 | 187513646 | 187503647 | 187503770 | 187520541 | 187520613 | 0 | 1 | 0 | 2 | 0 | 0 | 3 | 2 | 0 | 0 | 0 | 3 | 1 | 5 | 0 | 7 | 3 | 6 | 1 | 0 | 180 | 149 | 1 | 1 | NA | 0.216 | 0 | 1 | NA | 0 | 0.453 | 0.121 | 0.805 | NA | 0.061 |
| SE | 83287 | ENSG00000003436 | TFPI | chr2 | - | 187513615 | 187513646 | 187503647 | 187503770 | 187529363 | 187529485 | 0 | 1 | 0 | 0 | 0 | 1 | 1 | 0 | 2 | 0 | 0 | 5 | 1 | 4 | 0 | 0 | 0 | 7 | 1 | 4 | 180 | 149 | 1 | 1 | 0 | 0.453 | NA | 0 | NA | NA | 1 | 0.106 | 0.768 | 0 | -0.318 |
| SE | 83289 | ENSG00000003436 | TFPI | chr2 | - | 187520541 | 187520613 | 187503647 | 187503770 | 187544503 | 187544637 | 0 | 2 | 2 | 0 | 0 | 0 | 1 | 2 | 0 | 1 | 18 | 3 | 4 | 1 | 0 | 0 | 0 | 0 | 0 | 0 | 221 | 149 | 1 | 1 | NA | 0.574 | 0.403 | NA | 0 | 1 | 1 | 1 | 1 | NA | -0.674 |
| SE | 83291 | ENSG00000003436 | TFPI | chr2 | - | 187520541 | 187520613 | 187513615 | 187513646 | 187554199 | 187554351 | 0 | 0 | 0 | 0 | 0 | 2 | 2 | 3 | 5 | 1 | 0 | 0 | 0 | 1 | 0 | 18 | 4 | 3 | 4 | 6 | 221 | 149 | 1 | 1 | 0 | 0 | 0 | 0 | 0 | 0 | 0 | 0 | 0.144 | 0 | -0.029 |
| SE | 83296 | ENSG00000100528 | CNIH1 | chr14 | - | 54430260 | 54430404 | 54423560 | 54427841 | 54432107 | 54432220 | 248 | 136 | 199 | 223 | 215 | 0 | 0 | 1 | 0 | 0 | 128 | 168 | 120 | 97 | 75 | 0 | 0 | 0 | 0 | 0 | 293 | 149 | 1 | 1 | 1 | 1 | 0.99 | 1 | 1 | 1 | 1 | 1 | 1 | 1 | -0.002 |
| SE | 83297 | ENSG00000100528 | CNIH1 | chr14 | - | 54430260 | 54430404 | 54423560 | 54427841 | 54436368 | 54436437 | 119 | 65 | 120 | 135 | 113 | 1 | 0 | 0 | 0 | 0 | 25 | 71 | 89 | 58 | 56 | 0 | 0 | 0 | 0 | 0 | 293 | 149 | 1 | 1 | 0.984 | 1 | 1 | 1 | 1 | 1 | 1 | 1 | 1 | 1 | -0.003 |
| SE | 83298 | ENSG00000100528 | CNIH1 | chr14 | - | 54432107 | 54432220 | 54426935 | 54427841 | 54436368 | 54436437 | 87 | 47 | 61 | 66 | 48 | 1 | 0 | 0 | 0 | 0 | 42 | 70 | 49 | 28 | 24 | 0 | 0 | 0 | 0 | 0 | 262 | 149 | 1 | 1 | 0.98 | 1 | 1 | 1 | 1 | 1 | 1 | 1 | 1 | 1 | -0.004 |
| SE | 83300 | ENSG00000100528 | CNIH1 | chr14 | - | 54436078 | 54436126 | 54432107 | 54432220 | 54436368 | 54436437 | 1 | 0 | 0 | 0 | 4 | 147 | 70 | 93 | 98 | 72 | 0 | 0 | 0 | 0 | 0 | 79 | 130 | 75 | 40 | 45 | 197 | 149 | 1 | 1 | 0.005 | 0 | 0 | 0 | 0.04 | 0 | 0 | 0 | 0 | 0 | 0.009 |
| SE | 83302 | ENSG00000100528 | CNIH1 | chr14 | - | 54436669 | 54436785 | 54436368 | 54436437 | 54441246 | 54441344 | 0 | 0 | 2 | 0 | 0 | 74 | 46 | 67 | 66 | 48 | 0 | 0 | 0 | 0 | 0 | 47 | 85 | 54 | 20 | 37 | 265 | 149 | 1 | 1 | 0 | 0 | 0.017 | 0 | 0 | 0 | 0 | 0 | 0 | 0 | 0.003 |
| SE | 83303 | ENSG00000168078 | PBK | chr8 | - | 27810957 | 27811134 | 27809623 | 27810501 | 27820564 | 27820694 | 0 | 0 | 15 | 10 | 19 | 0 | 0 | 0 | 0 | 0 | 0 | 8 | 11 | 0 | 0 | 0 | 0 | 1 | 0 | 0 | 298 | 149 | 1 | 1 | NA | NA | 1 | 1 | 1 | NA | 1 | 0.846 | NA | NA | 0.077 |
| SE | 83304 | ENSG00000168078 | PBK | chr8 | - | 27810957 | 27811167 | 27810200 | 27810501 | 27820564 | 27820694 | 0 | 0 | 9 | 3 | 12 | 0 | 0 | 0 | 0 | 0 | 0 | 5 | 4 | 0 | 0 | 0 | 0 | 1 | 0 | 0 | 298 | 149 | 1 | 1 | NA | NA | 1 | 1 | 1 | NA | 1 | 0.667 | NA | NA | 0.166 |
| SE | 83306 | ENSG00000229021 | NA | chr1 | + | 152016899 | 152018528 | 152016005 | 152016075 | 152018626 | 152018837 | 0 | 0 | 0 | 0 | 2 | 1 | 1 | 0 | 0 | 0 | 0 | 0 | 0 | 0 | 3 | 0 | 0 | 0 | 0 | 0 | 298 | 149 | 1 | 1 | 0 | 0 | NA | NA | 1 | NA | NA | NA | NA | 1 | -0.667 |
| SE | 83307 | ENSG00000182487 | NCF1B | chr7 | + | 73225710 | 73225786 | 73223877 | 73223956 | 73225885 | 73226051 | 165 | 137 | 137 | 75 | 266 | 0 | 0 | 1 | 0 | 0 | 1043 | 769 | 348 | 754 | 1631 | 1 | 0 | 0 | 0 | 4 | 225 | 149 | 1 | 1 | 1 | 1 | 0.989 | 1 | 1 | 0.999 | 1 | 1 | 1 | 0.996 | -0.001 |
| SE | 83312 | ENSG00000161010 | MRNIP | chr5 | - | 179844151 | 179844227 | 179841906 | 179842064 | 179851242 | 179851337 | 12 | 14 | 13 | 12 | 22 | 0 | 0 | 1 | 0 | 0 | 1 | 18 | 13 | 8 | 44 | 0 | 0 | 1 | 0 | 0 | 225 | 149 | 1 | 1 | 1 | 1 | 0.896 | 1 | 1 | 1 | 1 | 0.896 | 1 | 1 | 0 |
| SE | 83313 | ENSG00000161010 | MRNIP | chr5 | - | 179844151 | 179844227 | 179841906 | 179842064 | 179853196 | 179853276 | 11 | 15 | 13 | 12 | 22 | 0 | 0 | 2 | 0 | 0 | 1 | 21 | 13 | 8 | 56 | 0 | 0 | 0 | 3 | 0 | 225 | 149 | 1 | 1 | 1 | 1 | 0.811 | 1 | 1 | 1 | 1 | 1 | 0.638 | 1 | 0.035 |
| SE | 83315 | ENSG00000161010 | MRNIP | chr5 | - | 179847977 | 179848066 | 179841906 | 179842064 | 179851242 | 179851337 | 0 | 1 | 5 | 5 | 0 | 0 | 0 | 1 | 0 | 0 | 2 | 7 | 11 | 6 | 4 | 0 | 0 | 1 | 0 | 0 | 238 | 149 | 1 | 1 | NA | 1 | 0.758 | 1 | NA | 1 | 1 | 0.873 | 1 | 1 | -0.055 |

| | | | | | | | | | | | | | | | | | | | | | | | | | | | | | | | | | | | | | | | | | | | | | | |
|---|---|---|---|---|---|---|---|---|---|---|---|---|---|---|---|---|---|---|---|---|---|---|---|---|---|---|---|---|---|---|---|---|---|---|---|---|---|---|---|---|---|---|---|---|---|---|
| SE | 83316 | ENSG00000161010 | MRNIP | chr5 | - | 179847977 | 179848066 | 179841906 | 179842064 | 179853196 | 179853276 | 6 | 3 | 5 | 7 | 5 | 0 | 0 | 2 | 0 | 0 | 14 | 9 | 15 | 13 | 27 | 0 | 0 | 0 | 3 | 0 | 238 | 149 | 1 | 1 | 1 | 1 | 0.61 | 1 | 1 | 1 | 1 | 1 | 0.731 | 1 | -0.024 |
| SE | 83318 | ENSG00000161010 | MRNIP | chr5 | - | 179847977 | 179848066 | 179844151 | 179844227 | 179851242 | 179851337 | 21 | 24 | 27 | 40 | 22 | 1 | 0 | 0 | 0 | 0 | 43 | 37 | 38 | 30 | 55 | 0 | 0 | 0 | 0 | 0 | 238 | 149 | 1 | 1 | 0.929 | 1 | 1 | 1 | 1 | 1 | 1 | 1 | 1 | 1 | -0.014 |
| SE | 83321 | ENSG00000161010 | MRNIP | chr5 | - | 179851242 | 179851337 | 179841907 | 179842064 | 179853196 | 179853437 | 0 | 1 | 1 | 0 | 0 | 0 | 0 | 2 | 0 | 0 | 0 | 0 | 0 | 0 | 0 | 0 | 0 | 0 | 3 | 0 | 244 | 149 | 1 | 1 | NA | 1 | 0.234 | NA | NA | NA | NA | NA | 0 | NA | 0.617 |
| SE | 83323 | ENSG00000161010 | MRNIP | chr5 | - | 179851242 | 179851337 | 179844151 | 179844227 | 179853196 | 179853437 | 1 | 1 | 0 | 0 | 0 | 0 | 1 | 0 | 0 | 0 | 0 | 0 | 0 | 0 | 0 | 0 | 5 | 0 | 0 | 12 | 244 | 149 | 1 | 1 | 1 | 0.379 | NA | NA | NA | NA | 0 | NA | NA | 0 | 0.69 |
| SE | 83327 | ENSG00000161010 | MRNIP | chr5 | - | 179851242 | 179851424 | 179842046 | 179842064 | 179853196 | 179853276 | 0 | 1 | 1 | 5 | 1 | 0 | 0 | 2 | 0 | 0 | 2 | 4 | 5 | 18 | 16 | 0 | 0 | 0 | 3 | 0 | 298 | 149 | 1 | 1 | NA | 1 | 0.2 | 1 | 1 | 1 | 1 | 1 | 0.75 | 1 | -0.15 |
| SE | 83328 | ENSG00000161010 | MRNIP | chr5 | - | 179851242 | 179851424 | 179842046 | 179842064 | 179853377 | 179853437 | 0 | 0 | 1 | 0 | 0 | 0 | 1 | 3 | 0 | 1 | 0 | 0 | 1 | 0 | 0 | 0 | 0 | 0 | 0 | 0 | 298 | 149 | 1 | 1 | NA | 0 | 0.143 | NA | 0 | NA | NA | 1 | NA | NA | -0.952 |
| SE | 83329 | ENSG00000161010 | MRNIP | chr5 | - | 179851242 | 179851424 | 179844151 | 179844227 | 179853196 | 179853276 | 1 | 1 | 0 | 5 | 1 | 0 | 1 | 0 | 0 | 0 | 2 | 4 | 4 | 18 | 16 | 0 | 5 | 0 | 0 | 12 | 298 | 149 | 1 | 1 | 1 | 0.333 | NA | 1 | 1 | 1 | 0.286 | 1 | 1 | 0.4 | 0.096 |
| SE | 83333 | ENSG00000161010 | MRNIP | chr5 | - | 179853196 | 179853276 | 179841906 | 179842064 | 179853377 | 179853437 | 0 | 0 | 6 | 4 | 6 | 0 | 1 | 3 | 0 | 1 | 0 | 5 | 3 | 9 | 6 | 0 | 0 | 0 | 0 | 0 | 229 | 149 | 1 | 1 | NA | 0 | 0.565 | 1 | 0.796 | NA | 1 | 1 | 1 | 1 | -0.41 |
| SE | 83334 | ENSG00000161010 | MRNIP | chr5 | - | 179853196 | 179853276 | 179844151 | 179844227 | 179853377 | 179853437 | 0 | 0 | 4 | 4 | 6 | 1 | 1 | 2 | 1 | 0 | 0 | 10 | 3 | 6 | 6 | 0 | 0 | 0 | 0 | 0 | 229 | 149 | 1 | 1 | 0 | 0 | 0.565 | 0.722 | 1 | NA | 1 | 1 | 1 | 1 | -0.543 |
| SE | 83341 | ENSG00000087274 | ADD1 | chr4 | + | 2853623 | 2853698 | 2843952 | 2844024 | 2875895 | 2876405 | 0 | 0 | 0 | 0 | 0 | 147 | 97 | 115 | 142 | 73 | 0 | 0 | 0 | 0 | 3 | 176 | 166 | 84 | 160 | 273 | 224 | 149 | 1 | 1 | 0 | 0 | 0 | 0 | 0 | 0 | 0 | 0 | 0 | 0.007 | -0.001 |
| SE | 83342 | ENSG00000087274 | ADD1 | chr4 | + | 2853623 | 2853698 | 2853133 | 2853203 | 2875895 | 2876405 | 0 | 0 | 0 | 0 | 0 | 0 | 0 | 0 | 0 | 1 | 0 | 0 | 0 | 0 | 3 | 0 | 0 | 0 | 0 | 0 | 224 | 149 | 1 | 1 | NA | NA | NA | NA | 0 | NA | NA | NA | NA | 1 | -1 |
| SE | 83343 | ENSG00000087274 | ADD1 | chr4 | + | 2854853 | 2855029 | 2853623 | 2853698 | 2875895 | 2876405 | 0 | 0 | 0 | 0 | 1 | 0 | 0 | 0 | 0 | 0 | 0 | 0 | 0 | 3 | 0 | 0 | 0 | 0 | 0 | 3 | 298 | 149 | 1 | 1 | NA | NA | NA | NA | 1 | NA | NA | NA | 1 | 0 | 0.5 |
| SE | 83345 | ENSG00000087274 | ADD1 | chr4 | + | 2867971 | 2868129 | 2844149 | 2844834 | 2875895 | 2876110 | 16 | 6 | 9 | 5 | 5 | 0 | 1 | 1 | 3 | 1 | 34 | 0 | 8 | 0 | 0 | 0 | 0 | 0 | 0 | 0 | 298 | 149 | 1 | 1 | 1 | 0.75 | 0.818 | 0.455 | 0.714 | 1 | NA | 1 | NA | NA | -0.253 |
| SE | 83346 | ENSG00000087274 | ADD1 | chr4 | + | 2873014 | 2873135 | 2844400 | 2844834 | 2875895 | 2876110 | 2 | 0 | 0 | 1 | 0 | 0 | 1 | 1 | 3 | 1 | 0 | 1 | 7 | 0 | 0 | 0 | 0 | 0 | 0 | 0 | 270 | 149 | 1 | 1 | 1 | 0 | 0 | 0.155 | 0 | NA | 1 | 1 | NA | NA | -0.769 |
| SE | 83347 | ENSG00000087274 | ADD1 | chr4 | + | 2881897 | 2882060 | 2875895 | 2876110 | 2884514 | 2884569 | 1271 | 814 | 1045 | 1316 | 1496 | 0 | 0 | 3 | 3 | 1 | 1996 | 1444 | 952 | 1406 | 2509 | 0 | 5 | 0 | 0 | 5 | 298 | 149 | 1 | 1 | 1 | 1 | 0.994 | 0.995 | 0.999 | 1 | 0.993 | 1 | 1 | 0.996 | 0 |
| SE | 83348 | ENSG00000087274 | ADD1 | chr4 | + | 2884514 | 2884666 | 2875895 | 2876110 | 2894012 | 2894093 | 478 | 259 | 386 | 436 | 556 | 0 | 2 | 1 | 1 | 3 | 374 | 424 | 304 | 367 | 684 | 0 | 0 | 2 | 1 | 0 | 298 | 149 | 1 | 1 | 1 | 0.985 | 0.995 | 0.995 | 0.989 | 1 | 1 | 0.987 | 0.995 | 1 | -0.004 |
| SE | 83349 | ENSG00000087274 | ADD1 | chr4 | + | 2894012 | 2894093 | 2884514 | 2884666 | 2894581 | 2894682 | 973 | 514 | 747 | 900 | 1040 | 6 | 0 | 11 | 4 | 0 | 808 | 925 | 634 | 735 | 1491 | 3 | 8 | 3 | 3 | 0 | 230 | 149 | 1 | 1 | 0.991 | 1 | 0.978 | 0.993 | 1 | 0.994 | 0.987 | 0.993 | 0.994 | 1 | -0.001 |
| SE | 83350 | ENSG00000087274 | ADD1 | chr4 | + | 2898432 | 2898531 | 2898183 | 2898327 | 2899258 | 2899435 | 400 | 255 | 410 | 419 | 651 | 4 | 0 | 0 | 0 | 0 | 549 | 444 | 257 | 409 | 642 | 0 | 0 | 0 | 1 | 0 | 248 | 149 | 1 | 1 | 0.984 | 1 | 1 | 1 | 1 | 1 | 1 | 1 | 0.996 | 1 | -0.002 |
| SE | 83351 | ENSG00000087274 | ADD1 | chr4 | + | 2899258 | 2899435 | 2898183 | 2898327 | 2904763 | 2904777 | 400 | 249 | 406 | 381 | 576 | 0 | 0 | 0 | 0 | 0 | 645 | 421 | 334 | 405 | 637 | 6 | 0 | 3 | 0 | 0 | 298 | 149 | 1 | 1 | 1 | 1 | 1 | 1 | 1 | 0.982 | 1 | 0.982 | 1 | 1 | 0.007 |
| SE | 83352 | ENSG00000087274 | ADD1 | chr4 | + | 2904763 | 2905015 | 2899258 | 2899435 | 2907742 | 2907844 | 1253 | 738 | 1033 | 1176 | 1363 | 0 | 0 | 0 | 0 | 1 | 2267 | 1598 | 918 | 1336 | 2878 | 0 | 0 | 0 | 0 | 10 | 298 | 149 | 1 | 1 | 1 | 1 | 1 | 1 | 0.999 | 1 | 1 | 1 | 1 | 0.993 | 0.001 |
| SE | 83353 | ENSG00000087274 | ADD1 | chr4 | + | 2904763 | 2905108 | 2899258 | 2899435 | 2907742 | 2907844 | 448 | 282 | 454 | 422 | 644 | 0 | 0 | 0 | 0 | 1 | 773 | 465 | 396 | 478 | 725 | 0 | 0 | 0 | 0 | 10 | 298 | 149 | 1 | 1 | 1 | 1 | 1 | 1 | 0.997 | 1 | 1 | 1 | 1 | 0.973 | 0.005 |
| SE | 83354 | ENSG00000087274 | ADD1 | chr4 | + | 2907742 | 2907844 | 2904763 | 2905015 | 2914883 | 2915040 | 391 | 193 | 243 | 370 | 255 | 1 | 0 | 1 | 1 | 0 | 764 | 464 | 231 | 373 | 1042 | 0 | 0 | 1 | 2 | 0 | 251 | 149 | 1 | 1 | 0.996 | 1 | 0.993 | 0.995 | 1 | 1 | 1 | 0.993 | 0.991 | 1 | 0 |
| SE | 83355 | ENSG00000087274 | ADD1 | chr4 | + | 2908514 | 2908604 | 2907203 | 2907844 | 2914883 | 2915040 | 1153 | 735 | 953 | 1180 | 1150 | 4 | 5 | 5 | 8 | 1 | 2007 | 1673 | 902 | 1415 | 2803 | 0 | 7 | 5 | 7 | 8 | 239 | 149 | 1 | 1 | 0.994 | 0.989 | 0.992 | 0.989 | 0.999 | 1 | 0.993 | 0.991 | 0.992 | 0.995 | -0.002 |
| SE | 83358 | ENSG00000087274 | ADD1 | chr4 | + | 2914883 | 2915040 | 2908514 | 2908604 | 2926013 | 2926100 | 788 | 464 | 678 | 872 | 689 | 0 | 0 | 0 | 1 | 4 | 1140 | 1060 | 574 | 878 | 1777 | 0 | 6 | 6 | 4 | 8 | 298 | 149 | 1 | 1 | 1 | 1 | 1 | 0.998 | 0.989 | 1 | 0.989 | 0.98 | 0.991 | 0.991 | 0.007 |
| SE | 83359 | ENSG00000087274 | ADD1 | chr4 | + | 2914883 | 2915040 | 2908514 | 2908604 | 2928170 | 2928431 | 540 | 329 | 447 | 617 | 489 | 0 | 0 | 0 | 0 | 0 | 846 | 749 | 412 | 651 | 1335 | 0 | 0 | 0 | 3 | 0 | 298 | 149 | 1 | 1 | 1 | 1 | 1 | 1 | 1 | 1 | 1 | 1 | 0.991 | 1 | 0.002 |
| SE | 83360 | ENSG00000087274 | ADD1 | chr4 | + | 2926013 | 2926112 | 2908514 | 2908604 | 2928170 | 2928431 | 277 | 137 | 262 | 234 | 294 | 0 | 0 | 0 | 0 | 0 | 398 | 374 | 219 | 239 | 665 | 0 | 0 | 0 | 3 | 0 | 248 | 149 | 1 | 1 | 1 | 1 | 1 | 1 | 1 | 1 | 1 | 1 | 0.98 | 1 | 0.004 |
| SE | 83361 | ENSG00000087274 | ADD1 | chr4 | + | 2926013 | 2926112 | 2914883 | 2915040 | 2928170 | 2928431 | 524 | 273 | 492 | 487 | 489 | 1 | 0 | 0 | 0 | 0 | 686 | 672 | 372 | 462 | 1080 | 0 | 0 | 1 | 0 | 0 | 248 | 149 | 1 | 1 | 0.997 | 1 | 1 | 1 | 1 | 1 | 1 | 0.996 | 1 | 1 | 0 |
| SE | 83367 | ENSG00000166716 | ZNF592 | chr15 | + | 84790704 | 84790872 | 84782656 | 84784895 | 84797868 | 84798045 | 109 | 60 | 71 | 101 | 47 | 0 | 4 | 1 | 0 | 0 | 134 | 70 | 43 | 107 | 89 | 0 | 0 | 0 | 0 | 0 | 298 | 149 | 1 | 1 | 1 | 0.882 | 0.973 | 1 | 1 | 1 | 1 | 1 | 1 | 1 | -0.029 |
| SE | 83368 | ENSG00000166716 | ZNF592 | chr15 | + | 84790704 | 84790883 | 84782653 | 84784895 | 84797868 | 84798045 | 147 | 80 | 99 | 140 | 62 | 0 | 4 | 1 | 0 | 0 | 167 | 100 | 59 | 159 | 108 | 0 | 0 | 0 | 0 | 0 | 298 | 149 | 1 | 1 | 1 | 0.909 | 0.98 | 1 | 1 | 1 | 1 | 1 | 1 | 1 | -0.022 |
| SE | 83371 | ENSG00000274523 | RCC1L | chr7 | - | 75052710 | 75052796 | 75042182 | 75043109 | 75055900 | 75056074 | 111 | 50 | 108 | 116 | 101 | 0 | 0 | 0 | 0 | 0 | 105 | 125 | 43 | 58 | 71 | 0 | 0 | 0 | 0 | 1 | 235 | 149 | 1 | 1 | 1 | 1 | 1 | 1 | 1 | 1 | 1 | 1 | 1 | 0.978 | 0.004 |
| SE | 83374 | ENSG00000274523 | RCC1L | chr7 | - | 75055900 | 75056074 | 75052710 | 75052796 | 75057528 | 75057616 | 189 | 66 | 162 | 161 | 162 | 0 | 1 | 0 | 2 | 0 | 182 | 184 | 62 | 107 | 171 | 0 | 0 | 1 | 0 | 4 | 298 | 149 | 1 | 1 | 1 | 0.971 | 1 | 0.976 | 1 | 1 | 1 | 0.969 | 1 | 0.955 | 0.005 |
| SE | 83375 | ENSG00000274523 | RCC1L | chr7 | - | 75055900 | 75056074 | 75052710 | 75052796 | 75058587 | 75058769 | 104 | 47 | 99 | 94 | 90 | 0 | 1 | 0 | 1 | 0 | 100 | 97 | 40 | 52 | 71 | 0 | 0 | 0 | 0 | 0 | 298 | 149 | 1 | 1 | 1 | 0.959 | 1 | 0.979 | 1 | 1 | 1 | 1 | 1 | 1 | -0.012 |
| SE | 83376 | ENSG00000274523 | RCC1L | chr7 | - | 75057528 | 75057616 | 75052710 | 75052796 | 75058587 | 75058769 | 34 | 8 | 28 | 22 | 25 | 0 | 1 | 0 | 1 | 0 | 67 | 41 | 18 | 20 | 43 | 0 | 0 | 0 | 0 | 0 | 237 | 149 | 1 | 1 | 1 | 0.834 | 1 | 0.933 | 1 | 1 | 1 | 1 | 1 | 1 | -0.047 |
| SE | 83377 | ENSG00000274523 | RCC1L | chr7 | - | 75057528 | 75057616 | 75055574 | 75056074 | 75058587 | 75058769 | 118 | 25 | 91 | 85 | 97 | 0 | 0 | 2 | 1 | 0 | 149 | 128 | 39 | 75 | 139 | 0 | 0 | 0 | 0 | 0 | 237 | 149 | 1 | 1 | 1 | 1 | 0.966 | 0.982 | 1 | 1 | 1 | 1 | 1 | 1 | -0.01 |
| SE | 83378 | ENSG00000274523 | RCC1L | chr7 | - | 75061206 | 75061291 | 75058587 | 75058769 | 75063291 | 75063343 | 64 | 34 | 113 | 77 | 132 | 0 | 0 | 0 | 0 | 2 | 110 | 104 | 56 | 29 | 112 | 0 | 0 | 0 | 0 | 0 | 234 | 149 | 1 | 1 | 1 | 1 | 1 | 1 | 0.977 | 1 | 1 | 1 | 1 | 1 | -0.005 |
| SE | 83379 | ENSG00000274523 | RCC1L | chr7 | - | 75063291 | 75063343 | 75061206 | 75061291 | 75064581 | 75064648 | 60 | 27 | 89 | 56 | 107 | 0 | 1 | 0 | 2 | 0 | 94 | 91 | 38 | 28 | 69 | 0 | 0 | 0 | 1 | 0 | 201 | 149 | 1 | 1 | 1 | 0.952 | 1 | 0.954 | 1 | 1 | 1 | 1 | 0.954 | 1 | -0.01 |
| SE | 83383 | ENSG00000162959 | MEMO1 | chr2 | - | 32010186 | 32010264 | 31943301 | 31943383 | 32010993 | 32011230 | 48 | 13 | 25 | 25 | 22 | 0 | 0 | 0 | 0 | 1 | 24 | 19 | 34 | 38 | 50 | 0 | 0 | 0 | 3 | 0 | 227 | 149 | 1 | 1 | 1 | 1 | 1 | 1 | 0.935 | 1 | 1 | 1 | 0.893 | 1 | 0.008 |
| SE | 83384 | ENSG00000174428 | GTF2IRD2B | chr7 | + | 75093850 | 75095029 | 75092572 | 75092765 | 75108959 | 75109063 | 0 | 1 | 3 | 3 | 0 | 23 | 9 | 22 | 15 | 0 | 0 | 0 | 0 | 0 | 2 | 38 | 26 | 13 | 11 | 33 | 298 | 149 | 1 | 1 | 0 | 0.053 | 0.064 | 0.091 | NA | 0 | 0 | 0 | 0 | 0.029 | 0.046 |
| SE | 83386 | ENSG00000174428 | GTF2IRD2B | chr7 | + | 75123135 | 75123319 | 75120890 | 75121010 | 75123487 | 75123516 | 42 | 22 | 33 | 27 | 36 | 0 | 0 | 1 | 0 | 0 | 55 | 34 | 13 | 30 | 13 | 0 | 0 | 0 | 0 | 0 | 298 | 149 | 1 | 1 | 1 | 1 | 0.943 | 1 | 1 | 1 | 1 | 1 | 1 | 1 | -0.011 |
| SE | 83388 | ENSG00000174428 | GTF2IRD2B | chr7 | + | 75123487 | 75123516 | 75120890 | 75121010 | 75125915 | 75125970 | 0 | 0 | 1 | 0 | 0 | 0 | 3 | 1 | 4 | 4 | 0 | 0 | 0 | 0 | 0 | 0 | 1 | 4 | 0 | 7 | 178 | 149 | 1 | 1 | NA | 0 | 0.456 | 0 | 0 | NA | 0 | 0 | NA | 0 | 0.114 |
| SE | 83389 | ENSG00000174428 | GTF2IRD2B | chr7 | + | 75123487 | 75123516 | 75123135 | 75123319 | 75125915 | 75125970 | 11 | 3 | 6 | 1 | 3 | 0 | 1 | 3 | 4 | 1 | 0 | 5 | 3 | 6 | 0 | 17 | 7 | 8 | 8 | 0 | 178 | 149 | 1 | 1 | 1 | 0.715 | 0.626 | 0.173 | 0.715 | 0 | 0.374 | 0.239 | 0.386 | NA | 0.396 |
| SE | 83391 | ENSG00000174428 | GTF2IRD2B | chr7 | + | 75133134 | 75133212 | 75126341 | 75126385 | 75135000 | 75135057 | 0 | 1 | 1 | 3 | 7 | 0 | 0 | 1 | 0 | 0 | 0 | 5 | 2 | 3 | 0 | 0 | 0 | 0 | 0 | 0 | 227 | 149 | 1 | 1 | NA | 1 | 0.396 | 1 | 1 | NA | 1 | 1 | 1 | NA | -0.151 |
| SE | 83393 | ENSG00000091073 | DTX2 | chr7 | + | 76463444 | 76463709 | 76461696 | 76461836 | 76480420 | 76480584 | 7 | 8 | 12 | 11 | 15 | 1 | 0 | 0 | 0 | 0 | 0 | 14 | 6 | 14 | 40 | 0 | 0 | 0 | 0 | 0 | 298 | 149 | 1 | 1 | 0.778 | 1 | 1 | 1 | 1 | NA | 1 | 1 | 1 | 1 | -0.044 |
| SE | 83394 | ENSG00000091073 | DTX2 | chr7 | + | 76463544 | 76463709 | 76461700 | 76461836 | 76476942 | 76477178 | 1 | 3 | 0 | 1 | 1 | 0 | 0 | 0 | 0 | 0 | 0 | 7 | 0 | 0 | 20 | 0 | 0 | 5 | 0 | 0 | 298 | 149 | 1 | 1 | 1 | 1 | NA | 1 | 1 | NA | 0.412 | NA | NA | 1 | 0.294 |
| SE | 83396 | ENSG00000091073 | DTX2 | chr7 | + | 76463544 | 76463709 | 76461700 | 76461836 | 76480420 | 76480777 | 8 | 10 | 11 | 12 | 16 | 1 | 0 | 0 | 0 | 0 | 0 | 19 | 6 | 13 | 50 | 0 | 0 | 0 | 0 | 0 | 298 | 149 | 1 | 1 | 0.8 | 1 | 1 | 1 | 1 | NA | 1 | 1 | 1 | 1 | -0.04 |
| SE | 83398 | ENSG00000091073 | DTX2 | chr7 | + | 76470299 | 76470419 | 76463544 | 76463709 | 76477113 | 76477178 | 0 | 1 | 1 | 0 | 0 | 1 | 1 | 0 | 1 | 0 | 0 | 0 | 0 | 7 | 0 | 12 | 0 | 5 | 1 | 0 | 269 | 149 | 1 | 1 | 0 | 0.356 | 1 | 0 | NA | 0 | NA | 0 | 0.795 | NA | 0.074 |
| SE | 83399 | ENSG00000091073 | DTX2 | chr7 | + | 76470299 | 76470419 | 76463544 | 76463709 | 76480420 | 76480551 | 0 | 1 | 2 | 0 | 1 | 7 | 8 | 11 | 11 | 15 | 0 | 0 | 0 | 5 | 0 | 0 | 12 | 6 | 13 | 40 | 269 | 149 | 1 | 1 | 0 | 0.065 | 0.091 | 0 | 0.036 | NA | 0 | 0 | 0.176 | 0 | -0.006 |
| SE | 83400 | ENSG00000091073 | DTX2 | chr7 | + | 76476942 | 76477178 | 76461696 | 76461836 | 76480420 | 76480447 | 0 | 0 | 0 | 0 | 0 | 1 | 0 | 0 | 0 | 0 | 0 | 5 | 3 | 0 | 0 | 0 | 0 | 0 | 0 | 0 | 298 | 149 | 1 | 1 | 0 | NA | NA | NA | NA | NA | 1 | 1 | NA | NA | -1 |
| SE | 83401 | ENSG00000091073 | DTX2 | chr7 | + | 76476942 | 76477178 | 76463544 | 76463709 | 76480420 | 76480447 | 0 | 1 | 0 | 0 | 0 | 7 | 8 | 11 | 11 | 15 | 0 | 0 | 3 | 0 | 10 | 0 | 12 | 6 | 13 | 40 | 298 | 149 | 1 | 1 | 0 | 0.059 | 0 | 0 | 0 | NA | 0 | 0.2 | 0 | 0.111 | -0.066 |
| SE | 83402 | ENSG00000091073 | DTX2 | chr7 | + | 76477113 | 76477178 | 76461719 | 76461836 | 76480420 | 76480695 | 0 | 0 | 0 | 0 | 0 | 1 | 0 | 0 | 0 | 0 | 0 | 0 | 3 | 0 | 0 | 0 | 0 | 0 | 0 | 0 | 214 | 149 | 1 | 1 | 0 | NA | NA | NA | NA | NA | NA | 1 | NA | NA | -1 |
| SE | 83403 | ENSG00000091073 | DTX2 | chr7 | + | 76477113 | 76477178 | 76463544 | 76463709 | 76480420 | 76480695 | 1 | 1 | 0 | 1 | 0 | 7 | 8 | 11 | 11 | 15 | 12 | 0 | 8 | 1 | 0 | 0 | 12 | 6 | 13 | 40 | 214 | 149 | 1 | 1 | 0.09 | 0.08 | 0 | 0.06 | 0 | 1 | 0 | 0.481 | 0.051 | 0 | -0.26 |
| SE | 83404 | ENSG00000091073 | DTX2 | chr7 | + | 76477113 | 76477178 | 76470299 | 76470419 | 76480420 | 76480695 | 0 | 0 | 0 | 0 | 0 | 0 | 1 | 1 | 0 | 1 | 0 | 0 | 3 | 7 | 0 | 0 | 0 | 0 | 5 | 0 | 214 | 149 | 1 | 1 | NA | 0 | 0 | NA | 0 | NA | NA | 1 | 0.494 | NA | -0.747 |
| SE | 83405 | ENSG00000091073 | DTX2 | chr7 | + | 76480420 | 76480777 | 76463544 | 76463709 | 76482507 | 76482563 | 12 | 11 | 20 | 22 | 21 | 0 | 0 | 0 | 0 | 0 | 30 | 32 | 20 | 24 | 63 | 0 | 0 | 0 | 0 | 2 | 298 | 149 | 1 | 1 | 1 | 1 | 1 | 1 | 1 | 1 | 1 | 1 | 1 | 0.94 | 0.012 |
| SE | 83410 | ENSG00000146707 | POMZP3 | chr7 | - | 76618182 | 76618615 | 76611721 | 76611813 | 76625521 | 76625683 | 8 | 10 | 7 | 7 | 11 | 1 | 0 | 0 | 0 | 0 | 0 | 24 | 10 | 5 | 7 | 0 | 0 | 0 | 0 | 7 | 298 | 149 | 1 | 1 | 0.8 | 1 | 1 | 1 | 1 | NA | 1 | 1 | 1 | 0.333 | 0.127 |
| SE | 83415 | ENSG00000175274 | TP53I11 | chr11 | - | 44937108 | 44937352 | 44936803 | 44936899 | 44937554 | 44937613 | 2 | 0 | 2 | 1 | 3 | 0 | 0 | 0 | 0 | 0 | 1 | 12 | 2 | 11 | 0 | 0 | 0 | 0 | 1 | 0 | 298 | 149 | 1 | 1 | 1 | NA | 1 | 1 | 1 | 1 | 1 | 1 | 0.846 | NA | 0.038 |
| SE | 83416 | ENSG00000175274 | TP53I11 | chr11 | - | 44937303 | 44937352 | 44936802 | 44936899 | 44937554 | 44937613 | 21 | 15 | 16 | 20 | 12 | 0 | 0 | 0 | 0 | 0 | 5 | 43 | 6 | 36 | 12 | 0 | 0 | 0 | 1 | 0 | 198 | 149 | 1 | 1 | 1 | 1 | 1 | 1 | 1 | 1 | 1 | 1 | 0.964 | 1 | 0.007 |
| SE | 83418 | ENSG00000175274 | TP53I11 | chr11 | - | 44939113 | 44939190 | 44938206 | 44938366 | 44950676 | 44950752 | 3 | 0 | 0 | 0 | 0 | 2 | 0 | 3 | 2 | 0 | 0 | 0 | 0 | 0 | 0 | 0 | 0 | 0 | 7 | 0 | 226 | 149 | 1 | 1 | 0.497 | NA | 0 | 0 | NA | NA | NA | NA | 0 | NA | 0.166 |
| SE | 83421 | ENSG00000175274 | TP53I11 | chr11 | - | 44939165 | 44939281 | 44938206 | 44938366 | 44950676 | 44950894 | 3 | 2 | 0 | 0 | 0 | 2 | 0 | 3 | 2 | 0 | 0 | 0 | 0 | 8 | 0 | 0 | 0 | 0 | 7 | 0 | 265 | 149 | 1 | 1 | 0.458 | 1 | 0 | 0 | NA | NA | NA | | 0.391 | NA | -0.027 |
| SE | 83422 | ENSG00000175274 | TP53I11 | chr11 | - | 44939165 | 44939281 | 44938206 | 44938366 | 44950876 | 44951042 | 3 | 1 | 0 | 0 | 0 | 0 | 1 | 0 | 0 | 0 | 0 | 0 | 0 | 6 | 0 | 0 | 2 | 0 | 0 | 0 | 265 | 149 | 1 | 1 | 1 | 0.36 | NA | NA | NA | NA | 0 | NA | 1 | NA | 0.18 |
| SE | 83426 | ENSG00000175274 | TP53I11 | chr11 | - | 44941002 | 44941077 | 44939165 | 44939281 | 44950676 | 44950845 | 0 | 0 | 0 | 0 | 0 | 0 | 1 | 0 | 0 | 0 | 14 | 0 | 0 | 0 | 0 | 0 | 0 | 0 | 2 | 0 | 224 | 149 | 1 | 1 | NA | 0 | NA | NA | | 1 | NA | NA | 0 | NA | -0.5 |
| SE | 83427 | ENSG00000122545 | SEPTIN7 | chr7 | + | 35831352 | 35831496 | 35801217 | 35801270 | 35832797 | 35832900 | 12 | 8 | 15 | 10 | 20 | 10 | 13 | 9 | 13 | 14 | 0 | 4 | 6 | 0 | 0 | 0 | 2 | 9 | 1 | 0 | 293 | 149 | 1 | 1 | 0.379 | 0.238 | 0.459 | 0.281 | 0.421 | NA | 0.504 | 0.253 | 0 | NA | 0.103 |
| SE | 83428 | ENSG00000122545 | SEPTIN7 | chr7 | + | 35831424 | 35831496 | 35801218 | 35801270 | 35832797 | 35832900 | 11 | 7 | 12 | 11 | 12 | 10 | 13 | 9 | 13 | 14 | 0 | 10 | 2 | 0 | 0 | 0 | 2 | 9 | 1 | 0 | 221 | 149 | 1 | 1 | 0.426 | 0.266 | 0.473 | 0.363 | 0.366 | NA | 0.771 | 0.13 | 0 | NA | 0.078 |
| SE | 83429 | ENSG00000122545 | SEPTIN7 | chr7 | + | 35831491 | 35831496 | 35800931 | 35801270 | 35832797 | 35832900 | 114 | 84 | 86 | 109 | 56 | 10 | 13 | 9 | 13 | 14 | 12 | 55 | 11 | 13 | 0 | 0 | 2 | 9 | 1 | 0 | 154 | 149 | 1 | 1 | 0.917 | 0.862 | 0.902 | 0.89 | 0.795 | 1 | 0.964 | 0.542 | 0.926 | NA | 0.015 |
| SE | 83430 | ENSG00000122545 | SEPTIN7 | chr7 | + | 35831491 | 35831496 | 35800931 | 35801270 | 35832803 | 35832900 | 14 | 6 | 11 | 5 | 1 | 0 | 0 | 0 | 1 | 0 | 0 | 2 | 0 | 0 | 0 | 0 | 0 | 0 | 0 | 0 | 154 | 149 | 1 | 1 | 1 | 1 | 1 | 0.829 | 1 | NA | 1 | NA | NA | NA | -0.034 |
| SE | 83431 | ENSG00000122545 | SEPTIN7 | chr7 | + | 35831491 | 35832034 | 35801218 | 35801270 | 35832797 | 35832845 | 7 | 1 | 8 | 2 | 6 | 10 | 13 | 9 | 13 | 14 | 3 | 13 | 2 | 5 | 0 | 0 | 2 | 9 | 1 | 0 | 298 | 149 | 1 | 1 | 0.259 | 0.037 | 0.308 | 0.071 | 0.176 | 1 | 0.765 | 0.1 | 0.714 | NA | -0.475 |
| SE | 83432 | ENSG00000122545 | SEPTIN7 | chr7 | + | 35832792 | 35832900 | 35800985 | 35801270 | 35863551 | 35863658 | 534 | 402 | 342 | 545 | 763 | 4 | 2 | 3 | 6 | 1 | 43 | 218 | 96 | 32 | 27 | 0 | 0 | 0 | 0 | 0 | 257 | 149 | 1 | 1 | 0.987 | 0.991 | 0.985 | 0.981 | 0.998 | 1 | 1 | 1 | 1 | 1 | -0.012 |
| SE | 83433 | ENSG00000122545 | SEPTIN7 | chr7 | + | 35832797 | 35832893 | 35801218 | 35801270 | 35863551 | 35863658 | 0 | 0 | 0 | 0 | 4 | 4 | 2 | 3 | 6 | 1 | 0 | 0 | 6 | 1 | 0 | 0 | 0 | 0 | 0 | 0 | 245 | 149 | 1 | 1 | 0 | 0 | 0 | 0 | 0.709 | NA | NA | 1 | 1 | NA | -0.858 |
| SE | 83434 | ENSG00000122545 | SEPTIN7 | chr7 | + | 35832797 | 35832900 | 35800931 | 35801270 | 35863551 | 35863613 | 544 | 414 | 351 | 558 | 773 | 4 | 2 | 3 | 6 | 1 | 43 | 220 | 105 | 33 | 27 | 0 | 0 | 0 | 0 | 0 | 252 | 149 | 1 | 1 | 0.988 | 0.992 | 0.986 | 0.982 | 0.998 | 1 | 1 | 1 | 1 | 1 | -0.011 |
| SE | 83435 | ENSG00000122545 | SEPTIN7 | chr7 | + | 35832803 | 35832900 | 35801215 | 35801270 | 35863551 | 35863658 | 499 | 371 | 321 | 516 | 701 | 4 | 2 | 3 | 6 | 1 | 43 | 203 | 91 | 32 | 27 | 0 | 0 | 0 | 0 | 0 | 246 | 149 | 1 | 1 | 0.987 | 0.991 | 0.985 | 0.981 | 0.998 | 1 | 1 | 1 | 1 | 1 | -0.012 |
| SE | 83439 | ENSG00000122545 | SEPTIN7 | chr7 | + | 35873640 | 35873775 | 35872665 | 35872766 | 35879822 | 35879872 | 2596 | 2102 | 1931 | 2737 | 4900 | 5 | 1 | 0 | 0 | 0 | 106 | 957 | 638 | 188 | 113 | 0 | 0 | 0 | 0 | 0 | 284 | 149 | 1 | 1 | 0.996 | 0.999 | 1 | 1 | 1 | 1 | 1 | 1 | 1 | 1 | -0.001 |
| SE | 83441 | ENSG00000128944 | KNSTRN | chr15 | + | 40383227 | 40383322 | 40382720 | 40383044 | 40386404 | 40386494 | 6 | 1 | 1 | 0 | 2 | 0 | 0 | 0 | 0 | 0 | 0 | 0 | 4 | 1 | 3 | 0 | 0 | 1 | 0 | 0 | 244 | 149 | 1 | 1 | 1 | 1 | 1 | NA | 1 | NA | NA | 0.71 | 1 | 1 | 0.097 |
| SE | 83443 | ENSG00000128944 | KNSTRN | chr15 | + | 40384456 | 40384519 | 40383227 | 40383322 | 40386361 | 40386494 | 1 | 0 | 0 | 1 | 0 | 13 | 2 | 1 | 12 | 2 | 0 | 4 | 0 | 0 | 0 | 0 | 0 | 3 | 1 | 0 | 212 | 149 | 1 | 1 | 0.051 | 0 | 0 | 0.055 | 0 | NA | 1 | 0 | 0 | NA | -0.312 |
| SE | 83445 | ENSG00000128944 | KNSTRN | chr15 | + | 40391492 | 40391554 | 40389835 | 40389929 | 40391948 | 40392023 | 20 | 9 | 16 | 19 | 29 | 0 | 0 | 1 | 0 | 2 | 6 | 22 | 25 | 20 | 24 | 0 | 0 | 0 | 0 | 0 | 211 | 149 | 1 | 1 | 1 | 1 | 0.919 | 1 | 0.911 | 1 | 1 | 1 | 1 | 1 | -0.034 |
| SE | 83447 | ENSG00000219545 | UMAD1 | chr7 | + | 7641503 | 7641705 | 7640710 | 7640821 | 7673308 | 7673453 | 1 | 2 | 5 | 1 | 0 | 0 | 1 | 9 | 0 | 1 | 0 | 0 | 2 | 0 | 0 | 0 | 0 | 5 | 0 | 0 | 298 | 149 | 1 | 1 | 1 | 0.5 | 0.217 | 1 | 0 | NA | NA | 0.167 | NA | NA | 0.376 |
| SE | 83448 | ENSG00000219545 | UMAD1 | chr7 | + | 7676101 | 7676208 | 7673308 | 7673453 | 7689334 | 7689460 | 6 | 2 | 5 | 1 | 0 | 0 | 0 | 2 | 2 | 0 | 0 | 3 | 6 | 0 | 0 | 0 | 5 | 5 | 1 | 0 | 256 | 149 | 1 | 1 | 1 | 1 | 0.593 | 0.225 | NA | NA | 0.259 | 0.411 | 0 | NA | 0.481 |
| SE | 83450 | ENSG00000219545 | UMAD1 | chr7 | + | 7689334 | 7689460 | 7676101 | 7676208 | 7692347 | 7692508 | 1 | 0 | 1 | 1 | 0 | 0 | 0 | 0 | 0 | 0 | 0 | 8 | 8 | 1 | 0 | 8 | 0 | 0 | 0 | 0 | 275 | 149 | 1 | 1 | 1 | NA | 1 | 1 | NA | 0 | 1 | 1 | 1 | NA | 0.25 |
| SE | 83452 | ENSG00000219545 | UMAD1 | chr7 | + | 7868329 | 7868533 | 7801669 | 7801743 | 7877280 | 7879205 | 4 | 1 | 7 | 0 | 10 | 17 | 13 | 25 | 18 | 30 | 0 | 6 | 0 | 0 | 0 | 3 | 20 | 16 | 8 | 0 | 298 | 149 | 1 | 1 | 0.105 | 0.037 | 0.123 | 0 | 0.143 | 0 | 0 | 0.158 | 0 | NA | 0.042 |
| SE | 83453 | ENSG00000219545 | UMAD1 | chr7 | + | 7868329 | 7868608 | 7801669 | 7801743 | 7877280 | 7877661 | 5 | 1 | 6 | 0 | 10 | 17 | 13 | 25 | 18 | 30 | 0 | 2 | 9 | 0 | 0 | 3 | 20 | 16 | 8 | 0 | 298 | 149 | 1 | 1 | 0.128 | 0.037 | 0.107 | 0 | 0.143 | 0 | 0.048 | 0.22 | 0 | NA | 0.016 |
| SE | 83454 | ENSG00000156831 | NSMCE2 | chr8 | + | 125094467 | 125094533 | 125091863 | 125091958 | 125102013 | 125102146 | 0 | 0 | 0 | 0 | 2 | 14 | 6 | 7 | 1 | 1 | 0 | 0 | 0 | 0 | 0 | 8 | 4 | 7 | 19 | 4 | 215 | 149 | 1 | 1 | 0 | 0 | 0 | 0 | 0.581 | 0 | 0 | 0 | 0 | 0 | 0.116 |
| SE | 83455 | ENSG00000156831 | NSMCE2 | chr8 | + | 125094467 | 125094533 | 125091863 | 125091958 | 125102050 | 125102146 | 0 | 0 | 0 | 0 | 0 | 7 | 7 | 6 | 8 | 4 | 0 | 0 | 0 | 2 | 0 | 0 | 4 | 7 | 1 | 0 | 215 | 149 | 1 | 1 | 0 | 0 | 0 | 0 | 0 | NA | 0 | 0 | 0.581 | NA | -0.194 |
| SE | 83456 | ENSG00000156831 | NSMCE2 | chr8 | + | 125094467 | 125094533 | 125091863 | 125091958 | 125102316 | 125102464 | 0 | 0 | 0 | 0 | 0 | 4 | 2 | 2 | 8 | 1 | 0 | 0 | 2 | 0 | 0 | 0 | 12 | 14 | 2 | 11 | 215 | 149 | 1 | 1 | 0 | 0 | 0 | 0 | 0 | NA | 0 | 0.09 | 0 | 0 | -0.022 |
| SE | 83457 | ENSG00000156831 | NSMCE2 | chr8 | + | 125102013 | 125102146 | 125091678 | 125091900 | 125102316 | 125102487 | 15 | 14 | 19 | 19 | 12 | 0 | 0 | 1 | 0 | 0 | 22 | 15 | 22 | 27 | 8 | 0 | 0 | 1 | 0 | 0 | 282 | 149 | 1 | 1 | 1 | 1 | 0.909 | 1 | 1 | 1 | 1 | 0.921 | 1 | 1 | -0.002 |
| SE | 83459 | ENSG00000156831 | NSMCE2 | chr8 | + | 125102013 | 125102146 | 125094467 | 125094533 | 125102316 | 125102487 | 15 | 14 | 19 | 19 | 14 | 0 | 0 | 0 | 0 | 0 | 22 | 14 | 22 | 27 | 8 | 0 | 0 | 2 | 0 | 0 | 282 | 149 | 1 | 1 | 1 | 1 | 1 | 1 | 1 | 1 | 1 | 0.853 | 1 | 1 | 0.029 |
| SE | 83460 | ENSG00000156831 | NSMCE2 | chr8 | + | 125102050 | 125102146 | 125091843 | 125091900 | 125102316 | 125102464 | 14 | 13 | 20 | 16 | 17 | 0 | 0 | 1 | 0 | 0 | 22 | 12 | 19 | 24 | 13 | 0 | 0 | 1 | 0 | 0 | 245 | 149 | 1 | 1 | 1 | 1 | 0.924 | 1 | 1 | 1 | 1 | 0.92 | 1 | 1 | 0.001 |
| SE | 83462 | ENSG00000156831 | NSMCE2 | chr8 | + | 125102050 | 125102146 | 125094467 | 125094533 | 125102316 | 125102464 | 14 | 13 | 19 | 15 | 12 | 0 | 0 | 0 | 0 | 0 | 22 | 12 | 18 | 26 | 8 | 0 | 0 | 2 | 0 | 0 | 245 | 149 | 1 | 1 | 1 | 1 | 1 | 1 | 1 | 1 | 1 | 0.846 | 1 | 1 | 0.031 |
| SE | 83463 | ENSG00000156831 | NSMCE2 | chr8 | + | 125102316 | 125102487 | 125091869 | 125091958 | 125151170 | 125151277 | 61 | 53 | 70 | 75 | 118 | 0 | 1 | 0 | 0 | 0 | 126 | 92 | 109 | 81 | 74 | 0 | 0 | 0 | 0 | 0 | 298 | 149 | 1 | 1 | 1 | 0.964 | 1 | 1 | 1 | 1 | 1 | 1 | 1 | 1 | -0.007 |
| SE | 83464 | ENSG00000156831 | NSMCE2 | chr8 | + | 125126965 | 125127067 | 125102357 | 125102487 | 125151170 | 125151277 | 0 | 0 | 0 | 0 | 0 | 57 | 51 | 68 | 67 | 117 | 0 | 1 | 0 | 0 | 0 | 126 | 80 | 95 | 79 | 63 | 251 | 149 | 1 | 1 | 0 | 0 | 0 | 0 | 0 | 0 | 0.007 | 0 | 0 | 0 | -0.001 |
| SE | 83465 | ENSG00000156831 | NSMCE2 | chr8 | + | 125130212 | 125130354 | 125102316 | 125102487 | 125151170 | 125151277 | 1 | 2 | 0 | 0 | 0 | 57 | 51 | 68 | 67 | 117 | 0 | 0 | 2 | 0 | 6 | 126 | 80 | 95 | 79 | 63 | 291 | 149 | 1 | 1 | 0.009 | 0.02 | 0 | 0 | 0 | 0 | 0 | 0.011 | 0 | 0.046 | -0.006 |
| SE | 83466 | ENSG00000156831 | NSMCE2 | chr8 | + | 125151170 | 125151277 | 125102316 | 125102487 | 125182102 | 125182256 | 197 | 172 | 205 | 195 | 424 | 0 | 0 | 0 | 0 | 3 | 377 | 242 | 243 | 269 | 195 | 0 | 0 | 0 | 0 | 0 | 256 | 149 | 1 | 1 | 1 | 1 | 1 | 1 | 0.988 | 1 | 1 | 1 | 1 | 1 | -0.002 |
| SE | 83468 | ENSG00000156831 | NSMCE2 | chr8 | + | 125357711 | 125357818 | 125357218 | 125357319 | 125366767 | 125367075 | 304 | 222 | 258 | 253 | 587 | 1 | 0 | 0 | 1 | 1 | 492 | 304 | 339 | 342 | 260 | 0 | 0 | 0 | 0 | 0 | 256 | 149 | 1 | 1 | 0.994 | 1 | 1 | 0.993 | 0.997 | 1 | 1 | 1 | 1 | 1 | -0.003 |
| SE | 83470 | ENSG00000106603 | COA1 | chr7 | - | 43648599 | 43648652 | 43646471 | 43647634 | 43650492 | 43650712 | 40 | 36 | 59 | 69 | 39 | 0 | 0 | 0 | 0 | 0 | 53 | 49 | 28 | 35 | 37 | 0 | 0 | 2 | 0 | 0 | 202 | 149 | 1 | 1 | 1 | 1 | 1 | 1 | 1 | 1 | 0.948 | 1 | 1 | 1 | 0.01 |
| SE | 83472 | ENSG00000106603 | COA1 | chr7 | - | 43650611 | 43650712 | 43647227 | 43648652 | 43656032 | 43656153 | 11 | 18 | 18 | 12 | 9 | 0 | 3 | 4 | 0 | 0 | 0 | 10 | 12 | 13 | 8 | 0 | 1 | 0 | 5 | 0 | 250 | 149 | 1 | 1 | 1 | 0.781 | 0.728 | 1 | 1 | NA | 0.856 | 1 | 0.608 | 1 | 0.036 |
| SE | 83473 | ENSG00000106603 | COA1 | chr7 | - | 43650611 | 43650712 | 43647227 | 43648652 | 43657215 | 43657269 | 14 | 13 | 15 | 13 | 9 | 1 | 0 | 0 | 1 | 0 | 7 | 0 | 12 | 13 | 13 | 0 | 0 | 0 | 0 | 0 | 250 | 149 | 1 | 1 | 0.893 | 1 | 1 | 0.886 | 1 | 1 | NA | 1 | 1 | 1 | -0.044 |
| SE | 83475 | ENSG00000106603 | COA1 | chr7 | - | 43650611 | 43650712 | 43647227 | 43648652 | 43711399 | 43711449 | 11 | 13 | 15 | 15 | 11 | 0 | 0 | 0 | 0 | 0 | 0 | 0 | 12 | 13 | 8 | 1 | 0 | 0 | 0 | 0 | 250 | 149 | 1 | 1 | 1 | 1 | 1 | 1 | 1 | 0 | NA | 1 | 1 | 1 | 0.25 |
| SE | 83477 | ENSG00000106603 | COA1 | chr7 | - | 43656032 | 43656153 | 43648599 | 43648652 | 43657215 | 43657269 | 0 | 5 | 9 | 0 | 0 | 1 | 0 | 0 | 1 | 0 | 0 | 4 | 0 | 5 | 0 | 0 | 0 | 0 | 0 | 0 | 270 | 149 | 1 | 1 | 0 | 1 | 1 | 0 | NA | NA | 1 | NA | 1 | NA | -0.5 |
| SE | 83478 | ENSG00000106603 | COA1 | chr7 | - | 43656032 | 43656153 | 43648599 | 43648652 | 43665657 | 43665722 | 0 | 3 | 4 | 0 | 0 | 9 | 4 | 4 | 11 | 20 | 0 | 0 | 0 | 5 | 0 | 0 | 2 | 16 | 6 | 4 | 270 | 149 | 1 | 1 | 0 | 0.293 | 0.356 | 0 | 0 | NA | 0 | 0 | 0.315 | 0 | 0.051 |
| SE | 83480 | ENSG00000106603 | COA1 | chr7 | - | 43656032 | 43656153 | 43650611 | 43650712 | 43657215 | 43657269 | 0 | 9 | 10 | 0 | 0 | 3 | 0 | 0 | 1 | 0 | 0 | 14 | 0 | 0 | 0 | 7 | 0 | 0 | 0 | 5 | 270 | 149 | 1 | 1 | 0 | 1 | 1 | 0 | NA | 0 | 1 | NA | NA | 0 | 0.167 |
| SE | 83481 | ENSG00000106603 | COA1 | chr7 | - | 43656032 | 43656153 | 43650611 | 43650712 | 43665657 | 43665722 | 0 | 7 | 6 | 0 | 0 | 4 | 0 | 1 | 1 | 1 | 0 | 10 | 0 | 0 | 0 | 14 | 0 | 0 | 0 | 0 | 270 | 149 | 1 | 1 | 0 | 1 | 0.768 | 0 | 0 | 0 | 1 | NA | NA | NA | -0.146 |
| SE | 83483 | ENSG00000106603 | COA1 | chr7 | - | 43657215 | 43657269 | 43648599 | 43648652 | 43665657 | 43665722 | 3 | 2 | 2 | 1 | 1 | 9 | 4 | 4 | 11 | 20 | 0 | 4 | 0 | 0 | 0 | 0 | 2 | 16 | 6 | 4 | 203 | 149 | 1 | 1 | 0.197 | 0.268 | 0.268 | 0.063 | 0.035 | NA | 0.595 | 0 | 0 | 0 | 0.017 |
| SE | 83484 | ENSG00000106603 | COA1 | chr7 | - | 43657215 | 43657269 | 43648599 | 43648652 | 43729428 | 43729505 | 0 | 0 | 0 | 1 | 0 | 37 | 19 | 14 | 30 | 44 | 0 | 0 | 0 | 0 | 0 | 42 | 21 | 24 | 16 | 22 | 203 | 149 | 1 | 1 | 0 | 0 | 0 | 0.024 | 0 | 0 | 0 | 0 | 0 | 0 | 0.005 |
| SE | 83485 | ENSG00000106603 | COA1 | chr7 | - | 43657215 | 43657269 | 43650611 | 43650712 | 43665657 | 43665722 | 2 | 2 | 2 | 1 | 1 | 4 | 0 | 1 | 1 | 1 | 7 | 4 | 0 | 0 | 5 | 14 | 0 | 0 | 0 | 0 | 203 | 149 | 1 | 1 | 0.268 | 1 | 0.595 | 0.423 | 0.423 | 0.268 | 1 | NA | NA | 1 | -0.214 |
| SE | 83487 | ENSG00000106603 | COA1 | chr7 | - | 43657215 | 43657269 | 43656032 | 43656153 | 43665657 | 43665722 | 2 | 4 | 8 | 1 | 1 | 0 | 0 | 1 | 0 | 0 | 0 | 8 | 0 | 0 | 0 | 0 | 0 | 0 | 0 | 0 | 203 | 149 | 1 | 1 | 1 | 1 | 0.854 | 1 | 1 | NA | 1 | NA | NA | NA | -0.029 |
| SE | 83488 | ENSG00000106603 | COA1 | chr7 | - | 43657215 | 43657269 | 43656032 | 43656153 | 43729428 | 43729505 | 0 | 0 | 4 | 0 | 0 | 0 | 4 | 7 | 0 | 0 | 0 | 4 | 0 | 0 | 0 | 0 | 13 | 0 | 10 | 0 | 203 | 149 | 1 | 1 | NA | 0 | 0.295 | NA | NA | NA | 0.184 | NA | 0 | NA | 0.055 |
| SE | 83489 | ENSG00000106603 | COA1 | chr7 | - | 43658711 | 43659144 | 43657221 | 43657269 | 43665657 | 43665722 | 1 | 0 | 0 | 0 | 0 | 3 | 4 | 4 | 1 | 1 | 0 | 0 | 0 | 0 | 0 | 0 | 4 | 0 | 0 | 5 | 298 | 149 | 1 | 1 | 0.143 | 0 | 0 | 0 | 0 | NA | 0 | NA | NA | 0 | 0.029 |
| SE | 83490 | ENSG00000106603 | COA1 | chr7 | - | 43658711 | 43659144 | 43657221 | 43657269 | 43729428 | 43729499 | 0 | 0 | 0 | 1 | 0 | 0 | 0 | 0 | 1 | 0 | 0 | 1 | 1 | 0 | 0 | 0 | 0 | 0 | 0 | 0 | 298 | 149 | 1 | 1 | NA | NA | NA | 0 | NA | NA | 1 | 1 | NA | NA | -1 |
| SE | 83491 | ENSG00000106603 | COA1 | chr7 | - | 43665657 | 43665722 | 43648599 | 43648652 | 43711399 | 43711449 | 0 | 1 | 1 | 5 | 4 | 0 | 0 | 0 | 0 | 0 | 0 | 6 | 1 | 0 | 0 | 1 | 0 | 0 | 0 | 0 | 214 | 149 | 1 | 1 | NA | 1 | 1 | 1 | 1 | 0 | 1 | 1 | NA | NA | 0.333 |
| SE | 83493 | ENSG00000106603 | COA1 | chr7 | - | 43665657 | 43665722 | 43650611 | 43650712 | 43711399 | 43711449 | 0 | 0 | 0 | 3 | 0 | 0 | 0 | 0 | 3 | 2 | 0 | 6 | 1 | 0 | 0 | 0 | 0 | 0 | 0 | 0 | 214 | 149 | 1 | 1 | NA | NA | NA | 0.41 | 0 | NA | 1 | 1 | NA | NA | -0.795 |
| SE | 83495 | ENSG00000106603 | COA1 | chr7 | - | 43665657 | 43665722 | 43656032 | 43656153 | 43729428 | 43729505 | 1 | 0 | 1 | 0 | 0 | 0 | 4 | 7 | 0 | 0 | 0 | 0 | 0 | 0 | 0 | 0 | 13 | 0 | 10 | 0 | 214 | 149 | 1 | 1 | 1 | 0 | 0.09 | NA | NA | NA | 0 | NA | 0 | NA | 0.363 |
| SE | 83496 | ENSG00000106603 | COA1 | chr7 | - | 43665657 | 43665722 | 43657215 | 43657269 | 43729428 | 43729505 | 4 | 4 | 4 | 1 | 1 | 0 | 0 | 0 | 1 | 0 | 0 | 4 | 0 | 0 | 5 | 0 | 0 | 0 | 0 | 0 | 214 | 149 | 1 | 1 | 1 | 1 | 1 | 0.41 | 1 | NA | 1 | NA | NA | 1 | -0.118 |
| SE | 83497 | ENSG00000106603 | COA1 | chr7 | - | 43665657 | 43665722 | 43658711 | 43659144 | 43711399 | 43711449 | 1 | 0 | 0 | 3 | 0 | 0 | 0 | 0 | 0 | 0 | 0 | 6 | 1 | 0 | 0 | 0 | 0 | 2 | 0 | 0 | 214 | 149 | 1 | 1 | 1 | NA | NA | 1 | NA | NA | 1 | 0.258 | NA | NA | 0.371 |
| SE | 83498 | ENSG00000106603 | COA1 | chr7 | - | 43665657 | 43665722 | 43658711 | 43659144 | 43729428 | 43729505 | 2 | 0 | 0 | 0 | 0 | 0 | 0 | 0 | 0 | 0 | 0 | 0 | 0 | 0 | 0 | 0 | 1 | 1 | 0 | 0 | 214 | 149 | 1 | 1 | 1 | NA | NA | NA | NA | NA | 0 | 0 | NA | NA | 1 |
| SE | 83499 | ENSG00000106603 | COA1 | chr7 | - | 43711399 | 43711449 | 43648599 | 43648652 | 43729428 | 43729523 | 0 | 0 | 0 | 0 | 0 | 37 | 19 | 14 | 30 | 44 | 1 | 0 | 0 | 0 | 0 | 42 | 21 | 24 | 16 | 22 | 199 | 149 | 1 | 1 | 0 | 0 | 0 | 0 | 0 | 0.018 | 0 | 0 | 0 | 0 | -0.004 |
| SE | 83500 | ENSG00000106603 | COA1 | chr7 | - | 43711399 | 43711449 | 43650611 | 43650712 | 43729428 | 43729523 | 0 | 0 | 0 | 3 | 2 | 15 | 11 | 20 | 17 | 19 | 0 | 0 | 0 | 0 | 0 | 10 | 11 | 3 | 2 | 13 | 199 | 149 | 1 | 1 | 0 | 0 | 0 | 0.117 | 0.073 | 0 | 0 | 0 | 0 | 0 | 0.038 |
| SE | 83504 | ENSG00000213719 | CLIC1 | chr6 | - | 31732216 | 31732398 | 31730580 | 31731003 | 31733565 | 31733672 | 4431 | 2716 | 3106 | 3954 | 4756 | 0 | 1 | 0 | 0 | 1 | 10657 | 5449 | 4034 | 6622 | 10243 | 0 | 4 | 0 | 4 | 0 | 298 | 149 | 1 | 1 | 1 | 0.999 | 1 | 1 | 1 | 1 | 0.999 | 1 | 0.999 | 1 | 0 |
| SE | 83505 | ENSG00000213719 | CLIC1 | chr6 | - | 31733565 | 31733672 | 31732216 | 31732398 | 31733835 | 31733961 | 3255 | 1763 | 2521 | 2934 | 3447 | 0 | 0 | 0 | 0 | 0 | 6666 | 3912 | 2856 | 4230 | 7032 | 12 | 4 | 1 | 2 | 0 | 256 | 149 | 1 | 1 | 1 | 1 | 1 | 1 | 1 | 0.997 | 0.998 | 0.999 | 0.999 | 1 | 0.001 |
| SE | 83506 | ENSG00000213719 | CLIC1 | chr6 | - | 31736261 | 31736350 | 31734153 | 31734263 | 31736482 | 31736545 | 1426 | 543 | 1120 | 1358 | 1736 | 0 | 0 | 0 | 0 | 0 | 2384 | 1290 | 1222 | 1737 | 2857 | 0 | 0 | 1 | 0 | 0 | 238 | 149 | 1 | 1 | 1 | 1 | 1 | 1 | 1 | 1 | 1 | 0.999 | 1 | 1 | 0 |
| SE | 83507 | ENSG00000234444 | ZNF736 | chr7 | + | 64312807 | 64312851 | 64308460 | 64308625 | 64313872 | 64314017 | 1 | 0 | 0 | 2 | 0 | 0 | 0 | 1 | 0 | 2 | 0 | 0 | 0 | 0 | 0 | 0 | 0 | 2 | 0 | 0 | 193 | 149 | 1 | 1 | 1 | NA | 0 | 1 | 0 | NA | NA | 0 | NA | NA | 0.5 |
| SE | 83513 | ENSG00000196313 | POM121 | chr7 | + | 72940413 | 72940516 | 72929939 | 72930111 | 72941836 | 72943522 | 7 | 8 | 8 | 2 | 20 | 0 | 0 | 0 | 0 | 0 | 2 | 2 | 5 | 9 | 4 | 0 | 0 | 2 | 0 | 0 | 252 | 149 | 1 | 1 | 1 | 1 | 1 | 1 | 1 | 1 | 1 | 0.596 | 1 | 1 | 0.081 |
| SE | 83514 | ENSG00000196313 | POM121 | chr7 | + | 72940413 | 72940516 | 72939846 | 72939968 | 72940816 | 72940993 | 35 | 18 | 43 | 33 | 55 | 0 | 0 | 0 | 0 | 0 | 70 | 61 | 33 | 55 | 75 | 0 | 0 | 0 | 0 | 1 | 252 | 149 | 1 | 1 | 1 | 1 | 1 | 1 | 1 | 1 | 1 | 1 | 1 | 0.978 | 0.004 |
| SE | 83516 | ENSG00000161203 | AP2M1 | chr3 | + | 184176950 | 184177067 | 184174688 | 184174959 | 184178856 | 184179122 | 542 | 269 | 377 | 529 | 340 | 2 | 2 | 0 | 1 | 0 | 760 | 691 | 354 | 508 | 1853 | 0 | 11 | 0 | 0 | 0 | 266 | 149 | 1 | 1 | 0.993 | 0.987 | 1 | 0.997 | 1 | 1 | 0.972 | 1 | 1 | 1 | 0.001 |
| SE | 83521 | ENSG00000161203 | AP2M1 | chr3 | + | 184178856 | 184179122 | 184176777 | 184177067 | 184180168 | 184180251 | 1186 | 681 | 818 | 1159 | 794 | 0 | 0 | 5 | 1 | 0 | 1864 | 1360 | 831 | 1089 | 3784 | 6 | 2 | 2 | 2 | 27 | 298 | 149 | 1 | 1 | 1 | 1 | 0.988 | 0.998 | 1 | 0.994 | 0.997 | 0.995 | 0.996 | 0.986 | 0.004 |
| SE | 83522 | ENSG00000161203 | AP2M1 | chr3 | + | 184178856 | 184179122 | 184176777 | 184177067 | 184180848 | 184180881 | 454 | 231 | 293 | 436 | 289 | 0 | 0 | 0 | 0 | 0 | 630 | 573 | 290 | 395 | 1453 | 0 | 0 | 0 | 1 | 0 | 298 | 149 | 1 | 1 | 1 | 1 | 1 | 1 | 1 | 1 | 1 | 1 | 0.995 | 1 | 0.001 |
| SE | 83524 | ENSG00000161203 | AP2M1 | chr3 | + | 184180168 | 184180251 | 184176777 | 184177067 | 184180848 | 184180984 | 34 | 24 | 32 | 46 | 23 | 0 | 0 | 0 | 0 | 0 | 110 | 74 | 50 | 55 | 187 | 0 | 0 | 0 | 1 | 0 | 232 | 149 | 1 | 1 | 1 | 1 | 1 | 1 | 1 | 1 | 1 | 1 | 0.972 | 1 | 0.006 |
| SE | 83525 | ENSG00000161203 | AP2M1 | chr3 | + | 184180168 | 184180251 | 184178856 | 184179122 | 184180848 | 184180984 | 651 | 384 | 434 | 638 | 440 | 0 | 0 | 0 | 0 | 0 | 1138 | 693 | 486 | 587 | 1965 | 0 | 0 | 0 | 3 | 0 | 232 | 149 | 1 | 1 | 1 | 1 | 1 | 1 | 1 | 1 | 1 | 1 | 0.992 | 1 | 0.002 |
| SE | 83527 | ENSG00000161203 | AP2M1 | chr3 | + | 184180644 | 184180741 | 184180168 | 184180251 | 184180848 | 184180915 | 0 | 5 | 7 | 1 | 0 | 220 | 102 | 165 | 180 | 153 | 0 | 5 | 0 | 7 | 9 | 361 | 233 | 168 | 176 | 572 | 246 | 149 | 1 | 1 | 0 | 0.029 | 0.025 | 0.003 | 0 | 0 | 0.013 | 0 | 0.024 | 0.009 | 0.002 |
| SE | 83529 | ENSG00000161203 | AP2M1 | chr3 | + | 184181084 | 184181226 | 184180848 | 184180984 | 184181695 | 184181793 | 755 | 443 | 587 | 800 | 692 | 0 | 0 | 0 | 0 | 0 | 1127 | 902 | 566 | 841 | 2204 | 0 | 0 | 1 | 0 | 0 | 291 | 149 | 1 | 1 | 1 | 1 | 1 | 1 | 1 | 1 | 1 | 0.997 | 1 | 1 | 0.001 |
| SE | 83530 | ENSG00000161203 | AP2M1 | chr3 | + | 184181084 | 184181226 | 184180848 | 184180984 | 184181911 | 184182047 | 404 | 219 | 285 | 415 | 320 | 0 | 0 | 2 | 0 | 0 | 650 | 485 | 348 | 454 | 1176 | 0 | 0 | 0 | 0 | 0 | 291 | 149 | 1 | 1 | 1 | 1 | 0.986 | 1 | 1 | 1 | 1 | 1 | 1 | 1 | -0.003 |
| SE | 83531 | ENSG00000161203 | AP2M1 | chr3 | + | 184181695 | 184181815 | 184180848 | 184180984 | 184181911 | 184182047 | 237 | 142 | 225 | 293 | 320 | 0 | 0 | 2 | 0 | 0 | 396 | 288 | 140 | 372 | 973 | 0 | 0 | 0 | 0 | 0 | 269 | 149 | 1 | 1 | 1 | 1 | 0.984 | 1 | 1 | 1 | 1 | 1 | 1 | 1 | -0.003 |
| SE | 83532 | ENSG00000161203 | AP2M1 | chr3 | + | 184181695 | 184181815 | 184181084 | 184181226 | 184181911 | 184182047 | 591 | 367 | 527 | 680 | 683 | 2 | 1 | 3 | 0 | 5 | 908 | 729 | 357 | 752 | 1968 | 0 | 5 | 0 | 0 | 0 | 269 | 149 | 1 | 1 | 0.994 | 0.995 | 0.99 | 1 | 0.987 | 1 | 0.988 | 1 | 1 | 1 | -0.004 |
| SE | 83533 | ENSG00000161203 | AP2M1 | chr3 | + | 184182150 | 184182248 | 184181911 | 184182047 | 184182756 | 184182868 | 793 | 423 | 629 | 832 | 1160 | 0 | 0 | 1 | 0 | 0 | 1141 | 917 | 504 | 851 | 2500 | 0 | 0 | 0 | 0 | 0 | 247 | 149 | 1 | 1 | 1 | 1 | 0.997 | 1 | 1 | 1 | 1 | 1 | 1 | 1 | -0.001 |
| SE | 83534 | ENSG00000171115 | GIMAP8 | chr7 | + | 150470828 | 150470874 | 150466670 | 150467334 | 150474011 | 150474638 | 434 | 331 | 316 | 260 | 409 | 1 | 0 | 1 | 0 | 4 | 167 | 157 | 50 | 80 | 129 | 0 | 0 | 0 | 0 | 0 | 195 | 149 | 1 | 1 | 0.997 | 1 | 0.996 | 1 | 0.987 | 1 | 1 | 1 | 1 | 1 | -0.004 |
| SE | 83535 | ENSG00000171115 | GIMAP8 | chr7 | + | 150470828 | 150470874 | 150466670 | 150467334 | 150477091 | 150479393 | 226 | 171 | 163 | 137 | 178 | 0 | 0 | 2 | 0 | 2 | 86 | 76 | 30 | 50 | 66 | 0 | 0 | 0 | 0 | 0 | 195 | 149 | 1 | 1 | 1 | 1 | 0.984 | 1 | 0.986 | 1 | 1 | 1 | 1 | 1 | -0.006 |
| SE | 83536 | ENSG00000171115 | GIMAP8 | chr7 | + | 150474011 | 150474638 | 150466670 | 150467334 | 150477091 | 150479393 | 132 | 108 | 115 | 74 | 132 | 0 | 0 | 2 | 0 | 2 | 79 | 63 | 16 | 27 | 24 | 0 | 0 | 0 | 0 | 0 | 298 | 149 | 1 | 1 | 1 | 1 | 0.966 | 1 | 0.971 | 1 | 1 | 1 | 1 | 1 | -0.013 |
| SE | 83538 | ENSG00000106526 | ACTR3C | chr7 | - | 150289449 | 150289593 | 150286366 | 150286540 | 150293311 | 150293419 | 12 | 16 | 12 | 20 | 6 | 0 | 0 | 0 | 0 | 0 | 16 | 16 | 8 | 10 | 17 | 0 | 1 | 0 | 0 | 0 | 293 | 149 | 1 | 1 | 1 | 1 | 1 | 1 | 1 | 1 | 0.891 | 1 | 1 | 1 | 0.022 |
| SE | 83547 | ENSG00000259024 | TVP23C-CDRT4 | chr17 | - | 15440207 | 15440285 | 15436020 | 15438200 | 15545784 | 15545916 | 6 | 3 | 8 | 3 | 12 | 1 | 1 | 0 | 0 | 1 | 0 | 1 | 2 | 0 | 0 | 0 | 0 | 0 | 0 | 0 | 227 | 149 | 1 | 1 | 0.798 | 0.663 | 1 | 1 | 0.887 | NA | 1 | 1 | NA | NA | -0.13 |
| SE | 83548 | ENSG00000259024 | TVP23C-CDRT4 | chr17 | - | 15503097 | 15503232 | 15437887 | 15438200 | 15540432 | 15540561 | 3 | 0 | 1 | 4 | 0 | 0 | 0 | 0 | 0 | 0 | 0 | 0 | 3 | 0 | 0 | 0 | 0 | 0 | 4 | 0 | 284 | 149 | 1 | 1 | 1 | NA | 1 | 1 | NA | NA | NA | 1 | 0 | NA | 0.5 |
| SE | 83549 | ENSG00000259024 | TVP23C-CDRT4 | chr17 | - | 15503097 | 15503232 | 15437887 | 15438200 | 15545784 | 15545916 | 3 | 0 | 8 | 1 | 1 | 1 | 1 | 0 | 0 | 1 | 0 | 2 | 3 | 0 | 4 | 0 | 0 | 0 | 0 | 0 | 284 | 149 | 1 | 1 | 0.611 | 0 | 1 | 1 | 0.344 | NA | 1 | 1 | NA | 1 | -0.409 |
| SE | 83550 | ENSG00000259024 | TVP23C-CDRT4 | chr17 | - | 15503097 | 15503232 | 15440207 | 15440285 | 15545784 | 15545916 | 8 | 4 | 9 | 3 | 9 | 2 | 0 | 5 | 0 | 6 | 0 | 3 | 11 | 0 | 17 | 0 | 1 | 0 | 0 | 0 | 284 | 149 | 1 | 1 | 0.677 | 1 | 0.486 | 1 | 0.44 | NA | 0.611 | 1 | NA | 1 | -0.15 |
| SE | 83551 | ENSG00000259024 | TVP23C-CDRT4 | chr17 | - | 15540432 | 15540561 | 15437887 | 15438200 | 15545784 | 15545916 | 14 | 9 | 0 | 34 | 0 | 1 | 1 | 0 | 0 | 1 | 5 | 6 | 14 | 42 | 0 | 0 | 0 | 0 | 0 | 0 | 278 | 149 | 1 | 1 | 0.882 | 0.828 | NA | 1 | 0 | 1 | 1 | 1 | 1 | NA | -0.323 |
| SE | 83558 | ENSG00000124222 | STX16 | chr20 | + | 58659622 | 58659634 | 58651282 | 58652138 | 58667200 | 58667318 | 0 | 1 | 3 | 4 | 0 | 0 | 0 | 0 | 0 | 0 | 0 | 0 | 4 | 9 | 0 | 0 | 1 | 3 | 0 | 13 | 161 | 149 | 1 | 1 | NA | 1 | 1 | 1 | NA | NA | 0 | 0.552 | 1 | 0 | 0.612 |
| SE | 83563 | ENSG00000124222 | STX16 | chr20 | + | 58667486 | 58667597 | 58652072 | 58652138 | 58667986 | 58668127 | 336 | 296 | 413 | 391 | 350 | 0 | 0 | 1 | 0 | 0 | 400 | 358 | 383 | 404 | 293 | 0 | 1 | 0 | 0 | 0 | 260 | 149 | 1 | 1 | 1 | 1 | 0.996 | 1 | 1 | 1 | 0.995 | 1 | 1 | 1 | 0 |
| SE | 83564 | ENSG00000124222 | STX16 | chr20 | + | 58667489 | 58667597 | 58651604 | 58652087 | 58667986 | 58668127 | 546 | 503 | 636 | 636 | 533 | 0 | 0 | 5 | 0 | 0 | 696 | 636 | 647 | 675 | 503 | 0 | 0 | 2 | 0 | 0 | 257 | 149 | 1 | 1 | 1 | 1 | 0.987 | 1 | 1 | 1 | 1 | 0.995 | 1 | 1 | -0.002 |
| SE | 83565 | ENSG00000124222 | STX16 | chr20 | + | 58667489 | 58667597 | 58651604 | 58652087 | 58670511 | 58670603 | 171 | 168 | 170 | 202 | 133 | 0 | 0 | 1 | 0 | 0 | 227 | 217 | 219 | 210 | 191 | 0 | 0 | 0 | 5 | 0 | 257 | 149 | 1 | 1 | 1 | 1 | 0.99 | 1 | 1 | 1 | 1 | 1 | 0.961 | 1 | 0.006 |
| SE | 83566 | ENSG00000124222 | STX16 | chr20 | + | 58667489 | 58667597 | 58651745 | 58652138 | 58667986 | 58668127 | 396 | 362 | 501 | 487 | 415 | 0 | 0 | 1 | 0 | 0 | 471 | 451 | 450 | 488 | 349 | 0 | 1 | 0 | 0 | 0 | 257 | 149 | 1 | 1 | 1 | 1 | 0.997 | 1 | 1 | 1 | 0.996 | 1 | 1 | 1 | 0 |
| SE | 83567 | ENSG00000124222 | STX16 | chr20 | + | 58667986 | 58668127 | 58651271 | 58652087 | 58670511 | 58670547 | 70 | 41 | 64 | 46 | 44 | 0 | 0 | 1 | 0 | 0 | 68 | 52 | 91 | 63 | 79 | 0 | 0 | 0 | 5 | 0 | 290 | 149 | 1 | 1 | 1 | 1 | 0.97 | 1 | 1 | 1 | 1 | 1 | 0.866 | 1 | 0.021 |
| SE | 83568 | ENSG00000124222 | STX16 | chr20 | + | 58667986 | 58668127 | 58667486 | 58667597 | 58670511 | 58670547 | 463 | 392 | 545 | 499 | 465 | 0 | 3 | 0 | 1 | 0 | 533 | 486 | 527 | 534 | 415 | 0 | 0 | 1 | 0 | 0 | 290 | 149 | 1 | 1 | 1 | 0.985 | 1 | 0.996 | 1 | 1 | 1 | 0.996 | 1 | 1 | -0.003 |
| SE | 83573 | ENSG00000006757 | PNPLA4 | chrX | - | 7912027 | 7912093 | 7901988 | 7902141 | 7921712 | 7921848 | 36 | 23 | 34 | 21 | 44 | 1 | 0 | 0 | 0 | 0 | 24 | 18 | 3 | 14 | 10 | 0 | 0 | 0 | 0 | 0 | 215 | 149 | 1 | 1 | 0.961 | 1 | 1 | 1 | 1 | 1 | 1 | 1 | 1 | 1 | -0.008 |
| SE | 83574 | ENSG00000006757 | PNPLA4 | chrX | - | 7922003 | 7922098 | 7921712 | 7921848 | 7925939 | 7926132 | 81 | 42 | 88 | 56 | 145 | 0 | 1 | 0 | 0 | 0 | 90 | 28 | 20 | 62 | 53 | 0 | 0 | 0 | 0 | 0 | 244 | 149 | 1 | 1 | 1 | 0.962 | 1 | 1 | 1 | 1 | 1 | 1 | 1 | 1 | -0.008 |
| SE | 83575 | ENSG00000006757 | PNPLA4 | chrX | - | 7925939 | 7926132 | 7922003 | 7922098 | 7927670 | 7927739 | 39 | 21 | 47 | 24 | 60 | 0 | 0 | 0 | 0 | 3 | 62 | 12 | 8 | 27 | 27 | 0 | 3 | 0 | 0 | 0 | 298 | 149 | 1 | 1 | 1 | 1 | 1 | 1 | 0.909 | 1 | 0.667 | 1 | 1 | 1 | 0.048 |
| SE | 83576 | ENSG00000006757 | PNPLA4 | chrX | - | 7925939 | 7926132 | 7922003 | 7922098 | 7927685 | 7927739 | 37 | 20 | 38 | 23 | 54 | 2 | 0 | 0 | 0 | 0 | 52 | 10 | 8 | 31 | 27 | 0 | 0 | 0 | 0 | 0 | 298 | 149 | 1 | 1 | 0.902 | 1 | 1 | 1 | 1 | 1 | 1 | 1 | 1 | 1 | -0.02 |
| SE | 83580 | ENSG00000130649 | CYP2E1 | chr10 | + | 133533755 | 133533897 | 133532691 | 133532868 | 133537062 | 133537250 | 4 | 6 | 6 | 4 | 3 | 0 | 0 | 0 | 0 | 0 | 5 | 4 | 2 | 0 | 6 | 0 | 0 | 1 | 0 | 0 | 291 | 149 | 1 | 1 | 1 | 1 | 1 | 1 | 1 | 1 | 1 | 0.506 | NA | 1 | 0.123 |
| SE | 83583 | ENSG00000107897 | ACBD5 | chr10 | - | 27208245 | 27208445 | 27205197 | 27205248 | 27210813 | 27211081 | 95 | 80 | 108 | 135 | 105 | 1 | 0 | 0 | 0 | 0 | 21 | 82 | 79 | 76 | 16 | 0 | 0 | 1 | 0 | 0 | 298 | 149 | 1 | 1 | 0.979 | 1 | 1 | 1 | 1 | 1 | 1 | 0.975 | 1 | 1 | 0.001 |
| SE | 83586 | ENSG00000107897 | ACBD5 | chr10 | - | 27215534 | 27215641 | 27210813 | 27211081 | 27217979 | 27218183 | 289 | 219 | 201 | 241 | 322 | 0 | 4 | 0 | 0 | 0 | 128 | 215 | 288 | 159 | 15 | 0 | 0 | 0 | 0 | 0 | 256 | 149 | 1 | 1 | 1 | 0.97 | 1 | 1 | 1 | 1 | 1 | 1 | 1 | 1 | -0.006 |
| SE | 83588 | ENSG00000107897 | ACBD5 | chr10 | - | 27220372 | 27220464 | 27219722 | 27219803 | 27223206 | 27223239 | 0 | 0 | 4 | 1 | 0 | 0 | 0 | 0 | 1 | 0 | 0 | 0 | 0 | 0 | 0 | 0 | 0 | 0 | 1 | 0 | 241 | 149 | 1 | 1 | NA | NA | 1 | 0.382 | NA | NA | NA | NA | 0 | NA | 0.691 |
| SE | 83590 | ENSG00000107897 | ACBD5 | chr10 | - | 27220372 | 27220464 | 27219722 | 27219857 | 27223206 | 27223239 | 2 | 0 | 7 | 10 | 2 | 34 | 26 | 24 | 34 | 30 | 0 | 0 | 0 | 0 | 0 | 13 | 24 | 20 | 1 | 0 | 241 | 149 | 1 | 1 | 0.035 | 0 | 0.153 | 0.154 | 0.04 | 0 | 0 | 0 | 0 | NA | 0.076 |
| SE | 83595 | ENSG00000107897 | ACBD5 | chr10 | - | 27223206 | 27223239 | 27220372 | 27220464 | 27223337 | 27223452 | 10 | 6 | 9 | 11 | 5 | 0 | 0 | 2 | 0 | 3 | 0 | 3 | 1 | 0 | 0 | 0 | 0 | 0 | 0 | 0 | 182 | 149 | 1 | 1 | 1 | 1 | 0.787 | 1 | 0.577 | NA | 1 | 1 | NA | NA | -0.127 |
| SE | 83596 | ENSG00000107897 | ACBD5 | chr10 | - | 27231747 | 27231820 | 27223337 | 27223452 | 27240318 | 27240484 | 49 | 34 | 36 | 64 | 51 | 0 | 0 | 0 | 0 | 0 | 42 | 52 | 36 | 20 | 4 | 0 | 0 | 1 | 0 | 0 | 222 | 149 | 1 | 1 | 1 | 1 | 1 | 1 | 1 | 1 | 1 | 0.96 | 1 | 1 | 0.008 |
| SE | 83597 | ENSG00000107897 | ACBD5 | chr10 | - | 27234357 | 27234488 | 27231747 | 27231820 | 27235091 | 27235212 | 0 | 0 | 5 | 0 | 0 | 62 | 50 | 53 | 96 | 72 | 0 | 0 | 0 | 0 | 0 | 49 | 84 | 63 | 48 | 13 | 280 | 149 | 1 | 1 | 0 | 0 | 0.048 | 0 | 0 | 0 | 0 | 0 | 0 | 0 | 0.01 |
| SE | 83598 | ENSG00000107897 | ACBD5 | chr10 | - | 27235091 | 27235212 | 27230230 | 27231820 | 27240318 | 27240474 | 95 | 73 | 81 | 133 | 93 | 0 | 1 | 0 | 1 | 0 | 49 | 108 | 100 | 61 | 19 | 0 | 0 | 1 | 4 | 0 | 270 | 149 | 1 | 1 | 1 | 0.976 | 1 | 0.987 | 1 | 1 | 1 | 0.982 | 0.894 | 1 | 0.017 |
| SE | 83603 | ENSG00000234129 | NA | chrX | - | 11055935 | 11056036 | 11038538 | 11038655 | 11106900 | 11106990 | 0 | 6 | 0 | 1 | 1 | 1 | 0 | 0 | 1 | 0 | 4 | 0 | 0 | 0 | 0 | 0 | 0 | 4 | 0 | 0 | 250 | 149 | 1 | 1 | 0 | 1 | NA | 0.373 | 1 | 1 | NA | 0 | NA | NA | 0.093 |
| SE | 83605 | ENSG00000169100 | SLC25A6 | chrX | - | 1387278 | 1387419 | 1386151 | 1386759 | 1389240 | 1389727 | 1600 | 715 | 1591 | 1568 | 1913 | 1 | 1 | 0 | 2 | 0 | 2231 | 1578 | 1331 | 1115 | 3836 | 0 | 2 | 1 | 7 | 0 | 290 | 149 | 1 | 1 | 0.999 | 0.997 | 1 | 0.998 | 1 | 1 | 0.998 | 0.999 | 0.988 | 1 | 0.002 |
| SE | 83608 | ENSG00000169084 | DHRSX | chrX | - | 2243022 | 2243230 | 2219505 | 2221229 | 2266739 | 2266947 | 73 | 38 | 81 | 55 | 131 | 0 | 0 | 0 | 0 | 0 | 123 | 69 | 76 | 45 | 105 | 0 | 0 | 2 | 0 | 0 | 298 | 149 | 1 | 1 | 1 | 1 | 1 | 1 | 1 | 1 | 1 | 0.95 | 1 | 1 | 0.01 |
| SE | 83612 | ENSG00000169084 | DHRSX | chrX | - | 2408744 | 2408813 | 2291501 | 2291603 | 2425196 | 2425304 | 26 | 20 | 15 | 21 | 33 | 0 | 0 | 0 | 0 | 0 | 23 | 30 | 27 | 35 | 68 | 0 | 0 | 0 | 1 | 0 | 218 | 149 | 1 | 1 | 1 | 1 | 1 | 1 | 1 | 1 | 1 | 1 | 0.96 | 1 | 0.008 |
| SE | 83613 | ENSG00000130066 | SAT1 | chrX | + | 23783799 | 23783883 | 23783657 | 23783709 | 23784293 | 23784403 | 670 | 626 | 451 | 812 | 380 | 1 | 0 | 0 | 0 | 0 | 897 | 715 | 1240 | 1150 | 806 | 0 | 0 | 0 | 0 | 0 | 233 | 149 | 1 | 1 | 0.998 | 1 | 1 | 1 | 1 | 1 | 1 | 1 | 1 | 1 | 0 |
| SE | 83618 | ENSG00000032389 | EIPR1 | chr2 | - | 3257298 | 3257455 | 3214148 | 3214248 | 3261714 | 3261827 | 27 | 21 | 32 | 39 | 18 | 0 | 0 | 0 | 0 | 2 | 65 | 52 | 46 | 47 | 28 | 0 | 3 | 0 | 0 | 0 | 298 | 149 | 1 | 1 | 1 | 1 | 1 | 1 | 0.818 | 1 | 0.897 | 1 | 1 | 1 | -0.016 |
| SE | 83619 | ENSG00000032389 | EIPR1 | chr2 | - | 3257298 | 3257455 | 3214148 | 3214248 | 3274338 | 3274419 | 27 | 19 | 37 | 39 | 20 | 0 | 0 | 1 | 0 | 3 | 65 | 52 | 46 | 47 | 28 | 0 | 0 | 0 | 0 | 0 | 298 | 149 | 1 | 1 | 1 | 1 | 0.949 | 1 | 0.769 | 1 | 1 | 1 | 1 | 1 | -0.056 |
| SE | 83621 | ENSG00000032389 | EIPR1 | chr2 | - | 3261714 | 3261827 | 3214148 | 3214248 | 3338016 | 3338149 | 2 | 0 | 0 | 0 | 4 | 10 | 6 | 8 | 2 | 4 | 0 | 3 | 0 | 0 | 0 | 4 | 12 | 8 | 8 | 0 | 262 | 149 | 1 | 1 | 0.102 | 0 | 0 | 0 | 0.363 | 0 | 0.124 | 0 | 0 | NA | 0.062 |
| SE | 83622 | ENSG00000032389 | EIPR1 | chr2 | - | 3261714 | 3261827 | 3257298 | 3257455 | 3338016 | 3338149 | 2 | 2 | 0 | 1 | 2 | 45 | 32 | 42 | 42 | 38 | 0 | 0 | 0 | 0 | 0 | 79 | 30 | 39 | 56 | 54 | 262 | 149 | 1 | 1 | 0.025 | 0.034 | 0 | 0.013 | 0.029 | 0 | 0 | 0 | 0 | 0 | 0.02 |
| SE | 83623 | ENSG00000032389 | EIPR1 | chr2 | - | 3274338 | 3274419 | 3214148 | 3214248 | 3338016 | 3338149 | 0 | 0 | 2 | 0 | 5 | 10 | 6 | 8 | 2 | 4 | 0 | 0 | 1 | 0 | 0 | 4 | 12 | 8 | 8 | 0 | 230 | 149 | 1 | 1 | 0 | 0 | 0.139 | 0 | 0.447 | 0 | 0 | 0.075 | 0 | NA | 0.098 |
| SE | 83626 | ENSG00000032389 | EIPR1 | chr2 | - | 3318856 | 3318958 | 3257350 | 3257455 | 3338016 | 3338149 | 0 | 3 | 0 | 0 | 0 | 45 | 32 | 42 | 42 | 38 | 0 | 0 | 0 | 0 | 7 | 79 | 30 | 39 | 56 | 54 | 251 | 149 | 1 | 1 | 0 | 0.053 | 0 | 0 | 0 | 0 | 0 | 0 | 0 | 0.071 | -0.004 |
| SE | 83628 | ENSG00000032389 | EIPR1 | chr2 | - | 3353205 | 3353323 | 3338016 | 3338149 | 3354549 | 3354633 | 0 | 0 | 0 | 0 | 0 | 52 | 39 | 54 | 43 | 28 | 0 | 4 | 0 | 0 | 0 | 42 | 36 | 12 | 43 | 45 | 267 | 149 | 1 | 1 | 0 | 0 | 0 | 0 | 0 | 0 | 0.058 | 0 | 0 | 0 | -0.012 |
| SE | 83629 | ENSG00000152082 | MZT2B | chr2 | + | 130182626 | 130182775 | 130181736 | 130181797 | 130190468 | 130190695 | 27 | 18 | 25 | 24 | 8 | 1 | 0 | 0 | 0 | 0 | 31 | 18 | 24 | 16 | 12 | 0 | 0 | 0 | 0 | 0 | 298 | 149 | 1 | 1 | 0.931 | 1 | 1 | 1 | 1 | 1 | 1 | 1 | 1 | 1 | -0.014 |
| SE | 83630 | ENSG00000152082 | MZT2B | chr2 | + | 130182626 | 130182775 | 130181771 | 130181830 | 130190468 | 130190695 | 27 | 18 | 25 | 26 | 8 | 0 | 0 | 1 | 0 | 0 | 23 | 18 | 26 | 16 | 12 | 0 | 0 | 0 | 0 | 0 | 298 | 149 | 1 | 1 | 1 | 1 | 0.926 | 1 | 1 | 1 | 1 | 1 | 1 | 1 | -0.015 |
| SE | 83633 | ENSG00000169093 | ASMTL | chrX | - | 1417972 | 1418116 | 1412731 | 1412854 | 1421657 | 1421842 | 29 | 5 | 25 | 47 | 51 | 0 | 1 | 0 | 0 | 0 | 10 | 51 | 20 | 13 | 80 | 0 | 0 | 0 | 0 | 0 | 293 | 149 | 1 | 1 | 1 | 0.718 | 1 | 1 | 1 | 1 | 1 | 1 | 1 | 1 | -0.056 |
| SE | 83636 | ENSG00000223511 | NA | chrX | + | 1733949 | 1734196 | 1732583 | 1732723 | 1741930 | 1741988 | 2 | 7 | 7 | 3 | 7 | 0 | 1 | 0 | 0 | 0 | 25 | 9 | 8 | 9 | 4 | 0 | 0 | 0 | 0 | 0 | 298 | 149 | 1 | 1 | 1 | 0.778 | 1 | 1 | 1 | 1 | 1 | 1 | 1 | 1 | -0.044 |
| SE | 83637 | ENSG00000166579 | NDEL1 | chr17 | + | 8431201 | 8431359 | 8413131 | 8413269 | 8444259 | 8444357 | 0 | 0 | 0 | 0 | 0 | 15 | 8 | 21 | 36 | 32 | 0 | 0 | 5 | 0 | 0 | 36 | 41 | 114 | 55 | 19 | 298 | 149 | 1 | 1 | 0 | 0 | 0 | 0 | 0 | 0 | 0 | 0.021 | 0 | 0 | -0.004 |
| SE | 83638 | ENSG00000166579 | NDEL1 | chr17 | + | 8443926 | 8444176 | 8435921 | 8436045 | 8444259 | 8444357 | 2 | 0 | 0 | 3 | 0 | 179 | 111 | 137 | 120 | 74 | 0 | 0 | 0 | 1 | 0 | 238 | 183 | 189 | 190 | 138 | 298 | 149 | 1 | 1 | 0.006 | 0 | 0 | 0.012 | 0 | 0 | 0 | 0 | 0.003 | 0 | 0.003 |
| SE | 83639 | ENSG00000166579 | NDEL1 | chr17 | + | 8444623 | 8444742 | 8444259 | 8444357 | 8445710 | 8445864 | 1 | 4 | 3 | 3 | 0 | 517 | 430 | 345 | 485 | 407 | 14 | 0 | 1 | 0 | 0 | 968 | 699 | 807 | 908 | 628 | 268 | 149 | 1 | 1 | 0.001 | 0.005 | 0.005 | 0.003 | 0 | 0.008 | 0 | 0.001 | 0 | 0 | 0.001 |
| SE | 83642 | ENSG00000166579 | NDEL1 | chr17 | + | 8450779 | 8450953 | 8448549 | 8448686 | 8454795 | 8454887 | 569 | 449 | 411 | 676 | 599 | 0 | 0 | 0 | 0 | 0 | 983 | 598 | 1176 | 942 | 813 | 0 | 0 | 1 | 0 | 0 | 298 | 149 | 1 | 1 | 1 | 1 | 1 | 1 | 1 | 1 | 0 | 0.998 | 1 | 1 | 0 |
| SE | 83643 | ENSG00000166579 | NDEL1 | chr17 | + | 8454795 | 8454887 | 8448549 | 8448686 | 8466929 | 8468160 | 1 | 2 | 0 | 4 | 3 | 0 | 0 | 0 | 1 | 0 | 0 | 1 | 8 | 3 | 0 | 0 | 0 | 0 | 0 | 0 | 241 | 149 | 1 | 1 | 1 | 1 | NA | 0.712 | 1 | NA | 1 | 1 | 1 | NA | -0.072 |
| SE | 83646 | ENSG00000166579 | NDEL1 | chr17 | + | 8460008 | 8460160 | 8454795 | 8454887 | 8490246 | 8490411 | 310 | 230 | 245 | 316 | 221 | 0 | 0 | 0 | 0 | 0 | 395 | 286 | 463 | 388 | 353 | 0 | 0 | 2 | 2 | 0 | 298 | 149 | 1 | 1 | 1 | 1 | 1 | 1 | 1 | 1 | 1 | 0.991 | 0.99 | 1 | 0.004 |
| SE | 83648 | ENSG00000004961 | HCCS | chrX | + | 11112018 | 11112160 | 11111331 | 11111518 | 11114834 | 11114986 | 97 | 46 | 81 | 62 | 61 | 0 | 0 | 0 | 0 | 0 | 60 | 85 | 64 | 59 | 48 | 0 | 0 | 2 | 0 | 0 | 291 | 149 | 1 | 1 | 1 | 1 | 1 | 1 | 1 | 1 | 1 | 0.942 | 1 | 1 | 0.012 |
| SE | 83649 | ENSG00000004961 | HCCS | chrX | + | 11114834 | 11114986 | 11112018 | 11112160 | 11117266 | 11117415 | 128 | 71 | 119 | 78 | 88 | 0 | 0 | 0 | 0 | 0 | 178 | 114 | 98 | 84 | 146 | 0 | 0 | 3 | 0 | 0 | 298 | 149 | 1 | 1 | 1 | 1 | 1 | 1 | 1 | 1 | 1 | 0.942 | 1 | 1 | 0.012 |
| SE | 83655 | ENSG00000050405 | LIMA1 | chr12 | - | 50192451 | 50192561 | 50181903 | 50182037 | 50195829 | 50195887 | 58 | 67 | 67 | 103 | 126 | 0 | 0 | 1 | 1 | 0 | 30 | 113 | 69 | 26 | 6 | 0 | 0 | 0 | 0 | 0 | 259 | 149 | 1 | 1 | 1 | 1 | 0.975 | 0.983 | 1 | 1 | 1 | 1 | 1 | 1 | -0.008 |
| SE | 83656 | ENSG00000050405 | LIMA1 | chr12 | - | 50192451 | 50192564 | 50181903 | 50182037 | 50195829 | 50195887 | 47 | 57 | 65 | 76 | 91 | 0 | 0 | 1 | 1 | 0 | 21 | 86 | 47 | 24 | 6 | 0 | 0 | 0 | 0 | 0 | 262 | 149 | 1 | 1 | 1 | 1 | 0.974 | 0.977 | 1 | 1 | 1 | 1 | 1 | 1 | -0.01 |
| SE | 83657 | ENSG00000050405 | LIMA1 | chr12 | - | 50201302 | 50201324 | 50200776 | 50200884 | 50204551 | 50204700 | 0 | 0 | 0 | 0 | 2 | 45 | 50 | 48 | 71 | 96 | 0 | 0 | 0 | 0 | 0 | 16 | 92 | 30 | 31 | 0 | 171 | 149 | 1 | 1 | 0 | 0 | 0 | 0 | 0.018 | 0 | 0 | 0 | 0 | NA | 0.004 |
| SE | 83658 | ENSG00000050405 | LIMA1 | chr12 | - | 50204551 | 50204700 | 50200776 | 50200884 | 50205983 | 50206068 | 83 | 91 | 101 | 125 | 204 | 0 | 0 | 0 | 0 | 0 | 23 | 169 | 46 | 59 | 0 | 0 | 0 | 1 | 0 | 0 | 298 | 149 | 1 | 1 | 1 | 1 | 1 | 1 | 1 | 1 | 1 | 0.958 | 1 | NA | 0.01 |
| SE | 83663 | ENSG00000029363 | BCLAF1 | chr6 | - | 136270322 | 136270383 | 136269261 | 136269612 | 136271994 | 136272079 | 37 | 41 | 39 | 77 | 92 | 208 | 216 | 240 | 300 | 324 | 4 | 25 | 17 | 7 | 0 | 30 | 137 | 221 | 30 | 0 | 210 | 149 | 1 | 1 | 0.112 | 0.119 | 0.103 | 0.154 | 0.168 | 0.086 | 0.115 | 0.052 | 0.142 | NA | 0.032 |
| [illegible] | | | | | | | | | | | | | | | | | | | | | | | | | | | | | | | | | | | | | | | | | | | | | | |

| | | | | | | | | | | | | | | | | | | | | | | | | | | | | | | | | | | | | | | | | | | | | | | |
|---|---|---|---|---|---|---|---|---|---|---|---|---|---|---|---|---|---|---|---|---|---|---|---|---|---|---|---|---|---|---|---|---|---|---|---|---|---|---|---|---|---|---|---|---|---|---|
| SE | 83698 | ENSG00000177324 | BEND2 | chrX | - | 18180509 | 18180650 | 18177568 | 18177769 | 18201814 | 18201940 | 32 | 32 | 6 | 51 | 68 | 0 | 0 | 0 | 0 | 0 | 14 | 19 | 23 | 46 | 25 | 0 | 0 | 0 | 1 | 0 | 290 | 149 | 1 | 1 | 1 | 1 | 1 | 1 | 1 | 1 | 1 | 1 | 0.959 | 1 | 0.008 |
| SE | 83699 | ENSG00000177324 | BEND2 | chrX | - | 18191000 | 18191108 | 18177568 | 18177769 | 18201814 | 18201940 | 2 | 4 | 1 | 1 | 2 | 0 | 0 | 0 | 0 | 0 | 0 | 0 | 1 | 4 | 0 | 0 | 0 | 0 | 1 | 0 | 257 | 149 | 1 | 1 | 1 | 1 | 1 | 1 | 1 | NA | NA | 1 | 0.699 | NA | 0.151 |
| SE | 83705 | ENSG00000130150 | MOSPD2 | chrX | + | 14900574 | 14900635 | 14897083 | 14897238 | 14902965 | 14903004 | 222 | 121 | 157 | 147 | 298 | 1 | 0 | 0 | 0 | 4 | 24 | 90 | 19 | 17 | 0 | 0 | 0 | 0 | 0 | 0 | 210 | 149 | 1 | 1 | 0.994 | 1 | 1 | 1 | 0.981 | 1 | 1 | 1 | 1 | NA | -0.005 |
| SE | 83706 | ENSG00000130150 | MOSPD2 | chrX | + | 14902965 | 14903004 | 14873688 | 14873758 | 14911236 | 14911413 | 0 | 1 | 1 | 0 | 0 | 0 | 0 | 0 | 0 | 0 | 0 | 0 | 1 | 0 | 0 | 0 | 0 | 1 | 0 | 0 | 188 | 149 | 1 | 1 | NA | 1 | 1 | NA | NA | NA | NA | 0.442 | NA | NA | 0.558 |
| SE | 83707 | ENSG00000130150 | MOSPD2 | chrX | + | 14902965 | 14903004 | 14897083 | 14897238 | 14908859 | 14908984 | 39 | 30 | 22 | 36 | 85 | 0 | 0 | 0 | 1 | 0 | 11 | 8 | 5 | 6 | 0 | 0 | 0 | 0 | 0 | 0 | 188 | 149 | 1 | 1 | 1 | 1 | 1 | 0.966 | 1 | 1 | 1 | 1 | 1 | NA | -0.007 |
| SE | 83708 | ENSG00000130150 | MOSPD2 | chrX | + | 14908859 | 14908984 | 14902965 | 14903004 | 14911236 | 14911413 | 322 | 171 | 232 | 235 | 443 | 0 | 3 | 1 | 0 | 0 | 52 | 141 | 30 | 30 | 0 | 0 | 0 | 1 | 0 | 0 | 274 | 149 | 1 | 1 | 1 | 0.969 | 0.992 | 1 | 1 | 1 | 1 | 0.942 | 1 | NA | 0.007 |
| SE | 83709 | ENSG00000130150 | MOSPD2 | chrX | + | 14915667 | 14915764 | 14914502 | 14914599 | 14916196 | 14916326 | 153 | 69 | 98 | 90 | 186 | 4 | 6 | 7 | 2 | 4 | 15 | 70 | 27 | 8 | 0 | 0 | 6 | 1 | 0 | 0 | 246 | 149 | 1 | 1 | 0.959 | 0.874 | 0.895 | 0.965 | 0.966 | 1 | 0.876 | 0.942 | 1 | NA | -0.023 |
| SE | 83710 | ENSG00000170293 | CMTM8 | chr3 | + | 32357372 | 32357546 | 32238679 | 32239119 | 32367871 | 32367988 | 5 | 1 | 11 | 0 | 8 | 0 | 0 | 0 | 0 | 0 | 0 | 5 | 1 | 0 | 0 | 0 | 0 | 6 | 0 | 0 | 298 | 149 | 1 | 1 | 1 | 1 | 1 | NA | 1 | NA | 1 | 0.077 | NA | NA | 0.462 |
| SE | 83715 | ENSG00000196998 | WDR45 | chrX | - | 49076429 | 49076524 | 49075865 | 49075945 | 49077836 | 49077911 | 79 | 34 | 57 | 82 | 31 | 0 | 0 | 0 | 0 | 0 | 105 | 99 | 72 | 96 | 156 | 0 | 0 | 2 | 0 | 0 | 244 | 149 | 1 | 1 | 1 | 1 | 1 | 1 | 1 | 1 | 1 | 0.956 | 1 | 1 | 0.009 |
| SE | 83717 | ENSG00000196998 | WDR45 | chrX | - | 49076644 | 49076750 | 49076405 | 49076524 | 49077836 | 49077911 | 117 | 27 | 107 | 112 | 55 | 0 | 0 | 0 | 0 | 0 | 151 | 198 | 116 | 93 | 210 | 0 | 2 | 0 | 0 | 0 | 255 | 149 | 1 | 1 | 1 | 1 | 1 | 1 | 1 | 1 | 0.983 | 1 | 1 | 1 | 0.003 |
| SE | 83719 | ENSG00000196998 | WDR45 | chrX | - | 49076644 | 49076753 | 49076429 | 49076524 | 49077836 | 49077911 | 113 | 28 | 98 | 107 | 56 | 0 | 0 | 0 | 0 | 0 | 142 | 178 | 99 | 88 | 199 | 0 | 2 | 0 | 0 | 0 | 258 | 149 | 1 | 1 | 1 | 1 | 1 | 1 | 1 | 1 | 0.981 | 1 | 1 | 1 | 0.004 |
| SE | 83722 | ENSG00000196998 | WDR45 | chrX | - | 49077642 | 49077747 | 49075865 | 49075945 | 49077836 | 49077911 | 27 | 11 | 28 | 34 | 21 | 0 | 0 | 0 | 0 | 0 | 31 | 75 | 33 | 41 | 132 | 0 | 0 | 2 | 0 | 0 | 254 | 149 | 1 | 1 | 1 | 1 | 1 | 1 | 1 | 1 | 1 | 0.906 | 1 | 1 | 0.019 |
| SE | 83723 | ENSG00000196998 | WDR45 | chrX | - | 49077642 | 49077747 | 49076429 | 49076524 | 49077836 | 49077911 | 27 | 11 | 28 | 34 | 21 | 0 | 0 | 0 | 0 | 0 | 31 | 75 | 33 | 41 | 132 | 0 | 2 | 0 | 0 | 0 | 254 | 149 | 1 | 1 | 1 | 1 | 1 | 1 | 1 | 1 | 0.957 | 1 | 1 | 1 | 0.009 |
| SE | 83728 | ENSG00000118513 | MYB | chr6 | + | 135187833 | 135187905 | 135185876 | 135186020 | 135189790 | 135189883 | 24 | 19 | 33 | 29 | 25 | 0 | 0 | 0 | 0 | 0 | 30 | 31 | 41 | 101 | 22 | 0 | 0 | 0 | 0 | 2 | 221 | 149 | 1 | 1 | 1 | 1 | 1 | 1 | 1 | 1 | 1 | 1 | 1 | 0.881 | 0.024 |
| SE | 83730 | ENSG00000118513 | MYB | chr6 | + | 135194355 | 135194451 | 135193837 | 135193918 | 135195747 | 135195917 | 5 | 9 | 18 | 5 | 12 | 0 | 0 | 0 | 0 | 0 | 6 | 0 | 7 | 22 | 0 | 0 | 0 | 0 | 5 | 0 | 245 | 149 | 1 | 1 | 1 | 1 | 1 | 1 | 1 | 1 | NA | 1 | 0.728 | NA | 0.091 |
| SE | 83732 | ENSG00000118513 | MYB | chr6 | + | 135194959 | 135195081 | 135194355 | 135194451 | 135195747 | 135195917 | 1 | 1 | 0 | 0 | 0 | 1 | 0 | 5 | 0 | 1 | 0 | 0 | 0 | 10 | 0 | 6 | 0 | 0 | 9 | 0 | 271 | 149 | 1 | 1 | 0.355 | 1 | 0 | NA | 0 | 0 | NA | NA | 0.379 | NA | 0.149 |
| SE | 83740 | ENSG00000118513 | MYB | chr6 | + | 135196725 | 135197323 | 135195747 | 135195917 | 135198907 | 135199050 | 0 | 0 | 0 | 0 | 7 | 1 | 0 | 0 | 0 | 0 | 0 | 0 | 1 | 14 | 0 | 0 | 0 | 0 | 0 | 0 | 298 | 149 | 1 | 1 | 0 | NA | NA | NA | 1 | NA | NA | 1 | 1 | NA | -0.5 |
| SE | 83742 | ENSG00000118513 | MYB | chr6 | + | 135196960 | 135197323 | 135195747 | 135195917 | 135198907 | 135199050 | 0 | 0 | 1 | 0 | 7 | 1 | 0 | 0 | 0 | 0 | 0 | 0 | 1 | 14 | 0 | 0 | 0 | 0 | 0 | 0 | 298 | 149 | 1 | 1 | 0 | NA | 1 | NA | 1 | NA | NA | 1 | 1 | NA | -0.333 |
| SE | 83745 | ENSG00000118513 | MYB | chr6 | + | 135199525 | 135199581 | 135198907 | 135199050 | 135200084 | 135200199 | 1 | 0 | 5 | 1 | 4 | 3 | 16 | 7 | 10 | 27 | 0 | 5 | 0 | 8 | 0 | 0 | 9 | 22 | 25 | 2 | 205 | 149 | 1 | 1 | 0.195 | 0 | 0.342 | 0.068 | 0.097 | NA | 0.288 | 0 | 0.189 | 0 | 0.021 |
| SE | 83751 | ENSG00000149970 | CNKSR2 | chrX | + | 21468594 | 21468674 | 21440693 | 21440781 | 21470765 | 21470807 | 0 | 0 | 0 | 1 | 0 | 14 | 2 | 13 | 8 | 26 | 0 | 0 | 0 | 0 | 0 | 0 | 3 | 6 | 4 | 5 | 229 | 149 | 1 | 1 | 0 | 0 | 0 | 0.075 | 0 | NA | 0 | 0 | 0 | 0 | 0.015 |
| SE | 83752 | ENSG00000149970 | CNKSR2 | chrX | + | 21516484 | 21516631 | 21501474 | 21501588 | 21526866 | 21527000 | 6 | 5 | 5 | 4 | 8 | 0 | 1 | 4 | 2 | 12 | 0 | 1 | 14 | 2 | 0 | 0 | 1 | 0 | 0 | 0 | 296 | 149 | 1 | 1 | 1 | 0.716 | 0.386 | 0.502 | 0.251 | NA | 0.335 | 1 | 1 | NA | -0.207 |
| SE | 83754 | ENSG00000149970 | CNKSR2 | chrX | + | 21561470 | 21561560 | 21531855 | 21532067 | 21563237 | 21563452 | 13 | 9 | 11 | 14 | 15 | 9 | 2 | 9 | 14 | 26 | 0 | 12 | 0 | 4 | 13 | 0 | 1 | 0 | 7 | 0 | 239 | 149 | 1 | 1 | 0.474 | 0.737 | 0.432 | 0.384 | 0.265 | NA | 0.882 | NA | 0.263 | 1 | -0.257 |
| SE | 83758 | ENSG00000149970 | CNKSR2 | chrX | + | 21609070 | 21609617 | 21606778 | 21606879 | 21648830 | 21649027 | 9 | 5 | 30 | 7 | 19 | 0 | 0 | 0 | 3 | 0 | 0 | 21 | 10 | 2 | 3 | 0 | 0 | 0 | 0 | 0 | 298 | 149 | 1 | 1 | 1 | 1 | 1 | 0.538 | 1 | NA | 1 | 1 | 1 | 1 | -0.092 |
| SE | 83760 | ENSG00000149970 | CNKSR2 | chrX | + | 21637273 | 21637414 | 21609070 | 21609617 | 21648830 | 21649027 | 1 | 0 | 0 | 1 | 0 | 4 | 2 | 9 | 1 | 15 | 0 | 0 | 0 | 0 | 0 | 0 | 7 | 1 | 0 | 0 | 290 | 149 | 1 | 1 | 0.114 | 0 | 0 | 0.339 | 0 | NA | 0 | 0 | NA | NA | 0.091 |
| SE | 83763 | ENSG00000196696 | PDXDC2P-NPIPB14P | chr16 | - | 69986380 | 69986509 | 69982657 | 69982757 | 69990596 | 69990683 | 132 | 65 | 71 | 120 | 104 | 0 | 0 | 0 | 1 | 0 | 182 | 110 | 118 | 122 | 15 | 0 | 0 | 0 | 0 | 0 | 278 | 149 | 1 | 1 | 1 | 1 | 1 | 0.985 | 1 | 1 | 1 | 1 | 1 | 1 | -0.003 |
| SE | 83764 | ENSG00000196696 | PDXDC2P-NPIPB14P | chr16 | - | 69989955 | 69990208 | 69986380 | 69986509 | 69990596 | 69990683 | 0 | 1 | 0 | 0 | 4 | 52 | 16 | 29 | 51 | 34 | 0 | 0 | 0 | 0 | 0 | 85 | 23 | 34 | 49 | 2 | 298 | 149 | 1 | 1 | 0 | 0.03 | 0 | 0 | 0.056 | 0 | 0 | 0 | 0 | 0 | 0.017 |
| SE | 83770 | ENSG00000196696 | PDXDC2P-NPIPB14P | chr16 | - | 70011840 | 70011929 | 69996389 | 69996468 | 70018046 | 70018097 | 0 | 1 | 0 | 4 | 2 | 1 | 0 | 0 | 0 | 2 | 0 | 2 | 0 | 5 | 0 | 0 | 1 | 0 | 0 | 0 | 238 | 149 | 1 | 1 | 0 | 1 | NA | 1 | 0.385 | NA | 0.556 | NA | 1 | NA | -0.182 |
| SE | 83771 | ENSG00000196696 | PDXDC2P-NPIPB14P | chr16 | - | 70038904 | 70039100 | 70036431 | 70036578 | 70042350 | 70042416 | 72 | 84 | 51 | 84 | 56 | 0 | 0 | 0 | 2 | 0 | 85 | 112 | 109 | 129 | 59 | 0 | 0 | 2 | 0 | 0 | 298 | 149 | 1 | 1 | 1 | 1 | 1 | 0.955 | 1 | 1 | 1 | 0.965 | 1 | 1 | -0.002 |
| SE | 83772 | ENSG00000196696 | PDXDC2P-NPIPB14P | chr16 | - | 70042350 | 70042416 | 70038904 | 70039100 | 70042929 | 70043003 | 44 | 47 | 25 | 60 | 32 | 0 | 0 | 0 | 0 | 0 | 75 | 63 | 101 | 97 | 31 | 0 | 0 | 0 | 3 | 0 | 215 | 149 | 1 | 1 | 1 | 1 | 1 | 1 | 1 | 1 | 1 | 1 | 0.957 | 1 | 0.009 |
| SE | 83779 | ENSG00000111412 | SPRING1 | chr12 | - | 116719762 | 116719876 | 116710170 | 116717893 | 116737789 | 116738069 | 25 | 9 | 16 | 15 | 14 | 0 | 0 | 1 | 0 | 0 | 6 | 22 | 2 | 12 | 24 | 0 | 0 | 0 | 0 | 0 | 263 | 149 | 1 | 1 | 1 | 1 | 0.901 | 1 | 1 | 1 | 1 | 1 | 1 | 1 | -0.02 |
| SE | 83781 | ENSG00000111412 | SPRING1 | chr12 | - | 116720295 | 116720447 | 116719762 | 116719876 | 116737789 | 116738070 | 149 | 72 | 96 | 147 | 133 | 4 | 0 | 0 | 0 | 0 | 41 | 159 | 55 | 54 | 98 | 0 | 0 | 0 | 2 | 0 | 298 | 149 | 1 | 1 | 0.949 | 1 | 1 | 1 | 1 | 1 | 1 | 1 | 0.931 | 1 | 0.004 |
| SE | 83783 | ENSG00000111412 | SPRING1 | chr12 | - | 116723066 | 116723223 | 116720295 | 116720447 | 116737789 | 116738069 | 62 | 29 | 60 | 59 | 33 | 1 | 0 | 1 | 1 | 0 | 35 | 62 | 19 | 22 | 67 | 0 | 6 | 1 | 0 | 3 | 298 | 149 | 1 | 1 | 0.969 | 1 | 0.968 | 0.967 | 1 | 1 | 0.838 | 0.905 | 1 | 0.918 | 0.049 |
| SE | 83787 | ENSG00000078804 | TP53INP2 | chr20 | + | 34705357 | 34705474 | 34704343 | 34704512 | 34708690 | 34708863 | 0 | 0 | 0 | 2 | 0 | 0 | 0 | 0 | 0 | 0 | 0 | 6 | 0 | 3 | 19 | 0 | 0 | 0 | 1 | 4 | 266 | 149 | 1 | 1 | NA | NA | NA | 1 | NA | NA | 1 | NA | 0.627 | 0.727 | 0.215 |
| SE | 83788 | ENSG00000078804 | TP53INP2 | chr20 | + | 34709235 | 34709524 | 34708690 | 34708863 | 34710057 | 34713436 | 4 | 0 | 2 | 0 | 0 | 0 | 1 | 0 | 0 | 0 | 4 | 4 | 4 | 8 | 35 | 0 | 0 | 0 | 0 | 0 | 298 | 149 | 1 | 1 | 1 | 0 | 1 | NA | NA | 1 | 1 | 1 | 1 | 1 | -0.333 |
| SE | 83789 | ENSG00000173674 | EIF1AX | chrX | - | 20132181 | 20132263 | 20130515 | 20130607 | 20133956 | 20134007 | 340 | 305 | 398 | 312 | 631 | 3 | 0 | 0 | 1 | 0 | 155 | 78 | 144 | 184 | 24 | 0 | 0 | 0 | 0 | 0 | 231 | 149 | 1 | 1 | 0.987 | 1 | 1 | 0.995 | 1 | 1 | 1 | 1 | 1 | 1 | -0.004 |
| SE | 83790 | ENSG00000173674 | EIF1AX | chrX | - | 20133956 | 20134007 | 20132181 | 20132263 | 20135737 | 20135841 | 348 | 291 | 370 | 299 | 570 | 2 | 2 | 0 | 0 | 0 | 145 | 103 | 135 | 152 | 22 | 0 | 0 | 0 | 3 | 0 | 200 | 149 | 1 | 1 | 0.992 | 0.991 | 1 | 1 | 1 | 1 | 1 | 1 | 0.974 | 1 | 0.002 |
| SE | 83791 | ENSG00000173674 | EIF1AX | chrX | - | 20138538 | 20138622 | 20135737 | 20135841 | 20141624 | 20141838 | 299 | 201 | 286 | 237 | 341 | 0 | 0 | 2 | 0 | 0 | 105 | 90 | 94 | 100 | 55 | 0 | 0 | 0 | 3 | 0 | 233 | 149 | 1 | 1 | 1 | 1 | 0.989 | 1 | 1 | 1 | 1 | 1 | 1 | 1 | -0.002 |
| SE | 83792 | ENSG00000257924 | LINC02416 | chr12 | + | 47367291 | 47367424 | 47353753 | 47353822 | 47369657 | 47369935 | 5 | 0 | 1 | 0 | 4 | 1 | 0 | 0 | 0 | 0 | 8 | 3 | 0 | 1 | 0 | 0 | 0 | 0 | 3 | 0 | 282 | 149 | 1 | 1 | 0.725 | NA | 1 | NA | 1 | 1 | 1 | NA | 0.15 | NA | 0.192 |
| SE | 83793 | ENSG00000165168 | CYBB | chrX | + | 37782087 | 37782183 | 37780058 | 37780122 | 37783489 | 37783600 | 1772 | 1179 | 1146 | 1365 | 1204 | 2 | 2 | 0 | 0 | 0 | 1802 | 1687 | 828 | 2859 | 856 | 0 | 4 | 0 | 6 | 0 | 245 | 149 | 1 | 1 | 0.998 | 0.997 | 1 | 1 | 1 | 1 | 0.996 | 1 | 0.997 | 1 | 0 |
| SE | 83795 | ENSG00000165168 | CYBB | chrX | + | 37783489 | 37783600 | 37780058 | 37780122 | 37791974 | 37792059 | 829 | 575 | 450 | 569 | 379 | 1 | 0 | 1 | 2 | 7 | 796 | 906 | 428 | 1315 | 362 | 0 | 0 | 2 | 5 | 0 | 260 | 149 | 1 | 1 | 0.998 | 1 | 0.996 | 0.994 | 0.969 | 1 | 1 | 0.992 | 0.993 | 1 | -0.006 |
| SE | 83796 | ENSG00000165168 | CYBB | chrX | + | 37783489 | 37783600 | 37782087 | 37782183 | 37791974 | 37792059 | 2477 | 1662 | 1524 | 1830 | 1513 | 0 | 0 | 0 | 2 | 0 | 2407 | 2386 | 1180 | 3958 | 1190 | 0 | 2 | 2 | 0 | 0 | 260 | 149 | 1 | 1 | 1 | 1 | 1 | 0.998 | 1 | 1 | 0.999 | 0.997 | 1 | 1 | 0 |
| SE | 83797 | ENSG00000165168 | CYBB | chrX | + | 37791974 | 37792059 | 37780058 | 37780122 | 37803876 | 37804130 | 1 | 0 | 1 | 0 | 0 | 0 | 0 | 0 | 0 | 0 | 0 | 0 | 1 | 0 | 0 | 0 | 0 | 3 | 0 | 0 | 234 | 149 | 1 | 1 | 1 | NA | 1 | NA | NA | NA | NA | 0.175 | NA | NA | 0.825 |
| SE | 83800 | ENSG00000165168 | CYBB | chrX | + | 37793664 | 37793810 | 37791974 | 37792059 | 37801255 | 37801348 | 1196 | 857 | 789 | 996 | 1371 | 0 | 0 | 0 | 0 | 2 | 1340 | 1205 | 641 | 2155 | 651 | 0 | 0 | 0 | 0 | 0 | 295 | 149 | 1 | 1 | 1 | 1 | 1 | 1 | 0.997 | 1 | 1 | 1 | 1 | 1 | -0.001 |
| SE | 83801 | ENSG00000165168 | CYBB | chrX | + | 37793664 | 37793810 | 37791974 | 37792059 | 37803876 | 37804130 | 1196 | 857 | 789 | 997 | 1371 | 0 | 0 | 0 | 0 | 0 | 1340 | 1205 | 642 | 2157 | 651 | 0 | 0 | 1 | 0 | 0 | 295 | 149 | 1 | 1 | 1 | 1 | 1 | 1 | 1 | 1 | 1 | 0.997 | 1 | 1 | 0.001 |
| SE | 83802 | ENSG00000165168 | CYBB | chrX | + | 37795950 | 37796141 | 37791974 | 37792059 | 37798954 | 37799084 | 3009 | 2861 | 1927 | 2475 | 2993 | 15 | 7 | 9 | 4 | 7 | 5451 | 4048 | 2208 | 7625 | 2072 | 4 | 8 | 6 | 6 | 13 | 298 | 149 | 1 | 1 | 0.99 | 0.995 | 0.991 | 0.997 | 0.995 | 0.999 | 0.996 | 0.995 | 0.998 | 0.988 | -0.002 |
| SE | 83803 | ENSG00000165168 | CYBB | chrX | + | 37795950 | 37796141 | 37793664 | 37793810 | 37798954 | 37799084 | 5236 | 4996 | 3406 | 4386 | 6165 | 8 | 10 | 7 | 6 | 3 | 9289 | 7079 | 3748 | 12946 | 3620 | 2 | 9 | 2 | 8 | 12 | 298 | 149 | 1 | 1 | 0.997 | 0.996 | 0.996 | 0.997 | 0.999 | 1 | 0.997 | 0.999 | 0.999 | 0.993 | -0.001 |
| SE | 83804 | ENSG00000165168 | CYBB | chrX | + | 37798954 | 37799084 | 37791974 | 37792059 | 37801255 | 37801348 | 909 | 534 | 584 | 613 | 976 | 0 | 0 | 0 | 0 | 2 | 611 | 737 | 357 | 1186 | 355 | 0 | 0 | 0 | 0 | 0 | 279 | 149 | 1 | 1 | 1 | 1 | 1 | 1 | 0.996 | 1 | 1 | 1 | 1 | 1 | -0.001 |
| SE | 83805 | ENSG00000165168 | CYBB | chrX | + | 37798954 | 37799084 | 37791974 | 37792059 | 37803876 | 37804079 | 15 | 7 | 10 | 4 | 7 | 0 | 0 | 0 | 0 | 0 | 4 | 8 | 6 | 8 | 13 | 0 | 0 | 1 | 0 | 0 | 279 | 149 | 1 | 1 | 1 | 1 | 1 | 1 | 1 | 1 | 1 | 0.762 | 1 | 1 | 0.048 |
| SE | 83806 | ENSG00000165168 | CYBB | chrX | + | 37798954 | 37799084 | 37793664 | 37793810 | 37801255 | 37801348 | 902 | 537 | 582 | 615 | 972 | 1 | 0 | 0 | 0 | 0 | 609 | 738 | 353 | 1188 | 354 | 0 | 0 | 0 | 0 | 0 | 279 | 149 | 1 | 1 | 0.998 | 1 | 1 | 1 | 1 | 1 | 1 | 1 | 1 | 1 | 0 |
| SE | 83807 | ENSG00000165168 | CYBB | chrX | + | 37798954 | 37799084 | 37793664 | 37793810 | 37803876 | 37804079 | 8 | 10 | 8 | 6 | 3 | 1 | 0 | 0 | 1 | 0 | 2 | 9 | 2 | 10 | 12 | 0 | 0 | 1 | 2 | 0 | 279 | 149 | 1 | 1 | 0.81 | 1 | 1 | 0.762 | 1 | 1 | 1 | 0.516 | 0.728 | 1 | 0.066 |
| SE | 83808 | ENSG00000165168 | CYBB | chrX | + | 37801255 | 37801348 | 37791974 | 37792059 | 37803876 | 37804079 | 1409 | 892 | 797 | 963 | 1284 | 0 | 0 | 0 | 0 | 0 | 1108 | 1125 | 581 | 2052 | 442 | 0 | 0 | 1 | 0 | 0 | 242 | 149 | 1 | 1 | 1 | 1 | 1 | 1 | 1 | 1 | 1 | 0.997 | 1 | 1 | 0.001 |
| SE | 83809 | ENSG00000165168 | CYBB | chrX | + | 37801255 | 37801348 | 37793664 | 37793810 | 37803876 | 37804079 | 1410 | 892 | 797 | 963 | 1284 | 1 | 0 | 0 | 1 | 0 | 1108 | 1125 | 581 | 2052 | 442 | 0 | 0 | 1 | 2 | 0 | 242 | 149 | 1 | 1 | 0.999 | 1 | 1 | 0.998 | 1 | 1 | 1 | 0.997 | 0.998 | 1 | 0 |
| SE | 83810 | ENSG00000165168 | CYBB | chrX | + | 37801255 | 37801348 | 37798954 | 37799084 | 37803876 | 37804079 | 2316 | 1422 | 1390 | 1587 | 2294 | 0 | 0 | 0 | 1 | 0 | 1711 | 1900 | 944 | 3246 | 795 | 0 | 0 | 0 | 2 | 0 | 242 | 149 | 1 | 1 | 1 | 1 | 0.999 | 1 | 1 | 1 | 1 | 1 | 0.999 | 1 | 0 |
| SE | 83811 | ENSG00000165168 | CYBB | chrX | + | 37803876 | 37804130 | 37801255 | 37801348 | 37805005 | 37805168 | 3396 | 2217 | 1999 | 2465 | 3020 | 0 | 0 | 1 | 0 | 1 | 2693 | 2949 | 1445 | 5086 | 1118 | 0 | 0 | 0 | 1 | 0 | 298 | 149 | 1 | 1 | 1 | 1 | 1 | 0.999 | 1 | 1 | 1 | 1 | 1 | 1 | 0 |
| SE | 83812 | ENSG00000135070 | ISCA1 | chr9 | - | 86272006 | 86272112 | 86264545 | 86266191 | 86282377 | 86282538 | 158 | 95 | 105 | 145 | 174 | 0 | 0 | 0 | 1 | 0 | 105 | 129 | 154 | 164 | 175 | 0 | 0 | 0 | 0 | 0 | 255 | 149 | 1 | 1 | 1 | 1 | 1 | 0.988 | 1 | 1 | 1 | 1 | 1 | 1 | -0.002 |
| SE | 83813 | ENSG00000135070 | ISCA1 | chr9 | - | 86274188 | 86274242 | 86272006 | 86272112 | 86282377 | 86282538 | 253 | 167 | 170 | 244 | 170 | 0 | 0 | 0 | 1 | 0 | 248 | 344 | 246 | 212 | 378 | 0 | 0 | 0 | 4 | 0 | 203 | 149 | 1 | 1 | 1 | 1 | 1 | 1 | 1 | 1 | 1 | 1 | 0.975 | 1 | 0.005 |
| SE | 83815 | ENSG00000167615 | LENG8 | chr19 | + | 54451289 | 54451382 | 54449189 | 54449306 | 54452092 | 54452267 | 174 | 116 | 189 | 186 | 106 | 1 | 1 | 1 | 1 | 3 | 330 | 234 | 120 | 194 | 330 | 0 | 0 | 1 | 0 | 0 | 242 | 149 | 1 | 1 | 0.991 | 0.986 | 0.991 | 0.991 | 0.956 | 1 | 1 | 0.987 | 1 | 1 | -0.014 |
| SE | 83816 | ENSG00000167615 | LENG8 | chr19 | + | 54452650 | 54452752 | 54451991 | 54452267 | 54454429 | 54454682 | 180 | 97 | 223 | 251 | 171 | 0 | 0 | 0 | 0 | 0 | 323 | 297 | 181 | 328 | 532 | 0 | 0 | 0 | 1 | 3 | 251 | 149 | 1 | 1 | 1 | 1 | 1 | 1 | 1 | 1 | 1 | 1 | 0.995 | 0.991 | 0.003 |
| SE | 83821 | ENSG00000169891 | REPS2 | chrX | + | 17006220 | 17006344 | 16946657 | 16947134 | 17022122 | 17022271 | 10 | 8 | 4 | 23 | 35 | 0 | 0 | 0 | 2 | 0 | 0 | 25 | 27 | 17 | 86 | 0 | 0 | 0 | 0 | 0 | 273 | 149 | 1 | 1 | 1 | 1 | 1 | 0.863 | 1 | NA | 1 | 1 | 1 | 1 | -0.027 |
| SE | 83824 | ENSG00000169891 | REPS2 | chrX | + | 17054807 | 17054950 | 17052381 | 17052445 | 17062437 | 17062532 | 37 | 31 | 11 | 19 | 10 | 0 | 0 | 1 | 0 | 0 | 27 | 39 | 59 | 62 | 94 | 0 | 0 | 0 | 2 | 0 | 292 | 149 | 1 | 1 | 1 | 1 | 0.849 | 1 | 1 | 1 | 1 | 1 | 0.941 | 1 | -0.018 |
| SE | 83825 | ENSG00000169891 | REPS2 | chrX | + | 17062437 | 17062532 | 17054807 | 17054950 | 17068401 | 17068471 | 33 | 35 | 9 | 29 | 51 | 0 | 0 | 0 | 0 | 0 | 57 | 43 | 54 | 67 | 92 | 0 | 0 | 1 | 0 | 0 | 244 | 149 | 1 | 1 | 1 | 1 | 1 | 1 | 1 | 1 | 1 | 0.971 | 1 | 1 | 0.006 |
| SE | 83826 | ENSG00000169891 | REPS2 | chrX | + | 17068401 | 17068471 | 17062437 | 17062532 | 17103717 | 17103779 | 4 | 10 | 6 | 10 | 8 | 0 | 0 | 0 | 0 | 0 | 0 | 10 | 18 | 18 | 24 | 0 | 0 | 1 | 0 | 0 | 219 | 149 | 1 | 1 | 1 | 1 | 1 | 1 | 1 | NA | 1 | 0.925 | 1 | 1 | 0.019 |
| SE | 83827 | ENSG00000169891 | REPS2 | chrX | + | 17069939 | 17069993 | 17068401 | 17068471 | 17103717 | 17103779 | 24 | 32 | 6 | 26 | 65 | 0 | 0 | 0 | 2 | 0 | 74 | 29 | 37 | 47 | 117 | 0 | 0 | 0 | 0 | 0 | 203 | 149 | 1 | 1 | 1 | 1 | 1 | 0.905 | 1 | 1 | 1 | 1 | 1 | 1 | -0.019 |
| SE | 83829 | ENSG00000169891 | REPS2 | chrX | + | 17077270 | 17077407 | 17074113 | 17074159 | 17103717 | 17103779 | 17 | 15 | 9 | 14 | 50 | 0 | 0 | 0 | 0 | 0 | 39 | 19 | 76 | 15 | 29 | 0 | 2 | 2 | 0 | 0 | 286 | 149 | 1 | 1 | 1 | 1 | 1 | 1 | 1 | 1 | 0.832 | 0.952 | 1 | 1 | 0.043 |
| SE | 83831 | ENSG00000248309 | MEF2C-AS1 | chr5 | + | 88892166 | 88892277 | 88883416 | 88883485 | 88905208 | 88905270 | 2 | 1 | 0 | 0 | 7 | 0 | 4 | 3 | 5 | 4 | 6 | 0 | 2 | 0 | 0 | 4 | 15 | 1 | 0 | 0 | 260 | 149 | 1 | 1 | 1 | 0.125 | 0 | 0 | 0.501 | 0.462 | 0 | 0.534 | NA | NA | -0.007 |
| SE | 83832 | ENSG00000248309 | MEF2C-AS1 | chr5 | + | 88892166 | 88892277 | 88889453 | 88889480 | 88905208 | 88905270 | 0 | 1 | 0 | 0 | 4 | 0 | 0 | 1 | 0 | 0 | 0 | 0 | 1 | 0 | 0 | 0 | 4 | 0 | 0 | 0 | 260 | 149 | 1 | 1 | NA | 1 | 0 | NA | 1 | NA | 0 | 1 | NA | NA | 0.167 |
| SE | 83833 | ENSG00000248309 | MEF2C-AS1 | chr5 | + | 88903937 | 88904160 | 88883398 | 88883485 | 88905208 | 88905270 | 0 | 0 | 0 | 2 | 0 | 0 | 4 | 3 | 5 | 4 | 0 | 0 | 0 | 0 | 0 | 4 | 15 | 1 | 0 | 0 | 298 | 149 | 1 | 1 | NA | 0 | 0 | 0.167 | 0 | 0 | 0 | 0 | NA | NA | 0.042 |
| SE | 83834 | ENSG00000248309 | MEF2C-AS1 | chr5 | + | 88903937 | 88904160 | 88892256 | 88892277 | 88905208 | 88905270 | 0 | 0 | 1 | 0 | 0 | 2 | 1 | 0 | 0 | 4 | 0 | 0 | 0 | 0 | 0 | 0 | 0 | 1 | 0 | 0 | 298 | 149 | 1 | 1 | 0 | 0 | 1 | NA | 0 | NA | NA | 0 | NA | NA | 0.25 |
| SE | 83835 | ENSG00000248309 | MEF2C-AS1 | chr5 | + | 88903937 | 88904203 | 88889450 | 88889480 | 88905208 | 88905270 | 0 | 0 | 0 | 0 | 0 | 0 | 0 | 1 | 0 | 0 | 0 | 0 | 0 | 0 | 8 | 0 | 4 | 0 | 0 | 0 | 298 | 149 | 1 | 1 | NA | NA | 0 | NA | NA | NA | 0 | NA | NA | 1 | -0.5 |
| SE | 83836 | ENSG00000248309 | MEF2C-AS1 | chr5 | + | 88905701 | 88905773 | 88905208 | 88905270 | 88943015 | 88944993 | 1 | 0 | 1 | 0 | 7 | 3 | 4 | 4 | 1 | 6 | 0 | 0 | 0 | 0 | 0 | 16 | 20 | 2 | 0 | 19 | 221 | 149 | 1 | 1 | 0.183 | 0 | 0.144 | 0 | 0.44 | 0 | 0 | 0 | NA | 0 | 0.153 |
| SE | 83837 | ENSG00000248309 | MEF2C-AS1 | chr5 | + | 88905701 | 88905773 | 88905208 | 88905270 | 88965840 | 88967279 | 1 | 0 | 1 | 0 | 0 | 2 | 1 | 1 | 5 | 0 | 0 | 3 | 0 | 0 | 4 | 15 | 3 | 0 | 3 | 0 | 221 | 149 | 1 | 1 | 0.252 | 0 | 0.403 | 0 | NA | 0 | 0.403 | NA | 0 | 1 | -0.187 |
| SE | 83839 | ENSG00000248309 | MEF2C-AS1 | chr5 | + | 88948135 | 88948215 | 88943015 | 88943122 | 88965840 | 88965872 | 1 | 0 | 3 | 2 | 0 | 1 | 1 | 0 | 5 | 2 | 13 | 3 | 1 | 0 | 0 | 0 | 13 | 2 | 0 | 0 | 229 | 149 | 1 | 1 | 0.394 | 0 | 1 | 0.207 | 0 | 1 | 0.131 | 0.245 | NA | NA | -0.138 |
| SE | 83840 | ENSG00000248309 | MEF2C-AS1 | chr5 | + | 88965840 | 88965915 | 88905208 | 88905270 | 89094958 | 89095029 | 4 | 1 | 1 | 5 | 0 | 2 | 1 | 0 | 0 | 4 | 19 | 5 | 0 | 0 | 5 | 0 | 0 | 0 | 0 | 0 | 224 | 149 | 1 | 1 | 0.571 | 0.399 | 1 | 1 | 0 | 1 | 1 | NA | | 1 | -0.406 |
| SE | 83841 | ENSG00000248309 | MEF2C-AS1 | chr5 | + | 89094958 | 89095029 | 88965840 | 88965915 | 89150532 | 89150595 | 4 | 1 | 0 | 6 | 9 | 0 | 1 | 0 | 0 | 0 | 23 | 12 | 2 | 0 | 5 | 0 | 0 | 0 | 0 | 0 | 220 | 149 | 1 | 1 | 1 | 0.404 | NA | 1 | 1 | 1 | 1 | 1 | NA | 1 | -0.149 |
| SE | 83842 | ENSG00000248309 | MEF2C-AS1 | chr5 | + | 89150532 | 89150595 | 89094958 | 89095029 | 89168249 | 89168668 | 5 | 1 | 0 | 4 | 7 | 2 | 0 | 0 | 1 | 0 | 3 | 6 | 0 | 0 | 6 | 0 | 0 | 0 | 0 | 0 | 212 | 149 | 1 | 1 | 0.637 | 1 | NA | 0.738 | 1 | 1 | 1 | NA | NA | 1 | -0.156 |
| SE | 83843 | ENSG00000248309 | MEF2C-AS1 | chr5 | + | 89150723 | 89150777 | 89150532 | 89150595 | 89168249 | 89168668 | 1 | 5 | 0 | 2 | 15 | 2 | 1 | 0 | 2 | 7 | 4 | 0 | 0 | 0 | 0 | 3 | 4 | 0 | 0 | 1 | 203 | 149 | 1 | 1 | 0.268 | 0.786 | NA | 0.423 | 0.611 | 0.495 | 0 | NA | NA | 0 | 0.357 |
| SE | 83845 | ENSG00000158526 | TSR2 | chrX | + | 54443399 | 54443491 | 54440689 | 54440780 | 54444007 | 54444184 | 187 | 110 | 163 | 164 | 199 | 0 | 0 | 0 | 0 | 0 | 352 | 195 | 219 | 162 | 264 | 0 | 0 | 0 | 4 | 0 | 241 | 149 | 1 | 1 | 1 | 1 | 1 | 1 | 1 | 1 | 1 | 1 | 0.962 | 1 | 0.008 |
| SE | 83847 | ENSG00000204837 | FGF7P3 | chr9 | - | 39830012 | 39830110 | 39818820 | 39818924 | 39863832 | 39863942 | 1 | 1 | 7 | 0 | 0 | 0 | 0 | 1 | 0 | 0 | 0 | 0 | 0 | 0 | 0 | 0 | 3 | 2 | 0 | 0 | 247 | 149 | 1 | 1 | 1 | 1 | 0.809 | NA | NA | NA | 0 | 0 | NA | NA | 0.936 |
| SE | 83848 | ENSG00000204837 | FGF7P3 | chr9 | - | 39830012 | 39830110 | 39818820 | 39818924 | 39873909 | 39874206 | 0 | 0 | 6 | 0 | 0 | 4 | 0 | 0 | 3 | 0 | 0 | 0 | 2 | 0 | 0 | 0 | 0 | 0 | 0 | 0 | 247 | 149 | 1 | 1 | 0 | NA | 1 | 0 | NA | NA | NA | 1 | NA | NA | -0.667 |
| SE | 83849 | ENSG00000204837 | FGF7P3 | chr9 | - | 39863832 | 39863942 | 39818820 | 39818924 | 39873909 | 39874206 | 0 | 0 | 1 | 0 | 0 | 4 | 0 | 0 | 3 | 0 | 0 | 3 | 2 | 0 | 0 | 0 | 0 | 0 | 0 | 0 | 259 | 149 | 1 | 1 | 0 | NA | 1 | 0 | NA | NA | 1 | 1 | NA | NA | -0.667 |
| SE | 83850 | ENSG00000204837 | FGF7P3 | chr9 | - | 39863832 | 39863942 | 39830012 | 39830110 | 39873909 | 39874206 | 1 | 1 | 1 | 0 | 0 | 0 | 0 | 0 | 0 | 0 | 0 | 0 | 0 | 0 | 0 | 0 | 0 | 2 | 0 | 0 | 259 | 149 | 1 | 1 | 1 | 1 | 1 | NA | NA | NA | NA | 0 | NA | NA | 1 |
| SE | 83851 | ENSG00000130741 | EIF2S3 | chrX | + | 24055614 | 24055678 | 24054945 | 24055037 | 24057420 | 24057548 | 810 | 433 | 777 | 693 | 604 | 0 | 0 | 0 | 0 | 0 | 555 | 284 | 466 | 381 | 156 | 0 | 2 | 0 | 0 | 0 | 213 | 149 | 1 | 1 | 1 | 1 | 1 | 1 | 1 | 1 | 0.99 | 1 | 1 | 1 | 0.002 |
| SE | 83853 | ENSG00000130741 | EIF2S3 | chrX | + | 24060087 | 24060182 | 24057632 | 24057754 | 24064200 | 24064335 | 498 | 361 | 556 | 433 | 633 | 0 | 0 | 0 | 0 | 0 | 328 | 205 | 303 | 373 | 149 | 0 | 8 | 0 | 0 | 0 | 244 | 149 | 1 | 1 | 1 | 1 | 1 | 1 | 1 | 1 | 0.94 | 1 | 1 | 1 | 0.012 |
| SE | 83854 | ENSG00000130741 | EIF2S3 | chrX | + | 24062415 | 24062574 | 24057632 | 24057754 | 24064200 | 24064335 | 1420 | 1147 | 1542 | 1340 | 1752 | 0 | 0 | 0 | 0 | 0 | 1403 | 725 | 915 | 1230 | 321 | 0 | 8 | 0 | 0 | 0 | 298 | 149 | 1 | 1 | 1 | 1 | 1 | 1 | 1 | 1 | 0.978 | 1 | 1 | 1 | 0.004 |
| SE | 83855 | ENSG00000130741 | EIF2S3 | chrX | + | 24062415 | 24062574 | 24060087 | 24060182 | 24064200 | 24064335 | 2788 | 2126 | 3072 | 2702 | 3296 | 1 | 0 | 2 | 0 | 0 | 2356 | 1313 | 1738 | 2327 | 647 | 0 | 0 | 0 | 0 | 0 | 298 | 149 | 1 | 1 | 0.999 | 1 | 0.999 | 1 | 1 | 1 | 1 | 1 | 1 | 1 | 0 |
| SE | 83856 | ENSG00000130741 | EIF2S3 | chrX | + | 24064200 | 24064335 | 24062415 | 24062574 | 24071557 | 24071727 | 1394 | 1124 | 1516 | 1305 | 1699 | 0 | 1 | 0 | 0 | 0 | 1374 | 708 | 893 | 1196 | 321 | 0 | 0 | 0 | 0 | 0 | 284 | 149 | 1 | 1 | 1 | 0.998 | 1 | 1 | 1 | 1 | 1 | 1 | 1 | 1 | 0 |
| SE | 83857 | ENSG00000130741 | EIF2S3 | chrX | + | 24065997 | 24066092 | 24064200 | 24064335 | 24071557 | 24071727 | 613 | 494 | 691 | 582 | 726 | 0 | 0 | 0 | 0 | 1 | 490 | 328 | 359 | 485 | 161 | 0 | 0 | 0 | 1 | 0 | 244 | 149 | 1 | 1 | 1 | 1 | 1 | 1 | 0.998 | 1 | 1 | 1 | 0.997 | 1 | 0 |
| SE | 83858 | ENSG00000130741 | EIF2S3 | chrX | + | 24067963 | 24068108 | 24065997 | 24066092 | 24071557 | 24071727 | 2680 | 2104 | 2740 | 2518 | 2779 | 0 | 0 | 0 | 5 | 0 | 2328 | 1373 | 1614 | 2239 | 682 | 2 | 0 | 1 | 3 | 0 | 294 | 149 | 1 | 1 | 1 | 1 | 1 | 0.996 | 1 | 0.998 | 1 | 0.999 | 0.997 | 1 | 0 |
| SE | 83859 | ENSG00000130741 | EIF2S3 | chrX | + | 24067963 | 24068108 | 24065997 | 24066092 | 24073090 | 24073263 | 1578 | 1289 | 1638 | 1456 | 1606 | 0 | 0 | 1 | 0 | 0 | 1564 | 880 | 1019 | 1288 | 353 | 0 | 0 | 0 | 0 | 0 | 294 | 149 | 1 | 1 | 1 | 1 | 0.999 | 1 | 1 | 1 | 1 | 1 | 1 | 1 | 0 |
| SE | 83860 | ENSG00000130741 | EIF2S3 | chrX | + | 24067963 | 24068108 | 24065997 | 24066092 | 24076721 | 24078810 | 1578 | 1287 | 1638 | 1455 | 1607 | 1 | 0 | 0 | 0 | 0 | 1564 | 880 | 1023 | 1284 | 353 | 0 | 0 | 0 | 0 | 0 | 294 | 149 | 1 | 1 | 0.999 | 1 | 1 | 1 | 1 | 1 | 1 | 1 | 1 | 1 | 0 |
| SE | 83861 | ENSG00000130741 | EIF2S3 | chrX | + | 24071557 | 24071727 | 24065997 | 24066092 | 24073090 | 24073263 | 1111 | 990 | 1297 | 1222 | 1389 | 0 | 0 | 1 | 0 | 0 | 1036 | 591 | 933 | 1215 | 413 | 0 | 0 | 0 | 0 | 0 | 298 | 149 | 1 | 1 | 1 | 1 | 0.998 | 1 | 1 | 1 | 1 | 1 | 1 | 1 | 0 |
| SE | 83862 | ENSG00000130741 | EIF2S3 | chrX | + | 24071557 | 24071727 | 24065997 | 24066092 | 24076721 | 24078810 | 22 | 5 | 26 | 20 | 22 | 1 | 0 | 0 | 0 | 0 | 33 | 6 | 10 | 29 | 0 | 0 | 0 | 0 | 0 | 0 | 298 | 149 | 1 | 1 | 0.917 | 1 | 1 | 1 | 1 | 1 | 1 | 1 | 1 | NA | -0.017 |
| SE | 83863 | ENSG00000130741 | EIF2S3 | chrX | + | 24071557 | 24071727 | 24067963 | 24068108 | 24073090 | 24073263 | 2215 | 1811 | 2408 | 2288 | 2572 | 0 | 3 | 0 | 1 | 0 | 1798 | 1088 | 1531 | 2168 | 744 | 0 | 0 | 0 | 4 | 0 | 298 | 149 | 1 | 1 | 1 | 0.997 | 1 | 0.999 | 1 | 1 | 1 | 1 | 0.996 | 1 | 0 |
| SE | 83864 | ENSG00000130741 | EIF2S3 | chrX | + | 24071557 | 24071727 | 24067963 | 24068108 | 24076721 | 24078810 | 1126 | 826 | 1137 | 1086 | 1205 | 0 | 1 | 0 | 0 | 1 | 795 | 503 | 608 | 982 | 331 | 0 | 0 | 4 | 0 | 0 | 298 | 149 | 1 | 1 | 1 | 0.998 | 1 | 1 | 0.998 | 1 | 1 | 0.987 | 1 | 1 | 0.002 |
| SE | 83865 | ENSG00000130741 | EIF2S3 | chrX | + | 24073090 | 24073263 | 24065997 | 24066092 | 24076721 | 24078810 | 1677 | 1335 | 1597 | 1404 | 2258 | 1 | 0 | 0 | 0 | 0 | 960 | 492 | 850 | 1203 | 341 | 0 | 0 | 0 | 0 | 0 | 298 | 149 | 1 | 1 | 0.999 | 1 | 1 | 1 | 1 | 1 | 1 | 1 | 1 | 1 | 0 |
| SE | 83866 | ENSG00000130741 | EIF2S3 | chrX | + | 24073090 | 24073263 | 24067963 | 24068108 | 24076721 | 24078810 | 1677 | 1338 | 1596 | 1405 | 2258 | 0 | 1 | 0 | 0 | 1 | 960 | 492 | 850 | 1207 | 341 | 0 | 0 | 4 | 0 | 0 | 298 | 149 | 1 | 1 | 1 | 0.999 | 1 | 1 | 0.999 | 1 | 1 | 0.991 | 1 | 1 | 0.001 |
| SE | 83868 | ENSG00000116106 | EPHA4 | chr2 | - | 221421979 | 221422070 | 221420444 | 221420552 | 221425208 | 221425540 | 0 | 0 | 1 | 0 | 0 | 21 | 18 | 43 | 30 | 34 | 0 | 0 | 0 | 0 | 0 | 0 | 0 | 14 | 8 | 8 | 240 | 149 | 1 | 1 | 0 | 0 | 0.014 | 0 | 0 | NA | NA | 0 | 0 | 0 | 0.003 |
| SE | 83869 | ENSG00000116106 | EPHA4 | chr2 | - | 221429957 | 221430151 | 221426434 | 221426619 | 221434141 | 221434291 | 35 | 20 | 68 | 46 | 51 | 0 | 0 | 1 | 0 | 3 | 0 | 8 | 13 | 27 | 16 | 0 | 0 | 0 | 0 | 0 | 298 | 149 | 1 | 1 | 1 | 1 | 0.971 | 1 | 0.895 | NA | 1 | 1 | 1 | 1 | -0.027 |
| SE | 83870 | ENSG00000116106 | EPHA4 | chr2 | - | 221434141 | 221434291 | 221429957 | 221430151 | 221436398 | 221436608 | 85 | 31 | 94 | 66 | 70 | 0 | 0 | 0 | 1 | 0 | 17 | 39 | 34 | 18 | 24 | 0 | 0 | 0 | 0 | 0 | 298 | 149 | 1 | 1 | 1 | 1 | 1 | 0.971 | 1 | 1 | 1 | 1 | 1 | 1 | -0.006 |
| SE | 83871 | ENSG00000116106 | EPHA4 | chr2 | - | 221437060 | 221437122 | 221436398 | 221436608 | 221442828 | 221443014 | 35 | 20 | 53 | 39 | 26 | 4 | 0 | 0 | 0 | 0 | 9 | 17 | 11 | 20 | 0 | 0 | 0 | 0 | 0 | 0 | 211 | 149 | 1 | 1 | 0.861 | 1 | 1 | 1 | 1 | 1 | 1 | 1 | 1 | NA | -0.028 |
| SE | 83872 | ENSG00000116106 | EPHA4 | chr2 | - | 221455546 | 221455658 | 221446122 | 221446181 | 221456612 | 221456772 | 54 | 23 | 57 | 26 | 71 | 1 | 0 | 5 | 2 | 0 | 0 | 4 | 15 | 12 | 8 | 0 | 4 | 0 | 0 | 0 | 261 | 149 | 1 | 1 | 0.969 | 1 | 0.867 | 0.881 | 1 | NA | 0.363 | 1 | 1 | 1 | 0.103 |
| SE | 83873 | ENSG00000116106 | EPHA4 | chr2 | - | 221456612 | 221456772 | 221455546 | 221455658 | 221457865 | 221457990 | 64 | 26 | 74 | 39 | 86 | 0 | 0 | 0 | 0 | 0 | 0 | 14 | 25 | 18 | 8 | 0 | 0 | 1 | 0 | 0 | 298 | 149 | 1 | 1 | 1 | 1 | 1 | 1 | 1 | NA | 1 | 0.926 | 1 | 1 | 0.018 |
| SE | 83874 | ENSG00000116106 | EPHA4 | chr2 | - | 221568717 | 221568785 | 221563730 | 221564394 | 221572157 | 221572281 | 54 | 39 | 98 | 91 | 62 | 3 | 1 | 2 | 2 | 0 | 0 | 17 | 59 | 13 | 12 | 0 | 9 | 3 | 0 | 0 | 217 | 149 | 1 | 1 | 0.925 | 0.964 | 0.971 | 0.969 | 1 | NA | 0.565 | 0.931 | 1 | 1 | 0.092 |
| SE | 83881 | ENSG00000157168 | NRG1 | chr8 | + | 32742007 | 32742075 | 32727948 | 32728078 | 32754371 | 32754474 | 36 | 23 | 14 | 9 | 53 | 0 | 0 | 1 | 0 | 0 | 56 | 6 | 22 | 13 | 36 | 0 | 0 | 0 | 0 | 0 | 217 | 149 | 1 | 1 | 1 | 1 | 0.906 | 1 | 1 | 1 | 1 | 1 | 1 | 1 | -0.019 |
| SE | 83882 | ENSG00000157168 | NRG1 | chr8 | + | 32742674 | 32742733 | 32727948 | 32728078 | 32754371 | 32754474 | 12 | 7 | 3 | 3 | 7 | 0 | 0 | 1 | 0 | 0 | 27 | 4 | 7 | 4 | 2 | 0 | 0 | 0 | 0 | 0 | 208 | 149 | 1 | 1 | 1 | 1 | 0.682 | 1 | 1 | 1 | 1 | 1 | 1 | 1 | -0.064 |
| SE | 83885 | ENSG00000157168 | NRG1 | chr8 | + | 32763217 | 32763359 | 32760199 | 32760406 | 32763747 | 32763998 | 4 | 2 | 1 | 2 | 11 | 14 | 6 | 15 | 3 | 14 | 0 | 0 | 1 | 2 | 6 | 27 | 0 | 6 | 11 | 28 | 291 | 149 | 1 | 1 | 0.128 | 0.146 | 0.033 | 0.254 | 0.287 | 0 | NA | 0.079 | 0.085 | 0.099 | 0.104 |
| SE | 83887 | ENSG00000168807 | SNTB2 | chr16 | + | 69260049 | 69260127 | 69245601 | 69245815 | 69270142 | 69270200 | 38 | 15 | 18 | 26 | 23 | 0 | 0 | 0 | 0 | 1 | 20 | 10 | 23 | 12 | 9 | 1 | 0 | 0 | 0 | 0 | 227 | 149 | 1 | 1 | 1 | 1 | 1 | 1 | 0.938 | 0.929 | 1 | 1 | 1 | 1 | 0.002 |
| SE | 83888 | ENSG00000168807 | SNTB2 | chr16 | + | 69260049 | 69260260 | 69245601 | 69245815 | 69270142 | 69270285 | 115 | 46 | 76 | 77 | 63 | 0 | 0 | 0 | 0 | 1 | 47 | 77 | 53 | 44 | 31 | 1 | 0 | 0 | 0 | 0 | 298 | 149 | 1 | 1 | 1 | 1 | 1 | 1 | 0.969 | 0.959 | 1 | 1 | 1 | 1 | 0.002 |
| SE | 83889 | ENSG00000168807 | SNTB2 | chr16 | + | 69260170 | 69260260 | 69245601 | 69245815 | 69270142 | 69270181 | 24 | 6 | 18 | 21 | 4 | 0 | 0 | 0 | 0 | 1 | 3 | 31 | 4 | 12 | 8 | 1 | 0 | 0 | 0 | 0 | 239 | 149 | 1 | 1 | 1 | 1 | 1 | 1 | 0.714 | 0.652 | 1 | 1 | 1 | 1 | 0.012 |
| SE | 83890 | ENSG00000168807 | SNTB2 | chr16 | + | 69284047 | 69284244 | 69270142 | 69270285 | 69299589 | 69299774 | 56 | 34 | 55 | 69 | 48 | 1 | 1 | 3 | 0 | 0 | 10 | 75 | 46 | 35 | 43 | 0 | 0 | 0 | 0 | 0 | 298 | 149 | 1 | 1 | 0.966 | 0.944 | 0.902 | 1 | 1 | 1 | 1 | 1 | 1 | 1 | -0.038 |
| SE | 83891 | ENSG00000171365 | CLCN5 | chrX | + | 50050282 | 50050413 | 50042337 | 50042462 | 50054439 | 50054633 | 1 | 0 | 0 | 0 | 1 | 1 | 0 | 2 | 0 | 0 | 0 | 0 | 0 | 0 | 0 | 0 | 0 | 6 | 0 | 0 | 280 | 149 | 1 | 1 | 0.347 | NA | 0 | NA | 1 | NA | NA | 0 | NA | NA | 0.449 |
| SE | 83892 | ENSG00000171365 | CLCN5 | chrX | + | 50054439 | 50054633 | 50042337 | 50042462 | 50069878 | 50070030 | 1 | 0 | 2 | 0 | 0 | 5 | 10 | 11 | 10 | 14 | 18 | 0 | 7 | 0 | 0 | 6 | 0 | 0 | 7 | 0 | 298 | 149 | 1 | 1 | 0.091 | 0 | 0.083 | 0 | 0 | 0.6 | NA | 1 | 0 | NA | -0.499 |
| SE | 83904 | ENSG00000130985 | UBA1 | chrX | + | 47200893 | 47201000 | 47199479 | 47199614 | 47201275 | 47201366 | 380 | 160 | 331 | 341 | 247 | 3 | 2 | 1 | 1 | 4 | 557 | 352 | 206 | 327 | 608 | 0 | 0 | 0 | 0 | 0 | 256 | 149 | 1 | 1 | 0.987 | 0.979 | 0.995 | 0.995 | 0.973 | 1 | 1 | 1 | 1 | 1 | -0.014 |
| SE | 83905 | ENSG00000130985 | UBA1 | chrX | + | 47203540 | 47203696 | 47203133 | 47203214 | 47205947 | 47206113 | 526 | 289 | 440 | 528 | 708 | 0 | 0 | 0 | 0 | 0 | 585 | 447 | 214 | 480 | 995 | 0 | 0 | 0 | 4 | 0 | 298 | 149 | 1 | 1 | 1 | 1 | 1 | 1 | 1 | 1 | 1 | 1 | 0.984 | 1 | 0.003 |
| SE | 83906 | ENSG00000130985 | UBA1 | chrX | + | 47205947 | 47206113 | 47203540 | 47203696 | 47206247 | 47206444 | 815 | 431 | 729 | 858 | 1012 | 0 | 0 | 0 | 0 | 3 | 823 | 763 | 361 | 727 | 1400 | 0 | 0 | 0 | 0 | 0 | 298 | 149 | 1 | 1 | 1 | 1 | 1 | 1 | 0.994 | 1 | 1 | 1 | 1 | 1 | -0.001 |
| SE | 83907 | ENSG00000130985 | UBA1 | chrX | + | 47206077 | 47206113 | 47203540 | 47203696 | 47206247 | 47206664 | 145 | 81 | 126 | 185 | 79 | 0 | 0 | 0 | 0 | 3 | 118 | 148 | 73 | 139 | 295 | 0 | 0 | 0 | 0 | 0 | 185 | 149 | 1 | 1 | 1 | 1 | 1 | 1 | 0.955 | 1 | 1 | 1 | 1 | 1 | -0.009 |
| SE | 83908 | ENSG00000130985 | UBA1 | chrX | + | 47206247 | 47206444 | 47203540 | 47203696 | 47209622 | 47209687 | 261 | 132 | 252 | 262 | 304 | 1 | 0 | 0 | 0 | 4 | 311 | 264 | 97 | 212 | 339 | 0 | 2 | 0 | 0 | 0 | 298 | 149 | 1 | 1 | 0.992 | 1 | 1 | 1 | 0.974 | 1 | 0.985 | 1 | 1 | 1 | -0.004 |
| SE | 83909 | ENSG00000130985 | UBA1 | chrX | + | 47211035 | 47211225 | 47210841 | 47210916 | 47212423 | 47212512 | 380 | 181 | 303 | 347 | 284 | 0 | 0 | 1 | 0 | 0 | 425 | 328 | 186 | 364 | 746 | 0 | 0 | 0 | 0 | 0 | 298 | 149 | 1 | 1 | 1 | 1 | 0.993 | 1 | 1 | 1 | 1 | 1 | 1 | 1 | -0.001 |
| SE | 83910 | ENSG00000130985 | UBA1 | chrX | + | 47214536 | 47214637 | 47214326 | 47214428 | 47214793 | 47215126 | 1251 | 741 | 1058 | 1179 | 1818 | 6 | 1 | 2 | 3 | 4 | 2112 | 1236 | 813 | 1252 | 1985 | 8 | 3 | 0 | 4 | 0 | 250 | 149 | 1 | 1 | 0.992 | 0.998 | 0.997 | 0.996 | 0.996 | 0.994 | 0.996 | 1 | 0.995 | 1 | -0.001 |
| SE | 83915 | ENSG00000167778 | SPRYD3 | chr12 | - | 53074648 | 53075219 | 53073285 | 53073471 | 53077114 | 53077261 | 111 | 42 | 80 | 96 | 46 | 2 | 2 | 0 | 0 | 0 | 69 | 26 | 29 | 58 | 95 | 0 | 0 | 0 | 0 | 0 | 298 | 149 | 1 | 1 | 0.965 | 0.913 | 1 | 1 | 1 | 1 | 1 | 1 | 1 | 1 | -0.024 |
| SE | 83916 | ENSG00000167778 | SPRYD3 | chr12 | - | 53075094 | 53075219 | 53073285 | 53073471 | 53077114 | 53077261 | 2 | 2 | 1 | 1 | 4 | 2 | 2 | 0 | 0 | 0 | 4 | 12 | 3 | 7 | 0 | 0 | 0 | 0 | 0 | 0 | 274 | 149 | 1 | 1 | 0.352 | 0.352 | 1 | 1 | 1 | 1 | 1 | 1 | 1 | NA | -0.259 |
| SE | 83922 | ENSG00000112561 | TFEB | chr6 | - | 41686089 | 41686237 | 41683977 | 41685078 | 41687093 | 41687169 | 223 | 82 | 113 | 110 | 144 | 1 | 0 | 0 | 0 | 0 | 225 | 218 | 102 | 114 | 297 | 0 | 0 | 0 | 0 | 0 | 297 | 149 | 1 | 1 | 0.991 | 1 | 1 | 1 | 1 | 1 | 1 | 1 | 1 | 1 | -0.002 |
| SE | 83923 | ENSG00000112561 | TFEB | chr6 | - | 41686089 | 41686356 | 41683993 | 41685078 | 41687093 | 41687169 | 81 | 23 | 41 | 32 | 43 | 1 | 0 | 0 | 0 | 0 | 85 | 76 | 36 | 35 | 70 | 0 | 0 | 0 | 0 | 0 | 298 | 149 | 1 | 1 | 0.976 | 1 | 1 | 1 | 1 | 1 | 1 | 1 | 1 | 1 | -0.005 |
| SE | 83925 | ENSG00000112561 | TFEB | chr6 | - | 41690662 | 41690917 | 41689730 | 41689811 | 41691000 | 41691235 | 117 | 27 | 58 | 73 | 47 | 0 | 0 | 1 | 0 | 1 | 50 | 151 | 61 | 48 | 188 | 0 | 0 | 2 | 0 | 3 | 298 | 149 | 1 | 1 | 1 | 1 | 0.967 | 1 | 0.959 | 1 | 1 | 0.938 | 1 | 0.969 | 0.004 |
| SE | 83926 | ENSG00000112561 | TFEB | chr6 | - | 41705646 | 41705752 | 41691000 | 41691235 | 41735349 | 41735608 | 3 | 0 | 4 | 0 | 0 | 17 | 3 | 9 | 13 | 3 | 0 | 0 | 0 | 0 | 0 | 18 | 11 | 0 | 9 | 25 | 255 | 149 | 1 | 1 | 0.093 | 0 | 0.206 | 0 | 0 | 0 | 0 | NA | 0 | 0 | 0.06 |
| SE | 83927 | ENSG00000112561 | TFEB | chr6 | - | 41732648 | 41732739 | 41691000 | 41691235 | 41733663 | 41733690 | 20 | 5 | 17 | 6 | 21 | 1 | 0 | 0 | 0 | 0 | 14 | 3 | 7 | 8 | 5 | 0 | 0 | 0 | 0 | 0 | 240 | 149 | 1 | 1 | 0.925 | 1 | 1 | 1 | 1 | 1 | 1 | 1 | 1 | 1 | -0.015 |
| SE | 83934 | ENSG00000251595 | ABCA11P | chr4 | - | 437445 | 437511 | 425814 | 426973 | 472574 | 472701 | 7 | 3 | 10 | 9 | 2 | 2 | 0 | 4 | 0 | 3 | 12 | 4 | 0 | 3 | 1 | 0 | 0 | 0 | 0 | 0 | 215 | 149 | 1 | 1 | 0.708 | 1 | 0.634 | 1 | 0.316 | 1 | 1 | NA | 1 | 1 | -0.268 |
| SE | 83936 | ENSG00000147421 | HMBOX1 | chr8 | + | 28891759 | 28891790 | 28890403 | 28891019 | 28963810 | 28963890 | 0 | 0 | 0 | 1 | 0 | 3 | 6 | 1 | 3 | 3 | 0 | 0 | 0 | 0 | 0 | 13 | 5 | 3 | 0 | 0 | 180 | 149 | 1 | 1 | 0 | 0 | 0 | 0.216 | 0 | 0 | 0 | 0 | NA | NA | 0.043 |
| SE | 83937 | ENSG00000147421 | HMBOX1 | chr8 | + | 28891759 | 28891790 | 28890624 | 28890678 | 28963810 | 28963890 | 1 | 0 | 0 | 2 | 0 | 28 | 31 | 24 | 47 | 19 | 0 | 0 | 0 | 0 | 0 | 28 | 38 | 34 | 37 | 18 | 180 | 149 | 1 | 1 | 0.029 | 0 | 0 | 0.034 | 0 | 0 | 0 | 0 | 0 | 0 | 0.013 |
| SE | 83941 | ENSG00000147421 | HMBOX1 | chr8 | + | 29042638 | 29042718 | 29018759 | 29018913 | 29045360 | 29045443 | 1 | 2 | 1 | 5 | 0 | 91 | 67 | 87 | 90 | 73 | 6 | 0 | 0 | 0 | 1 | 42 | 79 | 67 | 59 | 29 | 229 | 149 | 1 | 1 | 0.007 | 0.019 | 0.007 | 0.035 | 0 | 0.085 | 0 | 0 | 0 | 0.022 | -0.008 |
| SE | 83942 | ENSG00000147421 | HMBOX1 | chr8 | + | 29045360 | 29045443 | 29018759 | 29018913 | 29047357 | 29047453 | 152 | 132 | 144 | 146 | 139 | 0 | 0 | 0 | 1 | 0 | 78 | 120 | 115 | 117 | 38 | 0 | 0 | 0 | 0 | 0 | 232 | 149 | 1 | 1 | 1 | 1 | 1 | 0.989 | 1 | 1 | 1 | 1 | 1 | 1 | -0.002 |
| SE | 83943 | ENSG00000147421 | HMBOX1 | chr8 | + | 29047357 | 29047453 | 29045360 | 29045443 | 29048953 | 29049048 | 102 | 93 | 96 | 91 | 87 | 2 | 0 | 2 | 0 | 0 | 54 | 64 | 70 | 79 | 14 | 0 | 0 | 0 | 0 | 0 | 245 | 149 | 1 | 1 | 0.969 | 1 | 0.967 | 1 | 1 | 1 | 1 | 1 | 1 | 1 | -0.013 |
| SE | 83948 | ENSG00000139218 | SCAF11 | chr12 | - | 45922462 | 45922582 | 45919130 | 45919309 | 45926141 | 45928859 | 5 | 13 | 5 | 6 | 5 | 0 | 0 | 0 | 1 | 0 | 0 | 0 | 0 | 1 | 0 | 0 | 0 | 0 | 0 | 0 | 269 | 149 | 1 | 1 | 1 | 1 | 1 | 0.769 | 1 | NA | NA | NA | 1 | NA | -0.046 |
| SE | 83950 | ENSG00000139218 | SCAF11 | chr12 | - | 45922935 | 45923154 | 45922355 | 45922582 | 45926141 | 45928859 | 226 | 230 | 170 | 225 | 242 | 0 | 0 | 0 | 2 | 3 | 0 | 21 | 31 | 9 | 0 | 0 | 0 | 0 | 0 | 0 | 298 | 149 | 1 | 1 | 1 | 1 | 1 | 0.983 | 0.976 | NA | 1 | 1 | 1 | NA | -0.008 |
| SE | 83951 | ENSG00000139218 | SCAF11 | chr12 | - | 45924727 | 45925074 | 45922462 | 45922582 | 45926141 | 45928859 | 277 | 225 | 211 | 220 | 597 | 0 | 0 | 0 | 2 | 3 | 1 | 51 | 96 | 17 | 0 | 0 | 0 | 0 | 0 | 0 | 298 | 149 | 1 | 1 | 1 | 1 | 1 | 0.982 | 0.99 | 1 | 1 | 1 | 1 | NA | -0.006 |
| SE | 83952 | ENSG00000139218 | SCAF11 | chr12 | - | 45924727 | 45925074 | 45922935 | 45923154 | 45926141 | 45928859 | 669 | 621 | 528 | 599 | 1107 | 0 | 1 | 1 | 0 | 1 | 46 | 135 | 189 | 27 | 3 | 0 | 0 | 0 | 0 | 0 | 298 | 149 | 1 | 1 | 1 | 0.997 | 0.996 | 1 | 0.998 | 1 | 1 | 1 | 1 | 1 | -0.002 |
| SE | 83953 | ENSG00000139218 | SCAF11 | chr12 | - | 45934175 | 45934285 | 45933130 | 45933232 | 45934446 | 45934505 | 502 | 395 | 399 | 444 | 901 | 0 | 1 | 0 | 0 | 0 | 7 | 153 | 127 | 63 | 44 | 0 | 0 | 0 | 0 | 0 | 259 | 149 | 1 | 1 | 1 | 0.996 | 1 | 1 | 1 | 1 | 1 | 1 | 1 | 1 | -0.001 |
| SE | 83954 | ENSG00000139218 | SCAF11 | chr12 | - | 45951649 | 45951727 | 45948436 | 45948537 | 45961699 | 45961857 | 472 | 488 | 417 | 486 | 1031 | 0 | 0 | 0 | 0 | 2 | 132 | 182 | 188 | 102 | 160 | 0 | 0 | 0 | 0 | 0 | 227 | 149 | 1 | 1 | 1 | 1 | 1 | 1 | 0.997 | 1 | 1 | 1 | 1 | 1 | -0.001 |
| SE | 83955 | ENSG00000139218 | SCAF11 | chr12 | - | 45951649 | 45951727 | 45948436 | 45948537 | 45964106 | 45964188 | 127 | 138 | 138 | 137 | 309 | 0 | 0 | 0 | 1 | 0 | 42 | 70 | 81 | 42 | 86 | 0 | 5 | 0 | 0 | 0 | 227 | 149 | 1 | 1 | 1 | 1 | 1 | 0.989 | 1 | 1 | 0.902 | 1 | 1 | 1 | 0.017 |
| SE | 83958 | ENSG00000139218 | SCAF11 | chr12 | - | 45961699 | 45961857 | 45960454 | 45961322 | 45964106 | 45964188 | 193 | 187 | 137 | 215 | 345 | 0 | 0 | 0 | 0 | 0 | 20 | 57 | 76 | 14 | 25 | 0 | 0 | 0 | 2 | 0 | 298 | 149 | 1 | 1 | 1 | 1 | 1 | 1 | 1 | 1 | 1 | 1 | 0.778 | 1 | 0.044 |
| SE | 83959 | ENSG00000139218 | SCAF11 | chr12 | - | 45964106 | 45964188 | 45961406 | 45961857 | 45990352 | 45990574 | 207 | 191 | 149 | 230 | 347 | 0 | 0 | 0 | 1 | 0 | 20 | 76 | 77 | 12 | 31 | 0 | 0 | 0 | 0 | 0 | 231 | 149 | 1 | 1 | 1 | 1 | 1 | 0.993 | 1 | 1 | 1 | 1 | 1 | 1 | -0.001 |
| SE | 83961 | ENSG00000068400 | GRIPAP1 | chrX | - | 48975154 | 48975309 | 48973722 | 48974285 | 48976017 | 48976111 | 56 | 44 | 54 | 86 | 45 | 0 | 0 | 1 | 0 | 0 | 58 | 82 | 49 | 90 | 140 | 0 | 0 | 0 | 0 | 0 | 298 | 149 | 1 | 1 | 1 | 1 | 0.964 | 1 | 1 | 1 | 1 | 1 | 1 | 1 | -0.007 |
| SE | 83963 | ENSG00000068400 | GRIPAP1 | chrX | - | 48981794 | 48981872 | 48981595 | 48981691 | 48983227 | 48983440 | 38 | 21 | 15 | 30 | 19 | 0 | 0 | 0 | 0 | 0 | 13 | 48 | 61 | 51 | 86 | 0 | 0 | 0 | 3 | 0 | 227 | 149 | 1 | 1 | 1 | 1 | 1 | 1 | 1 | 1 | 1 | 1 | 0.918 | 1 | 0.016 |
| SE | 83964 | ENSG00000068400 | GRIPAP1 | chrX | - | 48982978 | 48983092 | 48981595 | 48981691 | 48983227 | 48983440 | 100 | 48 | 63 | 109 | 65 | 0 | 0 | 0 | 0 | 0 | 205 | 160 | 86 | 194 | 211 | 0 | 0 | 0 | 3 | 0 | 263 | 149 | 1 | 1 | 1 | 1 | 1 | 1 | 1 | 1 | 1 | 1 | 0.973 | 1 | 0.005 |
| SE | 83965 | ENSG00000068400 | GRIPAP1 | chrX | - | 48982978 | 48983092 | 48981794 | 48981872 | 48983227 | 48983440 | 199 | 116 | 143 | 202 | 102 | 0 | 0 | 0 | 0 | 0 | 454 | 305 | 198 | 399 | 419 | 0 | 0 | 2 | 0 | 0 | 263 | 149 | 1 | 1 | 1 | 1 | 1 | 1 | 1 | 1 | 1 | 0.982 | 1 | 1 | 0.004 |
| SE | 83966 | ENSG00000068400 | GRIPAP1 | chrX | - | 48987784 | 48987877 | 48985267 | 48985402 | 48988120 | 48988198 | 70 | 63 | 57 | 73 | 103 | 0 | 0 | 0 | 0 | 0 | 95 | 113 | 81 | 167 | 103 | 0 | 1 | 0 | 0 | 0 | 242 | 149 | 1 | 1 | 1 | 1 | 1 | 1 | 1 | 1 | 0.986 | 1 | 1 | 1 | 0.003 |
| SE | 83968 | ENSG00000068400 | GRIPAP1 | chrX | - | 48990925 | 48991110 | 48990684 | 48990732 | 48993427 | 48993578 | 125 | 77 | 96 | 97 | 67 | 0 | 0 | 0 | 0 | 0 | 214 | 149 | 149 | 275 | 217 | 0 | 0 | 0 | 2 | 0 | 298 | 149 | 1 | 1 | 1 | 1 | 1 | 1 | 1 | 1 | 1 | 1 | 0.986 | 1 | 0.003 |
| SE | 83989 | ENSG00000149809 | TM7SF2 | chr11 | + | 65115313 | 65115394 | 65114912 | 65115081 | 65115475 | 65115514 | 20 | 8 | 21 | 13 | 55 | 1 | 0 | 1 | 1 | 0 | 0 | 32 | 11 | 5 | 47 | 0 | 0 | 0 | 0 | 0 | 230 | 149 | 1 | 1 | 0.928 | 1 | 0.932 | 0.894 | 1 | NA | 1 | 1 | 1 | 1 | -0.049 |
| SE | 83991 | ENSG00000149809 | TM7SF2 | chr11 | + | 65115774 | 65115822 | 65115475 | 65115598 | 65115892 | 65115957 | 0 | 0 | 1 | 0 | 0 | 25 | 7 | 15 | 15 | 27 | 0 | 0 | 0 | 0 | 0 | 0 | 13 | 12 | 2 | 20 | 197 | 149 | 1 | 1 | 0 | 0 | 0.048 | 0 | 0 | NA | 0 | 0 | 0 | 0 | 0.01 |
| SE | 83994 | ENSG00000112715 | VEGFA | chr6 | + | 43780731 | 43780785 | 43778888 | 43778918 | 43781955 | 43782087 | 0 | 2 | 0 | 1 | 0 | 10 | 3 | 6 | 9 | 0 | 0 | 1 | 0 | 2 | 0 | 13 | 19 | 4 | 15 | 12 | 203 | 149 | 1 | 1 | 0 | 0.329 | 0 | 0.075 | NA | 0 | 0.037 | 0 | 0.089 | 0 | 0.076 |
| SE | 83995 | ENSG00000112715 | VEGFA | chr6 | + | 43780731 | 43780803 | 43778888 | 43778918 | 43781955 | 43782087 | 0 | 2 | 1 | 5 | 0 | 10 | 3 | 6 | 9 | 0 | 5 | 3 | 1 | 5 | 0 | 13 | 19 | 4 | 15 | 12 | 221 | 149 | 1 | 1 | 0 | 0.31 | 0.101 | 0.272 | NA | 0.206 | 0.096 | 0.144 | 0.183 | 0 | 0.045 |
| SE | 83996 | ENSG00000112715 | VEGFA | chr6 | + | 43780731 | 43780803 | 43778888 | 43778918 | 43784540 | 43784562 | 0 | 2 | 0 | 1 | 0 | 5 | 8 | 5 | 4 | 7 | 0 | 1 | 0 | 2 | 0 | 17 | 28 | 9 | 48 | 15 | 221 | 149 | 1 | 1 | 0 | 0.144 | 0 | 0.144 | 0 | 0 | 0.024 | 0 | 0.027 | 0 | 0.047 |
| SE | 83997 | ENSG00000112715 | VEGFA | chr6 | + | 43780731 | 43780854 | 43778888 | 43778918 | 43781955 | 43782087 | 0 | 2 | 0 | 1 | 0 | 10 | 3 | 6 | 9 | 0 | 0 | 1 | 0 | 2 | 0 | 13 | 19 | 4 | 15 | 12 | 272 | 149 | 1 | 1 | 0 | 0.268 | 0 | 0.057 | NA | 0 | 0.028 | 0 | 0.068 | 0 | 0.062 |
| SE | 84002 | ENSG00000156697 | UTP14A | chrX | + | 129907366 | 129907442 | 129906120 | 129906236 | 129908059 | 129908130 | 43 | 14 | 26 | 29 | 14 | 0 | 0 | 0 | 0 | 0 | 46 | 20 | 34 | 11 | 10 | 0 | 0 | 3 | 0 | 0 | 225 | 149 | 1 | 1 | 1 | 1 | 1 | 1 | 1 | 1 | 1 | 0.882 | 1 | 1 | 0.024 |
| SE | 84004 | ENSG00000156697 | UTP14A | chrX | + | 129920674 | 129920752 | 129920456 | 129920580 | 129921193 | 129921587 | 100 | 93 | 84 | 96 | 136 | 0 | 0 | 0 | 0 | 0 | 156 | 70 | 46 | 89 | 89 | 0 | 3 | 0 | 0 | 0 | 227 | 149 | 1 | 1 | 1 | 1 | 1 | 1 | 1 | 1 | 0.939 | 1 | 1 | 1 | 0.012 |
| SE | 84005 | ENSG00000261644 | CYLD-AS1 | chr16 | - | 50729842 | 50729953 | 50729012 | 50729150 | 50732217 | 50732360 | 0 | 0 | 5 | 0 | 5 | 0 | 0 | 0 | 2 | 0 | 0 | 0 | 0 | 2 | 0 | 0 | 0 | 0 | 0 | 0 | 260 | 149 | 1 | 1 | NA | NA | 1 | 0 | 1 | NA | NA | NA | 1 | NA | -0.333 |
| SE | 84007 | ENSG00000102145 | GATA1 | chrX | + | 48791090 | 48791329 | 48786572 | 48786645 | 48791843 | 48792221 | 3 | 5 | 0 | 1 | 3 | 0 | 1 | 0 | 0 | 0 | 22 | 0 | 8 | 10 | 0 | 0 | 0 | 0 | 0 | 0 | 298 | 149 | 1 | 1 | 1 | 0.714 | NA | 1 | 1 | 1 | NA | 1 | 1 | NA | -0.072 |
| SE | 84009 | ENSG00000244055 | NA | chr7 | + | 92462957 | 92463026 | 92461318 | 92461410 | 92491242 | 92491610 | 21 | 14 | 17 | 9 | 35 | 8 | 0 | 1 | 6 | 1 | 0 | 20 | 18 | 12 | 34 | 0 | 6 | 9 | 0 | 0 | 218 | 149 | 1 | 1 | 0.642 | 1 | 0.921 | 0.506 | 0.96 | NA | 0.695 | 0.578 | 1 | 1 | -0.012 |
| SE | 84011 | ENSG00000184293 | CLECL1 | chr12 | - | 9718640 | 9718774 | 9715859 | 9716776 | 9722643 | 9722661 | 5 | 2 | 0 | 0 | 0 | 7 | 10 | 2 | 8 | 6 | 0 | 0 | 0 | 0 | 0 | 0 | 11 | 0 | 0 | 0 | 283 | 149 | 1 | 1 | 0.273 | 0.095 | 0 | 0 | 0 | NA | 0 | NA | NA | NA | 0.074 |
| SE | 84014 | ENSG00000111581 | NUP107 | chr12 | + | 68691967 | 68692112 | 68690467 | 68690746 | 68696850 | 68696922 | 79 | 55 | 49 | 76 | 61 | 0 | 0 | 0 | 0 | 0 | 23 | 38 | 25 | 9 | 18 | 0 | 0 | 1 | 0 | 0 | 294 | 149 | 1 | 1 | 1 | 1 | 1 | 1 | 1 | 1 | 1 | 0.927 | 1 | 1 | 0.015 |
| SE | 84015 | ENSG00000111581 | NUP107 | chr12 | + | 68696818 | 68696922 | 68691967 | 68692112 | 68700725 | 68700853 | 145 | 79 | 103 | 140 | 141 | 0 | 0 | 0 | 0 | 0 | 21 | 76 | 28 | 18 | 18 | 0 | 0 | 1 | 0 | 0 | 253 | 149 | 1 | 1 | 1 | 1 | 1 | 1 | 1 | 1 | 1 | 0.943 | 1 | 1 | 0.011 |
| SE | 84016 | ENSG00000111581 | NUP107 | chr12 | + | 68696850 | 68696922 | 68691967 | 68692112 | 68700725 | 68700814 | 35 | 34 | 23 | 39 | 23 | 0 | 0 | 0 | 0 | 0 | 10 | 8 | 12 | 4 | 9 | 0 | 0 | 1 | 0 | 0 | 221 | 149 | 1 | 1 | 1 | 1 | 1 | 1 | 1 | 1 | 1 | 0.89 | 1 | 1 | 0.022 |
| SE | 84017 | ENSG00000111581 | NUP107 | chr12 | + | 68700725 | 68700853 | 68696818 | 68696922 | 68702735 | 68702784 | 175 | 118 | 140 | 183 | 212 | 0 | 0 | 1 | 0 | 0 | 38 | 100 | 62 | 38 | 39 | 0 | 0 | 0 | 0 | 0 | 277 | 149 | 1 | 1 | 1 | 1 | 0.987 | 1 | 1 | 1 | 1 | 1 | 1 | 1 | -0.003 |
| SE | 84018 | ENSG00000111581 | NUP107 | chr12 | + | 68700725 | 68700853 | 68696818 | 68696922 | 68709237 | 68709309 | 94 | 52 | 66 | 95 | 81 | 0 | 0 | 1 | 0 | 0 | 21 | 49 | 20 | 17 | 21 | 0 | 0 | 0 | 0 | 0 | 277 | 149 | 1 | 1 | 1 | 1 | 0.973 | 1 | 1 | 1 | 1 | 1 | 1 | 1 | -0.005 |
| SE | 84019 | ENSG00000111581 | NUP107 | chr12 | + | 68702735 | 68702784 | 68696818 | 68696922 | 68709237 | 68709265 | 33 | 18 | 29 | 34 | 40 | 0 | 0 | 1 | 0 | 0 | 0 | 3 | 6 | 12 | 0 | 0 | 0 | 0 | 0 | 0 | 198 | 149 | 1 | 1 | 1 | 1 | 0.956 | 1 | 1 | NA | 1 | 1 | 1 | NA | -0.009 |
| SE | 84020 | ENSG00000111581 | NUP107 | chr12 | + | 68702735 | 68702784 | 68700725 | 68700853 | 68709237 | 68709265 | 118 | 86 | 105 | 122 | 165 | 2 | 0 | 1 | 0 | 0 | 17 | 59 | 47 | 33 | 18 | 0 | 0 | 0 | 0 | 0 | 198 | 149 | 1 | 1 | 0.978 | 1 | 0.988 | 1 | 1 | 1 | 1 | 1 | 1 | 1 | -0.007 |
| SE | 84021 | ENSG00000111581 | NUP107 | chr12 | + | 68709237 | 68709309 | 68702735 | 68702784 | 68710004 | 68710093 | 138 | 91 | 129 | 152 | 206 | 0 | 0 | 1 | 0 | 0 | 21 | 56 | 44 | 36 | 27 | 0 | 0 | 0 | 0 | 0 | 221 | 149 | 1 | 1 | 1 | 1 | 0.989 | 1 | 1 | 1 | 1 | 1 | 1 | 1 | -0.002 |
| SE | 84023 | ENSG00000111581 | NUP107 | chr12 | + | 68721117 | 68721177 | 68719577 | 68719654 | 68721840 | 68721986 | 75 | 50 | 55 | 110 | 78 | 1 | 0 | 2 | 1 | 0 | 21 | 49 | 44 | 10 | 45 | 0 | 0 | 2 | 0 | 3 | 209 | 149 | 1 | 1 | 0.982 | 1 | 0.951 | 0.987 | 1 | 1 | 1 | 0.94 | 1 | 0.914 | 0.013 |
| SE | 84024 | ENSG00000111581 | NUP107 | chr12 | + | 68725726 | 68725796 | 68722103 | 68722152 | 68726498 | 68726617 | 175 | 141 | 134 | 139 | 250 | 0 | 0 | 0 | 0 | 0 | 63 | 59 | 83 | 12 | 39 | 0 | 0 | 1 | 0 | 0 | 219 | 149 | 1 | 1 | 1 | 1 | 1 | 1 | 1 | 1 | 1 | 0.983 | 1 | 1 | 0.003 |
| SE | 84025 | ENSG00000111581 | NUP107 | chr12 | + | 68731109 | 68731260 | 68727350 | 68727389 | 68731606 | 68731719 | 255 | 250 | 263 | 239 | 284 | 1 | 0 | 0 | 2 | 0 | 30 | 182 | 138 | 94 | 73 | 0 | 2 | 2 | 0 | 0 | 298 | 149 | 1 | 1 | 0.992 | 1 | 1 | 0.984 | 1 | 1 | 0.978 | 0.972 | 1 | 1 | 0.005 |
| SE | 84026 | ENSG00000111581 | NUP107 | chr12 | + | 68731606 | 68731719 | 68727350 | 68727389 | 68732636 | 68732739 | 65 | 65 | 38 | 82 | 44 | 1 | 1 | 1 | 1 | 0 | 0 | 23 | 15 | 26 | 8 | 0 | 2 | 4 | 2 | 0 | 262 | 149 | 1 | 1 | 0.974 | 0.974 | 0.956 | 0.979 | 1 | NA | 0.867 | 0.681 | 0.881 | 1 | 0.119 |
| SE | 84027 | ENSG00000111581 | NUP107 | chr12 | + | 68735230 | 68735344 | 68734707 | 68734833 | 68741812 | 68741980 | 164 | 137 | 96 | 182 | 151 | 0 | 0 | 0 | 0 | 0 | 27 | 79 | 43 | 35 | 5 | 0 | 0 | 1 | 0 | 0 | 263 | 149 | 1 | 1 | 1 | 1 | 1 | 1 | 1 | 1 | 1 | 0.961 | 1 | 1 | 0.008 |
| SE | 84028 | ENSG00000110848 | CD69 | chr12 | - | 9754586 | 9754690 | 9752485 | 9753589 | 9755061 | 9755261 | 651 | 310 | 693 | 964 | 1739 | 1 | 0 | 0 | 0 | 0 | 8 | 592 | 185 | 41 | 13 | 0 | 0 | 0 | 0 | 0 | 253 | 149 | 1 | 1 | 0.997 | 1 | 1 | 1 | 1 | 1 | 1 | 1 | 1 | 1 | -0.001 |
| SE | 84029 | ENSG00000110848 | CD69 | chr12 | - | 9755061 | 9755261 | 9754138 | 9754690 | 9756296 | 9756419 | 977 | 435 | 958 | 1409 | 2082 | 2 | 1 | 0 | 5 | 0 | 36 | 956 | 268 | 75 | 32 | 0 | 0 | 0 | 0 | 0 | 298 | 149 | 1 | 1 | 0.996 | 0.995 | 1 | 0.993 | 1 | 1 | 1 | 1 | 1 | 1 | -0.003 |
| SE | 84030 | ENSG00000110848 | CD69 | chr12 | - | 9756296 | 9756419 | 9754567 | 9755261 | 9760756 | 9760901 | 815 | 356 | 790 | 1093 | 1483 | 0 | 0 | 0 | 1 | 0 | 30 | 795 | 205 | 51 | 46 | 0 | 0 | 0 | 0 | 0 | 272 | 149 | 1 | 1 | 1 | 1 | 1 | 0.998 | 1 | 1 | 1 | 1 | 1 | 1 | 0 |
| SE | 84032 | ENSG00000178761 | FAM219B | chr15 | - | 74901168 | 74901256 | 74899991 | 74900137 | 74901949 | 74902063 | 0 | 0 | 1 | 0 | 0 | 2 | 3 | 3 | 1 | 4 | 0 | 0 | 0 | 0 | 0 | 0 | 9 | 3 | 6 | 28 | 237 | 149 | 1 | 1 | 0 | 0 | 0.173 | 0 | 0 | NA | 0 | 0 | 0 | 0 | 0.035 |
| SE | 84033 | ENSG00000178761 | FAM219B | chr15 | - | 74901949 | 74902063 | 74899991 | 74900137 | 74902603 | 74902786 | 6 | 8 | 8 | 5 | 6 | 0 | 0 | 0 | 0 | 0 | 0 | 18 | 6 | 11 | 34 | 0 | 0 | 1 | 4 | 4 | 263 | 149 | 1 | 1 | 1 | 1 | 1 | 1 | 1 | NA | 1 | 0.773 | 0.609 | 0.828 | 0.198 |
| SE | 84038 | ENSG00000178761 | FAM219B | chr15 | - | 74906323 | 74906365 | 74905153 | 74905231 | 74906586 | 74906848 | 3 | 1 | 2 | 3 | 1 | 5 | 8 | 3 | 5 | 0 | 0 | 3 | 0 | 0 | 8 | 0 | 6 | 0 | 2 | 0 | 191 | 149 | 1 | 1 | 0.319 | 0.089 | 0.342 | 0.319 | 1 | NA | 0.281 | NA | 0 | 1 | -0.013 |
| SE | 84039 | ENSG00000182220 | ATP6AP2 | chrX | + | 40588985 | 40589116 | 40580893 | 40581102 | 40591233 | 40591365 | 334 | 164 | 239 | 273 | 337 | 3 | 0 | 2 | 1 | 0 | 261 | 250 | 238 | 269 | 138 | 0 | 0 | 8 | 0 | 0 | 280 | 149 | 1 | 1 | 0.983 | 1 | 0.985 | 0.993 | 1 | 1 | 1 | 0.941 | 1 | 1 | 0.004 |
| SE | 84040 | ENSG00000182220 | ATP6AP2 | chrX | + | 40591233 | 40591365 | 40588985 | 40589116 | 40597248 | 40597344 | 565 | 294 | 419 | 438 | 664 | 0 | 0 | 0 | 2 | 0 | 338 | 415 | 424 | 448 | 238 | 0 | 0 | 1 | 0 | 0 | 281 | 149 | 1 | 1 | 1 | 1 | 1 | 0.991 | 1 | 1 | 1 | 0.996 | 1 | 1 | -0.001 |
| SE | 84041 | ENSG00000182220 | ATP6AP2 | chrX | + | 40597248 | 40597344 | 40591233 | 40591365 | 40597526 | 40597661 | 627 | 336 | 501 | 528 | 887 | 4 | 2 | 2 | 1 | 4 | 411 | 543 | 475 | 507 | 270 | 0 | 3 | 2 | 0 | 6 | 245 | 149 | 1 | 1 | 0.99 | 0.99 | 0.993 | 0.997 | 0.993 | 1 | 0.991 | 0.993 | 1 | 0.965 | 0.003 |
| SE | 84045 | ENSG00000182220 | ATP6AP2 | chrX | + | 40598651 | 40598734 | 40597526 | 40597664 | 40599591 | 40599741 | 80 | 53 | 50 | 76 | 108 | 2 | 0 | 0 | 0 | 0 | 64 | 59 | 77 | 100 | 19 | 0 | 0 | 0 | 0 | 0 | 232 | 149 | 1 | 1 | 0.963 | 1 | 1 | 1 | 1 | 1 | 1 | 1 | 1 | 1 | -0.007 |
| SE | 84047 | ENSG00000182220 | ATP6AP2 | chrX | + | 40598680 | 40598734 | 40597526 | 40597664 | 40599591 | 40599741 | 465 | 321 | 324 | 389 | 651 | 2 | 0 | 0 | 0 | 0 | 227 | 385 | 367 | 384 | 136 | 0 | 0 | 0 | 0 | 0 | 203 | 149 | 1 | 1 | 0.994 | 1 | 1 | 1 | 1 | 1 | 1 | 1 | 1 | 1 | -0.001 |
| SE | 84048 | ENSG00000182220 | ATP6AP2 | chrX | + | 40598935 | 40599018 | 40598680 | 40598734 | 40599591 | 40599741 | 6 | 2 | 6 | 3 | 0 | 383 | 266 | 253 | 326 | 535 | 0 | 2 | 11 | 0 | 0 | 199 | 314 | 298 | 353 | 104 | 232 | 149 | 1 | 1 | 0.01 | 0.005 | 0.015 | 0.006 | 0 | 0 | 0.004 | 0.023 | 0 | 0 | 0.002 |
| SE | 84049 | ENSG00000182220 | ATP6AP2 | chrX | + | 40600761 | 40600881 | 40599591 | 40599741 | 40605560 | 40605594 | 1037 | 612 | 630 | 718 | 927 | 1 | 0 | 0 | 3 | 0 | 674 | 846 | 855 | 1037 | 345 | 0 | 0 | 3 | 6 | 0 | 269 | 149 | 1 | 1 | 0.998 | 1 | 1 | 0.993 | 1 | 1 | 1 | 0.994 | 0.99 | 1 | 0.001 |
| SE | 84052 | ENSG00000182220 | ATP6AP2 | chrX | + | 40603336 | 40603439 | 40600777 | 40600881 | 40605560 | 40605887 | 1 | 0 | 3 | 0 | 0 | 579 | 315 | 370 | 391 | 544 | 0 | 0 | 0 | 0 | 0 | 302 | 512 | 436 | 550 | 183 | 252 | 149 | 1 | 1 | 0.001 | 0 | 0.005 | 0 | 0 | 0 | 0 | 0 | 0 | 0 | 0.001 |
| SE | 84053 | ENSG00000197157 | SND1 | chr7 | + | 127686612 | 127686762 | 127652193 | 127652451 | 127694827 | 127694948 | 415 | 185 | 298 | 301 | 145 | 1 | 0 | 0 | 0 | 0 | 464 | 315 | 295 | 322 | 442 | 0 | 0 | 0 | 0 | 0 | 298 | 149 | 1 | 1 | 0.995 | 1 | 1 | 1 | 1 | 1 | 1 | 1 | 1 | 1 | -0.001 |
| SE | 84054 | ENSG00000197157 | SND1 | chr7 | + | 127694827 | 127694948 | 127686612 | 127686762 | 127698874 | 127698937 | 452 | 215 | 345 | 389 | 277 | 0 | 0 | 0 | 1 | 0 | 704 | 361 | 357 | 454 | 562 | 0 | 4 | 0 | 4 | 0 | 270 | 149 | 1 | 1 | 1 | 1 | 1 | 0.995 | 1 | 1 | 0.98 | 1 | 0.984 | 1 | 0.006 |
| SE | 84055 | ENSG00000197157 | SND1 | chr7 | + | 127702434 | 127702526 | 127701162 | 127701323 | 127703164 | 127703191 | 464 | 276 | 488 | 445 | 725 | 0 | 0 | 0 | 0 | 1 | 724 | 473 | 435 | 383 | 683 | 0 | 1 | 0 | 0 | 0 | 241 | 149 | 1 | 1 | 1 | 1 | 1 | 1 | 0.998 | 1 | 0.997 | 1 | 1 | 1 | 0 |
| SE | 84056 | ENSG00000197157 | SND1 | chr7 | + | 127703164 | 127703323 | 127701162 | 127701323 | 127704838 | 127704945 | 457 | 304 | 463 | 497 | 407 | 0 | 0 | 0 | 0 | 0 | 762 | 552 | 471 | 410 | 935 | 0 | 0 | 1 | 0 | 0 | 298 | 149 | 1 | 1 | 1 | 1 | 1 | 1 | 1 | 1 | 1 | 0.996 | 1 | 1 | 0.001 |
| SE | 84057 | ENSG00000197157 | SND1 | chr7 | + | 127707556 | 127707647 | 127704838 | 127704945 | 127721286 | 127721400 | 561 | 434 | 638 | 641 | 480 | 0 | 0 | 1 | 0 | 0 | 1123 | 690 | 663 | 714 | 1160 | 0 | 0 | 3 | 0 | 0 | 240 | 149 | 1 | 1 | 1 | 1 | 0.997 | 1 | 1 | 1 | 1 | 0.993 | 1 | 1 | 0.001 |
| SE | 84058 | ENSG00000197157 | SND1 | chr7 | + | 127721286 | 127721400 | 127707556 | 127707647 | 127807483 | 127807573 | 704 | 511 | 801 | 803 | 682 | 0 | 0 | 0 | 2 | 0 | 1248 | 789 | 711 | 800 | 1464 | 0 | 0 | 0 | 0 | 0 | 263 | 149 | 1 | 1 | 1 | 1 | 1 | 0.996 | 1 | 1 | 1 | 1 | 1 | 1 | -0.001 |
| SE | 84059 | ENSG00000197157 | SND1 | chr7 | + | 127807483 | 127807573 | 127721286 | 127721400 | 127844323 | 127844424 | 610 | 470 | 671 | 728 | 713 | 1 | 0 | 0 | 0 | 0 | 1058 | 687 | 636 | 697 | 1254 | 0 | 2 | 0 | 0 | 0 | 239 | 149 | 1 | 1 | 0.997 | 1 | 1 | 1 | 1 | 1 | 0.995 | 1 | 1 | 1 | 0 |
| SE | 84060 | ENSG00000197157 | SND1 | chr7 | + | 127844323 | 127844424 | 127807483 | 127807573 | 127887901 | 127887949 | 434 | 298 | 496 | 538 | 563 | 0 | 0 | 0 | 0 | 0 | 587 | 435 | 413 | 459 | 753 | 0 | 0 | 3 | 3 | 0 | 250 | 149 | 1 | 1 | 1 | 1 | 1 | 1 | 1 | 1 | 1 | 0.988 | 0.989 | 1 | 0.005 |
| SE | 84062 | ENSG00000197157 | SND1 | chr7 | + | 127990946 | 127991056 | 127929187 | 127929329 | 128008033 | 128008144 | 173 | 129 | 188 | 171 | 247 | 0 | 0 | 0 | 1 | 0 | 291 | 176 | 190 | 173 | 227 | 0 | 0 | 0 | 0 | 0 | 259 | 149 | 1 | 1 | 1 | 1 | 1 | 0.99 | 1 | 1 | 1 | 1 | 1 | 1 | -0.002 |
| SE | 84063 | ENSG00000197157 | SND1 | chr7 | + | 127990946 | 127991056 | 127929187 | 127929329 | 128074501 | 128074690 | 312 | 209 | 307 | 309 | 341 | 1 | 1 | 0 | 0 | 0 | 457 | 331 | 342 | 310 | 435 | 0 | 0 | 2 | 0 | 0 | 259 | 149 | 1 | 1 | 0.994 | 0.992 | 1 | 1 | 1 | 1 | 1 | 0.99 | 1 | 1 | -0.001 |
| SE | 84064 | ENSG00000197157 | SND1 | chr7 | + | 128008033 | 128008144 | 127929187 | 127929329 | 128074501 | 128074690 | 1 | 0 | 0 | 2 | 0 | 1 | 1 | 0 | 0 | 0 | 0 | 0 | 0 | 1 | 0 | 0 | 0 | 2 | 0 | 0 | 260 | 149 | 1 | 1 | 0.364 | 0 | NA | 1 | NA | NA | NA | 0 | 1 | NA | -0.045 |
| SE | 84067 | ENSG00000197157 | SND1 | chr7 | + | 128081359 | 128081501 | 127990946 | 127991056 | 128091836 | 128091881 | 1 | 4 | 3 | 0 | 3 | 0 | 0 | 0 | 0 | 0 | 12 | 8 | 0 | 0 | 0 | 0 | 0 | 0 | 1 | 0 | 291 | 149 | 1 | 1 | 1 | 1 | 1 | NA | 1 | 1 | 1 | NA | 0 | NA | 0.333 |
| SE | 84068 | ENSG00000197157 | SND1 | chr7 | + | 128084723 | 128084847 | 128081359 | 128081501 | 128085710 | 128085780 | 401 | 228 | 420 | 320 | 450 | 0 | 1 | 0 | 0 | 0 | 514 | 345 | 351 | 346 | 656 | 8 | 0 | 0 | 0 | 4 | 273 | 149 | 1 | 1 | 1 | 0.992 | 1 | 1 | 1 | 0.972 | 1 | 1 | 1 | 0.989 | 0.006 |
| SE | 84069 | ENSG00000197157 | SND1 | chr7 | + | 128084723 | 128084847 | 128081359 | 128081501 | 128086937 | 128087051 | 185 | 102 | 212 | 166 | 176 | 0 | 0 | 0 | 0 | 0 | 300 | 163 | 159 | 199 | 332 | 0 | 0 | 0 | 2 | 0 | 273 | 149 | 1 | 1 | 1 | 1 | 1 | 1 | 1 | 1 | 1 | 1 | 0.982 | 1 | 0.004 |
| SE | 84070 | ENSG00000197157 | SND1 | chr7 | + | 128085710 | 128085780 | 128081359 | 128081501 | 128086937 | 128087051 | 140 | 66 | 169 | 123 | 169 | 0 | 0 | 0 | 0 | 0 | 244 | 81 | 106 | 129 | 235 | 0 | 0 | 0 | 2 | 0 | 219 | 149 | 1 | 1 | 1 | 1 | 1 | 1 | 1 | 1 | 1 | 1 | 0.978 | 1 | 0.004 |
| SE | 84071 | ENSG00000197157 | SND1 | chr7 | + | 128085710 | 128085780 | 128084723 | 128084847 | 128086937 | 128087051 | 356 | 191 | 377 | 277 | 443 | 0 | 0 | 0 | 0 | 0 | 450 | 263 | 300 | 276 | 549 | 0 | 0 | 2 | 0 | 0 | 219 | 149 | 1 | 1 | 1 | 1 | 1 | 1 | 1 | 1 | 1 | 0.99 | 1 | 1 | 0.002 |
| SE | 84072 | ENSG00000197157 | SND1 | chr7 | + | 128086937 | 128087051 | 128081359 | 128081501 | 128091836 | 128091881 | 0 | 0 | 0 | 0 | 0 | 0 | 0 | 1 | 0 | 0 | 0 | 2 | 0 | 11 | 0 | 0 | 0 | 0 | 0 | 0 | 263 | 149 | 1 | 1 | NA | NA | 0 | NA | NA | NA | 1 | NA | 1 | NA | -1 |
| SE | 84074 | ENSG00000102316 | MAGED2 | chrX | + | 54809302 | 54809376 | 54807598 | 54807802 | 54809721 | 54809808 | 142 | 82 | 88 | 112 | 102 | 2 | 0 | 0 | 0 | 0 | 75 | 63 | 44 | 46 | 230 | 0 | 0 | 0 | 0 | 0 | 223 | 149 | 1 | 1 | 0.979 | 1 | 1 | 1 | 1 | 1 | 1 | 1 | 1 | 1 | -0.004 |
| SE | 84075 | ENSG00000102316 | MAGED2 | chrX | + | 54809721 | 54810213 | 54809302 | 54809376 | 54810820 | 54811129 | 268 | 158 | 188 | 204 | 238 | 0 | 3 | 0 | 0 | 0 | 208 | 171 | 119 | 118 | 522 | 0 | 0 | 0 | 0 | 0 | 298 | 149 | 1 | 1 | 1 | 0.963 | 1 | 1 | 1 | 1 | 1 | 1 | 1 | 1 | -0.007 |
| SE | 84076 | ENSG00000102316 | MAGED2 | chrX | + | 54810117 | 54810213 | 54809302 | 54809376 | 54810820 | 54811129 | 79 | 55 | 60 | 61 | 106 | 0 | 3 | 0 | 0 | 0 | 66 | 52 | 44 | 37 | 155 | 0 | 0 | 0 | 0 | 0 | 245 | 149 | 1 | 1 | 1 | 0.918 | 1 | 1 | 1 | 1 | 1 | 1 | 1 | 1 | -0.016 |
| SE | 84077 | ENSG00000102316 | MAGED2 | chrX | + | 54810820 | 54811129 | 54809721 | 54810213 | 54812944 | 54813024 | 108 | 72 | 76 | 79 | 122 | 0 | 0 | 0 | 0 | 0 | 123 | 67 | 50 | 62 | 245 | 0 | 0 | 1 | 0 | 0 | 298 | 149 | 1 | 1 | 1 | 1 | 1 | 1 | 1 | 1 | 1 | 0.962 | 1 | 1 | 0.008 |
| SE | 84078 | ENSG00000102316 | MAGED2 | chrX | + | 54811249 | 54811313 | 54810820 | 54811129 | 54811573 | 54811644 | 103 | 77 | 105 | 111 | 138 | 0 | 0 | 0 | 2 | 0 | 106 | 75 | 47 | 73 | 251 | 0 | 0 | 2 | 3 | 0 | 213 | 149 | 1 | 1 | 1 | 1 | 1 | 0.975 | 1 | 1 | 1 | 0.943 | 0.945 | 1 | 0.017 |
| SE | 84080 | ENSG00000102316 | MAGED2 | chrX | + | 54813117 | 54813160 | 54812944 | 54813024 | 54813487 | 54813550 | 190 | 101 | 161 | 182 | 268 | 1 | 0 | 0 | 1 | 0 | 111 | 228 | 78 | 113 | 459 | 0 | 0 | 0 | 0 | 0 | 192 | 149 | 1 | 1 | 0.993 | 1 | 1 | 0.993 | 1 | 1 | 1 | 1 | 1 | 1 | -0.003 |

| | | | | | | | | | | | | | |
|---|---|---|---|---|---|---|---|---|---|---|---|---|---|
| SE | 84081 | ENSG00000102316 | MAGED2 | chrX | + | 54814660 | 54814775 | 54813487 | 54813550 | 54815247 | 54815690 | 262 | 126 |
| SE | 84083 | ENSG00000197892 | KIF13B | chr8 | - | 29075280 | 29075343 | 29071619 | 29072316 | 29099132 | 29099241 | 14 | 6 |
| SE | 84084 | ENSG00000197892 | KIF13B | chr8 | - | 29092744 | 29092878 | 29071619 | 29072316 | 29099132 | 29099241 | 50 | 35 |
| SE | 84086 | ENSG00000197892 | KIF13B | chr8 | - | 29109917 | 29110070 | 29109433 | 29109511 | 29113462 | 29113555 | 123 | 86 |
| SE | 84087 | ENSG00000197892 | KIF13B | chr8 | - | 29116830 | 29117007 | 29113462 | 29113555 | 29118867 | 29118992 | 90 | 66 |
| SE | 84088 | ENSG00000197892 | KIF13B | chr8 | - | 29124023 | 29124123 | 29123365 | 29123492 | 29126481 | 29126511 | 266 | 219 |
| SE | 84090 | ENSG00000197892 | KIF13B | chr8 | - | 29132307 | 29132465 | 29130532 | 29130665 | 29140062 | 29140191 | 95 | 85 |
| SE | 84092 | ENSG00000197892 | KIF13B | chr8 | - | 29146377 | 29146540 | 29142156 | 29142303 | 29147391 | 29147602 | 140 | 86 |
| SE | 84093 | ENSG00000197892 | KIF13B | chr8 | - | 29148576 | 29148767 | 29147391 | 29147602 | 29150296 | 29150383 | 199 | 243 |
| SE | 84094 | ENSG00000197892 | KIF13B | chr8 | - | 29150296 | 29150383 | 29148576 | 29148767 | 29155725 | 29155856 | 160 | 174 |
| SE | 84096 | ENSG00000197892 | KIF13B | chr8 | - | 29186291 | 29186472 | 29181918 | 29182006 | 29188514 | 29188607 | 256 | 216 |
| SE | 84098 | ENSG00000197892 | KIF13B | chr8 | - | 29249984 | 29250084 | 29245345 | 29245439 | 29262979 | 29263062 | 0 | 2 |
| SE | 84118 | ENSG00000203965 | EFCAB7 | chr1 | + | 63531819 | 63532031 | 63525571 | 63525759 | 63532669 | 63532696 | 53 | 52 |
| SE | 84119 | ENSG00000203965 | EFCAB7 | chr1 | + | 63555357 | 63555515 | 63551724 | 63551834 | 63557113 | 63557247 | 31 | 23 |
| SE | 84121 | ENSG00000140543 | DET1 | chr15 | - | 88530622 | 88531473 | 88527598 | 88527786 | 88531612 | 88531715 | 29 | 13 |
| SE | 84123 | ENSG00000140543 | DET1 | chr15 | - | 88536311 | 88536381 | 88530622 | 88531715 | 88546539 | 88546653 | 0 | 1 |
| SE | 84124 | ENSG00000094841 | UPRT | chrX | + | 75291392 | 75291560 | 75274126 | 75274640 | 75293471 | 75293514 | 2 | 0 |
| SE | 84125 | ENSG00000094841 | UPRT | chrX | + | 75291674 | 75291747 | 75291392 | 75291560 | 75293471 | 75293514 | 2 | 0 |
| SE | 84127 | ENSG00000094841 | UPRT | chrX | + | 75293471 | 75293514 | 75274727 | 75274758 | 75296341 | 75296411 | 21 | 13 |
| SE | 84129 | ENSG00000094841 | UPRT | chrX | + | 75296341 | 75296411 | 75293471 | 75293514 | 75297490 | 75297553 | 68 | 49 |
| SE | 84133 | ENSG00000229151 | NA | chrX | - | 51340072 | 51340132 | 51329423 | 51329565 | 51356943 | 51357027 | 6 | 4 |
| SE | 84134 | ENSG00000229151 | NA | chrX | - | 51340283 | 51340402 | 51340072 | 51340132 | 51356943 | 51357027 | 6 | 3 |
| SE | 84135 | ENSG00000115816 | CEBPZ | chr2 | - | 37210998 | 37211082 | 37202949 | 37203008 | 37211842 | 37212039 | 143 | 91 |
| SE | 84136 | ENSG00000115816 | CEBPZ | chr2 | - | 37216983 | 37217037 | 37216315 | 37216418 | 37220384 | 37220473 | 261 | 143 |
| SE | 84137 | ENSG00000115816 | CEBPZ | chr2 | - | 37222379 | 37222563 | 37220384 | 37220473 | 37223169 | 37223401 | 324 | 204 |
| SE | 84138 | ENSG00000115816 | CEBPZ | chr2 | - | 37223169 | 37223401 | 37220384 | 37220473 | 37227543 | 37229036 | 174 | 109 |
| SE | 84139 | ENSG00000165259 | HDX | chrX | - | 84440531 | 84440585 | 84361465 | 84361612 | 84468471 | 84469575 | 3 | 8 |
| SE | 84140 | ENSG00000165259 | HDX | chrX | - | 84468471 | 84469575 | 84440531 | 84440585 | 84475250 | 84475397 | 23 | 21 |
| SE | 84141 | ENSG00000165259 | HDX | chrX | - | 84468471 | 84469575 | 84440531 | 84440585 | 84488023 | 84488132 | 7 | 9 |
| SE | 84142 | ENSG00000165259 | HDX | chrX | - | 84475250 | 84475397 | 84440531 | 84440585 | 84488023 | 84488132 | 2 | 3 |
| SE | 84143 | ENSG00000165259 | HDX | chrX | - | 84475250 | 84475397 | 84468471 | 84469575 | 84488023 | 84488132 | 18 | 15 |
| SE | 84144 | ENSG00000165259 | HDX | chrX | - | 84475250 | 84475397 | 84468471 | 84469575 | 84502341 | 84502453 | 20 | 15 |
| SE | 84145 | ENSG00000165259 | HDX | chrX | - | 84488023 | 84488132 | 84468471 | 84469575 | 84502341 | 84502453 | 0 | 0 |
| SE | 84146 | ENSG00000165259 | HDX | chrX | - | 84488023 | 84488132 | 84475250 | 84475397 | 84501428 | 84501511 | 2 | 3 |
| SE | 84147 | ENSG00000165259 | HDX | chrX | - | 84488023 | 84488132 | 84475250 | 84475397 | 84502341 | 84502453 | 2 | 3 |
| SE | 84148 | ENSG00000165259 | HDX | chrX | - | 84501428 | 84501511 | 84475250 | 84475397 | 84502341 | 84502453 | 2 | 0 |
| SE | 84149 | ENSG00000165259 | HDX | chrX | - | 84501428 | 84501511 | 84488023 | 84488132 | 84502341 | 84502453 | 2 | 0 |
| SE | 84150 | ENSG00000140511 | HAPLN3 | chr15 | - | 88878966 | 88879269 | 88877293 | 88878256 | 88881356 | 88881725 | 18 | 10 |
| SE | 84151 | ENSG00000140511 | HAPLN3 | chr15 | - | 88879417 | 88879452 | 88879118 | 88879269 | 88881356 | 88881725 | 1 | 2 |
| SE | 84158 | ENSG00000124333 | VAMP7 | chrX | + | 155895622 | 155895680 | 155889457 | 155889608 | 155898111 | 155898249 | 83 | 48 |
| SE | 84160 | ENSG00000124333 | VAMP7 | chrX | + | 155895622 | 155895680 | 155889457 | 155889633 | 155898111 | 155898249 | 103 | 63 |
| SE | 84161 | ENSG00000124333 | VAMP7 | chrX | + | 155900496 | 155900587 | 155898111 | 155898249 | 155919812 | 155919880 | 436 | 318 |
| SE | 84162 | ENSG00000124333 | VAMP7 | chrX | + | 155900496 | 155900587 | 155898111 | 155898249 | 155939700 | 155939793 | 221 | 190 |
| SE | 84166 | ENSG00000137642 | SORL1 | chr11 | + | 121478117 | 121478243 | 121470006 | 121470123 | 121488031 | 121488193 | 1185 | 617 |
| SE | 84167 | ENSG00000137642 | SORL1 | chr11 | + | 121488031 | 121488193 | 121478117 | 121478243 | 121490042 | 121490110 | 918 | 624 |
| SE | 84168 | ENSG00000137642 | SORL1 | chr11 | + | 121488031 | 121488193 | 121478117 | 121478243 | 121496868 | 121497049 | 538 | 356 |
| SE | 84169 | ENSG00000137642 | SORL1 | chr11 | + | 121490042 | 121490110 | 121478117 | 121478243 | 121496868 | 121497049 | 211 | 83 |
| SE | 84170 | ENSG00000137642 | SORL1 | chr11 | + | 121490042 | 121490110 | 121488031 | 121488193 | 121496868 | 121497049 | 589 | 351 |
| SE | 84171 | ENSG00000137642 | SORL1 | chr11 | + | 121496868 | 121497049 | 121478117 | 121478243 | 121513002 | 121513104 | 907 | 543 |
| SE | 84172 | ENSG00000137642 | SORL1 | chr11 | + | 121513002 | 121513104 | 121496868 | 121497049 | 121514151 | 121514321 | 1679 | 1054 |
| SE | 84174 | ENSG00000137642 | SORL1 | chr11 | + | 121520656 | 121520849 | 121496868 | 121497049 | 121532463 | 121532552 | 2 | 0 |
| SE | 84175 | ENSG00000137642 | SORL1 | chr11 | + | 121522585 | 121522703 | 121520656 | 121520849 | 121522915 | 121522989 | 437 | 191 |
| SE | 84176 | ENSG00000137642 | SORL1 | chr11 | + | 121522915 | 121522989 | 121520656 | 121520849 | 121532463 | 121532552 | 327 | 163 |
| SE | 84177 | ENSG00000137642 | SORL1 | chr11 | + | 121532463 | 121532552 | 121522915 | 121522989 | 121543547 | 121543726 | 844 | 476 |
| SE | 84178 | ENSG00000137642 | SORL1 | chr11 | + | 121543547 | 121543726 | 121532463 | 121532552 | 121545242 | 121545429 | 1808 | 1060 |
| SE | 84179 | ENSG00000137642 | SORL1 | chr11 | + | 121543547 | 121543726 | 121532463 | 121532552 | 121549959 | 121550088 | 766 | 438 |
| SE | 84181 | ENSG00000137642 | SORL1 | chr11 | + | 121545242 | 121545429 | 121543547 | 121543726 | 121549959 | 121550088 | 1385 | 818 |
| SE | 84183 | ENSG00000137642 | SORL1 | chr11 | + | 121550584 | 121550670 | 121545242 | 121545429 | 121553936 | 121554109 | 178 | 78 |
| SE | 84185 | ENSG00000137642 | SORL1 | chr11 | + | 121559518 | 121559657 | 121558590 | 121558837 | 121566939 | 121567113 | 1345 | 655 |
| SE | 84187 | ENSG00000137642 | SORL1 | chr11 | + | 121577280 | 121577400 | 121574240 | 121574363 | 121583457 | 121583583 | 891 | 506 |
| SE | 84188 | ENSG00000137642 | SORL1 | chr11 | + | 121604192 | 121604324 | 121595622 | 121595772 | 121605112 | 121605239 | 694 | 300 |
| SE | 84194 | ENSG00000137642 | SORL1 | chr11 | + | 121625084 | 121625277 | 121622161 | 121622268 | 121627554 | 121627767 | 1312 | 775 |
| SE | 84195 | ENSG00000137642 | SORL1 | chr11 | + | 121625084 | 121625277 | 121622161 | 121622268 | 121629495 | 121629678 | 658 | 388 |
| SE | 84196 | ENSG00000137642 | SORL1 | chr11 | + | 121627554 | 121627767 | 121622161 | 121622268 | 121629495 | 121629678 | 423 | 225 |
| SE | 84197 | ENSG00000137642 | SORL1 | chr11 | + | 121627554 | 121627767 | 121625084 | 121625277 | 121629495 | 121629678 | 1078 | 613 |
| SE | 84206 | ENSG00000068354 | TBC1D25 | chrX | + | 48541332 | 48541442 | 48539713 | 48539920 | 48558896 | 48559024 | 14 | 0 |
| SE | 84208 | ENSG00000068354 | TBC1D25 | chrX | + | 48558896 | 48559024 | 48544868 | 48545023 | 48559157 | 48559284 | 82 | 28 |
| SE | 84209 | ENSG00000273398 | NA | chr2 | - | 68141693 | 68141795 | 68138741 | 68138857 | 68144584 | 68144715 | 83 | 93 |
| SE | 84211 | ENSG00000214783 | POLR2J4 | chr7 | - | 43986600 | 43986677 | 43973209 | 43973323 | 44013592 | 44013679 | 1 | 1 |
| SE | 84212 | ENSG00000214783 | POLR2J4 | chr7 | - | 43987677 | 43987888 | 43986600 | 43986677 | 44014605 | 44014783 | 7 | 9 |
| SE | 84213 | ENSG00000214783 | POLR2J4 | chr7 | - | 43988085 | 43988247 | 43986600 | 43986677 | 44013592 | 44013679 | 16 | 10 |
| SE | 84214 | ENSG00000214783 | POLR2J4 | chr7 | - | 43988085 | 43988247 | 43986600 | 43986677 | 44014605 | 44014783 | 0 | 0 |
| SE | 84216 | ENSG00000214783 | POLR2J4 | chr7 | - | 44013592 | 44013679 | 43986600 | 43986677 | 44014605 | 44014783 | 2 | 4 |
| SE | 84217 | ENSG00000214783 | POLR2J4 | chr7 | - | 44013592 | 44013679 | 43988085 | 43988247 | 44014605 | 44014783 | 12 | 10 |
| SE | 84223 | ENSG00000158813 | EDA | chrX | + | 70033397 | 70033528 | 70030468 | 70030520 | 70035357 | 70039469 | 1 | 1 |
| SE | 84225 | ENSG00000188419 | CHM | chrX | - | 85878964 | 85879063 | 85873051 | 85873212 | 85894187 | 85894284 | 71 | 49 |
| SE | 84226 | ENSG00000188419 | CHM | chrX | - | 85978766 | 85978891 | 85963664 | 85964052 | 86027490 | 86027557 | 62 | 76 |
| SE | 84227 | ENSG00000188419 | CHM | chrX | - | 85981736 | 85981809 | 85978655 | 85978891 | 86027490 | 86027557 | 59 | 48 |
| SE | 84229 | ENSG00000260302 | LINC01882 | chr18 | - | 12765276 | 12765455 | 12749293 | 12749533 | 12775668 | 12775821 | 7 | 1 |
| SE | 84235 | ENSG00000112305 | SMAP1 | chr6 | + | 70798656 | 70798737 | 70773349 | 70773425 | 70836940 | 70837028 | 58 | 52 |
| SE | 84236 | ENSG00000112305 | SMAP1 | chr6 | + | 70856858 | 70857030 | 70852539 | 70852664 | 70860199 | 70861166 | 34 | 33 |
| SE | 84238 | ENSG00000112305 | SMAP1 | chr6 | + | 70859328 | 70859416 | 70857921 | 70858229 | 70860199 | 70861166 | 1 | 0 |
| SE | 84242 | ENSG00000184867 | ARMCX2 | chrX | - | 101659075 | 101659136 | 101658460 | 101658525 | 101659757 | 101659844 | 3 | 1 |
| SE | 84244 | ENSG00000184867 | ARMCX2 | chrX | - | 101659075 | 101659148 | 101658460 | 101658525 | 101659757 | 101659834 | 0 | 1 |
| SE | 84248 | ENSG00000115241 | PPM1G | chr2 | - | 27382125 | 27382228 | 27381194 | 27381805 | 27382475 | 27382605 | 718 | 478 |
| SE | 84249 | ENSG00000115241 | PPM1G | chr2 | - | 27382475 | 27382605 | 27382125 | 27382228 | 27383365 | 27383600 | 598 | 381 |
| SE | 84250 | ENSG00000115241 | PPM1G | chr2 | - | 27382475 | 27382605 | 27382125 | 27382228 | 27383951 | 27384092 | 416 | 289 |
| SE | 84251 | ENSG00000115241 | PPM1G | chr2 | - | 27383365 | 27383600 | 27382125 | 27382228 | 27383951 | 27384092 | 318 | 194 |
| SE | 84253 | ENSG00000115241 | PPM1G | chr2 | - | 27383951 | 27384092 | 27383365 | 27383600 | 27384672 | 27384767 | 658 | 375 |
| SE | 84254 | ENSG00000115241 | PPM1G | chr2 | - | 27384672 | 27384767 | 27383951 | 27384092 | 27384974 | 27387158 | 215 | 122 |
| SE | 84255 | ENSG00000115241 | PPM1G | chr2 | - | 27384672 | 27385088 | 27383951 | 27384092 | 27385746 | 27385879 | 435 | 259 |
| SE | 84257 | ENSG00000172943 | PHF8 | chrX | - | 53985815 | 53985949 | 53984913 | 53985227 | 53987077 | 53987163 | 208 | 147 |
| SE | 84260 | ENSG00000172943 | PHF8 | chrX | - | 53995130 | 53995205 | 53992793 | 53992839 | 53995692 | 53995782 | 0 | 0 |
| SE | 84261 | ENSG00000172943 | PHF8 | chrX | - | 54011121 | 54011284 | 54002594 | 54002682 | 54014376 | 54014449 | 195 | 102 |
| SE | 84262 | ENSG00000172943 | PHF8 | chrX | - | 54014376 | 54014563 | 54011121 | 54011284 | 54016594 | 54016736 | 254 | 137 |
| SE | 84263 | ENSG00000172943 | PHF8 | chrX | - | 54016594 | 54016736 | 54014376 | 54014563 | 54017660 | 54017821 | 221 | 124 |
| SE | 84264 | ENSG00000172943 | PHF8 | chrX | - | 54016594 | 54016736 | 54014376 | 54014563 | 54022258 | 54022262 | 124 | 69 |
| SE | 84265 | ENSG00000172943 | PHF8 | chrX | - | 54017660 | 54017821 | 54014376 | 54014563 | 54022258 | 54022262 | 86 | 49 |
| SE | 84266 | ENSG00000172943 | PHF8 | chrX | - | 54017660 | 54017821 | 54016594 | 54016736 | 54022258 | 54022262 | 183 | 105 |
| SE | 84267 | ENSG00000198182 | ZNF607 | chr19 | - | 37706381 | 37706583 | 37699324 | 37699895 | 37707913 | 37708009 | 0 | 2 |
| SE | 84268 | ENSG00000198182 | ZNF607 | chr19 | - | 37711609 | 37711692 | 37709695 | 37709822 | 37719268 | 37719723 | 10 | 13 |
| SE | 84271 | ENSG00000255730 | NA | chr19 | + | 41378280 | 41378519 | 41376756 | 41376854 | 41410636 | 41410816 | 14 | 5 |
| SE | 84273 | ENSG00000075218 | GTSE1 | chr22 | + | 46308149 | 46308207 | 46297379 | 46297479 | 46308318 | 46308943 | 2 | 0 |
| SE | 84276 | ENSG00000109736 | MFSD10 | chr4 | - | 2931554 | 2931663 | 2931374 | 2931465 | 2931824 | 2931936 | 31 | 4 |
| SE | 84277 | ENSG00000109736 | MFSD10 | chr4 | - | 2932350 | 2932409 | 2932044 | 2932153 | 2932599 | 2932758 | 6 | 0 |
| SE | 84278 | ENSG00000109736 | MFSD10 | chr4 | - | 2932350 | 2932512 | 2931824 | 2932153 | 2932599 | 2932690 | 33 | 6 |
| SE | 84279 | ENSG00000105327 | BBC3 | chr19 | - | 47226563 | 47226754 | 47220823 | 47221918 | 47230928 | 47231199 | 2 | 5 |
| SE | 84283 | ENSG00000169598 | DFFB | chr1 | + | 3860419 | 3860506 | 3858717 | 3858844 | 3865811 | 3866000 | 0 | 1 |
| SE | 84285 | ENSG00000169598 | DFFB | chr1 | + | 3872471 | 3872572 | 3869604 | 3869775 | 3883506 | 3885420 | 24 | 16 |
| SE | 84287 | ENSG00000165813 | CCDC186 | chr10 | - | 114125885 | 114126105 | 114120861 | 114125226 | 114127460 | 114127671 | 324 | 464 |
| SE | 84289 | ENSG00000165813 | CCDC186 | chr10 | - | 114162636 | 114163329 | 114157553 | 114157680 | 114173140 | 114173265 | 89 | 91 |
| SE | 84291 | ENSG00000165813 | CCDC186 | chr10 | - | 114173140 | 114173265 | 114157553 | 114157680 | 114174014 | 114174202 | 18 | 17 |
| SE | 84297 | ENSG00000172613 | RAD9A | chr11 | + | 67393690 | 67393790 | 67392881 | 67393610 | 67395715 | 67395825 | 16 | 6 |
| SE | 84298 | ENSG00000172613 | RAD9A | chr11 | + | 67393722 | 67393790 | 67393495 | 67393610 | 67395715 | 67395807 | 3 | 2 |
| SE | 84300 | ENSG00000147202 | DIAPH2 | chrX | + | 96735757 | 96735790 | 96684662 | 96685190 | 96738585 | 96738762 | 12 | 26 |
| SE | 84301 | ENSG00000147202 | DIAPH2 | chrX | + | 96758153 | 96758258 | 96738585 | 96738762 | 96763099 | 96763120 | 203 | 204 |
| SE | 84303 | ENSG00000147202 | DIAPH2 | chrX | + | 96763099 | 96763120 | 96738585 | 96738762 | 96881578 | 96881718 | 4 | 7 |
| SE | 84306 | ENSG00000147202 | DIAPH2 | chrX | + | 96948934 | 96949039 | 96945526 | 96945591 | 96957827 | 96958148 | 300 | 364 |
| SE | 84307 | ENSG00000181220 | ZNF746 | chr7 | - | 149477563 | 149477755 | 149476921 | 149477047 | 149492858 | 149492972 | 161 | 74 |
| SE | 84311 | ENSG00000239407 | NA | chrX | + | 102901022 | 102901133 | 102900749 | 102900832 | 102901630 | 102901771 | 16 | 22 |
| SE | 84314 | ENSG00000164062 | APEH | chr3 | + | 49675890 | 49675966 | 49675182 | 49675787 | 49676608 | 49676700 | 117 | 55 |
| SE | 84316 | ENSG00000164062 | APEH | chr3 | + | 49676055 | 49676219 | 49675890 | 49675966 | 49676608 | 49676700 | 315 | 180 |
| SE | 84317 | ENSG00000164062 | APEH | chr3 | + | 49676055 | 49676515 | 49675890 | 49675966 | 49676608 | 49676700 | 363 | 192 |
| SE | 84320 | ENSG00000164062 | APEH | chr3 | + | 49676776 | 49676817 | 49676608 | 49676700 | 49676902 | 49676995 | 114 | 42 |
| SE | 84322 | ENSG00000164062 | APEH | chr3 | + | 49681100 | 49681239 | 49680540 | 49680629 | 49681721 | 49681805 | 114 | 47 |
| SE | 84323 | ENSG00000164062 | APEH | chr3 | + | 49681721 | 49681805 | 49681100 | 49681239 | 49681886 | 49681967 | 65 | 28 |
| SE | 84325 | ENSG00000164062 | APEH | chr3 | + | 49681886 | 49681967 | 49681721 | 49681805 | 49682347 | 49682436 | 42 | 22 |
| SE | 84326 | ENSG00000164062 | APEH | chr3 | + | 49682347 | 49682436 | 49681886 | 49681967 | 49682545 | 49682736 | 132 | 52 |
| SE | 84328 | ENSG00000008256 | CYTH3 | chr7 | - | 6188704 | 6188889 | 6187656 | 6187721 | 6190448 | 6190531 | 0 | 0 |
| SE | 84329 | ENSG00000138593 | SECISBP2L | chr15 | - | 48999832 | 48999987 | 48996366 | 48996586 | 49000876 | 49001097 | 572 | 543 |
| SE | 84330 | ENSG00000138593 | SECISBP2L | chr15 | - | 49009215 | 49009378 | 49000876 | 49001097 | 49011730 | 49011863 | 229 | 150 |
| SE | 84331 | ENSG00000138593 | SECISBP2L | chr15 | - | 49011730 | 49011863 | 49009215 | 49009378 | 49012647 | 49012817 | 237 | 165 |
| SE | 84332 | ENSG00000138593 | SECISBP2L | chr15 | - | 49016559 | 49016701 | 49012647 | 49012817 | 49017547 | 49017628 | 78 | 75 |
| SE | 84334 | ENSG00000138593 | SECISBP2L | chr15 | - | 49017547 | 49017628 | 49016847 | 49017015 | 49019417 | 49019552 | 312 | 227 |
| SE | 84335 | ENSG00000138593 | SECISBP2L | chr15 | - | 49017547 | 49017628 | 49016847 | 49017015 | 49027364 | 49027480 | 223 | 147 |
| SE | 84336 | ENSG00000138593 | SECISBP2L | chr15 | - | 49019417 | 49019552 | 49016847 | 49017015 | 49027364 | 49027480 | 121 | 107 |
| SE | 84338 | ENSG00000138593 | SECISBP2L | chr15 | - | 49028143 | 49028168 | 49027364 | 49027480 | 49028452 | 49028487 | 108 | 72 |
| SE | 84339 | ENSG00000138593 | SECISBP2L | chr15 | - | 49032964 | 49033100 | 49028452 | 49028682 | 49035333 | 49035456 | 232 | 145 |
| SE | 84340 | ENSG00000159256 | MORC3 | chr21 | + | 36333645 | 36333718 | 36320188 | 36320303 | 36336873 | 36337006 | 127 | 96 |
| SE | 84341 | ENSG00000159256 | MORC3 | chr21 | + | 36341398 | 36341546 | 36338773 | 36338921 | 36344578 | 36344707 | 2092 | 1508 |
| SE | 84342 | ENSG00000159256 | MORC3 | chr21 | + | 36349310 | 36349408 | 36344911 | 36345031 | 36356619 | 36356724 | 365 | 317 |
| SE | 84343 | ENSG00000159256 | MORC3 | chr21 | + | 36349310 | 36349408 | 36344911 | 36345031 | 36359954 | 36360049 | 148 | 118 |
| SE | 84344 | ENSG00000159256 | MORC3 | chr21 | + | 36356619 | 36356724 | 36344911 | 36345031 | 36359954 | 36360049 | 98 | 130 |
| SE | 84345 | ENSG00000159256 | MORC3 | chr21 | + | 36356619 | 36356724 | 36349310 | 36349408 | 36359954 | 36360049 | 312 | 318 |
| SE | 84346 | ENSG00000159256 | MORC3 | chr21 | + | 36362182 | 36362228 | 36360183 | 36360258 | 36364092 | 36364259 | 264 | 212 |
| SE | 84354 | ENSG00000230590 | FTX | chrX | - | 74247331 | 74247424 | 74196350 | 74203909 | 74274341 | 74274410 | 1 | 0 |
| SE | 84355 | ENSG00000230590 | FTX | chrX | - | 74247331 | 74247424 | 74217687 | 74218725 | 74274341 | 74274410 | 1 | 0 |
| SE | 84359 | ENSG00000230590 | FTX | chrX | - | 74274341 | 74274410 | 74068088 | 74069584 | 74280393 | 74280476 | 1 | 3 |
| SE | 84360 | ENSG00000230590 | FTX | chrX | - | 74274341 | 74274410 | 74202514 | 74203953 | 74280393 | 74280476 | 10 | 12 |
| SE | 84361 | ENSG00000230590 | FTX | chrX | - | 74274341 | 74274410 | 74202650 | 74203909 | 74280393 | 74280476 | 5 | 6 |
| SE | 84362 | ENSG00000230590 | FTX | chrX | - | 74274341 | 74274457 | 74202674 | 74203953 | 74280393 | 74280494 | 4 | 7 |
| SE | 84363 | ENSG00000230590 | FTX | chrX | - | 74280376 | 74280494 | 74274341 | 74274410 | 74280930 | 74281001 | 98 | 115 |
| SE | 84364 | ENSG00000230590 | FTX | chrX | - | 74280393 | 74280494 | 74274259 | 74274410 | 74280930 | 74281001 | 97 | 111 |
| SE | 84365 | ENSG00000230590 | FTX | chrX | - | 74280930 | 74281001 | 74280376 | 74280494 | 74281701 | 74281848 | 104 | 116 |
| SE | 84366 | ENSG00000230590 | FTX | chrX | - | 74280930 | 74281085 | 74278125 | 74280494 | 74281701 | 74281816 | 237 | 261 |
| SE | 84367 | ENSG00000230590 | FTX | chrX | - | 74284742 | 74284944 | 74281701 | 74281848 | 74285914 | 74286326 | 3 | 7 |
| SE | 84368 | ENSG00000230590 | FTX | chrX | - | 74284742 | 74284944 | 74281701 | 74281848 | 74291757 | 74292027 | 3 | 5 |
| SE | 84371 | ENSG00000230590 | FTX | chrX | - | 74284742 | 74284948 | 74281701 | 74281848 | 74285914 | 74286170 | 3 | 7 |
| SE | 84372 | ENSG00000230590 | FTX | chrX | - | 74284742 | 74284948 | 74281701 | 74281848 | 74291757 | 74292064 | 3 | 5 |
| SE | 84378 | ENSG00000230590 | FTX | chrX | - | 74285914 | 74286170 | 74284742 | 74284948 | 74291757 | 74292079 | 0 | 5 |
| SE | 84379 | ENSG00000230590 | FTX | chrX | - | 74285914 | 74286170 | 74284742 | 74284948 | 74292168 | 74292349 | 1 | 4 |
| SE | 84381 | ENSG00000230590 | FTX | chrX | - | 74290394 | 74290430 | 74281701 | 74281848 | 74292426 | 74292620 | 0 | 0 |
| SE | 84383 | ENSG00000136143 | SUCLA2 | chr13 | - | 47956839 | 47956878 | 47954395 | 47954557 | 47968594 | 47968733 | 5 | 10 |
| SE | 84384 | ENSG00000136143 | SUCLA2 | chr13 | - | 47965577 | 47965640 | 47954395 | 47954557 | 47968594 | 47968733 | 1 | 5 |
| SE | 84385 | ENSG00000136143 | SUCLA2 | chr13 | - | 47973263 | 47973392 | 47968594 | 47968733 | 47988540 | 47988548 | 178 | 142 |
| SE | 84389 | ENSG00000136143 | SUCLA2 | chr13 | - | 47994799 | 47994896 | 47991502 | 47991678 | 47996842 | 47997023 | 5 | 2 |
| SE | 84390 | ENSG00000136143 | SUCLA2 | chr13 | - | 47996842 | 47997023 | 47988881 | 47988981 | 48001179 | 48001266 | 86 | 76 |
| SE | 84392 | ENSG00000188917 | TRMT2B | chrX | - | 101020486 | 101020588 | 101019283 | 101019403 | 101021100 | 101021315 | 111 | 65 |
| SE | 84393 | ENSG00000188917 | TRMT2B | chrX | - | 101020486 | 101020588 | 101019283 | 101019403 | 101021967 | 101022062 | 59 | 28 |
| SE | 84394 | ENSG00000188917 | TRMT2B | chrX | - | 101021100 | 101021315 | 101019283 | 101019403 | 101021967 | 101022062 | 34 | 18 |
| SE | 84395 | ENSG00000188917 | TRMT2B | chrX | - | 101021100 | 101021315 | 101019283 | 101019403 | 101035612 | 101035683 | 2 | 0 |
| SE | 84396 | ENSG00000188917 | TRMT2B | chrX | - | 101021100 | 101021315 | 101020486 | 101020588 | 101021967 | 101022062 | 87 | 55 |
| SE | 84397 | ENSG00000188917 | TRMT2B | chrX | - | 101021967 | 101022062 | 101019283 | 101019403 | 101035612 | 101035683 | 0 | 0 |
| SE | 84398 | ENSG00000188917 | TRMT2B | chrX | - | 101021967 | 101022062 | 101020486 | 101020588 | 101023469 | 101023616 | 34 | 14 |
| SE | 84399 | ENSG00000188917 | TRMT2B | chrX | - | 101021967 | 101022062 | 101021100 | 101021315 | 101035612 | 101035683 | 13 | 5 |
| SE | 84400 | ENSG00000188917 | TRMT2B | chrX | - | 101023469 | 101023616 | 101021967 | 101022062 | 101035612 | 101035683 | 111 | 52 |
| SE | 84401 | ENSG00000188917 | TRMT2B | chrX | - | 101035612 | 101035683 | 101023469 | 101023616 | 101041316 | 101041371 | 38 | 18 |
| SE | 84402 | ENSG00000188917 | TRMT2B | chrX | - | 101035612 | 101035683 | 101023469 | 101023616 | 101051816 | 101052054 | 38 | 18 |
| SE | 84403 | ENSG00000188917 | TRMT2B | chrX | - | 101036973 | 101037073 | 101035612 | 101035683 | 101041316 | 101041371 | 80 | 47 |
| SE | 84404 | ENSG00000188917 | TRMT2B | chrX | - | 101036973 | 101037073 | 101035612 | 101035683 | 101051816 | 101052054 | 59 | 29 |
| SE | 84409 | ENSG00000188917 | TRMT2B | chrX | - | 101041316 | 101041371 | 101023469 | 101023616 | 101042232 | 101042312 | 8 | 3 |
| SE | 84410 | ENSG00000188917 | TRMT2B | chrX | - | 101041316 | 101041371 | 101023469 | 101023616 | 101051816 | 101052054 | 1 | 0 |
| SE | 84411 | ENSG00000188917 | TRMT2B | chrX | - | 101041316 | 101041371 | 101035612 | 101035683 | 101051783 | 101052082 | 1 | 0 |
| SE | 84412 | ENSG00000188917 | TRMT2B | chrX | - | 101041316 | 101041371 | 101035612 | 101035683 | 101051816 | 101052054 | 1 | 0 |
| SE | 84413 | ENSG00000188917 | TRMT2B | chrX | - | 101041316 | 101041371 | 101036973 | 101037073 | 101042041 | 101042312 | 37 | 25 |
| SE | 84419 | ENSG00000188917 | TRMT2B | chrX | - | 101041316 | 101041371 | 101037916 | 101038051 | 101051783 | 101052082 | 28 | 11 |
| SE | 84420 | ENSG00000188917 | TRMT2B | chrX | - | 101041316 | 101041371 | 101037916 | 101038051 | 101051816 | 101052054 | 28 | 11 |
| SE | 84422 | ENSG00000188917 | TRMT2B | chrX | - | 101042041 | 101042312 | 101036973 | 101037073 | 101051816 | 101052054 | 2 | 4 |
| SE | 84423 | ENSG00000188917 | TRMT2B | chrX | - | 101042041 | 101042312 | 101041316 | 101041371 | 101051250 | 101051366 | 78 | 39 |
| SE | 84424 | ENSG00000188917 | TRMT2B | chrX | - | 101042041 | 101042312 | 101041316 | 101041371 | 101051643 | 101051733 | 57 | 24 |
| SE | 84425 | ENSG00000188917 | TRMT2B | chrX | - | 101042041 | 101042312 | 101041316 | 101041371 | 101051783 | 101052116 | 69 | 29 |
| SE | 84427 | ENSG00000188917 | TRMT2B | chrX | - | 101042232 | 101042312 | 101037916 | 101038051 | 101051783 | 101052082 | 12 | 3 |
| SE | 84428 | ENSG00000188917 | TRMT2B | chrX | - | 101042232 | 101042312 | 101041316 | 101041371 | 101051783 | 101052082 | 22 | 6 |
| SE | 84429 | ENSG00000188917 | TRMT2B | chrX | - | 101051250 | 101051366 | 101036973 | 101037073 | 101051816 | 101052054 | 1 | 2 |
| SE | 84430 | ENSG00000188917 | TRMT2B | chrX | - | 101051250 | 101051366 | 101041316 | 101041371 | 101051643 | 101051733 | 2 | 3 |
| SE | 84431 | ENSG00000188917 | TRMT2B | chrX | - | 101051250 | 101051366 | 101041316 | 101041371 | 101051783 | 101052116 | 0 | 2 |
| SE | 84432 | ENSG00000188917 | TRMT2B | chrX | - | 101051250 | 101051366 | 101041316 | 101041371 | 101051816 | 101052054 | 1 | 2 |
| SE | 84437 | ENSG00000188917 | TRMT2B | chrX | - | 101051250 | 101051733 | 101037916 | 101038051 | 101051816 | 101052111 | 21 | 14 |
| SE | 84445 | ENSG00000102245 | CD40LG | chrX | + | 136654372 | 136654430 | 136650265 | 136650397 | 136656355 | 136656418 | 247 | 182 |
| SE | 84446 | ENSG00000102245 | CD40LG | chrX | + | 136656355 | 136656418 | 136654372 | 136654430 | 136659038 | 136660390 | 166 | 127 |
| SE | 84454 | ENSG00000137522 | RNF121 | chr11 | + | 71962151 | 71962289 | 71960749 | 71960891 | 71982760 | 71982915 | 2 | 0 |
| SE | 84456 | ENSG00000137522 | RNF121 | chr11 | + | 71978085 | 71978235 | 71960749 | 71960891 | 71987003 | 71987028 | 0 | 0 |
| SE | 84457 | ENSG00000137522 | RNF121 | chr11 | + | 71982760 | 71982915 | 71960749 | 71960891 | 71987003 | 71987028 | 35 | 17 |
| SE | 84458 | ENSG00000137522 | RNF121 | chr11 | + | 71982760 | 71982915 | 71960749 | 71960891 | 71990596 | 71990717 | 12 | 11 |
| SE | 84459 | ENSG00000137522 | RNF121 | chr11 | + | 71982760 | 71982915 | 71978085 | 71978235 | 71987003 | 71987028 | 32 | 7 |
| SE | 84460 | ENSG00000137522 | RNF121 | chr11 | + | 71982806 | 71982915 | 71960749 | 71960891 | 71987003 | 71987111 | 20 | 5 |
| SE | 84461 | ENSG00000137522 | RNF121 | chr11 | + | 71982806 | 71982915 | 71960749 | 71960891 | 71990596 | 71990663 | 2 | 0 |
| SE | 84464 | ENSG00000137522 | RNF121 | chr11 | + | 71990596 | 71990717 | 71960749 | 71960891 | 71994718 | 71994777 | 40 | 13 |
| SE | 84465 | ENSG00000137522 | RNF121 | chr11 | + | 71995449 | 71995551 | 71994718 | 71994852 | 71996194 | 71996275 | 70 | 34 |

| | | | | | | | | | | | | | | | | | | | | | | | | | | | | | | | | | | | | | | | | | | | | | | |
|---|---|---|---|---|---|---|---|---|---|---|---|---|---|---|---|---|---|---|---|---|---|---|---|---|---|---|---|---|---|---|---|---|---|---|---|---|---|---|---|---|---|---|---|---|---|---|
| SE | 84467 | ENSG00000126947 | ARMCX1 | chrX | + | 101550970 | 101551024 | 101550546 | 101550660 | 101551568 | 101551633 | 7 | 8 | 10 | 1 | 1 | 0 | 0 | 0 | 1 | 0 | 5 | 4 | 5 | 5 | 1 | 0 | 0 | 0 | 0 | 0 | 203 | 149 | 1 | 1 | 1 | 1 | 1 | 0.423 | 1 | 1 | 1 | 1 | 1 | 1 | -0.115 |
| SE | 84470 | ENSG00000017797 | RALBP1 | chr18 | + | 9512990 | 9513289 | 9475512 | 9475691 | 9516844 | 9517158 | 96 | 82 | 81 | 116 | 123 | 0 | 2 | 0 | 0 | 0 | 203 | 94 | 92 | 173 | 139 | 0 | 0 | 0 | 0 | 0 | 298 | 149 | 1 | 1 | 1 | 0.953 | 1 | 1 | 1 | 1 | 1 | 1 | 1 | 1 | -0.009 |
| SE | 84471 | ENSG00000017797 | RALBP1 | chr18 | + | 9522159 | 9522509 | 9516844 | 9517303 | 9524593 | 9524755 | 519 | 355 | 397 | 452 | 651 | 1 | 0 | 1 | 4 | 2 | 819 | 400 | 384 | 625 | 459 | 0 | 0 | 0 | 0 | 0 | 298 | 149 | 1 | 1 | 0.996 | 1 | 0.995 | 0.983 | 0.994 | 1 | 1 | 1 | 1 | 1 | -0.006 |
| SE | 84472 | ENSG00000017797 | RALBP1 | chr18 | + | 9524593 | 9524755 | 9522159 | 9522509 | 9525719 | 9525851 | 321 | 207 | 238 | 300 | 503 | 0 | 0 | 0 | 0 | 0 | 447 | 320 | 258 | 361 | 347 | 0 | 6 | 0 | 0 | 0 | 298 | 149 | 1 | 1 | 1 | 1 | 1 | 1 | 1 | 1 | 0.964 | 1 | 1 | 1 | 0.007 |
| SE | 84473 | ENSG00000017797 | RALBP1 | chr18 | + | 9524593 | 9524755 | 9522159 | 9522509 | 9530833 | 9530941 | 156 | 73 | 104 | 126 | 214 | 0 | 0 | 0 | 0 | 0 | 104 | 106 | 87 | 121 | 78 | 0 | 3 | 0 | 0 | 0 | 298 | 149 | 1 | 1 | 1 | 1 | 1 | 1 | 1 | 1 | 0.946 | 1 | 1 | 1 | 0.011 |
| SE | 84474 | ENSG00000017797 | RALBP1 | chr18 | + | 9525719 | 9525851 | 9522159 | 9522509 | 9530833 | 9530941 | 150 | 128 | 129 | 169 | 207 | 0 | 0 | 0 | 0 | 0 | 251 | 147 | 165 | 193 | 144 | 0 | 3 | 0 | 0 | 0 | 281 | 149 | 1 | 1 | 1 | 1 | 1 | 1 | 1 | 1 | 0.963 | 1 | 1 | 1 | 0.007 |
| SE | 84475 | ENSG00000017797 | RALBP1 | chr18 | + | 9525719 | 9525851 | 9524593 | 9524755 | 9530833 | 9530941 | 309 | 254 | 257 | 332 | 477 | 0 | 0 | 0 | 1 | 0 | 587 | 354 | 334 | 422 | 408 | 0 | 0 | 0 | 0 | 0 | 281 | 149 | 1 | 1 | 1 | 1 | 1 | 0.994 | 1 | 1 | 1 | 1 | 1 | 1 | -0.001 |
| SE | 84476 | ENSG00000017797 | RALBP1 | chr18 | + | 9533334 | 9533456 | 9530833 | 9530941 | 9533702 | 9533826 | 570 | 460 | 456 | 583 | 785 | 0 | 0 | 0 | 0 | 0 | 1017 | 490 | 459 | 837 | 679 | 1 | 0 | 0 | 0 | 0 | 271 | 149 | 1 | 1 | 1 | 1 | 1 | 1 | 1 | 0.998 | 1 | 1 | 1 | 1 | 0 |
| SE | 84477 | ENSG00000148985 | PGAP2 | chr11 | + | 3811249 | 3811424 | 3808289 | 3808315 | 3822908 | 3822993 | 11 | 1 | 8 | 10 | 0 | 0 | 3 | 0 | 0 | 0 | 0 | 11 | 2 | 0 | 15 | 0 | 0 | 0 | 4 | 0 | 298 | 149 | 1 | 1 | 1 | 0.143 | 1 | 1 | NA | NA | 1 | 1 | 0 | 1 | 0.036 |
| SE | 84478 | ENSG00000148985 | PGAP2 | chr11 | + | 3811249 | 3811424 | 3808289 | 3808315 | 3823612 | 3823740 | 10 | 1 | 8 | 10 | 0 | 2 | 0 | 0 | 0 | 0 | 0 | 9 | 2 | 0 | 8 | 0 | 0 | 0 | 0 | 0 | 298 | 149 | 1 | 1 | 0.714 | 1 | 1 | 1 | NA | NA | 1 | 1 | NA | 1 | -0.072 |
| SE | 84479 | ENSG00000148985 | PGAP2 | chr11 | + | 3811249 | 3811424 | 3808289 | 3808315 | 3823882 | 3824135 | 14 | 2 | 12 | 16 | 4 | 0 | 0 | 2 | 0 | 0 | 0 | 11 | 2 | 1 | 8 | 0 | 0 | 0 | 0 | 0 | 298 | 149 | 1 | 1 | 1 | 1 | 0.75 | 1 | 1 | NA | 1 | 1 | 1 | 1 | -0.05 |
| SE | 84480 | ENSG00000148985 | PGAP2 | chr11 | + | 3811249 | 3811424 | 3808289 | 3808315 | 3824269 | 3824364 | 11 | 1 | 8 | 12 | 0 | 0 | 0 | 2 | 0 | 0 | 0 | 9 | 2 | 0 | 8 | 0 | 4 | 2 | 0 | 0 | 298 | 149 | 1 | 1 | 1 | 1 | 0.667 | 1 | NA | NA | 0.529 | 0.333 | NA | 1 | 0.296 |
| SE | 84483 | ENSG00000148985 | PGAP2 | chr11 | + | 3817352 | 3817535 | 3811249 | 3811424 | 3823882 | 3824135 | 0 | 0 | 0 | 2 | 0 | 4 | 1 | 4 | 6 | 4 | 0 | 2 | 0 | 0 | 0 | 0 | 2 | 0 | 1 | 0 | 298 | 149 | 1 | 1 | 0 | 0 | 0 | 0.143 | 0 | NA | 0.333 | NA | 0 | NA | -0.138 |
| SE | 84484 | ENSG00000148985 | PGAP2 | chr11 | + | 3817431 | 3817535 | 3811249 | 3811424 | 3823882 | 3824056 | 0 | 0 | 0 | 0 | 0 | 4 | 1 | 4 | 6 | 4 | 0 | 1 | 0 | 0 | 0 | 0 | 2 | 0 | 1 | 0 | 253 | 149 | 1 | 1 | 0 | 0 | 0 | 0 | 0 | NA | 0.227 | NA | 0 | NA | -0.114 |
| SE | 84486 | ENSG00000148985 | PGAP2 | chr11 | + | 3822908 | 3822993 | 3808293 | 3808315 | 3823882 | 3824135 | 0 | 0 | 2 | 0 | 0 | 0 | 0 | 2 | 0 | 0 | 0 | 0 | 2 | 8 | 0 | 0 | 0 | 0 | 0 | 0 | 234 | 149 | 1 | 1 | NA | NA | 0.389 | NA | NA | NA | NA | 1 | 1 | NA | -0.611 |
| SE | 84488 | ENSG00000148985 | PGAP2 | chr11 | + | 3822908 | 3822993 | 3811249 | 3811424 | 3823882 | 3824135 | 1 | 0 | 2 | 0 | 0 | 4 | 1 | 4 | 6 | 4 | 0 | 2 | 2 | 4 | 7 | 0 | 2 | 0 | 1 | 0 | 234 | 149 | 1 | 1 | 0.137 | 0 | 0.241 | 0 | 0 | NA | 0.389 | 1 | 0.718 | 1 | -0.701 |
| SE | 84491 | ENSG00000148985 | PGAP2 | chr11 | + | 3823612 | 3823740 | 3811249 | 3811424 | 3823882 | 3824135 | 0 | 2 | 0 | 0 | 0 | 4 | 1 | 4 | 6 | 4 | 0 | 0 | 0 | 0 | 0 | 0 | 2 | 0 | 1 | 0 | 277 | 149 | 1 | 1 | 0 | 0.518 | 0 | 0 | 0 | NA | 0 | NA | 0 | NA | 0.104 |
| SE | 84492 | ENSG00000148985 | PGAP2 | chr11 | + | 3823612 | 3823740 | 3822908 | 3822993 | 3823882 | 3824135 | 0 | 5 | 0 | 0 | 0 | 0 | 0 | 2 | 0 | 0 | 0 | 0 | 0 | 0 | 0 | 0 | 2 | 2 | 8 | 0 | 277 | 149 | 1 | 1 | NA | 1 | 0 | NA | NA | NA | 0 | 0 | 0 | NA | 0.5 |
| SE | 84493 | ENSG00000148985 | PGAP2 | chr11 | + | 3823616 | 3823740 | 3822908 | 3822993 | 3823882 | 3823915 | 0 | 2 | 0 | 0 | 0 | 0 | 0 | 2 | 0 | 0 | 0 | 0 | 0 | 0 | 0 | 0 | 2 | 2 | 8 | 0 | 273 | 149 | 1 | 1 | NA | 1 | 0 | NA | NA | NA | 0 | 0 | 0 | NA | 0.5 |
| SE | 84494 | ENSG00000148985 | PGAP2 | chr11 | + | 3823882 | 3824135 | 3808293 | 3808315 | 3824269 | 3824343 | 5 | 2 | 15 | 4 | 5 | 0 | 0 | 2 | 0 | 0 | 0 | 2 | 2 | 2 | 0 | 0 | 4 | 2 | 0 | 0 | 298 | 149 | 1 | 1 | 1 | 1 | 0.789 | 1 | 1 | NA | 0.2 | 0.333 | 1 | NA | 0.447 |
| SE | 84495 | ENSG00000148985 | PGAP2 | chr11 | + | 3823882 | 3824135 | 3811249 | 3811424 | 3824269 | 3824343 | 9 | 3 | 17 | 10 | 9 | 1 | 0 | 0 | 2 | 0 | 0 | 4 | 2 | 3 | 0 | 0 | 0 | 0 | 0 | 0 | 298 | 149 | 1 | 1 | 0.818 | 1 | 1 | 0.714 | 1 | NA | 1 | 1 | 1 | NA | -0.094 |
| SE | 84496 | ENSG00000128731 | HERC2 | chr15 | - | 28130162 | 28130302 | 28125005 | 28125193 | 28130502 | 28130594 | 103 | 62 | 147 | 104 | 83 | 0 | 0 | 0 | 0 | 0 | 131 | 64 | 66 | 91 | 97 | 0 | 0 | 0 | 3 | 0 | 289 | 149 | 1 | 1 | 1 | 1 | 1 | 1 | 1 | 1 | 1 | 1 | 0.94 | 1 | 0.012 |
| SE | 84498 | ENSG00000128731 | HERC2 | chr15 | - | 28141431 | 28141630 | 28135477 | 28135692 | 28141730 | 28141846 | 57 | 43 | 102 | 82 | 71 | 0 | 0 | 1 | 0 | 0 | 38 | 78 | 46 | 75 | 38 | 0 | 0 | 0 | 0 | 0 | 298 | 149 | 1 | 1 | 1 | 1 | 0.981 | 1 | 1 | 1 | 1 | 1 | 1 | 1 | -0.004 |
| SE | 84499 | ENSG00000128731 | HERC2 | chr15 | - | 28142237 | 28142393 | 28141730 | 28141846 | 28142826 | 28142952 | 98 | 40 | 116 | 85 | 114 | 0 | 0 | 0 | 0 | 0 | 64 | 75 | 79 | 84 | 84 | 0 | 1 | 0 | 1 | 0 | 298 | 149 | 1 | 1 | 1 | 1 | 1 | 1 | 1 | 1 | 0.974 | 1 | 0.977 | 1 | 0.01 |
| SE | 84500 | ENSG00000128731 | HERC2 | chr15 | - | 28146236 | 28146344 | 28144672 | 28144804 | 28152676 | 28152830 | 28 | 6 | 28 | 28 | 6 | 0 | 0 | 2 | 0 | 0 | 10 | 33 | 27 | 13 | 0 | 0 | 0 | 0 | 1 | 0 | 257 | 149 | 1 | 1 | 1 | 1 | 0.89 | 1 | 1 | 1 | 1 | 1 | 0.883 | NA | 0.007 |
| SE | 84504 | ENSG00000128731 | HERC2 | chr15 | - | 28182400 | 28182512 | 28179141 | 28179223 | 28186576 | 28186752 | 68 | 50 | 90 | 83 | 55 | 0 | 0 | 0 | 0 | 0 | 38 | 82 | 51 | 129 | 110 | 0 | 0 | 1 | 0 | 0 | 261 | 149 | 1 | 1 | 1 | 1 | 1 | 1 | 1 | 1 | 1 | 0.967 | 1 | 1 | 0.007 |
| SE | 84505 | ENSG00000128731 | HERC2 | chr15 | - | 28182400 | 28182512 | 28179141 | 28179223 | 28190964 | 28191056 | 32 | 28 | 42 | 28 | 21 | 0 | 0 | 0 | 0 | 0 | 0 | 33 | 34 | 50 | 57 | 0 | 0 | 0 | 2 | 0 | 261 | 149 | 1 | 1 | 1 | 1 | 1 | 1 | 1 | NA | 1 | 1 | 0.935 | 1 | 0.016 |
| SE | 84506 | ENSG00000128731 | HERC2 | chr15 | - | 28186576 | 28186752 | 28179141 | 28179223 | 28190964 | 28191056 | 37 | 33 | 40 | 62 | 49 | 0 | 0 | 0 | 0 | 0 | 79 | 70 | 21 | 33 | 29 | 0 | 0 | 0 | 2 | 0 | 298 | 149 | 1 | 1 | 1 | 1 | 1 | 1 | 1 | 1 | 1 | 1 | 0.892 | 1 | 0.022 |
| SE | 84507 | ENSG00000128731 | HERC2 | chr15 | - | 28186576 | 28186752 | 28182400 | 28182512 | 28190964 | 28191056 | 73 | 55 | 88 | 117 | 83 | 0 | 0 | 0 | 0 | 0 | 117 | 119 | 37 | 113 | 82 | 0 | 0 | 0 | 1 | 0 | 298 | 149 | 1 | 1 | 1 | 1 | 1 | 1 | 1 | 1 | 1 | 1 | 0.983 | 1 | 0.003 |
| SE | 84509 | ENSG00000128731 | HERC2 | chr15 | - | 28213741 | 28213972 | 28212444 | 28212583 | 28214075 | 28214272 | 51 | 21 | 77 | 92 | 39 | 0 | 0 | 0 | 3 | 0 | 3 | 43 | 19 | 45 | 55 | 0 | 0 | 0 | 0 | 0 | 298 | 149 | 1 | 1 | 1 | 1 | 1 | 0.939 | 1 | 1 | 1 | 1 | 1 | 1 | -0.012 |
| SE | 84510 | ENSG00000128731 | HERC2 | chr15 | - | 28228217 | 28228409 | 28222027 | 28222215 | 28229194 | 28229346 | 64 | 26 | 49 | 78 | 83 | 0 | 0 | 0 | 0 | 1 | 28 | 32 | 48 | 78 | 60 | 0 | 0 | 0 | 0 | 0 | 298 | 149 | 1 | 1 | 1 | 1 | 1 | 1 | 0.976 | 1 | 1 | 1 | 1 | 1 | -0.005 |
| SE | 84512 | ENSG00000128731 | HERC2 | chr15 | - | 28245880 | 28246066 | 28238601 | 28238772 | 28248551 | 28248736 | 53 | 63 | 51 | 65 | 57 | 0 | 0 | 0 | 0 | 0 | 104 | 65 | 44 | 73 | 54 | 0 | 0 | 1 | 0 | 0 | 298 | 149 | 1 | 1 | 1 | 1 | 1 | 1 | 1 | 1 | 1 | 0.957 | 1 | 1 | 0.009 |
| SE | 84513 | ENSG00000128731 | HERC2 | chr15 | - | 28246741 | 28246897 | 28245880 | 28246066 | 28248551 | 28248736 | 86 | 55 | 80 | 80 | 91 | 1 | 0 | 0 | 2 | 0 | 80 | 95 | 84 | 53 | 79 | 0 | 0 | 0 | 0 | 0 | 298 | 149 | 1 | 1 | 0.977 | 1 | 1 | 0.952 | 1 | 1 | 1 | 1 | 1 | 1 | -0.014 |
| SE | 84514 | ENSG00000128731 | HERC2 | chr15 | - | 28274290 | 28274447 | 28272893 | 28273004 | 28274904 | 28275005 | 29 | 5 | 27 | 28 | 10 | 1 | 0 | 0 | 0 | 0 | 12 | 21 | 15 | 21 | 18 | 0 | 0 | 0 | 0 | 0 | 298 | 149 | 1 | 1 | 0.935 | 1 | 1 | 1 | 1 | 1 | 1 | 1 | 1 | 1 | -0.013 |
| SE | 84515 | ENSG00000128731 | HERC2 | chr15 | - | 28274904 | 28275005 | 28272893 | 28273004 | 28280067 | 28280287 | 22 | 13 | 37 | 25 | 30 | 4 | 1 | 2 | 4 | 7 | 0 | 44 | 29 | 15 | 19 | 0 | 7 | 3 | 5 | 0 | 250 | 149 | 1 | 1 | 0.766 | 0.886 | 0.917 | 0.788 | 0.719 | NA | 0.789 | 0.852 | 0.641 | 1 | -0.005 |
| SE | 84516 | ENSG00000128731 | HERC2 | chr15 | - | 28292887 | 28293022 | 28280033 | 28280287 | 28299401 | 28299483 | 74 | 37 | 86 | 76 | 80 | 0 | 0 | 1 | 0 | 0 | 69 | 119 | 71 | 32 | 73 | 0 | 0 | 0 | 0 | 0 | 284 | 149 | 1 | 1 | 1 | 1 | 0.978 | 1 | 1 | 1 | 1 | 1 | 1 | 1 | -0.004 |
| SE | 84518 | ENSG00000128731 | HERC2 | chr15 | - | 28299401 | 28299516 | 28292887 | 28293022 | 28321361 | 28321464 | 39 | 11 | 32 | 23 | 38 | 2 | 0 | 0 | 0 | 0 | 59 | 55 | 29 | 28 | 26 | 0 | 0 | 0 | 0 | 0 | 264 | 149 | 1 | 1 | 0.917 | 1 | 1 | 1 | 1 | 1 | 1 | 1 | 1 | 1 | -0.017 |
| SE | 84521 | ENSG00000198960 | ARMCX6 | chrX | - | 101617502 | 101617584 | 101615124 | 101616766 | 101617884 | 101617910 | 203 | 152 | 120 | 187 | 413 | 1 | 2 | 0 | 1 | 2 | 326 | 189 | 91 | 191 | 706 | 0 | 5 | 2 | 2 | 0 | 231 | 149 | 1 | 1 | 0.992 | 0.98 | 1 | 0.992 | 0.993 | 1 | 0.961 | 0.967 | 0.984 | 1 | 0.009 |
| SE | 84522 | ENSG00000273674 | NA | chr15 | - | 50890855 | 50891077 | 50874571 | 50874726 | 50908551 | 50908572 | 0 | 0 | 0 | 2 | 0 | 5 | 0 | 0 | 0 | 0 | 0 | 0 | 1 | 0 | 0 | 0 | 0 | 0 | 0 | 0 | 298 | 149 | 1 | 1 | 0 | NA | NA | 1 | NA | NA | NA | 1 | NA | NA | -0.5 |
| SE | 84524 | ENSG00000177853 | ZNF518A | chr10 | + | 96135169 | 96135289 | 96133582 | 96133648 | 96151415 | 96151566 | 0 | 0 | 2 | 0 | 0 | 14 | 6 | 8 | 4 | 11 | 0 | 0 | 0 | 0 | 0 | 0 | 0 | 0 | 2 | 0 | 269 | 149 | 1 | 1 | 0 | 0 | 0.122 | 0 | 0 | NA | NA | NA | 0 | NA | 0.024 |
| SE | 84525 | ENSG00000177853 | ZNF518A | chr10 | + | 96135169 | 96135289 | 96133582 | 96133648 | 96155325 | 96155398 | 0 | 0 | 2 | 2 | 1 | 26 | 17 | 11 | 40 | 32 | 0 | 0 | 0 | 0 | 0 | 0 | 8 | 1 | 0 | 2 | 269 | 149 | 1 | 1 | 0 | 0 | 0.091 | 0.027 | 0.017 | NA | 0 | 0 | NA | 0 | 0.027 |
| SE | 84526 | ENSG00000177853 | ZNF518A | chr10 | + | 96151415 | 96151566 | 96133555 | 96133648 | 96155325 | 96155398 | 30 | 7 | 14 | 7 | 36 | 26 | 17 | 11 | 40 | 32 | 0 | 0 | 0 | 2 | 0 | 0 | 8 | 1 | 0 | 2 | 298 | 149 | 1 | 1 | 0.366 | 0.171 | 0.389 | 0.08 | 0.36 | NA | 0 | 0 | 1 | 0 | 0.023 |
| SE | 84527 | ENSG00000177853 | ZNF518A | chr10 | + | 96151415 | 96151566 | 96133555 | 96133648 | 96155923 | 96156047 | 14 | 6 | 8 | 4 | 11 | 7 | 6 | 3 | 6 | 10 | 0 | 0 | 0 | 2 | 0 | 0 | 0 | 0 | 0 | 0 | 298 | 149 | 1 | 1 | 0.5 | 0.333 | 0.571 | 0.25 | 0.355 | NA | NA | NA | 1 | NA | -0.598 |
| SE | 84529 | ENSG00000177853 | ZNF518A | chr10 | + | 96155325 | 96155398 | 96133555 | 96133648 | 96155923 | 96156019 | 7 | 7 | 6 | 10 | 9 | 7 | 6 | 3 | 6 | 10 | 0 | 1 | 0 | 0 | 0 | 0 | 0 | 0 | 0 | 0 | 222 | 149 | 1 | 1 | 0.402 | 0.439 | 0.573 | 0.528 | 0.377 | NA | 1 | NA | NA | NA | -0.536 |
| SE | 84531 | ENSG00000102401 | ARMCX3 | chrX | + | 101623813 | 101623893 | 101623127 | 101623276 | 101624479 | 101624550 | 11 | 10 | 10 | 6 | 9 | 0 | 0 | 0 | 0 | 0 | 12 | 0 | 8 | 3 | 25 | 0 | 0 | 1 | 0 | 0 | 229 | 149 | 1 | 1 | 1 | 1 | 1 | 1 | 1 | 1 | NA | 0.839 | 1 | 1 | 0.04 |
| SE | 84542 | ENSG00000069509 | FUNDC1 | chrX | - | 44527236 | 44527365 | 44523638 | 44524275 | 44538466 | 44538542 | 101 | 75 | 69 | 118 | 164 | 0 | 0 | 0 | 0 | 0 | 71 | 48 | 53 | 74 | 18 | 0 | 3 | 0 | 2 | 0 | 278 | 149 | 1 | 1 | 1 | 1 | 1 | 1 | 1 | 1 | 0.896 | 1 | 0.952 | 1 | 0.03 |
| SE | 84543 | ENSG00000069509 | FUNDC1 | chrX | - | 44538466 | 44538542 | 44527236 | 44527365 | 44541944 | 44542101 | 82 | 84 | 88 | 90 | 193 | 3 | 0 | 3 | 0 | 1 | 8 | 61 | 38 | 44 | 0 | 0 | 0 | 0 | 0 | 0 | 225 | 149 | 1 | 1 | 0.948 | 1 | 0.951 | 1 | 0.992 | 1 | 1 | 1 | 1 | NA | -0.022 |
| SE | 84544 | ENSG00000156170 | NDUFAF6 | chr8 | + | 94945482 | 94945619 | 94895812 | 94895927 | 94957994 | 94958179 | 3 | 0 | 0 | 0 | 0 | 0 | 1 | 0 | 0 | 0 | 0 | 0 | 0 | 2 | 0 | 0 | 0 | 0 | 5 | 0 | 286 | 149 | 1 | 1 | 1 | 0 | NA | NA | NA | NA | NA | NA | 0.172 | NA | 0.328 |
| SE | 84545 | ENSG00000156170 | NDUFAF6 | chr8 | + | 94975317 | 94975415 | 94957994 | 94958179 | 94975816 | 94975983 | 7 | 2 | 0 | 3 | 3 | 1 | 2 | 0 | 3 | 0 | 0 | 11 | 4 | 0 | 0 | 0 | 3 | 0 | 2 | 0 | 247 | 149 | 1 | 1 | 0.809 | 0.376 | NA | 0.376 | 1 | NA | 0.689 | 1 | 0 | NA | 0.077 |
| SE | 84546 | ENSG00000156170 | NDUFAF6 | chr8 | + | 94975317 | 94975415 | 94957994 | 94958179 | 94980854 | 94980973 | 0 | 2 | 1 | 2 | 2 | 2 | 0 | 1 | 0 | 5 | 0 | 0 | 1 | 0 | 0 | 0 | 1 | 0 | 3 | 0 | 247 | 149 | 1 | 1 | 0 | 1 | 0.376 | 1 | 0.194 | NA | 0 | 1 | 0 | NA | 0.181 |
| SE | 84551 | ENSG00000156170 | NDUFAF6 | chr8 | + | 95031994 | 95032094 | 95024988 | 95025205 | 95035453 | 95035576 | 8 | 4 | 5 | 17 | 14 | 0 | 0 | 0 | 0 | 0 | 4 | 11 | 9 | 16 | 2 | 0 | 0 | 2 | 0 | 0 | 249 | 149 | 1 | 1 | 1 | 1 | 1 | 1 | 1 | 1 | 1 | 0.729 | 1 | 1 | 0.054 |
| SE | 84552 | ENSG00000156170 | NDUFAF6 | chr8 | + | 95031994 | 95032098 | 94957994 | 94958179 | 95035453 | 95035576 | 9 | 3 | 11 | 8 | 19 | 0 | 0 | 0 | 1 | 0 | 4 | 14 | 3 | 4 | 1 | 0 | 0 | 0 | 0 | 0 | 253 | 149 | 1 | 1 | 1 | 1 | 1 | 0.825 | 1 | 1 | 1 | 1 | 1 | 1 | -0.035 |
| SE | 84553 | ENSG00000156170 | NDUFAF6 | chr8 | + | 95031994 | 95032098 | 95024993 | 95025205 | 95035453 | 95035576 | 11 | 3 | 12 | 13 | 19 | 0 | 0 | 0 | 0 | 0 | 8 | 14 | 4 | 4 | 1 | 0 | 0 | 2 | 0 | 0 | 253 | 149 | 1 | 1 | 1 | 1 | 1 | 1 | 1 | 1 | 1 | 0.541 | 1 | 1 | 0.092 |
| SE | 84557 | ENSG00000156170 | NDUFAF6 | chr8 | + | 95040705 | 95040927 | 95035453 | 95035576 | 95041569 | 95041626 | 0 | 0 | 4 | 0 | 0 | 35 | 29 | 29 | 45 | 93 | 0 | 0 | 0 | 0 | 0 | 24 | 27 | 34 | 28 | 46 | 298 | 149 | 1 | 1 | 0 | 0 | 0.065 | 0 | 0 | 0 | 0 | 0 | 0 | 0 | 0.013 |
| SE | 84560 | ENSG00000156170 | NDUFAF6 | chr8 | + | 95052173 | 95052230 | 95048456 | 95048558 | 95075632 | 95076168 | 3 | 0 | 0 | 0 | 2 | 0 | 0 | 1 | 0 | 0 | 8 | 0 | 1 | 1 | 8 | 0 | 0 | 1 | 1 | 0 | 206 | 149 | 1 | 1 | 1 | NA | 0 | NA | 1 | 1 | NA | 0.42 | 0.42 | 1 | -0.043 |
| SE | 84561 | ENSG00000166432 | ZMAT1 | chrX | - | 101887197 | 101887304 | 101886631 | 101886731 | 101892704 | 101892754 | 5 | 10 | 7 | 16 | 0 | 1 | 6 | 1 | 6 | 4 | 0 | 1 | 17 | 0 | 2 | 0 | 0 | 0 | 0 | 0 | 256 | 149 | 1 | 1 | 0.744 | 0.492 | 0.803 | 0.608 | 0 | NA | 1 | 1 | NA | 1 | -0.471 |
| SE | 84565 | ENSG00000166432 | ZMAT1 | chrX | - | 101918512 | 101918633 | 101904223 | 101904330 | 101931716 | 101931764 | 4 | 0 | 0 | 0 | 0 | 16 | 11 | 16 | 23 | 12 | 0 | 0 | 0 | 0 | 0 | 0 | 4 | 3 | 2 | 0 | 270 | 149 | 1 | 1 | 0.121 | 0 | 0 | 0 | 0 | NA | 0 | 0 | 0 | NA | 0.024 |
| SE | 84580 | ENSG00000134668 | SPOCD1 | chr1 | - | 31798206 | 31798323 | 31794017 | 31794235 | 31798441 | 31798601 | 11 | 0 | 3 | 11 | 5 | 0 | 0 | 0 | 2 | 0 | 33 | 14 | 5 | 4 | 31 | 0 | 0 | 0 | 0 | 0 | 266 | 149 | 1 | 1 | 1 | NA | 1 | 0.755 | 1 | 1 | 1 | 1 | 1 | 1 | -0.061 |
| SE | 84590 | ENSG00000132356 | PRKAA1 | chr5 | - | 40769415 | 40769503 | 40767465 | 40767690 | 40775409 | 40775503 | 101 | 72 | 92 | 121 | 188 | 0 | 0 | 2 | 0 | 0 | 27 | 42 | 68 | 87 | 33 | 0 | 0 | 0 | 0 | 0 | 237 | 149 | 1 | 1 | 1 | 1 | 0.967 | 1 | 1 | 1 | 1 | 1 | 1 | 1 | -0.007 |
| SE | 84591 | ENSG00000132356 | PRKAA1 | chr5 | - | 40771718 | 40771863 | 40767506 | 40767690 | 40775409 | 40775503 | 172 | 128 | 144 | 202 | 158 | 0 | 0 | 2 | 0 | 0 | 32 | 141 | 142 | 69 | 18 | 0 | 0 | 0 | 0 | 0 | 294 | 149 | 1 | 1 | 1 | 1 | 0.973 | 1 | 1 | 1 | 1 | 1 | 1 | 1 | -0.005 |
| SE | 84592 | ENSG00000132356 | PRKAA1 | chr5 | - | 40771718 | 40771863 | 40769415 | 40769503 | 40775409 | 40775503 | 535 | 392 | 411 | 589 | 502 | 0 | 0 | 1 | 0 | 0 | 127 | 314 | 381 | 199 | 44 | 0 | 0 | 2 | 0 | 0 | 294 | 149 | 1 | 1 | 1 | 1 | 0.995 | 1 | 1 | 1 | 1 | 0.99 | 1 | 1 | 0.001 |
| SE | 84594 | ENSG00000132356 | PRKAA1 | chr5 | - | 40775409 | 40775503 | 40771718 | 40771863 | 40777444 | 40777586 | 363 | 260 | 321 | 352 | 342 | 1 | 0 | 0 | 0 | 0 | 55 | 279 | 227 | 185 | 18 | 0 | 0 | 0 | 0 | 0 | 243 | 149 | 1 | 1 | 0.996 | 1 | 1 | 1 | 1 | 1 | 1 | 1 | 1 | 1 | -0.001 |
| SE | 84597 | ENSG00000153130 | SCOC | chr4 | + | 140379568 | 140379652 | 140379120 | 140379192 | 140380961 | 140381126 | 104 | 66 | 99 | 71 | 182 | 5 | 0 | 2 | 1 | 7 | 0 | 32 | 2 | 18 | 9 | 0 | 0 | 8 | 0 | 0 | 233 | 149 | 1 | 1 | 0.93 | 1 | 0.969 | 0.978 | 0.943 | NA | 1 | 0.138 | 1 | 1 | 0.179 |
| SE | 84598 | ENSG00000245958 | SEPTIN7P14 | chr4 | + | 119457970 | 119458077 | 119454895 | 119455133 | 119460182 | 119460242 | 0 | 4 | 0 | 3 | 0 | 14 | 5 | 8 | 1 | 4 | 0 | 0 | 0 | 0 | 0 | 5 | 5 | 12 | 6 | 0 | 256 | 149 | 1 | 1 | 0 | 0.318 | 0 | 0.636 | 0 | 0 | 0 | 0 | 0 | NA | 0.191 |
| SE | 84599 | ENSG00000245958 | SEPTIN7P14 | chr4 | + | 119457970 | 119458170 | 119454790 | 119455133 | 119460182 | 119460244 | 0 | 3 | 0 | 3 | 0 | 14 | 5 | 8 | 1 | 4 | 0 | 0 | 0 | 0 | 0 | 5 | 5 | 12 | 6 | 0 | 298 | 149 | 1 | 1 | 0 | 0.231 | 0 | 0.6 | 0 | 0 | 0 | 0 | 0 | NA | 0.166 |
| SE | 84600 | ENSG00000245958 | SEPTIN7P14 | chr4 | + | 119460182 | 119460242 | 119454819 | 119455133 | 119460730 | 119460827 | 20 | 7 | 19 | 12 | 17 | 0 | 0 | 0 | 0 | 0 | 5 | 8 | 22 | 10 | 7 | 0 | 0 | 4 | 0 | 0 | 209 | 149 | 1 | 1 | 1 | 1 | 1 | 1 | 1 | 1 | 1 | 0.797 | 1 | 1 | 0.041 |
| SE | 84601 | ENSG00000245958 | SEPTIN7P14 | chr4 | + | 119461716 | 119461851 | 119460730 | 119460827 | 119462895 | 119462979 | 58 | 59 | 73 | 101 | 117 | 0 | 0 | 2 | 0 | 0 | 23 | 48 | 105 | 32 | 51 | 0 | 0 | 1 | 0 | 0 | 284 | 149 | 1 | 1 | 1 | 1 | 0.95 | 1 | 1 | 1 | 1 | 1 | 1 | 1 | -0.01 |
| SE | 84604 | ENSG00000245958 | SEPTIN7P14 | chr4 | + | 119493332 | 119493437 | 119462895 | 119462979 | 119494432 | 119494523 | 83 | 70 | 77 | 79 | 120 | 0 | 0 | 1 | 2 | 0 | 49 | 61 | 89 | 22 | 74 | 0 | 2 | 0 | 0 | 0 | 254 | 149 | 1 | 1 | 1 | 1 | 0.978 | 0.959 | 1 | 1 | 0.947 | 1 | 1 | 1 | -0.002 |
| SE | 84607 | ENSG00000245958 | SEPTIN7P14 | chr4 | + | 119528739 | 119528789 | 119512812 | 119512970 | 119550378 | 119550443 | 6 | 9 | 4 | 26 | 30 | 2 | 2 | 0 | 5 | 1 | 0 | 1 | 4 | 3 | 5 | 0 | 1 | 0 | 0 | 0 | 199 | 149 | 1 | 1 | 0.692 | 0.771 | 1 | 0.796 | 0.957 | NA | 1 | 1 | 1 | 1 | -0.157 |
| SE | 84608 | ENSG00000245958 | SEPTIN7P14 | chr4 | + | 119543752 | 119543845 | 119528739 | 119528789 | 119550378 | 119550443 | 1 | 0 | 0 | 2 | 9 | 6 | 7 | 2 | 20 | 23 | 6 | 0 | 2 | 0 | 0 | 0 | 1 | 3 | 0 | 5 | 242 | 149 | 1 | 1 | 0.093 | 0 | 0 | 0.058 | 0.194 | 1 | 0 | 0.291 | NA | 0 | -0.254 |
| SE | 84609 | ENSG00000015285 | WAS | chrX | + | 48684282 | 48684423 | 48683798 | 48683985 | 48685546 | 48685633 | 603 | 301 | 408 | 432 | 251 | 0 | 0 | 0 | 0 | 0 | 588 | 474 | 340 | 534 | 778 | 0 | 0 | 1 | 0 | 0 | 290 | 149 | 1 | 1 | 1 | 1 | 1 | 1 | 1 | 1 | 1 | 0.994 | 1 | 1 | 0.001 |
| SE | 84610 | ENSG00000015285 | WAS | chrX | + | 48685546 | 48685633 | 48684282 | 48684423 | 48685733 | 48685836 | 429 | 215 | 299 | 317 | 188 | 7 | 2 | 1 | 0 | 0 | 495 | 412 | 277 | 417 | 571 | 0 | 2 | 0 | 6 | 7 | 236 | 149 | 1 | 1 | 0.975 | 0.985 | 0.995 | 1 | 1 | 1 | 0.992 | 1 | 0.978 | 0.981 | 0.001 |
| SE | 84611 | ENSG00000015285 | WAS | chrX | + | 48686780 | 48686955 | 48686080 | 48686134 | 48688053 | 48688096 | 653 | 368 | 497 | 614 | 591 | 0 | 0 | 0 | 0 | 1 | 1249 | 866 | 535 | 904 | 1521 | 0 | 0 | 0 | 0 | 1 | 298 | 149 | 1 | 1 | 1 | 1 | 1 | 1 | 0.997 | 1 | 1 | 1 | 1 | 0.999 | 0 |
| SE | 84612 | ENSG00000015285 | WAS | chrX | + | 48688053 | 48688096 | 48686780 | 48686955 | 48688299 | 48688453 | 380 | 170 | 292 | 319 | 321 | 0 | 0 | 0 | 0 | 0 | 612 | 438 | 264 | 428 | 851 | 5 | 0 | 0 | 0 | 0 | 192 | 149 | 1 | 1 | 1 | 1 | 1 | 1 | 1 | 0.99 | 1 | 1 | 1 | 1 | 0.002 |
| SE | 84613 | ENSG00000015285 | WAS | chrX | + | 48688659 | 48689066 | 48688299 | 48688453 | 48689319 | 48689434 | 338 | 163 | 233 | 292 | 240 | 0 | 0 | 0 | 2 | 0 | 468 | 415 | 228 | 462 | 583 | 4 | 0 | 4 | 0 | 0 | 298 | 149 | 1 | 1 | 1 | 1 | 1 | 0.986 | 1 | 0.983 | 1 | 0.966 | 1 | 1 | 0.007 |
| SE | 84614 | ENSG00000015285 | WAS | chrX | + | 48689319 | 48689434 | 48688659 | 48689066 | 48691106 | 48691427 | 322 | 143 | 196 | 229 | 202 | 0 | 0 | 0 | 0 | 0 | 511 | 354 | 159 | 365 | 504 | 0 | 0 | 1 | 0 | 0 | 264 | 149 | 1 | 1 | 1 | 1 | 1 | 1 | 1 | 1 | 1 | 0.989 | 1 | 1 | 0.002 |
| SE | 84631 | ENSG00000090238 | YPEL3 | chr16 | - | 30094997 | 30095146 | 30094852 | 30094897 | 30095251 | 30095434 | 22 | 12 | 26 | 31 | 5 | 0 | 0 | 0 | 0 | 0 | 52 | 60 | 34 | 31 | 38 | 0 | 2 | 0 | 0 | 0 | 298 | 149 | 1 | 1 | 1 | 1 | 1 | 1 | 1 | 1 | 0.938 | 1 | 1 | 1 | 0.012 |
| SE | 84632 | ENSG00000090238 | YPEL3 | chr16 | - | 30095102 | 30095146 | 30093761 | 30094897 | 30095251 | 30095335 | 138 | 53 | 83 | 170 | 49 | 0 | 0 | 0 | 0 | 0 | 207 | 167 | 159 | 103 | 321 | 0 | 2 | 0 | 0 | 0 | 193 | 149 | 1 | 1 | 1 | 1 | 1 | 1 | 1 | 1 | 0.985 | 1 | 1 | 1 | 0.003 |
| SE | 84633 | ENSG00000262919 | CCNQ | chrX | - | 153592505 | 153592733 | 153587924 | 153588454 | 153594546 | 153594679 | 21 | 15 | 16 | 16 | 5 | 0 | 1 | 0 | 1 | 0 | 0 | 2 | 3 | 15 | 17 | 0 | 2 | 1 | 0 | 0 | 298 | 149 | 1 | 1 | 1 | 0.882 | 1 | 0.889 | 1 | NA | 0.333 | 0.6 | 1 | 1 | 0.221 |
| SE | 84634 | ENSG00000262919 | CCNQ | chrX | - | 153592505 | 153592733 | 153587931 | 153588394 | 153594546 | 153594679 | 22 | 12 | 18 | 22 | 6 | 0 | 0 | 0 | 0 | 1 | 0 | 5 | 17 | 22 | 17 | 0 | 0 | 3 | 0 | 0 | 298 | 149 | 1 | 1 | 1 | 1 | 1 | 1 | 0.75 | NA | 1 | 0.739 | 1 | 1 | 0.015 |
| SE | 84644 | ENSG00000131196 | NFATC1 | chr18 | + | 79461310 | 79461366 | 79433578 | 79433738 | 79467449 | 79467582 | 22 | 13 | 20 | 26 | 43 | 2 | 0 | 0 | 0 | 0 | 0 | 19 | 8 | 6 | 6 | 0 | 0 | 0 | 0 | 0 | 205 | 149 | 1 | 1 | 0.889 | 1 | 1 | 1 | 1 | NA | 1 | 1 | 1 | 1 | -0.022 |
| SE | 84645 | ENSG00000131196 | NFATC1 | chr18 | + | 79461310 | 79461366 | 79451675 | 79451816 | 79467449 | 79467582 | 83 | 41 | 56 | 76 | 87 | 0 | 0 | 0 | 0 | 0 | 17 | 53 | 32 | 31 | 32 | 0 | 0 | 1 | 0 | 0 | 205 | 149 | 1 | 1 | 1 | 1 | 1 | 1 | 1 | 1 | 1 | 0.959 | 1 | 1 | 0.008 |
| SE | 84646 | ENSG00000131196 | NFATC1 | chr18 | + | 79461310 | 79461366 | 79451675 | 79451816 | 79486247 | 79486630 | 10 | 2 | 16 | 12 | 13 | 0 | 0 | 0 | 0 | 0 | 6 | 5 | 1 | 7 | 0 | 0 | 3 | 0 | 0 | 0 | 205 | 149 | 1 | 1 | 1 | 1 | 1 | 1 | 1 | 1 | 0.548 | 1 | 1 | NA | 0.113 |
| SE | 84647 | ENSG00000131196 | NFATC1 | chr18 | + | 79467449 | 79467582 | 79451675 | 79451816 | 79486247 | 79486630 | 75 | 35 | 48 | 60 | 117 | 0 | 0 | 0 | 0 | 0 | 33 | 57 | 18 | 62 | 59 | 0 | 3 | 0 | 0 | 0 | 282 | 149 | 1 | 1 | 1 | 1 | 1 | 1 | 1 | 1 | 0.909 | 1 | 1 | 1 | 0.018 |
| SE | 84648 | ENSG00000131196 | NFATC1 | chr18 | + | 79467449 | 79467582 | 79451675 | 79451816 | 79527527 | 79527577 | 1 | 0 | 3 | 0 | 2 | 0 | 0 | 3 | 0 | 0 | 0 | 0 | 2 | 0 | 2 | 0 | 0 | 0 | 0 | 0 | 282 | 149 | 1 | 1 | 1 | NA | 0.346 | NA | 1 | NA | NA | 1 | NA | 1 | -0.218 |
| SE | 84649 | ENSG00000131196 | NFATC1 | chr18 | + | 79467449 | 79467582 | 79461310 | 79461366 | 79486247 | 79486630 | 146 | 74 | 87 | 123 | 190 | 0 | 0 | 0 | 0 | 0 | 44 | 113 | 48 | 86 | 99 | 0 | 1 | 0 | 0 | 0 | 282 | 149 | 1 | 1 | 1 | 1 | 1 | 1 | 1 | 1 | 0.984 | 1 | 1 | 1 | 0.003 |
| SE | 84650 | ENSG00000131196 | NFATC1 | chr18 | + | 79486247 | 79486630 | 79451675 | 79451816 | 79527527 | 79529323 | 8 | 4 | 4 | 7 | 0 | 0 | 0 | 3 | 0 | 0 | 0 | 14 | 2 | 2 | 1 | 0 | 0 | 0 | 0 | 0 | 298 | 149 | 1 | 1 | 1 | 1 | 0.4 | 1 | NA | NA | 1 | 1 | 1 | 1 | -0.15 |
| SE | 84652 | ENSG00000131196 | NFATC1 | chr18 | + | 79486247 | 79486937 | 79451675 | 79451816 | 79527527 | 79527741 | 18 | 11 | 23 | 32 | 17 | 0 | 0 | 3 | 0 | 0 | 49 | 15 | 13 | 32 | 54 | 0 | 0 | 0 | 0 | 0 | 298 | 149 | 1 | 1 | 1 | 1 | 0.793 | 1 | 1 | 1 | 1 | 1 | 1 | 1 | -0.041 |
| SE | 84664 | ENSG00000142528 | ZNF473 | chr19 | + | 50030891 | 50031091 | 50025892 | 50026122 | 50039160 | 50039287 | 16 | 1 | 13 | 4 | 0 | 7 | 1 | 0 | 3 | 0 | 8 | 29 | 2 | 3 | 0 | 2 | 4 | 0 | 0 | 0 | 298 | 149 | 1 | 1 | 0.533 | 0.333 | 1 | 0.4 | NA | 0.667 | 0.784 | 1 | 1 | NA | -0.296 |
| SE | 84667 | ENSG00000152465 | NMT2 | chr10 | - | 15119342 | 15119513 | 15112795 | 15112963 | 15128349 | 15128458 | 229 | 133 | 369 | 187 | 404 | 0 | 0 | 1 | 0 | 0 | 35 | 83 | 88 | 59 | 9 | 0 | 0 | 0 | 0 | 0 | 298 | 149 | 1 | 1 | 1 | 1 | 0.995 | 1 | 1 | 1 | 1 | 1 | 1 | 1 | -0.001 |
| SE | 84670 | ENSG00000224016 | NA | chr7 | - | 149892273 | 149892398 | 149891190 | 149891301 | 149909649 | 149909704 | 1 | 2 | 2 | 1 | 7 | 0 | 0 | 1 | 0 | 0 | 32 | 5 | 2 | 1 | 28 | 0 | 0 | 0 | 0 | 0 | 274 | 149 | 1 | 1 | 1 | 1 | 0.521 | 1 | 1 | 1 | 1 | 1 | 1 | 1 | -0.096 |
| SE | 84687 | ENSG00000258867 | LINC01146 | chr14 | + | 88044614 | 88044848 | 88025406 | 88025481 | 88079103 | 88079368 | 2 | 0 | 3 | 1 | 0 | 0 | 0 | 1 | 0 | 0 | 0 | 0 | 2 | 0 | 0 | 0 | 0 | 0 | 0 | 0 | 298 | 149 | 1 | 1 | 1 | NA | 0.6 | 1 | NA | NA | NA | 1 | NA | NA | -0.133 |
| SE | 84696 | ENSG00000258867 | LINC01146 | chr14 | + | 88044685 | 88044848 | 88025406 | 88025481 | 88079103 | 88079368 | 2 | 0 | 4 | 1 | 1 | 0 | 0 | 1 | 0 | 0 | 0 | 0 | 2 | 0 | 0 | 0 | 0 | 0 | 0 | 0 | 298 | 149 | 1 | 1 | 1 | NA | 0.667 | 1 | 1 | NA | NA | 1 | NA | NA | -0.083 |
| SE | 84707 | ENSG00000258867 | LINC01146 | chr14 | + | 88079103 | 88079368 | 88044496 | 88044848 | 88084036 | 88084155 | 7 | 0 | 3 | 1 | 6 | 1 | 0 | 4 | 2 | 1 | 0 | 1 | 2 | 0 | 0 | 0 | 0 | 0 | 0 | 0 | 298 | 149 | 1 | 1 | 0.778 | NA | 0.273 | 0.2 | 0.75 | NA | 1 | 1 | NA | NA | -0.5 |
| SE | 84708 | ENSG00000258867 | LINC01146 | chr14 | + | 88079103 | 88079368 | 88044496 | 88044848 | 88086827 | 88087110 | 2 | 0 | 3 | 1 | 0 | 0 | 0 | 0 | 1 | 0 | 0 | 0 | 2 | 0 | 0 | 0 | 0 | 0 | 0 | 0 | 298 | 149 | 1 | 1 | 1 | NA | 1 | 0.333 | NA | NA | NA | 1 | NA | NA | -0.222 |
| SE | 84711 | ENSG00000258867 | LINC01146 | chr14 | + | 88079103 | 88079368 | 88044496 | 88044848 | 88096498 | 88097629 | 2 | 0 | 3 | 1 | 0 | 1 | 0 | 4 | 0 | 0 | 0 | 0 | 2 | 0 | 0 | 0 | 0 | 0 | 0 | 0 | 298 | 149 | 1 | 1 | 0.5 | NA | 0.273 | 1 | NA | NA | NA | 1 | NA | NA | -0.409 |
| SE | 84718 | ENSG00000258867 | LINC01146 | chr14 | + | 88084036 | 88084155 | 88044511 | 88044848 | 88096498 | 88096583 | 6 | 1 | 8 | 3 | 1 | 1 | 0 | 4 | 0 | 0 | 0 | 2 | 1 | 0 | 0 | 0 | 0 | 0 | 0 | 0 | 268 | 149 | 1 | 1 | 0.769 | 1 | 0.527 | 1 | 1 | NA | 1 | 1 | NA | NA | -0.141 |
| SE | 84723 | ENSG00000258867 | LINC01146 | chr14 | + | 88084983 | 88085036 | 88084036 | 88084155 | 88086827 | 88087023 | 0 | 0 | 0 | 0 | 0 | 3 | 1 | 0 | 2 | 1 | 0 | 1 | 0 | 0 | 0 | 0 | 0 | 0 | 0 | 0 | 202 | 149 | 1 | 1 | 0 | 0 | NA | 0 | 0 | NA | 1 | NA | NA | NA | -1 |
| SE | 84729 | ENSG00000258867 | LINC01146 | chr14 | + | 88086827 | 88087023 | 88084036 | 88084155 | 88096498 | 88097625 | 3 | 1 | 2 | 2 | 1 | 9 | 1 | 4 | 1 | 0 | 0 | 0 | 0 | 0 | 0 | 0 | 2 | 1 | 0 | 0 | 298 | 149 | 1 | 1 | 0.143 | 0.333 | 0.2 | 0.5 | 1 | NA | 0 | 0 | NA | NA | 0.435 |
| SE | 84733 | ENSG00000258867 | LINC01146 | chr14 | + | 88086827 | 88087037 | 88084036 | 88084155 | 88096498 | 88096594 | 3 | 1 | 0 | 2 | 1 | 9 | 1 | 4 | 1 | 0 | 0 | 0 | 0 | 0 | 0 | 0 | 2 | 1 | 0 | 0 | 298 | 149 | 1 | 1 | 0.143 | 0.333 | 0 | 0.5 | 1 | NA | 0 | 0 | NA | NA | 0.395 |
| SE | 84739 | ENSG00000258867 | LINC01146 | chr14 | + | 88090022 | 88090210 | 88084036 | 88084155 | 88096498 | 88097608 | 0 | 1 | 0 | 0 | 0 | 9 | 1 | 4 | 1 | 0 | 0 | 0 | 0 | 0 | 0 | 0 | 2 | 1 | 0 | 0 | 298 | 149 | 1 | 1 | 0 | 0.333 | 0 | 0 | NA | NA | 0 | 0 | NA | NA | 0.083 |
| SE | 84744 | ENSG00000258867 | LINC01146 | chr14 | + | 88091335 | 88091449 | 88090022 | 88090210 | 88096498 | 88097623 | 0 | 0 | 1 | 2 | 4 | 0 | 1 | 0 | 0 | 0 | 0 | 4 | 1 | 2 | 2 | 0 | 0 | 0 | 0 | 0 | 263 | 149 | 1 | 1 | NA | 0 | 1 | 1 | 1 | NA | 1 | 1 | 1 | 1 | -0.25 |
| SE | 84745 | ENSG00000258867 | LINC01146 | chr14 | + | 88091335 | 88091652 | 88090022 | 88090210 | 88096498 | 88097622 | 1 | 1 | 6 | 3 | 8 | 0 | 1 | 0 | 0 | 0 | 0 | 10 | 1 | 4 | 2 | 0 | 0 | 0 | 0 | 0 | 298 | 149 | 1 | 1 | 1 | 0.333 | 1 | 1 | 1 | NA | 1 | 1 | 1 | 1 | -0.133 |
| SE | 84746 | ENSG00000286163 | NA | chrX | + | 121151610 | 121151724 | 121093391 | 121093477 | 121164058 | 121164157 | 0 | 0 | 0 | 0 | 0 | 2 | 0 | 0 | 0 | 0 | 0 | 0 | 3 | 0 | 0 | 0 | 0 | 0 | 0 | 0 | 263 | 149 | 1 | 1 | 0 | NA | NA | NA | NA | NA | NA | 1 | NA | NA | -1 |
| SE | 84747 | ENSG00000187676 | B3GLCT | chr13 | + | 31269213 | 31269277 | 31260945 | 31261082 | 31274508 | 31274628 | 41 | 16 | 19 | 30 | 49 | 0 | 0 | 1 | 0 | 0 | 66 | 10 | 9 | 27 | 4 | 0 | 0 | 0 | 2 | 0 | 213 | 149 | 1 | 1 | 1 | 1 | 0.93 | 1 | 1 | 1 | 1 | 1 | 0.904 | 1 | 0.005 |
| SE | 84748 | ENSG00000187676 | B3GLCT | chr13 | + | 31274508 | 31274628 | 31269213 | 31269277 | 31276701 | 31276771 | 39 | 11 | 20 | 26 | 38 | 0 | 1 | 0 | 1 | 0 | 35 | 19 | 14 | 31 | 4 | 0 | 0 | 0 | 0 | 0 | 269 | 149 | 1 | 1 | 1 | 0.859 | 1 | 0.935 | 1 | 1 | 1 | 1 | 1 | 1 | -0.041 |
| SE | 84749 | ENSG00000187676 | B3GLCT | chr13 | + | 31276701 | 31276771 | 31274508 | 31274628 | 31284647 | 31284761 | 44 | 12 | 26 | 24 | 34 | 0 | 0 | 1 | 0 | 0 | 9 | 22 | 20 | 27 | 0 | 0 | 0 | 0 | 0 | 0 | 219 | 149 | 1 | 1 | 1 | 1 | 0.946 | 1 | 1 | 1 | 1 | 1 | 1 | NA | -0.011 |
| SE | 84751 | ENSG00000187676 | B3GLCT | chr13 | + | 31317565 | 31317685 | 31286719 | 31286819 | 31323750 | 31323895 | 55 | 20 | 31 | 37 | 49 | 2 | 0 | 1 | 2 | 0 | 16 | 20 | 31 | 38 | 0 | 0 | 0 | 2 | 0 | 0 | 269 | 149 | 1 | 1 | 0.938 | 1 | 0.945 | 0.911 | 1 | 1 | 1 | 0.896 | 1 | NA | -0.015 |
| SE | 84752 | ENSG00000130734 | ATG4D | chr19 | + | 10546838 | 10547071 | 10544956 | 10545130 | 10547188 | 10547253 | 22 | 4 | 20 | 8 | 4 | 0 | 0 | 0 | 0 | 0 | 0 | 1 | 9 | 11 | 17 | 0 | 0 | 1 | 0 | 0 | 298 | 149 | 1 | 1 | 1 | 1 | 1 | 1 | 1 | NA | 1 | 0.818 | 1 | 1 | 0.045 |
| SE | 84753 | ENSG00000130734 | ATG4D | chr19 | + | 10546838 | 10547075 | 10544782 | 10545130 | 10547188 | 10547253 | 21 | 4 | 16 | 7 | 4 | 0 | 0 | 0 | 0 | 0 | 0 | 1 | 9 | 14 | 30 | 0 | 0 | 1 | 0 | 0 | 298 | 149 | 1 | 1 | 1 | 1 | 1 | 1 | 1 | NA | 1 | 0.818 | 1 | 1 | 0.045 |
| SE | 84754 | ENSG00000130734 | ATG4D | chr19 | + | 10546838 | 10547082 | 10545119 | 10545130 | 10547188 | 10547253 | 21 | 4 | 16 | 7 | 4 | 0 | 0 | 0 | 0 | 0 | 0 | 1 | 9 | 11 | 17 | 0 | 0 | 1 | 0 | 0 | 298 | 149 | 1 | 1 | 1 | 1 | 1 | 1 | 1 | NA | 1 | 0.818 | 1 | 1 | 0.045 |
| SE | 84755 | ENSG00000130734 | ATG4D | chr19 | + | 10546838 | 10547115 | 10544956 | 10545130 | 10547188 | 10547253 | 31 | 9 | 26 | 12 | 12 | 0 | 0 | 0 | 0 | 0 | 11 | 6 | 10 | 15 | 36 | 0 | 0 | 1 | 0 | 0 | 298 | 149 | 1 | 1 | 1 | 1 | 1 | 1 | 1 | 1 | 1 | 0.833 | 1 | 1 | 0.033 |
| SE | 84756 | ENSG00000130734 | ATG4D | chr19 | + | 10546838 | 10547115 | 10544956 | 10545130 | 10548903 | 10549034 | 21 | 4 | 18 | 7 | 4 | 0 | 1 | 0 | 1 | 0 | 0 | 1 | 12 | 11 | 17 | 0 | 15 | 0 | 3 | 4 | 298 | 149 | 1 | 1 | 1 | 0.667 | 1 | 0.778 | 1 | NA | 0.032 | 1 | 0.647 | 0.68 | 0.299 |
| SE | 84757 | ENSG00000130734 | ATG4D | chr19 | + | 10546838 | 10547253 | 10544956 | 10545130 | 10548903 | 10549034 | 30 | 5 | 26 | 9 | 10 | 0 | 1 | 0 | 1 | 0 | 0 | 16 | 23 | 13 | 21 | 0 | 15 | 0 | 3 | 4 | 298 | 149 | 1 | 1 | 1 | 0.714 | 1 | 0.818 | 1 | NA | 0.348 | 1 | 0.684 | 0.724 | 0.217 |
| SE | 84758 | ENSG00000130734 | ATG4D | chr19 | + | 10547188 | 10547253 | 10544956 | 10545130 | 10548903 | 10549034 | 9 | 1 | 8 | 2 | 6 | 0 | 1 | 0 | 1 | 0 | 0 | 15 | 14 | 2 | 4 | 0 | 15 | 0 | 3 | 4 | 214 | 149 | 1 | 1 | 1 | 0.41 | 1 | 0.582 | 1 | NA | 0.41 | 1 | 0.317 | 0.41 | 0.264 |
| SE | 84759 | ENSG00000130734 | ATG4D | chr19 | + | 10547188 | 10547253 | 10546838 | 10547115 | 10548903 | 10549034 | 19 | 6 | 18 | 7 | 14 | 0 | 0 | 2 | 0 | 0 | 0 | 20 | 14 | 6 | 17 | 0 | 0 | 3 | 0 | 0 | 214 | 149 | 1 | 1 | 1 | 1 | 0.862 | 1 | 1 | NA | 1 | 0.765 | 1 | 1 | 0.031 |
| SE | 84761 | ENSG00000130734 | ATG4D | chr19 | + | 10551896 | 10551975 | 10548903 | 10549034 | 10552044 | 10552121 | 25 | 6 | 25 | 7 | 11 | 1 | 0 | 0 | 1 | 0 | 8 | 24 | 32 | 14 | 23 | 0 | 0 | 0 | 0 | 0 | 228 | 149 | 1 | 1 | 0.942 | 1 | 1 | 0.821 | 1 | 1 | 1 | 1 | 1 | 1 | -0.047 |
| SE | 84763 | ENSG00000158201 | ABHD3 | chr18 | - | 21656860 | 21657026 | 21650900 | 21651763 | 21657103 | 21657152 | 145 | 82 | 124 | 104 | 222 | 0 | 0 | 0 | 0 | 0 | 51 | 74 | 37 | 33 | 21 | 0 | 2 | 0 | 0 | 0 | 298 | 149 | 1 | 1 | 1 | 1 | 1 | 1 | 1 | 1 | 0.949 | 1 | 1 | 1 | 0.01 |
| SE | 84764 | ENSG00000158201 | ABHD3 | chr18 | - | 21657103 | 21657152 | 21656860 | 21657026 | 21659169 | 21659343 | 140 | 88 | 127 | 143 | 211 | 1 | 0 | 0 | 0 | 0 | 56 | 71 | 41 | 48 | 21 | 0 | 0 | 0 | 0 | 0 | 198 | 149 | 1 | 1 | 0.991 | 1 | 1 | 1 | 1 | 1 | 1 | 1 | 1 | 1 | -0.002 |
| SE | 84765 | ENSG00000158201 | ABHD3 | chr18 | - | 21663677 | 21664230 | 21659169 | 21659343 | 21683919 | 21683965 | 60 | 56 | 70 | 75 | 119 | 0 | 0 | 1 | 0 | 1 | 20 | 49 | 20 | 28 | 34 | 0 | 0 | 0 | 0 | 0 | 298 | 149 | 1 | 1 | 1 | 1 | 0.972 | 1 | 0.983 | 1 | 1 | 1 | 1 | 1 | -0.009 |
| SE | 84766 | ENSG00000158201 | ABHD3 | chr18 | - | 21664117 | 21664230 | 21659169 | 21659343 | 21683919 | 21683965 | 62 | 47 | 57 | 73 | 75 | 0 | 0 | 1 | 0 | 1 | 30 | 58 | 15 | 32 | 39 | 0 | 0 | 0 | 0 | 0 | 262 | 149 | 1 | 1 | 1 | 1 | 0.97 | 1 | 0.977 | 1 | 1 | 1 | 1 | 1 | -0.011 |
| SE | 84769 | ENSG00000101856 | PGRMC1 | chrX | + | 119240308 | 119240464 | 119236284 | 119236691 | 119243150 | 119244466 | 267 | 244 | 93 | 396 | 173 | 1 | 1 | 0 | 0 | 0 | 634 | 399 | 348 | 460 | 1201 | 0 | 0 | 0 | 0 | 0 | 298 | 149 | 1 | 1 | 0.993 | 0.992 | 1 | 1 | 1 | 1 | 1 | 1 | 1 | 1 | -0.003 |
| SE | 84774 | ENSG00000125676 | THOC2 | chrX | - | 123625911 | 123626069 | 123624540 | 123624669 | 123626520 | 123626636 | 361 | 269 | 310 | 359 | 470 | 1 | 0 | 0 | 1 | 0 | 24 | 121 | 84 | 16 | 34 | 0 | 0 | 0 | 0 | 0 | 298 | 149 | 1 | 1 | 0.994 | 1 | 1 | 0.994 | 1 | 1 | 1 | 1 | 1 | 1 | -0.002 |
| SE | 84775 | ENSG00000125676 | THOC2 | chrX | - | 123632860 | 123633040 | 123631687 | 123631852 | 123633952 | 123634070 | 278 | 164 | 204 | 274 | 264 | 0 | 2 | 0 | 0 | 0 | 13 | 93 | 62 | 32 | 27 | 0 | 3 | 0 | 0 | 0 | 298 | 149 | 1 | 1 | 1 | 0.976 | 1 | 1 | 1 | 1 | 0.939 | 1 | 1 | 1 | 0.007 |
| SE | 84776 | ENSG00000125676 | THOC2 | chrX | - | 123668158 | 123668314 | 123667105 | 123667278 | 123671668 | 123671761 | 304 | 327 | 239 | 358 | 387 | 0 | 0 | 1 | 0 | 0 | 91 | 196 | 100 | 62 | 123 | 0 | 0 | 0 | 0 | 0 | 298 | 149 | 1 | 1 | 1 | 1 | 0.992 | 1 | 1 | 1 | 1 | 1 | 1 | 1 | -0.002 |
| SE | 84777 | ENSG00000125676 | THOC2 | chrX | - | 123686547 | 123686714 | 123671668 | 123671761 | 123696020 | 123696154 | 422 | 389 | 367 | 417 | 611 | 0 | 0 | 0 | 0 | 1 | 53 | 144 | 111 | 52 | 66 | 0 | 0 | 0 | 0 | 0 | 298 | 149 | 1 | 1 | 1 | 1 | 1 | 1 | 0.997 | 1 | 1 | 1 | 1 | 1 | -0.001 |
| SE | 84778 | ENSG00000125676 | THOC2 | chrX | - | 123696020 | 123696154 | 123671668 | 123671761 | 123696720 | 123696835 | 205 | 133 | 150 | 221 | 245 | 0 | 0 | 0 | 1 | 0 | 37 | 90 | 58 | 20 | 28 | 0 | 0 | 0 | 0 | 0 | 283 | 149 | 1 | 1 | 1 | 1 | 1 | 0.991 | 1 | 1 | 1 | 1 | 1 | 1 | -0.002 |
| SE | 84779 | ENSG00000125676 | THOC2 | chrX | - | 123703453 | 123703505 | 123697680 | 123697751 | 123706857 | 123706949 | 193 | 170 | 150 | 204 | 274 | 1 | 0 | 0 | 0 | 0 | 16 | 100 | 66 | 38 | 12 | 0 | 5 | 0 | 0 | 0 | 201 | 149 | 1 | 1 | 0.993 | 1 | 1 | 1 | 1 | 1 | 0.937 | 1 | 1 | 1 | 0.011 |
| SE | 84785 | ENSG00000204271 | SPIN3 | chrX | - | 56993786 | 56993888 | 56990955 | 56992583 | 56994722 | 56994949 | 3 | 1 | 4 | 1 | 0 | 0 | 1 | 0 | 0 | 0 | 0 | 0 | 4 | 0 | 0 | 0 | 0 | 0 | 0 | 0 | 251 | 149 | 1 | 1 | 1 | 0.372 | 1 | 1 | NA | NA | NA | 1 | NA | NA | -0.157 |
| SE | 84787 | ENSG00000183401 | CCDC159 | chr19 | + | 11349653 | 11349687 | 11346532 | 11346624 | 11349937 | 11350026 | 19 | 10 | 9 | 7 | 10 | 1 | 0 | 0 | 0 | 0 | 44 | 19 | 17 | 29 | 31 | 0 | 0 | 0 | 0 | 0 | 183 | 149 | 1 | 1 | 0.939 | 1 | 1 | 1 | 1 | 1 | 1 | 1 | 1 | 1 | -0.012 |
| SE | 84793 | ENSG00000173681 | BCLAF3 | chrX | - | 19923157 | 19923407 | 19916254 | 19917334 | 19929784 | 19929940 | 0 | 0 | 1 | 0 | 0 | 22 | 23 | 19 | 15 | 20 | 0 | 0 | 0 | 0 | 0 | 0 | 10 | 16 | 0 | 0 | 298 | 149 | 1 | 1 | 0 | 0 | 0.026 | 0 | 0 | NA | 0 | 0 | NA | NA | 0.005 |
| SE | 84794 | ENSG00000173681 | BCLAF3 | chrX | - | 19929784 | 19929940 | 19912859 | 19917334 | 19935808 | 19935898 | 43 | 40 | 39 | 39 | 51 | 0 | 5 | 0 | 0 | 0 | 12 | 45 | 42 | 0 | 9 | 0 | 8 | 0 | 0 | 0 | 298 | 149 | 1 | 1 | 1 | 0.8 | 1 | 1 | 1 | 1 | 0.738 | 1 | NA | 1 | 0.025 |
| SE | 84795 | ENSG00000173681 | BCLAF3 | chrX | - | 19929784 | 19929940 | 19912859 | 19917334 | 19937504 | 19937532 | 23 | 29 | 21 | 15 | 28 | 0 | 1 | 0 | 1 | 1 | 0 | 12 | 27 | 0 | 0 | 0 | 0 | 3 | 0 | 0 | 298 | 149 | 1 | 1 | 1 | 0.935 | 1 | 0.882 | 0.933 | NA | 1 | 0.818 | NA | NA | 0.041 |
| SE | 84796 | ENSG00000173681 | BCLAF3 | chrX | - | 19935808 | 19935898 | 19912859 | 19917334 | 19937504 | 19937532 | 17 | 26 | 32 | 29 | 32 | 0 | 1 | 0 | 1 | 1 | 4 | 28 | 24 | 7 | 0 | 0 | 0 | 3 | 0 | 0 | 239 | 149 | 1 | 1 | 1 | 0.942 | 1 | 0.948 | 0.952 | 1 | 1 | 0.833 | 1 | NA | 0.01 |
| SE | 84799 | ENSG00000173681 | BCLAF3 | chrX | - | 19937417 | 19937532 | 19935808 | 19935898 | 19952987 | 19953051 | 8 | 2 | 4 | 11 | 5 | 0 | 0 | 0 | 0 | 3 | 0 | 0 | 1 | 2 | 0 | 0 | 0 | 0 | 0 | 0 | 264 | 149 | 1 | 1 | 1 | 1 | 1 | 1 | 0.485 | NA | NA | 1 | 1 | NA | -0.103 |
| SE | 84800 | ENSG00000173681 | BCLAF3 | chrX | - | 19937504 | 19937532 | 19935808 | 19935898 | 19952987 | 19953051 | 0 | 4 | 4 | 1 | 2 | 0 | 0 | 0 | 0 | 3 | 0 | 1 | 1 | 0 | 0 | 0 | 0 | 0 | 0 | 0 | 177 | 149 | 1 | 1 | NA | 1 | 1 | 1 | 0.359 | NA | 1 | 1 | NA | NA | -0.16 |
| SE | 84801 | ENSG00000173681 | BCLAF3 | chrX | - | 19950752 | 19950868 | 19937417 | 19937532 | 19952987 | 19953051 | 83 | 96 | 86 | 102 | 148 | 0 | 1 | 1 | 0 | 1 | 47 | 86 | 101 | 22 | 8 | 0 | 0 | 0 | 0 | 0 | 265 | 149 | 1 | 1 | 1 | 0.982 | 0.98 | 1 | 0.988 | 1 | 1 | 1 | 1 | 1 | -0.01 |
| SE | 84803 | ENSG00000137801 | THBS1 | chr15 | + | 39588041 | 39588218 | 39587346 | 39587520 | 39588525 | 39588699 | 160 | 123 | 59 | 254 | 234 | 0 | 0 | 0 | 0 | 0 | 441 | 444 | 532 | 205 | 1878 | 0 | 1 | 0 | 0 | 0 | 298 | 149 | 1 | 1 | 1 | 1 | 1 | 1 | 1 | 1 | 0.996 | 1 | 1 | 1 | 0.001 |
| SE | 84804 | ENSG00000137801 | THBS1 | chr15 | + | 39588041 | 39588218 | 39587346 | 39587520 | 39588958 | 39589086 | 94 | 66 | 30 | 129 | 109 | 0 | 0 | 0 | 0 | 0 | 253 | 294 | 331 | 107 | 955 | 0 | 3 | 0 | 0 | 0 | 298 | 149 | 1 | 1 | 1 | 1 | 1 | 1 | 1 | 1 | 0.98 | 1 | 1 | 1 | 0.004 |
| SE | 84805 | ENSG00000137801 | THBS1 | chr15 | + | 39588525 | 39588699 | 39587346 | 39587520 | 39588958 | 39589086 | 119 | 55 | 28 | 147 | 170 | 0 | 0 | 0 | 0 | 0 | 255 | 218 | 227 | 65 | 924 | 0 | 3 | 0 | 0 | 0 | 298 | 149 | 1 | 1 | 1 | 1 | 1 | 1 | 1 | 1 | 0.973 | 1 | 1 | 1 | 0.005 |
| SE | 84807 | ENSG00000137801 | THBS1 | chr15 | + | 39589804 | 39590023 | 39589201 | 39589354 | 39590515 | 39590623 | 205 | 145 | 61 | 307 | 135 | 0 | 0 | 0 | 0 | 6 | 862 | 578 | 550 | 142 | 2642 | 0 | 0 | 0 | 0 | 0 | 298 | 149 | 1 | 1 | 1 | 1 | 1 | 1 | 0.918 | 1 | 1 | 1 | 1 | 1 | -0.016 |
| SE | 84808 | ENSG00000137801 | THBS1 | chr15 | + | 39590515 | 39590623 | 39589804 | 39590023 | 39591190 | 39591350 | 247 | 270 | 91 | 426 | 242 | 0 | 0 | 4 | 0 | 0 | 1314 | 634 | 884 | 326 | 3745 | 0 | 0 | 0 | 0 | 0 | 257 | 149 | 1 | 1 | 1 | 1 | 0.93 | 1 | 1 | 1 | 1 | 1 | 1 | 1 | -0.014 |
| SE | 84809 | ENSG00000139508 | SLC46A3 | chr13 | - | 28703942 | 28704099 | 28700079 | 28701581 | 28710759 | 28710843 | 222 | 140 | 200 | 227 | 257 | 0 | 0 | 0 | 1 | 0 | 229 | 127 | 94 | 53 | 26 | 0 | 0 | 0 | 0 | 0 | 298 | 149 | 1 | 1 | 1 | 1 | 1 | 0.991 | 1 | 1 | 1 | 1 | 1 | 1 | -0.002 |
| SE | 84810 | ENSG00000142675 | CNKSR1 | chr1 | + | 26184070 | 26184141 | 26183728 | 26183830 | 26184213 | 26184287 | 1 | 0 | 0 | 0 | 2 | 0 | 0 | 0 | 0 | 2 | 19 | 0 | 0 | 6 | 0 | 0 | 0 | 0 | 0 | 0 | 220 | 149 | 1 | 1 | 1 | NA | NA | NA | 0.404 | 1 | NA | 1 | NA | 1 | -0.298 |
| SE | 84811 | ENSG00000142675 | CNKSR1 | chr1 | + | 26185013 | 26185186 | 26184584 | 26184612 | 26187167 | 26187241 | 4 | 0 | 2 | 0 | 3 | 0 | 0 | 1 | 0 | 0 | 0 | 4 | 0 | 0 | 0 | 0 | 0 | 0 | 0 | 0 | 298 | 149 | 1 | 1 | 1 | NA | 0.5 | NA | 1 | NA | 1 | NA | NA | NA | -0.167 |
| SE | 84819 | ENSG00000107185 | RGP1 | chr9 | + | 35751626 | 35751754 | 35751265 | 35751412 | 35751955 | 35752145 | 125 | 72 | 122 | 126 | 134 | 0 | 0 | 0 | 0 | 0 | 108 | 115 | 64 | 108 | 211 | 0 | 6 | 0 | 0 | 0 | 277 | 149 | 1 | 1 | 1 | 1 | 1 | 1 | 1 | 1 | 0.912 | 1 | 1 | 1 | 0.018 |
| SE | 84824 | ENSG00000121390 | PSPC1 | chr13 | - | 19705661 | 19705831 | 19677723 | 19677823 | 19730238 | 19730344 | 0 | 0 | 2 | 3 | 5 | 60 | 28 | 42 | 20 | 60 | 0 | 4 | 1 | 3 | 0 | 6 | 23 | 27 | 14 | 0 | 298 | 149 | 1 | 1 | 0 | 0 | 0.023 | 0.07 | 0.04 | 0 | 0.08 | 0.018 | 0.097 | NA | -0.022 |
| SE | 84825 | ENSG00000121390 | PSPC1 | chr13 | - | 19709541 | 19709599 | 19705661 | 19705831 | 19730238 | 19730344 | 118 | 77 | 105 | 103 | 173 | 0 | 0 | 0 | 3 | 0 | 149 | 96 | 144 | 198 | 123 | 0 | 4 | 1 | 3 | 0 | 207 | 149 | 1 | 1 | 1 | 1 | 1 | 0.961 | 1 | 1 | 0.945 | 0.99 | 0.979 | 1 | 0.009 |
| SE | 84826 | ENSG00000121390 | PSPC1 | chr13 | - | 19741564 | 19741649 | 19730238 | 19730344 | 19751270 | 19751467 | 388 | 315 | 323 | 406 | 435 | 0 | 1 | 0 | 0 | 0 | 660 | 555 | 497 | 635 | 503 | 0 | 0 | 0 | 0 | 0 | 234 | 149 | 1 | 1 | 1 | 0.995 | 1 | 1 | 1 | 1 | 1 | 1 | 1 | 1 | -0.001 |
| SE | 84827 | ENSG00000121390 | PSPC1 | chr13 | - | 19751270 | 19751467 | 19741564 | 19741649 | 19759322 | 19759418 | 532 | 478 | 474 | 556 | 540 | 0 | 0 | 0 | 0 | 0 | 855 | 776 | 761 | 847 | 690 | 0 | 0 | 0 | 7 | 0 | 298 | 149 | 1 | 1 | 1 | 1 | 1 | 1 | 1 | 1 | 1 | 1 | 0.984 | 1 | 0.003 |
| SE | 84833 | ENSG00000094631 | HDAC6 | chrX | + | 48802662 | 48802785 | 48802015 | 48802272 | 48802870 | 48802930 | 42 | 39 | 36 | 46 | 39 | 0 | 0 | 0 | 0 | 0 | 44 | 45 | 25 | 57 | 47 | 0 | 3 | 0 | 0 | 0 | 272 | 149 | 1 | 1 | 1 | 1 | 1 | 1 | 1 | 1 | 0.892 | 1 | 1 | 1 | 0.022 |
| SE | 84835 | ENSG00000094631 | HDAC6 | chrX | + | 48808257 | 48808326 | 48808032 | 48808137 | 48814439 | 48814566 | 22 | 18 | 29 | 29 | 47 | 0 | 0 | 0 | 0 | 0 | 59 | 21 | 26 | 25 | 88 | 0 | 1 | 0 | 0 | 4 | 218 | 149 | 1 | 1 | 1 | 1 | 1 | 1 | 1 | 1 | 0.935 | 1 | 1 | 0.938 | 0.025 |
| SE | 84837 | ENSG00000094631 | HDAC6 | chrX | + | 48811002 | 48811072 | 48808257 | 48808326 | 48814439 | 48814566 | 0 | 0 | 4 | 0 | 0 | 14 | 18 | 27 | 23 | 38 | 0 | 0 | 0 | 0 | 0 | 45 | 11 | 21 | 25 | 67 | 219 | 149 | 1 | 1 | 0 | 0 | 0.092 | 0 | 0 | 0 | 0 | 0 | 0 | 0 | 0.018 |
| SE | 84838 | ENSG00000094631 | HDAC6 | chrX | + | 48817325 | 48817459 | 48816464 | 48816633 | 48818040 | 48818109 | 25 | 6 | 20 | 7 | 6 | 0 | 0 | 1 | 0 | 0 | 5 | 28 | 13 | 22 | 29 | 0 | 0 | 0 | 0 | 0 | 283 | 149 | 1 | 1 | 1 | 1 | 0.913 | 1 | 1 | 1 | 1 | 1 | 1 | 1 | -0.017 |
| SE | 84839 | ENSG00000094631 | HDAC6 | chrX | + | 48817359 | 48817459 | 48816472 | 48816633 | 48818040 | 48818109 | 5 | 2 | 11 | 6 | 5 | 0 | 0 | 1 | 0 | 0 | 0 | 10 | 3 | 11 | 19 | 0 | 0 | 0 | 0 | 0 | 249 | 149 | 1 | 1 | 1 | 1 | 0.868 | 1 | 1 | NA | 1 | 1 | 1 | 1 | -0.026 |
| SE | 84840 | ENSG00000094631 | HDAC6 | chrX | + | 48818219 | 48818412 | 48818040 | 48818109 | 48820105 | 48820180 | 21 | 7 | 30 | 25 | 6 | 1 | 0 | 1 | 0 | 0 | 20 | 27 | 14 | 13 | 49 | 0 | 3 | 0 | 0 | 0 | 298 | 149 | 1 | 1 | 0.913 | 1 | 0.938 | 1 | 1 | 1 | 0.818 | 1 | 1 | 1 | 0.007 |
| SE | 84845 | ENSG00000094631 | HDAC6 | chrX | + | 48824165 | 48824294 | 48823921 | 48824068 | 48824543 | 48824860 | 76 | 53 | 93 | 97 | 82 | 0 | 1 | 0 | 0 | 0 | 162 | 137 | 83 | 43 | 258 | 0 | 3 | 0 | 0 | 0 | 278 | 149 | 1 | 1 | 1 | 0.966 | 1 | 1 | 1 | 1 | 0.961 | 1 | 1 | 1 | 0.001 |
| SE | 84847 | ENSG00000077514 | POLD3 | chr11 | + | 74612877 | 74613010 | 74611498 | 74611538 | 74618536 | 74618594 | 127 | 90 | 83 | 99 | 138 | 0 | 0 | 0 | 0 | 0 | 162 | 78 | 56 | 72 | 29 | 0 | 0 | 0 | 1 | 0 | 282 | 149 | 1 | 1 | 1 | 1 | 1 | 1 | 1 | 1 | 1 | 1 | 0.974 | 1 | 0.005 |
| SE | 84848 | ENSG00000077514 | POLD3 | chr11 | + | 74620016 | 74620089 | 74618536 | 74618804 | 74625407 | 74625573 | 121 | 54 | 99 | 73 | 203 | 0 | 0 | 0 | 0 | 0 | 182 | 159 | 72 | 170 | 66 | 0 | 0 | 1 | 0 | 0 | 222 | 149 | 1 | 1 | 1 | 1 | 1 | 1 | 1 | 1 | 1 | 0.98 | 1 | 1 | 0.004 |
| SE | 84850 | ENSG00000269556 | TMEM185A | chrX | - | 149603986 | 149604070 | 149600302 | 149600479 | 149608626 | 149608834 | 50 | 31 | 38 | 67 | 130 | 0 | 0 | 0 | 0 | 0 | 66 | 34 | 23 | 32 | 394 | 0 | 0 | 0 | 0 | 3 | 233 | 149 | 1 | 1 | 1 | 1 | 1 | 1 | 1 | 1 | 1 | 1 | 1 | 0.988 | 0.002 |
| SE | 84853 | ENSG00000269556 | TMEM185A | chrX | - | 149608626 | 149608834 | 149603708 | 149604070 | 149611286 | 149611463 | 52 | 33 | 33 | 62 | 92 | 0 | 0 | 0 | 1 | 0 | 55 | 18 | 24 | 23 | 332 | 0 | 0 | 0 | 0 | 0 | 298 | 149 | 1 | 1 | 1 | 1 | 1 | 0.969 | 1 | 1 | 1 | 1 | 1 | 1 | -0.006 |
| SE | 84855 | ENSG00000269556 | TMEM185A | chrX | - | 149611286 | 149611463 | 149603986 | 149604070 | 149631542 | 149631698 | 9 | 3 | 8 | 8 | 8 | 4 | 0 | 0 | 2 | 0 | 2 | 0 | 2 | 7 | 62 | 0 | 0 | 0 | 0 | 0 | 298 | 149 | 1 | 1 | 0.529 | 1 | 1 | 0.667 | 1 | 1 | NA | 1 | 1 | 1 | -0.161 |
| SE | 84856 | ENSG00000269556 | TMEM185A | chrX | - | 149611286 | 149611463 | 149608626 | 149608829 | 149631542 | 149631698 | 12 | 3 | 9 | 12 | 8 | 0 | 0 | 1 | 0 | 0 | 2 | 0 | 2 | 7 | 76 | 0 | 0 | 0 | 0 | 0 | 298 | 149 | 1 | 1 | 1 | 1 | 0.818 | 1 | 1 | 1 | NA | 1 | 1 | 1 | -0.036 |
| SE | 84861 | ENSG00000164074 | ABHD18 | chr4 | + | 127982938 | 127983047 | 127965305 | 127965606 | 127984338 | 127984423 | 88 | 78 | 112 | 104 | 86 | 0 | 0 | 0 | 0 | 0 | 113 | 57 | 111 | 91 | 96 | 0 | 4 | 0 | 0 | 0 | 258 | 149 | 1 | 1 | 1 | 1 | 1 | 1 | 1 | 1 | 0.892 | 1 | 1 | 1 | 0.022 |
| SE | 84869 | ENSG00000164074 | ABHD18 | chr4 | + | 128020079 | 128020169 | 128011672 | 128011700 | 128028474 | 128028853 | 3 | 1 | 2 | 3 | 3 | 0 | 0 | 0 | 0 | 1 | 0 | 2 | 4 | 0 | 0 | 0 | 0 | 0 | 0 | 0 | 239 | 149 | 1 | 1 | 1 | 1 | 1 | 1 | 0.652 | NA | 1 | 1 | NA | NA | -0.07 |
| SE | 84871 | ENSG00000164074 | ABHD18 | chr4 | + | 128028474 | 128028853 | 128021136 | 128021238 | 128030509 | 128030672 | 154 | 103 | 115 | 151 | 211 | 0 | 0 | 1 | 0 | 2 | 95 | 109 | 141 | 126 | 49 | 0 | 0 | 1 | 0 | 0 | 298 | 149 | 1 | 1 | 1 | 1 | 0.983 | 1 | 0.981 | 1 | 1 | 0.986 | 1 | 1 | -0.004 |
| SE | 84872 | ENSG00000164074 | ABHD18 | chr4 | + | 128030509 | 128030672 | 128028474 | 128028853 | 128035761 | 128036286 | 182 | 142 | 154 | 136 | 261 | 2 | 0 | 0 | 0 | 4 | 82 | 102 | 151 | 151 | 12 | 0 | 0 | 0 | 0 | 0 | 298 | 149 | 1 | 1 | 0.978 | 1 | 1 | 1 | 0.97 | 1 | 1 | 1 | 1 | 1 | -0.01 |
| SE | 84875 | ENSG00000156970 | BUB1B | chr15 | + | 40185164 | 40185379 | 40183713 | 40183883 | 40185550 | 40185642 | 13 | 4 | 25 | 14 | 4 | 0 | 0 | 0 | 0 | 0 | 28 | 13 | 32 | 27 | 10 | 0 | 0 | 1 | 2 | 0 | 298 | 149 | 1 | 1 | 1 | 1 | 1 | 1 | 1 | 1 | 1 | 0.941 | 0.871 | 1 | 0.038 |
| SE | 84878 | ENSG00000156970 | BUB1B | chr15 | + | 40199614 | 40199727 | 40196544 | 40196774 | 40200243 | 40200289 | 2 | 6 | 15 | 16 | 9 | 0 | 0 | 0 | 2 | 0 | 4 | 29 | 60 | 34 | 8 | 0 | 0 | 0 | 3 | 0 | 262 | 149 | 1 | 1 | 1 | 1 | 1 | 0.82 | 1 | 1 | 1 | 1 | 0.866 | 1 | -0.009 |
| SE | 84879 | ENSG00000102241 | HTATSF1 | chrX | + | 136500157 | 136500205 | 136499597 | 136499778 | 136500663 | 136500818 | 237 | 227 | 231 | 237 | 319 | 1 | 0 | 0 | 0 | 0 | 166 | 242 | 275 | 334 | 37 | 0 | 1 | 0 | 0 | 0 | 197 | 149 | 1 | 1 | 0.994 | 1 | 1 | 1 | 1 | 1 | 0.995 | 1 | 1 | 1 | 0 |
| SE | 84880 | ENSG00000157881 | PANK4 | chr1 | - | 2510677 | 2510748 | 2510056 | 2510157 | 2511337 | 2511387 | 14 | 9 | 28 | 29 | 27 | 0 | 0 | 0 | 0 | 1 | 22 | 5 | 26 | 3 | 27 | 0 | 0 | 0 | 0 | 0 | 220 | 149 | 1 | 1 | 1 | 1 | 1 | 1 | 0.948 | 1 | 1 | 1 | 1 | 1 | -0.01 |
| SE | 84881 | ENSG00000157881 | PANK4 | chr1 | - | 2510677 | 2510782 | 2510056 | 2510157 | 2511337 | 2511387 | 28 | 8 | 29 | 14 | 24 | 0 | 0 | 0 | 0 | 1 | 27 | 2 | 14 | 5 | 30 | 0 | 0 | 0 | 0 | 0 | 254 | 149 | 1 | 1 | 1 | 1 | 1 | 1 | 0.934 | 1 | 1 | 1 | 1 | 1 | -0.013 |
| SE | 84885 | ENSG00000157881 | PANK4 | chr1 | - | 2515561 | 2515717 | 2514353 | 2514466 | 2519142 | 2519295 | 8 | 5 | 8 | 21 | 12 | 0 | 0 | 0 | 1 | 0 | 0 | 7 | 2 | 11 | 6 | 0 | 2 | 0 | 0 | 0 | 298 | 149 | 1 | 1 | 1 | 1 | 1 | 0.913 | 1 | NA | 0.636 | 1 | 1 | 1 | 0.074 |
| SE | 84886 | ENSG00000157881 | PANK4 | chr1 | - | 2518163 | 2518264 | 2514353 | 2514466 | 2519142 | 2519295 | 3 | 2 | 0 | 3 | 1 | 0 | 0 | 0 | 1 | 0 | 10 | 0 | 2 | 3 | 5 | 0 | 2 | 0 | 0 | 0 | 250 | 149 | 1 | 1 | 1 | 1 | NA | 0.641 | 1 | 1 | 0 | 1 | 1 | 1 | 0.11 |
| SE | 84887 | ENSG00000157881 | PANK4 | chr1 | - | 2518163 | 2518264 | 2515561 | 2515717 | 2518515 | 2518597 | 33 | 5 | 24 | 33 | 20 | 0 | 2 | 0 | 0 | 0 | 37 | 6 | 18 | 29 | 0 | 0 | 0 | 0 | 0 | 0 | 250 | 149 | 1 | 1 | 1 | 0.598 | 1 | 1 | 1 | 1 | 1 | 1 | 1 | NA | -0.08 |
| SE | 84888 | ENSG00000157881 | PANK4 | chr1 | - | 2518163 | 2518264 | 2515561 | 2515717 | 2519142 | 2519295 | 14 | 3 | 9 | 19 | 6 | 0 | 0 | 0 | 1 | 0 | 25 | 4 | 10 | 14 | 0 | 0 | 0 | 0 | 0 | 0 | 250 | 149 | 1 | 1 | 1 | 1 | 1 | 0.919 | 1 | 1 | 1 | 1 | 1 | NA | -0.016 |
| SE | 84889 | ENSG00000157881 | PANK4 | chr1 | - | 2518515 | 2518597 | 2515561 | 2515717 | 2519142 | 2519295 | 16 | 4 | 10 | 17 | 8 | 0 | 0 | 0 | 1 | 0 | 13 | 18 | 7 | 0 | 39 | 0 | 0 | 0 | 0 | 0 | 231 | 149 | 1 | 1 | 1 | 1 | 1 | 0.916 | 1 | 1 | 1 | 1 | NA | 1 | -0.017 |
| SE | 84892 | ENSG00000063127 | SLC6A16 | chr19 | - | 49290604 | 49290767 | 49289637 | 49290392 | 49293222 | 49293382 | 7 | 2 | 20 | 2 | 8 | 0 | 0 | 1 | 0 | 0 | 0 | 16 | 7 | 6 | 0 | 0 | 3 | 0 | 4 | 0 | 298 | 149 | 1 | 1 | 1 | 1 | 0.909 | 1 | 1 | NA | 0.727 | 1 | 0.429 | NA | 0.263 |
| SE | 84894 | ENSG00000063127 | SLC6A16 | chr19 | - | 49309650 | 49309826 | 49294366 | 49294553 | 49310039 | 49310146 | 1 | 0 | 0 | 1 | 0 | 0 | 0 | 0 | 0 | 0 | 0 | 0 | 2 | 0 | 5 | 9 | 1 | 0 | 0 | 0 | 298 | 149 | 1 | 1 | 1 | NA | NA | 1 | NA | 0 | 0 | 1 | NA | 1 | 0.5 |
| SE | 84895 | ENSG00000100883 | SRP54 | chr14 | + | 34983129 | 34983215 | 34982681 | 34983003 | 34996676 | 34996787 | 46 | 29 | 40 | 63 | 53 | 0 | 0 | 0 | 2 | 0 | 0 | 61 | 47 | 29 | 7 | 0 | 0 | 0 | 0 | 0 | 235 | 149 | 1 | 1 | 1 | 1 | 1 | 0.952 | 1 | NA | 1 | 1 | 1 | 1 | -0.01 |
| SE | 84897 | ENSG00000100883 | SRP54 | chr14 | + | 34999557 | 34999649 | 34996676 | 34996787 | 35000935 | 35001020 | 188 | 137 | 198 | 218 | 274 | 0 | 0 | 0 | 2 | 0 | 14 | 221 | 150 | 156 | 16 | 0 | 0 | 0 | 0 | 0 | 241 | 149 | 1 | 1 | 1 | 1 | 1 | 0.985 | 1 | 1 | 1 | 1 | 1 | 1 | -0.003 |
| SE | 84902 | ENSG00000100883 | SRP54 | chr14 | + | 35013801 | 35013902 | 35013345 | 35013494 | 35014743 | 35014830 | 434 | 351 | 437 | 424 | 370 | 0 | 0 | 0 | 0 | 0 | 149 | 436 | 461 | 371 | 90 | 0 | 0 | 1 | 0 | 0 | 250 | 149 | 1 | 1 | 1 | 1 | 1 | 1 | 1 | 1 | 1 | 0.996 | 1 | 1 | 0.001 |
| SE | 84904 | ENSG00000100883 | SRP54 | chr14 | + | 35018965 | 35019074 | 35014743 | 35014830 | 35028087 | 35028183 | 0 | 0 | 1 | 0 | 0 | 0 | 4 | 0 | 0 | 0 | 0 | 4 | 0 | 0 | 0 | 0 | 0 | 0 | 0 | 0 | 258 | 149 | 1 | 1 | NA | 0 | 1 | NA | NA | NA | 1 | NA | NA | NA | -0.5 |
| SE | 84912 | ENSG00000162398 | LEXM | chr1 | + | 54807888 | 54808011 | 54807525 | 54807690 | 54813813 | 54813975 | 26 | 10 | 8 | 19 | 4 | 0 | 0 | 0 | 0 | 0 | 14 | 8 | 4 | 0 | 3 | 0 | 2 | 1 | 0 | 0 | 272 | 149 | 1 | 1 | 1 | 1 | 1 | 1 | 1 | 1 | 0.687 | 0.687 | NA | 1 | 0.156 |
| SE | 84913 | ENSG00000162398 | LEXM | chr1 | + | 54811793 | 54811931 | 54807888 | 54808011 | 54812045 | 54812189 | 27 | 8 | 12 | 21 | 9 | 2 | 0 | 0 | 0 | 0 | 14 | 0 | 12 | 0 | 11 | 0 | 0 | 0 | 0 | 0 | 287 | 149 | 1 | 1 | 0.875 | 1 | 1 | 1 | 1 | 1 | NA | 1 | NA | 1 | -0.025 |
| SE | 84914 | ENSG00000162398 | LEXM | chr1 | + | 54812045 | 54812189 | 54807888 | 54808011 | 54813813 | 54813975 | 10 | 4 | 9 | 14 | 8 | 0 | 2 | 0 | 1 | 0 | 0 | 0 | 1 | 0 | 4 | 0 | 1 | 1 | 0 | 0 | 293 | 149 | 1 | 1 | 1 | 0.504 | 1 | 0.877 | 1 | NA | 0 | 0.337 | NA | 1 | 0.431 |
| SE | 84915 | ENSG00000162398 | LEXM | chr1 | + | 54841615 | 54841702 | 54816988 | 54817122 | 54841800 | 54842252 | 3 | 1 | 0 | 0 | 0 | 1 | 0 | 0 | 4 | 0 | 0 | 0 | 2 | 0 | 0 | 0 | 0 | 0 | 0 | 0 | 236 | 149 | 1 | 1 | 0.654 | 1 | NA | 0 | NA | NA | NA | 1 | NA | NA | -0.449 |
| SE | 84931 | ENSG00000179195 | ZNF664 | chr12 | + | 124003422 | 124003540 | 123988042 | 123988138 | 124004877 | 124004935 | 5 | 2 | 2 | 2 | 0 | 20 | 12 | 13 | 14 | 3 | 17 | 0 | 3 | 3 | 0 | 0 | 17 | 16 | 7 | 0 | 267 | 149 | 1 | 1 | 0.122 | 0.085 | 0.079 | 0.074 | 0 | 1 | 0 | 0.095 | 0.193 | NA | -0.25 |
| SE | 84935 | ENSG00000170734 | POLH | chr6 | + | 43587271 | 43587489 | 43583006 | 43583141 | 43597695 | 43597865 | 64 | 78 | 93 | 110 | 157 | 1 | 0 | 1 | 0 | 0 | 104 | 60 | 33 | 57 | 179 | 0 | 0 | 0 | 0 | 0 | 298 | 149 | 1 | 1 | 0.97 | 1 | 0.979 | 1 | 1 | 1 | 1 | 1 | 1 | 1 | -0.01 |
| SE | 84940 | ENSG00000259736 | CRTC3-AS1 | chr15 | - | 90714920 | 90715042 | 90712992 | 90713299 | 90716980 | 90717141 | 3 | 0 | 4 | 0 | 4 | 0 | 1 | 0 | 0 | 0 | 0 | 9 | 0 | 6 | 24 | 0 | 0 | 0 | 5 | 0 | 271 | 149 | 1 | 1 | 1 | 0 | 1 | NA | 1 | NA | 1 | NA | 0.398 | 1 | -0.049 |
| SE | 84941 | ENSG00000139436 | GIT2 | chr12 | - | 109934021 | 109934085 | 109929801 | 109933190 | 109938379 | 109938568 | 141 | 104 | 112 | 128 | 190 | 0 | 1 | 0 | 0 | 0 | 134 | 152 | 117 | 93 | 119 | 0 | 1 | 4 | 2 | 0 | 213 | 149 | 1 | 1 | 1 | 0.986 | 1 | 1 | 1 | 1 | 0.991 | 0.953 | 0.97 | 1 | 0.014 |
| SE | 84950 | ENSG00000139436 | GIT2 | chr12 | - | 109947255 | 109947504 | 109945259 | 109945349 | 109951211 | 109951316 | 67 | 32 | 39 | 71 | 101 | 0 | 0 | 0 | 0 | 2 | 70 | 79 | 37 | 57 | 19 | 0 | 0 | 3 | 0 | 0 | 298 | 149 | 1 | 1 | 1 | 1 | 1 | 1 | 0.962 | 1 | 1 | 0.86 | 1 | 1 | 0.02 |
| SE | 84951 | ENSG00000139436 | GIT2 | chr12 | - | 109947255 | 109947504 | 109945259 | 109945349 | 109953091 | 109953234 | 74 | 39 | 49 | 70 | 116 | 1 | 0 | 0 | 1 | 2 | 119 | 105 | 51 | 73 | 36 | 0 | 0 | 2 | 0 | 0 | 298 | 149 | 1 | 1 | 0.974 | 1 | 1 | 0.972 | 0.967 | 1 | 1 | 0.927 | 1 | 1 | -0.003 |
| SE | 84954 | ENSG00000139436 | GIT2 | chr12 | - | 109951166 | 109951316 | 109945259 | 109945349 | 109953091 | 109953234 | 119 | 94 | 95 | 110 | 111 | 1 | 0 | 0 | 1 | 2 | 110 | 111 | 115 | 168 | 125 | 0 | 0 | 2 | 0 | 0 | 298 | 149 | 1 | 1 | 0.983 | 1 | 1 | 0.982 | 0.965 | 1 | 1 | 0.966 | 1 | 1 | -0.007 |
| SE | 84958 | ENSG00000139436 | GIT2 | chr12 | - | 109951211 | 109951316 | 109945259 | 109945349 | 109953091 | 109953234 | 85 | 76 | 76 | 89 | 96 | 1 | 0 | 0 | 1 | 2 | 96 | 94 | 96 | 137 | 101 | 0 | 0 | 2 | 0 | 0 | 254 | 149 | 1 | 1 | 0.98 | 1 | 1 | 0.981 | 0.966 | 1 | 1 | 0.966 | 1 | 1 | -0.008 |
| SE | 84965 | ENSG00000139436 | GIT2 | chr12 | - | 109967457 | 109967503 | 109965525 | 109965577 | 109980951 | 109981046 | 489 | 410 | 435 | 451 | 459 | 0 | 0 | 1 | 0 | 0 | 875 | 581 | 488 | 650 | 439 | 0 | 0 | 0 | 0 | 0 | 195 | 149 | 1 | 1 | 1 | 1 | 0.997 | 1 | 1 | 1 | 1 | 1 | 1 | 1 | -0.001 |
| SE | 84967 | ENSG00000139436 | GIT2 | chr12 | - | 109988962 | 109989068 | 109983607 | 109983694 | 109989689 | 109989802 | 662 | 368 | 506 | 623 | 753 | 0 | 0 | 2 | 3 | 0 | 849 | 602 | 602 | 880 | 470 | 0 | 4 | 2 | 0 | 0 | 255 | 149 | 1 | 1 | 1 | 1 | 0.993 | 0.992 | 1 | 1 | 0.989 | 0.994 | 1 | 1 | 0 |
| SE | 84968 | ENSG00000139436 | GIT2 | chr12 | - | 109989689 | 109989802 | 109983607 | 109983694 | 109991626 | 109991760 | 115 | 58 | 95 | 127 | 46 | 0 | 0 | 0 | 0 | 0 | 104 | 104 | 124 | 130 | 68 | 0 | 0 | 1 | 0 | 0 | 262 | 149 | 1 | 1 | 1 | 1 | 1 | 1 | 1 | 1 | 1 | 0.986 | 1 | 1 | 0.003 |
| SE | 84969 | ENSG00000139436 | GIT2 | chr12 | - | 109989689 | 109989802 | 109983607 | 109983694 | 109996172 | 109996224 | 0 | 0 | 2 | 2 | 0 | 6 | 0 | 0 | 0 | 0 | 0 | 4 | 2 | 0 | 0 | 0 | 0 | 0 | 0 | 0 | 262 | 149 | 1 | 1 | 0 | NA | 1 | 1 | NA | NA | 1 | 1 | NA | NA | -0.333 |
| SE | 84971 | ENSG00000139436 | GIT2 | chr12 | - | 109991626 | 109991760 | 109989689 | 109989802 | 109996172 | 109996224 | 231 | 120 | 220 | 264 | 75 | 0 | 0 | 0 | 0 | 1 | 212 | 254 | 226 | 255 | 144 | 0 | 0 | 0 | 0 | 0 | 283 | 149 | 1 | 1 | 1 | 1 | 1 | 1 | 0.975 | 1 | 1 | 1 | 1 | 1 | -0.005 |
| SE | 84973 | ENSG00000170832 | USP32 | chr17 | - | 60205446 | 60205658 | 60198259 | 60198444 | 60207020 | 60207132 | 329 | 310 | 220 | 264 | 593 | 1 | 4 | 3 | 2 | 0 | 272 | 344 | 449 | 392 | 44 | 0 | 3 | 2 | 0 | 0 | 298 | 149 | 1 | 1 | 0.994 | 0.975 | 0.973 | 0.985 | 1 | 1 | 0.983 | 0.991 | 1 | 1 | -0.009 |
| SE | 84974 | ENSG00000170832 | USP32 | chr17 | - | 60205446 | 60205658 | 60198259 | 60198444 | 60208058 | 60208210 | 203 | 200 | 151 | 170 | 349 | 0 | 2 | 0 | 0 | 0 | 129 | 241 | 270 | 246 | 15 | 0 | 0 | 0 | 0 | 0 | 298 | 149 | 1 | 1 | 1 | 0.98 | 1 | 1 | 1 | 1 | 1 | 1 | 1 | 1 | -0.004 |
| SE | 84975 | ENSG00000170832 | USP32 | chr17 | - | 60207020 | 60207132 | 60198259 | 60198444 | 60208058 | 60208210 | 63 | 61 | 42 | 39 | 122 | 0 | 2 | 0 | 0 | 0 | 56 | 69 | 66 | 71 | 12 | 0 | 0 | 0 | 0 | 0 | 261 | 149 | 1 | 1 | 1 | 0.946 | 1 | 1 | 1 | 1 | 1 | 1 | 1 | 1 | -0.011 |
| SE | 84977 | ENSG00000170832 | USP32 | chr17 | - | 60208653 | 60208828 | 60208058 | 60208210 | 60209369 | 60209543 | 102 | 104 | 86 | 128 | 222 | 0 | 0 | 2 | 1 | 0 | 27 | 82 | 176 | 108 | 37 | 0 | 1 | 0 | 0 | 0 | 298 | 149 | 1 | 1 | 1 | 1 | 0.956 | 0.985 | 1 | 1 | 0.976 | 1 | 1 | 1 | -0.007 |
| SE | 84983 | ENSG00000170832 | USP32 | chr17 | - | 60223410 | 60223586 | 60222408 | 60222549 | 60226038 | 60226063 | 291 | 305 | 220 | 294 | 353 | 5 | 0 | 1 | 0 | 0 | 307 | 327 | 396 | 330 | 24 | 0 | 4 | 3 | 0 | 0 | 298 | 149 | 1 | 1 | 0.967 | 1 | 0.991 | 1 | 1 | 1 | 0.976 | 0.985 | 1 | 1 | -0.001 |
| SE | 84984 | ENSG00000170832 | USP32 | chr17 | - | 60255174 | 60255258 | 60252380 | 60252442 | 60265411 | 60265474 | 159 | 212 | 154 | 172 | 272 | 2 | 1 | 0 | 1 | 2 | 183 | 320 | 325 | 300 | 40 | 3 | 0 | 0 | 0 | 0 | 233 | 149 | 1 | 1 | 0.981 | 0.993 | 1 | 0.991 | 0.989 | 0.975 | 1 | 1 | 1 | 1 | -0.004 |
| SE | 84985 | ENSG00000170832 | USP32 | chr17 | - | 60269449 | 60269557 | 60265975 | 60266091 | 60271349 | 60271481 | 316 | 329 | 232 | 345 | 356 | 0 | 0 | 0 | 0 | 0 | 338 | 444 | 476 | 495 | 258 | 0 | 2 | 0 | 4 | 0 | 257 | 149 | 1 | 1 | 1 | 1 | 1 | 1 | 1 | 1 | 0.992 | 1 | 0.986 | 1 | 0.004 |
| SE | 84987 | ENSG00000170832 | USP32 | chr17 | - | 60271349 | 60271481 | 60269449 | 60269557 | 60294682 | 60294801 | 218 | 241 | 156 | 240 | 226 | 0 | 0 | 1 | 1 | 0 | 234 | 310 | 314 | 355 | 183 | 0 | 8 | 0 | 0 | 0 | 281 | 149 | 1 | 1 | 1 | 1 | 0.988 | 0.992 | 1 | 1 | 0.954 | 1 | 1 | 1 | 0.005 |
| SE | 84992 | ENSG00000253540 | FAM86HP | chr3 | - | 130105529 | 130105631 | 130104165 | 130104297 | 130111278 | 130111433 | 0 | 0 | 3 | 0 | 0 | 1 | 0 | 0 | 0 | 0 | 0 | 2 | 0 | 0 | 0 | 0 | 0 | 0 | 0 | 2 | 251 | 149 | 1 | 1 | 0 | NA | 1 | NA | NA | NA | 1 | NA | NA | 0 | 0 |

| | | | | | | | | | | | | | |
|---|---|---|---|---|---|---|---|---|---|---|---|---|---|
| SE | 84993 | ENSG00000112984 | KIF20A | chr5 | + | 138184511 | 138184676 | 138184238 | 138184404 | 138184806 | 138184946 | 0 | 2 |
| SE | 84994 | ENSG00000112984 | KIF20A | chr5 | + | 138185511 | 138185710 | 138185094 | 138185197 | 138185960 | 138186052 | 0 | 8 |
| SE | 84996 | ENSG00000101901 | ALG13 | chrX | + | 111686136 | 111686174 | 111684964 | 111685103 | 111689086 | 111689197 | 11 | 9 |
| SE | 84998 | ENSG00000101901 | ALG13 | chrX | + | 111689086 | 111689197 | 111684964 | 111685103 | 111711674 | 111711725 | 1 | 5 |
| SE | 85000 | ENSG00000101901 | ALG13 | chrX | + | 111689086 | 111689197 | 111684964 | 111685103 | 111717845 | 111717927 | 1 | 4 |
| SE | 85003 | ENSG00000101901 | ALG13 | chrX | + | 111708964 | 111709048 | 111684964 | 111685103 | 111711674 | 111711725 | 8 | 6 |
| SE | 85006 | ENSG00000101901 | ALG13 | chrX | + | 111711674 | 111711725 | 111689086 | 111689197 | 111712483 | 111712493 | 14 | 19 |
| SE | 85007 | ENSG00000101901 | ALG13 | chrX | + | 111712483 | 111712530 | 111689086 | 111689197 | 111713224 | 111713297 | 16 | 14 |
| SE | 85011 | ENSG00000101901 | ALG13 | chrX | + | 111717845 | 111717927 | 111684964 | 111685103 | 111718111 | 111718274 | 45 | 81 |
| SE | 85015 | ENSG00000101901 | ALG13 | chrX | + | 111721602 | 111721711 | 111720094 | 111720170 | 111724933 | 111725061 | 36 | 48 |
| SE | 85018 | ENSG00000101901 | ALG13 | chrX | + | 111723797 | 111723898 | 111720094 | 111720170 | 111724933 | 111725061 | 49 | 80 |
| SE | 85019 | ENSG00000101901 | ALG13 | chrX | + | 111727613 | 111727770 | 111727331 | 111727445 | 111728184 | 111728270 | 125 | 129 |
| SE | 85021 | ENSG00000101901 | ALG13 | chrX | + | 111730394 | 111730427 | 111727613 | 111727770 | 111744667 | 111744904 | 0 | 0 |
| SE | 85023 | ENSG00000101901 | ALG13 | chrX | + | 111730524 | 111730580 | 111730394 | 111730427 | 111744667 | 111744904 | 4 | 4 |
| SE | 85024 | ENSG00000101901 | ALG13 | chrX | + | 111736713 | 111736879 | 111727613 | 111727770 | 111744667 | 111744904 | 35 | 37 |
| SE | 85031 | ENSG00000204632 | HLA-G | chr6 | + | 29829815 | 29829932 | 29829417 | 29829693 | 29830377 | 29830410 | 0 | 0 |
| SE | 85036 | ENSG00000081386 | ZNF510 | chr9 | - | 96774787 | 96774846 | 96763505 | 96763632 | 96775999 | 96776245 | 47 | 37 |
| SE | 85053 | ENSG00000147459 | DOCK5 | chr8 | + | 25268844 | 25268885 | 25243673 | 25243757 | 25275385 | 25275441 | 74 | 56 |
| SE | 85054 | ENSG00000147459 | DOCK5 | chr8 | + | 25268844 | 25268885 | 25243673 | 25243757 | 25292023 | 25292172 | 12 | 5 |
| SE | 85056 | ENSG00000147459 | DOCK5 | chr8 | + | 25275385 | 25275441 | 25268844 | 25268885 | 25292023 | 25292172 | 42 | 33 |
| SE | 85057 | ENSG00000147459 | DOCK5 | chr8 | + | 25278568 | 25278665 | 25275385 | 25275441 | 25292023 | 25292172 | 83 | 72 |
| SE | 85058 | ENSG00000147459 | DOCK5 | chr8 | + | 25296512 | 25296648 | 25292023 | 25292172 | 25298943 | 25299101 | 166 | 101 |
| SE | 85059 | ENSG00000147459 | DOCK5 | chr8 | + | 25300575 | 25300657 | 25298943 | 25299101 | 25302324 | 25302454 | 100 | 73 |
| SE | 85060 | ENSG00000147459 | DOCK5 | chr8 | + | 25317006 | 25317131 | 25310406 | 25310532 | 25319577 | 25319676 | 287 | 251 |
| SE | 85062 | ENSG00000147459 | DOCK5 | chr8 | + | 25341738 | 25341809 | 25336238 | 25336373 | 25342400 | 25342507 | 6 | 15 |
| SE | 85063 | ENSG00000147459 | DOCK5 | chr8 | + | 25342400 | 25342507 | 25336238 | 25336373 | 25345474 | 25345611 | 33 | 27 |
| SE | 85068 | ENSG00000147459 | DOCK5 | chr8 | + | 25368570 | 25368687 | 25368191 | 25368250 | 25369555 | 25369641 | 69 | 35 |
| SE | 85070 | ENSG00000147459 | DOCK5 | chr8 | + | 25368570 | 25368725 | 25368191 | 25368250 | 25369555 | 25369641 | 240 | 176 |
| SE | 85071 | ENSG00000147459 | DOCK5 | chr8 | + | 25374563 | 25374654 | 25373617 | 25373658 | 25377304 | 25377424 | 88 | 60 |
| SE | 85073 | ENSG00000147459 | DOCK5 | chr8 | + | 25395542 | 25395719 | 25392795 | 25392882 | 25399910 | 25399994 | 123 | 84 |
| SE | 85075 | ENSG00000147459 | DOCK5 | chr8 | + | 25408801 | 25408940 | 25403557 | 25403724 | 25410098 | 25410202 | 21 | 13 |
| SE | 85078 | ENSG00000103037 | SETD6 | chr16 | + | 58516477 | 58516672 | 58516201 | 58516343 | 58518050 | 58518231 | 4 | 1 |
| SE | 85079 | ENSG00000103037 | SETD6 | chr16 | + | 58516612 | 58516672 | 58516201 | 58516343 | 58518050 | 58518231 | 0 | 1 |
| SE | 85080 | ENSG00000103037 | SETD6 | chr16 | + | 58516807 | 58516928 | 58516477 | 58516672 | 58518019 | 58518231 | 21 | 14 |
| SE | 85085 | ENSG00000164162 | ANAPC10 | chr4 | - | 145064571 | 145064692 | 144994574 | 144995603 | 145095984 | 145096111 | 37 | 23 |
| SE | 85086 | ENSG00000164162 | ANAPC10 | chr4 | - | 145081659 | 145081750 | 144994924 | 144995603 | 145095984 | 145096111 | 31 | 17 |
| SE | 85088 | ENSG00000164162 | ANAPC10 | chr4 | - | 145097494 | 145097562 | 145095984 | 145096111 | 145097687 | 145097760 | 0 | 0 |
| SE | 85089 | ENSG00000164162 | ANAPC10 | chr4 | - | 145097687 | 145097760 | 145095984 | 145096111 | 145098119 | 145098180 | 3 | 0 |
| SE | 85093 | ENSG00000139146 | SINHCAF | chr12 | - | 31287633 | 31287784 | 31280583 | 31282871 | 31293804 | 31293931 | 367 | 254 |
| SE | 85096 | ENSG00000165775 | FUNDC2 | chrX | + | 155046508 | 155046584 | 155033402 | 155033553 | 155051669 | 155051732 | 325 | 266 |
| SE | 85097 | ENSG00000165775 | FUNDC2 | chrX | + | 155047368 | 155047391 | 155046508 | 155046584 | 155051669 | 155051801 | 3 | 0 |
| SE | 85098 | ENSG00000165775 | FUNDC2 | chrX | + | 155051669 | 155051801 | 155046508 | 155046584 | 155054594 | 155054613 | 453 | 302 |
| SE | 85104 | ENSG00000147586 | MRPS28 | chr8 | - | 80028694 | 80028796 | 80002998 | 80003120 | 80030035 | 80030289 | 0 | 3 |
| SE | 85105 | ENSG00000147586 | MRPS28 | chr8 | - | 80028694 | 80028796 | 80002998 | 80003120 | 80030053 | 80030232 | 0 | 2 |
| SE | 85106 | ENSG00000147586 | MRPS28 | chr8 | - | 80028694 | 80028796 | 80002998 | 80003180 | 80029760 | 80029912 | 3 | 3 |
| SE | 85107 | ENSG00000147586 | MRPS28 | chr8 | - | 80028694 | 80028796 | 80002998 | 80003180 | 80029904 | 80029974 | 6 | 4 |
| SE | 85115 | ENSG00000107371 | EXOSC3 | chr9 | - | 37783913 | 37784063 | 37780274 | 37780880 | 37784720 | 37784745 | 21 | 29 |
| SE | 85121 | ENSG00000214827 | MTCP1 | chrX | - | 155065905 | 155066057 | 155065618 | 155065789 | 155070693 | 155071136 | 3 | 6 |
| SE | 85126 | ENSG00000285171 | NA | chrX | - | 71106353 | 71106426 | 71105999 | 71106137 | 71106531 | 71106590 | 1 | 4 |
| SE | 85127 | ENSG00000285171 | NA | chrX | - | 71106531 | 71106590 | 71105999 | 71106137 | 71108276 | 71108346 | 0 | 1 |
| SE | 85130 | ENSG00000285171 | NA | chrX | - | 71106531 | 71106770 | 71105999 | 71106137 | 71108276 | 71108346 | 1 | 0 |
| SE | 85134 | ENSG00000285171 | NA | chrX | - | 71109227 | 71109390 | 71108276 | 71108346 | 71110155 | 71110295 | 1389 | 656 |
| SE | 85137 | ENSG00000285171 | NA | chrX | - | 71110896 | 71111050 | 71110503 | 71110688 | 71111424 | 71111571 | 1612 | 680 |
| SE | 85141 | ENSG00000152766 | ANKRD22 | chr10 | - | 88823279 | 88823378 | 88819895 | 88823018 | 88826037 | 88826115 | 2 | 8 |
| SE | 85142 | ENSG00000196152 | ZNF79 | chr9 | + | 127428551 | 127428660 | 127424373 | 127424803 | 127428831 | 127428920 | 3 | 4 |
| SE | 85143 | ENSG00000196152 | ZNF79 | chr9 | + | 127428551 | 127428660 | 127424373 | 127424910 | 127428831 | 127428920 | 3 | 1 |
| SE | 85144 | ENSG00000136504 | KAT7 | chr17 | + | 49791885 | 49792033 | 49788647 | 49788849 | 49796749 | 49796926 | 284 | 154 |
| SE | 85146 | ENSG00000136504 | KAT7 | chr17 | + | 49796749 | 49796926 | 49791885 | 49792033 | 49805362 | 49805445 | 195 | 101 |
| SE | 85147 | ENSG00000136504 | KAT7 | chr17 | + | 49796749 | 49796926 | 49791885 | 49792033 | 49811475 | 49811574 | 198 | 101 |
| SE | 85148 | ENSG00000136504 | KAT7 | chr17 | + | 49798318 | 49798558 | 49791885 | 49792033 | 49805362 | 49805445 | 273 | 155 |
| SE | 85149 | ENSG00000136504 | KAT7 | chr17 | + | 49798318 | 49798558 | 49791885 | 49792033 | 49809118 | 49809208 | 8 | 3 |
| SE | 85150 | ENSG00000136504 | KAT7 | chr17 | + | 49798318 | 49798558 | 49791885 | 49792033 | 49811475 | 49811574 | 2 | 2 |
| SE | 85151 | ENSG00000136504 | KAT7 | chr17 | + | 49798318 | 49798558 | 49796749 | 49796926 | 49805362 | 49805445 | 537 | 288 |
| SE | 85152 | ENSG00000136504 | KAT7 | chr17 | + | 49798318 | 49798558 | 49796749 | 49796926 | 49811475 | 49811574 | 266 | 135 |
| SE | 85153 | ENSG00000136504 | KAT7 | chr17 | + | 49805362 | 49805445 | 49791885 | 49792033 | 49809118 | 49809208 | 152 | 68 |
| SE | 85154 | ENSG00000136504 | KAT7 | chr17 | + | 49805362 | 49805445 | 49791885 | 49792033 | 49811475 | 49811574 | 43 | 25 |
| SE | 85155 | ENSG00000136504 | KAT7 | chr17 | + | 49805362 | 49805445 | 49796749 | 49796926 | 49811475 | 49811574 | 43 | 25 |
| SE | 85159 | ENSG00000136504 | KAT7 | chr17 | + | 49809118 | 49809208 | 49791885 | 49792033 | 49811475 | 49811574 | 219 | 139 |
| SE | 85162 | ENSG00000136504 | KAT7 | chr17 | + | 49811475 | 49811574 | 49796749 | 49796926 | 49815802 | 49815913 | 191 | 159 |
| SE | 85163 | ENSG00000136504 | KAT7 | chr17 | + | 49811475 | 49811574 | 49805362 | 49805445 | 49815802 | 49815913 | 248 | 204 |
| SE | 85164 | ENSG00000136504 | KAT7 | chr17 | + | 49811565 | 49811574 | 49796749 | 49796926 | 49815802 | 49815896 | 69 | 61 |
| SE | 85165 | ENSG00000136504 | KAT7 | chr17 | + | 49815802 | 49815913 | 49811475 | 49811574 | 49821336 | 49821426 | 299 | 206 |
| SE | 85170 | ENSG00000241489 | NA | chrX | - | 149529512 | 149529593 | 149529366 | 149529428 | 149532881 | 149532968 | 28 | 21 |
| SE | 85171 | ENSG00000241489 | NA | chrX | - | 149536173 | 149536213 | 149532881 | 149534374 | 149538114 | 149538172 | 1 | 1 |
| SE | 85172 | ENSG00000241769 | LINC00893 | chrX | - | 149538723 | 149538906 | 149538114 | 149538172 | 149540694 | 149540813 | 10 | 7 |
| SE | 85175 | ENSG00000162669 | HFM1 | chr1 | - | 91316390 | 91316476 | 91316100 | 91316184 | 91319077 | 91319209 | 1 | 1 |
| SE | 85180 | ENSG00000162669 | HFM1 | chr1 | - | 91350737 | 91350871 | 91347428 | 91347476 | 91352505 | 91352651 | 0 | 2 |
| SE | 85181 | ENSG00000162669 | HFM1 | chr1 | - | 91351548 | 91351643 | 91350737 | 91350871 | 91352505 | 91352651 | 1 | 4 |
| SE | 85184 | ENSG00000140332 | TLE3 | chr15 | - | 70051390 | 70051467 | 70047789 | 70050204 | 70052373 | 70052524 | 100 | 64 |
| SE | 85187 | ENSG00000140332 | TLE3 | chr15 | - | 70055048 | 70055296 | 70053226 | 70053374 | 70056297 | 70056374 | 1 | 0 |
| SE | 85188 | ENSG00000140332 | TLE3 | chr15 | - | 70055048 | 70055298 | 70053226 | 70053374 | 70056297 | 70056374 | 86 | 20 |
| SE | 85189 | ENSG00000140332 | TLE3 | chr15 | - | 70056297 | 70056374 | 70055048 | 70055298 | 70057458 | 70057658 | 92 | 24 |
| SE | 85190 | ENSG00000140332 | TLE3 | chr15 | - | 70056297 | 70056374 | 70055048 | 70055298 | 70057473 | 70057519 | 85 | 22 |
| SE | 85191 | ENSG00000140332 | TLE3 | chr15 | - | 70059409 | 70059460 | 70058662 | 70058815 | 70060529 | 70060649 | 286 | 212 |
| SE | 85197 | ENSG00000140332 | TLE3 | chr15 | - | 70066013 | 70066215 | 70064453 | 70064470 | 70074532 | 70074607 | 185 | 112 |
| SE | 85200 | ENSG00000140332 | TLE3 | chr15 | - | 70066013 | 70066218 | 70064453 | 70064470 | 70074502 | 70074607 | 200 | 121 |
| SE | 85201 | ENSG00000140332 | TLE3 | chr15 | - | 70066013 | 70066218 | 70064453 | 70064470 | 70074532 | 70074607 | 279 | 160 |
| SE | 85204 | ENSG00000233382 | NKAPP1 | chrX | - | 120240226 | 120240355 | 120137041 | 120137189 | 120244763 | 120245233 | 15 | 6 |
| SE | 85207 | ENSG00000110583 | NAA40 | chr11 | + | 63946950 | 63947003 | 63945839 | 63945935 | 63952237 | 63952333 | 51 | 26 |
| SE | 85210 | ENSG00000099204 | ABLIM1 | chr10 | - | 114447879 | 114448020 | 114445311 | 114445403 | 114465697 | 114465827 | 70 | 59 |
| SE | 85212 | ENSG00000099204 | ABLIM1 | chr10 | - | 114447879 | 114448026 | 114445311 | 114445403 | 114465697 | 114465827 | 72 | 57 |
| SE | 85213 | ENSG00000099204 | ABLIM1 | chr10 | - | 114451623 | 114451671 | 114445311 | 114445403 | 114465697 | 114465827 | 102 | 65 |
| SE | 85214 | ENSG00000099204 | ABLIM1 | chr10 | - | 114451623 | 114451671 | 114447879 | 114448020 | 114465697 | 114465827 | 146 | 94 |
| SE | 85215 | ENSG00000099204 | ABLIM1 | chr10 | - | 114451623 | 114451671 | 114447879 | 114448026 | 114465697 | 114465827 | 105 | 70 |
| SE | 85218 | ENSG00000099204 | ABLIM1 | chr10 | - | 114465697 | 114465827 | 114451623 | 114451671 | 114468180 | 114468216 | 485 | 373 |
| SE | 85220 | ENSG00000099204 | ABLIM1 | chr10 | - | 114468180 | 114468216 | 114465697 | 114465827 | 114472976 | 114473132 | 406 | 303 |
| SE | 85221 | ENSG00000099204 | ABLIM1 | chr10 | - | 114472976 | 114473132 | 114468180 | 114468216 | 114473878 | 114473956 | 571 | 478 |
| SE | 85222 | ENSG00000099204 | ABLIM1 | chr10 | - | 114472976 | 114473132 | 114468180 | 114468216 | 114487957 | 114488016 | 376 | 283 |
| SE | 85223 | ENSG00000099204 | ABLIM1 | chr10 | - | 114473878 | 114473956 | 114468180 | 114468216 | 114487957 | 114487993 | 91 | 74 |
| SE | 85226 | ENSG00000099204 | ABLIM1 | chr10 | - | 114485296 | 114485380 | 114473878 | 114473956 | 114487957 | 114488016 | 0 | 0 |
| SE | 85227 | ENSG00000099204 | ABLIM1 | chr10 | - | 114571296 | 114571406 | 114547490 | 114547776 | 114575415 | 114575537 | 15 | 28 |
| SE | 85229 | ENSG00000009694 | TENM1 | chrX | - | 124382669 | 124382812 | 124375902 | 124381294 | 124383633 | 124384854 | 38 | 43 |
| SE | 85230 | ENSG00000009694 | TENM1 | chrX | - | 124487208 | 124487229 | 124481731 | 124481964 | 124497015 | 124497143 | 0 | 5 |
| SE | 85231 | ENSG00000009694 | TENM1 | chrX | - | 124546873 | 124547090 | 124529863 | 124529983 | 124561670 | 124561817 | 31 | 9 |
| SE | 85232 | ENSG00000009694 | TENM1 | chrX | - | 124563748 | 124563772 | 124561670 | 124561817 | 124565374 | 124565560 | 0 | 0 |
| SE | 85236 | ENSG00000183337 | BCOR | chrX | - | 40062138 | 40062393 | 40057154 | 40057321 | 40062745 | 40063071 | 133 | 102 |
| SE | 85239 | ENSG00000147905 | ZCCHC7 | chr9 | + | 37121599 | 37121672 | 37120604 | 37120776 | 37126311 | 37126942 | 2 | 0 |
| SE | 85240 | ENSG00000147905 | ZCCHC7 | chr9 | + | 37126311 | 37126942 | 37120573 | 37120623 | 37302187 | 37302231 | 178 | 141 |
| SE | 85244 | ENSG00000147905 | ZCCHC7 | chr9 | + | 37302187 | 37302231 | 37126311 | 37126942 | 37305543 | 37305714 | 56 | 42 |
| SE | 85246 | ENSG00000147905 | ZCCHC7 | chr9 | + | 37304187 | 37304313 | 37302187 | 37302231 | 37305543 | 37305714 | 444 | 316 |
| SE | 85248 | ENSG00000147905 | ZCCHC7 | chr9 | + | 37349356 | 37349452 | 37327798 | 37327834 | 37354709 | 37354802 | 152 | 85 |
| SE | 85249 | ENSG00000147905 | ZCCHC7 | chr9 | + | 37354709 | 37354824 | 37349356 | 37349452 | 37356834 | 37356980 | 122 | 74 |
| SE | 85252 | ENSG00000241769 | LINC00893 | chrX | - | 149535946 | 149535982 | 149534203 | 149534374 | 149536173 | 149536213 | 1 | 1 |
| SE | 85254 | ENSG00000241769 | LINC00893 | chrX | - | 149536173 | 149536255 | 149533629 | 149534374 | 149538114 | 149538172 | 2 | 5 |
| SE | 85255 | ENSG00000241769 | LINC00893 | chrX | - | 149537632 | 149537742 | 149535935 | 149536213 | 149538114 | 149538172 | 2 | 2 |
| SE | 85256 | ENSG00000241769 | LINC00893 | chrX | - | 149537632 | 149537778 | 149533629 | 149534374 | 149538114 | 149538172 | 17 | 2 |
| SE | 85257 | ENSG00000241769 | LINC00893 | chrX | - | 149537632 | 149537778 | 149536172 | 149536255 | 149538114 | 149538172 | 16 | 3 |
| SE | 85267 | ENSG00000176124 | DLEU1 | chr13 | + | 50390197 | 50390291 | 50083141 | 50083194 | 50415088 | 50415202 | 1 | 2 |
| SE | 85268 | ENSG00000176124 | DLEU1 | chr13 | + | 50390197 | 50390291 | 50185879 | 50186070 | 50415088 | 50415202 | 6 | 5 |
| SE | 85270 | ENSG00000176124 | DLEU1 | chr13 | + | 50390197 | 50390291 | 50310161 | 50310341 | 50415088 | 50415202 | 1 | 2 |
| SE | 85272 | ENSG00000176124 | DLEU1 | chr13 | + | 50415088 | 50415202 | 50390197 | 50390291 | 50433412 | 50433550 | 7 | 8 |
| SE | 85274 | ENSG00000176124 | DLEU1 | chr13 | + | 50433412 | 50433550 | 50415088 | 50415202 | 50492650 | 50492761 | 4 | 7 |
| SE | 85281 | ENSG00000176124 | DLEU1 | chr13 | + | 50492650 | 50492761 | 50433412 | 50433550 | 50648589 | 50648642 | 1 | 4 |
| SE | 85286 | ENSG00000176124 | DLEU1 | chr13 | + | 50589474 | 50589576 | 50433412 | 50433550 | 50648589 | 50648642 | 0 | 0 |
| SE | 85291 | ENSG00000176124 | DLEU1 | chr13 | + | 50719646 | 50719710 | 50433412 | 50433550 | 50757762 | 50757965 | 0 | 1 |
| SE | 85292 | ENSG00000176124 | DLEU1 | chr13 | + | 50719646 | 50719710 | 50712537 | 50715887 | 50757762 | 50757965 | 0 | 2 |
| SE | 85293 | ENSG00000176124 | DLEU1 | chr13 | + | 50723138 | 50723236 | 50719646 | 50719710 | 50725991 | 50726168 | 3 | 2 |
| SE | 85294 | ENSG00000176124 | DLEU1 | chr13 | + | 50725991 | 50726168 | 50719646 | 50719710 | 50757762 | 50757965 | 0 | 1 |
| SE | 85298 | ENSG00000117143 | UAP1 | chr1 | + | 162590322 | 162590511 | 162588692 | 162588833 | 162597791 | 162597858 | 93 | 41 |
| SE | 85299 | ENSG00000117143 | UAP1 | chr1 | + | 162592731 | 162592782 | 162590322 | 162590511 | 162597791 | 162597858 | 3 | 2 |
| SE | 85300 | ENSG00000117143 | UAP1 | chr1 | + | 162592734 | 162592782 | 162590322 | 162590511 | 162597791 | 162597858 | 0 | 0 |
| SE | 85304 | ENSG00000054983 | GALC | chr14 | - | 87981413 | 87981447 | 87976357 | 87976488 | 87982204 | 87982243 | 2 | 1 |
| SE | 85305 | ENSG00000054983 | GALC | chr14 | - | 87982204 | 87982243 | 87976357 | 87976488 | 87984393 | 87984533 | 225 | 116 |
| SE | 85306 | ENSG00000054983 | GALC | chr14 | - | 87984393 | 87984533 | 87982204 | 87982243 | 87986488 | 87986602 | 264 | 164 |
| SE | 85308 | ENSG00000054983 | GALC | chr14 | - | 87988454 | 87988523 | 87987751 | 87988207 | 87992303 | 87992405 | 49 | 29 |
| SE | 85309 | ENSG00000054983 | GALC | chr14 | - | 87988454 | 87988523 | 87987751 | 87988207 | 87992969 | 87993154 | 60 | 35 |
| SE | 85311 | ENSG00000054983 | GALC | chr14 | - | 87992303 | 87992405 | 87988454 | 87988523 | 87992969 | 87993122 | 3 | 0 |
| SE | 85313 | ENSG00000147394 | ZNF185 | chrX | + | 152920327 | 152920411 | 152918982 | 152919081 | 152920709 | 152920748 | 39 | 38 |
| SE | 85314 | ENSG00000147394 | ZNF185 | chrX | + | 152920706 | 152920748 | 152920327 | 152920411 | 152922172 | 152922256 | 68 | 69 |
| SE | 85315 | ENSG00000147394 | ZNF185 | chrX | + | 152920709 | 152920748 | 152920327 | 152920411 | 152922172 | 152922256 | 36 | 39 |
| SE | 85317 | ENSG00000147394 | ZNF185 | chrX | + | 152922172 | 152922256 | 152920706 | 152920748 | 152931671 | 152931776 | 50 | 55 |
| SE | 85320 | ENSG00000147394 | ZNF185 | chrX | + | 152928574 | 152928661 | 152920706 | 152920748 | 152931671 | 152931776 | 13 | 22 |
| SE | 85325 | ENSG00000147394 | ZNF185 | chrX | + | 152932872 | 152932971 | 152931671 | 152931776 | 152945266 | 152945464 | 43 | 38 |
| SE | 85327 | ENSG00000147394 | ZNF185 | chrX | + | 152938073 | 152938163 | 152931671 | 152931776 | 152945266 | 152945464 | 50 | 61 |
| SE | 85332 | ENSG00000147394 | ZNF185 | chrX | + | 152969381 | 152969484 | 152967166 | 152967238 | 152970442 | 152970541 | 188 | 158 |
| SE | 85333 | ENSG00000147394 | ZNF185 | chrX | + | 152970442 | 152970541 | 152969381 | 152969484 | 152971273 | 152973471 | 182 | 115 |
| SE | 85335 | ENSG00000014257 | ACP3 | chr3 | + | 132332191 | 132332344 | 132331646 | 132331733 | 132337455 | 132337554 | 66 | 82 |
| SE | 85337 | ENSG00000014257 | ACP3 | chr3 | + | 132352719 | 132352823 | 132349919 | 132350002 | 132367703 | 132368298 | 18 | 38 |
| SE | 85344 | ENSG00000102181 | CD99L2 | chrX | - | 150828305 | 150829604 | 150816006 | 150816078 | 150831230 | 150831293 | 1 | 1 |
| SE | 85345 | ENSG00000102181 | CD99L2 | chrX | - | 150828305 | 150829604 | 150816278 | 150816455 | 150831230 | 150831293 | 0 | 1 |
| SE | 85346 | ENSG00000185010 | F8 | chrX | - | 154902050 | 154902167 | 154901370 | 154901442 | 154903905 | 154904088 | 0 | 1 |
| SE | 85350 | ENSG00000137337 | MDC1 | chr6 | - | 30703620 | 30706098 | 30703417 | 30703537 | 30707383 | 30707454 | 41 | 45 |
| SE | 85352 | ENSG00000137337 | MDC1 | chr6 | - | 30711411 | 30711504 | 30707383 | 30707454 | 30711666 | 30711726 | 32 | 16 |
| SE | 85354 | ENSG00000155966 | AFF2 | chrX | + | 148837646 | 148837733 | 148809875 | 148809920 | 148842965 | 148843002 | 2 | 1 |
| SE | 85355 | ENSG00000155966 | AFF2 | chrX | + | 148842965 | 148843002 | 148809875 | 148809920 | 148843381 | 148843415 | 5 | 1 |
| SE | 85357 | ENSG00000155966 | AFF2 | chrX | + | 148973470 | 148973607 | 148967628 | 148967692 | 148977932 | 148978004 | 1 | 2 |
| SE | 85361 | ENSG00000235437 | LINC01278 | chrX | - | 63253448 | 63253539 | 63239786 | 63239830 | 63259667 | 63259817 | 0 | 0 |
| SE | 85374 | ENSG00000235437 | LINC01278 | chrX | - | 63432363 | 63432491 | 63396557 | 63399903 | 63509943 | 63510078 | 11 | 6 |
| SE | 85376 | ENSG00000235437 | LINC01278 | chrX | - | 63432363 | 63432491 | 63396557 | 63399903 | 63560831 | 63561052 | 17 | 8 |
| SE | 85378 | ENSG00000235437 | LINC01278 | chrX | - | 63509943 | 63510079 | 63399428 | 63399903 | 63560831 | 63560948 | 3 | 1 |
| SE | 85381 | ENSG00000235437 | LINC01278 | chrX | - | 63516909 | 63517082 | 63397669 | 63399903 | 63560831 | 63560915 | 2 | 1 |
| SE | 85386 | ENSG00000198157 | HMGN5 | chrX | - | 81119787 | 81119817 | 81118729 | 81118759 | 81121534 | 81121672 | 10 | 1 |
| SE | 85387 | ENSG00000198157 | HMGN5 | chrX | - | 81195117 | 81195297 | 81121534 | 81121672 | 81201736 | 81201886 | 6 | 6 |
| SE | 85393 | ENSG00000188211 | NCR3LG1 | chr11 | + | 17356650 | 17357001 | 17351799 | 17352039 | 17367008 | 17367347 | 26 | 12 |
| SE | 85400 | ENSG00000155085 | AK9 | chr6 | - | 109495337 | 109495440 | 109493980 | 109494095 | 109497464 | 109497563 | 1 | 0 |
| SE | 85401 | ENSG00000155085 | AK9 | chr6 | - | 109514223 | 109514437 | 109509178 | 109509380 | 109515856 | 109516075 | 3 | 2 |
| SE | 85403 | ENSG00000155085 | AK9 | chr6 | - | 109529010 | 109529073 | 109516429 | 109516642 | 109533250 | 109533333 | 2 | 11 |
| SE | 85404 | ENSG00000155085 | AK9 | chr6 | - | 109533250 | 109533470 | 109527628 | 109529073 | 109542046 | 109542171 | 10 | 25 |
| SE | 85405 | ENSG00000155085 | AK9 | chr6 | - | 109564079 | 109564280 | 109563596 | 109563712 | 109564755 | 109564845 | 6 | 24 |
| SE | 85406 | ENSG00000155085 | AK9 | chr6 | - | 109585122 | 109585237 | 109579344 | 109579626 | 109610364 | 109610513 | 0 | 1 |
| SE | 85407 | ENSG00000155085 | AK9 | chr6 | - | 109585915 | 109586072 | 109585122 | 109585237 | 109610364 | 109610513 | 10 | 9 |
| SE | 85408 | ENSG00000155085 | AK9 | chr6 | - | 109612009 | 109612093 | 109610364 | 109610513 | 109614182 | 109614296 | 8 | 1 |
| SE | 85409 | ENSG00000155085 | AK9 | chr6 | - | 109619091 | 109619236 | 109614384 | 109614480 | 109632922 | 109633103 | 23 | 16 |
| SE | 85410 | ENSG00000155085 | AK9 | chr6 | - | 109619091 | 109619236 | 109614384 | 109614480 | 109633183 | 109633323 | 17 | 11 |
| SE | 85411 | ENSG00000155085 | AK9 | chr6 | - | 109619091 | 109619236 | 109614384 | 109614480 | 109641517 | 109641616 | 20 | 18 |
| SE | 85412 | ENSG00000155085 | AK9 | chr6 | - | 109632922 | 109633103 | 109614384 | 109614480 | 109633183 | 109633323 | 18 | 19 |
| SE | 85413 | ENSG00000155085 | AK9 | chr6 | - | 109632922 | 109633103 | 109619091 | 109619236 | 109633183 | 109633323 | 23 | 24 |
| SE | 85414 | ENSG00000155085 | AK9 | chr6 | - | 109633183 | 109633323 | 109614384 | 109614480 | 109641517 | 109641616 | 24 | 42 |
| SE | 85415 | ENSG00000155085 | AK9 | chr6 | - | 109633183 | 109633323 | 109619091 | 109619236 | 109641517 | 109641616 | 24 | 42 |
| SE | 85418 | ENSG00000155085 | AK9 | chr6 | - | 109659227 | 109659413 | 109644613 | 109644688 | 109662550 | 109662657 | 54 | 58 |
| SE | 85419 | ENSG00000155085 | AK9 | chr6 | - | 109662550 | 109662663 | 109659227 | 109659413 | 109671918 | 109672015 | 102 | 76 |
| SE | 85420 | ENSG00000155085 | AK9 | chr6 | - | 109675628 | 109675756 | 109674197 | 109674261 | 109690471 | 109690670 | 19 | 5 |
| SE | 85421 | ENSG00000101928 | MOSPD1 | chrX | - | 134891478 | 134891640 | 134887631 | 134889192 | 134896816 | 134897034 | 16 | 16 |
| SE | 85422 | ENSG00000101928 | MOSPD1 | chrX | - | 134899089 | 134899165 | 134896813 | 134897034 | 134899279 | 134899534 | 20 | 11 |
| SE | 85423 | ENSG00000101928 | MOSPD1 | chrX | - | 134899279 | 134899534 | 134896816 | 134897034 | 134915181 | 134915227 | 14 | 7 |
| SE | 85426 | ENSG00000122126 | OCRL | chrX | + | 129557860 | 129557950 | 129557324 | 129557435 | 129558632 | 129558753 | 37 | 23 |
| SE | 85427 | ENSG00000122126 | OCRL | chrX | + | 129560549 | 129560651 | 129558839 | 129559001 | 129561178 | 129561293 | 42 | 29 |
| SE | 85430 | ENSG00000122126 | OCRL | chrX | + | 129584343 | 129584367 | 129576316 | 129576552 | 129587001 | 129587118 | 4 | 0 |
| SE | 85431 | ENSG00000122126 | OCRL | chrX | + | 129589236 | 129589364 | 129588885 | 129589013 | 129589844 | 129589956 | 2 | 2 |
| SE | 85433 | ENSG00000137216 | TMEM63B | chr6 | + | 44134560 | 44134743 | 44127719 | 44127866 | 44136348 | 44136439 | 0 | 0 |
| SE | 85436 | ENSG00000137216 | TMEM63B | chr6 | + | 44150223 | 44150310 | 44149858 | 44149965 | 44150563 | 44150629 | 19 | 15 |
| SE | 85437 | ENSG00000137216 | TMEM63B | chr6 | + | 44150563 | 44150629 | 44150223 | 44150310 | 44151845 | 44152008 | 17 | 18 |
| SE | 85446 | ENSG00000179241 | LDLRAD3 | chr11 | + | 36227084 | 36227430 | 36098326 | 36098461 | 36229159 | 36229211 | 8 | 11 |
| SE | 85455 | ENSG00000177485 | ZBTB33 | chrX | + | 120252668 | 120252770 | 120250811 | 120250977 | 120253413 | 120258398 | 6 | 4 |
| SE | 85457 | ENSG00000116791 | CRYZ | chr1 | - | 74706898 | 74706994 | 74705481 | 74706457 | 74707102 | 74707204 | 69 | 98 |
| SE | 85458 | ENSG00000116791 | CRYZ | chr1 | - | 74706898 | 74706994 | 74705481 | 74706457 | 74710097 | 74710247 | 21 | 23 |
| SE | 85459 | ENSG00000116791 | CRYZ | chr1 | - | 74707102 | 74707204 | 74705481 | 74706457 | 74710097 | 74710247 | 11 | 42 |
| SE | 85460 | ENSG00000116791 | CRYZ | chr1 | - | 74707102 | 74707204 | 74706898 | 74706994 | 74710097 | 74710247 | 63 | 119 |
| SE | 85461 | ENSG00000116791 | CRYZ | chr1 | - | 74710097 | 74710247 | 74707102 | 74707204 | 74723117 | 74723270 | 43 | 84 |
| SE | 85462 | ENSG00000116791 | CRYZ | chr1 | - | 74714578 | 74714630 | 74710097 | 74710247 | 74723117 | 74723270 | 49 | 54 |
| SE | 85463 | ENSG00000116791 | CRYZ | chr1 | - | 74719208 | 74719372 | 74714578 | 74714630 | 74723117 | 74723270 | 122 | 107 |
| SE | 85464 | ENSG00000116791 | CRYZ | chr1 | - | 74724710 | 74724834 | 74723117 | 74723270 | 74730381 | 74730420 | 43 | 18 |
| SE | 85465 | ENSG00000116791 | CRYZ | chr1 | - | 74724710 | 74724834 | 74723117 | 74723270 | 74732955 | 74733022 | 54 | 28 |
| SE | 85466 | ENSG00000116791 | CRYZ | chr1 | - | 74730381 | 74730420 | 74723117 | 74723270 | 74732955 | 74733061 | 0 | 0 |
| SE | 85468 | ENSG00000116791 | CRYZ | chr1 | - | 74730381 | 74730420 | 74724710 | 74724834 | 74732955 | 74733061 | 2 | 1 |
| SE | 85473 | ENSG00000133703 | KRAS | chr12 | - | 25225613 | 25225773 | 25205245 | 25209911 | 25245273 | 25245395 | 84 | 60 |
| SE | 85475 | ENSG00000133703 | KRAS | chr12 | - | 25245273 | 25245395 | 25227233 | 25227412 | 25250750 | 25250929 | 213 | 166 |
| SE | 85478 | ENSG00000131724 | IL13RA1 | chrX | + | 118758054 | 118758242 | 118749657 | 118749778 | 118761137 | 118761289 | 401 | 363 |
| SE | 85480 | ENSG00000131724 | IL13RA1 | chrX | + | 118766529 | 118766577 | 118761137 | 118761289 | 118773878 | 118773975 | 51 | 35 |

| | | | | | | | | | | | | | | | | | | | | | | | | | | | | | | | | | | | | | | | | | | | | | | |
|---|---|---|---|---|---|---|---|---|---|---|---|---|---|---|---|---|---|---|---|---|---|---|---|---|---|---|---|---|---|---|---|---|---|---|---|---|---|---|---|---|---|---|---|---|---|---|
|SE|85481|ENSG00000131724|IL13RA1|chrX|+|118766843|118766976|118766529|118766577|118773878|118773975|358|339|222|414|729|1|0|0|0|0|667|324|637|456|137|0|0|0|0|0|282|149|1|1|0.995|1|1|1|1|1|1|1|1|1|-0.001|
|SE|85485|ENSG00000167548|KMT2D|chr12|-|49024577|49024708|49022589|49022875|49024809|49024946|154|97|196|156|128|0|0|0|1|0|127|223|156|242|419|0|0|0|0|0|280|149|1|1|1|1|1|0.988|1|1|1|1|1|1|-0.002|
|SE|85492|ENSG00000167548|KMT2D|chr12|-|49042088|49042330|49041916|49041990|49042560|49042645|189|100|173|162|126|0|0|0|2|0|240|160|122|199|264|0|0|0|2|0|298|149|1|1|1|1|1|0.976|1|1|1|1|0.98|1|-0.001|
|SE|85493|ENSG00000167548|KMT2D|chr12|-|49043634|49043782|49043362|49043428|49043867|49043998|248|142|214|254|202|0|0|0|0|0|430|243|282|298|246|0|0|1|6|3|297|149|1|1|1|1|1|1|1|1|1|0.993|0.961|0.976|0.014|
|SE|85494|ENSG00000167548|KMT2D|chr12|-|49044190|49044304|49043867|49043998|49044402|49044522|215|136|173|197|106|0|0|0|0|0|368|241|163|273|282|0|0|3|0|0|263|149|1|1|1|1|1|1|1|1|1|0.969|1|1|0.006|
|SE|85495|ENSG00000167548|KMT2D|chr12|-|49044199|49044304|49043867|49043998|49044402|49044522|237|129|201|195|130|0|0|0|0|0|475|266|233|262|378|0|0|3|0|0|254|149|1|1|1|1|1|1|1|1|1|0.979|1|1|0.004|
|SE|85496|ENSG00000167548|KMT2D|chr12|-|49046608|49046790|49046259|49046424|49047964|49048069|178|62|125|110|145|0|0|1|0|0|112|125|107|114|233|0|0|1|3|0|298|149|1|1|1|1|0.984|1|1|1|1|0.982|0.95|1|0.01|
|SE|85498|ENSG00000167548|KMT2D|chr12|-|49048658|49048769|49046259|49046424|49049104|49049218|29|12|27|12|35|0|1|0|0|0|16|28|20|39|35|0|0|0|0|0|260|149|1|1|1|0.873|1|1|1|1|1|1|1|1|-0.025|
|SE|85500|ENSG00000167548|KMT2D|chr12|-|49054306|49054416|49053977|49054140|49054899|49055026|12|12|10|16|5|0|0|0|0|0|54|27|14|35|38|0|0|2|0|0|259|149|1|1|1|1|1|1|1|1|1|0.801|1|1|0.04|
|SE|85501|ENSG00000167548|KMT2D|chr12|-|49054527|49054751|49053977|49054140|49054899|49055026|48|22|36|44|26|0|0|0|0|0|63|60|73|74|83|0|0|2|0|0|298|149|1|1|1|1|1|1|1|1|1|0.948|1|1|0.01|
|SE|85502|ENSG00000167548|KMT2D|chr12|-|49054527|49054751|49054306|49054416|49054899|49055026|117|66|78|111|55|0|0|0|0|0|204|152|146|178|330|6|0|0|0|0|298|149|1|1|1|1|1|1|1|0.944|1|1|1|1|0.011|
|SE|85506|ENSG00000151778|SERP2|chr13|+|44390356|44390578|44379640|44379713|44397271|44397710|2|0|2|1|12|0|0|0|0|0|0|1|4|0|3|0|5|2|0|0|298|149|1|1|1|NA|1|1|1|NA|0.091|0.5|NA|1|0.47|
|SE|85512|ENSG00000163697|APBB2|chr4|-|41033235|41033304|41013582|41014398|41065575|41065673|0|0|0|1|0|0|0|0|0|0|0|0|0|6|53|0|0|0|6|4|218|149|1|1|NA|NA|NA|1|NA|NA|NA|NA|0.406|0.901|0.347|
|SE|85516|ENSG00000163697|APBB2|chr4|-|41100638|41100750|41065575|41065673|41142986|41143142|0|0|0|1|0|0|0|0|0|0|0|0|0|6|47|0|0|0|0|16|261|149|1|1|NA|NA|NA|1|NA|NA|NA|NA|1|0.626|0.187|
|SE|85520|ENSG00000139083|ETV6|chr12|+|11752449|11752579|11649673|11650160|11825643|11825713|81|63|85|91|150|0|0|0|0|0|192|117|136|124|135|0|0|1|0|0|279|149|1|1|1|1|1|1|1|1|1|0.986|1|1|0.003|
|SE|85524|ENSG00000139083|ETV6|chr12|+|11839139|11839304|11752449|11752579|11869423|11869969|123|93|121|140|185|1|0|0|0|0|336|162|118|187|252|0|0|0|0|0|298|149|1|1|0.984|1|1|1|1|1|1|1|1|1|-0.003|
|SE|85525|ENSG00000139083|ETV6|chr12|+|11853426|11853561|11752449|11752579|11869423|11869969|76|97|70|97|143|1|0|0|0|0|217|102|142|175|253|0|0|0|0|0|284|149|1|1|0.976|1|1|1|1|1|1|1|1|1|-0.005|
|SE|85529|ENSG00000100802|C14orf93|chr14|-|22987902|22988015|22985893|22987634|22989741|22989845|68|52|56|69|87|2|0|0|0|0|141|76|92|111|41|0|0|2|0|0|262|149|1|1|0.951|1|1|1|1|1|1|0.963|1|1|-0.002|
|SE|85530|ENSG00000100802|C14orf93|chr14|-|22989741|22989845|22987902|22988015|22990065|22990127|68|63|55|66|71|0|0|0|1|0|175|61|71|93|32|0|0|0|0|0|253|149|1|1|1|1|1|0.975|1|1|1|1|1|1|-0.005|
|SE|85531|ENSG00000100802|C14orf93|chr14|-|22999956|23000117|22998772|22999156|23000958|23001060|4|0|0|0|6|6|2|5|2|7|0|0|0|0|0|13|14|2|0|14|298|149|1|1|0.25|0|0|0|0.3|0|0|0|NA|0|0.11|
|SE|85532|ENSG00000100802|C14orf93|chr14|-|22999956|23000117|22998772|22999156|23010016|23010166|4|4|0|0|0|1|1|1|0|3|0|0|0|0|0|0|1|1|5|5|298|149|1|1|0.667|0.667|0|NA|0|NA|0|0|0|0|0.334|
|SE|85533|ENSG00000100802|C14orf93|chr14|-|22999956|23000117|22998772|22999156|23010100|23010143|4|0|0|0|0|2|1|1|1|1|0|0|0|0|0|8|0|3|0|0|298|149|1|1|0.5|0|0|0|0|0|NA|0|NA|NA|0.1|
|SE|85534|ENSG00000100802|C14orf93|chr14|-|22999956|23000117|22999044|22999362|23000958|23001060|0|4|2|0|6|4|0|2|1|2|0|0|0|0|0|0|0|0|2|0|298|149|1|1|0|1|0.333|0|0.6|NA|NA|NA|0|NA|0.387|
|SE|85535|ENSG00000100802|C14orf93|chr14|-|22999956|23000117|22999044|22999362|23010016|23010166|0|8|2|0|0|0|0|2|0|0|0|0|0|0|0|0|0|0|0|3|298|149|1|1|NA|1|0.333|NA|NA|NA|NA|NA|NA|0|0.666|
|SE|85536|ENSG00000100802|C14orf93|chr14|-|22999956|23000117|22999044|22999362|23010100|23010143|0|4|2|0|0|0|0|0|0|1|0|0|0|0|0|0|0|2|0|7|298|149|1|1|NA|1|1|NA|0|NA|NA|0|NA|0|0.667|
|SE|85537|ENSG00000100802|C14orf93|chr14|-|22999956|23000117|22999120|22999402|23000958|23001060|1|0|0|0|6|0|0|0|0|0|0|0|0|0|0|0|0|0|4|0|298|149|1|1|1|NA|NA|NA|1|NA|NA|NA|0|NA|1|
|SE|85538|ENSG00000100802|C14orf93|chr14|-|22999956|23000117|22999120|22999402|23010100|23010143|1|0|0|0|0|9|6|11|9|20|0|0|0|0|0|10|6|5|7|12|298|149|1|1|0.053|0|0|0|0|0|0|0|0|0|0.011|
|SE|85539|ENSG00000100802|C14orf93|chr14|-|23000958|23001031|22998704|22999156|23010100|23010143|3|1|3|1|3|2|1|1|1|1|0|9|1|0|14|8|0|3|0|0|222|149|1|1|0.502|0.402|0.668|0.402|0.668|0|1|0.183|NA|1|-0.017|
|SE|85540|ENSG00000100802|C14orf93|chr14|-|23000958|23001037|22998426|22999156|23010016|23010143|3|1|3|1|3|1|1|1|0|3|0|7|4|0|1|0|1|1|5|5|228|149|1|1|0.662|0.395|0.662|1|0.395|NA|0.821|0.723|0|0.116|0.208|
|SE|85541|ENSG00000100802|C14orf93|chr14|-|23000958|23001037|22998426|22999156|23010100|23010128|4|2|5|2|6|2|1|1|1|1|6|7|2|0|1|8|0|3|0|0|228|149|1|1|0.567|0.567|0.766|0.567|0.797|0.329|1|0.303|NA|1|-0.005|
|SE|85542|ENSG00000100802|C14orf93|chr14|-|23000958|23001037|22999044|22999362|23010016|23010143|2|0|0|1|0|0|0|2|0|0|0|0|3|0|0|0|0|0|0|3|228|149|1|1|1|NA|0|1|NA|NA|NA|1|NA|0|0.167|
|SE|85543|ENSG00000100802|C14orf93|chr14|-|23000958|23001037|22999044|22999362|23010100|23010128|2|0|1|1|0|0|0|0|0|1|0|0|0|2|0|0|0|2|0|7|228|149|1|1|1|NA|1|1|0|NA|NA|0|1|0|0.417|
|SE|85546|ENSG00000100802|C14orf93|chr14|-|23005064|23005130|23000958|23001037|23010016|23010143|2|3|0|0|3|0|0|0|0|0|7|0|0|8|1|0|0|3|0|0|215|149|1|1|1|1|NA|NA|1|1|NA|0|1|1|0.25|
|SE|85548|ENSG00000100532|CGRRF1|chr14|+|54522453|54522593|54509811|54510063|54530048|54530226|92|38|43|71|109|1|0|0|0|0|31|86|42|28|83|0|0|0|0|0|289|149|1|1|0.979|1|1|1|1|1|1|1|1|1|-0.004|
|SE|85553|ENSG00000008294|SPAG9|chr17|-|50977107|50977221|50974770|50974947|50982523|50982672|65|38|50|55|67|0|0|1|0|0|8|47|40|62|19|0|0|0|0|0|263|149|1|1|1|1|0.966|1|1|1|1|1|1|1|-0.007|
|SE|85554|ENSG00000008294|SPAG9|chr17|-|50979745|50979917|50976599|50977221|50982523|50982672|129|106|148|139|158|0|0|0|0|0|38|137|139|102|57|0|0|2|0|0|298|149|1|1|1|1|1|1|1|1|1|0.972|1|1|0.006|
|SE|85556|ENSG00000008294|SPAG9|chr17|-|50984922|50984990|50982523|50982672|50985697|50985778|112|70|105|109|143|0|0|0|0|0|66|75|114|69|94|0|4|2|0|0|217|149|1|1|1|1|1|1|1|1|0.928|0.975|1|1|0.019|
|SE|85557|ENSG00000008294|SPAG9|chr17|-|50985697|50985778|50982523|50982672|50987111|50987237|45|23|45|49|65|0|0|0|0|1|21|29|65|34|8|0|0|0|0|0|230|149|1|1|1|1|1|1|0.977|1|1|1|1|1|-0.005|
|SE|85558|ENSG00000008294|SPAG9|chr17|-|50989676|50989827|50987111|50987237|50990449|50990668|82|41|68|72|100|0|0|0|0|1|12|63|86|35|33|0|2|0|0|0|298|149|1|1|1|1|1|1|0.98|1|0.94|1|1|1|0.008|
|SE|85559|ENSG00000008294|SPAG9|chr17|-|50989676|50989872|50987111|50987237|50990449|50990585|123|70|98|113|160|0|0|0|0|1|33|96|167|70|42|0|2|0|0|0|298|149|1|1|1|1|1|1|0.988|1|0.96|1|1|1|0.006|
|SE|85560|ENSG00000008294|SPAG9|chr17|-|50995056|50995224|50993763|50993935|50995443|50995533|288|235|207|236|288|0|2|0|0|0|129|162|313|132|24|0|5|0|1|0|298|149|1|1|1|0.983|1|1|1|1|0.942|1|0.985|1|0.011|
|SE|85561|ENSG00000008294|SPAG9|chr17|-|50995443|50995533|50993763|50993935|50996564|50996694|8|10|8|5|7|0|0|0|0|0|9|10|5|3|0|0|0|1|0|0|239|149|1|1|1|1|1|1|1|1|1|0.757|1|NA|0.061|
|SE|85564|ENSG00000008294|SPAG9|chr17|-|51007836|51007851|51007268|51007326|51014231|51014353|0|4|0|0|2|160|212|182|219|272|0|1|0|0|0|84|259|476|253|55|164|149|1|1|0|0.017|0|0|0.007|0|0.003|0|0|0|0.004|
|SE|85568|ENSG00000008294|SPAG9|chr17|-|51047374|51047469|51041500|51041651|51056411|51056424|235|202|179|245|393|0|3|0|1|4|78|316|345|268|87|0|0|0|0|0|244|149|1|1|1|0.976|1|0.993|0.984|1|1|1|1|1|-0.009|
|SE|85569|ENSG00000008294|SPAG9|chr17|-|51047374|51047469|51041500|51041651|51079583|51079622|75|66|67|97|163|0|0|0|0|0|39|130|135|89|15|0|0|2|0|0|244|149|1|1|1|1|1|1|1|1|1|0.976|1|1|0.005|
|SE|85570|ENSG00000008294|SPAG9|chr17|-|51056411|51056482|51041500|51041651|51079583|51079622|50|38|43|44|73|0|0|0|0|0|50|54|67|59|7|0|0|2|0|0|220|149|1|1|1|1|1|1|1|1|1|0.958|1|1|0.008|
|SE|85571|ENSG00000008294|SPAG9|chr17|-|51056411|51056482|51047374|51047469|51079583|51079622|206|164|148|188|283|0|0|1|0|0|89|223|277|237|72|0|0|2|2|0|220|149|1|1|1|1|0.99|1|1|1|1|0.989|0.988|1|0.003|
|SE|85572|ENSG00000008294|SPAG9|chr17|-|51079583|51079704|51056411|51056482|51120353|51120734|173|140|133|166|187|1|0|0|0|0|101|196|259|188|36|0|0|0|0|0|270|149|1|1|0.99|1|1|1|1|1|1|1|1|1|-0.002|
|SE|85574|ENSG00000236287|ZBED5|chr11|-|10838268|10838573|10836983|10837067|10856143|10856218|1|1|2|2|0|1|0|0|0|0|0|0|3|0|0|0|0|0|0|0|298|149|1|1|0.333|1|1|1|NA|NA|NA|1|NA|NA|-0.167|
|SE|85583|ENSG00000104870|FCGRT|chr19|+|49513881|49514133|49513153|49513473|49524506|49524776|102|40|54|64|19|1|0|0|0|1|111|78|75|109|95|0|0|0|0|0|298|149|1|1|0.981|1|1|1|0.905|1|1|1|1|1|-0.023|
|SE|85585|ENSG00000104870|FCGRT|chr19|+|49515176|49515269|49514210|49514486|49521694|49521813|0|0|0|0|0|0|1|1|5|0|0|3|0|0|0|12|2|0|0|11|242|149|1|1|NA|0|0|0|NA|0|0.48|NA|NA|0|-0.16|
|SE|85587|ENSG00000104870|FCGRT|chr19|+|49516113|49516199|49514210|49514486|49524506|49524664|1|0|0|0|0|109|48|56|92|81|0|0|0|2|0|74|57|39|67|156|235|149|1|1|0.006|0|0|0|0|0|0|0|0.019|0|-0.003|
|SE|85588|ENSG00000104870|FCGRT|chr19|+|49520668|49521201|49514210|49514486|49524506|49525263|0|0|0|0|0|109|48|56|92|81|0|0|1|3|0|74|57|39|67|156|298|149|1|1|0|0|0|0|0|0|0|0.013|0.022|0|-0.007|
|SE|85593|ENSG00000104870|FCGRT|chr19|+|49524506|49525573|49513194|49514486|49526009|49526331|500|247|307|355|428|2|1|0|4|0|589|351|204|341|720|0|0|3|4|0|298|149|1|1|0.992|0.992|1|0.978|1|1|1|0.971|0.977|1|0.003|
|SE|85594|ENSG00000104870|FCGRT|chr19|+|49525456|49525573|49513254|49513473|49526009|49526329|390|199|244|256|343|0|0|0|0|1|508|294|165|267|564|0|0|0|0|0|266|149|1|1|1|1|1|1|0.995|1|1|1|1|1|-0.001|
|SE|85595|ENSG00000104870|FCGRT|chr19|+|49525456|49525573|49514210|49514486|49526009|49526329|390|199|245|256|343|2|1|0|4|0|508|294|165|269|575|0|0|3|4|0|266|149|1|1|0.991|0.991|1|0.973|1|1|1|0.969|0.974|1|0.002|
|SE|85598|ENSG00000226521|NA|chr17|+|20722034|20722135|20721210|20721467|20724595|20724650|0|0|0|0|0|0|0|0|0|3|0|0|0|0|7|0|0|0|0|0|250|149|1|1|NA|NA|NA|NA|0|NA|NA|NA|NA|1|-1|
|SE|85601|ENSG00000163389|POGLUT1|chr3|+|119479907|119480172|119477312|119477448|119485327|119485387|39|21|50|57|32|0|0|0|0|0|27|19|33|53|0|0|0|2|0|0|298|149|1|1|1|1|1|1|1|1|1|0.892|1|NA|0.027|
|SE|85602|ENSG00000163389|POGLUT1|chr3|+|119480050|119480172|119477312|119477448|119485327|119485357|70|35|78|86|67|0|0|0|0|0|37|29|47|75|5|0|0|2|0|0|271|149|1|1|1|1|1|1|1|1|1|0.928|1|1|0.014|
|SE|85603|ENSG00000163389|POGLUT1|chr3|+|119488928|119488987|119486832|119486932|119490550|119490667|66|47|32|51|91|0|0|0|0|2|42|50|26|25|28|0|0|0|0|0|208|149|1|1|1|1|1|1|0.97|1|1|1|1|1|-0.006|
|SE|85604|ENSG00000163389|POGLUT1|chr3|+|119490550|119490718|119486832|119486932|119491517|119491574|32|31|36|48|57|0|0|0|0|0|12|35|22|13|7|0|0|0|1|0|298|149|1|1|1|1|1|1|1|1|1|1|0.867|1|0.027|
|SE|85607|ENSG00000147155|EBP|chrX|+|48523698|48524072|48521807|48521907|48526988|48527025|108|43|71|70|31|0|0|0|0|0|133|127|99|95|170|0|0|0|3|0|298|149|1|1|1|1|1|1|1|1|1|1|0.941|1|0.012|
|SE|85608|ENSG00000147155|EBP|chrX|+|48523753|48524072|48521804|48521907|48526988|48527025|108|44|71|70|31|0|0|0|0|0|133|127|100|98|183|0|0|0|3|0|298|149|1|1|1|1|1|1|1|1|1|1|0.942|1|0.012|
|SE|85609|ENSG00000147155|EBP|chrX|+|48523922|48524072|48521798|48521907|48526988|48527025|111|46|71|70|31|0|0|0|0|0|133|127|99|95|170|0|0|0|3|0|298|149|1|1|1|1|1|1|1|1|1|1|0.941|1|0.012|
|SE|85610|ENSG00000147010|SH3KBP1|chrX|-|19537716|19537780|19533976|19536458|19541924|19542193|418|261|331|389|637|0|0|0|2|0|458|313|263|296|387|0|1|0|0|0|213|149|1|1|1|1|1|0.993|1|1|0.995|1|1|1|0|
|SE|85612|ENSG00000147010|SH3KBP1|chrX|-|19549973|19550083|19541924|19542193|19569102|19569188|173|86|148|158|125|0|0|0|0|0|245|152|162|177|160|0|2|0|0|0|259|149|1|1|1|1|1|1|1|1|0.978|1|1|1|0.004|
|SE|85613|ENSG00000147010|SH3KBP1|chrX|-|19594948|19595000|19592066|19592147|19607937|19608045|522|381|479|564|911|0|0|0|0|0|1224|736|685|815|720|0|0|0|8|7|201|149|1|1|1|1|1|1|1|1|1|1|0.987|0.987|0.005|
|SE|85615|ENSG00000147010|SH3KBP1|chrX|-|19631863|19631958|19607937|19608045|19645399|19645475|1070|791|1017|1028|1458|0|0|0|1|0|1591|1166|1075|1283|1116|0|3|0|0|0|244|149|1|1|1|1|1|0.998|1|1|0.996|1|1|1|0|
|SE|85616|ENSG00000147010|SH3KBP1|chrX|-|19645399|19645475|19631863|19631958|19683822|19684028|1001|716|954|926|1146|0|0|0|4|0|1405|1012|870|1091|1008|0|0|0|0|0|225|149|1|1|1|1|1|0.994|1|1|1|1|1|1|-0.001|
|SE|85617|ENSG00000147010|SH3KBP1|chrX|-|19645399|19645475|19631863|19631958|19695611|19695741|225|173|249|245|309|0|0|1|0|0|428|270|200|273|201|5|0|2|1|0|225|149|1|1|1|1|0.994|1|1|0.983|1|0.985|0.994|1|0.006|
|SE|85618|ENSG00000147010|SH3KBP1|chrX|-|19683822|19684028|19631863|19631958|19695611|19695741|557|274|403|492|506|0|0|1|0|0|987|519|463|594|595|5|0|2|1|0|298|149|1|1|1|1|0.995|1|1|0.99|1|0.991|0.997|1|0.003|
|SE|85620|ENSG00000147010|SH3KBP1|chrX|-|19687616|19687691|19683822|19684028|19694994|19695051|3|2|1|0|0|3|6|1|3|8|0|6|3|8|9|0|4|6|6|0|224|149|1|1|0.399|0.181|0.399|0|0|NA|0.499|0.25|0.47|1|-0.359|
|SE|85621|ENSG00000147010|SH3KBP1|chrX|-|19687616|19687691|19683822|19684028|19695611|19695741|5|2|2|1|2|557|274|403|488|506|0|11|8|10|5|987|519|463|594|595|224|149|1|1|0.006|0.005|0.003|0.001|0.003|0|0.014|0.011|0.011|0.006|-0.005|
|SE|85623|ENSG00000147010|SH3KBP1|chrX|-|19694994|19695051|19687616|19687691|19695611|19695741|6|0|1|4|6|3|0|2|1|2|0|0|1|2|23|0|5|5|2|5|206|149|1|1|0.591|NA|0.266|0.743|0.685|NA|0|0.126|0.42|0.769|0.243|
|SE|85624|ENSG00000147010|SH3KBP1|chrX|-|19695611|19695741|19645399|19645475|19706880|19706984|298|161|222|244|244|0|0|2|0|0|443|389|218|251|324|0|0|0|0|0|279|149|1|1|1|1|0.983|1|1|1|1|1|1|1|-0.003|
|SE|85625|ENSG00000147010|SH3KBP1|chrX|-|19695611|19695741|19645399|19645475|19799633|19799751|10|3|2|7|1|0|0|2|0|0|8|8|6|5|14|0|0|0|0|0|279|149|1|1|1|1|0.348|1|1|1|1|1|1|1|-0.13|
|SE|85626|ENSG00000147010|SH3KBP1|chrX|-|19706880|19706984|19645399|19645475|19836124|19836282|0|0|1|0|0|0|0|0|0|0|0|0|3|0|0|0|0|0|1|0|253|149|1|1|NA|NA|1|NA|NA|NA|NA|1|0|NA|0.5|
|SE|85627|ENSG00000147010|SH3KBP1|chrX|-|19706880|19706984|19695611|19695741|19746317|19746411|628|362|497|551|556|0|0|2|0|0|1052|746|525|644|702|0|0|0|0|0|253|149|1|1|1|1|0.993|1|1|1|1|1|1|1|-0.001|
|SE|85628|ENSG00000147010|SH3KBP1|chrX|-|19746317|19746441|19695611|19695741|19799633|19799751|114|77|132|153|109|0|0|0|1|0|45|148|79|55|69|0|0|0|0|0|273|149|1|1|1|1|1|0.988|1|1|1|1|1|1|-0.002|
|SE|85629|ENSG00000147010|SH3KBP1|chrX|-|19746317|19746441|19706880|19706984|19836124|19836282|422|276|367|417|368|0|0|0|0|0|893|556|397|601|602|0|0|3|0|0|273|149|1|1|1|1|1|1|1|1|1|0.986|1|1|0.003|
|SE|85630|ENSG00000108679|LGALS3BP|chr17|-|78972969|78973222|78971254|78972704|78974687|78974772|71|20|50|67|46|0|1|1|0|0|196|244|36|329|172|0|0|0|0|0|298|149|1|1|1|0.909|0.962|1|1|1|1|1|1|1|-0.026|
|SE|85631|ENSG00000108679|LGALS3BP|chr17|-|78972969|78973343|78972469|78972704|78974687|78974819|47|8|30|38|35|0|1|1|0|0|103|148|19|173|70|0|0|0|0|0|298|149|1|1|1|0.8|0.938|1|1|1|1|1|1|1|-0.052|
|SE|85634|ENSG00000108679|LGALS3BP|chr17|-|78977403|78977473|78977139|78977214|78979823|78979843|0|0|0|3|0|50|20|37|56|13|0|0|0|0|0|123|146|47|249|105|219|149|1|1|0|0|0|0.035|0|0|0|0|0|0|0.007|
|SE|85639|ENSG00000173660|UQCRH|chr1|+|46310154|46310316|46309100|46309127|46316551|46316776|375|225|375|399|352|2|0|1|4|2|464|307|315|428|701|0|0|0|0|0|298|149|1|1|0.989|1|0.995|0.98|0.989|1|1|1|1|1|-0.009|
|SE|85646|ENSG00000166881|NEMP1|chr12|-|57064070|57064185|57063118|57063344|57064645|57064739|63|34|54|41|80|6|4|4|0|1|30|30|14|8|0|0|5|3|0|0|264|149|1|1|0.856|0.828|0.884|1|0.978|1|0.772|0.725|1|NA|0.035|
|SE|85647|ENSG00000166881|NEMP1|chr12|-|57069233|57069306|57064645|57064739|57070673|57070893|17|16|13|14|49|0|2|4|0|1|24|13|6|5|0|0|1|0|0|0|222|149|1|1|1|0.843|0.686|1|0.97|1|0.897|1|1|NA|-0.074|
|SE|85648|ENSG00000166881|NEMP1|chr12|-|57069233|57069306|57064645|57064739|57070819|57070893|12|4|8|3|20|2|1|9|3|7|8|7|2|3|0|0|1|0|0|0|222|149|1|1|0.801|0.729|0.374|0.402|0.657|1|0.825|1|1|NA|-0.364|
|SE|85650|ENSG00000101199|ARFGAP1|chr20|+|63276090|63276200|63275576|63275640|63276479|63276547|47|11|59|57|41|0|0|0|0|0|29|57|51|43|95|0|0|0|3|0|259|149|1|1|1|1|1|1|1|1|1|1|0.892|1|0.022|
|SE|85651|ENSG00000101199|ARFGAP1|chr20|+|63277204|63277305|63276479|63276651|63278116|63278203|106|40|91|92|60|1|0|0|4|0|150|99|55|102|127|3|2|1|0|0|250|149|1|1|0.984|1|1|0.932|1|0.968|0.967|0.97|1|1|0.002|
|SE|85653|ENSG00000101199|ARFGAP1|chr20|+|63282818|63282851|63281290|63281347|63284865|63284922|24|25|29|37|20|3|0|0|1|0|73|37|24|27|51|0|0|0|0|0|182|149|1|1|0.868|1|1|0.968|1|1|1|1|1|1|-0.033|
|SE|85655|ENSG00000101199|ARFGAP1|chr20|+|63285979|63286218|63285653|63285713|63286365|63286442|0|0|0|0|0|60|30|46|52|27|0|0|0|3|0|102|44|30|57|87|298|149|1|1|0|0|0|0|0|0|0|0|0.026|0|-0.005|
|SE|85656|ENSG00000119760|SUPT7L|chr2|-|27655364|27655602|27650808|27653747|27657344|27657669|195|125|146|211|113|0|0|0|1|0|141|187|174|159|125|0|0|0|0|0|298|149|1|1|1|1|1|0.991|1|1|1|1|1|1|-0.002|
|SE|85658|ENSG00000119760|SUPT7L|chr2|-|27660983|27661388|27657344|27657669|27662178|27662281|125|89|123|142|140|0|0|0|0|0|140|112|130|137|135|0|0|0|4|0|298|149|1|1|1|1|1|1|1|1|1|1|0.945|1|0.011|
|SE|85659|ENSG00000119760|SUPT7L|chr2|-|27660983|27661601|27657344|27657669|27662178|27662281|91|64|82|93|102|0|0|0|0|0|90|96|103|66|76|0|0|0|4|0|298|149|1|1|1|1|1|1|1|1|1|1|0.892|1|0.022|
|SE|85662|ENSG00000161791|FMNL3|chr12|-|49651932|49652212|49651381|49651450|49653225|49653327|190|114|139|206|200|0|0|1|0|0|95|106|72|85|197|0|0|0|0|0|298|149|1|1|1|1|0.986|1|1|1|1|1|1|1|-0.003|
|SE|85663|ENSG00000161791|FMNL3|chr12|-|49653225|49653327|49651932|49652212|49653724|49653874|195|141|169|232|207|0|0|0|0|0|170|138|98|103|276|0|1|0|0|0|251|149|1|1|1|1|1|1|1|1|0.988|1|1|1|0.002|
|SE|85664|ENSG00000161791|FMNL3|chr12|-|49654909|49654984|49654191|49654302|49656403|49656497|220|140|188|247|191|0|0|0|0|0|287|117|84|140|226|0|0|0|2|0|224|149|1|1|1|1|1|1|1|1|1|1|0.979|1|0.004|
|SE|85666|ENSG00000177646|ACAD9|chr3|+|128884652|128884746|128879595|128879841|128893554|128893656|105|90|77|87|163|0|2|2|2|0|247|134|105|189|231|0|2|0|0|0|243|149|1|1|1|0.965|0.959|0.964|1|1|0.976|1|1|1|-0.018|
|SE|85667|ENSG00000177646|ACAD9|chr3|+|128884652|128884746|128879622|128880017|128893554|128893656|86|70|65|73|146|0|0|2|0|2|195|123|105|157|183|0|0|0|0|0|243|149|1|1|1|1|0.952|1|0.978|1|1|1|1|1|-0.014|
|SE|85669|ENSG00000177646|ACAD9|chr3|+|128884656|128884746|128879603|128879841|128893554|128893656|76|58|58|59|133|0|2|2|2|0|168|81|87|129|153|0|2|0|0|0|239|149|1|1|1|0.948|0.948|0.948|1|1|0.962|1|1|1|-0.024|
|SE|85670|ENSG00000177646|ACAD9|chr3|+|128884656|128884746|128879628|128880017|128893554|128893656|76|58|55|59|133|0|0|2|0|2|168|81|87|129|153|0|0|0|0|0|239|149|1|1|1|1|0.945|1|0.976|1|1|1|1|1|-0.016|
|SE|85673|ENSG00000177646|ACAD9|chr3|+|128884674|128884746|128879642|128880017|128893554|128893656|67|48|50|52|94|0|0|2|0|2|144|67|71|122|149|0|0|0|0|0|221|149|1|1|1|1|0.944|1|0.969|1|1|1|1|1|-0.017|
|SE|85674|ENSG00000177646|ACAD9|chr3|+|128895309|128895416|128893518|128893656|128896435|128896466|201|149|144|139|390|0|0|0|0|0|383|226|183|278|393|0|0|1|0|0|256|149|1|1|1|1|1|1|1|1|1|0.991|1|1|0.002|
|SE|85675|ENSG00000177646|ACAD9|chr3|+|128895309|128895416|128893518|128893656|128897631|128897710|95|80|72|81|178|0|0|0|0|0|213|106|74|143|172|0|0|0|3|0|256|149|1|1|1|1|1|1|1|1|1|1|0.965|1|0.007|
|SE|85676|ENSG00000177646|ACAD9|chr3|+|128896435|128896536|128893554|128893656|128897631|128897710|24|10|13|26|48|0|0|0|0|0|27|26|26|24|20|0|0|0|3|0|250|149|1|1|1|1|1|1|1|1|1|1|0.827|1|0.035|
|SE|85677|ENSG00000177646|ACAD9|chr3|+|128896435|128896536|128895309|128895416|128897631|128897710|130|79|86|86|260|0|0|1|1|0|197|145|134|159|241|0|0|0|0|0|250|149|1|1|1|1|0.981|0.981|1|1|1|1|1|1|-0.008|
|SE|85678|ENSG00000177646|ACAD9|chr3|+|128898430|128898506|128897631|128897710|128899286|128899461|0|0|2|1|0|27|17|44|40|34|0|0|0|1|1|44|25|32|52|58|225|149|1|1|0|0|0.029|0.016|0|0|0|0|0.013|0.011|0.004|
|SE|85679|ENSG00000177646|ACAD9|chr3|+|128907907|128908264|128906120|128906249|128908972|128909099|12|9|13|15|16|0|1|0|0|0|5|11|2|12|20|0|0|0|0|0|298|149|1|1|1|0.818|1|1|1|1|1|1|1|1|-0.036|
|SE|85680|ENSG00000177646|ACAD9|chr3|+|128908184|128908264|128906120|128906249|128908972|128909099|27|13|27|24|25|0|1|0|0|0|30|38|8|22|50|0|0|0|0|0|229|149|1|1|1|0.894|1|1|1|1|1|1|1|1|-0.021|
|SE|85681|ENSG00000177646|ACAD9|chr3|+|128910390|128910539|128910020|128910149|128910740|128910813|2|0|0|0|3|24|17|29|19|5|0|0|0|0|0|19|13|7|28|33|298|149|1|1|0.04|0|0|0|0.231|0|0|0|0|0|0.054|
|SE|85685|ENSG00000177981|ASB8|chr12|-|48150086|48150170|48149838|48149998|48151200|48151305|2|0|1|0|4|105|26|68|63|123|2|0|0|1|0|128|99|68|54|109|233|149|1|1|0.012|0|0.009|0|0.02|0.01|0|0|0.012|0|0.004|
|SE|85691|ENSG00000177981|ASB8|chr12|-|48157121|48157257|48153367|48153529|48157458|48157517|0|0|0|0|0|31|10|24|13|21|4|0|0|0|0|27|16|31|15|18|285|149|1|1|0|0|0|0|0|0.072|0|0|0|0|-0.014|
|SE|85696|ENSG00000175463|TBC1D10C|chr11|+|67405084|67405184|67404082|67404354|67405398|67405506|285|77|237|194|130|0|0|1|0|0|111|226|134|131|378|0|0|0|2|0|249|149|1|1|1|1|0.993|1|1|1|1|1|0.975|1|0.004|
|SE|85700|ENSG00000175463|TBC1D10C|chr11|+|67405594|67406692|67405594|67405701|67408978|67409133|4|0|3|2|0|0|0|0|1|0|0|7|0|0|0|0|0|0|0|0|215|149|1|1|1|NA|1|0.581|NA|NA|1|NA|NA|NA|-0.14|
|SE|85701|ENSG00000175463|TBC1D10C|chr11|+|67406626|67406692|67405902|67406017|67406826|67407016|129|36|118|93|39|0|0|0|0|0|10|94|44|30|55|0|0|4|0|0|215|149|1|1|1|1|1|1|1|1|1|0.884|1|1|0.023|
|SE|85702|ENSG00000175463|TBC1D10C|chr11|+|67406626|67406738|67405902|67406017|67406826|67407016|75|21|61|60|31|0|0|0|0|0|10|66|55|24|113|0|0|4|0|0|261|149|1|1|1|1|1|1|1|1|1|0.887|1|1|0.023|
|SE|85708|ENSG00000072778|ACADVL|chr17|+|7220923|7221058|7220757|7220830|7221951|7221998|123|52|85|94|128|0|0|0|0|0|147|153|83|115|234|0|0|1|0|0|284|149|1|1|1|1|1|1|1|1|1|0.978|1|1|0.004|
|SE|85709|ENSG00000072778|ACADVL|chr17|+|7221518|7221682|7220923|7221058|7221951|7222081|275|164|191|242|241|0|0|0|0|2|375|301|204|246|656|1|0|3|1|0|298|149|1|1|1|1|1|1|0.984|0.995|1|0.971|0.992|1|0.005|
|SE|85710|ENSG00000072778|ACADVL|chr17|+|7221537|7221682|7220765|7220830|7221951|7221956|273|162|193|239|241|0|0|0|0|0|386|302|209|246|656|0|0|1|0|0|294|149|1|1|1|1|1|1|1|1|1|0.991|1|1|0.002|
|SE|85711|ENSG00000072778|ACADVL|chr17|+|7221537|7221682|7220923|7221058|7221951|7221956|434|255|294|425|367|0|0|0|0|2|506|473|316|368|1048|1|0|3|1|0|294|149|1|1|1|1|1|1|0.989|0.996|1|0.982|0.995|1|0.003|
|SE|85712|ENSG00000072778|ACADVL|chr17|+|7221951|7222081|7221518|7221682|7222176|7222302|344|195|230|312|294|0|0|0|0|0|467|385|242|330|878|0|0|0|1|0|279|149|1|1|1|1|1|1|1|1|1|1|0.994|1|0.001|
|SE|85714|ENSG00000072778|ACADVL|chr17|+|7223132|7223237|7222666|7222865|7223643|7223730|771|582|488|735|950|0|0|0|0|0|1917|1294|898|1530|2948|0|0|2|0|0|254|149|1|1|1|1|1|1|1|1|1|0.996|1|1|0.001|
|SE|85715|ENSG00000072778|ACADVL|chr17|+|7223421|7223730|7223106|7223237|7223812|7223830|251|144|175|243|244|2|2|0|0|0|438|408|253|373|789|0|0|0|0|0|298|149|1|1|0.984|0.973|1|1|1|1|1|1|1|1|-0.009|
|SE|85716|ENSG00000072778|ACADVL|chr17|+|7223643|7223730|7222639|7223237|7223812|7223847|866|624|575|849|1042|2|2|0|0|0|1947|1382|956|1600|3024|0|0|0|0|0|236|149|1|1|0.996|0.995|1|1|1|1|1|1|1|1|-0.002|
|SE|85718|ENSG00000104356|POP1|chr8|+|98123335|98123479|98117292|98117390|98127594|98127762|11|11|4|14|8|0|2|0|0|0|26|11|13|16|11|0|0|0|0|0|293|149|1|1|1|0.737|1|1|1|1|1|1|1|1|-0.053|
|SE|85722|ENSG00000168994|PXDC1|chr6|-|3739057|3739161|3738061|3738148|3751275|3751529|0|1|0|0|0|2|3|2|3|0|0|0|0|0|2|0|0|0|5|20|253|149|1|1|0|0.164|0|0|NA|NA|NA|NA|0|0.056|0.013|
|SE|85731|ENSG00000134294|SLC38A2|chr12|-|46362493|46362638|46362283|46362381|46363020|46363145|696|438|612|684|606|0|0|0|0|0|70|326|388|181|49|0|5|0|0|0|294|149|1|1|1|1|1|1|1|1|0.971|1|1|1|0.006|
|SE|85735|ENSG00000134294|SLC38A2|chr12|-|46371177|46371379|46370775|46370857|46372508|46372771|420|285|347|435|433|0|1|0|0|0|99|288|230|122|48|0|0|0|0|0|298|149|1|1|1|0.993|1|1|1|1|1|1|1|1|-0.001|
|SE|85737|ENSG00000146701|MDH2|chr7|+|76054829|76054998|76048105|76048226|76057409|76057447|117|45|131|120|79|0|0|0|0|0|380|276|155|135|304|0|0|3|5|0|298|149|1|1|1|1|1|1|1|1|1|0.963|0.931|1|0.021|
|SE|85738|ENSG00000146701|MDH2|chr7|+|76060372|76060498|76057968|76058078|76063514|76063592|550|263|579|491|725|0|2|0|0|0|775|799|478|492|1048|0|6|0|0|0|275|149|1|1|1|0.986|1|1|1|1|0.986|1|1|1|0|
|SE|85741|ENSG00000151413|NUBPL|chr14|+|31667820|31667979|31599288|31599379|31673354|31673394|0|1|0|0|0|7|7|16|18|21|0|0|0|0|0|21|13|7|22|8|298|149|1|1|0|0.067|0|0|0|0|0|0|0|0|0.013|
|SE|85743|ENSG00000151413|NUBPL|chr14|+|31673483|31673574|31599288|31599379|31787779|31787793|2|0|10|11|9|0|0|3|2|0|3|0|0|1|0|0|0|0|0|0|240|149|1|1|1|NA|0.674|0.773|1|1|NA|NA|1|NA|-0.138|
|SE|85744|ENSG00000151413|NUBPL|chr14|+|31673483|31673574|31599288|31599379|31826628|31826714|0|0|4|2|0|0|0|0|1|0|0|0|0|1|0|0|0|0|0|0|240|149|1|1|NA|NA|1|0.554|NA|NA|NA|NA|1|NA|-0.223|
|SE|85745|ENSG00000151413|NUBPL|chr14|+|31787779|31787873|31599288|31599379|31826628|31826714|11|6|7|7|9|0|0|0|1|0|8|1|3|1|0|0|0|0|0|0|243|149|1|1|1|1|1|0.811|1|1|1|1|1|NA|-0.038|
|SE|85747|ENSG00000075539|FRYL|chr4|-|48500029|48500220|48497356|48499680|48501622|48501733|363|255|278|280|480|0|0|0|1|1|9|124|93|20|6|0|0|0|0|0|298|149|1|1|1|1|1|0.993|0.996|1|1|1|1|1|-0.002|
|SE|85748|ENSG00000075539|FRYL|chr4|-|48501622|48501733|48500029|48500220|48505546|48505615|213|131|179|168|233|0|1|0|0|0|4|70|70|7|0|0|0|3|0|0|260|149|1|1|1|0.987|1|1|1|1|1|0.93|1|NA|0.015|
|SE|85750|ENSG00000075539|FRYL|chr4|-|48505546|48505615|48501622|48501733|48510058|48510157|178|121|125|136|152|0|0|0|0|0|12|40|58|6|0|0|3|0|0|0|218|149|1|1|1|1|1|1|1|1|0.901|1|1|NA|0.025|
|SE|85751|ENSG00000075539|FRYL|chr4|-|48510058|48510157|48505546|48505615|48510834|48510984|243|149|155|184|180|0|0|0|2|0|8|54|60|9|0|0|0|6|0|0|248|149|1|1|1|1|1|0.982|1|1|1|0.857|1|NA|0.032|
|SE|85753|ENSG00000075539|FRYL|chr4|-|48527476|48527653|48522900|48523104|48527970|48528045|320|276|247|359|392|0|0|0|0|0|30|163|94|26|6|0|0|3|0|0|298|149|1|1|1|1|1|1|1|1|1|0.94|1|1|0.012|
|SE|85754|ENSG00000075539|FRYL|chr4|-|48531155|48531353|48528174|48528336|48534544|48534685|154|95|149|131|182|0|1|1|7|3|0|60|61|8|0|0|3|0|0|0|298|149|1|1|1|0.979|0.987|0.903|0.968|NA|0.909|1|1|NA|-0.002|
|SE|85756|ENSG00000075539|FRYL|chr4|-|48540352|48540960|48539970|48540068|48542026|48542121|314|275|275|323|382|0|0|0|1|0|24|138|106|51|4|0|0|0|0|0|298|149|1|1|1|1|1|0.994|1|1|1|1|1|1|-0.001|
|SE|85757|ENSG00000075539|FRYL|chr4|-|48542026|48542121|48540352|48540960|48543806|48543997|250|221|191|219|226|2|0|3|5|0|29|104|93|44|0|0|1|0|0|0|244|149|1|1|0.987|1|0.975|0.964|1|1|0.984|1|1|NA|-0.011|
|SE|85758|ENSG00000075539|FRYL|chr4|-|48544782|48544904|48543806|48543997|48546066|48546268|196|144|144|167|249|0|0|0|2|0|19|102|45|15|0|0|0|0|0|0|271|149|1|1|1|1|1|0.979|1|1|1|1|1|NA|-0.004|
|SE|85759|ENSG00000075539|FRYL|chr4|-|48547583|48547769|48546066|48546271|48548689|48548793|210|141|139|226|149|0|0|2|0|0|23|91|81|8|10|0|0|0|0|0|298|149|1|1|1|1|0.972|1|1|1|1|1|1|1|-0.006|
|SE|85762|ENSG00000075539|FRYL|chr4|-|48561467|48561636|48557452|48557712|48562888|48562988|151|142|193|196|197|0|0|0|0|0|10|75|81|21|39|0|3|0|0|0|298|149|1|1|1|1|1|1|1|1|0.926|1|1|1|0.015|
|SE|85763|ENSG00000075539|FRYL|chr4|-|48575116|48575241|48573185|48573243|48576029|48576222|200|131|188|195|229|0|0|0|0|0|26|108|97|59|11|0|0|0|4|0|274|149|1|1|1|1|1|1|1|1|1|1|0.889|1|0.022|
|SE|85764|ENSG00000075539|FRYL|chr4|-|48605740|48605833|48603289|48603388|48606437|48606606|129|109|121|109|247|0|2|0|0|0|0|73|52|16|38|0|0|0|0|0|242|149|1|1|1|0.971|1|1|1|NA|1|1|1|1|-0.006|
|SE|85765|ENSG00000075539|FRYL|chr4|-|48608986|48609067|48606437|48606606|48619273|48619370|14|14|26|22|34|0|0|0|2|2|0|21|8|1|0|0|0|0|0|0|230|149|1|1|1|1|1|0.877|0.917|NA|1|1|1|NA|-0.041|
|SE|85766|ENSG00000075539|FRYL|chr4|-|48609743|48609823|48608986|48609067|48619273|48619370|210|220|217|284|332|3|0|5|0|2|24|107|73|44|0|0|9|0|0|0|229|149|1|1|0.979|1|0.966|1|0.991|1|0.886|1|1|NA|0.016|
|SE|85767|ENSG00000075539|FRYL|chr4|-|48634290|48634490|48623125|48623179|48684672|48684795|455|419|379|480|329|0|0|1|0|0|92|262|182|109|95|0|2|0|0|0|298|149|1|1|1|1|0.995|1|1|1|0.985|1|1|1|0.002|
|SE|85769|ENSG00000075539|FRYL|chr4|-|48684672|48684795|48634290|48634490|48780077|48780271|170|168|168|195|114|0|0|0|2|0|16|91|61|46|19|0|0|0|1|0|272|149|1|1|1|1|1|0.982|1|1|1|1|0.962|1|0.004|
|SE|85770|ENSG00000075539|FRYL|chr4|-|48710518|48710698|48634417|48634490|48780077|48780248|45|33|30|47|22|0|0|0|2|0|7|18|13|20|7|0|0|0|1|0|298|149|1|1|1|1|1|0.922|1|1|1|1|0.909|1|0.003|
|SE|85772|ENSG00000075539|FRYL|chr4|-|48727430|48727491|48710518|48710698|48780077|48780129|0|0|0|2|0|30|17|25|41|17|0|0|0|0|0|7|11|10|20|7|210|149|1|1|0|0|0|0.033|0|0|0|0|0|0|0.007|
|SE|85776|ENSG00000256861|NA|chr12|-|122249870|122250045|122242381|122242508|122261260|122261447|3|0|1|0|0|0|0|0|4|0|0|0|5|0|11|0|2|0|0|0|298|149|1|1|1|NA|1|0|NA|NA|0|1|NA|1|0|
|SE|85777|ENSG00000162065|TBC1D24|chr16|+|2493596|2493706|2475117|2475170|2496033|2497113|0|2|0|0|0|6|4|6|1|0|0|0|0|0|0|0|0|1|5|0|259|149|1|1|0|0.223|0|0|NA|NA|NA|0|0|NA|0.056|
|SE|85786|ENSG00000258529|NA|chr11|-|111809642|111809773|111786285|111786520|111836164|111836294|81|69|113|78|107|0|0|3|0|0|121|165|113|142|94|0|0|0|0|0|280|149|1|1|1|1|0.952|1|1|1|1|1|1|1|-0.01|
|SE|85787|ENSG00000258529|NA|chr11|-|111836164|111836294|111809642|111809773|111837467|111837615|146|90|151|104|138|0|1|0|2|0|157|161|141|135|116|0|0|1|1|0|279|149|1|1|1|0.98|1|0.965|1|1|1|0.987|0.986|1|-0.006|
|SE|85788|ENSG00000258529|NA|chr11|-|111837467|111837615|111809642|111809773|111844600|111844723|0|4|1|4|0|0|0|0|0|0|0|0|14|9|0|3|0|0|0|0|297|149|1|1|NA|1|1|1|NA|0|NA|1|1|NA|0.333|
|SE|85789|ENSG00000258529|NA|chr11|-|111838248|111838378|111837467|111837615|111840654|111840809|86|53|89|83|124|1|0|0|0|0|68|54|92|64|86|0|0|0|0|0|279|149|1|1|0.979|1|1|1|1|1|1|1|1|1|-0.004|
|SE|85794|ENSG00000086848|ALG9|chr11|-|111865180|111865251|111860546|111860635|111868601|111868736|55|40|34|86|53|0|0|0|0|1|31|48|88|47|15|0|0|2|0|0|220|149|1|1|1|1|1|1|0.973|1|1|0.968|1|1|0.001|
|SE|85797|ENSG00000258529|NA|chr11|-|111876480|111876639|111875697|111876104|111876807|111876844|19|17|26|12|31|3|0|1|0|0|0|14|0|3|0|0|0|6|0|0|298|149|1|1|0.76|1|0.929|1|1|NA|1|0|1|NA|0.271|
|SE|85802|ENSG00000126756|UXT|chrX|-|47652080|47652144|47651795|47651895|47657182|47657290|805|468|843|825|1747|0|0|1|0|0|576|592|415|544|793|0|0|0|0|0|213|149|1|1|1|1|0.998|1|1|1|1|1|1|1|0|
|SE|85803|ENSG00000126756|UXT|chrX|-|47657182|47657290|47652080|47652144|47657571|47657639|1191|700|1243|1198|2501|0|0|2|6|2|880|809|576|715|1165|0|1|3|1|14|257|149|1|1|1|1|0.997|0.991|0.999|1|0.998|0.991|0.998|0.98|0.004|
|SE|85807|ENSG00000163605|PPP4R2|chr3|+|73063672|73063747|73061022|73061060|73064064|73064146|115|77|98|111|182|0|0|2|0|0|31|57|113|32|19|0|0|1|0|0|224|149|1|1|1|1|0.97|1|1|1|1|0.987|1|1|-0.003|
|SE|85809|ENSG00000163605|PPP4R2|chr3|+|73064064|73064146|73063672|73063747|73064851|73064982|77|56|82|76|159|1|0|2|1|1|11|40|42|14|0|0|0|7|0|0|231|149|1|1|0.98|1|0.964|0.98|0.99|1|1|0.795|1|NA|0.034|
|SE|85811|ENSG00000118922|KLF12|chr13|-|73764937|73765000|73715367|73715525|73813151|73813287|420|358|397|352|374|0|0|0|2|0|43|265|240|53|14|0|0|0|0|0|212|149|1|1|1|1|1|0.992|1|1|1|1|1|1|-0.002|
|SE|85812|ENSG00000118922|KLF12|chr13|-|73813151|73813287|73764937|73765000|73845826|73846373|311|288|285|283|274|1|0|1|0|0|62|196|191|60|43|0|0|0|0|0|285|149|1|1|0.994|1|0.993|1|1|1|1|1|1|1|-0.003|
|SE|85813|ENSG00000118922|KLF12|chr13|-|73813151|73813287|73764937|73765000|73943980|73944070|270|240|246|235|259|0|0|3|0|0|36|167|158|42|4|0|0|0|0|0|285|149|1|1|1|1|0.977|1|1|1|1|1|1|1|-0.005|
|SE|85814|ENSG00000118922|KLF12|chr13|-|73845826|73846373|73764937|73765000|73943980|73944070|360|247|298|266|485|0|0|3|0|0|156|154|206|63|89|0|0|0|0|0|298|149|1|1|1|1|0.98|1|1|1|1|1|1|1|-0.004|
|SE|85815|ENSG00000118922|KLF12|chr13|-|73845826|73846373|73764937|73765000|73994989|73995053|3|8|7|2|7|2|0|0|0|0|0|1|7|0|0|0|0|0|0|0|298|149|1|1|0.429|1|1|1|1|NA|1|1|NA|NA|-0.114|
|SE|85816|ENSG00000118922|KLF12|chr13|-|73845826|73846373|73812872|73813287|73943980|73944070|402|298|340|322|503|2|2|1|2|3|182|183|239|81|128|0|0|0|0|0|298|149|1|1|0.99|0.987|0.994|0.988|0.988|1|1|1|1|1|-0.011|
|SE|85817|ENSG00000118922|KLF12|chr13|-|73943980|73944070|73764937|73765000|73994989|73995053|167|108|125|122|162|2|0|0|0|0|34|68|57|25|57|0|0|0|0|0|239|149|1|1|0.981|1|1|1|1|1|1|1|1|1|-0.004|
|SE|85819|ENSG00000118922|KLF12|chr13|-|73943980|73944070|73845826|73846373|74133738|74133929|230|174|183|178|358|0|0|1|0|0|100|132|129|48|57|0|0|0|0|0|239|149|1|1|1|1|0.991|1|1|1|1|1|1|1|-0.002|
|SE|85820|ENSG00000118922|KLF12|chr13|-|73994989|73995053|73845826|73846373|74133738|74133929|3|6|9|1|10|0|0|1|0|0|0|1|0|0|0|0|0|0|0|0|213|149|1|1|1|1|0.863|1|1|NA|1|NA|NA|NA|-0.027|
|SE|85821|ENSG00000118922|KLF12|chr13|-|73994989|73995053|73943980|73944070|74133738|74133929|232|132|196|154|249|1|0|0|0|1|70|79|105|24|68|0|0|0|0|0|213|149|1|1|0.994|1|1|1|0.994|1|1|1|1|1|-0.002|
|SE|85823|ENSG00000138594|TMOD3|chr15|+|51869216|51869373|51862810|51863010|51887588|51887711|618|619|553|759|899|0|1|0|0|3|397|590|657|813|382|0|5|3|0|0|298|149|1|1|1|0.997|1|1|0.993|1|0.983|0.991|1|1|0.003|
|SE|85824|ENSG00000138594|TMOD3|chr15|+|51893814|51893945|51862810|51863010|51900154|51900298|5|1|1|2|0|0|0|0|0|0|1|3|0|5|5|0|0|0|0|2|280|149|1|1|1|1|1|1|NA|1|1|NA|1|0.571|0.107|
|SE|85825|ENSG00000138594|TMOD3|chr15|+|51896418|51896526|51893814|51893945|51900154|51900298|554|401|389|561|696|5|0|1|2|0|344|426|490|540|243|1|3|0|5|0|257|149|1|1|0.985|1|0.996|0.994|1|0.995|0.988|1|0.984|1|0.002|
|SE|85826|ENSG00000138594|TMOD3|chr15|+|51911013|51911152|51908775|51908862|51916014|51916133|4|2|9|1|9|0|0|0|2|0|0|7|9|3|0|0|0|0|0|0|288|149|1|1|1|1|1|0.206|1|NA|1|1|1|NA|-0.159|
|SE|85827|ENSG00000138594|TMOD3|chr15|+|51921495|51921616|51916014|51916133|51929432|51929454|0|0|2|6|1|0|0|0|1|0|0|0|1|18|0|0|0|0|0|0|270|149|1|1|NA|NA|1|0.768|1|NA|NA|1|1|NA|-0.077|
|SE|85828|ENSG00000138594|TMOD3|chr15|+|51934249|51934354|51916014|51916133|51946989|51947295|4|5|4|1|16|1|0|1|1|0|0|0|12|9|0|0|0|5|4|0|254|149|1|1|0.701|1|0.701|0.37|1|NA|NA|0.585|0.569|NA|0.177|
|SE|85829|ENSG00000138594|TMOD3|chr15|+|51934249|51934354|51921493|51921616|51946989|51947295|3|3|3|3|14|0|1|0|0|0|0|0|11|5|0|0|0|0|0|0|254|149|1|1|1|0.638|1|1|1|NA|NA|1|1|NA|-0.072|
|SE|85830|ENSG00000147130|ZMYM3|chrX|-|71242169|71242424|71241226|71241344|71242969|71243084|27|19|51|47|88|0|0|1|0|0|25|43|19|15|45|0|0|0|0|0|298|149|1|1|1|1|0.962|1|1|1|1|1|1|1|-0.008|
|SE|85834|ENSG00000147130|ZMYM3|chrX|-|71252588|71253274|71251557|71251601|71254028|71254127|27|13|48|50|30|0|0|0|0|0|32|68|16|32|46|0|0|0|4|0|298|149|1|1|1|1|1|1|1|1|1|1|0.8|1|0.04|
|SE|85845|ENSG00000196776|CD47|chr3|-|108059451|108059544|108058336|108058429|108060744|108060852|859|610|725|838|1141|0|1|0|0|0|454|492|459|508|306|0|0|0|0|0|242|149|1|1|1|0.997|1|1|1|1|1|1|1|1|-0.001|
|SE|85846|ENSG00000196776|CD47|chr3|-|108060744|108060852|108059451|108059544|108071092|108071182|1418|965|1089|1314|1329|0|3|2|0|0|636|839|734|741|479|0|0|1|0|0|257|149|1|1|1|0.995|0.997|1|1|1|1|0.998|1|1|-0.001|
|SE|85847|ENSG00000196776|CD47|chr3|-|108071092|108071182|108060744|108060852|108079990|108080344|1483|1085|1153|1361|1360|0|0|0|0|3|665|992|793|805|544|0|0|0|0|0|239|149|1|1|1|1|1|1|0.996|1|1|1|1|1|-0.001|
|SE|85848|ENSG00000196776|CD47|chr3|-|108079990|108080344|108071092|108071182|108090862|108091025|993|710|782|936|929|3|0|0|0|0|515|743|527|549|387|0|0|0|0|0|298|149|1|1|0.994|1|1|1|1|1|1|1|1|1|-0.001|
|SE|85850|ENSG00000185973|TMLHE|chrX|-|155506897|155507134|155492356|155492495|155511672|155511792|137|78|96|91|145|3|0|0|0|0|77|81|49|43|75|0|0|0|0|0|298|149|1|1|0.958|1|1|1|1|1|1|1|1|1|-0.008|
|SE|85851|ENSG00000185973|TMLHE|chrX|-|155511672|155511792|155506897|155507134|155513985|155514265|126|90|88|81|161|0|0|0|0|0|112|104|61|46|97|0|0|1|0|0|269|149|1|1|1|1|1|1|1|1|1|0.971|1|1|0.006|
|SE|85852|ENSG00000185973|TMLHE|chrX|-|155513985|155514265|155506897|155507134|155524455|155524632|51|44|41|32|57|1|0|0|0|0|22|46|32|43|49|0|0|0|0|0|298|149|1|1|0.962|1|1|1|1|1|1|1|1|1|-0.008|
|SE|85855|ENSG00000223959|AFG3L1P|chr16|+|89977604|89977802|89972616|89972765|89978809|89978878|12|14|12|7|9|0|2|1|0|0|0|10|10|41|10|0|0|0|0|5|298|149|1|1|1|0.778|0.857|1|1|NA|1|1|1|0.5|0.052|
|SE|85856|ENSG00000223959|AFG3L1P|chr16|+|89977638|89977802|89972585|89972765|89978809|89978878|16|17|17|16|9|0|2|1|0|0|0|16|13|50|16|0|0|0|0|5|298|149|1|1|1|0.81|0.895|1|1|NA|1|1|1|0.615|0.037|
|SE|85857|ENSG00000223959|AFG3L1P|chr16|+|89977638|89977802|89972585|89972765|89980241|89980336|5|3|5|15|0|1|0|0|0|0|0|6|6|5|11|6|0|0|0|0|298|149|1|1|0.714|1|1|1|NA|NA|1|1|1|1|-0.072|
|SE|85858|ENSG00000223959|AFG3L1P|chr16|+|89977677|89977802|89972592|89972765|89978809|89978878|12|15|11|10|9|0|2|1|0|0|0|10|12|36|15|0|0|0|0|5|274|149|1|1|1|0.803|0.857|1|1|NA|1|1|1|0.62|0.027|
|SE|85859|ENSG00000223959|AFG3L1P|chr16|+|89978809|89978878|89972585|89972765|89980241|89980336|12|10|11|17|8|1|0|0|0|0|44|16|18|17|28|0|0|0|0|0|218|149|1|1|0.891|1|1|1|1|1|1|1|1|1|-0.022|
|SE|85860|ENSG00000223959|AFG3L1P|chr16|+|89978809|89978878|89977604|89977802|89980241|89980336|26|23|22|24|17|1|0|0|6|0|44|26|28|56|33|0|0|2|2|0|218|149|1|1|0.947|1|1|0.732|1|1|1|0.905|0.95|1|-0.035|
|SE|85861|ENSG00000223959|AFG3L1P|chr16|+|89985445|89985570|89984530|89984599|89988681|89988951|0|0|1|2|0|0|0|0|0|0|0|0|1|1|0|0|2|0|0|0|274|149|1|1|NA|NA|1|1|NA|NA|0|1|1|NA|0.333|
|SE|85866|ENSG00000223959|AFG3L1P|chr16|+|89990865|89991003|89984530|89984599|89992720|89992828|9|7|12|11|9|0|0|0|0|1|22|13|14|12|33|0|0|0|0|0|287|149|1|1|1|1|1|1|0.824|1|1|1|1|1|-0.035|
|SE|85870|ENSG00000041988|THAP3|chr1|+|6629642|6629724|6628498|6628691|6630287|6630353|0|0|0|4|0|20|4|8|8|3|0|0|0|0|0|15|5|3|0|11|231|149|1|1|0|0|0|0.244|0|0|0|0|NA|0|0.049|
|SE|85897|ENSG00000013563|DNASE1L1|chrX|-|154409417|154409675|154409111|154409196|154411890|154412090|6|5|9|3|3|0|0|2|0|0|16|8|6|12|0|0|0|0|0|0|298|149|1|1|1|1|0.692|1|1|1|1|1|1|NA|-0.062|
|SE|85898|ENSG00000013563|DNASE1L1|chrX|-|154409417|154409813|154409111|154409196|154411890|154412089|3|4|3|1|3|0|0|2|0|0|10|1|6|12|0|0|0|0|0|0|298|149|1|1|1|1|0.429|1|1|1|1|1|1|NA|-0.114|
|SE|85900|ENSG00000126903|SLC10A3|chrX|-|154489941|154490027|154487310|154489082|154490593|154490628|0|2|0|0|0|0|0|0|0|0|0|2|0|0|0|8|0|0|0|0|235|149|1|1|NA|1|NA|NA|NA|0|1|NA|NA|NA|0.5|
|SE|85901|ENSG00000126903|SLC10A3|chrX|-|154490306|154490358|154487342|154489082|154490593|154490614|8|1|7|6|3|0|0|0|0|0|12|10|0|2|25|8|0|0|0|0|201|149|1|1|1|1|1|1|1|0.527|1|NA|1|1|0.118|
|SE|85902|ENSG00000076924|XAB2|chr19|-|7626135|7626270|7625879|7626044|7627242|7627440|124|32|90|85|34|0|0|0|0|0|101|143|52|109|127|0|0|2|0|0|284|149|1|1|1|1|1|1|1|1|1|0.932|1|1|0.014|
|SE|85921|ENSG00000106976|DNM1|chr9|+|128247923|128247935|128247374|128247486|128248582|128248753|2|2|4|5|0|0|0|0|2|0|30|11|7|7|61|3|0|1|0|0|161|149|1|1|1|1|1|0.698|NA|0.902|1|0.866|1|1|-0.029|
|SE|85938|ENSG00000005302|MSL3|chrX|+|11759792|11759875|11759603|11759712|11760402|11760498|171|195|133|203|236|0|0|0|0|0|302|231|213|224|113|0|3|0|0|0|232|149|1|1|1|1|1|1|1|1|0.98|1|1|1|0.004|
|SE|85939|ENSG00000005302|MSL3|chrX|+|11759792|11759875|11759603|11759712|11760836|11760937|2|4|0|1|3|0|0|0|0|0|0|0|2|1|0|0|0|1|0|0|232|149|1|1|1|1|NA|1|1|NA|NA|0.562|1|NA|0.219|
|SE|85940|ENSG00000005302|MSL3|chrX|+|11760402|11760498|11759401|11759875|11760836|11760937|375|374|359|417|538|2|4|0|0|0|552|470|383|455|180|0|0|0|1|0|245|149|1|1|0.991|0.983|1|1|1|1|1|1|0.996|1|-0.004|

| | | | | | | | | | | | | | | | | | | | | | | | | | | | | | | | | | | | | | | | | | | | | | | |
|---|---|---|---|---|---|---|---|---|---|---|---|---|---|---|---|---|---|---|---|---|---|---|---|---|---|---|---|---|---|---|---|---|---|---|---|---|---|---|---|---|---|---|---|---|---|---|
| SE | 85941 | ENSG00000005302 | MSL3 | chrX | + | 11760402 | 11760498 | 11759603 | 11759712 | 11760836 | 11760937 | 135 | 112 | 166 | 152 | 246 | 0 | 0 | 0 | 0 | 0 | 116 | 199 | 120 | 158 | 44 | 0 | 0 | 1 | 0 | 0 | 245 | 149 | 1 | 1 | 1 | 1 | 1 | 1 | 1 | 1 | 1 | 0.986 | 1 | 1 | 0.003 |
| SE | 85943 | ENSG00000005302 | MSL3 | chrX | + | 11762836 | 11762997 | 11760836 | 11762252 | 11763779 | 11763810 | 547 | 494 | 587 | 609 | 1302 | 0 | 1 | 0 | 0 | 0 | 609 | 548 | 398 | 507 | 91 | 0 | 7 | 0 | 0 | 0 | 298 | 149 | 1 | 1 | 1 | 0.996 | 1 | 1 | 1 | 1 | 0.975 | | 1 | 1 | 0.004 |
| SE | 85944 | ENSG00000005302 | MSL3 | chrX | + | 11766960 | 11767046 | 11765537 | 11765729 | 11767144 | 11767286 | 0 | 1 | 0 | 0 | 0 | 0 | 2 | 0 | 0 | 0 | 0 | 0 | 0 | 0 | 0 | 0 | 2 | 4 | 1 | 0 | 235 | 149 | 1 | 1 | NA | 0.241 | NA | NA | NA | NA | 0 | 0 | 0 | NA | 0.241 |
| SE | 85945 | ENSG00000005302 | MSL3 | chrX | + | 11766960 | 11767052 | 11765537 | 11765729 | 11767144 | 11767286 | 1 | 5 | 2 | 0 | 0 | 0 | 2 | 0 | 0 | 0 | 0 | 4 | 0 | 0 | 0 | 0 | 2 | 4 | 1 | 0 | 241 | 149 | 1 | 1 | 1 | 0.607 | 1 | NA | NA | NA | 0.553 | 0 | 0 | NA | 0.685 |
| SE | 85947 | ENSG00000005302 | MSL3 | chrX | + | 11766960 | 11767169 | 11765537 | 11765729 | 11768621 | 11768682 | 0 | 1 | 0 | 0 | 0 | 0 | 0 | 0 | 0 | 0 | 0 | 0 | 0 | 0 | 0 | 0 | 0 | 0 | 1 | 0 | 298 | 149 | 1 | 1 | NA | 1 | NA | NA | NA | NA | NA | NA | 0 | NA | 1 |
| SE | 85953 | ENSG00000005302 | MSL3 | chrX | + | 11772155 | 11772255 | 11763779 | 11768682 | 11772620 | 11772705 | 283 | 167 | 244 | 236 | 308 | 0 | 0 | 0 | 0 | 4 | 275 | 232 | 200 | 229 | 124 | 0 | 0 | 0 | 0 | 0 | 249 | 149 | 1 | 1 | 1 | 1 | 1 | 1 | 0.979 | 1 | 1 | 1 | 1 | 1 | -0.004 |
| SE | 85958 | ENSG00000111203 | ITFG2 | chr12 | + | 2820723 | 2820872 | 2819657 | 2820225 | 2821691 | 2821792 | 37 | 13 | 50 | 40 | 28 | 0 | 0 | 1 | 0 | 0 | 84 | 45 | 31 | 45 | 54 | 0 | 0 | 0 | 0 | 0 | 298 | 149 | 1 | 1 | 1 | 1 | 0.962 | 1 | 1 | 1 | 1 | 1 | 1 | 1 | -0.008 |
| SE | 85959 | ENSG00000111203 | ITFG2 | chr12 | + | 2821261 | 2821359 | 2820723 | 2820872 | 2821542 | 2821596 | 89 | 70 | 101 | 91 | 126 | 0 | 0 | 0 | 0 | 0 | 57 | 130 | 58 | 86 | 196 | 0 | 0 | 2 | 0 | 0 | 247 | 149 | 1 | 1 | 1 | 1 | 1 | 1 | 1 | 1 | 1 | 0.946 | 1 | 1 | 0.011 |
| SE | 85963 | ENSG00000111203 | ITFG2 | chr12 | + | 2821691 | 2821792 | 2821261 | 2821359 | 2822793 | 2822911 | 53 | 40 | 71 | 44 | 52 | 0 | 0 | 1 | 0 | 0 | 84 | 75 | 34 | 53 | 68 | 0 | 0 | 2 | 0 | 0 | 250 | 149 | 1 | 1 | 1 | 1 | 0.977 | 1 | 1 | 1 | 1 | 0.91 | 1 | 1 | 0.013 |
| SE | 85965 | ENSG00000111203 | ITFG2 | chr12 | + | 2824089 | 2824252 | 2823769 | 2823943 | 2836744 | 2836956 | 87 | 40 | 83 | 53 | 132 | 0 | 0 | 0 | 0 | 0 | 92 | 57 | 55 | 44 | 113 | 0 | 0 | 0 | 4 | 0 | 298 | 149 | 1 | 1 | 1 | 1 | 1 | 1 | 1 | 1 | 1 | 1 | 0.846 | 1 | 0.031 |
| SE | 85966 | ENSG00000111203 | ITFG2 | chr12 | + | 2830216 | 2830885 | 2824089 | 2824252 | 2836744 | 2836956 | 15 | 9 | 25 | 13 | 10 | 0 | 2 | 0 | 0 | 0 | 38 | 25 | 10 | 22 | 28 | 0 | 2 | 0 | 0 | 0 | 298 | 149 | 1 | 1 | 1 | 0.692 | 1 | 1 | 1 | 1 | 0.862 | 1 | 1 | 1 | -0.034 |
| SE | 85968 | ENSG00000111203 | ITFG2 | chr12 | + | 2830833 | 2830885 | 2824089 | 2824252 | 2836744 | 2836956 | 11 | 6 | 23 | 1 | 9 | 0 | 2 | 0 | 0 | 0 | 28 | 18 | 9 | 11 | 20 | 0 | 2 | 0 | 0 | 0 | 201 | 149 | 1 | 1 | 1 | 0.69 | 1 | 1 | 1 | 1 | 0.87 | 1 | 1 | 1 | -0.036 |
| SE | 85970 | ENSG00000111203 | ITFG2 | chr12 | + | 2836744 | 2836956 | 2830216 | 2830885 | 2858011 | 2858331 | 17 | 14 | 33 | 20 | 21 | 0 | 1 | 0 | 0 | 0 | 59 | 27 | 22 | 30 | 31 | 0 | 0 | 0 | 0 | 0 | 298 | 149 | 1 | 1 | 1 | 0.875 | 1 | 1 | 1 | 1 | 1 | 1 | 1 | 1 | -0.025 |
| SE | 85972 | ENSG00000111203 | ITFG2 | chr12 | + | 2858011 | 2858331 | 2840841 | 2840995 | 2859533 | 2859790 | 25 | 10 | 27 | 15 | 13 | 0 | 0 | 0 | 0 | 0 | 21 | 19 | 14 | 22 | 16 | 0 | 0 | 0 | 2 | 0 | 298 | 149 | 1 | 1 | 1 | 1 | 1 | 1 | 1 | 1 | 1 | 1 | 0.846 | 1 | 0.031 |
| SE | 85973 | ENSG00000144161 | ZC3H8 | chr2 | - | 112240367 | 112240492 | 112238314 | 112238528 | 112250190 | 112250272 | 2 | 0 | 2 | 0 | 0 | 23 | 19 | 21 | 36 | 48 | 0 | 1 | 1 | 1 | 2 | 0 | 16 | 20 | 8 | 7 | 274 | 149 | 1 | 1 | 0.045 | 0 | 0.049 | 0 | 0 | NA | 0.033 | 0.026 | 0.064 | 0.134 | -0.045 |
| SE | 85979 | ENSG00000165609 | NUDT5 | chr10 | - | 12170716 | 12170770 | 12165329 | 12167811 | 12170899 | 12170908 | 259 | 166 | 260 | 251 | 282 | 1 | 0 | 0 | 1 | 0 | 226 | 201 | 199 | 228 | 168 | 0 | 0 | 0 | 2 | 0 | 203 | 149 | 1 | 1 | 0.995 | 1 | 1 | 0.995 | 1 | 1 | 1 | 1 | 0.988 | 1 | 0 |
| SE | 85980 | ENSG00000165609 | NUDT5 | chr10 | - | 12170899 | 12170908 | 12170716 | 12170770 | 12172764 | 12172866 | 129 | 74 | 131 | 131 | 168 | 1 | 0 | 0 | 0 | 1 | 116 | 98 | 94 | 110 | 84 | 0 | 1 | 1 | 2 | 0 | 158 | 149 | 1 | 1 | 0.992 | 1 | 1 | 1 | 0.994 | 1 | 0.989 | 0.989 | 0.981 | 1 | 0.005 |
| SE | 85982 | ENSG00000165609 | NUDT5 | chr10 | - | 12180394 | 12180489 | 12179082 | 12179132 | 12184888 | 12184956 | 1 | 3 | 0 | 4 | 0 | 216 | 166 | 166 | 160 | 264 | 0 | 2 | 1 | 0 | 0 | 135 | 106 | 88 | 136 | 134 | 244 | 149 | 1 | 1 | 0.003 | 0.011 | 0 | 0.015 | 0 | 0 | 0.011 | 0.007 | 0 | 0 | 0.002 |
| SE | 85986 | ENSG00000149600 | COMMD7 | chr20 | - | 32742475 | 32742565 | 32728088 | 32728142 | 32743307 | 32743445 | 0 | 0 | 2 | 0 | 0 | 14 | 10 | 20 | 15 | 7 | 0 | 0 | 0 | 0 | 0 | 24 | 3 | 15 | 4 | 21 | 239 | 149 | 1 | 1 | 0 | 0 | 0.059 | 0 | 0 | 0 | 0 | 0 | 0 | 0 | 0.012 |
| SE | 85989 | ENSG00000204241 | LINC02731 | chr11 | + | 134044107 | 134044802 | 134039761 | 134040080 | 134046760 | 134046945 | 0 | 0 | 3 | 0 | 1 | 0 | 0 | 0 | 0 | 2 | 0 | 0 | 0 | 0 | 0 | 0 | 3 | 0 | 0 | 0 | 298 | 149 | 1 | 1 | NA | NA | 1 | NA | 0.2 | NA | 0 | NA | NA | NA | 0.6 |
| SE | 85990 | ENSG00000244754 | N4BP2L2 | chr13 | - | 32446358 | 32446480 | 32442387 | 32444126 | 32477886 | 32478048 | 2 | 0 | 2 | 6 | 5 | 2 | 1 | 0 | 2 | 0 | 0 | 4 | 0 | 0 | 0 | 0 | 0 | 0 | 0 | 0 | 271 | 149 | 1 | 1 | 0.355 | 0 | 1 | 0.623 | 1 | NA | 1 | NA | NA | NA | -0.404 |
| SE | 85991 | ENSG00000244754 | N4BP2L2 | chr13 | - | 32446358 | 32446480 | 32442387 | 32444126 | 32480589 | 32480638 | 3 | 5 | 4 | 9 | 6 | 21 | 17 | 8 | 15 | 22 | 0 | 4 | 0 | 0 | 0 | 0 | 5 | 6 | 8 | 0 | 271 | 149 | 1 | 1 | 0.073 | 0.139 | 0.216 | 0.248 | 0.13 | NA | 0.305 | 0 | 0 | NA | 0.06 |
| SE | 85992 | ENSG00000244754 | N4BP2L2 | chr13 | - | 32446358 | 32446480 | 32442387 | 32444126 | 32517856 | 32518003 | 3 | 0 | 2 | 11 | 5 | 1 | 3 | 5 | 0 | 14 | 0 | 4 | 3 | 0 | 0 | 0 | 0 | 1 | 0 | 0 | 271 | 149 | 1 | 1 | 0.623 | 0 | 0.18 | 1 | 0.164 | NA | 1 | 0.623 | NA | NA | -0.418 |
| SE | 85993 | ENSG00000244754 | N4BP2L2 | chr13 | - | 32477886 | 32478048 | 32442387 | 32444126 | 32478227 | 32478275 | 2 | 1 | 0 | 2 | 0 | 0 | 2 | 0 | 0 | 2 | 0 | 0 | 1 | 0 | 0 | 0 | 0 | 0 | 0 | 0 | 298 | 149 | 1 | 1 | 1 | 0.2 | NA | 1 | 0 | NA | NA | 1 | NA | NA | -0.45 |
| SE | 85994 | ENSG00000244754 | N4BP2L2 | chr13 | - | 32477886 | 32478048 | 32442387 | 32444126 | 32480589 | 32480637 | 4 | 3 | 2 | 12 | 6 | 21 | 17 | 8 | 15 | 22 | 0 | 0 | 12 | 2 | 4 | 0 | 5 | 6 | 8 | 0 | 298 | 149 | 1 | 1 | 0.087 | 0.081 | 0.111 | 0.286 | 0.12 | NA | 0 | 0.5 | 0.111 | 1 | -0.266 |
| SE | 85995 | ENSG00000244754 | N4BP2L2 | chr13 | - | 32477886 | 32478048 | 32446358 | 32446480 | 32480589 | 32480637 | 2 | 2 | 2 | 10 | 6 | 1 | 5 | 2 | 3 | 1 | 0 | 0 | 12 | 2 | 4 | 0 | 0 | 0 | 0 | 0 | 298 | 149 | 1 | 1 | 0.5 | 0.167 | 0.333 | 0.625 | 0.75 | NA | NA | 1 | 1 | 1 | -0.525 |
| SE | 85996 | ENSG00000244754 | N4BP2L2 | chr13 | - | 32478227 | 32478275 | 32442387 | 32444126 | 32480589 | 32480638 | 0 | 2 | 0 | 0 | 2 | 21 | 17 | 8 | 15 | 22 | 0 | 0 | 0 | 0 | 0 | 0 | 5 | 6 | 8 | 0 | 197 | 149 | 1 | 1 | 0 | 0.082 | 0 | 0 | 0.064 | NA | 0 | 0 | 0 | NA | 0.029 |
| SE | 85997 | ENSG00000244754 | N4BP2L2 | chr13 | - | 32478227 | 32478275 | 32477886 | 32478048 | 32480589 | 32480638 | 0 | 0 | 0 | 0 | 0 | 2 | 2 | 2 | 10 | 6 | 0 | 0 | 1 | 0 | 0 | 0 | 0 | 12 | 2 | 4 | 197 | 149 | 1 | 1 | 0 | 0 | 0 | 0 | 0 | NA | NA | 0.059 | 0 | 0 | -0.02 |
| SE | 85998 | ENSG00000244754 | N4BP2L2 | chr13 | - | 32480589 | 32480637 | 32443064 | 32444126 | 32517856 | 32517920 | 23 | 25 | 15 | 25 | 32 | 1 | 3 | 5 | 0 | 14 | 9 | 10 | 9 | 4 | 0 | 0 | 0 | 1 | 0 | 0 | 197 | 149 | 1 | 1 | 0.946 | 0.863 | 0.694 | 1 | 0.634 | 1 | 1 | 0.872 | 1 | NA | -0.141 |
| SE | 85999 | ENSG00000244754 | N4BP2L2 | chr13 | - | 32480589 | 32480637 | 32443064 | 32444126 | 32535768 | 32537027 | 14 | 18 | 4 | 8 | 6 | 1 | 0 | 0 | 0 | 0 | 0 | 0 | 0 | 3 | 0 | 0 | 0 | 0 | 0 | 0 | 197 | 149 | 1 | 1 | 0.914 | 1 | 1 | 1 | 1 | NA | NA | NA | 1 | NA | -0.017 |
| SE | 86001 | ENSG00000244754 | N4BP2L2 | chr13 | - | 32521372 | 32521449 | 32510291 | 32518003 | 32527407 | 32527532 | 634 | 538 | 448 | 556 | 1119 | 4 | 2 | 0 | 3 | 0 | 332 | 279 | 271 | 207 | 341 | 0 | 0 | 0 | 6 | 0 | 226 | 149 | 1 | 1 | 0.991 | 0.994 | 1 | 0.992 | 1 | 1 | 1 | 1 | 0.958 | 1 | 0.004 |
| SE | 86002 | ENSG00000244754 | N4BP2L2 | chr13 | - | 32521372 | 32521449 | 32510291 | 32518003 | 32535768 | 32537027 | 635 | 538 | 450 | 554 | 1121 | 3 | 2 | 5 | 2 | 3 | 332 | 279 | 271 | 207 | 341 | 0 | 0 | 0 | 0 | 0 | 226 | 149 | 1 | 1 | 0.993 | 0.994 | 0.983 | 0.995 | 0.996 | 1 | 1 | 1 | 1 | 1 | -0.008 |
| SE | 86004 | ENSG00000244754 | N4BP2L2 | chr13 | - | 32522181 | 32522270 | 32517874 | 32518003 | 32527457 | 32527532 | 19 | 8 | 10 | 14 | 3 | 1 | 0 | 0 | 1 | 0 | 37 | 11 | 10 | 2 | 19 | 0 | 0 | 0 | 0 | 0 | 238 | 149 | 1 | 1 | 0.922 | 1 | 1 | 0.898 | 1 | 1 | 1 | 1 | 1 | 1 | -0.036 |
| SE | 86005 | ENSG00000244754 | N4BP2L2 | chr13 | - | 32522181 | 32522270 | 32517874 | 32518003 | 32535768 | 32537027 | 10 | 4 | 7 | 7 | 22 | 3 | 2 | 5 | 2 | 3 | 0 | 3 | 0 | 5 | 0 | 0 | 0 | 0 | 0 | 0 | 238 | 149 | 1 | 1 | 0.676 | 0.556 | 0.467 | 0.687 | 0.821 | NA | 1 | NA | 1 | NA | -0.359 |
| SE | 86006 | ENSG00000244754 | N4BP2L2 | chr13 | - | 32522181 | 32522270 | 32521372 | 32521449 | 32527407 | 32527532 | 882 | 621 | 672 | 875 | 1740 | 0 | 0 | 0 | 3 | 1 | 440 | 483 | 400 | 277 | 657 | 0 | 0 | 0 | 0 | 0 | 238 | 149 | 1 | 1 | 1 | 1 | 1 | 0.995 | 0.999 | 1 | 1 | 1 | 1 | 1 | -0.001 |
| SE | 86007 | ENSG00000244754 | N4BP2L2 | chr13 | - | 32522181 | 32522270 | 32521372 | 32521449 | 32535768 | 32537027 | 502 | 362 | 391 | 512 | 861 | 1 | 0 | 2 | 1 | 3 | 324 | 290 | 241 | 188 | 378 | 0 | 0 | 0 | 0 | 0 | 238 | 149 | 1 | 1 | 0.997 | 1 | 0.992 | 0.997 | 0.994 | 1 | 1 | 1 | 1 | 1 | -0.004 |
| SE | 86009 | ENSG00000244754 | N4BP2L2 | chr13 | - | 32527407 | 32527532 | 32521372 | 32521449 | 32535768 | 32535869 | 235 | 136 | 194 | 216 | 423 | 1 | 0 | 2 | 1 | 3 | 9 | 54 | 73 | 25 | 4 | 0 | 0 | 0 | 0 | 0 | 274 | 149 | 1 | 1 | 0.992 | 1 | 0.981 | 0.992 | 0.987 | 1 | 1 | 1 | 1 | 1 | -0.01 |
| SE | 86011 | ENSG00000244754 | N4BP2L2 | chr13 | - | 32527407 | 32527532 | 32522181 | 32522270 | 32536932 | 32537027 | 300 | 206 | 214 | 287 | 777 | 0 | 1 | 2 | 0 | 0 | 106 | 147 | 116 | 83 | 251 | 0 | 0 | 0 | 0 | 0 | 274 | 149 | 1 | 1 | 1 | 0.991 | 0.983 | 1 | 1 | 1 | 1 | 1 | 1 | 1 | -0.005 |
| SE | 86012 | ENSG00000244754 | N4BP2L2 | chr13 | - | 32527407 | 32527532 | 32522181 | 32522270 | 32537005 | 32537027 | 305 | 210 | 216 | 287 | 777 | 0 | 0 | 0 | 0 | 3 | 106 | 147 | 116 | 83 | 251 | 0 | 0 | 0 | 0 | 0 | 274 | 149 | 1 | 1 | 1 | 1 | 1 | 1 | 0.993 | 1 | 1 | 1 | 1 | 1 | -0.001 |
| SE | 86015 | ENSG00000244754 | N4BP2L2 | chr13 | - | 32527457 | 32527532 | 32522181 | 32522270 | 32537005 | 32537027 | 44 | 20 | 21 | 33 | 87 | 0 | 0 | 0 | 0 | 3 | 31 | 32 | 25 | 19 | 72 | 0 | 0 | 0 | 0 | 0 | 224 | 149 | 1 | 1 | 1 | 1 | 1 | 1 | 0.951 | 1 | 1 | 1 | 1 | 1 | -0.01 |
| SE | 86017 | ENSG00000244754 | N4BP2L2 | chr13 | - | 32535768 | 32537027 | 32480589 | 32480637 | 32538682 | 32538802 | 168 | 152 | 146 | 153 | 142 | 0 | 0 | 0 | 0 | 2 | 18 | 124 | 100 | 20 | 23 | 0 | 0 | 0 | 0 | 0 | 298 | 149 | 1 | 1 | 1 | 1 | 1 | 1 | 0.973 | 1 | 1 | 1 | 1 | 1 | -0.005 |
| SE | 86018 | ENSG00000244754 | N4BP2L2 | chr13 | - | 32535768 | 32537027 | 32480589 | 32480637 | 32538777 | 32538825 | 20 | 18 | 12 | 25 | 23 | 1 | 0 | 0 | 0 | 0 | 0 | 21 | 6 | 0 | 0 | 0 | 0 | 0 | 0 | 0 | 298 | 149 | 1 | 1 | 0.909 | 1 | 1 | 1 | 1 | NA | 1 | 1 | NA | NA | -0.018 |
| SE | 86021 | ENSG00000244754 | N4BP2L2 | chr13 | - | 32535768 | 32537027 | 32526818 | 32527532 | 32538777 | 32538825 | 353 | 218 | 294 | 311 | 593 | 0 | 0 | 0 | 0 | 0 | 14 | 115 | 107 | 32 | 4 | 0 | 0 | 0 | 1 | 0 | 298 | 149 | 1 | 1 | 1 | 1 | 1 | 1 | 1 | 1 | 1 | 1 | 0.941 | 1 | 0.012 |
| SE | 86022 | ENSG00000244754 | N4BP2L2 | chr13 | - | 32536932 | 32537027 | 32527407 | 32527532 | 32538617 | 32538684 | 160 | 161 | 137 | 152 | 142 | 11 | 5 | 5 | 4 | 3 | 29 | 76 | 65 | 53 | 6 | 0 | 1 | 3 | 0 | 0 | 244 | 149 | 1 | 1 | 0.899 | 0.952 | 0.944 | 0.959 | 0.967 | 1 | 0.979 | 0.93 | 1 | 1 | -0.038 |
| SE | 86025 | ENSG00000244754 | N4BP2L2 | chr13 | - | 32537005 | 32537027 | 32527407 | 32527532 | 32538777 | 32538813 | 11 | 5 | 8 | 0 | 0 | 0 | 0 | 0 | 0 | 0 | 0 | 0 | 2 | 1 | 6 | 0 | 0 | 0 | 1 | 0 | 171 | 149 | 1 | 1 | 1 | 1 | 1 | NA | NA | NA | NA | 1 | 0.466 | 1 | 0.178 |
| SE | 86026 | ENSG00000244754 | N4BP2L2 | chr13 | - | 32538617 | 32538684 | 32527457 | 32527532 | 32538777 | 32538792 | 10 | 2 | 3 | 4 | 2 | 0 | 0 | 0 | 0 | 0 | 0 | 1 | 3 | 0 | 0 | 0 | 0 | 0 | 1 | 0 | 216 | 149 | 1 | 1 | 1 | 1 | 1 | 1 | 1 | NA | 1 | 1 | 0 | NA | 0.333 |
| SE | 86029 | ENSG00000156313 | RPGR | chrX | - | 38275088 | 38275146 | 38273385 | 38273477 | 38276586 | 38276772 | 77 | 61 | 42 | 47 | 184 | 0 | 0 | 0 | 1 | 0 | 3 | 23 | 18 | 7 | 16 | 0 | 0 | 0 | 0 | 0 | 207 | 149 | 1 | 1 | 1 | 1 | 1 | 0.971 | 1 | 1 | 1 | 1 | 1 | 1 | -0.006 |
| SE | 86031 | ENSG00000156313 | RPGR | chrX | - | 38287093 | 38287245 | 38276586 | 38276772 | 38287860 | 38288041 | 38 | 26 | 27 | 46 | 44 | 0 | 3 | 0 | 0 | 2 | 0 | 24 | 17 | 17 | 0 | 0 | 0 | 0 | 0 | 0 | 298 | 149 | 1 | 1 | 1 | 0.812 | 1 | 1 | 0.917 | NA | 1 | 1 | 1 | NA | -0.054 |
| SE | 86033 | ENSG00000156313 | RPGR | chrX | - | 38287093 | 38287245 | 38276586 | 38276772 | 38290958 | 38291024 | 19 | 3 | 11 | 17 | 20 | 5 | 4 | 4 | 2 | 4 | 0 | 20 | 5 | 0 | 0 | 0 | 0 | 1 | 0 | 4 | 298 | 149 | 1 | 1 | 0.655 | 0.273 | 0.579 | 0.81 | 0.714 | NA | 1 | 0.714 | NA | 0 | 0.035 |
| SE | 86039 | ENSG00000156313 | RPGR | chrX | - | 38291392 | 38291484 | 38290958 | 38291024 | 38297283 | 38297452 | 38 | 16 | 30 | 26 | 48 | 2 | 2 | 0 | 0 | 0 | 5 | 5 | 23 | 11 | 4 | 0 | 0 | 2 | 0 | 0 | 241 | 149 | 1 | 1 | 0.922 | 0.832 | 1 | 1 | 1 | 1 | 1 | 0.877 | 1 | 1 | -0.025 |
| SE | 86040 | ENSG00000156313 | RPGR | chrX | - | 38295984 | 38296144 | 38291392 | 38291484 | 38297283 | 38297452 | 0 | 0 | 0 | 1 | 0 | 26 | 6 | 18 | 14 | 36 | 0 | 0 | 0 | 0 | 0 | 0 | 1 | 11 | 6 | 0 | 298 | 149 | 1 | 1 | 0 | 0 | 0 | 0.034 | 0 | NA | 0 | 0 | 0 | NA | 0.007 |
| SE | 86041 | ENSG00000156313 | RPGR | chrX | - | 38298955 | 38299141 | 38297283 | 38297452 | 38301246 | 38301371 | 15 | 15 | 12 | 31 | 41 | 0 | 1 | 0 | 2 | 0 | 14 | 6 | 6 | 0 | 6 | 0 | 0 | 0 | 0 | 0 | 298 | 149 | 1 | 1 | 1 | 0.882 | 1 | 0.886 | 1 | 1 | 1 | 1 | NA | 1 | -0.046 |
| SE | 86044 | ENSG00000156313 | RPGR | chrX | - | 38310614 | 38310773 | 38304634 | 38304790 | 38317315 | 38317465 | 42 | 27 | 33 | 35 | 67 | 0 | 0 | 0 | 1 | 0 | 14 | 2 | 21 | 8 | 26 | 0 | 0 | 0 | 0 | 0 | 298 | 149 | 1 | 1 | 1 | 1 | 1 | 0.946 | 1 | 1 | 1 | 1 | 1 | 1 | -0.011 |
| SE | 86046 | ENSG00000156313 | RPGR | chrX | - | 38318828 | 38318987 | 38317315 | 38317465 | 38322852 | 38322945 | 17 | 34 | 40 | 30 | 20 | 1 | 0 | 0 | 0 | 0 | 21 | 11 | 37 | 7 | 7 | 0 | 0 | 0 | 0 | 0 | 298 | 149 | 1 | 1 | 0.895 | 1 | 1 | 1 | 1 | 1 | 1 | 1 | 1 | 1 | -0.021 |
| SE | 86047 | ENSG00000156313 | RPGR | chrX | - | 38321026 | 38321089 | 38318828 | 38318987 | 38322852 | 38322945 | 36 | 37 | 38 | 36 | 16 | 0 | 0 | 0 | 4 | 0 | 31 | 13 | 22 | 9 | 20 | 0 | 3 | 1 | 0 | 0 | 212 | 149 | 1 | 1 | 1 | 1 | 1 | 0.863 | 1 | 1 | 0.753 | 0.939 | 1 | 1 | 0.034 |
| SE | 86053 | ENSG00000085415 | SEH1L | chr18 | + | 12978751 | 12978892 | 12971152 | 12971251 | 12982517 | 12982675 | 143 | 82 | 143 | 157 | 161 | 1 | 1 | 0 | 0 | 0 | 104 | 69 | 65 | 91 | 17 | 0 | 0 | 0 | 0 | 0 | 290 | 149 | 1 | 1 | 0.987 | 0.977 | 1 | | 1 | 1 | 1 | 1 | 1 | 1 | -0.007 |
| SE | 86054 | ENSG00000137310 | TCF19 | chr6 | + | 31166279 | 31166365 | 31161448 | 31162005 | 31166997 | 31167159 | 0 | 0 | 0 | 0 | 0 | 0 | 0 | 0 | 1 | 0 | 2 | 0 | 0 | 0 | 0 | 0 | 0 | 0 | 0 | 0 | 235 | 149 | 1 | 1 | NA | NA | NA | 0 | NA | 1 | NA | NA | NA | NA | -1 |
| SE | 86055 | ENSG00000164031 | DNAJB14 | chr4 | - | 99905596 | 99905706 | 99903725 | 99903898 | 99906037 | 99906153 | 226 | 195 | 181 | 240 | 255 | 0 | 2 | 0 | 0 | 0 | 34 | 71 | 29 | 20 | 9 | 0 | 0 | 0 | 0 | 0 | 259 | 149 | 1 | 1 | 1 | 0.982 | 1 | | 1 | 1 | 1 | 1 | 1 | 1 | -0.004 |
| SE | 86056 | ENSG00000164031 | DNAJB14 | chr4 | - | 99906037 | 99906153 | 99905596 | 99905706 | 99906516 | 99906597 | 9 | 5 | 15 | 18 | 6 | 218 | 206 | 155 | 219 | 328 | 0 | 4 | 1 | 0 | 0 | 31 | 44 | 35 | 16 | 0 | 265 | 149 | 1 | 1 | 0.023 | 0.013 | 0.052 | 0.044 | 0.01 | 0 | 0.049 | 0.016 | 0 | NA | 0.012 |
| SE | 86057 | ENSG00000164031 | DNAJB14 | chr4 | - | 99908710 | 99908896 | 99906516 | 99906611 | 99923039 | 99923185 | 695 | 544 | 576 | 656 | 973 | 1 | 0 | 0 | 1 | 0 | 80 | 173 | 130 | 75 | 67 | 0 | 0 | 0 | 0 | 0 | 298 | 149 | 1 | 1 | 0.997 | 1 | 1 | 0.997 | 1 | 1 | 1 | 1 | 1 | 1 | -0.001 |
| SE | 86058 | ENSG00000164031 | DNAJB14 | chr4 | - | 99910348 | 99910375 | 99908710 | 99908896 | 99923039 | 99923185 | 1 | 5 | 1 | 7 | 4 | 400 | 279 | 308 | 367 | 504 | 0 | 4 | 0 | 0 | 0 | 56 | 113 | 84 | 47 | 29 | 176 | 149 | 1 | 1 | 0.002 | 0.015 | 0.003 | 0.016 | 0.007 | 0 | 0.029 | 0 | 0 | 0 | 0.003 |
| SE | 86059 | ENSG00000164031 | DNAJB14 | chr4 | - | 99923039 | 99923185 | 99908710 | 99908896 | 99930449 | 99930621 | 629 | 452 | 465 | 571 | 786 | 2 | 0 | 1 | 0 | 0 | 104 | 171 | 131 | 62 | 37 | 0 | 0 | 0 | 0 | 0 | 295 | 149 | 1 | 1 | 0.994 | 1 | 0.996 | 1 | 1 | 1 | 1 | 1 | 1 | 1 | -0.002 |
| SE | 86060 | ENSG00000164031 | DNAJB14 | chr4 | - | 99924759 | 99924815 | 99923039 | 99923185 | 99927716 | 99927838 | 5 | 12 | 5 | 1 | 3 | 1 | 3 | 1 | 6 | 20 | 0 | 0 | 0 | 0 | 0 | 0 | 0 | 0 | 2 | 0 | 205 | 149 | 1 | 1 | 0.784 | 0.744 | 0.784 | 0.108 | 0.098 | NA | NA | NA | 0 | NA | 0.504 |
| SE | 86061 | ENSG00000164031 | DNAJB14 | chr4 | - | 99924759 | 99924815 | 99923039 | 99923185 | 99928541 | 99928617 | 3 | 7 | 4 | 1 | 11 | 1 | 1 | 3 | 1 | 0 | 0 | 0 | 0 | 0 | 0 | 0 | 0 | 2 | 2 | 0 | 205 | 149 | 1 | 1 | 0.686 | 0.836 | 0.492 | 0.421 | 1 | NA | NA | 0 | 0 | NA | 0.687 |
| SE | 86063 | ENSG00000164031 | DNAJB14 | chr4 | - | 99924759 | 99924815 | 99923039 | 99923185 | 99946438 | 99946558 | 4 | 7 | 4 | 1 | 3 | 4 | 4 | 2 | 0 | 1 | 0 | 0 | 0 | 0 | 0 | 0 | 3 | 0 | 0 | 0 | 205 | 149 | 1 | 1 | 0.421 | 0.56 | 0.592 | 1 | 0.686 | NA | 0 | NA | NA | NA | 0.652 |
| SE | 86064 | ENSG00000164031 | DNAJB14 | chr4 | - | 99927716 | 99927838 | 99923039 | 99923185 | 99928541 | 99928617 | 5 | 11 | 6 | 15 | 26 | 1 | 1 | 3 | 1 | 0 | 0 | 2 | 1 | 14 | 5 | 0 | 0 | 2 | 2 | 0 | 271 | 149 | 1 | 1 | 0.733 | 0.858 | 0.524 | 0.892 | 1 | NA | 1 | 0.216 | 0.794 | 1 | 0.049 |
| SE | 86066 | ENSG00000164031 | DNAJB14 | chr4 | - | 99927716 | 99927838 | 99924759 | 99924815 | 99928541 | 99928617 | 6 | 13 | 6 | 9 | 6 | 0 | 0 | 0 | 0 | 8 | 0 | 2 | 1 | 12 | 5 | 0 | 0 | 0 | 0 | 0 | 271 | 149 | 1 | 1 | 1 | 1 | 1 | 1 | 0.292 | NA | 1 | 1 | 1 | 1 | -0.142 |
| SE | 86067 | ENSG00000164031 | DNAJB14 | chr4 | - | 99927716 | 99927838 | 99924759 | 99924815 | 99930449 | 99930589 | 2 | 5 | 1 | 0 | 0 | 1 | 2 | 10 | 5 | 3 | 0 | 0 | 0 | 0 | 0 | 6 | 3 | 1 | 0 | 4 | 271 | 149 | 1 | 1 | 0.524 | 0.579 | 0.052 | 0 | 0 | 0 | 0 | 0 | NA | 0 | 0.231 |
| SE | 86069 | ENSG00000164031 | DNAJB14 | chr4 | - | 99928541 | 99928617 | 99924759 | 99924815 | 99930449 | 99930534 | 1 | 11 | 2 | 4 | 11 | 1 | 2 | 10 | 5 | 3 | 0 | 0 | 1 | 0 | 0 | 6 | 3 | 1 | 0 | 4 | 225 | 149 | 1 | 1 | 0.398 | 0.785 | 0.117 | 0.346 | 0.708 | 0 | 0 | 0.398 | NA | 0 | 0.371 |
| SE | 86072 | ENSG00000164031 | DNAJB14 | chr4 | - | 99930449 | 99930621 | 99924759 | 99924815 | 99946438 | 99946558 | 55 | 26 | 43 | 62 | 43 | 1 | 0 | 0 | 0 | 0 | 6 | 12 | 3 | 1 | 4 | 0 | 0 | 0 | 0 | 0 | 298 | 149 | 1 | 1 | 0.965 | 1 | 1 | 1 | 1 | 1 | 1 | 1 | 1 | 1 | -0.007 |
| SE | 86073 | ENSG00000156531 | PHF6 | chrX | + | 134377571 | 134377755 | 134373311 | 134373467 | 134393500 | 134393634 | 74 | 40 | 47 | 44 | 37 | 0 | 0 | 0 | 0 | 0 | 9 | 30 | 23 | 1 | 0 | 0 | 5 | 3 | 0 | 0 | 298 | 149 | 1 | 1 | 1 | 1 | 1 | 1 | 1 | 1 | 0.75 | 0.793 | 1 | NA | 0.114 |
| SE | 86074 | ENSG00000156531 | PHF6 | chrX | + | 134378004 | 134378106 | 134377571 | 134377755 | 134393500 | 134393634 | 127 | 130 | 117 | 132 | 269 | 1 | 1 | 0 | 0 | 8 | 0 | 102 | 102 | 15 | 17 | 0 | 0 | 0 | 0 | 0 | 251 | 149 | 1 | 1 | 0.987 | 0.987 | 1 | 1 | 0.952 | NA | 1 | 1 | 1 | 1 | -0.015 |
| SE | 86077 | ENSG00000169359 | SLC33A1 | chr3 | - | 155829687 | 155829755 | 155827855 | 155828377 | 155829855 | 155829903 | 20 | 17 | 20 | 27 | 30 | 0 | 0 | 1 | 0 | 0 | 7 | 14 | 7 | 3 | 12 | 0 | 0 | 0 | 0 | 0 | 217 | 149 | 1 | 1 | 1 | 1 | 0.932 | 1 | 1 | 1 | 1 | 1 | 1 | 1 | -0.014 |
| SE | 86079 | ENSG00000169359 | SLC33A1 | chr3 | - | 155833467 | 155834041 | 155827773 | 155828377 | 155842431 | 155842619 | 89 | 52 | 98 | 64 | 185 | 0 | 0 | 0 | 0 | 2 | 67 | 74 | 25 | 41 | 91 | 0 | 0 | 0 | 0 | 0 | 298 | 149 | 1 | 1 | 1 | 1 | 1 | 1 | 0.979 | 1 | 1 | 1 | 1 | 1 | -0.004 |
| SE | 86080 | ENSG00000169359 | SLC33A1 | chr3 | - | 155833467 | 155834041 | 155829687 | 155829903 | 155842431 | 155842619 | 156 | 104 | 146 | 122 | 277 | 7 | 3 | 4 | 2 | 26 | 67 | 106 | 45 | 71 | 166 | 0 | 3 | 2 | 0 | 0 | 298 | 149 | 1 | 1 | 0.918 | 0.945 | 0.948 | 0.968 | 0.842 | 1 | 0.946 | 0.918 | 1 | 1 | -0.049 |
| SE | 86081 | ENSG00000169359 | SLC33A1 | chr3 | - | 155833856 | 155834041 | 155833467 | 155833585 | 155853107 | 155854456 | 32 | 37 | 27 | 29 | 58 | 0 | 1 | 0 | 0 | 0 | 9 | 29 | 7 | 33 | 43 | 0 | 0 | 0 | 0 | 0 | 298 | 149 | 1 | 1 | 1 | 1 | 0.931 | 1 | 1 | 1 | 1 | 1 | 1 | 1 | -0.014 |
| SE | 86082 | ENSG00000169359 | SLC33A1 | chr3 | - | 155842431 | 155842619 | 155833467 | 155833585 | 155853107 | 155854456 | 4 | 11 | 13 | 5 | 14 | 0 | 0 | 1 | 0 | 0 | 3 | 12 | 13 | 15 | 7 | 0 | 0 | 0 | 0 | 0 | 298 | 149 | 1 | 1 | 1 | 1 | 0.867 | 1 | 1 | 1 | 1 | 1 | 1 | 1 | -0.027 |
| SE | 86089 | ENSG00000102054 | RBBP7 | chrX | - | 16849243 | 16849301 | 16845827 | 16845938 | 16852045 | 16852065 | 339 | 220 | 259 | 337 | 423 | 0 | 0 | 0 | 0 | 3 | 197 | 366 | 278 | 340 | 151 | 0 | 0 | 0 | 0 | 0 | 207 | 149 | 1 | 1 | 1 | 1 | 1 | 1 | 0.99 | 1 | 1 | 1 | 1 | 1 | -0.002 |
| SE | 86090 | ENSG00000102054 | RBBP7 | chrX | - | 16852045 | 16852122 | 16848990 | 16849301 | 16853681 | 16853842 | 143 | 91 | 107 | 116 | 170 | 0 | 0 | 0 | 1 | 0 | 77 | 121 | 105 | 132 | 55 | 0 | 1 | 0 | 0 | 0 | 226 | 149 | 1 | 1 | 1 | 1 | 1 | 0.987 | 1 | 1 | 0.988 | 1 | 1 | 1 | 0 |
| SE | 86091 | ENSG00000102054 | RBBP7 | chrX | - | 16852415 | 16852550 | 16852045 | 16852122 | 16852748 | 16852875 | 5 | 2 | 3 | 4 | 0 | 0 | 0 | 0 | 3 | 0 | 32 | 0 | 1 | 4 | 0 | 0 | 3 | 7 | 4 | 0 | 284 | 149 | 1 | 1 | 1 | 1 | 1 | 0.412 | NA | 1 | 0 | 0.07 | 0.344 | NA | 0.499 |
| SE | 86093 | ENSG00000102054 | RBBP7 | chrX | - | 16852748 | 16852875 | 16852045 | 16852122 | 16853681 | 16853842 | 212 | 130 | 178 | 200 | 248 | 0 | 0 | 0 | 0 | 0 | 135 | 108 | 140 | 135 | 41 | 0 | 0 | 0 | 4 | 0 | 276 | 149 | 1 | 1 | 1 | 1 | 1 | 1 | 1 | 1 | 1 | 1 | 0.948 | 1 | 0.01 |
| SE | 86094 | ENSG00000102054 | RBBP7 | chrX | - | 16857593 | 16857709 | 16852551 | 16853842 | 16858675 | 16858803 | 464 | 262 | 389 | 390 | 592 | 0 | 2 | 0 | 0 | 0 | 280 | 311 | 308 | 273 | 133 | 0 | 0 | 0 | 0 | 0 | 265 | 149 | 1 | 1 | 1 | 0.987 | 1 | 1 | 1 | 1 | 1 | 1 | 1 | 1 | -0.003 |
| SE | 86095 | ENSG00000102054 | RBBP7 | chrX | - | 16862954 | 16863100 | 16858675 | 16858849 | 16869075 | 16869220 | 783 | 553 | 811 | 812 | 1310 | 2 | 1 | 1 | 7 | 4 | 741 | 790 | 719 | 767 | 466 | 0 | 0 | 0 | 0 | 0 | 295 | 149 | 1 | 1 | 0.995 | 0.996 | 0.998 | 0.983 | 0.994 | 1 | 1 | 1 | 1 | 1 | -0.007 |
| SE | 86096 | ENSG00000102054 | RBBP7 | chrX | - | 16869075 | 16869220 | 16849243 | 16849301 | 16870037 | 16870361 | 138 | 105 | 124 | 142 | 133 | 0 | 0 | 1 | 0 | 0 | 96 | 71 | 95 | 100 | 39 | 0 | 0 | 0 | 0 | 0 | 294 | 149 | 1 | 1 | 1 | 1 | 0.984 | 1 | 1 | 1 | 1 | 1 | 1 | 1 | -0.003 |
| SE | 86101 | ENSG00000120860 | WASHC3 | chr12 | - | 102025973 | 102026038 | 102012839 | 102013192 | 102039867 | 102039975 | 385 | 280 | 277 | 346 | 553 | 0 | 0 | 0 | 0 | 0 | 372 | 303 | 229 | 371 | 177 | 0 | 8 | 0 | 0 | 0 | 214 | 149 | 1 | 1 | 1 | 1 | 1 | 1 | 1 | 1 | 0.963 | 1 | 1 | 1 | 0.007 |
| SE | 86102 | ENSG00000120860 | WASHC3 | chr12 | - | 102039867 | 102039975 | 102025973 | 102026038 | 102044104 | 102044212 | 271 | 199 | 197 | 223 | 393 | 0 | 0 | 0 | 1 | 0 | 221 | 190 | 152 | 179 | 94 | 0 | 0 | 2 | 0 | 0 | 257 | 149 | 1 | 1 | 1 | 1 | 1 | 0.992 | 1 | 1 | 1 | 0.978 | 1 | 1 | 0.003 |
| SE | 86103 | ENSG00000120860 | WASHC3 | chr12 | - | 102039867 | 102039978 | 102025920 | 102026038 | 102044104 | 102044212 | 392 | 294 | 302 | 337 | 625 | 0 | 0 | 0 | 1 | 0 | 301 | 280 | 237 | 297 | 138 | 0 | 0 | 2 | 0 | 0 | 260 | 149 | 1 | 1 | 1 | 1 | 1 | 0.995 | 1 | 1 | 1 | 0.985 | 1 | 1 | 0.002 |
| SE | 86112 | ENSG00000047644 | WWC3 | chrX | + | 10079724 | 10079804 | 10078798 | 10078875 | 10090770 | 10090900 | 73 | 60 | 69 | 49 | 47 | 0 | 0 | 0 | 0 | 0 | 139 | 115 | 54 | 83 | 266 | 0 | 0 | 0 | 5 | 0 | 229 | 149 | 1 | 1 | 1 | 1 | 1 | 1 | 1 | 1 | 1 | 1 | 0.915 | 1 | 0.017 |
| SE | 86113 | ENSG00000047644 | WWC3 | chrX | + | 10109777 | 10110020 | 10098505 | 10098579 | 10116417 | 10116507 | 113 | 73 | 75 | 96 | 61 | 2 | 0 | 0 | 0 | 0 | 207 | 88 | 63 | 115 | 242 | 0 | 0 | 2 | 2 | 0 | 298 | 149 | 1 | 1 | 0.966 | 1 | 1 | 1 | 1 | 1 | 1 | 0.94 | 0.966 | 1 | 0.012 |
| SE | 86114 | ENSG00000047644 | WWC3 | chrX | + | 10117123 | 10117689 | 10116417 | 10116507 | 10122618 | 10122739 | 40 | 19 | 31 | 33 | 33 | 0 | 0 | 0 | 4 | 0 | 14 | 45 | 28 | 37 | 67 | 0 | 0 | 2 | 0 | 0 | 298 | 149 | 1 | 1 | 1 | 1 | 1 | 0.805 | 1 | 1 | 1 | 0.875 | 1 | 1 | -0.014 |
| SE | 86115 | ENSG00000047644 | WWC3 | chrX | + | 10125023 | 10125110 | 10124264 | 10124339 | 10128556 | 10128708 | 16 | 6 | 15 | 3 | 10 | 0 | 0 | 0 | 1 | 0 | 35 | 6 | 18 | 34 | 7 | 0 | 0 | 0 | 0 | 0 | 236 | 149 | 1 | 1 | 1 | 1 | 1 | 0.654 | 1 | 1 | 1 | 1 | 1 | 1 | -0.069 |
| SE | 86116 | ENSG00000047644 | WWC3 | chrX | + | 10126113 | 10126306 | 10125023 | 10125110 | 10127987 | 10128161 | 77 | 27 | 47 | 61 | 32 | 0 | 0 | 0 | 0 | 0 | 74 | 64 | 64 | 35 | 147 | 0 | 0 | 0 | 4 | 0 | 298 | 149 | 1 | 1 | 1 | 1 | 1 | 1 | 1 | 1 | 1 | 1 | 0.814 | 1 | 0.037 |
| SE | 86120 | ENSG00000164754 | RAD21 | chr8 | - | 116848945 | 116849029 | 116845933 | 116847691 | 116850617 | 116850767 | 448 | 305 | 334 | 444 | 768 | 0 | 2 | 0 | 0 | 0 | 66 | 188 | 228 | 89 | 46 | 0 | 1 | 0 | 0 | 0 | 233 | 149 | 1 | 1 | 1 | 0.99 | 1 | 1 | 1 | 1 | 0.992 | 1 | 1 | 1 | 0 |
| SE | 86121 | ENSG00000164754 | RAD21 | chr8 | - | 116852548 | 116852708 | 116851947 | 116852096 | 116854244 | 116854468 | 1077 | 693 | 1033 | 1131 | 1712 | 0 | 2 | 0 | 0 | 0 | 302 | 761 | 931 | 449 | 205 | 0 | 0 | 0 | 0 | 0 | 298 | 149 | 1 | 1 | 1 | 0.994 | 1 | 1 | 1 | 1 | 1 | 1 | 1 | 1 | -0.001 |
| SE | 86122 | ENSG00000164754 | RAD21 | chr8 | - | 116854244 | 116854468 | 116852548 | 116852708 | 116856165 | 116856288 | 1009 | 673 | 897 | 1044 | 1603 | 0 | 0 | 2 | 0 | 0 | 299 | 680 | 915 | 309 | 176 | 0 | 0 | 4 | 4 | 0 | 298 | 149 | 1 | 1 | 1 | 1 | 0.996 | 1 | 1 | 1 | 1 | 0.991 | 0.975 | 1 | 0.006 |
| SE | 86123 | ENSG00000164754 | RAD21 | chr8 | - | 116856645 | 116856771 | 116856165 | 116856288 | 116857266 | 116857473 | 1407 | 1038 | 1324 | 1423 | 2556 | 0 | 1 | 0 | 0 | 0 | 362 | 838 | 1080 | 357 | 98 | 0 | 0 | 0 | 0 | 0 | 275 | 149 | 1 | 1 | 1 | 0.998 | 1 | 1 | 1 | 1 | 1 | 1 | 1 | 1 | 0 |
| SE | 86124 | ENSG00000164754 | RAD21 | chr8 | - | 116858351 | 116858458 | 116857266 | 116857473 | 116861840 | 116861940 | 900 | 681 | 809 | 953 | 1360 | 1 | 1 | 1 | 0 | 0 | 233 | 527 | 611 | 245 | 68 | 0 | 0 | 0 | 0 | 0 | 256 | 149 | 1 | 1 | 0.998 | 0.997 | 0.998 | 1 | 1 | 1 | 1 | 1 | 1 | 1 | -0.001 |
| SE | 86125 | ENSG00000101916 | TLR8 | chrX | + | 12910305 | 12910442 | 12906638 | 12906709 | 12919043 | 12922202 | 21 | 14 | 16 | 6 | 26 | 220 | 148 | 107 | 121 | 360 | 0 | 13 | 17 | 14 | 0 | 0 | 201 | 126 | 83 | 12 | 286 | 149 | 1 | 1 | 0.047 | 0.047 | 0.072 | 0.025 | 0.036 | NA | 0.033 | 0.066 | 0.081 | 0 | 0 |
| SE | 86127 | ENSG00000090520 | DNAJB11 | chr3 | + | 186577667 | 186577800 | 186575839 | 186575937 | 186581994 | 186582077 | 68 | 38 | 66 | 55 | 40 | 0 | 0 | 0 | 0 | 0 | 79 | 61 | 62 | 93 | 45 | 0 | 0 | 2 | 0 | 0 | 282 | 149 | 1 | 1 | 1 | 1 | 1 | 1 | 1 | 1 | 1 | 0.942 | 1 | 1 | 0.012 |
| SE | 86128 | ENSG00000090520 | DNAJB11 | chr3 | + | 186577667 | 186581513 | 186575839 | 186575937 | 186581994 | 186582077 | 133 | 81 | 143 | 126 | 88 | 0 | 0 | 0 | 0 | 0 | 145 | 137 | 181 | 166 | 82 | 0 | 0 | 2 | 0 | 0 | 298 | 149 | 1 | 1 | 1 | 1 | 1 | 1 | 1 | 1 | 1 | 0.978 | 1 | 1 | 0.004 |
| SE | 86129 | ENSG00000090520 | DNAJB11 | chr3 | + | 186581370 | 186581513 | 186575869 | 186575937 | 186581994 | 186582077 | 64 | 41 | 73 | 70 | 44 | 0 | 0 | 0 | 0 | 0 | 66 | 76 | 118 | 78 | 37 | 0 | 0 | 2 | 0 | 0 | 292 | 149 | 1 | 1 | 1 | 1 | 1 | 1 | 1 | 1 | 1 | 0.968 | 1 | 1 | 0.006 |
| SE | 86130 | ENSG00000090520 | DNAJB11 | chr3 | + | 186581370 | 186581513 | 186577667 | 186577800 | 186581994 | 186582077 | 118 | 70 | 130 | 131 | 86 | 0 | 0 | 0 | 0 | 0 | 178 | 159 | 183 | 140 | 72 | 0 | 0 | 0 | 5 | 0 | 292 | 149 | 1 | 1 | 1 | 1 | 1 | 1 | 1 | 1 | | 1 | 0.935 | 1 | 0.013 |
| SE | 86134 | ENSG00000197713 | RPE | chr2 | + | 210003437 | 210003468 | 210002627 | 210002783 | 210015972 | 210016112 | 3 | 4 | 3 | 8 | 2 | 8 | 7 | 11 | 6 | 0 | 0 | 5 | 1 | 7 | 0 | 0 | 2 | 0 | 9 | 17 | 180 | 149 | 1 | 1 | 0.237 | 0.321 | 0.184 | 0.525 | 1 | NA | 0.674 | 1 | 0.392 | 0 | -0.063 |
| SE | 86141 | ENSG00000197713 | RPE | chr2 | + | 210017789 | 210017887 | 210017472 | 210017559 | 210018152 | 210018221 | 0 | 1 | 1 | 0 | 0 | 9 | 1 | 6 | 7 | 4 | 0 | 0 | 0 | 0 | 0 | 6 | 0 | 0 | 11 | 0 | 247 | 149 | 1 | 1 | 0 | 0.376 | 0.091 | 0 | 0 | 0 | NA | NA | 0 | NA | 0.093 |
| SE | 86142 | ENSG00000197713 | RPE | chr2 | + | 210017789 | 210017887 | 210017472 | 210017559 | 210019668 | 210019735 | 1 | 0 | 6 | 1 | 0 | 73 | 45 | 53 | 79 | 65 | 0 | 0 | 0 | 0 | 0 | 22 | 66 | 37 | 36 | 33 | 247 | 149 | 1 | 1 | 0.008 | 0 | 0.064 | 0.008 | 0 | 0 | 0 | 0 | 0 | 0 | 0.016 |
| SE | 86144 | ENSG00000197713 | RPE | chr2 | + | 210018152 | 210018221 | 210017789 | 210017887 | 210019668 | 210019940 | 4 | 3 | 2 | 2 | 1 | 1 | 0 | 5 | 1 | 0 | 0 | 3 | 2 | 0 | 0 | 0 | 0 | 0 | 0 | 0 | 218 | 149 | 1 | 1 | 0.732 | 1 | 0.215 | 0.578 | 1 | NA | 1 | 1 | NA | NA | -0.295 |
| SE | 86148 | ENSG00000017483 | SLC38A5 | chrX | - | 48466229 | 48466322 | 48466014 | 48466093 | 48466798 | 48466872 | 53 | 20 | 68 | 27 | 11 | 0 | 0 | 0 | 1 | 0 | 36 | 72 | 72 | 10 | 54 | 0 | 0 | 0 | 0 | 0 | 242 | 149 | 1 | 1 | 1 | 1 | 1 | 0.943 | 1 | 1 | 1 | 1 | 1 | 1 | -0.011 |
| SE | 86149 | ENSG00000068438 | FTSJ1 | chrX | + | 48476562 | 48476668 | 48476160 | 48476396 | 48477960 | 48478168 | 4 | 0 | 2 | 1 | 2 | 127 | 51 | 100 | 107 | 49 | 0 | 0 | 1 | 4 | 0 | 88 | 115 | 86 | 115 | 212 | 255 | 149 | 1 | 1 | 0.018 | 0 | 0.012 | 0.005 | 0.023 | 0 | 0 | 0.007 | 0.02 | 0 | 0.006 |
| SE | 86151 | ENSG00000068438 | FTSJ1 | chrX | + | 48478448 | 48478518 | 48477960 | 48478168 | 48478616 | 48478707 | 145 | 70 | 108 | 138 | 55 | 0 | 1 | 0 | 0 | 0 | 92 | 90 | 68 | 136 | 138 | 0 | 0 | 0 | 0 | 0 | 219 | 149 | 1 | 1 | 1 | 0.979 | 1 | 1 | 1 | 1 | 1 | 1 | 1 | 1 | -0.004 |
| SE | 86152 | ENSG00000068438 | FTSJ1 | chrX | + | 48478616 | 48478680 | 48477960 | 48478518 | 48479037 | 48479116 | 25 | 6 | 15 | 38 | 19 | 0 | 0 | 0 | 0 | 0 | 0 | 17 | 6 | 6 | 15 | 0 | 0 | 4 | 0 | 0 | 213 | 149 | 1 | 1 | 1 | 1 | 1 | 1 | 1 | NA | 1 | 0.512 | 1 | 1 | 0.122 |
| SE | 86153 | ENSG00000068438 | FTSJ1 | chrX | + | 48478616 | 48478707 | 48478448 | 48478518 | 48479037 | 48479070 | 103 | 54 | 109 | 127 | 47 | 0 | 0 | 0 | 0 | 0 | 73 | 74 | 67 | 73 | 83 | 0 | 0 | 4 | 0 | 0 | 240 | 149 | 1 | 1 | 1 | 1 | 1 | 1 | 1 | 1 | 1 | 0.912 | 1 | 1 | 0.018 |
| SE | 86156 | ENSG00000068438 | FTSJ1 | chrX | + | 48483240 | 48483341 | 48482985 | 48483027 | 48485735 | 48485984 | 0 | 0 | 6 | 1 | 1 | 97 | 38 | 67 | 74 | 77 | 0 | 1 | 2 | 0 | 0 | 47 | 114 | 52 | 76 | 122 | 250 | 149 | 1 | 1 | 0 | 0 | 0.051 | 0.008 | 0.008 | 0 | 0.005 | 0.022 | 0 | 0 | 0.008 |
| SE | 86164 | ENSG00000113845 | TIMMDC1 | chr3 | + | 119503953 | 119504021 | 119498580 | 119498927 | 119517204 | 119517315 | 3 | 0 | 0 | 0 | 1 | 0 | 0 | 0 | 0 | 0 | 0 | 0 | 0 | 0 | 0 | 0 | 0 | 1 | 8 | 5 | 217 | 149 | 1 | 1 | 1 | NA | NA | NA | 1 | NA | NA | 0 | 0 | 0 | 1 |
| SE | 86169 | ENSG00000113845 | TIMMDC1 | chr3 | + | 119513640 | 119513719 | 119503953 | 119504021 | 119517204 | 119517315 | 137 | 110 | 132 | 149 | 193 | 3 | 0 | 0 | 1 | 1 | 318 | 112 | 59 | 231 | 263 | 0 | 0 | 0 | 0 | 0 | 228 | 149 | 1 | 1 | 0.968 | 1 | 1 | 0.99 | 0.992 | 1 | 1 | 1 | 1 | 1 | -0.01 |
| SE | 86178 | ENSG00000115275 | MOGS | chr2 | - | 74464495 | 74464722 | 74461183 | 74463386 | 74464895 | 74464987 | 58 | 14 | 30 | 45 | 18 | 2 | 0 | 1 | 1 | 0 | 19 | 40 | 31 | 21 | 17 | 0 | 0 | 0 | 4 | 0 | 298 | 149 | 1 | 1 | 0.935 | 1 | 0.938 | 0.957 | 1 | 1 | 1 | 1 | 0.724 | 1 | 0.021 |
| SE | 86180 | ENSG00000115275 | MOGS | chr2 | - | 74464495 | 74464722 | 74463189 | 74463331 | 74464895 | 74464987 | 18 | 5 | 9 | 15 | 0 | 0 | 0 | 1 | 0 | 0 | 9 | 15 | 14 | 6 | 13 | 0 | 0 | 0 | 0 | 0 | 298 | 149 | 1 | 1 | 1 | 1 | 0.818 | 1 | NA | 1 | 1 | 1 | 1 | 1 | -0.045 |
| SE | 86182 | ENSG00000115275 | MOGS | chr2 | - | 74464665 | 74464722 | 74463189 | 74463331 | 74464895 | 74465294 | 3 | 0 | 1 | 2 | 0 | 0 | 0 | 1 | 0 | 0 | 0 | 6 | 5 | 2 | 0 | 0 | 0 | 0 | 0 | 0 | 206 | 149 | 1 | 1 | 1 | NA | 0.42 | 1 | NA | NA | 1 | 1 | 1 | NA | -0.193 |
| SE | 86187 | ENSG00000102030 | NAA10 | chrX | - | 153932070 | 153932115 | 153930762 | 153930847 | 153932315 | 153932431 | 209 | 115 | 164 | 170 | 242 | 1 | 4 | 1 | 2 | 0 | 177 | 231 | 115 | 164 | 358 | 0 | 0 | 1 | 3 | 0 | 194 | 149 | 1 | 1 | 0.994 | 0.957 | 0.992 | 0.985 | 1 | 1 | 1 | 0.989 | 0.977 | 1 | -0.008 |
| SE | 86188 | ENSG00000102030 | NAA10 | chrX | - | 153933960 | 153934001 | 153932538 | 153932584 | 153934376 | 153934475 | 16 | 12 | 15 | 8 | 15 | 0 | 0 | 0 | 1 | 0 | 6 | 2 | 12 | 17 | 22 | 0 | 0 | 0 | 0 | 0 | 190 | 149 | 1 | 1 | 1 | 1 | 1 | 0.863 | 1 | 1 | 1 | 1 | 1 | 1 | -0.027 |
| SE | 86189 | ENSG00000102030 | NAA10 | chrX | - | 153933960 | 153934001 | 153932538 | 153932584 | 153934883 | 153935015 | 1 | 1 | 0 | 1 | 0 | 1 | 0 | 0 | 0 | 0 | 0 | 0 | 2 | 4 | 0 | 0 | 0 | 0 | 0 | 0 | 190 | 149 | 1 | 1 | 0.44 | 1 | NA | 1 | NA | NA | NA | 1 | 1 | NA | -0.187 |
| SE | 86192 | ENSG00000102032 | RENBP | chrX | - | 153941949 | 153942031 | 153941477 | 153941653 | 153942854 | 153943079 | 54 | 24 | 29 | 33 | 31 | 3 | 0 | 0 | 0 | 0 | 60 | 37 | 25 | 27 | 48 | 0 | 0 | 0 | 0 | 0 | 231 | 149 | 1 | 1 | 0.921 | 1 | 1 | | 1 | 1 | 1 | 1 | 1 | 1 | -0.016 |
| SE | 86197 | ENSG00000171311 | EXOSC1 | chr10 | - | 97438662 | 97438703 | 97437699 | 97437750 | 97441170 | 97441259 | 32 | 24 | 30 | 43 | 28 | 0 | 1 | 0 | 0 | 0 | 37 | 52 | 12 | 44 | 27 | 0 | 0 | 0 | 0 | 0 | 190 | 149 | 1 | 1 | 1 | 0.95 | 1 | 1 | 1 | 1 | 1 | 1 | 1 | 1 | -0.01 |
| SE | 86198 | ENSG00000171311 | EXOSC1 | chr10 | - | 97438669 | 97438703 | 97437699 | 97437750 | 97441170 | 97441259 | 116 | 80 | 126 | 145 | 105 | 0 | 1 | 0 | 0 | 0 | 109 | 103 | 47 | 135 | 127 | 0 | 0 | 0 | 0 | 0 | 183 | 149 | 1 | 1 | 1 | 0.985 | 1 | 1 | 1 | 1 | 1 | 1 | 1 | 1 | -0.003 |
| SE | 86200 | ENSG00000171311 | EXOSC1 | chr10 | - | 97441170 | 97441259 | 97438662 | 97438703 | 97445731 | 97445847 | 79 | 44 | 75 | 87 | 48 | 0 | 0 | 0 | 0 | 0 | 34 | 32 | 30 | 73 | 83 | 0 | 2 | 0 | 0 | 0 | 238 | 149 | 1 | 1 | 1 | 1 | 1 | 1 | 1 | 1 | 0.909 | 1 | 1 | 1 | 0.018 |
| SE | 86203 | ENSG00000172534 | HCFC1 | chrX | - | 153950242 | 153950543 | 153949552 | 153949616 | 153951349 | 153951487 | 107 | 61 | 86 | 67 | 109 | 0 | 0 | 0 | 0 | 0 | 95 | 88 | 50 | 137 | 146 | 0 | 0 | 0 | 1 | 0 | 298 | 149 | 1 | 1 | 1 | 1 | 1 | 1 | 1 | 1 | 1 | 1 | 0.986 | 1 | 0.003 |
| SE | 86205 | ENSG00000172534 | HCFC1 | chrX | - | 153951840 | 153952158 | 153951588 | 153951707 | 153952513 | 153952958 | 153 | 50 | 129 | 128 | 91 | 0 | 0 | 0 | 1 | 0 | 73 | 144 | 56 | 143 | 162 | 0 | 0 | 0 | 0 | 0 | 298 | 149 | 1 | 1 | 1 | 1 | 1 | 0.985 | 1 | 1 | 1 | 1 | 1 | 1 | -0.003 |
| SE | 86206 | ENSG00000172534 | HCFC1 | chrX | - | 153951840 | 153952161 | 153951588 | 153951707 | 153952513 | 153952958 | 100 | 36 | 95 | 93 | 45 | 0 | 0 | 0 | 1 | 0 | 57 | 120 | 55 | 114 | 122 | 0 | 0 | 0 | 0 | 0 | 298 | 149 | 1 | 1 | 1 | 1 | 1 | 0.979 | 1 | 1 | 1 | 1 | 1 | 1 | -0.004 |
| SE | 86210 | ENSG00000172534 | HCFC1 | chrX | - | 153959801 | 153960161 | 153959330 | 153959491 | 153960234 | 153960414 | 258 | 146 | 235 | 285 | 286 | 4 | 0 | 1 | 0 | 4 | 369 | 344 | 189 | 281 | 482 | 0 | 0 | 0 | 2 | 0 | 298 | 149 | 1 | 1 | 0.97 | 1 | 0.992 | 1 | 0.973 | 1 | 1 | 1 | 0.986 | 1 | -0.01 |
| SE | 86212 | ENSG00000172534 | HCFC1 | chrX | - | 153963224 | 153963433 | 153962221 | 153962306 | 153964123 | 153964284 | 363 | 147 | 287 | 345 | 411 | 0 | 0 | 0 | 0 | 0 | 321 | 355 | 234 | 369 | 493 | 1 | 0 | 0 | 2 | 0 | 298 | 149 | 1 | 1 | 1 | 1 | 1 | 1 | 1 | 0.994 | 1 | 1 | 0.989 | 1 | 0.003 |
| SE | 86213 | ENSG00000172534 | HCFC1 | chrX | - | 153963224 | 153963433 | 153962221 | 153962306 | 153964577 | 153964726 | 208 | 88 | 182 | 196 | 287 | 0 | 0 | 0 | 0 | 0 | 195 | 214 | 131 | 174 | 311 | 0 | 0 | 0 | 2 | 0 | 298 | 149 | 1 | 1 | 1 | 1 | 1 | 1 | 1 | 1 | 1 | 1 | 0.978 | 1 | 0.004 |
| SE | 86214 | ENSG00000172534 | HCFC1 | chrX | - | 153964123 | 153964284 | 153962221 | 153962306 | 153964577 | 153964726 | 70 | 20 | 63 | 57 | 36 | 0 | 0 | 0 | 0 | 0 | 64 | 74 | 44 | 38 | 127 | 0 | 0 | 0 | 2 | 0 | 298 | 149 | 1 | 1 | 1 | 1 | 1 | 1 | 1 | 1 | 1 | 1 | 0.905 | 1 | 0.019 |
| SE | 86215 | ENSG00000172534 | HCFC1 | chrX | - | 153964123 | 153964284 | 153963224 | 153963433 | 153964577 | 153964726 | 225 | 79 | 168 | 207 | 160 | 0 | 0 | 0 | 1 | 0 | 189 | 215 | 147 | 231 | 309 | 0 | 0 | 0 | 0 | 0 | 298 | 149 | 1 | 1 | 1 | 1 | 1 | 0.99 | 1 | 1 | 1 | 1 | 1 | 1 | -0.002 |
| SE | 86217 | ENSG00000126953 | TIMM8A | chrX | - | 101348060 | 101348401 | 101345660 | 101346660 | 101348532 | 101348685 | 1 | 0 | 0 | 2 | 0 | 13 | 9 | 4 | 7 | 5 | 0 | 6 | 5 | 0 | 0 | 0 | 0 | 2 | 1 | 5 | 298 | 149 | 1 | 1 | 0.037 | 0 | 0 | 0.125 | 0 | NA | 1 | 0.556 | 0 | 0 | -0.357 |
| SE | 86218 | ENSG00000010671 | BTK | chrX | - | 101354629 | 101354694 | 101353869 | 101353988 | 101356051 | 101356268 | 79 | 63 | 75 | 111 | 201 | 0 | 0 | 0 | 2 | 2 | 229 | 134 | 50 | 157 | 150 | 0 | 0 | 2 | 0 | 0 | 214 | 149 | 1 | 1 | 1 | 1 | 1 | 0.975 | 0.986 | 1 | 1 | 0.946 | 1 | 1 | 0.003 |
| SE | 86220 | ENSG00000010671 | BTK | chrX | - | 101360087 | 101360150 | 101359292 | 101359347 | 101360567 | 101360755 | 795 | 835 | 473 | 846 | 862 | 0 | 0 | 1 | 1 | 0 | 2525 | 1010 | 702 | 1535 | 1921 | 0 | 0 | 0 | 0 | 0 | 212 | 149 | 1 | 1 | 1 | 1 | 0.997 | 0.998 | 1 | 1 | 1 | 1 | 1 | 1 | -0.001 |
| SE | 86221 | ENSG00000010671 | BTK | chrX | - | 101371632 | 101371701 | 101369997 | 101370079 | 101374535 | 101374634 | 316 | 262 | 222 | 303 | 467 | 0 | 0 | 0 | 1 | 0 | 503 | 329 | 193 | 333 | 569 | 0 | 0 | 1 | 0 | 0 | 218 | 149 | 1 | 1 | 1 | 1 | 1 | 1 | 1 | 1 | 1 | 0.992 | 1 | 1 | 0.002 |
| SE | 86222 | ENSG00000010671 | BTK | chrX | - | 101374535 | 101374634 | 101371632 | 101371701 | 101375143 | 101375284 | 222 | 213 | 176 | 235 | 380 | 0 | 0 | 1 | 0 | 0 | 459 | 270 | 143 | 277 | 387 | 0 | 0 | 0 | 0 | 0 | 248 | 149 | 1 | 1 | 1 | 1 | 0.991 | 1 | 1 | 1 | 1 | 1 | 1 | 1 | -0.002 |
| SE | 86223 | ENSG00000010671 | BTK | chrX | - | 101375143 | 101375314 | 101371632 | 101371701 | 101386061 | 101386148 | 111 | 93 | 78 | 114 | 191 | 0 | 0 | 0 | 0 | 0 | 192 | 142 | 56 | 133 | 202 | 0 | 0 | 3 | 0 | 0 | 298 | 149 | 1 | 1 | 1 | 1 | 1 | 1 | 1 | 1 | 1 | 0.903 | 1 | 1 | 0.019 |
| SE | 86224 | ENSG00000010671 | BTK | chrX | - | 101377858 | 101378023 | 101375143 | 101375314 | 101386061 | 101386224 | 2 | 4 | 0 | 0 | 3 | 111 | 93 | 77 | 114 | 191 | 0 | 0 | 0 | 0 | 0 | 192 | 142 | 56 | 133 | 202 | 298 | 149 | 1 | 1 | 0.009 | 0.021 | 0 | 0 | 0.008 | 0 | 0 | 0 | 0 | 0 | 0.008 |
| SE | 86225 | ENSG00000241343 | RPL36A | chrX | + | 101392049 | 101392095 | 101391754 | 101391822 | 101395334 | 101395457 | 2 | 5 | 0 | 0 | 5 | 5633 | 4223 | 5532 | 5575 | 5605 | 3 | 9 | 5 | 7 | 32 | 9119 | 3799 | 6011 | 5670 | 6800 | 195 | 149 | 1 | 1 | 0 | 0.001 | 0 | 0 | 0.001 | 0 | 0.002 | 0.001 | 0.001 | 0.004 | -0.001 |
| SE | 86226 | ENSG00000241343 | RPL36A | chrX | + | 101395334 | 101395457 | 101391754 | 101391822 | 101395727 | 101395786 | 5598 | 4120 | 5538 | 5498 | 5989 | 0 | 1 | 2 | 0 | 0 | 8684 | 3693 | 5761 | 5511 | 6596 | 0 | 0 | 0 | 0 | 0 | 272 | 149 | 1 | 1 | 1 | 1 | 0.999 | 1 | 1 | 1 | 1 | 1 | 1 | 1 | 0 |
| SE | 86227 | ENSG00000257529 | RPL36A-HNRNPH2 | chrX | + | 101395334 | 101395457 | 101391754 | 101391822 | 101411935 | 101412165 | 4600 | 3398 | 4568 | 4532 | 4890 | 0 | 0 | 0 | 1 | 0 | 6789 | 3033 | 4406 | 4347 | 5140 | 0 | 0 | 0 | 0 | 0 | 272 | 149 | 1 | 1 | 1 | 1 | 1 | 1 | 1 | 1 | 1 | 1 | 1 | 1 | 0 |
| SE | 86228 | ENSG00000177042 | TMEM80 | chr11 | + | 700614 | 700707 | 698868 | 700235 | 702944 | 703146 | 122 | 45 | 58 | 52 | 88 | 0 | 0 | 0 | 1 | 0 | 55 | 92 | 35 | 60 | 65 | 0 | 0 | 0 | 0 | 0 | 242 | 149 | 1 | 1 | 1 | 1 | 1 | 0.97 | 1 | 1 | 1 | 1 | 1 | 1 | -0.006 |
| SE | 86230 | ENSG00000184634 | MED12 | chrX | + | 71120970 | 71121152 | 71120013 | 71120170 | 71121326 | 71121437 | 199 | 153 | 175 | 203 | 303 | 0 | 0 | 0 | 0 | 0 | 315 | 241 | 115 | 226 | 375 | 0 | 0 | 5 | 0 | 0 | 298 | 149 | 1 | 1 | 1 | 1 | 1 | 1 | 1 | 1 | 1 | 0.92 | 1 | 1 | 0.016 |
| SE | 86231 | ENSG00000184634 | MED12 | chrX | + | 71122507 | 71122607 | 71122199 | 71122346 | 71122737 | 71122874 | 61 | 44 | 69 | 66 | 55 | 0 | 0 | 0 | 0 | 0 | 101 | 68 | 48 | 65 | 143 | 0 | 0 | 0 | 0 | 6 | 249 | 149 | 1 | 1 | 1 | 1 | 1 | 1 | 1 | 1 | 1 | 1 | 1 | 0.934 | 0.013 |
| SE | 86234 | ENSG00000184634 | MED12 | chrX | + | 71125662 | 71125713 | 71125350 | 71125495 | 71126035 | 71126154 | 36 | 24 | 38 | 45 | 63 | 1 | 0 | 0 | 1 | 3 | 54 | 49 | 23 | 25 | 23 | 0 | 0 | 0 | 0 | 0 | 200 | 149 | 1 | 1 | 0.964 | 1 | 1 | 0.971 | 0.94 | 1 | 1 | 1 | 1 | 1 | -0.025 |
| SE | 86235 | ENSG00000184634 | MED12 | chrX | + | 71126968 | 71127132 | 71126260 | 71126484 | 71127335 | 71127386 | 138 | 79 | 131 | 148 | 102 | 0 | 0 | 0 | 1 | 0 | 92 | 94 | 55 | 95 | 221 | 0 | 0 | 1 | 4 | 0 | 298 | 149 | 1 | 1 | 1 | 1 | 1 | 0.987 | 1 | 1 | 1 | 0.965 | 0.922 | 1 | 0.02 |
| SE | 86236 | ENSG00000184634 | MED12 | chrX | + | 71132072 | 71132206 | 71131549 | 71131621 | 71132376 | 71132538 | 111 | 33 | 106 | 97 | 116 | 0 | 0 | 0 | 4 | 0 | 65 | 58 | 33 | 98 | 60 | 0 | 0 | 0 | 0 | 0 | 283 | 149 | 1 | 1 | 1 | 1 | 1 | 0.927 | 1 | 1 | 1 | 1 | 1 | 1 | -0.015 |
| SE | 86239 | ENSG00000184634 | MED12 | chrX | + | 71137186 | 71137383 | 71136878 | 71137029 | 71137557 | 71137635 | 42 | 8 | 34 | 17 | 30 | 0 | 0 | 0 | 2 | 0 | 0 | 17 | 4 | 29 | 47 | 0 | 2 | 0 | 0 | 6 | 298 | 149 | 1 | 1 | 1 | 1 | 1 | 0.81 | 1 | NA | 0.81 | 1 | 1 | 0.797 | 0.06 |
| SE | 86241 | ENSG00000134516 | DOCK2 | chr5 | + | 169669287 | 169669328 | 169654402 | 169654486 | 169670541 | 169670597 | 277 | 140 | 174 | 176 | 174 | 1 | 0 | 0 | 0 | 4 | 314 | 261 | 242 | 254 | 195 | 0 | 0 | 5 | 0 | 0 | 190 | 149 | 1 | 1 | 0.995 | 1 | 1 | 1 | 0.972 | 1 | 1 | 0.974 | 1 | 1 | -0.001 |
| SE | 86243 | ENSG00000134516 | DOCK2 | chr5 | + | 169669287 | 169669328 | 169654402 | 169654486 | 169674296 | 169674445 | 62 | 21 | 41 | 27 | 5 | 0 | 0 | 0 | 0 | 0 | 12 | 56 | 18 | 55 | 45 | 0 | 3 | 0 | 0 | 0 | 190 | 149 | 1 | 1 | 1 | 1 | 1 | 1 | 1 | 1 | 0.936 | 1 | 1 | 1 | 0.013 |
| SE | 86246 | ENSG00000134516 | DOCK2 | chr5 | + | 169671077 | 169671174 | 169654402 | 169654486 | 169674296 | 169674445 | 295 | 230 | 214 | 252 | 347 | 0 | 0 | 0 | 0 | 0 | 505 | 396 | 331 | 358 | 224 | 0 | 3 | 0 | 0 | 0 | 246 | 149 | 1 | 1 | 1 | 1 | 1 | 1 | 1 | 1 | 0.988 | 1 | 1 | 1 | 0.002 |
| SE | 86247 | ENSG00000134516 | DOCK2 | chr5 | + | 169671077 | 169671174 | 169654402 | 169654486 | 169681743 | 169681879 | 0 | 3 | 0 | 1 | 0 | 1 | 0 | 0 | 0 | 0 | 0 | 0 | 6 | 7 | 4 | 0 | 4 | 0 | 0 | 0 | 246 | 149 | 1 | 1 | 0 | 1 | NA | 1 | NA | NA | 0 | 1 | 1 | 1 | -0.083 |
| SE | 86248 | ENSG00000134516 | DOCK2 | chr5 | + | 169671077 | 169671174 | 169669287 | 169669328 | 169674296 | 169674445 | 296 | 230 | 215 | 252 | 341 | 2 | 1 | 0 | 0 | 0 | 507 | 404 | 328 | 356 | 220 | 0 | 0 | 1 | 1 | 0 | 246 | 149 | 1 | 1 | 0.989 | 0.993 | 1 | 1 | 1 | 1 | 1 | 0.995 | 0.995 | 1 | -0.002 |
| SE | 86249 | ENSG00000134516 | DOCK2 | chr5 | + | 169674296 | 169674445 | 169654402 | 169654486 | 169681743 | 169681879 | 601 | 565 | 514 | 527 | 653 | 1 | 0 | 0 | 0 | 0 | 1202 | 791 | 752 | 800 | 498 | 0 | 4 | 0 | 0 | 0 | 298 | 149 | 1 | 1 | 0.997 | 1 | 1 | 1 | 1 | 1 | 0.99 | 1 | 1 | 1 | 0.001 |
| SE | 86250 | ENSG00000134516 | DOCK2 | chr5 | + | 169674296 | 169674445 | 169671077 | 169671174 | 169681743 | 169681879 | 1150 | 961 | 873 | 946 | 1195 | 0 | 4 | 0 | 0 | 0 | 2009 | 1410 | 1347 | 1431 | 861 | 0 | 0 | 0 | 0 | 0 | 298 | 149 | 1 | 1 | 1 | 0.992 | 1 | 1 | 1 | 1 | 1 | 1 | 1 | 1 | -0.002 |
| SE | 86252 | ENSG00000134516 | DOCK2 | chr5 | + | 169684195 | 169684350 | 169681743 | 169681879 | 169695802 | 169695938 | 767 | 536 | 649 | 625 | 813 | 0 | 0 | 0 | 1 | 0 | 1006 | 746 | 643 | 777 | 434 | 4 | 6 | 1 | 0 | 0 | 298 | 149 | 1 | 1 | 1 | 1 | 1 | 0.997 | 1 | 0.992 | 0.984 | 0.997 | 1 | 1 | 0.005 |
| SE | 86253 | ENSG00000134516 | DOCK2 | chr5 | + | 169689251 | 169689333 | 169681743 | 169681879 | 169695802 | 169695938 | 364 | 244 | 291 | 300 | 358 | 0 | 0 | 0 | 1 | 0 | 452 | 392 | 428 | 466 | 276 | 4 | 6 | 1 | 0 | 0 | 231 | 149 | 1 | 1 | 1 | 1 | 1 | 0.995 | 1 | 0.986 | 0.977 | 0.996 | 1 | 1 | 0.007 |
| SE | 86254 | ENSG00000134516 | DOCK2 | chr5 | + | 169689251 | 169689333 | 169684195 | 169684350 | 169695802 | 169695938 | 947 | 648 | 826 | 791 | 830 | 2 | 3 | 0 | 2 | 2 | 1247 | 905 | 969 | 1261 | 629 | 4 | 5 | 6 | 12 | 0 | 231 | 149 | 1 | 1 | 0.997 | 0.993 | 1 | 0.996 | 0.996 | 0.995 | 0.992 | 0.99 | 0.985 | 1 | 0.004 |
| SE | 86255 | ENSG00000134516 | DOCK2 | chr5 | + | 169695802 | 169695938 | 169689251 | 169689333 | 169698373 | 169698449 | 1690 | 1177 | 1319 | 1433 | 1804 | 0 | 0 | 0 | 1 | 0 | 2286 | 1793 | 1726 | 1925 | 1076 | 0 | 0 | 0 | 0 | 0 | 285 | 149 | 1 | 1 | 1 | 1 | 1 | 0.999 | 1 | 1 | 1 | 1 | 1 | 1 | 0 |
| SE | 86256 | ENSG00000134516 | DOCK2 | chr5 | + | 169699381 | 169699458 | 169698373 | 169698449 | 169700013 | 169700039 | 1617 | 1108 | 1047 | 1370 | 1472 | 0 | 0 | 0 | 0 | 0 | 2473 | 1836 | 1635 | 1884 | 992 | 0 | 0 | 0 | 3 | 0 | 226 | 149 | 1 | 1 | 1 | 1 | 1 | 1 | 1 | 1 | 1 | 1 | 0.998 | 1 | 0 |
| SE | 86257 | ENSG00000134516 | DOCK2 | chr5 | + | 169702302 | 169702427 | 169700013 | 169700139 | 169708168 | 169708267 | 1335 | 894 | 927 | 1168 | 1088 | 1 | 1 | 0 | 0 | 0 | 2145 | 1570 | 1288 | 1670 | 879 | 0 | 0 | 0 | 0 | 0 | 274 | 149 | 1 | 1 | 0.999 | 0.998 | 1 | 1 | 1 | 1 | 1 | 1 | 1 | 1 | -0.001 |
| SE | 86260 | ENSG00000134516 | DOCK2 | chr5 | + | 169714027 | 169714211 | 169712119 | 169712223 | 169714359 | 169714403 | 2050 | 1487 | 1512 | 1815 | 2237 | 0 | 0 | 0 | 1 | 0 | 3682 | 2508 | 2444 | 2802 | 1779 | 0 | 0 | 0 | 1 | 0 | 298 | 149 | 1 | 1 | 1 | 1 | 1 | 0.999 | 1 | 1 | 1 | 1 | 0.999 | 1 | 0 |
| SE | 86261 | ENSG00000134516 | DOCK2 | chr5 | + | 169714359 | 169714457 | 169714027 | 169714211 | 169716212 | 169716302 | 1103 | 890 | 764 | 938 | 1080 | 0 | 1 | 0 | 3 | 0 | 2234 | 1363 | 1430 | 1565 | 1060 | 0 | 2 | 1 | 3 | 0 | 247 | 149 | 1 | 1 | 1 | 0.998 | 1 | 0.995 | 1 | 1 | 0.998 | 0.999 | 0.997 | 1 | 0 |
| SE | 86262 | ENSG00000134516 | DOCK2 | chr5 | + | 169717383 | 169717484 | 169716212 | 169716302 | 169718656 | 169718791 | 1036 | 849 | 676 | 829 | 1047 | 0 | 0 | 0 | 0 | 0 | 1934 | 1170 | 1219 | 1407 | 902 | 0 | 0 | 0 | 2 | 0 | 250 | 149 | 1 | 1 | 1 | 1 | 1 | 1 | 1 | 1 | 1 | 1 | 0.998 | 1 | 0 |
| SE | 86263 | ENSG00000134516 | DOCK2 | chr5 | + | 169759704 | 169759775 | 169747395 | 169747504 | 169761518 | 169761625 | 973 | 606 | 592 | 858 | 1629 | 0 | 1 | 0 | 0 | 0 | 1729 | 979 | 1060 | 1163 | 716 | 0 | 0 | 1 | 0 | 8 | 220 | 149 | 1 | 1 | 1 | 0.998 | 1 | 1 | 1 | 1 | 1 | 0.999 | 1 | 0.984 | 0.003 |
| SE | 86267 | ENSG00000134516 | DOCK2 | chr5 | + | 169840756 | 169840852 | 169803057 | 169803206 | 169983067 | 169983166 | 565 | 345 | 496 | 445 | 516 | 0 | 3 | 0 | 0 | 0 | 720 | 444 | 531 | 606 | 282 | 0 | 3 | 0 | 0 | 0 | 245 | 149 | 1 | 1 | 1 | 0.986 | 1 | 1 | 1 | 1 | 0.989 | 1 | 1 | 1 | -0.001 |
| SE | 86268 | ENSG00000134516 | DOCK2 | chr5 | + | 169983067 | 169983166 | 169840756 | 169840852 | 169985827 | 169985922 | 569 | 334 | 467 | 452 | 538 | 0 | 0 | 0 | 1 | 0 | 944 | 556 | 583 | 751 | 445 | 0 | 1 | 0 | 3 | 0 | 248 | 149 | 1 | 1 | 1 | 1 | 1 | 0.996 | 1 | 1 | 0.997 | 1 | 0.993 | 1 | 0.001 |
| SE | 86269 | ENSG00000134516 | DOCK2 | chr5 | + | 169997973 | 169998024 | 169996085 | 169996164 | 170008496 | 170008597 | 0 | 0 | 0 | 0 | 0 | 755 | 409 | 509 | 567 | 740 | 0 | 1 | 2 | 0 | 0 | 1235 | 758 | 737 | 1010 | 481 | 200 | 149 | 1 | 1 | 0 | 0 | 0 | 0 | 0 | 0 | 0.001 | 0.002 | 0 | 0 | -0.001 |
| SE | 86272 | ENSG00000134516 | DOCK2 | chr5 | + | 170034398 | 170034555 | 170027862 | 170027948 | 170036514 | 170036555 | 837 | 551 | 657 | 843 | 846 | 0 | 0 | 1 | 1 | 0 | 1586 | 792 | 845 | 1036 | 494 | 0 | 6 | 0 | 0 | 0 | 298 | 149 | 1 | 1 | 1 | 1 | 0.997 | 0.998 | 1 | 1 | 0.985 | 1 | 1 | 1 | 0.002 |
| SE | 86273 | ENSG00000134516 | DOCK2 | chr5 | + | 170034398 | 170034555 | 170027862 | 170027948 | 170041054 | 170041145 | 551 | 411 | 483 | 570 | 659 | 0 | 0 | 0 | 1 | 0 | 1185 | 561 | 637 | 751 | 369 | 0 | 0 | 0 | 0 | 0 | 298 | 149 | 1 | 1 | 1 | 1 | 1 | 0.997 | 1 | 1 | 1 | 1 | 1 | 1 | -0.001 |
| SE | 86274 | ENSG00000134516 | DOCK2 | chr5 | + | 170036514 | 170036555 | 170027862 | 170027948 | 170041054 | 170041145 | 95 | 80 | 76 | 98 | 175 | 0 | 0 | 0 | 1 | 0 | 148 | 148 | 108 | 133 | 96 | 0 | 0 | 0 | 0 | 0 | 190 | 149 | 1 | 1 | 1 | 1 | 1 | 0.987 | 1 | 1 | 1 | 1 | 1 | 1 | -0.003 |
| SE | 86275 | ENSG00000134516 | DOCK2 | chr5 | + | 170036514 | 170036555 | 170034398 | 170034555 | 170041054 | 170041145 | 375 | 222 | 238 | 359 | 351 | 0 | 4 | 0 | 0 | 0 | 548 | 366 | 315 | 414 | 218 | 0 | 0 | 0 | 0 | 0 | 190 | 149 | 1 | 1 | 1 | 0.978 | 1 | 1 | 1 | 1 | 1 | 1 | 1 | 1 | -0.004 |
| SE | 86276 | ENSG00000134516 | DOCK2 | chr5 | + | 170045815 | 170045905 | 170042012 | 170042132 | 170047509 | 170047614 | 460 | 252 | 269 | 352 | 385 | 0 | 4 | 1 | 0 | 0 | 550 | 388 | 385 | 400 | 333 | 0 | 0 | 2 | 0 | 0 | 239 | 149 | 1 | 1 | 1 | 0.975 | 0.994 | 1 | 1 | 1 | 1 | 0.992 | 1 | 1 | -0.005 |
| SE | 86277 | ENSG00000134516 | DOCK2 | chr5 | + | 170067509 | 170067686 | 170056437 | 170057666 | 170069136 | 170069220 | 652 | 330 | 486 | 593 | 754 | 0 | 0 | 0 | 0 | 0 | 838 | 568 | 496 | 638 | 269 | 0 | 0 | 1 | 0 | 0 | 298 | 149 | 1 | 1 | 1 | 1 | 1 | 1 | 1 | 1 | 1 | 0.996 | 1 | 1 | 0.001 |
| SE | 86278 | ENSG00000134516 | DOCK2 | chr5 | + | 170069136 | 170069220 | 170067509 | 170067686 | 170075946 | 170076084 | 345 | 169 | 235 | 270 | 304 | 5 | 0 | 3 | 4 | 2 | 425 | 282 | 301 | 352 | 240 | 0 | 8 | 5 | 3 | 0 | 233 | 149 | 1 | 1 | 0.978 | 1 | 0.98 | 0.977 | 0.99 | 1 | 0.958 | 0.975 | 0.987 | 1 | 0.001 |
| SE | 86281 | ENSG00000134516 | DOCK2 | chr5 | + | 170075946 | 170076084 | 170069136 | 170069220 | 170077709 | 170077837 | 626 | 280 | 372 | 456 | 620 | 0 | 0 | 0 | 0 | 0 | 764 | 474 | 471 | 604 | 398 | 0 | 1 | 7 | 5 | 0 | 287 | 149 | 1 | 1 | 1 | 1 | 1 | 1 | 1 | 1 | 0.996 | 0.972 | 0.984 | 1 | 0.01 |
| SE | 86282 | ENSG00000134516 | DOCK2 | chr5 | + | 170077709 | 170077837 | 170069136 | 170069220 | 170078974 | 170079146 | 225 | 112 | 207 | 177 | 236 | 1 | 0 | 1 | 0 | 0 | 396 | 207 | 253 | 294 | 144 | 5 | 0 | 1 | 0 | 0 | 277 | 149 | 1 | 1 | 0.992 | 1 | 0.991 | 1 | 1 | 0.977 | 1 | 0.993 | 1 | 1 | 0.003 |
| SE | 86283 | ENSG00000134516 | DOCK2 | chr5 | + | 170080162 | 170080283 | 170078974 | 170079146 | 170081841 | 170081984 | 435 | 193 | 336 | 320 | 208 | 2 | 0 | 0 | 4 | 0 | 450 | 287 | 275 | 297 | 214 | 0 | 2 | 2 | 0 | 0 | 270 | 149 | 1 | 1 | 0.992 | 1 | 1 | 0.978 | 1 | 1 | 0.988 | 0.987 | 1 | 1 | -0.001 |
| SE | 86285 | ENSG00000243364 | EFNA4 | chr1 | + | 155067371 | 155067440 | 155066729 | 155067016 | 155069018 | 155069538 | 10 | 1 | 7 | 1 | 0 | 2 | 0 | 0 | 0 | 0 | 0 | 10 | 8 | 5 | 20 | 0 | 0 | 0 | 0 | 0 | 218 | 149 | 1 | 1 | 0.774 | 1 | 1 | 1 | NA | NA | 1 | 1 | 1 | 1 | -0.056 |
| SE | 86299 | ENSG00000230162 | CT45A11P | chrX | - | 135944795 | 135945034 | 135942477 | 135943103 | 135946964 | 135947139 | 2 | 2 | 3 | 0 | 1 | 0 | 0 | 1 | 0 | 0 | 0 | 1 | 0 | 5 | 32 | 0 | 0 | 0 | 0 | 0 | 298 | 149 | 1 | 1 | 1 | 1 | 0.6 | NA | 1 | NA | 1 | NA | 1 | 1 | -0.1 |
| SE | 86300 | ENSG00000147224 | PRPS1 | chrX | + | 107636587 | 107636679 | 107628484 | 107628750 | 107639294 | 107639478 | 1 | 0 | 0 | 0 | 0 | 155 | 55 | 163 | 176 | 120 | 0 | 0 | 0 | 0 | 0 | 112 | 165 | 115 | 138 | 139 | 241 | 149 | 1 | 1 | 0.004 | 0 | 0 | 0 | 0 | 0 | 0 | 0 | 0 | 0 | 0.001 |
| SE | 86302 | ENSG00000147224 | PRPS1 | chrX | + | 107639294 | 107639478 | 107628427 | 107628750 | 107640901 | 107640911 | 272 | 106 | 294 | 353 | 265 | 1 | 0 | 0 | 1 | 0 | 214 | 272 | 222 | 233 | 263 | 0 | 0 | 0 | 0 | 0 | 298 | 149 | 1 | 1 | 0.993 | 1 | 1 | 0.994 | 1 | 1 | 1 | 1 | 1 | 1 | -0.003 |
| SE | 86303 | ENSG00000147224 | PRPS1 | chrX | + | 107639294 | 107639478 | 107628427 | 107628750 | 107645176 | 107645350 | 155 | 55 | 163 | 176 | 120 | 0 | 0 | 0 | 0 | 0 | 112 | 165 | 115 | 138 | 139 | 0 | 0 | 0 | 4 | 0 | 298 | 149 | 1 | 1 | 1 | 1 | 1 | | 1 | 1 | 1 | 1 | 0.945 | 1 | 0.011 |
| SE | 86304 | ENSG00000147224 | PRPS1 | chrX | + | 107640901 | 107641000 | 107628515 | 107628750 | 107645176 | 107645569 | 0 | 0 | 0 | 2 | 1 | 0 | 0 | 0 | 0 | 0 | 0 | 0 | 0 | 4 | 0 | 0 | 0 | 0 | 4 | 0 | 248 | 149 | 1 | 1 | NA | NA | NA | 1 | 1 | NA | NA | NA | 0.375 | NA | 0.625 |
| SE | 86305 | ENSG00000147224 | PRPS1 | chrX | + | 107640901 | 107641000 | 107639294 | 107639478 | 107642365 | 107642490 | 170 | 105 | 201 | 224 | 186 | 1 | 0 | 0 | 0 | 0 | 131 | 166 | 167 | 130 | 152 | 0 | 0 | 0 | 0 | 6 | 248 | 149 | 1 | 1 | 0.99 | 1 | 1 | 1 | 1 | 1 | 1 | 1 | 1 | 0.938 | 0.01 |
| SE | 86309 | ENSG00000147224 | PRPS1 | chrX | + | 107642365 | 107642490 | 107640901 | 107641000 | 107645176 | 107645350 | 186 | 139 | 253 | 227 | 281 | 0 | 0 | 1 | 1 | 1 | 126 | 192 | 169 | 135 | 134 | 0 | 0 | 0 | 4 | 0 | 274 | 149 | 1 | 1 | 1 | 1 | 0.993 | 0.992 | 0.993 | 1 | 1 | 1 | 0.948 | 1 | 0.006 |
| SE | 86310 | ENSG00000147224 | PRPS1 | chrX | + | 107643222 | 107643286 | 107642365 | 107642490 | 107645176 | 107645350 | 1 | 0 | 0 | 0 | 0 | 111 | 76 | 143 | 144 | 200 | 0 | 0 | 0 | 0 | 0 | 83 | 119 | 102 | 80 | 63 | 213 | 149 | 1 | 1 | 0.006 | 0 | 0 | 0 | 0 | 0 | 0 | 0 | 0 | 0 | 0.001 |
| SE | 86316 | ENSG00000176092 | CRYBG2 | chr1 | - | 26338011 | 26338047 | 26337537 | 26337674 | 26338350 | 26338439 | 0 | 0 | 0 | 3 | 0 | 0 | 0 | 0 | 0 | 0 | 0 | 2 | 0 | 0 | 0 | 0 | 3 | 0 | 0 | 0 | 185 | 149 | 1 | 1 | NA | NA | NA | 1 | NA | NA | 0.349 | NA | NA | NA | 0.651 |
| SE | 86318 | ENSG00000213965 | NUDT19 | chr19 | + | 32709184 | 32709392 | 32691820 | 32692674 | 32711751 | 32713792 | 41 | 24 | 36 | 33 | 48 | 0 | 1 | 0 | 0 | 0 | 50 | 24 | 24 | 69 | 56 | 0 | 0 | 0 | 0 | 0 | 298 | 149 | 1 | 1 | 1 | 0.923 | 1 | 1 | 1 | 1 | 1 | 1 | 1 | 1 | -0.015 |
| SE | 86320 | ENSG00000135048 | CEMIP2 | chr9 | - | 71685742 | 71685846 | 71683365 | 71685393 | 71694508 | 71694607 | 239 | 122 | 89 | 165 | 94 | 0 | 0 | 2 | 0 | 0 | 30 | 153 | 237 | 127 | 30 | 0 | 0 | 3 | 0 | 0 | 253 | 149 | 1 | 1 | 1 | 1 | 0.963 | 1 | 1 | 1 | 1 | 0.979 | 1 | 1 | -0.003 |
| SE | 86321 | ENSG00000135048 | CEMIP2 | chr9 | - | 71690091 | 71690246 | 71683365 | 71685393 | 71694508 | 71694607 | 54 | 25 | 34 | 61 | 32 | 0 | 0 | 2 | 0 | 0 | 41 | 50 | 53 | 71 | 11 | 0 | 0 | 3 | 0 | 0 | 298 | 149 | 1 | 1 | 1 | 1 | 0.895 | 1 | 1 | 1 | 1 | 0.898 | 1 | 1 | -0.001 |
| SE | 86323 | ENSG00000135048 | CEMIP2 | chr9 | - | 71697984 | 71698204 | 71694508 | 71694607 | 71700641 | 71700824 | 247 | 114 | 128 | 129 | 202 | 0 | 0 | 0 | 0 | 0 | 146 | 168 | 370 | 205 | 26 | 0 | 6 | 1 | 0 | 0 | 298 | 149 | 1 | 1 | 1 | 1 | 1 | 1 | 1 | 1 | 0.933 | 0.995 | 1 | 1 | 0.014 |
| SE | 86324 | ENSG00000135048 | CEMIP2 | chr9 | - | 71717947 | 71718079 | 71716516 | 71716552 | 71722426 | 71722515 | 442 | 207 | 255 | 307 | 290 | 0 | 0 | 0 | 0 | 0 | 205 | 534 | 896 | 236 | 22 | 0 | 1 | 0 | 0 | 0 | 281 | 149 | 1 | 1 | 1 | 1 | 1 | 1 | 1 | 1 | 0.996 | 1 | 1 | 1 | 0.001 |
| SE | 86325 | ENSG00000135048 | CEMIP2 | chr9 | - | 71729844 | 71729914 | 71725580 | 71725709 | 71730047 | 71730253 | 656 | 304 | 443 | 479 | 413 | 0 | 0 | 0 | 0 | 0 | 362 | 657 | 1316 | 470 | 73 | 0 | 0 | 2 | 0 | 0 | 219 | 149 | 1 | 1 | 1 | 1 | 1 | 1 | 1 | 1 | 1 | 0.998 | 1 | 1 | 0 |
| SE | 86326 | ENSG00000135048 | CEMIP2 | chr9 | - | 71730047 | 71730253 | 71725580 | 71725709 | 71730704 | 71730914 | 272 | 107 | 159 | 202 | 182 | 0 | 0 | 0 | 0 | 0 | 60 | 285 | 459 | 166 | 28 | 0 | 1 | 1 | 0 | 0 | 298 | 149 | 1 | 1 | 1 | 1 | 1 | 1 | 1 | 1 | 0.993 | 0.996 | 1 | 1 | 0.002 |
| SE | 86331 | ENSG00000135048 | CEMIP2 | chr9 | - | 71740062 | 71740232 | 71732350 | 71732520 | 71768356 | 71768386 | 23 | 16 | 9 | 20 | 23 | 0 | 0 | 0 | 0 | 0 | 0 | 79 | 133 | 25 | 29 | 0 | 2 | 2 | 0 | 0 | 298 | 149 | 1 | 1 | 1 | 1 | 1 | 1 | 1 | NA | 0.952 | 0.971 | 1 | 1 | 0.019 |
| SE | 86333 | ENSG00000135048 | CEMIP2 | chr9 | - | 71745017 | 71745579 | 71732350 | 71732520 | 71750042 | 71750385 | 0 | 0 | 1 | 0 | 0 | 0 | 0 | 0 | 0 | 6 | 0 | 8 | 14 | 7 | 0 | 11 | 4 | 4 | 0 | 0 | 298 | 149 | 1 | 1 | NA | NA | 1 | NA | 0 | 0 | 0.5 | 0.636 | 1 | NA | -0.034 |
| SE | 86334 | ENSG00000135048 | CEMIP2 | chr9 | - | 71745017 | 71745579 | 71740062 | 71740232 | 71746200 | 71746341 | 456 | 199 | 243 | 286 | 199 | 0 | 2 | 0 | 0 | 0 | 130 | 493 | 515 | 233 | 47 | 0 | 1 | 7 | 0 | 0 | 298 | 149 | 1 | 1 | 1 | 0.98 | 1 | 1 | 1 | 1 | 0.996 | 0.974 | 1 | 1 | 0.002 |
| SE | 86337 | ENSG00000135048 | CEMIP2 | chr9 | - | 71746200 | 71746341 | 71745017 | 71745579 | 71750042 | 71750385 | 545 | 264 | 257 | 316 | 307 | 0 | 0 | 0 | 0 | 0 | 304 | 450 | 456 | 293 | 105 | 0 | 8 | 0 | 0 | 0 | 290 | 149 | 1 | 1 | 1 | 1 | 1 | 1 | 1 | 1 | 0.967 | 1 | 1 | 1 | 0.007 |
| SE | 86338 | ENSG00000135048 | CEMIP2 | chr9 | - | 71750042 | 71750385 | 71732350 | 71732520 | 71768356 | 71768386 | 55 | 13 | 33 | 30 | 35 | 0 | 0 | 0 | 0 | 0 | 22 | 64 | 66 | 43 | 25 | 0 | 2 | 2 | 0 | 0 | 298 | 149 | 1 | 1 | 1 | 1 | 1 | 1 | 1 | 1 | 0.941 | 0.943 | 1 | 1 | 0.023 |
| SE | 86344 | ENSG00000074966 | TXK | chr4 | - | 48076401 | 48076466 | 48071516 | 48071674 | 48079911 | 48080066 | 83 | 41 | 50 | 52 | 54 | 0 | 0 | 0 | 1 | 0 | 0 | 2 | 32 | 4 | 0 | 0 | 0 | 2 | 0 | 0 | 214 | 149 | 1 | 1 | 1 | 1 | 1 | 0.973 | 1 | NA | 1 | 0.918 | 1 | NA | 0.022 |
| SE | 86345 | ENSG00000074966 | TXK | chr4 | - | 48089749 | 48089824 | 48086465 | 48086637 | 48104900 | 48104955 | 57 | 30 | 47 | 40 | 36 | 2 | 0 | 0 | 0 | 0 | 0 | 17 | 34 | 12 | 0 | 0 | 0 | 0 | 0 | 0 | 224 | 149 | 1 | 1 | 0.95 | 1 | 1 | 1 | 1 | NA | 1 | 1 | 1 | NA | -0.01 |
| SE | 86347 | ENSG00000074966 | TXK | chr4 | - | 48095142 | 48095222 | 48089384 | 48089824 | 48104900 | 48104955 | 142 | 123 | 170 | 90 | 148 | 1 | 0 | 0 | 0 | 0 | 38 | 64 | 122 | 35 | 24 | 0 | 0 | 2 | 0 | 0 | 229 | 149 | 1 | 1 | 0.989 | 1 | 1 | 1 | 1 | 1 | 1 | 0.975 | 1 | 1 | 0.003 |
| SE | 86350 | ENSG00000074966 | TXK | chr4 | - | 48112306 | 48112512 | 48110537 | 48110603 | 48113206 | 48113309 | 573 | 354 | 451 | 382 | 422 | 0 | 0 | 0 | 1 | 0 | 138 | 119 | 409 | 106 | 100 | 5 | 0 | 0 | 0 | 0 | 298 | 149 | 1 | 1 | 1 | 1 | 1 | 0.995 | 1 | 0.932 | 1 | 1 | 1 | 1 | 0.013 |
| SE | 86351 | ENSG00000074966 | TXK | chr4 | - | 48114347 | 48114402 | 48113206 | 48113309 | 48134154 | 48134229 | 147 | 140 | 127 | 78 | 156 | 2 | 2 | 6 | 2 | 5 | 19 | 32 | 92 | 46 | 0 | 10 | 4 | 12 | 1 | 0 | 204 | 149 | 1 | 1 | 0.982 | 0.981 | 0.939 | 0.966 | 0.958 | 0.581 | 0.854 | 0.848 | 0.971 | NA | 0.152 |
| SE | 86353 | ENSG00000185591 | SP1 | chr12 | + | 53406584 | 53406753 | 53382109 | 53383622 | 53409361 | 53409561 | 376 | 237 | 328 | 350 | 352 | 0 | 0 | 0 | 0 | 0 | 534 | 372 | 323 | 519 | 209 | 0 | 0 | 1 | 0 | 0 | 298 | 149 | 1 | 1 | 1 | 1 | 1 | 1 | 1 | 1 | 1 | 0.994 | 1 | 1 | 0.001 |
| SE | 86354 | ENSG00000185591 | SP1 | chr12 | + | 53409361 | 53409561 | 53406584 | 53406753 | 53410926 | 53416446 | 514 | 346 | 461 | 553 | 654 | 2 | 0 | 0 | 0 | 0 | 890 | 628 | 489 | 800 | 385 | 0 | 0 | 0 | 0 | 0 | 298 | 149 | 1 | 1 | 0.992 | 1 | 1 | 1 | 1 | 1 | 1 | 1 | 1 | 1 | -0.002 |
| SE | 86357 | ENSG00000136541 | ERMN | chr2 | - | 157324117 | 157324269 | 157321738 | 157321791 | 157324669 | 157324762 | 0 | 0 | 0 | 0 | 0 | 11 | 17 | 9 | 8 | 11 | 0 | 0 | 2 | 0 | 0 | 0 | 16 | 40 | 10 | 20 | 298 | 149 | 1 | 1 | 0 | 0 | 0 | 0 | 0 | NA | 0 | 0.024 | 0 | 0 | -0.006 |

|||||||||||||||||||||||||||||||||||||||||||||||
|---|---|---|---|---|---|---|---|---|---|---|---|---|---|---|---|---|---|---|---|---|---|---|---|---|---|---|---|---|---|---|---|---|---|---|---|---|---|---|---|---|---|---|---|---|---|---|
|SE|86358|ENSG00000136541|ERMN|chr2|-|157324146|157324269|157321684|157321791|157324669|157324762|0|0|0|0|0|11|17|9|8|11|0|0|2|0|0|0|16|40|10|20|272|149|1|1|0|0|0|0|0|NA|0|0.027|0|0|-0.007|
|SE|86360|ENSG00000035681|NSMAF|chr8|-|58585897|58586000|58585651|58585761|58586457|58586608|228|209|225|171|241|0|0|4|2|0|162|166|172|155|135|0|2|1|0|0|252|149|1|1|1|1|0.971|0.981|1|1|0.98|0.99|1|1|-0.004|
|SE|86363|ENSG00000035681|NSMAF|chr8|-|58603209|58603386|58602057|58602137|58605926|58606035|249|165|315|248|343|0|0|2|0|0|322|208|146|171|95|0|0|0|0|0|298|149|1|1|1|1|0.987|1|1|1|1|1|1|1|-0.003|
|SE|86365|ENSG00000035681|NSMAF|chr8|-|58607768|58607840|58603209|58603386|58609603|58609733|21|17|28|28|58|0|0|1|0|0|11|14|11|14|1|0|0|0|0|0|221|149|1|1|1|1|0.95|1|1|1|1|1|1|1|-0.01|
|SE|86366|ENSG00000035681|NSMAF|chr8|-|58607768|58607840|58605926|58606035|58623219|58623272|86|60|88|76|159|0|0|0|1|0|118|52|37|76|13|0|0|0|0|0|221|149|1|1|1|1|1|0.981|1|1|1|1|1|1|-0.004|
|SE|86367|ENSG00000035681|NSMAF|chr8|-|58609603|58609733|58607768|58607840|58623219|58623272|181|117|184|184|300|1|0|0|0|0|136|124|74|112|18|0|0|0|0|0|279|149|1|1|0.99|1|1|1|1|1|1|1|1|1|-0.002|
|SE|86368|ENSG00000035681|NSMAF|chr8|-|58631495|58631546|58623708|58623780|58635304|58635372|50|32|35|48|73|3|0|2|0|0|34|9|24|19|9|0|0|0|0|2|200|149|1|1|0.925|1|0.929|1|1|1|1|1|1|0.77|0.017|
|SE|86369|ENSG00000035681|NSMAF|chr8|-|58635188|58635225|58631364|58631546|58635304|58635372|173|78|109|126|203|0|0|0|0|0|121|101|80|47|21|0|0|1|0|0|186|149|1|1|1|1|1|1|1|1|1|0.985|1|1|0.003|
|SE|86370|ENSG00000035681|NSMAF|chr8|-|58637233|58637389|58635467|58635546|58639967|58640102|7|2|4|4|0|0|0|0|0|0|0|0|5|4|14|0|1|0|0|0|298|149|1|1|1|1|1|1|NA|NA|0|1|1|1|0.25|
|SE|86372|ENSG00000035681|NSMAF|chr8|-|58639967|58640102|58635467|58635546|58642983|58643073|0|0|0|0|0|124|47|82|89|64|0|1|0|0|0|83|53|58|74|22|284|149|1|1|0|0|0|0|0|0|0.01|0|0|0|-0.002|
|SE|86373|ENSG00000035681|NSMAF|chr8|-|58639967|58640102|58637233|58637389|58642983|58643073|1|0|0|2|0|10|3|1|3|1|0|0|0|3|0|0|0|3|1|0|284|149|1|1|0.05|0|0|0.259|0|NA|NA|0|0.611|NA|-0.244|
|SE|86379|ENSG00000123562|MORF4L2|chrX|-|103678498|103678600|103675495|103677051|103684680|103684729|292|253|314|340|551|2|1|0|2|3|388|213|315|267|83|0|0|0|0|0|251|149|1|1|0.989|0.993|1|0.99|0.991|1|1|1|1|1|-0.007|
|SE|86381|ENSG00000123562|MORF4L2|chrX|-|103678498|103678620|103676825|103677051|103684680|103684729|282|245|309|317|500|2|1|0|2|3|390|187|304|247|82|0|0|0|0|0|271|149|1|1|0.987|0.993|1|0.989|0.989|1|1|1|1|1|-0.008|
|SE|86383|ENSG00000123562|MORF4L2|chrX|-|103678498|103678651|103675497|103677051|103684680|103684729|302|272|338|339|574|2|1|0|2|3|408|210|331|265|82|0|0|0|0|0|298|149|1|1|0.987|0.993|1|0.988|0.99|1|1|1|1|1|-0.008|
|SE|86391|ENSG00000278175|GLIDR|chr9|-|39809485|39809615|39803495|39803859|39809730|39809946|4|2|5|7|8|1|1|1|0|2|0|9|0|1|30|9|5|0|5|13|279|149|1|1|0.681|0.516|0.728|1|0.681|0|0.49|NA|0.097|0.552|0.436|
|SE|86393|ENSG00000102007|PLP2|chrX|+|49173387|49173483|49173128|49173281|49174334|49174425|558|348|515|488|622|0|0|0|0|0|1272|784|604|1212|1734|0|0|0|0|8|245|149|1|1|1|1|1|1|1|1|1|1|1|0.992|0.002|
|SE|86397|ENSG00000102001|CACNA1F|chrX|-|49217897|49218005|49217754|49217807|49218454|49218542|1|0|3|2|7|1|0|0|0|2|0|4|0|8|8|0|0|0|0|0|257|149|1|1|0.367|NA|1|1|0.67|NA|1|NA|1|1|-0.241|
|SE|86398|ENSG00000102001|CACNA1F|chrX|-|49224760|49224986|49222928|49223136|49225908|49226069|2|0|1|0|0|0|0|1|0|0|0|2|0|0|14|0|2|0|0|7|298|149|1|1|1|NA|0.333|NA|NA|NA|0.333|NA|NA|0.5|0.25|
|SE|86399|ENSG00000102001|CACNA1F|chrX|-|49225908|49226069|49222928|49223136|49226216|49226243|0|0|2|0|0|0|0|0|0|0|0|2|0|0|16|0|0|2|0|0|298|149|1|1|NA|NA|1|NA|NA|NA|1|0|NA|1|0.333|
|SE|86401|ENSG00000131381|RBSN|chr3|-|15075046|15075116|15074904|15074930|15075289|15075417|5|0|3|8|0|0|0|0|0|1|12|16|1|7|0|0|0|0|0|0|219|149|1|1|1|NA|1|1|0|1|1|1|1|NA|-0.25|
|SE|86403|ENSG00000131381|RBSN|chr3|-|15075289|15075417|15074904|15074930|15075605|15075710|0|0|0|0|1|21|9|34|24|21|0|0|0|0|0|32|17|19|33|17|277|149|1|1|0|0|0|0|0.025|0|0|0|0|0|0.005|
|SE|86406|ENSG00000131381|RBSN|chr3|-|15077061|15077164|15075605|15075710|15078074|15078161|76|18|51|88|102|0|0|0|0|0|122|77|86|138|165|0|0|0|0|5|252|149|1|1|1|1|1|1|1|1|1|1|1|0.951|0.01|
|SE|86407|ENSG00000131381|RBSN|chr3|-|15078074|15078161|15077061|15077164|15080731|15080802|58|13|39|59|85|0|0|0|0|0|105|62|65|88|130|0|0|0|1|0|236|149|1|1|1|1|1|1|1|1|1|1|0.982|1|0.004|
|SE|86409|ENSG00000131381|RBSN|chr3|-|15090398|15090539|15085860|15085961|15095972|15096288|67|37|94|83|109|1|0|1|2|1|98|110|90|113|109|8|4|0|2|2|290|149|1|1|0.972|1|0.98|0.955|0.982|0.863|0.934|1|0.967|0.966|0.032|
|SE|86411|ENSG00000131381|RBSN|chr3|-|15096534|15096710|15095670|15096288|15099034|15099143|24|10|24|33|49|0|0|0|4|0|17|37|29|24|32|0|0|3|0|0|298|149|1|1|1|1|1|0.805|1|1|1|0.829|1|1|-0.005|
|SE|86419|ENSG00000196092|PAX5|chr9|-|36882003|36882105|36846842|36846929|36966548|36966724|8|1|2|10|7|0|0|1|0|0|0|15|0|1|12|0|0|1|0|0|251|149|1|1|1|1|0.543|1|1|NA|1|0|1|1|0.159|
|SE|86421|ENSG00000196092|PAX5|chr9|-|36923354|36923484|36846842|36846929|36966548|36966724|25|7|9|28|16|0|0|1|0|0|2|32|13|17|44|0|0|1|0|0|279|149|1|1|1|1|0.828|1|1|1|1|0.874|1|1|-0.009|
|SE|86423|ENSG00000196092|PAX5|chr9|-|36923354|36923484|36882003|36882105|36966548|36966724|35|17|19|43|23|2|0|0|0|0|15|52|17|34|61|0|0|0|0|0|279|149|1|1|0.903|1|1|1|1|1|1|1|1|1|-0.019|
|SE|86429|ENSG00000196092|PAX5|chr9|-|37014996|37015194|37006472|37006537|37020635|37020801|144|67|105|129|174|0|0|0|0|0|200|327|56|136|270|0|0|0|4|0|298|149|1|1|1|1|1|1|1|1|1|1|0.944|1|0.011|
|SE|86432|ENSG00000023839|ABCC2|chr10|+|99796390|99796853|99794412|99794468|99797096|99797331|0|0|0|1|0|1|5|2|7|0|0|0|0|0|0|2|8|4|0|0|298|149|1|1|0|0|0|0.067|NA|0|0|0|NA|NA|0.017|
|SE|86433|ENSG00000023839|ABCC2|chr10|+|99797096|99797331|99794412|99794468|99799206|99799220|1|5|3|8|0|0|0|0|0|0|2|8|4|0|0|0|0|2|0|0|298|149|1|1|1|1|1|1|NA|1|1|0.5|NA|NA|0.167|
|SE|86434|ENSG00000023839|ABCC2|chr10|+|99819088|99819269|99818789|99818957|99830306|99830433|15|4|4|10|5|0|0|0|0|0|0|6|6|4|10|0|0|4|0|0|298|149|1|1|1|1|1|1|1|NA|1|0.429|1|1|0.143|
|SE|86435|ENSG00000023839|ABCC2|chr10|+|99830715|99830851|99830306|99830433|99831610|99831830|16|1|9|6|10|0|0|0|0|0|3|13|6|0|4|0|4|0|0|0|285|149|1|1|1|1|1|1|1|1|0.63|1|NA|1|0.093|
|SE|86436|ENSG00000023839|ABCC2|chr10|+|99841966|99842093|99836090|99836290|99843798|99843900|4|1|3|5|2|1|0|0|0|0|0|0|3|2|0|0|0|0|0|0|276|149|1|1|0.683|1|1|1|1|NA|NA|1|1|NA|-0.063|
|SE|86439|ENSG00000049769|PPP1R3F|chrX|+|49285833|49287727|49281980|49282063|49299275|49299435|6|7|14|3|13|4|1|1|7|7|0|5|0|3|7|1|0|0|0|0|298|149|1|1|0.429|0.778|0.875|0.176|0.481|0|1|NA|1|1|-0.202|
|SE|86441|ENSG00000147118|ZNF182|chrX|-|47988523|47988617|47983284|47983411|48002594|48002653|4|8|15|17|6|15|2|11|7|11|0|0|8|0|0|0|1|13|0|0|243|149|1|1|0.141|0.71|0.455|0.598|0.251|NA|0|0.274|NA|NA|0.294|
|SE|86445|ENSG00000104921|FCER2|chr19|-|7697012|7697075|7696824|7696914|7697235|7697298|141|49|102|55|125|1|1|0|0|2|129|54|13|61|206|0|0|0|0|0|212|149|1|1|0.99|0.972|1|1|0.978|1|1|1|1|1|-0.012|
|SE|86447|ENSG00000102103|PQBP1|chrX|+|48898491|48898576|48897480|48898093|48901189|48901301|112|45|105|107|114|0|0|0|1|0|184|153|65|82|256|0|0|0|0|0|234|149|1|1|1|1|1|0.986|1|1|1|1|1|1|-0.003|
|SE|86448|ENSG00000102103|PQBP1|chrX|+|48898491|48898576|48897911|48898082|48901189|48901301|174|104|169|167|224|2|0|0|0|1|255|228|123|104|350|0|0|0|0|0|234|149|1|1|0.982|1|1|1|0.993|1|1|1|1|1|-0.005|
|SE|86449|ENSG00000102103|PQBP1|chrX|+|48898491|48898576|48897939|48898014|48901189|48901301|138|60|108|135|161|1|0|0|0|1|212|140|76|76|197|3|0|1|0|0|234|149|1|1|0.989|1|1|1|0.99|0.978|1|0.98|1|1|0.004|
|SE|86454|ENSG00000102103|PQBP1|chrX|+|48902232|48902795|48901929|48902042|48902927|48903084|236|146|167|208|177|0|0|0|0|0|318|142|167|138|250|1|0|0|0|0|298|149|1|1|1|1|1|1|1|0.994|1|1|1|1|0.001|
|SE|86455|ENSG00000102103|PQBP1|chrX|+|48902731|48902795|48901929|48902042|48902927|48903056|55|34|47|43|31|0|0|0|0|0|90|27|34|12|28|1|0|0|0|0|213|149|1|1|1|1|1|1|1|0.984|1|1|1|1|0.003|
|SE|86458|ENSG00000067221|STOML1|chr15|-|73989029|73989257|73988665|73988802|73990350|73990457|6|9|2|0|2|0|0|0|1|0|0|1|0|6|0|0|0|0|0|0|298|149|1|1|1|1|1|0|1|NA|1|NA|1|NA|-0.2|
|SE|86459|ENSG00000067221|STOML1|chr15|-|73989107|73989257|73988598|73988802|73990350|73990457|9|10|2|0|2|0|0|0|1|0|0|3|0|10|0|0|0|0|0|0|298|149|1|1|1|1|1|0|1|NA|1|NA|1|NA|-0.2|
|SE|86460|ENSG00000067221|STOML1|chr15|-|73990814|73990945|73990350|73990457|73992090|73992230|0|1|2|0|0|1|4|1|1|0|0|0|0|0|0|0|0|0|3|0|280|149|1|1|0|0.117|0.516|0|NA|NA|NA|NA|0|NA|0.158|
|SE|86464|ENSG00000215021|PHB2|chr12|-|6967170|6967335|6966423|6966500|6967895|6967962|193|117|198|190|450|0|0|1|1|0|296|128|135|185|231|0|0|0|0|0|298|149|1|1|1|1|0.99|0.99|1|1|1|1|1|1|-0.004|
|SE|86473|ENSG00000101966|XIAP|chrX|+|123860166|123860293|123859723|123859905|123885630|123885711|18|4|10|10|0|2|0|3|1|1|0|2|9|2|0|0|3|1|1|0|276|149|1|1|0.829|1|0.643|0.844|0|NA|0.265|0.829|0.519|NA|0.126|
|SE|86474|ENSG00000101966|XIAP|chrX|+|123860166|123860293|123859723|123859905|123888618|123888718|38|18|14|40|1|0|0|0|10|5|0|3|1|12|0|0|10|7|0|0|276|149|1|1|1|1|1|0.683|0.097|NA|0.139|0.072|1|NA|0.352|
|SE|86478|ENSG00000101966|XIAP|chrX|+|123891237|123891309|123888618|123888718|123892730|123892773|195|157|137|204|165|0|0|2|2|0|63|148|139|97|4|0|0|0|3|0|221|149|1|1|1|1|0.979|0.986|1|1|1|1|0.956|1|0.002|
|SE|86479|ENSG00000101966|XIAP|chrX|+|123891237|123891316|123888618|123888718|123892730|123892773|426|331|270|381|453|0|0|2|2|0|63|230|298|180|7|0|0|0|3|0|228|149|1|1|1|1|0.989|0.992|1|1|1|1|0.975|1|0.001|
|SE|86480|ENSG00000101966|XIAP|chrX|+|123892730|123892773|123891237|123891316|123900492|123900566|246|182|144|185|306|3|0|0|0|0|11|84|164|90|3|0|0|0|0|0|192|149|1|1|0.985|1|1|1|1|1|1|1|1|1|-0.003|
|SE|86486|ENSG00000071889|FAM3A|chrX|-|154508473|154508597|154508288|154508347|154511847|154511871|66|24|87|37|20|0|0|0|0|1|71|38|19|8|67|0|0|0|0|0|273|149|1|1|1|1|1|1|0.916|1|1|1|1|1|-0.017|
|SE|86487|ENSG00000071889|FAM3A|chrX|-|154508473|154508652|154508288|154508347|154511847|154511871|16|10|39|29|21|0|0|0|0|1|40|27|13|4|40|0|0|0|0|0|298|149|1|1|1|1|1|1|0.913|1|1|1|1|1|-0.017|
|SE|86488|ENSG00000071889|FAM3A|chrX|-|154508473|154508811|154508288|154508347|154511847|154511871|21|11|37|27|13|0|0|0|0|1|40|28|12|4|40|0|0|0|0|0|298|149|1|1|1|1|1|1|0.867|1|1|1|1|1|-0.027|
|SE|86492|ENSG00000185485|SDHAP1|chr3|-|195960299|195960423|195959747|195960086|195963299|195963421|0|2|0|0|2|33|28|24|27|32|0|0|0|0|0|20|16|31|9|41|273|149|1|1|0|0.038|0|0|0.033|0|0|0|0|0|0.014|
|SE|86496|ENSG00000185485|SDHAP1|chr3|-|195984111|195984276|195982161|195982310|195985626|195985789|25|38|80|63|41|0|0|0|0|0|79|71|61|76|80|0|0|0|0|2|298|149|1|1|1|1|1|1|1|1|1|1|1|0.952|0.01|
|SE|86497|ENSG00000185485|SDHAP1|chr3|-|195984111|195984276|195982161|195982310|195990189|195990258|14|19|49|42|26|0|0|1|0|0|18|46|33|29|27|0|0|0|0|0|298|149|1|1|1|1|0.961|1|1|1|1|1|1|1|-0.008|
|SE|86500|ENSG00000067992|PDK3|chrX|+|24494741|24494883|24465269|24465561|24498828|24498900|115|107|87|101|167|1|0|0|1|0|128|93|88|90|273|0|0|2|0|0|291|149|1|1|0.983|1|1|0.981|1|1|1|0.957|1|1|0.001|
|SE|86501|ENSG00000067992|PDK3|chrX|+|24518932|24519010|24498828|24498900|24526197|24526274|35|24|20|40|53|0|0|0|0|0|8|19|15|12|26|0|1|0|0|0|227|149|1|1|1|1|1|1|1|1|0.926|1|1|1|0.015|
|SE|86512|ENSG00000189221|MAOA|chrX|+|43657861|43657969|43654906|43656414|43683512|43683607|0|0|0|0|0|0|0|0|0|4|0|0|12|0|0|0|26|33|2|0|257|149|1|1|NA|NA|NA|NA|0|NA|0|0.174|0|NA|-0.058|
|SE|86513|ENSG00000189221|MAOA|chrX|+|43711871|43711976|43693290|43693428|43712704|43712796|0|3|0|0|0|0|0|0|0|0|16|13|77|6|0|0|0|2|0|0|254|149|1|1|NA|1|NA|NA|NA|1|1|0.958|1|NA|0.01|
|SE|86514|ENSG00000189221|MAOA|chrX|+|43728172|43728314|43712704|43712796|43731240|43731382|0|1|0|0|0|0|0|0|0|0|0|0|67|6|0|0|0|3|3|0|291|149|1|1|NA|1|NA|NA|NA|NA|NA|0.92|0.506|NA|0.287|
|SE|86517|ENSG00000085224|ATRX|chrX|-|77589833|77589940|77574249|77574358|77593695|77593849|321|288|250|320|249|0|0|0|3|0|22|43|31|4|40|0|0|0|0|0|256|149|1|1|1|1|1|0.984|1|1|1|1|1|1|-0.003|
|SE|86518|ENSG00000085224|ATRX|chrX|-|77593695|77593849|77574249|77574358|77599410|77599580|221|177|156|198|212|0|0|1|0|0|8|46|21|2|22|0|0|0|0|0|298|149|1|1|1|1|0.987|1|1|1|1|1|1|1|-0.003|
|SE|86520|ENSG00000085224|ATRX|chrX|-|77618805|77618981|77613260|77616730|77620394|77620532|519|354|400|451|431|0|1|0|4|1|21|79|37|29|9|0|0|0|0|0|298|149|1|1|1|0.994|1|0.983|0.995|1|1|1|1|1|-0.006|
|SE|86521|ENSG00000085224|ATRX|chrX|-|77663381|77663558|77656559|77656653|77664644|77664778|237|262|161|234|333|0|0|0|3|3|19|95|39|39|54|0|0|0|0|0|298|149|1|1|1|1|1|0.975|0.982|1|1|1|1|1|-0.009|
|SE|86523|ENSG00000085224|ATRX|chrX|-|77681519|77684593|77664644|77664778|77688817|77688927|0|5|1|0|2|0|6|0|0|0|4|3|0|3|0|0|0|0|0|0|298|149|1|1|NA|0.294|1|NA|1|1|1|NA|1|NA|-0.235|
|SE|86524|ENSG00000085224|ATRX|chrX|-|77684938|77685006|77681519|77684593|77688817|77688924|155|97|109|130|233|0|0|1|0|0|16|23|7|7|7|0|0|0|0|0|217|149|1|1|1|1|0.987|1|1|1|1|1|1|1|-0.003|
|SE|86525|ENSG00000085224|ATRX|chrX|-|77690802|77690925|77688817|77688927|77693823|77693937|1|5|1|0|2|65|37|36|61|61|0|0|0|0|0|30|9|9|7|0|272|149|1|1|0.008|0.069|0.015|0|0.018|0|0|0|0|NA|0.022|
|SE|86528|ENSG00000085224|ATRX|chrX|-|77693823|77693937|77688817|77688927|77696576|77696704|293|306|185|297|310|24|23|20|25|36|70|66|35|27|0|16|13|3|0|0|263|149|1|1|0.874|0.883|0.84|0.871|0.83|0.713|0.742|0.869|1|NA|0.029|
|SE|86533|ENSG00000083750|RRAGB|chrX|+|55722185|55722285|55719313|55719347|55729293|55729360|133|115|120|178|110|1|0|0|0|0|47|100|85|142|63|0|0|0|0|0|249|149|1|1|0.988|1|1|1|1|1|1|1|1|1|-0.002|
|SE|86535|ENSG00000083750|RRAGB|chrX|+|55755840|55755932|55753391|55753514|55757215|55757331|26|16|10|26|49|0|0|1|1|0|0|9|0|3|5|0|0|0|0|0|241|149|1|1|1|1|0.861|0.941|1|NA|1|NA|1|1|-0.04|
|SE|86536|ENSG00000083750|RRAGB|chrX|+|55757215|55757331|55753391|55753514|55758245|55758348|13|5|14|19|15|0|0|0|0|0|0|6|0|3|5|0|0|0|3|0|265|149|1|1|1|1|1|1|1|NA|1|NA|0.36|1|0.213|
|SE|86543|ENSG00000136938|ANP32B|chr9|+|98004963|98005153|97998555|97998678|98011270|98011389|1691|1412|1866|2123|3291|0|0|0|0|0|5050|1691|2402|3164|6734|0|0|2|6|0|298|149|1|1|1|1|1|1|1|1|1|0.998|0.996|1|0.001|
|SE|86544|ENSG00000136938|ANP32B|chr9|+|98011270|98011389|98004963|98005153|98012420|98012472|595|581|822|979|1354|0|0|0|0|0|3190|859|1396|2032|3914|0|0|0|4|0|268|149|1|1|1|1|1|1|1|1|1|1|0.996|1|0.001|
|SE|86545|ENSG00000136938|ANP32B|chr9|+|98012420|98012472|98004963|98005153|98015363|98015943|23|16|13|19|27|0|1|0|0|0|73|11|25|18|47|0|0|0|0|0|201|149|1|1|1|0.922|1|1|1|1|1|1|1|1|-0.016|
|SE|86547|ENSG00000109111|SUPT6H|chr17|+|28673370|28673510|28670907|28671231|28674282|28674323|124|104|111|167|145|0|0|0|0|0|337|360|410|326|272|0|1|0|0|0|289|149|1|1|1|1|1|1|1|1|0.995|1|1|1|0.001|
|SE|86549|ENSG00000109111|SUPT6H|chr17|+|28678544|28678634|28678075|28678192|28678820|28678963|185|95|206|216|184|0|0|0|0|0|370|268|257|220|221|0|0|0|2|0|239|149|1|1|1|1|1|1|1|1|1|1|0.986|1|0.003|
|SE|86551|ENSG00000109111|SUPT6H|chr17|+|28682941|28683092|28682726|28682856|28683267|28683422|197|145|168|198|174|0|0|1|0|0|249|200|219|179|230|0|0|0|0|0|298|149|1|1|1|1|0.988|1|1|1|1|1|1|1|-0.002|
|SE|86552|ENSG00000109111|SUPT6H|chr17|+|28683267|28683422|28682726|28682856|28683620|28683816|178|125|155|121|173|0|0|0|0|0|293|212|177|220|277|0|8|0|0|0|298|149|1|1|1|1|1|1|1|1|0.93|1|1|1|0.014|
|SE|86555|ENSG00000109111|SUPT6H|chr17|+|28690920|28691063|28690081|28690229|28695351|28695547|142|76|132|113|114|0|0|0|1|1|129|144|143|162|126|0|0|2|4|0|292|149|1|1|1|1|1|0.983|0.983|1|1|0.973|0.954|1|0.008|
|SE|86556|ENSG00000109111|SUPT6H|chr17|+|28693695|28693836|28690081|28690229|28695351|28695547|151|105|127|147|216|0|0|0|1|1|249|216|196|248|240|0|0|2|4|0|290|149|1|1|1|1|1|0.987|0.991|1|1|0.981|0.97|1|0.005|
|SE|86557|ENSG00000109111|SUPT6H|chr17|+|28693695|28693836|28690920|28691063|28695351|28695547|301|200|271|295|326|0|0|0|0|0|596|468|410|438|433|0|0|5|2|2|290|149|1|1|1|1|1|1|1|1|1|0.977|0.991|0.991|0.008|
|SE|86558|ENSG00000109111|SUPT6H|chr17|+|28695351|28695547|28690920|28691063|28696843|28697082|139|103|110|121|153|0|0|0|0|0|128|105|174|158|230|0|0|5|0|0|298|149|1|1|1|1|1|1|1|1|1|0.946|1|1|0.011|
|SE|86560|ENSG00000126759|CFP|chrX|-|47628475|47628553|47628121|47628277|47629523|47629674|0|1|0|1|0|200|81|98|114|51|0|0|0|4|0|196|149|81|133|348|227|149|1|1|0|0.008|0|0.006|0|0|0|0|0.019|0|-0.001|
|SE|86561|ENSG00000126759|CFP|chrX|-|47629523|47629645|47628101|47628277|47629768|47629895|195|72|93|113|50|1|0|2|0|0|200|144|80|135|319|0|0|2|3|0|271|149|1|1|0.991|1|0.962|1|1|1|1|0.957|0.961|1|0.007|
|SE|86562|ENSG00000126759|CFP|chrX|-|47629523|47629674|47628101|47628277|47629768|47629909|370|145|183|196|98|1|0|2|0|0|306|234|125|267|588|0|0|2|3|0|298|149|1|1|0.995|1|0.979|1|1|1|1|0.969|0.978|1|0.005|
|SE|86565|ENSG00000136908|DPM2|chr9|-|127936552|127936655|127935098|127935780|127937433|127937523|50|18|24|23|11|0|0|0|0|0|0|84|9|12|39|0|0|1|0|0|252|149|1|1|1|1|1|1|1|NA|1|0.842|1|1|0.039|
|SE|86566|ENSG00000136908|DPM2|chr9|-|127937433|127937523|127936033|127936655|127937817|127937843|26|13|8|17|3|0|0|1|0|0|0|41|5|1|17|0|0|0|0|0|239|149|1|1|1|1|0.833|1|1|NA|1|1|1|1|-0.033|
|SE|86569|ENSG00000134575|ACP2|chr11|-|47244734|47244867|47243238|47243321|47245304|47245394|87|30|46|49|28|0|0|1|1|1|83|105|65|41|123|0|0|0|9|3|282|149|1|1|1|1|0.96|0.963|0.937|1|1|1|0.706|0.956|0.04|
|SE|86570|ENSG00000134575|ACP2|chr11|-|47244734|47244867|47243238|47243321|47245681|47245834|49|11|15|24|13|0|1|0|0|0|47|50|23|15|36|0|0|0|0|0|282|149|1|1|1|0.853|1|1|1|1|1|1|1|1|-0.029|
|SE|86571|ENSG00000134575|ACP2|chr11|-|47245304|47245394|47243238|47243321|47245681|47245834|0|0|1|1|1|0|1|0|0|0|0|0|0|0|3|0|0|0|0|0|239|149|1|1|NA|0|1|1|1|NA|NA|NA|NA|1|-0.25|
|SE|86573|ENSG00000134575|ACP2|chr11|-|47245304|47245422|47243238|47243321|47245681|47245834|0|0|1|1|1|0|1|0|0|0|0|0|0|0|3|0|0|0|0|0|267|149|1|1|NA|0|1|1|1|NA|NA|NA|NA|1|-0.25|
|SE|86575|ENSG00000134575|ACP2|chr11|-|47245304|47245427|47243238|47243321|47245681|47245834|0|0|1|1|1|0|1|0|0|0|0|0|0|0|3|0|0|0|0|0|272|149|1|1|NA|0|1|1|1|NA|NA|NA|NA|1|-0.25|
|SE|86578|ENSG00000134575|ACP2|chr11|-|47248037|47248133|47247392|47247727|47248484|47248628|18|9|2|10|10|0|0|0|0|0|10|25|8|7|5|0|0|0|1|0|245|149|1|1|1|1|1|1|1|1|1|1|0.81|1|0.038|
|SE|86579|ENSG00000134575|ACP2|chr11|-|47248037|47248133|47247392|47247727|47248675|47248802|47|13|22|24|11|1|0|0|0|0|74|77|26|35|45|0|0|0|0|0|245|149|1|1|0.966|1|1|1|1|1|1|1|1|1|-0.007|
|SE|86581|ENSG00000020181|ADGRA2|chr8|+|37833009|37833208|37831422|37831587|37833687|37833837|14|3|4|10|6|1|0|0|1|0|4|4|0|16|32|0|0|0|0|0|298|149|1|1|0.875|1|1|0.833|1|1|1|NA|1|1|-0.058|
|SE|86582|ENSG00000020181|ADGRA2|chr8|+|37835173|37835398|37833966|37834128|37835553|37835770|14|2|4|5|11|0|0|0|0|1|5|7|1|9|23|0|0|0|0|0|298|149|1|1|1|1|1|1|0.846|1|1|1|1|1|-0.031|
|SE|86583|ENSG00000020181|ADGRA2|chr8|+|37840759|37840849|37840120|37840266|37841085|37844896|2|1|2|2|1|0|0|0|2|0|17|0|10|0|15|0|0|0|0|0|239|149|1|1|1|1|1|0.384|1|1|NA|1|NA|1|-0.123|
|SE|86594|ENSG00000159374|M1AP|chr2|-|74559697|74559709|74557882|74558874|74560150|74560291|1|0|0|1|0|0|0|0|1|0|0|0|0|9|0|0|0|0|0|1|161|149|1|1|1|NA|NA|0.481|NA|NA|NA|NA|1|0|0.24|
|SE|86596|ENSG00000159374|M1AP|chr2|-|74562216|74562423|74560150|74560291|74575437|74575579|2|0|2|3|0|0|0|0|0|0|14|5|2|36|7|0|0|0|3|0|298|149|1|1|1|NA|1|1|NA|1|1|1|0.857|1|0.029|
|SE|86599|ENSG00000107951|MTPAP|chr10|-|30336802|30337027|30326423|30326635|30340225|30340450|73|44|95|54|154|0|0|1|0|0|37|33|30|46|26|0|0|0|0|0|298|149|1|1|1|1|0.979|1|1|1|1|1|1|1|-0.004|
|SE|86607|ENSG00000148143|ZNF462|chr9|+|107003293|107003426|106984185|106984409|107009544|107009668|0|0|1|2|0|0|0|0|0|0|0|0|6|1|0|0|1|0|0|0|282|149|1|1|NA|NA|1|1|NA|NA|0|1|1|NA|0.333|
|SE|86609|ENSG00000012048|BRCA1|chr17|-|43063873|43063951|43063332|43063373|43067607|43067695|21|9|12|12|11|0|0|0|0|0|41|12|7|3|7|0|0|0|0|1|227|149|1|1|1|1|1|1|1|1|1|1|1|0.821|0.036|
|SE|86611|ENSG00000012048|BRCA1|chr17|-|43079333|43079399|43076487|43076614|43082403|43082575|5|1|2|0|0|9|11|7|2|2|0|0|0|0|0|14|11|4|6|0|215|149|1|1|0.278|0.059|0.165|0|0|0|0|0|0|NA|0.1|
|SE|86612|ENSG00000012048|BRCA1|chr17|-|43091434|43094860|43090943|43091032|43095845|43095922|14|15|11|14|27|0|0|0|2|0|7|10|8|18|0|0|0|0|0|0|298|149|1|1|1|1|1|0.778|1|1|1|1|1|NA|-0.044|
|SE|86613|ENSG00000012048|BRCA1|chr17|-|43094743|43094860|43090943|43091032|43095845|43095922|7|8|8|6|14|0|0|0|2|0|10|4|0|6|0|0|0|0|0|0|266|149|1|1|1|1|1|0.627|1|1|1|NA|1|NA|-0.075|
|SE|86614|ENSG00000012048|BRCA1|chr17|-|43095845|43095922|43091097|43094860|43097243|43097289|11|16|15|15|25|1|0|0|2|0|1|15|11|13|6|0|0|0|0|0|226|149|1|1|0.879|1|1|0.832|1|1|1|1|1|1|-0.058|
|SE|86615|ENSG00000012048|BRCA1|chr17|-|43095845|43095922|43091097|43094860|43099774|43099877|3|3|4|5|7|1|0|2|0|0|0|3|4|6|0|0|0|1|0|0|226|149|1|1|0.664|1|0.569|1|1|NA|1|0.725|1|NA|-0.062|
|SE|86616|ENSG00000012048|BRCA1|chr17|-|43097243|43097289|43091097|43094860|43099774|43099877|5|8|7|7|0|1|0|2|0|0|0|0|1|2|6|0|0|1|0|0|195|149|1|1|0.793|1|0.728|1|NA|NA|NA|0.433|1|1|0.069|
|SE|86617|ENSG00000012048|BRCA1|chr17|-|43097243|43097289|43095845|43095922|43099774|43099877|12|21|18|18|18|0|0|0|3|0|1|12|8|9|12|0|0|0|0|0|195|149|1|1|1|1|1|0.821|1|1|1|1|1|1|-0.036|
|SE|86618|ENSG00000012048|BRCA1|chr17|-|43106455|43106533|43104867|43104956|43115725|43115779|24|10|15|14|31|0|0|0|0|0|32|19|13|24|30|0|0|0|4|0|227|149|1|1|1|1|1|1|1|1|1|1|0.798|1|0.04|
|SE|86619|ENSG00000012048|BRCA1|chr17|-|43106477|43106533|43104867|43104956|43115725|43115779|9|4|1|1|4|0|0|0|0|0|0|8|5|11|0|0|0|0|4|0|205|149|1|1|1|1|1|1|1|NA|1|1|0.667|NA|0.111|
|SE|86622|ENSG00000012048|BRCA1|chr17|-|43115725|43115779|43106455|43106533|43124016|43124115|24|13|14|16|21|0|0|0|0|0|35|21|17|22|9|0|0|0|3|0|203|149|1|1|1|1|1|1|1|1|1|1|0.843|1|0.031|
|SE|86624|ENSG00000012048|BRCA1|chr17|-|43121557|43121676|43115725|43115779|43124016|43124115|0|0|0|0|1|22|6|11|15|12|8|0|0|0|0|35|9|16|20|0|268|149|1|1|0|0|0|0|0.044|0.113|0|0|0|NA|-0.019|
|SE|86625|ENSG00000012048|BRCA1|chr17|-|43124016|43124115|43115725|43115779|43125276|43125343|21|5|7|10|13|3|0|0|0|0|20|9|13|17|0|0|0|0|0|0|248|149|1|1|0.808|1|1|1|1|1|1|1|1|NA|-0.038|
|SE|86627|ENSG00000186312|CA5BP1|chrX|+|15688660|15688858|15675689|15675778|15697239|15697460|9|9|11|27|20|1|0|0|0|0|54|9|7|12|13|0|0|0|0|0|298|149|1|1|0.818|1|1|1|1|1|1|1|1|1|-0.036|
|SE|86631|ENSG00000102265|TIMP1|chrX|+|47583407|47583536|47582407|47582474|47584935|47585015|97|33|53|121|21|0|0|0|1|0|210|254|135|219|1003|0|0|2|0|0|278|149|1|1|1|1|1|0.985|1|1|1|0.973|1|1|0.002|
|SE|86632|ENSG00000102265|TIMP1|chrX|+|47585204|47585331|47584935|47585015|47585542|47585667|412|155|164|389|387|1|0|0|0|0|768|717|369|429|3206|0|0|0|0|0|276|149|1|1|0.996|1|1|1|1|1|1|1|1|1|-0.001|
|SE|86633|ENSG00000185015|CA13|chr8|+|85224954|85225295|85220926|85221125|85230084|85230200|7|10|11|11|6|4|2|2|2|0|0|22|3|4|0|0|5|0|6|0|298|149|1|1|0.467|0.714|0.733|0.733|1|NA|0.688|1|0.25|NA|0.083|
|SE|86634|ENSG00000185015|CA13|chr8|+|85230084|85230200|85224954|85225295|85250739|85250937|13|6|12|13|7|0|0|0|2|0|0|22|0|0|6|0|0|0|0|0|265|149|1|1|1|1|1|0.785|1|NA|1|NA|NA|1|-0.043|
|SE|86635|ENSG00000185015|CA13|chr8|+|85237718|85237870|85230084|85230200|85250739|85250797|0|0|0|1|2|10|5|11|6|6|0|0|0|0|0|0|0|0|0|6|298|149|1|1|0|0|0|0.077|0.143|NA|NA|NA|NA|0|0.044|
|SE|86638|ENSG00000169249|ZRSR2|chrX|+|15803687|15803796|15799871|15799953|15804110|15804197|59|39|45|39|44|0|0|0|1|0|10|13|46|64|19|0|0|0|0|0|258|149|1|1|1|1|1|0.957|1|1|1|1|1|1|-0.009|
|SE|86639|ENSG00000169249|ZRSR2|chrX|+|15808232|15808271|15804110|15804197|15809199|15809318|74|55|89|89|59|4|2|5|1|0|50|42|62|96|0|0|4|0|8|0|188|149|1|1|0.936|0.956|0.934|0.986|1|1|0.893|1|0.905|NA|0.013|
|SE|86640|ENSG00000169249|ZRSR2|chrX|+|15820206|15820316|15818586|15818642|15822730|15823260|99|58|115|107|119|0|0|0|0|0|83|68|60|66|3|0|0|1|0|0|259|149|1|1|1|1|1|1|1|1|1|0.972|1|1|0.006|
|SE|86644|ENSG00000158578|ALAS2|chrX|-|55023756|55023867|55021051|55021274|55024717|55024840|12|3|10|5|3|0|0|2|0|0|0|5|2|135|0|0|4|0|82|4|260|149|1|1|1|1|0.741|1|1|NA|0.417|1|0.485|0|0.473|
|SE|86645|ENSG00000158578|ALAS2|chrX|-|55024717|55024840|55021051|55021274|55025819|55026015|3|0|6|5|5|3|0|0|0|0|0|11|2|200|9|0|0|0|0|0|272|149|1|1|0.354|NA|1|1|1|NA|1|1|1|1|-0.161|
|SE|86647|ENSG00000158578|ALAS2|chrX|-|55027750|55027857|55025819|55026015|55030941|55030977|3|0|1|0|3|8|0|0|2|1|0|0|0|16|0|0|0|1|22|5|256|149|1|1|0.179|NA|1|0|0.636|NA|NA|0|0.297|0|0.355|
|SE|86650|ENSG00000196924|FLNA|chrX|-|154350908|154351041|154350030|154350207|154351580|154351696|5023|2178|2647|4020|1729|0|0|5|0|0|6908|4801|2724|3764|23291|0|0|0|0|0|282|149|1|1|1|1|0.996|1|1|1|1|1|1|1|-0.001|
|SE|86651|ENSG00000196924|FLNA|chrX|-|154351883|154352021|154350030|154351696|154352180|154352447|3365|1220|1889|2595|1193|0|0|1|0|0|3385|3145|1656|2004|13111|0|0|0|0|0|287|149|1|1|1|1|0.999|1|1|1|1|1|1|1|0|
|SE|86652|ENSG00000196924|FLNA|chrX|-|154352180|154352447|154351727|154352021|154352771|154352924|1834|688|1065|1403|761|0|0|0|0|0|1943|1680|918|1155|7006|9|4|0|11|0|298|149|1|1|1|1|1|1|1|0.991|0.995|1|0.981|1|0.007|
|SE|86655|ENSG00000196924|FLNA|chrX|-|154353914|154354043|154353553|154353727|154354150|154354291|3751|1616|2026|3116|2528|0|0|0|0|0|5586|3698|2293|2884|18151|0|3|0|0|0|278|149|1|1|1|1|1|1|1|1|0.998|1|1|1|0|
|SE|86658|ENSG00000196924|FLNA|chrX|-|154359989|154360587|154359731|154359905|154361307|154361570|2523|775|1277|1691|745|0|0|0|0|1|2258|2165|976|1262|9637|0|0|0|0|0|298|149|1|1|1|1|1|1|0.997|1|1|1|1|1|-0.001|
|SE|86659|ENSG00000196924|FLNA|chrX|-|154361307|154361570|154359989|154360587|154361669|154361787|2001|657|971|1315|751|0|0|0|0|0|1665|1677|764|1017|7391|0|5|2|0|0|298|149|1|1|1|1|1|1|1|1|0.994|0.995|1|1|0.002|
|SE|86660|ENSG00000196924|FLNA|chrX|-|154361669|154361787|154361307|154361570|154361978|154362148|1714|579|881|1211|595|0|0|1|0|0|1530|1276|710|930|6029|0|0|0|3|0|267|149|1|1|1|1|0.998|1|1|1|1|1|0.994|1|0.001|
|SE|86661|ENSG00000196924|FLNA|chrX|-|154362417|154362578|154362241|154362332|154362660|154362784|3312|1278|1831|2569|1830|1|0|0|1|0|3321|2748|1520|2290|12077|0|0|0|0|0|298|149|1|1|0.999|1|1|0.999|1|1|1|1|1|1|0|
|SE|86662|ENSG00000196924|FLNA|chrX|-|154365348|154365486|154365135|154365259|154366023|154366224|4166|1691|2185|3564|1246|0|0|1|0|0|6313|4199|2473|3363|20213|0|0|0|0|0|287|149|1|1|1|1|0.999|1|1|1|1|1|1|1|0|
|SE|86663|ENSG00000196924|FLNA|chrX|-|154365744|154365801|154365348|154365486|154366023|154366224|0|0|1|0|0|1882|700|982|1543|446|0|4|2|10|0|2582|1900|1144|1414|9433|206|149|1|1|0|0|0.001|0|0|0|0.002|0.001|0.005|0|-0.001|
|SE|86666|ENSG00000196924|FLNA|chrX|-|154367396|154367544|154366731|154366850|154370872|154371203|2110|921|1117|1711|935|0|0|0|0|0|3020|1926|1093|1802|9416|0|0|2|0|0|297|149|1|1|1|1|1|1|1|1|1|0.996|1|1|0.001|
|SE|86667|ENSG00000196924|FLNA|chrX|-|154367640|154367738|154367396|154367544|154367841|154368090|1795|552|910|1269|545|0|0|0|1|0|1977|1499|807|1071|7505|0|0|0|0|0|247|149|1|1|1|1|1|0.999|1|1|1|1|1|1|0|
|SE|86668|ENSG00000196924|FLNA|chrX|-|154367841|154368090|154367396|154367544|154370872|154371203|1531|589|808|1151|526|0|0|0|2|0|1329|1063|654|770|5146|8|0|0|8|0|298|149|1|1|1|1|1|0.997|1|0.988|1|1|0.98|1|0.006|
|SE|86669|ENSG00000196924|FLNA|chrX|-|154370872|154371361|154367841|154368090|154374505|154374634|1551|589|811|1158|526|0|0|0|0|0|1329|1073|659|779|5244|0|0|1|0|0|298|149|1|1|1|1|1|1|1|1|1|0.997|1|1|0.001|
|SE|86672|ENSG00000269335|IKBKG|chrX|+|154556164|154556376|154551987|154552189|154560406|154560511|99|65|64|94|36|0|0|0|0|0|149|109|62|56|433|0|0|0|0|1|298|149|1|1|1|1|1|1|1|1|1|1|1|0.995|0.001|
|SE|86673|ENSG00000269335|IKBKG|chrX|+|154556164|154556376|154551987|154552189|154562809|154562953|98|65|64|94|36|0|0|0|0|0|149|110|59|51|433|0|0|1|0|0|298|149|1|1|1|1|1|1|1|1|1|0.967|1|1|0.007|
|SE|86674|ENSG00000269335|IKBKG|chrX|+|154558531|154558650|154556164|154556376|154560406|154560511|21|15|4|8|4|1|0|0|0|0|0|5|3|3|48|0|0|3|5|0|268|149|1|1|0.921|1|1|1|1|NA|1|0.357|0.25|1|0.332|
|SE|86675|ENSG00000269335|IKBKG|chrX|+|154558534|154558650|154556164|154556376|154560406|154560559|16|10|3|5|1|1|0|0|0|0|17|0|2|8|27|0|0|3|5|0|265|149|1|1|0.9|1|1|1|1|1|NA|0.273|0.474|1|0.293|
|SE|86679|ENSG00000165175|MID1IP1|chrX|+|38802534|38804413|38801431|38801487|38804682|38806528|94|30|63|67|46|2|0|0|0|0|80|44|54|117|116|0|0|0|0|0|298|149|1|1|0.959|1|1|1|1|1|1|1|1|1|-0.008|
|SE|86680|ENSG00000165175|MID1IP1|chrX|+|38804309|38804413|38801453|38801487|38804682|38806433|71|14|49|54|41|2|0|0|0|0|49|36|30|67|77|0|0|0|0|0|253|149|1|1|0.954|1|1|1|1|1|1|1|1|1|-0.009|
|SE|86681|ENSG00000254685|FPGT|chr1|+|74200691|74200818|74199663|74199831|74201317|74201410|0|0|0|0|1|67|31|37|32|64|0|0|0|0|0|0|12|7|4|4|276|149|1|1|0|0|0|0|0.008|NA|0|0|0|0|0.002|
|SE|86686|ENSG00000059728|MXD1|chr2|+|69921735|69921765|69916120|69916220|69935350|69935465|134|76|117|132|118|0|0|1|0|0|180|179|597|424|104|0|3|0|0|0|179|149|1|1|1|1|0.99|1|1|1|0.98|1|1|1|0.002|
|SE|86687|ENSG00000059728|MXD1|chr2|+|69937018|69937394|69935350|69935465|69938096|69938192|117|92|98|138|136|1|0|0|0|0|189|222|660|454|137|0|0|0|0|0|298|149|1|1|0.983|1|1|1|1|1|1|1|1|1|-0.003|
|SE|86688|ENSG00000059728|MXD1|chr2|+|69937234|69937394|69935350|69935465|69938096|69938259|265|200|225|280|338|1|0|0|0|0|471|439|1376|914|267|0|0|0|0|0|298|149|1|1|0.993|1|1|1|1|1|1|1|1|1|-0.001|
|SE|86689|ENSG00000147140|NONO|chrX|+|71284396|71284453|71283589|71283721|71284983|71285097|5|1|6|5|4|0|0|0|0|0|0|6|17|8|0|0|0|2|0|0|206|149|1|1|1|1|1|1|1|NA|1|0.86|1|NA|0.047|
|SE|86692|ENSG00000147140|NONO|chrX|+|71284983|71285097|71284396|71284453|71285364|71286174|4|0|2|2|2|0|9|5|3|4|0|6|11|8|0|2|3|1|1|0|263|149|1|1|1|0|0.185|0.274|0.221|0|0.531|0.862|0.819|NA|-0.217|
|SE|86694|ENSG00000147140|NONO|chrX|+|71288426|71288534|71284396|71284453|71290628|71290791|4|0|2|0|0|305|192|311|251|239|0|0|0|10|0|204|185|137|150|331|257|149|1|1|0.008|0|0.004|0|0|0|0|0|0.037|0|-0.005|
|SE|86697|ENSG00000147140|NONO|chrX|+|71294226|71294528|71290628|71291972|71296564|71296660|750|539|773|713|1106|0|0|3|6|1|1049|761|637|637|1052|0|7|5|4|0|298|149|1|1|1|1|0.992|0.983|0.998|1|0.982|0.985|0.988|1|0.004|
|SE|86699|ENSG00000147140|NONO|chrX|+|71296564|71296660|71294226|71294348|71296850|71297047|106|112|153|117|149|0|0|0|0|0|178|139|122|141|184|0|0|2|0|0|245|149|1|1|1|1|1|1|1|1|1|0.974|1|1|0.005|
|SE|86700|ENSG00000147140|NONO|chrX|+|71297587|71297677|71297376|71297461|71297835|71297938|0|0|0|0|0|230|154|258|252|218|0|0|0|1|0|497|290|192|235|342|239|149|1|1|0|0|0|0|0|0|0|0|0.003|0|-0.001|
|SE|86703|ENSG00000147140|NONO|chrX|+|71298468|71298508|71297376|71297938|71298706|71298816|312|166|335|327|317|0|0|0|0|0|418|291|261|291|416|0|0|0|3|0|189|149|1|1|1|1|1|1|1|1|1|1|0.987|1|0.003|
|SE|86704|ENSG00000147140|NONO|chrX|+|71298477|71298508|71297835|71297938|71298706|71298816|36|23|57|42|62|0|0|0|0|0|51|27|45|44|35|0|0|0|3|0|180|149|1|1|1|1|1|1|1|1|1|1|0.924|1|0.015|
|SE|86706|ENSG00000128654|MTX2|chr2|+|176270215|176270419|176269394|176269669|176296859|176296907|4|7|12|8|12|9|3|4|4|1|0|3|7|7|0|0|2|9|3|3|298|149|1|1|0.182|0.538|0.6|0.5|0.857|NA|0.429|0.28|0.538|0|0.224|
|SE|86707|ENSG00000128654|MTX2|chr2|+|176304240|176304365|176297848|176297895|176323391|176323464|6|7|6|4|11|19|10|24|14|17|0|0|0|0|14|0|9|11|9|13|274|149|1|1|0.147|0.276|0.12|0.134|0.26|NA|0|0|0|0.369|0.095|
|SE|86708|ENSG00000128654|MTX2|chr2|+|176323391|176323464|176297848|176297895|176326824|176326901|51|30|65|35|38|0|0|0|6|5|0|21|36|20|50|0|0|3|0|3|222|149|1|1|1|1|1|0.797|0.836|NA|1|0.89|1|0.918|-0.025|
|SE|86712|ENSG00000128654|MTX2|chr2|+|176329369|176329426|176328873|176328912|176337492|176337630|0|0|0|1|1|2|0|0|0|3|0|0|0|0|0|0|0|0|0|5|206|149|1|1|0|NA|NA|1|0.194|NA|NA|NA|NA|0|0.398|
|SE|86714|ENSG00000128654|MTX2|chr2|+|176330583|176330660|176329300|176329426|176337492|176337630|32|32|27|52|50|1|2|0|0|1|37|14|7|29|12|0|0|0|0|0|226|149|1|1|0.955|0.913|1|1|0.971|1|1|1|1|1|-0.032|
|SE|86715|ENSG00000198814|GK|chrX|+|30665510|30665584|30653358|30653615|30668011|30668118|68|56|72|70|81|1|0|0|0|0|49|43|114|79|63|0|0|0|0|0|223|149|1|1|0.978|1|1|1|1|1|1|1|1|1|-0.004|
|SE|86716|ENSG00000198814|GK|chrX|+|30665510|30665584|30653358|30653615|30677374|30677452|19|16|27|23|26|0|0|0|0|0|10|11|34|22|20|0|0|1|0|0|223|149|1|1|1|1|1|1|1|1|1|0.958|1|1|0.008|
|SE|86717|ENSG00000198814|GK|chrX|+|30668011|30668118|30653358|30653615|30677374|30677452|50|21|27|27|36|0|0|0|0|0|70|28|22|25|12|0|0|1|0|0|256|149|1|1|1|1|1|1|1|1|1|0.928|1|1|0.014|
|SE|86720|ENSG00000198814|GK|chrX|+|30691122|30691199|30677374|30677452|30694399|30694537|69|38|32|54|79|0|0|2|0|0|96|30|42|36|21|0|0|4|0|0|226|149|1|1|1|1|0.913|1|1|1|1|0.874|1|1|0.008|
|SE|86721|ENSG00000198814|GK|chrX|+|30691122|30691199|30677374|30677452|30696041|30696103|30|14|17|28|28|0|0|0|0|4|72|25|27|19|11|1|0|2|0|0|226|149|1|1|1|1|1|1|0.822|0.979|1|0.899|1|1|-0.011|
|SE|86722|ENSG00000198814|GK|chrX|+|30694399|30694537|30677374|30677452|30696041|30696062|52|43|35|58|85|0|0|0|0|4|64|38|35|28|24|1|0|2|0|0|287|149|1|1|1|1|1|1|0.917|0.971|1|0.901|1|1|0.009|
|SE|86725|ENSG00000198814|GK|chrX|+|30695082|30695132|30691122|30691199|30696041|30696151|0|0|0|0|0|0|1|1|7|0|0|0|4|0|0|0|6|5|4|0|199|149|1|1|NA|0|0|0|NA|NA|0|0.375|0|NA|-0.125|
|SE|86726|ENSG00000198814|GK|chrX|+|30695082|30695132|30694399|30694537|30696041|30696151|0|0|0|0|3|52|43|34|60|87|0|0|2|1|0|64|38|31|28|24|199|149|1|1|0|0|0|0|0.025|0|0|0.046|0.026|0|-0.009|
|SE|86728|ENSG00000198814|GK|chrX|+|30700413|30700449|30696616|30696683|30707555|30707598|19|10|9|15|16|1|1|1|1|0|14|15|10|12|11|14|1|2|0|0|185|149|1|1|0.939|0.89|0.879|0.924|1|0.446|0.924|0.801|1|1|0.092|
|SE|86729|ENSG00000198814|GK|chrX|+|30700837|30700905|30700413|30700449|30707555|30707598|146|88|71|107|149|0|0|0|0|2|112|90|75|70|62|0|0|0|0|0|217|149|1|1|1|1|1|1|0.981|1|1|1|1|1|-0.004|
|SE|86730|ENSG00000198814|GK|chrX|+|30720010|30720095|30719418|30719515|30720620|30720741|76|60|54|87|137|0|0|0|0|0|121|80|64|31|43|0|0|3|1|0|234|149|1|1|1|1|1|1|1|1|1|0.931|0.952|1|0.023|
|SE|86735|ENSG00000102230|PCYT1B|chrX|-|24618984|24619084|24607744|24607861|24672569|24672677|25|9|4|13|41|0|1|0|0|3|67|56|4|15|255|0|0|0|0|0|249|149|1|1|1|0.843|1|1|0.891|1|1|1|1|1|-0.053|
|SE|86739|ENSG00000101868|POLA1|chrX|+|24733754|24733816|24732369|24732454|24735398|24735488|36|14|15|20|42|2|0|0|0|0|0|23|17|6|0|0|0|0|0|0|211|149|1|1|0.927|1|1|1|1|NA|1|1|1|NA|-0.015|
|SE|86740|ENSG00000101868|POLA1|chrX|+|24737624|24737741|24735398|24735488|24739374|24739550|35|26|25|29|36|0|0|0|0|0|0|22|19|6|10|0|0|1|0|0|266|149|1|1|1|1|1|1|1|NA|1|0.914|1|1|0.021|
|SE|86744|ENSG00000101868|POLA1|chrX|+|24743229|24743329|24742001|24742121|24748310|24748460|43|27|34|71|65|1|0|0|0|0|8|35|46|35|0|0|0|1|0|0|249|149|1|1|0.963|1|1|1|1|1|1|0.965|1|NA|0.001|
|SE|86746|ENSG00000101868|POLA1|chrX|+|24745417|24745542|24743229|24743329|24748310|24748460|66|87|61|102|141|1|0|0|2|0|0|68|76|56|2|0|0|4|2|0|274|149|1|1|0.973|1|1|0.965|1|NA|1|0.912|0.938|1|0.025|
|SE|86747|ENSG00000101868|POLA1|chrX|+|24748869|24748992|24748310|24748460|24809897|24809930|82|97|84|121|148|0|0|0|1|0|0|75|81|89|2|0|0|0|0|0|272|149|1|1|1|1|1|0.985|1|NA|1|1|1|1|-0.003|
|SE|86748|ENSG00000101868|POLA1|chrX|+|24775196|24775356|24748869|24748992|24809897|24809930|0|0|1|0|1|16|17|19|42|49|0|0|0|0|0|0|11|7|7|0|298|149|1|1|0|0|0.026|0|0.01|NA|0|0|0|NA|0.007|
|SE|86749|ENSG00000101868|POLA1|chrX|+|24888005|24888122|24843545|24843677|24930452|24930549|48|18|54|23|76|0|1|1|0|0|26|11|40|21|26|0|0|0|0|0|266|149|1|1|1|0.91|0.968|1|1|1|1|1|1|1|-0.024|
|SE|86751|ENSG00000124275|MTRR|chr5|+|7870769|7870923|7869103|7869189|7875257|7875375|135|96|142|109|116|0|1|0|2|0|82|135|141|120|13|0|0|0|0|0|298|149|1|1|1|0.98|1|0.965|1|1|1|1|1|1|-0.011|
|SE|86753|ENSG00000124275|MTRR|chr5|+|7872229|7872305|7870769|7870923|7873372|7873526|0|0|0|0|1|20|22|32|29|31|0|0|0|0|0|18|23|11|6|0|225|149|1|1|0|0|0|0|0.021|0|0|0|0|NA|0.004|
|SE|86757|ENSG00000124275|MTRR|chr5|+|7883154|7883277|7877943|7878322|7885700|7885854|162|117|212|171|280|0|0|0|0|0|49|157|171|86|46|0|0|1|0|0|272|149|1|1|1|1|1|1|1|1|1|0.989|1|1|0.002|
|SE|86761|ENSG00000124275|MTRR|chr5|+|7889094|7889275|7886578|7886703|7891371|7891388|133|113|154|131|208|0|0|1|0|0|97|113|100|75|27|0|0|0|0|0|298|149|1|1|1|1|0.987|1|1|1|1|1|1|1|-0.003|
|SE|86762|ENSG00000124275|MTRR|chr5|+|7895733|7895852|7892726|7892913|7896863|7896956|78|68|136|109|124|0|0|0|1|0|48|66|118|51|41|0|0|0|0|0|268|149|1|1|1|1|1|0.984|1|1|1|1|1|1|-0.003|
|SE|86763|ENSG00000256294|ZNF225|chr19|+|44114048|44114213|44113414|44113569|44115759|44115842|0|1|0|1|1|3|1|4|13|0|0|0|0|0|0|0|1|0|1|0|298|149|1|1|0|0.333|0|0.037|1|NA|0|NA|0|NA|0.274|
|SE|86764|ENSG00000256294|ZNF225|chr19|+|44129041|44129098|44118481|44118574|44130849|44132874|0|0|0|0|2|7|7|11|3|15|0|0|0|0|0|0|0|1|0|0|206|149|1|1|0|0|0|0|0.088|NA|NA|0|NA|NA|0.018|
|SE|86765|ENSG00000182484|WASH6P|chrX|+|156022313|156022834|156021998|156022145|156023011|156023213|42|19|25|27|111|0|0|0|1|0|55|44|14|38|54|0|0|0|0|1|298|149|1|1|1|1|1|0.931|1|1|1|1|1|0.964|-0.007|
|SE|86766|ENSG00000182484|WASH6P|chrX|+|156022322|156022459|156021998|156022145|156023011|156023207|5|4|1|4|11|0|0|0|1|0|11|7|8|1|2|0|0|0|0|1|286|149|1|1|1|1|1|0.676|1|1|1|1|0.51|[illegible]|[illegible]|
|[illegible]|[illegible]|[illegible]|[illegible]|[illegible]|[illegible]|[illegible]|[illegible]|[illegible]|[illegible]|[illegible]|[illegible]|[illegible]|[illegible]|[illegible]|[illegible]|[illegible]|[illegible]|[illegible]|[illegible]|[illegible]|[illegible]|[illegible]|[illegible]|[illegible]|[illegible]|[illegible]|[illegible]|[illegible]|[illegible]|[illegible]|[illegible]|[illegible]|[illegible]|[illegible]|[illegible]|[illegible]|[illegible]|[illegible]|[illegible]|[illegible]|[illegible]|[illegible]|[illegible]|[illegible]|[illegible]|[illegible]|

| | | | | | | | | | | | | | | | | | | | | | | | | | | | | | | | | | | |
|---|---|---|---|---|---|---|---|---|---|---|---|---|---|---|---|---|---|---|---|---|---|---|---|---|---|---|---|---|---|---|---|---|---|---|
| SE | 86805 | ENSG00000166707 | ZCCHC18 | chrX | + | 104113433 | 104113573 | 104113194 | 104113246 | 104115251 | 104115842 | 6 | 14 | 23 | 7 | 20 | 0 | 0 | 1 | 0 | 1 | 33 | 8 | 2 | 1 | 2 | 0 | 0 | 0 | 0 | 0 | 289 | 149 | [illegible] |
| SE | 86807 | ENSG00000068308 | OTUD5 | chrX | - | 48923632 | 48923746 | 48922023 | 48923295 | 48923850 | 48924052 | 191 | 88 | 164 | 193 | 150 | 0 | 0 | 0 | 0 | 0 | 254 | 241 | 150 | 173 | 350 | 0 | 2 | 2 | 1 | 0 | 263 | 149 | [illegible] |
| SE | 86808 | ENSG00000068308 | OTUD5 | chrX | - | 48923850 | 48924052 | 48922023 | 48923295 | 48925846 | 48926050 | 186 | 115 | 180 | 176 | 185 | 2 | 0 | 0 | 0 | 0 | 131 | 121 | 123 | 97 | 205 | 0 | 0 | 0 | 0 | 0 | 298 | 149 | [illegible] |
| SE | 86810 | ENSG00000163682 | RPL9 | chr4 | - | 39454863 | 39454929 | 39454561 | 39454649 | 39456405 | 39456538 | 2239 | 1331 | 2495 | 1909 | 5723 | 2 | 1 | 0 | 0 | 0 | 1225 | 1115 | 1094 | 793 | 1509 | 0 | 0 | 0 | 0 | 6 | 215 | 149 | [illegible] |
| SE | 86811 | ENSG00000163682 | RPL9 | chr4 | - | 39454863 | 39454944 | 39454532 | 39454649 | 39456405 | 39456538 | 5110 | 2767 | 3280 | 2514 | 6791 | 2 | 1 | 0 | 0 | 0 | 1361 | 1289 | 2093 | 1784 | 3042 | 0 | 0 | 0 | 0 | 6 | 230 | 149 | [illegible] |
| SE | 86812 | ENSG00000163682 | RPL9 | chr4 | - | 39454863 | 39454944 | 39454532 | 39454649 | 39457585 | 39457681 | 1803 | 995 | 1594 | 1174 | 4375 | 0 | 0 | 0 | 0 | 0 | 462 | 639 | 768 | 650 | 972 | 0 | 0 | 3 | 0 | 0 | 230 | 149 | [illegible] |
| SE | 86813 | ENSG00000163682 | RPL9 | chr4 | - | 39456405 | 39456538 | 39454532 | 39454649 | 39457585 | 39457681 | 4120 | 2469 | 2958 | 2355 | 6472 | 0 | 0 | 0 | 0 | 0 | 1642 | 1364 | 1901 | 1536 | 3188 | 0 | 0 | 3 | 0 | 0 | 282 | 149 | [illegible] |
| SE | 86814 | ENSG00000163682 | RPL9 | chr4 | - | 39456405 | 39456538 | 39454863 | 39454944 | 39457585 | 39457681 | 7427 | 4241 | 4646 | 3700 | 8895 | 1 | 1 | 3 | 5 | 7 | 2541 | 2024 | 3238 | 2678 | 5256 | 0 | 16 | 12 | 12 | 4 | 282 | 149 | [illegible] |
| SE | 86815 | ENSG00000163682 | RPL9 | chr4 | - | 39457585 | 39457681 | 39454863 | 39454944 | 39458193 | 39458309 | 830 | 424 | 605 | 550 | 815 | 1 | 0 | 0 | 0 | 0 | 397 | 226 | 435 | 360 | 719 | 0 | 0 | 0 | 0 | 0 | 245 | 149 | [illegible] |
| SE | 86816 | ENSG00000163682 | RPL9 | chr4 | - | 39458393 | 39458440 | 39457804 | 39458309 | 39458890 | 39458915 | 745 | 371 | 775 | 686 | 1016 | 0 | 0 | 0 | 3 | 0 | 425 | 296 | 369 | 295 | 546 | 0 | 0 | 0 | 3 | 0 | 196 | 149 | [illegible] |
| SE | 86817 | ENSG00000163682 | RPL9 | chr4 | - | 39458393 | 39458441 | 39458193 | 39458309 | 39458890 | 39458897 | 474 | 246 | 472 | 414 | 804 | 0 | 0 | 0 | 3 | 0 | 333 | 190 | 251 | 185 | 466 | 0 | 0 | 0 | 3 | 0 | 197 | 149 | [illegible] |
| SE | 86818 | ENSG00000163682 | RPL9 | chr4 | - | 39458393 | 39458598 | 39458193 | 39458309 | 39458890 | 39458923 | 524 | 254 | 535 | 477 | 834 | 0 | 0 | 0 | 3 | 0 | 386 | 233 | 264 | 211 | 513 | 0 | 0 | 0 | 3 | 0 | 298 | 149 | [illegible] |
| SE | 86819 | ENSG00000005889 | ZFX | chrX | + | 24151690 | 24151800 | 24149728 | 24149794 | 24152719 | 24152830 | 23 | 10 | 14 | 27 | 14 | 2 | 0 | 2 | 3 | 0 | 0 | 5 | 1 | 20 | 0 | 0 | 0 | 0 | 0 | 0 | 259 | 149 | [illegible] |
| SE | 86820 | ENSG00000005889 | ZFX | chrX | + | 24152719 | 24152830 | 24149728 | 24149794 | 24172714 | 24172800 | 30 | 16 | 17 | 21 | 31 | 6 | 4 | 11 | 14 | 10 | 0 | 9 | 11 | 1 | 0 | 0 | 8 | 11 | 7 | 0 | 260 | 149 | [illegible] |
| SE | 86821 | ENSG00000005889 | ZFX | chrX | + | 24161714 | 24161842 | 24152719 | 24152830 | 24172714 | 24172800 | 25 | 16 | 25 | 55 | 14 | 41 | 20 | 23 | 29 | 31 | 0 | 10 | 3 | 1 | 6 | 0 | 9 | 11 | 1 | 0 | 277 | 149 | [illegible] |
| SE | 86822 | ENSG00000005889 | ZFX | chrX | + | 24172714 | 24172800 | 24149628 | 24149794 | 24179182 | 24179219 | 76 | 37 | 73 | 85 | 76 | 0 | 0 | 1 | 0 | 0 | 38 | 22 | 31 | 26 | 0 | 0 | 0 | 0 | 0 | 0 | 235 | 149 | [illegible] |
| SE | 86826 | ENSG00000005889 | ZFX | chrX | + | 24175388 | 24175443 | 24172755 | 24172800 | 24179182 | 24179550 | 5 | 3 | 4 | 10 | 16 | 98 | 45 | 90 | 98 | 86 | 0 | 0 | 2 | 0 | 0 | 38 | 31 | 33 | 29 | 0 | 204 | 149 | [illegible] |
| SE | 86827 | ENSG00000005889 | ZFX | chrX | + | 24175388 | 24175443 | 24173606 | 24173723 | 24177683 | 24177866 | 0 | 0 | 0 | 6 | 0 | 1 | 0 | 0 | 0 | 0 | 6 | 2 | 0 | 0 | 0 | 0 | 0 | 0 | 0 | 0 | 204 | 149 | [illegible] |
| SE | 86829 | ENSG00000005889 | ZFX | chrX | + | 24177683 | 24177866 | 24173606 | 24173723 | 24179182 | 24179550 | 4 | 1 | 1 | 1 | 0 | 23 | 8 | 10 | 7 | 10 | 0 | 4 | 0 | 0 | 0 | 0 | 7 | 1 | 4 | 17 | 298 | 149 | [illegible] |
| SE | 86830 | ENSG00000005889 | ZFX | chrX | + | 24177683 | 24177866 | 24175388 | 24175443 | 24179182 | 24179550 | 3 | 1 | 1 | 2 | 0 | 5 | 3 | 4 | 5 | 16 | 0 | 4 | 0 | 0 | 0 | 0 | 0 | 3 | 0 | 0 | 298 | 149 | [illegible] |

| | | | | | | | | | | | | | | | | | | | | | | | | | | | | | | | | | | | | | | | | | | | | | | |
|---|---|---|---|---|---|---|---|---|---|---|---|---|---|---|---|---|---|---|---|---|---|---|---|---|---|---|---|---|---|---|---|---|---|---|---|---|---|---|---|---|---|---|---|---|---|---|
| SE | 87249 | ENSG00000046651 | OFD1 | chrX | + | 13753367 | 13753441 | 13751248 | 13751368 | 13755150 | 13755242 | 101 | 62 | 99 | 78 | 170 | 0 | 2 | 1 | 0 | 0 | 24 | 61 | 69 | 23 | 3 | 0 | 0 | 0 | 0 | 0 | 223 | 149 | 1 | 1 | 1 | 0.954 | 0.985 | 1 | 1 | 1 | 1 | 1 | 1 | 1 | -0.012 |
| SE | 87250 | ENSG00000046651 | OFD1 | chrX | + | 13753367 | 13753441 | 13751248 | 13751368 | 13756577 | 13756767 | 32 | 27 | 45 | 29 | 103 | 2 | 0 | 0 | 1 | 5 | 4 | 34 | 33 | 12 | 0 | 11 | 6 | 1 | 0 | 0 | 223 | 149 | 1 | 1 | 0.914 | 1 | 1 | 0.951 | 0.932 | 0.195 | 0.791 | 0.957 | 1 | NA | 0.224 |
| SE | 87253 | ENSG00000046651 | OFD1 | chrX | + | 13755150 | 13755242 | 13752703 | 13753441 | 13756577 | 13756767 | 253 | 184 | 185 | 220 | 310 | 1 | 1 | 0 | 0 | 0 | 81 | 118 | 156 | 80 | 26 | 0 | 0 | 0 | 0 | 0 | 241 | 149 | 1 | 1 | 0.994 | 0.991 | 1 | 1 | 1 | 1 | 1 | 1 | 1 | 1 | -0.003 |
| SE | 87255 | ENSG00000046651 | OFD1 | chrX | + | 13758336 | 13758448 | 13757659 | 13757790 | 13768053 | 13768224 | 23 | 29 | 26 | 33 | 28 | 0 | 0 | 0 | 0 | 0 | 10 | 24 | 18 | 14 | 0 | 0 | 0 | 1 | 0 | 0 | 261 | 149 | 1 | 1 | 1 | 1 | 1 | 1 | 1 | 1 | 1 | 0.911 | 1 | NA | 0.022 |
| SE | 87257 | ENSG00000046651 | OFD1 | chrX | + | 13763744 | 13763855 | 13762343 | 13762444 | 13768053 | 13768224 | 86 | 94 | 111 | 83 | 79 | 2 | 3 | 2 | 2 | 0 | 51 | 44 | 39 | 35 | 0 | 18 | 1 | 7 | 0 | 0 | 260 | 149 | 1 | 1 | 0.961 | 0.947 | 0.97 | 0.96 | 1 | 0.619 | 0.962 | 0.761 | 1 | NA | 0.132 |
| SE | 87258 | ENSG00000046651 | OFD1 | chrX | + | 13767126 | 13767284 | 13762343 | 13762444 | 13768053 | 13768224 | 77 | 81 | 65 | 50 | 79 | 2 | 3 | 2 | 2 | 0 | 10 | 56 | 61 | 39 | 0 | 18 | 1 | 7 | 0 | 0 | 298 | 149 | 1 | 1 | 0.951 | 0.931 | 0.942 | 0.926 | 1 | 0.217 | 0.966 | 0.813 | 1 | NA | 0.201 |
| SE | 87262 | ENSG00000147133 | TAF1 | chrX | + | 71382760 | 71382868 | 71382535 | 71382663 | 71383961 | 71384135 | 106 | 86 | 91 | 95 | 175 | 0 | 0 | 0 | 0 | 0 | 62 | 91 | 105 | 68 | 46 | 0 | 0 | 2 | 0 | 0 | 257 | 149 | 1 | 1 | 1 | 1 | 1 | 1 | 1 | 1 | 1 | 0.968 | 1 | 1 | 0.006 |
| SE | 87263 | ENSG00000147133 | TAF1 | chrX | + | 71382990 | 71383164 | 71382760 | 71382868 | 71383961 | 71384135 | 235 | 183 | 201 | 238 | 341 | 0 | 0 | 0 | 0 | 0 | 122 | 263 | 214 | 117 | 79 | 0 | 0 | 1 | 0 | 0 | 298 | 149 | 1 | 1 | 1 | 1 | 1 | 1 | 1 | 1 | 1 | 0.991 | 1 | 1 | 0.002 |
| SE | 87265 | ENSG00000147133 | TAF1 | chrX | + | 71388236 | 71388378 | 71387260 | 71387461 | 71389584 | 71389665 | 99 | 79 | 76 | 76 | 57 | 0 | 0 | 0 | 0 | 0 | 87 | 51 | 102 | 72 | 17 | 0 | 1 | 0 | 0 | 0 | 291 | 149 | 1 | 1 | 1 | 1 | 1 | 1 | 1 | 1 | 0.963 | 1 | 1 | 1 | 0.007 |
| SE | 87266 | ENSG00000147133 | TAF1 | chrX | + | 71388737 | 71388868 | 71387260 | 71387461 | 71389584 | 71389665 | 102 | 89 | 76 | 95 | 144 | 0 | 0 | 0 | 0 | 0 | 119 | 90 | 86 | 85 | 13 | 0 | 1 | 0 | 0 | 0 | 280 | 149 | 1 | 1 | 1 | 1 | 1 | 1 | 1 | 1 | 0.98 | 1 | 1 | 1 | 0.004 |
| SE | 87268 | ENSG00000147133 | TAF1 | chrX | + | 71392870 | 71392994 | 71392568 | 71392718 | 71393300 | 71393476 | 111 | 117 | 88 | 106 | 123 | 0 | 0 | 0 | 0 | 0 | 131 | 67 | 130 | 150 | 7 | 0 | 0 | 0 | 2 | 0 | 273 | 149 | 1 | 1 | 1 | 1 | 1 | 1 | 1 | 1 | 1 | 1 | 0.976 | 1 | 0.005 |
| SE | 87269 | ENSG00000147133 | TAF1 | chrX | + | 71392874 | 71392994 | 71392568 | 71392718 | 71393300 | 71393476 | 202 | 219 | 158 | 187 | 203 | 0 | 0 | 0 | 0 | 0 | 214 | 132 | 217 | 244 | 7 | 0 | 0 | 0 | 2 | 0 | 269 | 149 | 1 | 1 | 1 | 1 | 1 | 1 | 1 | 1 | 1 | 1 | 0.985 | 1 | 0.003 |
| SE | 87271 | ENSG00000147133 | TAF1 | chrX | + | 71397252 | 71397466 | 71394066 | 71394245 | 71398571 | 71398737 | 298 | 248 | 289 | 285 | 314 | 0 | 0 | 0 | 0 | 0 | 262 | 288 | 299 | 220 | 72 | 0 | 0 | 2 | 0 | 5 | 298 | 149 | 1 | 1 | 1 | 1 | 1 | 1 | 1 | 1 | 1 | 0.987 | 1 | 0.878 | 0.027 |
| SE | 87273 | ENSG00000147133 | TAF1 | chrX | + | 71423116 | 71423239 | 71421308 | 71421376 | 71423973 | 71424066 | 252 | 156 | 172 | 214 | 294 | 0 | 0 | 0 | 1 | 0 | 92 | 113 | 114 | 160 | 38 | 0 | 0 | 1 | 0 | 0 | 272 | 149 | 1 | 1 | 1 | 1 | 1 | 0.992 | 1 | 1 | 1 | 0.984 | 1 | 1 | 0.002 |
| SE | 87274 | ENSG00000147133 | TAF1 | chrX | + | 71423116 | 71423264 | 71421308 | 71421376 | 71423973 | 71424066 | 95 | 53 | 73 | 79 | 91 | 0 | 0 | 0 | 1 | 0 | 46 | 52 | 65 | 81 | 21 | 0 | 0 | 1 | 0 | 0 | 297 | 149 | 1 | 1 | 1 | 1 | 1 | 0.975 | 1 | 1 | 1 | 0.97 | 1 | 1 | 0.001 |
| SE | 87275 | ENSG00000147133 | TAF1 | chrX | + | 71424153 | 71424238 | 71423116 | 71424066 | 71454169 | 71454237 | 232 | 208 | 178 | 201 | 304 | 0 | 0 | 0 | 1 | 0 | 84 | 131 | 129 | 180 | 19 | 0 | 0 | 0 | 0 | 0 | 234 | 149 | 1 | 1 | 1 | 1 | 1 | 0.992 | 1 | 1 | 1 | 1 | 1 | 1 | -0.002 |
| SE | 87276 | ENSG00000147133 | TAF1 | chrX | + | 71441699 | 71441721 | 71424153 | 71424238 | 71454169 | 71454237 | 0 | 0 | 0 | 0 | 0 | 87 | 106 | 87 | 88 | 171 | 0 | 0 | 0 | 1 | 0 | 8 | 70 | 56 | 77 | 6 | 171 | 149 | 1 | 1 | 0 | 0 | 0 | 0 | 0 | 0 | 0 | 0 | 0.011 | 0 | -0.002 |
| SE | 87279 | ENSG00000147133 | TAF1 | chrX | + | 71459149 | 71459251 | 71458240 | 71458357 | 71459551 | 71459708 | 13 | 15 | 12 | 19 | 25 | 1 | 0 | 0 | 2 | 7 | 0 | 19 | 16 | 9 | 8 | 0 | 1 | 0 | 6 | 0 | 251 | 149 | 1 | 1 | 0.885 | 1 | 1 | 0.849 | 0.679 | NA | 0.919 | 1 | 0.471 | 1 | 0.035 |
| SE | 87282 | ENSG00000147144 | CCDC120 | chrX | + | 49064369 | 49064664 | 49063860 | 49064001 | 49065453 | 49065628 | 8 | 0 | 0 | 0 | 2 | 0 | 0 | 0 | 0 | 0 | 0 | 0 | 2 | 0 | 2 | 0 | 0 | 2 | 0 | 0 | 298 | 149 | 1 | 1 | 1 | NA | NA | NA | 1 | NA | NA | 0.333 | NA | 1 | 0.334 |
| SE | 87283 | ENSG00000147144 | CCDC120 | chrX | + | 49065035 | 49065098 | 49064369 | 49064664 | 49065453 | 49065628 | 1 | 0 | 0 | 2 | 0 | 2 | 0 | 0 | 0 | 0 | 14 | 6 | 6 | 0 | 0 | 0 | 0 | 0 | 0 | 0 | 212 | 149 | 1 | 1 | 0.26 | NA | NA | 1 | NA | 1 | 1 | 1 | NA | NA | -0.37 |
| SE | 87285 | ENSG00000172732 | MUS81 | chr11 | + | 65862443 | 65862529 | 65862210 | 65862279 | 65863064 | 65863089 | 70 | 27 | 64 | 68 | 26 | 0 | 0 | 0 | 0 | 0 | 83 | 70 | 28 | 61 | 90 | 0 | 5 | 0 | 0 | 0 | 235 | 149 | 1 | 1 | 1 | 1 | 1 | 1 | 1 | 1 | 0.899 | 1 | 1 | 1 | 0.02 |
| SE | 87286 | ENSG00000243279 | PRAF2 | chrX | - | 49072432 | 49072650 | 49071160 | 49072008 | 49073808 | 49074002 | 38 | 10 | 20 | 15 | 11 | 0 | 1 | 0 | 0 | 0 | 17 | 28 | 8 | 7 | 118 | 0 | 0 | 0 | 0 | 0 | 298 | 149 | 1 | 1 | 1 | 0.833 | 1 | 1 | 1 | 1 | 1 | 1 | 1 | 1 | -0.033 |
| SE | 87290 | ENSG00000123643 | SLC36A1 | chr5 | + | 151464513 | 151464602 | 151463552 | 151463643 | 151465073 | 151465149 | 53 | 33 | 40 | 29 | 35 | 0 | 0 | 0 | 0 | 0 | 102 | 71 | 56 | 48 | 112 | 0 | 2 | 2 | 0 | 0 | 238 | 149 | 1 | 1 | 1 | 1 | 1 | 1 | 1 | 1 | 0.957 | 0.946 | 1 | 1 | 0.019 |
| SE | 87291 | ENSG00000123643 | SLC36A1 | chr5 | + | 151469245 | 151469271 | 151467706 | 151467925 | 151473672 | 151473771 | 1 | 0 | 0 | 1 | 0 | 21 | 12 | 14 | 14 | 13 | 0 | 0 | 4 | 0 | 0 | 23 | 45 | 26 | 37 | 37 | 175 | 149 | 1 | 1 | 0.039 | 0 | 0 | 0.057 | 0 | 0 | 0 | 0.116 | 0 | 0 | -0.004 |
| SE | 87293 | ENSG00000123643 | SLC36A1 | chr5 | + | 151479319 | 151479489 | 151476589 | 151476756 | 151487982 | 151488269 | 83 | 44 | 45 | 49 | 58 | 0 | 0 | 0 | 0 | 0 | 97 | 91 | 62 | 101 | 102 | 0 | 0 | 3 | 0 | 0 | 298 | 149 | 1 | 1 | 1 | 1 | 1 | 1 | 1 | 1 | 1 | 0.912 | 1 | 1 | 0.018 |
| SE | 87295 | ENSG00000083457 | ITGAE | chr17 | - | 3720306 | 3720402 | 3716687 | 3716798 | 3723687 | 3723837 | 81 | 28 | 67 | 94 | 75 | 0 | 0 | 1 | 0 | 0 | 59 | 54 | 78 | 50 | 111 | 0 | 5 | 0 | 0 | 0 | 245 | 149 | 1 | 1 | 1 | 1 | 0.976 | 1 | 1 | 1 | 0.868 | 1 | 1 | 1 | 0.022 |
| SE | 87297 | ENSG00000083457 | ITGAE | chr17 | - | 3729477 | 3729555 | 3728104 | 3728168 | 3731103 | 3731183 | 2 | 0 | 2 | 2 | 0 | 0 | 0 | 0 | 0 | 0 | 13 | 0 | 0 | 4 | 0 | 0 | 0 | 0 | 3 | 0 | 227 | 149 | 1 | 1 | 1 | NA | 1 | 1 | NA | 1 | NA | NA | 0.467 | NA | 0.266 |
| SE | 87299 | ENSG00000083457 | ITGAE | chr17 | - | 3739804 | 3739878 | 3734816 | 3734949 | 3743488 | 3743540 | 0 | 2 | 0 | 0 | 0 | 0 | 0 | 0 | 0 | 0 | 2 | 0 | 0 | 0 | 0 | 0 | 0 | 5 | 0 | 0 | 223 | 149 | 1 | 1 | NA | 1 | NA | NA | NA | 1 | NA | 0 | NA | NA | 0.5 |
| SE | 87304 | ENSG00000215301 | DDX3X | chrX | + | 41336082 | 41336159 | 41334605 | 41334769 | 41337407 | 41337465 | 2 | 1 | 2 | 2 | 0 | 1 | 0 | 2 | 2 | 1 | 10 | 0 | 0 | 0 | 0 | 0 | 0 | 1 | 3 | 0 | 226 | 149 | 1 | 1 | 0.569 | 1 | 0.397 | 0.397 | 0 | 1 | NA | 0 | 0 | NA | 0.139 |
| SE | 87305 | ENSG00000215301 | DDX3X | chrX | + | 41336082 | 41336159 | 41334609 | 41334815 | 41337407 | 41337465 | 2 | 1 | 2 | 2 | 0 | 0 | 0 | 3 | 0 | 0 | 10 | 0 | 0 | 0 | 0 | 0 | 0 | 0 | 0 | 0 | 226 | 149 | 1 | 1 | 1 | 1 | 0.305 | 1 | NA | 1 | NA | NA | NA | NA | -0.174 |
| SE | 87306 | ENSG00000215301 | DDX3X | chrX | + | 41337407 | 41337465 | 41333347 | 41334297 | 41341483 | 41341616 | 1146 | 578 | 939 | 1130 | 372 | 1 | 1 | 2 | 2 | 3 | 264 | 524 | 677 | 428 | 170 | 0 | 0 | 0 | 0 | 0 | 207 | 149 | 1 | 1 | 0.999 | 0.998 | 0.997 | 0.998 | 0.989 | 1 | 1 | 1 | 1 | 1 | -0.004 |
| SE | 87307 | ENSG00000215301 | DDX3X | chrX | + | 41337407 | 41339083 | 41334161 | 41334297 | 41341483 | 41341616 | 1393 | 735 | 1177 | 1379 | 536 | 1 | 1 | 2 | 2 | 3 | 283 | 612 | 881 | 516 | 200 | 0 | 0 | 0 | 0 | 0 | 298 | 149 | 1 | 1 | 0.999 | 0.997 | 0.997 | 0.997 | 0.989 | 1 | 1 | 1 | 1 | 1 | -0.004 |
| SE | 87308 | ENSG00000215301 | DDX3X | chrX | + | 41337690 | 41338912 | 41337407 | 41337465 | 41339035 | 41339083 | 3 | 2 | 3 | 4 | 7 | 1127 | 591 | 871 | 1034 | 437 | 0 | 0 | 0 | 0 | 0 | 177 | 530 | 614 | 413 | 178 | 298 | 149 | 1 | 1 | 0.001 | 0.002 | 0.002 | 0.002 | 0.008 | 0 | 0 | 0 | 0 | 0 | 0.003 |
| SE | 87310 | ENSG00000215301 | DDX3X | chrX | + | 41339035 | 41339083 | 41337407 | 41337465 | 41340635 | 41340814 | 180 | 120 | 148 | 196 | 80 | 0 | 0 | 2 | 0 | 0 | 23 | 123 | 134 | 98 | 67 | 0 | 2 | 0 | 0 | 0 | 197 | 149 | 1 | 1 | 1 | 1 | 0.982 | 1 | 1 | 1 | 0.979 | 1 | 1 | 1 | 0.001 |
| SE | 87313 | ENSG00000215301 | DDX3X | chrX | + | 41340635 | 41340814 | 41337407 | 41337465 | 41341483 | 41341616 | 8 | 19 | 14 | 26 | 16 | 23 | 7 | 9 | 32 | 30 | 7 | 9 | 17 | 13 | 0 | 19 | 13 | 8 | 24 | 0 | 298 | 149 | 1 | 1 | 0.148 | 0.576 | 0.437 | 0.289 | 0.211 | 0.156 | 0.257 | 0.515 | 0.213 | NA | 0.047 |
| SE | 87315 | ENSG00000215301 | DDX3X | chrX | + | 41341948 | 41342046 | 41341483 | 41341616 | 41342494 | 41342506 | 4 | 7 | 8 | 4 | 1 | 1238 | 857 | 1263 | 1522 | 992 | 0 | 2 | 10 | 4 | 0 | 439 | 776 | 1089 | 606 | 137 | 247 | 149 | 1 | 1 | 0.002 | 0.005 | 0.004 | 0.002 | 0.001 | 0 | 0.002 | 0.006 | 0.004 | 0 | 0 |
| SE | 87316 | ENSG00000215301 | DDX3X | chrX | + | 41342736 | 41342836 | 41340011 | 41342653 | 41343215 | 41343351 | 1434 | 972 | 1207 | 1605 | 1989 | 0 | 0 | 0 | 1 | 2 | 350 | 627 | 1015 | 687 | 139 | 0 | 0 | 2 | 0 | 0 | 249 | 149 | 1 | 1 | 1 | 1 | 1 | 0.999 | 0.998 | 1 | 1 | 0.997 | 1 | 1 | 0 |
| SE | 87317 | ENSG00000215301 | DDX3X | chrX | + | 41342736 | 41342836 | 41340011 | 41342653 | 41343736 | 41343822 | 654 | 415 | 552 | 688 | 923 | 1 | 0 | 0 | 0 | 0 | 158 | 302 | 354 | 300 | 38 | 0 | 0 | 0 | 0 | 0 | 249 | 149 | 1 | 1 | 0.997 | 1 | 1 | 1 | 1 | 1 | 1 | 1 | 1 | 1 | -0.001 |
| SE | 87318 | ENSG00000215301 | DDX3X | chrX | + | 41342736 | 41344128 | 41342494 | 41342653 | 41345179 | 41345324 | 645 | 408 | 541 | 684 | 908 | 2 | 0 | 0 | 1 | 0 | 158 | 276 | 348 | 307 | 38 | 0 | 0 | 0 | 0 | 0 | 298 | 149 | 1 | 1 | 0.994 | 1 | 1 | 0.997 | 1 | 1 | 1 | 1 | 1 | 1 | -0.002 |
| SE | 87320 | ENSG00000215301 | DDX3X | chrX | + | 41343215 | 41343351 | 41340005 | 41342836 | 41344029 | 41344128 | 849 | 617 | 713 | 1005 | 1092 | 1 | 0 | 3 | 3 | 6 | 256 | 452 | 712 | 451 | 118 | 0 | 0 | 0 | 1 | 0 | 285 | 149 | 1 | 1 | 0.998 | 1 | 0.992 | 0.994 | 0.99 | 1 | 1 | 1 | 0.996 | 1 | -0.004 |
| SE | 87321 | ENSG00000215301 | DDX3X | chrX | + | 41343215 | 41343351 | 41340011 | 41342653 | 41343736 | 41343822 | 380 | 289 | 345 | 474 | 578 | 1 | 0 | 0 | 0 | 0 | 67 | 202 | 328 | 217 | 30 | 0 | 0 | 0 | 0 | 0 | 285 | 149 | 1 | 1 | 0.995 | 1 | 1 | 1 | 1 | 1 | 1 | 1 | 1 | 1 | -0.001 |
| SE | 87325 | ENSG00000215301 | DDX3X | chrX | + | 41343736 | 41343822 | 41342639 | 41343351 | 41344238 | 41344399 | 188 | 124 | 164 | 235 | 245 | 1 | 0 | 0 | 2 | 0 | 23 | 68 | 173 | 101 | 5 | 0 | 0 | 0 | 0 | 0 | 235 | 149 | 1 | 1 | 0.992 | 1 | 1 | 0.987 | 1 | 1 | 1 | 1 | 1 | 1 | -0.004 |
| SE | 87328 | ENSG00000215301 | DDX3X | chrX | + | 41343736 | 41344128 | 41343215 | 41343351 | 41344238 | 41344399 | 773 | 514 | 702 | 957 | 884 | 1 | 0 | 0 | 2 | 0 | 126 | 379 | 599 | 391 | 46 | 0 | 0 | 0 | 0 | 0 | 298 | 149 | 1 | 1 | 0.997 | 1 | 1 | 0.996 | 1 | 1 | 1 | 1 | 1 | 1 | -0.001 |
| SE | 87329 | ENSG00000215301 | DDX3X | chrX | + | 41344029 | 41344128 | 41343215 | 41343351 | 41344238 | 41344399 | 574 | 380 | 525 | 692 | 613 | 1 | 0 | 0 | 2 | 0 | 97 | 314 | 420 | 290 | 51 | 0 | 0 | 0 | 0 | 0 | 248 | 149 | 1 | 1 | 0.997 | 1 | 1 | 0.995 | 1 | 1 | 1 | 1 | 1 | 1 | -0.002 |
| SE | 87331 | ENSG00000215301 | DDX3X | chrX | + | 41344238 | 41344399 | 41339942 | 41344128 | 41345179 | 41345324 | 1335 | 878 | 1100 | 1503 | 1413 | 1 | 0 | 0 | 0 | 0 | 161 | 666 | 864 | 510 | 74 | 0 | 0 | 0 | 0 | 0 | 298 | 149 | 1 | 1 | 0.999 | 1 | 1 | 1 | 1 | 1 | 1 | 1 | 1 | 1 | 0 |
| SE | 87332 | ENSG00000215301 | DDX3X | chrX | + | 41344238 | 41344564 | 41344029 | 41344128 | 41345179 | 41345324 | 692 | 447 | 601 | 809 | 783 | 1 | 0 | 0 | 0 | 0 | 117 | 341 | 482 | 330 | 48 | 0 | 0 | 0 | 0 | 0 | 298 | 149 | 1 | 1 | 0.997 | 1 | 1 | 1 | 1 | 1 | 1 | 1 | 1 | 1 | -0.001 |
| SE | 87333 | ENSG00000215301 | DDX3X | chrX | + | 41345179 | 41345324 | 41344190 | 41344399 | 41345403 | 41345548 | 1747 | 1181 | 1546 | 1934 | 2240 | 0 | 0 | 2 | 0 | 0 | 175 | 785 | 1145 | 487 | 116 | 0 | 0 | 0 | 0 | 0 | 294 | 149 | 1 | 1 | 1 | 1 | 0.997 | 1 | 1 | 1 | 1 | 1 | 1 | 1 | -0.001 |
| SE | 87340 | ENSG00000215301 | DDX3X | chrX | + | 41347311 | 41347451 | 41346858 | 41347012 | 41364273 | 41364472 | 627 | 450 | 476 | 631 | 853 | 0 | 0 | 0 | 0 | 1 | 65 | 228 | 418 | 223 | 40 | 0 | 0 | 0 | 0 | 0 | 289 | 149 | 1 | 1 | 1 | 1 | 1 | 1 | 0.998 | 1 | 1 | 1 | 1 | 1 | 0 |
| SE | 87343 | ENSG00000215301 | DDX3X | chrX | + | 41347314 | 41347451 | 41346858 | 41347012 | 41364273 | 41364463 | 297 | 187 | 283 | 290 | 407 | 0 | 0 | 0 | 0 | 1 | 9 | 168 | 243 | 119 | 36 | 0 | 0 | 0 | 0 | 0 | 286 | 149 | 1 | 1 | 1 | 1 | 1 | 1 | 0.995 | 1 | 1 | 1 | 1 | 1 | -0.001 |
| SE | 87346 | ENSG00000215301 | DDX3X | chrX | + | 41347639 | 41347915 | 41346858 | 41347012 | 41364273 | 41364472 | 39 | 22 | 19 | 58 | 50 | 0 | 0 | 0 | 0 | 1 | 12 | 54 | 56 | 33 | 0 | 0 | 0 | 0 | 0 | 0 | 298 | 149 | 1 | 1 | 1 | 1 | 1 | 1 | 0.962 | 1 | 1 | 1 | 1 | NA | -0.008 |
| SE | 87354 | ENSG00000147677 | EIF3H | chr8 | - | 116648805 | 116648926 | 116646470 | 116646603 | 116655855 | 116656005 | 2352 | 1807 | 2358 | 2356 | 4904 | 0 | 0 | 1 | 0 | 0 | 1874 | 1569 | 2040 | 1683 | 823 | 0 | 0 | 7 | 0 | 0 | 270 | 149 | 1 | 1 | 1 | 1 | 0.999 | 1 | 1 | 1 | 1 | 0.994 | 1 | 1 | 0.001 |
| SE | 87355 | ENSG00000147677 | EIF3H | chr8 | - | 116655855 | 116656005 | 116648805 | 116648926 | 116657214 | 116657314 | 3224 | 2436 | 3098 | 3063 | 6561 | 3 | 0 | 0 | 0 | 7 | 2380 | 1864 | 2479 | 2074 | 1174 | 0 | 0 | 0 | 5 | 0 | 298 | 149 | 1 | 1 | 0.998 | 1 | 1 | 1 | 0.998 | 1 | 1 | 1 | 0.995 | 1 | 0 |
| SE | 87356 | ENSG00000147677 | EIF3H | chr8 | - | 116655855 | 116656005 | 116648805 | 116648926 | 116726015 | 116726160 | 1814 | 1362 | 1708 | 1788 | 3794 | 2 | 0 | 2 | 3 | 0 | 1472 | 1058 | 1440 | 1227 | 665 | 0 | 0 | 0 | 0 | 0 | 298 | 149 | 1 | 1 | 0.998 | 1 | 0.998 | 0.997 | 1 | 1 | 1 | 1 | 1 | 1 | -0.001 |
| SE | 87357 | ENSG00000147677 | EIF3H | chr8 | - | 116655855 | 116656005 | 116648805 | 116648926 | 116755665 | 116755784 | 1819 | 1362 | 1707 | 1788 | 3791 | 1 | 0 | 1 | 0 | 0 | 1472 | 1058 | 1440 | 1227 | 665 | 0 | 0 | 1 | 0 | 0 | 298 | 149 | 1 | 1 | 0.999 | 1 | 0.999 | 1 | 1 | 1 | 1 | 0.999 | 1 | 1 | 0 |
| SE | 87358 | ENSG00000147677 | EIF3H | chr8 | - | 116657214 | 116657314 | 116655855 | 116656005 | 116658812 | 116658980 | 2147 | 1504 | 2220 | 1959 | 4325 | 1 | 0 | 0 | 0 | 3 | 1265 | 1219 | 1458 | 1182 | 776 | 0 | 1 | 1 | 0 | 0 | 249 | 149 | 1 | 1 | 0.999 | 1 | 1 | 1 | 0.999 | 1 | 0.999 | 0.999 | 1 | 1 | 0 |
| SE | 87359 | ENSG00000147677 | EIF3H | chr8 | - | 116658812 | 116658980 | 116655855 | 116656005 | 116726015 | 116726160 | 1101 | 678 | 1399 | 1131 | 2022 | 0 | 2 | 3 | 0 | 3 | 403 | 664 | 597 | 533 | 238 | 0 | 0 | 0 | 0 | 0 | 298 | 149 | 1 | 1 | 1 | 0.994 | 0.996 | 1 | 0.997 | 1 | 1 | 1 | 1 | 1 | -0.003 |
| SE | 87360 | ENSG00000147677 | EIF3H | chr8 | - | 116689068 | 116689140 | 116658831 | 116658980 | 116726015 | 116726172 | 2 | 0 | 4 | 1 | 2 | 1100 | 678 | 1399 | 1131 | 2019 | 0 | 0 | 0 | 0 | 0 | 403 | 663 | 596 | 533 | 238 | 221 | 149 | 1 | 1 | 0.001 | 0 | 0.002 | 0.001 | 0.001 | 0 | 0 | 0 | 0 | 0 | 0.001 |
| SE | 87361 | ENSG00000147677 | EIF3H | chr8 | - | 116697052 | 116697237 | 116658812 | 116658980 | 116726015 | 116726172 | 0 | 0 | 0 | 1 | 0 | 1100 | 678 | 1399 | 1131 | 2019 | 0 | 0 | 0 | 0 | 0 | 403 | 663 | 596 | 533 | 238 | 298 | 149 | 1 | 1 | 0 | 0 | 0 | 0 | 0 | 0 | 0 | 0 | 0 | 0 | 0 |
| SE | 87363 | ENSG00000147677 | EIF3H | chr8 | - | 116726015 | 116726172 | 116648805 | 116648926 | 116755665 | 116755797 | 768 | 477 | 1036 | 885 | 1055 | 1 | 0 | 1 | 0 | 0 | 471 | 538 | 470 | 428 | 297 | 0 | 0 | 1 | 0 | 0 | 298 | 149 | 1 | 1 | 0.997 | 1 | 0.998 | 1 | 1 | 1 | 1 | 0.996 | 1 | 1 | 0 |
| SE | 87364 | ENSG00000147677 | EIF3H | chr8 | - | 116726015 | 116726172 | 116655855 | 116656005 | 116755665 | 116755797 | 766 | 479 | 1037 | 882 | 1058 | 5 | 2 | 2 | 0 | 0 | 471 | 538 | 470 | 428 | 297 | 0 | 0 | 0 | 0 | 0 | 298 | 149 | 1 | 1 | 0.987 | 0.992 | 0.996 | 1 | 1 | 1 | 1 | 1 | 1 | 1 | -0.005 |
| SE | 87365 | ENSG00000147677 | EIF3H | chr8 | - | 116726015 | 116726172 | 116658248 | 116658980 | 116737236 | 116737287 | 1100 | 681 | 1401 | 1133 | 2022 | 0 | 0 | 1 | 4 | 0 | 405 | 663 | 598 | 535 | 238 | 0 | 0 | 0 | 0 | 0 | 298 | 149 | 1 | 1 | 1 | 1 | 0.999 | 0.993 | 1 | 1 | 1 | 1 | 1 | 1 | -0.002 |
| SE | 87368 | ENSG00000147677 | EIF3H | chr8 | - | 116734276 | 116734324 | 116726015 | 116726172 | 116755665 | 116755822 | 0 | 3 | 3 | 0 | 0 | 766 | 477 | 1034 | 882 | 1055 | 0 | 2 | 0 | 0 | 0 | 471 | 538 | 470 | 428 | 297 | 197 | 149 | 1 | 1 | 0 | 0.005 | 0.002 | 0 | 0 | 0 | 0.003 | 0 | 0 | 0 | 0.001 |
| SE | 87369 | ENSG00000147677 | EIF3H | chr8 | - | 116734276 | 116734354 | 116726015 | 116726172 | 116755665 | 116755823 | 0 | 3 | 2 | 0 | 0 | 766 | 477 | 1034 | 882 | 1055 | 0 | 0 | 0 | 0 | 0 | 471 | 538 | 470 | 428 | 297 | 227 | 149 | 1 | 1 | 0 | 0.004 | 0.001 | 0 | 0 | 0 | 0 | 0 | 0 | 0 | 0.001 |
| SE | 87370 | ENSG00000147677 | EIF3H | chr8 | - | 116737236 | 116737287 | 116658812 | 116658980 | 116755665 | 116755829 | 0 | 1 | 1 | 4 | 0 | 2 | 8 | 18 | 12 | 6 | 0 | 0 | 0 | 2 | 0 | 4 | 13 | 27 | 0 | 0 | 200 | 149 | 1 | 1 | 0 | 0.085 | 0.04 | 0.199 | 0 | 0 | 0 | 0 | 1 | NA | -0.185 |
| SE | 87371 | ENSG00000147677 | EIF3H | chr8 | - | 116737236 | 116737287 | 116726015 | 116726172 | 116755665 | 116755829 | 0 | 4 | 2 | 2 | 3 | 766 | 477 | 1034 | 882 | 1055 | 2 | 0 | 2 | 4 | 0 | 471 | 538 | 470 | 428 | 297 | 200 | 149 | 1 | 1 | 0 | 0.006 | 0.001 | 0.002 | 0.002 | 0.003 | 0 | 0.003 | 0.007 | 0 | 0 |
| SE | 87372 | ENSG00000134982 | APC | chr5 | + | 112738352 | 112738460 | 112737894 | 112737925 | 112754872 | 112755025 | 0 | 0 | 3 | 0 | 0 | 1 | 0 | 4 | 4 | 0 | 0 | 0 | 0 | 0 | 0 | 0 | 0 | 2 | 0 | 0 | 257 | 149 | 1 | 1 | 0 | NA | 0.303 | 0 | NA | NA | NA | 0 | NA | NA | 0.101 |
| SE | 87374 | ENSG00000134982 | APC | chr5 | + | 112754872 | 112755025 | 112737884 | 112737925 | 112766325 | 112766383 | 73 | 51 | 66 | 71 | 33 | 0 | 0 | 1 | 0 | 0 | 41 | 23 | 15 | 17 | 12 | 0 | 0 | 2 | 0 | 0 | 298 | 149 | 1 | 1 | 1 | 1 | 0.971 | 1 | 1 | 1 | 1 | 1 | 1 | 1 | -0.006 |
| SE | 87375 | ENSG00000134982 | APC | chr5 | + | 112766325 | 112766410 | 112754872 | 112755025 | 112767188 | 112767322 | 120 | 74 | 94 | 104 | 97 | 0 | 0 | 1 | 3 | 0 | 49 | 40 | 31 | 28 | 27 | 0 | 1 | 0 | 0 | 0 | 234 | 149 | 1 | 1 | 1 | 1 | 0.984 | 0.957 | 1 | 1 | 0.962 | 1 | 1 | 1 | -0.004 |
| SE | 87376 | ENSG00000134982 | APC | chr5 | + | 112775628 | 112775737 | 112767188 | 112767390 | 112780789 | 112780903 | 206 | 171 | 144 | 208 | 229 | 0 | 0 | 0 | 1 | 0 | 25 | 87 | 70 | 64 | 87 | 0 | 0 | 2 | 0 | 0 | 258 | 149 | 1 | 1 | 1 | 1 | 1 | 0.992 | 1 | 1 | 1 | 0.953 | 1 | 1 | 0.008 |
| SE | 87377 | ENSG00000134982 | APC | chr5 | + | 112792445 | 112792529 | 112780789 | 112780903 | 112801278 | 112801383 | 132 | 126 | 101 | 102 | 147 | 0 | 0 | 1 | 0 | 0 | 4 | 44 | 57 | 35 | 64 | 0 | 0 | 0 | 0 | 0 | 233 | 149 | 1 | 1 | 1 | 1 | 0.985 | 1 | 1 | 1 | 1 | 1 | 1 | 1 | -0.003 |
| SE | 87378 | ENSG00000134982 | APC | chr5 | + | 112810125 | 112810311 | 112801278 | 112801383 | 112815494 | 112815593 | 1 | 0 | 1 | 1 | 1 | 76 | 58 | 73 | 69 | 116 | 0 | 0 | 3 | 0 | 0 | 0 | 30 | 28 | 20 | 13 | 298 | 149 | 1 | 1 | 0.007 | 0 | 0.007 | 0.007 | 0.004 | NA | 0 | 0.051 | 0 | 0 | -0.008 |
| SE | 87379 | ENSG00000134982 | APC | chr5 | + | 112815494 | 112815593 | 112801278 | 112801383 | 112818965 | 112819344 | 99 | 80 | 94 | 92 | 133 | 0 | 0 | 0 | 1 | 0 | 0 | 41 | 33 | 24 | 13 | 0 | 0 | 0 | 0 | 0 | 248 | 149 | 1 | 1 | 1 | 1 | 1 | 0.982 | 1 | NA | 1 | 1 | 1 | 1 | -0.004 |
| SE | 87381 | ENSG00000134982 | APC | chr5 | + | 112821895 | 112821991 | 112818965 | 112819344 | 112827107 | 112827247 | 105 | 112 | 65 | 132 | 130 | 0 | 0 | 0 | 0 | 2 | 21 | 60 | 31 | 41 | 10 | 0 | 0 | 0 | 0 | 0 | 245 | 149 | 1 | 1 | 1 | 1 | 1 | 1 | 0.975 | 1 | 1 | 1 | 1 | 1 | -0.005 |
| SE | 87383 | ENSG00000134982 | APC | chr5 | + | 112827107 | 112827247 | 112821895 | 112821991 | 112827928 | 112828006 | 148 | 126 | 96 | 143 | 123 | 0 | 0 | 0 | 0 | 0 | 1 | 68 | 37 | 39 | 15 | 0 | 0 | 2 | 0 | 0 | 289 | 149 | 1 | 1 | 1 | 1 | 1 | 1 | 1 | 1 | 1 | 0.905 | 1 | 1 | 0.019 |
| SE | 87385 | ENSG00000115128 | SF3B6 | chr2 | - | 24074075 | 24074194 | 24068320 | 24068459 | 24076199 | 24076326 | 807 | 457 | 616 | 649 | 802 | 5 | 0 | 5 | 3 | 0 | 782 | 620 | 583 | 975 | 407 | 0 | 4 | 1 | 6 | 0 | 268 | 149 | 1 | 1 | 0.989 | 1 | 0.986 | 0.992 | 1 | 1 | 0.989 | 0.997 | 0.989 | 1 | -0.002 |
| SE | 87386 | ENSG00000117724 | CENPF | chr1 | + | 214644556 | 214647400 | 214639920 | 214643324 | 214648674 | 214648827 | 20 | 7 | 25 | 32 | 14 | 2 | 0 | 0 | 0 | 0 | 0 | 0 | 1 | 11 | 0 | 0 | 0 | 0 | 0 | 0 | 298 | 149 | 1 | 1 | 0.833 | 1 | 1 | 1 | 1 | NA | NA | 1 | 1 | NA | -0.033 |
| SE | 87395 | ENSG00000234665 | LERFS | chr9 | - | 62877148 | 62877428 | 62869076 | 62869304 | 62897418 | 62898095 | 0 | 0 | 0 | 0 | 0 | 16 | 0 | 0 | 8 | 0 | 0 | 0 | 1 | 0 | 0 | 0 | 1 | 2 | 0 | 0 | 298 | 149 | 1 | 1 | 0 | NA | NA | 0 | NA | NA | 0 | 0.2 | NA | NA | -0.1 |
| SE | 87396 | ENSG00000234665 | LERFS | chr9 | - | 62880522 | 62880630 | 62869189 | 62869304 | 62897418 | 62897707 | 0 | 0 | 0 | 6 | 10 | 16 | 0 | 0 | 8 | 0 | 0 | 0 | 0 | 0 | 0 | 0 | 1 | 2 | 0 | 0 | 257 | 149 | 1 | 1 | 0 | NA | NA | 0.303 | 1 | NA | 0 | 0 | NA | NA | 0.434 |
| SE | 87397 | ENSG00000234665 | LERFS | chr9 | - | 62880522 | 62880630 | 62869189 | 62869304 | 62897528 | 62898095 | 0 | 0 | 0 | 6 | 10 | 1 | 0 | 0 | 1 | 0 | 0 | 3 | 0 | 0 | 0 | 0 | 0 | 0 | 0 | 0 | 257 | 149 | 1 | 1 | 0 | NA | NA | 0.777 | 1 | NA | 1 | NA | NA | NA | -0.408 |
| SE | 87398 | ENSG00000186376 | ZNF75D | chrX | - | 135291008 | 135291135 | 135285790 | 135287846 | 135295767 | 135296033 | 35 | 15 | 27 | 29 | 49 | 0 | 0 | 0 | 0 | 0 | 0 | 4 | 9 | 14 | 5 | 0 | 0 | 1 | 0 | 0 | 276 | 149 | 1 | 1 | 1 | 1 | 1 | 1 | 1 | NA | 1 | 0.829 | 1 | 1 | 0.043 |
| SE | 87400 | ENSG00000186376 | ZNF75D | chrX | - | 135291471 | 135291563 | 135291008 | 135291135 | 135292280 | 135292473 | 23 | 6 | 20 | 11 | 32 | 0 | 0 | 2 | 0 | 0 | 0 | 8 | 9 | 28 | 0 | 0 | 0 | 0 | 0 | 0 | 241 | 149 | 1 | 1 | 1 | 1 | 0.861 | 1 | 1 | NA | 1 | 1 | 1 | NA | -0.028 |
| SE | 87401 | ENSG00000186376 | ZNF75D | chrX | - | 135291471 | 135291563 | 135291008 | 135291135 | 135341767 | 135344109 | 12 | 1 | 14 | 5 | 16 | 0 | 0 | 0 | 1 | 2 | 0 | 3 | 5 | 5 | 0 | 19 | 0 | 0 | 2 | 0 | 241 | 149 | 1 | 1 | 1 | 1 | 1 | 0.756 | 0.832 | 0 | 1 | 1 | 0.607 | NA | 0.266 |
| SE | 87403 | ENSG00000186376 | ZNF75D | chrX | - | 135292280 | 135292473 | 135291008 | 135291135 | 135295767 | 135296039 | 3 | 1 | 5 | 0 | 3 | 1 | 2 | 0 | 0 | 3 | 0 | 5 | 3 | 1 | 0 | 0 | 0 | 0 | 0 | 0 | 298 | 149 | 1 | 1 | 0.6 | 0.2 | 1 | NA | 0.333 | NA | 1 | 1 | 1 | NA | -0.467 |
| SE | 87404 | ENSG00000186376 | ZNF75D | chrX | - | 135292280 | 135292473 | 135291008 | 135291135 | 135341767 | 135344109 | 0 | 0 | 4 | 4 | 0 | 0 | 0 | 0 | 1 | 2 | 0 | 0 | 0 | 0 | 0 | 19 | 0 | 0 | 2 | 0 | 298 | 149 | 1 | 1 | NA | NA | 1 | 0.667 | 0 | 0 | NA | NA | 0 | NA | 0.556 |
| SE | 87405 | ENSG00000186376 | ZNF75D | chrX | - | 135292280 | 135292473 | 135291008 | 135291135 | 135343260 | 135344087 | 1 | 0 | 2 | 2 | 1 | 2 | 2 | 1 | 0 | 0 | 0 | 0 | 0 | 5 | 0 | 0 | 0 | 0 | 0 | 0 | 298 | 149 | 1 | 1 | 0.2 | 0 | 0.5 | 1 | 1 | NA | NA | NA | 1 | NA | -0.46 |
| SE | 87406 | ENSG00000186376 | ZNF75D | chrX | - | 135292280 | 135292473 | 135291471 | 135291563 | 135341767 | 135344109 | 11 | 5 | 8 | 10 | 16 | 0 | 0 | 0 | 0 | 0 | 0 | 5 | 5 | 23 | 0 | 0 | 0 | 1 | 0 | 0 | 298 | 149 | 1 | 1 | 1 | 1 | 1 | 1 | 1 | NA | 1 | 0.714 | 1 | NA | 0.095 |
| SE | 87407 | ENSG00000186376 | ZNF75D | chrX | - | 135293729 | 135294258 | 135291008 | 135291135 | 135295767 | 135296033 | 21 | 11 | 12 | 13 | 13 | 1 | 2 | 0 | 0 | 3 | 0 | 6 | 0 | 3 | 0 | 0 | 0 | 0 | 0 | 0 | 298 | 149 | 1 | 1 | 0.913 | 0.733 | 1 | 1 | 0.684 | NA | 1 | NA | 1 | NA | -0.134 |
| SE | 87408 | ENSG00000186376 | ZNF75D | chrX | - | 135293729 | 135294258 | 135291008 | 135291135 | 135341767 | 135344109 | 0 | 0 | 1 | 1 | 0 | 0 | 0 | 0 | 1 | 2 | 0 | 0 | 0 | 0 | 0 | 19 | 0 | 0 | 2 | 0 | 298 | 149 | 1 | 1 | NA | NA | 1 | 0.333 | 0 | 0 | NA | NA | 0 | NA | 0.444 |
| SE | 87410 | ENSG00000186376 | ZNF75D | chrX | - | 135293729 | 135294258 | 135291471 | 135291563 | 135341767 | 135344109 | 24 | 16 | 9 | 29 | 15 | 0 | 0 | 2 | 4 | 0 | 4 | 4 | 0 | 7 | 6 | 0 | 0 | 0 | 0 | 0 | 298 | 149 | 1 | 1 | 1 | 1 | 0.692 | 0.784 | 1 | 1 | 1 | NA | 1 | 1 | -0.105 |
| SE | 87411 | ENSG00000186376 | ZNF75D | chrX | - | 135295767 | 135296039 | 135285790 | 135287846 | 135341767 | 135344109 | 11 | 2 | 4 | 14 | 10 | 2 | 0 | 1 | 0 | 0 | 0 | 0 | 4 | 5 | 5 | 0 | 1 | 3 | 0 | 0 | 298 | 149 | 1 | 1 | 0.733 | 1 | 0.667 | 1 | 1 | NA | 0 | 0.4 | 1 | 1 | 0.28 |
| SE | 87412 | ENSG00000186376 | ZNF75D | chrX | - | 135295767 | 135296039 | 135291008 | 135291135 | 135341767 | 135344109 | 12 | 4 | 4 | 14 | 13 | 0 | 0 | 0 | 1 | 2 | 0 | 0 | 3 | 5 | 5 | 19 | 0 | 0 | 2 | 0 | 298 | 149 | 1 | 1 | 1 | 1 | 1 | 0.875 | 0.765 | 0 | NA | 1 | 0.556 | 1 | 0.289 |
| SE | 87413 | ENSG00000186376 | ZNF75D | chrX | - | 135295767 | 135296039 | 135291008 | 135291135 | 135343260 | 135344087 | 3 | 2 | 2 | 3 | 4 | 2 | 2 | 1 | 0 | 0 | 0 | 0 | 1 | 0 | 0 | 0 | 0 | 0 | 0 | 0 | 298 | 149 | 1 | 1 | 0.429 | 0.333 | 0.5 | 1 | 1 | NA | NA | 1 | NA | NA | -0.348 |
| SE | 87414 | ENSG00000186376 | ZNF75D | chrX | - | 135295767 | 135296039 | 135292280 | 135292473 | 135341767 | 135344109 | 14 | 3 | 7 | 14 | 13 | 0 | 0 | 2 | 4 | 0 | 0 | 5 | 6 | 6 | 5 | 0 | 0 | 0 | 0 | 0 | 298 | 149 | 1 | 1 | 1 | 1 | 0.636 | 0.636 | 1 | NA | 1 | 1 | 1 | 1 | -0.146 |
| SE | 87415 | ENSG00000186376 | ZNF75D | chrX | - | 135295767 | 135296039 | 135292280 | 135292473 | 135343260 | 135344087 | 5 | 1 | 5 | 3 | 4 | 1 | 0 | 0 | 2 | 1 | 0 | 5 | 4 | 1 | 0 | 0 | 0 | 0 | 5 | 0 | 298 | 149 | 1 | 1 | 0.714 | 1 | 1 | 0.429 | 0.667 | NA | 1 | 1 | 0.091 | NA | 0.065 |
| SE | 87416 | ENSG00000186376 | ZNF75D | chrX | - | 135295767 | 135296039 | 135293729 | 135294258 | 135341767 | 135344109 | 32 | 13 | 16 | 27 | 23 | 0 | 0 | 1 | 1 | 0 | 0 | 6 | 3 | 8 | 5 | 0 | 0 | 0 | 0 | 0 | 298 | 149 | 1 | 1 | 1 | 1 | 0.889 | 0.931 | 1 | NA | 1 | 1 | 1 | 1 | -0.036 |
| SE | 87418 | ENSG00000021762 | OSBPL5 | chr11 | - | 3093745 | 3093835 | 3093384 | 3093663 | 3094236 | 3094334 | 388 | 135 | 156 | 216 | 115 | 0 | 0 | 0 | 1 | 0 | 290 | 195 | 136 | 111 | 333 | 0 | 0 | 0 | 0 | 0 | 239 | 149 | 1 | 1 | 1 | 1 | 1 | 0.993 | 1 | 1 | 1 | 1 | 1 | 1 | -0.001 |
| SE | 87421 | ENSG00000021762 | OSBPL5 | chr11 | - | 3107262 | 3107455 | 3104192 | 3104377 | 3107770 | 3107945 | 144 | 30 | 44 | 61 | 11 | 0 | 0 | 0 | 0 | 0 | 54 | 49 | 22 | 37 | 101 | 0 | 0 | 1 | 0 | 0 | 298 | 149 | 1 | 1 | 1 | 1 | 1 | 1 | 1 | 1 | 1 | 0.917 | 1 | 1 | 0.017 |
| SE | 87424 | ENSG00000021762 | OSBPL5 | chr11 | - | 3121996 | 3122098 | 3119546 | 3119631 | 3122347 | 3122428 | 67 | 14 | 20 | 32 | 16 | 0 | 0 | 0 | 1 | 0 | 62 | 37 | 40 | 27 | 24 | 0 | 0 | 0 | 0 | 0 | 251 | 149 | 1 | 1 | 1 | 1 | 1 | 0.95 | 1 | 1 | 1 | 1 | 1 | 1 | -0.01 |
| SE | 87426 | ENSG00000125351 | UPF3B | chrX | - | 119840645 | 119840684 | 119838366 | 119838527 | 119841075 | 119841258 | 0 | 1 | 3 | 2 | 2 | 24 | 24 | 8 | 7 | 18 | 0 | 3 | 0 | 2 | 0 | 5 | 14 | 27 | 31 | 0 | 188 | 149 | 1 | 1 | 0 | 0.032 | 0.229 | 0.185 | 0.081 | 0 | 0.145 | 0 | 0.049 | NA | 0.057 |
| SE | 87428 | ENSG00000125351 | UPF3B | chrX | - | 119845197 | 119845296 | 119843190 | 119843301 | 119851494 | 119851601 | 82 | 65 | 68 | 90 | 111 | 0 | 2 | 0 | 1 | 2 | 13 | 99 | 68 | 50 | 4 | 0 | 3 | 0 | 0 | 0 | 248 | 149 | 1 | 1 | 1 | 0.951 | 1 | 0.982 | 0.971 | 1 | 0.952 | 1 | 1 | 1 | -0.01 |
| SE | 87429 | ENSG00000183354 | KIAA2026 | chr9 | - | 5910614 | 5910707 | 5881595 | 5881675 | 5913857 | 5913860 | 35 | 14 | 21 | 28 | 43 | 0 | 0 | 4 | 1 | 0 | 0 | 32 | 19 | 11 | 0 | 0 | 0 | 3 | 1 | 0 | 242 | 149 | 1 | 1 | 1 | 1 | 0.764 | 0.945 | 1 | NA | 1 | 0.796 | 0.871 | NA | 0.053 |
| SE | 87430 | ENSG00000183354 | KIAA2026 | chr9 | - | 5913857 | 5914062 | 5881595 | 5881675 | 5920543 | 5921097 | 28 | 16 | 21 | 21 | 30 | 1 | 0 | 2 | 0 | 0 | 4 | 23 | 29 | 9 | 8 | 0 | 0 | 0 | 0 | 0 | 298 | 149 | 1 | 1 | 0.933 | 1 | 0.84 | 1 | 1 | 1 | 1 | 1 | 1 | 1 | -0.045 |
| SE | 87431 | ENSG00000183354 | KIAA2026 | chr9 | - | 5913857 | 5914062 | 5881595 | 5881675 | 5933750 | 5933881 | 0 | 2 | 4 | 1 | 0 | 0 | 2 | 0 | 0 | 0 | 4 | 0 | 3 | 1 | 0 | 0 | 0 | 0 | 0 | 0 | 298 | 149 | 1 | 1 | NA | 0.333 | 1 | 1 | NA | 1 | NA | 1 | 1 | NA | -0.222 |
| SE | 87432 | ENSG00000183354 | KIAA2026 | chr9 | - | 5913857 | 5914062 | 5910614 | 5910707 | 5924657 | 5924845 | 20 | 9 | 21 | 24 | 30 | 0 | 0 | 0 | 0 | 0 | 0 | 24 | 16 | 7 | 0 | 0 | 0 | 1 | 0 | 0 | 298 | 149 | 1 | 1 | 1 | 1 | 1 | 1 | 1 | NA | 1 | 0.889 | 1 | NA | 0.037 |
| SE | 87433 | ENSG00000183354 | KIAA2026 | chr9 | - | 5933750 | 5933881 | 5913857 | 5914062 | 5944870 | 5944960 | 72 | 46 | 55 | 75 | 184 | 0 | 0 | 0 | 0 | 0 | 15 | 67 | 55 | 7 | 25 | 0 | 0 | 0 | 1 | 0 | 280 | 149 | 1 | 1 | 1 | 1 | 1 | 1 | 1 | 1 | 1 | 1 | 0.788 | 1 | 0.042 |
| SE | 87435 | ENSG00000183354 | KIAA2026 | chr9 | - | 5933750 | 5933881 | 5929043 | 5929159 | 5954015 | 5954095 | 18 | 13 | 22 | 18 | 22 | 29 | 22 | 11 | 21 | 29 | 8 | 19 | 15 | 0 | 0 | 0 | 22 | 13 | 13 | 0 | 280 | 149 | 1 | 1 | 0.248 | 0.239 | 0.516 | 0.313 | 0.288 | 1 | 0.315 | 0.38 | 0 | NA | -0.103 |
| SE | 87436 | ENSG00000183354 | KIAA2026 | chr9 | - | 5933750 | 5933881 | 5929043 | 5929159 | 5968018 | 5968081 | 10 | 8 | 16 | 17 | 7 | 6 | 0 | 2 | 4 | 5 | 8 | 2 | 11 | 0 | 0 | 0 | 0 | 2 | 6 | 0 | 280 | 149 | 1 | 1 | 0.47 | 1 | 0.81 | 0.693 | 0.427 | 1 | 1 | 0.745 | 0 | NA | -0.006 |
| SE | 87440 | ENSG00000183354 | KIAA2026 | chr9 | - | 5944870 | 5944960 | 5933750 | 5933881 | 5968018 | 5968081 | 26 | 19 | 26 | 48 | 76 | 0 | 0 | 0 | 0 | 0 | 4 | 36 | 45 | 6 | 21 | 0 | 0 | 2 | 0 | 0 | 239 | 149 | 1 | 1 | 1 | 1 | 1 | 1 | 1 | 1 | 1 | 0.933 | 1 | 1 | 0.013 |
| SE | 87442 | ENSG00000183354 | KIAA2026 | chr9 | - | 5954015 | 5954095 | 5933750 | 5933881 | 5968018 | 5968081 | 146 | 129 | 118 | 121 | 300 | 0 | 0 | 0 | 0 | 0 | 12 | 139 | 101 | 66 | 5 | 0 | 0 | 2 | 0 | 0 | 229 | 149 | 1 | 1 | 1 | 1 | 1 | 1 | 1 | 1 | 1 | 0.97 | 1 | 1 | 0.006 |
| SE | 87444 | ENSG00000183354 | KIAA2026 | chr9 | - | 5954015 | 5954095 | 5944870 | 5944960 | 5988344 | 5988545 | 112 | 100 | 96 | 90 | 198 | 0 | 0 | 2 | 0 | 0 | 21 | 125 | 72 | 26 | 9 | 0 | 0 | 2 | 0 | 0 | 229 | 149 | 1 | 1 | 1 | 1 | 0.969 | 1 | 1 | 1 | 1 | 1 | 1 | 1 | -0.006 |
| SE | 87445 | ENSG00000183354 | KIAA2026 | chr9 | - | 5968018 | 5969436 | 5944870 | 5944960 | 5988344 | 5988545 | 73 | 61 | 54 | 102 | 132 | 0 | 0 | 2 | 0 | 0 | 52 | 69 | 79 | 67 | 14 | 0 | 0 | 0 | 0 | 0 | 298 | 149 | 1 | 1 | 1 | 1 | 0.931 | 1 | 1 | 1 | 1 | 1 | 1 | 1 | -0.014 |
| SE | 87446 | ENSG00000183354 | KIAA2026 | chr9 | - | 5968018 | 5969436 | 5954015 | 5954095 | 5988344 | 5988545 | 275 | 266 | 212 | 288 | 555 | 0 | 0 | 0 | 0 | 1 | 75 | 255 | 238 | 165 | 25 | 0 | 0 | 0 | 0 | 0 | 298 | 149 | 1 | 1 | 1 | 1 | 1 | 1 | 0.996 | 1 | 1 | 1 | 1 | 1 | -0.001 |
| SE | 87447 | ENSG00000183354 | KIAA2026 | chr9 | - | 5988344 | 5988545 | 5968018 | 5969436 | 6007194 | 6007787 | 155 | 139 | 128 | 192 | 168 | 0 | 0 | 0 | 2 | 0 | 84 | 133 | 144 | 132 | 23 | 0 | 0 | 0 | 0 | 0 | 298 | 149 | 1 | 1 | 1 | 1 | 1 | 0.98 | 1 | 1 | 1 | 1 | 1 | 1 | -0.004 |
| SE | 87452 | ENSG00000100483 | VCPKMT | chr14 | - | 50113369 | 50113512 | 50108631 | 50109713 | 50114284 | 50114404 | 1 | 0 | 2 | 7 | 6 | 7 | 2 | 6 | 2 | 4 | 0 | 0 | 1 | 0 | 0 | 0 | 8 | 5 | 0 | 10 | 292 | 149 | 1 | 1 | 0.068 | 0 | 0.145 | 0.641 | 0.434 | NA | 0 | 0.093 | NA | 0 | 0.227 |
| SE | 87453 | ENSG00000100483 | VCPKMT | chr14 | - | 50113369 | 50113512 | 50112614 | 50112719 | 50114284 | 50114404 | 1 | 0 | 3 | 7 | 8 | 11 | 15 | 8 | 17 | 40 | 0 | 2 | 1 | 0 | 0 | 9 | 18 | 33 | 11 | 0 | 292 | 149 | 1 | 1 | 0.044 | 0 | 0.161 | 0.174 | 0.093 | 0 | 0.054 | 0.015 | 0 | NA | 0.077 |
| SE | 87454 | ENSG00000100483 | VCPKMT | chr14 | - | 50114284 | 50114404 | 50109536 | 50109713 | 50115838 | 50115911 | 48 | 28 | 62 | 56 | 66 | 1 | 0 | 0 | 0 | 0 | 14 | 34 | 65 | 15 | 18 | 0 | 0 | 0 | 0 | 0 | 269 | 149 | 1 | 1 | 0.964 | 1 | 1 | 1 | 1 | 1 | 1 | 1 | 1 | 1 | -0.007 |
| SE | 87456 | ENSG00000257122 | RRN3P3 | chr16 | - | 22426212 | 22426366 | 22418671 | 22420594 | 22429795 | 22429948 | 8 | 3 | 7 | 6 | 6 | 1 | 0 | 1 | 5 | 2 | 0 | 15 | 9 | 6 | 32 | 0 | 3 | 0 | 5 | 0 | 298 | 149 | 1 | 1 | 0.8 | 1 | 0.778 | 0.375 | 0.6 | NA | 0.714 | 1 | 0.375 | 1 | -0.062 |
| SE | 87459 | ENSG00000102172 | SMS | chrX | + | 21972506 | 21972571 | 21967195 | 21967316 | 21977060 | 21977236 | 308 | 208 | 210 | 246 | 207 | 2 | 0 | 1 | 1 | 0 | 398 | 282 | 231 | 316 | 240 | 0 | 0 | 0 | 0 | 0 | 214 | 149 | 1 | 1 | 0.991 | 1 | 0.993 | 0.994 | 1 | 1 | 1 | 1 | 1 | 1 | -0.004 |
| SE | 87460 | ENSG00000271147 | ARMCX5-GPRASP2 | chrX | + | 102601463 | 102601509 | 102599525 | 102599847 | 102605481 | 102605653 | 25 | 10 | 11 | 20 | 16 | 9 | 6 | 4 | 13 | 12 | 0 | 7 | 15 | 2 | 6 | 0 | 2 | 1 | 6 | 0 | 195 | 149 | 1 | 1 | 0.68 | 0.56 | 0.678 | 0.54 | 0.505 | NA | 0.728 | 0.92 | 0.203 | 1 | -0.12 |
| SE | 87461 | ENSG00000271147 | ARMCX5-GPRASP2 | chrX | + | 102601463 | 102601509 | 102599705 | 102599859 | 102605481 | 102605653 | 8 | 5 | 4 | 3 | 7 | 1 | 0 | 1 | 0 | 2 | 0 | 0 | 5 | 0 | 15 | 0 | 0 | 3 | 0 | 0 | 195 | 149 | 1 | 1 | 0.859 | 1 | 0.753 | 1 | 0.728 | NA | NA | 0.56 | NA | 1 | 0.088 |
| SE | 87462 | ENSG00000271147 | ARMCX5-GPRASP2 | chrX | + | 102601879 | 102601977 | 102599525 | 102599847 | 102605481 | 102605653 | 1 | 1 | 4 | 0 | 5 | 9 | 6 | 4 | 13 | 12 | 4 | 0 | 0 | 0 | 0 | 0 | 2 | 1 | 6 | 0 | 247 | 149 | 1 | 1 | 0.063 | 0.091 | 0.376 | 0 | 0.201 | 1 | 0 | 0 | 0 | NA | -0.104 |
| SE | 87464 | ENSG00000271147 | ARMCX5-GPRASP2 | chrX | + | 102605481 | 102605653 | 102599525 | 102599847 | 102639399 | 102639527 | 29 | 18 | 18 | 35 | 27 | 2 | 0 | 1 | 0 | 0 | 14 | 6 | 10 | 12 | 5 | 0 | 0 | 0 | 0 | 0 | 298 | 149 | 1 | 1 | 0.879 | 1 | 0.9 | 1 | 1 | 1 | 1 | 1 | 1 | 1 | -0.044 |
| SE | 87465 | ENSG00000271147 | ARMCX5-GPRASP2 | chrX | + | 102637995 | 102638045 | 102605481 | 102605653 | 102639399 | 102639624 | 1 | 0 | 0 | 0 | 0 | 20 | 12 | 14 | 22 | 15 | 0 | 0 | 0 | 0 | 0 | 14 | 4 | 9 | 6 | 5 | 199 | 149 | 1 | 1 | 0.036 | 0 | 0 | 0 | 0 | 0 | 0 | 0 | 0 | 0 | 0.007 |
| SE | 87466 | ENSG00000271147 | ARMCX5-GPRASP2 | chrX | + | 102639399 | 102639533 | 102605481 | 102605653 | 102649953 | 102650145 | 21 | 12 | 14 | 25 | 22 | 0 | 1 | 1 | 0 | 0 | 14 | 4 | 9 | 6 | 5 | 0 | 0 | 0 | 0 | 0 | 283 | 149 | 1 | 1 | 1 | 0.863 | 0.881 | 1 | 1 | 1 | 1 | 1 | 1 | 1 | -0.051 |
| SE | 87468 | ENSG00000271147 | ARMCX5-GPRASP2 | chrX | + | 102713781 | 102713849 | 102605481 | 102605653 | 102714188 | 102714199 | 4 | 0 | 1 | 1 | 3 | 0 | 0 | 0 | 0 | 0 | 0 | 0 | 6 | 4 | 0 | 0 | 0 | 0 | 8 | 0 | 217 | 149 | 1 | 1 | 1 | NA | 1 | 1 | 1 | NA | NA | 1 | 0.256 | NA | 0.372 |
| SE | 87470 | ENSG00000068366 | ACSL4 | chrX | - | 109683135 | 109683372 | 109682718 | 109682896 | 109696143 | 109696196 | 189 | 166 | 126 | 175 | 173 | 0 | 0 | 0 | 0 | 0 | 88 | 156 | 119 | 47 | 94 | 0 | 0 | 2 | 0 | 0 | 298 | 149 | 1 | 1 | 1 | 1 | 1 | 1 | 1 | 1 | 1 | 0.967 | 1 | 1 | 0.007 |
| SE | 87471 | ENSG00000068366 | ACSL4 | chrX | - | 109683135 | 109683375 | 109682718 | 109682896 | 109696143 | 109696196 | 191 | 161 | 136 | 191 | 180 | 0 | 0 | 0 | 0 | 0 | 88 | 162 | 119 | 47 | 93 | 0 | 0 | 2 | 0 | 0 | 298 | 149 | 1 | 1 | 1 | 1 | 1 | 1 | 1 | 1 | 1 | 0.967 | 1 | 1 | 0.007 |
| SE | 87472 | ENSG00000068366 | ACSL4 | chrX | - | 109683135 | 109683552 | 109682718 | 109682896 | 109696143 | 109696196 | 185 | 155 | 123 | 173 | 174 | 0 | 0 | 0 | 0 | 0 | 88 | 156 | 119 | 47 | 84 | 0 | 0 | 2 | 0 | 0 | 298 | 149 | 1 | 1 | 1 | 1 | 1 | 1 | 1 | 1 | 1 | 0.967 | 1 | 1 | 0.007 |
| SE | 87473 | ENSG00000068366 | ACSL4 | chrX | - | 109683135 | 109683666 | 109682718 | 109682896 | 109696143 | 109696196 | 190 | 155 | 124 | 172 | 173 | 0 | 0 | 0 | 0 | 0 | 88 | 156 | 120 | 47 | 84 | 0 | 0 | 2 | 0 | 0 | 298 | 149 | 1 | 1 | 1 | 1 | 1 | 1 | 1 | 1 | 1 | 0.968 | 1 | 1 | 0.006 |
| SE | 87475 | ENSG00000068366 | ACSL4 | chrX | - | 109691002 | 109691131 | 109683135 | 109683372 | 109696143 | 109696196 | 0 | 4 | 0 | 2 | 1 | 4 | 11 | 3 | 3 | 3 | 0 | 0 | 0 | 0 | 0 | 0 | 0 | 0 | 0 | 10 | 278 | 149 | 1 | 1 | 0 | 0.163 | 0 | 0.263 | 0.152 | NA | NA | NA | NA | 0 | 0.116 |
| SE | 87476 | ENSG00000068366 | ACSL4 | chrX | - | 109691002 | 109691131 | 109683135 | 109683375 | 109696143 | 109696196 | 0 | 7 | 0 | 5 | 2 | 6 | 6 | 13 | 19 | 10 | 0 | 0 | 0 | 0 | 0 | 0 | 6 | 0 | 0 | 9 | 278 | 149 | 1 | 1 | 0 | 0.385 | 0 | 0.124 | 0.097 | NA | 0 | NA | NA | 0 | 0.121 |
| SE | 87477 | ENSG00000068366 | ACSL4 | chrX | - | 109696143 | 109696196 | 109683135 | 109683372 | 109733138 | 109733198 | 7 | 12 | 3 | 4 | 4 | 0 | 0 | 1 | 0 | 0 | 0 | 0 | 0 | 0 | 10 | 0 | 0 | 0 | 0 | 0 | 202 | 149 | 1 | 1 | 1 | 1 | 0.689 | 1 | 1 | NA | NA | NA | NA | 1 | -0.062 |
| SE | 87478 | ENSG00000068366 | ACSL4 | chrX | - | 109696143 | 109696196 | 109683135 | 109683375 | 109733138 | 109733198 | 9 | 7 | 13 | 20 | 11 | 0 | 1 | 0 | 1 | 0 | 0 | 6 | 0 | 0 | 9 | 0 | 2 | 0 | 0 | 0 | 202 | 149 | 1 | 1 | 1 | 0.838 | 1 | 0.937 | 1 | NA | 0.689 | NA | NA | 1 | 0.111 |
| SE | 87480 | ENSG00000068366 | ACSL4 | chrX | - | 109696143 | 109696196 | 109683135 | 109683666 | 109733138 | 109733198 | 8 | 1 | 1 | 1 | 4 | 3 | 1 | 0 | 0 | 0 | 0 | 0 | 1 | 0 | 0 | 0 | 2 | 0 | 0 | 0 | 202 | 149 | 1 | 1 | 0.663 | 0.425 | 1 | 1 | 1 | NA | NA | 1 | NA | NA | -0.182 |
| SE | 87497 | ENSG00000198954 | KIFBP | chr10 | + | 69006841 | 69006929 | 69005731 | 69005915 | 69007626 | 69007751 | 2 | 0 | 0 | 0 | 2 | 0 | 2 | 8 | 6 | 7 | 0 | 1 | 0 | 0 | 0 | 4 | 7 | 1 | 7 | 0 | 237 | 149 | 1 | 1 | 1 | 0 | 0 | 0 | 0.152 | 0 | 0.082 | 0 | 0 | NA | 0.21 |
| SE | 87498 | ENSG00000198954 | KIFBP | chr10 | + | 69006841 | 69006929 | 69005731 | 69005915 | 69008840 | 69008925 | 1 | 0 | 0 | 0 | 0 | 10 | 14 | 8 | 19 | 9 | 0 | 1 | 0 | 0 | 0 | 16 | 3 | 1 | 1 | 0 | 237 | 149 | 1 | 1 | 0.059 | 0 | 0 | 0 | 0 | 0 | 0.173 | 0 | 0 | NA | -0.031 |
| SE | 87499 | ENSG00000198954 | KIFBP | chr10 | + | 69007626 | 69007751 | 69005731 | 69005915 | 69008840 | 69008925 | 18 | 6 | 14 | 19 | 24 | 10 | 14 | 8 | 19 | 9 | 15 | 18 | 12 | 10 | 0 | 16 | 3 | 1 | 1 | 0 | 274 | 149 | 1 | 1 | 0.495 | 0.189 | 0.488 | 0.352 | 0.592 | 0.338 | 0.765 | 0.867 | 0.845 | NA | -0.281 |
| SE | 87500 | ENSG00000198954 | KIFBP | chr10 | + | 69007626 | 69007751 | 69005731 | 69005915 | 69010899 | 69011015 | 0 | 2 | 6 | 6 | 3 | 1 | 0 | 0 | 1 | 0 | 0 | 7 | 0 | 0 | 0 | 0 | 0 | 0 | 0 | 0 | 274 | 149 | 1 | 1 | 0 | 1 | 1 | 0.765 | 1 | NA | 1 | NA | NA | NA | -0.247 |
| SE | 87502 | ENSG00000198954 | KIFBP | chr10 | + | 69008840 | 69008925 | 69005731 | 69005915 | 69010899 | 69011015 | 16 | 29 | 22 | 37 | 17 | 1 | 0 | 0 | 1 | 0 | 16 | 8 | 6 | 4 | 0 | 0 | 0 | 0 | 0 | 0 | 234 | 149 | 1 | 1 | 0.911 | 1 | 1 | 0.959 | 1 | 1 | 1 | 1 | 1 | NA | -0.026 |
| SE | 87503 | ENSG00000198954 | KIFBP | chr10 | + | 69008840 | 69008925 | 69005731 | 69007751 | 69010899 | 69011015 | 24 | 19 | 22 | 32 | 29 | 0 | 0 | 0 | 1 | 0 | 23 | 16 | 17 | 13 | 0 | 0 | 0 | 0 | 0 | 0 | 234 | 149 | 1 | 1 | 1 | 1 | 1 | 0.953 | 1 | 1 | 1 | 1 | 1 | NA | -0.009 |
| SE | 87506 | ENSG00000198954 | KIFBP | chr10 | + | 69015357 | 69015436 | 69010899 | 69011015 | 69015540 | 69016978 | 0 | 0 | 0 | 1 | 0 | 15 | 20 | 20 | 26 | 21 | 0 | 0 | 0 | 0 | 0 | 0 | 4 | 12 | 6 | 0 | 228 | 149 | 1 | 1 | 0 | 0 | 0 | 0.025 | 0 | NA | 0 | 0 | 0 | NA | 0.005 |
| SE | 87515 | ENSG00000239306 | RBM14 | chr11 | + | 66624213 | 66624324 | 66616625 | 66617057 | 66626460 | 66627343 | 40 | 16 | 38 | 31 | 17 | 0 | 0 | 2 | 0 | 0 | 36 | 30 | 6 | 29 | 48 | 0 | 0 | 0 | 1 | 0 | 260 | 149 | 1 | 1 | 1 | 1 | 0.916 | 1 | 1 | 1 | 1 | 1 | 0.943 | 1 | -0.005 |
| SE | 87516 | ENSG00000239306 | RBM14 | chr11 | + | 66624213 | 66625678 | 66616629 | 66617057 | 66626460 | 66629934 | 71 | 21 | 69 | 54 | 40 | 0 | 0 | 2 | 0 | 0 | 46 | 42 | 14 | 46 | 89 | 0 | 0 | 0 | 1 | 0 | 298 | 149 | 1 | 1 | 1 | 1 | 0.945 | 1 | 1 | 1 | 1 | 1 | 0.958 | 1 | -0.003 |
| SE | 87527 | ENSG00000147050 | KDM6A | chrX | + | 44961283 | 44961392 | 44873714 | 44873987 | 44974665 | 44974715 | 188 | 116 | 144 | 124 | 96 | 0 | 0 | 2 | 0 | 2 | 119 | 81 | 160 | 119 | 54 | 0 | 0 | 2 | 0 | 0 | 258 | 149 | 1 | 1 | 1 | 1 | 0.977 | 1 | 0.965 | 1 | 1 | 0.979 | 1 | 1 | -0.007 |
| SE | 87528 | ENSG00000147050 | KDM6A | chrX | + | 44961283 | 44961392 | 44873714 | 44873987 | 45010960 | 45011019 | 51 | 23 | 51 | 32 | 8 | 0 | 0 | 0 | 3 | 0 | 50 | 21 | 37 | 32 | 24 | 0 | 0 | 1 | 3 | 0 | 258 | 149 | 1 | 1 | 1 | 1 | 1 | 0.86 | 1 | 1 | 1 | 0.955 | 0.86 | 1 | 0.009 |
| SE | 87529 | ENSG00000147050 | KDM6A | chrX | + | 44974665 | 44974715 | 44873923 | 44873987 | 45010960 | 45011019 | 78 | 40 | 51 | 39 | 40 | 0 | 0 | 0 | 3 | 0 | 42 | 18 | 48 | 38 | 9 | 0 | 0 | 1 | 3 | 0 | 199 | 149 | 1 | 1 | 1 | 1 | 1 | 0.907 | 1 | 1 | 1 | 0.973 | 0.905 | 1 | 0.006 |
| SE | 87532 | ENSG00000147050 | KDM6A | chrX | + | 45010960 | 45011019 | 44974665 | 44974715 | 45020609 | 45020730 | 191 | 125 | 137 | 134 | 139 | 0 | 0 | 0 | 0 | 0 | 115 | 73 | 148 | 117 | 23 | 0 | 0 | 1 | 5 | 0 | 208 | 149 | 1 | 1 | 1 | 1 | 1 | 1 | 1 | 1 | 1 | 0.991 | 0.944 | 1 | 0.013 |
| SE | 87533 | ENSG00000147050 | KDM6A | chrX | + | 45010960 | 45011019 | 44974665 | 44974715 | 45034930 | 45034985 | 66 | 53 | 50 | 49 | 54 | 0 | 0 | 0 | 0 | 2 | 23 | 18 | 72 | 45 | 24 | 0 | 0 | 0 | 2 | 0 | 208 | 149 | 1 | 1 | 1 | 1 | 1 | 1 | 0.951 | 1 | 1 | 1 | 0.942 | 1 | 0.002 |
| SE | 87534 | ENSG00000147050 | KDM6A | chrX | + | 45020609 | 45020730 | 44974665 | 44974715 | 45034930 | 45034985 | 129 | 100 | 95 | 100 | 169 | 0 | 0 | 0 | 0 | 2 | 63 | 71 | 57 | 90 | 22 | 0 | 0 | 0 | 2 | 0 | 270 | 149 | 1 | 1 | 1 | 1 | 1 | 1 | 0.979 | 1 | 1 | 1 | 0.961 | 1 | 0.004 |
| SE | 87536 | ENSG00000147050 | KDM6A | chrX | + | 45051708 | 45051802 | 45037654 | 45037689 | 45053828 | 45053955 | 196 | 130 | 185 | 155 | 289 | 0 | 0 | 0 | 0 | 2 | 74 | 130 | 82 | 124 | 39 | 0 | 0 | 0 | 0 | 0 | 243 | 149 | 1 | 1 | 1 | 1 | 1 | 1 | 0.989 | 1 | 1 | 1 | 1 | 1 | -0.002 |
| SE | 87537 | ENSG00000147050 | KDM6A | chrX | + | 45053828 | 45053955 | 45051708 | 45051802 | 45059005 | 45059104 | 282 | 173 | 240 | 212 | 322 | 0 | 1 | 2 | 0 | 3 | 128 | 157 | 144 | 174 | 49 | 0 | 0 | 0 | 6 | 0 | 276 | 149 | 1 | 1 | 1 | 0.989 | 0.985 | 1 | 0.983 | 1 | 1 | 1 | 0.94 | 1 | 0.003 |
| SE | 87538 | ENSG00000147050 | KDM6A | chrX | + | 45059005 | 45059104 | 45053828 | 45053955 | 45059246 | 45059466 | 205 | 118 | 156 | 189 | 218 | 1 | 0 | 0 | 2 | 0 | 98 | 92 | 102 | 150 | 31 | 0 | 0 | 0 | 0 | 0 | 248 | 149 | 1 | 1 | 0.992 | 1 | 1 | 0.983 | 1 | 1 | 1 | 1 | 1 | 1 | -0.005 |
| SE | 87543 | ENSG00000147050 | KDM6A | chrX | + | 45061323 | 45061419 | 45059246 | 45059466 | 45063421 | 45063817 | 26 | 25 | 32 | 26 | 55 | 0 | 0 | 1 | 0 | 0 | 0 | 20 | 20 | 28 | 19 | 0 | 0 | 0 | 0 | 0 | 245 | 149 | 1 | 1 | 1 | 1 | 0.951 | 1 | 1 | NA | 1 | 1 | 1 | 1 | -0.01 |
| SE | 87546 | ENSG00000147050 | KDM6A | chrX | + | 45069578 | 45070357 | 45062646 | 45063817 | 45076696 | 45076826 | 232 | 205 | 253 | 265 | 324 | 0 | 1 | 0 | 0 | 0 | 144 | 211 | 225 | 275 | 30 | 0 | 0 | 0 | 0 | 0 | 298 | 149 | 1 | 1 | 1 | 0.99 | 1 | 1 | 1 | 1 | 1 | 1 | 1 | 1 | -0.002 |
| SE | 87547 | ENSG00000147050 | KDM6A | chrX | + | 45078399 | 45078505 | 45069578 | 45076826 | 45079145 | 45079351 | 243 | 218 | 173 | 271 | 262 | 1 | 0 | 0 | 0 | 0 | 184 | 196 | 196 | 189 | 20 | 0 | 0 | 0 | 0 | 0 | 255 | 149 | 1 | 1 | 0.993 | 1 | 1 | 1 | 1 | 1 | 1 | 1 | 1 | 1 | -0.001 |
| SE | 87548 | ENSG00000147050 | KDM6A | chrX | + | 45078399 | 45078505 | 45069578 | 45076826 | 45082575 | 45082640 | 104 | 100 | 64 | 111 | 125 | 0 | 1 | 0 | 0 | 0 | 73 | 95 | 103 | 77 | 12 | 0 | 0 | 0 | 0 | 0 | 255 | 149 | 1 | 1 | 1 | 0.983 | 1 | 1 | 1 | 1 | 1 | 1 | 1 | 1 | -0.003 |
| SE | 87549 | ENSG00000147050 | KDM6A | chrX | + | 45079145 | 45079351 | 45069578 | 45076826 | 45082575 | 45082640 | 188 | 160 | 165 | 164 | 340 | 0 | 1 | 0 | 0 | 0 | 124 | 110 | 205 | 202 | 13 | 0 | 0 | 0 | 0 | 0 | 298 | 149 | 1 | 1 | 1 | 0.988 | 1 | 1 | 1 | 1 | 1 | 1 | 1 | 1 | -0.002 |
| SE | 87550 | ENSG00000147050 | KDM6A | chrX | + | 45079145 | 45079351 | 45078399 | 45078505 | 45082575 | 45082640 | 331 | 282 | 278 | 327 | 478 | 1 | 3 | 2 | 2 | 1 | 243 | 211 | 304 | 314 | 21 | 0 | 0 | 0 | 0 | 0 | 298 | 149 | 1 | 1 | 0.994 | 0.979 | 0.986 | 0.988 | 0.996 | 1 | 1 | 1 | 1 | 1 | -0.011 |
| SE | 87551 | ENSG00000147050 | KDM6A | chrX | + | 45082575 | 45082640 | 45079145 | 45079351 | 45082714 | 45082789 | 242 | 220 | 205 | 220 | 533 | 0 | 0 | 0 | 1 | 0 | 145 | 135 | 288 | 310 | 56 | 0 | 0 | 0 | 0 | 0 | 214 | 149 | 1 | 1 | 1 | 1 | 1 | 0.994 | 1 | 1 | 1 | 1 | 1 | 1 | -0.001 |
| SE | 87552 | ENSG00000147050 | KDM6A | chrX | + | 45082714 | 45082789 | 45082397 | 45082640 | 45083459 | 45083523 | 293 | 275 | 244 | 262 | 560 | 0 | 0 | 1 | 0 | 0 | 164 | 158 | 300 | 366 | 50 | 0 | 0 | 0 | 0 | 0 | 224 | 149 | 1 | 1 | 1 | 1 | 0.994 | 1 | 1 | 1 | 1 | 1 | 1 | 1 | -0.001 |
| SE | 87553 | ENSG00000147050 | KDM6A | chrX | + | 45082714 | 45082789 | 45082397 | 45082640 | 45083518 | 45083608 | 94 | 83 | 95 | 91 | 158 | 0 | 0 | 0 | 0 | 0 | 42 | 63 | 97 | 90 | 6 | 0 | 0 | 1 | 0 | 0 | 224 | 149 | 1 | 1 | 1 | 1 | 1 | 1 | 1 | 1 | 1 | 0.985 | 1 | 1 | 0.003 |
| SE | 87555 | ENSG00000147050 | KDM6A | chrX | + | 45090722 | 45090832 | 45089742 | 45089930 | 45107409 | 45107536 | 111 | 100 | 95 | 85 | 132 | 0 | 0 | 2 | 0 | 3 | 19 | 75 | 62 | 65 | 12 | 0 | 4 | 0 | 0 | 0 | 259 | 149 | 1 | 1 | 1 | 1 | 0.965 | 1 | 0.962 | 1 | 0.915 | 1 | 1 | 1 | 0.002 |
| SE | 87556 | ENSG00000147050 | KDM6A | chrX | + | 45090722 | 45090864 | 45089742 | 45089930 | 45107409 | 45107536 | 229 | 186 | 226 | 193 | 263 | 0 | 0 | 2 | 0 | 3 | 64 | 127 | 169 | 162 | 16 | 0 | 4 | 0 | 0 | 0 | 291 | 149 | 1 | 1 | 1 | 1 | 0.983 | 1 | 0.978 | 1 | 0.942 | 1 | 1 | 1 | 0.004 |
| SE | 87559 | ENSG00000147050 | KDM6A | chrX | + | 45110078 | 45110249 | 45104223 | 45107536 | 45111381 | 45112158 | 139 | 113 | 212 | 152 | 250 | 1 | 0 | 1 | 1 | 0 | 72 | 100 | 150 | 132 | 27 | 0 | 0 | 0 | 0 | 0 | 298 | 149 | 1 | 1 | 0.986 | 1 | 0.991 | 0.987 | 1 | 1 | 1 | 1 | 1 | 1 | -0.007 |
| SE | 87561 | ENSG00000112159 | MDN1 | chr6 | - | 89656699 | 89656801 | 89655763 | 89655968 | 89658208 | 89658370 | 118 | 129 | 127 | 164 | 164 | 0 | 0 | 0 | 0 | 0 | 91 | 103 | 104 | 130 | 5 | 0 | 0 | 1 | 0 | 0 | 251 | 149 | 1 | 1 | 1 | 1 | 1 | 1 | 1 | 1 | 1 | 0.984 | 1 | 1 | 0.003 |
| SE | 87563 | ENSG00000112159 | MDN1 | chr6 | - | 89670918 | 89671080 | 89668013 | 89668151 | 89672199 | 89672363 | 72 | 46 | 74 | 68 | 39 | 0 | 0 | 1 | 1 | 0 | 37 | 67 | 40 | 46 | 0 | 0 | 0 | 0 | 0 | 0 | 298 | 149 | 1 | 1 | 1 | 1 | 0.974 | 0.971 | 1 | 1 | 1 | 1 | 1 | NA | -0.011 |
| SE | 87564 | ENSG00000112159 | MDN1 | chr6 | - | 89684875 | 89684985 | 89683830 | 89683904 | 89685826 | 89685973 | 104 | 57 | 91 | 69 | 101 | 0 | 1 | 0 | 0 | 3 | 89 | 83 | 50 | 81 | 27 | 0 | 0 | 0 | 0 | 0 | 259 | 149 | 1 | 1 | 1 | 0.97 | 1 | 1 | 0.951 | 1 | 1 | 1 | 1 | 1 | -0.016 |
| SE | 87565 | ENSG00000112159 | MDN1 | chr6 | - | 89685826 | 89685973 | 89684875 | 89684985 | 89686901 | 89687023 | 113 | 53 | 118 | 73 | 111 | 0 | 0 | 0 | 0 | 0 | 105 | 75 | 51 | 87 | 36 | 0 | 0 | 0 | 3 | 0 | 296 | 149 | 1 | 1 | 1 | 1 | 1 | 1 | 1 | 1 | 1 | 1 | 0.936 | 1 | 0.013 |
| SE | 87567 | ENSG00000112159 | MDN1 | chr6 | - | 89689869 | 89690143 | 89688077 | 89688173 | 89690672 | 89690834 | 48 | 42 | 63 | 42 | 85 | 0 | 0 | 0 | 4 | 0 | 12 | 43 | 40 | 34 | 21 | 0 | 0 | 0 | 0 | 0 | 298 | 149 | 1 | 1 | 1 | 1 | 1 | 0.84 | 1 | 1 | 1 | 1 | 1 | 1 | -0.032 |
| SE | 87568 | ENSG00000112159 | MDN1 | chr6 | - | 89689869 | 89690143 | 89688077 | 89688173 | 89692442 | 89693148 | 2 | 1 | 2 | 0 | 0 | 0 | 0 | 0 | 0 | 0 | 0 | 1 | 4 | 4 | 2 | 0 | 0 | 6 | 0 | 0 | 298 | 149 | 1 | 1 | 1 | 1 | 1 | NA | NA | NA | 1 | 0.25 | 1 | 1 | 0.188 |
| SE | 87570 | ENSG00000112159 | MDN1 | chr6 | - | 89690672 | 89690834 | 89689869 | 89690143 | 89692442 | 89693148 | 131 | 72 | 126 | 115 | 133 | 1 | 0 | 0 | 0 | 0 | 101 | 79 | 93 | 85 | 73 | 0 | 0 | 0 | 0 | 0 | 298 | 149 | 1 | 1 | 0.985 | 1 | 1 | 1 | 1 | 1 | 1 | 1 | 1 | 1 | -0.003 |
| SE | 87571 | ENSG00000112159 | MDN1 | chr6 | - | 89696359 | 89696574 | 89695604 | 89695992 | 89698864 | 89699035 | 46 | 41 | 70 | 83 | 83 | 0 | 0 | 0 | 0 | 0 | 14 | 71 | 60 | 59 | 60 | 0 | 4 | 0 | 0 | 0 | 298 | 149 | 1 | 1 | 1 | 1 | 1 | 1 | 1 | 1 | 0.899 | 1 | 1 | 1 | 0.02 |
| SE | 87572 | ENSG00000112159 | MDN1 | chr6 | - | 89700645 | 89700856 | 89700062 | 89700294 | 89701557 | 89701678 | 114 | 106 | 139 | 106 | 183 | 0 | 0 | 0 | 0 | 0 | 74 | 90 | 110 | 62 | 37 | 0 | 0 | 0 | 3 | 0 | 298 | 149 | 1 | 1 | 1 | 1 | 1 | 1 | 1 | 1 | 1 | 1 | 0.912 | 1 | 0.018 |
| SE | 87573 | ENSG00000112159 | MDN1 | chr6 | - | 89710680 | 89710794 | 89708495 | 89708628 | 89712035 | 89712256 | 116 | 55 | 112 | 79 | 129 | 0 | 0 | 0 | 0 | 0 | 21 | 93 | 68 | 82 | 8 | 0 | 0 | 1 | 0 | 0 | 263 | 149 | 1 | 1 | 1 | 1 | 1 | 1 | 1 | 1 | 1 | 0.975 | 1 | 1 | 0.005 |
| SE | 87574 | ENSG00000112159 | MDN1 | chr6 | - | 89710680 | 89710794 | 89708495 | 89708628 | 89712574 | 89712786 | 66 | 30 | 67 | 41 | 75 | 0 | 0 | 0 | 0 | 0 | 21 | 46 | 33 | 47 | 0 | 10 | 0 | 0 | 0 | 0 | 263 | 149 | 1 | 1 | 1 | 1 | 1 | 1 | 1 | 0.543 | 1 | 1 | 1 | NA | 0.114 |
| SE | 87577 | ENSG00000112159 | MDN1 | chr6 | - | 89713147 | 89713296 | 89712574 | 89712786 | 89714542 | 89714751 | 100 | 61 | 135 | 134 | 79 | 1 | 0 | 0 | 1 | 0 | 109 | 154 | 106 | 114 | 17 | 0 | 4 | 1 | 0 | 0 | 298 | 149 | 1 | 1 | 0.98 | 1 | 1 | 0.985 | 1 | 1 | 0.951 | 0.981 | 1 | 1 | 0.007 |
| SE | 87578 | ENSG00000112159 | MDN1 | chr6 | - | 89728930 | 89729139 | 89727832 | 89727955 | 89730725 | 89730923 | 184 | 127 | 203 | 205 | 293 | 0 | 0 | 0 | 0 | 0 | 200 | 148 | 121 | 150 | 52 | 0 | 0 | 0 | 4 | 0 | 298 | 149 | 1 | 1 | 1 | 1 | 1 | 1 | 1 | 1 | 1 | 1 | 0.949 | 1 | 0.01 |
| SE | 87579 | ENSG00000112159 | MDN1 | chr6 | - | 89738325 | 89738455 | 89732556 | 89732775 | 89743149 | 89743280 | 64 | 60 | 89 | 73 | 91 | 0 | 0 | 0 | 0 | 0 | 75 | 93 | 58 | 85 | 31 | 0 | 0 | 0 | 2 | 0 | 279 | 149 | 1 | 1 | 1 | 1 | 1 | 1 | 1 | 1 | 1 | 1 | 0.958 | 1 | 0.008 |
| SE | 87580 | ENSG00000112159 | MDN1 | chr6 | - | 89740233 | 89740378 | 89738325 | 89738455 | 89743149 | 89743280 | 104 | 66 | 122 | 123 | 134 | 0 | 0 | 0 | 0 | 0 | 122 | 97 | 81 | 90 | 39 | 0 | 0 | 1 | 0 | 0 | 294 | 149 | 1 | 1 | 1 | 1 | 1 | 1 | 1 | 1 | 1 | 0.976 | 1 | 1 | 0.005 |
| SE | 87584 | ENSG00000112159 | MDN1 | chr6 | - | 89751430 | 89751582 | 89750353 | 89750532 | 89753511 | 89753622 | 165 | 101 | 133 | 156 | 141 | 1 | 0 | 2 | 0 | 0 | 118 | 157 | 150 | 156 | 111 | 0 | 0 | 0 | 3 | 0 | 298 | 149 | 1 | 1 | 0.988 | 1 | 0.971 | 1 | 1 | 1 | 1 | 1 | 0.963 | 1 | -0.001 |
| SE | 87585 | ENSG00000112159 | MDN1 | chr6 | - | 89771560 | 89771621 | 89762318 | 89762530 | 89772572 | 89772721 | 101 | 96 | 146 | 113 | 115 | 3 | 0 | 0 | 0 | 0 | 131 | 147 | 115 | 83 | 85 | 0 | 0 | 4 | 0 | 0 | 210 | 149 | 1 | 1 | 0.96 | 1 | 1 | 1 | 1 | 1 | 1 | 0.953 | 1 | 1 | 0.001 |
| SE | 87587 | ENSG00000112159 | MDN1 | chr6 | - | 89776599 | 89776695 | 89774620 | 89774733 | 89781398 | 89781592 | 101 | 114 | 124 | 105 | 119 | 0 | 1 | 0 | 0 | 0 | 87 | 149 | 152 | 118 | 82 | 0 | 0 | 0 | 0 | 0 | 245 | 149 | 1 | 1 | 1 | 0.986 | 1 | 1 | 1 | 1 | 1 | 1 | 1 | 1 | -0.003 |
| SE | 87588 | ENSG00000112159 | MDN1 | chr6 | - | 89780211 | 89780293 | 89774620 | 89774733 | 89781398 | 89781592 | 29 | 33 | 35 | 63 | 96 | 0 | 1 | 0 | 0 | 0 | 32 | 64 | 56 | 23 | 13 | 0 | 0 | 0 | 0 | 0 | 231 | 149 | 1 | 1 | 1 | 0.955 | 1 | 1 | 1 | 1 | 1 | 1 | 1 | 1 | -0.009 |
| SE | 87590 | ENSG00000112159 | MDN1 | chr6 | - | 89790158 | 89790401 | 89789779 | 89789911 | 89793761 | 89793954 | 89 | 53 | 81 | 100 | 59 | 2 | 2 | 1 | 0 | 0 | 103 | 121 | 66 | 68 | 75 | 0 | 0 | 1 | 2 | 0 | 298 | 149 | 1 | 1 | 0.957 | 0.93 | 0.976 | 1 | 1 | 1 | 1 | 0.971 | 0.944 | 1 | -0.01 |
| SE | 87591 | ENSG00000112159 | MDN1 | chr6 | - | 89793761 | 89793954 | 89789779 | 89789911 | 89794099 | 89794207 | 59 | 81 | 71 | 71 | 29 | 0 | 0 | 0 | 0 | 0 | 72 | 90 | 51 | 54 | 28 | 4 | 0 | 0 | 0 | 0 | 298 | 149 | 1 | 1 | 1 | 1 | 1 | 1 | 1 | 0.9 | 1 | 1 | 1 | 1 | 0.02 |
| SE | 87592 | ENSG00000166788 | SAAL1 | chr11 | - | 18083534 | 18083709 | 18081410 | 18081503 | 18086865 | 18086960 | 35 | 18 | 35 | 25 | 20 | 0 | 0 | 0 | 0 | 0 | 20 | 11 | 21 | 12 | 20 | 0 | 0 | 2 | 0 | 0 | 298 | 149 | 1 | 1 | 1 | 1 | 1 | 1 | 1 | 1 | 1 | 0.84 | 1 | 1 | 0.032 |
| SE | 87593 | ENSG00000166788 | SAAL1 | chr11 | - | 18083534 | 18083728 | 18081410 | 18081503 | 18086865 | 18087054 | 42 | 30 | 32 | 32 | 27 | 0 | 0 | 0 | 0 | 0 | 20 | 18 | 20 | 15 | 23 | 0 | 0 | 2 | 0 | 0 | 298 | 149 | 1 | 1 | 1 | 1 | 1 | 1 | 1 | 1 | 1 | 0.833 | 1 | 1 | 0.033 |
| SE | 87594 | ENSG00000166788 | SAAL1 | chr11 | - | 18083534 | 18083731 | 18081410 | 18081503 | 18086865 | 18087054 | 69 | 43 | 56 | 58 | 67 | 0 | 0 | 0 | 0 | 0 | 75 | 53 | 54 | 25 | 49 | 0 | 0 | 2 | 0 | 0 | 298 | 149 | 1 | 1 | 1 | 1 | 1 | 1 | 1 | 1 | 1 | 0.931 | 1 | 1 | 0.014 |
| SE | 87597 | ENSG00000177112 | IRAG1-AS1 | chr11 | + | 10589011 | 10589150 | 10541260 | 10541542 | 10594116 | 10594228 | 0 | 0 | 0 | 0 | 0 | 7 | 0 | 0 | 1 | 0 | 0 | 6 | 0 | 0 | 0 | 6 | 8 | 3 | 0 | 0 | 288 | 149 | 1 | 1 | 0 | NA | NA | 0 | NA | 0 | 0.28 | 0 | NA | NA | -0.093 |
| SE | 87599 | ENSG00000227953 | LINC01341 | chr1 | + | 246781478 | 246781627 | 246780383 | 246780583 | 246789507 | 246789698 | 0 | 1 | 0 | 0 | 0 | 0 | 0 | 0 | 0 | 0 | 0 | 0 | 0 | 0 | 0 | 0 | 0 | 0 | 6 | 0 | 298 | 149 | 1 | 1 | NA | 1 | NA | NA | NA | NA | NA | NA | 0 | NA | 1 |
| SE | 87606 | ENSG00000102078 | SLC25A14 | chrX | + | 130340106 | 130340344 | 130339887 | 130339976 | 130345181 | 130345275 | 0 | 0 | 0 | 2 | 0 | 4 | 2 | 4 | 4 | 4 | 0 | 0 | 0 | 0 | 0 | 0 | 0 | 0 | 2 | 0 | 298 | 149 | 1 | 1 | 0 | 0 | 0 | 0.2 | 0 | NA | NA | NA | 0 | NA | 0.04 |
| SE | 87607 | ENSG00000102078 | SLC25A14 | chrX | + | 130340106 | 130340353 | 130339912 | 130339976 | 130345181 | 130345275 | 0 | 0 | 1 | 2 | 0 | 4 | 2 | 4 | 4 | 4 | 0 | 0 | 0 | 4 | 0 | 0 | 0 | 0 | 2 | 0 | 298 | 149 | 1 | 1 | 0 | 0 | 0.111 | 0.2 | 0 | NA | NA | NA | 0.5 | NA | -0.438 |
| SE | 87608 | ENSG00000102078 | SLC25A14 | chrX | + | 130345181 | 130345275 | 130339887 | 130339976 | 130346543 | 130346691 | 7 | 9 | 9 | 14 | 5 | 0 | 0 | 0 | 0 | 0 | 0 | 8 | 7 | 5 | 1 | 0 | 0 | 4 | 0 | 0 | 243 | 149 | 1 | 1 | 1 | 1 | 1 | 1 | 1 | NA | 1 | 0.518 | 1 | 1 | 0.121 |
| SE | 87609 | ENSG00000102078 | SLC25A14 | chrX | + | 130346543 | 130346691 | 130339887 | 130339976 | 130349250 | 130349345 | 21 | 22 | 17 | 30 | 20 | 1 | 1 | 1 | 0 | 0 | 40 | 18 | 31 | 20 | 3 | 0 | 0 | 0 | 0 | 6 | 297 | 149 | 1 | 1 | 0.913 | 0.917 | 0.895 | 1 | 1 | 1 | 1 | 1 | 1 | 0.201 | 0.105 |
| SE | 87614 | ENSG00000134597 | RBMX2 | chrX | + | 130402254 | 130402370 | 130401986 | 130402037 | 130403801 | 130403853 | 44 | 24 | 26 | 29 | 30 | 0 | 0 | 2 | 0 | 0 | 77 | 38 | 64 | 34 | 47 | 0 | 0 | 0 | 0 | 0 | 265 | 149 | 1 | 1 | 1 | 1 | 0.88 | 1 | 1 | 1 | 1 | 1 | 1 | 1 | -0.024 |
| SE | 87615 | ENSG00000134597 | RBMX2 | chrX | + | 130403801 | 130403853 | 130402000 | 130402370 | 130409256 | 130409386 | 44 | 27 | 33 | 32 | 42 | 0 | 0 | 1 | 2 | 0 | 74 | 42 | 62 | 40 | 50 | 0 | 0 | 0 | 0 | 0 | 201 | 149 | 1 | 1 | 1 | 1 | 0.961 | 0.922 | 1 | 1 | 1 | 1 | 1 | 1 | -0.023 |
| SE | 87616 | ENSG00000008838 | MED24 | chr17 | - | 40027382 | 40027465 | 40026855 | 40027034 | 40027908 | 40027946 | 76 | 46 | 78 | 73 | 132 | 0 | 0 | 0 | 2 | 0 | 103 | 125 | 51 | 68 | 167 | 0 | 1 | 0 | 0 | 0 | 232 | 149 | 1 | 1 | 1 | 1 | 1 | 0.959 | 1 | 1 | 0.988 | 1 | 1 | 1 | -0.006 |
| SE | 87619 | ENSG00000008838 | MED24 | chr17 | - | 40036115 | 40036154 | 40035721 | 40035795 | 40053297 | 40053380 | 83 | 50 | 92 | 110 | 105 | 1 | 1 | 0 | 2 | 1 | 180 | 142 | 50 | 62 | 189 | 2 | 0 | 0 | 0 | 0 | 188 | 149 | 1 | 1 | 0.985 | 0.975 | 1 | 0.978 | 0.988 | 0.986 | 1 | 1 | 1 | 1 | -0.012 |
| SE | 87625 | ENSG00000140678 | ITGAX | chr16 | + | 31355892 | 31355998 | 31355173 | 31355291 | 31356624 | 31356728 | 156 | 95 | 82 | 94 | 41 | 3 | 0 | 0 | 0 | 0 | 155 | 141 | 89 | 92 | 295 | 0 | 0 | 0 | 0 | 0 | 255 | 149 | 1 | 1 | 0.968 | 1 | 1 | 1 | 1 | 1 | 1 | 1 | 1 | 1 | -0.006 |
| SE | 87626 | ENSG00000140678 | ITGAX | chr16 | + | 31357252 | 31357364 | 31357030 | 31357101 | 31359699 | 31359830 | 264 | 237 | 144 | 182 | 148 | 4 | 3 | 1 | 0 | 0 | 342 | 339 | 179 | 190 | 594 | 0 | 4 | 0 | 0 | 0 | 261 | 149 | 1 | 1 | 0.974 | 0.978 | 0.988 | 1 | 1 | 1 | 0.98 | 1 | 1 | 1 | -0.008 |
| SE | 87627 | ENSG00000140678 | ITGAX | chr16 | + | 31359919 | 31360065 | 31359699 | 31359830 | 31361062 | 31361213 | 126 | 127 | 100 | 108 | 117 | 0 | 0 | 0 | 0 | 0 | 298 | 176 | 111 | 123 | 289 | 0 | 4 | 2 | 0 | 0 | 295 | 149 | 1 | 1 | 1 | 1 | 1 | 1 | 1 | 1 | 0.957 | 0.966 | 1 | 1 | 0.015 |
| SE | 87632 | ENSG00000140678 | ITGAX | chr16 | + | 31372377 | 31372509 | 31371473 | 31371784 | 31372596 | 31372654 | 330 | 248 | 224 | 250 | 223 | 0 | 0 | 0 | 1 | 0 | 717 | 462 | 282 | 264 | 654 | 0 | 0 | 0 | 0 | 0 | 281 | 149 | 1 | 1 | 1 | 1 | 1 | 0.993 | 1 | 1 | 1 | 1 | 1 | 1 | -0.001 |
| SE | 87634 | ENSG00000140678 | ITGAX | chr16 | + | 31379756 | 31379864 | 31379567 | 31379646 | 31379981 | 31380065 | 336 | 260 | 279 | 333 | 254 | 0 | 2 | 0 | 0 | 0 | 760 | 365 | 255 | 305 | 601 | 0 | 0 | 0 | 0 | 0 | 257 | 149 | 1 | 1 | 1 | 0.987 | 1 | 1 | 1 | 1 | 1 | 1 | 1 | 1 | -0.003 |
| SE | 87635 | ENSG00000102174 | PHEX | chrX | + | 22111466 | 22111560 | 22099005 | 22099151 | 22114457 | 22114571 | 9 | 9 | 17 | 13 | 17 | 0 | 0 | 0 | 1 | 0 | 0 | 10 | 6 | 0 | 0 | 0 | 0 | 0 | 0 | 0 | 243 | 149 | 1 | 1 | 1 | 1 | 1 | 0.889 | 1 | NA | 1 | 1 | NA | NA | -0.022 |
| SE | 87636 | ENSG00000102174 | PHEX | chrX | + | 22140245 | 22140406 | 22133522 | 22133624 | 22168311 | 22168389 | 0 | 3 | 0 | 0 | 0 | 2 | 2 | 0 | 2 | 0 | 0 | 0 | 0 | 0 | 0 | 0 | 0 | 3 | 0 | 0 | 298 | 149 | 1 | 1 | 0 | 0.429 | NA | 0 | NA | NA | NA | 0 | NA | NA | 0.143 |
| SE | 87637 | ENSG00000102174 | PHEX | chrX | + | 22178272 | 22178376 | 22168311 | 22168389 | 22190443 | 22190502 | 3 | 9 | 15 | 5 | 12 | 0 | 1 | 0 | 0 | 0 | 0 | 0 | 5 | 0 | 0 | 0 | 0 | 0 | 0 | 0 | 253 | 149 | 1 | 1 | 1 | 0.841 | 1 | 1 | 1 | NA | NA | 1 | NA | NA | -0.032 |
| SE | 87642 | ENSG00000117676 | RPS6KA1 | chr1 | + | 26546866 | 26546983 | 26545841 | 26546035 | 26547188 | 26547270 | 126 | 36 | 84 | 73 | 35 | 2 | 0 | 1 | 2 | 0 | 158 | 128 | 51 | 99 | 160 | 0 | 1 | 1 | 6 | 0 | 266 | 149 | 1 | 1 | 0.972 | 1 | 0.979 | 0.953 | 1 | 1 | 0.986 | 0.966 | 0.902 | 1 | 0.01 |
| SE | 87645 | ENSG00000117676 | RPS6KA1 | chr1 | + | 26554595 | 26554738 | 26554213 | 26554251 | 26555150 | 26555221 | 425 | 240 | 261 | 327 | 404 | 0 | 0 | 1 | 4 | 2 | 473 | 491 | 260 | 348 | 793 | 0 | 0 | 2 | 1 | 0 | 292 | 149 | 1 | 1 | 1 | 1 | 0.993 | 0.977 | 0.99 | 1 | 1 | 0.985 | 0.994 | 1 | -0.004 |
| SE | 87648 | ENSG00000117676 | RPS6KA1 | chr1 | + | 26556075 | 26556163 | 26555536 | 26555625 | 26556653 | 26556718 | 0 | 0 | 1 | 0 | 0 | 200 | 94 | 124 | 100 | 148 | 1 | 0 | 1 | 0 | 2 | 194 | 126 | 138 | 84 | 332 | 237 | 149 | 1 | 1 | 0 | 0 | 0.005 | 0 | 0 | 0.003 | 0 | 0.005 | 0 | 0.004 | -0.001 |

| | | | | | | | | | | | | | | | | | | | | | | | | | | | | | | | | | | | | | | | | | | | | | | |
|---|---|---|---|---|---|---|---|---|---|---|---|---|---|---|---|---|---|---|---|---|---|---|---|---|---|---|---|---|---|---|---|---|---|---|---|---|---|---|---|---|---|---|---|---|---|---|
| SE | 87650 | ENSG00000117676 | RPS6KA1 | chr1 | + | 26556653 | 26556718 | 26555536 | 26555625 | 26556997 | 26557036 | 323 | 145 | 191 | 178 | 207 | 1 | 0 | 0 | 0 | 0 | 300 | 252 | 211 | 162 | 431 | 0 | 0 | 0 | 0 | 0 | 214 | 149 | 1 | 1 | 0.996 | 1 | 1 | 1 | 1 | 1 | 1 | 1 | 1 | 1 | -0.001 |
| SE | 87651 | ENSG00000117676 | RPS6KA1 | chr1 | + | 26560725 | 26560851 | 26558806 | 26558937 | 26561044 | 26561134 | 368 | 164 | 235 | 260 | 120 | 0 | 0 | 0 | 0 | 0 | 458 | 502 | 154 | 317 | 409 | 0 | 0 | 0 | 2 | 0 | 275 | 149 | 1 | 1 | 1 | 1 | 1 | 1 | 1 | 1 | 1 | 1 | 0.988 | 1 | 0.002 |
| SE | 87653 | ENSG00000117676 | RPS6KA1 | chr1 | + | 26571448 | 26571610 | 26561504 | 26561663 | 26571848 | 26571925 | 545 | 224 | 317 | 347 | 340 | 2 | 0 | 1 | 0 | 4 | 541 | 441 | 268 | 331 | 835 | 0 | 1 | 0 | 0 | 0 | 298 | 149 | 1 | 1 | 0.993 | 1 | 0.994 | 1 | 0.977 | 1 | 0.995 | 1 | 1 | 1 | -0.006 |
| SE | 87658 | ENSG00000143569 | UBAP2L | chr1 | + | 154225083 | 154225213 | 154220848 | 154220975 | 154227281 | 154227359 | 268 | 240 | 240 | 342 | 234 | 0 | 0 | 0 | 0 | 0 | 893 | 335 | 264 | 464 | 561 | 4 | 2 | 1 | 6 | 0 | 279 | 149 | 1 | 1 | 1 | 1 | 1 | 1 | 1 | 0.992 | 0.989 | 0.993 | 0.976 | 1 | 0.01 |
| SE | 87659 | ENSG00000143569 | UBAP2L | chr1 | + | 154227281 | 154227359 | 154225081 | 154225213 | 154228614 | 154228725 | 516 | 526 | 482 | 656 | 651 | 0 | 0 | 0 | 2 | 0 | 1626 | 753 | 505 | 905 | 1054 | 0 | 0 | 0 | 0 | 0 | 227 | 149 | 1 | 1 | 1 | 1 | 1 | 0.995 | 1 | 1 | 1 | 1 | 1 | 1 | -0.001 |
| SE | 87660 | ENSG00000143569 | UBAP2L | chr1 | + | 154228614 | 154228725 | 154227281 | 154227359 | 154234590 | 154234738 | 941 | 939 | 842 | 1144 | 1295 | 0 | 0 | 0 | 0 | 0 | 2724 | 1271 | 810 | 1402 | 1991 | 15 | 0 | 0 | 3 | 0 | 260 | 149 | 1 | 1 | 1 | 1 | 1 | 1 | 1 | 0.99 | 1 | 1 | 0.996 | 1 | 0.003 |
| SE | 87667 | ENSG00000143569 | UBAP2L | chr1 | + | 154235195 | 154235291 | 154234590 | 154234759 | 154236565 | 154236611 | 444 | 335 | 470 | 509 | 732 | 1 | 0 | 0 | 0 | 0 | 675 | 511 | 353 | 530 | 656 | 0 | 2 | 0 | 0 | 0 | 245 | 149 | 1 | 1 | 0.996 | 1 | 1 | 1 | 1 | 1 | 0.994 | 1 | 1 | 1 | 0 |
| SE | 87668 | ENSG00000143569 | UBAP2L | chr1 | + | 154235195 | 154235291 | 154234590 | 154234792 | 154236565 | 154236611 | 238 | 192 | 236 | 257 | 379 | 0 | 0 | 0 | 0 | 1 | 445 | 267 | 208 | 293 | 321 | 0 | 0 | 0 | 0 | 0 | 245 | 149 | 1 | 1 | 1 | 1 | 1 | 1 | 0.996 | 1 | 1 | 1 | 1 | 1 | -0.001 |
| SE | 87669 | ENSG00000143569 | UBAP2L | chr1 | + | 154236565 | 154236611 | 154235195 | 154235291 | 154237023 | 154237136 | 580 | 495 | 545 | 614 | 789 | 0 | 0 | 0 | 0 | 0 | 1077 | 647 | 550 | 684 | 920 | 0 | 0 | 0 | 6 | 0 | 195 | 149 | 1 | 1 | 1 | 1 | 1 | 1 | 1 | 1 | 1 | 1 | 0.989 | 1 | 0.002 |
| SE | 87671 | ENSG00000143569 | UBAP2L | chr1 | + | 154246203 | 154246375 | 154242929 | 154243302 | 154249238 | 154249437 | 393 | 219 | 406 | 319 | 695 | 3 | 0 | 0 | 1 | 0 | 406 | 404 | 262 | 439 | 408 | 0 | 0 | 0 | 0 | 0 | 298 | 149 | 1 | 1 | 0.985 | 1 | 1 | 0.994 | 1 | 1 | 1 | 1 | 1 | 1 | -0.004 |
| SE | 87676 | ENSG00000143569 | UBAP2L | chr1 | + | 154268756 | 154268954 | 154266500 | 154266568 | 154270199 | 154270847 | 72 | 33 | 71 | 77 | 61 | 0 | 0 | 0 | 0 | 0 | 128 | 40 | 63 | 71 | 141 | 0 | 1 | 0 | 0 | 0 | 298 | 149 | 1 | 1 | 1 | 1 | 1 | 1 | 1 | 1 | 0.952 | 1 | 1 | 1 | 0.01 |
| SE | 87681 | ENSG00000102119 | EMD | chrX | + | 154380233 | 154380367 | 154379941 | 154380019 | 154380752 | 154380802 | 126 | 73 | 83 | 88 | 81 | 0 | 0 | 0 | 0 | 0 | 161 | 98 | 83 | 123 | 304 | 0 | 4 | 1 | 0 | 0 | 283 | 149 | 1 | 1 | 1 | 1 | 1 | 1 | 1 | 1 | 0.928 | 0.978 | 1 | 1 | 0.019 |
| SE | 87683 | ENSG00000204116 | CHIC1 | chrX | + | 73577406 | 73577461 | 73563199 | 73563580 | 73584416 | 73584572 | 46 | 13 | 30 | 32 | 37 | 1 | 0 | 0 | 3 | 0 | 6 | 7 | 10 | 0 | 3 | 0 | 0 | 0 | 0 | 0 | 204 | 149 | 1 | 1 | 0.971 | 1 | 1 | 0.886 | 1 | 1 | 1 | 1 | NA | 1 | -0.029 |
| SE | 87684 | ENSG00000204116 | CHIC1 | chrX | + | 73578703 | 73578785 | 73577406 | 73577461 | 73584416 | 73584572 | 3 | 1 | 3 | 1 | 0 | 37 | 8 | 22 | 29 | 22 | 0 | 0 | 0 | 0 | 0 | 6 | 6 | 9 | 0 | 0 | 231 | 149 | 1 | 1 | 0.05 | 0.075 | 0.081 | 0.022 | 0 | 0 | 0 | 0 | NA | NA | 0.046 |
| SE | 87685 | ENSG00000204116 | CHIC1 | chrX | + | 73679653 | 73679713 | 73679325 | 73679382 | 73680954 | 73681116 | 32 | 15 | 27 | 45 | 50 | 0 | 0 | 0 | 4 | 0 | 0 | 18 | 7 | 0 | 6 | 0 | 0 | 0 | 0 | 0 | 209 | 149 | 1 | 1 | 1 | 1 | 1 | 0.889 | 1 | NA | 1 | 1 | NA | 1 | -0.022 |
| SE | 87686 | ENSG00000204116 | CHIC1 | chrX | + | 73679653 | 73679713 | 73679325 | 73679382 | 73686595 | 73686773 | 20 | 11 | 12 | 30 | 13 | 1 | 0 | 0 | 0 | 0 | 0 | 7 | 2 | 0 | 0 | 0 | 0 | 0 | 0 | 0 | 209 | 149 | 1 | 1 | 0.934 | 1 | 1 | 1 | 1 | NA | 1 | 1 | NA | NA | -0.013 |
| SE | 87687 | ENSG00000204116 | CHIC1 | chrX | + | 73680954 | 73681116 | 73679653 | 73679713 | 73686595 | 73686773 | 12 | 4 | 15 | 16 | 39 | 0 | 0 | 0 | 0 | 0 | 0 | 15 | 6 | 0 | 12 | 0 | 4 | 0 | 0 | 0 | 298 | 149 | 1 | 1 | 1 | 1 | 1 | 1 | 1 | NA | 0.652 | 1 | NA | 1 | 0.116 |
| SE | 87688 | ENSG00000229807 | XIST | chrX | - | 73822066 | 73822216 | 73820716 | 73821029 | 73826114 | 73827984 | 31 | 24 | 20 | 27 | 61 | 34 | 19 | 33 | 30 | 36 | 0 | 0 | 0 | 3 | 0 | 13 | 0 | 0 | 15 | 0 | 298 | 149 | 1 | 1 | 0.313 | 0.387 | 0.233 | 0.31 | 0.459 | 0 | NA | NA | 0.091 | NA | 0.295 |
| SE | 87689 | ENSG00000229807 | XIST | chrX | - | 73822070 | 73822216 | 73820650 | 73821029 | 73826114 | 73826347 | 100 | 70 | 80 | 84 | 107 | 34 | 19 | 33 | 30 | 36 | 17 | 0 | 0 | 13 | 0 | 13 | 0 | 0 | 15 | 0 | 295 | 149 | 1 | 1 | 0.598 | 0.65 | 0.55 | 0.586 | 0.6 | 0.398 | NA | NA | 0.304 | NA | 0.246 |
| SE | 87690 | ENSG00000229807 | XIST | chrX | - | 73822070 | 73822216 | 73820650 | 73821029 | 73826737 | 73826889 | 69 | 46 | 61 | 57 | 46 | 3 | 4 | 7 | 5 | 10 | 17 | 0 | 0 | 10 | 0 | 0 | 0 | 0 | 0 | 0 | 295 | 149 | 1 | 1 | 0.921 | 0.853 | 0.815 | 0.852 | 0.699 | 1 | NA | NA | 1 | NA | -0.172 |
| SE | 87691 | ENSG00000229807 | XIST | chrX | - | 73822070 | 73822233 | 73820655 | 73821029 | 73826114 | 73826301 | 70 | 46 | 64 | 57 | 46 | 34 | 19 | 33 | 30 | 36 | 17 | 0 | 0 | 10 | 0 | 13 | 0 | 0 | 15 | 0 | 298 | 149 | 1 | 1 | 0.507 | 0.548 | 0.492 | 0.487 | 0.39 | 0.395 | NA | NA | 0.25 | NA | 0.162 |
| SE | 87692 | ENSG00000229807 | XIST | chrX | - | 73826114 | 73826347 | 73820655 | 73821029 | 73826737 | 73826792 | 35 | 19 | 35 | 30 | 38 | 3 | 4 | 7 | 5 | 10 | 13 | 0 | 0 | 15 | 0 | 0 | 0 | 0 | 0 | 0 | 298 | 149 | 1 | 1 | 0.854 | 0.704 | 0.714 | 0.75 | 0.655 | 1 | NA | NA | 1 | NA | -0.265 |
| SE | 87693 | ENSG00000229807 | XIST | chrX | - | 73826114 | 73826347 | 73822070 | 73822216 | 73826737 | 73826792 | 32 | 24 | 22 | 27 | 63 | 0 | 0 | 1 | 0 | 0 | 0 | 0 | 0 | 3 | 0 | 0 | 0 | 0 | 0 | 0 | 298 | 149 | 1 | 1 | 1 | 1 | 0.917 | 1 | 1 | NA | NA | NA | 1 | NA | -0.017 |
| SE | 87694 | ENSG00000229807 | XIST | chrX | - | 73830622 | 73830790 | 73829067 | 73829231 | 73831065 | 73831274 | 0 | 11 | 1 | 1 | 5 | 384 | 314 | 491 | 383 | 384 | 0 | 0 | 0 | 0 | 0 | 88 | 0 | 0 | 53 | 0 | 298 | 149 | 1 | 1 | 0 | 0.017 | 0.001 | 0.001 | 0.006 | 0 | NA | NA | 0 | NA | 0.005 |
| SE | 87695 | ENSG00000229807 | XIST | chrX | - | 73831065 | 73831206 | 73829067 | 73829231 | 73833237 | 73833374 | 400 | 329 | 506 | 397 | 405 | 4 | 0 | 0 | 0 | 0 | 107 | 0 | 0 | 55 | 0 | 0 | 0 | 0 | 0 | 0 | 290 | 149 | 1 | 1 | 0.981 | 1 | 1 | 1 | 1 | 1 | NA | NA | 1 | NA | -0.004 |
| SE | 87696 | ENSG00000229807 | XIST | chrX | - | 73831065 | 73831206 | 73829067 | 73829231 | 73837439 | 73837503 | 383 | 318 | 485 | 381 | 373 | 4 | 2 | 1 | 2 | 0 | 84 | 0 | 0 | 52 | 0 | 0 | 0 | 0 | 0 | 0 | 290 | 149 | 1 | 1 | 0.98 | 0.988 | 0.996 | 0.99 | 1 | 1 | NA | NA | 1 | NA | -0.009 |
| SE | 87697 | ENSG00000229807 | XIST | chrX | - | 73831065 | 73831274 | 73829067 | 73829231 | 73833237 | 73833374 | 1265 | 1282 | 1289 | 1147 | 1227 | 4 | 0 | 0 | 0 | 0 | 297 | 1 | 0 | 296 | 0 | 0 | 0 | 0 | 0 | 0 | 298 | 149 | 1 | 1 | 0.994 | 1 | 1 | 1 | 1 | 1 | 1 | NA | 1 | NA | -0.001 |
| SE | 87698 | ENSG00000229807 | XIST | chrX | - | 73831065 | 73831274 | 73829067 | 73829231 | 73837439 | 73837503 | 561 | 526 | 682 | 549 | 570 | 4 | 2 | 1 | 2 | 0 | 154 | 0 | 0 | 106 | 0 | 0 | 0 | 0 | 0 | 0 | 298 | 149 | 1 | 1 | 0.986 | 0.992 | 0.997 | 0.993 | 1 | 1 | NA | NA | 1 | NA | -0.006 |
| SE | 87699 | ENSG00000229807 | XIST | chrX | - | 73833237 | 73833374 | 73829067 | 73829231 | 73837439 | 73837503 | 708 | 614 | 606 | 621 | 430 | 4 | 2 | 1 | 2 | 0 | 93 | 0 | 0 | 161 | 0 | 0 | 0 | 0 | 0 | 0 | 286 | 149 | 1 | 1 | 0.989 | 0.994 | 0.997 | 0.994 | 1 | 1 | NA | NA | 1 | NA | -0.005 |
| SE | 87700 | ENSG00000229807 | XIST | chrX | - | 73833237 | 73833374 | 73831065 | 73831206 | 73837439 | 73837503 | 722 | 630 | 629 | 639 | 463 | 1 | 5 | 2 | 2 | 1 | 116 | 0 | 0 | 164 | 0 | 0 | 0 | 0 | 0 | 0 | 286 | 149 | 1 | 1 | 0.997 | 0.985 | 0.994 | 0.994 | 0.996 | 1 | NA | NA | 1 | NA | -0.007 |
| SE | 87701 | ENSG00000229807 | XIST | chrX | - | 73833237 | 73833374 | 73831065 | 73831206 | 73841381 | 73841473 | 28 | 31 | 28 | 32 | 45 | 0 | 0 | 1 | 0 | 0 | 23 | 0 | 0 | 3 | 0 | 0 | 0 | 0 | 0 | 0 | 286 | 149 | 1 | 1 | 1 | 1 | 0.936 | 1 | 1 | 1 | NA | NA | 1 | NA | -0.013 |
| SE | 87702 | ENSG00000229807 | XIST | chrX | - | 73833237 | 73833374 | 73831065 | 73831274 | 73837439 | 73837503 | 1578 | 1578 | 1400 | 1386 | 1265 | 177 | 212 | 192 | 167 | 188 | 302 | 1 | 0 | 404 | 0 | 66 | 0 | 0 | 53 | 0 | 286 | 149 | 1 | 1 | 0.823 | 0.795 | 0.792 | 0.812 | 0.778 | 0.704 | 1 | NA | 0.799 | NA | -0.034 |
| SE | 87703 | ENSG00000229807 | XIST | chrX | - | 73833237 | 73833374 | 73831065 | 73831274 | 73841381 | 73841473 | 867 | 958 | 767 | 759 | 839 | 18 | 12 | 6 | 3 | 4 | 209 | 1 | 0 | 242 | 0 | 0 | 0 | 0 | 0 | 0 | 286 | 149 | 1 | 1 | 0.962 | 0.977 | 0.985 | 0.992 | 0.991 | 1 | 1 | NA | 1 | NA | -0.019 |
| SE | 87704 | ENSG00000229807 | XIST | chrX | - | 73837439 | 73837503 | 73831065 | 73831206 | 73841381 | 73841473 | 292 | 301 | 298 | 244 | 392 | 0 | 0 | 1 | 0 | 0 | 87 | 0 | 0 | 72 | 0 | 0 | 0 | 0 | 0 | 0 | 213 | 149 | 1 | 1 | 1 | 1 | 0.995 | 1 | 1 | 1 | NA | NA | 1 | NA | -0.001 |
| SE | 87705 | ENSG00000229807 | XIST | chrX | - | 73837439 | 73837503 | 73831065 | 73831274 | 73841381 | 73841473 | 468 | 508 | 480 | 406 | 575 | 18 | 12 | 6 | 3 | 4 | 153 | 0 | 0 | 125 | 0 | 0 | 0 | 0 | 0 | 0 | 213 | 149 | 1 | 1 | 0.948 | 0.967 | 0.982 | 0.99 | 0.99 | 1 | NA | NA | 1 | NA | -0.025 |
| SE | 87706 | ENSG00000229807 | XIST | chrX | - | 73837439 | 73837503 | 73833237 | 73833374 | 73841381 | 73841473 | 1008 | 922 | 919 | 878 | 819 | 11 | 15 | 6 | 14 | 12 | 179 | 0 | 0 | 230 | 0 | 0 | 0 | 0 | 0 | 0 | 213 | 149 | 1 | 1 | 0.985 | 0.977 | 0.991 | 0.978 | 0.979 | 1 | NA | NA | 1 | NA | -0.018 |
| SE | 87708 | ENSG00000229807 | XIST | chrX | - | 73841381 | 73841473 | 73837439 | 73837503 | 73850501 | 73851715 | 630 | 600 | 583 | 560 | 433 | 0 | 0 | 1 | 0 | 0 | 103 | 0 | 0 | 162 | 0 | 0 | 0 | 0 | 0 | 0 | 241 | 149 | 1 | 1 | 1 | 1 | 0.997 | 1 | 1 | 1 | NA | NA | 1 | NA | -0.001 |
| SE | 87713 | ENSG00000242574 | HLA-DMB | chr6 | - | 32937171 | 32937456 | 32935535 | 32935652 | 32938683 | 32938688 | 521 | 294 | 438 | 488 | 594 | 0 | 0 | 1 | 0 | 0 | 948 | 357 | 200 | 556 | 1184 | 0 | 3 | 0 | 0 | 4 | 298 | 149 | 1 | 1 | 1 | 1 | 0.995 | 1 | 1 | 1 | 0.983 | 1 | 1 | 0.993 | 0.004 |
| SE | 87714 | ENSG00000115896 | PLCL1 | chr2 | + | 198001951 | 198002008 | 197804701 | 197805339 | 198083757 | 198086232 | 2 | 4 | 5 | 1 | 7 | 2 | 0 | 4 | 9 | 1 | 0 | 0 | 2 | 2 | 0 | 0 | 0 | 4 | 1 | 0 | 206 | 149 | 1 | 1 | 0.42 | 1 | 0.475 | 0.074 | 0.835 | NA | NA | 0.266 | 0.591 | NA | 0.132 |
| SE | 87715 | ENSG00000115896 | PLCL1 | chr2 | + | 198101284 | 198101360 | 198088857 | 198089061 | 198146779 | 198146932 | 28 | 39 | 30 | 22 | 42 | 0 | 0 | 4 | 0 | 0 | 0 | 25 | 19 | 17 | 0 | 0 | 0 | 0 | 0 | 0 | 225 | 149 | 1 | 1 | 1 | 1 | 0.832 | 1 | 1 | NA | 1 | 1 | 1 | NA | -0.034 |
| SE | 87716 | ENSG00000115896 | PLCL1 | chr2 | + | 198103826 | 198103936 | 198101284 | 198101360 | 198146779 | 198146932 | 54 | 82 | 74 | 82 | 124 | 1 | 0 | 0 | 0 | 0 | 26 | 55 | 64 | 32 | 3 | 0 | 0 | 0 | 0 | 0 | 259 | 149 | 1 | 1 | 0.969 | 1 | 1 | 1 | 1 | 1 | 1 | 1 | 1 | 1 | -0.006 |
| SE | 87717 | ENSG00000115896 | PLCL1 | chr2 | + | 198387852 | 198388005 | 198355294 | 198355407 | 198571353 | 198571477 | 0 | 0 | 0 | 4 | 2 | 0 | 3 | 0 | 0 | 0 | 0 | 0 | 0 | 0 | 0 | 0 | 0 | 0 | 1 | 0 | 298 | 149 | 1 | 1 | NA | 0 | NA | 1 | 1 | NA | NA | NA | 0 | NA | 0.667 |
| SE | 87718 | ENSG00000115896 | PLCL1 | chr2 | + | 198571353 | 198571477 | 198387852 | 198388005 | 198571855 | 198572581 | 0 | 4 | 0 | 2 | 2 | 0 | 0 | 0 | 2 | 0 | 4 | 0 | 2 | 1 | 0 | 0 | 0 | 0 | 0 | 0 | 273 | 149 | 1 | 1 | NA | 1 | NA | 0.353 | 1 | 1 | NA | 1 | 1 | NA | -0.216 |
| SE | 87721 | ENSG00000107815 | TWNK | chr10 | + | 100990435 | 100990543 | 100989643 | 100989884 | 100990868 | 100991010 | 38 | 21 | 35 | 24 | 58 | 0 | 0 | 1 | 0 | 0 | 0 | 19 | 10 | 10 | 3 | 0 | 0 | 0 | 0 | 0 | 257 | 149 | 1 | 1 | 1 | 1 | 0.953 | 1 | 1 | NA | 1 | 1 | 1 | 1 | -0.009 |
| SE | 87722 | ENSG00000107815 | TWNK | chr10 | + | 100990435 | 100990663 | 100989643 | 100989884 | 100990868 | 100991010 | 14 | 12 | 19 | 10 | 28 | 0 | 0 | 1 | 0 | 0 | 0 | 8 | 10 | 12 | 0 | 0 | 0 | 0 | 0 | 0 | 298 | 149 | 1 | 1 | 1 | 1 | 0.905 | 1 | 1 | NA | 1 | 1 | 1 | NA | -0.019 |
| SE | 87723 | ENSG00000107815 | TWNK | chr10 | + | 100990868 | 100991010 | 100989643 | 100989884 | 100993189 | 100993580 | 19 | 13 | 16 | 13 | 19 | 2 | 0 | 0 | 0 | 0 | 0 | 18 | 10 | 8 | 3 | 0 | 0 | 0 | 0 | 0 | 291 | 149 | 1 | 1 | 0.829 | 1 | 1 | 1 | 1 | NA | 1 | 1 | 1 | 1 | -0.034 |
| SE | 87727 | ENSG00000198947 | DMD | chrX | - | 31172347 | 31172413 | 31169442 | 31169601 | 31173538 | 31173604 | 2 | 0 | 4 | 3 | 0 | 1 | 0 | 0 | 0 | 2 | 0 | 0 | 4 | 6 | 0 | 0 | 0 | 0 | 0 | 0 | 215 | 149 | 1 | 1 | 0.581 | NA | 1 | 1 | 0 | NA | NA | 1 | 1 | NA | -0.355 |
| SE | 87728 | ENSG00000198947 | DMD | chrX | - | 31173538 | 31173604 | 31172347 | 31172413 | 31178668 | 31178805 | 1 | 0 | 2 | 0 | 0 | 0 | 0 | 0 | 1 | 0 | 0 | 0 | 2 | 0 | 0 | 0 | 0 | 0 | 0 | 0 | 215 | 149 | 1 | 1 | 1 | NA | 1 | 0 | NA | NA | NA | 1 | NA | NA | -0.333 |
| SE | 87729 | ENSG00000198947 | DMD | chrX | - | 31177931 | 31177970 | 31173538 | 31173604 | 31178668 | 31178805 | 1 | 0 | 2 | 1 | 2 | 0 | 0 | 2 | 0 | 0 | 0 | 0 | 0 | 3 | 0 | 0 | 0 | 2 | 0 | 0 | 188 | 149 | 1 | 1 | 1 | NA | 0.442 | 1 | 1 | NA | NA | 0 | 1 | NA | 0.361 |
| SE | 87731 | ENSG00000198947 | DMD | chrX | - | 31247693 | 31247768 | 31223046 | 31223121 | 31260954 | 31261016 | 0 | 0 | 0 | 0 | 0 | 0 | 0 | 1 | 0 | 0 | 0 | 6 | 1 | 0 | 0 | 0 | 0 | 0 | 0 | 0 | 224 | 149 | 1 | 1 | NA | NA | 0 | NA | NA | NA | 1 | 1 | NA | NA | -1 |
| SE | 87740 | ENSG00000171100 | MTM1 | chrX | + | 150598591 | 150598686 | 150596497 | 150596570 | 150614588 | 150614699 | 77 | 67 | 93 | 104 | 135 | 1 | 1 | 0 | 1 | 0 | 57 | 74 | 88 | 38 | 22 | 0 | 0 | 0 | 0 | 0 | 244 | 149 | 1 | 1 | 0.979 | 0.976 | 1 | 0.984 | 1 | 1 | 1 | 1 | 1 | 1 | -0.012 |
| SE | 87741 | ENSG00000171100 | MTM1 | chrX | + | 150614588 | 150614699 | 150598591 | 150598686 | 150619037 | 150619139 | 74 | 81 | 102 | 116 | 175 | 1 | 0 | 0 | 0 | 0 | 75 | 89 | 71 | 49 | 13 | 0 | 2 | 1 | 0 | 0 | 260 | 149 | 1 | 1 | 0.977 | 1 | 1 | 1 | 1 | 1 | 0.962 | 0.976 | 1 | 1 | 0.008 |
| SE | 87742 | ENSG00000171100 | MTM1 | chrX | + | 150614588 | 150614699 | 150598591 | 150598686 | 150638942 | 150639026 | 20 | 24 | 33 | 32 | 59 | 0 | 0 | 2 | 2 | 0 | 34 | 21 | 25 | 22 | 5 | 0 | 0 | 0 | 0 | 0 | 260 | 149 | 1 | 1 | 1 | 1 | 0.904 | 0.902 | 1 | 1 | 1 | 1 | 1 | 1 | -0.039 |
| SE | 87743 | ENSG00000171100 | MTM1 | chrX | + | 150619037 | 150619139 | 150598591 | 150598686 | 150638942 | 150639026 | 61 | 53 | 49 | 59 | 120 | 0 | 0 | 2 | 2 | 0 | 8 | 76 | 69 | 27 | 4 | 0 | 0 | 0 | 0 | 0 | 251 | 149 | 1 | 1 | 1 | 1 | 0.936 | 0.946 | 1 | 1 | 1 | 1 | 1 | 1 | -0.024 |
| SE | 87744 | ENSG00000171100 | MTM1 | chrX | + | 150619037 | 150619139 | 150614588 | 150614699 | 150638942 | 150639026 | 113 | 111 | 122 | 148 | 243 | 0 | 0 | 0 | 1 | 3 | 49 | 142 | 117 | 52 | 12 | 0 | 0 | 0 | 0 | 0 | 251 | 149 | 1 | 1 | 1 | 1 | 1 | 0.989 | 0.98 | 1 | 1 | 1 | 1 | 1 | -0.006 |
| SE | 87745 | ENSG00000171100 | MTM1 | chrX | + | 150645682 | 150645871 | 150641268 | 150641418 | 150649715 | 150649901 | 240 | 288 | 283 | 292 | 295 | 2 | 0 | 1 | 0 | 0 | 141 | 277 | 274 | 126 | 113 | 0 | 0 | 0 | 0 | 0 | 298 | 149 | 1 | 1 | 0.984 | 1 | 0.993 | 1 | 1 | 1 | 1 | 1 | 1 | 1 | -0.005 |
| SE | 87751 | ENSG00000063601 | MTMR1 | chrX | + | 150712943 | 150712970 | 150712341 | 150712365 | 150718624 | 150718700 | 0 | 0 | 0 | 0 | 1 | 66 | 55 | 98 | 84 | 115 | 0 | 0 | 1 | 0 | 0 | 68 | 68 | 45 | 55 | 34 | 176 | 149 | 1 | 1 | 0 | 0 | 0 | 0 | 0.007 | 0 | 0 | 0.018 | 0 | 0 | -0.002 |
| SE | 87752 | ENSG00000063601 | MTMR1 | chrX | + | 150714478 | 150714529 | 150699194 | 150699300 | 150718624 | 150718700 | 0 | 0 | 0 | 0 | 0 | 3 | 0 | 1 | 4 | 0 | 0 | 0 | 5 | 0 | 0 | 0 | 4 | 7 | 4 | 0 | 200 | 149 | 1 | 1 | 0 | NA | 0 | 0 | NA | NA | 0 | 0.347 | 0 | NA | -0.116 |
| SE | 87753 | ENSG00000063601 | MTMR1 | chrX | + | 150714478 | 150714529 | 150712341 | 150712365 | 150718624 | 150718700 | 0 | 0 | 0 | 0 | 0 | 66 | 55 | 98 | 84 | 115 | 0 | 0 | 5 | 0 | 0 | 68 | 68 | 45 | 55 | 34 | 200 | 149 | 1 | 1 | 0 | 0 | 0 | 0 | 0 | 0 | 0 | 0.076 | 0 | 0 | -0.015 |
| SE | 87754 | ENSG00000063601 | MTMR1 | chrX | + | 150714478 | 150714529 | 150712943 | 150712970 | 150718624 | 150718700 | 0 | 0 | 0 | 0 | 0 | 0 | 0 | 0 | 0 | 1 | 0 | 0 | 5 | 0 | 0 | 0 | 0 | 1 | 0 | 0 | 200 | 149 | 1 | 1 | NA | NA | NA | NA | 0 | NA | NA | 0.788 | NA | NA | -0.788 |
| SE | 87757 | ENSG00000063601 | MTMR1 | chrX | + | 150730108 | 150730210 | 150727683 | 150727791 | 150730524 | 150730565 | 156 | 108 | 153 | 156 | 274 | 1 | 0 | 0 | 0 | 0 | 65 | 174 | 120 | 143 | 109 | 0 | 0 | 1 | 0 | 0 | 251 | 149 | 1 | 1 | 0.989 | 1 | 1 | 1 | 1 | 1 | 1 | 0.986 | 1 | 1 | 0.001 |
| SE | 87759 | ENSG00000063601 | MTMR1 | chrX | + | 150750729 | 150750843 | 150744360 | 150744453 | 150755688 | 150755865 | 110 | 70 | 109 | 95 | 99 | 1 | 1 | 0 | 0 | 3 | 131 | 72 | 62 | 51 | 89 | 0 | 0 | 0 | 0 | 0 | 263 | 149 | 1 | 1 | 0.984 | 0.975 | 1 | 1 | 0.949 | 1 | 1 | 1 | 1 | 1 | -0.018 |
| SE | 87760 | ENSG00000063601 | MTMR1 | chrX | + | 150750729 | 150750843 | 150744360 | 150744453 | 150762564 | 150762737 | 50 | 32 | 40 | 40 | 41 | 1 | 0 | 0 | 0 | 0 | 47 | 25 | 33 | 27 | 58 | 0 | 0 | 0 | 0 | 0 | 263 | 149 | 1 | 1 | 0.966 | 1 | 1 | 1 | 1 | 1 | 1 | 1 | 1 | 1 | -0.007 |
| SE | 87762 | ENSG00000063601 | MTMR1 | chrX | + | 150755688 | 150755865 | 150744360 | 150744453 | 150762564 | 150762737 | 34 | 38 | 45 | 43 | 51 | 1 | 0 | 0 | 0 | 0 | 57 | 26 | 20 | 19 | 22 | 0 | 0 | 0 | 0 | 0 | 298 | 149 | 1 | 1 | 0.944 | 1 | 1 | 1 | 1 | 1 | 1 | 1 | 1 | 1 | -0.011 |
| SE | 87764 | ENSG00000063601 | MTMR1 | chrX | + | 150762564 | 150762737 | 150755688 | 150755865 | 150764511 | 150765102 | 33 | 37 | 47 | 43 | 48 | 0 | 0 | 0 | 0 | 0 | 57 | 26 | 20 | 19 | 22 | 0 | 0 | 1 | 0 | 0 | 298 | 149 | 1 | 1 | 1 | 1 | 1 | 1 | 1 | 1 | 1 | 0.909 | 1 | 1 | 0.018 |
| SE | 87768 | ENSG00000196459 | TRAPPC2 | chrX | - | 13734043 | 13734185 | 13720427 | 13720497 | 13734524 | 13734620 | 27 | 11 | 18 | 30 | 5 | 1 | 1 | 0 | 1 | 0 | 18 | 51 | 14 | 45 | 29 | 0 | 0 | 1 | 0 | 0 | 291 | 149 | 1 | 1 | 0.933 | 0.849 | 1 | 0.939 | 1 | 1 | 1 | 0.878 | 1 | 1 | -0.031 |
| SE | 87770 | ENSG00000102158 | MAGT1 | chrX | - | 77830804 | 77830895 | 77825746 | 77829235 | 77841245 | 77841320 | 226 | 165 | 186 | 207 | 323 | 4 | 0 | 0 | 2 | 0 | 82 | 119 | 162 | 156 | 46 | 0 | 0 | 3 | 0 | 0 | 240 | 149 | 1 | 1 | 0.972 | 1 | 1 | 0.985 | 1 | 1 | 1 | 0.971 | 1 | 1 | -0.003 |
| SE | 87771 | ENSG00000102158 | MAGT1 | chrX | - | 77855500 | 77855590 | 77853900 | 77853964 | 77856732 | 77856873 | 172 | 112 | 155 | 178 | 151 | 0 | 0 | 0 | 0 | 2 | 123 | 69 | 167 | 112 | 71 | 0 | 0 | 3 | 0 | 0 | 239 | 149 | 1 | 1 | 1 | 1 | 1 | 1 | 0.979 | 1 | 1 | 0.972 | 1 | 1 | 0.001 |
| SE | 87772 | ENSG00000165675 | ENOX2 | chrX | - | 130627957 | 130628043 | 130622324 | 130625445 | 130631467 | 130631576 | 19 | 18 | 10 | 22 | 23 | 0 | 0 | 1 | 0 | 0 | 10 | 27 | 9 | 11 | 7 | 0 | 0 | 0 | 0 | 0 | 235 | 149 | 1 | 1 | 1 | 1 | 0.864 | 1 | 1 | 1 | 1 | 1 | 1 | 1 | -0.027 |
| SE | 87773 | ENSG00000165675 | ENOX2 | chrX | - | 130631467 | 130631576 | 130627957 | 130628043 | 130634983 | 130635091 | 18 | 23 | 16 | 33 | 35 | 0 | 0 | 0 | 1 | 0 | 12 | 21 | 10 | 8 | 14 | 0 | 0 | 0 | 0 | 0 | 258 | 149 | 1 | 1 | 1 | 1 | 1 | 0.95 | 1 | 1 | 1 | 1 | 1 | 1 | -0.01 |
| SE | 87774 | ENSG00000165675 | ENOX2 | chrX | - | 130669964 | 130670198 | 130667529 | 130667742 | 130679541 | 130679748 | 83 | 41 | 60 | 52 | 53 | 0 | 0 | 2 | 0 | 0 | 30 | 35 | 55 | 35 | 5 | 0 | 0 | 0 | 0 | 0 | 298 | 149 | 1 | 1 | 1 | 1 | 0.938 | 1 | 1 | 1 | 1 | 1 | 1 | 1 | -0.012 |
| SE | 87776 | ENSG00000165675 | ENOX2 | chrX | - | 130688862 | 130689018 | 130679541 | 130679748 | 130703119 | 130703254 | 61 | 50 | 38 | 37 | 41 | 0 | 0 | 0 | 0 | 1 | 11 | 25 | 21 | 36 | 38 | 0 | 0 | 0 | 0 | 0 | 298 | 149 | 1 | 1 | 1 | 1 | 1 | 1 | 0.953 | 1 | 1 | 1 | 1 | 1 | -0.009 |
| SE | 87778 | ENSG00000165675 | ENOX2 | chrX | - | 130709242 | 130709331 | 130703119 | 130703254 | 130901683 | 130901731 | 4 | 2 | 7 | 1 | 3 | 0 | 1 | 6 | 0 | 4 | 9 | 7 | 0 | 5 | 1 | 0 | 2 | 0 | 0 | 0 | 238 | 149 | 1 | 1 | 1 | 0.556 | 0.422 | 1 | 0.32 | 1 | 0.687 | NA | 1 | 1 | -0.262 |
| SE | 87780 | ENSG00000165675 | ENOX2 | chrX | - | 130783546 | 130783690 | 130709242 | 130709331 | 130901683 | 130901731 | 23 | 16 | 26 | 30 | 8 | 0 | 1 | 0 | 0 | 0 | 30 | 15 | 13 | 15 | 0 | 0 | 2 | 0 | 0 | 0 | 293 | 149 | 1 | 1 | 1 | 0.891 | 1 | 1 | 1 | 1 | 0.792 | 1 | 1 | NA | 0.03 |
| SE | 87782 | ENSG00000157514 | TSC22D3 | chrX | - | 107715898 | 107715950 | 107713220 | 107714749 | 107716689 | 107717135 | 1438 | 780 | 1408 | 1496 | 1536 | 0 | 0 | 0 | 0 | 0 | 882 | 2459 | 2276 | 1327 | 3656 | 0 | 0 | 0 | 1 | 0 | 201 | 149 | 1 | 1 | 1 | 1 | 1 | 1 | 1 | 1 | 1 | 1 | 0.999 | 1 | 0 |
| SE | 87783 | ENSG00000157514 | TSC22D3 | chrX | - | 107715898 | 107715950 | 107713220 | 107714749 | 107775099 | 107775294 | 626 | 348 | 640 | 731 | 692 | 0 | 0 | 0 | 1 | 0 | 400 | 1046 | 1022 | 645 | 1996 | 0 | 0 | 0 | 0 | 0 | 201 | 149 | 1 | 1 | 1 | 1 | 1 | 0.998 | 1 | 1 | 1 | 1 | 1 | 1 | 0 |
| SE | 87788 | ENSG00000230074 | NA | chr9 | + | 34678095 | 34678268 | 34665606 | 34665767 | 34680905 | 34681981 | 0 | 4 | 1 | 6 | 0 | 0 | 0 | 0 | 2 | 0 | 0 | 10 | 8 | 0 | 0 | 0 | 0 | 0 | 0 | 5 | 298 | 149 | 1 | 1 | NA | 1 | 1 | 0.6 | NA | NA | 1 | 1 | NA | 0 | 0.2 |
| SE | 87789 | ENSG00000230074 | NA | chr9 | + | 34678099 | 34678268 | 34665664 | 34665767 | 34680905 | 34681298 | 0 | 3 | 0 | 6 | 0 | 0 | 0 | 0 | 2 | 0 | 0 | 10 | 8 | 0 | 0 | 0 | 0 | 0 | 0 | 5 | 298 | 149 | 1 | 1 | NA | 1 | NA | 0.6 | NA | NA | 1 | 1 | NA | 0 | 0.133 |
| SE | 87800 | ENSG00000111237 | VPS29 | chr12 | - | 110496011 | 110496203 | 110492920 | 110493231 | 110502048 | 110502108 | 840 | 551 | 686 | 713 | 464 | 0 | 0 | 0 | 2 | 0 | 647 | 588 | 486 | 731 | 525 | 0 | 0 | 0 | 0 | 0 | 298 | 149 | 1 | 1 | 1 | 1 | 1 | 0.994 | 1 | 1 | 1 | 1 | 1 | 1 | -0.001 |
| SE | 87802 | ENSG00000111237 | VPS29 | chr12 | - | 110496822 | 110497002 | 110496045 | 110496203 | 110499456 | 110499546 | 1 | 2 | 0 | 0 | 5 | 8 | 6 | 9 | 6 | 11 | 0 | 0 | 0 | 2 | 0 | 0 | 7 | 2 | 5 | 12 | 298 | 149 | 1 | 1 | 0.059 | 0.143 | 0 | 0 | 0.185 | NA | 0 | 0 | 0.167 | 0 | 0.036 |
| SE | 87803 | ENSG00000111237 | VPS29 | chr12 | - | 110496822 | 110497002 | 110496045 | 110496203 | 110499534 | 110499546 | 1 | 0 | 0 | 1 | 7 | 101 | 61 | 80 | 102 | 34 | 0 | 0 | 0 | 2 | 0 | 97 | 58 | 49 | 62 | 28 | 298 | 149 | 1 | 1 | 0.005 | 0 | 0 | 0.005 | 0.093 | 0 | 0 | 0 | 0.016 | 0 | 0.017 |
| SE | 87807 | ENSG00000111237 | VPS29 | chr12 | - | 110498819 | 110498859 | 110496822 | 110497002 | 110499456 | 110499546 | 2 | 0 | 1 | 4 | 3 | 0 | 2 | 0 | 0 | 0 | 4 | 0 | 0 | 0 | 2 | 0 | 0 | 0 | 0 | 0 | 189 | 149 | 1 | 1 | 1 | 0 | 1 | 1 | 1 | 1 | NA | NA | NA | 1 | -0.2 |
| SE | 87808 | ENSG00000111237 | VPS29 | chr12 | - | 110498819 | 110498859 | 110496822 | 110497002 | 110499534 | 110499546 | 2 | 0 | 3 | 9 | 5 | 1 | 0 | 0 | 1 | 2 | 4 | 3 | 0 | 0 | 1 | 0 | 0 | 0 | 0 | 0 | 189 | 149 | 1 | 1 | 0.612 | NA | 1 | 0.876 | 0.663 | 1 | 1 | NA | NA | 1 | -0.212 |
| SE | 87809 | ENSG00000111237 | VPS29 | chr12 | - | 110498819 | 110498859 | 110496822 | 110497002 | 110502048 | 110502078 | 2 | 0 | 1 | 4 | 3 | 1 | 1 | 0 | 0 | 0 | 4 | 0 | 0 | 0 | 1 | 0 | 0 | 3 | 0 | 0 | 189 | 149 | 1 | 1 | 0.612 | 0 | 1 | 1 | 1 | 1 | NA | 0 | NA | 1 | 0.056 |
| SE | 87815 | ENSG00000111237 | VPS29 | chr12 | - | 110499534 | 110499546 | 110496822 | 110497002 | 110502048 | 110502099 | 1 | 0 | 0 | 1 | 2 | 1 | 1 | 0 | 0 | 0 | 0 | 0 | 0 | 0 | 0 | 0 | 0 | 3 | 0 | 0 | 161 | 149 | 1 | 1 | 0.481 | 0 | NA | 1 | 1 | NA | NA | 0 | NA | NA | 0.62 |
| SE | 87816 | ENSG00000111237 | VPS29 | chr12 | - | 110499534 | 110499546 | 110498819 | 110498859 | 110502048 | 110502099 | 15 | 7 | 11 | 13 | 21 | 0 | 1 | 0 | 0 | 0 | 21 | 10 | 28 | 19 | 4 | 0 | 0 | 3 | 0 | 0 | 161 | 149 | 1 | 1 | 1 | 0.866 | 1 | 1 | 1 | 1 | 1 | 1 | 1 | 1 | -0.027 |
| SE | 87817 | ENSG00000111237 | VPS29 | chr12 | - | 110501416 | 110501559 | 110496822 | 110497002 | 110502048 | 110502092 | 0 | 0 | 0 | 2 | 0 | 1 | 1 | 0 | 0 | 0 | 0 | 0 | 0 | 0 | 0 | 0 | 0 | 3 | 0 | 0 | 292 | 149 | 1 | 1 | 0 | 0 | NA | 1 | NA | NA | NA | 0 | NA | NA | 0.333 |
| SE | 87818 | ENSG00000111237 | VPS29 | chr12 | - | 110501416 | 110501559 | 110499534 | 110499546 | 110502048 | 110502092 | 1 | 3 | 3 | 3 | 1 | 111 | 62 | 93 | 95 | 45 | 0 | 2 | 0 | 0 | 0 | 73 | 64 | 61 | 79 | 49 | 292 | 149 | 1 | 1 | 0.005 | 0.024 | 0.016 | 0.016 | 0.011 | 0 | 0.016 | 0 | 0 | 0 | 0.011 |
| SE | 87830 | ENSG00000072506 | HSD17B10 | chrX | - | 53431797 | 53431906 | 53431260 | 53431594 | 53431987 | 53432116 | 214 | 94 | 149 | 175 | 109 | 0 | 0 | 2 | 0 | 0 | 347 | 257 | 138 | 180 | 304 | 0 | 0 | 0 | 0 | 0 | 258 | 149 | 1 | 1 | 1 | 1 | 0.977 | 1 | 1 | 1 | 1 | 1 | 1 | 1 | -0.005 |
| SE | 87831 | ENSG00000072506 | HSD17B10 | chrX | - | 53431824 | 53431906 | 53431257 | 53431594 | 53431987 | 53432116 | 114 | 67 | 66 | 103 | 51 | 0 | 0 | 2 | 0 | 0 | 243 | 149 | 107 | 126 | 242 | 0 | 0 | 0 | 0 | 0 | 231 | 149 | 1 | 1 | 1 | 1 | 0.955 | 1 | 1 | 1 | 1 | 1 | 1 | 1 | -0.009 |
| SE | 87832 | ENSG00000101152 | DNAJC5 | chr20 | + | 63929311 | 63929525 | 63928334 | 63928452 | 63930850 | 63931022 | 257 | 164 | 189 | 228 | 228 | 0 | 0 | 0 | 0 | 0 | 551 | 296 | 303 | 398 | 562 | 0 | 0 | 2 | 0 | 0 | 298 | 149 | 1 | 1 | 1 | 1 | 1 | 1 | 1 | 1 | 1 | 0.987 | 1 | 1 | 0.003 |
| SE | 87834 | ENSG00000189420 | ZFP92 | chrX | + | 153411964 | 153412013 | 153411471 | 153411703 | 153418304 | 153418355 | 1 | 0 | 3 | 0 | 0 | 0 | 0 | 1 | 1 | 0 | 0 | 0 | 5 | 0 | 0 | 0 | 0 | 0 | 2 | 0 | 198 | 149 | 1 | 1 | 1 | NA | 0.693 | 0 | NA | NA | NA | 1 | 0 | NA | 0.064 |
| SE | 87847 | ENSG00000101752 | MIB1 | chr18 | + | 21773623 | 21773728 | 21768622 | 21768752 | 21778102 | 21778169 | 160 | 88 | 83 | 115 | 134 | 1 | 0 | 0 | 0 | 0 | 35 | 53 | 34 | 56 | 89 | 0 | 0 | 0 | 0 | 0 | 254 | 149 | 1 | 1 | 0.989 | 1 | 1 | 1 | 1 | 1 | 1 | 1 | 1 | 1 | -0.002 |
| SE | 87850 | ENSG00000101752 | MIB1 | chr18 | + | 21846943 | 21847125 | 21844091 | 21844253 | 21849195 | 21849388 | 121 | 42 | 87 | 120 | 110 | 0 | 0 | 1 | 0 | 0 | 34 | 88 | 64 | 13 | 0 | 0 | 0 | 0 | 0 | 0 | 298 | 149 | 1 | 1 | 1 | 1 | 0.978 | 1 | 1 | 1 | 1 | 1 | 1 | NA | -0.004 |
| SE | 87852 | ENSG00000172725 | CORO1B | chr11 | - | 67440701 | 67440836 | 67440334 | 67440439 | 67441124 | 67441244 | 2 | 0 | 0 | 1 | 0 | 31 | 12 | 16 | 13 | 4 | 0 | 2 | 0 | 0 | 0 | 15 | 34 | 8 | 16 | 72 | 284 | 149 | 1 | 1 | 0.033 | 0 | 0 | 0.039 | 0 | 0 | 0.03 | 0 | 0 | 0 | 0.008 |
| SE | 87853 | ENSG00000067840 | PDZD4 | chrX | - | 153807278 | 153807369 | 153806741 | 153806840 | 153808341 | 153808595 | 51 | 10 | 37 | 47 | 10 | 0 | 0 | 0 | 2 | 0 | 14 | 19 | 2 | 4 | 19 | 0 | 0 | 0 | 0 | 0 | 240 | 149 | 1 | 1 | 1 | 1 | 1 | 0.936 | 1 | 1 | 1 | 1 | 1 | 1 | -0.013 |
| SE | 87854 | ENSG00000067840 | PDZD4 | chrX | - | 153807896 | 153808139 | 153807278 | 153807369 | 153808341 | 153808595 | 0 | 1 | 4 | 0 | 5 | 24 | 0 | 10 | 19 | 4 | 0 | 1 | 0 | 0 | 0 | 0 | 3 | 1 | 2 | 2 | 298 | 149 | 1 | 1 | 0 | 1 | 0.167 | 0 | 0.385 | NA | 0.143 | 0 | 0 | 0 | 0.275 |
| SE | 87855 | ENSG00000067840 | PDZD4 | chrX | - | 153808341 | 153808595 | 153806741 | 153806840 | 153830238 | 153830364 | 25 | 3 | 10 | 23 | 4 | 1 | 0 | 1 | 1 | 1 | 24 | 17 | 3 | 0 | 10 | 0 | 0 | 0 | 0 | 3 | 298 | 149 | 1 | 1 | 0.926 | 1 | 0.833 | 0.92 | 0.667 | 1 | 1 | 1 | NA | 0.625 | -0.037 |
| SE | 87856 | ENSG00000198932 | GPRASP1 | chrX | + | 102651686 | 102651733 | 102651482 | 102651595 | 102652174 | 102652265 | 0 | 5 | 19 | 1 | 7 | 0 | 1 | 7 | 0 | 0 | 0 | 6 | 4 | 0 | 0 | 0 | 0 | 0 | 0 | 0 | 196 | 149 | 1 | 1 | NA | 0.792 | 0.674 | 1 | 1 | NA | 1 | 1 | NA | NA | -0.133 |
| SE | 87857 | ENSG00000198932 | GPRASP1 | chrX | + | 102651686 | 102651908 | 102651091 | 102651595 | 102652174 | 102652265 | 10 | 4 | 19 | 7 | 1 | 0 | 1 | 7 | 0 | 0 | 0 | 13 | 6 | 0 | 0 | 0 | 0 | 0 | 0 | 0 | 298 | 149 | 1 | 1 | 1 | 0.667 | 0.576 | 1 | 1 | NA | 1 | 1 | NA | NA | -0.151 |
| SE | 87858 | ENSG00000198932 | GPRASP1 | chrX | + | 102652174 | 102652265 | 102651365 | 102651595 | 102652830 | 102652898 | 9 | 4 | 17 | 2 | 5 | 2 | 0 | 0 | 0 | 0 | 0 | 0 | 6 | 5 | 0 | 0 | 0 | 0 | 0 | 0 | 240 | 149 | 1 | 1 | 0.736 | 1 | 1 | 1 | 1 | NA | NA | 1 | 1 | NA | -0.053 |
| SE | 87859 | ENSG00000198932 | GPRASP1 | chrX | + | 102652174 | 102652265 | 102651365 | 102651595 | 102653220 | 102653298 | 0 | 1 | 3 | 0 | 0 | 0 | 0 | 0 | 0 | 0 | 0 | 0 | 1 | 0 | 0 | 0 | 0 | 3 | 0 | 0 | 240 | 149 | 1 | 1 | NA | 1 | 1 | NA | NA | NA | NA | 0.171 | NA | NA | 0.829 |
| SE | 87860 | ENSG00000198932 | GPRASP1 | chrX | + | 102652830 | 102652898 | 102651365 | 102651595 | 102653220 | 102653298 | 6 | 0 | 9 | 2 | 0 | 0 | 0 | 0 | 0 | 0 | 0 | 0 | 0 | 5 | 0 | 0 | 0 | 3 | 0 | 0 | 217 | 149 | 1 | 1 | 1 | NA | 1 | 1 | NA | NA | NA | 0 | 1 | NA | 0.5 |
| SE | 87861 | ENSG00000198932 | GPRASP1 | chrX | + | 102652830 | 102652898 | 102652174 | 102652265 | 102653220 | 102653298 | 19 | 8 | 27 | 6 | 5 | 0 | 1 | 1 | 0 | 2 | 0 | 7 | 6 | 10 | 0 | 0 | 0 | 2 | 0 | 0 | 217 | 149 | 1 | 1 | 1 | 0.846 | 0.949 | 1 | 0.632 | NA | 1 | 0.673 | 1 | NA | -0.006 |
| SE | 87862 | ENSG00000198932 | GPRASP1 | chrX | + | 102653220 | 102653298 | 102652174 | 102652265 | 102653579 | 102659074 | 9 | 16 | 22 | 16 | 15 | 0 | 0 | 0 | 0 | 0 | 7 | 4 | 9 | 1 | 3 | 0 | 0 | 3 | 0 | 0 | 227 | 149 | 1 | 1 | 1 | 1 | 1 | 1 | 1 | 1 | 1 | 0.663 | 1 | 1 | 0.067 |
| SE | 87864 | ENSG00000156504 | PABIR2 | chrX | - | 134785885 | 134785947 | 134781820 | 134781917 | 134787471 | 134787533 | 109 | 47 | 65 | 77 | 119 | 2 | 0 | 0 | 0 | 0 | 90 | 46 | 61 | 48 | 106 | 0 | 0 | 1 | 0 | 0 | 211 | 149 | 1 | 1 | 0.975 | 1 | 1 | 1 | 1 | 1 | 1 | 0.977 | 1 | 1 | 0 |
| SE | 87865 | ENSG00000156504 | PABIR2 | chrX | - | 134785885 | 134785950 | 134781820 | 134781917 | 134787471 | 134787533 | 128 | 98 | 122 | 159 | 136 | 2 | 0 | 0 | 0 | 0 | 142 | 114 | 86 | 120 | 269 | 0 | 0 | 1 | 0 | 0 | 214 | 149 | 1 | 1 | 0.978 | 1 | 1 | 1 | 1 | 1 | 1 | 0.984 | 1 | 1 | -0.001 |
| SE | 87866 | ENSG00000156504 | PABIR2 | chrX | - | 134787471 | 134787533 | 134785885 | 134785950 | 134788729 | 134788831 | 134 | 98 | 129 | 183 | 146 | 0 | 0 | 0 | 1 | 0 | 130 | 116 | 91 | 153 | 295 | 0 | 0 | 0 | 0 | 0 | 211 | 149 | 1 | 1 | 1 | 1 | 1 | 0.992 | 1 | 1 | 1 | 1 | 1 | 1 | -0.002 |
| SE | 87868 | ENSG00000156504 | PABIR2 | chrX | - | 134796107 | 134796355 | 134793814 | 134793893 | 134797120 | 134797187 | 60 | 51 | 73 | 70 | 41 | 0 | 0 | 0 | 0 | 0 | 89 | 86 | 61 | 95 | 167 | 0 | 0 | 1 | 0 | 0 | 298 | 149 | 1 | 1 | 1 | 1 | 1 | 1 | 1 | 1 | 1 | 0.968 | 1 | 1 | 0.006 |
| SE | 87869 | ENSG00000247746 | USP51 | chrX | - | 55489168 | 55489365 | 55484615 | 55488988 | 55489775 | 55489848 | 0 | 0 | 0 | 2 | 0 | 0 | 0 | 0 | 0 | 0 | 0 | 0 | 0 | 0 | 0 | 0 | 0 | 1 | 0 | 0 | 298 | 149 | 1 | 1 | NA | NA | NA | 1 | NA | NA | NA | 0 | NA | NA | 1 |
| SE | 87870 | ENSG00000233672 | RNASEH2B-AS1 | chr13 | - | 50886057 | 50886127 | 50882510 | 50882645 | 50889608 | 50889995 | 0 | 0 | 1 | 0 | 6 | 5 | 0 | 3 | 6 | 5 | 0 | 0 | 0 | 0 | 0 | 0 | 6 | 0 | 1 | 0 | 219 | 149 | 1 | 1 | 0 | NA | 0.185 | 0 | 0.449 | NA | 0 | NA | 0 | NA | 0.159 |
| SE | 87872 | ENSG00000233672 | RNASEH2B-AS1 | chr13 | - | 50891303 | 50891504 | 50889608 | 50889735 | 50904720 | 50904863 | 0 | 4 | 0 | 0 | 3 | 3 | 0 | 0 | 1 | 9 | 0 | 2 | 0 | 1 | 0 | 0 | 0 | 0 | 0 | 0 | 298 | 149 | 1 | 1 | 0 | 1 | NA | 0 | 0.143 | NA | 1 | NA | 1 | NA | -0.714 |
| SE | 87873 | ENSG00000233672 | RNASEH2B-AS1 | chr13 | - | 50904720 | 50904863 | 50889608 | 50889735 | 50910482 | 50910578 | 4 | 0 | 0 | 1 | 9 | 1 | 0 | 1 | 3 | 0 | 0 | 4 | 0 | 0 | 0 | 0 | 0 | 0 | 0 | 0 | 292 | 149 | 1 | 1 | 0.671 | NA | 0 | 0.145 | 1 | NA | 1 | NA | NA | NA | -0.546 |
| SE | 87874 | ENSG00000182774 | RPS17 | chr15 | - | 82537921 | 82538042 | 82536755 | 82536881 | 82538305 | 82538371 | 0 | 1 | 0 | 0 | 2 | 16555 | 10024 | 16622 | 15880 | 15755 | 0 | 0 | 2 | 1 | 0 | 18562 | 10889 | 11021 | 9958 | 18827 | 270 | 149 | 1 | 1 | 0 | 0 | 0 | 0 | 0 | 0 | 0 | 0 | 0 | 0 | 0 |
| SE | 87875 | ENSG00000182774 | RPS17 | chr15 | - | 82539299 | 82539375 | 82538879 | 82538985 | 82539980 | 82540131 | 0 | 0 | 0 | 2 | 0 | 8410 | 4079 | 9401 | 8587 | 4313 | 0 | 0 | 0 | 0 | 0 | 8982 | 6182 | 5898 | 5048 | 9179 | 225 | 149 | 1 | 1 | 0 | 0 | 0 | 0 | 0 | 0 | 0 | 0 | 0 | 0 | 0 |
| SE | 87876 | ENSG00000101974 | ATP11C | chrX | - | 139731650 | 139731755 | 139726347 | 139728962 | 139732434 | 139732525 | 9 | 3 | 6 | 7 | 13 | 0 | 0 | 2 | 0 | 5 | 0 | 6 | 9 | 1 | 0 | 0 | 0 | 0 | 0 | 0 | 254 | 149 | 1 | 1 | 1 | 1 | 0.638 | 1 | 0.604 | NA | 1 | 1 | 1 | NA | -0.152 |
| SE | 87877 | ENSG00000101974 | ATP11C | chrX | - | 139731650 | 139731755 | 139726347 | 139728962 | 139737730 | 139737796 | 9 | 3 | 6 | 10 | 13 | 0 | 6 | 0 | 1 | 0 | 0 | 6 | 8 | 1 | 0 | 0 | 0 | 3 | 0 | 0 | 254 | 149 | 1 | 1 | 1 | 0.227 | 1 | 0.854 | 1 | NA | 1 | 0.61 | 1 | NA | -0.054 |
| SE | 87878 | ENSG00000101974 | ATP11C | chrX | - | 139731650 | 139731755 | 139726347 | 139728962 | 139737915 | 139738069 | 32 | 15 | 31 | 18 | 45 | 47 | 15 | 32 | 29 | 88 | 0 | 30 | 19 | 1 | 0 | 0 | 17 | 9 | 8 | 0 | 254 | 149 | 1 | 1 | 0.285 | 0.37 | 0.362 | 0.267 | 0.231 | NA | 0.509 | 0.553 | 0.068 | NA | -0.074 |
| SE | 87879 | ENSG00000101974 | ATP11C | chrX | - | 139732434 | 139732525 | 139728785 | 139728962 | 139737915 | 139738069 | 0 | 0 | 2 | 0 | 5 | 47 | 15 | 32 | 29 | 88 | 0 | 4 | 0 | 0 | 0 | 0 | 17 | 9 | 8 | 0 | 240 | 149 | 1 | 1 | 0 | 0 | 0.037 | 0 | 0.034 | NA | 0.127 | 0 | 0 | NA | -0.028 |
| SE | 87880 | ENSG00000101974 | ATP11C | chrX | - | 139732434 | 139732525 | 139731650 | 139731755 | 139737915 | 139738069 | 0 | 0 | 1 | 0 | 0 | 23 | 12 | 25 | 11 | 32 | 0 | 4 | 0 | 0 | 0 | 0 | 24 | 11 | 0 | 0 | 240 | 149 | 1 | 1 | 0 | 0 | 0.024 | 0 | 0 | NA | 0.094 | 0 | NA | NA | -0.042 |
| SE | 87881 | ENSG00000101974 | ATP11C | chrX | - | 139737730 | 139737796 | 139726352 | 139728962 | 139737915 | 139738069 | 0 | 11 | 0 | 1 | 0 | 47 | 15 | 32 | 29 | 88 | 0 | 0 | 3 | 0 | 0 | 0 | 17 | 9 | 8 | 0 | 215 | 149 | 1 | 1 | 0 | 0.337 | 0 | 0.023 | 0 | NA | 0 | 0.188 | 0 | NA | 0.009 |
| SE | 87882 | ENSG00000101974 | ATP11C | chrX | - | 139737730 | 139737796 | 139731650 | 139731755 | 139737915 | 139738069 | 0 | 5 | 0 | 3 | 0 | 23 | 12 | 25 | 11 | 32 | 0 | 0 | 0 | 0 | 0 | 0 | 24 | 11 | 0 | 0 | 215 | 149 | 1 | 1 | 0 | 0.224 | 0 | 0.159 | 0 | NA | 0 | 0 | NA | NA | 0.077 |
| SE | 87883 | ENSG00000101974 | ATP11C | chrX | - | 139743558 | 139743624 | 139740990 | 139741094 | 139745721 | 139745791 | 48 | 38 | 52 | 43 | 62 | 0 | 0 | 2 | 0 | 0 | 0 | 36 | 19 | 16 | 0 | 0 | 0 | 0 | 0 | 0 | 215 | 149 | 1 | 1 | 1 | 1 | 0.947 | 1 | 1 | NA | 1 | 1 | 1 | NA | -0.011 |
| SE | 87884 | ENSG00000101974 | ATP11C | chrX | - | 139745721 | 139745857 | 139743558 | 139743624 | 139750024 | 139750053 | 57 | 35 | 50 | 48 | 73 | 0 | 0 | 0 | 1 | 0 | 0 | 39 | 14 | 9 | 0 | 0 | 0 | 0 | 0 | 0 | 285 | 149 | 1 | 1 | 1 | 1 | 1 | 0.962 | 1 | NA | 1 | 1 | 1 | NA | -0.008 |
| SE | 87885 | ENSG00000101974 | ATP11C | chrX | - | 139763315 | 139763418 | 139761960 | 139762106 | 139768259 | 139768434 | 136 | 64 | 103 | 111 | 136 | 1 | 2 | 0 | 0 | 0 | 0 | 50 | 39 | 29 | 13 | 0 | 0 | 3 | 0 | 2 | 252 | 149 | 1 | 1 | 0.988 | 0.95 | 1 | 1 | 1 | NA | 1 | 0.885 | 1 | 0.794 | 0.068 |
| SE | 87886 | ENSG00000101974 | ATP11C | chrX | - | 139774689 | 139774953 | 139768259 | 139768434 | 139782546 | 139782728 | 148 | 159 | 141 | 179 | 208 | 3 | 0 | 0 | 0 | 0 | 20 | 90 | 82 | 28 | 23 | 0 | 0 | 0 | 0 | 0 | 298 | 149 | 1 | 1 | 0.961 | 1 | 1 | 1 | 1 | 1 | 1 | 1 | 1 | 1 | -0.008 |
| SE | 87888 | ENSG00000101974 | ATP11C | chrX | - | 139785225 | 139785299 | 139783163 | 139783267 | 139788191 | 139788343 | 58 | 54 | 49 | 62 | 100 | 0 | 0 | 0 | 0 | 1 | 5 | 22 | 28 | 1 | 2 | 0 | 0 | 0 | 0 | 0 | 223 | 149 | 1 | 1 | 1 | 1 | 1 | 1 | 0.985 | 1 | 1 | 1 | 1 | 1 | -0.003 |
| SE | 87889 | ENSG00000101974 | ATP11C | chrX | - | 139787172 | 139787244 | 139785225 | 139785299 | 139788191 | 139788343 | 149 | 131 | 142 | 144 | 189 | 6 | 2 | 3 | 8 | 0 | 0 | 79 | 105 | 39 | 3 | 0 | 5 | 7 | 0 | 0 | 221 | 149 | 1 | 1 | 0.944 | 0.978 | 0.97 | 0.924 | 1 | NA | 0.914 | 0.91 | 1 | 1 | 0.007 |
| SE | 87890 | ENSG00000101974 | ATP11C | chrX | - | 139796272 | 139796470 | 139789326 | 139789488 | 139798272 | 139798354 | 49 | 54 | 43 | 45 | 108 | 0 | 0 | 0 | 0 | 0 | 12 | 34 | 36 | 10 | 0 | 0 | 0 | 1 | 0 | 0 | 298 | 149 | 1 | 1 | 1 | 1 | 1 | 1 | 1 | 1 | 1 | 0.947 | 1 | NA | 0.013 |
| SE | 87891 | ENSG00000101974 | ATP11C | chrX | - | 139797175 | 139797326 | 139796272 | 139796470 | 139798272 | 139798354 | 191 | 143 | 191 | 153 | 140 | 0 | 0 | 0 | 4 | 0 | 20 | 49 | 100 | 32 | 16 | 0 | 0 | 0 | 0 | 0 | 298 | 149 | 1 | 1 | 1 | 1 | 1 | 0.95 | 1 | 1 | 1 | 1 | 1 | 1 | -0.01 |
| SE | 87892 | ENSG00000101974 | ATP11C | chrX | - | 139814877 | 139814985 | 139804470 | 139804599 | 139816862 | 139816943 | 90 | 90 | 64 | 73 | 144 | 0 | 1 | 1 | 1 | 0 | 23 | 48 | 37 | 27 | 19 | 0 | 0 | 1 | 0 | 0 | 257 | 149 | 1 | 1 | 1 | 0.981 | 0.974 | 0.977 | 1 | 1 | 1 | 0.955 | 1 | 1 | -0.005 |
| SE | 87895 | ENSG00000167862 | MRPL58 | chr17 | + | 75018585 | 75018694 | 75017077 | 75017114 | 75019699 | 75019759 | 0 | 1 | 3 | 1 | 0 | 28 | 13 | 34 | 32 | 35 | 2 | 1 | 2 | 10 | 14 | 28 | 18 | 23 | 23 | 56 | 258 | 149 | 1 | 1 | 0 | 0.043 | 0.048 | 0.018 | 0 | 0.04 | 0.031 | 0.048 | 0.201 | 0.126 | -0.067 |
| SE | 87898 | ENSG00000119328 | ABITRAM | chr9 | + | 108935637 | 108935689 | 108934399 | 108934565 | 108950506 | 108950744 | 0 | 6 | 6 | 4 | 0 | 0 | 0 | 1 | 0 | 0 | 0 | 0 | 0 | 0 | 4 | 0 | 0 | 0 | 0 | 0 | 201 | 149 | 1 | 1 | NA | 1 | 0.816 | 1 | NA | NA | NA | NA | NA | 1 | -0.061 |
| SE | 87899 | ENSG00000119328 | ABITRAM | chr9 | + | 108936307 | 108936437 | 108934539 | 108934565 | 108939195 | 108939272 | 19 | 5 | 16 | 16 | 23 | 2 | 0 | 0 | 0 | 0 | 0 | 1 | 6 | 5 | 0 | 0 | 0 | 0 | 0 | 0 | 279 | 149 | 1 | 1 | 0.835 | 1 | 1 | 1 | 1 | NA | 1 | 1 | 1 | NA | -0.033 |
| SE | 87901 | ENSG00000119328 | ABITRAM | chr9 | + | 108936307 | 108936437 | 108935637 | 108935689 | 108939195 | 108939272 | 50 | 12 | 42 | 52 | 41 | 0 | 1 | 0 | 0 | 3 | 12 | 9 | 8 | 14 | 0 | 0 | 0 | 0 | 0 | 0 | 279 | 149 | 1 | 1 | 1 | 0.865 | 1 | 1 | 0.879 | 1 | 1 | 1 | 1 | NA | -0.051 |
| SE | 87910 | ENSG00000215374 | FAM66B | chr8 | - | 7324821 | 7324889 | 7324296 | 7324364 | 7325136 | 7325198 | 10 | 0 | 11 | 3 | 9 | 0 | 0 | 0 | 0 | 0 | 0 | 5 | 2 | 0 | 0 | 0 | 0 | 0 | 0 | 4 | 217 | 149 | 1 | 1 | 1 | NA | 1 | 1 | 1 | NA | 1 | 1 | NA | 0 | 0.333 |
| SE | 87911 | ENSG00000215374 | FAM66B | chr8 | - | 7324821 | 7324960 | 7322689 | 7324364 | 7325136 | 7325198 | 2 | 0 | 3 | 1 | 3 | 0 | 0 | 0 | 0 | 0 | 0 | 5 | 1 | 0 | 0 | 0 | 0 | 0 | 0 | 4 | 288 | 149 | 1 | 1 | 1 | NA | 1 | 1 | 1 | NA | 1 | 1 | NA | 0 | 0.333 |
| SE | 87912 | ENSG00000215374 | FAM66B | chr8 | - | 7324821 | 7324977 | 7322669 | 7324364 | 7325136 | 7325198 | 2 | 0 | 3 | 1 | 3 | 0 | 0 | 0 | 0 | 0 | 0 | 5 | 0 | 0 | 0 | 0 | 0 | 0 | 0 | 4 | 298 | 149 | 1 | 1 | 1 | NA | 1 | 1 | 1 | NA | 1 | NA | NA | 0 | 0.5 |
| SE | 87913 | ENSG00000173273 | TNKS | chr8 | + | 9704662 | 9704757 | 9680724 | 9680800 | 9706186 | 9706253 | 37 | 15 | 16 | 35 | 47 | 0 | 0 | 0 | 0 | 0 | 0 | 14 | 8 | 0 | 3 | 0 | 0 | 1 | 1 | 0 | 244 | 149 | 1 | 1 | 1 | 1 | 1 | 1 | 1 | NA | 1 | 0.83 | 0 | 1 | 0.292 |
| SE | 87914 | ENSG00000173273 | TNKS | chr8 | + | 9708370 | 9708492 | 9706810 | 9706997 | 9709954 | 9710046 | 117 | 116 | 142 | 125 | 192 | 0 | 0 | 1 | 0 | 11 | 175 | 120 | 125 | 113 | 98 | 5 | 0 | 1 | 0 | 3 | 271 | 149 | 1 | 1 | 1 | 1 | 0.987 | 1 | 0.906 | 0.951 | 1 | 0.986 | 1 | 0.947 | 0.002 |
| SE | 87915 | ENSG00000173273 | TNKS | chr8 | + | 9720373 | 9720545 | 9710141 | 9710220 | 9726640 | 9726720 | 69 | 71 | 102 | 110 | 89 | 0 | 0 | 0 | 0 | 0 | 24 | 116 | 71 | 39 | 30 | 0 | 1 | 0 | 0 | 0 | 298 | 149 | 1 | 1 | 1 | 1 | 1 | 1 | 1 | 1 | 0.983 | 1 | 1 | 1 | 0.003 |
| SE | 87917 | ENSG00000173273 | TNKS | chr8 | + | 9730889 | 9731035 | 9726640 | 9726720 | 9733278 | 9733444 | 47 | 28 | 101 | 61 | 43 | 2 | 1 | 0 | 1 | 1 | 66 | 69 | 59 | 35 | 24 | 0 | 3 | 1 | 0 | 0 | 295 | 149 | 1 | 1 | 0.922 | 0.934 | 1 | 0.969 | 0.956 | 1 | 0.921 | 0.968 | 1 | 1 | -0.022 |
| SE | 87920 | ENSG00000173273 | TNKS | chr8 | + | 9733278 | 9733444 | 9730889 | 9731035 | 9734864 | 9735084 | 65 | 42 | 112 | 56 | 97 | 0 | 0 | 0 | 0 | 0 | 100 | 62 | 71 | 67 | 36 | 0 | 3 | 0 | 0 | 0 | 298 | 149 | 1 | 1 | 1 | 1 | 1 | 1 | 1 | 1 | 0.912 | 1 | 1 | 1 | 0.018 |
| SE | 87924 | ENSG00000173273 | TNKS | chr8 | + | 9763146 | 9763244 | 9761515 | 9761636 | 9764715 | 9764790 | 128 | 88 | 90 | 63 | 127 | 0 | 0 | 0 | 0 | 0 | 38 | 87 | 63 | 91 | 20 | 0 | 1 | 0 | 0 | 0 | 247 | 149 | 1 | 1 | 1 | 1 | 1 | 1 | 1 | 1 | 0.981 | 1 | 1 | 1 | 0.004 |
| SE | 87925 | ENSG00000173273 | TNKS | chr8 | + | 9766238 | 9766425 | 9765691 | 9765797 | 9770105 | 9770262 | 152 | 157 | 161 | 116 | 238 | 1 | 0 | 0 | 0 | 3 | 77 | 121 | 142 | 119 | 77 | 0 | 0 | 0 | 0 | 0 | 298 | 149 | 1 | 1 | 0.987 | 1 | 1 | 1 | 0.975 | 1 | 1 | 1 | 1 | 1 | -0.008 |
| SE | 87927 | ENSG00000113384 | GOLPH3 | chr5 | - | 32143748 | 32143880 | 32135571 | 32135686 | 32173809 | 32174319 | 712 | 534 | 521 | 564 | 969 | 2 | 1 | 5 | 1 | 2 | 996 | 664 | 487 | 634 | 618 | 0 | 0 | 4 | 2 | 2 | 281 | 149 | 1 | 1 | 0.995 | 0.996 | 0.982 | 0.997 | 0.996 | 1 | 1 | 0.985 | 0.994 | 0.994 | -0.001 |
| SE | 87928 | ENSG00000147408 | CSGALNACT1 | chr8 | - | 19418655 | 19418750 | 19408612 | 19408694 | 19420339 | 19420518 | 107 | 137 | 238 | 143 | 322 | 0 | 0 | 1 | 1 | 0 | 120 | 158 | 212 | 150 | 17 | 0 | 0 | 0 | 3 | 0 | 244 | 149 | 1 | 1 | 1 | 1 | 0.993 | 0.989 | 1 | 1 | 1 | 1 | 0.968 | 1 | 0.003 |
| SE | 87929 | ENSG00000147408 | CSGALNACT1 | chr8 | - | 19439829 | 19439931 | 19420339 | 19420518 | 19458425 | 19458642 | 54 | 72 | 120 | 84 | 149 | 0 | 0 | 0 | 0 | 1 | 69 | 57 | 111 | 91 | 4 | 0 | 0 | 0 | 0 | 0 | 251 | 149 | 1 | 1 | 1 | 1 | 1 | 1 | 0.989 | 1 | 1 | 1 | 1 | 1 | -0.002 |
| SE | 87930 | ENSG00000147408 | CSGALNACT1 | chr8 | - | 19457529 | 19457816 | 19439829 | 19439931 | 19458425 | 19458642 | 10 | 12 | 26 | 4 | 41 | 15 | 12 | 30 | 15 | 46 | 0 | 1 | 9 | 4 | 0 | 12 | 13 | 41 | 32 | 0 | 298 | 149 | 1 | 1 | 0.25 | 0.333 | 0.302 | 0.118 | 0.308 | 0 | 0.037 | 0.099 | 0.059 | NA | 0.213 |
| SE | 87931 | ENSG00000147408 | CSGALNACT1 | chr8 | - | 19458425 | 19458642 | 19439829 | 19439931 | 19601770 | 19601865 | 15 | 16 | 32 | 15 | 46 | 0 | 0 | 0 | 0 | 0 | 12 | 13 | 41 | 32 | 0 | 0 | 0 | 2 | 0 | 0 | 298 | 149 | 1 | 1 | 1 | 1 | 1 | 1 | 1 | 1 | 1 | 0.911 | 1 | NA | 0.022 |
| SE | 87934 | ENSG00000147408 | CSGALNACT1 | chr8 | - | 19591159 | 19591278 | 19505200 | 19506130 | 19601770 | 19601865 | 2 | 0 | 0 | 0 | 2 | 3 | 5 | 12 | 3 | 3 | 0 | 1 | 0 | 0 | 0 | 16 | 0 | 10 | 19 | 3 | 268 | 149 | 1 | 1 | 0.27 | 0 | 0 | 0 | 0.27 | 0 | 1 | 0 | 0 | 0 | -0.092 |
| SE | 87938 | ENSG00000147408 | CSGALNACT1 | chr8 | - | 19732455 | 19732566 | 19601770 | 19601865 | 19757185 | 19757217 | 0 | 1 | 1 | 0 | 1 | 2 | 0 | 2 | 3 | 0 | 0 | 2 | 0 | 2 | 0 | 0 | 0 | 0 | 0 | 0 | 260 | 149 | 1 | 1 | 0 | 1 | 0.223 | 0 | 1 | NA | 1 | NA | 1 | NA | -0.555 |
| SE | 87939 | ENSG00000104361 | NIPAL2 | chr8 | - | 98193384 | 98193412 | 98189832 | 98193090 | 98194727 | 98194822 | 0 | 0 | 0 | 2 | 0 | 47 | 34 | 17 | 14 | 60 | 0 | 0 | 0 | 4 | 0 | 7 | 26 | 20 | 13 | 6 | 177 | 149 | 1 | 1 | 0 | 0 | 0 | 0.107 | 0 | 0 | 0 | 0 | 0.206 | 0 | -0.02 |
| SE | 87941 | ENSG00000104361 | NIPAL2 | chr8 | - | 98194727 | 98194822 | 98189825 | 98193090 | 98203107 | 98203196 | 25 | 25 | 14 | 12 | 38 | 0 | 0 | 0 | 0 | 4 | 7 | 16 | 19 | 0 | 6 | 0 | 0 | 0 | 0 | 0 | 244 | 149 | 1 | 1 | 1 | 1 | 1 | 1 | 0.853 | 1 | 1 | 1 | NA | 1 | -0.029 |
| SE | 87942 | ENSG00000104361 | NIPAL2 | chr8 | - | 98195941 | 98196005 | 98189825 | 98193090 | 98203107 | 98203196 | 31 | 13 | 4 | 12 | 37 | 0 | 0 | 0 | 0 | 4 | 6 | 7 | 8 | 5 | 0 | 0 | 0 | 0 | 0 | 0 | 213 | 149 | 1 | 1 | 1 | 1 | 1 | 1 | 0.866 | 1 | 1 | 1 | 1 | NA | -0.027 |
| SE | 87943 | ENSG00000104361 | NIPAL2 | chr8 | - | 98195941 | 98196005 | 98194727 | 98194822 | 98203107 | 98203196 | 60 | 28 | 13 | 30 | 83 | 0 | 2 | 3 | 3 | 4 | 6 | 14 | 14 | 27 | 0 | 0 | 0 | 0 | 0 | 0 | 213 | 149 | 1 | 1 | 1 | 0.907 | 0.752 | 0.875 | 0.936 | 1 | 1 | 1 | 1 | NA | -0.106 |
| SE | 87945 | ENSG00000104361 | NIPAL2 | chr8 | - | 98236154 | 98236214 | 98222478 | 98222600 | 98252462 | 98252601 | 45 | 23 | 38 | 44 | 29 | 0 | 0 | 0 | 0 | 0 | 7 | 22 | 24 | 39 | 0 | 0 | 0 | 2 | 0 | 0 | 209 | 149 | 1 | 1 | 1 | 1 | 1 | 1 | 1 | 1 | 1 | 0.895 | 1 | NA | 0.026 |
| SE | 87952 | ENSG00000120699 | EXOSC8 | chr13 | + | 37001748 | 37001801 | 37000785 | 37000822 | 37002272 | 37002309 | 2 | 1 | 8 | 7 | 8 | 40 | 19 | 31 | 27 | 40 | 0 | 3 | 2 | 11 | 6 | 0 | 15 | 17 | 2 | 0 | 202 | 149 | 1 | 1 | 0.036 | 0.037 | 0.16 | 0.161 | 0.129 | NA | 0.129 | 0.08 | 0.802 | 1 | -0.398 |
| SE | 87954 | ENSG00000120699 | EXOSC8 | chr13 | + | 37002272 | 37002309 | 37000785 | 37000822 | 37002487 | 37002551 | 87 | 49 | 66 | 67 | 99 | 3 | 1 | 0 | 1 | 0 | 0 | 18 | 35 | 13 | 5 | 0 | 0 | 0 | 0 | 0 | 186 | 149 | 1 | 1 | 0.959 | 0.975 | 1 | 0.982 | 1 | NA | 1 | 1 | 1 | 1 | -0.017 |
| SE | 87955 | ENSG00000120699 | EXOSC8 | chr13 | + | 37002272 | 37002309 | 37001473 | 37001801 | 37002487 | 37002551 | 49 | 32 | 48 | 58 | 76 | 0 | 0 | 0 | 2 | 0 | 10 | 6 | 25 | 26 | 11 | 0 | 0 | 0 | 0 | 0 | 186 | 149 | 1 | 1 | 1 | 1 | 1 | 0.959 | 1 | 1 | 1 | 1 | 1 | 1 | -0.008 |
| SE | 87956 | ENSG00000120699 | EXOSC8 | chr13 | + | 37002933 | 37003007 | 37002487 | 37002551 | 37003494 | 37003517 | 85 | 50 | 76 | 84 | 98 | 0 | 0 | 0 | 0 | 5 | 0 | 18 | 46 | 24 | 11 | 0 | 0 | 0 | 0 | 0 | 223 | 149 | 1 | 1 | 1 | 1 | 1 | 1 | 0.929 | NA | 1 | 1 | 1 | 1 | -0.014 |
| SE | 87958 | ENSG00000120699 | EXOSC8 | chr13 | + | 37003494 | 37003517 | 37002933 | 37003007 | 37004515 | 37004525 | 4 | 1 | 7 | 6 | 12 | 65 | 51 | 75 | 90 | 101 | 0 | 0 | 7 | 0 | 0 | 13 | 28 | 36 | 3 | 0 | 172 | 149 | 1 | 1 | 0.051 | 0.017 | 0.075 | 0.055 | 0.093 | 0 | 0 | 0.144 | 0 | NA | 0.022 |
| SE | 87966 | ENSG00000168066 | SF1 | chr11 | - | 64765950 | 64766155 | 64764605 | 64765518 | 64767191 | 64767251 | 1 | 2 | 7 | 4 | 0 | 2 | 1 | 5 | 7 | 0 | 10 | 0 | 0 | 0 | 0 | 0 | 3 | 6 | 2 | 0 | 298 | 149 | 1 | 1 | 0.2 | 0.5 | 0.412 | 0.222 | NA | 1 | 0 | 0 | 0 | NA | 0.083 |
| SE | 87968 | ENSG00000168066 | SF1 | chr11 | - | 64766899 | 64767079 | 64764605 | 64766155 | 64767191 | 64767246 | 167 | 115 | 184 | 196 | 82 | 1 | 0 | 2 | 1 | 0 | 362 | 317 | 191 | 384 | 218 | 0 | 0 | 0 | 0 | 0 | 298 | 149 | 1 | 1 | 0.988 | 1 | 0.979 | 0.99 | 1 | 1 | 1 | 1 | 1 | 1 | -0.009 |
| SE | 87969 | ENSG00000168066 | SF1 | chr11 | - | 64769963 | 64770053 | 64769425 | 64769609 | 64776497 | 64776626 | 247 | 157 | 273 | 303 | 300 | 0 | 0 | 0 | 0 | 0 | 435 | 300 | 324 | 521 | 500 | 0 | 0 | 0 | 0 | 4 | 239 | 149 | 1 | 1 | 1 | 1 | 1 | 1 | 1 | 1 | 1 | 1 | 1 | 0.987 | 0.003 |
| SE | 87970 | ENSG00000168066 | SF1 | chr11 | - | 64770255 | 64770408 | 64769963 | 64770053 | 64776497 | 64776626 | 783 | 423 | 695 | 872 | 745 | 0 | 0 | 0 | 0 | 0 | 1479 | 1077 | 964 | 1438 | 1256 | 0 | 0 | 0 | 3 | 0 | 298 | 149 | 1 | 1 | 1 | 1 | 1 | 1 | 1 | 1 | 1 | 1 | 0.996 | 1 | 0.001 |
| SE | 87971 | ENSG00000168066 | SF1 | chr11 | - | 64771463 | 64771537 | 64770277 | 64770408 | 64773429 | 64773505 | 8 | 3 | 2 | 8 | 7 | 453 | 253 | 423 | 476 | 551 | 0 | 6 | 10 | 6 | 9 | 818 | 485 | 432 | 641 | 1049 | 223 | 149 | 1 | 1 | 0.012 | 0.008 | 0.003 | 0.011 | 0.008 | 0 | 0.008 | 0.015 | 0.006 | 0.006 | 0.001 |
| SE | 87977 | ENSG00000198373 | WWP2 | chr16 | + | 69775078 | 69775140 | 69762361 | 69762391 | 69776188 | 69776241 | 0 | 0 | 0 | 1 | 2 | 19 | 9 | 17 | 20 | 12 | 3 | 1 | 0 | 0 | 0 | 19 | 67 | 16 | 40 | 59 | 211 | 149 | 1 | 1 | 0 | 0 | 0 | 0.034 | 0.105 | 0.1 | 0.01 | 0 | 0 | 0 | 0.006 |
| SE | 87978 | ENSG00000198373 | WWP2 | chr16 | + | 69776188 | 69776241 | 69762327 | 69762391 | 69783031 | 69783152 | 3 | 0 | 2 | 2 | 0 | 0 | 0 | 1 | 0 | 0 | 0 | 0 | 0 | 0 | 7 | 0 | 0 | 0 | 0 | 0 | 202 | 149 | 1 | 1 | 1 | NA | 0.596 | 1 | NA | NA | NA | NA | NA | 1 | -0.135 |
| SE | 87981 | ENSG00000198373 | WWP2 | chr16 | + | 69786995 | 69787080 | 69762331 | 69762391 | 69798681 | 69798796 | 141 | 75 | 99 | 108 | 111 | 0 | 0 | 0 | 0 | 0 | 418 | 204 | 163 | 260 | 242 | 0 | 0 | 1 | 0 | 0 | 234 | 149 | 1 | 1 | 1 | 1 | 1 | 1 | 1 | 1 | 1 | 0.99 | 1 | 1 | 0.002 |
| SE | 87984 | ENSG00000198373 | WWP2 | chr16 | + | 69840125 | 69840263 | 69799045 | 69799295 | 69871803 | 69871865 | 134 | 82 | 100 | 91 | 168 | 0 | 0 | 0 | 0 | 0 | 313 | 195 | 123 | 205 | 250 | 0 | 2 | 0 | 0 | 0 | 287 | 149 | 1 | 1 | 1 | 1 | 1 | 1 | 1 | 1 | 0.981 | 1 | 1 | 1 | 0.004 |
| SE | 87985 | ENSG00000198373 | WWP2 | chr16 | + | 69842023 | 69842120 | 69840125 | 69840263 | 69871803 | 69871865 | 266 | 154 | 194 | 227 | 294 | 0 | 1 | 0 | 0 | 0 | 505 | 310 | 221 | 305 | 434 | 0 | 2 | 0 | 0 | 0 | 246 | 149 | 1 | 1 | 1 | 0.989 | 1 | 1 | 1 | 1 | 0.989 | 1 | 1 | 1 | 0 |
| SE | 87987 | ENSG00000198373 | WWP2 | chr16 | + | 69917708 | 69917883 | 69908760 | 69908850 | 69925429 | 69925484 | 86 | 57 | 75 | 109 | 60 | 0 | 0 | 0 | 0 | 0 | 106 | 171 | 60 | 120 | 150 | 0 | 3 | 2 | 0 | 0 | 298 | 149 | 1 | 1 | 1 | 1 | 1 | 1 | 1 | 1 | 0.966 | 0.938 | 1 | 1 | 0.019 |
| SE | 87989 | ENSG00000198373 | WWP2 | chr16 | + | 69931151 | 69931227 | 69930129 | 69930258 | 69931508 | 69931580 | 217 | 93 | 180 | 189 | 237 | 0 | 0 | 0 | 4 | 0 | 230 | 172 | 115 | 189 | 297 | 0 | 0 | 0 | 0 | 0 | 225 | 149 | 1 | 1 | 1 | 1 | 1 | 0.969 | 1 | 1 | 1 | 1 | 1 | 1 | -0.006 |
| SE | 87990 | ENSG00000198373 | WWP2 | chr16 | + | 69933969 | 69934129 | 69931801 | 69931890 | 69935852 | 69935986 | 158 | 105 | 108 | 129 | 215 | 0 | 0 | 0 | 0 | 0 | 291 | 200 | 139 | 192 | 383 | 0 | 0 | 0 | 1 | 0 | 298 | 149 | 1 | 1 | 1 | 1 | 1 | 1 | 1 | 1 | 1 | 1 | 0.99 | 1 | 0.002 |
| SE | 87993 | ENSG00000139233 | LLPH | chr12 | - | 66128895 | 66129113 | 66116554 | 66124018 | 66130704 | 66130750 | 91 | 91 | 89 | 129 | 86 | 0 | 0 | 0 | 1 | 0 | 31 | 68 | 178 | 127 | 29 | 0 | 0 | 0 | 0 | 0 | 298 | 149 | 1 | 1 | 1 | 1 | 1 | 0.985 | 1 | 1 | 1 | 1 | 1 | 1 | -0.003 |
| SE | 87997 | ENSG00000219481 | NBPF1 | chr1 | - | 16583593 | 16583696 | 16582543 | 16582758 | 16585488 | 16585698 | 44 | 34 | 20 | 59 | 74 | 0 | 0 | 2 | 0 | 0 | 29 | 6 | 15 | 17 | 4 | 0 | 0 | 0 | 0 | 0 | 252 | 149 | 1 | 1 | 1 | 1 | 0.855 | 1 | 1 | 1 | 1 | 1 | 1 | 1 | -0.029 |
| SE | 87999 | ENSG00000219481 | NBPF1 | chr1 | - | 16594930 | 16595013 | 16594591 | 16594661 | 16608507 | 16608650 | 33 | 38 | 30 | 36 | 28 | 0 | 0 | 0 | 2 | 0 | 9 | 1 | 10 | 6 | 27 | 0 | 2 | 0 | 0 | 0 | 232 | 149 | 1 | 1 | 1 | 1 | 1 | 0.92 | 1 | 1 | 0.243 | 1 | 1 | 1 | 0.135 |
| SE | 88001 | ENSG00000234424 | NA | chr9 | + | 85764707 | 85764925 | 85753305 | 85753419 | 85770283 | 85770490 | 8 | 1 | 3 | 6 | 7 | 0 | 0 | 0 | 0 | 0 | 0 | 6 | 5 | 0 | 0 | 0 | 0 | 5 | 0 | 0 | 298 | 149 | 1 | 1 | 1 | 1 | 1 | 1 | 1 | NA | 1 | 0.333 | NA | NA | 0.334 |
| SE | 88002 | ENSG00000234424 | NA | chr9 | + | 85770283 | 85770490 | 85753305 | 85753419 | 85773837 | 85774036 | 5 | 4 | 1 | 1 | 0 | 0 | 0 | 0 | 0 | 0 | 0 | 5 | 6 | 0 | 0 | 0 | 0 | 1 | 0 | 0 | 298 | 149 | 1 | 1 | 1 | 1 | 1 | 1 | NA | NA | 1 | 0.75 | NA | NA | 0.125 |
| SE | 88008 | ENSG00000258388 | PPT2-EGFL8 | chr6 | + | 32164578 | 32164657 | 32162567 | 32162622 | 32166137 | 32166232 | 3 | 0 | 2 | 6 | 0 | 0 | 1 | 0 | 0 | 0 | 0 | 0 | 3 | 0 | 0 | 0 | 0 | 0 | 0 | 0 | 228 | 149 | 1 | 1 | 1 | 0 | 1 | 1 | NA | NA | NA | 1 | NA | NA | -0.25 |
| SE | 88009 | ENSG00000258388 | PPT2-EGFL8 | chr6 | + | 32164578 | 32164681 | 32162567 | 32162622 | 32166137 | 32166620 | 3 | 1 | 7 | 5 | 0 | 0 | 1 | 0 | 0 | 0 | 0 | 2 | 5 | 0 | 0 | 0 | 0 | 0 | 0 | 0 | 252 | 149 | 1 | 1 | 1 | 0.372 | 1 | 1 | NA | NA | 1 | 1 | NA | NA | -0.157 |
| SE | 88012 | ENSG00000147439 | BIN3 | chr8 | - | 22624221 | 22624363 | 22623914 | 22624049 | 22629963 | 22630004 | 347 | 213 | 271 | 260 | 301 | 0 | 0 | 0 | 0 | 0 | 328 | 273 | 353 | 482 | 394 | 0 | 0 | 0 | 3 | 0 | 291 | 149 | 1 | 1 | 1 | 1 | 1 | 1 | 1 | 1 | 1 | 1 | 0.988 | 1 | 0.002 |
| SE | 88014 | ENSG00000147439 | BIN3 | chr8 | - | 22630441 | 22630578 | 22629703 | 22630004 | 22636921 | 22636962 | 172 | 107 | 154 | 148 | 178 | 4 | 0 | 0 | 0 | 0 | 256 | 177 | 260 | 302 | 361 | 0 | 0 | 4 | 1 | 0 | 286 | 149 | 1 | 1 | 0.957 | 1 | 1 | 1 | 1 | 1 | 1 | 0.971 | 0.994 | 1 | -0.002 |
| SE | 88017 | ENSG00000147439 | BIN3 | chr8 | - | 22636524 | 22636586 | 22629703 | 22630004 | 22636921 | 22636962 | 15 | 11 | 17 | 12 | 5 | 4 | 0 | 0 | 0 | 0 | 14 | 23 | 16 | 26 | 18 | 0 | 0 | 4 | 1 | 0 | 211 | 149 | 1 | 1 | 0.726 | 1 | 1 | 1 | 1 | 1 | 1 | 0.739 | 0.948 | 1 | 0.008 |
| SE | 88020 | ENSG00000147439 | BIN3 | chr8 | - | 22636921 | 22636962 | 22630441 | 22630578 | 22644754 | 22644803 | 14 | 10 | 20 | 21 | 24 | 0 | 0 | 0 | 0 | 4 | 33 | 29 | 23 | 36 | 55 | 0 | 0 | 2 | 2 | 0 | 190 | 149 | 1 | 1 | 1 | 1 | 1 | 1 | 0.825 | 1 | 1 | 0.9 | 0.934 | 1 | -0.002 |
| SE | 88022 | ENSG00000249709 | ZNF564 | chr19 | - | 12541424 | 12541537 | 12528569 | 12528696 | 12551329 | 12551542 | 0 | 0 | 2 | 0 | 0 | 46 | 32 | 31 | 25 | 12 | 0 | 0 | 0 | 0 | 0 | 17 | 58 | 58 | 36 | 0 | 262 | 149 | 1 | 1 | 0 | 0 | 0.035 | 0 | 0 | 0 | 0 | 0 | 0 | NA | 0.007 |
| SE | 88023 | ENSG00000249709 | ZNF564 | chr19 | - | 12550462 | 12550652 | 12528569 | 12528696 | 12551329 | 12551467 | 0 | 0 | 0 | 2 | 5 | 46 | 32 | 31 | 25 | 12 | 0 | 0 | 0 | 0 | 0 | 17 | 58 | 58 | 36 | 0 | 298 | 149 | 1 | 1 | 0 | 0 | 0 | 0.038 | 0.172 | 0 | 0 | 0 | 0 | NA | 0.042 |
| SE | 88033 | ENSG00000156171 | DRAM2 | chr1 | - | 111120515 | 111120693 | 111119876 | 111119959 | 111124741 | 111124881 | 498 | 326 | 443 | 577 | 907 | 1 | 2 | 2 | 4 | 5 | 353 | 349 | 307 | 246 | 149 | 0 | 0 | 0 | 4 | 0 | 298 | 149 | 1 | 1 | 0.996 | 0.988 | 0.991 | 0.986 | 0.989 | 1 | 1 | 1 | 0.969 | 1 | -0.004 |
| SE | 88038 | ENSG00000156171 | DRAM2 | chr1 | - | 111124741 | 111124881 | 111119876 | 111119959 | 111131423 | 111131568 | 1 | 2 | 2 | 6 | 5 | 0 | 0 | 1 | 0 | 0 | 0 | 0 | 2 | 4 | 0 | 0 | 7 | 7 | 0 | 0 | 289 | 149 | 1 | 1 | 1 | 1 | 0.508 | 1 | 1 | NA | 0 | 0.128 | 1 | NA | 0.526 |
| SE | 88042 | ENSG00000156171 | DRAM2 | chr1 | - | 111126226 | 111126294 | 111124741 | 111124881 | 111131423 | 111131568 | 57 | 40 | 37 | 46 | 59 | 0 | 0 | 0 | 2 | 0 | 46 | 40 | 20 | 17 | 16 | 0 | 0 | 2 | 0 | 0 | 217 | 149 | 1 | 1 | 1 | 1 | 1 | 0.94 | 1 | 1 | 1 | 0.873 | 1 | 1 | 0.013 |
| SE | 88044 | ENSG00000156171 | DRAM2 | chr1 | - | 111139500 | 111139666 | 111137522 | 111137586 | 111140037 | 111140090 | 90 | 76 | 69 | 1 | 61 | 5 | 0 | 6 | 14 | 4 | 71 | 0 | 25 | 89 | 14 | 0 | 0 | 19 | 0 | 0 | 298 | 149 | 1 | 1 | 0.9 | 1 | 0.852 | 0.034 | 0.884 | 1 | NA | 0.397 | 1 | 1 | -0.115 |
| SE | 88045 | ENSG00000156171 | DRAM2 | chr1 | - | 111139587 | 111139711 | 111137522 | 111137586 | 111140037 | 111140062 | 5 | 4 | 12 | 22 | 2 | 5 | 0 | 6 | 14 | 4 | 0 | 11 | 0 | 0 | 3 | 0 | 0 | 19 | 0 | 0 | 273 | 149 | 1 | 1 | 0.353 | 1 | 0.522 | 0.462 | 0.214 | NA | 1 | 0 | NA | 1 | -0.156 |
| SE | 88049 | ENSG00000239213 | NCK1-DT | chr3 | - | 136846527 | 136846586 | 136841024 | 136842569 | 136861746 | 136862031 | 12 | 6 | 9 | 14 | 13 | 0 | 0 | 0 | 1 | 0 | 7 | 13 | 8 | 8 | 38 | 0 | 0 | 0 | 0 | 0 | 208 | 149 | 1 | 1 | 1 | 1 | 1 | 0.909 | 1 | 1 | 1 | 1 | 1 | 1 | -0.018 |
| SE | 88050 | ENSG00000239213 | NCK1-DT | chr3 | - | 136846527 | 136846586 | 136841662 | 136842293 | 136861746 | 136862031 | 4 | 1 | 3 | 6 | 7 | 0 | 0 | 0 | 1 | 0 | 4 | 4 | 5 | 4 | 0 | 0 | 0 | 0 | 0 | 0 | 208 | 149 | 1 | 1 | 1 | 1 | 1 | 0.811 | 1 | 1 | 1 | 1 | 1 | NA | -0.038 |
| SE | 88054 | ENSG00000180263 | FGD6 | chr12 | - | 95084497 | 95084646 | 95076748 | 95081556 | 95085779 | 95085908 | 41 | 34 | 13 | 26 | 27 | 0 | 0 | 0 | 0 | 3 | 7 | 28 | 30 | 10 | 1 | 0 | 0 | 0 | 0 | 0 | 298 | 149 | 1 | 1 | 1 | 1 | 1 | 1 | 0.818 | 1 | 1 | 1 | 1 | 1 | -0.036 |
| SE | 88055 | ENSG00000180263 | FGD6 | chr12 | - | 95105006 | 95105086 | 95094591 | 95094694 | 95106953 | 95107037 | 37 | 19 | 16 | 24 | 27 | 0 | 0 | 0 | 0 | 0 | 21 | 13 | 38 | 30 | 0 | 0 | 0 | 0 | 7 | 0 | 229 | 149 | 1 | 1 | 1 | 1 | 1 | 1 | 1 | 1 | 1 | 1 | 0.736 | NA | 0.066 |
| SE | 88056 | ENSG00000180263 | FGD6 | chr12 | - | 95107562 | 95107631 | 95106953 | 95107037 | 95108347 | 95108419 | 11 | 13 | 7 | 20 | 17 | 0 | 0 | 2 | 0 | 0 | 7 | 17 | 40 | 12 | 0 | 0 | 0 | 0 | 0 | 0 | 218 | 149 | 1 | 1 | 1 | 1 | 0.705 | 1 | 1 | 1 | 1 | 1 | 1 | NA | -0.059 |
| SE | 88057 | ENSG00000180263 | FGD6 | chr12 | - | 95134738 | 95134826 | 95113650 | 95113701 | 95137521 | 95137678 | 19 | 28 | 22 | 21 | 32 | 1 | 0 | 0 | 0 | 1 | 15 | 18 | 28 | 12 | 30 | 0 | 0 | 0 | 0 | 0 | 237 | 149 | 1 | 1 | 0.923 | 1 | 1 | 1 | 0.953 | 1 | 1 | 1 | 1 | 1 | -0.025 |
| SE | 88058 | ENSG00000180263 | FGD6 | chr12 | - | 95141387 | 95141539 | 95137521 | 95137678 | 95152810 | 95152841 | 50 | 49 | 26 | 54 | 64 | 0 | 0 | 0 | 0 | 0 | 21 | 46 | 53 | 49 | 8 | 0 | 0 | 0 | 4 | 0 | 298 | 149 | 1 | 1 | 1 | 1 | 1 | 1 | 1 | 1 | 1 | 1 | 0.86 | 1 | 0.028 |
| SE | 88059 | ENSG00000180263 | FGD6 | chr12 | - | 95208842 | 95211267 | 95172599 | 95172744 | 95217224 | 95217379 | 22 | 22 | 11 | 19 | 20 | 0 | 1 | 1 | 1 | 0 | 19 | 39 | 32 | 16 | 0 | 0 | 2 | 0 | 2 | 0 | 298 | 149 | 1 | 1 | 1 | 0.917 | 0.846 | 0.905 | 1 | 1 | 0.907 | 1 | 0.8 | NA | 0.007 |
| SE | 88066 | ENSG00000126432 | PRDX5 | chr11 | + | 64319733 | 64319868 | 64318120 | 64318386 | 64320660 | 64320792 | 292 | 119 | 188 | 205 | 89 | 0 | 5 | 2 | 0 | 0 | 304 | 296 | 176 | 476 | 572 | 0 | 0 | 1 | 0 | 0 | 284 | 149 | 1 | 1 | 1 | 0.926 | 0.98 | 1 | 1 | 1 | 1 | 0.989 | 1 | 1 | -0.017 |
| SE | 88068 | ENSG00000126432 | PRDX5 | chr11 | + | 64320660 | 64320792 | 64318120 | 64318386 | 64320864 | 64320903 | 83 | 62 | 73 | 68 | 72 | 1 | 0 | 2 | 2 | 0 | 134 | 101 | 66 | 174 | 168 | 12 | 6 | 0 | 0 | 0 | 281 | 149 | 1 | 1 | 0.978 | 1 | 0.951 | 0.947 | 1 | 0.856 | 0.899 | 1 | 1 | 1 | 0.024 |
| SE | 88071 | ENSG00000198440 | ZNF583 | chr19 | + | 56413958 | 56414085 | 56407025 | 56407123 | 56422890 | 56423047 | 17 | 19 | 12 | 17 | 31 | 0 | 0 | 1 | 0 | 0 | 9 | 13 | 12 | 5 | 3 | 0 | 0 | 0 | 0 | 0 | 276 | 149 | 1 | 1 | 1 | 1 | 0.866 | 1 | 1 | 1 | 1 | 1 | 1 | 1 | -0.027 |
| SE | 88072 | ENSG00000198440 | ZNF583 | chr19 | + | 56414344 | 56414440 | 56413958 | 56414085 | 56422890 | 56423047 | 39 | 20 | 24 | 22 | 56 | 2 | 0 | 0 | 0 | 9 | 9 | 12 | 19 | 7 | 0 | 0 | 0 | 0 | 0 | 0 | 245 | 149 | 1 | 1 | 0.922 | 1 | 1 | 1 | 0.791 | 1 | 1 | 1 | 1 | NA | -0.057 |
| SE | 88075 | ENSG00000196663 | TECPR2 | chr14 | + | 102376649 | 102376940 | 102362940 | 102363116 | 102407337 | 102407466 | 32 | 21 | 25 | 25 | 37 | 0 | 0 | 0 | 0 | 0 | 26 | 46 | 52 | 38 | 48 | 3 | 0 | 0 | 0 | 0 | 298 | 149 | 1 | 1 | 1 | 1 | 1 | 1 | 1 | 0.812 | 1 | 1 | 1 | 1 | 0.038 |
| SE | 88076 | ENSG00000196663 | TECPR2 | chr14 | + | 102424978 | 102425291 | 102414635 | 102414793 | 102428249 | 102428382 | 90 | 49 | 73 | 80 | 102 | 0 | 0 | 0 | 0 | 0 | 110 | 84 | 96 | 81 | 64 | 0 | 0 | 1 | 0 | 0 | 298 | 149 | 1 | 1 | 1 | 1 | 1 | 1 | 1 | 1 | 1 | 0.98 | 1 | 1 | 0.004 |
| SE | 88077 | ENSG00000196663 | TECPR2 | chr14 | + | 102428249 | 102428382 | 102414635 | 102414793 | 102431795 | 102432128 | 11 | 10 | 18 | 24 | 49 | 0 | 0 | 0 | 0 | 0 | 29 | 31 | 35 | 43 | 62 | 0 | 0 | 1 | 0 | 0 | 282 | 149 | 1 | 1 | 1 | 1 | 1 | 1 | 1 | 1 | 1 | 0.949 | 1 | 1 | 0.01 |
| SE | 88078 | ENSG00000196663 | TECPR2 | chr14 | + | 102428249 | 102428382 | 102414635 | 102414793 | 102434234 | 102435211 | 2 | 2 | 5 | 6 | 9 | 0 | 0 | 0 | 0 | 0 | 0 | 0 | 7 | 1 | 0 | 0 | 0 | 0 | 0 | 7 | 282 | 149 | 1 | 1 | 1 | 1 | 1 | 1 | 1 | NA | NA | 1 | 1 | 0 | 0.333 |

| | | | | | | | |
|---|---|---|---|---|---|---|---|
| SE | 88082 | ENSG00000142046 | TMEM91 | chr19 | + | 41378280 | 41378519 |
| SE | 88085 | ENSG00000142046 | TMEM91 | chr19 | + | 41383560 | 41383632 |
| SE | 88086 | ENSG00000142046 | TMEM91 | chr19 | + | 41383560 | 41383632 |
| SE | 88087 | ENSG00000241852 | C8orf58 | chr8 | + | 22602536 | 22602643 |
| SE | 88088 | ENSG00000241852 | C8orf58 | chr8 | + | 22602560 | 22602643 |
| SE | 88090 | ENSG00000139197 | PEX5 | chr12 | + | 7190021 | 7190216 |
| SE | 88098 | ENSG00000139197 | PEX5 | chr12 | + | 7202611 | 7202704 |
| SE | 88099 | ENSG00000139197 | PEX5 | chr12 | + | 7202611 | 7202704 |
| SE | 88104 | ENSG00000197265 | GTF2E2 | chr8 | - | 30607056 | 30607150 |
| SE | 88105 | ENSG00000197265 | GTF2E2 | chr8 | - | 30612298 | 30612481 |
| SE | 88106 | ENSG00000197265 | GTF2E2 | chr8 | - | 30635031 | 30635123 |
| SE | 88107 | ENSG00000197265 | GTF2E2 | chr8 | - | 30636929 | 30636960 |
| SE | 88109 | ENSG00000197265 | GTF2E2 | chr8 | - | 30637244 | 30637274 |
| SE | 88112 | ENSG00000104228 | TRIM35 | chr8 | - | 27298463 | 27298559 |
| SE | 88116 | ENSG00000100522 | GNPNAT1 | chr14 | - | 52787832 | 52787871 |
| SE | 88117 | ENSG00000158483 | FAM86C1P | chr11 | + | 71791735 | 71791819 |
| SE | 88118 | ENSG00000158483 | FAM86C1P | chr11 | + | 71791735 | 71791819 |
| SE | 88119 | ENSG00000158483 | FAM86C1P | chr11 | + | 71791738 | 71791819 |
| SE | 88122 | ENSG00000158483 | FAM86C1P | chr11 | + | 71793379 | 71793481 |
| SE | 88123 | ENSG00000158483 | FAM86C1P | chr11 | + | 71796016 | 71796166 |
| SE | 88124 | ENSG00000185495 | SEPTIN7P13 | chr1 | + | 224002521 | 224002622 |
| SE | 88125 | ENSG00000185495 | SEPTIN7P13 | chr1 | + | 224007293 | 224007417 |
| SE | 88126 | ENSG00000111364 | DDX55 | chr12 | + | 123605930 | 123606159 |
| SE | 88127 | ENSG00000111364 | DDX55 | chr12 | + | 123607431 | 123607523 |
| SE | 88134 | ENSG00000111364 | DDX55 | chr12 | + | 123618668 | 123618837 |
| SE | 88135 | ENSG00000104689 | TNFRSF10A | chr8 | - | 23199265 | 23199448 |
| SE | 88136 | ENSG00000104689 | TNFRSF10A | chr8 | - | 23200504 | 23200600 |
| SE | 88137 | ENSG00000104689 | TNFRSF10A | chr8 | - | 23200686 | 23200760 |
| SE | 88138 | ENSG00000104689 | TNFRSF10A | chr8 | - | 23212115 | 23212212 |
| SE | 88139 | ENSG00000198042 | MAK16 | chr8 | + | 33488987 | 33489139 |
| SE | 88140 | ENSG00000198042 | MAK16 | chr8 | + | 33490284 | 33490339 |
| SE | 88141 | ENSG00000198042 | MAK16 | chr8 | + | 33497231 | 33497297 |
| SE | 88142 | ENSG00000036565 | SLC18A1 | chr8 | - | 20147257 | 20147391 |
| SE | 88143 | ENSG00000036565 | SLC18A1 | chr8 | - | 20148006 | 20148070 |
| SE | 88149 | ENSG00000172728 | FUT10 | chr8 | - | 33394217 | 33394255 |
| SE | 88150 | ENSG00000172728 | FUT10 | chr8 | - | 33460711 | 33460771 |
| SE | 88155 | ENSG00000130227 | XPO7 | chr8 | + | 21969482 | 21969576 |
| SE | 88158 | ENSG00000130227 | XPO7 | chr8 | + | 21977769 | 21977843 |
| SE | 88159 | ENSG00000130227 | XPO7 | chr8 | + | 21987140 | 21987276 |
| SE | 88161 | ENSG00000147475 | ERLIN2 | chr8 | + | 37737907 | 37738029 |
| SE | 88162 | ENSG00000147475 | ERLIN2 | chr8 | + | 37740364 | 37740446 |
| SE | 88164 | ENSG00000147475 | ERLIN2 | chr8 | + | 37744354 | 37744416 |
| SE | 88165 | ENSG00000092964 | DPYSL2 | chr8 | + | 26627214 | 26627295 |
| SE | 88166 | ENSG00000092964 | DPYSL2 | chr8 | + | 26643438 | 26643595 |
| SE | 88169 | ENSG00000227678 | NA | chr6 | - | 130142683 | 130142813 |
| SE | 88170 | ENSG00000227678 | NA | chr6 | - | 130142683 | 130142813 |
| SE | 88171 | ENSG00000227678 | NA | chr6 | - | 130143009 | 130143116 |
| SE | 88173 | ENSG00000227678 | NA | chr6 | - | 130143749 | 130144051 |
| SE | 88174 | ENSG00000227678 | NA | chr6 | - | 130144433 | 130144804 |
| SE | 88187 | ENSG00000136279 | DBNL | chr7 | + | 44050224 | 44050280 |
| SE | 88189 | ENSG00000136279 | DBNL | chr7 | + | 44051829 | 44051942 |
| SE | 88191 | ENSG00000136279 | DBNL | chr7 | + | 44051829 | 44051942 |
| SE | 88199 | ENSG00000136279 | DBNL | chr7 | + | 44059353 | 44059449 |
| SE | 88201 | ENSG00000262580 | NA | chr17 | - | 80204498 | 80204596 |
| SE | 88202 | ENSG00000165949 | IFI27 | chr14 | + | 94110925 | 94110978 |
| SE | 88203 | ENSG00000165949 | IFI27 | chr14 | + | 94111624 | 94111773 |
| SE | 88204 | ENSG00000165949 | IFI27 | chr14 | + | 94111624 | 94111773 |
| SE | 88205 | ENSG00000165949 | IFI27 | chr14 | + | 94111658 | 94111773 |
| SE | 88206 | ENSG00000165949 | IFI27 | chr14 | + | 94111658 | 94111773 |
| SE | 88207 | ENSG00000165949 | IFI27 | chr14 | + | 94114850 | 94114880 |
| SE | 88208 | ENSG00000165949 | IFI27 | chr14 | + | 94114850 | 94114880 |
| SE | 88211 | ENSG00000104626 | ERI1 | chr8 | + | 9009031 | 9009138 |
| SE | 88212 | ENSG00000104626 | ERI1 | chr8 | + | 9018296 | 9018406 |
| SE | 88213 | ENSG00000104626 | ERI1 | chr8 | + | 9029791 | 9030081 |
| SE | 88215 | ENSG00000104626 | ERI1 | chr8 | + | 9088459 | 9088569 |
| SE | 88216 | ENSG00000104626 | ERI1 | chr8 | + | 9096742 | 9096860 |
| SE | 88223 | ENSG00000134056 | MRPS36 | chr5 | + | 69226337 | 69226401 |
| SE | 88224 | ENSG00000134056 | MRPS36 | chr5 | + | 69226337 | 69226401 |
| SE | 88225 | ENSG00000134056 | MRPS36 | chr5 | + | 69226341 | 69226401 |
| SE | 88227 | ENSG00000134056 | MRPS36 | chr5 | + | 69228199 | 69228387 |
| SE | 88228 | ENSG00000134056 | MRPS36 | chr5 | + | 69228199 | 69228387 |
| SE | 88232 | ENSG00000163428 | LRRC58 | chr3 | - | 120335824 | 120335953 |
| SE | 88233 | ENSG00000104660 | LEPROTL1 | chr8 | + | 30101897 | 30101973 |
| SE | 88234 | ENSG00000104660 | LEPROTL1 | chr8 | + | 30104299 | 30104486 |
| SE | 88236 | ENSG00000104660 | LEPROTL1 | chr8 | + | 30104733 | 30104991 |
| SE | 88237 | ENSG00000104660 | LEPROTL1 | chr8 | + | 30132374 | 30132489 |
| SE | 88241 | ENSG00000147416 | ATP6V1B2 | chr8 | + | 20203958 | 20204083 |
| SE | 88243 | ENSG00000147416 | ATP6V1B2 | chr8 | + | 20210568 | 20210646 |
| SE | 88245 | ENSG00000147416 | ATP6V1B2 | chr8 | + | 20216412 | 20216495 |
| SE | 88246 | ENSG00000254362 | NA | chr8 | + | 26441171 | 26441251 |
| SE | 88249 | ENSG00000068383 | INPP5A | chr10 | + | 132607914 | 132607956 |
| SE | 88250 | ENSG00000068383 | INPP5A | chr10 | + | 132645867 | 132645968 |
| SE | 88251 | ENSG00000068383 | INPP5A | chr10 | + | 132650417 | 132650505 |
| SE | 88252 | ENSG00000068383 | INPP5A | chr10 | + | 132690391 | 132690455 |
| SE | 88253 | ENSG00000068383 | INPP5A | chr10 | + | 132697815 | 132697919 |
| SE | 88254 | ENSG00000068383 | INPP5A | chr10 | + | 132726820 | 132726905 |
| SE | 88255 | ENSG00000068383 | INPP5A | chr10 | + | 132726820 | 132726905 |
| SE | 88256 | ENSG00000068383 | INPP5A | chr10 | + | 132726820 | 132726905 |
| SE | 88257 | ENSG00000068383 | INPP5A | chr10 | + | 132765772 | 132765846 |
| SE | 88258 | ENSG00000068383 | INPP5A | chr10 | + | 132765772 | 132765846 |
| SE | 88261 | ENSG00000104756 | KCTD9 | chr8 | - | 25432503 | 25432637 |
| SE | 88263 | ENSG00000104756 | KCTD9 | chr8 | - | 25446079 | 25446250 |
| SE | 88264 | ENSG00000104756 | KCTD9 | chr8 | - | 25446128 | 25446250 |
| SE | 88267 | ENSG00000275023 | MLLT6 | chr17 | + | 38706949 | 38707029 |
| SE | 88269 | ENSG00000275023 | MLLT6 | chr17 | + | 38716366 | 38716981 |
| SE | 88274 | ENSG00000147471 | PLPBP | chr8 | + | 37775338 | 37775481 |
| SE | 88289 | ENSG00000197467 | COL13A1 | chr10 | + | 69925803 | 69925872 |
| SE | 88290 | ENSG00000197467 | COL13A1 | chr10 | + | 69930399 | 69930552 |
| SE | 88305 | ENSG00000134013 | LOXL2 | chr8 | - | 23298835 | 23298947 |
| SE | 88306 | ENSG00000134013 | LOXL2 | chr8 | - | 23328381 | 23328565 |
| SE | 88307 | ENSG00000134013 | LOXL2 | chr8 | - | 23328381 | 23328565 |
| SE | 88308 | ENSG00000134013 | LOXL2 | chr8 | - | 23328381 | 23328586 |
| SE | 88309 | ENSG00000134013 | LOXL2 | chr8 | - | 23333400 | 23333623 |
| SE | 88310 | ENSG00000134013 | LOXL2 | chr8 | - | 23333400 | 23333623 |
| SE | 88311 | ENSG00000134013 | LOXL2 | chr8 | - | 23370642 | 23370762 |
| SE | 88324 | ENSG00000145868 | FBXO38 | chr5 | + | 148401981 | 148402145 |
| SE | 88325 | ENSG00000145868 | FBXO38 | chr5 | + | 148402347 | 148402513 |
| SE | 88326 | ENSG00000145868 | FBXO38 | chr5 | + | 148427212 | 148427722 |
| SE | 88327 | ENSG00000145868 | FBXO38 | chr5 | + | 148427212 | 148427947 |
| SE | 88329 | ENSG00000145868 | FBXO38 | chr5 | + | 148438331 | 148438498 |
| SE | 88330 | ENSG00000145868 | FBXO38 | chr5 | + | 148439646 | 148439792 |
| SE | 88331 | ENSG00000145868 | FBXO38 | chr5 | + | 148439646 | 148439792 |
| SE | 88335 | ENSG00000104219 | ZDHHC2 | chr8 | + | 17197581 | 17197651 |
| SE | 88340 | ENSG00000104219 | ZDHHC2 | chr8 | + | 17210387 | 17210480 |
| SE | 88341 | ENSG00000104219 | ZDHHC2 | chr8 | + | 17215236 | 17215349 |
| SE | 88342 | ENSG00000104219 | ZDHHC2 | chr8 | + | 17217171 | 17217246 |
| SE | 88344 | ENSG00000188938 | FAM120AOS | chr9 | - | 93447257 | 93447697 |
| SE | 88345 | ENSG00000188938 | FAM120AOS | chr9 | - | 93448368 | 93448489 |
| SE | 88347 | ENSG00000215548 | FRG1JP | chr9 | - | 63837730 | 63837832 |
| SE | 88351 | ENSG00000215548 | FRG1JP | chr9 | - | 63842823 | 63842949 |
| SE | 88354 | ENSG00000166167 | BTRC | chr10 | + | 101430344 | 101430452 |
| SE | 88355 | ENSG00000166167 | BTRC | chr10 | + | 101453500 | 101453556 |
| SE | 88356 | ENSG00000166167 | BTRC | chr10 | + | 101453500 | 101453556 |
| SE | 88360 | ENSG00000166167 | BTRC | chr10 | + | 101534660 | 101534910 |
| SE | 88361 | ENSG00000166167 | BTRC | chr10 | + | 101538292 | 101538371 |
| SE | 88363 | ENSG00000168300 | PCMTD1 | chr8 | - | 51845660 | 51845763 |
| SE | 88364 | ENSG00000168300 | PCMTD1 | chr8 | - | 51845660 | 51845763 |
| SE | 88370 | ENSG00000168300 | PCMTD1 | chr8 | - | 51870247 | 51870390 |
| SE | 88372 | ENSG00000087088 | BAX | chr19 | + | 48955547 | 48955599 |
| SE | 88373 | ENSG00000087088 | BAX | chr19 | + | 48955547 | 48955833 |
| SE | 88374 | ENSG00000087088 | BAX | chr19 | + | 48955686 | 48955833 |
| SE | 88375 | ENSG00000087088 | BAX | chr19 | + | 48955686 | 48955833 |
| SE | 88376 | ENSG00000087088 | BAX | chr19 | + | 48960238 | 48960336 |
| SE | 88377 | ENSG00000096968 | JAK2 | chr9 | + | 4985939 | 4986022 |
| SE | 88378 | ENSG00000096968 | JAK2 | chr9 | + | 5029782 | 5029906 |
| SE | 88379 | ENSG00000096968 | JAK2 | chr9 | + | 5050685 | 5050831 |
| SE | 88380 | ENSG00000096968 | JAK2 | chr9 | + | 5066677 | 5066789 |
| SE | 88381 | ENSG00000096968 | JAK2 | chr9 | + | 5073697 | 5073785 |
| SE | 88382 | ENSG00000096968 | JAK2 | chr9 | + | 5090738 | 5090911 |
| SE | 88383 | ENSG00000096968 | JAK2 | chr9 | + | 5126332 | 5126446 |
| SE | 88384 | ENSG00000053108 | FSTL4 | chr5 | - | 133224189 | 133224216 |
| SE | 88385 | ENSG00000121316 | PLBD1 | chr12 | - | 14540763 | 14540902 |
| SE | 88386 | ENSG00000121316 | PLBD1 | chr12 | - | 14542207 | 14542291 |
| SE | 88387 | ENSG00000121316 | PLBD1 | chr12 | - | 14553192 | 14553412 |
| SE | 88390 | ENSG00000145041 | DCAF1 | chr3 | - | 51403142 | 51403395 |
| SE | 88391 | ENSG00000145041 | DCAF1 | chr3 | - | 51412378 | 51412480 |
| SE | 88392 | ENSG00000145041 | DCAF1 | chr3 | - | 51427371 | 51427541 |
| SE | 88393 | ENSG00000145041 | DCAF1 | chr3 | - | 51429260 | 51429470 |
| SE | 88396 | ENSG00000145041 | DCAF1 | chr3 | - | 51441384 | 51441897 |
| SE | 88397 | ENSG00000145041 | DCAF1 | chr3 | - | 51443765 | 51443903 |
| SE | 88398 | ENSG00000145041 | DCAF1 | chr3 | - | 51443765 | 51443903 |
| SE | 88401 | ENSG00000145041 | DCAF1 | chr3 | - | 51470928 | 51471005 |
| SE | 88402 | ENSG00000145041 | DCAF1 | chr3 | - | 51483718 | 51483836 |
| SE | 88403 | ENSG00000145041 | DCAF1 | chr3 | - | 51483718 | 51483836 |
| SE | 88404 | ENSG00000145041 | DCAF1 | chr3 | - | 51496733 | 51496780 |
| SE | 88407 | ENSG00000168172 | HOOK3 | chr8 | + | 42906172 | 42906258 |
| SE | 88408 | ENSG00000168172 | HOOK3 | chr8 | + | 42906172 | 42906258 |
| SE | 88410 | ENSG00000168172 | HOOK3 | chr8 | + | 42925556 | 42925629 |
| SE | 88411 | ENSG00000168172 | HOOK3 | chr8 | + | 42930121 | 42930172 |
| SE | 88412 | ENSG00000168172 | HOOK3 | chr8 | + | 42950387 | 42950455 |
| SE | 88413 | ENSG00000168172 | HOOK3 | chr8 | + | 42966472 | 42966613 |
| SE | 88417 | ENSG00000168172 | HOOK3 | chr8 | + | 42986654 | 42986795 |
| SE | 88418 | ENSG00000168172 | HOOK3 | chr8 | + | 42986654 | 42986795 |
| SE | 88419 | ENSG00000168172 | HOOK3 | chr8 | + | 42994433 | 42994492 |
| SE | 88420 | ENSG00000168172 | HOOK3 | chr8 | + | 42997549 | 42997637 |
| SE | 88425 | ENSG00000168172 | HOOK3 | chr8 | + | 43002106 | 43002141 |
| SE | 88426 | ENSG00000168172 | HOOK3 | chr8 | + | 43002106 | 43002141 |
| SE | 88427 | ENSG00000168172 | HOOK3 | chr8 | + | 43007846 | 43007929 |
| SE | 88428 | ENSG00000168172 | HOOK3 | chr8 | + | 43007846 | 43007929 |
| SE | 88429 | ENSG00000221874 | ZNF816-ZNF321P | chr19 | - | 52952750 | 52952877 |
| SE | 88433 | ENSG00000179715 | PCED1B | chr12 | + | 47104112 | 47104195 |
| SE | 88434 | ENSG00000179715 | PCED1B | chr12 | + | 47216221 | 47216689 |
| SE | 88438 | ENSG00000221914 | PPP2R2A | chr8 | + | 26293665 | 26293740 |
| SE | 88439 | ENSG00000221914 | PPP2R2A | chr8 | + | 26293665 | 26293860 |
| SE | 88441 | ENSG00000221914 | PPP2R2A | chr8 | + | 26333485 | 26333570 |
| SE | 88442 | ENSG00000221914 | PPP2R2A | chr8 | + | 26338889 | 26338987 |
| SE | 88443 | ENSG00000221914 | PPP2R2A | chr8 | + | 26338889 | 26338987 |
| SE | 88444 | ENSG00000221914 | PPP2R2A | chr8 | + | 26339902 | 26339973 |
| SE | 88448 | ENSG00000221914 | PPP2R2A | chr8 | + | 26354467 | 26354633 |
| SE | 88449 | ENSG00000221914 | PPP2R2A | chr8 | + | 26358906 | 26359032 |
| SE | 88450 | ENSG00000221914 | PPP2R2A | chr8 | + | 26358911 | 26359032 |
| SE | 88451 | ENSG00000221914 | PPP2R2A | chr8 | + | 26360168 | 26360281 |
| SE | 88455 | ENSG00000110025 | SNX15 | chr11 | + | 65032167 | 65032203 |
| SE | 88456 | ENSG00000110025 | SNX15 | chr11 | + | 65038571 | 65038829 |
| SE | 88458 | ENSG00000152601 | MBNL1 | chr3 | + | 152299404 | 152300367 |
| SE | 88463 | ENSG00000152601 | MBNL1 | chr3 | + | 152335110 | 152335230 |
| SE | 88466 | ENSG00000152601 | MBNL1 | chr3 | + | 152335110 | 152335230 |
| SE | 88468 | ENSG00000152601 | MBNL1 | chr3 | + | 152396166 | 152396224 |
| SE | 88469 | ENSG00000152601 | MBNL1 | chr3 | + | 152396166 | 152396224 |
| SE | 88471 | ENSG00000152601 | MBNL1 | chr3 | + | 152414940 | 152415111 |
| SE | 88472 | ENSG00000152601 | MBNL1 | chr3 | + | 152414940 | 152415111 |
| SE | 88474 | ENSG00000152601 | MBNL1 | chr3 | + | 152414940 | 152415111 |
| SE | 88475 | ENSG00000152601 | MBNL1 | chr3 | + | 152414940 | 152415111 |
| SE | 88477 | ENSG00000152601 | MBNL1 | chr3 | + | 152432716 | 152432920 |
| SE | 88479 | ENSG00000152601 | MBNL1 | chr3 | + | 152445281 | 152445539 |
| SE | 88480 | ENSG00000152601 | MBNL1 | chr3 | + | 152445281 | 152445539 |
| SE | 88481 | ENSG00000152601 | MBNL1 | chr3 | + | 152445281 | 152445539 |
| SE | 88484 | ENSG00000152601 | MBNL1 | chr3 | + | 152447619 | 152447773 |
| SE | 88485 | ENSG00000152601 | MBNL1 | chr3 | + | 152447619 | 152447773 |
| SE | 88486 | ENSG00000152601 | MBNL1 | chr3 | + | 152447619 | 152447773 |
| SE | 88488 | ENSG00000152601 | MBNL1 | chr3 | + | 152455541 | 152455577 |
| SE | 88489 | ENSG00000152601 | MBNL1 | chr3 | + | 152455541 | 152455577 |
| SE | 88493 | ENSG00000152601 | MBNL1 | chr3 | + | 152456266 | 152456361 |
| SE | 88496 | ENSG00000152601 | MBNL1 | chr3 | + | 152458129 | 152458193 |
| SE | 88498 | ENSG00000256150 | ITFG2-AS1 | chr12 | - | 2787341 | 2787487 |
| SE | 88499 | ENSG00000197217 | ENTPD4 | chr8 | - | 23403953 | 23404073 |
| SE | 88500 | ENSG00000197217 | ENTPD4 | chr8 | - | 23435391 | 23435477 |
| SE | 88501 | ENSG00000197217 | ENTPD4 | chr8 | - | 23435391 | 23435477 |
| SE | 88505 | ENSG00000197217 | ENTPD4 | chr8 | - | 23449892 | 23449997 |

| | | | | | | | | | | | | | | | |
|---|---|---|---|---|---|---|---|---|---|---|---|---|---|---|---|
| SE | 88514 | ENSG00000188385 | JAKMIP3 | chr10 | + | 132166982 | 132167055 | 132164669 | 132164735 | 132167952 | 132169033 | 0 | 0 | 0 | 0 |
| SE | 88516 | ENSG00000188385 | JAKMIP3 | chr10 | + | 132167013 | 132167055 | 132164669 | 132164735 | 132167952 | 132168271 | 0 | 0 | 0 | 0 |
| SE | 88522 | ENSG00000138378 | STAT4 | chr2 | - | 191030971 | 191031080 | 191029575 | 191029866 | 191031449 | 191031516 | 455 | 248 | 308 | 349 |
| SE | 88524 | ENSG00000138378 | STAT4 | chr2 | - | 191039198 | 191039297 | 191036163 | 191036299 | 191041064 | 191041148 | 602 | 396 | 504 | 514 |
| SE | 88526 | ENSG00000138378 | STAT4 | chr2 | - | 191058017 | 191058111 | 191054489 | 191054534 | 191058201 | 191058219 | 574 | 336 | 468 | 480 |
| SE | 88527 | ENSG00000138378 | STAT4 | chr2 | - | 191058201 | 191058219 | 191058017 | 191058111 | 191058709 | 191058769 | 247 | 127 | 177 | 180 |
| SE | 88529 | ENSG00000138378 | STAT4 | chr2 | - | 191066429 | 191066515 | 191064806 | 191064958 | 191068059 | 191068594 | 207 | 126 | 185 | 185 |
| SE | 88530 | ENSG00000138378 | STAT4 | chr2 | - | 191066429 | 191066515 | 191064806 | 191064958 | 191069692 | 191069771 | 501 | 330 | 419 | 452 |
| SE | 88531 | ENSG00000138378 | STAT4 | chr2 | - | 191068059 | 191068594 | 191064806 | 191064958 | 191069692 | 191069771 | 1 | 1 | 5 | 5 |
| SE | 88533 | ENSG00000138378 | STAT4 | chr2 | - | 191073097 | 191073190 | 191069692 | 191069771 | 191076226 | 191076325 | 348 | 227 | 317 | 306 |
| SE | 88534 | ENSG00000138378 | STAT4 | chr2 | - | 191076226 | 191076325 | 191073097 | 191073190 | 191146612 | 191146757 | 359 | 227 | 307 | 267 |
| SE | 88587 | ENSG00000120915 | EPHX2 | chr8 | + | 27500925 | 27501010 | 27491124 | 27491309 | 27503603 | 27503759 | 28 | 18 | 35 | 19 |
| SE | 88588 | ENSG00000120915 | EPHX2 | chr8 | + | 27500927 | 27501010 | 27491194 | 27491309 | 27503603 | 27503763 | 19 | 7 | 27 | 15 |
| SE | 88589 | ENSG00000120915 | EPHX2 | chr8 | + | 27500963 | 27501010 | 27491165 | 27491309 | 27503603 | 27503763 | 19 | 7 | 24 | 15 |
| SE | 88590 | ENSG00000120915 | EPHX2 | chr8 | + | 27506871 | 27506994 | 27504955 | 27505146 | 27511835 | 27511910 | 54 | 29 | 64 | 57 |
| SE | 88592 | ENSG00000120915 | EPHX2 | chr8 | + | 27516319 | 27516398 | 27511835 | 27511910 | 27518037 | 27518072 | 11 | 10 | 13 | 15 |
| SE | 88593 | ENSG00000120915 | EPHX2 | chr8 | + | 27522422 | 27522508 | 27520882 | 27520909 | 27538658 | 27538692 | 16 | 9 | 14 | 15 |
| SE | 88594 | ENSG00000120915 | EPHX2 | chr8 | + | 27525361 | 27525473 | 27522422 | 27522508 | 27538658 | 27538692 | 10 | 17 | 27 | 17 |
| SE | 88595 | ENSG00000120915 | EPHX2 | chr8 | + | 27536783 | 27536855 | 27525361 | 27525473 | 27538658 | 27538692 | 19 | 24 | 37 | 38 |
| SE | 88598 | ENSG00000254979 | NA | chr11 | - | 57409174 | 57409298 | 57399495 | 57399566 | 57414614 | 57414731 | 0 | 1 | 0 | 0 |
| SE | 88599 | ENSG00000134014 | ELP3 | chr8 | + | 28097218 | 28097318 | 28093066 | 28093233 | 28099827 | 28099966 | 92 | 60 | 92 | 90 |
| SE | 88600 | ENSG00000134014 | ELP3 | chr8 | + | 28099827 | 28099966 | 28097218 | 28097318 | 28106712 | 28106783 | 262 | 133 | 231 | 249 |
| SE | 88603 | ENSG00000134014 | ELP3 | chr8 | + | 28113018 | 28113173 | 28110369 | 28110438 | 28129501 | 28129663 | 346 | 271 | 299 | 286 |
| SE | 88606 | ENSG00000134014 | ELP3 | chr8 | + | 28137697 | 28137891 | 28132277 | 28132404 | 28155941 | 28156032 | 211 | 119 | 180 | 142 |
| SE | 88607 | ENSG00000134014 | ELP3 | chr8 | + | 28155941 | 28156032 | 28137697 | 28137891 | 28158567 | 28158625 | 222 | 122 | 160 | 139 |
| SE | 88608 | ENSG00000134014 | ELP3 | chr8 | + | 28160228 | 28160456 | 28158567 | 28158633 | 28161996 | 28162078 | 280 | 204 | 210 | 255 |
| SE | 88609 | ENSG00000134014 | ELP3 | chr8 | + | 28161996 | 28162078 | 28160228 | 28160456 | 28189648 | 28189992 | 149 | 110 | 140 | 165 |
| SE | 88612 | ENSG00000169139 | UBE2V2 | chr8 | + | 48034865 | 48035061 | 48008418 | 48008470 | 48043032 | 48043181 | 2 | 0 | 4 | 4 |
| SE | 88615 | ENSG00000169139 | UBE2V2 | chr8 | + | 48049852 | 48049978 | 48041746 | 48043181 | 48060681 | 48060897 | 412 | 267 | 315 | 293 |
| SE | 88618 | ENSG00000168575 | SLC20A2 | chr8 | - | 42439449 | 42439653 | 42436988 | 42437577 | 42444645 | 42444762 | 41 | 29 | 52 | 63 |
| SE | 88619 | ENSG00000168575 | SLC20A2 | chr8 | - | 42444645 | 42444762 | 42439449 | 42439653 | 42459895 | 42459992 | 55 | 39 | 75 | 86 |
| SE | 88620 | ENSG00000168575 | SLC20A2 | chr8 | - | 42459895 | 42459992 | 42444645 | 42444762 | 42463004 | 42463090 | 53 | 43 | 59 | 80 |
| SE | 88621 | ENSG00000168575 | SLC20A2 | chr8 | - | 42463004 | 42463090 | 42444645 | 42444762 | 42472101 | 42472654 | 1 | 1 | 0 | 0 |
| SE | 88622 | ENSG00000168575 | SLC20A2 | chr8 | - | 42465776 | 42465917 | 42463004 | 42463090 | 42472101 | 42472654 | 37 | 22 | 32 | 35 |
| SE | 88634 | ENSG00000178234 | GALNT11 | chr7 | + | 152027596 | 152027718 | 152025713 | 152025884 | 152079286 | 152079408 | 0 | 0 | 1 | 0 |
| SE | 88640 | ENSG00000178234 | GALNT11 | chr7 | + | 152103111 | 152103278 | 152094189 | 152094522 | 152105244 | 152105370 | 77 | 29 | 90 | 96 |
| SE | 88641 | ENSG00000178234 | GALNT11 | chr7 | + | 152105244 | 152105370 | 152103111 | 152103278 | 152108037 | 152108287 | 102 | 39 | 117 | 145 |
| SE | 88642 | ENSG00000178234 | GALNT11 | chr7 | + | 152110527 | 152110645 | 152107249 | 152108287 | 152113245 | 152113373 | 76 | 40 | 71 | 92 |
| SE | 88644 | ENSG00000178234 | GALNT11 | chr7 | + | 152117156 | 152117375 | 152113245 | 152113398 | 152118677 | 152118769 | 209 | 112 | 150 | 245 |
| SE | 88645 | ENSG00000178234 | GALNT11 | chr7 | + | 152118677 | 152118782 | 152113245 | 152113398 | 152120830 | 152120968 | 75 | 51 | 49 | 47 |
| SE | 88646 | ENSG00000187735 | TCEA1 | chr8 | - | 53970391 | 53970463 | 53966551 | 53968112 | 53984362 | 53984517 | 60 | 23 | 45 | 52 |
| SE | 88647 | ENSG00000187735 | TCEA1 | chr8 | - | 53978522 | 53978697 | 53970391 | 53970463 | 53979024 | 53979171 | 0 | 0 | 2 | 1 |
| SE | 88648 | ENSG00000187735 | TCEA1 | chr8 | - | 53979024 | 53979171 | 53970391 | 53970463 | 53984362 | 53984517 | 893 | 632 | 729 | 769 |
| SE | 88651 | ENSG00000187735 | TCEA1 | chr8 | - | 53993667 | 53993755 | 53988113 | 53988259 | 53999944 | 54000050 | 1035 | 859 | 869 | 1074 |
| SE | 88652 | ENSG00000187735 | TCEA1 | chr8 | - | 53999944 | 54000050 | 53993375 | 53993755 | 54010429 | 54010492 | 951 | 740 | 799 | 967 |
| SE | 88653 | ENSG00000187735 | TCEA1 | chr8 | - | 53999944 | 54000050 | 53993375 | 53993755 | 54022062 | 54022448 | 323 | 270 | 303 | 386 |
| SE | 88655 | ENSG00000187735 | TCEA1 | chr8 | - | 54002891 | 54003086 | 54000009 | 54000050 | 54010429 | 54010492 | 3 | 2 | 1 | 1 |
| SE | 88656 | ENSG00000187735 | TCEA1 | chr8 | - | 54002891 | 54003092 | 53999944 | 54000050 | 54010429 | 54010492 | 3 | 2 | 1 | 1 |
| SE | 88657 | ENSG00000187735 | TCEA1 | chr8 | - | 54010429 | 54010492 | 53993375 | 53993755 | 54022062 | 54022125 | 25 | 9 | 16 | 21 |
| SE | 88659 | ENSG00000187735 | TCEA1 | chr8 | - | 54021525 | 54021586 | 54010429 | 54010492 | 54022062 | 54022310 | 1 | 3 | 0 | 1 |
| SE | 88661 | ENSG00000151090 | THRB | chr3 | - | 24237990 | 24238081 | 24228937 | 24229001 | 24297225 | 24297371 | 0 | 0 | 0 | 1 |
| SE | 88662 | ENSG00000151090 | THRB | chr3 | - | 24237990 | 24238081 | 24228937 | 24229001 | 24337299 | 24337371 | 0 | 0 | 0 | 1 |
| SE | 88666 | ENSG00000151090 | THRB | chr3 | - | 24318357 | 24318494 | 24228937 | 24229001 | 24337299 | 24337371 | 2 | 0 | 0 | 0 |
| SE | 88672 | ENSG00000126216 | TUBGCP3 | chr13 | - | 112504031 | 112504163 | 112499044 | 112499185 | 112504625 | 112504714 | 149 | 102 | 99 | 116 |
| SE | 88673 | ENSG00000126216 | TUBGCP3 | chr13 | - | 112504625 | 112504714 | 112499044 | 112499185 | 112516439 | 112516575 | 12 | 5 | 10 | 5 |
| SE | 88674 | ENSG00000126216 | TUBGCP3 | chr13 | - | 112519885 | 112520021 | 112518974 | 112519043 | 112522319 | 112522509 | 151 | 73 | 96 | 127 |
| SE | 88676 | ENSG00000126216 | TUBGCP3 | chr13 | - | 112548107 | 112548176 | 112527373 | 112527484 | 112554056 | 112554182 | 29 | 21 | 15 | 13 |
| SE | 88677 | ENSG00000126216 | TUBGCP3 | chr13 | - | 112554886 | 112555005 | 112554056 | 112554182 | 112556051 | 112556224 | 128 | 102 | 82 | 104 |
| SE | 88678 | ENSG00000126216 | TUBGCP3 | chr13 | - | 112558195 | 112558413 | 112556051 | 112556224 | 112559321 | 112559399 | 205 | 139 | 174 | 175 |
| SE | 88679 | ENSG00000126216 | TUBGCP3 | chr13 | - | 112584038 | 112584161 | 112569151 | 112569259 | 112587904 | 112588205 | 0 | 1 | 2 | 2 |
| SE | 88691 | ENSG00000183077 | AFMID | chr17 | + | 78190969 | 78191060 | 78187316 | 78187433 | 78197145 | 78197560 | 9 | 5 | 2 | 6 |
| SE | 88694 | ENSG00000183077 | AFMID | chr17 | + | 78202702 | 78202751 | 78190969 | 78191060 | 78205945 | 78206050 | 4 | 8 | 4 | 2 |
| SE | 88695 | ENSG00000183077 | AFMID | chr17 | + | 78202702 | 78202751 | 78190969 | 78191060 | 78206910 | 78207187 | 1 | 3 | 1 | 0 |
| SE | 88697 | ENSG00000183077 | AFMID | chr17 | + | 78204827 | 78204900 | 78202702 | 78202751 | 78205439 | 78205518 | 1 | 1 | 1 | 1 |
| SE | 88699 | ENSG00000183077 | AFMID | chr17 | + | 78205092 | 78205190 | 78204827 | 78204900 | 78205439 | 78205518 | 6 | 3 | 3 | 0 |
| SE | 88700 | ENSG00000183077 | AFMID | chr17 | + | 78205439 | 78205518 | 78202702 | 78202751 | 78205602 | 78205738 | 1 | 1 | 3 | 0 |
| SE | 88702 | ENSG00000183077 | AFMID | chr17 | + | 78205439 | 78205518 | 78204827 | 78204900 | 78205945 | 78206050 | 3 | 0 | 0 | 0 |
| SE | 88706 | ENSG00000183077 | AFMID | chr17 | + | 78205602 | 78205753 | 78205439 | 78205518 | 78205945 | 78206050 | 1 | 1 | 2 | 0 |
| SE | 88711 | ENSG00000104738 | MCM4 | chr8 | + | 47966186 | 47966275 | 47964591 | 47964712 | 47967364 | 47967485 | 20 | 22 | 26 | 23 |
| SE | 88712 | ENSG00000104738 | MCM4 | chr8 | + | 47966186 | 47966407 | 47964573 | 47964712 | 47967364 | 47967485 | 41 | 44 | 80 | 92 |
| SE | 88714 | ENSG00000104738 | MCM4 | chr8 | + | 47969278 | 47969401 | 47967364 | 47967485 | 47969797 | 47970057 | 0 | 0 | 0 | 0 |
| SE | 88717 | ENSG00000225921 | NOL7 | chr6 | + | 13617769 | 13617801 | 13616462 | 13616521 | 13618057 | 13618139 | 224 | 144 | 225 | 196 |
| SE | 88718 | ENSG00000225921 | NOL7 | chr6 | + | 13620753 | 13620856 | 13620407 | 13620485 | 13632392 | 13632739 | 114 | 93 | 97 | 99 |
| SE | 88721 | ENSG00000172954 | LCLAT1 | chr2 | + | 30459551 | 30459722 | 30447242 | 30447383 | 30476576 | 30476635 | 3 | 5 | 2 | 9 |
| SE | 88723 | ENSG00000172954 | LCLAT1 | chr2 | + | 30459551 | 30459722 | 30447242 | 30447383 | 30525586 | 30525755 | 8 | 7 | 2 | 13 |
| SE | 88724 | ENSG00000172954 | LCLAT1 | chr2 | + | 30459551 | 30459722 | 30447256 | 30447404 | 30525586 | 30525755 | 6 | 3 | 5 | 7 |
| SE | 88727 | ENSG00000172954 | LCLAT1 | chr2 | + | 30476401 | 30476635 | 30459551 | 30459722 | 30525586 | 30525755 | 2 | 2 | 1 | 4 |
| SE | 88728 | ENSG00000172954 | LCLAT1 | chr2 | + | 30476576 | 30476635 | 30447255 | 30447383 | 30525586 | 30525755 | 6 | 3 | 1 | 5 |
| SE | 88729 | ENSG00000172954 | LCLAT1 | chr2 | + | 30476576 | 30476635 | 30459551 | 30459722 | 30525586 | 30525755 | 3 | 3 | 3 | 7 |
| SE | 88730 | ENSG00000172954 | LCLAT1 | chr2 | + | 30489159 | 30489240 | 30447248 | 30447383 | 30525586 | 30525755 | 0 | 0 | 0 | 0 |
| SE | 88731 | ENSG00000172954 | LCLAT1 | chr2 | + | 30489159 | 30489240 | 30459551 | 30459722 | 30525586 | 30525755 | 0 | 1 | 0 | 0 |
| SE | 88734 | ENSG00000172954 | LCLAT1 | chr2 | + | 30552492 | 30552649 | 30533115 | 30533314 | 30562145 | 30562292 | 21 | 38 | 20 | 29 |
| SE | 88735 | ENSG00000172954 | LCLAT1 | chr2 | + | 30562145 | 30562292 | 30533115 | 30533314 | 30568059 | 30568176 | 96 | 75 | 71 | 85 |
| SE | 88736 | ENSG00000258659 | TRIM34 | chr11 | + | 5632254 | 5632754 | 5625000 | 5625060 | 5633803 | 5633899 | 118 | 95 | 102 | 142 |
| SE | 88737 | ENSG00000258659 | TRIM34 | chr11 | + | 5633803 | 5633899 | 5625000 | 5625060 | 5634630 | 5634861 | 70 | 55 | 71 | 64 |
| SE | 88738 | ENSG00000258659 | TRIM34 | chr11 | + | 5642816 | 5642843 | 5642405 | 5642506 | 5643143 | 5644398 | 58 | 21 | 26 | 27 |
| SE | 88739 | ENSG00000258659 | TRIM34 | chr11 | + | 5642816 | 5642853 | 5642405 | 5642506 | 5643143 | 5643809 | 11 | 1 | 5 | 6 |
| SE | 88746 | ENSG00000205581 | HMGN1 | chr21 | - | 39345829 | 39345958 | 39345145 | 39345274 | 39347915 | 39347952 | 236 | 304 | 338 | 322 |
| SE | 88749 | ENSG00000205581 | HMGN1 | chr21 | - | 39347113 | 39347292 | 39345829 | 39345958 | 39347378 | 39347483 | 23 | 6 | 9 | 11 |
| SE | 88751 | ENSG00000205581 | HMGN1 | chr21 | - | 39347246 | 39347292 | 39345145 | 39345274 | 39348291 | 39348339 | 0 | 0 | 1 | 0 |
| SE | 88752 | ENSG00000205581 | HMGN1 | chr21 | - | 39347246 | 39347292 | 39345829 | 39345958 | 39347378 | 39347483 | 5 | 6 | 9 | 4 |
| SE | 88754 | ENSG00000205581 | HMGN1 | chr21 | - | 39347378 | 39347478 | 39345145 | 39345274 | 39347915 | 39347952 | 7 | 3 | 6 | 4 |
| SE | 88756 | ENSG00000205581 | HMGN1 | chr21 | - | 39347378 | 39347478 | 39345829 | 39345958 | 39347915 | 39347952 | 0 | 1 | 10 | 8 |
| SE | 88760 | ENSG00000205581 | HMGN1 | chr21 | - | 39347378 | 39347483 | 39345829 | 39345958 | 39347915 | 39347952 | 0 | 2 | 10 | 8 |
| SE | 88763 | ENSG00000205581 | HMGN1 | chr21 | - | 39347915 | 39347952 | 39345145 | 39345274 | 39348291 | 39348339 | 0 | 4 | 4 | 3 |
| SE | 88765 | ENSG00000205581 | HMGN1 | chr21 | - | 39347915 | 39347952 | 39347378 | 39347478 | 39348291 | 39348339 | 0 | 0 | 2 | 3 |
| SE | 88767 | ENSG00000205581 | HMGN1 | chr21 | - | 39348421 | 39348451 | 39345145 | 39348339 | 39348544 | 39348577 | 286 | 233 | 339 | 359 |
| SE | 88781 | ENSG00000140632 | GLYR1 | chr16 | - | 4811622 | 4811802 | 4811169 | 4811294 | 4812085 | 4812248 | 285 | 145 | 241 | 206 |
| SE | 88782 | ENSG00000140632 | GLYR1 | chr16 | - | 4812085 | 4812248 | 4811622 | 4811802 | 4813736 | 4813838 | 174 | 120 | 172 | 153 |
| SE | 88788 | ENSG00000140632 | GLYR1 | chr16 | - | 4822874 | 4822931 | 4817597 | 4817697 | 4823820 | 4823907 | 51 | 41 | 49 | 59 |
| SE | 88789 | ENSG00000140632 | GLYR1 | chr16 | - | 4822874 | 4822931 | 4821546 | 4821597 | 4823820 | 4823907 | 260 | 184 | 242 | 252 |
| SE | 88801 | ENSG00000165061 | ZMAT4 | chr8 | - | 40674703 | 40674931 | 40581164 | 40581261 | 40697244 | 40697401 | 2 | 0 | 1 | 8 |
| SE | 88802 | ENSG00000165061 | ZMAT4 | chr8 | - | 40786688 | 40786782 | 40767640 | 40767730 | 40808508 | 40808571 | 1 | 1 | 0 | 0 |
| SE | 88804 | ENSG00000165061 | ZMAT4 | chr8 | - | 40808508 | 40808571 | 40767640 | 40767730 | 40825574 | 40825680 | 4 | 2 | 1 | 6 |
| SE | 88806 | ENSG00000104679 | R3HCC1 | chr8 | + | 23293302 | 23293373 | 23290975 | 23291533 | 23294768 | 23294864 | 85 | 45 | 86 | 82 |
| SE | 88809 | ENSG00000174197 | MGA | chr15 | + | 41668827 | 41669958 | 41660396 | 41660525 | 41696074 | 41697023 | 111 | 81 | 95 | 116 |
| SE | 88810 | ENSG00000174197 | MGA | chr15 | + | 41708103 | 41708208 | 41707727 | 41707859 | 41710690 | 41711349 | 109 | 67 | 93 | 93 |
| SE | 88811 | ENSG00000174197 | MGA | chr15 | + | 41727179 | 41727406 | 41713150 | 41713496 | 41729163 | 41729349 | 136 | 104 | 168 | 128 |
| SE | 88813 | ENSG00000174197 | MGA | chr15 | + | 41740052 | 41740203 | 41736180 | 41736368 | 41748636 | 41748927 | 17 | 8 | 3 | 20 |
| SE | 88814 | ENSG00000174197 | MGA | chr15 | + | 41740052 | 41740203 | 41736180 | 41736698 | 41748636 | 41748927 | 45 | 24 | 29 | 47 |
| SE | 88816 | ENSG00000174197 | MGA | chr15 | + | 41748636 | 41748927 | 41736180 | 41736368 | 41754436 | 41754567 | 0 | 5 | 1 | 0 |
| SE | 88817 | ENSG00000174197 | MGA | chr15 | + | 41748636 | 41748927 | 41736180 | 41736698 | 41754436 | 41754567 | 0 | 2 | 0 | 0 |
| SE | 88818 | ENSG00000174197 | MGA | chr15 | + | 41749110 | 41750615 | 41748636 | 41748927 | 41754436 | 41754567 | 161 | 122 | 161 | 131 |
| SE | 88819 | ENSG00000174197 | MGA | chr15 | + | 41760322 | 41760529 | 41757787 | 41757839 | 41761738 | 41761769 | 207 | 167 | 156 | 153 |
| SE | 88821 | ENSG00000174197 | MGA | chr15 | + | 41762128 | 41762362 | 41761738 | 41761850 | 41764885 | 41765062 | 271 | 183 | 209 | 180 |
| SE | 88822 | ENSG00000167904 | TMEM68 | chr8 | - | 55743480 | 55743620 | 55738743 | 55740218 | 55745060 | 55745121 | 61 | 86 | 65 | 92 |
| SE | 88823 | ENSG00000167904 | TMEM68 | chr8 | - | 55743480 | 55743620 | 55738743 | 55740218 | 55750963 | 55751157 | 24 | 29 | 27 | 43 |
| SE | 88824 | ENSG00000167904 | TMEM68 | chr8 | - | 55745060 | 55745121 | 55740171 | 55740218 | 55750963 | 55751157 | 26 | 12 | 11 | 30 |
| SE | 88825 | ENSG00000167904 | TMEM68 | chr8 | - | 55745060 | 55745121 | 55743480 | 55743620 | 55750963 | 55751157 | 59 | 70 | 46 | 75 |
| SE | 88830 | ENSG00000167904 | TMEM68 | chr8 | - | 55763636 | 55763701 | 55762634 | 55763028 | 55763935 | 55763980 | 1 | 0 | 0 | 0 |
| SE | 88838 | ENSG00000104412 | EMC2 | chr8 | + | 108446161 | 108446301 | 108443600 | 108443698 | 108449822 | 108449936 | 2 | 0 | 2 | 0 |
| SE | 88839 | ENSG00000104412 | EMC2 | chr8 | + | 108450427 | 108450492 | 108449822 | 108449936 | 108453061 | 108453147 | 218 | 154 | 115 | 170 |
| SE | 88841 | ENSG00000104412 | EMC2 | chr8 | + | 108475881 | 108475963 | 108470061 | 108470121 | 108476781 | 108476892 | 179 | 110 | 121 | 148 |
| SE | 88842 | ENSG00000104412 | EMC2 | chr8 | + | 108476781 | 108476892 | 108470061 | 108470121 | 108486511 | 108486636 | 0 | 1 | 0 | 4 |
| SE | 88843 | ENSG00000104412 | EMC2 | chr8 | + | 108479005 | 108479110 | 108476781 | 108476892 | 108486511 | 108486636 | 246 | 223 | 181 | 215 |
| SE | 88852 | ENSG00000116824 | CD2 | chr1 | + | 116764483 | 116764606 | 116760401 | 116760632 | 116768463 | 116769229 | 1191 | 617 | 848 | 827 |
| SE | 88853 | ENSG00000228801 | NA | chr8 | + | 51913056 | 51913180 | 51899324 | 51899846 | 51946435 | 51949874 | 3 | 3 | 11 | 0 |
| SE | 88864 | ENSG00000104442 | ARMC1 | chr8 | - | 65622262 | 65622354 | 65613243 | 65613433 | 65627215 | 65627433 | 110 | 71 | 109 | 89 |
| SE | 88865 | ENSG00000104442 | ARMC1 | chr8 | - | 65632897 | 65632997 | 65627215 | 65627433 | 65633997 | 65634177 | 12 | 0 | 4 | 1 |
| SE | 88868 | ENSG00000183255 | PTTG1IP | chr21 | - | 44855209 | 44855256 | 44849597 | 44851627 | 44856192 | 44856364 | 343 | 228 | 213 | 267 |
| SE | 88871 | ENSG00000183255 | PTTG1IP | chr21 | - | 44856192 | 44856364 | 44849597 | 44851627 | 44873501 | 44873690 | 0 | 0 | 0 | 0 |
| SE | 88873 | ENSG00000183255 | PTTG1IP | chr21 | - | 44861162 | 44861271 | 44849584 | 44851627 | 44873501 | 44873690 | 1 | 1 | 0 | 0 |
| SE | 88875 | ENSG00000183255 | PTTG1IP | chr21 | - | 44861162 | 44861271 | 44856192 | 44856364 | 44873501 | 44873690 | 211 | 157 | 111 | 145 |
| SE | 88878 | ENSG00000183255 | PTTG1IP | chr21 | - | 44865394 | 44865447 | 44861162 | 44861271 | 44872853 | 44873032 | 105 | 105 | 86 | 112 |
| SE | 88880 | ENSG00000183255 | PTTG1IP | chr21 | - | 44865570 | 44865650 | 44865394 | 44865447 | 44872853 | 44873032 | 1 | 0 | 1 | 1 |
| SE | 88882 | ENSG00000167302 | TEPSIN | chr17 | - | 81232314 | 81232518 | 81231846 | 81232021 | 81233431 | 81233503 | 42 | 12 | 19 | 25 |
| SE | 88883 | ENSG00000167302 | TEPSIN | chr17 | - | 81237550 | 81237660 | 81237386 | 81237459 | 81238513 | 81238873 | 6 | 4 | 6 | 6 |
| SE | 88909 | ENSG00000158669 | GPAT4 | chr8 | + | 41611902 | 41611992 | 41609654 | 41609955 | 41612179 | 41612236 | 129 | 60 | 106 | 115 |
| SE | 88910 | ENSG00000158669 | GPAT4 | chr8 | + | 41611902 | 41611992 | 41609654 | 41609955 | 41614385 | 41614441 | 1 | 0 | 2 | 3 |
| SE | 88911 | ENSG00000158669 | GPAT4 | chr8 | + | 41612179 | 41612273 | 41609654 | 41609955 | 41614385 | 41614441 | 1 | 0 | 4 | 1 |
| SE | 88912 | ENSG00000158669 | GPAT4 | chr8 | + | 41612179 | 41612273 | 41611902 | 41611992 | 41612844 | 41612960 | 256 | 133 | 235 | 233 |
| SE | 88913 | ENSG00000158669 | GPAT4 | chr8 | + | 41612179 | 41612273 | 41611902 | 41611992 | 41614385 | 41614441 | 86 | 38 | 96 | 76 |
| SE | 88915 | ENSG00000158669 | GPAT4 | chr8 | + | 41612844 | 41612960 | 41611902 | 41611992 | 41614385 | 41614441 | 178 | 116 | 139 | 197 |
| SE | 88924 | ENSG00000164823 | OSGIN2 | chr8 | + | 89914076 | 89914213 | 89909566 | 89909721 | 89914554 | 89914746 | 235 | 122 | 152 | 197 |
| SE | 88928 | ENSG00000198919 | DZIP3 | chr3 | + | 108603018 | 108603234 | 108602832 | 108602907 | 108605334 | 108605438 | 0 | 0 | 2 | 0 |
| SE | 88929 | ENSG00000198919 | DZIP3 | chr3 | + | 108605334 | 108605438 | 108589704 | 108589839 | 108608088 | 108608158 | 28 | 16 | 21 | 24 |
| SE | 88930 | ENSG00000198919 | DZIP3 | chr3 | + | 108636615 | 108636708 | 108634870 | 108634972 | 108637495 | 108637548 | 84 | 94 | 106 | 108 |
| SE | 88931 | ENSG00000198919 | DZIP3 | chr3 | + | 108644163 | 108644781 | 108642437 | 108642514 | 108646616 | 108646649 | 59 | 61 | 71 | 81 |
| SE | 88932 | ENSG00000198919 | DZIP3 | chr3 | + | 108646616 | 108646649 | 108644163 | 108644781 | 108647942 | 108648112 | 43 | 48 | 66 | 57 |
| SE | 88933 | ENSG00000198919 | DZIP3 | chr3 | + | 108647942 | 108648112 | 108646616 | 108646649 | 108648917 | 108648962 | 42 | 48 | 62 | 50 |
| SE | 88934 | ENSG00000198919 | DZIP3 | chr3 | + | 108669680 | 108669749 | 108662129 | 108662257 | 108672559 | 108672656 | 54 | 83 | 49 | 52 |
| SE | 88935 | ENSG00000198919 | DZIP3 | chr3 | + | 108686444 | 108686584 | 108684215 | 108684341 | 108687975 | 108688096 | 38 | 11 | 26 | 24 |
| SE | 88936 | ENSG00000253738 | OTUD6B-AS1 | chr8 | - | 91068935 | 91069822 | 91062909 | 91063746 | 91069930 | 91070583 | 103 | 109 | 119 | 136 |
| SE | 88937 | ENSG00000253738 | OTUD6B-AS1 | chr8 | - | 91069578 | 91069822 | 91059317 | 91063746 | 91069930 | 91070149 | 119 | 113 | 133 | 165 |
| SE | 88955 | ENSG00000148516 | ZEB1 | chr10 | + | 31355143 | 31355192 | 31319221 | 31319292 | 31373017 | 31373173 | 1 | 0 | 1 | 2 |
| SE | 88956 | ENSG00000148516 | ZEB1 | chr10 | + | 31355143 | 31355192 | 31319221 | 31319292 | 31387123 | 31387266 | 0 | 4 | 0 | 0 |
| SE | 88958 | ENSG00000148516 | ZEB1 | chr10 | + | 31355146 | 31355192 | 31319211 | 31319292 | 31461036 | 31461237 | 0 | 1 | 0 | 1 |
| SE | 88959 | ENSG00000148516 | ZEB1 | chr10 | + | 31373017 | 31373173 | 31318782 | 31319133 | 31387123 | 31387266 | 41 | 32 | 40 | 33 |
| SE | 88962 | ENSG00000148516 | ZEB1 | chr10 | + | 31373017 | 31373173 | 31355143 | 31355192 | 31387123 | 31387266 | 42 | 32 | 41 | 35 |
| SE | 88966 | ENSG00000148516 | ZEB1 | chr10 | + | 31387123 | 31387266 | 31320605 | 31320698 | 31461036 | 31461237 | 31 | 37 | 33 | 32 |
| SE | 88967 | ENSG00000148516 | ZEB1 | chr10 | + | 31387123 | 31387266 | 31321473 | 31321532 | 31461036 | 31461237 | 31 | 37 | 33 | 28 |
| SE | 88970 | ENSG00000148516 | ZEB1 | chr10 | + | 31389604 | 31389660 | 31387719 | 31387798 | 31461036 | 31461236 | 0 | 0 | 0 | 1 |
| SE | 88971 | ENSG00000148516 | ZEB1 | chr10 | + | 31389604 | 31389695 | 31387719 | 31387798 | 31461036 | 31461180 | 2 | 0 | 0 | 3 |
| SE | 88974 | ENSG00000148516 | ZEB1 | chr10 | + | 31495775 | 31495838 | 31461036 | 31461237 | 31502347 | 31502423 | 28 | 37 | 41 | 37 |
| SE | 88975 | ENSG00000148516 | ZEB1 | chr10 | + | 31495778 | 31495838 | 31461036 | 31461237 | 31502347 | 31502406 | 29 | 42 | 41 | 47 |
| SE | 88976 | ENSG00000148516 | ZEB1 | chr10 | + | 31502347 | 31502509 | 31461036 | 31461237 | 31510672 | 31510875 | 23 | 13 | 30 | 30 |
| SE | 88978 | ENSG00000165806 | CASP7 | chr10 | + | 113691996 | 113692070 | 113679668 | 113679978 | 113697493 | 113697603 | 0 | 1 | 1 | 0 |
| SE | 88982 | ENSG00000214954 | LRRC69 | chr8 | + | 91124492 | 91124619 | 91102661 | 91102844 | 91127087 | 91127160 | 0 | 1 | 7 | 5 |
| SE | 88983 | ENSG00000214954 | LRRC69 | chr8 | + | 91127087 | 91127160 | 91102618 | 91102844 | 91133109 | 91133305 | 3 | 4 | 0 | 8 |
| SE | 88984 | ENSG00000214954 | LRRC69 | chr8 | + | 91130298 | 91130417 | 91127087 | 91127160 | 91133109 | 91133305 | 1 | 2 | 0 | 0 |
| SE | 88985 | ENSG00000214954 | LRRC69 | chr8 | + | 91130758 | 91130836 | 91130298 | 91130417 | 91133109 | 91133305 | 0 | 0 | 0 | 2 |
| SE | 88987 | ENSG00000214954 | LRRC69 | chr8 | + | 91135667 | 91135739 | 91133109 | 91133305 | 91189521 | 91189623 | 1 | 0 | 1 | 1 |
| SE | 88988 | ENSG00000214954 | LRRC69 | chr8 | + | 91135667 | 91135739 | 91133109 | 91133305 | 91200612 | 91200792 | 1 | 0 | 1 | 1 |
| SE | 88991 | ENSG00000214954 | LRRC69 | chr8 | + | 91200612 | 91200792 | 91189521 | 91189623 | 91218889 | 91219000 | 0 | 0 | 2 | 4 |
| SE | 88993 | ENSG00000177565 | TBL1XR1 | chr3 | - | 177032970 | 177033010 | 177026372 | 177026474 | 177034197 | 177034203 | 16 | 17 | 16 | 19 |
| SE | 88994 | ENSG00000177565 | TBL1XR1 | chr3 | - | 177032970 | 177033136 | 177026372 | 177026474 | 177034197 | 177034325 | 619 | 522 | 575 | 594 |
| SE | 88995 | ENSG00000177565 | TBL1XR1 | chr3 | - | 177037636 | 177037842 | 177034197 | 177034325 | 177038097 | 177038172 | 0 | 0 | 0 | 0 |
| SE | 88996 | ENSG00000177565 | TBL1XR1 | chr3 | - | 177046128 | 177046189 | 177038312 | 177038434 | 177047299 | 177047397 | 606 | 522 | 607 | 598 |
| SE | 88997 | ENSG00000177565 | TBL1XR1 | chr3 | - | 177051503 | 177051726 | 177050477 | 177050610 | 177053772 | 177053918 | 459 | 393 | 428 | 459 |
| SE | 88999 | ENSG00000177565 | TBL1XR1 | chr3 | - | 177064919 | 177065022 | 177053772 | 177053918 | 177079826 | 177079945 | 159 | 147 | 156 | 179 |
| SE | 89002 | ENSG00000177565 | TBL1XR1 | chr3 | - | 177064919 | 177065022 | 177053772 | 177053918 | 177197120 | 177197252 | 140 | 136 | 157 | 166 |
| SE | 89004 | ENSG00000177565 | TBL1XR1 | chr3 | - | 177079826 | 177079945 | 177053772 | 177053918 | 177098465 | 177098541 | 9 | 3 | 7 | 8 |
| SE | 89006 | ENSG00000177565 | TBL1XR1 | chr3 | - | 177079826 | 177079945 | 177053772 | 177053918 | 177197120 | 177197184 | 5 | 0 | 0 | 1 |
| SE | 89008 | ENSG00000177565 | TBL1XR1 | chr3 | - | 177079826 | 177079945 | 177064919 | 177065022 | 177196339 | 177196435 | 24 | 9 | 7 | 14 |
| SE | 89010 | ENSG00000177565 | TBL1XR1 | chr3 | - | 177086964 | 177087041 | 177064919 | 177065022 | 177091150 | 177091180 | 3 | 0 | 3 | 2 |
| SE | 89012 | ENSG00000177565 | TBL1XR1 | chr3 | - | 177086964 | 177087041 | 177079886 | 177079945 | 177098465 | 177098541 | 1 | 0 | 0 | 0 |
| SE | 89013 | ENSG00000177565 | TBL1XR1 | chr3 | - | 177091150 | 177091180 | 177064919 | 177065022 | 177098465 | 177098541 | 0 | 0 | 1 | 0 |
| SE | 89014 | ENSG00000177565 | TBL1XR1 | chr3 | - | 177091150 | 177091180 | 177086964 | 177087041 | 177098465 | 177098541 | 1 | 0 | 0 | 0 |
| SE | 89016 | ENSG00000177565 | TBL1XR1 | chr3 | - | 177098465 | 177098541 | 177053772 | 177053918 | 177197120 | 177197192 | 11 | 6 | 12 | 12 |
| SE | 89017 | ENSG00000177565 | TBL1XR1 | chr3 | - | 177098465 | 177098541 | 177064919 | 177065022 | 177131362 | 177131449 | 33 | 35 | 57 | 37 |
| SE | 89018 | ENSG00000177565 | TBL1XR1 | chr3 | - | 177098465 | 177098541 | 177064919 | 177065022 | 177160870 | 177160988 | 48 | 37 | 71 | 47 |
| SE | 89019 | ENSG00000177565 | TBL1XR1 | chr3 | - | 177098465 | 177098541 | 177064919 | 177065022 | 177196339 | 177196353 | 35 | 35 | 58 | 48 |
| SE | 89021 | ENSG00000177565 | TBL1XR1 | chr3 | - | 177098465 | 177098541 | 177079453 | 177079945 | 177131362 | 177131449 | 12 | 9 | 7 | 8 |
| SE | 89022 | ENSG00000177565 | TBL1XR1 | chr3 | - | 177098465 | 177098541 | 177079453 | 177079945 | 177160870 | 177160988 | 22 | 10 | 12 | 14 |
| SE | 89023 | ENSG00000177565 | TBL1XR1 | chr3 | - | 177098465 | 177098541 | 177079453 | 177079945 | 177196339 | 177196353 | 13 | 10 | 5 | 12 |
| SE | 89024 | ENSG00000177565 | TBL1XR1 | chr3 | - | 177098465 | 177098541 | 177079453 | 177079945 | 177197120 | 177197192 | 20 | 12 | 19 | 19 |
| SE | 89025 | ENSG00000177565 | TBL1XR1 | chr3 | - | 177110808 | 177110887 | 177064919 | 177065022 | 177197120 | 177197227 | 0 | 2 | 4 | 2 |
| SE | 89028 | ENSG00000177565 | TBL1XR1 | chr3 | - | 177131362 | 177131449 | 177079886 | 177079945 | 177197120 | 177197163 | 0 | 0 | 0 | 0 |
| SE | 89030 | ENSG00000177565 | TBL1XR1 | chr3 | - | 177160870 | 177160988 | 177064919 | 177065022 | 177163977 | 177164001 | 3 | 0 | 2 | 6 |
| SE | 89031 | ENSG00000177565 | TBL1XR1 | chr3 | - | 177160870 | 177160988 | 177064919 | 177065022 | 177196339 | 177196484 | 3 | 0 | 3 | 4 |
| SE | 89032 | ENSG00000177565 | TBL1XR1 | chr3 | - | 177160870 | 177160988 | 177064919 | 177065022 | 177197120 | 177197225 | 5 | 5 | 3 | 5 |
| SE | 89033 | ENSG00000177565 | TBL1XR1 | chr3 | - | 177160870 | 177160988 | 177079886 | 177079945 | 177196339 | 177196484 | 3 | 0 | 2 | 1 |
| SE | 89034 | ENSG00000177565 | TBL1XR1 | chr3 | - | 177160870 | 177160988 | 177079886 | 177079945 | 177197120 | 177197225 | 5 | 5 | 2 | 2 |
| SE | 89035 | ENSG00000177565 | TBL1XR1 | chr3 | - | 177160870 | 177160988 | 177098465 | 177098541 | 177163977 | 177164001 | 14 | 5 | 13 | 14 |
| SE | 89036 | ENSG00000177565 | TBL1XR1 | chr3 | - | 177160870 | 177160988 | 177098465 | 177098541 | 177196339 | 177196484 | 14 | 5 | 14 | 12 |
| SE | 89039 | ENSG00000177565 | TBL1XR1 | chr3 | - | 177160870 | 177160988 | 177131377 | 177131449 | 177196339 | 177196484 | 1 | 0 | 2 | 1 |
| SE | 89040 | ENSG00000177565 | TBL1XR1 | chr3 | - | 177160870 | 177160988 | 177131377 | 177131449 | 177197120 | 177197225 | 3 | 5 | 2 | 2 |
| SE | 89042 | ENSG00000177565 | TBL1XR1 | chr3 | - | 177163977 | 177164001 | 177064978 | 177065022 | 177197120 | 177197194 | 0 | 0 | 0 | 0 |
| SE | 89044 | ENSG00000177565 | TBL1XR1 | chr3 | - | 177163977 | 177164001 | 177098486 | 177098541 | 177197120 | 177197194 | 0 | 0 | 0 | 0 |

| | | | | | | | | | | | | | | | | | | | | | |
|---|---|---|---|---|---|---|---|---|---|---|---|---|---|---|---|---|---|---|---|---|---|
| SE | 89047 | ENSG00000177565 | TBL1XR1 | chr3 | - | 177163977 | 177164001 | 177160870 | 177160988 | 177196339 | 177196484 | 1 | 0 | 0 | 0 | 0 | 1 | 0 | 2 | 1 | 2 |
| SE | 89048 | ENSG00000177565 | TBL1XR1 | chr3 | - | 177163977 | 177164001 | 177160870 | 177160988 | 177197120 | 177197194 | 0 | 0 | 1 | 3 | 0 | 10 | 6 | 6 | 2 | 0 |
| SE | 89049 | ENSG00000067601 | PMS2P4 | chr7 | - | 67292462 | 67292549 | 67287540 | 67287719 | 67295617 | 67295801 | 7 | 10 | 6 | 0 | 8 | 0 | 0 | 0 | 1 | 0 |
| SE | 89051 | ENSG00000067601 | PMS2P4 | chr7 | - | 67297216 | 67297303 | 67295617 | 67295801 | 67299284 | 67299424 | 37 | 29 | 17 | 24 | 22 | 0 | 0 | 0 | 3 | 0 |
| SE | 89052 | ENSG00000120832 | MTERF2 | chr12 | - | 106980620 | 106980713 | 106978455 | 106978771 | 106985114 | 106985286 | 3 | 1 | 0 | 0 | 6 | 13 | 11 | 15 | 17 | 36 |
| SE | 89053 | ENSG00000120832 | MTERF2 | chr12 | - | 106985114 | 106985215 | 106977290 | 106978771 | 106986968 | 106987149 | 11 | 2 | 14 | 16 | 33 | 0 | 5 | 0 | 0 | 0 |
| SE | 89054 | ENSG00000120832 | MTERF2 | chr12 | - | 106985114 | 106985225 | 106977276 | 106978771 | 106986968 | 106987122 | 12 | 10 | 21 | 19 | 35 | 0 | 5 | 0 | 0 | 0 |
| SE | 89058 | ENSG00000104081 | BMF | chr15 | - | 40105794 | 40106091 | 40091733 | 40091888 | 40108772 | 40108879 | 4 | 2 | 4 | 5 | 0 | 0 | 0 | 0 | 0 | 0 |
| SE | 89060 | ENSG00000104081 | BMF | chr15 | - | 40108258 | 40108357 | 40105794 | 40106091 | 40108772 | 40108892 | 9 | 3 | 2 | 5 | 0 | 1 | 0 | 1 | 0 | 0 |
| SE | 89061 | ENSG00000104081 | BMF | chr15 | - | 40108258 | 40108386 | 40105794 | 40106091 | 40108772 | 40108879 | 13 | 4 | 6 | 6 | 0 | 1 | 0 | 1 | 0 | 0 |
| SE | 89062 | ENSG00000171988 | JMJD1C | chr10 | - | 63168434 | 63168566 | 63167220 | 63168134 | 63176296 | 63176473 | 669 | 641 | 641 | 675 | 1266 | 1 | 0 | 0 | 1 | 0 |
| SE | 89063 | ENSG00000171988 | JMJD1C | chr10 | - | 63183446 | 63183569 | 63176799 | 63177856 | 63184607 | 63184738 | 532 | 465 | 464 | 589 | 894 | 8 | 0 | 1 | 3 | 2 |
| SE | 89065 | ENSG00000171988 | JMJD1C | chr10 | - | 63185562 | 63185653 | 63184607 | 63184738 | 63186214 | 63186383 | 331 | 349 | 279 | 375 | 479 | 0 | 0 | 0 | 0 | 0 |
| SE | 89067 | ENSG00000171988 | JMJD1C | chr10 | - | 63186214 | 63186383 | 63185562 | 63185653 | 63190893 | 63191108 | 234 | 231 | 202 | 285 | 375 | 0 | 0 | 1 | 0 | 0 |
| SE | 89068 | ENSG00000171988 | JMJD1C | chr10 | - | 63189167 | 63189446 | 63185562 | 63185653 | 63190893 | 63191108 | 327 | 272 | 320 | 343 | 523 | 0 | 0 | 1 | 0 | 0 |
| SE | 89069 | ENSG00000171988 | JMJD1C | chr10 | - | 63189167 | 63189446 | 63186214 | 63186383 | 63190893 | 63191108 | 504 | 427 | 443 | 526 | 747 | 0 | 3 | 0 | 0 | 5 |
| SE | 89070 | ENSG00000171988 | JMJD1C | chr10 | - | 63198512 | 63198727 | 63197410 | 63197563 | 63200475 | 63200677 | 489 | 466 | 459 | 577 | 627 | 1 | 1 | 0 | 1 | 0 |
| SE | 89074 | ENSG00000171988 | JMJD1C | chr10 | - | 63209062 | 63209235 | 63206594 | 63208801 | 63213472 | 63215151 | 377 | 345 | 340 | 481 | 277 | 0 | 1 | 0 | 3 | 0 |
| SE | 89075 | ENSG00000171988 | JMJD1C | chr10 | - | 63213472 | 63215151 | 63209062 | 63209235 | 63217206 | 63217331 | 177 | 150 | 176 | 230 | 159 | 6 | 0 | 0 | 1 | 2 |
| SE | 89076 | ENSG00000171988 | JMJD1C | chr10 | - | 63215262 | 63215455 | 63213472 | 63215151 | 63217206 | 63217331 | 435 | 463 | 370 | 508 | 969 | 1 | 0 | 0 | 0 | 0 |
| SE | 89077 | ENSG00000171988 | JMJD1C | chr10 | - | 63215552 | 63215617 | 63215437 | 63215455 | 63217206 | 63217331 | 46 | 56 | 43 | 61 | 99 | 3 | 0 | 2 | 4 | 8 |
| SE | 89079 | ENSG00000171988 | JMJD1C | chr10 | - | 63215552 | 63215696 | 63215262 | 63215455 | 63264650 | 63264764 | 200 | 216 | 185 | 263 | 506 | 0 | 0 | 0 | 0 | 2 |
| SE | 89080 | ENSG00000171988 | JMJD1C | chr10 | - | 63217206 | 63217331 | 63215262 | 63215455 | 63264650 | 63264764 | 110 | 97 | 94 | 133 | 276 | 0 | 0 | 0 | 0 | 2 |
| SE | 89081 | ENSG00000171988 | JMJD1C | chr10 | - | 63217206 | 63217331 | 63215552 | 63215696 | 63264650 | 63264764 | 276 | 241 | 223 | 315 | 648 | 0 | 0 | 0 | 0 | 0 |
| SE | 89082 | ENSG00000171988 | JMJD1C | chr10 | - | 63217206 | 63217331 | 63215552 | 63215696 | 63268662 | 63269187 | 164 | 147 | 125 | 171 | 349 | 0 | 0 | 0 | 0 | 0 |
| SE | 89086 | ENSG00000171988 | JMJD1C | chr10 | - | 63264650 | 63264764 | 63215552 | 63215696 | 63268662 | 63268875 | 133 | 143 | 123 | 176 | 286 | 0 | 0 | 0 | 0 | 0 |
| SE | 89090 | ENSG00000171988 | JMJD1C | chr10 | - | 63264650 | 63264764 | 63219876 | 63219983 | 63380317 | 63380482 | 78 | 108 | 85 | 90 | 173 | 0 | 3 | 0 | 2 | 0 |
| SE | 89091 | ENSG00000155100 | OTUD6B | chr8 | + | 91073830 | 91073911 | 91071137 | 91071289 | 91078355 | 91078668 | 53 | 48 | 45 | 59 | 61 | 0 | 1 | 0 | 0 | 0 |
| SE | 89092 | ENSG00000155100 | OTUD6B | chr8 | + | 91076591 | 91076731 | 91073830 | 91073911 | 91078355 | 91078668 | 17 | 18 | 25 | 46 | 45 | 45 | 31 | 28 | 29 | 37 |
| SE | 89093 | ENSG00000160588 | MPZL3 | chr11 | - | 118235423 | 118235589 | 118233459 | 118233523 | 118237049 | 118237260 | 77 | 52 | 66 | 76 | 45 | 2 | 0 | 0 | 0 | 0 |
| SE | 89094 | ENSG00000160588 | MPZL3 | chr11 | - | 118235423 | 118235589 | 118233459 | 118233523 | 118240210 | 118240377 | 43 | 33 | 38 | 40 | 36 | 0 | 0 | 2 | 2 | 0 |
| SE | 89095 | ENSG00000160588 | MPZL3 | chr11 | - | 118237049 | 118237260 | 118233459 | 118233523 | 118240210 | 118240341 | 36 | 34 | 49 | 48 | 49 | 0 | 0 | 2 | 2 | 0 |
| SE | 89099 | ENSG00000064115 | TM7SF3 | chr12 | - | 26975495 | 26975658 | 26971578 | 26974227 | 26976259 | 26976357 | 260 | 134 | 161 | 190 | 238 | 0 | 0 | 0 | 0 | 0 |
| SE | 89100 | ENSG00000064115 | TM7SF3 | chr12 | - | 26975495 | 26975658 | 26971578 | 26974227 | 26979783 | 26979936 | 159 | 82 | 101 | 125 | 155 | 0 | 0 | 0 | 1 | 0 |
| SE | 89102 | ENSG00000064115 | TM7SF3 | chr12 | - | 26975495 | 26975658 | 26971578 | 26974227 | 26995236 | 26995268 | 153 | 80 | 99 | 121 | 154 | 0 | 0 | 0 | 0 | 3 |
| SE | 89103 | ENSG00000064115 | TM7SF3 | chr12 | - | 26976259 | 26976357 | 26971578 | 26974227 | 26979783 | 26979936 | 24 | 7 | 10 | 23 | 17 | 0 | 0 | 0 | 1 | 0 |
| SE | 89104 | ENSG00000064115 | TM7SF3 | chr12 | - | 26976259 | 26976357 | 26971578 | 26974227 | 26980565 | 26980608 | 2 | 1 | 2 | 0 | 0 | 0 | 0 | 0 | 1 | 0 |
| SE | 89111 | ENSG00000064115 | TM7SF3 | chr12 | - | 26980565 | 26980646 | 26971578 | 26974227 | 26995236 | 26995268 | 0 | 0 | 0 | 1 | 0 | 0 | 0 | 0 | 0 | 3 |
| SE | 89112 | ENSG00000064115 | TM7SF3 | chr12 | - | 26980565 | 26980646 | 26975495 | 26975658 | 26995236 | 26995268 | 2 | 4 | 1 | 0 | 1 | 1 | 0 | 0 | 0 | 0 |
| SE | 89115 | ENSG00000064115 | TM7SF3 | chr12 | - | 26982772 | 26982859 | 26980565 | 26980646 | 26990449 | 26990627 | 119 | 55 | 65 | 68 | 108 | 0 | 0 | 0 | 3 | 0 |
| SE | 89116 | ENSG00000064115 | TM7SF3 | chr12 | - | 26982772 | 26982859 | 26980565 | 26980646 | 26995236 | 26995268 | 46 | 22 | 23 | 30 | 56 | 0 | 0 | 0 | 0 | 0 |
| SE | 89122 | ENSG00000064115 | TM7SF3 | chr12 | - | 26995236 | 26995408 | 26983492 | 26983596 | 27014077 | 27014347 | 2 | 0 | 4 | 5 | 4 | 0 | 1 | 0 | 0 | 0 |
| SE | 89123 | ENSG00000064115 | TM7SF3 | chr12 | - | 26995236 | 26995408 | 26990449 | 26990627 | 26996741 | 26996785 | 202 | 87 | 161 | 150 | 293 | 2 | 0 | 1 | 0 | 0 |
| SE | 89124 | ENSG00000064115 | TM7SF3 | chr12 | - | 26996741 | 26996862 | 26990449 | 26990627 | 26999525 | 26999648 | 77 | 34 | 55 | 50 | 89 | 0 | 0 | 1 | 0 | 0 |
| SE | 89128 | ENSG00000064115 | TM7SF3 | chr12 | - | 26999525 | 26999648 | 26996741 | 26996862 | 27014077 | 27014398 | 129 | 61 | 98 | 99 | 232 | 1 | 0 | 0 | 0 | 0 |
| SE | 89130 | ENSG00000064115 | TM7SF3 | chr12 | - | 26999525 | 26999676 | 26996741 | 26996862 | 27014077 | 27014211 | 138 | 69 | 109 | 101 | 244 | 1 | 0 | 0 | 0 | 0 |
| SE | 89133 | ENSG00000064115 | TM7SF3 | chr12 | - | 27003235 | 27003390 | 26999525 | 26999648 | 27014077 | 27014384 | 15 | 8 | 10 | 13 | 12 | 1 | 0 | 0 | 0 | 0 |
| SE | 89134 | ENSG00000064115 | TM7SF3 | chr12 | - | 27003235 | 27003390 | 26999525 | 26999676 | 27014077 | 27014384 | 127 | 69 | 95 | 88 | 210 | 0 | 0 | 2 | 0 | 0 |
| SE | 89135 | ENSG00000064115 | TM7SF3 | chr12 | - | 27003235 | 27003405 | 26999588 | 26999676 | 27014077 | 27014398 | 112 | 64 | 85 | 76 | 198 | 0 | 0 | 2 | 0 | 0 |
| SE | 89139 | ENSG00000154589 | LY96 | chr8 | + | 74010000 | 74010129 | 73991425 | 73991554 | 74026788 | 74026841 | 54 | 20 | 28 | 44 | 141 | 2 | 0 | 0 | 0 | 0 |
| SE | 89140 | ENSG00000154589 | LY96 | chr8 | + | 74026788 | 74026841 | 74010000 | 74010129 | 74028955 | 74029074 | 35 | 8 | 16 | 21 | 85 | 0 | 0 | 0 | 0 | 3 |
| SE | 89141 | ENSG00000111276 | CDKN1B | chr12 | + | 12718824 | 12718954 | 12717367 | 12718314 | 12721035 | 12721204 | 1303 | 981 | 1480 | 1375 | 2407 | 0 | 0 | 0 | 1 | 0 |
| SE | 89144 | ENSG00000235531 | MSC-AS1 | chr8 | + | 71844228 | 71844422 | 71843122 | 71843704 | 71962922 | 71963048 | 8 | 3 | 3 | 1 | 4 | 0 | 0 | 0 | 0 | 0 |
| SE | 89146 | ENSG00000235531 | MSC-AS1 | chr8 | + | 71965408 | 71965534 | 71962922 | 71963048 | 71966741 | 71966779 | 3 | 11 | 9 | 11 | 9 | 0 | 2 | 0 | 0 | 0 |
| SE | 89147 | ENSG00000235531 | MSC-AS1 | chr8 | + | 71965408 | 71965534 | 71962922 | 71963048 | 71973537 | 71973659 | 3 | 11 | 7 | 10 | 9 | 0 | 0 | 0 | 1 | 0 |
| SE | 89150 | ENSG00000235531 | MSC-AS1 | chr8 | + | 71973537 | 71973659 | 71965408 | 71965534 | 72025361 | 72025434 | 0 | 0 | 0 | 3 | 0 | 0 | 0 | 0 | 0 | 7 |
| SE | 89151 | ENSG00000235531 | MSC-AS1 | chr8 | + | 71973537 | 71973659 | 71965408 | 71965534 | 72052538 | 72054918 | 0 | 0 | 0 | 2 | 0 | 5 | 8 | 10 | 6 | 3 |
| SE | 89154 | ENSG00000235531 | MSC-AS1 | chr8 | + | 72055464 | 72055582 | 72052538 | 72052616 | 72060362 | 72060441 | 3 | 0 | 0 | 1 | 1 | 1 | 1 | 0 | 0 | 0 |
| SE | 89156 | ENSG00000143321 | HDGF | chr1 | - | 156744162 | 156744348 | 156743651 | 156743878 | 156745007 | 156745146 | 1898 | 1396 | 1504 | 1932 | 2112 | 0 | 0 | 0 | 1 | 0 |
| SE | 89158 | ENSG00000143321 | HDGF | chr1 | - | 156745007 | 156745146 | 156744162 | 156744348 | 156745296 | 156745373 | 1809 | 1399 | 1425 | 1876 | 2034 | 1 | 0 | 0 | 2 | 4 |
| SE | 89159 | ENSG00000143321 | HDGF | chr1 | - | 156745007 | 156745373 | 156744162 | 156744348 | 156751342 | 156751496 | 1127 | 870 | 855 | 1062 | 1207 | 0 | 0 | 0 | 0 | 0 |
| SE | 89160 | ENSG00000143321 | HDGF | chr1 | - | 156745296 | 156745373 | 156744162 | 156744348 | 156751342 | 156751544 | 18 | 6 | 10 | 14 | 4 | 0 | 0 | 0 | 0 | 0 |
| SE | 89161 | ENSG00000075856 | SART3 | chr12 | - | 108525456 | 108525609 | 108523868 | 108524506 | 108526098 | 108526185 | 187 | 112 | 149 | 142 | 157 | 0 | 0 | 4 | 0 | 0 |
| SE | 89162 | ENSG00000075856 | SART3 | chr12 | - | 108531203 | 108531280 | 108530141 | 108530310 | 108535358 | 108535468 | 19 | 21 | 27 | 20 | 19 | 0 | 0 | 0 | 1 | 0 |
| SE | 89171 | ENSG00000075856 | SART3 | chr12 | - | 108538064 | 108538203 | 108537487 | 108537595 | 108543027 | 108543152 | 140 | 63 | 118 | 99 | 102 | 0 | 1 | 0 | 0 | 0 |
| SE | 89172 | ENSG00000075856 | SART3 | chr12 | - | 108538933 | 108539089 | 108537487 | 108537595 | 108543027 | 108543152 | 60 | 33 | 54 | 38 | 60 | 0 | 1 | 0 | 0 | 0 |
| SE | 89173 | ENSG00000075856 | SART3 | chr12 | - | 108538933 | 108539089 | 108538064 | 108538203 | 108543027 | 108543152 | 189 | 95 | 168 | 134 | 130 | 0 | 0 | 0 | 0 | 0 |
| SE | 89176 | ENSG00000075856 | SART3 | chr12 | - | 108543027 | 108543152 | 108538933 | 108539089 | 108544426 | 108544478 | 188 | 92 | 160 | 182 | 218 | 1 | 0 | 0 | 0 | 0 |
| SE | 89182 | ENSG00000162676 | GFI1 | chr1 | - | 92483372 | 92483586 | 92482863 | 92483046 | 92486013 | 92486071 | 47 | 16 | 25 | 29 | 23 | 1 | 0 | 0 | 0 | 0 |
| SE | 89183 | ENSG00000147649 | MTDH | chr8 | + | 97661071 | 97661173 | 97644183 | 97644887 | 97686667 | 97686752 | 219 | 198 | 239 | 229 | 348 | 2 | 0 | 0 | 0 | 0 |
| SE | 89184 | ENSG00000147649 | MTDH | chr8 | + | 97686667 | 97686752 | 97661071 | 97661173 | 97689037 | 97689103 | 97 | 92 | 104 | 104 | 165 | 2 | 0 | 0 | 0 | 0 |
| SE | 89185 | ENSG00000147649 | MTDH | chr8 | + | 97686667 | 97686752 | 97661071 | 97661173 | 97690951 | 97691164 | 95 | 93 | 103 | 95 | 161 | 0 | 0 | 0 | 1 | 0 |
| SE | 89187 | ENSG00000147649 | MTDH | chr8 | + | 97687428 | 97687605 | 97686667 | 97686752 | 97690951 | 97691164 | 267 | 208 | 281 | 257 | 508 | 0 | 1 | 0 | 0 | 2 |
| SE | 89188 | ENSG00000147649 | MTDH | chr8 | + | 97689037 | 97689103 | 97661071 | 97661173 | 97690951 | 97691188 | 134 | 101 | 136 | 88 | 238 | 0 | 0 | 0 | 1 | 0 |
| SE | 89189 | ENSG00000147649 | MTDH | chr8 | + | 97689037 | 97689103 | 97686667 | 97686752 | 97690951 | 97691188 | 134 | 101 | 137 | 97 | 244 | 0 | 1 | 0 | 0 | 2 |
| SE | 89192 | ENSG00000147649 | MTDH | chr8 | + | 97699753 | 97699852 | 97690951 | 97691188 | 97713661 | 97713769 | 204 | 183 | 219 | 214 | 372 | 1 | 2 | 2 | 0 | 0 |
| SE | 89193 | ENSG00000147649 | MTDH | chr8 | + | 97706625 | 97706750 | 97690951 | 97691188 | 97713661 | 97713769 | 552 | 456 | 554 | 655 | 856 | 1 | 2 | 2 | 0 | 0 |
| SE | 89195 | ENSG00000147649 | MTDH | chr8 | + | 97713661 | 97713769 | 97706411 | 97706750 | 97719048 | 97719087 | 697 | 623 | 713 | 837 | 1255 | 0 | 0 | 0 | 0 | 0 |
| SE | 89197 | ENSG00000147649 | MTDH | chr8 | + | 97722878 | 97723035 | 97719048 | 97719189 | 97724599 | 97724617 | 707 | 588 | 685 | 716 | 1207 | 0 | 0 | 0 | 0 | 0 |
| SE | 89200 | ENSG00000165660 | ABRAXAS2 | chr10 | + | 124816575 | 124816612 | 124806830 | 124806921 | 124819383 | 124819450 | 84 | 72 | 68 | 77 | 84 | 0 | 0 | 0 | 0 | 0 |
| SE | 89201 | ENSG00000165660 | ABRAXAS2 | chr10 | + | 124816575 | 124816612 | 124806830 | 124806921 | 124826594 | 124826785 | 12 | 12 | 20 | 14 | 1 | 0 | 0 | 2 | 0 | 0 |
| SE | 89202 | ENSG00000165660 | ABRAXAS2 | chr10 | + | 124819383 | 124819450 | 124806830 | 124806921 | 124826594 | 124826785 | 30 | 16 | 37 | 32 | 48 | 0 | 0 | 2 | 0 | 0 |
| SE | 89203 | ENSG00000165660 | ABRAXAS2 | chr10 | + | 124819383 | 124819450 | 124816575 | 124816612 | 124826594 | 124826785 | 108 | 80 | 91 | 99 | 142 | 0 | 4 | 2 | 0 | 1 |
| SE | 89204 | ENSG00000273066 | NA | chr9 | + | 136809254 | 136809429 | 136807739 | 136808326 | 136809687 | 136810042 | 0 | 0 | 1 | 2 | 0 | 0 | 0 | 0 | 1 | 0 |
| SE | 89208 | ENSG00000100403 | ZC3H7B | chr22 | + | 41320654 | 41320713 | 41301524 | 41301772 | 41325563 | 41325597 | 76 | 61 | 49 | 72 | 47 | 0 | 0 | 0 | 0 | 0 |
| SE | 89209 | ENSG00000100403 | ZC3H7B | chr22 | + | 41325720 | 41325918 | 41325563 | 41325597 | 41330022 | 41330103 | 47 | 51 | 48 | 67 | 62 | 0 | 1 | 0 | 0 | 0 |
| SE | 89210 | ENSG00000100403 | ZC3H7B | chr22 | + | 41327205 | 41327364 | 41325720 | 41325918 | 41330022 | 41330103 | 125 | 94 | 95 | 149 | 105 | 0 | 0 | 0 | 0 | 0 |
| SE | 89212 | ENSG00000100403 | ZC3H7B | chr22 | + | 41341087 | 41341146 | 41339815 | 41340137 | 41342528 | 41342628 | 39 | 16 | 38 | 27 | 18 | 1 | 0 | 0 | 0 | 0 |
| SE | 89213 | ENSG00000100403 | ZC3H7B | chr22 | + | 41342528 | 41342628 | 41339815 | 41340137 | 41343414 | 41343576 | 18 | 5 | 10 | 14 | 12 | 0 | 0 | 0 | 1 | 0 |
| SE | 89214 | ENSG00000100403 | ZC3H7B | chr22 | + | 41348066 | 41348167 | 41346002 | 41346208 | 41349119 | 41349301 | 91 | 37 | 61 | 67 | 51 | 1 | 0 | 0 | 0 | 0 |
| SE | 89215 | ENSG00000100403 | ZC3H7B | chr22 | + | 41351560 | 41351646 | 41349119 | 41349301 | 41355468 | 41355602 | 56 | 43 | 52 | 58 | 62 | 0 | 0 | 0 | 0 | 0 |
| SE | 89216 | ENSG00000164024 | METAP1 | chr4 | + | 99005768 | 99005866 | 98995720 | 98995867 | 99028866 | 99028918 | 1 | 0 | 0 | 3 | 0 | 40 | 27 | 24 | 23 | 9 |
| SE | 89217 | ENSG00000164024 | METAP1 | chr4 | + | 99005768 | 99005870 | 98995765 | 98995867 | 99028866 | 99028918 | 0 | 0 | 0 | 3 | 0 | 40 | 27 | 24 | 23 | 9 |
| SE | 89218 | ENSG00000164024 | METAP1 | chr4 | + | 99031507 | 99031662 | 99028866 | 99028918 | 99034229 | 99034342 | 0 | 0 | 0 | 0 | 0 | 66 | 46 | 60 | 57 | 33 |
| SE | 89219 | ENSG00000164024 | METAP1 | chr4 | + | 99035399 | 99035460 | 99034229 | 99034342 | 99039373 | 99039465 | 123 | 99 | 124 | 145 | 145 | 0 | 0 | 0 | 0 | 0 |
| SE | 89220 | ENSG00000164024 | METAP1 | chr4 | + | 99035953 | 99036011 | 99035399 | 99035460 | 99039373 | 99039465 | 0 | 1 | 1 | 0 | 2 | 76 | 70 | 83 | 94 | 110 |
| SE | 89221 | ENSG00000164024 | METAP1 | chr4 | + | 99036042 | 99036089 | 99035399 | 99035460 | 99037766 | 99037941 | 2 | 2 | 0 | 0 | 0 | 1 | 1 | 0 | 0 | 0 |
| SE | 89224 | ENSG00000164024 | METAP1 | chr4 | + | 99037766 | 99037941 | 99036042 | 99036089 | 99039373 | 99039465 | 1 | 3 | 1 | 2 | 5 | 0 | 4 | 2 | 0 | 2 |
| SE | 89228 | ENSG00000164024 | METAP1 | chr4 | + | 99057752 | 99057818 | 99048732 | 99048876 | 99061153 | 99061519 | 172 | 117 | 131 | 147 | 146 | 1 | 0 | 0 | 0 | 0 |
| SE | 89229 | ENSG00000137571 | SLCO5A1 | chr8 | - | 69676608 | 69676673 | 69667045 | 69673326 | 69679377 | 69679619 | 0 | 0 | 0 | 1 | 0 | 0 | 0 | 1 | 0 | 0 |
| SE | 89230 | ENSG00000137571 | SLCO5A1 | chr8 | - | 69738039 | 69738204 | 69705030 | 69705229 | 69755423 | 69755641 | 0 | 0 | 0 | 4 | 4 | 0 | 0 | 1 | 0 | 0 |
| SE | 89231 | ENSG00000094914 | AAAS | chr12 | - | 53307844 | 53307929 | 53307456 | 53307713 | 53308051 | 53308133 | 85 | 30 | 92 | 89 | 115 | 3 | 0 | 0 | 1 | 0 |
| SE | 89232 | ENSG00000094914 | AAAS | chr12 | - | 53308724 | 53309020 | 53308434 | 53308528 | 53309156 | 53309281 | 38 | 16 | 27 | 31 | 33 | 0 | 1 | 0 | 0 | 0 |
| SE | 89247 | ENSG00000137574 | TGS1 | chr8 | + | 55795977 | 55796152 | 55792697 | 55792784 | 55798913 | 55798998 | 139 | 85 | 143 | 117 | 349 | 1 | 0 | 0 | 0 | 0 |
| SE | 89248 | ENSG00000137574 | TGS1 | chr8 | + | 55798913 | 55799220 | 55795977 | 55796152 | 55802456 | 55802606 | 118 | 97 | 140 | 143 | 333 | 0 | 0 | 1 | 0 | 0 |
| SE | 89249 | ENSG00000137574 | TGS1 | chr8 | + | 55813054 | 55813118 | 55804892 | 55805036 | 55824580 | 55825448 | 45 | 37 | 40 | 46 | 20 | 2 | 0 | 0 | 0 | 0 |
| SE | 89250 | ENSG00000175606 | TMEM70 | chr8 | + | 73978755 | 73978861 | 73976194 | 73976491 | 73981154 | 73982783 | 47 | 40 | 52 | 58 | 94 | 0 | 0 | 0 | 0 | 0 |
| SE | 89251 | ENSG00000175606 | TMEM70 | chr8 | + | 73978755 | 73978875 | 73976206 | 73976491 | 73981154 | 73981366 | 11 | 16 | 11 | 24 | 15 | 0 | 0 | 0 | 0 | 0 |
| SE | 89254 | ENSG00000104325 | DECR1 | chr8 | + | 90005291 | 90005369 | 90001475 | 90001561 | 90015540 | 90015666 | 0 | 1 | 0 | 0 | 0 | 0 | 4 | 4 | 4 | 0 |
| SE | 89255 | ENSG00000104325 | DECR1 | chr8 | + | 90005291 | 90005369 | 90001475 | 90001561 | 90017123 | 90017326 | 2 | 3 | 0 | 0 | 0 | 90 | 52 | 76 | 74 | 68 |
| SE | 89256 | ENSG00000104325 | DECR1 | chr8 | + | 90005447 | 90005731 | 90001476 | 90001561 | 90006035 | 90006233 | 4 | 2 | 6 | 2 | 10 | 4 | 0 | 0 | 0 | 0 |
| SE | 89258 | ENSG00000104325 | DECR1 | chr8 | + | 90005447 | 90005845 | 90000141 | 90001561 | 90006035 | 90006277 | 5 | 9 | 10 | 2 | 11 | 4 | 0 | 0 | 0 | 0 |
| SE | 89260 | ENSG00000104325 | DECR1 | chr8 | + | 90005447 | 90005845 | 90001476 | 90001561 | 90017202 | 90017326 | 4 | 2 | 6 | 2 | 10 | 1 | 0 | 0 | 0 | 0 |
| SE | 89261 | ENSG00000104325 | DECR1 | chr8 | + | 90005447 | 90005845 | 90001476 | 90001561 | 90018908 | 90018966 | 5 | 2 | 6 | 2 | 10 | 25 | 5 | 15 | 19 | 1 |
| SE | 89262 | ENSG00000104325 | DECR1 | chr8 | + | 90005447 | 90005845 | 90005291 | 90005369 | 90017123 | 90017326 | 1 | 1 | 2 | 1 | 0 | 2 | 2 | 0 | 0 | 0 |
| SE | 89264 | ENSG00000104325 | DECR1 | chr8 | + | 90006035 | 90006277 | 90005447 | 90005845 | 90017123 | 90017326 | 4 | 8 | 6 | 1 | 5 | 1 | 0 | 0 | 1 | 0 |
| SE | 89266 | ENSG00000104325 | DECR1 | chr8 | + | 90015540 | 90015666 | 90005291 | 90005369 | 90017123 | 90017326 | 1 | 3 | 1 | 5 | 0 | 2 | 2 | 0 | 0 | 0 |
| SE | 89268 | ENSG00000104325 | DECR1 | chr8 | + | 90017123 | 90017326 | 90005447 | 90005845 | 90018908 | 90018966 | 193 | 134 | 196 | 177 | 208 | 1 | 0 | 0 | 0 | 0 |
| SE | 89270 | ENSG00000104325 | DECR1 | chr8 | + | 90017202 | 90017326 | 90005447 | 90005845 | 90018908 | 90018966 | 171 | 120 | 172 | 164 | 180 | 1 | 0 | 0 | 0 | 0 |
| SE | 89272 | ENSG00000104325 | DECR1 | chr8 | + | 90036840 | 90036940 | 90020908 | 90021056 | 90042727 | 90042748 | 664 | 464 | 488 | 614 | 914 | 7 | 5 | 2 | 4 | 1 |
| SE | 89274 | ENSG00000231205 | ZNF826P | chr19 | - | 20417665 | 20417783 | 20408949 | 20409045 | 20424785 | 20424956 | 2 | 0 | 4 | 2 | 0 | 0 | 0 | 2 | 0 | 0 |
| SE | 89279 | ENSG00000166169 | POLL | chr10 | - | 101582762 | 101582891 | 101580247 | 101580416 | 101583507 | 101583681 | 50 | 26 | 55 | 56 | 26 | 0 | 0 | 0 | 0 | 0 |
| SE | 89282 | ENSG00000166169 | POLL | chr10 | - | 101583507 | 101583681 | 101580247 | 101580416 | 101584601 | 101584720 | 31 | 11 | 16 | 27 | 25 | 4 | 0 | 0 | 0 | 0 |
| SE | 89283 | ENSG00000166169 | POLL | chr10 | - | 101583507 | 101583681 | 101580247 | 101580416 | 101587245 | 101587406 | 3 | 1 | 0 | 0 | 1 | 2 | 1 | 4 | 3 | 0 |
| SE | 89284 | ENSG00000166169 | POLL | chr10 | - | 101583507 | 101583681 | 101580247 | 101580416 | 101588203 | 101588228 | 0 | 1 | 0 | 0 | 0 | 1 | 0 | 0 | 0 | 0 |
| SE | 89289 | ENSG00000166169 | POLL | chr10 | - | 101584601 | 101584720 | 101582762 | 101582891 | 101584877 | 101584919 | 2 | 0 | 2 | 3 | 0 | 1 | 0 | 0 | 0 | 0 |
| SE | 89291 | ENSG00000166169 | POLL | chr10 | - | 101584601 | 101584919 | 101580247 | 101580416 | 101587245 | 101587406 | 6 | 1 | 1 | 0 | 0 | 2 | 1 | 4 | 3 | 0 |
| SE | 89293 | ENSG00000166169 | POLL | chr10 | - | 101584601 | 101584919 | 101582762 | 101582891 | 101587245 | 101587406 | 4 | 1 | 3 | 4 | 0 | 10 | 2 | 6 | 3 | 0 |
| SE | 89296 | ENSG00000166169 | POLL | chr10 | - | 101584601 | 101584919 | 101583507 | 101583681 | 101588203 | 101588248 | 29 | 11 | 16 | 26 | 25 | 0 | 1 | 0 | 0 | 0 |
| SE | 89297 | ENSG00000166169 | POLL | chr10 | - | 101584877 | 101584919 | 101582762 | 101582891 | 101587245 | 101587406 | 0 | 0 | 0 | 0 | 0 | 10 | 2 | 6 | 3 | 0 |
| SE | 89298 | ENSG00000166169 | POLL | chr10 | - | 101584877 | 101584919 | 101583507 | 101583681 | 101585315 | 101585478 | 7 | 7 | 4 | 4 | 8 | 1 | 0 | 0 | 3 | 0 |
| SE | 89299 | ENSG00000166169 | POLL | chr10 | - | 101584877 | 101584919 | 101583507 | 101583681 | 101587245 | 101587406 | 3 | 0 | 5 | 3 | 8 | 3 | 1 | 0 | 0 | 1 |
| SE | 89301 | ENSG00000166169 | POLL | chr10 | - | 101585315 | 101585478 | 101583507 | 101583681 | 101587245 | 101587382 | 2 | 0 | 0 | 3 | 2 | 3 | 1 | 0 | 0 | 1 |
| SE | 89304 | ENSG00000166169 | POLL | chr10 | - | 101585861 | 101586156 | 101583507 | 101583681 | 101587121 | 101587200 | 0 | 2 | 1 | 0 | 2 | 0 | 1 | 0 | 0 | 1 |
| SE | 89306 | ENSG00000166169 | POLL | chr10 | - | 101585861 | 101586156 | 101583507 | 101583681 | 101588203 | 101588228 | 4 | 2 | 1 | 0 | 0 | 0 | 1 | 0 | 0 | 0 |
| SE | 89309 | ENSG00000166169 | POLL | chr10 | - | 101585861 | 101586156 | 101585315 | 101585478 | 101587245 | 101587382 | 33 | 12 | 34 | 37 | 36 | 1 | 0 | 0 | 0 | 2 |
| SE | 89310 | ENSG00000166169 | POLL | chr10 | - | 101587121 | 101587200 | 101583507 | 101583681 | 101588203 | 101588248 | 0 | 1 | 0 | 0 | 0 | 0 | 1 | 0 | 0 | 0 |
| SE | 89311 | ENSG00000166169 | POLL | chr10 | - | 101587121 | 101587200 | 101585861 | 101586156 | 101588203 | 101588248 | 0 | 2 | 1 | 0 | 2 | 4 | 2 | 1 | 0 | 0 |
| SE | 89312 | ENSG00000166169 | POLL | chr10 | - | 101587245 | 101587382 | 101584686 | 101584919 | 101588203 | 101588230 | 2 | 1 | 3 | 2 | 0 | 1 | 0 | 0 | 0 | 0 |
| SE | 89313 | ENSG00000166169 | POLL | chr10 | - | 101587245 | 101587382 | 101585861 | 101586156 | 101588203 | 101588230 | 12 | 9 | 9 | 18 | 23 | 4 | 2 | 1 | 0 | 0 |
| SE | 89314 | ENSG00000166169 | POLL | chr10 | - | 101587245 | 101587406 | 101580247 | 101580416 | 101588203 | 101588232 | 6 | 2 | 7 | 3 | 3 | 1 | 0 | 0 | 0 | 0 |
| SE | 89315 | ENSG00000166169 | POLL | chr10 | - | 101587245 | 101587406 | 101583507 | 101583681 | 101588203 | 101588232 | 7 | 2 | 3 | 0 | 4 | 0 | 1 | 0 | 0 | 0 |
| SE | 89316 | ENSG00000166169 | POLL | chr10 | - | 101587245 | 101587406 | 101584601 | 101584919 | 101588203 | 101588232 | 6 | 2 | 6 | 2 | 3 | 1 | 0 | 0 | 0 | 0 |
| SE | 89319 | ENSG00000120963 | ZNF706 | chr8 | - | 101199989 | 101200097 | 101177877 | 101178387 | 101201606 | 101201743 | 357 | 263 | 338 | 359 | 224 | 0 | 2 | 0 | 0 | 0 |
| SE | 89328 | ENSG00000120963 | ZNF706 | chr8 | - | 101204649 | 101204693 | 101202332 | 101202447 | 101205434 | 101205682 | 0 | 3 | 5 | 1 | 0 | 0 | 0 | 0 | 2 | 0 |
| SE | 89329 | ENSG00000120963 | ZNF706 | chr8 | - | 101204649 | 101204779 | 101202332 | 101202447 | 101205434 | 101205732 | 0 | 3 | 2 | 2 | 0 | 0 | 0 | 0 | 2 | 0 |
| SE | 89330 | ENSG00000120963 | ZNF706 | chr8 | - | 101204649 | 101204779 | 101202332 | 101202452 | 101205434 | 101205732 | 1 | 1 | 0 | 1 | 0 | 0 | 0 | 0 | 0 | 0 |
| SE | 89331 | ENSG00000165392 | WRN | chr8 | + | 31058371 | 31058543 | 31033787 | 31033973 | 31080866 | 31081296 | 12 | 23 | 26 | 10 | 6 | 1 | 0 | 0 | 0 | 0 |
| SE | 89333 | ENSG00000165392 | WRN | chr8 | + | 31067032 | 31067182 | 31064914 | 31065063 | 31068257 | 31068327 | 64 | 54 | 73 | 63 | 68 | 2 | 0 | 0 | 0 | 0 |
| SE | 89335 | ENSG00000165392 | WRN | chr8 | + | 31068257 | 31068327 | 31067032 | 31067182 | 31076172 | 31076287 | 80 | 69 | 52 | 62 | 71 | 0 | 0 | 0 | 1 | 0 |
| SE | 89336 | ENSG00000165392 | WRN | chr8 | + | 31076172 | 31076287 | 31068257 | 31068327 | 31080866 | 31081296 | 162 | 151 | 117 | 135 | 187 | 1 | 0 | 0 | 2 | 1 |
| SE | 89337 | ENSG00000165392 | WRN | chr8 | + | 31083698 | 31083779 | 31080866 | 31081296 | 31085165 | 31085246 | 63 | 43 | 46 | 67 | 84 | 0 | 0 | 2 | 0 | 1 |
| SE | 89339 | ENSG00000165392 | WRN | chr8 | + | 31090833 | 31090942 | 31090464 | 31090532 | 31091829 | 31091898 | 37 | 36 | 54 | 48 | 80 | 2 | 0 | 0 | 0 | 0 |
| SE | 89340 | ENSG00000165392 | WRN | chr8 | + | 31120242 | 31120424 | 31116353 | 31116528 | 31124521 | 31124623 | 69 | 40 | 54 | 51 | 103 | 0 | 0 | 0 | 1 | 0 |
| SE | 89341 | ENSG00000165392 | WRN | chr8 | + | 31124521 | 31124623 | 31119955 | 31120424 | 31124907 | 31124965 | 68 | 54 | 55 | 70 | 91 | 1 | 0 | 0 | 0 | 0 |
| SE | 89352 | ENSG00000115419 | GLS | chr2 | + | 190895151 | 190895248 | 190880820 | 190881470 | 190900563 | 190900693 | 19 | 10 | 18 | 18 | 7 | 0 | 0 | 3 | 0 | 0 |
| SE | 89353 | ENSG00000115419 | GLS | chr2 | + | 190895603 | 190895725 | 190895151 | 190895248 | 190900563 | 190900693 | 642 | 423 | 497 | 622 | 734 | 0 | 1 | 1 | 0 | 0 |
| SE | 89358 | ENSG00000115419 | GLS | chr2 | + | 190921023 | 190921056 | 190910262 | 190910321 | 190921144 | 190921203 | 552 | 474 | 488 | 599 | 639 | 0 | 0 | 0 | 0 | 0 |
| SE | 89359 | ENSG00000115419 | GLS | chr2 | + | 190924542 | 190924593 | 190923916 | 190923983 | 190927305 | 190927393 | 256 | 225 | 223 | 273 | 347 | 0 | 1 | 0 | 0 | 0 |
| SE | 89361 | ENSG00000115419 | GLS | chr2 | + | 190931544 | 190931637 | 190930436 | 190930568 | 190932715 | 190933008 | 658 | 605 | 598 | 773 | 745 | 2 | 2 | 0 | 0 | 0 |
| SE | 89362 | ENSG00000115419 | GLS | chr2 | + | 190954754 | 190954818 | 190954583 | 190954660 | 190962829 | 190963158 | 115 | 41 | 86 | 87 | 197 | 1 | 0 | 0 | 2 | 0 |
| SE | 89363 | ENSG00000103266 | STUB1 | chr16 | + | 681062 | 681350 | 680223 | 680684 | 681437 | 681603 | 145 | 37 | 106 | 103 | 42 | 2 | 0 | 0 | 0 | 0 |
| SE | 89364 | ENSG00000103266 | STUB1 | chr16 | + | 681151 | 681350 | 680409 | 680684 | 681437 | 681603 | 255 | 72 | 193 | 179 | 78 | 2 | 0 | 0 | 0 | 0 |
| SE | 89365 | ENSG00000103266 | STUB1 | chr16 | + | 681208 | 681350 | 680423 | 680684 | 681437 | 681603 | 144 | 37 | 102 | 106 | 42 | 2 | 0 | 0 | 0 | 0 |
| SE | 89366 | ENSG00000103266 | STUB1 | chr16 | + | 681437 | 681603 | 680223 | 680684 | 681792 | 681880 | 105 | 29 | 70 | 101 | 39 | 2 | 2 | 0 | 0 | 0 |
| SE | 89367 | ENSG00000103266 | STUB1 | chr16 | + | 681792 | 681880 | 680223 | 680684 | 682019 | 682076 | 52 | 31 | 45 | 49 | 68 | 0 | 0 | 0 | 0 | 2 |
| SE | 89369 | ENSG00000155099 | PIP4P2 | chr8 | - | 91020156 | 91020263 | 91018389 | 91018513 | 91021255 | 91021404 | 126 | 119 | 87 | 170 | 231 | 0 | 1 | 0 | 0 | 0 |
| SE | 89376 | ENSG00000108946 | PRKAR1A | chr17 | + | 68527839 | 68527900 | 68525753 | 68525912 | 68528869 | 68528991 | 2001 | 1684 | 1360 | 2292 | 2271 | 0 | 0 | 0 | 2 | 0 |
| SE | 89377 | ENSG00000108946 | PRKAR1A | chr17 | + | 68528630 | 68528719 | 68527839 | 68527900 | 68528869 | 68528991 | 0 | 3 | 7 | 9 | 8 | 1802 | 1563 | 1143 | 1981 | 1974 |
| SE | 89385 | ENSG00000132541 | RIDA | chr8 | - | 98108645 | 98108713 | 98106271 | 98106326 | 98117031 | 98117159 | 4 | 8 | 5 | 9 | 0 | 1 | 1 | 0 | 0 | 0 |
| SE | 89386 | ENSG00000132541 | RIDA | chr8 | - | 98108645 | 98108751 | 98106271 | 98106326 | 98117031 | 98117171 | 16 | 15 | 12 | 23 | 27 | 1 | 1 | 0 | 0 | 0 |
| SE | 89387 | ENSG00000132541 | RIDA | chr8 | - | 98108645 | 98108768 | 98106271 | 98106326 | 98117031 | 98117110 | 4 | 11 | 5 | 10 | 3 | 1 | 1 | 0 | 0 | 0 |
| SE | 89388 | ENSG00000132541 | RIDA | chr8 | - | 98113637 | 98113825 | 98108645 | 98108751 | 98117031 | 98117114 | 0 | 0 | 0 | 0 | 0 | 12 | 5 | 7 | 13 | 24 |
| SE | 89390 | ENSG00000123124 | WWP1 | chr8 | + | 86374029 | 86374120 | 86342546 | 86342930 | 86380725 | 86380864 | 87 | 71 | 87 | 109 | 147 | 0 | 1 | 0 | 0 | 0 |
| SE | 89391 | ENSG00000123124 | WWP1 | chr8 | + | 86380725 | 86380864 | 86374029 | 86374120 | 86398341 | 86398479 | 141 | 95 | 113 | 159 | 173 | 0 | 0 | 0 | 0 | 2 |
| SE | 89392 | ENSG00000123124 | WWP1 | chr8 | + | 86381504 | 86381629 | 86380725 | 86380864 | 86398341 | 86398479 | 311 | 292 | 367 | 355 | 754 | 2 | 2 | 0 | 0 | 0 |
| SE | 89393 | ENSG00000123124 | WWP1 | chr8 | + | 86398341 | 86398479 | 86381504 | 86381629 | 86398571 | 86398638 | 341 | 353 | 382 | 400 | 998 | 0 | 0 | 0 | 1 | 0 |
| SE | 89397 | ENSG00000123124 | WWP1 | chr8 | + | 86438584 | 86438673 | 86435610 | 86435704 | 86442618 | 86442778 | 264 | 185 | 254 | 221 | 341 | 0 | 0 | 0 | 1 | 0 |
| SE | 89398 | ENSG00000123124 | WWP1 | chr8 | + | 86440625 | 86440733 | 86438584 | 86438673 | 86442618 | 86442778 | 0 | 0 | 0 | 1 | 0 | 118 | 63 | 90 | 84 | 207 |
| SE | 89399 | ENSG00000123124 | WWP1 | chr8 | + | 86442618 | 86442778 | 86438584 | 86438673 | 86448147 | 86448281 | 304 | 217 | 231 | 239 | 546 | 0 | 0 | 0 | 0 | 0 |
| SE | 89400 | ENSG00000123124 | WWP1 | chr8 | + | 86448147 | 86448281 | 86438584 | 86438673 | 86448372 | 86448435 | 237 | 186 | 191 | 203 | 372 | 0 | 0 | 5 | 0 | 0 |
| SE | 89401 | ENSG00000123124 | WWP1 | chr8 | + | 86452558 | 86452679 | 86448372 | 86448513 | 86457920 | 86458025 | 436 | 376 | 383 | 484 | 804 | 5 | 2 | 1 | 2 | 0 |
| SE | 89416 | ENSG00000253394 | LINC00534 | chr8 | + | 90236024 | 90236219 | 90221499 | 90221746 | 90246179 | 90246270 | 0 | 0 | 0 | 0 | 0 | 6 | 8 | 3 | 25 | 26 |
| SE | 89419 | ENSG00000253394 | LINC00534 | chr8 | + | 90351540 | 90351666 | 90246179 | 90246270 | 90582503 | 90582644 | 2 | 0 | 0 | 3 | 9 | 0 | 2 | 0 | 0 | 0 |
| SE | 89420 | ENSG00000253394 | LINC00534 | chr8 | + | 90387566 | 90387684 | 90246179 | 90246270 | 90430367 | 90431344 | 2 | 1 | 0 | 1 | 7 | 0 | 0 | 0 | 3 | 0 |
| SE | 89421 | ENSG00000253394 | LINC00534 | chr8 | + | 90454408 | 90454523 | 90246179 | 90246270 | 90539337 | 90539474 | 5 | 10 | 3 | 11 | 15 | 0 | 0 | 0 | 2 | 0 |
| SE | 89423 | ENSG00000253394 | LINC00534 | chr8 | + | 90454408 | 90454523 | 90246179 | 90246270 | 90569134 | 90569318 | 4 | 9 | 3 | 11 | 15 | 0 | 0 | 0 | 0 | 2 |
| SE | 89424 | ENSG00000253394 | LINC00534 | chr8 | + | 90454408 | 90454523 | 90246179 | 90246270 | 90582503 | 90582644 | 4 | 11 | 3 | 13 | 15 | 0 | 2 | 0 | 0 | 0 |
| SE | 89425 | ENSG00000253394 | LINC00534 | chr8 | + | 90454408 | 90454523 | 90246179 | 90246270 | 90669039 | 90669133 | 4 | 9 | 3 | 11 | 15 | 0 | 0 | 0 | 2 | 0 |
| SE | 89426 | ENSG00000253394 | LINC00534 | chr8 | + | 90454408 | 90454523 | 90351540 | 90351666 | 90582503 | 90582644 | 0 | 4 | 1 | 3 | 0 | 0 | 0 | 0 | 1 | 0 |
| SE | 89427 | ENSG00000253394 | LINC00534 | chr8 | + | 90508558 | 90508681 | 90454408 | 90454523 | 90509989 | 90510479 | 0 | 0 | 0 | 0 | 0 | 0 | 0 | 0 | 2 | 0 |
| SE | 89428 | ENSG00000253394 | LINC00534 | chr8 | + | 90566488 | 90566549 | 90454408 | 90454523 | 90569134 | 90570420 | 0 | 0 | 0 | 0 | 1 | 0 | 0 | 0 | 0 | 0 |
| SE | 89429 | ENSG00000253394 | LINC00534 | chr8 | + | 90569134 | 90569318 | 90454408 | 90454523 | 90582503 | 90582644 | 0 | 1 | 0 | 0 | 5 | 0 | 2 | 0 | 2 | 0 |
| SE | 89430 | ENSG00000253394 | LINC00534 | chr8 | + | 90582503 | 90582644 | 90569134 | 90569318 | 90594124 | 90595421 | 0 | 3 | 0 | 0 | 7 | 0 | 0 | 0 | 3 | 0 |
| SE | 89431 | ENSG00000253394 | LINC00534 | chr8 | + | 90669039 | 90669133 | 90454408 | 90454523 | 90687360 | 90687863 | 0 | 0 | 0 | 0 | 0 | 2 | 0 | 0 | 0 | 0 |
| SE | 89433 | ENSG00000121879 | PIK3CA | chr3 | + | 179210411 | 179210565 | 179210185 | 179210338 | 179218209 | 179218334 | 87 | 55 | 73 | 89 | 147 | 0 | 1 | 0 | 1 | 0 |
| SE | 89434 | ENSG00000121879 | PIK3CA | chr3 | + | 179210430 | 179210565 | 179210185 | 179210338 | 179218209 | 179218334 | 294 | 204 | 245 | 277 | 448 | 0 | 1 | 0 | 1 | 0 |
| SE | 89435 | ENSG00000121879 | PIK3CA | chr3 | + | 179219195 | 179219277 | 179218208 | 179218334 | 179219570 | 179219735 | 86 | 60 | 77 | 96 | 174 | 0 | 0 | 0 | 1 | 1 |
| SE | 89437 | ENSG00000121879 | PIK3CA | chr3 | + | 179224699 | 179224821 | 179224080 | 179224187 | 179225961 | 179226040 | 271 | 225 | 261 | 304 | 326 | 1 | 1 | 0 | 0 | 0 |
| SE | 89439 | ENSG00000121879 | PIK3CA | chr3 | + | 179233288 | 179233371 | 179230224 | 179230376 | 179234093 | 179235098 | 0 | 7 | 3 | 5 | 5 | 114 | 92 | 87 | 105 | 159 |
| SE | 89442 | ENSG00000166946 | CCNDBP1 | chr15 | + | 43185819 | 43186233 | 43185117 | 43185607 | 43189198 | 43189280 | 122 | 33 | 90 | 84 | 60 | 0 | 0 | 1 | 0 | 0 |
| SE | 89445 | ENSG00000166946 | CCNDBP1 | chr15 | + | 43191394 | 43191675 | 43190965 | 43191042 | 43192742 | 43192803 | 678 | 327 | 558 | 700 | 958 | 0 | 1 | 0 | 0 | 0 |
| SE | 89446 | ENSG00000166946 | CCNDBP1 | chr15 | + | 43192742 | 43192803 | 43191394 | 43191675 | 43194414 | 43194461 | 471 | 246 | 373 | 460 | 719 | 0 | 0 | 0 | 0 | 1 |
| SE | 89452 | ENSG00000133739 | LRRCC1 | chr8 | + | 85109594 | 85109800 | 85107237 | 85107399 | 85110114 | 85110180 | 7 | 3 | 5 | 9 | 16 | 2 | 1 | 2 | 4 | 8 |
| SE | 89453 | ENSG00000133739 | LRRCC1 | chr8 | + | 85115099 | 85115275 | 85112931 | 85113099 | 85123412 | 85123606 | 8 | 6 | 9 | 3 | 21 | 0 | 0 | 0 | 2 | 0 |
| SE | 89454 | ENSG00000133739 | LRRCC1 | chr8 | + | 85115099 | 85115275 | 85112931 | 85113099 | 85124791 | 85124939 | 8 | 7 | 9 | 3 | 21 | 0 | 0 | 1 | 0 | 0 |
| SE | 89455 | ENSG00000133739 | LRRCC1 | chr8 | + | 85115374 | 85115584 | 85115099 | 85115275 | 85123412 | 85123606 | 39 | 26 | 22 | 23 | 36 | 0 | 0 | 0 | 0 | 0 |
| SE | 89456 | ENSG00000133739 | LRRCC1 | chr8 | + | 85115374 | 85115584 | 85115099 | 85115275 | 85124791 | 85124939 | 30 | 16 | 9 | 18 | 14 | 0 | 1 | 0 | 0 | 0 |
| SE | 89457 | ENSG00000133739 | LRRCC1 | chr8 | + | 85123412 | 85123606 | 85112931 | 85113099 | 85124791 | 85124939 | 3 | 11 | 6 | 5 | 10 | 0 | 0 | 1 | 0 | 0 |

| | | | | | | | | | | | | | | | | | | | | | | | | | |
|---|---|---|---|---|---|---|---|---|---|---|---|---|---|---|---|---|---|---|---|---|---|---|---|---|---|
| 89047 | 0 | 0 | 0 | 0 | 0 | 0 | 2 | 0 | 0 | 0 | 173 | 149 | 1 | 1 | 0.463 | NA | 0 | 0 | 0 | NA | 0 | NA | NA | NA | 0.116 |
| 89048 | 0 | 0 | 0 | 0 | 0 | 0 | 4 | 5 | 1 | 0 | 173 | 149 | 1 | 1 | 0 | 0 | 0.126 | 0.564 | NA | NA | 0 | 0 | 0 | NA | 0.172 |
| 89049 | 22 | 5 | 6 | 0 | 1 | 0 | 0 | 0 | 0 | 0 | 236 | 149 | 1 | 1 | 1 | 1 | 1 | 0 | 1 | 1 | 1 | 1 | NA | 1 | -0.2 |
| 89051 | 28 | 48 | 25 | 21 | 28 | 0 | 0 | 4 | 0 | 0 | 236 | 149 | 1 | 1 | 1 | 1 | 1 | 0.835 | 1 | 1 | 1 | 0.798 | 1 | 1 | 0.007 |
| 89052 | 0 | 3 | 3 | 0 | 0 | 0 | 4 | 9 | 0 | 0 | 242 | 149 | 1 | 1 | 0.124 | 0.053 | 0 | 0 | 0.093 | NA | 0.316 | 0.17 | NA | NA | -0.189 |
| 89053 | 0 | 4 | 3 | 0 | 0 | 0 | 0 | 0 | 0 | 0 | 250 | 149 | 1 | 1 | 1 | 0.193 | 1 | 1 | 1 | NA | 1 | 1 | NA | NA | -0.161 |
| 89054 | 0 | 4 | 3 | 0 | 0 | 0 | 0 | 0 | 0 | 0 | 260 | 149 | 1 | 1 | 1 | 0.534 | 1 | 1 | 1 | NA | 1 | 1 | NA | NA | -0.093 |
| 89058 | 33 | 16 | 5 | 9 | 35 | 0 | 0 | 0 | 0 | 2 | 298 | 149 | 1 | 1 | 1 | 1 | 1 | 1 | NA | 1 | 1 | 1 | 1 | 0.897 | 0.021 |
| 89060 | 6 | 5 | 8 | 22 | 13 | 0 | 0 | 0 | 0 | 0 | 248 | 149 | 1 | 1 | 0.844 | 1 | 0.546 | 1 | NA | 1 | 1 | 1 | 1 | 1 | -0.153 |
| 89061 | 6 | 19 | 17 | 28 | 39 | 0 | 0 | 0 | 0 | 0 | 277 | 149 | 1 | 1 | 0.875 | 1 | 0.763 | 1 | NA | 1 | 1 | 1 | 1 | 1 | -0.091 |
| 89062 | 86 | 165 | 75 | 43 | 74 | 0 | 0 | 1 | 0 | 0 | 281 | 149 | 1 | 1 | 0.997 | 1 | 1 | 0.997 | 1 | 1 | 1 | 0.975 | 1 | 1 | 0.004 |
| 89063 | 98 | 116 | 98 | 16 | 21 | 0 | 0 | 2 | 0 | 0 | 272 | 149 | 1 | 1 | 0.973 | 1 | 0.996 | 0.991 | 0.996 | 1 | 1 | 0.964 | 1 | 1 | -0.002 |
| 89065 | 54 | 61 | 84 | 15 | 62 | 0 | 0 | 4 | 0 | 0 | 240 | 149 | 1 | 1 | 1 | 1 | 1 | 1 | 1 | 1 | 1 | 0.929 | 1 | 1 | 0.014 |
| 89067 | 42 | 33 | 48 | 7 | 35 | 0 | 0 | 0 | 0 | 0 | 298 | 149 | 1 | 1 | 1 | 1 | 0.99 | 1 | 1 | 1 | 1 | 1 | 1 | 1 | -0.002 |
| 89068 | 24 | 95 | 46 | 29 | 23 | 0 | 0 | 0 | 0 | 0 | 298 | 149 | 1 | 1 | 1 | 1 | 0.994 | 1 | 1 | 1 | 1 | 1 | 1 | 1 | -0.001 |
| 89069 | 38 | 117 | 80 | 41 | 40 | 0 | 0 | 0 | 0 | 0 | 298 | 149 | 1 | 1 | 1 | 0.986 | 1 | 1 | 0.987 | 1 | 1 | 1 | 1 | 1 | -0.005 |
| 89070 | 72 | 145 | 133 | 5 | 4 | 0 | 0 | 0 | 0 | 0 | 298 | 149 | 1 | 1 | 0.996 | 0.996 | 1 | 0.997 | 1 | 1 | 1 | 1 | 1 | 1 | -0.002 |
| 89074 | 0 | 190 | 154 | 24 | 18 | 0 | 4 | 0 | 0 | 0 | 298 | 149 | 1 | 1 | 1 | 0.994 | 1 | 0.988 | 1 | NA | 0.96 | 1 | 1 | 1 | 0.006 |
| 89075 | 0 | 121 | 65 | 8 | 18 | 0 | 0 | 0 | 0 | 0 | 298 | 149 | 1 | 1 | 0.937 | 1 | 1 | 0.991 | 0.975 | NA | 1 | 1 | 1 | 1 | -0.019 |
| 89076 | 12 | 211 | 205 | 39 | 33 | 0 | 2 | 0 | 0 | 0 | 298 | 149 | 1 | 1 | 0.995 | 1 | 1 | 1 | 1 | 1 | 0.981 | 1 | 1 | 1 | 0.003 |
| 89077 | 3 | 25 | 11 | 2 | 0 | 0 | 6 | 5 | 2 | 0 | 214 | 149 | 1 | 1 | 0.914 | 1 | 0.937 | 0.914 | 0.896 | 1 | 0.744 | 0.605 | 0.41 | NA | 0.242 |
| 89079 | 46 | 144 | 92 | 15 | 17 | 0 | 0 | 0 | 2 | 0 | 293 | 149 | 1 | 1 | 1 | 1 | 1 | 1 | 0.992 | 1 | 1 | 1 | 0.792 | 1 | 0.04 |
| 89080 | 57 | 42 | 42 | 24 | 12 | 0 | 0 | 0 | 2 | 0 | 274 | 149 | 1 | 1 | 1 | 1 | 1 | 1 | 0.987 | 1 | 1 | 1 | 0.867 | 1 | 0.024 |
| 89081 | 104 | 139 | 97 | 36 | 35 | 0 | 0 | 1 | 0 | 0 | 274 | 149 | 1 | 1 | 1 | 1 | 1 | 1 | 1 | 1 | 1 | 0.981 | 1 | 1 | 0.004 |
| 89082 | 47 | 106 | 50 | 14 | 23 | 0 | 0 | 0 | 1 | 0 | 274 | 149 | 1 | 1 | 1 | 1 | 1 | 1 | 1 | 1 | 1 | 1 | 1 | 0.926 | 0.015 |
| 89086 | 29 | 69 | 59 | 53 | 24 | 0 | 0 | 0 | 1 | 0 | 263 | 149 | 1 | 1 | 1 | 1 | 1 | 1 | 1 | 1 | 1 | 1 | 1 | 0.931 | 0.014 |
| 89090 | 33 | 29 | 40 | 8 | 23 | 0 | 3 | 0 | 0 | 0 | 263 | 149 | 1 | 1 | 1 | 0.953 | 1 | 0.962 | 1 | 1 | 0.846 | 1 | 1 | 1 | 0.014 |
| 89091 | 0 | 13 | 11 | 3 | 3 | 0 | 0 | 0 | 0 | 0 | 230 | 149 | 1 | 1 | 1 | 0.969 | 1 | 1 | 1 | NA | 1 | 1 | 1 | 1 | -0.006 |
| 89092 | 4 | 7 | 3 | 4 | 0 | 0 | 11 | 10 | 3 | 0 | 289 | 149 | 1 | 1 | 0.163 | 0.23 | 0.315 | 0.45 | 0.385 | 1 | 0.247 | 0.134 | 0.407 | NA | -0.138 |
| 89093 | 110 | 10 | 123 | 94 | 76 | 0 | 1 | 0 | 0 | 0 | 298 | 149 | 1 | 1 | 0.951 | 1 | 1 | 1 | 1 | 1 | 0.833 | 1 | 1 | 1 | 0.024 |
| 89094 | 71 | 12 | 67 | 56 | 33 | 0 | 0 | 1 | 2 | 0 | 298 | 149 | 1 | 1 | 1 | 1 | 0.905 | 0.909 | 1 | 1 | 1 | 0.971 | 0.933 | 1 | -0.018 |
| 89095 | 62 | 17 | 63 | 72 | 28 | 0 | 0 | 1 | 2 | 0 | 298 | 149 | 1 | 1 | 1 | 1 | 0.925 | 0.923 | 1 | 1 | 1 | 0.969 | 0.947 | 1 | -0.014 |
| 89099 | 225 | 185 | 142 | 234 | 390 | 0 | 0 | 1 | 0 | 0 | 298 | 149 | 1 | 1 | 1 | 1 | 1 | 1 | 1 | 1 | 1 | 0.986 | 1 | 1 | 0.003 |
| 89100 | 137 | 170 | 99 | 179 | 238 | 0 | 0 | 0 | 0 | 0 | 298 | 149 | 1 | 1 | 1 | 1 | 1 | 0.984 | 1 | 1 | 1 | 1 | 1 | 1 | -0.003 |
| 89102 | 132 | 141 | 103 | 162 | 231 | 0 | 0 | 0 | 0 | 0 | 298 | 149 | 1 | 1 | 1 | 1 | 1 | 1 | 0.963 | 1 | 1 | 1 | 1 | 1 | -0.007 |
| 89103 | 66 | 15 | 12 | 12 | 21 | 0 | 0 | 0 | 0 | 0 | 247 | 149 | 1 | 1 | 1 | 1 | 1 | 0.933 | 1 | 1 | 1 | 1 | 1 | 1 | -0.013 |
| 89104 | 0 | 0 | 5 | 0 | 0 | 0 | 4 | 0 | 0 | 10 | 247 | 149 | 1 | 1 | 1 | 1 | 1 | 0 | NA | NA | 0 | 1 | NA | 0 | 0.417 |
| 89111 | 0 | 0 | 1 | 0 | 0 | 0 | 0 | 0 | 0 | 0 | 230 | 149 | 1 | 1 | NA | NA | NA | 1 | 0 | NA | NA | 1 | NA | NA | -0.5 |
| 89112 | 0 | 0 | 2 | 3 | 4 | 0 | 0 | 7 | 0 | 6 | 230 | 149 | 1 | 1 | 0.564 | 1 | 1 | NA | 1 | NA | NA | 0.156 | 1 | 0.302 | 0.405 |
| 89115 | 86 | 82 | 53 | 66 | 185 | 0 | 0 | 0 | 0 | 0 | 236 | 149 | 1 | 1 | 1 | 1 | 1 | 0.935 | 1 | 1 | 1 | 1 | 1 | 1 | -0.013 |
| 89116 | 51 | 31 | 28 | 12 | 68 | 0 | 0 | 1 | 0 | 0 | 236 | 149 | 1 | 1 | 1 | 1 | 1 | 1 | 1 | 1 | 1 | 0.946 | 1 | 1 | 0.011 |
| 89122 | 18 | 3 | 0 | 3 | 12 | 0 | 0 | 0 | 0 | 0 | 298 | 149 | 1 | 1 | 1 | 0 | 1 | 1 | 1 | 1 | 1 | NA | 1 | 1 | -0.2 |
| 89123 | 208 | 170 | 81 | 154 | 190 | 0 | 0 | 0 | 0 | 0 | 298 | 149 | 1 | 1 | 0.981 | 1 | 0.988 | 1 | 1 | 1 | 1 | 1 | 1 | 1 | -0.006 |
| 89124 | 37 | 56 | 30 | 20 | 62 | 0 | 0 | 0 | 0 | 0 | 270 | 149 | 1 | 1 | 1 | 1 | 0.968 | 1 | 1 | 1 | 1 | 1 | 1 | 1 | -0.006 |
| 89128 | 96 | 115 | 61 | 61 | 111 | 3 | 0 | 2 | 0 | 0 | 272 | 149 | 1 | 1 | 0.986 | 1 | 1 | 1 | 1 | 0.946 | 1 | 0.944 | 1 | 1 | 0.019 |
| 89130 | 96 | 134 | 66 | 65 | 119 | 3 | 0 | 2 | 0 | 0 | 298 | 149 | 1 | 1 | 0.986 | 1 | 1 | 1 | 1 | 0.941 | 1 | 0.943 | 1 | 1 | 0.02 |
| 89133 | 12 | 3 | 5 | 14 | 10 | 0 | 0 | 1 | 0 | 0 | 298 | 149 | 1 | 1 | 0.882 | 1 | 1 | 1 | 1 | 1 | 1 | 0.714 | 1 | 1 | 0.034 |
| 89134 | 116 | 87 | 43 | 71 | 113 | 0 | 0 | 0 | 3 | 1 | 298 | 149 | 1 | 1 | 1 | 1 | 0.96 | 1 | 1 | 1 | 1 | 1 | 0.922 | 0.983 | 0.011 |
| 89135 | 104 | 84 | 38 | 59 | 103 | 0 | 0 | 0 | 3 | 1 | 298 | 149 | 1 | 1 | 1 | 1 | 0.955 | 1 | 1 | 1 | 1 | 1 | 0.908 | 0.981 | 0.013 |
| 89139 | 74 | 41 | 42 | 56 | 34 | 0 | 0 | 0 | 0 | 0 | 278 | 149 | 1 | 1 | 0.935 | 1 | 1 | 1 | 1 | 1 | 1 | 1 | 1 | 1 | -0.013 |
| 89140 | 54 | 19 | 30 | 36 | 20 | 0 | 0 | 0 | 0 | 0 | 202 | 149 | 1 | 1 | 1 | 1 | 1 | 1 | 0.954 | 1 | 1 | 1 | 1 | 1 | -0.009 |
| 89141 | 892 | 1043 | 1271 | 832 | 680 | 0 | 0 | 0 | 0 | 0 | 279 | 149 | 1 | 1 | 1 | 1 | 1 | 0.999 | 1 | 1 | 1 | 1 | 1 | 1 | 0 |
| 89144 | 0 | 12 | 1 | 0 | 18 | 0 | 0 | 0 | 2 | 0 | 298 | 149 | 1 | 1 | 1 | 1 | 1 | 1 | 1 | NA | 1 | 1 | 0 | 1 | 0.25 |
| 89146 | 7 | 1 | 15 | 1 | 63 | 0 | 0 | 0 | 0 | 0 | 275 | 149 | 1 | 1 | 1 | 0.749 | 1 | 1 | 1 | 1 | 1 | 1 | 1 | 1 | -0.05 |
| 89147 | 7 | 1 | 15 | 1 | 41 | 0 | 0 | 0 | 0 | 0 | 275 | 149 | 1 | 1 | 1 | 1 | 1 | 0.844 | 1 | 1 | 1 | 1 | 1 | 1 | -0.031 |
| 89150 | 0 | 5 | 0 | 0 | 0 | 0 | 0 | 2 | 0 | 0 | 271 | 149 | 1 | 1 | NA | NA | NA | 1 | 0 | NA | 1 | 0 | NA | NA | 0 |
| 89151 | 0 | 0 | 0 | 0 | 0 | 29 | 5 | 3 | 3 | 58 | 271 | 149 | 1 | 1 | 0 | 0 | 0 | 0.155 | 0 | 0 | 0 | 0 | 0 | 0 | 0.031 |
| 89154 | 16 | 0 | 0 | 8 | 27 | 0 | 2 | 0 | 0 | 7 | 267 | 149 | 1 | 1 | 0.626 | 0 | NA | 1 | 1 | 1 | 0 | NA | 1 | 0.683 | -0.014 |
| 89156 | 7841 | 2651 | 2256 | 5499 | 7219 | 0 | 0 | 4 | 0 | 0 | 298 | 149 | 1 | 1 | 1 | 1 | 1 | 0.999 | 1 | 1 | 1 | 0.996 | 1 | 1 | 0.001 |
| 89158 | 8405 | 2732 | 2462 | 5711 | 7132 | 6 | 2 | 2 | 17 | 0 | 288 | 149 | 1 | 1 | 0.999 | 1 | 1 | 0.998 | 0.996 | 0.999 | 0.999 | 0.998 | 0.994 | 1 | 0.001 |
| 89159 | 4698 | 1527 | 1325 | 3351 | 3981 | 0 | 0 | 0 | 1 | 0 | 298 | 149 | 1 | 1 | 1 | 1 | 1 | 1 | 1 | 1 | 1 | 1 | 0.999 | 1 | 0 |
| 89160 | 58 | 20 | 14 | 60 | 54 | 0 | 0 | 0 | 1 | 0 | 226 | 149 | 1 | 1 | 1 | 1 | 1 | 1 | 1 | 1 | 1 | 1 | 0.975 | 1 | 0.005 |
| 89161 | 91 | 146 | 154 | 178 | 153 | 0 | 0 | 0 | 0 | 0 | 298 | 149 | 1 | 1 | 1 | 1 | 0.949 | 1 | 1 | 1 | 1 | 1 | 1 | 1 | -0.01 |
| 89162 | 41 | 19 | 52 | 29 | 8 | 0 | 0 | 0 | 0 | 0 | 226 | 149 | 1 | 1 | 1 | 1 | 1 | 0.93 | 1 | 1 | 1 | 1 | 1 | 1 | -0.014 |
| 89171 | 75 | 98 | 78 | 84 | 75 | 0 | 0 | 0 | 0 | 0 | 288 | 149 | 1 | 1 | 1 | 0.97 | 1 | 1 | 1 | 1 | 1 | 1 | 1 | 1 | -0.006 |
| 89172 | 24 | 39 | 35 | 31 | 22 | 0 | 0 | 0 | 0 | 0 | 298 | 149 | 1 | 1 | 1 | 0.943 | 1 | 1 | 1 | 1 | 1 | 1 | 1 | 1 | -0.011 |
| 89173 | 79 | 106 | 80 | 98 | 90 | 0 | 4 | 0 | 1 | 0 | 298 | 149 | 1 | 1 | 1 | 1 | 1 | 1 | 1 | 1 | 0.93 | 1 | 0.98 | 1 | 0.018 |
| 89176 | 152 | 130 | 136 | 82 | 123 | 0 | 0 | 0 | 0 | 0 | 274 | 149 | 1 | 1 | 0.99 | 1 | 1 | 1 | 1 | 1 | 1 | 1 | 1 | 1 | -0.002 |
| 89182 | 0 | 63 | 20 | 13 | 6 | 0 | 0 | 3 | 0 | 0 | 298 | 149 | 1 | 1 | 0.959 | 1 | 1 | 1 | 1 | NA | 1 | 1 | 1 | 1 | -0.008 |
| 89183 | 125 | 191 | 277 | 283 | 66 | 0 | 0 | 3 | 0 | 0 | 251 | 149 | 1 | 1 | 0.985 | 1 | 1 | 1 | 1 | 1 | 1 | 0.982 | 1 | 1 | 0.001 |
| 89184 | 100 | 103 | 131 | 109 | 30 | 0 | 0 | 0 | 0 | 0 | 234 | 149 | 1 | 1 | 0.969 | 1 | 1 | 1 | 1 | 1 | 1 | 1 | 1 | 1 | -0.006 |
| 89185 | 81 | 89 | 107 | 102 | 26 | 0 | 1 | 0 | 0 | 0 | 234 | 149 | 1 | 1 | 1 | 1 | 1 | 0.984 | 1 | 1 | 0.983 | 1 | 1 | 1 | 0 |
| 89187 | 77 | 249 | 245 | 269 | 63 | 0 | 3 | 0 | 0 | 0 | 298 | 149 | 1 | 1 | 1 | 0.99 | 1 | 1 | 0.992 | 1 | 0.976 | 1 | 1 | 1 | 0.001 |
| 89188 | 136 | 102 | 141 | 173 | 52 | 0 | 1 | 0 | 0 | 0 | 215 | 149 | 1 | 1 | 1 | 1 | 1 | 0.984 | 1 | 1 | 0.986 | 1 | 1 | 1 | 0 |
| 89189 | 155 | 116 | 164 | 180 | 56 | 0 | 3 | 0 | 0 | 0 | 215 | 149 | 1 | 1 | 1 | 0.986 | 1 | 1 | 0.988 | 1 | 0.964 | 1 | 1 | 1 | 0.002 |
| 89192 | 67 | 270 | 261 | 216 | 42 | 0 | 1 | 6 | 1 | 0 | 248 | 149 | 1 | 1 | 0.992 | 0.982 | 0.985 | 1 | 1 | 1 | 0.994 | 0.963 | 0.992 | 1 | 0.002 |
| 89193 | 190 | 502 | 753 | 521 | 57 | 0 | 1 | 6 | 1 | 0 | 274 | 149 | 1 | 1 | 0.997 | 0.992 | 0.993 | 1 | 1 | 1 | 0.996 | 0.986 | 0.996 | 1 | 0.001 |
| 89195 | 244 | 699 | 973 | 689 | 92 | 0 | 0 | 0 | 5 | 0 | 257 | 149 | 1 | 1 | 1 | 1 | 1 | 1 | 1 | 1 | 1 | 1 | 0.988 | 1 | 0.002 |
| 89197 | 219 | 658 | 866 | 723 | 44 | 0 | 0 | 1 | 0 | 0 | 298 | 149 | 1 | 1 | 1 | 1 | 1 | 1 | 1 | 1 | 1 | 0.998 | 1 | 1 | 0 |
| 89200 | 31 | 54 | 89 | 53 | 10 | 0 | 0 | 1 | 0 | 0 | 186 | 149 | 1 | 1 | 1 | 1 | 1 | 1 | 1 | 1 | 1 | 0.986 | 1 | 1 | 0.003 |
| 89201 | 0 | 7 | 19 | 15 | 0 | 0 | 0 | 0 | 0 | 0 | 186 | 149 | 1 | 1 | 1 | 1 | 0.889 | 1 | 1 | NA | 1 | 1 | 1 | NA | -0.022 |
| 89202 | 29 | 14 | 28 | 7 | 5 | 0 | 0 | 0 | 0 | 0 | 216 | 149 | 1 | 1 | 1 | 1 | 0.927 | 1 | 1 | 1 | 1 | 1 | 1 | 1 | -0.015 |
| 89203 | 63 | 72 | 101 | 51 | 13 | 0 | 4 | 1 | 0 | 0 | 216 | 149 | 1 | 1 | 1 | 0.932 | 0.969 | 1 | 0.99 | 1 | 0.925 | 0.986 | 1 | 1 | -0.004 |
| 89204 | 0 | 0 | 3 | 0 | 0 | 0 | 0 | 0 | 0 | 0 | 298 | 149 | 1 | 1 | NA | NA | 1 | 0.5 | NA | NA | NA | 1 | NA | NA | -0.25 |
| 89208 | 88 | 129 | 45 | 102 | 131 | 0 | 0 | 1 | 0 | 0 | 208 | 149 | 1 | 1 | 1 | 1 | 1 | 1 | 1 | 1 | 1 | 0.97 | 1 | 1 | 0.006 |
| 89209 | 110 | 117 | 31 | 66 | 71 | 0 | 0 | 0 | 0 | 0 | 298 | 149 | 1 | 1 | 1 | 0.962 | 1 | 1 | 1 | 1 | 1 | 1 | 1 | 1 | -0.008 |
| 89210 | 185 | 166 | 50 | 126 | 207 | 0 | 5 | 0 | 0 | 0 | 298 | 149 | 1 | 1 | 1 | 1 | 1 | 1 | 1 | 1 | 0.943 | 1 | 1 | 1 | 0.011 |
| 89212 | 23 | 81 | 16 | 39 | 107 | 0 | 0 | 0 | 0 | 0 | 208 | 149 | 1 | 1 | 0.965 | 1 | 1 | 1 | 1 | 1 | 1 | 1 | 1 | 1 | -0.007 |
| 89213 | 38 | 25 | 21 | 13 | 27 | 0 | 0 | 0 | 0 | 0 | 249 | 149 | 1 | 1 | 1 | 1 | 1 | 0.893 | 1 | 1 | 1 | 1 | 1 | 1 | -0.021 |
| 89214 | 48 | 38 | 39 | 35 | 81 | 0 | 0 | 0 | 0 | 0 | 250 | 149 | 1 | 1 | 0.982 | 1 | 1 | 1 | 1 | 1 | 1 | 1 | 1 | 1 | -0.004 |
| 89215 | 47 | 87 | 46 | 76 | 124 | 0 | 5 | 0 | 0 | 0 | 235 | 149 | 1 | 1 | 1 | 1 | 1 | 1 | 1 | 1 | 0.917 | 1 | 1 | 1 | 0.017 |
| 89216 | 0 | 0 | 0 | 0 | 0 | 36 | 27 | 24 | 19 | 0 | 247 | 149 | 1 | 1 | 0.015 | 0 | 0 | 0.073 | 0 | 0 | 0 | 0 | 0 | NA | 0.018 |
| 89217 | 0 | 0 | 0 | 0 | 0 | 36 | 27 | 24 | 19 | 0 | 251 | 149 | 1 | 1 | 0 | 0 | 0 | 0.072 | 0 | 0 | 0 | 0 | 0 | NA | 0.014 |
| 89218 | 0 | 0 | 2 | 0 | 0 | 36 | 45 | 36 | 53 | 5 | 298 | 149 | 1 | 1 | 0 | 0 | 0 | 0 | 0 | 0 | 0 | 0.027 | 0 | 0 | -0.005 |
| 89219 | 56 | 114 | 64 | 99 | 6 | 0 | 4 | 0 | 0 | 0 | 210 | 149 | 1 | 1 | 1 | 1 | 1 | 1 | 1 | 1 | 0.953 | 1 | 1 | 1 | 0.009 |
| 89220 | 0 | 2 | 0 | 0 | 0 | 43 | 86 | 45 | 72 | 6 | 207 | 149 | 1 | 1 | 0 | 0.01 | 0.009 | 0 | 0.013 | 0 | 0.016 | 0 | 0 | 0 | 0.003 |
| 89221 | 0 | 1 | 0 | 0 | 0 | 0 | 0 | 0 | 0 | 0 | 196 | 149 | 1 | 1 | 0.603 | 0.603 | NA | NA | NA | NA | 1 | NA | NA | NA | -0.397 |
| 89224 | 0 | 1 | 0 | 0 | 0 | 7 | 1 | 4 | 0 | 0 | 298 | 149 | 1 | 1 | 1 | 0.273 | 0.2 | 1 | 0.556 | 0 | 0.333 | 0 | NA | NA | 0.495 |
| 89228 | 37 | 136 | 75 | 56 | 8 | 0 | 0 | 0 | 0 | 0 | 215 | 149 | 1 | 1 | 0.992 | 1 | 1 | 1 | 1 | 1 | 1 | 1 | 1 | 1 | -0.002 |
| 89229 | 0 | 2 | 0 | 0 | 0 | 0 | 0 | 0 | 0 | 0 | 214 | 149 | 1 | 1 | NA | NA | 0 | 1 | NA | NA | 1 | NA | NA | NA | -0.5 |
| 89230 | 0 | 10 | 5 | 10 | 11 | 0 | 0 | 0 | 0 | 0 | 298 | 149 | 1 | 1 | NA | NA | 0 | 1 | 1 | NA | 1 | 1 | 1 | 1 | -0.333 |
| 89231 | 113 | 98 | 61 | 93 | 125 | 0 | 0 | 0 | 0 | 0 | 234 | 149 | 1 | 1 | 0.947 | 1 | 1 | 1 | 1 | 1 | 1 | 1 | 1 | 1 | -0.011 |
| 89232 | 57 | 54 | 30 | 37 | 45 | 0 | 0 | 0 | 0 | 0 | 298 | 149 | 1 | 1 | 1 | 0.889 | 1 | 1 | 1 | 1 | 1 | 1 | 1 | 1 | -0.022 |
| 89247 | 4 | 98 | 121 | 55 | 25 | 0 | 0 | 0 | 0 | 0 | 298 | 149 | 1 | 1 | 0.986 | 1 | 1 | 1 | 1 | 1 | 1 | 1 | 1 | 1 | -0.003 |
| 89248 | 28 | 110 | 134 | 62 | 17 | 0 | 2 | 0 | 0 | 0 | 298 | 149 | 1 | 1 | 1 | 1 | 0.986 | 1 | 1 | 1 | 0.965 | 1 | 1 | 1 | 0.004 |
| 89249 | 17 | 28 | 35 | 32 | 6 | 0 | 2 | 0 | 0 | 2 | 213 | 149 | 1 | 1 | 0.94 | 1 | 1 | 1 | 1 | 1 | 0.907 | 1 | 1 | 0.677 | 0.071 |
| 89250 | 56 | 51 | 65 | 61 | 25 | 0 | 0 | 0 | 0 | 2 | 255 | 149 | 1 | 1 | 1 | 1 | 1 | 1 | 1 | 1 | 1 | 1 | 1 | 0.88 | 0.024 |
| 89251 | 14 | 22 | 10 | 17 | 7 | 0 | 0 | 0 | 0 | 2 | 269 | 149 | 1 | 1 | 1 | 1 | 1 | 1 | 1 | 1 | 1 | 1 | 1 | 0.66 | 0.068 |
| 89254 | 0 | 3 | 0 | 0 | 4 | 0 | 3 | 2 | 19 | 0 | 227 | 149 | 1 | 1 | NA | 0.141 | 0 | 0 | NA | NA | 0.396 | 0 | 0 | 1 | -0.302 |
| 89255 | 0 | 2 | 0 | 5 | 2 | 168 | 95 | 114 | 111 | 51 | 227 | 149 | 1 | 1 | 0.014 | 0.036 | 0 | 0 | 0 | 0 | 0.014 | 0 | 0.029 | 0.025 | -0.004 |
| 89256 | 12 | 9 | 7 | 16 | 0 | 0 | 0 | 0 | 0 | 0 | 298 | 149 | 1 | 1 | 0.333 | 1 | 1 | 1 | 1 | 1 | 1 | 1 | 1 | NA | -0.133 |
| 89258 | 11 | 12 | 11 | 4 | 0 | 0 | 0 | 0 | 0 | 0 | 298 | 149 | 1 | 1 | 0.385 | 1 | 1 | 1 | 1 | 1 | 1 | 1 | 1 | NA | -0.123 |
| 89260 | 11 | 9 | 8 | 4 | 0 | 0 | 7 | 0 | 0 | 0 | 298 | 149 | 1 | 1 | 0.667 | 1 | 1 | 1 | 1 | 1 | 0.391 | 1 | 1 | NA | 0.086 |
| 89261 | 11 | 9 | 7 | 4 | 0 | 0 | 4 | 11 | 4 | 0 | 298 | 149 | 1 | 1 | 0.091 | 0.167 | 0.167 | 0.05 | 0.833 | 1 | 0.529 | 0.241 | 0.333 | NA | -0.264 |
| 89262 | 0 | 4 | 2 | 2 | 0 | 0 | 2 | 0 | 5 | 0 | 298 | 149 | 1 | 1 | 0.2 | 0.2 | 1 | 1 | NA | NA | 0.5 | 1 | 0.167 | NA | 0.044 |
| 89264 | 0 | 3 | 9 | 0 | 0 | 0 | 4 | 0 | 2 | 0 | 298 | 149 | 1 | 1 | 0.667 | 1 | 1 | 0.333 | 1 | NA | 0.273 | 1 | 0 | NA | 0.376 |
| 89266 | 0 | 8 | 4 | 4 | 2 | 0 | 2 | 0 | 5 | 0 | 275 | 149 | 1 | 1 | 0.213 | 0.448 | 1 | 1 | NA | NA | 0.684 | 1 | 0.302 | 1 | -0.081 |
| 89268 | 313 | 240 | 216 | 233 | 61 | 0 | 0 | 0 | 0 | 0 | 298 | 149 | 1 | 1 | 0.99 | 1 | 1 | 1 | 1 | 1 | 1 | 1 | 1 | 1 | -0.002 |
| 89270 | 306 | 212 | 198 | 206 | 61 | 0 | 0 | 0 | 0 | 0 | 273 | 149 | 1 | 1 | 0.989 | 1 | 1 | 1 | 1 | 1 | 1 | 1 | 1 | 1 | -0.002 |
| 89272 | 686 | 535 | 459 | 464 | 153 | 12 | 0 | 1 | 0 | 0 | 249 | 149 | 1 | 1 | 0.983 | 0.982 | 0.993 | 0.989 | 0.998 | 0.972 | 1 | 0.996 | 1 | 1 | -0.005 |
| 89274 | 0 | 0 | 0 | 0 | 0 | 0 | 0 | 0 | 1 | 0 | 267 | 149 | 1 | 1 | 1 | NA | 0.527 | 1 | NA | NA | NA | NA | 0 | NA | 0.842 |
| 89279 | 46 | 52 | 34 | 29 | 67 | 0 | 0 | 0 | 4 | 0 | 278 | 149 | 1 | 1 | 1 | 1 | 1 | 1 | 1 | 1 | 1 | 1 | 0.795 | 1 | 0.041 |
| 89282 | 10 | 17 | 15 | 28 | 42 | 0 | 0 | 0 | 0 | 0 | 298 | 149 | 1 | 1 | 0.795 | 1 | 1 | 1 | 1 | 1 | 1 | 1 | 1 | 1 | -0.041 |
| 89283 | 11 | 0 | 0 | 4 | 7 | 14 | 0 | 9 | 6 | 6 | 298 | 149 | 1 | 1 | 0.429 | 0.333 | 0 | 0 | 1 | 0.282 | NA | 0 | 0.25 | 0.368 | 0.127 |
| 89284 | 0 | 0 | 0 | 4 | 0 | 0 | 0 | 0 | 0 | 0 | 298 | 149 | 1 | 1 | 0 | 1 | NA | NA | NA | NA | NA | NA | 1 | NA | -0.5 |
| 89289 | 0 | 0 | 0 | 0 | 10 | 0 | 0 | 0 | 0 | 10 | 268 | 149 | 1 | 1 | 1 | NA | 1 | 1 | NA | NA | NA | NA | NA | 0.357 | 0.643 |
| 89291 | 0 | 0 | 0 | 1 | 0 | 14 | 0 | 9 | 6 | 6 | 298 | 149 | 1 | 1 | 0.6 | 0.333 | 0.111 | 0 | NA | 0 | NA | 0 | 0.077 | 0 | 0.242 |
| 89293 | 0 | 0 | 0 | 1 | 10 | 4 | 14 | 1 | 4 | 2 | 298 | 149 | 1 | 1 | 0.167 | 0.2 | 0.2 | 0.4 | NA | 0 | 0 | 0 | 0.111 | 0.714 | 0.077 |
| 89296 | 10 | 16 | 15 | 24 | 42 | 0 | 0 | 0 | 0 | 0 | 298 | 149 | 1 | 1 | 1 | 0.846 | 1 | 1 | 1 | 1 | 1 | 1 | 1 | 1 | -0.031 |
| 89297 | 0 | 0 | 0 | 1 | 10 | 4 | 14 | 1 | 4 | 2 | 191 | 149 | 1 | 1 | 0 | 0 | 0 | 0 | NA | 0 | 0 | 0 | 0.163 | 0.796 | -0.192 |
| 89298 | 22 | 6 | 0 | 3 | 24 | 13 | 5 | 0 | 5 | 0 | 191 | 149 | 1 | 1 | 0.845 | 1 | 1 | 0.51 | 1 | 0.569 | 0.484 | NA | 0.319 | 1 | 0.278 |
| 89299 | 15 | 2 | 0 | 2 | 24 | 11 | 0 | 0 | 0 | 7 | 191 | 149 | 1 | 1 | 0.438 | 0 | 1 | 1 | 0.862 | 0.515 | 1 | NA | 1 | 0.728 | -0.151 |
| 89301 | 13 | 7 | 0 | 5 | 0 | 11 | 0 | 0 | 0 | 7 | 298 | 149 | 1 | 1 | 0.25 | 0 | NA | 1 | 0.5 | 0.371 | 1 | NA | 1 | 0 | -0.155 |
| 89304 | 0 | 0 | 6 | 2 | 4 | 0 | 2 | 0 | 0 | 0 | 298 | 149 | 1 | 1 | NA | 0.5 | 1 | NA | 0.5 | NA | 0 | 1 | 1 | 1 | -0.083 |
| 89306 | 0 | 0 | 0 | 0 | 4 | 0 | 0 | 0 | 0 | 0 | 298 | 149 | 1 | 1 | 1 | 0.5 | 1 | NA | NA | NA | NA | NA | NA | 1 | -0.167 |
| 89309 | 26 | 32 | 10 | 21 | 14 | 0 | 2 | 0 | 0 | 0 | 298 | 149 | 1 | 1 | 0.943 | 1 | 1 | 1 | 0.9 | 1 | 0.889 | 1 | 1 | 1 | -0.009 |
| 89310 | 0 | 2 | 0 | 0 | 0 | 0 | 0 | 0 | 0 | 0 | 228 | 149 | 1 | 1 | NA | 0.395 | NA | NA | NA | NA | 1 | NA | NA | NA | -0.605 |
| 89311 | 0 | 0 | 6 | 2 | 0 | 0 | 0 | 0 | 0 | 0 | 228 | 149 | 1 | 1 | 0 | 0.395 | 0.395 | NA | 1 | NA | NA | 1 | 1 | NA | -0.552 |
| 89312 | 0 | 0 | 0 | 1 | 0 | 0 | 0 | 0 | 0 | 0 | 286 | 149 | 1 | 1 | 0.51 | 1 | 1 | 1 | NA | NA | NA | NA | 1 | NA | -0.123 |
| 89313 | 19 | 10 | 2 | 0 | 3 | 0 | 0 | 0 | 0 | 0 | 286 | 149 | 1 | 1 | 0.61 | 0.701 | 0.824 | 1 | 1 | 1 | 1 | 1 | NA | 1 | -0.173 |
| 89314 | 18 | 1 | 23 | 6 | 15 | 0 | 0 | 0 | 0 | 0 | 298 | 149 | 1 | 1 | 0.75 | 1 | 1 | 1 | 1 | 1 | 1 | 1 | 1 | 1 | -0.05 |
| 89315 | 15 | 1 | 14 | 0 | 16 | 0 | 0 | 0 | 0 | 0 | 298 | 149 | 1 | 1 | 1 | 0.5 | 1 | NA | 1 | 1 | 1 | 1 | NA | 1 | -0.125 |
| 89316 | 4 | 1 | 14 | 1 | 9 | 0 | 0 | 0 | 0 | 0 | 298 | 149 | 1 | 1 | 0.75 | 1 | 1 | 1 | 1 | 1 | 1 | 1 | 1 | 1 | -0.05 |
| 89319 | 462 | 286 | 227 | 222 | 310 | 0 | 0 | 0 | 0 | 0 | 257 | 149 | 1 | 1 | 1 | 0.987 | 1 | 1 | 1 | 1 | 1 | 1 | 1 | 1 | -0.003 |
| 89328 | 4 | 4 | 4 | 2 | 0 | 0 | 5 | 2 | 0 | 7 | 193 | 149 | 1 | 1 | NA | 1 | 1 | 0.279 | NA | 1 | 0.382 | 0.607 | 1 | 0 | 0.162 |
| 89329 | 4 | 4 | 3 | 0 | 0 | 0 | 5 | 2 | 0 | 7 | 279 | 149 | 1 | 1 | NA | 1 | 1 | 0.348 | NA | 1 | 0.299 | 0.445 | NA | 0 | 0.347 |
| 89330 | 0 | 0 | 0 | 0 | 0 | 0 | 0 | 3 | 0 | 0 | 279 | 149 | 1 | 1 | 1 | 1 | NA | 1 | NA | NA | NA | 0 | NA | NA | 1 |
| 89331 | 6 | 1 | 10 | 4 | 0 | 0 | 0 | 0 | 0 | 0 | 298 | 149 | 1 | 1 | 0.857 | 1 | 1 | 1 | 1 | 1 | 1 | 1 | 1 | NA | -0.029 |
| 89333 | 4 | 28 | 9 | 14 | 7 | 0 | 0 | 0 | 0 | 0 | 298 | 149 | 1 | 1 | 0.941 | 1 | 1 | 1 | 1 | 1 | 1 | 1 | 1 | 1 | -0.012 |
| 89335 | 10 | 24 | 18 | 19 | 7 | 0 | 2 | 0 | 0 | 0 | 219 | 149 | 1 | 1 | 1 | 1 | 1 | 0.977 | 1 | 1 | 0.891 | 1 | 1 | 1 | 0.017 |
| 89336 | 22 | 61 | 75 | 48 | 45 | 0 | 4 | 0 | 0 | 0 | 264 | 149 | 1 | 1 | 0.989 | 1 | 1 | 0.974 | 0.991 | 1 | 0.896 | 1 | 1 | 1 | 0.012 |
| 89337 | 20 | 9 | 35 | 11 | 17 | 0 | 0 | 0 | 0 | 0 | 230 | 149 | 1 | 1 | 1 | 1 | 0.937 | 1 | 0.982 | 1 | 1 | 1 | 1 | 1 | -0.016 |
| 89339 | 5 | 13 | 17 | 8 | 0 | 0 | 0 | 0 | 0 | 0 | 258 | 149 | 1 | 1 | 0.914 | 1 | 1 | 1 | 1 | 1 | 1 | 1 | 1 | NA | -0.017 |
| 89340 | 16 | 3 | 22 | 7 | 0 | 0 | 0 | 0 | 0 | 0 | 298 | 149 | 1 | 1 | 1 | 1 | 1 | 0.962 | 1 | 1 | 1 | 1 | 1 | NA | -0.008 |
| 89341 | 16 | 17 | 31 | 7 | 0 | 0 | 0 | 0 | 0 | 0 | 251 | 149 | 1 | 1 | 0.976 | 1 | 1 | 1 | 1 | 1 | 1 | 1 | 1 | NA | -0.005 |
| 89352 | 0 | 2 | 11 | 0 | 0 | 0 | 0 | 0 | 0 | 0 | 246 | 149 | 1 | 1 | 1 | 1 | 0.784 | 1 | 1 | NA | 1 | 1 | NA | NA | -0.043 |
| 89353 | 172 | 189 | 159 | 90 | 40 | 0 | 0 | 0 | 0 | 0 | 271 | 149 | 1 | 1 | 1 | 0.996 | 0.996 | 1 | 1 | 1 | 1 | 1 | 1 | 1 | -0.002 |
| 89358 | 253 | 205 | 173 | 127 | 214 | 0 | 0 | 0 | 0 | 3 | 182 | 149 | 1 | 1 | 1 | 1 | 1 | 1 | 1 | 1 | 1 | 1 | 1 | 0.983 | 0.003 |
| 89359 | 54 | 65 | 76 | 57 | 78 | 0 | 0 | 0 | 0 | 0 | 200 | 149 | 1 | 1 | 1 | 0.994 | 1 | 1 | 1 | 1 | 1 | 1 | 1 | 1 | -0.001 |
| 89361 | 24 | 196 | 169 | 44 | 28 | 0 | 0 | 2 | 2 | 0 | 242 | 149 | 1 | 1 | 0.995 | 0.995 | 1 | 1 | 1 | 1 | 1 | 0.981 | 0.931 | 1 | 0.016 |
| 89362 | 0 | 32 | 22 | 7 | 0 | 0 | 0 | 0 | 0 | 0 | 213 | 149 | 1 | 1 | 0.988 | 1 | 1 | 0.968 | 1 | NA | 1 | 1 | 1 | NA | -0.009 |
| 89363 | 145 | 122 | 81 | 68 | 242 | 0 | 0 | 0 | 0 | 0 | 298 | 149 | 1 | 1 | 0.973 | 1 | 1 | 1 | 1 | 1 | 1 | 1 | 1 | 1 | -0.005 |
| 89364 | 238 | 188 | 132 | 101 | 422 | 0 | 0 | 0 | 0 | 0 | 298 | 149 | 1 | 1 | 0.985 | 1 | 1 | 1 | 1 | 1 | 1 | 1 | 1 | 1 | -0.003 |
| 89365 | 145 | 118 | 78 | 68 | 259 | 0 | 0 | 0 | 0 | 0 | 291 | 149 | 1 | 1 | 0.974 | 1 | 1 | 1 | 1 | 1 | 1 | 1 | 1 | 1 | -0.005 |
| 89366 | 42 | 114 | 66 | 67 | 133 | 0 | 0 | 0 | 0 | 0 | 298 | 149 | 1 | 1 | 0.963 | 0.879 | 1 | 1 | 1 | 1 | 1 | 1 | 1 | 1 | -0.032 |
| 89367 | 34 | 55 | 47 | 34 | 67 | 0 | 0 | 0 | 0 | 0 | 237 | 149 | 1 | 1 | 1 | 1 | 1 | 1 | 0.955 | 1 | 1 | 1 | 1 | 1 | -0.009 |
| 89369 | 308 | 287 | 190 | 399 | 460 | 0 | 0 | 0 | 0 | 0 | 256 | 149 | 1 | 1 | 1 | 0.986 | 1 | 1 | 1 | 1 | 1 | 1 | 1 | 1 | -0.003 |
| 89376 | 1584 | 1714 | 1669 | 2190 | 1161 | 0 | 0 | 0 | 0 | 0 | 210 | 149 | 1 | 1 | 1 | 1 | 1 | 0.999 | 1 | 1 | 1 | 1 | 1 | 1 | 0 |
| 89377 | 0 | 0 | 0 | 0 | 0 | 1362 | 1542 | 1428 | 1890 | 1030 | 238 | 149 | 1 | 1 | 0 | 0.001 | 0.004 | 0.003 | 0.003 | 0 | 0 | 0 | 0 | 0 | 0.002 |
| 89385 | 0 | 0 | 0 | 11 | 2 | 0 | 0 | 0 | 0 | 0 | 217 | 149 | 1 | 1 | 0.733 | 0.846 | 1 | 1 | NA | NA | NA | NA | 1 | 1 | -0.105 |
| 89386 | 0 | 1 | 1 | 11 | 2 | 0 | 0 | 0 | 0 | 0 | 255 | 149 | 1 | 1 | 0.903 | 0.898 | 1 | 1 | 1 | NA | 1 | 1 | 1 | 1 | -0.04 |
| 89387 | 0 | 1 | 0 | 11 | 2 | 0 | 0 | 0 | 0 | 0 | 272 | 149 | 1 | 1 | 0.687 | 0.858 | 1 | 1 | 1 | NA | 1 | NA | 1 | 1 | -0.091 |
| 89388 | 0 | 0 | 1 | 0 | 0 | 0 | 0 | 1 | 0 | 0 | 298 | 149 | 1 | 1 | 0 | 0 | 0 | 0 | 0 | NA | NA | 0.333 | NA | NA | -0.333 |
| 89390 | 6 | 50 | 52 | 11 | 2 | 0 | 0 | 0 | 0 | 0 | 240 | 149 | 1 | 1 | 1 | 0.978 | 1 | 1 | 1 | 1 | 1 | 1 | 1 | 1 | -0.004 |
| 89391 | 3 | 75 | 66 | 33 | 15 | 0 | 0 | 0 | 0 | 0 | 288 | 149 | 1 | 1 | 1 | 1 | 1 | 1 | 0.978 | 1 | 1 | 1 | 1 | 1 | -0.004 |
| 89392 | 48 | 155 | 218 | 103 | 49 | 0 | 0 | 0 | 0 | 0 | 274 | 149 | 1 | 1 | 0.988 | 0.988 | 1 | 1 | 1 | 1 | 1 | 1 | 1 | 1 | -0.005 |
| 89393 | 85 | 158 | 227 | 119 | 64 | 0 | 0 | 0 | 0 | 0 | 287 | 149 | 1 | 1 | 1 | 1 | 1 | 0.995 | 1 | 1 | 1 | 1 | 1 | 1 | -0.001 |
| 89397 | 20 | 141 | 175 | 61 | 19 | 0 | 0 | 0 | 0 | 0 | 238 | 149 | 1 | 1 | 1 | 1 | 1 | 0.993 | 1 | 1 | 1 | 1 | 1 | 1 | -0.001 |
| 89398 | 0 | 3 | 0 | 0 | 0 | 4 | 32 | 44 | 20 | 0 | 257 | 149 | 1 | 1 | 0 | 0 | 0 | 0.007 | 0 | 0 | 0.052 | 0 | 0 | NA | -0.012 |
| 89399 | 14 | 110 | 122 | 62 | 21 | 0 | 0 | 1 | 0 | 0 | 298 | 149 | 1 | 1 | 1 | 1 | 1 | 1 | 1 | 1 | 1 | 0.984 | 1 | 1 | 0.003 |
| 89400 | 15 | 115 | 146 | 61 | 15 | 0 | 0 | 0 | 0 | 0 | 283 | 149 | 1 | 1 | 1 | 1 | 0.953 | 1 | 1 | 1 | 1 | 1 | 1 | 1 | -0.009 |
| 89401 | 36 | 233 | 299 | 73 | 30 | 0 | 0 | 0 | 0 | 0 | 270 | 149 | 1 | 1 | 0.98 | 0.99 | 0.995 | 0.993 | 1 | 1 | 1 | 1 | 1 | 1 | -0.008 |
| 89416 | 0 | 0 | 0 | 0 | 4 | 37 | 73 | 21 | 8 | 156 | 298 | 149 | 1 | 1 | 0 | 0 | 0 | 0 | 0 | 0 | 0 | 0 | 0 | 0.013 | -0.003 |
| 89419 | 11 | 5 | 5 | 0 | 14 | 0 | 0 | 0 | 0 | 0 | 275 | 149 | 1 | 1 | 1 | 0 | NA | 1 | 1 | 1 | 1 | 1 | NA | 1 | -0.25 |
| 89420 | 0 | 28 | 0 | 2 | 36 | 0 | 5 | 0 | 0 | 0 | 267 | 149 | 1 | 1 | 1 | 1 | NA | 0.157 | 1 | NA | 0.758 | NA | 1 | 1 | -0.13 |
| 89421 | 15 | 23 | 10 | 4 | 143 | 0 | 0 | 0 | 0 | 0 | 264 | 149 | 1 | 1 | 1 | 1 | 1 | 0.756 | 1 | 1 | 1 | 1 | 1 | 1 | -0.049 |
| 89423 | 15 | 23 | 10 | 4 | 149 | 0 | 0 | 0 | 0 | 0 | 264 | 149 | 1 | 1 | 1 | 1 | 1 | 1 | 0.809 | 1 | 1 | 1 | 1 | 1 | -0.038 |
| 89424 | 15 | 36 | 12 | 7 | 143 | 0 | 0 | 0 | 0 | 0 | 264 | 149 | 1 | 1 | 1 | 0.756 | 1 | 1 | 1 | 1 | 1 | 1 | 1 | 1 | -0.049 |
| 89425 | 23 | 23 | 10 | 4 | 143 | 0 | 0 | 0 | 0 | 0 | 264 | 149 | 1 | 1 | 1 | 1 | 1 | 0.756 | 1 | 1 | 1 | 1 | 1 | 1 | -0.049 |
| 89426 | 0 | 7 | 2 | 3 | 0 | 0 | 0 | 0 | 0 | 0 | 264 | 149 | 1 | 1 | NA | 1 | 1 | 0.629 | NA | NA | 1 | 1 | 1 | NA | -0.124 |
| 89427 | 0 | 0 | 0 | 0 | 7 | 0 | 0 | 0 | 0 | 0 | 272 | 149 | 1 | 1 | NA | NA | NA | 0 | NA | NA | NA | NA | NA | 1 | -1 |
| 89428 | 0 | 0 | 0 | 0 | 2 | 0 | 0 | 0 | 0 | 6 | 210 | 149 | 1 | 1 | NA | NA | NA | NA | 1 | NA | NA | NA | NA | 0.191 | 0.809 |
| 89429 | 0 | 0 | 0 | 0 | 6 | 0 | 13 | 2 | 3 | 0 | 298 | 149 | 1 | 1 | NA | 0.2 | NA | 0 | 1 | NA | 0 | 0 | 0 | 1 | 0.15 |
| 89430 | 0 | 5 | 2 | 0 | 0 | 0 | 0 | 0 | 0 | 0 | 290 | 149 | 1 | 1 | NA | 1 | NA | 0 | 1 | NA | 1 | 1 | NA | NA | -0.333 |
| 89431 | 0 | 0 | 0 | 0 | 2 | 0 | 0 | 0 | 0 | 0 | 243 | 149 | 1 | 1 | 0 | NA | NA | NA | NA | NA | NA | NA | NA | 1 | -1 |
| 89433 | 11 | 35 | 25 | 5 | 6 | 0 | 0 | 0 | 0 | 0 | 298 | 149 | 1 | 1 | 1 | 0.965 | 1 | 0.978 | 1 | 1 | 1 | 1 | 1 | 1 | -0.011 |
| 89434 | 62 | 123 | 111 | 10 | 5 | 0 | 0 | 0 | 0 | 0 | 284 | 149 | 1 | 1 | 1 | 0.991 | 1 | 0.993 | 1 | 1 | 1 | 1 | 1 | 1 | -0.003 |
| 89435 | 35 | 23 | 46 | 0 | 14 | 0 | 0 | 0 | 0 | 0 | 231 | 149 | 1 | 1 | 1 | 1 | 1 | 0.984 | 0.991 | 1 | 1 | 1 | NA | 1 | -0.005 |
| 89437 | 0 | 65 | 177 | 16 | 10 | 0 | 2 | 0 | 1 | 0 | 271 | 149 | 1 | 1 | 0.993 | 0.992 | 1 | 1 | 1 | NA | 0.947 | 1 | 0.898 | 1 | 0.036 |
| 89439 | 0 | 0 | 2 | 0 | 0 | 0 | 36 | 44 | 5 | 9 | 232 | 149 | 1 | 1 | 0 | 0.047 | 0.022 | 0.03 | 0.02 | NA | 0 | 0.028 | 0 | 0 | 0.017 |
| 89442 | 52 | 96 | 80 | 106 | 158 | 0 | 0 | 0 | 5 | 0 | 298 | 149 | 1 | 1 | 1 | 1 | 0.978 | 1 | 1 | 1 | 1 | 1 | 0.914 | 1 | 0.013 |
| 89445 | 795 | 702 | 523 | 515 | 691 | 0 | 0 | 0 | 0 | 0 | 298 | 149 | 1 | 1 | 1 | 0.994 | 1 | 1 | 1 | 1 | 1 | 1 | 1 | 1 | -0.001 |
| 89446 | 615 | 442 | 317 | 303 | 521 | 0 | 0 | 0 | 0 | 0 | 210 | 149 | 1 | 1 | 1 | 1 | 1 | 1 | 0.998 | 1 | 1 | 1 | 1 | 1 | 0 |
| 89452 | 0 | 1 | 1 | 0 | 0 | 0 | 1 | 1 | 0 | 0 | 298 | 149 | 1 | 1 | 0.636 | 0.6 | 0.556 | 0.529 | 0.5 | NA | 0.333 | 0.333 | NA | NA | 0.231 |
| 89453 | 0 | 1 | 1 | 5 | 0 | 0 | 0 | 0 | 0 | 0 | 298 | 149 | 1 | 1 | 1 | 1 | 1 | 0.429 | 1 | NA | 1 | 1 | 1 | NA | -0.114 |
| 89454 | 0 | 1 | 1 | 2 | 0 | 0 | 0 | 0 | 0 | 0 | 298 | 149 | 1 | 1 | 1 | 1 | 0.818 | 1 | 1 | NA | 1 | 1 | 1 | NA | -0.036 |
| 89455 | 0 | 12 | 4 | 2 | 0 | 0 | 0 | 0 | 3 | 0 | 298 | 149 | 1 | 1 | 1 | 1 | 1 | 1 | 1 | NA | 1 | 1 | 0.25 | NA | 0.25 |
| 89456 | 0 | 12 | 4 | 2 | 0 | 0 | 0 | 0 | 0 | 0 | 298 | 149 | 1 | 1 | 1 | 0.889 | 1 | 1 | 1 | NA | 1 | 1 | 1 | NA | -0.022 |
| 89457 | 0 | 10 | 0 | 2 | 0 | 0 | 0 | 0 | 0 | 0 | 298 | 149 | 1 | 1 | 1 | 1 | 0.75 | 1 | 1 | NA | 1 | NA | 1 | NA | -0.05 |

| | | | | | | | | | | | | | | | | | | | | | | | | | | | | | | | | | | | | | | | | | | | | | | |
|---|---|---|---|---|---|---|---|---|---|---|---|---|---|---|---|---|---|---|---|---|---|---|---|---|---|---|---|---|---|---|---|---|---|---|---|---|---|---|---|---|---|---|---|---|---|---|
| SE | 89458 | ENSG00000133739 | LRRCC1 | chr8 | + | 85123412 | 85123606 | 85115099 | 85115275 | 85124791 | 85124939 | 3 | 11 | 6 | 3 | 10 | 0 | 1 | 0 | 0 | 0 | 0 | 10 | 0 | 5 | 0 | 0 | 0 | 0 | 0 | 0 | 298 | 149 | 1 | 1 | 1 | 0.846 | 1 | 1 | 1 | NA | 1 | NA | 1 | NA | -0.031 |
| SE | 89459 | ENSG00000133739 | LRRCC1 | chr8 | + | 85123412 | 85123606 | 85115374 | 85115584 | 85124791 | 85124939 | 16 | 23 | 21 | 8 | 32 | 4 | 2 | 2 | 0 | 0 | 0 | 10 | 0 | 2 | 0 | 0 | 0 | 0 | 0 | 0 | 298 | 149 | 1 | 1 | 0.667 | 0.852 | 0.84 | 1 | 1 | NA | 1 | NA | 1 | NA | -0.128 |
| SE | 89460 | ENSG00000133739 | LRRCC1 | chr8 | + | 85129174 | 85129379 | 85126688 | 85126837 | 85129918 | 85130058 | 31 | 20 | 11 | 5 | 25 | 0 | 0 | 0 | 1 | 0 | 0 | 0 | 0 | 0 | 0 | 0 | 1 | 0 | 0 | 0 | 298 | 149 | 1 | 1 | 1 | 1 | 1 | 0.714 | 1 | NA | 0 | NA | NA | NA | 0.943 |
| SE | 89461 | ENSG00000133739 | LRRCC1 | chr8 | + | 85134846 | 85135032 | 85131759 | 85131961 | 85135788 | 85135963 | 48 | 28 | 29 | 47 | 59 | 0 | 0 | 0 | 0 | 4 | 0 | 19 | 0 | 0 | 0 | 3 | 0 | 0 | 0 | 0 | 298 | 149 | 1 | 1 | 1 | 1 | 1 | 1 | 0.881 | 0 | 1 | NA | NA | NA | 0.476 |
| SE | 89462 | ENSG00000226091 | LINC00937 | chr12 | - | 8370212 | 8370429 | 8356962 | 8357141 | 8390269 | 8390969 | 1 | 0 | 0 | 5 | 2 | 2 | 0 | 1 | 0 | 0 | 0 | 0 | 0 | 4 | 0 | 0 | 0 | 1 | 0 | 0 | 298 | 149 | 1 | 1 | 0.2 | NA | 0 | 1 | 1 | NA | NA | 0 | 1 | NA | 0.05 |
| SE | 89464 | ENSG00000226091 | LINC00937 | chr12 | - | 8385143 | 8385299 | 8382843 | 8383075 | 8385687 | 8385846 | 0 | 7 | 4 | 2 | 2 | 0 | 0 | 1 | 0 | 0 | 6 | 0 | 1 | 6 | 0 | 0 | 0 | 0 | 3 | 0 | 298 | 149 | 1 | 1 | NA | 1 | 0.667 | 1 | 1 | 1 | NA | 1 | 0.5 | NA | 0.083 |
| SE | 89465 | ENSG00000226091 | LINC00937 | chr12 | - | 8385143 | 8385299 | 8384104 | 8384421 | 8385687 | 8385846 | 0 | 7 | 6 | 5 | 1 | 0 | 1 | 1 | 1 | 0 | 15 | 0 | 10 | 19 | 9 | 0 | 0 | 0 | 0 | 0 | 298 | 149 | 1 | 1 | NA | 0.778 | 0.75 | 0.714 | 1 | 1 | NA | 1 | 1 | 1 | -0.19 |
| SE | 89466 | ENSG00000226091 | LINC00937 | chr12 | - | 8390901 | 8390969 | 8390269 | 8390511 | 8396422 | 8396495 | 4 | 2 | 2 | 2 | 0 | 0 | 0 | 2 | 0 | 0 | 10 | 3 | 3 | 4 | 0 | 0 | 0 | 0 | 0 | 0 | 217 | 149 | 1 | 1 | 1 | 1 | 0.407 | 1 | NA | 1 | 1 | 1 | 1 | NA | -0.148 |
| SE | 89467 | ENSG00000145779 | TNFAIP8 | chr5 | + | 119316184 | 119316302 | 119268753 | 119268907 | 119392815 | 119393084 | 1 | 0 | 3 | 1 | 3 | 210 | 74 | 240 | 294 | 32 | 0 | 0 | 0 | 0 | 0 | 58 | 135 | 118 | 155 | 76 | 267 | 149 | 1 | 1 | 0.003 | 0 | 0.007 | 0.002 | 0.05 | 0 | 0 | 0 | 0 | 0 | 0.012 |
| SE | 89470 | ENSG00000145779 | TNFAIP8 | chr5 | + | 119391364 | 119391483 | 119268753 | 119268907 | 119392815 | 119394599 | 0 | 2 | 0 | 0 | 0 | 210 | 74 | 240 | 294 | 32 | 0 | 8 | 0 | 0 | 0 | 58 | 135 | 118 | 155 | 76 | 268 | 149 | 1 | 1 | 0 | 0.015 | 0 | 0 | 0 | 0 | 0.032 | 0 | 0 | 0 | -0.003 |
| SE | 89471 | ENSG00000145779 | TNFAIP8 | chr5 | + | 119391364 | 119391483 | 119356034 | 119356121 | 119392815 | 119394599 | 0 | 2 | 0 | 0 | 0 | 66 | 50 | 41 | 35 | 32 | 0 | 4 | 0 | 0 | 0 | 13 | 13 | 37 | 11 | 0 | 268 | 149 | 1 | 1 | 0 | 0.022 | 0 | 0 | 0 | 0 | 0.146 | 0 | 0 | NA | -0.032 |
| SE | 89472 | ENSG00000064313 | TAF2 | chr8 | - | 119744880 | 119745036 | 119744287 | 119744393 | 119746704 | 119746934 | 6 | 4 | 0 | 10 | 10 | 42 | 34 | 22 | 41 | 65 | 0 | 0 | 1 | 0 | 0 | 0 | 39 | 16 | 11 | 6 | 298 | 149 | 1 | 1 | 0.067 | 0.056 | 0 | 0.109 | 0.071 | NA | 0 | 0.03 | 0 | 0 | 0.053 |
| SE | 89473 | ENSG00000064313 | TAF2 | chr8 | - | 119744880 | 119745036 | 119744287 | 119744393 | 119756005 | 119756115 | 2 | 2 | 0 | 8 | 2 | 3 | 1 | 0 | 2 | 0 | 0 | 0 | 2 | 0 | 0 | 0 | 0 | 2 | 0 | 0 | 298 | 149 | 1 | 1 | 0.25 | 0.5 | NA | 0.667 | 1 | NA | NA | 0.333 | NA | NA | 0.271 |
| SE | 89474 | ENSG00000064313 | TAF2 | chr8 | - | 119746704 | 119746934 | 119744287 | 119744393 | 119756005 | 119756115 | 94 | 79 | 50 | 78 | 135 | 3 | 1 | 0 | 2 | 0 | 0 | 63 | 47 | 19 | 12 | 0 | 0 | 2 | 0 | 0 | 298 | 149 | 1 | 1 | 0.94 | 0.975 | 1 | 0.951 | 1 | NA | 1 | 0.922 | 1 | 1 | -0.007 |
| SE | 89475 | ENSG00000064313 | TAF2 | chr8 | - | 119746704 | 119746934 | 119744880 | 119745036 | 119756005 | 119756115 | 56 | 47 | 28 | 39 | 78 | 0 | 0 | 0 | 0 | 0 | 0 | 24 | 31 | 8 | 6 | 0 | 0 | 1 | 0 | 0 | 298 | 149 | 1 | 1 | 1 | 1 | 1 | 1 | 1 | NA | 1 | 0.939 | 1 | 1 | 0.015 |
| SE | 89478 | ENSG00000064313 | TAF2 | chr8 | - | 119783380 | 119783633 | 119781052 | 119781193 | 119785200 | 119785266 | 156 | 136 | 128 | 114 | 252 | 2 | 0 | 0 | 0 | 0 | 65 | 115 | 58 | 33 | 44 | 0 | 2 | 0 | 0 | 0 | 298 | 149 | 1 | 1 | 0.975 | 1 | 1 | 1 | 1 | 1 | 0.966 | 1 | 1 | 1 | 0.002 |
| SE | 89479 | ENSG00000064313 | TAF2 | chr8 | - | 119788337 | 119788447 | 119785200 | 119785266 | 119788789 | 119788904 | 135 | 121 | 91 | 114 | 164 | 0 | 0 | 0 | 0 | 0 | 63 | 137 | 113 | 24 | 32 | 0 | 0 | 0 | 2 | 0 | 259 | 149 | 1 | 1 | 1 | 1 | 1 | 1 | 1 | 1 | 1 | 1 | 0.873 | 1 | 0.025 |
| SE | 89480 | ENSG00000064313 | TAF2 | chr8 | - | 119791323 | 119791459 | 119789591 | 119789746 | 119793365 | 119793451 | 206 | 149 | 188 | 205 | 170 | 1 | 0 | 0 | 0 | 0 | 111 | 151 | 106 | 66 | 104 | 0 | 0 | 0 | 0 | 0 | 285 | 149 | 1 | 1 | 0.991 | 1 | 1 | 1 | 1 | 1 | 1 | 1 | 1 | 1 | -0.002 |
| SE | 89481 | ENSG00000064313 | TAF2 | chr8 | - | 119791323 | 119791459 | 119789591 | 119789746 | 119795531 | 119795631 | 149 | 126 | 144 | 157 | 110 | 0 | 0 | 0 | 0 | 0 | 105 | 118 | 119 | 59 | 98 | 0 | 0 | 6 | 0 | 0 | 285 | 149 | 1 | 1 | 1 | 1 | 1 | 1 | 1 | 1 | 1 | 0.912 | 1 | 1 | 0.018 |
| SE | 89485 | ENSG00000064313 | TAF2 | chr8 | - | 119801793 | 119802025 | 119797661 | 119797846 | 119803877 | 119804019 | 186 | 69 | 113 | 115 | 323 | 0 | 3 | 0 | 1 | 0 | 0 | 39 | 44 | 12 | 53 | 0 | 7 | 10 | 1 | 0 | 298 | 149 | 1 | 1 | 1 | 0.92 | 1 | 0.983 | 1 | NA | 0.736 | 0.688 | 0.857 | 1 | 0.16 |
| SE | 89486 | ENSG00000064313 | TAF2 | chr8 | - | 119803877 | 119804019 | 119801793 | 119802025 | 119806282 | 119806401 | 120 | 62 | 88 | 140 | 210 | 0 | 0 | 0 | 1 | 5 | 18 | 41 | 51 | 29 | 12 | 0 | 0 | 0 | 0 | 0 | 291 | 149 | 1 | 1 | 1 | 1 | 1 | 0.986 | 0.956 | 1 | 1 | 1 | 1 | 1 | -0.012 |
| SE | 89496 | ENSG00000156162 | DPY19L4 | chr8 | + | 94726330 | 94726441 | 94719702 | 94720014 | 94738368 | 94738459 | 23 | 28 | 21 | 18 | 20 | 0 | 1 | 0 | 0 | 0 | 0 | 19 | 9 | 17 | 6 | 0 | 0 | 0 | 0 | 0 | 260 | 149 | 1 | 1 | 1 | 0.941 | 1 | 1 | 1 | NA | 1 | 1 | 1 | 1 | -0.012 |
| SE | 89497 | ENSG00000156162 | DPY19L4 | chr8 | + | 94734629 | 94734754 | 94726330 | 94726441 | 94738368 | 94738459 | 58 | 58 | 75 | 91 | 144 | 14 | 19 | 19 | 9 | 18 | 0 | 14 | 21 | 40 | 25 | 0 | 17 | 7 | 8 | 0 | 274 | 149 | 1 | 1 | 0.693 | 0.624 | 0.682 | 0.846 | 0.813 | NA | 0.309 | 0.62 | 0.731 | 1 | 0.067 |
| SE | 89498 | ENSG00000156162 | DPY19L4 | chr8 | + | 94734629 | 94734754 | 94726330 | 94726441 | 94739412 | 94739534 | 29 | 25 | 35 | 45 | 74 | 0 | 1 | 0 | 0 | 3 | 0 | 4 | 10 | 26 | 14 | 0 | 0 | 0 | 0 | 0 | 274 | 149 | 1 | 1 | 1 | 0.931 | 1 | 1 | 0.931 | NA | 1 | 1 | 1 | 1 | -0.028 |
| SE | 89499 | ENSG00000156162 | DPY19L4 | chr8 | + | 94738368 | 94738459 | 94726330 | 94726441 | 94739412 | 94739534 | 32 | 25 | 37 | 21 | 78 | 0 | 1 | 0 | 0 | 3 | 0 | 24 | 19 | 14 | 9 | 0 | 0 | 0 | 0 | 0 | 240 | 149 | 1 | 1 | 1 | 0.939 | 1 | 1 | 0.942 | NA | 1 | 1 | 1 | 1 | -0.024 |
| SE | 89500 | ENSG00000156162 | DPY19L4 | chr8 | + | 94738368 | 94738459 | 94734629 | 94734754 | 94739412 | 94739534 | 48 | 40 | 61 | 62 | 146 | 1 | 1 | 1 | 6 | 12 | 0 | 17 | 24 | 24 | 20 | 0 | 0 | 1 | 4 | 0 | 240 | 149 | 1 | 1 | 0.968 | 0.961 | 0.974 | 0.865 | 0.883 | NA | 1 | 0.937 | 0.788 | 1 | -0.001 |
| SE | 89501 | ENSG00000156162 | DPY19L4 | chr8 | + | 94739412 | 94739534 | 94738368 | 94738459 | 94739644 | 94739725 | 97 | 94 | 106 | 100 | 157 | 0 | 0 | 2 | 1 | 0 | 9 | 39 | 43 | 48 | 43 | 0 | 0 | 0 | 0 | 0 | 271 | 149 | 1 | 1 | 1 | 1 | 0.967 | 0.982 | 1 | 1 | 1 | 1 | 1 | 1 | -0.01 |
| SE | 89503 | ENSG00000156162 | DPY19L4 | chr8 | + | 94761699 | 94761834 | 94739644 | 94739790 | 94766611 | 94766685 | 4 | 3 | 10 | 7 | 1 | 0 | 0 | 0 | 2 | 0 | 0 | 4 | 1 | 0 | 18 | 0 | 0 | 0 | 0 | 0 | 284 | 149 | 1 | 1 | 1 | 1 | 1 | 0.647 | 1 | NA | 1 | 1 | NA | 1 | -0.071 |
| SE | 89504 | ENSG00000156162 | DPY19L4 | chr8 | + | 94761699 | 94761834 | 94756035 | 94756159 | 94765182 | 94765314 | 52 | 38 | 38 | 34 | 39 | 0 | 0 | 0 | 0 | 2 | 18 | 6 | 14 | 11 | 22 | 0 | 0 | 0 | 0 | 0 | 284 | 149 | 1 | 1 | 1 | 1 | 1 | 1 | 0.911 | 1 | 1 | 1 | 1 | 1 | -0.018 |
| SE | 89506 | ENSG00000156162 | DPY19L4 | chr8 | + | 94765182 | 94765314 | 94761699 | 94761834 | 94766611 | 94766685 | 13 | 14 | 13 | 10 | 23 | 0 | 0 | 2 | 0 | 0 | 5 | 0 | 3 | 4 | 17 | 0 | 0 | 0 | 0 | 0 | 281 | 149 | 1 | 1 | 1 | 1 | 0.775 | 1 | 1 | 1 | NA | 1 | 1 | 1 | -0.045 |
| SE | 89507 | ENSG00000156162 | DPY19L4 | chr8 | + | 94765710 | 94765809 | 94761699 | 94761834 | 94766611 | 94766685 | 30 | 18 | 36 | 10 | 16 | 0 | 0 | 2 | 0 | 0 | 8 | 8 | 10 | 17 | 10 | 0 | 0 | 0 | 0 | 0 | 248 | 149 | 1 | 1 | 1 | 1 | 0.915 | 1 | 1 | 1 | 1 | 1 | 1 | 1 | -0.017 |
| SE | 89508 | ENSG00000156162 | DPY19L4 | chr8 | + | 94765710 | 94765809 | 94765182 | 94765314 | 94766611 | 94766685 | 64 | 51 | 79 | 41 | 52 | 1 | 0 | 0 | 0 | 0 | 13 | 11 | 18 | 23 | 30 | 0 | 0 | 0 | 0 | 0 | 248 | 149 | 1 | 1 | 0.975 | 1 | 1 | 1 | 1 | 1 | 1 | 1 | 1 | 1 | -0.005 |
| SE | 89509 | ENSG00000156162 | DPY19L4 | chr8 | + | 94781083 | 94781166 | 94780358 | 94780415 | 94783669 | 94783802 | 41 | 32 | 39 | 39 | 106 | 1 | 0 | 0 | 0 | 1 | 0 | 22 | 19 | 7 | 0 | 0 | 0 | 0 | 1 | 0 | 232 | 149 | 1 | 1 | 0.963 | 1 | 1 | 1 | 0.986 | NA | 1 | 1 | 0.818 | NA | 0.05 |
| SE | 89510 | ENSG00000108239 | TBC1D12 | chr10 | + | 94497054 | 94497172 | 94493364 | 94493447 | 94500220 | 94500327 | 24 | 47 | 22 | 36 | 89 | 0 | 0 | 0 | 1 | 0 | 32 | 0 | 40 | 53 | 1 | 0 | 0 | 0 | 0 | 0 | 267 | 149 | 1 | 1 | 1 | 1 | 1 | 0.953 | 1 | 1 | NA | 1 | 1 | 1 | -0.009 |
| SE | 89511 | ENSG00000108239 | TBC1D12 | chr10 | + | 94500220 | 94500327 | 94497054 | 94497172 | 94507266 | 94507347 | 16 | 30 | 11 | 20 | 59 | 0 | 0 | 1 | 0 | 0 | 18 | 6 | 14 | 16 | 0 | 0 | 0 | 0 | 0 | 0 | 256 | 149 | 1 | 1 | 1 | 1 | 0.865 | 1 | 1 | 1 | 1 | 1 | 1 | NA | -0.027 |
| SE | 89512 | ENSG00000108239 | TBC1D12 | chr10 | + | 94500220 | 94500327 | 94497054 | 94497172 | 94510090 | 94510179 | 7 | 26 | 8 | 16 | 41 | 0 | 0 | 0 | 0 | 0 | 6 | 0 | 8 | 10 | 0 | 0 | 0 | 1 | 0 | 0 | 256 | 149 | 1 | 1 | 1 | 1 | 1 | 1 | 1 | 1 | NA | 0.823 | 1 | NA | 0.059 |
| SE | 89513 | ENSG00000108239 | TBC1D12 | chr10 | + | 94507266 | 94507347 | 94497054 | 94497172 | 94510090 | 94510179 | 4 | 8 | 8 | 17 | 14 | 0 | 0 | 0 | 0 | 0 | 20 | 4 | 15 | 5 | 0 | 0 | 0 | 1 | 0 | 0 | 230 | 149 | 1 | 1 | 1 | 1 | 1 | 1 | 1 | 1 | 1 | 0.907 | 1 | NA | 0.023 |
| SE | 89514 | ENSG00000108239 | TBC1D12 | chr10 | + | 94507266 | 94507347 | 94500220 | 94500327 | 94510090 | 94510179 | 13 | 15 | 10 | 24 | 31 | 0 | 3 | 0 | 2 | 0 | 32 | 7 | 21 | 9 | 0 | 0 | 0 | 0 | 0 | 0 | 230 | 149 | 1 | 1 | 1 | 0.764 | 1 | 0.886 | 1 | 1 | 1 | 1 | 1 | NA | -0.07 |
| SE | 89515 | ENSG00000108239 | TBC1D12 | chr10 | + | 94510090 | 94510179 | 94500220 | 94500327 | 94511582 | 94511654 | 1 | 9 | 1 | 5 | 8 | 0 | 1 | 0 | 0 | 0 | 0 | 0 | 1 | 0 | 0 | 0 | 0 | 0 | 0 | 0 | 238 | 149 | 1 | 1 | 1 | 0.849 | 1 | 1 | 1 | NA | NA | 1 | NA | NA | -0.03 |
| SE | 89516 | ENSG00000108239 | TBC1D12 | chr10 | + | 94521954 | 94522083 | 94511582 | 94511654 | 94531201 | 94531460 | 2 | 3 | 1 | 4 | 15 | 1 | 0 | 0 | 0 | 0 | 0 | 0 | 1 | 0 | 0 | 0 | 0 | 1 | 0 | 0 | 278 | 149 | 1 | 1 | 0.517 | 1 | 1 | 1 | 1 | NA | NA | 0.349 | NA | NA | 0.554 |
| SE | 89517 | ENSG00000108239 | TBC1D12 | chr10 | + | 94522343 | 94522453 | 94521954 | 94522083 | 94531201 | 94531460 | 20 | 17 | 10 | 40 | 28 | 1 | 0 | 0 | 1 | 0 | 0 | 0 | 13 | 17 | 0 | 0 | 0 | 0 | 0 | 0 | 259 | 149 | 1 | 1 | 0.92 | 1 | 1 | 0.958 | 1 | NA | NA | 1 | 1 | NA | -0.024 |
| SE | 89518 | ENSG00000108239 | TBC1D12 | chr10 | + | 94531201 | 94531460 | 94522343 | 94522453 | 94533027 | 94533297 | 25 | 21 | 21 | 46 | 39 | 0 | 0 | 0 | 0 | 5 | 0 | 0 | 31 | 36 | 0 | 0 | 0 | 0 | 0 | 0 | 298 | 149 | 1 | 1 | 1 | 1 | 1 | 1 | 0.796 | NA | NA | 1 | 1 | NA | -0.041 |
| SE | 89529 | ENSG00000110074 | FOXRED1 | chr11 | + | 126271219 | 126271657 | 126269200 | 126269291 | 126272968 | 126273079 | 20 | 9 | 10 | 18 | 10 | 0 | 0 | 2 | 0 | 0 | 24 | 5 | 17 | 15 | 15 | 0 | 0 | 0 | 0 | 0 | 298 | 149 | 1 | 1 | 1 | 1 | 0.714 | 1 | 1 | 1 | 1 | 1 | 1 | 1 | -0.057 |
| SE | 89530 | ENSG00000110074 | FOXRED1 | chr11 | + | 126271436 | 126271657 | 126269153 | 126269291 | 126272968 | 126273079 | 33 | 20 | 18 | 24 | 12 | 0 | 0 | 2 | 0 | 0 | 59 | 20 | 21 | 23 | 26 | 0 | 0 | 0 | 0 | 0 | 298 | 149 | 1 | 1 | 1 | 1 | 0.818 | 1 | 1 | 1 | 1 | 1 | 1 | 1 | -0.036 |
| SE | 89535 | ENSG00000110074 | FOXRED1 | chr11 | + | 126274926 | 126275021 | 126273335 | 126273454 | 126275793 | 126275867 | 9 | 4 | 9 | 4 | 6 | 0 | 1 | 0 | 1 | 0 | 25 | 0 | 7 | 8 | 11 | 0 | 0 | 1 | 2 | 0 | 244 | 149 | 1 | 1 | 1 | 0.71 | 1 | 0.71 | 1 | 1 | NA | 0.81 | 0.71 | 1 | 0.004 |
| SE | 89537 | ENSG00000110074 | FOXRED1 | chr11 | + | 126274930 | 126275021 | 126273335 | 126273454 | 126275793 | 126275870 | 0 | 0 | 1 | 0 | 0 | 0 | 1 | 0 | 1 | 0 | 0 | 0 | 0 | 0 | 0 | 0 | 0 | 1 | 2 | 0 | 240 | 149 | 1 | 1 | NA | 0 | 1 | 0 | NA | NA | NA | 0 | 0 | NA | 0.333 |

[illegible]

| | | | | | | | | | | | | | | | | | | | | | | | | | | | | | | | | | | | | | | | | | | | | | | |
|---|---|---|---|---|---|---|---|---|---|---|---|---|---|---|---|---|---|---|---|---|---|---|---|---|---|---|---|---|---|---|---|---|---|---|---|---|---|---|---|---|---|---|---|---|---|---|
| SE | 89847 | ENSG00000137770 | CTDSPL2 | chr15 | + | 44490783 | 44490999 | 44486550 | 44486700 | 44496379 | 44496458 | 480 | 330 | 363 | 401 | 763 | 1 | 0 | 0 | 0 | 0 | 99 | 296 | 357 | 86 | 96 | 0 | 0 | 0 | 0 | 0 | 298 | 149 | 1 | 1 | 0.996 | 1 | 1 | 1 | 1 | 1 | 1 | 1 | 1 | 1 | -0.001 |
| SE | 89848 | ENSG00000137770 | CTDSPL2 | chr15 | + | 44496379 | 44496458 | 44490783 | 44490999 | 44497026 | 44497138 | 207 | 167 | 202 | 159 | 333 | 0 | 0 | 0 | 0 | 2 | 25 | 107 | 140 | 39 | 39 | 0 | 0 | 0 | 0 | 0 | 228 | 149 | 1 | 1 | 1 | 1 | 1 | 1 | 0.991 | 1 | 1 | 1 | 1 | 1 | -0.002 |
| SE | 89849 | ENSG00000137770 | CTDSPL2 | chr15 | + | 44497026 | 44497138 | 44496379 | 44496458 | 44499726 | 44499813 | 282 | 214 | 240 | 247 | 508 | 0 | 0 | 0 | 0 | 0 | 23 | 144 | 169 | 69 | 28 | 0 | 0 | 1 | 0 | 0 | 261 | 149 | 1 | 1 | 1 | 1 | 1 | 1 | 1 | 1 | 1 | 0.99 | 1 | 1 | 0.002 |
| SE | 89850 | ENSG00000137770 | CTDSPL2 | chr15 | + | 44501961 | 44502048 | 44499726 | 44499813 | 44503423 | 44503879 | 8 | 4 | 11 | 7 | 13 | 6 | 5 | 8 | 2 | 20 | 0 | 13 | 4 | 11 | 0 | 0 | 3 | 4 | 5 | 11 | 236 | 149 | 1 | 1 | 0.457 | 0.336 | 0.465 | 0.688 | 0.291 | NA | 0.732 | 0.387 | 0.581 | 0 | 0.022 |
| SE | 89852 | ENSG00000137770 | CTDSPL2 | chr15 | + | 44518653 | 44518864 | 44514764 | 44514844 | 44519168 | 44519295 | 4 | 1 | 0 | 0 | 8 | 188 | 120 | 150 | 182 | 382 | 0 | 0 | 0 | 4 | 0 | 0 | 131 | 158 | 26 | 5 | 298 | 149 | 1 | 1 | 0.011 | 0.004 | 0 | 0 | 0.01 | NA | 0 | 0 | 0.071 | 0 | -0.013 |
| SE | 89853 | ENSG00000137770 | CTDSPL2 | chr15 | + | 44521263 | 44521406 | 44519223 | 44519295 | 44524108 | 44524436 | 58 | 48 | 55 | 49 | 129 | 0 | 0 | 1 | 2 | 4 | 0 | 51 | 43 | 11 | 7 | 0 | 0 | 0 | 0 | 0 | 292 | 149 | 1 | 1 | 1 | 1 | 0.966 | 0.926 | 0.943 | NA | 1 | 1 | 1 | 1 | -0.033 |
| SE | 89854 | ENSG00000137770 | CTDSPL2 | chr15 | + | 44521310 | 44521406 | 44519168 | 44519295 | 44524108 | 44524543 | 208 | 140 | 183 | 186 | 502 | 0 | 0 | 1 | 2 | 4 | 0 | 67 | 108 | 16 | 11 | 0 | 0 | 0 | 0 | 0 | 245 | 149 | 1 | 1 | 1 | 1 | 0.991 | 0.983 | 0.987 | NA | 1 | 1 | 1 | 1 | -0.008 |
| SE | 89860 | ENSG00000137492 | THAP12 | chr11 | - | 76355559 | 76355654 | 76350060 | 76352794 | 76360955 | 76361063 | 107 | 75 | 53 | 92 | 223 | 3 | 1 | 0 | 0 | 0 | 18 | 78 | 79 | 62 | 0 | 0 | 1 | 2 | 0 | 0 | 244 | 149 | 1 | 1 | 0.956 | 0.979 | 1 | 1 | 1 | 1 | 0.979 | 0.96 | 1 | NA | 0.002 |
| SE | 89861 | ENSG00000137492 | THAP12 | chr11 | - | 76355617 | 76355654 | 76349897 | 76352794 | 76360955 | 76361063 | 384 | 285 | 300 | 359 | 1048 | 3 | 1 | 0 | 0 | 0 | 38 | 315 | 309 | 193 | 12 | 0 | 1 | 2 | 0 | 0 | 186 | 149 | 1 | 1 | 0.99 | 0.996 | 1 | 1 | 1 | 1 | 0.996 | 0.992 | 1 | 1 | 0 |
| SE | 89862 | ENSG00000137492 | THAP12 | chr11 | - | 76360955 | 76361063 | 76355559 | 76355654 | 76365851 | 76365972 | 471 | 331 | 363 | 410 | 1326 | 2 | 0 | 0 | 0 | 0 | 41 | 344 | 334 | 158 | 37 | 0 | 0 | 3 | 0 | 0 | 257 | 149 | 1 | 1 | 0.993 | 1 | 1 | 1 | 1 | 1 | 1 | 0.985 | 1 | 1 | 0.002 |
| SE | 89865 | ENSG00000137492 | THAP12 | chr11 | - | 76364337 | 76364432 | 76360955 | 76361063 | 76365779 | 76365972 | 52 | 54 | 69 | 47 | 112 | 1 | 0 | 0 | 0 | 3 | 26 | 89 | 52 | 53 | 19 | 0 | 0 | 4 | 0 | 0 | 244 | 149 | 1 | 1 | 0.969 | 1 | 1 | 1 | 0.958 | 1 | 1 | 0.888 | 1 | 1 | 0.008 |
| SE | 89867 | ENSG00000137492 | THAP12 | chr11 | - | 76364640 | 76364720 | 76360955 | 76361063 | 76365851 | 76365972 | 1 | 2 | 2 | 2 | 4 | 241 | 180 | 175 | 196 | 689 | 0 | 0 | 2 | 0 | 0 | 11 | 168 | 157 | 70 | 25 | 229 | 149 | 1 | 1 | 0.003 | 0.007 | 0.007 | 0.007 | 0.004 | 0 | 0 | 0.008 | 0 | 0 | 0.004 |
| SE | 89868 | ENSG00000137492 | THAP12 | chr11 | - | 76364640 | 76364720 | 76364337 | 76364432 | 76365851 | 76365972 | 0 | 0 | 1 | 1 | 0 | 43 | 63 | 76 | 65 | 100 | 0 | 0 | 4 | 0 | 0 | 13 | 79 | 70 | 47 | 9 | 229 | 149 | 1 | 1 | 0 | 0 | 0.008 | 0.01 | 0 | 0 | 0 | 0.036 | 0 | 0 | -0.004 |
| SE | 89872 | ENSG00000034677 | RNF19A | chr8 | - | 100259853 | 100259997 | 100257059 | 100259246 | 100268784 | 100268920 | 308 | 206 | 214 | 260 | 430 | 1 | 0 | 0 | 0 | 0 | 56 | 367 | 636 | 173 | 35 | 0 | 3 | 3 | 0 | 0 | 293 | 149 | 1 | 1 | 0.994 | 1 | 1 | 1 | 1 | 1 | 0.984 | 0.991 | 1 | 1 | 0.004 |
| SE | 89874 | ENSG00000034677 | RNF19A | chr8 | - | 100261541 | 100261755 | 100257059 | 100259246 | 100264670 | 100264785 | 32 | 18 | 19 | 25 | 8 | 1 | 3 | 2 | 5 | 2 | 13 | 75 | 139 | 50 | 0 | 0 | 19 | 12 | 4 | 0 | 298 | 149 | 1 | 1 | 0.941 | 0.75 | 0.826 | 0.714 | 0.667 | 1 | 0.664 | 0.853 | 0.862 | NA | -0.065 |
| SE | 89875 | ENSG00000034677 | RNF19A | chr8 | - | 100261541 | 100261755 | 100257059 | 100259246 | 100268784 | 100268920 | 8 | 2 | 1 | 4 | 1 | 1 | 0 | 0 | 0 | 0 | 5 | 4 | 7 | 0 | 0 | 0 | 3 | 3 | 0 | 0 | 298 | 149 | 1 | 1 | 0.8 | 1 | 1 | 1 | 1 | 1 | 0.4 | 0.538 | NA | NA | 0.314 |
| SE | 89878 | ENSG00000034677 | RNF19A | chr8 | - | 100261541 | 100261755 | 100259853 | 100259997 | 100268784 | 100268920 | 248 | 193 | 162 | 179 | 188 | 2 | 0 | 0 | 1 | 0 | 40 | 199 | 378 | 74 | 18 | 0 | 0 | 4 | 0 | 0 | 298 | 149 | 1 | 1 | 0.984 | 1 | 1 | 0.989 | 1 | 1 | 1 | 0.979 | 1 | 1 | -0.001 |
| SE | 89881 | ENSG00000034677 | RNF19A | chr8 | - | 100264670 | 100264785 | 100257059 | 100259246 | 100268784 | 100268920 | 75 | 48 | 64 | 72 | 70 | 1 | 0 | 0 | 0 | 0 | 13 | 168 | 219 | 22 | 38 | 0 | 3 | 3 | 0 | 0 | 264 | 149 | 1 | 1 | 0.977 | 1 | 1 | 1 | 1 | 1 | 0.969 | 0.976 | 1 | 1 | 0.006 |
| SE | 89882 | ENSG00000034677 | RNF19A | chr8 | - | 100264670 | 100264785 | 100259853 | 100259997 | 100268784 | 100268920 | 78 | 47 | 68 | 78 | 75 | 2 | 0 | 0 | 1 | 0 | 13 | 175 | 218 | 23 | 38 | 0 | 0 | 4 | 0 | 0 | 264 | 149 | 1 | 1 | 0.957 | 1 | 1 | 0.978 | 1 | 1 | 1 | 0.969 | 1 | 1 | -0.007 |
| SE | 89883 | ENSG00000034677 | RNF19A | chr8 | - | 100264670 | 100264785 | 100261541 | 100261755 | 100268784 | 100268920 | 98 | 59 | 80 | 90 | 75 | 0 | 0 | 0 | 2 | 0 | 21 | 221 | 333 | 68 | 38 | 0 | 0 | 2 | 0 | 0 | 264 | 149 | 1 | 1 | 1 | 1 | 1 | 0.962 | 1 | 1 | 1 | 0.989 | 1 | 1 | -0.005 |
| SE | 89884 | ENSG00000034677 | RNF19A | chr8 | - | 100274952 | 100275161 | 100269868 | 100270013 | 100287500 | 100288267 | 384 | 220 | 230 | 311 | 304 | 16 | 6 | 10 | 11 | 9 | 73 | 405 | 824 | 199 | 77 | 0 | 35 | 22 | 4 | 0 | 298 | 149 | 1 | 1 | 0.923 | 0.948 | 0.92 | 0.934 | 0.944 | 1 | 0.853 | 0.949 | 0.961 | 1 | -0.019 |
| SE | 89885 | ENSG00000034677 | RNF19A | chr8 | - | 100300527 | 100300651 | 100287500 | 100288267 | 100303159 | 100303259 | 19 | 16 | 7 | 14 | 8 | 10 | 9 | 7 | 19 | 4 | 0 | 0 | 6 | 2 | 0 | 10 | 10 | 4 | 1 | 0 | 273 | 149 | 1 | 1 | 0.509 | 0.492 | 0.353 | 0.287 | 0.522 | 0 | 0 | 0.45 | 0.522 | NA | 0.19 |
| SE | 89886 | ENSG00000034677 | RNF19A | chr8 | - | 100312343 | 100312410 | 100287677 | 100288267 | 100313290 | 100313372 | 0 | 0 | 2 | 3 | 2 | 1 | 2 | 1 | 0 | 3 | 0 | 4 | 4 | 0 | 0 | 0 | 0 | 6 | 1 | 0 | 216 | 149 | 1 | 1 | 0 | 0 | 0.58 | 1 | 0.315 | NA | 1 | 0.315 | 0 | NA | -0.059 |
| SE | 89888 | ENSG00000147576 | ADHFE1 | chr8 | + | 66440161 | 66440199 | 66432497 | 66432575 | 66442797 | 66442844 | 8 | 7 | 6 | 6 | 8 | 0 | 0 | 2 | 0 | 0 | 7 | 2 | 8 | 0 | 3 | 0 | 4 | 0 | 0 | 0 | 187 | 149 | 1 | 1 | 1 | 1 | 0.705 | 1 | 1 | 1 | 0.285 | 1 | NA | 1 | 0.12 |
| SE | 89889 | ENSG00000147576 | ADHFE1 | chr8 | + | 66440161 | 66440199 | 66432497 | 66432575 | 66444366 | 66444420 | 6 | 1 | 7 | 1 | 3 | 1 | 4 | 9 | 3 | 0 | 0 | 5 | 0 | 0 | 0 | 0 | 3 | 0 | 2 | 0 | 187 | 149 | 1 | 1 | 0.827 | 0.166 | 0.383 | 0.21 | 1 | NA | 0.57 | NA | 0 | NA | 0.232 |
| SE | 89890 | ENSG00000147576 | ADHFE1 | chr8 | + | 66440161 | 66440199 | 66432509 | 66432891 | 66442797 | 66442844 | 4 | 2 | 2 | 2 | 9 | 0 | 0 | 0 | 0 | 0 | 2 | 2 | 1 | 0 | 3 | 0 | 4 | 0 | 0 | 0 | 187 | 149 | 1 | 1 | 1 | 1 | 1 | 1 | 1 | 1 | 0.285 | 1 | NA | 1 | 0.179 |
| SE | 89891 | ENSG00000147576 | ADHFE1 | chr8 | + | 66440161 | 66440199 | 66432509 | 66432891 | 66444366 | 66444420 | 4 | 0 | 3 | 0 | 1 | 0 | 0 | 0 | 0 | 0 | 0 | 0 | 0 | 0 | 0 | 0 | 0 | 2 | 0 | 0 | 187 | 149 | 1 | 1 | 1 | NA | 1 | NA | 1 | NA | NA | 0 | NA | NA | 1 |
| SE | 89892 | ENSG00000147576 | ADHFE1 | chr8 | + | 66442797 | 66442844 | 66432504 | 66432575 | 66444366 | 66444420 | 10 | 1 | 4 | 6 | 6 | 1 | 4 | 9 | 3 | 0 | 0 | 9 | 6 | 2 | 0 | 0 | 3 | 0 | 2 | 0 | 196 | 149 | 1 | 1 | 0.884 | 0.16 | 0.253 | 0.603 | 1 | NA | 0.695 | 1 | 0.432 | NA | -0.129 |
| SE | 89893 | ENSG00000147576 | ADHFE1 | chr8 | + | 66442797 | 66442844 | 66432509 | 66432891 | 66444366 | 66444420 | 10 | 1 | 2 | 6 | 6 | 0 | 0 | 0 | 0 | 0 | 0 | 9 | 6 | 2 | 0 | 0 | 0 | 2 | 0 | 0 | 196 | 149 | 1 | 1 | 1 | 1 | 1 | 1 | 1 | NA | 1 | 0.695 | 1 | NA | 0.102 |
| SE | 89897 | ENSG00000120903 | CHRNA2 | chr8 | - | 27475212 | 27475272 | 27471001 | 27471194 | 27478823 | 27479059 | 0 | 0 | 0 | 0 | 0 | 0 | 0 | 0 | 1 | 2 | 0 | 0 | 0 | 0 | 2 | 0 | 1 | 1 | 2 | 0 | 209 | 149 | 1 | 1 | NA | NA | NA | 0 | 0 | NA | 0 | 0 | 0 | 1 | -0.25 |
| SE | 89899 | ENSG00000120903 | CHRNA2 | chr8 | - | 27478823 | 27478894 | 27471188 | 27471194 | 27479019 | 27479042 | 0 | 0 | 0 | 1 | 0 | 0 | 0 | 0 | 0 | 0 | 0 | 0 | 0 | 1 | 0 | 10 | 4 | 0 | 0 | 5 | 220 | 149 | 1 | 1 | NA | NA | NA | 1 | NA | 0 | 0 | NA | 1 | 0 | 0.75 |
| SE | 89900 | ENSG00000198088 | NUP62CL | chrX | - | 107150855 | 107150967 | 107147742 | 107147809 | 107153171 | 107153306 | 3 | 2 | 4 | 1 | 2 | 0 | 1 | 8 | 3 | 2 | 0 | 0 | 3 | 0 | 0 | 0 | 0 | 0 | 0 | 0 | 261 | 149 | 1 | 1 | 1 | 0.533 | 0.222 | 0.16 | 0.363 | NA | NA | 1 | NA | NA | -0.544 |
| SE | 89906 | ENSG00000104375 | STK3 | chr8 | - | 98721073 | 98721160 | 98707146 | 98707311 | 98722742 | 98722969 | 13 | 7 | 4 | 13 | 8 | 0 | 3 | 0 | 0 | 1 | 23 | 12 | 10 | 23 | 43 | 0 | 0 | 0 | 0 | 0 | 236 | 149 | 1 | 1 | 1 | 0.596 | 1 | 1 | 0.835 | 1 | 1 | 1 | 1 | 1 | -0.114 |
| SE | 89911 | ENSG00000108576 | SLC6A4 | chr17 | - | 30203171 | 30203339 | 30194318 | 30198530 | 30207731 | 30207832 | 2 | 1 | 0 | 9 | 12 | 0 | 0 | 0 | 0 | 0 | 58 | 5 | 9 | 16 | 55 | 0 | 0 | 2 | 0 | 0 | 298 | 149 | 1 | 1 | 1 | 1 | NA | 1 | 1 | 1 | 1 | 0.692 | 1 | 1 | 0.062 |
| SE | 89912 | ENSG00000108576 | SLC6A4 | chr17 | - | 30209142 | 30209242 | 30207731 | 30207832 | 30210514 | 30210646 | 1 | 1 | 1 | 4 | 1 | 0 | 0 | 0 | 0 | 0 | 34 | 11 | 10 | 7 | 15 | 0 | 0 | 2 | 2 | 0 | 249 | 149 | 1 | 1 | 1 | 1 | 1 | 1 | 1 | 1 | 1 | 0.749 | 0.677 | 1 | 0.115 |
| SE | 89913 | ENSG00000108576 | SLC6A4 | chr17 | - | 30222818 | 30222915 | 30221615 | 30222081 | 30235612 | 30235697 | 2 | 1 | 0 | 3 | 1 | 1 | 0 | 0 | 2 | 0 | 18 | 6 | 2 | 9 | 21 | 0 | 0 | 2 | 0 | 0 | 246 | 149 | 1 | 1 | 0.548 | 1 | NA | 0.476 | 1 | 1 | 1 | 0.377 | 1 | 1 | -0.119 |
| SE | 89953 | ENSG00000147679 | UTP23 | chr8 | + | 116770191 | 116770366 | 116766507 | 116766791 | 116779482 | 116779586 | 32 | 19 | 40 | 25 | 29 | 1 | 2 | 0 | 0 | 0 | 5 | 7 | 14 | 12 | 3 | 0 | 0 | 0 | 0 | 0 | 298 | 149 | 1 | 1 | 0.941 | 0.826 | 1 | 1 | 1 | 1 | 1 | 1 | 1 | 1 | -0.047 |
| SE | 89958 | ENSG00000254275 | LINC00824 | chr8 | - | 128560239 | 128560548 | 128559022 | 128559576 | 128561069 | 128561126 | 1 | 0 | 1 | 3 | 0 | 0 | 0 | 4 | 0 | 0 | 0 | 0 | 1 | 1 | 0 | 0 | 1 | 0 | 2 | 0 | 298 | 149 | 1 | 1 | 1 | NA | 0.111 | 1 | NA | NA | 0 | 1 | 0.2 | NA | 0.304 |
| SE | 89959 | ENSG00000067533 | RRP15 | chr1 | + | 218305027 | 218305125 | 218302293 | 218302559 | 218307430 | 218307632 | 38 | 40 | 66 | 53 | 57 | 0 | 1 | 0 | 0 | 0 | 0 | 10 | 18 | 21 | 1 | 0 | 0 | 0 | 0 | 0 | 247 | 149 | 1 | 1 | 1 | 0.96 | 1 | 1 | 1 | NA | 1 | 1 | 1 | 1 | -0.008 |
| SE | 89960 | ENSG00000067533 | RRP15 | chr1 | + | 218307430 | 218307632 | 218302293 | 218302559 | 218330947 | 218337983 | 30 | 38 | 43 | 26 | 24 | 0 | 0 | 0 | 0 | 0 | 7 | 6 | 7 | 2 | 4 | 0 | 0 | 2 | 0 | 0 | 298 | 149 | 1 | 1 | 1 | 1 | 1 | 1 | 1 | 1 | 1 | 0.636 | 1 | 1 | 0.073 |
| SE | 89961 | ENSG00000100380 | ST13 | chr22 | - | 40829625 | 40829674 | 40827095 | 40827229 | 40830839 | 40830956 | 387 | 281 | 485 | 439 | 657 | 1 | 0 | 0 | 0 | 0 | 207 | 203 | 280 | 179 | 143 | 0 | 0 | 0 | 0 | 0 | 198 | 149 | 1 | 1 | 0.997 | 1 | 1 | 1 | 1 | 1 | 1 | 1 | 1 | 1 | -0.001 |
| SE | 89963 | ENSG00000100380 | ST13 | chr22 | - | 40830839 | 40830956 | 40829625 | 40829674 | 40835559 | 40835905 | 178 | 143 | 219 | 219 | 267 | 0 | 0 | 0 | 0 | 0 | 96 | 86 | 94 | 66 | 79 | 0 | 1 | 0 | 0 | 0 | 266 | 149 | 1 | 1 | 1 | 1 | 1 | 1 | 1 | 1 | 0.98 | 1 | 1 | 1 | 0.004 |
| SE | 89964 | ENSG00000100380 | ST13 | chr22 | - | 40832086 | 40832216 | 40830874 | 40830956 | 40832568 | 40832671 | 0 | 0 | 0 | 0 | 1 | 553 | 459 | 648 | 748 | 647 | 0 | 0 | 0 | 0 | 0 | 565 | 311 | 405 | 512 | 319 | 279 | 149 | 1 | 1 | 0 | 0 | 0 | 0 | 0.001 | 0 | 0 | 0 | 0 | 0 | 0 |
| SE | 89965 | ENSG00000100380 | ST13 | chr22 | - | 40832568 | 40832671 | 40829625 | 40829674 | 40835559 | 40835670 | 461 | 292 | 478 | 480 | 735 | 0 | 0 | 0 | 0 | 0 | 342 | 230 | 285 | 257 | 110 | 0 | 1 | 0 | 0 | 0 | 252 | 149 | 1 | 1 | 1 | 1 | 1 | 1 | 1 | 1 | 0.993 | 1 | 1 | 1 | 0.001 |
| SE | 89966 | ENSG00000100380 | ST13 | chr22 | - | 40832568 | 40832671 | 40830839 | 40830956 | 40835559 | 40835670 | 1007 | 748 | 1117 | 1217 | 1356 | 1 | 4 | 0 | 0 | 3 | 907 | 532 | 677 | 768 | 429 | 0 | 0 | 0 | 0 | 0 | 252 | 149 | 1 | 1 | 0.998 | 0.991 | 1 | 1 | 0.996 | 1 | 1 | 1 | 1 | 1 | -0.003 |
| SE | 89967 | ENSG00000100380 | ST13 | chr22 | - | 40835802 | 40835887 | 40835559 | 40835670 | 40840625 | 40840692 | 1542 | 1109 | 1727 | 1575 | 2608 | 0 | 0 | 0 | 1 | 0 | 1154 | 829 | 920 | 884 | 552 | 0 | 3 | 0 | 0 | 0 | 234 | 149 | 1 | 1 | 1 | 1 | 1 | 0.999 | 1 | 1 | 0.994 | 1 | 1 | 1 | 0.001 |
| SE | 89968 | ENSG00000100380 | ST13 | chr22 | - | 40848293 | 40848369 | 40844838 | 40844909 | 40850822 | 40850880 | 330 | 252 | 371 | 293 | 463 | 0 | 2 | 2 | 2 | 0 | 165 | 166 | 234 | 192 | 115 | 0 | 5 | 3 | 0 | 0 | 225 | 149 | 1 | 1 | 1 | 0.988 | 0.992 | 0.99 | 1 | 1 | 0.956 | 0.981 | 1 | 1 | 0.007 |
| SE | 89969 | ENSG00000100380 | ST13 | chr22 | - | 40850822 | 40850880 | 40844838 | 40844909 | 40856430 | 40856565 | 36 | 32 | 34 | 34 | 35 | 0 | 0 | 0 | 1 | 1 | 17 | 17 | 13 | 27 | 7 | 0 | 0 | 0 | 0 | 0 | 207 | 149 | 1 | 1 | 1 | 1 | 1 | 0.961 | 0.962 | 1 | 1 | 1 | 1 | 1 | -0.015 |
| SE | 89970 | ENSG00000285655 | NA | chr8 | + | 66442797 | 66442844 | 66430184 | 66430378 | 66444366 | 66444420 | 12 | 1 | 2 | 6 | 6 | 0 | 0 | 0 | 0 | 0 | 0 | 5 | 6 | 2 | 0 | 0 | 1 | 4 | 0 | 0 | 196 | 149 | 1 | 1 | 1 | 1 | 1 | 1 | 1 | NA | 0.792 | 0.533 | 1 | NA | 0.225 |
| SE | 89972 | ENSG00000153317 | ASAP1 | chr8 | - | 130057953 | 130058076 | 130052103 | 130054805 | 130060578 | 130061069 | 154 | 173 | 118 | 198 | 113 | 0 | 0 | 3 | 0 | 0 | 336 | 279 | 255 | 425 | 281 | 0 | 0 | 0 | 1 | 0 | 272 | 149 | 1 | 1 | 1 | 1 | 0.956 | 1 | 1 | 1 | 1 | 1 | 0.996 | 1 | -0.008 |
| SE | 89973 | ENSG00000153317 | ASAP1 | chr8 | - | 130079901 | 130079971 | 130076347 | 130076406 | 130091972 | 130092143 | 63 | 84 | 81 | 64 | 127 | 0 | 0 | 0 | 0 | 0 | 243 | 178 | 121 | 262 | 245 | 0 | 0 | 0 | 3 | 0 | 219 | 149 | 1 | 1 | 1 | 1 | 1 | 1 | 1 | 1 | 1 | 1 | 0.983 | 1 | 0.003 |
| SE | 89974 | ENSG00000153317 | ASAP1 | chr8 | - | 130079901 | 130079971 | 130076347 | 130076406 | 130112093 | 130112322 | 49 | 51 | 39 | 28 | 64 | 0 | 0 | 0 | 0 | 0 | 117 | 109 | 99 | 131 | 95 | 0 | 0 | 0 | 3 | 0 | 219 | 149 | 1 | 1 | 1 | 1 | 1 | 1 | 1 | 1 | 1 | 1 | 0.967 | 1 | 0.007 |
| SE | 89975 | ENSG00000153317 | ASAP1 | chr8 | - | 130091972 | 130092143 | 130076347 | 130076406 | 130112093 | 130112322 | 19 | 27 | 25 | 29 | 59 | 0 | 0 | 0 | 0 | 0 | 77 | 55 | 32 | 45 | 104 | 0 | 0 | 0 | 3 | 0 | 298 | 149 | 1 | 1 | 1 | 1 | 1 | 1 | 1 | 1 | 1 | 1 | 0.882 | 1 | 0.024 |
| SE | 89978 | ENSG00000153317 | ASAP1 | chr8 | - | 130112093 | 130112322 | 130079901 | 130079971 | 130115627 | 130115735 | 256 | 235 | 204 | 200 | 409 | 0 | 0 | 0 | 0 | 0 | 673 | 389 | 393 | 651 | 555 | 0 | 0 | 7 | 0 | 0 | 298 | 149 | 1 | 1 | 1 | 1 | 1 | 1 | 1 | 1 | 1 | 0.966 | 1 | 1 | 0.007 |
| SE | 89979 | ENSG00000153317 | ASAP1 | chr8 | - | 130112093 | 130112322 | 130091972 | 130092143 | 130115627 | 130115735 | 259 | 255 | 222 | 223 | 463 | 0 | 0 | 0 | 0 | 0 | 711 | 424 | 375 | 665 | 622 | 0 | 2 | 2 | 0 | 0 | 298 | 149 | 1 | 1 | 1 | 1 | 1 | 1 | 1 | 1 | 0.991 | 0.989 | 1 | 1 | 0.004 |
| SE | 89980 | ENSG00000153317 | ASAP1 | chr8 | - | 130124012 | 130124104 | 130118488 | 130118675 | 130125955 | 130126089 | 292 | 380 | 230 | 344 | 420 | 0 | 0 | 0 | 0 | 0 | 790 | 497 | 464 | 832 | 1019 | 0 | 0 | 0 | 1 | 0 | 241 | 149 | 1 | 1 | 1 | 1 | 1 | 1 | 1 | 1 | 1 | 1 | 0.998 | 1 | 0 |
| SE | 89981 | ENSG00000153317 | ASAP1 | chr8 | - | 130134295 | 130134344 | 130127926 | 130128090 | 130136950 | 130137038 | 186 | 289 | 118 | 205 | 337 | 0 | 0 | 1 | 0 | 0 | 417 | 234 | 245 | 522 | 504 | 0 | 0 | 0 | 2 | 0 | 198 | 149 | 1 | 1 | 1 | 1 | 0.989 | 1 | 1 | 1 | 1 | 1 | 0.995 | 1 | -0.001 |
| SE | 89982 | ENSG00000153317 | ASAP1 | chr8 | - | 130160784 | 130160793 | 130159863 | 130159964 | 130167535 | 130167622 | 1 | 2 | 2 | 3 | 0 | 464 | 592 | 329 | 566 | 763 | 0 | 8 | 6 | 1 | 0 | 1873 | 822 | 782 | 1481 | 2083 | 158 | 149 | 1 | 1 | 0.002 | 0.003 | 0.006 | 0.005 | 0 | 0 | 0.009 | 0.007 | 0.001 | 0 | 0 |
| SE | 89984 | ENSG00000153317 | ASAP1 | chr8 | - | 130180750 | 130180880 | 130168991 | 130169067 | 130187235 | 130187285 | 236 | 291 | 202 | 245 | 331 | 0 | 2 | 0 | 0 | 3 | 1188 | 557 | 434 | 985 | 944 | 0 | 0 | 0 | 0 | 0 | 279 | 149 | 1 | 1 | 1 | 0.987 | 1 | 1 | 0.983 | 1 | 1 | 1 | 1 | 1 | -0.006 |
| SE | 89986 | ENSG00000153317 | ASAP1 | chr8 | - | 130188108 | 130188183 | 130180750 | 130180880 | 130214555 | 130214701 | 79 | 122 | 88 | 108 | 158 | 0 | 0 | 0 | 0 | 1 | 398 | 199 | 161 | 364 | 306 | 0 | 1 | 0 | 0 | 0 | 224 | 149 | 1 | 1 | 1 | 1 | 1 | 1 | 0.991 | 1 | 0.993 | 1 | 1 | 1 | 0 |
| SE | 89987 | ENSG00000153317 | ASAP1 | chr8 | - | 130208629 | 130208746 | 130188108 | 130188183 | 130214555 | 130214624 | 0 | 0 | 1 | 1 | 0 | 169 | 208 | 152 | 184 | 257 | 0 | 1 | 6 | 12 | 1 | 734 | 402 | 271 | 675 | 531 | 266 | 149 | 1 | 1 | 0 | 0 | 0.004 | 0.003 | 0 | 0 | 0.001 | 0.012 | 0.01 | 0.001 | -0.003 |
| SE | 89989 | ENSG00000153317 | ASAP1 | chr8 | - | 130236921 | 130236994 | 130214555 | 130214701 | 130358016 | 130358143 | 220 | 209 | 151 | 226 | 186 | 0 | 0 | 0 | 0 | 0 | 584 | 409 | 366 | 615 | 584 | 0 | 0 | 0 | 4 | 0 | 222 | 149 | 1 | 1 | 1 | 1 | 1 | 1 | 1 | 1 | 1 | 1 | 0.99 | 1 | 0.002 |
| SE | 89990 | ENSG00000153317 | ASAP1 | chr8 | - | 130361669 | 130361771 | 130358016 | 130358143 | 130401884 | 130401915 | 2 | 1 | 1 | 4 | 8 | 3 | 2 | 1 | 0 | 0 | 0 | 1 | 1 | 0 | 0 | 0 | 0 | 1 | 0 | 0 | 251 | 149 | 1 | 1 | 0.284 | 0.229 | 0.372 | 1 | 1 | NA | 1 | 0.372 | NA | NA | -0.109 |
| SE | 89994 | ENSG00000147654 | EBAG9 | chr8 | + | 109554728 | 109554887 | 109553864 | 109553943 | 109556934 | 109557027 | 337 | 249 | 297 | 225 | 709 | 0 | 0 | 0 | 0 | 0 | 100 | 208 | 172 | 104 | 21 | 0 | 0 | 4 | 0 | 0 | 298 | 149 | 1 | 1 | 1 | 1 | 1 | 1 | 1 | 1 | 1 | 0.956 | 1 | 1 | 0.009 |
| SE | 89995 | ENSG00000147654 | EBAG9 | chr8 | + | 109556934 | 109557042 | 109554728 | 109554887 | 109560837 | 109560929 | 286 | 260 | 314 | 212 | 634 | 0 | 0 | 1 | 0 | 0 | 59 | 241 | 177 | 110 | 24 | 0 | 0 | 3 | 0 | 0 | 257 | 149 | 1 | 1 | 1 | 1 | 0.995 | 1 | 1 | 1 | 1 | 0.972 | 1 | 1 | 0.005 |
| SE | 89996 | ENSG00000147654 | EBAG9 | chr8 | + | 109557730 | 109557919 | 109556934 | 109557042 | 109560837 | 109560866 | 0 | 0 | 0 | 0 | 2 | 151 | 148 | 182 | 136 | 366 | 0 | 2 | 1 | 0 | 0 | 26 | 141 | 92 | 56 | 16 | 298 | 149 | 1 | 1 | 0 | 0 | 0 | 0 | 0.003 | 0 | 0.007 | 0.005 | 0 | 0 | -0.002 |
| SE | 89997 | ENSG00000147654 | EBAG9 | chr8 | + | 109560837 | 109560929 | 109556934 | 109557042 | 109564438 | 109564508 | 220 | 213 | 235 | 188 | 513 | 0 | 0 | 0 | 0 | 1 | 26 | 175 | 129 | 72 | 16 | 0 | 0 | 0 | 0 | 0 | 241 | 149 | 1 | 1 | 1 | 1 | 1 | 1 | 0.997 | 1 | 1 | 1 | 1 | 1 | -0.001 |
| SE | 89998 | ENSG00000147654 | EBAG9 | chr8 | + | 109563396 | 109563535 | 109560837 | 109560929 | 109564438 | 109565996 | 2 | 0 | 0 | 0 | 0 | 152 | 147 | 154 | 118 | 340 | 0 | 0 | 2 | 0 | 0 | 0 | 113 | 91 | 55 | 10 | 288 | 149 | 1 | 1 | 0.007 | 0 | 0 | 0 | 0 | NA | 0 | 0.011 | 0 | 0 | -0.001 |
| SE | 89999 | ENSG00000147654 | EBAG9 | chr8 | + | 109563396 | 109563535 | 109560837 | 109560929 | 109564442 | 109564559 | 0 | 0 | 0 | 0 | 0 | 0 | 0 | 4 | 0 | 0 | 0 | 0 | 2 | 0 | 0 | 0 | 0 | 0 | 0 | 0 | 288 | 149 | 1 | 1 | NA | NA | 0 | NA | NA | NA | NA | 1 | NA | NA | -1 |
| SE | 90002 | ENSG00000104783 | KCNN4 | chr19 | - | 43776540 | 43776636 | 43774191 | 43774619 | 43780702 | 43780973 | 6 | 0 | 11 | 16 | 4 | 0 | 0 | 1 | 0 | 0 | 53 | 3 | 0 | 2 | 0 | 0 | 0 | 0 | 0 | 0 | 245 | 149 | 1 | 1 | 1 | NA | 0.87 | 1 | 1 | 1 | 1 | NA | 1 | NA | -0.032 |
| SE | 90004 | ENSG00000078487 | ZCWPW1 | chr7 | - | 100401888 | 100402041 | 100400852 | 100401336 | 100402515 | 100402576 | 3 | 7 | 18 | 10 | 1 | 0 | 0 | 0 | 0 | 0 | 15 | 6 | 2 | 0 | 13 | 0 | 1 | 0 | 0 | 0 | 298 | 149 | 1 | 1 | 1 | 1 | 1 | 1 | 1 | 1 | 0.75 | 1 | NA | 1 | 0.062 |
| SE | 90005 | ENSG00000078487 | ZCWPW1 | chr7 | - | 100402515 | 100402576 | 100400854 | 100401336 | 100403693 | 100403785 | 0 | 0 | 0 | 0 | 0 | 0 | 1 | 3 | 2 | 1 | 0 | 1 | 0 | 2 | 0 | 0 | 0 | 4 | 0 | 14 | 210 | 149 | 1 | 1 | NA | 0 | 0 | 0 | 0 | NA | 1 | 0 | 1 | 0 | -0.5 |
| SE | 90006 | ENSG00000078487 | ZCWPW1 | chr7 | - | 100405012 | 100405093 | 100404177 | 100404244 | 100406693 | 100406798 | 10 | 9 | 12 | 14 | 30 | 1 | 0 | 0 | 0 | 0 | 32 | 6 | 10 | 21 | 2 | 0 | 0 | 0 | 2 | 0 | 230 | 149 | 1 | 1 | 0.866 | 1 | 1 | 1 | 1 | 1 | 1 | 1 | 0.872 | 1 | -0.001 |
| SE | 90008 | ENSG00000078487 | ZCWPW1 | chr7 | - | 100407227 | 100407303 | 100406693 | 100406798 | 100408538 | 100408659 | 7 | 5 | 13 | 6 | 11 | 2 | 3 | 0 | 0 | 2 | 12 | 3 | 0 | 11 | 11 | 0 | 0 | 0 | 5 | 0 | 225 | 149 | 1 | 1 | 0.699 | 0.525 | 1 | 1 | 0.785 | 1 | 1 | NA | 0.593 | 1 | -0.096 |
| SE | 90009 | ENSG00000078487 | ZCWPW1 | chr7 | - | 100415974 | 100416097 | 100409427 | 100409544 | 100425029 | 100425136 | 4 | 3 | 10 | 7 | 29 | 0 | 0 | 0 | 2 | 3 | 10 | 0 | 8 | 9 | 0 | 1 | 0 | 0 | 6 | 0 | 272 | 149 | 1 | 1 | 1 | 1 | 1 | 0.657 | 0.841 | 0.846 | NA | 1 | 1 | NA | -0.049 |
| SE | 90010 | ENSG00000078487 | ZCWPW1 | chr7 | - | 100416304 | 100416456 | 100415974 | 100416097 | 100417065 | 100417183 | 9 | 17 | 20 | 17 | 29 | 0 | 0 | 0 | 0 | 0 | 14 | 15 | 32 | 21 | 0 | 0 | 0 | 0 | 0 | 0 | 298 | 149 | 1 | 1 | 1 | 1 | 1 | 1 | 1 | 1 | 1 | 1 | 0.636 | NA | 0.091 |
| SE | 90011 | ENSG00000078487 | ZCWPW1 | chr7 | - | 100416304 | 100416456 | 100415974 | 100416097 | 100425029 | 100425136 | 5 | 12 | 11 | 4 | 8 | 0 | 0 | 0 | 0 | 0 | 5 | 5 | 22 | 14 | 0 | 0 | 0 | 1 | 0 | 0 | 298 | 149 | 1 | 1 | 1 | 1 | 1 | 1 | 1 | 1 | 1 | 0.917 | 1 | NA | 0.021 |
| SE | 90012 | ENSG00000078487 | ZCWPW1 | chr7 | - | 100420621 | 100420678 | 100419629 | 100419883 | 100425029 | 100425136 | 13 | 11 | 40 | 9 | 12 | 0 | 0 | 0 | 0 | 0 | 24 | 7 | 7 | 8 | 14 | 0 | 0 | 3 | 0 | 0 | 206 | 149 | 1 | 1 | 1 | 1 | 1 | 1 | 1 | 1 | 1 | 0.628 | 1 | 1 | 0.074 |
| SE | 90016 | ENSG00000155926 | SLA | chr8 | - | 133039997 | 133040130 | 133036727 | 133038737 | 133044983 | 133045115 | 476 | 217 | 357 | 304 | 229 | 0 | 0 | 0 | 0 | 0 | 219 | 364 | 711 | 289 | 185 | 0 | 0 | 0 | 0 | 3 | 282 | 149 | 1 | 1 | 1 | 1 | 1 | 1 | 1 | 1 | 1 | 1 | 1 | 0.97 | 0.006 |
| SE | 90021 | ENSG00000155926 | SLA | chr8 | - | 133063579 | 133063728 | 133060117 | 133060200 | 133074852 | 133075130 | 0 | 0 | 7 | 1 | 0 | 13 | 15 | 8 | 11 | 2 | 0 | 6 | 12 | 0 | 0 | 0 | 3 | 32 | 14 | 6 | 298 | 149 | 1 | 1 | 0 | 0 | 0.304 | 0.043 | 0 | NA | 0.5 | 0.158 | 0 | 0 | -0.095 |
| SE | 90033 | ENSG00000144713 | RPL32 | chr3 | - | 12840141 | 12840242 | 12839050 | 12839530 | 12841493 | 12841543 | 10766 | 5539 | 13555 | 9406 | 21234 | 5 | 0 | 5 | 4 | 6 | 6432 | 6999 | 5307 | 4121 | 9178 | 0 | 0 | 2 | 0 | 0 | 250 | 149 | 1 | 1 | 0.999 | 1 | 0.999 | 0.999 | 1 | 1 | 1 | 0.999 | 1 | 1 | 0 |
| SE | 90034 | ENSG00000144713 | RPL32 | chr3 | - | 12840141 | 12840341 | 12839348 | 12839530 | 12841493 | 12841544 | 7257 | 4120 | 9049 | 6434 | 19169 | 5 | 0 | 5 | 4 | 6 | 4610 | 4353 | 3567 | 2884 | 6234 | 0 | 0 | 2 | 0 | 0 | 298 | 149 | 1 | 1 | 0.999 | 1 | 0.999 | 0.999 | 0.999 | 1 | 1 | 0.999 | 1 | 1 | -0.001 |
| SE | 90035 | ENSG00000144713 | RPL32 | chr3 | - | 12840141 | 12840353 | 12839348 | 12839530 | 12841493 | 12841545 | 7242 | 4116 | 9029 | 6424 | 19163 | 5 | 0 | 5 | 4 | 6 | 4606 | 4340 | 3557 | 2858 | 6209 | 0 | 0 | 2 | 0 | 0 | 298 | 149 | 1 | 1 | 0.999 | 1 | 0.999 | 0.999 | 0.999 | 1 | 1 | 0.999 | 1 | 1 | -0.001 |
| SE | 90039 | ENSG00000247081 | BAALC-AS1 | chr8 | - | 103268691 | 103268843 | 103268390 | 103268472 | 103285837 | 103285902 | 1 | 0 | 0 | 0 | 0 | 0 | 0 | 0 | 1 | 0 | 0 | 0 | 0 | 0 | 0 | 0 | 0 | 1 | 0 | 0 | 298 | 149 | 1 | 1 | 1 | NA | NA | 0 | NA | NA | NA | 0 | NA | NA | 0.5 |
| SE | 90044 | ENSG00000140525 | FANCI | chr15 | + | 89245606 | 89245708 | 89244003 | 89244033 | 89247628 | 89247731 | 0 | 0 | 1 | 0 | 0 | 4 | 2 | 6 | 2 | 2 | 0 | 0 | 0 | 0 | 0 | 0 | 0 | 0 | 5 | 1 | 251 | 149 | 1 | 1 | 0 | 0 | 0.09 | 0 | 0 | NA | NA | NA | 0 | 0 | 0.018 |
| SE | 90045 | ENSG00000140525 | FANCI | chr15 | + | 89245606 | 89245708 | 89244033 | 89244067 | 89247628 | 89247731 | 0 | 0 | 0 | 0 | 0 | 0 | 3 | 0 | 0 | 0 | 0 | 0 | 0 | 3 | 0 | 0 | 0 | 4 | 19 | 0 | 251 | 149 | 1 | 1 | NA | 0 | NA | NA | NA | NA | NA | 0 | 0.086 | NA | -0.043 |
| SE | 90046 | ENSG00000140525 | FANCI | chr15 | + | 89245755 | 89245839 | 89243989 | 89244044 | 89247628 | 89247731 | 0 | 0 | 1 | 1 | 1 | 5 | 0 | 0 | 0 | 0 | 0 | 2 | 4 | 4 | 0 | 0 | 3 | 4 | 0 | 0 | 233 | 149 | 1 | 1 | 0 | NA | 1 | 1 | 1 | NA | 0.299 | 0.39 | 1 | NA | 0.187 |
| SE | 90047 | ENSG00000140525 | FANCI | chr15 | + | 89245755 | 89245839 | 89244003 | 89244033 | 89247628 | 89247731 | 0 | 0 | 1 | 2 | 1 | 4 | 2 | 6 | 2 | 2 | 0 | 2 | 4 | 4 | 0 | 0 | 0 | 0 | 5 | 1 | 233 | 149 | 1 | 1 | 0 | 0 | 0.096 | 0.39 | 0.242 | NA | 1 | 1 | 0.338 | 0 | -0.439 |
| SE | 90048 | ENSG00000140525 | FANCI | chr15 | + | 89245755 | 89245839 | 89244016 | 89244067 | 89247628 | 89247731 | 0 | 0 | 1 | 1 | 1 | 0 | 3 | 0 | 0 | 0 | 0 | 3 | 4 | 4 | 0 | 0 | 0 | 4 | 19 | 0 | 233 | 149 | 1 | 1 | NA | 0 | 1 | 1 | 1 | NA | 1 | 0.39 | 0.119 | NA | 0.247 |
| SE | 90051 | ENSG00000140525 | FANCI | chr15 | + | 89273376 | 89273469 | 89268398 | 89268525 | 89274167 | 89274304 | 23 | 9 | 27 | 37 | 37 | 0 | 0 | 0 | 0 | 0 | 5 | 21 | 34 | 44 | 18 | 0 | 1 | 0 | 0 | 0 | 242 | 149 | 1 | 1 | 1 | 1 | 1 | 1 | 1 | 1 | 0.928 | 1 | 1 | 1 | 0.014 |
| SE | 90052 | ENSG00000140525 | FANCI | chr15 | + | 89278686 | 89278774 | 89276710 | 89276891 | 89281169 | 89281300 | 29 | 10 | 26 | 52 | 55 | 1 | 0 | 0 | 0 | 0 | 0 | 27 | 49 | 43 | 2 | 0 | 0 | 1 | 0 | 0 | 237 | 149 | 1 | 1 | 0.948 | 1 | 1 | 1 | 1 | NA | 1 | 0.969 | 1 | 1 | -0.003 |
| SE | 90053 | ENSG00000140525 | FANCI | chr15 | + | 89281764 | 89281835 | 89281169 | 89281300 | 89285095 | 89285218 | 12 | 1 | 10 | 5 | 15 | 0 | 0 | 0 | 0 | 0 | 14 | 21 | 14 | 15 | 0 | 0 | 0 | 3 | 0 | 0 | 220 | 149 | 1 | 1 | 1 | 1 | 1 | 1 | 1 | 1 | 1 | 0.76 | 1 | NA | 0.06 |
| SE | 90054 | ENSG00000140525 | FANCI | chr15 | + | 89281764 | 89283250 | 89281169 | 89281300 | 89285095 | 89285218 | 18 | 2 | 22 | 16 | 30 | 0 | 0 | 0 | 0 | 0 | 23 | 36 | 30 | 27 | 6 | 0 | 0 | 3 | 0 | 0 | 298 | 149 | 1 | 1 | 1 | 1 | 1 | 1 | 1 | 1 | 1 | 0.833 | 1 | 1 | 0.033 |
| SE | 90055 | ENSG00000140525 | FANCI | chr15 | + | 89283135 | 89283250 | 89281764 | 89281835 | 89285095 | 89285218 | 8 | 1 | 14 | 17 | 15 | 1 | 0 | 0 | 0 | 0 | 9 | 22 | 7 | 18 | 0 | 0 | 0 | 0 | 0 | 0 | 264 | 149 | 1 | 1 | 0.819 | 1 | 1 | 1 | 1 | 1 | 1 | 1 | 1 | NA | -0.036 |
| SE | 90056 | ENSG00000140525 | FANCI | chr15 | + | 89292941 | 89293063 | 89292687 | 89292864 | 89299799 | 89299966 | 12 | 4 | 22 | 16 | 12 | 0 | 0 | 0 | 0 | 0 | 0 | 20 | 14 | 26 | 8 | 0 | 2 | 1 | 2 | 0 | 271 | 149 | 1 | 1 | 1 | 1 | 1 | 1 | 1 | NA | 0.846 | 0.885 | 0.877 | 1 | 0.098 |
| SE | 90057 | ENSG00000140525 | FANCI | chr15 | + | 89293832 | 89293997 | 89292941 | 89293063 | 89299799 | 89299966 | 23 | 8 | 39 | 19 | 24 | 0 | 0 | 0 | 2 | 0 | 0 | 27 | 21 | 31 | 0 | 0 | 0 | 5 | 0 | 0 | 298 | 149 | 1 | 1 | 1 | 1 | 1 | 0.826 | 1 | NA | 1 | 0.677 | 1 | NA | 0.073 |
| SE | 90058 | ENSG00000140525 | FANCI | chr15 | + | 89294914 | 89295094 | 89293832 | 89293997 | 89299799 | 89299966 | 32 | 23 | 37 | 38 | 50 | 2 | 1 | 7 | 1 | 4 | 0 | 20 | 41 | 56 | 0 | 0 | 8 | 0 | 0 | 0 | 298 | 149 | 1 | 1 | 0.889 | 0.92 | 0.725 | 0.95 | 0.862 | NA | 0.556 | 1 | 1 | NA | 0.017 |
| SE | 90059 | ENSG00000140525 | FANCI | chr15 | + | 89294956 | 89295094 | 89293832 | 89293997 | 89299799 | 89299966 | 9 | 13 | 19 | 21 | 29 | 2 | 1 | 7 | 1 | 4 | 0 | 22 | 13 | 23 | 0 | 0 | 8 | 0 | 0 | 0 | 287 | 149 | 1 | 1 | 0.7 | 0.871 | 0.585 | 0.916 | 0.79 | NA | 0.588 | 1 | 1 | NA | -0.09 |
| SE | 90060 | ENSG00000140525 | FANCI | chr15 | + | 89301325 | 89301442 | 89300299 | 89300385 | 89303863 | 89303915 | 48 | 17 | 59 | 48 | 32 | 0 | 0 | 0 | 0 | 0 | 5 | 21 | 47 | 53 | 0 | 0 | 0 | 1 | 0 | 0 | 266 | 149 | 1 | 1 | 1 | 1 | 1 | 1 | 1 | 1 | 1 | 0.963 | 1 | NA | 0.009 |
| SE | 90061 | ENSG00000140525 | FANCI | chr15 | + | 89306006 | 89306194 | 89305604 | 89305698 | 89307475 | 89307529 | 43 | 12 | 37 | 44 | 27 | 0 | 0 | 1 | 0 | 0 | 0 | 60 | 50 | 89 | 15 | 0 | 0 | 1 | 0 | 0 | 298 | 149 | 1 | 1 | 1 | 1 | 0.949 | 1 | 1 | NA | 1 | 0.962 | 1 | 1 | -0.001 |
| SE | 90062 | ENSG00000140525 | FANCI | chr15 | + | 89306009 | 89306194 | 89305604 | 89305698 | 89307475 | 89307529 | 27 | 7 | 28 | 27 | 14 | 0 | 0 | 1 | 0 | 0 | 0 | 40 | 31 | 51 | 24 | 0 | 0 | 1 | 0 | 0 | 298 | 149 | 1 | 1 | 1 | 1 | 0.933 | 1 | 1 | NA | 1 | 0.939 | 1 | 1 | 0.002 |
| SE | 90063 | ENSG00000140525 | FANCI | chr15 | + | 89307612 | 89307672 | 89307475 | 89307529 | 89312903 | 89312972 | 22 | 15 | 28 | 34 | 27 | 0 | 0 | 0 | 0 | 0 | 0 | 38 | 34 | 66 | 12 | 0 | 0 | 5 | 0 | 0 | 209 | 149 | 1 | 1 | 1 | 1 | 1 | 1 | 1 | NA | 1 | 0.829 | 1 | 1 | 0.043 |
| SE | 90067 | ENSG00000156787 | TBC1D31 | chr8 | + | 123105287 | 123105464 | 123100806 | 123101007 | 123109316 | 123109396 | 351 | 189 | 213 | 271 | 481 | 2 | 0 | 0 | 2 | 0 | 137 | 184 | 108 | 90 | 156 | 0 | 2 | 0 | 0 | 0 | 298 | 149 | 1 | 1 | 0.989 | 1 | 1 | 0.985 | 1 | 1 | 0.979 | 1 | 1 | 1 | -0.001 |
| SE | 90068 | ENSG00000156787 | TBC1D31 | chr8 | + | 123120054 | 123120188 | 123109473 | 123109620 | 123126055 | 123126189 | 129 | 77 | 118 | 130 | 389 | 3 | 2 | 3 | 3 | 11 | 0 | 42 | 32 | 23 | 14 | 0 | 6 | 0 | 0 | 0 | 283 | 149 | 1 | 1 | 0.958 | 0.953 | 0.954 | 0.958 | 0.949 | NA | 0.787 | 1 | 1 | 1 | 0.008 |
| SE | 90069 | ENSG00000156787 | TBC1D31 | chr8 | + | 123128280 | 123128513 | 123126507 | 123126687 | 123129065 | 123129172 | 216 | 193 | 215 | 246 | 304 | 0 | 0 | 2 | 2 | 4 | 33 | 119 | 65 | 14 | 38 | 0 | 0 | 0 | 0 | 0 | 298 | 149 | 1 | 1 | 1 | 1 | 0.982 | 0.984 | 0.974 | 1 | 1 | 1 | 1 | 1 | -0.012 |
| SE | 90070 | ENSG00000156787 | TBC1D31 | chr8 | + | 123128280 | 123128513 | 123126507 | 123126687 | 123129069 | 123129218 | 76 | 54 | 80 | 92 | 87 | 8 | 10 | 14 | 11 | 7 | 0 | 34 | 20 | 6 | 3 | 0 | 0 | 4 | 0 | 0 | 298 | 149 | 1 | 1 | 0.826 | 0.73 | 0.741 | 0.807 | 0.861 | NA | 1 | 0.714 | 1 | 1 | -0.136 |
| SE | 90071 | ENSG00000156787 | TBC1D31 | chr8 | + | 123130197 | 123130333 | 123129065 | 123129218 | 123140760 | 123140901 | 124 | 96 | 90 | 113 | 170 | 0 | 1 | 0 | 0 | 0 | 5 | 63 | 40 | 4 | 20 | 0 | 0 | 0 | 0 | 0 | 285 | 149 | 1 | 1 | 1 | 0.98 | 1 | 1 | 1 | 1 | 1 | 1 | 1 | 1 | -0.004 |
| SE | 90073 | ENSG00000156787 | TBC1D31 | chr8 | + | 123140760 | 123140901 | 123134113 | 123134206 | 123144716 | 123144855 | 102 | 68 | 73 | 86 | 94 | 0 | 0 | 1 | 1 | 4 | 0 | 23 | 15 | 3 | 1 | 0 | 0 | 0 | 0 | 0 | 290 | 149 | 1 | 1 | 1 | 1 | 1 | 0.978 | 0.924 | NA | 1 | 1 | 1 | 1 | -0.02 |
| SE | 90074 | ENSG00000156787 | TBC1D31 | chr8 | + | 123142261 | 123142456 | 123140760 | 123140901 | 123144716 | 123144855 | 64 | 47 | 104 | 139 | 219 | 41 | 25 | 7 | 12 | 11 | 23 | 17 | 9 | 4 | 0 | 0 | 15 | 7 | 2 | 0 | 298 | 149 | 1 | 1 | 0.438 | 0.485 | 0.881 | 0.853 | 0.909 | 1 | 0.362 | 0.391 | 0.5 | NA | 0.15 |
| SE | 90075 | ENSG00000156787 | TBC1D31 | chr8 | + | 123150035 | 123150128 | 123144716 | 123144855 | 123151805 | 123151954 | 89 | 66 | 68 | 91 | 134 | 2 | 3 | 1 | 0 | 0 | 0 | 23 | 6 | 0 | 0 | 0 | 1 | 0 | 0 | 0 | 242 | 149 | 1 | 1 | 0.965 | 0.931 | 0.977 | 1 | 1 | NA | 0.934 | 1 | NA | NA | 0.008 |
| SE | 90076 | ENSG00000165156 | ZHX1 | chr8 | - | 123253321 | 123256171 | 123248450 | 123250320 | 123267272 | 123267386 | 113 | 87 | 102 | 112 | 150 | 5 | 0 | 3 | 7 | 11 | 11 | 60 | 32 | 5 | 0 | 0 | 0 | 0 | 0 | 0 | 298 | 149 | 1 | 1 | 0.919 | 1 | 0.944 | 0.889 | 0.872 | 1 | 1 | 1 | 1 | NA | -0.075 |
| SE | 90077 | ENSG00000165156 | ZHX1 | chr8 | - | 123257032 | 123257217 | 123256119 | 123256171 | 123267272 | 123267386 | 0 | 0 | 3 | 0 | 4 | 38 | 30 | 46 | 44 | 52 | 0 | 0 | 0 | 5 | 0 | 11 | 25 | 13 | 4 | 0 | 298 | 149 | 1 | 1 | 0 | 0 | 0.032 | 0 | 0.037 | 0 | 0 | 0 | 0.385 | NA | -0.082 |
| SE | 90101 | ENSG00000104549 | SQLE | chr8 | + | 125020783 | 125020871 | 125018630 | 125018727 | 125021752 | 125022281 | 53 | 35 | 28 | 23 | 47 | 0 | 1 | 0 | 0 | 0 | 22 | 20 | 32 | 37 | 11 | 0 | 3 | 0 | 0 | 0 | 237 | 149 | 1 | 1 | 1 | 0.957 | 1 | 1 | 1 | 1 | 0.807 | 1 | 1 | 1 | 0.03 |
| SE | 90109 | ENSG00000158106 | RHPN1 | chr8 | + | 143378695 | 143378820 | 143378268 | 143378346 | 143378911 | 143379078 | 3 | 1 | 3 | 0 | 2 | 0 | 0 | 0 | 0 | 0 | 0 | 3 | 0 | 7 | 0 | 3 | 0 | 0 | 0 | 0 | 274 | 149 | 1 | 1 | 1 | 1 | 1 | NA | 1 | 0 | 1 | NA | 1 | NA | 0.333 |
| SE | 90114 | ENSG00000253819 | LINC01151 | chr8 | - | 122562537 | 122562711 | 122559822 | 122561919 | 122670359 | 122670500 | 0 | 0 | 0 | 0 | 0 | 1 | 0 | 0 | 1 | 0 | 6 | 0 | 2 | 0 | 0 | 0 | 0 | 0 | 0 | 0 | 298 | 149 | 1 | 1 | NA | NA | NA | NA | 0 | 1 | NA | 1 | NA | NA | -1 |
| SE | 90118 | ENSG00000147592 | LACTB2 | chr8 | - | 70640901 | 70641050 | 70638547 | 70638629 | 70644064 | 70644243 | 152 | 96 | 83 | 110 | 249 | 2 | 3 | 0 | 0 | 2 | 52 | 59 | 38 | 47 | 29 | 0 | 0 | 0 | 0 | 0 | 298 | 149 | 1 | 1 | 0.974 | 0.941 | 1 | 1 | 0.984 | 1 | 1 | 1 | 1 | 1 | -0.02 |
| SE | 90119 | ENSG00000160932 | LY6E | chr8 | + | 143018832 | 143019119 | 143018527 | 143018586 | 143020261 | 143020377 | 0 | 0 | 0 | 1 | 0 | 1 | 1 | 1 | 3 | 0 | 0 | 3 | 0 | 8 | 0 | 9 | 3 | 0 | 4 | 0 | 298 | 149 | 1 | 1 | 0 | 0 | 0 | 0.143 | NA | 0 | 0.333 | NA | 0.5 | NA | -0.242 |
| SE | 90122 | ENSG00000160932 | LY6E | chr8 | + | 143020261 | 143020377 | 143018516 | 143018586 | 143020882 | 143020991 | 1 | 1 | 1 | 3 | 2 | 119 | 76 | 89 | 131 | 28 | 9 | 9 | 0 | 7 | 0 | 435 | 423 | 165 | 376 | 166 | 265 | 149 | 1 | 1 | 0.005 | 0.007 | 0.006 | 0.013 | 0.039 | 0.011 | 0.012 | 0 | 0.01 | 0 | 0.007 |
| SE | 90124 | ENSG00000160932 | LY6E | chr8 | + | 143020261 | 143020377 | 143018832 | 143019119 | 143020882 | 143020991 | 0 | 0 | 0 | 0 | 2 | 1 | 2 | 0 | 3 | 0 | 0 | 7 | 0 | 3 | 0 | 0 | 2 | 1 | 15 | 0 | 265 | 149 | 1 | 1 | 0 | 0 | NA | 0 | 1 | NA | 0.663 | 0 | 0.101 | NA | -0.005 |
| SE | 90125 | ENSG00000160932 | LY6E | chr8 | + | 143020261 | 143020377 | 143018832 | 143019119 | 143020887 | 143020991 | 0 | 1 | 1 | 0 | 0 | 1 | 1 | 0 | 1 | 0 | 13 | 1 | 1 | 7 | 0 | 0 | 4 | 0 | 5 | 0 | 265 | 149 | 1 | 1 | 0 | 0.36 | 1 | 0 | NA | 1 | 0.123 | 1 | 0.44 | NA | -0.301 |
| SE | 90126 | ENSG00000160932 | LY6E | chr8 | + | 143020882 | 143020991 | 143018528 | 143018586 | 143021313 | 143021433 | 424 | 257 | 318 | 460 | 267 | 0 | 0 | 0 | 1 | 0 | 1471 | 1367 | 608 | 1216 | 477 | 0 | 0 | 0 | 0 | 0 | 258 | 149 | 1 | 1 | 1 | 1 | 1 | 0.996 | 1 | 1 | 1 | 1 | 1 | 1 | -0.001 |
| SE | 90127 | ENSG00000160932 | LY6E | chr8 | + | 143020887 | 143020991 | 143018482 | 143018586 | 143021313 | 143021433 | 337 | 203 | 252 | 351 | 236 | 0 | 0 | 0 | 1 | 0 | 1171 | 1076 | 466 | 909 | 367 | 0 | 0 | 0 | 0 | 0 | 253 | 149 | 1 | 1 | 1 | 1 | 1 | 0.995 | 1 | 1 | 1 | 1 | 1 | 1 | -0.001 |
| SE | 90128 | ENSG00000160932 | LY6E | chr8 | + | 143020916 | 143020991 | 143018541 | 143018586 | 143021313 | 143021433 | 271 | 153 | 187 | 289 | 228 | 0 | 0 | 0 | 1 | 0 | 900 | 825 | 399 | 705 | 256 | 0 | 0 | 0 | 0 | 0 | 224 | 149 | 1 | 1 | 1 | 1 | 1 | 0.995 | 1 | 1 | 1 | 1 | 1 | 1 | -0.001 |
| SE | 90144 | ENSG00000185730 | ZNF696 | chr8 | + | 143295378 | 143295470 | 143292971 | 143293065 | 143295739 | 143295812 | 2 | 0 | 0 | 0 | 1 | 4 | 0 | 6 | 1 | 0 | 0 | 0 | 0 | 0 | 0 | 0 | 2 | 0 | 0 | 0 | 241 | 149 | 1 | 1 | 0.236 | NA | 0 | 0 | 1 | NA | 0 | NA | NA | NA | 0.309 |
| SE | 90148 | ENSG00000241560 | ZBTB20-AS1 | chr3 | + | 114367131 | 114367221 | 114366966 | 114367028 | 114387422 | 114387579 | 3 | 1 | 0 | 0 | 0 | 2 | 0 | 0 | 1 | 0 | 0 | 3 | 0 | 0 | 0 | 0 | 0 | 0 | 0 | 0 | 239 | 149 | 1 | 1 | 0.483 | 1 | NA | NA | NA | NA | 1 | NA | NA | NA | -0.258 |
| SE | 90149 | ENSG00000241560 | ZBTB20-AS1 | chr3 | + | 114367134 | 114367221 | 114366966 | 114367028 | 114387422 | 114387579 | 3 | 1 | 0 | 0 | 0 | 2 | 0 | 0 | 1 | 0 | 0 | 3 | 0 | 0 | 0 | 0 | 0 | 0 | 0 | 0 | 236 | 149 | 1 | 1 | 0.486 | 1 | NA | NA | NA | NA | 1 | NA | NA | NA | -0.257 |
| SE | 90151 | ENSG00000241560 | ZBTB20-AS1 | chr3 | + | 114375793 | 114375836 | 114366966 | 114367221 | 114387422 | 114387579 | 1 | 0 | 0 | 0 | 0 | 3 | 3 | 0 | 0 | 0 | 0 | 0 | 0 | 0 | 0 | 0 | 3 | 0 | 0 | 0 | 192 | 149 | 1 | 1 | 0.206 | 0 | NA | NA | NA | NA | 0 | NA | NA | NA | 0.103 |
| SE | 90152 | ENSG00000241560 | ZBTB20-AS1 | chr3 | + | 114387670 | 114387775 | 114387422 | 114387579 | 114388395 | 114388931 | 6 | 0 | 2 | 0 | 0 | 1 | 0 | 0 | 1 | 0 | 0 | 2 | 0 | 0 | 0 | 0 | 0 | 0 | 0 | 0 | 254 | 149 | 1 | 1 | 0.779 | NA | 1 | 0 | NA | NA | 1 | NA | NA | NA | -0.407 |
| SE | 90155 | ENSG00000035928 | RFC1 | chr4 | - | 39303059 | 39303151 | 39302736 | 39302874 | 39304813 | 39304928 | 182 | 146 | 180 | 212 | 161 | 0 | 0 | 1 | 0 | 0 | 31 | 128 | 208 | 48 | 28 | 0 | 0 | 0 | 0 | 0 | 241 | 149 | 1 | 1 | 1 | 1 | 0.991 | 1 | 1 | 1 | 1 | 1 | 1 | 1 | -0.002 |
| SE | 90156 | ENSG00000035928 | RFC1 | chr4 | - | 39306591 | 39306701 | 39304813 | 39304928 | 39307375 | 39307471 | 93 | 65 | 98 | 96 | 65 | 0 | 0 | 0 | 0 | 0 | 3 | 53 | 82 | 12 | 0 | 0 | 2 | 0 | 0 | 0 | 259 | 149 | 1 | 1 | 1 | 1 | 1 | 1 | 1 | 1 | 0.938 | 1 | 1 | NA | 0.016 |
| SE | 90158 | ENSG00000035928 | RFC1 | chr4 | - | 39306591 | 39306704 | 39304813 | 39304928 | 39308635 | 39309032 | 130 | 84 | 130 | 123 | 94 | 0 | 2 | 2 | 4 | 9 | 3 | 64 | 92 | 17 | 0 | 0 | 4 | 6 | 0 | 0 | 262 | 149 | 1 | 1 | 1 | 0.96 | 0.974 | 0.946 | 0.856 | 1 | 0.901 | 0.897 | 1 | NA | -0.002 |
| SE | 90159 | ENSG00000035928 | RFC1 | chr4 | - | 39307375 | 39307471 | 39304813 | 39304928 | 39308635 | 39309032 | 0 | 0 | 0 | 1 | 1 | 0 | 2 | 2 | 4 | 9 | 0 | 4 | 0 | 0 | 0 | 0 | 4 | 6 | 0 | 0 | 245 | 149 | 1 | 1 | NA | 0 | 0 | 0.132 | 0.063 | NA | 0.378 | 0 | NA | NA | -0.14 |
| SE | 90160 | ENSG00000035928 | RFC1 | chr4 | - | 39307375 | 39307471 | 39306591 | 39306701 | 39308635 | 39309032 | 0 | 0 | 0 | 2 | 1 | 139 | 96 | 143 | 136 | 128 | 0 | 2 | 0 | 0 | 0 | 0 | 65 | 95 | 19 | 1 | 245 | 149 | 1 | 1 | 0 | 0 | 0 | 0.009 | 0.005 | NA | 0.018 | 0 | 0 | 0 | -0.002 |
| SE | 90161 | ENSG00000035928 | RFC1 | chr4 | - | 39307375 | 39307471 | 39306633 | 39306704 | 39308635 | 39309032 | 0 | 0 | 0 | 3 | 1 | 37 | 21 | 32 | 27 | 29 | 0 | 3 | 3 | 0 | 0 | 0 | 11 | 10 | 11 | 0 | 245 | 149 | 1 | 1 | 0 | 0 | 0 | 0.063 | 0.021 | NA | 0.142 | 0.154 | 0 | NA | -0.082 |
| SE | 90162 | ENSG00000035928 | RFC1 | chr4 | - | 39308635 | 39309032 | 39304813 | 39304928 | 39311444 | 39311549 | 46 | 66 | 59 | 86 | 76 | 3 | 0 | 1 | 3 | 7 | 12 | 70 | 106 | 37 | 0 | 0 | 9 | 10 | 2 | 0 | 298 | 149 | 1 | 1 | 0.885 | 1 | 0.967 | 0.935 | 0.844 | 1 | 0.795 | 0.841 | 0.902 | NA | 0.042 |
| SE | 90164 | ENSG00000035928 | RFC1 | chr4 | - | 39320382 | 39320669 | 39316914 | 39317022 | 39327523 | 39327756 | 173 | 130 | 172 | 134 | 303 | 0 | 0 | 0 | 2 | 1 | 4 | 160 | 148 | 87 | 3 | 0 | 0 | 0 | 0 | 0 | 298 | 149 | 1 | 1 | 1 | 1 | 1 | 0.971 | 0.993 | 1 | 1 | 1 | 1 | 1 | -0.007 |
| SE | 90166 | ENSG00000035928 | RFC1 | chr4 | - | 39321286 | 39321374 | 39320382 | 39320669 | 39326562 | 39326640 | 75 | 60 | 100 | 94 | 179 | 1 | 2 | 0 | 2 | 0 | 0 | 64 | 72 | 25 | 12 | 0 | 12 | 3 | 0 | 0 | 237 | 149 | 1 | 1 | 0.979 | 0.95 | 1 | 0.967 | 1 | NA | 0.77 | 0.938 | 1 | 1 | 0.052 |
| SE | 90167 | ENSG00000035928 | RFC1 | chr4 | - | 39321286 | 39321374 | 39320382 | 39320669 | 39327523 | 39327756 | 56 | 48 | 86 | 68 | 166 | 1 | 0 | 0 | 1 | 1 | 0 | 40 | 38 | 17 | 9 | 0 | 0 | 0 | 0 | 0 | 237 | 149 | 1 | 1 | 1 | 1 | 1 | 0.977 | 0.991 | NA | 1 | 1 | 1 | 1 | -0.006 |
| SE | 90168 | ENSG00000035928 | RFC1 | chr4 | - | 39323339 | 39323417 | 39320382 | 39320669 | 39326562 | 39326640 | 100 | 86 | 108 | 109 | 189 | 1 | 2 | 0 | 2 | 0 | 0 | 87 | 67 | 58 | 2 | 0 | 12 | 3 | 0 | 0 | 227 | 149 | 1 | 1 | 0.985 | 0.966 | 1 | 0.973 | 1 | NA | 0.826 | 0.936 | 1 | 1 | 0.044 |
| SE | 90170 | ENSG00000035928 | RFC1 | chr4 | - | 39326562 | 39326640 | 39320382 | 39320669 | 39327523 | 39327756 | 61 | 85 | 70 | 87 | 123 | 0 | 0 | 0 | 1 | 1 | 6 | 111 | 101 | 35 | 22 | 0 | 0 | 0 | 0 | 0 | 227 | 149 | 1 | 1 | 1 | 1 | 1 | 0.983 | 0.988 | 1 | 1 | 1 | 1 | 1 | -0.006 |
| SE | 90171 | ENSG00000035928 | RFC1 | chr4 | - | 39326562 | 39326640 | 39321286 | 39321374 | 39327523 | 39327756 | 79 | 95 | 85 | 111 | 136 | 0 | 0 | 1 | 0 | 0 | 6 | 123 | 137 | 43 | 25 | 0 | 0 | 1 | 0 | 0 | 227 | 149 | 1 | 1 | 1 | 1 | 0.982 | 1 | 1 | 1 | 1 | 0.989 | 1 | 1 | -0.001 |
| SE | 90172 | ENSG00000035928 | RFC1 | chr4 | - | 39326562 | 39326640 | 39323339 | 39323417 | 39327523 | 39327756 | 203 | 208 | 216 | 255 | 417 | 2 | 1 | 0 | 1 | 0 | 13 | 230 | 213 | 107 | 24 | 0 | 0 | 1 | 2 | 0 | 227 | 149 | 1 | 1 | 0.985 | 0.993 | 1 | 0.994 | 1 | 1 | 1 | 0.993 | 0.972 | 1 | 0.001 |
| SE | 90173 | ENSG00000035928 | RFC1 | chr4 | - | 39341486 | 39341665 | 39327523 | 39327756 | 39342344 | 39342467 | 0 | 0 | 0 | 0 | 0 | 103 | 66 | 97 | 106 | 219 | 0 | 0 | 2 | 0 | 0 | 2 | 128 | 117 | 80 | 0 | 298 | 149 | 1 | 1 | 0 | 0 | 0 | 0 | 0 | 0 | 0 | 0.008 | 0 | NA | -0.002 |
| SE | 90174 | ENSG00000035928 | RFC1 | chr4 | - | 39355876 | 39355997 | 39351347 | 39351476 | 39366238 | 39366362 | 0 | 0 | 4 | 2 | 0 | 40 | 35 | 42 | 52 | 36 | 0 | 0 | 0 | 0 | 0 | 17 | 80 | 44 | 14 | 0 | 270 | 149 | 1 | 1 | 0 | 0 | 0.05 | 0.021 | 0 | 0 | 0 | 0 | 0 | NA | 0.014 |
| SE | 90175 | ENSG00000198876 | DCAF12 | chr9 | - | 34089411 | 34089590 | 34086386 | 34088508 | 34093285 | 34093448 | 207 | 139 | 162 | 165 | 194 | 0 | 0 | 0 | 0 | 0 | 255 | 209 | 112 | 246 | 293 | 4 | 0 | 0 | 0 | 0 | 298 | 149 | 1 | 1 | 1 | 1 | 1 | 1 | 1 | 0.97 | 1 | 1 | 1 | 1 | 0.006 |
| SE | 90177 | ENSG00000198876 | DCAF12 | chr9 | - | 34096715 | 34096781 | 34093285 | 34093448 | 34098323 | 34098517 | 134 | 85 | 120 | 140 | 161 | 1 | 0 | 0 | 3 | 0 | 116 | 103 | 85 | 173 | 218 | 0 | 0 | 3 | 0 | 0 | 215 | 149 | 1 | 1 | 0.989 | 1 | 1 | 0.97 | 1 | 1 | 1 | 0.952 | 1 | 1 | 0.001 |
| SE | 90178 | ENSG00000198876 | DCAF12 | chr9 | - | 34106433 | 34106494 | 34098323 | 34098517 | 34107358 | 34107565 | 228 | 104 | 154 | 178 | 257 | 2 | 0 | 1 | 0 | 0 | 215 | 128 | 77 | 246 | 276 | 0 | 0 | 1 | 0 | 0 | 210 | 149 | 1 | 1 | 0.988 | 1 | 0.991 | 1 | 1 | 1 | 1 | 0.982 | 1 | 1 | -0.001 |
| SE | 90181 | ENSG00000171045 | TSNARE1 | chr8 | - | 142284412 | 142284485 | 142274780 | 142274863 | 142300485 | 142300644 | 25 | 16 | 16 | 25 | 31 | 0 | 0 | 0 | 0 | 0 | 0 | 39 | 19 | 2 | 78 | 0 | 0 | 5 | 0 | 0 | 222 | 149 | 1 | 1 | 1 | 1 | 1 | 1 | 1 | NA | 1 | 0.718 | 1 | 1 | 0.071 |
| SE | 90182 | ENSG00000171045 | TSNARE1 | chr8 | - | 142318543 | 142318634 | 142315002 | 142315089 | 142330900 | 142330970 | 17 | 6 | 21 | 15 | 3 | 0 | 0 | 2 | 0 | 0 | 14 | 8 | 3 | 9 | 46 | 0 | 0 | 0 | 0 | 0 | 240 | 149 | 1 | 1 | 1 | 1 | 0.867 | 1 | 1 | 1 | 1 | 1 | 1 | 1 | -0.027 |
| SE | 90184 | ENSG00000171045 | TSNARE1 | chr8 | - | 142343965 | 142344472 | 142331753 | 142331831 | 142354636 | 142354724 | 9 | 0 | 12 | 6 | 4 | 0 | 0 | 1 | 0 | 0 | 0 | 5 | 2 | 3 | 3 | 3 | 0 | 0 | 2 | 14 | 298 | 149 | 1 | 1 | 1 | NA | 0.857 | 1 | 1 | 0 | 1 | 1 | 0.429 | 0.097 | 0.459 |
| SE | 90185 | ENSG00000171045 | TSNARE1 | chr8 | - | 142345742 | 142345892 | 142343965 | 142344472 | 142354636 | 142354724 | 18 | 8 | 31 | 33 | 28 | 0 | 0 | 0 | 1 | 2 | 10 | 5 | 15 | 36 | 65 | 0 | 0 | 0 | 0 | 0 | 298 | 149 | 1 | 1 | 1 | 1 | 1 | 0.943 | 0.875 | 1 | 1 | 1 | 1 | 1 | -0.036 |
| SE | 90186 | ENSG00000171045 | TSNARE1 | chr8 | - | 142354636 | 142354763 | 142345742 | 142345892 | 142403103 | 142403168 | 9 | 6 | 14 | 17 | 8 | 0 | 0 | 0 | 0 | 0 | 0 | 4 | 6 | 15 | 31 | 0 | 0 | 0 | 3 | 0 | 276 | 149 | 1 | 1 | 1 | 1 | 1 | 1 | 1 | NA | 1 | 1 | 0.73 | 1 | 0.068 |
| SE | 90187 | ENSG00000171045 | TSNARE1 | chr8 | - | 142365910 | 142366017 | 142354636 | 142354763 | 142403103 | 142403182 | 0 | 0 | 0 | 1 | 6 | 4 | 3 | 6 | 5 | 0 | 0 | 1 | 0 | 0 | 8 | 0 | 4 | 0 | 5 | 0 | 256 | 149 | 1 | 1 | 0 | 0 | 0 | 0.104 | 1 | NA | 0.127 | NA | 0 | 1 | -0.155 |
| SE | 90190 | ENSG00000198169 | ZNF251 | chr8 | - | 144754191 | 144754321 | 144753453 | 144753796 | 144754695 | 144754803 | 13 | 6 | 9 | 9 | 23 | 3 | 0 | 0 | 0 | 0 | 7 | 12 | 9 | 17 | 23 | 0 | 0 | 0 | 0 | 0 | 279 | 149 | 1 | 1 | 0.698 | 1 | 1 | 1 | 1 | 1 |  | 1 | 1 | 1 | -0.06 |
| SE | 90192 | ENSG00000177335 | C8orf31 | chr8 | + | 143042957 | 143043087 | 143039260 | 143039569 | 143043177 | 143043262 | 0 | 0 | 0 | 0 | 0 | 1 | 0 | 0 | 0 | 0 | 0 | 3 | 0 | 0 | 0 | 0 | 0 | 0 | 0 | 0 | 279 | 149 | 1 | 1 | 0 | NA | NA | NA | NA | NA | 1 | NA | NA | NA | -1 |
| SE | 90207 | ENSG00000104324 | CPQ | chr8 | + | 96834972 | 96835180 | 96784863 | 96785330 | 96879797 | 96880005 | 389 | 200 | 152 | 267 | 219 | 0 | 0 | 0 | 0 | 0 | 581 | 336 | 363 | 219 | 663 | 0 | 0 | 9 | 0 | 0 | 298 | 149 | 1 | 1 | 1 | 1 | 1 | 1 | 1 | 1 | 1 | 0.953 | 1 | 1 | 0.009 |
| SE | 90208 | ENSG00000104324 | CPQ | chr8 | + | 96879797 | 96880005 | 96834972 | 96835180 | 96965934 | 96966046 | 220 | 107 | 123 | 176 | 125 | 1 | 0 | 0 | 0 | 0 | 234 | 252 | 176 | 145 | 391 | 0 | 0 | 0 | 0 | 0 | 298 | 149 | 1 | 1 | 0.991 | 1 | 1 | 1 | 1 | 1 | 1 | 1 | 1 | 1 | -0.002 |
| SE | 90209 | ENSG00000104324 | CPQ | chr8 | + | 96965934 | 96966046 | 96879797 | 96880005 | 97029402 | 97029494 | 178 | 69 | 92 | 148 | 105 | 0 | 0 | 0 | 0 | 0 | 243 | 147 | 126 | 125 | 250 | 0 | 2 | 1 | 0 | 0 | 261 | 149 | 1 | 1 | 1 | 1 | 1 | 1 | 1 | 1 | 0.977 | 0.986 | 1 | 1 | 0.007 |
| SE | 90210 | ENSG00000104324 | CPQ | chr8 | + | 97056405 | 97056545 | 97055526 | 97055645 | 97066008 | 97066210 | 1 | 8 | 1 | 1 | 0 | 1 | 1 | 2 | 0 | 5 | 15 | 0 | 10 | 0 | 2 | 0 | 0 | 2 | 3 | 4 | 289 | 149 | 1 | 1 | 0.34 | 0.805 | 0.205 | 1 | 0 | 1 | NA | 0.721 | 0 | 0.205 | -0.012 |
| SE | 90211 | ENSG00000104324 | CPQ | chr8 | + | 97066008 | 97066210 | 97029402 | 97029494 | 97143019 | 97143188 | 166 | 57 | 108 | 96 | 114 | 0 | 0 | 0 | 0 | 0 | 139 | 112 | 117 | 104 | 281 | 0 | 5 | 0 | 0 | 0 | 298 | 149 | 1 | 1 | 1 | 1 | 1 | 1 | 1 | 1 | 0.918 | 1 | 1 | 1 | 0.016 |
| SE | 90212 | ENSG00000104324 | CPQ | chr8 | + | 97066008 | 97066210 | 97055526 | 97055645 | 97143019 | 97143188 | 62 | 24 | 61 | 39 | 63 | 0 | 0 | 1 | 0 | 0 | 87 | 40 | 41 | 23 | 108 | 0 | 0 | 0 | 0 | 0 | 298 | 149 | 1 | 1 | 1 | 1 | 0.968 | 1 | 1 | 1 | 1 | 1 | 1 | 1 | -0.006 |
| SE | 90213 | ENSG00000132294 | EFR3A | chr8 | + | 131949968 | 131950090 | 131946482 | 131946633 | 131953817 | 131953836 | 287 | 169 | 185 | 230 | 398 | 0 | 4 | 0 | 0 | 0 | 80 | 106 | 88 | 98 | 67 | 0 | 0 | 0 | 0 | 0 | 271 | 149 | 1 | 1 | 1 | 0.959 | 1 | 1 | 1 | 1 | 1 | 1 | 1 | 1 | -0.008 |
| SE | 90214 | ENSG00000132294 | EFR3A | chr8 | + | 131959584 | 131959663 | 131955767 | 131955905 | 131968294 | 131968430 | 381 | 285 | 276 | 370 | 506 | 0 | 0 | 0 | 2 | 0 | 112 | 231 | 156 | 135 | 95 | 0 | 0 | 0 | 0 | 0 | 228 | 149 | 1 | 1 | 1 | 1 | 1 | 0.992 | 1 | 1 | 1 | 1 | 1 | 1 | -0.002 |
| SE | 90215 | ENSG00000132294 | EFR3A | chr8 | + | 131976026 | 131976141 | 131970475 | 131970643 | 131977040 | 131977092 | 478 | 374 | 361 | 380 | 392 | 0 | 0 | 0 | 0 | 0 | 99 | 283 | 208 | 236 | 45 | 0 | 0 | 5 | 0 | 0 | 264 | 149 | 1 | 1 | 1 | 1 | 1 | 1 | 1 | 1 | 1 | 0.959 | 1 | 1 | 0.008 |
| SE | 90228 | ENSG00000172269 | DPAGT1 | chr11 | - | 119100629 | 119100750 | 119100261 | 119100408 | 119101017 | 119101138 | 12 | 10 | 10 | 31 | 7 | 0 | 0 | 1 | 0 | 0 | 9 | 26 | 7 | 6 | 47 | 0 | 0 | 0 | 0 | 0 | 270 | 149 | 1 | 1 | 1 | 1 | 0.847 | 1 | 1 | 1 | 1 | 1 | 1 | 1 | -0.031 |
| SE | 90229 | ENSG00000172269 | DPAGT1 | chr11 | - | 119100629 | 119100827 | 119100261 | 119100408 | 119101017 | 119101138 | 12 | 10 | 14 | 33 | 8 | 0 | 0 | 1 | 0 | 0 | 31 | 26 | 9 | 8 | 54 | 0 | 0 | 0 | 0 | 0 | 298 | 149 | 1 | 1 | 1 | 1 | 0.875 | 1 | 1 | 1 | 1 | 1 | 1 | 1 | -0.025 |
| SE | 90231 | ENSG00000172269 | DPAGT1 | chr11 | - | 119100629 | 119100843 | 119100261 | 119100408 | 119101017 | 119101047 | 21 | 18 | 15 | 49 | 14 | 0 | 0 | 1 | 0 | 0 | 47 | 33 | 13 | 13 | 69 | 0 | 0 | 0 | 0 | 0 | 298 | 149 | 1 | 1 | 1 | 1 | 0.882 | 1 | 1 | 1 | 1 | 1 | 1 | 1 | -0.024 |
| SE | 90233 | ENSG00000172269 | DPAGT1 | chr11 | - | 119100629 | 119100877 | 119100261 | 119100408 | 119101017 | 119101138 | 12 | 10 | 14 | 33 | 8 | 0 | 0 | 1 | 0 | 0 | 31 | 26 | 9 | 8 | 54 | 0 | 0 | 0 | 0 | 0 | 298 | 149 | 1 | 1 | 1 | 1 | 0.875 | 1 | 1 | 1 | 1 | 1 | 1 | 1 | -0.025 |
| SE | 90235 | ENSG00000172269 | DPAGT1 | chr11 | - | 119100629 | 119101138 | 119100261 | 119100408 | 119101494 | 119101878 | 47 | 22 | 25 | 57 | 13 | 0 | 0 | 0 | 1 | 0 | 62 | 48 | 35 | 23 | 85 | 0 | 0 | 0 | 0 | 5 | 298 | 149 | 1 | 1 | 1 | 1 | 1 | 0.966 | 1 | 1 | 1 | 1 | 1 | 0.895 | 0.014 |
| SE | 90238 | ENSG00000172269 | DPAGT1 | chr11 | - | 119101017 | 119101138 | 119100629 | 119100827 | 119101494 | 119101777 | 35 | 12 | 11 | 24 | 5 | 1 | 0 | 1 | 0 | 0 | 31 | 22 | 26 | 15 | 31 | 0 | 0 | 0 | 0 | 0 | 270 | 149 | 1 | 1 | 0.951 | 1 | 0.859 | 1 | 1 | 1 | 1 | 1 | 1 | 1 | -0.038 |
| SE | 90241 | ENSG00000066855 | MTFR1 | chr8 | + | 65682043 | 65682080 | 65669870 | 65670018 | 65682352 | 65682451 | 0 | 0 | 0 | 0 | 0 | 17 | 11 | 12 | 16 | 25 | 0 | 4 | 1 | 0 | 0 | 6 | 4 | 11 | 14 | 0 | 186 | 149 | 1 | 1 | 0 | 0 | 0 | 0 | 0 | 0 | 0.445 | 0.068 | 0 | NA | -0.128 |
| SE | 90245 | ENSG00000066855 | MTFR1 | chr8 | + | 65693643 | 65693759 | 65682327 | 65682451 | 65704693 | 65704747 | 44 | 38 | 26 | 46 | 94 | 0 | 0 | 1 | 0 | 0 | 21 | 24 | 34 | 23 | 19 | 0 | 0 | 0 | 0 | 0 | 265 | 149 | 1 | 1 | 1 | 1 | 0.936 | 1 | 1 | 1 | 1 | 1 | 1 | 1 | -0.013 |
| SE | 90246 | ENSG00000066855 | MTFR1 | chr8 | + | 65719379 | 65719481 | 65707842 | 65708011 | 65728360 | 65728603 | 6 | 4 | 6 | 5 | 8 | 0 | 0 | 0 | 0 | 0 | 0 | 0 | 9 | 4 | 0 | 0 | 0 | 0 | 2 | 0 | 251 | 149 | 1 | 1 | 1 | 1 | 1 | 1 | 1 | NA | NA | 1 | 0.543 | NA | 0.228 |
| SE | 90247 | ENSG00000066855 | MTFR1 | chr8 | + | 65719379 | 65719481 | 65707842 | 65708011 | 65770946 | 65771261 | 4 | 3 | 6 | 5 | 7 | 0 | 0 | 0 | 0 | 0 | 0 | 0 | 6 | 4 | 0 | 0 | 0 | 2 | 0 | 0 | 251 | 149 | 1 | 1 | 1 | 1 | 1 | 1 | 1 | NA | NA | 0.64 | 1 | NA | 0.18 |
| SE | 90257 | ENSG00000249859 | PVT1 | chr8 | + | 127795893 | 127796008 | 127794541 | 127794734 | 127855154 | 127855319 | 3 | 0 | 0 | 0 | 4 | 0 | 0 | 0 | 2 | 0 | 1 | 3 | 2 | 4 | 13 | 9 | 0 | 0 | 0 | 5 | 264 | 149 | 1 | 1 | 1 | NA | NA | 0 | 1 | 0.059 | 1 | 1 | 1 | 0.595 | -0.064 |
| SE | 90261 | ENSG00000249859 | PVT1 | chr8 | + | 127795893 | 127796071 | 127794530 | 127794734 | 127855154 | 127855319 | 1 | 0 | 0 | 0 | 0 | 0 | 0 | 0 | 2 | 0 | 1 | 7 | 2 | 4 | 13 | 9 | 0 | 0 | 0 | 5 | 298 | 149 | 1 | 1 | 1 | NA | NA | 0 | NA | 0.053 | 1 | 1 | 1 | 0.565 | -0.224 |
| SE | 90271 | ENSG00000249859 | PVT1 | chr8 | + | 127818899 | 127818998 | 127794699 | 127794734 | 127819908 | 127820382 | 0 | 0 | 1 | 0 | 0 | 1 | 0 | 0 | 0 | 0 | 0 | 0 | 0 | 2 | 0 | 0 | 0 | 0 | 0 | 0 | 248 | 149 | 1 | 1 | 0 | NA | 1 | NA | NA | NA | NA | NA | 1 | NA | -0.5 |
| SE | 90274 | ENSG00000249859 | PVT1 | chr8 | + | 127855154 | 127855319 | 127795220 | 127796008 | 127890588 | 127890998 | 4 | 1 | 1 | 2 | 9 | 6 | 7 | 7 | 6 | 7 | 0 | 0 | 2 | 1 | 0 | 12 | 20 | 11 | 5 | 0 | 298 | 149 | 1 | 1 | 0.25 | 0.067 | 0.067 | 0.143 | 0.391 | 0 | 0 | 0.083 | 0.091 | NA | 0.14 |
| SE | 90275 | ENSG00000249859 | PVT1 | chr8 | + | 127855154 | 127855319 | 127795357 | 127796071 | 127890588 | 127890998 | 2 | 1 | 1 | 2 | 4 | 2 | 1 | 1 | 8 | 4 | 0 | 0 | 1 | 1 | 0 | 0 | 8 | 10 | 12 | 13 | 298 | 149 | 1 | 1 | 0.333 | 0.333 | 0.333 | 0.111 | 0.333 | NA | 0 | 0.048 | 0.04 | 0 | 0.267 |
| SE | 90287 | ENSG00000249859 | PVT1 | chr8 | + | 127932027 | 127932198 | 127890588 | 127890998 | 127932464 | 127932701 | 1 | 0 | 0 | 1 | 0 | 4 | 8 | 12 | 10 | 23 | 0 | 0 | 0 | 0 | 0 | 26 | 34 | 15 | 10 | 14 | 298 | 149 | 1 | 1 | 0.111 | 0 | 0 | 0.048 | 0 | 0 | 0 | 0 | 0 | 0 | 0.032 |
| SE | 90288 | ENSG00000249859 | PVT1 | chr8 | + | 127932052 | 127932198 | 127890588 | 127890998 | 127932464 | 127932700 | 1 | 0 | 0 | 1 | 0 | 4 | 8 | 12 | 10 | 23 | 0 | 0 | 0 | 0 | 0 | 26 | 34 | 15 | 10 | 14 | 295 | 149 | 1 | 1 | 0.112 | 0 | 0 | 0.048 | 0 | 0 | 0 | 0 | 0 | 0 | 0.032 |
| SE | 90289 | ENSG00000249859 | PVT1 | chr8 | + | 127932464 | 127932474 | 127890588 | 127890998 | 127939507 | 127939676 | 1 | 0 | 3 | 0 | 0 | 8 | 6 | 12 | 9 | 7 | 0 | 0 | 0 | 0 | 0 | 6 | 23 | 9 | 4 | 5 | 159 | 149 | 1 | 1 | 0.105 | 0 | 0.19 | 0 | 0 | 0 | 0 | 0 | 0 | 0 | 0.059 |
| SE | 90291 | ENSG00000249859 | PVT1 | chr8 | + | 127939507 | 127939676 | 127890586 | 127890998 | 127969415 | 127969537 | 8 | 6 | 13 | 9 | 7 | 0 | 0 | 0 | 1 | 0 | 6 | 23 | 9 | 4 | 9 | 0 | 0 | 0 | 0 | 0 | 298 | 149 | 1 | 1 | 1 | 1 | 1 | 0.818 | 1 | 1 | 1 | 1 | 1 | 1 | -0.036 |
| SE | 90292 | ENSG00000249859 | PVT1 | chr8 | + | 127939507 | 127939676 | 127890586 | 127890998 | 127983903 | 127984204 | 8 | 6 | 13 | 10 | 8 | 0 | 0 | 0 | 0 | 0 | 6 | 23 | 9 | 4 | 5 | 0 | 0 | 2 | 0 | 0 | 298 | 149 | 1 | 1 | 1 | 1 | 1 | 1 | 1 | 1 | 1 | 0.692 | 1 | 1 | 0.062 |
| SE | 90294 | ENSG00000249859 | PVT1 | chr8 | + | 127969415 | 127969537 | 127890588 | 127890998 | 127989161 | 127989291 | 1 | 0 | 1 | 1 | 0 | 1 | 1 | 2 | 3 | 1 | 0 | 0 | 6 | 2 | 0 | 0 | 0 | 3 | 3 | 0 | 271 | 149 | 1 | 1 | 0.355 | 0 | 0.216 | 0.155 | 0 | NA | NA | 0.524 | 0.268 | NA | -0.251 |
| SE | 90295 | ENSG00000249859 | PVT1 | chr8 | + | 127969415 | 127969537 | 127939507 | 127939676 | 127982359 | 127982479 | 0 | 0 | 1 | 0 | 0 | 1 | 0 | 0 | 0 | 0 | 0 | 0 | 0 | 0 | 4 | 0 | 0 | 0 | 0 | 0 | 271 | 149 | 1 | 1 | 0 | NA | 1 | NA | NA | NA | NA | NA | NA | 1 | -0.5 |
| SE | 90297 | ENSG00000249859 | PVT1 | chr8 | + | 127982359 | 127982479 | 127939507 | 127939676 | 127983903 | 127984204 | 1 | 1 | 6 | 4 | 0 | 0 | 0 | 1 | 1 | 1 | 0 | 2 | 0 | 0 | 4 | 0 | 0 | 0 | 0 | 0 | 269 | 149 | 1 | 1 | 1 | 1 | 0.769 | 0.689 | 0 | NA | 1 | NA | NA | 1 | -0.308 |
| SE | 90298 | ENSG00000249859 | PVT1 | chr8 | + | 127982359 | 127982479 | 127939507 | 127939676 | 127989161 | 127989291 | 1 | 0 | 2 | 0 | 0 | 6 | 2 | 11 | 9 | 2 | 0 | 0 | 0 | 0 | 0 | 25 | 7 | 9 | 6 | 6 | 269 | 149 | 1 | 1 | 0.085 | 0 | 0.091 | 0 | 0 | 0 | 0 | 0 | 0 | 0 | 0.035 |
| SE | 90299 | ENSG00000249859 | PVT1 | chr8 | + | 127982359 | 127982479 | 127969415 | 127969537 | 127989161 | 127989291 | 0 | 0 | 2 | 0 | 0 | 1 | 0 | 1 | 0 | 0 | 0 | 0 | 0 | 0 | 4 | 0 | 0 | 6 | 2 | 0 | 269 | 149 | 1 | 1 | 0 | NA | 0.526 | NA | NA | NA | NA | 0 | 0 | 1 | -0.07 |
| SE | 90302 | ENSG00000249859 | PVT1 | chr8 | + | 127983903 | 127984204 | 127982359 | 127982479 | 127989161 | 127989291 | 10 | 3 | 18 | 15 | 16 | 0 | 0 | 2 | 0 | 0 | 20 | 6 | 3 | 13 | 8 | 0 | 0 | 0 | 0 | 0 | 298 | 149 | 1 | 1 | 1 | 1 | 0.818 | 1 | 1 | 1 | 1 | 1 | 1 | 1 | -0.036 |
| SE | 90303 | ENSG00000249859 | PVT1 | chr8 | + | 127989161 | 127989291 | 127890588 | 127890998 | 128009589 | 128009718 | 22 | 4 | 21 | 17 | 45 | 0 | 0 | 0 | 0 | 0 | 22 | 38 | 15 | 21 | 30 | 0 | 0 | 2 | 0 | 0 | 279 | 149 | 1 | 1 | 1 | 1 | 1 | 1 | 1 | 1 | 1 | 0.8 | 1 | 1 | 0.04 |
| SE | 90305 | ENSG00000249859 | PVT1 | chr8 | + | 127994882 | 127995093 | 127989161 | 127989291 | 127996560 | 127996598 | 0 | 0 | 0 | 3 | 0 | 11 | 4 | 29 | 8 | 29 | 6 | 0 | 2 | 7 | 0 | 13 | 6 | 14 | 4 | 0 | 298 | 149 | 1 | 1 | 0 | 0 | 0 | 0.158 | 0 | 0.188 | 0 | 0.067 | 0.467 | NA | -0.149 |
| SE | 90306 | ENSG00000249859 | PVT1 | chr8 | + | 127996560 | 127996598 | 127989161 | 127989291 | 127998199 | 127998359 | 1 | 0 | 16 | 5 | 5 | 0 | 0 | 0 | 0 | 0 | 13 | 5 | 9 | 0 | 0 | 0 | 0 | 1 | 0 | 0 | 187 | 149 | 1 | 1 | 1 | NA | 1 | 1 | 1 | 1 | 1 | 0.878 | NA | NA | 0.041 |
| SE | 90307 | ENSG00000249859 | PVT1 | chr8 | + | 127996560 | 127996598 | 127989161 | 127989291 | 127998911 | 127999334 | 1 | 0 | 19 | 5 | 6 | 0 | 0 | 1 | 1 | 0 | 13 | 3 | 6 | 0 | 0 | 0 | 0 | 0 | 0 | 7 | 187 | 149 | 1 | 1 | 1 | NA | 0.938 | 0.799 | 1 | 1 | 1 | 1 | NA | 0 | 0.184 |
| SE | 90310 | ENSG00000249859 | PVT1 | chr8 | + | 127998199 | 127998359 | 127996560 | 127996598 | 128009589 | 128009718 | 9 | 2 | 5 | 1 | 2 | 0 | 0 | 1 | 0 | 0 | 3 | 10 | 9 | 4 | 11 | 0 | 0 | 0 | 0 | 0 | 298 | 149 | 1 | 1 | 1 | 1 | 0.714 | 1 | 1 | 1 | 1 | 1 | 1 | 1 | -0.057 |
| SE | 90314 | ENSG00000249859 | PVT1 | chr8 | + | 128010317 | 128010444 | 128008878 | 128009718 | 128070159 | 128070272 | 52 | 37 | 52 | 20 | 79 | 0 | 0 | 0 | 0 | 1 | 67 | 64 | 34 | 20 | 45 | 0 | 0 | 1 | 0 | 0 | 276 | 149 | 1 | 1 | 1 | 1 | 1 | 1 | 0.977 | 1 | 1 | 0.948 | 1 | 1 | 0.006 |
| SE | 90316 | ENSG00000249859 | PVT1 | chr8 | + | 128031958 | 128032272 | 128010317 | 128010444 | 128070159 | 128070272 | 0 | 0 | 0 | 0 | 0 | 19 | 9 | 18 | 5 | 37 | 0 | 0 | 0 | 0 | 9 | 26 | 15 | 7 | 0 | 13 | 298 | 149 | 1 | 1 | 0 | 0 | 0 | 0 | 0 | 0 | 0 | 0 | NA | 0.257 | -0.064 |
| SE | 90318 | ENSG00000249859 | PVT1 | chr8 | + | 128038916 | 128038980 | 127989161 | 127989291 | 128070159 | 128071539 | 0 | 0 | 1 | 0 | 0 | 14 | 1 | 6 | 4 | 7 | 0 | 0 | 0 | 0 | 0 | 0 | 0 | 4 | 0 | 10 | 213 | 149 | 1 | 1 | 0 | 0 | 0.104 | 0 | 0 | NA | NA | 0 | NA | 0 | 0.021 |
| SE | 90320 | ENSG00000249859 | PVT1 | chr8 | + | 128038916 | 128038980 | 128010317 | 128010444 | 128070159 | 128071539 | 0 | 0 | 1 | 0 | 0 | 19 | 9 | 18 | 5 | 37 | 0 | 0 | 4 | 0 | 0 | 26 | 15 | 7 | 0 | 13 | 213 | 149 | 1 | 1 | 0 | 0 | 0.037 | 0 | 0 | 0 | 0 | 0.286 | NA | 0 | -0.064 |
| SE | 90321 | ENSG00000249859 | PVT1 | chr8 | + | 128038916 | 128038980 | 128010317 | 128010444 | 128096517 | 128096763 | 0 | 0 | 0 | 0 | 0 | 5 | 5 | 12 | 10 | 15 | 0 | 0 | 7 | 0 | 0 | 10 | 4 | 7 | 3 | 0 | 213 | 149 | 1 | 1 | 0 | 0 | 0 | 0 | 0 | 0 | 0 | 0.412 | 0 | NA | -0.103 |
| SE | 90322 | ENSG00000249859 | PVT1 | chr8 | + | 128048447 | 128048609 | 127989161 | 127989291 | 128070159 | 128070272 | 2 | 1 | 0 | 0 | 0 | 14 | 1 | 6 | 4 | 7 | 0 | 0 | 0 | 0 | 0 | 0 | 0 | 4 | 0 | 10 | 298 | 149 | 1 | 1 | 0.067 | 0.333 | 0 | 0 | 0 | NA | NA | 0 | NA | 0 | 0.08 |
| SE | 90324 | ENSG00000249859 | PVT1 | chr8 | + | 128048447 | 128048609 | 128010317 | 128010444 | 128070159 | 128070272 | 4 | 3 | 0 | 2 | 0 | 19 | 9 | 18 | 5 | 37 | 0 | 0 | 0 | 0 | 0 | 26 | 15 | 7 | 0 | 13 | 298 | 149 | 1 | 1 | 0.095 | 0.143 | 0 | 0.167 | 0 | 0 | 0 | 0 | NA | 0 | 0.081 |
| SE | 90325 | ENSG00000249859 | PVT1 | chr8 | + | 128048447 | 128048609 | 128010317 | 128010444 | 128096517 | 128096654 | 2 | 2 | 0 | 2 | 0 | 5 | 5 | 12 | 10 | 15 | 0 | 0 | 0 | 0 | 0 | 10 | 4 | 7 | 3 | 0 | 298 | 149 | 1 | 1 | 0.167 | 0.167 | 0 | 0.091 | 0 | 0 | 0 | 0 | 0 | NA | 0.085 |
| SE | 90330 | ENSG00000249859 | PVT1 | chr8 | + | 128049584 | 128049748 | 128049151 | 128049441 | 128070159 | 128070272 | 1 | 0 | 0 | 0 | 0 | 0 | 0 | 0 | 0 | 0 | 0 | 0 | 0 | 0 | 0 | 12 | 0 | 0 | 0 | 0 | 298 | 149 | 1 | 1 | 1 | NA | NA | NA | NA | 0 | NA | NA | NA | NA | 1 |
| SE | 90333 | ENSG00000249859 | PVT1 | chr8 | + | 128070159 | 128070272 | 127989161 | 127989291 | 128082752 | 128082848 | 15 | 2 | 8 | 4 | 2 | 0 | 0 | 2 | 0 | 0 | 12 | 0 | 7 | 0 | 10 | 0 | 0 | 0 | 0 | 0 | 262 | 149 | 1 | 1 | 1 | 1 | 0.695 | 1 | 1 | 1 | NA | 1 | NA | 1 | -0.061 |
| SE | 90335 | ENSG00000249859 | PVT1 | chr8 | + | 128070159 | 128070272 | 127989161 | 127989291 | 128096520 | 128096578 | 15 | 1 | 7 | 5 | 2 | 0 | 0 | 0 | 0 | 0 | 0 | 0 | 2 | 0 | 10 | 0 | 0 | 2 | 0 | 0 | 262 | 149 | 1 | 1 | 1 | 1 | 1 | 1 | 1 | NA | NA | 0.363 | NA | 1 | 0.319 |
| SE | 90339 | ENSG00000249859 | PVT1 | chr8 | + | 128070159 | 128070272 | 128010317 | 128010444 | 128096520 | 128096578 | 18 | 7 | 15 | 6 | 34 | 3 | 0 | 0 | 0 | 2 | 15 | 15 | 6 | 0 | 13 | 0 | 0 | 0 | 0 | 0 | 262 | 149 | 1 | 1 | 0.773 | 1 | 1 | 1 | 0.906 | 1 | 1 | 1 | NA | 1 | -0.064 |
| SE | 90340 | ENSG00000249859 | PVT1 | chr8 | + | 128070159 | 128070272 | 128031958 | 128032272 | 128096517 | 128096654 | 7 | 8 | 12 | 4 | 2 | 0 | 0 | 0 | 0 | 0 | 12 | 4 | 6 | 1 | 0 | 0 | 0 | 1 | 0 | 0 | 262 | 149 | 1 | 1 | 1 | 1 | 1 | 1 | 1 | 1 | 1 | 0.773 | 1 | NA | 0.057 |
| SE | 90348 | ENSG00000249859 | PVT1 | chr8 | + | 128070159 | 128070361 | 127989161 | 127989291 | 128082752 | 128082848 | 14 | 1 | 6 | 4 | 6 | 0 | 0 | 2 | 0 | 0 | 0 | 0 | 2 | 0 | 10 | 0 | 0 | 0 | 0 | 0 | 298 | 149 | 1 | 1 | 1 | 1 | 0.6 | 1 | 1 | NA | NA | 1 | NA | 1 | -0.08 |

| | | | | | | | | | | | | | | | | | | | | | | | | | | | | | | | | | | | | | | | | | | | | | | |
|---|---|---|---|---|---|---|---|---|---|---|---|---|---|---|---|---|---|---|---|---|---|---|---|---|---|---|---|---|---|---|---|---|---|---|---|---|---|---|---|---|---|---|---|---|---|---|
| SE | 90349 | ENSG00000249859 | PVT1 | chr8 | + | 128070159 | 128070361 | 127989161 | 127989291 | 128096517 | 128096654 | 25 | 2 | 16 | 6 | 7 | 4 | 0 | 5 | 3 | 13 | 0 | 7 | 2 | 0 | 17 | 0 | 0 | 9 | 0 | 0 | 298 | 149 | 1 | 1 | 0.758 | 1 | 0.615 | 0.5 | 0.212 | NA | 1 | 0.1 | NA | 1 | -0.083 |
| SE | 90350 | ENSG00000249859 | PVT1 | chr8 | + | 128070159 | 128070361 | 127989161 | 127989291 | 128096520 | 128096654 | 14 | 1 | 6 | 4 | 4 | 0 | 0 | 0 | 0 | 0 | 0 | 0 | 2 | 0 | 10 | 0 | 0 | 2 | 0 | 0 | 298 | 149 | 1 | 1 | 1 | 1 | 1 | 1 | 1 | NA | NA | 0.333 | NA | 1 | 0.334 |
| SE | 90354 | ENSG00000249859 | PVT1 | chr8 | + | 128070159 | 128070361 | 128010317 | 128010444 | 128096520 | 128096654 | 18 | 9 | 16 | 5 | 36 | 3 | 0 | 0 | 0 | 2 | 15 | 15 | 7 | 0 | 13 | 0 | 0 | 0 | 0 | 0 | 298 | 149 | 1 | 1 | 0.75 | 1 | 1 | 1 | 0.9 | 1 | 1 | 1 | NA | 1 | -0.07 |
| SE | 90360 | ENSG00000249859 | PVT1 | chr8 | + | 128070159 | 128070361 | 128049408 | 128049748 | 128096517 | 128096654 | 12 | 1 | 10 | 2 | 3 | 0 | 0 | 2 | 1 | 3 | 0 | 7 | 0 | 0 | 7 | 0 | 3 | 0 | 0 | 4 | 298 | 149 | 1 | 1 | 1 | 1 | 0.714 | 0.5 | 0.333 | NA | 0.538 | NA | NA | 0.467 | 0.207 |
| SE | 90362 | ENSG00000249859 | PVT1 | chr8 | + | 128070161 | 128070272 | 127989161 | 127989291 | 128096517 | 128096654 | 7 | 8 | 12 | 4 | 2 | 4 | 0 | 5 | 3 | 13 | 12 | 4 | 6 | 1 | 0 | 0 | 0 | 9 | 0 | 0 | 260 | 149 | 1 | 1 | 0.501 | 1 | 0.579 | 0.433 | 0.081 | 1 | 1 | 0.276 | 1 | NA | -0.3 |
| SE | 90363 | ENSG00000249859 | PVT1 | chr8 | + | 128074069 | 128074291 | 128070159 | 128070361 | 128096517 | 128096654 | 0 | 0 | 0 | 0 | 0 | 11 | 1 | 10 | 2 | 3 | 0 | 0 | 1 | 0 | 0 | 0 | 7 | 0 | 0 | 7 | 298 | 149 | 1 | 1 | 0 | 0 | 0 | 0 | 0 | NA | 0 | 1 | NA | 0 | -0.333 |
| SE | 90364 | ENSG00000249859 | PVT1 | chr8 | + | 128074069 | 128074291 | 128070161 | 128070272 | 128096517 | 128096654 | 0 | 0 | 0 | 0 | 0 | 12 | 8 | 14 | 4 | 6 | 0 | 0 | 2 | 0 | 0 | 35 | 4 | 6 | 1 | 0 | 298 | 149 | 1 | 1 | 0 | 0 | 0 | 0 | 0 | 0 | 0 | 0.143 | 0 | NA | -0.036 |
| SE | 90365 | ENSG00000249859 | PVT1 | chr8 | + | 128074072 | 128074291 | 128070159 | 128070272 | 128080302 | 128080430 | 0 | 0 | 0 | 0 | 0 | 1 | 0 | 0 | 0 | 0 | 0 | 0 | 1 | 0 | 0 | 0 | 5 | 0 | 0 | 6 | 298 | 149 | 1 | 1 | 0 | NA | NA | NA | NA | NA | 0 | 1 | NA | 0 | -0.333 |
| SE | 90366 | ENSG00000249859 | PVT1 | chr8 | + | 128074072 | 128074291 | 128070159 | 128070272 | 128096517 | 128096654 | 0 | 0 | 0 | 0 | 0 | 12 | 8 | 14 | 4 | 6 | 0 | 0 | 1 | 0 | 0 | 35 | 4 | 6 | 1 | 0 | 298 | 149 | 1 | 1 | 0 | 0 | 0 | 0 | 0 | 0 | 0 | 0.077 | 0 | NA | -0.019 |
| SE | 90367 | ENSG00000249859 | PVT1 | chr8 | + | 128080302 | 128080430 | 128070159 | 128070361 | 128082752 | 128083366 | 0 | 0 | 0 | 0 | 0 | 0 | 0 | 0 | 0 | 2 | 0 | 0 | 2 | 0 | 0 | 0 | 0 | 0 | 0 | 0 | 277 | 149 | 1 | 1 | NA | NA | NA | NA | 0 | NA | NA | 1 | NA | NA | -1 |
| SE | 90368 | ENSG00000249859 | PVT1 | chr8 | + | 128080302 | 128080430 | 128070159 | 128070361 | 128096517 | 128096654 | 0 | 0 | 0 | 0 | 0 | 11 | 1 | 10 | 2 | 3 | 0 | 0 | 3 | 0 | 0 | 0 | 7 | 0 | 0 | 7 | 277 | 149 | 1 | 1 | 0 | 0 | 0 | 0 | 0 | NA | 0 | 1 | NA | 0 | -0.333 |
| SE | 90370 | ENSG00000249859 | PVT1 | chr8 | + | 128080302 | 128080430 | 128070161 | 128070272 | 128082752 | 128083366 | 1 | 0 | 0 | 0 | 0 | 1 | 1 | 2 | 0 | 0 | 0 | 5 | 2 | 0 | 6 | 12 | 0 | 5 | 0 | 0 | 277 | 149 | 1 | 1 | 0.35 | 0 | 0 | NA | NA | 0 | 1 | 0.177 | NA | 1 | -0.428 |
| SE | 90372 | ENSG00000249859 | PVT1 | chr8 | + | 128080302 | 128080430 | 128070161 | 128070272 | 128096520 | 128096654 | 1 | 0 | 0 | 1 | 0 | 1 | 0 | 1 | 1 | 0 | 0 | 5 | 0 | 0 | 6 | 0 | 0 | 0 | 0 | 0 | 277 | 149 | 1 | 1 | 0.35 | NA | 0 | 0.35 | NA | NA | 1 | NA | NA | 1 | -0.767 |
| SE | 90374 | ENSG00000249859 | PVT1 | chr8 | + | 128082752 | 128082848 | 127989161 | 127989291 | 128096517 | 128096654 | 0 | 2 | 5 | 0 | 0 | 4 | 0 | 5 | 3 | 13 | 0 | 0 | 2 | 0 | 0 | 0 | 0 | 9 | 0 | 0 | 245 | 149 | 1 | 1 | 0 | 1 | 0.378 | 0 | 0 | NA | NA | 0.119 | NA | NA | 0.157 |
| SE | 90375 | ENSG00000249859 | PVT1 | chr8 | + | 128082752 | 128082848 | 127989161 | 127989291 | 128096520 | 128096654 | 0 | 0 | 1 | 0 | 0 | 0 | 0 | 0 | 0 | 0 | 0 | 0 | 0 | 0 | 0 | 0 | 0 | 2 | 0 | 0 | 245 | 149 | 1 | 1 | NA | NA | 1 | NA | NA | NA | NA | 0 | NA | NA | 1 |
| SE | 90377 | ENSG00000249859 | PVT1 | chr8 | + | 128082752 | 128082848 | 128070159 | 128070272 | 128096517 | 128096654 | 0 | 3 | 5 | 0 | 0 | 12 | 8 | 14 | 4 | 6 | 12 | 0 | 7 | 0 | 0 | 35 | 4 | 6 | 1 | 0 | 245 | 149 | 1 | 1 | 0 | 0.186 | 0.178 | 0 | 0 | 0.173 | 0 | 0.415 | 0 | NA | -0.074 |
| SE | 90378 | ENSG00000249859 | PVT1 | chr8 | + | 128082752 | 128082848 | 128070159 | 128070272 | 128096520 | 128096654 | 1 | 0 | 1 | 0 | 0 | 1 | 0 | 1 | 1 | 0 | 0 | 0 | 2 | 0 | 0 | 0 | 0 | 0 | 0 | 0 | 245 | 149 | 1 | 1 | 0.378 | NA | 0.378 | 0 | NA | NA | NA | 1 | NA | NA | -0.748 |
| SE | 90379 | ENSG00000249859 | PVT1 | chr8 | + | 128082752 | 128082848 | 128070159 | 128070361 | 128096517 | 128096654 | 0 | 2 | 3 | 0 | 2 | 11 | 1 | 10 | 2 | 3 | 0 | 0 | 2 | 0 | 0 | 0 | 7 | 0 | 0 | 7 | 245 | 149 | 1 | 1 | 0 | 0.549 | 0.154 | 0 | 0.288 | NA | 0 | 1 | NA | 0 | -0.135 |
| SE | 90381 | ENSG00000249859 | PVT1 | chr8 | + | 128084542 | 128084621 | 128070159 | 128070361 | 128096517 | 128096654 | 0 | 1 | 0 | 1 | 0 | 11 | 1 | 10 | 2 | 3 | 0 | 0 | 0 | 4 | 0 | 0 | 7 | 0 | 0 | 7 | 228 | 149 | 1 | 1 | 0 | 0.395 | 0 | 0.246 | 0 | NA | 0 | NA | 1 | 0 | -0.205 |
| SE | 90382 | ENSG00000249859 | PVT1 | chr8 | + | 128084542 | 128084621 | 128082377 | 128082848 | 128096517 | 128096654 | 2 | 1 | 0 | 1 | 0 | 0 | 7 | 5 | 0 | 2 | 0 | 0 | 0 | 4 | 0 | 14 | 0 | 8 | 4 | 0 | 228 | 149 | 1 | 1 | 1 | 0.085 | 0 | 1 | 0 | 0 | NA | 0 | 0.395 | NA | 0.285 |
| SE | 90383 | ENSG00000249859 | PVT1 | chr8 | + | 128090247 | 128090288 | 128070159 | 128070272 | 128096517 | 128096654 | 0 | 0 | 0 | 0 | 0 | 12 | 8 | 14 | 4 | 6 | 0 | 0 | 2 | 0 | 0 | 35 | 4 | 6 | 1 | 0 | 190 | 149 | 1 | 1 | 0 | 0 | 0 | 0 | 0 | 0 | 0 | 0.207 | 0 | NA | -0.052 |
| SE | 90384 | ENSG00000249859 | PVT1 | chr8 | + | 128090247 | 128090288 | 128084542 | 128084621 | 128096517 | 128096654 | 0 | 0 | 0 | 0 | 0 | 2 | 1 | 0 | 1 | 0 | 0 | 0 | 2 | 0 | 0 | 0 | 0 | 0 | 4 | 0 | 190 | 149 | 1 | 1 | 0 | 0 | NA | 0 | NA | NA | NA | 1 | 0 | NA | -0.5 |
| SE | 90386 | ENSG00000249859 | PVT1 | chr8 | + | 128091246 | 128091353 | 128010317 | 128010444 | 128096517 | 128096654 | 0 | 0 | 0 | 0 | 4 | 5 | 5 | 12 | 10 | 15 | 0 | 0 | 0 | 0 | 0 | 10 | 4 | 7 | 3 | 0 | 256 | 149 | 1 | 1 | 0 | 0 | 0 | 0 | 0.134 | 0 | 0 | 0 | 0 | NA | 0.027 |
| SE | 90387 | ENSG00000249859 | PVT1 | chr8 | + | 128091246 | 128091353 | 128090247 | 128090288 | 128096517 | 128096654 | 0 | 0 | 0 | 0 | 3 | 0 | 0 | 0 | 0 | 0 | 0 | 0 | 0 | 0 | 0 | 0 | 0 | 4 | 0 | 0 | 256 | 149 | 1 | 1 | NA | NA | NA | NA | 1 | NA | NA | 0 | NA | NA | 1 |
| SE | 90394 | ENSG00000170633 | RNF34 | chr12 | + | 121402741 | 121402807 | 121400118 | 121400218 | 121420241 | 121420334 | 2 | 3 | 1 | 2 | 0 | 0 | 0 | 0 | 0 | 0 | 0 | 0 | 0 | 0 | 0 | 0 | 2 | 0 | 0 | 34 | 215 | 149 | 1 | 1 | 1 | 1 | 1 | 1 | NA | NA | 0 | NA | NA | 0 | 1 |
| SE | 90396 | ENSG00000170633 | RNF34 | chr12 | + | 121416158 | 121416377 | 121400040 | 121400218 | 121417503 | 121417521 | 184 | 102 | 129 | 190 | 193 | 2 | 0 | 0 | 0 | 0 | 75 | 92 | 109 | 83 | 84 | 0 | 0 | 2 | 0 | 0 | 298 | 149 | 1 | 1 | 0.979 | 1 | 1 | 1 | 1 | 1 | 1 | 0.965 | 1 | 1 | 0.003 |
| SE | 90399 | ENSG00000170633 | RNF34 | chr12 | + | 121417503 | 121417911 | 121400040 | 121400218 | 121423385 | 121423517 | 5 | 0 | 2 | 3 | 3 | 0 | 1 | 2 | 1 | 0 | 8 | 6 | 5 | 4 | 0 | 0 | 0 | 0 | 0 | 0 | 298 | 149 | 1 | 1 | 1 | 0 | 0.333 | 0.6 | 1 | 1 | 1 | 1 | 1 | NA | -0.413 |
| SE | 90401 | ENSG00000170633 | RNF34 | chr12 | + | 121417860 | 121417911 | 121400082 | 121400218 | 121423385 | 121424352 | 1 | 0 | 1 | 3 | 0 | 0 | 1 | 2 | 1 | 0 | 0 | 4 | 3 | 2 | 0 | 0 | 0 | 0 | 0 | 0 | 200 | 149 | 1 | 1 | 1 | 0 | 0.271 | 0.691 | NA | NA | 1 | 1 | 1 | NA | -0.51 |
| SE | 90402 | ENSG00000170633 | RNF34 | chr12 | + | 121420241 | 121420334 | 121400040 | 121400218 | 121423385 | 121423517 | 1 | 0 | 0 | 0 | 2 | 0 | 1 | 2 | 1 | 0 | 0 | 2 | 0 | 2 | 28 | 0 | 0 | 0 | 0 | 0 | 242 | 149 | 1 | 1 | 1 | 0 | 0 | 0 | 1 | NA | 1 | NA | 1 | 1 | -0.6 |
| SE | 90404 | ENSG00000170633 | RNF34 | chr12 | + | 121420576 | 121420778 | 121419897 | 121420334 | 121423385 | 121423517 | 273 | 133 | 242 | 214 | 370 | 2 | 0 | 1 | 0 | 2 | 147 | 187 | 160 | 146 | 241 | 0 | 2 | 0 | 2 | 0 | 298 | 149 | 1 | 1 | 0.986 | 1 | 0.992 | 1 | 0.989 | 1 | 0.979 | 1 | 0.973 | 1 | 0.003 |
| SE | 90407 | ENSG00000255325 | NA | chr8 | - | 122486432 | 122486611 | 122485514 | 122485695 | 122487386 | 122487466 | 3 | 2 | 0 | 3 | 2 | 5 | 4 | 0 | 10 | 2 | 0 | 10 | 2 | 7 | 32 | 5 | 16 | 4 | 3 | 75 | 298 | 149 | 1 | 1 | 0.231 | 0.2 | NA | 0.13 | 0.333 | 0 | 0.238 | 0.2 | 0.538 | 0.176 | -0.007 |
| SE | 90408 | ENSG00000047249 | ATP6V1H | chr8 | - | 53743576 | 53743690 | 53715542 | 53716024 | 53756554 | 53756656 | 171 | 93 | 128 | 112 | 144 | 1 | 0 | 0 | 3 | 0 | 87 | 107 | 57 | 77 | 96 | 0 | 0 | 0 | 0 | 0 | 263 | 149 | 1 | 1 | 0.99 | 1 | 1 | 0.955 | 1 | 1 | 1 | 1 | 1 | 1 | -0.011 |
| SE | 90410 | ENSG00000047249 | ATP6V1H | chr8 | - | 53756554 | 53756656 | 53743576 | 53743690 | 53769617 | 53769743 | 166 | 103 | 113 | 105 | 209 | 0 | 0 | 0 | 0 | 3 | 131 | 138 | 75 | 121 | 89 | 0 | 0 | 0 | 0 | 0 | 251 | 149 | 1 | 1 | 1 | 1 | 1 | 1 | 0.976 | 1 | 1 | 1 | 1 | 1 | -0.005 |
| SE | 90411 | ENSG00000047249 | ATP6V1H | chr8 | - | 53769617 | 53769743 | 53756261 | 53756656 | 53795646 | 53795839 | 125 | 83 | 97 | 67 | 193 | 0 | 0 | 3 | 0 | 0 | 111 | 115 | 56 | 102 | 66 | 0 | 0 | 0 | 0 | 0 | 275 | 149 | 1 | 1 | 1 | 1 | 0.946 | 1 | 1 | 1 | 1 | 1 | 1 | 1 | -0.011 |
| SE | 90415 | ENSG00000047249 | ATP6V1H | chr8 | - | 53795646 | 53795839 | 53771988 | 53772167 | 53801798 | 53801868 | 108 | 94 | 77 | 100 | 110 | 0 | 0 | 0 | 0 | 0 | 140 | 85 | 77 | 79 | 51 | 0 | 0 | 2 | 0 | 0 | 298 | 149 | 1 | 1 | 1 | 1 | 1 | 1 | 1 | 1 | 1 | 0.951 | 1 | 1 | 0.01 |
| SE | 90416 | ENSG00000047249 | ATP6V1H | chr8 | - | 53801798 | 53801896 | 53771988 | 53772167 | 53811163 | 53811217 | 34 | 35 | 32 | 28 | 17 | 0 | 0 | 0 | 0 | 0 | 31 | 15 | 35 | 30 | 5 | 0 | 0 | 3 | 0 | 0 | 247 | 149 | 1 | 1 | 1 | 1 | 1 | 1 | 1 | 1 | 1 | 0.876 | 1 | 1 | 0.025 |
| SE | 90420 | ENSG00000047249 | ATP6V1H | chr8 | - | 53829443 | 53829533 | 53817416 | 53817530 | 53832983 | 53833009 | 124 | 85 | 116 | 143 | 203 | 0 | 0 | 0 | 1 | 0 | 98 | 84 | 111 | 104 | 94 | 0 | 0 | 0 | 0 | 0 | 239 | 149 | 1 | 1 | 1 | 1 | 1 | 0.989 | 1 | 1 | 1 | 1 | 1 | 1 | -0.002 |
| SE | 90422 | ENSG00000123908 | AGO2 | chr8 | - | 140547467 | 140547627 | 140544212 | 140544303 | 140549113 | 140549298 | 107 | 46 | 57 | 53 | 44 | 1 | 0 | 0 | 1 | 0 | 48 | 62 | 76 | 68 | 100 | 0 | 0 | 0 | 0 | 2 | 298 | 149 | 1 | 1 | 0.982 | 1 | 1 | 1 | 1 | 1 | 1 | 1 | 1 | 0.962 | 0.004 |
| SE | 90423 | ENSG00000123908 | AGO2 | chr8 | - | 140549113 | 140549298 | 140547467 | 140547627 | 140551302 | 140551436 | 206 | 103 | 139 | 152 | 131 | 0 | 0 | 0 | 1 | 0 | 133 | 154 | 166 | 107 | 247 | 0 | 0 | 0 | 0 | 0 | 298 | 149 | 1 | 1 | 1 | 1 | 1 | 0.987 | 1 | 1 | 1 | 1 | 1 | 1 | -0.003 |
| SE | 90424 | ENSG00000123908 | AGO2 | chr8 | - | 140558484 | 140558572 | 140557088 | 140557236 | 140559394 | 140559529 | 203 | 127 | 192 | 219 | 308 | 1 | 0 | 1 | 1 | 0 | 167 | 251 | 238 | 225 | 408 | 0 | 0 | 0 | 0 | 0 | 237 | 149 | 1 | 1 | 0.992 | 1 | 0.992 | 0.993 | 1 | 1 | 1 | 1 | 1 | 1 | -0.005 |
| SE | 90425 | ENSG00000123908 | AGO2 | chr8 | - | 140559394 | 140559529 | 140558484 | 140558572 | 140560373 | 140560510 | 178 | 103 | 167 | 197 | 236 | 0 | 0 | 0 | 0 | 0 | 142 | 200 | 230 | 165 | 343 | 0 | 0 | 0 | 0 | 3 | 284 | 149 | 1 | 1 | 1 | 1 | 1 | 1 | 1 | 1 | 1 | 1 | 1 | 0.984 | 0.003 |
| SE | 90427 | ENSG00000165923 | AGBL2 | chr11 | - | 47666955 | 47667063 | 47663025 | 47663112 | 47667570 | 47667696 | 1 | 2 | 4 | 1 | 6 | 0 | 0 | 1 | 0 | 0 | 2 | 4 | 0 | 0 | 0 | 0 | 0 | 0 | 0 | 0 | 257 | 149 | 1 | 1 | 1 | 1 | 0.699 | 1 | 1 | 1 | 1 | NA | NA | NA | -0.06 |
| SE | 90428 | ENSG00000165923 | AGBL2 | chr11 | - | 47677270 | 47677401 | 47668840 | 47668907 | 47679972 | 47680073 | 2 | 2 | 2 | 1 | 6 | 2 | 0 | 5 | 1 | 0 | 0 | 5 | 0 | 0 | 0 | 0 | 0 | 2 | 7 | 0 | 280 | 149 | 1 | 1 | 0.347 | 1 | 0.176 | 0.347 | 1 | NA | 1 | 0 | 0 | NA | 0.241 |
| SE | 90429 | ENSG00000165923 | AGBL2 | chr11 | - | 47679972 | 47680073 | 47668840 | 47668907 | 47681968 | 47682095 | 5 | 0 | 5 | 5 | 3 | 0 | 0 | 2 | 0 | 0 | 0 | 7 | 2 | 9 | 0 | 0 | 0 | 0 | 0 | 0 | 250 | 149 | 1 | 1 | 1 | NA | 0.598 | 1 | 1 | NA | 1 | 1 | 1 | NA | -0.101 |
| SE | 90430 | ENSG00000165923 | AGBL2 | chr11 | - | 47679972 | 47680073 | 47668840 | 47668907 | 47690075 | 47690370 | 2 | 0 | 5 | 1 | 1 | 0 | 2 | 5 | 1 | 0 | 0 | 0 | 2 | 7 | 0 | 0 | 0 | 0 | 0 | 0 | 250 | 149 | 1 | 1 | 1 | 0 | 0.373 | 0.373 | 1 | NA | NA | 1 | 1 | NA | -0.451 |
| SE | 90432 | ENSG00000165923 | AGBL2 | chr11 | - | 47681968 | 47682095 | 47679972 | 47680073 | 47690075 | 47690370 | 6 | 0 | 0 | 3 | 1 | 1 | 0 | 0 | 0 | 1 | 0 | 0 | 0 | 2 | 0 | 0 | 0 | 0 | 0 | 0 | 276 | 149 | 1 | 1 | 0.764 | NA | NA | 1 | 0.351 | NA | NA | NA | 1 | NA | -0.295 |
| SE | 90433 | ENSG00000165923 | AGBL2 | chr11 | - | 47685892 | 47686049 | 47681968 | 47682095 | 47690075 | 47690858 | 6 | 0 | 5 | 11 | 14 | 4 | 0 | 0 | 2 | 0 | 0 | 16 | 3 | 0 | 0 | 0 | 0 | 0 | 2 | 0 | 298 | 149 | 1 | 1 | 0.429 | NA | 1 | 0.733 | 1 | NA | 1 | 1 | 0 | NA | 0.124 |
| SE | 90436 | ENSG00000165923 | AGBL2 | chr11 | - | 47705863 | 47705917 | 47704542 | 47704728 | 47710376 | 47710511 | 6 | 5 | 0 | 6 | 0 | 1 | 0 | 0 | 3 | 0 | 0 | 7 | 2 | 3 | 0 | 0 | 0 | 2 | 2 | 0 | 203 | 149 | 1 | 1 | 0.815 | 1 | NA | 0.595 | NA | NA | 1 | 0.423 | 0.524 | NA | 0.154 |
| SE | 90437 | ENSG00000165923 | AGBL2 | chr11 | - | 47710376 | 47710511 | 47705863 | 47705917 | 47714283 | 47714347 | 9 | 13 | 2 | 3 | 0 | 0 | 0 | 1 | 0 | 0 | 0 | 8 | 4 | 3 | 0 | 0 | 0 | 0 | 0 | 0 | 284 | 149 | 1 | 1 | 1 | 1 | 0.512 | 1 | NA | NA | 1 | 1 | 1 | NA | -0.122 |
| SE | 90438 | ENSG00000165923 | AGBL2 | chr11 | - | 47710697 | 47710902 | 47710376 | 47710511 | 47714074 | 47714146 | 3 | 0 | 0 | 2 | 0 | 0 | 2 | 0 | 0 | 0 | 0 | 0 | 0 | 0 | 0 | 0 | 3 | 0 | 0 | 0 | 298 | 149 | 1 | 1 | 1 | 0 | NA | 1 | NA | NA | 0 | NA | NA | NA | 0.667 |
| SE | 90439 | ENSG00000165923 | AGBL2 | chr11 | - | 47710697 | 47710902 | 47710376 | 47710511 | 47714283 | 47714347 | 3 | 0 | 0 | 2 | 0 | 3 | 10 | 2 | 1 | 0 | 0 | 0 | 0 | 0 | 0 | 0 | 1 | 2 | 0 | 0 | 298 | 149 | 1 | 1 | 0.333 | 0 | 0 | 0.5 | NA | NA | 0 | 0 | NA | NA | 0.208 |
| SE | 90440 | ENSG00000165923 | AGBL2 | chr11 | - | 47714074 | 47714146 | 47710383 | 47710511 | 47714283 | 47714347 | 0 | 2 | 0 | 0 | 0 | 3 | 10 | 2 | 1 | 0 | 0 | 3 | 0 | 0 | 0 | 0 | 1 | 2 | 0 | 0 | 221 | 149 | 1 | 1 | 0 | 0.119 | 0 | 0 | NA | NA | 0.669 | 0 | NA | NA | -0.305 |
| SE | 90448 | ENSG00000070081 | NUCB2 | chr11 | + | 17282788 | 17282943 | 17276733 | 17276848 | 17295323 | 17295424 | 118 | 78 | 164 | 122 | 170 | 0 | 0 | 0 | 0 | 0 | 14 | 35 | 29 | 32 | 7 | 0 | 0 | 0 | 1 | 0 | 298 | 149 | 1 | 1 | 1 | 1 | 1 | 1 | 1 | 1 | 1 | 1 | 0.941 | 1 | 0.012 |
| SE | 90450 | ENSG00000070081 | NUCB2 | chr11 | + | 17282788 | 17282943 | 17276983 | 17277095 | 17295323 | 17295424 | 121 | 77 | 171 | 122 | 168 | 0 | 0 | 1 | 0 | 0 | 14 | 34 | 30 | 41 | 7 | 0 | 0 | 0 | 0 | 0 | 298 | 149 | 1 | 1 | 1 | 1 | 0.988 | 1 | 1 | 1 | 1 | 1 | 1 | 1 | -0.002 |
| SE | 90452 | ENSG00000070081 | NUCB2 | chr11 | + | 17286633 | 17286713 | 17282788 | 17282943 | 17291239 | 17291365 | 0 | 10 | 7 | 3 | 9 | 0 | 0 | 4 | 5 | 1 | 0 | 0 | 1 | 2 | 0 | 0 | 0 | 5 | 0 | 0 | 229 | 149 | 1 | 1 | NA | 1 | 0.532 | 0.281 | 0.854 | NA | NA | 0.115 | 1 | NA | 0.109 |
| SE | 90454 | ENSG00000070081 | NUCB2 | chr11 | + | 17286633 | 17286717 | 17282788 | 17282943 | 17291239 | 17291365 | 0 | 10 | 7 | 4 | 9 | 0 | 0 | 4 | 5 | 1 | 0 | 0 | 1 | 4 | 0 | 0 | 0 | 5 | 0 | 0 | 233 | 149 | 1 | 1 | NA | 1 | 0.528 | 0.338 | 0.852 | NA | NA | 0.113 | 1 | NA | 0.123 |
| SE | 90457 | ENSG00000070081 | NUCB2 | chr11 | + | 17291239 | 17291365 | 17286633 | 17286713 | 17295323 | 17295441 | 1 | 2 | 4 | 12 | 3 | 0 | 3 | 10 | 7 | 13 | 15 | 4 | 6 | 0 | 0 | 0 | 4 | 1 | 3 | 0 | 275 | 149 | 1 | 1 | 1 | 0.265 | 0.178 | 0.482 | 0.111 | 1 | 0.351 | 0.765 | 0 | NA | -0.122 |
| SE | 90458 | ENSG00000070081 | NUCB2 | chr11 | + | 17291239 | 17291365 | 17286633 | 17286717 | 17295323 | 17295441 | 1 | 2 | 4 | 12 | 3 | 17 | 11 | 21 | 19 | 27 | 15 | 4 | 6 | 2 | 0 | 0 | 2 | 12 | 14 | 0 | 275 | 149 | 1 | 1 | 0.031 | 0.09 | 0.094 | 0.255 | 0.057 | 1 | 0.52 | 0.213 | 0.072 | NA | -0.346 |
| SE | 90460 | ENSG00000070081 | NUCB2 | chr11 | + | 17311184 | 17311283 | 17310824 | 17311010 | 17311871 | 17311930 | 139 | 251 | 221 | 254 | 393 | 0 | 0 | 1 | 0 | 0 | 75 | 76 | 104 | 105 | 21 | 0 | 0 | 0 | 0 | 0 | 248 | 149 | 1 | 1 | 1 | 1 | 0.993 | 1 | 1 | 1 | 1 | 1 | 1 | 1 | -0.001 |
| SE | 90461 | ENSG00000070081 | NUCB2 | chr11 | + | 17311192 | 17311283 | 17310824 | 17311010 | 17311871 | 17311930 | 400 | 625 | 569 | 623 | 1088 | 0 | 0 | 1 | 0 | 0 | 100 | 206 | 233 | 255 | 60 | 0 | 0 | 0 | 0 | 0 | 240 | 149 | 1 | 1 | 1 | 1 | 0.997 | 1 | 1 | 1 | 1 | 1 | 1 | 1 | -0.001 |
| SE | 90464 | ENSG00000070081 | NUCB2 | chr11 | + | 17330126 | 17330297 | 17315385 | 17315475 | 17330901 | 17330983 | 497 | 484 | 579 | 646 | 1197 | 0 | 0 | 1 | 1 | 0 | 20 | 102 | 131 | 73 | 26 | 0 | 0 | 0 | 0 | 0 | 298 | 149 | 1 | 1 | 1 | 1 | 0.997 | 0.997 | 1 | 1 | 1 | 1 | 1 | 1 | -0.001 |
| SE | 90465 | ENSG00000070081 | NUCB2 | chr11 | + | 17330901 | 17330983 | 17330126 | 17330297 | 17337347 | 17337534 | 218 | 215 | 212 | 289 | 409 | 2 | 0 | 0 | 0 | 0 | 0 | 25 | 46 | 22 | 6 | 0 | 0 | 0 | 0 | 0 | 231 | 149 | 1 | 1 | 0.986 | 1 | 1 | 1 | 1 | NA | 1 | 1 | 1 | 1 | -0.003 |
| SE | 90466 | ENSG00000070081 | NUCB2 | chr11 | + | 17337347 | 17337534 | 17330901 | 17330983 | 17349344 | 17349980 | 5 | 14 | 17 | 8 | 20 | 2 | 4 | 2 | 6 | 0 | 0 | 12 | 8 | 3 | 0 | 0 | 0 | 0 | 0 | 0 | 298 | 149 | 1 | 1 | 0.556 | 0.636 | 0.81 | 0.4 | 1 | NA | 1 | 1 | 1 | NA | -0.32 |
| SE | 90467 | ENSG00000151461 | UPF2 | chr10 | - | 11952065 | 11952249 | 11948368 | 11948508 | 11955231 | 11955507 | 226 | 248 | 291 | 281 | 474 | 0 | 0 | 2 | 2 | 1 | 62 | 138 | 208 | 53 | 17 | 0 | 2 | 1 | 0 | 0 | 298 | 149 | 1 | 1 | 1 | 1 | 0.986 | 0.986 | 0.996 | 1 | 0.972 | 0.99 | 1 | 1 | 0.001 |
| SE | 90468 | ENSG00000151461 | UPF2 | chr10 | - | 11952065 | 11952249 | 11948368 | 11948508 | 11956319 | 11956523 | 170 | 190 | 196 | 202 | 335 | 0 | 0 | 2 | 0 | 0 | 49 | 98 | 152 | 45 | 17 | 0 | 0 | 0 | 0 | 0 | 298 | 149 | 1 | 1 | 1 | 1 | 0.98 | 1 | 1 | 1 | 1 | 1 | 1 | 1 | -0.004 |

| | | | | | | | |
|---|---|---|---|---|---|---|---|
| SE | 90813 | ENSG00000173064 | HECTD4 | chr12 | - | 112210014 | [illegible] |
| SE | 90814 | ENSG00000173064 | HECTD4 | chr12 | - | 112212486 | [illegible] |
| SE | 90815 | ENSG00000173064 | HECTD4 | chr12 | - | 112228088 | [illegible] |
| SE | 90816 | ENSG00000173064 | HECTD4 | chr12 | - | 112229697 | [illegible] |
| SE | 90817 | ENSG00000173064 | HECTD4 | chr12 | - | 112230686 | [illegible] |
| SE | 90818 | ENSG00000173064 | HECTD4 | chr12 | - | 112231512 | [illegible] |
| SE | 90821 | ENSG00000173064 | HECTD4 | chr12 | - | 112239051 | [illegible] |
| SE | 90824 | ENSG00000173064 | HECTD4 | chr12 | - | 112250143 | [illegible] |
| SE | 90825 | ENSG00000173064 | HECTD4 | chr12 | - | 112250970 | [illegible] |
| SE | 90826 | ENSG00000173064 | HECTD4 | chr12 | - | 112252423 | [illegible] |
| SE | 90827 | ENSG00000173064 | HECTD4 | chr12 | - | 112252423 | [illegible] |
| SE | 90828 | ENSG00000173064 | HECTD4 | chr12 | - | 112258495 | [illegible] |
| SE | 90829 | ENSG00000173064 | HECTD4 | chr12 | - | 112259111 | [illegible] |
| SE | 90831 | ENSG00000173064 | HECTD4 | chr12 | - | 112270226 | [illegible] |
| SE | 90834 | ENSG00000173064 | HECTD4 | chr12 | - | 112279227 | [illegible] |
| SE | 90836 | ENSG00000173064 | HECTD4 | chr12 | - | 112308752 | [illegible] |
| SE | 90837 | ENSG00000173064 | HECTD4 | chr12 | - | 112309560 | [illegible] |
| SE | 90841 | ENSG00000129696 | TTI2 | chr8 | - | 33500327 | [illegible] |
| SE | 90842 | ENSG00000129696 | TTI2 | chr8 | - | 33503428 | [illegible] |
| SE | 90849 | ENSG00000166689 | PLEKHA7 | chr11 | - | 16787109 | [illegible] |
| SE | 90850 | ENSG00000166689 | PLEKHA7 | chr11 | - | 16790797 | [illegible] |
| SE | 90851 | ENSG00000166689 | PLEKHA7 | chr11 | - | 16807143 | [illegible] |
| SE | 90858 | ENSG00000254054 | NA | chr8 | + | 17907192 | [illegible] |
| SE | 90859 | ENSG00000082516 | GEMIN5 | chr5 | - | 154898439 | [illegible] |
| SE | 90860 | ENSG00000082516 | GEMIN5 | chr5 | - | 154901338 | [illegible] |
| SE | 90861 | ENSG00000082516 | GEMIN5 | chr5 | - | 154904506 | [illegible] |
| SE | 90862 | ENSG00000082516 | GEMIN5 | chr5 | - | 154911726 | [illegible] |
| SE | 90864 | ENSG00000082516 | GEMIN5 | chr5 | - | 154932098 | [illegible] |
| SE | 90865 | ENSG00000104763 | ASAH1 | chr8 | - | 18059340 | [illegible] |
| SE | 90866 | ENSG00000104763 | ASAH1 | chr8 | - | 18062247 | [illegible] |
| SE | 90867 | ENSG00000104763 | ASAH1 | chr8 | - | 18062278 | [illegible] |
| SE | 90869 | ENSG00000104763 | ASAH1 | chr8 | - | 18064122 | [illegible] |
| SE | 90870 | ENSG00000104763 | ASAH1 | chr8 | - | 18064456 | [illegible] |
| SE | 90871 | ENSG00000104763 | ASAH1 | chr8 | - | 18064456 | [illegible] |
| SE | 90872 | ENSG00000104763 | ASAH1 | chr8 | - | 18064456 | [illegible] |
| SE | 90873 | ENSG00000104763 | ASAH1 | chr8 | - | 18064456 | [illegible] |
| SE | 90874 | ENSG00000104763 | ASAH1 | chr8 | - | 18064456 | [illegible] |
| SE | 90875 | ENSG00000104763 | ASAH1 | chr8 | - | 18067219 | [illegible] |
| SE | 90876 | ENSG00000104763 | ASAH1 | chr8 | - | 18069791 | [illegible] |
| SE | 90878 | ENSG00000104763 | ASAH1 | chr8 | - | 18069791 | [illegible] |
| SE | 90881 | ENSG00000104763 | ASAH1 | chr8 | - | 18071299 | [illegible] |
| SE | 90883 | ENSG00000104763 | ASAH1 | chr8 | - | 18071299 | [illegible] |
| SE | 90885 | ENSG00000104763 | ASAH1 | chr8 | - | 18073255 | [illegible] |
| SE | 90889 | ENSG00000245281 | NA | chr8 | + | 18085241 | [illegible] |
| SE | 90890 | ENSG00000245281 | NA | chr8 | + | 18087024 | [illegible] |
| SE | 90891 | ENSG00000258311 | NA | chr12 | + | 55716932 | [illegible] |
| SE | 90893 | ENSG00000185728 | YTHDF3 | chr8 | + | 63169386 | [illegible] |
| SE | 90896 | ENSG00000185728 | YTHDF3 | chr8 | + | 63172652 | [illegible] |
| SE | 90897 | ENSG00000185728 | YTHDF3 | chr8 | + | 63173650 | [illegible] |
| SE | 90898 | ENSG00000185728 | YTHDF3 | chr8 | + | 63173650 | [illegible] |
| SE | 90900 | ENSG00000185728 | YTHDF3 | chr8 | + | 63175330 | [illegible] |
| SE | 90902 | ENSG00000185728 | YTHDF3 | chr8 | + | 63186146 | [illegible] |
| SE | 90903 | ENSG00000185728 | YTHDF3 | chr8 | + | 63186146 | [illegible] |
| SE | 90906 | ENSG00000107020 | PLGRKT | chr9 | - | 5381862 | [illegible] |
| SE | 90907 | ENSG00000107020 | PLGRKT | chr9 | - | 5431896 | [illegible] |
| SE | 90908 | ENSG00000107020 | PLGRKT | chr9 | - | 5431896 | [illegible] |
| SE | 90909 | ENSG00000107020 | PLGRKT | chr9 | - | 5436568 | [illegible] |
| SE | 90914 | ENSG00000185122 | HSF1 | chr8 | + | 144311504 | [illegible] |
| SE | 90917 | ENSG00000185122 | HSF1 | chr8 | + | 144313845 | [illegible] |
| SE | 90919 | ENSG00000182325 | FBXL6 | chr8 | - | 144357438 | [illegible] |
| SE | 90921 | ENSG00000181396 | OGFOD3 | chr17 | - | 82394330 | [illegible] |
| SE | 90922 | ENSG00000181396 | OGFOD3 | chr17 | - | 82406417 | [illegible] |
| SE | 90927 | ENSG00000147804 | SLC39A4 | chr8 | - | 144413236 | [illegible] |
| SE | 90930 | ENSG00000160948 | VPS28 | chr8 | - | 144424717 | [illegible] |
| SE | 90931 | ENSG00000160948 | VPS28 | chr8 | - | 144424945 | [illegible] |
| SE | 90932 | ENSG00000160948 | VPS28 | chr8 | - | 144424945 | [illegible] |
| SE | 90933 | ENSG00000160948 | VPS28 | chr8 | - | 144425874 | [illegible] |
| SE | 90934 | ENSG00000160948 | VPS28 | chr8 | - | 144425921 | [illegible] |
| SE | 90935 | ENSG00000160948 | VPS28 | chr8 | - | 144426025 | [illegible] |
| SE | 90937 | ENSG00000185803 | SLC52A2 | chr8 | + | 144359622 | [illegible] |
| SE | 90941 | ENSG00000185803 | SLC52A2 | chr8 | + | 144360114 | [illegible] |
| SE | 90943 | ENSG00000185803 | SLC52A2 | chr8 | + | 144360171 | [illegible] |
| SE | 90945 | ENSG00000185803 | SLC52A2 | chr8 | + | 144360600 | [illegible] |
| SE | 90946 | ENSG00000181826 | RELL1 | chr4 | - | 37631384 | [illegible] |
| SE | 90947 | ENSG00000171428 | NAT1 | chr8 | + | 18211192 | [illegible] |
| SE | 90948 | ENSG00000171428 | NAT1 | chr8 | + | 18212390 | [illegible] |
| SE | 90950 | ENSG00000171428 | NAT1 | chr8 | + | 18212390 | [illegible] |
| SE | 90951 | ENSG00000171428 | NAT1 | chr8 | + | 18216876 | [illegible] |
| SE | 90952 | ENSG00000171428 | NAT1 | chr8 | + | 18216876 | [illegible] |
| SE | 90955 | ENSG00000166225 | FRS2 | chr12 | + | 69480298 | [illegible] |
| SE | 90956 | ENSG00000166225 | FRS2 | chr12 | + | 69486201 | [illegible] |
| SE | 90957 | ENSG00000166225 | FRS2 | chr12 | + | 69491537 | [illegible] |
| SE | 90958 | ENSG00000166225 | FRS2 | chr12 | + | 69537915 | [illegible] |
| SE | 90959 | ENSG00000166225 | FRS2 | chr12 | + | 69537915 | [illegible] |
| SE | 90960 | ENSG00000166225 | FRS2 | chr12 | + | 69557799 | [illegible] |
| SE | 90961 | ENSG00000166225 | FRS2 | chr12 | + | 69557799 | [illegible] |
| SE | 90962 | ENSG00000166225 | FRS2 | chr12 | + | 69562179 | [illegible] |
| SE | 90963 | ENSG00000166225 | FRS2 | chr12 | + | 69569004 | [illegible] |
| SE | 90964 | ENSG00000166225 | FRS2 | chr12 | + | 69570330 | [illegible] |
| SE | 90965 | ENSG00000166225 | FRS2 | chr12 | + | 69572117 | [illegible] |
| SE | 90966 | ENSG00000170379 | TCAF2 | chr7 | + | 143702980 | [illegible] |
| SE | 90967 | ENSG00000170379 | TCAF2 | chr7 | + | 143702983 | [illegible] |
| SE | 90968 | ENSG00000170379 | TCAF2 | chr7 | + | 143715845 | [illegible] |
| SE | 90970 | ENSG00000181690 | PLAG1 | chr8 | - | 56168027 | [illegible] |
| SE | 90971 | ENSG00000181690 | PLAG1 | chr8 | - | 56179408 | [illegible] |
| SE | 90981 | ENSG00000132669 | RIN2 | chr20 | + | 19974653 | [illegible] |
| SE | 90982 | ENSG00000132669 | RIN2 | chr20 | + | 19990005 | [illegible] |
| SE | 90984 | ENSG00000132669 | RIN2 | chr20 | + | 19992167 | [illegible] |
| SE | 90985 | ENSG00000132669 | RIN2 | chr20 | + | 19996678 | [illegible] |
| SE | 90987 | ENSG00000166866 | MYO1A | chr12 | - | 57043855 | [illegible] |
| SE | 90988 | ENSG00000166866 | MYO1A | chr12 | - | 57046551 | [illegible] |
| SE | 90989 | ENSG00000166866 | MYO1A | chr12 | - | 57047060 | [illegible] |
| SE | 90996 | ENSG00000133742 | CA1 | chr8 | - | 85341753 | [illegible] |
| SE | 91000 | ENSG00000133742 | CA1 | chr8 | - | 85342748 | [illegible] |
| SE | 91006 | ENSG00000176209 | SMIM19 | chr8 | + | 42546468 | [illegible] |
| SE | 91008 | ENSG00000176209 | SMIM19 | chr8 | + | 42546468 | [illegible] |
| SE | 91009 | ENSG00000154582 | ELOC | chr8 | - | 73955910 | [illegible] |
| SE | 91010 | ENSG00000154582 | ELOC | chr8 | - | 73959764 | [illegible] |
| SE | 91011 | ENSG00000154582 | ELOC | chr8 | - | 73959764 | [illegible] |
| SE | 91012 | ENSG00000154582 | ELOC | chr8 | - | 73959764 | [illegible] |
| SE | 91013 | ENSG00000154582 | ELOC | chr8 | - | 73964486 | [illegible] |
| SE | 91014 | ENSG00000154582 | ELOC | chr8 | - | 73964486 | [illegible] |
| SE | 91017 | ENSG00000154582 | ELOC | chr8 | - | 73969559 | [illegible] |
| SE | 91018 | ENSG00000154582 | ELOC | chr8 | - | 73969559 | [illegible] |
| SE | 91021 | ENSG00000154582 | ELOC | chr8 | - | 73969559 | [illegible] |
| SE | 91022 | ENSG00000154582 | ELOC | chr8 | - | 73970578 | [illegible] |
| SE | 91024 | ENSG00000120910 | PPP3CC | chr8 | + | 22498000 | [illegible] |
| SE | 91025 | ENSG00000120910 | PPP3CC | chr8 | + | 22498000 | [illegible] |
| SE | 91026 | ENSG00000120910 | PPP3CC | chr8 | + | 22511085 | [illegible] |
| SE | 91031 | ENSG00000120910 | PPP3CC | chr8 | + | 22527391 | [illegible] |
| SE | 91034 | ENSG00000120910 | PPP3CC | chr8 | + | 22528505 | [illegible] |
| SE | 91037 | ENSG00000254673 | NA | chr8 | + | 43069554 | [illegible] |
| SE | 91038 | ENSG00000254673 | NA | chr8 | + | 43069554 | [illegible] |
| SE | 91039 | ENSG00000254673 | NA | chr8 | + | 43072180 | [illegible] |
| SE | 91042 | ENSG00000168522 | FNTA | chr8 | + | 43064100 | [illegible] |
| SE | 91043 | ENSG00000168522 | FNTA | chr8 | + | 43064100 | [illegible] |
| SE | 91044 | ENSG00000168522 | FNTA | chr8 | + | 43069554 | [illegible] |
| SE | 91045 | ENSG00000168522 | FNTA | chr8 | + | 43072180 | [illegible] |
| SE | 91046 | ENSG00000168522 | FNTA | chr8 | + | 43077215 | [illegible] |
| SE | 91047 | ENSG00000168522 | FNTA | chr8 | + | 43077215 | [illegible] |
| SE | 91048 | ENSG00000168522 | FNTA | chr8 | + | 43077215 | [illegible] |
| SE | 91049 | ENSG00000168522 | FNTA | chr8 | + | 43083117 | [illegible] |
| SE | 91050 | ENSG00000168522 | FNTA | chr8 | + | 43083117 | [illegible] |
| SE | 91051 | ENSG00000168522 | FNTA | chr8 | + | 43084709 | [illegible] |
| SE | 91052 | ENSG00000104164 | BLOC1S6 | chr15 | + | 45591457 | [illegible] |
| SE | 91056 | ENSG00000104164 | BLOC1S6 | chr15 | + | 45592134 | [illegible] |
| SE | 91058 | ENSG00000104164 | BLOC1S6 | chr15 | + | 45592134 | [illegible] |
| SE | 91060 | ENSG00000104164 | BLOC1S6 | chr15 | + | 45592134 | [illegible] |
| SE | 91061 | ENSG00000104164 | BLOC1S6 | chr15 | + | 45592134 | [illegible] |
| SE | 91068 | ENSG00000104164 | BLOC1S6 | chr15 | + | 45602356 | [illegible] |
| SE | 91069 | ENSG00000104164 | BLOC1S6 | chr15 | + | 45602356 | [illegible] |
| SE | 91071 | ENSG00000104164 | BLOC1S6 | chr15 | + | 45603099 | [illegible] |
| SE | 91074 | ENSG00000104164 | BLOC1S6 | chr15 | + | 45603099 | [illegible] |
| SE | 91077 | ENSG00000104164 | BLOC1S6 | chr15 | + | 45605427 | [illegible] |
| SE | 91081 | ENSG00000165102 | HGSNAT | chr8 | + | 43167943 | [illegible] |
| SE | 91085 | ENSG00000165102 | HGSNAT | chr8 | + | 43178073 | [illegible] |
| SE | 91087 | ENSG00000165102 | HGSNAT | chr8 | + | 43192303 | [illegible] |
| SE | 91094 | ENSG00000185900 | POMK | chr8 | + | 43097523 | [illegible] |
| SE | 91097 | ENSG00000185900 | POMK | chr8 | + | 43102504 | [illegible] |
| SE | 91099 | ENSG00000185900 | POMK | chr8 | + | 43109563 | [illegible] |
| SE | 91105 | ENSG00000107864 | CPEB3 | chr10 | - | 92144944 | [illegible] |
| SE | 91106 | ENSG00000107864 | CPEB3 | chr10 | - | 92144944 | [illegible] |
| SE | 91108 | ENSG00000107864 | CPEB3 | chr10 | - | 92144944 | [illegible] |
| SE | 91110 | ENSG00000173848 | NET1 | chr10 | + | 5451829 | [illegible] |
| SE | 91112 | ENSG00000173848 | NET1 | chr10 | + | 5453249 | [illegible] |
| SE | 91113 | ENSG00000173848 | NET1 | chr10 | + | 5456086 | [illegible] |
| SE | 91117 | ENSG00000121022 | COPS5 | chr8 | - | 67045811 | [illegible] |
| SE | 91120 | ENSG00000121022 | COPS5 | chr8 | - | 67058082 | [illegible] |
| SE | 91123 | ENSG00000121022 | COPS5 | chr8 | - | 67061291 | [illegible] |
| SE | 91127 | ENSG00000180900 | SCRIB | chr8 | - | 143793256 | [illegible] |
| SE | 91128 | ENSG00000180900 | SCRIB | chr8 | - | 143803382 | [illegible] |
| SE | 91138 | ENSG00000116852 | KIF21B | chr1 | - | 200990916 | [illegible] |
| SE | 91141 | ENSG00000116852 | KIF21B | chr1 | - | 201002160 | [illegible] |
| SE | 91142 | ENSG00000116852 | KIF21B | chr1 | - | 201003585 | [illegible] |
| SE | 91143 | ENSG00000116852 | KIF21B | chr1 | - | 201004339 | [illegible] |
| SE | 91149 | ENSG00000179091 | CYC1 | chr8 | + | 144095832 | [illegible] |
| SE | 91152 | ENSG00000183049 | CAMK1D | chr10 | + | 12666735 | [illegible] |
| SE | 91154 | ENSG00000183049 | CAMK1D | chr10 | + | 12814194 | [illegible] |
| SE | 91156 | ENSG00000164941 | INTS8 | chr8 | + | 94824892 | [illegible] |
| SE | 91159 | ENSG00000164941 | INTS8 | chr8 | + | 94827262 | [illegible] |
| SE | 91160 | ENSG00000164941 | INTS8 | chr8 | + | 94827721 | [illegible] |
| SE | 91161 | ENSG00000164941 | INTS8 | chr8 | + | 94828974 | [illegible] |
| SE | 91162 | ENSG00000164941 | INTS8 | chr8 | + | 94831991 | [illegible] |
| SE | 91165 | ENSG00000164941 | INTS8 | chr8 | + | 94838462 | [illegible] |
| SE | 91167 | ENSG00000164941 | INTS8 | chr8 | + | 94838493 | [illegible] |
| SE | 91168 | ENSG00000164941 | INTS8 | chr8 | + | 94841490 | [illegible] |
| SE | 91169 | ENSG00000164941 | INTS8 | chr8 | + | 94841490 | [illegible] |
| SE | 91170 | ENSG00000164941 | INTS8 | chr8 | + | 94842346 | [illegible] |
| SE | 91172 | ENSG00000164941 | INTS8 | chr8 | + | 94849915 | [illegible] |
| SE | 91173 | ENSG00000164941 | INTS8 | chr8 | + | 94849915 | [illegible] |
| SE | 91174 | ENSG00000164941 | INTS8 | chr8 | + | 94851552 | [illegible] |
| SE | 91175 | ENSG00000164941 | INTS8 | chr8 | + | 94851552 | [illegible] |
| SE | 91177 | ENSG00000164941 | INTS8 | chr8 | + | 94859510 | [illegible] |
| SE | 91178 | ENSG00000164941 | INTS8 | chr8 | + | 94865505 | [illegible] |
| SE | 91180 | ENSG00000164941 | INTS8 | chr8 | + | 94865505 | [illegible] |
| SE | 91182 | ENSG00000164941 | INTS8 | chr8 | + | 94866157 | [illegible] |
| SE | 91183 | ENSG00000164941 | INTS8 | chr8 | + | 94866311 | [illegible] |
| SE | 91186 | ENSG00000164941 | INTS8 | chr8 | + | 94871883 | [illegible] |
| SE | 91188 | ENSG00000164941 | INTS8 | chr8 | + | 94876073 | [illegible] |
| SE | 91189 | ENSG00000175073 | VCPIP1 | chr8 | - | 66651457 | [illegible] |
| SE | 91190 | ENSG00000288596 | C8orf44 | chr8 | + | 66676744 | [illegible] |
| SE | 91191 | ENSG00000288596 | C8orf44 | chr8 | + | 66676744 | [illegible] |
| SE | 91192 | ENSG00000288596 | C8orf44 | chr8 | + | 66676744 | [illegible] |
| SE | 91194 | ENSG00000288596 | C8orf44 | chr8 | + | 66677641 | [illegible] |
| SE | 91195 | ENSG00000288596 | C8orf44 | chr8 | + | 66680392 | [illegible] |
| SE | 91196 | ENSG00000116604 | MEF2D | chr1 | - | 156467992 | [illegible] |
| SE | 91203 | ENSG00000116604 | MEF2D | chr1 | - | 156499431 | [illegible] |
| SE | 91204 | ENSG00000288602 | C8orf44-SGK3 | chr8 | + | 66690704 | [illegible] |
| SE | 91206 | ENSG00000288602 | C8orf44-SGK3 | chr8 | + | 66828653 | [illegible] |
| SE | 91207 | ENSG00000288602 | C8orf44-SGK3 | chr8 | + | 66831253 | [illegible] |
| SE | 91208 | ENSG00000288602 | C8orf44-SGK3 | chr8 | + | 66840002 | [illegible] |
| SE | 91209 | ENSG00000288602 | C8orf44-SGK3 | chr8 | + | 66840210 | [illegible] |
| SE | 91211 | ENSG00000104205 | SGK3 | chr8 | + | 66843451 | [illegible] |
| SE | 91213 | ENSG00000181894 | ZNF329 | chr19 | - | 58142556 | [illegible] |
| SE | 91214 | ENSG00000181894 | ZNF329 | chr19 | - | 58143104 | [illegible] |
| SE | 91215 | ENSG00000164989 | CCDC171 | chr9 | + | 15594040 | [illegible] |
| SE | 91216 | ENSG00000164989 | CCDC171 | chr9 | + | 15678757 | [illegible] |
| SE | 91218 | ENSG00000164989 | CCDC171 | chr9 | + | 15724775 | [illegible] |
| SE | 91222 | ENSG00000104205 | SGK3 | chr8 | + | 66790722 | [illegible] |

| | | | | | | | | | | | | | | | | | | |
|---|---|---|---|---|---|---|---|---|---|---|---|---|---|---|---|---|---|---|
| SE | 91230 | ENSG00000147364 | FBXO25 | chr8 | + | 413072 | 413213 | 406807 | 407066 | 431340 | 431444 | 83 | 79 | 71 | 59 | 49 | 0 | 0 |
| SE | 91232 | ENSG00000147364 | FBXO25 | chr8 | + | 435614 | 435707 | 431340 | 431444 | 449989 | 450083 | 90 | 73 | 72 | 53 | 162 | 0 | 0 |
| SE | 91237 | ENSG00000171055 | FEZ2 | chr2 | - | 36555976 | 36556060 | 36555682 | 36555748 | 36558437 | 36558513 | 0 | 0 | 0 | 0 | 1 | 187 | 159 |
| SE | 91241 | ENSG00000171055 | FEZ2 | chr2 | - | 36590902 | 36591011 | 36583352 | 36583469 | 36597876 | 36598166 | 65 | 44 | 50 | 40 | 24 | 0 | 1 |
| SE | 91242 | ENSG00000085719 | CPNE3 | chr8 | + | 86515461 | 86515499 | 86514434 | 86514541 | 86528535 | 86528677 | 50 | 21 | 41 | 36 | 59 | 2 | 1 |
| SE | 91243 | ENSG00000085719 | CPNE3 | chr8 | + | 86521603 | 86521714 | 86515461 | 86515499 | 86528535 | 86528677 | 0 | 0 | 0 | 0 | 0 | 52 | 22 |
| SE | 91245 | ENSG00000085719 | CPNE3 | chr8 | + | 86524876 | 86524997 | 86515461 | 86515499 | 86528535 | 86528677 | 0 | 0 | 0 | 8 | 1 | 52 | 22 |
| SE | 91246 | ENSG00000085719 | CPNE3 | chr8 | + | 86532508 | 86532580 | 86531154 | 86531229 | 86537562 | 86537646 | 365 | 239 | 263 | 296 | 621 | 0 | 0 |
| SE | 91247 | ENSG00000085719 | CPNE3 | chr8 | + | 86537562 | 86537646 | 86532508 | 86532580 | 86540244 | 86540334 | 331 | 251 | 240 | 284 | 542 | 0 | 0 |
| SE | 91248 | ENSG00000085719 | CPNE3 | chr8 | + | 86546594 | 86546681 | 86544739 | 86544838 | 86547710 | 86547770 | 245 | 148 | 174 | 233 | 600 | 0 | 3 |
| SE | 91249 | ENSG00000085719 | CPNE3 | chr8 | + | 86554850 | 86554984 | 86551182 | 86551234 | 86558287 | 86561411 | 121 | 76 | 134 | 117 | 167 | 0 | 0 |
| SE | 91251 | ENSG00000117560 | FASLG | chr1 | + | 172660094 | 172660140 | 172659102 | 172659549 | 172664333 | 172664390 | 44 | 17 | 16 | 38 | 9 | 1 | 0 |
| SE | 91252 | ENSG00000117560 | FASLG | chr1 | + | 172664333 | 172664390 | 172660094 | 172660140 | 172665621 | 172666075 | 52 | 14 | 32 | 52 | 12 | 0 | 0 |
| SE | 91255 | ENSG00000154114 | TBCEL | chr11 | + | 121035732 | 121035944 | 121024071 | 121024154 | 121045673 | 121045823 | 0 | 1 | 0 | 0 | 0 | 0 | 0 |
| SE | 91256 | ENSG00000154114 | TBCEL | chr11 | + | 121035732 | 121035944 | 121024120 | 121024291 | 121036504 | 121036612 | 13 | 12 | 15 | 8 | 15 | 2 | 0 |
| SE | 91257 | ENSG00000154114 | TBCEL | chr11 | + | 121035732 | 121035944 | 121024120 | 121024291 | 121045673 | 121045823 | 0 | 1 | 4 | 0 | 0 | 2 | 2 |
| SE | 91260 | ENSG00000154114 | TBCEL | chr11 | + | 121036504 | 121036612 | 121035732 | 121035944 | 121045673 | 121045823 | 17 | 18 | 18 | 17 | 31 | 0 | 1 |
| SE | 91262 | ENSG00000154114 | TBCEL | chr11 | + | 121053550 | 121053732 | 121047527 | 121047667 | 121086777 | 121087096 | 61 | 36 | 48 | 82 | 114 | 2 | 2 |
| SE | 91264 | ENSG00000154114 | TBCEL | chr11 | + | 121053629 | 121053732 | 121047527 | 121047667 | 121057507 | 121057614 | 25 | 7 | 13 | 21 | 32 | 1 | 0 |
| SE | 91266 | ENSG00000154114 | TBCEL | chr11 | + | 121053629 | 121053732 | 121047527 | 121047667 | 121086777 | 121086849 | 24 | 7 | 12 | 21 | 32 | 2 | 2 |
| SE | 91267 | ENSG00000154114 | TBCEL | chr11 | + | 121055051 | 121055308 | 121047527 | 121047667 | 121057507 | 121057614 | 2 | 5 | 4 | 9 | 5 | 1 | 0 |
| SE | 91269 | ENSG00000154114 | TBCEL | chr11 | + | 121055051 | 121055308 | 121047527 | 121047667 | 121086777 | 121086849 | 2 | 5 | 3 | 9 | 5 | 2 | 2 |
| SE | 91272 | ENSG00000154114 | TBCEL | chr11 | + | 121055051 | 121055308 | 121053550 | 121053732 | 121086777 | 121086849 | 59 | 21 | 44 | 49 | 51 | 0 | 0 |
| SE | 91273 | ENSG00000154114 | TBCEL | chr11 | + | 121057507 | 121057614 | 121047527 | 121047667 | 121058344 | 121058471 | 3 | 3 | 2 | 0 | 0 | 11 | 4 |
| SE | 91276 | ENSG00000120526 | NUDCD1 | chr8 | - | 109293343 | 109293524 | 109289750 | 109289933 | 109296383 | 109296569 | 133 | 97 | 115 | 119 | 218 | 0 | 1 |
| SE | 91277 | ENSG00000120526 | NUDCD1 | chr8 | - | 109296383 | 109296569 | 109289750 | 109293524 | 109322308 | 109322463 | 111 | 71 | 109 | 140 | 176 | 1 | 0 |
| SE | 91278 | ENSG00000120526 | NUDCD1 | chr8 | - | 109313796 | 109313888 | 109296383 | 109296569 | 109322308 | 109322463 | 3 | 2 | 2 | 3 | 14 | 57 | 38 |
| SE | 91280 | ENSG00000155265 | GOLGA7B | chr10 | + | 97859457 | 97859583 | 97849842 | 97850315 | 97863929 | 97864082 | 11 | 0 | 7 | 5 | 2 | 0 | 0 |
| SE | 91282 | ENSG00000111530 | CAND1 | chr12 | + | 67297406 | 67297663 | 67294954 | 67295156 | 67297747 | 67297848 | 336 | 254 | 286 | 282 | 441 | 0 | 0 |
| SE | 91283 | ENSG00000111530 | CAND1 | chr12 | + | 67304604 | 67304746 | 67302322 | 67302615 | 67305103 | 67305312 | 443 | 289 | 373 | 408 | 705 | 1 | 1 |
| SE | 91284 | ENSG00000111530 | CAND1 | chr12 | + | 67305103 | 67305312 | 67302322 | 67302615 | 67307396 | 67307492 | 2 | 2 | 4 | 13 | 6 | 0 | 0 |
| SE | 91285 | ENSG00000111530 | CAND1 | chr12 | + | 67305103 | 67306597 | 67302322 | 67302615 | 67307396 | 67307492 | 349 | 252 | 240 | 292 | 521 | 0 | 0 |
| SE | 91286 | ENSG00000089234 | BRAP | chr12 | - | 111649938 | 111650042 | 111642145 | 111644562 | 111655565 | 111655655 | 135 | 65 | 109 | 93 | 139 | 0 | 0 |
| SE | 91287 | ENSG00000089234 | BRAP | chr12 | - | 111649938 | 111650042 | 111642145 | 111644562 | 111658735 | 111658845 | 61 | 34 | 51 | 28 | 80 | 0 | 0 |
| SE | 91288 | ENSG00000089234 | BRAP | chr12 | - | 111655565 | 111655655 | 111642145 | 111644562 | 111658735 | 111658845 | 56 | 32 | 85 | 53 | 132 | 0 | 0 |
| SE | 91291 | ENSG00000089234 | BRAP | chr12 | - | 111658735 | 111658845 | 111655565 | 111655655 | 111665638 | 111665787 | 95 | 45 | 96 | 67 | 159 | 2 | 0 |
| SE | 91292 | ENSG00000089234 | BRAP | chr12 | - | 111659206 | 111659345 | 111655565 | 111655655 | 111660599 | 111660675 | 126 | 138 | 146 | 182 | 304 | 2 | 0 |
| SE | 91293 | ENSG00000089234 | BRAP | chr12 | - | 111659206 | 111659345 | 111655565 | 111655655 | 111665638 | 111665787 | 0 | 0 | 4 | 1 | 0 | 2 | 0 |
| SE | 91294 | ENSG00000089234 | BRAP | chr12 | - | 111659206 | 111659345 | 111655565 | 111655655 | 111672660 | 111672774 | 0 | 0 | 4 | 1 | 0 | 0 | 0 |
| SE | 91295 | ENSG00000089234 | BRAP | chr12 | - | 111659206 | 111659345 | 111658735 | 111658845 | 111665638 | 111665787 | 169 | 168 | 163 | 210 | 343 | 0 | 0 |
| SE | 91296 | ENSG00000089234 | BRAP | chr12 | - | 111660599 | 111660675 | 111655565 | 111655655 | 111665638 | 111665787 | 46 | 25 | 39 | 49 | 51 | 2 | 0 |
| SE | 91297 | ENSG00000089234 | BRAP | chr12 | - | 111660599 | 111660675 | 111655565 | 111655655 | 111672660 | 111672774 | 8 | 1 | 2 | 4 | 2 | 0 | 0 |
| SE | 91298 | ENSG00000089234 | BRAP | chr12 | - | 111660599 | 111660675 | 111655565 | 111655655 | 111683145 | 111683307 | 2 | 0 | 2 | 1 | 0 | 0 | 0 |
| SE | 91299 | ENSG00000089234 | BRAP | chr12 | - | 111660599 | 111660675 | 111659206 | 111659345 | 111665638 | 111665787 | 179 | 171 | 192 | 237 | 379 | 0 | 0 |
| SE | 91300 | ENSG00000089234 | BRAP | chr12 | - | 111660599 | 111660675 | 111659206 | 111659345 | 111672660 | 111672774 | 112 | 106 | 124 | 154 | 291 | 0 | 0 |
| SE | 91301 | ENSG00000089234 | BRAP | chr12 | - | 111665638 | 111665787 | 111655565 | 111655655 | 111672660 | 111672774 | 58 | 22 | 42 | 66 | 60 | 0 | 0 |
| SE | 91302 | ENSG00000089234 | BRAP | chr12 | - | 111665638 | 111665787 | 111659206 | 111659345 | 111672660 | 111672774 | 56 | 22 | 42 | 66 | 60 | 0 | 0 |
| SE | 91304 | ENSG00000089234 | BRAP | chr12 | - | 111665638 | 111665787 | 111660599 | 111660675 | 111679150 | 111679340 | 73 | 67 | 69 | 89 | 94 | 0 | 0 |
| SE | 91305 | ENSG00000089234 | BRAP | chr12 | - | 111672660 | 111672774 | 111655565 | 111655655 | 111683145 | 111683307 | 0 | 0 | 0 | 1 | 2 | 0 | 0 |
| SE | 91306 | ENSG00000089234 | BRAP | chr12 | - | 111672660 | 111672774 | 111660599 | 111660675 | 111679150 | 111679340 | 62 | 34 | 45 | 47 | 60 | 0 | 0 |
| SE | 91307 | ENSG00000089234 | BRAP | chr12 | - | 111672660 | 111672774 | 111660599 | 111660675 | 111683145 | 111683307 | 6 | 0 | 2 | 5 | 4 | 0 | 0 |
| SE | 91308 | ENSG00000089234 | BRAP | chr12 | - | 111672660 | 111672774 | 111665638 | 111665787 | 111679150 | 111679340 | 112 | 53 | 84 | 107 | 116 | 0 | 0 |
| SE | 91309 | ENSG00000089234 | BRAP | chr12 | - | 111679150 | 111679340 | 111660599 | 111660675 | 111683145 | 111683307 | 0 | 2 | 0 | 0 | 5 | 0 | 0 |
| SE | 91310 | ENSG00000089234 | BRAP | chr12 | - | 111679150 | 111679340 | 111672660 | 111672774 | 111683145 | 111683307 | 70 | 45 | 60 | 63 | 72 | 0 | 0 |
| SE | 91312 | ENSG00000089234 | BRAP | chr12 | - | 111681636 | 111681835 | 111679150 | 111679340 | 111685710 | 111685956 | 143 | 123 | 124 | 134 | 265 | 0 | 0 |
| SE | 91313 | ENSG00000089234 | BRAP | chr12 | - | 111683145 | 111683307 | 111679150 | 111679340 | 111685710 | 111685956 | 32 | 12 | 18 | 32 | 30 | 0 | 0 |
| SE | 91316 | ENSG00000186106 | ANKRD46 | chr8 | - | 100532355 | 100532460 | 100529522 | 100529860 | 100533208 | 100533311 | 5 | 0 | 0 | 3 | 0 | 29 | 12 |
| SE | 91322 | ENSG00000136156 | ITM2B | chr13 | + | 48233929 | 48234071 | 48233218 | 48233477 | 48253807 | 48253936 | 0 | 0 | 0 | 0 | 0 | 677 | 311 |
| SE | 91324 | ENSG00000136156 | ITM2B | chr13 | + | 48239225 | 48239332 | 48233205 | 48233477 | 48253807 | 48253936 | 8 | 5 | 0 | 7 | 12 | 677 | 311 |
| SE | 91327 | ENSG00000136156 | ITM2B | chr13 | + | 48256176 | 48256383 | 48253807 | 48253936 | 48258796 | 48258878 | 2087 | 1325 | 1574 | 1935 | 3145 | 2 | 3 |
| SE | 91328 | ENSG00000136156 | ITM2B | chr13 | + | 48258125 | 48258236 | 48256176 | 48256383 | 48258796 | 48258878 | 7446 | 5425 | 5713 | 7902 | 19199 | 5 | 1 |
| SE | 91329 | ENSG00000136156 | ITM2B | chr13 | + | 48258796 | 48258947 | 48258125 | 48258236 | 48261138 | 48261388 | 11064 | 8308 | 8364 | 11728 | 29357 | 0 | 0 |
| SE | 91330 | ENSG00000120868 | APAF1 | chr12 | + | 98659159 | 98659343 | 98649486 | 98649684 | 98662455 | 98662568 | 408 | 392 | 226 | 427 | 360 | 5 | 0 |
| SE | 91332 | ENSG00000120868 | APAF1 | chr12 | + | 98671534 | 98671719 | 98670972 | 98671086 | 98677424 | 98677551 | 311 | 215 | 152 | 245 | 377 | 1 | 1 |
| SE | 91335 | ENSG00000120868 | APAF1 | chr12 | + | 98708584 | 98708704 | 98706484 | 98706610 | 98712318 | 98712435 | 212 | 132 | 129 | 203 | 314 | 1 | 0 |
| SE | 91336 | ENSG00000120868 | APAF1 | chr12 | + | 98715426 | 98715552 | 98712318 | 98712435 | 98723192 | 98723312 | 201 | 120 | 121 | 172 | 281 | 0 | 0 |
| SE | 91338 | ENSG00000235173 | HGH1 | chr8 | + | 144138506 | 144138607 | 144137773 | 144138428 | 144138714 | 144138813 | 10 | 0 | 5 | 10 | 0 | 0 | 0 |
| SE | 91342 | ENSG00000167632 | TRAPPC9 | chr8 | - | 140252776 | 140252929 | 140221458 | 140221583 | 140275657 | 140275821 | 230 | 153 | 222 | 233 | 422 | 0 | 0 |
| SE | 91343 | ENSG00000167632 | TRAPPC9 | chr8 | - | 140283888 | 140284021 | 140275657 | 140275821 | 140287607 | 140287734 | 96 | 27 | 96 | 94 | 103 | 0 | 0 |
| SE | 91344 | ENSG00000167632 | TRAPPC9 | chr8 | - | 140287607 | 140287734 | 140283888 | 140284021 | 140290992 | 140291078 | 98 | 52 | 107 | 104 | 116 | 1 | 0 |
| SE | 91346 | ENSG00000167632 | TRAPPC9 | chr8 | - | 140300468 | 140300614 | 140290992 | 140291078 | 140311247 | 140311374 | 162 | 107 | 166 | 221 | 233 | 0 | 0 |
| SE | 91347 | ENSG00000167632 | TRAPPC9 | chr8 | - | 140311247 | 140311374 | 140300468 | 140300614 | 140360049 | 140360193 | 160 | 75 | 174 | 166 | 244 | 0 | 0 |
| SE | 91348 | ENSG00000167632 | TRAPPC9 | chr8 | - | 140360049 | 140360193 | 140311247 | 140311374 | 140370963 | 140371180 | 79 | 34 | 101 | 78 | 112 | 0 | 1 |
| SE | 91349 | ENSG00000167632 | TRAPPC9 | chr8 | - | 140370963 | 140371180 | 140360049 | 140360193 | 140397619 | 140397745 | 110 | 63 | 105 | 82 | 97 | 0 | 0 |
| SE | 91351 | ENSG00000167632 | TRAPPC9 | chr8 | - | 140426614 | 140426641 | 140405576 | 140405698 | 140439051 | 140439197 | 12 | 15 | 11 | 10 | 9 | 0 | 0 |
| SE | 91352 | ENSG00000167632 | TRAPPC9 | chr8 | - | 140435111 | 140435240 | 140405576 | 140405698 | 140439051 | 140439197 | 86 | 40 | 72 | 105 | 50 | 0 | 0 |
| SE | 91353 | ENSG00000167632 | TRAPPC9 | chr8 | - | 140435111 | 140435240 | 140426614 | 140426641 | 140439051 | 140439197 | 138 | 68 | 103 | 149 | 58 | 3 | 0 |
| SE | 91354 | ENSG00000167632 | TRAPPC9 | chr8 | - | 140439051 | 140439197 | 140435111 | 140435240 | 140450789 | 140450930 | 124 | 67 | 100 | 160 | 100 | 0 | 1 |
| SE | 91355 | ENSG00000238083 | LRRC37A2 | chr17 | + | 46512315 | 46515321 | 46511510 | 46511608 | 46517361 | 46517433 | 11 | 2 | 15 | 12 | 6 | 8 | 1 |
| SE | 91358 | ENSG00000238083 | LRRC37A2 | chr17 | + | 46523803 | 46523884 | 46520211 | 46520283 | 46540175 | 46540247 | 10 | 2 | 7 | 21 | 16 | 2 | 2 |
| SE | 91360 | ENSG00000238083 | LRRC37A2 | chr17 | + | 46546254 | 46546373 | 46520211 | 46520283 | 46548311 | 46549843 | 4 | 1 | 6 | 5 | 7 | 0 | 0 |
| SE | 91361 | ENSG00000238083 | LRRC37A2 | chr17 | + | 46546254 | 46546373 | 46540806 | 46540881 | 46548311 | 46549843 | 10 | 3 | 6 | 13 | 23 | 0 | 0 |
| SE | 91362 | ENSG00000189298 | ZKSCAN3 | chr6 | + | 28350271 | 28350388 | 28349949 | 28350067 | 28359524 | 28359988 | 0 | 0 | 1 | 3 | 0 | 6 | 6 |
| SE | 91363 | ENSG00000189298 | ZKSCAN3 | chr6 | + | 28350271 | 28350388 | 28349949 | 28350067 | 28361323 | 28361471 | 0 | 0 | 0 | 3 | 0 | 2 | 0 |
| SE | 91364 | ENSG00000189298 | ZKSCAN3 | chr6 | + | 28359524 | 28359988 | 28349946 | 28350067 | 28361323 | 28361471 | 6 | 8 | 12 | 1 | 9 | 2 | 0 |
| SE | 91365 | ENSG00000189298 | ZKSCAN3 | chr6 | + | 28359524 | 28359988 | 28350271 | 28350388 | 28361323 | 28361471 | 0 | 2 | 2 | 1 | 0 | 0 | 0 |
| SE | 91366 | ENSG00000189298 | ZKSCAN3 | chr6 | + | 28361323 | 28361471 | 28350271 | 28350388 | 28363302 | 28363385 | 17 | 12 | 22 | 8 | 11 | 2 | 0 |
| SE | 91367 | ENSG00000189298 | ZKSCAN3 | chr6 | + | 28363302 | 28363385 | 28361323 | 28361471 | 28365425 | 28366746 | 14 | 13 | 19 | 8 | 7 | 0 | 1 |
| SE | 91368 | ENSG00000189298 | ZKSCAN3 | chr6 | + | 28363691 | 28363815 | 28363302 | 28363385 | 28365425 | 28366746 | 29 | 16 | 31 | 24 | 34 | 3 | 4 |
| SE | 91369 | ENSG00000100288 | CHKB | chr22 | - | 50579726 | 50579802 | 50579425 | 50579507 | 50579973 | 50580082 | 182 | 76 | 104 | 114 | 158 | 2 | 0 |
| SE | 91371 | ENSG00000100288 | CHKB | chr22 | - | 50579973 | 50580082 | 50579726 | 50579802 | 50580189 | 50580253 | 196 | 115 | 149 | 192 | 155 | 0 | 0 |
| SE | 91372 | ENSG00000100288 | CHKB | chr22 | - | 50580056 | 50580082 | 50579726 | 50579802 | 50580189 | 50580271 | 34 | 18 | 27 | 34 | 31 | 0 | 0 |
| SE | 91374 | ENSG00000100288 | CHKB | chr22 | - | 50582028 | 50582095 | 50581748 | 50581862 | 50582248 | 50582357 | 0 | 0 | 0 | 0 | 0 | 7 | 1 |
| SE | 91375 | ENSG00000242779 | ZNF702P | chr19 | - | 52974031 | 52974153 | 52967085 | 52970842 | 52980891 | 52980972 | 0 | 1 | 1 | 0 | 0 | 7 | 8 |
| SE | 91379 | ENSG00000104408 | EIF3E | chr8 | - | 108216411 | 108216513 | 108214606 | 108214716 | 108217333 | 108217460 | 3521 | 2281 | 3584 | 2825 | 6239 | 0 | 1 |
| SE | 91380 | ENSG00000104408 | EIF3E | chr8 | - | 108217333 | 108217460 | 108214606 | 108214716 | 108228266 | 108228391 | 1681 | 1044 | 1711 | 1318 | 1648 | 0 | 0 |
| SE | 91381 | ENSG00000104408 | EIF3E | chr8 | - | 108219749 | 108219830 | 108217333 | 108217460 | 108228266 | 108228391 | 0 | 0 | 0 | 2 | 0 | 1783 | 1117 |
| SE | 91382 | ENSG00000104408 | EIF3E | chr8 | - | 108225711 | 108225833 | 108217333 | 108217460 | 108228266 | 108228391 | 1 | 0 | 0 | 3 | 0 | 1783 | 1117 |
| SE | 91383 | ENSG00000104408 | EIF3E | chr8 | - | 108239957 | 108240075 | 108236160 | 108236203 | 108241798 | 108241913 | 6966 | 4951 | 7068 | 6002 | 7086 | 0 | 0 |
| SE | 91384 | ENSG00000104408 | EIF3E | chr8 | - | 108239957 | 108240075 | 108236160 | 108236203 | 108248612 | 108248730 | 4465 | 3169 | 4566 | 3884 | 3917 | 0 | 0 |
| SE | 91385 | ENSG00000104408 | EIF3E | chr8 | - | 108239957 | 108240098 | 108236160 | 108236203 | 108241798 | 108241913 | 4503 | 3198 | 4616 | 3910 | 3954 | 0 | 0 |
| SE | 91387 | ENSG00000104408 | EIF3E | chr8 | - | 108241798 | 108241913 | 108236160 | 108236203 | 108248612 | 108248705 | 2031 | 1107 | 2093 | 1698 | 1867 | 0 | 0 |
| SE | 91388 | ENSG00000104408 | EIF3E | chr8 | - | 108241798 | 108241913 | 108239957 | 108240075 | 108242111 | 108242256 | 1837 | 1390 | 1831 | 1570 | 2579 | 2 | 2 |
| SE | 91389 | ENSG00000104408 | EIF3E | chr8 | - | 108241798 | 108241913 | 108239957 | 108240075 | 108248612 | 108248705 | 4569 | 2917 | 4637 | 3851 | 5097 | 4 | 3 |
| SE | 91391 | ENSG00000104408 | EIF3E | chr8 | - | 108241885 | 108241913 | 108239957 | 108240075 | 108248612 | 108248719 | 4 | 3 | 4 | 5 | 4 | 4 | 3 |
| SE | 91392 | ENSG00000104408 | EIF3E | chr8 | - | 108242111 | 108242256 | 108241798 | 108241913 | 108248612 | 108248717 | 7 | 12 | 9 | 11 | 24 | 2744 | 1545 |
| SE | 91393 | ENSG00000104408 | EIF3E | chr8 | - | 108242111 | 108242346 | 108241798 | 108241913 | 108248612 | 108248716 | 8 | 12 | 9 | 11 | 24 | 2744 | 1545 |
| SE | 91394 | ENSG00000104408 | EIF3E | chr8 | - | 108243292 | 108243463 | 108241798 | 108241913 | 108248612 | 108248716 | 2 | 1 | 0 | 3 | 6 | 2744 | 1545 |
| SE | 91402 | ENSG00000158473 | CD1D | chr1 | + | 158183935 | 158184035 | 158182877 | 158183156 | 158184128 | 158186427 | 109 | 50 | 54 | 93 | 33 | 0 | 0 |
| SE | 91403 | ENSG00000008988 | RPS20 | chr8 | - | 56073694 | 56073768 | 56069592 | 56073272 | 56074059 | 56074144 | 15513 | 10805 | 17006 | 15379 | 26187 | 10 | 2 |
| SE | 91404 | ENSG00000008988 | RPS20 | chr8 | - | 56073694 | 56073768 | 56069592 | 56073272 | 56074380 | 56074510 | 5150 | 3686 | 5634 | 5016 | 9291 | 0 | 0 |
| SE | 91405 | ENSG00000008988 | RPS20 | chr8 | - | 56073694 | 56074159 | 56073126 | 56073272 | 56074380 | 56074506 | 9426 | 6096 | 11099 | 9376 | 13352 | 0 | 0 |
| SE | 91406 | ENSG00000008988 | RPS20 | chr8 | - | 56074059 | 56074150 | 56073053 | 56073272 | 56074380 | 56074472 | 7 | 1 | 4 | 12 | 5 | 0 | 0 |
| SE | 91408 | ENSG00000008988 | RPS20 | chr8 | - | 56074059 | 56074159 | 56073074 | 56073272 | 56074380 | 56074498 | 4256 | 2382 | 5449 | 4335 | 4017 | 0 | 0 |
| SE | 91409 | ENSG00000164733 | CTSB | chr8 | - | 11845081 | 11845222 | 11843009 | 11844241 | 11845660 | 11845789 | 1306 | 916 | 903 | 1138 | 890 | 0 | 0 |
| SE | 91410 | ENSG00000164733 | CTSB | chr8 | - | 11846561 | 11846616 | 11845660 | 11845789 | 11847051 | 11847168 | 0 | 0 | 2 | 0 | 1 | 1246 | 941 |
| SE | 91414 | ENSG00000164733 | CTSB | chr8 | - | 11849045 | 11849164 | 11848066 | 11848152 | 11850865 | 11850980 | 684 | 329 | 386 | 687 | 584 | 3 | 1 |
| SE | 91415 | ENSG00000164733 | CTSB | chr8 | - | 11849045 | 11849164 | 11848066 | 11848152 | 11852609 | 11852695 | 382 | 163 | 206 | 368 | 293 | 0 | 0 |
| SE | 91416 | ENSG00000164733 | CTSB | chr8 | - | 11849045 | 11849215 | 11848066 | 11848152 | 11850865 | 11850980 | 385 | 163 | 206 | 368 | 293 | 3 | 1 |
| SE | 91417 | ENSG00000164733 | CTSB | chr8 | - | 11849045 | 11849228 | 11848066 | 11848152 | 11850865 | 11850980 | 382 | 164 | 206 | 368 | 294 | 3 | 1 |
| SE | 91418 | ENSG00000164733 | CTSB | chr8 | - | 11849997 | 11850047 | 11849045 | 11849164 | 11850865 | 11850980 | 0 | 0 | 0 | 0 | 0 | 303 | 166 |
| SE | 91419 | ENSG00000164733 | CTSB | chr8 | - | 11850865 | 11850980 | 11847051 | 11849164 | 11852609 | 11852695 | 546 | 274 | 318 | 540 | 502 | 0 | 0 |
| SE | 91420 | ENSG00000164733 | CTSB | chr8 | - | 11850865 | 11850980 | 11848066 | 11848152 | 11852609 | 11852695 | 246 | 113 | 139 | 223 | 215 | 0 | 0 |
| SE | 91421 | ENSG00000164733 | CTSB | chr8 | - | 11854679 | 11854914 | 11853328 | 11853479 | 11864375 | 11864463 | 0 | 0 | 0 | 0 | 0 | 0 | 1 |
| SE | 91422 | ENSG00000164733 | CTSB | chr8 | - | 11854679 | 11854914 | 11853328 | 11853479 | 11868000 | 11868122 | 0 | 0 | 0 | 0 | 0 | 0 | 0 |
| SE | 91424 | ENSG00000164733 | CTSB | chr8 | - | 11857857 | 11857940 | 11853328 | 11853479 | 11864375 | 11864463 | 0 | 0 | 2 | 2 | 0 | 0 | 1 |
| SE | 91425 | ENSG00000164733 | CTSB | chr8 | - | 11857857 | 11857940 | 11853328 | 11853479 | 11867253 | 11868076 | 0 | 0 | 2 | 2 | 0 | 0 | 0 |
| SE | 91426 | ENSG00000164733 | CTSB | chr8 | - | 11857857 | 11857940 | 11853328 | 11853479 | 11868000 | 11868114 | 0 | 0 | 2 | 3 | 0 | 0 | 0 |
| SE | 91429 | ENSG00000164733 | CTSB | chr8 | - | 11861360 | 11861479 | 11853328 | 11853479 | 11862344 | 11862478 | 0 | 1 | 0 | 4 | 1 | 0 | 0 |
| SE | 91430 | ENSG00000164733 | CTSB | chr8 | - | 11861360 | 11861479 | 11853328 | 11853479 | 11868000 | 11868090 | 0 | 0 | 1 | 4 | 0 | 0 | 0 |
| SE | 91431 | ENSG00000164733 | CTSB | chr8 | - | 11861405 | 11861479 | 11853328 | 11853479 | 11864375 | 11864463 | 0 | 0 | 1 | 1 | 3 | 0 | 1 |
| SE | 91432 | ENSG00000164733 | CTSB | chr8 | - | 11861405 | 11861479 | 11853328 | 11853479 | 11868000 | 11868114 | 0 | 0 | 1 | 1 | 3 | 0 | 0 |
| SE | 91433 | ENSG00000164733 | CTSB | chr8 | - | 11862344 | 11862478 | 11853328 | 11853479 | 11864375 | 11864463 | 0 | 0 | 2 | 3 | 0 | 0 | 1 |
| SE | 91434 | ENSG00000164733 | CTSB | chr8 | - | 11862344 | 11862478 | 11853328 | 11853479 | 11867969 | 11868052 | 0 | 0 | 3 | 3 | 0 | 0 | 0 |
| SE | 91435 | ENSG00000164733 | CTSB | chr8 | - | 11862344 | 11862478 | 11853328 | 11853479 | 11868000 | 11868090 | 0 | 0 | 2 | 4 | 0 | 0 | 0 |
| SE | 91436 | ENSG00000164733 | CTSB | chr8 | - | 11862344 | 11862478 | 11861360 | 11861479 | 11864375 | 11864463 | 0 | 0 | 0 | 0 | 1 | 0 | 0 |
| SE | 91437 | ENSG00000164733 | CTSB | chr8 | - | 11862344 | 11862478 | 11861360 | 11861479 | 11868000 | 11868090 | 0 | 0 | 0 | 1 | 1 | 0 | 0 |
| SE | 91438 | ENSG00000164733 | CTSB | chr8 | - | 11864375 | 11864463 | 11853328 | 11853479 | 11865463 | 11865560 | 0 | 1 | 5 | 7 | 3 | 0 | 0 |
| SE | 91440 | ENSG00000164733 | CTSB | chr8 | - | 11864375 | 11864463 | 11853328 | 11853479 | 11867253 | 11867391 | 0 | 1 | 5 | 8 | 3 | 0 | 0 |
| SE | 91441 | ENSG00000164733 | CTSB | chr8 | - | 11864375 | 11864463 | 11853328 | 11853479 | 11868000 | 11868040 | 0 | 1 | 6 | 9 | 3 | 0 | 0 |
| SE | 91443 | ENSG00000164733 | CTSB | chr8 | - | 11864375 | 11864463 | 11857857 | 11857940 | 11867253 | 11867391 | 0 | 0 | 0 | 1 | 0 | 0 | 0 |
| SE | 91444 | ENSG00000164733 | CTSB | chr8 | - | 11864375 | 11864463 | 11857857 | 11857940 | 11868000 | 11868040 | 0 | 0 | 1 | 1 | 0 | 0 | 0 |
| SE | 91445 | ENSG00000164733 | CTSB | chr8 | - | 11864375 | 11864463 | 11861299 | 11861479 | 11868000 | 11868040 | 0 | 0 | 1 | 1 | 0 | 0 | 0 |
| SE | 91446 | ENSG00000164733 | CTSB | chr8 | - | 11864375 | 11864463 | 11862344 | 11862478 | 11868000 | 11868040 | 0 | 0 | 1 | 1 | 0 | 0 | 0 |
| SE | 91447 | ENSG00000164733 | CTSB | chr8 | - | 11864375 | 11864483 | 11853328 | 11853479 | 11867253 | 11867530 | 0 | 1 | 5 | 7 | 3 | 0 | 0 |
| SE | 91448 | ENSG00000164733 | CTSB | chr8 | - | 11864375 | 11864483 | 11853328 | 11853479 | 11868000 | 11868112 | 0 | 1 | 5 | 7 | 3 | 0 | 0 |
| SE | 91449 | ENSG00000164733 | CTSB | chr8 | - | 11864375 | 11864483 | 11861405 | 11861479 | 11868000 | 11868112 | 0 | 0 | 0 | 0 | 0 | 0 | 0 |
| SE | 91452 | ENSG00000164733 | CTSB | chr8 | - | 11865463 | 11865584 | 11853328 | 11853479 | 11867253 | 11867391 | 0 | 0 | 0 | 0 | 0 | 0 | 0 |
| SE | 91453 | ENSG00000164733 | CTSB | chr8 | - | 11865463 | 11865584 | 11853328 | 11853479 | 11868000 | 11868126 | 0 | 0 | 0 | 0 | 0 | 0 | 0 |
| SE | 91458 | ENSG00000164733 | CTSB | chr8 | - | 11867253 | 11867391 | 11853328 | 11853479 | 11868000 | 11868089 | 0 | 0 | 1 | 2 | 0 | 0 | 0 |
| SE | 91459 | ENSG00000164733 | CTSB | chr8 | - | 11867253 | 11867391 | 11864375 | 11864463 | 11868000 | 11868089 | 0 | 0 | 0 | 1 | 0 | 0 | 0 |
| SE | 91462 | ENSG00000204685 | STARD7-AS1 | chr2 | + | 96237904 | 96238099 | 96208406 | 96208979 | 96239712 | 96240688 | 5 | 5 | 6 | 3 | 9 | 0 | 3 |
| SE | 91468 | ENSG00000197140 | ADAM32 | chr8 | + | 39169915 | 39169997 | 39165029 | 39165196 | 39186908 | 39187045 | 0 | 6 | 0 | 0 | 0 | 0 | 0 |
| SE | 91470 | ENSG00000197140 | ADAM32 | chr8 | + | 39186908 | 39187045 | 39165029 | 39165196 | 39211143 | 39211324 | 0 | 0 | 0 | 1 | 0 | 0 | 0 |
| SE | 91471 | ENSG00000197140 | ADAM32 | chr8 | + | 39186908 | 39187045 | 39169915 | 39169997 | 39211143 | 39211324 | 0 | 6 | 1 | 1 | 0 | 0 | 1 |
| SE | 91474 | ENSG00000197140 | ADAM32 | chr8 | + | 39211143 | 39211324 | 39186908 | 39187045 | 39221609 | 39221702 | 0 | 0 | 0 | 2 | 0 | 0 | 0 |
| SE | 91480 | ENSG00000147601 | TERF1 | chr8 | + | 73012855 | 73012931 | 73008863 | 73009205 | 73013894 | 73013990 | 0 | 0 | 0 | 0 | 0 | 5 | 4 |
| SE | 91482 | ENSG00000147601 | TERF1 | chr8 | + | 73022215 | 73022302 | 73020683 | 73020805 | 73024821 | 73024971 | 155 | 124 | 160 | 168 | 254 | 0 | 0 |
| SE | 91485 | ENSG00000147601 | TERF1 | chr8 | + | 73030794 | 73030955 | 73026939 | 73027052 | 73032041 | 73032133 | 3 | 0 | 2 | 0 | 0 | 54 | 42 |
| SE | 91486 | ENSG00000147601 | TERF1 | chr8 | + | 73032041 | 73032133 | 73026939 | 73027052 | 73039115 | 73039219 | 83 | 60 | 98 | 100 | 110 | 1 | 0 |
| SE | 91487 | ENSG00000147601 | TERF1 | chr8 | + | 73038705 | 73038795 | 73032041 | 73032133 | 73039115 | 73039219 | 8 | 11 | 9 | 22 | 14 | 67 | 40 |
| SE | 91489 | ENSG00000147601 | TERF1 | chr8 | + | 73039544 | 73039619 | 73039115 | 73039219 | 73045960 | 73047479 | 3 | 1 | 1 | 4 | 10 | 69 | 49 |
| SE | 91490 | ENSG00000224597 | SVIL-AS1 | chr10 | + | 29418181 | 29418304 | 29415356 | 29415412 | 29420490 | 29420524 | 0 | 1 | 0 | 0 | 0 | 23 | 29 |
| SE | 91491 | ENSG00000224597 | SVIL-AS1 | chr10 | + | 29458273 | 29458520 | 29415356 | 29415412 | 29467239 | 29467278 | 0 | 2 | 2 | 2 | 0 | 1 | 0 |
| SE | 91492 | ENSG00000224597 | SVIL-AS1 | chr10 | + | 29458273 | 29458520 | 29415356 | 29415412 | 29482832 | 29483432 | 0 | 2 | 2 | 2 | 0 | 0 | 2 |
| SE | 91503 | ENSG00000135679 | MDM2 | chr12 | + | 68813553 | 68813628 | 68809207 | 68809292 | 68816811 | 68816945 | 328 | 239 | 252 | 255 | 199 | 0 | 0 |
| SE | 91520 | ENSG00000135679 | MDM2 | chr12 | + | 68820884 | 68820910 | 68820324 | 68820374 | 68824362 | 68824430 | 0 | 0 | 1 | 0 | 0 | 217 | 161 |
| SE | 91521 | ENSG00000135679 | MDM2 | chr12 | + | 68823414 | 68823514 | 68820324 | 68820374 | 68824362 | 68824430 | 0 | 2 | 0 | 0 | 3 | 217 | 161 |
| SE | 91524 | ENSG00000135679 | MDM2 | chr12 | + | 68824362 | 68824430 | 68820324 | 68820374 | 68828770 | 68828931 | 122 | 81 | 102 | 101 | 134 | 3 | 0 |
| SE | 91527 | ENSG00000135679 | MDM2 | chr12 | + | 68824554 | 68824651 | 68824362 | 68824430 | 68828770 | 68828931 | 468 | 418 | 400 | 397 | 557 | 2 | 4 |
| SE | 91533 | ENSG00000135679 | MDM2 | chr12 | + | 68835828 | 68835843 | 68828770 | 68828931 | 68836671 | 68836749 | 26 | 18 | 22 | 20 | 43 | 0 | 0 |
| SE | 91537 | ENSG00000135679 | MDM2 | chr12 | + | 68835828 | 68835984 | 68828770 | 68828931 | 68836671 | 68836749 | 565 | 456 | 449 | 438 | 738 | 0 | 0 |
| SE | 91538 | ENSG00000135679 | MDM2 | chr12 | + | 68835828 | 68835984 | 68828770 | 68828931 | 68839273 | 68839314 | 250 | 188 | 188 | 186 | 273 | 0 | 0 |
| SE | 91539 | ENSG00000135679 | MDM2 | chr12 | + | 68836671 | 68836749 | 68828770 | 68828931 | 68839273 | 68839282 | 246 | 290 | 237 | 258 | 456 | 0 | 0 |
| SE | 91541 | ENSG00000139546 | TARBP2 | chr12 | + | 53502014 | 53502184 | 53500920 | 53501461 | 53503026 | 53503129 | 26 | 6 | 34 | 51 | 20 | 3 | 0 |
| SE | 91543 | ENSG00000139546 | TARBP2 | chr12 | + | 53502014 | 53502184 | 53500920 | 53501461 | 53503729 | 53503808 | 7 | 4 | 9 | 9 | 3 | 0 | 2 |
| SE | 91545 | ENSG00000139546 | TARBP2 | chr12 | + | 53502014 | 53502184 | 53501590 | 53501708 | 53503729 | 53503808 | 0 | 0 | 8 | 8 | 0 | 0 | 0 |
| SE | 91546 | ENSG00000139546 | TARBP2 | chr12 | + | 53502019 | 53502184 | 53501365 | 53501461 | 53503026 | 53503129 | 19 | 2 | 30 | 42 | 17 | 3 | 0 |
| SE | 91549 | ENSG00000139546 | TARBP2 | chr12 | + | 53502059 | 53502184 | 53501305 | 53501461 | 53503026 | 53503212 | 18 | 1 | 23 | 34 | 17 | 3 | 0 |
| SE | 91552 | ENSG00000139546 | TARBP2 | chr12 | + | 53503026 | 53503129 | 53501365 | 53501461 | 53503729 | 53503808 | 5 | 0 | 0 | 0 | 0 | 0 | 2 |
| SE | 91553 | ENSG00000139546 | TARBP2 | chr12 | + | 53503026 | 53503129 | 53501596 | 53501708 | 53503729 | 53503808 | 2 | 0 | 0 | 0 | 0 | 0 | 0 |
| SE | 91554 | ENSG00000139546 | TARBP2 | chr12 | + | 53503026 | 53503129 | 53502014 | 53502184 | 53503712 | 53503808 | 26 | 3 | 32 | 40 | 23 | 0 | 1 |
| SE | 91556 | ENSG00000139546 | TARBP2 | chr12 | + | 53503026 | 53503212 | 53502014 | 53502184 | 53503712 | 53503808 | 16 | 2 | 24 | 37 | 17 | 0 | 1 |
| SE | 91558 | ENSG00000139546 | TARBP2 | chr12 | + | 53505134 | 53505262 | 53504611 | 53504815 | 53505648 | 53505686 | 27 | 8 | 13 | 6 | 11 | 3 | 0 |
| SE | 91560 | ENSG00000139546 | TARBP2 | chr12 | + | 53505648 | 53505850 | 53504697 | 53504815 | 53505990 | 53506229 | 20 | 5 | 5 | 6 | 4 | 0 | 0 |
| SE | 91566 | ENSG00000070501 | POLB | chr8 | + | 42350006 | 42350065 | 42349015 | 42349090 | 42355515 | 42355567 | 17 | 14 | 19 | 16 | 13 | 1 | 0 |
| SE | 91567 | ENSG00000070501 | POLB | chr8 | + | 42352518 | 42352568 | 42349015 | 42349090 | 42355515 | 42355567 | 9 | 5 | 26 | 16 | 22 | 1 | 0 |
| SE | 91568 | ENSG00000070501 | POLB | chr8 | + | 42352518 | 42352568 | 42350006 | 42350065 | 42355515 | 42355567 | 75 | 56 | 98 | 96 | 129 | 0 | 0 |
| SE | 91572 | ENSG00000070501 | POLB | chr8 | + | 42362611 | 42362698 | 42361294 | 42361365 | 42369848 | 42369988 | 21 | 12 | 30 | 32 | 51 | 0 | 0 |
| SE | 91574 | ENSG00000070501 | POLB | chr8 | + | 42369270 | 42369335 | 42361294 | 42361365 | 42369848 | 42369988 | 65 | 47 | 50 | 66 | 117 | 0 | 0 |
| SE | 91575 | ENSG00000070501 | POLB | chr8 | + | 42369270 | 42369335 | 42361294 | 42361365 | 42371562 | 42371752 | 5 | 7 | 1 | 5 | 0 | 0 | 1 |
| SE | 91576 | ENSG00000070501 | POLB | chr8 | + | 42369270 | 42369335 | 42362611 | 42362698 | 42369848 | 42369988 | 121 | 83 | 113 | 129 | 247 | 0 | 1 |
| SE | 91578 | ENSG00000070501 | POLB | chr8 | + | 42369848 | 42369988 | 42361294 | 42361365 | 42371562 | 42371765 | 82 | 67 | 64 | 82 | 76 | 0 | 1 |
| SE | 91582 | ENSG00000147687 | TATDN1 | chr8 | - | 124503839 | 124503951 | 124495471 | 124495542 | 124504270 | 124504347 | 11 | 1 | 1 | 3 | 10 | 79 | 42 |
| SE | 91583 | ENSG00000147687 | TATDN1 | chr8 | - | 124503839 | 124503951 | 124495471 | 124495542 | 124504274 | 124504347 | 12 | 0 | 1 | 6 | 16 | 3 | 3 |
| SE | 91584 | ENSG00000147687 | TATDN1 | chr8 | - | 124518741 | 124518881 | 124515886 | 124516030 | 124522936 | 124523002 | 27 | 24 | 24 | 32 | 40 | 0 | 0 |
| SE | 91585 | ENSG00000147687 | TATDN1 | chr8 | - | 124518817 | 124518881 | 124515886 | 124516030 | 124522936 | 124523002 | 80 | 77 | 78 | 92 | 87 | 0 | 0 |
| SE | 91587 | ENSG00000147687 | TATDN1 | chr8 | - | 124522150 | 124522200 | 124518741 | 124518881 | 124522936 | 124523002 | 45 | 46 | 60 | 60 | 57 | 7 | 4 |
| SE | 91591 | ENSG00000147687 | TATDN1 | chr8 | - | 124522936 | 124523002 | 124521636 | 124522200 | 124539024 | 124539070 | 19 | 23 | 27 | 30 | 17 | 0 | 0 |
| SE | 91594 | ENSG00000104687 | GSR | chr8 | - | 30681929 | 30682061 | 30680903 | 30681037 | 30684087 | 30684199 | 682 | 450 | 511 | 657 | 859 | 0 | 0 |
| SE | 91595 | ENSG00000104687 | GSR | chr8 | - | 30687522 | 30687639 | 30684087 | 30684199 | 30689160 | 30689319 | 0 | 0 | 2 | 0 | 0 | 218 | 139 |
| SE | 91597 | ENSG00000104687 | GSR | chr8 | - | 30689160 | 30689319 | 30684087 | 30684199 | 30696379 | 30696479 | 219 | 142 | 180 | 210 | 248 | 0 | 0 |
| SE | 91598 | ENSG00000104687 | GSR | chr8 | - | 30692968 | 30693055 | 30684087 | 30684199 | 30696379 | 30696479 | 70 | 41 | 66 | 72 | 46 | 0 | 0 |
| SE | 91600 | ENSG00000104687 | GSR | chr8 | - | 30700080 | 30700135 | 30696379 | 30696479 | 30703092 | 30703240 | 265 | 155 | 186 | 235 | 180 | 3 | 0 |
| SE | 91601 | ENSG00000104687 | GSR | chr8 | - | 30700080 | 30700135 | 30696379 | 30696479 | 30708071 | 30708141 | 36 | 19 | 34 | 36 | 33 | 0 | 0 |
| SE | 91602 | ENSG00000104687 | GSR | chr8 | - | 30703092 | 30703240 | 30696379 | 30696479 | 30708071 | 30708141 | 316 | 150 | 204 | 266 | 299 | 0 | 0 |
| SE | 91603 | ENSG00000104687 | GSR | chr8 | - | 30703092 | 30703240 | 30700080 | 30700135 | 30708071 | 30708141 | 544 | 286 | 356 | 467 | 455 | 0 | 0 |
| SE | 91604 | ENSG00000104687 | GSR | chr8 | - | 30704646 | 30704864 | 30703092 | 30703240 | 30708071 | 30708141 | 0 | 1 | 0 | 1 | 0 | 313 | 151 |
| SE | 91605 | ENSG00000104687 | GSR | chr8 | - | 30706989 | 30707104 | 30703092 | 30703240 | 30708071 | 30708141 | 4 | 2 | 0 | 4 | 2 | 313 | 151 |

| | | | | | | | | | | | | | | | | | | | | | | | | | | | | | | | | | | | | | | | | | | | | | | |
|---|---|---|---|---|---|---|---|---|---|---|---|---|---|---|---|---|---|---|---|---|---|---|---|---|---|---|---|---|---|---|---|---|---|---|---|---|---|---|---|---|---|---|---|---|---|---|
| SE | 91606 | ENSG00000104687 | GSR | chr8 | - | 30708071 | 30708141 | 30703092 | 30703240 | 30709813 | 30709902 | 428 | 194 | 255 | 373 | 411 | 0 | 0 | 0 | 0 | 0 | 620 | 647 | 266 | 413 | 835 | 0 | 4 | 1 | 2 | 0 | 219 | 149 | 1 | 1 | 1 | 1 | 1 | 1 | 1 | 1 | 0.991 | 0.995 | 0.993 | 1 | 0.004 |
| SE | 91607 | ENSG00000104691 | UBXN8 | chr8 | + | 30751395 | 30751518 | 30744163 | 30744277 | 30753034 | 30753105 | 31 | 40 | 30 | 43 | 34 | 0 | 1 | 0 | 0 | 0 | 2 | 10 | 3 | 18 | 5 | 0 | 0 | 0 | 0 | 0 | 272 | 149 | 1 | 1 | 1 | 0.956 | 1 | 1 | 1 | 1 | 1 | 1 | 1 | 1 | -0.009 |
| SE | 91608 | ENSG00000104691 | UBXN8 | chr8 | + | 30751395 | 30751518 | 30744173 | 30744460 | 30753034 | 30753105 | 30 | 40 | 28 | 44 | 43 | 1 | 0 | 0 | 0 | 0 | 1 | 5 | 3 | 18 | 5 | 0 | 0 | 0 | 0 | 0 | 272 | 149 | 1 | 1 | 0.943 | 1 | 1 | 1 | 1 | 1 | 1 | 1 | 1 | 1 | -0.011 |
| SE | 91609 | ENSG00000104691 | UBXN8 | chr8 | + | 30751506 | 30751518 | 30744189 | 30744277 | 30753034 | 30753105 | 1 | 2 | 5 | 5 | 3 | 0 | 1 | 0 | 0 | 0 | 0 | 0 | 0 | 4 | 0 | 0 | 0 | 0 | 0 | 0 | 161 | 149 | 1 | 1 | 1 | 0.649 | 1 | 1 | 1 | NA | NA | NA | 1 | NA | -0.07 |
| SE | 91610 | ENSG00000104691 | UBXN8 | chr8 | + | 30754664 | 30754749 | 30753034 | 30753105 | 30756764 | 30756887 | 24 | 29 | 17 | 25 | 32 | 0 | 0 | 0 | 0 | 3 | 1 | 4 | 4 | 14 | 5 | 0 | 0 | 0 | 0 | 0 | 234 | 149 | 1 | 1 | 1 | 1 | 1 | 1 | 0.872 | 1 | 1 | 1 | 1 | 1 | -0.026 |
| SE | 91611 | ENSG00000104691 | UBXN8 | chr8 | + | 30754664 | 30754787 | 30753034 | 30753105 | 30756764 | 30756887 | 92 | 71 | 53 | 67 | 82 | 0 | 0 | 0 | 0 | 3 | 1 | 31 | 14 | 29 | 26 | 0 | 0 | 0 | 0 | 0 | 272 | 149 | 1 | 1 | 1 | 1 | 1 | 1 | 0.937 | 1 | 1 | 1 | 1 | 1 | -0.013 |
| SE | 91612 | ENSG00000184445 | KNTC1 | chr12 | + | 122529990 | 122530192 | 122527248 | 122527351 | 122534663 | 122534784 | 57 | 70 | 68 | 59 | 86 | 5 | 8 | 5 | 4 | 1 | 16 | 55 | 51 | 46 | 0 | 0 | 1 | 7 | 6 | 0 | 298 | 149 | 1 | 1 | 0.851 | 0.814 | 0.872 | 0.881 | 0.977 | 1 | 0.965 | 0.785 | 0.793 | NA | -0.007 |
| SE | 91613 | ENSG00000184445 | KNTC1 | chr12 | + | 122530021 | 122530192 | 122527247 | 122527351 | 122534663 | 122534784 | 104 | 113 | 116 | 99 | 122 | 5 | 8 | 5 | 4 | 1 | 44 | 104 | 93 | 74 | 0 | 0 | 1 | 7 | 6 | 0 | 298 | 149 | 1 | 1 | 0.912 | 0.876 | 0.921 | 0.925 | 0.984 | 1 | 0.981 | 0.869 | 0.86 | NA | -0.004 |
| SE | 91614 | ENSG00000184445 | KNTC1 | chr12 | + | 122534663 | 122534784 | 122527247 | 122527351 | 122538338 | 122538454 | 51 | 40 | 39 | 29 | 50 | 0 | 0 | 0 | 0 | 0 | 18 | 33 | 19 | 19 | 0 | 0 | 0 | 2 | 0 | 0 | 270 | 149 | 1 | 1 | 1 | 1 | 1 | 1 | 1 | 1 | 1 | 0.84 | 1 | NA | 0.04 |
| SE | 91615 | ENSG00000184445 | KNTC1 | chr12 | + | 122543599 | 122543634 | 122542049 | 122542127 | 122544158 | 122544269 | 49 | 39 | 35 | 46 | 84 | 0 | 0 | 0 | 0 | 0 | 15 | 35 | 31 | 6 | 0 | 13 | 0 | 0 | 0 | 0 | 184 | 149 | 1 | 1 | 1 | 1 | 1 | 1 | 1 | 0.483 | 1 | 1 | 1 | NA | 0.129 |
| SE | 91617 | ENSG00000184445 | KNTC1 | chr12 | + | 122549765 | 122549864 | 122547914 | 122547969 | 122551318 | 122551362 | 47 | 48 | 44 | 57 | 146 | 0 | 0 | 0 | 0 | 0 | 15 | 74 | 38 | 37 | 8 | 0 | 0 | 1 | 0 | 0 | 248 | 149 | 1 | 1 | 1 | 1 | 1 | 1 | 1 | 1 | 1 | 0.958 | 1 | 1 | 0.008 |
| SE | 91618 | ENSG00000184445 | KNTC1 | chr12 | + | 122551318 | 122551362 | 122549765 | 122549864 | 122551457 | 122551523 | 46 | 49 | 46 | 59 | 113 | 1 | 0 | 0 | 0 | 0 | 15 | 50 | 25 | 25 | 7 | 0 | 0 | 0 | 0 | 0 | 193 | 149 | 1 | 1 | 0.973 | 1 | 1 | 1 | 1 | 1 | 1 | 1 | 1 | 1 | -0.005 |
| SE | 91619 | ENSG00000184445 | KNTC1 | chr12 | + | 122568260 | 122568372 | 122562637 | 122562699 | 122569680 | 122569824 | 59 | 33 | 66 | 51 | 49 | 1 | 0 | 0 | 1 | 0 | 18 | 30 | 19 | 24 | 6 | 0 | 0 | 0 | 0 | 0 | 261 | 149 | 1 | 1 | 0.971 | 1 | 1 | 0.967 | 1 | 1 | 1 | 1 | 1 | 1 | -0.012 |
| SE | 91621 | ENSG00000184445 | KNTC1 | chr12 | + | 122586700 | 122586757 | 122585635 | 122585774 | 122587710 | 122587874 | 24 | 15 | 10 | 18 | 28 | 0 | 0 | 0 | 0 | 0 | 13 | 22 | 22 | 17 | 0 | 0 | 4 | 0 | 0 | 0 | 206 | 149 | 1 | 1 | 1 | 1 | 1 | 1 | 1 | 1 | 0.799 | 1 | 1 | NA | 0.05 |
| SE | 91622 | ENSG00000184445 | KNTC1 | chr12 | + | 122597730 | 122597938 | 122593415 | 122594385 | 122601535 | 122601607 | 35 | 39 | 40 | 40 | 54 | 1 | 1 | 6 | 4 | 1 | 7 | 35 | 36 | 28 | 0 | 0 | 0 | 0 | 1 | 0 | 298 | 149 | 1 | 1 | 0.946 | 0.951 | 0.769 | 0.833 | 0.964 | 1 | 1 | 1 | 0.933 | NA | -0.091 |
| SE | 91623 | ENSG00000184445 | KNTC1 | chr12 | + | 122597730 | 122597938 | 122593415 | 122594385 | 122602568 | 122602740 | 19 | 6 | 15 | 15 | 18 | 0 | 0 | 0 | 0 | 2 | 0 | 11 | 1 | 12 | 0 | 0 | 0 | 0 | 0 | 0 | 298 | 149 | 1 | 1 | 1 | 1 | 1 | 1 | 0.818 | NA | 1 | 1 | 1 | NA | -0.036 |
| SE | 91624 | ENSG00000184445 | KNTC1 | chr12 | + | 122601535 | 122601625 | 122594275 | 122594385 | 122602568 | 122602740 | 17 | 8 | 27 | 16 | 30 | 0 | 0 | 0 | 0 | 2 | 0 | 23 | 7 | 20 | 0 | 0 | 0 | 0 | 0 | 0 | 239 | 149 | 1 | 1 | 1 | 1 | 1 | 1 | 0.903 | NA | 1 | 1 | 1 | NA | -0.019 |
| SE | 91626 | ENSG00000184445 | KNTC1 | chr12 | + | 122601535 | 122601625 | 122597242 | 122597938 | 122602568 | 122602740 | 34 | 40 | 46 | 37 | 64 | 2 | 0 | 0 | 0 | 0 | 7 | 49 | 42 | 35 | 0 | 0 | 2 | 0 | 0 | 0 | 239 | 149 | 1 | 1 | 0.914 | 1 | 1 | 1 | 1 | 1 | 0.939 | 1 | 1 | NA | -0.002 |
| SE | 91627 | ENSG00000184445 | KNTC1 | chr12 | + | 122602828 | 122602887 | 122602396 | 122602740 | 122603026 | 122603243 | 42 | 49 | 52 | 52 | 111 | 0 | 0 | 0 | 0 | 2 | 3 | 55 | 59 | 17 | 2 | 0 | 0 | 0 | 0 | 0 | 208 | 149 | 1 | 1 | 1 | 1 | 1 | 1 | 0.975 | 1 | 1 | 1 | 1 | 1 | -0.005 |
| SE | 91628 | ENSG00000184445 | KNTC1 | chr12 | + | 122613111 | 122613230 | 122610821 | 122610900 | 122613625 | 122613669 | 90 | 58 | 105 | 68 | 175 | 0 | 0 | 1 | 0 | 0 | 30 | 31 | 44 | 10 | 0 | 0 | 0 | 0 | 4 | 0 | 268 | 149 | 1 | 1 | 1 | 1 | 0.983 | 1 | 1 | 1 | 1 | 1 | 0.582 | NA | 0.101 |
| SE | 91629 | ENSG00000184445 | KNTC1 | chr12 | + | 122615469 | 122615526 | 122614990 | 122615086 | 122618342 | 122618397 | 34 | 28 | 29 | 35 | 32 | 0 | 0 | 0 | 1 | 0 | 3 | 18 | 30 | 5 | 5 | 0 | 0 | 0 | 0 | 0 | 206 | 149 | 1 | 1 | 1 | 1 | 1 | 0.962 | 1 | 1 | 1 | 1 | 1 | 1 | -0.008 |
| SE | 91631 | ENSG00000184445 | KNTC1 | chr12 | + | 122618342 | 122618397 | 122615469 | 122615526 | 122618481 | 122618537 | 36 | 25 | 38 | 41 | 15 | 0 | 0 | 1 | 0 | 0 | 3 | 25 | 25 | 6 | 5 | 0 | 0 | 0 | 0 | 0 | 204 | 149 | 1 | 1 | 1 | 1 | 0.965 | 1 | 1 | 1 | 1 | 1 | 1 | 1 | -0.007 |
| SE | 91632 | ENSG00000184445 | KNTC1 | chr12 | + | 122620478 | 122620608 | 122618481 | 122618545 | 122621880 | 122621970 | 67 | 44 | 60 | 88 | 27 | 0 | 0 | 0 | 0 | 0 | 4 | 73 | 59 | 10 | 19 | 0 | 0 | 0 | 1 | 0 | 279 | 149 | 1 | 1 | 1 | 1 | 1 | 1 | 1 | 1 | 1 | 1 | 0.842 | 1 | 0.032 |
| SE | 91633 | ENSG00000184445 | KNTC1 | chr12 | + | 122624597 | 122624688 | 122622461 | 122622607 | 122626204 | 122626332 | 37 | 44 | 47 | 44 | 60 | 0 | 0 | 0 | 0 | 0 | 23 | 44 | 52 | 32 | 0 | 0 | 0 | 2 | 0 | 0 | 240 | 149 | 1 | 1 | 1 | 1 | 1 | 1 | 1 | 1 | 1 | 0.942 | 1 | NA | 0.014 |
| SE | 91634 | ENSG00000104695 | PPP2CB | chr8 | - | 30797580 | 30797754 | 30794191 | 30794281 | 30799545 | 30799648 | 255 | 167 | 151 | 221 | 473 | 0 | 1 | 0 | 0 | 0 | 219 | 240 | 112 | 187 | 288 | 0 | 0 | 0 | 0 | 0 | 298 | 149 | 1 | 1 | 1 | 0.988 | 1 | 1 | 1 | 1 | 1 | 1 | 1 | 1 | -0.002 |
| SE | 91635 | ENSG00000104695 | PPP2CB | chr8 | - | 30799545 | 30799755 | 30797580 | 30797754 | 30812319 | 30812818 | 207 | 164 | 144 | 178 | 356 | 0 | 0 | 0 | 0 | 0 | 238 | 194 | 99 | 147 | 268 | 0 | 2 | 0 | 0 | 0 | 298 | 149 | 1 | 1 | 1 | 1 | 1 | 1 | 1 | 1 | 0.98 | 1 | 1 | 1 | 0.004 |
| SE | 91640 | ENSG00000105339 | DENND3 | chr8 | + | 141144147 | 141144259 | 141141202 | 141141324 | 141150833 | 141150953 | 136 | 45 | 99 | 87 | 220 | 0 | 0 | 0 | 0 | 0 | 250 | 229 | 219 | 254 | 264 | 0 | 0 | 1 | 0 | 0 | 261 | 149 | 1 | 1 | 1 | 1 | 1 | 1 | 1 | 1 | 1 | 0.992 | 1 | 1 | 0.002 |
| SE | 91641 | ENSG00000105339 | DENND3 | chr8 | + | 141150285 | 141150348 | 141144147 | 141144259 | 141150833 | 141150908 | 0 | 1 | 0 | 0 | 2 | 82 | 26 | 58 | 56 | 143 | 0 | 0 | 0 | 4 | 0 | 134 | 136 | 114 | 142 | 102 | 212 | 149 | 1 | 1 | 0 | 0.026 | 0 | 0 | 0.01 | 0 | 0 | 0 | 0.019 | 0 | 0.003 |
| SE | 91643 | ENSG00000105339 | DENND3 | chr8 | + | 141151618 | 141151837 | 141150833 | 141150953 | 141160631 | 141160787 | 49 | 26 | 13 | 37 | 56 | 0 | 0 | 1 | 0 | 0 | 35 | 84 | 97 | 95 | 123 | 0 | 0 | 0 | 0 | 0 | 298 | 149 | 1 | 1 | 1 | 1 | 0.867 | 1 | 1 | 1 | 1 | 1 | 1 | 1 | -0.027 |
| SE | 91644 | ENSG00000105339 | DENND3 | chr8 | + | 141155848 | 141155970 | 141151618 | 141151837 | 141160631 | 141160787 | 108 | 54 | 84 | 100 | 41 | 0 | 0 | 0 | 0 | 0 | 262 | 180 | 225 | 301 | 337 | 0 | 1 | 0 | 0 | 0 | 271 | 149 | 1 | 1 | 1 | 1 | 1 | 1 | 1 | 1 | 0.99 | 1 | 1 | 1 | 0.002 |
| SE | 91647 | ENSG00000105339 | DENND3 | chr8 | + | 141160631 | 141160787 | 141155848 | 141155970 | 141166189 | 141166233 | 74 | 39 | 69 | 62 | 34 | 1 | 1 | 0 | 0 | 0 | 189 | 102 | 165 | 222 | 272 | 0 | 0 | 0 | 1 | 0 | 298 | 149 | 1 | 1 | 0.974 | 0.951 | 1 | 1 | 1 | 1 | 1 | 1 | 0.991 | 1 | -0.013 |
| SE | 91648 | ENSG00000105339 | DENND3 | chr8 | + | 141160631 | 141160787 | 141157394 | 141158287 | 141163332 | 141163429 | 42 | 27 | 43 | 26 | 42 | 0 | 0 | 2 | 0 | 0 | 120 | 87 | 119 | 134 | 165 | 0 | 0 | 0 | 0 | 0 | 298 | 149 | 1 | 1 | 1 | 1 | 0.915 | 1 | 1 | 1 | 1 | 1 | 1 | 1 | -0.017 |
| SE | 91650 | ENSG00000105339 | DENND3 | chr8 | + | 141163332 | 141163429 | 141155848 | 141155970 | 141166189 | 141166389 | 6 | 7 | 3 | 7 | 4 | 1 | 1 | 0 | 0 | 0 | 12 | 12 | 9 | 14 | 11 | 0 | 0 | 0 | 1 | 0 | 246 | 149 | 1 | 1 | 0.784 | 0.809 | 1 | 1 | 1 | 1 | 1 | 1 | 0.895 | 1 | -0.06 |
| SE | 91653 | ENSG00000105339 | DENND3 | chr8 | + | 141165185 | 141165289 | 141155848 | 141155970 | 141166189 | 141166233 | 63 | 49 | 42 | 50 | 48 | 1 | 1 | 0 | 0 | 0 | 130 | 83 | 124 | 145 | 200 | 0 | 0 | 0 | 1 | 0 | 253 | 149 | 1 | 1 | 0.974 | 0.967 | 1 | 1 | 1 | 1 | 1 | 1 | 0.988 | 1 | -0.009 |
| SE | 91655 | ENSG00000105339 | DENND3 | chr8 | + | 141165185 | 141165289 | 141163332 | 141163429 | 141166189 | 141166233 | 114 | 97 | 89 | 99 | 119 | 0 | 0 | 0 | 0 | 0 | 252 | 199 | 225 | 306 | 360 | 0 | 0 | 1 | 3 | 0 | 253 | 149 | 1 | 1 | 1 | 1 | 1 | 1 | 1 | 1 | 1 | 0.993 | 0.984 | 1 | 0.005 |
| SE | 91657 | ENSG00000105339 | DENND3 | chr8 | + | 141175199 | 141175459 | 141168003 | 141168525 | 141176590 | 141176652 | 136 | 90 | 76 | 102 | 138 | 0 | 0 | 0 | 0 | 0 | 309 | 175 | 331 | 282 | 600 | 0 | 0 | 2 | 0 | 7 | 298 | 149 | 1 | 1 | 1 | 1 | 1 | 1 | 1 | 1 | 1 | 0.988 | 1 | 0.977 | 0.007 |
| SE | 91658 | ENSG00000105339 | DENND3 | chr8 | + | 141190283 | 141190417 | 141188348 | 141189146 | 141192330 | 141192449 | 54 | 39 | 26 | 65 | 42 | 1 | 0 | 0 | 0 | 0 | 106 | 76 | 135 | 129 | 145 | 0 | 0 | 1 | 0 | 0 | 283 | 149 | 1 | 1 | 0.966 | 1 | 1 | 1 | 1 | 1 | 1 | 0.986 | 1 | 1 | -0.004 |
| SE | 91663 | ENSG00000177570 | SAMD12 | chr8 | - | 118239714 | 118239975 | 118197305 | 118197732 | 118379559 | 118379700 | 0 | 0 | 0 | 0 | 3 | 0 | 2 | 0 | 4 | 1 | 28 | 0 | 0 | 0 | 0 | 0 | 0 | 10 | 0 | 0 | 298 | 149 | 1 | 1 | NA | 0 | NA | 0 | 0.6 | 1 | NA | 0 | NA | NA | -0.3 |
| SE | 91665 | ENSG00000177570 | SAMD12 | chr8 | - | 118439831 | 118439961 | 118377983 | 118379700 | 118580714 | 118580893 | 5 | 34 | 7 | 25 | 23 | 0 | 1 | 1 | 0 | 2 | 8 | 4 | 34 | 0 | 7 | 0 | 0 | 0 | 0 | 0 | 279 | 149 | 1 | 1 | 1 | 0.948 | 0.789 | 1 | 0.86 | 1 | 1 | 1 | NA | 1 | -0.081 |
| SE | 91666 | ENSG00000177570 | SAMD12 | chr8 | - | 118439861 | 118439961 | 118379559 | 118379700 | 118580714 | 118580893 | 0 | 10 | 3 | 10 | 6 | 0 | 1 | 1 | 0 | 2 | 0 | 0 | 11 | 2 | 7 | 0 | 0 | 0 | 0 | 0 | 249 | 149 | 1 | 1 | NA | 0.857 | 0.642 | 1 | 0.642 | NA | NA | 1 | 1 | 1 | -0.215 |
| SE | 91667 | ENSG00000177570 | SAMD12 | chr8 | - | 118492915 | 118493004 | 118439861 | 118439961 | 118580714 | 118580893 | 0 | 2 | 0 | 0 | 0 | 2 | 19 | 5 | 14 | 11 | 0 | 0 | 0 | 0 | 0 | 0 | 1 | 19 | 2 | 7 | 238 | 149 | 1 | 1 | 0 | 0.062 | 0 | 0 | 0 | NA | 0 | 0 | 0 | 0 | 0.012 |
| SE | 91670 | ENSG00000257303 | NA | chr12 | + | 56312406 | 56312499 | 56300075 | 56300187 | 56314528 | 56314802 | 1 | 5 | 8 | 3 | 0 | 3 | 1 | 0 | 0 | 0 | 0 | 0 | 5 | 0 | 0 | 0 | 8 | 0 | 0 | 0 | 242 | 149 | 1 | 1 | 0.17 | 0.755 | 1 | 1 | NA | NA | 0 | 1 | NA | NA | 0.231 |
| SE | 91671 | ENSG00000187954 | CYHR1 | chr8 | - | 144464459 | 144464612 | 144453221 | 144453460 | 144464808 | 144465035 | 8 | 4 | 6 | 8 | 12 | 0 | 0 | 0 | 0 | 0 | 0 | 4 | 2 | 1 | 10 | 0 | 0 | 0 | 4 | 0 | 298 | 149 | 1 | 1 | 1 | 1 | 1 | 1 | 1 | NA | 1 | 1 | 0.111 | 1 | 0.222 |
| SE | 91672 | ENSG00000092094 | OSGEP | chr14 | - | 20447903 | 20447994 | 20447614 | 20447690 | 20448105 | 20448171 | 239 | 254 | 177 | 199 | 236 | 0 | 0 | 0 | 3 | 0 | 382 | 168 | 156 | 286 | 302 | 0 | 0 | 0 | 8 | 0 | 240 | 149 | 1 | 1 | 1 | 1 | 1 | 0.976 | 1 | 1 | 1 | 1 | 0.957 | 1 | 0.004 |
| SE | 91673 | ENSG00000092094 | OSGEP | chr14 | - | 20448732 | 20448811 | 20447614 | 20448171 | 20448963 | 20449013 | 245 | 223 | 197 | 212 | 274 | 0 | 0 | 1 | 0 | 0 | 386 | 160 | 157 | 300 | 396 | 0 | 0 | 0 | 0 | 0 | 228 | 149 | 1 | 1 | 1 | 1 | 0.992 | 1 | 1 | 1 | 1 | 1 | 1 | 1 | -0.002 |
| SE | 91674 | ENSG00000092094 | OSGEP | chr14 | - | 20451256 | 20451452 | 20449170 | 20449301 | 20451973 | 20452149 | 6 | 5 | 7 | 5 | 5 | 0 | 0 | 0 | 0 | 0 | 0 | 0 | 0 | 0 | 0 | 0 | 0 | 0 | 3 | 0 | 298 | 149 | 1 | 1 | 1 | 1 | 1 | 1 | 1 | NA | NA | NA | 0 | NA | 1 |
| SE | 91678 | ENSG00000178209 | PLEC | chr8 | - | 143931936 | 143932032 | 143931533 | 143931659 | 143932129 | 143932234 | 469 | 255 | 391 | 438 | 273 | 0 | 0 | 1 | 1 | 0 | 550 | 412 | 268 | 398 | 675 | 0 | 0 | 0 | 1 | 0 | 245 | 149 | 1 | 1 | 1 | 1 | 0.996 | 0.996 | 1 | 1 | 1 | 1 | 0.996 | 1 | -0.001 |
| SE | 91679 | ENSG00000178209 | PLEC | chr8 | - | 143932634 | 143932712 | 143932399 | 143932561 | 143932792 | 143933111 | 437 | 219 | 400 | 385 | 164 | 0 | 0 | 1 | 0 | 0 | 797 | 568 | 352 | 498 | 954 | 6 | 0 | 0 | 0 | 0 | 227 | 149 | 1 | 1 | 1 | 1 | 0.996 | 1 | 1 | 0.989 | 1 | 1 | 1 | 1 | 0.001 |
| SE | 91680 | ENSG00000178209 | PLEC | chr8 | - | 143937748 | 143937784 | 143937164 | 143937242 | 143938150 | 143938240 | 0 | 0 | 0 | 0 | 0 | 218 | 99 | 187 | 184 | 190 | 0 | 0 | 0 | 4 | 0 | 188 | 190 | 154 | 213 | 355 | 185 | 149 | 1 | 1 | 0 | 0 | 0 | 0 | 0 | 0 | 0 | 0 | 0.015 | 0 | -0.003 |
| SE | 91688 | ENSG00000168476 | REEP4 | chr8 | - | 22138638 | 22138793 | 22138019 | 22138552 | 22138925 | 22139061 | 43 | 22 | 12 | 34 | 9 | 0 | 0 | 0 | 0 | 0 | 16 | 44 | 36 | 31 | 35 | 0 | 0 | 0 | 0 | 4 | 298 | 149 | 1 | 1 | 1 | 1 | 1 | 1 | 1 | 1 | 1 | 1 | 1 | 0.814 | 0.037 |
| SE | 91690 | ENSG00000168476 | REEP4 | chr8 | - | 22140606 | 22140697 | 22140205 | 22140248 | 22141450 | 22141951 | 18 | 6 | 14 | 10 | 13 | 0 | 0 | 2 | 0 | 0 | 28 | 26 | 11 | 0 | 20 | 0 | 0 | 0 | 0 | 0 | 240 | 149 | 1 | 1 | 1 | 1 | 0.813 | 1 | 1 | 1 | 1 | 1 | NA | 1 | -0.037 |
| SE | 91691 | ENSG00000168476 | REEP4 | chr8 | - | 22140624 | 22140697 | 22140171 | 22140248 | 22141450 | 22141540 | 50 | 29 | 29 | 24 | 35 | 0 | 0 | 2 | 0 | 0 | 69 | 45 | 48 | 20 | 65 | 0 | 0 | 0 | 0 | 0 | 222 | 149 | 1 | 1 | 1 | 1 | 0.907 | 1 | 1 | 1 | 1 | 1 | 1 | 1 | -0.019 |
| SE | 91693 | ENSG00000197181 | PIWIL2 | chr8 | + | 22281376 | 22281515 | 22281119 | 22281207 | 22283033 | 22283240 | 0 | 0 | 2 | 1 | 0 | 0 | 0 | 0 | 0 | 0 | 4 | 0 | 4 | 0 | 0 | 0 | 4 | 0 | 0 | 0 | 288 | 149 | 1 | 1 | NA | NA | 1 | 1 | NA | 1 | 0 | 1 | NA | NA | 0.333 |
| SE | 91697 | ENSG00000285130 | NA | chr9 | + | 69053139 | 69053260 | 69046384 | 69046482 | 69064937 | 69065035 | 75 | 39 | 87 | 58 | 78 | 0 | 1 | 0 | 0 | 0 | 102 | 66 | 27 | 53 | 42 | 0 | 0 | 0 | 0 | 0 | 270 | 149 | 1 | 1 | 1 | 0.956 | 1 | 1 | 1 | 1 | 1 | 1 | 1 | 1 | -0.009 |
| SE | 91699 | ENSG00000285130 | NA | chr9 | + | 69216338 | 69216463 | 69212547 | 69212601 | 69218256 | 69218359 | 38 | 48 | 20 | 62 | 21 | 0 | 0 | 0 | 0 | 2 | 99 | 42 | 29 | 95 | 123 | 0 | 0 | 0 | 0 | 0 | 274 | 149 | 1 | 1 | 1 | 1 | 1 | 1 | 0.851 | 1 | 1 | 1 | 1 | 1 | -0.03 |
| SE | 91700 | ENSG00000285130 | NA | chr9 | + | 69220886 | 69221496 | 69218256 | 69218359 | 69225303 | 69225407 | 42 | 40 | 22 | 63 | 19 | 1 | 0 | 0 | 0 | 0 | 79 | 25 | 12 | 56 | 117 | 0 | 0 | 0 | 0 | 0 | 298 | 149 | 1 | 1 | 0.955 | 1 | 1 | 1 | 1 | 1 | 1 | 1 | 1 | 1 | -0.009 |
| SE | 91701 | ENSG00000285130 | NA | chr9 | + | 69226021 | 69226175 | 69225303 | 69225407 | 69227764 | 69227873 | 134 | 149 | 85 | 170 | 164 | 0 | 0 | 1 | 3 | 0 | 419 | 106 | 92 | 174 | 598 | 0 | 1 | 4 | 1 | 0 | 298 | 149 | 1 | 1 | 1 | 1 | 0.977 | 0.966 | 1 | 1 | 0.981 | 0.92 | 0.989 | 1 | 0.011 |
| SE | 91708 | ENSG00000149084 | HSD17B12 | chr11 | + | 43750910 | 43750957 | 43680737 | 43680987 | 43798319 | 43798427 | 1 | 3 | 1 | 2 | 1 | 0 | 0 | 0 | 0 | 0 | 0 | 2 | 1 | 3 | 7 | 0 | 0 | 0 | 0 | 5 | 196 | 149 | 1 | 1 | 1 | 1 | 1 | 1 | 1 | NA | 1 | 1 | 1 | 0.516 | 0.121 |
| SE | 91709 | ENSG00000149084 | HSD17B12 | chr11 | + | 43754045 | 43754121 | 43750910 | 43750957 | 43798319 | 43798427 | 166 | 157 | 147 | 160 | 196 | 0 | 2 | 0 | 0 | 1 | 121 | 108 | 95 | 146 | 400 | 0 | 0 | 0 | 0 | 7 | 225 | 149 | 1 | 1 | 1 | 0.981 | 1 | 1 | 0.992 | 1 | 1 | 1 | 1 | 0.974 | 0 |
| SE | 91710 | ENSG00000149084 | HSD17B12 | chr11 | + | 43798319 | 43798427 | 43754043 | 43754121 | 43815436 | 43815501 | 241 | 218 | 200 | 223 | 353 | 0 | 0 | 1 | 0 | 0 | 136 | 156 | 121 | 195 | 558 | 0 | 0 | 0 | 0 | 0 | 257 | 149 | 1 | 1 | 1 | 1 | 0.991 | 1 | 1 | 1 | 1 | 1 | 1 | 1 | -0.002 |
| SE | 91711 | ENSG00000149084 | HSD17B12 | chr11 | + | 43830975 | 43831010 | 43816346 | 43816391 | 43838316 | 43838374 | 229 | 172 | 209 | 221 | 310 | 0 | 0 | 1 | 0 | 0 | 118 | 188 | 136 | 229 | 479 | 0 | 0 | 0 | 0 | 0 | 184 | 149 | 1 | 1 | 1 | 1 | 0.994 | 1 | 1 | 1 | 1 | 1 | 1 | 1 | -0.001 |
| SE | 91715 | ENSG00000134539 | KLRD1 | chr12 | + | 10311463 | 10311615 | 10307977 | 10309688 | 10313406 | 10313513 | 704 | 218 | 247 | 333 | 561 | 3 | 1 | 1 | 2 | 3 | 52 | 215 | 201 | 69 | 29 | 0 | 0 | 1 | 2 | 0 | 298 | 149 | 1 | 1 | 0.992 | 0.991 | 0.992 | 0.988 | 0.989 | 1 | 1 | 0.99 | 0.945 | 1 | 0.003 |
| SE | 91716 | ENSG00000134539 | KLRD1 | chr12 | + | 10311463 | 10311615 | 10307977 | 10309688 | 10313409 | 10313513 | 1653 | 544 | 565 | 860 | 928 | 2 | 0 | 0 | 1 | 0 | 112 | 638 | 618 | 228 | 85 | 0 | 0 | 3 | 1 | 0 | 298 | 149 | 1 | 1 | 0.998 | 1 | 1 | 0.998 | 1 | 1 | 1 | 0.99 | 0.991 | 1 | 0.003 |
| SE | 91717 | ENSG00000134539 | KLRD1 | chr12 | + | 10311463 | 10311619 | 10309625 | 10309688 | 10313406 | 10313513 | 450 | 160 | 184 | 235 | 453 | 3 | 1 | 1 | 2 | 3 | 33 | 154 | 146 | 46 | 13 | 0 | 0 | 1 | 2 | 0 | 298 | 149 | 1 | 1 | 0.987 | 0.988 | 0.989 | 0.983 | 0.987 | 1 | 1 | 0.986 | 0.92 | 1 | 0.006 |
| SE | 91718 | ENSG00000134539 | KLRD1 | chr12 | + | 10311463 | 10311619 | 10309625 | 10309688 | 10313409 | 10313513 | 455 | 163 | 188 | 238 | 453 | 2 | 0 | 0 | 1 | 0 | 33 | 157 | 144 | 46 | 13 | 0 | 0 | 3 | 1 | 0 | 298 | 149 | 1 | 1 | 0.991 | 1 | 1 | 0.992 | 1 | 1 | 1 | 0.96 | 0.958 | 1 | 0.013 |
| SE | 91719 | ENSG00000134539 | KLRD1 | chr12 | + | 10313406 | 10313513 | 10311463 | 10311615 | 10314672 | 10314977 | 1534 | 418 | 442 | 747 | 384 | 0 | 0 | 1 | 3 | 1 | 105 | 639 | 650 | 162 | 93 | 0 | 4 | 0 | 0 | 0 | 256 | 149 | 1 | 1 | 1 | 1 | 0.996 | 0.993 | 0.996 | 1 | 0.989 | 1 | 1 | 1 | -0.001 |
| SE | 91720 | ENSG00000134539 | KLRD1 | chr12 | + | 10313409 | 10313513 | 10311463 | 10311615 | 10314672 | 10314818 | 2458 | 726 | 748 | 1265 | 720 | 0 | 0 | 1 | 3 | 1 | 165 | 1044 | 1053 | 314 | 144 | 0 | 4 | 0 | 0 | 0 | 253 | 149 | 1 | 1 | 1 | 1 | 0.998 | 0.996 | 0.998 | 1 | 0.994 | 1 | 1 | 1 | 0 |
| SE | 91721 | ENSG00000176623 | RMDN1 | chr8 | - | 86470217 | 86470410 | 86468267 | 86468423 | 86474819 | 86474914 | 21 | 6 | 4 | 24 | 24 | 0 | 0 | 1 | 0 | 0 | 0 | 1 | 23 | 4 | 7 | 0 | 0 | 0 | 0 | 0 | 298 | 149 | 1 | 1 | 1 | 1 | 0.667 | 1 | 1 | NA | 1 | 1 | 1 | 1 | -0.067 |
| SE | 91722 | ENSG00000176623 | RMDN1 | chr8 | - | 86474819 | 86474914 | 86470217 | 86470410 | 86478922 | 86479010 | 10 | 4 | 3 | 13 | 14 | 0 | 0 | 0 | 1 | 4 | 0 | 0 | 8 | 4 | 6 | 0 | 0 | 0 | 0 | 0 | 244 | 149 | 1 | 1 | 1 | 1 | 1 | 0.888 | 0.681 | NA | NA | 1 | 1 | 1 | -0.086 |
| SE | 91724 | ENSG00000176623 | RMDN1 | chr8 | - | 86474819 | 86474953 | 86470217 | 86470410 | 86478922 | 86479010 | 10 | 4 | 3 | 16 | 25 | 0 | 0 | 0 | 1 | 4 | 0 | 1 | 21 | 4 | 7 | 0 | 0 | 0 | 0 | 0 | 283 | 149 | 1 | 1 | 1 | 1 | 1 | 0.894 | 0.767 | NA | 1 | 1 | 1 | 1 | -0.068 |
| SE | 91725 | ENSG00000176623 | RMDN1 | chr8 | - | 86474819 | 86474953 | 86472348 | 86472479 | 86477293 | 86477324 | 166 | 151 | 120 | 162 | 342 | 2 | 0 | 0 | 0 | 0 | 114 | 121 | 120 | 177 | 112 | 0 | 4 | 0 | 0 | 0 | 283 | 149 | 1 | 1 | 0.978 | 1 | 1 | 1 | 1 | 1 | 0.941 | 1 | 1 | 1 | 0.007 |
| SE | 91726 | ENSG00000176623 | RMDN1 | chr8 | - | 86477293 | 86477324 | 86474819 | 86474914 | 86478922 | 86479010 | 41 | 30 | 23 | 30 | 62 | 0 | 0 | 0 | 1 | 0 | 38 | 32 | 32 | 38 | 6 | 0 | 0 | 1 | 0 | 0 | 180 | 149 | 1 | 1 | 1 | 1 | 1 | 0.961 | 1 | 1 | 1 | 0.964 | 1 | 1 | -0.001 |
| SE | 91729 | ENSG00000176623 | RMDN1 | chr8 | - | 86494828 | 86494966 | 86488551 | 86488639 | 86505299 | 86505379 | 9 | 1 | 4 | 2 | 9 | 0 | 0 | 2 | 0 | 0 | 0 | 4 | 1 | 2 | 0 | 0 | 0 | 0 | 4 | 0 | 287 | 149 | 1 | 1 | 1 | 1 | 0.509 | 1 | 1 | NA | 1 | 1 | 0.206 | NA | 0.166 |
| SE | 91732 | ENSG00000176623 | RMDN1 | chr8 | - | 86505299 | 86505379 | 86494828 | 86494966 | 86506994 | 86507112 | 0 | 0 | 0 | 1 | 0 | 3 | 2 | 1 | 3 | 11 | 0 | 0 | 0 | 4 | 0 | 0 | 3 | 1 | 0 | 0 | 229 | 149 | 1 | 1 | 0 | 0 | 0 | 0.178 | 0 | NA | 0 | 0 | 1 | NA | -0.298 |
| SE | 91733 | ENSG00000176623 | RMDN1 | chr8 | - | 86506994 | 86507112 | 86488551 | 86488639 | 86508491 | 86508619 | 79 | 49 | 49 | 77 | 20 | 1 | 0 | 0 | 0 | 0 | 35 | 80 | 22 | 51 | 18 | 0 | 0 | 0 | 0 | 0 | 267 | 149 | 1 | 1 | 0.978 | 1 | 1 | 1 | 1 | 1 | 1 | 1 | 1 | 1 | -0.004 |
| SE | 91735 | ENSG00000182986 | ZNF320 | chr19 | - | 52891228 | 52891346 | 52890240 | 52890328 | 52897519 | 52897693 | 11 | 24 | 17 | 9 | 4 | 0 | 0 | 0 | 0 | 0 | 11 | 15 | 10 | 16 | 4 | 0 | 0 | 0 | 2 | 0 | 267 | 149 | 1 | 1 | 1 | 1 | 1 | 1 | 1 | 1 | 1 | 1 | 0.817 | 1 | 0.037 |
| SE | 91736 | ENSG00000182986 | ZNF320 | chr19 | - | 52891228 | 52891346 | 52890240 | 52890328 | 52897647 | 52897684 | 12 | 23 | 13 | 9 | 4 | 0 | 0 | 2 | 2 | 0 | 11 | 15 | 9 | 19 | 4 | 0 | 0 | 0 | 1 | 0 | 267 | 149 | 1 | 1 | 1 | 1 | 0.784 | 0.715 | 1 | 1 | 1 | 1 | 0.914 | 1 | -0.083 |
| SE | 91737 | ENSG00000182986 | ZNF320 | chr19 | - | 52893383 | 52893619 | 52891228 | 52891346 | 52893778 | 52893856 | 5 | 7 | 3 | 6 | 2 | 3 | 1 | 6 | 3 | 2 | 0 | 3 | 0 | 0 | 0 | 0 | 3 | 3 | 4 | 0 | 298 | 149 | 1 | 1 | 0.455 | 0.778 | 0.2 | 0.5 | 0.333 | NA | 0.333 | 0 | 0 | NA | 0.342 |
| SE | 91738 | ENSG00000182986 | ZNF320 | chr19 | - | 52893383 | 52893619 | 52891228 | 52891346 | 52897509 | 52897650 | 2 | 6 | 1 | 4 | 0 | 0 | 0 | 1 | 1 | 0 | 0 | 3 | 0 | 0 | 0 | 0 | 2 | 0 | 0 | 0 | 298 | 149 | 1 | 1 | 1 | 1 | 0.333 | 0.667 | NA | NA | 0.429 | NA | NA | NA | 0.321 |
| SE | 91739 | ENSG00000182986 | ZNF320 | chr19 | - | 52893383 | 52893619 | 52891228 | 52891346 | 52897519 | 52897655 | 3 | 5 | 1 | 4 | 0 | 0 | 1 | 4 | 0 | 0 | 0 | 9 | 0 | 0 | 0 | 0 | 0 | 1 | 0 | 0 | 298 | 149 | 1 | 1 | 1 | 0.714 | 0.111 | 1 | NA | NA | 1 | 0 | NA | NA | 0.206 |
| SE | 91740 | ENSG00000182986 | ZNF320 | chr19 | - | 52893778 | 52893856 | 52890240 | 52890328 | 52897519 | 52897635 | 2 | 0 | 1 | 1 | 0 | 0 | 0 | 0 | 0 | 0 | 0 | 7 | 0 | 0 | 0 | 0 | 0 | 0 | 2 | 0 | 227 | 149 | 1 | 1 | 1 | NA | 1 | 1 | NA | NA | 1 | NA | 0 | NA | 0.5 |
| SE | 91741 | ENSG00000182986 | ZNF320 | chr19 | - | 52893778 | 52893856 | 52890240 | 52890328 | 52897647 | 52897686 | 2 | 0 | 0 | 0 | 0 | 0 | 0 | 2 | 2 | 0 | 0 | 0 | 0 | 0 | 0 | 0 | 0 | 0 | 1 | 0 | 227 | 149 | 1 | 1 | 1 | NA | 0 | 0 | NA | NA | NA | NA | 0 | NA | 0.333 |
| SE | 91742 | ENSG00000182986 | ZNF320 | chr19 | - | 52893778 | 52893856 | 52891228 | 52891346 | 52897519 | 52897635 | 2 | 1 | 5 | 3 | 1 | 0 | 1 | 4 | 0 | 0 | 0 | 10 | 0 | 4 | 0 | 0 | 0 | 1 | 0 | 0 | 227 | 149 | 1 | 1 | 1 | 0.396 | 0.451 | 1 | 1 | NA | 1 | 0 | 1 | NA | 0.103 |
| SE | 91743 | ENSG00000182986 | ZNF320 | chr19 | - | 52893778 | 52893856 | 52891228 | 52891346 | 52897647 | 52897686 | 3 | 1 | 5 | 2 | 1 | 1 | 0 | 0 | 0 | 0 | 0 | 3 | 0 | 4 | 0 | 0 | 0 | 0 | 3 | 0 | 227 | 149 | 1 | 1 | 0.663 | 1 | 1 | 1 | 1 | NA | 1 | NA | 0.467 | NA | 0.199 |
| SE | 91744 | ENSG00000182986 | ZNF320 | chr19 | - | 52893778 | 52893856 | 52893383 | 52893619 | 52897519 | 52897635 | 1 | 0 | 3 | 2 | 0 | 1 | 0 | 0 | 0 | 0 | 0 | 7 | 0 | 0 | 0 | 0 | 6 | 0 | 0 | 0 | 227 | 149 | 1 | 1 | 0.396 | NA | 1 | 1 | NA | NA | 0.434 | NA | NA | NA | 0.365 |
| SE | 91745 | ENSG00000182986 | ZNF320 | chr19 | - | 52893778 | 52893927 | 52890240 | 52890328 | 52897519 | 52897654 | 2 | 0 | 0 | 0 | 0 | 0 | 0 | 0 | 0 | 0 | 0 | 4 | 0 | 0 | 0 | 0 | 0 | 0 | 2 | 0 | 298 | 149 | 1 | 1 | 1 | NA | NA | NA | NA | NA | 1 | NA | 0 | NA | 0.5 |
| SE | 91746 | ENSG00000182986 | ZNF320 | chr19 | - | 52893778 | 52893927 | 52891228 | 52891346 | 52897519 | 52897654 | 2 | 1 | 3 | 3 | 2 | 0 | 1 | 4 | 0 | 0 | 0 | 3 | 3 | 4 | 0 | 0 | 0 | 1 | 0 | 0 | 298 | 149 | 1 | 1 | 1 | 0.333 | 0.273 | 1 | 1 | NA | 1 | 0.6 | 1 | NA | -0.145 |
| SE | 91748 | ENSG00000182986 | ZNF320 | chr19 | - | 52895338 | 52895435 | 52893778 | 52893856 | 52897519 | 52897560 | 0 | 1 | 0 | 0 | 0 | 0 | 0 | 2 | 1 | 0 | 0 | 0 | 0 | 0 | 0 | 0 | 7 | 0 | 0 | 0 | 246 | 149 | 1 | 1 | NA | 1 | 0 | 0 | NA | NA | 0 | NA | NA | NA | 0.333 |
| SE | 91750 | ENSG00000173295 | FAM86B3P | chr8 | + | 8236031 | 8236299 | 8235815 | 8235949 | 8237125 | 8237275 | 2 | 0 | 0 | 9 | 0 | 0 | 0 | 0 | 0 | 0 | 0 | 1 | 0 | 0 | 0 | 8 | 0 | 0 | 0 | 0 | 298 | 149 | 1 | 1 | 1 | NA | NA | 1 | NA | 0 | 1 | NA | NA | NA | 0.5 |
| SE | 91752 | ENSG00000143748 | NVL | chr1 | - | 224232088 | 224232182 | 224231225 | 224231296 | 224233200 | 224233289 | 2 | 0 | 1 | 1 | 4 | 49 | 32 | 45 | 47 | 119 | 0 | 2 | 0 | 0 | 0 | 21 | 37 | 25 | 18 | 9 | 243 | 149 | 1 | 1 | 0.024 | 0 | 0.013 | 0.013 | 0.02 | 0 | 0.032 | 0 | 0 | 0 | 0.008 |
| SE | 91754 | ENSG00000143748 | NVL | chr1 | - | 224233200 | 224233289 | 224231225 | 224231296 | 224236505 | 224236582 | 69 | 45 | 70 | 65 | 165 | 2 | 0 | 1 | 0 | 0 | 35 | 58 | 33 | 30 | 25 | 0 | 0 | 0 | 0 | 0 | 238 | 149 | 1 | 1 | 0.956 | 1 | 0.978 | 1 | 1 | 1 | 1 | 1 | 1 | 1 | -0.013 |
| SE | 91759 | ENSG00000143748 | NVL | chr1 | - | 224304735 | 224304812 | 224303722 | 224303857 | 224305033 | 224305166 | 79 | 59 | 47 | 61 | 123 | 0 | 0 | 1 | 0 | 0 | 64 | 53 | 45 | 54 | 32 | 0 | 0 | 0 | 0 | 0 | 226 | 149 | 1 | 1 | 1 | 1 | 0.969 | 1 | 1 | 1 | 1 | 1 | 1 | 1 | -0.006 |
| SE | 91760 | ENSG00000143748 | NVL | chr1 | - | 224307990 | 224308263 | 224305033 | 224305166 | 224311799 | 224311857 | 105 | 117 | 121 | 123 | 215 | 1 | 0 | 1 | 1 | 0 | 85 | 116 | 133 | 126 | 22 | 0 | 1 | 0 | 0 | 0 | 298 | 149 | 1 | 1 | 0.981 | 1 | 0.984 | 0.984 | 1 | 1 | 0.983 | 1 | 1 | 1 | -0.007 |
| SE | 91766 | ENSG00000143748 | NVL | chr1 | - | 224317693 | 224317793 | 224312726 | 224313151 | 224317877 | 224317925 | 64 | 42 | 57 | 53 | 100 | 1 | 0 | 0 | 0 | 0 | 58 | 51 | 85 | 47 | 74 | 0 | 0 | 0 | 0 | 0 | 249 | 149 | 1 | 1 | 0.975 | 1 | 1 | 1 | 1 | 1 | 1 | 1 | 1 | 1 | -0.005 |
| SE | 91767 | ENSG00000143748 | NVL | chr1 | - | 224326390 | 224326464 | 224317877 | 224317925 | 224330070 | 224330137 | 7 | 7 | 6 | 6 | 2 | 0 | 0 | 0 | 0 | 1 | 17 | 0 | 3 | 5 | 20 | 0 | 0 | 0 | 0 | 0 | 223 | 149 | 1 | 1 | 1 | 1 | 1 | 1 | 0.572 | 1 | NA | 1 | 1 | 1 | -0.086 |
| SE | 91773 | ENSG00000110841 | PPFIBP1 | chr12 | + | 27674191 | 27674221 | 27673766 | 27673827 | 27676427 | 27676599 | 11 | 3 | 4 | 6 | 5 | 1 | 1 | 0 | 0 | 6 | 0 | 1 | 5 | 12 | 5 | 0 | 1 | 0 | 0 | 9 | 179 | 149 | 1 | 1 | 0.902 | 0.714 | 1 | 1 | 0.41 | NA | 0.454 | 1 | 1 | 0.316 | 0.113 |
| SE | 91774 | ENSG00000110841 | PPFIBP1 | chr12 | + | 27677063 | 27677096 | 27676427 | 27676599 | 27679488 | 27679639 | 1 | 0 | 0 | 0 | 0 | 3 | 4 | 2 | 4 | 4 | 0 | 0 | 0 | 0 | 6 | 0 | 1 | 2 | 12 | 0 | 182 | 149 | 1 | 1 | 0.214 | 0 | 0 | 0 | 0 | NA | 0 | 0 | 0 | 1 | -0.207 |
| SE | 91776 | ENSG00000253981 | ALG1L13P | chr8 | - | 8242786 | 8242926 | 8241321 | 8241400 | 8244459 | 8244667 | 14 | 1 | 8 | 6 | 18 | 0 | 0 | 0 | 0 | 2 | 20 | 13 | 40 | 22 | 0 | 0 | 0 | 1 | 0 | 0 | 289 | 149 | 1 | 1 | 1 | 1 | 1 | 1 | 0.823 | 1 | 1 | 0.954 | 1 | NA | -0.024 |
| SE | 91779 | ENSG00000133794 | ARNTL | chr11 | + | 13309986 | 13310059 | 13277751 | 13277923 | 13349944 | 13350070 | 36 | 27 | 34 | 34 | 22 | 0 | 0 | 0 | 0 | 0 | 6 | 28 | 52 | 28 | 18 | 0 | 3 | 0 | 0 | 0 | 222 | 149 | 1 | 1 | 1 | 1 | 1 | 1 | 1 | 1 | 0.862 | 1 | 1 | 1 | 0.028 |
| SE | 91780 | ENSG00000133794 | ARNTL | chr11 | + | 13321820 | 13321919 | 13309986 | 13310059 | 13326412 | 13326467 | 0 | 0 | 0 | 0 | 0 | 22 | 3 | 7 | 17 | 19 | 0 | 0 | 4 | 0 | 0 | 6 | 23 | 25 | 20 | 8 | 248 | 149 | 1 | 1 | 0 | 0 | 0 | 0 | 0 | 0 | 0 | 0.088 | 0 | 0 | -0.018 |
| SE | 91782 | ENSG00000133794 | ARNTL | chr11 | + | 13326412 | 13326467 | 13309986 | 13310059 | 13354300 | 13354448 | 9 | 0 | 2 | 5 | 0 | 1 | 1 | 0 | 0 | 0 | 1 | 6 | 10 | 2 | 8 | 0 | 0 | 0 | 0 | 0 | 204 | 149 | 1 | 1 | 0.868 | 0 | 1 | 1 | NA | 1 | 1 | 1 | 1 | 1 | -0.283 |
| SE | 91783 | ENSG00000133794 | ARNTL | chr11 | + | 13349944 | 13350070 | 13309985 | 13310059 | 13354300 | 13354448 | 118 | 62 | 96 | 93 | 73 | 1 | 1 | 0 | 0 | 0 | 51 | 79 | 168 | 102 | 82 | 0 | 0 | 0 | 0 | 0 | 275 | 149 | 1 | 1 | 0.985 | 0.971 | 1 | 1 | 1 | 1 | 1 | 1 | 1 | 1 | -0.009 |
| SE | 91784 | ENSG00000133794 | ARNTL | chr11 | + | 13349944 | 13350070 | 13326412 | 13326467 | 13354300 | 13354448 | 107 | 45 | 84 | 85 | 80 | 2 | 0 | 0 | 0 | 0 | 48 | 80 | 173 | 111 | 72 | 0 | 0 | 3 | 0 | 0 | 275 | 149 | 1 | 1 | 0.967 | 1 | 1 | 1 | 1 | 1 | 1 | 0.969 | 1 | 1 | 0 |
| SE | 91785 | ENSG00000133794 | ARNTL | chr11 | + | 13354300 | 13354448 | 13349944 | 13350070 | 13356738 | 13356777 | 142 | 77 | 108 | 117 | 104 | 0 | 0 | 0 | 0 | 0 | 75 | 105 | 211 | 160 | 88 | 0 | 0 | 2 | 0 | 0 | 297 | 149 | 1 | 1 | 1 | 1 | 1 | 1 | 1 | 1 | 1 | 0.981 | 1 | 1 | 0.004 |
| SE | 91790 | ENSG00000133794 | ARNTL | chr11 | + | 13355225 | 13355296 | 13354989 | 13355010 | 13356738 | 13356777 | 10 | 2 | 4 | 3 | 9 | 1 | 1 | 1 | 2 | 1 | 0 | 3 | 4 | 0 | 0 | 0 | 0 | 2 | 0 | 0 | 220 | 149 | 1 | 1 | 0.871 | 0.575 | 0.73 | 0.504 | 0.859 | NA | 1 | 0.575 | NA | NA | -0.08 |
| SE | 91793 | ENSG00000133794 | ARNTL | chr11 | + | 13358433 | 13358591 | 13356877 | 13357096 | 13360344 | 13360424 | 197 | 116 | 116 | 141 | 215 | 0 | 0 | 2 | 0 | 3 | 52 | 195 | 268 | 197 | 42 | 0 | 0 | 0 | 4 | 0 | 298 | 149 | 1 | 1 | 1 | 1 | 0.967 | 1 | 0.973 | 1 | 1 | 1 | 0.961 | 1 | -0.004 |
| SE | 91796 | ENSG00000133794 | ARNTL | chr11 | + | 13369617 | 13369767 | 13366664 | 13366782 | 13372165 | 13372409 | 146 | 66 | 88 | 153 | 182 | 0 | 0 | 0 | 1 | 0 | 30 | 59 | 130 | 48 | 15 | 0 | 0 | 0 | 0 | 0 | 298 | 149 | 1 | 1 | 1 | 1 | 1 | 0.987 | 1 | 1 | 1 | 1 | 1 | 1 | -0.003 |
| SE | 91797 | ENSG00000133794 | ARNTL | chr11 | + | 13369696 | 13369767 | 13366664 | 13366782 | 13372165 | 13372409 | 26 | 16 | 22 | 31 | 15 | 0 | 0 | 0 | 1 | 0 | 0 | 16 | 30 | 4 | 0 | 0 | 0 | 0 | 0 | 0 | 220 | 149 | 1 | 1 | 1 | 1 | 1 | 0.955 | 1 | NA | 1 | 1 | 1 | NA | -0.009 |
| SE | 91800 | ENSG00000133794 | ARNTL | chr11 | + | 13381160 | 13381254 | 13378321 | 13378445 | 13385688 | 13385791 | 170 | 68 | 147 | 165 | 213 | 0 | 0 | 2 | 0 | 0 | 68 | 223 | 265 | 148 | 46 | 0 | 0 | 1 | 1 | 0 | 243 | 149 | 1 | 1 | 1 | 1 | 0.978 | 1 | 1 | 1 | 1 | 0.994 | 0.989 | 1 | -0.001 |
| SE | 91801 | ENSG00000133794 | ARNTL | chr11 | + | 13385688 | 13385791 | 13378336 | 13381254 | 13386595 | 13387180 | 128 | 52 | 110 | 129 | 177 | 0 | 0 | 0 | 2 | 0 | 63 | 167 | 202 | 122 | 23 | 0 | 0 | 0 | 0 | 0 | 252 | 149 | 1 | 1 | 1 | 1 | 1 | 0.974 | 1 | 1 | 1 | 1 | 1 | 1 | -0.005 |
| SE | 91806 | ENSG00000156738 | MS4A1 | chr11 | + | 60458048 | 60458208 | 60455845 | 60455945 | 60461071 | 60461160 | 0 | 1 | 0 | 0 | 0 | 257 | 155 | 223 | 244 | 319 | 0 | 0 | 0 | 0 | 0 | 37 | 142 | 18 | 4 | 35 | 298 | 149 | 1 | 1 | 0 | 0.003 | 0 | 0 | 0 | 0 | 0 | 0 | 0 | 0 | 0.001 |
| SE | 91809 | ENSG00000156738 | MS4A1 | chr11 | + | 60462417 | 60462533 | 60462184 | 60462298 | 60463001 | 60463121 | 481 | 341 | 408 | 539 | 1059 | 0 | 0 | 0 | 0 | 0 | 139 | 427 | 78 | 133 | 98 | 0 | 3 | 0 | 0 | 0 | 265 | 149 | 1 | 1 | 1 | 1 | 1 | 1 | 1 | 1 | 0.988 | 1 | 1 | 1 | 0.002 |
| SE | 91810 | ENSG00000156738 | MS4A1 | chr11 | + | 60465920 | 60466157 | 60464287 | 60464344 | 60466958 | 60467060 | 1175 | 981 | 1023 | 1438 | 2488 | 4 | 0 | 0 | 0 | 0 | 390 | 1141 | 199 | 274 | 247 | 0 | 0 | 1 | 3 | 0 | 298 | 149 | 1 | 1 | 0.993 | 1 | 1 | 1 | 1 | 1 | 1 | 0.99 | 0.979 | 1 | 0.005 |
| SE | 91811 | ENSG00000156738 | MS4A1 | chr11 | + | 60465920 | 60466157 | 60464287 | 60464344 | 60466997 | 60467060 | 734 | 613 | 618 | 907 | 1678 | 1 | 1 | 3 | 4 | 0 | 206 | 673 | 92 | 164 | 138 | 0 | 1 | 0 | 0 | 0 | 298 | 149 | 1 | 1 | 0.997 | 0.997 | 0.99 | 0.991 | 1 | 1 | 0.997 | 1 | 1 | 1 | -0.004 |
| SE | 91812 | ENSG00000104290 | FZD3 | chr8 | + | 28499932 | 28499978 | 28494211 | 28494343 | 28502669 | 28503202 | 7 | 2 | 3 | 1 | 0 | 3 | 1 | 0 | 0 | 0 | 0 | 10 | 1 | 0 | 0 | 0 | 0 | 0 | 0 | 0 | 195 | 149 | 1 | 1 | 0.641 | 0.604 | 1 | 1 | NA | NA | 1 | 1 | NA | NA | -0.189 |
| SE | 91813 | ENSG00000104290 | FZD3 | chr8 | + | 28551602 | 28551751 | 28527146 | 28528164 | 28555737 | 28555971 | 3 | 9 | 7 | 11 | 5 | 0 | 0 | 0 | 0 | 0 | 0 | 2 | 2 | 0 | 0 | 0 | 0 | 3 | 0 | 0 | 298 | 149 | 1 | 1 | 1 | 1 | 1 | 1 | 1 | NA | 1 | 0.25 | NA | NA | 0.375 |
| SE | 91821 | ENSG00000198626 | RYR2 | chr1 | + | 237806136 | 237806283 | 237801855 | 237801916 | 237808900 | 237809035 | 1 | 1 | 3 | 2 | 8 | 0 | 0 | 0 | 1 | 0 | 0 | 0 | 8 | 9 | 5 | 0 | 0 | 0 | 3 | 0 | 296 | 149 | 1 | 1 | 1 | 1 | 1 | 0.502 | 1 | NA | NA | 1 | 0.602 | 1 | 0.033 |
| SE | 91822 | ENSG00000198626 | RYR2 | chr1 | + | 237808900 | 237809035 | 237801855 | 237801916 | 237819035 | 237819192 | 1 | 0 | 3 | 4 | 8 | 0 | 0 | 0 | 0 | 0 | 0 | 3 | 6 | 8 | 5 | 0 | 0 | 0 | 1 | 0 | 284 | 149 | 1 | 1 | 1 | NA | 1 | 1 | 1 | NA | 1 | 1 | 0.808 | 1 | 0.048 |
| SE | 91827 | ENSG00000104365 | IKBKB | chr8 | + | 42293442 | 42293512 | 42290155 | 42290273 | 42305186 | 42305236 | 65 | 28 | 44 | 64 | 28 | 0 | 0 | 0 | 2 | 0 | 162 | 68 | 62 | 48 | 95 | 0 | 0 | 0 | 0 | 0 | 219 | 149 | 1 | 1 | 1 | 1 | 1 | 0.956 | 1 | 1 | 1 | 1 | 1 | 1 | -0.009 |
| SE | 91828 | ENSG00000104365 | IKBKB | chr8 | + | 42293442 | 42293568 | 42290155 | 42290273 | 42305186 | 42305275 | 29 | 16 | 33 | 15 | 3 | 0 | 0 | 0 | 2 | 0 | 31 | 26 | 17 | 14 | 27 | 0 | 0 | 0 | 0 | 0 | 275 | 149 | 1 | 1 | 1 | 1 | 1 | 0.803 | 1 | 1 | 1 | 1 | 1 | 1 | -0.039 |
| SE | 91829 | ENSG00000104365 | IKBKB | chr8 | + | 42306342 | 42306432 | 42305186 | 42305275 | 42308900 | 42309025 | 103 | 47 | 73 | 97 | 58 | 0 | 0 | 1 | 0 | 0 | 216 | 81 | 57 | 92 | 138 | 0 | 0 | 0 | 0 | 0 | 239 | 149 | 1 | 1 | 1 | 1 | 0.978 | 1 | 1 | 1 | 1 | 1 | 1 | 1 | -0.004 |
| SE | 91830 | ENSG00000104365 | IKBKB | chr8 | + | 42306367 | 42306432 | 42305186 | 42305275 | 42308900 | 42308932 | 17 | 10 | 17 | 24 | 12 | 0 | 0 | 1 | 0 | 0 | 25 | 28 | 20 | 34 | 32 | 0 | 0 | 0 | 0 | 0 | 214 | 149 | 1 | 1 | 1 | 1 | 0.922 | 1 | 1 | 1 | 1 | 1 | 1 | 1 | -0.016 |
| SE | 91832 | ENSG00000104365 | IKBKB | chr8 | + | 42314321 | 42314429 | 42308900 | 42309025 | 42316209 | 42316339 | 155 | 62 | 104 | 152 | 152 | 0 | 1 | 0 | 0 | 0 | 201 | 162 | 112 | 171 | 141 | 0 | 0 | 0 | 0 | 0 | 257 | 149 | 1 | 1 | 1 | 0.973 | 1 | 1 | 1 | 1 | 1 | 1 | 1 | 1 | -0.005 |
| SE | 91833 | ENSG00000104365 | IKBKB | chr8 | + | 42317656 | 42317771 | 42316709 | 42316904 | 42318551 | 42318675 | 310 | 156 | 187 | 246 | 277 | 1 | 0 | 0 | 0 | 0 | 306 | 219 | 165 | 242 | 319 | 0 | 0 | 0 | 2 | 0 | 264 | 149 | 1 | 1 | 0.994 | 1 | 1 | 1 | 1 | 1 | 1 | 1 | 0.986 | 1 | 0.002 |
| SE | 91834 | ENSG00000104365 | IKBKB | chr8 | + | 42317709 | 42317771 | 42316709 | 42316904 | 42318551 | 42318675 | 91 | 50 | 62 | 83 | 90 | 1 | 0 | 0 | 0 | 0 | 101 | 92 | 81 | 65 | 100 | 0 | 0 | 0 | 2 | 0 | 211 | 149 | 1 | 1 | 0.985 | 1 | 1 | 1 | 1 | 1 | 1 | 1 | 0.958 | 1 | 0.005 |
| SE | 91835 | ENSG00000104365 | IKBKB | chr8 | + | 42319584 | 42319646 | 42319269 | 42319421 | 42320674 | 42320844 | 80 | 39 | 41 | 63 | 53 | 0 | 0 | 1 | 0 | 0 | 82 | 69 | 35 | 72 | 98 | 0 | 0 | 0 | 0 | 0 | 211 | 149 | 1 | 1 | 1 | 1 | 0.967 | 1 | 1 | 1 | 1 | 1 | 1 | 1 | -0.007 |
| SE | 91837 | ENSG00000104365 | IKBKB | chr8 | + | 42325969 | 42326097 | 42322053 | 42322494 | 42330913 | 42331880 | 120 | 74 | 87 | 94 | 158 | 0 | 0 | 0 | 0 | 0 | 66 | 86 | 46 | 84 | 101 | 0 | 0 | 1 | 0 | 0 | 277 | 149 | 1 | 1 | 1 | 1 | 1 | 1 | 1 | 1 | 1 | 0.961 | 1 | 1 | 0.008 |
| SE | 91838 | ENSG00000104365 | IKBKB | chr8 | + | 42329123 | 42329214 | 42325609 | 42326097 | 42330913 | 42331067 | 178 | 90 | 115 | 174 | 195 | 0 | 0 | 1 | 0 | 0 | 174 | 160 | 80 | 147 | 216 | 0 | 0 | 0 | 0 | 0 | 240 | 149 | 1 | 1 | 1 | 1 | 0.986 | 1 | 1 | 1 | 1 | 1 | 1 | 1 | -0.003 |
| SE | 91839 | ENSG00000104365 | IKBKB | chr8 | + | 42329367 | 42329499 | 42329123 | 42329214 | 42330913 | 42331527 | 0 | 0 | 1 | 0 | 0 | 103 | 49 | 67 | 117 | 133 | 0 | 0 | 0 | 0 | 0 | 80 | 111 | 53 | 79 | 110 | 281 | 149 | 1 | 1 | 0 | 0 | 0.008 | 0 | 0 | 0 | 0 | 0 | 0 | 0 | 0.002 |
| SE | 91842 | ENSG00000211772 | TRBC2 | chr7 | + | 142801943 | 142801961 | 142801040 | 142801427 | 142802104 | 142802211 | 1214 | 817 | 1256 | 1767 | 1212 | 10 | 0 | 15 | 14 | 3 | 501 | 1257 | 696 | 515 | 1379 | 0 | 0 | 2 | 0 | 0 | 167 | 149 | 1 | 1 | 0.991 | 1 | 0.987 | 0.991 | 0.997 | 1 | 1 | 0.997 | 1 | 1 | -0.006 |
| SE | 91844 | ENSG00000164830 | OXR1 | chr8 | + | 106683198 | 106683306 | 106679209 | 106679292 | 106684245 | 106684340 | 443 | 327 | 284 | 291 | 1100 | 0 | 0 | 1 | 0 | 0 | 50 | 194 | 198 | 77 | 50 | 0 | 0 | 0 | 0 | 0 | 257 | 149 | 1 | 1 | 1 | 1 | 0.994 | 1 | 1 | 1 | 1 | 1 | 1 | 1 | -0.001 |
| SE | 91845 | ENSG00000164830 | OXR1 | chr8 | + | 106692727 | 106692877 | 106684245 | 106684359 | 106702905 | 106703087 | 265 | 193 | 253 | 261 | 485 | 0 | 2 | 0 | 0 | 3 | 32 | 186 | 118 | 50 | 25 | 0 | 0 | 2 | 0 | 0 | 298 | 149 | 1 | 1 | 1 | 0.98 | 1 | 1 | 0.988 | 1 | 1 | 0.967 | 1 | 1 | 0 |
| SE | 91846 | ENSG00000164830 | OXR1 | chr8 | + | 106692727 | 106692877 | 106684245 | 106684359 | 106706381 | 106706982 | 176 | 134 | 167 | 144 | 321 | 0 | 0 | 2 | 0 | 0 | 22 | 108 | 77 | 27 | 17 | 0 | 0 | 0 | 0 | 0 | 298 | 149 | 1 | 1 | 1 | 1 | 1 | 0.973 | 1 | 1 | 1 | 1 | 1 | 1 | -0.005 |
| SE | 91847 | ENSG00000164830 | OXR1 | chr8 | + | 106702905 | 106703090 | 106684245 | 106684359 | 106706381 | 106706982 | 74 | 102 | 75 | 93 | 112 | 0 | 0 | 2 | 0 | 0 | 21 | 43 | 90 | 26 | 10 | 0 | 0 | 0 | 0 | 0 | 298 | 149 | 1 | 1 | 1 | 1 | 1 | 0.959 | 1 | 1 | 1 | 1 | 1 | 1 | -0.008 |
| SE | 91849 | ENSG00000164830 | OXR1 | chr8 | + | 106713822 | 106713985 | 106710621 | 106710790 | 106739457 | 106739583 | 338 | 212 | 207 | 268 | 370 | 2 | 1 | 1 | 0 | 0 | 50 | 130 | 97 | 51 | 12 | 0 | 2 | 0 | 0 | 0 | 298 | 149 | 1 | 1 | 0.988 | 0.991 | 0.99 | 1 | 1 | 1 | 0.97 | 1 | 1 | 1 | 0 |
| SE | 91851 | ENSG00000164830 | OXR1 | chr8 | + | 106737519 | 106737600 | 106726106 | 106726309 | 106739457 | 106739583 | 23 | 15 | 18 | 12 | 23 | 5 | 4 | 2 | 1 | 6 | 0 | 29 | 13 | 1 | 4 | 0 | 0 | 0 | 0 | 0 | 230 | 149 | 1 | 1 | 0.749 | 0.708 | 0.854 | 0.886 | 0.713 | NA | 1 | 1 | 1 | 1 | -0.218 |
| SE | 91854 | ENSG00000164830 | OXR1 | chr8 | + | 106740342 | 106740495 | 106713822 | 106713985 | 106742221 | 106742317 | 213 | 175 | 220 | 203 | 207 | 2 | 2 | 0 | 2 | 0 | 23 | 131 | 95 | 50 | 2 | 0 | 0 | 0 | 0 | 0 | 298 | 149 | 1 | 1 | 0.982 | 0.978 | 1 | 0.981 | 1 | 1 | 1 | 1 | 1 | 1 | -0.012 |
| SE | 91855 | ENSG00000164830 | OXR1 | chr8 | + | 106740342 | 106740495 | 106739457 | 106739583 | 106742221 | 106742317 | 421 | 337 | 427 | 400 | 494 | 0 | 4 | 0 | 3 | 0 | 30 | 227 | 174 | 85 | 17 | 0 | 0 | 0 | 0 | 0 | 298 | 149 | 1 | 1 | 1 | 0.977 | 1 | 0.985 | 1 | 1 | 1 | 1 | 1 | 1 | -0.008 |
| SE | 91856 | ENSG00000164830 | OXR1 | chr8 | + | 106745788 | 106745862 | 106742221 | 106742317 | 106750805 | 106751173 | 111 | 88 | 107 | 104 | 132 | 0 | 1 | 3 | 2 | 1 | 0 | 48 | 44 | 21 | 0 | 0 | 1 | 0 | 3 | 0 | 223 | 149 | 1 | 1 | 1 | 0.983 | 0.96 | 0.972 | 0.989 | NA | 0.97 | 1 | 0.824 | NA | 0.049 |
| SE | 91858 | ENSG00000154188 | ANGPT1 | chr8 | - | 107303239 | 107303367 | 107293935 | 107294037 | 107321895 | 107322128 | 41 | 49 | 27 | 77 | 68 | 0 | 2 | 0 | 0 | 0 | 179 | 31 | 24 | 54 | 156 | 0 | 4 | 0 | 0 | 0 | 277 | 149 | 1 | 1 | 1 | 0.929 | 1 | 1 | 1 | 1 | 0.807 | 1 | 1 | 1 | 0.024 |
| SE | 91860 | ENSG00000154188 | ANGPT1 | chr8 | - | 107321895 | 107322128 | 107303239 | 107303367 | 107336149 | 107336271 | 40 | 60 | 11 | 105 | 130 | 2 | 0 | 0 | 0 | 0 | 283 | 55 | 64 | 93 | 247 | 0 | 0 | 0 | 0 | 0 | 298 | 149 | 1 | 1 | 0.909 | 1 | 1 | 1 | 1 | 1 | 1 | 1 | 1 | 1 | -0.018 |
| SE | 91861 | ENSG00000154188 | ANGPT1 | chr8 | - | 107321898 | 107322128 | 107303239 | 107303367 | 107336149 | 107336271 | 29 | 56 | 12 | 93 | 113 | 2 | 0 | 0 | 0 | 0 | 204 | 48 | 63 | 76 | 184 | 0 | 0 | 0 | 0 | 0 | 298 | 149 | 1 | 1 | 0.879 | 1 | 1 | 1 | 1 | 1 | 1 | 1 | 1 | 1 | -0.024 |
| SE | 91868 | ENSG00000170786 | SDR16C5 | chr8 | - | 56312156 | 56312288 | 56308927 | 56309027 | 56316014 | 56316361 | 11 | 0 | 0 | 5 | 11 | 0 | 0 | 0 | 0 | 1 | 0 | 0 | 0 | 12 | 0 | 0 | 0 | 0 | 0 | 0 | 281 | 149 | 1 | 1 | 1 | NA | NA | 1 | 0.854 | NA | NA | NA | 1 | NA | -0.049 |
| SE | 91870 | ENSG00000113494 | PRLR | chr5 | - | 35068778 | 35068878 | 35068215 | 35068285 | 35070123 | 35070265 | 3 | 11 | 1 | 1 | 2 | 0 | 0 | 0 | 0 | 0 | 0 | 0 | 1 | 0 | 0 | 0 | 0 | 3 | 0 | 0 | 249 | 149 | 1 | 1 | 1 | 1 | 1 | 1 | 1 | NA | NA | 0.166 | NA | NA | 0.834 |
| SE | 91878 | ENSG00000104518 | GSDMD | chr8 | + | 143555933 | 143556030 | 143555058 | 143555196 | 143558377 | 143558451 | 0 | 2 | 2 | 5 | 1 | 0 | 0 | 0 | 3 | 0 | 0 | 3 | 0 | 0 | 0 | 0 | 0 | 0 | 0 | 0 | 246 | 149 | 1 | 1 | NA | 1 | 1 | 0.502 | 1 | NA | 1 | NA | NA | NA | -0.125 |
| SE | 91880 | ENSG00000104518 | GSDMD | chr8 | + | 143561958 | 143562131 | 143561729 | 143561828 | 143562208 | 143562350 | 134 | 40 | 127 | 107 | 28 | 0 | 0 | 0 | 0 | 0 | 102 | 81 | 32 | 79 | 132 | 0 | 0 | 1 | 0 | 0 | 298 | 149 | 1 | 1 | 1 | 1 | 1 | 1 | 1 | 1 | 1 | 0.941 | 1 | 1 | 0.012 |
| SE | 91882 | ENSG00000204839 | MROH6 | chr8 | - | 143568141 | 143568261 | 143567785 | 143567888 | 143568551 | 143568719 | 1 | 0 | 1 | 0 | 0 | 2 | 0 | 0 | 0 | 0 | 0 | 1 | 0 | 9 | 2 | 0 | 0 | 0 | 0 | 0 | 269 | 149 | 1 | 1 | 0.217 | NA | 1 | NA | NA | NA | 1 | NA | 1 | 1 | -0.391 |
| SE | 91884 | ENSG00000254995 | STX16-NPEPL1 | chr20 | + | 58670511 | 58670603 | 58667489 | 58667597 | 58691723 | 58691780 | 0 | 3 | 0 | 1 | 0 | 0 | 0 | 0 | 0 | 0 | 0 | 0 | 2 | 0 | 0 | 9 | 0 | 0 | 0 | 0 | 241 | 149 | 1 | 1 | NA | 1 | NA | 1 | NA | 0 | NA | 1 | NA | NA | 0.5 |
| SE | 91885 | ENSG00000254995 | STX16-NPEPL1 | chr20 | + | 58671153 | 58671297 | 58670511 | 58670603 | 58691723 | 58691780 | 250 | 183 | 216 | 229 | 143 | 0 | 0 | 0 | 0 | 0 | 405 | 227 | 227 | 261 | 265 | 0 | 0 | 1 | 0 | 0 | 293 | 149 | 1 | 1 | 1 | 1 | 1 | 1 | 1 | 1 | 1 | 0.991 | 1 | 1 | 0.002 |
| SE | 91890 | ENSG00000104529 | EEF1D | chr8 | - | 143580011 | 143580206 | 143579696 | 143579830 | 143580505 | 143580632 | 2316 | 1102 | 2563 | 2150 | 1951 | 0 | 0 | 0 | 0 | 0 | 2116 | 1782 | 1405 | 1391 | 2565 | 0 | 0 | 0 | 1 | 0 | 298 | 149 | 1 | 1 | 1 | 1 | 1 | 1 | 1 | 1 | 1 | 1 | 0.999 | 1 | 0 |
| SE | 91891 | ENSG00000104529 | EEF1D | chr8 | - | 143580505 | 143580632 | 143580009 | 143580206 | 143581053 | 143581154 | 1377 | 670 | 1485 | 1258 | 1373 | 0 | 0 | 0 | 2 | 0 | 1224 | 1079 | 790 | 852 | 1476 | 0 | 0 | 0 | 0 | 0 | 276 | 149 | 1 | 1 | 1 | 1 | 1 | 0.997 | 1 | 1 | 1 | 1 | 1 | 1 | -0.001 |
| SE | 91892 | ENSG00000104529 | EEF1D | chr8 | - | 143580505 | 143580727 | 143580011 | 143580206 | 143581053 | 143581154 | 3957 | 1972 | 3966 | 3517 | 4014 | 0 | 0 | 0 | 2 | 0 | 3644 | 3206 | 2351 | 2253 | 4270 | 0 | 0 | 0 | 0 | 0 | 298 | 149 | 1 | 1 | 1 | 1 | 1 | 0.999 | 1 | 1 | 1 | 1 | 1 | 1 | 0 |
| SE | 91893 | ENSG00000104529 | EEF1D | chr8 | - | 143581228 | 143581328 | 143581053 | 143581154 | 143586218 | 143586290 | 2294 | 1114 | 2294 | 1937 | 2559 | 0 | 0 | 1 | 0 | 0 | 2092 | 1812 | 1359 | 1264 | 2392 | 0 | 0 | 0 | 2 | 0 | 249 | 149 | 1 | 1 | 1 | 1 | 0.999 | 1 | 1 | 1 | 1 | 1 | 0.997 | 1 | 0 |
| SE | 91894 | ENSG00000104529 | EEF1D | chr8 | - | 143581228 | 143581328 | 143581053 | 143581154 | 143586728 | 143586849 | 1615 | 876 | 1449 | 1327 | 2011 | 0 | 0 | 0 | 4 | 3 | 1472 | 1114 | 943 | 818 | 1807 | 0 | 0 | 1 | 5 | 0 | 249 | 149 | 1 | 1 | 1 | 1 | 1 | 0.995 | 0.998 | 1 | 1 | 0.998 | 0.99 | 1 | 0.001 |
| SE | 91895 | ENSG00000104529 | EEF1D | chr8 | - | 143581228 | 143581328 | 143581053 | 143581154 | 143597347 | 143597362 | 1223 | 681 | 1136 | 1039 | 1401 | 0 | 0 | 0 | 2 | 0 | 1160 | 882 | 735 | 656 | 1353 | 0 | 0 | 0 | 0 | 0 | 249 | 149 | 1 | 1 | 1 | 1 | 1 | 0.997 | 1 | 1 | 1 | 1 | 1 | 1 | -0.001 |
| SE | 91896 | ENSG00000104529 | EEF1D | chr8 | - | 143581285 | 143581328 | 143581053 | 143581154 | 143586218 | 143586290 | 376 | 146 | 399 | 306 | 536 | 0 | 0 | 1 | 0 | 0 | 251 | 255 | 197 | 208 | 225 | 0 | 0 | 0 | 2 | 0 | 192 | 149 | 1 | 1 | 1 | 1 | 0.997 | 1 | 1 | 1 | 1 | 1 | 0.988 | 1 | 0.002 |
| SE | 91898 | ENSG00000104529 | EEF1D | chr8 | - | 143581285 | 143581328 | 143581053 | 143581154 | 143597347 | 143597397 | 14 | 10 | 10 | 9 | 17 | 0 | 0 | 0 | 2 | 0 | 0 | 0 | 3 | 10 | 0 | 0 | 0 | 0 | 0 | 0 | 192 | 149 | 1 | 1 | 1 | 1 | 1 | 0.777 | 1 | NA | NA | 1 | 1 | NA | -0.045 |
| SE | 91899 | ENSG00000104529 | EEF1D | chr8 | - | 143581285 | 143581706 | 143581053 | 143581154 | 143586218 | 143586290 | 14 | 10 | 10 | 9 | 17 | 0 | 0 | 1 | 0 | 0 | 0 | 0 | 3 | 10 | 0 | 0 | 0 | 0 | 2 | 0 | 298 | 149 | 1 | 1 | 1 | 1 | 0.833 | 1 | 1 | NA | NA | 1 | 0.714 | NA | 0.11 |
| SE | 91900 | ENSG00000104529 | EEF1D | chr8 | - | 143586218 | 143586290 | 143581053 | 143581154 | 143586728 | 143586849 | 248 | 101 | 293 | 214 | 368 | 0 | 0 | 0 | 4 | 3 | 136 | 184 | 157 | 118 | 286 | 0 | 0 | 1 | 5 | 0 | 221 | 149 | 1 | 1 | 1 | 1 | 1 | 0.973 | 0.988 | 1 | 1 | 0.991 | 0.941 | 1 | 0.006 |
| SE | 91902 | ENSG00000104529 | EEF1D | chr8 | - | 143586728 | 143586849 | 143581053 | 143581154 | 143597347 | 143597384 | 37 | 14 | 41 | 31 | 12 | 0 | 0 | 0 | 2 | 0 | 10 | 27 | 19 | 14 | 40 | 0 | 0 | 0 | 0 | 0 | 270 | 149 | 1 | 1 | 1 | 1 | 1 | 0.895 | 1 | 1 | 1 | 1 | 1 | 1 | -0.021 |
| SE | 91903 | ENSG00000104529 | EEF1D | chr8 | - | 143586728 | 143586849 | 143581228 | 143581328 | 143597347 | 143597384 | 425 | 214 | 341 | 308 | 618 | 0 | 0 | 0 | 0 | 0 | 321 | 244 | 220 | 174 | 433 | 0 | 0 | 0 | 0 | 4 | 270 | 149 | 1 | 1 | 1 | 1 | 1 | 1 | 1 | 1 | 1 | 1 | 1 | 0.984 | 0.003 |
| SE | 91904 | ENSG00000104529 | EEF1D | chr8 | - | 143586728 | 143586852 | 143581053 | 143581154 | 143597347 | 143597387 | 230 | 65 | 279 | 167 | 134 | 0 | 0 | 0 | 2 | 0 | 136 | 157 | 124 | 125 | 72 | 0 | 0 | 0 | 0 | 0 | 273 | 149 | 1 | 1 | 1 | 1 | 1 | 0.979 | 1 | 1 | 1 | 1 | 1 | 1 | -0.004 |
| SE | 91905 | ENSG00000104529 | EEF1D | chr8 | - | 143586728 | 143586852 | 143581228 | 143581328 | 143597347 | 143597387 | 629 | 267 | 593 | 458 | 754 | 0 | 0 | 0 | 0 | 0 | 460 | 384 | 328 | 285 | 499 | 0 | 0 | 0 | 0 | 4 | 273 | 149 | 1 | 1 | 1 | 1 | 1 | 1 | 1 | 1 | 1 | 1 | 1 | 0.986 | 0.003 |
| SE | 91908 | ENSG00000104529 | EEF1D | chr8 | - | 143590607 | 143590729 | 143589510 | 143590081 | 143597347 | 143597415 | 12 | 5 | 9 | 16 | 4 | 3 | 0 | 10 | 2 | 0 | 0 | 1 | 0 | 0 | 17 | 0 | 0 | 0 | 0 | 0 | 271 | 149 | 1 | 1 | 0.687 | 1 | 0.331 | 0.815 | 1 | NA | 1 | NA | NA | 1 | -0.233 |
| SE | 91909 | ENSG00000104529 | EEF1D | chr8 | - | 143590607 | 143590735 | 143589694 | 143590081 | 143592646 | 143592893 | 13 | 5 | 9 | 19 | 4 | 1 | 1 | 1 | 2 | 1 | 0 | 1 | 0 | 0 | 21 | 0 | 0 | 2 | 1 | 2 | 277 | 149 | 1 | 1 | 0.875 | 0.729 | 0.829 | 0.836 | 0.683 | NA | 1 | 0 | 0 | 0.85 | 0.328 |
| SE | 91910 | ENSG00000104529 | EEF1D | chr8 | - | 143590607 | 143590735 | 143589694 | 143590081 | 143597347 | 143597415 | 12 | 5 | 9 | 21 | 4 | 3 | 0 | 10 | 2 | 0 | 0 | 1 | 0 | 0 | 21 | 0 | 0 | 0 | 0 | 0 | 277 | 149 | 1 | 1 | 0.683 | 1 | 0.326 | 0.85 | 1 | NA | 1 | NA | NA | 1 | -0.228 |
| SE | 91911 | ENSG00000104529 | EEF1D | chr8 | - | 143590607 | 143590738 | 143588990 | 143590081 | 143592646 | 143592797 | 12 | 5 | 11 | 23 | 4 | 1 | 1 | 1 | 2 | 1 | 0 | 1 | 0 | 0 | 21 | 0 | 0 | 2 | 1 | 2 | 280 | 149 | 1 | 1 | 0.865 | 0.727 | 0.854 | 0.86 | 0.68 | NA | 1 | 0 | 0 | 0.848 | 0.335 |
| SE | 91912 | ENSG00000104529 | EEF1D | chr8 | - | 143590607 | 143590738 | 143588990 | 143590081 | 143592714 | 143592893 | 12 | 5 | 9 | 19 | 4 | 2 | 0 | 0 | 0 | 0 | 0 | 1 | 0 | 0 | 21 | 0 | 0 | 0 | 4 | 0 | 280 | 149 | 1 | 1 | 0.761 | 1 | 1 | 1 | 1 | NA | 1 | NA | 0 | 1 | 0.286 |
| SE | 91914 | ENSG00000104529 | EEF1D | chr8 | - | 143590607 | 143590738 | 143588990 | 143590081 | 143596878 | 143597105 | 12 | 5 | 9 | 19 | 4 | 0 | 0 | 0 | 1 | 0 | 0 | 1 | 0 | 0 | 21 | 0 | 0 | 0 | 0 | 0 | 280 | 149 | 1 | 1 | 1 | 1 | 1 | 0.91 | 1 | NA | 1 | NA | NA | 1 | -0.018 |
| SE | 91915 | ENSG00000104529 | EEF1D | chr8 | - | 143590607 | 143590738 | 143588990 | 143590081 | 143597347 | 143597415 | 15 | 9 | 12 | 21 | 4 | 3 | 0 | 10 | 2 | 0 | 0 | 2 | 0 | 3 | 22 | 0 | 0 | 0 | 0 | 0 | 280 | 149 | 1 | 1 | 0.727 | 1 | 0.39 | 0.848 | 1 | NA | 1 | NA | 1 | 1 | -0.207 |
| SE | 91917 | ENSG00000104529 | EEF1D | chr8 | - | 143590644 | 143590738 | 143589643 | 143590081 | 143597347 | 143597415 | 0 | 0 | 0 | 0 | 0 | 3 | 0 | 10 | 2 | 0 | 0 | 0 | 0 | 3 | 0 | 0 | 0 | 0 | 0 | 0 | 243 | 149 | 1 | 1 | 0 | NA | 0 | 0 | NA | NA | NA | NA | 1 | NA | -1 |
| SE | 91920 | ENSG00000104529 | EEF1D | chr8 | - | 143592646 | 143592797 | 143590607 | 143590738 | 143597347 | 143597380 | 0 | 0 | 2 | 4 | 0 | 3 | 4 | 3 | 2 | 0 | 0 | 0 | 0 | 0 | 0 | 0 | 1 | 0 | 3 | 1 | 298 | 149 | 1 | 1 | 0 | 0 | 0.25 | 0.5 | NA | NA | 0 | NA | 0 | 0 | 0.188 |
| SE | 91921 | ENSG00000104529 | EEF1D | chr8 | - | 143592646 | 143592893 | 143589837 | 143590081 | 143597347 | 143597413 | 3 | 1 | 1 | 2 | 1 | 3 | 0 | 10 | 2 | 0 | 0 | 0 | 2 | 1 | 2 | 0 | 0 | 0 | 0 | 0 | 298 | 149 | 1 | 1 | 0.333 | 1 | 0.048 | 0.333 | 1 | NA | NA | 1 | 1 | 1 | -0.457 |
| SE | 91924 | ENSG00000104529 | EEF1D | chr8 | - | 143592646 | 143592893 | 143590607 | 143590738 | 143597347 | 143597413 | 2 | 0 | 2 | 4 | 0 | 3 | 4 | 3 | 2 | 0 | 0 | 0 | 0 | 0 | 0 | 0 | 1 | 0 | 3 | 1 | 298 | 149 | 1 | 1 | 0.25 | 0 | 0.25 | 0.5 | NA | NA | 0 | NA | 0 | 0 | 0.25 |
| SE | 91925 | ENSG00000104529 | EEF1D | chr8 | - | 143592646 | 143592942 | 143589829 | 143590081 | 143597347 | 143597415 | 2 | 1 | 1 | 3 | 1 | 3 | 0 | 10 | 2 | 0 | 0 | 0 | 2 | 1 | 2 | 0 | 0 | 0 | 0 | 0 | 298 | 149 | 1 | 1 | 0.25 | 1 | 0.048 | 0.429 | 1 | NA | NA | 1 | 1 | 1 | -0.455 |
| SE | 91926 | ENSG00000104529 | EEF1D | chr8 | - | 143592714 | 143592893 | 143586728 | 143586852 | 143593892 | 143594071 | 0 | 0 | 0 | 0 | 1 | 0 | 0 | 0 | 0 | 1 | 0 | 0 | 0 | 0 | 0 | 0 | 0 | 0 | 1 | 0 | 298 | 149 | 1 | 1 | NA | NA | NA | NA | 0.333 | NA | NA | NA | 0 | NA | 0.333 |
| SE | 91927 | ENSG00000104529 | EEF1D | chr8 | - | 143592714 | 143592893 | 143586728 | 143586852 | 143597347 | 143597415 | 2 | 0 | 0 | 0 | 0 | 255 | 68 | 312 | 182 | 140 | 0 | 0 | 0 | 0 | 0 | 153 | 177 | 144 | 130 | 92 | 298 | 149 | 1 | 1 | 0.004 | 0 | 0 | 0 | 0 | 0 | 0 | 0 | 0 | 0 | 0.001 |
| SE | 91928 | ENSG00000104529 | EEF1D | chr8 | - | 143592714 | 143592893 | 143588990 | 143590081 | 143597347 | 143597415 | 4 | 0 | 0 | 0 | 0 | 3 | 0 | 10 | 2 | 0 | 0 | 0 | 0 | 4 | 0 | 0 | 0 | 0 | 0 | 0 | 298 | 149 | 1 | 1 | 0.4 | NA | 0 | 0 | NA | NA | NA | NA | 1 | NA | -0.867 |
| SE | 91930 | ENSG00000104529 | EEF1D | chr8 | - | 143592714 | 143592893 | 143590607 | 143590738 | 143597347 | 143597415 | 2 | 0 | 0 | 0 | 0 | 3 | 4 | 3 | 2 | 0 | 0 | 0 | 0 | 0 | 0 | 0 | 1 | 0 | 3 | 1 | 298 | 149 | 1 | 1 | 0.25 | 0 | 0 | 0 | NA | NA | 0 | NA | 0 | 0 | 0.062 |
| SE | 91931 | ENSG00000104529 | EEF1D | chr8 | - | 143592714 | 143592942 | 143586728 | 143586852 | 143597347 | 143597365 | 1 | 0 | 0 | 1 | 0 | 255 | 68 | 312 | 182 | 140 | 0 | 0 | 0 | 0 | 0 | 153 | 177 | 144 | 130 | 92 | 298 | 149 | 1 | 1 | 0.002 | 0 | 0 | 0.003 | 0 | 0 | 0 | 0 | 0 | 0 | 0.001 |
| SE | 91933 | ENSG00000104529 | EEF1D | chr8 | - | 143593874 | 143594071 | 143592714 | 143592893 | 143597347 | 143597362 | 0 | 0 | 2 | 0 | 4 | 2 | 0 | 0 | 0 | 0 | 0 | 0 | 0 | 0 | 3 | 0 | 0 | 0 | 0 | 0 | 298 | 149 | 1 | 1 | 0 | NA | 1 | NA | 1 | NA | NA | NA | NA | 1 | -0.333 |
| SE | 91934 | ENSG00000104529 | EEF1D | chr8 | - | 143593874 | 143594071 | 143592714 | 143592942 | 143597347 | 143597362 | 0 | 0 | 2 | 0 | 0 | 1 | 0 | 0 | 1 | 0 | 0 | 0 | 0 | 0 | 3 | 0 | 0 | 0 | 0 | 0 | 298 | 149 | 1 | 1 | 0 | NA | 1 | 0 | NA | NA | NA | NA | NA | 1 | -0.667 |
| SE | 91935 | ENSG00000104529 | EEF1D | chr8 | - | 143593892 | 143594071 | 143586728 | 143586849 | 143597347 | 143597394 | 0 | 0 | 2 | 0 | 0 | 44 | 18 | 45 | 32 | 10 | 0 | 0 | 0 | 0 | 3 | 10 | 28 | 26 | 9 | 40 | 298 | 149 | 1 | 1 | 0 | 0 | 0.022 | 0 | 0 | 0 | 0 | 0 | 0 | 0.036 | -0.003 |
| SE | 91936 | ENSG00000104529 | EEF1D | chr8 | - | 143593892 | 143594071 | 143586728 | 143586852 | 143597347 | 143597394 | 0 | 0 | 2 | 0 | 1 | 255 | 68 | 312 | 182 | 140 | 0 | 0 | 0 | 1 | 3 | 153 | 177 | 144 | 130 | 92 | 298 | 149 | 1 | 1 | 0 | 0 | 0.003 | 0 | 0.004 | 0 | 0 | 0 | 0.004 | 0.016 | -0.003 |
| SE | 91937 | ENSG00000104529 | EEF1D | chr8 | - | 143593892 | 143594071 | 143590607 | 143590738 | 143597347 | 143597394 | 0 | 0 | 3 | 1 | 0 | 3 | 4 | 3 | 2 | 0 | 0 | 0 | 0 | 0 | 3 | 0 | 1 | 0 | 3 | 1 | 298 | 149 | 1 | 1 | 0 | 0 | 0.333 | 0.2 | NA | NA | 0 | NA | 0 | 0.6 | -0.067 |
| SE | 91938 | ENSG00000104529 | EEF1D | chr8 | - | 143593892 | 143594071 | 143592646 | 143592893 | 143597347 | 143597394 | 0 | 0 | 2 | 0 | 1 | 2 | 0 | 0 | 0 | 0 | 0 | 0 | 0 | 0 | 3 | 0 | 0 | 0 | 0 | 0 | 298 | 149 | 1 | 1 | 0 | NA | 1 | NA | 1 | NA | NA | NA | NA | 1 | -0.333 |
| SE | 91939 | ENSG00000104529 | EEF1D | chr8 | - | 143594415 | 143594518 | 143593874 | 143594071 | 143597347 | 143597393 | 0 | 0 | 3 | 0 | 0 | 0 | 0 | 2 | 0 | 0 | 0 | 0 | 0 | 3 | 0 | 0 | 0 | 0 | 0 | 3 | 252 | 149 | 1 | 1 | NA | NA | 0.47 | NA | NA | NA | NA | NA | 1 | 0 | -0.03 |
| SE | 91941 | ENSG00000104522 | GFUS | chr8 | - | 143614163 | 143614228 | 143613750 | 143613817 | 143614319 | 143614453 | 67 | 16 | 59 | 62 | 15 | 0 | 0 | 0 | 0 | 0 | 88 | 77 | 20 | 59 | 130 | 0 | 0 | 1 | 0 | 0 | 214 | 149 | 1 | 1 | 1 | 1 | 1 | 1 | 1 | 1 | 1 | 0.933 | 1 | 1 | 0.013 |
| SE | 91942 | ENSG00000104522 | GFUS | chr8 | - | 143614623 | 143614697 | 143614319 | 143614453 | 143614786 | 143614915 | 64 | 24 | 60 | 68 | 76 | 0 | 0 | 2 | 0 | 0 | 54 | 107 | 26 | 53 | 129 | 0 | 0 | 0 | 0 | 0 | 223 | 149 | 1 | 1 | 1 | 1 | 0.952 | 1 | 1 | 1 | 1 | 1 | 1 | 1 | -0.01 |
| SE | 91960 | ENSG00000253320 | MAILR | chr8 | + | 102941920 | 102942053 | 102937623 | 102937757 | 102977586 | 102977738 | 2 | 3 | 0 | 2 | 4 | 0 | 0 | 0 | 0 | 0 | 0 | 4 | 0 | 0 | 1 | 7 | 0 | 0 | 0 | 0 | 282 | 149 | 1 | 1 | 1 | 1 | NA | 1 | 1 | 0 | 1 | NA | NA | 1 | 0.333 |
| SE | 91961 | ENSG00000253320 | MAILR | chr8 | + | 102967596 | 102967673 | 102879894 | 102880058 | 102977586 | 102978955 | 0 | 0 | 0 | 2 | 0 | 0 | 0 | 0 | 0 | 0 | 0 | 0 | 0 | 0 | 0 | 6 | 0 | 0 | 0 | 0 | 226 | 149 | 1 | 1 | NA | NA | NA | 1 | NA | NA | 0 | NA | NA | NA | 1 |
| SE | 91962 | ENSG00000253320 | MAILR | chr8 | + | 102992249 | 102992407 | 102978784 | 102978955 | 102995624 | 102995682 | 0 | 2 | 0 | 0 | 1 | 2 | 0 | 1 | 2 | 2 | 0 | 0 | 0 | 0 | 0 | 7 | 3 | 1 | 4 | 0 | 298 | 149 | 1 | 1 | 0 | 1 | 0 | 0 | 0.2 | 0 | 0 | 0 | 0 | NA | 0.24 |
| SE | 91963 | ENSG00000253320 | MAILR | chr8 | + | 102995624 | 102995682 | 102977586 | 102978955 | 103000008 | 103000184 | 2 | 0 | 1 | 2 | 2 | 7 | 10 | 7 | 3 | 1 | 7 | 3 | 1 | 4 | 0 | 7 | 2 | 9 | 0 | 0 | 207 | 149 | 1 | 1 | 0.171 | 0 | 0.093 | 0.324 | 0.59 | 0.419 | 0.519 | 0.074 | 1 | NA | -0.267 |
| SE | 91965 | ENSG00000183808 | RBM12B | chr8 | - | 93737306 | 93737355 | 93728154 | 93736438 | 93740880 | 93741017 | 152 | 199 | 142 | 174 | 121 | 2 | 0 | 2 | 0 | 0 | 93 | 116 | 74 | 136 | 28 | 0 | 3 | 2 | 0 | 0 | 198 | 149 | 1 | 1 | 0.983 | 1 | 0.982 | 1 | 1 | 1 | 0.967 | 0.965 | 1 | 1 | 0.007 |
| SE | 91966 | ENSG00000183808 | RBM12B | chr8 | - | 93740628 | 93740696 | 93728154 | 93736438 | 93740880 | 93741009 | 8 | 5 | 12 | 19 | 4 | 2 | 0 | 2 | 0 | 0 | 3 | 14 | 11 | 20 | 0 | 0 | 3 | 2 | 0 | 0 | 217 | 149 | 1 | 1 | 0.733 | 1 | 0.805 | 1 | 1 | 1 | 0.762 | 0.791 | 1 | NA | 0.019 |
| SE | 91967 | ENSG00000183808 | RBM12B | chr8 | - | 93740628 | 93740696 | 93737306 | 93737355 | 93740862 | 93740989 | 49 | 38 | 37 | 42 | 69 | 2 | 1 | 2 | 2 | 0 | 0 | 27 | 24 | 30 | 7 | 0 | 0 | 0 | 0 | 0 | 217 | 149 | 1 | 1 | 0.944 | 0.963 | 0.927 | 0.935 | 1 | NA | 1 | 1 | 1 | 1 | -0.046 |
| SE | 91971 | ENSG00000119397 | CNTRL | chr9 | + | 121080305 | 121080478 | 121074954 | 121075067 | 121088295 | 121088543 | 75 | 87 | 126 | 108 | 69 | 49 | 31 | 44 | 64 | 21 | 0 | 59 | 19 | 35 | 4 | 0 | 52 | 40 | 19 | 20 | 298 | 149 | 1 | 1 | 0.434 | 0.584 | 0.589 | 0.458 | 0.622 | NA | 0.362 | 0.192 | 0.479 | 0.091 | 0.256 |
| SE | 91972 | ENSG00000119397 | CNTRL | chr9 | + | 121080305 | 121080478 | 121074954 | 121075067 | 121090274 | 121090405 | 22 | 29 | 46 | 24 | 24 | 0 | 0 | 0 | 0 | 2 | 0 | 16 | 7 | 3 | 0 | 0 | 0 | 0 | 4 | 0 | 298 | 149 | 1 | 1 | 1 | 1 | 1 | 1 | 0.857 | NA | 1 | 1 | 0.273 | NA | 0.214 |
| SE | 91973 | ENSG00000119397 | CNTRL | chr9 | + | 121088295 | 121088543 | 121074752 | 121075067 | 121090274 | 121090405 | 330 | 315 | 311 | 407 | 219 | 0 | 0 | 0 | 0 | 2 | 68 | 245 | 202 | 115 | 76 | 0 | 0 | 0 | 4 | 0 | 298 | 149 | 1 | 1 | 1 | 1 | 1 | 1 | 0.982 | 1 | 1 | 1 | 0.935 | 1 | 0.009 |
| SE | 91974 | ENSG00000119397 | CNTRL | chr9 | + | 121088295 | 121088543 | 121080305 | 121080478 | 121090274 | 121090405 | 334 | 342 | 348 | 428 | 243 | 0 | 0 | 1 | 1 | 0 | 68 | 236 | 176 | 128 | 60 | 0 | 0 | 2 | 0 | 0 | 298 | 149 | 1 | 1 | 1 | 1 | 0.994 | 0.995 | 1 | 1 | 1 | 0.978 | 1 | 1 | 0.002 |
| SE | 91975 | ENSG00000119397 | CNTRL | chr9 | + | 121090274 | 121090405 | 121080305 | 121080478 | 121096421 | 121096563 | 8 | 2 | 1 | 4 | 2 | 0 | 0 | 0 | 1 | 0 | 0 | 5 | 2 | 0 | 0 | 0 | 0 | 0 | 0 | 0 | 280 | 149 | 1 | 1 | 1 | 1 | 1 | 0.68 | 1 | NA | 1 | 1 | NA | NA | -0.064 |
| SE | 91977 | ENSG00000119397 | CNTRL | chr9 | + | 121096421 | 121096563 | 121094887 | 121095018 | 121098385 | 121098572 | 267 | 303 | 234 | 269 | 437 | 0 | 0 | 0 | 0 | 0 | 32 | 134 | 111 | 25 | 26 | 0 | 0 | 4 | 0 | 0 | 291 | 149 | 1 | 1 | 1 | 1 | 1 | 1 | 1 | 1 | 1 | 0.934 | 1 | 1 | 0.013 |
| SE | 91980 | ENSG00000119397 | CNTRL | chr9 | + | 121113501 | 121113724 | 121107801 | 121107995 | 121115090 | 121115200 | 108 | 137 | 119 | 110 | 123 | 0 | 1 | 0 | 1 | 0 | 0 | 95 | 76 | 11 | 14 | 0 | 0 | 0 | 0 | 0 | 298 | 149 | 1 | 1 | 1 | 0.986 | 1 | 0.982 | 1 | NA | 1 | 1 | 1 | 1 | -0.006 |
| SE | 91981 | ENSG00000119397 | CNTRL | chr9 | + | 121118345 | 121118540 | 121115090 | 121115200 | 121123930 | 121124084 | 176 | 202 | 108 | 170 | 193 | 0 | 2 | 2 | 0 | 0 | 15 | 124 | 64 | 33 | 22 | 0 | 0 | 0 | 0 | 0 | 298 | 149 | 1 | 1 | 1 | 0.981 | 0.964 | 1 | 1 | 1 | 1 | 1 | 1 | 1 | -0.011 |
| SE | 91982 | ENSG00000119397 | CNTRL | chr9 | + | 121118345 | 121118540 | 121115090 | 121115200 | 121138544 | 121138679 | 100 | 128 | 74 | 107 | 101 | 0 | 0 | 0 | 0 | 3 | 6 | 59 | 44 | 16 | 14 | 0 | 0 | 0 | 0 | 0 | 298 | 149 | 1 | 1 | 1 | 1 | 1 | 1 | 0.944 | 1 | 1 | 1 | 1 | 1 | -0.011 |
| SE | 91983 | ENSG00000119397 | CNTRL | chr9 | + | 121123930 | 121124084 | 121115090 | 121115200 | 121138544 | 121138679 | 1 | 2 | 8 | 6 | 6 | 0 | 0 | 0 | 0 | 3 | 9 | 0 | 6 | 5 | 0 | 0 | 0 | 0 | 0 | 0 | 298 | 149 | 1 | 1 | 1 | 1 | 1 | 1 | 0.5 | 1 | NA | 1 | 1 | NA | -0.1 |
| SE | 91984 | ENSG00000119397 | CNTRL | chr9 | + | 121123930 | 121124084 | 121118345 | 121118540 | 121138544 | 121138679 | 77 | 74 | 40 | 74 | 106 | 0 | 0 | 0 | 2 | 0 | 18 | 65 | 27 | 22 | 8 | 0 | 0 | 1 | 0 | 0 | 298 | 149 | 1 | 1 | 1 | 1 | 1 | 0.949 | 1 | 1 | 1 | 0.931 | 1 | 1 | 0.004 |
| SE | 91985 | ENSG00000119397 | CNTRL | chr9 | + | 121125715 | 121125936 | 121123930 | 121124084 | 121135805 | 121135982 | 254 | 310 | 253 | 301 | 216 | 0 | 1 | 1 | 0 | 1 | 20 | 135 | 114 | 70 | 17 | 0 | 0 | 0 | 0 | 0 | 298 | 149 | 1 | 1 | 1 | 0.994 | 0.992 | 1 | 0.991 | 1 | 1 | 1 | 1 | 1 | -0.005 |
| SE | 91987 | ENSG00000119397 | CNTRL | chr9 | + | 121135805 | 121135982 | 121123930 | 121124084 | 121138544 | 121138679 | 119 | 82 | 100 | 113 | 100 | 1 | 0 | 6 | 6 | 6 | 42 | 46 | 27 | 29 | 0 | 9 | 0 | 6 | 5 | 0 | 298 | 149 | 1 | 1 | 0.983 | 1 | 0.893 | 0.904 | 0.893 | 0.7 | 1 | 0.692 | 0.744 | NA | 0.151 |

| | | | | | | | | | | | | | | | | | | | | | | | | | | | | | | | | | | | | | | | | | | | | | | |
|---|---|---|---|---|---|---|---|---|---|---|---|---|---|---|---|---|---|---|---|---|---|---|---|---|---|---|---|---|---|---|---|---|---|---|---|---|---|---|---|---|---|---|---|---|---|---|
| SE | 91988 | ENSG00000119397 | CNTRL | chr9 | + | 121135805 | 121135982 | 121125715 | 121125936 | 121138544 | 121138679 | 201 | 173 | 187 | 219 | 184 | 12 | 11 | 15 | 8 | 11 | 57 | 120 | 46 | 56 | 0 | 5 | 17 | 9 | 2 | 0 | 298 | 149 | 1 | 1 | 0.893 | 0.887 | 0.862 | 0.932 | 0.893 | 0.851 | 0.779 | 0.719 | 0.933 | NA | 0.073 |
| SE | 91989 | ENSG00000119397 | CNTRL | chr9 | + | 121138544 | 121138679 | 121115090 | 121115200 | 121142090 | 121142270 | 2 | 0 | 0 | 0 | 4 | 0 | 0 | 1 | 0 | 0 | 0 | 7 | 6 | 0 | 10 | 0 | 0 | 0 | 0 | 0 | 284 | 149 | 1 | 1 | 1 | NA | 0 | NA | 1 | NA | 1 | 1 | NA | 1 | -0.333 |
| SE | 91992 | ENSG00000119397 | CNTRL | chr9 | + | 121144842 | 121144959 | 121143902 | 121144082 | 121145243 | 121145385 | 262 | 293 | 310 | 321 | 355 | 1 | 0 | 0 | 3 | 1 | 89 | 105 | 108 | 69 | 16 | 0 | 2 | 2 | 0 | 0 | 266 | 149 | 1 | 1 | 0.993 | 1 | 1 | 0.984 | 0.995 | 1 | 0.967 | 0.968 | 1 | 1 | 0.007 |
| SE | 91994 | ENSG00000119397 | CNTRL | chr9 | + | 121150169 | 121150305 | 121148671 | 121148861 | 121152484 | 121152693 | 146 | 166 | 176 | 196 | 227 | 0 | 2 | 0 | 1 | 0 | 62 | 101 | 66 | 30 | 6 | 0 | 0 | 0 | 0 | 0 | 285 | 149 | 1 | 1 | 1 | 0.977 | 1 | 0.99 | 1 | 1 | 1 | 1 | 1 | 1 | -0.007 |
| SE | 91995 | ENSG00000119397 | CNTRL | chr9 | + | 121150169 | 121150483 | 121148671 | 121148861 | 121152484 | 121152693 | 168 | 202 | 214 | 243 | 246 | 0 | 2 | 0 | 1 | 0 | 68 | 118 | 75 | 38 | 10 | 0 | 0 | 0 | 0 | 0 | 298 | 149 | 1 | 1 | 1 | 0.981 | 1 | 0.992 | 1 | 1 | 1 | 1 | 1 | 1 | -0.005 |
| SE | 91996 | ENSG00000119397 | CNTRL | chr9 | + | 121150169 | 121150483 | 121148671 | 121148861 | 121154720 | 121154913 | 146 | 169 | 180 | 206 | 231 | 0 | 0 | 0 | 0 | 0 | 64 | 101 | 69 | 30 | 6 | 0 | 0 | 0 | 3 | 0 | 298 | 149 | 1 | 1 | 1 | 1 | 1 | 1 | 1 | 1 | 1 | 1 | 0.833 | 1 | 0.033 |
| SE | 91998 | ENSG00000119397 | CNTRL | chr9 | + | 121152484 | 121152693 | 121150169 | 121150483 | 121154720 | 121154913 | 98 | 151 | 137 | 130 | 138 | 0 | 2 | 0 | 3 | 1 | 21 | 49 | 60 | 11 | 16 | 2 | 0 | 3 | 0 | 0 | 298 | 149 | 1 | 1 | 1 | 0.974 | 1 | 0.956 | 0.986 | 0.84 | 1 | 0.909 | 1 | 1 | 0.033 |
| SE | 92001 | ENSG00000119397 | CNTRL | chr9 | + | 121161855 | 121161971 | 121160142 | 121160302 | 121164942 | 121165100 | 125 | 172 | 121 | 160 | 154 | 4 | 1 | 0 | 1 | 1 | 27 | 89 | 31 | 11 | 8 | 0 | 0 | 0 | 0 | 0 | 265 | 149 | 1 | 1 | 0.946 | 0.99 | 1 | 0.989 | 0.989 | 1 | 1 | 1 | 1 | 1 | -0.017 |
| SE | 92002 | ENSG00000119397 | CNTRL | chr9 | + | 121161855 | 121161971 | 121160142 | 121160302 | 121167488 | 121167677 | 124 | 169 | 119 | 160 | 155 | 2 | 0 | 0 | 0 | 0 | 27 | 86 | 29 | 11 | 8 | 0 | 0 | 0 | 0 | 0 | 265 | 149 | 1 | 1 | 0.972 | 1 | 1 | 1 | 1 | 1 | 1 | 1 | 1 | 1 | -0.006 |
| SE | 92003 | ENSG00000119397 | CNTRL | chr9 | + | 121162053 | 121162271 | 121161855 | 121161971 | 121164942 | 121165100 | 307 | 336 | 294 | 326 | 448 | 1 | 3 | 2 | 0 | 0 | 55 | 135 | 84 | 52 | 53 | 0 | 3 | 2 | 0 | 0 | 298 | 149 | 1 | 1 | 0.994 | 0.982 | 0.987 | 1 | 1 | 1 | 0.957 | 0.955 | 1 | 1 | 0.01 |
| SE | 92004 | ENSG00000119397 | CNTRL | chr9 | + | 121162053 | 121162271 | 121161855 | 121161971 | 121167488 | 121167677 | 174 | 182 | 151 | 196 | 201 | 0 | 0 | 0 | 0 | 1 | 22 | 62 | 46 | 18 | 22 | 0 | 0 | 0 | 0 | 0 | 298 | 149 | 1 | 1 | 1 | 1 | 1 | 1 | 0.99 | 1 | 1 | 1 | 1 | 1 | -0.002 |
| SE | 92005 | ENSG00000119397 | CNTRL | chr9 | + | 121164942 | 121165100 | 121162053 | 121162271 | 121166106 | 121166180 | 284 | 297 | 285 | 257 | 538 | 0 | 0 | 0 | 2 | 0 | 49 | 149 | 88 | 65 | 56 | 0 | 0 | 0 | 0 | 0 | 298 | 149 | 1 | 1 | 1 | 1 | 1 | 0.985 | 1 | 1 | 1 | 1 | 1 | 1 | -0.003 |
| SE | 92006 | ENSG00000119397 | CNTRL | chr9 | + | 121166106 | 121166180 | 121162053 | 121162271 | 121167488 | 121167677 | 42 | 47 | 55 | 64 | 68 | 0 | 0 | 0 | 0 | 0 | 0 | 14 | 12 | 7 | 0 | 0 | 0 | 4 | 0 | 0 | 223 | 149 | 1 | 1 | 1 | 1 | 1 | 1 | 1 | NA | 1 | 0.667 | 1 | NA | 0.111 |
| SE | 92007 | ENSG00000119397 | CNTRL | chr9 | + | 121167488 | 121167677 | 121162053 | 121162271 | 121168095 | 121168321 | 41 | 55 | 50 | 46 | 23 | 0 | 2 | 0 | 0 | 0 | 0 | 4 | 27 | 4 | 6 | 0 | 0 | 0 | 0 | 0 | 298 | 149 | 1 | 1 | 1 | 0.932 | 1 | 1 | 1 | NA | 1 | 1 | 1 | 1 | -0.014 |
| SE | 92009 | ENSG00000119397 | CNTRL | chr9 | + | 121175017 | 121175224 | 121173674 | 121173737 | 121177162 | 121177610 | 135 | 129 | 158 | 168 | 275 | 0 | 2 | 1 | 0 | 1 | 0 | 44 | 32 | 12 | 16 | 0 | 0 | 0 | 0 | 0 | 298 | 149 | 1 | 1 | 1 | 0.97 | 0.988 | 1 | 0.993 | NA | 1 | 1 | 1 | 1 | -0.01 |
| SE | 92011 | ENSG00000147789 | ZNF7 | chr8 | + | 144828728 | 144828950 | 144827578 | 144827609 | 144829042 | 144829090 | 2 | 0 | 0 | 0 | 0 | 3 | 1 | 6 | 5 | 0 | 0 | 0 | 0 | 0 | 0 | 6 | 6 | 0 | 13 | 0 | 298 | 149 | 1 | 1 | 0.25 | 0 | 0 | 0 | NA | 0 | 0 | NA | 0 | NA | 0.062 |
| SE | 92012 | ENSG00000147789 | ZNF7 | chr8 | + | 144829042 | 144829090 | 144827517 | 144827609 | 144837390 | 144837507 | 0 | 0 | 2 | 3 | 0 | 0 | 0 | 2 | 1 | 0 | 0 | 0 | 0 | 1 | 0 | 0 | 0 | 0 | 0 | 0 | 197 | 149 | 1 | 1 | NA | NA | 0.431 | 0.694 | NA | NA | NA | NA | 1 | NA | -0.438 |
| SE | 92013 | ENSG00000147789 | ZNF7 | chr8 | + | 144829042 | 144829604 | 144827554 | 144827609 | 144837390 | 144837507 | 6 | 4 | 17 | 5 | 20 | 0 | 0 | 2 | 1 | 0 | 19 | 27 | 8 | 15 | 0 | 0 | 0 | 0 | 0 | 0 | 298 | 149 | 1 | 1 | 1 | 1 | 0.81 | 0.714 | 1 | 1 | 1 | 1 | 1 | NA | -0.095 |
| SE | 92014 | ENSG00000147789 | ZNF7 | chr8 | + | 144829426 | 144829604 | 144829042 | 144829090 | 144837390 | 144837507 | 6 | 3 | 23 | 3 | 21 | 0 | 0 | 3 | 3 | 0 | 19 | 32 | 8 | 3 | 0 | 0 | 0 | 0 | 1 | 0 | 298 | 149 | 1 | 1 | 1 | 1 | 0.793 | 0.333 | 1 | 1 | 1 | 1 | 0.6 | NA | -0.075 |
| SE | 92015 | ENSG00000147789 | ZNF7 | chr8 | + | 144829444 | 144829604 | 144829042 | 144829090 | 144837390 | 144837507 | 6 | 3 | 17 | 4 | 20 | 0 | 0 | 3 | 3 | 0 | 26 | 29 | 10 | 3 | 0 | 0 | 0 | 0 | 1 | 0 | 298 | 149 | 1 | 1 | 1 | 1 | 0.739 | 0.4 | 1 | 1 | 1 | 1 | 0.6 | NA | -0.072 |
| SE | 92016 | ENSG00000147789 | ZNF7 | chr8 | + | 144829477 | 144829604 | 144829041 | 144829090 | 144837390 | 144837507 | 11 | 3 | 21 | 5 | 24 | 0 | 0 | 3 | 3 | 0 | 25 | 33 | 8 | 3 | 0 | 0 | 0 | 0 | 1 | 0 | 276 | 149 | 1 | 1 | 1 | 1 | 0.791 | 0.474 | 1 | 1 | 1 | 1 | 0.618 | NA | -0.051 |
| SE | 92017 | ENSG00000147789 | ZNF7 | chr8 | + | 144829522 | 144829604 | 144829042 | 144829090 | 144837390 | 144837507 | 6 | 1 | 12 | 3 | 13 | 0 | 0 | 3 | 3 | 0 | 0 | 19 | 3 | 3 | 0 | 0 | 0 | 0 | 1 | 0 | 231 | 149 | 1 | 1 | 1 | 1 | 0.721 | 0.392 | 1 | NA | 1 | 1 | 0.659 | NA | -0.064 |
| SE | 92019 | ENSG00000135315 | CEP162 | chr6 | - | 84149561 | 84149703 | 84146686 | 84146785 | 84152544 | 84153179 | 50 | 52 | 46 | 55 | 92 | 5 | 4 | 0 | 6 | 2 | 24 | 25 | 7 | 10 | 0 | 0 | 0 | 0 | 0 | 0 | 291 | 149 | 1 | 1 | 0.837 | 0.869 | 1 | 0.824 | 0.959 | 1 | 1 | 1 | 1 | NA | -0.102 |
| SE | 92020 | ENSG00000135315 | CEP162 | chr6 | - | 84169327 | 84169433 | 84163143 | 84163270 | 84171605 | 84171718 | 65 | 81 | 57 | 87 | 118 | 1 | 1 | 1 | 1 | 13 | 27 | 19 | 2 | 0 | 3 | 0 | 0 | 0 | 0 | 1 | 255 | 149 | 1 | 1 | 0.974 | 0.979 | 0.971 | 0.981 | 0.841 | 1 | 1 | 1 | NA | 0.637 | 0.04 |
| SE | 92021 | ENSG00000135315 | CEP162 | chr6 | - | 84171605 | 84171718 | 84163143 | 84163270 | 84174047 | 84174188 | 34 | 30 | 29 | 40 | 63 | 0 | 0 | 0 | 4 | 0 | 31 | 7 | 10 | 17 | 17 | 0 | 0 | 0 | 0 | 0 | 262 | 149 | 1 | 1 | 1 | 1 | 1 | 0.85 | 1 | 1 | 1 | 1 | 1 | 1 | -0.03 |
| SE | 92022 | ENSG00000135315 | CEP162 | chr6 | - | 84175209 | 84175347 | 84174726 | 84174954 | 84185186 | 84185448 | 33 | 23 | 12 | 40 | 37 | 0 | 1 | 0 | 1 | 0 | 11 | 16 | 2 | 4 | 0 | 0 | 0 | 0 | 0 | 0 | 287 | 149 | 1 | 1 | 1 | 0.923 | 1 | 0.954 | 1 | 1 | 1 | 1 | 1 | NA | -0.025 |
| SE | 92023 | ENSG00000135315 | CEP162 | chr6 | - | 84175213 | 84175347 | 84174726 | 84174954 | 84185186 | 84185448 | 87 | 79 | 33 | 81 | 80 | 0 | 1 | 0 | 1 | 0 | 22 | 24 | 10 | 28 | 0 | 0 | 0 | 0 | 0 | 0 | 283 | 149 | 1 | 1 | 1 | 0.977 | 1 | 0.977 | 1 | 1 | 1 | 1 | 1 | NA | -0.009 |
| SE | 92024 | ENSG00000135315 | CEP162 | chr6 | - | 84201736 | 84201767 | 84200788 | 84200905 | 84203980 | 84204096 | 33 | 29 | 17 | 44 | 83 | 0 | 1 | 0 | 0 | 2 | 35 | 19 | 11 | 9 | 31 | 0 | 0 | 0 | 2 | 0 | 180 | 149 | 1 | 1 | 1 | 0.96 | 1 | 1 | 0.972 | 1 | 1 | 1 | 0.788 | 1 | 0.029 |
| SE | 92025 | ENSG00000135315 | CEP162 | chr6 | - | 84203980 | 84204096 | 84200788 | 84200905 | 84212956 | 84213024 | 22 | 14 | 11 | 17 | 33 | 0 | 0 | 0 | 2 | 0 | 0 | 16 | 2 | 5 | 3 | 0 | 0 | 0 | 0 | 0 | 265 | 149 | 1 | 1 | 1 | 1 | 1 | 0.827 | 1 | NA | 1 | 1 | 1 | 1 | -0.035 |
| SE | 92027 | ENSG00000135315 | CEP162 | chr6 | - | 84217798 | 84217860 | 84215775 | 84215922 | 84221056 | 84221171 | 0 | 0 | 1 | 0 | 1 | 23 | 29 | 12 | 19 | 23 | 0 | 0 | 0 | 0 | 0 | 0 | 6 | 0 | 0 | 3 | 211 | 149 | 1 | 1 | 0 | 0 | 0.056 | 0 | 0.03 | NA | 0 | NA | NA | 0 | 0.017 |
| SE | 92029 | ENSG00000135315 | CEP162 | chr6 | - | 84219102 | 84219214 | 84217798 | 84217860 | 84221056 | 84221171 | 9 | 5 | 9 | 5 | 10 | 0 | 0 | 1 | 0 | 0 | 10 | 8 | 0 | 4 | 0 | 0 | 0 | 0 | 0 | 0 | 261 | 149 | 1 | 1 | 1 | 1 | 0.837 | 1 | 1 | 1 | 1 | NA | 1 | NA | -0.033 |
| SE | 92030 | ENSG00000135315 | CEP162 | chr6 | - | 84226336 | 84226452 | 84221056 | 84221171 | 84227579 | 84227608 | 22 | 10 | 7 | 9 | 34 | 6 | 7 | 3 | 2 | 3 | 20 | 4 | 8 | 5 | 0 | 0 | 0 | 0 | 0 | 0 | 265 | 149 | 1 | 1 | 0.673 | 0.445 | 0.567 | 0.717 | 0.864 | 1 | 1 | 1 | 1 | NA | -0.347 |
| SE | 92031 | ENSG00000196150 | ZNF250 | chr8 | - | 144886839 | 144886902 | 144876950 | 144882836 | 144889580 | 144889694 | 16 | 13 | 3 | 13 | 22 | 0 | 1 | 0 | 0 | 0 | 11 | 0 | 6 | 0 | 7 | 0 | 0 | 0 | 0 | 0 | 212 | 149 | 1 | 1 | 1 | 0.901 | 1 | 1 | 1 | 1 | NA | 1 | NA | 1 | -0.02 |
| SE | 92032 | ENSG00000196150 | ZNF250 | chr8 | - | 144886839 | 144886902 | 144880364 | 144880458 | 144889580 | 144889694 | 4 | 10 | 4 | 6 | 4 | 0 | 0 | 0 | 0 | 0 | 0 | 4 | 1 | 4 | 10 | 0 | 0 | 0 | 4 | 0 | 212 | 149 | 1 | 1 | 1 | 1 | 1 | 1 | 1 | NA | 1 | 1 | 0.413 | 1 | 0.147 |
| SE | 92033 | ENSG00000196150 | ZNF250 | chr8 | - | 144889580 | 144889694 | 144886839 | 144886902 | 144889932 | 144890059 | 18 | 26 | 11 | 8 | 19 | 1 | 0 | 0 | 0 | 0 | 29 | 0 | 24 | 12 | 17 | 0 | 0 | 2 | 1 | 4 | 263 | 149 | 1 | 1 | 0.911 | 1 | 1 | 1 | 1 | 1 | NA | 0.872 | 0.872 | 0.707 | 0.119 |
| SE | 92035 | ENSG00000258526 | NA | chr14 | + | 39433487 | 39433729 | 39432598 | 39432675 | 39470317 | 39470400 | 0 | 0 | 0 | 1 | 0 | 1 | 0 | 3 | 1 | 0 | 0 | 0 | 0 | 0 | 0 | 0 | 0 | 5 | 0 | 0 | 298 | 149 | 1 | 1 | 0 | NA | 0 | 0.333 | NA | NA | NA | 0 | NA | NA | 0.111 |
| SE | 92042 | ENSG00000258526 | NA | chr14 | + | 40264814 | 40264929 | 40249687 | 40249753 | 40386251 | 40386443 | 1 | 0 | 0 | 0 | 5 | 0 | 0 | 0 | 0 | 0 | 0 | 8 | 0 | 0 | 7 | 0 | 0 | 1 | 0 | 0 | 264 | 149 | 1 | 1 | 1 | NA | NA | NA | 1 | NA | 1 | 0 | NA | 1 | 0.333 |
| SE | 92043 | ENSG00000258526 | NA | chr14 | + | 40318435 | 40318556 | 40264814 | 40264929 | 40371520 | 40371578 | 2 | 0 | 2 | 4 | 0 | 0 | 1 | 0 | 0 | 0 | 0 | 0 | 0 | 0 | 7 | 0 | 0 | 0 | 0 | 0 | 270 | 149 | 1 | 1 | 1 | 0 | 1 | 1 | NA | NA | NA | NA | NA | 1 | -0.25 |
| SE | 92044 | ENSG00000258526 | NA | chr14 | + | 40318435 | 40318556 | 40264814 | 40264929 | 40386251 | 40386443 | 2 | 1 | 2 | 6 | 0 | 0 | 0 | 0 | 0 | 3 | 0 | 0 | 0 | 0 | 7 | 0 | 4 | 0 | 0 | 0 | 270 | 149 | 1 | 1 | 1 | 1 | 1 | 1 | 0 | NA | 0 | NA | NA | 1 | 0.3 |
| SE | 92045 | ENSG00000258526 | NA | chr14 | + | 40371520 | 40371578 | 40264814 | 40264929 | 40386251 | 40388040 | 0 | 1 | 0 | 0 | 0 | 0 | 0 | 0 | 0 | 3 | 0 | 0 | 0 | 0 | 0 | 0 | 4 | 0 | 0 | 0 | 207 | 149 | 1 | 1 | NA | 1 | NA | NA | 0 | NA | 0 | NA | NA | NA | 0.5 |
| SE | 92047 | ENSG00000182197 | EXT1 | chr8 | - | 117812871 | 117812961 | 117807216 | 117807377 | 117818434 | 117818530 | 28 | 13 | 26 | 33 | 44 | 0 | 0 | 0 | 0 | 0 | 94 | 26 | 29 | 34 | 26 | 0 | 0 | 0 | 1 | 0 | 239 | 149 | 1 | 1 | 1 | 1 | 1 | 1 | 1 | 1 | 1 | 1 | 0.955 | 1 | 0.009 |
| SE | 92049 | ENSG00000182197 | EXT1 | chr8 | - | 117819675 | 117819794 | 117818434 | 117818530 | 117822464 | 117822597 | 46 | 31 | 30 | 38 | 67 | 0 | 0 | 0 | 0 | 0 | 36 | 10 | 38 | 37 | 86 | 3 | 0 | 0 | 1 | 0 | 268 | 149 | 1 | 1 | 1 | 1 | 1 | 1 | 1 | 0.87 | 1 | 1 | 0.954 | 1 | 0.035 |
| SE | 92050 | ENSG00000182197 | EXT1 | chr8 | - | 117822464 | 117822597 | 117818434 | 117818530 | 117830229 | 117830349 | 26 | 24 | 23 | 16 | 32 | 0 | 0 | 1 | 0 | 0 | 41 | 14 | 24 | 63 | 21 | 0 | 0 | 0 | 0 | 0 | 282 | 149 | 1 | 1 | 1 | 1 | 0.924 | 1 | 1 | 1 | 1 | 1 | 1 | 1 | -0.015 |
| SE | 92051 | ENSG00000182197 | EXT1 | chr8 | - | 117822464 | 117822597 | 117819675 | 117819794 | 117835443 | 117835551 | 7 | 7 | 5 | 3 | 29 | 0 | 0 | 0 | 0 | 0 | 0 | 2 | 3 | 16 | 17 | 0 | 0 | 0 | 1 | 0 | 282 | 149 | 1 | 1 | 1 | 1 | 1 | 1 | 1 | NA | 1 | 1 | 0.894 | 1 | 0.026 |
| SE | 92052 | ENSG00000182197 | EXT1 | chr8 | - | 117830229 | 117830349 | 117822464 | 117822597 | 117835443 | 117835551 | 73 | 61 | 57 | 57 | 63 | 0 | 0 | 0 | 0 | 0 | 89 | 29 | 64 | 126 | 82 | 0 | 0 | 2 | 0 | 0 | 269 | 149 | 1 | 1 | 1 | 1 | 1 | 1 | 1 | 1 | 1 | 0.947 | 1 | 1 | 0.011 |
| SE | 92054 | ENSG00000182197 | EXT1 | chr8 | - | 117835443 | 117835551 | 117830229 | 117830349 | 117837107 | 117837201 | 67 | 54 | 57 | 54 | 50 | 1 | 0 | 0 | 0 | 0 | 73 | 35 | 38 | 114 | 49 | 0 | 0 | 0 | 0 | 0 | 257 | 149 | 1 | 1 | 0.975 | 1 | 1 | 1 | 1 | 1 | 1 | 1 | 1 | 1 | -0.005 |
| SE | 92060 | ENSG00000136573 | BLK | chr8 | + | 11555331 | 11555484 | 11554742 | 11554889 | 11556657 | 11556788 | 108 | 54 | 93 | 106 | 66 | 0 | 0 | 0 | 0 | 0 | 148 | 144 | 72 | 43 | 191 | 0 | 0 | 0 | 2 | 0 | 298 | 149 | 1 | 1 | 1 | 1 | 1 | 1 | 1 | 1 | 1 | 1 | 0.915 | 1 | 0.017 |
| SE | 92062 | ENSG00000165934 | CPSF2 | chr14 | + | 92126121 | 92126180 | 92121968 | 92122128 | 92130950 | 92131133 | 77 | 45 | 42 | 39 | 85 | 0 | 0 | 0 | 0 | 0 | 4 | 60 | 28 | 12 | 14 | 0 | 0 | 1 | 0 | 0 | 208 | 149 | 1 | 1 | 1 | 1 | 1 | 1 | 1 | 1 | 1 | 0.953 | 1 | 1 | 0.009 |
| SE | 92064 | ENSG00000165934 | CPSF2 | chr14 | + | 92142163 | 92142351 | 92138231 | 92138347 | 92143003 | 92143294 | 254 | 228 | 196 | 215 | 358 | 0 | 0 | 0 | 0 | 1 | 68 | 194 | 97 | 124 | 61 | 0 | 2 | 0 | 0 | 0 | 298 | 149 | 1 | 1 | 1 | 1 | 1 | 1 | 0.994 | 1 | 0.98 | 1 | 1 | 1 | 0.003 |
| SE | 92065 | ENSG00000165934 | CPSF2 | chr14 | + | 92156478 | 92156631 | 92155122 | 92155323 | 92157658 | 92157884 | 483 | 371 | 362 | 432 | 654 | 1 | 0 | 0 | 0 | 2 | 119 | 340 | 279 | 248 | 19 | 0 | 1 | 0 | 0 | 0 | 298 | 149 | 1 | 1 | 0.996 | 1 | 1 | 1 | 0.994 | 1 | 0.994 | 1 | 1 | 1 | -0.001 |
| SE | 92068 | ENSG00000108010 | GLRX3 | chr10 | + | 130169432 | 130169490 | 130166918 | 130166980 | 130171583 | 130171636 | 236 | 158 | 235 | 267 | 361 | 0 | 0 | 0 | 0 | 2 | 445 | 260 | 255 | 280 | 342 | 0 | 0 | 1 | 0 | 0 | 207 | 149 | 1 | 1 | 1 | 1 | 1 | 1 | 0.992 | 1 | 1 | 0.995 | 1 | 1 | -0.001 |
| SE | 92069 | ENSG00000108010 | GLRX3 | chr10 | + | 130174866 | 130174906 | 130171583 | 130171636 | 130179341 | 130179420 | 67 | 62 | 79 | 90 | 113 | 0 | 0 | 2 | 0 | 0 | 152 | 90 | 108 | 125 | 127 | 0 | 0 | 0 | 0 | 0 | 189 | 149 | 1 | 1 | 1 | 1 | 0.969 | 1 | 1 | 1 | 1 | 1 | 1 | 1 | -0.006 |
| SE | 92071 | ENSG00000108010 | GLRX3 | chr10 | + | 130179341 | 130179420 | 130174996 | 130175089 | 130180112 | 130180321 | 125 | 75 | 101 | 106 | 200 | 0 | 0 | 0 | 0 | 1 | 288 | 125 | 156 | 148 | 150 | 0 | 0 | 0 | 0 | 0 | 228 | 149 | 1 | 1 | 1 | 1 | 1 | 1 | 0.992 | 1 | 1 | 1 | 1 | 1 | -0.002 |
| SE | 92072 | ENSG00000226752 | CUTALP | chr9 | + | 120844992 | 120845196 | 120843099 | 120843439 | 120847293 | 120847353 | 6 | 15 | 5 | 7 | 47 | 0 | 0 | 0 | 0 | 3 | 7 | 23 | 14 | 4 | 29 | 0 | 0 | 0 | 0 | 0 | 298 | 149 | 1 | 1 | 1 | 1 | 1 | 1 | 0.887 | 1 | 1 | 1 | 1 | 1 | -0.023 |
| SE | 92074 | ENSG00000226752 | CUTALP | chr9 | + | 120844992 | 120845196 | 120844712 | 120844899 | 120851237 | 120851355 | 12 | 37 | 9 | 15 | 68 | 0 | 0 | 1 | 0 | 0 | 17 | 21 | 51 | 54 | 25 | 0 | 0 | 0 | 0 | 0 | 298 | 149 | 1 | 1 | 1 | 1 | 0.818 | 1 | 1 | 1 | 1 | 1 | 1 | 1 | -0.036 |
| SE | 92076 | ENSG00000226752 | CUTALP | chr9 | + | 120847293 | 120847353 | 120844712 | 120844899 | 120851237 | 120851355 | 13 | 25 | 10 | 5 | 47 | 0 | 0 | 1 | 0 | 0 | 20 | 16 | 45 | 24 | 14 | 0 | 0 | 0 | 0 | 0 | 209 | 149 | 1 | 1 | 1 | 1 | 0.877 | 1 | 1 | 1 | 1 | 1 | 1 | 1 | -0.025 |
| SE | 92077 | ENSG00000226752 | CUTALP | chr9 | + | 120847293 | 120847353 | 120844873 | 120845196 | 120851237 | 120851355 | 19 | 37 | 14 | 7 | 86 | 0 | 0 | 0 | 0 | 0 | 27 | 38 | 56 | 27 | 24 | 0 | 0 | 0 | 2 | 0 | 209 | 149 | 1 | 1 | 1 | 1 | 1 | 1 | 1 | 1 | 1 | 1 | 0.906 | 1 | 0.019 |
| SE | 92080 | ENSG00000226752 | CUTALP | chr9 | + | 120850362 | 120850657 | 120847293 | 120847353 | 120851237 | 120851524 | 0 | 1 | 0 | 0 | 0 | 17 | 35 | 22 | 10 | 92 | 0 | 0 | 1 | 0 | 0 | 27 | 28 | 61 | 35 | 14 | 298 | 149 | 1 | 1 | 0 | 0.014 | 0 | 0 | 0 | 0 | 0 | 0.008 | 0 | 0 | 0.001 |
| SE | 92084 | ENSG00000167325 | RRM1 | chr11 | + | 4105605 | 4105691 | 4101992 | 4102081 | 4106045 | 4106223 | 0 | 0 | 0 | 0 | 0 | 73 | 32 | 55 | 55 | 59 | 0 | 0 | 2 | 0 | 0 | 54 | 31 | 53 | 69 | 8 | 235 | 149 | 1 | 1 | 0 | 0 | 0 | 0 | 0 | 0 | 0 | 0.023 | 0 | 0 | -0.005 |
| SE | 92086 | ENSG00000167325 | RRM1 | chr11 | + | 4118319 | 4118461 | 4111899 | 4112062 | 4119844 | 4119928 | 113 | 54 | 174 | 121 | 202 | 0 | 3 | 0 | 0 | 0 | 57 | 50 | 90 | 132 | 24 | 0 | 0 | 0 | 0 | 0 | 291 | 149 | 1 | 1 | 1 | 0.902 | 1 | 1 | 1 | 1 | 1 | 1 | 1 | 1 | -0.02 |
| SE | 92088 | ENSG00000167325 | RRM1 | chr11 | + | 4132285 | 4132421 | 4129073 | 4129150 | 4133562 | 4133658 | 215 | 106 | 254 | 206 | 226 | 0 | 0 | 0 | 0 | 2 | 74 | 134 | 195 | 243 | 4 | 0 | 0 | 1 | 5 | 0 | 285 | 149 | 1 | 1 | 1 | 1 | 1 | 1 | 0.983 | 1 | 1 | 0.99 | 0.962 | 1 | 0.006 |
| SE | 92093 | ENSG00000036448 | MYOM2 | chr8 | + | 2129126 | 2129232 | 2124178 | 2124217 | 2140722 | 2140818 | 290 | 963 | 959 | 1537 | 12 | 0 | 0 | 0 | 1 | 0 | 23 | 19 | 10 | 98 | 0 | 0 | 0 | 0 | 0 | 0 | 255 | 149 | 1 | 1 | 1 | 1 | 1 | 0.999 | 1 | 1 | 1 | 1 | 1 | NA | 0 |
| SE | 92094 | ENSG00000036448 | MYOM2 | chr8 | + | 2141140 | 2141177 | 2140722 | 2140886 | 2142374 | 2142397 | 177 | 343 | 545 | 672 | 3 | 0 | 1 | 0 | 3 | 0 | 0 | 12 | 2 | 10 | 0 | 0 | 0 | 0 | 0 | 0 | 186 | 149 | 1 | 1 | 1 | 0.996 | 1 | 0.994 | 1 | NA | 1 | 1 | 1 | NA | -0.002 |
| SE | 92095 | ENSG00000036448 | MYOM2 | chr8 | + | 2141140 | 2141177 | 2140722 | 2140886 | 2143400 | 2143456 | 34 | 48 | 104 | 147 | 1 | 9 | 41 | 45 | 54 | 2 | 0 | 1 | 8 | 2 | 0 | 0 | 7 | 0 | 0 | 0 | 186 | 149 | 1 | 1 | 0.752 | 0.484 | 0.649 | 0.686 | 0.286 | NA | 0.103 | 1 | 1 | NA | -0.13 |
| SE | 92096 | ENSG00000036448 | MYOM2 | chr8 | + | 2142374 | 2142397 | 2140722 | 2140886 | 2143400 | 2143456 | 16 | 38 | 24 | 56 | 0 | 9 | 41 | 45 | 54 | 2 | 0 | 0 | 0 | 0 | 0 | 0 | 7 | 0 | 0 | 0 | 172 | 149 | 1 | 1 | 0.606 | 0.445 | 0.316 | 0.473 | 0 | NA | 0 | NA | NA | NA | 0.368 |
| SE | 92097 | ENSG00000036448 | MYOM2 | chr8 | + | 2142374 | 2142397 | 2141140 | 2141177 | 2143400 | 2143456 | 177 | 363 | 511 | 662 | 2 | 14 | 13 | 36 | 70 | 0 | 0 | 11 | 2 | 10 | 0 | 0 | 0 | 8 | 2 | 0 | 172 | 149 | 1 | 1 | 0.916 | 0.96 | 0.925 | 0.891 | 1 | NA | 1 | 0.178 | 0.812 | NA | 0.275 |
| SE | 92098 | ENSG00000076554 | TPD52 | chr8 | - | 80044166 | 80044208 | 80042619 | 80042668 | 80050444 | 80050471 | 0 | 0 | 0 | 2 | 0 | 3 | 0 | 1 | 0 | 0 | 0 | 0 | 0 | 0 | 0 | 0 | 1 | 4 | 0 | 0 | 191 | 149 | 1 | 1 | 0 | NA | 0 | 1 | NA | NA | 0 | 0 | NA | NA | 0.333 |
| SE | 92099 | ENSG00000076554 | TPD52 | chr8 | - | 80044166 | 80044208 | 80042619 | 80042668 | 80051526 | 80051628 | 0 | 0 | 0 | 4 | 0 | 215 | 59 | 130 | 118 | 168 | 0 | 0 | 0 | 0 | 0 | 44 | 147 | 104 | 40 | 55 | 191 | 149 | 1 | 1 | 0 | 0 | 0 | 0.026 | 0 | 0 | 0 | 0 | 0 | 0 | 0.005 |
| SE | 92100 | ENSG00000076554 | TPD52 | chr8 | - | 80050345 | 80050471 | 80044166 | 80044208 | 80051526 | 80051628 | 0 | 0 | 1 | 1 | 0 | 0 | 0 | 0 | 2 | 0 | 0 | 1 | 0 | 0 | 0 | 0 | 0 | 0 | 0 | 0 | 275 | 149 | 1 | 1 | NA | NA | 1 | 0.213 | NA | NA | 1 | NA | NA | NA | -0.393 |
| SE | 92102 | ENSG00000076554 | TPD52 | chr8 | - | 80050444 | 80050471 | 80042619 | 80042668 | 80051526 | 80051628 | 3 | 0 | 2 | 1 | 0 | 215 | 59 | 130 | 118 | 168 | 0 | 2 | 4 | 0 | 0 | 44 | 147 | 104 | 40 | 55 | 176 | 149 | 1 | 1 | 0.012 | 0 | 0.013 | 0.007 | 0 | 0 | 0.011 | 0.032 | 0 | 0 | -0.002 |
| SE | 92103 | ENSG00000076554 | TPD52 | chr8 | - | 80050444 | 80050471 | 80044166 | 80044208 | 80051526 | 80051628 | 0 | 0 | 1 | 1 | 0 | 0 | 0 | 0 | 2 | 0 | 0 | 1 | 0 | 0 | 0 | 0 | 0 | 0 | 0 | 0 | 176 | 149 | 1 | 1 | NA | NA | 1 | 0.297 | NA | NA | 1 | NA | NA | NA | -0.352 |
| SE | 92104 | ENSG00000076554 | TPD52 | chr8 | - | 80052605 | 80052675 | 80051526 | 80051628 | 80053281 | 80053353 | 2 | 0 | 3 | 5 | 0 | 178 | 60 | 123 | 136 | 190 | 0 | 0 | 0 | 0 | 0 | 47 | 99 | 83 | 31 | 63 | 219 | 149 | 1 | 1 | 0.008 | 0 | 0.016 | 0.024 | 0 | 0 | 0 | 0 | 0 | 0 | 0.01 |
| SE | 92105 | ENSG00000076554 | TPD52 | chr8 | - | 80064477 | 80064593 | 80053278 | 80053430 | 80119814 | 80119884 | 40 | 16 | 40 | 35 | 59 | 1 | 0 | 0 | 0 | 0 | 24 | 14 | 22 | 7 | 21 | 0 | 0 | 0 | 0 | 0 | 265 | 149 | 1 | 1 | 0.957 | 1 | 1 | 1 | 1 | 1 | 1 | 1 | 1 | 1 | -0.009 |
| SE | 92106 | ENSG00000076554 | TPD52 | chr8 | - | 80064477 | 80064593 | 80053278 | 80053430 | 80171424 | 80171558 | 51 | 20 | 50 | 51 | 60 | 0 | 0 | 1 | 0 | 0 | 24 | 18 | 24 | 10 | 22 | 0 | 0 | 0 | 0 | 0 | 265 | 149 | 1 | 1 | 1 | 1 | 0.966 | 1 | 1 | 1 | 1 | 1 | 1 | 1 | -0.007 |
| SE | 92108 | ENSG00000076554 | TPD52 | chr8 | - | 80119814 | 80119884 | 80064477 | 80064593 | 80171424 | 80171471 | 0 | 0 | 1 | 0 | 0 | 11 | 4 | 11 | 16 | 1 | 0 | 0 | 0 | 0 | 0 | 0 | 4 | 2 | 3 | 1 | 219 | 149 | 1 | 1 | 0 | 0 | 0.058 | 0 | 0 | NA | 0 | 0 | 0 | 0 | 0.012 |
| SE | 92109 | ENSG00000120162 | MOB3B | chr9 | - | 27359033 | 27359236 | 27325208 | 27330616 | 27455132 | 27455748 | 26 | 14 | 24 | 29 | 43 | 0 | 0 | 0 | 1 | 0 | 39 | 58 | 16 | 19 | 36 | 0 | 0 | 0 | 0 | 0 | 298 | 149 | 1 | 1 | 1 | 1 | 1 | 0.935 | 1 | 1 | 1 | 1 | 1 | 1 | -0.013 |
| SE | 92115 | ENSG00000066117 | SMARCD1 | chr12 | + | 50088520 | 50088637 | 50086755 | 50086878 | 50089883 | 50089985 | 213 | 150 | 178 | 282 | 308 | 0 | 0 | 0 | 1 | 0 | 398 | 424 | 310 | 320 | 427 | 0 | 0 | 0 | 1 | 0 | 266 | 149 | 1 | 1 | 1 | 1 | 1 | 0.994 | 1 | 1 | 1 | 1 | 0.994 | 1 | 0 |
| SE | 92116 | ENSG00000066117 | SMARCD1 | chr12 | + | 50089883 | 50089985 | 50088520 | 50088637 | 50090240 | 50090326 | 298 | 198 | 245 | 369 | 450 | 0 | 0 | 0 | 1 | 0 | 445 | 522 | 354 | 350 | 484 | 2 | 0 | 0 | 0 | 8 | 251 | 149 | 1 | 1 | 1 | 1 | 1 | 1 | 1 | 0.992 | 1 | 1 | 1 | 0.973 | 0.007 |
| SE | 92127 | ENSG00000124380 | SNRNP27 | chr2 | + | 69904511 | 69904633 | 69904253 | 69904425 | 69905265 | 69905575 | 37 | 19 | 13 | 12 | 16 | 0 | 2 | 0 | 1 | 0 | 0 | 17 | 13 | 13 | 7 | 0 | 0 | 0 | 0 | 0 | 271 | 149 | 1 | 1 | 1 | 0.839 | 1 | 0.868 | 1 | NA | 1 | 1 | 1 | 1 | -0.059 |
| SE | 92128 | ENSG00000164935 | DCSTAMP | chr8 | + | 104340286 | 104340440 | 104339825 | 104339862 | 104348540 | 104348931 | 0 | 2 | 0 | 0 | 0 | 0 | 0 | 0 | 1 | 0 | 16 | 0 | 0 | 0 | 0 | 0 | 0 | 7 | 0 | 0 | 298 | 149 | 1 | 1 | NA | 1 | NA | 0 | NA | 1 | NA | 0 | NA | NA | 0 |
| SE | 92133 | ENSG00000108666 | C17orf75 | chr17 | - | 32333416 | 32333520 | 32324564 | 32324774 | 32334468 | 32334605 | 37 | 27 | 47 | 43 | 52 | 0 | 0 | 2 | 0 | 0 | 22 | 12 | 26 | 16 | 2 | 0 | 0 | 0 | 0 | 0 | 253 | 149 | 1 | 1 | 1 | 1 | 0.933 | 1 | 1 | 1 | 1 | 1 | 1 | 1 | -0.013 |
| SE | 92139 | ENSG00000249898 | MCPH1-AS1 | chr8 | - | 6634594 | 6635419 | 6627021 | 6627310 | 6669988 | 6670240 | 2 | 16 | 0 | 7 | 21 | 0 | 0 | 0 | 0 | 0 | 0 | 7 | 1 | 3 | 5 | 0 | 0 | 0 | 1 | 0 | 298 | 149 | 1 | 1 | 1 | 1 | NA | 1 | 1 | NA | 1 | 1 | 0.6 | 1 | 0.1 |
| SE | 92141 | ENSG00000249898 | MCPH1-AS1 | chr8 | - | 6669988 | 6670240 | 6627150 | 6627310 | 6708140 | 6708179 | 1 | 1 | 4 | 6 | 3 | 0 | 0 | 0 | 1 | 0 | 0 | 5 | 5 | 3 | 28 | 0 | 5 | 1 | 0 | 15 | 298 | 149 | 1 | 1 | 1 | 1 | 1 | 0.75 | 1 | NA | 0.333 | 0.714 | 1 | 0.483 | 0.318 |
| SE | 92145 | ENSG00000232104 | RFX3-AS1 | chr9 | + | 3568793 | 3568889 | 3546777 | 3546912 | 3606692 | 3606738 | 3 | 5 | 1 | 1 | 2 | 0 | 0 | 0 | 0 | 0 | 0 | 5 | 6 | 0 | 3 | 0 | 0 | 8 | 0 | 0 | 245 | 149 | 1 | 1 | 1 | 1 | 1 | 1 | 1 | NA | 1 | 0.313 | NA | 1 | 0.229 |
| SE | 92146 | ENSG00000232104 | RFX3-AS1 | chr9 | + | 3602522 | 3602584 | 3546777 | 3546912 | 3606692 | 3606738 | 20 | 7 | 10 | 12 | 7 | 0 | 0 | 0 | 0 | 0 | 0 | 12 | 0 | 8 | 0 | 0 | 0 | 8 | 0 | 0 | 211 | 149 | 1 | 1 | 1 | 1 | 1 | 1 | 1 | NA | 1 | 0 | 1 | NA | 0.333 |
| SE | 92147 | ENSG00000232104 | RFX3-AS1 | chr9 | + | 3602522 | 3602584 | 3568793 | 3568889 | 3606692 | 3606738 | 16 | 8 | 8 | 18 | 6 | 0 | 1 | 0 | 0 | 0 | 0 | 13 | 0 | 3 | 0 | 0 | 0 | 0 | 0 | 0 | 211 | 149 | 1 | 1 | 1 | 0.85 | 1 | 1 | 1 | NA | 1 | NA | 1 | NA | -0.03 |
| SE | 92148 | ENSG00000232104 | RFX3-AS1 | chr9 | + | 3671473 | 3671645 | 3668164 | 3668408 | 3674410 | 3674648 | 32 | 40 | 13 | 27 | 23 | 1 | 0 | 0 | 0 | 0 | 3 | 10 | 3 | 17 | 0 | 0 | 0 | 0 | 0 | 0 | 298 | 149 | 1 | 1 | 0.941 | 1 | 1 | 1 | 1 | 1 | 1 | 1 | 1 | NA | -0.012 |
| SE | 92149 | ENSG00000232104 | RFX3-AS1 | chr9 | + | 3674410 | 3674648 | 3668164 | 3668408 | 3690836 | 3690959 | 5 | 7 | 7 | 3 | 5 | 0 | 1 | 0 | 0 | 0 | 6 | 2 | 6 | 0 | 0 | 0 | 0 | 0 | 0 | 0 | 298 | 149 | 1 | 1 | 1 | 0.778 | 1 | 1 | 1 | 1 | 1 | 1 | NA | NA | -0.044 |
| SE | 92150 | ENSG00000177599 | ZNF491 | chr19 | + | 11804541 | 11804667 | 11798522 | 11798727 | 11805946 | 11808503 | 1 | 2 | 4 | 1 | 10 | 0 | 0 | 0 | 4 | 0 | 0 | 0 | 0 | 2 | 0 | 0 | 0 | 0 | 0 | 0 | 275 | 149 | 1 | 1 | 1 | 1 | 1 | 0.119 | 1 | NA | NA | NA | 1 | NA | -0.176 |
| SE | 92151 | ENSG00000035499 | DEPDC1B | chr5 | - | 60599074 | 60599260 | 60596911 | 60597914 | 60603390 | 60603567 | 8 | 0 | 8 | 15 | 15 | 0 | 0 | 0 | 2 | 0 | 0 | 4 | 16 | 37 | 0 | 0 | 0 | 0 | 0 | 0 | 298 | 149 | 1 | 1 | 1 | NA | 1 | 0.789 | 1 | NA | 1 | 1 | 1 | NA | -0.053 |
| SE | 92152 | ENSG00000035499 | DEPDC1B | chr5 | - | 60605689 | 60605856 | 60603390 | 60603567 | 60638749 | 60638890 | 3 | 0 | 8 | 0 | 11 | 0 | 0 | 0 | 0 | 0 | 0 | 6 | 26 | 15 | 0 | 0 | 5 | 0 | 0 | 0 | 298 | 149 | 1 | 1 | 1 | NA | 1 | NA | 1 | NA | 0.375 | 1 | 1 | NA | 0.208 |
| SE | 92153 | ENSG00000035499 | DEPDC1B | chr5 | - | 60687724 | 60687838 | 60686961 | 60687227 | 60700045 | 60700190 | 0 | 0 | 0 | 0 | 0 | 7 | 0 | 3 | 4 | 0 | 10 | 0 | 0 | 7 | 4 | 0 | 3 | 6 | 3 | 3 | 263 | 149 | 1 | 1 | 0 | NA | 0 | 0 | NA | 1 | 0 | 0 | 0.569 | 0.43 | -0.4 |
| SE | 92155 | ENSG00000090612 | ZNF268 | chr12 | + | 133187871 | 133188072 | 133181945 | 133182030 | 133191488 | 133191615 | 34 | 36 | 24 | 35 | 27 | 0 | 0 | 0 | 0 | 2 | 0 | 32 | 14 | 19 | 10 | 0 | 0 | 0 | 0 | 0 | 298 | 149 | 1 | 1 | 1 | 1 | 1 | 1 | 0.871 | NA | 1 | 1 | 1 | 1 | -0.026 |
| SE | 92156 | ENSG00000090612 | ZNF268 | chr12 | + | 133187871 | 133188072 | 133181945 | 133182030 | 133191907 | 133192003 | 22 | 14 | 14 | 21 | 14 | 0 | 0 | 0 | 0 | 0 | 0 | 23 | 7 | 13 | 6 | 0 | 2 | 0 | 5 | 0 | 298 | 149 | 1 | 1 | 1 | 1 | 1 | 1 | 1 | NA | 0.852 | 1 | 0.565 | 1 | 0.146 |
| SE | 92157 | ENSG00000090612 | ZNF268 | chr12 | + | 133191488 | 133191615 | 133181945 | 133182030 | 133191907 | 133192003 | 13 | 16 | 11 | 17 | 14 | 0 | 0 | 0 | 0 | 0 | 5 | 5 | 7 | 0 | 0 | 0 | 2 | 0 | 5 | 0 | 276 | 149 | 1 | 1 | 1 | 1 | 1 | 1 | 1 | 1 | 0.574 | 1 | 0 | NA | 0.357 |
| SE | 92158 | ENSG00000090612 | ZNF268 | chr12 | + | 133191488 | 133191615 | 133187864 | 133188072 | 133191907 | 133192003 | 26 | 38 | 21 | 31 | 26 | 1 | 0 | 0 | 0 | 1 | 5 | 14 | 14 | 6 | 4 | 0 | 0 | 0 | 0 | 0 | 276 | 149 | 1 | 1 | 0.933 | 1 | 1 | 1 | 0.933 | 1 | 1 | 1 | | 1 | -0.027 |
| SE | 92159 | ENSG00000090612 | ZNF268 | chr12 | + | 133191907 | 133192003 | 133181945 | 133182030 | 133202143 | 133202219 | 9 | 15 | 12 | 13 | 14 | 1 | 1 | 2 | 0 | 0 | 0 | 13 | 6 | 6 | 4 | 0 | 0 | 0 | 2 | 0 | 245 | 149 | 1 | 1 | 0.846 | 0.901 | 0.785 | 1 | 1 | NA | 1 | 1 | 0.646 | 1 | -0.005 |
| SE | 92163 | ENSG00000147862 | NFIB | chr9 | - | 14102421 | 14102510 | 14081842 | 14088326 | 14120439 | 14120624 | 0 | 1 | 0 | 0 | 3 | 0 | 4 | 0 | 1 | 3 | 14 | 0 | 0 | 4 | 0 | 6 | 0 | 3 | 8 | 0 | 238 | 149 | 1 | 1 | NA | 0.135 | NA | 0 | 0.385 | 0.594 | NA | 0 | 0.238 | NA | -0.104 |
| SE | 92165 | ENSG00000099219 | ERMP1 | chr9 | - | 5798805 | 5799008 | 5797816 | 5797932 | 5801175 | 5801328 | 180 | 132 | 152 | 182 | 241 | 0 | 2 | 0 | 0 | 0 | 54 | 44 | 90 | 58 | 39 | 0 | 0 | 0 | 0 | 0 | 298 | 149 | 1 | 1 | 1 | 0.971 | 1 | 1 | 1 | 1 | 1 | 1 | 1 | 1 | -0.006 |
| SE | 92166 | ENSG00000099219 | ERMP1 | chr9 | - | 5805026 | 5805217 | 5801175 | 5801328 | 5805610 | 5805785 | 69 | 37 | 58 | 66 | 101 | 16 | 2 | 7 | 8 | 9 | 38 | 30 | 20 | 19 | 0 | 0 | 6 | 7 | 0 | 0 | 298 | 149 | 1 | 1 | 0.683 | 0.902 | 0.806 | 0.805 | 0.849 | 1 | 0.714 | 0.588 | 1 | NA | -0.017 |
| SE | 92168 | ENSG00000099219 | ERMP1 | chr9 | - | 5812888 | 5813035 | 5812124 | 5812217 | 5823895 | 5824001 | 132 | 63 | 73 | 100 | 62 | 0 | 0 | 0 | 0 | 0 | 57 | 32 | 59 | 40 | 8 | 0 | 0 | 0 | 2 | 0 | 296 | 149 | 1 | 1 | 1 | 1 | 1 | 1 | 1 | 1 | 1 | 1 | 0.91 | 1 | 0.018 |
| SE | 92169 | ENSG00000099219 | ERMP1 | chr9 | - | 5823895 | 5824001 | 5812736 | 5813035 | 5830726 | 5831028 | 38 | 24 | 37 | 30 | 21 | 3 | 0 | 0 | 0 | 0 | 11 | 10 | 29 | 11 | 3 | 0 | 0 | 0 | 0 | 0 | 255 | 149 | 1 | 1 | 0.881 | 1 | 1 | 1 | 1 | 1 | 1 | 1 | 1 | 1 | -0.024 |
| SE | 92174 | ENSG00000153786 | ZDHHC7 | chr16 | - | 84981869 | 84981994 | 84979188 | 84979285 | 84990303 | 84990635 | 254 | 138 | 160 | 161 | 148 | 0 | 0 | 0 | 2 | 0 | 455 | 157 | 163 | 278 | 773 | 0 | 0 | 0 | 3 | 0 | 274 | 149 | 1 | 1 | 1 | 1 | 1 | 0.978 | 1 | 1 | 1 | 1 | 0.981 | 1 | -0.001 |
| SE | 92176 | ENSG00000122692 | SMU1 | chr9 | - | 33053122 | 33053290 | 33048105 | 33048258 | 33056112 | 33056239 | 462 | 282 | 391 | 444 | 756 | 3 | 3 | 1 | 0 | 8 | 689 | 501 | 319 | 485 | 268 | 10 | 1 | 1 | 1 | 0 | 298 | 149 | 1 | 1 | 0.987 | 0.979 | 0.995 | 1 | 0.979 | 0.972 | 0.996 | 0.994 | 0.996 | 1 | -0.004 |
| SE | 92177 | ENSG00000122692 | SMU1 | chr9 | - | 33053122 | 33053290 | 33048105 | 33048258 | 33056836 | 33056964 | 164 | 104 | 149 | 191 | 283 | 0 | 0 | 0 | 0 | 2 | 276 | 162 | 109 | 190 | 90 | 0 | 0 | 0 | 0 | 0 | 298 | 149 | 1 | 1 | 1 | 1 | 1 | 1 | 0.986 | 1 | 1 | 1 | 1 | 1 | -0.003 |
| SE | 92178 | ENSG00000122692 | SMU1 | chr9 | - | 33056112 | 33056239 | 33048105 | 33048258 | 33056836 | 33056964 | 437 | 334 | 339 | 372 | 643 | 0 | 0 | 0 | 0 | 2 | 752 | 529 | 378 | 613 | 228 | 0 | 0 | 0 | 0 | 0 | 276 | 149 | 1 | 1 | 1 | 1 | 1 | 1 | 0.994 | 1 | 1 | 1 | 1 | 1 | -0.001 |
| SE | 92179 | ENSG00000122692 | SMU1 | chr9 | - | 33056112 | 33056239 | 33053122 | 33053290 | 33056836 | 33056964 | 725 | 505 | 577 | 618 | 1093 | 0 | 0 | 0 | 0 | 0 | 1151 | 865 | 585 | 905 | 404 | 0 | 0 | 0 | 1 | 0 | 276 | 149 | 1 | 1 | 1 | 1 | 1 | 1 | 1 | 1 | 1 | 1 | 0.998 | 1 | 0 |
| SE | 92180 | ENSG00000122692 | SMU1 | chr9 | - | 33056836 | 33056964 | 33056112 | 33056239 | 33057597 | 33057714 | 775 | 533 | 627 | 704 | 1061 | 2 | 0 | 0 | 0 | 0 | 1033 | 924 | 707 | 1135 | 367 | 0 | 2 | 0 | 3 | 0 | 277 | 149 | 1 | 1 | 0.995 | 1 | 1 | 1 | 1 | 1 | 0.996 | 1 | 0.995 | 1 | 0.001 |
| SE | 92182 | ENSG00000122692 | SMU1 | chr9 | - | 33060464 | 33060584 | 33057597 | 33057714 | 33068823 | 33068934 | 515 | 394 | 416 | 492 | 476 | 0 | 0 | 0 | 0 | 0 | 688 | 595 | 516 | 790 | 352 | 3 | 0 | 0 | 0 | 0 | 269 | 149 | 1 | 1 | 1 | 1 | 1 | 1 | 1 | 0.992 | 1 | 1 | 1 | 1 | 0.002 |
| SE | 92183 | ENSG00000122692 | SMU1 | chr9 | - | 33062048 | 33062177 | 33057597 | 33057714 | 33068823 | 33068934 | 157 | 90 | 109 | 161 | 100 | 0 | 0 | 0 | 0 | 0 | 153 | 198 | 167 | 244 | 58 | 3 | 0 | 0 | 0 | 0 | 278 | 149 | 1 | 1 | 1 | 1 | 1 | 1 | 1 | 0.965 | 1 | 1 | 1 | 1 | 0.007 |
| SE | 92184 | ENSG00000122692 | SMU1 | chr9 | - | 33062048 | 33062177 | 33060464 | 33060584 | 33068823 | 33068934 | 596 | 352 | 439 | 551 | 467 | 0 | 0 | 0 | 0 | 0 | 680 | 606 | 580 | 708 | 350 | 0 | 0 | 0 | 1 | 0 | 278 | 149 | 1 | 1 | 1 | 1 | 1 | 1 | 1 | 1 | 1 | 1 | 0.997 | 1 | 0.001 |
| SE | 92185 | ENSG00000122692 | SMU1 | chr9 | - | 33068823 | 33068934 | 33062048 | 33062177 | 33071739 | 33071892 | 313 | 199 | 262 | 365 | 446 | 0 | 0 | 1 | 0 | 0 | 392 | 346 | 262 | 423 | 154 | 0 | 0 | 0 | 0 | 0 | 260 | 149 | 1 | 1 | 1 | 1 | 0.993 | 1 | 1 | 1 | 1 | 1 | 1 | 1 | -0.001 |
| SE | 92190 | ENSG00000099810 | MTAP | chr9 | + | 21808806 | 21809000 | 21802641 | 21802950 | 21815432 | 21815519 | 0 | 1 | 0 | 0 | 0 | 1 | 0 | 0 | 0 | 0 | 0 | 1 | 0 | 1 | 0 | 0 | 0 | 1 | 0 | 0 | 298 | 149 | 1 | 1 | 0 | 1 | NA | NA | NA | NA | 1 | 0 | 1 | NA | -0.167 |
| SE | 92192 | ENSG00000099810 | MTAP | chr9 | + | 21808806 | 21809000 | 21803277 | 21803381 | 21815432 | 21815519 | 0 | 0 | 0 | 0 | 0 | 0 | 0 | 2 | 0 | 0 | 0 | 1 | 0 | 1 | 0 | 0 | 0 | 0 | 0 | 0 | 298 | 149 | 1 | 1 | NA | NA | 0 | NA | NA | NA | 1 | NA | 1 | NA | -1 |
| SE | 92193 | ENSG00000099810 | MTAP | chr9 | + | 21813765 | 21813886 | 21808806 | 21809000 | 21815432 | 21815519 | 0 | 1 | 3 | 0 | 0 | 0 | 0 | 0 | 0 | 0 | 0 | 0 | 0 | 0 | 0 | 0 | 1 | 0 | 1 | 0 | 270 | 149 | 1 | 1 | NA | 1 | 1 | NA | NA | NA | 0 | NA | 0 | NA | 1 |
| SE | 92194 | ENSG00000099810 | MTAP | chr9 | + | 21816713 | 21816772 | 21815432 | 21815519 | 21818034 | 21818045 | 67 | 37 | 80 | 102 | 78 | 1 | 0 | 0 | 0 | 0 | 58 | 27 | 31 | 41 | 23 | 0 | 0 | 0 | 0 | 0 | 208 | 149 | 1 | 1 | 0.98 | 1 | 1 | 1 | 1 | 1 | 1 | 1 | 1 | 1 | -0.004 |
| SE | 92195 | ENSG00000099810 | MTAP | chr9 | + | 21837907 | 21838010 | 21818034 | 21818202 | 21854630 | 21854870 | 142 | 129 | 137 | 108 | 106 | 1 | 1 | 1 | 1 | 0 | 64 | 96 | 92 | 78 | 21 | 0 | 1 | 0 | 0 | 0 | 252 | 149 | 1 | 1 | 0.988 | 0.987 | 0.988 | 0.985 | 1 | 1 | 0.983 | 1 | 1 | 1 | -0.007 |
| SE | 92196 | ENSG00000099810 | MTAP | chr9 | + | 21859302 | 21859425 | 21854630 | 21854870 | 21931007 | 21937651 | 42 | 36 | 36 | 46 | 15 | 1 | 2 | 9 | 3 | 3 | 0 | 48 | 38 | 20 | 10 | 0 | 5 | 3 | 8 | 0 | 272 | 149 | 1 | 1 | 0.958 | 0.908 | 0.687 | 0.894 | 0.733 | NA | 0.84 | 0.874 | 0.578 | 1 | 0.013 |
| SE | 92198 | ENSG00000099810 | MTAP | chr9 | + | 21861766 | 21861817 | 21859322 | 21859425 | 21931007 | 21937651 | 2 | 0 | 0 | 1 | 0 | 7 | 1 | 2 | 4 | 3 | 0 | 0 | 0 | 0 | 0 | 0 | 13 | 7 | 4 | 0 | 200 | 149 | 1 | 1 | 0.176 | 0 | 0 | 0.157 | 0 | NA | 0 | 0 | 0 | NA | 0.067 |
| SE | 92201 | ENSG00000164904 | ALDH7A1 | chr5 | - | 126568258 | 126568356 | 126561082 | 126561124 | 126570781 | 126570859 | 6 | 3 | 5 | 3 | 0 | 0 | 0 | 0 | 2 | 0 | 0 | 4 | 1 | 1 | 0 | 0 | 0 | 0 | 0 | 0 | 247 | 149 | 1 | 1 | 1 | 1 | 1 | 0.475 | NA | NA | 1 | 1 | 1 | NA | -0.131 |
| SE | 92218 | ENSG00000143198 | MGST3 | chr1 | + | 165649754 | 165649964 | 165631238 | 165631293 | 165651013 | 165651087 | 205 | 130 | 165 | 154 | 185 | 0 | 0 | 0 | 0 | 0 | 357 | 170 | 98 | 260 | 294 | 0 | 4 | 0 | 0 | 0 | 298 | 149 | 1 | 1 | 1 | 1 | 1 | 1 | 1 | 1 | 0.955 | 1 | 1 | 1 | 0.009 |
| SE | 92219 | ENSG00000143198 | MGST3 | chr1 | + | 165649840 | 165649964 | 165631212 | 165631293 | 165651013 | 165651087 | 299 | 191 | 265 | 242 | 236 | 0 | 0 | 0 | 0 | 0 | 510 | 283 | 169 | 404 | 683 | 0 | 4 | 0 | 0 | 0 | 273 | 149 | 1 | 1 | 1 | 1 | 1 | 1 | 1 | 1 | 0.975 | 1 | 1 | 1 | 0.005 |
| SE | 92220 | ENSG00000143198 | MGST3 | chr1 | + | 165651013 | 165651087 | 165649754 | 165649964 | 165651977 | 165652035 | 507 | 355 | 467 | 427 | 400 | 5 | 1 | 1 | 2 | 0 | 957 | 514 | 294 | 776 | 1070 | 0 | 2 | 0 | 7 | 0 | 223 | 149 | 1 | 1 | 0.985 | 0.996 | 0.997 | 0.993 | 1 | 1 | 0.994 | 1 | 0.987 | 1 | -0.002 |
| SE | 92221 | ENSG00000143198 | MGST3 | chr1 | + | 165651013 | 165651087 | 165649754 | 165649964 | 165654278 | 165654351 | 197 | 127 | 155 | 153 | 171 | 0 | 0 | 0 | 0 | 0 | 323 | 168 | 114 | 254 | 355 | 0 | 0 | 0 | 4 | 0 | 223 | 149 | 1 | 1 | 1 | 1 | 1 | 1 | 1 | 1 | 1 | 1 | 0.977 | 1 | 0.005 |
| SE | 92222 | ENSG00000143198 | MGST3 | chr1 | + | 165651013 | 165651087 | 165649754 | 165649964 | 165661428 | 165661796 | 197 | 127 | 155 | 153 | 171 | 0 | 1 | 0 | 0 | 0 | 313 | 164 | 107 | 250 | 345 | 0 | 0 | 0 | 0 | 0 | 223 | 149 | 1 | 1 | 1 | 0.988 | 1 | 1 | 1 | 1 | 1 | 1 | 1 | 1 | -0.002 |
| SE | 92223 | ENSG00000143198 | MGST3 | chr1 | + | 165651977 | 165652035 | 165649754 | 165649964 | 165654278 | 165654351 | 45 | 15 | 27 | 23 | 14 | 0 | 0 | 0 | 0 | 0 | 38 | 20 | 23 | 41 | 32 | 0 | 0 | 0 | 4 | 0 | 207 | 149 | 1 | 1 | 1 | 1 | 1 | 1 | 1 | 1 | 1 | 1 | 0.881 | 1 | 0.024 |
| SE | 92225 | ENSG00000143198 | MGST3 | chr1 | + | 165654278 | 165654351 | 165649754 | 165649964 | 165661428 | 165661796 | 0 | 0 | 0 | 0 | 0 | 0 | 1 | 0 | 0 | 0 | 0 | 0 | 0 | 4 | 0 | 0 | 0 | 0 | 0 | 0 | 222 | 149 | 1 | 1 | NA | 0 | NA | NA | NA | NA | NA | NA | 1 | NA | -1 |
| SE | 92226 | ENSG00000143198 | MGST3 | chr1 | + | 165654278 | 165654351 | 165651013 | 165651087 | 165661428 | 165661796 | 0 | 0 | 1 | 0 | 0 | 0 | 0 | 0 | 0 | 3 | 10 | 4 | 7 | 4 | 0 | 0 | 0 | 0 | 0 | 0 | 222 | 149 | 1 | 1 | NA | NA | 1 | NA | 0 | 1 | 1 | 1 | 1 | NA | -0.5 |
| SE | 92230 | ENSG00000134531 | EMP1 | chr12 | + | 13211468 | 13211588 | 13196725 | 13196872 | 13213478 | 13213575 | 21 | 10 | 7 | 27 | 13 | 0 | 0 | 0 | 0 | 0 | 83 | 7 | 14 | 125 | 0 | 6 | 0 | 0 | 0 | 0 | 269 | 149 | 1 | 1 | 1 | 1 | 1 | 1 | 1 | 0.885 | 1 | 1 | 1 | NA | 0.029 |
| SE | 92236 | ENSG00000114209 | PDCD10 | chr3 | - | 167704841 | 167704895 | 167697008 | 167697126 | 167720061 | 167720098 | 447 | 382 | 305 | 443 | 481 | 0 | 0 | 0 | 0 | 0 | 202 | 438 | 167 | 140 | 100 | 0 | 0 | 1 | 0 | 0 | 203 | 149 | 1 | 1 | 1 | 1 | 1 | 1 | 1 | 1 | 1 | 0.992 | 1 | 1 | 0.002 |
| SE | 92237 | ENSG00000114209 | PDCD10 | chr3 | - | 167704841 | 167704895 | 167697008 | 167697126 | 167734805 | 167734832 | 135 | 124 | 90 | 149 | 137 | 0 | 0 | 0 | 0 | 0 | 61 | 118 | 56 | 52 | 35 | 0 | 3 | 0 | 0 | 0 | 203 | 149 | 1 | 1 | 1 | 1 | 1 | 1 | 1 | 1 | 0.967 | 1 | 1 | 1 | 0.007 |
| SE | 92238 | ENSG00000114209 | PDCD10 | chr3 | - | 167720061 | 167720273 | 167697008 | 167697126 | 167734805 | 167734929 | 180 | 106 | 105 | 153 | 154 | 0 | 0 | 0 | 0 | 0 | 57 | 160 | 69 | 37 | 14 | 0 | 3 | 0 | 0 | 0 | 298 | 149 | 1 | 1 | 1 | 1 | 1 | 1 | 1 | 1 | 0.964 | 1 | 1 | 1 | 0.007 |
| SE | 92240 | ENSG00000114209 | PDCD10 | chr3 | - | 167720061 | 167720273 | 167704557 | 167704895 | 167734805 | 167734929 | 509 | 373 | 332 | 462 | 504 | 0 | 0 | 1 | 3 | 0 | 198 | 485 | 194 | 131 | 79 | 0 | 0 | 0 | 0 | 0 | 298 | 149 | 1 | 1 | 1 | 1 | 0.994 | 0.987 | 1 | 1 | 1 | 1 | 1 | 1 | -0.004 |
| SE | 92247 | ENSG00000114209 | PDCD10 | chr3 | - | 167734213 | 167734345 | 167725400 | 167725473 | 167734805 | 167734832 | 6 | 2 | 2 | 1 | 3 | 59 | 45 | 40 | 62 | 77 | 0 | 6 | 0 | 0 | 0 | 0 | 43 | 9 | 9 | 0 | 281 | 149 | 1 | 1 | 0.051 | 0.023 | 0.026 | 0.008 | 0.02 | NA | 0.069 | 0 | 0 | NA | 0.003 |
| SE | 92249 | ENSG00000074071 | MRPS34 | chr16 | - | 1772603 | 1772646 | 1771894 | 1772513 | 1772798 | 1773134 | 121 | 35 | 66 | 88 | 20 | 0 | 0 | 0 | 2 | 0 | 108 | 59 | 53 | 56 | 175 | 0 | 0 | 1 | 0 | 0 | 192 | 149 | 1 | 1 | 1 | 1 | 1 | 0.972 | 1 | 1 | 1 | 0.976 | 1 | 1 | -0.001 |
| SE | 92250 | ENSG00000177076 | ACER2 | chr9 | + | 19434946 | 19435084 | 19424699 | 19424841 | 19446280 | 19446418 | 19 | 15 | 17 | 34 | 60 | 0 | 0 | 0 | 0 | 0 | 45 | 89 | 22 | 41 | 174 | 0 | 4 | 0 | 0 | 0 | 287 | 149 | 1 | 1 | 1 | 1 | 1 | 1 | 1 | 1 | 0.92 | 1 | 1 | 1 | 0.016 |
| SE | 92253 | ENSG00000233554 | B4GALT1-AS1 | chr9 | + | 33178895 | 33178957 | 33177419 | 33177469 | 33179324 | 33179984 | 5 | 4 | 1 | 0 | 0 | 0 | 0 | 0 | 1 | 0 | 0 | 2 | 0 | 1 | 11 | 0 | 0 | 0 | 0 | 0 | 211 | 149 | 1 | 1 | 1 | 1 | 1 | 0 | NA | NA | 1 | NA | 1 | 1 | -0.25 |
| SE | 92254 | ENSG00000115091 | ACTR3 | chr2 | + | 113913171 | 113913227 | 113890088 | 113890323 | 113916883 | 113917008 | 1095 | 882 | 803 | 1026 | 725 | 0 | 2 | 0 | 0 | 0 | 627 | 843 | 563 | 826 | 217 | 0 | 0 | 2 | 0 | 0 | 205 | 149 | 1 | 1 | 1 | 0.997 | 1 | 1 | 1 | 1 | 1 | 0.995 | 1 | 1 | 0 |
| SE | 92255 | ENSG00000115091 | ACTR3 | chr2 | + | 113916883 | 113917008 | 113913171 | 113913227 | 113927344 | 113927455 | 2510 | 2057 | 1904 | 2432 | 1988 | 0 | 0 | 2 | 0 | 0 | 1358 | 1893 | 1403 | 1907 | 484 | 0 | 0 | 0 | 0 | 0 | 274 | 149 | 1 | 1 | 1 | 1 | 0.998 | 1 | 1 | 1 | 1 | 1 | 1 | 1 | 0 |
| SE | 92257 | ENSG00000115091 | ACTR3 | chr2 | + | 113916883 | 113917008 | 113913171 | 113913227 | 113951478 | 113951571 | 867 | 740 | 646 | 838 | 605 | 0 | 0 | 0 | 0 | 0 | 459 | 692 | 477 | 698 | 155 | 0 | 3 | 0 | 0 | 0 | 274 | 149 | 1 | 1 | 1 | 1 | 1 | 1 | 1 | 1 | 0.992 | 1 | 1 | 1 | 0.002 |
| SE | 92259 | ENSG00000115091 | ACTR3 | chr2 | + | 113927344 | 113927455 | 113913171 | 113913227 | 113951478 | 113951571 | 2 | 0 | 2 | 0 | 0 | 0 | 0 | 0 | 0 | 0 | 0 | 0 | 0 | 0 | 0 | 0 | 3 | 0 | 0 | 0 | 260 | 149 | 1 | 1 | 1 | NA | 1 | NA | NA | NA | 0 | NA | NA | NA | 1 |
| SE | 92260 | ENSG00000115091 | ACTR3 | chr2 | + | 113927344 | 113927455 | 113916883 | 113917008 | 113931300 | 113931369 | 2765 | 2125 | 2156 | 2579 | 2811 | 0 | 0 | 0 | 2 | 0 | 1541 | 1947 | 1628 | 2088 | 545 | 0 | 2 | 0 | 0 | 0 | 260 | 149 | 1 | 1 | 1 | 1 | 1 | 0.999 | 1 | 1 | 0.998 | 1 | 1 | 1 | 0 |
| SE | 92261 | ENSG00000115091 | ACTR3 | chr2 | + | 113927344 | 113927455 | 113916883 | 113917008 | 113951478 | 113951571 | 1206 | 970 | 870 | 1120 | 913 | 0 | 0 | 0 | 0 | 2 | 662 | 918 | 639 | 816 | 231 | 0 | 0 | 0 | 0 | 0 | 260 | 149 | 1 | 1 | 1 | 1 | 1 | 1 | 0.996 | 1 | 1 | 1 | 1 | 1 | -0.001 |
| SE | 92263 | ENSG00000115091 | ACTR3 | chr2 | + | 113931300 | 113931396 | 113916883 | 113917008 | 113951478 | 113951571 | 0 | 0 | 0 | 0 | 0 | 0 | 0 | 0 | 0 | 2 | 0 | 5 | 1 | 0 | 0 | 0 | 0 | 0 | 0 | 0 | 245 | 149 | 1 | 1 | NA | NA | NA | NA | 0 | NA | 1 | 1 | NA | NA | -1 |
| SE | 92264 | ENSG00000115091 | ACTR3 | chr2 | + | 113931300 | 113931396 | 113927344 | 113927455 | 113951478 | 113951571 | 661 | 529 | 595 | 665 | 783 | 2 | 0 | 0 | 0 | 0 | 366 | 406 | 420 | 517 | 146 | 0 | 0 | 0 | 0 | 0 | 245 | 149 | 1 | 1 | 0.995 | 1 | 1 | 1 | 1 | 1 | 1 | 1 | 1 | 1 | -0.001 |
| SE | 92265 | ENSG00000115091 | ACTR3 | chr2 | + | 113934278 | 113934386 | 113931300 | 113931396 | 113939958 | 113940102 | 3972 | 2997 | 3147 | 3550 | 4780 | 0 | 0 | 0 | 0 | 4 | 2662 | 2604 | 1847 | 3372 | 752 | 0 | 0 | 0 | 4 | 0 | 257 | 149 | 1 | 1 | 1 | 1 | 1 | 1 | 0.999 | 1 | 1 | 1 | 0.998 | 1 | 0 |
| SE | 92266 | ENSG00000115091 | ACTR3 | chr2 | + | 113934278 | 113934386 | 113931300 | 113931396 | 113951478 | 113951571 | 2193 | 1597 | 1722 | 1931 | 2559 | 0 | 0 | 0 | 0 | 0 | 1406 | 1384 | 969 | 1812 | 428 | 0 | 3 | 1 | 0 | 0 | 257 | 149 | 1 | 1 | 1 | 1 | 1 | 1 | 1 | 1 | 0.996 | 0.998 | 1 | 1 | 0.001 |
| SE | 92267 | ENSG00000115091 | ACTR3 | chr2 | + | 113939958 | 113940102 | 113931300 | 113931396 | 113951478 | 113951571 | 0 | 1 | 0 | 0 | 4 | 0 | 0 | 0 | 0 | 0 | 0 | 0 | 0 | 4 | 0 | 0 | 3 | 1 | 0 | 0 | 293 | 149 | 1 | 1 | NA | 1 | NA | NA | 1 | NA | 0 | 0 | 1 | NA | 0.667 |
| SE | 92268 | ENSG00000115091 | ACTR3 | chr2 | + | 113939958 | 113940102 | 113934278 | 113934386 | 113951478 | 113951571 | 1799 | 1416 | 1429 | 1646 | 2241 | 0 | 0 | 0 | 0 | 2 | 1264 | 1240 | 905 | 1584 | 299 | 6 | 0 | 0 | 0 | 0 | 293 | 149 | 1 | 1 | 1 | 1 | 1 | 1 | 0.998 | 0.991 | 1 | 1 | 1 | 1 | 0.001 |
| SE | 92269 | ENSG00000115091 | ACTR3 | chr2 | + | 113942185 | 113942359 | 113939958 | 113940102 | 113951478 | 113951571 | 2405 | 1994 | 1865 | 2278 | 3006 | 0 | 1 | 0 | 0 | 0 | 1472 | 1597 | 1423 | 1903 | 595 | 0 | 0 | 0 | 0 | 0 | 298 | 149 | 1 | 1 | 1 | 0.999 | 1 | 1 | 1 | 1 | 1 | 1 | 1 | 1 | 0 |
| SE | 92274 | ENSG00000132326 | PER2 | chr2 | - | 238261728 | 238261837 | 238260827 | 238260953 | 238262190 | 238262344 | 26 | 30 | 18 | 28 | 17 | 0 | 0 | 0 | 1 | 0 | 14 | 26 | 15 | 1 | 0 | 0 | 0 | 0 | 3 | 0 | 258 | 149 | 1 | 1 | 1 | 1 | 1 | 0.942 | 1 | 1 | 1 | 1 | 0.161 | NA | 0.198 |
| SE | 92275 | ENSG00000132326 | PER2 | chr2 | - | 238262190 | 238262344 | 238261440 | 238261837 | 238262951 | 238262975 | 53 | 29 | 37 | 57 | 40 | 0 | 0 | 0 | 0 | 0 | 71 | 54 | 24 | 27 | 66 | 0 | 0 | 1 | 0 | 0 | 298 | 149 | 1 | 1 | 1 | 1 | 1 | 1 | 1 | 1 | 1 | 0.923 | 1 | 1 | 0.015 |
| SE | 92277 | ENSG00000132326 | PER2 | chr2 | - | 238268055 | 238268198 | 238265511 | 238265590 | 238268922 | 238268974 | 82 | 52 | 96 | 82 | 81 | 0 | 1 | 0 | 0 | 0 | 136 | 48 | 65 | 67 | 96 | 0 | 0 | 0 | 0 | 0 | 292 | 149 | 1 | 1 | 1 | 0.964 | 1 | 1 | 1 | 1 | 1 | 1 | 1 | 1 | -0.007 |
| SE | 92291 | ENSG00000125743 | SNRPD2 | chr19 | - | 45690571 | 45690667 | 45688386 | 45688566 | 45691886 | 45691940 | 0 | 4 | 0 | 2 | 1 | 170 | 71 | 188 | 129 | 176 | 0 | 0 | 0 | 0 | 0 | 120 | 107 | 53 | 73 | 117 | 245 | 149 | 1 | 1 | 0 | 0.033 | 0 | 0.009 | 0.003 | 0 | 0 | 0 | 0 | 0 | 0.009 |
| SE | 92292 | ENSG00000157045 | NTAN1 | chr16 | - | 15041622 | 15041676 | 15041067 | 15041121 | 15044333 | 15044407 | 274 | 205 | 238 | 349 | 450 | 0 | 0 | 1 | 0 | 0 | 594 | 436 | 297 | 482 | 1132 | 0 | 0 | 0 | 0 | 0 | 203 | 149 | 1 | 1 | 1 | 1 | 0.994 | 1 | 1 | 1 | 1 | 1 | 1 | 1 | -0.001 |
| SE | 92293 | ENSG00000157045 | NTAN1 | chr16 | - | 15044333 | 15044407 | 15041622 | 15041676 | 15047417 | 15047550 | 93 | 60 | 92 | 102 | 143 | 0 | 0 | 1 | 0 | 2 | 180 | 164 | 90 | 132 | 317 | 0 | 2 | 0 | 3 | 0 | 223 | 149 | 1 | 1 | 1 | 1 | 1 | 1 | 0.979 | 1 | 0.982 | 1 | 0.967 | 1 | 0.006 |
| SE | 92294 | ENSG00000157045 | NTAN1 | chr16 | - | 15044333 | 15044407 | 15041622 | 15041676 | 15047441 | 15047550 | 220 | 158 | 179 | 257 | 317 | 1 | 0 | 0 | 0 | 2 | 500 | 281 | 240 | 382 | 855 | 0 | 0 | 0 | 3 | 3 | 223 | 149 | 1 | 1 | 0.993 | 1 | 1 | 1 | 0.991 | 1 | 1 | 1 | 0.988 | 0.995 | 0 |
| SE | 92295 | ENSG00000157045 | NTAN1 | chr16 | - | 15047854 | 15047920 | 15047215 | 15047550 | 15047996 | 15048099 | 24 | 20 | 18 | 16 | 21 | 0 | 0 | 0 | 0 | 0 | 22 | 27 | 50 | 35 | 74 | 0 | 0 | 2 | 0 | 0 | 215 | 149 | 1 | 1 | 1 | 1 | 1 | 1 | 1 | 1 | 1 | 0.945 | 1 | 1 | 0.011 |
| SE | 92297 | ENSG00000157045 | NTAN1 | chr16 | - | 15047996 | 15048099 | 15047427 | 15047550 | 15055890 | 15056074 | 6 | 1 | 0 | 0 | 0 | 0 | 0 | 0 | 1 | 0 | 0 | 0 | 9 | 0 | 6 | 4 | 2 | 0 | 4 | 0 | 252 | 149 | 1 | 1 | 1 | 1 | NA | 0 | NA | 0 | 0 | 1 | 0 | 1 | 0.267 |
| SE | 92299 | ENSG00000157045 | NTAN1 | chr16 | - | 15048077 | 15048099 | 15047854 | 15047920 | 15055890 | 15056039 | 3 | 0 | 0 | 0 | 0 | 1 | 1 | 0 | 2 | 0 | 0 | 0 | 0 | 0 | 0 | 0 | 0 | 1 | 3 | 8 | 171 | 149 | 1 | 1 | 0.723 | 0 | NA | 0 | NA | NA | NA | 0 | 0 | 0 | 0.241 |
| SE | 92301 | ENSG00000204130 | RUFY2 | chr10 | - | 68345701 | 68345911 | 68341106 | 68341288 | 68346006 | 68346084 | 27 | 22 | 24 | 21 | 32 | 5 | 0 | 0 | 0 | 2 | 0 | 17 | 10 | 1 | 0 | 0 | 0 | 0 | 0 | 0 | 298 | 149 | 1 | 1 | 0.73 | 1 | 1 | 1 | 0.889 | NA | 1 | 1 | 1 | NA | -0.076 |
| SE | 92302 | ENSG00000204130 | RUFY2 | chr10 | - | 68345830 | 68345911 | 68341129 | 68341288 | 68346006 | 68346084 | 9 | 11 | 10 | 9 | 19 | 5 | 0 | 0 | 0 | 2 | 0 | 10 | 4 | 0 | 0 | 0 | 0 | 0 | 0 | 0 | 230 | 149 | 1 | 1 | 0.538 | 1 | 1 | 1 | 0.86 | NA | 1 | 1 | NA | NA | -0.12 |
| SE | 92305 | ENSG00000204130 | RUFY2 | chr10 | - | 68393137 | 68393203 | 68386058 | 68386128 | 68394074 | 68394136 | 82 | 85 | 77 | 55 | 132 | 0 | 2 | 1 | 1 | 0 | 41 | 55 | 53 | 51 | 20 | 0 | 0 | 1 | 0 | 0 | 215 | 149 | 1 | 1 | 1 | 0.967 | 0.982 | 0.974 | 1 | 1 | 1 | 0.973 | 1 | 1 | -0.01 |
| SE | 92306 | ENSG00000204130 | RUFY2 | chr10 | - | 68394074 | 68394136 | 68386058 | 68386128 | 68401619 | 68401737 | 0 | 2 | 1 | 1 | 0 | 0 | 0 | 0 | 0 | 0 | 0 | 0 | 0 | 0 | 0 | 0 | 0 | 0 | 2 | 0 | 211 | 149 | 1 | 1 | NA | 1 | 1 | 1 | NA | NA | NA | NA | 0 | NA | 1 |
| SE | 92309 | ENSG00000204130 | RUFY2 | chr10 | - | 68404670 | 68404844 | 68401619 | 68401737 | 68407185 | 68407236 | 73 | 56 | 74 | 73 | 68 | 1 | 0 | 2 | 0 | 0 | 105 | 58 | 84 | 104 | 53 | 0 | 2 | 0 | 0 | 0 | 298 | 149 | 1 | 1 | 0.973 | 1 | 0.949 | 1 | 1 | 1 | 0.935 | 1 | 1 | 1 | -0.003 |
| SE | 92310 | ENSG00000160867 | FGFR4 | chr5 | + | 177088238 | 177088514 | 177086915 | 177087033 | 177089549 | 177089693 | 0 | 0 | 0 | 0 | 4 | 0 | 0 | 0 | 0 | 0 | 0 | 0 | 0 | 0 | 0 | 12 | 0 | 0 | 0 | 0 | 298 | 149 | 1 | 1 | NA | NA | NA | NA | 1 | 0 | NA | NA | NA | NA | 1 |
| SE | 92313 | ENSG00000188921 | HACD4 | chr9 | - | 21011588 | 21011695 | 21008020 | 21008146 | 21015897 | 21016010 | 359 | 290 | 204 | 313 | 792 | 0 | 1 | 0 | 0 | 0 | 393 | 251 | 186 | 405 | 511 | 0 | 0 | 0 | 0 | 0 | 256 | 149 | 1 | 1 | 1 | 0.994 | 1 | 1 | 1 | 1 | 1 | 1 | 1 | 1 | -0.001 |
| SE | 92314 | ENSG00000188921 | HACD4 | chr9 | - | 21011588 | 21011695 | 21008020 | 21008146 | 21026595 | 21026723 | 132 | 123 | 77 | 91 | 261 | 0 | 3 | 1 | 0 | 0 | 184 | 90 | 71 | 151 | 193 | 0 | 1 | 0 | 0 | 0 | 256 | 149 | 1 | 1 | 1 | 0.96 | 0.978 | 1 | 1 | 1 | 0.981 | 1 | 1 | 1 | -0.009 |
| SE | 92315 | ENSG00000188921 | HACD4 | chr9 | - | 21015897 | 21016010 | 21008020 | 21008146 | 21026595 | 21026723 | 87 | 80 | 54 | 96 | 135 | 0 | 3 | 1 | 0 | 0 | 46 | 91 | 41 | 77 | 142 | 0 | 1 | 0 | 0 | 0 | 262 | 149 | 1 | 1 | 1 | 0.938 | 0.968 | 1 | 1 | 1 | 0.981 | 1 | 1 | 1 | -0.015 |
| SE | 92316 | ENSG00000188921 | HACD4 | chr9 | - | 21015897 | 21016010 | 21011588 | 21011695 | 21026595 | 21026723 | 356 | 284 | 205 | 366 | 739 | 1 | 0 | 0 | 0 | 0 | 324 | 323 | 199 | 360 | 515 | 4 | 0 | 0 | 0 | 0 | 262 | 149 | 1 | 1 | 0.995 | 1 | 1 | 1 | 1 | 0.979 | 1 | 1 | 1 | 1 | 0.003 |
| SE | 92318 | ENSG00000137509 | PRCP | chr11 | - | 82825450 | 82825599 | 82824399 | 82825122 | 82838386 | 82838574 | 2 | 0 | 1 | 0 | 3 | 280 | 167 | 194 | 236 | 543 | 10 | 0 | 0 | 7 | 0 | 691 | 193 | 120 | 318 | 431 | 298 | 149 | 1 | 1 | 0.004 | 0 | 0.003 | 0 | 0.003 | 0.007 | 0 | 0 | 0.011 | 0 | -0.002 |
| SE | 92319 | ENSG00000137509 | PRCP | chr11 | - | 82838386 | 82838574 | 82822935 | 82825122 | 82849048 | 82849218 | 282 | 167 | 194 | 239 | 543 | 0 | 0 | 2 | 0 | 0 | 691 | 193 | 120 | 318 | 431 | 0 | 0 | 0 | 0 | 0 | 298 | 149 | 1 | 1 | 1 | 1 | 0.98 | 1 | 1 | 1 | 1 | 1 | 1 | 1 | -0.004 |
| SE | 92320 | ENSG00000137509 | PRCP | chr11 | - | 82839260 | 82839425 | 82836559 | 82838574 | 82843183 | 82843304 | 143 | 121 | 107 | 136 | 187 | 0 | 0 | 0 | 0 | 0 | 226 | 66 | 76 | 163 | 246 | 0 | 0 | 0 | 2 | 0 | 298 | 149 | 1 | 1 | 1 | 1 | 1 | 1 | 1 | 1 | 1 | 1 | 0.976 | 1 | 0.005 |
| SE | 92321 | ENSG00000137509 | PRCP | chr11 | - | 82839260 | 82839425 | 82836559 | 82838574 | 82849048 | 82849218 | 214 | 183 | 175 | 190 | 310 | 2 | 0 | 0 | 3 | 0 | 403 | 97 | 123 | 289 | 408 | 0 | 0 | 0 | 0 | 0 | 298 | 149 | 1 | 1 | 0.982 | 1 | 1 | 0.969 | 1 | 1 | 1 | 1 | 1 | 1 | -0.01 |
| SE | 92322 | ENSG00000137509 | PRCP | chr11 | - | 82839260 | 82839425 | 82836559 | 82838574 | 82849913 | 82850071 | 144 | 121 | 108 | 136 | 187 | 0 | 0 | 0 | 0 | 0 | 226 | 64 | 76 | 163 | 246 | 0 | 0 | 1 | 0 | 0 | 298 | 149 | 1 | 1 | 1 | 1 | 1 | 1 | 1 | 1 | 1 | 0.974 | 1 | 1 | 0.005 |
| SE | 92323 | ENSG00000137509 | PRCP | chr11 | - | 82839260 | 82839425 | 82836559 | 82838574 | 82859976 | 82860117 | 143 | 121 | 107 | 135 | 187 | 0 | 1 | 0 | 0 | 0 | 226 | 64 | 76 | 163 | 246 | 0 | 0 | 0 | 0 | 0 | 298 | 149 | 1 | 1 | 1 | 0.984 | 1 | 1 | 1 | 1 | 1 | 1 | 1 | 1 | -0.003 |
| SE | 92324 | ENSG00000137509 | PRCP | chr11 | - | 82843183 | 82843304 | 82838386 | 82838574 | 82849048 | 82849218 | 0 | 0 | 0 | 1 | 3 | 2 | 0 | 0 | 3 | 0 | 0 | 0 | 1 | 2 | 0 | 0 | 0 | 0 | 0 | 0 | 270 | 149 | 1 | 1 | 0 | NA | NA | 0.155 | 1 | NA | NA | 1 | 1 | NA | -0.615 |
| SE | 92325 | ENSG00000137509 | PRCP | chr11 | - | 82843183 | 82843304 | 82838386 | 82838574 | 82849913 | 82850071 | 0 | 1 | 0 | 0 | 0 | 0 | 0 | 0 | 0 | 0 | 0 | 0 | 2 | 2 | 9 | 0 | 0 | 1 | 0 | 0 | 270 | 149 | 1 | 1 | NA | 1 | NA | NA | NA | NA | NA | 0.525 | 1 | 1 | 0.158 |
| SE | 92328 | ENSG00000137509 | PRCP | chr11 | - | 82843183 | 82843304 | 82839260 | 82839425 | 82849048 | 82849218 | 0 | 0 | 0 | 2 | 3 | 71 | 62 | 68 | 55 | 123 | 0 | 0 | 1 | 0 | 0 | 177 | 33 | 47 | 126 | 162 | 270 | 149 | 1 | 1 | 0 | 0 | 0 | 0.02 | 0.013 | 0 | 0 | 0.012 | 0 | 0 | 0.004 |
| SE | 92329 | ENSG00000137509 | PRCP | chr11 | - | 82843183 | 82843304 | 82839260 | 82839425 | 82849913 | 82850071 | 0 | 1 | 0 | 1 | 0 | 1 | 0 | 1 | 1 | 0 | 0 | 2 | 2 | 0 | 9 | 0 | 0 | 0 | 0 | 0 | 270 | 149 | 1 | 1 | 0 | 1 | 0 | 0.356 | NA | NA | 1 | 1 | NA | 1 | -0.661 |
| SE | 92330 | ENSG00000137509 | PRCP | chr11 | - | 82849048 | 82849218 | 82836559 | 82838574 | 82849913 | 82850071 | 107 | 67 | 93 | 86 | 152 | 0 | 0 | 0 | 0 | 0 | 78 | 76 | 34 | 97 | 137 | 0 | 0 | 1 | 0 | 0 | 298 | 149 | 1 | 1 | 1 | 1 | 1 | 1 | 1 | 1 | 1 | 0.944 | 1 | 1 | 0.011 |
| SE | 92331 | ENSG00000137509 | PRCP | chr11 | - | 82849048 | 82849218 | 82839260 | 82839425 | 82849913 | 82850071 | 176 | 129 | 161 | 138 | 275 | 1 | 0 | 1 | 1 | 0 | 255 | 109 | 81 | 223 | 299 | 0 | 0 | 0 | 0 | 0 | 298 | 149 | 1 | 1 | 0.989 | 1 | 0.988 | 0.986 | 1 | 1 | 1 | 1 | 1 | 1 | -0.007 |
| SE | 92338 | ENSG00000027697 | IFNGR1 | chr6 | - | 137204331 | 137204504 | 137203498 | 137203685 | 137206135 | 137206206 | 1264 | 1038 | 748 | 873 | 1554 | 0 | 1 | 0 | 0 | 0 | 318 | 1196 | 1866 | 525 | 185 | 0 | 0 | 0 | 0 | 0 | 298 | 149 | 1 | 1 | 1 | 0.998 | 1 | 1 | 1 | 1 | 1 | 1 | 1 | 1 | 0 |
| SE | 92341 | ENSG00000027697 | IFNGR1 | chr6 | - | 137209849 | 137209972 | 137206962 | 137207077 | 137219242 | 137219313 | 4 | 0 | 0 | 4 | 0 | 267 | 167 | 135 | 156 | 41 | 0 | 2 | 0 | 1 | 0 | 63 | 346 | 397 | 85 | 29 | 272 | 149 | 1 | 1 | 0.008 | 0 | 0 | 0.014 | 0 | 0 | 0.003 | 0 | 0.006 | 0 | 0.003 |
| SE | 92343 | ENSG00000027697 | IFNGR1 | chr6 | - | 137209849 | 137209984 | 137206962 | 137207077 | 137219242 | 137219312 | 4 | 0 | 3 | 6 | 0 | 267 | 167 | 135 | 156 | 41 | 0 | 2 | 0 | 1 | 0 | 63 | 346 | 397 | 85 | 29 | 284 | 149 | 1 | 1 | 0.008 | 0 | 0.012 | 0.02 | 0 | 0 | 0.003 | 0 | 0.006 | 0 | 0.006 |
| SE | 92347 | ENSG00000177888 | ZBTB41 | chr1 | - | 197175930 | 197176014 | 197175009 | 197175115 | 197176563 | 197176670 | 9 | 9 | 5 | 8 | 18 | 27 | 36 | 26 | 16 | 86 | 10 | 10 | 0 | 0 | 0 | 0 | 1 | 6 | 0 | 7 | 233 | 149 | 1 | 1 | 0.176 | 0.138 | 0.11 | 0.242 | 0.118 | 1 | 0.865 | 0 | NA | 0 | -0.309 |
| SE | 92348 | ENSG00000165271 | NOL6 | chr9 | - | 33466070 | 33466225 | 33465733 | 33465897 | 33466307 | 33466425 | 17 | 20 | 22 | 15 | 28 | 0 | 0 | 0 | 1 | 0 | 44 | 51 | 9 | 10 | 69 | 0 | 0 | 0 | 0 | 0 | 298 | 149 | 1 | 1 | 1 | 1 | 1 | 0.882 | 1 | 1 | 1 | 1 | 1 | 1 | -0.024 |
| SE | 92349 | ENSG00000165271 | NOL6 | chr9 | - | 33467393 | 33467516 | 33467113 | 33467262 | 33467690 | 33467868 | 27 | 19 | 32 | 24 | 19 | 0 | 0 | 0 | 0 | 0 | 0 | 13 | 11 | 1 | 24 | 0 | 0 | 1 | 0 | 0 | 272 | 149 | 1 | 1 | 1 | 1 | 1 | 1 | 1 | NA | 1 | 0.858 | 1 | 1 | 0.035 |
| SE | 92350 | ENSG00000165271 | NOL6 | chr9 | - | 33467690 | 33467868 | 33467393 | 33467516 | 33468029 | 33468145 | 41 | 23 | 30 | 9 | 39 | 0 | 0 | 0 | 0 | 0 | 0 | 30 | 13 | 6 | 14 | 0 | 0 | 1 | 0 | 8 | 298 | 149 | 1 | 1 | 1 | 1 | 1 | 1 | 1 | NA | 1 | 0.867 | 1 | 0.467 | 0.166 |
| SE | 92354 | ENSG00000143437 | ARNT | chr1 | - | 150814076 | 150814239 | 150813171 | 150813338 | 150816258 | 150816400 | 89 | 36 | 87 | 103 | 62 | 1 | 0 | 0 | 0 | 0 | 82 | 113 | 73 | 76 | 72 | 0 | 0 | 0 | 0 | 0 | 298 | 149 | 1 | 1 | 0.978 | 1 | 1 | 1 | 1 | 1 | 1 | 1 | 1 | 1 | -0.004 |
| SE | 92355 | ENSG00000143437 | ARNT | chr1 | - | 150816787 | 150816890 | 150816258 | 150816406 | 150817081 | 150817202 | 205 | 138 | 155 | 223 | 424 | 0 | 0 | 0 | 0 | 0 | 332 | 241 | 132 | 192 | 155 | 0 | 0 | 0 | 4 | 0 | 252 | 149 | 1 | 1 | 1 | 1 | 1 | 1 | 1 | 1 | 1 | 1 | 0.966 | 1 | 0.007 |
| SE | 92358 | ENSG00000143437 | ARNT | chr1 | - | 150834537 | 150834640 | 150832333 | 150832399 | 150836279 | 150836493 | 395 | 392 | 355 | 498 | 622 | 0 | 1 | 0 | 0 | 0 | 1137 | 531 | 537 | 602 | 689 | 0 | 0 | 0 | 0 | 0 | 252 | 149 | 1 | 1 | 1 | 0.996 | 1 | 1 | 1 | 1 | 1 | 1 | 1 | 1 | -0.001 |
| SE | 92359 | ENSG00000143437 | ARNT | chr1 | - | 150839440 | 150839654 | 150836279 | 150836493 | 150842423 | 150842468 | 161 | 103 | 111 | 130 | 126 | 0 | 0 | 0 | 0 | 0 | 206 | 168 | 135 | 120 | 244 | 0 | 0 | 1 | 5 | 0 | 298 | 149 | 1 | 1 | 1 | 1 | 1 | 1 | 1 | 1 | 1 | 0.985 | 0.923 | 1 | 0.018 |
| SE | 92363 | ENSG00000143437 | ARNT | chr1 | - | 150853155 | 150853278 | 150852761 | 150852806 | 150858348 | 150858460 | 0 | 3 | 0 | 0 | 0 | 43 | 32 | 33 | 73 | 19 | 0 | 0 | 0 | 0 | 0 | 46 | 30 | 65 | 30 | 61 | 272 | 149 | 1 | 1 | 0 | 0.049 | 0 | 0 | 0 | 0 | 0 | 0 | 0 | 0 | 0.01 |
| SE | 92365 | ENSG00000080503 | SMARCA2 | chr9 | + | 2028986 | 2029247 | 2015185 | 2015404 | 2032951 | 2033081 | 288 | 163 | 293 | 264 | 163 | 0 | 2 | 0 | 0 | 0 | 215 | 245 | 229 | 146 | 130 | 0 | 0 | 0 | 0 | 4 | 298 | 149 | 1 | 1 | 1 | 0.976 | 1 | 1 | 1 | 1 | 1 | 1 | 1 | 0.942 | 0.007 |
| SE | 92366 | ENSG00000080503 | SMARCA2 | chr9 | + | 2028986 | 2029247 | 2015970 | 2016143 | 2032951 | 2033081 | 243 | 129 | 239 | 221 | 125 | 0 | 0 | 0 | 1 | 0 | 139 | 213 | 180 | 110 | 119 | 0 | 0 | 0 | 0 | 0 | 298 | 149 | 1 | 1 | 1 | 1 | 1 | 0.991 | 1 | 1 | 1 | 1 | 1 | 1 | -0.002 |
| SE | 92367 | ENSG00000080503 | SMARCA2 | chr9 | + | 2028986 | 2029247 | 2017086 | 2017207 | 2032951 | 2033081 | 255 | 142 | 247 | 228 | 131 | 0 | 1 | 0 | 0 | 0 | 152 | 205 | 193 | 122 | 120 | 0 | 0 | 0 | 0 | 0 | 298 | 149 | 1 | 1 | 1 | 0.986 | 1 | 1 | 1 | 1 | 1 | 1 | 1 | 1 | -0.003 |
| SE | 92368 | ENSG00000080503 | SMARCA2 | chr9 | + | 2028986 | 2029247 | 2017487 | 2017799 | 2032951 | 2033081 | 253 | 133 | 255 | 231 | 124 | 0 | 1 | 0 | 1 | 0 | 151 | 207 | 194 | 118 | 131 | 0 | 0 | 0 | 0 | 0 | 298 | 149 | 1 | 1 | 1 | 0.985 | 1 | 0.991 | 1 | 1 | 1 | 1 | 1 | 1 | -0.005 |
| SE | 92369 | ENSG00000080503 | SMARCA2 | chr9 | + | 2047228 | 2047484 | 2039465 | 2039900 | 2054596 | 2054723 | 86 | 54 | 64 | 102 | 60 | 3 | 0 | 0 | 1 | 0 | 85 | 61 | 52 | 53 | 61 | 0 | 0 | 1 | 0 | 0 | 298 | 149 | 1 | 1 | 0.935 | 1 | 1 | 0.981 | 1 | 1 | 1 | 0.963 | 1 | 1 | -0.009 |
| SE | 92370 | ENSG00000080503 | SMARCA2 | chr9 | + | 2054596 | 2054723 | 2039465 | 2039900 | 2056671 | 2056845 | 231 | 144 | 181 | 328 | 276 | 0 | 0 | 0 | 0 | 0 | 205 | 190 | 133 | 141 | 136 | 0 | 0 | 3 | 0 | 0 | 276 | 149 | 1 | 1 | 1 | 1 | 1 | 1 | 1 | 1 | 1 | 0.96 | 1 | 1 | 0.008 |
| SE | 92371 | ENSG00000080503 | SMARCA2 | chr9 | + | 2056671 | 2056845 | 2054596 | 2054723 | 2058290 | 2058464 | 491 | 369 | 454 | 720 | 594 | 0 | 0 | 0 | 0 | 0 | 430 | 376 | 404 | 395 | 323 | 0 | 0 | 0 | 2 | 0 | 298 | 149 | 1 | 1 | 1 | 1 | 1 | 1 | 1 | 1 | 1 | 1 | 0.99 | 1 | 0.002 |
| SE | 92372 | ENSG00000080503 | SMARCA2 | chr9 | + | 2060815 | 2060986 | 2058290 | 2058464 | 2070417 | 2070471 | 418 | 392 | 434 | 631 | 455 | 0 | 0 | 0 | 0 | 0 | 594 | 371 | 408 | 396 | 327 | 0 | 0 | 0 | 0 | 5 | 298 | 149 | 1 | 1 | 1 | 1 | 1 | 1 | 1 | 1 | 1 | 1 | 1 | 0.97 | 0.006 |
| SE | 92373 | ENSG00000080503 | SMARCA2 | chr9 | + | 2072031 | 2072067 | 2070432 | 2070471 | 2073211 | 2073247 | 0 | 0 | 1 | 2 | 1 | 173 | 117 | 163 | 238 | 197 | 0 | 0 | 0 | 1 | 0 | 146 | 138 | 120 | 139 | 116 | 185 | 149 | 1 | 1 | 0 | 0 | 0.005 | 0.007 | 0.004 | 0 | 0 | 0 | 0.006 | 0 | 0.002 |
| SE | 92374 | ENSG00000080503 | SMARCA2 | chr9 | + | 2073565 | 2073623 | 2073096 | 2073342 | 2076228 | 2076329 | 366 | 238 | 288 | 389 | 352 | 0 | 0 | 0 | 1 | 0 | 240 | 213 | 283 | 216 | 222 | 0 | 0 | 2 | 0 | 0 | 207 | 149 | 1 | 1 | 1 | 1 | 1 | 0.996 | 1 | 1 | 1 | 0.99 | 1 | 1 | 0.001 |
| SE | 92379 | ENSG00000080503 | SMARCA2 | chr9 | + | 2104002 | 2104169 | 2101569 | 2101616 | 2110253 | 2110360 | 337 | 186 | 329 | 298 | 334 | 0 | 0 | 0 | 0 | 0 | 165 | 207 | 142 | 121 | 132 | 0 | 1 | 0 | 0 | 0 | 298 | 149 | 1 | 1 | 1 | 1 | 1 | 1 | 1 | 1 | 0.99 | 1 | 1 | 1 | 0.002 |
| SE | 92380 | ENSG00000080503 | SMARCA2 | chr9 | + | 2110253 | 2110417 | 2104002 | 2104169 | 2115821 | 2116049 | 337 | 175 | 347 | 335 | 292 | 0 | 0 | 0 | 0 | 0 | 199 | 229 | 137 | 168 | 88 | 0 | 0 | 2 | 0 | 2 | 298 | 149 | 1 | 1 | 1 | 1 | 1 | 1 | 1 | 1 | 1 | 0.972 | 1 | 0.957 | 0.014 |
| SE | 92381 | ENSG00000080503 | SMARCA2 | chr9 | + | 2115821 | 2116049 | 2104002 | 2104169 | 2119457 | 2119535 | 153 | 120 | 141 | 177 | 175 | 0 | 0 | 0 | 1 | 2 | 104 | 154 | 97 | 134 | 63 | 0 | 0 | 0 | 0 | 0 | 298 | 149 | 1 | 1 | 1 | 1 | 1 | 0.989 | 0.978 | 1 | 1 | 1 | 1 | 1 | -0.007 |
| SE | 92383 | ENSG00000080503 | SMARCA2 | chr9 | + | 2123718 | 2123937 | 2119457 | 2119535 | 2161685 | 2161795 | 353 | 310 | 286 | 400 | 398 | 0 | 0 | 0 | 0 | 0 | 303 | 294 | 303 | 369 | 225 | 0 | 0 | 1 | 0 | 0 | 298 | 149 | 1 | 1 | 1 | 1 | 1 | 1 | 1 | 1 | 1 | 0.993 | 1 | 1 | 0.001 |
| SE | 92386 | ENSG00000080503 | SMARCA2 | chr9 | + | 2159813 | 2159909 | 2158451 | 2158553 | 2161685 | 2161903 | 6 | 1 | 0 | 0 | 4 | 0 | 0 | 0 | 1 | 0 | 0 | 3 | 1 | 1 | 0 | 0 | 0 | 0 | 0 | 0 | 245 | 149 | 1 | 1 | 1 | 1 | NA | 0 | 1 | NA | 1 | 1 | 1 | NA | -0.25 |

| | | | | | | | | | | | | | |
|---|---|---|---|---|---|---|---|---|---|---|---|---|---|
| SE | 92392 | ENSG00000080503 | SMARCA2 | chr9 | + | 2161000 | 2161035 | 2159813 | 2159909 | 2161685 | 2161843 | 2 | 0 |
| SE | 92404 | ENSG00000010610 | CD4 | chr12 | + | 6799537 | 6799623 | 6789513 | 6789662 | 6800071 | 6800187 | 2 | 2 |
| SE | 92407 | ENSG00000010610 | CD4 | chr12 | + | 6800306 | 6800471 | 6800071 | 6800187 | 6814141 | 6814233 | 821 | 445 |
| SE | 92408 | ENSG00000010610 | CD4 | chr12 | + | 6814758 | 6814992 | 6814141 | 6814300 | 6816055 | 6816204 | 489 | 292 |
| SE | 92409 | ENSG00000010610 | CD4 | chr12 | + | 6816055 | 6816403 | 6814141 | 6814300 | 6817129 | 6817330 | 479 | 201 |
| SE | 92410 | ENSG00000104853 | CLPTM1 | chr19 | + | 44961962 | 44962075 | 44955379 | 44955467 | 44973086 | 44973210 | 128 | 64 |
| SE | 92412 | ENSG00000104853 | CLPTM1 | chr19 | + | 44990849 | 44990945 | 44989983 | 44990585 | 44992232 | 44992400 | 54 | 15 |
| SE | 92413 | ENSG00000104853 | CLPTM1 | chr19 | + | 44991237 | 44991373 | 44990849 | 44990945 | 44992232 | 44992400 | 194 | 83 |
| SE | 92443 | ENSG00000204778 | CBWD4P | chr9 | - | 65289826 | 65289895 | 65289657 | 65289723 | 65301424 | 65301476 | 0 | 0 |
| SE | 92444 | ENSG00000136950 | ARPC5L | chr9 | + | 124874974 | 124875151 | 124872588 | 124873764 | 124876877 | 124877401 | 382 | 254 |
| SE | 92446 | ENSG00000165092 | ALDH1A1 | chr9 | - | 72911957 | 72912122 | 72909601 | 72909759 | 72916919 | 72917104 | 344 | 268 |
| SE | 92447 | ENSG00000165092 | ALDH1A1 | chr9 | - | 72924018 | 72924132 | 72918719 | 72918822 | 72925483 | 72925612 | 207 | 148 |
| SE | 92451 | ENSG00000163378 | EOGT | chr3 | - | 68998010 | 68998114 | 68988505 | 68989017 | 69001607 | 69001641 | 34 | 40 |
| SE | 92452 | ENSG00000163378 | EOGT | chr3 | - | 68998010 | 68998114 | 68988505 | 68989017 | 69004377 | 69004482 | 16 | 14 |
| SE | 92454 | ENSG00000163378 | EOGT | chr3 | - | 69001607 | 69001714 | 68998010 | 68998114 | 69004377 | 69004482 | 37 | 37 |
| SE | 92458 | ENSG00000100815 | TRIP11 | chr14 | - | 91974626 | 91974743 | 91972716 | 91972861 | 91975171 | 91975286 | 110 | 111 |
| SE | 92459 | ENSG00000100815 | TRIP11 | chr14 | - | 91974626 | 91974743 | 91972716 | 91972861 | 91976107 | 91976189 | 61 | 61 |
| SE | 92460 | ENSG00000100815 | TRIP11 | chr14 | - | 91975171 | 91975286 | 91972716 | 91972861 | 91976107 | 91976189 | 132 | 143 |
| SE | 92462 | ENSG00000100815 | TRIP11 | chr14 | - | 91976107 | 91976189 | 91974626 | 91974743 | 91977125 | 91977248 | 25 | 16 |
| SE | 92463 | ENSG00000100815 | TRIP11 | chr14 | - | 91976107 | 91976189 | 91975171 | 91975286 | 91988283 | 91988383 | 206 | 204 |
| SE | 92464 | ENSG00000100815 | TRIP11 | chr14 | - | 91977125 | 91977248 | 91976107 | 91976189 | 91988283 | 91988383 | 38 | 29 |
| SE | 92466 | ENSG00000100815 | TRIP11 | chr14 | - | 91995351 | 91995515 | 91993808 | 91993912 | 91999967 | 92000108 | 78 | 74 |
| SE | 92467 | ENSG00000100815 | TRIP11 | chr14 | - | 91999239 | 91999433 | 91993808 | 91993912 | 91999967 | 92000108 | 112 | 82 |
| SE | 92468 | ENSG00000100815 | TRIP11 | chr14 | - | 91999239 | 91999433 | 91995351 | 91995515 | 91999967 | 92000108 | 180 | 137 |
| SE | 92469 | ENSG00000100815 | TRIP11 | chr14 | - | 92003418 | 92006448 | 91999967 | 92000108 | 92007639 | 92007852 | 139 | 182 |
| SE | 92470 | ENSG00000100815 | TRIP11 | chr14 | - | 92007639 | 92007852 | 91999967 | 92000108 | 92010985 | 92011072 | 143 | 172 |
| SE | 92473 | ENSG00000100815 | TRIP11 | chr14 | - | 92011754 | 92011795 | 92007639 | 92007852 | 92014214 | 92014577 | 26 | 53 |
| SE | 92475 | ENSG00000100815 | TRIP11 | chr14 | - | 92014214 | 92014577 | 92011754 | 92011795 | 92015695 | 92015751 | 197 | 243 |
| SE | 92476 | ENSG00000100815 | TRIP11 | chr14 | - | 92021555 | 92021831 | 92017681 | 92017750 | 92025309 | 92025420 | 274 | 181 |

| | | | | | | | | | | | | |
|---|---|---|---|---|---|---|---|---|---|---|---|---|
| SE | 92836 | ENSG00000130958 | SLC35D2 | chr9 | - | 96343903 | 96343996 | 96336716 | 96336784 | 96351102 | 96351171 | 58 | 58 | [illegible] |
| SE | 92837 | ENSG00000130958 | SLC35D2 | chr9 | - | 96343903 | 96343996 | 96336716 | 96336784 | 96360153 | 96360192 | 55 | 55 | [illegible] |
| SE | 92838 | ENSG00000130958 | SLC35D2 | chr9 | - | 96343903 | 96343996 | 96336716 | 96336784 | 96364463 | 96364537 | 56 | 59 | [illegible] |
| SE | 92839 | ENSG00000130958 | SLC35D2 | chr9 | - | 96345298 | 96345401 | 96336716 | 96336784 | 96351102 | 96351171 | 76 | 39 | [illegible] |
| SE | 92840 | ENSG00000130958 | SLC35D2 | chr9 | - | 96345298 | 96345401 | 96336716 | 96336784 | 96360153 | 96360192 | 0 | 2 | [illegible] |
| SE | 92841 | ENSG00000130958 | SLC35D2 | chr9 | - | 96345298 | 96345401 | 96336716 | 96336784 | 96364463 | 96364537 | 0 | 2 | [illegible] |
| SE | 92845 | ENSG00000130958 | SLC35D2 | chr9 | - | 96351102 | 96351171 | 96324090 | 96324169 | 96360153 | 96360192 | 2 | 0 | [illegible] |
| SE | 92847 | ENSG00000130958 | SLC35D2 | chr9 | - | 96351102 | 96351171 | 96343903 | 96343996 | 96360153 | 96360192 | 1 | 2 | [illegible] |
| SE | 92849 | ENSG00000130958 | SLC35D2 | chr9 | - | 96352037 | 96352109 | 96350748 | 96351171 | 96360153 | 96360192 | 130 | 124 | [illegible] |
| SE | 92854 | ENSG00000130958 | SLC35D2 | chr9 | - | 96360153 | 96360221 | 96345298 | 96345401 | 96364463 | 96364550 | 23 | 23 | [illegible] |
| SE | 92858 | ENSG00000130958 | SLC35D2 | chr9 | - | 96373999 | 96374082 | 96368271 | 96368305 | 96383476 | 96383680 | 0 | 0 | [illegible] |
| SE | 92859 | ENSG00000147872 | PLIN2 | chr9 | - | 19116358 | 19116649 | 19108374 | 19108747 | 19118320 | 19118446 | 239 | 104 | [illegible] |
| SE | 92869 | ENSG00000135018 | UBQLN1 | chr9 | - | 83665029 | 83665145 | 83663874 | 83664043 | 83666349 | 83666394 | 706 | 410 | [illegible] |
| SE | 92871 | ENSG00000135018 | UBQLN1 | chr9 | - | 83666349 | 83666394 | 83663874 | 83664043 | 83669184 | 83669291 | 1 | 0 | [illegible] |
| SE | 92875 | ENSG00000135018 | UBQLN1 | chr9 | - | 83669184 | 83669327 | 83665029 | 83665145 | 83707499 | 83707934 | 27 | 17 | [illegible] |
| SE | 92876 | ENSG00000135018 | UBQLN1 | chr9 | - | 83677726 | 83677961 | 83666349 | 83669327 | 83678440 | 83678599 | 640 | 401 | [illegible] |
| SE | 92877 | ENSG00000135018 | UBQLN1 | chr9 | - | 83677726 | 83677961 | 83666349 | 83669327 | 83682950 | 83683066 | 360 | 249 | [illegible] |
| SE | 92880 | ENSG00000135018 | UBQLN1 | chr9 | - | 83678440 | 83678599 | 83666349 | 83669327 | 83707499 | 83707601 | 8 | 3 | [illegible] |
| SE | 92881 | ENSG00000135018 | UBQLN1 | chr9 | - | 83678440 | 83678599 | 83677726 | 83677961 | 83682950 | 83683066 | 285 | 163 | [illegible] |
| SE | 92882 | ENSG00000135018 | UBQLN1 | chr9 | - | 83679774 | 83680037 | 83666349 | 83669327 | 83682950 | 83683066 | 355 | 258 | [illegible] |
| SE | 92884 | ENSG00000135018 | UBQLN1 | chr9 | - | 83679774 | 83680037 | 83678440 | 83678599 | 83686003 | 83686053 | 481 | 319 | [illegible] |
| SE | 92885 | ENSG00000135018 | UBQLN1 | chr9 | - | 83682950 | 83683066 | 83666349 | 83669327 | 83707499 | 83707934 | 3 | 1 | [illegible] |
| SE | 92886 | ENSG00000135018 | UBQLN1 | chr9 | - | 83682950 | 83683066 | 83678440 | 83678599 | 83686003 | 83686053 | 214 | 145 | [illegible] |
| SE | 92888 | ENSG00000135018 | UBQLN1 | chr9 | - | 83682950 | 83683066 | 83679774 | 83680037 | 83686003 | 83686053 | 562 | 391 | [illegible] |
| SE | 92889 | ENSG00000135018 | UBQLN1 | chr9 | - | 83686003 | 83686155 | 83678440 | 83678599 | 83707499 | 83707934 | 124 | 72 | [illegible] |
| SE | 92890 | ENSG00000135018 | UBQLN1 | chr9 | - | 83686003 | 83686155 | 83682950 | 83683066 | 83707499 | 83707934 | 370 | 247 | [illegible] |
| SE | 92893 | ENSG00000136874 | STX17 | chr9 | + | 99928777 | 99928843 | 99915177 | 99915362 | 99951059 | 99951184 | 127 | 150 | [illegible] |
| SE | 92894 | ENSG00000136874 | STX17 | chr9 | + | 99945718 | 99945850 | 99928777 | 99928843 | 99951059 | 99951285 | 0 | 0 | [illegible] |
| SE | 92895 | ENSG00000136874 | STX17 | chr9 | + | 99959916 | 99960032 | 99951059 | 99951285 | 99960104 | 99960155 | 129 | 88 | [illegible] |
| SE | 92896 | ENSG00000136874 | STX17 | chr9 | + | 99960104 | 99960155 | 99959916 | 99960032 | 99967652 | 99967739 | 184 | 123 | [illegible] |
| SE | 92897 | ENSG00000100353 | EIF3D | chr22 | - | 36517300 | 36517431 | 36516704 | 36516790 | 36518762 | 36518910 | 549 | 269 | [illegible] |
| SE | 92898 | ENSG00000100353 | EIF3D | chr22 | - | 36517300 | 36517493 | 36516704 | 36516790 | 36518762 | 36518910 | 215 | 108 | [illegible] |
| SE | 92899 | ENSG00000100353 | EIF3D | chr22 | - | 36520575 | 36520688 | 36519404 | 36519537 | 36523208 | 36523281 | 2086 | 1510 | [illegible] |
| SE | 92900 | ENSG00000100353 | EIF3D | chr22 | - | 36526819 | 36526905 | 36525998 | 36526131 | 36529075 | 36529166 | 0 | 2 | [illegible] |
| SE | 92901 | ENSG00000006607 | FARP2 | chr2 | + | 241373083 | 241373290 | 241356284 | 241356388 | 241403827 | 241403932 | 14 | 2 | [illegible] |
| SE | 92905 | ENSG00000006607 | FARP2 | chr2 | + | 241404798 | 241404841 | 241403827 | 241403932 | 241407536 | 241407608 | 28 | 23 | [illegible] |
| SE | 92906 | ENSG00000006607 | FARP2 | chr2 | + | 241404798 | 241404841 | 241403827 | 241403932 | 241411032 | 241411130 | 6 | 3 | [illegible] |
| SE | 92907 | ENSG00000006607 | FARP2 | chr2 | + | 241405255 | 241405402 | 241404798 | 241404841 | 241407536 | 241407615 | 0 | 0 | [illegible] |
| SE | 92908 | ENSG00000006607 | FARP2 | chr2 | + | 241407536 | 241407615 | 241403827 | 241403932 | 241411032 | 241411130 | 13 | 11 | [illegible] |
| SE | 92909 | ENSG00000006607 | FARP2 | chr2 | + | 241407536 | 241407615 | 241404798 | 241404841 | 241411032 | 241411130 | 36 | 32 | [illegible] |
| SE | 92910 | ENSG00000006607 | FARP2 | chr2 | + | 241434157 | 241434321 | 241431678 | 241431774 | 241434961 | 241435030 | 82 | 62 | [illegible] |
| SE | 92911 | ENSG00000006607 | FARP2 | chr2 | + | 241436480 | 241436538 | 241434961 | 241435030 | 241441303 | 241441556 | 42 | 28 | [illegible] |
| SE | 92916 | ENSG00000131089 | ARHGEF9 | chrX | - | 63643979 | 63644048 | 63634966 | 63638209 | 63665885 | 63666017 | 13 | 8 | [illegible] |
| SE | 92918 | ENSG00000131089 | ARHGEF9 | chrX | - | 63655493 | 63655737 | 63643979 | 63644048 | 63674037 | 63674167 | 71 | 32 | [illegible] |
| SE | 92921 | ENSG00000131089 | ARHGEF9 | chrX | - | 63665885 | 63666017 | 63643979 | 63644048 | 63674037 | 63674138 | 62 | 42 | [illegible] |
| SE | 92923 | ENSG00000131089 | ARHGEF9 | chrX | - | 63701489 | 63701642 | 63697124 | 63697304 | 63703169 | 63703284 | 0 | 0 | [illegible] |
| SE | 92925 | ENSG00000131089 | ARHGEF9 | chrX | - | 63706257 | 63706449 | 63697124 | 63697304 | 63754314 | 63755063 | 60 | 26 | [illegible] |
| SE | 92927 | ENSG00000131089 | ARHGEF9 | chrX | - | 63715987 | 63716119 | 63706257 | 63706449 | 63724531 | 63724711 | 0 | 0 | [illegible] |
| SE | 92928 | ENSG00000131089 | ARHGEF9 | chrX | - | 63719673 | 63719856 | 63706257 | 63706449 | 63754314 | 63754676 | 1 | 0 | [illegible] |
| SE | 92930 | ENSG00000131089 | ARHGEF9 | chrX | - | 63724531 | 63724711 | 63706257 | 63706449 | 63743368 | 63743446 | 40 | 21 | [illegible] |
| SE | 92936 | ENSG00000131089 | ARHGEF9 | chrX | - | 63724795 | 63724846 | 63724531 | 63724711 | 63754314 | 63754676 | 2 | 0 | [illegible] |
| SE | 92938 | ENSG00000131089 | ARHGEF9 | chrX | - | 63725682 | 63725735 | 63724531 | 63724711 | 63754314 | 63755124 | 1 | 0 | [illegible] |
| SE | 92939 | ENSG00000131089 | ARHGEF9 | chrX | - | 63726645 | 63726693 | 63724531 | 63724711 | 63754303 | 63754747 | 0 | 0 | [illegible] |
| SE | 92940 | ENSG00000131089 | ARHGEF9 | chrX | - | 63727190 | 63727419 | 63724531 | 63724711 | 63736989 | 63737118 | 5 | 5 | [illegible] |
| SE | 92942 | ENSG00000131089 | ARHGEF9 | chrX | - | 63736989 | 63737118 | 63724531 | 63724711 | 63743368 | 63743446 | 0 | 1 | [illegible] |
| SE | 92943 | ENSG00000131089 | ARHGEF9 | chrX | - | 63736989 | 63737118 | 63724531 | 63724711 | 63754303 | 63754676 | 0 | 1 | [illegible] |
| SE | 92944 | ENSG00000131089 | ARHGEF9 | chrX | - | 63736989 | 63737118 | 63724531 | 63724711 | 63754314 | 63755124 | 0 | 2 | [illegible] |
| SE | 92945 | ENSG00000131089 | ARHGEF9 | chrX | - | 63743368 | 63743446 | 63706257 | 63706449 | 63754303 | 63755113 | 0 | 0 | [illegible] |
| SE | 92948 | ENSG00000131089 | ARHGEF9 | chrX | - | 63743368 | 63743446 | 63724531 | 63724711 | 63754303 | 63755113 | 1 | 0 | [illegible] |
| SE | 92949 | ENSG00000131089 | ARHGEF9 | chrX | - | 63743368 | 63743446 | 63724531 | 63724711 | 63754307 | 63754676 | 1 | 0 | [illegible] |
| SE | 92950 | ENSG00000131089 | ARHGEF9 | chrX | - | 63743368 | 63743446 | 63724531 | 63724711 | 63754314 | 63754740 | 1 | 0 | [illegible] |
| SE | 92953 | ENSG00000131089 | ARHGEF9 | chrX | - | 63754303 | 63754676 | 63724531 | 63724711 | 63754920 | 63755078 | 8 | 10 | [illegible] |
| SE | 92962 | ENSG00000108669 | CYTH1 | chr17 | - | 78692416 | 78692493 | 78680970 | 78681042 | 78698268 | 78698380 | 602 | 396 | [illegible] |
| SE | 92963 | ENSG00000108669 | CYTH1 | chr17 | - | 78692416 | 78692493 | 78680970 | 78681042 | 78698819 | 78698968 | 598 | 395 | [illegible] |
| SE | 92964 | ENSG00000108669 | CYTH1 | chr17 | - | 78692416 | 78692493 | 78680970 | 78681042 | 78700330 | 78700443 | 598 | 395 | [illegible] |
| SE | 92966 | ENSG00000108669 | CYTH1 | chr17 | - | 78696006 | 78696009 | 78692416 | 78692493 | 78698819 | 78698968 | 0 | 0 | [illegible] |
| SE | 92967 | ENSG00000108669 | CYTH1 | chr17 | - | 78696006 | 78696009 | 78692416 | 78692493 | 78700330 | 78700443 | 0 | 0 | [illegible] |
| SE | 92969 | ENSG00000108669 | CYTH1 | chr17 | - | 78698268 | 78698380 | 78680970 | 78681042 | 78700330 | 78700443 | 0 | 1 | [illegible] |
| SE | 92970 | ENSG00000108669 | CYTH1 | chr17 | - | 78698268 | 78698380 | 78692416 | 78692493 | 78698819 | 78698968 | 431 | 217 | [illegible] |
| SE | 92971 | ENSG00000108669 | CYTH1 | chr17 | - | 78698268 | 78698380 | 78692416 | 78692493 | 78700330 | 78700443 | 6 | 1 | [illegible] |
| SE | 92972 | ENSG00000108669 | CYTH1 | chr17 | - | 78698268 | 78698380 | 78696006 | 78696009 | 78698819 | 78698968 | 1038 | 555 | [illegible] |
| SE | 92973 | ENSG00000108669 | CYTH1 | chr17 | - | 78698268 | 78698380 | 78696006 | 78696009 | 78700330 | 78700443 | 522 | 290 | [illegible] |
| SE | 92974 | ENSG00000108669 | CYTH1 | chr17 | - | 78698819 | 78698968 | 78680970 | 78681042 | 78700330 | 78700443 | 835 | 440 | [illegible] |
| SE | 92975 | ENSG00000108669 | CYTH1 | chr17 | - | 78698819 | 78698968 | 78692416 | 78692493 | 78700330 | 78700443 | 835 | 440 | [illegible] |
| SE | 92976 | ENSG00000108669 | CYTH1 | chr17 | - | 78698819 | 78698968 | 78696006 | 78696009 | 78700330 | 78700443 | 838 | 440 | [illegible] |
| SE | 92977 | ENSG00000108669 | CYTH1 | chr17 | - | 78698819 | 78698968 | 78698267 | 78698380 | 78700330 | 78700443 | 1389 | 730 | [illegible] |
| SE | 92979 | ENSG00000108669 | CYTH1 | chr17 | - | 78708196 | 78708261 | 78702121 | 78702240 | 78709649 | 78709732 | 145 | 111 | [illegible] |
| SE | 92981 | ENSG00000148082 | SHC3 | chr9 | - | 89037992 | 89038288 | 89005770 | 89013575 | 89042025 | 89042137 | 0 | 1 | [illegible] |
| SE | 92991 | ENSG00000139726 | DENR | chr12 | + | 122768781 | 122768921 | 122767487 | 122767604 | 122769033 | 122771064 | 766 | 582 | [illegible] |
| SE | 92992 | ENSG00000095209 | TMEM38B | chr9 | + | 105702706 | 105702822 | 105694543 | 105694772 | 105705596 | 105705753 | 0 | 0 | [illegible] |
| SE | 92993 | ENSG00000095209 | TMEM38B | chr9 | + | 105721536 | 105721721 | 105705596 | 105705753 | 105722533 | 105722621 | 27 | 21 | [illegible] |
| SE | 92994 | ENSG00000095209 | TMEM38B | chr9 | + | 105722533 | 105722621 | 105705596 | 105705753 | 105748072 | 105748190 | 52 | 29 | [illegible] |
| SE | 92995 | ENSG00000095209 | TMEM38B | chr9 | + | 105748072 | 105748190 | 105722533 | 105722621 | 105773864 | 105776612 | 93 | 47 | [illegible] |
| SE | 92996 | ENSG00000095209 | TMEM38B | chr9 | + | 105749075 | 105749111 | 105748072 | 105748190 | 105773864 | 105774079 | 0 | 2 | [illegible] |
| SE | 93011 | ENSG00000147162 | OGT | chrX | + | 71536177 | 71536358 | 71533103 | 71533336 | 71537828 | 71538072 | 1881 | 1430 | [illegible] |
| SE | 93012 | ENSG00000147162 | OGT | chrX | + | 71536207 | 71536358 | 71533103 | 71533336 | 71537828 | 71538072 | 1059 | 883 | [illegible] |
| SE | 93013 | ENSG00000147162 | OGT | chrX | + | 71537828 | 71538072 | 71533103 | 71533336 | 71544566 | 71544624 | 508 | 377 | [illegible] |
| SE | 93014 | ENSG00000147162 | OGT | chrX | + | 71537828 | 71538072 | 71533103 | 71533336 | 71547906 | 71548023 | 9 | 10 | [illegible] |
| SE | 93015 | ENSG00000147162 | OGT | chrX | + | 71537828 | 71538072 | 71533103 | 71533336 | 71554512 | 71554592 | 2 | 8 | [illegible] |
| SE | 93016 | ENSG00000147162 | OGT | chrX | + | 71544566 | 71544635 | 71533103 | 71533336 | 71547906 | 71548023 | 73 | 30 | [illegible] |
| SE | 93018 | ENSG00000147162 | OGT | chrX | + | 71544566 | 71548023 | 71533103 | 71533336 | 71554512 | 71554592 | 600 | 492 | [illegible] |
| SE | 93019 | ENSG00000147162 | OGT | chrX | + | 71544566 | 71548023 | 71537828 | 71538072 | 71554512 | 71554592 | 1071 | 849 | [illegible] |
| SE | 93020 | ENSG00000147162 | OGT | chrX | + | 71544566 | 71548023 | 71537828 | 71538072 | 71555953 | 71556094 | 473 | 358 | [illegible] |
| SE | 93021 | ENSG00000147162 | OGT | chrX | + | 71547906 | 71548023 | 71533103 | 71533336 | 71554512 | 71554592 | 491 | 400 | [illegible] |
| SE | 93023 | ENSG00000147162 | OGT | chrX | + | 71547906 | 71548023 | 71537828 | 71538072 | 71555953 | 71556094 | 8 | 3 | [illegible] |
| SE | 93024 | ENSG00000147162 | OGT | chrX | + | 71554512 | 71554592 | 71537828 | 71538072 | 71555953 | 71556094 | 260 | 204 | [illegible] |
| SE | 93025 | ENSG00000147162 | OGT | chrX | + | 71554512 | 71554592 | 71544566 | 71548023 | 71555189 | 71555214 | 1065 | 847 | [illegible] |
| SE | 93026 | ENSG00000147162 | OGT | chrX | + | 71554512 | 71554592 | 71544566 | 71548023 | 71555953 | 71556094 | 759 | 611 | [illegible] |
| SE | 93027 | ENSG00000147162 | OGT | chrX | + | 71555189 | 71555385 | 71547906 | 71548023 | 71555953 | 71556094 | 632 | 589 | [illegible] |
| SE | 93029 | ENSG00000147162 | OGT | chrX | + | 71555953 | 71556094 | 71554512 | 71554592 | 71556679 | 71556778 | 996 | 789 | [illegible] |
| SE | 93030 | ENSG00000147162 | OGT | chrX | + | 71556951 | 71557105 | 71556679 | 71556780 | 71557194 | 71557296 | 1074 | 818 | [illegible] |
| SE | 93032 | ENSG00000147162 | OGT | chrX | + | 71561774 | 71561900 | 71559587 | 71559677 | 71562846 | 71563017 | 961 | 755 | [illegible] |
| SE | 93033 | ENSG00000147162 | OGT | chrX | + | 71563129 | 71563246 | 71562846 | 71563017 | 71563328 | 71563499 | 2482 | 2783 | [illegible] |
| SE | 93034 | ENSG00000107242 | PIP5K1B | chr9 | + | 68742500 | 68742657 | 68705239 | 68705762 | 68786082 | 68786160 | 3 | 1 | [illegible] |
| SE | 93038 | ENSG00000107242 | PIP5K1B | chr9 | + | 68863836 | 68863967 | 68822614 | 68822683 | 68888980 | 68889133 | 23 | 8 | [illegible] |
| SE | 93039 | ENSG00000107242 | PIP5K1B | chr9 | + | 68876676 | 68876794 | 68822614 | 68822683 | 68888980 | 68889133 | 33 | 48 | [illegible] |
| SE | 93042 | ENSG00000107242 | PIP5K1B | chr9 | + | 68934889 | 68935045 | 68923301 | 68923386 | 68940645 | 68940790 | 85 | 89 | [illegible] |
| SE | 93043 | ENSG00000107242 | PIP5K1B | chr9 | + | 68940645 | 68940790 | 68934889 | 68935045 | 68991139 | 68991257 | 52 | 58 | [illegible] |
| SE | 93044 | ENSG00000107242 | PIP5K1B | chr9 | + | 68991139 | 68991257 | 68940645 | 68940790 | 69008446 | 69009176 | 23 | 17 | [illegible] |
| SE | 93047 | ENSG00000165138 | ANKS6 | chr9 | - | 98768080 | 98768250 | 98756419 | 98756603 | 98770895 | 98771046 | 5 | 0 | [illegible] |
| SE | 93048 | ENSG00000165138 | ANKS6 | chr9 | - | 98770895 | 98771046 | 98756419 | 98756603 | 98773876 | 98774080 | 2 | 0 | [illegible] |
| SE | 93050 | ENSG00000165138 | ANKS6 | chr9 | - | 98782466 | 98782573 | 98780188 | 98780337 | 98783952 | 98784157 | 18 | 10 | [illegible] |
| SE | 93053 | ENSG00000136052 | SLC41A2 | chr12 | - | 104931631 | 104931791 | 104928299 | 104928694 | 104958189 | 104958319 | 4 | 1 | [illegible] |
| SE | 93057 | ENSG00000163820 | FYCO1 | chr3 | - | 45923655 | 45923765 | 45917898 | 45921840 | 45931070 | 45931281 | 122 | 42 | [illegible] |
| SE | 93058 | ENSG00000163820 | FYCO1 | chr3 | - | 45931070 | 45931281 | 45923655 | 45923765 | 45936447 | 45936543 | 138 | 54 | [illegible] |
| SE | 93062 | ENSG00000163820 | FYCO1 | chr3 | - | 45965032 | 45965125 | 45962224 | 45962392 | 45966276 | 45968703 | 23 | 25 | [illegible] |
| SE | 93064 | ENSG00000163820 | FYCO1 | chr3 | - | 45973087 | 45973231 | 45969674 | 45969765 | 45975238 | 45975345 | 213 | 125 | [illegible] |
| SE | 93068 | ENSG00000163820 | FYCO1 | chr3 | - | 45979704 | 45979830 | 45975238 | 45975345 | 45981569 | 45981676 | 135 | 68 | [illegible] |
| SE | 93069 | ENSG00000285269 | NA | chr9 | - | 96249909 | 96250052 | 96249750 | 96249786 | 96251417 | 96251485 | 1 | 0 | [illegible] |
| SE | 93070 | ENSG00000285269 | NA | chr9 | - | 96249909 | 96250485 | 96249750 | 96249786 | 96251417 | 96251485 | 0 | 0 | [illegible] |
| SE | 93071 | ENSG00000285269 | NA | chr9 | - | 96250295 | 96250485 | 96249909 | 96250052 | 96251417 | 96251485 | 0 | 0 | [illegible] |
| SE | 93075 | ENSG00000154277 | UCHL1 | chr4 | + | 41261875 | 41261923 | 41261714 | 41261800 | 41263224 | 41263291 | 0 | 0 | [illegible] |
| SE | 93082 | ENSG00000175938 | ORAI3 | chr16 | + | 30953184 | 30953712 | 30949122 | 30949517 | 30956199 | 30956461 | 6 | 7 | [illegible] |
| SE | 93083 | ENSG00000113924 | HGD | chr3 | - | 120633146 | 120633328 | 120628171 | 120628529 | 120641588 | 120641693 | 6 | 15 | [illegible] |
| SE | 93085 | ENSG00000113924 | HGD | chr3 | - | 120638454 | 120638581 | 120633146 | 120633328 | 120646972 | 120647052 | 3 | 7 | [illegible] |
| SE | 93087 | ENSG00000113924 | HGD | chr3 | - | 120641588 | 120641693 | 120633146 | 120633328 | 120646972 | 120647052 | 13 | 25 | [illegible] |
| SE | 93088 | ENSG00000113924 | HGD | chr3 | - | 120641588 | 120641693 | 120638454 | 120638581 | 120646972 | 120647052 | 22 | 24 | [illegible] |
| SE | 93093 | ENSG00000113924 | HGD | chr3 | - | 120646266 | 120646366 | 120644318 | 120644443 | 120646972 | 120647052 | 0 | 2 | [illegible] |
| SE | 93095 | ENSG00000113924 | HGD | chr3 | - | 120677903 | 120677991 | 120675791 | 120675863 | 120682096 | 120682131 | 0 | 0 | [illegible] |
| SE | 93096 | ENSG00000113924 | HGD | chr3 | - | 120677903 | 120678029 | 120675791 | 120675863 | 120682096 | 120682243 | 0 | 0 | [illegible] |
| SE | 93100 | ENSG00000105879 | CBLL1 | chr7 | + | 107755417 | 107755491 | 107753894 | 107753978 | 107758142 | 107758180 | 406 | 320 | [illegible] |
| SE | 93122 | ENSG00000079387 | SENP1 | chr12 | - | 48066926 | 48066965 | 48065595 | 48065680 | 48071666 | 48071721 | 86 | 81 | [illegible] |
| SE | 93123 | ENSG00000079387 | SENP1 | chr12 | - | 48083590 | 48083762 | 48074689 | 48074793 | 48088800 | 48088960 | 151 | 182 | [illegible] |
| SE | 93125 | ENSG00000079387 | SENP1 | chr12 | - | 48096342 | 48096427 | 48088552 | 48088960 | 48097993 | 48098034 | 94 | 70 | [illegible] |
| SE | 93126 | ENSG00000079387 | SENP1 | chr12 | - | 48097993 | 48098124 | 48096342 | 48096427 | 48101468 | 48101472 | 113 | 55 | [illegible] |
| SE | 93128 | ENSG00000079387 | SENP1 | chr12 | - | 48099061 | 48099154 | 48097993 | 48098124 | 48106027 | 48106052 | 2 | 2 | [illegible] |
| SE | 93129 | ENSG00000079387 | SENP1 | chr12 | - | 48101468 | 48101516 | 48097736 | 48098124 | 48105401 | 48105834 | 46 | 13 | [illegible] |
| SE | 93132 | ENSG00000283486 | FAM95C | chr9 | - | 38541288 | 38541562 | 38540682 | 38540849 | 38542311 | 38543156 | 2 | 0 | [illegible] |
| SE | 93134 | ENSG00000283486 | FAM95C | chr9 | - | 38573083 | 38573129 | 38572005 | 38572059 | 38575475 | 38575698 | 7 | 2 | [illegible] |
| SE | 93135 | ENSG00000164975 | SNAPC3 | chr9 | + | 15447094 | 15447244 | 15444601 | 15444706 | 15451319 | 15451402 | 162 | 230 | [illegible] |
| SE | 93137 | ENSG00000164975 | SNAPC3 | chr9 | + | 15453040 | 15453205 | 15451319 | 15451402 | 15457959 | 15458067 | 195 | 217 | [illegible] |
| SE | 93138 | ENSG00000164975 | SNAPC3 | chr9 | + | 15459718 | 15460313 | 15457959 | 15458067 | 15461630 | 15461805 | 182 | 173 | [illegible] |
| SE | 93139 | ENSG00000164975 | SNAPC3 | chr9 | + | 15462864 | 15462990 | 15461630 | 15461805 | 15465509 | 15465832 | 3 | 1 | [illegible] |
| SE | 93140 | ENSG00000148158 | SNX30 | chr9 | + | 112817704 | 112817815 | 112804775 | 112804967 | 112830724 | 112830883 | 216 | 130 | [illegible] |
| SE | 93141 | ENSG00000148158 | SNX30 | chr9 | + | 112836213 | 112836409 | 112830724 | 112830883 | 112838497 | 112838697 | 175 | 182 | [illegible] |
| SE | 93142 | ENSG00000148158 | SNX30 | chr9 | + | 112838497 | 112838697 | 112830724 | 112830883 | 112850858 | 112850945 | 116 | 92 | [illegible] |
| SE | 93144 | ENSG00000148158 | SNX30 | chr9 | + | 112875168 | 112875291 | 112864246 | 112864399 | 112875831 | 112875929 | 22 | 5 | [illegible] |
| SE | 93147 | ENSG00000077721 | UBE2A | chrX | + | 119581506 | 119581596 | 119575374 | 119575400 | 119582596 | 119582676 | 26 | 13 | [illegible] |
| SE | 93148 | ENSG00000197969 | VPS13A | chr9 | + | 77205312 | 77205408 | 77201364 | 77201407 | 77205977 | 77206079 | 231 | 252 | [illegible] |
| SE | 93149 | ENSG00000197969 | VPS13A | chr9 | + | 77209422 | 77209532 | 77205977 | 77206079 | 77210615 | 77210675 | 219 | 265 | [illegible] |
| SE | 93150 | ENSG00000197969 | VPS13A | chr9 | + | 77214328 | 77214386 | 77213233 | 77213314 | 77219953 | 77220081 | 229 | 234 | [illegible] |
| SE | 93151 | ENSG00000197969 | VPS13A | chr9 | + | 77219953 | 77220081 | 77214328 | 77214386 | 77220276 | 77220383 | 264 | 279 | [illegible] |
| SE | 93153 | ENSG00000197969 | VPS13A | chr9 | + | 77247258 | 77247395 | 77238271 | 77238386 | 77250096 | 77250229 | 133 | 78 | [illegible] |
| SE | 93156 | ENSG00000197969 | VPS13A | chr9 | + | 77307944 | 77308098 | 77302914 | 77303062 | 77313991 | 77314119 | 112 | 124 | [illegible] |
| SE | 93157 | ENSG00000197969 | VPS13A | chr9 | + | 77353408 | 77353641 | 77351316 | 77351446 | 77356713 | 77356867 | 195 | 226 | [illegible] |
| SE | 93158 | ENSG00000197969 | VPS13A | chr9 | + | 77356713 | 77356867 | 77353408 | 77353641 | 77357691 | 77357838 | 147 | 187 | [illegible] |
| SE | 93159 | ENSG00000197969 | VPS13A | chr9 | + | 77381975 | 77382087 | 77371025 | 77371149 | 77403235 | 77403321 | 187 | 204 | [illegible] |
| SE | 93163 | ENSG00000137757 | CASP5 | chr11 | - | 104997382 | 104997492 | 104995739 | 104995842 | 104998884 | 104999028 | 34 | 33 | [illegible] |
| SE | 93165 | ENSG00000137757 | CASP5 | chr11 | - | 105003273 | 105003383 | 105002027 | 105002201 | 105023129 | 105023136 | 7 | 13 | [illegible] |
| SE | 93167 | ENSG00000137757 | CASP5 | chr11 | - | 105007082 | 105007334 | 105003273 | 105003383 | 105008806 | 105008980 | 14 | 19 | [illegible] |
| SE | 93168 | ENSG00000137757 | CASP5 | chr11 | - | 105007082 | 105007334 | 105003273 | 105003383 | 105023129 | 105023136 | 10 | 14 | [illegible] |
| SE | 93169 | ENSG00000137757 | CASP5 | chr11 | - | 105008806 | 105008980 | 105003273 | 105003383 | 105023129 | 105023168 | 1 | 5 | [illegible] |
| SE | 93170 | ENSG00000137757 | CASP5 | chr11 | - | 105008806 | 105008980 | 105007082 | 105007334 | 105023129 | 105023168 | 7 | 9 | [illegible] |
| SE | 93171 | ENSG00000137757 | CASP5 | chr11 | - | 105008806 | 105009019 | 105003273 | 105003383 | 105023129 | 105023136 | 0 | 2 | [illegible] |
| SE | 93172 | ENSG00000137757 | CASP5 | chr11 | - | 105008806 | 105009019 | 105007082 | 105007334 | 105023129 | 105023136 | 6 | 6 | [illegible] |
| SE | 93173 | ENSG00000137757 | CASP5 | chr11 | - | 105008806 | 105009053 | 105003273 | 105003383 | 105023129 | 105023143 | 0 | 2 | [illegible] |
| SE | 93174 | ENSG00000137757 | CASP5 | chr11 | - | 105008806 | 105009053 | 105007082 | 105007334 | 105023129 | 105023143 | 6 | 6 | [illegible] |
| SE | 93177 | ENSG00000106785 | TRIM14 | chr9 | - | 98099930 | 98100164 | 98094866 | 98095029 | 98109888 | 98109984 | 267 | 157 | [illegible] |
| SE | 93181 | ENSG00000163558 | PRKCI | chr3 | + | 170263378 | 170263429 | 170259968 | 170260058 | 170267914 | 170268000 | 63 | 76 | [illegible] |
| SE | 93184 | ENSG00000163558 | PRKCI | chr3 | + | 170291853 | 170291941 | 170284460 | 170284596 | 170293382 | 170293508 | 24 | 23 | [illegible] |
| SE | 93185 | ENSG00000163558 | PRKCI | chr3 | + | 170293382 | 170293508 | 170291641 | 170291941 | 170295910 | 170295990 | 30 | 48 | [illegible] |
| SE | 93188 | ENSG00000139112 | GABARAPL1 | chr12 | + | 10213835 | 10214187 | 10212892 | 10213219 | 10214668 | 10214779 | 4 | 5 | [illegible] |
| SE | 93189 | ENSG00000139112 | GABARAPL1 | chr12 | + | 10213835 | 10214187 | 10212892 | 10213219 | 10218062 | 10218141 | 0 | 2 | [illegible] |
| SE | 93193 | ENSG00000139112 | GABARAPL1 | chr12 | + | 10218062 | 10218141 | 10212835 | 10213219 | 10220439 | 10220481 | 250 | 125 | [illegible] |
| SE | 93194 | ENSG00000139112 | GABARAPL1 | chr12 | + | 10218062 | 10218141 | 10213105 | 10214779 | 10220439 | 10220481 | 119 | 68 | [illegible] |
| SE | 93196 | ENSG00000136848 | DAB2IP | chr9 | + | 121759884 | 121760439 | 121758897 | 121758996 | 121763504 | 121763649 | 2 | 4 | [illegible] |
| SE | 93198 | ENSG00000137207 | YIPF3 | chr6 | - | 43512439 | 43512563 | 43511831 | 43512315 | 43512760 | 43512874 | 262 | 94 | [illegible] |
| SE | 93202 | ENSG00000137207 | YIPF3 | chr6 | - | 43515888 | 43516046 | 43515594 | 43515701 | 43516726 | 43516919 | 109 | 53 | [illegible] |
| SE | 93203 | ENSG00000137207 | YIPF3 | chr6 | - | 43515888 | 43516095 | 43515536 | 43515701 | 43516726 | 43516926 | 258 | 132 | [illegible] |
| SE | 93204 | ENSG00000137207 | YIPF3 | chr6 | - | 43515888 | 43516113 | 43515594 | 43515701 | 43516726 | 43516941 | 108 | 53 | [illegible] |
| SE | 93205 | ENSG00000137207 | YIPF3 | chr6 | - | 43515888 | 43516187 | 43515594 | 43515701 | 43516726 | 43516933 | 112 | 59 | [illegible] |
| SE | 93206 | ENSG00000137207 | YIPF3 | chr6 | - | 43515888 | 43516190 | 43515594 | 43515701 | 43516726 | 43516972 | 117 | 57 | [illegible] |
| SE | 93210 | ENSG00000182158 | CREB3L2 | chr7 | - | 137928149 | 137928366 | 137915836 | 137916012 | 138001603 | 138001827 | 36 | 14 | [illegible] |
| SE | 93211 | ENSG00000113328 | CCNG1 | chr5 | + | 163439256 | 163439346 | 163437613 | 163437804 | 163441077 | 163441331 | 13 | 3 | [illegible] |
| SE | 93212 | ENSG00000113328 | CCNG1 | chr5 | + | 163439256 | 163439520 | 163437594 | 163437804 | 163441077 | 163441157 | 156 | 70 | [illegible] |
| SE | 93213 | ENSG00000113328 | CCNG1 | chr5 | + | 163441885 | 163441964 | 163441077 | 163441331 | 163442373 | 163442568 | 115 | 84 | [illegible] |
| SE | 93215 | ENSG00000113328 | CCNG1 | chr5 | + | 163442373 | 163442568 | 163442044 | 163442143 | 163443673 | 163443729 | 648 | 457 | [illegible] |
| SE | 93217 | ENSG00000166507 | NDST2 | chr10 | - | 73804772 | 73804869 | 73803892 | 73804016 | 73805586 | 73805769 | 164 | 96 | [illegible] |
| SE | 93218 | ENSG00000166507 | NDST2 | chr10 | - | 73805899 | 73806028 | 73805586 | 73805769 | 73807107 | 73807195 | 87 | 47 | [illegible] |
| SE | 93219 | ENSG00000166507 | NDST2 | chr10 | - | 73806288 | 73806474 | 73805899 | 73806028 | 73806656 | 73806811 | 102 | 73 | [illegible] |
| SE | 93223 | ENSG00000166507 | NDST2 | chr10 | - | 73807107 | 73807195 | 73805899 | 73806028 | 73807383 | 73808729 | 50 | 23 | [illegible] |
| SE | 93225 | ENSG00000110455 | ACCS | chr11 | + | 44071255 | 44071315 | 44067627 | 44067915 | 44073446 | 44073517 | 44 | 10 | [illegible] |
| SE | 93230 | ENSG00000110455 | ACCS | chr11 | + | 44073446 | 44073535 | 44067627 | 44067915 | 44074611 | 44074681 | 0 | 0 | [illegible] |
| SE | 93232 | ENSG00000110455 | ACCS | chr11 | + | 44077844 | 44077922 | 44077278 | 44077376 | 44078683 | 44078784 | 20 | 7 | [illegible] |
| SE | 93233 | ENSG00000110455 | ACCS | chr11 | + | 44081178 | 44081320 | 44081017 | 44081065 | 44083168 | 44083311 | 58 | 14 | [illegible] |
| SE | 93235 | ENSG00000110455 | ACCS | chr11 | + | 44083168 | 44083311 | 44081178 | 44081320 | 44083694 | 44083893 | 32 | 6 | [illegible] |
| SE | 93236 | ENSG00000110455 | ACCS | chr11 | + | 44083423 | 44083577 | 44083168 | 44083311 | 44083694 | 44084222 | 37 | 7 | [illegible] |
| SE | 93237 | ENSG00000124702 | KLHDC3 | chr6 | + | 43017133 | 43017346 | 43014102 | 43014348 | 43017852 | 43017968 | 40 | 13 | [illegible] |
| SE | 93239 | ENSG00000124702 | KLHDC3 | chr6 | + | 43017518 | 43017695 | 43017133 | 43017346 | 43018144 | 43018216 | 128 | 56 | [illegible] |
| SE | 93240 | ENSG00000124702 | KLHDC3 | chr6 | + | 43017852 | 43017968 | 43017133 | 43017346 | 43018144 | 43018216 | 115 | 59 | [illegible] |
| SE | 93242 | ENSG00000124702 | KLHDC3 | chr6 | + | 43019287 | 43019366 | 43019091 | 43019165 | 43020666 | 43020725 | 438 | 177 | [illegible] |
| SE | 93244 | ENSG00000136937 | NCBP1 | chr9 | + | 97643203 | 97643360 | 97641561 | 97641662 | 97645116 | 97645224 | 157 | 126 | [illegible] |
| SE | 93245 | ENSG00000136937 | NCBP1 | chr9 | + | 97643203 | 97643360 | 97641561 | 97641662 | 97647491 | 97647561 | 49 | 40 | [illegible] |
| SE | 93246 | ENSG00000136937 | NCBP1 | chr9 | + | 97645116 | 97645224 | 97641561 | 97641662 | 97647491 | 97647561 | 6 | 4 | [illegible] |
| SE | 93248 | ENSG00000136937 | NCBP1 | chr9 | + | 97645116 | 97645224 | 97643203 | 97643360 | 97647491 | 97647561 | 98 | 70 | [illegible] |
| SE | 93249 | ENSG00000136937 | NCBP1 | chr9 | + | 97645610 | 97645732 | 97643203 | 97643360 | 97647491 | 97647561 | 64 | 58 | [illegible] |
| SE | 93253 | ENSG00000136937 | NCBP1 | chr9 | + | 97662953 | 97663047 | 97662041 | 97662144 | 97664339 | 97664443 | 196 | 171 | [illegible] |
| SE | 93255 | ENSG00000256448 | NA | chr11 | + | 73406442 | 73406501 | 73405296 | 73405384 | 73410485 | 73410672 | 2 | 0 | [illegible] |
| SE | 93256 | ENSG00000225032 | NA | chr9 | + | 127817758 | 127817857 | 127816065 | 127817343 | 127818310 | 127818389 | 2 | 6 | [illegible] |

| | | | | | | | | | | | | |
|---|---|---|---|---|---|---|---|---|---|---|---|---|
| SE | 93259 | ENSG00000148908 | RGS10 | chr10 | - | 119526031 | 119526118 | 119515508 | 119515652 | 119527305 | 119527424 | 462 | 
| SE | 93260 | ENSG00000148908 | RGS10 | chr10 | - | 119527305 | 119527424 | 119515508 | 119515652 | 119542589 | 119542719 | 12 |
| SE | 93264 | ENSG00000132334 | PTPRE | chr10 | + | 127982273 | 127982296 | 127907102 | 127907309 | 128040874 | 128040990 | 1 |
| SE | 93265 | ENSG00000132334 | PTPRE | chr10 | + | 128056122 | 128056213 | 128049529 | 128049666 | 128060938 | 128061015 | 399 |
| SE | 93267 | ENSG00000132334 | PTPRE | chr10 | + | 128066074 | 128066194 | 128062918 | 128063180 | 128068122 | 128068286 | 385 |
| SE | 93268 | ENSG00000132334 | PTPRE | chr10 | + | 128068122 | 128068286 | 128062918 | 128063180 | 128070807 | 128070841 | 0 |
| SE | 93269 | ENSG00000132334 | PTPRE | chr10 | + | 128072137 | 128072214 | 128070807 | 128070901 | 128073336 | 128073471 | 350 |
| SE | 93270 | ENSG00000132334 | PTPRE | chr10 | + | 128073336 | 128073471 | 128072137 | 128072214 | 128076602 | 128076728 | 729 |
| SE | 93272 | ENSG00000132334 | PTPRE | chr10 | + | 128079559 | 128079695 | 128077616 | 128077783 | 128082831 | 128085855 | 338 |
| SE | 93274 | ENSG00000056586 | RC3H2 | chr9 | - | 122855731 | 122855878 | 122855183 | 122855397 | 122857922 | 122858093 | 322 |
| SE | 93277 | ENSG00000056586 | RC3H2 | chr9 | - | 122859916 | 122860131 | 122858668 | 122859102 | 122877470 | 122877583 | 60 |
| SE | 93278 | ENSG00000056586 | RC3H2 | chr9 | - | 122865348 | 122865541 | 122858668 | 122859102 | 122877470 | 122877583 | 7 |
| SE | 93279 | ENSG00000056586 | RC3H2 | chr9 | - | 122865348 | 122865541 | 122859916 | 122860131 | 122865623 | 122865657 | 105 |
| SE | 93281 | ENSG00000056586 | RC3H2 | chr9 | - | 122865348 | 122865657 | 122858668 | 122859102 | 122877470 | 122877583 | 105 |
| SE | 93285 | ENSG00000056586 | RC3H2 | chr9 | - | 122883203 | 122883379 | 122880593 | 122880794 | 122890311 | 122890545 | 170 |
| SE | 93289 | ENSG00000090432 | MUL1 | chr1 | - | 20502068 | 20502189 | 20499447 | 20501419 | 20503221 | 20503309 | 159 |
| SE | 93304 | ENSG00000099139 | PCSK5 | chr9 | + | 76023737 | 76023881 | 75986131 | 75986245 | 76026960 | 76027037 | 46 |
| SE | 93307 | ENSG00000099139 | PCSK5 | chr9 | + | 76157044 | 76157162 | 76107250 | 76107351 | 76158982 | 76159171 | 17 |
| SE | 93308 | ENSG00000099139 | PCSK5 | chr9 | + | 76179595 | 76179698 | 76174985 | 76175129 | 76181397 | 76181591 | 22 |
| SE | 93314 | ENSG00000196781 | TLE1 | chr9 | - | 81612291 | 81612393 | 81611805 | 81611959 | 81613421 | 81613521 | 3 |
| SE | 93315 | ENSG00000196781 | TLE1 | chr9 | - | 81612291 | 81612393 | 81611842 | 81611956 | 81613376 | 81613521 | 12 |
| SE | 93316 | ENSG00000196781 | TLE1 | chr9 | - | 81612291 | 81612393 | 81611842 | 81611956 | 81613421 | 81613521 | 9 |
| SE | 93317 | ENSG00000196781 | TLE1 | chr9 | - | 81633347 | 81633364 | 81620440 | 81620557 | 81634096 | 81634301 | 37 |
| SE | 93318 | ENSG00000196781 | TLE1 | chr9 | - | 81633347 | 81633364 | 81620440 | 81620557 | 81652183 | 81652288 | 3 |
| SE | 93319 | ENSG00000196781 | TLE1 | chr9 | - | 81633347 | 81633364 | 81620440 | 81620557 | 81652213 | 81652288 | 4 |
| SE | 93320 | ENSG00000196781 | TLE1 | chr9 | - | 81634096 | 81634301 | 81620440 | 81620557 | 81652183 | 81652288 | 12 |
| SE | 93322 | ENSG00000196781 | TLE1 | chr9 | - | 81634096 | 81634301 | 81633347 | 81633364 | 81652183 | 81652288 | 47 |
| SE | 93323 | ENSG00000196781 | TLE1 | chr9 | - | 81634096 | 81634301 | 81633347 | 81633364 | 81652213 | 81652288 | 86 |
| SE | 93324 | ENSG00000196781 | TLE1 | chr9 | - | 81652183 | 81652288 | 81634096 | 81634301 | 81653973 | 81654036 | 48 |
| [illegible] | | | | | | | | | | | | |

| | | | | | | | | | | | | | | |
|---|---|---|---|---|---|---|---|---|---|---|---|---|---|---|
| SE | 93651 | ENSG00000152763 | DNAI4 | chr1 | - | 66893228 | 66893413 | 66890182 | 66891266 | 66905200 | 66905303 | 56 | 34 | [illegible] |
| SE | 93654 | ENSG00000165304 | MELK | chr9 | + | 36581643 | 36581739 | 36572861 | 36573007 | 36589535 | 36589652 | 2 | 0 | [illegible] |
| SE | 93655 | ENSG00000165304 | MELK | chr9 | + | 36583626 | 36583712 | 36581643 | 36581739 | 36589535 | 36589652 | 3 | 10 | [illegible] |
| SE | 93658 | ENSG00000165304 | MELK | chr9 | + | 36589535 | 36589652 | 36583626 | 36583712 | 36594627 | 36594771 | 4 | 10 | [illegible] |
| SE | 93659 | ENSG00000165304 | MELK | chr9 | + | 36594627 | 36594771 | 36589535 | 36589652 | 36597221 | 36597290 | 4 | 2 | [illegible] |
| SE | 93660 | ENSG00000165304 | MELK | chr9 | + | 36597221 | 36597290 | 36589535 | 36589652 | 36599393 | 36599486 | 2 | 1 | [illegible] |
| SE | 93663 | ENSG00000165304 | MELK | chr9 | + | 36642996 | 36643083 | 36633101 | 36633200 | 36651745 | 36651877 | 0 | 0 | [illegible] |
| SE | 93665 | ENSG00000165304 | MELK | chr9 | + | 36674833 | 36674937 | 36670997 | 36671166 | 36677159 | 36677417 | 1 | 0 | [illegible] |
| SE | 93667 | ENSG00000148153 | INIP | chr9 | - | 112694130 | 112694233 | 112687131 | 112687633 | 112716460 | 112716541 | 33 | 18 | [illegible] |
| SE | 93668 | ENSG00000148153 | INIP | chr9 | - | 112694130 | 112694233 | 112689526 | 112689617 | 112701831 | 112701880 | 55 | 50 | [illegible] |
| SE | 93672 | ENSG00000148153 | INIP | chr9 | - | 112701831 | 112701880 | 112694130 | 112694233 | 112717986 | 112718089 | 3 | 0 | [illegible] |
| SE | 93674 | ENSG00000148153 | INIP | chr9 | - | 112716460 | 112716541 | 112701787 | 112701880 | 112717986 | 112718086 | 27 | 19 | [illegible] |
| SE | 93676 | ENSG00000170264 | FAM161A | chr2 | - | 61838537 | 61838705 | 61836009 | 61836109 | 61839420 | 61840581 | 0 | 0 | [illegible] |
| SE | 93679 | ENSG00000158467 | AHCYL2 | chr7 | + | 129400289 | 129400384 | 129397221 | 129397324 | 129403378 | 129403485 | 117 | 89 | [illegible] |
| SE | 93680 | ENSG00000158467 | AHCYL2 | chr7 | + | 129405835 | 129405899 | 129405096 | 129405213 | 129406377 | 129406466 | 84 | 79 | [illegible] |
| SE | 93681 | ENSG00000158467 | AHCYL2 | chr7 | + | 129406377 | 129406466 | 129405835 | 129405899 | 129409475 | 129409546 | 83 | 76 | [illegible] |
| SE | 93682 | ENSG00000158467 | AHCYL2 | chr7 | + | 129426442 | 129426563 | 129425062 | 129425141 | 129427038 | 129427175 | 36 | 24 | [illegible] |
| SE | 93683 | ENSG00000125482 | TTF1 | chr9 | - | 132401454 | 132402828 | 132400034 | 132400258 | 132406789 | 132406840 | 28 | 16 | [illegible] |
| SE | 93684 | ENSG00000198780 | FAM169A | chr5 | - | 74813839 | 74814019 | 74805155 | 74805284 | 74834425 | 74834597 | 76 | 42 | [illegible] |
| SE | 93685 | ENSG00000198780 | FAM169A | chr5 | - | 74813839 | 74814019 | 74805155 | 74805284 | 74838964 | 74839050 | 35 | 24 | [illegible] |
| SE | 93688 | ENSG00000198780 | FAM169A | chr5 | - | 74834425 | 74834597 | 74813839 | 74814019 | 74840073 | 74840173 | 49 | 26 | [illegible] |
| SE | 93695 | ENSG00000136828 | RALGPS1 | chr9 | + | 126965843 | 126965951 | 126962224 | 126962346 | 126977694 | 126977745 | 26 | 24 | [illegible] |
| SE | 93696 | ENSG00000136828 | RALGPS1 | chr9 | + | 127034430 | 127034514 | 126977694 | 126977745 | 127050042 | 127050132 | 19 | 15 | [illegible] |
| SE | 93697 | ENSG00000136828 | RALGPS1 | chr9 | + | 127034430 | 127034514 | 126977694 | 126977745 | 127052846 | 127052939 | 13 | 8 | [illegible] |
| SE | 93698 | ENSG00000136828 | RALGPS1 | chr9 | + | 127050042 | 127050132 | 126977694 | 126977745 | 127052846 | 127052939 | 15 | 12 | [illegible] |
| SE | 93700 | ENSG00000136828 | RALGPS1 | chr9 | + | 127174714 | 127174782 | 127168678 | 127168772 | 127195090 | 127195217 | 26 | 20 | [illegible] |
| SE | 93701 | ENSG00000136828 | RALGPS1 | chr9 | + | 127196473 | 127196631 | 127195090 | 127195217 | 127210694 | 127210778 | 12 | 4 | [illegible] |
| SE | 93702 | ENSG00000136828 | RALGPS1 | chr9 | + | 127199014 | 127199066 | 127196473 | 127196631 | 127210694 | 127210778 | 4 | 0 | [illegible] |
| SE | 93703 | ENSG00000136828 | RALGPS1 | chr9 | + | 127199014 | 127199066 | 127196473 | 127196631 | 127212130 | 127212236 | 0 | 0 | [illegible] |
| SE | 93704 | ENSG00000136828 | RALGPS1 | chr9 | + | 127210694 | 127210778 | 127196473 | 127196631 | 127212130 | 127212236 | 3 | 2 | [illegible] |
| SE | 93708 | ENSG00000197329 | PELI1 | chr2 | - | 64100397 | 64100499 | 64096412 | 64096610 | 64104700 | 64104830 | 203 | 192 | [illegible] |
| SE | 93709 | ENSG00000197329 | PELI1 | chr2 | - | 64100397 | 64100499 | 64096412 | 64096610 | 64108239 | 64108379 | 62 | 52 | [illegible] |
| SE | 93711 | ENSG00000197329 | PELI1 | chr2 | - | 64104700 | 64104830 | 64096412 | 64096610 | 64108239 | 64108379 | 94 | 97 | [illegible] |
| SE | 93713 | ENSG00000197329 | PELI1 | chr2 | - | 64108239 | 64108379 | 64104700 | 64104830 | 64144080 | 64144362 | 124 | 124 | [illegible] |
| SE | 93729 | ENSG00000270820 | USP34-DT | chr2 | + | 61480324 | 61480583 | 61471187 | 61471379 | 61482626 | 61483913 | 4 | 0 | [illegible] |
| SE | 93732 | ENSG00000168003 | SLC3A2 | chr11 | + | 62870744 | 62870795 | 62856110 | 62856381 | 62871576 | 62871669 | 0 | 0 | [illegible] |
| SE | 93733 | ENSG00000168003 | SLC3A2 | chr11 | + | 62870744 | 62870837 | 62856101 | 62856381 | 62871576 | 62871669 | 2 | 1 | [illegible] |
| SE | 93735 | ENSG00000168003 | SLC3A2 | chr11 | + | 62871576 | 62871669 | 62856003 | 62856381 | 62881018 | 62881447 | 0 | 0 | [illegible] |
| SE | 93736 | ENSG00000168003 | SLC3A2 | chr11 | + | 62876783 | 62876879 | 62871576 | 62871669 | 62881018 | 62881447 | 2 | 0 | [illegible] |
| SE | 93737 | ENSG00000168003 | SLC3A2 | chr11 | + | 62876786 | 62876879 | 62871576 | 62871669 | 62881018 | 62881447 | 2 | 0 | [illegible] |
| SE | 93738 | ENSG00000168003 | SLC3A2 | chr11 | + | 62882504 | 62882618 | 62881698 | 62882066 | 62882907 | 62882999 | 2 | 0 | [illegible] |
| SE | 93739 | ENSG00000168003 | SLC3A2 | chr11 | + | 62882529 | 62882618 | 62881721 | 62882066 | 62882907 | 62882999 | 1 | 0 | [illegible] |
| SE | 93740 | ENSG00000168003 | SLC3A2 | chr11 | + | 62882666 | 62882807 | 62881698 | 62882066 | 62882907 | 62882999 | 0 | 0 | [illegible] |
| SE | 93741 | ENSG00000168003 | SLC3A2 | chr11 | + | 62882907 | 62882999 | 62881698 | 62882066 | 62884456 | 62884512 | 220 | 115 | [illegible] |
| SE | 93742 | ENSG00000168003 | SLC3A2 | chr11 | + | 62883988 | 62884112 | 62882907 | 62882999 | 62884456 | 62884525 | 0 | 1 | [illegible] |
| SE | 93743 | ENSG00000168003 | SLC3A2 | chr11 | + | 62886436 | 62886568 | 62885464 | 62885608 | 62888134 | 62888218 | 2 | 0 | [illegible] |
| SE | 93744 | ENSG00000168003 | SLC3A2 | chr11 | + | 62886436 | 62886568 | 62885464 | 62885608 | 62888330 | 62888880 | 0 | 0 | [illegible] |
| SE | 93745 | ENSG00000168003 | SLC3A2 | chr11 | + | 62886440 | 62886568 | 62885464 | 62885608 | 62888134 | 62888218 | 2 | 0 | [illegible] |
| SE | 93746 | ENSG00000168003 | SLC3A2 | chr11 | + | 62886440 | 62886568 | 62885464 | 62885608 | 62888330 | 62888849 | 0 | 0 | [illegible] |
| SE | 93750 | ENSG00000168003 | SLC3A2 | chr11 | + | 62888134 | 62888218 | 62886436 | 62886568 | 62888330 | 62888368 | 56 | 20 | [illegible] |
| SE | 93755 | ENSG00000143409 | MINDY1 | chr1 | - | 151002164 | 151002170 | 151001724 | 151001782 | 151006311 | 151006871 | 1 | 0 | [illegible] |
| SE | 93760 | ENSG00000070950 | RAD18 | chr3 | - | 8902379 | 8902520 | 8898893 | 8899047 | 8912311 | 8912372 | 80 | 45 | [illegible] |
| SE | 93761 | ENSG00000070950 | RAD18 | chr3 | - | 8913643 | 8913720 | 8912311 | 8912372 | 8935870 | 8936055 | 39 | 42 | [illegible] |
| SE | 93762 | ENSG00000070950 | RAD18 | chr3 | - | 8947219 | 8947290 | 8941466 | 8941804 | 8958919 | 8959001 | 24 | 11 | [illegible] |
| SE | 93763 | ENSG00000070950 | RAD18 | chr3 | - | 8948508 | 8948570 | 8946951 | 8947290 | 8958919 | 8959001 | 27 | 14 | [illegible] |
| SE | 93765 | ENSG00000148484 | RSU1 | chr10 | - | 16695022 | 16695155 | 16590610 | 16593496 | 16752538 | 16752653 | 828 | 973 | [illegible] |
| SE | 93766 | ENSG00000148484 | RSU1 | chr10 | - | 16727030 | 16727157 | 16695022 | 16695155 | 16752538 | 16752653 | 1 | 4 | [illegible] |
| SE | 93767 | ENSG00000148484 | RSU1 | chr10 | - | 16752917 | 16753000 | 16752538 | 16752653 | 16754870 | 16754989 | 996 | 1173 | [illegible] |
| SE | 93768 | ENSG00000148484 | RSU1 | chr10 | - | 16764389 | 16764510 | 16695022 | 16695155 | 16782033 | 16782084 | 368 | 447 | [illegible] |
| SE | 93769 | ENSG00000148484 | RSU1 | chr10 | - | 16764389 | 16764510 | 16695022 | 16695155 | 16816972 | 16817084 | 4 | 1 | [illegible] |
| SE | 93770 | ENSG00000148484 | RSU1 | chr10 | - | 16764389 | 16764510 | 16754870 | 16754989 | 16816972 | 16817084 | 546 | 703 | [illegible] |
| SE | 93771 | ENSG00000148484 | RSU1 | chr10 | - | 16782033 | 16782084 | 16695022 | 16695155 | 16816972 | 16817084 | 85 | 134 | [illegible] |
| SE | 93778 | ENSG00000116701 | NCF2 | chr1 | - | 183560095 | 183560273 | 183555561 | 183556230 | 183563194 | 183563306 | 1619 | 1093 | [illegible] |
| SE | 93779 | ENSG00000116701 | NCF2 | chr1 | - | 183563433 | 183563585 | 183563194 | 183563306 | 183564004 | 183564030 | 1490 | 1024 | [illegible] |
| SE | 93780 | ENSG00000116701 | NCF2 | chr1 | - | 183564004 | 183564030 | 183563433 | 183563585 | 183565703 | 183565779 | 729 | 539 | [illegible] |
| SE | 93781 | ENSG00000116701 | NCF2 | chr1 | - | 183565703 | 183565779 | 183564004 | 183564030 | 183566919 | 183566988 | 1351 | 811 | [illegible] |
| SE | 93782 | ENSG00000116701 | NCF2 | chr1 | - | 183565703 | 183565779 | 183564004 | 183564030 | 183569141 | 183569225 | 201 | 152 | [illegible] |
| SE | 93783 | ENSG00000116701 | NCF2 | chr1 | - | 183565703 | 183565779 | 183564004 | 183564030 | 183570779 | 183570839 | 200 | 152 | [illegible] |
| SE | 93784 | ENSG00000116701 | NCF2 | chr1 | - | 183566919 | 183566988 | 183564004 | 183564030 | 183569141 | 183569185 | 1 | 2 | [illegible] |
| SE | 93785 | ENSG00000116701 | NCF2 | chr1 | - | 183566919 | 183566988 | 183564004 | 183564030 | 183570779 | 183570839 | 1 | 2 | [illegible] |
| SE | 93786 | ENSG00000116701 | NCF2 | chr1 | - | 183566919 | 183566988 | 183565703 | 183565779 | 183567203 | 183567345 | 1649 | 901 | [illegible] |
| SE | 93787 | ENSG00000116701 | NCF2 | chr1 | - | 183566919 | 183566988 | 183565703 | 183565779 | 183569141 | 183569185 | 396 | 297 | [illegible] |
| SE | 93788 | ENSG00000116701 | NCF2 | chr1 | - | 183566919 | 183566988 | 183565703 | 183565779 | 183570779 | 183570839 | 396 | 297 | [illegible] |
| SE | 93789 | ENSG00000116701 | NCF2 | chr1 | - | 183567203 | 183567345 | 183565703 | 183565779 | 183569141 | 183569185 | 954 | 469 | [illegible] |
| SE | 93790 | ENSG00000116701 | NCF2 | chr1 | - | 183567203 | 183567345 | 183566919 | 183566988 | 183569141 | 183569185 | 2208 | 1076 | [illegible] |
| SE | 93791 | ENSG00000116701 | NCF2 | chr1 | - | 183569141 | 183569185 | 183564004 | 183564030 | 183570779 | 183570839 | 117 | 103 | [illegible] |
| SE | 93793 | ENSG00000116701 | NCF2 | chr1 | - | 183570779 | 183570839 | 183564004 | 183564030 | 183573184 | 183573292 | 101 | 53 | [illegible] |
| SE | 93794 | ENSG00000116701 | NCF2 | chr1 | - | 183570779 | 183570839 | 183565703 | 183565779 | 183577598 | 183577707 | 0 | 1 | [illegible] |
| SE | 93795 | ENSG00000116701 | NCF2 | chr1 | - | 183570779 | 183570839 | 183569141 | 183569185 | 183574486 | 183574621 | 386 | 221 | [illegible] |
| SE | 93798 | ENSG00000116701 | NCF2 | chr1 | - | 183573184 | 183573292 | 183570779 | 183570839 | 183586894 | 183586977 | 542 | 297 | [illegible] |
| SE | 93801 | ENSG00000116701 | NCF2 | chr1 | - | 183577598 | 183577707 | 183570779 | 183570839 | 183586894 | 183586977 | 355 | 266 | [illegible] |
| SE | 93802 | ENSG00000116701 | NCF2 | chr1 | - | 183577598 | 183577707 | 183573184 | 183573292 | 183586894 | 183586977 | 363 | 268 | [illegible] |
| SE | 93803 | ENSG00000116701 | NCF2 | chr1 | - | 183586894 | 183586977 | 183577598 | 183577707 | 183590155 | 183590359 | 1624 | 991 | [illegible] |
| SE | 93814 | ENSG00000091164 | TXNL1 | chr18 | - | 56616244 | 56616314 | 56614423 | 56614596 | 56618003 | 56618126 | 504 | 342 | [illegible] |
| SE | 93816 | ENSG00000091164 | TXNL1 | chr18 | - | 56638342 | 56638505 | 56626360 | 56626457 | 56647448 | 56647535 | 71 | 47 | [illegible] |
| SE | 93817 | ENSG00000091164 | TXNL1 | chr18 | - | 56638342 | 56639039 | 56626360 | 56626457 | 56651016 | 56651274 | 71 | 47 | [illegible] |
| SE | 93818 | ENSG00000091164 | TXNL1 | chr18 | - | 56638961 | 56639039 | 56638358 | 56638505 | 56647448 | 56647535 | 7 | 1 | [illegible] |
| SE | 93819 | ENSG00000091164 | TXNL1 | chr18 | - | 56647448 | 56647535 | 56626360 | 56626457 | 56651016 | 56651128 | 2 | 0 | [illegible] |
| SE | 93820 | ENSG00000091164 | TXNL1 | chr18 | - | 56647448 | 56647535 | 56638961 | 56639039 | 56651016 | 56651128 | 8 | 1 | [illegible] |
| SE | 93821 | ENSG00000129675 | ARHGEF6 | chrX | - | 136672019 | 136672119 | 136669481 | 136669536 | 136675006 | 136675096 | 655 | 607 | [illegible] |
| SE | 93823 | ENSG00000129675 | ARHGEF6 | chrX | - | 136677935 | 136677956 | 136676623 | 136676717 | 136680730 | 136680876 | 18 | 39 | [illegible] |
| SE | 93824 | ENSG00000129675 | ARHGEF6 | chrX | - | 136679534 | 136679660 | 136676623 | 136676717 | 136680730 | 136680876 | 444 | 369 | [illegible] |
| SE | 93825 | ENSG00000129675 | ARHGEF6 | chrX | - | 136679534 | 136679660 | 136677935 | 136677956 | 136680730 | 136680876 | 604 | 492 | [illegible] |
| SE | 93827 | ENSG00000129675 | ARHGEF6 | chrX | - | 136779413 | 136779497 | 136747507 | 136747592 | 136780717 | 136780932 | 425 | 336 | [illegible] |
| SE | 93832 | ENSG00000156042 | CFAP70 | chr10 | - | 73277239 | 73277361 | 73275445 | 73275598 | 73278178 | 73278337 | 1 | 1 | [illegible] |
| SE | 93834 | ENSG00000156042 | CFAP70 | chr10 | - | 73298906 | 73299097 | 73297041 | 73297173 | 73299604 | 73299665 | 8 | 2 | [illegible] |
| SE | 93835 | ENSG00000156042 | CFAP70 | chr10 | - | 73298906 | 73299101 | 73293262 | 73297173 | 73299604 | 73299665 | 14 | 4 | [illegible] |
| SE | 93836 | ENSG00000156042 | CFAP70 | chr10 | - | 73348155 | 73348235 | 73345064 | 73345244 | 73348422 | 73348521 | 0 | 0 | [illegible] |
| SE | 93840 | ENSG00000116580 | GON4L | chr1 | - | 155751766 | 155751869 | 155749658 | 155750733 | 155754374 | 155754488 | 27 | 29 | [illegible] |
| SE | 93841 | ENSG00000116580 | GON4L | chr1 | - | 155751959 | 155752587 | 155749716 | 155750733 | 155754374 | 155754488 | 6 | 2 | [illegible] |
| SE | 93842 | ENSG00000116580 | GON4L | chr1 | - | 155751959 | 155752587 | 155749716 | 155750733 | 155756957 | 155757077 | 0 | 0 | [illegible] |
| SE | 93843 | ENSG00000116580 | GON4L | chr1 | - | 155751959 | 155752587 | 155751766 | 155751869 | 155754374 | 155754488 | 32 | 27 | [illegible] |
| SE | 93844 | ENSG00000116580 | GON4L | chr1 | - | 155751959 | 155752590 | 155749658 | 155750733 | 155754374 | 155754488 | 2 | 0 | [illegible] |
| SE | 93845 | ENSG00000116580 | GON4L | chr1 | - | 155751959 | 155752590 | 155751766 | 155751869 | 155754374 | 155754488 | 28 | 25 | [illegible] |
| SE | 93847 | ENSG00000116580 | GON4L | chr1 | - | 155753203 | 155753414 | 155751959 | 155752587 | 155756957 | 155757077 | 44 | 22 | [illegible] |
| SE | 93848 | ENSG00000116580 | GON4L | chr1 | - | 155753203 | 155753414 | 155751959 | 155752590 | 155754374 | 155754488 | 80 | 43 | [illegible] |
| SE | 93850 | ENSG00000116580 | GON4L | chr1 | - | 155754374 | 155754488 | 155751959 | 155752587 | 155756957 | 155757077 | 54 | 21 | [illegible] |
| SE | 93851 | ENSG00000116580 | GON4L | chr1 | - | 155754374 | 155754488 | 155753203 | 155753414 | 155756957 | 155757077 | 126 | 60 | [illegible] |
| SE | 93852 | ENSG00000116580 | GON4L | chr1 | - | 155762189 | 155762374 | 155760443 | 155760641 | 155763311 | 155763564 | 160 | 129 | [illegible] |
| SE | 93854 | ENSG00000116580 | GON4L | chr1 | - | 155764999 | 155766709 | 155763311 | 155763564 | 155767424 | 155767541 | 177 | 139 | [illegible] |
| SE | 93856 | ENSG00000116580 | GON4L | chr1 | - | 155771066 | 155771217 | 155767424 | 155767541 | 155773065 | 155773210 | 99 | 86 | [illegible] |
| SE | 93857 | ENSG00000116580 | GON4L | chr1 | - | 155773065 | 155773210 | 155771066 | 155771217 | 155774962 | 155775173 | 102 | 83 | [illegible] |
| SE | 93858 | ENSG00000116580 | GON4L | chr1 | - | 155773065 | 155773210 | 155771066 | 155771217 | 155775001 | 155775173 | 180 | 135 | [illegible] |
| SE | 93860 | ENSG00000116580 | GON4L | chr1 | - | 155777621 | 155777820 | 155776394 | 155776481 | 155783985 | 155784089 | 128 | 113 | [illegible] |
| SE | 93862 | ENSG00000116580 | GON4L | chr1 | - | 155813633 | 155813804 | 155804948 | 155805141 | 155815804 | 155815900 | 96 | 77 | [illegible] |
| SE | 93864 | ENSG00000116580 | GON4L | chr1 | - | 155820605 | 155820656 | 155815804 | 155816262 | 155821473 | 155821520 | 152 | 124 | [illegible] |
| SE | 93865 | ENSG00000116580 | GON4L | chr1 | - | 155821473 | 155821548 | 155820605 | 155820656 | 155822285 | 155822476 | 204 | 152 | [illegible] |
| SE | 93866 | ENSG00000116580 | GON4L | chr1 | - | 155826836 | 155827028 | 155822285 | 155822476 | 155853275 | 155853806 | 223 | 235 | [illegible] |
| SE | 93871 | ENSG00000116580 | GON4L | chr1 | - | 155853275 | 155853806 | 155826836 | 155827028 | 155857146 | 155857214 | 167 | 162 | [illegible] |
| SE | 93872 | ENSG00000116580 | GON4L | chr1 | - | 155853275 | 155853806 | 155826836 | 155827028 | 155857158 | 155857204 | 150 | 156 | [illegible] |
| SE | 93875 | ENSG00000107872 | FBXL15 | chr10 | + | 102421786 | 102422292 | 102421353 | 102421507 | 102422803 | 102423136 | 42 | 6 | [illegible] |
| SE | 93877 | ENSG00000089053 | ANAPC5 | chr12 | - | 121318276 | 121318404 | 121309700 | 121309863 | 121318500 | 121318608 | 143 | 84 | [illegible] |
| SE | 93878 | ENSG00000089053 | ANAPC5 | chr12 | - | 121318276 | 121318424 | 121309700 | 121309863 | 121318500 | 121318608 | 279 | 177 | [illegible] |
| SE | 93879 | ENSG00000089053 | ANAPC5 | chr12 | - | 121320384 | 121320459 | 121319696 | 121319818 | 121327095 | 121327231 | 250 | 157 | [illegible] |
| SE | 93880 | ENSG00000089053 | ANAPC5 | chr12 | - | 121327095 | 121327231 | 121320384 | 121320459 | 121328315 | 121328458 | 309 | 166 | [illegible] |
| SE | 93882 | ENSG00000089053 | ANAPC5 | chr12 | - | 121330582 | 121330672 | 121328315 | 121328458 | 121335532 | 121335682 | 54 | 43 | [illegible] |
| SE | 93885 | ENSG00000089053 | ANAPC5 | chr12 | - | 121337290 | 121337392 | 121335516 | 121335723 | 121342002 | 121342069 | 413 | 332 | [illegible] |
| SE | 93886 | ENSG00000089053 | ANAPC5 | chr12 | - | 121342002 | 121342069 | 121337290 | 121337392 | 121345838 | 121346031 | 437 | 359 | [illegible] |
| SE | 93888 | ENSG00000089053 | ANAPC5 | chr12 | - | 121347801 | 121347881 | 121345838 | 121347005 | 121352133 | 121352304 | 148 | 79 | [illegible] |
| SE | 93892 | ENSG00000184465 | WDR27 | chr6 | - | 169572418 | 169572540 | 169558320 | 169559927 | 169582835 | 169582934 | 0 | 0 | [illegible] |
| SE | 93893 | ENSG00000184465 | WDR27 | chr6 | - | 169636370 | 169636504 | 169634427 | 169634525 | 169638538 | 169638660 | 0 | 2 | [illegible] |
| SE | 93894 | ENSG00000184465 | WDR27 | chr6 | - | 169638538 | 169638660 | 169634427 | 169634525 | 169643696 | 169643786 | 7 | 7 | [illegible] |
| SE | 93895 | ENSG00000184465 | WDR27 | chr6 | - | 169638538 | 169638660 | 169634427 | 169634525 | 169658275 | 169658358 | 2 | 2 | [illegible] |
| SE | 93896 | ENSG00000184465 | WDR27 | chr6 | - | 169643696 | 169643786 | 169634427 | 169634525 | 169658275 | 169658358 | 1 | 2 | [illegible] |
| SE | 93897 | ENSG00000184465 | WDR27 | chr6 | - | 169643696 | 169643786 | 169638538 | 169638660 | 169647772 | 169647870 | 7 | 12 | [illegible] |
| SE | 93898 | ENSG00000184465 | WDR27 | chr6 | - | 169643696 | 169643786 | 169638538 | 169638660 | 169658275 | 169658358 | 4 | 1 | [illegible] |
| SE | 93899 | ENSG00000184465 | WDR27 | chr6 | - | 169651929 | 169652008 | 169649197 | 169649275 | 169658275 | 169658358 | 4 | 4 | [illegible] |
| SE | 93900 | ENSG00000184465 | WDR27 | chr6 | - | 169659085 | 169659207 | 169658275 | 169658358 | 169659450 | 169659518 | 8 | 11 | [illegible] |
| SE | 93901 | ENSG00000184465 | WDR27 | chr6 | - | 169659450 | 169659518 | 169659085 | 169659207 | 169660662 | 169660692 | 11 | 6 | [illegible] |
| SE | 93903 | ENSG00000184465 | WDR27 | chr6 | - | 169663562 | 169663649 | 169662303 | 169662424 | 169664165 | 169664286 | 0 | 4 | [illegible] |
| SE | 93904 | ENSG00000184465 | WDR27 | chr6 | - | 169663562 | 169663706 | 169662303 | 169662424 | 169664165 | 169664632 | 0 | 2 | [illegible] |
| SE | 93905 | ENSG00000184465 | WDR27 | chr6 | - | 169663562 | 169663878 | 169662303 | 169662424 | 169664165 | 169664286 | 0 | 6 | [illegible] |
| SE | 93906 | ENSG00000184465 | WDR27 | chr6 | - | 169664165 | 169664286 | 169662303 | 169662424 | 169672254 | 169672396 | 9 | 2 | [illegible] |
| SE | 93907 | ENSG00000184465 | WDR27 | chr6 | - | 169665485 | 169665556 | 169664165 | 169664286 | 169667135 | 169667187 | 1 | 3 | [illegible] |
| SE | 93911 | ENSG00000184465 | WDR27 | chr6 | - | 169667135 | 169667187 | 169664285 | 169665556 | 169672254 | 169672396 | 2 | 0 | [illegible] |
| SE | 93914 | ENSG00000184465 | WDR27 | chr6 | - | 169667135 | 169668776 | 169665134 | 169665556 | 169672254 | 169672396 | 2 | 0 | [illegible] |
| SE | 93916 | ENSG00000184465 | WDR27 | chr6 | - | 169667981 | 169668185 | 169667135 | 169667187 | 169670568 | 169670693 | 1 | 4 | [illegible] |
| SE | 93918 | ENSG00000184465 | WDR27 | chr6 | - | 169670568 | 169670693 | 169665485 | 169665556 | 169672254 | 169672396 | 5 | 6 | [illegible] |
| SE | 93919 | ENSG00000184465 | WDR27 | chr6 | - | 169670568 | 169670693 | 169666917 | 169668776 | 169672254 | 169672396 | 8 | 7 | [illegible] |
| SE | 93920 | ENSG00000184465 | WDR27 | chr6 | - | 169670568 | 169670693 | 169667135 | 169667187 | 169672254 | 169672396 | 5 | 6 | [illegible] |
| SE | 93924 | ENSG00000184465 | WDR27 | chr6 | - | 169694462 | 169695065 | 169689126 | 169689308 | 169701550 | 169701854 | 3 | 2 | [illegible] |
| SE | 93926 | ENSG00000144524 | COPS7B | chr2 | + | 231788554 | 231788732 | 231781670 | 231781883 | 231794262 | 231794351 | 23 | 17 | [illegible] |
| SE | 93931 | ENSG00000144524 | COPS7B | chr2 | + | 231791732 | 231791808 | 231781677 | 231781883 | 231794262 | 231794351 | 22 | 11 | [illegible] |
| SE | 93942 | ENSG00000196116 | TDRD7 | chr9 | + | 97460177 | 97460764 | 97439244 | 97439318 | 97464841 | 97465028 | 58 | 62 | [illegible] |
| SE | 93944 | ENSG00000196116 | TDRD7 | chr9 | + | 97460177 | 97460764 | 97441657 | 97441875 | 97472292 | 97472495 | 51 | 28 | [illegible] |
| SE | 93945 | ENSG00000196116 | TDRD7 | chr9 | + | 97464841 | 97465028 | 97441657 | 97441875 | 97472292 | 97472495 | 12 | 6 | [illegible] |
| SE | 93946 | ENSG00000196116 | TDRD7 | chr9 | + | 97464841 | 97465028 | 97460177 | 97460764 | 97472292 | 97472495 | 65 | 66 | [illegible] |
| SE | 93950 | ENSG00000196116 | TDRD7 | chr9 | + | 97480827 | 97480938 | 97478438 | 97478573 | 97482848 | 97483351 | 56 | 45 | [illegible] |
| SE | 93951 | ENSG00000196116 | TDRD7 | chr9 | + | 97487171 | 97487332 | 97482848 | 97483351 | 97495662 | 97496125 | 95 | 110 | [illegible] |
| SE | 93953 | ENSG00000114030 | KPNA1 | chr3 | - | 122433660 | 122433788 | 122427537 | 122427716 | 122437169 | 122437295 | 146 | 85 | [illegible] |
| SE | 93954 | ENSG00000114030 | KPNA1 | chr3 | - | 122437169 | 122437295 | 122433660 | 122433788 | 122442037 | 122442116 | 119 | 76 | [illegible] |
| SE | 93955 | ENSG00000114030 | KPNA1 | chr3 | - | 122451533 | 122451633 | 122449573 | 122449737 | 122453869 | 122454001 | 190 | 144 | [illegible] |
| SE | 93956 | ENSG00000114030 | KPNA1 | chr3 | - | 122451975 | 122452064 | 122451533 | 122451633 | 122453869 | 122454001 | 126 | 62 | [illegible] |
| SE | 93957 | ENSG00000114030 | KPNA1 | chr3 | - | 122453869 | 122454001 | 122449573 | 122449737 | 122461223 | 122461318 | 68 | 21 | [illegible] |
| SE | 93958 | ENSG00000114030 | KPNA1 | chr3 | - | 122453869 | 122454001 | 122451975 | 122452064 | 122461223 | 122461318 | 163 | 59 | [illegible] |
| SE | 93959 | ENSG00000114030 | KPNA1 | chr3 | - | 122457725 | 122457848 | 122453869 | 122454001 | 122459438 | 122459681 | 17 | 13 | [illegible] |
| SE | 93960 | ENSG00000114030 | KPNA1 | chr3 | - | 122457725 | 122457848 | 122453869 | 122454001 | 122459773 | 122460657 | 12 | 8 | [illegible] |
| SE | 93962 | ENSG00000114030 | KPNA1 | chr3 | - | 122459438 | 122459681 | 122453869 | 122454001 | 122459773 | 122460657 | 2 | 5 | [illegible] |
| SE | 93963 | ENSG00000114030 | KPNA1 | chr3 | - | 122459438 | 122459681 | 122453869 | 122454001 | 122461223 | 122461318 | 0 | 0 | [illegible] |
| SE | 93964 | ENSG00000114030 | KPNA1 | chr3 | - | 122459438 | 122459681 | 122457725 | 122457848 | 122459773 | 122460657 | 8 | 13 | [illegible] |
| SE | 93965 | ENSG00000114030 | KPNA1 | chr3 | - | 122459438 | 122459681 | 122457725 | 122457848 | 122461223 | 122461318 | 6 | 8 | [illegible] |
| SE | 93966 | ENSG00000114030 | KPNA1 | chr3 | - | 122461223 | 122461318 | 122457725 | 122457848 | 122463941 | 122464041 | 122 | 72 | [illegible] |
| SE | 93970 | ENSG00000149177 | PTPRJ | chr11 | + | 48130458 | 48130716 | 48127779 | 48128043 | 48136038 | 48136296 | 944 | 621 | [illegible] |
| SE | 93971 | ENSG00000149177 | PTPRJ | chr11 | + | 48144674 | 48144885 | 48142918 | 48143050 | 48146875 | 48146963 | 421 | 169 | [illegible] |
| SE | 93972 | ENSG00000149177 | PTPRJ | chr11 | + | 48144674 | 48144885 | 48142918 | 48143050 | 48149446 | 48149486 | 421 | 169 | [illegible] |
| SE | 93973 | ENSG00000149177 | PTPRJ | chr11 | + | 48144999 | 48145124 | 48144674 | 48144885 | 48146875 | 48146963 | 470 | 260 | [illegible] |
| SE | 93974 | ENSG00000149177 | PTPRJ | chr11 | + | 48144999 | 48145124 | 48144674 | 48144885 | 48149446 | 48149486 | 285 | 168 | [illegible] |
| SE | 93975 | ENSG00000149177 | PTPRJ | chr11 | + | 48146875 | 48146963 | 48142918 | 48143050 | 48149446 | 48149486 | 304 | 186 | [illegible] |
| SE | 93976 | ENSG00000149177 | PTPRJ | chr11 | + | 48146875 | 48146963 | 48144674 | 48144885 | 48149446 | 48149486 | 301 | 186 | [illegible] |
| SE | 93978 | ENSG00000149177 | PTPRJ | chr11 | + | 48146875 | 48146963 | 48144997 | 48145124 | 48150095 | 48150183 | 139 | 69 | [illegible] |
| SE | 93979 | ENSG00000149177 | PTPRJ | chr11 | + | 48149446 | 48149488 | 48144999 | 48145124 | 48150095 | 48150183 | 0 | 0 | [illegible] |
| SE | 93980 | ENSG00000149177 | PTPRJ | chr11 | + | 48149446 | 48149488 | 48146875 | 48146963 | 48150095 | 48150183 | 34 | 23 | [illegible] |
| SE | 93981 | ENSG00000149177 | PTPRJ | chr11 | + | 48149989 | 48149998 | 48149446 | 48149488 | 48150095 | 48150183 | 409 | 276 | [illegible] |
| SE | 93983 | ENSG00000254659 | LINC02715 | chr11 | - | 109746553 | 109746619 | 109744951 | 109745023 | 109780981 | 109781036 | 0 | 0 | [illegible] |
| SE | 93985 | ENSG00000213625 | LEPROT | chr1 | + | 65429861 | 65430048 | 65425302 | 65425378 | 65431802 | 65436007 | 204 | 136 | [illegible] |
| SE | 93986 | ENSG00000213625 | LEPROT | chr1 | + | 65429930 | 65430048 | 65425302 | 65425378 | 65431802 | 65431920 | 129 | 90 | [illegible] |
| SE | 93988 | ENSG00000080802 | CNOT4 | chr7 | - | 135398168 | 135398226 | 135395633 | 135395883 | 135410514 | 135410648 | 92 | 58 | [illegible] |
| SE | 93989 | ENSG00000080802 | CNOT4 | chr7 | - | 135398168 | 135398226 | 135395633 | 135395883 | 135410523 | 135410648 | 148 | 74 | [illegible] |
| SE | 93990 | ENSG00000080802 | CNOT4 | chr7 | - | 135413487 | 135413613 | 135410514 | 135410648 | 135422155 | 135422353 | 60 | 28 | [illegible] |
| SE | 93992 | ENSG00000080802 | CNOT4 | chr7 | - | 135414330 | 135414432 | 135413487 | 135413613 | 135422155 | 135422353 | 142 | 175 | [illegible] |
| SE | 93993 | ENSG00000080802 | CNOT4 | chr7 | - | 135415175 | 135415262 | 135413487 | 135413613 | 135422155 | 135422353 | 87 | 87 | [illegible] |
| SE | 93994 | ENSG00000080802 | CNOT4 | chr7 | - | 135415175 | 135415262 | 135414330 | 135414432 | 135422155 | 135422353 | 242 | 216 | [illegible] |
| SE | 93996 | ENSG00000080802 | CNOT4 | chr7 | - | 135509070 | 135509157 | 135438157 | 135438423 | 135509888 | 135509938 | 0 | 0 | [illegible] |
| SE | 93997 | ENSG00000140988 | RPS2 | chr16 | - | 1962496 | 1962656 | 1962057 | 1962270 | 1962735 | 1962909 | 5615 | 2162 | [illegible] |
| SE | 93998 | ENSG00000140988 | RPS2 | chr16 | - | 1962496 | 1962656 | 1962057 | 1962270 | 1963148 | 1963256 | 2895 | 1088 | [illegible] |
| SE | 93999 | ENSG00000140988 | RPS2 | chr16 | - | 1962496 | 1962909 | 1962060 | 1962270 | 1963148 | 1963256 | 9781 | 5104 | [illegible] |
| SE | 94000 | ENSG00000140988 | RPS2 | chr16 | - | 1962735 | 1962909 | 1962060 | 1962270 | 1963148 | 1963256 | 6887 | 4018 | [illegible] |
| SE | 94001 | ENSG00000140988 | RPS2 | chr16 | - | 1962735 | 1962909 | 1962060 | 1962656 | 1963148 | 1963256 | 9606 | 5094 | [illegible] |
| SE | 94002 | ENSG00000140988 | RPS2 | chr16 | - | 1963148 | 1963256 | 1962373 | 1962909 | 1964275 | 1964365 | 13638 | 7681 | [illegible] |
| SE | 94003 | ENSG00000140988 | RPS2 | chr16 | - | 1963148 | 1963256 | 1962373 | 1962909 | 1964448 | 1964812 | 6667 | 3877 | [illegible] |
| SE | 94006 | ENSG00000140988 | RPS2 | chr16 | - | 1964275 | 1964365 | 1962496 | 1963256 | 1964448 | 1964581 | 7264 | 4043 | [illegible] |
| SE | 94009 | ENSG00000130827 | PLXNA3 | chrX | + | 154469687 | 154469784 | 154469378 | 154469482 | 154469976 | 154470167 | 39 | 14 | [illegible] |
| SE | 94010 | ENSG00000238164 | TNFRSF14-AS1 | chr1 | - | 2550572 | 2550731 | 2549919 | 2550274 | 2550991 | 2552106 | 2 | 0 | [illegible] |
| SE | 94011 | ENSG00000238164 | TNFRSF14-AS1 | chr1 | - | 2553625 | 2553814 | 2551774 | 2552703 | 2554100 | 2554296 | 1 | 1 | [illegible] |
| SE | 94012 | ENSG00000099381 | SETD1A | chr16 | + | 30963432 | 30963554 | 30961266 | 30961537 | 30964093 | 30964323 | 108 | 54 | [illegible] |
| SE | 94013 | ENSG00000099381 | SETD1A | chr16 | + | 30964093 | 30964323 | 30963432 | 30963554 | 30964611 | 30965461 | 76 | 25 | [illegible] |
| SE | 94016 | ENSG00000099381 | SETD1A | chr16 | + | 30967500 | 30967588 | 30966883 | 30967060 | 30969304 | 30969462 | 147 | 92 | [illegible] |
| SE | 94017 | ENSG00000099381 | SETD1A | chr16 | + | 30969304 | 30969462 | 30967500 | 30967588 | 30969601 | 30969689 | 194 | 115 | [illegible] |
| SE | 94020 | ENSG00000136856 | SLC2A8 | chr9 | + | 127402556 | 127402753 | 127399906 | 127400006 | 127403659 | 127403803 | 0 | 2 | [illegible] |

| | | | | | | | | | | | | | | | | | | | | | | | | | | | | | | | | | | | | | | | | | | | | | | |
|---|---|---|---|---|---|---|---|---|---|---|---|---|---|---|---|---|---|---|---|---|---|---|---|---|---|---|---|---|---|---|---|---|---|---|---|---|---|---|---|---|---|---|---|---|---|---|
| SE | 94029 | ENSG00000110955 | ATP5F1B | chr12 | - | 56642672 | 56642684 | 56642457 | 56642580 | 56643402 | 56643573 | 38 | 6 | 27 | 17 | 34 | 0 | 0 | 1 | 3 | 0 | 39 | 35 | 18 | 36 | 63 | 2 | 0 | 3 | 0 | 0 | 161 | 149 | 1 | 1 | 1 | 1 | 0.962 | 0.84 | 1 | 0.947 | 1 | 0.847 | 1 | 1 | 0.002 |
| SE | 94030 | ENSG00000110955 | ATP5F1B | chr12 | - | 56642672 | 56642831 | 56642457 | 56642580 | 56643402 | 56643587 | 1399 | 731 | 1225 | 1229 | 1687 | 0 | 0 | 1 | 3 | 0 | 1509 | 1368 | 1006 | 1175 | 2080 | 2 | 0 | 3 | 0 | 0 | 298 | 149 | 1 | 1 | 1 | 1 | 0.998 | 0.995 | 1 | 0.997 | 1 | 0.994 | 1 | 1 | 0 |
| SE | 94031 | ENSG00000110955 | ATP5F1B | chr12 | - | 56643836 | 56643958 | 56643402 | 56643587 | 56645170 | 56645353 | 710 | 401 | 586 | 657 | 834 | 0 | 0 | 0 | 0 | 0 | 1037 | 897 | 573 | 637 | 1030 | 0 | 1 | 0 | 0 | 0 | 271 | 149 | 1 | 1 | 1 | 1 | 1 | 1 | 1 | 1 | 0.998 | 1 | 1 | 1 | 0 |
| SE | 94032 | ENSG00000110955 | ATP5F1B | chr12 | - | 56644780 | 56644955 | 56643836 | 56643958 | 56645170 | 56645353 | 1020 | 568 | 867 | 927 | 940 | 0 | 0 | 2 | 0 | 0 | 1843 | 1231 | 827 | 1189 | 1587 | 0 | 0 | 0 | 0 | 0 | 298 | 149 | 1 | 1 | 1 | 1 | 0.995 | 1 | 1 | 1 | 1 | 1 | 1 | 1 | -0.001 |
| SE | 94034 | ENSG00000167094 | TTC16 | chr9 | + | 127724755 | 127724897 | 127724119 | 127724364 | 127726238 | 127726404 | 5 | 0 | 1 | 0 | 2 | 2 | 4 | 6 | 4 | 5 | 0 | 5 | 0 | 0 | 0 | 0 | 5 | 0 | 0 | 3 | 291 | 149 | 1 | 1 | 0.561 | 0 | 0.079 | 0 | 0.17 | NA | 0.339 | NA | NA | 0 | -0.008 |
| SE | 94035 | ENSG00000167094 | TTC16 | chr9 | + | 127724755 | 127724897 | 127724119 | 127724364 | 127727269 | 127727465 | 0 | 0 | 0 | 0 | 2 | 1 | 0 | 0 | 0 | 0 | 0 | 4 | 0 | 0 | 0 | 0 | 0 | 2 | 0 | 0 | 291 | 149 | 1 | 1 | 0 | NA | NA | NA | 1 | NA | 1 | 0 | NA | NA | 0 |
| SE | 94037 | ENSG00000167094 | TTC16 | chr9 | + | 127726238 | 127726404 | 127724119 | 127724897 | 127727269 | 127727465 | 17 | 4 | 3 | 15 | 10 | 0 | 0 | 0 | 0 | 0 | 32 | 21 | 0 | 8 | 14 | 0 | 4 | 0 | 0 | 0 | 298 | 149 | 1 | 1 | 1 | 1 | 1 | 1 | 1 | 1 | 0.724 | NA | 1 | 1 | 0.069 |
| SE | 94039 | ENSG00000167094 | TTC16 | chr9 | + | 127729580 | 127729668 | 127727269 | 127727465 | 127730635 | 127731589 | 34 | 5 | 2 | 15 | 8 | 0 | 1 | 1 | 2 | 0 | 17 | 36 | 11 | 9 | 12 | 0 | 1 | 0 | 0 | 0 | 237 | 149 | 1 | 1 | 1 | 0.759 | 0.557 | 0.825 | 1 | 1 | 0.958 | 1 | 1 | 1 | -0.163 |
| SE | 94040 | ENSG00000100296 | THOC5 | chr22 | - | 29525837 | 29525946 | 29520997 | 29521099 | 29528077 | 29528177 | 83 | 44 | 86 | 63 | 74 | 0 | 0 | 0 | 0 | 0 | 118 | 92 | 36 | 64 | 177 | 0 | 1 | 0 | 0 | 0 | 258 | 149 | 1 | 1 | 1 | 1 | 1 | 1 | 1 | 1 | 0.982 | 1 | 1 | 1 | 0.004 |
| SE | 94044 | ENSG00000100296 | THOC5 | chr22 | - | 29542858 | 29542956 | 29539329 | 29539476 | 29543428 | 29543542 | 120 | 98 | 125 | 137 | 193 | 0 | 0 | 0 | 1 | 2 | 268 | 174 | 154 | 132 | 282 | 0 | 0 | 0 | 0 | 4 | 247 | 149 | 1 | 1 | 1 | 1 | 1 | 0.988 | 0.983 | 1 | 1 | 1 | 1 | 0.977 | -0.001 |
| SE | 94047 | ENSG00000148346 | LCN2 | chr9 | + | 128150237 | 128150374 | 128149452 | 128149663 | 128151637 | 128151717 | 32 | 51 | 55 | 263 | 74 | 0 | 0 | 0 | 0 | 0 | 359 | 286 | 119 | 6871 | 1097 | 0 | 0 | 0 | 7 | 8 | 286 | 149 | 1 | 1 | 1 | 1 | 1 | 1 | 1 | 1 | 1 | 1 | 0.998 | 0.986 | 0.003 |
| SE | 94048 | ENSG00000148346 | LCN2 | chr9 | + | 128150374 | 128150556 | 128149440 | 128149663 | 128151637 | 128151717 | 0 | 0 | 1 | 2 | 0 | 0 | 0 | 0 | 0 | 0 | 0 | 0 | 0 | 60 | 32 | 0 | 0 | 0 | 7 | 8 | 298 | 149 | 1 | 1 | NA | NA | 1 | 1 | NA | NA | NA | NA | 0.811 | 0.667 | 0.261 |
| SE | 94050 | ENSG00000148346 | LCN2 | chr9 | + | 128152182 | 128152284 | 128151905 | 128152025 | 128153099 | 128153126 | 85 | 119 | 73 | 290 | 159 | 0 | 0 | 0 | 0 | 0 | 677 | 680 | 191 | 9867 | 3167 | 0 | 0 | 0 | 0 | 5 | 251 | 149 | 1 | 1 | 1 | 1 | 1 | 1 | 1 | 1 | 1 | 1 | 1 | 0.997 | 0.001 |
| SE | 94053 | ENSG00000148337 | CIZ1 | chr9 | - | 128178368 | 128178490 | 128177565 | 128177763 | 128179072 | 128179415 | 151 | 59 | 124 | 149 | 146 | 0 | 0 | 0 | 1 | 0 | 172 | 141 | 85 | 103 | 255 | 0 | 3 | 0 | 0 | 0 | 271 | 149 | 1 | 1 | 1 | 1 | 1 | 0.988 | 1 | 1 | 0.963 | 1 | 1 | 1 | 0.005 |
| SE | 94054 | ENSG00000148337 | CIZ1 | chr9 | - | 128178708 | 128178904 | 128177565 | 128177763 | 128179072 | 128179415 | 16 | 6 | 17 | 10 | 1 | 0 | 0 | 0 | 1 | 0 | 2 | 10 | 3 | 15 | 24 | 0 | 3 | 0 | 0 | 0 | 298 | 149 | 1 | 1 | 1 | 1 | 1 | 0.833 | 1 | 1 | 0.625 | 1 | 1 | 1 | 0.042 |
| SE | 94055 | ENSG00000148337 | CIZ1 | chr9 | - | 128178708 | 128178904 | 128177565 | 128177763 | 128179243 | 128179415 | 6 | 0 | 0 | 3 | 0 | 0 | 0 | 1 | 0 | 0 | 2 | 0 | 1 | 9 | 9 | 0 | 0 | 0 | 0 | 0 | 298 | 149 | 1 | 1 | 1 | NA | 0 | 1 | NA | 1 | NA | 1 | 1 | 1 | -0.333 |
| SE | 94057 | ENSG00000148337 | CIZ1 | chr9 | - | 128178988 | 128179415 | 128178708 | 128178904 | 128180414 | 128180523 | 86 | 43 | 100 | 70 | 34 | 0 | 0 | 1 | 0 | 0 | 110 | 101 | 74 | 165 | 212 | 0 | 0 | 0 | 0 | 0 | 298 | 149 | 1 | 1 | 1 | 1 | 0.98 | 1 | 1 | 1 | 1 | 1 | 1 | 1 | -0.004 |
| SE | 94058 | ENSG00000148337 | CIZ1 | chr9 | - | 128179072 | 128179415 | 128178708 | 128178904 | 128180414 | 128180523 | 96 | 49 | 116 | 77 | 35 | 0 | 0 | 1 | 0 | 0 | 110 | 111 | 76 | 171 | 227 | 0 | 0 | 0 | 0 | 0 | 298 | 149 | 1 | 1 | 1 | 1 | 0.983 | 1 | 1 | 1 | 1 | 1 | 1 | 1 | -0.003 |
| SE | 94059 | ENSG00000148337 | CIZ1 | chr9 | - | 128179243 | 128179415 | 128178708 | 128178904 | 128180414 | 128180523 | 86 | 43 | 99 | 70 | 34 | 0 | 0 | 1 | 0 | 0 | 110 | 101 | 74 | 165 | 212 | 0 | 0 | 0 | 0 | 0 | 298 | 149 | 1 | 1 | 1 | 1 | 0.98 | 1 | 1 | 1 | 1 | 1 | 1 | 1 | -0.004 |
| SE | 94066 | ENSG00000167985 | SDHAF2 | chr11 | + | 61437624 | 61437848 | 61430123 | 61430182 | 61438003 | 61438113 | 512 | 266 | 336 | 369 | 620 | 3 | 0 | 5 | 1 | 4 | 365 | 428 | 247 | 296 | 403 | 0 | 3 | 3 | 0 | 4 | 298 | 149 | 1 | 1 | 0.988 | 1 | 0.971 | 0.995 | 0.987 | 1 | 0.986 | 0.976 | 1 | 0.981 | 0 |
| SE | 94070 | ENSG00000167985 | SDHAF2 | chr11 | + | 61438244 | 61438342 | 61438003 | 61438113 | 61445940 | 61446005 | 0 | 0 | 1 | 0 | 1 | 146 | 90 | 114 | 105 | 196 | 0 | 2 | 2 | 0 | 6 | 135 | 132 | 90 | 91 | 143 | 247 | 149 | 1 | 1 | 0 | 0 | 0.005 | 0 | 0.003 | 0 | 0.009 | 0.013 | 0 | 0.025 | -0.008 |
| SE | 94071 | ENSG00000167985 | SDHAF2 | chr11 | + | 61438294 | 61438342 | 61438003 | 61438113 | 61445940 | 61446059 | 1 | 3 | 1 | 0 | 0 | 146 | 90 | 114 | 105 | 196 | 0 | 0 | 4 | 0 | 0 | 135 | 132 | 90 | 91 | 143 | 197 | 149 | 1 | 1 | 0.005 | 0.025 | 0.007 | 0 | 0 | 0 | 0 | 0.033 | 0 | 0 | 0.001 |
| SE | 94074 | ENSG00000175854 | SWI5 | chr9 | + | 128313829 | 128313905 | 128285938 | 128286033 | 128315423 | 128316123 | 1 | 0 | 5 | 2 | 0 | 1 | 1 | 0 | 0 | 0 | 0 | 6 | 6 | 6 | 0 | 0 | 3 | 4 | 0 | 0 | 225 | 149 | 1 | 1 | 0.398 | 0 | 1 | 1 | NA | NA | 0.57 | 0.498 | 1 | NA | -0.09 |
| SE | 94075 | ENSG00000175854 | SWI5 | chr9 | + | 128314097 | 128314247 | 128313829 | 128313905 | 128315423 | 128316123 | 1 | 0 | 3 | 0 | 0 | 1 | 0 | 2 | 1 | 0 | 0 | 10 | 0 | 0 | 0 | 0 | 3 | 3 | 3 | 0 | 298 | 149 | 1 | 1 | 0.333 | NA | 0.429 | 0 | NA | NA | 0.625 | 0 | 0 | NA | 0.046 |
| SE | 94076 | ENSG00000167112 | TRUB2 | chr9 | - | 128315566 | 128315628 | 128313807 | 128313889 | 128315821 | 128315896 | 13 | 2 | 13 | 7 | 6 | 0 | 0 | 0 | 1 | 0 | 23 | 6 | 13 | 1 | 31 | 0 | 0 | 0 | 0 | 0 | 211 | 149 | 1 | 1 | 1 | 1 | 1 | 0.832 | 1 | 1 | 1 | 1 | 1 | 1 | -0.034 |
| SE | 94079 | ENSG00000167113 | COQ4 | chr9 | + | 128332152 | 128332282 | 128325778 | 128325881 | 128332849 | 128332943 | 34 | 15 | 35 | 29 | 20 | 0 | 0 | 0 | 1 | 0 | 43 | 26 | 19 | 24 | 56 | 0 | 0 | 0 | 0 | 0 | 279 | 149 | 1 | 1 | 1 | 1 | 1 | 0.939 | 1 | 1 | 1 | 1 | 1 | 1 | -0.012 |
| SE | 94080 | ENSG00000167114 | SLC27A4 | chr9 | + | 128353024 | 128353234 | 128352637 | 128352747 | 128353414 | 128353541 | 34 | 13 | 32 | 48 | 9 | 0 | 0 | 0 | 3 | 0 | 27 | 31 | 13 | 13 | 68 | 0 | 0 | 0 | 0 | 0 | 298 | 149 | 1 | 1 | 1 | 1 | 1 | 0.889 | 1 | 1 | 1 | 1 | 1 | 1 | -0.022 |
| SE | 94089 | ENSG00000136811 | ODF2 | chr9 | + | 128460157 | 128460299 | 128459566 | 128459657 | 128460512 | 128460666 | 2 | 0 | 0 | 1 | 2 | 58 | 15 | 22 | 44 | 26 | 0 | 4 | 0 | 0 | 0 | 34 | 30 | 33 | 37 | 67 | 291 | 149 | 1 | 1 | 0.017 | 0 | 0 | 0.012 | 0.038 | 0 | 0.064 | 0 | 0 | 0 | 0.001 |
| SE | 94090 | ENSG00000136811 | ODF2 | chr9 | + | 128460157 | 128460299 | 128459566 | 128459657 | 128460549 | 128460666 | 0 | 0 | 0 | 1 | 1 | 5 | 4 | 6 | 11 | 0 | 0 | 4 | 0 | 0 | 0 | 0 | 6 | 3 | 16 | 23 | 291 | 149 | 1 | 1 | 0 | 0 | 0 | 0.044 | 1 | NA | 0.254 | 0 | 0 | 0 | 0.145 |
| SE | 94091 | ENSG00000136811 | ODF2 | chr9 | + | 128460512 | 128460666 | 128459566 | 128459657 | 128460941 | 128461010 | 114 | 55 | 51 | 95 | 38 | 0 | 2 | 0 | 3 | 0 | 70 | 89 | 52 | 95 | 126 | 0 | 0 | 1 | 4 | 0 | 298 | 149 | 1 | 1 | 1 | 0.932 | 1 | 0.941 | 1 | 1 | 1 | 0.963 | 0.922 | 1 | -0.002 |
| SE | 94093 | ENSG00000136811 | ODF2 | chr9 | + | 128460941 | 128461010 | 128460512 | 128460666 | 128469182 | 128469353 | 31 | 39 | 30 | 42 | 18 | 0 | 0 | 0 | 2 | 0 | 52 | 63 | 23 | 82 | 85 | 0 | 0 | 0 | 0 | 0 | 218 | 149 | 1 | 1 | 1 | 1 | 1 | 0.935 | 1 | 1 | 1 | 1 | 1 | 1 | -0.013 |
| SE | 94094 | ENSG00000136811 | ODF2 | chr9 | + | 128460941 | 128461067 | 128460512 | 128460666 | 128469182 | 128469234 | 98 | 59 | 33 | 87 | 44 | 0 | 0 | 0 | 2 | 0 | 48 | 69 | 30 | 88 | 171 | 0 | 0 | 0 | 0 | 0 | 275 | 149 | 1 | 1 | 1 | 1 | 1 | 0.959 | 1 | 1 | 1 | 1 | 1 | 1 | -0.008 |
| SE | 94096 | ENSG00000136811 | ODF2 | chr9 | + | 128469182 | 128469353 | 128460941 | 128461010 | 128472912 | 128473042 | 14 | 16 | 20 | 18 | 14 | 0 | 0 | 0 | 0 | 0 | 44 | 27 | 13 | 48 | 27 | 4 | 0 | 0 | 0 | 0 | 298 | 149 | 1 | 1 | 1 | 1 | 1 | 1 | 1 | 0.846 | 1 | 1 | 1 | 1 | 0.031 |
| SE | 94099 | ENSG00000136811 | ODF2 | chr9 | + | 128471307 | 128471468 | 128460941 | 128461010 | 128472912 | 128473042 | 65 | 52 | 60 | 101 | 118 | 0 | 0 | 0 | 0 | 0 | 149 | 66 | 88 | 118 | 183 | 4 | 0 | 0 | 0 | 0 | 298 | 149 | 1 | 1 | 1 | 1 | 1 | 1 | 1 | 0.949 | 1 | 1 | 1 | 1 | 0.01 |
| SE | 94101 | ENSG00000136811 | ODF2 | chr9 | + | 128471307 | 128471468 | 128469182 | 128469353 | 128472912 | 128473042 | 164 | 107 | 108 | 201 | 205 | 2 | 0 | 1 | 0 | 0 | 373 | 122 | 152 | 180 | 367 | 0 | 0 | 2 | 1 | 0 | 298 | 149 | 1 | 1 | 0.976 | 1 | 0.982 | 1 | 1 | 1 | 1 | 0.974 | 0.989 | 1 | -0.001 |
| SE | 94102 | ENSG00000136811 | ODF2 | chr9 | + | 128482815 | 128482887 | 128481579 | 128481651 | 128483937 | 128484054 | 114 | 90 | 107 | 156 | 93 | 1 | 0 | 0 | 2 | 0 | 197 | 100 | 123 | 136 | 328 | 0 | 0 | 0 | 0 | 0 | 221 | 149 | 1 | 1 | 0.987 | 1 | 1 | 0.981 | 1 | 1 | 1 | 1 | 1 | 1 | -0.006 |
| SE | 94103 | ENSG00000136811 | ODF2 | chr9 | + | 128483937 | 128484054 | 128482815 | 128482887 | 128484700 | 128484886 | 185 | 191 | 201 | 239 | 149 | 0 | 0 | 0 | 2 | 0 | 439 | 184 | 211 | 254 | 595 | 0 | 0 | 0 | 0 | 0 | 266 | 149 | 1 | 1 | 1 | 1 | 1 | 0.985 | 1 | 1 | 1 | 1 | 1 | 1 | -0.003 |
| SE | 94104 | ENSG00000136811 | ODF2 | chr9 | + | 128496040 | 128496141 | 128494509 | 128494668 | 128498412 | 128498575 | 144 | 84 | 97 | 102 | 94 | 2 | 0 | 0 | 0 | 0 | 300 | 189 | 79 | 150 | 222 | 0 | 0 | 3 | 0 | 0 | 250 | 149 | 1 | 1 | 0.977 | 1 | 1 | 1 | 1 | 1 | 1 | 0.94 | 1 | 1 | 0.007 |
| SE | 94105 | ENSG00000136811 | ODF2 | chr9 | + | 128496040 | 128496297 | 128494509 | 128494668 | 128498412 | 128498575 | 49 | 32 | 36 | 39 | 29 | 2 | 0 | 0 | 0 | 0 | 100 | 67 | 47 | 71 | 77 | 0 | 0 | 3 | 0 | 0 | 298 | 149 | 1 | 1 | 0.925 | 1 | 1 | 1 | 1 | 1 | 1 | 0.887 | 1 | 1 | 0.008 |
| SE | 94107 | ENSG00000113734 | BNIP1 | chr5 | + | 173158743 | 173158845 | 173154321 | 173154413 | 173159932 | 173160051 | 87 | 44 | 55 | 68 | 87 | 0 | 0 | 1 | 2 | 0 | 289 | 106 | 142 | 83 | 143 | 3 | 0 | 0 | 0 | 0 | 251 | 149 | 1 | 1 | 1 | 1 | 0.97 | 0.953 | 1 | 0.983 | 1 | 1 | 1 | 1 | -0.012 |
| SE | 94108 | ENSG00000113734 | BNIP1 | chr5 | + | 173159932 | 173160051 | 173158743 | 173158845 | 173163724 | 173164057 | 78 | 45 | 62 | 68 | 79 | 1 | 0 | 0 | 1 | 0 | 304 | 98 | 115 | 64 | 122 | 0 | 0 | 0 | 0 | 0 | 268 | 149 | 1 | 1 | 0.977 | 1 | 1 | 0.974 | 1 | 1 | 1 | 1 | 1 | 1 | -0.01 |
| SE | 94109 | ENSG00000198018 | ENTPD7 | chr10 | + | 99661445 | 99661628 | 99659861 | 99659964 | 99679260 | 99679466 | 25 | 16 | 28 | 25 | 21 | 0 | 2 | 0 | 0 | 0 | 21 | 29 | 15 | 45 | 7 | 0 | 0 | 0 | 3 | 0 | 298 | 149 | 1 | 1 | 1 | 0.8 | 1 | 1 | 1 | 1 | 1 | 1 | 0.882 | 1 | -0.016 |
| SE | 94110 | ENSG00000198018 | ENTPD7 | chr10 | + | 99691384 | 99691518 | 99688693 | 99688750 | 99695955 | 99696122 | 28 | 28 | 31 | 22 | 28 | 0 | 1 | 0 | 0 | 0 | 0 | 11 | 27 | 66 | 0 | 0 | 0 | 0 | 0 | 0 | 283 | 149 | 1 | 1 | 1 | 0.936 | 1 | 1 | 1 | NA | 1 | 1 | 1 | NA | -0.013 |
| SE | 94111 | ENSG00000198018 | ENTPD7 | chr10 | + | 99698533 | 99698858 | 99695955 | 99696122 | 99700972 | 99701058 | 17 | 11 | 27 | 31 | 33 | 1 | 0 | 0 | 4 | 0 | 0 | 25 | 18 | 44 | 0 | 0 | 3 | 0 | 0 | 0 | 298 | 149 | 1 | 1 | 0.895 | 1 | 1 | 0.795 | 1 | NA | 0.806 | 1 | 1 | NA | 0.003 |
| SE | 94112 | ENSG00000122435 | TRMT13 | chr1 | + | 100140181 | 100140251 | 100139648 | 100139711 | 100140407 | 100140514 | 73 | 87 | 87 | 81 | 203 | 18 | 6 | 10 | 11 | 13 | 0 | 22 | 11 | 12 | 0 | 0 | 2 | 4 | 6 | 0 | 219 | 149 | 1 | 1 | 0.734 | 0.908 | 0.855 | 0.834 | 0.914 | NA | 0.882 | 0.652 | 0.576 | NA | 0.146 |
| SE | 94113 | ENSG00000122435 | TRMT13 | chr1 | + | 100140181 | 100140251 | 100139648 | 100139711 | 100140449 | 100140812 | 40 | 46 | 32 | 30 | 63 | 10 | 7 | 6 | 2 | 0 | 0 | 15 | 12 | 4 | 0 | 0 | 7 | 2 | 2 | 0 | 219 | 149 | 1 | 1 | 0.731 | 0.817 | 0.784 | 0.911 | 1 | NA | 0.593 | 0.803 | 0.576 | NA | 0.191 |
| SE | 94114 | ENSG00000122435 | TRMT13 | chr1 | + | 100147893 | 100148326 | 100144068 | 100144143 | 100148624 | 100150498 | 34 | 30 | 45 | 28 | 110 | 3 | 0 | 1 | 3 | 6 | 0 | 13 | 6 | 0 | 0 | 0 | 0 | 0 | 0 | 0 | 298 | 149 | 1 | 1 | 0.85 | 1 | 0.957 | 0.824 | 0.902 | NA | 1 | 1 | NA | NA | -0.093 |
| SE | 94116 | ENSG00000139644 | TMBIM6 | chr12 | + | 49741956 | 49742115 | 49741548 | 49741611 | 49742437 | 49742607 | 11 | 5 | 5 | 5 | 2 | 0 | 0 | 0 | 1 | 0 | 13 | 1 | 7 | 12 | 0 | 0 | 0 | 0 | 0 | 0 | 298 | 149 | 1 | 1 | 1 | 1 | 1 | 0.714 | 1 | 1 | 1 | 1 | 1 | NA | -0.057 |
| SE | 94117 | ENSG00000139644 | TMBIM6 | chr12 | + | 49741956 | 49742115 | 49741548 | 49741611 | 49743416 | 49743565 | 12 | 5 | 2 | 5 | 2 | 0 | 0 | 0 | 2 | 0 | 13 | 1 | 7 | 12 | 0 | 0 | 0 | 1 | 0 | 0 | 298 | 149 | 1 | 1 | 1 | 1 | 1 | 0.556 | 1 | 1 | 1 | 0.778 | 1 | NA | -0.033 |
| SE | 94118 | ENSG00000139644 | TMBIM6 | chr12 | + | 49741956 | 49742115 | 49741548 | 49741611 | 49744412 | 49744542 | 11 | 6 | 2 | 8 | 2 | 4 | 0 | 2 | 2 | 0 | 13 | 1 | 7 | 12 | 0 | 0 | 0 | 0 | 0 | 0 | 298 | 149 | 1 | 1 | 0.579 | 1 | 0.333 | 0.667 | 1 | 1 | 1 | 1 | 1 | NA | -0.284 |
| SE | 94119 | ENSG00000139644 | TMBIM6 | chr12 | + | 49741956 | 49742115 | 49741548 | 49741611 | 49749617 | 49749856 | 11 | 5 | 2 | 5 | 2 | 0 | 0 | 0 | 6 | 0 | 13 | 1 | 7 | 12 | 0 | 0 | 0 | 0 | 0 | 0 | 298 | 149 | 1 | 1 | 1 | 1 | 1 | 0.294 | 1 | 1 | 1 | 1 | 1 | NA | -0.141 |
| SE | 94124 | ENSG00000139644 | TMBIM6 | chr12 | + | 49742437 | 49742607 | 49741570 | 49741611 | 49743416 | 49743565 | 2 | 0 | 1 | 3 | 0 | 0 | 0 | 0 | 2 | 0 | 0 | 8 | 1 | 0 | 0 | 0 | 0 | 1 | 0 | 0 | 298 | 149 | 1 | 1 | 1 | NA | 1 | 0.429 | NA | NA | 1 | 0.333 | NA | NA | 0.143 |
| SE | 94125 | ENSG00000139644 | TMBIM6 | chr12 | + | 49742437 | 49742607 | 49741570 | 49741611 | 49752463 | 49752524 | 2 | 2 | 6 | 5 | 0 | 320 | 116 | 272 | 243 | 214 | 5 | 4 | 5 | 0 | 8 | 259 | 232 | 143 | 129 | 396 | 298 | 149 | 1 | 1 | 0.003 | 0.009 | 0.011 | 0.01 | 0 | 0.01 | 0.009 | 0.017 | 0 | 0.01 | -0.003 |
| SE | 94126 | ENSG00000139644 | TMBIM6 | chr12 | + | 49742437 | 49742607 | 49741607 | 49741768 | 49752463 | 49752524 | 4 | 2 | 7 | 4 | 0 | 3 | 1 | 4 | 1 | 1 | 5 | 4 | 5 | 0 | 8 | 2 | 5 | 0 | 0 | 0 | 298 | 149 | 1 | 1 | 0.4 | 0.5 | 0.467 | 0.667 | 0 | 0.556 | 0.286 | 1 | NA | 1 | -0.304 |
| SE | 94127 | ENSG00000139644 | TMBIM6 | chr12 | + | 49742437 | 49742607 | 49741956 | 49742115 | 49743416 | 49743565 | 2 | 0 | 4 | 2 | 0 | 1 | 0 | 0 | 0 | 0 | 0 | 8 | 1 | 0 | 0 | 0 | 0 | 0 | 0 | 0 | 298 | 149 | 1 | 1 | 0.5 | NA | 1 | 1 | NA | NA | 1 | 1 | NA | NA | -0.167 |
| SE | 94129 | ENSG00000139644 | TMBIM6 | chr12 | + | 49743416 | 49743565 | 49741556 | 49741611 | 49744412 | 49744542 | 0 | 0 | 0 | 4 | 0 | 4 | 0 | 2 | 2 | 0 | 0 | 0 | 1 | 0 | 0 | 0 | 0 | 0 | 0 | 0 | 298 | 149 | 1 | 1 | 0 | NA | 0 | 0.5 | NA | NA | NA | 1 | NA | NA | -0.833 |
| SE | 94133 | ENSG00000139644 | TMBIM6 | chr12 | + | 49743416 | 49743565 | 49742437 | 49742607 | 49752463 | 49752549 | 6 | 1 | 1 | 9 | 0 | 2 | 2 | 6 | 4 | 0 | 0 | 28 | 6 | 7 | 0 | 5 | 4 | 5 | 0 | 8 | 298 | 149 | 1 | 1 | 0.6 | 0.2 | 0.077 | 0.529 | NA | 0 | 0.778 | 0.375 | 1 | 0 | -0.079 |
| SE | 94136 | ENSG00000139644 | TMBIM6 | chr12 | + | 49744412 | 49744542 | 49743416 | 49743565 | 49752463 | 49752549 | 6 | 1 | 1 | 8 | 7 | 4 | 1 | 0 | 7 | 0 | 6 | 0 | 4 | 0 | 0 | 0 | 20 | 5 | 7 | 0 | 279 | 149 | 1 | 1 | 0.445 | 0.348 | 1 | 0.379 | 1 | 1 | 0 | 0.299 | 0 | NA | 0.31 |
| SE | 94137 | ENSG00000139644 | TMBIM6 | chr12 | + | 49749617 | 49749734 | 49741543 | 49741611 | 49752463 | 49752549 | 0 | 0 | 0 | 12 | 0 | 320 | 116 | 272 | 243 | 214 | 0 | 0 | 2 | 0 | 0 | 259 | 232 | 143 | 129 | 396 | 266 | 149 | 1 | 1 | 0 | 0 | 0 | 0.027 | 0 | 0 | 0 | 0.008 | 0 | 0 | 0.004 |
| SE | 94138 | ENSG00000139644 | TMBIM6 | chr12 | + | 49749617 | 49749856 | 49741556 | 49741611 | 49752463 | 49752542 | 0 | 0 | 0 | 8 | 0 | 320 | 116 | 272 | 243 | 214 | 0 | 0 | 0 | 0 | 0 | 259 | 232 | 143 | 129 | 396 | 298 | 149 | 1 | 1 | 0 | 0 | 0 | 0.016 | 0 | 0 | 0 | 0 | 0 | 0 | 0.003 |
| SE | 94139 | ENSG00000139644 | TMBIM6 | chr12 | + | 49749617 | 49749856 | 49741956 | 49742115 | 49752463 | 49752542 | 0 | 0 | 0 | 2 | 0 | 26 | 7 | 4 | 14 | 7 | 0 | 0 | 0 | 0 | 0 | 8 | 11 | 7 | 11 | 32 | 298 | 149 | 1 | 1 | 0 | 0 | 0 | 0.067 | 0 | 0 | 0 | 0 | 0 | 0 | 0.013 |
| SE | 94140 | ENSG00000139644 | TMBIM6 | chr12 | + | 49752972 | 49753081 | 49752455 | 49752549 | 49755634 | 49755736 | 2026 | 846 | 1682 | 1755 | 1902 | 0 | 0 | 0 | 0 | 3 | 1517 | 1744 | 987 | 1182 | 3135 | 0 | 0 | 0 | 0 | 0 | 258 | 149 | 1 | 1 | 1 | 1 | 1 | 1 | 0.997 | 1 | 1 | 1 | 1 | 1 | -0.001 |
| SE | 94142 | ENSG00000139644 | TMBIM6 | chr12 | + | 49758382 | 49758480 | 49758206 | 49758275 | 49758682 | 49758686 | 1916 | 821 | 1571 | 1690 | 1662 | 0 | 0 | 0 | 0 | 0 | 1738 | 1879 | 1138 | 1453 | 3388 | 13 | 0 | 0 | 0 | 5 | 247 | 149 | 1 | 1 | 1 | 1 | 1 | 1 | 1 | 0.988 | 1 | 1 | 1 | 0.998 | 0.003 |
| SE | 94144 | ENSG00000090054 | SPTLC1 | chr9 | - | 92047612 | 92047708 | 92047171 | 92047268 | 92049959 | 92050067 | 418 | 230 | 227 | 317 | 364 | 0 | 1 | 0 | 0 | 0 | 164 | 286 | 180 | 270 | 78 | 0 | 0 | 0 | 4 | 0 | 245 | 149 | 1 | 1 | 1 | 0.993 | 1 | 1 | 1 | 1 | 1 | 1 | 0.976 | 1 | 0.003 |
| SE | 94145 | ENSG00000090054 | SPTLC1 | chr9 | - | 92050295 | 92050491 | 92049959 | 92050067 | 92055404 | 92055494 | 0 | 0 | 0 | 1 | 0 | 192 | 124 | 136 | 163 | 226 | 0 | 0 | 0 | 0 | 0 | 101 | 142 | 76 | 103 | 27 | 298 | 149 | 1 | 1 | 0 | 0 | 0 | 0.003 | 0 | 0 | 0 | 0 | 0 | 0 | 0.001 |
| SE | 94146 | ENSG00000090054 | SPTLC1 | chr9 | - | 92067965 | 92068098 | 92059178 | 92059308 | 92080015 | 92080088 | 300 | 230 | 223 | 326 | 225 | 0 | 1 | 0 | 0 | 0 | 233 | 254 | 209 | 292 | 72 | 0 | 0 | 0 | 0 | 0 | 282 | 149 | 1 | 1 | 1 | 0.992 | 1 | 1 | 1 | 1 | 1 | 1 | 1 | 1 | -0.002 |
| SE | 94147 | ENSG00000090054 | SPTLC1 | chr9 | - | 92070123 | 92070296 | 92067965 | 92068098 | 92076797 | 92077035 | 3 | 1 | 3 | 6 | 3 | 0 | 0 | 10 | 6 | 1 | 20 | 9 | 12 | 6 | 0 | 0 | 0 | 6 | 6 | 0 | 298 | 149 | 1 | 1 | 1 | 1 | 0.13 | 0.333 | 0.6 | 1 | 1 | 0.5 | 0.333 | NA | -0.096 |
| SE | 94150 | ENSG00000090054 | SPTLC1 | chr9 | - | 92076797 | 92077035 | 92067965 | 92068098 | 92080869 | 92080963 | 0 | 0 | 10 | 6 | 1 | 2 | 0 | 0 | 0 | 0 | 0 | 0 | 6 | 8 | 0 | 0 | 0 | 0 | 0 | 0 | 298 | 149 | 1 | 1 | 0 | NA | 1 | 1 | 1 | NA | NA | 1 | 1 | NA | -0.25 |
| SE | 94151 | ENSG00000090054 | SPTLC1 | chr9 | - | 92076797 | 92077035 | 92070123 | 92070296 | 92080015 | 92080076 | 0 | 1 | 1 | 2 | 5 | 0 | 1 | 0 | 0 | 0 | 0 | 6 | 8 | 2 | 15 | 0 | 0 | 3 | 3 | 0 | 298 | 149 | 1 | 1 | NA | 0.333 | 1 | 1 | 1 | NA | 1 | 0.571 | 0.25 | 1 | 0.128 |
| SE | 94153 | ENSG00000090054 | SPTLC1 | chr9 | - | 92080015 | 92080088 | 92059178 | 92059308 | 92108739 | 92108834 | 0 | 3 | 1 | 0 | 0 | 0 | 0 | 0 | 0 | 0 | 0 | 0 | 0 | 0 | 0 | 0 | 5 | 0 | 0 | 0 | 222 | 149 | 1 | 1 | NA | 1 | 1 | NA | NA | NA | 0 | NA | NA | NA | 1 |
| SE | 94154 | ENSG00000090054 | SPTLC1 | chr9 | - | 92080015 | 92080088 | 92067965 | 92068098 | 92080869 | 92080963 | 362 | 267 | 283 | 394 | 217 | 2 | 0 | 0 | 0 | 0 | 303 | 289 | 320 | 371 | 113 | 0 | 0 | 0 | 0 | 0 | 222 | 149 | 1 | 1 | 0.992 | 1 | 1 | 1 | 1 | 1 | 1 | 1 | 1 | 1 | -0.002 |
| SE | 94155 | ENSG00000090054 | SPTLC1 | chr9 | - | 92080015 | 92080088 | 92076797 | 92077035 | 92080869 | 92080963 | 133 | 100 | 107 | 141 | 60 | 0 | 0 | 0 | 0 | 0 | 90 | 114 | 139 | 119 | 62 | 0 | 0 | 0 | 2 | 0 | 222 | 149 | 1 | 1 | 1 | 1 | 1 | 1 | 1 | 1 | 1 | 1 | 0.976 | 1 | 0.005 |
| SE | 94156 | ENSG00000090054 | SPTLC1 | chr9 | - | 92080869 | 92080963 | 92079852 | 92080088 | 92108739 | 92108768 | 346 | 237 | 278 | 379 | 301 | 0 | 3 | 1 | 0 | 0 | 347 | 271 | 295 | 350 | 164 | 0 | 0 | 0 | 0 | 0 | 243 | 149 | 1 | 1 | 1 | 0.98 | 0.994 | 1 | 1 | 1 | 1 | 1 | 1 | 1 | -0.005 |
| SE | 94161 | ENSG00000198000 | NOL8 | chr9 | - | 92319942 | 92320137 | 92319220 | 92319356 | 92321667 | 92321746 | 18 | 5 | 13 | 5 | 13 | 37 | 43 | 26 | 49 | 40 | 0 | 2 | 4 | 4 | 0 | 0 | 26 | 24 | 3 | 24 | 298 | 149 | 1 | 1 | 0.196 | 0.055 | 0.2 | 0.049 | 0.14 | NA | 0.037 | 0.077 | 0.4 | 0 | -0.001 |
| SE | 94162 | ENSG00000198000 | NOL8 | chr9 | - | 92319942 | 92320379 | 92319220 | 92319356 | 92321667 | 92321746 | 19 | 7 | 12 | 14 | 14 | 37 | 43 | 26 | 49 | 40 | 0 | 8 | 16 | 7 | 0 | 0 | 26 | 24 | 3 | 24 | 298 | 149 | 1 | 1 | 0.204 | 0.075 | 0.188 | 0.125 | 0.149 | NA | 0.133 | 0.25 | 0.538 | 0 | -0.082 |
| SE | 94163 | ENSG00000198000 | NOL8 | chr9 | - | 92321667 | 92321746 | 92319987 | 92320382 | 92323440 | 92323503 | 36 | 27 | 27 | 33 | 36 | 0 | 0 | 0 | 0 | 1 | 0 | 27 | 21 | 0 | 0 | 0 | 0 | 0 | 0 | 0 | 228 | 149 | 1 | 1 | 1 | 1 | 1 | 1 | 0.959 | NA | 1 | 1 | NA | NA | -0.008 |
| SE | 94164 | ENSG00000198000 | NOL8 | chr9 | - | 92323235 | 92323351 | 92321667 | 92321746 | 92323440 | 92323503 | 29 | 22 | 26 | 37 | 34 | 57 | 53 | 43 | 64 | 55 | 0 | 14 | 13 | 8 | 9 | 0 | 33 | 34 | 1 | 18 | 265 | 149 | 1 | 1 | 0.222 | 0.189 | 0.254 | 0.245 | 0.258 | NA | 0.193 | 0.177 | 0.818 | 0.219 | -0.118 |
| SE | 94165 | ENSG00000198000 | NOL8 | chr9 | - | 92323440 | 92323503 | 92321667 | 92321746 | 92324022 | 92324080 | 93 | 86 | 83 | 123 | 95 | 0 | 1 | 0 | 0 | 0 | 4 | 62 | 62 | 2 | 28 | 0 | 0 | 1 | 0 | 0 | 212 | 149 | 1 | 1 | 1 | 0.984 | 1 | 1 | 1 | 1 | 1 | 0.978 | 1 | 1 | 0.001 |
| SE | 94175 | ENSG00000152795 | HNRNPDL | chr4 | - | 82427189 | 82427304 | 82426036 | 82426129 | 82428017 | 82428179 | 48 | 33 | 33 | 61 | 71 | 1 | 0 | 0 | 0 | 0 | 65 | 58 | 83 | 71 | 19 | 0 | 0 | 0 | 0 | 0 | 264 | 149 | 1 | 1 | 0.964 | 1 | 1 | 1 | 1 | 1 | 1 | 1 | 1 | 1 | -0.007 |
| SE | 94176 | ENSG00000152795 | HNRNPDL | chr4 | - | 82427432 | 82427564 | 82427189 | 82427304 | 82428017 | 82428179 | 4872 | 4182 | 4588 | 5065 | 9344 | 0 | 0 | 0 | 3 | 0 | 1638 | 3993 | 4211 | 2638 | 863 | 0 | 0 | 0 | 0 | 0 | 281 | 149 | 1 | 1 | 1 | 1 | 1 | 0.999 | 1 | 1 | 1 | 1 | 1 | 1 | 0 |
| SE | 94177 | ENSG00000152795 | HNRNPDL | chr4 | - | 82428017 | 82428179 | 82427432 | 82427564 | 82429247 | 82429397 | 2273 | 1896 | 2032 | 2277 | 4551 | 0 | 1 | 0 | 1 | 6 | 704 | 1756 | 1840 | 1121 | 383 | 0 | 1 | 0 | 0 | 0 | 298 | 149 | 1 | 1 | 1 | 0.999 | 1 | 0.999 | 0.997 | 1 | 0.999 | 1 | 1 | 1 | -0.001 |
| SE | 94178 | ENSG00000152795 | HNRNPDL | chr4 | - | 82428277 | 82428446 | 82428017 | 82428179 | 82429247 | 82429397 | 2011 | 1509 | 1799 | 1851 | 3345 | 0 | 0 | 0 | 0 | 0 | 555 | 1472 | 1451 | 1036 | 277 | 0 | 2 | 0 | 0 | 0 | 298 | 149 | 1 | 1 | 1 | 1 | 1 | 1 | 1 | 1 | 0.997 | 1 | 1 | 1 | 0.001 |
| SE | 94179 | ENSG00000127080 | IPPK | chr9 | - | 92619485 | 92619565 | 92613182 | 92616057 | 92634385 | 92634488 | 24 | 27 | 31 | 32 | 12 | 0 | 0 | 0 | 0 | 0 | 39 | 28 | 19 | 20 | 18 | 0 | 0 | 0 | 1 | 0 | 229 | 149 | 1 | 1 | 1 | 1 | 1 | 1 | 1 | 1 | 1 | 1 | 0.929 | 1 | 0.014 |
| SE | 94186 | ENSG00000167272 | POP5 | chr12 | - | 120579513 | 120579597 | 120578763 | 120579412 | 120579773 | 120579923 | 102 | 66 | 92 | 131 | 190 | 0 | 0 | 5 | 0 | 0 | 95 | 55 | 27 | 91 | 64 | 0 | 0 | 0 | 0 | 0 | 233 | 149 | 1 | 1 | 1 | 1 | 0.922 | 1 | 1 | 1 | 1 | 1 | 1 | 1 | -0.016 |
| SE | 94187 | ENSG00000167272 | POP5 | chr12 | - | 120579513 | 120579597 | 120578763 | 120579412 | 120581114 | 120581257 | 49 | 44 | 54 | 71 | 98 | 0 | 0 | 0 | 0 | 0 | 71 | 61 | 13 | 46 | 52 | 0 | 1 | 0 | 0 | 0 | 233 | 149 | 1 | 1 | 1 | 1 | 1 | 1 | 1 | 1 | 0.975 | 1 | 1 | 1 | 0.005 |
| SE | 94188 | ENSG00000167272 | POP5 | chr12 | - | 120579513 | 120579923 | 120579351 | 120579412 | 120581114 | 120581251 | 79 | 62 | 96 | 98 | 103 | 0 | 0 | 0 | 0 | 0 | 92 | 50 | 30 | 65 | 64 | 0 | 1 | 0 | 0 | 0 | 298 | 149 | 1 | 1 | 1 | 1 | 1 | 1 | 1 | 1 | 0.962 | 1 | 1 | 1 | 0.008 |
| SE | 94189 | ENSG00000167272 | POP5 | chr12 | - | 120579773 | 120579923 | 120578763 | 120579412 | 120580995 | 120581257 | 1 | 0 | 5 | 5 | 0 | 0 | 0 | 0 | 0 | 0 | 0 | 0 | 0 | 0 | 0 | 0 | 0 | 1 | 0 | 0 | 298 | 149 | 1 | 1 | 1 | NA | 1 | 1 | NA | NA | NA | 0 | NA | NA | 1 |
| SE | 94190 | ENSG00000167272 | POP5 | chr12 | - | 120579773 | 120579923 | 120578763 | 120579412 | 120581114 | 120581257 | 37 | 18 | 38 | 22 | 4 | 0 | 0 | 0 | 0 | 0 | 34 | 16 | 18 | 28 | 24 | 0 | 1 | 0 | 0 | 0 | 298 | 149 | 1 | 1 | 1 | 1 | 1 | 1 | 1 | 1 | 0.889 | 1 | 1 | 1 | 0.022 |
| SE | 94193 | ENSG00000137078 | SIT1 | chr9 | - | 35650356 | 35650415 | 35650183 | 35650245 | 35650753 | 35650910 | 20 | 7 | 17 | 10 | 6 | 0 | 0 | 0 | 0 | 1 | 0 | 39 | 3 | 0 | 0 | 0 | 0 | 0 | 0 | 7 | 208 | 149 | 1 | 1 | 1 | 1 | 1 | 1 | 0.811 | NA | 1 | 1 | NA | 0 | 0.296 |
| SE | 94195 | ENSG00000159884 | CCDC107 | chr9 | + | 35660745 | 35660858 | 35660556 | 35660647 | 35660939 | 35661499 | 104 | 62 | 57 | 78 | 24 | 0 | 0 | 0 | 0 | 0 | 40 | 108 | 47 | 69 | 154 | 0 | 0 | 0 | 1 | 0 | 262 | 149 | 1 | 1 | 1 | 1 | 1 | 1 | 1 | 1 | 1 | 1 | 0.975 | 1 | 0.005 |
| SE | 94196 | ENSG00000159884 | CCDC107 | chr9 | + | 35660765 | 35660858 | 35660556 | 35660647 | 35660939 | 35661177 | 39 | 22 | 18 | 29 | 7 | 0 | 0 | 0 | 0 | 0 | 10 | 42 | 18 | 19 | 50 | 0 | 0 | 0 | 1 | 0 | 242 | 149 | 1 | 1 | 1 | 1 | 1 | 1 | 1 | 1 | 1 | 1 | 0.921 | 1 | 0.016 |
| SE | 94197 | ENSG00000198467 | TPM2 | chr9 | - | 35684731 | 35684807 | 35684487 | 35684550 | 35685063 | 35685139 | 22 | 28 | 18 | 34 | 5 | 4 | 1 | 5 | 2 | 1 | 0 | 6 | 7 | 7 | 46 | 0 | 2 | 7 | 6 | 27 | 225 | 149 | 1 | 1 | 0.785 | 0.949 | 0.704 | 0.918 | 0.768 | NA | 0.665 | 0.398 | 0.436 | 0.53 | 0.318 |
| SE | 94202 | ENSG00000123179 | EBPL | chr13 | - | 49669776 | 49669846 | 49660722 | 49661208 | 49686474 | 49686595 | 1 | 0 | 1 | 1 | 4 | 0 | 0 | 0 | 0 | 2 | 0 | 2 | 0 | 0 | 0 | 0 | 0 | 0 | 0 | 0 | 219 | 149 | 1 | 1 | 1 | NA | 1 | 1 | 0.576 | NA | 1 | NA | NA | NA | -0.106 |
| SE | 94203 | ENSG00000123179 | EBPL | chr13 | - | 49669776 | 49669846 | 49660722 | 49661208 | 49691253 | 49691463 | 1 | 0 | 2 | 0 | 4 | 2 | 0 | 0 | 2 | 0 | 0 | 2 | 0 | 0 | 0 | 0 | 4 | 0 | 1 | 0 | 219 | 149 | 1 | 1 | 0.254 | NA | 1 | 0 | 1 | NA | 0.254 | NA | 0 | NA | 0.436 |
| SE | 94204 | ENSG00000123179 | EBPL | chr13 | - | 49669776 | 49669846 | 49663056 | 49663195 | 49691253 | 49691463 | 2 | 4 | 4 | 2 | 10 | 0 | 0 | 1 | 5 | 0 | 0 | 17 | 5 | 5 | 13 | 0 | 0 | 2 | 0 | 0 | 219 | 149 | 1 | 1 | 1 | 1 | 0.731 | 0.214 | 1 | NA | 1 | 0.63 | 1 | 1 | -0.119 |
| SE | 94205 | ENSG00000123179 | EBPL | chr13 | - | 49686474 | 49686595 | 49660722 | 49661208 | 49691253 | 49691463 | 0 | 0 | 0 | 0 | 2 | 2 | 0 | 0 | 2 | 0 | 0 | 0 | 0 | 0 | 0 | 0 | 4 | 0 | 1 | 0 | 270 | 149 | 1 | 1 | 0 | NA | NA | 0 | 1 | NA | 0 | NA | 0 | NA | 0.333 |
| SE | 94206 | ENSG00000123179 | EBPL | chr13 | - | 49686474 | 49686595 | 49669776 | 49669846 | 49691253 | 49691463 | 0 | 0 | 0 | 1 | 0 | 0 | 1 | 2 | 0 | 0 | 0 | 0 | 0 | 0 | 0 | 0 | 0 | 0 | 3 | 0 | 270 | 149 | 1 | 1 | NA | 0 | 0 | 1 | NA | NA | NA | NA | 0 | NA | 0.333 |
| SE | 94211 | ENSG00000077463 | SIRT6 | chr19 | - | 4177078 | 4177138 | 4175027 | 4175937 | 4180781 | 4180909 | 18 | 13 | 15 | 14 | 30 | 1 | 0 | 0 | 0 | 0 | 0 | 36 | 20 | 21 | 59 | 0 | 0 | 0 | 0 | 0 | 209 | 149 | 1 | 1 | 0.928 | 1 | 1 | 1 | 1 | NA | 1 | 1 | 1 | 1 | -0.014 |
| SE | 94217 | ENSG00000137103 | TMEM8B | chr9 | + | 35835010 | 35835218 | 35834460 | 35834650 | 35841133 | 35841267 | 17 | 4 | 11 | 12 | 7 | 0 | 1 | 0 | 0 | 0 | 32 | 0 | 0 | 1 | 0 | 0 | 1 | 0 | 0 | 0 | 298 | 149 | 1 | 1 | 1 | 0.667 | 1 | 1 | 1 | 1 | 0 | NA | 1 | NA | 0.267 |
| SE | 94219 | ENSG00000137103 | TMEM8B | chr9 | + | 35842391 | 35842717 | 35841525 | 35841794 | 35845974 | 35846068 | 8 | 4 | 2 | 15 | 5 | 1 | 0 | 0 | 0 | 0 | 3 | 9 | 8 | 9 | 3 | 0 | 0 | 0 | 0 | 0 | 298 | 149 | 1 | 1 | 0.8 | 1 | 1 | 1 | 1 | 1 | 1 | 1 | 1 | 1 | -0.04 |
| SE | 94224 | ENSG00000171843 | MLLT3 | chr9 | - | 20354807 | 20354879 | 20345076 | 20346574 | 20360741 | 20360841 | 9 | 11 | 10 | 7 | 11 | 0 | 4 | 0 | 0 | 0 | 0 | 3 | 5 | 0 | 0 | 0 | 5 | 0 | 0 | 0 | 221 | 149 | 1 | 1 | 1 | 0.65 | 1 | 1 | 1 | NA | 0.288 | 1 | NA | NA | 0.286 |
| SE | 94227 | ENSG00000171843 | MLLT3 | chr9 | - | 20380206 | 20380327 | 20365668 | 20365744 | 20382380 | 20382452 | 1 | 2 | 0 | 0 | 0 | 3 | 0 | 0 | 0 | 0 | 0 | 0 | 2 | 0 | 0 | 0 | 22 | 0 | 3 | 13 | 270 | 149 | 1 | 1 | 0.155 | 1 | NA | NA | NA | NA | 0 | 1 | 0 | 0 | 0.328 |
| SE | 94228 | ENSG00000171843 | MLLT3 | chr9 | - | 20413720 | 20414425 | 20365668 | 20365744 | 20448122 | 20448266 | 197 | 172 | 225 | 248 | 823 | 0 | 0 | 1 | 1 | 0 | 57 | 122 | 184 | 102 | 36 | 0 | 0 | 0 | 0 | 0 | 298 | 149 | 1 | 1 | 1 | 1 | 0.991 | 0.992 | 1 | 1 | 1 | 1 | 1 | 1 | -0.003 |
| SE | 94229 | ENSG00000171843 | MLLT3 | chr9 | - | 20448122 | 20448266 | 20413720 | 20414425 | 20620653 | 20620834 | 27 | 30 | 43 | 39 | 43 | 0 | 1 | 0 | 0 | 0 | 0 | 9 | 12 | 15 | 0 | 0 | 0 | 0 | 0 | 0 | 293 | 149 | 1 | 1 | 1 | 0.938 | 1 | 1 | 1 | NA | 1 | 1 | 1 | NA | -0.012 |
| SE | 94230 | ENSG00000171843 | MLLT3 | chr9 | - | 20456703 | 20456786 | 20448122 | 20448266 | 20620653 | 20620834 | 42 | 42 | 72 | 69 | 132 | 0 | 0 | 0 | 0 | 1 | 34 | 28 | 24 | 51 | 50 | 0 | 0 | 0 | 0 | 0 | 232 | 149 | 1 | 1 | 1 | 1 | 1 | 1 | 0.988 | 1 | 1 | 1 | 1 | 1 | -0.002 |
| SE | 94231 | ENSG00000213722 | DDAH2 | chr6 | - | 31728172 | 31728292 | 31727542 | 31727643 | 31728450 | 31728524 | 16 | 3 | 8 | 14 | 1 | 1 | 0 | 1 | 0 | 0 | 21 | 8 | 8 | 4 | 11 | 0 | 0 | 0 | 0 | 0 | 269 | 149 | 1 | 1 | 0.899 | 1 | 0.816 | 1 | 1 | 1 | 1 | 1 | 1 | 1 | -0.057 |
| SE | 94232 | ENSG00000213722 | DDAH2 | chr6 | - | 31728172 | 31728292 | 31727542 | 31727692 | 31728450 | 31728524 | 35 | 12 | 18 | 31 | 2 | 0 | 0 | 0 | 2 | 0 | 26 | 20 | 9 | 17 | 29 | 0 | 0 | 0 | 0 | 0 | 269 | 149 | 1 | 1 | 1 | 1 | 1 | 0.896 | 1 | 1 | 1 | 1 | 1 | 1 | -0.021 |
| SE | 94233 | ENSG00000213722 | DDAH2 | chr6 | - | 31728450 | 31728524 | 31728172 | 31728292 | 31728645 | 31728745 | 29 | 20 | 27 | 33 | 17 | 0 | 0 | 0 | 2 | 0 | 50 | 25 | 29 | 29 | 57 | 0 | 0 | 0 | 0 | 0 | 223 | 149 | 1 | 1 | 1 | 1 | 1 | 0.917 | 1 | 1 | 1 | 1 | 1 | 1 | -0.017 |
| SE | 94237 | ENSG00000130653 | PNPLA7 | chr9 | - | 137547308 | 137547396 | 137546829 | 137546909 | 137550167 | 137550384 | 1 | 0 | 2 | 1 | 1 | 0 | 0 | 0 | 0 | 0 | 8 | 5 | 2 | 0 | 5 | 0 | 3 | 0 | 0 | 0 | 237 | 149 | 1 | 1 | 1 | NA | 1 | 1 | 1 | 1 | 0.512 | 1 | NA | 1 | 0.122 |
| SE | 94239 | ENSG00000197070 | ARRDC1 | chr9 | + | 137613412 | 137613510 | 137612895 | 137613006 | 137613614 | 137613769 | 29 | 13 | 23 | 23 | 15 | 1 | 0 | 0 | 0 | 0 | 36 | 45 | 9 | 26 | 44 | 0 | 0 | 0 | 0 | 0 | 247 | 149 | 1 | 1 | 0.946 | 1 | 1 | 1 | 1 | 1 | 1 | 1 | 1 | 1 | -0.011 |
| SE | 94240 | ENSG00000197070 | ARRDC1 | chr9 | + | 137613422 | 137613510 | 137612895 | 137613006 | 137613614 | 137613769 | 30 | 13 | 22 | 23 | 15 | 1 | 0 | 0 | 0 | 0 | 36 | 45 | 8 | 26 | 44 | 0 | 0 | 0 | 0 | 0 | 237 | 149 | 1 | 1 | 0.95 | 1 | 1 | 1 | 1 | 1 | 1 | 1 | 1 | 1 | -0.01 |
| SE | 94241 | ENSG00000197070 | ARRDC1 | chr9 | + | 137613459 | 137613510 | 137611450 | 137613006 | 137613614 | 137613769 | 62 | 21 | 50 | 44 | 51 | 1 | 0 | 0 | 0 | 0 | 69 | 113 | 25 | 43 | 88 | 0 | 0 | 0 | 0 | 0 | 200 | 149 | 1 | 1 | 0.979 | 1 | 1 | 1 | 1 | 1 | 1 | 1 | 1 | 1 | -0.004 |
| SE | 94254 | ENSG00000138663 | COPS4 | chr4 | + | 83049165 | 83049317 | 83045596 | 83045705 | 83049880 | 83049959 | 224 | 131 | 162 | 164 | 312 | 0 | 0 | 0 | 2 | 0 | 23 | 193 | 159 | 108 | 34 | 0 | 0 | 0 | 0 | 0 | 298 | 149 | 1 | 1 | 1 | 1 | 1 | 0.976 | 1 | 1 | 1 | 1 | 1 | 1 | -0.005 |
| SE | 94255 | ENSG00000138663 | COPS4 | chr4 | + | 83057257 | 83057408 | 83056925 | 83057079 | 83063075 | 83063246 | 198 | 102 | 115 | 142 | 110 | 0 | 0 | 0 | 2 | 0 | 69 | 158 | 91 | 75 | 52 | 0 | 0 | 0 | 0 | 0 | 298 | 149 | 1 | 1 | 1 | 1 | 1 | 0.973 | 1 | 1 | 1 | 1 | 1 | 1 | -0.005 |
| SE | 94257 | ENSG00000138663 | COPS4 | chr4 | + | 83066437 | 83066553 | 83062974 | 83063246 | 83068437 | 83068506 | 126 | 84 | 120 | 99 | 182 | 4 | 0 | 0 | 0 | 2 | 48 | 139 | 77 | 118 | 41 | 0 | 0 | 0 | 0 | 0 | 265 | 149 | 1 | 1 | 0.947 | 1 | 1 | 1 | 0.981 | 1 | 1 | 1 | 1 | 1 | -0.014 |
| SE | 94258 | ENSG00000138663 | COPS4 | chr4 | + | 83068437 | 83068522 | 83066437 | 83066553 | 83075296 | 83075371 | 189 | 149 | 134 | 123 | 339 | 0 | 0 | 0 | 0 | 1 | 65 | 161 | 93 | 117 | 30 | 0 | 0 | 0 | 0 | 0 | 234 | 149 | 1 | 1 | 1 | 1 | 1 | 1 | 0.995 | 1 | 1 | 1 | 1 | 1 | -0.001 |
| SE | 94259 | ENSG00000138663 | COPS4 | chr4 | + | 83073205 | 83073353 | 83068437 | 83068522 | 83075296 | 83075756 | 10 | 8 | 5 | 8 | 15 | 133 | 112 | 73 | 77 | 280 | 0 | 2 | 0 | 0 | 0 | 27 | 86 | 42 | 59 | 0 | 297 | 149 | 1 | 1 | 0.036 | 0.035 | 0.033 | 0.05 | 0.026 | 0 | 0.012 | 0 | 0 | NA | 0.033 |
| SE | 94264 | ENSG00000157954 | WIPI2 | chr7 | + | 5214534 | 5214704 | 5199575 | 5199658 | 5216562 | 5216659 | 295 | 150 | 269 | 246 | 311 | 0 | 0 | 0 | 0 | 0 | 331 | 384 | 190 | 225 | 330 | 0 | 0 | 0 | 0 | 6 | 298 | 149 | 1 | 1 | 1 | 1 | 1 | 1 | 1 | 1 | 1 | 1 | 1 | 0.965 | 0.007 |
| SE | 94265 | ENSG00000157954 | WIPI2 | chr7 | + | 5216748 | 5216863 | 5216590 | 5216659 | 5217089 | 5217187 | 0 | 1 | 2 | 1 | 0 | 77 | 44 | 78 | 77 | 85 | 0 | 0 | 2 | 0 | 8 | 95 | 157 | 65 | 89 | 134 | 264 | 149 | 1 | 1 | 0 | 0.013 | 0.014 | 0.007 | 0 | 0 | 0 | 0.017 | 0 | 0.033 | -0.003 |
| SE | 94266 | ENSG00000157954 | WIPI2 | chr7 | + | 5217089 | 5217187 | 5216562 | 5216659 | 5217921 | 5218013 | 166 | 74 | 191 | 156 | 150 | 0 | 0 | 0 | 1 | 0 | 216 | 312 | 148 | 196 | 289 | 0 | 0 | 0 | 0 | 0 | 247 | 149 | 1 | 1 | 1 | 1 | 1 | 0.989 | 1 | 1 | 1 | 1 | 1 | 1 | -0.002 |
| SE | 94267 | ENSG00000048828 | FAM120A | chr9 | + | 93471140 | 93471387 | 93451684 | 93452389 | 93476255 | 93476338 | 764 | 390 | 529 | 575 | 841 | 0 | 0 | 2 | 0 | 0 | 834 | 621 | 454 | 571 | 699 | 0 | 0 | 0 | 0 | 0 | 298 | 149 | 1 | 1 | 1 | 1 | 0.992 | 1 | 1 | 1 | 1 | 1 | 1 | 1 | -0.002 |
| SE | 94268 | ENSG00000048828 | FAM120A | chr9 | + | 93476255 | 93476338 | 93471140 | 93471387 | 93497470 | 93497599 | 935 | 542 | 688 | 750 | 1320 | 0 | 0 | 0 | 3 | 0 | 1308 | 755 | 558 | 753 | 818 | 0 | 0 | 3 | 0 | 6 | 232 | 149 | 1 | 1 | 1 | 1 | 1 | 0.994 | 1 | 1 | 1 | 0.992 | 1 | 0.989 | 0.003 |
| SE | 94269 | ENSG00000048828 | FAM120A | chr9 | + | 93497470 | 93497599 | 93476255 | 93476338 | 93498789 | 93498886 | 1162 | 760 | 917 | 1077 | 1754 | 0 | 0 | 0 | 0 | 0 | 1812 | 1072 | 801 | 1114 | 1161 | 0 | 2 | 0 | 0 | 0 | 278 | 149 | 1 | 1 | 1 | 1 | 1 | 1 | 1 | 1 | 0.997 | 1 | 1 | 1 | 0.001 |
| SE | 94271 | ENSG00000048828 | FAM120A | chr9 | + | 93529352 | 93529580 | 93527154 | 93527242 | 93532154 | 93532329 | 623 | 394 | 522 | 647 | 469 | 0 | 2 | 0 | 0 | 0 | 1038 | 579 | 474 | 741 | 798 | 0 | 0 | 0 | 0 | 0 | 298 | 149 | 1 | 1 | 1 | 0.99 | 1 | 1 | 1 | 1 | 1 | 1 | 1 | 1 | -0.002 |
| SE | 94274 | ENSG00000048828 | FAM120A | chr9 | + | 93556381 | 93556591 | 93550576 | 93550691 | 93557826 | 93558010 | 648 | 327 | 461 | 461 | 760 | 2 | 0 | 0 | 0 | 0 | 551 | 468 | 352 | 414 | 489 | 0 | 0 | 2 | 0 | 0 | 298 | 149 | 1 | 1 | 0.994 | 1 | 1 | 1 | 1 | 1 | 1 | 0.989 | 1 | 1 | 0.001 |
| SE | 94276 | ENSG00000048828 | FAM120A | chr9 | + | 93561108 | 93561250 | 93558580 | 93558718 | 93562207 | 93562304 | 221 | 98 | 138 | 208 | 56 | 0 | 1 | 0 | 3 | 0 | 239 | 176 | 181 | 184 | 320 | 0 | 0 | 0 | 2 | 2 | 291 | 149 | 1 | 1 | 1 | 0.98 | 1 | 0.973 | 1 | 1 | 1 | 1 | 0.979 | 0.988 | -0.003 |
| SE | 94277 | ENSG00000048828 | FAM120A | chr9 | + | 93561111 | 93561250 | 93558580 | 93558718 | 93562207 | 93562304 | 171 | 89 | 113 | 164 | 54 | 0 | 1 | 0 | 3 | 0 | 209 | 159 | 146 | 145 | 243 | 0 | 0 | 0 | 2 | 2 | 288 | 149 | 1 | 1 | 1 | 0.979 | 1 | 0.966 | 1 | 1 | 1 | 1 | 0.974 | 0.984 | -0.003 |
| SE | 94279 | ENSG00000107104 | KANK1 | chr9 | + | 549106 | 549721 | 540507 | 540667 | 676889 | 677009 | 1 | 0 | 0 | 0 | 0 | 0 | 1 | 3 | 0 | 3 | 0 | 0 | 0 | 0 | 0 | 15 | 0 | 3 | 0 | 0 | 298 | 149 | 1 | 1 | 1 | 0 | 0 | NA | 0 | 0 | NA | 0 | NA | NA | 0.25 |
| SE | 94280 | ENSG00000107104 | KANK1 | chr9 | + | 730050 | 730248 | 710803 | 713464 | 731157 | 731266 | 14 | 5 | 12 | 3 | 7 | 0 | 0 | 2 | 0 | 0 | 0 | 13 | 20 | 4 | 33 | 0 | 0 | 0 | 0 | 0 | 298 | 149 | 1 | 1 | 1 | 1 | 0.75 | 1 | 1 | NA | 1 | 1 | 1 | 1 | -0.05 |
| SE | 94281 | ENSG00000107104 | KANK1 | chr9 | + | 731157 | 731266 | 710803 | 713464 | 734747 | 734835 | 0 | 0 | 4 | 0 | 6 | 0 | 0 | 0 | 0 | 0 | 0 | 0 | 0 | 0 | 2 | 0 | 0 | 0 | 2 | 0 | 258 | 149 | 1 | 1 | NA | NA | 1 | NA | 1 | NA | NA | NA | 0 | 1 | 0.5 |
| SE | 94283 | ENSG00000107104 | KANK1 | chr9 | + | 735711 | 735780 | 734747 | 734835 | 738284 | 738504 | 1 | 0 | 0 | 0 | 0 | 14 | 5 | 28 | 1 | 50 | 0 | 0 | 0 | 0 | 0 | 0 | 2 | 1 | 6 | 5 | 218 | 149 | 1 | 1 | 0.047 | 0 | 0 | 0 | 0 | NA | 0 | 0 | 0 | 0 | 0.009 |
| SE | 94284 | ENSG00000107104 | KANK1 | chr9 | + | 742204 | 742405 | 740791 | 740934 | 744490 | 744589 | 14 | 1 | 37 | 13 | 10 | 0 | 0 | 3 | 0 | 0 | 9 | 4 | 14 | 0 | 0 | 0 | 0 | 0 | 0 | 0 | 298 | 149 | 1 | 1 | 1 | 1 | 0.86 | 1 | 1 | 1 | 1 | 1 | NA | NA | -0.028 |
| SE | 94285 | ENSG00000147684 | NDUFB9 | chr8 | + | 124543086 | 124543279 | 124539100 | 124539287 | 124546999 | 124547113 | 327 | 166 | 265 | 255 | 183 | 2 | 0 | 0 | 0 | 0 | 363 | 351 | 226 | 364 | 483 | 0 | 0 | 0 | 0 | 0 | 298 | 149 | 1 | 1 | 0.988 | 1 | 1 | 1 | 1 | 1 | 1 | 1 | 1 | 1 | -0.002 |
| SE | 94286 | ENSG00000147684 | NDUFB9 | chr8 | + | 124543088 | 124543279 | 124539177 | 124539287 | 124546999 | 124547113 | 231 | 123 | 200 | 181 | 149 | 2 | 0 | 0 | 0 | 0 | 283 | 234 | 179 | 265 | 357 | 0 | 0 | 0 | 0 | 0 | 298 | 149 | 1 | 1 | 0.983 | 1 | 1 | 1 | 1 | 1 | 1 | 1 | 1 | 1 | -0.003 |
| SE | 94287 | ENSG00000147684 | NDUFB9 | chr8 | + | 124543119 | 124543279 | 124539143 | 124539287 | 124546999 | 124547113 | 231 | 123 | 201 | 183 | 148 | 2 | 0 | 0 | 0 | 0 | 283 | 234 | 183 | 267 | 357 | 0 | 0 | 0 | 0 | 0 | 298 | 149 | 1 | 1 | 0.983 | 1 | 1 | 1 | 1 | 1 | 1 | 1 | 1 | 1 | -0.003 |
| SE | 94290 | ENSG00000147684 | NDUFB9 | chr8 | + | 124550443 | 124550501 | 124549773 | 124549833 | 124552116 | 124552318 | 1 | 2 | 2 | 0 | 14 | 0 | 0 | 0 | 0 | 0 | 0 | 2 | 4 | 0 | 0 | 0 | 0 | 0 | 1 | 0 | 207 | 149 | 1 | 1 | 1 | 1 | 1 | NA | 1 | NA | 1 | 1 | 0 | NA | 0.333 |
| SE | 94292 | ENSG00000132740 | IGHMBP2 | chr11 | + | 68906068 | 68906238 | 68903862 | 68904038 | 68908144 | 68908337 | 38 | 12 | 23 | 19 | 9 | 0 | 0 | 0 | 0 | 0 | 45 | 77 | 38 | 43 | 12 | 0 | 0 | 0 | 2 | 0 | 298 | 149 | 1 | 1 | 1 | 1 | 1 | 1 | 1 | 1 | 1 | 1 | 0.915 | 1 | 0.017 |
| SE | 94293 | ENSG00000132740 | IGHMBP2 | chr11 | + | 68906068 | 68906242 | 68903912 | 68904038 | 68908144 | 68908337 | 8 | 1 | 6 | 5 | 6 | 0 | 0 | 0 | 0 | 0 | 4 | 25 | 18 | 4 | 0 | 0 | 0 | 0 | 2 | 0 | 298 | 149 | 1 | 1 | 1 | 1 | 1 | 1 | 1 | 1 | 1 | 1 | 0.5 | NA | 0.125 |
| SE | 94294 | ENSG00000132740 | IGHMBP2 | chr11 | + | 68906068 | 68906384 | 68903925 | 68904038 | 68908144 | 68908192 | 7 | 3 | 6 | 4 | 5 | 0 | 0 | 0 | 0 | 0 | 4 | 25 | 23 | 4 | 0 | 0 | 0 | 0 | 2 | 0 | 298 | 149 | 1 | 1 | 1 | 1 | 1 | 1 | 1 | 1 | 1 | 1 | 0.5 | NA | 0.125 |
| SE | 94297 | ENSG00000132740 | IGHMBP2 | chr11 | + | 68914184 | 68914350 | 68911439 | 68911603 | 68914822 | 68915023 | 0 | 1 | 0 | 1 | 2 | 32 | 9 | 47 | 17 | 44 | 0 | 0 | 0 | 0 | 0 | 29 | 51 | 22 | 59 | 79 | 298 | 149 | 1 | 1 | 0 | 0.053 | 0 | 0.029 | 0.022 | 0 | 0 | 0 | 0 | 0 | 0.021 |
| SE | 94298 | ENSG00000132740 | IGHMBP2 | chr11 | + | 68917160 | 68917301 | 68914822 | 68915023 | 68917735 | 68917883 | 0 | 0 | 0 | 0 | 0 | 20 | 26 | 18 | 18 | 14 | 0 | 0 | 0 | 0 | 4 | 62 | 22 | 23 | 14 | 20 | 290 | 149 | 1 | 1 | 0 | 0 | 0 | 0 | 0 | 0 | 0 | 0 | 0 | 0.093 | -0.019 |
| SE | 94307 | ENSG00000132740 | IGHMBP2 | chr11 | + | 68932926 | 68933123 | 68930322 | 68930434 | 68933298 | 68933481 | 0 | 0 | 0 | 2 | 0 | 4 | 6 | 3 | 4 | 0 | 0 | 0 | 0 | 0 | 0 | 0 | 0 | 1 | 0 | 6 | 298 | 149 | 1 | 1 | 0 | 0 | 0 | 0.2 | NA | NA | NA | 0 | NA | 0 | 0.05 |
| SE | 94321 | ENSG00000179630 | LACC1 | chr13 | + | 43882184 | 43882363 | 43880951 | 43881547 | 43883770 | 43883936 | 19 | 18 | 21 | 14 | 28 | 0 | 1 | 1 | 1 | 2 | 0 | 0 | 1 | 0 | 0 | 0 | 0 | 0 | 0 | 0 | 298 | 149 | 1 | 1 | 1 | 0.9 | 0.913 | 0.875 | 0.875 | NA | NA | 1 | NA | NA | -0.087 |
| SE | 94322 | ENSG00000179630 | LACC1 | chr13 | + | 43890113 | 43890274 | 43888756 | 43888982 | 43891448 | 43893931 | 16 | 25 | 3 | 11 | 34 | 4 | 3 | 1 | 0 | 2 | 0 | 2 | 0 | 0 | 0 | 0 | 0 | 0 | 0 | 0 | 298 | 149 | 1 | 1 | 0.667 | 0.806 | 0.6 | 1 | 0.895 | NA | 1 | NA | NA | NA | -0.206 |
| SE | 94329 | ENSG00000178445 | GLDC | chr9 | - | 6610191 | 6610356 | 6606591 | 6606669 | 6620183 | 6620319 | 4 | 4 | 49 | 17 | 6 | 0 | 0 | 0 | 0 | 0 | 0 | 8 | 24 | 26 | 0 | 0 | 0 | 0 | 3 | 0 | 298 | 149 | 1 | 1 | 1 | 1 | 1 | 1 | 1 | NA | 1 | 1 | 0.812 | NA | 0.063 |
| SE | 94330 | ENSG00000178445 | GLDC | chr9 | - | 6620183 | 6620319 | 6610191 | 6610356 | 6644613 | 6644692 | 9 | 7 | 58 | 19 | 7 | 0 | 0 | 1 | 0 | 0 | 0 | 5 | 28 | 11 | 0 | 0 | 0 | 0 | 0 | 0 | 285 | 149 | 1 | 1 | 1 | 1 | 0.968 | 1 | 1 | NA | 1 | 1 | 1 | NA | -0.006 |
| SE | 94331 | ENSG00000136936 | XPA | chr9 | - | 97682199 | 97682430 | 97674908 | 97675587 | 97684922 | 97685040 | 11 | 10 | 10 | 12 | 1 | 27 | 15 | 12 | 26 | 24 | 0 | 6 | 1 | 0 | 0 | 36 | 19 | 13 | 8 | 0 | 298 | 149 | 1 | 1 | 0.169 | 0.25 | 0.294 | 0.188 | 0.02 | 0 | 0.136 | 0.037 | 0 | NA | 0.141 |
| SE | 94332 | ENSG00000136936 | XPA | chr9 | - | 97684922 | 97685040 | 97674908 | 97675587 | 97693648 | 97693759 | 24 | 12 | 10 | 22 | 18 | 0 | 0 | 0 | 0 | 0 | 25 | 12 | 9 | 8 | 0 | 0 | 7 | 0 | 0 | 0 | 267 | 149 | 1 | 1 | 1 | 1 | 1 | 1 | 1 | 1 | 0.489 | 1 | 1 | NA | 0.128 |
| SE | 94333 | ENSG00000136936 | XPA | chr9 | - | 97687095 | 97687261 | 97684922 | 97685040 | 97693648 | 97693759 | 261 | 236 | 267 | 265 | 594 | 1 | 0 | 0 | 0 | 0 | 180 | 254 | 242 | 149 | 115 | 0 | 2 | 0 | 0 | 0 | 298 | 149 | 1 | 1 | 0.992 | 1 | 1 | 1 | 1 | 1 | 0.984 | 1 | 1 | 1 | 0.002 |
| SE | 94339 | ENSG00000165802 | NSMF | chr9 | - | 137455238 | 137455307 | 137454390 | 137454443 | 137457406 | 137457901 | 39 | 22 | 20 | 29 | 32 | 0 | 0 | 0 | 0 | 0 | 13 | 33 | 11 | 5 | 69 | 0 | 0 | 0 | 1 | 0 | 218 | 149 | 1 | 1 | 1 | 1 | 1 | 1 | 1 | 1 | 1 | 1 | 0.774 | 1 | 0.045 |
| SE | 94341 | ENSG00000165802 | NSMF | chr9 | - | 137455628 | 137455634 | 137455238 | 137455307 | 137456410 | 137456486 | 4 | 0 | 0 | 0 | 0 | 18 | 18 | 17 | 25 | 28 | 2 | 0 | 0 | 0 | 0 | 26 | 22 | 5 | 32 | 11 | 155 | 149 | 1 | 1 | 0.176 | 0 | 0 | 0 | 0 | 0.069 | 0 | 0 | 0 | 0 | 0.021 |
| SE | 94342 | ENSG00000165802 | NSMF | chr9 | - | 137456410 | 137456486 | 137454390 | 137454443 | 137457406 | 137457901 | 18 | 7 | 13 | 16 | 3 | 0 | 0 | 0 | 0 | 0 | 20 | 20 | 17 | 5 | 33 | 0 | 0 | 0 | 1 | 0 | 225 | 149 | 1 | 1 | 1 | 1 | 1 | 1 | 1 | 1 | 1 | 1 | 0.768 | 1 | 0.046 |
| SE | 94343 | ENSG00000165802 | NSMF | chr9 | - | 137456410 | 137456486 | 137455238 | 137455307 | 137457406 | 137457901 | 36 | 20 | 30 | 38 | 31 | 0 | 0 | 0 | 0 | 0 | 41 | 41 | 19 | 37 | 41 | 0 | 2 | 0 | 0 | 0 | 225 | 149 | 1 | 1 | 1 | 1 | 1 | 1 | 1 | 1 | 0.931 | 1 | 1 | 1 | 0.014 |
| SE | 94344 | ENSG00000173611 | SCAI | chr9 | - | 124974174 | 124974276 | 124971670 | 124971844 | 124976113 | 124976186 | 1 | 0 | 0 | 0 | 0 | 25 | 10 | 12 | 7 | 20 | 0 | 0 | 0 | 0 | 0 | 0 | 0 | 18 | 3 | 0 | 251 | 149 | 1 | 1 | 0.023 | 0 | 0 | 0 | 0 | NA | NA | 0 | 0 | NA | 0.005 |
| SE | 94346 | ENSG00000173611 | SCAI | chr9 | - | 124975246 | 124975389 | 124971670 | 124971844 | 124976113 | 124976186 | 8 | 5 | 2 | 5 | 1 | 25 | 10 | 12 | 7 | 20 | 0 | 0 | 0 | 2 | 2 | 0 | 0 | 18 | 3 | 0 | 292 | 149 | 1 | 1 | 0.14 | 0.203 | 0.078 | 0.267 | 0.025 | NA | NA | 0 | 0.254 | 1 | -0.275 |
| SE | 94347 | ENSG00000173611 | SCAI | chr9 | - | 125065989 | 125066058 | 125055875 | 125056007 | 125142632 | 125142677 | 14 | 18 | 16 | 11 | 33 | 0 | 0 | 7 | 2 | 2 | 0 | 3 | 10 | 13 | 0 | 0 | 3 | 0 | 0 | 0 | 218 | 149 | 1 | 1 | 1 | 1 | 0.61 | 0.79 | 0.919 | NA | 0.406 | 1 | 1 | NA | 0.062 |
| SE | 94358 | ENSG00000021776 | AQR | chr15 | - | 34875934 | 34876006 | 34873827 | 34874864 | 34882501 | 34882639 | 176 | 101 | 119 | 155 | 307 | 0 | 1 | 0 | 0 | 0 | 19 | 79 | 70 | 41 | 0 | 0 | 0 | 0 | 0 | 0 | 221 | 149 | 1 | 1 | 1 | 0.986 | 1 | 1 | 1 | 1 | 1 | 1 | 1 | NA | -0.003 |
| SE | 94359 | ENSG00000021776 | AQR | chr15 | - | 34897558 | 34897705 | 34896896 | 34896966 | 34900621 | 34900863 | 269 | 203 | 241 | 284 | 352 | 0 | 0 | 1 | 5 | 0 | 14 | 159 | 181 | 100 | 39 | 0 | 0 | 2 | 0 | 0 | 296 | 149 | 1 | 1 | 1 | 1 | 0.992 | 0.966 | 1 | 1 | 1 | 0.979 | 1 | 1 | -0.004 |
| SE | 94360 | ENSG00000021776 | AQR | chr15 | - | 34900621 | 34900863 | 34897558 | 34897705 | 34904335 | 34904505 | 421 | 312 | 340 | 367 | 445 | 0 | 1 | 0 | 0 | 0 | 11 | 260 | 205 | 135 | 49 | 0 | 0 | 0 | 0 | 0 | 298 | 149 | 1 | 1 | 1 | 0.994 | 1 | 1 | 1 | 1 | 1 | 1 | 1 | 1 | -0.001 |
| SE | 94361 | ENSG00000021776 | AQR | chr15 | - | 34904335 | 34904505 | 34900621 | 34900863 | 34906544 | 34906712 | 492 | 445 | 349 | 468 | 500 | 0 | 0 | 0 | 0 | 0 | 90 | 371 | 280 | 246 | 20 | 0 | 0 | 2 | 0 | 0 | 298 | 149 | 1 | 1 | 1 | 1 | 1 | 1 | 1 | 1 | 1 | 0.986 | 1 | 1 | 0.003 |
| SE | 94362 | ENSG00000021776 | AQR | chr15 | - | 34915037 | 34915179 | 34910134 | 34910313 | 34918257 | 34918378 | 213 | 113 | 206 | 200 | 306 | 4 | 0 | 1 | 0 | 0 | 44 | 150 | 115 | 70 | 42 | 0 | 3 | 0 | 0 | 0 | 291 | 149 | 1 | 1 | 0.965 | 1 | 0.991 | 1 | 1 | 1 | 0.962 | 1 | 1 | 1 | -0.001 |
| SE | 94364 | ENSG00000021776 | AQR | chr15 | - | 34934570 | 34934635 | 34932317 | 34932434 | 34938736 | 34938813 | 175 | 111 | 187 | 137 | 359 | 0 | 7 | 0 | 0 | 0 | 15 | 111 | 93 | 90 | 8 | 0 | 0 | 0 | 0 | 0 | 214 | 149 | 1 | 1 | 1 | 0.917 | 1 | 1 | 1 | 1 | 1 | 1 | 1 | 1 | -0.017 |
| SE | 94365 | ENSG00000021776 | AQR | chr15 | - | 34938736 | 34938813 | 34934570 | 34934635 | 34940898 | 34940999 | 299 | 235 | 288 | 264 | 625 | 2 | 1 | 1 | 1 | 0 | 37 | 183 | 180 | 154 | 14 | 0 | 0 | 0 | 0 | 0 | 226 | 149 | 1 | 1 | 0.99 | 0.994 | 0.995 | 0.994 | 1 | 1 | 1 | 1 | 1 | 1 | -0.005 |
| SE | 94366 | ENSG00000021776 | AQR | chr15 | - | 34944287 | 34944428 | 34942011 | 34942080 | 34948263 | 34948384 | 358 | 343 | 323 | 356 | 669 | 1 | 0 | 0 | 0 | 0 | 110 | 283 | 255 | 189 | 52 | 0 | 0 | 0 | 0 | 0 | 290 | 149 | 1 | 1 | 0.995 | 1 | 1 | 1 | 1 | 1 | 1 | 1 | 1 | 1 | -0.001 |
| SE | 94369 | ENSG00000021776 | AQR | chr15 | - | 34964233 | 34964290 | 34960773 | 34960814 | 34969538 | 34969733 | 154 | 145 | 133 | 155 | 177 | 0 | 3 | 0 | 0 | 0 | 98 | 131 | 92 | 75 | 27 | 0 | 0 | 0 | 0 | 0 | 206 | 149 | 1 | 1 | 1 | 0.972 | 1 | 1 | 1 | 1 | 1 | 1 | 1 | 1 | -0.006 |
| SE | 94370 | ENSG00000149308 | NPAT | chr11 | - | 108169927 | 108170043 | 108169743 | 108169852 | 108172198 | 108173851 | 216 | 182 | 230 | 219 | 411 | 1 | 0 | 0 | 1 | 1 | 8 | 73 | 53 | 13 | 6 | 0 | 4 | 0 | 0 | 0 | 265 | 149 | 1 | 1 | 0.992 | 1 | 1 | 0.992 | 0.996 | 1 | 0.911 | 1 | 1 | 1 | 0.014 |
| SE | 94371 | ENSG00000149308 | NPAT | chr11 | - | 108172198 | 108173851 | 108169927 | 108170043 | 108176245 | 108176374 | 304 | 253 | 285 | 290 | 534 | 1 | 1 | 0 | 0 | 0 | 0 | 135 | 80 | 10 | 0 | 0 | 0 | 0 | 0 | 0 | 298 | 149 | 1 | 1 | 0.993 | 0.992 | 1 | 1 | 1 | NA | 1 | 1 | 1 | NA | -0.003 |
| SE | 94372 | ENSG00000149308 | NPAT | chr11 | - | 108176245 | 108176374 | 108171131 | 108173851 | 108176993 | 108177090 | 338 | 263 | 306 | 316 | 570 | 1 | 1 | 0 | 0 | 0 | 12 | 150 | 71 | 14 | 8 | 0 | 2 | 1 | 0 | 0 | 278 | 149 | 1 | 1 | 0.995 | 0.993 | 1 | 1 | 1 | 1 | 0.976 | 0.974 | 1 | 1 | 0.008 |
| SE | 94373 | ENSG00000149308 | NPAT | chr11 | - | 108176993 | 108177090 | 108171131 | 108173851 | 108185231 | 108185319 | 78 | 76 | 83 | 85 | 183 | 0 | 0 | 1 | 1 | 0 | 0 | 16 | 19 | 4 | 0 | 0 | 0 | 1 | 0 | 0 | 246 | 149 | 1 | 1 | 1 | 1 | 0.98 | 0.981 | 1 | NA | 1 | 0.92 | 1 | NA | 0.019 |
| SE | 94375 | ENSG00000149308 | NPAT | chr11 | - | 108197301 | 108197420 | 108193956 | 108194017 | 108222499 | 108222636 | 384 | 273 | 396 | 450 | 365 | 0 | 0 | 0 | 0 | 0 | 23 | 173 | 95 | 31 | 16 | 0 | 0 | 1 | 0 | 0 | 268 | 149 | 1 | 1 | 1 | 1 | 1 | 1 | 1 | 1 | 1 | 0.981 | 1 | 1 | 0.004 |
| SE | 94380 | ENSG00000196502 | SULT1A1 | chr16 | - | 28608477 | 28608603 | 28608290 | 28608388 | 28608707 | 28608859 | 135 | 39 | 127 | 106 | 111 | 0 | 0 | 1 | 0 | 0 | 165 | 289 | 76 | 191 | 340 | 0 | 6 | 0 | 0 | 0 | 275 | 149 | 1 | 1 | 1 | 1 | 0.986 | 1 | 1 | 1 | 0.963 | 1 | 1 | 1 | 0.005 |
| SE | 94381 | ENSG00000196502 | SULT1A1 | chr16 | - | 28609058 | 28610191 | 28608707 | 28608859 | 28620062 | 28620133 | 1 | 7 | 1 | 6 | 10 | 0 | 0 | 1 | 0 | 0 | 9 | 1 | 3 | 0 | 22 | 0 | 0 | 4 | 0 | 0 | 298 | 149 | 1 | 1 | 1 | 1 | 0.333 | 1 | 1 | 1 | 1 | 0.273 | NA | 1 | 0.048 |
| SE | 94382 | ENSG00000196502 | SULT1A1 | chr16 | - | 28609362 | 28609435 | 28608707 | 28608859 | 28609930 | 28610191 | 3 | 0 | 0 | 1 | 0 | 78 | 37 | 98 | 59 | 96 | 0 | 0 | 2 | 0 | 0 | 118 | 294 | 51 | 176 | 258 | 222 | 149 | 1 | 1 | 0.025 | 0 | 0 | 0.011 | 0 | 0 | 0 | 0.026 | 0 | 0 | 0.002 |
| SE | 94383 | ENSG00000196502 | SULT1A1 | chr16 | - | 28609930 | 28610191 | 28608707 | 28608859 | 28620062 | 28620133 | 79 | 42 | 88 | 54 | 98 | 0 | 0 | 1 | 0 | 0 | 127 | 288 | 49 | 174 | 270 | 0 | 0 | 4 | 0 | 0 | 298 | 149 | 1 | 1 | 1 | 1 | 0.978 | 1 | 1 | 1 | 1 | 0.86 | 1 | 1 | 0.024 |
| SE | 94384 | ENSG00000106868 | SUSD1 | chr9 | - | 112042100 | 112042165 | 112041866 | 112041960 | 112052154 | 112052349 | 9 | 6 | 13 | 4 | 5 | 0 | 5 | 0 | 1 | 1 | 0 | 10 | 9 | 36 | 35 | 0 | 0 | 2 | 10 | 0 | 214 | 149 | 1 | 1 | 1 | 0.455 | 1 | 0.736 | 0.777 | NA | 1 | 0.758 | 0.715 | 1 | -0.075 |
| SE | 94388 | ENSG00000106868 | SUSD1 | chr9 | - | 112052154 | 112052349 | 112042100 | 112042165 | 112058427 | 112058471 | 14 | 13 | 16 | 5 | 3 | 0 | 1 | 0 | 1 | 0 | 12 | 22 | 5 | 44 | 14 | 0 | 0 | 2 | 0 | 0 | 298 | 149 | 1 | 1 | 1 | 0.867 | 1 | 0.714 | 1 | 1 | 1 | 0.556 | 1 | 1 | 0.005 |
| SE | 94390 | ENSG00000106868 | SUSD1 | chr9 | - | 112052398 | 112052438 | 112042100 | 112042165 | 112058427 | 112058686 | 18 | 8 | 16 | 23 | 22 | 0 | 1 | 0 | 1 | 0 | 18 | 19 | 1 | 14 | 22 | 0 | 0 | 2 | 0 | 0 | 189 | 149 | 1 | 1 | 1 | 0.863 | 1 | 0.948 | 1 | 1 | 1 | 0.283 | 1 | 1 | 0.106 |
| SE | 94391 | ENSG00000106868 | SUSD1 | chr9 | - | 112058427 | 112058686 | 112052398 | 112052438 | 112078537 | 112078724 | 62 | 44 | 58 | 61 | 91 | 0 | 0 | 0 | 2 | 0 | 78 | 72 | 22 | 69 | 197 | 0 | 0 | 0 | 0 | 0 | 298 | 149 | 1 | 1 | 1 | 1 | 1 | 0.938 | 1 | 1 | 1 | 1 | 1 | 1 | -0.012 |
| SE | 94392 | ENSG00000106868 | SUSD1 | chr9 | - | 112062936 | 112063033 | 112058427 | 112058686 | 112078537 | 112078724 | 212 | 149 | 120 | 219 | 197 | 0 | 0 | 0 | 0 | 0 | 467 | 311 | 166 | 255 | 785 | 0 | 0 | 0 | 1 | 0 | 246 | 149 | 1 | 1 | 1 | 1 | 1 | 1 | 1 | 1 | 1 | 1 | 0.994 | 1 | 0.001 |
| SE | 94394 | ENSG00000106868 | SUSD1 | chr9 | - | 112078537 | 112078724 | 112062936 | 112063033 | 112098469 | 112098662 | 148 | 117 | 96 | 169 | 157 | 1 | 0 | 0 | 0 | 0 | 329 | 256 | 123 | 208 | 660 | 0 | 0 | 2 | 3 | 0 | 298 | 149 | 1 | 1 | 0.987 | 1 | 1 | 1 | 1 | 1 | 1 | 0.969 | 0.972 | 1 | 0.009 |
| SE | 94395 | ENSG00000106868 | SUSD1 | chr9 | - | 112080073 | 112080165 | 112062936 | 112063033 | 112098469 | 112098662 | 136 | 118 | 59 | 126 | 132 | 1 | 0 | 0 | 0 | 0 | 258 | 217 | 111 | 190 | 547 | 0 | 0 | 2 | 3 | 0 | 241 | 149 | 1 | 1 | 0.988 | 1 | 1 | 1 | 1 | 1 | 1 | 0.972 | 0.975 | 1 | 0.008 |
| SE | 94396 | ENSG00000106868 | SUSD1 | chr9 | - | 112080073 | 112080165 | 112062936 | 112063033 | 112111653 | 112111840 | 4 | 10 | 2 | 9 | 1 | 0 | 0 | 0 | 0 | 2 | 10 | 3 | 0 | 10 | 4 | 0 | 0 | 0 | 0 | 0 | 241 | 149 | 1 | 1 | 1 | 1 | 1 | 1 | 0.236 | 1 | 1 | NA | 1 | 1 | -0.153 |
| SE | 94397 | ENSG00000106868 | SUSD1 | chr9 | - | 112080073 | 112080165 | 112078537 | 112078724 | 112098469 | 112098662 | 315 | 224 | 157 | 285 | 281 | 0 | 0 | 0 | 1 | 3 | 693 | 393 | 217 | 376 | 1114 | 0 | 0 | 0 | 3 | 0 | 241 | 149 | 1 | 1 | 1 | 1 | 1 | 0.994 | 0.983 | 1 | 1 | 1 | 0.987 | 1 | -0.002 |
| SE | 94398 | ENSG00000106868 | SUSD1 | chr9 | - | 112098469 | 112098662 | 112062936 | 112063033 | 112111653 | 112111840 | 1 | 0 | 0 | 2 | 0 | 0 | 0 | 0 | 0 | 2 | 0 | 0 | 7 | 3 | 0 | 0 | 0 | 0 | 0 | 0 | 298 | 149 | 1 | 1 | 1 | NA | NA | 1 | 0 | NA | NA | 1 | 1 | NA | -0.333 |
| SE | 94399 | ENSG00000106868 | SUSD1 | chr9 | - | 112098469 | 112098662 | 112080073 | 112080165 | 112102175 | 112102285 | 416 | 359 | 216 | 434 | 516 | 1 | 1 | 0 | 0 | 0 | 974 | 712 | 324 | 543 | 1701 | 0 | 4 | 1 | 0 | 0 | 298 | 149 | 1 | 1 | 0.995 | 0.994 | 1 | 1 | 1 | 1 | 0.989 | 0.994 | 1 | 1 | 0.001 |
| SE | 94402 | ENSG00000106868 | SUSD1 | chr9 | - | 112102175 | 112102285 | 112098469 | 112098662 | 112111653 | 112111840 | 301 | 320 | 181 | 371 | 400 | 0 | 0 | 0 | 2 | 0 | 827 | 641 | 288 | 444 | 1384 | 0 | 0 | 5 | 0 | 0 | 259 | 149 | 1 | 1 | 1 | 1 | 1 | 0.991 | 1 | 1 | 1 | 0.971 | 1 | 1 | 0.004 |
| SE | 94404 | ENSG00000106868 | SUSD1 | chr9 | - | 112142319 | 112142499 | 112124256 | 112124436 | 112143470 | 112143623 | 222 | 185 | 164 | 232 | 210 | 1 | 0 | 0 | 0 | 0 | 627 | 389 | 244 | 342 | 887 | 0 | 0 | 3 | 0 | 0 | 298 | 149 | 1 | 1 | 0.991 | 1 | 1 | 1 | 1 | 1 | 1 | 0.976 | 1 | 1 | 0.003 |
| SE | 94405 | ENSG00000106868 | SUSD1 | chr9 | - | 112149243 | 112149399 | 112143470 | 112143623 | 112157499 | 112157613 | 264 | 167 | 138 | 230 | 265 | 0 | 0 | 0 | 1 | 0 | 524 | 319 | 137 | 340 | 936 | 0 | 7 | 0 | 2 | 1 | 298 | 149 | 1 | 1 | 1 | 1 | 1 | 0.991 | 1 | 1 | 0.958 | 1 | 0.988 | 0.998 | 0.009 |
| SE | 94406 | ENSG00000106868 | SUSD1 | chr9 | - | 112157499 | 112157613 | 112149243 | 112149399 | 112175132 | 112175185 | 194 | 119 | 82 | 143 | 133 | 2 | 0 | 0 | 0 | 0 | 326 | 223 | 110 | 241 | 654 | 0 | 0 | 0 | 0 | 0 | 263 | 149 | 1 | 1 | 0.982 | 1 | 1 | 1 | 1 | 1 | 1 | 1 | 1 | 1 | -0.004 |
| SE | 94410 | ENSG00000107331 | ABCA2 | chr9 | - | 137023837 | 137023840 | 137022940 | 137023052 | 137024142 | 137024236 | 1 | 6 | 1 | 2 | 0 | 1 | 0 | 0 | 8 | 0 | 0 | 1 | 0 | 8 | 0 | 0 | 1 | 1 | 0 | 0 | 152 | 149 | 1 | 1 | 0.495 | 1 | 1 | 0.197 | NA | NA | 0.495 | 0 | 1 | NA | 0.175 |
| SE | 94412 | ENSG00000169519 | METTL15 | chr11 | + | 28122141 | 28122180 | 28113317 | 28113604 | 28123861 | 28123922 | 14 | 7 | 1 | 2 | 2 | 1 | 0 | 0 | 0 | 0 | 5 | 6 | 0 | 0 | 0 | 0 | 0 | 0 | 1 | 0 | 188 | 149 | 1 | 1 | 0.917 | 1 | 1 | 1 | 1 | 1 | 1 | NA | 0 | NA | 0.317 |
| SE | 94413 | ENSG00000169519 | METTL15 | chr11 | + | 28122141 | 28122180 | 28113317 | 28113604 | 28211061 | 28211198 | 7 | 9 | 1 | 2 | 0 | 15 | 21 | 31 | 25 | 24 | 0 | 6 | 0 | 0 | 0 | 0 | 8 | 16 | 2 | 12 | 188 | 149 | 1 | 1 | 0.27 | 0.254 | 0.025 | 0.06 | 0 | NA | 0.373 | 0 | 0 | 0 | 0.029 |
| SE | 94414 | ENSG00000169519 | METTL15 | chr11 | + | 28123861 | 28123922 | 28113317 | 28113604 | 28125509 | 28125587 | 2 | 5 | 5 | 0 | 1 | 1 | 0 | 0 | 0 | 0 | 5 | 0 | 0 | 1 | 0 | 0 | 0 | 0 | 0 | 0 | 210 | 149 | 1 | 1 | 0.587 | 1 | 1 | NA | 1 | 1 | NA | NA | 1 | NA | -0.103 |
| SE | 94418 | ENSG00000169519 | METTL15 | chr11 | + | 28290205 | 28290397 | 28211061 | 28211198 | 28296752 | 28296931 | 19 | 16 | 34 | 28 | 67 | 0 | 2 | 0 | 0 | 0 | 39 | 18 | 38 | 7 | 12 | 0 | 0 | 2 | 0 | 0 | 298 | 149 | 1 | 1 | 1 | 0.8 | 1 | 1 | 1 | 1 | 1 | 0.905 | 1 | 1 | -0.021 |
| SE | 94419 | ENSG00000169519 | METTL15 | chr11 | + | 28290205 | 28290397 | 28211061 | 28211198 | 28296796 | 28296931 | 22 | 12 | 36 | 21 | 61 | 0 | 1 | 1 | 2 | 0 | 31 | 20 | 37 | 6 | 9 | 0 | 0 | 0 | 0 | 0 | 298 | 149 | 1 | 1 | 1 | 0.857 | 0.947 | 0.84 | 1 | 1 | 1 | 1 | 1 | 1 | -0.071 |
| SE | 94420 | ENSG00000169519 | METTL15 | chr11 | + | 28296752 | 28296931 | 28290205 | 28290397 | 28328094 | 28328165 | 6 | 6 | 8 | 16 | 25 | 0 | 1 | 0 | 0 | 0 | 12 | 6 | 10 | 1 | 7 | 0 | 0 | 0 | 0 | 0 | 298 | 149 | 1 | 1 | 1 | 0.75 | 1 | 1 | 1 | 1 | 1 | 1 | 1 | 1 | -0.05 |
| SE | 94423 | ENSG00000169519 | METTL15 | chr11 | + | 28296796 | 28296931 | 28290205 | 28290397 | 28330395 | 28333507 | 17 | 4 | 13 | 8 | 20 | 0 | 2 | 3 | 0 | 3 | 4 | 13 | 15 | 0 | 4 | 0 | 0 | 2 | 0 | 0 | 284 | 149 | 1 | 1 | 1 | 0.512 | 0.695 | 1 | 0.778 | 1 | 1 | 0.797 | NA | 1 | -0.152 |
| SE | 94424 | ENSG00000169519 | METTL15 | chr11 | + | 28328094 | 28328165 | 28290205 | 28290397 | 28330395 | 28333507 | 0 | 2 | 0 | 2 | 0 | 0 | 2 | 3 | 0 | 3 | 0 | 0 | 1 | 0 | 0 | 0 | 0 | 2 | 0 | 0 | 220 | 149 | 1 | 1 | NA | 0.404 | 0 | 1 | 0 | NA | NA | 0.253 | NA | NA | 0.098 |
| SE | 94425 | ENSG00000169519 | METTL15 | chr11 | + | 28328094 | 28328165 | 28296752 | 28296931 | 28330395 | 28333507 | 0 | 1 | 1 | 4 | 0 | 8 | 3 | 4 | 2 | 1 | 0 | 0 | 1 | 0 | 0 | 0 | 5 | 6 | 0 | 0 | 220 | 149 | 1 | 1 | 0 | 0.184 | 0.145 | 0.575 | 0 | NA | 0 | 0.101 | NA | NA | 0.13 |
| SE | 94427 | ENSG00000169519 | METTL15 | chr11 | + | 28330395 | 28330760 | 28296752 | 28296931 | 28352089 | 28352158 | 8 | 3 | 4 | 2 | 1 | 2 | 1 | 0 | 0 | 0 | 0 | 6 | 6 | 0 | 0 | 0 | 4 | 2 | 0 | 10 | 298 | 149 | 1 | 1 | 0.667 | 0.6 | 1 | 1 | 1 | NA | 0.429 | 0.6 | NA | 0 | 0.51 |
| SE | 94431 | ENSG00000204147 | ASAH2B | chr10 | + | 50745127 | 50745274 | 50742914 | 50743010 | 50749342 | 50749402 | 0 | 0 | 4 | 0 | 0 | 0 | 0 | 2 | 3 | 0 | 0 | 0 | 0 | 0 | 0 | 0 | 0 | 1 | 0 | 0 | 296 | 149 | 1 | 1 | NA | NA | 0.502 | 0 | NA | NA | NA | 0 | NA | NA | 0.251 |
| SE | 94432 | ENSG00000204147 | ASAH2B | chr10 | + | 50745201 | 50745274 | 50742914 | 50743010 | 50749342 | 50749402 | 1 | 2 | 3 | 0 | 0 | 0 | 0 | 2 | 3 | 0 | 0 | 0 | 0 | 0 | 0 | 0 | 0 | 1 | 0 | 0 | 222 | 149 | 1 | 1 | 1 | 1 | 0.502 | 0 | NA | NA | NA | 0 | NA | NA | 0.625 |
| SE | 94433 | ENSG00000176248 | ANAPC2 | chr9 | - | 137186223 | 137186356 | 137184912 | 137185087 | 137187480 | 137188103 | 79 | 23 | 32 | 65 | 45 | 1 | 0 | 0 | 0 | 0 | 20 | 43 | 18 | 9 | 34 | 0 | 0 | 0 | 0 | 0 | 282 | 149 | 1 | 1 | 0.977 | 1 | 1 | 1 | 1 | 1 | 1 | 1 | 1 | 1 | -0.005 |
| SE | 94444 | ENSG00000184384 | MAML2 | chr11 | - | 95991519 | 95991723 | 95985530 | 95985642 | 96091891 | 96093517 | 183 | 151 | 280 | 294 | 365 | 0 | 0 | 2 | 0 | 0 | 62 | 189 | 183 | 82 | 17 | 0 | 0 | 0 | 0 | 0 | 298 | 149 | 1 | 1 | 1 | 1 | 0.986 | 1 | 1 | 1 | 1 | 1 | 1 | 1 | -0.003 |
| SE | 94445 | ENSG00000099958 | DERL3 | chr22 | - | 23838351 | 23838445 | 23837658 | 23837854 | 23838563 | 23838637 | 1 | 2 | 28 | 13 | 2 | 0 | 0 | 1 | 0 | 0 | 0 | 19 | 11 | 12 | 2 | 0 | 0 | 0 | 0 | 0 | 243 | 149 | 1 | 1 | 1 | 1 | 0.945 | 1 | 1 | NA | 1 | 1 | 1 | 1 | -0.011 |
| SE | 94446 | ENSG00000122490 | SLC66A2 | chr18 | - | 79919183 | 79919400 | 79902419 | 79904183 | 79930227 | 79930311 | 12 | 1 | 11 | 8 | 1 | 1 | 0 | 0 | 0 | 0 | 0 | 15 | 0 | 1 | 14 | 0 | 0 | 0 | 0 | 0 | 298 | 149 | 1 | 1 | 0.857 | 1 | 1 | 1 | 1 | NA | 1 | NA | 1 | 1 | -0.029 |
| SE | 94447 | ENSG00000122490 | SLC66A2 | chr18 | - | 79919183 | 79919400 | 79902419 | 79904183 | 79933968 | 79934022 | 22 | 1 | 14 | 10 | 0 | 1 | 0 | 0 | 4 | 0 | 8 | 21 | 1 | 8 | 18 | 0 | 0 | 13 | 5 | 3 | 298 | 149 | 1 | 1 | 0.917 | 1 | 1 | 0.556 | NA | 1 | 1 | 0.037 | 0.444 | 0.75 | 0.222 |
| SE | 94448 | ENSG00000122490 | SLC66A2 | chr18 | - | 79919183 | 79919400 | 79902419 | 79904183 | 79943328 | 79943462 | 19 | 5 | 11 | 12 | 19 | 0 | 4 | 3 | 6 | 8 | 0 | 28 | 9 | 17 | 30 | 0 | 17 | 1 | 0 | 2 | 298 | 149 | 1 | 1 | 1 | 0.385 | 0.647 | 0.5 | 0.543 | NA | 0.452 | 0.818 | 1 | 0.882 | -0.173 |
| SE | 94449 | ENSG00000122490 | SLC66A2 | chr18 | - | 79930227 | 79930311 | 79903577 | 79904183 | 79933968 | 79934022 | 2 | 0 | 0 | 3 | 1 | 1 | 0 | 0 | 4 | 0 | 0 | 0 | 0 | 0 | 0 | 0 | 0 | 13 | 5 | 3 | 233 | 149 | 1 | 1 | 0.561 | NA | NA | 0.324 | 1 | NA | NA | 0 | 0 | 0 | 0.628 |

| | | | | | | | | | | | | | | | | | | | | | | | | | | | | | | | | | | | | | | | | | | | | | | |
|---|---|---|---|---|---|---|---|---|---|---|---|---|---|---|---|---|---|---|---|---|---|---|---|---|---|---|---|---|---|---|---|---|---|---|---|---|---|---|---|---|---|---|---|---|---|---|
| SE | 94450 | ENSG00000122490 | SLC66A2 | chr18 | - | 79930227 | 79930311 | 79903577 | 79904183 | 79943328 | 79943462 | 0 | 0 | 0 | 0 | 0 | 0 | 4 | 3 | 6 | 8 | 0 | 0 | 2 | 0 | 0 | 0 | 17 | 1 | 0 | 2 | 233 | 149 | 1 | 1 | NA | 0 | 0 | 0 | 0 | NA | 0 | 0.561 | NA | 0 | -0.187 |
| SE | 94451 | ENSG00000122490 | SLC66A2 | chr18 | - | 79930227 | 79930311 | 79919183 | 79919400 | 79933968 | 79934022 | 3 | 0 | 3 | 7 | 2 | 12 | 0 | 7 | 6 | 0 | 0 | 7 | 0 | 1 | 10 | 8 | 13 | 1 | 8 | 14 | 233 | 149 | 1 | 1 | 0.138 | NA | 0.215 | 0.427 | 1 | 0 | 0.256 | 0 | 0.074 | 0.314 | 0.316 |
| SE | 94452 | ENSG00000122490 | SLC66A2 | chr18 | - | 79930227 | 79930311 | 79919183 | 79919400 | 79943328 | 79943462 | 1 | 0 | 2 | 4 | 1 | 9 | 4 | 4 | 8 | 19 | 0 | 7 | 2 | 0 | 10 | 0 | 20 | 9 | 17 | 26 | 233 | 149 | 1 | 1 | 0.066 | 0 | 0.242 | 0.242 | 0.033 | NA | 0.183 | 0.124 | 0 | 0.197 | -0.009 |
| SE | 94455 | ENSG00000122490 | SLC66A2 | chr18 | - | 79933968 | 79934022 | 79930227 | 79930311 | 79943328 | 79943462 | 4 | 5 | 4 | 5 | 2 | 0 | 0 | 1 | 0 | 0 | 3 | 5 | 4 | 9 | 19 | 0 | 0 | 2 | 0 | 0 | 203 | 149 | 1 | 1 | 1 | 1 | 0.746 | 1 | 1 | 1 | 1 | 0.595 | 1 | 1 | 0.03 |
| SE | 94456 | ENSG00000122490 | SLC66A2 | chr18 | - | 79947266 | 79947341 | 79943328 | 79943462 | 79950723 | 79950866 | 0 | 0 | 0 | 1 | 0 | 16 | 8 | 11 | 13 | 19 | 0 | 0 | 0 | 0 | 0 | 12 | 35 | 9 | 35 | 32 | 224 | 149 | 1 | 1 | 0 | 0 | 0 | 0.049 | 0 | 0 | 0 | 0 | 0 | 0 | 0.01 |
| SE | 94460 | ENSG00000175611 | LINC00476 | chr9 | - | 95850226 | 95850292 | 95813169 | 95813922 | 95874979 | 95875587 | 23 | 0 | 14 | 10 | 23 | 0 | 3 | 0 | 0 | 0 | 0 | 5 | 8 | 0 | 18 | 0 | 0 | 0 | 0 | 0 | 215 | 149 | 1 | 1 | 1 | 0 | 1 | 1 | 1 | NA | 1 | 1 | NA | 1 | -0.2 |
| SE | 94463 | ENSG00000175611 | LINC00476 | chr9 | - | 95851823 | 95852029 | 95850226 | 95850292 | 95875864 | 95876049 | 1 | 0 | 0 | 0 | 1 | 15 | 0 | 10 | 3 | 3 | 0 | 0 | 0 | 0 | 0 | 0 | 7 | 7 | 0 | 20 | 298 | 149 | 1 | 1 | 0.032 | NA | 0 | 0 | 0.143 | NA | 0 | 0 | NA | 0 | 0.044 |
| SE | 94464 | ENSG00000175611 | LINC00476 | chr9 | - | 95874508 | 95874548 | 95870282 | 95870441 | 95874979 | 95875070 | 4 | 1 | 3 | 0 | 5 | 1 | 2 | 6 | 2 | 18 | 6 | 8 | 1 | 0 | 0 | 6 | 0 | 0 | 0 | 0 | 189 | 149 | 1 | 1 | 0.759 | 0.283 | 0.283 | 0 | 0.18 | 0.441 | 1 | 1 | NA | NA | -0.513 |
| SE | 94465 | ENSG00000175611 | LINC00476 | chr9 | - | 95874508 | 95874622 | 95870283 | 95870441 | 95874979 | 95875976 | 4 | 5 | 0 | 0 | 0 | 1 | 2 | 6 | 2 | 18 | 6 | 0 | 3 | 0 | 0 | 6 | 0 | 0 | 0 | 0 | 263 | 149 | 1 | 1 | 0.694 | 0.586 | 0 | 0 | 0 | 0.362 | NA | 1 | NA | NA | -0.425 |
| SE | 94469 | ENSG00000188649 | CC2D2B | chr10 | + | 95937990 | 95938189 | 95927236 | 95927332 | 95949895 | 95950105 | 9 | 3 | 1 | 1 | 7 | 0 | 4 | 0 | 0 | 0 | 0 | 4 | 0 | 0 | 0 | 0 | 0 | 0 | 0 | 0 | 298 | 149 | 1 | 1 | 1 | 0.273 | 1 | 1 | 1 | NA | 1 | NA | NA | NA | -0.145 |
| SE | 94470 | ENSG00000188649 | CC2D2B | chr10 | + | 95952322 | 95952425 | 95949967 | 95950105 | 95955393 | 95955491 | 8 | 4 | 1 | 1 | 1 | 3 | 0 | 6 | 1 | 4 | 0 | 7 | 3 | 0 | 3 | 0 | 0 | 2 | 0 | 0 | 252 | 149 | 1 | 1 | 0.612 | 1 | 0.09 | 0.372 | 0.129 | NA | 1 | 0.47 | NA | 1 | -0.383 |
| SE | 94471 | ENSG00000188649 | CC2D2B | chr10 | + | 95952322 | 95952425 | 95949967 | 95950105 | 95961828 | 95961969 | 4 | 2 | 1 | 1 | 1 | 1 | 0 | 0 | 0 | 0 | 0 | 7 | 2 | 0 | 3 | 0 | 0 | 1 | 0 | 0 | 252 | 149 | 1 | 1 | 0.703 | 1 | 1 | 1 | 1 | NA | 1 | 0.542 | NA | 1 | 0.093 |
| SE | 94472 | ENSG00000188649 | CC2D2B | chr10 | + | 95955393 | 95955491 | 95949895 | 95950105 | 95961828 | 95961969 | 7 | 6 | 9 | 4 | 7 | 1 | 0 | 0 | 0 | 0 | 0 | 0 | 8 | 0 | 0 | 0 | 0 | 1 | 0 | 0 | 247 | 149 | 1 | 1 | 0.809 | 1 | 1 | 1 | 1 | NA | NA | 0.828 | NA | NA | 0.134 |
| SE | 94473 | ENSG00000188649 | CC2D2B | chr10 | + | 95955393 | 95955491 | 95952322 | 95952425 | 95961828 | 95961969 | 8 | 8 | 3 | 4 | 3 | 0 | 0 | 0 | 1 | 0 | 0 | 0 | 6 | 0 | 0 | 0 | 0 | 0 | 0 | 0 | 247 | 149 | 1 | 1 | 1 | 1 | 1 | 0.707 | 1 | NA | NA | 1 | NA | NA | -0.059 |
| SE | 94474 | ENSG00000188649 | CC2D2B | chr10 | + | 95968723 | 95968901 | 95966189 | 95966302 | 95972065 | 95972216 | 0 | 2 | 0 | 0 | 6 | 0 | 1 | 0 | 0 | 0 | 0 | 0 | 5 | 0 | 0 | 0 | 0 | 0 | 3 | 0 | 298 | 149 | 1 | 1 | NA | 0.5 | NA | NA | 1 | NA | NA | 1 | 0 | NA | 0.25 |
| SE | 94475 | ENSG00000188649 | CC2D2B | chr10 | + | 95968723 | 95968901 | 95966189 | 95966302 | 95974008 | 95974156 | 0 | 3 | 0 | 0 | 6 | 1 | 0 | 0 | 0 | 0 | 0 | 0 | 6 | 0 | 0 | 0 | 0 | 0 | 0 | 0 | 298 | 149 | 1 | 1 | 0 | 1 | NA | NA | 1 | NA | NA | 1 | NA | NA | -0.333 |
| SE | 94476 | ENSG00000188649 | CC2D2B | chr10 | + | 95968723 | 95968901 | 95966189 | 95966302 | 95981974 | 95982113 | 0 | 2 | 1 | 0 | 6 | 0 | 1 | 0 | 5 | 4 | 0 | 0 | 5 | 0 | 0 | 6 | 0 | 0 | 0 | 0 | 298 | 149 | 1 | 1 | NA | 0.5 | 1 | 0 | 0.429 | 0 | NA | 1 | NA | NA | -0.018 |
| SE | 94477 | ENSG00000188649 | CC2D2B | chr10 | + | 95972065 | 95972216 | 95966189 | 95966302 | 95974008 | 95974156 | 0 | 1 | 0 | 0 | 0 | 1 | 0 | 0 | 0 | 0 | 0 | 4 | 0 | 5 | 0 | 0 | 0 | 0 | 0 | 0 | 298 | 149 | 1 | 1 | 0 | 1 | NA | NA | NA | NA | 1 | NA | 1 | NA | -0.5 |
| SE | 94478 | ENSG00000188649 | CC2D2B | chr10 | + | 95972065 | 95972216 | 95968723 | 95968901 | 95974008 | 95974156 | 0 | 0 | 0 | 0 | 0 | 0 | 1 | 0 | 0 | 0 | 0 | 4 | 0 | 2 | 0 | 0 | 0 | 1 | 0 | 0 | 298 | 149 | 1 | 1 | NA | 0 | NA | NA | NA | NA | 1 | 0 | 1 | NA | -0.667 |
| SE | 94479 | ENSG00000188649 | CC2D2B | chr10 | + | 95974008 | 95974156 | 95966189 | 95966302 | 95981974 | 95982113 | 1 | 1 | 0 | 0 | 1 | 0 | 1 | 0 | 5 | 4 | 2 | 0 | 0 | 5 | 0 | 6 | 0 | 0 | 0 | 0 | 297 | 149 | 1 | 1 | 1 | 0.334 | NA | 0 | 0.111 | 0.143 | NA | NA | 1 | NA | -0.21 |
| SE | 94480 | ENSG00000188649 | CC2D2B | chr10 | + | 95974008 | 95974156 | 95968723 | 95968901 | 95981974 | 95982113 | 0 | 2 | 0 | 0 | 1 | 0 | 0 | 1 | 0 | 0 | 2 | 0 | 1 | 5 | 0 | 0 | 0 | 0 | 0 | 0 | 297 | 149 | 1 | 1 | NA | 1 | 0 | NA | 1 | 1 | NA | 1 | 1 | NA | -0.333 |
| SE | 94481 | ENSG00000188649 | CC2D2B | chr10 | + | 95983605 | 95983809 | 95981974 | 95982113 | 95988249 | 95988342 | 4 | 1 | 1 | 7 | 2 | 0 | 0 | 0 | 0 | 1 | 6 | 0 | 1 | 0 | 0 | 0 | 0 | 0 | 0 | 0 | 298 | 149 | 1 | 1 | 1 | 1 | 1 | 1 | 0.5 | 1 | NA | 1 | NA | NA | -0.1 |
| SE | 94482 | ENSG00000188649 | CC2D2B | chr10 | + | 95983605 | 95983809 | 95981974 | 95982113 | 95991369 | 95991461 | 1 | 0 | 1 | 2 | 2 | 0 | 4 | 0 | 0 | 0 | 6 | 0 | 0 | 0 | 0 | 0 | 0 | 0 | 0 | 0 | 298 | 149 | 1 | 1 | 1 | 0 | 1 | 1 | 1 | 1 | NA | NA | NA | NA | -0.2 |
| SE | 94484 | ENSG00000188649 | CC2D2B | chr10 | + | 95988249 | 95988342 | 95981974 | 95982113 | 95991369 | 95991461 | 2 | 0 | 0 | 1 | 1 | 0 | 4 | 0 | 0 | 0 | 0 | 4 | 0 | 0 | 0 | 0 | 0 | 0 | 0 | 0 | 242 | 149 | 1 | 1 | 1 | 0 | NA | 1 | 1 | NA | 1 | NA | NA | NA | -0.25 |
| SE | 94486 | ENSG00000188649 | CC2D2B | chr10 | + | 95991369 | 95991461 | 95983605 | 95983809 | 95992526 | 95992697 | 2 | 6 | 1 | 2 | 1 | 0 | 2 | 0 | 0 | 0 | 0 | 0 | 7 | 0 | 0 | 0 | 0 | 0 | 0 | 0 | 241 | 149 | 1 | 1 | 1 | 0.65 | 1 | 1 | 1 | NA | NA | 1 | NA | NA | -0.07 |
| SE | 94487 | ENSG00000188649 | CC2D2B | chr10 | + | 95993008 | 95993112 | 95992526 | 95992697 | 95995268 | 95995365 | 0 | 4 | 0 | 0 | 4 | 0 | 1 | 0 | 0 | 0 | 0 | 2 | 6 | 0 | 0 | 0 | 0 | 0 | 0 | 0 | 253 | 149 | 1 | 1 | NA | 0.702 | NA | NA | 1 | NA | 1 | 1 | NA | NA | -0.149 |
| SE | 94489 | ENSG00000188649 | CC2D2B | chr10 | + | 96004151 | 96004248 | 95996142 | 95996252 | 96009824 | 96009923 | 5 | 0 | 0 | 0 | 1 | 0 | 0 | 2 | 0 | 0 | 0 | 0 | 0 | 0 | 0 | 0 | 2 | 0 | 0 | 0 | 246 | 149 | 1 | 1 | 1 | NA | 0 | NA | 1 | NA | 0 | NA | NA | NA | 0.667 |
| SE | 94496 | ENSG00000109689 | STIM2 | chr4 | + | 26999231 | 26999347 | 26995378 | 26995490 | 27002216 | 27002228 | 284 | 209 | 259 | 283 | 404 | 1 | 0 | 0 | 2 | 2 | 195 | 279 | 212 | 207 | 88 | 0 | 0 | 0 | 0 | 0 | 265 | 149 | 1 | 1 | 0.994 | 1 | 1 | 0.988 | 0.991 | 1 | 1 | 1 | 1 | 1 | -0.005 |
| SE | 94500 | ENSG00000109689 | STIM2 | chr4 | + | 27021000 | 27021068 | 27017710 | 27017984 | 27022518 | 27024187 | 1 | 0 | 0 | 0 | 0 | 54 | 44 | 73 | 64 | 103 | 0 | 0 | 0 | 0 | 0 | 13 | 62 | 61 | 24 | 35 | 217 | 149 | 1 | 1 | 0.013 | 0 | 0 | 0 | 0 | 0 | 0 | 0 | 0 | 0 | 0.003 |
| SE | 94506 | ENSG00000070061 | ELP1 | chr9 | - | 108874894 | 108874963 | 108868650 | 108869182 | 108877994 | 108878149 | 58 | 56 | 56 | 46 | 84 | 0 | 0 | 1 | 0 | 0 | 13 | 20 | 45 | 20 | 1 | 0 | 0 | 0 | 0 | 0 | 218 | 149 | 1 | 1 | 1 | 1 | 0.975 | 1 | 1 | 1 | 1 | 1 | 1 | 1 | -0.005 |
| SE | 94507 | ENSG00000070061 | ELP1 | chr9 | - | 108874894 | 108874970 | 108866897 | 108869182 | 108877994 | 108878149 | 165 | 121 | 152 | 142 | 287 | 0 | 0 | 1 | 0 | 0 | 19 | 61 | 97 | 65 | 18 | 0 | 0 | 0 | 0 | 0 | 225 | 149 | 1 | 1 | 1 | 1 | 0.99 | 1 | 1 | 1 | 1 | 1 | 1 | 1 | -0.002 |
| SE | 94509 | ENSG00000070061 | ELP1 | chr9 | - | 108877612 | 108877800 | 108874894 | 108874970 | 108877994 | 108878149 | 1 | 0 | 0 | 0 | 0 | 112 | 64 | 97 | 98 | 209 | 0 | 0 | 0 | 0 | 0 | 6 | 41 | 56 | 45 | 17 | 298 | 149 | 1 | 1 | 0.004 | 0 | 0 | 0 | 0 | 0 | 0 | 0 | 0 | 0 | 0.001 |
| SE | 94510 | ENSG00000070061 | ELP1 | chr9 | - | 108879445 | 108879557 | 108878622 | 108878750 | 108880051 | 108880165 | 117 | 92 | 113 | 87 | 151 | 0 | 0 | 0 | 1 | 0 | 23 | 89 | 73 | 32 | 0 | 0 | 0 | 1 | 9 | 0 | 261 | 149 | 1 | 1 | 1 | 1 | 1 | 0.98 | 1 | 1 | 1 | 0.977 | 0.67 | NA | 0.084 |
| SE | 94511 | ENSG00000070061 | ELP1 | chr9 | - | 108884795 | 108884912 | 108882124 | 108882187 | 108889331 | 108889393 | 0 | 0 | 1 | 0 | 0 | 53 | 49 | 45 | 55 | 43 | 0 | 0 | 0 | 0 | 0 | 0 | 44 | 45 | 51 | 11 | 266 | 149 | 1 | 1 | 0 | 0 | 0.012 | 0 | 0 | NA | 0 | 0 | 0 | 0 | 0.002 |
| SE | 94514 | ENSG00000070061 | ELP1 | chr9 | - | 108916203 | 108916297 | 108912263 | 108912494 | 108922841 | 108922927 | 57 | 35 | 41 | 60 | 70 | 0 | 0 | 0 | 1 | 0 | 5 | 41 | 40 | 17 | 11 | 0 | 0 | 0 | 0 | 0 | 243 | 149 | 1 | 1 | 1 | 1 | 1 | 0.974 | 1 | 1 | 1 | 1 | 1 | 1 | -0.005 |
| SE | 94515 | ENSG00000070061 | ELP1 | chr9 | - | 108916203 | 108919349 | 108912263 | 108912494 | 108922841 | 108922927 | 110 | 74 | 95 | 123 | 127 | 0 | 0 | 0 | 1 | 0 | 26 | 74 | 78 | 32 | 27 | 0 | 0 | 0 | 0 | 0 | 298 | 149 | 1 | 1 | 1 | 1 | 1 | 0.984 | 1 | 1 | 1 | 1 | 1 | 1 | -0.003 |
| SE | 94516 | ENSG00000070061 | ELP1 | chr9 | - | 108917546 | 108917670 | 108916203 | 108916297 | 108922841 | 108922927 | 40 | 31 | 38 | 38 | 46 | 0 | 0 | 0 | 0 | 3 | 18 | 21 | 36 | 11 | 9 | 0 | 0 | 0 | 0 | 0 | 273 | 149 | 1 | 1 | 1 | 1 | 1 | 1 | 0.893 | 1 | 1 | 1 | 1 | 1 | -0.021 |
| SE | 94517 | ENSG00000070061 | ELP1 | chr9 | - | 108917546 | 108919349 | 108916203 | 108916297 | 108922841 | 108922927 | 93 | 66 | 89 | 104 | 103 | 0 | 0 | 0 | 0 | 3 | 32 | 53 | 71 | 26 | 25 | 0 | 0 | 0 | 0 | 0 | 298 | 149 | 1 | 1 | 1 | 1 | 1 | 1 | 0.945 | 1 | 1 | 1 | 1 | 1 | -0.011 |
| SE | 94518 | ENSG00000070061 | ELP1 | chr9 | - | 108918810 | 108918901 | 108917546 | 108917670 | 108919252 | 108919349 | 40 | 15 | 50 | 35 | 39 | 0 | 0 | 0 | 0 | 0 | 0 | 30 | 12 | 11 | 5 | 0 | 0 | 4 | 0 | 0 | 240 | 149 | 1 | 1 | 1 | 1 | 1 | 1 | 1 | NA | 1 | 0.651 | 1 | 1 | 0.087 |
| SE | 94519 | ENSG00000070061 | ELP1 | chr9 | - | 108918810 | 108918901 | 108917546 | 108917670 | 108922841 | 108922927 | 19 | 11 | 22 | 15 | 26 | 0 | 3 | 1 | 2 | 0 | 0 | 25 | 11 | 5 | 1 | 0 | 0 | 3 | 0 | 0 | 240 | 149 | 1 | 1 | 1 | 0.695 | 0.932 | 0.823 | 1 | NA | 1 | 0.695 | 1 | 1 | -0.034 |
| SE | 94521 | ENSG00000070061 | ELP1 | chr9 | - | 108919252 | 108919349 | 108917546 | 108917670 | 108926522 | 108926603 | 0 | 0 | 0 | 0 | 0 | 0 | 0 | 0 | 1 | 0 | 0 | 0 | 6 | 0 | 0 | 0 | 0 | 0 | 0 | 0 | 246 | 149 | 1 | 1 | NA | NA | NA | 0 | NA | NA | NA | 1 | NA | NA | -1 |
| SE | 94522 | ENSG00000070061 | ELP1 | chr9 | - | 108919252 | 108919349 | 108918810 | 108918901 | 108922841 | 108922927 | 73 | 35 | 75 | 72 | 63 | 0 | 0 | 0 | 0 | 0 | 14 | 37 | 38 | 23 | 20 | 0 | 0 | 0 | 2 | 0 | 246 | 149 | 1 | 1 | 1 | 1 | 1 | 1 | 1 | 1 | 1 | 1 | 0.874 | 1 | 0.025 |
| SE | 94524 | ENSG00000070061 | ELP1 | chr9 | - | 108922841 | 108922927 | 108916203 | 108919349 | 108926522 | 108926595 | 97 | 78 | 86 | 110 | 98 | 0 | 0 | 0 | 1 | 0 | 49 | 55 | 61 | 47 | 23 | 0 | 0 | 2 | 0 | 0 | 235 | 149 | 1 | 1 | 1 | 1 | 1 | 0.986 | 1 | 1 | 1 | 0.951 | 1 | 1 | 0.007 |
| SE | 94525 | ENSG00000070061 | ELP1 | chr9 | - | 108922841 | 108922927 | 108917546 | 108917670 | 108926522 | 108926595 | 34 | 41 | 31 | 42 | 39 | 0 | 0 | 0 | 1 | 0 | 35 | 23 | 26 | 29 | 7 | 0 | 0 | 0 | 0 | 0 | 235 | 149 | 1 | 1 | 1 | 1 | 1 | 0.964 | 1 | 1 | 1 | 1 | 1 | 1 | -0.007 |
| SE | 94526 | ENSG00000070061 | ELP1 | chr9 | - | 108930996 | 108931056 | 108929768 | 108929921 | 108933863 | 108933943 | 27 | 21 | 26 | 43 | 19 | 2 | 0 | 0 | 0 | 2 | 5 | 19 | 34 | 23 | 3 | 0 | 0 | 0 | 0 | 0 | 209 | 149 | 1 | 1 | 0.906 | 1 | 1 | 1 | 0.871 | 1 | 1 | 1 | 1 | 1 | -0.045 |
| SE | 94527 | ENSG00000070061 | ELP1 | chr9 | - | 108930996 | 108931199 | 108929768 | 108929921 | 108933863 | 108934113 | 37 | 36 | 37 | 44 | 51 | 2 | 0 | 0 | 0 | 2 | 12 | 34 | 45 | 43 | 6 | 0 | 0 | 0 | 0 | 0 | 298 | 149 | 1 | 1 | 0.902 | 1 | 1 | 1 | 0.927 | 1 | 1 | 1 | 1 | 1 | -0.034 |
| SE | 94528 | ENSG00000070061 | ELP1 | chr9 | - | 108930996 | 108931201 | 108929768 | 108929921 | 108933863 | 108934113 | 41 | 45 | 45 | 55 | 53 | 2 | 0 | 0 | 0 | 2 | 26 | 37 | 47 | 45 | 16 | 0 | 0 | 0 | 0 | 0 | 298 | 149 | 1 | 1 | 0.911 | 1 | 1 | 1 | 0.93 | 1 | 1 | 1 | 1 | 1 | -0.032 |
| SE | 94530 | ENSG00000107164 | FUBP3 | chr9 | + | 130595482 | 130595588 | 130579576 | 130579764 | 130609953 | 130609987 | 107 | 97 | 90 | 74 | 61 | 1 | 0 | 0 | 0 | 0 | 118 | 131 | 55 | 47 | 40 | 0 | 0 | 0 | 0 | 0 | 255 | 149 | 1 | 1 | 0.984 | 1 | 1 | 1 | 1 | 1 | 1 | 1 | 1 | 1 | -0.003 |
| SE | 94532 | ENSG00000107164 | FUBP3 | chr9 | + | 130614287 | 130614345 | 130612955 | 130613027 | 130616354 | 130616517 | 195 | 133 | 203 | 205 | 224 | 2 | 1 | 3 | 3 | 0 | 110 | 164 | 170 | 207 | 12 | 0 | 0 | 0 | 0 | 0 | 207 | 149 | 1 | 1 | 0.986 | 0.99 | 0.98 | 0.98 | 1 | 1 | 1 | 1 | 1 | 1 | -0.013 |
| SE | 94533 | ENSG00000107164 | FUBP3 | chr9 | + | 130616354 | 130616517 | 130614287 | 130614345 | 130617796 | 130617895 | 261 | 193 | 272 | 289 | 268 | 4 | 0 | 0 | 0 | 0 | 143 | 244 | 225 | 256 | 38 | 0 | 0 | 2 | 0 | 0 | 298 | 149 | 1 | 1 | 0.97 | 1 | 1 | 1 | 1 | 1 | 1 | 0.983 | 1 | 1 | -0.003 |
| SE | 94534 | ENSG00000107164 | FUBP3 | chr9 | + | 130631160 | 130631293 | 130630627 | 130630788 | 130631556 | 130631630 | 0 | 0 | 9 | 6 | 7 | 24 | 8 | 22 | 29 | 6 | 0 | 2 | 0 | 6 | 6 | 0 | 10 | 8 | 23 | 12 | 282 | 149 | 1 | 1 | 0 | 0 | 0.178 | 0.099 | 0.381 | NA | 0.096 | 0 | 0.121 | 0.209 | 0.025 |
| SE | 94537 | ENSG00000097007 | ABL1 | chr9 | + | 130874867 | 130875052 | 130872859 | 130873037 | 130878414 | 130878567 | 115 | 46 | 80 | 79 | 80 | 1 | 0 | 1 | 0 | 0 | 160 | 110 | 99 | 113 | 233 | 0 | 0 | 0 | 0 | 1 | 298 | 149 | 1 | 1 | 0.983 | 1 | 0.976 | 1 | 1 | 1 | 1 | 1 | 1 | 0.991 | -0.006 |
| SE | 94538 | ENSG00000179163 | FUCA1 | chr1 | - | 23846073 | 23846173 | 23845076 | 23845855 | 23848648 | 23848839 | 133 | 132 | 151 | 84 | 262 | 1 | 0 | 0 | 0 | 0 | 154 | 117 | 119 | 177 | 181 | 0 | 0 | 0 | 0 | 0 | 249 | 149 | 1 | 1 | 0.988 | 1 | 1 | 1 | 1 | 1 | 1 | 1 | 1 | 1 | -0.002 |
| SE | 94539 | ENSG00000179163 | FUCA1 | chr1 | - | 23859797 | 23859903 | 23854359 | 23854560 | 23863133 | 23863271 | 181 | 123 | 169 | 140 | 219 | 0 | 1 | 2 | 0 | 0 | 397 | 237 | 156 | 218 | 224 | 0 | 0 | 0 | 0 | 0 | 255 | 149 | 1 | 1 | 1 | 0.986 | 0.98 | 1 | 1 | 1 | 1 | 1 | 1 | 1 | -0.007 |
| SE | 94542 | ENSG00000233901 | LINC01503 | chr9 | + | 129341735 | 129342043 | 129338568 | 129338917 | 129355457 | 129355598 | 8 | 4 | 8 | 3 | 17 | 0 | 1 | 0 | 0 | 0 | 7 | 3 | 2 | 4 | 15 | 0 | 0 | 0 | 0 | 0 | 298 | 149 | 1 | 1 | 1 | 0.667 | 1 | 1 | 1 | 1 | 1 | 1 | 1 | 1 | -0.067 |
| SE | 94545 | ENSG00000233901 | LINC01503 | chr9 | + | 129341831 | 129342043 | 129338562 | 129338917 | 129355457 | 129355598 | 1 | 3 | 0 | 1 | 0 | 0 | 1 | 0 | 0 | 0 | 0 | 0 | 0 | 4 | 0 | 0 | 0 | 0 | 0 | 0 | 298 | 149 | 1 | 1 | 1 | 0.6 | NA | 1 | NA | NA | NA | NA | 1 | NA | -0.133 |
| SE | 94547 | ENSG00000171488 | LRRC8C | chr1 | + | 89686469 | 89686611 | 89633071 | 89633322 | 89712708 | 89719533 | 192 | 88 | 169 | 170 | 238 | 0 | 2 | 0 | 0 | 0 | 19 | 139 | 116 | 18 | 73 | 0 | 0 | 2 | 0 | 0 | 291 | 149 | 1 | 1 | 1 | 0.957 | 1 | 1 | 1 | 1 | 1 | 0.967 | 1 | 1 | -0.002 |
| SE | 94548 | ENSG00000171488 | LRRC8C | chr1 | + | 89706301 | 89706400 | 89686469 | 89686611 | 89712708 | 89719533 | 2 | 0 | 1 | 3 | 1 | 167 | 72 | 155 | 125 | 238 | 0 | 0 | 0 | 0 | 0 | 16 | 122 | 90 | 15 | 66 | 248 | 149 | 1 | 1 | 0.007 | 0 | 0.004 | 0.014 | 0.003 | 0 | 0 | 0 | 0 | 0 | 0.006 |
| SE | 94552 | ENSG00000132950 | ZMYM5 | chr13 | - | 19838699 | 19838985 | 19835476 | 19835689 | 19851688 | 19852190 | 11 | 14 | 9 | 8 | 20 | 0 | 2 | 1 | 1 | 2 | 17 | 13 | 11 | 6 | 0 | 0 | 0 | 0 | 0 | 0 | 298 | 149 | 1 | 1 | 1 | 0.778 | 0.818 | 0.8 | 0.833 | 1 | 1 | 1 | 1 | NA | -0.154 |
| SE | 94558 | ENSG00000141665 | FBXO15 | chr18 | - | 74129404 | 74129614 | 74125974 | 74126101 | 74130415 | 74130658 | 4 | 0 | 1 | 4 | 12 | 0 | 0 | 0 | 2 | 0 | 0 | 6 | 0 | 0 | 0 | 0 | 0 | 0 | 0 | 0 | 298 | 149 | 1 | 1 | 1 | NA | 1 | 0.5 | 1 | NA | 1 | NA | NA | NA | -0.125 |
| SE | 94563 | ENSG00000198771 | RCSD1 | chr1 | + | 167633648 | 167633806 | 167630265 | 167630429 | 167638389 | 167638442 | 0 | 0 | 0 | 0 | 0 | 3 | 2 | 1 | 2 | 4 | 0 | 0 | 0 | 2 | 0 | 5 | 10 | 3 | 7 | 4 | 298 | 149 | 1 | 1 | 0 | 0 | 0 | 0 | 0 | 0 | 0 | 0 | 0.125 | 0 | -0.025 |
| SE | 94564 | ENSG00000198771 | RCSD1 | chr1 | + | 167638389 | 167638442 | 167630265 | 167630429 | 167640410 | 167640474 | 6 | 1 | 0 | 6 | 5 | 0 | 0 | 0 | 0 | 0 | 5 | 24 | 1 | 11 | 20 | 0 | 0 | 0 | 0 | 3 | 202 | 149 | 1 | 1 | 1 | 1 | NA | 1 | 1 | 1 | 1 | 1 | 1 | 0.831 | 0.034 |
| SE | 94565 | ENSG00000198771 | RCSD1 | chr1 | + | 167638389 | 167638442 | 167630265 | 167630429 | 167640570 | 167640712 | 0 | 1 | 0 | 0 | 0 | 0 | 0 | 0 | 0 | 0 | 0 | 0 | 0 | 11 | 0 | 0 | 6 | 4 | 0 | 9 | 202 | 149 | 1 | 1 | NA | 1 | NA | NA | NA | NA | 0 | 0 | 1 | 0 | 0.75 |
| SE | 94566 | ENSG00000198771 | RCSD1 | chr1 | + | 167638389 | 167638442 | 167630265 | 167630429 | 167641390 | 167641741 | 0 | 1 | 0 | 1 | 0 | 0 | 0 | 0 | 1 | 0 | 4 | 1 | 0 | 2 | 0 | 0 | 0 | 0 | 0 | 0 | 202 | 149 | 1 | 1 | NA | 1 | NA | 0.425 | NA | 1 | 1 | NA | 1 | NA | -0.287 |
| SE | 94569 | ENSG00000198771 | RCSD1 | chr1 | + | 167640410 | 167640474 | 167630265 | 167630429 | 167641390 | 167641741 | 0 | 0 | 0 | 0 | 0 | 0 | 0 | 0 | 1 | 0 | 0 | 0 | 0 | 0 | 3 | 0 | 0 | 0 | 0 | 0 | 213 | 149 | 1 | 1 | NA | NA | NA | 0 | NA | NA | NA | NA | NA | 1 | -1 |
| SE | 94571 | ENSG00000198771 | RCSD1 | chr1 | + | 167640410 | 167640474 | 167638389 | 167638442 | 167641390 | 167641741 | 3 | 0 | 0 | 2 | 4 | 0 | 0 | 0 | 1 | 0 | 5 | 5 | 0 | 0 | 4 | 4 | 1 | 0 | 0 | 9 | 213 | 149 | 1 | 1 | 1 | NA | NA | 0.583 | 1 | 0.466 | 0.778 | NA | NA | 0.237 | 0.367 |
| SE | 94572 | ENSG00000198771 | RCSD1 | chr1 | + | 167640570 | 167640712 | 167630248 | 167630429 | 167641390 | 167641741 | 6 | 3 | 6 | 7 | 2 | 0 | 0 | 0 | 1 | 0 | 0 | 38 | 8 | 22 | 27 | 0 | 0 | 0 | 0 | 0 | 291 | 149 | 1 | 1 | 1 | 1 | 1 | 0.782 | 1 | NA | 1 | 1 | 1 | 1 | -0.044 |
| SE | 94575 | ENSG00000198771 | RCSD1 | chr1 | + | 167640570 | 167640712 | 167638389 | 167638442 | 167683899 | 167684001 | 2 | 0 | 0 | 0 | 0 | 0 | 2 | 0 | 0 | 0 | 0 | 0 | 0 | 13 | 0 | 0 | 3 | 4 | 2 | 0 | 291 | 149 | 1 | 1 | 1 | 0 | NA | NA | NA | NA | 0 | 0 | 0.769 | NA | 0.244 |
| SE | 94576 | ENSG00000198771 | RCSD1 | chr1 | + | 167640570 | 167640712 | 167640410 | 167640474 | 167641390 | 167641741 | 13 | 9 | 10 | 25 | 12 | 1 | 0 | 0 | 0 | 0 | 0 | 74 | 6 | 41 | 69 | 0 | 2 | 0 | 0 | 0 | 291 | 149 | 1 | 1 | 0.869 | 1 | 1 | 1 | 1 | NA | 0.95 | 1 | 1 | 1 | -0.014 |
| SE | 94578 | ENSG00000198771 | RCSD1 | chr1 | + | 167685420 | 167685510 | 167683899 | 167684001 | 167694098 | 167694302 | 51 | 34 | 56 | 47 | 52 | 0 | 0 | 0 | 0 | 0 | 77 | 69 | 15 | 38 | 17 | 0 | 1 | 0 | 0 | 0 | 239 | 149 | 1 | 1 | 1 | 1 | 1 | 1 | 1 | 1 | 0.977 | 1 | 1 | 1 | 0.005 |
| SE | 94579 | ENSG00000198771 | RCSD1 | chr1 | + | 167690048 | 167690120 | 167683899 | 167684001 | 167694098 | 167694302 | 318 | 159 | 271 | 219 | 227 | 0 | 0 | 0 | 0 | 0 | 259 | 237 | 137 | 127 | 320 | 0 | 1 | 0 | 0 | 0 | 221 | 149 | 1 | 1 | 1 | 1 | 1 | 1 | 1 | 1 | 0.994 | 1 | 1 | 1 | 0.001 |
| SE | 94581 | ENSG00000198771 | RCSD1 | chr1 | + | 167694098 | 167694302 | 167690048 | 167690120 | 167697098 | 167697287 | 1139 | 605 | 905 | 822 | 789 | 5 | 4 | 4 | 1 | 2 | 1036 | 1050 | 525 | 610 | 1290 | 7 | 0 | 0 | 0 | 0 | 298 | 149 | 1 | 1 | 0.991 | 0.987 | 0.991 | 0.998 | 0.995 | 0.987 | 1 | 1 | 1 | 1 | -0.005 |
| SE | 94584 | ENSG00000198771 | RCSD1 | chr1 | + | 167697098 | 167697842 | 167694098 | 167694302 | 167704663 | 167706248 | 867 | 508 | 715 | 682 | 518 | 0 | 0 | 1 | 1 | 0 | 863 | 958 | 440 | 626 | 1150 | 0 | 2 | 1 | 0 | 0 | 298 | 149 | 1 | 1 | 1 | 1 | 0.997 | 0.997 | 1 | 1 | 0.996 | 0.995 | 1 | 1 | 0.001 |
| SE | 94588 | ENSG00000106829 | TLE4 | chr9 | + | 79601414 | 79601509 | 79576132 | 79576177 | 79612655 | 79612718 | 0 | 0 | 1 | 0 | 0 | 223 | 176 | 164 | 201 | 227 | 0 | 0 | 0 | 1 | 0 | 208 | 187 | 225 | 203 | 232 | 244 | 149 | 1 | 1 | 0 | 0 | 0.004 | 0 | 0 | 0 | 0 | 0 | 0.003 | 0 | 0 |
| SE | 94590 | ENSG00000106829 | TLE4 | chr9 | + | 79612655 | 79612718 | 79576132 | 79576177 | 79649819 | 79649858 | 66 | 60 | 75 | 64 | 108 | 0 | 0 | 0 | 0 | 0 | 92 | 77 | 85 | 65 | 69 | 0 | 0 | 1 | 0 | 0 | 212 | 149 | 1 | 1 | 1 | 1 | 1 | 1 | 1 | 1 | 1 | 0.984 | 1 | 1 | 0.003 |
| SE | 94592 | ENSG00000106829 | TLE4 | chr9 | + | 79627373 | 79627448 | 79576132 | 79576177 | 79649819 | 79649858 | 9 | 4 | 12 | 4 | 9 | 0 | 0 | 0 | 0 | 0 | 8 | 17 | 38 | 9 | 17 | 0 | 0 | 1 | 0 | 0 | 224 | 149 | 1 | 1 | 1 | 1 | 1 | 1 | 1 | 1 | 1 | 0.962 | 1 | 1 | 0.008 |
| SE | 94594 | ENSG00000106829 | TLE4 | chr9 | + | 79627373 | 79627448 | 79612655 | 79612718 | 79649480 | 79649724 | 129 | 100 | 76 | 112 | 106 | 0 | 0 | 0 | 0 | 0 | 124 | 70 | 94 | 93 | 137 | 0 | 0 | 0 | 2 | 0 | 224 | 149 | 1 | 1 | 1 | 1 | 1 | 1 | 1 | 1 | 1 | 1 | 0.969 | 1 | 0.006 |
| SE | 94598 | ENSG00000106829 | TLE4 | chr9 | + | 79649819 | 79649858 | 79576132 | 79576177 | 79652589 | 79652798 | 0 | 2 | 1 | 0 | 3 | 1 | 0 | 5 | 3 | 0 | 0 | 8 | 8 | 0 | 4 | 0 | 0 | 0 | 4 | 0 | 188 | 149 | 1 | 1 | 0 | 1 | 0.137 | 0 | 1 | NA | 1 | 1 | 0 | 1 | -0.323 |
| SE | 94599 | ENSG00000106829 | TLE4 | chr9 | + | 79649819 | 79649858 | 79612655 | 79612718 | 79652589 | 79652798 | 1 | 2 | 2 | 0 | 7 | 7 | 9 | 5 | 8 | 0 | 0 | 8 | 11 | 0 | 4 | 0 | 1 | 1 | 18 | 0 | 188 | 149 | 1 | 1 | 0.102 | 0.15 | 0.241 | 0 | 1 | NA | 0.864 | 0.897 | 0 | 1 | -0.392 |
| SE | 94600 | ENSG00000106829 | TLE4 | chr9 | + | 79649819 | 79649858 | 79612655 | 79612718 | 79652592 | 79652799 | 2 | 0 | 0 | 2 | 2 | 11 | 1 | 2 | 6 | 10 | 18 | 0 | 0 | 4 | 0 | 0 | 5 | 0 | 0 | 0 | 188 | 149 | 1 | 1 | 0.126 | 0 | 0 | 0.209 | 0.137 | 1 | 0 | NA | 1 | NA | -0.572 |
| SE | 94603 | ENSG00000106829 | TLE4 | chr9 | + | 79652592 | 79652794 | 79627373 | 79627448 | 79704782 | 79704902 | 43 | 32 | 57 | 46 | 53 | 0 | 0 | 2 | 1 | 0 | 43 | 45 | 22 | 50 | 35 | 6 | 0 | 0 | 7 | 13 | 298 | 149 | 1 | 1 | 1 | 1 | 0.934 | 0.958 | 1 | 0.782 | 1 | 1 | 0.781 | 0.574 | 0.151 |
| SE | 94605 | ENSG00000106829 | TLE4 | chr9 | + | 79671208 | 79671338 | 79654058 | 79654075 | 79704782 | 79704902 | 0 | 2 | 0 | 0 | 1 | 259 | 132 | 185 | 183 | 334 | 0 | 0 | 0 | 0 | 0 | 282 | 180 | 219 | 172 | 400 | 279 | 149 | 1 | 1 | 0 | 0.008 | 0 | 0 | 0.002 | 0 | 0 | 0 | 0 | 0 | 0.002 |
| SE | 94607 | ENSG00000106829 | TLE4 | chr9 | + | 79706746 | 79706899 | 79705888 | 79705942 | 79707122 | 79707218 | 296 | 176 | 229 | 229 | 328 | 4 | 2 | 0 | 0 | 0 | 285 | 304 | 248 | 222 | 348 | 0 | 0 | 0 | 0 | 0 | 298 | 149 | 1 | 1 | 0.974 | 0.978 | 1 | 1 | 1 | 1 | 1 | 1 | 1 | 1 | -0.01 |
| SE | 94611 | ENSG00000106829 | TLE4 | chr9 | + | 79708592 | 79708786 | 79708117 | 79708250 | 79709622 | 79709672 | 222 | 106 | 176 | 177 | 152 | 0 | 0 | 1 | 1 | 0 | 115 | 180 | 145 | 151 | 198 | 0 | 0 | 0 | 0 | 0 | 298 | 149 | 1 | 1 | 1 | 1 | 0.989 | 0.989 | 1 | 1 | 1 | 1 | 1 | 1 | -0.004 |
| SE | 94613 | ENSG00000106829 | TLE4 | chr9 | + | 79722450 | 79722601 | 79721740 | 79721888 | 79722958 | 79723035 | 187 | 82 | 132 | 141 | 194 | 0 | 0 | 0 | 0 | 0 | 178 | 162 | 151 | 166 | 154 | 4 | 0 | 0 | 0 | 0 | 298 | 149 | 1 | 1 | 1 | 1 | 1 | 1 | 1 | 0.957 | 1 | 1 | 1 | 1 | 0.009 |
| SE | 94615 | ENSG00000204394 | VARS1 | chr6 | - | 31781690 | 31781952 | 31781357 | 31781606 | 31782086 | 31782420 | 37 | 23 | 41 | 42 | 20 | 0 | 0 | 0 | 0 | 0 | 57 | 34 | 33 | 33 | 69 | 0 | 3 | 0 | 0 | 0 | 298 | 149 | 1 | 1 | 1 | 1 | 1 | 1 | 1 | 1 | 0.85 | 1 | 1 | 1 | 0.03 |
| SE | 94616 | ENSG00000204394 | VARS1 | chr6 | - | 31783095 | 31783186 | 31782411 | 31782845 | 31784213 | 31784308 | 63 | 22 | 44 | 65 | 51 | 0 | 0 | 0 | 0 | 0 | 114 | 51 | 52 | 37 | 69 | 0 | 0 | 0 | 4 | 0 | 240 | 149 | 1 | 1 | 1 | 1 | 1 | 1 | 1 | 1 | 1 | 1 | 0.852 | 1 | 0.03 |
| SE | 94618 | ENSG00000204394 | VARS1 | chr6 | - | 31785245 | 31785327 | 31784594 | 31784714 | 31785568 | 31785733 | 54 | 34 | 57 | 67 | 49 | 0 | 1 | 0 | 0 | 0 | 41 | 38 | 54 | 44 | 98 | 0 | 0 | 0 | 3 | 0 | 231 | 149 | 1 | 1 | 1 | 0.956 | 1 | 1 | 1 | 1 | 1 | 1 | 0.904 | 1 | 0.01 |
| SE | 94630 | ENSG00000248167 | TRIM39-RPP21 | chr6 | + | 30341711 | 30341896 | 30340504 | 30340620 | 30345297 | 30345398 | 44 | 39 | 51 | 57 | 55 | 0 | 0 | 0 | 0 | 1 | 66 | 62 | 69 | 67 | 41 | 0 | 0 | 1 | 0 | 0 | 298 | 149 | 1 | 1 | 1 | 1 | 1 | 1 | 0.965 | 1 | 1 | 0.972 | 1 | 1 | -0.001 |
| SE | 94631 | ENSG00000241370 | RPP21 | chr6 | + | 30345490 | 30345573 | 30345158 | 30345398 | 30346431 | 30346557 | 48 | 26 | 15 | 29 | 18 | 0 | 0 | 0 | 0 | 0 | 12 | 39 | 13 | 19 | 53 | 2 | 0 | 0 | 0 | 0 | 232 | 149 | 1 | 1 | 1 | 1 | 1 | 1 | 1 | 0.794 | 1 | 1 | 1 | 1 | 0.041 |
| SE | 94632 | ENSG00000136867 | SLC31A2 | chr9 | + | 113157726 | 113157793 | 113151006 | 113151080 | 113161508 | 113161698 | 110 | 20 | 18 | 46 | 6 | 0 | 0 | 0 | 0 | 0 | 38 | 44 | 45 | 65 | 31 | 0 | 0 | 0 | 2 | 0 | 216 | 149 | 1 | 1 | 1 | 1 | 1 | 1 | 1 | 1 | 1 | 1 | 0.957 | 1 | 0.009 |
| SE | 94633 | ENSG00000136867 | SLC31A2 | chr9 | + | 113158043 | 113158096 | 113157726 | 113157793 | 113161508 | 113161698 | 0 | 1 | 0 | 2 | 0 | 115 | 22 | 20 | 49 | 7 | 0 | 6 | 0 | 0 | 0 | 39 | 51 | 45 | 74 | 31 | 202 | 149 | 1 | 1 | 0 | 0.032 | 0 | 0.029 | 0 | 0 | 0.08 | 0 | 0 | 0 | -0.004 |
| SE | 94634 | ENSG00000136867 | SLC31A2 | chr9 | + | 113161508 | 113161698 | 113157726 | 113157793 | 113162748 | 113164137 | 179 | 41 | 55 | 86 | 36 | 0 | 0 | 0 | 0 | 0 | 89 | 83 | 55 | 100 | 67 | 0 | 3 | 0 | 0 | 0 | 298 | 149 | 1 | 1 | 1 | 1 | 1 | 1 | 1 | 1 | 0.933 | 1 | 1 | 1 | 0.013 |
| SE | 94635 | ENSG00000011454 | RABGAP1 | chr9 | + | 122945331 | 122945432 | 122941032 | 122941088 | 122957010 | 122957209 | 2 | 1 | 2 | 1 | 1 | 19 | 3 | 11 | 2 | 5 | 9 | 7 | 0 | 1 | 0 | 5 | 8 | 5 | 2 | 0 | 250 | 149 | 1 | 1 | 0.059 | 0.166 | 0.098 | 0.23 | 0.107 | 0.518 | 0.343 | 0 | 0.23 | NA | -0.141 |
| SE | 94636 | ENSG00000011454 | RABGAP1 | chr9 | + | 122957010 | 122957209 | 122941019 | 122941093 | 122986214 | 122986419 | 16 | 10 | 15 | 32 | 18 | 0 | 0 | 1 | 0 | 0 | 25 | 20 | 16 | 20 | 3 | 0 | 0 | 0 | 0 | 0 | 298 | 149 | 1 | 1 | 1 | 1 | 0.882 | 1 | 1 | 1 | 1 | 1 | 1 | 1 | -0.024 |
| SE | 94639 | ENSG00000011454 | RABGAP1 | chr9 | + | 122989296 | 122989471 | 122986214 | 122986419 | 122996040 | 122996151 | 88 | 81 | 93 | 106 | 199 | 0 | 1 | 0 | 0 | 0 | 31 | 104 | 55 | 51 | 51 | 0 | 0 | 0 | 0 | 0 | 298 | 149 | 1 | 1 | 1 | 0.976 | 1 | 1 | 1 | 1 | 1 | 1 | 1 | 1 | -0.005 |
| SE | 94641 | ENSG00000011454 | RABGAP1 | chr9 | + | 122997258 | 122997361 | 122996538 | 122996605 | 122998596 | 122998766 | 243 | 203 | 206 | 264 | 351 | 0 | 0 | 0 | 0 | 0 | 134 | 342 | 214 | 206 | 74 | 0 | 3 | 0 | 0 | 0 | 252 | 149 | 1 | 1 | 1 | 1 | 1 | 1 | 1 | 1 | 0.985 | 1 | 1 | 1 | 0.003 |
| SE | 94642 | ENSG00000011454 | RABGAP1 | chr9 | + | 122998596 | 122998766 | 122997258 | 122997361 | 123010353 | 123010528 | 323 | 259 | 275 | 414 | 558 | 2 | 1 | 0 | 0 | 0 | 209 | 518 | 332 | 362 | 108 | 0 | 1 | 0 | 0 | 5 | 298 | 149 | 1 | 1 | 0.988 | 0.992 | 1 | 1 | 1 | 1 | 0.996 | 1 | 1 | 0.915 | 0.014 |
| SE | 94647 | ENSG00000011454 | RABGAP1 | chr9 | + | 123071411 | 123071501 | 123070349 | 123070424 | 123076244 | 123076286 | 0 | 7 | 3 | 1 | 0 | 0 | 2 | 0 | 0 | 0 | 0 | 7 | 1 | 0 | 0 | 0 | 0 | 2 | 4 | 8 | 239 | 149 | 1 | 1 | NA | 0.686 | 1 | 1 | NA | NA | 1 | 0.238 | 0 | 0 | 0.586 |
| SE | 94648 | ENSG00000011454 | RABGAP1 | chr9 | + | 123073551 | 123073677 | 123065347 | 123065461 | 123076244 | 123076286 | 1 | 0 | 1 | 1 | 0 | 0 | 0 | 0 | 0 | 0 | 9 | 3 | 5 | 6 | 0 | 0 | 0 | 6 | 0 | 0 | 275 | 149 | 1 | 1 | 1 | NA | 1 | 1 | NA | 1 | 1 | 0.311 | 1 | NA | 0.172 |
| SE | 94649 | ENSG00000011454 | RABGAP1 | chr9 | + | 123073551 | 123073677 | 123070349 | 123070424 | 123074284 | 123074310 | 147 | 109 | 107 | 148 | 228 | 0 | 0 | 0 | 0 | 0 | 39 | 109 | 103 | 109 | 84 | 0 | 0 | 0 | 0 | 4 | 275 | 149 | 1 | 1 | 1 | 1 | 1 | 1 | 1 | 1 | 1 | 1 | 1 | 0.919 | 0.016 |
| SE | 94651 | ENSG00000011454 | RABGAP1 | chr9 | + | 123073551 | 123073677 | 123071411 | 123071501 | 123076244 | 123076286 | 5 | 0 | 8 | 2 | 2 | 0 | 2 | 0 | 0 | 0 | 0 | 3 | 5 | 9 | 0 | 0 | 0 | 0 | 0 | 0 | 275 | 149 | 1 | 1 | 1 | 0 | 1 | 1 | 1 | NA | 1 | 1 | 1 | NA | -0.2 |
| SE | 94656 | ENSG00000165209 | STRBP | chr9 | - | 123115928 | 123116098 | 123109556 | 123109753 | 123122192 | 123122326 | 0 | 0 | 0 | 0 | 0 | 0 | 2 | 0 | 0 | 0 | 0 | 2 | 0 | 0 | 0 | 0 | 0 | 0 | 0 | 0 | 298 | 149 | 1 | 1 | NA | 0 | NA | NA | NA | NA | 1 | NA | NA | NA | -1 |
| SE | 94658 | ENSG00000165209 | STRBP | chr9 | - | 123128213 | 123128258 | 123121616 | 123125673 | 123132844 | 123132967 | 81 | 73 | 93 | 96 | 220 | 0 | 0 | 0 | 0 | 0 | 21 | 47 | 54 | 28 | 0 | 0 | 0 | 2 | 0 | 0 | 194 | 149 | 1 | 1 | 1 | 1 | 1 | 1 | 1 | 1 | 1 | 0.954 | 1 | NA | 0.012 |
| SE | 94659 | ENSG00000165209 | STRBP | chr9 | - | 123136040 | 123136181 | 123132844 | 123132968 | 123136380 | 123136515 | 145 | 116 | 168 | 154 | 215 | 0 | 0 | 0 | 0 | 0 | 33 | 133 | 101 | 52 | 0 | 0 | 0 | 2 | 0 | 0 | 290 | 149 | 1 | 1 | 1 | 1 | 1 | 1 | 1 | 1 | 1 | 0.963 | 1 | NA | 0.009 |
| SE | 94660 | ENSG00000165209 | STRBP | chr9 | - | 123139528 | 123139687 | 123136380 | 123136515 | 123146854 | 123147054 | 162 | 120 | 192 | 214 | 187 | 1 | 1 | 1 | 1 | 2 | 62 | 175 | 136 | 96 | 46 | 0 | 0 | 0 | 4 | 0 | 298 | 149 | 1 | 1 | 0.988 | 0.984 | 0.99 | 0.991 | 0.979 | 1 | 1 | 1 | 0.923 | 1 | 0.002 |
| SE | 94661 | ENSG00000165209 | STRBP | chr9 | - | 123158011 | 123158123 | 123147777 | 123147870 | 123158331 | 123158429 | 8 | 3 | 78 | 92 | 86 | 2 | 0 | 0 | 0 | 0 | 17 | 52 | 29 | 21 | 7 | 0 | 0 | 0 | 0 | 0 | 261 | 149 | 1 | 1 | 0.695 | 1 | 1 | 1 | 1 | 1 | 1 | 1 | 1 | 1 | -0.061 |
| SE | 94662 | ENSG00000165209 | STRBP | chr9 | - | 123159095 | 123159207 | 123158331 | 123158429 | 123160366 | 123160462 | 135 | 108 | 157 | 208 | 248 | 0 | 0 | 1 | 0 | 0 | 5 | 135 | 80 | 76 | 32 | 0 | 0 | 0 | 0 | 0 | 261 | 149 | 1 | 1 | 1 | 1 | 0.989 | 1 | 1 | 1 | 1 | 1 | 1 | 1 | -0.002 |
| SE | 94663 | ENSG00000165209 | STRBP | chr9 | - | 123159095 | 123159207 | 123158331 | 123158429 | 123160976 | 123161068 | 39 | 30 | 66 | 64 | 94 | 0 | 0 | 0 | 4 | 0 | 0 | 51 | 43 | 19 | 12 | 0 | 0 | 2 | 0 | 0 | 261 | 149 | 1 | 1 | 1 | 1 | 1 | 0.901 | 1 | NA | 1 | 0.925 | 1 | 1 | -0.001 |
| SE | 94664 | ENSG00000165209 | STRBP | chr9 | - | 123160366 | 123160462 | 123158331 | 123158429 | 123160976 | 123161068 | 81 | 63 | 86 | 98 | 138 | 0 | 0 | 0 | 4 | 0 | 3 | 57 | 39 | 38 | 15 | 0 | 0 | 2 | 0 | 0 | 245 | 149 | 1 | 1 | 1 | 1 | 1 | 0.937 | 1 | 1 | 1 | 0.922 | 1 | 1 | 0.003 |
| SE | 94666 | ENSG00000165209 | STRBP | chr9 | - | 123160976 | 123161068 | 123160366 | 123160462 | 123169901 | 123170046 | 117 | 106 | 142 | 185 | 190 | 0 | 0 | 1 | 1 | 5 | 12 | 97 | 62 | 73 | 31 | 0 | 0 | 0 | 0 | 0 | 241 | 149 | 1 | 1 | 1 | 1 | 0.989 | 0.991 | 0.959 | 1 | 1 | 1 | 1 | 1 | -0.012 |
| SE | 94672 | ENSG00000165209 | STRBP | chr9 | - | 123236829 | 123236966 | 123184131 | 123184298 | 123268435 | 123268586 | 9 | 2 | 8 | 2 | 0 | 5 | 0 | 5 | 4 | 1 | 0 | 11 | 0 | 7 | 0 | 0 | 8 | 2 | 5 | 0 | 286 | 149 | 1 | 1 | 0.484 | 1 | 0.455 | 0.207 | 0 | NA | 0.417 | 0 | 0.422 | NA | 0.15 |
| SE | 94674 | ENSG00000165060 | FXN | chr9 | + | 69064937 | 69065043 | 69053139 | 69053260 | 69072611 | 69073022 | 37 | 18 | 32 | 26 | 23 | 1 | 0 | 1 | 1 | 0 | 56 | 50 | 10 | 23 | 11 | 9 | 0 | 0 | 0 | 0 | 255 | 149 | 1 | 1 | 0.956 | 1 | 0.949 | 0.938 | 1 | 0.784 | 1 | 1 | 1 | 1 | 0.012 |
| SE | 94679 | ENSG00000284052 | NA | chr2 | - | 190629750 | 190629825 | 190628068 | 190628715 | 190648219 | 190648474 | 0 | 0 | 0 | 0 | 0 | 0 | 0 | 0 | 1 | 0 | 0 | 0 | 0 | 10 | 0 | 0 | 0 | 2 | 0 | 0 | 224 | 149 | 1 | 1 | NA | NA | NA | 0 | NA | NA | NA | 0 | 1 | NA | -0.5 |
| SE | 94680 | ENSG00000284052 | NA | chr2 | - | 190640950 | 190641462 | 190627877 | 190628715 | 190648219 | 190648336 | 0 | 0 | 1 | 0 | 0 | 0 | 0 | 0 | 1 | 0 | 0 | 0 | 0 | 0 | 0 | 0 | 0 | 2 | 0 | 0 | 298 | 149 | 1 | 1 | NA | NA | 1 | 0 | NA | NA | NA | 0 | NA | NA | 0.5 |
| SE | 94684 | ENSG00000172889 | EGFL7 | chr9 | + | 136669605 | 136669721 | 136668556 | 136668673 | 136669913 | 136670009 | 2 | 1 | 0 | 3 | 0 | 3 | 0 | 0 | 0 | 0 | 0 | 0 | 0 | 12 | 51 | 0 | 0 | 0 | 0 | 0 | 265 | 149 | 1 | 1 | 0.273 | 1 | NA | 1 | NA | NA | NA | NA | 1 | 1 | -0.242 |
| SE | 94686 | ENSG00000204003 | NA | chr9 | - | 136740835 | 136740943 | 136739949 | 136740048 | 136741880 | 136742020 | 7 | 11 | 1 | 9 | 8 | 3 | 6 | 8 | 11 | 4 | 42 | 3 | 0 | 8 | 0 | 0 | 6 | 0 | 2 | 0 | 257 | 149 | 1 | 1 | 0.575 | 0.515 | 0.068 | 0.322 | 0.537 | 1 | 0.225 | NA | 0.699 | NA | -0.238 |
| SE | 94691 | ENSG00000198382 | UVRAG | chr11 | + | 75879878 | 75880040 | 75861745 | 75861780 | 75888828 | 75888903 | 143 | 121 | 118 | 106 | 159 | 0 | 2 | 0 | 0 | 0 | 134 | 190 | 125 | 79 | 70 | 0 | 0 | 0 | 0 | 0 | 298 | 149 | 1 | 1 | 1 | 0.968 | 1 | 1 | 1 | 1 | 1 | 1 | 1 | 1 | -0.006 |
| SE | 94692 | ENSG00000198382 | UVRAG | chr11 | + | 75888828 | 75888903 | 75879878 | 75880040 | 75911953 | 75912039 | 146 | 134 | 133 | 134 | 166 | 3 | 0 | 0 | 0 | 0 | 123 | 222 | 140 | 105 | 80 | 0 | 0 | 0 | 0 | 0 | 224 | 149 | 1 | 1 | 0.97 | 1 | 1 | 1 | 1 | 1 | 1 | 1 | 1 | 1 | -0.006 |
| SE | 94693 | ENSG00000198382 | UVRAG | chr11 | + | 75961443 | 75961549 | 75911953 | 75912039 | 75983386 | 75983513 | 232 | 179 | 143 | 177 | 232 | 0 | 0 | 2 | 0 | 2 | 187 | 327 | 197 | 169 | 111 | 0 | 0 | 1 | 0 | 0 | 255 | 149 | 1 | 1 | 1 | 1 | 0.977 | 1 | 0.985 | 1 | 1 | 0.991 | 1 | 1 | -0.006 |
| SE | 94694 | ENSG00000198382 | UVRAG | chr11 | + | 76004004 | 76004089 | 75983386 | 75983513 | 76007533 | 76007621 | 149 | 127 | 117 | 140 | 239 | 0 | 0 | 0 | 0 | 0 | 201 | 164 | 83 | 179 | 77 | 0 | 1 | 0 | 0 | 0 | 234 | 149 | 1 | 1 | 1 | 1 | 1 | 1 | 1 | 1 | 0.991 | 1 | 1 | 1 | 0.002 |
| SE | 94696 | ENSG00000198382 | UVRAG | chr11 | + | 76016814 | 76016980 | 76008806 | 76008867 | 76065709 | 76065788 | 225 | 214 | 185 | 204 | 390 | 0 | 2 | 0 | 0 | 0 | 307 | 348 | 260 | 246 | 150 | 0 | 0 | 2 | 2 | 0 | 298 | 149 | 1 | 1 | 1 | 0.982 | 1 | 1 | 1 | 1 | 1 | 0.985 | 0.984 | 1 | 0.003 |
| SE | 94697 | ENSG00000198382 | UVRAG | chr11 | + | 76115923 | 76116015 | 76065709 | 76065788 | 76140710 | 76141500 | 159 | 106 | 107 | 166 | 179 | 0 | 0 | 0 | 0 | 0 | 227 | 158 | 188 | 185 | 105 | 0 | 0 | 1 | 0 | 0 | 241 | 149 | 1 | 1 | 1 | 1 | 1 | 1 | 1 | 1 | 1 | 0.991 | 1 | 1 | 0.002 |
| SE | 94698 | ENSG00000198382 | UVRAG | chr11 | + | 76137193 | 76137452 | 76115899 | 76116015 | 76140710 | 76141660 | 0 | 2 | 0 | 1 | 0 | 93 | 64 | 64 | 98 | 98 | 0 | 0 | 0 | 0 | 0 | 100 | 76 | 99 | 125 | 59 | 298 | 149 | 1 | 1 | 0 | 0.015 | 0 | 0.005 | 0 | 0 | 0 | 0 | 0 | 0 | 0.004 |
| SE | 94699 | ENSG00000213213 | CCDC183 | chr9 | + | 136799712 | 136799790 | 136799101 | 136799223 | 136800001 | 136800169 | 3 | 1 | 4 | 1 | 0 | 0 | 0 | 0 | 0 | 0 | 0 | 3 | 0 | 0 | 0 | 0 | 10 | 0 | 0 | 0 | 227 | 149 | 1 | 1 | 1 | 1 | 1 | 1 | NA | NA | 0.165 | NA | NA | NA | 0.835 |
| SE | 94700 | ENSG00000213213 | CCDC183 | chr9 | + | 136805356 | 136805457 | 136804761 | 136804816 | 136806077 | 136806238 | 2 | 6 | 1 | 2 | 8 | 0 | 0 | 1 | 0 | 0 | 0 | 8 | 0 | 6 | 0 | 0 | 0 | 0 | 0 | 0 | 250 | 149 | 1 | 1 | 1 | 1 | 0.373 | 1 | 1 | NA | 1 | NA | 1 | NA | -0.125 |
| SE | 94701 | ENSG00000213213 | CCDC183 | chr9 | + | 136807280 | 136807352 | 136806969 | 136807066 | 136807571 | 136807741 | 0 | 0 | 3 | 1 | 0 | 3 | 0 | 1 | 0 | 0 | 0 | 0 | 0 | 0 | 0 | 0 | 0 | 2 | 0 | 0 | 221 | 149 | 1 | 1 | 0 | NA | 0.669 | 1 | NA | NA | NA | 0 | NA | NA | 0.556 |
| SE | 94703 | ENSG00000165688 | PMPCA | chr9 | + | 136412809 | 136412892 | 136412489 | 136412569 | 136414552 | 136414647 | 223 | 147 | 197 | 230 | 202 | 1 | 0 | 0 | 0 | 0 | 302 | 171 | 201 | 318 | 396 | 0 | 0 | 0 | 0 | 0 | 232 | 149 | 1 | 1 | 0.993 | 1 | 1 | 1 | 1 | 1 | 1 | 1 | 1 | 1 | -0.001 |
| SE | 94704 | ENSG00000165688 | PMPCA | chr9 | + | 136412809 | 136412892 | 136412489 | 136412569 | 136416290 | 136416391 | 139 | 77 | 106 | 151 | 97 | 2 | 0 | 1 | 0 | 0 | 224 | 127 | 146 | 283 | 375 | 0 | 0 | 0 | 0 | 0 | 232 | 149 | 1 | 1 | 0.978 | 1 | 0.986 | 1 | 1 | 1 | 1 | 1 | 1 | 1 | -0.007 |
| SE | 94705 | ENSG00000165688 | PMPCA | chr9 | + | 136414552 | 136414647 | 136412489 | 136412569 | 136416290 | 136416297 | 43 | 33 | 66 | 88 | 87 | 2 | 0 | 1 | 0 | 0 | 92 | 48 | 80 | 76 | 60 | 0 | 0 | 0 | 0 | 0 | 244 | 149 | 1 | 1 | 0.929 | 1 | 0.976 | 1 | 1 | 1 | 1 | 1 | 1 | 1 | -0.019 |
| SE | 94707 | ENSG00000165688 | PMPCA | chr9 | + | 136416290 | 136416391 | 136414552 | 136414647 | 136416999 | 136417195 | 76 | 46 | 70 | 76 | 86 | 2 | 0 | 0 | 0 | 0 | 87 | 55 | 48 | 70 | 95 | 0 | 0 | 0 | 0 | 0 | 250 | 149 | 1 | 1 | 0.958 | 1 | 1 | 1 | 1 | 1 | 1 | 1 | 1 | 1 | -0.008 |
| SE | 94708 | ENSG00000184939 | ZFP90 | chr16 | + | 68539757 | 68539825 | 68533697 | 68533855 | 68557997 | 68558101 | 2 | 2 | 4 | 11 | 0 | 0 | 4 | 0 | 4 | 0 | 0 | 13 | 4 | 14 | 0 | 0 | 0 | 0 | 0 | 0 | 217 | 149 | 1 | 1 | 1 | 0.256 | 1 | 0.654 | NA | NA | 1 | 1 | 1 | NA | -0.272 |
| SE | 94710 | ENSG00000106701 | FSD1L | chr9 | + | 105461519 | 105461615 | 105447795 | 105448235 | 105464235 | 105464331 | 5 | 0 | 0 | 2 | 2 | 2 | 3 | 3 | 7 | 6 | 0 | 0 | 0 | 0 | 0 | 0 | 3 | 0 | 0 | 4 | 245 | 149 | 1 | 1 | 0.603 | 0 | 0 | 0.148 | 0.169 | NA | 0 | NA | NA | 0 | 0.184 |
| SE | 94711 | ENSG00000106701 | FSD1L | chr9 | + | 105471903 | 105472005 | 105468192 | 105468324 | 105484380 | 105484502 | 17 | 6 | 8 | 13 | 17 | 1 | 0 | 0 | 0 | 0 | 0 | 2 | 0 | 0 | 0 | 0 | 0 | 1 | 0 | 0 | 251 | 149 | 1 | 1 | 0.91 | 1 | 1 | 1 | 1 | NA | 1 | 0 | NA | NA | 0.482 |
| SE | 94712 | ENSG00000106701 | FSD1L | chr9 | + | 105471903 | 105472005 | 105468192 | 105468324 | 105506398 | 105506475 | 12 | 5 | 7 | 11 | 17 | 1 | 0 | 0 | 1 | 0 | 0 | 2 | 0 | 0 | 0 | 0 | 0 | 0 | 0 | 0 | 251 | 149 | 1 | 1 | 0.877 | 1 | 1 | 0.867 | 1 | NA | 1 | NA | NA | NA | -0.051 |
| SE | 94713 | ENSG00000106701 | FSD1L | chr9 | + | 105479353 | 105479376 | 105471903 | 105472005 | 105484380 | 105484502 | 6 | 6 | 8 | 5 | 14 | 5 | 1 | 2 | 3 | 0 | 0 | 9 | 0 | 0 | 5 | 0 | 0 | 0 | 0 | 0 | 172 | 149 | 1 | 1 | 0.51 | 0.839 | 0.776 | 0.591 | 1 | NA | 1 | NA | NA | 1 | -0.257 |
| SE | 94714 | ENSG00000106701 | FSD1L | chr9 | + | 105484380 | 105484502 | 105468192 | 105468324 | 105506398 | 105506475 | 6 | 6 | 12 | 4 | 10 | 1 | 0 | 0 | 1 | 0 | 0 | 9 | 5 | 0 | 0 | 0 | 0 | 0 | 0 | 0 | 271 | 149 | 1 | 1 | 0.767 | 1 | 1 | 0.687 | 1 | NA | 1 | 1 | NA | NA | -0.109 |
| SE | 94715 | ENSG00000106701 | FSD1L | chr9 | + | 105484380 | 105484502 | 105471903 | 105472005 | 105506398 | 105506475 | 10 | 7 | 14 | 7 | 10 | 0 | 0 | 1 | 1 | 0 | 0 | 9 | 4 | 0 | 0 | 0 | 0 | 0 | 0 | 0 | 271 | 149 | 1 | 1 | 1 | 1 | 0.885 | 0.794 | 1 | NA | 1 | 1 | NA | NA | -0.064 |
| SE | 94716 | ENSG00000106701 | FSD1L | chr9 | + | 105506616 | 105508715 | 105506398 | 105506608 | 105512806 | 105512936 | 27 | 35 | 18 | 24 | 38 | 0 | 2 | 0 | 0 | 0 | 0 | 24 | 0 | 10 | 0 | 0 | 2 | 0 | 0 | 0 | 248 | 149 | 1 | 1 | 1 | 0.913 | 1 | 1 | 1 | NA | 0.878 | NA | 1 | NA | 0.044 |
| SE | 94717 | ENSG00000106701 | FSD1L | chr9 | + | 105513598 | 105513631 | 105512806 | 105512936 | 105534492 | 105534593 | 1 | 0 | 0 | 1 | 1 | 9 | 16 | 3 | 6 | 15 | 0 | 0 | 0 | 12 | 0 | 0 | 14 | 0 | 24 | 0 | 182 | 149 | 1 | 1 | 0.083 | 0 | 0 | 0.12 | 0.052 | NA | 0 | NA | 0.29 | NA | -0.094 |
| SE | 94718 | ENSG00000106701 | FSD1L | chr9 | + | 105535066 | 105535318 | 105534492 | 105534593 | 105539262 | 105539351 | 33 | 17 | 22 | 16 | 29 | 3 | 0 | 0 | 1 | 0 | 0 | 17 | 9 | 35 | 4 | 0 | 0 | 1 | 0 | 0 | 298 | 149 | 1 | 1 | 0.846 | 1 | 1 | 0.889 | 1 | NA | 1 | 0.818 | 1 | 1 | -0.007 |
| SE | 94719 | ENSG00000106701 | FSD1L | chr9 | + | 105539262 | 105539351 | 105534492 | 105534593 | 105546357 | 105546602 | 6 | 2 | 3 | 2 | 9 | 0 | 0 | 1 | 0 | 0 | 0 | 0 | 1 | 0 | 0 | 0 | 0 | 0 | 0 | 0 | 238 | 149 | 1 | 1 | 1 | 1 | 0.653 | 1 | 1 | NA | NA | 1 | NA | NA | -0.069 |
| SE | 94721 | ENSG00000106948 | AKNA | chr9 | - | 114341532 | 114341725 | 114334155 | 114337306 | 114342008 | 114342125 | 1101 | 461 | 818 | 764 | 586 | 0 | 0 | 0 | 4 | 0 | 576 | 1081 | 586 | 517 | 1131 | 0 | 0 | 0 | 0 | 0 | 298 | 149 | 1 | 1 | 1 | 1 | 1 | 0.99 | 1 | 1 | 1 | 1 | 1 | 1 | -0.002 |
| SE | 94724 | ENSG00000106948 | AKNA | chr9 | - | 114343707 | 114346009 | 114342008 | 114342125 | 114346668 | 114346774 | 951 | 404 | 696 | 739 | 436 | 0 | 0 | 0 | 0 | 0 | 549 | 858 | 490 | 527 | 879 | 0 | 8 | 0 | 0 | 0 | 298 | 149 | 1 | 1 | 1 | 1 | 1 | 1 | 1 | 1 | 0.982 | 1 | 1 | 1 | 0.004 |
| SE | 94725 | ENSG00000106948 | AKNA | chr9 | - | 114345862 | 114346009 | 114342008 | 114342125 | 114346668 | 114346784 | 442 | 208 | 343 | 379 | 202 | 0 | 0 | 0 | 0 | 0 | 386 | 631 | 441 | 362 | 637 | 0 | 8 | 0 | 0 | 0 | 296 | 149 | 1 | 1 | 1 | 1 | 1 | 1 | 1 | 1 | 0.975 | 1 | 1 | 1 | 0.005 |
| SE | 94726 | ENSG00000106948 | AKNA | chr9 | - | 114345862 | 114346009 | 114343707 | 114343803 | 114347723 | 114347900 | 761 | 386 | 631 | 642 | 548 | 7 | 0 | 2 | 0 | 0 | 313 | 739 | 256 | 263 | 512 | 0 | 0 | 0 | 0 | 0 | 296 | 149 | 1 | 1 | 0.982 | 1 | 0.994 | 1 | 1 | 1 | 1 | 1 | 1 | 1 | -0.005 |
| SE | 94727 | ENSG00000106948 | AKNA | chr9 | - | 114346668 | 114346784 | 114345862 | 114346009 | 114347723 | 114347900 | 567 | 245 | 448 | 463 | 193 | 0 | 3 | 0 | 2 | 0 | 329 | 591 | 331 | 311 | 557 | 0 | 0 | 0 | 1 | 0 | 265 | 149 | 1 | 1 | 1 | 0.979 | 1 | 0.992 | 1 | 1 | 1 | 1 | 0.994 | 1 | -0.005 |
| SE | 94728 | ENSG00000106948 | AKNA | chr9 | - | 114350858 | 114351021 | 114347723 | 114347900 | 114355924 | 114356136 | 620 | 269 | 398 | 516 | 293 | 0 | 0 | 0 | 0 | 0 | 583 | 637 | 424 | 306 | 675 | 0 | 0 | 3 | 0 | 0 | 298 | 149 | 1 | 1 | 1 | 1 | 1 | 1 | 1 | 1 | 1 | 0.986 | 1 | 1 | 0.003 |
| SE | 94731 | ENSG00000106948 | AKNA | chr9 | - | 114357920 | 114358167 | 114355924 | 114356136 | 114359593 | 114359794 | 551 | 200 | 456 | 388 | 445 | 0 | 0 | 0 | 0 | 0 | 274 | 480 | 246 | 219 | 388 | 0 | 0 | 0 | 1 | 0 | 298 | 149 | 1 | 1 | 1 | 1 | 1 | 1 | 1 | 1 | 1 | 1 | 0.991 | 1 | 0.002 |
| SE | 94733 | ENSG00000106948 | AKNA | chr9 | - | 114359895 | 114360062 | 114357920 | 114358167 | 114361703 | 114361911 | 363 | 147 | 238 | 267 | 165 | 0 | 1 | 0 | 0 | 0 | 227 | 285 | 198 | 221 | 342 | 0 | 0 | 0 | 0 | 0 | 298 | 149 | 1 | 1 | 1 | 0.987 | 1 | 1 | 1 | 1 | 1 | 1 | 1 | 1 | -0.003 |
| SE | 94734 | ENSG00000106948 | AKNA | chr9 | - | 114367542 | 114367697 | 114364559 | 114364619 | 114368438 | 114368595 | 519 | 181 | 309 | 345 | 178 | 0 | 0 | 2 | 0 | 0 | 209 | 454 | 229 | 253 | 444 | 0 | 0 | 0 | 0 | 0 | 298 | 149 | 1 | 1 | 1 | 1 | 0.987 | 1 | 1 | 1 | 1 | 1 | 1 | 1 | -0.003 |
| SE | 94735 | ENSG00000106948 | AKNA | chr9 | - | 114368199 | 114368246 | 114367542 | 114367697 | 114368438 | 114368595 | 0 | 1 | 2 | 0 | 0 | 193 | 67 | 112 | 122 | 42 | 0 | 0 | 0 | 0 | 0 | 50 | 127 | 74 | 81 | 125 | 196 | 149 | 1 | 1 | 0 | 0.011 | 0.013 | 0 | 0 | 0 | 0 | 0 | 0 | 0 | 0.005 |
| SE | 94736 | ENSG00000106948 | AKNA | chr9 | - | 114376465 | 114377532 | 114374092 | 114374167 | 114381059 | 114381446 | 1371 | 559 | 992 | 1072 | 830 | 0 | 1 | 1 | 0 | 0 | 1033 | 1409 | 889 | 717 | 1607 | 0 | 0 | 10 | 2 | 0 | 298 | 149 | 1 | 1 | 1 | 0.996 | 0.998 | 1 | 1 | 1 | 1 | 0.978 | 0.994 | 1 | 0.004 |
| SE | 94737 | ENSG00000106948 | AKNA | chr9 | - | 114376465 | 114377532 | 114374092 | 114374167 | 114394356 | 114394405 | 845 | 351 | 561 | 611 | 566 | 0 | 2 | 0 | 0 | 0 | 594 | 754 | 507 | 441 | 985 | 0 | 0 | 0 | 0 | 0 | 298 | 149 | 1 | 1 | 1 | 0.989 | 1 | 1 | 1 | 1 | 1 | 1 | 1 | 1 | -0.002 |
| SE | 94738 | ENSG00000106948 | AKNA | chr9 | - | 114381059 | 114381446 | 114374092 | 114374167 | 114394356 | 114394405 | 83 | 58 | 74 | 86 | 45 | 0 | 2 | 0 | 0 | 0 | 79 | 83 | 118 | 93 | 158 | 0 | 0 | 0 | 0 | 0 | 298 | 149 | 1 | 1 | 1 | 0.935 | 1 | 1 | 1 | 1 | 1 | 1 | 1 | 1 | -0.013 |
| SE | 94741 | ENSG00000119408 | NEK6 | chr9 | + | 124292848 | 124293013 | 124257962 | 124258085 | 124301935 | 124302054 | 4 | 2 | 2 | 0 | 2 | 12 | 0 | 3 | 11 | 1 | 1 | 0 | 1 | 5 | 0 | 7 | 0 | 0 | 4 | 0 | 298 | 149 | 1 | 1 | 0.143 | 1 | 0.25 | 0 | 0.5 | 0.067 | NA | 1 | 0.385 | NA | -0.105 |
| SE | 94742 | ENSG00000119408 | NEK6 | chr9 | + | 124292848 | 124293013 | 124286439 | 124286490 | 124301935 | 124302054 | 4 | 2 | 2 | 0 | 2 | 0 | 0 | 0 | 1 | 0 | 1 | 0 | 1 | 5 | 0 | 0 | 0 | 0 | 0 | 0 | 298 | 149 | 1 | 1 | 1 | 1 | 1 | 0 | 1 | 1 | NA | 1 | 1 | NA | -0.2 |
| SE | 94743 | ENSG00000119408 | NEK6 | chr9 | + | 124292848 | 124293013 | 124291978 | 124292088 | 124301935 | 124302054 | 4 | 2 | 2 | 0 | 2 | 2 | 0 | 0 | 0 | 0 | 19 | 0 | 1 | 5 | 0 | 0 | 0 | 0 | 0 | 0 | 298 | 149 | 1 | 1 | 0.5 | 1 | 1 | NA | 1 | 1 | NA | 1 | 1 | NA | -0.125 |
| SE | 94744 | ENSG00000119408 | NEK6 | chr9 | + | 124301935 | 124302054 | 124257948 | 124258085 | 124312508 | 124312649 | 45 | 12 | 26 | 36 | 12 | 0 | 0 | 0 | 1 | 0 | 179 | 45 | 26 | 65 | 150 | 0 | 0 | 0 | 0 | 0 | 268 | 149 | 1 | 1 | 1 | 1 | 1 | 0.952 | 1 | 1 | 1 | 1 | 1 | 1 | -0.01 |
| SE | 94745 | ENSG00000119408 | NEK6 | chr9 | + | 124327337 | 124327445 | 124326329 | 124326438 | 124339570 | 124339607 | 149 | 60 | 119 | 214 | 156 | 0 | 0 | 0 | 0 | 0 | 345 | 100 | 96 | 122 | 322 | 0 | 0 | 0 | 4 | 0 | 257 | 149 | 1 | 1 | 1 | 1 | 1 | 1 | 1 | 1 | 1 | 1 | 0.946 | 1 | 0.011 |
| SE | 94746 | ENSG00000119408 | NEK6 | chr9 | + | 124339570 | 124339665 | 124327337 | 124327445 | 124347708 | 124347797 | 110 | 51 | 108 | 161 | 163 | 0 | 0 | 3 | 1 | 0 | 295 | 84 | 73 | 113 | 257 | 0 | 0 | 0 | 0 | 2 | 244 | 149 | 1 | 1 | 1 | 1 | 0.956 | 0.99 | 1 | 1 | 1 | 1 | 1 | 0.987 | -0.008 |
| SE | 94749 | ENSG00000107077 | KDM4C | chr9 | + | 6814630 | 6814745 | 6805598 | 6805774 | 6849506 | 6849700 | 513 | 437 | 576 | 575 | 383 | 0 | 0 | 0 | 2 | 2 | 558 | 436 | 552 | 558 | 286 | 0 | 0 | 3 | 0 | 0 | 264 | 149 | 1 | 1 | 1 | 1 | 1 | 0.994 | 0.991 | 1 | 1 | 0.99 | 1 | 1 | -0.001 |
| SE | 94750 | ENSG00000107077 | KDM4C | chr9 | + | 6814630 | 6814745 | 6805598 | 6805774 | 6880011 | 6880061 | 375 | 305 | 376 | 411 | 218 | 0 | 0 | 1 | 0 | 0 | 477 | 325 | 446 | 472 | 233 | 0 | 0 | 0 | 0 | 0 | 264 | 149 | 1 | 1 | 1 | 1 | 0.995 | 1 | 1 | 1 | 1 | 1 | 1 | 1 | -0.001 |
| SE | 94751 | ENSG00000107077 | KDM4C | chr9 | + | 6849506 | 6849700 | 6805598 | 6805774 | 6880011 | 6880061 | 189 | 154 | 165 | 199 | 180 | 0 | 0 | 1 | 0 | 0 | 198 | 105 | 137 | 140 | 110 | 0 | 0 | 0 | 0 | 0 | 298 | 149 | 1 | 1 | 1 | 1 | 0.988 | 1 | 1 | 1 | 1 | 1 | 1 | 1 | -0.002 |
| SE | 94755 | ENSG00000107077 | KDM4C | chr9 | + | 6990415 | 6990524 | 6986343 | 6986666 | 7011697 | 7011879 | 142 | 109 | 180 | 114 | 207 | 0 | 2 | 0 | 0 | 0 | 64 | 50 | 80 | 129 | 54 | 0 | 0 | 0 | 0 | 0 | 258 | 149 | 1 | 1 | 1 | 0.969 | 1 | 1 | 1 | 1 | 1 | 1 | 1 | 1 | -0.006 |
| SE | 94756 | ENSG00000107077 | KDM4C | chr9 | + | 7011697 | 7011879 | 6990415 | 6990524 | 7015852 | 7015929 | 91 | 71 | 111 | 63 | 123 | 2 | 0 | 1 | 0 | 0 | 37 | 33 | 39 | 58 | 40 | 0 | 0 | 0 | 0 | 0 | 298 | 149 | 1 | 1 | 0.958 | 1 | 0.982 | 1 | 1 | 1 | 1 | 1 | 1 | 1 | -0.012 |
| SE | 94757 | ENSG00000107077 | KDM4C | chr9 | + | 7011697 | 7011879 | 6990415 | 6990524 | 7046861 | 7046917 | 92 | 71 | 110 | 64 | 125 | 0 | 0 | 0 | 0 | 1 | 37 | 33 | 39 | 58 | 40 | 0 | 0 | 0 | 0 | 0 | 298 | 149 | 1 | 1 | 1 | 1 | 1 | 1 | 0.984 | 1 | 1 | 1 | 1 | 1 | -0.003 |
| SE | 94758 | ENSG00000107077 | KDM4C | chr9 | + | 7013787 | 7014001 | 7011697 | 7011879 | 7015852 | 7015929 | 288 | 234 | 291 | 287 | 233 | 1 | 0 | 1 | 0 | 0 | 260 | 186 | 178 | 203 | 175 | 0 | 0 | 0 | 0 | 0 | 298 | 149 | 1 | 1 | 0.993 | 1 | 0.993 | 1 | 1 | 1 | 1 | 1 | 1 | 1 | -0.003 |
| SE | 94759 | ENSG00000107077 | KDM4C | chr9 | + | 7013787 | 7014001 | 7011697 | 7011879 | 7046861 | 7046917 | 151 | 109 | 158 | 115 | 109 | 2 | 0 | 0 | 1 | 2 | 107 | 92 | 67 | 84 | 77 | 0 | 0 | 0 | 0 | 0 | 298 | 149 | 1 | 1 | 0.974 | 1 | 1 | 0.983 | 0.965 | 1 | 1 | 1 | 1 | 1 | -0.016 |
| SE | 94760 | ENSG00000107077 | KDM4C | chr9 | + | 7015852 | 7015929 | 6990415 | 6990524 | 7046861 | 7046917 | 99 | 59 | 82 | 76 | 88 | 0 | 0 | 0 | 0 | 1 | 133 | 56 | 48 | 42 | 40 | 0 | 0 | 0 | 0 | 0 | 226 | 149 | 1 | 1 | 1 | 1 | 1 | 1 | 0.983 | 1 | 1 | 1 | 1 | 1 | -0.003 |
| SE | 94761 | ENSG00000107077 | KDM4C | chr9 | + | 7015852 | 7015929 | 7011697 | 7011879 | 7046861 | 7046917 | 98 | 59 | 82 | 76 | 88 | 2 | 0 | 0 | 1 | 2 | 133 | 56 | 48 | 42 | 40 | 0 | 0 | 0 | 0 | 0 | 226 | 149 | 1 | 1 | 0.97 | 1 | 1 | 0.98 | 0.967 | 1 | 1 | 1 | 1 | 1 | -0.017 |
| SE | 94763 | ENSG00000107077 | KDM4C | chr9 | + | 7015852 | 7015929 | 7013787 | 7014001 | 7128065 | 7128236 | 43 | 56 | 45 | 60 | 41 | 0 | 0 | 0 | 0 | 0 | 52 | 55 | 48 | 35 | 27 | 0 | 0 | 0 | 2 | 0 | 226 | 149 | 1 | 1 | 1 | 1 | 1 | 1 | 1 | 1 | 1 | 1 | 0.92 | 1 | 0.016 |
| SE | 94766 | ENSG00000107077 | KDM4C | chr9 | + | 7103684 | 7103870 | 7049091 | 7049200 | 7128065 | 7128236 | 232 | 118 | 212 | 203 | 229 | 0 | 0 | 0 | 0 | 0 | 189 | 166 | 104 | 132 | 86 | 3 | 0 | 2 | 0 | 0 | 298 | 149 | 1 | 1 | 1 | 1 | 1 | 1 | 1 | 0.969 | 1 | 0.963 | 1 | 1 | 0.014 |
| SE | 94768 | ENSG00000107077 | KDM4C | chr9 | + | 7105401 | 7105500 | 7103684 | 7103870 | 7165237 | 7165357 | 4 | 5 | 0 | 2 | 1 | 3 | 1 | 1 | 1 | 9 | 0 | 4 | 2 | 0 | 0 | 0 | 0 | 3 | 1 | 0 | 248 | 149 | 1 | 1 | 0.445 | 0.75 | 0 | 0.546 | 0.063 | NA | 1 | 0.286 | 0 | NA | -0.068 |
| SE | 94769 | ENSG00000107077 | KDM4C | chr9 | + | 7128065 | 7128236 | 6893094 | 6893232 | 7165237 | 7165357 | 153 | 126 | 157 | 180 | 194 | 0 | 0 | 0 | 0 | 0 | 156 | 147 | 76 | 100 | 41 | 0 | 2 | 1 | 0 | 0 | 298 | 149 | 1 | 1 | 1 | 1 | 1 | 1 | 1 | 1 | 0.974 | 0.974 | 1 | 1 | 0.01 |
| SE | 94772 | ENSG00000107077 | KDM4C | chr9 | + | 7128065 | 7128236 | 7105401 | 7105500 | 7165237 | 7165357 | 174 | 135 | 171 | 200 | 209 | 2 | 0 | 0 | 0 | 0 | 155 | 165 | 75 | 108 | 42 | 0 | 0 | 0 | 0 | 0 | 298 | 149 | 1 | 1 | 0.978 | 1 | 1 | 1 | 1 | 1 | 1 | 1 | 1 | 1 | -0.004 |
| SE | 94775 | ENSG00000119314 | PTBP3 | chr9 | - | 112224132 | 112224210 | 112217715 | 112223983 | 112227410 | 112227627 | 768 | 477 | 520 | 608 | 1811 | 1 | 2 | 0 | 3 | 4 | 111 | 450 | 633 | 215 | 53 | 0 | 0 | 4 | 0 | 0 | 227 | 149 | 1 | 1 | 0.998 | 0.994 | 1 | 0.993 | 0.997 | 1 | 1 | 0.99 | 1 | 1 | -0.002 |
| SE | 94776 | ENSG00000119314 | PTBP3 | chr9 | - | 112231379 | 112231413 | 112228379 | 112228472 | 112232098 | 112232238 | 655 | 394 | 501 | 544 | 862 | 1 | 0 | 0 | 1 | 0 | 160 | 548 | 595 | 383 | 166 | 0 | 0 | 0 | 0 | 8 | 183 | 149 | 1 | 1 | 0.998 | 1 | 1 | 0.998 | 1 | 1 | 1 | 1 | 1 | 0.944 | 0.01 |
| SE | 94778 | ENSG00000119314 | PTBP3 | chr9 | - | 112252677 | 112252788 | 112234819 | 112234897 | 112268048 | 112268195 | 14 | 11 | 7 | 6 | 5 | 1 | 0 | 0 | 0 | 0 | 0 | 1 | 1 | 1 | 0 | 0 | 0 | 0 | 0 | 0 | 260 | 149 | 1 | 1 | 0.889 | 1 | 1 | 1 | 1 | NA | 1 | 1 | 1 | NA | -0.022 |
| SE | 94779 | ENSG00000119314 | PTBP3 | chr9 | - | 112262434 | 112262599 | 112252677 | 112252788 | 112268048 | 112268195 | 1158 | 702 | 885 | 863 | 1614 | 3 | 3 | 5 | 2 | 1 | 183 | 793 | 826 | 433 | 380 | 0 | 1 | 4 | 1 | 0 | 298 | 149 | 1 | 1 | 0.995 | 0.992 | 0.989 | 0.995 | 0.999 | 1 | 0.997 | 0.99 | 0.995 | 1 | -0.002 |
| SE | 94780 | ENSG00000119314 | PTBP3 | chr9 | - | 112268048 | 112268195 | 112234819 | 112234897 | 112297831 | 112297916 | 38 | 20 | 20 | 30 | 49 | 0 | 0 | 0 | 0 | 0 | 8 | 19 | 17 | 19 | 2 | 0 | 0 | 0 | 2 | 0 | 296 | 149 | 1 | 1 | 1 | 1 | 1 | 1 | 1 | 1 | 1 | 1 | 0.827 | 1 | 0.035 |
| SE | 94781 | ENSG00000119314 | PTBP3 | chr9 | - | 112268048 | 112268195 | 112262434 | 112262599 | 112275843 | 112276013 | 1236 | 741 | 935 | 976 | 1717 | 0 | 1 | 0 | 1 | 0 | 331 | 902 | 944 | 545 | 512 | 0 | 0 | 1 | 0 | 0 | 296 | 149 | 1 | 1 | 1 | 0.997 | 1 | 0.998 | 1 | 1 | 1 | 0.998 | 1 | 1 | -0.001 |
| SE | 94788 | ENSG00000101464 | PIGU | chr20 | - | 34581547 | 34581672 | 34575103 | 34575246 | 34588452 | 34588607 | 7 | 2 | 15 | 6 | 37 | 0 | 0 | 2 | 0 | 0 | 6 | 7 | 11 | 12 | 24 | 0 | 0 | 0 | 0 | 0 | 274 | 149 | 1 | 1 | 1 | 1 | 0.803 | 1 | 1 | 1 | 1 | 1 | 1 | 1 | -0.039 |
| SE | 94789 | ENSG00000101464 | PIGU | chr20 | - | 34585436 | 34585580 | 34575103 | 34575246 | 34588452 | 34588607 | 24 | 15 | 29 | 16 | 39 | 0 | 0 | 2 | 0 | 0 | 22 | 10 | 18 | 24 | 4 | 0 | 0 | 0 | 0 | 0 | 293 | 149 | 1 | 1 | 1 | 1 | 0.881 | 1 | 1 | 1 | 1 | 1 | 1 | 1 | -0.024 |
| SE | 94790 | ENSG00000101464 | PIGU | chr20 | - | 34585436 | 34585580 | 34581547 | 34581672 | 34588452 | 34588607 | 32 | 17 | 46 | 18 | 57 | 0 | 0 | 0 | 0 | 4 | 38 | 19 | 26 | 29 | 13 | 0 | 0 | 0 | 0 | 0 | 293 | 149 | 1 | 1 | 1 | 1 | 1 | 1 | 0.879 | 1 | 1 | 1 | 1 | 1 | -0.024 |
| SE | 94792 | ENSG00000101464 | PIGU | chr20 | - | 34634614 | 34634715 | 34616041 | 34616139 | 34637875 | 34637985 | 66 | 28 | 56 | 56 | 70 | 0 | 0 | 0 | 0 | 2 | 37 | 24 | 23 | 30 | 30 | 0 | 0 | 2 | 0 | 0 | 250 | 149 | 1 | 1 | 1 | 1 | 1 | 1 | 0.954 | 1 | 1 | 0.873 | 1 | 1 | 0.016 |
| SE | 94793 | ENSG00000101464 | PIGU | chr20 | - | 34645274 | 34645334 | 34643566 | 34644226 | 34657179 | 34657244 | 29 | 18 | 33 | 41 | 42 | 2 | 0 | 0 | 0 | 0 | 75 | 50 | 22 | 21 | 73 | 0 | 0 | 0 | 0 | 0 | 209 | 149 | 1 | 1 | 0.912 | 1 | 1 | 1 | 1 | 1 | 1 | 1 | 1 | 1 | -0.018 |
| SE | 94797 | ENSG00000165275 | TRMT10B | chr9 | + | 37763628 | 37763753 | 37761902 | 37762117 | 37768075 | 37768228 | 7 | 2 | 12 | 10 | 16 | 0 | 0 | 0 | 0 | 0 | 4 | 9 | 0 | 7 | 7 | 0 | 0 | 0 | 0 | 6 | 274 | 149 | 1 | 1 | 1 | 1 | 1 | 1 | 1 | 1 | 1 | NA | 1 | 0.388 | 0.153 |
| SE | 94799 | ENSG00000165275 | TRMT10B | chr9 | + | 37763628 | 37763865 | 37761902 | 37762117 | 37768075 | 37768163 | 10 | 11 | 17 | 14 | 7 | 0 | 0 | 0 | 0 | 0 | 0 | 9 | 10 | 7 | 0 | 0 | 0 | 0 | 0 | 6 | 298 | 149 | 1 | 1 | 1 | 1 | 1 | 1 | 1 | NA | 1 | 1 | 1 | 0 | 0.25 |
| SE | 94800 | ENSG00000165275 | TRMT10B | chr9 | + | 37763628 | 37763865 | 37762576 | 37762685 | 37768075 | 37768163 | 28 | 26 | 34 | 32 | 29 | 0 | 0 | 0 | 5 | 0 | 0 | 18 | 25 | 45 | 16 | 0 | 0 | 0 | 0 | 0 | 298 | 149 | 1 | 1 | 1 | 1 | 1 | 0.762 | 1 | NA | 1 | 1 | 1 | 1 | -0.048 |
| SE | 94801 | ENSG00000165275 | TRMT10B | chr9 | + | 37768075 | 37768228 | 37763628 | 37763753 | 37769940 | 37770019 | 43 | 44 | 33 | 32 | 79 | 0 | 2 | 0 | 0 | 0 | 42 | 26 | 10 | 39 | 18 | 0 | 0 | 0 | 0 | 0 | 298 | 149 | 1 | 1 | 1 | 0.917 | 1 | 1 | 1 | 1 | 1 | 1 | 1 | 1 | -0.017 |
| SE | 94802 | ENSG00000165275 | TRMT10B | chr9 | + | 37768075 | 37768228 | 37763628 | 37763865 | 37770671 | 37770739 | 10 | 14 | 25 | 16 | 10 | 2 | 0 | 0 | 0 | 0 | 0 | 16 | 14 | 13 | 6 | 0 | 0 | 0 | 0 | 0 | 298 | 149 | 1 | 1 | 0.714 | 1 | 1 | 1 | 1 | NA | 1 | 1 | 1 | 1 | -0.057 |
| SE | 94804 | ENSG00000165275 | TRMT10B | chr9 | + | 37776281 | 37776405 | 37770671 | 37770739 | 37777600 | 37778022 | 50 | 50 | 54 | 45 | 78 | 0 | 1 | 0 | 0 | 0 | 44 | 22 | 25 | 50 | 9 | 0 | 0 | 0 | 0 | 0 | 273 | 149 | 1 | 1 | 1 | 0.965 | 1 | 1 | 1 | 1 | 1 | 1 | 1 | 1 | -0.007 |
| SE | 94805 | ENSG00000122696 | SLC25A51 | chr9 | - | 37899828 | 37899950 | 37887603 | 37888592 | 37903530 | 37904110 | 74 | 49 | 62 | 71 | 113 | 0 | 0 | 0 | 0 | 0 | 26 | 28 | 38 | 94 | 24 | 0 | 0 | 1 | 0 | 0 | 271 | 149 | 1 | 1 | 1 | 1 | 1 | 1 | 1 | 1 | 1 | 0.954 | 1 | 1 | 0.009 |
| SE | 94806 | ENSG00000122696 | SLC25A51 | chr9 | - | 37899828 | 37899950 | 37887603 | 37888592 | 37904067 | 37904127 | 90 | 43 | 72 | 63 | 91 | 0 | 0 | 1 | 2 | 0 | 14 | 14 | 47 | 84 | 12 | 0 | 0 | 0 | 0 | 0 | 271 | 149 | 1 | 1 | 1 | 1 | 0.975 | 0.945 | 1 | 1 | 1 | 1 | 1 | 1 | -0.016 |
| SE | 94807 | ENSG00000163527 | STT3B | chr3 | + | 31576395 | 31576504 | 31532637 | 31533312 | 31579808 | 31580096 | 474 | 334 | 458 | 394 | 375 | 0 | 0 | 0 | 2 | 3 | 156 | 249 | 290 | 188 | 226 | 0 | 0 | 0 | 0 | 0 | 258 | 149 | 1 | 1 | 1 | 1 | 1 | 0.991 | 0.986 | 1 | 1 | 1 | 1 | 1 | -0.005 |
| SE | 94808 | ENSG00000163527 | STT3B | chr3 | + | 31576395 | 31576504 | 31532637 | 31533312 | 31596797 | 31596863 | 77 | 36 | 52 | 50 | 1 | 0 | 0 | 1 | 0 | 0 | 20 | 31 | 44 | 16 | 23 | 0 | 0 | 0 | 0 | 0 | 258 | 149 | 1 | 1 | 1 | 1 | 0.968 | 1 | 1 | 1 | 1 | 1 | 1 | 1 | -0.006 |
| SE | 94809 | ENSG00000163527 | STT3B | chr3 | + | 31579808 | 31580096 | 31532637 | 31533312 | 31596797 | 31596863 | 162 | 147 | 211 | 184 | 188 | 0 | 0 | 1 | 0 | 0 | 42 | 74 | 65 | 37 | 28 | 0 | 0 | 0 | 0 | 0 | 298 | 149 | 1 | 1 | 1 | 1 | 0.991 | 1 | 1 | 1 | 1 | 1 | 1 | 1 | -0.002 |
| SE | 94810 | ENSG00000163527 | STT3B | chr3 | + | 31579808 | 31580096 | 31576395 | 31576504 | 31596797 | 31596863 | 561 | 446 | 618 | 529 | 562 | 0 | 0 | 0 | 0 | 1 | 178 | 292 | 313 | 209 | 231 | 0 | 0 | 0 | 0 | 0 | 298 | 149 | 1 | 1 | 1 | 1 | 1 | 1 | 0.996 | 1 | 1 | 1 | 1 | 1 | -0.001 |
| SE | 94812 | ENSG00000163527 | STT3B | chr3 | + | 31619675 | 31619830 | 31617939 | 31617988 | 31622096 | 31622308 | 1014 | 626 | 948 | 853 | 804 | 3 | 0 | 0 | 1 | 0 | 164 | 699 | 800 | 378 | 55 | 0 | 0 | 4 | 0 | 0 | 298 | 149 | 1 | 1 | 0.994 | 1 | 1 | 0.998 | 1 | 1 | 1 | 0.99 | 1 | 1 | 0 |
| SE | 94813 | ENSG00000163527 | STT3B | chr3 | + | 31619675 | 31619830 | 31617939 | 31617988 | 31623673 | 31623861 | 376 | 181 | 378 | 315 | 219 | 1 | 0 | 0 | 0 | 0 | 66 | 240 | 304 | 112 | 29 | 0 | 0 | 0 | 0 | 0 | 298 | 149 | 1 | 1 | 0.995 | 1 | 1 | 1 | 1 | 1 | 1 | 1 | 1 | 1 | -0.001 |
| SE | 94814 | ENSG00000163527 | STT3B | chr3 | + | 31622096 | 31622308 | 31617939 | 31617988 | 31623673 | 31623861 | 420 | 301 | 392 | 365 | 504 | 1 | 0 | 0 | 0 | 0 | 26 | 209 | 253 | 107 | 16 | 0 | 0 | 0 | 0 | 0 | 298 | 149 | 1 | 1 | 0.995 | 1 | 1 | 1 | 1 | 1 | 1 | 1 | 1 | 1 | -0.001 |
| SE | 94815 | ENSG00000163527 | STT3B | chr3 | + | 31622096 | 31622308 | 31619675 | 31619830 | 31623673 | 31623861 | 1055 | 746 | 962 | 902 | 1089 | 0 | 0 | 0 | 0 | 0 | 124 | 668 | 746 | 373 | 42 | 0 | 0 | 1 | 0 | 0 | 298 | 149 | 1 | 1 | 1 | 1 | 1 | 1 | 1 | 1 | 1 | 0.997 | 1 | 1 | 0.001 |
| SE | 94817 | ENSG00000163527 | STT3B | chr3 | + | 31624913 | 31625085 | 31623673 | 31623861 | 31629297 | 31629411 | 497 | 402 | 500 | 481 | 583 | 0 | 0 | 0 | 0 | 0 | 40 | 271 | 265 | 186 | 19 | 0 | 0 | 4 | 0 | 0 | 298 | 149 | 1 | 1 | 1 | 1 | 1 | 1 | 1 | 1 | 1 | 0.971 | 1 | 1 | 0.006 |
| SE | 94818 | ENSG00000163527 | STT3B | chr3 | + | 31625953 | 31626127 | 31623673 | 31623861 | 31629297 | 31629411 | 612 | 465 | 693 | 648 | 1337 | 0 | 0 | 0 | 0 | 0 | 118 | 442 | 539 | 195 | 31 | 0 | 0 | 4 | 0 | 0 | 298 | 149 | 1 | 1 | 1 | 1 | 1 | 1 | 1 | 1 | 1 | 0.985 | 1 | 1 | 0.003 |
| SE | 94819 | ENSG00000163527 | STT3B | chr3 | + | 31625953 | 31626127 | 31624913 | 31625085 | 31629297 | 31629411 | 1066 | 826 | 1154 | 1038 | 1927 | 2 | 0 | 0 | 0 | 0 | 189 | 630 | 821 | 308 | 47 | 0 | 0 | 0 | 0 | 0 | 298 | 149 | 1 | 1 | 0.996 | 1 | 1 | 1 | 1 | 1 | 1 | 1 | 1 | 1 | -0.001 |
| SE | 94820 | ENSG00000163527 | STT3B | chr3 | + | 31629297 | 31629411 | 31625953 | 31626127 | 31632934 | 31633147 | 954 | 747 | 1075 | 1027 | 2070 | 0 | 1 | 1 | 0 | 1 | 182 | 666 | 790 | 293 | 62 | 0 | 0 | 0 | 0 | 0 | 263 | 149 | 1 | 1 | 1 | 0.998 | 0.998 | 1 | 0.999 | 1 | 1 | 1 | 1 | 1 | -0.001 |
| SE | 94821 | ENSG00000163527 | STT3B | chr3 | + | 31632934 | 31633147 | 31629250 | 31629411 | 31635983 | 31636045 | 922 | 751 | 1009 | 925 | 1706 | 0 | 0 | 0 | 3 | 0 | 127 | 526 | 649 | 276 | 73 | 0 | 0 | 2 | 0 | 0 | 298 | 149 | 1 | 1 | 1 | 1 | 1 | 0.994 | 1 | 1 | 1 | 0.994 | 1 | 1 | 0 |
| SE | 94822 | ENSG00000163527 | STT3B | chr3 | + | 31632934 | 31633184 | 31629297 | 31629411 | 31635983 | 31636464 | 594 | 482 | 670 | 618 | 1195 | 0 | 0 | 0 | 3 | 0 | 96 | 387 | 469 | 177 | 41 | 0 | 0 | 2 | 0 | 0 | 298 | 149 | 1 | 1 | 1 | 1 | 1 | 0.99 | 1 | 1 | 1 | 0.992 | 1 | 1 | 0 |

| | | | | | | | | | | | | | | | | | | | | | | | | | | | | | | | | | | | | | | | | | | | | | | |
|---|---|---|---|---|---|---|---|---|---|---|---|---|---|---|---|---|---|---|---|---|---|---|---|---|---|---|---|---|---|---|---|---|---|---|---|---|---|---|---|---|---|---|---|---|---|---|
| SE | 94832 | ENSG00000126882 | FAM78A | chr9 | - | 131262690 | 131262737 | 131258075 | 131261350 | 131263477 | 131264715 | 5 | 1 | 6 | 8 | 1 | 1 | 3 | 12 | 3 | 4 | 0 | 6 | 1 | 15 | 35 | 0 | 6 | 0 | 6 | 0 | 196 | 149 | 1 | 1 | 0.792 | 0.202 | 0.275 | 0.67 | 0.16 | NA | 0.432 | 1 | 0.655 | 1 | -0.352 |
| SE | 94836 | ENSG00000126882 | FAM78A | chr9 | - | 131264583 | 131264627 | 131258075 | 131261350 | 131270459 | 131270523 | 4 | 0 | 2 | 2 | 6 | 2 | 1 | 2 | 3 | 4 | 5 | 0 | 3 | 0 | 9 | 3 | 0 | 0 | 2 | 0 | 193 | 149 | 1 | 1 | 0.607 | 0 | 0.436 | 0.34 | 0.537 | 0.563 | NA | 1 | 0 | 1 | -0.257 |
| SE | 94838 | ENSG00000124145 | SDC4 | chr20 | - | 45330365 | 45330564 | 45325287 | 45327415 | 45333022 | 45333069 | 40 | 29 | 18 | 40 | 50 | 0 | 0 | 0 | 0 | 0 | 212 | 70 | 159 | 112 | 179 | 0 | 0 | 0 | 2 | 0 | 298 | 149 | 1 | 1 | 1 | 1 | 1 | 1 | 1 | 1 | 1 | 1 | 0.966 | 1 | 0.007 |
| SE | 94842 | ENSG00000257524 | NA | chr9 | - | 127878386 | 127878423 | 127874610 | 127874649 | 127885185 | 127885589 | 0 | 0 | 0 | 0 | 3 | 38 | 17 | 35 | 32 | 18 | 0 | 0 | 0 | 0 | 0 | 40 | 41 | 30 | 31 | 29 | 186 | 149 | 1 | 1 | 0 | 0 | 0 | 0 | 0.118 | 0 | 0 | 0 | 0 | 0 | 0.024 |
| SE | 94843 | ENSG00000257524 | NA | chr9 | - | 127878386 | 127878479 | 127874610 | 127874649 | 127879513 | 127879624 | 0 | 0 | 0 | 1 | 0 | 0 | 0 | 5 | 0 | 0 | 0 | 0 | 0 | 0 | 0 | 0 | 0 | 3 | 0 | 0 | 242 | 149 | 1 | 1 | NA | NA | 0 | 1 | NA | NA | NA | 0 | NA | NA | 0.5 |
| SE | 94844 | ENSG00000257524 | NA | chr9 | - | 127879513 | 127879624 | 127874610 | 127874649 | 127885185 | 127885542 | 0 | 0 | 6 | 0 | 0 | 38 | 17 | 35 | 32 | 18 | 0 | 0 | 6 | 0 | 0 | 40 | 41 | 30 | 31 | 29 | 260 | 149 | 1 | 1 | 0 | 0 | 0.089 | 0 | 0 | 0 | 0 | 0.103 | 0 | 0 | -0.003 |
| SE | 94845 | ENSG00000257524 | NA | chr9 | - | 127879513 | 127879624 | 127878386 | 127878423 | 127885185 | 127885542 | 0 | 0 | 1 | 0 | 0 | 0 | 0 | 0 | 0 | 3 | 0 | 0 | 3 | 0 | 0 | 0 | 0 | 0 | 0 | 0 | 260 | 149 | 1 | 1 | NA | NA | 1 | NA | 0 | NA | NA | 1 | NA | NA | -0.5 |
| SE | 94846 | ENSG00000257524 | NA | chr9 | - | 127886635 | 127886788 | 127885185 | 127885343 | 127887483 | 127887591 | 53 | 4 | 46 | 41 | 16 | 0 | 0 | 0 | 0 | 0 | 0 | 32 | 27 | 17 | 52 | 0 | 0 | 0 | 5 | 0 | 298 | 149 | 1 | 1 | 1 | 1 | 1 | 1 | 1 | NA | 1 | 1 | 0.63 | 1 | 0.093 |
| SE | 94852 | ENSG00000136840 | ST6GALNAC4 | chr9 | - | 127916407 | 127916494 | 127914655 | 127914841 | 127916825 | 127917018 | 8 | 2 | 10 | 4 | 0 | 0 | 0 | 2 | 3 | 0 | 0 | 0 | 2 | 3 | 22 | 0 | 0 | 0 | 0 | 6 | 236 | 149 | 1 | 1 | 1 | 1 | 0.759 | 0.457 | NA | NA | NA | 1 | 1 | 0.698 | -0.095 |
| SE | 94854 | ENSG00000167106 | FAM102A | chr9 | - | 127945366 | 127945562 | 127944776 | 127944898 | 127945648 | 127945753 | 215 | 91 | 303 | 211 | 101 | 0 | 0 | 0 | 0 | 0 | 8 | 128 | 121 | 24 | 63 | 0 | 2 | 0 | 0 | 0 | 298 | 149 | 1 | 1 | 1 | 1 | 1 | 1 | 1 | 1 | 0.97 | 1 | 1 | 1 | 0.006 |
| SE | 94855 | ENSG00000167106 | FAM102A | chr9 | - | 127948070 | 127948225 | 127945648 | 127945753 | 127948367 | 127948414 | 323 | 167 | 623 | 321 | 464 | 0 | 0 | 1 | 0 | 0 | 31 | 315 | 160 | 116 | 204 | 0 | 0 | 0 | 0 | 0 | 298 | 149 | 1 | 1 | 1 | 1 | 0.997 | 1 | 1 | 1 | | 1 | 1 | 1 | -0.001 |
| SE | 94856 | ENSG00000167106 | FAM102A | chr9 | - | 127948367 | 127948414 | 127945648 | 127945753 | 127950422 | 127950539 | 29 | 20 | 53 | 28 | 63 | 0 | 0 | 0 | 0 | 0 | 0 | 12 | 15 | 1 | 36 | 0 | 0 | 2 | 0 | 6 | 196 | 149 | 1 | 1 | 1 | 1 | 1 | 1 | 1 | NA | 1 | 0.851 | 1 | 0.82 | 0.082 |
| SE | 94857 | ENSG00000167106 | FAM102A | chr9 | - | 127948367 | 127948414 | 127945648 | 127945753 | 127953574 | 127953604 | 0 | 0 | 1 | 0 | 3 | 1 | 0 | 2 | 1 | 0 | 0 | 4 | 0 | 0 | 0 | 0 | 0 | 0 | 0 | 0 | 196 | 149 | 1 | 1 | 0 | NA | 0.275 | 0 | 1 | NA | 1 | NA | NA | NA | -0.681 |
| SE | 94860 | ENSG00000285283 | NA | chr11 | + | 32100547 | 32100608 | 32098349 | 32098528 | 32103280 | 32103480 | 163 | 111 | 99 | 174 | 165 | 0 | 0 | 0 | 0 | 0 | 390 | 86 | 88 | 129 | 117 | 0 | 0 | 0 | 1 | 0 | 210 | 149 | 1 | 1 | 1 | 1 | 1 | 1 | 1 | 1 | 1 | | 0.989 | 1 | 0.002 |
| SE | 94862 | ENSG00000196597 | ZNF782 | chr9 | - | 96827079 | 96827181 | 96816268 | 96819778 | 96844889 | 96845016 | 25 | 39 | 23 | 43 | 57 | 0 | 0 | 0 | 0 | 0 | 28 | 24 | 35 | 0 | 19 | 0 | 5 | 1 | 0 | 0 | 251 | 149 | 1 | 1 | 1 | 1 | 1 | 1 | 1 | 1 | 0.74 | 0.954 | NA | 1 | 0.077 |
| SE | 94863 | ENSG00000196597 | ZNF782 | chr9 | - | 96827079 | 96827181 | 96816268 | 96819778 | 96851946 | 96852005 | 15 | 36 | 11 | 26 | 39 | 0 | 0 | 1 | 0 | 0 | 15 | 13 | 27 | 0 | 12 | 0 | 0 | 0 | 0 | 0 | 251 | 149 | 1 | 1 | 1 | 1 | 0.867 | 1 | 1 | 1 | 1 | 1 | NA | 1 | -0.027 |
| SE | 94864 | ENSG00000196597 | ZNF782 | chr9 | - | 96844889 | 96845016 | 96816268 | 96819778 | 96851946 | 96852005 | 14 | 14 | 11 | 14 | 15 | 0 | 0 | 1 | 0 | 0 | 13 | 17 | 24 | 7 | 9 | 0 | 0 | 0 | 0 | 0 | 276 | 149 | 1 | 1 | 1 | 1 | 0.856 | 1 | 1 | 1 | 1 | 1 | 1 | 1 | -0.029 |
| SE | 94866 | ENSG00000196597 | ZNF782 | chr9 | - | 96844889 | 96845016 | 96827079 | 96827181 | 96854087 | 96854360 | 15 | 13 | 14 | 18 | 20 | 0 | 0 | 0 | 1 | 0 | 13 | 18 | 8 | 0 | 7 | 0 | 0 | 0 | 0 | 0 | 276 | 149 | 1 | 1 | 1 | 1 | 1 | 0.907 | 1 | 1 | 1 | 1 | NA | 1 | -0.019 |
| SE | 94867 | ENSG00000196597 | ZNF782 | chr9 | - | 96851946 | 96852005 | 96827079 | 96827181 | 96854087 | 96854183 | 4 | 7 | 3 | 1 | 6 | 0 | 0 | 0 | 1 | 0 | 0 | 0 | 1 | 0 | 0 | 0 | 0 | 0 | 0 | 0 | 208 | 149 | 1 | 1 | 1 | 1 | 1 | 0.417 | 1 | NA | NA | 1 | NA | NA | -0.117 |
| SE | 94868 | ENSG00000196597 | ZNF782 | chr9 | - | 96851946 | 96852005 | 96844889 | 96845016 | 96854087 | 96854183 | 7 | 9 | 10 | 9 | 12 | 0 | 1 | 0 | 1 | 0 | 5 | 10 | 8 | 0 | 7 | 0 | 0 | 0 | 0 | 0 | 208 | 149 | 1 | 1 | 1 | 0.866 | 1 | 0.866 | 1 | 1 | 1 | 1 | NA | 1 | -0.054 |
| SE | 94870 | ENSG00000254093 | PINX1 | chr8 | - | 10787623 | 10787825 | 10765854 | 10765916 | 10826151 | 10826244 | 2 | 0 | 0 | 0 | 4 | 1 | 3 | 7 | 5 | 12 | 0 | 0 | 12 | 2 | 0 | 0 | 0 | 0 | 1 | 0 | 298 | 149 | 1 | 1 | 0.5 | 0 | 0 | 0 | 0.143 | NA | NA | 1 | 0.5 | NA | -0.621 |
| SE | 94871 | ENSG00000254093 | PINX1 | chr8 | - | 10820192 | 10820269 | 10764960 | 10765916 | 10826151 | 10826244 | 7 | 2 | 5 | 16 | 12 | 1 | 3 | 7 | 5 | 12 | 0 | 7 | 4 | 11 | 0 | 0 | 0 | 0 | 1 | 0 | 226 | 149 | 1 | 1 | 0.822 | 0.305 | 0.32 | 0.678 | 0.397 | NA | 1 | 1 | 0.879 | NA | -0.455 |
| SE | 94872 | ENSG00000254093 | PINX1 | chr8 | - | 10820192 | 10820310 | 10787521 | 10787825 | 10826151 | 10826169 | 0 | 0 | 0 | 1 | 0 | 2 | 0 | 0 | 0 | 4 | 0 | 0 | 1 | 0 | 0 | 0 | 0 | 12 | 2 | 0 | 267 | 149 | 1 | 1 | 0 | NA | NA | 1 | 0 | NA | NA | 0.044 | 0 | NA | 0.311 |
| SE | 94874 | ENSG00000136144 | RCBTB1 | chr13 | - | 49537869 | 49538083 | 49533917 | 49534262 | 49540875 | 49541006 | 6 | 3 | 2 | 8 | 6 | 30 | 12 | 20 | 25 | 35 | 0 | 5 | 2 | 0 | 0 | 45 | 18 | 15 | 13 | 0 | 298 | 149 | 1 | 1 | 0.091 | 0.111 | 0.048 | 0.138 | 0.079 | 0 | 0.122 | 0.062 | 0 | NA | 0.047 |
| SE | 94875 | ENSG00000136144 | RCBTB1 | chr13 | - | 49551325 | 49551468 | 49549457 | 49549648 | 49552177 | 49552285 | 70 | 66 | 83 | 81 | 73 | 0 | 0 | 0 | 2 | 0 | 67 | 112 | 64 | 59 | 44 | 0 | 0 | 0 | 0 | 0 | 292 | 149 | 1 | 1 | 1 | 1 | 1 | 0.954 | 1 | 1 | | 1 | 1 | 1 | -0.009 |
| SE | 94877 | ENSG00000188283 | ZNF383 | chr19 | + | 37219598 | 37219749 | 37217925 | 37218274 | 37224817 | 37224879 | 0 | 0 | 1 | 2 | 4 | 0 | 0 | 0 | 3 | 10 | 0 | 9 | 1 | 13 | 0 | 0 | 0 | 0 | 0 | 0 | 298 | 149 | 1 | 1 | NA | NA | 1 | 0.25 | 0.167 | NA | 1 | 1 | 1 | NA | -0.528 |
| SE | 94878 | ENSG00000188283 | ZNF383 | chr19 | + | 37219598 | 37219749 | 37217925 | 37218274 | 37230408 | 37230462 | 0 | 0 | 2 | 0 | 0 | 2 | 5 | 1 | 2 | 2 | 0 | 0 | 5 | 5 | 0 | 0 | 0 | 0 | 0 | 0 | 298 | 149 | 1 | 1 | 0 | 0 | 0.5 | 0 | 0 | NA | NA | 1 | 1 | NA | -0.9 |
| SE | 94879 | ENSG00000188283 | ZNF383 | chr19 | + | 37224817 | 37224939 | 37217925 | 37218274 | 37230408 | 37230462 | 3 | 0 | 7 | 7 | 12 | 2 | 5 | 1 | 2 | 2 | 9 | 14 | 3 | 5 | 0 | 0 | 0 | 0 | 0 | 0 | 271 | 149 | 1 | 1 | 0.452 | 0 | 0.794 | 0.658 | 0.767 | 1 | 1 | 1 | 1 | NA | -0.466 |
| SE | 94880 | ENSG00000188283 | ZNF383 | chr19 | + | 37224817 | 37224939 | 37219598 | 37219749 | 37230408 | 37230462 | 3 | 0 | 8 | 6 | 6 | 0 | 0 | 4 | 0 | 0 | 9 | 14 | 3 | 8 | 0 | 0 | 0 | 4 | 0 | 0 | 271 | 149 | 1 | 1 | 1 | NA | 0.524 | 1 | 1 | 1 | 1 | 0.292 | 1 | NA | 0.058 |
| SE | 94881 | ENSG00000188283 | ZNF383 | chr19 | + | 37235978 | 37236074 | 37235548 | 37235675 | 37242468 | 37242579 | 71 | 61 | 66 | 102 | 76 | 2 | 2 | 0 | 0 | 0 | 9 | 47 | 66 | 25 | 14 | 0 | 0 | 0 | 0 | 0 | 245 | 149 | 1 | 1 | 0.956 | 0.949 | 1 | 1 | 1 | 1 | 1 | 1 | 1 | 1 | -0.019 |
| SE | 94889 | ENSG00000119446 | RBM18 | chr9 | - | 122254273 | 122254367 | 122251846 | 122251973 | 122261379 | 122261508 | 9 | 5 | 5 | 10 | 3 | 24 | 14 | 18 | 26 | 29 | 0 | 7 | 8 | 2 | 0 | 0 | 14 | 19 | 10 | 4 | 243 | 149 | 1 | 1 | 0.187 | 0.18 | 0.146 | 0.191 | 0.06 | NA | 0.235 | 0.205 | 0.109 | 0 | 0.016 |
| SE | 94917 | ENSG00000095397 | WHRN | chr9 | - | 114466266 | 114466392 | 114424987 | 114426413 | 114478552 | 114478572 | 12 | 0 | 6 | 3 | 0 | 0 | 0 | 0 | 0 | 0 | 0 | 0 | 2 | 1 | 20 | 0 | 0 | 0 | 1 | 0 | 275 | 149 | 1 | 1 | 1 | NA | 1 | 1 | NA | NA | NA | 1 | 0.351 | 1 | 0.216 |
| SE | 94918 | ENSG00000157693 | TMEM268 | chr9 | + | 114626895 | 114627006 | 114624349 | 114624459 | 114628100 | 114628250 | 16 | 19 | 16 | 18 | 6 | 0 | 1 | 0 | 0 | 0 | 45 | 25 | 12 | 29 | 14 | 4 | 0 | 0 | 0 | 0 | 260 | 149 | 1 | 1 | 1 | 0.916 | 1 | 1 | 1 | 0.866 | 1 | 1 | 1 | 1 | 0.01 |
| SE | 94919 | ENSG00000157693 | TMEM268 | chr9 | + | 114626898 | 114627006 | 114624135 | 114624459 | 114628100 | 114628225 | 48 | 34 | 33 | 43 | 19 | 0 | 1 | 0 | 0 | 0 | 102 | 30 | 28 | 55 | 35 | 4 | 0 | 0 | 0 | 0 | 257 | 149 | 1 | 1 | 1 | 0.952 | 1 | 1 | 1 | 0.937 | | 1 | 1 | 1 | 0.003 |
| SE | 94920 | ENSG00000157693 | TMEM268 | chr9 | + | 114633767 | 114633878 | 114617117 | 114617301 | 114636989 | 114637070 | 27 | 17 | 36 | 36 | 23 | 0 | 0 | 1 | 0 | 0 | 22 | 33 | 38 | 49 | 46 | 0 | 2 | 0 | 0 | 0 | 260 | 149 | 1 | 1 | 1 | 1 | 0.954 | 1 | 1 | 1 | 0.904 | 1 | 1 | 1 | 0.01 |
| SE | 94923 | ENSG00000148655 | LRMDA | chr10 | + | 76058665 | 76058783 | 76047163 | 76047303 | 76324400 | 76324485 | 29 | 30 | 22 | 20 | 26 | 1 | 0 | 0 | 0 | 0 | 105 | 50 | 31 | 105 | 130 | 0 | 3 | 0 | 0 | 0 | 267 | 149 | 1 | 1 | 0.942 | 1 | 1 | 1 | 1 | 1 | 0.903 | 1 | 1 | 1 | 0.008 |
| SE | 94926 | ENSG00000148655 | LRMDA | chr10 | + | 76535797 | 76535987 | 76324400 | 76324485 | 76557208 | 76557366 | 0 | 1 | 0 | 0 | 3 | 3 | 14 | 5 | 7 | 19 | 0 | 0 | 0 | 0 | 4 | 35 | 7 | 13 | 6 | 41 | 298 | 149 | 1 | 1 | 0 | 0.034 | 0 | 0 | 0.073 | 0 | 0 | 0 | 0 | 0.047 | 0.012 |
| SE | 94927 | ENSG00000136888 | ATP6V1G1 | chr9 | + | 114592551 | 114592652 | 114587768 | 114587920 | 114597569 | 114598496 | 339 | 219 | 306 | 278 | 321 | 0 | 0 | 1 | 0 | 0 | 265 | 273 | 202 | 352 | 224 | 0 | 1 | 0 | 0 | 0 | 250 | 149 | 1 | 1 | 1 | 1 | 0.995 | 1 | 1 | 1 | 0.994 | 1 | 1 | 1 | 0 |
| SE | 94928 | ENSG00000136888 | ATP6V1G1 | chr9 | + | 114592551 | 114592657 | 114587705 | 114587920 | 114597569 | 114598384 | 117 | 49 | 98 | 72 | 55 | 0 | 0 | 1 | 0 | 0 | 82 | 99 | 65 | 129 | 77 | 0 | 1 | 0 | 0 | 0 | 255 | 149 | 1 | 1 | 1 | 1 | 0.983 | 1 | 1 | 1 | 0.983 | 1 | 1 | 1 | 0 |
| SE | 94929 | ENSG00000136888 | ATP6V1G1 | chr9 | + | 114596506 | 114596588 | 114592551 | 114592652 | 114597569 | 114597743 | 0 | 0 | 0 | 0 | 0 | 224 | 170 | 208 | 209 | 268 | 0 | 2 | 0 | 0 | 0 | 190 | 174 | 137 | 225 | 147 | 231 | 149 | 1 | 1 | 0 | 0 | 0 | 0 | 0 | 0 | 0.007 | 0 | 0 | 0 | -0.001 |
| SE | 94932 | ENSG00000119403 | PHF19 | chr9 | - | 120869181 | 120869330 | 120866869 | 120866965 | 120869844 | 120869945 | 34 | 9 | 31 | 34 | 22 | 2 | 0 | 0 | 0 | 0 | 0 | 19 | 11 | 13 | 29 | 0 | 0 | 0 | 0 | 0 | 298 | 149 | 1 | 1 | 0.895 | 1 | 1 | 1 | 1 | NA | 1 | 1 | 1 | 1 | -0.021 |
| SE | 94935 | ENSG00000148180 | GSN | chr9 | + | 121281469 | 121281562 | 121265660 | 121265754 | 121301962 | 121302167 | 13 | 2 | 5 | 19 | 7 | 1 | 0 | 0 | 0 | 1 | 25 | 24 | 19 | 43 | 151 | 0 | 0 | 0 | 0 | 0 | 242 | 149 | 1 | 1 | 0.889 | 1 | | 1 | 0.812 | 1 | 1 | 1 | 1 | 1 | -0.06 |
| SE | 94939 | ENSG00000148180 | GSN | chr9 | + | 121297765 | 121297842 | 121283318 | 121283392 | 121301962 | 121302149 | 0 | 0 | 0 | 1 | 0 | 0 | 0 | 0 | 0 | 0 | 0 | 0 | 0 | 0 | 0 | 0 | 0 | 0 | 8 | 0 | 226 | 149 | 1 | 1 | NA | NA | NA | 1 | NA | NA | NA | NA | 0 | NA | 1 |
| SE | 94941 | ENSG00000148180 | GSN | chr9 | + | 121300055 | 121300126 | 121286122 | 121286185 | 121301962 | 121302149 | 0 | 0 | 0 | 0 | 0 | 0 | 1 | 0 | 1 | 3 | 0 | 7 | 0 | 0 | 7 | 0 | 0 | 1 | 0 | 24 | 220 | 149 | 1 | 1 | NA | 0 | NA | 0 | 0 | NA | 1 | 0 | NA | 0.165 | -0.388 |
| SE | 94942 | ENSG00000148180 | GSN | chr9 | + | 121300055 | 121300126 | 121297765 | 121297842 | 121301962 | 121302149 | 0 | 0 | 0 | 0 | 0 | 0 | 0 | 0 | 1 | 0 | 0 | 7 | 0 | 0 | 7 | 0 | 0 | 0 | 0 | 0 | 220 | 149 | 1 | 1 | NA | NA | NA | 0 | NA | NA | 1 | NA | NA | 1 | -1 |
| SE | 94943 | ENSG00000148180 | GSN | chr9 | + | 121312338 | 121312488 | 121310683 | 121310845 | 121313933 | 121314023 | 172 | 107 | 155 | 240 | 137 | 0 | 0 | 1 | 2 | 0 | 503 | 337 | 151 | 656 | 1363 | 1 | 2 | 1 | 3 | 0 | 298 | 149 | 1 | 1 | 1 | 1 | 0.987 | 0.984 | 1 | 0.996 | 0.988 | 0.987 | 0.991 | 1 | 0.002 |
| SE | 94944 | ENSG00000148180 | GSN | chr9 | + | 121321267 | 121321401 | 121318664 | 121318880 | 121324553 | 121324644 | 172 | 102 | 158 | 216 | 123 | 0 | 0 | 0 | 0 | 0 | 361 | 323 | 160 | 563 | 1288 | 0 | 0 | 3 | 0 | 0 | 283 | 149 | 1 | 1 | 1 | 1 | 1 | 1 | 1 | 1 | 1 | 0.966 | 1 | 1 | 0.007 |
| SE | 94945 | ENSG00000148180 | GSN | chr9 | + | 121327307 | 121327482 | 121326511 | 121326682 | 121328890 | 121329015 | 204 | 80 | 110 | 208 | 107 | 0 | 0 | 0 | 0 | 0 | 258 | 195 | 97 | 488 | 976 | 0 | 0 | 0 | 0 | 6 | 298 | 149 | 1 | 1 | 1 | 1 | 1 | 1 | 1 | 1 | 1 | 1 | 1 | 0.988 | 0.002 |
| SE | 94946 | ENSG00000148180 | GSN | chr9 | + | 121331387 | 121331448 | 121328852 | 121329315 | 121332433 | 121332589 | 226 | 140 | 191 | 227 | 257 | 0 | 0 | 0 | 0 | 0 | 531 | 302 | 166 | 836 | 1517 | 0 | 0 | 0 | 2 | 0 | 210 | 149 | 1 | 1 | 1 | 1 | 1 | 1 | 1 | 1 | 1 | 1 | 0.997 | 1 | 0.001 |
| SE | 94948 | ENSG00000198242 | RPL23A | chr17 | + | 28722722 | 28722899 | 28720568 | 28720890 | 28723570 | 28723640 | 11059 | 5903 | 11578 | 10068 | 14278 | 0 | 0 | 0 | 0 | 0 | 5187 | 6330 | 5292 | 4488 | 4687 | 0 | 0 | 1 | 0 | 0 | 298 | 149 | 1 | 1 | 1 | 1 | 1 | 1 | 1 | 1 | 1 | 1 | 1 | 1 | 0 |
| SE | 94958 | ENSG00000266714 | MYO15B | chr17 | + | 75590625 | 75590689 | 75590340 | 75590385 | 75590906 | 75591016 | 1 | 1 | 5 | 6 | 2 | 1 | 0 | 0 | 0 | 0 | 0 | 0 | 0 | 5 | 5 | 0 | 0 | 0 | 0 | 0 | 213 | 149 | 1 | 1 | 0.412 | 1 | 1 | | 1 | NA | NA | NA | 1 | 1 | -0.118 |
| SE | 94959 | ENSG00000266714 | MYO15B | chr17 | + | 75590625 | 75590689 | 75590340 | 75590385 | 75591171 | 75591246 | 1 | 4 | 6 | 9 | 4 | 2 | 0 | 0 | 1 | 0 | 19 | 9 | 0 | 1 | 0 | 0 | 0 | 0 | 0 | 0 | 213 | 149 | 1 | 1 | 0.259 | 1 | 1 | 0.863 | 1 | 1 | 1 | NA | 1 | NA | -0.176 |
| SE | 94960 | ENSG00000266714 | MYO15B | chr17 | + | 75590906 | 75591016 | 75588057 | 75590385 | 75591171 | 75591246 | 2 | 1 | 2 | 0 | 0 | 2 | 0 | 0 | 1 | 0 | 0 | 2 | 1 | 6 | 10 | 0 | 0 | 0 | 0 | 0 | 259 | 149 | 1 | 1 | 0.365 | 1 | 1 | 0 | NA | NA | 1 | 1 | 1 | 1 | -0.409 |
| SE | 94965 | ENSG00000266714 | MYO15B | chr17 | + | 75594474 | 75594588 | 75592678 | 75592840 | 75594700 | 75594759 | 10 | 8 | 16 | 28 | 11 | 0 | 2 | 0 | 0 | 0 | 22 | 28 | 8 | 51 | 33 | 0 | 0 | 0 | 0 | 0 | 263 | 149 | 1 | 1 | 1 | 0.694 | 1 | 1 | 1 | 1 | 1 | 1 | 1 | 1 | -0.061 |
| SE | 94967 | ENSG00000266714 | MYO15B | chr17 | + | 75594700 | 75594759 | 75592678 | 75592840 | 75594839 | 75594972 | 0 | 1 | 2 | 1 | 0 | 1 | 0 | 0 | 0 | 2 | 0 | 13 | 5 | 10 | 18 | 0 | 0 | 9 | 15 | 0 | 208 | 149 | 1 | 1 | 0 | 1 | 1 | 1 | 0 | NA | 1 | 0.285 | 0.323 | 1 | -0.052 |
| SE | 94968 | ENSG00000266714 | MYO15B | chr17 | + | 75594700 | 75594759 | 75594474 | 75594588 | 75594839 | 75594972 | 3 | 5 | 11 | 10 | 4 | 0 | 0 | 0 | 0 | 0 | 0 | 29 | 5 | 19 | 41 | 0 | 0 | 0 | 3 | 0 | 208 | 149 | 1 | 1 | 1 | 1 | 1 | 1 | 1 | NA | 1 | 1 | 0.819 | 1 | 0.045 |
| SE | 94969 | ENSG00000266714 | MYO15B | chr17 | + | 75594839 | 75594972 | 75592678 | 75592840 | 75596459 | 75596555 | 4 | 5 | 4 | 9 | 2 | 0 | 0 | 0 | 0 | 0 | 0 | 7 | 10 | 25 | 0 | 0 | 0 | 0 | 2 | 0 | 282 | 149 | 1 | 1 | 1 | 1 | 1 | 1 | 1 | NA | 1 | 1 | 0.869 | NA | 0.044 |
| SE | 94971 | ENSG00000266714 | MYO15B | chr17 | + | 75600630 | 75600680 | 75596767 | 75596899 | 75601437 | 75601563 | 0 | 0 | 0 | 2 | 0 | 28 | 15 | 24 | 19 | 20 | 0 | 0 | 0 | 0 | 0 | 13 | 35 | 9 | 33 | 68 | 199 | 149 | 1 | 1 | 0 | 0 | 0 | 0.073 | 0 | 0 | 0 | 0 | 0 | 0 | 0.015 |
| SE | 94972 | ENSG00000266714 | MYO15B | chr17 | + | 75601437 | 75601563 | 75596767 | 75596899 | 75602516 | 75602594 | 40 | 26 | 36 | 34 | 29 | 0 | 1 | 0 | 2 | 0 | 39 | 57 | 29 | 67 | 77 | 0 | 0 | 0 | 2 | 0 | 275 | 149 | 1 | 1 | 1 | 0.934 | 1 | 0.902 | 1 | 1 | 1 | 1 | 0.948 | 1 | -0.022 |
| SE | 94973 | ENSG00000266714 | MYO15B | chr17 | + | 75603187 | 75603312 | 75603031 | 75603077 | 75605503 | 75605621 | 16 | 12 | 15 | 14 | 2 | 1 | 0 | 0 | 0 | 0 | 6 | 16 | 13 | 38 | 35 | 0 | 0 | 0 | 0 | 0 | 274 | 149 | 1 | 1 | 0.897 | 1 | 1 | 1 | 1 | 1 | 1 | 1 | 1 | 1 | -0.021 |
| SE | 94974 | ENSG00000266714 | MYO15B | chr17 | + | 75603187 | 75603312 | 75603031 | 75603077 | 75610165 | 75610259 | 9 | 8 | 12 | 11 | 1 | 1 | 0 | 0 | 0 | 0 | 6 | 16 | 13 | 21 | 17 | 0 | 1 | 0 | 0 | 0 | 274 | 149 | 1 | 1 | 0.83 | 1 | 1 | 1 | 1 | 1 | 0.897 | 1 | 1 | 1 | -0.013 |
| SE | 94976 | ENSG00000266714 | MYO15B | chr17 | + | 75605863 | 75606021 | 75603187 | 75603312 | 75610165 | 75610259 | 2 | 2 | 3 | 0 | 3 | 0 | 0 | 2 | 0 | 0 | 12 | 4 | 4 | 12 | 3 | 0 | 0 | 0 | 2 | 0 | 298 | 149 | 1 | 1 | 1 | 1 | 0.429 | NA | 1 | | 1 | 1 | 0.75 | 1 | -0.093 |
| SE | 94979 | ENSG00000266714 | MYO15B | chr17 | + | 75613980 | 75614033 | 75613704 | 75613777 | 75614198 | 75614360 | 4 | 2 | 4 | 6 | 0 | 2 | 4 | 1 | 1 | 0 | 13 | 0 | 1 | 5 | 30 | 5 | 0 | 5 | 12 | 9 | 202 | 149 | 1 | 1 | 0.596 | 0.269 | 0.747 | 0.816 | NA | 0.657 | NA | 0.129 | 0.235 | 0.711 | 0.174 |
| SE | 94980 | ENSG00000266714 | MYO15B | chr17 | + | 75616068 | 75616147 | 75615692 | 75615884 | 75616523 | 75616785 | 6 | 4 | 3 | 3 | 0 | 0 | 0 | 0 | 0 | 0 | 0 | 7 | 3 | 20 | 1 | 0 | 0 | 1 | 0 | 0 | 228 | 149 | 1 | 1 | 1 | 1 | 1 | 1 | NA | NA | 1 | 0.662 | 1 | 1 | 0.085 |
| SE | 94984 | ENSG00000266714 | MYO15B | chr17 | + | 75623780 | 75623854 | 75621881 | 75622080 | 75623948 | 75624058 | 39 | 39 | 37 | 42 | 22 | 0 | 0 | 1 | 0 | 0 | 59 | 42 | 35 | 95 | 91 | 0 | 0 | 0 | 0 | 0 | 223 | 149 | 1 | 1 | 1 | 1 | 0.961 | 1 | 1 | 1 | 1 | 1 | 1 | 1 | -0.008 |
| SE | 94996 | ENSG00000096996 | IL12RB1 | chr19 | - | 18063875 | 18064010 | 18062180 | 18062277 | 18066541 | 18066697 | 45 | 18 | 44 | 43 | 27 | 0 | 0 | 0 | 0 | 0 | 30 | 61 | 16 | 26 | 53 | 0 | 0 | 0 | 3 | 0 | 284 | 149 | 1 | 1 | 1 | 1 | 1 | 1 | 1 | 1 | 1 | 1 | 0.82 | 1 | 0.036 |
| SE | 94997 | ENSG00000096996 | IL12RB1 | chr19 | - | 18072111 | 18072349 | 18069545 | 18069713 | 18073516 | 18073599 | 83 | 33 | 64 | 69 | 35 | 0 | 0 | 0 | 0 | 0 | 47 | 78 | 44 | 34 | 96 | 0 | 0 | 0 | 2 | 0 | 298 | 149 | 1 | 1 | 1 | 1 | 1 | 1 | 1 | 1 | 1 | 1 | 0.895 | 1 | 0.021 |
| SE | 94999 | ENSG00000096996 | IL12RB1 | chr19 | - | 18073516 | 18073599 | 18069545 | 18069713 | 18075748 | 18075868 | 45 | 13 | 36 | 36 | 27 | 1 | 0 | 1 | 0 | 0 | 37 | 48 | 12 | 6 | 24 | 0 | 0 | 0 | 0 | 0 | 232 | 149 | 1 | 1 | 0.967 | 1 | 0.959 | 1 | 1 | 1 | 1 | 1 | 1 | 1 | -0.015 |
| SE | 95008 | ENSG00000135047 | CTSL | chr9 | + | 87727360 | 87727468 | 87726121 | 87726253 | 87727593 | 87727729 | 0 | 0 | 0 | 0 | 0 | 10 | 10 | 11 | 29 | 9 | 0 | 1 | 0 | 0 | 0 | 23 | 5 | 18 | 19 | 41 | 257 | 149 | 1 | 1 | 0 | 0 | 0 | 0 | 0 | 0 | 0.104 | 0 | 0 | 0 | -0.021 |
| SE | 95009 | ENSG00000135047 | CTSL | chr9 | + | 87727360 | 87727468 | 87726121 | 87726308 | 87727593 | 87727729 | 0 | 0 | 0 | 0 | 0 | 4 | 4 | 0 | 6 | 1 | 0 | 1 | 0 | 0 | 0 | 12 | 5 | 3 | 12 | 0 | 257 | 149 | 1 | 1 | 0 | 0 | NA | 0 | 0 | 0 | 0.104 | 0 | 0 | NA | -0.026 |
| SE | 95012 | ENSG00000135047 | CTSL | chr9 | + | 87728249 | 87728396 | 87726838 | 87728149 | 87729572 | 87729735 | 16 | 35 | 5 | 14 | 40 | 0 | 0 | 0 | 1 | 0 | 58 | 17 | 25 | 42 | 29 | 2 | 0 | 0 | 0 | 0 | 296 | 149 | 1 | 1 | 1 | | 1 | 0.876 | | 0.936 | 1 | 1 | 1 | 1 | -0.012 |
| SE | 95016 | ENSG00000172270 | BSG | chr19 | + | 577773 | 578121 | 572595 | 572701 | 579499 | 579629 | 1 | 1 | 3 | 1 | 1 | 617 | 240 | 299 | 510 | 313 | 0 | 4 | 3 | 8 | 0 | 717 | 666 | 458 | 752 | 1196 | 298 | 149 | 1 | 1 | 0.001 | 0.002 | 0.005 | 0.001 | 0.002 | 0 | 0.003 | 0.003 | 0.005 | 0 | 0 |
| SE | 95019 | ENSG00000172270 | BSG | chr19 | + | 579499 | 579656 | 572528 | 572701 | 580378 | 580461 | 1315 | 580 | 733 | 1160 | 816 | 0 | 0 | 0 | 0 | 1 | 1671 | 1593 | 1137 | 1908 | 2876 | 0 | 0 | 0 | 0 | 0 | 298 | 149 | 1 | 1 | 1 | 1 | 1 | 1 | 0.998 | 1 | 1 | 1 | 1 | 1 | 0 |
| SE | 95021 | ENSG00000172270 | BSG | chr19 | + | 579508 | 579656 | 572609 | 572701 | 580378 | 580461 | 698 | 340 | 434 | 650 | 503 | 0 | 0 | 0 | 0 | 1 | 954 | 927 | 679 | 1156 | 1680 | 0 | 0 | 0 | 0 | 0 | 297 | 149 | 1 | 1 | 1 | 1 | 1 | 1 | 0.996 | 1 | 1 | 1 | 1 | 1 | -0.001 |
| SE | 95022 | ENSG00000172270 | BSG | chr19 | + | 579523 | 579656 | 572571 | 572701 | 580378 | 580461 | 690 | 336 | 425 | 637 | 495 | 0 | 0 | 0 | 0 | 1 | 935 | 895 | 664 | 1130 | 1604 | 0 | 0 | 0 | 0 | 0 | 282 | 149 | 1 | 1 | 1 | 1 | 1 | 1 | 0.996 | 1 | 1 | 1 | 1 | 1 | -0.001 |
| SE | 95024 | ENSG00000060237 | WNK1 | chr12 | + | 813641 | 813814 | 752578 | 754324 | 827041 | 827262 | 997 | 795 | 870 | 1017 | 1397 | 0 | 1 | 0 | 0 | 0 | 778 | 1027 | 930 | 975 | 668 | 8 | 0 | 0 | 0 | 0 | 298 | 149 | 1 | 1 | 1 | 0.997 | 1 | 1 | 1 | 0.98 | 1 | 1 | 1 | 1 | 0.003 |
| SE | 95027 | ENSG00000060237 | WNK1 | chr12 | + | 862082 | 862270 | 859244 | 859464 | 878211 | 878361 | 64 | 25 | 65 | 58 | 29 | 0 | 0 | 0 | 0 | 0 | 23 | 35 | 72 | 70 | 26 | 0 | 2 | 0 | 0 | 0 | 298 | 149 | 1 | 1 | 1 | 1 | 1 | 1 | 1 | 1 | 0.897 | 1 | 1 | 1 | 0.021 |
| SE | 95032 | ENSG00000060237 | WNK1 | chr12 | + | 871264 | 871348 | 867865 | 869104 | 878214 | 878361 | 10 | 3 | 10 | 6 | 4 | 0 | 0 | 0 | 0 | 0 | 13 | 11 | 7 | 7 | 0 | 0 | 0 | 1 | 0 | 0 | 233 | 149 | 1 | 1 | 1 | 1 | 1 | 1 | 1 | 1 | 1 | 0.817 | 1 | NA | 0.046 |
| SE | 95038 | ENSG00000060237 | WNK1 | chr12 | + | 890452 | 890510 | 889139 | 889223 | 894561 | 894635 | 60 | 52 | 45 | 54 | 80 | 2 | 0 | 2 | 0 | 1 | 102 | 61 | 55 | 45 | 19 | 0 | 0 | 1 | 0 | 0 | 207 | 149 | 1 | 1 | 0.956 | 1 | 0.942 | 1 | 0.983 | 1 | 1 | 0.975 | 1 | 1 | -0.019 |
| SE | 95039 | ENSG00000060237 | WNK1 | chr12 | + | 890452 | 890513 | 889139 | 889223 | 894561 | 894635 | 314 | 264 | 259 | 325 | 378 | 2 | 0 | 2 | 0 | 1 | 264 | 347 | 379 | 290 | 180 | 0 | 0 | 1 | 0 | 0 | 210 | 149 | 1 | 1 | 0.991 | 1 | 0.989 | 1 | 0.996 | 1 | 1 | 0.996 | 1 | 1 | -0.004 |
| SE | 95040 | ENSG00000060237 | WNK1 | chr12 | + | 894561 | 894635 | 890452 | 890513 | 896070 | 896732 | 460 | 394 | 445 | 496 | 596 | 0 | 0 | 0 | 1 | 0 | 366 | 510 | 508 | 477 | 260 | 0 | 0 | 0 | 1 | 0 | 223 | 149 | 1 | 1 | 1 | 1 | 1 | 0.997 | 1 | 1 | 1 | 1 | 0.997 | 1 | 0 |
| SE | 95048 | ENSG00000148334 | PTGES2 | chr9 | - | 128125895 | 128126260 | 128125243 | 128125441 | 128127438 | 128127726 | 0 | 0 | 1 | 0 | 0 | 5 | 4 | 3 | 4 | 3 | 0 | 0 | 0 | 0 | 0 | 0 | 4 | 1 | 10 | 4 | 298 | 149 | 1 | 1 | 0 | 0 | 0.143 | 0 | 0 | NA | 0 | 0 | 0 | 0 | 0.029 |
| SE | 95049 | ENSG00000198887 | SMC5 | chr9 | + | 70267922 | 70267975 | 70264303 | 70264445 | 70277309 | 70277472 | 91 | 90 | 80 | 114 | 131 | 3 | 0 | 0 | 0 | 0 | 6 | 77 | 43 | 24 | 18 | 0 | 0 | 0 | 0 | 0 | 202 | 149 | 1 | 1 | 0.957 | 1 | 1 | 1 | 1 | 1 | 1 | 1 | 1 | 1 | -0.009 |
| SE | 95050 | ENSG00000198887 | SMC5 | chr9 | + | 70267922 | 70267975 | 70264303 | 70264445 | 70280758 | 70280899 | 36 | 37 | 26 | 43 | 43 | 0 | 0 | 0 | 1 | 1 | 6 | 46 | 13 | 22 | 8 | 0 | 0 | 0 | 0 | 0 | 202 | 149 | 1 | 1 | 1 | 1 | 1 | 0.969 | 0.969 | 1 | 1 | 1 | 1 | 1 | -0.012 |
| SE | 95051 | ENSG00000198887 | SMC5 | chr9 | + | 70277309 | 70277472 | 70264303 | 70264445 | 70280758 | 70280899 | 3 | 0 | 0 | 3 | 0 | 0 | 0 | 0 | 1 | 1 | 0 | 0 | 1 | 1 | 0 | 0 | 0 | 0 | 0 | 0 | 298 | 149 | 1 | 1 | 1 | NA | NA | 0.6 | 0 | NA | NA | 1 | 1 | NA | -0.467 |
| SE | 95052 | ENSG00000198887 | SMC5 | chr9 | + | 70277309 | 70277472 | 70267922 | 70267975 | 70280758 | 70280899 | 59 | 59 | 55 | 75 | 93 | 4 | 6 | 1 | 0 | 2 | 0 | 33 | 31 | 3 | 10 | 0 | 2 | 0 | 0 | 0 | 298 | 149 | 1 | 1 | 0.881 | 0.831 | 0.965 | 1 | 0.959 | NA | 0.892 | 1 | 1 | 1 | -0.046 |
| SE | 95053 | ENSG00000198887 | SMC5 | chr9 | + | 70278490 | 70278625 | 70277309 | 70277472 | 70280758 | 70280899 | 278 | 301 | 259 | 301 | 262 | 0 | 0 | 1 | 3 | 0 | 50 | 201 | 274 | 75 | 59 | 0 | 0 | 1 | 1 | 0 | 284 | 149 | 1 | 1 | 1 | 1 | 1 | 0.981 | 1 | 1 | 1 | 0.993 | 0.975 | 1 | 0.003 |
| SE | 95056 | ENSG00000198887 | SMC5 | chr9 | + | 70300045 | 70300200 | 70297965 | 70298221 | 70314741 | 70314836 | 132 | 192 | 103 | 157 | 264 | 0 | 0 | 0 | 0 | 0 | 0 | 93 | 114 | 16 | 19 | 0 | 0 | 3 | 0 | 0 | 298 | 149 | 1 | 1 | 1 | 1 | 1 | 1 | 1 | NA | 1 | 0.95 | 1 | 1 | 0.012 |
| SE | 95058 | ENSG00000198887 | SMC5 | chr9 | + | 70344143 | 70344269 | 70324020 | 70324143 | 70346604 | 70346649 | 198 | 185 | 188 | 269 | 354 | 0 | 1 | 0 | 0 | 0 | 0 | 63 | 57 | 0 | 0 | 0 | 0 | 0 | 0 | 0 | 275 | 149 | 1 | 1 | 1 | 0.99 | 1 | 1 | 1 | NA | 1 | 1 | NA | NA | -0.002 |
| SE | 95059 | ENSG00000198887 | SMC5 | chr9 | + | 70344143 | 70344269 | 70324020 | 70324143 | 70347065 | 70347161 | 195 | 181 | 179 | 250 | 311 | 4 | 2 | 6 | 1 | 17 | 0 | 37 | 49 | 0 | 1 | 0 | 0 | 0 | 0 | 0 | 275 | 149 | 1 | 1 | 0.964 | 0.98 | 0.942 | 0.993 | 0.908 | NA | 1 | 1 | NA | 1 | -0.043 |
| SE | 95060 | ENSG00000198887 | SMC5 | chr9 | + | 70346604 | 70346649 | 70324020 | 70324143 | 70347065 | 70347161 | 17 | 21 | 24 | 27 | 38 | 4 | 2 | 6 | 1 | 17 | 0 | 6 | 10 | 0 | 0 | 0 | 0 | 0 | 0 | 0 | 194 | 149 | 1 | 1 | 0.765 | 0.89 | 0.754 | 0.954 | 0.632 | NA | 1 | 1 | NA | NA | -0.201 |
| SE | 95061 | ENSG00000198887 | SMC5 | chr9 | + | 70346604 | 70346649 | 70344143 | 70344269 | 70347065 | 70347161 | 69 | 59 | 72 | 89 | 138 | 48 | 36 | 39 | 43 | 57 | 0 | 35 | 25 | 0 | 0 | 0 | 3 | 7 | 0 | 1 | 194 | 149 | 1 | 1 | 0.525 | 0.557 | 0.586 | 0.614 | 0.65 | NA | 0.9 | 0.733 | NA | 0 | 0.042 |
| SE | 95062 | ENSG00000198887 | SMC5 | chr9 | + | 70347612 | 70347717 | 70347065 | 70347161 | 70347918 | 70348038 | 210 | 191 | 162 | 195 | 296 | 0 | 2 | 2 | 1 | 5 | 29 | 59 | 55 | 5 | 2 | 0 | 1 | 3 | 0 | 0 | 254 | 149 | 1 | 1 | 1 | 0.982 | 0.979 | 0.991 | 0.972 | 1 | 0.972 | 0.915 | 1 | 1 | 0.007 |
| SE | 95063 | ENSG00000198887 | SMC5 | chr9 | + | 70347612 | 70347717 | 70347065 | 70347161 | 70350113 | 70350293 | 124 | 132 | 96 | 123 | 170 | 0 | 0 | 0 | 3 | 0 | 29 | 41 | 30 | 1 | 1 | 0 | 0 | 0 | 0 | 0 | 254 | 149 | 1 | 1 | 1 | 1 | 1 | 0.96 | 1 | 1 | 1 | 1 | 1 | 1 | -0.008 |
| SE | 95064 | ENSG00000198887 | SMC5 | chr9 | + | 70347918 | 70348038 | 70347065 | 70347161 | 70350113 | 70350293 | 88 | 59 | 63 | 91 | 175 | 0 | 0 | 0 | 3 | 0 | 0 | 15 | 21 | 2 | 0 | 0 | 0 | 0 | 0 | 0 | 269 | 149 | 1 | 1 | 1 | 1 | 1 | 0.944 | 1 | NA | 1 | 1 | 1 | NA | -0.011 |
| SE | 95065 | ENSG00000198887 | SMC5 | chr9 | + | 70347918 | 70348038 | 70347612 | 70347717 | 70350113 | 70350293 | 173 | 116 | 128 | 163 | 299 | 0 | 0 | 1 | 1 | 3 | 0 | 28 | 42 | 6 | 1 | 0 | 0 | 0 | 0 | 0 | 269 | 149 | 1 | 1 | 1 | 1 | 0.986 | 0.989 | 0.982 | NA | 1 | 1 | 1 | 1 | -0.009 |
| SE | 95066 | ENSG00000198887 | SMC5 | chr9 | + | 70350113 | 70350293 | 70347918 | 70348038 | 70350375 | 70350471 | 221 | 164 | 172 | 198 | 458 | 0 | 0 | 0 | 2 | 0 | 8 | 40 | 54 | 4 | 12 | 0 | 0 | 0 | 0 | 0 | 298 | 149 | 1 | 1 | 1 | 1 | 1 | 0.98 | 1 | 1 | 1 | 1 | 1 | 1 | -0.004 |
| SE | 95067 | ENSG00000198887 | SMC5 | chr9 | + | 70350113 | 70350293 | 70347918 | 70348038 | 70352190 | 70352245 | 112 | 77 | 83 | 109 | 217 | 0 | 1 | 0 | 3 | 0 | 0 | 18 | 23 | 2 | 0 | 0 | 0 | 0 | 0 | 0 | 298 | 149 | 1 | 1 | 1 | 0.975 | 1 | 0.948 | 1 | NA | 1 | 1 | 1 | NA | -0.015 |
| SE | 95068 | ENSG00000198887 | SMC5 | chr9 | + | 70350375 | 70350471 | 70347918 | 70348038 | 70352190 | 70352245 | 107 | 94 | 97 | 103 | 199 | 0 | 1 | 0 | 3 | 0 | 0 | 31 | 19 | 1 | 12 | 0 | 0 | 0 | 0 | 0 | 245 | 149 | 1 | 1 | 1 | 0.983 | 1 | 0.954 | 1 | NA | 1 | 1 | 1 | 1 | -0.013 |
| SE | 95069 | ENSG00000198887 | SMC5 | chr9 | + | 70350375 | 70350471 | 70350113 | 70350293 | 70352190 | 70352245 | 216 | 181 | 186 | 190 | 440 | 1 | 0 | 0 | 1 | 0 | 8 | 53 | 50 | 3 | 24 | 0 | 0 | 0 | 0 | 0 | 245 | 149 | 1 | 1 | 0.992 | 1 | 1 | 0.991 | 1 | 1 | 1 | 1 | 1 | 1 | -0.003 |
| SE | 95071 | ENSG00000166348 | USP54 | chr10 | - | 73504849 | 73504990 | 73500654 | 73500838 | 73505307 | 73505426 | 2 | 0 | 9 | 2 | 10 | 1 | 0 | 1 | 6 | 7 | 0 | 0 | 0 | 0 | 0 | 0 | 4 | 0 | 4 | 0 | 290 | 149 | 1 | 1 | 0.507 | NA | 0.822 | 0.146 | 0.423 | NA | 0 | NA | 0 | NA | 0.474 |
| SE | 95075 | ENSG00000166348 | USP54 | chr10 | - | 73520907 | 73521027 | 73518721 | 73519992 | 73523582 | 73523750 | 9 | 13 | 7 | 22 | 11 | 0 | 0 | 0 | 2 | 0 | 0 | 3 | 12 | 13 | 1 | 0 | 0 | 0 | 0 | 0 | 269 | 149 | 1 | 1 | 1 | 1 | 1 | 0.859 | 1 | NA | 1 | 1 | 1 | 1 | -0.028 |
| SE | 95077 | ENSG00000166348 | USP54 | chr10 | - | 73530703 | 73530835 | 73530142 | 73530523 | 73536268 | 73536437 | 6 | 13 | 14 | 10 | 13 | 0 | 0 | 0 | 0 | 0 | 0 | 15 | 2 | 6 | 2 | 0 | 0 | 2 | 0 | 0 | 281 | 149 | 1 | 1 | 1 | 1 | 1 | 1 | 1 | NA | 1 | 0.347 | 1 | 1 | 0.163 |
| SE | 95078 | ENSG00000166348 | USP54 | chr10 | - | 73534599 | 73534770 | 73530142 | 73530523 | 73536268 | 73536437 | 0 | 2 | 0 | 3 | 0 | 0 | 0 | 0 | 0 | 0 | 0 | 0 | 0 | 0 | 0 | 0 | 0 | 2 | 0 | 0 | 298 | 149 | 1 | 1 | NA | 1 | NA | 1 | NA | NA | NA | 0 | NA | NA | 1 |
| SE | 95079 | ENSG00000166348 | USP54 | chr10 | - | 73534599 | 73534770 | 73530703 | 73530835 | 73536268 | 73536437 | 1 | 2 | 0 | 3 | 0 | 4 | 4 | 6 | 0 | 4 | 0 | 2 | 0 | 0 | 0 | 0 | 3 | 0 | 0 | 0 | 298 | 149 | 1 | 1 | 0.111 | 0.2 | 0 | 1 | 0 | NA | NA | 0.25 | NA | NA | 0.012 |
| SE | 95080 | ENSG00000166348 | USP54 | chr10 | - | 73576135 | 73576361 | 73575511 | 73575675 | 73625936 | 73625953 | 0 | 1 | 0 | 2 | 4 | 0 | 0 | 1 | 0 | 0 | 0 | 0 | 1 | 0 | 5 | 0 | 0 | 0 | 0 | 0 | 298 | 149 | 1 | 1 | NA | 1 | 0 | 1 | 1 | NA | NA | 1 | NA | 1 | -0.25 |
| SE | 95082 | ENSG00000165028 | NIPSNAP3B | chr9 | + | 104766324 | 104766535 | 104764128 | 104764300 | 104768862 | 104769021 | 21 | 13 | 12 | 12 | 28 | 3 | 0 | 4 | 1 | 0 | 0 | 11 | 1 | 4 | 16 | 0 | 2 | 0 | 0 | 0 | 298 | 149 | 1 | 1 | 0.778 | 1 | 0.6 | 0.857 | 1 | NA | 0.733 | 1 | 1 | 1 | -0.086 |
| SE | 95083 | ENSG00000165028 | NIPSNAP3B | chr9 | + | 104770848 | 104770998 | 104768862 | 104769021 | 104772821 | 104772908 | 6 | 4 | 8 | 6 | 6 | 0 | 1 | 0 | 2 | 1 | 0 | 0 | 0 | 6 | 0 | 16 | 0 | 1 | 0 | 5 | 298 | 149 | 1 | 1 | 1 | 0.667 | 1 | 0.6 | 0.75 | 0 | NA | 0 | 1 | 0 | 0.553 |
| SE | 95084 | ENSG00000165028 | NIPSNAP3B | chr9 | + | 104771233 | 104771316 | 104770848 | 104770998 | 104772821 | 104772908 | 0 | 1 | 0 | 7 | 0 | 1 | 0 | 1 | 1 | 2 | 0 | 0 | 2 | 0 | 0 | 0 | 0 | 0 | 4 | 0 | 232 | 149 | 1 | 1 | 0 | 1 | 0 | 0.818 | 0 | NA | NA | 1 | 0 | NA | -0.136 |
| SE | 95085 | ENSG00000165028 | NIPSNAP3B | chr9 | + | 104772996 | 104773176 | 104772821 | 104772908 | 104777127 | 104777457 | 4 | 2 | 3 | 0 | 1 | 0 | 0 | 1 | 0 | 2 | 0 | 0 | 0 | 8 | 0 | 0 | 0 | 0 | 0 | 0 | 298 | 149 | 1 | 1 | 1 | 1 | 0.6 | NA | 0.2 | NA | NA | NA | 1 | NA | -0.3 |
| SE | 95088 | ENSG00000135119 | RNFT2 | chr12 | + | 116833791 | 116833941 | 116779194 | 116779348 | 116849313 | 116849484 | 2 | 3 | 4 | 3 | 10 | 0 | 0 | 0 | 0 | 2 | 0 | 3 | 2 | 3 | 9 | 0 | 0 | 0 | 0 | 1 | 298 | 149 | 1 | 1 | 1 | 1 | 1 | 1 | 1 | NA | 1 | 1 | 1 | 0.818 | 0.045 |
| SE | 95089 | ENSG00000135119 | RNFT2 | chr12 | + | 116835959 | 116836025 | 116833791 | 116833941 | 116836180 | 116836282 | 6 | 3 | 13 | 3 | 6 | 0 | 0 | 0 | 0 | 1 | 0 | 4 | 0 | 2 | 0 | 0 | 0 | 0 | 0 | 0 | 215 | 149 | 1 | 1 | 1 | 1 | 1 | 1 | 0.806 | NA | 1 | NA | 1 | NA | -0.039 |
| SE | 95090 | ENSG00000135119 | RNFT2 | chr12 | + | 116836180 | 116836282 | 116833791 | 116833941 | 116849313 | 116849484 | 0 | 1 | 7 | 1 | 4 | 0 | 0 | 0 | 1 | 0 | 0 | 3 | 11 | 0 | 6 | 0 | 0 | 0 | 0 | 0 | 251 | 149 | 1 | 1 | NA | 1 | 1 | 0.372 | 1 | NA | 1 | 1 | NA | 1 | -0.157 |
| SE | 95092 | ENSG00000148358 | GPR107 | chr9 | + | 130075635 | 130075749 | 130053425 | 130054073 | 130076411 | 130076462 | 98 | 60 | 66 | 103 | 89 | 0 | 0 | 0 | 0 | 0 | 154 | 142 | 113 | 172 | 176 | 0 | 0 | 1 | 6 | 0 | 263 | 149 | 1 | 1 | 1 | 1 | 1 | 1 | 1 | 1 | 1 | 0.985 | 0.942 | 1 | 0.015 |
| SE | 95095 | ENSG00000148358 | GPR107 | chr9 | + | 130086419 | 130086476 | 130083301 | 130083602 | 130090875 | 130090906 | 156 | 92 | 115 | 156 | 268 | 2 | 2 | 1 | 0 | 0 | 337 | 178 | 167 | 227 | 284 | 0 | 6 | 0 | 0 | 0 | 206 | 149 | 1 | 1 | 0.983 | 0.971 | 0.988 | 1 | 1 | 1 | 0.955 | 1 | 1 | 1 | -0.003 |
| SE | 95097 | ENSG00000148358 | GPR107 | chr9 | + | 130092247 | 130092381 | 130090875 | 130090983 | 130099456 | 130099532 | 272 | 138 | 154 | 217 | 401 | 3 | 0 | 2 | 1 | 0 | 464 | 293 | 281 | 350 | 403 | 0 | 0 | 0 | 0 | 0 | 283 | 149 | 1 | 1 | 0.979 | 1 | 0.976 | 0.991 | 1 | 1 | 1 | 1 | 1 | 1 | -0.011 |
| SE | 95099 | ENSG00000148358 | GPR107 | chr9 | + | 130099456 | 130099532 | 130092247 | 130092381 | 130107495 | 130107539 | 57 | 23 | 28 | 24 | 40 | 0 | 0 | 0 | 0 | 0 | 57 | 43 | 41 | 40 | 55 | 0 | 0 | 0 | 2 | 0 | 225 | 149 | 1 | 1 | 1 | 1 | 1 | 1 | 1 | 1 | 1 | 1 | 0.93 | 1 | 0.014 |
| SE | 95101 | ENSG00000148358 | GPR107 | chr9 | + | 130101105 | 130101223 | 130099456 | 130099532 | 130107495 | 130107539 | 1 | 0 | 8 | 6 | 12 | 0 | 0 | 0 | 0 | 0 | 30 | 16 | 11 | 9 | 13 | 0 | 0 | 2 | 4 | 0 | 267 | 149 | 1 | 1 | 1 | NA | 1 | 1 | 1 | 1 | 1 | 0.754 | 0.557 | 1 | 0.138 |
| SE | 95104 | ENSG00000148358 | GPR107 | chr9 | + | 130110369 | 130110426 | 130107495 | 130107539 | 130127482 | 130127531 | 4 | 2 | 4 | 2 | 4 | 0 | 0 | 0 | 0 | 0 | 0 | 0 | 2 | 10 | 0 | 0 | 0 | 0 | 8 | 0 | 206 | 149 | 1 | 1 | 1 | 1 | 1 | 1 | 1 | NA | NA | 1 | 0.475 | NA | 0.262 |
| SE | 95107 | ENSG00000148358 | GPR107 | chr9 | + | 130124914 | 130124964 | 130110369 | 130110426 | 130127482 | 130127531 | 59 | 28 | 29 | 35 | 24 | 0 | 0 | 2 | 0 | 0 | 61 | 34 | 31 | 42 | 49 | 0 | 0 | 0 | 0 | 0 | 199 | 149 | 1 | 1 | 1 | 1 | 0.916 | 1 | 1 | 1 | 1 | 1 | 1 | 1 | -0.017 |
| SE | 95108 | ENSG00000148358 | GPR107 | chr9 | + | 130128639 | 130128761 | 130127482 | 130127566 | 130135024 | 130136123 | 113 | 63 | 96 | 113 | 94 | 0 | 3 | 0 | 0 | 0 | 200 | 97 | 110 | 129 | 133 | 0 | 0 | 0 | 0 | 0 | 271 | 149 | 1 | 1 | 1 | 0.92 | 1 | 1 | 1 | 1 | 1 | 1 | 1 | 1 | -0.016 |
| SE | 95110 | ENSG00000096872 | IFT74 | chr9 | + | 26992382 | 26992677 | 26990133 | 26990195 | 27009019 | 27009158 | 0 | 3 | 1 | 0 | 4 | 19 | 16 | 22 | 25 | 48 | 0 | 0 | 0 | 0 | 0 | 3 | 20 | 6 | 6 | 0 | 298 | 149 | 1 | 1 | 0 | 0.086 | 0.022 | 0 | 0.04 | 0 | 0 | 0 | 0 | NA | 0.03 |
| SE | 95111 | ENSG00000096872 | IFT74 | chr9 | + | 27009019 | 27009158 | 26990133 | 26990195 | 27011905 | 27011968 | 45 | 68 | 55 | 66 | 115 | 0 | 0 | 2 | 0 | 0 | 41 | 38 | 38 | 19 | 30 | 0 | 1 | 0 | 1 | 0 | 288 | 149 | 1 | 1 | 1 | 1 | 0.934 | 1 | 1 | 1 | 0.952 | 1 | 0.908 | 1 | 0.015 |
| SE | 95113 | ENSG00000175893 | ZDHHC21 | chr9 | - | 14639895 | 14640012 | 14619638 | 14619682 | 14658748 | 14658887 | 117 | 97 | 65 | 151 | 156 | 0 | 0 | 1 | 0 | 0 | 20 | 50 | 23 | 23 | 15 | 0 | 0 | 0 | 0 | 0 | 266 | 149 | 1 | 1 | 1 | 1 | 0.973 | 1 | 1 | 1 | 1 | 1 | 1 | 1 | -0.005 |
| SE | 95114 | ENSG00000175893 | ZDHHC21 | chr9 | - | 14658748 | 14658887 | 14639895 | 14640012 | 14662214 | 14662326 | 152 | 79 | 76 | 115 | 206 | 0 | 0 | 0 | 1 | 2 | 28 | 62 | 27 | 30 | 57 | 0 | 2 | 0 | 0 | 0 | 288 | 149 | 1 | 1 | 1 | 1 | 1 | 0.983 | 0.982 | 1 | 0.941 | 1 | 1 | 1 | 0.005 |
| SE | 95115 | ENSG00000175893 | ZDHHC21 | chr9 | - | 14662214 | 14662326 | 14639895 | 14640012 | 14690336 | 14690385 | 0 | 0 | 2 | 1 | 2 | 0 | 0 | 0 | 0 | 0 | 0 | 4 | 0 | 0 | 0 | 0 | 2 | 0 | 0 | 0 | 261 | 149 | 1 | 1 | NA | NA | 1 | 1 | 1 | NA | 0.533 | NA | NA | NA | 0.467 |
| SE | 95116 | ENSG00000175893 | ZDHHC21 | chr9 | - | 14680032 | 14680162 | 14639895 | 14640012 | 14690336 | 14690385 | 18 | 23 | 21 | 19 | 12 | 0 | 0 | 0 | 0 | 0 | 3 | 8 | 2 | 0 | 2 | 0 | 2 | 0 | 0 | 0 | 279 | 149 | 1 | 1 | 1 | 1 | 1 | 1 | 1 | 1 | 0.681 | 1 | NA | 1 | 0.08 |
| SE | 95117 | ENSG00000175893 | ZDHHC21 | chr9 | - | 14683570 | 14683729 | 14680032 | 14680162 | 14690336 | 14690385 | 13 | 5 | 14 | 21 | 2 | 22 | 22 | 22 | 21 | 12 | 7 | 5 | 1 | 0 | 0 | 3 | 8 | 2 | 0 | 2 | 298 | 149 | 1 | 1 | 0.228 | 0.102 | 0.241 | 0.333 | 0.077 | 0.538 | 0.238 | 0.2 | NA | 0 | -0.048 |
| SE | 95119 | ENSG00000136854 | STXBP1 | chr9 | + | 127642468 | 127642563 | 127612276 | 127612440 | 127651602 | 127651652 | 0 | 0 | 0 | 0 | 0 | 1 | 0 | 3 | 2 | 6 | 0 | 0 | 0 | 0 | 2 | 0 | 0 | 0 | 0 | 20 | 244 | 149 | 1 | 1 | 0 | NA | 0 | 0 | 0 | NA | NA | NA | NA | 0.058 | -0.058 |
| SE | 95129 | ENSG00000165282 | PIGO | chr9 | - | 35091239 | 35091291 | 35090465 | 35090672 | 35092542 | 35092767 | 9 | 2 | 1 | 4 | 1 | 1 | 0 | 0 | 0 | 0 | 0 | 4 | 6 | 1 | 0 | 0 | 0 | 0 | 0 | 0 | 201 | 149 | 1 | 1 | 0.87 | 1 | | | 1 | NA | 1 | 1 | 1 | NA | -0.026 |
| SE | 95130 | ENSG00000165282 | PIGO | chr9 | - | 35091239 | 35092767 | 35090465 | 35090672 | 35095054 | 35095566 | 6 | 2 | 4 | 5 | 3 | 0 | 1 | 0 | 0 | 0 | 0 | 2 | 9 | 0 | 2 | 0 | 0 | 0 | 0 | 0 | 298 | 149 | 1 | 1 | 1 | 0.5 | 1 | 1 | 1 | NA | 1 | 1 | NA | 1 | -0.1 |
| SE | 95131 | ENSG00000165282 | PIGO | chr9 | - | 35091239 | 35094359 | 35090465 | 35090672 | 35095054 | 35095566 | 33 | 18 | 53 | 27 | 20 | 0 | 1 | 0 | 0 | 0 | 36 | 16 | 24 | 6 | 24 | 0 | 0 | 0 | 0 | 0 | 298 | 149 | 1 | 1 | 1 | 0.9 | 1 | 1 | 1 | 1 | 1 | 1 | 1 | 1 | -0.02 |
| SE | 95132 | ENSG00000165282 | PIGO | chr9 | - | 35092542 | 35092767 | 35090465 | 35090672 | 35095054 | 35095566 | 1 | 0 | 0 | 0 | 0 | 0 | 1 | 0 | 0 | 0 | 0 | 2 | 0 | 0 | 0 | 0 | 0 | 0 | 0 | 0 | 298 | 149 | 1 | 1 | 1 | 0 | NA | NA | NA | NA | 1 | NA | NA | NA | -0.5 |
| SE | 95133 | ENSG00000165282 | PIGO | chr9 | - | 35092542 | 35092767 | 35091239 | 35091291 | 35093029 | 35093209 | 37 | 12 | 16 | 19 | 19 | 0 | 0 | 1 | 0 | 0 | 6 | 23 | 14 | 5 | 13 | 0 | 0 | 0 | 0 | 0 | 298 | 149 | 1 | 1 | 1 | 1 | 0.889 | 1 | 1 | 1 | 1 | 1 | 1 | 1 | -0.022 |
| SE | 95134 | ENSG00000165282 | PIGO | chr9 | - | 35092542 | 35092767 | 35091239 | 35091291 | 35093420 | 35093580 | 8 | 2 | 2 | 4 | 0 | 0 | 0 | 1 | 0 | 0 | 0 | 4 | 3 | 1 | 0 | 0 | 0 | 0 | 0 | 0 | 298 | 149 | 1 | 1 | 1 | 1 | 0.5 | 1 | NA | NA | 1 | 1 | 1 | NA | -0.125 |
| SE | 95135 | ENSG00000165282 | PIGO | chr9 | - | 35093029 | 35093209 | 35091239 | 35091291 | 35093420 | 35093580 | 18 | 16 | 15 | 16 | 18 | 0 | 0 | 1 | 0 | 0 | 3 | 13 | 6 | 4 | 8 | 0 | 0 | 0 | 0 | 0 | 298 | 149 | 1 | 1 | 1 | 1 | 0.882 | 1 | 1 | 1 | 1 | 1 | 1 | 1 | -0.024 |
| SE | 95136 | ENSG00000165282 | PIGO | chr9 | - | 35093029 | 35093209 | 35091239 | 35091291 | 35093900 | 35094024 | 0 | 0 | 3 | 0 | 0 | 0 | 0 | 0 | 0 | 0 | 0 | 0 | 0 | 0 | 5 | 0 | 0 | 0 | 3 | 0 | 298 | 149 | 1 | 1 | NA | NA | 1 | NA | | NA | NA | NA | 0 | 1 | 0.5 |
| SE | 95137 | ENSG00000165282 | PIGO | chr9 | - | 35093029 | 35093209 | 35091239 | 35092767 | 35093420 | 35093580 | 47 | 26 | 29 | 32 | 37 | 0 | 0 | 1 | 1 | 0 | 9 | 32 | 17 | 8 | 21 | 0 | 0 | 0 | 0 | 0 | 298 | 149 | 1 | 1 | 1 | 1 | 0.935 | 0.941 | 1 | 1 | 1 | 1 | 1 | 1 | -0.025 |
| SE | 95138 | ENSG00000165282 | PIGO | chr9 | - | 35093420 | 35093580 | 35091239 | 35091291 | 35093900 | 35094024 | 28 | 21 | 30 | 36 | 40 | 0 | 0 | 0 | 0 | 0 | 26 | 46 | 3 | 11 | 59 | 0 | 0 | 0 | 3 | 0 | 298 | 149 | 1 | 1 | 1 | 1 | 1 | 1 | 1 | 1 | 1 | 1 | 0.647 | 1 | 0.071 |
| SE | 95141 | ENSG00000165282 | PIGO | chr9 | - | 35093900 | 35094024 | 35088689 | 35093580 | 35094215 | 35094359 | 41 | 34 | 43 | 45 | 59 | 0 | 1 | 0 | 0 | 0 | 42 | 57 | 16 | 30 | 76 | 0 | 0 | 0 | 1 | 0 | 273 | 149 | 1 | 1 | 1 | 0.949 | 1 | 1 | 1 | 1 | 1 | 1 | 0.942 | 1 | 0.001 |
| SE | 95144 | ENSG00000165282 | PIGO | chr9 | - | 35096154 | 35096480 | 35095143 | 35095566 | 35096560 | 35096594 | 0 | 0 | 1 | 0 | 0 | 5 | 0 | 4 | 0 | 0 | 0 | 0 | 0 | 0 | 0 | 0 | 4 | 6 | 0 | 10 | 298 | 149 | 1 | 1 | 0 | NA | 0.111 | NA | | NA | 0 | 0 | NA | 0 | 0.056 |
| SE | 95145 | ENSG00000165283 | STOML2 | chr9 | - | 35101134 | 35101279 | 35100597 | 35101011 | 35101709 | 35101811 | 102 | 51 | 86 | 84 | 183 | 0 | 0 | 0 | 0 | 0 | 68 | 98 | 46 | 43 | 56 | 0 | 0 | 0 | 2 | 0 | 294 | 149 | 1 | 1 | 1 | 1 | 1 | 1 | 1 | 1 | 1 | 1 | 0.916 | 1 | 0.017 |
| SE | 95147 | ENSG00000165283 | STOML2 | chr9 | - | 35102094 | 35102194 | 35101903 | 35101962 | 35102685 | 35102823 | 111 | 55 | 92 | 80 | 99 | 0 | 0 | 1 | 0 | 0 | 57 | 87 | 59 | 62 | 167 | 0 | 0 | 0 | 0 | 0 | 249 | 149 | 1 | 1 | 1 | 1 | 0.982 | 1 | 1 | 1 | 1 | 1 | 1 | 1 | -0.004 |
| SE | 95148 | ENSG00000165283 | STOML2 | chr9 | - | 35102685 | 35102823 | 35102094 | 35102194 | 35103049 | 35103131 | 62 | 27 | 29 | 24 | 11 | 0 | 0 | 0 | 3 | 0 | 0 | 27 | 14 | 30 | 42 | 0 | 0 | 0 | 0 | 0 | 287 | 149 | 1 | 1 | 1 | 1 | 1 | 0.806 | 1 | NA | 1 | 1 | 1 | 1 | -0.039 |
| SE | 95149 | ENSG00000005238 | FAM214B | chr9 | - | 35105681 | 35105857 | 35104111 | 35105374 | 35105940 | 35106009 | 51 | 40 | 38 | 41 | 50 | 0 | 0 | 1 | 0 | 0 | 54 | 57 | 68 | 76 | 213 | 0 | 0 | 0 | 1 | 0 | 298 | 149 | 1 | 1 | 1 | 1 | 0.95 | 1 | 1 | 1 | 1 | 1 | 0.974 | 1 | -0.005 |
| SE | 95150 | ENSG00000005238 | FAM214B | chr9 | - | 35105681 | 35105857 | 35104111 | 35105374 | 35106251 | 35106410 | 15 | 22 | 17 | 23 | 16 | 0 | 0 | 0 | 1 | 0 | 43 | 36 | 37 | 59 | 132 | 3 | 0 | 0 | 0 | 0 | 298 | 149 | 1 | 1 | 1 | 1 | 1 | 0.92 | 1 | 0.878 | 1 | 1 | 1 | 1 | 0.008 |
| SE | 95153 | ENSG00000005238 | FAM214B | chr9 | - | 35106537 | 35106643 | 35106251 | 35106410 | 35106832 | 35106891 | 64 | 27 | 46 | 49 | 13 | 0 | 0 | 0 | 0 | 0 | 70 | 114 | 58 | 68 | 176 | 0 | 2 | 0 | 0 | 2 | 255 | 149 | 1 | 1 | 1 | 1 | 1 | 1 | 1 | 1 | 0.971 | 1 | 1 | 0.981 | 0.01 |
| SE | 95155 | ENSG00000198722 | UNC13B | chr9 | + | 35236468 | 35236586 | 35231119 | 35231219 | 35237702 | 35237826 | 8 | 4 | 4 | 3 | 7 | 0 | 0 | 2 | 0 | 0 | 13 | 7 | 2 | 11 | 95 | 0 | 0 | 0 | 0 | 0 | 267 | 149 | 1 | 1 | 1 | 1 | 0.527 | 1 | 1 | 1 | 1 | 1 | 1 | 1 | -0.095 |
| SE | 95157 | ENSG00000198722 | UNC13B | chr9 | + | 35300165 | 35308412 | 35295695 | 35295930 | 35310466 | 35310781 | 1 | 0 | 0 | 0 | 0 | 0 | 0 | 0 | 0 | 1 | 0 | 0 | 0 | 0 | 0 | 13 | 2 | 5 | 5 | 44 | 298 | 149 | 1 | 1 | 1 | NA | NA | NA | 0 | 0 | 0 | 0 | 0 | 0 | 0.5 |
| SE | 95160 | ENSG00000198722 | UNC13B | chr9 | + | 35375126 | 35375201 | 35370317 | 35370396 | 35376027 | 35376247 | 1 | 2 | 1 | 12 | 0 | 0 | 0 | 0 | 0 | 0 | 16 | 1 | 20 | 0 | 45 | 0 | 0 | 0 | 0 | 3 | 224 | 149 | 1 | 1 | 1 | 1 | 1 | 1 | NA | 1 | 1 | 1 | NA | 0.909 | 0.023 |
| SE | 95161 | ENSG00000198722 | UNC13B | chr9 | + | 35381099 | 35381215 | 35380469 | 35380639 | 35381555 | 35381719 | 1 | 1 | 0 | 0 | 6 | 0 | 0 | 0 | 0 | 0 | 11 | 0 | 7 | 9 | 79 | 0 | 0 | 1 | 4 | 0 | 265 | 149 | 1 | 1 | 1 | 1 | NA | NA | 1 | 1 | NA | 0.797 | 0.559 | 1 | 0.161 |
| SE | 95170 | ENSG00000213930 | GALT | chr9 | + | 34647388 | 34647567 | 34647122 | 34647258 | 34647831 | 34647961 | 0 | 0 | 0 | 0 | 1 | 3 | 0 | 4 | 0 | 0 | 0 | 0 | 0 | 0 | 0 | 10 | 1 | 0 | 0 | 0 | 298 | 149 | 1 | 1 | 0 | NA | 0 | NA | 1 | 0 | 0 | NA | NA | NA | 0.333 |
| SE | 95171 | ENSG00000213930 | GALT | chr9 | + | 34647474 | 34647567 | 34646899 | 34647258 | 34647831 | 34647961 | 4 | 0 | 3 | 4 | 5 | 3 | 0 | 4 | 0 | 0 | 0 | 0 | 1 | 5 | 0 | 10 | 1 | 0 | 0 | 0 | 242 | 149 | 1 | 1 | 0.451 | NA | 0.316 | 1 | 1 | 0 | 0 | 1 | 1 | NA | 0.192 |
| SE | 95178 | ENSG00000213930 | GALT | chr9 | + | 34647540 | 34647567 | 34647088 | 34647258 | 34647831 | 34647905 | 0 | 0 | 3 | 0 | 1 | 3 | 0 | 4 | 0 | 0 | 0 | 0 | 2 | 0 | 0 | 10 | 1 | 0 | 0 | 0 | 176 | 149 | 1 | 1 | 0 | NA | 0.388 | NA | 1 | 0 | 0 | 1 | NA | NA | 0.129 |
| SE | 95180 | ENSG00000213930 | GALT | chr9 | + | 34647656 | 34647705 | 34647388 | 34647567 | 34647831 | 34647905 | 54 | 41 | 71 | 61 | 48 | 2 | 0 | 0 | 0 | 1 | 59 | 60 | 47 | 41 | 71 | 0 | 0 | 0 | 0 | 0 | 198 | 149 | 1 | 1 | 0.953 | 1 | 1 | 1 | 0.973 | 1 | 1 | 1 | 1 | 1 | -0.015 |
| SE | 95181 | ENSG00000213930 | GALT | chr9 | + | 34647831 | 34647961 | 34647088 | 34647258 | 34648047 | 34648171 | 10 | 0 | 10 | 4 | 2 | 0 | 0 | 1 | 0 | 0 | 22 | 1 | 10 | 11 | 14 | 0 | 0 | 0 | 0 | 0 | 279 | 149 | 1 | 1 | 1 | NA | 0.842 | 1 | 1 | 1 | 1 | 1 | 1 | 1 | -0.039 |
| SE | 95183 | ENSG00000213930 | GALT | chr9 | + | 34647831 | 34647961 | 34647088 | 34647258 | 34648333 | 34648375 | 4 | 0 | 4 | 1 | 0 | 0 | 0 | 0 | 0 | 0 | 10 | 1 | 0 | 0 | 0 | 0 | 0 | 0 | 0 | 5 | 279 | 149 | 1 | 1 | 1 | NA | 1 | 1 | NA | 1 | 1 | NA | NA | 0 | 0.333 |
| SE | 95184 | ENSG00000213930 | GALT | chr9 | + | 34647831 | 34647961 | 34647491 | 34647705 | 34648114 | 34648171 | 138 | 78 | 171 | 103 | 97 | 0 | 0 | 0 | 0 | 1 | 164 | 169 | 87 | 113 | 173 | 0 | 0 | 2 | 0 | 0 | 279 | 149 | 1 | 1 | 1 | 1 | 1 | 1 | 0.981 | 1 | 1 | 0.959 | 1 | 1 | 0.004 |
| SE | 95185 | ENSG00000213930 | GALT | chr9 | + | 34647831 | 34647961 | 34647491 | 34647705 | 34648333 | 34648375 | 45 | 22 | 45 | 36 | 44 | 0 | 0 | 0 | 0 | 0 | 54 | 41 | 31 | 43 | 39 | 0 | 1 | 0 | 0 | 0 | 279 | 149 | 1 | 1 | 1 | 1 | 1 | 1 | 1 | 1 | 0.956 | 1 | 1 | 1 | 0.009 |
| SE | 95187 | ENSG00000213930 | GALT | chr9 | + | 34648047 | 34648171 | 34647656 | 34647961 | 34648333 | 34648375 | 33 | 29 | 34 | 22 | 11 | 2 | 0 | 0 | 1 | 0 | 34 | 6 | 18 | 28 | 36 | 0 | 0 | 0 | 0 | 0 | 273 | 149 | 1 | 1 | 0.9 | 1 | 1 | 0.923 | 1 | 1 | 1 | 1 | 1 | 1 | -0.035 |
| SE | 95188 | ENSG00000213930 | GALT | chr9 | + | 34648091 | 34648171 | 34647831 | 34647961 | 34648333 | 34648456 | 26 | 27 | 27 | 19 | 14 | 2 | 0 | 0 | 1 | 0 | 22 | 6 | 12 | 17 | 22 | 0 | 0 | 0 | 0 | 0 | 229 | 149 | 1 | 1 | 0.894 | 1 | 1 | 0.925 | 1 | 1 | 1 | 1 | 1 | 1 | -0.036 |
| SE | 95189 | ENSG00000213930 | GALT | chr9 | + | 34648114 | 34648171 | 34646892 | 34647961 | 34648333 | 34648456 | 118 | 82 | 149 | 84 | 61 | 2 | 0 | 0 | 1 | 0 | 132 | 138 | 64 | 82 | 163 | 0 | 0 | 0 | 0 | 0 | 206 | 149 | 1 | 1 | 0.977 | 1 | 1 | 0.984 | 1 | 1 | 1 | 1 | 1 | 1 | -0.008 |
| SE | 95190 | ENSG00000213930 | GALT | chr9 | + | 34648114 | 34648171 | 34647491 | 34647705 | 34648333 | 34648456 | 20 | 25 | 26 | 16 | 9 | 0 | 0 | 0 | 0 | 0 | 22 | 6 | 10 | 12 | 22 | 0 | 1 | 0 | 0 | 0 | 206 | 149 | 1 | 1 | 1 | 1 | 1 | 1 | 1 | 1 | 0.813 | 1 | 1 | 1 | 0.037 |
| SE | 95192 | ENSG00000213930 | GALT | chr9 | + | 34648761 | 34648894 | 34648333 | 34648456 | 34649409 | 34649564 | 42 | 25 | 45 | 28 | 20 | 3 | 0 | 0 | 0 | 0 | 26 | 21 | 22 | 1 | 70 | 0 | 0 | 0 | 0 | 0 | 282 | 149 | 1 | 1 | 0.881 | 1 | 1 | 1 | 1 | 1 | 1 | 1 | 1 | 1 | -0.024 |
| SE | 95193 | ENSG00000213930 | GALT | chr9 | + | 34648761 | 34649081 | 34648424 | 34648456 | 34649409 | 34649564 | 66 | 41 | 92 | 66 | 46 | 3 | 0 | 0 | 0 | 0 | 44 | 48 | 28 | 18 | 141 | 0 | 0 | 0 | 0 | 0 | 298 | 149 | 1 | 1 | 0.917 | 1 | 1 | 1 | 1 | 1 | 1 | 1 | 1 | 1 | -0.017 |
| SE | 95197 | ENSG00000148308 | GTF3C5 | chr9 | + | 133042086 | 133042306 | 133030710 | 133030910 | 133043727 | 133043926 | 20 | 10 | 20 | 17 | 12 | 0 | 1 | 0 | 0 | 0 | 22 | 37 | 12 | 29 | 34 | 0 | 0 | 0 | 0 | 0 | 298 | 149 | 1 | 1 | 1 | 0.833 | 1 | 1 | 1 | 1 | 1 | 1 | 1 | 1 | -0.033 |
| SE | 95198 | ENSG00000148308 | GTF3C5 | chr9 | + | 133042086 | 133042306 | 133030710 | 133030910 | 133050782 | 133050896 | 0 | 0 | 0 | 0 | 0 | 0 | 0 | 0 | 1 | 0 | 0 | 7 | 0 | 0 | 0 | 0 | 0 | 0 | 0 | 0 | 298 | 149 | 1 | 1 | NA | NA | NA | 0 | NA | NA | 1 | NA | NA | NA | -1 |
| SE | 95200 | ENSG00000148308 | GTF3C5 | chr9 | + | 133042086 | 133042306 | 133030996 | 133031164 | 133050782 | 133050896 | 16 | 9 | 9 | 10 | 5 | 0 | 0 | 0 | 1 | 0 | 20 | 8 | 12 | 20 | 29 | 0 | 0 | 0 | 0 | 0 | 298 | 149 | 1 | 1 | 1 | 1 | 1 | 0.833 | 1 | 1 | 1 | 1 | 1 | 1 | -0.033 |
| SE | 95201 | ENSG00000148308 | GTF3C5 | chr9 | + | 133043727 | 133043926 | 133030710 | 133030910 | 133050782 | 133050896 | 14 | 7 | 11 | 21 | 10 | 0 | 0 | 0 | 1 | 0 | 41 | 34 | 14 | 28 | 41 | 0 | 0 | 0 | 0 | 0 | 298 | 149 | 1 | 1 | 1 | 1 | 1 | 0.913 | 1 | 1 | 1 | 1 | 1 | 1 | -0.017 |
| SE | 95202 | ENSG00000148308 | GTF3C5 | chr9 | + | 133043727 | 133043926 | 133031016 | 133031164 | 133050782 | 133050896 | 15 | 8 | 11 | 24 | 10 | 0 | 0 | 0 | 1 | 0 | 41 | 34 | 14 | 28 | 41 | 0 | 0 | 0 | 0 | 0 | 298 | 149 | 1 | 1 | 1 | 1 | 1 | 0.923 | 1 | 1 | 1 | 1 | 1 | 1 | -0.015 |
| SE | 95203 | ENSG00000148308 | GTF3C5 | chr9 | + | 133043727 | 133043926 | 133042086 | 133042306 | 133050782 | 133050896 | 34 | 16 | 31 | 38 | 22 | 0 | 0 | 0 | 0 | 0 | 63 | 67 | 26 | 57 | 75 | 0 | 3 | 0 | 0 | 0 | 298 | 149 | 1 | 1 | 1 | 1 | 1 | 1 | 1 | 1 | 0.918 | 1 | 1 | 1 | 0.016 |
| SE | 95216 | ENSG00000148300 | REXO4 | chr9 | - | 133407806 | 133407881 | 133406058 | 133407072 | 133408767 | 133408842 | 85 | 41 | 101 | 95 | 173 | 0 | 0 | 0 | 0 | 0 | 77 | 84 | 50 | 54 | 42 | 0 | 0 | 0 | 1 | 0 | 224 | 149 | 1 | 1 | 1 | 1 | 1 | 0.984 | 1 | 1 | 1 | 1 | 0.973 | 1 | 0.002 |
| SE | 95217 | ENSG00000148300 | REXO4 | chr9 | - | 133408767 | 133408842 | 133407806 | 133407881 | 133410984 | 133411073 | 102 | 52 | 103 | 110 | 164 | 2 | 0 | 0 | 0 | 0 | 94 | 104 | 51 | 64 | 52 | 0 | 0 | 0 | 0 | 0 | 224 | 149 | 1 | 1 | 0.971 | 1 | 1 | 1 | 1 | 1 | 1 | 1 | 1 | 1 | -0.006 |
| SE | 95226 | ENSG00000167553 | TUBA1C | chr12 | + | 49269827 | 49269976 | 49269464 | 49269687 | 49272252 | 49273328 | 190 | 95 | 139 | 280 | 156 | 0 | 0 | 0 | 0 | 1 | 234 | 253 | 153 | 241 | 1002 | 0 | 0 | 0 | 0 | 0 | 298 | 149 | 1 | 1 | 1 | 1 | 1 | 1 | 0.987 | 1 | 1 | 1 | 1 | 1 | -0.003 |
| SE | 95227 | ENSG00000160323 | ADAMTS13 | chr9 | + | 133425528 | 133425612 | 133424320 | 133424478 | 133425937 | 133426062 | 0 | 0 | 1 | 0 | 0 | 0 | 0 | 0 | 1 | 0 | 0 | 0 | 6 | 0 | 0 | 0 | 0 | 0 | 0 | 0 | 233 | 149 | 1 | 1 | NA | NA | 1 | 0 | NA | NA | NA | 1 | NA | NA | -0.5 |
| SE | 95229 | ENSG00000160323 | ADAMTS13 | chr9 | + | 133438245 | 133438366 | 133437748 | 133437897 | 133439365 | 133439446 | 0 | 2 | 0 | 0 | 0 | 0 | 0 | 0 | 0 | 0 | 0 | 0 | 0 | 0 | 0 | 0 | 5 | 3 | 0 | 0 | 270 | 149 | 1 | 1 | NA | 1 | NA | NA | NA | NA | 0 | 0 | NA | NA | 1 |
| SE | 95232 | ENSG00000160325 | CACFD1 | chr9 | + | 133465034 | 133465158 | 133463482 | 133463555 | 133465321 | 133465447 | 0 | 0 | 1 | 0 | 0 | 1 | 2 | 2 | 0 | 0 | 0 | 0 | 0 | 0 | 0 | 0 | 0 | 0 | 2 | 5 | 273 | 149 | 1 | 1 | 0 | 0 | 0.214 | NA | NA | NA | NA | NA | 0 | 0 | 0.071 |
| SE | 95233 | ENSG00000160325 | CACFD1 | chr9 | + | 133465321 | 133465447 | 133463482 | 133463555 | 133467920 | 133468028 | 1 | 2 | 2 | 0 | 4 | 0 | 1 | 0 | 0 | 0 | 0 | 2 | 3 | 5 | 5 | 0 | 0 | 0 | 0 | 0 | 275 | 149 | 1 | 1 | 1 | 0.52 | 1 | NA | 1 | NA | 1 | 1 | 1 | 1 | -0.12 |
| SE | 95234 | ENSG00000160325 | CACFD1 | chr9 | + | 133468339 | 133468464 | 133467920 | 133468028 | 133468562 | 133470787 | 0 | 0 | 1 | 0 | 3 | 1 | 0 | 2 | 2 | 0 | 0 | 0 | 0 | 0 | 0 | 0 | 3 | 4 | 0 | 5 | 274 | 149 | 1 | 1 | 0 | NA | 0.214 | 0 | 1 | NA | 0 | 0 | NA | 0 | 0.303 |
| SE | 95235 | ENSG00000122565 | CBX3 | chr7 | + | 26202970 | 26203022 | 26201161 | 26201561 | 26206367 | 26206510 | 118 | 90 | 97 | 115 | 160 | 0 | 0 | 0 | 2 | 0 | 3 | 72 | 58 | 39 | 10 | 0 | 0 | 0 | 0 | 0 | 201 | 149 | 1 | 1 | 1 | 1 | 1 | 0.977 | 1 | 1 | 1 | 1 | 1 | 1 | -0.005 |
| SE | 95236 | ENSG00000122565 | CBX3 | chr7 | + | 26208392 | 26208555 | 26206367 | 26206510 | 26211661 | 26211717 | 1393 | 984 | 1302 | 1311 | 1903 | 2 | 0 | 0 | 2 | 0 | 302 | 713 | 580 | 452 | 168 | 0 | 0 | 0 | 0 | 0 | 298 | 149 | 1 | 1 | 0.997 | 1 | 1 | 0.997 | 1 | 1 | 1 | 1 | 1 | 1 | -0.001 |
| SE | 95237 | ENSG00000122565 | CBX3 | chr7 | + | 26208465 | 26208555 | 26206367 | 26206510 | 26211661 | 26211756 | 513 | 357 | 494 | 465 | 655 | 2 | 0 | 0 | 2 | 0 | 135 | 340 | 267 | 199 | 91 | 0 | 0 | 0 | 0 | 0 | 239 | 149 | 1 | 1 | 0.994 | 1 | 1 | 0.993 | 1 | 1 | 1 | 1 | 1 | 1 | -0.003 |
| SE | 95239 | ENSG00000159712 | ANKRD18CP | chr9 | - | 97179996 | 97180916 | 97178263 | 97178413 | 97185441 | 97185515 | 0 | 2 | 0 | 0 | 3 | 0 | 0 | 1 | 0 | 0 | 0 | 6 | 0 | 0 | 0 | 6 | 1 | 5 | 3 | 0 | 298 | 149 | 1 | 1 | NA | | 0 | NA | 1 | 0 | 0.75 | 0 | 0 | NA | 0.479 |
| SE | 95240 | ENSG00000159712 | ANKRD18CP | chr9 | - | 97200458 | 97200565 | 97194702 | 97194840 | 97211372 | 97211487 | 1 | 1 | 0 | 0 | 0 | 0 | 1 | 0 | 0 | 0 | 0 | 0 | 0 | 0 | 0 | 0 | 1 | 0 | 0 | 0 | 256 | 149 | 1 | 1 | 1 | 0.368 | NA | NA | NA | NA | 0 | NA | NA | NA | 0.684 |
| SE | 95250 | ENSG00000113356 | POLR3G | chr5 | + | 90497655 | 90497706 | 90495676 | 90495733 | 90501905 | 90501988 | 11 | 14 | 10 | 10 | 4 | 0 | 0 | 0 | 0 | 1 | 24 | 4 | 3 | 23 | 18 | 0 | 0 | 0 | 0 | 0 | 200 | 149 | 1 | 1 | 1 | 1 | 1 | 1 | 0.749 | 1 | 1 | 1 | 1 | 1 | -0.05 |
| SE | 95251 | ENSG00000113356 | POLR3G | chr5 | + | 90497655 | 90497735 | 90495676 | 90495733 | 90501905 | 90501988 | 5 | 4 | 2 | 5 | 10 | 0 | 0 | 0 | 0 | 1 | 0 | 1 | 6 | 6 | 6 | 0 | 0 | 0 | 0 | 0 | 229 | 149 | 1 | 1 | 1 | 1 | 1 | 1 | 0.867 | NA | 1 | 1 | 1 | 1 | -0.027 |
| SE | 95253 | ENSG00000165219 | GAPVD1 | chr9 | + | 125263550 | 125263700 | 125261830 | 125261959 | 125295457 | 125295522 | 9 | 1 | 1 | 1 | 0 | 5 | 2 | 1 | 2 | 1 | 12 | 4 | 0 | 0 | 0 | 0 | 3 | 0 | 0 | 2 | 298 | 149 | 1 | 1 | 0.474 | 0.2 | 0.333 | 0.2 | 0 | 1 | 0.4 | NA | NA | 0 | -0.225 |
| SE | 95255 | ENSG00000165219 | GAPVD1 | chr9 | + | 125268935 | 125268984 | 125261809 | 125261959 | 125295457 | 125295574 | 1 | 1 | 3 | 4 | 2 | 5 | 2 | 1 | 2 | 1 | 0 | 0 | 0 | 5 | 0 | 0 | 3 | 0 | 0 | 2 | 198 | 149 | 1 | 1 | 0.131 | 0.273 | 0.693 | 0.601 | 0.601 | NA | 0 | NA | 1 | 0 | 0.126 |
| SE | 95258 | ENSG00000165219 | GAPVD1 | chr9 | + | 125287761 | 125287883 | 125263550 | 125263700 | 125298889 | 125299106 | 2 | 2 | 0 | 2 | 0 | 1 | 0 | 0 | 1 | 0 | 0 | 11 | 0 | 2 | 9 | 0 | 0 | 0 | 0 | 0 | 271 | 149 | 1 | 1 | 0.524 | 1 | NA | 0.524 | NA | NA | 1 | NA | 1 | 1 | -0.317 |
| SE | 95259 | ENSG00000165219 | GAPVD1 | chr9 | + | 125287761 | 125287883 | 125268935 | 125268984 | 125295457 | 125295522 | 5 | 1 | 1 | 3 | 0 | 2 | 0 | 3 | 4 | 2 | 0 | 2 | 0 | 0 | 0 | 0 | 0 | 0 | 5 | 0 | 271 | 149 | 1 | 1 | 0.579 | 1 | 0.155 | 0.292 | 0 | NA | 1 | NA | 0 | NA | -0.095 |
| SE | 95263 | ENSG00000165219 | GAPVD1 | chr9 | + | 125298889 | 125299106 | 125261793 | 125261959 | 125301982 | 125302148 | 120 | 109 | 113 | 126 | 179 | 1 | 0 | 0 | 0 | 0 | 201 | 160 | 128 | 226 | 70 | 0 | 0 | 0 | 0 | 0 | 298 | 149 | 1 | 1 | 0.984 | 1 | 1 | 1 | 1 | 1 | 1 | 1 | 1 | 1 | -0.003 |
| SE | 95264 | ENSG00000165219 | GAPVD1 | chr9 | + | 125301982 | 125302826 | 125298885 | 125299106 | 125305062 | 125305149 | 163 | 153 | 134 | 182 | 302 | 0 | 0 | 0 | 1 | 0 | 326 | 205 | 164 | 315 | 72 | 0 | 0 | 0 | 0 | 0 | 298 | 149 | 1 | 1 | 1 | 1 | 1 | 0.989 | 1 | 1 | 1 | 1 | 1 | 1 | -0.002 |
| SE | 95267 | ENSG00000165219 | GAPVD1 | chr9 | + | 125323797 | 125323923 | 125321432 | 125321499 | 125326415 | 125326589 | 289 | 179 | 189 | 272 | 311 | 1 | 1 | 0 | 1 | 0 | 282 | 281 | 241 | 298 | 157 | 0 | 0 | 0 | 1 | 0 | 275 | 149 | 1 | 1 | 0.994 | 0.99 | 1 | 0.993 | 1 | 1 | 1 | 1 | 0.994 | 1 | -0.003 |
| SE | 95268 | ENSG00000165219 | GAPVD1 | chr9 | + | 125323797 | 125323923 | 125321432 | 125321562 | 125326415 | 125326589 | 350 | 228 | 228 | 300 | 319 | 0 | 0 | 1 | 2 | 3 | 331 | 326 | 269 | 394 | 154 | 0 | 1 | 0 | 0 | 4 | 275 | 149 | 1 | 1 | 1 | 1 | 0.992 | 0.988 | 0.983 | 1 | 0.994 | 1 | 1 | 0.954 | 0.003 |
| SE | 95269 | ENSG00000165219 | GAPVD1 | chr9 | + | 125326415 | 125326589 | 125323797 | 125323923 | 125330077 | 125330218 | 387 | 240 | 242 | 308 | 414 | 0 | 0 | 1 | 0 | 0 | 410 | 351 | 339 | 409 | 235 | 0 | 0 | 0 | 0 | 0 | 298 | 149 | 1 | 1 | 1 | 1 | 0.992 | 1 | 1 | 1 | 1 | 1 | 1 | 1 | -0.002 |
| SE | 95270 | ENSG00000165219 | GAPVD1 | chr9 | + | 125330077 | 125330218 | 125323797 | 125323923 | 125331925 | 125331972 | 80 | 37 | 54 | 71 | 86 | 0 | 0 | 0 | 0 | 0 | 70 | 77 | 68 | 62 | 23 | 2 | 0 | 0 | 0 | 0 | 290 | 149 | 1 | 1 | 1 | 1 | 1 | 1 | 1 | 0.947 | 1 | 1 | 1 | 1 | 0.011 |
| SE | 95272 | ENSG00000165219 | GAPVD1 | chr9 | + | 125337017 | 125337095 | 125332509 | 125332629 | 125337220 | 125337591 | 112 | 62 | 96 | 79 | 113 | 0 | 0 | 0 | 0 | 2 | 114 | 101 | 66 | 92 | 102 | 0 | 0 | 0 | 0 | 0 | 227 | 149 | 1 | 1 | 1 | 1 | 1 | 1 | 0.974 | 1 | 1 | 1 | 1 | 1 | -0.005 |
| SE | 95277 | ENSG00000122729 | ACO1 | chr9 | + | 32386370 | 32386438 | 32384619 | 32384735 | 32405484 | 32405603 | 0 | 0 | 0 | 0 | 0 | 2 | 1 | 3 | 2 | 7 | 0 | 3 | 0 | 4 | 0 | 12 | 2 | 1 | 4 | 0 | 217 | 149 | 1 | 1 | 0 | 0 | 0 | 0 | 0 | 0 | 0.507 | 0 | 0.407 | NA | -0.228 |
| SE | 95279 | ENSG00000122729 | ACO1 | chr9 | + | 32418327 | 32418511 | 32418127 | 32418197 | 32419037 | 32419177 | 24 | 12 | 13 | 33 | 27 | 0 | 0 | 0 | 1 | 1 | 32 | 9 | 17 | 35 | 34 | 0 | 8 | 0 | 0 | 0 | 298 | 149 | 1 | 1 | 1 | 1 | 1 | 0.943 | 0.931 | 1 | 0.36 | 1 | 1 | 1 | 0.103 |
| SE | 95280 | ENSG00000172716 | SLFN11 | chr17 | - | 35360242 | 35360371 | 35353335 | 35354059 | 35362738 | 35363826 | 48 | 24 | 104 | 149 | 170 | 2 | 0 | 3 | 1 | 0 | 170 | 163 | 164 | 124 | 69 | 5 | 1 | 0 | 0 | 3 | 278 | 149 | 1 | 1 | 0.928 | 1 | 0.949 | 0.988 | 1 | 0.948 | 0.989 | 1 | 1 | 0.925 | 0.001 |
| SE | 95289 | ENSG00000172716 | SLFN11 | chr17 | - | 35367540 | 35367700 | 35363536 | 35363826 | 35373473 | 35373620 | 10 | 5 | 7 | 16 | 0 | 8 | 4 | 7 | 1 | 0 | 22 | 11 | 5 | 8 | 6 | 50 | 8 | 11 | 7 | 0 | 298 | 149 | 1 | 1 | 0.385 | 0.385 | 0.333 | 0.889 | NA | 0.18 | 0.407 | 0.185 | 0.364 | 1 | 0.071 |
| SE | 95290 | ENSG00000172716 | SLFN11 | chr17 | - | 35367540 | 35367700 | 35363536 | 35363826 | 35373581 | 35373620 | 2 | 1 | 1 | 0 | 0 | 5 | 1 | 3 | 3 | 0 | 0 | 1 | 2 | 4 | 6 | 8 | 1 | 6 | 1 | 10 | 298 | 149 | 1 | 1 | 0.167 | 0.333 | 0.143 | 0 | NA | 0 | 0.333 | 0.143 | 0.667 | 0 | -0.068 |
| SE | 95292 | ENSG00000172716 | SLFN11 | chr17 | - | 35367540 | 35367700 | 35366946 | 35367043 | 35373581 | 35373620 | 2 | 1 | 1 | 1 | 0 | 0 | 0 | 0 | 0 | 0 | 0 | 1 | 0 | 1 | 0 | 14 | 0 | 1 | 0 | 7 | 298 | 149 | 1 | 1 | 1 | 1 | 1 | 1 | NA | 0 | 1 | 0 | 1 | 0 | 0.6 |
| SE | 95294 | ENSG00000172716 | SLFN11 | chr17 | - | 35367540 | 35367700 | 35366946 | 35367063 | 35373581 | 35373620 | 7 | 8 | 3 | 0 | 6 | 0 | 1 | 3 | 0 | 2 | 11 | 1 | 0 | 4 | 0 | 0 | 3 | 0 | 0 | 0 | 298 | 149 | 1 | 1 | 1 | 0.8 | 0.333 | NA | 0.6 | 1 | 0.143 | NA | 1 | NA | -0.031 |
| SE | 95295 | ENSG00000172716 | SLFN11 | chr17 | - | 35367602 | 35367700 | 35362738 | 35363826 | 35373473 | 35373695 | 8 | 6 | 8 | 23 | 0 | 8 | 4 | 7 | 1 | 0 | 32 | 20 | 14 | 12 | 16 | 50 | 8 | 11 | 7 | 0 | 247 | 149 | 1 | 1 | 0.376 | 0.475 | 0.408 | 0.933 | NA | 0.279 | 0.601 | 0.434 | 0.508 | 1 | -0.016 |
| SE | 95296 | ENSG00000172716 | SLFN11 | chr17 | - | 35367602 | 35367700 | 35362738 | 35363826 | 35373581 | 35373620 | 2 | 1 | 1 | 5 | 0 | 5 | 1 | 3 | 3 | 0 | 10 | 9 | 12 | 10 | 10 | 8 | 1 | 6 | 1 | 10 | 247 | 149 | 1 | 1 | 0.194 | 0.376 | 0.167 | 0.501 | NA | 0.43 | 0.844 | 0.547 | 0.858 | 0.376 | -0.301 |
| SE | 95298 | ENSG00000172716 | SLFN11 | chr17 | - | 35367602 | 35367700 | 35366946 | 35367043 | 35373581 | 35373620 | 8 | 4 | 3 | 3 | 4 | 0 | 0 | 0 | 0 | 0 | 0 | 0 | 1 | 9 | 0 | 14 | 0 | 1 | 0 | 7 | 247 | 149 | 1 | 1 | 1 | 1 | 1 | 1 | 1 | 0 | NA | 0.376 | 1 | 0 | 0.656 |
| SE | 95302 | ENSG00000197579 | TOPORS | chr9 | - | 32550773 | 32550968 | 32540543 | 32544326 | 32552433 | 32552586 | 33 | 20 | 50 | 39 | 48 | 3 | 4 | 1 | 8 | 5 | 8 | 15 | 40 | 2 | 0 | 0 | 2 | 1 | 2 | 0 | 298 | 149 | 1 | 1 | 0.846 | 0.714 | 0.962 | 0.709 | 0.828 | 1 | 0.789 | 0.952 | 0.333 | NA | 0.043 |
| SE | 95305 | ENSG00000148400 | NOTCH1 | chr9 | - | 136515980 | 136516094 | 136515482 | 136515716 | 136517271 | 136517385 | 74 | 20 | 58 | 81 | 32 | 0 | 0 | 0 | 0 | 0 | 41 | 51 | 63 | 36 | 111 | 0 | 0 | 2 | 0 | 0 | 263 | 149 | 1 | 1 | 1 | 1 | 1 | 1 | 1 | 1 | 1 | 0.947 | 1 | 1 | 0.011 |

| | | | | | | | | | | | | | | |
|---|---|---|---|---|---|---|---|---|---|---|---|---|---|---|
| SE | 95308 | ENSG00000148400 | NOTCH1 | chr9 | - | 136517271 | 136517385 | 136515980 | 136516094 | 136517751 | 136517937 | 60 | 14 | [illegible] |
| SE | 95309 | ENSG00000148400 | NOTCH1 | chr9 | - | 136518136 | 136518292 | 136517751 | 136517937 | 136518590 | 136518824 | 197 | 59 | [illegible] |
| SE | 95310 | ENSG00000148400 | NOTCH1 | chr9 | - | 136519442 | 136519565 | 136518590 | 136518824 | 136544023 | 136544102 | 147 | 52 | [illegible] |
| SE | 95311 | ENSG00000148400 | NOTCH1 | chr9 | - | 136523716 | 136523979 | 136518590 | 136518824 | 136544023 | 136544102 | 36 | 9 | [illegible] |
| SE | 95314 | ENSG00000165699 | TSC1 | chr9 | - | 132900714 | 132900837 | 132897183 | 132897610 | 132901630 | 132901699 | 33 | 26 | [illegible] |
| SE | 95315 | ENSG00000165699 | TSC1 | chr9 | - | 132902604 | 132902783 | 132901588 | 132901699 | 132903650 | 132903817 | 35 | 25 | [illegible] |
| SE | 95316 | ENSG00000165699 | TSC1 | chr9 | - | 132902604 | 132902787 | 132901588 | 132901699 | 132903650 | 132903802 | 66 | 45 | [illegible] |
| SE | 95319 | ENSG00000165699 | TSC1 | chr9 | - | 132904410 | 132904454 | 132903650 | 132903817 | 132905580 | 132905621 | 32 | 23 | [illegible] |
| SE | 95320 | ENSG00000165699 | TSC1 | chr9 | - | 132909806 | 132910692 | 132907300 | 132907370 | 132911001 | 132911113 | 29 | 6 | [illegible] |
| SE | 95321 | ENSG00000165699 | TSC1 | chr9 | - | 132909806 | 132910692 | 132907300 | 132907370 | 132911452 | 132911568 | 2 | 0 | [illegible] |
| SE | 95323 | ENSG00000165699 | TSC1 | chr9 | - | 132910570 | 132910689 | 132907300 | 132907370 | 132911452 | 132911568 | 14 | 19 | [illegible] |
| SE | 95324 | ENSG00000165699 | TSC1 | chr9 | - | 132910570 | 132910692 | 132907300 | 132907370 | 132911001 | 132911113 | 42 | 25 | [illegible] |
| SE | 95325 | ENSG00000165699 | TSC1 | chr9 | - | 132910570 | 132910692 | 132907300 | 132907370 | 132911452 | 132911568 | 14 | 19 | [illegible] |
| SE | 95326 | ENSG00000165699 | TSC1 | chr9 | - | 132911001 | 132911113 | 132907300 | 132907370 | 132911452 | 132911568 | 15 | 12 | [illegible] |
| SE | 95327 | ENSG00000165699 | TSC1 | chr9 | - | 132911001 | 132911113 | 132909751 | 132910692 | 132911409 | 132911568 | 20 | 6 | [illegible] |
| SE | 95330 | ENSG00000165699 | TSC1 | chr9 | - | 132921362 | 132921436 | 132912281 | 132912457 | 132921818 | 132921973 | 104 | 75 | [illegible] |
| SE | 95335 | ENSG00000165699 | TSC1 | chr9 | - | 132925586 | 132925739 | 132923347 | 132923492 | 132928766 | 132928952 | 20 | 9 | [illegible] |
| SE | 95336 | ENSG00000165699 | TSC1 | chr9 | - | 132925586 | 132927304 | 132923347 | 132923492 | 132928766 | 132928952 | 98 | 70 | [illegible] |
| SE | 95337 | ENSG00000165699 | TSC1 | chr9 | - | 132927200 | 132927304 | 132923347 | 132923492 | 132928766 | 132928952 | 79 | 62 | [illegible] |
| SE | 95338 | ENSG00000165699 | TSC1 | chr9 | - | 132927200 | 132927304 | 132925586 | 132925739 | 132928766 | 132928952 | 100 | 82 | [illegible] |
| SE | 95342 | ENSG00000165699 | TSC1 | chr9 | - | 132934465 | 132934540 | 132931208 | 132931427 | 132935032 | 132935095 | 0 | 0 | [illegible] |
| SE | 95343 | ENSG00000165699 | TSC1 | chr9 | - | 132935032 | 132935095 | 132928766 | 132928952 | 132944542 | 132944572 | 54 | 37 | [illegible] |
| SE | 95345 | ENSG00000165699 | TSC1 | chr9 | - | 132943467 | 132943706 | 132940734 | 132942426 | 132944542 | 132944616 | 0 | 0 | [illegible] |
| SE | 95346 | ENSG00000132481 | TRIM47 | chr17 | - | 75875900 | 75876099 | 75875399 | 75875474 | 75876261 | 75876492 | 1 | 0 | [illegible] |
| SE | 95348 | ENSG00000115459 | ELMOD3 | chr2 | + | 85355075 | 85355170 | 85354393 | 85354829 | 85356601 | 85356667 | 7 | 2 | [illegible] |
| SE | 95351 | ENSG00000115459 | ELMOD3 | chr2 | + | 85355554 | 85355598 | 85354719 | 85354829 | 85356966 | 85357252 | 11 | 2 | [illegible] |
| SE | 95352 | ENSG00000115459 | ELMOD3 | chr2 | + | 85355554 | 85355598 | 85355075 | 85355170 | 85356601 | 85356667 | 1 | 2 | [illegible] |
| SE | 95353 | ENSG00000115459 | ELMOD3 | chr2 | + | 85355554 | 85355598 | 85355075 | 85355170 | 85356966 | 85357252 | 12 | 4 | [illegible] |
| SE | 95354 | ENSG00000115459 | ELMOD3 | chr2 | + | 85355554 | 85355716 | 85354725 | 85354829 | 85356601 | 85356667 | 2 | 5 | [illegible] |
| SE | 95356 | ENSG00000115459 | ELMOD3 | chr2 | + | 85355554 | 85355716 | 85355075 | 85355170 | 85356601 | 85356667 | 6 | 3 | [illegible] |
| SE | 95359 | ENSG00000115459 | ELMOD3 | chr2 | + | 85355554 | 85355896 | 85354805 | 85354829 | 85356601 | 85356667 | 2 | 5 | [illegible] |
| SE | 95361 | ENSG00000115459 | ELMOD3 | chr2 | + | 85356601 | 85356667 | 85354752 | 85354829 | 85356966 | 85357252 | 0 | 0 | [illegible] |
| SE | 95364 | ENSG00000115459 | ELMOD3 | chr2 | + | 85356601 | 85356667 | 85355554 | 85355716 | 85356966 | 85357252 | 0 | 0 | [illegible] |
| SE | 95365 | ENSG00000115459 | ELMOD3 | chr2 | + | 85356601 | 85356667 | 85355554 | 85355896 | 85356966 | 85357252 | 0 | 0 | [illegible] |
| SE | 95367 | ENSG00000115459 | ELMOD3 | chr2 | + | 85356966 | 85357252 | 85355916 | 85356533 | 85362185 | 85362260 | 73 | 53 | [illegible] |
| SE | 95371 | ENSG00000115459 | ELMOD3 | chr2 | + | 85371085 | 85371209 | 85368685 | 85368754 | 85377343 | 85377474 | 10 | 0 | [illegible] |
| SE | 95372 | ENSG00000115459 | ELMOD3 | chr2 | + | 85371085 | 85371562 | 85368685 | 85368754 | 85377343 | 85377474 | 17 | 9 | [illegible] |
| SE | 95376 | ENSG00000181404 | WASHC1 | chr9 | - | 15080 | 15149 | 14474 | 14940 | 15908 | 16061 | 15 | 1 | [illegible] |
| SE | 95377 | ENSG00000181404 | WASHC1 | chr9 | - | 17343 | 17479 | 16964 | 17166 | 17718 | 17855 | 46 | 6 | [illegible] |
| SE | 95378 | ENSG00000181404 | WASHC1 | chr9 | - | 17718 | 17855 | 17343 | 17479 | 18027 | 18174 | 48 | 11 | [illegible] |
| SE | 95380 | ENSG00000198431 | TXNRD1 | chr12 | + | 104288930 | 104289040 | 104258018 | 104258079 | 104311289 | 104311412 | 95 | 66 | [illegible] |
| SE | 95384 | ENSG00000198431 | TXNRD1 | chr12 | + | 104289465 | 104289542 | 104288894 | 104289040 | 104311289 | 104311412 | 0 | 0 | [illegible] |
| SE | 95386 | ENSG00000198431 | TXNRD1 | chr12 | + | 104290992 | 104291104 | 104289465 | 104289542 | 104311289 | 104311412 | 2 | 0 | [illegible] |
| SE | 95389 | ENSG00000198431 | TXNRD1 | chr12 | + | 104311289 | 104311412 | 104288717 | 104289040 | 104313244 | 104313316 | 192 | 125 | [illegible] |
| SE | 95390 | ENSG00000198431 | TXNRD1 | chr12 | + | 104313244 | 104313317 | 104311289 | 104311412 | 104315776 | 104315826 | 254 | 149 | [illegible] |
| SE | 95391 | ENSG00000198431 | TXNRD1 | chr12 | + | 104319469 | 104319585 | 104318912 | 104319055 | 104321090 | 104321316 | 235 | 171 | [illegible] |
| SE | 95392 | ENSG00000198431 | TXNRD1 | chr12 | + | 104321090 | 104321316 | 104319469 | 104319585 | 104325336 | 104325429 | 313 | 238 | [illegible] |
| SE | 95393 | ENSG00000198431 | TXNRD1 | chr12 | + | 104326346 | 104326423 | 104325336 | 104325429 | 104327514 | 104327574 | 319 | 231 | [illegible] |
| SE | 95394 | ENSG00000198431 | TXNRD1 | chr12 | + | 104346018 | 104346155 | 104339138 | 104339273 | 104348352 | 104350283 | 0 | 0 | [illegible] |
| SE | 95397 | ENSG00000151239 | TWF1 | chr12 | - | 43797707 | 43797833 | 43797301 | 43797452 | 43799397 | 43799502 | 269 | 208 | [illegible] |
| SE | 95401 | ENSG00000151239 | TWF1 | chr12 | - | 43805523 | 43805637 | 43804494 | 43804572 | 43806220 | 43806296 | 0 | 0 | [illegible] |
| SE | 95402 | ENSG00000151239 | TWF1 | chr12 | - | 43805830 | 43805885 | 43804494 | 43804572 | 43806220 | 43806287 | 1 | 1 | [illegible] |
| SE | 95403 | ENSG00000151239 | TWF1 | chr12 | - | 43805830 | 43806002 | 43804494 | 43804572 | 43806220 | 43806282 | 2 | 1 | [illegible] |
| SE | 95409 | ENSG00000148356 | LRSAM1 | chr9 | + | 127460876 | 127460998 | 127459002 | 127459071 | 127461172 | 127461257 | 0 | 0 | [illegible] |
| SE | 95411 | ENSG00000148356 | LRSAM1 | chr9 | + | 127473800 | 127473931 | 127462251 | 127462373 | 127478933 | 127478963 | 39 | 21 | [illegible] |
| SE | 95412 | ENSG00000148356 | LRSAM1 | chr9 | + | 127482732 | 127482813 | 127481182 | 127481227 | 127482949 | 127483020 | 3 | 0 | [illegible] |
| SE | 95413 | ENSG00000148356 | LRSAM1 | chr9 | + | 127482949 | 127483020 | 127481182 | 127481227 | 127485735 | 127485765 | 42 | 23 | [illegible] |
| SE | 95414 | ENSG00000148356 | LRSAM1 | chr9 | + | 127486404 | 127486694 | 127485735 | 127485835 | 127487494 | 127487553 | 0 | 0 | [illegible] |
| SE | 95427 | ENSG00000234705 | HMGA1P4 | chr9 | - | 128663511 | 128663576 | 128663133 | 128663400 | 128684365 | 128684477 | 0 | 3 | [illegible] |
| SE | 95429 | ENSG00000123353 | ORMDL2 | chr12 | + | 55818998 | 55819173 | 55818040 | 55818112 | 55819341 | 55819493 | 147 | 44 | [illegible] |
| SE | 95430 | ENSG00000123353 | ORMDL2 | chr12 | + | 55819090 | 55819173 | 55818040 | 55818112 | 55819341 | 55819493 | 48 | 15 | [illegible] |
| SE | 95432 | ENSG00000167130 | DOLPP1 | chr9 | + | 129085206 | 129085306 | 129084667 | 129084768 | 129085517 | 129085616 | 21 | 4 | [illegible] |
| SE | 95438 | ENSG00000119383 | PTPA | chr9 | + | 129123051 | 129123138 | 129120512 | 129120610 | 129131521 | 129131639 | 37 | 36 | [illegible] |
| SE | 95440 | ENSG00000119383 | PTPA | chr9 | + | 129128984 | 129129110 | 129120512 | 129120610 | 129131521 | 129131639 | 85 | 32 | [illegible] |
| SE | 95441 | ENSG00000119383 | PTPA | chr9 | + | 129128984 | 129129110 | 129123051 | 129123138 | 129131521 | 129131639 | 138 | 78 | [illegible] |
| SE | 95442 | ENSG00000119383 | PTPA | chr9 | + | 129128984 | 129129110 | 129123051 | 129123138 | 129136470 | 129136556 | 47 | 36 | [illegible] |
| SE | 95443 | ENSG00000119383 | PTPA | chr9 | + | 129131521 | 129131639 | 129123051 | 129123138 | 129136470 | 129136556 | 4 | 2 | [illegible] |
| SE | 95444 | ENSG00000119383 | PTPA | chr9 | + | 129131521 | 129131639 | 129128984 | 129129110 | 129134794 | 129134827 | 169 | 79 | [illegible] |
| SE | 95445 | ENSG00000119383 | PTPA | chr9 | + | 129131521 | 129131639 | 129128984 | 129129110 | 129136470 | 129136556 | 87 | 41 | [illegible] |
| SE | 95448 | ENSG00000119383 | PTPA | chr9 | + | 129136470 | 129136595 | 129134794 | 129134894 | 129137591 | 129137692 | 216 | 130 | [illegible] |
| SE | 95449 | ENSG00000160447 | PKN3 | chr9 | + | 128715368 | 128715460 | 128715171 | 128715235 | 128716746 | 128716923 | 2 | 0 | [illegible] |
| SE | 95450 | ENSG00000160445 | ZER1 | chr9 | - | 128735333 | 128735431 | 128733425 | 128733528 | 128739930 | 128740119 | 161 | 93 | [illegible] |
| SE | 95451 | ENSG00000160445 | ZER1 | chr9 | - | 128735333 | 128735431 | 128733425 | 128733528 | 128740771 | 128740887 | 70 | 37 | [illegible] |
| SE | 95452 | ENSG00000160445 | ZER1 | chr9 | - | 128739930 | 128740119 | 128733425 | 128733528 | 128740771 | 128740887 | 82 | 25 | [illegible] |
| SE | 95455 | ENSG00000180773 | SLC36A4 | chr11 | - | 93167943 | 93168171 | 93165917 | 93166016 | 93180796 | 93180881 | 121 | 76 | [illegible] |
| SE | 95457 | ENSG00000180773 | SLC36A4 | chr11 | - | 93180796 | 93180881 | 93170153 | 93170221 | 93181690 | 93181786 | 15 | 8 | [illegible] |
| SE | 95461 | ENSG00000171097 | KYAT1 | chr9 | - | 128846679 | 128846859 | 128845352 | 128845411 | 128881896 | 128881931 | 0 | 4 | [illegible] |
| SE | 95463 | ENSG00000136802 | LRRC8A | chr9 | + | 128886014 | 128886121 | 128882132 | 128882250 | 128907156 | 128909321 | 55 | 11 | [illegible] |
| SE | 95468 | ENSG00000095319 | NUP188 | chr9 | + | 128958009 | 128958054 | 128956951 | 128957032 | 128958801 | 128958894 | 141 | 81 | [illegible] |
| SE | 95469 | ENSG00000095319 | NUP188 | chr9 | + | 128968505 | 128968717 | 128959014 | 128959134 | 128969399 | 128969514 | 259 | 163 | [illegible] |
| SE | 95470 | ENSG00000095319 | NUP188 | chr9 | + | 128968505 | 128968717 | 128959014 | 128959134 | 128970757 | 128970958 | 178 | 104 | [illegible] |
| SE | 95471 | ENSG00000095319 | NUP188 | chr9 | + | 128969399 | 128969514 | 128959014 | 128959134 | 128970757 | 128970958 | 71 | 26 | [illegible] |
| SE | 95473 | ENSG00000095319 | NUP188 | chr9 | + | 128970757 | 128970958 | 128959014 | 128959134 | 128973159 | 128973249 | 89 | 40 | [illegible] |
| SE | 95474 | ENSG00000095319 | NUP188 | chr9 | + | 128973159 | 128973249 | 128970757 | 128970958 | 128979261 | 128979327 | 128 | 53 | [illegible] |
| SE | 95475 | ENSG00000095319 | NUP188 | chr9 | + | 128990119 | 128990226 | 128988046 | 128988186 | 128993196 | 128993403 | 56 | 28 | [illegible] |
| SE | 95476 | ENSG00000095319 | NUP188 | chr9 | + | 128990152 | 128990226 | 128988046 | 128988186 | 128993196 | 128993230 | 21 | 9 | [illegible] |
| SE | 95478 | ENSG00000095319 | NUP188 | chr9 | + | 128998537 | 128998623 | 128998150 | 128998228 | 128999623 | 128999805 | 9 | 7 | [illegible] |
| SE | 95481 | ENSG00000108424 | KPNB1 | chr17 | + | 47656859 | 47657060 | 47652693 | 47652876 | 47658507 | 47658641 | 874 | 572 | [illegible] |
| SE | 95482 | ENSG00000108424 | KPNB1 | chr17 | + | 47663088 | 47663178 | 47660881 | 47661178 | 47668185 | 47668410 | 246 | 131 | [illegible] |
| SE | 95483 | ENSG00000108424 | KPNB1 | chr17 | + | 47664158 | 47664269 | 47663088 | 47663178 | 47665056 | 47665158 | 839 | 592 | [illegible] |
| SE | 95484 | ENSG00000108424 | KPNB1 | chr17 | + | 47665056 | 47665158 | 47663088 | 47663178 | 47668185 | 47668410 | 336 | 229 | [illegible] |
| SE | 95488 | ENSG00000108424 | KPNB1 | chr17 | + | 47676408 | 47676491 | 47674637 | 47674782 | 47677019 | 47677127 | 399 | 279 | [illegible] |
| SE | 95489 | ENSG00000108424 | KPNB1 | chr17 | + | 47677019 | 47677127 | 47674637 | 47674782 | 47678045 | 47678189 | 239 | 146 | [illegible] |
| SE | 95490 | ENSG00000108424 | KPNB1 | chr17 | + | 47680019 | 47680134 | 47678307 | 47678413 | 47680507 | 47680669 | 1155 | 927 | [illegible] |
| SE | 95492 | ENSG00000114999 | TTL | chr2 | + | 112494142 | 112494375 | 112485916 | 112485995 | 112501205 | 112501341 | 112 | 109 | [illegible] |
| SE | 95493 | ENSG00000114999 | TTL | chr2 | + | 112520281 | 112520425 | 112502911 | 112503181 | 112528680 | 112541739 | 53 | 46 | [illegible] |
| SE | 95495 | ENSG00000025423 | HSD17B6 | chr12 | + | 56763197 | 56763414 | 56752160 | 56752318 | 56773833 | 56774078 | 1 | 0 | [illegible] |
| SE | 95499 | ENSG00000025423 | HSD17B6 | chr12 | + | 56763217 | 56763414 | 56752160 | 56752318 | 56773833 | 56774165 | 1 | 0 | [illegible] |
| SE | 95501 | ENSG00000025423 | HSD17B6 | chr12 | + | 56785901 | 56786014 | 56784938 | 56785016 | 56787124 | 56787345 | 0 | 0 | [illegible] |
| SE | 95505 | ENSG00000119333 | DYNC2I2 | chr9 | - | 128635657 | 128635767 | 128635091 | 128635259 | 128636280 | 128636438 | 6 | 1 | [illegible] |
| SE | 95507 | ENSG00000137145 | DENND4C | chr9 | + | 19288595 | 19288665 | 19286768 | 19287021 | 19290703 | 19290876 | 263 | 164 | [illegible] |
| SE | 95512 | ENSG00000137145 | DENND4C | chr9 | + | 19326063 | 19326194 | 19325938 | 19325974 | 19328029 | 19328162 | 402 | 324 | [illegible] |
| SE | 95513 | ENSG00000137145 | DENND4C | chr9 | + | 19328029 | 19328162 | 19326063 | 19326194 | 19331977 | 19332184 | 409 | 308 | [illegible] |
| SE | 95515 | ENSG00000137145 | DENND4C | chr9 | + | 19331977 | 19332188 | 19328029 | 19328162 | 19334976 | 19335105 | 208 | 155 | [illegible] |
| SE | 95518 | ENSG00000137145 | DENND4C | chr9 | + | 19342632 | 19342779 | 19340991 | 19341114 | 19345920 | 19347086 | 262 | 161 | [illegible] |
| SE | 95519 | ENSG00000137145 | DENND4C | chr9 | + | 19350701 | 19350879 | 19345920 | 19347086 | 19352072 | 19352182 | 182 | 124 | [illegible] |
| SE | 95520 | ENSG00000137145 | DENND4C | chr9 | + | 19352489 | 19352665 | 19352072 | 19352182 | 19361845 | 19361963 | 48 | 37 | [illegible] |
| SE | 95522 | ENSG00000137145 | DENND4C | chr9 | + | 19361845 | 19361963 | 19360243 | 19360489 | 19369836 | 19369987 | 389 | 331 | [illegible] |
| SE | 95523 | ENSG00000137145 | DENND4C | chr9 | + | 19369836 | 19369987 | 19361845 | 19361963 | 19371755 | 19371820 | 328 | 282 | [illegible] |
| SE | 95524 | ENSG00000137145 | DENND4C | chr9 | + | 19369836 | 19369987 | 19361845 | 19361963 | 19372036 | 19372880 | 255 | 231 | [illegible] |
| SE | 95525 | ENSG00000137145 | DENND4C | chr9 | + | 19371755 | 19371820 | 19361845 | 19361963 | 19372036 | 19372594 | 34 | 26 | [illegible] |
| SE | 95526 | ENSG00000137145 | DENND4C | chr9 | + | 19371755 | 19371820 | 19369836 | 19369987 | 19372036 | 19372594 | 116 | 87 | [illegible] |
| SE | 95527 | ENSG00000187764 | SEMA4D | chr9 | - | 89376832 | 89377051 | 89363740 | 89363950 | 89381054 | 89381098 | 2 | 0 | [illegible] |
| SE | 95528 | ENSG00000187764 | SEMA4D | chr9 | - | 89381054 | 89381098 | 89363740 | 89363950 | 89386366 | 89386482 | 0 | 0 | [illegible] |
| SE | 95529 | ENSG00000187764 | SEMA4D | chr9 | - | 89381054 | 89381098 | 89376385 | 89379629 | 89381173 | 89381346 | 262 | 134 | [illegible] |
| SE | 95530 | ENSG00000187764 | SEMA4D | chr9 | - | 89381173 | 89381346 | 89381054 | 89381098 | 89386366 | 89386482 | 357 | 180 | [illegible] |
| SE | 95531 | ENSG00000187764 | SEMA4D | chr9 | - | 89381173 | 89381346 | 89381054 | 89381098 | 89455887 | 89455953 | 174 | 81 | [illegible] |
| SE | 95532 | ENSG00000187764 | SEMA4D | chr9 | - | 89386366 | 89386482 | 89363740 | 89363950 | 89387385 | 89387608 | 259 | 123 | [illegible] |
| SE | 95533 | ENSG00000187764 | SEMA4D | chr9 | - | 89386366 | 89386482 | 89381173 | 89381346 | 89387385 | 89387608 | 441 | 222 | [illegible] |
| SE | 95536 | ENSG00000187764 | SEMA4D | chr9 | - | 89391263 | 89391415 | 89387385 | 89387608 | 89392422 | 89392536 | 618 | 452 | [illegible] |
| SE | 95537 | ENSG00000187764 | SEMA4D | chr9 | - | 89392422 | 89392536 | 89391263 | 89391415 | 89393561 | 89393655 | 1070 | 735 | [illegible] |
| SE | 95539 | ENSG00000187764 | SEMA4D | chr9 | - | 89418055 | 89418118 | 89405350 | 89405699 | 89418362 | 89418437 | 145 | 121 | [illegible] |
| SE | 95546 | ENSG00000187764 | SEMA4D | chr9 | - | 89455887 | 89455953 | 89418055 | 89418118 | 89497918 | 89498113 | 70 | 50 | [illegible] |
| SE | 95547 | ENSG00000148219 | ASTN2 | chr9 | - | 116487358 | 116487500 | 116442452 | 116442553 | 116618323 | 116618472 | 16 | 2 | [illegible] |
| SE | 95549 | ENSG00000196363 | WDR5 | chr9 | + | 134135678 | 134135842 | 134135376 | 134135441 | 134139819 | 134139958 | 0 | 1 | [illegible] |
| SE | 95552 | ENSG00000204136 | GGTA1 | chr9 | - | 121452143 | 121452281 | 121444990 | 121447597 | 121455654 | 121455841 | 2 | 0 | [illegible] |
| SE | 95556 | ENSG00000165948 | IFI27L1 | chr14 | + | 94094749 | 94094899 | 94081395 | 94081449 | 94096855 | 94096965 | 0 | 0 | [illegible] |
| SE | 95558 | ENSG00000165948 | IFI27L1 | chr14 | + | 94097491 | 94097678 | 94096886 | 94096965 | 94100738 | 94100771 | 0 | 0 | [illegible] |
| SE | 95563 | ENSG00000180104 | EXOC3 | chr5 | + | 444925 | 445064 | 443267 | 443290 | 446149 | 446349 | 5 | 3 | [illegible] |
| SE | 95565 | ENSG00000180104 | EXOC3 | chr5 | + | 457899 | 458025 | 453369 | 454051 | 461959 | 462070 | 0 | 1 | [illegible] |
| SE | 95567 | ENSG00000106546 | AHR | chr7 | + | 17338985 | 17340228 | 17335644 | 17335786 | 17342920 | 17343853 | 474 | 600 | [illegible] |
| SE | 95568 | ENSG00000106952 | TNFSF8 | chr9 | - | 114905827 | 114905899 | 114901144 | 114904325 | 114918095 | 114918138 | 123 | 89 | [illegible] |
| SE | 95569 | ENSG00000165684 | SNAPC4 | chr9 | - | 136383952 | 136384032 | 136383185 | 136383668 | 136384719 | 136384814 | 44 | 57 | [illegible] |
| SE | 95570 | ENSG00000165684 | SNAPC4 | chr9 | - | 136396976 | 136397023 | 136395602 | 136395770 | 136398298 | 136398437 | 8 | 7 | [illegible] |
| SE | 95572 | ENSG00000109220 | CHIC2 | chr4 | - | 54048954 | 54049110 | 54014062 | 54014119 | 54049250 | 54049305 | 481 | 279 | [illegible] |
| SE | 95573 | ENSG00000109220 | CHIC2 | chr4 | - | 54049038 | 54049110 | 54014062 | 54014119 | 54049250 | 54049305 | 123 | 59 | [illegible] |
| SE | 95574 | ENSG00000109220 | CHIC2 | chr4 | - | 54049250 | 54049305 | 54014062 | 54014119 | 54064181 | 54064298 | 42 | 20 | [illegible] |
| SE | 95590 | ENSG00000130714 | POMT1 | chr9 | + | 131504453 | 131504649 | 131504188 | 131504340 | 131506402 | 131506453 | 0 | 0 | [illegible] |
| SE | 95595 | ENSG00000130714 | POMT1 | chr9 | + | 131506182 | 131506220 | 131504188 | 131504340 | 131506402 | 131506453 | 3 | 1 | [illegible] |
| SE | 95601 | ENSG00000130714 | POMT1 | chr9 | + | 131507350 | 131507514 | 131506402 | 131506453 | 131508910 | 131509022 | 5 | 4 | [illegible] |
| SE | 95603 | ENSG00000130714 | POMT1 | chr9 | + | 131507367 | 131507514 | 131504997 | 131506453 | 131508910 | 131508959 | 21 | 7 | [illegible] |
| SE | 95604 | ENSG00000130714 | POMT1 | chr9 | + | 131507367 | 131507514 | 131504997 | 131506453 | 131509742 | 131509808 | 16 | 3 | [illegible] |
| SE | 95605 | ENSG00000130714 | POMT1 | chr9 | + | 131507367 | 131509022 | 131506402 | 131506453 | 131509742 | 131509808 | 28 | 12 | [illegible] |
| SE | 95608 | ENSG00000130714 | POMT1 | chr9 | + | 131508910 | 131509022 | 131506402 | 131506453 | 131509742 | 131509808 | 12 | 7 | [illegible] |
| SE | 95610 | ENSG00000130714 | POMT1 | chr9 | + | 131508910 | 131509022 | 131507350 | 131507514 | 131509742 | 131509808 | 17 | 11 | [illegible] |
| SE | 95613 | ENSG00000130714 | POMT1 | chr9 | + | 131509742 | 131509808 | 131506402 | 131509022 | 131510259 | 131510415 | 9 | 7 | [illegible] |
| SE | 95614 | ENSG00000130714 | POMT1 | chr9 | + | 131509902 | 131509996 | 131508910 | 131509022 | 131510259 | 131510415 | 7 | 1 | [illegible] |
| SE | 95623 | ENSG00000142396 | ERVK3-1 | chr19 | + | 58311198 | 58311380 | 58306087 | 58306216 | 58312020 | 58312061 | 0 | 0 | [illegible] |
| SE | 95625 | ENSG00000142396 | ERVK3-1 | chr19 | + | 58312027 | 58312462 | 58306087 | 58306216 | 58314747 | 58314781 | 115 | 113 | [illegible] |
| SE | 95626 | ENSG00000142396 | ERVK3-1 | chr19 | + | 58312196 | 58312462 | 58306087 | 58306216 | 58314747 | 58315202 | 122 | 115 | [illegible] |
| SE | 95632 | ENSG00000166341 | DCHS1 | chr11 | - | 6633788 | 6634020 | 6633411 | 6633648 | 6634117 | 6634306 | 4 | 4 | [illegible] |
| SE | 95633 | ENSG00000137073 | UBAP2 | chr9 | - | 33927796 | 33927992 | 33926988 | 33927080 | 33932561 | 33932628 | 14 | 6 | [illegible] |
| SE | 95635 | ENSG00000137073 | UBAP2 | chr9 | - | 33932561 | 33932628 | 33927796 | 33927992 | 33933489 | 33933628 | 20 | 20 | [illegible] |
| SE | 95636 | ENSG00000137073 | UBAP2 | chr9 | - | 33943419 | 33943589 | 33941648 | 33941862 | 33944364 | 33944639 | 60 | 35 | [illegible] |
| SE | 95637 | ENSG00000137073 | UBAP2 | chr9 | - | 33943419 | 33943589 | 33941648 | 33941862 | 33948373 | 33948587 | 32 | 28 | [illegible] |
| SE | 95638 | ENSG00000137073 | UBAP2 | chr9 | - | 33944364 | 33944639 | 33941648 | 33941862 | 33948373 | 33948587 | 59 | 30 | [illegible] |
| SE | 95641 | ENSG00000137073 | UBAP2 | chr9 | - | 33948373 | 33948587 | 33943419 | 33943589 | 33953284 | 33953474 | 78 | 60 | [illegible] |
| SE | 95643 | ENSG00000137073 | UBAP2 | chr9 | - | 33948373 | 33948587 | 33944364 | 33944639 | 33960825 | 33960878 | 62 | 31 | [illegible] |
| SE | 95644 | ENSG00000137073 | UBAP2 | chr9 | - | 33953284 | 33953474 | 33944364 | 33944639 | 33960825 | 33960878 | 59 | 27 | [illegible] |
| SE | 95645 | ENSG00000137073 | UBAP2 | chr9 | - | 33953284 | 33953474 | 33948373 | 33948587 | 33956078 | 33956146 | 390 | 255 | [illegible] |
| SE | 95651 | ENSG00000137073 | UBAP2 | chr9 | - | 33956078 | 33956146 | 33953284 | 33953474 | 33971650 | 33971737 | 188 | 117 | [illegible] |
| SE | 95653 | ENSG00000137073 | UBAP2 | chr9 | - | 33960825 | 33960878 | 33953284 | 33953474 | 33971650 | 33971754 | 44 | 18 | [illegible] |
| SE | 95654 | ENSG00000137073 | UBAP2 | chr9 | - | 33960825 | 33960878 | 33953284 | 33953474 | 33986759 | 33986837 | 41 | 16 | [illegible] |
| SE | 95655 | ENSG00000137073 | UBAP2 | chr9 | - | 33960825 | 33960878 | 33956078 | 33956146 | 33971650 | 33971754 | 55 | 44 | [illegible] |
| SE | 95656 | ENSG00000137073 | UBAP2 | chr9 | - | 33963725 | 33963791 | 33960825 | 33960878 | 33971650 | 33971737 | 202 | 142 | [illegible] |
| SE | 95657 | ENSG00000137073 | UBAP2 | chr9 | - | 33963725 | 33963791 | 33960825 | 33960878 | 33986759 | 33986837 | 113 | 63 | [illegible] |
| SE | 95661 | ENSG00000137073 | UBAP2 | chr9 | - | 33996222 | 33996333 | 33988972 | 33989126 | 33998786 | 33998864 | 156 | 135 | [illegible] |
| SE | 95662 | ENSG00000137073 | UBAP2 | chr9 | - | 33996222 | 33996333 | 33988972 | 33989126 | 34011626 | 34011664 | 70 | 68 | [illegible] |
| SE | 95663 | ENSG00000137073 | UBAP2 | chr9 | - | 33996222 | 33997036 | 33988972 | 33989126 | 33998786 | 33998864 | 70 | 68 | [illegible] |
| SE | 95664 | ENSG00000137073 | UBAP2 | chr9 | - | 33998786 | 33998864 | 33988972 | 33989126 | 34011626 | 34011664 | 0 | 0 | [illegible] |
| SE | 95665 | ENSG00000137073 | UBAP2 | chr9 | - | 33998786 | 33998864 | 33996222 | 33996333 | 34011626 | 34011664 | 25 | 11 | [illegible] |
| SE | 95666 | ENSG00000137073 | UBAP2 | chr9 | - | 34011626 | 34011664 | 33996477 | 33998864 | 34014503 | 34014625 | 6 | 2 | [illegible] |
| SE | 95668 | ENSG00000137073 | UBAP2 | chr9 | - | 34014503 | 34014625 | 33998786 | 33998864 | 34017049 | 34017189 | 0 | 0 | [illegible] |
| SE | 95669 | ENSG00000137073 | UBAP2 | chr9 | - | 34014503 | 34014625 | 34011626 | 34011664 | 34017049 | 34017189 | 0 | 0 | [illegible] |
| SE | 95670 | ENSG00000137073 | UBAP2 | chr9 | - | 34017049 | 34017189 | 33994480 | 33998864 | 34048824 | 34048868 | 95 | 84 | [illegible] |
| SE | 95682 | ENSG00000107249 | GLIS3 | chr9 | - | 4125733 | 4125941 | 4117767 | 4118881 | 4286037 | 4286523 | 2 | 1 | [illegible] |
| SE | 95688 | ENSG00000132589 | FLOT2 | chr17 | - | 28881191 | 28881375 | 28880712 | 28880862 | 28881813 | 28882028 | 871 | 542 | [illegible] |
| SE | 95689 | ENSG00000132589 | FLOT2 | chr17 | - | 28884224 | 28884315 | 28882978 | 28883231 | 28885611 | 28885685 | 139 | 45 | [illegible] |
| SE | 95693 | ENSG00000132589 | FLOT2 | chr17 | - | 28885611 | 28885685 | 28883107 | 28883231 | 28885856 | 28885947 | 0 | 0 | [illegible] |
| SE | 95694 | ENSG00000132589 | FLOT2 | chr17 | - | 28885611 | 28885685 | 28884224 | 28884315 | 28885856 | 28885947 | 0 | 0 | [illegible] |
| SE | 95696 | ENSG00000132589 | FLOT2 | chr17 | - | 28885856 | 28885947 | 28883107 | 28883231 | 28897525 | 28897652 | 4 | 0 | [illegible] |
| SE | 95698 | ENSG00000132589 | FLOT2 | chr17 | - | 28885856 | 28885947 | 28884224 | 28884315 | 28897525 | 28897652 | 4 | 0 | [illegible] |
| SE | 95704 | ENSG00000103187 | COTL1 | chr16 | - | 84590104 | 84590262 | 84565595 | 84566955 | 84617500 | 84617583 | 2639 | 1355 | [illegible] |
| SE | 95705 | ENSG00000103187 | COTL1 | chr16 | - | 84617500 | 84617583 | 84590104 | 84590262 | 84617837 | 84618078 | 1128 | 489 | [illegible] |
| SE | 95707 | ENSG00000106397 | PLOD3 | chr7 | - | 101212529 | 101212655 | 101212252 | 101212374 | 101212841 | 101212943 | 146 | 75 | [illegible] |
| SE | 95710 | ENSG00000165140 | FBP1 | chr9 | - | 94606814 | 94606952 | 94605456 | 94605576 | 94609920 | 94610061 | 283 | 137 | [illegible] |
| SE | 95711 | ENSG00000165140 | FBP1 | chr9 | - | 94606814 | 94607112 | 94605456 | 94605576 | 94609920 | 94610061 | 121 | 51 | [illegible] |
| SE | 95712 | ENSG00000165140 | FBP1 | chr9 | - | 94609920 | 94610061 | 94605456 | 94605576 | 94617767 | 94617860 | 103 | 66 | [illegible] |
| SE | 95713 | ENSG00000165140 | FBP1 | chr9 | - | 94609920 | 94610061 | 94606814 | 94606952 | 94620328 | 94620491 | 159 | 93 | [illegible] |
| SE | 95716 | ENSG00000185920 | PTCH1 | chr9 | - | 95449840 | 95449940 | 95449068 | 95449323 | 95453477 | 95453620 | 46 | 12 | [illegible] |
| SE | 95717 | ENSG00000185920 | PTCH1 | chr9 | - | 95449840 | 95453620 | 95449068 | 95449323 | 95456275 | 95456413 | 46 | 10 | [illegible] |
| SE | 95718 | ENSG00000185920 | PTCH1 | chr9 | - | 95453477 | 95453620 | 95449068 | 95449323 | 95456275 | 95456413 | 22 | 9 | [illegible] |
| SE | 95719 | ENSG00000185920 | PTCH1 | chr9 | - | 95456275 | 95456413 | 95449840 | 95453620 | 95458012 | 95458293 | 25 | 9 | [illegible] |
| SE | 95720 | ENSG00000185920 | PTCH1 | chr9 | - | 95469812 | 95469931 | 95468750 | 95469153 | 95476033 | 95476159 | 67 | 18 | [illegible] |
| SE | 95721 | ENSG00000185920 | PTCH1 | chr9 | - | 95474011 | 95474143 | 95469812 | 95469931 | 95476033 | 95476159 | 3 | 3 | [illegible] |
| SE | 95723 | ENSG00000185920 | PTCH1 | chr9 | - | 95477546 | 95477702 | 95476758 | 95476857 | 95478054 | 95478186 | 48 | 10 | [illegible] |
| SE | 95724 | ENSG00000185920 | PTCH1 | chr9 | - | 95478054 | 95478186 | 95476758 | 95476857 | 95478999 | 95479147 | 77 | 27 | [illegible] |
| SE | 95739 | ENSG00000164978 | NUDT2 | chr9 | + | 34336228 | 34336341 | 34329568 | 34329599 | 34338712 | 34338847 | 5 | 1 | [illegible] |
| SE | 95740 | ENSG00000164978 | NUDT2 | chr9 | + | 34336299 | 34336341 | 34329505 | 34329599 | 34338712 | 34338847 | 4 | 0 | [illegible] |
| SE | 95742 | ENSG00000164978 | NUDT2 | chr9 | + | 34339023 | 34339166 | 34338712 | 34338847 | 34343123 | 34343699 | 103 | 112 | [illegible] |
| SE | 95743 | ENSG00000173258 | ZNF483 | chr9 | + | 111527267 | 111527807 | 111525158 | 111525262 | 111530874 | 111530963 | 21 | 18 | [illegible] |
| SE | 95744 | ENSG00000173258 | ZNF483 | chr9 | + | 111530874 | 111530963 | 111527267 | 111527807 | 111533738 | 111533865 | 24 | 21 | [illegible] |
| SE | 95745 | ENSG00000173258 | ZNF483 | chr9 | + | 111534260 | 111534353 | 111533738 | 111533865 | 111541656 | 111555417 | 32 | 21 | [illegible] |
| SE | 95746 | ENSG00000173258 | ZNF483 | chr9 | + | 111534260 | 111534353 | 111533738 | 111533865 | 111576364 | 111577844 | 10 | 6 | [illegible] |
| SE | 95747 | ENSG00000070476 | ZXDC | chr3 | - | 126441764 | 126441946 | 126439631 | 126439727 | 126459652 | 126459737 | 23 | 12 | [illegible] |
| SE | 95752 | ENSG00000070476 | ZXDC | chr3 | - | 126466154 | 126466325 | 126461082 | 126462220 | 126470894 | 126471025 | 199 | 141 | [illegible] |
| SE | 95753 | ENSG00000070476 | ZXDC | chr3 | - | 126472152 | 126472305 | 126471972 | 126472051 | 126474958 | 126474987 | 184 | 99 | [illegible] |
| SE | 95755 | ENSG00000178177 | LCORL | chr4 | - | 17866998 | 17867025 | 17841772 | 17845901 | 17886067 | 17886161 | 2 | 0 | [illegible] |
| SE | 95756 | ENSG00000178177 | LCORL | chr4 | - | 17873387 | 17878213 | 17841198 | 17845901 | 17886067 | 17886161 | 29 | 5 | [illegible] |

| | | | | | | | | | | | | | | |
|---|---|---|---|---|---|---|---|---|---|---|---|---|---|---|
| SE | 95757 | ENSG00000178177 | LCORL | chr4 | - | 17873387 | 17878213 | 17841198 | 17845901 | 17909093 | 17909345 | 13 | 2 | [illegible] |
| SE | 95758 | ENSG00000178177 | LCORL | chr4 | - | 17884274 | 17884752 | 17866998 | 17867025 | 17886067 | 17886161 | 36 | 26 | [illegible] |
| SE | 95759 | ENSG00000178177 | LCORL | chr4 | - | 17886067 | 17886161 | 17841198 | 17845901 | 17909093 | 17909345 | 35 | 39 | [illegible] |
| SE | 95760 | ENSG00000178177 | LCORL | chr4 | - | 17886067 | 17886161 | 17841198 | 17845901 | 17961902 | 17962032 | 6 | 10 | [illegible] |
| SE | 95761 | ENSG00000178177 | LCORL | chr4 | - | 17886067 | 17886161 | 17873387 | 17878213 | 17909093 | 17909345 | 47 | 33 | [illegible] |
| SE | 95762 | ENSG00000178177 | LCORL | chr4 | - | 17909093 | 17909345 | 17845496 | 17845901 | 17961902 | 17962032 | 53 | 40 | [illegible] |
| SE | 95763 | ENSG00000178177 | LCORL | chr4 | - | 17909093 | 17909345 | 17886067 | 17886161 | 17960264 | 17960311 | 54 | 43 | [illegible] |
| SE | 95764 | ENSG00000178177 | LCORL | chr4 | - | 17909093 | 17909345 | 17886067 | 17886161 | 17961902 | 17962032 | 96 | 81 | [illegible] |
| SE | 95765 | ENSG00000178177 | LCORL | chr4 | - | 17960264 | 17960311 | 17886067 | 17886161 | 17961902 | 17962032 | 1 | 3 | [illegible] |
| SE | 95767 | ENSG00000178177 | LCORL | chr4 | - | 17962969 | 17963049 | 17961902 | 17962032 | 17972819 | 17972885 | 94 | 56 | [illegible] |
| SE | 95770 | ENSG00000125944 | HNRNPR | chr1 | - | 23321527 | 23321663 | 23313552 | 23313702 | 23323555 | 23323732 | 581 | 431 | [illegible] |
| SE | 95775 | ENSG00000125944 | HNRNPR | chr1 | - | 23338062 | 23338163 | 23337753 | 23337861 | 23338489 | 23338608 | 1 | 1 | [illegible] |
| SE | 95783 | ENSG00000148120 | AOPEP | chr9 | + | 94792764 | 94792918 | 94759648 | 94760580 | 94800756 | 94801002 | 10 | 6 | [illegible] |
| SE | 95786 | ENSG00000148120 | AOPEP | chr9 | + | 94967757 | 94967801 | 94955907 | 94956015 | 94979366 | 94979427 | 9 | 5 | [illegible] |
| SE | 95787 | ENSG00000148120 | AOPEP | chr9 | + | 95019449 | 95020233 | 95005541 | 95005616 | 95060693 | 95060987 | 0 | 0 | [illegible] |
| SE | 95788 | ENSG00000148120 | AOPEP | chr9 | + | 95060693 | 95060810 | 95005541 | 95005616 | 95080693 | 95080756 | 5 | 5 | [illegible] |
| SE | 95789 | ENSG00000148120 | AOPEP | chr9 | + | 95082574 | 95082719 | 95080693 | 95080780 | 95086681 | 95086918 | 5 | 1 | [illegible] |
| SE | 95791 | ENSG00000143614 | GATAD2B | chr1 | - | 153818039 | 153818171 | 153817371 | 153817542 | 153818790 | 153818922 | 206 | 133 | [illegible] |
| SE | 95792 | ENSG00000143614 | GATAD2B | chr1 | - | 153818790 | 153818922 | 153817371 | 153817542 | 153819605 | 153819735 | 156 | 70 | [illegible] |
| SE | 95795 | ENSG00000143614 | GATAD2B | chr1 | - | 153914673 | 153914740 | 153828012 | 153828348 | 153922732 | 153922968 | 0 | 0 | [illegible] |
| SE | 95796 | ENSG00000175764 | TTLL11 | chr9 | - | 121860336 | 121860443 | 121815673 | 121822879 | 121870496 | 121870748 | 2 | 2 | [illegible] |
| SE | 95797 | ENSG00000175764 | TTLL11 | chr9 | - | 121989194 | 121989770 | 121974883 | 121974979 | 122031722 | 122031856 | 13 | 12 | [illegible] |
| SE | 95798 | ENSG00000175764 | TTLL11 | chr9 | - | 121989194 | 121989770 | 121974883 | 121974979 | 122039271 | 122039368 | 11 | 13 | [illegible] |
| SE | 95799 | ENSG00000175764 | TTLL11 | chr9 | - | 122031722 | 122031856 | 121974883 | 121974979 | 122039271 | 122039368 | 2 | 1 | [illegible] |
| SE | 95802 | ENSG00000167721 | TSR1 | chr17 | - | 2330514 | 2330625 | 2329342 | 2329475 | 2330946 | 2330966 | 175 | 114 | [illegible] |
| SE | 95804 | ENSG00000167721 | TSR1 | chr17 | - | 2332960 | 2333124 | 2332168 | 2332359 | 2333556 | 2333716 | 131 | 172 | [illegible] |
| SE | 95812 | ENSG00000215126 | CBWD6 | chr9 | - | 41160804 | 41161021 | 41156382 | 41156439 | 41162332 | 41162377 | 0 | 1 | [illegible] |
| SE | 95813 | ENSG00000215126 | CBWD6 | chr9 | - | 41160937 | 41161021 | 41156382 | 41156439 | 41162332 | 41162377 | 0 | 1 | [illegible] |
| SE | 95814 | ENSG00000215126 | CBWD6 | chr9 | - | 41161600 | 41161723 | 41156382 | 41156439 | 41162332 | 41162377 | 4 | 5 | [illegible] |
| SE | 95816 | ENSG00000215126 | CBWD6 | chr9 | - | 41170278 | 41170423 | 41166783 | 41166830 | 41172711 | 41172749 | 1 | 0 | [illegible] |
| SE | 95818 | ENSG00000215126 | CBWD6 | chr9 | - | 41170283 | 41170423 | 41166783 | 41166830 | 41172711 | 41172749 | 2 | 0 | [illegible] |
| SE | 95819 | ENSG00000215126 | CBWD6 | chr9 | - | 41171846 | 41171934 | 41166783 | 41166830 | 41172711 | 41172749 | 14 | 7 | [illegible] |
| SE | 95821 | ENSG00000215126 | CBWD6 | chr9 | - | 41174292 | 41174352 | 41172711 | 41172749 | 41182374 | 41182395 | 2 | 0 | [illegible] |
| SE | 95829 | ENSG00000080608 | PUM3 | chr9 | - | 2824716 | 2824815 | 2823780 | 2823834 | 2827072 | 2827151 | 92 | 64 | [illegible] |
| SE | 95830 | ENSG00000080608 | PUM3 | chr9 | - | 2824716 | 2824815 | 2823780 | 2823834 | 2828674 | 2828778 | 39 | 25 | [illegible] |
| SE | 95831 | ENSG00000080608 | PUM3 | chr9 | - | 2827072 | 2827151 | 2823780 | 2823834 | 2828674 | 2828778 | 32 | 24 | [illegible] |
| SE | 95832 | ENSG00000080608 | PUM3 | chr9 | - | 2827072 | 2827151 | 2824716 | 2824815 | 2828674 | 2828778 | 98 | 68 | [illegible] |
| SE | 95833 | ENSG00000080608 | PUM3 | chr9 | - | 2838425 | 2838517 | 2837179 | 2837401 | 2844044 | 2844095 | 159 | 198 | [illegible] |
| SE | 95836 | ENSG00000141551 | CSNK1D | chr17 | - | 82252433 | 82252527 | 82251378 | 82251527 | 82265685 | 82265796 | 202 | 137 | [illegible] |
| SE | 95837 | ENSG00000141551 | CSNK1D | chr17 | - | 82252433 | 82252604 | 82251378 | 82251527 | 82265685 | 82265796 | 259 | 170 | [illegible] |
| SE | 95838 | ENSG00000141551 | CSNK1D | chr17 | - | 82253015 | 82253244 | 82252433 | 82252527 | 82265685 | 82265796 | 10 | 2 | [illegible] |
| SE | 95839 | ENSG00000141551 | CSNK1D | chr17 | - | 82253015 | 82253244 | 82252433 | 82252604 | 82255428 | 82255516 | 665 | 438 | [illegible] |
| SE | 95840 | ENSG00000141551 | CSNK1D | chr17 | - | 82253015 | 82253244 | 82252433 | 82252604 | 82265685 | 82265796 | 475 | 308 | [illegible] |
| SE | 95843 | ENSG00000141551 | CSNK1D | chr17 | - | 82255428 | 82255577 | 82253015 | 82253244 | 82273305 | 82273431 | 192 | 130 | [illegible] |
| SE | 95844 | ENSG00000141551 | CSNK1D | chr17 | - | 82265685 | 82265796 | 82253015 | 82253244 | 82273305 | 82273369 | 87 | 31 | [illegible] |
| SE | 95846 | ENSG00000141551 | CSNK1D | chr17 | - | 82266724 | 82266894 | 82265685 | 82265796 | 82273305 | 82273457 | 1 | 0 | [illegible] |
| SE | 95847 | ENSG00000141551 | CSNK1D | chr17 | - | 82272069 | 82272316 | 82265685 | 82265796 | 82273305 | 82273731 | 0 | 1 | [illegible] |
| SE | 95850 | ENSG00000107263 | RAPGEF1 | chr9 | - | 131598198 | 131598310 | 131588800 | 131588986 | 131621795 | 131621838 | 1 | 9 | [illegible] |
| SE | 95852 | ENSG00000107263 | RAPGEF1 | chr9 | - | 131603960 | 131604053 | 131602060 | 131602149 | 131604930 | 131605188 | 5 | 0 | [illegible] |
| SE | 95854 | ENSG00000107263 | RAPGEF1 | chr9 | - | 131625921 | 131626422 | 131621795 | 131621998 | 131643238 | 131643417 | 251 | 184 | [illegible] |
| SE | 95856 | ENSG00000107263 | RAPGEF1 | chr9 | - | 131628548 | 131628672 | 131627912 | 131628096 | 131629101 | 131629254 | 462 | 167 | [illegible] |
| SE | 95857 | ENSG00000107263 | RAPGEF1 | chr9 | - | 131629101 | 131629254 | 131625921 | 131626422 | 131643238 | 131643391 | 1 | 0 | [illegible] |
| SE | 95861 | ENSG00000107263 | RAPGEF1 | chr9 | - | 131638634 | 131638791 | 131630235 | 131630324 | 131643238 | 131643391 | 989 | 474 | [illegible] |
| SE | 95862 | ENSG00000107263 | RAPGEF1 | chr9 | - | 131643238 | 131643417 | 131625921 | 131626422 | 131650128 | 131650242 | 210 | 102 | [illegible] |
| SE | 95863 | ENSG00000107263 | RAPGEF1 | chr9 | - | 131643238 | 131643417 | 131625921 | 131626422 | 131739769 | 131739944 | 2 | 0 | [illegible] |
| SE | 95865 | ENSG00000107263 | RAPGEF1 | chr9 | - | 131643238 | 131643417 | 131638634 | 131638791 | 131650809 | 131650949 | 623 | 313 | [illegible] |
| SE | 95868 | ENSG00000107263 | RAPGEF1 | chr9 | - | 131650809 | 131650949 | 131643238 | 131643417 | 131739769 | 131739944 | 135 | 91 | [illegible] |
| SE | 95869 | ENSG00000148396 | SEC16A | chr9 | - | 136443822 | 136443900 | 136440096 | 136441823 | 136445644 | 136445719 | 79 | 45 | [illegible] |
| SE | 95876 | ENSG00000148396 | SEC16A | chr9 | - | 136451255 | 136451408 | 136448083 | 136448161 | 136453427 | 136453510 | 190 | 97 | [illegible] |
| SE | 95879 | ENSG00000148396 | SEC16A | chr9 | - | 136460041 | 136460123 | 136459756 | 136459874 | 136461176 | 136461274 | 210 | 79 | [illegible] |
| SE | 95884 | ENSG00000148396 | SEC16A | chr9 | - | 136465961 | 136466136 | 136464419 | 136464562 | 136466263 | 136466462 | 123 | 48 | [illegible] |
| SE | 95886 | ENSG00000080298 | RFX3 | chr9 | - | 3262934 | 3263084 | 3256990 | 3257199 | 3266207 | 3266305 | 50 | 34 | [illegible] |
| SE | 95890 | ENSG00000080298 | RFX3 | chr9 | - | 3288130 | 3288250 | 3277339 | 3277461 | 3293076 | 3293258 | 62 | 27 | [illegible] |
| SE | 95892 | ENSG00000080298 | RFX3 | chr9 | - | 3346666 | 3346764 | 3330258 | 3330517 | 3395471 | 3395588 | 93 | 64 | [illegible] |
| SE | 95893 | ENSG00000080298 | RFX3 | chr9 | - | 3488775 | 3488895 | 3395471 | 3395596 | 3490278 | 3490345 | 4 | 0 | [illegible] |
| SE | 95894 | ENSG00000080298 | RFX3 | chr9 | - | 3490278 | 3490345 | 3395471 | 3395596 | 3525746 | 3525983 | 0 | 0 | [illegible] |
| SE | 95897 | ENSG00000188566 | NDOR1 | chr9 | + | 137212501 | 137212599 | 137206231 | 137206309 | 137213779 | 137213878 | 33 | 11 | [illegible] |
| SE | 95899 | ENSG00000188566 | NDOR1 | chr9 | + | 137215406 | 137215521 | 137215094 | 137215202 | 137215658 | 137215805 | 7 | 2 | [illegible] |
| SE | 95900 | ENSG00000188566 | NDOR1 | chr9 | + | 137215427 | 137215521 | 137215094 | 137215202 | 137215658 | 137215805 | 5 | 1 | [illegible] |
| SE | 95901 | ENSG00000188566 | NDOR1 | chr9 | + | 137215658 | 137215805 | 137215094 | 137215202 | 137215898 | 137216017 | 9 | 1 | [illegible] |
| SE | 95902 | ENSG00000188566 | NDOR1 | chr9 | + | 137216066 | 137216188 | 137215898 | 137216017 | 137216271 | 137217009 | 14 | 3 | [illegible] |
| SE | 95903 | ENSG00000188566 | NDOR1 | chr9 | + | 137216093 | 137216166 | 137215898 | 137216017 | 137216271 | 137216989 | 6 | 0 | [illegible] |
| SE | 95904 | ENSG00000188566 | NDOR1 | chr9 | + | 137216093 | 137216188 | 137215898 | 137216017 | 137216271 | 137216444 | 24 | 5 | [illegible] |
| SE | 95907 | ENSG00000198121 | LPAR1 | chr9 | - | 111036121 | 111036201 | 110973480 | 110973558 | 111038166 | 111038470 | 1 | 1 | [illegible] |
| SE | 95909 | ENSG00000148343 | MIGA2 | chr9 | + | 129047104 | 129047224 | 129042303 | 129042514 | 129048426 | 129048539 | 1 | 0 | [illegible] |
| SE | 95914 | ENSG00000188352 | FOCAD | chr9 | + | 20694519 | 20694564 | 20658690 | 20658826 | 20715321 | 20715410 | 0 | 0 | [illegible] |
| SE | 95916 | ENSG00000188352 | FOCAD | chr9 | + | 20770031 | 20770238 | 20764868 | 20765073 | 20778680 | 20778768 | 50 | 57 | [illegible] |
| SE | 95917 | ENSG00000188352 | FOCAD | chr9 | + | 20819795 | 20819900 | 20789350 | 20789608 | 20820323 | 20820425 | 39 | 38 | [illegible] |
| SE | 95919 | ENSG00000188352 | FOCAD | chr9 | + | 20907149 | 20907242 | 20885108 | 20885230 | 20912865 | 20912908 | 51 | 35 | [illegible] |
| SE | 95920 | ENSG00000188352 | FOCAD | chr9 | + | 20923659 | 20923768 | 20916892 | 20916937 | 20926300 | 20926417 | 37 | 25 | [illegible] |
| SE | 95922 | ENSG00000188352 | FOCAD | chr9 | + | 20944626 | 20944774 | 20933013 | 20933103 | 20948270 | 20948393 | 22 | 30 | [illegible] |
| SE | 95923 | ENSG00000188352 | FOCAD | chr9 | + | 20946700 | 20946820 | 20944626 | 20944774 | 20948270 | 20948393 | 31 | 21 | [illegible] |
| SE | 95924 | ENSG00000188352 | FOCAD | chr9 | + | 20957501 | 20957617 | 20952984 | 20953065 | 20976419 | 20976548 | 0 | 0 | [illegible] |
| SE | 95926 | ENSG00000188352 | FOCAD | chr9 | + | 20982356 | 20982446 | 20981425 | 20981686 | 20986287 | 20986465 | 35 | 19 | [illegible] |
| SE | 95928 | ENSG00000166199 | ALKBH3 | chr11 | + | 43892040 | 43892129 | 43889724 | 43889828 | 43897194 | 43897354 | 62 | 19 | [illegible] |
| SE | 95935 | ENSG00000111249 | CUX2 | chr12 | + | 111217889 | 111217937 | 111214199 | 111214310 | 111263760 | 111263839 | 8 | 4 | [illegible] |
| SE | 95938 | ENSG00000136868 | SLC31A1 | chr9 | + | 113258693 | 113258862 | 113257112 | 113257185 | 113260271 | 113264492 | 82 | 61 | [illegible] |
| SE | 95939 | ENSG00000176386 | CDC26 | chr9 | - | 113272426 | 113272548 | 113266991 | 113267439 | 113273328 | 113273438 | 426 | 321 | [illegible] |
| SE | 95951 | ENSG00000172340 | SUCLG2 | chr3 | - | 67508806 | 67508903 | 67498133 | 67498295 | 67518246 | 67518336 | 286 | 210 | [illegible] |
| SE | 95952 | ENSG00000172340 | SUCLG2 | chr3 | - | 67520481 | 67520634 | 67518246 | 67518336 | 67528131 | 67528222 | 353 | 261 | [illegible] |
| SE | 95953 | ENSG00000172340 | SUCLG2 | chr3 | - | 67528131 | 67528222 | 67520481 | 67520634 | 67609454 | 67609596 | 165 | 143 | [illegible] |
| SE | 95954 | ENSG00000172340 | SUCLG2 | chr3 | - | 67529086 | 67529186 | 67528131 | 67528222 | 67609454 | 67609596 | 261 | 222 | [illegible] |
| SE | 95955 | ENSG00000172340 | SUCLG2 | chr3 | - | 67609454 | 67609596 | 67529086 | 67529186 | 67654502 | 67654607 | 205 | 188 | [illegible] |
| SE | 95960 | ENSG00000198208 | RPS6KL1 | chr14 | - | 74911793 | 74911841 | 74910889 | 74911380 | 74919844 | 74919969 | 0 | 0 | [illegible] |
| SE | 95967 | ENSG00000260691 | ANKRD20A1 | chr9 | + | 67877759 | 67877790 | 67871156 | 67871212 | 67880475 | 67880545 | 1 | 0 | [illegible] |
| SE | 95999 | ENSG00000256591 | NA | chr11 | + | 61437624 | 61437848 | 61430055 | 61430182 | 61468663 | 61468714 | 196 | 84 | [illegible] |
| SE | 96000 | ENSG00000256591 | NA | chr11 | + | 61438003 | 61438113 | 61430120 | 61430182 | 61468663 | 61468716 | 3 | 2 | [illegible] |
| SE | 96001 | ENSG00000256591 | NA | chr11 | + | 61438003 | 61438113 | 61430120 | 61430182 | 61481781 | 61481879 | 3 | 0 | [illegible] |
| SE | 96002 | ENSG00000256591 | NA | chr11 | + | 61438003 | 61438113 | 61430120 | 61430182 | 61484687 | 61484816 | 5 | 0 | [illegible] |
| SE | 96003 | ENSG00000256591 | NA | chr11 | + | 61438003 | 61438113 | 61437624 | 61437848 | 61468663 | 61468716 | 212 | 107 | [illegible] |
| SE | 96005 | ENSG00000256591 | NA | chr11 | + | 61438003 | 61438120 | 61430055 | 61430182 | 61468663 | 61468714 | 3 | 0 | [illegible] |
| SE | 96007 | ENSG00000256591 | NA | chr11 | + | 61438003 | 61438120 | 61437624 | 61437848 | 61468663 | 61468714 | 208 | 103 | [illegible] |
| SE | 96010 | ENSG00000256591 | NA | chr11 | + | 61445940 | 61446231 | 61437624 | 61437848 | 61468663 | 61468716 | 5 | 3 | [illegible] |
| SE | 96012 | ENSG00000256591 | NA | chr11 | + | 61445940 | 61446231 | 61438003 | 61438113 | 61481781 | 61481918 | 146 | 90 | [illegible] |
| SE | 96016 | ENSG00000256591 | NA | chr11 | + | 61481781 | 61481918 | 61430124 | 61430182 | 61484687 | 61484727 | 0 | 0 | [illegible] |
| SE | 96017 | ENSG00000256591 | NA | chr11 | + | 61481781 | 61481918 | 61438003 | 61438113 | 61484687 | 61484727 | 0 | 0 | [illegible] |
| SE | 96018 | ENSG00000256591 | NA | chr11 | + | 61482654 | 61482808 | 61430125 | 61430182 | 61484687 | 61484762 | 4 | 0 | [illegible] |
| SE | 96019 | ENSG00000074416 | MGLL | chr3 | - | 127692965 | 127693201 | 127692131 | 127692323 | 127694974 | 127695190 | 0 | 1 | [illegible] |
| SE | 96020 | ENSG00000074416 | MGLL | chr3 | - | 127694974 | 127695190 | 127689061 | 127692323 | 127710575 | 127710665 | 12 | 19 | [illegible] |
| SE | 96023 | ENSG00000074416 | MGLL | chr3 | - | 127721052 | 127721163 | 127710575 | 127710665 | 127722429 | 127722566 | 43 | 34 | [illegible] |
| SE | 96024 | ENSG00000170909 | OSCAR | chr19 | - | 54099589 | 54099601 | 54096861 | 54097164 | 54099747 | 54099780 | 1 | 0 | [illegible] |
| SE | 96025 | ENSG00000170909 | OSCAR | chr19 | - | 54099747 | 54099780 | 54096861 | 54097164 | 54100755 | 54100792 | 4 | 1 | [illegible] |
| SE | 96027 | ENSG00000078902 | TOLLIP | chr11 | - | 1286001 | 1286092 | 1274370 | 1277253 | 1288623 | 1288776 | 107 | 38 | [illegible] |
| SE | 96030 | ENSG00000078902 | TOLLIP | chr11 | - | 1295644 | 1295794 | 1290094 | 1290409 | 1309465 | 1309590 | 41 | 16 | [illegible] |
| SE | 96031 | ENSG00000188868 | ZNF563 | chr19 | - | 12319429 | 12319833 | 12319186 | 12319311 | 12321271 | 12321332 | 2 | 12 | [illegible] |
| SE | 96032 | ENSG00000188868 | ZNF563 | chr19 | - | 12321271 | 12321332 | 12317476 | 12319833 | 12322584 | 12322711 | 0 | 9 | [illegible] |
| SE | 96069 | ENSG00000132109 | TRIM21 | chr11 | - | 4385677 | 4385853 | 4384896 | 4385394 | 4386156 | 4386188 | 229 | 135 | [illegible] |
| SE | 96071 | ENSG00000132109 | TRIM21 | chr11 | - | 4386967 | 4386990 | 4384896 | 4385853 | 4388299 | 4388530 | 29 | 8 | [illegible] |
| SE | 96073 | ENSG00000132109 | TRIM21 | chr11 | - | 4389653 | 4389749 | 4388299 | 4388530 | 4390001 | 4390458 | 428 | 300 | [illegible] |
| SE | 96074 | ENSG00000166313 | APBB1 | chr11 | - | 6401976 | 6401982 | 6401051 | 6401688 | 6402081 | 6402209 | 0 | 0 | [illegible] |
| SE | 96079 | ENSG00000174607 | UGT8 | chr4 | + | 114619422 | 114619525 | 114598769 | 114598974 | 114622878 | 114623305 | 0 | 2 | [illegible] |
| SE | 96080 | ENSG00000174607 | UGT8 | chr4 | + | 114665679 | 114665756 | 114663994 | 114664137 | 114668084 | 114668304 | 15 | 3 | [illegible] |
| SE | 96081 | ENSG00000159445 | THEM4 | chr1 | - | 151888272 | 151888383 | 151877000 | 151877125 | 151889213 | 151889373 | 199 | 178 | [illegible] |
| SE | 96082 | ENSG00000159445 | THEM4 | chr1 | - | 151889213 | 151889373 | 151888272 | 151888383 | 151895007 | 151895194 | 257 | 196 | [illegible] |
| SE | 96087 | ENSG00000083799 | CYLD | chr16 | + | 50744729 | 50744866 | 50742761 | 50742822 | 50749575 | 50749798 | 84 | 49 | [illegible] |
| SE | 96090 | ENSG00000083799 | CYLD | chr16 | + | 50751603 | 50751906 | 50749548 | 50750202 | 50754318 | 50754424 | 847 | 772 | [illegible] |
| SE | 96091 | ENSG00000083799 | CYLD | chr16 | + | 50754318 | 50754424 | 50751603 | 50751906 | 50776178 | 50776277 | 572 | 500 | [illegible] |
| SE | 96093 | ENSG00000083799 | CYLD | chr16 | + | 50775165 | 50775174 | 50754318 | 50754424 | 50776178 | 50776277 | 0 | 0 | [illegible] |
| SE | 96094 | ENSG00000083799 | CYLD | chr16 | + | 50776178 | 50776277 | 50751603 | 50751906 | 50777824 | 50777941 | 294 | 276 | [illegible] |
| SE | 96096 | ENSG00000083799 | CYLD | chr16 | + | 50777824 | 50777941 | 50754318 | 50754424 | 50779664 | 50780044 | 452 | 430 | [illegible] |
| SE | 96097 | ENSG00000083799 | CYLD | chr16 | + | 50779664 | 50780044 | 50777824 | 50777941 | 50782324 | 50782466 | 509 | 482 | [illegible] |
| SE | 96099 | ENSG00000083799 | CYLD | chr16 | + | 50786854 | 50786946 | 50784328 | 50784451 | 50787785 | 50787852 | 586 | 500 | [illegible] |
| SE | 96100 | ENSG00000083799 | CYLD | chr16 | + | 50786854 | 50786946 | 50784328 | 50784451 | 50791557 | 50791690 | 293 | 264 | [illegible] |
| SE | 96101 | ENSG00000083799 | CYLD | chr16 | + | 50787785 | 50787852 | 50784328 | 50784451 | 50791557 | 50791690 | 75 | 105 | [illegible] |
| SE | 96102 | ENSG00000083799 | CYLD | chr16 | + | 50787785 | 50787852 | 50786854 | 50786946 | 50791557 | 50791690 | 352 | 328 | [illegible] |
| SE | 96103 | ENSG00000083799 | CYLD | chr16 | + | 50794211 | 50794428 | 50793545 | 50793664 | 50796323 | 50796711 | 421 | 369 | [illegible] |
| SE | 96104 | ENSG00000083799 | CYLD | chr16 | + | 50796323 | 50796711 | 50794211 | 50794428 | 50800819 | 50800872 | 169 | 138 | [illegible] |
| SE | 96105 | ENSG00000123360 | PDE1B | chr12 | + | 54554216 | 54554377 | 54549859 | 54549985 | 54566973 | 54567087 | 0 | 6 | [illegible] |
| SE | 96106 | ENSG00000123360 | PDE1B | chr12 | + | 54575550 | 54575632 | 54574756 | 54575218 | 54575991 | 54576100 | 11 | 5 | [illegible] |
| SE | 96117 | ENSG00000166405 | RIC3 | chr11 | - | 8126658 | 8126804 | 8108941 | 8111137 | 8139966 | 8140193 | 19 | 21 | [illegible] |
| SE | 96121 | ENSG00000166405 | RIC3 | chr11 | - | 8128218 | 8128354 | 8126658 | 8126804 | 8137377 | 8137471 | 5 | 0 | [illegible] |
| SE | 96122 | ENSG00000166405 | RIC3 | chr11 | - | 8128218 | 8128354 | 8126658 | 8126807 | 8137377 | 8137471 | 1 | 0 | [illegible] |
| SE | 96123 | ENSG00000166405 | RIC3 | chr11 | - | 8128218 | 8128354 | 8126658 | 8126891 | 8137377 | 8137471 | 1 | 1 | [illegible] |
| SE | 96124 | ENSG00000166405 | RIC3 | chr11 | - | 8137377 | 8137471 | 8106055 | 8111137 | 8138271 | 8138347 | 15 | 20 | [illegible] |
| SE | 96125 | ENSG00000166405 | RIC3 | chr11 | - | 8137377 | 8137471 | 8135581 | 8135844 | 8138271 | 8138347 | 15 | 22 | [illegible] |
| SE | 96126 | ENSG00000166405 | RIC3 | chr11 | - | 8138271 | 8138347 | 8110697 | 8111137 | 8139966 | 8140186 | 0 | 0 | [illegible] |
| SE | 96129 | ENSG00000166405 | RIC3 | chr11 | - | 8153374 | 8153423 | 8139966 | 8140193 | 8168865 | 8169008 | 0 | 8 | [illegible] |
| SE | 96130 | ENSG00000149294 | NCAM1 | chr11 | + | 113202378 | 113202453 | 112961274 | 112961664 | 113204285 | 113204504 | 151 | 52 | [illegible] |
| SE | 96131 | ENSG00000149294 | NCAM1 | chr11 | + | 113221295 | 113221325 | 113214368 | 113214511 | 113231195 | 113231273 | 1 | 2 | [illegible] |
| SE | 96132 | ENSG00000149294 | NCAM1 | chr11 | + | 113221295 | 113221325 | 113214368 | 113214511 | 113231644 | 113231795 | 14 | 10 | [illegible] |
| SE | 96133 | ENSG00000149294 | NCAM1 | chr11 | + | 113231195 | 113231273 | 113214368 | 113214511 | 113231644 | 113231795 | 1 | 0 | [illegible] |
| SE | 96138 | ENSG00000149294 | NCAM1 | chr11 | + | 113246367 | 113246370 | 113235032 | 113235164 | 113255876 | 113256001 | 1 | 0 | [illegible] |
| SE | 96141 | ENSG00000149294 | NCAM1 | chr11 | + | 113272932 | 113273748 | 113271759 | 113271876 | 113275266 | 113275740 | 0 | 0 | [illegible] |
| SE | 96142 | ENSG00000089351 | GRAMD1A | chr19 | + | 35009422 | 35009443 | 35009118 | 35009329 | 35009887 | 35009972 | 52 | 18 | [illegible] |
| SE | 96144 | ENSG00000089351 | GRAMD1A | chr19 | + | 35013540 | 35013691 | 35011473 | 35011554 | 35015823 | 35015967 | 13 | 13 | [illegible] |
| SE | 96145 | ENSG00000089351 | GRAMD1A | chr19 | + | 35014188 | 35014387 | 35013540 | 35013691 | 35015823 | 35015967 | 112 | 48 | [illegible] |
| SE | 96146 | ENSG00000089351 | GRAMD1A | chr19 | + | 35021950 | 35022038 | 35021690 | 35021864 | 35023235 | 35023343 | 79 | 46 | [illegible] |
| SE | 96148 | ENSG00000089351 | GRAMD1A | chr19 | + | 35023426 | 35023547 | 35023235 | 35023343 | 35026048 | 35026469 | 151 | 117 | [illegible] |
| SE | 96149 | ENSG00000115084 | SLC35F5 | chr2 | - | 113717752 | 113717848 | 113714646 | 113715195 | 113719153 | 113719244 | 14 | 15 | [illegible] |
| SE | 96150 | ENSG00000115084 | SLC35F5 | chr2 | - | 113717752 | 113717850 | 113705010 | 113705548 | 113719153 | 113719280 | 42 | 43 | [illegible] |
| SE | 96152 | ENSG00000115084 | SLC35F5 | chr2 | - | 113729400 | 113729505 | 113725377 | 113725537 | 113731583 | 113731648 | 81 | 54 | [illegible] |
| SE | 96154 | ENSG00000115084 | SLC35F5 | chr2 | - | 113734585 | 113734673 | 113731583 | 113731648 | 113735776 | 113735856 | 71 | 61 | [illegible] |
| SE | 96155 | ENSG00000115084 | SLC35F5 | chr2 | - | 113746276 | 113746339 | 113743712 | 113743794 | 113750424 | 113750568 | 76 | 64 | [illegible] |
| SE | 96156 | ENSG00000115084 | SLC35F5 | chr2 | - | 113746276 | 113746339 | 113743712 | 113743794 | 113755164 | 113755306 | 27 | 14 | [illegible] |
| SE | 96157 | ENSG00000115084 | SLC35F5 | chr2 | - | 113750424 | 113750568 | 113743712 | 113743794 | 113755164 | 113755306 | 48 | 40 | [illegible] |
| SE | 96158 | ENSG00000115084 | SLC35F5 | chr2 | - | 113750424 | 113750568 | 113746276 | 113746339 | 113755164 | 113755306 | 96 | 91 | [illegible] |
| SE | 96159 | ENSG00000115084 | SLC35F5 | chr2 | - | 113750424 | 113750568 | 113746276 | 113746339 | 113755453 | 113755544 | 49 | 51 | [illegible] |
| SE | 96160 | ENSG00000115084 | SLC35F5 | chr2 | - | 113755164 | 113755306 | 113746276 | 113746339 | 113755453 | 113755544 | 44 | 13 | [illegible] |
| SE | 96161 | ENSG00000115084 | SLC35F5 | chr2 | - | 113755164 | 113755306 | 113750424 | 113750568 | 113755453 | 113755544 | 91 | 52 | [illegible] |
| SE | 96164 | ENSG00000186868 | MAPT | chr17 | + | 45971858 | 45971945 | 45962320 | 45962470 | 45978374 | 45978425 | 2 | 2 | [illegible] |
| SE | 96174 | ENSG00000110756 | HPS5 | chr11 | - | 18280508 | 18280642 | 18279617 | 18279942 | 18281949 | 18282220 | 1 | 0 | [illegible] |
| SE | 96175 | ENSG00000110756 | HPS5 | chr11 | - | 18281949 | 18282220 | 18278667 | 18279942 | 18283794 | 18283901 | 44 | 35 | [illegible] |
| SE | 96176 | ENSG00000110756 | HPS5 | chr11 | - | 18283794 | 18283901 | 18281913 | 18282220 | 18285345 | 18285459 | 51 | 32 | [illegible] |
| SE | 96177 | ENSG00000110756 | HPS5 | chr11 | - | 18285345 | 18285459 | 18281949 | 18282220 | 18287534 | 18287690 | 0 | 1 | [illegible] |
| SE | 96179 | ENSG00000110756 | HPS5 | chr11 | - | 18286590 | 18286710 | 18285345 | 18285459 | 18287534 | 18287690 | 78 | 49 | [illegible] |
| SE | 96184 | ENSG00000110756 | HPS5 | chr11 | - | 18295019 | 18295169 | 18292898 | 18292976 | 18295998 | 18296045 | 98 | 66 | [illegible] |
| SE | 96185 | ENSG00000110756 | HPS5 | chr11 | - | 18298791 | 18298970 | 18297558 | 18297717 | 18300827 | 18300916 | 23 | 30 | [illegible] |
| SE | 96186 | ENSG00000110756 | HPS5 | chr11 | - | 18306134 | 18306347 | 18305421 | 18305493 | 18308945 | 18309079 | 68 | 44 | [illegible] |
| SE | 96187 | ENSG00000110756 | HPS5 | chr11 | - | 18310740 | 18310933 | 18308945 | 18309079 | 18311386 | 18311451 | 75 | 48 | [illegible] |
| SE | 96188 | ENSG00000110756 | HPS5 | chr11 | - | 18311913 | 18312024 | 18311386 | 18311451 | 18317750 | 18317907 | 53 | 37 | [illegible] |
| SE | 96189 | ENSG00000110756 | HPS5 | chr11 | - | 18311913 | 18312024 | 18311386 | 18311451 | 18321945 | 18322174 | 38 | 28 | [illegible] |
| SE | 96190 | ENSG00000110756 | HPS5 | chr11 | - | 18311913 | 18312024 | 18311386 | 18311451 | 18322076 | 18322198 | 33 | 21 | [illegible] |
| SE | 96192 | ENSG00000110756 | HPS5 | chr11 | - | 18314403 | 18314519 | 18311386 | 18311451 | 18322076 | 18322198 | 0 | 0 | [illegible] |
| SE | 96195 | ENSG00000110756 | HPS5 | chr11 | - | 18317750 | 18317907 | 18311386 | 18311451 | 18321945 | 18322088 | 13 | 6 | [illegible] |
| SE | 96196 | ENSG00000110756 | HPS5 | chr11 | - | 18317750 | 18317907 | 18311386 | 18311451 | 18322076 | 18322198 | 4 | 4 | [illegible] |
| SE | 96200 | ENSG00000110756 | HPS5 | chr11 | - | 18317750 | 18317907 | 18314403 | 18314519 | 18322076 | 18322198 | 2 | 4 | [illegible] |
| SE | 96201 | ENSG00000151117 | TMEM86A | chr11 | + | 18699810 | 18700049 | 18698778 | 18698907 | 18700932 | 18701115 | 2 | 0 | [illegible] |
| SE | 96202 | ENSG00000151117 | TMEM86A | chr11 | + | 18699817 | 18700049 | 18698803 | 18698907 | 18700932 | 18701154 | 2 | 0 | [illegible] |
| SE | 96203 | ENSG00000117751 | PPP1R8 | chr1 | + | 27832755 | 27832816 | 27830781 | 27830891 | 27838698 | 27838852 | 101 | 55 | [illegible] |
| SE | 96204 | ENSG00000117751 | PPP1R8 | chr1 | + | 27838698 | 27838852 | 27832755 | 27832816 | 27841013 | 27841234 | 234 | 131 | [illegible] |
| SE | 96206 | ENSG00000117751 | PPP1R8 | chr1 | + | 27841013 | 27841234 | 27832755 | 27832816 | 27843185 | 27843330 | 192 | 129 | [illegible] |
| SE | 96208 | ENSG00000117751 | PPP1R8 | chr1 | + | 27843185 | 27843330 | 27838698 | 27838852 | 27847027 | 27847092 | 232 | 153 | [illegible] |
| SE | 96209 | ENSG00000117751 | PPP1R8 | chr1 | + | 27847027 | 27847092 | 27843185 | 27843330 | 27850092 | 27850432 | 240 | 159 | [illegible] |
| SE | 96210 | ENSG00000260220 | CCDC187 | chr9 | - | 136274657 | 136274749 | 136268048 | 136268125 | 136274882 | 136275007 | 0 | 0 | [illegible] |
| SE | 96211 | ENSG00000260220 | CCDC187 | chr9 | - | 136300219 | 136300318 | 136297713 | 136297821 | 136302811 | 136303293 | 0 | 3 | [illegible] |
| SE | 96212 | ENSG00000110768 | GTF2H1 | chr11 | + | 18324235 | 18324296 | 18322546 | 18322740 | 18325946 | 18326153 | 0 | 0 | [illegible] |
| SE | 96213 | ENSG00000110768 | GTF2H1 | chr11 | + | 18324235 | 18324296 | 18322546 | 18322740 | 18333059 | 18333228 | 3 | 0 | [illegible] |
| SE | 96214 | ENSG00000110768 | GTF2H1 | chr11 | + | 18325946 | 18326019 | 18322568 | 18322740 | 18333059 | 18333228 | 0 | 0 | [illegible] |
| SE | 96216 | ENSG00000110768 | GTF2H1 | chr11 | + | 18325946 | 18326153 | 18324235 | 18324296 | 18333059 | 18333183 | 1 | 0 | [illegible] |
| SE | 96217 | ENSG00000110768 | GTF2H1 | chr11 | + | 18335753 | 18335946 | 18332661 | 18333228 | 18338108 | 18338274 | 257 | 148 | [illegible] |
| SE | 96218 | ENSG00000110768 | GTF2H1 | chr11 | + | 18338108 | 18338274 | 18333059 | 18333228 | 18341260 | 18341410 | 0 | 1 | [illegible] |
| SE | 96219 | ENSG00000110768 | GTF2H1 | chr11 | + | 18338108 | 18338274 | 18335753 | 18335946 | 18339563 | 18339657 | 323 | 207 | [illegible] |
| SE | 96224 | ENSG00000133943 | DGLUCY | chr14 | + | 91128777 | 91128912 | 91114685 | 91114994 | 91145220 | 91145313 | 0 | 0 | [illegible] |
| SE | 96225 | ENSG00000133943 | DGLUCY | chr14 | + | 91145220 | 91145313 | 91114062 | 91114283 | 91147780 | 91147858 | 64 | 52 | [illegible] |
| SE | 96227 | ENSG00000133943 | DGLUCY | chr14 | + | 91145220 | 91145313 | 91114369 | 91114528 | 91147780 | 91147858 | 42 | 26 | [illegible] |
| SE | 96237 | ENSG00000133943 | DGLUCY | chr14 | + | 91157638 | 91157690 | 91114072 | 91114283 | 91160265 | 91160369 | 90 | 60 | [illegible] |
| SE | 96238 | ENSG00000133943 | DGLUCY | chr14 | + | 91158855 | 91158972 | 91157638 | 91157690 | 91160265 | 91160397 | 0 | 0 | [illegible] |
| SE | 96239 | ENSG00000133943 | DGLUCY | chr14 | + | 91165246 | 91165337 | 91160265 | 91160397 | 91167224 | 91167378 | 0 | 0 | [illegible] |
| SE | 96240 | ENSG00000133943 | DGLUCY | chr14 | + | 91167224 | 91167378 | 91160265 | 91160397 | 91173288 | 91173439 | 114 | 92 | [illegible] |

| | | | | | | | | | | | | | | | | | | | | | | | | | | | | | |
|---|---|---|---|---|---|---|---|---|---|---|---|---|---|---|---|---|---|---|---|---|---|---|---|---|---|---|---|---|---|
| SE | 96242 | ENSG00000133943 | DGLUCY | chr14 | + | 91170002 | 91170201 | 91167224 | 91167378 | 91173288 | 91173436 | 69 | 47 | 79 | 70 | 53 | 0 | 1 | 0 | 1 | 3 | 85 | 52 | 32 | 63 | 135 | 0 | 0 | 1 |
| SE | 96243 | ENSG00000133943 | DGLUCY | chr14 | + | 91173288 | 91173403 | 91160265 | 91160397 | 91175933 | 91176056 | 1 | 0 | 5 | 5 | 0 | 0 | 1 | 0 | 0 | 0 | 2 | 25 | 0 | 3 | 4 | 0 | 0 | 0 |
| SE | 96244 | ENSG00000133943 | DGLUCY | chr14 | + | 91173288 | 91173439 | 91160265 | 91160397 | 91175933 | 91176056 | 19 | 9 | 31 | 48 | 6 | 0 | 1 | 0 | 0 | 0 | 43 | 22 | 14 | 18 | 46 | 0 | 0 | 0 |
| SE | 96246 | ENSG00000133943 | DGLUCY | chr14 | + | 91173288 | 91173439 | 91170002 | 91170201 | 91175933 | 91176056 | 42 | 23 | 62 | 81 | 24 | 0 | 0 | 3 | 0 | 0 | 58 | 34 | 15 | 29 | 105 | 0 | 0 | 0 |
| SE | 96248 | ENSG00000133943 | DGLUCY | chr14 | + | 91175933 | 91176056 | 91173288 | 91173403 | 91181185 | 91181389 | 47 | 28 | 80 | 70 | 44 | 0 | 0 | 0 | 0 | 0 | 36 | 80 | 30 | 79 | 201 | 0 | 4 | 0 |
| SE | 96250 | ENSG00000133943 | DGLUCY | chr14 | + | 91181185 | 91181389 | 91173288 | 91173439 | 91188909 | 91189170 | 75 | 69 | 74 | 95 | 91 | 0 | 0 | 0 | 0 | 0 | 139 | 52 | 98 | 105 | 217 | 0 | 0 | 0 |
| SE | 96255 | ENSG00000133943 | DGLUCY | chr14 | + | 91223678 | 91223795 | 91215404 | 91215793 | 91224683 | 91224822 | 23 | 4 | 11 | 7 | 9 | 4 | 1 | 0 | 0 | 0 | 0 | 3 | 26 | 11 | 20 | 0 | 9 | 6 |
| SE | 96256 | ENSG00000133943 | DGLUCY | chr14 | + | 91223678 | 91223795 | 91215856 | 91216460 | 91224683 | 91224822 | 23 | 4 | 11 | 7 | 9 | 1 | 0 | 0 | 0 | 1 | 0 | 3 | 24 | 11 | 20 | 0 | 0 | 0 |
| SE | 96257 | ENSG00000110696 | C11orf58 | chr11 | + | 16744600 | 16744684 | 16738646 | 16738841 | 16748096 | 16748157 | 443 | 314 | 352 | 400 | 326 | 0 | 0 | 1 | 1 | 0 | 180 | 346 | 383 | 385 | 48 | 0 | 0 | 5 |
| SE | 96261 | ENSG00000110696 | C11orf58 | chr11 | + | 16752784 | 16752894 | 16746703 | 16748157 | 16754870 | 16754906 | 865 | 700 | 678 | 788 | 1127 | 2 | 1 | 3 | 1 | 14 | 300 | 724 | 765 | 650 | 85 | 0 | 0 | 3 |
| SE | 96265 | ENSG00000185345 | PRKN | chr6 | - | 162054090 | 162054174 | 161972945 | 161973417 | 162201130 | 162201252 | 3 | 2 | 2 | 10 | 0 | 0 | 0 | 1 | 0 | 0 | 0 | 0 | 2 | 0 | 1 | 0 | 0 | 0 |
| SE | 96267 | ENSG00000185345 | PRKN | chr6 | - | 162201130 | 162201252 | 162054090 | 162054174 | 162262524 | 162262765 | 5 | 1 | 5 | 7 | 0 | 0 | 0 | 1 | 0 | 0 | 0 | 0 | 5 | 0 | 1 | 0 | 0 | 0 |
| SE | 96273 | ENSG00000139324 | TMTC3 | chr12 | + | 88172965 | 88173089 | 88172596 | 88172745 | 88174606 | 88174727 | 4 | 0 | 1 | 6 | 5 | 15 | 14 | 18 | 16 | 28 | 0 | 0 | 0 | 0 | 6 | 0 | 0 | 3 |
| SE | 96274 | ENSG00000139324 | TMTC3 | chr12 | + | 88174606 | 88174727 | 88172596 | 88172745 | 88176207 | 88176319 | 27 | 20 | 40 | 48 | 67 | 0 | 0 | 2 | 2 | 0 | 0 | 0 | 8 | 9 | 13 | 0 | 0 | 0 |
| SE | 96275 | ENSG00000139324 | TMTC3 | chr12 | + | 88192603 | 88192830 | 88190452 | 88190622 | 88194837 | 88199887 | 85 | 59 | 75 | 79 | 105 | 2 | 0 | 0 | 0 | 3 | 9 | 21 | 17 | 0 | 0 | 0 | 0 | 0 |
| SE | 96278 | ENSG00000029153 | ARNTL2 | chr12 | + | 27370128 | 27370230 | 27368261 | 27368412 | 27376345 | 27376387 | 1 | 0 | 4 | 0 | 0 | 5 | 0 | 0 | 0 | 0 | 8 | 0 | 0 | 0 | 0 | 0 | 0 | 0 |
| SE | 96279 | ENSG00000029153 | ARNTL2 | chr12 | + | 27370128 | 27370230 | 27368261 | 27368412 | 27380246 | 27380404 | 1 | 0 | 4 | 2 | 0 | 2 | 2 | 2 | 1 | 0 | 0 | 0 | 0 | 0 | 0 | 0 | 0 | 0 |
| SE | 96282 | ENSG00000029153 | ARNTL2 | chr12 | + | 27376345 | 27376387 | 27370128 | 27370230 | 27380246 | 27380404 | 0 | 0 | 0 | 0 | 0 | 0 | 0 | 0 | 2 | 0 | 5 | 0 | 0 | 0 | 0 | 0 | 0 | 0 |
| SE | 96283 | ENSG00000029153 | ARNTL2 | chr12 | + | 27387227 | 27387320 | 27385480 | 27385560 | 27389178 | 27389296 | 7 | 17 | 2 | 6 | 18 | 0 | 0 | 0 | 0 | 0 | 2 | 5 | 2 | 17 | 0 | 0 | 0 | 0 |
| SE | 96284 | ENSG00000029153 | ARNTL2 | chr12 | + | 27389178 | 27389296 | 27385480 | 27385560 | 27390095 | 27390239 | 6 | 5 | 3 | 5 | 7 | 0 | 0 | 1 | 0 | 0 | 3 | 6 | 1 | 7 | 0 | 0 | 0 | 0 |
| SE | 96287 | ENSG00000125611 | CHCHD5 | chr2 | + | 112585973 | 112586114 | 112584239 | 112584486 | 112586199 | 112586365 | 24 | 0 | 11 | 7 | 5 | 1 | 0 | 0 | 0 | 0 | 25 | 12 | 4 | 4 | 22 | 0 | 0 | 0 |
| SE | 96288 | ENSG00000125611 | CHCHD5 | chr2 | + | 112585973 | 112586114 | 112584440 | 112584649 | 112586199 | 112586365 | 49 | 10 | 36 | 34 | 37 | 1 | 0 | 1 | 0 | 0 | 60 | 48 | 14 | 20 | 56 | 0 | 0 | 0 |
| SE | 96290 | ENSG00000125611 | CHCHD5 | chr2 | + | 112586199 | 112586365 | 112584440 | 112584649 | 112588865 | 112589040 | 48 | 16 | 29 | 26 | 21 | 5 | 0 | 0 | 0 | 4 | 0 | 41 | 22 | 25 | 48 | 0 | 0 | 0 |
| SE | 96295 | ENSG00000167333 | TRIM68 | chr11 | - | 4601026 | 4601127 | 4598671 | 4600826 | 4601663 | 4601686 | 37 | 8 | 46 | 25 | 28 | 0 | 0 | 0 | 1 | 0 | 0 | 25 | 17 | 3 | 38 | 0 | 0 | 2 |
| SE | 96297 | ENSG00000167333 | TRIM68 | chr11 | - | 4605078 | 4605561 | 4603240 | 4603340 | 4608026 | 4608138 | 16 | 9 | 24 | 19 | 11 | 3 | 0 | 0 | 1 | 0 | 18 | 5 | 4 | 9 | 25 | 0 | 0 | 0 |
| SE | 96301 | ENSG00000152219 | ARL14EP | chr11 | + | 30332865 | 30332993 | 30330885 | 30331374 | 30336566 | 30338223 | 296 | 418 | 421 | 309 | 635 | 0 | 0 | 5 | 0 | 2 | 119 | 173 | 309 | 84 | 18 | 0 | 0 | 0 |
| SE | 96304 | ENSG00000110422 | HIPK3 | chr11 | + | 33286412 | 33287511 | 33258330 | 33258690 | 33328509 | 33328633 | 440 | 402 | 296 | 461 | 609 | 0 | 0 | 0 | 0 | 0 | 196 | 383 | 366 | 163 | 86 | 0 | 0 | 3 |
| SE | 96305 | ENSG00000110422 | HIPK3 | chr11 | + | 33328509 | 33328633 | 33286412 | 33287511 | 33337074 | 33337194 | 862 | 746 | 585 | 897 | 1093 | 0 | 1 | 2 | 2 | 0 | 364 | 736 | 691 | 321 | 144 | 0 | 0 | 0 |
| SE | 96306 | ENSG00000110422 | HIPK3 | chr11 | + | 33328509 | 33328633 | 33286412 | 33287511 | 33338756 | 33338843 | 390 | 332 | 248 | 398 | 539 | 0 | 0 | 0 | 0 | 2 | 166 | 340 | 329 | 148 | 76 | 0 | 0 | 0 |
| SE | 96308 | ENSG00000110422 | HIPK3 | chr11 | + | 33335657 | 33335766 | 33328599 | 33328633 | 33337074 | 33337194 | 4 | 6 | 0 | 1 | 6 | 486 | 429 | 343 | 514 | 575 | 0 | 0 | 2 | 0 | 0 | 210 | 409 | 385 |
| SE | 96309 | ENSG00000110422 | HIPK3 | chr11 | + | 33337074 | 33337194 | 33286412 | 33287511 | 33338756 | 33338842 | 290 | 209 | 158 | 236 | 427 | 0 | 0 | 0 | 0 | 2 | 73 | 174 | 166 | 97 | 86 | 0 | 0 | 0 |
| SE | 96311 | ENSG00000110422 | HIPK3 | chr11 | + | 33347851 | 33348013 | 33347628 | 33347753 | 33348521 | 33348818 | 342 | 327 | 283 | 343 | 496 | 1 | 0 | 0 | 0 | 0 | 160 | 199 | 278 | 117 | 38 | 0 | 0 | 0 |
| SE | 96314 | ENSG00000110422 | HIPK3 | chr11 | + | 33348521 | 33348818 | 33347851 | 33348013 | 33349146 | 33349287 | 570 | 475 | 397 | 491 | 996 | 2 | 7 | 2 | 1 | 0 | 235 | 387 | 379 | 253 | 66 | 0 | 1 | 2 |
| SE | 96316 | ENSG00000110422 | HIPK3 | chr11 | + | 33349146 | 33349287 | 33348521 | 33348818 | 33351607 | 33351843 | 759 | 554 | 520 | 587 | 1147 | 0 | 1 | 0 | 0 | 0 | 167 | 521 | 435 | 273 | 81 | 0 | 0 | 0 |
| SE | 96319 | ENSG00000110717 | NDUFS8 | chr11 | + | 68035627 | 68035788 | 68033140 | 68033283 | 68036252 | 68036381 | 1 | 0 | 1 | 0 | 5 | 89 | 24 | 52 | 68 | 41 | 0 | 1 | 0 | 0 | 0 | 76 | 48 | 36 |
| [illegible] | | | | | | | | | | | | | | | | | | | | | | | | | | | | | |

| | | | | | | | | | | | | | |
|---|---|---|---|---|---|---|---|---|---|---|---|---|---|
| SE | 96735 | ENSG00000162599 | NFIA | chr1 | + | 61385870 | 61385924 | 61383236 | 61383365 | 61404103 | 61404226 | 0 | 0 |
| SE | 96737 | ENSG00000162599 | NFIA | chr1 | + | 61426464 | 61426556 | 61406561 | 61406727 | 61455302 | 61455487 | 8 | 0 |
| SE | 96739 | ENSG00000162599 | NFIA | chr1 | + | 61443522 | 61443577 | 61426464 | 61426556 | 61455302 | 61456742 | 4 | 0 |
| SE | 96740 | ENSG00000185238 | PRMT3 | chr11 | + | 20388018 | 20388154 | 20387715 | 20387774 | 20389743 | 20389826 | 23 | 16 |
| SE | 96742 | ENSG00000185238 | PRMT3 | chr11 | + | 20392896 | 20392999 | 20389743 | 20389826 | 20395802 | 20395962 | 43 | 35 |
| SE | 96744 | ENSG00000185238 | PRMT3 | chr11 | + | 20395802 | 20395962 | 20389743 | 20389826 | 20402918 | 20402984 | 1 | 2 |
| SE | 96746 | ENSG00000185238 | PRMT3 | chr11 | + | 20426765 | 20426865 | 20405445 | 20408032 | 20452129 | 20452208 | 76 | 55 |
| SE | 96747 | ENSG00000185238 | PRMT3 | chr11 | + | 20452129 | 20452208 | 20426765 | 20426865 | 20461979 | 20462167 | 65 | 48 |
| SE | 96748 | ENSG00000183431 | SF3A3 | chr1 | - | 37960119 | 37960175 | 37956974 | 37958263 | 37968043 | 37968134 | 359 | 259 |
| SE | 96749 | ENSG00000183431 | SF3A3 | chr1 | - | 37968043 | 37968134 | 37960119 | 37960175 | 37969353 | 37969464 | 478 | 369 |
| SE | 96750 | ENSG00000183431 | SF3A3 | chr1 | - | 37969353 | 37969464 | 37968043 | 37968134 | 37969570 | 37969735 | 328 | 277 |
| SE | 96751 | ENSG00000183431 | SF3A3 | chr1 | - | 37976883 | 37976953 | 37969570 | 37969735 | 37978719 | 37978827 | 181 | 151 |
| SE | 96753 | ENSG00000183431 | SF3A3 | chr1 | - | 37978719 | 37978827 | 37969570 | 37969735 | 37979464 | 37979533 | 2 | 3 |
| SE | 96758 | ENSG00000183431 | SF3A3 | chr1 | - | 37984168 | 37984260 | 37981728 | 37981811 | 37984706 | 37984779 | 290 | 262 |
| SE | 96761 | ENSG00000183431 | SF3A3 | chr1 | - | 37987783 | 37987836 | 37987572 | 37987678 | 37989547 | 37989595 | 239 | 151 |
| SE | 96762 | ENSG00000183431 | SF3A3 | chr1 | - | 37989547 | 37989595 | 37987783 | 37987836 | 37989869 | 37989973 | 273 | 174 |
| SE | 96763 | ENSG00000052841 | TTC17 | chr11 | + | 43370078 | 43370385 | 43358953 | 43359113 | 43379232 | 43379322 | 0 | 3 |
| SE | 96764 | ENSG00000052841 | TTC17 | chr11 | + | 43389651 | 43389821 | 43379232 | 43379322 | 43391464 | 43391576 | 121 | 100 |
| SE | 96765 | ENSG00000052841 | TTC17 | chr11 | + | 43389651 | 43389821 | 43379232 | 43379322 | 43391820 | 43391870 | 86 | 75 |
| SE | 96766 | ENSG00000052841 | TTC17 | chr11 | + | 43391464 | 43391576 | 43379232 | 43379322 | 43391820 | 43391870 | 178 | 119 |
| SE | 96767 | ENSG00000052841 | TTC17 | chr11 | + | 43391464 | 43391576 | 43389651 | 43389821 | 43391820 | 43391870 | 212 | 142 |
| SE | 96769 | ENSG00000052841 | TTC17 | chr11 | + | 43414589 | 43414776 | 43407352 | 43407577 | 43436163 | 43436334 | 280 | 229 |
| SE | 96771 | ENSG00000052841 | TTC17 | chr11 | + | 43436163 | 43436334 | 43407352 | 43407577 | 43443324 | 43443584 | 337 | 274 |
| SE | 96777 | ENSG00000109911 | ELP4 | chr11 | + | 31580682 | 31580800 | 31539661 | 31539783 | 31594769 | 31594901 | 0 | 3 |
| SE | 96780 | ENSG00000109911 | ELP4 | chr11 | + | 31628238 | 31628419 | 31627109 | 31627194 | 31632216 | 31632405 | 2 | 0 |
| SE | 96786 | ENSG00000109911 | ELP4 | chr11 | + | 31779712 | 31779945 | 31650180 | 31650221 | 31783392 | 31783757 | 1 | 0 |
| SE | 96789 | ENSG00000259719 | LINC02284 | chr14 | + | 56317891 | 56318086 | 56310879 | 56311102 | 56321485 | 56321611 | 7 | 11 |
| SE | 96792 | ENSG00000259719 | LINC02284 | chr14 | + | 56390575 | 56390660 | 56344973 | 56345070 | 56426187 | 56427032 | 0 | 0 |
| SE | 96793 | ENSG00000151773 | CCDC122 | chr13 | - | 43868693 | 43868803 | 43859671 | 43860070 | 43869330 | 43869489 | 2 | 6 |
| SE | 96795 | ENSG00000151773 | CCDC122 | chr13 | - | 43869330 | 43869489 | 43859671 | 43860070 | 43874841 | 43874927 | 3 | 5 |
| SE | 96796 | ENSG00000151773 | CCDC122 | chr13 | - | 43869330 | 43869489 | 43868693 | 43868803 | 43874841 | 43874927 | 3 | 8 |
| SE | 96802 | ENSG00000175220 | ARHGAP1 | chr11 | - | 46679647 | 46679776 | 46679364 | 46679468 | 46680204 | 46680282 | 286 | 84 |
| SE | 96803 | ENSG00000175220 | ARHGAP1 | chr11 | - | 46679647 | 46679831 | 46679364 | 46679468 | 46680204 | 46680282 | 137 | 35 |
| SE | 96806 | ENSG00000205220 | PSMB10 | chr16 | - | 67936397 | 67936485 | 67936214 | 67936312 | 67936693 | 67936847 | 218 | 77 |
| SE | 96807 | ENSG00000166889 | PATL1 | chr11 | - | 59647753 | 59647913 | 59642879 | 59643035 | 59649461 | 59649610 | 203 | 200 |
| SE | 96808 | ENSG00000166889 | PATL1 | chr11 | - | 59651543 | 59651641 | 59650753 | 59650813 | 59652463 | 59652587 | 301 | 202 |
| SE | 96810 | ENSG00000166889 | PATL1 | chr11 | - | 59656498 | 59656600 | 59655522 | 59655740 | 59658865 | 59658946 | 15 | 7 |
| SE | 96811 | ENSG00000166889 | PATL1 | chr11 | - | 59657529 | 59657724 | 59656498 | 59656600 | 59658865 | 59658946 | 178 | 109 |
| SE | 96812 | ENSG00000166889 | PATL1 | chr11 | - | 59659251 | 59659469 | 59658865 | 59658946 | 59666852 | 59666964 | 583 | 510 |
| SE | 96816 | ENSG00000147874 | HAUS6 | chr9 | - | 19076601 | 19076704 | 19070218 | 19070300 | 19080478 | 19080672 | 59 | 36 |
| SE | 96817 | ENSG00000147874 | HAUS6 | chr9 | - | 19078175 | 19078302 | 19070218 | 19070300 | 19080478 | 19080672 | 34 | 36 |
| SE | 96818 | ENSG00000147874 | HAUS6 | chr9 | - | 19078175 | 19078302 | 19076601 | 19076704 | 19080478 | 19080672 | 103 | 90 |
| SE | 96819 | ENSG00000147874 | HAUS6 | chr9 | - | 19080478 | 19080672 | 19078175 | 19078302 | 19082872 | 19083043 | 103 | 91 |
| SE | 96820 | ENSG00000147874 | HAUS6 | chr9 | - | 19082872 | 19083043 | 19080478 | 19080672 | 19086733 | 19086782 | 155 | 111 |
| SE | 96827 | ENSG00000152270 | PDE3B | chr11 | + | 14786436 | 14786685 | 14771936 | 14771987 | 14789105 | 14789242 | 382 | 368 |
| SE | 96828 | ENSG00000152270 | PDE3B | chr11 | + | 14803943 | 14804050 | 14643803 | 14645053 | 14818182 | 14818393 | 169 | 118 |
| SE | 96829 | ENSG00000152270 | PDE3B | chr11 | + | 14803943 | 14804050 | 14788938 | 14789242 | 14818182 | 14818393 | 378 | 294 |
| SE | 96831 | ENSG00000152270 | PDE3B | chr11 | + | 14843826 | 14844026 | 14834981 | 14835095 | 14859042 | 14859246 | 209 | 202 |
| SE | 96832 | ENSG00000152270 | PDE3B | chr11 | + | 14859042 | 14859246 | 14843826 | 14844026 | 14861204 | 14861366 | 250 | 229 |
| SE | 96833 | ENSG00000152270 | PDE3B | chr11 | + | 14861204 | 14861366 | 14843826 | 14844026 | 14867505 | 14867758 | 90 | 74 |
| SE | 96835 | ENSG00000149516 | MS4A3 | chr11 | + | 60061145 | 60061316 | 60056627 | 60056740 | 60066950 | 60067112 | 2 | 4 |
| SE | 96837 | ENSG00000149516 | MS4A3 | chr11 | + | 60061489 | 60061554 | 60061145 | 60061316 | 60064261 | 60064318 | 0 | 0 |
| SE | 96840 | ENSG00000149516 | MS4A3 | chr11 | + | 60062467 | 60062605 | 60061145 | 60061316 | 60066950 | 60067112 | 5 | 5 |
| SE | 96841 | ENSG00000149516 | MS4A3 | chr11 | + | 60064261 | 60064318 | 60056586 | 60056740 | 60066950 | 60067112 | 22 | 11 |
| SE | 96842 | ENSG00000149516 | MS4A3 | chr11 | + | 60064261 | 60064318 | 60061145 | 60061316 | 60066950 | 60067112 | 12 | 12 |
| SE | 96843 | ENSG00000149516 | MS4A3 | chr11 | + | 60064261 | 60064318 | 60061489 | 60061554 | 60066950 | 60067112 | 12 | 8 |
| SE | 96844 | ENSG00000149516 | MS4A3 | chr11 | + | 60064261 | 60064318 | 60062467 | 60062605 | 60066950 | 60067112 | 29 | 15 |
| SE | 96845 | ENSG00000149516 | MS4A3 | chr11 | + | 60066950 | 60067112 | 60064261 | 60064318 | 60069573 | 60069675 | 54 | 34 |
| SE | 96846 | ENSG00000149516 | MS4A3 | chr11 | + | 60069573 | 60069675 | 60066950 | 60067112 | 60070203 | 60070233 | 35 | 32 |
| SE | 96848 | ENSG00000083535 | PIBF1 | chr13 | + | 72821848 | 72821982 | 72797906 | 72798026 | 72827009 | 72827118 | 186 | 179 |
| SE | 96849 | ENSG00000083535 | PIBF1 | chr13 | + | 72827732 | 72827914 | 72827009 | 72827118 | 72835242 | 72835368 | 183 | 174 |
| SE | 96850 | ENSG00000083535 | PIBF1 | chr13 | + | 72854056 | 72854155 | 72835242 | 72835368 | 72893783 | 72893949 | 73 | 84 |
| SE | 96854 | ENSG00000083535 | PIBF1 | chr13 | + | 72917075 | 72917166 | 72908530 | 72908681 | 72931164 | 72931267 | 122 | 129 |
| SE | 96855 | ENSG00000083535 | PIBF1 | chr13 | + | 72965273 | 72965404 | 72931164 | 72931267 | 72973590 | 72973675 | 93 | 78 |
| SE | 96856 | ENSG00000083535 | PIBF1 | chr13 | + | 72965273 | 72965404 | 72931164 | 72931267 | 72998821 | 72998995 | 37 | 36 |
| SE | 96857 | ENSG00000083535 | PIBF1 | chr13 | + | 72973590 | 72973675 | 72931164 | 72931267 | 72998821 | 72998995 | 41 | 31 |
| SE | 96858 | ENSG00000083535 | PIBF1 | chr13 | + | 72973590 | 72973675 | 72965273 | 72965404 | 72998821 | 72998995 | 101 | 77 |
| SE | 96859 | ENSG00000083535 | PIBF1 | chr13 | + | 72998821 | 72998995 | 72973590 | 72973675 | 73015868 | 73015980 | 79 | 74 |
| SE | 96864 | ENSG00000126934 | MAP2K2 | chr19 | - | 4097278 | 4097343 | 4094452 | 4095449 | 4101018 | 4101047 | 86 | 21 |
| SE | 96865 | ENSG00000126934 | MAP2K2 | chr19 | - | 4099200 | 4099414 | 4097278 | 4097343 | 4101018 | 4101143 | 301 | 85 |
| SE | 96866 | ENSG00000126934 | MAP2K2 | chr19 | - | 4103411 | 4103484 | 4102375 | 4102453 | 4110508 | 4110655 | 0 | 0 |
| SE | 96867 | ENSG00000126934 | MAP2K2 | chr19 | - | 4117418 | 4117629 | 4110508 | 4110655 | 4123783 | 4123815 | 220 | 105 |
| SE | 96868 | ENSG00000136816 | TOR1B | chr9 | + | 129807187 | 129807363 | 129804072 | 129804338 | 129808904 | 129809032 | 235 | 146 |
| SE | 96869 | ENSG00000136816 | TOR1B | chr9 | + | 129807187 | 129807363 | 129804357 | 129804479 | 129808904 | 129809032 | 122 | 70 |
| SE | 96870 | ENSG00000136816 | TOR1B | chr9 | + | 129808904 | 129809032 | 129804072 | 129804338 | 129810226 | 129810316 | 3 | 0 |
| SE | 96882 | ENSG00000100731 | PCNX1 | chr14 | + | 70962225 | 70962331 | 70946881 | 70947123 | 70976941 | 70978648 | 229 | 161 |
| SE | 96883 | ENSG00000100731 | PCNX1 | chr14 | + | 70969858 | 70969999 | 70969020 | 70969110 | 70976941 | 70977151 | 0 | 4 |
| SE | 96886 | ENSG00000100731 | PCNX1 | chr14 | + | 70988566 | 70988699 | 70976941 | 70978648 | 71011491 | 71011549 | 48 | 20 |
| SE | 96891 | ENSG00000100731 | PCNX1 | chr14 | + | 70995740 | 70995925 | 70976941 | 70978648 | 71012999 | 71013202 | 9 | 7 |
| SE | 96893 | ENSG00000100731 | PCNX1 | chr14 | + | 70995740 | 70995925 | 70988566 | 70988699 | 71012999 | 71013202 | 15 | 7 |
| SE | 96897 | ENSG00000100731 | PCNX1 | chr14 | + | 71009633 | 71009724 | 70988566 | 70988699 | 71012999 | 71013202 | 0 | 1 |
| SE | 96898 | ENSG00000100731 | PCNX1 | chr14 | + | 71009633 | 71009724 | 70995740 | 70995925 | 71012999 | 71013202 | 20 | 8 |
| SE | 96899 | ENSG00000100731 | PCNX1 | chr14 | + | 71011491 | 71011549 | 70976941 | 70978648 | 71012999 | 71013202 | 9 | 5 |
| SE | 96900 | ENSG00000100731 | PCNX1 | chr14 | + | 71011491 | 71011549 | 70988566 | 70988699 | 71012999 | 71013202 | 10 | 5 |
| SE | 96902 | ENSG00000100731 | PCNX1 | chr14 | + | 71019008 | 71019162 | 71012984 | 71013202 | 71023199 | 71023232 | 229 | 143 |
| SE | 96903 | ENSG00000100731 | PCNX1 | chr14 | + | 71026116 | 71026288 | 71023199 | 71023232 | 71026771 | 71026882 | 213 | 121 |
| SE | 96904 | ENSG00000100731 | PCNX1 | chr14 | + | 71036064 | 71036157 | 71033930 | 71034036 | 71045132 | 71045283 | 107 | 64 |
| SE | 96908 | ENSG00000100731 | PCNX1 | chr14 | + | 71051882 | 71052012 | 71047806 | 71047984 | 71073544 | 71073759 | 0 | 0 |
| SE | 96909 | ENSG00000100731 | PCNX1 | chr14 | + | 71053296 | 71053412 | 71051882 | 71052012 | 71055503 | 71055562 | 1 | 1 |
| SE | 96910 | ENSG00000100731 | PCNX1 | chr14 | + | 71053296 | 71053412 | 71051882 | 71052012 | 71073544 | 71073759 | 1 | 0 |
| SE | 96911 | ENSG00000100731 | PCNX1 | chr14 | + | 71055503 | 71055562 | 71047806 | 71047984 | 71073544 | 71073798 | 6 | 0 |
| SE | 96912 | ENSG00000100731 | PCNX1 | chr14 | + | 71055503 | 71055562 | 71051882 | 71052012 | 71073544 | 71073798 | 67 | 45 |
| SE | 96915 | ENSG00000100731 | PCNX1 | chr14 | + | 71101989 | 71102220 | 71089191 | 71089342 | 71103394 | 71103669 | 221 | 141 |
| SE | 96916 | ENSG00000100731 | PCNX1 | chr14 | + | 71103394 | 71103669 | 71101989 | 71102220 | 71105234 | 71105440 | 310 | 131 |
| SE | 96917 | ENSG00000100731 | PCNX1 | chr14 | + | 71108603 | 71109046 | 71105234 | 71105440 | 71109451 | 71109592 | 464 | 359 |
| SE | 96918 | ENSG00000239445 | ST3GAL6-AS1 | chr3 | - | 98714407 | 98714498 | 98706274 | 98708316 | 98715375 | 98715521 | 4 | 1 |
| SE | 96919 | ENSG00000239445 | ST3GAL6-AS1 | chr3 | - | 98714407 | 98714498 | 98709698 | 98710162 | 98715375 | 98715521 | 4 | 0 |
| SE | 96920 | ENSG00000239445 | ST3GAL6-AS1 | chr3 | - | 98715375 | 98715521 | 98706235 | 98708316 | 98718418 | 98718521 | 0 | 0 |
| SE | 96921 | ENSG00000239445 | ST3GAL6-AS1 | chr3 | - | 98715375 | 98715521 | 98706235 | 98708316 | 98718423 | 98718479 | 0 | 0 |
| SE | 96922 | ENSG00000239445 | ST3GAL6-AS1 | chr3 | - | 98715649 | 98715777 | 98711284 | 98715521 | 98718418 | 98718521 | 6 | 0 |
| SE | 96923 | ENSG00000239445 | ST3GAL6-AS1 | chr3 | - | 98715649 | 98715777 | 98711284 | 98715521 | 98718423 | 98718499 | 3 | 0 |
| SE | 96924 | ENSG00000239445 | ST3GAL6-AS1 | chr3 | - | 98718418 | 98718521 | 98715649 | 98715777 | 98732425 | 98732455 | 2 | 0 |
| SE | 96925 | ENSG00000239445 | ST3GAL6-AS1 | chr3 | - | 98718423 | 98718521 | 98713315 | 98715777 | 98732425 | 98732490 | 0 | 0 |
| SE | 96928 | ENSG00000089159 | PXN | chr12 | - | 120215929 | 120216090 | 120215559 | 120215661 | 120216272 | 120217116 | 2 | 0 |
| SE | 96932 | ENSG00000089159 | PXN | chr12 | - | 120216840 | 120217116 | 120216272 | 120216581 | 120221622 | 120221758 | 0 | 0 |
| SE | 96934 | ENSG00000089159 | PXN | chr12 | - | 120219206 | 120220091 | 120216272 | 120217116 | 120221622 | 120221758 | 0 | 0 |
| SE | 96936 | ENSG00000089159 | PXN | chr12 | - | 120223717 | 120223833 | 120222862 | 120222999 | 120224150 | 120224377 | 532 | 202 |
| SE | 96937 | ENSG00000089159 | PXN | chr12 | - | 120224642 | 120224727 | 120224150 | 120224377 | 120225908 | 120226055 | 5 | 0 |
| SE | 96944 | ENSG00000089159 | PXN | chr12 | - | 120264231 | 120264316 | 120224150 | 120224377 | 120265616 | 120265706 | 0 | 0 |
| SE | 96946 | ENSG00000062822 | POLD1 | chr19 | + | 50387659 | 50387804 | 50384372 | 50384390 | 50398850 | 50399053 | 0 | 0 |
| SE | 96948 | ENSG00000062822 | POLD1 | chr19 | + | 50395024 | 50395150 | 50384341 | 50384390 | 50398847 | 50399053 | 0 | 0 |
| SE | 96950 | ENSG00000062822 | POLD1 | chr19 | + | 50395024 | 50395150 | 50393974 | 50394063 | 50398850 | 50399053 | 5 | 1 |
| SE | 96954 | ENSG00000156689 | GLYATL2 | chr11 | - | 58839534 | 58839652 | 58838260 | 58838368 | 58844433 | 58844704 | 0 | 0 |
| SE | 96957 | ENSG00000183597 | TANGO2 | chr22 | + | 20036759 | 20036854 | 20021068 | 20021246 | 20043354 | 20043443 | 33 | 11 |
| SE | 96958 | ENSG00000183597 | TANGO2 | chr22 | + | 20036759 | 20037094 | 20021138 | 20021246 | 20043354 | 20043443 | 17 | 13 |
| SE | 96959 | ENSG00000183597 | TANGO2 | chr22 | + | 20047969 | 20048328 | 20043354 | 20043443 | 20052464 | 20052584 | 0 | 0 |
| SE | 96963 | ENSG00000183597 | TANGO2 | chr22 | + | 20053436 | 20053551 | 20052464 | 20052584 | 20063337 | 20063442 | 33 | 18 |
| SE | 96966 | ENSG00000183597 | TANGO2 | chr22 | + | 20055942 | 20056013 | 20052464 | 20052584 | 20064541 | 20065925 | 28 | 22 |
| SE | 96969 | ENSG00000183597 | TANGO2 | chr22 | + | 20061529 | 20061681 | 20053436 | 20053551 | 20063337 | 20063383 | 1 | 2 |
| SE | 96970 | ENSG00000183597 | TANGO2 | chr22 | + | 20061529 | 20061683 | 20052464 | 20052584 | 20063337 | 20063411 | 30 | 21 |
| SE | 96973 | ENSG00000183597 | TANGO2 | chr22 | + | 20063337 | 20063442 | 20052464 | 20052584 | 20064541 | 20064907 | 17 | 21 |
| SE | 96974 | ENSG00000183597 | TANGO2 | chr22 | + | 20063337 | 20063442 | 20055942 | 20056013 | 20064541 | 20064907 | 18 | 21 |
| SE | 96976 | ENSG00000136270 | TBRG4 | chr7 | - | 45104098 | 45104256 | 45103030 | 45103443 | 45105440 | 45105669 | 28 | 12 |
| SE | 96978 | ENSG00000136270 | TBRG4 | chr7 | - | 45104537 | 45104709 | 45103631 | 45104256 | 45105440 | 45105478 | 81 | 29 |
| SE | 96979 | ENSG00000136270 | TBRG4 | chr7 | - | 45110744 | 45110877 | 45108927 | 45109287 | 45111642 | 45111697 | 0 | 0 |
| SE | 96988 | ENSG00000254772 | EEF1G | chr11 | - | 62570964 | 62571108 | 62567398 | 62567528 | 62571539 | 62571682 | 8518 | 5036 |
| SE | 96989 | ENSG00000254772 | EEF1G | chr11 | - | 62571539 | 62571682 | 62570964 | 62571108 | 62571837 | 62571901 | 11558 | 6904 |
| SE | 96990 | ENSG00000254772 | EEF1G | chr11 | - | 62571623 | 62571682 | 62570964 | 62571108 | 62571837 | 62571901 | 2695 | 1593 |
| SE | 96992 | ENSG00000254772 | EEF1G | chr11 | - | 62572583 | 62572742 | 62571837 | 62571901 | 62573830 | 62573891 | 6650 | 3665 |
| SE | 96995 | ENSG00000153339 | TRAPPC8 | chr18 | - | 31832083 | 31832173 | 31829196 | 31830989 | 31846715 | 31846817 | 77 | 54 |
| SE | 96997 | ENSG00000153339 | TRAPPC8 | chr18 | - | 31839311 | 31839457 | 31832083 | 31832173 | 31846715 | 31846817 | 432 | 345 |
| SE | 96998 | ENSG00000153339 | TRAPPC8 | chr18 | - | 31849565 | 31849739 | 31846715 | 31846817 | 31852445 | 31852504 | 537 | 418 |
| SE | 96999 | ENSG00000153339 | TRAPPC8 | chr18 | - | 31853848 | 31853945 | 31852594 | 31852663 | 31855659 | 31855807 | 308 | 223 |
| SE | 97000 | ENSG00000153339 | TRAPPC8 | chr18 | - | 31855659 | 31855807 | 31853848 | 31853945 | 31857539 | 31857720 | 319 | 263 |
| SE | 97001 | ENSG00000153339 | TRAPPC8 | chr18 | - | 31857539 | 31857720 | 31855659 | 31855807 | 31864626 | 31864781 | 141 | 108 |
| SE | 97002 | ENSG00000153339 | TRAPPC8 | chr18 | - | 31857539 | 31857982 | 31855659 | 31855807 | 31864626 | 31864781 | 204 | 146 |
| SE | 97003 | ENSG00000153339 | TRAPPC8 | chr18 | - | 31866848 | 31866975 | 31864626 | 31864781 | 31867401 | 31867476 | 223 | 149 |
| SE | 97004 | ENSG00000153339 | TRAPPC8 | chr18 | - | 31874479 | 31874704 | 31873429 | 31873538 | 31890734 | 31890866 | 281 | 234 |
| SE | 97005 | ENSG00000153339 | TRAPPC8 | chr18 | - | 31890734 | 31890866 | 31873569 | 31874704 | 31897785 | 31897891 | 126 | 126 |
| SE | 97007 | ENSG00000153339 | TRAPPC8 | chr18 | - | 31913368 | 31913522 | 31909666 | 31909760 | 31931328 | 31931523 | 215 | 138 |
| SE | 97011 | ENSG00000148950 | IMMP1L | chr11 | - | 31447982 | 31449091 | 31433459 | 31433570 | 31456259 | 31456386 | 0 | 0 |
| SE | 97012 | ENSG00000148950 | IMMP1L | chr11 | - | 31456259 | 31456386 | 31433459 | 31433570 | 31460625 | 31460714 | 33 | 19 |
| SE | 97014 | ENSG00000148950 | IMMP1L | chr11 | - | 31460625 | 31460714 | 31433459 | 31433570 | 31463171 | 31463305 | 12 | 3 |
| SE | 97015 | ENSG00000148950 | IMMP1L | chr11 | - | 31460625 | 31460714 | 31433459 | 31433570 | 31509518 | 31509592 | 1 | 0 |
| SE | 97017 | ENSG00000148950 | IMMP1L | chr11 | - | 31460625 | 31460714 | 31456259 | 31456386 | 31509518 | 31509592 | 9 | 12 |
| SE | 97019 | ENSG00000148950 | IMMP1L | chr11 | - | 31463171 | 31463305 | 31456312 | 31456386 | 31469739 | 31469797 | 0 | 0 |
| SE | 97020 | ENSG00000148950 | IMMP1L | chr11 | - | 31463171 | 31463305 | 31456312 | 31456386 | 31509518 | 31509574 | 5 | 0 |
| SE | 97021 | ENSG00000148950 | IMMP1L | chr11 | - | 31463171 | 31463305 | 31460625 | 31460714 | 31509518 | 31509574 | 27 | 7 |
| SE | 97022 | ENSG00000148950 | IMMP1L | chr11 | - | 31469739 | 31469797 | 31463171 | 31463305 | 31473723 | 31473786 | 0 | 0 |
| SE | 97025 | ENSG00000148950 | IMMP1L | chr11 | - | 31473723 | 31473786 | 31463171 | 31463305 | 31509518 | 31509594 | 2 | 0 |
| SE | 97027 | ENSG00000148950 | IMMP1L | chr11 | - | 31477500 | 31477759 | 31433459 | 31433570 | 31509518 | 31509592 | 0 | 0 |
| SE | 97033 | ENSG00000251562 | MALAT1 | chr11 | + | 65498968 | 65503905 | 65498681 | 65498734 | 65504132 | 65504206 | 0 | 0 |
| SE | 97035 | ENSG00000251562 | MALAT1 | chr11 | + | 65505503 | 65505662 | 65504518 | 65505019 | 65506385 | 65506469 | 10 | 2 |
| SE | 97036 | ENSG00000251562 | MALAT1 | chr11 | + | 65505590 | 65505662 | 65504518 | 65505019 | 65506385 | 65506465 | 0 | 5 |
| SE | 97040 | ENSG00000166444 | DENND2B | chr11 | - | 8713997 | 8714042 | 8712550 | 8712735 | 8714609 | 8714706 | 2 | 0 |
| SE | 97060 | ENSG00000175216 | CKAP5 | chr11 | - | 46753309 | 46753497 | 46751345 | 46751534 | 46758922 | 46759043 | 2 | 5 |
| SE | 97061 | ENSG00000175216 | CKAP5 | chr11 | - | 46754887 | 46754983 | 46753453 | 46753497 | 46758922 | 46759043 | 30 | 21 |
| SE | 97062 | ENSG00000175216 | CKAP5 | chr11 | - | 46754887 | 46755067 | 46753309 | 46753497 | 46758922 | 46759043 | 106 | 86 |
| SE | 97063 | ENSG00000175216 | CKAP5 | chr11 | - | 46759268 | 46759442 | 46758922 | 46759043 | 46760611 | 46760784 | 145 | 117 |
| SE | 97065 | ENSG00000175216 | CKAP5 | chr11 | - | 46763480 | 46763630 | 46762975 | 46763179 | 46765130 | 46765256 | 180 | 61 |
| SE | 97066 | ENSG00000175216 | CKAP5 | chr11 | - | 46767574 | 46767663 | 46765130 | 46765256 | 46769962 | 46770098 | 126 | 65 |
| SE | 97067 | ENSG00000175216 | CKAP5 | chr11 | - | 46777438 | 46777552 | 46776254 | 46776383 | 46778138 | 46778313 | 180 | 114 |
| SE | 97068 | ENSG00000175216 | CKAP5 | chr11 | - | 46783273 | 46783368 | 46780427 | 46780485 | 46784487 | 46784673 | 138 | 87 |
| SE | 97069 | ENSG00000175216 | CKAP5 | chr11 | - | 46784487 | 46784673 | 46783273 | 46783368 | 46788680 | 46788773 | 123 | 90 |
| SE | 97071 | ENSG00000175216 | CKAP5 | chr11 | - | 46818309 | 46818503 | 46816197 | 46816404 | 46821174 | 46821257 | 345 | 255 |
| SE | 97075 | ENSG00000173599 | PC | chr11 | - | 66954248 | 66954436 | 66952295 | 66952468 | 66958321 | 66958383 | 1 | 0 |
| SE | 97076 | ENSG00000173599 | PC | chr11 | - | 66957739 | 66957996 | 66952429 | 66952468 | 66958321 | 66958372 | 1 | 1 |
| SE | 97077 | ENSG00000173599 | PC | chr11 | - | 66957868 | 66957996 | 66952429 | 66952468 | 66958321 | 66958344 | 1 | 0 |
| SE | 97085 | ENSG00000154258 | ABCA9 | chr17 | - | 68989812 | 68989930 | 68989026 | 68989118 | 68990836 | 68990957 | 1 | 0 |
| SE | 97087 | ENSG00000154258 | ABCA9 | chr17 | - | 69035659 | 69035801 | 69035245 | 69035431 | 69043488 | 69043715 | 0 | 0 |
| SE | 97088 | ENSG00000154258 | ABCA9 | chr17 | - | 69049282 | 69049490 | 69045171 | 69045336 | 69051030 | 69051139 | 0 | 0 |
| SE | 97097 | ENSG00000151465 | CDC123 | chr10 | + | 12210289 | 12210322 | 12209966 | 12210024 | 12215739 | 12215835 | 209 | 105 |
| SE | 97099 | ENSG00000151465 | CDC123 | chr10 | + | 12217360 | 12217467 | 12198704 | 12198776 | 12230947 | 12230978 | 218 | 126 |
| SE | 97100 | ENSG00000151465 | CDC123 | chr10 | + | 12237143 | 12237266 | 12235047 | 12235123 | 12238456 | 12238485 | 593 | 408 |
| SE | 97102 | ENSG00000151465 | CDC123 | chr10 | + | 12238456 | 12238485 | 12237143 | 12237266 | 12246148 | 12246277 | 416 | 313 |
| SE | 97106 | ENSG00000149806 | FAU | chr11 | - | 65120980 | 65121036 | 65120629 | 65120806 | 65121499 | 65121539 | 3854 | 2020 |
| SE | 97107 | ENSG00000110025 | SNX15 | chr11 | + | 65032167 | 65032271 | 65027471 | 65027636 | 65032430 | 65032551 | 6 | 1 |
| SE | 97110 | ENSG00000205639 | MFSD2B | chr2 | + | 24017285 | 24017364 | 24016844 | 24016968 | 24017457 | 24017588 | 5 | 0 |
| SE | 97112 | ENSG00000052126 | PLEKHA5 | chr12 | + | 19243179 | 19243443 | 19132392 | 19132450 | 19253939 | 19254009 | 0 | 0 |
| SE | 97114 | ENSG00000052126 | PLEKHA5 | chr12 | + | 19255044 | 19255165 | 19253939 | 19254023 | 19257432 | 19257443 | 70 | 19 |
| SE | 97115 | ENSG00000052126 | PLEKHA5 | chr12 | + | 19260948 | 19261021 | 19257432 | 19257537 | 19265749 | 19265817 | 57 | 10 |
| SE | 97116 | ENSG00000052126 | PLEKHA5 | chr12 | + | 19265749 | 19265850 | 19260948 | 19261021 | 19269769 | 19269885 | 71 | 14 |
| SE | 97117 | ENSG00000052126 | PLEKHA5 | chr12 | + | 19269769 | 19269885 | 19265749 | 19265850 | 19270187 | 19270205 | 37 | 11 |
| SE | 97118 | ENSG00000052126 | PLEKHA5 | chr12 | + | 19269769 | 19269885 | 19265749 | 19265850 | 19274515 | 19274983 | 86 | 28 |
| SE | 97119 | ENSG00000052126 | PLEKHA5 | chr12 | + | 19270187 | 19270205 | 19265749 | 19265850 | 19274515 | 19274983 | 0 | 0 |
| SE | 97120 | ENSG00000052126 | PLEKHA5 | chr12 | + | 19270187 | 19270205 | 19269769 | 19269885 | 19274515 | 19274983 | 1 | 0 |
| SE | 97121 | ENSG00000052126 | PLEKHA5 | chr12 | + | 19287472 | 19287556 | 19283279 | 19283745 | 19320561 | 19320624 | 26 | 5 |
| SE | 97126 | ENSG00000052126 | PLEKHA5 | chr12 | + | 19346993 | 19347182 | 19345841 | 19345888 | 19348398 | 19348519 | 0 | 2 |
| SE | 97127 | ENSG00000052126 | PLEKHA5 | chr12 | + | 19348398 | 19348519 | 19345841 | 19345888 | 19353883 | 19354002 | 48 | 9 |
| SE | 97128 | ENSG00000052126 | PLEKHA5 | chr12 | + | 19369692 | 19369798 | 19365963 | 19366109 | 19375530 | 19375592 | 21 | 5 |
| SE | 97137 | ENSG00000146112 | PPP1R18 | chr6 | - | 30679178 | 30679389 | 30676388 | 30677288 | 30684407 | 30684544 | 645 | 317 |
| SE | 97141 | ENSG00000162139 | NEU3 | chr11 | + | 75004304 | 75004405 | 74994508 | 74994720 | 75005412 | 75005469 | 0 | 0 |
| SE | 97145 | ENSG00000153827 | TRIP12 | chr2 | - | 229768615 | 229768719 | 229763836 | 229767750 | 229769230 | 229769325 | 739 | 616 |
| SE | 97146 | ENSG00000153827 | TRIP12 | chr2 | - | 229769230 | 229769325 | 229763836 | 229767750 | 229774096 | 229774261 | 3 | 3 |
| SE | 97148 | ENSG00000153827 | TRIP12 | chr2 | - | 229774096 | 229774261 | 229771518 | 229771632 | 229777314 | 229777479 | 536 | 463 |
| SE | 97149 | ENSG00000153827 | TRIP12 | chr2 | - | 229777314 | 229777479 | 229773560 | 229774261 | 229778432 | 229778465 | 698 | 595 |
| SE | 97150 | ENSG00000153827 | TRIP12 | chr2 | - | 229777314 | 229777479 | 229773560 | 229774261 | 229785756 | 229785855 | 287 | 281 |
| SE | 97151 | ENSG00000153827 | TRIP12 | chr2 | - | 229778432 | 229778587 | 229774096 | 229774261 | 229785756 | 229785855 | 3 | 5 |
| SE | 97152 | ENSG00000153827 | TRIP12 | chr2 | - | 229778432 | 229778587 | 229777314 | 229777479 | 229785756 | 229785855 | 412 | 315 |
| SE | 97153 | ENSG00000153827 | TRIP12 | chr2 | - | 229778875 | 229778990 | 229778432 | 229778587 | 229785756 | 229785855 | 470 | 305 |
| SE | 97154 | ENSG00000153827 | TRIP12 | chr2 | - | 229778875 | 229778990 | 229778432 | 229778587 | 229789610 | 229789762 | 249 | 152 |
| SE | 97155 | ENSG00000153827 | TRIP12 | chr2 | - | 229785756 | 229785855 | 229778432 | 229778587 | 229787504 | 229787661 | 172 | 154 |
| SE | 97156 | ENSG00000153827 | TRIP12 | chr2 | - | 229785756 | 229785855 | 229778875 | 229778990 | 229788797 | 229788940 | 129 | 115 |
| SE | 97157 | ENSG00000153827 | TRIP12 | chr2 | - | 229787504 | 229787661 | 229785756 | 229785855 | 229788797 | 229788940 | 529 | 401 |
| SE | 97158 | ENSG00000153827 | TRIP12 | chr2 | - | 229788797 | 229788940 | 229778875 | 229778990 | 229789610 | 229789762 | 339 | 235 |
| SE | 97159 | ENSG00000153827 | TRIP12 | chr2 | - | 229789610 | 229789762 | 229788797 | 229788940 | 229791123 | 229791251 | 601 | 425 |
| SE | 97160 | ENSG00000153827 | TRIP12 | chr2 | - | 229791865 | 229792065 | 229791123 | 229791251 | 229792152 | 229792226 | 666 | 564 |
| SE | 97161 | ENSG00000153827 | TRIP12 | chr2 | - | 229796590 | 229796782 | 229795178 | 229795330 | 229797689 | 229797831 | 507 | 308 |
| SE | 97162 | ENSG00000153827 | TRIP12 | chr2 | - | 229803570 | 229803689 | 229802250 | 229802459 | 229803998 | 229804227 | 551 | 387 |

| | | | | | | | | | | |
|---|---|---|---|---|---|---|---|---|---|---|
| SE | 97163 | ENSG00000153827 | TRIP12 | chr2 | - | 229803998 | 229804227 | 229802250 | 229802459 | 229805729 | 229805802 | [illegible] |
| SE | 97164 | ENSG00000153827 | TRIP12 | chr2 | - | 229808251 | 229808369 | 229807318 | 229807864 | 229810879 | 229811045 | [illegible] |
| SE | 97165 | ENSG00000153827 | TRIP12 | chr2 | - | 229813869 | 229814031 | 229811135 | 229811204 | 229814232 | 229814325 | [illegible] |
| SE | 97167 | ENSG00000153827 | TRIP12 | chr2 | - | 229830755 | 229830821 | 229829192 | 229829288 | 229836847 | 229836984 | [illegible] |
| SE | 97168 | ENSG00000153827 | TRIP12 | chr2 | - | 229830755 | 229830839 | 229829192 | 229829288 | 229836847 | 229836984 | [illegible] |
| SE | 97170 | ENSG00000153827 | TRIP12 | chr2 | - | 229858771 | 229859574 | 229840821 | 229840927 | 229879981 | 229880128 | [illegible] |
| SE | 97172 | ENSG00000153827 | TRIP12 | chr2 | - | 229879981 | 229880128 | 229858771 | 229859574 | 229908964 | 229909086 | [illegible] |
| SE | 97173 | ENSG00000153827 | TRIP12 | chr2 | - | 229879981 | 229880128 | 229858771 | 229859574 | 229921879 | 229921903 | [illegible] |
| SE | 97174 | ENSG00000153827 | TRIP12 | chr2 | - | 229900736 | 229900858 | 229879981 | 229880128 | 229921879 | 229921994 | [illegible] |
| SE | 97175 | ENSG00000153827 | TRIP12 | chr2 | - | 229908964 | 229909086 | 229858771 | 229859574 | 229921879 | 229922009 | [illegible] |
| SE | 97176 | ENSG00000153827 | TRIP12 | chr2 | - | 229908964 | 229909086 | 229879981 | 229880128 | 229921270 | 229921371 | [illegible] |
| SE | 97180 | ENSG00000106290 | TAF6 | chr7 | - | 100108366 | 100108540 | 100107925 | 100108123 | 100109947 | 100110073 | [illegible] |
| SE | 97182 | ENSG00000106290 | TAF6 | chr7 | - | 100113348 | 100113405 | 100112797 | 100112917 | 100113867 | 100113954 | [illegible] |
| SE | 97183 | ENSG00000106290 | TAF6 | chr7 | - | 100113348 | 100113769 | 100112797 | 100112917 | 100113867 | 100113954 | [illegible] |
| SE | 97184 | ENSG00000106290 | TAF6 | chr7 | - | 100113615 | 100113769 | 100112797 | 100112917 | 100113867 | 100113954 | [illegible] |
| SE | 97187 | ENSG00000106290 | TAF6 | chr7 | - | 100116513 | 100116612 | 100114181 | 100114268 | 100118517 | 100118649 | [illegible] |
| SE | 97188 | ENSG00000106290 | TAF6 | chr7 | - | 100116513 | 100116633 | 100114168 | 100114268 | 100119203 | 100119304 | [illegible] |
| SE | 97190 | ENSG00000106290 | TAF6 | chr7 | - | 100118517 | 100118649 | 100114053 | 100114268 | 100119203 | 100119339 | [illegible] |
| SE | 97191 | ENSG00000106290 | TAF6 | chr7 | - | 100118517 | 100118649 | 100114053 | 100114268 | 100119231 | 100119348 | [illegible] |
| SE | 97192 | ENSG00000273045 | C2orf15 | chr2 | + | 99147026 | 99147122 | 99142284 | 99142401 | 99147401 | 99147493 | [illegible] |
| SE | 97196 | ENSG00000147316 | MCPH1 | chr8 | + | 6414764 | 6414883 | 6409278 | 6409370 | 6431498 | 6431586 | [illegible] |
| SE | 97198 | ENSG00000147316 | MCPH1 | chr8 | + | 6436047 | 6436162 | 6431498 | 6431586 | 6442066 | 6442156 | [illegible] |
| SE | 97199 | ENSG00000147316 | MCPH1 | chr8 | + | 6438952 | 6439096 | 6431498 | 6431586 | 6442066 | 6442156 | [illegible] |
| SE | 97200 | ENSG00000147316 | MCPH1 | chr8 | + | 6438952 | 6439096 | 6436047 | 6436162 | 6442066 | 6442156 | [illegible] |
| SE | 97201 | ENSG00000147316 | MCPH1 | chr8 | + | 6442066 | 6442156 | 6438952 | 6439096 | 6444392 | 6445547 | [illegible] |
| SE | 97202 | ENSG00000147316 | MCPH1 | chr8 | + | 6442066 | 6442156 | 6438952 | 6439096 | 6455142 | 6455252 | [illegible] |
| SE | 97203 | ENSG00000147316 | MCPH1 | chr8 | + | 6444392 | 6445547 | 6438952 | 6439096 | 6455142 | 6455252 | [illegible] |
| SE | 97204 | ENSG00000147316 | MCPH1 | chr8 | + | 6444392 | 6445547 | 6442066 | 6442156 | 6455142 | 6455252 | [illegible] |
| SE | 97205 | ENSG00000147316 | MCPH1 | chr8 | + | 6621453 | 6621691 | 6499851 | 6499929 | 6642993 | 6648508 | [illegible] |
| SE | 97206 | ENSG00000147316 | MCPH1 | chr8 | + | 6642679 | 6642839 | 6621451 | 6621691 | 6642993 | 6643370 | [illegible] |
| SE | 97207 | ENSG00000285816 | NA | chr11 | + | 65318913 | 65318983 | 65295863 | 65295990 | 65320543 | 65322561 | [illegible] |
| SE | 97208 | ENSG00000173653 | RCE1 | chr11 | + | 66844285 | 66844364 | 66843955 | 66844039 | 66844868 | 66845008 | [illegible] |
| SE | 97211 | ENSG00000119392 | GLE1 | chr9 | + | 128508875 | 128509097 | 128504654 | 128504904 | 128515528 | 128515639 | [illegible] |
| SE | 97214 | ENSG00000119392 | GLE1 | chr9 | + | 128515528 | 128515639 | 128508875 | 128509097 | 128522667 | 128522816 | [illegible] |
| SE | 97215 | ENSG00000119392 | GLE1 | chr9 | + | 128522667 | 128522816 | 128515528 | 128515639 | 128523279 | 128523340 | [illegible] |
| SE | 97219 | ENSG00000119392 | GLE1 | chr9 | + | 128536354 | 128536484 | 128533760 | 128533951 | 128537985 | 128538090 | [illegible] |
| SE | 97220 | ENSG00000119392 | GLE1 | chr9 | + | 128537985 | 128538090 | 128536070 | 128536484 | 128539615 | 128539698 | [illegible] |
| SE | 97221 | ENSG00000213465 | ARL2 | chr11 | + | 65020418 | 65020499 | 65018570 | 65018733 | 65021720 | 65021888 | [illegible] |
| SE | 97222 | ENSG00000254536 | NA | chr10 | + | 133399169 | 133399226 | 133398434 | 133398515 | 133402148 | 133402245 | [illegible] |
| SE | 97229 | ENSG00000073921 | PICALM | chr11 | - | 85981128 | 85981228 | 85974707 | 85974812 | 85981744 | 85981775 | [illegible] |
| SE | 97230 | ENSG00000073921 | PICALM | chr11 | - | 85981871 | 85981988 | 85981744 | 85981775 | 85983865 | 85983900 | [illegible] |
| SE | 97231 | ENSG00000073921 | PICALM | chr11 | - | 85981871 | 85982003 | 85981744 | 85981775 | 85983865 | 85983973 | [illegible] |
| SE | 97232 | ENSG00000073921 | PICALM | chr11 | - | 85983865 | 85983973 | 85981871 | 85981988 | 85996825 | 85996929 | [illegible] |
| SE | 97233 | ENSG00000073921 | PICALM | chr11 | - | 85983865 | 85983973 | 85981871 | 85982003 | 85996825 | 85996929 | [illegible] |
| SE | 97236 | ENSG00000073921 | PICALM | chr11 | - | 85996825 | 85996929 | 85990249 | 85990399 | 86000642 | 86000779 | [illegible] |
| SE | 97240 | ENSG00000073921 | PICALM | chr11 | - | 86001034 | 86001158 | 86000642 | 86000779 | 86003365 | 86003451 | [illegible] |
| SE | 97241 | ENSG00000073921 | PICALM | chr11 | - | 86007541 | 86007583 | 86003365 | 86003451 | 86011029 | 86011136 | [illegible] |
| SE | 97242 | ENSG00000073921 | PICALM | chr11 | - | 86012280 | 86012392 | 86011029 | 86011136 | 86014869 | 86014963 | [illegible] |
| SE | 97248 | ENSG00000073921 | PICALM | chr11 | - | 86026291 | 86026367 | 86014869 | 86014963 | 86068650 | 86068733 | [illegible] |
| SE | 97250 | ENSG00000073921 | PICALM | chr11 | - | 86026291 | 86026367 | 86023415 | 86023541 | 86031468 | 86031611 | [illegible] |
| SE | 97251 | ENSG00000073921 | PICALM | chr11 | - | 86031468 | 86031611 | 86014869 | 86014963 | 86068650 | 86068733 | [illegible] |
| SE | 97252 | ENSG00000073921 | PICALM | chr11 | - | 86031468 | 86031611 | 86026291 | 86026367 | 86068650 | 86068733 | [illegible] |
| SE | 97253 | ENSG00000073921 | PICALM | chr11 | - | 86068650 | 86068733 | 86031468 | 86031611 | 86069719 | 86069811 | [illegible] |
| SE | 97256 | ENSG00000134627 | PIWIL4 | chr11 | + | 94607438 | 94607639 | 94603983 | 94604056 | 94616492 | 94616564 | [illegible] |
| SE | 97259 | ENSG00000134627 | PIWIL4 | chr11 | + | 94617953 | 94618107 | 94616492 | 94616563 | 94619759 | 94619885 | [illegible] |
| SE | 97260 | ENSG00000156110 | ADK | chr10 | + | 74174415 | 74174618 | 74151234 | 74151343 | 74200763 | 74200838 | [illegible] |
| SE | 97261 | ENSG00000156110 | ADK | chr10 | + | 74200763 | 74200838 | 74151201 | 74151343 | 74224537 | 74224591 | [illegible] |
| SE | 97263 | ENSG00000156110 | ADK | chr10 | + | 74242737 | 74243033 | 74224537 | 74224591 | 74314666 | 74314745 | [illegible] |
| SE | 97266 | ENSG00000156110 | ADK | chr10 | + | 74525255 | 74525426 | 74398470 | 74398579 | 74600378 | 74600493 | [illegible] |
| SE | 97268 | ENSG00000156110 | ADK | chr10 | + | 74589281 | 74589317 | 74398470 | 74398579 | 74600378 | 74600493 | [illegible] |
| SE | 97269 | ENSG00000156110 | ADK | chr10 | + | 74589281 | 74589317 | 74398470 | 74398579 | 74670182 | 74670269 | [illegible] |
| SE | 97270 | ENSG00000156110 | ADK | chr10 | + | 74589281 | 74589317 | 74525255 | 74525426 | 74600378 | 74600493 | [illegible] |
| SE | 97271 | ENSG00000156110 | ADK | chr10 | + | 74600378 | 74600493 | 74398470 | 74398579 | 74670182 | 74670269 | [illegible] |
| SE | 97274 | ENSG00000156110 | ADK | chr10 | + | 74680411 | 74680514 | 74670182 | 74670269 | 74708320 | 74708820 | [illegible] |
| SE | 97275 | ENSG00000156110 | ADK | chr10 | + | 74686698 | 74686820 | 74670182 | 74670269 | 74708320 | 74708934 | [illegible] |
| SE | 97282 | ENSG00000081026 | MAGI3 | chr1 | + | 113580541 | 113580661 | 113549514 | 113549631 | 113585386 | 113585596 | [illegible] |
| SE | 97284 | ENSG00000081026 | MAGI3 | chr1 | + | 113672614 | 113672741 | 113671733 | 113671836 | 113673321 | 113673465 | [illegible] |
| SE | 97287 | ENSG00000153107 | ANAPC1 | chr2 | - | 111780298 | 111780385 | 111778674 | 111778770 | 111782368 | 111782507 | [illegible] |
| SE | 97289 | ENSG00000153107 | ANAPC1 | chr2 | - | 111805801 | 111805893 | 111803704 | 111803839 | 111808946 | 111809181 | [illegible] |
| SE | 97290 | ENSG00000153107 | ANAPC1 | chr2 | - | 111824965 | 111825036 | 111822421 | 111822600 | 111825130 | 111825167 | [illegible] |
| SE | 97291 | ENSG00000153107 | ANAPC1 | chr2 | - | 111825130 | 111825167 | 111822421 | 111822600 | 111825776 | 111825855 | [illegible] |
| SE | 97292 | ENSG00000153107 | ANAPC1 | chr2 | - | 111863674 | 111863895 | 111862388 | 111862598 | 111864805 | 111864951 | [illegible] |
| SE | 97294 | ENSG00000153107 | ANAPC1 | chr2 | - | 111864805 | 111864951 | 111863674 | 111863895 | 111872629 | 111872712 | [illegible] |
| SE | 97295 | ENSG00000153107 | ANAPC1 | chr2 | - | 111868022 | 111868096 | 111863674 | 111863895 | 111872629 | 111872712 | [illegible] |
| SE | 97298 | ENSG00000153107 | ANAPC1 | chr2 | - | 111873307 | 111873408 | 111872629 | 111872712 | 111878809 | 111878971 | [illegible] |
| SE | 97299 | ENSG00000153107 | ANAPC1 | chr2 | - | 111873612 | 111873664 | 111872629 | 111872712 | 111878809 | 111878971 | [illegible] |
| SE | 97301 | ENSG00000153107 | ANAPC1 | chr2 | - | 111878809 | 111878971 | 111872629 | 111872712 | 111880612 | 111880825 | [illegible] |
| SE | 97305 | ENSG00000139626 | ITGB7 | chr12 | - | 53193139 | 53193363 | 53192690 | 53192910 | 53193707 | 53193901 | [illegible] |
| SE | 97306 | ENSG00000139626 | ITGB7 | chr12 | - | 53193707 | 53193901 | 53192329 | 53192538 | 53194197 | 53194258 | [illegible] |
| SE | 97309 | ENSG00000139626 | ITGB7 | chr12 | - | 53201071 | 53201194 | 53200242 | 53200446 | 53206673 | 53206804 | [illegible] |
| SE | 97314 | ENSG00000139626 | ITGB7 | chr12 | - | 53206673 | 53206804 | 53201071 | 53201194 | 53207201 | 53207254 | [illegible] |
| SE | 97320 | ENSG00000070718 | AP3M2 | chr8 | + | 42162280 | 42162418 | 42157940 | 42158112 | 42165070 | 42165133 | [illegible] |
| SE | 97321 | ENSG00000070718 | AP3M2 | chr8 | + | 42165070 | 42165156 | 42162068 | 42162418 | 42165426 | 42165560 | [illegible] |
| SE | 97322 | ENSG00000070718 | AP3M2 | chr8 | + | 42165426 | 42165560 | 42165070 | 42165156 | 42167163 | 42167193 | [illegible] |
| SE | 97323 | ENSG00000070718 | AP3M2 | chr8 | + | 42165426 | 42165560 | 42165070 | 42165156 | 42167665 | 42167810 | [illegible] |
| SE | 97324 | ENSG00000070718 | AP3M2 | chr8 | + | 42167163 | 42167371 | 42165070 | 42165156 | 42167665 | 42167810 | [illegible] |
| SE | 97326 | ENSG00000070718 | AP3M2 | chr8 | + | 42168196 | 42168249 | 42167665 | 42167810 | 42168960 | 42169141 | [illegible] |
| SE | 97338 | ENSG00000116747 | RO60 | chr1 | + | 193069033 | 193069634 | 193060033 | 193060222 | 193075819 | 193076040 | [illegible] |
| SE | 97341 | ENSG00000116747 | RO60 | chr1 | + | 193075819 | 193076040 | 193069015 | 193069634 | 193076500 | 193076647 | [illegible] |
| SE | 97345 | ENSG00000229124 | VIM-AS1 | chr10 | - | 17217601 | 17217675 | 17216166 | 17216274 | 17226855 | 17226898 | [illegible] |
| SE | 97346 | ENSG00000229124 | VIM-AS1 | chr10 | - | 17217601 | 17217675 | 17216166 | 17216274 | 17229344 | 17229985 | [illegible] |
| SE | 97347 | ENSG00000204577 | LILRB3 | chr19 | - | 54221082 | 54221379 | 54220527 | 54220830 | 54221913 | 54222130 | [illegible] |
| SE | 97348 | ENSG00000204577 | LILRB3 | chr19 | - | 54221617 | 54221786 | 54221082 | 54221379 | 54221913 | 54222130 | [illegible] |
| SE | 97349 | ENSG00000204577 | LILRB3 | chr19 | - | 54222746 | 54222782 | 54222277 | 54222562 | 54222942 | 54222978 | [illegible] |
| SE | 97350 | ENSG00000261645 | DISC1FP1 | chr11 | + | 90361558 | 90361622 | 90251203 | 90251317 | 90546753 | 90546895 | [illegible] |
| SE | 97352 | ENSG00000261645 | DISC1FP1 | chr11 | + | 90602804 | 90602908 | 90546753 | 90546895 | 90801765 | 90801862 | [illegible] |
| SE | 97355 | ENSG00000261645 | DISC1FP1 | chr11 | + | 90801765 | 90801862 | 90602804 | 90602908 | 90844879 | 90844949 | [illegible] |
| SE | 97358 | ENSG00000151532 | VTI1A | chr10 | + | 112448381 | 112448489 | 112446997 | 112447467 | 112448881 | 112450726 | [illegible] |
| SE | 97359 | ENSG00000151532 | VTI1A | chr10 | + | 112527086 | 112527164 | 112464546 | 112464657 | 112538245 | 112538330 | [illegible] |
| SE | 97360 | ENSG00000151532 | VTI1A | chr10 | + | 112533529 | 112533550 | 112527086 | 112527164 | 112538245 | 112538301 | [illegible] |
| SE | 97364 | ENSG00000110218 | PANX1 | chr11 | + | 94179601 | 94180257 | 94153490 | 94153630 | 94180789 | 94181968 | [illegible] |
| SE | 97365 | ENSG00000110218 | PANX1 | chr11 | + | 94179601 | 94180257 | 94178368 | 94178592 | 94180789 | 94181968 | [illegible] |
| SE | 97368 | ENSG00000126391 | FRMD8 | chr11 | + | 65389360 | 65389528 | 65386927 | 65387121 | 65393572 | 65393590 | [illegible] |
| SE | 97370 | ENSG00000164169 | PRMT9 | chr4 | - | 147638959 | 147639082 | 147637784 | 147638747 | 147642786 | 147642940 | [illegible] |
| SE | 97371 | ENSG00000164169 | PRMT9 | chr4 | - | 147657791 | 147657975 | 147653851 | 147654566 | 147660845 | 147661038 | [illegible] |
| SE | 97372 | ENSG00000164169 | PRMT9 | chr4 | - | 147670640 | 147670743 | 147668538 | 147668645 | 147672958 | 147673126 | [illegible] |
| SE | 97373 | ENSG00000164169 | PRMT9 | chr4 | - | 147673637 | 147673874 | 147672958 | 147673126 | 147680322 | 147680471 | [illegible] |
| SE | 97374 | ENSG00000164169 | PRMT9 | chr4 | - | 147680322 | 147680471 | 147673637 | 147673874 | 147683798 | 147684163 | [illegible] |
| SE | 97379 | ENSG00000113163 | CERT1 | chr5 | - | 75381071 | 75381201 | 75377774 | 75379473 | 75381948 | 75382077 | [illegible] |
| SE | 97380 | ENSG00000113163 | CERT1 | chr5 | - | 75384641 | 75384712 | 75381948 | 75382077 | 75385901 | 75385923 | [illegible] |
| SE | 97383 | ENSG00000113163 | CERT1 | chr5 | - | 75389591 | 75389687 | 75385901 | 75386034 | 75400204 | 75400262 | [illegible] |
| SE | 97387 | ENSG00000113163 | CERT1 | chr5 | - | 75400204 | 75400297 | 75389591 | 75389687 | 75411010 | 75411103 | [illegible] |
| SE | 97388 | ENSG00000113163 | CERT1 | chr5 | - | 75400204 | 75403058 | 75389591 | 75389687 | 75411010 | 75411103 | [illegible] |
| SE | 97389 | ENSG00000113163 | CERT1 | chr5 | - | 75402674 | 75402792 | 75400204 | 75400297 | 75402971 | 75403058 | [illegible] |
| SE | 97391 | ENSG00000113163 | CERT1 | chr5 | - | 75402971 | 75403058 | 75400204 | 75400297 | 75411010 | 75411103 | [illegible] |
| SE | 97392 | ENSG00000113163 | CERT1 | chr5 | - | 75411010 | 75411103 | 75400204 | 75403058 | 75416875 | 75417033 | [illegible] |
| SE | 97393 | ENSG00000113163 | CERT1 | chr5 | - | 75419340 | 75419424 | 75416875 | 75417033 | 75425360 | 75425473 | [illegible] |
| SE | 97394 | ENSG00000113163 | CERT1 | chr5 | - | 75425269 | 75425499 | 75419340 | 75419424 | 75426370 | 75426438 | [illegible] |
| SE | 97395 | ENSG00000113163 | CERT1 | chr5 | - | 75425291 | 75425499 | 75419340 | 75419424 | 75426370 | 75426478 | [illegible] |
| SE | 97396 | ENSG00000113163 | CERT1 | chr5 | - | 75425360 | 75425499 | 75419340 | 75419424 | 75426370 | 75426478 | [illegible] |
| SE | 97398 | ENSG00000113163 | CERT1 | chr5 | - | 75511111 | 75511505 | 75505981 | 75506116 | 75511758 | 75512138 | [illegible] |
| SE | 97403 | ENSG00000070018 | LRP6 | chr12 | - | 12164272 | 12164562 | 12162192 | 12162419 | 12165078 | 12165295 | [illegible] |
| SE | 97408 | ENSG00000066294 | CD84 | chr1 | - | 160550937 | 160551035 | 160549916 | 160549979 | 160552704 | 160552737 | [illegible] |
| SE | 97409 | ENSG00000066294 | CD84 | chr1 | - | 160550937 | 160551035 | 160549916 | 160549979 | 160553359 | 160553497 | [illegible] |
| SE | 97410 | ENSG00000066294 | CD84 | chr1 | - | 160550937 | 160551035 | 160549916 | 160549979 | 160553377 | 160553497 | [illegible] |
| SE | 97412 | ENSG00000066294 | CD84 | chr1 | - | 160552704 | 160552737 | 160549916 | 160549979 | 160553359 | 160553497 | [illegible] |
| SE | 97413 | ENSG00000066294 | CD84 | chr1 | - | 160552704 | 160552737 | 160549916 | 160549979 | 160553377 | 160553497 | [illegible] |
| SE | 97414 | ENSG00000066294 | CD84 | chr1 | - | 160552704 | 160552737 | 160550937 | 160551035 | 160553359 | 160553497 | [illegible] |
| SE | 97417 | ENSG00000066294 | CD84 | chr1 | - | 160553359 | 160553497 | 160550937 | 160551035 | 160553894 | 160554146 | [illegible] |
| SE | 97419 | ENSG00000066294 | CD84 | chr1 | - | 160553377 | 160553497 | 160550937 | 160551035 | 160553894 | 160554146 | [illegible] |
| SE | 97422 | ENSG00000118113 | MMP8 | chr11 | - | 102722990 | 102723078 | 102722428 | 102722673 | 102723479 | 102723546 | [illegible] |
| SE | 97423 | ENSG00000118113 | MMP8 | chr11 | - | 102722990 | 102723078 | 102722428 | 102722673 | 102724753 | 102725003 | [illegible] |
| SE | 97424 | ENSG00000118113 | MMP8 | chr11 | - | 102723479 | 102723546 | 102722428 | 102722673 | 102724753 | 102724941 | [illegible] |
| SE | 97425 | ENSG00000118113 | MMP8 | chr11 | - | 102723479 | 102723546 | 102722990 | 102723078 | 102724753 | 102724941 | [illegible] |
| SE | 97430 | ENSG00000165895 | ARHGAP42 | chr11 | + | 100816714 | 100816856 | 100795104 | 100795166 | 100859553 | 100859625 | [illegible] |
| SE | 97432 | ENSG00000165895 | ARHGAP42 | chr11 | + | 100973174 | 100973334 | 100965670 | 100965776 | 100974458 | 100974603 | [illegible] |
| SE | 97433 | ENSG00000165895 | ARHGAP42 | chr11 | + | 100986073 | 100986352 | 100978986 | 100979049 | 100987512 | 100987592 | [illegible] |
| SE | 97435 | ENSG00000168876 | ANKRD49 | chr11 | + | 94497901 | 94497964 | 94496603 | 94497118 | 94498070 | 94498214 | [illegible] |
| SE | 97436 | ENSG00000168876 | ANKRD49 | chr11 | + | 94497901 | 94497964 | 94496754 | 94497261 | 94498070 | 94498214 | [illegible] |
| SE | 97440 | ENSG00000023445 | BIRC3 | chr11 | + | 102319876 | 102320006 | 102317483 | 102317571 | 102325465 | 102325565 | [illegible] |
| SE | 97443 | ENSG00000023445 | BIRC3 | chr11 | + | 102321836 | 102325362 | 102319876 | 102320006 | 102325465 | 102325565 | [illegible] |
| SE | 97444 | ENSG00000023445 | BIRC3 | chr11 | + | 102326728 | 102326851 | 102325465 | 102325565 | 102328051 | 102328130 | [illegible] |
| SE | 97445 | ENSG00000023445 | BIRC3 | chr11 | + | 102328051 | 102328130 | 102325465 | 102325565 | 102328896 | 102328945 | [illegible] |
| SE | 97446 | ENSG00000023445 | BIRC3 | chr11 | + | 102335965 | 102336220 | 102330998 | 102331241 | 102336759 | 102336801 | [illegible] |
| SE | 97447 | ENSG00000149289 | ZC3H12C | chr11 | + | 110159255 | 110159490 | 110152918 | 110153058 | 110163272 | 110163379 | [illegible] |
| SE | 97448 | ENSG00000149289 | ZC3H12C | chr11 | + | 110163272 | 110163379 | 110159255 | 110159490 | 110164340 | 110165934 | [illegible] |
| SE | 97450 | ENSG00000142185 | TRPM2 | chr21 | + | 44364113 | 44364282 | 44354647 | 44354736 | 44366753 | 44366761 | [illegible] |
| SE | 97451 | ENSG00000142185 | TRPM2 | chr21 | + | 44364113 | 44364282 | 44354647 | 44354736 | 44377711 | 44377773 | [illegible] |
| SE | 97454 | ENSG00000142185 | TRPM2 | chr21 | + | 44427009 | 44427111 | 44426659 | 44426736 | 44435130 | 44435217 | [illegible] |
| SE | 97455 | ENSG00000110777 | POU2AF1 | chr11 | - | 111357794 | 111357837 | 111357444 | 111357710 | 111358787 | 111358918 | [illegible] |
| SE | 97456 | ENSG00000110777 | POU2AF1 | chr11 | - | 111358787 | 111358918 | 111357444 | 111357710 | 111379161 | 111379275 | [illegible] |
| SE | 97458 | ENSG00000110777 | POU2AF1 | chr11 | - | 111377773 | 111377852 | 111363330 | 111363434 | 111379161 | 111379275 | [illegible] |
| SE | 97475 | ENSG00000166575 | TMEM135 | chr11 | + | 87071522 | 87071615 | 87067693 | 87067821 | 87091361 | 87091395 | [illegible] |
| SE | 97478 | ENSG00000166575 | TMEM135 | chr11 | + | 87295781 | 87295823 | 87236637 | 87236684 | 87302295 | 87302442 | [illegible] |
| SE | 97479 | ENSG00000166575 | TMEM135 | chr11 | + | 87305935 | 87306005 | 87302295 | 87302442 | 87309504 | 87309672 | [illegible] |
| SE | 97480 | ENSG00000166575 | TMEM135 | chr11 | + | 87314470 | 87314547 | 87313424 | 87313488 | 87318136 | 87318235 | [illegible] |
| SE | 97481 | ENSG00000076984 | MAP2K7 | chr19 | + | 7904450 | 7904548 | 7903906 | 7904068 | 7905807 | 7907066 | [illegible] |
| SE | 97483 | ENSG00000137672 | TRPC6 | chr11 | - | 101469426 | 101469501 | 101455017 | 101455101 | 101471182 | 101471386 | [illegible] |
| SE | 97491 | ENSG00000173120 | KDM2A | chr11 | + | 67143203 | 67143371 | 67121233 | 67121358 | 67149054 | 67149207 | [illegible] |
| SE | 97492 | ENSG00000173120 | KDM2A | chr11 | + | 67143203 | 67143371 | 67121233 | 67121358 | 67180078 | 67180453 | [illegible] |
| SE | 97493 | ENSG00000173120 | KDM2A | chr11 | + | 67180078 | 67180217 | 67121233 | 67121358 | 67207509 | 67207688 | [illegible] |
| SE | 97495 | ENSG00000173120 | KDM2A | chr11 | + | 67181845 | 67181892 | 67180078 | 67180217 | 67207509 | 67207688 | [illegible] |
| SE | 97499 | ENSG00000173120 | KDM2A | chr11 | + | 67215339 | 67215446 | 67207509 | 67207688 | 67217730 | 67217884 | [illegible] |
| SE | 97500 | ENSG00000173120 | KDM2A | chr11 | + | 67215339 | 67215446 | 67207509 | 67207688 | 67217755 | 67217884 | [illegible] |
| SE | 97503 | ENSG00000173120 | KDM2A | chr11 | + | 67217730 | 67217884 | 67207509 | 67207688 | 67228036 | 67228163 | [illegible] |
| SE | 97504 | ENSG00000173120 | KDM2A | chr11 | + | 67217755 | 67217884 | 67207601 | 67207688 | 67228036 | 67228163 | [illegible] |
| SE | 97505 | ENSG00000173120 | KDM2A | chr11 | + | 67219287 | 67219403 | 67217730 | 67217884 | 67228036 | 67228163 | [illegible] |
| SE | 97506 | ENSG00000173120 | KDM2A | chr11 | + | 67228036 | 67228163 | 67207509 | 67207688 | 67243008 | 67243092 | [illegible] |
| SE | 97507 | ENSG00000173120 | KDM2A | chr11 | + | 67231565 | 67231960 | 67228036 | 67228163 | 67243008 | 67243092 | [illegible] |
| SE | 97508 | ENSG00000173120 | KDM2A | chr11 | + | 67243008 | 67243092 | 67228036 | 67228163 | 67245188 | 67245458 | [illegible] |
| SE | 97510 | ENSG00000173120 | KDM2A | chr11 | + | 67243008 | 67243092 | 67240046 | 67240369 | 67243506 | 67243816 | [illegible] |
| SE | 97511 | ENSG00000173120 | KDM2A | chr11 | + | 67243008 | 67243092 | 67240046 | 67240369 | 67245188 | 67245458 | [illegible] |
| SE | 97512 | ENSG00000173120 | KDM2A | chr11 | + | 67245188 | 67245458 | 67243008 | 67243092 | 67245984 | 67246116 | [illegible] |
| SE | 97513 | ENSG00000173120 | KDM2A | chr11 | + | 67245984 | 67246116 | 67243008 | 67243092 | 67248280 | 67248370 | [illegible] |
| SE | 97514 | ENSG00000173120 | KDM2A | chr11 | + | 67248280 | 67248370 | 67245984 | 67246116 | 67250085 | 67250346 | [illegible] |
| SE | 97515 | ENSG00000152404 | CWF19L2 | chr11 | - | 107345413 | 107345490 | 107336557 | 107336713 | 107348936 | 107349053 | [illegible] |
| SE | 97516 | ENSG00000152404 | CWF19L2 | chr11 | - | 107390073 | 107390211 | 107353523 | 107353736 | 107392778 | 107392895 | [illegible] |
| SE | 97518 | ENSG00000152404 | CWF19L2 | chr11 | - | 107418193 | 107418287 | 107392778 | 107392895 | 107428798 | 107429451 | [illegible] |
| SE | 97520 | ENSG00000152404 | CWF19L2 | chr11 | - | 107439089 | 107439183 | 107433633 | 107433749 | 107441502 | 107441622 | [illegible] |
| SE | 97521 | ENSG00000152404 | CWF19L2 | chr11 | - | 107455665 | 107455776 | 107454449 | 107454572 | 107457711 | 107457844 | [illegible] |
| SE | 97526 | ENSG00000164414 | SLC35A1 | chr6 | + | 87477361 | 87477539 | 87472924 | 87473019 | 87501157 | 87501310 | [illegible] |
| SE | 97527 | ENSG00000164414 | SLC35A1 | chr6 | + | 87500507 | 87500667 | 87472924 | 87473019 | 87501157 | 87501310 | [illegible] |
| SE | 97531 | ENSG00000149761 | NUDT22 | chr11 | + | 64227400 | 64227493 | 64226608 | 64227132 | 64229246 | 64229344 | [illegible] |
| SE | 97532 | ENSG00000149761 | NUDT22 | chr11 | + | 64227567 | 64227666 | 64226634 | 64227132 | 64229246 | 64229344 | [illegible] |
| SE | 97534 | ENSG00000149761 | NUDT22 | chr11 | + | 64227567 | 64227666 | 64227400 | 64227493 | 64229246 | 64229344 | [illegible] |
| SE | 97540 | ENSG00000109452 | INPP4B | chr4 | - | 142086143 | 142086256 | 142082030 | 142082185 | 142108092 | 142108190 | [illegible] |
| SE | 97542 | ENSG00000109452 | INPP4B | chr4 | - | 142122127 | 142122245 | 142112541 | 142112682 | 142123291 | 142123415 | [illegible] |
| SE | 97543 | ENSG00000109452 | INPP4B | chr4 | - | 142123291 | 142123415 | 142112541 | 142112682 | 142124587 | 142124760 | [illegible] |
| SE | 97545 | ENSG00000109452 | INPP4B | chr4 | - | 142193086 | 142193195 | 142173631 | 142173809 | 142208424 | 142208529 | [illegible] |
| SE | 97546 | ENSG00000109452 | INPP4B | chr4 | - | 142193086 | 142193195 | 142173631 | 142173809 | 142208446 | 142208529 | [illegible] |
| SE | 97547 | ENSG00000109452 | INPP4B | chr4 | - | 142208424 | 142208529 | 142193086 | 142193195 | 142208895 | 142209026 | [illegible] |
| SE | 97548 | ENSG00000109452 | INPP4B | chr4 | - | 142208446 | 142208529 | 142193086 | 142193195 | 142208895 | 142209026 | [illegible] |
| SE | 97549 | ENSG00000109452 | INPP4B | chr4 | - | 142260491 | 142260564 | 142237863 | 142238011 | 142270662 | 142270774 | [illegible] |
| SE | 97550 | ENSG00000109452 | INPP4B | chr4 | - | 142270662 | 142270774 | 142260491 | 142260564 | 142305457 | 142305537 | [illegible] |
| SE | 97551 | ENSG00000109452 | INPP4B | chr4 | - | 142305457 | 142305537 | 142270662 | 142270774 | 142402937 | 142403009 | [illegible] |
| SE | 97553 | ENSG00000109452 | INPP4B | chr4 | - | 142347373 | 142347556 | 142270662 | 142270774 | 142402937 | 142403009 | [illegible] |
| SE | 97554 | ENSG00000109452 | INPP4B | chr4 | - | 142347373 | 142347556 | 142314711 | 142314762 | 142402937 | 142403009 | [illegible] |
| SE | 97555 | ENSG00000109452 | INPP4B | chr4 | - | 142383961 | 142384116 | 142314711 | 142314762 | 142402937 | 142403054 | [illegible] |
| SE | 97558 | ENSG00000109452 | INPP4B | chr4 | - | 142425206 | 142425263 | 142423779 | 142423817 | 142429172 | 142429217 | [illegible] |
| SE | 97560 | ENSG00000109452 | INPP4B | chr4 | - | 142467956 | 142468055 | 142462662 | 142462726 | 142725838 | 142725901 | [illegible] |
| SE | 97563 | ENSG00000109452 | INPP4B | chr4 | - | 142660776 | 142660896 | 142654511 | 142654680 | 142725838 | 142725901 | [illegible] |
| SE | 97566 | ENSG00000166266 | CUL5 | chr11 | + | 108033801 | 108033911 | 108008897 | 108009372 | 108046269 | 108046369 | [illegible] |
| SE | 97569 | ENSG00000166266 | CUL5 | chr11 | + | 108052659 | 108052801 | 108046269 | 108046369 | 108054646 | 108054792 | [illegible] |
| SE | 97571 | ENSG00000166266 | CUL5 | chr11 | + | 108078175 | 108078240 | 108073389 | 108073497 | 108088526 | 108088659 | [illegible] |
| SE | 97572 | ENSG00000166266 | CUL5 | chr11 | + | 108094390 | 108094514 | 108089491 | 108089623 | 108094811 | 108094987 | [illegible] |
| SE | 97573 | ENSG00000166266 | CUL5 | chr11 | + | 108097635 | 108097754 | 108095529 | 108095691 | 108104189 | 108104902 | [illegible] |
| SE | 97574 | ENSG00000166266 | CUL5 | chr11 | + | 108098405 | 108098529 | 108097635 | 108097754 | 108104189 | 108104902 | [illegible] |
| SE | 97578 | ENSG00000142186 | SCYL1 | chr11 | + | 65536258 | 65536334 | 65535952 | 65536141 | 65536585 | 65536626 | [illegible] |
| SE | 97583 | ENSG00000174928 | C3orf33 | chr3 | - | 155805596 | 155805723 | 155802531 | 155802591 | 155806138 | 155806251 | [illegible] |

| | | | | | | | | | | | | | | |
|---|---|---|---|---|---|---|---|---|---|---|---|---|---|---|
| SE | 97584 | ENSG00000255666 | LINC02700 | chr11 | + | 94643969 | 94644134 | 94641999 | 94642157 | 94647334 | 94647578 | 1 | 3 | [illegible] |
| SE | 97588 | ENSG00000110721 | CHKA | chr11 | - | 68064524 | 68064631 | 68061952 | 68062034 | 68065785 | 68065894 | 27 | 29 | [illegible] |
| SE | 97589 | ENSG00000110721 | CHKA | chr11 | - | 68074716 | 68074830 | 68070723 | 68070857 | 68081403 | 68081457 | 29 | 33 | [illegible] |
| SE | 97593 | ENSG00000170209 | ANKK1 | chr11 | + | 113394727 | 113394782 | 113393632 | 113393775 | 113394928 | 113395080 | 0 | 0 | [illegible] |
| SE | 97594 | ENSG00000139722 | VPS37B | chr12 | - | 122868479 | 122868562 | 122865329 | 122867607 | 122870889 | 122871055 | 335 | 196 | [illegible] |
| SE | 97596 | ENSG00000139722 | VPS37B | chr12 | - | 122870889 | 122871055 | 122867153 | 122867607 | 122895951 | 122896127 | 0 | 0 | [illegible] |
| SE | 97607 | ENSG00000179832 | MROH1 | chr8 | + | 144243862 | 144243942 | 144243493 | 144243616 | 144244221 | 144244336 | 72 | 79 | [illegible] |
| SE | 97609 | ENSG00000179832 | MROH1 | chr8 | + | 144260185 | 144260374 | 144259910 | 144260057 | 144260676 | 144260832 | 12 | 6 | [illegible] |
| SE | 97610 | ENSG00000179832 | MROH1 | chr8 | + | 144261283 | 144261349 | 144261113 | 144261216 | 144261654 | 144261809 | 9 | 6 | [illegible] |
| SE | 97611 | ENSG00000118363 | SPCS2 | chr11 | + | 74963594 | 74963687 | 74949264 | 74949399 | 74965033 | 74965117 | 0 | 0 | [illegible] |
| SE | 97612 | ENSG00000118363 | SPCS2 | chr11 | + | 74965033 | 74965117 | 74949246 | 74949399 | 74969639 | 74969699 | 68 | 30 | [illegible] |
| SE | 97613 | ENSG00000118363 | SPCS2 | chr11 | + | 74965033 | 74965117 | 74949246 | 74949399 | 74976856 | 74977087 | 60 | 25 | [illegible] |
| SE | 97615 | ENSG00000118363 | SPCS2 | chr11 | + | 74969639 | 74969699 | 74949263 | 74949399 | 74976856 | 74977348 | 69 | 51 | [illegible] |
| SE | 97618 | ENSG00000189164 | ZNF527 | chr19 | + | 37380047 | 37380372 | 37379119 | 37379246 | 37388305 | 37393066 | 9 | 6 | [illegible] |
| SE | 97619 | ENSG00000189164 | ZNF527 | chr19 | + | 37380276 | 37380372 | 37379119 | 37379236 | 37388305 | 37388783 | 10 | 5 | [illegible] |
| SE | 97620 | ENSG00000189164 | ZNF527 | chr19 | + | 37380276 | 37380372 | 37379119 | 37379246 | 37388305 | 37388783 | 18 | 14 | [illegible] |
| SE | 97629 | ENSG00000152558 | TMEM123 | chr11 | - | 102401538 | 102401692 | 102396331 | 102398891 | 102401915 | 102402206 | 2495 | 2772 | [illegible] |
| SE | 97633 | ENSG00000198879 | SFMBT2 | chr10 | - | 7227854 | 7227937 | 7220410 | 7220537 | 7243557 | 7243705 | 67 | 38 | [illegible] |
| SE | 97634 | ENSG00000198879 | SFMBT2 | chr10 | - | 7276891 | 7276989 | 7248547 | 7248649 | 7283903 | 7283959 | 99 | 71 | [illegible] |
| SE | 97635 | ENSG00000198879 | SFMBT2 | chr10 | - | 7276891 | 7276989 | 7248547 | 7248649 | 7285865 | 7285954 | 34 | 27 | [illegible] |
| SE | 97637 | ENSG00000198879 | SFMBT2 | chr10 | - | 7283903 | [illegible] | [illegible] | [illegible] | [illegible] | [illegible] | [illegible] | [illegible] | [illegible] |
| SE | 97638 | ENSG00000198879 | SFMBT2 | chr10 | - | 7283903 | [illegible] | [illegible] | [illegible] | [illegible] | [illegible] | [illegible] | [illegible] | [illegible] |
| SE | 97639 | ENSG00000198879 | SFMBT2 | chr10 | - | 7283903 | [illegible] | [illegible] | [illegible] | [illegible] | [illegible] | [illegible] | [illegible] | [illegible] |
| SE | 97641 | ENSG00000198879 | SFMBT2 | chr10 | - | 7285865 | [illegible] | [illegible] | [illegible] | [illegible] | [illegible] | [illegible] | [illegible] | [illegible] |
| SE | 97642 | ENSG00000198879 | SFMBT2 | chr10 | - | 7285865 | [illegible] | [illegible] | [illegible] | [illegible] | [illegible] | [illegible] | [illegible] | [illegible] |
| SE | 97643 | ENSG00000198879 | SFMBT2 | chr10 | - | 7365223 | [illegible] | [illegible] | [illegible] | [illegible] | [illegible] | [illegible] | [illegible] | [illegible] |
| SE | 97647 | ENSG00000149823 | VPS51 | chr11 | + | 65111160 | [illegible] | [illegible] | [illegible] | [illegible] | [illegible] | [illegible] | [illegible] | [illegible] |
| SE | 97651 | ENSG00000147457 | CHMP7 | chr8 | + | 23247986 | [illegible] | [illegible] | [illegible] | [illegible] | [illegible] | [illegible] | [illegible] | [illegible] |
| SE | 97652 | ENSG00000147457 | CHMP7 | chr8 | + | 23247986 | [illegible] | [illegible] | [illegible] | [illegible] | [illegible] | [illegible] | [illegible] | [illegible] |
| SE | 97653 | ENSG00000147457 | CHMP7 | chr8 | + | 23249209 | [illegible] | [illegible] | [illegible] | [illegible] | [illegible] | [illegible] | [illegible] | [illegible] |
| SE | 97654 | ENSG00000147457 | CHMP7 | chr8 | + | 23249209 | [illegible] | [illegible] | [illegible] | [illegible] | [illegible] | [illegible] | [illegible] | [illegible] |
| SE | 97658 | ENSG00000147457 | CHMP7 | chr8 | + | 23255246 | [illegible] | [illegible] | [illegible] | [illegible] | [illegible] | [illegible] | [illegible] | [illegible] |
| SE | 97659 | ENSG00000147457 | CHMP7 | chr8 | + | 23256459 | [illegible] | [illegible] | [illegible] | [illegible] | [illegible] | [illegible] | [illegible] | [illegible] |
| SE | 97662 | ENSG00000212127 | TAS2R14 | chr12 | - | 10973654 | [illegible] | [illegible] | [illegible] | [illegible] | [illegible] | [illegible] | [illegible] | [illegible] |
| SE | 97663 | ENSG00000212127 | TAS2R14 | chr12 | - | 10973654 | [illegible] | [illegible] | [illegible] | [illegible] | [illegible] | [illegible] | [illegible] | [illegible] |
| SE | 97664 | ENSG00000212127 | TAS2R14 | chr12 | - | 11034781 | [illegible] | [illegible] | [illegible] | [illegible] | [illegible] | [illegible] | [illegible] | [illegible] |
| SE | 97665 | ENSG00000150687 | PRSS23 | chr11 | + | 86939019 | [illegible] | [illegible] | [illegible] | [illegible] | [illegible] | [illegible] | [illegible] | [illegible] |
| SE | 97667 | ENSG00000110367 | DDX6 | chr11 | - | 118754704 | [illegible] | [illegible] | [illegible] | [illegible] | [illegible] | [illegible] | [illegible] | [illegible] |
| SE | 97668 | ENSG00000110367 | DDX6 | chr11 | - | 118755401 | [illegible] | [illegible] | [illegible] | [illegible] | [illegible] | [illegible] | [illegible] | [illegible] |
| SE | 97670 | ENSG00000110367 | DDX6 | chr11 | - | 118759921 | [illegible] | [illegible] | [illegible] | [illegible] | [illegible] | [illegible] | [illegible] | [illegible] |
| SE | 97671 | ENSG00000110367 | DDX6 | chr11 | - | 118763211 | [illegible] | [illegible] | [illegible] | [illegible] | [illegible] | [illegible] | [illegible] | [illegible] |
| SE | 97673 | ENSG00000110367 | DDX6 | chr11 | - | 118768222 | [illegible] | [illegible] | [illegible] | [illegible] | [illegible] | [illegible] | [illegible] | [illegible] |
| SE | 97674 | ENSG00000110367 | DDX6 | chr11 | - | 118779631 | [illegible] | [illegible] | [illegible] | [illegible] | [illegible] | [illegible] | [illegible] | [illegible] |
| SE | 97675 | ENSG00000110367 | DDX6 | chr11 | - | 118779631 | [illegible] | [illegible] | [illegible] | [illegible] | [illegible] | [illegible] | [illegible] | [illegible] |
| SE | 97676 | ENSG00000110367 | DDX6 | chr11 | - | 118781120 | [illegible] | [illegible] | [illegible] | [illegible] | [illegible] | [illegible] | [illegible] | [illegible] |
| SE | 97677 | ENSG00000110367 | DDX6 | chr11 | - | 118781120 | [illegible] | [illegible] | [illegible] | [illegible] | [illegible] | [illegible] | [illegible] | [illegible] |
| SE | 97678 | ENSG00000110367 | DDX6 | chr11 | - | 118786051 | [illegible] | [illegible] | [illegible] | [illegible] | [illegible] | [illegible] | [illegible] | [illegible] |
| SE | 97681 | ENSG00000122218 | COPA | chr1 | - | 160294767 | [illegible] | [illegible] | [illegible] | [illegible] | [illegible] | [illegible] | [illegible] | [illegible] |
| SE | 97682 | ENSG00000122218 | COPA | chr1 | - | 160298844 | [illegible] | [illegible] | [illegible] | [illegible] | [illegible] | [illegible] | [illegible] | [illegible] |
| SE | 97683 | ENSG00000122218 | COPA | chr1 | - | 160298844 | [illegible] | [illegible] | [illegible] | [illegible] | [illegible] | [illegible] | [illegible] | [illegible] |
| SE | 97687 | ENSG00000122218 | COPA | chr1 | - | 160307162 | [illegible] | [illegible] | [illegible] | [illegible] | [illegible] | [illegible] | [illegible] | [illegible] |
| SE | 97689 | ENSG00000122218 | COPA | chr1 | - | 160325542 | [illegible] | [illegible] | [illegible] | [illegible] | [illegible] | [illegible] | [illegible] | [illegible] |
| SE | 97690 | ENSG00000122218 | COPA | chr1 | - | 160325542 | [illegible] | [illegible] | [illegible] | [illegible] | [illegible] | [illegible] | [illegible] | [illegible] |
| SE | 97691 | ENSG00000122218 | COPA | chr1 | - | 160332447 | [illegible] | [illegible] | [illegible] | [illegible] | [illegible] | [illegible] | [illegible] | [illegible] |
| SE | 97692 | ENSG00000122218 | COPA | chr1 | - | 160332447 | [illegible] | [illegible] | [illegible] | [illegible] | [illegible] | [illegible] | [illegible] | [illegible] |
| SE | 97693 | ENSG00000122218 | COPA | chr1 | - | 160332447 | [illegible] | [illegible] | [illegible] | [illegible] | [illegible] | [illegible] | [illegible] | [illegible] |
| SE | 97694 | ENSG00000122218 | COPA | chr1 | - | 160333602 | [illegible] | [illegible] | [illegible] | [illegible] | [illegible] | [illegible] | [illegible] | [illegible] |
| SE | 97695 | ENSG00000122218 | COPA | chr1 | - | 160333602 | [illegible] | [illegible] | [illegible] | [illegible] | [illegible] | [illegible] | [illegible] | [illegible] |
| SE | 97698 | ENSG00000122218 | COPA | chr1 | - | 160335241 | [illegible] | [illegible] | [illegible] | [illegible] | [illegible] | [illegible] | [illegible] | [illegible] |
| SE | 97699 | ENSG00000122218 | COPA | chr1 | - | 160339908 | [illegible] | [illegible] | [illegible] | [illegible] | [illegible] | [illegible] | [illegible] | [illegible] |
| SE | 97700 | ENSG00000122218 | COPA | chr1 | - | 160340180 | [illegible] | [illegible] | [illegible] | [illegible] | [illegible] | [illegible] | [illegible] | [illegible] |
| SE | 97704 | ENSG00000170145 | SIK2 | chr11 | + | 111703202 | [illegible] | [illegible] | [illegible] | [illegible] | [illegible] | [illegible] | [illegible] | [illegible] |
| SE | 97705 | ENSG00000119318 | RAD23B | chr9 | + | 107283526 | [illegible] | [illegible] | [illegible] | [illegible] | [illegible] | [illegible] | [illegible] | [illegible] |
| SE | 97706 | ENSG00000119318 | RAD23B | chr9 | + | 107300140 | [illegible] | [illegible] | [illegible] | [illegible] | [illegible] | [illegible] | [illegible] | [illegible] |
| SE | 97708 | ENSG00000119318 | RAD23B | chr9 | + | 107302034 | [illegible] | [illegible] | [illegible] | [illegible] | [illegible] | [illegible] | [illegible] | [illegible] |
| SE | 97709 | ENSG00000119318 | RAD23B | chr9 | + | 107302034 | [illegible] | [illegible] | [illegible] | [illegible] | [illegible] | [illegible] | [illegible] | [illegible] |
| SE | 97710 | ENSG00000119318 | RAD23B | chr9 | + | 107324833 | [illegible] | [illegible] | [illegible] | [illegible] | [illegible] | [illegible] | [illegible] | [illegible] |
| SE | 97712 | ENSG00000110711 | AIP | chr11 | + | 67487005 | [illegible] | [illegible] | [illegible] | [illegible] | [illegible] | [illegible] | [illegible] | [illegible] |
| SE | 97713 | ENSG00000110711 | AIP | chr11 | + | 67489266 | [illegible] | [illegible] | [illegible] | [illegible] | [illegible] | [illegible] | [illegible] | [illegible] |
| SE | 97717 | ENSG00000149212 | SESN3 | chr11 | - | 95185255 | [illegible] | [illegible] | [illegible] | [illegible] | [illegible] | [illegible] | [illegible] | [illegible] |
| SE | 97718 | ENSG00000149212 | SESN3 | chr11 | - | 95189778 | [illegible] | [illegible] | [illegible] | [illegible] | [illegible] | [illegible] | [illegible] | [illegible] |
| SE | 97719 | ENSG00000149212 | SESN3 | chr11 | - | 95189778 | [illegible] | [illegible] | [illegible] | [illegible] | [illegible] | [illegible] | [illegible] | [illegible] |
| SE | 97721 | ENSG00000149212 | SESN3 | chr11 | - | 95201236 | [illegible] | [illegible] | [illegible] | [illegible] | [illegible] | [illegible] | [illegible] | [illegible] |
| SE | 97728 | ENSG00000166947 | EPB42 | chr15 | - | 43216267 | [illegible] | [illegible] | [illegible] | [illegible] | [illegible] | [illegible] | [illegible] | [illegible] |
| SE | 97732 | ENSG00000095139 | ARCN1 | chr11 | + | 118573619 | [illegible] | [illegible] | [illegible] | [illegible] | [illegible] | [illegible] | [illegible] | [illegible] |
| SE | 97734 | ENSG00000095139 | ARCN1 | chr11 | + | 118583178 | [illegible] | [illegible] | [illegible] | [illegible] | [illegible] | [illegible] | [illegible] | [illegible] |
| SE | 97735 | ENSG00000095139 | ARCN1 | chr11 | + | 118583808 | [illegible] | [illegible] | [illegible] | [illegible] | [illegible] | [illegible] | [illegible] | [illegible] |
| SE | 97736 | ENSG00000095139 | ARCN1 | chr11 | + | 118584479 | [illegible] | [illegible] | [illegible] | [illegible] | [illegible] | [illegible] | [illegible] | [illegible] |
| SE | 97737 | ENSG00000095139 | ARCN1 | chr11 | + | 118592708 | [illegible] | [illegible] | [illegible] | [illegible] | [illegible] | [illegible] | [illegible] | [illegible] |
| SE | 97738 | ENSG00000095139 | ARCN1 | chr11 | + | 118592708 | [illegible] | [illegible] | [illegible] | [illegible] | [illegible] | [illegible] | [illegible] | [illegible] |
| SE | 97739 | ENSG00000095139 | ARCN1 | chr11 | + | 118593589 | [illegible] | [illegible] | [illegible] | [illegible] | [illegible] | [illegible] | [illegible] | [illegible] |
| SE | 97740 | ENSG00000095139 | ARCN1 | chr11 | + | 118593589 | [illegible] | [illegible] | [illegible] | [illegible] | [illegible] | [illegible] | [illegible] | [illegible] |
| SE | 97747 | ENSG00000146223 | RPL7L1 | chr6 | + | 42883857 | [illegible] | [illegible] | [illegible] | [illegible] | [illegible] | [illegible] | [illegible] | [illegible] |
| SE | 97748 | ENSG00000146223 | RPL7L1 | chr6 | + | 42884612 | [illegible] | [illegible] | [illegible] | [illegible] | [illegible] | [illegible] | [illegible] | [illegible] |
| SE | 97750 | ENSG00000146223 | RPL7L1 | chr6 | + | 42886255 | [illegible] | [illegible] | [illegible] | [illegible] | [illegible] | [illegible] | [illegible] | [illegible] |
| SE | 97752 | ENSG00000166900 | STX3 | chr11 | + | 59788872 | [illegible] | [illegible] | [illegible] | [illegible] | [illegible] | [illegible] | [illegible] | [illegible] |
| SE | 97753 | ENSG00000166900 | STX3 | chr11 | + | 59793379 | [illegible] | [illegible] | [illegible] | [illegible] | [illegible] | [illegible] | [illegible] | [illegible] |
| SE | 97758 | ENSG00000166900 | STX3 | chr11 | + | 59797282 | [illegible] | [illegible] | [illegible] | [illegible] | [illegible] | [illegible] | [illegible] | [illegible] |
| SE | 97762 | ENSG00000137714 | FDX1 | chr11 | + | 110456917 | [illegible] | [illegible] | [illegible] | [illegible] | [illegible] | [illegible] | [illegible] | [illegible] |
| SE | 97763 | ENSG00000125245 | GPR18 | chr13 | - | 99256327 | [illegible] | [illegible] | [illegible] | [illegible] | [illegible] | [illegible] | [illegible] | [illegible] |
| SE | 97766 | ENSG00000155158 | TTC39B | chr9 | - | 15182306 | [illegible] | [illegible] | [illegible] | [illegible] | [illegible] | [illegible] | [illegible] | [illegible] |
| SE | 97768 | ENSG00000155158 | TTC39B | chr9 | - | 15185279 | [illegible] | [illegible] | [illegible] | [illegible] | [illegible] | [illegible] | [illegible] | [illegible] |
| SE | 97769 | ENSG00000155158 | TTC39B | chr9 | - | 15186943 | [illegible] | [illegible] | [illegible] | [illegible] | [illegible] | [illegible] | [illegible] | [illegible] |
| SE | 97770 | ENSG00000155158 | TTC39B | chr9 | - | 15199860 | [illegible] | [illegible] | [illegible] | [illegible] | [illegible] | [illegible] | [illegible] | [illegible] |
| SE | 97771 | ENSG00000155158 | TTC39B | chr9 | - | 15203822 | [illegible] | [illegible] | [illegible] | [illegible] | [illegible] | [illegible] | [illegible] | [illegible] |
| SE | 97772 | ENSG00000155158 | TTC39B | chr9 | - | 15210087 | [illegible] | [illegible] | [illegible] | [illegible] | [illegible] | [illegible] | [illegible] | [illegible] |
| SE | 97773 | ENSG00000155158 | TTC39B | chr9 | - | 15211265 | [illegible] | [illegible] | [illegible] | [illegible] | [illegible] | [illegible] | [illegible] | [illegible] |
| SE | 97774 | ENSG00000155158 | TTC39B | chr9 | - | 15211265 | [illegible] | [illegible] | [illegible] | [illegible] | [illegible] | [illegible] | [illegible] | [illegible] |
| SE | 97775 | ENSG00000155158 | TTC39B | chr9 | - | 15214138 | [illegible] | [illegible] | [illegible] | [illegible] | [illegible] | [illegible] | [illegible] | [illegible] |
| SE | 97776 | ENSG00000155158 | TTC39B | chr9 | - | 15214138 | [illegible] | [illegible] | [illegible] | [illegible] | [illegible] | [illegible] | [illegible] | [illegible] |
| SE | 97787 | ENSG00000166452 | AKIP1 | chr11 | + | 8912452 | [illegible] | [illegible] | [illegible] | [illegible] | [illegible] | [illegible] | [illegible] | [illegible] |
| SE | 97788 | ENSG00000166452 | AKIP1 | chr11 | + | 8914825 | [illegible] | [illegible] | [illegible] | [illegible] | [illegible] | [illegible] | [illegible] | [illegible] |
| SE | 97789 | ENSG00000166452 | AKIP1 | chr11 | + | 8914825 | [illegible] | [illegible] | [illegible] | [illegible] | [illegible] | [illegible] | [illegible] | [illegible] |
| SE | 97794 | ENSG00000196914 | ARHGEF12 | chr11 | + | 120406117 | [illegible] | [illegible] | [illegible] | [illegible] | [illegible] | [illegible] | [illegible] | [illegible] |
| SE | 97795 | ENSG00000196914 | ARHGEF12 | chr11 | + | 120407737 | [illegible] | [illegible] | [illegible] | [illegible] | [illegible] | [illegible] | [illegible] | [illegible] |
| SE | 97798 | ENSG00000196914 | ARHGEF12 | chr11 | + | 120437307 | [illegible] | [illegible] | [illegible] | [illegible] | [illegible] | [illegible] | [illegible] | [illegible] |
| SE | 97799 | ENSG00000196914 | ARHGEF12 | chr11 | + | 120442103 | [illegible] | [illegible] | [illegible] | [illegible] | [illegible] | [illegible] | [illegible] | [illegible] |
| SE | 97800 | ENSG00000196914 | ARHGEF12 | chr11 | + | 120465236 | [illegible] | [illegible] | [illegible] | [illegible] | [illegible] | [illegible] | [illegible] | [illegible] |
| SE | 97801 | ENSG00000011114 | BTBD7 | chr14 | - | 93248475 | [illegible] | [illegible] | [illegible] | [illegible] | [illegible] | [illegible] | [illegible] | [illegible] |
| SE | 97803 | ENSG00000011114 | BTBD7 | chr14 | - | 93261601 | [illegible] | [illegible] | [illegible] | [illegible] | [illegible] | [illegible] | [illegible] | [illegible] |
| SE | 97807 | ENSG00000011114 | BTBD7 | chr14 | - | 93293857 | [illegible] | [illegible] | [illegible] | [illegible] | [illegible] | [illegible] | [illegible] | [illegible] |
| SE | 97808 | ENSG00000011114 | BTBD7 | chr14 | - | 93295969 | [illegible] | [illegible] | [illegible] | [illegible] | [illegible] | [illegible] | [illegible] | [illegible] |
| SE | 97809 | ENSG00000255422 | NA | chr11 | + | 118704607 | [illegible] | [illegible] | [illegible] | [illegible] | [illegible] | [illegible] | [illegible] | [illegible] |
| SE | 97813 | ENSG00000196323 | ZBTB44 | chr11 | - | 130231638 | [illegible] | [illegible] | [illegible] | [illegible] | [illegible] | [illegible] | [illegible] | [illegible] |
| SE | 97814 | ENSG00000196323 | ZBTB44 | chr11 | - | 130234155 | [illegible] | [illegible] | [illegible] | [illegible] | [illegible] | [illegible] | [illegible] | [illegible] |
| SE | 97815 | ENSG00000196323 | ZBTB44 | chr11 | - | 130234155 | [illegible] | [illegible] | [illegible] | [illegible] | [illegible] | [illegible] | [illegible] | [illegible] |
| SE | 97816 | ENSG00000196323 | ZBTB44 | chr11 | - | 130236109 | [illegible] | [illegible] | [illegible] | [illegible] | [illegible] | [illegible] | [illegible] | [illegible] |
| SE | 97817 | ENSG00000196323 | ZBTB44 | chr11 | - | 130236109 | [illegible] | [illegible] | [illegible] | [illegible] | [illegible] | [illegible] | [illegible] | [illegible] |
| SE | 97820 | ENSG00000196323 | ZBTB44 | chr11 | - | 130238443 | [illegible] | [illegible] | [illegible] | [illegible] | [illegible] | [illegible] | [illegible] | [illegible] |
| SE | 97821 | ENSG00000196323 | ZBTB44 | chr11 | - | 130239811 | [illegible] | [illegible] | [illegible] | [illegible] | [illegible] | [illegible] | [illegible] | [illegible] |
| SE | 97822 | ENSG00000196323 | ZBTB44 | chr11 | - | 130260855 | [illegible] | [illegible] | [illegible] | [illegible] | [illegible] | [illegible] | [illegible] | [illegible] |
| SE | 97842 | ENSG00000142669 | SH3BGRL3 | chr1 | + | 26280764 | [illegible] | [illegible] | [illegible] | [illegible] | [illegible] | [illegible] | [illegible] | [illegible] |
| SE | 97843 | ENSG00000109971 | HSPA8 | chr11 | - | 123058631 | [illegible] | [illegible] | [illegible] | [illegible] | [illegible] | [illegible] | [illegible] | [illegible] |
| SE | 97844 | ENSG00000109971 | HSPA8 | chr11 | - | 123060115 | [illegible] | [illegible] | [illegible] | [illegible] | [illegible] | [illegible] | [illegible] | [illegible] |
| SE | 97845 | ENSG00000109971 | HSPA8 | chr11 | - | 123060115 | [illegible] | [illegible] | [illegible] | [illegible] | [illegible] | [illegible] | [illegible] | [illegible] |
| SE | 97848 | ENSG00000109971 | HSPA8 | chr11 | - | 123060592 | [illegible] | [illegible] | [illegible] | [illegible] | [illegible] | [illegible] | [illegible] | [illegible] |
| SE | 97850 | ENSG00000109971 | HSPA8 | chr11 | - | 123061119 | [illegible] | [illegible] | [illegible] | [illegible] | [illegible] | [illegible] | [illegible] | [illegible] |
| SE | 97851 | ENSG00000109971 | HSPA8 | chr11 | - | 123061119 | [illegible] | [illegible] | [illegible] | [illegible] | [illegible] | [illegible] | [illegible] | [illegible] |
| SE | 97852 | ENSG00000116786 | PLEKHM2 | chr1 | + | 15716706 | [illegible] | [illegible] | [illegible] | [illegible] | [illegible] | [illegible] | [illegible] | [illegible] |
| SE | 97853 | ENSG00000116786 | PLEKHM2 | chr1 | + | 15719733 | [illegible] | [illegible] | [illegible] | [illegible] | [illegible] | [illegible] | [illegible] | [illegible] |
| SE | 97855 | ENSG00000138092 | CENPO | chr2 | + | 24793851 | [illegible] | [illegible] | [illegible] | [illegible] | [illegible] | [illegible] | [illegible] | [illegible] |
| SE | 97856 | ENSG00000138092 | CENPO | chr2 | + | 24793851 | [illegible] | [illegible] | [illegible] | [illegible] | [illegible] | [illegible] | [illegible] | [illegible] |
| SE | 97857 | ENSG00000138092 | CENPO | chr2 | + | 24799674 | [illegible] | [illegible] | [illegible] | [illegible] | [illegible] | [illegible] | [illegible] | [illegible] |
| SE | 97858 | ENSG00000138092 | CENPO | chr2 | + | 24815496 | [illegible] | [illegible] | [illegible] | [illegible] | [illegible] | [illegible] | [illegible] | [illegible] |
| SE | 97859 | ENSG00000138092 | CENPO | chr2 | + | 24815597 | [illegible] | [illegible] | [illegible] | [illegible] | [illegible] | [illegible] | [illegible] | [illegible] |
| SE | 97862 | ENSG00000149557 | FEZ1 | chr11 | - | 125448501 | [illegible] | [illegible] | [illegible] | [illegible] | [illegible] | [illegible] | [illegible] | [illegible] |
| SE | 97876 | ENSG00000151500 | THYN1 | chr11 | - | 134249827 | [illegible] | [illegible] | [illegible] | [illegible] | [illegible] | [illegible] | [illegible] | [illegible] |
| SE | 97877 | ENSG00000151500 | THYN1 | chr11 | - | 134251748 | [illegible] | [illegible] | [illegible] | [illegible] | [illegible] | [illegible] | [illegible] | [illegible] |
| SE | 97886 | ENSG00000149548 | CCDC15 | chr11 | + | 124975095 | [illegible] | [illegible] | [illegible] | [illegible] | [illegible] | [illegible] | [illegible] | [illegible] |
| SE | 97887 | ENSG00000149548 | CCDC15 | chr11 | + | 124977477 | [illegible] | [illegible] | [illegible] | [illegible] | [illegible] | [illegible] | [illegible] | [illegible] |
| SE | 97888 | ENSG00000149548 | CCDC15 | chr11 | + | 124991460 | [illegible] | [illegible] | [illegible] | [illegible] | [illegible] | [illegible] | [illegible] | [illegible] |
| SE | 97889 | ENSG00000149548 | CCDC15 | chr11 | + | 124992579 | [illegible] | [illegible] | [illegible] | [illegible] | [illegible] | [illegible] | [illegible] | [illegible] |
| SE | 97891 | ENSG00000149548 | CCDC15 | chr11 | + | 124993168 | [illegible] | [illegible] | [illegible] | [illegible] | [illegible] | [illegible] | [illegible] | [illegible] |
| SE | 97892 | ENSG00000149548 | CCDC15 | chr11 | + | 125003866 | [illegible] | [illegible] | [illegible] | [illegible] | [illegible] | [illegible] | [illegible] | [illegible] |
| SE | 97893 | ENSG00000149548 | CCDC15 | chr11 | + | 125003866 | [illegible] | [illegible] | [illegible] | [illegible] | [illegible] | [illegible] | [illegible] | [illegible] |
| SE | 97894 | ENSG00000145555 | MYO10 | chr5 | - | 16670525 | [illegible] | [illegible] | [illegible] | [illegible] | [illegible] | [illegible] | [illegible] | [illegible] |
| SE | 97897 | ENSG00000197860 | SGTB | chr5 | - | 65672243 | [illegible] | [illegible] | [illegible] | [illegible] | [illegible] | [illegible] | [illegible] | [illegible] |
| SE | 97899 | ENSG00000197860 | SGTB | chr5 | - | 65704278 | [illegible] | [illegible] | [illegible] | [illegible] | [illegible] | [illegible] | [illegible] | [illegible] |
| SE | 97901 | ENSG00000197860 | SGTB | chr5 | - | 65712960 | [illegible] | [illegible] | [illegible] | [illegible] | [illegible] | [illegible] | [illegible] | [illegible] |
| SE | 97908 | ENSG00000134910 | STT3A | chr11 | + | 125593637 | [illegible] | [illegible] | [illegible] | [illegible] | [illegible] | [illegible] | [illegible] | [illegible] |
| SE | 97909 | ENSG00000134910 | STT3A | chr11 | + | 125594049 | [illegible] | [illegible] | [illegible] | [illegible] | [illegible] | [illegible] | [illegible] | [illegible] |
| SE | 97911 | ENSG00000134910 | STT3A | chr11 | + | 125597058 | [illegible] | [illegible] | [illegible] | [illegible] | [illegible] | [illegible] | [illegible] | [illegible] |
| SE | 97914 | ENSG00000134910 | STT3A | chr11 | + | 125611427 | [illegible] | [illegible] | [illegible] | [illegible] | [illegible] | [illegible] | [illegible] | [illegible] |
| SE | 97915 | ENSG00000134910 | STT3A | chr11 | + | 125612591 | [illegible] | [illegible] | [illegible] | [illegible] | [illegible] | [illegible] | [illegible] | [illegible] |
| SE | 97916 | ENSG00000134910 | STT3A | chr11 | + | 125614086 | [illegible] | [illegible] | [illegible] | [illegible] | [illegible] | [illegible] | [illegible] | [illegible] |
| SE | 97917 | ENSG00000109929 | SC5D | chr11 | + | 121306385 | [illegible] | [illegible] | [illegible] | [illegible] | [illegible] | [illegible] | [illegible] | [illegible] |
| SE | 97918 | ENSG00000109929 | SC5D | chr11 | + | 121306458 | [illegible] | [illegible] | [illegible] | [illegible] | [illegible] | [illegible] | [illegible] | [illegible] |
| SE | 97919 | ENSG00000095370 | SH2D3C | chr9 | - | 127742848 | [illegible] | [illegible] | [illegible] | [illegible] | [illegible] | [illegible] | [illegible] | [illegible] |
| SE | 97922 | ENSG00000095370 | SH2D3C | chr9 | - | 127761610 | [illegible] | [illegible] | [illegible] | [illegible] | [illegible] | [illegible] | [illegible] | [illegible] |
| SE | 97924 | ENSG00000095370 | SH2D3C | chr9 | - | 127766900 | [illegible] | [illegible] | [illegible] | [illegible] | [illegible] | [illegible] | [illegible] | [illegible] |
| SE | 97926 | ENSG00000129255 | MPDU1 | chr17 | + | 7585731 | [illegible] | [illegible] | [illegible] | [illegible] | [illegible] | [illegible] | [illegible] | [illegible] |
| SE | 97927 | ENSG00000129255 | MPDU1 | chr17 | + | 7585945 | [illegible] | [illegible] | [illegible] | [illegible] | [illegible] | [illegible] | [illegible] | [illegible] |
| SE | 97928 | ENSG00000129255 | MPDU1 | chr17 | + | 7585950 | [illegible] | [illegible] | [illegible] | [illegible] | [illegible] | [illegible] | [illegible] | [illegible] |
| SE | 97934 | ENSG00000129255 | MPDU1 | chr17 | + | 7586898 | [illegible] | [illegible] | [illegible] | [illegible] | [illegible] | [illegible] | [illegible] | [illegible] |
| SE | 97941 | ENSG00000149547 | EI24 | chr11 | + | 125575802 | [illegible] | [illegible] | [illegible] | [illegible] | [illegible] | [illegible] | [illegible] | [illegible] |
| SE | 97944 | ENSG00000149547 | EI24 | chr11 | + | 125578948 | [illegible] | [illegible] | [illegible] | [illegible] | [illegible] | [illegible] | [illegible] | [illegible] |
| SE | 97945 | ENSG00000149547 | EI24 | chr11 | + | 125578948 | [illegible] | [illegible] | [illegible] | [illegible] | [illegible] | [illegible] | [illegible] | [illegible] |
| SE | 97946 | ENSG00000149547 | EI24 | chr11 | + | 125580092 | [illegible] | [illegible] | [illegible] | [illegible] | [illegible] | [illegible] | [illegible] | [illegible] |
| SE | 97948 | ENSG00000149547 | EI24 | chr11 | + | 125580092 | [illegible] | [illegible] | [illegible] | [illegible] | [illegible] | [illegible] | [illegible] | [illegible] |
| SE | 97949 | ENSG00000149547 | EI24 | chr11 | + | 125581210 | [illegible] | [illegible] | [illegible] | [illegible] | [illegible] | [illegible] | [illegible] | [illegible] |
| SE | 97950 | ENSG00000149547 | EI24 | chr11 | + | 125581210 | [illegible] | [illegible] | [illegible] | [illegible] | [illegible] | [illegible] | [illegible] | [illegible] |
| SE | 97951 | ENSG00000149547 | EI24 | chr11 | + | 125582345 | [illegible] | [illegible] | [illegible] | [illegible] | [illegible] | [illegible] | [illegible] | [illegible] |
| SE | 97954 | ENSG00000137500 | CCDC90B | chr11 | - | 83273646 | [illegible] | [illegible] | [illegible] | [illegible] | [illegible] | [illegible] | [illegible] | [illegible] |
| SE | 97961 | ENSG00000137500 | CCDC90B | chr11 | - | 83278725 | [illegible] | [illegible] | [illegible] | [illegible] | [illegible] | [illegible] | [illegible] | [illegible] |
| SE | 97963 | ENSG00000137500 | CCDC90B | chr11 | - | 83280140 | [illegible] | [illegible] | [illegible] | [illegible] | [illegible] | [illegible] | [illegible] | [illegible] |
| SE | 97964 | ENSG00000137500 | CCDC90B | chr11 | - | 83280140 | [illegible] | [illegible] | [illegible] | [illegible] | [illegible] | [illegible] | [illegible] | [illegible] |
| SE | 97965 | ENSG00000137500 | CCDC90B | chr11 | - | 83280140 | [illegible] | [illegible] | [illegible] | [illegible] | [illegible] | [illegible] | [illegible] | [illegible] |
| SE | 97966 | ENSG00000137500 | CCDC90B | chr11 | - | 83280140 | [illegible] | [illegible] | [illegible] | [illegible] | [illegible] | [illegible] | [illegible] | [illegible] |
| SE | 97967 | ENSG00000137500 | CCDC90B | chr11 | - | 83285126 | [illegible] | [illegible] | [illegible] | [illegible] | [illegible] | [illegible] | [illegible] | [illegible] |
| SE | 97968 | ENSG00000137500 | CCDC90B | chr11 | - | 83285126 | [illegible] | [illegible] | [illegible] | [illegible] | [illegible] | [illegible] | [illegible] | [illegible] |
| SE | 97971 | ENSG00000099251 | HSD17B7P2 | chr10 | + | 38369123 | [illegible] | [illegible] | [illegible] | [illegible] | [illegible] | [illegible] | [illegible] | [illegible] |
| SE | 97979 | ENSG00000273415 | LINC02725 | chr11 | + | 128167171 | [illegible] | [illegible] | [illegible] | [illegible] | [illegible] | [illegible] | [illegible] | [illegible] |
| SE | 97984 | ENSG00000197798 | FAM118B | chr11 | + | 126233585 | [illegible] | [illegible] | [illegible] | [illegible] | [illegible] | [illegible] | [illegible] | [illegible] |
| SE | 97985 | ENSG00000197798 | FAM118B | chr11 | + | 126234994 | [illegible] | [illegible] | [illegible] | [illegible] | [illegible] | [illegible] | [illegible] | [illegible] |
| SE | 97986 | ENSG00000197798 | FAM118B | chr11 | + | 126234994 | [illegible] | [illegible] | [illegible] | [illegible] | [illegible] | [illegible] | [illegible] | [illegible] |
| SE | 97987 | ENSG00000197798 | FAM118B | chr11 | + | 126240791 | [illegible] | [illegible] | [illegible] | [illegible] | [illegible] | [illegible] | [illegible] | [illegible] |
| SE | 97989 | ENSG00000197798 | FAM118B | chr11 | + | 126250505 | [illegible] | [illegible] | [illegible] | [illegible] | [illegible] | [illegible] | [illegible] | [illegible] |
| SE | 97990 | ENSG00000197798 | FAM118B | chr11 | + | 126250505 | [illegible] | [illegible] | [illegible] | [illegible] | [illegible] | [illegible] | [illegible] | [illegible] |
| SE | 97991 | ENSG00000197798 | FAM118B | chr11 | + | 126254304 | [illegible] | [illegible] | [illegible] | [illegible] | [illegible] | [illegible] | [illegible] | [illegible] |
| SE | 97992 | ENSG00000197798 | FAM118B | chr11 | + | 126254304 | [illegible] | [illegible] | [illegible] | [illegible] | [illegible] | [illegible] | [illegible] | [illegible] |
| SE | 97993 | ENSG00000197798 | FAM118B | chr11 | + | 126261424 | [illegible] | [illegible] | [illegible] | [illegible] | [illegible] | [illegible] | [illegible] | [illegible] |
| SE | 97994 | ENSG00000197798 | FAM118B | chr11 | + | 126261427 | [illegible] | [illegible] | [illegible] | [illegible] | [illegible] | [illegible] | [illegible] | [illegible] |
| SE | 97995 | ENSG00000205758 | CRYZL1 | chr21 | - | 33591161 | [illegible] | [illegible] | [illegible] | [illegible] | [illegible] | [illegible] | [illegible] | [illegible] |
| SE | 97996 | ENSG00000205758 | CRYZL1 | chr21 | - | 33597279 | [illegible] | [illegible] | [illegible] | [illegible] | [illegible] | [illegible] | [illegible] | [illegible] |
| SE | 97998 | ENSG00000205758 | CRYZL1 | chr21 | - | 33597279 | [illegible] | [illegible] | [illegible] | [illegible] | [illegible] | [illegible] | [illegible] | [illegible] |
| SE | 97999 | ENSG00000205758 | CRYZL1 | chr21 | - | 33613537 | [illegible] | [illegible] | [illegible] | [illegible] | [illegible] | [illegible] | [illegible] | [illegible] |
| SE | 98000 | ENSG00000205758 | CRYZL1 | chr21 | - | 33616705 | [illegible] | [illegible] | [illegible] | [illegible] | [illegible] | [illegible] | [illegible] | [illegible] |
| SE | 98004 | ENSG00000205758 | CRYZL1 | chr21 | - | 33641178 | [illegible] | [illegible] | [illegible] | [illegible] | [illegible] | [illegible] | [illegible] | [illegible] |
| SE | 98011 | ENSG00000188997 | KCTD21 | chr11 | - | 78176887 | [illegible] | [illegible] | [illegible] | [illegible] | [illegible] | [illegible] | [illegible] | [illegible] |
| SE | 98013 | ENSG00000188997 | KCTD21 | chr11 | - | 78186680 | [illegible] | [illegible] | [illegible] | [illegible] | [illegible] | [illegible] | [illegible] | [illegible] |
| SE | 98028 | ENSG00000183648 | NDUFB1 | chr14 | - | 92118848 | [illegible] | [illegible] | [illegible] | [illegible] | [illegible] | [illegible] | [illegible] | [illegible] |
| SE | 98031 | ENSG00000085117 | CD82 | chr11 | + | 44566272 | [illegible] | [illegible] | [illegible] | [illegible] | [illegible] | [illegible] | [illegible] | [illegible] |
| SE | 98032 | ENSG00000085117 | CD82 | chr11 | + | 44585142 | [illegible] | [illegible] | [illegible] | [illegible] | [illegible] | [illegible] | [illegible] | [illegible] |
| SE | 98034 | ENSG00000085117 | CD82 | chr11 | + | 44594642 | [illegible] | [illegible] | [illegible] | [illegible] | [illegible] | [illegible] | [illegible] | [illegible] |
| SE | 98035 | ENSG00000085117 | CD82 | chr11 | + | 44605354 | [illegible] | [illegible] | [illegible] | [illegible] | [illegible] | [illegible] | [illegible] | [illegible] |
| SE | 98036 | ENSG00000085117 | CD82 | chr11 | + | 44618639 | [illegible] | [illegible] | [illegible] | [illegible] | [illegible] | [illegible] | [illegible] | [illegible] |
| SE | 98038 | ENSG00000139697 | SBNO1 | chr12 | - | 123304581 | [illegible] | [illegible] | [illegible] | [illegible] | [illegible] | [illegible] | [illegible] | [illegible] |
| SE | 98039 | ENSG00000139697 | SBNO1 | chr12 | - | 123309309 | [illegible] | [illegible] | [illegible] | [illegible] | [illegible] | [illegible] | [illegible] | [illegible] |
| SE | 98040 | ENSG00000139697 | SBNO1 | chr12 | - | 123311054 | [illegible] | [illegible] | [illegible] | [illegible] | [illegible] | [illegible] | [illegible] | [illegible] |

| | | | | | | | | | | | | | | | | | | | | | | | | | | | | | | | | | | | | | | | | | | | | | | |
|---|---|---|---|---|---|---|---|---|---|---|---|---|---|---|---|---|---|---|---|---|---|---|---|---|---|---|---|---|---|---|---|---|---|---|---|---|---|---|---|---|---|---|---|---|---|---|
| SE | 98041 | ENSG00000139697 | SBNO1 | chr12 | - | 123313619 | 123313719 | 123311054 | 123311129 | 123315372 | 123315444 | 253 | 214 | 252 | 245 | 394 | 0 | 0 | 0 | 3 | 3 | 36 | 200 | 177 | 101 | 0 | 0 | 1 | 0 | 0 | 0 | 249 | 149 | 1 | 1 | 1 | 1 | 1 | 0.98 | 0.987 | 1 | 0.992 | 1 | 1 | NA | -0.005 |
| SE | 98042 | ENSG00000139697 | SBNO1 | chr12 | - | 123317220 | 123317356 | 123315547 | 123315660 | 123319899 | 123320031 | 544 | 531 | 475 | 672 | 478 | 9 | 3 | 0 | 0 | 0 | 165 | 531 | 830 | 415 | 42 | 4 | 3 | 1 | 4 | 0 | 285 | 149 | 1 | 1 | 0.969 | 0.989 | 1 | 1 | 1 | 0.956 | 0.989 | 0.998 | 0.982 | 1 | 0.007 |
| SE | 98043 | ENSG00000139697 | SBNO1 | chr12 | - | 123334052 | 123334213 | 123331241 | 123331375 | 123336394 | 123336491 | 399 | 349 | 362 | 343 | 482 | 0 | 0 | 0 | 1 | 0 | 221 | 367 | 542 | 280 | 73 | 0 | 0 | 0 | 0 | 0 | 298 | 149 | 1 | 1 | 1 | 1 | 1 | 0.994 | 1 | 1 | 1 | 1 | 1 | 1 | -0.001 |
| SE | 98044 | ENSG00000139697 | SBNO1 | chr12 | - | 123345257 | 123345567 | 123340987 | 123341088 | 123348028 | 123348133 | 428 | 351 | 333 | 496 | 454 | 5 | 0 | 2 | 0 | 0 | 364 | 517 | 405 | 314 | 129 | 0 | 0 | 0 | 1 | 0 | 298 | 149 | 1 | 1 | 0.977 | 1 | 0.988 | 1 | 1 | 1 | 1 | 1 | 0.994 | 1 | -0.006 |
| SE | 98045 | ENSG00000139697 | SBNO1 | chr12 | - | 123345257 | 123345570 | 123340987 | 123341088 | 123348028 | 123348133 | 413 | 363 | 339 | 462 | 490 | 5 | 0 | 2 | 0 | 0 | 337 | 511 | 373 | 263 | 113 | 0 | 0 | 0 | 1 | 0 | 298 | 149 | 1 | 1 | 0.976 | 1 | 0.988 | 1 | 1 | 1 | 1 | 1 | 0.992 | 1 | -0.006 |
| SE | 98046 | ENSG00000139697 | SBNO1 | chr12 | - | 123348028 | 123348133 | 123345257 | 123345567 | 123350309 | 123350441 | 173 | 116 | 167 | 194 | 274 | 2 | 4 | 0 | 0 | 0 | 104 | 119 | 149 | 160 | 74 | 0 | 0 | 1 | 0 | 0 | 254 | 149 | 1 | 1 | 0.981 | 0.944 | 1 | 1 | 1 | 1 | 1 | 0.989 | 1 | 1 | -0.013 |
| SE | 98048 | ENSG00000139697 | SBNO1 | chr12 | - | 123350309 | 123350441 | 123348028 | 123348133 | 123364700 | 123364843 | 162 | 105 | 155 | 176 | 232 | 1 | 3 | 0 | 0 | 0 | 79 | 107 | 115 | 119 | 59 | 0 | 0 | 0 | 1 | 0 | 281 | 149 | 1 | 1 | 0.988 | 0.949 | 1 | 1 | 1 | 1 | 1 | 1 | 0.984 | 1 | -0.009 |
| SE | 98049 | ENSG00000109881 | CCDC34 | chr11 | - | 27350331 | 27350439 | 27341391 | 27341550 | 27357402 | 27357541 | 1 | 1 | 4 | 2 | 2 | 0 | 0 | 0 | 0 | 2 | 0 | 0 | 2 | 21 | 1 | 0 | 0 | 0 | 0 | 0 | 257 | 149 | 1 | 1 | 1 | 1 | 1 | 1 | 0.367 | NA | NA | 1 | 1 | 1 | -0.127 |
| SE | 98050 | ENSG00000110318 | CEP126 | chr11 | + | 101955791 | 101956943 | 101948030 | 101948142 | 101958167 | 101958366 | 12 | 12 | 12 | 12 | 14 | 7 | 1 | 7 | 13 | 4 | 0 | 2 | 0 | 0 | 0 | 0 | 3 | 0 | 0 | 0 | 298 | 149 | 1 | 1 | 0.462 | 0.857 | 0.462 | 0.316 | 0.636 | NA | 0.25 | NA | NA | NA | 0.297 |
| SE | 98051 | ENSG00000110318 | CEP126 | chr11 | + | 101956261 | 101956943 | 101948030 | 101948142 | 101958167 | 101958366 | 13 | 12 | 11 | 13 | 10 | 7 | 1 | 7 | 13 | 4 | 0 | 4 | 0 | 0 | 0 | 0 | 3 | 0 | 0 | 0 | 298 | 149 | 1 | 1 | 0.481 | 0.857 | 0.44 | 0.333 | 0.556 | NA | 0.4 | NA | NA | NA | 0.133 |
| SE | 98052 | ENSG00000110318 | CEP126 | chr11 | + | 101956867 | 101956943 | 101948030 | 101948142 | 101958167 | 101958366 | 6 | 4 | 2 | 7 | 4 | 7 | 1 | 7 | 13 | 4 | 0 | 2 | 0 | 0 | 0 | 0 | 3 | 0 | 0 | 0 | 225 | 149 | 1 | 1 | 0.362 | 0.726 | 0.159 | 0.263 | 0.398 | NA | 0.306 | NA | NA | NA | 0.076 |
| SE | 98053 | ENSG00000110318 | CEP126 | chr11 | + | 101958167 | 101958366 | 101948030 | 101948142 | 101978346 | 101978459 | 8 | 1 | 9 | 13 | 4 | 0 | 0 | 2 | 0 | 0 | 0 | 3 | 0 | 0 | 0 | 0 | 0 | 0 | 0 | 0 | 298 | 149 | 1 | 1 | 1 | 1 | 0.692 | 1 | 1 | NA | 1 | NA | NA | NA | -0.062 |
| SE | 98054 | ENSG00000110318 | CEP126 | chr11 | + | 101961740 | 101963880 | 101958167 | 101958366 | 101978346 | 101978459 | 48 | 33 | 24 | 28 | 29 | 1 | 0 | 2 | 0 | 0 | 0 | 5 | 0 | 3 | 0 | 0 | 0 | 0 | 0 | 0 | 298 | 149 | 1 | 1 | 0.96 | 1 | 0.857 | 1 | 1 | NA | 1 | NA | 1 | NA | -0.037 |
| SE | 98055 | ENSG00000110318 | CEP126 | chr11 | + | 101978346 | 101978459 | 101961740 | 101963880 | 101981888 | 101981964 | 37 | 33 | 18 | 19 | 21 | 5 | 0 | 0 | 6 | 10 | 0 | 0 | 0 | 6 | 0 | 0 | 0 | 0 | 0 | 0 | 262 | 149 | 1 | 1 | 0.808 | 1 | 1 | 0.643 | 0.544 | NA | NA | NA | 1 | NA | -0.201 |
| SE | 98057 | ENSG00000186314 | PRELID2 | chr5 | - | 145764930 | 145765000 | 145703903 | 145704295 | 145796441 | 145796531 | 7 | 1 | 2 | 4 | 10 | 0 | 0 | 0 | 0 | 1 | 14 | 6 | 0 | 6 | 19 | 0 | 0 | 0 | 1 | 0 | 219 | 149 | 1 | 1 | 1 | 1 | 1 | 1 | 0.872 | 1 | 1 | NA | 0.803 | 1 | 0.024 |
| SE | 98060 | ENSG00000186314 | PRELID2 | chr5 | - | 145825997 | 145826200 | 145823076 | 145823134 | 145835176 | 145835336 | 11 | 8 | 1 | 21 | 11 | 0 | 0 | 1 | 2 | 0 | 79 | 50 | 9 | 22 | 150 | 0 | 0 | 0 | 0 | 0 | 298 | 149 | 1 | 1 | 1 | 1 | 0.333 | 0.84 | 1 | 1 | 1 | 1 | 1 | 1 | -0.165 |
| SE | 98062 | ENSG00000182240 | BACE2 | chr21 | + | 41250751 | 41250901 | 41245961 | 41246063 | 41275370 | 41276597 | 5 | 4 | 7 | 4 | 10 | 0 | 0 | 0 | 0 | 0 | 0 | 3 | 1 | 13 | 14 | 0 | 1 | 0 | 1 | 0 | 298 | 149 | 1 | 1 | 1 | 1 | 1 | 1 | 1 | NA | 0.6 | 1 | 0.867 | 1 | 0.133 |
| SE | 98063 | ENSG00000182240 | BACE2 | chr21 | + | 41257157 | 41257326 | 41245961 | 41246063 | 41275370 | 41275381 | 4 | 3 | 1 | 2 | 1 | 0 | 0 | 0 | 0 | 0 | 0 | 6 | 1 | 0 | 1 | 0 | 1 | 0 | 1 | 0 | 298 | 149 | 1 | 1 | 1 | 1 | 1 | 1 | 1 | NA | 0.75 | 1 | 0 | 1 | 0.312 |
| SE | 98064 | ENSG00000182240 | BACE2 | chr21 | + | 41257157 | 41257326 | 41250751 | 41250901 | 41275370 | 41275381 | 14 | 5 | 13 | 8 | 9 | 1 | 0 | 0 | 0 | 0 | 0 | 17 | 5 | 0 | 14 | 0 | 2 | 0 | 0 | 0 | 298 | 149 | 1 | 1 | 0.875 | 1 | 1 | 1 | 1 | NA | 0.81 | 1 | NA | 1 | 0.038 |
| SE | 98065 | ENSG00000144785 | NA | chr12 | - | 56309965 | 56310029 | 56300159 | 56300220 | 56310957 | 56311054 | 1 | 4 | 0 | 0 | 1 | 0 | 0 | 0 | 0 | 0 | 6 | 0 | 1 | 12 | 0 | 5 | 0 | 0 | 5 | 0 | 213 | 149 | 1 | 1 | 1 | 1 | NA | NA | 1 | 0.456 | NA | 1 | 0.627 | NA | 0.306 |
| SE | 98067 | ENSG00000144785 | NA | chr12 | - | 56311210 | 56311414 | 56310957 | 56311054 | 56314850 | 56314966 | 242 | 126 | 217 | 208 | 181 | 0 | 0 | 0 | 1 | 0 | 402 | 188 | 180 | 277 | 273 | 0 | 0 | 0 | 0 | 0 | 298 | 149 | 1 | 1 | 1 | 1 | 1 | 0.99 | 1 | 1 | 1 | 1 | 1 | 1 | -0.002 |
| SE | 98070 | ENSG00000066926 | FECH | chr18 | - | 57554259 | 57554424 | 57551174 | 57551374 | 57554844 | 57554952 | 73 | 34 | 75 | 44 | 119 | 0 | 0 | 0 | 0 | 0 | 56 | 39 | 43 | 48 | 161 | 0 | 0 | 0 | 3 | 0 | 298 | 149 | 1 | 1 | 1 | 1 | 1 | 1 | 1 | 1 | 1 | 1 | 0.889 | 1 | 0.022 |
| SE | 98076 | ENSG00000126705 | AHDC1 | chr1 | - | 27547260 | 27552189 | 27534034 | 27534916 | 27552891 | 27552929 | 25 | 3 | 14 | 18 | 13 | 0 | 0 | 0 | 0 | 0 | 12 | 7 | 2 | 6 | 28 | 0 | 0 | 2 | 0 | 0 | 298 | 149 | 1 | 1 | 1 | 1 | 1 | 1 | 1 | 1 | 1 | 0.333 | 1 | 1 | 0.133 |
| SE | 98084 | ENSG00000176273 | SLC35G1 | chr10 | + | 93912666 | 93912799 | 93906251 | 93906385 | 93954008 | 93954168 | 2 | 2 | 2 | 0 | 0 | 0 | 1 | 0 | 1 | 0 | 0 | 1 | 0 | 0 | 0 | 0 | 0 | 0 | 0 | 0 | 282 | 149 | 1 | 1 | 1 | 0.514 | 1 | 0 | NA | NA | 1 | NA | NA | NA | -0.371 |
| SE | 98085 | ENSG00000239523 | MYLK-AS1 | chr3 | + | 123611234 | 123611355 | 123585641 | 123585801 | 123629490 | 123629681 | 0 | 2 | 2 | 0 | 4 | 1 | 0 | 0 | 0 | 0 | 0 | 0 | 0 | 0 | 13 | 0 | 2 | 0 | 0 | 8 | 270 | 149 | 1 | 1 | 0 | 1 | 1 | NA | 1 | NA | 0 | NA | NA | 0.473 | 0.514 |
| SE | 98086 | ENSG00000239523 | MYLK-AS1 | chr3 | + | 123611234 | 123611355 | 123585641 | 123585801 | 123629493 | 123629822 | 4 | 5 | 1 | 0 | 0 | 1 | 0 | 0 | 0 | 1 | 0 | 0 | 0 | 0 | 13 | 0 | 1 | 0 | 0 | 6 | 270 | 149 | 1 | 1 | 0.688 | 1 | 1 | NA | 0 | NA | 0 | NA | NA | 0.545 | 0.399 |
| SE | 98087 | ENSG00000239523 | MYLK-AS1 | chr3 | + | 123611234 | 123611355 | 123585646 | 123585794 | 123612238 | 123612579 | 4 | 2 | 0 | 5 | 4 | 4 | 0 | 0 | 0 | 0 | 0 | 0 | 0 | 5 | 0 | 0 | 0 | 0 | 0 | 0 | 270 | 149 | 1 | 1 | 0.356 | 1 | NA | 1 | 1 | NA | NA | NA | 1 | NA | -0.161 |
| SE | 98088 | ENSG00000239523 | MYLK-AS1 | chr3 | + | 123611234 | 123611355 | 123585646 | 123585794 | 123629490 | 123629681 | 1 | 0 | 2 | 1 | 4 | 0 | 0 | 0 | 0 | 0 | 0 | 0 | 0 | 0 | 0 | 0 | 3 | 0 | 0 | 0 | 270 | 149 | 1 | 1 | 1 | NA | 1 | 1 | 1 | NA | 0 | NA | NA | NA | 1 |

[illegible]

| | | | | | | | | | | | | | | | | | | | | | | | | | | | | | | | | | | | | | | | | | | | | | | |
|---|---|---|---|---|---|---|---|---|---|---|---|---|---|---|---|---|---|---|---|---|---|---|---|---|---|---|---|---|---|---|---|---|---|---|---|---|---|---|---|---|---|---|---|---|---|---|
| SE | 98530 | ENSG00000130935 | NOL11 | chr17 | + | 67719110 | 67719167 | 67717930 | 67718088 | 67719673 | 67719787 | 0 | 4 | 0 | 0 | 0 | 97 | 104 | 109 | 108 | 91 | 0 | 0 | 0 | 0 | 0 | 10 | 99 | 85 | 48 | 19 | 206 | 149 | 1 | 1 | 0 | 0.027 | 0 | 0 | 0 | 0 | 0 | 0 | 0 | 0 | 0.005 |
| SE | 98531 | ENSG00000130935 | NOL11 | chr17 | + | 67720346 | 67720444 | 67719905 | 67719962 | 67721377 | 67721440 | 1 | 2 | 0 | 0 | 0 | 185 | 132 | 195 | 160 | 203 | 5 | 1 | 0 | 0 | 0 | 78 | 135 | 105 | 51 | 31 | 247 | 149 | 1 | 1 | 0.003 | 0.009 | 0 | 0 | 0 | 0.037 | 0.004 | 0 | 0 | 0 | -0.006 |
| SE | 98532 | ENSG00000130935 | NOL11 | chr17 | + | 67722579 | 67722637 | 67721377 | 67721526 | 67724048 | 67724193 | 267 | 151 | 238 | 200 | 522 | 0 | 1 | 0 | 4 | 0 | 35 | 102 | 112 | 39 | 10 | 0 | 0 | 0 | 0 | 0 | 207 | 149 | 1 | 1 | 1 | 0.991 | 1 | 0.973 | 1 | 1 | 1 | 1 | 1 | 1 | -0.007 |
| SE | 98533 | ENSG00000130935 | NOL11 | chr17 | + | 67726459 | 67726648 | 67724048 | 67724193 | 67734362 | 67734439 | 343 | 217 | 350 | 292 | 720 | 0 | 0 | 0 | 0 | 1 | 30 | 157 | 192 | 60 | 28 | 0 | 0 | 0 | 0 | 0 | 298 | 149 | 1 | 1 | 1 | 1 | 1 | 1 | 0.997 | 1 | 1 | 1 | 1 | 1 | -0.001 |
| SE | 98534 | ENSG00000130935 | NOL11 | chr17 | + | 67734362 | 67734439 | 67726459 | 67726648 | 67735899 | 67736023 | 121 | 73 | 136 | 94 | 313 | 2 | 0 | 0 | 0 | 1 | 18 | 63 | 79 | 25 | 40 | 0 | 0 | 0 | 0 | 0 | 226 | 149 | 1 | 1 | 0.976 | 1 | 1 | 1 | 0.995 | 1 | 1 | 1 | 1 | 1 | -0.006 |
| SE | 98536 | ENSG00000130935 | NOL11 | chr17 | + | 67735899 | 67736023 | 67734362 | 67734439 | 67737070 | 67737145 | 66 | 55 | 69 | 69 | 153 | 0 | 0 | 0 | 0 | 0 | 66 | 79 | 46 | 32 | 40 | 0 | 3 | 0 | 0 | 0 | 273 | 149 | 1 | 1 | 1 | 1 | 1 | 1 | 1 | 1 | 0.935 | 1 | 1 | 1 | 0.013 |
| SE | 98538 | ENSG00000130935 | NOL11 | chr17 | + | 67737070 | 67737145 | 67736579 | 67736754 | 67737507 | 67737689 | 378 | 211 | 347 | 307 | 522 | 1 | 0 | 1 | 0 | 0 | 35 | 145 | 192 | 82 | 11 | 0 | 0 | 0 | 0 | 0 | 224 | 149 | 1 | 1 | 0.996 | 1 | 0.996 | 1 | 1 | 1 | 1 | 1 | 1 | 1 | -0.002 |
| SE | 98539 | ENSG00000118096 | IFT46 | chr11 | - | 118545408 | 118545494 | 118544527 | 118545011 | 118545792 | 118545853 | 21 | 5 | 35 | 23 | 37 | 0 | 1 | 0 | 0 | 0 | 46 | 12 | 12 | 6 | 4 | 0 | 0 | 0 | 0 | 0 | 235 | 149 | 1 | 1 | 1 | 0.76 | 1 | 1 | 1 | 1 | 1 | 1 | 1 | 1 | -0.048 |
| SE | 98541 | ENSG00000118096 | IFT46 | chr11 | - | 118564964 | 118565061 | 118559784 | 118559864 | 118565789 | 118565810 | 36 | 25 | 31 | 44 | 29 | 0 | 0 | 3 | 0 | 0 | 11 | 20 | 22 | 28 | 43 | 0 | 0 | 0 | 0 | 0 | 246 | 149 | 1 | 1 | 1 | 1 | 0.862 | 1 | 1 | 1 | 1 | 1 | 1 | 1 | -0.028 |
| SE | 98542 | ENSG00000118096 | IFT46 | chr11 | - | 118564964 | 118565091 | 118559784 | 118559864 | 118565789 | 118566011 | 10 | 16 | 14 | 24 | 12 | 0 | 0 | 3 | 0 | 0 | 11 | 7 | 4 | 13 | 23 | 0 | 0 | 0 | 0 | 0 | 276 | 149 | 1 | 1 | 1 | 1 | 0.716 | 1 | 1 | 1 | 1 | 1 | 1 | 1 | -0.057 |
| SE | 98549 | ENSG00000067715 | SYT1 | chr12 | + | 78995758 | 78995847 | 78977784 | 78977931 | 79047296 | 79047362 | 0 | 0 | 2 | 0 | 0 | 0 | 0 | 0 | 0 | 0 | 0 | 0 | 0 | 0 | 0 | 0 | 0 | 2 | 0 | 0 | 238 | 149 | 1 | 1 | NA | NA | 1 | NA | NA | NA | NA | 0 | NA | NA | 1 |
| SE | 98551 | ENSG00000067715 | SYT1 | chr12 | + | 79103013 | 79103055 | 79047296 | 79047362 | 79217502 | 79217685 | 0 | 0 | 2 | 0 | 0 | 0 | 0 | 0 | 2 | 1 | 0 | 0 | 0 | 0 | 1 | 0 | 0 | 2 | 0 | 0 | 191 | 149 | 1 | 1 | NA | NA | 1 | 0 | 0 | NA | NA | 0 | NA | 1 | -0.167 |
| SE | 98552 | ENSG00000051009 | FAM160A2 | chr11 | - | 6214732 | 6214911 | 6214410 | 6214573 | 6217370 | 6218150 | 51 | 18 | 34 | 37 | 24 | 1 | 0 | 0 | 1 | 0 | 16 | 60 | 24 | 32 | 24 | 0 | 2 | 3 | 0 | 0 | 298 | 149 | 1 | 1 | 0.962 | 1 | 1 | 0.949 | 1 | 1 | 0.938 | 0.8 | 1 | 1 | 0.035 |
| SE | 98553 | ENSG00000051009 | FAM160A2 | chr11 | - | 6218557 | 6218763 | 6217370 | 6218150 | 6222441 | 6222609 | 8 | 1 | 7 | 14 | 8 | 0 | 0 | 2 | 0 | 0 | 0 | 0 | 0 | 0 | 13 | 0 | 0 | 0 | 0 | 0 | 298 | 149 | 1 | 1 | 1 | 1 | 0.636 | 1 | 1 | NA | NA | NA | NA | 1 | -0.073 |
| SE | 98554 | ENSG00000051009 | FAM160A2 | chr11 | - | 6218599 | 6218763 | 6216569 | 6218150 | 6222441 | 6222609 | 70 | 16 | 29 | 55 | 20 | 0 | 0 | 2 | 0 | 0 | 28 | 42 | 28 | 25 | 57 | 0 | 0 | 0 | 0 | 0 | 298 | 149 | 1 | 1 | 1 | 1 | 0.879 | 1 | 1 | 1 | 1 | 1 | 1 | 1 | -0.024 |
| SE | 98555 | ENSG00000051009 | FAM160A2 | chr11 | - | 6218970 | 6219050 | 6218557 | 6218763 | 6222441 | 6222609 | 126 | 38 | 71 | 101 | 53 | 0 | 0 | 0 | 2 | 0 | 76 | 93 | 46 | 100 | 191 | 0 | 0 | 0 | 0 | 0 | 229 | 149 | 1 | 1 | 1 | 1 | 1 | 0.97 | 1 | 1 | 1 | 1 | 1 | 1 | -0.006 |
| SE | 98557 | ENSG00000051009 | FAM160A2 | chr11 | - | 6223079 | 6223238 | 6222810 | 6222897 | 6223609 | 6224248 | 57 | 17 | 25 | 31 | 31 | 0 | 0 | 0 | 0 | 0 | 48 | 59 | 24 | 44 | 50 | 0 | 0 | 0 | 2 | 0 | 298 | 149 | 1 | 1 | 1 | 1 | 1 | 1 | 1 | 1 | 1 | 1 | 0.917 | 1 | 0.017 |
| SE | 98558 | ENSG00000051009 | FAM160A2 | chr11 | - | 6224378 | 6224707 | 6223609 | 6224248 | 6234543 | 6234636 | 85 | 36 | 67 | 58 | 83 | 0 | 0 | 0 | 0 | 0 | 51 | 52 | 20 | 45 | 92 | 0 | 0 | 0 | 4 | 0 | 298 | 149 | 1 | 1 | 1 | 1 | 1 | 1 | 1 | 1 | 1 | 1 | 0.849 | 1 | 0.03 |
| SE | 98562 | ENSG00000134955 | SLC37A2 | chr11 | + | 125077229 | 125077323 | 125076756 | 125076838 | 125077449 | 125077528 | 33 | 19 | 9 | 54 | 20 | 1 | 0 | 1 | 1 | 1 | 60 | 15 | 20 | 11 | 38 | 0 | 0 | 0 | 0 | 6 | 243 | 149 | 1 | 1 | 0.953 | 1 | 0.847 | 0.971 | 0.925 | 1 | 1 | 1 | 1 | 0.795 | -0.02 |
| SE | 98564 | ENSG00000134955 | SLC37A2 | chr11 | + | 125079683 | 125079760 | 125077449 | 125077528 | 125080613 | 125080647 | 11 | 5 | 9 | 16 | 3 | 1 | 0 | 0 | 0 | 0 | 8 | 11 | 2 | 30 | 35 | 0 | 0 | 0 | 0 | 0 | 226 | 149 | 1 | 1 | 0.879 | 1 | 1 | 1 | 1 | 1 | 1 | 1 | 1 | 1 | -0.024 |
| SE | 98565 | ENSG00000134955 | SLC37A2 | chr11 | + | 125081753 | 125081906 | 125081420 | 125081458 | 125082243 | 125082334 | 31 | 10 | 11 | 40 | 10 | 0 | 0 | 0 | 0 | 0 | 47 | 25 | 5 | 46 | 60 | 0 | 0 | 1 | 0 | 0 | 298 | 149 | 1 | 1 | 1 | 1 | 1 | 1 | 1 | 1 | 1 | 0.714 | 1 | 1 | 0.057 |
| SE | 98569 | ENSG00000025796 | SEC63 | chr6 | - | 107872807 | 107872912 | 107867755 | 107871847 | 107876563 | 107876662 | 679 | 752 | 582 | 843 | 866 | 0 | 0 | 0 | 2 | 0 | 158 | 543 | 1115 | 394 | 42 | 0 | 0 | 0 | 0 | 0 | 254 | 149 | 1 | 1 | 1 | 1 | 1 | 0.996 | 1 | 1 | 1 | 1 | 1 | 1 | -0.001 |
| SE | 98571 | ENSG00000025796 | SEC63 | chr6 | - | 107881148 | 107881250 | 107876563 | 107876662 | 107882987 | 107883146 | 405 | 431 | 371 | 498 | 494 | 0 | 0 | 0 | 0 | 0 | 135 | 345 | 667 | 235 | 17 | 0 | 0 | 0 | 3 | 0 | 251 | 149 | 1 | 1 | 1 | 1 | 1 | 1 | 1 | 1 | 1 | 1 | 0.979 | 1 | 0.004 |
| SE | 98572 | ENSG00000025796 | SEC63 | chr6 | - | 107882987 | 107883146 | 107876563 | 107876662 | 107893481 | 107893655 | 46 | 56 | 58 | 56 | 76 | 0 | 0 | 0 | 1 | 1 | 8 | 51 | 75 | 44 | 9 | 0 | 0 | 0 | 0 | 0 | 298 | 149 | 1 | 1 | 1 | 1 | 1 | 0.966 | 0.974 | 1 | 1 | 1 | 1 | 1 | -0.012 |
| SE | 98573 | ENSG00000025796 | SEC63 | chr6 | - | 107897648 | 107897731 | 107893579 | 107893897 | 107901369 | 107901517 | 139 | 127 | 128 | 120 | 207 | 0 | 2 | 0 | 0 | 0 | 37 | 119 | 174 | 101 | 22 | 0 | 0 | 0 | 0 | 0 | 232 | 149 | 1 | 1 | 1 | 0.976 | 1 | 1 | 1 | 1 | 1 | 1 | 1 | 1 | -0.005 |
| SE | 98574 | ENSG00000025796 | SEC63 | chr6 | - | 107908926 | 107909035 | 107906682 | 107906777 | 107911345 | 107911396 | 425 | 316 | 382 | 352 | 922 | 0 | 0 | 0 | 0 | 0 | 163 | 256 | 257 | 166 | 85 | 0 | 7 | 0 | 0 | 0 | 258 | 149 | 1 | 1 | 1 | 1 | 1 | 1 | 1 | 1 | 0.955 | 1 | 1 | 1 | 0.009 |
| SE | 98575 | ENSG00000025796 | SEC63 | chr6 | - | 107911345 | 107911396 | 107908926 | 107909035 | 107912715 | 107912774 | 335 | 265 | 296 | 310 | 826 | 0 | 0 | 0 | 1 | 0 | 126 | 199 | 190 | 145 | 77 | 0 | 0 | 0 | 0 | 0 | 200 | 149 | 1 | 1 | 1 | 1 | 1 | 0.996 | 1 | 1 | 1 | 1 | 1 | 1 | -0.001 |
| SE | 98576 | ENSG00000025796 | SEC63 | chr6 | - | 107924817 | 107924932 | 107921796 | 107921909 | 107929414 | 107929514 | 325 | 259 | 332 | 296 | 538 | 1 | 0 | 0 | 1 | 0 | 296 | 187 | 297 | 242 | 133 | 2 | 0 | 0 | 0 | 0 | 264 | 149 | 1 | 1 | 0.995 | 1 | 1 | 0.994 | 1 | 0.988 | 1 | 1 | 1 | 1 | 0 |
| SE | 98577 | ENSG00000025796 | SEC63 | chr6 | - | 107929414 | 107929514 | 107924817 | 107924932 | 107957885 | 107957931 | 241 | 195 | 274 | 216 | 271 | 0 | 1 | 0 | 2 | 0 | 219 | 195 | 303 | 259 | 90 | 0 | 0 | 0 | 0 | 0 | 249 | 149 | 1 | 1 | 1 | 0.992 | 1 | 0.985 | 1 | 1 | 1 | 1 | 1 | 1 | -0.005 |
| SE | 98578 | ENSG00000025796 | SEC63 | chr6 | - | 107955964 | 107956106 | 107913365 | 107913427 | 107957885 | 107958042 | 0 | 3 | 0 | 10 | 3 | 1 | 0 | 4 | 0 | 0 | 0 | 0 | 1 | 2 | 0 | 0 | 0 | 0 | 0 | 0 | 291 | 149 | 1 | 1 | 0 | 1 | 0 | 1 | 1 | NA | NA | 1 | 1 | NA | -0.4 |
| SE | 98582 | ENSG00000149269 | PAK1 | chr11 | - | 77325327 | 77325376 | 77322947 | 77323360 | 77336085 | 77336282 | 6 | 1 | 1 | 0 | 2 | 6 | 2 | 0 | 0 | 0 | 10 | 1 | 1 | 4 | 11 | 4 | 0 | 2 | 1 | 8 | 198 | 149 | 1 | 1 | 0.429 | 0.273 | 1 | NA | 1 | 0.653 | 1 | 0.273 | 0.751 | 0.509 | 0.038 |
| SE | 98583 | ENSG00000149269 | PAK1 | chr11 | - | 77332729 | 77332867 | 77322016 | 77323360 | 77336085 | 77336282 | 315 | 324 | 285 | 340 | 605 | 6 | 2 | 0 | 0 | 0 | 768 | 222 | 270 | 450 | 621 | 4 | 0 | 2 | 1 | 8 | 287 | 149 | 1 | 1 | 0.965 | 0.988 | 1 | 1 | 1 | 0.99 | 1 | 0.986 | 0.996 | 0.976 | 0.001 |
| SE | 98584 | ENSG00000149269 | PAK1 | chr11 | - | 77332729 | 77332867 | 77325327 | 77325376 | 77336085 | 77336282 | 174 | 194 | 161 | 203 | 307 | 0 | 0 | 1 | 0 | 0 | 457 | 126 | 154 | 273 | 397 | 0 | 0 | 0 | 0 | 0 | 287 | 149 | 1 | 1 | 1 | 1 | 0.988 | 1 | 1 | 1 | 1 | 1 | 1 | 1 | -0.002 |
| SE | 98585 | ENSG00000149269 | PAK1 | chr11 | - | 77337323 | 77337423 | 77336085 | 77336282 | 77340645 | 77340763 | 276 | 205 | 212 | 264 | 411 | 1 | 0 | 1 | 0 | 0 | 456 | 157 | 170 | 370 | 290 | 0 | 3 | 5 | 0 | 0 | 249 | 149 | 1 | 1 | 0.994 | 1 | 0.992 | 1 | 1 | 1 | 0.969 | 0.953 | 1 | 1 | 0.013 |
| SE | 98586 | ENSG00000149269 | PAK1 | chr11 | - | 77343818 | 77343931 | 77340645 | 77340763 | 77349238 | 77349287 | 246 | 212 | 190 | 215 | 395 | 0 | 0 | 0 | 0 | 0 | 600 | 148 | 146 | 320 | 477 | 0 | 0 | 0 | 0 | 3 | 262 | 149 | 1 | 1 | 1 | 1 | 1 | 1 | 1 | 1 | 1 | 1 | 1 | 0.989 | 0.002 |
| SE | 98587 | ENSG00000149269 | PAK1 | chr11 | - | 77349238 | 77349287 | 77343818 | 77343931 | 77358897 | 77358953 | 15 | 10 | 14 | 8 | 12 | 0 | 0 | 1 | 0 | 0 | 19 | 8 | 10 | 28 | 28 | 0 | 0 | 0 | 4 | 0 | 198 | 149 | 1 | 1 | 1 | 1 | 0.913 | 1 | 1 | 1 | 1 | 1 | 0.84 | 1 | 0.015 |
| SE | 98588 | ENSG00000149269 | PAK1 | chr11 | - | 77353535 | 77353599 | 77349238 | 77349287 | 77358897 | 77359017 | 88 | 72 | 48 | 55 | 138 | 0 | 0 | 1 | 0 | 0 | 205 | 33 | 45 | 108 | 226 | 0 | 0 | 0 | 0 | 0 | 213 | 149 | 1 | 1 | 1 | 1 | 0.971 | 1 | 1 | 1 | 1 | 1 | 1 | 1 | -0.006 |
| SE | 98589 | ENSG00000149269 | PAK1 | chr11 | - | 77355667 | 77355842 | 77353535 | 77353599 | 77358897 | 77359017 | 222 | 200 | 206 | 211 | 581 | 0 | 0 | 0 | 2 | 0 | 707 | 209 | 194 | 479 | 627 | 0 | 0 | 0 | 0 | 0 | 298 | 149 | 1 | 1 | 1 | 1 | 1 | 0.981 | 1 | 1 | 1 | 1 | 1 | 1 | -0.004 |
| SE | 98590 | ENSG00000149269 | PAK1 | chr11 | - | 77392330 | 77392538 | 77379893 | 77379994 | 77411773 | 77411944 | 124 | 128 | 102 | 124 | 178 | 0 | 0 | 0 | 0 | 0 | 496 | 163 | 122 | 302 | 320 | 0 | 0 | 0 | 1 | 7 | 298 | 149 | 1 | 1 | 1 | 1 | 1 | 1 | 1 | 1 | 1 | 1 | 0.993 | 0.958 | 0.01 |
| SE | 98591 | ENSG00000149269 | PAK1 | chr11 | - | 77392330 | 77392541 | 77379893 | 77379994 | 77411773 | 77412064 | 162 | 158 | 129 | 138 | 213 | 0 | 0 | 0 | 0 | 0 | 576 | 197 | 152 | 389 | 414 | 0 | 0 | 0 | 1 | 7 | 298 | 149 | 1 | 1 | 1 | 1 | 1 | 1 | 1 | 1 | 1 | 1 | 0.995 | 0.967 | 0.008 |
| SE | 98595 | ENSG00000149269 | PAK1 | chr11 | - | 77408153 | 77408232 | 77392330 | 77392538 | 77411773 | 77411933 | 0 | 0 | 1 | 0 | 3 | 6 | 4 | 1 | 3 | 5 | 0 | 0 | 0 | 0 | 10 | 6 | 7 | 4 | 0 | 0 | 228 | 149 | 1 | 1 | 0 | 0 | 0.395 | 0 | 0.282 | 0 | 0 | 0 | NA | 1 | -0.115 |
| SE | 98597 | ENSG00000149269 | PAK1 | chr11 | - | 77455880 | 77455937 | 77392330 | 77392541 | 77473551 | 77473735 | 0 | 0 | 0 | 0 | 0 | 1 | 3 | 6 | 2 | 7 | 0 | 0 | 1 | 0 | 0 | 0 | 0 | 0 | 12 | 0 | 206 | 149 | 1 | 1 | 0 | 0 | 0 | 0 | 0 | NA | NA | 1 | 0 | NA | -0.5 |
| SE | 98598 | ENSG00000136527 | TRA2B | chr3 | - | 185919436 | 185919496 | 185918364 | 185918438 | 185921103 | 185921187 | 210 | 192 | 214 | 276 | 271 | 4 | 1 | 0 | 3 | 3 | 242 | 365 | 247 | 327 | 129 | 0 | 0 | 0 | 0 | 0 | 209 | 149 | 1 | 1 | 0.974 | 0.993 | 1 | 0.985 | 0.985 | 1 | 1 | 1 | 1 | 1 | -0.013 |
| SE | 98599 | ENSG00000136527 | TRA2B | chr3 | - | 185921103 | 185921187 | 185919436 | 185919496 | 185922010 | 185922107 | 715 | 600 | 709 | 906 | 956 | 1 | 0 | 0 | 0 | 0 | 891 | 1048 | 801 | 1162 | 451 | 0 | 0 | 0 | 0 | 0 | 233 | 149 | 1 | 1 | 0.998 | 1 | 1 | 1 | 1 | 1 | 1 | 1 | 1 | 1 | 0 |
| SE | 98600 | ENSG00000136527 | TRA2B | chr3 | - | 185922010 | 185922126 | 185921103 | 185921187 | 185923795 | 185923893 | 1549 | 1262 | 1418 | 1809 | 2172 | 0 | 1 | 0 | 0 | 0 | 2136 | 2176 | 1648 | 2562 | 1018 | 0 | 0 | 0 | 0 | 0 | 265 | 149 | 1 | 1 | 1 | 0.999 | 1 | 1 | 1 | 1 | 1 | 1 | 1 | 1 | 0 |
| SE | 98603 | ENSG00000136527 | TRA2B | chr3 | - | 185925463 | 185925626 | 185923795 | 185923984 | 185931576 | 185931852 | 699 | 445 | 702 | 772 | 774 | 0 | 0 | 0 | 2 | 0 | 739 | 861 | 680 | 923 | 476 | 0 | 0 | 0 | 0 | 0 | 298 | 149 | 1 | 1 | 1 | 1 | 1 | 0.995 | 1 | 1 | 1 | 1 | 1 | 1 | -0.001 |
| SE | 98608 | ENSG00000136527 | TRA2B | chr3 | - | 185934428 | 185934505 | 185931576 | 185931852 | 185936237 | 185936600 | 3 | 3 | 3 | 2 | 5 | 0 | 0 | 0 | 0 | 1 | 0 | 3 | 6 | 0 | 0 | 0 | 0 | 0 | 0 | 0 | 226 | 149 | 1 | 1 | 1 | 1 | 1 | 1 | 0.767 | NA | 1 | 1 | NA | NA | -0.047 |
| SE | 98609 | ENSG00000049449 | RCN1 | chr11 | + | 32097143 | 32097337 | 32091073 | 32091450 | 32098349 | 32098458 | 139 | 91 | 112 | 117 | 149 | 0 | 1 | 2 | 3 | 0 | 283 | 79 | 73 | 100 | 101 | 0 | 0 | 0 | 1 | 0 | 298 | 149 | 1 | 1 | 1 | 0.978 | 0.966 | 0.951 | 1 | 1 | 1 | 1 | 0.98 | 1 | -0.017 |
| SE | 98614 | ENSG00000137710 | RDX | chr11 | - | 110247702 | 110247833 | 110237491 | 110237652 | 110253945 | 110254109 | 199 | 220 | 152 | 223 | 376 | 2 | 0 | 0 | 0 | 0 | 37 | 162 | 180 | 58 | 29 | 0 | 0 | 0 | 0 | 0 | 280 | 149 | 1 | 1 | 0.981 | 1 | 1 | 1 | 1 | 1 | 1 | 1 | 1 | 1 | -0.004 |
| SE | 98617 | ENSG00000137710 | RDX | chr11 | - | 110264778 | 110264874 | 110263959 | 110264234 | 110272535 | 110272619 | 43 | 40 | 28 | 45 | 56 | 2 | 0 | 1 | 0 | 0 | 2 | 25 | 27 | 7 | 2 | 0 | 1 | 0 | 0 | 0 | 245 | 149 | 1 | 1 | 0.929 | 1 | 0.945 | 1 | 1 | 1 | 0.938 | 1 | 1 | 1 | -0.013 |
| SE | 98626 | ENSG00000151498 | ACAD8 | chr11 | + | 134256547 | 134256648 | 134253557 | 134253709 | 134257087 | 134257257 | 19 | 11 | 10 | 23 | 4 | 0 | 0 | 0 | 1 | 0 | 36 | 43 | 7 | 34 | 46 | 0 | 0 | 0 | 0 | 0 | 250 | 149 | 1 | 1 | 1 | 1 | 1 | 0.932 | 1 | 1 | 1 | 1 | 1 | 1 | -0.014 |
| SE | 98631 | ENSG00000151498 | ACAD8 | chr11 | + | 134258514 | 134258624 | 134253568 | 134253709 | 134259007 | 134259084 | 49 | 21 | 29 | 40 | 16 | 1 | 0 | 0 | 0 | 0 | 94 | 36 | 26 | 40 | 88 | 0 | 0 | 0 | 0 | 0 | 259 | 149 | 1 | 1 | 0.966 | 1 | 1 | 1 | 1 | 1 | 1 | 1 | 1 | 1 | -0.007 |
| SE | 98632 | ENSG00000151498 | ACAD8 | chr11 | + | 134258514 | 134258624 | 134257087 | 134257257 | 134259007 | 134259084 | 75 | 33 | 55 | 62 | 35 | 0 | 0 | 0 | 2 | 0 | 132 | 56 | 53 | 62 | 142 | 0 | 0 | 0 | 0 | 0 | 259 | 149 | 1 | 1 | 1 | 1 | 1 | 0.947 | 1 | 1 | 1 | 1 | 1 | 1 | -0.011 |
| SE | 98634 | ENSG00000149179 | C11orf49 | chr11 | + | 46948025 | 46948164 | 46936778 | 46936851 | 46987207 | 46987310 | 0 | 0 | 2 | 0 | 0 | 10 | 16 | 7 | 12 | 7 | 0 | 0 | 1 | 0 | 0 | 18 | 15 | 3 | 5 | 23 | 288 | 149 | 1 | 1 | 0 | 0 | 0.129 | 0 | 0 | 0 | 0 | 0.147 | 0 | 0 | -0.004 |
| SE | 98635 | ENSG00000149179 | C11orf49 | chr11 | + | 46987207 | 46987310 | 46936696 | 46936851 | 47052387 | 47052518 | 20 | 26 | 14 | 13 | 21 | 1 | 0 | 0 | 0 | 0 | 35 | 21 | 9 | 13 | 28 | 0 | 0 | 0 | 1 | 3 | 252 | 149 | 1 | 1 | 0.922 | 1 | 1 | 1 | 1 | 1 | 1 | 1 | 0.885 | 0.847 | 0.038 |
| SE | 98639 | ENSG00000149179 | C11orf49 | chr11 | + | 47052387 | 47052518 | 46987207 | 46987310 | 47137644 | 47137729 | 20 | 19 | 21 | 24 | 39 | 0 | 0 | 1 | 0 | 0 | 17 | 18 | 17 | 24 | 33 | 0 | 0 | 0 | 0 | 0 | 280 | 149 | 1 | 1 | 1 | 1 | 0.918 | 1 | 1 | 1 | 1 | 1 | 1 | 1 | -0.016 |
| SE | 98645 | ENSG00000107036 | RIC1 | chr9 | + | 5689958 | 5690038 | 5656582 | 5656690 | 5713895 | 5714003 | 112 | 96 | 102 | 100 | 116 | 5 | 0 | 0 | 0 | 0 | 110 | 167 | 91 | 95 | 12 | 0 | 0 | 2 | 3 | 0 | 229 | 149 | 1 | 1 | 0.936 | 1 | 1 | 1 | 1 | 1 | 1 | 0.967 | 0.954 | 1 | 0.003 |
| SE | 98650 | ENSG00000107036 | RIC1 | chr9 | + | 5742868 | 5743013 | 5738449 | 5738538 | 5743688 | 5743737 | 215 | 162 | 188 | 212 | 317 | 0 | 0 | 0 | 0 | 0 | 118 | 200 | 167 | 166 | 62 | 0 | 5 | 0 | 2 | 0 | 294 | 149 | 1 | 1 | 1 | 1 | 1 | 1 | 1 | 1 | 0.953 | 1 | 0.977 | 1 | 0.014 |
| SE | 98652 | ENSG00000107036 | RIC1 | chr9 | + | 5748584 | 5748714 | 5747301 | 5747505 | 5753199 | 5753238 | 2 | 3 | 3 | 1 | 2 | 157 | 126 | 142 | 165 | 178 | 0 | 5 | 5 | 0 | 0 | 121 | 144 | 144 | 171 | 31 | 279 | 149 | 1 | 1 | 0.007 | 0.013 | 0.011 | 0.003 | 0.006 | 0 | 0.018 | 0.018 | 0 | 0 | 0.001 |
| SE | 98653 | ENSG00000107036 | RIC1 | chr9 | + | 5748584 | 5748714 | 5747301 | 5747505 | 5754840 | 5754930 | 1 | 2 | 2 | 1 | 2 | 0 | 0 | 0 | 0 | 0 | 0 | 5 | 4 | 0 | 0 | 0 | 0 | 3 | 0 | 0 | 279 | 149 | 1 | 1 | 1 | 1 | 1 | 1 | 1 | NA | 1 | 0.416 | NA | NA | 0.292 |
| SE | 98654 | ENSG00000107036 | RIC1 | chr9 | + | 5753199 | 5753238 | 5747301 | 5747505 | 5753535 | 5753646 | 294 | 235 | 241 | 293 | 290 | 0 | 0 | 0 | 0 | 1 | 254 | 271 | 249 | 309 | 66 | 0 | 0 | 0 | 0 | 0 | 188 | 149 | 1 | 1 | 1 | 1 | 1 | 1 | 0.996 | 1 | 1 | 1 | 1 | 1 | -0.001 |
| SE | 98655 | ENSG00000107036 | RIC1 | chr9 | + | 5753199 | 5753238 | 5747301 | 5747505 | 5754840 | 5754930 | 56 | 50 | 53 | 77 | 85 | 0 | 0 | 0 | 0 | 0 | 74 | 75 | 79 | 52 | 8 | 0 | 0 | 3 | 0 | 0 | 188 | 149 | 1 | 1 | 1 | 1 | 1 | 1 | 1 | 1 | 1 | 0.954 | 1 | 1 | 0.009 |
| SE | 98657 | ENSG00000107036 | RIC1 | chr9 | + | 5753535 | 5753646 | 5747301 | 5747505 | 5754840 | 5754930 | 27 | 13 | 13 | 20 | 36 | 0 | 0 | 0 | 0 | 0 | 11 | 24 | 9 | 9 | 0 | 0 | 0 | 3 | 0 | 0 | 260 | 149 | 1 | 1 | 1 | 1 | 1 | 1 | 1 | 1 | 1 | 0.632 | 1 | NA | 0.092 |
| SE | 98659 | ENSG00000107036 | RIC1 | chr9 | + | 5754840 | 5754930 | 5753535 | 5753646 | 5756211 | 5756372 | 224 | 141 | 155 | 163 | 229 | 0 | 0 | 0 | 0 | 0 | 154 | 167 | 152 | 121 | 37 | 0 | 0 | 1 | 0 | 0 | 239 | 149 | 1 | 1 | 1 | 1 | 1 | 1 | 1 | 1 | 1 | 0.99 | 1 | 1 | 0.002 |
| SE | 98664 | ENSG00000107036 | RIC1 | chr9 | + | 5765661 | 5765798 | 5765413 | 5765572 | 5768969 | 5769106 | 122 | 95 | 104 | 160 | 191 | 0 | 0 | 0 | 1 | 0 | 91 | 106 | 89 | 69 | 41 | 0 | 0 | 0 | 0 | 0 | 286 | 149 | 1 | 1 | 1 | 1 | 1 | 0.988 | 1 | 1 | 1 | 1 | 1 | 1 | -0.002 |
| SE | 98667 | ENSG00000130592 | LSP1 | chr11 | + | 1880086 | 1880224 | 1853083 | 1853197 | 1883931 | 1883997 | 981 | 562 | 647 | 769 | 329 | 0 | 0 | 0 | 0 | 0 | 1645 | 997 | 878 | 1322 | 2441 | 0 | 0 | 2 | 0 | 0 | 287 | 149 | 1 | 1 | 1 | 1 | 1 | 1 | 1 | 1 | 1 | 0.996 | 1 | 1 | 0.001 |
| SE | 98671 | ENSG00000130592 | LSP1 | chr11 | + | 1883418 | 1883560 | 1853083 | 1853197 | 1883931 | 1883997 | 1438 | 646 | 902 | 842 | 1079 | 0 | 0 | 0 | 0 | 0 | 1105 | 839 | 706 | 909 | 1669 | 0 | 0 | 2 | 0 | 0 | 291 | 149 | 1 | 1 | 1 | 1 | 1 | 1 | 1 | 1 | 1 | 0.994 | 1 | 1 | 0.001 |
| SE | 98672 | ENSG00000130592 | LSP1 | chr11 | + | 1883418 | 1883560 | 1880086 | 1880224 | 1883931 | 1883997 | 1453 | 651 | 909 | 861 | 1085 | 0 | 0 | 2 | 0 | 0 | 1113 | 874 | 717 | 948 | 1738 | 0 | 0 | 0 | 0 | 0 | 291 | 149 | 1 | 1 | 1 | 1 | 0.996 | 1 | 1 | 1 | 1 | 1 | 1 | 1 | -0.001 |
| SE | 98673 | ENSG00000130592 | LSP1 | chr11 | + | 1884279 | 1884323 | 1883931 | 1884024 | 1884499 | 1884581 | 1612 | 932 | 1145 | 1258 | 1706 | 0 | 4 | 1 | 0 | 2 | 1703 | 1335 | 983 | 1388 | 2485 | 0 | 0 | 1 | 0 | 0 | 193 | 149 | 1 | 1 | 1 | 0.994 | 0.999 | 1 | 0.998 | 1 | 1 | 0.999 | 1 | 1 | -0.002 |
| SE | 98674 | ENSG00000130592 | LSP1 | chr11 | + | 1887236 | 1887314 | 1886731 | 1886866 | 1887473 | 1887502 | 2348 | 1190 | 1635 | 1605 | 2585 | 1 | 0 | 0 | 1 | 7 | 1249 | 1982 | 1515 | 1613 | 2559 | 0 | 0 | 3 | 1 | 9 | 227 | 149 | 1 | 1 | 0.999 | 1 | 1 | 0.999 | 0.996 | 1 | 1 | 0.997 | 0.999 | 0.995 | 0.001 |
| SE | 98675 | ENSG00000130592 | LSP1 | chr11 | + | 1887473 | 1887576 | 1887236 | 1887314 | 1891772 | 1892263 | 3535 | 1877 | 2352 | 2240 | 5154 | 0 | 0 | 3 | 0 | 0 | 1449 | 2699 | 2093 | 2343 | 3262 | 0 | 0 | 0 | 0 | 0 | 252 | 149 | 1 | 1 | 1 | 1 | 0.998 | 1 | 1 | 1 | 1 | 1 | 1 | 1 | 0 |
| SE | 98677 | ENSG00000205531 | NAP1L4 | chr11 | - | 2950968 | 2951016 | 2949226 | 2949264 | 2951258 | 2951295 | 0 | 0 | 2 | 0 | 1 | 96 | 47 | 93 | 96 | 150 | 0 | 3 | 4 | 0 | 0 | 58 | 44 | 56 | 72 | 77 | 197 | 149 | 1 | 1 | 0 | 0 | 0.016 | 0 | 0.005 | 0 | 0.049 | 0.051 | 0 | 0 | -0.016 |
| SE | 98679 | ENSG00000205531 | NAP1L4 | chr11 | - | 2958398 | 2958544 | 2955743 | 2955766 | 2959769 | 2959909 | 1111 | 849 | 1114 | 1168 | 1598 | 0 | 0 | 0 | 0 | 0 | 1068 | 956 | 626 | 654 | 1384 | 0 | 0 | 0 | 0 | 3 | 295 | 149 | 1 | 1 | 1 | 1 | 1 | 1 | 1 | 1 | 1 | 1 | 1 | 0.996 | 0.001 |
| SE | 98680 | ENSG00000205531 | NAP1L4 | chr11 | - | 2958398 | 2958544 | 2955743 | 2955766 | 2964679 | 2964712 | 441 | 364 | 387 | 421 | 669 | 0 | 0 | 0 | 0 | 3 | 529 | 328 | 202 | 267 | 665 | 0 | 0 | 0 | 0 | 0 | 295 | 149 | 1 | 1 | 1 | 1 | 1 | 1 | 0.991 | 1 | 1 | 1 | 1 | 1 | -0.002 |
| SE | 98681 | ENSG00000205531 | NAP1L4 | chr11 | - | 2959769 | 2959909 | 2955743 | 2955766 | 2964679 | 2964712 | 551 | 372 | 556 | 558 | 1004 | 0 | 0 | 0 | 0 | 3 | 327 | 394 | 345 | 313 | 516 | 0 | 0 | 0 | 0 | 0 | 289 | 149 | 1 | 1 | 1 | 1 | 1 | 1 | 0.994 | 1 | 1 | 1 | 1 | 1 | -0.001 |
| SE | 98682 | ENSG00000205531 | NAP1L4 | chr11 | - | 2959769 | 2959909 | 2958398 | 2958544 | 2964679 | 2964712 | 1219 | 855 | 1280 | 1305 | 1933 | 0 | 2 | 0 | 0 | 0 | 866 | 1024 | 770 | 700 | 1226 | 0 | 0 | 0 | 0 | 0 | 289 | 149 | 1 | 1 | 1 | 0.995 | 1 | 1 | 1 | 1 | 1 | 1 | 1 | 1 | -0.001 |
| SE | 98686 | ENSG00000205531 | NAP1L4 | chr11 | - | 2983745 | 2983813 | 2979206 | 2979237 | 2988076 | 2988126 | 4 | 0 | 0 | 0 | 0 | 1 | 1 | 1 | 4 | 2 | 0 | 0 | 0 | 0 | 0 | 0 | 10 | 9 | 3 | 0 | 217 | 149 | 1 | 1 | 0.733 | 0 | 0 | 0 | 0 | NA | 0 | 0 | 0 | NA | 0.147 |
| SE | 98688 | ENSG00000205531 | NAP1L4 | chr11 | - | 2988076 | 2988126 | 2979206 | 2979237 | 2989128 | 2989197 | 5 | 2 | 5 | 5 | 4 | 10 | 8 | 8 | 11 | 13 | 0 | 10 | 9 | 3 | 0 | 0 | 12 | 4 | 12 | 3 | 199 | 149 | 1 | 1 | 0.272 | 0.158 | 0.319 | 0.254 | 0.187 | NA | 0.384 | 0.628 | 0.158 | 0 | -0.054 |
| SE | 98690 | ENSG00000205531 | NAP1L4 | chr11 | - | 2989128 | 2989271 | 2979206 | 2979237 | 2990908 | 2990992 | 10 | 3 | 5 | 10 | 11 | 4 | 2 | 5 | 8 | 3 | 0 | 12 | 3 | 12 | 5 | 0 | 0 | 0 | 3 | 25 | 292 | 149 | 1 | 1 | 0.561 | 0.434 | 0.338 | 0.389 | 0.652 | NA | 1 | 1 | 0.671 | 0.093 | -0.216 |
| SE | 98692 | ENSG00000205531 | NAP1L4 | chr11 | - | 2989128 | 2989271 | 2979206 | 2979237 | 2992228 | 2992258 | 10 | 4 | 4 | 10 | 11 | 2 | 1 | 0 | 0 | 0 | 0 | 12 | 3 | 12 | 3 | 0 | 3 | 0 | 0 | 0 | 292 | 149 | 1 | 1 | 0.718 | 0.671 | 1 | 1 | 1 | NA | 0.671 | 1 | 1 | 1 | -0.04 |
| SE | 98694 | ENSG00000205531 | NAP1L4 | chr11 | - | 2990908 | 2990992 | 2979206 | 2979237 | 2992228 | 2992259 | 2 | 0 | 4 | 2 | 2 | 2 | 1 | 0 | 0 | 0 | 0 | 0 | 0 | 3 | 6 | 0 | 3 | 0 | 0 | 0 | 233 | 149 | 1 | 1 | 0.39 | 0 | 1 | 1 | 1 | NA | 0 | NA | 1 | 1 | 0.011 |
| SE | 98696 | ENSG00000205531 | NAP1L4 | chr11 | - | 2990908 | 2990992 | 2989128 | 2989271 | 2992228 | 2992259 | 0 | 0 | 0 | 0 | 0 | 0 | 1 | 0 | 0 | 0 | 0 | 0 | 0 | 0 | 2 | 0 | 0 | 0 | 0 | 0 | 233 | 149 | 1 | 1 | NA | 0 | NA | NA | NA | NA | NA | NA | NA | 1 | -1 |
| SE | 98697 | ENSG00000123810 | B9D2 | chr19 | - | 41357896 | 41358022 | 41354416 | 41355013 | 41363957 | 41364149 | 1 | 2 | 0 | 3 | 3 | 0 | 0 | 0 | 0 | 0 | 0 | 0 | 3 | 0 | 6 | 0 | 0 | 0 | 1 | 0 | 275 | 149 | 1 | 1 | 1 | 1 | NA | 1 | 1 | NA | NA | 1 | 0 | 1 | 0.333 |
| SE | 98698 | ENSG00000123810 | B9D2 | chr19 | - | 41362982 | 41363077 | 41357896 | 41358022 | 41363431 | 41363523 | 0 | 1 | 0 | 2 | 0 | 9 | 5 | 3 | 13 | 1 | 0 | 0 | 0 | 0 | 0 | 0 | 23 | 3 | 14 | 11 | 244 | 149 | 1 | 1 | 0 | 0.109 | 0 | 0.086 | 0 | NA | 0 | 0 | 0 | 0 | 0.039 |
| SE | 98699 | ENSG00000123810 | B9D2 | chr19 | - | 41363431 | 41363523 | 41357896 | 41358022 | 41363957 | 41364092 | 14 | 13 | 10 | 14 | 7 | 0 | 0 | 0 | 1 | 0 | 13 | 35 | 9 | 33 | 21 | 0 | 0 | 0 | 0 | 0 | 241 | 149 | 1 | 1 | 1 | 1 | 1 | 0.896 | 1 | 1 | 1 | 1 | 1 | 1 | -0.021 |
| SE | 98700 | ENSG00000140299 | BNIP2 | chr15 | - | 59665349 | 59665442 | 59662892 | 59664120 | 59668102 | 59668138 | 2 | 1 | 0 | 3 | 0 | 83 | 44 | 41 | 67 | 144 | 0 | 0 | 0 | 0 | 0 | 11 | 25 | 26 | 13 | 3 | 242 | 149 | 1 | 1 | 0.015 | 0.014 | 0 | 0.027 | 0 | 0 | 0 | 0 | 0 | 0 | 0.011 |
| SE | 98701 | ENSG00000140299 | BNIP2 | chr15 | - | 59665349 | 59665442 | 59662892 | 59664120 | 59668891 | 59668949 | 0 | 6 | 0 | 2 | 4 | 138 | 119 | 120 | 171 | 287 | 0 | 0 | 0 | 0 | 6 | 0 | 49 | 82 | 58 | 16 | 242 | 149 | 1 | 1 | 0 | 0.03 | 0 | 0.007 | 0.009 | NA | 0 | 0 | 0 | 0.188 | -0.038 |
| SE | 98703 | ENSG00000140299 | BNIP2 | chr15 | - | 59668102 | 59668138 | 59665349 | 59665442 | 59668891 | 59668949 | 22 | 13 | 6 | 23 | 56 | 0 | 5 | 0 | 1 | 5 | 0 | 4 | 5 | 3 | 0 | 0 | 0 | 0 | 0 | 6 | 185 | 149 | 1 | 1 | 1 | 0.677 | 1 | 0.949 | 0.9 | NA | 1 | 1 | 1 | 0 | 0.155 |
| SE | 98704 | ENSG00000140299 | BNIP2 | chr15 | - | 59668891 | 59668990 | 59659145 | 59664120 | 59671182 | 59671314 | 84 | 84 | 74 | 109 | 115 | 0 | 0 | 0 | 0 | 2 | 0 | 38 | 51 | 43 | 16 | 0 | 0 | 0 | 0 | 0 | 248 | 149 | 1 | 1 | 1 | 1 | 1 | 1 | 0.972 | NA | 1 | 1 | 1 | 1 | -0.006 |
| SE | 98706 | ENSG00000140299 | BNIP2 | chr15 | - | 59671182 | 59671314 | 59669275 | 59669362 | 59672636 | 59672739 | 701 | 527 | 591 | 730 | 1565 | 1 | 0 | 2 | 1 | 1 | 100 | 364 | 457 | 300 | 118 | 0 | 0 | 0 | 1 | 0 | 281 | 149 | 1 | 1 | 0.997 | 1 | 0.994 | 0.997 | 0.999 | 1 | 1 | 1 | 0.994 | 1 | -0.001 |
| SE | 98707 | ENSG00000140299 | BNIP2 | chr15 | - | 59672636 | 59672739 | 59671180 | 59671314 | 59677910 | 59678016 | 690 | 502 | 586 | 773 | 1264 | 0 | 0 | 0 | 1 | 0 | 147 | 439 | 473 | 283 | 120 | 0 | 0 | 0 | 0 | 0 | 252 | 149 | 1 | 1 | 1 | 1 | 1 | 0.998 | 1 | 1 | 1 | 1 | 1 | 1 | 0 |
| SE | 98708 | ENSG00000140299 | BNIP2 | chr15 | - | 59678747 | 59678834 | 59677910 | 59678087 | 59679591 | 59679768 | 0 | 0 | 1 | 3 | 0 | 437 | 453 | 340 | 439 | 681 | 0 | 0 | 0 | 0 | 0 | 155 | 355 | 445 | 296 | 179 | 236 | 149 | 1 | 1 | 0 | 0 | 0.002 | 0.004 | 0 | 0 | 0 | 0 | 0 | 0 | 0.001 |
| SE | 98713 | ENSG00000140299 | BNIP2 | chr15 | - | 59680240 | 59680308 | 59679255 | 59679768 | 59682407 | 59682502 | 1204 | 1464 | 939 | 1320 | 909 | 0 | 0 | 0 | 1 | 3 | 835 | 1209 | 1785 | 1252 | 437 | 0 | 0 | 0 | 0 | 0 | 217 | 149 | 1 | 1 | 1 | 1 | 1 | 0.999 | 0.995 | 1 | 1 | 1 | 1 | 1 | -0.001 |
| SE | 98714 | ENSG00000140299 | BNIP2 | chr15 | - | 59682407 | 59682514 | 59680240 | 59680308 | 59689134 | 59689309 | 527 | 701 | 426 | 579 | 389 | 0 | 0 | 0 | 3 | 0 | 307 | 548 | 859 | 560 | 174 | 0 | 0 | 1 | 0 | 0 | 256 | 149 | 1 | 1 | 1 | 1 | 1 | 0.991 | 1 | 1 | 1 | 0.998 | 1 | 1 | -0.001 |
| SE | 98719 | ENSG00000107290 | SETX | chr9 | - | 132269402 | 132269489 | 132266126 | 132266249 | 132269614 | 132269702 | 71 | 37 | 38 | 90 | 105 | 6 | 5 | 1 | 0 | 5 | 23 | 34 | 41 | 29 | 0 | 0 | 0 | 0 | 1 | 0 | 236 | 149 | 1 | 1 | 0.882 | 0.824 | 0.96 | 1 | 0.93 | 1 | 1 | 1 | 0.948 | NA | -0.068 |
| SE | 98721 | ENSG00000107290 | SETX | chr9 | - | 132269614 | 132269702 | 132269402 | 132269489 | 132271709 | 132271808 | 301 | 238 | 202 | 283 | 467 | 1 | 0 | 0 | 0 | 0 | 94 | 203 | 131 | 82 | 7 | 0 | 0 | 0 | 0 | 0 | 237 | 149 | 1 | 1 | 0.995 | 1 | 1 | 1 | 1 | 1 | 1 | 1 | 1 | 1 | -0.001 |
| SE | 98722 | ENSG00000107290 | SETX | chr9 | - | 132275255 | 132275420 | 132271709 | 132271808 | 132277059 | 132277152 | 840 | 658 | 506 | 767 | 1037 | 1 | 0 | 0 | 0 | 0 | 169 | 504 | 409 | 164 | 116 | 0 | 0 | 0 | 0 | 0 | 298 | 149 | 1 | 1 | 0.998 | 1 | 1 | 1 | 1 | 1 | 1 | 1 | 1 | 1 | 0 |
| SE | 98723 | ENSG00000107290 | SETX | chr9 | - | 132278069 | 132278257 | 132277059 | 132277152 | 132281466 | 132281574 | 799 | 629 | 589 | 796 | 743 | 4 | 1 | 0 | 5 | 0 | 124 | 564 | 483 | 210 | 95 | 0 | 0 | 0 | 0 | 0 | 298 | 149 | 1 | 1 | 0.99 | 0.997 | 1 | 0.988 | 1 | 1 | 1 | 1 | 1 | 1 | -0.005 |
| SE | 98725 | ENSG00000107290 | SETX | chr9 | - | 132286422 | 132286494 | 132281466 | 132281574 | 132288235 | 132288351 | 96 | 55 | 65 | 76 | 146 | 0 | 0 | 2 | 0 | 0 | 33 | 57 | 60 | 27 | 11 | 0 | 0 | 0 | 0 | 0 | 221 | 149 | 1 | 1 | 1 | 1 | 0.956 | 1 | 1 | 1 | 1 | 1 | 1 | 1 | -0.009 |
| SE | 98726 | ENSG00000107290 | SETX | chr9 | - | 132288235 | 132288351 | 132286422 | 132286494 | 132288549 | 132288651 | 571 | 444 | 388 | 556 | 842 | 0 | 0 | 0 | 1 | 0 | 153 | 366 | 328 | 109 | 30 | 0 | 1 | 0 | 0 | 0 | 265 | 149 | 1 | 1 | 1 | 1 | 1 | 0.997 | 1 | 1 | 0.995 | 1 | 1 | 1 | 0 |
| SE | 98727 | ENSG00000107290 | SETX | chr9 | - | 132295871 | 132296028 | 132288549 | 132288651 | 132296886 | 132297054 | 822 | 703 | 603 | 801 | 1268 | 0 | 0 | 0 | 2 | 2 | 169 | 509 | 545 | 134 | 48 | 0 | 0 | 0 | 0 | 0 | 298 | 149 | 1 | 1 | 1 | 1 | 1 | 0.995 | 0.997 | 1 | 1 | 1 | 1 | 1 | -0.002 |
| SE | 98728 | ENSG00000107290 | SETX | chr9 | - | 132300629 | 132300803 | 132298079 | 132298312 | 132311756 | 132311856 | 849 | 544 | 628 | 752 | 926 | 0 | 1 | 0 | 0 | 1 | 92 | 613 | 493 | 132 | 82 | 0 | 0 | 0 | 0 | 0 | 298 | 149 | 1 | 1 | 1 | 0.996 | 1 | 1 | 0.998 | 1 | 1 | 1 | 1 | 1 | -0.001 |
| SE | 98729 | ENSG00000107290 | SETX | chr9 | - | 132311756 | 132311856 | 132300629 | 132300803 | 132326323 | 132330499 | 857 | 592 | 566 | 760 | 659 | 0 | 4 | 0 | 0 | 0 | 84 | 573 | 517 | 101 | 28 | 0 | 0 | 0 | 3 | 0 | 249 | 149 | 1 | 1 | 1 | 0.989 | 1 | 1 | 1 | 1 | 1 | 1 | 0.953 | 1 | 0.007 |
| SE | 98732 | ENSG00000107290 | SETX | chr9 | - | 132336295 | 132336515 | 132334607 | 132334727 | 132342689 | 132342799 | 711 | 469 | 462 | 598 | 804 | 0 | 3 | 5 | 3 | 0 | 100 | 401 | 407 | 120 | 23 | 0 | 0 | 0 | 0 | 0 | 298 | 149 | 1 | 1 | 1 | 0.987 | 0.979 | 0.99 | 1 | 1 | 1 | 1 | 1 | 1 | -0.009 |
| SE | 98734 | ENSG00000107290 | SETX | chr9 | - | 132342689 | 132342799 | 132336295 | 132336515 | 132349251 | 132349435 | 85 | 64 | 64 | 92 | 119 | 0 | 0 | 0 | 1 | 0 | 8 | 49 | 45 | 16 | 6 | 0 | 0 | 0 | 0 | 0 | 259 | 149 | 1 | 1 | 1 | 1 | 1 | 0.981 | 1 | 1 | 1 | 1 | 1 | 1 | -0.004 |
| SE | 98735 | ENSG00000107290 | SETX | chr9 | - | 132346260 | 132346471 | 132336295 | 132336515 | 132349251 | 132349435 | 272 | 199 | 227 | 283 | 189 | 0 | 0 | 0 | 1 | 0 | 61 | 240 | 170 | 54 | 43 | 0 | 0 | 0 | 0 | 0 | 298 | 149 | 1 | 1 | 1 | 1 | 1 | 0.993 | 1 | 1 | 1 | 1 | 1 | 1 | -0.001 |
| SE | 98736 | ENSG00000107290 | SETX | chr9 | - | 132346260 | 132346471 | 132342689 | 132342799 | 132349251 | 132349435 | 580 | 381 | 422 | 591 | 464 | 1 | 1 | 0 | 1 | 0 | 134 | 405 | 362 | 75 | 57 | 0 | 0 | 0 | 0 | 0 | 298 | 149 | 1 | 1 | 0.997 | 0.995 | 1 | 0.997 | 1 | 1 | 1 | 1 | 1 | 1 | -0.002 |
| SE | 98737 | ENSG00000107290 | SETX | chr9 | - | 132349251 | 132349435 | 132346260 | 132346471 | 132353648 | 132353755 | 426 | 313 | 361 | 428 | 287 | 0 | 0 | 1 | 0 | 0 | 89 | 348 | 267 | 63 | 59 | 0 | 4 | 0 | 0 | 0 | 298 | 149 | 1 | 1 | 1 | 1 | 0.994 | 1 | 1 | 1 | 0.978 | 1 | 1 | 1 | 0.003 |
| SE | 98740 | ENSG00000110628 | SLC22A18 | chr11 | + | 2918905 | 2919008 | 2917987 | 2918095 | 2919306 | 2919407 | 0 | 1 | 0 | 0 | 0 | 3 | 4 | 2 | 2 | 0 | 0 | 0 | 0 | 0 | 0 | 0 | 5 | 0 | 6 | 10 | 252 | 149 | 1 | 1 | 0 | 0.129 | 0 | 0 | NA | NA | 0 | NA | 0 | 0 | 0.032 |
| SE | 98743 | ENSG00000246273 | SBF2-AS1 | chr11 | + | 9780375 | 9780575 | 9758295 | 9758503 | 9807847 | 9808038 | 0 | 0 | 0 | 1 | 0 | 3 | 1 | 1 | 0 | 1 | 0 | 0 | 0 | 0 | 0 | 4 | 0 | 0 | 0 | 0 | 298 | 149 | 1 | 1 | 0 | 0 | 0 | 1 | 0 | 0 | NA | NA | NA | NA | 0.2 |
| SE | 98745 | ENSG00000139211 | AMIGO2 | chr12 | - | 47079141 | 47079248 | 47076446 | 47079065 | 47079461 | 47079959 | 16 | 13 | 6 | 25 | 21 | 2 | 0 | 5 | 2 | 1 | 0 | 16 | 14 | 5 | 8 | 0 | 9 | 2 | 12 | 4 | 256 | 149 | 1 | 1 | 0.823 | 1 | 0.411 | 0.879 | 0.924 | NA | 0.509 | 0.803 | 0.195 | 0.538 | 0.296 |
| SE | 98746 | ENSG00000139211 | AMIGO2 | chr12 | - | 47079146 | 47079248 | 47076599 | 47079065 | 47079461 | 47079951 | 1 | 2 | 6 | 11 | 2 | 2 | 0 | 5 | 2 | 1 | 0 | 13 | 4 | 0 | 4 | 0 | 9 | 2 | 12 | 4 | 251 | 149 | 1 | 1 | 0.229 | 1 | 0.416 | 0.766 | 0.543 | NA | 0.462 | 0.543 | 0 | 0.372 | 0.247 |
| SE | 98748 | ENSG00000175166 | PSMD2 | chr3 | + | 184301770 | 184302071 | 184301536 | 184301658 | 184302369 | 184302422 | 109 | 59 | 63 | 87 | 82 | 0 | 0 | 0 | 1 | 0 | 143 | 104 | 44 | 79 | 162 | 0 | 0 | 3 | 0 | 0 | 298 | 149 | 1 | 1 | 1 | 1 | 1 | 0.978 | 1 | 1 | 1 | 0.88 | 1 | 1 | 0.02 |
| SE | 98749 | ENSG00000175166 | PSMD2 | chr3 | + | 184301793 | 184302071 | 184301536 | 184301658 | 184302369 | 184302528 | 109 | 57 | 61 | 80 | 81 | 0 | 0 | 0 | 1 | 0 | 138 | 97 | 43 | 75 | 162 | 0 | 0 | 3 | 0 | 0 | 298 | 149 | 1 | 1 | 1 | 1 | 1 | 0.976 | 1 | 1 | 1 | 0.878 | 1 | 1 | 0.02 |
| SE | 98751 | ENSG00000175166 | PSMD2 | chr3 | + | 184301846 | 184302071 | 184301536 | 184301658 | 184302369 | 184302528 | 400 | 271 | 336 | 388 | 389 | 0 | 0 | 0 | 1 | 0 | 682 | 551 | 303 | 383 | 834 | 0 | 0 | 3 | 0 | 0 | 298 | 149 | 1 | 1 | 1 | 1 | 1 | 0.995 | 1 | 1 | 1 | 0.981 | 1 | 1 | 0.003 |
| SE | 98752 | ENSG00000175166 | PSMD2 | chr3 | + | 184301846 | 184302071 | 184301536 | 184301658 | 184302678 | 184302823 | 291 | 214 | 275 | 308 | 308 | 0 | 0 | 0 | 0 | 0 | 548 | 455 | 260 | 308 | 672 | 4 | 0 | 1 | 0 | 0 | 298 | 149 | 1 | 1 | 1 | 1 | 1 | 1 | 1 | 0.986 | 1 | 0.992 | 1 | 1 | 0.004 |
| SE | 98753 | ENSG00000175166 | PSMD2 | chr3 | + | 184301857 | 184302071 | 184301536 | 184301658 | 184302369 | 184302528 | 109 | 57 | 61 | 80 | 81 | 0 | 0 | 0 | 1 | 0 | 138 | 97 | 43 | 75 | 162 | 0 | 0 | 3 | 0 | 0 | 298 | 149 | 1 | 1 | 1 | 1 | 1 | 0.976 | 1 | 1 | 1 | 0.878 | 1 | 1 | 0.02 |
| SE | 98755 | ENSG00000175166 | PSMD2 | chr3 | + | 184302369 | 184302528 | 184301536 | 184301658 | 184302678 | 184302692 | 115 | 72 | 110 | 124 | 152 | 0 | 0 | 0 | 0 | 0 | 211 | 172 | 115 | 163 | 262 | 4 | 0 | 1 | 0 | 0 | 298 | 149 | 1 | 1 | 1 | 1 | 1 | 1 | 1 | 0.963 | 1 | 0.983 | 1 | 1 | 0.011 |
| SE | 98756 | ENSG00000175166 | PSMD2 | chr3 | + | 184302369 | 184302528 | 184301721 | 184302071 | 184302678 | 184302692 | 224 | 129 | 171 | 203 | 233 | 0 | 0 | 0 | 0 | 0 | 349 | 269 | 155 | 238 | 424 | 4 | 1 | 0 | 0 | 0 | 298 | 149 | 1 | 1 | 1 | 1 | 1 | 1 | 1 | 0.978 | 0.993 | 1 | 1 | 1 | 0.006 |
| SE | 98757 | ENSG00000175166 | PSMD2 | chr3 | + | 184303946 | 184304074 | 184303642 | 184303749 | 184304303 | 184304391 | 245 | 142 | 168 | 178 | 253 | 0 | 0 | 0 | 0 | 0 | 249 | 220 | 146 | 132 | 383 | 0 | 0 | 0 | 0 | 7 | 277 | 149 | 1 | 1 | 1 | 1 | 1 | 1 | 1 | 1 | 1 | 1 | 1 | 0.967 | 0.007 |
| SE | 98760 | ENSG00000102781 | KATNAL1 | chr13 | - | 30240459 | 30240565 | 30231313 | 30231472 | 30240958 | 30241086 | 62 | 33 | 27 | 50 | 50 | 0 | 0 | 0 | 0 | 0 | 19 | 56 | 40 | 43 | 66 | 0 | 1 | 0 | 0 | 0 | 255 | 149 | 1 | 1 | 1 | 1 | 1 | 1 | 1 | 1 | 0.97 | 1 | 1 | 1 | 0.006 |
| SE | 98768 | ENSG00000157600 | TMEM164 | chrX | + | 110067346 | 110067396 | 110002634 | 110002772 | 110109079 | 110109146 | 31 | 5 | 17 | 27 | 8 | 0 | 0 | 0 | 0 | 0 | 44 | 22 | 4 | 22 | 92 | 0 | 3 | 0 | 2 | 0 | 199 | 149 | 1 | 1 | 1 | 1 | 1 | 1 | 1 | 1 | 0.846 | 1 | 0.892 | 1 | 0.052 |
| SE | 98769 | ENSG00000157600 | TMEM164 | chrX | + | 110067346 | 110067396 | 110002634 | 110002772 | 110144797 | 110144876 | 6 | 2 | 0 | 1 | 1 | 0 | 0 | 0 | 0 | 0 | 0 | 0 | 0 | 1 | 16 | 0 | 0 | 0 | 0 | 5 | 199 | 149 | 1 | 1 | 1 | 1 | NA | 1 | 1 | NA | NA | NA | 1 | 0.706 | 0.147 |
| SE | 98771 | ENSG00000157600 | TMEM164 | chrX | + | 110067346 | 110067396 | 110003500 | 110004164 | 110144797 | 110144876 | 2 | 2 | 2 | 6 | 2 | 0 | 0 | 0 | 0 | 0 | 0 | 0 | 3 | 0 | 29 | 0 | 0 | 0 | 0 | 5 | 199 | 149 | 1 | 1 | 1 | 1 | 1 | 1 | 1 | NA | NA | 1 | NA | 0.813 | 0.094 |
| SE | 98772 | ENSG00000157600 | TMEM164 | chrX | + | 110109079 | 110109146 | 110002634 | 110002772 | 110144797 | 110144876 | 30 | 22 | 15 | 44 | 10 | 0 | 0 | 0 | 0 | 0 | 46 | 44 | 21 | 36 | 123 | 0 | 0 | 0 | 0 | 5 | 216 | 149 | 1 | 1 | 1 | 1 | 1 | 1 | 1 | 1 | 1 | 1 | 1 | 0.944 | 0.011 |
| SE | 98773 | ENSG00000157600 | TMEM164 | chrX | + | 110109079 | 110109146 | 110003500 | 110004164 | 110144797 | 110144876 | 30 | 23 | 16 | 46 | 10 | 0 | 0 | 0 | 0 | 0 | 46 | 41 | 21 | 34 | 123 | 0 | 0 | 0 | 0 | 5 | 216 | 149 | 1 | 1 | 1 | 1 | 1 | 1 | 1 | 1 | 1 | 1 | 1 | 0.944 | 0.011 |
| SE | 98775 | ENSG00000157600 | TMEM164 | chrX | + | 110144797 | 110144876 | 110003500 | 110004164 | 110171419 | 110171511 | 22 | 13 | 20 | 28 | 23 | 0 | 1 | 0 | 0 | 0 | 12 | 22 | 11 | 18 | 85 | 0 | 0 | 0 | 0 | 0 | 228 | 149 | 1 | 1 | 1 | 0.895 | 1 | 1 | 1 | 1 | 1 | 1 | 1 | 1 | -0.021 |
| SE | 98776 | ENSG00000157600 | TMEM164 | chrX | + | 110171419 | 110171520 | 110144797 | 110144876 | 110173244 | 110173453 | 69 | 47 | 43 | 103 | 69 | 0 | 0 | 0 | 1 | 0 | 68 | 80 | 38 | 69 | 330 | 0 | 0 | 0 | 0 | 0 | 250 | 149 | 1 | 1 | 1 | 1 | 1 | 0.984 | 1 | 1 | 1 | 1 | 1 | 1 | -0.003 |
| SE | 98777 | ENSG00000110315 | RNF141 | chr11 | - | 10519033 | 10519141 | 10511672 | 10515066 | 10525191 | 10525373 | 200 | 129 | 160 | 185 | 402 | 0 | 0 | 0 | 1 | 0 | 172 | 157 | 209 | 141 | 129 | 0 | 0 | 0 | 0 | 0 | 257 | 149 | 1 | 1 | 1 | 1 | 1 | 0.991 | 1 | 1 | 1 | 1 | 1 | 1 | -0.002 |
| SE | 98778 | ENSG00000110315 | RNF141 | chr11 | - | 10531826 | 10532015 | 10530642 | 10530751 | 10534015 | 10534205 | 3 | 0 | 0 | 2 | 0 | 241 | 187 | 188 | 261 | 257 | 0 | 2 | 2 | 0 | 0 | 119 | 186 | 296 | 247 | 172 | 298 | 149 | 1 | 1 | 0.006 | 0 | 0 | 0.004 | 0 | 0 | 0.005 | 0.003 | 0 | 0 | 0 |
| SE | 98796 | ENSG00000121671 | CRY2 | chr11 | + | 45855981 | 45856090 | 45847405 | 45847705 | 45860847 | 45861032 | 16 | 16 | 22 | 12 | 7 | 0 | 0 | 0 | 2 | 0 | 17 | 26 | 19 | 6 | 15 | 0 | 0 | 0 | 0 | 0 | 258 | 149 | 1 | 1 | 1 | 1 | 1 | 0.776 | 1 | 1 | 1 | 1 | 1 | 1 | -0.045 |
| SE | 98797 | ENSG00000121671 | CRY2 | chr11 | + | 45858730 | 45858873 | 45855981 | 45856090 | 45860847 | 45861032 | 139 | 61 | 132 | 105 | 143 | 1 | 0 | 0 | 0 | 0 | 242 | 169 | 170 | 132 | 137 | 0 | 0 | 3 | 0 | 0 | 292 | 149 | 1 | 1 | 0.986 | 1 | 1 | 1 | 1 | 1 | 1 | 0.967 | 1 | 1 | 0.004 |
| SE | 98798 | ENSG00000121671 | CRY2 | chr11 | + | 45860847 | 45861032 | 45858730 | 45858873 | 45862059 | 45862148 | 112 | 53 | 101 | 78 | 111 | 2 | 0 | 0 | 0 | 0 | 175 | 103 | 113 | 92 | 117 | 0 | 0 | 0 | 0 | 0 | 298 | 149 | 1 | 1 | 0.966 | 1 | 1 | 1 | 1 | 1 | 1 | 1 | 1 | 1 | -0.007 |
| SE | 98800 | ENSG00000121671 | CRY2 | chr11 | + | 45870841 | 45870934 | 45870329 | 45870532 | 45872091 | 45872233 | 142 | 68 | 155 | 109 | 119 | 0 | 0 | 1 | 1 | 0 | 99 | 86 | 77 | 71 | 121 | 0 | 2 | 0 | 0 | 0 | 242 | 149 | 1 | 1 | 1 | 1 | 0.99 | 0.985 | 1 | 1 | 0.964 | 1 | 1 | 1 | 0.002 |
| SE | 98802 | ENSG00000165905 | LARGE2 | chr11 | + | 45926038 | 45926151 | 45924784 | 45924889 | 45926221 | 45926347 | 6 | 0 | 6 | 3 | 1 | 0 | 0 | 0 | 0 | 0 | 7 | 6 | 0 | 0 | 11 | 0 | 8 | 0 | 0 | 0 | 262 | 149 | 1 | 1 | 1 | NA | 1 | 1 | 1 | 1 | 0.299 | NA | NA | 1 | 0.234 |
| SE | 98803 | ENSG00000165905 | LARGE2 | chr11 | + | 45928176 | 45928372 | 45927748 | 45928069 | 45928629 | 45929082 | 4 | 3 | 3 | 8 | 10 | 1 | 0 | 0 | 1 | 0 | 0 | 2 | 1 | 8 | 0 | 0 | 1 | 0 | 0 | 0 | 298 | 149 | 1 | 1 | 0.667 | 1 | 1 | 0.8 | 1 | NA | 0.5 | 1 | 1 | NA | 0.06 |
| SE | 98804 | ENSG00000065717 | TLE2 | chr19 | - | 3000646 | 3000723 | 2997638 | 2997955 | 3002352 | 3002503 | 17 | 9 | 24 | 19 | 13 | 0 | 0 | 0 | 2 | 0 | 5 | 1 | 3 | 3 | 5 | 0 | 0 | 0 | 0 | 0 | 226 | 149 | 1 | 1 | 1 | 1 | 1 | 0.862 | 1 | 1 | 1 | 1 | 1 | 1 | -0.028 |
| SE | 98805 | ENSG00000065717 | TLE2 | chr19 | - | 3017839 | 3017859 | 3015652 | 3015760 | 3019282 | 3019463 | 2 | 0 | 9 | 0 | 0 | 3 | 0 | 0 | 0 | 0 | 0 | 0 | 2 | 3 | 0 | 0 | 0 | 0 | 0 | 0 | 169 | 149 | 1 | 1 | 0.37 | NA | 1 | NA | NA | NA | NA | 1 | 1 | NA | -0.315 |
| SE | 98806 | ENSG00000065717 | TLE2 | chr19 | - | 3019698 | 3019773 | 3015652 | 3015760 | 3025019 | 3025027 | 3 | 4 | 4 | 0 | 1 | 0 | 0 | 0 | 0 | 0 | 0 | 0 | 0 | 2 | 0 | 0 | 1 | 0 | 0 | 0 | 224 | 149 | 1 | 1 | 1 | 1 | 1 | NA | 1 | NA | 0 | NA | 1 | NA | 0.5 |
| SE | 98812 | ENSG00000149091 | DGKZ | chr11 | + | 46371312 | 46371384 | 46368001 | 46370009 | 46371486 | 46371603 | 484 | 213 | 350 | 402 | 361 | 6 | 0 | 1 | 2 | 0 | 414 | 426 | 234 | 368 | 605 | 0 | 0 | 1 | 0 | 0 | 221 | 149 | 1 | 1 | 0.982 | 1 | 0.996 | 0.993 | 1 | 1 | 1 | 0.994 | 1 | 1 | -0.005 |
| SE | 98814 | ENSG00000149091 | DGKZ | chr11 | + | 46372770 | 46372884 | 46372427 | 46372510 | 46372960 | 46373101 | 115 | 48 | 111 | 126 | 53 | 1 | 0 | 0 | 0 | 0 | 140 | 168 | 46 | 44 | 250 | 0 | 2 | 0 | 0 | 0 | 263 | 149 | 1 | 1 | 0.985 | 1 | 1 | 1 | 1 | 1 | 0.979 | 1 | 1 | 1 | 0.001 |
| SE | 98816 | ENSG00000149091 | DGKZ | chr11 | + | 46377072 | 46377212 | 46376523 | 46376564 | 46378197 | 46378229 | 156 | 62 | 150 | 158 | 124 | 0 | 0 | 0 | 0 | 0 | 123 | 231 | 98 | 116 | 374 | 0 | 0 | 1 | 0 | 6 | 289 | 149 | 1 | 1 | 1 | 1 | 1 | 1 | 1 | 1 | 1 | 0.981 | 1 | 0.97 | 0.01 |
| SE | 98817 | ENSG00000149091 | DGKZ | chr11 | + | 46378456 | 46378500 | 46378197 | 46378229 | 46378990 | 46379111 | 105 | 40 | 118 | 139 | 65 | 0 | 0 | 0 | 0 | 0 | 116 | 182 | 101 | 104 | 263 | 0 | 0 | 0 | 2 | 0 | 193 | 149 | 1 | 1 | 1 | 1 | 1 | 1 | 1 | 1 | 1 | 1 | 0.976 | 1 | 0.005 |
| SE | 98819 | ENSG00000149091 | DGKZ | chr11 | + | 46378990 | 46379111 | 46378456 | 46378500 | 46379202 | 46379236 | 254 | 88 | 201 | 250 | 121 | 0 | 0 | 1 | 0 | 0 | 264 | 282 | 150 | 189 | 463 | 7 | 0 | 2 | 0 | 0 | 270 | 149 | 1 | 1 | 1 | 1 | 0.991 | 1 | 1 | 0.954 | 1 | 0.976 | 1 | 1 | 0.012 |
| SE | 98820 | ENSG00000149091 | DGKZ | chr11 | + | 46379202 | 46379236 | 46378456 | 46378500 | 46379468 | 46379568 | 39 | 30 | 39 | 28 | 24 | 0 | 0 | 0 | 0 | 0 | 19 | 17 | 13 | 16 | 48 | 0 | 0 | 1 | 0 | 0 | 183 | 149 | 1 | 1 | 1 | 1 | 1 | 1 | 1 | 1 | 1 | 0.914 | 1 | 1 | 0.017 |
| SE | 98823 | ENSG00000135365 | PHF21A | chr11 | - | 45936493 | 45936569 | 45929322 | 45935739 | 45938156 | 45938264 | 106 | 52 | 87 | 105 | 138 | 0 | 0 | 1 | 0 | 3 | 143 | 155 | 129 | 215 | 184 | 0 | 0 | 0 | 0 | 0 | 225 | 149 | 1 | 1 | 1 | 1 | 0.983 | 1 | 0.968 | 1 | 1 | 1 | 1 | 1 | -0.01 |
| SE | 98827 | ENSG00000135365 | PHF21A | chr11 | - | 45946075 | 45946098 | 45938156 | 45938312 | 45948885 | 45948946 | 1 | 1 | 2 | 6 | 17 | 3 | 4 | 2 | 3 | 5 | 16 | 3 | 0 | 5 | 0 | 0 | 3 | 0 | 2 | 0 | 172 | 149 | 1 | 1 | 0.224 | 0.178 | 0.464 | 0.634 | 0.747 | 1 | 0.464 | NA | 0.684 | NA | -0.267 |
| SE | 98829 | ENSG00000135365 | PHF21A | chr11 | - | 45969814 | 45969901 | 45965314 | 45965608 | 45971115 | 45971367 | 25 | 15 | 20 | 22 | 25 | 1 | 0 | 0 | 0 | 0 | 65 | 36 | 62 | 40 | 86 | 0 | 0 | 5 | 0 | 0 | 236 | 149 | 1 | 1 | 0.94 | 1 | 1 | 1 | 1 | 1 | 1 | 0.887 | 1 | 1 | 0.011 |
| SE | 98830 | ENSG00000135365 | PHF21A | chr11 | - | 45969814 | 45969904 | 45965314 | 45965608 | 45971115 | 45971367 | 77 | 66 | 76 | 100 | 122 | 1 | 0 | 0 | 0 | 0 | 282 | 129 | 207 | 205 | 159 | 0 | 0 | 5 | 0 | 0 | 239 | 149 | 1 | 1 | 0.98 | 1 | 1 | 1 | 1 | 1 | 1 | 0.963 | 1 | 1 | 0.003 |
| SE | 98833 | ENSG00000135365 | PHF21A | chr11 | - | 45979759 | 45979966 | 45971115 | 45971367 | 46076753 | 46076819 | 286 | 191 | 219 | 244 | 521 | 0 | 0 | 0 | 0 | 9 | 383 | 372 | 339 | 485 | 285 | 0 | 0 | 0 | 0 | 0 | 298 | 149 | 1 | 1 | 1 | 1 | 1 | 1 | 0.967 | 1 | 1 | 1 | 1 | 1 | -0.007 |
| SE | 98835 | ENSG00000135365 | PHF21A | chr11 | - | 46076753 | 46076819 | 45979759 | 45979966 | 46084165 | 46084302 | 75 | 68 | 73 | 77 | 179 | 0 | 0 | 0 | 2 | 0 | 152 | 129 | 110 | 168 | 37 | 0 | 0 | 0 | 0 | 0 | 215 | 149 | 1 | 1 | 1 | 1 | 1 | 0.964 | 1 | 1 | 1 | 1 | 1 | 1 | -0.007 |
| SE | 98836 | ENSG00000135365 | PHF21A | chr11 | - | 46079133 | 46079166 | 46076753 | 46076819 | 46084165 | 46084302 | 351 | 320 | 257 | 338 | 784 | 0 | 0 | 0 | 0 | 0 | 638 | 445 | 381 | 601 | 427 | 0 | 3 | 0 | 0 | 0 | 182 | 149 | 1 | 1 | 1 | 1 | 1 | 1 | 1 | 1 | 0.992 | 1 | 1 | 1 | 0.002 |
| SE | 98837 | ENSG00000135365 | PHF21A | chr11 | - | 46083554 | 46083611 | 46079133 | 46079166 | 46084165 | 46084302 | 0 | 5 | 3 | 0 | 0 | 284 | 262 | 206 | 263 | 684 | 7 | 0 | 0 | 0 | 8 | 525 | 342 | 317 | 470 | 327 | 206 | 149 | 1 | 1 | 0 | 0.014 | 0.01 | 0 | 0 | 0.01 | 0 | 0 | 0 | 0.017 | -0.001 |
| SE | 98844 | ENSG00000135365 | PHF21A | chr11 | - | 46118741 | 46118800 | 46092182 | 46092223 | 46119723 | 46119787 | 4 | 0 | 2 | 4 | 2 | 0 | 2 | 0 | 1 | 0 | 19 | 12 | 0 | 7 | 0 | 0 | 5 | 0 | 0 | 4 | 208 | 149 | 1 | 1 | 1 | 0 | 1 | 0.741 | 1 | 1 | 0.632 | NA | 1 | 0 | 0.09 |
| SE | 98846 | ENSG00000135365 | PHF21A | chr11 | - | 46118741 | 46118800 | 46117921 | 46118114 | 46119723 | 46119787 | 7 | 1 | 2 | 4 | 0 | 1 | 0 | 0 | 0 | 0 | 3 | 8 | 0 | 4 | 0 | 0 | 0 | 0 | 0 | 0 | 208 | 149 | 1 | 1 | 0.834 | 1 | 1 | 1 | NA | 1 | 1 | NA | 1 | NA | -0.041 |
| SE | 98849 | ENSG00000077458 | FAM76B | chr11 | - | 95779606 | 95779687 | 95778821 | 95778957 | 95779878 | 95779926 | 193 | 174 | 237 | 229 | 445 | 0 | 0 | 0 | 0 | 1 | 13 | 92 | 73 | 50 | 13 | 0 | 0 | 0 | 0 | 0 | 230 | 149 | 1 | 1 | 1 | 1 | 1 | 1 | 0.997 | 1 | 1 | 1 | 1 | 1 | -0.001 |
| SE | 98850 | ENSG00000077458 | FAM76B | chr11 | - | 95788858 | 95788947 | 95788498 | 95788563 | 95789391 | 95789768 | 2 | 0 | 0 | 2 | 0 | 47 | 33 | 43 | 50 | 27 | 0 | 0 | 0 | 0 | 0 | 4 | 13 | 4 | 3 | 0 | 238 | 149 | 1 | 1 | 0.026 | 0 | 0 | 0.024 | 0 | 0 | 0 | 0 | 0 | NA | 0.01 |
| SE | 98853 | ENSG00000175592 | FOSL1 | chr11 | - | 65894013 | 65894116 | 65892644 | 65893296 | 65896808 | 65897006 | 2 | 2 | 0 | 4 | 3 | 0 | 0 | 0 | 0 | 0 | 0 | 1 | 1 | 26 | 30 | 0 | 0 | 0 | 2 | 0 | 252 | 149 | 1 | 1 | 1 | 1 | NA | 1 | 1 | NA | 1 | 1 | 0.885 | 1 | 0.029 |
| SE | 98854 | ENSG00000175592 | FOSL1 | chr11 | - | 65894013 | 65894116 | 65892644 | 65893296 | 65900240 | 65900419 | 2 | 2 | 0 | 4 | 3 | 4 | 0 | 0 | 0 | 0 | 0 | 1 | 1 | 28 | 30 | 0 | 0 | 0 | 0 | 0 | 252 | 149 | 1 | 1 | 0.228 | 1 | NA | 1 | 1 | NA | 1 | 1 | 1 | 1 | -0.193 |
| SE | 98855 | ENSG00000175592 | FOSL1 | chr11 | - | 65894013 | 65894121 | 65892048 | 65893296 | 65896808 | 65897006 | 2 | 4 | 0 | 5 | 3 | 0 | 0 | 0 | 0 | 0 | 11 | 1 | 3 | 36 | 30 | 0 | 0 | 0 | 2 | 0 | 257 | 149 | 1 | 1 | 1 | 1 | NA | 1 | 1 | 1 | 1 | 1 | 0.913 | 1 | 0.017 |
| SE | 98856 | ENSG00000175592 | FOSL1 | chr11 | - | 65894013 | 65894121 | 65892048 | 65893296 | 65900240 | 65900388 | 2 | 2 | 0 | 5 | 3 | 4 | 0 | 0 | 0 | 0 | 0 | 1 | 1 | 27 | 30 | 0 | 0 | 0 | 0 | 0 | 257 | 149 | 1 | 1 | 0.225 | 1 | NA | 1 | 1 | NA | 1 | 1 | 1 | 1 | -0.194 |
| SE | 98857 | ENSG00000175592 | FOSL1 | chr11 | - | 65894013 | 65894121 | 65892048 | 65893296 | 65900390 | 65900573 | 2 | 2 | 0 | 5 | 3 | 0 | 0 | 0 | 0 | 0 | 0 | 1 | 1 | 26 | 30 | 0 | 0 | 0 | 4 | 0 | 257 | 149 | 1 | 1 | 1 | 1 | NA | 1 | 1 | NA | 1 | 1 | 0.79 | 1 | 0.052 |
| SE | 98858 | ENSG00000175592 | FOSL1 | chr11 | - | 65896808 | 65897006 | 65892473 | 65893296 | 65900240 | 65900388 | 0 | 0 | 0 | 0 | 0 | 4 | 0 | 0 | 0 | 0 | 11 | 4 | 3 | 6 | 0 | 0 | 0 | 0 | 0 | 0 | 298 | 149 | 1 | 1 | 0 | NA | NA | NA | NA | 1 | 1 | 1 | 1 | NA | -1 |
| SE | 98860 | ENSG00000175592 | FOSL1 | chr11 | - | 65896808 | 65897006 | 65894013 | 65894121 | 65900240 | 65900388 | 0 | 2 | 0 | 0 | 0 | 0 | 0 | 0 | 0 | 0 | 22 | 4 | 5 | 14 | 0 | 0 | 0 | 0 | 1 | 0 | 298 | 149 | 1 | 1 | NA | 1 | NA | NA | NA | 1 | 1 | 1 | 0.875 | NA | 0.031 |
| SE | 98869 | ENSG00000175294 | CATSPER1 | chr11 | - | 66020139 | 66020200 | 66018787 | 66018902 | 66020810 | 66020954 | 9 | 6 | 4 | 1 | 7 | 0 | 0 | 0 | 1 | 0 | 21 | 14 | 10 | 5 | 6 | 0 | 0 | 0 | 0 | 0 | 210 | 149 | 1 | 1 | 1 | 1 | 1 | 0.415 | 1 | 1 | 1 | 1 | 1 | 1 | -0.117 |
| SE | 98870 | ENSG00000175294 | CATSPER1 | chr11 | - | 66020316 | 66020389 | 66020139 | 66020200 | 66020563 | 66020627 | 33 | 7 | 10 | 13 | 3 | 0 | 0 | 0 | 1 | 0 | 5 | 19 | 16 | 10 | 24 | 0 | 0 | 0 | 0 | 0 | 222 | 149 | 1 | 1 | 1 | 1 | 1 | 0.897 | 1 | 1 | 1 | 1 | 1 | 1 | -0.021 |
| SE | 98871 | ENSG00000175294 | CATSPER1 | chr11 | - | 66020316 | 66020389 | 66020139 | 66020200 | 66020810 | 66020954 | 26 | 13 | 10 | 16 | 1 | 0 | 0 | 0 | 0 | 0 | 9 | 17 | 12 | 8 | 7 | 0 | 0 | 0 | 3 | 0 | 222 | 149 | 1 | 1 | 1 | 1 | 1 | 1 | 1 | 1 | 1 | 1 | 0.642 | 1 | 0.072 |
| SE | 98872 | ENSG00000175294 | CATSPER1 | chr11 | - | 66020563 | 66020627 | 66020139 | 66020200 | 66020810 | 66020954 | 5 | 1 | 0 | 1 | 0 | 0 | 0 | 0 | 0 | 0 | 9 | 0 | 4 | 0 | 9 | 0 | 0 | 0 | 3 | 0 | 213 | 149 | 1 | 1 | 1 | 1 | NA | 1 | NA | 1 | NA | 1 | 0 | 1 | 0.25 |
| SE | 98874 | ENSG00000087365 | SF3B2 | chr11 | + | 66055075 | 66055315 | 66053026 | 66053104 | 66055534 | 66055585 | 300 | 140 | 299 | 242 | 154 | 0 | 0 | 0 | 2 | 0 | 248 | 294 | 193 | 196 | 323 | 0 | 0 | 0 | 3 | 0 | 298 | 149 | 1 | 1 | 1 | 1 | 1 | 0.984 | 1 | 1 | 1 | 1 | 0.97 | 1 | 0.003 |
| SE | 98875 | ENSG00000087365 | SF3B2 | chr11 | + | 66055075 | 66055320 | 66053026 | 66053104 | 66055534 | 66055586 | 205 | 85 | 204 | 160 | 125 | 0 | 0 | 0 | 2 | 0 | 134 | 167 | 108 | 128 | 163 | 0 | 0 | 0 | 3 | 0 | 298 | 149 | 1 | 1 | 1 | 1 | 1 | 0.976 | 1 | 1 | 1 | 1 | 0.955 | 1 | 0.004 |
| SE | 98876 | ENSG00000087365 | SF3B2 | chr11 | + | 66055534 | 66055585 | 66055075 | 66055315 | 66056837 | 66056955 | 193 | 91 | 166 | 145 | 80 | 0 | 1 | 0 | 0 | 0 | 217 | 184 | 162 | 153 | 374 | 0 | 1 | 0 | 0 | 8 | 200 | 149 | 1 | 1 | 1 | 0.985 | 1 | 1 | 1 | 1 | 0.993 | 1 | 1 | 0.972 | 0.004 |
| SE | 98878 | ENSG00000087365 | SF3B2 | chr11 | + | 66058313 | 66058405 | 66058053 | 66058150 | 66058829 | 66058930 | 184 | 125 | 155 | 234 | 159 | 0 | 0 | 0 | 0 | 0 | 332 | 256 | 232 | 339 | 350 | 0 | 0 | 0 | 3 | 0 | 241 | 149 | 1 | 1 | 1 | 1 | 1 | 1 | 1 | 1 | 1 | 1 | 0.986 | 1 | 0.003 |
| SE | 98879 | ENSG00000087365 | SF3B2 | chr11 | + | 66058829 | 66059045 | 66058313 | 66058405 | 66059200 | 66059338 | 401 | 271 | 323 | 492 | 307 | 0 | 0 | 0 | 0 | 3 | 628 | 340 | 503 | 629 | 619 | 0 | 0 | 0 | 5 | 7 | 298 | 149 | 1 | 1 | 1 | 1 | 1 | 1 | 0.981 | 1 | 1 | 1 | 0.984 | 0.978 | 0.004 |
| SE | 98880 | ENSG00000087365 | SF3B2 | chr11 | + | 66061685 | 66061775 | 66060581 | 66060731 | 66061890 | 66061998 | 400 | 263 | 398 | 386 | 653 | 0 | 0 | 0 | 0 | 0 | 720 | 468 | 386 | 455 | 617 | 0 | 0 | 0 | 10 | 0 | 239 | 149 | 1 | 1 | 1 | 1 | 1 | 1 | 1 | 1 | 1 | 1 | 0.966 | 1 | 0.007 |
| SE | 98881 | ENSG00000087365 | SF3B2 | chr11 | + | 66061890 | 66061998 | 66060581 | 66060731 | 66063008 | 66063116 | 201 | 126 | 206 | 215 | 252 | 0 | 0 | 0 | 2 | 0 | 381 | 212 | 210 | 307 | 486 | 0 | 0 | 1 | 0 | 0 | 257 | 149 | 1 | 1 | 1 | 1 | 1 | 0.984 | 1 | 1 | 1 | 0.992 | 1 | 1 | -0.002 |
| SE | 98882 | ENSG00000087365 | SF3B2 | chr11 | + | 66063399 | 66063542 | 66063008 | 66063116 | 66063627 | 66063729 | 540 | 388 | 528 | 564 | 658 | 0 | 0 | 0 | 0 | 0 | 1944 | 776 | 698 | 934 | 1601 | 0 | 0 | 0 | 2 | 0 | 292 | 149 | 1 | 1 | 1 | 1 | 1 | 1 | 1 | 1 | 1 | 1 | 0.996 | 1 | 0.001 |
| SE | 98886 | ENSG00000159352 | PSMD4 | chr1 | + | 151265393 | 151265609 | 151264831 | 151264918 | 151266003 | 151266112 | 296 | 160 | 239 | 221 | 320 | 0 | 1 | 0 | 0 | 0 | 459 | 249 | 143 | 277 | 443 | 8 | 0 | 0 | 0 | 3 | 298 | 149 | 1 | 1 | 1 | 0.988 | 1 | 1 | 1 | 0.966 | 1 | 1 | 1 | 0.987 | 0.007 |
| SE | 98888 | ENSG00000159352 | PSMD4 | chr1 | + | 151266003 | 151266112 | 151264831 | 151264918 | 151266307 | 151266439 | 138 | 51 | 122 | 117 | 101 | 0 | 0 | 0 | 0 | 0 | 97 | 185 | 68 | 123 | 219 | 0 | 0 | 0 | 6 | 0 | 258 | 149 | 1 | 1 | 1 | 1 | 1 | 1 | 1 | 1 | 1 | 1 | 0.922 | 1 | 0.016 |
| SE | 98892 | ENSG00000013523 | ANGEL1 | chr14 | - | 76804374 | 76804415 | 76803785 | 76803912 | 76806415 | 76806515 | 0 | 0 | 1 | 0 | 0 | 20 | 8 | 15 | 15 | 20 | 0 | 2 | 2 | 0 | 0 | 18 | 15 | 7 | 7 | 18 | 190 | 149 | 1 | 1 | 0 | 0 | 0.05 | 0 | 0 | 0 | 0.095 | 0.183 | 0 | 0 | -0.046 |
| SE | 98894 | ENSG00000013523 | ANGEL1 | chr14 | - | 76810088 | 76810254 | 76809180 | 76809643 | 76812763 | 76812861 | 0 | 1 | 2 | 4 | 0 | 12 | 3 | 8 | 12 | 0 | 0 | 0 | 0 | 0 | 0 | 0 | 17 | 2 | 12 | 4 | 298 | 149 | 1 | 1 | 0 | 0.143 | 0.111 | 0.143 | NA | NA | 0 | 0 | 0 | 0 | 0.099 |
| SE | 98899 | ENSG00000120756 | PLS1 | chr3 | + | 142645295 | 142645580 | 142596431 | 142596509 | 142664201 | 142664307 | 0 | 2 | 0 | 0 | 0 | 0 | 0 | 2 | 1 | 0 | 0 | 0 | 0 | 0 | 0 | 0 | 0 | 0 | 1 | 0 | 298 | 149 | 1 | 1 | NA | 1 | 0 | 0 | NA | NA | NA | NA | 0 | NA | 0.333 |
| SE | 98905 | ENSG00000174851 | YIF1A | chr11 | - | 66285700 | 66285758 | 66285380 | 66285536 | 66287597 | 66287676 | 27 | 9 | 13 | 10 | 6 | 0 | 0 | 1 | 0 | 0 | 11 | 16 | 2 | 8 | 24 | 0 | 0 | 0 | 0 | 0 | 207 | 149 | 1 | 1 | 1 | 1 | 0.903 | 1 | 1 | 1 | 1 | 1 | 1 | 1 | -0.019 |
| SE | 98912 | ENSG00000145649 | GZMA | chr5 | + | 55105473 | 55105618 | 55102645 | 55102752 | 55108124 | 55108394 | 1316 | 411 | 569 | 738 | 752 | 0 | 0 | 0 | 1 | 0 | 501 | 809 | 389 | 449 | 300 | 0 | 0 | 5 | 0 | 0 | 294 | 149 | 1 | 1 | 1 | 1 | 1 | 0.997 | 1 | 1 | 1 | 0.975 | 1 | 1 | 0.004 |
| SE | 98913 | ENSG00000145649 | GZMA | chr5 | + | 55107793 | 55107935 | 55102645 | 55102752 | 55108124 | 55108394 | 914 | 274 | 375 | 582 | 709 | 0 | 0 | 0 | 1 | 0 | 460 | 477 | 279 | 315 | 231 | 0 | 0 | 5 | 0 | 0 | 291 | 149 | 1 | 1 | 1 | 1 | 1 | 0.997 | 1 | 1 | 1 | 0.966 | 1 | 1 | 0.006 |
| SE | 98914 | ENSG00000145649 | GZMA | chr5 | + | 55107793 | 55107935 | 55105473 | 55105618 | 55108124 | 55108394 | 2317 | 791 | 986 | 1486 | 1713 | 0 | 1 | 4 | 0 | 0 | 1086 | 1259 | 650 | 821 | 452 | 0 | 0 | 1 | 0 | 0 | 291 | 149 | 1 | 1 | 1 | 0.998 | 0.992 | 1 | 1 | 1 | 1 | 0.997 | 1 | 1 | -0.001 |
| SE | 98918 | ENSG00000101158 | NELFCD | chr20 | + | 58989857 | 58989988 | 58989487 | 58989640 | 58990909 | 58991075 | 103 | 73 | 96 | 101 | 91 | 0 | 0 | 0 | 0 | 0 | 93 | 98 | 81 | 68 | 164 | 0 | 0 | 3 | 2 | 0 | 280 | 149 | 1 | 1 | 1 | 1 | 1 | 1 | 1 | 1 | 1 | 0.935 | 0.948 | 1 | 0.023 |
| SE | 98919 | ENSG00000101158 | NELFCD | chr20 | + | 58989857 | 58990232 | 58989487 | 58989640 | 58990909 | 58991075 | 57 | 42 | 46 | 53 | 57 | 0 | 0 | 0 | 0 | 0 | 54 | 54 | 38 | 44 | 103 | 0 | 0 | 3 | 2 | 0 | 298 | 149 | 1 | 1 | 1 | 1 | 1 | 1 | 1 | 1 | 1 | 0.864 | 0.917 | 1 | 0.044 |
| SE | 98920 | ENSG00000101158 | NELFCD | chr20 | + | 58993448 | 58993544 | 58992997 | 58993112 | 58994109 | 58994191 | 79 | 42 | 78 | 102 | 107 | 0 | 0 | 1 | 0 | 0 | 86 | 81 | 37 | 46 | 133 | 0 | 0 | 0 | 0 | 0 | 245 | 149 | 1 | 1 | 1 | 1 | 0.979 | 1 | 1 | 1 | 1 | 1 | 1 | 1 | -0.004 |
| SE | 98921 | ENSG00000101158 | NELFCD | chr20 | + | 58993623 | 58993675 | 58993448 | 58993544 | 58994109 | 58994191 | 26 | 16 | 36 | 27 | 40 | 2 | 0 | 0 | 0 | 0 | 42 | 39 | 20 | 16 | 66 | 0 | 0 | 0 | 0 | 0 | 201 | 149 | 1 | 1 | 0.906 | 1 | 1 | 1 | 1 | 1 | 1 | 1 | 1 | 1 | -0.019 |
| SE | 98922 | ENSG00000101158 | NELFCD | chr20 | + | 58993623 | 58993764 | 58993448 | 58993544 | 58994109 | 58994239 | 130 | 78 | 102 | 131 | 123 | 2 | 0 | 0 | 0 | 0 | 205 | 125 | 72 | 46 | 153 | 0 | 0 | 0 | 0 | 0 | 290 | 149 | 1 | 1 | 0.971 | 1 | 1 | 1 | 1 | 1 | 1 | 1 | 1 | 1 | -0.006 |
| SE | 98924 | ENSG00000102043 | MTMR8 | chrX | - | 64336078 | 64336128 | 64331421 | 64331757 | 64337267 | 64337393 | 12 | 10 | 20 | 4 | 15 | 0 | 0 | 2 | 0 | 0 | 7 | 19 | 4 | 5 | 0 | 0 | 0 | 0 | 0 | 0 | 199 | 149 | 1 | 1 | 1 | 1 | 1 | 0.6 | 1 | 1 | 1 | 1 | 1 | NA | -0.08 |
| SE | 98925 | ENSG00000102043 | MTMR8 | chrX | - | 64343610 | 64343720 | 64337267 | 64337393 | 64345044 | 64345158 | 4 | 3 | 19 | 8 | 9 | 0 | 1 | 4 | 3 | 6 | 3 | 23 | 2 | 4 | 0 | 0 | 5 | 0 | 0 | 0 | 259 | 149 | 1 | 1 | 1 | 0.633 | 0.732 | 0.605 | 0.463 | 1 | 0.726 | 1 | 1 | NA | -0.245 |
| SE | 98926 | ENSG00000102043 | MTMR8 | chrX | - | 64343610 | 64343720 | 64337267 | 64337393 | 64348659 | 64348794 | 3 | 2 | 7 | 4 | 5 | 1 | 0 | 1 | 1 | 0 | 0 | 17 | 1 | 4 | 0 | 0 | 0 | 0 | 0 | 0 | 259 | 149 | 1 | 1 | 0.633 | 1 | 0.801 | 0.697 | 1 | NA | 1 | 1 | 1 | NA | -0.174 |
| SE | 98927 | ENSG00000102043 | MTMR8 | chrX | - | 64345044 | 64345177 | 64337267 | 64337393 | 64348659 | 64348794 | 8 | 12 | 14 | 16 | 17 | 1 | 0 | 1 | 1 | 0 | 3 | 24 | 5 | 5 | 12 | 0 | 0 | 0 | 0 | 0 | 282 | 149 | 1 | 1 | 0.809 | 1 | 0.881 | 0.894 | 1 | 1 | 1 | 1 | 1 | 1 | -0.083 |
| SE | 98929 | ENSG00000102043 | MTMR8 | chrX | - | 64348659 | 64348794 | 64337267 | 64337393 | 64349941 | 64350040 | 14 | 18 | 19 | 18 | 31 | 0 | 0 | 0 | 0 | 1 | 0 | 13 | 6 | 9 | 1 | 0 | 0 | 0 | 0 | 0 | 284 | 149 | 1 | 1 | 1 | 1 | 1 | 1 | 0.942 | NA | 1 | 1 | 1 | 1 | -0.012 |
| SE | 98930 | ENSG00000102043 | MTMR8 | chrX | - | 64359404 | 64359527 | 64356175 | 64356338 | 64395339 | 64395452 | 16 | 13 | 8 | 6 | 7 | 1 | 0 | 1 | 2 | 0 | 0 | 10 | 13 | 11 | 0 | 0 | 0 | 6 | 0 | 0 | 272 | 149 | 1 | 1 | 0.898 | 1 | 0.814 | 0.622 | 1 | NA | 1 | 0.543 | 1 | NA | 0.019 |
| SE | 98932 | ENSG00000175115 | PACS1 | chr11 | + | 66219544 | 66219805 | 66216694 | 66216775 | 66220630 | 66220677 | 239 | 137 | 219 | 270 | 390 | 0 | 0 | 0 | 0 | 0 | 288 | 311 | 208 | 180 | 390 | 0 | 4 | 0 | 0 | 0 | 298 | 149 | 1 | 1 | 1 | 1 | 1 | 1 | 1 | 1 | 0.975 | 1 | 1 | 1 | 0.005 |
| SE | 98933 | ENSG00000175115 | PACS1 | chr11 | + | 66219745 | 66219805 | 66216694 | 66216775 | 66220630 | 66220791 | 577 | 344 | 532 | 605 | 876 | 0 | 0 | 0 | 0 | 0 | 683 | 773 | 506 | 474 | 1017 | 0 | 4 | 0 | 0 | 0 | 209 | 149 | 1 | 1 | 1 | 1 | 1 | 1 | 1 | 1 | 0.993 | 1 | 1 | 1 | 0.001 |
| SE | 98936 | ENSG00000175115 | PACS1 | chr11 | + | 66230804 | 66230940 | 66230379 | 66230663 | 66232171 | 66232276 | 538 | 271 | 494 | 529 | 675 | 0 | 0 | 0 | 2 | 0 | 274 | 539 | 357 | 270 | 781 | 0 | 0 | 3 | 0 | 0 | 285 | 149 | 1 | 1 | 1 | 1 | 1 | 0.993 | 1 | 1 | 1 | 0.984 | 1 | 1 | 0.002 |
| SE | 98938 | ENSG00000131981 | LGALS3 | chr14 | + | 55137369 | 55137391 | 55129251 | 55129300 | 55138044 | 55138292 | 104 | 35 | 53 | 56 | 37 | 1 | 0 | 0 | 3 | 0 | 117 | 56 | 30 | 45 | 90 | 0 | 0 | 0 | 4 | 0 | 171 | 149 | 1 | 1 | 0.989 | 1 | 1 | 0.942 | 1 | 1 | 1 | 1 | 0.907 | 1 | 0.005 |
| SE | 98940 | ENSG00000131981 | LGALS3 | chr14 | + | 55140274 | 55140363 | 55137187 | 55138368 | 55142583 | 55142678 | 402 | 277 | 246 | 318 | 167 | 0 | 1 | 0 | 0 | 0 | 990 | 302 | 222 | 525 | 799 | 5 | 1 | 0 | 2 | 0 | 238 | 149 | 1 | 1 | 1 | 0.994 | 1 | 1 | 1 | 0.992 | 0.995 | 1 | 0.994 | 1 | 0.003 |
| SE | 98944 | ENSG00000170291 | ELP5 | chr17 | + | 7258825 | 7258926 | 7258587 | 7258683 | 7259570 | 7259931 | 70 | 28 | 53 | 63 | 41 | 4 | 0 | 0 | 0 | 0 | 50 | 34 | 38 | 21 | 96 | 0 | 0 | 0 | 0 | 0 | 250 | 149 | 1 | 1 | 0.913 | 1 | 1 | 1 | 1 | 1 | 1 | 1 | 1 | 1 | -0.017 |
| SE | 98945 | ENSG00000134780 | DAGLA | chr11 | + | 61735560 | 61735644 | 61734848 | 61735002 | 61735738 | 61735816 | 9 | 7 | 5 | 9 | 2 | 1 | 0 | 0 | 0 | 0 | 2 | 10 | 7 | 8 | 0 | 0 | 0 | 0 | 0 | 0 | 233 | 149 | 1 | 1 | 0.852 | 1 | 1 | 1 | 1 | 1 | 1 | 1 | 1 | NA | -0.03 |
| SE | 98946 | ENSG00000134780 | DAGLA | chr11 | + | 61737181 | 61737324 | 61736269 | 61736350 | 61737686 | 61737755 | 6 | 8 | 12 | 9 | 2 | 0 | 0 | 0 | 0 | 0 | 0 | 6 | 0 | 4 | 33 | 0 | 0 | 0 | 3 | 0 | 292 | 149 | 1 | 1 | 1 | 1 | 1 | 1 | 1 | NA | 1 | NA | 0.405 | 1 | 0.198 |
| SE | 98947 | ENSG00000134780 | DAGLA | chr11 | + | 61738134 | 61738207 | 61737686 | 61737755 | 61739464 | 61739661 | 3 | 4 | 3 | 3 | 2 | 0 | 0 | 1 | 0 | 0 | 0 | 0 | 2 | 14 | 53 | 0 | 0 | 0 | 0 | 0 | 222 | 149 | 1 | 1 | 1 | 1 | 0.668 | 1 | 1 | NA | NA | 1 | 1 | 1 | -0.066 |
| SE | 98948 | ENSG00000221968 | FADS3 | chr11 | - | 61879311 | 61879509 | 61878640 | 61878847 | 61880040 | 61880151 | 17 | 2 | 28 | 26 | 10 | 1 | 0 | 0 | 0 | 0 | 15 | 42 | 19 | 3 | 41 | 0 | 0 | 0 | 0 | 0 | 298 | 149 | 1 | 1 | 0.895 | 1 | 1 | 1 | 1 | 1 | 1 | 1 | 1 | 1 | -0.021 |
| SE | 98949 | ENSG00000221968 | FADS3 | chr11 | - | 61879395 | 61879509 | 61878745 | 61878847 | 61880040 | 61880151 | 6 | 0 | 18 | 9 | 4 | 1 | 0 | 0 | 0 | 0 | 0 | 12 | 3 | 2 | 16 | 0 | 0 | 0 | 0 | 0 | 263 | 149 | 1 | 1 | 0.773 | NA | 1 | 1 | 1 | NA | 1 | 1 | 1 | 1 | -0.057 |
| SE | 98953 | ENSG00000177917 | ARL6IP6 | chr2 | + | 152719897 | 152720004 | 152717892 | 152717995 | 152720532 | 152720586 | 0 | 0 | 1 | 3 | 7 | 3 | 4 | 0 | 3 | 0 | 0 | 0 | 2 | 0 | 0 | 6 | 8 | 3 | 0 | 0 | 256 | 149 | 1 | 1 | 0 | 0 | 1 | 0.368 | 1 | 0 | 0 | 0.28 | NA | NA | 0.38 |
| SE | 98956 | ENSG00000177917 | ARL6IP6 | chr2 | + | 152734993 | 152735126 | 152720166 | 152720586 | 152759746 | 152759851 | 99 | 60 | 88 | 118 | 86 | 1 | 0 | 0 | 0 | 0 | 117 | 142 | 70 | 89 | 134 | 0 | 0 | 0 | 0 | 0 | 282 | 149 | 1 | 1 | 0.981 | 1 | 1 | 1 | 1 | 1 | 1 | 1 | 1 | 1 | -0.004 |

| | | | | | | | | | | | | | | | | | | | | |
|---|---|---|---|---|---|---|---|---|---|---|---|---|---|---|---|---|---|---|---|---|
| SE | 98960 | ENSG00000172932 | ANKRD13D | chr11 | + | 67299826 | 67299888 | 67297932 | 67299611 | 67299992 | 67300034 | 124 | 58 | [illegible] | [illegible] | [illegible] | [illegible] | [illegible] | [illegible] | [illegible] |
| SE | 98961 | ENSG00000075239 | ACAT1 | chr11 | + | 108127885 | 108127979 | 108121566 | 108121678 | 108128642 | 108129201 | 0 | 3 | [illegible] | [illegible] | [illegible] | [illegible] | [illegible] | [illegible] | [illegible] |
| SE | 98963 | ENSG00000075239 | ACAT1 | chr11 | + | 108128642 | 108129201 | 108121592 | 108121678 | 108131906 | 108131954 | 0 | 0 | [illegible] | [illegible] | [illegible] | [illegible] | [illegible] | [illegible] | [illegible] |
| SE | 98965 | ENSG00000075239 | ACAT1 | chr11 | + | 108128642 | 108129201 | 108127885 | 108127979 | 108131906 | 108131954 | 0 | 0 | 1 | 0 | 0 | 1 | 5 | 0 | 2 |
| SE | 98966 | ENSG00000075239 | ACAT1 | chr11 | + | 108129102 | 108129201 | 108127885 | 108127979 | 108131906 | 108131954 | 0 | 0 | 0 | 0 | 0 | 1 | 5 | 0 | 2 |
| SE | 98969 | ENSG00000075239 | ACAT1 | chr11 | + | 108134220 | 108134296 | 108133819 | 108133937 | 108135141 | 108135242 | 10 | 9 | 9 | 9 | 13 | 0 | 0 | 0 | 0 |
| SE | 98970 | ENSG00000075239 | ACAT1 | chr11 | + | 108134220 | 108134296 | 108133819 | 108133937 | 108138897 | 108139041 | 9 | 9 | 9 | 9 | 13 | 2 | 0 | 0 | 0 |
| SE | 98971 | ENSG00000075239 | ACAT1 | chr11 | + | 108134220 | 108134316 | 108133819 | 108133937 | 108135141 | 108135241 | 102 | 78 | 104 | 102 | 85 | 0 | 0 | 0 | 0 |
| SE | 98972 | ENSG00000075239 | ACAT1 | chr11 | + | 108134220 | 108134316 | 108133819 | 108133937 | 108138897 | 108139041 | 22 | 13 | 17 | 21 | 17 | 2 | 0 | 0 | 0 |
| SE | 98973 | ENSG00000075239 | ACAT1 | chr11 | + | 108135141 | 108135242 | 108133819 | 108133937 | 108138897 | 108139041 | 131 | 94 | 112 | 138 | 90 | 2 | 0 | 0 | 0 |
| SE | 98975 | ENSG00000075239 | ACAT1 | chr11 | + | 108135141 | 108135242 | 108134220 | 108134316 | 108138897 | 108139041 | 214 | 161 | 199 | 221 | 158 | 2 | 1 | 0 | 2 |
| SE | 98977 | ENSG00000075239 | ACAT1 | chr11 | + | 108143982 | 108144047 | 108142436 | 108142550 | 108146201 | 108146359 | 103 | 86 | 123 | 106 | 116 | 0 | 0 | 1 | 0 |
| SE | 98978 | ENSG00000076053 | RBM7 | chr11 | + | 114401697 | 114401860 | 114400029 | 114400767 | 114402827 | 114402915 | 270 | 188 | 294 | 418 | 597 | 3 | 1 | 2 | 1 |
| SE | 98981 | ENSG00000255663 | NA | chr11 | + | 114402827 | 114402915 | 114401697 | 114401860 | 114405705 | 114405799 | 193 | 114 | 204 | 271 | 417 | 0 | 1 | 0 | 0 |
| SE | 98982 | ENSG00000076043 | REXO2 | chr11 | + | 114440655 | 114440739 | 114439434 | 114439675 | 114443855 | 114443933 | 90 | 42 | 105 | 101 | 102 | 0 | 2 | 0 | 0 |
| SE | 98984 | ENSG00000076043 | REXO2 | chr11 | + | 114440655 | 114440739 | 114439434 | 114439675 | 114449845 | 114450279 | 19 | 2 | 20 | 13 | 3 | 0 | 0 | 0 | 0 |
| SE | 98985 | ENSG00000076043 | REXO2 | chr11 | + | 114443855 | 114443933 | 114439581 | 114439675 | 114445978 | 114446087 | 2 | 1 | 0 | 1 | 4 | 0 | 0 | 0 | 0 |
| SE | 98987 | ENSG00000076043 | REXO2 | chr11 | + | 114443855 | 114443933 | 114440655 | 114440739 | 114447825 | 114447852 | 39 | 20 | 39 | 66 | 57 | 1 | 0 | 0 | 0 |
| SE | 98992 | ENSG00000076043 | REXO2 | chr11 | + | 114445978 | 114446087 | 114440655 | 114440739 | 114447825 | 114447852 | 136 | 74 | 143 | 136 | 232 | 1 | 0 | 0 | 0 |
| SE | 98994 | ENSG00000076043 | REXO2 | chr11 | + | 114445978 | 114446087 | 114443855 | 114443933 | 114447825 | 114447852 | 139 | 70 | 144 | 130 | 242 | 0 | 0 | 0 | 2 |
| SE | 98996 | ENSG00000076043 | REXO2 | chr11 | + | 114447825 | 114447879 | 114445117 | 114446087 | 114449845 | 114449909 | 204 | 124 | 179 | 188 | 317 | 0 | 0 | 0 | 0 |
| SE | 98998 | ENSG00000136444 | RSAD1 | chr17 | + | 50479879 | 50480084 | 50479628 | 50479762 | 50482090 | 50482456 | 17 | 6 | 21 | 19 | 9 | 1 | 0 | 2 | 0 |
| SE | 99000 | ENSG00000136444 | RSAD1 | chr17 | + | 50483339 | 50483487 | 50482090 | 50482456 | 50484441 | 50484642 | 0 | 3 | 2 | 2 | 0 | 0 | 0 | 0 | 0 |
| SE | 99003 | ENSG00000136444 | RSAD1 | chr17 | + | 50484441 | 50484642 | 50482090 | 50482456 | 50484743 | 50484897 | 0 | 0 | 0 | 2 | 0 | 0 | 0 | 0 | 0 |
| SE | 99008 | ENSG00000110324 | IL10RA | chr11 | + | 117986606 | 117986813 | 117986393 | 117986534 | 117993240 | 117993410 | 2 | 0 | 1 | 0 | 0 | 0 | 0 | 0 | 0 |
| SE | 99011 | ENSG00000110324 | IL10RA | chr11 | + | 117988381 | 117988502 | 117986369 | 117986813 | 117993240 | 117993410 | 133 | 54 | 89 | 86 | 115 | 1 | 0 | 0 | 0 |
| SE | 99018 | ENSG00000160593 | JAML | chr11 | - | 118200473 | 118203665 | 118196734 | 118198091 | 118210486 | 118210712 | 863 | 647 | 596 | 632 | 1013 | 0 | 0 | 0 | 0 |
| SE | 99019 | ENSG00000160593 | JAML | chr11 | - | 118205881 | 118205991 | 118200473 | 118203665 | 118210486 | 118210712 | 800 | 736 | 556 | 614 | 772 | 0 | 0 | 2 | 3 |
| SE | 99021 | ENSG00000160593 | JAML | chr11 | - | 118210486 | 118210712 | 118205658 | 118205991 | 118212406 | 118212469 | 1626 | 1355 | 1204 | 1466 | 1515 | 0 | 0 | 0 | 0 |
| SE | 99022 | ENSG00000160593 | JAML | chr11 | - | 118214823 | 118214886 | 118212406 | 118212469 | 118224940 | 118224994 | 33 | 29 | 32 | 32 | 34 | 1 | 0 | 0 | 0 |
| SE | 99023 | ENSG00000160593 | JAML | chr11 | - | 118214823 | 118214886 | 118212406 | 118212561 | 118223998 | 118224087 | 182 | 152 | 115 | 136 | 188 | 0 | 0 | 0 | 3 |
| SE | 99028 | ENSG00000087053 | MTMR2 | chr11 | - | 95861991 | 95862102 | 95857551 | 95858632 | 95862271 | 95862366 | 111 | 75 | 66 | 107 | 99 | 0 | 0 | 0 | 0 |
| SE | 99032 | ENSG00000087053 | MTMR2 | chr11 | - | 95866402 | 95866486 | 95865600 | 95865676 | 95888155 | 95888261 | 0 | 0 | 0 | 0 | 1 | 30 | 12 | 13 | 16 |
| SE | 99036 | ENSG00000087053 | MTMR2 | chr11 | - | 95882373 | 95882496 | 95866402 | 95866486 | 95887611 | 95887684 | 2 | 0 | 1 | 1 | 4 | 0 | 0 | 0 | 2 |
| SE | 99037 | ENSG00000087053 | MTMR2 | chr11 | - | 95882373 | 95882496 | 95866402 | 95866486 | 95888155 | 95888261 | 0 | 0 | 1 | 3 | 3 | 0 | 0 | 0 | 1 |
| SE | 99038 | ENSG00000087053 | MTMR2 | chr11 | - | 95887325 | 95887373 | 95865600 | 95865676 | 95887611 | 95887684 | 0 | 0 | 2 | 0 | 0 | 16 | 10 | 12 | 22 |
| SE | 99039 | ENSG00000087053 | MTMR2 | chr11 | - | 95887325 | 95887373 | 95882373 | 95882496 | 95887611 | 95887684 | 0 | 0 | 3 | 0 | 0 | 3 | 0 | 1 | 1 |
| SE | 99041 | ENSG00000087053 | MTMR2 | chr11 | - | 95887611 | 95887684 | 95866402 | 95866486 | 95888155 | 95888261 | 5 | 6 | 6 | 4 | 17 | 0 | 0 | 0 | 1 |
| SE | 99046 | ENSG00000087053 | MTMR2 | chr11 | - | 95888155 | 95888261 | 95865439 | 95865676 | 95923874 | 95924100 | 31 | 11 | 16 | 22 | 5 | 0 | 0 | 0 | 0 |
| SE | 99049 | ENSG00000087053 | MTMR2 | chr11 | - | 95914241 | 95914312 | 95865600 | 95865676 | 95923874 | 95924000 | 11 | 5 | 7 | 8 | 2 | 0 | 0 | 0 | 0 |
| SE | 99057 | ENSG00000149577 | SIDT2 | chr11 | + | 117182518 | 117182620 | 117182059 | 117182105 | 117182722 | 117182806 | 206 | 135 | 92 | 283 | 242 | 0 | 0 | 0 | 1 |
| SE | 99058 | ENSG00000149577 | SIDT2 | chr11 | + | 117186129 | 117186223 | 117184073 | 117184139 | 117186583 | 117186636 | 85 | 75 | 47 | 157 | 48 | 1 | 0 | 0 | 0 |
| SE | 99061 | ENSG00000149577 | SIDT2 | chr11 | + | 117187891 | 117187954 | 117187627 | 117187699 | 117188707 | 117188826 | 0 | 0 | 0 | 0 | 0 | 137 | 104 | 66 | 250 |
| SE | 99062 | ENSG00000149577 | SIDT2 | chr11 | + | 117189951 | 117190025 | 117189334 | 117189401 | 117190165 | 117190282 | 152 | 90 | 71 | 206 | 75 | 0 | 0 | 1 | 0 |
| SE | 99063 | ENSG00000149577 | SIDT2 | chr11 | + | 117191877 | 117192014 | 117190622 | 117190740 | 117192253 | 117192362 | 281 | 190 | 101 | 413 | 173 | 0 | 0 | 0 | 0 |
| SE | 99064 | ENSG00000149577 | SIDT2 | chr11 | + | 117193152 | 117193258 | 117192819 | 117192866 | 117193852 | 117193963 | 120 | 47 | 29 | 169 | 19 | 0 | 0 | 0 | 0 |
| SE | 99067 | ENSG00000178105 | DDX10 | chr11 | + | 108715878 | 108715966 | 108706738 | 108706837 | 108719796 | 108719885 | 48 | 57 | 39 | 49 | 129 | 0 | 0 | 0 | 0 |
| SE | 99068 | ENSG00000178105 | DDX10 | chr11 | + | 108731537 | 108731679 | 108722996 | 108723462 | 108746245 | 108746295 | 2 | 0 | 1 | 1 | 2 | 0 | 2 | 0 | 2 |
| SE | 99069 | ENSG00000178105 | DDX10 | chr11 | + | 108731537 | 108731679 | 108722996 | 108723462 | 108767936 | 108768001 | 6 | 0 | 1 | 1 | 2 | 0 | 3 | 0 | 2 |
| SE | 99070 | ENSG00000178105 | DDX10 | chr11 | + | 108731537 | 108731679 | 108722996 | 108723462 | 108838445 | 108838565 | 4 | 1 | 1 | 2 | 6 | 45 | 95 | 67 | 56 |
| SE | 99072 | ENSG00000178105 | DDX10 | chr11 | + | 108746245 | 108746295 | 108722996 | 108723462 | 108838445 | 108838565 | 1 | 3 | 0 | 4 | 3 | 45 | 95 | 67 | 56 |
| SE | 99074 | ENSG00000178105 | DDX10 | chr11 | + | 108746245 | 108746295 | 108731537 | 108731679 | 108838445 | 108838565 | 1 | 3 | 0 | 4 | 3 | 2 | 1 | 0 | 1 |
| SE | 99076 | ENSG00000178105 | DDX10 | chr11 | + | 108767936 | 108768001 | 108731537 | 108731679 | 108838445 | 108838565 | 8 | 11 | 1 | 6 | 3 | 2 | 1 | 0 | 1 |
| SE | 99077 | ENSG00000178105 | DDX10 | chr11 | + | 108767936 | 108768001 | 108746245 | 108746295 | 108838445 | 108838565 | 4 | 13 | 1 | 8 | 3 | 1 | 4 | 0 | 4 |
| SE | 99078 | ENSG00000178105 | DDX10 | chr11 | + | 108917872 | 108918018 | 108852152 | 108852209 | 108940245 | 108940808 | 125 | 124 | 156 | 122 | 175 | 0 | 0 | 0 | 0 |
| SE | 99084 | ENSG00000110274 | CEP164 | chr11 | + | 117335604 | 117335680 | 117327853 | 117327904 | 117338565 | 117338668 | 18 | 7 | 19 | 25 | 11 | 0 | 0 | 0 | 1 |
| SE | 99095 | ENSG00000110274 | CEP164 | chr11 | + | 117373750 | 117373831 | 117371079 | 117371466 | 117380613 | 117380705 | 30 | 26 | 20 | 24 | 54 | 0 | 0 | 0 | 2 |
| SE | 99096 | ENSG00000110274 | CEP164 | chr11 | + | 117375707 | 117375791 | 117371079 | 117371466 | 117380613 | 117380705 | 20 | 7 | 13 | 18 | 3 | 0 | 0 | 0 | 2 |
| SE | 99097 | ENSG00000110274 | CEP164 | chr11 | + | 117375707 | 117375791 | 117373750 | 117373831 | 117380613 | 117380705 | 67 | 26 | 43 | 61 | 40 | 0 | 0 | 0 | 0 |
| SE | 99101 | ENSG00000110274 | CEP164 | chr11 | + | 117387202 | 117387412 | 117382795 | 117382942 | 117390776 | 117390908 | 40 | 35 | 25 | 52 | 33 | 0 | 0 | 0 | 1 |
| SE | 99103 | ENSG00000110274 | CEP164 | chr11 | + | 117390998 | 117391215 | 117390776 | 117390908 | 117392225 | 117392303 | 88 | 98 | 91 | 118 | 84 | 0 | 3 | 0 | 0 |
| SE | 99106 | ENSG00000110274 | CEP164 | chr11 | + | 117395122 | 117395191 | 117394919 | 117395003 | 117395546 | 117395722 | 30 | 14 | 29 | 31 | 23 | 0 | 0 | 0 | 0 |
| SE | 99112 | ENSG00000149136 | SSRP1 | chr11 | - | 57330854 | 57330927 | 57330290 | 57330429 | 57331667 | 57331889 | 201 | 142 | 213 | 186 | 216 | 0 | 0 | 0 | 1 |
| SE | 99113 | ENSG00000149136 | SSRP1 | chr11 | - | 57331667 | 57331889 | 57330290 | 57330927 | 57332151 | 57332280 | 276 | 131 | 287 | 268 | 289 | 1 | 0 | 0 | 2 |
| SE | 99116 | ENSG00000130764 | LRRC47 | chr1 | - | 3785086 | 3785203 | 3783995 | 3784111 | 3786848 | 3787310 | 93 | 30 | 72 | 104 | 68 | 2 | 0 | 0 | 0 |
| SE | 99127 | ENSG00000204946 | ZNF783 | chr7 | + | 149293750 | 149293842 | 149293376 | 149293492 | 149294379 | 149295185 | 2 | 5 | 7 | 2 | 4 | 0 | 0 | 2 | 0 |
| SE | 99129 | ENSG00000162341 | TPCN2 | chr11 | + | 69054032 | 69054097 | 69048931 | 69049106 | 69062883 | 69062990 | 1 | 0 | 0 | 3 | 0 | 0 | 0 | 0 | 0 |
| SE | 99130 | ENSG00000162341 | TPCN2 | chr11 | + | 69057577 | 69057694 | 69054032 | 69054097 | 69062883 | 69062990 | 21 | 2 | 15 | 4 | 4 | 0 | 0 | 0 | 0 |
| SE | 99131 | ENSG00000162341 | TPCN2 | chr11 | + | 69063894 | 69063967 | 69062883 | 69062990 | 69067502 | 69067605 | 43 | 12 | 22 | 19 | 10 | 0 | 0 | 1 | 0 |
| SE | 99132 | ENSG00000162341 | TPCN2 | chr11 | + | 69063894 | 69063967 | 69062883 | 69062990 | 69070429 | 69070495 | 7 | 0 | 4 | 4 | 2 | 1 | 2 | 0 | 0 |
| SE | 99133 | ENSG00000162341 | TPCN2 | chr11 | + | 69067502 | 69067605 | 69062883 | 69062990 | 69070429 | 69070495 | 11 | 6 | 9 | 4 | 5 | 1 | 2 | 0 | 0 |
| SE | 99138 | ENSG00000115414 | FN1 | chr2 | - | 215372108 | 215372300 | 215371908 | 215372015 | 215373321 | 215373411 | 0 | 5 | 3 | 0 | 0 | 0 | 9 | 2 | 0 |
| SE | 99139 | ENSG00000115414 | FN1 | chr2 | - | 215372108 | 215372375 | 215371908 | 215372015 | 215373321 | 215373411 | 0 | 19 | 2 | 2 | 0 | 0 | 9 | 2 | 0 |
| SE | 99140 | ENSG00000115414 | FN1 | chr2 | - | 215380810 | 215381080 | 215379129 | 215379317 | 215382211 | 215382325 | 0 | 23 | 4 | 5 | 0 | 0 | 45 | 3 | 7 |

| | | | | | | | | | | | | | | | | | | | | | | | | | | | | | | | | | | | | | | | | | | | | | | |
|---|---|---|---|---|---|---|---|---|---|---|---|---|---|---|---|---|---|---|---|---|---|---|---|---|---|---|---|---|---|---|---|---|---|---|---|---|---|---|---|---|---|---|---|---|---|---|
| SE | 99431 | ENSG00000110321 | EIF4G2 | chr11 | - | 10807254 | 10807319 | 10806819 | 10806885 | 10808704 | 10808923 | 470 | 336 | 391 | 550 | 498 | 1 | 0 | 0 | 1 | 0 | 287 | 479 | 325 | 519 | 269 | 0 | 0 | 9 | 2 | 0 | 214 | 149 | 1 | 1 | 0.997 | 1 | 1 | 0.997 | 1 | 1 | 1 | 0.962 | 0.994 | 1 | 0.008 |
| SE | 99433 | ENSG00000110321 | EIF4G2 | chr11 | - | 10807254 | 10807381 | 10806819 | 10806885 | 10808704 | 10808708 | 961 | 682 | 781 | 1105 | 1071 | 1 | 0 | 0 | 1 | 0 | 742 | 1010 | 731 | 1025 | 679 | 0 | 0 | 9 | 2 | 0 | 276 | 149 | 1 | 1 | 0.998 | 1 | 1 | 0.998 | 1 | 1 | 1 | 0.978 | 0.996 | 1 | 0.004 |
| SE | 99436 | ENSG00000247271 | ZBED5-AS1 | chr11 | + | 10865260 | 10865327 | 10858178 | 10858366 | 10906454 | 10906802 | 5 | 7 | 10 | 5 | 4 | 0 | 0 | 0 | 0 | 0 | 10 | 11 | 10 | 4 | 41 | 0 | 1 | 0 | 0 | 0 | 216 | 149 | 1 | 1 | 1 | 1 | 1 | 1 | 1 | 1 | 0.884 | 1 | 1 | 1 | 0.023 |
| SE | 99438 | ENSG00000109458 | GAB1 | chr4 | + | 143380611 | 143380739 | 143337177 | 143337260 | 143394816 | 143395720 | 1 | 0 | 0 | 0 | 0 | 0 | 0 | 0 | 0 | 0 | 0 | 0 | 1 | 0 | 0 | 0 | 0 | 4 | 3 | 0 | 277 | 149 | 1 | 1 | 1 | NA | NA | NA | NA | NA | NA | 0.119 | 0 | NA | 0.941 |
| SE | 99441 | ENSG00000109458 | GAB1 | chr4 | + | 143433490 | 143433716 | 143415476 | 143415771 | 143437998 | 143438600 | 57 | 51 | 52 | 32 | 100 | 0 | 0 | 0 | 0 | 3 | 47 | 67 | 54 | 14 | 18 | 0 | 0 | 4 | 0 | 0 | 298 | 149 | 1 | 1 | 1 | 1 | 1 | 1 | 0.943 | 1 | 1 | 0.871 | 1 | 1 | 0.014 |
| SE | 99443 | ENSG00000109458 | GAB1 | chr4 | + | 143457679 | 143457769 | 143440078 | 143440382 | 143459384 | 143459429 | 4 | 1 | 2 | 1 | 16 | 1 | 5 | 2 | 6 | 1 | 0 | 0 | 0 | 0 | 0 | 11 | 7 | 7 | 0 | 0 | 239 | 149 | 1 | 1 | 0.714 | 0.111 | 0.384 | 0.094 | 0.909 | 0 | 0 | 0 | NA | NA | 0.442 |
| SE | 99445 | ENSG00000105808 | RASA4 | chr7 | - | 102588233 | 102588445 | 102587446 | 102587770 | 102590158 | 102590296 | 0 | 4 | 3 | 8 | 2 | 0 | 0 | 1 | 0 | 0 | 14 | 0 | 0 | 0 | 0 | 0 | 0 | 0 | 0 | 0 | 298 | 149 | 1 | 1 | NA | 1 | 0.6 | 1 | 1 | 1 | NA | NA | NA | NA | -0.1 |
| SE | 99449 | ENSG00000285509 | TBCEL-TECTA | chr11 | + | 121105830 | 121105964 | 121102664 | 121102729 | 121109210 | 121109498 | 2 | 4 | 9 | 0 | 5 | 1 | 0 | 1 | 0 | 0 | 0 | 6 | 5 | 4 | 0 | 0 | 0 | 0 | 0 | 0 | 283 | 149 | 1 | 1 | 0.513 | 1 | 0.826 | NA | 1 | NA | 1 | 1 | 1 | NA | -0.165 |
| SE | 99450 | ENSG00000149743 | TRPT1 | chr11 | - | 64224099 | 64224162 | 64223829 | 64223967 | 64224284 | 64224341 | 3 | 2 | 2 | 0 | 1 | 1 | 0 | 0 | 1 | 0 | 0 | 5 | 0 | 0 | 0 | 0 | 0 | 0 | 3 | 0 | 212 | 149 | 1 | 1 | 0.678 | 1 | 1 | 0 | 1 | NA | 1 | NA | 0 | NA | 0.236 |
| SE | 99451 | ENSG00000149743 | TRPT1 | chr11 | - | 64224099 | 64224210 | 64223798 | 64223967 | 64224284 | 64224341 | 22 | 7 | 29 | 13 | 32 | 1 | 0 | 0 | 1 | 0 | 55 | 25 | 31 | 25 | 26 | 0 | 0 | 0 | 3 | 0 | 260 | 149 | 1 | 1 | 0.927 | 1 | 1 | 0.882 | 1 | 1 | 1 | 1 | 0.827 | 1 | -0.004 |
| SE | 99452 | ENSG00000149743 | TRPT1 | chr11 | - | 64224794 | 64224970 | 64224542 | 64224717 | 64225498 | 64225580 | 2 | 0 | 1 | 9 | 4 | 0 | 0 | 0 | 0 | 0 | 0 | 5 | 0 | 2 | 0 | 0 | 4 | 3 | 0 | 0 | 298 | 149 | 1 | 1 | 1 | NA | 1 | 1 | 1 | NA | 0.385 | 0 | 1 | NA | 0.538 |
| SE | 99453 | ENSG00000149743 | TRPT1 | chr11 | - | 64224800 | 64224970 | 64224542 | 64224717 | 64225498 | 64225580 | 18 | 5 | 16 | 22 | 10 | 0 | 0 | 0 | 0 | 0 | 0 | 17 | 4 | 8 | 26 | 0 | 4 | 3 | 0 | 0 | 298 | 149 | 1 | 1 | 1 | 1 | 1 | 1 | 1 | NA | 0.68 | 0.4 | 1 | 1 | 0.23 |
| SE | 99455 | ENSG00000149743 | TRPT1 | chr11 | - | 64225785 | 64225869 | 64224800 | 64225580 | 64226049 | 64226148 | 8 | 5 | 5 | 4 | 2 | 0 | 0 | 3 | 4 | 0 | 0 | 5 | 3 | 3 | 11 | 0 | 7 | 0 | 0 | 1 | 233 | 149 | 1 | 1 | 1 | 1 | 0.516 | 0.39 | 1 | NA | 0.314 | 1 | 1 | 0.876 | -0.016 |
| SE | 99468 | ENSG00000167283 | ATP5MG | chr11 | + | 118403923 | 118403985 | 118401618 | 118401717 | 118405656 | 118405709 | 0 | 0 | 0 | 0 | 0 | 1 | 1 | 1 | 6 | 1 | 0 | 3 | 0 | 2 | 0 | 0 | 5 | 2 | 4 | 10 | 211 | 149 | 1 | 1 | 0 | 0 | 0 | 0 | 0 | NA | 0.298 | 0 | 0.261 | 0 | -0.14 |
| SE | 99469 | ENSG00000167283 | ATP5MG | chr11 | + | 118403923 | 118403985 | 118401618 | 118401717 | 118406936 | 118407097 | 2 | 1 | 2 | 1 | 0 | 664 | 394 | 525 | 484 | 293 | 9 | 0 | 6 | 0 | 0 | 1462 | 534 | 570 | 760 | 1401 | 211 | 149 | 1 | 1 | 0.002 | 0.002 | 0.003 | 0.001 | 0 | 0.004 | 0 | 0.007 | 0 | 0 | -0.001 |
| SE | 99470 | ENSG00000167283 | ATP5MG | chr11 | + | 118405656 | 118405709 | 118401609 | 118401717 | 118406936 | 118407097 | 0 | 1 | 1 | 8 | 1 | 664 | 394 | 525 | 484 | 293 | 0 | 5 | 2 | 4 | 12 | 1462 | 534 | 570 | 760 | 1401 | 202 | 149 | 1 | 1 | 0 | 0.002 | 0.001 | 0.012 | 0.003 | 0 | 0.007 | 0.003 | 0.004 | 0.006 | 0 |
| SE | 99471 | ENSG00000167283 | ATP5MG | chr11 | + | 118405656 | 118405709 | 118401609 | 118401717 | 118408999 | 118409199 | 1 | 0 | 1 | 1 | 0 | 2 | 0 | 0 | 0 | 0 | 0 | 0 | 0 | 0 | 8 | 0 | 0 | 0 | 0 | 0 | 202 | 149 | 1 | 1 | 0.269 | NA | 1 | 1 | NA | NA | NA | NA | NA | 1 | -0.244 |
| SE | 99472 | ENSG00000167283 | ATP5MG | chr11 | + | 118405656 | 118405709 | 118403923 | 118403985 | 118406936 | 118407097 | 0 | 0 | 0 | 2 | 0 | 2 | 1 | 2 | 1 | 0 | 0 | 3 | 0 | 0 | 2 | 9 | 0 | 9 | 0 | 0 | 202 | 149 | 1 | 1 | 0 | 0 | 0 | 0.596 | NA | 0 | 1 | 0 | NA | 1 | -0.351 |
| SE | 99473 | ENSG00000167283 | ATP5MG | chr11 | + | 118406936 | 118406976 | 118401621 | 118401717 | 118408999 | 118409199 | 9 | 7 | 2 | 7 | 0 | 2 | 0 | 0 | 0 | 0 | 7 | 1 | 3 | 6 | 18 | 0 | 0 | 0 | 0 | 0 | 189 | 149 | 1 | 1 | 0.78 | 1 | 1 | 1 | NA | 1 | 1 | 1 | 1 | 1 | -0.055 |
| SE | 99474 | ENSG00000167283 | ATP5MG | chr11 | + | 118406936 | 118407097 | 118401605 | 118401717 | 118408999 | 118409199 | 1514 | 871 | 1175 | 1090 | 701 | 2 | 0 | 0 | 0 | 0 | 3037 | 1138 | 1232 | 1611 | 2943 | 0 | 0 | 0 | 0 | 0 | 298 | 149 | 1 | 1 | 0.997 | 1 | 1 | 1 | 1 | 1 | 1 | 1 | 1 | 1 | -0.001 |
| [illegible] | | | | | | | | | | | | | | | | | | | | | | | | | | | | | | | | | | | | | | | | | | | | | | |

| | | | | | | | | | | | | | |
|---|---|---|---|---|---|---|---|---|---|---|---|---|---|
| SE | 99799 | ENSG00000172543 | CTSW | chr11 | + | 65883265 | 65883424 | 65883068 | 65883133 | 65883507 | 65883741 | 1055 | 256 |
| SE | 99804 | ENSG00000115368 | WDR75 | chr2 | + | 189448800 | 189448834 | 189448378 | 189448508 | 189450902 | 189450922 | 0 | 0 |
| SE | 99805 | ENSG00000115368 | WDR75 | chr2 | + | 189457310 | 189457381 | 189455319 | 189455444 | 189458752 | 189458872 | 85 | 49 |
| SE | 99806 | ENSG00000115368 | WDR75 | chr2 | + | 189463693 | 189463753 | 189462483 | 189462642 | 189463845 | 189463961 | 97 | 80 |
| SE | 99807 | ENSG00000115368 | WDR75 | chr2 | + | 189465078 | 189465254 | 189463845 | 189463961 | 189466424 | 189466582 | 197 | 151 |
| SE | 99808 | ENSG00000115368 | WDR75 | chr2 | + | 189469343 | 189469439 | 189468474 | 189468569 | 189470075 | 189470245 | 123 | 86 |
| SE | 99809 | ENSG00000115368 | WDR75 | chr2 | + | 189469343 | 189469439 | 189468474 | 189468569 | 189470818 | 189470878 | 49 | 35 |
| SE | 99810 | ENSG00000115368 | WDR75 | chr2 | + | 189470075 | 189470245 | 189468474 | 189468569 | 189470818 | 189470878 | 48 | 41 |
| SE | 99814 | ENSG00000110060 | PUS3 | chr11 | - | 125895223 | 125895789 | 125893484 | 125894286 | 125895906 | 125896330 | 49 | 41 |
| SE | 99815 | ENSG00000110060 | PUS3 | chr11 | - | 125895906 | 125896330 | 125895223 | 125895789 | 125903169 | 125903206 | 37 | 27 |
| SE | 99820 | ENSG00000166483 | WEE1 | chr11 | + | 9581531 | 9581678 | 9577141 | 9577263 | 9585257 | 9585353 | 194 | 99 |
| SE | 99821 | ENSG00000087302 | RTRAF | chr14 | + | 51993722 | 51993822 | 51991316 | 51991441 | 51998493 | 51998580 | 640 | 596 |
| SE | 99822 | ENSG00000087302 | RTRAF | chr14 | + | 51998493 | 51998580 | 51993722 | 51993822 | 51999707 | 51999796 | 783 | 771 |
| SE | 99823 | ENSG00000087302 | RTRAF | chr14 | + | 52001797 | 52001866 | 51999707 | 51999796 | 52004193 | 52004242 | 799 | 718 |
| SE | 99824 | ENSG00000087302 | RTRAF | chr14 | + | 52001797 | 52001866 | 51999707 | 51999796 | 52004361 | 52004702 | 325 | 273 |
| SE | 99825 | ENSG00000087302 | RTRAF | chr14 | + | 52004193 | 52004242 | 51999707 | 51999796 | 52004361 | 52010694 | 565 | 437 |
| SE | 99826 | ENSG00000087302 | RTRAF | chr14 | + | 52004193 | 52004242 | 52001797 | 52001866 | 52004361 | 52010694 | 1044 | 888 |
| SE | 99827 | ENSG00000087302 | RTRAF | chr14 | + | 52004239 | 52004242 | 52001797 | 52001866 | 52004361 | 52004653 | 18 | 11 |
| SE | 99829 | ENSG00000013725 | CD6 | chr11 | + | 60987907 | 60988027 | 60971749 | 60971914 | 61006573 | 61006642 | 0 | 0 |
| SE | 99830 | ENSG00000013725 | CD6 | chr11 | + | 61006573 | 61006642 | 60971679 | 60971914 | 61007559 | 61007712 | 196 | 71 |

[illegible]

| | | | | | | | | | | | | | | | | | | | | | | | | | | | | | | | | | | | | | | | | | | | | | | |
|---|---|---|---|---|---|---|---|---|---|---|---|---|---|---|---|---|---|---|---|---|---|---|---|---|---|---|---|---|---|---|---|---|---|---|---|---|---|---|---|---|---|---|---|---|---|---|
| SE | 100174 | ENSG00000078124 | ACER3 | chr11 | + | 76994155 | 76994226 | 76985642 | 76985724 | 76998762 | 76998821 | 0 | 0 | 1 | 1 | 1 | 0 | 0 | 0 | 2 | 0 | 0 | 2 | 0 | 0 | 0 | 0 | 0 | 0 | 0 | 0 | 220 | 149 | 1 | 1 | NA | NA | 1 | 0.253 | 1 | NA | 1 | NA | NA | NA | -0.249 |
| SE | 100177 | ENSG00000078124 | ACER3 | chr11 | + | 76994155 | 76994226 | 76990538 | 76990574 | 77016674 | 77016689 | 4 | 3 | 0 | 7 | 4 | 0 | 0 | 0 | 0 | 0 | 17 | 11 | 2 | 9 | 5 | 0 | 0 | 0 | 0 | 11 | 220 | 149 | 1 | 1 | 1 | 1 | NA | 1 | 1 | 1 | 1 | 1 | 1 | 0.235 | 0.153 |
| SE | 100179 | ENSG00000078124 | ACER3 | chr11 | + | 76997205 | 76997273 | 76994155 | 76994226 | 76998762 | 76998821 | 3 | 5 | 3 | 7 | 1 | 0 | 0 | 2 | 1 | 1 | 15 | 2 | 5 | 0 | 9 | 0 | 2 | 0 | 0 | 0 | 217 | 149 | 1 | 1 | 1 | 1 | 0.507 | 0.828 | 0.407 | 1 | 0.407 | 1 | NA | 1 | -0.103 |
| SE | 100180 | ENSG00000078124 | ACER3 | chr11 | + | 76998762 | 76998821 | 76976288 | 76976341 | 77015015 | 77015117 | 15 | 25 | 13 | 15 | 55 | 2 | 0 | 0 | 0 | 0 | 7 | 28 | 16 | 23 | 20 | 0 | 0 | 0 | 0 | 0 | 208 | 149 | 1 | 1 | 0.843 | 1 | 1 | 1 | 1 | 1 | 1 | 1 | 1 | 1 | -0.031 |
| SE | 100181 | ENSG00000078124 | ACER3 | chr11 | + | 76998762 | 76998821 | 76976288 | 76976341 | 77016674 | 77016689 | 1 | 6 | 0 | 0 | 3 | 0 | 0 | 0 | 0 | 0 | 0 | 0 | 0 | 0 | 0 | 0 | 0 | 0 | 1 | 0 | 208 | 149 | 1 | 1 | 1 | 1 | NA | NA | 1 | NA | NA | NA | 0 | NA | 1 |
| SE | 100182 | ENSG00000078124 | ACER3 | chr11 | + | 76998762 | 76998821 | 76990538 | 76990574 | 77011329 | 77011375 | 11 | 10 | 2 | 17 | 15 | 0 | 1 | 0 | 1 | 1 | 26 | 14 | 11 | 12 | 3 | 0 | 2 | 4 | 0 | 0 | 208 | 149 | 1 | 1 | 1 | 0.878 | 1 | 0.924 | 0.915 | 1 | 0.834 | 0.663 | 1 | 1 | 0.044 |
| SE | 100184 | ENSG00000078124 | ACER3 | chr11 | + | 76998762 | 76998821 | 76994155 | 76994226 | 77016674 | 77016689 | 1 | 6 | 0 | 0 | 2 | 0 | 0 | 0 | 0 | 0 | 0 | 0 | 0 | 0 | 0 | 0 | 3 | 0 | 0 | 0 | 208 | 149 | 1 | 1 | 1 | 1 | NA | NA | 1 | NA | 0 | NA | NA | NA | 1 |
| SE | 100185 | ENSG00000078124 | ACER3 | chr11 | + | 77011329 | 77011375 | 76990538 | 76990574 | 77016674 | 77016779 | 0 | 0 | 1 | 1 | 2 | 0 | 0 | 0 | 0 | 0 | 0 | 0 | 3 | 0 | 0 | 0 | 0 | 0 | 0 | 11 | 195 | 149 | 1 | 1 | NA | NA | 1 | 1 | 1 | NA | NA | 1 | NA | 0 | 0.5 |
| SE | 100188 | ENSG00000078124 | ACER3 | chr11 | + | 77015015 | 77015117 | 76976288 | 76976341 | 77016674 | 77016689 | 39 | 66 | 17 | 39 | 132 | 0 | 0 | 0 | 0 | 0 | 8 | 58 | 34 | 40 | 37 | 0 | 0 | 0 | 1 | 0 | 251 | 149 | 1 | 1 | 1 | 1 | 1 | 1 | 1 | 1 | 1 | 1 | 0.96 | 1 | 0.008 |
| SE | 100190 | ENSG00000078124 | ACER3 | chr11 | + | 77015015 | 77015117 | 77011329 | 77011375 | 77016674 | 77016689 | 39 | 72 | 23 | 41 | 136 | 0 | 1 | 1 | 0 | 2 | 8 | 64 | 35 | 40 | 37 | 0 | 0 | 0 | 0 | 0 | 251 | 149 | 1 | 1 | 1 | 0.977 | 0.932 | 1 | 0.976 | 1 | 1 | 1 | 1 | 1 | -0.023 |
| SE | 100191 | ENSG00000078124 | ACER3 | chr11 | + | 77015015 | 77015659 | 76976288 | 76976341 | 77016674 | 77016779 | 13 | 2 | 2 | 4 | 3 | 0 | 0 | 0 | 0 | 0 | 8 | 0 | 5 | 0 | 0 | 0 | 0 | 0 | 1 | 0 | 298 | 149 | 1 | 1 | 1 | 1 | 1 | 1 | 1 | 1 | NA | 1 | 0 | NA | 0.333 |
| SE | 100193 | ENSG00000078124 | ACER3 | chr11 | + | 77015638 | 77015659 | 77015015 | 77015117 | 77016674 | 77016779 | 3 | 0 | 1 | 3 | 0 | 50 | 79 | 30 | 57 | 160 | 8 | 0 | 3 | 0 | 0 | 8 | 69 | 49 | 60 | 48 | 170 | 149 | 1 | 1 | 0.05 | 0 | 0.028 | 0.044 | 0 | 0.467 | 0 | 0.051 | 0 | 0 | -0.079 |
| SE | 100194 | ENSG00000078124 | ACER3 | chr11 | + | 77016674 | 77016779 | 77013434 | 77015117 | 77019730 | 77019767 | 101 | 162 | 68 | 120 | 284 | 0 | 0 | 0 | 0 | 0 | 61 | 125 | 105 | 132 | 79 | 0 | 0 | 0 | 1 | 0 | 254 | 149 | 1 | 1 | 1 | 1 | 1 | 1 | 1 | 1 | 1 | 1 | 0.987 | 1 | 0.003 |
| SE | 100197 | ENSG00000137474 | MYO7A | chr11 | + | 77177548 | 77177643 | 77175371 | 77175464 | 77179044 | 77179129 | 4 | 3 | 4 | 2 | 1 | 1 | 0 | 0 | 0 | 0 | 10 | 22 | 17 | 7 | 33 | 0 | 0 | 0 | 0 | 0 | 244 | 149 | 1 | 1 | 0.71 | 1 | 1 | 1 | 1 | 1 | 1 | 1 | 1 | 1 | -0.058 |
| SE | 100198 | ENSG00000137474 | MYO7A | chr11 | + | 77179734 | 77179953 | 77179044 | 77179129 | 77180373 | 77180481 | 5 | 5 | 3 | 4 | 7 | 1 | 0 | 0 | 0 | 0 | 30 | 67 | 19 | 17 | 51 | 0 | 0 | 0 | 2 | 0 | 298 | 149 | 1 | 1 | 0.714 | 1 | 1 | 1 | 1 | 1 | 1 | 1 | 0.81 | 1 | -0.019 |
| SE | 100199 | ENSG00000137474 | MYO7A | chr11 | + | 77182423 | 77182600 | 77181950 | 77182154 | 77184587 | 77184703 | 6 | 1 | 4 | 3 | 0 | 0 | 0 | 0 | 1 | 0 | 6 | 30 | 3 | 28 | 10 | 0 | 0 | 0 | 1 | 0 | 298 | 149 | 1 | 1 | 1 | 1 | 1 | 0.6 | NA | 1 | 1 | 1 | 0.933 | 1 | -0.087 |
| SE | 100200 | ENSG00000137474 | MYO7A | chr11 | + | 77183058 | 77183157 | 77182423 | 77182600 | 77184587 | 77184703 | 4 | 0 | 2 | 2 | 1 | 1 | 0 | 1 | 0 | 0 | 0 | 13 | 1 | 5 | 7 | 0 | 2 | 0 | 11 | 0 | 248 | 149 | 1 | 1 | 0.706 | NA | 0.546 | 1 | 1 | NA | 0.796 | 1 | 0.215 | 1 | 0.06 |
| SE | 100203 | ENSG00000137474 | MYO7A | chr11 | + | 77206995 | 77207402 | 77206096 | 77206202 | 77208429 | 77208517 | 5 | 0 | 0 | 2 | 0 | 0 | 0 | 1 | 0 | 0 | 10 | 28 | 9 | 9 | 17 | 0 | 0 | 0 | 0 | 0 | 298 | 149 | 1 | 1 | 1 | NA | 0 | 1 | NA | 1 | 1 | 1 | 1 | 1 | -0.333 |
| SE | 100204 | ENSG00000137474 | MYO7A | chr11 | + | 77207288 | 77207402 | 77206096 | 77206202 | 77208429 | 77208517 | 9 | 0 | 2 | 3 | 3 | 0 | 0 | 1 | 0 | 0 | 10 | 46 | 11 | 18 | 28 | 0 | 0 | 0 | 0 | 0 | 263 | 149 | 1 | 1 | 1 | NA | 0.531 | 1 | 1 | 1 | 1 | 1 | 1 | 1 | -0.117 |
| SE | 100220 | ENSG00000172183 | ISG20 | chr15 | + | 88638959 | 88639076 | 88635669 | 88635700 | 88652109 | 88652310 | 1 | 2 | 1 | 0 | 0 | 5 | 0 | 27 | 13 | 14 | 0 | 0 | 0 | 0 | 0 | 0 | 33 | 11 | 3 | 8 | 266 | 149 | 1 | 1 | 0.101 | 1 | 0.02 | 0 | 0 | NA | 0 | 0 | 0 | 0 | 0.224 |
| SE | 100222 | ENSG00000172183 | ISG20 | chr15 | + | 88638959 | 88639076 | 88636166 | 88636292 | 88652109 | 88652310 | 13 | 1 | 3 | 1 | 9 | 1 | 0 | 7 | 0 | 1 | 0 | 13 | 5 | 1 | 0 | 0 | 15 | 0 | 10 | 1 | 266 | 149 | 1 | 1 | 0.879 | 1 | 0.194 | 1 | 0.834 | NA | 0.327 | 1 | 0.053 | 0 | 0.436 |
| SE | 100223 | ENSG00000172183 | ISG20 | chr15 | + | 88639342 | 88639594 | 88635669 | 88635700 | 88650234 | 88651862 | 38 | 21 | 40 | 17 | 55 | 0 | 0 | 1 | 0 | 0 | 21 | 74 | 37 | 10 | 21 | 0 | 0 | 0 | 0 | 0 | 298 | 149 | 1 | 1 | 1 | 1 | 0.952 | 1 | 1 | 1 | 1 | 1 | 1 | 1 | -0.01 |
| SE | 100226 | ENSG00000172183 | ISG20 | chr15 | + | 88639342 | 88639594 | 88638952 | 88639076 | 88652109 | 88652310 | 377 | 169 | 387 | 316 | 218 | 1 | 0 | 0 | 0 | 0 | 222 | 587 | 456 | 283 | 223 | 0 | 0 | 0 | 0 | 0 | 298 | 149 | 1 | 1 | 0.995 | 1 | 1 | 1 | 1 | 1 | 1 | 1 | 1 | 1 | -0.001 |
| SE | 100227 | ENSG00000172183 | ISG20 | chr15 | + | 88650234 | 88652310 | 88639342 | 88639594 | 88655414 | 88655648 | 166 | 64 | 179 | 127 | 82 | 1 | 0 | 0 | 1 | 0 | 108 | 258 | 164 | 77 | 80 | 0 | 0 | 0 | 0 | 0 | 298 | 149 | 1 | 1 | 0.988 | 1 | 1 | 0.984 | 1 | 1 | 1 | 1 | 1 | 1 | -0.006 |
| SE | 100228 | ENSG00000172183 | ISG20 | chr15 | + | 88652109 | 88652310 | 88639342 | 88639594 | 88655414 | 88655621 | 360 | 162 | 403 | 295 | 153 | 1 | 0 | 0 | 1 | 0 | 185 | 603 | 398 | 215 | 189 | 0 | 0 | 0 | 0 | 0 | 298 | 149 | 1 | 1 | 0.994 | 1 | 1 | 0.993 | 1 | 1 | 1 | 1 | 1 | 1 | -0.003 |
| SE | 100229 | ENSG00000172183 | ISG20 | chr15 | + | 88652169 | 88652310 | 88639342 | 88639594 | 88655414 | 88655531 | 182 | 68 | 190 | 138 | 80 | 1 | 0 | 0 | 1 | 0 | 108 | 274 | 170 | 90 | 86 | 0 | 0 | 0 | 0 | 0 | 290 | 149 | 1 | 1 | 0.989 | 1 | 1 | 0.986 | 1 | 1 | 1 | 1 | 1 | 1 | -0.005 |
| SE | 100230 | ENSG00000287426 | NA | chr11 | - | 129783005 | 129783180 | 129767312 | 129767439 | 129813967 | 129814690 | 0 | 0 | 0 | 0 | 2 | 2 | 2 | 1 | 1 | 0 | 0 | 4 | 0 | 5 | 0 | 1 | 2 | 4 | 7 | 0 | 298 | 149 | 1 | 1 | 0 | 0 | 0 | 0 | 1 | 0 | 0.5 | 0 | 0.263 | NA | 0.009 |
| SE | 100231 | ENSG00000142583 | SLC2A5 | chr1 | - | 9069503 | 9069594 | 9047609 | 9047734 | 9085013 | 9085126 | 1 | 0 | 0 | 5 | 1 | 0 | 0 | 0 | 0 | 0 | 19 | 2 | 1 | 39 | 31 | 0 | 0 | 0 | 1 | 0 | 240 | 149 | 1 | 1 | 1 | NA | NA | 1 | 1 | 1 | 1 | 1 | 0.96 | 1 | 0.008 |
| SE | 100233 | ENSG00000142583 | SLC2A5 | chr1 | - | 9072303 | 9072385 | 9069503 | 9069594 | 9085013 | 9085166 | 0 | 0 | 0 | 0 | 0 | 0 | 0 | 0 | 5 | 1 | 0 | 0 | 0 | 3 | 7 | 19 | 2 | 1 | 53 | 31 | 231 | 149 | 1 | 1 | NA | NA | NA | 0 | 0 | 0 | 0 | 0 | 0.035 | 0.127 | -0.032 |
| SE | 100236 | ENSG00000084234 | APLP2 | chr11 | + | 130101925 | 130102000 | 130069893 | 130070082 | 130109428 | 130109602 | 0 | 1 | 0 | 0 | 0 | 215 | 88 | 96 | 97 | 31 | 0 | 0 | 0 | 0 | 0 | 409 | 329 | 175 | 313 | 381 | 224 | 149 | 1 | 1 | 0 | 0.008 | 0 | 0 | 0 | 0 | 0 | 0 | 0 | 0 | 0.002 |
| SE | 100237 | ENSG00000084234 | APLP2 | chr11 | + | 130109428 | 130109602 | 130069890 | 130070082 | 130110537 | 130110560 | 1175 | 713 | 658 | 721 | 1142 | 0 | 0 | 0 | 0 | 0 | 3507 | 2189 | 1197 | 2131 | 2547 | 0 | 9 | 1 | 0 | 0 | 298 | 149 | 1 | 1 | 1 | 1 | 1 | 1 | 1 | 1 | 0.992 | 0.998 | 1 | 1 | 0.002 |
| SE | 100239 | ENSG00000084234 | APLP2 | chr11 | + | 130110537 | 130110661 | 130069893 | 130070082 | 130120705 | 130120777 | 1161 | 770 | 812 | 969 | 1516 | 0 | 1 | 0 | 0 | 0 | 6845 | 2799 | 1774 | 3352 | 3986 | 0 | 0 | 1 | 0 | 0 | 273 | 149 | 1 | 1 | 1 | 0.998 | 1 | 1 | 1 | 1 | 1 | 0.999 | 1 | 1 | 0 |
| SE | 100240 | ENSG00000084234 | APLP2 | chr11 | + | 130120705 | 130120818 | 130110537 | 130110661 | 130121613 | 130121766 | 1843 | 1238 | 1306 | 1630 | 2804 | 0 | 0 | 0 | 1 | 0 | 11724 | 4642 | 2822 | 5725 | 6420 | 0 | 0 | 0 | 3 | 0 | 262 | 149 | 1 | 1 | 1 | 1 | 1 | 0.999 | 1 | 1 | 1 | 1 | 0.999 | 1 | 0 |
| SE | 100241 | ENSG00000084234 | APLP2 | chr11 | + | 130120705 | 130120818 | 130110537 | 130110661 | 130126699 | 130126830 | 1006 | 662 | 677 | 805 | 1407 | 0 | 0 | 0 | 0 | 0 | 5828 | 2402 | 1456 | 2908 | 3177 | 0 | 2 | 0 | 0 | 14 | 262 | 149 | 1 | 1 | 1 | 1 | 1 | 1 | 1 | 1 | 0.999 | 1 | 1 | 0.992 | 0.002 |
| SE | 100242 | ENSG00000084234 | APLP2 | chr11 | + | 130121613 | 130121810 | 130110537 | 130110661 | 130122304 | 130122513 | 628 | 492 | 289 | 509 | 665 | 0 | 0 | 0 | 0 | 0 | 3479 | 1373 | 909 | 1793 | 2113 | 0 | 0 | 0 | 1 | 0 | 298 | 149 | 1 | 1 | 1 | 1 | 1 | 1 | 1 | 1 | 1 | 1 | 0.999 | 1 | 0 |
| SE | 100243 | ENSG00000084234 | APLP2 | chr11 | + | 130121712 | 130121810 | 130110573 | 130110661 | 130122304 | 130122377 | 575 | 453 | 265 | 463 | 645 | 0 | 0 | 0 | 0 | 0 | 3132 | 1287 | 826 | 1599 | 1856 | 0 | 0 | 0 | 1 | 0 | 247 | 149 | 1 | 1 | 1 | 1 | 1 | 1 | 1 | 1 | 1 | 1 | 0.999 | 1 | 0 |
| SE | 100246 | ENSG00000084234 | APLP2 | chr11 | + | 130127765 | 130127840 | 130126699 | 130126830 | 130129047 | 130129206 | 1936 | 1119 | 1133 | 1276 | 1596 | 1 | 0 | 0 | 0 | 0 | 6066 | 3164 | 1815 | 3299 | 3665 | 0 | 0 | 0 | 0 | 0 | 224 | 149 | 1 | 1 | 0.999 | 1 | 1 | 1 | 1 | 1 | 1 | 1 | 1 | 1 | 0 |
| SE | 100254 | ENSG00000166471 | TMEM41B | chr11 | - | 9283370 | 9283593 | 9280653 | 9281041 | 9286454 | 9286593 | 82 | 79 | 109 | 93 | 112 | 0 | 0 | 0 | 0 | 0 | 63 | 120 | 127 | 102 | 4 | 0 | 0 | 0 | 2 | 0 | 298 | 149 | 1 | 1 | 1 | 1 | 1 | 1 | 1 | 1 | 1 | 1 | 0.962 | 1 | 0.008 |
| SE | 100256 | ENSG00000166471 | TMEM41B | chr11 | - | 9299583 | 9299701 | 9295095 | 9295387 | 9314320 | 9314567 | 38 | 26 | 59 | 45 | 77 | 0 | 0 | 0 | 1 | 0 | 0 | 31 | 23 | 23 | 20 | 0 | 0 | 0 | 0 | 0 | 267 | 149 | 1 | 1 | 1 | 1 | 1 | 0.962 | 1 | NA | 1 | 1 | 1 | 1 | -0.008 |
| SE | 100257 | ENSG00000110448 | CD5 | chr11 | + | 61115055 | 61115094 | 61102488 | 61102615 | 61118174 | 61118480 | 212 | 106 | 280 | 224 | 102 | 0 | 0 | 0 | 0 | 0 | 91 | 262 | 197 | 121 | 262 | 0 | 0 | 0 | 4 | 0 | 188 | 149 | 1 | 1 | 1 | 1 | 1 | 1 | 1 | 1 | 1 | 1 | 0.96 | 1 | 0.008 |
| SE | 100258 | ENSG00000110448 | CD5 | chr11 | + | 61121610 | 61121904 | 61119233 | 61119575 | 61122906 | 61123032 | 272 | 177 | 326 | 273 | 296 | 0 | 0 | 0 | 1 | 0 | 60 | 315 | 222 | 116 | 282 | 0 | 0 | 0 | 0 | 0 | 298 | 149 | 1 | 1 | 1 | 1 | 1 | 0.993 | 1 | 1 | 1 | 1 | 1 | 1 | -0.001 |
| SE | 100259 | ENSG00000110448 | CD5 | chr11 | + | 61121610 | 61121904 | 61119233 | 61119575 | 61123883 | 61123937 | 133 | 83 | 143 | 136 | 66 | 0 | 0 | 1 | 0 | 0 | 42 | 159 | 104 | 40 | 130 | 0 | 0 | 0 | 0 | 0 | 298 | 149 | 1 | 1 | 1 | 1 | 0.986 | 1 | 1 | 1 | 1 | 1 | 1 | 1 | -0.003 |
| SE | 100260 | ENSG00000110448 | CD5 | chr11 | + | 61122906 | 61123032 | 61119233 | 61119575 | 61123883 | 61123937 | 134 | 75 | 149 | 131 | 241 | 0 | 0 | 1 | 0 | 0 | 10 | 160 | 112 | 99 | 114 | 0 | 0 | 0 | 0 | 0 | 275 | 149 | 1 | 1 | 1 | 1 | 0.988 | 1 | 1 | 1 | 1 | 1 | 1 | 1 | -0.002 |
| SE | 100261 | ENSG00000110448 | CD5 | chr11 | + | 61122906 | 61123032 | 61121610 | 61121904 | 61123883 | 61123937 | 273 | 169 | 334 | 267 | 471 | 1 | 0 | 3 | 0 | 0 | 28 | 315 | 229 | 174 | 269 | 0 | 0 | 0 | 0 | 6 | 275 | 149 | 1 | 1 | 0.993 | 1 | 0.984 | 1 | 1 | 1 | 1 | 1 | 1 | 0.96 | 0.003 |
| SE | 100262 | ENSG00000110448 | CD5 | chr11 | + | 61125750 | 61125841 | 61125031 | 61125151 | 61126287 | 61127852 | 215 | 115 | 294 | 247 | 349 | 0 | 0 | 2 | 0 | 0 | 39 | 419 | 205 | 131 | 308 | 0 | 0 | 0 | 0 | 0 | 240 | 149 | 1 | 1 | 1 | 1 | 0.989 | 1 | 1 | 1 | 1 | 1 | 1 | 1 | -0.002 |
| SE | 100266 | ENSG00000171940 | ZNF217 | chr20 | - | 53597630 | 53597703 | 53582924 | 53583168 | 53609747 | 53609907 | 11 | 5 | 5 | 10 | 24 | 0 | 0 | 1 | 0 | 2 | 7 | 3 | 12 | 7 | 0 | 0 | 1 | 0 | 0 | 0 | 222 | 149 | 1 | 1 | 1 | 1 | 0.77 | 1 | 0.89 | 1 | 0.668 | 1 | 1 | NA | 0.015 |
| SE | 100267 | ENSG00000171940 | ZNF217 | chr20 | - | 53608353 | 53608444 | 53582541 | 53583168 | 53609747 | 53609907 | 2 | 0 | 2 | 1 | 1 | 0 | 0 | 1 | 0 | 2 | 0 | 3 | 0 | 0 | 0 | 0 | 1 | 0 | 0 | 0 | 240 | 149 | 1 | 1 | 1 | NA | 0.554 | 1 | 0.237 | NA | 0.651 | NA | NA | NA | 0.047 |
| SE | 100268 | ENSG00000171940 | ZNF217 | chr20 | - | 53608353 | 53608444 | 53597630 | 53597703 | 53609747 | 53609907 | 12 | 3 | 11 | 11 | 3 | 0 | 0 | 0 | 1 | 0 | 0 | 12 | 6 | 2 | 0 | 0 | 0 | 0 | 0 | 0 | 240 | 149 | 1 | 1 | 1 | 1 | 1 | 0.872 | 1 | NA | 1 | 1 | 1 | NA | -0.026 |
| SE | 100271 | ENSG00000167986 | DDB1 | chr11 | - | 61302581 | 61302751 | 61301711 | 61302359 | 61303045 | 61303155 | 462 | 253 | 431 | 402 | 611 | 0 | 0 | 0 | 2 | 0 | 580 | 361 | 201 | 236 | 446 | 8 | 1 | 0 | 0 | 2 | 298 | 149 | 1 | 1 | 1 | 1 | 1 | 0.99 | 1 | 0.973 | 0.994 | 1 | 1 | 0.991 | 0.006 |
| SE | 100272 | ENSG00000167986 | DDB1 | chr11 | - | 61308982 | 61309077 | 61303750 | 61304035 | 61310294 | 61310418 | 104 | 41 | 103 | 90 | 208 | 0 | 0 | 0 | 0 | 0 | 114 | 97 | 50 | 61 | 156 | 0 | 0 | 0 | 1 | 0 | 244 | 149 | 1 | 1 | 1 | 1 | 1 | 1 | 1 | 1 | 1 | 1 | 0.974 | 1 | 0.005 |
| SE | 100273 | ENSG00000167986 | DDB1 | chr11 | - | 61309795 | 61309861 | 61308982 | 61309077 | 61310294 | 61310418 | 106 | 63 | 91 | 100 | 90 | 0 | 0 | 0 | 1 | 0 | 92 | 89 | 57 | 101 | 99 | 0 | 0 | 0 | 0 | 0 | 215 | 149 | 1 | 1 | 1 | 1 | 1 | 0.986 | 1 | 1 | 1 | 1 | 1 | 1 | -0.003 |
| SE | 100274 | ENSG00000167986 | DDB1 | chr11 | - | 61309795 | 61309960 | 61308982 | 61309077 | 61310294 | 61310418 | 604 | 309 | 473 | 524 | 557 | 0 | 0 | 0 | 1 | 0 | 484 | 490 | 351 | 402 | 649 | 0 | 0 | 0 | 0 | 0 | 298 | 149 | 1 | 1 | 1 | 1 | 1 | 0.996 | 1 | 1 | 1 | 1 | 1 | 1 | -0.001 |
| SE | 100275 | ENSG00000167986 | DDB1 | chr11 | - | 61309795 | 61309960 | 61308982 | 61309077 | 61311783 | 61311895 | 308 | 164 | 236 | 285 | 276 | 0 | 0 | 0 | 3 | 0 | 281 | 251 | 172 | 234 | 375 | 0 | 1 | 0 | 0 | 0 | 298 | 149 | 1 | 1 | 1 | 1 | 1 | 0.979 | 1 | 1 | 0.992 | 1 | 1 | 1 | -0.003 |
| SE | 100276 | ENSG00000167986 | DDB1 | chr11 | - | 61309795 | 61309979 | 61308982 | 61309077 | 61310294 | 61310418 | 308 | 159 | 234 | 285 | 274 | 0 | 0 | 0 | 1 | 0 | 271 | 248 | 169 | 225 | 372 | 0 | 0 | 0 | 0 | 0 | 298 | 149 | 1 | 1 | 1 | 1 | 1 | 0.993 | 1 | 1 | 1 | 1 | 1 | 1 | -0.001 |
| SE | 100277 | ENSG00000167986 | DDB1 | chr11 | - | 61310294 | 61310418 | 61308982 | 61309077 | 61311783 | 61311895 | 262 | 116 | 181 | 237 | 329 | 0 | 0 | 0 | 3 | 0 | 240 | 237 | 170 | 183 | 396 | 0 | 1 | 0 | 0 | 0 | 273 | 149 | 1 | 1 | 1 | 1 | 1 | 0.977 | 1 | 1 | 0.992 | 1 | 1 | 1 | -0.003 |
| SE | 100279 | ENSG00000167986 | DDB1 | chr11 | - | 61313861 | 61313969 | 61313498 | 61313706 | 61314046 | 61314202 | 345 | 160 | 307 | 275 | 313 | 0 | 0 | 0 | 0 | 0 | 253 | 327 | 150 | 261 | 417 | 0 | 0 | 2 | 0 | 0 | 257 | 149 | 1 | 1 | 1 | 1 | 1 | 1 | 1 | 1 | 1 | 0.978 | 1 | 1 | 0.004 |
| SE | 100280 | ENSG00000167986 | DDB1 | chr11 | - | 61315331 | 61315523 | 61314452 | 61314486 | 61316284 | 61316393 | 0 | 0 | 0 | 1 | 0 | 269 | 137 | 240 | 221 | 509 | 0 | 0 | 0 | 0 | 0 | 173 | 203 | 128 | 146 | 306 | 298 | 149 | 1 | 1 | 0 | 0 | 0 | 0.002 | 0 | 0 | 0 | 0 | 0 | 0 | 0 |
| SE | 100282 | ENSG00000167986 | DDB1 | chr11 | - | 61316284 | 61316393 | 61314046 | 61314486 | 61321594 | 61321697 | 202 | 99 | 169 | 150 | 293 | 0 | 0 | 2 | 0 | 0 | 104 | 136 | 79 | 103 | 203 | 0 | 0 | 1 | 0 | 0 | 258 | 149 | 1 | 1 | 1 | 1 | 0.98 | 1 | 1 | 1 | 1 | 0.979 | 1 | 1 | 0 |
| SE | 100283 | ENSG00000167986 | DDB1 | chr11 | - | 61316491 | 61316567 | 61314883 | 61316393 | 61321594 | 61321697 | 384 | 208 | 336 | 346 | 502 | 0 | 1 | 0 | 0 | 1 | 384 | 312 | 253 | 283 | 483 | 0 | 0 | 0 | 0 | 0 | 225 | 149 | 1 | 1 | 1 | 0.993 | 1 | 1 | 0.997 | 1 | 1 | 1 | 1 | 1 | -0.002 |
| SE | 100284 | ENSG00000167986 | DDB1 | chr11 | - | 61321594 | 61321697 | 61316491 | 61316567 | 61322295 | 61322412 | 452 | 249 | 357 | 394 | 492 | 0 | 0 | 0 | 1 | 0 | 385 | 381 | 260 | 336 | 501 | 0 | 0 | 0 | 1 | 0 | 252 | 149 | 1 | 1 | 1 | 1 | 1 | 0.996 | 1 | 1 | 1 | 1 | 0.995 | 1 | 0 |
| SE | 100285 | ENSG00000167986 | DDB1 | chr11 | - | 61322295 | 61322412 | 61316491 | 61316567 | 61323978 | 61324137 | 0 | 0 | 0 | 1 | 0 | 1 | 0 | 0 | 0 | 0 | 18 | 0 | 9 | 5 | 4 | 0 | 0 | 1 | 0 | 0 | 266 | 149 | 1 | 1 | 0 | NA | NA | 1 | NA | 1 | NA | 0.834 | 1 | 1 | -0.459 |
| SE | 100286 | ENSG00000167986 | DDB1 | chr11 | - | 61323010 | 61323094 | 61314046 | 61314486 | 61323978 | 61324137 | 178 | 124 | 161 | 147 | 157 | 0 | 0 | 2 | 2 | 0 | 155 | 103 | 100 | 110 | 183 | 0 | 0 | 0 | 0 | 0 | 233 | 149 | 1 | 1 | 1 | 1 | 0.981 | 0.979 | 1 | 1 | 1 | 1 | 1 | 1 | -0.008 |
| SE | 100289 | ENSG00000167986 | DDB1 | chr11 | - | 61325610 | 61325708 | 61323660 | 61324137 | 61326778 | 61326893 | 257 | 125 | 215 | 207 | 292 | 0 | 0 | 1 | 2 | 0 | 456 | 281 | 148 | 197 | 399 | 0 | 0 | 0 | 0 | 0 | 247 | 149 | 1 | 1 | 1 | 1 | 0.992 | 0.984 | 1 | 1 | 1 | 1 | 1 | 1 | -0.005 |
| SE | 100291 | ENSG00000167986 | DDB1 | chr11 | - | 61326778 | 61326893 | 61323660 | 61324137 | 61331542 | 61331691 | 2 | 0 | 2 | 2 | 3 | 0 | 0 | 0 | 0 | 0 | 0 | 4 | 1 | 0 | 7 | 0 | 6 | 0 | 0 | 0 | 264 | 149 | 1 | 1 | 1 | NA | 1 | 1 | 1 | NA | 0.273 | 1 | NA | 1 | 0.242 |
| SE | 100295 | ENSG00000167986 | DDB1 | chr11 | - | 61331542 | 61331691 | 61326778 | 61326893 | 61332907 | 61333008 | 81 | 31 | 72 | 74 | 45 | 0 | 0 | 0 | 0 | 0 | 52 | 98 | 50 | 52 | 124 | 0 | 0 | 0 | 2 | 0 | 298 | 149 | 1 | 1 | 1 | 1 | 1 | 1 | 1 | 1 | 1 | 1 | 0.929 | 1 | 0.014 |
| SE | 100297 | ENSG00000162144 | CYB561A3 | chr11 | - | 61353783 | 61353823 | 61349646 | 61353139 | 61356529 | 61356728 | 5 | 2 | 7 | 3 | 6 | 0 | 0 | 0 | 1 | 0 | 0 | 21 | 2 | 4 | 19 | 0 | 0 | 0 | 0 | 0 | 189 | 149 | 1 | 1 | 1 | 1 | 1 | 0.703 | 1 | NA | 1 | 1 | 1 | 1 | -0.059 |
| SE | 100298 | ENSG00000162144 | CYB561A3 | chr11 | - | 61353783 | 61353992 | 61352955 | 61353139 | 61356529 | 61356728 | 121 | 29 | 63 | 76 | 80 | 0 | 0 | 0 | 1 | 0 | 61 | 148 | 23 | 84 | 286 | 0 | 0 | 0 | 0 | 0 | 298 | 149 | 1 | 1 | 1 | 1 | 1 | 0.974 | 1 | 1 | 1 | 1 | 1 | 1 | -0.005 |
| SE | 100299 | ENSG00000162144 | CYB561A3 | chr11 | - | 61356014 | 61356088 | 61353854 | 61353992 | 61356529 | 61356728 | 0 | 0 | 0 | 1 | 0 | 60 | 14 | 26 | 35 | 57 | 0 | 0 | 0 | 0 | 0 | 7 | 85 | 12 | 34 | 132 | 223 | 149 | 1 | 1 | 0 | 0 | 0 | 0.019 | 0 | 0 | 0 | 0 | 0 | 0 | 0.004 |
| SE | 100302 | ENSG00000162144 | CYB561A3 | chr11 | - | 61357732 | 61357828 | 61356410 | 61356728 | 61361732 | 61361819 | 31 | 16 | 43 | 35 | 52 | 1 | 0 | 1 | 0 | 0 | 13 | 40 | 10 | 26 | 64 | 0 | 0 | 0 | 0 | 0 | 245 | 149 | 1 | 1 | 0.95 | 1 | 0.963 | 1 | 1 | 1 | 1 | 1 | 1 | 1 | -0.017 |
| SE | 100303 | ENSG00000162144 | CYB561A3 | chr11 | - | 61359425 | 61359571 | 61356547 | 61356728 | 61361732 | 61361882 | 0 | 0 | 0 | 1 | 0 | 1 | 0 | 1 | 0 | 0 | 0 | 0 | 0 | 0 | 9 | 0 | 0 | 0 | 0 | 0 | 295 | 149 | 1 | 1 | 0 | NA | 0 | 1 | NA | NA | NA | NA | NA | 1 | -0.667 |
| SE | 100306 | ENSG00000149483 | TMEM138 | chr11 | + | 61367922 | 61367998 | 61365702 | 61366216 | 61368596 | 61368751 | 39 | 18 | 19 | 24 | 26 | 0 | 2 | 0 | 0 | 0 | 18 | 30 | 15 | 25 | 63 | 0 | 0 | 0 | 0 | 0 | 225 | 149 | 1 | 1 | 1 | 0.856 | 1 | 1 | 1 | 1 | 1 | 1 | 1 | 1 | -0.029 |
| SE | 100307 | ENSG00000187049 | TMEM216 | chr11 | + | 61393230 | 61393332 | 61392392 | 61392679 | 61393883 | 61393976 | 29 | 16 | 30 | 25 | 32 | 2 | 0 | 0 | 0 | 0 | 15 | 31 | 6 | 7 | 5 | 0 | 0 | 0 | 0 | 0 | 251 | 149 | 1 | 1 | 0.896 | 1 | 1 | 1 | 1 | 1 | 1 | 1 | 1 | 1 | -0.021 |
| SE | 100308 | ENSG00000118894 | EEF2KMT | chr16 | - | 5090083 | 5090349 | 5089106 | 5089256 | 5090431 | 5090565 | 10 | 1 | 2 | 13 | 4 | 1 | 0 | 0 | 0 | 0 | 7 | 8 | 16 | 8 | 12 | 0 | 0 | 0 | 2 | 0 | 298 | 149 | 1 | 1 | 0.833 | 1 | 1 | 1 | 1 | 1 | 1 | 1 | 0.667 | 1 | 0.033 |
| SE | 100311 | ENSG00000118894 | EEF2KMT | chr16 | - | 5091793 | 5091895 | 5090083 | 5090565 | 5093483 | 5093564 | 10 | 0 | 12 | 1 | 2 | 1 | 0 | 0 | 0 | 3 | 0 | 4 | 13 | 8 | 8 | 0 | 9 | 2 | 0 | 0 | 251 | 149 | 1 | 1 | 0.856 | NA | 1 | 1 | 0.284 | NA | 0.209 | 0.794 | 1 | 1 | 0.034 |
| SE | 100315 | ENSG00000118894 | EEF2KMT | chr16 | - | 5093483 | 5093564 | 5091793 | 5091895 | 5095451 | 5095514 | 11 | 0 | 13 | 2 | 4 | 0 | 0 | 0 | 0 | 2 | 0 | 2 | 4 | 4 | 0 | 0 | 0 | 0 | 0 | 8 | 230 | 149 | 1 | 1 | 1 | NA | 1 | 1 | 0.564 | NA | 1 | 1 | 1 | 0 | 0.141 |
| SE | 100316 | ENSG00000118894 | EEF2KMT | chr16 | - | 5097279 | 5097353 | 5095451 | 5095514 | 5097643 | 5097766 | 2 | 0 | 1 | 0 | 2 | 6 | 0 | 7 | 1 | 1 | 0 | 0 | 0 | 0 | 0 | 0 | 0 | 6 | 0 | 0 | 223 | 149 | 1 | 1 | 0.182 | NA | 0.087 | 0 | 0.572 | NA | NA | 0 | NA | NA | 0.21 |
| SE | 100317 | ENSG00000081237 | PTPRC | chr1 | + | 198692346 | 198692373 | 198639161 | 198639341 | 198696711 | 198696909 | 1155 | 722 | 922 | 1249 | 2326 | 1 | 0 | 0 | 0 | 0 | 203 | 695 | 358 | 120 | 66 | 0 | 0 | 0 | 0 | 0 | 176 | 149 | 1 | 1 | 0.999 | 1 | 1 | 1 | 1 | 1 | 1 | 1 | 1 | 1 | 0 |
| SE | 100318 | ENSG00000081237 | PTPRC | chr1 | + | 198692346 | 198692373 | 198639161 | 198639341 | 198703297 | 198703372 | 1043 | 721 | 691 | 1096 | 2361 | 0 | 1 | 0 | 0 | 0 | 103 | 499 | 278 | 77 | 75 | 0 | 0 | 0 | 0 | 0 | 176 | 149 | 1 | 1 | 1 | 0.998 | 1 | 1 | 1 | 1 | 1 | 1 | 1 | 1 | 0 |
| SE | 100327 | ENSG00000081237 | PTPRC | chr1 | + | 198696711 | 198696909 | 198694017 | 198694140 | 198703297 | 198703372 | 35 | 9 | 19 | 11 | 28 | 2 | 0 | 0 | 0 | 2 | 0 | 11 | 3 | 0 | 2 | 0 | 0 | 0 | 0 | 0 | 298 | 149 | 1 | 1 | 0.897 | 1 | 1 | 1 | 0.875 | NA | 1 | 1 | NA | 1 | -0.046 |
| SE | 100328 | ENSG00000081237 | PTPRC | chr1 | + | 198699563 | 198699704 | 198692346 | 198692373 | 198702386 | 198702530 | 1941 | 1280 | 1540 | 1880 | 3955 | 6 | 6 | 0 | 6 | 4 | 148 | 932 | 622 | 148 | 66 | 0 | 0 | 1 | 0 | 0 | 290 | 149 | 1 | 1 | 0.994 | 0.991 | 1 | 0.994 | 0.998 | 1 | 1 | 0.997 | 1 | 1 | -0.004 |
| SE | 100330 | ENSG00000081237 | PTPRC | chr1 | + | 198699563 | 198699704 | 198692346 | 198692373 | 198704471 | 198704485 | 994 | 886 | 975 | 1219 | 2979 | 1 | 0 | 0 | 0 | 0 | 53 | 566 | 372 | 93 | 43 | 0 | 0 | 0 | 0 | 0 | 290 | 149 | 1 | 1 | 0.998 | 1 | 1 | 1 | 1 | 1 | 1 | 1 | 1 | 1 | 0 |
| SE | 100331 | ENSG00000081237 | PTPRC | chr1 | + | 198699563 | 198699704 | 198694017 | 198694140 | 198703297 | 198703346 | 365 | 360 | 478 | 510 | 1188 | 2 | 0 | 0 | 0 | 2 | 39 | 321 | 155 | 38 | 22 | 0 | 0 | 0 | 0 | 0 | 290 | 149 | 1 | 1 | 0.989 | 1 | 1 | 1 | 0.997 | 1 | 1 | 1 | 1 | 1 | -0.003 |
| SE | 100332 | ENSG00000081237 | PTPRC | chr1 | + | 198699563 | 198699704 | 198696711 | 198696909 | 198702386 | 198702530 | 1874 | 805 | 1155 | 1324 | 2542 | 0 | 1 | 0 | 1 | 0 | 206 | 1000 | 500 | 133 | 107 | 0 | 0 | 2 | 0 | 0 | 290 | 149 | 1 | 1 | 1 | 0.998 | 1 | 0.999 | 1 | 1 | 1 | 0.992 | 1 | 1 | 0.001 |
| SE | 100333 | ENSG00000081237 | PTPRC | chr1 | + | 198699563 | 198699704 | 198696711 | 198696909 | 198703297 | 198703346 | 1280 | 759 | 1064 | 1166 | 2732 | 1 | 1 | 0 | 0 | 4 | 150 | 947 | 402 | 116 | 111 | 0 | 0 | 0 | 0 | 0 | 290 | 149 | 1 | 1 | 0.998 | 0.997 | 1 | 1 | 0.997 | 1 | 1 | 1 | 1 | 1 | -0.002 |
| SE | 100335 | ENSG00000081237 | PTPRC | chr1 | + | 198702386 | 198702530 | 198696711 | 198696909 | 198703297 | 198703346 | 480 | 285 | 404 | 474 | 350 | 1 | 1 | 0 | 0 | 4 | 25 | 212 | 130 | 26 | 0 | 0 | 0 | 0 | 0 | 0 | 293 | 149 | 1 | 1 | 0.996 | 0.993 | 1 | 1 | 0.978 | 1 | 1 | 1 | 1 | NA | -0.007 |
| SE | 100337 | ENSG00000081237 | PTPRC | chr1 | + | 198703297 | 198703372 | 198692346 | 198692373 | 198704471 | 198704485 | 1060 | 728 | 660 | 979 | 1851 | 1 | 0 | 0 | 0 | 0 | 82 | 397 | 232 | 75 | 52 | 0 | 0 | 0 | 0 | 0 | 224 | 149 | 1 | 1 | 0.999 | 1 | 1 | 1 | 1 | 1 | 1 | 1 | 1 | 1 | 0 |
| SE | 100338 | ENSG00000081237 | PTPRC | chr1 | + | 198703297 | 198703372 | 198699563 | 198699704 | 198704471 | 198704485 | 914 | 719 | 863 | 1020 | 1958 | 1 | 0 | 0 | 0 | 0 | 62 | 512 | 282 | 78 | 34 | 0 | 0 | 0 | 0 | 0 | 224 | 149 | 1 | 1 | 0.998 | 1 | 1 | 1 | 1 | 1 | 1 | 1 | 1 | 1 | 0 |
| SE | 100339 | ENSG00000081237 | PTPRC | chr1 | + | 198706733 | 198706952 | 198704471 | 198704498 | 198712952 | 198713072 | 1279 | 888 | 972 | 1243 | 2127 | 0 | 0 | 0 | 2 | 0 | 99 | 561 | 359 | 97 | 52 | 0 | 0 | 0 | 0 | 0 | 298 | 149 | 1 | 1 | 1 | 1 | 1 | 0.997 | 1 | 1 | 1 | 1 | 1 | 1 | -0.001 |
| SE | 100340 | ENSG00000081237 | PTPRC | chr1 | + | 198708132 | 198708261 | 198706733 | 198706952 | 198709686 | 198709824 | 7281 | 5289 | 5254 | 7095 | 21094 | 0 | 0 | 0 | 0 | 3 | 447 | 3461 | 2157 | 488 | 317 | 0 | 0 | 0 | 0 | 0 | 278 | 149 | 1 | 1 | 1 | 1 | 1 | 1 | 1 | 1 | 1 | 1 | 1 | 1 | 0 |
| SE | 100341 | ENSG00000081237 | PTPRC | chr1 | + | 198708132 | 198708261 | 198706733 | 198706952 | 198712952 | 198713072 | 2509 | 1677 | 1693 | 2260 | 7226 | 0 | 0 | 0 | 0 | 2 | 164 | 1074 | 671 | 156 | 106 | 0 | 3 | 1 | 0 | 0 | 278 | 149 | 1 | 1 | 1 | 1 | 1 | 1 | 0.999 | 1 | 0.995 | 0.997 | 1 | 1 | 0.001 |
| SE | 100342 | ENSG00000081237 | PTPRC | chr1 | + | 198709686 | 198709824 | 198706733 | 198706952 | 198712952 | 198713072 | 2800 | 1889 | 2028 | 2511 | 5905 | 0 | 0 | 0 | 0 | 2 | 106 | 1239 | 644 | 179 | 99 | 0 | 3 | 1 | 0 | 0 | 287 | 149 | 1 | 1 | 1 | 1 | 1 | 1 | 0.999 | 1 | 0.995 | 0.997 | 1 | 1 | 0.001 |
| SE | 100344 | ENSG00000081237 | PTPRC | chr1 | + | 198722415 | 198722476 | 198718093 | 198718302 | 198728339 | 198728448 | 2557 | 1855 | 1870 | 2488 | 3547 | 3 | 0 | 0 | 0 | 0 | 120 | 896 | 520 | 148 | 56 | 0 | 0 | 0 | 0 | 0 | 210 | 149 | 1 | 1 | 0.998 | 1 | 1 | 1 | 1 | 1 | 1 | 1 | 1 | 1 | 0 |
| SE | 100345 | ENSG00000081237 | PTPRC | chr1 | + | 198732299 | 198732390 | 198731616 | 198731726 | 198732479 | 198732556 | 8770 | 7194 | 6287 | 8661 | 11849 | 0 | 1 | 0 | 0 | 0 | 494 | 3746 | 2331 | 635 | 257 | 0 | 0 | 0 | 0 | 0 | 240 | 149 | 1 | 1 | 1 | 1 | 1 | 1 | 1 | 1 | 1 | 1 | 1 | 1 | 0 |
| SE | 100346 | ENSG00000081237 | PTPRC | chr1 | + | 198741868 | 198742026 | 198735126 | 198735252 | 198742231 | 198742367 | 6444 | 3899 | 4780 | 5467 | 7177 | 0 | 0 | 0 | 2 | 0 | 300 | 2488 | 1596 | 389 | 266 | 0 | 0 | 0 | 0 | 0 | 298 | 149 | 1 | 1 | 1 | 1 | 1 | 0.999 | 1 | 1 | 1 | 1 | 1 | 1 | 0 |
| SE | 100347 | ENSG00000081237 | PTPRC | chr1 | + | 198741868 | 198742026 | 198735126 | 198735252 | 198744053 | 198744203 | 3642 | 1820 | 2177 | 2564 | 3829 | 0 | 5 | 1 | 4 | 0 | 149 | 1208 | 880 | 210 | 133 | 0 | 0 | 0 | 1 | 6 | 298 | 149 | 1 | 1 | 1 | 0.995 | 0.999 | 0.997 | 1 | 1 | 1 | 1 | 0.991 | 0.917 | 0.017 |
| SE | 100349 | ENSG00000081237 | PTPRC | chr1 | + | 198742231 | 198742367 | 198741868 | 198742026 | 198744053 | 198744203 | 4127 | 2993 | 3507 | 4187 | 5429 | 1 | 3 | 0 | 4 | 2 | 253 | 1958 | 1203 | 262 | 234 | 0 | 0 | 0 | 0 | 0 | 285 | 149 | 1 | 1 | 1 | 0.998 | 1 | 0.998 | 0.999 | 1 | 1 | 1 | 1 | 1 | -0.001 |
| SE | 100350 | ENSG00000081237 | PTPRC | chr1 | + | 198748108 | 198748199 | 198744053 | 198744203 | 198750491 | 198750626 | 2771 | 2107 | 2007 | 2542 | 4200 | 0 | 0 | 2 | 0 | 0 | 106 | 1031 | 648 | 167 | 80 | 0 | 0 | 0 | 0 | 0 | 240 | 149 | 1 | 1 | 1 | 1 | 0.998 | 1 | 1 | 1 | 1 | 1 | 1 | 1 | 0 |
| SE | 100351 | ENSG00000081237 | PTPRC | chr1 | + | 198749415 | 198749549 | 198748108 | 198748199 | 198750491 | 198750626 | 9771 | 7761 | 6521 | 8837 | 11458 | 17 | 7 | 6 | 29 | 17 | 540 | 3967 | 2634 | 653 | 251 | 9 | 0 | 2 | 0 | 0 | 283 | 149 | 1 | 1 | 0.997 | 0.998 | 0.998 | 0.994 | 0.997 | 0.969 | 1 | 0.999 | 1 | 1 | 0.003 |
| SE | 100352 | ENSG00000174842 | GLMN | chr1 | - | 92264553 | 92264638 | 92263622 | 92263732 | 92266418 | 92266492 | 54 | 39 | 39 | 65 | 102 | 1 | 0 | 0 | 0 | 2 | 19 | 28 | 13 | 7 | 18 | 0 | 0 | 0 | 0 | 0 | 234 | 149 | 1 | 1 | 0.972 | 1 | 1 | 1 | 0.97 | 1 | 1 | 1 | 1 | 1 | -0.012 |
| SE | 100356 | ENSG00000174842 | GLMN | chr1 | - | 92297403 | 92297529 | 92291417 | 92291537 | 92297960 | 92298029 | 40 | 20 | 27 | 26 | 41 | 0 | 0 | 2 | 2 | 5 | 7 | 12 | 14 | 27 | 0 | 0 | 0 | 0 | 0 | 0 | 275 | 149 | 1 | 1 | 1 | 1 | 0.88 | 0.876 | 0.816 | 1 | 1 | 1 | 1 | NA | -0.086 |
| SE | 100358 | ENSG00000149541 | B3GAT3 | chr11 | - | 62621054 | 62621123 | 62620496 | 62620671 | 62621865 | 62622146 | 0 | 2 | 0 | 0 | 0 | 37 | 8 | 18 | 26 | 26 | 0 | 0 | 0 | 0 | 4 | 24 | 22 | 9 | 27 | 40 | 218 | 149 | 1 | 1 | 0 | 0.146 | 0 | 0 | 0 | 0 | 0 | 0 | 0 | 0.064 | 0.016 |
| SE | 100359 | ENSG00000149541 | B3GAT3 | chr11 | - | 62621054 | 62621293 | 62620496 | 62620671 | 62621865 | 62621984 | 0 | 2 | 0 | 0 | 0 | 37 | 8 | 18 | 26 | 26 | 0 | 0 | 0 | 0 | 4 | 24 | 22 | 9 | 27 | 40 | 298 | 149 | 1 | 1 | 0 | 0.111 | 0 | 0 | 0 | 0 | 0 | 0 | 0 | 0.048 | 0.013 |
| SE | 100361 | ENSG00000089597 | GANAB | chr11 | - | 62628768 | 62629012 | 62627288 | 62627353 | 62629193 | 62629295 | 617 | 291 | 543 | 526 | 365 | 0 | 0 | 0 | 0 | 0 | 635 | 600 | 345 | 385 | 787 | 0 | 0 | 2 | 0 | 0 | 298 | 149 | 1 | 1 | 1 | 1 | 1 | 1 | 1 | 1 | 1 | 0.989 | 1 | 1 | 0.002 |
| SE | 100363 | ENSG00000089597 | GANAB | chr11 | - | 62630196 | 62630276 | 62629813 | 62629957 | 62630378 | 62630505 | 533 | 208 | 446 | 420 | 250 | 0 | 0 | 0 | 0 | 0 | 410 | 511 | 281 | 314 | 584 | 4 | 0 | 1 | 0 | 0 | 229 | 149 | 1 | 1 | 1 | 1 | 1 | 1 | 1 | 0.985 | 1 | 0.995 | 1 | 1 | 0.004 |
| SE | 100364 | ENSG00000089597 | GANAB | chr11 | - | 62632564 | 62632745 | 62631029 | 62631183 | 62633004 | 62633101 | 384 | 158 | 380 | 361 | 358 | 0 | 0 | 0 | 0 | 0 | 371 | 381 | 228 | 201 | 470 | 0 | 6 | 0 | 1 | 0 | 298 | 149 | 1 | 1 | 1 | 1 | 1 | 1 | 1 | 1 | 0.969 | 1 | 0.99 | 1 | 0.008 |
| SE | 100366 | ENSG00000089597 | GANAB | chr11 | - | 62634309 | 62634375 | 62633440 | 62633567 | 62634820 | 62635000 | 19 | 3 | 4 | 10 | 16 | 0 | 1 | 0 | 8 | 3 | 0 | 13 | 0 | 12 | 22 | 0 | 0 | 4 | 2 | 8 | 215 | 149 | 1 | 1 | 1 | 0.675 | 1 | 0.464 | 0.787 | NA | 1 | 0 | 0.806 | 0.656 | 0.17 |
| SE | 100370 | ENSG00000089597 | GANAB | chr11 | - | 62639929 | 62640069 | 62639626 | 62639731 | 62646561 | 62646606 | 4 | 1 | 1 | 1 | 0 | 47 | 16 | 38 | 42 | 27 | 0 | 2 | 0 | 5 | 0 | 25 | 23 | 43 | 27 | 68 | 289 | 149 | 1 | 1 | 0.042 | 0.031 | 0.013 | 0.012 | 0 | 0 | 0.043 | 0 | 0.087 | 0 | -0.006 |
| SE | 100371 | ENSG00000162194 | LBHD1 | chr11 | - | 62669640 | 62669803 | 62667214 | 62667747 | 62669881 | 62670015 | 29 | 9 | 32 | 12 | 17 | 0 | 0 | 3 | 1 | 0 | 51 | 50 | 16 | 30 | 54 | 0 | 0 | 0 | 0 | 0 | 298 | 149 | 1 | 1 | 1 | 1 | 0.842 | 0.857 | 1 | 1 | 1 | 1 | 1 | 1 | -0.06 |
| SE | 100372 | ENSG00000162194 | LBHD1 | chr11 | - | 62671239 | 62671388 | 62669881 | 62670015 | 62671563 | 62671633 | 0 | 1 | 0 | 1 | 1 | 2 | 0 | 0 | 0 | 0 | 0 | 13 | 2 | 0 | 0 | 0 | 0 | 0 | 0 | 0 | 298 | 149 | 1 | 1 | 0 | 1 | NA | 1 | 1 | NA | 1 | 1 | NA | NA | -0.25 |
| SE | 100373 | ENSG00000099910 | KLHL22 | chr22 | - | 20457807 | 20458000 | 20446442 | 20446676 | 20464857 | 20465576 | 146 | 45 | 116 | 82 | 73 | 1 | 0 | 0 | 0 | 0 | 80 | 57 | 50 | 76 | 120 | 0 | 0 | 0 | 0 | 0 | 298 | 149 | 1 | 1 | 0.986 | 1 | 1 | 1 | 1 | 1 | 1 | 1 | 1 | 1 | -0.003 |
| SE | 100374 | ENSG00000099910 | KLHL22 | chr22 | - | 20471349 | 20471515 | 20464857 | 20465576 | 20488984 | 20489212 | 81 | 27 | 68 | 67 | 44 | 2 | 0 | 3 | 0 | 0 | 42 | 71 | 11 | 29 | 72 | 0 | 0 | 0 | 0 | 0 | 298 | 149 | 1 | 1 | 0.953 | 1 | 0.919 | 1 | 1 | 1 | 1 | 1 | 1 | 1 | -0.026 |
| SE | 100375 | ENSG00000099910 | KLHL22 | chr22 | - | 20471349 | 20471515 | 20465541 | 20465623 | 20488984 | 20489212 | 51 | 9 | 34 | 28 | 11 | 0 | 0 | 0 | 0 | 0 | 23 | 42 | 10 | 15 | 29 | 0 | 4 | 0 | 0 | 0 | 298 | 149 | 1 | 1 | 1 | 1 | 1 | 1 | 1 | 1 | 0.84 | 1 | 1 | 1 | 0.032 |
| SE | 100376 | ENSG00000099910 | KLHL22 | chr22 | - | 20488984 | 20489212 | 20471349 | 20471515 | 20495759 | 20495795 | 52 | 12 | 36 | 29 | 11 | 3 | 0 | 0 | 0 | 0 | 23 | 42 | 7 | 15 | 29 | 0 | 0 | 0 | 0 | 0 | 298 | 149 | 1 | 1 | 0.897 | 1 | 1 | 1 | 1 | 1 | 1 | 1 | 1 | 1 | -0.021 |
| SE | 100377 | ENSG00000099910 | KLHL22 | chr22 | - | 20488984 | 20489244 | 20471349 | 20471515 | 20495759 | 20495792 | 58 | 9 | 35 | 32 | 11 | 3 | 0 | 0 | 0 | 0 | 23 | 51 | 7 | 15 | 29 | 0 | 0 | 0 | 0 | 0 | 298 | 149 | 1 | 1 | 0.906 | 1 | 1 | 1 | 1 | 1 | 1 | 1 | 1 | 1 | -0.019 |
| SE | 100380 | ENSG00000133812 | SBF2 | chr11 | - | 9784350 | 9784438 | 9781506 | 9781638 | 9785124 | 9785318 | 62 | 76 | 63 | 53 | 104 | 0 | 0 | 2 | 0 | 0 | 8 | 90 | 73 | 79 | 27 | 0 | 0 | 0 | 0 | 0 | 237 | 149 | 1 | 1 | 1 | 1 | 0.952 | 1 | 1 | 1 | 1 | 1 | 1 | 1 | -0.01 |
| SE | 100383 | ENSG00000133812 | SBF2 | chr11 | - | 9807999 | 9808185 | 9790555 | 9790683 | 9808900 | 9809002 | 18 | 10 | 7 | 26 | 31 | 0 | 0 | 0 | 0 | 0 | 22 | 39 | 9 | 13 | 0 | 0 | 9 | 0 | 0 | 0 | 298 | 149 | 1 | 1 | 1 | 1 | 1 | 1 | 1 | 1 | 0.684 | 1 | 1 | NA | 0.079 |
| SE | 100386 | ENSG00000133812 | SBF2 | chr11 | - | 9819228 | 9819303 | 9816839 | 9817024 | 9829355 | 9829496 | 0 | 0 | 0 | 0 | 3 | 4 | 4 | 8 | 8 | 21 | 0 | 0 | 0 | 0 | 0 | 2 | 18 | 6 | 10 | 8 | 224 | 149 | 1 | 1 | 0 | 0 | 0 | 0 | 0.087 | 0 | 0 | 0 | 0 | 0 | 0.017 |
| SE | 100390 | ENSG00000133812 | SBF2 | chr11 | - | 9829355 | 9829496 | 9816839 | 9817024 | 9832223 | 9832420 | 7 | 16 | 16 | 13 | 40 | 1 | 0 | 3 | 0 | 0 | 2 | 23 | 12 | 19 | 13 | 0 | 0 | 1 | 0 | 0 | 290 | 149 | 1 | 1 | 0.782 | 1 | 0.733 | 1 | 1 | 1 | 1 | 0.86 | 1 | 1 | -0.069 |
| SE | 100391 | ENSG00000133812 | SBF2 | chr11 | - | 9845564 | 9845740 | 9842624 | 9842770 | 9846955 | 9847083 | 84 | 94 | 66 | 89 | 101 | 2 | 0 | 0 | 3 | 0 | 45 | 61 | 120 | 130 | 17 | 0 | 0 | 2 | 0 | 0 | 298 | 149 | 1 | 1 | 0.955 | 1 | 1 | 0.937 | 1 | 1 | 1 | 0.968 | 1 | 1 | -0.015 |
| SE | 100392 | ENSG00000133812 | SBF2 | chr11 | - | 9850022 | 9850218 | 9846955 | 9847083 | 9852675 | 9852749 | 63 | 49 | 43 | 56 | 87 | 0 | 1 | 0 | 0 | 0 | 8 | 45 | 54 | 49 | 0 | 0 | 0 | 0 | 0 | 0 | 298 | 149 | 1 | 1 | 1 | 0.961 | 1 | 1 | 1 | 1 | 1 | 1 | 1 | NA | -0.008 |
| SE | 100393 | ENSG00000133812 | SBF2 | chr11 | - | 9853539 | 9853712 | 9852675 | 9852749 | 9856457 | 9856720 | 145 | 109 | 80 | 110 | 96 | 0 | 1 | 0 | 0 | 0 | 33 | 89 | 108 | 90 | 8 | 0 | 0 | 0 | 0 | 0 | 298 | 149 | 1 | 1 | 1 | 0.982 | 1 | 1 | 1 | 1 | 1 | 1 | 1 | 1 | -0.004 |
| SE | 100394 | ENSG00000133812 | SBF2 | chr11 | - | 9961956 | 9962106 | 9895942 | 9896011 | 9963772 | 9963882 | 145 | 147 | 119 | 155 | 251 | 0 | 1 | 0 | 0 | 0 | 64 | 127 | 152 | 103 | 32 | 0 | 0 | 0 | 0 | 0 | 298 | 149 | 1 | 1 | 1 | 0.987 | 1 | 1 | 1 | 1 | 1 | 1 | 1 | 1 | -0.003 |
| SE | 100395 | ENSG00000133812 | SBF2 | chr11 | - | 9963772 | 9963882 | 9895942 | 9896011 | 9968340 | 9968356 | 46 | 37 | 33 | 41 | 41 | 0 | 0 | 0 | 0 | 0 | 16 | 51 | 50 | 40 | 9 | 0 | 0 | 0 | 2 | 0 | 259 | 149 | 1 | 1 | 1 | 1 | 1 | 1 | 1 | 1 | 1 | 1 | 0.92 | 1 | 0.016 |
| SE | 100396 | ENSG00000133812 | SBF2 | chr11 | - | 9968340 | 9968545 | 9963772 | 9963882 | 9989496 | 9989595 | 104 | 98 | 95 | 108 | 127 | 0 | 0 | 0 | 1 | 0 | 60 | 136 | 118 | 128 | 16 | 0 | 0 | 0 | 0 | 0 | 298 | 149 | 1 | 1 | 1 | 1 | 1 | 0.982 | 1 | 1 | 1 | 1 | 1 | 1 | -0.004 |
| SE | 100398 | ENSG00000133812 | SBF2 | chr11 | - | 9993920 | 9993998 | 9992989 | 9993103 | 9998265 | 9998379 | 77 | 90 | 60 | 62 | 113 | 0 | 0 | 0 | 1 | 0 | 39 | 77 | 136 | 70 | 16 | 0 | 0 | 0 | 0 | 0 | 227 | 149 | 1 | 1 | 1 | 1 | 1 | 0.976 | 1 | 1 | 1 | 1 | 1 | 1 | -0.005 |
| SE | 100399 | ENSG00000133812 | SBF2 | chr11 | - | 9998265 | 9998379 | 9993920 | 9993998 | 10000913 | 10001022 | 162 | 157 | 117 | 124 | 253 | 0 | 1 | 0 | 2 | 0 | 105 | 171 | 204 | 139 | 56 | 0 | 1 | 0 | 0 | 0 | 263 | 149 | 1 | 1 | 1 | 0.989 | 1 | 0.972 | 1 | 1 | 0.99 | 1 | 1 | 1 | -0.006 |
| SE | 100402 | ENSG00000133812 | SBF2 | chr11 | - | 10193901 | 10193987 | 10042843 | 10042981 | 10294014 | 10294207 | 66 | 65 | 54 | 60 | 38 | 0 | 0 | 0 | 2 | 0 | 67 | 60 | 61 | 86 | 20 | 0 | 0 | 0 | 0 | 0 | 235 | 149 | 1 | 1 | 1 | 1 | 1 | 0.95 | 1 | 1 | 1 | 1 | 1 | 1 | -0.01 |
| SE | 100403 | ENSG00000151376 | ME3 | chr11 | - | 86447064 | 86447207 | 86446313 | 86446487 | 86448149 | 86448255 | 12 | 10 | 14 | 3 | 14 | 0 | 1 | 0 | 0 | 0 | 0 | 1 | 12 | 6 | 0 | 0 | 0 | 5 | 0 | 0 | 292 | 149 | 1 | 1 | 1 | 0.836 | 1 | 1 | 1 | NA | 1 | 0.55 | 1 | NA | 0.117 |
| SE | 100404 | ENSG00000151376 | ME3 | chr11 | - | 86450300 | 86450398 | 86449888 | 86450002 | 86457365 | 86457759 | 9 | 13 | 19 | 11 | 14 | 0 | 1 | 0 | 0 | 0 | 27 | 18 | 11 | 8 | 11 | 0 | 0 | 0 | 0 | 0 | 247 | 149 | 1 | 1 | 1 | 0.887 | 1 | 1 | 1 | 1 | 1 | 1 | 1 | 1 | -0.023 |
| SE | 100405 | ENSG00000151376 | ME3 | chr11 | - | 86450300 | 86450398 | 86449888 | 86450002 | 86465090 | 86465200 | 6 | 11 | 18 | 5 | 7 | 0 | 0 | 0 | 0 | 0 | 20 | 8 | 6 | 0 | 6 | 0 | 1 | 0 | 0 | 0 | 247 | 149 | 1 | 1 | 1 | 1 | 1 | 1 | 1 | 1 | 0.828 | 1 | NA | 1 | 0.043 |
| SE | 100406 | ENSG00000151376 | ME3 | chr11 | - | 86457365 | 86457759 | 86449964 | 86450002 | 86465090 | 86465200 | 4 | 6 | 5 | 1 | 6 | 0 | 0 | 0 | 0 | 0 | 12 | 0 | 0 | 0 | 0 | 0 | 1 | 0 | 0 | 0 | 298 | 149 | 1 | 1 | 1 | 1 | 1 | 1 | 1 | 1 | 0 | NA | NA | NA | 0.5 |
| SE | 100409 | ENSG00000151376 | ME3 | chr11 | - | 86556552 | 86556702 | 86508791 | 86508867 | 86559689 | 86559823 | 23 | 18 | 31 | 6 | 4 | 0 | 0 | 0 | 0 | 0 | 0 | 17 | 14 | 12 | 22 | 0 | 1 | 0 | 0 | 0 | 298 | 149 | 1 | 1 | 1 | 1 | 1 | 1 | 1 | NA | 0.895 | 1 | 1 | 1 | 0.026 |
| SE | 100411 | ENSG00000167770 | OTUB1 | chr11 | + | 63988653 | 63988752 | 63988336 | 63988398 | 63996529 | 63996648 | 302 | 136 | 237 | 273 | 285 | 0 | 0 | 0 | 2 | 5 | 272 | 361 | 215 | 234 | 464 | 0 | 0 | 1 | 0 | 8 | 248 | 149 | 1 | 1 | 1 | 1 | 1 | 0.988 | 0.972 | 1 | 1 | 0.992 | 1 | 0.972 | -0.001 |
| SE | 100414 | ENSG00000167770 | OTUB1 | chr11 | + | 63996853 | 63996941 | 63996529 | 63996648 | 63997049 | 63997244 | 90 | 32 | 60 | 81 | 73 | 0 | 1 | 0 | 0 | 0 | 140 | 101 | 39 | 118 | 113 | 0 | 0 | 0 | 0 | 0 | 237 | 149 | 1 | 1 | 1 | 0.953 | 1 | 1 | 1 | 1 | 1 | 1 | 1 | 1 | -0.009 |
| SE | 100415 | ENSG00000167770 | OTUB1 | chr11 | + | 63996856 | 63996941 | 63996529 | 63996648 | 63997049 | 63997244 | 274 | 110 | 187 | 236 | 208 | 0 | 1 | 0 | 0 | 0 | 276 | 285 | 131 | 251 | 278 | 0 | 0 | 0 | 0 | 0 | 234 | 149 | 1 | 1 | 1 | 0.986 | 1 | 1 | 1 | 1 | 1 | 1 | 1 | 1 | -0.003 |
| SE | 100418 | ENSG00000072518 | MARK2 | chr11 | + | 63890221 | 63890295 | 63888695 | 63888964 | 63895158 | 63895226 | 3 | 9 | 3 | 2 | 8 | 2 | 0 | 4 | 2 | 0 | 0 | 0 | 1 | 0 | 4 | 0 | 0 | 0 | 1 | 3 | 223 | 149 | 1 | 1 | 0.501 | 1 | 0.334 | 0.401 | 1 | NA | NA | 1 | 0 | 0.471 | 0.157 |
| SE | 100420 | ENSG00000072518 | MARK2 | chr11 | + | 63900956 | 63901069 | 63900779 | 63900879 | 63902197 | 63902330 | 596 | 329 | 439 | 524 | 1084 | 0 | 0 | 0 | 0 | 0 | 611 | 575 | 311 | 552 | 564 | 0 | 0 | 2 | 6 | 0 | 262 | 149 | 1 | 1 | 1 | 1 | 1 | 1 | 1 | 1 | 1 | 0.989 | 0.981 | 1 | 0.006 |
| SE | 100421 | ENSG00000072518 | MARK2 | chr11 | + | 63902600 | 63902782 | 63902197 | 63902330 | 63903060 | 63903158 | 48 | 23 | 29 | 52 | 8 | 0 | 0 | 0 | 1 | 0 | 27 | 38 | 15 | 23 | 83 | 0 | 0 | 0 | 0 | 1 | 298 | 149 | 1 | 1 | 1 | 1 | 1 | 0.963 | 1 | 1 | 1 | 1 | 1 | 0.976 | -0.003 |
| SE | 100422 | ENSG00000072518 | MARK2 | chr11 | + | 63902603 | 63902782 | 63902197 | 63902330 | 63903060 | 63903132 | 103 | 51 | 55 | 116 | 71 | 0 | 0 | 0 | 1 | 0 | 108 | 94 | 42 | 98 | 179 | 0 | 0 | 0 | 0 | 1 | 298 | 149 | 1 | 1 | 1 | 1 | 1 | 0.983 | 1 | 1 | 1 | 1 | 1 | 0.989 | -0.001 |
| SE | 100424 | ENSG00000072518 | MARK2 | chr11 | + | 63903060 | 63903158 | 63902600 | 63902782 | 63904785 | 63905043 | 73 | 35 | 35 | 57 | 14 | 0 | 0 | 0 | 0 | 3 | 43 | 46 | 32 | 40 | 89 | 0 | 0 | 0 | 0 | 0 | 247 | 149 | 1 | 1 | 1 | 1 | 1 | 1 | 0.738 | 1 | 1 | 1 | 1 | 1 | -0.052 |
| SE | 100425 | ENSG00000072518 | MARK2 | chr11 | + | 63903985 | 63904147 | 63902600 | 63902782 | 63904785 | 63905043 | 59 | 35 | 33 | 43 | 25 | 0 | 0 | 0 | 0 | 3 | 44 | 48 | 42 | 87 | 69 | 0 | 0 | 0 | 0 | 0 | 298 | 149 | 1 | 1 | 1 | 1 | 1 | 1 | 0.806 | 1 | 1 | 1 | 1 | 1 | -0.039 |
| SE | 100427 | ENSG00000072518 | MARK2 | chr11 | + | 63903985 | 63904354 | 63902603 | 63902782 | 63904785 | 63905043 | 4 | 1 | 0 | 4 | 0 | 0 | 0 | 0 | 0 | 3 | 0 | 4 | 4 | 9 | 2 | 0 | 0 | 0 | 0 | 0 | 298 | 149 | 1 | 1 | 1 | 1 | NA | 1 | 0 | NA | 1 | 1 | 1 | 1 | -0.25 |
| SE | 100430 | ENSG00000072518 | MARK2 | chr11 | + | 63906087 | 63906114 | 63904785 | 63905043 | 63908876 | 63909539 | 1 | 0 | 5 | 0 | 0 | 3 | 1 | 0 | 0 | 3 | 0 | 0 | 2 | 0 | 0 | 0 | 7 | 0 | 0 | 12 | 176 | 149 | 1 | 1 | 0.22 | 0 | 1 | NA | NA | NA | 0 | 1 | NA | 0 | 0.073 |
| SE | 100431 | ENSG00000072518 | MARK2 | chr11 | + | 63906087 | 63907567 | 63904785 | 63905043 | 63908259 | 63908304 | 0 | 0 | 0 | 0 | 0 | 77 | 44 | 48 | 64 | 38 | 0 | 0 | 3 | 4 | 0 | 73 | 93 | 65 | 83 | 105 | 298 | 149 | 1 | 1 | 0 | 0 | 0 | 0 | 0 | 0 | 0 | 0.023 | 0.024 | 0 | -0.009 |
| SE | 100435 | ENSG00000132275 | RRP8 | chr11 | - | 6600925 | 6601602 | 6600668 | 6600775 | 6601851 | 6602215 | 101 | 56 | 107 | 66 | 166 | 0 | 0 | 1 | 0 | 0 | 67 | 58 | 52 | 47 | 93 | 0 | 0 | 0 | 0 | 0 | 298 | 149 | 1 | 1 | 1 | 1 | 0.982 | 1 | 1 | 1 | 1 | 1 | 1 | 1 | -0.004 |
| SE | 100437 | ENSG00000132275 | RRP8 | chr11 | - | 6601148 | 6601602 | 6600668 | 6600775 | 6601851 | 6602215 | 86 | 50 | 96 | 58 | 148 | 0 | 0 | 1 | 0 | 0 | 58 | 60 | 48 | 40 | 83 | 0 | 0 | 0 | 0 | 0 | 298 | 149 | 1 | 1 | 1 | 1 | 0.98 | 1 | 1 | 1 | 1 | 1 | 1 | 1 | -0.004 |
| SE | 100438 | ENSG00000132275 | RRP8 | chr11 | - | 6601148 | 6601602 | 6600668 | 6600775 | 6603403 | 6603616 | 0 | 0 | 0 | 0 | 0 | 0 | 0 | 1 | 0 | 0 | 0 | 4 | 0 | 1 | 0 | 0 | 0 | 6 | 9 | 6 | 298 | 149 | 1 | 1 | NA | NA | 0 | NA | NA | NA | 1 | 0 | 0.053 | 0 | -0.263 |
| SE | 100439 | ENSG00000132275 | RRP8 | chr11 | - | 6601851 | 6602215 | 6600668 | 6600775 | 6603403 | 6603604 | 19 | 15 | 23 | 13 | 11 | 0 | 0 | 1 | 0 | 0 | 0 | 1 | 6 | 4 | 3 | 0 | 0 | 6 | 9 | 6 | 298 | 149 | 1 | 1 | 1 | 1 | 0.92 | 1 | 1 | NA | 1 | 0.333 | 0.182 | 0.2 | 0.555 |
| SE | 100440 | ENSG00000132275 | RRP8 | chr11 | - | 6601851 | 6602215 | 6600925 | 6601602 | 6603403 | 6603604 | 105 | 65 | 118 | 71 | 159 | 0 | 0 | 0 | 0 | 0 | 58 | 57 | 54 | 44 | 86 | 0 | 0 | 0 | 1 | 0 | 298 | 149 | 1 | 1 | 1 | 1 | 1 | 1 | 1 | 1 | 1 | 1 | 0.957 | 1 | 0.009 |
| SE | 100441 | ENSG00000132275 | RRP8 | chr11 | - | 6601851 | 6602264 | 6601253 | 6601602 | 6603403 | 6603597 | 91 | 50 | 96 | 58 | 148 | 0 | 0 | 0 | 0 | 0 | 58 | 61 | 48 | 40 | 88 | 0 | 0 | 0 | 1 | 0 | 298 | 149 | 1 | 1 | 1 | 1 | 1 | 1 | 1 | 1 | 1 | 1 | 0.952 | 1 | 0.01 |
| SE | 100442 | ENSG00000165916 | PSMC3 | chr11 | - | 47420263 | 47420409 | 47419115 | 47419197 | 47422573 | 47422722 | 175 | 98 | 166 | 153 | 97 | 0 | 0 | 1 | 0 | 0 | 245 | 158 | 102 | 146 | 223 | 0 | 0 | 0 | 3 | 0 | 295 | 149 | 1 | 1 | 1 | 1 | 0.988 | 1 | 1 | 1 | 1 | 1 | 0.961 | 1 | 0.005 |
| SE | 100444 | ENSG00000165916 | PSMC3 | chr11 | - | 47422573 | 47422722 | 47420263 | 47420409 | 47422829 | 47422973 | 56 | 16 | 50 | 48 | 32 | 0 | 0 | 0 | 0 | 0 | 73 | 45 | 52 | 52 | 35 | 0 | 0 | 0 | 1 | 0 | 298 | 149 | 1 | 1 | 1 | 1 | 1 | 1 | 1 | 1 | 1 | 1 | 0.963 | 1 | 0.007 |
| SE | 100445 | ENSG00000165916 | PSMC3 | chr11 | - | 47424045 | 47424183 | 47422829 | 47422973 | 47424428 | 47424491 | 217 | 131 | 174 | 220 | 129 | 0 | 0 | 0 | 0 | 0 | 366 | 342 | 216 | 307 | 355 | 0 | 1 | 0 | 0 | 1 | 287 | 149 | 1 | 1 | 1 | 1 | 1 | 1 | 1 | 1 | 0.994 | 1 | 1 | 0.995 | 0.002 |
| SE | 100446 | ENSG00000165916 | PSMC3 | chr11 | - | 47424108 | 47424183 | 47422829 | 47422973 | 47424428 | 47424491 | 48 | 36 | 34 | 44 | 40 | 0 | 0 | 0 | 0 | 0 | 52 | 69 | 44 | 62 | 44 | 0 | 1 | 0 | 0 | 1 | 224 | 149 | 1 | 1 | 1 | 1 | 1 | 1 | 1 | 1 | 0.979 | 1 | 1 | 0.967 | 0.011 |
| SE | 100447 | ENSG00000165916 | PSMC3 | chr11 | - | 47424428 | 47424491 | 47424045 | 47424183 | 47424606 | 47424711 | 186 | 115 | 159 | 176 | 118 | 0 | 0 | 0 | 0 | 6 | 303 | 267 | 176 | 234 | 294 | 0 | 0 | 0 | 0 | 3 | 212 | 149 | 1 | 1 | 1 | 1 | 1 | 1 | 0.933 | 1 | 1 | 1 | 1 | 0.986 | -0.011 |
| SE | 100455 | ENSG00000103429 | BFAR | chr16 | + | 14649803 | 14649973 | 14648387 | 14648592 | 14661891 | 14662065 | 67 | 50 | 79 | 86 | 154 | 1 | 0 | 0 | 0 | 0 | 36 | 72 | 58 | 65 | 77 | 0 | 0 | 0 | 0 | 0 | 298 | 149 | 1 | 1 | 0.971 | 1 | 1 | 1 | 1 | 1 | 1 | 1 | 1 | 1 | -0.006 |
| SE | 100463 | ENSG00000204356 | NELFE | chr6 | - | 31957317 | 31957435 | 31956940 | 31957010 | 31958371 | 31958454 | 2 | 0 | 0 | 0 | 0 | 54 | 43 | 47 | 58 | 59 | 0 | 0 | 0 | 0 | 0 | 114 | 74 | 54 | 88 | 108 | 267 | 149 | 1 | 1 | 0.02 | 0 | 0 | 0 | 0 | 0 | 0 | 0 | 0 | 0 | 0.004 |
| SE | 100464 | ENSG00000204356 | NELFE | chr6 | - | 31957317 | 31957435 | 31956940 | 31957010 | 31958891 | 31958954 | 1 | 0 | 0 | 1 | 0 | 0 | 0 | 0 | 0 | 0 | 0 | 0 | 0 | 0 | 0 | 0 | 0 | 0 | 1 | 0 | 267 | 149 | 1 | 1 | 1 | NA | NA | 1 | NA | NA | NA | NA | 0 | NA | 1 |
| SE | 100465 | ENSG00000204356 | NELFE | chr6 | - | 31958307 | 31958454 | 31956940 | 31957010 | 31958891 | 31958968 | 0 | 4 | 2 | 5 | 3 | 0 | 0 | 0 | 0 | 0 | 0 | 3 | 1 | 4 | 0 | 0 | 0 | 0 | 1 | 0 | 296 | 149 | 1 | 1 | NA | 1 | 1 | 1 | 1 | NA | 1 | 1 | 0.668 | NA | 0.111 |
| SE | 100466 | ENSG00000204356 | NELFE | chr6 | - | 31958371 | 31958454 | 31956692 | 31957010 | 31958891 | 31958954 | 49 | 44 | 40 | 61 | 57 | 0 | 0 | 0 | 0 | 0 | 114 | 76 | 50 | 83 | 97 | 0 | 0 | 0 | 1 | 0 | 232 | 149 | 1 | 1 | 1 | 1 | 1 | 1 | 1 | 1 | 1 | 1 | 0.982 | 1 | 0.004 |
| SE | 100467 | ENSG00000204356 | NELFE | chr6 | - | 31958371 | 31958454 | 31957317 | 31957435 | 31958891 | 31958954 | 1 | 1 | 0 | 3 | 0 | 0 | 0 | 0 | 1 | 0 | 0 | 3 | 0 | 0 | 0 | 0 | 0 | 0 | 0 | 0 | 232 | 149 | 1 | 1 | 1 | 1 | NA | 0.658 | NA | NA | 1 | NA | NA | NA | -0.114 |
| SE | 100468 | ENSG00000235505 | CASP4LP | chr11 | - | 104905447 | 104905557 | 104903390 | 104903551 | 104908834 | 104909008 | 52 | 38 | 35 | 45 | 84 | 0 | 0 | 0 | 0 | 0 | 4 | 61 | 16 | 16 | 35 | 0 | 3 | 0 | 0 | 0 | 259 | 149 | 1 | 1 | 1 | 1 | 1 | 1 | 1 | 1 | 0.921 | 1 | 1 | 1 | 0.016 |
| SE | 100470 | ENSG00000235505 | CASP4LP | chr11 | - | 104906449 | 104907618 | 104905447 | 104905557 | 104918112 | 104918158 | 37 | 45 | 28 | 31 | 74 | 1 | 0 | 0 | 0 | 0 | 0 | 17 | 10 | 4 | 0 | 0 | 0 | 0 | 0 | 0 | 298 | 149 | 1 | 1 | 0.949 | 1 | 1 | 1 | 1 | NA | 1 | 1 | 1 | NA | -0.01 |
| SE | 100472 | ENSG00000235505 | CASP4LP | chr11 | - | 104908834 | 104909008 | 104905447 | 104905557 | 104918112 | 104918141 | 58 | 31 | 23 | 45 | 83 | 1 | 0 | 0 | 0 | 0 | 4 | 63 | 12 | 12 | 42 | 0 | 0 | 0 | 0 | 0 | 298 | 149 | 1 | 1 | 0.967 | 1 | 1 | 1 | 1 | 1 | 1 | 1 | 1 | 1 | -0.007 |
| SE | 100473 | ENSG00000235505 | CASP4LP | chr11 | - | 104908834 | 104909008 | 104906449 | 104907618 | 104918112 | 104918141 | 34 | 12 | 8 | 19 | 45 | 0 | 0 | 0 | 1 | 0 | 0 | 32 | 5 | 8 | 32 | 0 | 0 | 0 | 0 | 0 | 298 | 149 | 1 | 1 | 1 | 1 | 1 | 0.905 | 1 | NA | 1 | 1 | 1 | 1 | -0.019 |
| SE | 100475 | ENSG00000186166 | CENATAC | chr11 | + | 119001612 | 119001825 | 118999010 | 118999109 | 119010763 | 119010830 | 1 | 2 | 11 | 2 | 1 | 3 | 0 | 11 | 12 | 7 | 0 | 0 | 0 | 13 | 0 | 0 | 3 | 9 | 2 | 10 | 298 | 149 | 1 | 1 | 0.143 | 1 | 0.333 | 0.077 | 0.067 | NA | 0 | 0 | 0.765 | 0 | 0.133 |
| SE | 100476 | ENSG00000186166 | CENATAC | chr11 | + | 119012148 | 119012254 | 119011938 | 119012003 | 119013231 | 119013262 | 34 | 30 | 51 | 37 | 44 | 0 | 0 | 0 | 0 | 1 | 93 | 34 | 104 | 95 | 33 | 0 | 0 | 0 | 0 | 0 | 255 | 149 | 1 | 1 | 1 | 1 | 1 | 1 | 0.963 | 1 | 1 | 1 | 1 | 1 | -0.007 |
| SE | 100479 | ENSG00000137700 | SLC37A4 | chr11 | - | 119026505 | 119026688 | 119025966 | 119026080 | 119026936 | 119027095 | 2 | 0 | 8 | 2 | 8 | 1 | 0 | 0 | 0 | 0 | 0 | 7 | 0 | 0 | 0 | 0 | 0 | 0 | 0 | 0 | 298 | 149 | 1 | 1 | 0.5 | NA | 1 | 1 | 1 | NA | 1 | NA | NA | NA | -0.125 |
| SE | 100480 | ENSG00000137700 | SLC37A4 | chr11 | - | 119026570 | 119026688 | 119025966 | 119026080 | 119026936 | 119027095 | 2 | 0 | 8 | 2 | 8 | 1 | 0 | 0 | 0 | 0 | 0 | 7 | 0 | 0 | 0 | 0 | 0 | 0 | 0 | 0 | 267 | 149 | 1 | 1 | 0.527 | NA | 1 | 1 | 1 | NA | 1 | NA | NA | NA | -0.118 |
| SE | 100481 | ENSG00000137700 | SLC37A4 | chr11 | - | 119026602 | 119026688 | 119025966 | 119026080 | 119026936 | 119027095 | 5 | 1 | 15 | 8 | 16 | 1 | 0 | 0 | 0 | 0 | 0 | 14 | 0 | 7 | 0 | 0 | 0 | 0 | 0 | 0 | 235 | 149 | 1 | 1 | 0.76 | 1 | 1 | 1 | 1 | NA | 1 | NA | 1 | NA | -0.048 |
| SE | 100485 | ENSG00000137700 | SLC37A4 | chr11 | - | 119028734 | 119028834 | 119028193 | 119028426 | 119029221 | 119029564 | 0 | 0 | 0 | 0 | 2 | 1 | 5 | 7 | 10 | 8 | 0 | 0 | 0 | 0 | 0 | 0 | 4 | 2 | 6 | 0 | 249 | 149 | 1 | 1 | 0 | 0 | 0 | 0 | 0.13 | NA | 0 | 0 | 0 | NA | 0.026 |
| SE | 100486 | ENSG00000137700 | SLC37A4 | chr11 | - | 119030231 | 119030735 | 119029221 | 119029564 | 119030848 | 119030906 | 0 | 0 | 0 | 1 | 0 | 2 | 0 | 0 | 2 | 0 | 0 | 0 | 0 | 0 | 0 | 0 | 5 | 1 | 0 | 0 | 298 | 149 | 1 | 1 | 0 | NA | NA | 0.2 | NA | NA | 0 | 0 | NA | NA | 0.1 |
| SE | 100489 | ENSG00000213639 | PPP1CB | chr2 | + | 28776850 | 28776982 | 28751829 | 28752176 | 28781737 | 28781842 | 363 | 194 | 295 | 320 | 218 | 1 | 0 | 0 | 3 | 0 | 113 | 190 | 231 | 104 | 26 | 0 | 0 | 0 | 0 | 0 | 281 | 149 | 1 | 1 | 0.995 | 1 | 1 | 0.983 | 1 | 1 | 1 | 1 | 1 | 1 | -0.004 |
| SE | 100490 | ENSG00000213639 | PPP1CB | chr2 | + | 28778350 | 28778589 | 28776850 | 28776982 | 28778808 | 28778931 | 1 | 2 | 5 | 0 | 13 | 659 | 443 | 594 | 706 | 975 | 0 | 0 | 1 | 0 | 0 | 165 | 444 | 531 | 339 | 80 | 298 | 149 | 1 | 1 | 0.001 | 0.002 | 0.004 | 0 | 0.007 | 0 | 0 | 0.001 | 0 | 0 | 0.003 |
| SE | 100491 | ENSG00000213639 | PPP1CB | chr2 | + | 28778808 | 28779039 | 28776850 | 28776982 | 28781737 | 28781842 | 1241 | 907 | 1103 | 1266 | 2027 | 0 | 0 | 0 | 0 | 0 | 300 | 776 | 997 | 601 | 126 | 0 | 2 | 0 | 0 | 0 | 298 | 149 | 1 | 1 | 1 | 1 | 1 | 1 | 1 | 1 | 0.995 | 1 | 1 | 1 | 0.001 |
| SE | 100492 | ENSG00000213639 | PPP1CB | chr2 | + | 28778808 | 28779039 | 28776850 | 28776982 | 28783906 | 28783978 | 663 | 443 | 594 | 706 | 975 | 0 | 0 | 0 | 1 | 2 | 165 | 444 | 531 | 339 | 80 | 0 | 0 | 0 | 0 | 0 | 298 | 149 | 1 | 1 | 1 | 1 | 1 | 0.997 | 0.996 | 1 | 1 | 1 | 1 | 1 | -0.001 |
| SE | 100493 | ENSG00000213639 | PPP1CB | chr2 | + | 28781737 | 28781842 | 28776850 | 28776982 | 28783906 | 28783978 | 521 | 404 | 400 | 467 | 521 | 0 | 0 | 0 | 1 | 2 | 99 | 301 | 347 | 166 | 58 | 0 | 0 | 0 | 0 | 0 | 254 | 149 | 1 | 1 | 1 | 1 | 1 | 0.996 | 0.993 | 1 | 1 | 1 | 1 | 1 | -0.002 |
| SE | 100494 | ENSG00000213639 | PPP1CB | chr2 | + | 28781737 | 28781842 | 28778808 | 28779039 | 28783906 | 28783978 | 1089 | 849 | 892 | 1005 | 1462 | 4 | 0 | 0 | 0 | 0 | 234 | 614 | 802 | 413 | 104 | 0 | 0 | 0 | 0 | 0 | 254 | 149 | 1 | 1 | 0.994 | 1 | 1 | 1 | 1 | 1 | 1 | 1 | 1 | 1 | -0.001 |
| SE | 100498 | ENSG00000172375 | C2CD2L | chr11 | + | 119112699 | 119112874 | 119112481 | 119112609 | 119113610 | 119113712 | 18 | 7 | 15 | 25 | 11 | 2 | 0 | 0 | 0 | 0 | 13 | 25 | 30 | 25 | 62 | 0 | 0 | 0 | 0 | 0 | 298 | 149 | 1 | 1 | 0.818 | 1 | 1 | 1 | 1 | 1 | 1 | 1 | 1 | 1 | -0.036 |
| SE | 100502 | ENSG00000247708 | STX18-AS1 | chr4 | + | 4705672 | 4705910 | 4649372 | 4649542 | 4709515 | 4710938 | 0 | 0 | 6 | 2 | 0 | 0 | 0 | 4 | 0 | 4 | 0 | 2 | 0 | 0 | 0 | 0 | 0 | 0 | 0 | 0 | 298 | 149 | 1 | 1 | NA | NA | 0.429 | 1 | 0 | NA | 1 | NA | NA | NA | -0.524 |
| SE | 100503 | ENSG00000248712 | CCDC153 | chr11 | - | 119193150 | 119193243 | 119190709 | 119190840 | 119194922 | 119194994 | 5 | 5 | 2 | 1 | 1 | 0 | 0 | 0 | 0 | 0 | 0 | 3 | 0 | 0 | 36 | 0 | 0 | 0 | 0 | 4 | 242 | 149 | 1 | 1 | 1 | 1 | 1 | 1 | 1 | NA | 1 | NA | NA | 0.847 | 0.077 |
| SE | 100504 | ENSG00000248712 | CCDC153 | chr11 | - | 119193150 | 119193885 | 119190709 | 119190840 | 119194922 | 119194994 | 8 | 10 | 3 | 3 | 7 | 0 | 0 | 0 | 0 | 0 | 19 | 2 | 0 | 5 | 22 | 0 | 0 | 0 | 0 | 4 | 298 | 149 | 1 | 1 | 1 | 1 | 1 | 1 | 1 | 1 | 1 | NA | 1 | 0.733 | 0.067 |
| SE | 100505 | ENSG00000248712 | CCDC153 | chr11 | - | 119193765 | 119193885 | 119193150 | 119193243 | 119194922 | 119194994 | 7 | 6 | 1 | 2 | 6 | 1 | 0 | 0 | 0 | 0 | 22 | 0 | 0 | 7 | 4 | 0 | 1 | 0 | 0 | 18 | 269 | 149 | 1 | 1 | 0.795 | 1 | 1 | 1 | 1 | 1 | 0 | NA | 1 | 0.11 | 0.431 |
| SE | 100506 | ENSG00000054654 | SYNE2 | chr14 | + | 63941694 | 63941790 | 63940613 | 63940675 | 63941884 | 63941962 | 444 | 268 | 297 | 339 | 410 | 0 | 0 | 0 | 1 | 0 | 130 | 291 | 182 | 125 | 169 | 0 | 2 | 0 | 3 | 0 | 245 | 149 | 1 | 1 | 1 | 1 | 1 | 0.995 | 1 | 1 | 0.989 | 1 | 0.962 | 1 | 0.009 |
| SE | 100507 | ENSG00000054654 | SYNE2 | chr14 | + | 63949824 | 63950006 | 63942050 | 63942143 | 63954718 | 63954915 | 851 | 442 | 479 | 597 | 530 | 0 | 0 | 0 | 1 | 0 | 174 | 338 | 268 | 159 | 118 | 0 | 0 | 0 | 0 | 0 | 298 | 149 | 1 | 1 | 1 | 1 | 1 | 0.997 | 1 | 1 | 1 | 1 | 1 | 1 | -0.001 |
| SE | 100508 | ENSG00000054654 | SYNE2 | chr14 | + | 63963898 | 63964000 | 63961524 | 63961625 | 63967708 | 63967846 | 408 | 227 | 250 | 261 | 271 | 2 | 0 | 1 | 0 | 0 | 91 | 129 | 130 | 48 | 31 | 0 | 2 | 0 | 0 | 0 | 251 | 149 | 1 | 1 | 0.992 | 1 | 0.993 | 1 | 1 | 1 | 0.975 | 1 | 1 | 1 | 0.002 |
| SE | 100509 | ENSG00000054654 | SYNE2 | chr14 | + | 63967708 | 63967846 | 63961524 | 63961625 | 63976562 | 63976727 | 202 | 124 | 134 | 168 | 189 | 1 | 0 | 1 | 0 | 0 | 59 | 71 | 71 | 31 | 19 | 0 | 0 | 0 | 0 | 0 | 287 | 149 | 1 | 1 | 0.991 | 1 | 0.986 | 1 | 1 | 1 | 1 | 1 | 1 | 1 | -0.005 |
| SE | 100510 | ENSG00000054654 | SYNE2 | chr14 | + | 63976562 | 63976727 | 63967708 | 63967846 | 63977904 | 63978017 | 721 | 487 | 391 | 512 | 467 | 1 | 0 | 0 | 0 | 0 | 153 | 340 | 238 | 110 | 51 | 0 | 0 | 0 | 0 | 0 | 298 | 149 | 1 | 1 | 0.997 | 1 | 1 | 1 | 1 | 1 | 1 | 1 | 1 | 1 | -0.001 |
| SE | 100511 | ENSG00000054654 | SYNE2 | chr14 | + | 63982629 | 63982794 | 63980985 | 63981173 | 63983736 | 63983886 | 1142 | 800 | 616 | 780 | 1005 | 3 | 0 | 0 | 0 | 0 | 166 | 451 | 416 | 115 | 73 | 0 | 0 | 3 | 1 | 0 | 298 | 149 | 1 | 1 | 0.995 | 1 | 1 | 1 | 1 | 1 | 1 | 0.986 | 0.983 | 1 | 0.005 |
| SE | 100514 | ENSG00000054654 | SYNE2 | chr14 | + | 63997300 | 63997391 | 63996946 | 63997158 | 63998218 | 63998328 | 430 | 360 | 306 | 415 | 404 | 0 | 0 | 1 | 0 | 0 | 44 | 183 | 132 | 23 | 38 | 0 | 0 | 0 | 0 | 0 | 240 | 149 | 1 | 1 | 1 | 1 | 0.995 | 1 | 1 | 1 | 1 | 1 | 1 | 1 | -0.001 |
| SE | 100515 | ENSG00000054654 | SYNE2 | chr14 | + | 63997300 | 63997576 | 63996946 | 63997158 | 63998218 | 63998328 | 189 | 160 | 121 | 166 | 168 | 0 | 0 | 1 | 0 | 0 | 41 | 73 | 62 | 17 | 15 | 0 | 0 | 0 | 0 | 0 | 298 | 149 | 1 | 1 | 1 | 1 | 0.984 | 1 | 1 | 1 | 1 | 1 | 1 | 1 | -0.003 |
| SE | 100516 | ENSG00000054654 | SYNE2 | chr14 | + | 63998218 | 63998328 | 63997300 | 63997391 | 63998913 | 63999040 | 438 | 325 | 273 | 391 | 364 | 0 | 0 | 0 | 0 | 0 | 22 | 217 | 116 | 35 | 39 | 0 | 5 | 0 | 0 | 0 | 259 | 149 | 1 | 1 | 1 | 1 | 1 | 1 | 1 | 1 | 0.961 | 1 | 1 | 1 | 0.008 |
| SE | 100517 | ENSG00000054654 | SYNE2 | chr14 | + | 64002719 | 64003330 | 64001933 | 64002081 | 64016472 | 64016631 | 575 | 389 | 305 | 451 | 393 | 0 | 0 | 1 | 0 | 0 | 68 | 270 | 298 | 122 | 33 | 0 | 0 | 0 | 0 | 0 | 298 | 149 | 1 | 1 | 1 | 1 | 0.993 | 1 | 1 | 1 | 1 | 1 | 1 | 1 | -0.001 |
| [illegible] | [illegible] | [illegible] | [illegible] | [illegible] | [illegible] | [illegible] | [illegible] | [illegible] | [illegible] | [illegible] | [illegible] | [illegible] | [illegible] | [illegible] | [illegible] | [illegible] | [illegible] | [illegible] | [illegible] | [illegible] | [illegible] | [illegible] | [illegible] | [illegible] | [illegible] | [illegible] | [illegible] | [illegible] | [illegible] | [illegible] | [illegible] | [illegible] | [illegible] | [illegible] | [illegible] | [illegible] | [illegible] | [illegible] | [illegible] | [illegible] | [illegible] | [illegible] | [illegible] | [illegible] | [illegible] | [illegible] |

| | | | | | | | | | | | | | | | | | | | | | | | | | | | | | | | | | | | | | | | | | | | | | | |
|---|---|---|---|---|---|---|---|---|---|---|---|---|---|---|---|---|---|---|---|---|---|---|---|---|---|---|---|---|---|---|---|---|---|---|---|---|---|---|---|---|---|---|---|---|---|---|
| SE | 100546 | ENSG00000054654 | SYNE2 | chr14 | + | 64168876 | 64168971 | 64167494 | 64167639 | 64170227 | 64170321 | 358 | 228 | 183 | 267 | 212 | 0 | 0 | 0 | 0 | 0 | 67 | 60 | 142 | 44 | 38 | 0 | 3 | 0 | 0 | 0 | 244 | 149 | 1 | 1 | 1 | 1 | 1 | 1 | 1 | 1 | 0.924 | 1 | 1 | 1 | 0.015 |
| SE | 100549 | ENSG00000054654 | SYNE2 | chr14 | + | 64190070 | 64190237 | 64188549 | 64188708 | 64202800 | 64202963 | 377 | 213 | 196 | 282 | 293 | 0 | 0 | 0 | 0 | 4 | 30 | 109 | 115 | 43 | 35 | 0 | 0 | 0 | 0 | 0 | 298 | 149 | 1 | 1 | 1 | 1 | 1 | 1 | 0.973 | 1 | 1 | 1 | 1 | 1 | -0.005 |
| SE | 100552 | ENSG00000054654 | SYNE2 | chr14 | + | 64202800 | 64202963 | 64196783 | 64197104 | 64208757 | 64208945 | 280 | 123 | 162 | 228 | 157 | 0 | 0 | 0 | 0 | 0 | 34 | 78 | 89 | 44 | 22 | 0 | 5 | 0 | 0 | 0 | 298 | 149 | 1 | 1 | 1 | 1 | 1 | 1 | 1 | 1 | 0.886 | 1 | 1 | 1 | 0.023 |
| SE | 100555 | ENSG00000054654 | SYNE2 | chr14 | + | 64207957 | 64208192 | 64202800 | 64202963 | 64208757 | 64208945 | 0 | 2 | 1 | 0 | 0 | 280 | 122 | 161 | 229 | 164 | 0 | 0 | 1 | 0 | 0 | 34 | 79 | 89 | 44 | 18 | 298 | 149 | 1 | 1 | 0 | 0.008 | 0.003 | 0 | 0 | 0 | 0 | 0.006 | 0 | 0 | 0.001 |
| SE | 100556 | ENSG00000054654 | SYNE2 | chr14 | + | 64214193 | 64214470 | 64212810 | 64213005 | 64216247 | 64216387 | 261 | 105 | 148 | 209 | 146 | 0 | 0 | 2 | 0 | 0 | 113 | 116 | 98 | 27 | 71 | 0 | 0 | 0 | 0 | 0 | 298 | 149 | 1 | 1 | 1 | 1 | 0.974 | 1 | 1 | 1 | 1 | 1 | 1 | 1 | -0.005 |
| SE | 100558 | ENSG00000054654 | SYNE2 | chr14 | + | 64216247 | 64216387 | 64214143 | 64214470 | 64218397 | 64218512 | 171 | 60 | 104 | 162 | 102 | 0 | 0 | 0 | 0 | 0 | 18 | 40 | 70 | 29 | 80 | 0 | 4 | 0 | 0 | 0 | 289 | 149 | 1 | 1 | 1 | 1 | 1 | 1 | 1 | 1 | 0.838 | 1 | 1 | 1 | 0.032 |
| SE | 100566 | ENSG00000110011 | DNAJC4 | chr11 | + | 64231870 | 64231964 | 64230431 | 64230940 | 64233893 | 64233980 | 19 | 2 | 15 | 17 | 4 | 1 | 0 | 0 | 0 | 0 | 10 | 14 | 7 | 6 | 4 | 0 | 0 | 0 | 0 | 0 | 243 | 149 | 1 | 1 | 0.921 | 1 | 1 | 1 | 1 | 1 | 1 | 1 | 1 | 1 | -0.016 |
| SE | 100567 | ENSG00000084207 | GSTP1 | chr11 | + | 67584133 | 67584169 | 67583741 | 67583844 | 67584684 | 67584772 | 1 | 0 | 2 | 0 | 1 | 0 | 0 | 0 | 1 | 0 | 8 | 0 | 0 | 10 | 0 | 0 | 0 | 0 | 0 | 0 | 185 | 149 | 1 | 1 | 1 | NA | 1 | 0 | 1 | 1 | NA | NA | 1 | NA | -0.25 |
| SE | 100569 | ENSG00000084207 | GSTP1 | chr11 | + | 67584684 | 67584772 | 67584463 | 67584570 | 67585037 | 67585241 | 284 | 100 | 142 | 176 | 23 | 0 | 0 | 0 | 0 | 0 | 261 | 258 | 124 | 187 | 403 | 0 | 0 | 0 | 2 | 0 | 237 | 149 | 1 | 1 | 1 | 1 | 1 | 1 | 1 | 1 | 1 | 1 | 0.983 | 1 | 0.003 |
| SE | 100570 | ENSG00000084207 | GSTP1 | chr11 | + | 67586103 | 67586211 | 67584794 | 67585241 | 67586388 | 67586505 | 958 | 390 | 550 | 665 | 348 | 1 | 0 | 0 | 0 | 0 | 1012 | 877 | 336 | 723 | 1462 | 0 | 0 | 1 | 0 | 0 | 257 | 149 | 1 | 1 | 0.998 | 1 | 1 | 1 | 1 | 1 | 1 | 0.995 | 1 | 1 | 0.001 |
| SE | 100580 | ENSG00000184428 | TOP1MT | chr8 | - | 143327302 | 143327790 | 143326221 | 143326344 | 143329349 | 143329433 | 0 | 0 | 0 | 0 | 0 | 21 | 32 | 50 | 40 | 44 | 0 | 3 | 0 | 0 | 0 | 96 | 14 | 41 | 36 | 105 | 298 | 149 | 1 | 1 | 0 | 0 | 0 | 0 | 0 | 0 | 0.097 | 0 | 0 | 0 | -0.019 |
| SE | 100582 | ENSG00000184428 | TOP1MT | chr8 | - | 143332486 | 143332601 | 143331223 | 143331339 | 143334739 | 143334854 | 0 | 2 | 0 | 0 | 0 | 7 | 1 | 1 | 0 | 0 | 0 | 0 | 2 | 4 | 14 | 5 | 0 | 0 | 0 | 2 | 264 | 149 | 1 | 1 | 0 | 0.53 | 0 | NA | NA | 0 | NA | 1 | 1 | 0.798 | -0.523 |
| SE | 100583 | ENSG00000184428 | TOP1MT | chr8 | - | 143332486 | 143332657 | 143331223 | 143331339 | 143334739 | 143334866 | 0 | 2 | 0 | 0 | 0 | 7 | 1 | 1 | 0 | 0 | 0 | 0 | 2 | 4 | 14 | 5 | 0 | 0 | 0 | 2 | 298 | 149 | 1 | 1 | 0 | 0.5 | 0 | NA | NA | 0 | NA | 1 | 1 | 0.778 | -0.528 |
| SE | 100586 | ENSG00000184428 | TOP1MT | chr8 | - | 143334107 | 143334285 | 143331223 | 143331339 | 143334739 | 143334795 | 2 | 0 | 0 | 1 | 1 | 7 | 1 | 1 | 0 | 0 | 0 | 0 | 7 | 0 | 0 | 5 | 0 | 0 | 0 | 2 | 298 | 149 | 1 | 1 | 0.125 | 0 | 0 | 1 | 1 | 0 | NA | 1 | NA | 0 | 0.092 |
| SE | 100592 | ENSG00000102317 | RBM3 | chrX | + | 48575560 | 48575667 | 48574951 | 48575283 | 48577025 | 48577109 | 552 | 407 | 611 | 561 | 458 | 1 | 0 | 0 | 0 | 0 | 1551 | 785 | 918 | 827 | 1094 | 0 | 0 | 0 | 0 | 0 | 256 | 149 | 1 | 1 | 0.997 | 1 | 1 | 1 | 1 | 1 | 1 | 1 | 1 | 1 | -0.001 |
| SE | 100594 | ENSG00000102317 | RBM3 | chrX | + | 48575814 | 48576083 | 48575560 | 48575667 | 48576313 | 48576419 | 12 | 8 | 14 | 11 | 7 | 926 | 496 | 1165 | 836 | 906 | 28 | 8 | 6 | 10 | 23 | 1729 | 1243 | 1262 | 1175 | 1633 | 298 | 149 | 1 | 1 | 0.006 | 0.008 | 0.006 | 0.007 | 0.004 | 0.008 | 0.003 | 0.002 | 0.004 | 0.007 | 0.001 |
| SE | 100595 | ENSG00000102317 | RBM3 | chrX | + | 48575921 | 48576083 | 48575560 | 48575667 | 48576313 | 48576419 | 10 | 5 | 12 | 8 | 7 | 926 | 496 | 1165 | 836 | 906 | 0 | 6 | 1 | 5 | 29 | 1729 | 1243 | 1262 | 1175 | 1633 | 298 | 149 | 1 | 1 | 0.005 | 0.005 | 0.005 | 0.005 | 0.004 | 0 | 0.002 | 0 | 0.002 | 0.009 | 0.002 |
| SE | 100596 | ENSG00000102317 | RBM3 | chrX | + | 48576313 | 48576419 | 48575167 | 48575283 | 48577025 | 48577109 | 10 | 4 | 12 | 18 | 16 | 1 | 0 | 0 | 0 | 0 | 39 | 42 | 49 | 98 | 49 | 0 | 0 | 0 | 0 | 0 | 255 | 149 | 1 | 1 | 0.854 | 1 | 1 | 1 | 1 | 1 | 1 | 1 | 1 | 1 | -0.029 |
| SE | 100597 | ENSG00000102317 | RBM3 | chrX | + | 48576313 | 48576419 | 48575560 | 48575667 | 48577025 | 48577109 | 577 | 322 | 758 | 532 | 450 | 0 | 0 | 0 | 0 | 0 | 1089 | 770 | 773 | 696 | 1003 | 0 | 3 | 0 | 0 | 0 | 255 | 149 | 1 | 1 | 1 | 1 | 1 | 1 | 1 | 1 | 0.993 | 1 | 1 | 1 | 0.001 |
| SE | 100598 | ENSG00000102317 | RBM3 | chrX | + | 48576507 | 48576604 | 48575898 | 48576419 | 48577025 | 48577109 | 1948 | 1200 | 2257 | 1850 | 2011 | 0 | 0 | 0 | 0 | 0 | 3929 | 2409 | 2704 | 2653 | 3588 | 5 | 0 | 0 | 0 | 0 | 246 | 149 | 1 | 1 | 1 | 1 | 1 | 1 | 1 | 0.998 | 1 | 1 | 1 | 1 | 0 |
| SE | 100600 | ENSG00000144791 | LIMD1 | chr3 | + | 45634956 | 45635024 | 45612891 | 45612981 | 45636149 | 45636251 | 0 | 0 | 0 | 0 | 0 | 2 | 0 | 0 | 0 | 0 | 0 | 0 | 5 | 0 | 0 | 0 | 6 | 0 | 0 | 0 | 217 | 149 | 1 | 1 | 0 | NA | NA | NA | NA | NA | 0 | 1 | NA | NA | -0.5 |
| SE | 100601 | ENSG00000144791 | LIMD1 | chr3 | + | 45636149 | 45636251 | 45594323 | 45596287 | 45665649 | 45665717 | 58 | 45 | 46 | 61 | 29 | 0 | 1 | 0 | 0 | 0 | 62 | 42 | 32 | 29 | 53 | 0 | 0 | 0 | 0 | 0 | 251 | 149 | 1 | 1 | 1 | 0.964 | 1 | 1 | 1 | 1 | 1 | 1 | 1 | 1 | -0.007 |
| SE | 100602 | ENSG00000144791 | LIMD1 | chr3 | + | 45665649 | 45665717 | 45636149 | 45636251 | 45668293 | 45668356 | 66 | 63 | 61 | 71 | 53 | 0 | 0 | 0 | 1 | 0 | 67 | 53 | 49 | 55 | 83 | 0 | 0 | 0 | 3 | 0 | 217 | 149 | 1 | 1 | 1 | 1 | 1 | 0.98 | 1 | 1 | 1 | 1 | 0.926 | 1 | 0.011 |
| SE | 100604 | ENSG00000149292 | TTC12 | chr11 | + | 113315950 | 113316315 | 113314578 | 113314618 | 113323287 | 113323451 | 71 | 50 | 59 | 69 | 83 | 0 | 2 | 0 | 0 | 0 | 29 | 45 | 41 | 27 | 31 | 0 | 0 | 0 | 0 | 0 | 298 | 149 | 1 | 1 | 1 | 0.926 | 1 | 1 | 1 | 1 | 1 | 1 | 1 | 1 | -0.015 |
| SE | 100605 | ENSG00000149292 | TTC12 | chr11 | + | 113316242 | 113316315 | 113314582 | 113314618 | 113323287 | 113323451 | 94 | 50 | 80 | 76 | 142 | 0 | 2 | 0 | 0 | 0 | 76 | 51 | 69 | 35 | 44 | 0 | 0 | 0 | 0 | 0 | 222 | 149 | 1 | 1 | 1 | 0.944 | 1 | 1 | 1 | 1 | 1 | 1 | 1 | 1 | -0.011 |
| SE | 100606 | ENSG00000149292 | TTC12 | chr11 | + | 113324604 | 113324682 | 113323993 | 113324015 | 113325523 | 113325645 | 58 | 34 | 99 | 50 | 91 | 0 | 4 | 0 | 1 | 0 | 67 | 30 | 36 | 30 | 45 | 0 | 0 | 0 | 0 | 0 | 227 | 149 | 1 | 1 | 1 | 0.848 | 1 | 0.97 | 1 | 1 | 1 | 1 | 1 | 1 | -0.036 |
| SE | 100607 | ENSG00000149292 | TTC12 | chr11 | + | 113325523 | 113325645 | 113323993 | 113324015 | 113329919 | 113329977 | 23 | 8 | 39 | 18 | 25 | 0 | 0 | 0 | 0 | 0 | 8 | 7 | 4 | 11 | 14 | 0 | 0 | 0 | 2 | 0 | 271 | 149 | 1 | 1 | 1 | 1 | 1 | 1 | 1 | 1 | 1 | 1 | 0.751 | 1 | 0.05 |
| SE | 100609 | ENSG00000149292 | TTC12 | chr11 | + | 113329919 | 113329979 | 113325523 | 113325645 | 113334965 | 113335037 | 46 | 11 | 88 | 42 | 74 | 0 | 0 | 0 | 0 | 1 | 26 | 10 | 5 | 17 | 41 | 0 | 0 | 0 | 0 | 0 | 209 | 149 | 1 | 1 | 1 | 1 | 1 | 1 | 0.981 | 1 | 1 | 1 | 1 | 1 | -0.004 |
| SE | 100610 | ENSG00000149292 | TTC12 | chr11 | + | 113334965 | 113335037 | 113325523 | 113325645 | 113338773 | 113338785 | 11 | 7 | 16 | 16 | 20 | 0 | 0 | 0 | 0 | 1 | 6 | 4 | 8 | 4 | 14 | 0 | 0 | 0 | 0 | 0 | 221 | 149 | 1 | 1 | 1 | 1 | 1 | 1 | 0.931 | 1 | 1 | 1 | 1 | 1 | -0.014 |
| SE | 100611 | ENSG00000149292 | TTC12 | chr11 | + | 113338773 | 113338834 | 113334947 | 113335037 | 113339285 | 113339439 | 41 | 18 | 83 | 59 | 92 | 0 | 0 | 2 | 0 | 0 | 32 | 17 | 16 | 12 | 76 | 0 | 0 | 0 | 0 | 0 | 210 | 149 | 1 | 1 | 1 | 1 | 0.967 | 1 | 1 | 1 | 1 | 1 | 1 | 1 | -0.007 |
| SE | 100612 | ENSG00000149292 | TTC12 | chr11 | + | 113340663 | 113340733 | 113339285 | 113339474 | 113341836 | 113341925 | 41 | 32 | 77 | 40 | 78 | 2 | 1 | 1 | 1 | 0 | 34 | 38 | 26 | 17 | 19 | 0 | 2 | 5 | 3 | 0 | 219 | 149 | 1 | 1 | 0.933 | 0.956 | 0.981 | 0.965 | 1 | 1 | 0.928 | 0.78 | 0.794 | 1 | 0.067 |
| SE | 100614 | ENSG00000149292 | TTC12 | chr11 | + | 113364834 | 113365060 | 113363827 | 113363927 | 113368186 | 113368463 | 25 | 24 | 46 | 25 | 52 | 0 | 1 | 0 | 0 | 0 | 11 | 34 | 8 | 5 | 52 | 0 | 0 | 0 | 0 | 0 | 298 | 149 | 1 | 1 | 1 | 0.923 | 1 | 1 | 1 | 1 | 1 | 1 | 1 | 1 | -0.015 |
| SE | 100625 | ENSG00000283977 | NA | chr11 | + | 6502752 | 6502926 | 6482044 | 6482289 | 6508879 | 6508978 | 3 | 4 | 8 | 21 | 16 | 0 | 0 | 0 | 1 | 0 | 27 | 16 | 10 | 12 | 11 | 0 | 0 | 0 | 0 | 0 | 298 | 149 | 1 | 1 | 1 | 1 | 1 | 0.913 | 1 | 1 | 1 | 1 | 1 | 1 | -0.017 |
| SE | 100626 | ENSG00000179532 | DNHD1 | chr11 | + | 6497770 | 6497844 | 6497584 | 6497667 | 6498722 | 6498961 | 2 | 1 | 4 | 0 | 1 | 0 | 1 | 0 | 1 | 0 | 6 | 6 | 5 | 2 | 2 | 0 | 0 | 0 | 0 | 0 | 223 | 149 | 1 | 1 | 1 | 0.401 | 1 | 0 | 1 | 1 | 1 | 1 | 1 | 1 | -0.32 |
| SE | 100628 | ENSG00000179532 | DNHD1 | chr11 | + | 6533026 | 6533184 | 6528877 | 6529121 | 6533680 | 6533806 | 38 | 21 | 48 | 27 | 18 | 0 | 0 | 1 | 1 | 0 | 42 | 72 | 38 | 24 | 43 | 0 | 0 | 0 | 0 | 0 | 298 | 149 | 1 | 1 | 1 | 1 | 0.96 | 0.931 | 1 | 1 | 1 | 1 | 1 | 1 | -0.022 |
| SE | 100630 | ENSG00000179532 | DNHD1 | chr11 | + | 6538382 | 6538510 | 6533680 | 6534173 | 6538611 | 6538810 | 10 | 5 | 9 | 7 | 2 | 0 | 0 | 0 | 2 | 0 | 4 | 16 | 9 | 35 | 8 | 0 | 0 | 0 | 0 | 0 | 277 | 149 | 1 | 1 | 1 | 1 | 1 | 0.653 | 1 | 1 | 1 | 1 | 1 | 1 | -0.069 |
| SE | 100631 | ENSG00000179532 | DNHD1 | chr11 | + | 6538382 | 6538510 | 6533923 | 6534229 | 6538611 | 6538810 | 9 | 6 | 11 | 9 | 2 | 0 | 0 | 0 | 1 | 0 | 4 | 21 | 16 | 24 | 8 | 0 | 0 | 0 | 0 | 0 | 277 | 149 | 1 | 1 | 1 | 1 | 1 | 0.829 | 1 | 1 | 1 | 1 | 1 | 1 | -0.034 |
| SE | 100632 | ENSG00000179532 | DNHD1 | chr11 | + | 6539218 | 6539313 | 6538611 | 6538810 | 6539875 | 6540083 | 4 | 5 | 11 | 3 | 2 | 0 | 0 | 2 | 0 | 0 | 21 | 0 | 6 | 8 | 0 | 0 | 8 | 0 | 0 | 0 | 244 | 149 | 1 | 1 | 1 | 1 | 0.771 | 1 | 1 | 1 | 0 | 1 | 1 | NA | 0.204 |
| SE | 100633 | ENSG00000179532 | DNHD1 | chr11 | + | 6539875 | 6540083 | 6538611 | 6538810 | 6544120 | 6544246 | 3 | 1 | 7 | 6 | 4 | 0 | 0 | 0 | 0 | 0 | 7 | 14 | 0 | 1 | 0 | 0 | 0 | 2 | 0 | 0 | 298 | 149 | 1 | 1 | 1 | 1 | 1 | 1 | 1 | 1 | 1 | 0 | 1 | NA | 0.25 |
| SE | 100634 | ENSG00000179532 | DNHD1 | chr11 | + | 6544120 | 6544246 | 6538611 | 6538810 | 6544573 | 6544671 | 5 | 3 | 11 | 11 | 5 | 0 | 0 | 0 | 0 | 0 | 0 | 3 | 14 | 14 | 16 | 0 | 0 | 0 | 2 | 0 | 275 | 149 | 1 | 1 | 1 | 1 | 1 | 1 | 1 | NA | 1 | 1 | 0.791 | 1 | 0.052 |
| SE | 100635 | ENSG00000179532 | DNHD1 | chr11 | + | 6544120 | 6544246 | 6539875 | 6540083 | 6544791 | 6547666 | 1 | 1 | 4 | 6 | 4 | 0 | 0 | 0 | 0 | 0 | 0 | 6 | 0 | 1 | 0 | 0 | 0 | 0 | 4 | 0 | 275 | 149 | 1 | 1 | 1 | 1 | 1 | 1 | 1 | NA | 1 | NA | 0.119 | NA | 0.441 |
| SE | 100636 | ENSG00000179532 | DNHD1 | chr11 | + | 6544573 | 6544671 | 6538611 | 6538810 | 6544876 | 6547666 | 0 | 1 | 2 | 0 | 0 | 0 | 0 | 0 | 0 | 0 | 0 | 0 | 0 | 2 | 0 | 0 | 8 | 0 | 0 | 0 | 247 | 149 | 1 | 1 | NA | 1 | 1 | NA | NA | NA | 0 | NA | 1 | NA | 0.5 |
| SE | 100637 | ENSG00000179532 | DNHD1 | chr11 | + | 6544573 | 6544671 | 6544120 | 6544246 | 6544791 | 6547666 | 12 | 5 | 7 | 18 | 7 | 0 | 0 | 0 | 0 | 1 | 7 | 12 | 23 | 15 | 21 | 0 | 0 | 0 | 0 | 0 | 247 | 149 | 1 | 1 | 1 | 1 | 1 | 1 | 0.809 | 1 | 1 | 1 | 1 | 1 | -0.038 |
| SE | 100638 | ENSG00000179532 | DNHD1 | chr11 | + | 6547862 | 6548040 | 6544791 | 6547666 | 6548209 | 6548402 | 2 | 0 | 5 | 6 | 0 | 5 | 1 | 1 | 2 | 0 | 0 | 0 | 2 | 1 | 0 | 0 | 0 | 1 | 0 | 0 | 298 | 149 | 1 | 1 | 0.167 | 0 | 0.714 | 0.6 | NA | NA | NA | 0.5 | 1 | NA | -0.38 |
| SE | 100641 | ENSG00000179532 | DNHD1 | chr11 | + | 6570246 | 6570396 | 6570008 | 6570100 | 6570617 | 6571423 | 0 | 0 | 3 | 3 | 1 | 0 | 0 | 1 | 0 | 0 | 0 | 9 | 3 | 0 | 0 | 0 | 0 | 0 | 0 | 0 | 298 | 149 | 1 | 1 | NA | NA | 0.6 | 1 | 1 | NA | 1 | 1 | NA | NA | -0.133 |
| SE | 100644 | ENSG00000179532 | DNHD1 | chr11 | + | 6570617 | 6571423 | 6570008 | 6570396 | 6571635 | 6572019 | 1 | 1 | 6 | 6 | 4 | 0 | 0 | 0 | 0 | 0 | 3 | 9 | 4 | 0 | 1 | 0 | 3 | 0 | 0 | 0 | 298 | 149 | 1 | 1 | 1 | 1 | 1 | 1 | 1 | 1 | 0.6 | 1 | NA | 1 | 0.1 |
| SE | 100645 | ENSG00000166340 | TPP1 | chr11 | - | 6615170 | 6615329 | 6614865 | 6614991 | 6615441 | 6615545 | 695 | 358 | 454 | 530 | 582 | 0 | 0 | 1 | 0 | 0 | 882 | 719 | 422 | 793 | 2331 | 0 | 0 | 0 | 0 | 0 | 298 | 149 | 1 | 1 | 1 | 1 | 0.996 | 1 | 1 | 1 | 1 | 1 | 1 | 1 | -0.001 |
| SE | 100646 | ENSG00000166340 | TPP1 | chr11 | - | 6615764 | 6615848 | 6615441 | 6615562 | 6616004 | 6616074 | 1 | 0 | 1 | 0 | 1 | 122 | 62 | 94 | 102 | 76 | 0 | 0 | 0 | 7 | 0 | 158 | 131 | 75 | 153 | 395 | 233 | 149 | 1 | 1 | 0.005 | 0 | 0.007 | 0 | 0.008 | 0 | 0 | 0 | 0.028 | 0 | -0.002 |
| SE | 100647 | ENSG00000166340 | TPP1 | chr11 | - | 6615764 | 6615848 | 6615441 | 6615562 | 6616314 | 6616503 | 0 | 0 | 2 | 0 | 1 | 0 | 3 | 0 | 0 | 0 | 0 | 0 | 0 | 7 | 0 | 6 | 6 | 0 | 6 | 2 | 233 | 149 | 1 | 1 | NA | 0 | 1 | NA | 1 | 0 | 0 | NA | 0.427 | 0 | 0.56 |
| SE | 100649 | ENSG00000166340 | TPP1 | chr11 | - | 6616004 | 6616074 | 6615764 | 6615848 | 6616314 | 6616479 | 215 | 100 | 91 | 154 | 111 | 0 | 0 | 2 | 0 | 0 | 342 | 228 | 129 | 205 | 669 | 0 | 0 | 0 | 0 | 0 | 219 | 149 | 1 | 1 | 1 | 1 | 0.969 | 1 | 1 | 1 | 1 | 1 | 1 | 1 | -0.006 |
| SE | 100650 | ENSG00000166340 | TPP1 | chr11 | - | 6616314 | 6616503 | 6616004 | 6616074 | 6616660 | 6616859 | 590 | 269 | 323 | 424 | 323 | 0 | 3 | 0 | 0 | 0 | 796 | 610 | 286 | 542 | 1690 | 0 | 0 | 0 | 1 | 8 | 298 | 149 | 1 | 1 | 1 | 0.978 | 1 | 1 | 1 | 1 | 1 | 1 | 0.996 | 0.991 | -0.002 |
| SE | 100652 | ENSG00000166340 | TPP1 | chr11 | - | 6616660 | 6616859 | 6616314 | 6616503 | 6616974 | 6617153 | 700 | 308 | 358 | 458 | 284 | 1 | 0 | 1 | 0 | 0 | 810 | 624 | 309 | 571 | 1620 | 0 | 0 | 2 | 3 | 0 | 298 | 149 | 1 | 1 | 0.997 | 1 | 0.994 | 1 | 1 | 1 | 1 | 0.987 | 0.99 | 1 | 0.003 |
| SE | 100653 | ENSG00000166340 | TPP1 | chr11 | - | 6616974 | 6617153 | 6616314 | 6616503 | 6617300 | 6617428 | 286 | 141 | 192 | 208 | 272 | 0 | 0 | 0 | 0 | 0 | 376 | 319 | 166 | 315 | 792 | 0 | 0 | 0 | 0 | 9 | 298 | 149 | 1 | 1 | 1 | 1 | 1 | 1 | 1 | 1 | 1 | 1 | 1 | 0.978 | 0.004 |
| SE | 100655 | ENSG00000166340 | TPP1 | chr11 | - | 6616974 | 6617153 | 6616314 | 6616503 | 6617625 | 6617776 | 4 | 0 | 4 | 0 | 3 | 0 | 1 | 0 | 0 | 0 | 3 | 0 | 3 | 3 | 0 | 0 | 0 | 0 | 0 | 0 | 298 | 149 | 1 | 1 | 1 | 0 | 1 | NA | 1 | 1 | NA | 1 | 1 | NA | -0.25 |
| SE | 100656 | ENSG00000166340 | TPP1 | chr11 | - | 6616974 | 6617428 | 6616314 | 6616503 | 6617625 | 6617776 | 275 | 119 | 168 | 221 | 242 | 0 | 1 | 0 | 0 | 0 | 268 | 298 | 136 | 281 | 841 | 0 | 0 | 0 | 0 | 0 | 298 | 149 | 1 | 1 | 1 | 0.983 | 1 | 1 | 1 | 1 | 1 | 1 | 1 | 1 | -0.003 |
| SE | 100657 | ENSG00000166340 | TPP1 | chr11 | - | 6617300 | 6617428 | 6616314 | 6616503 | 6617625 | 6617776 | 274 | 119 | 166 | 221 | 242 | 0 | 1 | 0 | 0 | 0 | 268 | 298 | 134 | 278 | 848 | 0 | 0 | 0 | 0 | 0 | 277 | 149 | 1 | 1 | 1 | 0.985 | 1 | 1 | 1 | 1 | 1 | 1 | 1 | 1 | -0.003 |
| SE | 100658 | ENSG00000166340 | TPP1 | chr11 | - | 6617300 | 6617428 | 6616974 | 6617153 | 6617625 | 6617776 | 559 | 260 | 357 | 429 | 514 | 3 | 0 | 3 | 0 | 3 | 635 | 611 | 298 | 590 | 1626 | 3 | 0 | 1 | 0 | 0 | 277 | 149 | 1 | 1 | 0.99 | 1 | 0.985 | 1 | 0.989 | 0.991 | 1 | 0.994 | 1 | 1 | -0.004 |
| SE | 100660 | ENSG00000166340 | TPP1 | chr11 | - | 6617358 | 6617428 | 6616422 | 6616503 | 6617625 | 6617776 | 121 | 44 | 77 | 82 | 122 | 0 | 1 | 0 | 0 | 0 | 146 | 119 | 52 | 109 | 362 | 0 | 0 | 0 | 0 | 0 | 219 | 149 | 1 | 1 | 1 | 0.968 | 1 | 1 | 1 | 1 | 1 | 1 | 1 | 1 | -0.006 |
| SE | 100662 | ENSG00000166340 | TPP1 | chr11 | - | 6617625 | 6617776 | 6616974 | 6617153 | 6618775 | 6618808 | 315 | 137 | 175 | 226 | 184 | 1 | 0 | 0 | 0 | 0 | 302 | 282 | 130 | 349 | 911 | 0 | 0 | 0 | 0 | 0 | 298 | 149 | 1 | 1 | 0.994 | 1 | 1 | 1 | 1 | 1 | 1 | 1 | 1 | 1 | -0.001 |
| SE | 100663 | ENSG00000166340 | TPP1 | chr11 | - | 6618775 | 6618915 | 6616974 | 6617776 | 6619195 | 6619259 | 608 | 254 | 320 | 444 | 316 | 2 | 0 | 0 | 0 | 0 | 577 | 513 | 239 | 636 | 1788 | 3 | 1 | 2 | 0 | 0 | 289 | 149 | 1 | 1 | 0.994 | 1 | 1 | 1 | 1 | 0.99 | 0.996 | 0.984 | 1 | 1 | 0.005 |
| SE | 100670 | ENSG00000166387 | PPFIBP2 | chr11 | + | 7620934 | 7621027 | 7610290 | 7610422 | 7628284 | 7628346 | 22 | 10 | 10 | 22 | 19 | 0 | 0 | 0 | 2 | 0 | 45 | 26 | 27 | 11 | 31 | 0 | 0 | 0 | 0 | 0 | 242 | 149 | 1 | 1 | 1 | 1 | 1 | 0.871 | 1 | 1 | 1 | 1 | 1 | 1 | -0.026 |
| SE | 100672 | ENSG00000166387 | PPFIBP2 | chr11 | + | 7632866 | 7632934 | 7630922 | 7631028 | 7634494 | 7634552 | 41 | 29 | 41 | 26 | 138 | 0 | 0 | 0 | 3 | 0 | 66 | 36 | 31 | 27 | 43 | 0 | 0 | 0 | 0 | 0 | 217 | 149 | 1 | 1 | 1 | 1 | 1 | 0.856 | 1 | 1 | 1 | 1 | 1 | 1 | -0.029 |
| SE | 100674 | ENSG00000166387 | PPFIBP2 | chr11 | + | 7634494 | 7634552 | 7630922 | 7631028 | 7635551 | 7635593 | 10 | 12 | 9 | 14 | 40 | 0 | 0 | 0 | 0 | 1 | 18 | 0 | 7 | 9 | 0 | 0 | 0 | 0 | 0 | 5 | 207 | 149 | 1 | 1 | 1 | 1 | 1 | 1 | 0.966 | 1 | NA | 1 | 1 | 0 | 0.243 |
| SE | 100675 | ENSG00000166387 | PPFIBP2 | chr11 | + | 7634494 | 7634552 | 7630922 | 7631028 | 7639731 | 7639811 | 0 | 0 | 0 | 0 | 4 | 0 | 0 | 0 | 0 | 0 | 0 | 0 | 0 | 0 | 0 | 0 | 0 | 3 | 0 | 0 | 207 | 149 | 1 | 1 | NA | NA | NA | NA | 1 | NA | NA | 0 | NA | NA | 1 |
| SE | 100676 | ENSG00000166387 | PPFIBP2 | chr11 | + | 7634494 | 7634552 | 7632345 | 7632934 | 7635551 | 7635593 | 44 | 36 | 38 | 35 | 146 | 0 | 0 | 0 | 0 | 3 | 71 | 22 | 30 | 30 | 31 | 0 | 0 | 0 | 0 | 0 | 207 | 149 | 1 | 1 | 1 | 1 | 1 | 1 | 0.972 | 1 | 1 | 1 | 1 | 1 | -0.006 |
| SE | 100677 | ENSG00000166387 | PPFIBP2 | chr11 | + | 7635551 | 7635593 | 7630922 | 7631028 | 7639731 | 7639811 | 25 | 13 | 19 | 15 | 8 | 0 | 0 | 0 | 0 | 0 | 53 | 0 | 12 | 8 | 32 | 0 | 0 | 3 | 0 | 0 | 191 | 149 | 1 | 1 | 1 | 1 | 1 | 1 | 1 | 1 | NA | 0.757 | 1 | 1 | 0.061 |
| SE | 100678 | ENSG00000166387 | PPFIBP2 | chr11 | + | 7635551 | 7635593 | 7634494 | 7634552 | 7639731 | 7639811 | 62 | 42 | 47 | 41 | 119 | 0 | 0 | 0 | 0 | 4 | 115 | 10 | 38 | 31 | 58 | 0 | 0 | 0 | 0 | 0 | 191 | 149 | 1 | 1 | 1 | 1 | 1 | 1 | 0.959 | 1 | 1 | 1 | 1 | 1 | -0.008 |
| SE | 100683 | ENSG00000166387 | PPFIBP2 | chr11 | + | 7648799 | 7648911 | 7648107 | 7648537 | 7649146 | 7649172 | 59 | 27 | 63 | 45 | 125 | 0 | 1 | 0 | 0 | 0 | 114 | 73 | 46 | 47 | 72 | 0 | 0 | 0 | 0 | 0 | 261 | 149 | 1 | 1 | 1 | 0.939 | 1 | 1 | 1 | 1 | 1 | 1 | 1 | 1 | -0.012 |
| SE | 100684 | ENSG00000166387 | PPFIBP2 | chr11 | + | 7648799 | 7648911 | 7648107 | 7648537 | 7649531 | 7649654 | 19 | 10 | 25 | 16 | 41 | 0 | 0 | 0 | 0 | 0 | 80 | 33 | 22 | 26 | 48 | 0 | 0 | 0 | 2 | 0 | 261 | 149 | 1 | 1 | 1 | 1 | 1 | 1 | 1 | 1 | 1 | 1 | 0.881 | 1 | 0.024 |
| SE | 100685 | ENSG00000166387 | PPFIBP2 | chr11 | + | 7649146 | 7649235 | 7648386 | 7648537 | 7649531 | 7649654 | 17 | 12 | 27 | 20 | 82 | 0 | 0 | 0 | 0 | 0 | 11 | 21 | 11 | 5 | 16 | 0 | 0 | 0 | 2 | 0 | 238 | 149 | 1 | 1 | 1 | 1 | 1 | 1 | 1 | 1 | 1 | 1 | 0.61 | 1 | 0.078 |
| SE | 100686 | ENSG00000166387 | PPFIBP2 | chr11 | + | 7649146 | 7649235 | 7648799 | 7648911 | 7649531 | 7649654 | 58 | 28 | 67 | 50 | 172 | 0 | 0 | 0 | 0 | 1 | 51 | 61 | 35 | 26 | 40 | 0 | 0 | 0 | 0 | 0 | 238 | 149 | 1 | 1 | 1 | 1 | 1 | 1 | 0.991 | 1 | 1 | 1 | 1 | 1 | -0.002 |
| SE | 100689 | ENSG00000107372 | ZFAND5 | chr9 | - | 72359417 | 72359521 | 72356930 | 72357056 | 72360109 | 72360221 | 1215 | 934 | 872 | 1034 | 1611 | 1 | 5 | 2 | 9 | 4 | 983 | 1539 | 1394 | 1152 | 627 | 0 | 10 | 7 | 7 | 0 | 253 | 149 | 1 | 1 | 0.999 | 0.991 | 0.996 | 0.985 | 0.996 | 1 | 0.989 | 0.992 | 0.99 | 1 | -0.001 |
| SE | 100690 | ENSG00000107372 | ZFAND5 | chr9 | - | 72359417 | 72359521 | 72356930 | 72357056 | 72360627 | 72360784 | 199 | 161 | 172 | 197 | 360 | 0 | 0 | 0 | 0 | 0 | 128 | 196 | 216 | 164 | 80 | 0 | 5 | 0 | 0 | 0 | 253 | 149 | 1 | 1 | 1 | 1 | 1 | 1 | 1 | 1 | 0.958 | 1 | 1 | 1 | 0.008 |
| SE | 100691 | ENSG00000107372 | ZFAND5 | chr9 | - | 72360109 | 72360221 | 72356930 | 72357056 | 72360627 | 72360784 | 543 | 451 | 350 | 443 | 750 | 0 | 0 | 0 | 0 | 0 | 402 | 594 | 602 | 350 | 313 | 0 | 5 | 0 | 0 | 0 | 261 | 149 | 1 | 1 | 1 | 1 | 1 | 1 | 1 | 1 | 0.985 | 1 | 1 | 1 | 0.003 |
| SE | 100692 | ENSG00000107372 | ZFAND5 | chr9 | - | 72360109 | 72360221 | 72359417 | 72359521 | 72360627 | 72360784 | 1549 | 1210 | 1031 | 1260 | 1974 | 6 | 5 | 6 | 3 | 10 | 1257 | 1921 | 1751 | 1339 | 860 | 0 | 12 | 11 | 10 | 0 | 261 | 149 | 1 | 1 | 0.993 | 0.993 | 0.99 | 0.996 | 0.991 | 1 | 0.989 | 0.989 | 0.987 | 1 | 0 |
| SE | 100693 | ENSG00000107372 | ZFAND5 | chr9 | - | 72360627 | 72360784 | 72356930 | 72357056 | 72364695 | 72365235 | 2 | 0 | 1 | 3 | 0 | 0 | 0 | 0 | 0 | 0 | 10 | 9 | 2 | 1 | 0 | 0 | 0 | 1 | 0 | 0 | 298 | 149 | 1 | 1 | 1 | NA | 1 | 1 | NA | 1 | 1 | 0.5 | 1 | NA | 0.125 |
| SE | 100694 | ENSG00000107372 | ZFAND5 | chr9 | - | 72360627 | 72360787 | 72356930 | 72357056 | 72364695 | 72365235 | 49 | 18 | 33 | 33 | 12 | 0 | 0 | 0 | 0 | 0 | 70 | 67 | 61 | 77 | 64 | 0 | 0 | 1 | 0 | 0 | 298 | 149 | 1 | 1 | 1 | 1 | 1 | 1 | 1 | 1 | 1 | 0.968 | 1 | 1 | 0.006 |
| SE | 100695 | ENSG00000107372 | ZFAND5 | chr9 | - | 72363469 | 72363606 | 72360627 | 72360787 | 72364461 | 72364626 | 184 | 169 | 145 | 162 | 194 | 0 | 0 | 0 | 1 | 0 | 145 | 205 | 153 | 203 | 164 | 0 | 0 | 4 | 0 | 0 | 286 | 149 | 1 | 1 | 1 | 1 | 1 | 0.988 | 1 | 1 | 1 | 0.952 | 1 | 1 | 0.007 |
| SE | 100700 | ENSG00000109917 | ZPR1 | chr11 | - | 116782918 | 116783029 | 116782157 | 116782244 | 116783529 | 116783619 | 316 | 229 | 286 | 359 | 230 | 0 | 0 | 0 | 1 | 0 | 466 | 400 | 329 | 384 | 339 | 0 | 0 | 0 | 0 | 0 | 260 | 149 | 1 | 1 | 1 | 1 | 1 | 0.995 | 1 | 1 | 1 | 1 | 1 | 1 | -0.001 |
| SE | 100703 | ENSG00000109917 | ZPR1 | chr11 | - | 116787481 | 116787602 | 116786968 | 116787059 | 116787819 | 116787898 | 72 | 44 | 74 | 93 | 95 | 1 | 2 | 0 | 0 | 0 | 114 | 72 | 85 | 114 | 109 | 0 | 0 | 0 | 0 | 2 | 270 | 149 | 1 | 1 | 0.975 | 0.924 | 1 | 1 | 1 | 1 | 1 | 1 | 1 | 0.968 | -0.014 |
| SE | 100704 | ENSG00000109917 | ZPR1 | chr11 | - | 116787481 | 116787643 | 116786968 | 116787059 | 116787819 | 116787989 | 122 | 68 | 110 | 161 | 138 | 1 | 2 | 0 | 0 | 0 | 182 | 113 | 134 | 181 | 136 | 0 | 0 | 0 | 0 | 2 | 298 | 149 | 1 | 1 | 0.984 | 0.944 | 1 | 1 | 1 | 1 | 1 | 1 | 1 | 0.971 | -0.009 |
| SE | 100707 | ENSG00000187147 | RNF220 | chr1 | + | 44649882 | 44649957 | 44649660 | 44649769 | 44650703 | 44650808 | 392 | 156 | 363 | 355 | 439 | 1 | 2 | 0 | 1 | 1 | 371 | 338 | 251 | 163 | 484 | 0 | 0 | 1 | 3 | 0 | 224 | 149 | 1 | 1 | 0.996 | 0.981 | 1 | 0.996 | 0.997 | 1 | 1 | 0.994 | 0.973 | 1 | 0.001 |
| SE | 100708 | ENSG00000134954 | ETS1 | chr11 | - | 128463508 | 128463627 | 128458760 | 128462576 | 128480190 | 128480451 | 1438 | 797 | 1713 | 1282 | 1257 | 0 | 0 | 0 | 2 | 0 | 237 | 1008 | 1274 | 428 | 186 | 0 | 0 | 0 | 8 | 0 | 268 | 149 | 1 | 1 | 1 | 1 | 1 | 0.997 | 1 | 1 | 1 | 1 | 0.967 | 1 | 0.006 |
| SE | 100709 | ENSG00000134954 | ETS1 | chr11 | - | 128463508 | 128463627 | 128458760 | 128462576 | 128484822 | 128485071 | 716 | 389 | 872 | 679 | 713 | 0 | 0 | 0 | 0 | 0 | 182 | 617 | 693 | 231 | 108 | 0 | 0 | 1 | 0 | 0 | 268 | 149 | 1 | 1 | 1 | 1 | 1 | 1 | 1 | 1 | 1 | 0.997 | 1 | 1 | 0.001 |
| SE | 100710 | ENSG00000134954 | ETS1 | chr11 | - | 128480190 | 128480451 | 128458760 | 128462576 | 128484822 | 128485071 | 1595 | 695 | 1692 | 1294 | 1631 | 0 | 0 | 0 | 0 | 0 | 282 | 887 | 1121 | 405 | 225 | 0 | 0 | 1 | 0 | 0 | 298 | 149 | 1 | 1 | 1 | 1 | 1 | 1 | 1 | 1 | 1 | 0.998 | 1 | 1 | 0 |
| SE | 100712 | ENSG00000134954 | ETS1 | chr11 | - | 128480190 | 128480451 | 128463508 | 128463627 | 128521912 | 128522310 | 765 | 425 | 880 | 633 | 587 | 0 | 0 | 0 | 0 | 1 | 87 | 441 | 635 | 215 | 78 | 4 | 0 | 0 | 0 | 0 | 298 | 149 | 1 | 1 | 1 | 1 | 1 | 1 | 0.997 | 0.916 | 1 | 1 | 1 | 1 | 0.016 |
| SE | 100713 | ENSG00000134954 | ETS1 | chr11 | - | 128480400 | 128480451 | 128461493 | 128462576 | 128484822 | 128485071 | 369 | 137 | 395 | 311 | 386 | 0 | 0 | 0 | 0 | 0 | 24 | 202 | 227 | 78 | 25 | 0 | 0 | 1 | 0 | 0 | 200 | 149 | 1 | 1 | 1 | 1 | 1 | 1 | 1 | 1 | 1 | 0.994 | 1 | 1 | 0.001 |
| SE | 100714 | ENSG00000134954 | ETS1 | chr11 | - | 128486068 | 128486146 | 128484822 | 128485071 | 128490456 | 128490576 | 948 | 445 | 1059 | 839 | 1312 | 0 | 0 | 1 | 0 | 0 | 261 | 678 | 873 | 338 | 179 | 0 | 0 | 0 | 0 | 0 | 227 | 149 | 1 | 1 | 1 | 1 | 0.999 | 1 | 1 | 1 | 1 | 1 | 1 | 1 | 0 |
| SE | 100716 | ENSG00000134954 | ETS1 | chr11 | - | 128490456 | 128490576 | 128486068 | 128486146 | 128521912 | 128522201 | 849 | 459 | 1011 | 774 | 1568 | 0 | 0 | 1 | 1 | 1 | 93 | 573 | 830 | 316 | 101 | 0 | 3 | 0 | 0 | 0 | 269 | 149 | 1 | 1 | 1 | 1 | 0.998 | 0.998 | 0.999 | 1 | 0.991 | 1 | 1 | 1 | 0.001 |
| SE | 100721 | ENSG00000167286 | CD3D | chr11 | - | 118340374 | 118340593 | 118339774 | 118339906 | 118342552 | 118342631 | 1901 | 940 | 1734 | 1582 | 2499 | 1 | 0 | 1 | 0 | 0 | 619 | 1555 | 733 | 570 | 1118 | 0 | 0 | 1 | 0 | 0 | 298 | 149 | 1 | 1 | 0.999 | 1 | 0.999 | 1 | 1 | 1 | 1 | 0.997 | 1 | 1 | 0 |
| SE | 100725 | ENSG00000130830 | MPP1 | chrX | - | 154786203 | 154786340 | 154785050 | 154785157 | 154789953 | 154790022 | 173 | 173 | 110 | 334 | 349 | 0 | 0 | 0 | 0 | 4 | 712 | 314 | 269 | 352 | 2508 | 0 | 0 | 0 | 0 | 0 | 286 | 149 | 1 | 1 | 1 | 1 | 1 | 1 | 0.978 | 1 | 1 | 1 | 1 | 1 | -0.004 |
| SE | 100726 | ENSG00000130830 | MPP1 | chrX | - | 154786203 | 154786400 | 154784700 | 154785157 | 154789953 | 154790022 | 583 | 573 | 326 | 974 | 873 | 0 | 0 | 0 | 0 | 4 | 2292 | 1000 | 761 | 1222 | 7900 | 0 | 0 | 0 | 0 | 0 | 298 | 149 | 1 | 1 | 1 | 1 | 1 | 1 | 0.991 | 1 | 1 | 1 | 1 | 1 | -0.002 |
| SE | 100727 | ENSG00000130830 | MPP1 | chrX | - | 154786203 | 154786400 | 154784700 | 154785157 | 154805271 | 154805477 | 188 | 189 | 112 | 358 | 351 | 0 | 0 | 0 | 0 | 2 | 774 | 334 | 279 | 410 | 2727 | 0 | 0 | 0 | 0 | 17 | 298 | 149 | 1 | 1 | 1 | 1 | 1 | 1 | 0.989 | 1 | 1 | 1 | 1 | 0.988 | 0 |
| SE | 100728 | ENSG00000130830 | MPP1 | chrX | - | 154789953 | 154790022 | 154785050 | 154785157 | 154805271 | 154805485 | 0 | 0 | 0 | 0 | 0 | 0 | 0 | 0 | 0 | 2 | 10 | 0 | 0 | 0 | 0 | 0 | 0 | 0 | 0 | 17 | 218 | 149 | 1 | 1 | NA | NA | NA | NA | 0 | 1 | NA | NA | NA | 0 | -0.5 |
| SE | 100729 | ENSG00000130830 | MPP1 | chrX | - | 154789953 | 154790022 | 154786203 | 154786400 | 154790982 | 154791068 | 493 | 468 | 258 | 757 | 666 | 0 | 1 | 0 | 0 | 3 | 1942 | 883 | 614 | 996 | 6519 | 0 | 0 | 0 | 0 | 0 | 218 | 149 | 1 | 1 | 1 | 0.997 | 1 | 1 | 0.993 | 1 | 1 | 1 | 1 | 1 | -0.002 |
| SE | 100731 | ENSG00000130830 | MPP1 | chrX | - | 154790982 | 154791068 | 154789953 | 154790022 | 154791768 | 154791847 | 278 | 290 | 185 | 448 | 355 | 0 | 0 | 0 | 0 | 0 | 1167 | 560 | 334 | 542 | 3776 | 0 | 1 | 0 | 0 | 0 | 235 | 149 | 1 | 1 | 1 | 1 | 1 | 1 | 1 | 1 | 0.997 | 1 | 1 | 1 | 0.001 |
| SE | 100732 | ENSG00000130830 | MPP1 | chrX | - | 154799744 | 154799893 | 154792141 | 154792285 | 154804696 | 154804827 | 0 | 5 | 2 | 4 | 0 | 0 | 2 | 0 | 3 | 1 | 0 | 0 | 3 | 0 | 57 | 0 | 7 | 5 | 6 | 47 | 298 | 149 | 1 | 1 | NA | 0.556 | 1 | 0.4 | 0 | NA | 0 | 0.231 | 0 | 0.377 | 0.337 |
| SE | 100735 | ENSG00000130830 | MPP1 | chrX | - | 154804696 | 154804827 | 154799744 | 154799893 | 154805271 | 154805456 | 0 | 2 | 0 | 1 | 0 | 3 | 4 | 0 | 1 | 0 | 0 | 0 | 1 | 0 | 13 | 0 | 0 | 5 | 6 | 6 | 280 | 149 | 1 | 1 | 0 | 0.21 | NA | 0.347 | NA | NA | NA | 0.096 | 0 | 0.536 | -0.025 |
| SE | 100738 | ENSG00000143756 | FBXO28 | chr1 | + | 224134073 | 224134212 | 224130471 | 224130581 | 224157351 | 224158150 | 68 | 46 | 88 | 54 | 81 | 1 | 0 | 2 | 0 | 0 | 53 | 82 | 101 | 64 | 17 | 0 | 0 | 0 | 0 | 0 | 288 | 149 | 1 | 1 | 0.972 | 1 | 0.958 | 1 | 1 | 1 | 1 | 1 | 1 | 1 | -0.014 |
| SE | 100739 | ENSG00000143756 | FBXO28 | chr1 | + | 224153141 | 224153337 | 224130471 | 224130581 | 224157351 | 224158182 | 108 | 46 | 59 | 72 | 91 | 1 | 0 | 2 | 0 | 0 | 41 | 46 | 80 | 29 | 0 | 0 | 0 | 0 | 0 | 0 | 298 | 149 | 1 | 1 | 0.982 | 1 | 0.937 | 1 | 1 | 1 | 1 | 1 | 1 | NA | -0.016 |
| SE | 100743 | ENSG00000173992 | CCS | chr11 | + | 66600488 | 66600549 | 66599115 | 66599636 | 66605338 | 66605416 | 49 | 22 | 60 | 78 | 39 | 0 | 0 | 0 | 0 | 0 | 172 | 65 | 52 | 78 | 70 | 0 | 0 | 0 | 1 | 0 | 210 | 149 | 1 | 1 | 1 | 1 | 1 | 1 | 1 | 1 | 1 | 1 | 0.982 | 1 | 0.004 |
| SE | 100745 | ENSG00000248643 | RBM14-RBM4 | chr11 | + | 66639699 | 66640123 | 66616630 | 66617057 | 66646026 | 66646469 | 89 | 42 | 69 | 67 | 96 | 0 | 0 | 2 | 0 | 0 | 65 | 72 | 92 | 71 | 87 | 0 | 0 | 0 | 0 | 0 | 298 | 149 | 1 | 1 | 1 | 1 | 0.945 | 1 | 1 | 1 | 1 | 1 | 1 | 1 | -0.011 |
| SE | 100746 | ENSG00000248643 | RBM14-RBM4 | chr11 | + | 66643449 | 66644140 | 66616625 | 66617057 | 66646026 | 66646469 | 51 | 23 | 38 | 56 | 23 | 0 | 0 | 2 | 0 | 0 | 16 | 4 | 17 | 24 | 49 | 0 | 0 | 0 | 0 | 0 | 298 | 149 | 1 | 1 | 1 | 1 | 0.905 | 1 | 1 | 1 | 1 | 1 | 1 | 1 | -0.019 |
| SE | 100747 | ENSG00000106723 | SPIN1 | chr9 | + | 88415606 | 88415699 | 88388430 | 88388538 | 88418777 | 88418951 | 1 | 2 | 1 | 0 | 2 | 0 | 0 | 0 | 4 | 0 | 0 | 0 | 0 | 3 | 0 | 0 | 0 | 0 | 0 | 0 | 242 | 149 | 1 | 1 | 1 | 1 | 1 | 0 | 1 | NA | NA | NA | 1 | NA | -0.2 |
| SE | 100748 | ENSG00000106723 | SPIN1 | chr9 | + | 88415606 | 88415699 | 88388430 | 88388538 | 88418849 | 88418951 | 2 | 3 | 0 | 0 | 0 | 2 | 1 | 1 | 3 | 0 | 0 | 0 | 0 | 0 | 0 | 0 | 1 | 0 | 0 | 0 | 242 | 149 | 1 | 1 | 0.381 | 0.649 | 0 | 0 | NA | NA | 0 | NA | NA | NA | 0.258 |
| SE | 100749 | ENSG00000106723 | SPIN1 | chr9 | + | 88415606 | 88415699 | 88388430 | 88388538 | 88426381 | 88426591 | 1 | 1 | 1 | 2 | 10 | 38 | 14 | 20 | 25 | 19 | 0 | 0 | 0 | 8 | 0 | 0 | 15 | 22 | 20 | 18 | 242 | 149 | 1 | 1 | 0.016 | 0.042 | 0.03 | 0.047 | 0.245 | NA | 0 | 0 | 0.198 | 0 | 0.026 |
| SE | 100750 | ENSG00000106723 | SPIN1 | chr9 | + | 88416643 | 88416766 | 88415606 | 88415699 | 88418777 | 88418951 | 1 | 0 | 0 | 0 | 0 | 1 | 2 | 1 | 0 | 2 | 0 | 0 | 0 | 0 | 0 | 0 | 0 | 0 | 3 | 0 | 272 | 149 | 1 | 1 | 0.354 | 0 | 0 | NA | 0 | NA | NA | NA | 0 | NA | 0.088 |
| SE | 100751 | ENSG00000106723 | SPIN1 | chr9 | + | 88416643 | 88416766 | 88415606 | 88415699 | 88418849 | 88418951 | 1 | 0 | 0 | 0 | 0 | 2 | 3 | 0 | 0 | 0 | 0 | 0 | 1 | 0 | 0 | 0 | 0 | 0 | 0 | 0 | 272 | 149 | 1 | 1 | 0.215 | 0 | NA | NA | NA | NA | NA | 1 | NA | NA | -0.892 |
| SE | 100753 | ENSG00000106723 | SPIN1 | chr9 | + | 88418777 | 88418951 | 88415606 | 88415699 | 88426381 | 88426591 | 16 | 12 | 24 | 28 | 38 | 1 | 1 | 1 | 2 | 10 | 18 | 13 | 8 | 22 | 0 | 0 | 0 | 0 | 12 | 0 | 298 | 149 | 1 | 1 | 0.889 | 0.857 | 0.923 | 0.875 | 0.655 | 1 | 1 | 1 | 0.478 | NA | -0.03 |
| SE | 100755 | ENSG00000106723 | SPIN1 | chr9 | + | 88418849 | 88418951 | 88415606 | 88415699 | 88426381 | 88426591 | 16 | 12 | 13 | 26 | 26 | 1 | 1 | 1 | 2 | 10 | 18 | 11 | 4 | 16 | 0 | 0 | 0 | 0 | 12 | 0 | 251 | 149 | 1 | 1 | 0.905 | 0.877 | 0.885 | 0.885 | 0.607 | 1 | 1 | 1 | 0.442 | NA | -0.029 |
| SE | 100756 | ENSG00000106723 | SPIN1 | chr9 | + | 88426381 | 88426591 | 88388443 | 88388538 | 88448940 | 88448989 | 128 | 66 | 78 | 116 | 111 | 0 | 0 | 0 | 0 | 0 | 35 | 89 | 117 | 104 | 40 | 0 | 0 | 1 | 0 | 0 | 298 | 149 | 1 | 1 | 1 | 1 | 1 | 1 | 1 | 1 | 1 | 0.983 | 1 | 1 | 0.003 |
| SE | 100758 | ENSG00000106723 | SPIN1 | chr9 | + | 88468371 | 88468605 | 88448940 | 88448989 | 88475077 | 88475203 | 80 | 50 | 74 | 93 | 95 | 1 | 0 | 0 | 0 | 0 | 73 | 65 | 55 | 106 | 10 | 0 | 0 | 2 | 0 | 0 | 298 | 149 | 1 | 1 | 0.976 | 1 | 1 | 1 | 1 | 1 | 1 | 0.932 | 1 | 1 | 0.009 |
| SE | 100759 | ENSG00000173933 | RBM4 | chr11 | + | 66639699 | 66640123 | 66638666 | 66638752 | 66643449 | 66643528 | 354 | 203 | 279 | 331 | 271 | 1 | 0 | 0 | 0 | 0 | 489 | 358 | 266 | 373 | 372 | 0 | 0 | 0 | 0 | 0 | 298 | 149 | 1 | 1 | 0.994 | 1 | 1 | 1 | 1 | 1 | 1 | 1 | 1 | 1 | -0.001 |
| SE | 100760 | ENSG00000173933 | RBM4 | chr11 | + | 66643449 | 66643528 | 66639699 | 66640123 | 66643858 | 66644089 | 129 | 66 | 96 | 121 | 151 | 0 | 0 | 3 | 1 | 0 | 147 | 86 | 87 | 126 | 132 | 0 | 0 | 0 | 0 | 0 | 228 | 149 | 1 | 1 | 1 | 1 | 0.954 | 0.988 | 1 | 1 | 1 | 1 | 1 | 1 | -0.012 |
| SE | 100763 | ENSG00000173933 | RBM4 | chr11 | + | 66644633 | 66644743 | 66643749 | 66644140 | 66665855 | 66668374 | 1 | 3 | 2 | 1 | 5 | 3 | 5 | 3 | 7 | 3 | 0 | 2 | 0 | 0 | 6 | 32 | 4 | 5 | 0 | 9 | 259 | 149 | 1 | 1 | 0.161 | 0.257 | 0.277 | 0.076 | 0.489 | 0 | 0.223 | 0 | NA | 0.277 | 0.127 |
| SE | 100765 | ENSG00000173914 | RBM4B | chr11 | - | 66668614 | 66669291 | 66664997 | 66665578 | 66676667 | 66677091 | 28 | 17 | 36 | 37 | 26 | 0 | 0 | 3 | 2 | 0 | 57 | 18 | 13 | 24 | 44 | 0 | 0 | 0 | 0 | 0 | 298 | 149 | 1 | 1 | 1 | 1 | 0.857 | 0.902 | 1 | 1 | 1 | 1 | 1 | 1 | -0.048 |
| SE | 100767 | ENSG00000173542 | MOB1B | chr4 | + | 70942822 | 70942882 | 70902325 | 70902550 | 70950699 | 70950811 | 1 | 0 | 0 | 1 | 3 | 1 | 5 | 2 | 4 | 4 | 0 | 0 | 3 | 0 | 0 | 0 | 1 | 2 | 0 | 10 | 209 | 149 | 1 | 1 | 0.416 | 0 | 0 | 0.151 | 0.348 | NA | 0 | 0.517 | NA | 0 | 0.011 |
| SE | 100768 | ENSG00000173542 | MOB1B | chr4 | + | 70942822 | 70942882 | 70902325 | 70902550 | 70958873 | 70959040 | 0 | 0 | 0 | 1 | 0 | 25 | 51 | 24 | 72 | 35 | 0 | 0 | 0 | 0 | 0 | 51 | 70 | 26 | 14 | 90 | 209 | 149 | 1 | 1 | 0 | 0 | 0 | 0.01 | 0 | 0 | 0 | 0 | 0 | 0 | 0.002 |
| SE | 100770 | ENSG00000173542 | MOB1B | chr4 | + | 70950699 | 70950811 | 70942822 | 70942882 | 70958873 | 70959040 | 11 | 8 | 7 | 15 | 21 | 0 | 0 | 0 | 1 | 0 | 11 | 5 | 9 | 3 | 13 | 0 | 0 | 0 | 0 | 0 | 261 | 149 | 1 | 1 | 1 | 1 | 1 | 0.895 | 1 | 1 | 1 | 1 | 1 | 1 | -0.021 |
| SE | 100771 | ENSG00000173542 | MOB1B | chr4 | + | 70958873 | 70959040 | 70902346 | 70902550 | 70969930 | 70970024 | 127 | 248 | 117 | 349 | 229 | 1 | 0 | 0 | 0 | 0 | 309 | 249 | 110 | 114 | 378 | 0 | 0 | 0 | 0 | 0 | 298 | 149 | 1 | 1 | 0.984 | 1 | 1 | 1 | 1 | 1 | 1 | 1 | 1 | 1 | -0.003 |
| SE | 100772 | ENSG00000173542 | MOB1B | chr4 | + | 70969930 | 70970024 | 70958873 | 70959040 | 70975152 | 70975254 | 143 | 270 | 128 | 364 | 276 | 0 | 1 | 2 | 1 | 0 | 357 | 226 | 98 | 151 | 383 | 0 | 0 | 0 | 0 | 0 | 243 | 149 | 1 | 1 | 1 | 0.994 | 0.975 | 0.996 | 1 | 1 | 1 | 1 | 1 | 1 | -0.007 |
| SE | 100774 | ENSG00000173542 | MOB1B | chr4 | + | 70979127 | 70979291 | 70975152 | 70975286 | 70981979 | 70982242 | 318 | 662 | 201 | 765 | 897 | 1 | 0 | 2 | 7 | 0 | 363 | 439 | 184 | 327 | 546 | 0 | 0 | 0 | 2 | 9 | 298 | 149 | 1 | 1 | 0.994 | 1 | 0.98 | 0.982 | 1 | 1 | 1 | 1 | 0.988 | 0.968 | 0 |
| SE | 100779 | ENSG00000162408 | NOL9 | chr1 | - | 6531967 | 6532079 | 6528993 | 6529171 | 6532462 | 6532760 | 190 | 112 | 187 | 177 | 317 | 0 | 0 | 0 | 0 | 0 | 75 | 98 | 91 | 78 | 141 | 0 | 0 | 0 | 1 | 0 | 261 | 149 | 1 | 1 | 1 | 1 | 1 | 1 | 1 | 1 | 1 | 1 | 0.978 | 1 | 0.004 |
| SE | 100780 | ENSG00000162408 | NOL9 | chr1 | - | 6531967 | 6532079 | 6528993 | 6529171 | 6533279 | 6533441 | 59 | 41 | 65 | 60 | 83 | 0 | 1 | 0 | 0 | 0 | 42 | 51 | 26 | 42 | 41 | 0 | 0 | 0 | 0 | 0 | 261 | 149 | 1 | 1 | 1 | 0.959 | 1 | 1 | 1 | 1 | 1 | 1 | 1 | 1 | -0.008 |
| SE | 100781 | ENSG00000162408 | NOL9 | chr1 | - | 6532462 | 6532760 | 6528993 | 6529171 | 6533279 | 6533441 | 178 | 88 | 139 | 152 | 328 | 0 | 1 | 0 | 0 | 0 | 135 | 158 | 104 | 158 | 146 | 0 | 0 | 0 | 0 | 0 | 298 | 149 | 1 | 1 | 1 | 0.978 | 1 | 1 | 1 | 1 | 1 | 1 | 1 | 1 | -0.004 |
| SE | 100782 | ENSG00000162408 | NOL9 | chr1 | - | 6532462 | 6532760 | 6531967 | 6532079 | 6533279 | 6533441 | 309 | 159 | 261 | 269 | 562 | 0 | 0 | 0 | 0 | 0 | 168 | 205 | 169 | 193 | 246 | 0 | 0 | 1 | 0 | 0 | 298 | 149 | 1 | 1 | 1 | 1 | 1 | 1 | 1 | 1 | 1 | 0.988 | 1 | 1 | 0.002 |
| SE | 100783 | ENSG00000162408 | NOL9 | chr1 | - | 6533279 | 6533441 | 6532462 | 6532760 | 6544825 | 6544922 | 178 | 91 | 141 | 154 | 331 | 0 | 1 | 0 | 0 | 0 | 135 | 161 | 110 | 157 | 149 | 0 | 0 | 0 | 0 | 0 | 298 | 149 | 1 | 1 | 1 | 0.978 | 1 | 1 | 1 | 1 | 1 | 1 | 1 | 1 | -0.004 |
| SE | 100786 | ENSG00000162408 | NOL9 | chr1 | - | 6544825 | 6544922 | 6533279 | 6533441 | 6545044 | 6545180 | 65 | 43 | 70 | 62 | 90 | 3 | 0 | 0 | 0 | 0 | 67 | 64 | 33 | 54 | 57 | 0 | 0 | 0 | 0 | 0 | 246 | 149 | 1 | 1 | 0.929 | 1 | 1 | 1 | 1 | 1 | 1 | 1 | 1 | 1 | -0.014 |
| SE | 100788 | ENSG00000162408 | NOL9 | chr1 | - | 6549570 | 6549698 | 6533279 | 6533441 | 6550395 | 6550615 | 116 | 64 | 126 | 82 | 74 | 0 | 0 | 0 | 2 | 0 | 121 | 67 | 64 | 94 | 153 | 0 | 0 | 0 | 0 | 0 | 277 | 149 | 1 | 1 | 1 | 1 | 1 | 0.957 | 1 | 1 | 1 | 1 | 1 | 1 | -0.009 |
| SE | 100789 | ENSG00000162408 | NOL9 | chr1 | - | 6550395 | 6550615 | 6533279 | 6533441 | 6554106 | 6554513 | 26 | 14 | 30 | 24 | 5 | 0 | 0 | 0 | 1 | 0 | 55 | 15 | 26 | 21 | 29 | 0 | 0 | 0 | 0 | 0 | 298 | 149 | 1 | 1 | 1 | 1 | 1 | 0.923 | 1 | 1 | 1 | 1 | 1 | 1 | -0.015 |
| SE | 100790 | ENSG00000162408 | NOL9 | chr1 | - | 6550395 | 6550615 | 6549570 | 6549698 | 6554106 | 6554513 | 160 | 84 | 166 | 118 | 102 | 2 | 0 | 0 | 2 | 0 | 177 | 88 | 95 | 120 | 185 | 0 | 0 | 2 | 0 | 0 | 298 | 149 | 1 | 1 | 0.976 | 1 | 1 | 0.967 | 1 | 1 | 1 | 0.96 | 1 | 1 | -0.003 |
| SE | 100791 | ENSG00000135272 | MDFIC | chr7 | + | 114922926 | 114923127 | 114922358 | 114922636 | 114942274 | 114942397 | 37 | 9 | 14 | 12 | 4 | 0 | 0 | 0 | 1 | 0 | 0 | 5 | 7 | 3 | 7 | 0 | 0 | 0 | 0 | 0 | 298 | 149 | 1 | 1 | 1 | 1 | 1 | 0.857 | 1 | NA | 1 | 1 | 1 | 1 | -0.029 |
| SE | 100802 | ENSG00000149582 | TMEM25 | chr11 | + | 118533856 | 118533887 | 118532916 | 118533207 | 118534028 | 118534120 | 1 | 4 | 15 | 1 | 4 | 0 | 0 | 0 | 0 | 2 | 15 | 10 | 7 | 0 | 3 | 0 | 0 | 0 | 0 | 0 | 180 | 149 | 1 | 1 | 1 | 1 | 1 | 1 | 0.623 | 1 | 1 | 1 | NA | 1 | -0.075 |
| SE | 100838 | ENSG00000149311 | ATM | chr11 | + | 108223535 | 108223686 | 108223148 | 108223186 | 108224783 | 108224871 | 7 | 2 | 3 | 0 | 8 | 17 | 11 | 11 | 8 | 12 | 0 | 3 | 0 | 0 | 0 | 0 | 0 | 5 | 0 | 0 | 298 | 149 | 1 | 1 | 0.171 | 0.083 | 0.12 | 0 | 0.25 | NA | 1 | 0 | NA | NA | -0.375 |
| SE | 100840 | ENSG00000149311 | ATM | chr11 | + | 108223535 | 108223686 | 108223148 | 108223186 | 108227775 | 108227888 | 1 | 2 | 0 | 0 | 0 | 6 | 1 | 0 | 0 | 0 | 0 | 1 | 0 | 0 | 0 | 0 | 0 | 0 | 0 | 0 | 298 | 149 | 1 | 1 | 0.077 | 0.5 | NA | NA | NA | NA | 1 | NA | NA | NA | -0.712 |
| SE | 100849 | ENSG00000149311 | ATM | chr11 | + | 108226163 | 108226247 | 108224783 | 108224871 | 108227594 | 108227696 | 0 | 0 | 2 | 0 | 1 | 105 | 69 | 84 | 109 | 81 | 0 | 0 | 0 | 0 | 0 | 22 | 19 | 26 | 14 | 16 | 233 | 149 | 1 | 1 | 0 | 0 | 0.015 | 0 | 0.008 | 0 | 0 | 0 | 0 | 0 | 0.005 |
| SE | 100850 | ENSG00000149311 | ATM | chr11 | + | 108227594 | 108227696 | 108223066 | 108223186 | 108227775 | 108227888 | 668 | 464 | 630 | 592 | 981 | 6 | 1 | 0 | 0 | 0 | 135 | 200 | 116 | 87 | 57 | 0 | 0 | 0 | 0 | 0 | 251 | 149 | 1 | 1 | 0.985 | 0.996 | 1 | 1 | 1 | 1 | 1 | 1 | 1 | 1 | -0.004 |
| SE | 100851 | ENSG00000149311 | ATM | chr11 | + | 108227594 | 108227696 | 108223410 | 108223686 | 108227775 | 108227888 | 468 | 297 | 442 | 401 | 840 | 0 | 2 | 0 | 0 | 0 | 111 | 121 | 72 | 54 | 37 | 0 | 0 | 0 | 0 | 0 | 251 | 149 | 1 | 1 | 1 | 0.989 | 1 | 1 | 1 | 1 | 1 | 1 | 1 | 1 | -0.002 |
| SE | 100852 | ENSG00000149311 | ATM | chr11 | + | 108227598 | 108227696 | 108223173 | 108223186 | 108227775 | 108227888 | 406 | 256 | 394 | 332 | 719 | 6 | 1 | 0 | 0 | 0 | 111 | 104 | 63 | 54 | 37 | 0 | 0 | 0 | 0 | 0 | 247 | 149 | 1 | 1 | 0.976 | 0.994 | 1 | 1 | 1 | 1 | 1 | 1 | 1 | 1 | -0.006 |
| SE | 100854 | ENSG00000149311 | ATM | chr11 | + | 108235669 | 108235834 | 108229177 | 108229323 | 108246963 | 108247127 | 353 | 217 | 347 | 327 | 486 | 1 | 0 | 1 | 0 | 3 | 11 | 136 | 82 | 37 | 33 | 0 | 0 | 0 | 0 | 0 | 298 | 149 | 1 | 1 | 0.994 | 1 | 0.994 | 1 | 0.988 | 1 | 1 | 1 | 1 | 1 | -0.005 |
| SE | 100855 | ENSG00000149311 | ATM | chr11 | + | 108235669 | 108235834 | 108229177 | 108229323 | 108250700 | 108251072 | 353 | 217 | 345 | 322 | 482 | 0 | 0 | 0 | 2 | 0 | 11 | 136 | 80 | 37 | 33 | 0 | 0 | 0 | 0 | 0 | 298 | 149 | 1 | 1 | 1 | 1 | 1 | 0.988 | 1 | 1 | 1 | 1 | 1 | 1 | -0.002 |
| SE | 100856 | ENSG00000149311 | ATM | chr11 | + | 108243952 | 108244118 | 108229177 | 108229323 | 108246963 | 108247127 | 16 | 11 | 14 | 15 | 39 | 1 | 0 | 1 | 0 | 3 | 0 | 14 | 0 | 0 | 0 | 0 | 0 | 0 | 0 | 0 | 298 | 149 | 1 | 1 | 0.889 | 1 | 0.875 | 1 | 0.867 | NA | 1 | NA | NA | NA | -0.074 |
| SE | 100857 | ENSG00000149311 | ATM | chr11 | + | 108243952 | 108244118 | 108235669 | 108235834 | 108244787 | 108245019 | 560 | 409 | 658 | 535 | 1398 | 0 | 0 | 0 | 0 | 0 | 27 | 204 | 118 | 15 | 33 | 0 | 2 | 0 | 0 | 0 | 298 | 149 | 1 | 1 | 1 | 1 | 1 | 1 | 1 | 1 | 0.981 | 1 | 1 | 1 | 0.004 |
| SE | 100858 | ENSG00000149311 | ATM | chr11 | + | 108243952 | 108244118 | 108235669 | 108235834 | 108246963 | 108247127 | 212 | 144 | 276 | 190 | 451 | 1 | 0 | 3 | 5 | 4 | 0 | 64 | 32 | 6 | 11 | 0 | 0 | 2 | 0 | 0 | 298 | 149 | 1 | 1 | 0.991 | 1 | 0.979 | 0.95 | 0.983 | NA | 1 | 0.889 | 1 | 1 | 0.008 |
| SE | 100859 | ENSG00000149311 | ATM | chr11 | + | 108244787 | 108245026 | 108235669 | 108235834 | 108246963 | 108247127 | 359 | 352 | 384 | 413 | 841 | 1 | 0 | 3 | 5 | 4 | 36 | 142 | 88 | 21 | 13 | 0 | 0 | 2 | 0 | 0 | 298 | 149 | 1 | 1 | 0.994 | 1 | 0.985 | 0.976 | 0.991 | 1 | 1 | 0.957 | 1 | 1 | -0.002 |
| SE | 100860 | ENSG00000149311 | ATM | chr11 | + | 108244787 | 108245026 | 108243952 | 108244118 | 108246963 | 108247127 | 714 | 619 | 771 | 762 | 1799 | 7 | 1 | 5 | 4 | 9 | 63 | 285 | 174 | 30 | 35 | 0 | 5 | 0 | 0 | 0 | 298 | 149 | 1 | 1 | 0.981 | 0.997 | 0.987 | 0.99 | 0.99 | 1 | 0.966 | 1 | 1 | 1 | -0.004 |
| SE | 100861 | ENSG00000149311 | ATM | chr11 | + | 108246963 | 108247127 | 108229177 | 108229323 | 108248932 | 108249102 | 684 | 588 | 731 | 784 | 2056 | 0 | 0 | 0 | 0 | 0 | 31 | 183 | 147 | 35 | 67 | 0 | 0 | 1 | 0 | 0 | 298 | 149 | 1 | 1 | 1 | 1 | 1 | 1 | 1 | 1 | 1 | 0.987 | 1 | 1 | 0.003 |
| SE | 100862 | ENSG00000149311 | ATM | chr11 | + | 108246963 | 108247127 | 108229177 | 108229323 | 108250700 | 108251072 | 12 | 25 | 16 | 20 | 59 | 0 | 0 | 0 | 2 | 0 | 0 | 0 | 1 | 9 | 0 | 0 | 0 | 0 | 0 | 0 | 298 | 149 | 1 | 1 | 1 | 1 | 1 | 0.833 | 1 | NA | NA | 1 | 1 | NA | -0.033 |
| SE | 100863 | ENSG00000149311 | ATM | chr11 | + | 108246963 | 108247127 | 108235669 | 108235834 | 108250700 | 108251072 | 12 | 25 | 18 | 25 | 60 | 1 | 0 | 1 | 0 | 0 | 0 | 0 | 3 | 9 | 0 | 0 | 0 | 0 | 0 | 0 | 298 | 149 | 1 | 1 | 0.857 | 1 | 0.9 | 1 | 1 | NA | NA | 1 | 1 | NA | -0.049 |
| SE | 100864 | ENSG00000149311 | ATM | chr11 | + | 108246963 | 108247127 | 108244787 | 108245026 | 108248932 | 108249102 | 1042 | 940 | 1114 | 1197 | 2894 | 0 | 0 | 0 | 0 | 1 | 67 | 323 | 235 | 56 | 80 | 0 | 0 | 2 | 0 | 0 | 298 | 149 | 1 | 1 | 1 | 1 | 1 | 1 | 0.999 | 1 | 1 | 0.983 | 1 | 1 | 0.003 |
| SE | 100865 | ENSG00000149311 | ATM | chr11 | + | 108246963 | 108247136 | 108229177 | 108229323 | 108248932 | 108249102 | 3 | 0 | 1 | 0 | 3 | 0 | 0 | 0 | 0 | 0 | 0 | 0 | 0 | 0 | 0 | 0 | 0 | 1 | 0 | 0 | 298 | 149 | 1 | 1 | 1 | NA | 1 | NA | 1 | NA | NA | 0 | NA | NA | 1 |
| SE | 100866 | ENSG00000149311 | ATM | chr11 | + | 108246963 | 108247136 | 108244787 | 108245026 | 108248932 | 108249102 | 361 | 352 | 384 | 413 | 841 | 0 | 0 | 0 | 0 | 1 | 36 | 140 | 88 | 21 | 13 | 0 | 0 | 2 | 0 | 0 | 298 | 149 | 1 | 1 | 1 | 1 | 1 | 1 | 0.998 | 1 | 1 | 0.957 | 1 | 1 | 0.008 |
| SE | 100867 | ENSG00000149311 | ATM | chr11 | + | 108248932 | 108249102 | 108229177 | 108229323 | 108250700 | 108251072 | 558 | 509 | 595 | 639 | 1709 | 0 | 0 | 0 | 2 | 0 | 30 | 215 | 159 | 50 | 27 | 0 | 0 | 0 | 0 | 0 | 298 | 149 | 1 | 1 | 1 | 1 | 1 | 0.994 | 1 | 1 | 1 | 1 | 1 | 1 | -0.001 |
| SE | 100869 | ENSG00000149311 | ATM | chr11 | + | 108250700 | 108251072 | 108248932 | 108249102 | 108251836 | 108252031 | 1310 | 1253 | 1286 | 1431 | 3006 | 0 | 1 | 0 | 0 | 0 | 91 | 444 | 371 | 80 | 75 | 0 | 0 | 0 | 0 | 0 | 298 | 149 | 1 | 1 | 1 | 0.998 | 1 | 1 | 1 | 1 | 1 | 1 | 1 | 1 | 0 |
| SE | 100870 | ENSG00000149311 | ATM | chr11 | + | 108257480 | 108257606 | 108256214 | 108256340 | 108267170 | 108267342 | 231 | 175 | 219 | 197 | 434 | 0 | 1 | 2 | 2 | 5 | 25 | 53 | 61 | 10 | 25 | 0 | 1 | 2 | 0 | 0 | 275 | 149 | 1 | 1 | 1 | 0.99 | 0.983 | 0.982 | 0.979 | 1 | 0.966 | 0.943 | 1 | 1 | 0.005 |
| SE | 100871 | ENSG00000149311 | ATM | chr11 | + | 108258985 | 108259075 | 108257480 | 108257606 | 108267170 | 108267342 | 759 | 657 | 723 | 701 | 1431 | 3 | 3 | 2 | 2 | 5 | 69 | 161 | 184 | 24 | 56 | 0 | 0 | 0 | 0 | 0 | 239 | 149 | 1 | 1 | 0.994 | 0.993 | 0.996 | 0.995 | 0.994 | 1 | 1 | 1 | 1 | 1 | -0.006 |
| SE | 100872 | ENSG00000149311 | ATM | chr11 | + | 108267170 | 108267342 | 108258985 | 108259075 | 108268409 | 108268609 | 790 | 629 | 714 | 753 | 1198 | 1 | 2 | 1 | 5 | 1 | 111 | 147 | 138 | 36 | 55 | 0 | 0 | 0 | 0 | 0 | 298 | 149 | 1 | 1 | 0.997 | 0.994 | 0.997 | 0.987 | 0.998 | 1 | 1 | 1 | 1 | 1 | -0.005 |
| SE | 100873 | ENSG00000149311 | ATM | chr11 | + | 108267170 | 108267342 | 108258985 | 108259075 | 108271063 | 108271146 | 375 | 297 | 315 | 345 | 553 | 1 | 0 | 0 | 0 | 0 | 52 | 71 | 81 | 21 | 28 | 0 | 0 | 0 | 0 | 0 | 298 | 149 | 1 | 1 | 0.995 | 1 | 1 | 1 | 1 | 1 | 1 | 1 | 1 | 1 | -0.001 |
| SE | 100874 | ENSG00000149311 | ATM | chr11 | + | 108268409 | 108268609 | 108258985 | 108259075 | 108271063 | 108271146 | 349 | 243 | 316 | 310 | 706 | 1 | 0 | 0 | 0 | 0 | 20 | 46 | 91 | 14 | 14 | 0 | 0 | 0 | 0 | 0 | 298 | 149 | 1 | 1 | 0.994 | 1 | 1 | 1 | 1 | 1 | 1 | 1 | 1 | 1 | -0.001 |
| SE | 100876 | ENSG00000149311 | ATM | chr11 | + | 108271063 | 108271146 | 108268409 | 108268609 | 108271250 | 108271406 | 600 | 488 | 577 | 592 | 1168 | 3 | 1 | 1 | 4 | 0 | 33 | 112 | 162 | 18 | 23 | 0 | 0 | 0 | 0 | 0 | 232 | 149 | 1 | 1 | 0.992 | 0.997 | 0.997 | 0.99 | 1 | 1 | 1 | 1 | 1 | 1 | -0.005 |
| SE | 100877 | ENSG00000149311 | ATM | chr11 | + | 108272531 | 108272607 | 108271250 | 108271406 | 108272721 | 108272852 | 550 | 383 | 505 | 544 | 977 | 0 | 1 | 0 | 0 | 0 | 45 | 101 | 140 | 49 | 30 | 0 | 1 | 0 | 0 | 0 | 225 | 149 | 1 | 1 | 1 | 0.996 | 1 | 1 | 1 | 1 | 0.985 | 1 | 1 | 1 | 0.002 |
| SE | 100878 | ENSG00000149311 | ATM | chr11 | + | 108280994 | 108281168 | 108279490 | 108279608 | 108282709 | 108282879 | 496 | 360 | 420 | 402 | 1227 | 1 | 0 | 0 | 0 | 0 | 6 | 91 | 92 | 18 | 4 | 0 | 0 | 0 | 0 | 0 | 298 | 149 | 1 | 1 | 0.996 | 1 | 1 | 1 | 1 | 1 | 1 | 1 | 1 | 1 | -0.001 |
| SE | 100879 | ENSG00000149311 | ATM | chr11 | + | 108280994 | 108281168 | 108279490 | 108279608 | 108284226 | 108284473 | 164 | 115 | 148 | 128 | 369 | 0 | 0 | 0 | 2 | 0 | 0 | 24 | 27 | 1 | 0 | 0 | 0 | 0 | 0 | 0 | 298 | 149 | 1 | 1 | 1 | 1 | 1 | 0.97 | 1 | NA | 1 | 1 | 1 | NA | -0.006 |
| SE | 100880 | ENSG00000149311 | ATM | chr11 | + | 108282709 | 108282879 | 108279490 | 108279608 | 108284226 | 108284473 | 409 | 313 | 351 | 404 | 722 | 0 | 0 | 0 | 2 | 0 | 20 | 55 | 61 | 34 | 10 | 0 | 0 | 0 | 0 | 0 | 298 | 149 | 1 | 1 | 1 | 1 | 1 | 0.99 | 1 | 1 | 1 | 1 | 1 | 1 | -0.002 |
| SE | 100881 | ENSG00000149311 | ATM | chr11 | + | 108282709 | 108282879 | 108280994 | 108281168 | 108284226 | 108284473 | 740 | 560 | 625 | 679 | 1584 | 0 | 1 | 2 | 1 | 4 | 26 | 123 | 128 | 51 | 14 | 0 | 1 | 2 | 0 | 0 | 298 | 149 | 1 | 1 | 1 | 0.996 | 0.994 | 0.997 | 0.995 | 1 | 0.984 | 0.97 | 1 | 1 | 0.006 |
| SE | 100882 | ENSG00000149311 | ATM | chr11 | + | 108292618 | 108292793 | 108289601 | 108289801 | 108294926 | 108295059 | 277 | 256 | 322 | 327 | 690 | 0 | 1 | 0 | 1 | 0 | 14 | 65 | 63 | 15 | 10 | 0 | 0 | 0 | 0 | 0 | 298 | 149 | 1 | 1 | 1 | 0.992 | 1 | 0.994 | 1 | 1 | 1 | 1 | 1 | 1 | -0.003 |
| SE | 100883 | ENSG00000149311 | ATM | chr11 | + | 108292618 | 108292793 | 108289601 | 108289801 | 108297286 | 108297382 | 277 | 256 | 320 | 327 | 690 | 0 | 0 | 0 | 0 | 0 | 14 | 65 | 63 | 15 | 10 | 0 | 0 | 0 | 2 | 0 | 298 | 149 | 1 | 1 | 1 | 1 | 1 | 1 | 1 | 1 | 1 | 1 | 0.789 | 1 | 0.042 |
| SE | 100884 | ENSG00000149311 | ATM | chr11 | + | 108293312 | 108293477 | 108292618 | 108292793 | 108294926 | 108295059 | 467 | 398 | 509 | 493 | 714 | 0 | 0 | 2 | 0 | 0 | 39 | 131 | 95 | 30 | 20 | 0 | 0 | 0 | 0 | 0 | 298 | 149 | 1 | 1 | 1 | 1 | 0.992 | 1 | 1 | 1 | 1 | 1 | 1 | 1 | -0.002 |
| SE | 100889 | ENSG00000149311 | ATM | chr11 | + | 108294926 | 108296393 | 108293312 | 108293477 | 108297286 | 108297382 | 212 | 187 | 228 | 249 | 356 | 3 | 3 | 7 | 2 | 6 | 25 | 42 | 48 | 5 | 0 | 0 | 0 | 3 | 0 | 0 | 298 | 149 | 1 | 1 | 0.972 | 0.969 | 0.942 | 0.984 | 0.967 | 1 | 1 | 0.889 | 1 | NA | -0.005 |
| SE | 100890 | ENSG00000149311 | ATM | chr11 | + | 108297286 | 108297382 | 108293312 | 108293477 | 108299713 | 108299885 | 270 | 204 | 336 | 330 | 418 | 3 | 3 | 1 | 4 | 0 | 29 | 29 | 55 | 2 | 8 | 0 | 0 | 2 | 0 | 0 | 245 | 149 | 1 | 1 | 0.982 | 0.976 | 0.995 | 0.98 | 1 | 1 | 1 | 0.944 | 1 | 1 | -0.002 |
| SE | 100891 | ENSG00000149311 | ATM | chr11 | + | 108301647 | 108301789 | 108299713 | 108299885 | 108302852 | 108303029 | 786 | 718 | 821 | 917 | 1270 | 1 | 0 | 0 | 0 | 0 | 34 | 213 | 254 | 60 | 61 | 0 | 0 | 1 | 0 | 0 | 291 | 149 | 1 | 1 | 0.998 | 1 | 1 | 1 | 1 | 1 | 1 | 0.992 | 1 | 1 | 0.001 |
| SE | 100892 | ENSG00000149311 | ATM | chr11 | + | 108307896 | 108307984 | 108304674 | 108304852 | 108310159 | 108310315 | 1142 | 1049 | 1049 | 1201 | 1561 | 4 | 3 | 1 | 0 | 7 | 89 | 278 | 398 | 62 | 30 | 0 | 6 | 0 | 0 | 0 | 237 | 149 | 1 | 1 | 0.994 | 0.995 | 0.998 | 1 | 0.993 | 1 | 0.967 | 1 | 1 | 1 | 0.003 |
| SE | 100893 | ENSG00000149311 | ATM | chr11 | + | 108310159 | 108310315 | 108307803 | 108307984 | 108312410 | 108312498 | 2558 | 2615 | 2254 | 2774 | 3189 | 0 | 0 | 1 | 0 | 0 | 246 | 731 | 899 | 182 | 51 | 0 | 0 | 0 | 0 | 0 | 298 | 149 | 1 | 1 | 1 | 1 | 0.999 | 1 | 1 | 1 | 1 | 1 | 1 | 1 | 0 |
| SE | 100894 | ENSG00000149311 | ATM | chr11 | + | 108312410 | 108312498 | 108310159 | 108310315 | 108315822 | 108315894 | 1951 | 1970 | 1680 | 2087 | 2324 | 3 | 1 | 1 | 0 | 0 | 157 | 561 | 666 | 158 | 49 | 0 | 0 | 0 | 0 | 0 | 237 | 149 | 1 | 1 | 0.998 | 0.999 | 0.999 | 1 | 1 | 1 | 1 | 1 | 1 | 1 | -0.001 |
| SE | 100895 | ENSG00000149311 | ATM | chr11 | + | 108316010 | 108316113 | 108315822 | 108315911 | 108317372 | 108317467 | 308 | 207 | 258 | 300 | 339 | 0 | 0 | 0 | 0 | 1 | 5 | 80 | 71 | 10 | 11 | 0 | 0 | 0 | 0 | 0 | 252 | 149 | 1 | 1 | 1 | 1 | 1 | 1 | 0.995 | 1 | 1 | 1 | 1 | 1 | -0.001 |
| SE | 100896 | ENSG00000149311 | ATM | chr11 | + | 108319953 | 108320058 | 108317372 | 108317521 | 108321300 | 108321420 | 676 | 491 | 675 | 657 | 933 | 0 | 0 | 2 | 0 | 0 | 62 | 253 | 153 | 22 | 15 | 0 | 0 | 0 | 0 | 0 | 254 | 149 | 1 | 1 | 1 | 1 | 0.995 | 1 | 1 | 1 | 1 | 1 | 1 | 1 | -0.001 |
| SE | 100897 | ENSG00000149311 | ATM | chr11 | + | 108320709 | 108320780 | 108319953 | 108320058 | 108321300 | 108321420 | 0 | 0 | 0 | 1 | 0 | 436 | 308 | 478 | 456 | 669 | 0 | 0 | 0 | 0 | 0 | 46 | 158 | 101 | 6 | 11 | 220 | 149 | 1 | 1 | 0 | 0 | 0 | 0.001 | 0 | 0 | 0 | 0 | 0 | 0 | 0 |
| SE | 100898 | ENSG00000149311 | ATM | chr11 | + | 108327644 | 108327758 | 108326057 | 108326225 | 108329020 | 108329238 | 739 | 679 | 714 | 829 | 1439 | 0 | 0 | 1 | 2 | 1 | 16 | 178 | 184 | 19 | 14 | 0 | 5 | 0 | 0 | 0 | 263 | 149 | 1 | 1 | 1 | 1 | 0.998 | 0.996 | 0.999 | 1 | 0.953 | 1 | 1 | 1 | 0.008 |
| SE | 100899 | ENSG00000149311 | ATM | chr11 | + | 108331443 | 108331557 | 108330213 | 108330421 | 108331878 | 108332037 | 1438 | 1164 | 1521 | 1435 | 2319 | 0 | 0 | 0 | 0 | 0 | 104 | 461 | 518 | 114 | 81 | 0 | 0 | 1 | 0 | 0 | 263 | 149 | 1 | 1 | 1 | 1 | 1 | 1 | 1 | 1 | 1 | 0.997 | 1 | 1 | 0.001 |
| SE | 100900 | ENSG00000149311 | ATM | chr11 | + | 108331878 | 108332037 | 108331443 | 108331557 | 108332761 | 108332900 | 1699 | 1547 | 1658 | 1680 | 2458 | 2 | 4 | 1 | 5 | 7 | 107 | 479 | 512 | 110 | 103 | 0 | 2 | 1 | 0 | 0 | 298 | 149 | 1 | 1 | 0.998 | 0.995 | 0.999 | 0.994 | 0.994 | 1 | 0.992 | 0.996 | 1 | 1 | -0.002 |
| SE | 100901 | ENSG00000149311 | ATM | chr11 | + | 108331878 | 108332037 | 108331443 | 108331557 | 108333885 | 108333968 | 802 | 621 | 865 | 799 | 1302 | 4 | 1 | 2 | 0 | 0 | 53 | 238 | 240 | 37 | 35 | 0 | 0 | 0 | 0 | 0 | 298 | 149 | 1 | 1 | 0.99 | 0.997 | 0.995 | 1 | 1 | 1 | 1 | 1 | 1 | 1 | -0.004 |
| SE | 100902 | ENSG00000149311 | ATM | chr11 | + | 108332761 | 108332900 | 108331443 | 108331557 | 108333885 | 108333908 | 745 | 718 | 709 | 797 | 949 | 4 | 1 | 2 | 0 | 0 | 35 | 254 | 245 | 98 | 45 | 0 | 0 | 0 | 0 | 0 | 288 | 149 | 1 | 1 | 0.99 | 0.997 | 0.995 | 1 | 1 | 1 | 1 | 1 | 1 | 1 | -0.004 |
| SE | 100903 | ENSG00000149311 | ATM | chr11 | + | 108332761 | 108332900 | 108331878 | 108332037 | 108333885 | 108333908 | 1647 | 1637 | 1504 | 1683 | 2102 | 8 | 1 | 4 | 9 | 8 | 89 | 496 | 516 | 171 | 113 | 0 | 0 | 0 | 0 | 0 | 288 | 149 | 1 | 1 | 0.991 | 0.999 | 0.995 | 0.99 | 0.993 | 1 | 1 | 1 | 1 | 1 | -0.006 |
| SE | 100906 | ENSG00000149311 | ATM | chr11 | + | 108333885 | 108333968 | 108332499 | 108332900 | 108334968 | 108335109 | 1330 | 1309 | 1300 | 1482 | 1790 | 0 | 0 | 0 | 0 | 0 | 78 | 431 | 465 | 136 | 66 | 0 | 0 | 4 | 0 | 0 | 232 | 149 | 1 | 1 | 1 | 1 | 1 | 1 | 1 | 1 | 1 | 0.987 | 1 | 1 | 0.003 |
| SE | 100907 | ENSG00000149311 | ATM | chr11 | + | 108334968 | 108335109 | 108331443 | 108333968 | 108335844 | 108335961 | 1198 | 1108 | 1217 | 1395 | 1593 | 0 | 0 | 0 | 0 | 0 | 56 | 351 | 425 | 103 | 65 | 0 | 0 | 0 | 1 | 0 | 290 | 149 | 1 | 1 | 1 | 1 | 1 | 1 | 1 | 1 | 1 | 1 | 0.981 | 1 | 0.004 |
| SE | 100908 | ENSG00000149311 | ATM | chr11 | + | 108343221 | 108343371 | 108335844 | 108335961 | 108345742 | 108345908 | 800 | 670 | 765 | 806 | 932 | 1 | 0 | 0 | 2 | 0 | 12 | 179 | 175 | 32 | 24 | 0 | 0 | 0 | 0 | 0 | 298 | 149 | 1 | 1 | 0.998 | 1 | 1 | 0.995 | 1 | 1 | 1 | 1 | 1 | 1 | -0.001 |
| SE | 100909 | ENSG00000149311 | ATM | chr11 | + | 108347278 | 108347365 | 108345742 | 108345908 | 108353765 | 108353880 | 717 | 541 | 706 | 748 | 879 | 1 | 0 | 0 | 0 | 0 | 8 | 208 | 155 | 44 | 9 | 0 | 0 | 0 | 0 | 0 | 236 | 149 | 1 | 1 | 0.998 | 1 | 1 | 1 | 1 | 1 | 1 | 1 | 1 | 1 | 0 |
| SE | 100910 | ENSG00000149311 | ATM | chr11 | + | 108353765 | 108353880 | 108347278 | 108347365 | 108354810 | 108354874 | 741 | 579 | 769 | 722 | 906 | 1 | 1 | 1 | 1 | 0 | 41 | 177 | 162 | 38 | 6 | 0 | 0 | 0 | 0 | 0 | 264 | 149 | 1 | 1 | 0.998 | 0.997 | 0.998 | 0.998 | 1 | 1 | 1 | 1 | 1 | 1 | -0.002 |
| SE | 100915 | ENSG00000103479 | RBL2 | chr16 | + | 53442657 | 53442858 | 53439015 | 53439146 | 53447041 | 53447106 | 709 | 501 | 743 | 773 | 1422 | 3 | 0 | 0 | 4 | 12 | 185 | 483 | 396 | 316 | 107 | 0 | 0 | 0 | 0 | 0 | 298 | 149 | 1 | 1 | 0.992 | 1 | 1 | 0.99 | 0.983 | 1 | 1 | 1 | 1 | 1 | -0.007 |
| SE | 100916 | ENSG00000103479 | RBL2 | chr16 | + | 53442657 | 53442858 | 53439015 | 53439146 | 53451702 | 53451831 | 368 | 291 | 376 | 426 | 848 | 0 | 0 | 0 | 0 | 2 | 128 | 296 | 248 | 219 | 92 | 3 | 0 | 0 | 0 | 0 | 298 | 149 | 1 | 1 | 1 | 1 | 1 | 1 | 0.995 | 0.955 | 1 | 1 | 1 | 1 | 0.008 |
| SE | 100917 | ENSG00000103479 | RBL2 | chr16 | + | 53447041 | 53447106 | 53439015 | 53439146 | 53451702 | 53451831 | 118 | 77 | 153 | 150 | 376 | 0 | 0 | 0 | 0 | 2 | 46 | 66 | 53 | 49 | 30 | 3 | 0 | 0 | 0 | 0 | 214 | 149 | 1 | 1 | 1 | 1 | 1 | 1 | 0.992 | 0.914 | 1 | 1 | 1 | 1 | 0.016 |
| SE | 100919 | ENSG00000103479 | RBL2 | chr16 | + | 53451702 | 53451831 | 53447041 | 53447106 | 53453451 | 53453463 | 1156 | 838 | 1231 | 1277 | 2139 | 1 | 0 | 0 | 3 | 0 | 295 | 844 | 574 | 384 | 163 | 0 | 0 | 2 | 0 | 0 | 278 | 149 | 1 | 1 | 0.998 | 1 | 1 | 0.996 | 1 | 1 | 1 | 0.994 | 1 | 1 | 0 |
| SE | 100920 | ENSG00000103479 | RBL2 | chr16 | + | 53454225 | 53454347 | 53453704 | 53453769 | 53454655 | 53454842 | 2 | 2 | 1 | 2 | 11 | 768 | 543 | 695 | 792 | 1169 | 10 | 0 | 8 | 0 | 0 | 212 | 585 | 384 | 416 | 129 | 271 | 149 | 1 | 1 | 0.001 | 0.002 | 0.001 | 0.001 | 0.005 | 0.025 | 0 | 0.011 | 0 | 0 | -0.005 |
| SE | 100921 | ENSG00000103479 | RBL2 | chr16 | + | 53454655 | 53454842 | 53453704 | 53453769 | 53459450 | 53459617 | 1223 | 925 | 1202 | 1404 | 1812 | 1 | 0 | 0 | 0 | 0 | 326 | 877 | 613 | 568 | 163 | 0 | 0 | 0 | 0 | 0 | 298 | 149 | 1 | 1 | 0.998 | 1 | 1 | 1 | 1 | 1 | 1 | 1 | 1 | 1 | 0 |
| SE | 100922 | ENSG00000103479 | RBL2 | chr16 | + | 53459450 | 53459617 | 53454655 | 53454842 | 53461740 | 53461850 | 1140 | 970 | 1200 | 1405 | 1368 | 0 | 0 | 0 | 1 | 0 | 274 | 807 | 604 | 515 | 166 | 0 | 0 | 0 | 0 | 0 | 298 | 149 | 1 | 1 | 1 | 1 | 1 | 0.999 | 1 | 1 | 1 | 1 | 1 | 1 | 0 |
| SE | 100923 | ENSG00000103479 | RBL2 | chr16 | + | 53464225 | 53464363 | 53462551 | 53462655 | 53465437 | 53465602 | 1591 | 1176 | 1602 | 1728 | 2780 | 1 | 0 | 0 | 0 | 0 | 374 | 1236 | 840 | 771 | 278 | 0 | 0 | 0 | 0 | 0 | 287 | 149 | 1 | 1 | 0.999 | 1 | 1 | 1 | 1 | 1 | 1 | 1 | 1 | 1 | 0 |
| SE | 100924 | ENSG00000103479 | RBL2 | chr16 | + | 53465437 | 53465602 | 53462551 | 53462655 | 53467057 | 53467169 | 930 | 429 | 846 | 609 | 1873 | 1 | 0 | 1 | 0 | 0 | 291 | 386 | 380 | 435 | 102 | 0 | 0 | 0 | 0 | 0 | 298 | 149 | 1 | 1 | 0.998 | 1 | 0.998 | 1 | 1 | 1 | 1 | 1 | 1 | 1 | -0.001 |
| SE | 100926 | ENSG00000103479 | RBL2 | chr16 | + | 53479153 | 53479225 | 53470745 | 53470922 | 53479885 | 53479991 | 1406 | 1560 | 1507 | 1912 | 3358 | 4 | 0 | 6 | 0 | 5 | 339 | 1042 | 934 | 699 | 122 | 1 | 0 | 3 | 1 | 0 | 221 | 149 | 1 | 1 | 0.996 | 1 | 0.994 | 1 | 0.998 | 0.996 | 1 | 0.995 | 0.998 | 1 | 0 |
| SE | 100927 | ENSG00000103479 | RBL2 | chr16 | + | 53481670 | 53481835 | 53480566 | 53480769 | 53490129 | 53490301 | 1516 | 1347 | 1625 | 1780 | 3470 | 1 | 0 | 0 | 0 | 1 | 413 | 1204 | 956 | 754 | 157 | 0 | 0 | 0 | 0 | 0 | 298 | 149 | 1 | 1 | 0.999 | 1 | 1 | 1 | 0.999 | 1 | 1 | 1 | 1 | 1 | 0 |
| SE | 100931 | ENSG00000074319 | TSG101 | chr11 | - | 18479902 | 18480046 | 18469030 | 18469218 | 18480484 | 18480635 | 0 | 2 | 0 | 0 | 0 | 2 | 1 | 0 | 5 | 7 | 0 | 0 | 2 | 0 | 0 | 0 | 6 | 3 | 4 | 4 | 293 | 149 | 1 | 1 | 0 | 0.504 | NA | 0 | 0 | NA | 0 | 0.253 | 0 | 0 | 0.063 |
| SE | 100932 | ENSG00000074319 | TSG101 | chr11 | - | 18479902 | 18480046 | 18469030 | 18469218 | 18480507 | 18480635 | 0 | 0 | 0 | 0 | 0 | 0 | 0 | 0 | 0 | 2 | 0 | 0 | 2 | 0 | 0 | 6 | 0 | 0 | 0 | 0 | 293 | 149 | 1 | 1 | NA | NA | NA | NA | 0 | 0 | NA | 1 | NA | NA | -0.5 |
| SE | 100935 | ENSG00000074319 | TSG101 | chr11 | - | 18506856 | 18506923 | 18502484 | 18502577 | 18509541 | 18509665 | 254 | 158 | 184 | 228 | 361 | 0 | 0 | 0 | 1 | 2 | 319 | 200 | 241 | 241 | 298 | 0 | 0 | 3 | 6 | 0 | 216 | 149 | 1 | 1 | 1 | 1 | 1 | 0.994 | 0.992 | 1 | 1 | 0.982 | 0.965 | 1 | 0.008 |
| SE | 100941 | ENSG00000074319 | TSG101 | chr11 | - | 18516098 | 18516164 | 18514677 | 18514841 | 18519518 | 18519603 | 213 | 181 | 144 | 160 | 400 | 0 | 0 | 0 | 0 | 0 | 156 | 167 | 202 | 229 | 217 | 0 | 0 | 0 | 1 | 0 | 215 | 149 | 1 | 1 | 1 | 1 | 1 | 1 | 1 | 1 | 1 | 1 | 0.994 | 1 | 0.001 |
| SE | 100945 | ENSG00000116260 | QSOX1 | chr1 | + | 180166490 | 180166591 | 180154868 | 180155172 | 180175320 | 180175366 | 42 | 13 | 35 | 24 | 5 | 0 | 0 | 2 | 0 | 0 | 34 | 87 | 36 | 52 | 72 | 0 | 0 | 0 | 0 | 0 | 250 | 149 | 1 | 1 | 1 | 1 | 0.913 | 1 | 1 | 1 | 1 | 1 | 1 | 1 | -0.017 |
| SE | 100946 | ENSG00000116260 | QSOX1 | chr1 | + | 180175320 | 180175366 | 180166490 | 180166591 | 180178793 | 180178884 | 2 | 4 | 3 | 5 | 1 | 0 | 0 | 0 | 0 | 0 | 13 | 6 | 4 | 3 | 8 | 0 | 0 | 0 | 0 | 1 | 195 | 149 | 1 | 1 | 1 | 1 | 1 | 1 | 1 | 1 | 1 | 1 | 1 | 0.859 | 0.028 |
| SE | 100948 | ENSG00000116260 | QSOX1 | chr1 | + | 180190432 | 180190580 | 180189551 | 180189674 | 180196261 | 180196599 | 71 | 31 | 42 | 51 | 46 | 0 | 0 | 0 | 0 | 0 | 156 | 105 | 76 | 125 | 182 | 0 | 0 | 0 | 5 | 0 | 297 | 149 | 1 | 1 | 1 | 1 | 1 | 1 | 1 | 1 | 1 | 1 | 0.926 | 1 | 0.015 |
| SE | 100974 | ENSG00000151348 | EXT2 | chr11 | + | 44096224 | 44096321 | 44095822 | 44095852 | 44107682 | 44108248 | 1 | 0 | 0 | 0 | 0 | 17 | 12 | 13 | 13 | 9 | 0 | 0 | 0 | 0 | 0 | 15 | 13 | 6 | 2 | 10 | 246 | 149 | 1 | 1 | 0.034 | 0 | 0 | 0 | 0 | 0 | 0 | 0 | 0 | 0 | 0.007 |
| SE | 100975 | ENSG00000151348 | EXT2 | chr11 | + | 44103706 | 44103745 | 44095664 | 44095852 | 44107682 | 44108248 | 1 | 1 | 1 | 1 | 0 | 17 | 12 | 13 | 13 | 9 | 0 | 1 | 0 | 0 | 0 | 15 | 13 | 6 | 2 | 10 | 188 | 149 | 1 | 1 | 0.045 | 0.062 | 0.057 | 0.057 | 0 | 0 | 0.057 | 0 | 0 | 0 | 0.033 |
| SE | 100977 | ENSG00000151348 | EXT2 | chr11 | + | 44124788 | 44124984 | 44114184 | 44114301 | 44130044 | 44130138 | 32 | 18 | 37 | 46 | 15 | 1 | 0 | 0 | 0 | 0 | 53 | 15 | 24 | 11 | 38 | 0 | 0 | 0 | 0 | 0 | 298 | 149 | 1 | 1 | 0.941 | 1 | 1 | 1 | 1 | 1 | 1 | 1 | 1 | 1 | -0.012 |

| | | | | | | | | | | | | | | | | | | | | | | | | | | | | | | | | | | | | | | | | | | | | | | |
|---|---|---|---|---|---|---|---|---|---|---|---|---|---|---|---|---|---|---|---|---|---|---|---|---|---|---|---|---|---|---|---|---|---|---|---|---|---|---|---|---|---|---|---|---|---|---|
| SE | 100982 | ENSG00000151348 | EXT2 | chr11 | + | 44171610 | 44171742 | 44130042 | 44130138 | 44197828 | 44198018 | 92 | 38 | 50 | 100 | 78 | 0 | 0 | 0 | 0 | 0 | 52 | 90 | 45 | 30 | 53 | 0 | 0 | 1 | 0 | 0 | 281 | 149 | 1 | 1 | 1 | 1 | 1 | 1 | 1 | 1 | 1 | 0.96 | 1 | 1 | 0.008 |
| SE | 100985 | ENSG00000151348 | EXT2 | chr11 | + | 44192712 | 44192845 | 44171610 | 44171742 | 44197828 | 44198018 | 1 | 0 | 0 | 0 | 0 | 34 | 15 | 28 | 36 | 23 | 0 | 0 | 0 | 2 | 0 | 17 | 33 | 18 | 11 | 24 | 282 | 149 | 1 | 1 | 0.015 | 0 | 0 | 0 | 0 | 0 | 0 | 0 | 0.088 | 0 | -0.015 |
| SE | 100987 | ENSG00000151348 | EXT2 | chr11 | + | 44232352 | 44232496 | 44204982 | 44206959 | 44234114 | 44234243 | 184 | 92 | 100 | 174 | 170 | 0 | 0 | 2 | 0 | 4 | 192 | 139 | 84 | 148 | 145 | 0 | 0 | 5 | 0 | 0 | 293 | 149 | 1 | 1 | 1 | 1 | 0.962 | 1 | 0.956 | 1 | 1 | 0.895 | 1 | 1 | 0.005 |
| SE | 100988 | ENSG00000151348 | EXT2 | chr11 | + | 44234114 | 44234243 | 44204982 | 44206959 | 44244148 | 44244301 | 0 | 0 | 2 | 0 | 11 | 0 | 0 | 0 | 0 | 0 | 0 | 0 | 8 | 3 | 6 | 0 | 0 | 1 | 0 | 0 | 278 | 149 | 1 | 1 | NA | NA | 1 | NA | 1 | NA | NA | 0.811 | 1 | 1 | 0.063 |
| SE | 100993 | ENSG00000100600 | LGMN | chr14 | - | 92704639 | 92704707 | 92703808 | 92704361 | 92706482 | 92706653 | 23 | 12 | 23 | 36 | 2 | 1 | 0 | 0 | 0 | 0 | 12 | 25 | 15 | 7 | 74 | 0 | 0 | 0 | 0 | 0 | 217 | 149 | 1 | 1 | 0.94 | 1 | 1 | 1 | 1 | 1 | 1 | 1 | 1 | 1 | -0.012 |
| SE | 100997 | ENSG00000100600 | LGMN | chr14 | - | 92712722 | 92712871 | 92711836 | 92711955 | 92713822 | 92713885 | 10 | 7 | 12 | 39 | 53 | 0 | 0 | 0 | 0 | 0 | 43 | 16 | 8 | 9 | 44 | 0 | 0 | 1 | 0 | 0 | 298 | 149 | 1 | 1 | 1 | 1 | 1 | 1 | 1 | 1 | 1 | 0.8 | 1 | 1 | 0.04 |
| SE | 100998 | ENSG00000100600 | LGMN | chr14 | - | 92712804 | 92712871 | 92711836 | 92711955 | 92713822 | 92713885 | 43 | 41 | 57 | 112 | 138 | 0 | 0 | 0 | 0 | 0 | 80 | 57 | 36 | 36 | 189 | 0 | 0 | 1 | 0 | 0 | 216 | 149 | 1 | 1 | 1 | 1 | 1 | 1 | 1 | 1 | 1 | 0.961 | 1 | 1 | 0.008 |
| SE | 101008 | ENSG00000261008 | LINC01572 | chr16 | - | 72267551 | 72267654 | 72249162 | 72249236 | 72284571 | 72284728 | 0 | 2 | 0 | 0 | 0 | 1 | 0 | 1 | 0 | 2 | 0 | 0 | 0 | 0 | 0 | 0 | 0 | 2 | 0 | 0 | 252 | 149 | 1 | 1 | 0 | 1 | 0 | NA | 0 | NA | NA | 0 | NA | NA | 0.25 |
| SE | 101013 | ENSG00000261008 | LINC01572 | chr16 | - | 72522552 | 72522623 | 72426420 | 72427457 | 72523331 | 72523415 | 7 | 6 | 15 | 3 | 4 | 2 | 5 | 0 | 0 | 0 | 0 | 1 | 8 | 1 | 23 | 0 | 5 | 3 | 0 | 14 | 220 | 149 | 1 | 1 | 0.703 | 0.448 | 1 | 1 | 1 | NA | 0.119 | 0.644 | 1 | 0.527 | 0.258 |
| SE | 101014 | ENSG00000261008 | LINC01572 | chr16 | - | 72522552 | 72522623 | 72479750 | 72479790 | 72523331 | 72523415 | 2 | 3 | 2 | 0 | 0 | 3 | 0 | 0 | 0 | 0 | 0 | 0 | 2 | 2 | 16 | 0 | 0 | 0 | 0 | 0 | 220 | 149 | 1 | 1 | 0.311 | 1 | 1 | NA | NA | NA | NA | 1 | 1 | 1 | -0.23 |
| SE | 101016 | ENSG00000261008 | LINC01572 | chr16 | - | 72540577 | 72540611 | 72523331 | 72523415 | 72555264 | 72555326 | 6 | 13 | 7 | 1 | 2 | 0 | 0 | 0 | 1 | 0 | 0 | 12 | 7 | 0 | 23 | 0 | 0 | 0 | 1 | 7 | 183 | 149 | 1 | 1 | 1 | 1 | 1 | 0.449 | 1 | NA | 1 | 1 | 0 | 0.728 | 0.208 |
| SE | 101019 | ENSG00000261008 | LINC01572 | chr16 | - | 72555264 | 72555326 | 72540577 | 72540611 | 72561972 | 72562086 | 7 | 5 | 2 | 0 | 0 | 0 | 0 | 2 | 0 | 0 | 0 | 8 | 4 | 0 | 14 | 0 | 0 | 0 | 0 | 0 | 211 | 149 | 1 | 1 | 1 | 1 | 0.414 | NA | NA | NA | 1 | 1 | NA | 1 | -0.195 |
| SE | 101020 | ENSG00000261008 | LINC01572 | chr16 | - | 72555264 | 72555326 | 72540577 | 72540611 | 72570450 | 72570515 | 8 | 6 | 7 | 1 | 4 | 0 | 1 | 0 | 1 | 0 | 13 | 5 | 9 | 0 | 14 | 0 | 0 | 0 | 0 | 0 | 211 | 149 | 1 | 1 | 1 | 0.809 | 1 | 0.414 | 1 | 1 | 1 | 1 | NA | 1 | -0.155 |
| SE | 101021 | ENSG00000261008 | LINC01572 | chr16 | - | 72561972 | 72562086 | 72540577 | 72540611 | 72570450 | 72570515 | 0 | 0 | 3 | 0 | 0 | 0 | 1 | 0 | 1 | 0 | 0 | 2 | 0 | 0 | 0 | 0 | 0 | 0 | 0 | 0 | 263 | 149 | 1 | 1 | NA | 0 | 1 | 0 | NA | NA | 1 | NA | NA | NA | -0.667 |
| SE | 101022 | ENSG00000261008 | LINC01572 | chr16 | - | 72561972 | 72562086 | 72555264 | 72555326 | 72570450 | 72570515 | 0 | 0 | 1 | 0 | 0 | 1 | 1 | 5 | 2 | 4 | 0 | 5 | 0 | 0 | 0 | 13 | 0 | 5 | 0 | 0 | 263 | 149 | 1 | 1 | 0 | 0 | 0.102 | 0 | 0 | 0 | 1 | 0 | NA | NA | -0.313 |
| SE | 101023 | ENSG00000261008 | LINC01572 | chr16 | - | 72570434 | 72570515 | 72555264 | 72555326 | 72664891 | 72664947 | 2 | 5 | 2 | 0 | 0 | 2 | 0 | 0 | 0 | 0 | 0 | 8 | 3 | 0 | 14 | 0 | 0 | 0 | 0 | 0 | 230 | 149 | 1 | 1 | 0.393 | 1 | 1 | NA | NA | NA | 1 | 1 | NA | 1 | -0.202 |
| SE | 101024 | ENSG00000261008 | LINC01572 | chr16 | - | 72570450 | 72570515 | 72555264 | 72555326 | 72664891 | 72664970 | 1 | 1 | 5 | 2 | 4 | 2 | 0 | 0 | 0 | 0 | 0 | 0 | 5 | 0 | 14 | 0 | 0 | 0 | 0 | 0 | 214 | 149 | 1 | 1 | 0.258 | 1 | 1 | 1 | 1 | NA | NA | 1 | NA | 1 | -0.148 |
| SE | 101026 | ENSG00000261008 | LINC01572 | chr16 | - | 72617182 | 72617237 | 72570450 | 72570515 | 72664891 | 72665014 | 0 | 0 | 0 | 0 | 0 | 0 | 1 | 3 | 1 | 2 | 0 | 0 | 1 | 0 | 0 | 0 | 0 | 6 | 0 | 14 | 204 | 149 | 1 | 1 | NA | 0 | 0 | 0 | 0 | NA | NA | 0.109 | NA | 0 | -0.054 |
| SE | 101029 | ENSG00000187240 | DYNC2H1 | chr11 | + | 103125099 | 103125295 | 103122824 | 103123000 | 103128909 | 103129005 | 17 | 9 | 17 | 24 | 17 | 0 | 0 | 0 | 2 | 0 | 0 | 1 | 3 | 1 | 0 | 0 | 0 | 0 | 0 | 0 | 298 | 149 | 1 | 1 | 1 | 1 | 1 | 0.857 | 1 | NA | 1 | 1 | 1 | NA | -0.029 |
| SE | 101031 | ENSG00000187240 | DYNC2H1 | chr11 | + | 103191516 | 103191619 | 103189671 | 103189816 | 103192096 | 103192264 | 21 | 15 | 13 | 21 | 6 | 2 | 4 | 0 | 2 | 0 | 0 | 0 | 0 | 0 | 0 | 0 | 1 | 0 | 0 | 0 | 252 | 149 | 1 | 1 | 0.861 | 0.689 | 1 | 0.861 | 1 | NA | 0 | NA | NA | NA | 0.882 |
| SE | 101035 | ENSG00000118276 | B4GALT6 | chr18 | - | 31626988 | 31627121 | 31626282 | 31626384 | 31630958 | 31631146 | 39 | 29 | 19 | 31 | 25 | 1 | 0 | 0 | 0 | 0 | 0 | 25 | 5 | 14 | 9 | 0 | 0 | 0 | 0 | 0 | 282 | 149 | 1 | 1 | 0.954 | 1 | 1 | 1 | 1 | NA | 1 | 1 | 1 | 1 | -0.009 |
| SE | 101036 | ENSG00000118276 | B4GALT6 | chr18 | - | 31626998 | 31627121 | 31626282 | 31626384 | 31630958 | 31631146 | 72 | 48 | 37 | 67 | 36 | 1 | 0 | 0 | 0 | 0 | 0 | 30 | 17 | 14 | 17 | 0 | 0 | 0 | 0 | 0 | 272 | 149 | 1 | 1 | 0.975 | 1 | 1 | 1 | 1 | NA | 1 | 1 | 1 | 1 | -0.005 |
| SE | 101043 | ENSG00000118276 | B4GALT6 | chr18 | - | 31657975 | 31658089 | 31645354 | 31645479 | 31684311 | 31684571 | 40 | 21 | 12 | 31 | 28 | 1 | 0 | 0 | 0 | 0 | 40 | 5 | 5 | 10 | 27 | 0 | 0 | 0 | 0 | 0 | 263 | 149 | 1 | 1 | 0.958 | 1 | 1 | 1 | 1 | 1 | 1 | 1 | 1 | 1 | -0.008 |
| SE | 101044 | ENSG00000118276 | B4GALT6 | chr18 | - | 31666255 | 31666372 | 31645354 | 31645479 | 31684311 | 31684504 | 12 | 7 | 4 | 6 | 6 | 1 | 0 | 0 | 0 | 0 | 0 | 0 | 0 | 0 | 8 | 0 | 0 | 0 | 0 | 0 | 266 | 149 | 1 | 1 | 0.87 | 1 | 1 | 1 | 1 | NA | NA | NA | NA | 1 | -0.026 |
| SE | 101045 | ENSG00000118276 | B4GALT6 | chr18 | - | 31666255 | 31666372 | 31657974 | 31658089 | 31684311 | 31684504 | 62 | 37 | 10 | 43 | 59 | 0 | 0 | 1 | 1 | 0 | 26 | 13 | 18 | 39 | 39 | 0 | 0 | 0 | 0 | 0 | 266 | 149 | 1 | 1 | 1 | 1 | 0.849 | 0.96 | 1 | 1 | 1 | 1 | 1 | 1 | -0.038 |
| SE | 101049 | ENSG00000149089 | APIP | chr11 | - | 34883336 | 34883504 | 34882294 | 34882816 | 34888292 | 34888428 | 227 | 144 | 162 | 253 | 215 | 0 | 0 | 0 | 2 | 0 | 377 | 292 | 156 | 315 | 94 | 0 | 0 | 0 | 0 | 0 | 298 | 149 | 1 | 1 | 1 | 1 | 1 | 0.984 | 1 | 1 | 1 | 1 | 1 | 1 | -0.003 |
| SE | 101052 | ENSG00000110435 | PDHX | chr11 | + | 34947505 | 34947606 | 34931403 | 34931484 | 34984569 | 34984728 | 28 | 15 | 25 | 34 | 28 | 0 | 1 | 0 | 0 | 0 | 7 | 30 | 14 | 21 | 0 | 0 | 0 | 0 | 0 | 0 | 250 | 149 | 1 | 1 | 1 | 0.899 | 1 | 1 | 1 | 1 | 1 | 1 | 1 | NA | -0.02 |
| SE | 101057 | ENSG00000110330 | BIRC2 | chr11 | + | 102348280 | 102348407 | 102347210 | 102347376 | 102348597 | 102349013 | 6 | 0 | 0 | 0 | 0 | 37 | 21 | 34 | 35 | 12 | 0 | 0 | 0 | 0 | 0 | 1 | 29 | 10 | 5 | 18 | 276 | 149 | 1 | 1 | 0.08 | 0 | 0 | 0 | 0 | 0 | 0 | 0 | 0 | 0 | 0.016 |
| SE | 101058 | ENSG00000110330 | BIRC2 | chr11 | + | 102348597 | 102348719 | 102347213 | 102347376 | 102350345 | 102350565 | 35 | 17 | 34 | 33 | 8 | 0 | 0 | 1 | 0 | 0 | 1 | 29 | 10 | 2 | 18 | 0 | 0 | 0 | 0 | 0 | 271 | 149 | 1 | 1 | 1 | 1 | 0.949 | 1 | 1 | 1 | 1 | 1 | 1 | 1 | -0.01 |
| SE | 101061 | ENSG00000110330 | BIRC2 | chr11 | + | 102348597 | 102348764 | 102347216 | 102347376 | 102350345 | 102350565 | 37 | 21 | 34 | 35 | 18 | 0 | 0 | 1 | 0 | 0 | 1 | 29 | 10 | 5 | 18 | 0 | 0 | 0 | 0 | 0 | 298 | 149 | 1 | 1 | 1 | 1 | 0.944 | 1 | 1 | 1 | 1 | 1 | 1 | 1 | -0.011 |
| SE | 101062 | ENSG00000110330 | BIRC2 | chr11 | + | 102350843 | 102350943 | 102348597 | 102350749 | 102362895 | 102362974 | 215 | 194 | 235 | 249 | 308 | 1 | 0 | 0 | 3 | 0 | 0 | 87 | 87 | 31 | 19 | 0 | 0 | 2 | 2 | 0 | 249 | 149 | 1 | 1 | 0.992 | 1 | 1 | 0.98 | 1 | NA | 1 | 0.963 | 0.903 | 1 | 0.028 |
| SE | 101065 | ENSG00000187531 | SIRT7 | chr17 | - | 81913206 | 81913293 | 81911938 | 81912614 | 81913773 | 81913880 | 0 | 0 | 0 | 2 | 0 | 75 | 60 | 68 | 70 | 42 | 0 | 6 | 0 | 0 | 0 | 97 | 96 | 97 | 104 | 232 | 236 | 149 | 1 | 1 | 0 | 0 | 0 | 0.018 | 0 | 0 | 0.038 | 0 | 0 | 0 | -0.004 |
| SE | 101067 | ENSG00000187531 | SIRT7 | chr17 | - | 81917614 | 81917719 | 81915610 | 81915681 | 81917829 | 81917967 | 34 | 12 | 14 | 22 | 0 | 2 | 1 | 1 | 3 | 0 | 8 | 50 | 19 | 13 | 33 | 0 | 6 | 4 | 5 | 0 | 254 | 149 | 1 | 1 | 0.909 | 0.876 | 0.891 | 0.811 | NA | 1 | 0.83 | 0.736 | 0.604 | 1 | 0.038 |
| SE | 101069 | ENSG00000072952 | IRAG1 | chr11 | - | 10594145 | 10594195 | 10593491 | 10593599 | 10600917 | 10601059 | 13 | 9 | 6 | 27 | 12 | 0 | 0 | 0 | 0 | 0 | 9 | 11 | 24 | 12 | 99 | 0 | 0 | 1 | 0 | 0 | 199 | 149 | 1 | 1 | 1 | 1 | 1 | 1 | 1 | 1 | 1 | 0.947 | 1 | 1 | 0.011 |
| SE | 101070 | ENSG00000072952 | IRAG1 | chr11 | - | 10604404 | 10604545 | 10603119 | 10603251 | 10606741 | 10606772 | 22 | 19 | 9 | 32 | 25 | 0 | 0 | 0 | 0 | 0 | 45 | 44 | 19 | 7 | 239 | 0 | 2 | 0 | 0 | 0 | 290 | 149 | 1 | 1 | 1 | 1 | 1 | 1 | 1 | 1 | 0.919 | 1 | 1 | 1 | 0.016 |
| SE | 101077 | ENSG00000072952 | IRAG1 | chr11 | - | 10652024 | 10652182 | 10633967 | 10634071 | 10690264 | 10690369 | 0 | 0 | 0 | 0 | 0 | 0 | 0 | 0 | 0 | 5 | 0 | 0 | 0 | 2 | 19 | 0 | 0 | 0 | 1 | 12 | 298 | 149 | 1 | 1 | NA | NA | NA | NA | 0 | NA | NA | NA | 0.5 | 0.442 | -0.471 |
| SE | 101080 | ENSG00000072952 | IRAG1 | chr11 | - | 10687508 | 10687777 | 10633967 | 10634071 | 10690264 | 10690369 | 0 | 0 | 0 | 0 | 0 | 0 | 0 | 0 | 0 | 5 | 0 | 0 | 1 | 0 | 13 | 0 | 0 | 0 | 1 | 12 | 298 | 149 | 1 | 1 | NA | NA | NA | NA | 0 | NA | NA | 1 | 0 | 0.351 | -0.45 |
| SE | 101081 | ENSG00000072952 | IRAG1 | chr11 | - | 10687508 | 10687777 | 10652103 | 10652182 | 10690264 | 10690369 | 0 | 0 | 0 | 2 | 0 | 0 | 0 | 0 | 0 | 0 | 12 | 0 | 0 | 0 | 11 | 0 | 0 | 0 | 0 | 2 | 298 | 149 | 1 | 1 | NA | NA | NA | 1 | NA | 1 | NA | NA | NA | 0.733 | 0.133 |
| SE | 101082 | ENSG00000072952 | IRAG1 | chr11 | - | 10690264 | 10690369 | 10633967 | 10634071 | 10693535 | 10693576 | 0 | 0 | 0 | 0 | 5 | 0 | 0 | 0 | 0 | 0 | 0 | 0 | 0 | 1 | 16 | 0 | 8 | 0 | 0 | 13 | 254 | 149 | 1 | 1 | NA | NA | NA | NA | 1 | NA | 0 | NA | 1 | 0.419 | 0.527 |
| SE | 101085 | ENSG00000143429 | LSP1P4 | chr2 | - | 91654919 | 91654997 | 91636687 | 91637033 | 91655367 | 91655378 | 0 | 0 | 0 | 0 | 0 | 1 | 0 | 0 | 0 | 0 | 0 | 0 | 1 | 0 | 0 | 0 | 0 | 0 | 0 | 0 | 227 | 149 | 1 | 1 | 0 | NA | NA | NA | NA | NA | NA | 1 | NA | NA | -1 |
| SE | 101086 | ENSG00000164105 | SAP30 | chr4 | + | 173373938 | 173374037 | 173373389 | 173373515 | 173377204 | 173377532 | 137 | 102 | 82 | 82 | 195 | 0 | 0 | 0 | 0 | 0 | 118 | 382 | 535 | 110 | 229 | 0 | 2 | 0 | 0 | 0 | 248 | 149 | 1 | 1 | 1 | 1 | 1 | 1 | 1 | 1 | 0.991 | 1 | 1 | 1 | 0.002 |
| SE | 101090 | ENSG00000170959 | DCDC1 | chr11 | - | 31065018 | 31065153 | 31064468 | 31064626 | 31077864 | 31077925 | 0 | 0 | 0 | 1 | 0 | 0 | 0 | 0 | 0 | 2 | 0 | 0 | 0 | 4 | 0 | 0 | 0 | 0 | 0 | 0 | 284 | 149 | 1 | 1 | NA | NA | NA | 1 | 0 | NA | NA | NA | 1 | NA | -0.5 |
| SE | 101102 | ENSG00000103510 | KAT8 | chr16 | + | 31127188 | 31127353 | 31125583 | 31127088 | 31128049 | 31128139 | 138 | 70 | 136 | 128 | 195 | 0 | 0 | 1 | 0 | 0 | 87 | 144 | 70 | 101 | 274 | 0 | 0 | 0 | 0 | 0 | 298 | 149 | 1 | 1 | 1 | 1 | 0.986 | 1 | 1 | 1 | 1 | 1 | 1 | 1 | -0.003 |
| SE | 101104 | ENSG00000050165 | DKK3 | chr11 | - | 11998695 | 11998779 | 11968394 | 11968487 | 12002299 | 12002437 | 1 | 4 | 5 | 2 | 13 | 0 | 0 | 0 | 0 | 0 | 0 | 2 | 0 | 0 | 0 | 0 | 6 | 0 | 0 | 0 | 233 | 149 | 1 | 1 | 1 | 1 | 1 | 1 | 1 | 0 | 1 | NA | NA | NA | 0.5 |
| SE | 101106 | ENSG00000162241 | SLC25A45 | chr11 | - | 65377332 | 65377423 | 65376817 | 65377076 | 65379375 | 65379561 | 1 | 0 | 0 | 0 | 0 | 16 | 8 | 7 | 15 | 16 | 0 | 2 | 3 | 0 | 0 | 0 | 6 | 9 | 9 | 4 | 240 | 149 | 1 | 1 | 0.037 | 0 | 0 | 0 | 0 | NA | 0.171 | 0.171 | 0 | 0 | -0.078 |
| SE | 101107 | ENSG00000162241 | SLC25A45 | chr11 | - | 65377332 | 65377423 | 65376817 | 65377076 | 65379866 | 65379938 | 1 | 0 | 0 | 0 | 0 | 0 | 0 | 5 | 4 | 8 | 0 | 2 | 0 | 0 | 0 | 0 | 10 | 2 | 3 | 0 | 240 | 149 | 1 | 1 | 1 | NA | 0 | 0 | 0 | NA | 0.11 | 0 | 0 | NA | 0.213 |
| SE | 101109 | ENSG00000162241 | SLC25A45 | chr11 | - | 65379375 | 65379561 | 65377332 | 65377423 | 65379866 | 65379938 | 37 | 8 | 6 | 15 | 22 | 0 | 0 | 0 | 0 | 0 | 11 | 7 | 12 | 4 | 7 | 0 | 0 | 1 | 0 | 0 | 298 | 149 | 1 | 1 | 1 | 1 | 1 | 1 | 1 | 1 | 1 | 0.857 | 1 | 1 | 0.029 |
| SE | 101110 | ENSG00000162241 | SLC25A45 | chr11 | - | 65379866 | 65379938 | 65379375 | 65379561 | 65380131 | 65380175 | 46 | 24 | 13 | 29 | 32 | 0 | 0 | 0 | 0 | 0 | 23 | 21 | 22 | 5 | 43 | 0 | 2 | 0 | 0 | 0 | 221 | 149 | 1 | 1 | 1 | 1 | 1 | 1 | 1 | 1 | 0.876 | 1 | 1 | 1 | 0.025 |
| SE | 101115 | ENSG00000162241 | SLC25A45 | chr11 | - | 65380533 | 65380632 | 65380131 | 65380234 | 65381874 | 65381969 | 0 | 4 | 0 | 1 | 1 | 0 | 0 | 0 | 0 | 2 | 0 | 4 | 0 | 0 | 0 | 0 | 0 | 0 | 0 | 0 | 248 | 149 | 1 | 1 | NA | 1 | NA | 1 | 0.231 | NA | 1 | NA | NA | NA | -0.256 |
| SE | 101117 | ENSG00000162241 | SLC25A45 | chr11 | - | 65381874 | 65381969 | 65380131 | 65380175 | 65382485 | 65382571 | 9 | 1 | 9 | 8 | 22 | 3 | 1 | 2 | 5 | 2 | 0 | 26 | 6 | 3 | 23 | 0 | 7 | 5 | 0 | 19 | 244 | 149 | 1 | 1 | 0.647 | 0.379 | 0.733 | 0.494 | 0.87 | NA | 0.694 | 0.423 | 1 | 0.425 | -0.011 |
| SE | 101118 | ENSG00000162241 | SLC25A45 | chr11 | - | 65381874 | 65381969 | 65380131 | 65380234 | 65382485 | 65382571 | 9 | 1 | 8 | 6 | 22 | 0 | 0 | 0 | 1 | 0 | 0 | 20 | 6 | 3 | 23 | 0 | 0 | 0 | 0 | 0 | 244 | 149 | 1 | 1 | 1 | 1 | 1 | 0.786 | 1 | NA | 1 | 1 | 1 | 1 | -0.043 |
| SE | 101119 | ENSG00000162241 | SLC25A45 | chr11 | - | 65381914 | 65381969 | 65380131 | 65380175 | 65382485 | 65382646 | 16 | 2 | 9 | 17 | 21 | 3 | 1 | 2 | 5 | 2 | 0 | 25 | 18 | 5 | 46 | 0 | 7 | 5 | 0 | 19 | 204 | 149 | 1 | 1 | 0.796 | 0.594 | 0.767 | 0.713 | 0.885 | NA | 0.723 | 0.724 | 1 | 0.639 | -0.021 |
| SE | 101130 | ENSG00000175224 | ATG13 | chr11 | + | 46621041 | 46621157 | 46618259 | 46618375 | 46644278 | 46644331 | 0 | 0 | 0 | 1 | 0 | 10 | 4 | 12 | 6 | 14 | 0 | 0 | 0 | 0 | 0 | 9 | 10 | 0 | 4 | 10 | 265 | 149 | 1 | 1 | 0 | 0 | 0 | 0.086 | 0 | 0 | 0 | NA | 0 | 0 | 0.017 |
| SE | 101132 | ENSG00000175224 | ATG13 | chr11 | + | 46621059 | 46621157 | 46618324 | 46618375 | 46644278 | 46644360 | 2 | 0 | 0 | 0 | 0 | 10 | 4 | 12 | 6 | 14 | 0 | 0 | 0 | 0 | 0 | 9 | 10 | 0 | 4 | 10 | 247 | 149 | 1 | 1 | 0.108 | 0 | 0 | 0 | 0 | 0 | 0 | NA | 0 | 0 | 0.022 |
| SE | 101136 | ENSG00000175224 | ATG13 | chr11 | + | 46630896 | 46630991 | 46630044 | 46630100 | 46644278 | 46644360 | 1 | 0 | 0 | 1 | 0 | 45 | 29 | 40 | 37 | 27 | 0 | 0 | 0 | 1 | 0 | 33 | 47 | 23 | 16 | 59 | 244 | 149 | 1 | 1 | 0.013 | 0 | 0 | 0.016 | 0 | 0 | 0 | 0 | 0.037 | 0 | -0.002 |
| SE | 101139 | ENSG00000175224 | ATG13 | chr11 | + | 46644278 | 46644360 | 46617581 | 46617890 | 46645338 | 46645350 | 157 | 73 | 98 | 105 | 183 | 0 | 0 | 0 | 0 | 0 | 125 | 99 | 114 | 94 | 224 | 0 | 3 | 0 | 0 | 0 | 231 | 149 | 1 | 1 | 1 | 1 | 1 | 1 | 1 | 1 | 0.955 | 1 | 1 | 1 | 0.009 |
| SE | 101140 | ENSG00000175224 | ATG13 | chr11 | + | 46650176 | 46650317 | 46649136 | 46649183 | 46656232 | 46656273 | 559 | 333 | 481 | 485 | 745 | 0 | 0 | 0 | 0 | 0 | 698 | 488 | 460 | 497 | 729 | 0 | 0 | 0 | 4 | 0 | 290 | 149 | 1 | 1 | 1 | 1 | 1 | 1 | 1 | 1 | 1 | 1 | 0.985 | 1 | 0.003 |
| SE | 101141 | ENSG00000175224 | ATG13 | chr11 | + | 46657094 | 46657191 | 46656232 | 46656273 | 46657523 | 46657618 | 239 | 139 | 229 | 248 | 345 | 0 | 0 | 0 | 0 | 0 | 222 | 254 | 186 | 156 | 317 | 0 | 0 | 0 | 2 | 0 | 246 | 149 | 1 | 1 | 1 | 1 | 1 | 1 | 1 | 1 | 1 | 1 | 0.979 | 1 | 0.004 |
| SE | 101142 | ENSG00000175224 | ATG13 | chr11 | + | 46657523 | 46657622 | 46656950 | 46657191 | 46659391 | 46659485 | 193 | 112 | 178 | 187 | 297 | 0 | 0 | 0 | 0 | 0 | 238 | 175 | 181 | 161 | 303 | 0 | 0 | 0 | 1 | 0 | 248 | 149 | 1 | 1 | 1 | 1 | 1 | 1 | 1 | 1 | 1 | 1 | 0.99 | 1 | 0.002 |
| SE | 101146 | ENSG00000175224 | ATG13 | chr11 | + | 46665382 | 46665519 | 46659334 | 46659485 | 46667772 | 46667865 | 167 | 71 | 153 | 165 | 173 | 2 | 0 | 0 | 0 | 0 | 127 | 220 | 189 | 163 | 259 | 0 | 0 | 0 | 0 | 0 | 286 | 149 | 1 | 1 | 0.978 | 1 | 1 | 1 | 1 | 1 | 1 | 1 | 1 | 1 | -0.004 |
| SE | 101148 | ENSG00000175224 | ATG13 | chr11 | + | 46667772 | 46667887 | 46665382 | 46665519 | 46668498 | 46668576 | 166 | 66 | 135 | 144 | 142 | 0 | 0 | 0 | 3 | 0 | 151 | 191 | 168 | 119 | 209 | 0 | 0 | 2 | 0 | 0 | 264 | 149 | 1 | 1 | 1 | 1 | 1 | 0.964 | 1 | 1 | 1 | 0.979 | 1 | 1 | -0.003 |
| SE | 101151 | ENSG00000133816 | MICAL2 | chr11 | + | 12138389 | 12138460 | 12116916 | 12116916 | 12162078 | 12162419 | 18 | 24 | 19 | 21 | 19 | 0 | 0 | 0 | 0 | 0 | 27 | 15 | 16 | 17 | 50 | 0 | 0 | 1 | 0 | 0 | 220 | 149 | 1 | 1 | 1 | 1 | 1 | 1 | 1 | 1 | 1 | 0.916 | 1 | 1 | 0.017 |
| SE | 101155 | ENSG00000133816 | MICAL2 | chr11 | + | 12208022 | 12208139 | 12204249 | 12204457 | 12213254 | 12213410 | 103 | 56 | 52 | 80 | 82 | 0 | 0 | 0 | 0 | 0 | 181 | 26 | 48 | 92 | 100 | 0 | 0 | 0 | 0 | 4 | 266 | 149 | 1 | 1 | 1 | 1 | 1 | 1 | 1 | 1 | 1 | 1 | 1 | 0.933 | 0.013 |
| SE | 101157 | ENSG00000133816 | MICAL2 | chr11 | + | 12220200 | 12220458 | 12216218 | 12216319 | 12221643 | 12221759 | 83 | 33 | 67 | 87 | 95 | 3 | 1 | 0 | 0 | 0 | 135 | 57 | 38 | 81 | 113 | 0 | 0 | 0 | 0 | 8 | 298 | 149 | 1 | 1 | 0.933 | 0.943 | 1 | 1 | 1 | 1 | 1 | 1 | 1 | 0.876 | 0 |
| SE | 101158 | ENSG00000133816 | MICAL2 | chr11 | + | 12222616 | 12222743 | 12221643 | 12221759 | 12223410 | 12223501 | 133 | 60 | 77 | 110 | 71 | 0 | 0 | 0 | 0 | 0 | 192 | 64 | 31 | 89 | 204 | 0 | 0 | 0 | 2 | 0 | 276 | 149 | 1 | 1 | 1 | 1 | 1 | 1 | 1 | 1 | 1 | 1 | 0.96 | 1 | 0.008 |
| SE | 101163 | ENSG00000133816 | MICAL2 | chr11 | + | 12241039 | 12241162 | 12239435 | 12239585 | 12256784 | 12256971 | 0 | 1 | 0 | 2 | 0 | 85 | 49 | 63 | 100 | 61 | 0 | 0 | 0 | 0 | 0 | 201 | 56 | 30 | 65 | 102 | 272 | 149 | 1 | 1 | 0 | 0.011 | 0 | 0.011 | 0 | 0 | 0 | 0 | 0 | 0 | 0.004 |
| SE | 101165 | ENSG00000133816 | MICAL2 | chr11 | + | 12242670 | 12242772 | 12239435 | 12239585 | 12256784 | 12256971 | 4 | 0 | 0 | 2 | 0 | 85 | 49 | 63 | 100 | 61 | 0 | 0 | 0 | 0 | 4 | 201 | 56 | 30 | 65 | 102 | 251 | 149 | 1 | 1 | 0.027 | 0 | 0 | 0.012 | 0 | 0 | 0 | 0 | 0 | 0.023 | 0.003 |
| SE | 101167 | ENSG00000133816 | MICAL2 | chr11 | + | 12243986 | 12244112 | 12242670 | 12242772 | 12256784 | 12256971 | 0 | 0 | 0 | 0 | 0 | 4 | 0 | 0 | 0 | 0 | 0 | 5 | 0 | 0 | 0 | 0 | 0 | 0 | 0 | 4 | 275 | 149 | 1 | 1 | 0 | NA | NA | NA | NA | NA | 1 | NA | NA | 0 | -0.5 |
| SE | 101173 | ENSG00000133816 | MICAL2 | chr11 | + | 12255642 | 12255750 | 12243986 | 12244112 | 12256784 | 12256909 | 0 | 3 | 6 | 0 | 1 | 0 | 0 | 0 | 0 | 0 | 1 | 0 | 2 | 8 | 21 | 0 | 5 | 0 | 0 | 0 | 257 | 149 | 1 | 1 | NA | 1 | 1 | NA | 1 | 1 | 0 | 1 | 1 | 1 | 0.2 |
| SE | 101175 | ENSG00000133816 | MICAL2 | chr11 | + | 12256784 | 12256971 | 12239435 | 12239585 | 12258467 | 12258556 | 199 | 132 | 167 | 237 | 156 | 1 | 0 | 0 | 0 | 0 | 488 | 143 | 95 | 205 | 392 | 0 | 0 | 0 | 0 | 0 | 298 | 149 | 1 | 1 | 0.99 | 1 | 1 | 1 | 1 | 1 | 1 | 1 | 1 | 1 | -0.002 |
| SE | 101176 | ENSG00000133816 | MICAL2 | chr11 | + | 12258467 | 12258556 | 12253233 | 12256971 | 12259794 | 12259897 | 156 | 97 | 118 | 164 | 111 | 1 | 0 | 0 | 0 | 0 | 295 | 102 | 83 | 156 | 372 | 0 | 0 | 3 | 1 | 0 | 238 | 149 | 1 | 1 | 0.99 | 1 | 1 | 1 | 1 | 1 | 1 | 0.945 | 0.99 | 1 | 0.011 |
| SE | 101177 | ENSG00000133816 | MICAL2 | chr11 | + | 12259794 | 12259897 | 12258467 | 12258556 | 12275982 | 12276166 | 39 | 28 | 31 | 53 | 24 | 0 | 0 | 0 | 0 | 0 | 51 | 31 | 24 | 22 | 55 | 0 | 1 | 0 | 0 | 0 | 252 | 149 | 1 | 1 | 1 | 1 | 1 | 1 | 1 | 1 | 0.948 | 1 | 1 | 1 | 0.01 |
| SE | 101180 | ENSG00000133816 | MICAL2 | chr11 | + | 12276619 | 12276715 | 12276079 | 12276166 | 12280932 | 12281099 | 0 | 0 | 2 | 0 | 0 | 1 | 4 | 4 | 1 | 6 | 0 | 0 | 3 | 0 | 0 | 10 | 0 | 4 | 0 | 0 | 245 | 149 | 1 | 1 | 0 | 0 | 0.233 | 0 | 0 | 0 | NA | 0.313 | NA | NA | -0.11 |
| SE | 101181 | ENSG00000133816 | MICAL2 | chr11 | + | 12287086 | 12287164 | 12280932 | 12281099 | 12292142 | 12292297 | 8 | 11 | 4 | 8 | 10 | 1 | 0 | 0 | 0 | 0 | 6 | 1 | 18 | 4 | 12 | 0 | 0 | 0 | 0 | 0 | 227 | 149 | 1 | 1 | 0.84 | 1 | 1 | 1 | 1 | 1 | 1 | 1 | 1 | 1 | -0.032 |
| SE | 101182 | ENSG00000133816 | MICAL2 | chr11 | + | 12287089 | 12287164 | 12280944 | 12281099 | 12292142 | 12292255 | 3 | 13 | 3 | 10 | 10 | 1 | 0 | 0 | 0 | 0 | 11 | 1 | 16 | 8 | 3 | 0 | 0 | 0 | 0 | 0 | 224 | 149 | 1 | 1 | 0.666 | 1 | 1 | 1 | 1 | 1 | 1 | 1 | 1 | 1 | -0.067 |
| SE | 101183 | ENSG00000133816 | MICAL2 | chr11 | + | 12292142 | 12292297 | 12286899 | 12287164 | 12293560 | 12294745 | 14 | 25 | 8 | 15 | 36 | 0 | 0 | 0 | 0 | 0 | 3 | 3 | 25 | 20 | 14 | 0 | 0 | 2 | 0 | 0 | 298 | 149 | 1 | 1 | 1 | 1 | 1 | 1 | 1 | 1 | 1 | 0.862 | 1 | 1 | 0.028 |
| SE | 101184 | ENSG00000133816 | MICAL2 | chr11 | + | 12327172 | 12327266 | 12323977 | 12324070 | 12349837 | 12349937 | 8 | 7 | 6 | 21 | 5 | 0 | 0 | 0 | 0 | 0 | 0 | 14 | 4 | 16 | 0 | 0 | 3 | 3 | 0 | 0 | 243 | 149 | 1 | 1 | 1 | 1 | 1 | 1 | 1 | NA | 0.741 | 0.45 | 1 | NA | 0.27 |
| SE | 101194 | ENSG00000167535 | CACNB3 | chr12 | + | 48819635 | 48819942 | 48818908 | 48818974 | 48823343 | 48823466 | 0 | 1 | 2 | 0 | 0 | 0 | 0 | 0 | 0 | 1 | 0 | 0 | 0 | 3 | 0 | 0 | 0 | 0 | 0 | 0 | 298 | 149 | 1 | 1 | NA | 1 | 1 | NA | 0 | NA | NA | NA | 1 | NA | -0.333 |
| SE | 101199 | ENSG00000167535 | CACNB3 | chr12 | + | 48824948 | 48824968 | 48824668 | 48824733 | 48825162 | 48825243 | 2 | 3 | 5 | 3 | 0 | 0 | 2 | 0 | 0 | 0 | 0 | 0 | 0 | 3 | 6 | 6 | 0 | 5 | 2 | 0 | 169 | 149 | 1 | 1 | 1 | 0.569 | 1 | 1 | NA | 0 | NA | 0 | 0.569 | 1 | 0.5 |
| SE | 101201 | ENSG00000150433 | TMEM218 | chr11 | - | 125101443 | 125101521 | 125101200 | 125101303 | 125102131 | 125102317 | 0 | 0 | 0 | 2 | 0 | 4 | 3 | 4 | 9 | 11 | 0 | 1 | 0 | 0 | 0 | 3 | 2 | 4 | 1 | 21 | 227 | 149 | 1 | 1 | 0 | 0 | 0 | 0.127 | 0 | 0 | 0.247 | 0 | 0 | 0 | -0.024 |
| SE | 101202 | ENSG00000150433 | TMEM218 | chr11 | - | 125101443 | 125101521 | 125101200 | 125101303 | 125102636 | 125102809 | 0 | 1 | 0 | 1 | 0 | 1 | 1 | 1 | 2 | 1 | 0 | 0 | 0 | 0 | 0 | 0 | 2 | 0 | 0 | 1 | 227 | 149 | 1 | 1 | 0 | 0.396 | 0 | 0.247 | 0 | NA | 0 | NA | NA | 0 | 0.129 |
| SE | 101203 | ENSG00000150433 | TMEM218 | chr11 | - | 125101443 | 125101521 | 125101200 | 125101303 | 125102733 | 125102809 | 0 | 0 | 0 | 0 | 0 | 6 | 2 | 4 | 7 | 9 | 0 | 0 | 0 | 2 | 0 | 0 | 13 | 2 | 14 | 0 | 227 | 149 | 1 | 1 | 0 | 0 | 0 | 0 | 0 | NA | 0 | 0 | 0.086 | NA | -0.029 |
| SE | 101205 | ENSG00000150433 | TMEM218 | chr11 | - | 125102131 | 125102317 | 125101200 | 125101256 | 125102636 | 125102809 | 0 | 6 | 2 | 0 | 3 | 0 | 0 | 0 | 1 | 0 | 0 | 1 | 0 | 0 | 9 | 0 | 2 | 0 | 0 | 0 | 298 | 149 | 1 | 1 | NA | 1 | 1 | 0 | 1 | NA | 0.2 | NA | NA | 1 | 0.15 |
| SE | 101209 | ENSG00000150433 | TMEM218 | chr11 | - | 125102131 | 125102317 | 125101443 | 125101521 | 125102636 | 125102809 | 0 | 6 | 2 | 2 | 2 | 0 | 1 | 0 | 1 | 0 | 0 | 2 | 0 | 0 | 11 | 0 | 2 | 0 | 0 | 0 | 298 | 149 | 1 | 1 | NA | 0.75 | 1 | 0.5 | 1 | NA | 0.333 | NA | NA | 1 | 0.146 |
| SE | 101210 | ENSG00000150433 | TMEM218 | chr11 | - | 125102131 | 125102317 | 125101443 | 125101521 | 125102733 | 125102809 | 6 | 4 | 3 | 9 | 8 | 0 | 0 | 0 | 0 | 1 | 11 | 3 | 0 | 0 | 19 | 0 | 0 | 0 | 2 | 0 | 298 | 149 | 1 | 1 | 1 | 1 | 1 | 1 | 0.8 | 1 | 1 | NA | 0 | 1 | 0.21 |
| SE | 101211 | ENSG00000150433 | TMEM218 | chr11 | - | 125102131 | 125102351 | 125101200 | 125101256 | 125102636 | 125102809 | 1 | 0 | 2 | 0 | 1 | 0 | 0 | 0 | 1 | 0 | 0 | 0 | 0 | 0 | 0 | 0 | 2 | 0 | 0 | 0 | 298 | 149 | 1 | 1 | 1 | NA | 1 | 0 | 1 | NA | 0 | NA | NA | NA | 0.75 |
| SE | 101212 | ENSG00000150433 | TMEM218 | chr11 | - | 125102131 | 125102351 | 125101200 | 125101256 | 125102733 | 125102809 | 6 | 0 | 2 | 0 | 1 | 0 | 0 | 0 | 0 | 0 | 0 | 0 | 0 | 0 | 0 | 0 | 0 | 1 | 3 | 0 | 298 | 149 | 1 | 1 | 1 | NA | 1 | NA | 1 | NA | NA | 0 | 0 | NA | 1 |
| SE | 101215 | ENSG00000150433 | TMEM218 | chr11 | - | 125110490 | 125110549 | 125102733 | 125102809 | 125110643 | 125110746 | 4 | 0 | 2 | 3 | 3 | 4 | 4 | 5 | 0 | 0 | 0 | 1 | 0 | 0 | 0 | 0 | 2 | 0 | 2 | 0 | 208 | 149 | 1 | 1 | 0.417 | 0 | 0.223 | 1 | 1 | NA | 0.264 | NA | 0 | NA | 0.396 |
| SE | 101216 | ENSG00000150433 | TMEM218 | chr11 | - | 125110490 | 125110549 | 125102733 | 125102809 | 125111538 | 125111589 | 2 | 1 | 1 | 3 | 3 | 19 | 1 | 7 | 11 | 4 | 0 | 0 | 0 | 0 | 0 | 0 | 6 | 0 | 8 | 33 | 208 | 149 | 1 | 1 | 0.07 | 0.417 | 0.093 | 0.163 | 0.349 | NA | 0 | NA | 0 | 0 | 0.218 |
| SE | 101217 | ENSG00000150433 | TMEM218 | chr11 | - | 125110490 | 125110549 | 125102733 | 125102809 | 125111584 | 125111632 | 1 | 0 | 0 | 1 | 3 | 2 | 4 | 4 | 5 | 4 | 0 | 0 | 0 | 0 | 0 | 16 | 3 | 0 | 0 | 0 | 208 | 149 | 1 | 1 | 0.264 | 0 | 0 | 0.125 | 0.349 | 0 | 0 | NA | NA | NA | 0.148 |
| SE | 101218 | ENSG00000150433 | TMEM218 | chr11 | - | 125110643 | 125110746 | 125102131 | 125102809 | 125111538 | 125111579 | 8 | 4 | 5 | 0 | 0 | 19 | 1 | 7 | 11 | 4 | 0 | 2 | 0 | 2 | 0 | 0 | 6 | 0 | 8 | 33 | 252 | 149 | 1 | 1 | 0.199 | 0.703 | 0.297 | 0 | 0 | NA | 0.165 | NA | 0.129 | 0 | 0.142 |
| SE | 101219 | ENSG00000150433 | TMEM218 | chr11 | - | 125110643 | 125110746 | 125102131 | 125102809 | 125111584 | 125111611 | 4 | 4 | 3 | 1 | 0 | 2 | 4 | 4 | 5 | 4 | 0 | 0 | 0 | 2 | 0 | 16 | 3 | 0 | 0 | 0 | 252 | 149 | 1 | 1 | 0.542 | 0.372 | 0.307 | 0.106 | 0 | 0 | 0 | NA | 1 | NA | -0.068 |
| SE | 101220 | ENSG00000150433 | TMEM218 | chr11 | - | 125110643 | 125110746 | 125102131 | 125102809 | 125111586 | 125111613 | 4 | 4 | 3 | 0 | 0 | 1 | 5 | 2 | 2 | 1 | 0 | 0 | 0 | 2 | 0 | 0 | 0 | 2 | 8 | 0 | 252 | 149 | 1 | 1 | 0.703 | 0.321 | 0.47 | 0 | 0 | NA | NA | 0 | 0.129 | NA | 0.234 |
| SE | 101221 | ENSG00000150433 | TMEM218 | chr11 | - | 125110643 | 125110746 | 125110490 | 125110549 | 125111538 | 125111579 | 6 | 0 | 2 | 1 | 0 | 1 | 1 | 1 | 2 | 0 | 0 | 1 | 0 | 0 | 0 | 0 | 0 | 0 | 0 | 0 | 252 | 149 | 1 | 1 | 0.78 | 0 | 0.542 | 0.228 | NA | NA | 1 | NA | NA | NA | -0.613 |
| SE | 101222 | ENSG00000150433 | TMEM218 | chr11 | - | 125110643 | 125110746 | 125110490 | 125110549 | 125111584 | 125111611 | 2 | 0 | 0 | 3 | 0 | 1 | 0 | 0 | 0 | 0 | 0 | 1 | 0 | 0 | 0 | 0 | 0 | 0 | 0 | 0 | 252 | 149 | 1 | 1 | 0.542 | NA | NA | 1 | NA | NA | 1 | NA | NA | NA | -0.229 |
| SE | 101227 | ENSG00000019485 | PRDM11 | chr11 | + | 45224216 | 45224843 | 45219569 | 45219757 | 45225994 | 45235109 | 4 | 0 | 3 | 11 | 6 | 0 | 0 | 1 | 3 | 0 | 0 | 6 | 1 | 0 | 0 | 0 | 0 | 0 | 0 | 0 | 298 | 149 | 1 | 1 | 1 | NA | 0.6 | 0.647 | 1 | NA | 1 | 1 | NA | NA | -0.188 |
| SE | 101231 | ENSG00000160131 | VMA21 | chrX | + | 151396798 | 151397057 | 151396565 | 151396647 | 151403630 | 151403740 | 0 | 0 | 2 | 0 | 4 | 0 | 0 | 1 | 1 | 0 | 5 | 0 | 0 | 0 | 0 | 0 | 0 | 0 | 0 | 0 | 298 | 149 | 1 | 1 | NA | NA | 0.5 | 0 | 1 | 1 | NA | NA | NA | NA | -0.5 |
| SE | 101236 | ENSG00000144802 | NFKBIZ | chr3 | + | 101852737 | 101852768 | 101852084 | 101852224 | 101852885 | 101852989 | 369 | 356 | 321 | 449 | 800 | 0 | 0 | 5 | 0 | 0 | 948 | 405 | 435 | 750 | 416 | 0 | 1 | 0 | 0 | 0 | 180 | 149 | 1 | 1 | 1 | 1 | 0.982 | 1 | 1 | 1 | 0.997 | 1 | 1 | 1 | -0.003 |
| SE | 101238 | ENSG00000144802 | NFKBIZ | chr3 | + | 101855732 | 101855902 | 101855061 | 101855208 | 101857291 | 101857459 | 7 | 1 | 21 | 4 | 8 | 0 | 0 | 0 | 2 | 0 | 9 | 35 | 13 | 24 | 29 | 0 | 0 | 0 | 0 | 0 | 298 | 149 | 1 | 1 | 1 | 1 | 1 | 0.5 | 1 | 1 | 1 | 1 | 1 | 1 | -0.1 |
| SE | 101239 | ENSG00000144802 | NFKBIZ | chr3 | + | 101855732 | 101855902 | 101855394 | 101855458 | 101857072 | 101857183 | 276 | 207 | 259 | 348 | 419 | 0 | 4 | 0 | 0 | 0 | 306 | 367 | 309 | 276 | 170 | 0 | 0 | 0 | 0 | 3 | 298 | 149 | 1 | 1 | 1 | 0.963 | 1 | 1 | 1 | 1 | 1 | 1 | 1 | 0.966 | -0.001 |
| SE | 101241 | ENSG00000144802 | NFKBIZ | chr3 | + | 101857291 | 101857459 | 101855732 | 101855902 | 101859317 | 101859523 | 87 | 67 | 113 | 133 | 78 | 0 | 0 | 0 | 0 | 0 | 39 | 90 | 103 | 117 | 56 | 0 | 0 | 3 | 0 | 0 | 298 | 149 | 1 | 1 | 1 | 1 | 1 | 1 | 1 | 1 | 1 | 0.945 | 1 | 1 | 0.011 |
| SE | 101246 | ENSG00000166224 | SGPL1 | chr10 | + | 70844472 | 70844638 | 70816810 | 70816880 | 70851142 | 70851210 | 231 | 144 | 174 | 150 | 184 | 0 | 2 | 0 | 0 | 0 | 307 | 179 | 148 | 189 | 288 | 0 | 6 | 0 | 0 | 0 | 298 | 149 | 1 | 1 | 1 | 0.973 | 1 | 1 | 1 | 1 | 0.937 | 1 | 1 | 1 | 0.007 |
| SE | 101254 | ENSG00000166224 | SGPL1 | chr10 | + | 70871836 | 70871986 | 70869791 | 70869897 | 70875401 | 70875548 | 0 | 0 | 1 | 0 | 0 | 0 | 0 | 0 | 1 | 0 | 0 | 3 | 6 | 0 | 7 | 0 | 7 | 0 | 2 | 0 | 298 | 149 | 1 | 1 | NA | NA | 1 | 0 | NA | NA | 0.176 | 1 | 0 | 1 | -0.044 |
| SE | 101255 | ENSG00000166224 | SGPL1 | chr10 | + | 70871836 | 70871986 | 70871047 | 70871146 | 70873350 | 70873589 | 137 | 53 | 107 | 101 | 151 | 0 | 1 | 0 | 0 | 2 | 155 | 60 | 87 | 53 | 124 | 0 | 1 | 0 | 0 | 0 | 298 | 149 | 1 | 1 | 1 | 0.964 | 1 | 1 | 0.974 | 1 | 0.968 | 1 | 1 | 1 | -0.006 |
| SE | 101259 | ENSG00000166224 | SGPL1 | chr10 | + | 70873350 | 70873589 | 70871836 | 70871986 | 70875401 | 70875548 | 144 | 94 | 137 | 137 | 166 | 0 | 0 | 1 | 0 | 0 | 143 | 98 | 102 | 69 | 190 | 0 | 3 | 2 | 0 | 0 | 298 | 149 | 1 | 1 | 1 | 1 | 0.986 | 1 | 1 | 1 | 0.942 | 0.962 | 1 | 1 | 0.016 |
| SE | 101260 | ENSG00000166224 | SGPL1 | chr10 | + | 70875401 | 70875548 | 70868344 | 70868433 | 70876540 | 70876661 | 140 | 79 | 80 | 128 | 169 | 1 | 0 | 0 | 0 | 0 | 172 | 117 | 90 | 77 | 170 | 0 | 0 | 0 | 0 | 0 | 296 | 149 | 1 | 1 | 0.986 | 1 | 1 | 1 | 1 | 1 | 1 | 1 | 1 | 1 | -0.003 |
| SE | 101261 | ENSG00000166224 | SGPL1 | chr10 | + | 70875401 | 70875548 | 70871836 | 70871986 | 70876540 | 70876661 | 140 | 79 | 81 | 127 | 169 | 0 | 0 | 0 | 0 | 0 | 172 | 120 | 89 | 73 | 170 | 0 | 2 | 0 | 0 | 0 | 296 | 149 | 1 | 1 | 1 | 1 | 1 | 1 | 1 | 1 | 0.968 | 1 | 1 | 1 | 0.006 |
| SE | 101262 | ENSG00000166224 | SGPL1 | chr10 | + | 70876540 | 70876661 | 70875401 | 70875548 | 70877194 | 70881184 | 221 | 130 | 121 | 188 | 270 | 0 | 0 | 0 | 0 | 0 | 227 | 202 | 130 | 133 | 247 | 0 | 0 | 1 | 0 | 0 | 270 | 149 | 1 | 1 | 1 | 1 | 1 | 1 | 1 | 1 | 1 | 0.986 | 1 | 1 | 0.003 |
| SE | 101263 | ENSG00000086598 | TMED2 | chr12 | + | 123587610 | 123587631 | 123586746 | 123586939 | 123590341 | 123590446 | 0 | 0 | 0 | 0 | 0 | 915 | 707 | 887 | 970 | 1638 | 5 | 0 | 0 | 0 | 0 | 1430 | 917 | 1071 | 1221 | 406 | 170 | 149 | 1 | 1 | 0 | 0 | 0 | 0 | 0 | 0.003 | 0 | 0 | 0 | 0 | -0.001 |
| SE | 101264 | ENSG00000172465 | TCEAL1 | chrX | + | 103629493 | 103629561 | 103628703 | 103628793 | 103629884 | 103630447 | 5 | 1 | 6 | 1 | 2 | 0 | 0 | 0 | 0 | 0 | 12 | 6 | 1 | 2 | 10 | 0 | 0 | 0 | 2 | 0 | 217 | 149 | 1 | 1 | 1 | 1 | 1 | 1 | 1 | 1 | 1 | 1 | 0.407 | 1 | 0.119 |
| SE | 101265 | ENSG00000172465 | TCEAL1 | chrX | + | 103629493 | 103629561 | 103628998 | 103629025 | 103629884 | 103630447 | 6 | 3 | 8 | 2 | 2 | 1 | 0 | 0 | 0 | 0 | 12 | 8 | 1 | 2 | 7 | 0 | 0 | 0 | 0 | 0 | 217 | 149 | 1 | 1 | 0.805 | 1 | 1 | 1 | 1 | 1 | 1 | 1 | 1 | 1 | -0.039 |
| SE | 101266 | ENSG00000172465 | TCEAL1 | chrX | + | 103629493 | 103629573 | 103628974 | 103629025 | 103629884 | 103630953 | 4 | 1 | 1 | 3 | 3 | 1 | 0 | 0 | 0 | 0 | 0 | 0 | 0 | 0 | 2 | 0 | 0 | 0 | 0 | 0 | 229 | 149 | 1 | 1 | 0.722 | 1 | 1 | 1 | 1 | NA | NA | NA | NA | 1 | -0.056 |
| SE | 101267 | ENSG00000107485 | GATA3 | chr10 | + | 8063992 | 8064138 | 8058304 | 8058841 | 8069472 | 8069598 | 92 | 34 | 80 | 104 | 54 | 0 | 1 | 1 | 0 | 0 | 0 | 28 | 77 | 30 | 20 | 0 | 0 | 0 | 0 | 0 | 295 | 149 | 1 | 1 | 1 | 0.945 | 0.976 | 1 | 1 | NA | 1 | 1 | 1 | 1 | -0.016 |
| SE | 101268 | ENSG00000107485 | GATA3 | chr10 | + | 8063995 | 8064138 | 8058304 | 8058841 | 8069472 | 8069598 | 72 | 25 | 63 | 73 | 28 | 0 | 1 | 1 | 0 | 0 | 6 | 18 | 53 | 30 | 23 | 0 | 0 | 0 | 0 | 0 | 292 | 149 | 1 | 1 | 1 | 0.927 | 0.97 | 1 | 1 | 1 | 1 | 1 | 1 | 1 | -0.021 |
| SE | 101274 | ENSG00000165629 | ATP5F1C | chr10 | + | 7799771 | 7799915 | 7798989 | 7799194 | 7800026 | 7800052 | 1337 | 677 | 1305 | 1002 | 3019 | 2 | 0 | 11 | 1 | 3 | 1270 | 998 | 757 | 958 | 1020 | 0 | 0 | 2 | 2 | 0 | 293 | 149 | 1 | 1 | 0.997 | 1 | 0.984 | 0.998 | 0.998 | 1 | 1 | 0.995 | 0.996 | 1 | -0.003 |
| SE | 101278 | ENSG00000165629 | ATP5F1C | chr10 | + | 7806973 | 7807010 | 7802757 | 7802854 | 7807658 | 7807793 | 337 | 179 | 218 | 258 | 487 | 2 | 1 | 0 | 1 | 3 | 297 | 333 | 167 | 277 | 371 | 0 | 0 | 3 | 4 | 0 | 186 | 149 | 1 | 1 | 0.993 | 0.993 | 1 | 0.995 | 0.992 | 1 | 1 | 0.978 | 0.982 | 1 | 0.003 |
| SE | 101279 | ENSG00000215244 | LINC02649 | chr10 | + | 6327931 | 6329810 | 6326545 | 6326642 | 6331210 | 6331306 | 3 | 1 | 2 | 2 | 5 | 0 | 0 | 0 | 0 | 0 | 0 | 1 | 13 | 0 | 0 | 0 | 0 | 2 | 0 | 0 | 298 | 149 | 1 | 1 | 1 | 1 | 1 | 1 | 1 | NA | 1 | 0.765 | NA | NA | 0.117 |
| SE | 101280 | ENSG00000215244 | LINC02649 | chr10 | + | 6330952 | 6331016 | 6327931 | 6329810 | 6331210 | 6331306 | 1 | 1 | 0 | 0 | 1 | 0 | 0 | 2 | 0 | 0 | 0 | 0 | 0 | 0 | 0 | 0 | 0 | 2 | 0 | 0 | 213 | 149 | 1 | 1 | 1 | 1 | 0 | NA | 1 | NA | NA | 0 | NA | NA | 0.75 |
| SE | 101283 | ENSG00000065665 | SEC61A2 | chr10 | + | 12142432 | 12142565 | 12136104 | 12136170 | 12143116 | 12143195 | 0 | 0 | 1 | 0 | 0 | 16 | 14 | 15 | 28 | 12 | 0 | 8 | 1 | 0 | 0 | 0 | 31 | 25 | 24 | 4 | 282 | 149 | 1 | 1 | 0 | 0 | 0.034 | 0 | 0 | NA | 0.12 | 0.021 | 0 | 0 | -0.028 |
| SE | 101284 | ENSG00000065665 | SEC61A2 | chr10 | + | 12143116 | 12143195 | 12136104 | 12136170 | 12149594 | 12149726 | 26 | 21 | 19 | 39 | 31 | 0 | 0 | 0 | 2 | 0 | 0 | 39 | 39 | 39 | 8 | 0 | 0 | 0 | 0 | 0 | 228 | 149 | 1 | 1 | 1 | 1 | 1 | 0.927 | 1 | NA | 1 | 1 | 1 | 1 | -0.015 |
| SE | 101286 | ENSG00000065665 | SEC61A2 | chr10 | + | 12155777 | 12155931 | 12149851 | 12149961 | 12156906 | 12157067 | 3 | 1 | 0 | 0 | 2 | 3 | 0 | 2 | 6 | 0 | 0 | 1 | 0 | 0 | 0 | 0 | 11 | 11 | 3 | 0 | 298 | 149 | 1 | 1 | 0.333 | 1 | 0 | 0 | 1 | NA | 0.043 | 0 | 0 | NA | 0.452 |
| SE | 101290 | ENSG00000065665 | SEC61A2 | chr10 | + | 12167752 | 12167893 | 12162212 | 12162289 | 12169257 | 12169961 | 4 | 0 | 1 | 7 | 0 | 0 | 0 | 0 | 7 | 4 | 0 | 17 | 0 | 0 | 0 | 0 | 1 | 2 | 0 | 0 | 290 | 149 | 1 | 1 | 1 | NA | 1 | 0.339 | 0 | NA | 0.897 | 0 | NA | NA | 0.136 |
| SE | 101296 | ENSG00000136167 | LCP1 | chr13 | - | 46130813 | 46130938 | 46125919 | 46127723 | 46134126 | 46134250 | 9604 | 6697 | 6092 | 8330 | 14156 | 6 | 0 | 23 | 0 | 0 | 17078 | 8287 | 7291 | 13442 | 5187 | 0 | 0 | 1 | 0 | 0 | 274 | 149 | 1 | 1 | 0.999 | 1 | 0.993 | 1 | 1 | 1 | 1 | 1 | 1 | 1 | -0.002 |
| SE | 101297 | ENSG00000136167 | LCP1 | chr13 | - | 46134126 | 46134250 | 46130813 | 46130938 | 46142291 | 46142425 | 8271 | 5728 | 5298 | 7019 | 12081 | 0 | 1 | 1 | 0 | 0 | 12520 | 6478 | 5743 | 10301 | 4061 | 0 | 0 | 0 | 0 | 0 | 273 | 149 | 1 | 1 | 1 | 1 | 1 | 1 | 1 | 1 | 1 | 1 | 1 | 1 | 0 |
| SE | 101298 | ENSG00000136167 | LCP1 | chr13 | - | 46142291 | 46142425 | 46134126 | 46134250 | 46144441 | 46144520 | 4228 | 2999 | 2845 | 3772 | 5338 | 0 | 0 | 0 | 0 | 0 | 6139 | 3281 | 3004 | 5064 | 2017 | 0 | 5 | 0 | 0 | 0 | 283 | 149 | 1 | 1 | 1 | 1 | 1 | 1 | 1 | 1 | 0.997 | 1 | 1 | 1 | 0.001 |
| SE | 101299 | ENSG00000136167 | LCP1 | chr13 | - | 46143289 | 46143404 | 46142291 | 46142425 | 46144441 | 46144520 | 6769 | 4720 | 4688 | 6203 | 9394 | 0 | 0 | 0 | 0 | 1 | 13318 | 7345 | 6674 | 12045 | 4601 | 0 | 0 | 0 | 0 | 0 | 264 | 149 | 1 | 1 | 1 | 1 | 1 | 1 | 1 | 1 | 1 | 1 | 1 | 1 | 0 |
| SE | 101300 | ENSG00000136167 | LCP1 | chr13 | - | 46146907 | 46147103 | 46144441 | 46144520 | 46148351 | 46148447 | 8191 | 5344 | 5380 | 7109 | 8205 | 1 | 0 | 2 | 1 | 1 | 13638 | 7792 | 7129 | 12105 | 4867 | 0 | 2 | 0 | 7 | 5 | 298 | 149 | 1 | 1 | 1 | 1 | 0.999 | 1 | 1 | 1 | 0.999 | 1 | 0.999 | 0.998 | 0.001 |
| SE | 101301 | ENSG00000136167 | LCP1 | chr13 | - | 46148351 | 46148447 | 46146905 | 46147103 | 46150935 | 46151078 | 6243 | 3892 | 3916 | 5050 | 6158 | 0 | 0 | 2 | 0 | 0 | 8538 | 5058 | 4648 | 7694 | 3146 | 0 | 1 | 0 | 0 | 0 | 245 | 149 | 1 | 1 | 1 | 1 | 0.999 | 1 | 1 | 1 | 1 | 1 | 1 | 1 | 0 |
| SE | 101302 | ENSG00000136167 | LCP1 | chr13 | - | 46148781 | 46148808 | 46148351 | 46148447 | 46150935 | 46151078 | 0 | 0 | 1 | 2 | 0 | 4538 | 2827 | 2860 | 3689 | 4475 | 12 | 7 | 6 | 1 | 10 | 6632 | 3794 | 3455 | 5702 | 2273 | 176 | 149 | 1 | 1 | 0 | 0 | 0 | 0 | 0 | 0.002 | 0.002 | 0.001 | 0 | 0.004 | -0.002 |
| SE | 101303 | ENSG00000136167 | LCP1 | chr13 | - | 46150935 | 46151078 | 46148351 | 46148447 | 46152779 | 46152945 | 8529 | 5060 | 5627 | 7033 | 7961 | 3 | 3 | 2 | 1 | 0 | 11502 | 7125 | 6225 | 10110 | 4082 | 6 | 0 | 2 | 0 | 0 | 292 | 149 | 1 | 1 | 0.999 | 0.999 | 0.999 | 1 | 1 | 0.999 | 1 | 0.999 | 1 | 1 | 0 |
| SE | 101304 | ENSG00000136167 | LCP1 | chr13 | - | 46158521 | 46158651 | 46156437 | 46156570 | 46158825 | 46158989 | 7259 | 4507 | 4558 | 5868 | 14718 | 1 | 0 | 0 | 2 | 0 | 9304 | 5563 | 4635 | 8313 | 3605 | 0 | 0 | 6 | 3 | 0 | 279 | 149 | 1 | 1 | 1 | 1 | 1 | 0.999 | 1 | 1 | 1 | 0.998 | 0.999 | 1 | 0 |
| SE | 101305 | ENSG00000136167 | LCP1 | chr13 | - | 46158521 | 46158651 | 46156437 | 46156570 | 46159598 | 46159686 | 2048 | 1276 | 1302 | 1625 | 4932 | 0 | 0 | 0 | 0 | 3 | 2104 | 1470 | 1339 | 2234 | 1226 | 0 | 0 | 0 | 0 | 0 | 279 | 149 | 1 | 1 | 1 | 1 | 1 | 1 | 0.999 | 1 | 1 | 1 | 1 | 1 | 0 |
| SE | 101306 | ENSG00000136167 | LCP1 | chr13 | - | 46158825 | 46158989 | 46156437 | 46156570 | 46159598 | 46159686 | 5689 | 3513 | 3534 | 4632 | 7789 | 0 | 0 | 0 | 0 | 3 | 7428 | 4433 | 3478 | 6662 | 2431 | 0 | 0 | 0 | 0 | 0 | 298 | 149 | 1 | 1 | 1 | 1 | 1 | 1 | 0.999 | 1 | 1 | 1 | 1 | 1 | 0 |
| SE | 101307 | ENSG00000136167 | LCP1 | chr13 | - | 46158825 | 46158989 | 46158521 | 46158651 | 46159598 | 46159686 | 10902 | 6748 | 6793 | 8886 | 17590 | 3 | 4 | 3 | 13 | 15 | 14663 | 8539 | 6800 | 12813 | 4825 | 35 | 13 | 32 | 75 | 15 | 298 | 149 | 1 | 1 | 0.999 | 0.999 | 0.999 | 0.997 | 0.998 | 0.995 | 0.997 | 0.991 | 0.988 | 0.994 | 0.005 |
| SE | 101308 | ENSG00000136167 | LCP1 | chr13 | - | 46159598 | 46159686 | 46158825 | 46158989 | 46182110 | 46182177 | 5950 | 3580 | 3730 | 4796 | 7922 | 0 | 0 | 1 | 0 | 0 | 7618 | 4504 | 3568 | 6847 | 2544 | 0 | 0 | 0 | 0 | 0 | 237 | 149 | 1 | 1 | 1 | 1 | 1 | 1 | 1 | 1 | 1 | 1 | 1 | 1 | 0 |
| SE | 101312 | ENSG00000244627 | TPTEP2 | chr22 | - | 38347488 | 38347585 | 38336607 | 38336706 | 38361398 | 38361560 | 0 | 0 | 3 | 1 | 0 | 1 | 0 | 4 | 8 | 0 | 0 | 0 | 0 | 0 | 4 | 0 | 0 | 10 | 12 | 5 | 246 | 149 | 1 | 1 | 0 | NA | 0.312 | 0.07 | NA | NA | NA | 0 | 0 | 0.326 | 0.019 |
| SE | 101313 | ENSG00000244627 | TPTEP2 | chr22 | - | 38347488 | 38347585 | 38344623 | 38344741 | 38361398 | 38361560 | 0 | 0 | 1 | 1 | 0 | 2 | 1 | 1 | 1 | 3 | 0 | 0 | 0 | 0 | 8 | 0 | 1 | 2 | 0 | 13 | 246 | 149 | 1 | 1 | 0 | 0 | 0.377 | 0.377 | 0 | NA | 0 | 0 | NA | 0.272 | 0.06 |
| SE | 101314 | ENSG00000244627 | TPTEP2 | chr22 | - | 38363053 | 38363104 | 38361380 | 38361560 | 38369985 | 38370045 | 0 | 0 | 0 | 0 | 0 | 2 | 0 | 0 | 1 | 0 | 0 | 0 | 0 | 0 | 1 | 0 | 2 | 0 | 1 | 0 | 200 | 149 | 1 | 1 | 0 | NA | NA | 0 | NA | NA | 0 | NA | 0 | 1 | -0.333 |
| SE | 101315 | ENSG00000244627 | TPTEP2 | chr22 | - | 38363053 | 38363104 | 38361380 | 38361560 | 38380369 | 38380423 | 0 | 1 | 0 | 0 | 0 | 13 | 3 | 14 | 20 | 12 | 0 | 0 | 3 | 0 | 0 | 0 | 3 | 5 | 12 | 12 | 200 | 149 | 1 | 1 | 0 | 0.199 | 0 | 0 | 0 | NA | 0 | 0.309 | 0 | 0 | -0.037 |
| SE | 101316 | ENSG00000244627 | TPTEP2 | chr22 | - | 38369985 | 38370045 | 38361398 | 38361560 | 38380369 | 38380423 | 5 | 0 | 0 | 1 | 6 | 13 | 3 | 14 | 20 | 12 | 0 | 5 | 2 | 9 | 0 | 0 | 3 | 5 | 12 | 12 | 209 | 149 | 1 | 1 | 0.215 | 0 | 0 | 0.034 | 0.263 | NA | 0.543 | 0.222 | 0.348 | 0 | -0.176 |
| SE | 101317 | ENSG00000244627 | TPTEP2 | chr22 | - | 38369985 | 38370045 | 38363053 | 38363104 | 38380369 | 38380423 | 3 | 0 | 0 | 0 | 6 | 0 | 2 | 0 | 0 | 0 | 0 | 3 | 2 | 8 | 1 | 0 | 0 | 3 | 0 | 0 | 209 | 149 | 1 | 1 | 1 | 0 | NA | NA | 1 | NA | 1 | 0.322 | 1 | 1 | -0.164 |
| SE | 101318 | ENSG00000244627 | TPTEP2 | chr22 | - | 38381930 | 38381984 | 38380936 | 38380990 | 38384049 | 38384096 | 0 | 4 | 1 | 0 | 0 | 1 | 3 | 2 | 3 | 2 | 0 | 0 | 0 | 0 | 0 | 0 | 8 | 3 | 9 | 5 | 203 | 149 | 1 | 1 | 0 | 0.495 | 0.268 | 0 | 0 | NA | 0 | 0 | 0 | 0 | 0.153 |
| SE | 101335 | ENSG00000287277 | NA | chr10 | - | 6864086 | 6864134 | 6848869 | 6850849 | 6935112 | 6935408 | 0 | 0 | 0 | 0 | 0 | 2 | 0 | 0 | 0 | 0 | 0 | 0 | 1 | 0 | 0 | 0 | 0 | 0 | 0 | 0 | 197 | 149 | 1 | 1 | 0 | NA | NA | NA | NA | NA | NA | 1 | NA | NA | -1 |
| SE | 101336 | ENSG00000287277 | NA | chr10 | - | 6920428 | 6920538 | 6848869 | 6850849 | 6935112 | 6935408 | 0 | 0 | 0 | 0 | 0 | 2 | 0 | 0 | 0 | 0 | 0 | 17 | 0 | 0 | 0 | 0 | 0 | 0 | 0 | 0 | 259 | 149 | 1 | 1 | 0 | NA | NA | NA | NA | NA | 1 | NA | NA | NA | -1 |
| SE | 101338 | ENSG00000124795 | DEK | chr6 | - | 18226173 | 18226242 | 18223859 | 18225730 | 18236451 | 18236596 | 604 | 451 | 354 | 586 | 994 | 2 | 0 | 4 | 0 | 1 | 109 | 230 | 190 | 70 | 11 | 0 | 0 | 0 | 0 | 0 | 218 | 149 | 1 | 1 | 0.995 | 1 | 0.984 | 1 | 0.999 | 1 | 1 | 1 | 1 | 1 | -0.004 |
| SE | 101339 | ENSG00000124795 | DEK | chr6 | - | 18236451 | 18236596 | 18226173 | 18226242 | 18237380 | 18237516 | 556 | 424 | 338 | 532 | 829 | 4 | 1 | 1 | 0 | 1 | 109 | 194 | 177 | 62 | 11 | 0 | 0 | 0 | 0 | 0 | 294 | 149 | 1 | 1 | 0.986 | 0.995 | 0.994 | 1 | 0.998 | 1 | 1 | 1 | 1 | 1 | -0.005 |
| SE | 101340 | ENSG00000124795 | DEK | chr6 | - | 18236451 | 18236600 | 18224190 | 18226242 | 18237380 | 18237516 | 1177 | 901 | 750 | 1214 | 1701 | 4 | 1 | 1 | 0 | 1 | 294 | 487 | 397 | 179 | 115 | 0 | 0 | 0 | 0 | 0 | 298 | 149 | 1 | 1 | 0.993 | 0.998 | 0.997 | 1 | 0.999 | 1 | 1 | 1 | 1 | 1 | -0.003 |
| SE | 101341 | ENSG00000124795 | DEK | chr6 | - | 18237380 | 18237516 | 18224190 | 18226242 | 18249650 | 18249839 | 215 | 185 | 152 | 219 | 305 | 0 | 0 | 1 | 0 | 1 | 33 | 66 | 53 | 28 | 41 | 0 | 0 | 0 | 0 | 0 | 285 | 149 | 1 | 1 | 1 | 1 | 0.988 | 1 | 0.994 | 1 | 1 | 1 | 1 | 1 | -0.004 |
| SE | 101343 | ENSG00000124795 | DEK | chr6 | - | 18249650 | 18249839 | 18237049 | 18237516 | 18255730 | 18255851 | 789 | 735 | 549 | 810 | 1648 | 1 | 0 | 0 | 0 | 0 | 143 | 323 | 297 | 170 | 131 | 0 | 0 | 0 | 0 | 0 | 298 | 149 | 1 | 1 | 0.997 | 1 | 1 | 1 | 1 | 1 | 1 | 1 | 1 | 1 | -0.001 |
| SE | 101344 | ENSG00000124795 | DEK | chr6 | - | 18255730 | 18255851 | 18249650 | 18249839 | 18256360 | 18256374 | 922 | 893 | 631 | 963 | 1981 | 1 | 1 | 1 | 5 | 8 | 207 | 369 | 467 | 229 | 120 | 0 | 0 | 0 | 1 | 0 | 270 | 149 | 1 | 1 | 0.998 | 0.998 | 0.997 | 0.991 | 0.993 | 1 | 1 | 1 | 0.992 | 1 | -0.003 |
| SE | 101345 | ENSG00000124795 | DEK | chr6 | - | 18255730 | 18255851 | 18249650 | 18249839 | 18257952 | 18258031 | 513 | 491 | 352 | 492 | 1188 | 2 | 0 | 0 | 1 | 0 | 101 | 216 | 221 | 135 | 68 | 0 | 0 | 0 | 0 | 0 | 270 | 149 | 1 | 1 | 0.993 | 1 | 1 | 0.996 | 1 | 1 | 1 | 1 | 1 | 1 | -0.002 |
| SE | 101346 | ENSG00000124795 | DEK | chr6 | - | 18256360 | 18256455 | 18249650 | 18249839 | 18257952 | 18258031 | 422 | 456 | 282 | 398 | 788 | 2 | 0 | 0 | 1 | 0 | 91 | 183 | 241 | 121 | 104 | 0 | 0 | 0 | 0 | 0 | 244 | 149 | 1 | 1 | 0.992 | 1 | 1 | 0.996 | 1 | 1 | 1 | 1 | 1 | 1 | -0.002 |
| SE | 101347 | ENSG00000124795 | DEK | chr6 | - | 18256360 | 18256455 | 18255730 | 18255851 | 18257952 | 18258031 | 827 | 861 | 561 | 864 | 1574 | 0 | 0 | 0 | 0 | 0 | 197 | 338 | 495 | 218 | 156 | 0 | 0 | 4 | 0 | 0 | 244 | 149 | 1 | 1 | 1 | 1 | 1 | 1 | 1 | 1 | 1 | 0.987 | 1 | 1 | 0.003 |
| SE | 101348 | ENSG00000124795 | DEK | chr6 | - | 18257952 | 18258062 | 18256360 | 18256455 | 18258303 | 18258405 | 525 | 586 | 368 | 578 | 1081 | 1 | 0 | 0 | 0 | 0 | 144 | 258 | 383 | 185 | 144 | 0 | 0 | 0 | 0 | 0 | 259 | 149 | 1 | 1 | 0.997 | 1 | 1 | 1 | 1 | 1 | 1 | 1 | 1 | 1 | -0.001 |
| SE | 101349 | ENSG00000124795 | DEK | chr6 | - | 18257952 | 18258062 | 18256360 | 18256455 | 18263842 | 18263996 | 412 | 457 | 282 | 426 | 884 | 0 | 0 | 1 | 0 | 0 | 104 | 181 | 228 | 123 | 100 | 0 | 0 | 0 | 0 | 0 | 259 | 149 | 1 | 1 | 1 | 1 | 0.994 | 1 | 1 | 1 | 1 | 1 | 1 | 1 | -0.001 |
| SE | 101350 | ENSG00000124795 | DEK | chr6 | - | 18258303 | 18258405 | 18256360 | 18256455 | 18263842 | 18263996 | 83 | 61 | 46 | 51 | 61 | 0 | 0 | 1 | 0 | 0 | 2 | 37 | 26 | 19 | 40 | 0 | 0 | 0 | 0 | 0 | 251 | 149 | 1 | 1 | 1 | 1 | 0.965 | 1 | 1 | 1 | 1 | 1 | 1 | 1 | -0.007 |
| SE | 101358 | ENSG00000170525 | PFKFB3 | chr10 | + | 6219568 | 6219693 | 6217134 | 6217191 | 6220657 | 6220865 | 117 | 55 | 110 | 104 | 87 | 0 | 0 | 0 | 0 | 0 | 176 | 217 | 626 | 271 | 358 | 0 | 2 | 1 | 0 | 0 | 274 | 149 | 1 | 1 | 1 | 1 | 1 | 1 | 1 | 1 | 0.983 | 0.997 | 1 | 1 | 0.004 |
| SE | 101359 | ENSG00000170525 | PFKFB3 | chr10 | + | 6221640 | 6221745 | 6221380 | 6221527 | 6222854 | 6222984 | 56 | 18 | 45 | 71 | 37 | 0 | 0 | 0 | 0 | 0 | 284 | 120 | 348 | 270 | 251 | 0 | 0 | 0 | 4 | 0 | 254 | 149 | 1 | 1 | 1 | 1 | 1 | 1 | 1 | 1 | 1 | 1 | 0.975 | 1 | 0.005 |
| SE | 101361 | ENSG00000170525 | PFKFB3 | chr10 | + | 6224505 | 6224623 | 6224165 | 6224213 | 6226191 | 6226365 | 0 | 0 | 0 | 6 | 0 | 38 | 10 | 26 | 30 | 14 | 0 | 0 | 4 | 8 | 0 | 91 | 85 | 214 | 117 | 122 | 267 | 149 | 1 | 1 | 0 | 0 | 0 | 0.1 | 0 | 0 | 0 | 0.01 | 0.037 | 0 | 0.011 |
| SE | 101364 | ENSG00000170525 | PFKFB3 | chr10 | + | 6226191 | 6226365 | 6224148 | 6224213 | 6254177 | 6254644 | 38 | 10 | 26 | 30 | 14 | 0 | 0 | 0 | 0 | 0 | 91 | 85 | 216 | 117 | 122 | 0 | 0 | 1 | 0 | 0 | 298 | 149 | 1 | 1 | 1 | 1 | 1 | 1 | 1 | 1 | 1 | 0.991 | 1 | 1 | 0.002 |
| SE | 101368 | ENSG00000107537 | PHYH | chr10 | - | 13298186 | 13298245 | 13295227 | 13295606 | 13299419 | 13299549 | 9 | 9 | 4 | 4 | 14 | 1 | 0 | 0 | 0 | 0 | 13 | 0 | 10 | 12 | 0 | 0 | 0 | 0 | 0 | 0 | 208 | 149 | 1 | 1 | 0.866 | 1 | 1 | 1 | 1 | 1 | NA | 1 | 1 | NA | -0.027 |
| SE | 101379 | ENSG00000151657 | KIN | chr10 | - | 7753951 | 7754069 | 7753224 | 7753602 | 7759889 | 7759990 | 21 | 9 | 14 | 17 | 4 | 0 | 0 | 0 | 0 | 4 | 0 | 9 | 9 | 4 | 0 | 0 | 0 | 0 | 0 | 0 | 267 | 149 | 1 | 1 | 1 | 1 | 1 | 1 | 0.358 | NA | 1 | 1 | 1 | NA | -0.128 |
| SE | 101380 | ENSG00000151657 | KIN | chr10 | - | 7769215 | 7769345 | 7766052 | 7766103 | 7774830 | 7774878 | 133 | 104 | 122 | 99 | 199 | 1 | 0 | 0 | 0 | 0 | 10 | 77 | 104 | 80 | 18 | 0 | 0 | 0 | 0 | 0 | 279 | 149 | 1 | 1 | 0.986 | 1 | 1 | 1 | 1 | 1 | 1 | 1 | 1 | 1 | -0.003 |
| SE | 101382 | ENSG00000151657 | KIN | chr10 | - | 7775750 | 7775799 | 7774830 | 7774891 | 7778837 | 7779019 | 89 | 62 | 79 | 69 | 129 | 0 | 2 | 0 | 0 | 0 | 9 | 59 | 85 | 45 | 20 | 0 | 0 | 1 | 0 | 0 | 198 | 149 | 1 | 1 | 1 | 0.959 | 1 | 1 | 1 | 1 | 1 | 0.985 | 1 | 1 | -0.005 |
| SE | 101383 | ENSG00000151657 | KIN | chr10 | - | 7775750 | 7775799 | 7774830 | 7774891 | 7780055 | 7780178 | 18 | 10 | 17 | 20 | 44 | 0 | 0 | 0 | 0 | 0 | 0 | 21 | 27 | 6 | 0 | 0 | 0 | 0 | 1 | 0 | 198 | 149 | 1 | 1 | 1 | 1 | 1 | 1 | 1 | NA | 1 | 1 | 0.819 | NA | 0.06 |
| SE | 101384 | ENSG00000151657 | KIN | chr10 | - | 7778837 | 7779019 | 7774830 | 7774891 | 7780055 | 7780178 | 29 | 12 | 23 | 19 | 12 | 0 | 0 | 0 | 0 | 0 | 11 | 17 | 23 | 20 | 0 | 0 | 0 | 0 | 1 | 0 | 298 | 149 | 1 | 1 | 1 | 1 | 1 | 1 | 1 | 1 | 1 | 1 | 0.909 | NA | 0.023 |
| SE | 101385 | ENSG00000151657 | KIN | chr10 | - | 7778837 | 7779019 | 7775750 | 7775799 | 7780055 | 7780178 | 102 | 62 | 85 | 68 | 98 | 0 | 0 | 0 | 0 | 0 | 20 | 55 | 84 | 59 | 20 | 0 | 0 | 4 | 0 | 0 | 298 | 149 | 1 | 1 | 1 | 1 | 1 | 1 | 1 | 1 | 1 | 0.913 | 1 | 1 | 0.017 |
| SE | 101394 | ENSG00000131759 | RARA | chr17 | + | 40326474 | 40326551 | 40309191 | 40309286 | 40330856 | 40331396 | 1 | 4 | 1 | 3 | 3 | 1 | 0 | 2 | 1 | 3 | 5 | 0 | 0 | 6 | 0 | 0 | 0 | 0 | 2 | 0 | 226 | 149 | 1 | 1 | 0.397 | 1 | 0.248 | 0.664 | 0.397 | 1 | NA | NA | 0.664 | NA | -0.291 |
| SE | 101396 | ENSG00000131759 | RARA | chr17 | + | 40342795 | 40342867 | 40342398 | 40342706 | 40348315 | 40348464 | 1 | 2 | 11 | 10 | 11 | 0 | 0 | 0 | 0 | 0 | 0 | 0 | 4 | 3 | 9 | 2 | 0 | 0 | 0 | 0 | 221 | 149 | 1 | 1 | 1 | 1 | 1 | 1 | 1 | 0 | NA | 1 | 1 | 1 | 0.25 |
| SE | 101397 | ENSG00000134461 | ANKRD16 | chr10 | - | 5880297 | 5880376 | 5878086 | 5878287 | 5883005 | 5883167 | 23 | 7 | 35 | 17 | 20 | 2 | 0 | 0 | 0 | 0 | 36 | 36 | 8 | 46 | 18 | 0 | 0 | 0 | 0 | 0 | 228 | 149 | 1 | 1 | 0.883 | 1 | 1 | 1 | 1 | 1 | 1 | 1 | 1 | 1 | -0.023 |
| SE | 101400 | ENSG00000134470 | IL15RA | chr10 | - | 5960366 | 5960468 | 5959753 | 5959786 | 5963742 | 5963841 | 19 | 10 | 9 | 9 | 5 | 0 | 0 | 0 | 1 | 0 | 55 | 7 | 19 | 35 | 34 | 0 | 0 | 0 | 0 | 0 | 251 | 149 | 1 | 1 | 1 | 1 | 1 | 0.842 | 1 | 1 | 1 | 1 | 1 | 1 | -0.032 |
| SE | 101401 | ENSG00000134470 | IL15RA | chr10 | - | 5960366 | 5960567 | 5959753 | 5959786 | 5963742 | 5963832 | 29 | 15 | 13 | 9 | 5 | 0 | 0 | 0 | 1 | 0 | 55 | 7 | 33 | 39 | 35 | 0 | 0 | 0 | 0 | 0 | 298 | 149 | 1 | 1 | 1 | 1 | 1 | 0.818 | 1 | 1 | 1 | 1 | 1 | 1 | -0.036 |
| SE | 101402 | ENSG00000134470 | IL15RA | chr10 | - | 5963742 | 5963838 | 5960394 | 5960567 | 5966144 | 5966339 | 6 | 2 | 4 | 0 | 0 | 3 | 0 | 8 | 2 | 7 | 0 | 0 | 4 | 4 | 1 | 0 | 3 | 0 | 1 | 6 | 245 | 149 | 1 | 1 | 0.549 | 1 | 0.233 | 0 | 0 | NA | 0 | 1 | 0.709 | 0.092 | -0.094 |
| SE | 101405 | ENSG00000134470 | IL15RA | chr10 | - | 5966144 | 5966339 | 5960366 | 5960567 | 5977404 | 5977492 | 3 | 0 | 11 | 2 | 7 | 0 | 1 | 0 | 0 | 0 | 0 | 6 | 0 | 2 | 6 | 0 | 0 | 1 | 0 | 0 | 298 | 149 | 1 | 1 | 1 | 0 | 1 | 1 | 1 | NA | 1 | 0 | 1 | 1 | 0.05 |
| SE | 101406 | ENSG00000134470 | IL15RA | chr10 | - | 5966144 | 5966339 | 5960366 | 5960567 | 5977776 | 5978038 | 6 | 0 | 9 | 5 | 7 | 0 | 0 | 0 | 2 | 0 | 0 | 3 | 0 | 1 | 6 | 0 | 0 | 0 | 0 | 6 | 298 | 149 | 1 | 1 | 1 | NA | 1 | 0.556 | 1 | NA | 1 | NA | 1 | 0.333 | 0.111 |
| SE | 101407 | ENSG00000134470 | IL15RA | chr10 | - | 5968593 | 5968703 | 5966144 | 5966339 | 5968775 | 5968928 | 1 | 0 | 0 | 0 | 0 | 2 | 0 | 1 | 2 | 0 | 0 | 0 | 1 | 0 | 0 | 0 | 0 | 0 | 0 | 0 | 259 | 149 | 1 | 1 | 0.223 | NA | 0 | 0 | NA | NA | NA | 1 | NA | NA | -0.926 |
| SE | 101408 | ENSG00000134470 | IL15RA | chr10 | - | 5968775 | 5968928 | 5966144 | 5966339 | 5977404 | 5977611 | 2 | 0 | 1 | 2 | 0 | 0 | 0 | 3 | 0 | 0 | 0 | 0 | 0 | 0 | 0 | 0 | 3 | 0 | 1 | 0 | 298 | 149 | 1 | 1 | 1 | NA | 0.143 | 1 | NA | NA | 0 | NA | 0 | NA | 0.714 |
| SE | 101411 | ENSG00000152455 | SUV39H2 | chr10 | + | 14881499 | 14881645 | 14878862 | 14878919 | 14896845 | 14896977 | 1 | 1 | 4 | 0 | 7 | 0 | 0 | 0 | 1 | 5 | 15 | 0 | 0 | 0 | 0 | 0 | 0 | 6 | 0 | 0 | 295 | 149 | 1 | 1 | 1 | 1 | 1 | 0 | 0.414 | 1 | NA | 0 | NA | NA | 0.183 |
| SE | 101415 | ENSG00000204920 | ZNF155 | chr19 | + | 43990023 | 43990112 | 43988458 | 43988558 | 43991547 | 43991674 | 4 | 0 | 3 | 0 | 0 | 9 | 2 | 6 | 2 | 11 | 0 | 1 | 0 | 0 | 14 | 0 | 5 | 14 | 4 | 0 | 238 | 149 | 1 | 1 | 0.218 | 0 | 0.238 | 0 | 0 | NA | 0.111 | 0 | 0 | 1 | -0.187 |
| SE | 101416 | ENSG00000158545 | ZC3H18 | chr16 | + | 88577226 | 88586684 | 88577109 | 88577726 | 88622196 | 88622388 | 17 | 12 | 11 | 15 | 10 | 0 | 0 | 0 | 2 | 0 | 0 | 24 | 17 | 18 | 24 | 0 | 0 | 0 | 0 | 0 | 234 | 149 | 1 | 1 | 1 | 1 | 1 | 0.827 | 1 | NA | 1 | 1 | 1 | 1 | -0.035 |
| SE | 101418 | ENSG00000158545 | ZC3H18 | chr16 | + | 88598177 | 88598326 | 88586599 | 88586684 | 88598619 | 88598712 | 217 | 167 | 214 | 199 | 429 | 0 | 0 | 1 | 0 | 0 | 242 | 229 | 170 | 170 | 249 | 0 | 0 | 0 | 0 | 0 | 298 | 149 | 1 | 1 | 1 | 1 | 0.991 | 1 | 1 | 1 | 1 | 1 | 1 | 1 | -0.002 |
| SE | 101419 | ENSG00000158545 | ZC3H18 | chr16 | + | 88608933 | 88609051 | 88586599 | 88586684 | 88622196 | 88622388 | 8 | 5 | 0 | 4 | 0 | 0 | 0 | 0 | 0 | 0 | 64 | 53 | 49 | 67 | 43 | 0 | 0 | 0 | 0 | 7 | 267 | 149 | 1 | 1 | 1 | 1 | NA | 1 | NA | 1 | 1 | 1 | 1 | 0.774 | 0.045 |
| SE | 101420 | ENSG00000158545 | ZC3H18 | chr16 | + | 88608933 | 88609051 | 88599790 | 88599948 | 88611267 | 88611536 | 373 | 242 | 273 | 273 | 385 | 2 | 0 | 0 | 0 | 0 | 477 | 328 | 173 | 384 | 604 | 0 | 0 | 3 | 0 | 0 | 267 | 149 | 1 | 1 | 0.99 | 1 | 1 | 1 | 1 | 1 | 1 | 0.97 | 1 | 1 | 0.004 |
| SE | 101426 | ENSG00000152464 | RPP38 | chr10 | + | 15102217 | 15102336 | 15097179 | 15097409 | 15103304 | 15103713 | 8 | 1 | 8 | 4 | 2 | 11 | 6 | 3 | 5 | 5 | 0 | 3 | 7 | 0 | 10 | 0 | 7 | 4 | 9 | 0 | 268 | 149 | 1 | 1 | 0.288 | 0.085 | 0.597 | 0.308 | 0.182 | NA | 0.192 | 0.493 | 0 | 1 | -0.129 |
| SE | 101427 | ENSG00000152464 | RPP38 | chr10 | + | 15102217 | 15102336 | 15097354 | 15097766 | 15103304 | 15103713 | 5 | 4 | 4 | 6 | 3 | 5 | 6 | 3 | 2 | 4 | 0 | 0 | 8 | 0 | 9 | 0 | 1 | 3 | 8 | 7 | 268 | 149 | 1 | 1 | 0.357 | 0.27 | 0.426 | 0.625 | 0.294 | NA | 0 | 0.597 | 0 | 0.417 | 0.141 |
| SE | 101434 | ENSG00000232973 | CYP1B1-AS1 | chr2 | + | 38121984 | 38122225 | 38076265 | 38076560 | 38131510 | 38131561 | 2 | 1 | 0 | 4 | 2 | 0 | 0 | 0 | 0 | 0 | 0 | 9 | 2 | 4 | 0 | 0 | 3 | 0 | 0 | 0 | 298 | 149 | 1 | 1 | 1 | 1 | NA | 1 | 1 | NA | 0.6 | 1 | 1 | NA | 0.133 |
| SE | 101437 | ENSG00000264006 | AKR1C8P | chr10 | + | 5158599 | 5158709 | 5157634 | 5157800 | 5159882 | 5160005 | 0 | 9 | 2 | 0 | 11 | 0 | 0 | 1 | 0 | 0 | 0 | 6 | 0 | 0 | 6 | 0 | 0 | 0 | 0 | 0 | 259 | 149 | 1 | 1 | NA | 1 | 0.535 | NA | 1 | NA | 1 | NA | NA | 1 | -0.155 |
| SE | 101441 | ENSG00000165997 | ARL5B | chr10 | + | 18668529 | 18668677 | 18666574 | 18666635 | 18672621 | 18672705 | 88 | 88 | 73 | 102 | 210 | 0 | 0 | 0 | 0 | 1 | 58 | 33 | 56 | 34 | 27 | 0 | 0 | 1 | 0 | 0 | 297 | 149 | 1 | 1 | 1 | 1 | 1 | 1 | 0.991 | 1 | 1 | 0.966 | 1 | 1 | 0.005 |
| SE | 101446 | ENSG00000107611 | CUBN | chr10 | - | 16890370 | 16890527 | 16888416 | 16888566 | 16898995 | 16899183 | 23 | 8 | 6 | 13 | 60 | 1 | 0 | 0 | 2 | 0 | 0 | 3 | 8 | 4 | 0 | 0 | 0 | 0 | 0 | 0 | 298 | 149 | 1 | 1 | 0.92 | 1 | 1 | 0.765 | 1 | NA | 1 | 1 | 1 | NA | -0.063 |
| SE | 101447 | ENSG00000107611 | CUBN | chr10 | - | 16938962 | 16939147 | 16937591 | 16937784 | 16940031 | 16940237 | 6 | 5 | 9 | 9 | 11 | 0 | 0 | 0 | 0 | 0 | 24 | 19 | 4 | 1 | 0 | 0 | 0 | 0 | 2 | 0 | 298 | 149 | 1 | 1 | 1 | 1 | 1 | 1 | 1 | 1 | 1 | 1 | 0.2 | NA | 0.2 |
| SE | 101448 | ENSG00000107611 | CUBN | chr10 | - | 17019832 | 17019983 | 16990333 | 16990515 | 17041032 | 17041220 | 12 | 5 | 13 | 15 | 20 | 0 | 0 | 0 | 1 | 0 | 0 | 10 | 6 | 0 | 0 | 0 | 0 | 0 | 0 | 0 | 298 | 149 | 1 | 1 | 1 | 1 | 1 | 0.882 | 1 | NA | 1 | 1 | NA | NA | -0.024 |
| SE | 101449 | ENSG00000107611 | CUBN | chr10 | - | 17045006 | 17045188 | 17043826 | 17043983 | 17045933 | 17046094 | 27 | 11 | 23 | 19 | 17 | 0 | 2 | 0 | 0 | 0 | 0 | 8 | 8 | 0 | 0 | 0 | 0 | 0 | 0 | 0 | 298 | 149 | 1 | 1 | 1 | 0.733 | 1 | 1 | 1 | NA | 1 | 1 | NA | NA | -0.053 |

[illegible]

| | | | | | | | | | | | | | | |
|---|---|---|---|---|---|---|---|---|---|---|---|---|---|---|
| SE | 101476 | ENSG00000165309 | ARMC3 | chr10 | + | 22961883 | 22962078 | 22959398 | 22959574 | 22968305 | 22968489 | 7 | 7 | [illegible] |
| SE | 101477 | ENSG00000165309 | ARMC3 | chr10 | + | 22961883 | 22962078 | 22959398 | 22959574 | 22981339 | 22981492 | 7 | 6 | [illegible] |
| SE | 101478 | ENSG00000165309 | ARMC3 | chr10 | + | 22961938 | 22962078 | 22959398 | 22959574 | 22968305 | 22968489 | 0 | 2 | [illegible] |
| SE | 101479 | ENSG00000165309 | ARMC3 | chr10 | + | 22961938 | 22962078 | 22959398 | 22959574 | 22981339 | 22981492 | 0 | 1 | [illegible] |
| SE | 101480 | ENSG00000165309 | ARMC3 | chr10 | + | 22968305 | 22968489 | 22961883 | 22962078 | 22981339 | 22981453 | 1 | 12 | [illegible] |
| SE | 101481 | ENSG00000165309 | ARMC3 | chr10 | + | 22981339 | 22981492 | 22959398 | 22959574 | 22981594 | 22981608 | 2 | 10 | [illegible] |
| SE | 101482 | ENSG00000165309 | ARMC3 | chr10 | + | 22981339 | 22981492 | 22968305 | 22968489 | 22981594 | 22981608 | 3 | 20 | [illegible] |
| SE | 101483 | ENSG00000165309 | ARMC3 | chr10 | + | 23006883 | 23006981 | 23003245 | 23003414 | 23008275 | 23008374 | 1 | 6 | [illegible] |
| SE | 101485 | ENSG00000095794 | CREM | chr10 | + | 35137781 | 35137879 | 35126809 | 35127193 | 35148367 | 35148491 | 1 | 0 | [illegible] |
| SE | 101486 | ENSG00000095794 | CREM | chr10 | + | 35137781 | 35137879 | 35126809 | 35127193 | 35178888 | 35178986 | 0 | 2 | [illegible] |
| SE | 101487 | ENSG00000095794 | CREM | chr10 | + | 35137781 | 35137879 | 35127456 | 35127567 | 35148367 | 35148491 | 1 | 0 | [illegible] |
| SE | 101490 | ENSG00000095794 | CREM | chr10 | + | 35148367 | 35148491 | 35137781 | 35137879 | 35178888 | 35178986 | 12 | 14 | [illegible] |
| SE | 101493 | ENSG00000095794 | CREM | chr10 | + | 35178871 | 35178986 | 35148367 | 35148491 | 35179133 | 35179276 | 21 | 22 | [illegible] |
| SE | 101495 | ENSG00000095794 | CREM | chr10 | + | 35178888 | 35178986 | 35148367 | 35148491 | 35179133 | 35179276 | 33 | 36 | [illegible] |
| SE | 101497 | ENSG00000095794 | CREM | chr10 | + | 35179133 | 35179276 | 35148367 | 35148491 | 35188199 | 35188260 | 10 | 3 | [illegible] |
| SE | 101498 | ENSG00000095794 | CREM | chr10 | + | 35179133 | 35179276 | 35148367 | 35148491 | 35206894 | 35207051 | 14 | 17 | [illegible] |
| SE | 101500 | ENSG00000095794 | CREM | chr10 | + | 35188199 | 35188388 | 35148367 | 35148491 | 35206894 | 35207006 | 10 | 5 | [illegible] |
| SE | 101501 | ENSG00000095794 | CREM | chr10 | + | 35188199 | 35188388 | 35179133 | 35179276 | 35201450 | 35201486 | 13 | 3 | [illegible] |
| SE | 101505 | ENSG00000095794 | CREM | chr10 | + | 35201450 | 35201486 | 35188199 | 35188388 | 35206894 | 35207051 | 6 | 0 | [illegible] |
| SE | 101507 | ENSG00000095794 | CREM | chr10 | + | 35201450 | 35201486 | 35195864 | 35196142 | 35206894 | 35207051 | 4 | 2 | [illegible] |
| SE | 101508 | ENSG00000095794 | CREM | chr10 | + | 35201450 | 35201486 | 35195908 | 35196099 | 35206894 | 35207051 | 3 | 0 | [illegible] |
| SE | 101525 | ENSG00000241058 | NSUN6 | chr10 | - | 18609844 | 18609926 | 18596207 | 18596327 | 18614459 | 18614613 | 168 | 166 | [illegible] |
| SE | 101526 | ENSG00000241058 | NSUN6 | chr10 | - | 18609844 | 18609926 | 18596207 | 18596327 | 18616183 | 18616293 | 76 | 85 | [illegible] |
| SE | 101527 | ENSG00000241058 | NSUN6 | chr10 | - | 18614459 | 18614613 | 18596207 | 18596327 | 18616183 | 18616293 | 88 | 90 | [illegible] |
| SE | 101528 | ENSG00000241058 | NSUN6 | chr10 | - | 18614459 | 18614613 | 18609844 | 18609926 | 18616183 | 18616293 | 173 | 169 | [illegible] |
| SE | 101529 | ENSG00000241058 | NSUN6 | chr10 | - | 18616183 | 18616293 | 18596207 | 18596327 | 18648489 | 18648645 | 6 | 0 | [illegible] |
| SE | 101531 | ENSG00000241058 | NSUN6 | chr10 | - | 18648489 | 18648645 | 18616183 | 18616293 | 18651128 | 18651587 | 63 | 77 | [illegible] |
| SE | 101532 | ENSG00000241058 | NSUN6 | chr10 | - | 18658218 | 18658497 | 18654863 | 18655175 | 18659192 | 18659238 | 2 | 1 | [illegible] |
| SE | 101534 | ENSG00000233673 | ANAPC1P1 | chr2 | + | 86886583 | 86886729 | 86883355 | 86883429 | 86887772 | 86887993 | 3 | 22 | [illegible] |
| SE | 101535 | ENSG00000233673 | ANAPC1P1 | chr2 | + | 86893087 | 86893181 | 86889066 | 86889276 | 86893280 | 86893375 | 1 | 1 | [illegible] |
| SE | 101547 | ENSG00000088298 | EDEM2 | chr20 | - | 35118597 | 35118719 | 35115363 | 35115933 | 35126250 | 35126375 | 75 | 37 | [illegible] |
| SE | 101548 | ENSG00000088298 | EDEM2 | chr20 | - | 35123889 | 35124034 | 35115363 | 35115933 | 35126250 | 35126375 | 99 | 23 | [illegible] |
| SE | 101549 | ENSG00000088298 | EDEM2 | chr20 | - | 35123889 | 35124034 | 35118597 | 35118719 | 35126250 | 35126375 | 146 | 42 | [illegible] |
| SE | 101550 | ENSG00000088298 | EDEM2 | chr20 | - | 35126250 | 35126375 | 35123889 | 35124034 | 35131641 | 35131783 | 167 | 47 | [illegible] |
| SE | 101551 | ENSG00000088298 | EDEM2 | chr20 | - | 35126250 | 35126375 | 35123889 | 35124034 | 35137879 | 35138005 | 96 | 21 | [illegible] |
| SE | 101552 | ENSG00000088298 | EDEM2 | chr20 | - | 35131641 | 35131783 | 35123889 | 35124034 | 35137879 | 35138005 | 2 | 0 | [illegible] |
| SE | 101553 | ENSG00000088298 | EDEM2 | chr20 | - | 35131641 | 35131783 | 35126250 | 35126375 | 35134737 | 35134949 | 127 | 52 | [illegible] |
| SE | 101557 | ENSG00000088298 | EDEM2 | chr20 | - | 35134737 | 35134949 | 35126250 | 35126375 | 35142372 | 35142478 | 0 | 0 | [illegible] |
| SE | 101559 | ENSG00000088298 | EDEM2 | chr20 | - | 35134737 | 35134949 | 35131641 | 35131783 | 35142372 | 35142478 | 51 | 23 | [illegible] |
| SE | 101561 | ENSG00000088298 | EDEM2 | chr20 | - | 35137879 | 35138005 | 35131641 | 35131783 | 35142372 | 35142478 | 34 | 20 | [illegible] |
| SE | 101562 | ENSG00000088298 | EDEM2 | chr20 | - | 35137879 | 35138005 | 35134737 | 35134949 | 35142372 | 35142478 | 65 | 48 | [illegible] |
| SE | 101563 | ENSG00000088298 | EDEM2 | chr20 | - | 35142372 | 35142478 | 35137879 | 35138005 | 35144978 | 35145018 | 88 | 36 | [illegible] |
| SE | 101567 | ENSG00000088298 | EDEM2 | chr20 | - | 35146824 | 35146935 | 35144978 | 35145018 | 35147151 | 35147336 | 56 | 16 | [illegible] |
| SE | 101568 | ENSG00000233968 | NA | chr10 | - | 19724926 | 19724981 | 19722329 | 19722422 | 19728404 | 19728550 | 3 | 3 | [illegible] |
| SE | 101569 | ENSG00000233968 | NA | chr10 | - | 19724926 | 19724981 | 19722329 | 19722588 | 19728404 | 19728550 | 1 | 7 | [illegible] |
| SE | 101570 | ENSG00000133114 | GPALPP1 | chr13 | + | 45004304 | 45004437 | 44989537 | 44989742 | 45006201 | 45006303 | 162 | 128 | [illegible] |
| SE | 101571 | ENSG00000133114 | GPALPP1 | chr13 | + | 45004310 | 45004437 | 44989563 | 44989742 | 45006201 | 45006303 | 74 | 65 | [illegible] |
| SE | 101572 | ENSG00000133114 | GPALPP1 | chr13 | + | 45006201 | 45006303 | 44989537 | 44989742 | 45008794 | 45008879 | 84 | 61 | [illegible] |
| SE | 101573 | ENSG00000133114 | GPALPP1 | chr13 | + | 45029238 | 45032107 | 45027784 | 45028003 | 45033235 | 45033602 | 42 | 29 | [illegible] |
| SE | 101586 | ENSG00000171160 | MORN4 | chr10 | - | 97616677 | 97616787 | 97614554 | 97616411 | 97617207 | 97617322 | 0 | 0 | [illegible] |
| SE | 101588 | ENSG00000172671 | ZFAND4 | chr10 | - | 45624582 | 45624637 | 45618139 | 45618260 | 45625950 | 45627105 | 30 | 7 | [illegible] |
| SE | 101590 | ENSG00000172671 | ZFAND4 | chr10 | - | 45625950 | 45627105 | 45618139 | 45618260 | 45648293 | 45648534 | 7 | 2 | [illegible] |
| SE | 101591 | ENSG00000172671 | ZFAND4 | chr10 | - | 45639815 | 45639963 | 45618139 | 45618260 | 45648293 | 45648534 | 32 | 21 | [illegible] |
| SE | 101595 | ENSG00000172671 | ZFAND4 | chr10 | - | 45651965 | 45652033 | 45648293 | 45648534 | 45663541 | 45663842 | 34 | 13 | [illegible] |
| SE | 101596 | ENSG00000172671 | ZFAND4 | chr10 | - | 45651965 | 45652033 | 45650124 | 45651625 | 45652983 | 45653059 | 37 | 39 | [illegible] |
| SE | 101598 | ENSG00000172671 | ZFAND4 | chr10 | - | 45652983 | 45653059 | 45651965 | 45652033 | 45662579 | 45662681 | 29 | 14 | [illegible] |
| SE | 101602 | ENSG00000172671 | ZFAND4 | chr10 | - | 45663541 | 45663842 | 45652983 | 45653059 | 45672249 | 45672275 | 80 | 67 | [illegible] |
| SE | 101606 | ENSG00000011478 | QPCTL | chr19 | + | 45698546 | 45698699 | 45695400 | 45695718 | 45701797 | 45701914 | 14 | 4 | [illegible] |
| SE | 101607 | ENSG00000011478 | QPCTL | chr19 | + | 45698781 | 45698900 | 45698479 | 45698699 | 45701797 | 45701823 | 10 | 0 | [illegible] |
| SE | 101608 | ENSG00000011478 | QPCTL | chr19 | + | 45698800 | 45698900 | 45698546 | 45698699 | 45701797 | 45701914 | 12 | 1 | [illegible] |
| SE | 101609 | ENSG00000143702 | CEP170 | chr1 | - | 243126377 | 243126494 | 243125917 | 243126134 | 243126612 | 243126698 | 0 | 0 | [illegible] |
| SE | 101610 | ENSG00000143702 | CEP170 | chr1 | - | 243142285 | 243142463 | 243139936 | 243140107 | 243156220 | 243156347 | 46 | 32 | [illegible] |
| SE | 101611 | ENSG00000143702 | CEP170 | chr1 | - | 243142285 | 243142463 | 243139936 | 243140107 | 243164283 | 243166116 | 4 | 3 | [illegible] |
| SE | 101612 | ENSG00000143702 | CEP170 | chr1 | - | 243142315 | 243142463 | 243139936 | 243140107 | 243156220 | 243156455 | 56 | 42 | [illegible] |
| SE | 101613 | ENSG00000143702 | CEP170 | chr1 | - | 243142315 | 243142463 | 243139936 | 243140107 | 243164283 | 243166116 | 14 | 13 | [illegible] |
| SE | 101614 | ENSG00000143702 | CEP170 | chr1 | - | 243142366 | 243142463 | 243139936 | 243140107 | 243156220 | 243156347 | 39 | 18 | [illegible] |
| SE | 101615 | ENSG00000143702 | CEP170 | chr1 | - | 243142366 | 243142463 | 243139936 | 243140107 | 243164283 | 243164494 | 13 | 2 | [illegible] |
| SE | 101616 | ENSG00000143702 | CEP170 | chr1 | - | 243156220 | 243156347 | 243139936 | 243140107 | 243164283 | 243164494 | 10 | 9 | [illegible] |
| SE | 101617 | ENSG00000143702 | CEP170 | chr1 | - | 243156220 | 243156347 | 243142285 | 243142463 | 243164283 | 243164494 | 50 | 36 | [illegible] |
| SE | 101618 | ENSG00000143702 | CEP170 | chr1 | - | 243156220 | 243156455 | 243139936 | 243140107 | 243164283 | 243166116 | 58 | 71 | [illegible] |
| SE | 101619 | ENSG00000143702 | CEP170 | chr1 | - | 243156220 | 243156455 | 243142285 | 243142463 | 243164283 | 243166116 | 100 | 98 | [illegible] |
| SE | 101620 | ENSG00000143702 | CEP170 | chr1 | - | 243176701 | 243176816 | 243172696 | 243172846 | 243185778 | 243186072 | 0 | 3 | [illegible] |
| SE | 101621 | ENSG00000143702 | CEP170 | chr1 | - | 243185778 | 243186072 | 243172696 | 243172846 | 243186258 | 243186280 | 135 | 113 | [illegible] |
| SE | 101624 | ENSG00000142459 | EVI5L | chr19 | + | 7849255 | 7849330 | 7848909 | 7849145 | 7849996 | 7850122 | 31 | 3 | [illegible] |
| SE | 101645 | ENSG00000148450 | MSRB2 | chr10 | + | 23119045 | 23119134 | 23110241 | 23110318 | 23119303 | 23119451 | 0 | 0 | [illegible] |
| SE | 101653 | ENSG00000128245 | YWHAH | chr22 | + | 31950307 | 31950422 | 31944521 | 31944820 | 31956138 | 31957603 | 0 | 0 | [illegible] |
| SE | 101654 | ENSG00000128245 | YWHAH | chr22 | + | 31950307 | 31950422 | 31947421 | 31947485 | 31956138 | 31957603 | 0 | 0 | [illegible] |
| SE | 101656 | ENSG00000112425 | EPM2A | chr6 | - | 145671238 | 145671399 | 145635244 | 145635486 | 145686121 | 145686296 | 7 | 6 | [illegible] |
| SE | 101657 | ENSG00000112425 | EPM2A | chr6 | - | 145680082 | 145680130 | 145671304 | 145671399 | 145682408 | 145682659 | 0 | 0 | [illegible] |
| SE | 101658 | ENSG00000112425 | EPM2A | chr6 | - | 145682408 | 145682659 | 145671238 | 145671399 | 145686121 | 145686146 | 0 | 2 | [illegible] |
| SE | 101662 | ENSG00000099256 | PRTFDC1 | chr10 | - | 24851387 | 24851464 | 24848613 | 24849891 | 24855317 | 24855364 | 3 | 10 | [illegible] |
| SE | 101664 | ENSG00000204177 | BMS1P1 | chr10 | + | 46790961 | 46791041 | 46789496 | 46789686 | 46794726 | 46794915 | 15 | 16 | [illegible] |
| SE | 101666 | ENSG00000204177 | BMS1P1 | chr10 | + | 46795805 | 46795953 | 46790961 | 46791041 | 46796671 | 46796793 | 23 | 37 | [illegible] |
| SE | 101667 | ENSG00000204177 | BMS1P1 | chr10 | + | 46795805 | 46795953 | 46790961 | 46791041 | 46808995 | 46811989 | 4 | 3 | [illegible] |
| SE | 101668 | ENSG00000204177 | BMS1P1 | chr10 | + | 46796671 | 46796793 | 46790961 | 46791041 | 46798028 | 46798168 | 10 | 17 | [illegible] |
| SE | 101670 | ENSG00000204177 | BMS1P1 | chr10 | + | 46798028 | 46798168 | 46790961 | 46791041 | 46808995 | 46811989 | 1 | 3 | [illegible] |
| SE | 101671 | ENSG00000136935 | GOLGA1 | chr9 | - | 124908376 | 124908472 | 124900140 | 124900547 | 124911900 | 124911919 | 36 | 37 | [illegible] |
| SE | 101673 | ENSG00000136935 | GOLGA1 | chr9 | - | 124921128 | 124921240 | 124911900 | 124912026 | 124923094 | 124923223 | 23 | 34 | [illegible] |
| SE | 101674 | ENSG00000136935 | GOLGA1 | chr9 | - | 124921711 | 124921892 | 124911900 | 124912026 | 124923094 | 124923223 | 39 | 38 | [illegible] |
| SE | 101675 | ENSG00000136935 | GOLGA1 | chr9 | - | 124921711 | 124921892 | 124921128 | 124921240 | 124923094 | 124923223 | 40 | 38 | [illegible] |
| SE | 101676 | ENSG00000136935 | GOLGA1 | chr9 | - | 124921722 | 124921892 | 124921128 | 124921240 | 124923094 | 124923223 | 58 | 63 | [illegible] |
| SE | 101677 | ENSG00000136935 | GOLGA1 | chr9 | - | 124926708 | 124926741 | 124923094 | 124923223 | 124931315 | 124931406 | 5 | 3 | [illegible] |
| SE | 101679 | ENSG00000136935 | GOLGA1 | chr9 | - | 124928187 | 124928285 | 124926708 | 124926741 | 124931315 | 124931406 | 88 | 81 | [illegible] |
| SE | 101688 | ENSG00000180398 | MCFD2 | chr2 | - | 46909022 | 46909177 | 46907528 | 46907969 | 46915722 | 46915761 | 35 | 24 | [illegible] |
| SE | 101691 | ENSG00000180398 | MCFD2 | chr2 | - | 46910779 | 46910873 | 46909022 | 46909177 | 46915722 | 46915810 | 10 | 1 | [illegible] |
| SE | 101697 | ENSG00000176928 | GCNT4 | chr5 | - | 75047896 | 75048037 | 75025345 | 75030038 | 75052168 | 75052227 | 32 | 16 | [illegible] |
| SE | 101698 | ENSG00000176928 | GCNT4 | chr5 | - | 75052168 | 75052227 | 75047896 | 75048037 | 75052428 | 75052558 | 3 | 1 | [illegible] |
| SE | 101700 | ENSG00000136770 | DNAJC1 | chr10 | - | 21920797 | 21920963 | 21919831 | 21919929 | 21928505 | 21928552 | 91 | 46 | [illegible] |
| SE | 101701 | ENSG00000136770 | DNAJC1 | chr10 | - | 21928505 | 21928552 | 21920797 | 21920963 | 21929039 | 21929141 | 49 | 35 | [illegible] |
| SE | 101702 | ENSG00000155659 | VSIG4 | chrX | - | 66025477 | 66025618 | 66025024 | 66025129 | 66027448 | 66027526 | 0 | 0 | [illegible] |
| SE | 101704 | ENSG00000165322 | ARHGAP12 | chr10 | - | 31809229 | 31809307 | 31808993 | 31809128 | 31810648 | 31810747 | 98 | 60 | [illegible] |
| SE | 101705 | ENSG00000165322 | ARHGAP12 | chr10 | - | 31817787 | 31817886 | 31814258 | 31814361 | 31820386 | 31820488 | 139 | 109 | [illegible] |
| SE | 101706 | ENSG00000165322 | ARHGAP12 | chr10 | - | 31817787 | 31817886 | 31814258 | 31814361 | 31826303 | 31826385 | 14 | 13 | [illegible] |
| SE | 101707 | ENSG00000165322 | ARHGAP12 | chr10 | - | 31820386 | 31820488 | 31814258 | 31814361 | 31826303 | 31826385 | 80 | 78 | [illegible] |
| SE | 101708 | ENSG00000165322 | ARHGAP12 | chr10 | - | 31820386 | 31820488 | 31817787 | 31817886 | 31826303 | 31826385 | 209 | 183 | [illegible] |
| SE | 101709 | ENSG00000165322 | ARHGAP12 | chr10 | - | 31839304 | 31839319 | 31831738 | 31831800 | 31839636 | 31839711 | 9 | 7 | [illegible] |
| SE | 101710 | ENSG00000165322 | ARHGAP12 | chr10 | - | 31839636 | 31839711 | 31831738 | 31831800 | 31843460 | 31843586 | 21 | 20 | [illegible] |
| SE | 101711 | ENSG00000165322 | ARHGAP12 | chr10 | - | 31852516 | 31852597 | 31843460 | 31843586 | 31861394 | 31861658 | 45 | 39 | [illegible] |
| SE | 101713 | ENSG00000165322 | ARHGAP12 | chr10 | - | 31861394 | 31861658 | 31854065 | 31854206 | 31908171 | 31908874 | 132 | 88 | [illegible] |
| SE | 101735 | ENSG00000196693 | ZNF33B | chr10 | - | 42636919 | 42636972 | 42631928 | 42632024 | 42638473 | 42638521 | 3 | 0 | [illegible] |
| SE | 101736 | ENSG00000196693 | ZNF33B | chr10 | - | 42636919 | 42636972 | 42632294 | 42632439 | 42638473 | 42638521 | 58 | 54 | [illegible] |
| SE | 101737 | ENSG00000121903 | ZSCAN20 | chr1 | + | 33479178 | 33479705 | 33472644 | 33472691 | 33488464 | 33488651 | 2 | 1 | [illegible] |
| SE | 101740 | ENSG00000121903 | ZSCAN20 | chr1 | + | 33493186 | 33493615 | 33491024 | 33491702 | 33494217 | 33496507 | 5 | 2 | [illegible] |
| SE | 101743 | ENSG00000198056 | PRIM1 | chr12 | - | 56739293 | 56739363 | 56738433 | 56738525 | 56741434 | 56741576 | 110 | 79 | [illegible] |
| SE | 101750 | ENSG00000173210 | ABLIM3 | chr5 | + | 149237447 | 149237603 | 149233228 | 149233300 | 149239758 | 149239888 | 0 | 0 | [illegible] |
| SE | 101759 | ENSG00000176533 | GNG7 | chr19 | - | 2520607 | 2520725 | 2511218 | 2515147 | 2555148 | 2555188 | 27 | 10 | [illegible] |
| SE | 101760 | ENSG00000237840 | FAM21FP | chr10 | - | 45720993 | 45721064 | 45719836 | 45719955 | 45721698 | 45721786 | 3 | 0 | [illegible] |
| SE | 101761 | ENSG00000099942 | CRKL | chr22 | + | 20933778 | 20934244 | 20917406 | 20918245 | 20949710 | 20949927 | 415 | 222 | [illegible] |
| SE | 101764 | ENSG00000077327 | SPAG6 | chr10 | + | 22387822 | 22387996 | 22386753 | 22386959 | 22389159 | 22389312 | 0 | 0 | [illegible] |
| SE | 101765 | ENSG00000077327 | SPAG6 | chr10 | + | 22411030 | 22411176 | 22401160 | 22401277 | 22416618 | 22417609 | 0 | 0 | [illegible] |
| SE | 101771 | ENSG00000012779 | ALOX5 | chr10 | + | 45382482 | 45382681 | 45374175 | 45374429 | 45395854 | 45395936 | 474 | 290 | [illegible] |
| SE | 101773 | ENSG00000012779 | ALOX5 | chr10 | + | 45395854 | 45395936 | 45382482 | 45382681 | 45424040 | 45424147 | 153 | 95 | [illegible] |
| SE | 101774 | ENSG00000012779 | ALOX5 | chr10 | + | 45412188 | 45412313 | 45382482 | 45382681 | 45424040 | 45424147 | 401 | 258 | [illegible] |
| SE | 101775 | ENSG00000012779 | ALOX5 | chr10 | + | 45412190 | 45412313 | 45382482 | 45382681 | 45424040 | 45424147 | 402 | 259 | [illegible] |
| SE | 101776 | ENSG00000012779 | ALOX5 | chr10 | + | 45412190 | 45412313 | 45395854 | 45395936 | 45424040 | 45424147 | 755 | 509 | [illegible] |
| SE | 101777 | ENSG00000012779 | ALOX5 | chr10 | + | 45424959 | 45425132 | 45424040 | 45424147 | 45428617 | 45428764 | 636 | 344 | [illegible] |
| SE | 101778 | ENSG00000012779 | ALOX5 | chr10 | + | 45428617 | 45428764 | 45424959 | 45425132 | 45440429 | 45440633 | 374 | 169 | [illegible] |
| SE | 101782 | ENSG00000170759 | KIF5B | chr10 | - | 32028427 | 32028571 | 32022847 | 32023036 | 32031072 | 32031279 | 323 | 353 | [illegible] |
| SE | 101783 | ENSG00000170759 | KIF5B | chr10 | - | 32037253 | 32037378 | 32035889 | 32035994 | 32037519 | 32037607 | 798 | 662 | [illegible] |
| SE | 101785 | ENSG00000170759 | KIF5B | chr10 | - | 32038777 | 32038826 | 32038162 | 32038218 | 32040383 | 32040457 | 128 | 86 | [illegible] |
| SE | 101788 | ENSG00000170759 | KIF5B | chr10 | - | 32048463 | 32048551 | 32040383 | 32040457 | 32055847 | 32056425 | 399 | 236 | [illegible] |
| SE | 101794 | ENSG00000090621 | PABPC4 | chr1 | - | 39561684 | 39561787 | 39560708 | 39561122 | 39562072 | 39562203 | 282 | 146 | [illegible] |
| SE | 101795 | ENSG00000090621 | PABPC4 | chr1 | - | 39561727 | 39561787 | 39560815 | 39561122 | 39562072 | 39562203 | 64 | 31 | [illegible] |
| SE | 101796 | ENSG00000090621 | PABPC4 | chr1 | - | 39563613 | 39563741 | 39562322 | 39562416 | 39564470 | 39564542 | 134 | 61 | [illegible] |
| SE | 101802 | ENSG00000090621 | PABPC4 | chr1 | - | 39565105 | 39565378 | 39562322 | 39564773 | 39567750 | 39567846 | 639 | 448 | [illegible] |
| SE | 101803 | ENSG00000090621 | PABPC4 | chr1 | - | 39568801 | 39568939 | 39567750 | 39567846 | 39569594 | 39569689 | 1388 | 1100 | [illegible] |
| SE | 101809 | ENSG00000230155 | CEP250-AS1 | chr20 | - | 35477825 | 35477945 | 35476373 | 35476675 | 35481132 | 35481251 | 5 | 1 | [illegible] |
| SE | 101812 | ENSG00000230155 | CEP250-AS1 | chr20 | - | 35481132 | 35481251 | 35477825 | 35477945 | 35490855 | 35490917 | 0 | 6 | [illegible] |
| SE | 101813 | ENSG00000138286 | FAM149B1 | chr10 | + | 73177845 | 73177975 | 73174686 | 73174791 | 73192555 | 73192698 | 77 | 52 | [illegible] |
| SE | 101814 | ENSG00000138286 | FAM149B1 | chr10 | + | 73210250 | 73210438 | 73208618 | 73208786 | 73228059 | 73228160 | 71 | 58 | [illegible] |
| SE | 101815 | ENSG00000138286 | FAM149B1 | chr10 | + | 73210250 | 73210438 | 73208618 | 73208786 | 73234816 | 73234940 | 63 | 46 | [illegible] |
| SE | 101818 | ENSG00000138286 | FAM149B1 | chr10 | + | 73230421 | 73230525 | 73210250 | 73210438 | 73232938 | 73233163 | 39 | 28 | [illegible] |
| SE | 101819 | ENSG00000138286 | FAM149B1 | chr10 | + | 73232938 | 73233163 | 73210250 | 73210438 | 73234816 | 73234940 | 28 | 28 | [illegible] |
| SE | 101820 | ENSG00000138286 | FAM149B1 | chr10 | + | 73232938 | 73233197 | 73210266 | 73210438 | 73234816 | 73234940 | 3 | 0 | [illegible] |
| SE | 101824 | ENSG00000151150 | ANK3 | chr10 | - | 60044264 | 60044348 | 60042671 | 60042759 | 60055657 | 60055832 | 0 | 0 | [illegible] |
| SE | 101826 | ENSG00000151150 | ANK3 | chr10 | - | 60053676 | 60053727 | 60042671 | 60042759 | 60055657 | 60055832 | 0 | 0 | [illegible] |
| SE | 101828 | ENSG00000151150 | ANK3 | chr10 | - | 60059339 | 60059430 | 60055657 | 60056036 | 60059697 | 60060006 | 33 | 13 | [illegible] |
| SE | 101829 | ENSG00000151150 | ANK3 | chr10 | - | 60059697 | 60060006 | 60059339 | 60059430 | 60063110 | 60063254 | 43 | 16 | [illegible] |
| SE | 101830 | ENSG00000151150 | ANK3 | chr10 | - | 60059697 | 60060033 | 60059391 | 60059430 | 60063110 | 60063254 | 22 | 9 | [illegible] |
| SE | 101831 | ENSG00000151150 | ANK3 | chr10 | - | 60067934 | 60068009 | 60064156 | 60064288 | 60080536 | 60080618 | 42 | 30 | [illegible] |
| SE | 101834 | ENSG00000151150 | ANK3 | chr10 | - | 60073871 | 60076454 | 60067934 | 60068009 | 60080536 | 60080618 | 0 | 0 | [illegible] |
| SE | 101836 | ENSG00000151150 | ANK3 | chr10 | - | 60084601 | 60084830 | 60083491 | 60083617 | 60086676 | 60086884 | 26 | 22 | [illegible] |
| SE | 101837 | ENSG00000151150 | ANK3 | chr10 | - | 60085156 | 60085253 | 60083938 | 60084830 | 60086676 | 60086884 | 26 | 13 | [illegible] |
| SE | 101840 | ENSG00000151150 | ANK3 | chr10 | - | 60137341 | 60137353 | 60134270 | 60134358 | 60138963 | 60139087 | 0 | 0 | [illegible] |
| SE | 101845 | ENSG00000151150 | ANK3 | chr10 | - | 60166590 | 60166653 | 60145967 | 60146021 | 60166823 | 60166896 | 4 | 0 | [illegible] |
| SE | 101846 | ENSG00000151150 | ANK3 | chr10 | - | 60200128 | 60200227 | 60198339 | 60198537 | 60203001 | 60203100 | 19 | 3 | [illegible] |
| SE | 101847 | ENSG00000151150 | ANK3 | chr10 | - | 60200128 | 60200263 | 60198339 | 60198537 | 60203001 | 60203100 | 8 | 1 | [illegible] |
| SE | 101848 | ENSG00000151150 | ANK3 | chr10 | - | 60234687 | 60234786 | 60213411 | 60213510 | 60261858 | 60261957 | 33 | 12 | [illegible] |
| SE | 101849 | ENSG00000151150 | ANK3 | chr10 | - | 60615185 | 60615224 | 60279537 | 60279639 | 60733262 | 60733324 | 8 | 4 | [illegible] |
| SE | 101855 | ENSG00000144815 | NXPE3 | chr3 | + | 101779733 | 101779934 | 101779245 | 101779324 | 101782109 | 101782290 | 1 | 3 | [illegible] |
| SE | 101856 | ENSG00000144815 | NXPE3 | chr3 | + | 101782659 | 101782780 | 101781705 | 101782290 | 101785401 | 101785606 | 106 | 103 | [illegible] |
| SE | 101858 | ENSG00000144815 | NXPE3 | chr3 | + | 101801234 | 101801989 | 101785401 | 101785689 | 101807052 | 101807126 | 126 | 156 | [illegible] |
| SE | 101859 | ENSG00000144815 | NXPE3 | chr3 | + | 101801234 | 101801989 | 101785401 | 101785689 | 101816794 | 101817001 | 46 | 70 | [illegible] |
| SE | 101860 | ENSG00000144815 | NXPE3 | chr3 | + | 101807052 | 101807126 | 101785401 | 101785689 | 101816794 | 101816904 | 36 | 34 | [illegible] |
| SE | 101863 | ENSG00000169740 | ZNF32 | chr10 | - | 43646063 | 43646202 | 43643859 | 43644801 | 43648801 | 43648881 | 70 | 28 | [illegible] |
| SE | 101868 | ENSG00000026103 | FAS | chr10 | + | 89010752 | 89010815 | 89010538 | 89010600 | 89013342 | 89013367 | 17 | 24 | [illegible] |
| SE | 101869 | ENSG00000026103 | FAS | chr10 | + | 89011998 | 89012081 | 89010538 | 89010600 | 89013342 | 89013367 | 75 | 37 | [illegible] |
| SE | 101870 | ENSG00000026103 | FAS | chr10 | + | 89011998 | 89012081 | 89010538 | 89010600 | 89014118 | 89014450 | 11 | 3 | [illegible] |
| SE | 101871 | ENSG00000026103 | FAS | chr10 | + | 89011998 | 89012081 | 89010752 | 89010815 | 89013342 | 89013367 | 122 | 84 | [illegible] |
| SE | 101872 | ENSG00000026103 | FAS | chr10 | + | 89012198 | 89012335 | 89011998 | 89012081 | 89013342 | 89013367 | 1 | 0 | [illegible] |
| SE | 101873 | ENSG00000026103 | FAS | chr10 | + | 89013342 | 89013367 | 89010538 | 89010600 | 89014118 | 89014177 | 17 | 16 | [illegible] |
| SE | 101874 | ENSG00000026103 | FAS | chr10 | + | 89013342 | 89013367 | 89011998 | 89012081 | 89014118 | 89014177 | 121 | 86 | [illegible] |
| SE | 101876 | ENSG00000173915 | ATP5MK | chr10 | - | 103392190 | 103392283 | 103389040 | 103389166 | 103392370 | 103392466 | 1033 | 519 | [illegible] |
| SE | 101877 | ENSG00000173915 | ATP5MK | chr10 | - | 103392370 | 103392466 | 103392190 | 103392283 | 103396408 | 103396466 | 777 | 334 | [illegible] |
| SE | 101882 | ENSG00000164347 | GFM2 | chr5 | - | 74725639 | 74725755 | 74722378 | 74722561 | 74725940 | 74726126 | 73 | 38 | [illegible] |
| SE | 101883 | ENSG00000164347 | GFM2 | chr5 | - | 74730259 | 74730398 | 74725940 | 74726126 | 74730601 | 74730775 | 42 | 22 | [illegible] |
| SE | 101885 | ENSG00000164347 | GFM2 | chr5 | - | 74730601 | 74730775 | 74725940 | 74726126 | 74733021 | 74733098 | 1 | 0 | [illegible] |
| SE | 101886 | ENSG00000164347 | GFM2 | chr5 | - | 74730601 | 74730775 | 74730259 | 74730398 | 74733021 | 74733098 | 1 | 0 | [illegible] |
| SE | 101888 | ENSG00000164347 | GFM2 | chr5 | - | 74746104 | 74746165 | 74745677 | 74745857 | 74747691 | 74747780 | 28 | 35 | [illegible] |
| SE | 101889 | ENSG00000164347 | GFM2 | chr5 | - | 74750578 | 74750667 | 74747691 | 74747780 | 74751367 | 74751493 | 190 | 136 | [illegible] |
| SE | 101890 | ENSG00000164347 | GFM2 | chr5 | - | 74750578 | 74750667 | 74747691 | 74747780 | 74758848 | 74758946 | 89 | 61 | [illegible] |
| SE | 101891 | ENSG00000164347 | GFM2 | chr5 | - | 74751367 | 74751493 | 74747691 | 74747780 | 74758848 | 74758946 | 106 | 73 | [illegible] |
| SE | 101894 | ENSG00000164347 | GFM2 | chr5 | - | 74765034 | 74765152 | 74763679 | 74763766 | 74766937 | 74767105 | 0 | 1 | [illegible] |
| SE | 101896 | ENSG00000275004 | ZNF280B | chr22 | - | 22507809 | 22507870 | 22494062 | 22494180 | 22508658 | 22508733 | 0 | 2 | [illegible] |
| SE | 101913 | ENSG00000138326 | RPS24 | chr10 | + | 78035351 | 78035417 | 78033759 | 78033904 | 78035510 | 78035720 | 5770 | 3844 | [illegible] |
| SE | 101916 | ENSG00000138326 | RPS24 | chr10 | + | 78037438 | 78040225 | 78037193 | 78037304 | 78040614 | 78040716 | 2 | 2 | [illegible] |
| SE | 101920 | ENSG00000138326 | RPS24 | chr10 | + | 78040200 | 78040225 | 78037193 | 78037304 | 78040614 | 78040713 | 0 | 1 | [illegible] |
| SE | 101924 | ENSG00000138326 | RPS24 | chr10 | + | 78048980 | 78049225 | 78037193 | 78037304 | 78054530 | 78054676 | 1 | 2 | [illegible] |
| SE | 101939 | ENSG00000122033 | MTIF3 | chr13 | - | 27439988 | 27440449 | 27437115 | 27437273 | 27445087 | 27445137 | 176 | 127 | [illegible] |
| SE | 101941 | ENSG00000122033 | MTIF3 | chr13 | - | 27441016 | 27441077 | 27439988 | 27440449 | 27450508 | 27450564 | 58 | 29 | [illegible] |
| SE | 101942 | ENSG00000122033 | MTIF3 | chr13 | - | 27445087 | 27445137 | 27439988 | 27440449 | 27450141 | 27450196 | 44 | 33 | [illegible] |
| SE | 101943 | ENSG00000122033 | MTIF3 | chr13 | - | 27445087 | 27445137 | 27439988 | 27440449 | 27450508 | 27450587 | 44 | 33 | [illegible] |
| SE | 101945 | ENSG00000122033 | MTIF3 | chr13 | - | 27445087 | 27445156 | 27439988 | 27440449 | 27450141 | 27450199 | 59 | 37 | [illegible] |
| SE | 101946 | ENSG00000122033 | MTIF3 | chr13 | - | 27445087 | 27445156 | 27439988 | 27440449 | 27450508 | 27450564 | 92 | 68 | [illegible] |
| SE | 101948 | ENSG00000122033 | MTIF3 | chr13 | - | 27449851 | 27449918 | 27445087 | 27445137 | 27450063 | 27450143 | 0 | 0 | [illegible] |
| SE | 101949 | ENSG00000122033 | MTIF3 | chr13 | - | 27449851 | 27449918 | 27445087 | 27445137 | 27450141 | 27450199 | 0 | 0 | [illegible] |
| SE | 101950 | ENSG00000122033 | MTIF3 | chr13 | - | 27449851 | 27449918 | 27445087 | 27445137 | 27450508 | 27450564 | 0 | 0 | [illegible] |
| SE | 101951 | ENSG00000122033 | MTIF3 | chr13 | - | 27449851 | 27449918 | 27445087 | 27445156 | 27450063 | 27450143 | 4 | 2 | [illegible] |
| SE | 101955 | ENSG00000188404 | SELL | chr1 | - | 169696534 | 169696553 | 169690664 | 169691802 | 169701559 | 169701688 | 2207 | 1417 | [illegible] |
| SE | 101957 | ENSG00000188404 | SELL | chr1 | - | 169701559 | 169701688 | 169690664 | 169691802 | 169703254 | 169703440 | 1418 | 797 | [illegible] |
| SE | 101960 | ENSG00000188404 | SELL | chr1 | - | 169708416 | 169708803 | 169707341 | 169707449 | 169710435 | 169710517 | 3727 | 2140 | [illegible] |
| SE | 101962 | ENSG00000197771 | MCMBP | chr10 | - | 119836895 | 119837029 | 119835539 | 119835704 | 119838534 | 119838700 | 495 | 307 | [illegible] |
| SE | 101963 | ENSG00000197771 | MCMBP | chr10 | - | 119838534 | 119838700 | 119835539 | 119835704 | 119840842 | 119840960 | 450 | 280 | [illegible] |
| SE | 101964 | ENSG00000197771 | MCMBP | chr10 | - | 119840842 | 119840960 | 119838534 | 119838700 | 119842471 | 119842595 | 667 | 392 | [illegible] |
| SE | 101965 | ENSG00000197771 | MCMBP | chr10 | - | 119840842 | 119840960 | 119838534 | 119838700 | 119843253 | 119843426 | 418 | 256 | [illegible] |
| SE | 101966 | ENSG00000197771 | MCMBP | chr10 | - | 119842471 | 119842595 | 119838534 | 119838700 | 119843253 | 119843426 | 122 | 57 | [illegible] |
| SE | 101969 | ENSG00000119401 | TRIM32 | chr9 | + | 116697372 | 116697433 | 116687342 | 116687381 | 116697661 | 116698260 | 0 | 0 | [illegible] |
| SE | 101970 | ENSG00000119401 | TRIM32 | chr9 | + | 116697372 | 116697433 | 116687342 | 116687381 | 116697664 | 116701300 | 0 | 0 | [illegible] |
| SE | 101972 | ENSG00000107771 | CCSER2 | chr10 | + | 84425730 | 84425893 | 84373618 | 84373815 | 84463932 | 84464016 | 11 | 3 | [illegible] |

| | | | | | | | | | | | | | | | | | | | | | | | | | | | | | | | | | | | | | | | | | | | | | | |
|---|---|---|---|---|---|---|---|---|---|---|---|---|---|---|---|---|---|---|---|---|---|---|---|---|---|---|---|---|---|---|---|---|---|---|---|---|---|---|---|---|---|---|---|---|---|---|
| SE | 101973 | ENSG00000107771 | CCSER2 | chr10 | + | 84425730 | 84425893 | 84373618 | 84373815 | 84513448 | 84513661 | 4 | 4 | 2 | 3 | 12 | 1 | 0 | 0 | 0 | 0 | 0 | 11 | 6 | 3 | 0 | 0 | 0 | 0 | 0 | 0 | 298 | 149 | 1 | 1 | 0.667 | 1 | 1 | 1 | 1 | NA | 1 | 1 | 1 | NA | -0.067 |
| SE | 101975 | ENSG00000107771 | CCSER2 | chr10 | + | 84463932 | 84464016 | 84425730 | 84425893 | 84470371 | 84470458 | 74 | 39 | 69 | 44 | 166 | 0 | 0 | 1 | 0 | 0 | 28 | 26 | 35 | 26 | 11 | 0 | 0 | 0 | 0 | 0 | 233 | 149 | 1 | 1 | 1 | 1 | 0.978 | 1 | 1 | 1 | 1 | 1 | 1 | 1 | -0.004 |
| SE | 101977 | ENSG00000107771 | CCSER2 | chr10 | + | 84470371 | 84470458 | 84463932 | 84464016 | 84477574 | 84477664 | 168 | 104 | 183 | 138 | 463 | 1 | 0 | 0 | 0 | 0 | 64 | 77 | 137 | 43 | 36 | 0 | 0 | 0 | 0 | 0 | 236 | 149 | 1 | 1 | 0.991 | 1 | 1 | 1 | 1 | 1 | 1 | 1 | 1 | 1 | -0.002 |
| SE | 101979 | ENSG00000107771 | CCSER2 | chr10 | + | 84477574 | 84477664 | 84470371 | 84470458 | 84513448 | 84513661 | 116 | 81 | 141 | 87 | 286 | 0 | 0 | 0 | 0 | 0 | 38 | 62 | 86 | 49 | 17 | 0 | 0 | 4 | 0 | 0 | 239 | 149 | 1 | 1 | 1 | 1 | 1 | 1 | 1 | 1 | 1 | 0.931 | 1 | 1 | 0.014 |
| SE | 101983 | ENSG00000119541 | VPS4B | chr18 | - | 63393408 | 63393549 | 63389189 | 63391076 | 63397033 | 63397253 | 260 | 174 | 248 | 292 | 414 | 0 | 0 | 0 | 0 | 1 | 114 | 206 | 167 | 91 | 11 | 0 | 0 | 0 | 0 | 0 | 290 | 149 | 1 | 1 | 1 | 1 | 1 | 1 | 0.995 | 1 | 1 | 1 | 1 | 1 | -0.001 |
| SE | 101984 | ENSG00000119541 | VPS4B | chr18 | - | 63393408 | 63393549 | 63389189 | 63391076 | 63399241 | 63399323 | 159 | 104 | 136 | 152 | 210 | 0 | 0 | 0 | 0 | 1 | 66 | 116 | 89 | 66 | 6 | 0 | 0 | 0 | 0 | 0 | 290 | 149 | 1 | 1 | 1 | 1 | 1 | 1 | 0.991 | 1 | 1 | 1 | 1 | 1 | -0.002 |
| SE | 101985 | ENSG00000119541 | VPS4B | chr18 | - | 63393408 | 63393549 | 63389189 | 63391076 | 63400047 | 63400196 | 157 | 107 | 136 | 150 | 214 | 0 | 0 | 0 | 0 | 0 | 66 | 106 | 89 | 66 | 6 | 0 | 0 | 1 | 0 | 0 | 290 | 149 | 1 | 1 | 1 | 1 | 1 | 1 | 1 | 1 | 1 | 0.979 | 1 | 1 | 0.004 |
| SE | 101986 | ENSG00000119541 | VPS4B | chr18 | - | 63393408 | 63393566 | 63390862 | 63391076 | 63397033 | 63397124 | 158 | 104 | 134 | 151 | 211 | 0 | 0 | 0 | 0 | 1 | 66 | 106 | 86 | 67 | 6 | 0 | 0 | 0 | 0 | 0 | 298 | 149 | 1 | 1 | 1 | 1 | 1 | 1 | 0.991 | 1 | 1 | 1 | 1 | 1 | -0.002 |
| SE | 101987 | ENSG00000119541 | VPS4B | chr18 | - | 63397033 | 63397253 | 63389189 | 63391076 | 63399241 | 63399323 | 304 | 220 | 251 | 305 | 424 | 0 | 0 | 0 | 0 | 1 | 93 | 207 | 178 | 182 | 23 | 0 | 0 | 0 | 0 | 0 | 298 | 149 | 1 | 1 | 1 | 1 | 1 | 1 | 0.995 | 1 | 1 | 1 | 1 | 1 | -0.001 |
| SE | 101988 | ENSG00000119541 | VPS4B | chr18 | - | 63397033 | 63397253 | 63389189 | 63391076 | 63400047 | 63400196 | 1 | 0 | 0 | 0 | 1 | 0 | 0 | 0 | 0 | 0 | 0 | 0 | 0 | 0 | 0 | 0 | 0 | 1 | 0 | 0 | 298 | 149 | 1 | 1 | 1 | NA | NA | NA | 1 | NA | NA | 0 | NA | NA | 1 |
| SE | 101991 | ENSG00000119541 | VPS4B | chr18 | - | 63399241 | 63399323 | 63389189 | 63391076 | 63400047 | 63400196 | 163 | 89 | 109 | 115 | 195 | 0 | 0 | 0 | 0 | 0 | 31 | 86 | 110 | 72 | 20 | 0 | 0 | 1 | 0 | 0 | 231 | 149 | 1 | 1 | 1 | 1 | 1 | 1 | 1 | 1 | 1 | 0.986 | 1 | 1 | 0.003 |
| SE | 101993 | ENSG00000119541 | VPS4B | chr18 | - | 63399241 | 63399323 | 63396341 | 63397253 | 63400047 | 63400196 | 467 | 309 | 360 | 420 | 617 | 1 | 0 | 0 | 0 | 0 | 114 | 293 | 288 | 248 | 43 | 0 | 0 | 0 | 0 | 0 | 231 | 149 | 1 | 1 | 0.997 | 1 | 1 | 1 | 1 | 1 | 1 | 1 | 1 | 1 | -0.001 |
| SE | 101994 | ENSG00000119541 | VPS4B | chr18 | - | 63400047 | 63400196 | 63397033 | 63397253 | 63400546 | 63400703 | 258 | 181 | 213 | 209 | 398 | 0 | 0 | 0 | 0 | 0 | 102 | 148 | 175 | 99 | 39 | 0 | 4 | 0 | 0 | 0 | 298 | 149 | 1 | 1 | 1 | 1 | 1 | 1 | 1 | 1 | 0.949 | 1 | 1 | 1 | 0.01 |
| SE | 101995 | ENSG00000119541 | VPS4B | chr18 | - | 63407431 | 63407499 | 63403706 | 63403826 | 63410289 | 63410446 | 225 | 206 | 202 | 210 | 364 | 0 | 2 | 0 | 0 | 0 | 49 | 220 | 168 | 258 | 33 | 0 | 3 | 0 | 0 | 0 | 217 | 149 | 1 | 1 | 1 | 0.986 | 1 | 1 | 1 | 1 | 0.981 | 1 | 1 | 1 | 0.001 |
| SE | 101996 | ENSG00000119541 | VPS4B | chr18 | - | 63407431 | 63407499 | 63403706 | 63403826 | 63411466 | 63411578 | 92 | 75 | 70 | 84 | 173 | 0 | 0 | 0 | 1 | 0 | 29 | 82 | 77 | 77 | 19 | 0 | 0 | 0 | 0 | 0 | 217 | 149 | 1 | 1 | 1 | 1 | 1 | 0.983 | 1 | 1 | 1 | 1 | 1 | 1 | -0.003 |
| SE | 101997 | ENSG00000119541 | VPS4B | chr18 | - | 63410289 | 63410446 | 63403706 | 63403826 | 63411466 | 63411578 | 206 | 178 | 182 | 187 | 185 | 0 | 0 | 0 | 1 | 0 | 146 | 268 | 230 | 269 | 73 | 0 | 0 | 0 | 0 | 0 | 298 | 149 | 1 | 1 | 1 | 1 | 1 | 0.989 | 1 | 1 | 1 | 1 | 1 | 1 | -0.002 |
| SE | 102003 | ENSG00000176155 | CCDC57 | chr17 | - | 82134072 | 82134194 | 82128492 | 82128597 | 82151559 | 82151773 | 19 | 12 | 41 | 25 | 14 | 0 | 0 | 0 | 3 | 0 | 52 | 41 | 25 | 15 | 25 | 0 | 0 | 0 | 0 | 0 | 271 | 149 | 1 | 1 | 1 | 1 | 1 | 0.821 | 1 | 1 | 1 | 1 | 1 | 1 | -0.036 |
| SE | 102004 | ENSG00000176155 | CCDC57 | chr17 | - | 82134072 | 82134194 | 82128492 | 82128597 | 82157747 | 82157948 | 8 | 9 | 15 | 9 | 0 | 1 | 0 | 0 | 0 | 0 | 20 | 18 | 19 | 2 | 9 | 0 | 0 | 0 | 0 | 0 | 271 | 149 | 1 | 1 | 0.815 | 1 | 1 | 1 | NA | 1 | 1 | 1 | 1 | 1 | -0.046 |
| SE | 102006 | ENSG00000176155 | CCDC57 | chr17 | - | 82151559 | 82151773 | 82134072 | 82134194 | 82157747 | 82157948 | 12 | 3 | 26 | 17 | 14 | 0 | 0 | 0 | 0 | 0 | 32 | 23 | 8 | 13 | 16 | 0 | 0 | 2 | 0 | 0 | 298 | 149 | 1 | 1 | 1 | 1 | 1 | 1 | 1 | 1 | 1 | 0.667 | 1 | 1 | 0.067 |
| SE | 102009 | ENSG00000176155 | CCDC57 | chr17 | - | 82193981 | 82194139 | 82193755 | 82193830 | 82195262 | 82195364 | 23 | 31 | 38 | 54 | 29 | 0 | 0 | 0 | 0 | 0 | 48 | 40 | 23 | 42 | 78 | 0 | 0 | 2 | 0 | 0 | 298 | 149 | 1 | 1 | 1 | 1 | 1 | 1 | 1 | 1 | 1 | 0.852 | 1 | 1 | 0.03 |

[illegible]

| | | | | | | | | | | | | | | |
|---|---|---|---|---|---|---|---|---|---|---|---|---|---|---|
| SE | 102387 | ENSG00000065621 | GSTO2 | chr10 | + | 104279369 | 104279471 | 104277893 | 104278116 | 104297577 | 104297684 | 6 | 0 | [illegible] |
| SE | 102391 | ENSG00000069956 | MAPK6 | chr15 | + | 52027638 | 52027873 | 52024580 | 52024695 | 52045829 | 52046053 | 1 | 0 | [illegible] |
| SE | 102393 | ENSG00000069956 | MAPK6 | chr15 | + | 52042753 | 52042833 | 52019237 | 52019376 | 52045829 | 52046109 | 2 | 0 | [illegible] |
| SE | 102394 | ENSG00000069956 | MAPK6 | chr15 | + | 52045829 | 52047015 | 52019218 | 52019376 | 52049992 | 52050137 | 72 | 64 | [illegible] |
| SE | 102395 | ENSG00000069956 | MAPK6 | chr15 | + | 52049992 | 52050137 | 52045829 | 52047015 | 52058632 | 52058797 | 94 | 84 | [illegible] |
| SE | 102396 | ENSG00000069956 | MAPK6 | chr15 | + | 52058632 | 52058797 | 52049992 | 52050137 | 52061298 | 52061500 | 109 | 77 | [illegible] |
| SE | 102397 | ENSG00000069956 | MAPK6 | chr15 | + | 52061298 | 52061500 | 52058632 | 52058797 | 52063901 | 52067359 | 100 | 67 | [illegible] |
| SE | 102398 | ENSG00000156671 | SAMD8 | chr10 | + | 75164644 | 75164740 | 75150513 | 75151106 | 75168540 | 75168627 | 70 | 49 | [illegible] |
| SE | 102399 | ENSG00000156671 | SAMD8 | chr10 | + | 75164644 | 75164740 | 75150513 | 75151106 | 75176065 | 75176216 | 35 | 19 | [illegible] |
| SE | 102400 | ENSG00000156671 | SAMD8 | chr10 | + | 75168540 | 75168658 | 75150513 | 75151106 | 75176065 | 75176216 | 25 | 29 | [illegible] |
| SE | 102403 | ENSG00000255624 | C10orf88B | chr10 | + | 122883724 | 122883931 | 122881597 | 122881667 | 122884716 | 122884852 | 0 | 0 | [illegible] |
| SE | 102405 | ENSG00000255624 | C10orf88B | chr10 | + | 122884716 | 122884852 | 122881597 | 122881667 | 122888268 | 122888723 | 6 | 14 | [illegible] |
| SE | 102408 | ENSG00000138346 | DNA2 | chr10 | - | 68419802 | 68419892 | 68419033 | 68419213 | 68422224 | 68422429 | 13 | 5 | [illegible] |
| SE | 102410 | ENSG00000138346 | DNA2 | chr10 | - | 68437010 | 68437241 | 68432393 | 68432510 | 68442916 | 68443111 | 14 | 19 | [illegible] |
| SE | 102411 | ENSG00000138346 | DNA2 | chr10 | - | 68442916 | 68443111 | 68437010 | 68437241 | 68450027 | 68450247 | 12 | 17 | [illegible] |
| SE | 102412 | ENSG00000138346 | DNA2 | chr10 | - | 68444920 | 68445083 | 68442916 | 68443111 | 68450027 | 68450247 | 11 | 9 | [illegible] |
| SE | 102415 | ENSG00000138346 | DNA2 | chr10 | - | 68468122 | 68468306 | 68465666 | 68465812 | 68469980 | 68470163 | 59 | 42 | [illegible] |
| SE | 102416 | ENSG00000138346 | DNA2 | chr10 | - | 68470488 | 68470612 | 68469980 | 68470163 | 68471790 | 68471886 | 0 | 0 | [illegible] |
| SE | 102427 | ENSG00000166321 | NUDT13 | chr10 | + | 73125397 | 73125509 | 73125117 | 73125243 | 73126672 | 73126827 | 2 | 1 | [illegible] |
| SE | 102432 | ENSG00000227165 | WDR11-AS1 | chr10 | - | 120808232 | 120808341 | 120776734 | 120776884 | 120830218 | 120831161 | 0 | 0 | [illegible] |
| SE | 102441 | ENSG00000107593 | PKD2L1 | chr10 | - | 100288971 | 100289056 | 100288148 | 100288478 | 100290014 | 100290138 | 0 | 0 | [illegible] |
| SE | 102442 | ENSG00000129646 | QRICH2 | chr17 | - | 76277162 | 76277310 | 76276679 | 76276767 | 76277988 | 76278189 | 2 | 0 | [illegible] |
| SE | 102443 | ENSG00000129646 | QRICH2 | chr17 | - | 76289991 | 76290077 | 76287799 | 76287897 | 76291014 | 76291252 | 9 | 4 | [illegible] |
| SE | 102446 | ENSG00000137752 | CASP1 | chr11 | - | 105031164 | 105031280 | 105030329 | 105030503 | 105034207 | 105034474 | 802 | 615 | [illegible] |
| SE | 102451 | ENSG00000108100 | CCNY | chr10 | + | 35394815 | 35394878 | 35336873 | 35337207 | 35501500 | 35501535 | 0 | 1 | [illegible] |
| SE | 102452 | ENSG00000108100 | CCNY | chr10 | + | 35455356 | 35455463 | 35337051 | 35337207 | 35483403 | 35483478 | 1 | 1 | [illegible] |
| SE | 102453 | ENSG00000108100 | CCNY | chr10 | + | 35455356 | 35455463 | 35434112 | 35434220 | 35483403 | 35483478 | 1 | 1 | [illegible] |
| SE | 102455 | ENSG00000108100 | CCNY | chr10 | + | 35483403 | 35483478 | 35336508 | 35337207 | 35553018 | 35553183 | 159 | 80 | [illegible] |
| SE | 102456 | ENSG00000108100 | CCNY | chr10 | + | 35483403 | 35483478 | 35394815 | 35394878 | 35501500 | 35501535 | 228 | 148 | [illegible] |
| SE | 102457 | ENSG00000108100 | CCNY | chr10 | + | 35501500 | 35501535 | 35336873 | 35337207 | 35530123 | 35530243 | 0 | 0 | [illegible] |
| SE | 102458 | ENSG00000108100 | CCNY | chr10 | + | 35501500 | 35501535 | 35336873 | 35337207 | 35553018 | 35553183 | 0 | 0 | [illegible] |
| SE | 102459 | ENSG00000108100 | CCNY | chr10 | + | 35501500 | 35501535 | 35483403 | 35483478 | 35516522 | 35516575 | 546 | 377 | [illegible] |
| SE | 102460 | ENSG00000108100 | CCNY | chr10 | + | 35501500 | 35501535 | 35483403 | 35483478 | 35525963 | 35525999 | 129 | 70 | [illegible] |
| SE | 102461 | ENSG00000108100 | CCNY | chr10 | + | 35501500 | 35501535 | 35483403 | 35483478 | 35529972 | 35530030 | 113 | 69 | [illegible] |
| SE | 102462 | ENSG00000108100 | CCNY | chr10 | + | 35501500 | 35501535 | 35483403 | 35483478 | 35553018 | 35553183 | 112 | 66 | [illegible] |
| SE | 102463 | ENSG00000108100 | CCNY | chr10 | + | 35516522 | 35516623 | 35483403 | 35483478 | 35525963 | 35525999 | 296 | 230 | [illegible] |
| SE | 102464 | ENSG00000108100 | CCNY | chr10 | + | 35516522 | 35516623 | 35483403 | 35483478 | 35529972 | 35530030 | 2 | 3 | [illegible] |
| SE | 102467 | ENSG00000108100 | CCNY | chr10 | + | 35525963 | 35525999 | 35483403 | 35483478 | 35529972 | 35530030 | 26 | 21 | [illegible] |
| SE | 102469 | ENSG00000108100 | CCNY | chr10 | + | 35525963 | 35525999 | 35516522 | 35516623 | 35529972 | 35530030 | 451 | 353 | [illegible] |
| SE | 102470 | ENSG00000108100 | CCNY | chr10 | + | 35529972 | 35530030 | 35501500 | 35501535 | 35530123 | 35530243 | 52 | 24 | [illegible] |
| SE | 102471 | ENSG00000108100 | CCNY | chr10 | + | 35529972 | 35530030 | 35525963 | 35525999 | 35530123 | 35530243 | 414 | 294 | [illegible] |
| SE | 102472 | ENSG00000108100 | CCNY | chr10 | + | 35530123 | 35530243 | 35336508 | 35337207 | 35553018 | 35553183 | 272 | 186 | [illegible] |
| SE | 102473 | ENSG00000108100 | CCNY | chr10 | + | 35530123 | 35530243 | 35501500 | 35501535 | 35553018 | 35553183 | 272 | 186 | [illegible] |
| SE | 102474 | ENSG00000108100 | CCNY | chr10 | + | 35553018 | 35553185 | 35530123 | 35530243 | 35566022 | 35566128 | 599 | 347 | [illegible] |
| SE | 102475 | ENSG00000108100 | CCNY | chr10 | + | 35566022 | 35566185 | 35553018 | 35553185 | 35569053 | 35571924 | 660 | 320 | [illegible] |
| SE | 102476 | ENSG00000119979 | DENND10 | chr10 | + | 119104763 | 119105006 | 119104116 | 119104197 | 119107947 | 119108164 | 5 | 0 | [illegible] |
| SE | 102477 | ENSG00000119979 | DENND10 | chr10 | + | 119104763 | 119105006 | 119104116 | 119104197 | 119107967 | 119108164 | 5 | 2 | [illegible] |
| SE | 102478 | ENSG00000119979 | DENND10 | chr10 | + | 119104763 | 119105022 | 119104091 | 119104197 | 119107947 | 119108096 | 7 | 3 | [illegible] |
| SE | 102479 | ENSG00000119979 | DENND10 | chr10 | + | 119104763 | 119105022 | 119104091 | 119104197 | 119107967 | 119108164 | 5 | 1 | [illegible] |
| SE | 102480 | ENSG00000119979 | DENND10 | chr10 | + | 119105310 | 119105495 | 119104763 | 119105006 | 119107947 | 119108096 | 1 | 1 | [illegible] |
| SE | 102481 | ENSG00000119979 | DENND10 | chr10 | + | 119105310 | 119105495 | 119104763 | 119105006 | 119107967 | 119108164 | 7 | 9 | [illegible] |
| SE | 102482 | ENSG00000119979 | DENND10 | chr10 | + | 119105310 | 119105495 | 119104763 | 119105022 | 119107947 | 119108096 | 1 | 1 | [illegible] |
| SE | 102483 | ENSG00000119979 | DENND10 | chr10 | + | 119105310 | 119105495 | 119104763 | 119105022 | 119107967 | 119108164 | 7 | 9 | [illegible] |
| SE | 102485 | ENSG00000119979 | DENND10 | chr10 | + | 119111848 | 119111928 | 119107947 | 119108164 | 119117518 | 119117667 | 220 | 204 | [illegible] |
| SE | 102487 | ENSG00000119979 | DENND10 | chr10 | + | 119120340 | 119120452 | 119117518 | 119117667 | 119129514 | 119129546 | 200 | 162 | [illegible] |
| SE | 102488 | ENSG00000119979 | DENND10 | chr10 | + | 119123468 | 119123569 | 119117518 | 119117667 | 119129514 | 119129546 | 109 | 110 | [illegible] |
| SE | 102489 | ENSG00000119979 | DENND10 | chr10 | + | 119123468 | 119123569 | 119120340 | 119120452 | 119129514 | 119129546 | 369 | 365 | [illegible] |
| SE | 102490 | ENSG00000119979 | DENND10 | chr10 | + | 119123468 | 119123569 | 119120340 | 119120452 | 119132514 | 119132609 | 193 | 198 | [illegible] |
| SE | 102492 | ENSG00000119979 | DENND10 | chr10 | + | 119129514 | 119129622 | 119120340 | 119120452 | 119136470 | 119137826 | 1 | 2 | [illegible] |
| SE | 102493 | ENSG00000119979 | DENND10 | chr10 | + | 119129514 | 119129622 | 119123468 | 119123569 | 119132514 | 119132609 | 267 | 251 | [illegible] |
| SE | 102494 | ENSG00000119979 | DENND10 | chr10 | + | 119132514 | 119132609 | 119120340 | 119120452 | 119136470 | 119137826 | 47 | 51 | [illegible] |
| SE | 102495 | ENSG00000119979 | DENND10 | chr10 | + | 119132514 | 119132609 | 119129514 | 119129622 | 119136470 | 119137826 | 208 | 215 | [illegible] |
| SE | 102497 | ENSG00000107560 | RAB11FIP2 | chr10 | - | 118038971 | 118039440 | 118015064 | 118015110 | 118045810 | 118046592 | 52 | 48 | [illegible] |
| SE | 102498 | ENSG00000107560 | RAB11FIP2 | chr10 | - | 118040122 | 118040565 | 118038971 | 118039440 | 118045810 | 118046592 | 204 | 220 | [illegible] |
| SE | 102500 | ENSG00000169895 | SYAP1 | chrX | + | 16741715 | 16741789 | 16736165 | 16736232 | 16743700 | 16743840 | 118 | 92 | [illegible] |
| SE | 102501 | ENSG00000169895 | SYAP1 | chrX | + | 16754944 | 16755093 | 16743700 | 16743840 | 16756662 | 16756721 | 246 | 194 | [illegible] |
| SE | 102502 | ENSG00000169895 | SYAP1 | chrX | + | 16756662 | 16756721 | 16754944 | 16755093 | 16757161 | 16757309 | 239 | 164 | [illegible] |
| SE | 102503 | ENSG00000169895 | SYAP1 | chrX | + | 16757161 | 16757309 | 16756662 | 16756721 | 16760231 | 16765340 | 245 | 141 | [illegible] |
| SE | 102507 | ENSG00000146826 | TRAPPC14 | chr7 | - | 100155670 | 100155825 | 100155261 | 100155455 | 100156633 | 100156716 | 36 | 31 | [illegible] |
| SE | 102509 | ENSG00000146826 | TRAPPC14 | chr7 | - | 100156633 | 100156716 | 100155261 | 100155455 | 100156844 | 100156992 | 19 | 13 | [illegible] |
| SE | 102510 | ENSG00000146826 | TRAPPC14 | chr7 | - | 100156633 | 100156716 | 100156264 | 100156549 | 100156844 | 100156992 | 27 | 19 | [illegible] |
| SE | 102514 | ENSG00000166130 | IKBIP | chr12 | - | 98634295 | 98634413 | 98624248 | 98626766 | 98644522 | 98644788 | 61 | 37 | [illegible] |
| SE | 102516 | ENSG00000065154 | OAT | chr10 | - | 124403797 | 124403920 | 124402926 | 124403055 | 124405435 | 124405563 | 79 | 65 | [illegible] |
| SE | 102517 | ENSG00000065154 | OAT | chr10 | - | 124407302 | 124407476 | 124405435 | 124405563 | 124408541 | 124408637 | 1 | 0 | [illegible] |
| SE | 102518 | ENSG00000065154 | OAT | chr10 | - | 124411972 | 124412200 | 124408740 | 124408965 | 124418872 | 124418923 | 229 | 96 | [illegible] |
| SE | 102519 | ENSG00000065154 | OAT | chr10 | - | 124414345 | 124414432 | 124411972 | 124412200 | 124414952 | 124415072 | 2 | 0 | [illegible] |
| SE | 102521 | ENSG00000129450 | SIGLEC9 | chr19 | + | 51127029 | 51127296 | 51126080 | 51126128 | 51127948 | 51128039 | 58 | 29 | [illegible] |
| SE | 102522 | ENSG00000129450 | SIGLEC9 | chr19 | + | 51127948 | 51128039 | 51127029 | 51127296 | 51128413 | 51128510 | 117 | 54 | [illegible] |
| SE | 102524 | ENSG00000095574 | IKZF5 | chr10 | - | 122998492 | 122998658 | 122995993 | 122996176 | 123007025 | 123007176 | 85 | 57 | [illegible] |
| SE | 102525 | ENSG00000095574 | IKZF5 | chr10 | - | 122998492 | 122998671 | 122995993 | 122996176 | 123007025 | 123007176 | 158 | 89 | [illegible] |
| SE | 102527 | ENSG00000095574 | IKZF5 | chr10 | - | 123007025 | 123007176 | 122998492 | 122998658 | 123008693 | 123008795 | 70 | 44 | [illegible] |
| SE | 102529 | ENSG00000197857 | ZNF44 | chr19 | - | 12226371 | 12226527 | 12224685 | 12225279 | 12230459 | 12230515 | 0 | 0 | [illegible] |
| SE | 102530 | ENSG00000197857 | ZNF44 | chr19 | - | 12244580 | 12244673 | 12242706 | 12242844 | 12249977 | 12250038 | 0 | 1 | [illegible] |
| SE | 102531 | ENSG00000197857 | ZNF44 | chr19 | - | 12272486 | 12274063 | 12250241 | 12250368 | 12274972 | 12275033 | 38 | 31 | [illegible] |
| SE | 102532 | ENSG00000197857 | ZNF44 | chr19 | - | 12274972 | 12275033 | 12250241 | 12250368 | 12294691 | 12294794 | 4 | 2 | [illegible] |
| SE | 102533 | ENSG00000197857 | ZNF44 | chr19 | - | 12275955 | 12276082 | 12274972 | 12275033 | 12294691 | 12294834 | 41 | 49 | [illegible] |
| SE | 102537 | ENSG00000197857 | ZNF44 | chr19 | - | 12293231 | 12293375 | 12291027 | 12291250 | 12294691 | 12294794 | 6 | 2 | [illegible] |
| SE | 102539 | ENSG00000089876 | DHX32 | chr10 | - | 125839000 | 125839188 | 125838205 | 125838387 | 125840846 | 125840996 | 69 | 22 | [illegible] |
| SE | 102546 | ENSG00000186529 | CYP4F3 | chr19 | + | 15643906 | 15644051 | 15641308 | 15641613 | 15647051 | 15647105 | 1 | 0 | [illegible] |
| SE | 102548 | ENSG00000186529 | CYP4F3 | chr19 | + | 15645718 | 15645863 | 15643906 | 15644051 | 15647051 | 15647105 | 4 | 2 | [illegible] |
| SE | 102552 | ENSG00000203791 | EEF1AKMT2 | chr10 | - | 124789038 | 124789157 | 124774674 | 124774782 | 124791723 | 124791868 | 17 | 13 | [illegible] |
| SE | 102553 | ENSG00000077147 | TM9SF3 | chr10 | - | 96528030 | 96528177 | 96527412 | 96527496 | 96530539 | 96530608 | 481 | 256 | [illegible] |
| SE | 102554 | ENSG00000077147 | TM9SF3 | chr10 | - | 96533050 | 96533190 | 96530539 | 96530608 | 96544075 | 96544206 | 571 | 339 | [illegible] |
| SE | 102555 | ENSG00000077147 | TM9SF3 | chr10 | - | 96544075 | 96544206 | 96533050 | 96533190 | 96547894 | 96547989 | 862 | 534 | [illegible] |
| SE | 102556 | ENSG00000077147 | TM9SF3 | chr10 | - | 96559658 | 96559736 | 96552927 | 96553059 | 96561977 | 96562138 | 1128 | 947 | [illegible] |
| SE | 102557 | ENSG00000077147 | TM9SF3 | chr10 | - | 96576633 | 96576829 | 96565303 | 96565426 | 96586733 | 96587012 | 634 | 482 | [illegible] |
| SE | 102560 | ENSG00000171307 | ZDHHC16 | chr10 | + | 97450356 | 97450536 | 97446174 | 97446353 | 97451670 | 97451918 | 0 | 6 | [illegible] |
| SE | 102561 | ENSG00000171307 | ZDHHC16 | chr10 | + | 97451670 | 97451918 | 97446169 | 97446353 | 97452414 | 97452503 | 4 | 4 | [illegible] |
| SE | 102562 | ENSG00000171307 | ZDHHC16 | chr10 | + | 97452089 | 97452284 | 97451670 | 97451918 | 97452414 | 97452465 | 50 | 17 | [illegible] |
| SE | 102564 | ENSG00000171307 | ZDHHC16 | chr10 | + | 97455638 | 97455783 | 97454713 | 97454799 | 97455973 | 97456044 | 47 | 16 | [illegible] |
| SE | 102565 | ENSG00000171307 | ZDHHC16 | chr10 | + | 97455659 | 97455783 | 97454713 | 97454799 | 97455973 | 97456044 | 83 | 33 | [illegible] |
| SE | 102566 | ENSG00000171307 | ZDHHC16 | chr10 | + | 97455713 | 97455783 | 97454713 | 97454799 | 97455973 | 97456044 | 39 | 17 | [illegible] |
| SE | 102567 | ENSG00000171307 | ZDHHC16 | chr10 | + | 97455973 | 97456044 | 97455601 | 97455783 | 97456776 | 97456924 | 87 | 25 | [illegible] |
| SE | 102571 | ENSG00000183605 | SFXN4 | chr10 | - | 119155061 | 119155177 | 119147774 | 119147860 | 119156677 | 119156756 | 38 | 25 | [illegible] |
| SE | 102572 | ENSG00000183605 | SFXN4 | chr10 | - | 119160914 | 119160969 | 119159727 | 119159753 | 119162339 | 119162414 | 12 | 9 | [illegible] |
| SE | 102578 | ENSG00000120438 | TCP1 | chr6 | - | 159779894 | 159780087 | 159779626 | 159779790 | 159780442 | 159780566 | 922 | 698 | [illegible] |
| SE | 102586 | ENSG00000120438 | TCP1 | chr6 | - | 159788057 | 159788143 | 159787742 | 159787871 | 159789404 | 159789572 | 183 | 107 | [illegible] |
| SE | 102587 | ENSG00000120438 | TCP1 | chr6 | - | 159788609 | 159788696 | 159788057 | 159788143 | 159789404 | 159789672 | 0 | 1 | [illegible] |
| SE | 102591 | ENSG00000175470 | PPP2R2D | chr10 | + | 131940030 | 131940196 | 131901237 | 131901330 | 131955683 | 131956555 | 0 | 3 | [illegible] |
| SE | 102592 | ENSG00000175470 | PPP2R2D | chr10 | + | 131940581 | 131940694 | 131939775 | 131940196 | 131943967 | 131943997 | 188 | 133 | [illegible] |
| SE | 102595 | ENSG00000175470 | PPP2R2D | chr10 | + | 131940581 | 131940841 | 131940158 | 131940196 | 131943967 | 131944145 | 103 | 82 | [illegible] |
| SE | 102596 | ENSG00000175470 | PPP2R2D | chr10 | + | 131943967 | 131943997 | 131940162 | 131940196 | 131944094 | 131944145 | 0 | 0 | [illegible] |
| SE | 102602 | ENSG00000138161 | CUZD1 | chr10 | - | 122834705 | 122835097 | 122833671 | 122833940 | 122839016 | 122839231 | 7 | 5 | [illegible] |
| SE | 102603 | ENSG00000138161 | CUZD1 | chr10 | - | 122836177 | 122836350 | 122834705 | 122835097 | 122836830 | 122836899 | 9 | 3 | [illegible] |
| SE | 102606 | ENSG00000172845 | SP3 | chr2 | - | 173907885 | 173907965 | 173883275 | 173883488 | 173913069 | 173913266 | 5 | 6 | [illegible] |
| SE | 102609 | ENSG00000172845 | SP3 | chr2 | - | 173954872 | 173956232 | 173918592 | 173918785 | 173963760 | 173963883 | 809 | 562 | [illegible] |
| SE | 102610 | ENSG00000172845 | SP3 | chr2 | - | 173963760 | 173963883 | 173954872 | 173956232 | 173964404 | 173964553 | 95 | 44 | [illegible] |
| SE | 102611 | ENSG00000172845 | SP3 | chr2 | - | 173964404 | 173964553 | 173954872 | 173956232 | 173965164 | 173965373 | 14 | 5 | [illegible] |
| SE | 102612 | ENSG00000256377 | NA | chr12 | - | 27703460 | 27703620 | 27696900 | 27697298 | 27710598 | 27710731 | 0 | 3 | [illegible] |
| SE | 102613 | ENSG00000256377 | NA | chr12 | - | 27704737 | 27704924 | 27703460 | 27703620 | 27710598 | 27710770 | 0 | 19 | [illegible] |
| SE | 102619 | ENSG00000196177 | ACADSB | chr10 | + | 123040465 | 123040672 | 123037746 | 123037847 | 123041208 | 123041379 | 197 | 129 | [illegible] |
| SE | 102620 | ENSG00000196177 | ACADSB | chr10 | + | 123051048 | 123051186 | 123047208 | 123047298 | 123053060 | 123053160 | 86 | 93 | [illegible] |
| SE | 102621 | ENSG00000135972 | MRPS9 | chr2 | + | 105049170 | 105049350 | 105038068 | 105038227 | 105071312 | 105071375 | 184 | 148 | [illegible] |
| SE | 102624 | ENSG00000165646 | SLC18A2 | chr10 | + | 117255276 | 117255366 | 117254047 | 117254497 | 117255478 | 117255522 | 4 | 0 | [illegible] |
| SE | 102627 | ENSG00000248538 | NA | chr8 | + | 9154965 | 9155149 | 9151791 | 9151916 | 9156151 | 9156304 | 0 | 0 | [illegible] |
| SE | 102629 | ENSG00000248538 | NA | chr8 | + | 9189347 | 9189586 | 9156770 | 9156847 | 9198923 | 9198991 | 0 | 0 | [illegible] |
| SE | 102632 | ENSG00000248538 | NA | chr8 | + | 9189347 | 9189597 | 9156773 | 9156847 | 9198923 | 9198991 | 0 | 0 | [illegible] |
| SE | 102635 | ENSG00000248538 | NA | chr8 | + | 9193409 | 9194032 | 9189347 | 9189586 | 9198923 | 9198991 | 0 | 0 | [illegible] |
| SE | 102637 | ENSG00000248538 | NA | chr8 | + | 9198923 | 9198991 | 9189045 | 9189586 | 9202391 | 9202628 | 0 | 0 | [illegible] |
| SE | 102640 | ENSG00000248538 | NA | chr8 | + | 9198923 | 9198991 | 9189045 | 9189597 | 9202498 | 9202625 | 0 | 0 | [illegible] |
| SE | 102651 | ENSG00000165752 | STK32C | chr10 | - | 132209294 | 132209364 | 132209033 | 132209101 | 132222640 | 132222772 | 3 | 2 | [illegible] |
| SE | 102652 | ENSG00000165752 | STK32C | chr10 | - | 132222860 | 132222986 | 132222640 | 132222772 | 132224406 | 132224523 | 41 | 10 | [illegible] |
| SE | 102656 | ENSG00000185515 | BRCC3 | chrX | + | 155073373 | 155073431 | 155072326 | 155072343 | 155077169 | 155077283 | 16 | 19 | [illegible] |
| SE | 102657 | ENSG00000185515 | BRCC3 | chrX | + | 155073376 | 155073431 | 155072326 | 155072343 | 155077169 | 155077283 | 34 | 26 | [illegible] |
| SE | 102658 | ENSG00000185515 | BRCC3 | chrX | + | 155099314 | 155099389 | 155090783 | 155090839 | 155116056 | 155116183 | 30 | 3 | [illegible] |
| SE | 102659 | ENSG00000185515 | BRCC3 | chrX | + | 155099314 | 155099389 | 155090783 | 155090839 | 155116710 | 155116754 | 13 | 0 | [illegible] |
| SE | 102660 | ENSG00000185515 | BRCC3 | chrX | + | 155116056 | 155116188 | 155090783 | 155090839 | 155116710 | 155116754 | 71 | 78 | [illegible] |
| SE | 102662 | ENSG00000185515 | BRCC3 | chrX | + | 155116710 | 155116754 | 155116056 | 155116188 | 155119998 | 155120168 | 47 | 58 | [illegible] |
| SE | 102663 | ENSG00000175197 | DDIT3 | chr12 | - | 57517268 | 57517424 | 57516587 | 57517180 | 57517705 | 57517753 | 25 | 8 | [illegible] |
| SE | 102664 | ENSG00000175197 | DDIT3 | chr12 | - | 57517268 | 57517438 | 57516587 | 57517180 | 57517705 | 57517753 | 39 | 18 | [illegible] |
| SE | 102665 | ENSG00000175197 | DDIT3 | chr12 | - | 57517268 | 57517542 | 57516952 | 57517180 | 57517705 | 57517753 | 21 | 8 | [illegible] |
| SE | 102666 | ENSG00000175197 | DDIT3 | chr12 | - | 57517268 | 57517595 | 57516588 | 57517180 | 57517705 | 57517753 | 21 | 10 | [illegible] |
| SE | 102669 | ENSG00000175029 | CTBP2 | chr10 | - | 124993854 | 124993985 | 124993201 | 124993329 | 124994468 | 124994606 | 197 | 78 | [illegible] |
| SE | 102670 | ENSG00000175029 | CTBP2 | chr10 | - | 125038996 | 125039155 | 124994468 | 124994683 | 125110989 | 125111093 | 64 | 28 | [illegible] |
| SE | 102672 | ENSG00000175029 | CTBP2 | chr10 | - | 125038996 | 125039155 | 125003337 | 125003492 | 125158634 | 125158716 | 105 | 39 | [illegible] |
| SE | 102674 | ENSG00000175029 | CTBP2 | chr10 | - | 125110989 | 125111093 | 125003337 | 125003492 | 125158634 | 125158716 | 23 | 1 | [illegible] |
| SE | 102678 | ENSG00000175029 | CTBP2 | chr10 | - | 125122656 | 125122868 | 125110996 | 125111093 | 125159856 | 125159957 | 4 | 0 | [illegible] |
| SE | 102679 | ENSG00000175029 | CTBP2 | chr10 | - | 125122656 | 125122868 | 125110996 | 125111093 | 125160318 | 125160520 | 4 | 0 | [illegible] |
| SE | 102680 | ENSG00000175029 | CTBP2 | chr10 | - | 125133511 | 125133612 | 125038996 | 125039155 | 125158634 | 125158716 | 3 | 1 | [illegible] |
| SE | 102681 | ENSG00000175029 | CTBP2 | chr10 | - | 125133511 | 125133612 | 125038996 | 125039155 | 125160318 | 125160499 | 2 | 1 | [illegible] |
| SE | 102682 | ENSG00000175029 | CTBP2 | chr10 | - | 125133511 | 125133612 | 125110996 | 125111093 | 125138006 | 125138104 | 2 | 0 | [illegible] |
| SE | 102683 | ENSG00000175029 | CTBP2 | chr10 | - | 125133511 | 125133612 | 125110996 | 125111093 | 125158634 | 125158716 | 4 | 0 | [illegible] |
| SE | 102684 | ENSG00000175029 | CTBP2 | chr10 | - | 125133511 | 125133612 | 125110996 | 125111093 | 125160318 | 125160499 | 2 | 0 | [illegible] |
| SE | 102686 | ENSG00000107938 | EDRF1 | chr10 | + | 125723067 | 125723134 | 125721203 | 125721412 | 125723810 | 125723936 | 99 | 87 | [illegible] |
| SE | 102688 | ENSG00000107938 | EDRF1 | chr10 | + | 125729357 | 125729479 | 125725681 | 125725838 | 125730287 | 125730399 | 78 | 69 | [illegible] |
| SE | 102692 | ENSG00000107938 | EDRF1 | chr10 | + | 125734071 | 125734183 | 125733634 | 125733743 | 125735643 | 125735749 | 104 | 74 | [illegible] |
| SE | 102694 | ENSG00000107938 | EDRF1 | chr10 | + | 125742219 | 125742345 | 125741000 | 125741201 | 125743057 | 125743276 | 5 | 1 | [illegible] |
| SE | 102695 | ENSG00000107938 | EDRF1 | chr10 | + | 125752798 | 125752914 | 125749296 | 125749565 | 125753693 | 125753845 | 41 | 31 | [illegible] |
| SE | 102696 | ENSG00000107938 | EDRF1 | chr10 | + | 125753693 | 125753845 | 125749272 | 125749565 | 125763300 | 125763524 | 35 | 32 | [illegible] |
| SE | 102698 | ENSG00000074590 | NUAK1 | chr12 | - | 106086733 | 106086885 | 106083863 | 106083929 | 106106404 | 106106525 | 27 | 0 | [illegible] |
| SE | 102699 | ENSG00000180229 | HERC2P3 | chr15 | - | 20461191 | 20461342 | 20458231 | 20458447 | 20462342 | 20462446 | 11 | 5 | [illegible] |
| SE | 102701 | ENSG00000180229 | HERC2P3 | chr15 | - | 20466194 | 20466370 | 20462830 | 20463001 | 20505390 | 20505493 | 10 | 13 | [illegible] |
| SE | 102704 | ENSG00000180229 | HERC2P3 | chr15 | - | 20483402 | 20483517 | 20462830 | 20463001 | 20505390 | 20505462 | 16 | 8 | [illegible] |
| SE | 102705 | ENSG00000180229 | HERC2P3 | chr15 | - | 20483402 | 20483517 | 20466194 | 20466370 | 20505390 | 20505462 | 3 | 6 | [illegible] |
| SE | 102706 | ENSG00000180229 | HERC2P3 | chr15 | - | 20501545 | 20501714 | 20466194 | 20466370 | 20505390 | 20505485 | 0 | 0 | [illegible] |
| SE | 102719 | ENSG00000214279 | SCART1 | chr10 | + | 133464605 | 133465174 | 133459877 | 133460170 | 133467197 | 133467353 | 0 | 0 | [illegible] |
| SE | 102722 | ENSG00000214279 | SCART1 | chr10 | + | 133466234 | 133466381 | 133459486 | 133460170 | 133467197 | 133467353 | 3 | 4 | [illegible] |
| SE | 102723 | ENSG00000214279 | SCART1 | chr10 | + | 133493286 | 133493490 | 133491221 | 133491319 | 133497828 | 133498005 | 10 | 0 | [illegible] |
| SE | 102724 | ENSG00000235823 | OLMALINC | chr10 | + | 100380894 | 100381109 | 100373571 | 100374012 | 100381215 | 100381448 | 1 | 1 | [illegible] |
| SE | 102725 | ENSG00000235823 | OLMALINC | chr10 | + | 100380982 | 100381109 | 100373512 | 100374012 | 100381215 | 100381448 | 3 | 3 | [illegible] |
| SE | 102729 | ENSG00000111788 | NA | chr12 | + | 9294816 | 9294974 | 9294431 | 9294518 | 9297780 | 9297826 | 3 | 9 | [illegible] |
| SE | 102730 | ENSG00000111788 | NA | chr12 | + | 9302622 | 9302702 | 9302227 | 9302274 | 9303974 | 9304019 | 4 | 0 | [illegible] |
| SE | 102731 | ENSG00000111788 | NA | chr12 | + | 9309947 | 9310037 | 9307201 | 9307314 | 9310338 | 9310441 | 0 | 1 | [illegible] |
| SE | 102732 | ENSG00000111788 | NA | chr12 | + | 9310338 | 9310441 | 9307201 | 9307314 | 9311549 | 9311618 | 0 | 2 | [illegible] |
| SE | 102733 | ENSG00000111788 | NA | chr12 | + | 9311140 | 9311290 | 9310338 | 9310441 | 9311549 | 9311618 | 0 | 11 | [illegible] |
| SE | 102734 | ENSG00000111788 | NA | chr12 | + | 9312616 | 9312695 | 9311730 | 9311831 | 9312874 | 9313029 | 0 | 1 | [illegible] |
| SE | 102735 | ENSG00000111788 | NA | chr12 | + | 9312616 | 9312695 | 9312240 | 9312323 | 9312874 | 9313029 | 0 | 0 | [illegible] |
| SE | 102738 | ENSG00000040341 | STAU2 | chr8 | - | 73615674 | 73615782 | 73613743 | 73613956 | 73617291 | 73617451 | 203 | 151 | [illegible] |
| SE | 102739 | ENSG00000040341 | STAU2 | chr8 | - | 73615674 | 73615807 | 73613743 | 73613956 | 73617291 | 73617451 | 81 | 64 | [illegible] |
| SE | 102740 | ENSG00000040341 | STAU2 | chr8 | - | 73668980 | 73669110 | 73617291 | 73617451 | 73673106 | 73673242 | 0 | 0 | [illegible] |
| SE | 102742 | ENSG00000040341 | STAU2 | chr8 | - | 73673106 | 73673242 | 73617291 | 73617451 | 73688653 | 73688813 | 319 | 293 | [illegible] |
| SE | 102743 | ENSG00000040341 | STAU2 | chr8 | - | 73673106 | 73673242 | 73617291 | 73617451 | 73709031 | 73709177 | 109 | 85 | [illegible] |
| SE | 102744 | ENSG00000040341 | STAU2 | chr8 | - | 73673106 | 73673242 | 73617291 | 73617451 | 73738283 | 73738349 | 111 | 87 | [illegible] |
| SE | 102745 | ENSG00000040341 | STAU2 | chr8 | - | 73688653 | 73688813 | 73617291 | 73617451 | 73709031 | 73709162 | 23 | 23 | [illegible] |
| SE | 102746 | ENSG00000040341 | STAU2 | chr8 | - | 73688653 | 73688813 | 73617291 | 73617451 | 73738283 | 73738327 | 216 | 174 | [illegible] |
| SE | 102747 | ENSG00000040341 | STAU2 | chr8 | - | 73688653 | 73688813 | 73617291 | 73617451 | 73746782 | 73746840 | 5 | 1 | [illegible] |
| SE | 102749 | ENSG00000040341 | STAU2 | chr8 | - | 73688653 | 73688813 | 73672981 | 73673242 | 73709031 | 73709162 | 233 | 232 | [illegible] |
| SE | 102753 | ENSG00000040341 | STAU2 | chr8 | - | 73709031 | 73709162 | 73617291 | 73617451 | 73738283 | 73738344 | 25 | 31 | [illegible] |
| SE | 102759 | ENSG00000040341 | STAU2 | chr8 | - | 73738283 | 73738349 | 73617291 | 73617451 | 73746782 | 73746849 | 0 | 0 | [illegible] |
| SE | 102764 | ENSG00000040341 | STAU2 | chr8 | - | 73739755 | 73739868 | 73738283 | 73738349 | 73747386 | 73747473 | 141 | 127 | [illegible] |
| SE | 102765 | ENSG00000040341 | STAU2 | chr8 | - | 73742223 | 73742363 | 73739755 | 73739868 | 73746782 | 73746840 | 1 | 0 | [illegible] |
| SE | 102771 | ENSG00000105287 | PRKD2 | chr19 | - | 46678365 | 46678663 | 46675032 | 46675118 | 46681649 | 46681748 | 119 | 69 | [illegible] |
| SE | 102772 | ENSG00000105287 | PRKD2 | chr19 | - | 46678395 | 46678663 | 46675032 | 46675118 | 46681649 | 46681748 | 181 | 89 | [illegible] |
| SE | 102773 | ENSG00000105287 | PRKD2 | chr19 | - | 46681649 | 46681748 | 46678365 | 46678663 | 46689536 | 46689698 | 173 | 96 | [illegible] |
| SE | 102774 | ENSG00000105287 | PRKD2 | chr19 | - | 46685424 | 46685521 | 46681649 | 46681748 | 46689536 | 46689698 | 0 | 1 | [illegible] |
| SE | 102778 | ENSG00000198825 | INPP5F | chr10 | + | 119757891 | 119757977 | 119751075 | 119751156 | 119781634 | 119781771 | 0 | 0 | [illegible] |
| SE | 102779 | ENSG00000198825 | INPP5F | chr10 | + | 119757891 | 119757977 | 119751075 | 119751156 | 119791516 | 119791645 | 0 | 0 | [illegible] |
| SE | 102781 | ENSG00000198825 | INPP5F | chr10 | + | 119757894 | 119757977 | 119751075 | 119751156 | 119781634 | 119781771 | 0 | 0 | [illegible] |
| SE | 102782 | ENSG00000198825 | INPP5F | chr10 | + | 119757894 | 119757977 | 119751075 | 119751156 | 119791516 | 119791645 | 0 | 0 | [illegible] |
| SE | 102784 | ENSG00000198825 | INPP5F | chr10 | + | 119759856 | 119760068 | 119757891 | 119757977 | 119781634 | 119781771 | 0 | 0 | [illegible] |
| SE | 102785 | ENSG00000198825 | INPP5F | chr10 | + | 119781060 | 119781242 | 119751075 | 119751156 | 119781634 | 119781771 | 0 | 0 | [illegible] |
| SE | 102786 | ENSG00000198825 | INPP5F | chr10 | + | 119781060 | 119781242 | 119757894 | 119757977 | 119781634 | 119781771 | 0 | 0 | [illegible] |
| SE | 102788 | ENSG00000198825 | INPP5F | chr10 | + | 119781634 | 119781771 | 119750876 | 119751156 | 119791516 | 119791645 | 13 | 26 | [illegible] |
| SE | 102792 | ENSG00000198825 | INPP5F | chr10 | + | 119791516 | 119791645 | 119781634 | 119781771 | 119791868 | 119792036 | 21 | 32 | [illegible] |
| SE | 102795 | ENSG00000198825 | INPP5F | chr10 | + | 119798542 | 119798610 | 119797460 | 119797640 | 119804172 | 119804297 | 43 | 38 | [illegible] |
| SE | 102798 | ENSG00000198825 | INPP5F | chr10 | + | 119807867 | 119808060 | 119805383 | 119805461 | 119810099 | 119810217 | 24 | 13 | [illegible] |
| SE | 102799 | ENSG00000198825 | INPP5F | chr10 | + | 119807931 | 119808060 | 119805383 | 119805461 | 119810099 | 119810217 | 31 | 19 | [illegible] |
| SE | 102800 | ENSG00000198825 | INPP5F | chr10 | + | 119807931 | 119808060 | 119805383 | 119805461 | 119811809 | 119811955 | 8 | 10 | [illegible] |
| SE | 102801 | ENSG00000198825 | INPP5F | chr10 | + | 119810099 | 119810217 | 119805383 | 119805461 | 119811756 | 119811955 | 13 | 14 | [illegible] |

| | | | | | | | | | | | | | | | | | | | | | | | | | | | | | | | | | | | | | | | | | | | | | | | | |
|---|---|---|---|---|---|---|---|---|---|---|---|---|---|---|---|---|---|---|---|---|---|---|---|---|---|---|---|---|---|---|---|---|---|---|---|---|---|---|---|---|---|---|---|---|---|---|---|---|
| SE | 102802 | ENSG00000198825 | INPP5F | chr10 | + | 119810099 | 119810217 | 119805383 | 119805461 | 119811809 | 119811955 | 1 | 1 | 0 | 0 | 1 | 0 | 1 | 0 | 2 | 0 | 0 | 4 | 3 | 1 | 0 | 0 | 0 | 0 | 0 | 0 | 0 | 0 | 267 | 149 | 1 | 1 | 1 | 0.358 | NA | 0 | 1 | NA | 1 | 1 | 1 | NA | -0.41 |
| SE | 102803 | ENSG00000198825 | INPP5F | chr10 | + | 119810099 | 119810217 | 119807867 | 119808060 | 119811809 | 119811955 | 26 | 17 | 32 | 28 | 21 | 0 | 4 | 0 | 0 | 0 | 0 | 11 | 7 | 2 | 0 | 0 | 0 | 0 | 0 | 0 | 0 | 0 | 267 | 149 | 1 | 1 | 1 | 0.703 | 1 | 1 | 1 | NA | 1 | 1 | 1 | NA | -0.059 |
| SE | 102804 | ENSG00000198825 | INPP5F | chr10 | + | 119819766 | 119819834 | 119819446 | 119819539 | 119820845 | 119820917 | 0 | 0 | 1 | 0 | 2 | 0 | 0 | 0 | 1 | 0 | 0 | 3 | 0 | 0 | 0 | 0 | 0 | 0 | 0 | 0 | 0 | 0 | 217 | 149 | 1 | 1 | NA | NA | 1 | 0 | 1 | NA | 1 | NA | NA | NA | -0.333 |
| SE | 102805 | ENSG00000198825 | INPP5F | chr10 | + | 119820845 | 119820917 | 119811756 | 119811955 | 119822430 | 119822504 | 29 | 18 | 26 | 25 | 35 | 0 | 0 | 0 | 1 | 0 | 0 | 7 | 11 | 14 | 13 | 0 | 0 | 0 | 0 | 0 | 0 | 0 | 221 | 149 | 1 | 1 | 1 | 1 | 1 | 0.944 | 1 | NA | 1 | 1 | 1 | 1 | -0.011 |
| SE | 102806 | ENSG00000198825 | INPP5F | chr10 | + | 119823814 | 119823902 | 119823070 | 119823199 | 119826630 | 119826702 | 17 | 20 | 31 | 20 | 23 | 1 | 0 | 0 | 0 | 3 | 0 | 15 | 19 | 12 | 0 | 0 | 0 | 0 | 1 | 0 | 1 | 0 | 237 | 149 | 1 | 1 | 0.914 | 1 | 1 | 1 | 0.828 | NA | 1 | 1 | 0.883 | NA | -0.013 |
| SE | 102808 | ENSG00000165832 | TRUB1 | chr10 | + | 114942644 | 114942743 | 114938194 | 114938539 | 114951093 | 114951149 | 73 | 37 | 61 | 62 | 49 | 0 | 0 | 0 | 1 | 0 | 0 | 46 | 56 | 41 | 20 | 0 | 0 | 0 | 0 | 0 | 0 | 0 | 248 | 149 | 1 | 1 | 1 | 1 | 1 | 0.974 | 1 | NA | 1 | 1 | 1 | 1 | -0.005 |
| SE | 102809 | ENSG00000165832 | TRUB1 | chr10 | + | 114951093 | 114951149 | 114942644 | 114942743 | 114970367 | 114970440 | 20 | 12 | 15 | 17 | 7 | 0 | 0 | 0 | 0 | 2 | 0 | 10 | 13 | 10 | 0 | 0 | 0 | 0 | 0 | 0 | 0 | 0 | 205 | 149 | 1 | 1 | 1 | 1 | 1 | 1 | 0.718 | NA | 1 | 1 | 1 | NA | -0.056 |
| SE | 102813 | ENSG00000078668 | VDAC3 | chr8 | + | 42395083 | 42395133 | 42394209 | 42394278 | 42398711 | 42398864 | 998 | 775 | 733 | 1006 | 1381 | 0 | 0 | 2 | 4 | 4 | 1349 | 1260 | 761 | 996 | 1649 | 0 | 1 | 1 | 2 | 0 | 2 | 0 | 199 | 149 | 1 | 1 | 1 | 1 | 0.996 | 0.995 | 0.996 | 1 | 0.999 | 0.998 | 0.997 | 1 | -0.001 |
| SE | 102816 | ENSG00000078668 | VDAC3 | chr8 | + | 42398711 | 42398864 | 42395083 | 42395133 | 42399650 | 42399703 | 1276 | 1019 | 1049 | 1422 | 2014 | 0 | 0 | 0 | 0 | 0 | 1899 | 1630 | 1003 | 1375 | 2098 | 2 | 4 | 3 | 0 | 0 | 0 | 0 | 298 | 149 | 1 | 1 | 1 | 1 | 1 | 1 | 1 | 0.998 | 0.995 | 0.994 | 1 | 1 | 0.003 |
| SE | 102817 | ENSG00000078668 | VDAC3 | chr8 | + | 42399650 | 42399703 | 42398711 | 42398864 | 42401787 | 42401836 | 623 | 501 | 542 | 729 | 988 | 0 | 0 | 0 | 1 | 0 | 1118 | 854 | 503 | 701 | 1091 | 3 | 0 | 0 | 0 | 0 | 0 | 0 | 202 | 149 | 1 | 1 | 1 | 1 | 1 | 0.998 | 1 | 0.996 | 1 | 1 | 1 | 1 | 0 |
| SE | 102818 | ENSG00000078668 | VDAC3 | chr8 | + | 42404866 | 42404924 | 42403031 | 42403461 | 42405370 | 42405415 | 426 | 355 | 353 | 461 | 820 | 2 | 0 | 0 | 1 | 1 | 417 | 344 | 245 | 274 | 443 | 0 | 0 | 1 | 2 | 0 | 2 | 0 | 207 | 149 | 1 | 1 | 0.994 | 1 | 1 | 0.997 | 0.998 | 1 | 1 | 0.994 | 0.99 | 1 | 0.001 |
| SE | 102820 | ENSG00000148700 | ADD3 | chr10 | + | 110075617 | 110075709 | 110008230 | 110008299 | 110079239 | 110079297 | 0 | 1 | 2 | 0 | 0 | 3 | 3 | 5 | 11 | 3 | 0 | 0 | 4 | 0 | 0 | 0 | 2 | 16 | 0 | 0 | 0 | 0 | 241 | 149 | 1 | 1 | 0 | 0.171 | 0.198 | 0 | 0 | NA | 0 | 0.134 | NA | NA | 0.007 |
| SE | 102823 | ENSG00000148700 | ADD3 | chr10 | + | 110079239 | 110079297 | 110075617 | 110075709 | 110100624 | 110100704 | 47 | 38 | 43 | 27 | 66 | 0 | 0 | 1 | 3 | 4 | 17 | 31 | 34 | 11 | 0 | 0 | 12 | 7 | 0 | 0 | 0 | 0 | 207 | 149 | 1 | 1 | 1 | 1 | 0.969 | 0.866 | 0.922 | 1 | 0.65 | 0.778 | 1 | NA | 0.094 |
| SE | 102825 | ENSG00000148700 | ADD3 | chr10 | + | 110100624 | 110100848 | 110007963 | 110008299 | 110112776 | 110112906 | 941 | 661 | 971 | 971 | 1017 | 0 | 0 | 0 | 0 | 0 | 166 | 828 | 670 | 392 | 204 | 0 | 0 | 0 | 1 | 0 | 1 | 0 | 298 | 149 | 1 | 1 | 1 | 1 | 1 | 1 | 1 | 1 | 1 | 1 | 0.995 | 1 | 0.001 |
| SE | 102827 | ENSG00000148700 | ADD3 | chr10 | + | 110105999 | 110106053 | 110100624 | 110100848 | 110116258 | 110116410 | 15 | 8 | 15 | 8 | 14 | 0 | 1 | 0 | 0 | 0 | 0 | 17 | 4 | 1 | 0 | 0 | 0 | 2 | 0 | 0 | 0 | 0 | 203 | 149 | 1 | 1 | 1 | 0.854 | 1 | 1 | 1 | NA | 1 | 0.595 | 1 | NA | 0.106 |
| SE | 102829 | ENSG00000148700 | ADD3 | chr10 | + | 110112776 | 110112915 | 110100624 | 110100848 | 110116258 | 110116410 | 1392 | 1025 | 1421 | 1360 | 1631 | 0 | 1 | 0 | 0 | 0 | 266 | 1263 | 937 | 610 | 337 | 0 | 0 | 2 | 0 | 0 | 0 | 0 | 288 | 149 | 1 | 1 | 1 | 0.998 | 1 | 1 | 1 | 1 | 1 | 0.996 | 1 | 1 | 0 |
| SE | 102830 | ENSG00000148700 | ADD3 | chr10 | + | 110112776 | 110112915 | 110105999 | 110106053 | 110116258 | 110116410 | 648 | 478 | 654 | 608 | 870 | 0 | 0 | 0 | 0 | 0 | 129 | 621 | 411 | 303 | 170 | 0 | 0 | 1 | 0 | 0 | 0 | 0 | 288 | 149 | 1 | 1 | 1 | 1 | 1 | 1 | 1 | 1 | 1 | 0.995 | 1 | 1 | 0.001 |
| SE | 102833 | ENSG00000148700 | ADD3 | chr10 | + | 110125825 | 110125924 | 110124239 | 110124274 | 110126416 | 110126503 | 414 | 285 | 485 | 400 | 752 | 0 | 0 | 0 | 1 | 0 | 97 | 357 | 245 | 129 | 154 | 0 | 0 | 0 | 1 | 0 | 1 | 0 | 248 | 149 | 1 | 1 | 1 | 1 | 1 | 0.996 | 1 | 1 | 1 | 1 | 0.987 | 1 | 0.002 |
| SE | 102834 | ENSG00000148700 | ADD3 | chr10 | + | 110125825 | 110125945 | 110123836 | 110124274 | 110126416 | 110126503 | 1096 | 723 | 1180 | 1038 | 1751 | 0 | 0 | 0 | 1 | 0 | 151 | 815 | 622 | 355 | 343 | 0 | 0 | 0 | 1 | 0 | 1 | 0 | 269 | 149 | 1 | 1 | 1 | 1 | 1 | 0.998 | 1 | 1 | 1 | 1 | 0.995 | 1 | 0.001 |
| SE | 102842 | ENSG00000023191 | RNH1 | chr11 | - | 504823 | 504888 | 502061 | 502249 | 506608 | 506759 | 26 | 27 | 23 | 27 | 13 | 1 | 0 | 0 | 0 | 0 | 24 | 10 | 14 | 5 | 45 | 0 | 2 | 0 | 0 | 0 | 0 | 0 | 214 | 149 | 1 | 1 | 0.948 | 1 | 1 | 1 | 1 | 1 | 0.777 | 1 | 1 | 1 | 0.034 |
| SE | 102845 | ENSG00000023191 | RNH1 | chr11 | - | 504823 | 504996 | 502061 | 502249 | 506608 | 506800 | 52 | 36 | 57 | 75 | 38 | 1 | 0 | 0 | 0 | 0 | 77 | 10 | 22 | 17 | 52 | 0 | 2 | 0 | 0 | 0 | 0 | 0 | 298 | 149 | 1 | 1 | 0.963 | 1 | 1 | 1 | 1 | 1 | 0.714 | 1 | 1 | 1 | 0.05 |
| SE | 102849 | ENSG00000107554 | DNMBP | chr10 | - | 99888824 | 99888953 | 99886299 | 99886632 | 99894945 | 99895050 | 40 | 24 | 59 | 68 | 86 | 0 | 1 | 0 | 0 | 1 | 147 | 124 | 112 | 81 | 96 | 0 | 0 | 0 | 0 | 0 | 0 | 0 | 278 | 149 | 1 | 1 | 1 | 0.928 | 1 | 1 | 0.979 | 1 | 1 | 1 | 1 | 1 | -0.019 |
| SE | 102851 | ENSG00000107554 | DNMBP | chr10 | - | 99898085 | 99898285 | 99896266 | 99896397 | 99898742 | 99898760 | 38 | 22 | 14 | 43 | 29 | 0 | 0 | 0 | 0 | 1 | 48 | 48 | 45 | 38 | 9 | 0 | 0 | 0 | 0 | 0 | 0 | 0 | 298 | 149 | 1 | 1 | 1 | 1 | 1 | 1 | 0.935 | 1 | 1 | 1 | 1 | 1 | -0.013 |
| SE | 102852 | ENSG00000107554 | DNMBP | chr10 | - | 99899918 | 99900066 | 99898742 | 99898760 | 99907994 | 99908094 | 57 | 43 | 31 | 73 | 80 | 0 | 0 | 0 | 0 | 0 | 85 | 128 | 96 | 77 | 66 | 0 | 0 | 0 | 1 | 0 | 1 | 0 | 297 | 149 | 1 | 1 | 1 | 1 | 1 | 1 | 1 | 1 | 1 | 1 | 0.975 | 1 | 0.005 |
| SE | 102861 | ENSG00000182010 | RTKN2 | chr10 | - | 62204856 | 62205022 | 62199753 | 62199861 | 62217117 | 62217249 | 55 | 81 | 99 | 58 | 164 | 0 | 0 | 1 | 0 | 0 | 1 | 25 | 8 | 1 | 0 | 0 | 0 | 0 | 0 | 0 | 0 | 0 | 298 | 149 | 1 | 1 | 1 | 1 | 0.98 | 1 | 1 | 1 | 1 | 1 | 1 | NA | -0.004 |
| SE | 102862 | ENSG00000182010 | RTKN2 | chr10 | - | 62215028 | 62215091 | 62204856 | 62205022 | 62217117 | 62217249 | 2 | 0 | 0 | 0 | 0 | 19 | 23 | 29 | 16 | 43 | 0 | 0 | 3 | 0 | 0 | 0 | 9 | 0 | 0 | 0 | 0 | 0 | 212 | 149 | 1 | 1 | 0.069 | 0 | 0 | 0 | 0 | NA | 0 | 1 | NA | NA | -0.486 |
| SE | 102863 | ENSG00000182010 | RTKN2 | chr10 | - | 62217117 | 62217249 | 62204856 | 62205022 | 62223237 | 62223332 | 18 | 23 | 24 | 12 | 43 | 0 | 0 | 0 | 0 | 4 | 0 | 9 | 0 | 0 | 0 | 0 | 0 | 0 | 0 | 0 | 0 | 0 | 281 | 149 | 1 | 1 | 1 | 1 | 1 | 1 | 0.851 | NA | 1 | NA | NA | NA | -0.03 |
| SE | 102864 | ENSG00000182010 | RTKN2 | chr10 | - | 62218194 | 62218301 | 62217117 | 62217249 | 62223237 | 62223332 | 34 | 40 | 90 | 36 | 82 | 2 | 0 | 0 | 0 | 9 | 0 | 18 | 6 | 2 | 0 | 0 | 0 | 0 | 0 | 0 | 0 | 0 | 256 | 149 | 1 | 1 | 0.908 | 1 | 1 | 1 | 0.841 | NA | 1 | 1 | 1 | NA | -0.05 |
| SE | 102865 | ENSG00000182010 | RTKN2 | chr10 | - | 62223237 | 62223332 | 62204856 | 62205022 | 62236065 | 62236263 | 6 | 5 | 6 | 3 | 20 | 1 | 0 | 1 | 0 | 3 | 0 | 0 | 0 | 0 | 0 | 0 | 0 | 0 | 1 | 0 | 1 | 0 | 244 | 149 | 1 | 1 | 0.786 | 1 | 0.786 | 1 | 0.803 | NA | NA | NA | 0 | NA | 0.875 |
| SE | 102866 | ENSG00000182010 | RTKN2 | chr10 | - | 62223237 | 62223332 | 62218194 | 62218301 | 62236065 | 62236263 | 23 | 24 | 50 | 22 | 54 | 1 | 0 | 0 | 0 | 0 | 0 | 9 | 4 | 1 | 0 | 0 | 0 | 0 | 0 | 0 | 0 | 0 | 244 | 149 | 1 | 1 | 0.934 | 1 | 1 | 1 | 1 | NA | 1 | 1 | 1 | NA | -0.013 |
| SE | 102867 | ENSG00000163947 | ARHGEF3 | chr3 | - | 56732237 | 56732424 | 56727417 | 56729622 | 56737184 | 56737355 | 835 | 550 | 620 | 663 | 773 | 0 | 0 | 0 | 0 | 0 | 555 | 628 | 654 | 547 | 262 | 0 | 0 | 2 | 0 | 0 | 0 | 0 | 298 | 149 | 1 | 1 | 1 | 1 | 1 | 1 | 1 | 1 | 1 | 0.994 | 1 | 1 | 0.001 |
| SE | 102869 | ENSG00000163947 | ARHGEF3 | chr3 | - | 56751299 | 56751396 | 56745204 | 56745462 | 56753503 | 56753566 | 250 | 165 | 240 | 228 | 380 | 2 | 0 | 0 | 1 | 1 | 78 | 343 | 224 | 181 | 51 | 0 | 0 | 0 | 0 | 0 | 0 | 0 | 246 | 149 | 1 | 1 | 0.987 | 1 | 1 | 0.993 | 0.996 | 1 | 1 | 1 | 1 | 1 | -0.005 |
| SE | 102870 | ENSG00000163947 | ARHGEF3 | chr3 | - | 56753503 | 56753566 | 56751299 | 56751396 | 56754980 | 56755151 | 439 | 262 | 357 | 365 | 589 | 0 | 0 | 0 | 0 | 2 | 146 | 459 | 284 | 251 | 92 | 0 | 0 | 0 | 6 | 0 | 6 | 0 | 212 | 149 | 1 | 1 | 1 | 1 | 1 | 1 | 0.995 | 1 | 1 | 1 | 0.967 | 1 | 0.006 |
| SE | 102871 | ENSG00000163947 | ARHGEF3 | chr3 | - | 56753503 | 56753566 | 56751299 | 56751396 | 56773708 | 56773816 | 197 | 86 | 100 | 108 | 268 | 0 | 0 | 2 | 0 | 0 | 57 | 170 | 70 | 91 | 32 | 0 | 0 | 0 | 0 | 0 | 0 | 0 | 212 | 149 | 1 | 1 | 1 | 1 | 0.972 | 1 | 1 | 1 | 1 | 1 | 1 | 1 | -0.006 |
| SE | 102872 | ENSG00000163947 | ARHGEF3 | chr3 | - | 56754980 | 56755151 | 56751299 | 56751396 | 56773708 | 56773816 | 178 | 91 | 173 | 155 | 150 | 0 | 0 | 2 | 0 | 0 | 87 | 134 | 86 | 109 | 42 | 0 | 0 | 0 | 0 | 0 | 0 | 0 | 298 | 149 | 1 | 1 | 1 | 1 | 0.977 | 1 | 1 | 1 | 1 | 1 | 1 | 1 | -0.005 |

| | | | | | | | | | | | | | | | | | | | | | | | | | | | | | | | | | | | | | | | | | | | | | | |
|---|---|---|---|---|---|---|---|---|---|---|---|---|---|---|---|---|---|---|---|---|---|---|---|---|---|---|---|---|---|---|---|---|---|---|---|---|---|---|---|---|---|---|---|---|---|---|
| SE | 103237 | ENSG00000165732 | DDX21 | chr10 | + | 68968975 | 68969121 | 68967017 | 68967203 | 68970200 | 68970350 | 684 | 540 | 534 | 659 | 976 | 0 | 2 | 0 | 0 | 3 | 143 | 578 | 750 | 362 | 77 | 0 | 2 | 0 | 4 | 0 | 295 | 149 | 1 | 1 | 1 | 0.993 | 1 | 1 | 0.994 | 1 | 0.993 | 1 | 0.979 | 1 | 0.003 |
| SE | 103238 | ENSG00000165732 | DDX21 | chr10 | + | 68970200 | 68970350 | 68968975 | 68969121 | 68971890 | 68972052 | 687 | 518 | 547 | 634 | 774 | 3 | 1 | 2 | 0 | 0 | 134 | 478 | 612 | 260 | 33 | 0 | 0 | 2 | 0 | 0 | 298 | 149 | 1 | 1 | 0.991 | 0.996 | 0.993 | 1 | 1 | 1 | 1 | 0.994 | 1 | 1 | -0.003 |
| SE | 103239 | ENSG00000165732 | DDX21 | chr10 | + | 68970200 | 68970350 | 68968975 | 68969121 | 68973544 | 68973664 | 305 | 253 | 240 | 292 | 525 | 0 | 0 | 0 | 0 | 0 | 74 | 285 | 329 | 177 | 29 | 0 | 2 | 1 | 0 | 0 | 298 | 149 | 1 | 1 | 1 | 1 | 1 | 1 | 1 | 1 | 0.986 | 0.994 | 1 | 1 | 0.004 |
| SE | 103240 | ENSG00000165732 | DDX21 | chr10 | + | 68971890 | 68972052 | 68968975 | 68969121 | 68973544 | 68973664 | 472 | 361 | 364 | 450 | 304 | 0 | 0 | 0 | 0 | 0 | 78 | 351 | 415 | 141 | 4 | 0 | 2 | 1 | 0 | 0 | 298 | 149 | 1 | 1 | 1 | 1 | 1 | 1 | 1 | 1 | 0.989 | 0.995 | 1 | 1 | 0.003 |
| SE | 103242 | ENSG00000165732 | DDX21 | chr10 | + | 68973544 | 68973664 | 68970200 | 68970350 | 68977528 | 68977688 | 4 | 4 | 4 | 1 | 9 | 1 | 0 | 0 | 0 | 0 | 5 | 25 | 17 | 0 | 0 | 0 | 0 | 0 | 0 | 0 | 269 | 149 | 1 | 1 | 0.689 | 1 | 1 | 1 | 1 | 1 | 1 | 1 | NA | NA | -0.062 |
| SE | 103243 | ENSG00000165732 | DDX21 | chr10 | + | 68973544 | 68973664 | 68971890 | 68972052 | 68974669 | 68974743 | 519 | 407 | 405 | 500 | 339 | 0 | 0 | 0 | 0 | 0 | 90 | 390 | 469 | 150 | 4 | 0 | 3 | 0 | 0 | 0 | 269 | 149 | 1 | 1 | 1 | 1 | 1 | 1 | 1 | 1 | 0.986 | 1 | 1 | 1 | 0.003 |
| SE | 103244 | ENSG00000165732 | DDX21 | chr10 | + | 68974669 | 68974743 | 68973544 | 68973664 | 68977528 | 68977688 | 136 | 135 | 133 | 154 | 163 | 1 | 0 | 0 | 0 | 0 | 12 | 112 | 186 | 68 | 16 | 0 | 0 | 0 | 0 | 0 | 223 | 149 | 1 | 1 | 0.989 | 1 | 1 | 1 | 1 | 1 | 1 | 1 | 1 | 1 | -0.002 |
| SE | 103246 | ENSG00000165732 | DDX21 | chr10 | + | 68977528 | 68977688 | 68974669 | 68974743 | 68981536 | 68981581 | 128 | 128 | 129 | 157 | 149 | 0 | 0 | 0 | 0 | 2 | 12 | 104 | 175 | 79 | 16 | 0 | 0 | 0 | 0 | 0 | 298 | 149 | 1 | 1 | 1 | 1 | 1 | 1 | 0.974 | 1 | 1 | 1 | 1 | 1 | -0.005 |
| SE | 103247 | ENSG00000165732 | DDX21 | chr10 | + | 68978841 | 68978976 | 68974669 | 68974743 | 68981536 | 68981581 | 240 | 258 | 236 | 273 | 253 | 0 | 0 | 0 | 0 | 2 | 23 | 205 | 377 | 187 | 19 | 0 | 0 | 0 | 0 | 0 | 284 | 149 | 1 | 1 | 1 | 1 | 1 | 1 | 0.985 | 1 | 1 | 1 | 1 | 1 | -0.003 |
| SE | 103248 | ENSG00000165732 | DDX21 | chr10 | + | 68978841 | 68978976 | 68977528 | 68977688 | 68981536 | 68981581 | 376 | 401 | 372 | 404 | 368 | 0 | 1 | 0 | 2 | 0 | 44 | 350 | 569 | 283 | 31 | 0 | 0 | 0 | 0 | 0 | 284 | 149 | 1 | 1 | 1 | 0.995 | 1 | 0.991 | 1 | 1 | 1 | 1 | 1 | 1 | -0.003 |
| SE | 103249 | ENSG00000165732 | DDX21 | chr10 | + | 68981536 | 68981581 | 68978841 | 68978976 | 68982542 | 68985067 | 288 | 300 | 263 | 352 | 316 | 0 | 0 | 0 | 1 | 1 | 40 | 232 | 440 | 236 | 24 | 0 | 0 | 1 | 0 | 0 | 194 | 149 | 1 | 1 | 1 | 1 | 1 | 0.996 | 0.996 | 1 | 1 | 0.997 | 1 | 1 | -0.001 |
| SE | 103251 | ENSG00000172661 | WASHC2C | chr10 | + | 45728861 | 45729026 | 45727416 | 45727539 | 45737982 | 45738045 | 122 | 127 | 96 | 131 | 75 | 0 | 1 | 0 | 0 | 0 | 325 | 124 | 104 | 200 | 103 | 0 | 0 | 0 | 0 | 0 | 298 | 149 | 1 | 1 | 1 | 0.984 | 1 | 1 | 1 | 1 | 1 | 1 | 1 | 1 | -0.003 |
| SE | 103253 | ENSG00000172661 | WASHC2C | chr10 | + | 45737982 | 45738045 | 45728861 | 45729026 | 45743389 | 45743483 | 32 | 51 | 38 | 55 | 34 | 3 | 0 | 0 | 0 | 0 | 146 | 48 | 47 | 97 | 52 | 0 | 0 | 0 | 0 | 0 | 212 | 149 | 1 | 1 | 0.882 | 1 | 1 | 1 | 1 | 1 | 1 | 1 | 1 | 1 | -0.024 |
| SE | 103254 | ENSG00000172661 | WASHC2C | chr10 | + | 45740072 | 45740246 | 45728861 | 45729026 | 45743389 | 45743483 | 97 | 103 | 109 | 138 | 175 | 3 | 0 | 0 | 0 | 0 | 356 | 104 | 204 | 304 | 202 | 0 | 0 | 0 | 0 | 0 | 298 | 149 | 1 | 1 | 0.942 | 1 | 1 | 1 | 1 | 1 | 1 | 1 | 1 | 1 | -0.012 |
| SE | 103256 | ENSG00000172661 | WASHC2C | chr10 | + | 45746599 | 45746647 | 45744820 | 45744882 | 45750095 | 45750206 | 199 | 238 | 253 | 247 | 454 | 0 | 1 | 0 | 0 | 0 | 793 | 254 | 466 | 696 | 330 | 0 | 0 | 0 | 0 | 0 | 197 | 149 | 1 | 1 | 1 | 0.994 | 1 | 1 | 1 | 1 | 1 | 1 | 1 | 1 | -0.001 |
| SE | 103262 | ENSG00000172661 | WASHC2C | chr10 | + | 45769448 | 45769618 | 45765678 | 45765810 | 45773255 | 45773358 | 276 | 193 | 202 | 241 | 373 | 0 | 2 | 0 | 2 | 3 | 481 | 259 | 289 | 558 | 135 | 0 | 5 | 1 | 4 | 0 | 298 | 149 | 1 | 1 | 1 | 0.98 | 1 | 0.984 | 0.984 | 1 | 0.963 | 0.993 | 0.986 | 1 | 0.001 |
| SE | 103265 | ENSG00000172661 | WASHC2C | chr10 | + | 45784820 | 45784901 | 45784564 | 45784693 | 45788870 | 45789491 | 70 | 60 | 66 | 87 | 137 | 2 | 0 | 1 | 1 | 0 | 79 | 65 | 87 | 109 | 19 | 0 | 0 | 0 | 0 | 0 | 230 | 149 | 1 | 1 | 0.958 | 1 | 0.977 | 0.983 | 1 | 1 | 1 | 1 | 1 | 1 | -0.016 |
| SE | 103270 | ENSG00000172661 | WASHC2C | chr10 | + | 45786611 | 45786674 | 45784820 | 45784901 | 45788870 | 45789491 | 8 | 1 | 1 | 7 | 10 | 0 | 2 | 1 | 1 | 0 | 0 | 0 | 2 | 21 | 7 | 0 | 7 | 18 | 17 | 4 | 212 | 149 | 1 | 1 | 1 | 0.26 | 0.413 | 0.831 | 1 | NA | 0 | 0.072 | 0.465 | 0.552 | 0.429 |
| SE | 103272 | ENSG00000172661 | WASHC2C | chr10 | + | 45786611 | 45786674 | 45785508 | 45785631 | 45788870 | 45789491 | 58 | 41 | 67 | 58 | 126 | 0 | 2 | 0 | 0 | 1 | 11 | 34 | 47 | 56 | 18 | 0 | 0 | 1 | 2 | 0 | 212 | 149 | 1 | 1 | 1 | 0.935 | 1 | 1 | 0.989 | 1 | 1 | 0.971 | 0.952 | 1 | 0 |
| SE | 103273 | ENSG00000172661 | WASHC2C | chr10 | + | 45787034 | 45787247 | 45784820 | 45784901 | 45788870 | 45789491 | 59 | 34 | 57 | 55 | 23 | 0 | 2 | 1 | 1 | 0 | 92 | 52 | 52 | 68 | 60 | 0 | 7 | 18 | 17 | 4 | 298 | 149 | 1 | 1 | 1 | 0.895 | 0.966 | 0.965 | 1 | 1 | 0.788 | 0.591 | 0.667 | 0.882 | 0.18 |
| SE | 103274 | ENSG00000172661 | WASHC2C | chr10 | + | 45787034 | 45787247 | 45785508 | 45785631 | 45788870 | 45789491 | 57 | 37 | 54 | 60 | 18 | 0 | 2 | 0 | 0 | 1 | 87 | 53 | 55 | 55 | 60 | 0 | 0 | 1 | 2 | 0 | 298 | 149 | 1 | 1 | 1 | 0.902 | 1 | 1 | 0.9 | 1 | 1 | 0.965 | 0.932 | 1 | -0.019 |
| SE | 103276 | ENSG00000128805 | ARHGAP22 | chr10 | - | 48459683 | 48459891 | 48455001 | 48455134 | 48479635 | 48479764 | 2 | 0 | 3 | 0 | 4 | 3 | 0 | 0 | 0 | 3 | 0 | 0 | 0 | 0 | 6 | 0 | 0 | 0 | 0 | 0 | 298 | 149 | 1 | 1 | 0.25 | NA | 1 | NA | 0.4 | NA | NA | NA | NA | 1 | -0.45 |
| SE | 103278 | ENSG00000128805 | ARHGAP22 | chr10 | - | 48479587 | 48479764 | 48459683 | 48459891 | 48555462 | 48555550 | 1 | 0 | 1 | 0 | 2 | 0 | 0 | 1 | 0 | 0 | 0 | 10 | 0 | 0 | 0 | 0 | 0 | 0 | 0 | 0 | 298 | 149 | 1 | 1 | 1 | NA | 0.333 | NA | 1 | NA | 1 | NA | NA | NA | -0.222 |
| SE | 103280 | ENSG00000128805 | ARHGAP22 | chr10 | - | 48479635 | 48479764 | 48459683 | 48459891 | 48555462 | 48555550 | 1 | 0 | 1 | 0 | 4 | 0 | 0 | 1 | 0 | 0 | 0 | 10 | 0 | 0 | 0 | 0 | 0 | 0 | 0 | 0 | 278 | 149 | 1 | 1 | 1 | NA | 0.349 | NA | 1 | NA | 1 | NA | NA | NA | -0.217 |
| SE | 103285 | ENSG00000196233 | LCOR | chr10 | + | 96833405 | 96833479 | 96832905 | 96833114 | 96907264 | 96907330 | 16 | 11 | 13 | 22 | 37 | 0 | 0 | 0 | 0 | 0 | 9 | 11 | 24 | 31 | 26 | 0 | 1 | 0 | 0 | 0 | 223 | 149 | 1 | 1 | 1 | 1 | 1 | 1 | 1 | 1 | 0.88 | 1 | 1 | 1 | 0.024 |
| SE | 103288 | ENSG00000196233 | LCOR | chr10 | + | 96833776 | 96833841 | 96833405 | 96833479 | 96858291 | 96858416 | 1 | 0 | 0 | 0 | 2 | 1 | 2 | 1 | 0 | 0 | 0 | 6 | 2 | 0 | 0 | 0 | 0 | 1 | 6 | 2 | 214 | 149 | 1 | 1 | 0.41 | 0 | 0 | NA | 1 | NA | 1 | 0.582 | 0 | 0 | -0.043 |
| SE | 103291 | ENSG00000196233 | LCOR | chr10 | + | 96858291 | 96858416 | 96833776 | 96833841 | 96884402 | 96884708 | 1 | 0 | 0 | 0 | 2 | 0 | 1 | 0 | 2 | 0 | 0 | 12 | 0 | 0 | 0 | 0 | 3 | 0 | 0 | 0 | 274 | 149 | 1 | 1 | 1 | 0 | NA | 0 | 1 | NA | 0.685 | NA | NA | NA | -0.185 |
| SE | 103294 | ENSG00000196233 | LCOR | chr10 | + | 96944101 | 96944245 | 96907667 | 96907747 | 96952102 | 96952196 | 5 | 0 | 1 | 8 | 1 | 0 | 0 | 6 | 0 | 1 | 0 | 7 | 0 | 0 | 0 | 0 | 0 | 0 | 1 | 0 | 293 | 149 | 1 | 1 | 1 | NA | 0.078 | 1 | 0.337 | NA | 1 | NA | 0 | NA | 0.104 |
| SE | 103296 | ENSG00000196233 | LCOR | chr10 | + | 96944112 | 96944245 | 96907667 | 96907747 | 96952102 | 96952196 | 171 | 124 | 133 | 170 | 416 | 0 | 0 | 6 | 0 | 1 | 173 | 167 | 156 | 141 | 123 | 0 | 0 | 0 | 1 | 0 | 282 | 149 | 1 | 1 | 1 | 1 | 0.921 | 1 | 0.995 | 1 | 1 | 1 | 0.987 | 1 | -0.014 |

| | | | | | | | | | | | | | | | | | | | | | | | | | | | | | | | | | | | | | | | | | | | | | | |
|---|---|---|---|---|---|---|---|---|---|---|---|---|---|---|---|---|---|---|---|---|---|---|---|---|---|---|---|---|---|---|---|---|---|---|---|---|---|---|---|---|---|---|---|---|---|---|
| SE | 103648 | ENSG00000120616 | EPC1 | chr10 | - | 32273162 | 32273281 | 32272025 | 32272167 | 32273921 | 32274038 | 104 | 72 | 131 | 129 | 105 | 0 | 2 | 0 | 0 | 0 | 60 | 75 | 75 | 62 | 5 | 0 | 0 | 0 | 0 | 0 | 268 | 149 | 1 | 1 | 1 | 0.952 | 1 | 1 | 1 | 1 | 1 | 1 | 1 | 1 | -0.01 |
| SE | 103649 | ENSG00000120616 | EPC1 | chr10 | - | 32273162 | 32273281 | 32272025 | 32272167 | 32284697 | 32285050 | 360 | 290 | 464 | 484 | 913 | 2 | 2 | 0 | 0 | 1 | 202 | 389 | 354 | 233 | 43 | 0 | 5 | 0 | 0 | 0 | 268 | 149 | 1 | 1 | 0.99 | 0.988 | 1 | 1 | 0.998 | 1 | 0.977 | 1 | 1 | 1 | 0 |
| SE | 103650 | ENSG00000120616 | EPC1 | chr10 | - | 32273921 | 32274038 | 32272025 | 32272167 | 32284697 | 32285050 | 2 | 2 | 4 | 3 | 2 | 2 | 2 | 0 | 0 | 1 | 0 | 1 | 0 | 5 | 0 | 0 | 5 | 0 | 0 | 0 | 266 | 149 | 1 | 1 | 0.359 | 0.359 | 1 | 1 | 0.528 | NA | 0.101 | NA | 1 | NA | 0.099 |
| SE | 103651 | ENSG00000120616 | EPC1 | chr10 | - | 32273921 | 32274038 | 32273162 | 32273281 | 32284697 | 32285050 | 3 | 3 | 10 | 9 | 15 | 263 | 230 | 347 | 369 | 851 | 0 | 2 | 2 | 10 | 0 | 142 | 320 | 289 | 176 | 38 | 266 | 149 | 1 | 1 | 0.006 | 0.007 | 0.016 | 0.013 | 0.01 | 0 | 0.003 | 0.004 | 0.031 | 0 | 0.003 |
| SE | 103652 | ENSG00000120616 | EPC1 | chr10 | - | 32286925 | 32287015 | 32286693 | 32286842 | 32287097 | 32287274 | 710 | 608 | 616 | 831 | 758 | 0 | 2 | 0 | 0 | 0 | 640 | 815 | 889 | 642 | 183 | 0 | 0 | 0 | 0 | 0 | 239 | 149 | 1 | 1 | 1 | 0.995 | 1 | 1 | 1 | 1 | 1 | 1 | 1 | 1 | -0.001 |
| SE | 103654 | ENSG00000120616 | EPC1 | chr10 | - | 32305771 | 32305931 | 32293591 | 32293737 | 32378490 | 32378684 | 552 | 491 | 627 | 633 | 693 | 0 | 0 | 0 | 0 | 4 | 249 | 464 | 748 | 347 | 69 | 0 | 0 | 0 | 0 | 0 | 298 | 149 | 1 | 1 | 1 | 1 | 1 | 1 | 0.989 | 1 | 1 | 1 | 1 | 1 | -0.002 |
| SE | 103655 | ENSG00000185009 | AP3M1 | chr10 | - | 74124379 | 74124524 | 74120254 | 74123910 | 74126147 | 74126355 | 195 | 254 | 249 | 247 | 255 | 0 | 4 | 0 | 1 | 0 | 127 | 142 | 181 | 222 | 4 | 0 | 0 | 0 | 0 | 0 | 294 | 149 | 1 | 1 | 1 | 0.97 | 1 | 0.992 | 1 | 1 | 1 | 1 | 1 | 1 | -0.008 |
| SE | 103656 | ENSG00000185009 | AP3M1 | chr10 | - | 74126147 | 74126355 | 74124379 | 74124524 | 74129107 | 74129241 | 267 | 260 | 272 | 299 | 316 | 3 | 2 | 3 | 2 | 0 | 90 | 145 | 178 | 209 | 16 | 0 | 1 | 1 | 0 | 0 | 298 | 149 | 1 | 1 | 0.978 | 0.985 | 0.978 | 0.987 | 1 | 1 | 0.986 | 0.989 | 1 | 1 | -0.009 |
| SE | 103658 | ENSG00000185009 | AP3M1 | chr10 | - | 74138805 | 74138931 | 74138106 | 74138382 | 74150334 | 74150757 | 44 | 28 | 42 | 48 | 26 | 2 | 0 | 0 | 0 | 0 | 0 | 47 | 28 | 45 | 16 | 0 | 0 | 0 | 0 | 0 | 275 | 149 | 1 | 1 | 0.923 | 1 | 1 | 1 | 1 | NA | 1 | 1 | 1 | 1 | -0.015 |
| SE | 103660 | ENSG00000148634 | HERC4 | chr10 | - | 67936152 | 67936235 | 67932596 | 67932780 | 67939587 | 67939654 | 295 | 309 | 257 | 289 | 422 | 1 | 0 | 0 | 0 | 0 | 66 | 199 | 92 | 95 | 10 | 0 | 0 | 0 | 0 | 0 | 232 | 149 | 1 | 1 | 0.995 | 1 | 1 | 1 | 1 | 1 | 1 | 1 | 1 | 1 | -0.001 |
| SE | 103665 | ENSG00000148634 | HERC4 | chr10 | - | 67988662 | 67988835 | 67964791 | 67966802 | 67990210 | 67990241 | 477 | 359 | 389 | 342 | 335 | 1 | 1 | 1 | 2 | 0 | 125 | 301 | 198 | 185 | 206 | 0 | 0 | 0 | 0 | 0 | 298 | 149 | 1 | 1 | 0.996 | 0.994 | 0.995 | 0.988 | 1 | 1 | 1 | 1 | 1 | 1 | -0.005 |
| SE | 103670 | ENSG00000148634 | HERC4 | chr10 | - | 67990903 | 67991015 | 67966682 | 67966802 | 67992198 | 67992323 | 0 | 0 | 3 | 3 | 3 | 0 | 1 | 0 | 0 | 0 | 0 | 0 | 2 | 0 | 0 | 0 | 0 | 0 | 0 | 0 | 261 | 149 | 1 | 1 | NA | 0 | 1 | 1 | 1 | NA | NA | 1 | NA | NA | -0.25 |
| SE | 103671 | ENSG00000148634 | HERC4 | chr10 | - | 67990903 | 67991015 | 67990210 | 67990400 | 67992198 | 67992323 | 107 | 68 | 96 | 95 | 129 | 0 | 0 | 0 | 2 | 1 | 39 | 60 | 23 | 24 | 21 | 0 | 0 | 0 | 0 | 0 | 261 | 149 | 1 | 1 | 1 | 1 | 1 | 0.964 | 0.987 | 1 | 1 | 1 | 1 | 1 | -0.01 |
| SE | 103672 | ENSG00000148634 | HERC4 | chr10 | - | 67991139 | 67991199 | 67990903 | 67991015 | 67992198 | 67992323 | 192 | 150 | 176 | 183 | 364 | 0 | 0 | 1 | 0 | 0 | 49 | 89 | 58 | 64 | 75 | 0 | 0 | 0 | 0 | 0 | 209 | 149 | 1 | 1 | 1 | 1 | 0.992 | 1 | 1 | 1 | 1 | 1 | 1 | 1 | -0.002 |
| SE | 103673 | ENSG00000148634 | HERC4 | chr10 | - | 67996045 | 67996138 | 67992605 | 67992682 | 68014025 | 68014186 | 3 | 2 | 3 | 0 | 5 | 144 | 119 | 148 | 149 | 189 | 0 | 0 | 0 | 0 | 6 | 44 | 77 | 47 | 37 | 45 | 242 | 149 | 1 | 1 | 0.013 | 0.01 | 0.012 | 0 | 0.016 | 0 | 0 | 0 | 0 | 0.076 | -0.005 |
| SE | 103675 | ENSG00000148634 | HERC4 | chr10 | - | 68014025 | 68014186 | 67992605 | 67992682 | 68025545 | 68025676 | 251 | 210 | 253 | 244 | 292 | 0 | 0 | 0 | 0 | 0 | 60 | 124 | 73 | 54 | 54 | 0 | 0 | 2 | 0 | 0 | 298 | 149 | 1 | 1 | 1 | 1 | 1 | 1 | 1 | 1 | 1 | 0.948 | 1 | 1 | 0.01 |
| SE | 103676 | ENSG00000148634 | HERC4 | chr10 | - | 68025545 | 68025676 | 68014025 | 68014186 | 68032777 | 68032869 | 202 | 172 | 174 | 178 | 245 | 3 | 1 | 4 | 0 | 0 | 29 | 82 | 64 | 47 | 56 | 0 | 0 | 0 | 0 | 0 | 280 | 149 | 1 | 1 | 0.973 | 0.989 | 0.959 | 1 | 1 | 1 | 1 | 1 | 1 | 1 | -0.016 |
| SE | 103677 | ENSG00000148634 | HERC4 | chr10 | - | 68032777 | 68032869 | 68014025 | 68014186 | 68038092 | 68038169 | 0 | 1 | 3 | 0 | 0 | 0 | 0 | 1 | 0 | 1 | 0 | 0 | 0 | 0 | 0 | 0 | 2 | 0 | 0 | 0 | 241 | 149 | 1 | 1 | NA | 1 | 0.65 | NA | 0 | NA | 0 | NA | NA | NA | 0.55 |
| SE | 103679 | ENSG00000148634 | HERC4 | chr10 | - | 68033964 | 68034186 | 68032777 | 68032869 | 68038092 | 68038169 | 279 | 201 | 197 | 277 | 212 | 0 | 0 | 2 | 0 | 0 | 37 | 117 | 84 | 50 | 96 | 0 | 0 | 0 | 0 | 0 | 298 | 149 | 1 | 1 | 1 | 1 | 0.98 | 1 | 1 | 1 | 1 | 1 | 1 | 1 | -0.004 |
| SE | 103680 | ENSG00000148634 | HERC4 | chr10 | - | 68038092 | 68038169 | 68033964 | 68034186 | 68044403 | 68044563 | 181 | 129 | 145 | 202 | 135 | 0 | 0 | 0 | 0 | 0 | 45 | 70 | 56 | 37 | 55 | 0 | 0 | 2 | 0 | 0 | 226 | 149 | 1 | 1 | 1 | 1 | 1 | 1 | 1 | 1 | 1 | 0.949 | 1 | 1 | 0.01 |
| SE | 103681 | ENSG00000148634 | HERC4 | chr10 | - | 68038092 | 68038169 | 68033964 | 68034186 | 68072882 | 68073186 | 64 | 43 | 43 | 68 | 67 | 1 | 0 | 0 | 0 | 0 | 16 | 21 | 24 | 9 | 14 | 0 | 0 | 0 | 0 | 0 | 226 | 149 | 1 | 1 | 0.977 | 1 | 1 | 1 | 1 | 1 | 1 | 1 | 1 | 1 | -0.005 |
| SE | 103682 | ENSG00000148634 | HERC4 | chr10 | - | 68039416 | 68039488 | 68038092 | 68038169 | 68044403 | 68044563 | 2 | 1 | 1 | 0 | 0 | 119 | 88 | 105 | 134 | 71 | 0 | 0 | 4 | 0 | 0 | 29 | 56 | 32 | 28 | 51 | 221 | 149 | 1 | 1 | 0.011 | 0.008 | 0.006 | 0 | 0 | 0 | 0 | 0.078 | 0 | 0 | -0.011 |
| SE | 103683 | ENSG00000148634 | HERC4 | chr10 | - | 68044403 | 68044563 | 68033964 | 68034186 | 68072882 | 68073097 | 128 | 124 | 120 | 158 | 150 | 1 | 0 | 0 | 0 | 0 | 22 | 73 | 61 | 57 | 35 | 0 | 0 | 0 | 0 | 0 | 298 | 149 | 1 | 1 | 0.985 | 1 | 1 | 1 | 1 | 1 | 1 | 1 | 1 | 1 | -0.003 |
| SE | 103684 | ENSG00000148634 | HERC4 | chr10 | - | 68044403 | 68044563 | 68038092 | 68038169 | 68067084 | 68067217 | 135 | 96 | 123 | 152 | 78 | 0 | 0 | 2 | 0 | 1 | 29 | 70 | 43 | 44 | 61 | 0 | 2 | 0 | 2 | 0 | 298 | 149 | 1 | 1 | 1 | 1 | 0.969 | 1 | 0.975 | 1 | 0.946 | 1 | 0.917 | 1 | 0.016 |
| SE | 103685 | ENSG00000148634 | HERC4 | chr10 | - | 68044403 | 68044563 | 68038092 | 68038169 | 68072882 | 68073097 | 247 | 212 | 225 | 292 | 221 | 2 | 2 | 2 | 0 | 3 | 51 | 129 | 91 | 85 | 86 | 0 | 0 | 0 | 0 | 0 | 298 | 149 | 1 | 1 | 0.984 | 0.981 | 0.983 | 1 | 0.974 | 1 | 1 | 1 | 1 | 1 | -0.016 |
| SE | 103686 | ENSG00000148634 | HERC4 | chr10 | - | 68067084 | 68067217 | 68038092 | 68038169 | 68072882 | 68073186 | 32 | 19 | 25 | 42 | 32 | 2 | 2 | 2 | 0 | 3 | 0 | 24 | 16 | 2 | 11 | 0 | 0 | 0 | 0 | 0 | 282 | 149 | 1 | 1 | 0.894 | 0.834 | 0.869 | 1 | 0.849 | NA | 1 | 1 | 1 | 1 | -0.111 |
| SE | 103698 | ENSG00000125485 | DDX31 | chr9 | - | 132618329 | 132618441 | 132612086 | 132612255 | 132625663 | 132625745 | 49 | 17 | 37 | 23 | 47 | 0 | 0 | 1 | 0 | 0 | 10 | 9 | 27 | 29 | 8 | 0 | 0 | 0 | 0 | 0 | 261 | 149 | 1 | 1 | 1 | 1 | 0.955 | 1 | 1 | 1 | 1 | 1 | 1 | 1 | -0.009 |
| SE | 103700 | ENSG00000125485 | DDX31 | chr9 | - | 132625663 | 132625745 | 132612086 | 132612255 | 132630263 | 132630403 | 14 | 8 | 22 | 11 | 13 | 0 | 1 | 0 | 0 | 0 | 19 | 18 | 3 | 0 | 0 | 0 | 0 | 6 | 0 | 0 | 231 | 149 | 1 | 1 | 1 | 0.838 | 1 | 1 | 1 | 1 | 1 | 0.244 | NA | NA | 0.22 |
| SE | 103701 | ENSG00000125485 | DDX31 | chr9 | - | 132625663 | 132625745 | 132618329 | 132618441 | 132630263 | 132630403 | 36 | 19 | 43 | 22 | 40 | 0 | 0 | 0 | 0 | 0 | 19 | 22 | 12 | 7 | 0 | 7 | 0 | 0 | 1 | 0 | 231 | 149 | 1 | 1 | 1 | 1 | 1 | 1 | 1 | 0.636 | 1 | 1 | 0.819 | NA | 0.136 |
| SE | 103702 | ENSG00000125485 | DDX31 | chr9 | - | 132646822 | 132647058 | 132645894 | 132646071 | 132648188 | 132648295 | 50 | 43 | 71 | 40 | 95 | 1 | 0 | 0 | 0 | 0 | 22 | 37 | 25 | 17 | 30 | 0 | 0 | 0 | 0 | 0 | 298 | 149 | 1 | 1 | 0.962 | 1 | 1 | 1 | 1 | 1 | 1 | 1 | 1 | 1 | -0.008 |
| SE | 103703 | ENSG00000125485 | DDX31 | chr9 | - | 132651074 | 132651116 | 132650233 | 132650298 | 132652447 | 132652492 | 59 | 47 | 82 | 78 | 63 | 0 | 0 | 2 | 1 | 0 | 114 | 89 | 59 | 26 | 32 | 0 | 0 | 0 | 0 | 0 | 191 | 149 | 1 | 1 | 1 | 1 | 0.97 | 0.984 | 1 | 1 | 1 | 1 | 1 | 1 | -0.009 |
| SE | 103704 | ENSG00000168209 | DDIT4 | chr10 | + | 72274156 | 72274421 | 72273918 | 72274060 | 72274694 | 72275559 | 168 | 57 | 116 | 128 | 9 | 0 | 0 | 0 | 0 | 0 | 28 | 220 | 266 | 37 | 138 | 0 | 0 | 0 | 3 | 0 | 298 | 149 | 1 | 1 | 1 | 1 | 1 | 1 | 1 | 1 | 1 | 1 | 0.86 | 1 | 0.028 |
| SE | 103705 | ENSG00000168209 | DDIT4 | chr10 | + | 72274224 | 72274421 | 72273923 | 72274060 | 72274694 | 72276036 | 76 | 33 | 68 | 66 | 6 | 0 | 0 | 0 | 0 | 0 | 15 | 142 | 125 | 19 | 69 | 0 | 0 | 0 | 3 | 0 | 298 | 149 | 1 | 1 | 1 | 1 | 1 | 1 | 1 | 1 | 1 | 1 | 0.76 | 1 | 0.048 |
| SE | 103706 | ENSG00000139517 | LNX2 | chr13 | - | 27559841 | 27559985 | 27556235 | 27556413 | 27562412 | 27562781 | 43 | 28 | 42 | 39 | 55 | 0 | 0 | 3 | 0 | 0 | 12 | 40 | 28 | 36 | 7 | 3 | 0 | 0 | 0 | 0 | 293 | 149 | 1 | 1 | 1 | 0.826 | 1 | 1 | 1 | 0.67 | 1 | 1 | 1 | 1 | 0.031 |
| SE | 103712 | ENSG00000124587 | PEX6 | chr6 | - | 42966043 | 42966105 | 42965680 | 42965789 | 42966241 | 42966447 | 34 | 10 | 20 | 25 | 47 | 0 | 0 | 1 | 0 | 0 | 7 | 40 | 20 | 21 | 58 | 0 | 0 | 0 | 0 | 0 | 211 | 149 | 1 | 1 | 1 | 1 | 0.934 | 1 | 1 | 1 | 1 | 1 | 1 | 1 | -0.013 |
| SE | 103713 | ENSG00000124587 | PEX6 | chr6 | - | 42968873 | 42968985 | 42968289 | 42968498 | 42969667 | 42969801 | 35 | 15 | 12 | 30 | 35 | 0 | 1 | 1 | 2 | 0 | 22 | 9 | 4 | 0 | 33 | 0 | 4 | 0 | 0 | 0 | 261 | 149 | 1 | 1 | 1 | 0.895 | 0.873 | 0.895 | 1 | 1 | 0.562 | 1 | NA | 1 | 0.042 |
| SE | 103714 | ENSG00000124587 | PEX6 | chr6 | - | 42974002 | 42974086 | 42969884 | 42969987 | 42974874 | 42975038 | 73 | 24 | 35 | 38 | 72 | 0 | 0 | 0 | 1 | 0 | 0 | 28 | 9 | 29 | 20 | 0 | 0 | 0 | 0 | 0 | 233 | 149 | 1 | 1 | 1 | 1 | 1 | 0.96 | 1 | NA | 1 | | 1 | 1 | -0.008 |
| SE | 103720 | ENSG00000107863 | ARHGAP21 | chr10 | - | 24629995 | 24630050 | 24619472 | 24621369 | 24633401 | 24633480 | 65 | 103 | 27 | 123 | 232 | 0 | 0 | 0 | 0 | 1 | 44 | 86 | 59 | 98 | 83 | 0 | 0 | 3 | 3 | 0 | 204 | 149 | 1 | 1 | 1 | 1 | 1 | 1 | 0.994 | 1 | 1 | 0.935 | 0.96 | 1 | 0.02 |
| SE | 103721 | ENSG00000107863 | ARHGAP21 | chr10 | - | 24633401 | 24633480 | 24629013 | 24630050 | 24635010 | 24635103 | 71 | 94 | 34 | 136 | 223 | 0 | 1 | 0 | 0 | 0 | 85 | 97 | 84 | 94 | 118 | 0 | 0 | 0 | 0 | 0 | 228 | 149 | 1 | 1 | 1 | 0.984 | 1 | 1 | 1 | 1 | 1 | 1 | 1 | 1 | -0.003 |
| SE | 103722 | ENSG00000107863 | ARHGAP21 | chr10 | - | 24714296 | 24714378 | 24670338 | 24670397 | 24721836 | 24722279 | 0 | 0 | 0 | 0 | 0 | 22 | 15 | 9 | 34 | 39 | 0 | 0 | 0 | 3 | 0 | 24 | 28 | 29 | 23 | 38 | 231 | 149 | 1 | 1 | 0 | 0 | 0 | 0 | 0 | 0 | 0 | | 0.078 | 0 | -0.016 |
| SE | 103724 | ENSG00000136738 | STAM | chr10 | + | 17660463 | 17660548 | 17644150 | 17644379 | 17688026 | 17688173 | 16 | 16 | 19 | 21 | 16 | 0 | 0 | 0 | 0 | 0 | 8 | 8 | 27 | 9 | 0 | 0 | 1 | 2 | 0 | 0 | 234 | 149 | 1 | 1 | 1 | 1 | 1 | 1 | 1 | 1 | 0.836 | 0.896 | 1 | NA | 0.067 |
| SE | 103726 | ENSG00000136738 | STAM | chr10 | + | 17672974 | 17673078 | 17660463 | 17660548 | 17688026 | 17688173 | 1 | 0 | 0 | 3 | 0 | 6 | 6 | 3 | 3 | 9 | 0 | 0 | 1 | 3 | 0 | 0 | 2 | 9 | 2 | 0 | 253 | 149 | 1 | 1 | 0.089 | 0 | 0 | 0.371 | 0 | NA | 0 | 0.061 | 0.469 | NA | -0.085 |
| SE | 103727 | ENSG00000136738 | STAM | chr10 | + | 17684674 | 17684750 | 17644150 | 17644379 | 17688026 | 17688173 | 1 | 0 | 0 | 0 | 0 | 0 | 0 | 0 | 0 | 0 | 0 | 0 | 3 | 4 | 0 | 0 | 1 | 2 | 0 | 0 | 225 | 149 | 1 | 1 | 1 | NA | NA | NA | NA | NA | 0 | 0.498 | 1 | NA | 0.501 |
| SE | 103729 | ENSG00000136738 | STAM | chr10 | + | 17684674 | 17684750 | 17672974 | 17673078 | 17688026 | 17688173 | 0 | 0 | 1 | 0 | 8 | 0 | 0 | 0 | 3 | 0 | 0 | 0 | 0 | 2 | 0 | 0 | 1 | 0 | 0 | 0 | 225 | 149 | 1 | 1 | NA | NA | 1 | 0 | 1 | NA | NA | 0 | 1 | NA | 0.167 |
| SE | 103730 | ENSG00000136738 | STAM | chr10 | + | 17684831 | 17684927 | 17684674 | 17684750 | 17688026 | 17688173 | 141 | 97 | 107 | 162 | 298 | 0 | 0 | 0 | 0 | 0 | 44 | 110 | 112 | 111 | 28 | 0 | 0 | 1 | 1 | 0 | 245 | 149 | 1 | 1 | 1 | 1 | 1 | 1 | 1 | 1 | 1 | 0.986 | 0.985 | 1 | 0.006 |
| SE | 103731 | ENSG00000136738 | STAM | chr10 | + | 17688026 | 17688173 | 17660463 | 17660548 | 17693221 | 17693281 | 116 | 83 | 102 | 97 | 269 | 0 | 1 | 0 | 0 | 0 | 85 | 47 | 108 | 82 | 16 | 0 | 0 | 1 | 0 | 0 | 296 | 149 | 1 | 1 | 1 | 0.977 | 1 | 1 | 1 | 1 | 1 | 1 | 1 | 1 | -0.005 |
| SE | 103732 | ENSG00000136738 | STAM | chr10 | + | 17688026 | 17688173 | 17684831 | 17684927 | 17693221 | 17693281 | 205 | 145 | 170 | 205 | 475 | 0 | 0 | 0 | 1 | 0 | 125 | 115 | 178 | 149 | 34 | 0 | 0 | 0 | 0 | 0 | 296 | 149 | 1 | 1 | 1 | 1 | 1 | 0.99 | 1 | 1 | 1 | 1 | 1 | 1 | -0.002 |
| SE | 103733 | ENSG00000136738 | STAM | chr10 | + | 17693221 | 17693312 | 17688026 | 17688173 | 17695048 | 17695241 | 184 | 148 | 151 | 157 | 500 | 0 | 0 | 0 | 0 | 1 | 144 | 81 | 190 | 172 | 41 | 0 | 0 | 2 | 0 | 0 | 240 | 149 | 1 | 1 | 1 | 1 | 1 | 1 | 0.997 | 1 | 1 | 0.983 | 1 | 1 | 0.003 |
| SE | 103734 | ENSG00000136738 | STAM | chr10 | + | 17700190 | 17700279 | 17696774 | 17696869 | 17704430 | 17704518 | 205 | 209 | 185 | 234 | 417 | 0 | 0 | 0 | 0 | 0 | 112 | 242 | 269 | 244 | 119 | 0 | 2 | 0 | 0 | 0 | 238 | 149 | 1 | 1 | 1 | 1 | 1 | 1 | 1 | 1 | 0.987 | 1 | 1 | 1 | 0.003 |
| SE | 103735 | ENSG00000107937 | GTPBP4 | chr10 | + | 995928 | 996032 | 992488 | 992659 | 996105 | 996242 | 267 | 177 | 209 | 197 | 277 | 2 | 0 | 0 | 0 | 0 | 133 | 159 | 255 | 151 | 94 | 0 | 0 | 0 | 2 | 0 | 253 | 149 | 1 | 1 | 0.987 | 1 | 1 | 1 | 1 | 1 | 1 | 1 | 0.978 | 1 | 0.002 |
| SE | 103738 | ENSG00000107937 | GTPBP4 | chr10 | + | 1008957 | 1009035 | 1000947 | 1001013 | 1009528 | 1009580 | 32 | 16 | 22 | 24 | 26 | 0 | 0 | 0 | 0 | 0 | 4 | 10 | 47 | 11 | 6 | 0 | 0 | 1 | 0 | 0 | 227 | 149 | 1 | 1 | 1 | 1 | 1 | 1 | 1 | 1 | 1 | 0.969 | 1 | 1 | 0.006 |
| SE | 103739 | ENSG00000107937 | GTPBP4 | chr10 | + | 1012464 | 1012662 | 1010419 | 1010520 | 1014246 | 1014312 | 363 | 236 | 276 | 306 | 443 | 0 | 0 | 0 | 0 | 0 | 162 | 237 | 425 | 224 | 62 | 0 | 0 | 1 | 0 | 0 | 298 | 149 | 1 | 1 | 1 | 1 | 1 | 1 | 1 | 1 | 1 | 0.995 | 1 | 1 | 0.001 |
| SE | 103740 | ENSG00000153531 | ADPRHL1 | chr13 | - | 113424216 | 113424349 | 113421939 | 113422979 | 113425051 | 113425179 | 2 | 2 | 0 | 2 | 1 | 0 | 0 | 0 | 0 | 1 | 0 | 4 | 2 | 1 | 0 | 0 | 0 | 0 | 0 | 0 | 282 | 149 | 1 | 1 | 1 | 1 | NA | 1 | 0.346 | NA | 1 | 1 | 1 | NA | -0.163 |
| SE | 103744 | ENSG00000047056 | WDR37 | chr10 | + | 1072115 | 1072293 | 1056384 | 1056968 | 1077906 | 1078003 | 162 | 126 | 112 | 139 | 192 | 0 | 0 | 1 | 0 | 0 | 162 | 155 | 113 | 121 | 205 | 0 | 0 | 0 | 0 | 0 | 298 | 149 | 1 | 1 | 1 | 1 | 0.982 | 1 | 1 | 1 | 1 | 1 | 1 | 1 | -0.004 |
| SE | 103745 | ENSG00000047056 | WDR37 | chr10 | + | 1072115 | 1072293 | 1056403 | 1056564 | 1077906 | 1078003 | 150 | 121 | 102 | 142 | 187 | 0 | 0 | 0 | 1 | 0 | 190 | 177 | 121 | 143 | 228 | 0 | 0 | 0 | 0 | 0 | 298 | 149 | 1 | 1 | 1 | 1 | 1 | 0.986 | 1 | 1 | 1 | 1 | 1 | 1 | -0.003 |
| SE | 103747 | ENSG00000047056 | WDR37 | chr10 | + | 1077906 | 1078003 | 1072115 | 1072293 | 1080010 | 1080106 | 271 | 222 | 172 | 253 | 349 | 6 | 0 | 0 | 4 | 0 | 399 | 300 | 208 | 242 | 374 | 6 | 0 | 0 | 1 | 4 | 246 | 149 | 1 | 1 | 0.965 | 1 | 1 | 0.975 | 1 | 0.976 | 1 | 1 | 0.993 | 0.983 | -0.002 |
| SE | 103748 | ENSG00000047056 | WDR37 | chr10 | + | 1080010 | 1080106 | 1077906 | 1078003 | 1080411 | 1080476 | 290 | 213 | 235 | 285 | 338 | 0 | 0 | 0 | 0 | 0 | 462 | 341 | 200 | 278 | 489 | 0 | 0 | 0 | 1 | 0 | 245 | 149 | 1 | 1 | 1 | 1 | 1 | 1 | 1 | 1 | 1 | 1 | 0.994 | 1 | 0.001 |
| SE | 103749 | ENSG00000047056 | WDR37 | chr10 | + | 1080411 | 1080476 | 1077906 | 1078003 | 1084402 | 1084538 | 45 | 31 | 31 | 34 | 19 | 0 | 0 | 0 | 0 | 3 | 5 | 55 | 27 | 24 | 50 | 0 | 0 | 0 | 0 | 0 | 214 | 149 | 1 | 1 | 1 | 1 | 1 | 1 | 0.815 | 1 | 1 | 1 | 1 | 1 | -0.037 |
| SE | 103750 | ENSG00000047056 | WDR37 | chr10 | + | 1086285 | 1086357 | 1084402 | 1084538 | 1093451 | 1093496 | 179 | 116 | 138 | 164 | 124 | 0 | 0 | 0 | 0 | 0 | 169 | 138 | 104 | 124 | 254 | 0 | 0 | 0 | 0 | 1 | 221 | 149 | 1 | 1 | 1 | 1 | 1 | 1 | 1 | 1 | 1 | 1 | 1 | 0.994 | 0.001 |
| SE | 103751 | ENSG00000047056 | WDR37 | chr10 | + | 1103598 | 1103836 | 1096169 | 1096246 | 1105125 | 1105267 | 65 | 25 | 52 | 47 | 37 | 0 | 0 | 0 | 0 | 0 | 53 | 37 | 42 | 53 | 36 | 0 | 0 | 1 | 0 | 0 | 298 | 149 | 1 | 1 | 1 | 1 | 1 | 1 | 1 | 1 | 1 | 0.955 | 1 | 1 | 0.009 |
| SE | 103753 | ENSG00000047056 | WDR37 | chr10 | + | 1103601 | 1103836 | 1096169 | 1096246 | 1105125 | 1105267 | 45 | 29 | 39 | 38 | 18 | 0 | 0 | 0 | 0 | 0 | 43 | 26 | 33 | 50 | 34 | 0 | 0 | 1 | 0 | 0 | 298 | 149 | 1 | 1 | 1 | 1 | 1 | 1 | 1 | 1 | 1 | 0.943 | 1 | 1 | 0.011 |
| SE | 103756 | ENSG00000047056 | WDR37 | chr10 | + | 1105125 | 1105267 | 1103598 | 1103836 | 1124217 | 1124352 | 80 | 29 | 60 | 48 | 41 | 0 | 0 | 0 | 0 | 0 | 70 | 39 | 60 | 83 | 64 | 0 | 3 | 0 | 0 | 0 | 291 | 149 | 1 | 1 | 1 | 1 | 1 | 1 | 1 | 1 | 0.869 | 1 | 1 | 1 | 0.026 |
| SE | 103760 | ENSG00000239883 | PARGP1 | chr10 | - | 45963884 | 45964871 | 45962810 | 45962994 | 45971937 | 45972154 | 16 | 5 | 9 | 25 | 30 | 1 | 0 | 1 | 2 | 0 | 17 | 12 | 35 | 4 | 16 | 0 | 0 | 2 | 1 | 0 | 298 | 149 | 1 | 1 | 0.889 | 1 | 0.818 | 0.862 | 1 | 1 | 1 | 0.897 | 0.667 | 1 | 0.001 |
| SE | 103761 | ENSG00000118518 | RNF146 | chr6 | + | 127280515 | 127280604 | 127280230 | 127280340 | 127282286 | 127282395 | 9 | 2 | 23 | 7 | 9 | 3 | 0 | 0 | 1 | 0 | 0 | 12 | 0 | 0 | 0 | 0 | 0 | 1 | 0 | 0 | 238 | 149 | 1 | 1 | 0.653 | 1 | 1 | 0.814 | 1 | NA | 1 | 0 | NA | NA | 0.393 |
| SE | 103762 | ENSG00000118518 | RNF146 | chr6 | + | 127280515 | 127280604 | 127280230 | 127280340 | 127286049 | 127286177 | 11 | 3 | 24 | 8 | 26 | 28 | 9 | 4 | 15 | 12 | 0 | 17 | 2 | 0 | 0 | 0 | 2 | 11 | 2 | 0 | 238 | 149 | 1 | 1 | 0.197 | 0.173 | 0.79 | 0.25 | 0.576 | NA | 0.842 | 0.102 | 0 | NA | 0.083 |
| SE | 103763 | ENSG00000118518 | RNF146 | chr6 | + | 127282286 | 127282395 | 127280230 | 127280340 | 127286049 | 127286084 | 10 | 0 | 0 | 10 | 0 | 28 | 9 | 4 | 15 | 12 | 0 | 0 | 6 | 0 | 0 | 0 | 2 | 11 | 2 | 0 | 258 | 149 | 1 | 1 | 0.171 | 0 | 0 | 0.278 | 0 | NA | 0 | 0.24 | 0 | NA | 0.01 |
| SE | 103764 | ENSG00000118518 | RNF146 | chr6 | + | 127282286 | 127282395 | 127280230 | 127280340 | 127286615 | 127286849 | 2 | 1 | 0 | 0 | 0 | 19 | 4 | 14 | 13 | 20 | 0 | 0 | 1 | 0 | 0 | 11 | 7 | 7 | 22 | 0 | 258 | 149 | 1 | 1 | 0.057 | 0.126 | 0 | 0 | 0 | 0 | 0 | 0.076 | 0 | NA | 0.018 |
| SE | 103765 | ENSG00000118518 | RNF146 | chr6 | + | 127282286 | 127282395 | 127280515 | 127280604 | 127286049 | 127286084 | 7 | 0 | 3 | 9 | 0 | 4 | 1 | 4 | 1 | 17 | 0 | 3 | 5 | 0 | 0 | 0 | 8 | 2 | 0 | 0 | 258 | 149 | 1 | 1 | 0.503 | 0 | 0.302 | 0.839 | 0 | NA | 0.178 | 0.591 | NA | NA | -0.056 |
| SE | 103766 | ENSG00000118518 | RNF146 | chr6 | + | 127285207 | 127285348 | 127280230 | 127280340 | 127286049 | 127286097 | 8 | 2 | 10 | 3 | 0 | 28 | 9 | 4 | 15 | 12 | 0 | 3 | 0 | 16 | 0 | 0 | 2 | 11 | 2 | 0 | 290 | 149 | 1 | 1 | 0.128 | 0.102 | 0.562 | 0.093 | 0 | NA | 0.435 | 0 | 0.804 | NA | -0.236 |
| SE | 103767 | ENSG00000118518 | RNF146 | chr6 | + | 127286049 | 127286177 | 127280230 | 127280340 | 127286615 | 127286643 | 55 | 19 | 22 | 23 | 73 | 19 | 4 | 14 | 13 | 20 | 0 | 17 | 18 | 16 | 0 | 11 | 7 | 7 | 22 | 0 | 277 | 149 | 1 | 1 | 0.609 | 0.719 | 0.458 | 0.488 | 0.663 | 0 | 0.566 | 0.58 | 0.281 | NA | 0.231 |
| SE | 103768 | ENSG00000118518 | RNF146 | chr6 | + | 127286049 | 127286177 | 127282286 | 127282395 | 127286615 | 127286643 | 40 | 14 | 21 | 23 | 61 | 0 | 1 | 0 | 0 | 0 | 0 | 18 | 12 | 14 | 0 | 0 | 0 | 0 | 0 | 0 | 277 | 149 | 1 | 1 | 1 | 0.883 | 1 | 1 | 1 | NA | 1 | 1 | 1 | NA | -0.023 |
| SE | 103775 | ENSG00000107736 | CDH23 | chr10 | + | 71675111 | 71675176 | 71646458 | 71646617 | 71677455 | 71677693 | 12 | 11 | 0 | 5 | 7 | 0 | 0 | 1 | 0 | 0 | 20 | 29 | 12 | 0 | 9 | 0 | 0 | 0 | 0 | 0 | 214 | 149 | 1 | 1 | 1 | 1 | 0 | 1 | 1 | 1 | 1 | 1 | NA | 1 | -0.2 |
| SE | 103776 | ENSG00000107736 | CDH23 | chr10 | + | 71682444 | 71682572 | 71679386 | 71679492 | 71687646 | 71687719 | 4 | 6 | 0 | 4 | 3 | 0 | 0 | 0 | 0 | 0 | 3 | 6 | 8 | 6 | 6 | 10 | 0 | 2 | 0 | 0 | 277 | 149 | 1 | 1 | 1 | 1 | NA | 1 | 1 | 0.139 | 1 | 0.683 | 1 | 1 | 0.236 |
| SE | 103777 | ENSG00000107736 | CDH23 | chr10 | + | 71702021 | 71702211 | 71695417 | 71695525 | 71702548 | 71702694 | 19 | 14 | 17 | 10 | 29 | 0 | 0 | 0 | 0 | 0 | 12 | 26 | 5 | 24 | 36 | 0 | 0 | 0 | 2 | 0 | 298 | 149 | 1 | 1 | 1 | 1 | 1 | 1 | 1 | 1 | 1 | 1 | 0.857 | 1 | 0.029 |
| SE | 103779 | ENSG00000107736 | CDH23 | chr10 | + | 71730468 | 71730604 | 71724044 | 71724105 | 71731986 | 71732375 | 20 | 9 | 6 | 10 | 3 | 0 | 0 | 1 | 0 | 0 | 25 | 15 | 5 | 10 | 40 | 0 | 0 | 0 | 0 | 0 | 285 | 149 | 1 | 1 | 1 | 1 | 0.758 | 1 | 1 | 1 | 1 | 1 | 1 | 1 | -0.048 |
| SE | 103781 | ENSG00000107736 | CDH23 | chr10 | + | 71734655 | 71734658 | 71734239 | 71734341 | 71738497 | 71738647 | 2 | 1 | 1 | 2 | 0 | 8 | 11 | 1 | 11 | 0 | 0 | 0 | 0 | 1 | 15 | 0 | 3 | 2 | 0 | 34 | 152 | 149 | 1 | 1 | 0.197 | 0.082 | 0.495 | 0.151 | NA | NA | 0 | 0 | 1 | 0.302 | -0.094 |
| SE | 103782 | ENSG00000107736 | CDH23 | chr10 | + | 71738497 | 71738647 | 71734655 | 71734658 | 71739643 | 71739772 | 9 | 10 | 5 | 10 | 0 | 0 | 0 | 0 | 0 | 0 | 54 | 6 | 23 | 10 | 24 | 0 | 4 | 0 | 0 | 0 | 298 | 149 | 1 | 1 | 1 | 1 | 1 | 1 | NA | 1 | 0.429 | 1 | 1 | 1 | 0.114 |
| SE | 103783 | ENSG00000107736 | CDH23 | chr10 | + | 71791131 | 71791335 | 71790287 | 71790413 | 71793181 | 71793640 | 13 | 16 | 18 | 16 | 3 | 0 | 0 | 1 | 0 | 0 | 0 | 1 | 4 | 11 | 2 | 0 | 0 | 0 | 0 | 0 | 298 | 149 | 1 | 1 | 1 | 1 | 0.9 | 1 | 1 | NA | 1 | 1 | 1 | 1 | -0.02 |
| SE | 103788 | ENSG00000049323 | LTBP1 | chr2 | + | 33243661 | 33243784 | 33222079 | 33222151 | 33252676 | 33252844 | 35 | 42 | 14 | 92 | 114 | 0 | 0 | 0 | 1 | 0 | 67 | 28 | 46 | 56 | 607 | 0 | 0 | 0 | 0 | 2 | 272 | 149 | 1 | 1 | 1 | 1 | 1 | 0.981 | 1 | 1 | 1 | 1 | 1 | 0.994 | -0.003 |
| SE | 103793 | ENSG00000049323 | LTBP1 | chr2 | + | 33347366 | 33347510 | 33315143 | 33315269 | 33360596 | 33360779 | 33 | 41 | 11 | 110 | 112 | 0 | 0 | 0 | 1 | 0 | 102 | 19 | 54 | 47 | 1012 | 0 | 0 | 0 | 0 | 0 | 293 | 149 | 1 | 1 | 1 | 1 | 1 | 0.982 | 1 | 1 | 1 | 1 | 1 | 1 | -0.004 |
| SE | 103795 | ENSG00000108175 | ZMIZ1 | chr10 | + | 79208335 | 79208449 | 79201583 | 79201692 | 79216168 | 79216274 | 90 | 51 | 46 | 86 | 56 | 0 | 2 | 0 | 0 | 0 | 75 | 89 | 48 | 110 | 162 | 0 | 0 | 0 | 0 | 5 | 263 | 149 | 1 | 1 | 1 | 0.935 | 1 | 1 | 1 | 1 | 1 | 1 | 1 | 0.948 | -0.003 |
| SE | 103796 | ENSG00000108175 | ZMIZ1 | chr10 | + | 79277180 | 79277325 | 79216168 | 79216274 | 79289774 | 79289807 | 39 | 18 | 13 | 47 | 14 | 0 | 0 | 0 | 0 | 0 | 125 | 57 | 45 | 68 | 134 | 2 | 0 | 0 | 0 | 0 | 294 | 149 | 1 | 1 | 1 | 1 | 1 | 1 | 1 | 0.969 | 1 | 1 | 1 | 1 | 0.006 |
| SE | 103797 | ENSG00000108175 | ZMIZ1 | chr10 | + | 79293380 | 79293653 | 79292157 | 79292356 | 79296470 | 79296653 | 42 | 22 | 17 | 25 | 35 | 1 | 0 | 0 | 0 | 0 | 157 | 54 | 38 | 52 | 111 | 0 | 0 | 0 | 0 | 0 | 298 | 149 | 1 | 1 | 0.955 | 1 | 1 | 1 | 1 | 1 | 1 | 1 | 1 | 1 | -0.009 |
| SE | 103800 | ENSG00000139970 | RTN1 | chr14 | - | 59603211 | 59603258 | 59603064 | 59603123 | 59603851 | 59603921 | 93 | 84 | 73 | 93 | 110 | 0 | 0 | 0 | 3 | 0 | 258 | 46 | 28 | 44 | 638 | 0 | 0 | 1 | 1 | 0 | 196 | 149 | 1 | 1 | 1 | 1 | 1 | 0.959 | 1 | 1 | 1 | 0.955 | 0.971 | 1 | 0.007 |
| SE | 103801 | ENSG00000139970 | RTN1 | chr14 | - | 59605367 | 59605506 | 59603851 | 59603921 | 59607284 | 59607492 | 117 | 118 | 89 | 94 | 148 | 1 | 0 | 0 | 0 | 0 | 350 | 37 | 48 | 72 | 634 | 0 | 2 | 0 | 0 | 5 | 288 | 149 | 1 | 1 | 0.984 | 1 | 1 | 1 | 1 | 1 | 0.905 | 1 | 1 | 0.985 | 0.019 |
| SE | 103802 | ENSG00000139970 | RTN1 | chr14 | - | 59607284 | 59607492 | 59603851 | 59603921 | 59726918 | 59727668 | 37 | 40 | 34 | 14 | 40 | 0 | 0 | 0 | 1 | 0 | 81 | 11 | 15 | 47 | 155 | 0 | 0 | 0 | 0 | 0 | 298 | 149 | 1 | 1 | 1 | 1 | 1 | 0.875 | 1 | 1 | 1 | 1 | 1 | 1 | -0.025 |
| SE | 103804 | ENSG00000139970 | RTN1 | chr14 | - | 59610121 | 59610232 | 59607284 | 59607492 | 59726918 | 59727713 | 0 | 0 | 1 | 0 | 0 | 36 | 40 | 34 | 14 | 40 | 0 | 0 | 0 | 0 | 0 | 81 | 9 | 15 | 47 | 150 | 260 | 149 | 1 | 1 | 0 | 0 | 0.017 | 0 | 0 | 0 | 0 | 0 | 0 | 0 | 0.003 |
| SE | 103808 | ENSG00000225484 | NUTM2B-AS1 | chr10 | - | 79665496 | 79665594 | 79663087 | 79663234 | 79668385 | 79668452 | 0 | 0 | 1 | 0 | 0 | 0 | 0 | 0 | 3 | 0 | 7 | 21 | 0 | 0 | 0 | 0 | 0 | 0 | 0 | 0 | 247 | 149 | 1 | 1 | NA | NA | 1 | 0 | NA | 1 | 1 | NA | NA | NA | -0.5 |
| SE | 103811 | ENSG00000225484 | NUTM2B-AS1 | chr10 | - | 79723914 | 79724012 | 79716444 | 79723654 | 79766270 | 79766324 | 0 | 0 | 15 | 0 | 0 | 0 | 0 | 3 | 0 | 0 | 30 | 11 | 0 | 0 | 0 | 0 | 0 | 0 | 0 | 0 | 247 | 149 | 1 | 1 | NA | NA | 0.751 | NA | NA | 1 | 1 | NA | NA | NA | -0.249 |
| SE | 103814 | ENSG00000225484 | NUTM2B-AS1 | chr10 | - | 79806038 | 79806122 | 79804056 | 79804527 | 79808758 | 79808832 | 109 | 78 | 108 | 152 | 177 | 0 | 3 | 0 | 0 | 0 | 237 | 163 | 159 | 178 | 234 | 0 | 0 | 2 | 0 | 2 | 233 | 149 | 1 | 1 | 1 | 0.943 | 1 | 1 | 1 | 1 | 0.981 | 1 | 0.983 | 1 | -0.004 |
| SE | 103822 | ENSG00000062650 | WAPL | chr10 | - | 86446241 | 86446449 | 86443274 | 86443363 | 86451966 | 86452131 | 427 | 325 | 377 | 417 | 511 | 0 | 1 | 0 | 0 | 0 | 100 | 239 | 386 | 202 | 26 | 0 | 0 | 0 | 0 | 0 | 298 | 149 | 1 | 1 | 1 | 0.994 | 1 | 1 | 1 | 1 | 1 | 1 | 1 | 1 | -0.001 |
| SE | 103824 | ENSG00000062650 | WAPL | chr10 | - | 86460398 | 86460496 | 86458988 | 86459065 | 86461175 | 86461287 | 492 | 423 | 473 | 527 | 783 | 0 | 1 | 2 | 0 | 0 | 144 | 399 | 410 | 228 | 64 | 0 | 0 | 2 | 2 | 2 | 247 | 149 | 1 | 1 | 1 | 0.996 | 0.993 | 1 | 1 | 1 | 1 | 0.992 | 0.986 | 0.951 | 0.012 |
| SE | 103825 | ENSG00000062650 | WAPL | chr10 | - | 86461175 | 86461287 | 86460398 | 86460496 | 86467278 | 86467506 | 473 | 394 | 431 | 481 | 687 | 2 | 1 | 0 | 1 | 1 | 112 | 398 | 362 | 164 | 57 | 0 | 0 | 0 | 0 | 0 | 261 | 149 | 1 | 1 | 0.993 | 0.996 | 1 | 0.996 | 0.997 | 1 | 1 | 1 | 1 | 1 | -0.004 |
| SE | 103827 | ENSG00000062650 | WAPL | chr10 | - | 86470991 | 86471103 | 86467052 | 86467506 | 86472207 | 86472287 | 383 | 305 | 349 | 398 | 502 | 0 | 1 | 2 | 1 | 1 | 91 | 268 | 284 | 131 | 14 | 0 | 0 | 3 | 0 | 0 | 261 | 149 | 1 | 1 | 1 | 0.994 | 0.99 | 0.996 | 0.997 | 1 | 1 | 0.982 | 1 | 1 | -0.001 |
| SE | 103828 | ENSG00000062650 | WAPL | chr10 | - | 86472207 | 86472344 | 86467278 | 86467506 | 86472611 | 86472764 | 16 | 8 | 13 | 8 | 24 | 0 | 0 | 0 | 0 | 0 | 0 | 3 | 9 | 4 | 0 | 0 | 3 | 0 | 0 | 0 | 286 | 149 | 1 | 1 | 1 | 1 | 1 | 1 | 1 | NA | 0.343 | 1 | 1 | NA | 0.219 |
| SE | 103829 | ENSG00000062650 | WAPL | chr10 | - | 86472611 | 86472764 | 86472207 | 86472344 | 86473877 | 86473973 | 171 | 103 | 146 | 122 | 189 | 60 | 26 | 33 | 37 | 53 | 0 | 75 | 124 | 37 | 4 | 0 | 9 | 31 | 1 | 0 | 298 | 149 | 1 | 1 | 0.588 | 0.665 | 0.689 | 0.622 | 0.641 | NA | 0.806 | 0.667 | 0.949 | 1 | -0.215 |
| SE | 103830 | ENSG00000062650 | WAPL | chr10 | - | 86497200 | 86497319 | 86473877 | 86473973 | 86499717 | 86500743 | 193 | 140 | 154 | 183 | 288 | 0 | 0 | 0 | 4 | 0 | 20 | 162 | 172 | 88 | 6 | 0 | 0 | 5 | 0 | 0 | 268 | 149 | 1 | 1 | 1 | 1 | 1 | 0.962 | 1 | 1 | 1 | 0.95 | 1 | 1 | 0.002 |
| SE | 103831 | ENSG00000062650 | WAPL | chr10 | - | 86517570 | 86518091 | 86499717 | 86500743 | 86521364 | 86521792 | 171 | 108 | 138 | 171 | 190 | 3 | 0 | 0 | 0 | 0 | 10 | 125 | 160 | 73 | 5 | 0 | 0 | 0 | 0 | 0 | 298 | 149 | 1 | 1 | 0.966 | 1 | 1 | 1 | 1 | 1 | 1 | 1 | 1 | 1 | -0.007 |
| SE | 103833 | ENSG00000152582 | SPEF2 | chr5 | + | 35654539 | 35654726 | 35649360 | 35649425 | 35659018 | 35659207 | 49 | 39 | 43 | 44 | 61 | 0 | 0 | 0 | 0 | 0 | 5 | 3 | 6 | 8 | 8 | 0 | 0 | 2 | 0 | 0 | 298 | 149 | 1 | 1 | 1 | 1 | 1 | 1 | 1 | 1 | 1 | 0.6 | 1 | 1 | 0.08 |
| SE | 103835 | ENSG00000152582 | SPEF2 | chr5 | + | 35712811 | 35712886 | 35708947 | 35709121 | 35727674 | 35727823 | 0 | 1 | 2 | 2 | 3 | 0 | 2 | 0 | 1 | 0 | 0 | 0 | 1 | 0 | 0 | 0 | 0 | 0 | 0 | 0 | 224 | 149 | 1 | 1 | NA | 0.25 | 1 | 0.571 | 1 | NA | NA | 1 | NA | NA | -0.295 |
| SE | 103836 | ENSG00000152582 | SPEF2 | chr5 | + | 35759567 | 35759719 | 35753623 | 35753761 | 35763521 | 35763702 | 1 | 3 | 5 | 10 | 6 | 0 | 0 | 0 | 0 | 0 | 0 | 0 | 0 | 3 | 0 | 0 | 10 | 0 | 0 | 0 | 298 | 149 | 1 | 1 | 1 | 1 | 1 | 1 | 1 | NA | NA | 0 | 1 | NA | 0.5 |
| SE | 103840 | ENSG00000152582 | SPEF2 | chr5 | + | 35789157 | 35789375 | 35787865 | 35789031 | 35789496 | 35789705 | 5 | 4 | 4 | 7 | 4 | 0 | 0 | 0 | 0 | 1 | 0 | 0 | 0 | 3 | 0 | 0 | 0 | 0 | 0 | 0 | 298 | 149 | 1 | 1 | 1 | 1 | 1 | 1 | 0.667 | NA | NA | NA | 1 | NA | -0.067 |
| SE | 103848 | ENSG00000136807 | CDK9 | chr9 | + | 127786700 | 127786782 | 127785678 | 127786240 | 127787517 | 127787575 | 276 | 116 | 237 | 247 | 335 | 0 | 2 | 0 | 0 | 0 | 221 | 247 | 191 | 276 | 353 | 0 | 0 | 0 | 0 | 0 | 231 | 149 | 1 | 1 | 1 | 0.974 | 1 | 1 | 1 | 1 | 1 | 1 | 1 | 1 | -0.005 |
| SE | 103849 | ENSG00000136807 | CDK9 | chr9 | + | 127787946 | 127788113 | 127787499 | 127787608 | 127788213 | 127788385 | 326 | 191 | 330 | 327 | 509 | 0 | 0 | 0 | 0 | 0 | 262 | 338 | 247 | 239 | 479 | 0 | 0 | 1 | 0 | 0 | 298 | 149 | 1 | 1 | 1 | 1 | 1 | 1 | 1 | 1 | 1 | 0.992 | 1 | 1 | 0.002 |
| SE | 103855 | ENSG00000059573 | ALDH18A1 | chr10 | - | 95632958 | 95633049 | 95628367 | 95628492 | 95633490 | 95633649 | 128 | 47 | 135 | 132 | 121 | 0 | 0 | 0 | 1 | 4 | 154 | 133 | 66 | 81 | 106 | 0 | 0 | 1 | 0 | 0 | 240 | 149 | 1 | 1 | 1 | 1 | 1 | 0.988 | 0.949 | 1 | 1 | 0.976 | 1 | 1 | -0.008 |
| SE | 103856 | ENSG00000059573 | ALDH18A1 | chr10 | - | 95642991 | 95643206 | 95637286 | 95637436 | 95653289 | 95653405 | 95 | 40 | 85 | 79 | 40 | 0 | 0 | 0 | 1 | 0 | 32 | 63 | 31 | 31 | 4 | 0 | 0 | 0 | 0 | 0 | 298 | 149 | 1 | 1 | 1 | 1 | 1 | 0.975 | 1 | 1 | 1 | 1 | 1 | 1 | -0.005 |
| SE | 103857 | ENSG00000059573 | ALDH18A1 | chr10 | - | 95653289 | 95653405 | 95642991 | 95643206 | 95656596 | 95656706 | 90 | 38 | 68 | 67 | 34 | 0 | 0 | 1 | 2 | 0 | 25 | 36 | 26 | 32 | 9 | 0 | 0 | 0 | 0 | 0 | 265 | 149 | 1 | 1 | 1 | 1 | 0.975 | 0.95 | 1 | 1 | 1 | 1 | 1 | 1 | -0.015 |
| SE | 103860 | ENSG00000138185 | ENTPD1 | chr10 | + | 95839690 | 95839808 | 95823235 | 95823364 | 95844475 | 95844635 | 166 | 169 | 90 | 128 | 176 | 0 | 0 | 0 | 0 | 0 | 305 | 200 | 253 | 259 | 267 | 0 | 0 | 1 | 0 | 0 | 267 | 149 | 1 | 1 | 1 | 1 | 1 | 1 | 1 | 1 | 1 | 0.993 | 1 | 1 | 0.001 |
| SE | 103866 | ENSG00000119943 | PYROXD2 | chr10 | - | 98387864 | 98388084 | 98387200 | 98387307 | 98390597 | 98390754 | 1 | 6 | 2 | 0 | 4 | 3 | 2 | 0 | 1 | 6 | 0 | 0 | 0 | 0 | 0 | 0 | 0 | 0 | 0 | 7 | 298 | 149 | 1 | 1 | 0.143 | 0.6 | 1 | 0 | 0.25 | NA | NA | NA | NA | 0 | 0.399 |
| SE | 103867 | ENSG00000119943 | PYROXD2 | chr10 | - | 98388353 | 98388508 | 98387200 | 98387307 | 98390597 | 98390754 | 14 | 0 | 12 | 11 | 11 | 3 | 2 | 0 | 1 | 6 | 0 | 4 | 7 | 0 | 9 | 0 | 0 | 0 | 0 | 7 | 298 | 149 | 1 | 1 | 0.7 | 0 | 1 | 0.846 | 0.478 | NA | 1 | 1 | NA | 0.391 | -0.192 |
| SE | 103868 | ENSG00000119943 | PYROXD2 | chr10 | - | 98390597 | 98390754 | 98387200 | 98387307 | 98391009 | 98391082 | 30 | 9 | 13 | 11 | 42 | 3 | 0 | 1 | 0 | 0 | 0 | 12 | 10 | 1 | 7 | 0 | 0 | 0 | 0 | 0 | 298 | 149 | 1 | 1 | 0.833 | 1 | 0.867 | 1 | 1 | NA | 1 | 1 | 1 | 1 | -0.06 |
| SE | 103869 | ENSG00000119943 | PYROXD2 | chr10 | - | 98391009 | 98391082 | 98387200 | 98387307 | 98395195 | 98395293 | 3 | 0 | 0 | 0 | 0 | 0 | 0 | 1 | 0 | 0 | 0 | 0 | 0 | 0 | 5 | 0 | 0 | 0 | 0 | 0 | 222 | 149 | 1 | 1 | 1 | NA | 0 | NA | NA | NA | NA | NA | NA | 1 | -0.5 |
| SE | 103870 | ENSG00000119943 | PYROXD2 | chr10 | - | 98414731 | 98414834 | 98413947 | 98414295 | 98415008 | 98415058 | 5 | 1 | 9 | 3 | 3 | 0 | 2 | 1 | 1 | 3 | 0 | 13 | 0 | 12 | 0 | 6 | 2 | 0 | 10 | 1 | 252 | 149 | 1 | 1 | 1 | 0.228 | 0.842 | 0.639 | 0.372 | 0 | 0.794 | NA | 0.415 | 0 | 0.314 |
| SE | 103871 | ENSG00000107651 | SEC23IP | chr10 | + | 119898426 | 119898959 | 119892729 | 119892945 | 119902798 | 119903009 | 65 | 58 | 65 | 68 | 45 | 0 | 1 | [illegible] | 1 | 2 | 27 | 93 | 73 | 53 | 17 | 0 | 0 | 2 | 0 | 0 | 298 | 149 | 1 | 1 | 1 | 1 | 0.97 | 0.971 | 0.918 | 1 | 1 | 0.948 | 1 | 1 | -0.018 |
| SE | 103872 | ENSG00000107651 | SEC23IP | chr10 | + | 119904083 | 119904277 | 119902798 | 119903009 | 119909040 | 119909130 | 348 | 291 | 292 | 343 | 291 | 0 | 0 | 0 | 0 | 0 | 235 | 326 | 215 | 271 | 177 | 0 | 0 | 5 | 0 | 0 | 298 | 149 | 1 | 1 | 1 | 1 | 1 | 1 | 1 | 1 | 1 | 0.956 | 1 | 1 | 0.009 |
| SE | 103873 | ENSG00000107651 | SEC23IP | chr10 | + | 119909040 | 119909130 | 119904083 | 119904277 | 119912043 | 119912164 | 252 | 211 | 199 | 243 | 225 | 0 | 0 | 0 | 1 | 0 | 143 | 173 | 113 | 125 | 38 | 0 | 0 | 0 | 4 | 0 | 239 | 149 | 1 | 1 | 1 | 1 | 1 | 0.993 | 1 | 1 | 1 | 1 | 0.951 | 1 | 0.008 |
| SE | 103877 | ENSG00000107651 | SEC23IP | chr10 | + | 119930328 | 119930431 | 119926035 | 119926227 | 119932132 | 119932318 | 58 | 53 | 45 | 62 | 91 | 0 | 0 | 0 | 0 | 0 | 12 | 49 | 25 | 47 | 0 | 0 | 0 | 1 | 0 | 0 | 252 | 149 | 1 | 1 | 1 | 1 | 1 | 1 | 1 | 1 | 1 | 0.937 | 1 | NA | 0.016 |
| SE | 103878 | ENSG00000107651 | SEC23IP | chr10 | + | 119930328 | 119930431 | 119929606 | 119929762 | 119932132 | 119932318 | 186 | 132 | 150 | 170 | 250 | 1 | 1 | 2 | 0 | 8 | 50 | 93 | 63 | 87 | 0 | 0 | 0 | 0 | 0 | 0 | 252 | 149 | 1 | 1 | 0.991 | 0.987 | 0.978 | 1 | 0.949 | 1 | 1 | 1 | 1 | NA | -0.019 |
| SE | 103879 | ENSG00000107651 | SEC23IP | chr10 | + | 119933004 | 119933167 | 119932132 | 119932318 | 119933685 | 119933770 | 353 | 265 | 242 | 254 | 539 | 0 | 0 | 1 | 1 | 0 | 50 | 231 | 86 | 163 | 29 | 0 | 0 | 0 | 0 | 0 | 298 | 149 | 1 | 1 | 1 | 1 | 0.992 | 0.992 | 1 | 1 | 1 | 1 | 1 | 1 | -0.003 |
| SE | 103880 | ENSG00000119977 | TCTN3 | chr10 | - | 95669046 | 95669157 | 95663416 | 95664300 | 95680471 | 95680607 | 2 | 1 | 3 | 0 | 6 | 22 | 10 | 32 | 24 | 30 | 0 | 0 | 3 | 0 | 12 | 0 | 11 | 26 | 19 | 10 | 260 | 149 | 1 | 1 | 0.05 | 0.054 | 0.051 | 0 | 0.103 | NA | 0 | 0.062 | 0 | 0.407 | -0.066 |
| SE | 103881 | ENSG00000119977 | TCTN3 | chr10 | - | 95680471 | 95680609 | 95659822 | 95664300 | 95682650 | 95682804 | 46 | 23 | 59 | 44 | 56 | 1 | 0 | 0 | 3 | 0 | 0 | 22 | 38 | 22 | 14 | 0 | 0 | 0 | 0 | 0 | 287 | 149 | 1 | 1 | 0.96 | 1 | 1 | 0.884 | 1 | NA | 1 | 1 | 1 | 1 | -0.031 |
| SE | 103882 | ENSG00000119977 | TCTN3 | chr10 | - | 95680471 | 95680609 | 95659822 | 95664300 | 95683100 | 95683180 | 24 | 12 | 34 | 17 | 31 | 0 | 0 | 0 | 2 | 0 | 0 | 11 | 29 | 15 | 10 | 0 | 0 | 0 | 0 | 0 | 287 | 149 | 1 | 1 | 1 | 1 | 1 | 0.815 | 1 | NA | 1 | 1 | 1 | 1 | -0.037 |
| SE | 103883 | ENSG00000119977 | TCTN3 | chr10 | - | 95682650 | 95682804 | 95663400 | 95664300 | 95683100 | 95683180 | 46 | 35 | 49 | 45 | 74 | 0 | 0 | 0 | 2 | 0 | 49 | 45 | 6 | 17 | 21 | 0 | 0 | 0 | 0 | 0 | 298 | 149 | 1 | 1 | 1 | 1 | 1 | 0.918 | 1 | 1 | 1 | 1 | 1 | 1 | -0.016 |
| SE | 103889 | ENSG00000119977 | TCTN3 | chr10 | - | 95686494 | 95686530 | 95684498 | 95684624 | 95687043 | 95687159 | 11 | 7 | 13 | 14 | 40 | 0 | 0 | 1 | 0 | 0 | 13 | 6 | 12 | 2 | 14 | 0 | 0 | 0 | 0 | 0 | 185 | 149 | 1 | 1 | 1 | 1 | 0.913 | 1 | 1 | 1 | 1 | 1 | 1 | 1 | -0.017 |
| SE | 103890 | ENSG00000119977 | TCTN3 | chr10 | - | 95687043 | 95687159 | 95684532 | 95684624 | 95692919 | 95693038 | 0 | 0 | 1 | 0 | 0 | 0 | 0 | 0 | 0 | 0 | 0 | 0 | 0 | 0 | 0 | 0 | 1 | 0 | 0 | 0 | 265 | 149 | 1 | 1 | NA | NA | 1 | NA | NA | NA | 0 | NA | NA | NA | 1 |
| SE | 103895 | ENSG00000119977 | TCTN3 | chr10 | - | 95693352 | 95693476 | 95692863 | 95693038 | 95693643 | 95693899 | 46 | 25 | 63 | 54 | 50 | 0 | 0 | 0 | 1 | 0 | 54 | 51 | 69 | 23 | 52 | 0 | 0 | 3 | 0 | 0 | 273 | 149 | 1 | 1 | 1 | 1 | 1 | 0.967 | 1 | 1 | 1 | 0.926 | 1 | 1 | 0.008 |
| SE | 103896 | ENSG00000066084 | DIP2B | chr12 | + | 50640723 | 50640852 | 50625975 | 50626047 | 50660193 | 50660319 | 71 | 44 | 90 | 78 | 79 | 0 | 0 | 0 | 3 | 0 | 36 | 49 | 82 | 130 | 97 | 0 | 0 | 0 | 0 | 0 | 278 | 149 | 1 | 1 | 1 | 1 | 1 | 0.933 | 1 | 1 | 1 | 1 | 1 | 1 | -0.013 |
| SE | 103899 | ENSG00000066084 | DIP2B | chr12 | + | 50678678 | 50678876 | 50675328 | 50675448 | 50680671 | 50680763 | 128 | 65 | 175 | 142 | 191 | 0 | 0 | 1 | 0 | 0 | 167 | 139 | 147 | 270 | 129 | 0 | 3 | 0 | 1 | 0 | 298 | 149 | 1 | 1 | 1 | 1 | 0.989 | 1 | 1 | 1 | 0.959 | 1 | 0.993 | 1 | 0.007 |
| SE | 103906 | ENSG00000066084 | DIP2B | chr12 | + | 50714394 | 50714596 | 50708447 | 50708562 | 50718708 | 50718818 | 209 | 186 | 227 | 256 | 362 | 0 | 0 | 1 | 1 | 0 | 311 | 253 | 404 | 378 | 109 | 0 | 0 | 2 | 2 | 0 | 298 | 149 | 1 | 1 | 1 | 1 | 0.991 | 0.992 | 1 | 1 | 1 | 0.99 | 0.99 | 1 | 0.001 |
| SE | 103907 | ENSG00000066084 | DIP2B | chr12 | + | 50723201 | 50723323 | 50721272 | 50721396 | 50724774 | 50724886 | 59 | 47 | 83 | 68 | 105 | 0 | 0 | 0 | 0 | 0 | 79 | 76 | 69 | 79 | 66 | 0 | 0 | 1 | 0 | 0 | 271 | 149 | 1 | 1 | 1 | 1 | 1 | 1 | 1 | 1 | 1 | 0.974 | 1 | 1 | 0.005 |
| SE | 103909 | ENSG00000066084 | DIP2B | chr12 | + | 50727702 | 50727812 | 50724774 | 50724886 | 50728547 | 50728678 | 124 | 64 | 115 | 134 | 114 | 1 | 1 | 0 | 0 | 1 | 55 | 108 | 102 | 110 | 44 | 0 | 0 | 0 | 0 | 0 | 259 | 149 | 1 | 1 | 0.986 | 0.974 | 1 | 1 | 0.985 | 1 | 1 | 1 | 1 | 1 | -0.011 |
| SE | 103912 | ENSG00000066084 | DIP2B | chr12 | + | 50732365 | 50732536 | 50731368 | 50731537 | 50734134 | 50734196 | 135 | 93 | 115 | 156 | 91 | 1 | 1 | 0 | 0 | 0 | 149 | 158 | 144 | 177 | 22 | 0 | 0 | 0 | 0 | 0 | 298 | 149 | 1 | 1 | 0.985 | 0.979 | 1 | 1 | 1 | 1 | 1 | 1 | 1 | 1 | -0.007 |
| SE | 103913 | ENSG00000066084 | DIP2B | chr12 | + | 50735072 | 50735130 | 50734134 | 50734196 | 50739408 | 50739586 | 14 | 3 | 11 | 12 | 8 | 0 | 0 | 0 | 0 | 0 | 0 | 15 | 9 | 25 | 0 | 0 | 0 | 0 | 2 | 0 | 207 | 149 | 1 | 1 | 1 | 1 | 1 | 1 | 1 | NA | 1 | 1 | 0.9 | NA | 0.033 |
| SE | 103914 | ENSG00000066084 | DIP2B | chr12 | + | 50737035 | 50737110 | 50735072 | 50735130 | 50739408 | 50739586 | 69 | 57 | 92 | 89 | 98 | 0 | 0 | 0 | 0 | 1 | 61 | 78 | 48 | 120 | 26 | 0 | 0 | 0 | 0 | 0 | 224 | 149 | 1 | 1 | 1 | 1 | 1 | 1 | 0.985 | 1 | 1 | 1 | 1 | 1 | -0.003 |
| SE | 103915 | ENSG00000066084 | DIP2B | chr12 | + | 50741415 | 50741539 | 50739408 | 50739586 | 50744586 | 50745095 | 60 | 38 | 69 | 84 | 75 | 0 | 0 | 0 | 0 | 0 | 47 | 53 | 70 | 66 | 14 | 0 | 0 | 3 | 0 | 0 | 273 | 149 | 1 | 1 | 1 | 1 | 1 | 1 | 1 | 1 | 1 | 0.927 | 1 | 1 | 0.015 |
| SE | 103916 | ENSG00000184988 | TMEM106A | chr17 | + | 43212204 | 43212394 | 43211874 | 43211922 | 43213020 | 43213252 | 10 | 2 | 6 | 13 | 0 | 4 | 1 | 3 | 2 | 0 | 5 | 10 | 13 | 7 | 58 | 0 | 0 | 1 | 4 | 1 | 298 | 149 | 1 | 1 | 0.556 | 0.5 | 0.5 | 0.765 | NA | 1 | 1 | 0.867 | 0.467 | 0.967 | -0.28 |
| SE | 103917 | ENSG00000184988 | TMEM106A | chr17 | + | 43212226 | 43212394 | 43211874 | 43211922 | 43213020 | 43213045 | 13 | 2 | 6 | 23 | 0 | 4 | 1 | 3 | 2 | 0 | 5 | 16 | 15 | 9 | 73 | 0 | 0 | 1 | 4 | 1 | 298 | 149 | 1 | 1 | 0.619 | 0.5 | 0.5 | 0.852 | NA | 1 | 1 | 0.882 | 0.529 | 0.973 | -0.259 |
| SE | 103918 | ENSG00000184988 | TMEM106A | chr17 | + | 43212226 | 43212394 | 43211874 | 43211922 | 43213114 | 43213252 | 3 | 0 | 0 | 11 | 0 | 0 | 0 | 0 | 1 | 0 | 0 | 18 | 4 | 3 | 15 | 0 | 0 | 0 | 0 | 0 | 298 | 149 | 1 | 1 | 1 | NA | NA | 0.846 | NA | NA | 1 | 1 | 1 | 1 | -0.077 |
| SE | 103919 | ENSG00000184988 | TMEM106A | chr17 | + | 43212314 | 43212394 | 43211834 | 43211922 | 43213020 | 43213252 | 10 | 2 | 5 | 4 | 0 | 4 | 1 | 3 | 2 | 0 | 0 | 10 | 8 | 5 | 35 | 0 | 0 | 1 | 4 | 1 | 229 | 149 | 1 | 1 | 0.619 | 0.565 | 0.52 | 0.565 | NA | NA | 1 | 0.839 | 0.449 | 0.958 | -0.244 |
| SE | 103920 | ENSG00000184988 | TMEM106A | chr17 | + | 43215787 | 43215941 | 43213827 | 43213891 | 43216696 | 43216740 | 36 | 17 | 40 | 30 | 28 | 0 | 0 | 0 | 0 | 1 | 156 | 86 | 24 | 51 | 111 | 0 | 1 | 0 | 1 | 3 | 298 | 149 | 1 | 1 | 1 | 1 | 1 | 1 | 0.933 | 1 | 0.977 | 1 | 0.962 | 0.949 | 0.009 |
| SE | 103923 | ENSG00000188234 | AGAP4 | chr10 | - | 45834015 | 45834116 | 45831393 | 45831429 | 45844325 | 45844394 | 22 | 31 | 20 | 31 | 67 | 0 | 0 | 1 | 0 | 0 | 23 | 15 | 12 | 14 | 11 | 0 | 0 | 0 | 0 | 0 | 250 | 149 | 1 | 1 | 1 | 1 | 0.923 | 1 | 1 | 1 | 1 | 1 | 1 | 1 | -0.015 |
| SE | 103926 | ENSG00000188234 | AGAP4 | chr10 | - | 45847859 | 45847954 | 45847124 | 45847312 | 45848842 | 45848912 | 4 | 0 | 8 | 14 | 0 | 0 | 0 | 0 | 1 | 0 | 0 | 2 | 19 | 0 | 8 | 0 | 7 | 1 | 0 | 0 | 244 | 149 | 1 | 1 | 1 | NA | 1 | 0.895 | NA | NA | 0.149 | 0.921 | NA | 1 | 0.275 |
| SE | 103927 | ENSG00000188234 | AGAP4 | chr10 | - | 45847859 | 45847954 | 45847124 | 45847312 | 45853654 | 45853875 | 2 | 1 | 10 | 4 | 8 | 0 | 0 | 1 | 0 | 0 | 0 | 2 | 10 | 0 | 15 | 0 | 0 | 0 | 2 | 8 | 244 | 149 | 1 | 1 | 1 | 1 | 0.859 | 1 | 1 | NA | 1 | 1 | 0 | 0.534 | 0.338 |
| SE | 103929 | ENSG00000188234 | AGAP4 | chr10 | - | 45848842 | 45848912 | 45847859 | 45847954 | 45853654 | 45853875 | 6 | 2 | 3 | 15 | 7 | 0 | 1 | 3 | 0 | 8 | 5 | 1 | 13 | 0 | 0 | 0 | 0 | 0 | 0 | 7 | 219 | 149 | 1 | 1 | 1 | 0.576 | 0.405 | 1 | 0.373 | 1 | 1 | 1 | NA | 0 | -0.079 |
| SE | 103933 | ENSG00000152377 | SPOCK1 | chr5 | - | 136979331 | 136979469 | 136975297 | 136978844 | 136985139 | 136985202 | 1 | 3 | 1 | 9 | 4 | 0 | 1 | 0 | 0 | 0 | 20 | 20 | 21 | 16 | 0 | 4 | 1 | 0 | 0 | 0 | 287 | 149 | 1 | 1 | 1 | 0.609 | 1 | 1 | 1 | 0.722 | 0.912 | 1 | 1 | NA | 0.013 |
| SE | 103936 | ENSG00000171862 | PTEN | chr10 | + | 87925512 | 87925557 | 87894024 | 87894109 | 87931045 | 87931089 | 773 | 530 | 464 | 602 | 1270 | 0 | 3 | 2 | 1 | 8 | 338 | 452 | 628 | 333 | 173 | 0 | 0 | 0 | 3 | 0 | 194 | 149 | 1 | 1 | 1 | 0.993 | 0.994 | 0.998 | 0.992 | 1 | 1 | 1 | 0.988 | 1 | -0.002 |
| SE | 103938 | ENSG00000171862 | PTEN | chr10 | + | 87952117 | 87952259 | 87933012 | 87933251 | 87960893 | 87961118 | 556 | 355 | 367 | 574 | 754 | 1 | 0 | 0 | 0 | 0 | 138 | 443 | 526 | 284 | 90 | 0 | 0 | 0 | 0 | 0 | 291 | 149 | 1 | 1 | 0.996 | 1 | 1 | 1 | 1 | 1 | 1 | 1 | 1 | 1 | -0.001 |
| SE | 103939 | ENSG00000171862 | PTEN | chr10 | + | 87957852 | 87958019 | 87933012 | 87933251 | 87960893 | 87961118 | 1684 | 1398 | 1225 | 1774 | 3131 | 1 | 0 | 0 | 0 | 0 | 691 | 1701 | 2273 | 1002 | 266 | 0 | 0 | 0 | 0 | 0 | 298 | 149 | 1 | 1 | 0.999 | 1 | 1 | 1 | 1 | 1 | 1 | 1 | 1 | 1 | 0 |
| SE | 103940 | ENSG00000171862 | PTEN | chr10 | + | 87957852 | 87958019 | 87952117 | 87952259 | 87960893 | 87961118 | 2790 | 2400 | 2084 | 2934 | 5745 | 0 | 0 | 0 | 1 | 0 | 1146 | 2798 | 3819 | 1775 | 436 | 0 | 0 | 0 | 0 | 0 | 298 | 149 | 1 | 1 | 1 | 1 | 1 | 0.999 | 1 | 1 | 1 | 1 | 1 | 1 | 0 |
| SE | 103941 | ENSG00000150347 | ARID5B | chr10 | + | 61902158 | 61902413 | 61901683 | 61901730 | 61940182 | 61940408 | 211 | 93 | 137 | 151 | 132 | 1 | 0 | 1 | 0 | 0 | 132 | 221 | 230 | 60 | 21 | 0 | 0 | 0 | 0 | 0 | 298 | 149 | 1 | 1 | 0.991 | 1 | 0.986 | 1 | 1 | 1 | 1 | 1 | 1 | 1 | -0.005 |
| SE | 103943 | ENSG00000150347 | ARID5B | chr10 | + | 62050887 | 62051000 | 61940182 | 61940408 | 62057116 | 62057268 | 136 | 99 | 135 | 124 | 149 | 0 | 0 | 0 | 0 | 0 | 47 | 160 | 187 | 80 | 52 | 0 | 2 | 0 | 0 | 0 | 262 | 149 | 1 | 1 | 1 | 1 | 1 | 1 | 1 | 1 | 0.978 | 1 | 1 | 1 | 0.004 |
| SE | 103945 | ENSG00000150347 | ARID5B | chr10 | + | 62059242 | 62059295 | 62057116 | 62057318 | 62069699 | 62069797 | 180 | 132 | 180 | 150 | 245 | 0 | 1 | 1 | 0 | 0 | 101 | 240 | 253 | 171 | 31 | 0 | 3 | 0 | 0 | 0 | 202 | 149 | 1 | 1 | 1 | 0.99 | 0.993 | 1 | 1 | 1 | 0.983 | 1 | 1 | 1 | 0 |
| SE | 103946 | ENSG00000150347 | ARID5B | chr10 | + | 62059266 | 62059295 | 62057116 | 62057318 | 62069699 | 62069797 | 10 | 15 | 11 | 30 | 21 | 0 | 1 | 1 | 0 | 0 | 22 | 28 | 44 | 17 | 6 | 0 | 3 | 0 | 0 | 0 | 178 | 149 | 1 | 1 | 1 | 0.926 | 0.902 | 1 | 1 | 1 | 0.887 | 1 | 1 | 1 | -0.012 |
| SE | 103947 | ENSG00000150347 | ARID5B | chr10 | + | 62085701 | 62085900 | 62069699 | 62069797 | 62090861 | 62093064 | 111 | 111 | 145 | 120 | 222 | 1 | 0 | 0 | 0 | 0 | 59 | 196 | 171 | 117 | 16 | 0 | 0 | 0 | 0 | 0 | 298 | 149 | 1 | 1 | 0.982 | 1 | 1 | 1 | 1 | 1 | 1 | 1 | 1 | 1 | -0.004 |
| SE | 103952 | ENSG00000119906 | SLF2 | chr10 | + | 100945329 | 100945506 | 100944025 | 100944128 | 100947038 | 100947136 | 112 | 113 | 67 | 122 | 284 | 5 | 0 | 0 | 0 | 0 | 0 | 93 | 12 | 23 | 26 | 0 | 0 | 0 | 0 | 0 | 298 | 149 | 1 | 1 | 0.918 | 1 | 1 | 1 | 1 | NA | 1 | 1 | 1 | 1 | -0.016 |
| SE | 103953 | ENSG00000119906 | SLF2 | chr10 | + | 100950075 | 100950207 | 100947759 | 100947847 | 100950675 | 100950753 | 135 | 145 | 96 | 167 | 283 | 0 | 1 | 0 | 0 | 0 | 29 | 80 | 69 | 37 | 7 | 0 | 0 | 0 | 0 | 0 | 281 | 149 | 1 | 1 | 1 | 0.987 | 1 | 1 | 1 | 1 | 1 | 1 | 1 | 1 | -0.003 |
| SE | 103955 | ENSG00000119906 | SLF2 | chr10 | + | 100956450 | 100956537 | 100950675 | 100950753 | 100959427 | 100959496 | 125 | 106 | 75 | 116 | 223 | 0 | 0 | 0 | 1 | 0 | 0 | 72 | 90 | 24 | 0 | 0 | 0 | 0 | 0 | 0 | 236 | 149 | 1 | 1 | 1 | 1 | 1 | 0.987 | 1 | NA | 1 | 1 | 1 | NA | -0.003 |
| SE | 103956 | ENSG00000055950 | MRPL43 | chr10 | - | 100983947 | 100984074 | 100983754 | 100983817 | 100986748 | 100986975 | 0 | 0 | 0 | 0 | 0 | 0 | 1 | 0 | 0 | 0 | 0 | 13 | 1 | 5 | 0 | 0 | 1 | 0 | 0 | 0 | 276 | 149 | 1 | 1 | NA | 0 | NA | NA | NA | NA | 0.875 | 1 | 1 | NA | -0.958 |
| SE | 103957 | ENSG00000055950 | MRPL43 | chr10 | - | 100983947 | 100984074 | 100983754 | 100983817 | 100987089 | 100987270 | 0 | 0 | 0 | 0 | 0 | 0 | 0 | 1 | 0 | 0 | 0 | 6 | 0 | 0 | 0 | 0 | 0 | 0 | 0 | 0 | 276 | 149 | 1 | 1 | NA | NA | 0 | NA | NA | NA | 1 | NA | NA | NA | -1 |
| SE | 103960 | ENSG00000055950 | MRPL43 | chr10 | - | 100986686 | 100986975 | 100983754 | 100983817 | 100987089 | 100987196 | 34 | 22 | 32 | 21 | 47 | 0 | 0 | 1 | 0 | 0 | 59 | 16 | 22 | 52 | 36 | 0 | 0 | 0 | 0 | 0 | 298 | 149 | 1 | 1 | 1 | 1 | 0.941 | 1 | 1 | 1 | 1 | 1 | 1 | 1 | -0.012 |
| SE | 103962 | ENSG00000055950 | MRPL43 | chr10 | - | 100986748 | 100986975 | 100983754 | 100983817 | 100987089 | 100987196 | 32 | 23 | 32 | 21 | 47 | 0 | 0 | 1 | 0 | 0 | 59 | 17 | 22 | 52 | 36 | 0 | 0 | 0 | 0 | 0 | 298 | 149 | 1 | 1 | 1 | 1 | 0.941 | 1 | 1 | 1 | 1 | 1 | 1 | 1 | -0.012 |
| SE | 103964 | ENSG00000055950 | MRPL43 | chr10 | - | 100987089 | 100987196 | 100986275 | 100986975 | 100987312 | 100987433 | 43 | 24 | 36 | 25 | 46 | 2 | 0 | 0 | 0 | 0 | 93 | 29 | 25 | 59 | 45 | 0 | 0 | 0 | 0 | 0 | 256 | 149 | 1 | 1 | 0.926 | 1 | 1 | 1 | 1 | 1 | 1 | 1 | 1 | 1 | -0.015 |
| SE | 103984 | ENSG00000151651 | ADAM8 | chr10 | - | 133269444 | 133269529 | 133267928 | 133268118 | 133269896 | 133269974 | 63 | 27 | 32 | 33 | 8 | 0 | 0 | 0 | 1 | 0 | 38 | 43 | 21 | 29 | 90 | 0 | 0 | 0 | 2 | 0 | 234 | 149 | 1 | 1 | 1 | 1 | 1 | 0.955 | 1 | 1 | 1 | 1 | 0.902 | 1 | 0.011 |
| SE | 103986 | ENSG00000151651 | ADAM8 | chr10 | - | 133270880 | 133271070 | 133270735 | 133270805 | 133271199 | 133271289 | 165 | 69 | 72 | 79 | 28 | 1 | 0 | 0 | 0 | 0 | 103 | 104 | 86 | 64 | 198 | 0 | 0 | 0 | 0 | 0 | 298 | 149 | 1 | 1 | 0.988 | 1 | 1 | 1 | 1 | 1 | 1 | 1 | 1 | 1 | -0.002 |
| SE | 103987 | ENSG00000151651 | ADAM8 | chr10 | - | 133273761 | 133273838 | 133273253 | 133273443 | 133273950 | 133274029 | 275 | 87 | 136 | 149 | 43 | 0 | 0 | 0 | 0 | 0 | 255 | 157 | 96 | 74 | 220 | 0 | 2 | 0 | 0 | 0 | 226 | 149 | 1 | 1 | 1 | 1 | 1 | 1 | 1 | 1 | 0.981 | 1 | 1 | 1 | 0.004 |
| SE | 103991 | ENSG00000198546 | ZNF511 | chr10 | + | 133311715 | 133311841 | 133310163 | 133310288 | 133312787 | 133313117 | 114 | 66 | 105 | 94 | 84 | 0 | 0 | 0 | 0 | 3 | 106 | 117 | 93 | 164 | 162 | 0 | 0 | 1 | 0 | 0 | 275 | 149 | 1 | 1 | 1 | 1 | 1 | 1 | 0.938 | 1 | 1 | 0.981 | 1 | 1 | -0.009 |
| SE | 103992 | ENSG00000148803 | FUOM | chr10 | - | 133356639 | 133356738 | 133355519 | 133355811 | 133356942 | 133356996 | 33 | 16 | 22 | 33 | 13 | 0 | 0 | 0 | 2 | 0 | 33 | 51 | 22 | 15 | 48 | 0 | 0 | 0 | 0 | 0 | 248 | 149 | 1 | 1 | 1 | 1 | 1 | 0.908 | 1 | 1 | 1 | 1 | 1 | 1 | -0.018 |
| SE | 103993 | ENSG00000148803 | FUOM | chr10 | - | 133357186 | 133357255 | 133356942 | 133357013 | 133357922 | 133358019 | 17 | 8 | 15 | 12 | 4 | 0 | 0 | 0 | 2 | 0 | 35 | 35 | 15 | 5 | 42 | 0 | 0 | 0 | 0 | 0 | 218 | 149 | 1 | 1 | 1 | 1 | 1 | 0.804 | 1 | 1 | 1 | 1 | 1 | 1 | -0.039 |
| SE | 103994 | ENSG00000148803 | FUOM | chr10 | - | 133357186 | 133357462 | 133356942 | 133357013 | 133357922 | 133358001 | 10 | 4 | 7 | 3 | 6 | 0 | 0 | 0 | 2 | 0 | 23 | 11 | 7 | 1 | 31 | 0 | 0 | 0 | 0 | 0 | 298 | 149 | 1 | 1 | 1 | 1 | 1 | 0.429 | 1 | 1 | 1 | 1 | 1 | 1 | -0.114 |
| SE | 103995 | ENSG00000127884 | ECHS1 | chr10 | - | 133368922 | 133369022 | 133366888 | 133366993 | 133369903 | 133370031 | 163 | 40 | 153 | 122 | 130 | 0 | 0 | 0 | 1 | 0 | 58 | 76 | 31 | 72 | 170 | 0 | 0 | 0 | 0 | 0 | 249 | 149 | 1 | 1 | 1 | 1 | 1 | 0.986 | 1 | 1 | 1 | 1 | 1 | 1 | -0.003 |
| SE | 103996 | ENSG00000153832 | FBXO36 | chr2 | + | 229932770 | 229932886 | 229922301 | 229922609 | 229976240 | 229976349 | 0 | 0 | 1 | 6 | 0 | 3 | 0 | 0 | 4 | 8 | 0 | 0 | 0 | 0 | 0 | 0 | 4 | 7 | 0 | 8 | 265 | 149 | 1 | 1 | 0 | NA | 1 | 0.458 | 0 | NA | 0 | 0 | NA | 0 | 0.364 |
| SE | 104000 | ENSG00000266412 | NCOA4 | chr10 | - | 46009123 | 46009212 | 46006411 | 46006597 | 46009410 | 46009551 | 1 | 0 | 0 | 0 | 0 | 1477 | 960 | 765 | 1564 | 3296 | 0 | 0 | 0 | 0 | 3 | 987 | 680 | 581 | 1020 | 923 | 238 | 149 | 1 | 1 | 0 | 0 | 0 | 0 | 0 | 0 | 0 | 0 | 0 | 0.002 | 0 |
| SE | 104001 | ENSG00000266412 | NCOA4 | chr10 | - | 46009123 | 46009212 | 46006411 | 46006597 | 46010222 | 46011206 | 1 | 0 | 0 | 0 | 0 | 3 | 1 | 2 | 1 | 2 | 0 | 0 | 0 | 0 | 0 | 6 | 7 | 0 | 0 | 5 | 238 | 149 | 1 | 1 | 0.173 | 0 | 0 | 0 | 0 | 0 | 0 | NA | NA | 0 | 0.035 |
| SE | 104002 | ENSG00000266412 | NCOA4 | chr10 | - | 46009410 | 46009551 | 46005087 | 46006597 | 46010222 | 46011206 | 3048 | 2002 | 1588 | 2954 | 6369 | 3 | 1 | 2 | 1 | 2 | 2428 | 1717 | 1532 | 2351 | 1793 | 6 | 7 | 0 | 0 | 5 | 290 | 149 | 1 | 1 | 0.998 | 0.999 | 0.998 | 0.999 | 0.999 | 0.995 | 0.992 | 1 | 1 | 0.995 | 0.002 |
| SE | 104003 | ENSG00000266412 | NCOA4 | chr10 | - | 46009410 | 46009551 | 46009123 | 46009212 | 46010222 | 46011206 | 1610 | 1069 | 840 | 1433 | 3165 | 0 | 0 | 0 | 0 | 0 | 1484 | 1066 | 977 | 1355 | 909 | 0 | 3 | 0 | 0 | 0 | 290 | 149 | 1 | 1 | 1 | 1 | 1 | 1 | 1 | 1 | 0.995 | 1 | 1 | 1 | 0.001 |
| SE | 104004 | ENSG00000266412 | NCOA4 | chr10 | - | 46010222 | 46011206 | 46009410 | 46009551 | 46012882 | 46013026 | 2171 | 1375 | 1183 | 1978 | 3498 | 1 | 1 | 2 | 4 | 3 | 1902 | 1581 | 1451 | 1887 | 1180 | 2 | 3 | 1 | 0 | 0 | 298 | 149 | 1 | 1 | 0.999 | 0.999 | 0.997 | 0.996 | 0.998 | 0.998 | 0.996 | 0.999 | 1 | 1 | -0.001 |
| SE | 104005 | ENSG00000266412 | NCOA4 | chr10 | - | 46012882 | 46013026 | 46010222 | 46011206 | 46013549 | 46013639 | 1529 | 855 | 947 | 1379 | 1565 | 0 | 0 | 4 | 0 | 3 | 1486 | 1468 | 1330 | 1533 | 698 | 0 | 0 | 0 | 0 | 0 | 293 | 149 | 1 | 1 | 1 | 1 | 0.992 | 1 | 1 | 1 | 1 | 1 | 1 | 1 | -0.002 |
| SE | 104006 | ENSG00000266412 | NCOA4 | chr10 | - | 46013549 | 46013639 | 46012882 | 46013026 | 46014443 | 46014552 | 1709 | 994 | 1103 | 1466 | 1956 | 0 | 0 | 0 | 0 | 0 | 1929 | 1612 | 1454 | 1796 | 903 | 0 | 0 | 4 | 6 | 0 | 239 | 149 | 1 | 1 | 1 | 1 | 1 | 1 | 1 | 1 | 1 | 0.996 | 0.995 | 1 | 0.002 |
| SE | 104007 | ENSG00000266412 | NCOA4 | chr10 | - | 46014443 | 46014552 | 46013549 | 46013639 | 46014853 | 46014942 | 2027 | 1215 | 1334 | 1823 | 2254 | 2 | 0 | 0 | 0 | 0 | 2038 | 1806 | 1479 | 1915 | 1156 | 0 | 0 | 3 | 0 | 0 | 258 | 149 | 1 | 1 | 0.998 | 1 | 1 | 1 | 1 | 1 | 1 | 0.997 | 1 | 1 | 0 |
| SE | 104008 | ENSG00000266412 | NCOA4 | chr10 | - | 46014443 | 46014552 | 46013549 | 46013639 | 46015125 | 46015266 | 612 | 362 | 390 | 512 | 662 | 0 | 1 | 0 | 0 | 0 | 532 | 505 | 491 | 555 | 298 | 0 | 0 | 0 | 0 | 0 | 258 | 149 | 1 | 1 | 1 | 0.995 | 1 | 1 | 1 | 1 | 1 | 1 | 1 | 1 | -0.001 |
| SE | 104009 | ENSG00000266412 | NCOA4 | chr10 | - | 46014853 | 46014942 | 46013549 | 46013639 | 46015125 | 46015266 | 670 | 397 | 396 | 572 | 649 | 0 | 1 | 0 | 0 | 0 | 506 | 478 | 352 | 555 | 440 | 0 | 0 | 0 | 0 | 0 | 238 | 149 | 1 | 1 | 1 | 0.996 | 1 | 1 | 1 | 1 | 1 | 1 | 1 | 1 | -0.001 |
| SE | 104011 | ENSG00000266412 | NCOA4 | chr10 | - | 46015125 | 46015266 | 46014443 | 46014552 | 46016539 | 46016694 | 646 | 378 | 476 | 613 | 720 | 0 | 2 | 0 | 0 | 0 | 529 | 633 | 475 | 511 | 456 | 0 | 0 | 0 | 0 | 0 | 290 | 149 | 1 | 1 | 1 | 0.99 | 1 | 1 | 1 | 1 | 1 | 1 | 1 | 1 | -0.002 |
| SE | 104013 | ENSG00000266412 | NCOA4 | chr10 | - | 46016539 | 46016694 | 46014443 | 46014552 | 46030525 | 46030623 | 156 | 74 | 114 | 113 | 78 | 3 | 0 | 0 | 1 | 1 | 54 | 158 | 95 | 131 | 63 | 0 | 2 | 1 | 0 | 0 | 298 | 149 | 1 | 1 | 0.963 | 1 | 1 | 0.983 | 0.975 | 1 | 0.975 | 0.979 | 1 | 1 | -0.007 |
| SE | 104014 | ENSG00000266412 | NCOA4 | chr10 | - | 46016539 | 46016694 | 46015125 | 46015266 | 46023329 | 46023455 | 735 | 421 | 527 | 714 | 766 | 0 | 0 | 3 | 2 | 0 | 643 | 699 | 535 | 579 | 543 | 0 | 2 | 1 | 0 | 0 | 298 | 149 | 1 | 1 | 1 | 1 | 0.989 | 0.994 | 1 | 1 | 0.994 | 0.996 | 1 | 1 | -0.001 |
| SE | 104016 | ENSG00000266412 | NCOA4 | chr10 | - | 46027431 | 46027528 | 46016539 | 46016694 | 46030525 | 46030607 | 2 | 1 | 1 | 0 | 0 | 156 | 73 | 114 | 113 | 78 | 0 | 0 | 0 | 0 | 0 | 54 | 158 | 95 | 131 | 63 | 246 | 149 | 1 | 1 | 0.008 | 0.008 | 0.005 | 0 | 0 | 0 | 0 | 0 | 0 | 0 | 0.004 |
| SE | 104017 | ENSG00000170989 | S1PR1 | chr1 | + | 101241688 | 101241772 | 101236898 | 101237099 | 101242027 | 101243713 | 27 | 4 | 25 | 18 | 14 | 3 | 2 | 0 | 3 | 0 | 0 | 17 | 6 | 8 | 10 | 0 | 1 | 0 | 0 | 0 | 233 | 149 | 1 | 1 | 0.852 | 0.561 | 1 | 0.793 | 1 | NA | 0.916 | 1 | 1 | 1 | -0.138 |
| SE | 104019 | ENSG00000165983 | PTER | chr10 | + | 16437356 | 16437490 | 16436983 | 16437047 | 16486351 | 16486617 | 14 | 9 | 13 | 19 | 28 | 2 | 1 | 0 | 0 | 6 | 4 | 7 | 16 | 1 | 0 | 0 | 0 | 0 | 0 | 0 | 283 | 149 | 1 | 1 | 0.787 | 0.826 | 1 | 1 | 0.711 | 1 | 1 | 1 | 1 | NA | -0.135 |
| SE | 104021 | ENSG00000165983 | PTER | chr10 | + | 16484336 | 16484816 | 16437356 | 16437490 | 16486351 | 16486617 | 81 | 39 | 72 | 65 | 70 | 3 | 0 | 0 | 1 | 0 | 23 | 26 | 9 | 14 | 24 | 0 | 0 | 3 | 0 | 0 | 298 | 149 | 1 | 1 | 0.931 | 1 | 1 | 0.97 | 1 | 1 | 1 | 0.6 | 1 | 1 | 0.06 |
| SE | 104023 | ENSG00000139514 | SLC7A1 | chr13 | - | 29530537 | 29530712 | 29524131 | 29524253 | 29532823 | 29532982 | 67 | 39 | 51 | 35 | 69 | 0 | 0 | 0 | 0 | 3 | 41 | 59 | 89 | 35 | 29 | 0 | 0 | 1 | 0 | 0 | 298 | 149 | 1 | 1 | 1 | 1 | 1 | 1 | 0.92 | 1 | 1 | 0.978 | 1 | 1 | -0.012 |
| SE | 104025 | ENSG00000151923 | TIAL1 | chr10 | - | 119579934 | 119580010 | 119578725 | 119578834 | 119580470 | 119580538 | 204 | 134 | 158 | 171 | 352 | 0 | 0 | 0 | 3 | 0 | 78 | 118 | 131 | 134 | 132 | 0 | 0 | 0 | 0 | 0 | 225 | 149 | 1 | 1 | 1 | 1 | 1 | 0.974 | 1 | 1 | 1 | 1 | 1 | 1 | -0.005 |
| SE | 104028 | ENSG00000151923 | TIAL1 | chr10 | - | 119582458 | 119582557 | 119582168 | 119582223 | 119588151 | 119588210 | 448 | 303 | 454 | 390 | 506 | 0 | 0 | 0 | 0 | 0 | 268 | 291 | 233 | 296 | 211 | 0 | 6 | 0 | 0 | 0 | 248 | 149 | 1 | 1 | 1 | 1 | 1 | 1 | 1 | 1 | 0.967 | 1 | 1 | 1 | 0.007 |
| SE | 104029 | ENSG00000151923 | TIAL1 | chr10 | - | 119582458 | 119582608 | 119582168 | 119582223 | 119588151 | 119588248 | 482 | 329 | 399 | 432 | 547 | 0 | 0 | 0 | 0 | 0 | 160 | 334 | 221 | 237 | 254 | 0 | 6 | 0 | 0 | 0 | 298 | 149 | 1 | 1 | 1 | 1 | 1 | 1 | 1 | 1 | 0.965 | 1 | 1 | 1 | 0.007 |
| SE | 104030 | ENSG00000151923 | TIAL1 | chr10 | - | 119588151 | 119588248 | 119582458 | 119582557 | 119596433 | 119596495 | 345 | 223 | 340 | 309 | 218 | 1 | 0 | 0 | 0 | 0 | 180 | 254 | 168 | 242 | 153 | 0 | 0 | 0 | 0 | 0 | 246 | 149 | 1 | 1 | 0.995 | 1 | 1 | 1 | 1 | 1 | 1 | 1 | 1 | 1 | -0.001 |
| SE | 104031 | ENSG00000151923 | TIAL1 | chr10 | - | 119588151 | 119588248 | 119582458 | 119582608 | 119596433 | 119596495 | 256 | 180 | 226 | 250 | 171 | 0 | 0 | 0 | 2 | 0 | 43 | 223 | 120 | 134 | 139 | 0 | 0 | 0 | 0 | 0 | 246 | 149 | 1 | 1 | 1 | 1 | 1 | 0.987 | 1 | 1 | 1 | 1 | 1 | 1 | -0.003 |
| SE | 104032 | ENSG00000151923 | TIAL1 | chr10 | - | 119592254 | 119592296 | 119588151 | 119588248 | 119593451 | 119593494 | 0 | 1 | 0 | 0 | 0 | 66 | 32 | 47 | 43 | 53 | 0 | 0 | 1 | 0 | 0 | 68 | 47 | 48 | 37 | 15 | 191 | 149 | 1 | 1 | 0 | 0.024 | 0 | 0 | 0 | 0 | 0 | 0.016 | 0 | 0 | 0.002 |
| SE | 104033 | ENSG00000151923 | TIAL1 | chr10 | - | 119592254 | 119592296 | 119588151 | 119588248 | 119596433 | 119597029 | 2 | 0 | 0 | 0 | 0 | 204 | 114 | 195 | 184 | 94 | 0 | 0 | 1 | 0 | 0 | 59 | 151 | 76 | 119 | 85 | 191 | 149 | 1 | 1 | 0.008 | 0 | 0 | 0 | 0 | 0 | 0 | 0.01 | 0 | 0 | 0 |
| SE | 104034 | ENSG00000065809 | FAM107B | chr10 | - | 14527936 | 14528039 | 14521868 | 14522019 | 14530331 | 14530515 | 0 | 2 | 0 | 0 | 0 | 788 | 510 | 691 | 691 | 1451 | 0 | 0 | 0 | 0 | 0 | 261 | 609 | 942 | 779 | 97 | 252 | 149 | 1 | 1 | 0 | 0.002 | 0 | 0 | 0 | 0 | 0 | 0 | 0 | 0 | 0 |
| SE | 104036 | ENSG00000065809 | FAM107B | chr10 | - | 14542578 | 14542686 | 14530331 | 14530515 | 14571762 | 14572189 | 0 | 2 | 4 | 3 | 17 | 33 | 45 | 55 | 50 | 132 | 0 | 3 | 3 | 1 | 0 | 0 | 10 | 33 | 4 | 0 | 257 | 149 | 1 | 1 | 0 | 0.025 | 0.04 | 0.034 | 0.069 | NA | 0.148 | 0.05 | 0.127 | NA | -0.075 |
| SE | 104038 | ENSG00000065809 | FAM107B | chr10 | - | 14553321 | 14553387 | 14530331 | 14530515 | 14556361 | 14556405 | 271 | 138 | 249 | 186 | 440 | 1 | 1 | 1 | 0 | 0 | 120 | 254 | 372 | 292 | 35 | 0 | 0 | 0 | 0 | 0 | 215 | 149 | 1 | 1 | 0.995 | 0.99 | 0.994 | 1 | 1 | 1 | 1 | 1 | 1 | 1 | -0.004 |
| SE | 104041 | ENSG00000065809 | FAM107B | chr10 | - | 14553321 | 14553387 | 14530331 | 14530515 | 14571990 | 14572032 | 257 | 134 | 244 | 181 | 425 | 1 | 1 | 0 | 0 | 2 | 120 | 246 | 337 | 277 | 35 | 0 | 0 | 0 | 0 | 0 | 215 | 149 | 1 | 1 | 0.994 | 0.989 | 1 | 1 | 0.993 | 1 | 1 | 1 | 1 | 1 | -0.005 |
| SE | 104045 | ENSG00000065809 | FAM107B | chr10 | - | 14553321 | 14553387 | 14542578 | 14542686 | 14571762 | 14572155 | 17 | 10 | 21 | 17 | 31 | 0 | 0 | 2 | 0 | 0 | 0 | 12 | 44 | 2 | 0 | 0 | 0 | 0 | 0 | 0 | 215 | 149 | 1 | 1 | 1 | 1 | 0.879 | 1 | 1 | NA | 1 | 1 | 1 | NA | -0.024 |
| SE | 104046 | ENSG00000065809 | FAM107B | chr10 | - | 14556361 | 14556405 | 14553321 | 14553387 | 14571762 | 14572189 | 6 | 2 | 4 | 4 | 7 | 59 | 48 | 86 | 69 | 151 | 0 | 0 | 27 | 12 | 0 | 0 | 36 | 96 | 17 | 0 | 193 | 149 | 1 | 1 | 0.073 | 0.031 | 0.035 | 0.043 | 0.035 | NA | 0 | 0.178 | 0.353 | NA | -0.134 |
| SE | 104047 | ENSG00000065809 | FAM107B | chr10 | - | 14556361 | 14556405 | 14553321 | 14553387 | 14571923 | 14572030 | 4 | 1 | 4 | 2 | 8 | 7 | 6 | 14 | 7 | 23 | 0 | 6 | 17 | 9 | 0 | 0 | 4 | 3 | 3 | 0 | 193 | 149 | 1 | 1 | 0.306 | 0.114 | 0.181 | 0.181 | 0.212 | NA | 0.537 | 0.814 | 0.698 | NA | -0.484 |
| SE | 104048 | ENSG00000065809 | FAM107B | chr10 | - | 14556361 | 14556405 | 14553321 | 14553387 | 14572228 | 14572339 | 7 | 2 | 2 | 6 | 18 | 78 | 22 | 43 | 57 | 35 | 0 | 3 | 22 | 11 | 0 | 0 | 27 | 61 | 10 | 0 | 193 | 149 | 1 | 1 | 0.065 | 0.066 | 0.035 | 0.075 | 0.284 | NA | 0.079 | 0.218 | 0.459 | NA | -0.147 |
| SE | 104050 | ENSG00000120049 | KCNIP2 | chr10 | - | 101828150 | 101828258 | 101827888 | 101827993 | 101828388 | 101828459 | 5 | 4 | 13 | 5 | 9 | 0 | 0 | 0 | 1 | 0 | 0 | 12 | 1 | 13 | 18 | 0 | 0 | 0 | 0 | 0 | 257 | 149 | 1 | 1 | 1 | 1 | 1 | 0.744 | 1 | NA | 1 | 1 | 1 | 1 | -0.051 |
| SE | 104052 | ENSG00000120049 | KCNIP2 | chr10 | - | 101828818 | 101828846 | 101828626 | 101828696 | 101829074 | 101829199 | 1 | 0 | 0 | 0 | 0 | 1 | 2 | 3 | 2 | 1 | 0 | 0 | 0 | 0 | 0 | 0 | 2 | 1 | 1 | 0 | 177 | 149 | 1 | 1 | 0.457 | 0 | 0 | 0 | 0 | NA | 0 | 0 | 0 | NA | 0.091 |
| SE | 104053 | ENSG00000120049 | KCNIP2 | chr10 | - | 101828818 | 101828849 | 101828626 | 101828696 | 101829074 | 101829199 | 1 | 0 | 0 | 0 | 0 | 1 | 2 | 3 | 2 | 1 | 0 | 0 | 0 | 0 | 0 | 0 | 2 | 1 | 1 | 0 | 180 | 149 | 1 | 1 | 0.453 | 0 | 0 | 0 | 0 | NA | 0 | 0 | 0 | NA | 0.091 |
| SE | 104060 | ENSG00000171798 | KNDC1 | chr10 | + | 133206855 | 133206953 | 133206684 | 133206778 | 133207136 | 133207351 | 0 | 5 | 0 | 7 | 11 | 0 | 0 | 0 | 0 | 0 | 0 | 5 | 0 | 2 | 0 | 1 | 0 | 0 | 0 | 0 | 247 | 149 | 1 | 1 | NA | 1 | NA | 1 | 1 | 0 | 1 | NA | 1 | NA | 0.333 |
| SE | 104061 | ENSG00000171798 | KNDC1 | chr10 | + | 133209804 | 133209924 | 133207136 | 133207351 | 133210610 | 133210724 | 0 | 0 | 0 | 0 | 1 | 2 | 0 | 0 | 3 | 5 | 0 | 0 | 0 | 0 | 0 | 1 | 0 | 0 | 0 | 0 | 269 | 149 | 1 | 1 | 0 | NA | NA | 0 | 0.1 | 0 | NA | NA | NA | NA | 0.033 |

| | | | | | | | | | | | | | | |
|---|---|---|---|---|---|---|---|---|---|---|---|---|---|---|
| SE | 104065 | ENSG00000072694 | FCGR2B | chr1 | + | 161670251 | 161670272 | 161663146 | 161663353 | 161671391 | 161671649 | 1 | 6 | [illegible] |
| SE | 104069 | ENSG00000118432 | CNR1 | chr6 | - | 88164256 | 88164294 | 88147663 | 88147806 | 88165802 | 88166347 | 0 | 0 | [illegible] |
| SE | 104070 | ENSG00000257242 | LINC01619 | chr12 | - | 91987532 | 91987643 | 91985777 | 91986697 | 91989096 | 91989193 | 3 | 0 | [illegible] |
| SE | 104074 | ENSG00000257242 | LINC01619 | chr12 | - | 91989096 | 91989193 | 91987532 | 91987643 | 91993369 | 91993529 | 2 | 1 | [illegible] |
| SE | 104075 | ENSG00000257242 | LINC01619 | chr12 | - | 91990283 | 91990362 | 91989096 | 91989193 | 91993369 | 91993529 | 1 | 0 | [illegible] |
| SE | 104076 | ENSG00000257242 | LINC01619 | chr12 | - | 91990287 | 91990362 | 91989096 | 91989193 | 91993369 | 91993529 | 2 | 0 | [illegible] |
| SE | 104077 | ENSG00000257242 | LINC01619 | chr12 | - | 91993369 | 91993529 | 91989096 | 91989193 | 92142606 | 92142668 | 8 | 7 | [illegible] |
| SE | 104079 | ENSG00000139734 | DIAPH3 | chr13 | - | 60112009 | 60112186 | 60093627 | 60093732 | 60163586 | 60163766 | 0 | 0 | [illegible] |
| SE | 104080 | ENSG00000139734 | DIAPH3 | chr13 | - | 60132956 | 60132989 | 60112009 | 60112186 | 60163586 | 60163766 | 0 | 0 | [illegible] |
| SE | 104083 | ENSG00000122376 | SHLD2 | chr10 | + | 87096933 | 87096989 | 87094160 | 87094788 | 87151349 | 87151634 | 41 | 32 | [illegible] |
| SE | 104084 | ENSG00000122376 | SHLD2 | chr10 | + | 87151349 | 87152879 | 87096933 | 87096989 | 87158047 | 87158155 | 209 | 130 | [illegible] |
| SE | 104085 | ENSG00000122376 | SHLD2 | chr10 | + | 87151349 | 87152879 | 87096933 | 87096989 | 87170477 | 87170672 | 71 | 40 | [illegible] |
| SE | 104086 | ENSG00000122376 | SHLD2 | chr10 | + | 87158047 | 87158155 | 87096933 | 87096989 | 87170477 | 87170672 | 175 | 123 | [illegible] |
| SE | 104087 | ENSG00000122376 | SHLD2 | chr10 | + | 87158047 | 87158155 | 87151349 | 87152879 | 87170477 | 87170672 | 310 | 210 | [illegible] |
| SE | 104088 | ENSG00000122376 | SHLD2 | chr10 | + | 87170477 | 87170672 | 87158047 | 87158155 | 87170839 | 87170974 | 352 | 277 | [illegible] |
| SE | 104089 | ENSG00000122376 | SHLD2 | chr10 | + | 87175888 | 87176095 | 87170839 | 87170974 | 87180074 | 87180303 | 90 | 88 | [illegible] |
| SE | 104090 | ENSG00000122376 | SHLD2 | chr10 | + | 87180074 | 87180303 | 87170839 | 87170974 | 87187084 | 87187200 | 97 | 57 | [illegible] |
| SE | 104093 | ENSG00000099246 | RAB18 | chr10 | + | 27508901 | 27509033 | 27504303 | 27504437 | 27509874 | 27509930 | 1 | 1 | [illegible] |
| SE | 104096 | ENSG00000099246 | RAB18 | chr10 | + | 27513962 | 27514420 | 27509874 | 27509930 | 27526827 | 27526889 | 0 | 1 | [illegible] |
| SE | 104097 | ENSG00000099246 | RAB18 | chr10 | + | 27513965 | 27514420 | 27509874 | 27509930 | 27526827 | 27526889 | 0 | 1 | [illegible] |
| SE | 104102 | ENSG00000099246 | RAB18 | chr10 | + | 27526827 | 27526889 | 27509874 | 27509930 | 27532506 | 27532579 | 408 | 206 | [illegible] |
| SE | 104103 | ENSG00000099246 | RAB18 | chr10 | + | 27526827 | 27526889 | 27509874 | 27509930 | 27533734 | 27533853 | 172 | 75 | [illegible] |
| SE | 104108 | ENSG00000099246 | RAB18 | chr10 | + | 27526827 | 27532579 | 27509888 | 27509930 | 27533734 | 27533770 | 409 | 264 | [illegible] |
| SE | 104109 | ENSG00000099246 | RAB18 | chr10 | + | 27531528 | 27531615 | 27509874 | 27509930 | 27532506 | 27532579 | 6 | 7 | [illegible] |
| SE | 104111 | ENSG00000099246 | RAB18 | chr10 | + | 27532506 | 27532579 | 27509874 | 27509930 | 27533734 | 27533816 | 205 | 159 | [illegible] |
| SE | 104113 | ENSG00000099246 | RAB18 | chr10 | + | 27532506 | 27532579 | 27526827 | 27526889 | 27533734 | 27533816 | 443 | 295 | [illegible] |
| SE | 104116 | ENSG00000099246 | RAB18 | chr10 | + | 27533734 | 27533853 | 27526827 | 27526889 | 27533927 | 27533994 | 226 | 174 | [illegible] |
| SE | 104117 | ENSG00000099246 | RAB18 | chr10 | + | 27533927 | 27533994 | 27528515 | 27533853 | 27537875 | 27537917 | 593 | 476 | [illegible] |
| SE | 104118 | ENSG00000099246 | RAB18 | chr10 | + | 27533927 | 27533998 | 27533734 | 27533853 | 27537875 | 27538163 | 171 | 132 | [illegible] |
| SE | 104119 | ENSG00000099246 | RAB18 | chr10 | + | 27533940 | 27533994 | 27533734 | 27533853 | 27537875 | 27539185 | 239 | 202 | [illegible] |
| SE | 104126 | ENSG00000130783 | CCDC62 | chr12 | + | 122798084 | 122798200 | 122797306 | 122797395 | 122801123 | 122801852 | 6 | 2 | [illegible] |
| SE | 104129 | ENSG00000130640 | TUBGCP2 | chr10 | - | 133285084 | 133285213 | 133283881 | 133284002 | 133285455 | 133285595 | 168 | 76 | [illegible] |
| SE | 104130 | ENSG00000130640 | TUBGCP2 | chr10 | - | 133295321 | 133295439 | 133293561 | 133293769 | 133297951 | 133298111 | 0 | 1 | [illegible] |
| SE | 104131 | ENSG00000130640 | TUBGCP2 | chr10 | - | 133295523 | 133295911 | 133295363 | 133295439 | 133297951 | 133298023 | 2 | 0 | [illegible] |
| SE | 104132 | ENSG00000130640 | TUBGCP2 | chr10 | - | 133297320 | 133297404 | 133293561 | 133293769 | 133297951 | 133298111 | 0 | 0 | [illegible] |
| SE | 104133 | ENSG00000130640 | TUBGCP2 | chr10 | - | 133300451 | 133300547 | 133299984 | 133300113 | 133302791 | 133302980 | 1 | 0 | [illegible] |
| SE | 104138 | ENSG00000023041 | ZDHHC6 | chr10 | - | 112428381 | 112428438 | 112424427 | 112425593 | 112432239 | 112432286 | 5 | 4 | [illegible] |
| SE | 104139 | ENSG00000023041 | ZDHHC6 | chr10 | - | 112433239 | 112433281 | 112432375 | 112432521 | 112433540 | 112433570 | 146 | 168 | [illegible] |
| SE | 104143 | ENSG00000023041 | ZDHHC6 | chr10 | - | 112440533 | 112440695 | 112438335 | 112438389 | 112441728 | 112441812 | 178 | 186 | [illegible] |
| SE | 104145 | ENSG00000023041 | ZDHHC6 | chr10 | - | 112440533 | 112440695 | 112439505 | 112439626 | 112442191 | 112442259 | 96 | 70 | [illegible] |
| SE | 104149 | ENSG00000023041 | ZDHHC6 | chr10 | - | 112443514 | 112443606 | 112442191 | 112442259 | 112445169 | 112445650 | 63 | 41 | [illegible] |
| SE | 104151 | ENSG00000172731 | LRRC20 | chr10 | - | 70340552 | 70340702 | 70323862 | 70324030 | 70376451 | 70376596 | 4 | 0 | [illegible] |
| SE | 104164 | ENSG00000237489 | C10orf143 | chr10 | - | 130064861 | 130066105 | 130064339 | 130064383 | 130068275 | 130068608 | 3 | 0 | [illegible] |
| SE | 104166 | ENSG00000237489 | C10orf143 | chr10 | - | 130101093 | 130101211 | 130079736 | 130079901 | 130104082 | 130104230 | 7 | 0 | [illegible] |
| SE | 104172 | ENSG00000165626 | BEND7 | chr10 | - | 13518552 | 13518658 | 13499738 | 13500080 | 13526137 | 13526187 | 0 | 0 | [illegible] |
| SE | 104173 | ENSG00000165626 | BEND7 | chr10 | - | 13526137 | 13526221 | 13499777 | 13500080 | 13528472 | 13528974 | 1 | 2 | [illegible] |
| SE | 104175 | ENSG00000152782 | PANK1 | chr10 | - | 89611695 | 89612048 | 89599251 | 89599505 | 89644599 | 89645418 | 7 | 6 | [illegible] |
| SE | 104189 | ENSG00000198393 | ZNF26 | chr12 | + | 132987664 | 132987768 | 132986364 | 132986873 | 133007041 | 133007168 | 8 | 3 | [illegible] |
| SE | 104191 | ENSG00000198393 | ZNF26 | chr12 | + | 133001632 | 133001731 | 132987664 | 132987768 | 133007041 | 133007168 | 1 | 0 | [illegible] |
| SE | 104192 | ENSG00000198393 | ZNF26 | chr12 | + | 133007436 | 133007532 | 133004403 | 133007168 | 133010135 | 133010241 | 32 | 30 | [illegible] |
| SE | 104194 | ENSG00000141759 | TXNL4A | chr18 | - | 79977597 | 79977701 | 79970812 | 79973856 | 79986601 | 79986750 | 242 | 127 | [illegible] |
| SE | 104201 | ENSG00000148672 | GLUD1 | chr10 | - | 87053341 | 87053404 | 87050201 | 87051870 | 87057690 | 87057782 | 477 | 256 | [illegible] |
| SE | 104202 | ENSG00000148672 | GLUD1 | chr10 | - | 87060687 | 87060825 | 87060160 | 87060241 | 87060914 | 87061052 | 537 | 251 | [illegible] |
| SE | 104203 | ENSG00000148672 | GLUD1 | chr10 | - | 87060687 | 87060825 | 87060160 | 87060241 | 87062655 | 87062835 | 308 | 156 | [illegible] |
| SE | 104204 | ENSG00000148672 | GLUD1 | chr10 | - | 87060914 | 87061052 | 87060160 | 87060241 | 87062655 | 87062835 | 345 | 204 | [illegible] |
| SE | 104205 | ENSG00000148672 | GLUD1 | chr10 | - | 87060914 | 87061052 | 87060160 | 87060825 | 87062655 | 87062835 | 570 | 299 | [illegible] |
| SE | 104206 | ENSG00000148672 | GLUD1 | chr10 | - | 87067725 | 87067842 | 87062655 | 87062835 | 87068062 | 87068157 | 9 | 1 | [illegible] |
| SE | 104208 | ENSG00000148672 | GLUD1 | chr10 | - | 87078483 | 87078543 | 87068062 | 87068157 | 87094324 | 87094450 | 0 | 0 | [illegible] |
| SE | 104209 | ENSG00000148672 | GLUD1 | chr10 | - | 87078483 | 87078543 | 87076575 | 87076656 | 87089561 | 87089788 | 0 | 2 | [illegible] |
| SE | 104210 | ENSG00000148672 | GLUD1 | chr10 | - | 87078483 | 87078543 | 87076575 | 87076656 | 87094324 | 87094450 | 0 | 0 | [illegible] |
| SE | 104211 | ENSG00000148672 | GLUD1 | chr10 | - | 87089561 | 87089788 | 87076575 | 87076656 | 87091570 | 87091623 | 13 | 0 | [illegible] |
| SE | 104214 | ENSG00000148672 | GLUD1 | chr10 | - | 87089561 | 87089788 | 87078483 | 87078543 | 87094324 | 87094397 | 0 | 2 | [illegible] |
| SE | 104217 | ENSG00000148672 | GLUD1 | chr10 | - | 87091570 | 87091623 | 87076575 | 87076656 | 87094324 | 87094450 | 0 | 0 | [illegible] |
| SE | 104218 | ENSG00000148672 | GLUD1 | chr10 | - | 87091570 | 87091623 | 87089561 | 87089788 | 87092581 | 87092654 | 0 | 0 | [illegible] |
| SE | 104219 | ENSG00000148672 | GLUD1 | chr10 | - | 87091570 | 87091623 | 87089561 | 87089788 | 87094324 | 87094450 | 0 | 0 | [illegible] |
| SE | 104223 | ENSG00000148672 | GLUD1 | chr10 | - | 87092581 | 87092654 | 87089561 | 87089788 | 87094324 | 87094460 | 0 | 0 | [illegible] |
| SE | 104225 | ENSG00000148672 | GLUD1 | chr10 | - | 87092581 | 87092654 | 87091570 | 87091623 | 87094324 | 87094460 | 0 | 0 | [illegible] |
| SE | 104227 | ENSG00000139620 | KANSL2 | chr12 | - | 48654940 | 48655060 | 48653210 | 48654175 | 48660365 | 48660619 | 94 | 60 | [illegible] |
| SE | 104229 | ENSG00000139620 | KANSL2 | chr12 | - | 48661194 | 48661243 | 48660365 | 48660619 | 48662559 | 48662592 | 16 | 11 | [illegible] |
| SE | 104233 | ENSG00000139620 | KANSL2 | chr12 | - | 48679654 | 48679833 | 48679035 | 48679150 | 48681381 | 48681416 | 116 | 73 | [illegible] |
| SE | 104234 | ENSG00000139620 | KANSL2 | chr12 | - | 48681381 | 48681641 | 48679654 | 48679833 | 48682186 | 48682211 | 65 | 40 | [illegible] |
| SE | 104235 | ENSG00000139620 | KANSL2 | chr12 | - | 48681570 | 48681641 | 48679654 | 48679833 | 48682186 | 48682238 | 2 | 2 | [illegible] |
| SE | 104236 | ENSG00000139620 | KANSL2 | chr12 | - | 48681857 | 48682065 | 48681381 | 48681641 | 48682186 | 48682238 | 0 | 0 | [illegible] |
| SE | 104240 | ENSG00000128487 | SPECC1 | chr17 | + | 20017986 | 20018102 | 20010334 | 20010652 | 20057676 | 20057817 | 0 | 0 | [illegible] |
| SE | 104242 | ENSG00000128487 | SPECC1 | chr17 | + | 20057676 | 20057817 | 20017986 | 20018102 | 20096630 | 20096798 | 0 | 1 | [illegible] |
| SE | 104245 | ENSG00000128487 | SPECC1 | chr17 | + | 20124899 | 20124997 | 20110426 | 20110562 | 20227412 | 20227620 | 1 | 0 | [illegible] |
| SE | 104247 | ENSG00000128487 | SPECC1 | chr17 | + | 20204332 | 20205912 | 20110426 | 20110562 | 20227412 | 20227620 | 157 | 149 | [illegible] |
| SE | 104249 | ENSG00000128487 | SPECC1 | chr17 | + | 20204332 | 20205912 | 20155988 | 20156236 | 20227412 | 20227620 | 118 | 112 | [illegible] |
| SE | 104250 | ENSG00000128487 | SPECC1 | chr17 | + | 20232199 | 20232405 | 20231757 | 20231831 | 20245925 | 20246071 | 116 | 135 | [illegible] |
| SE | 104251 | ENSG00000128487 | SPECC1 | chr17 | + | 20232199 | 20232414 | 20231757 | 20231831 | 20245925 | 20246071 | 109 | 135 | [illegible] |
| SE | 104252 | ENSG00000128487 | SPECC1 | chr17 | + | 20245925 | 20246071 | 20232199 | 20232405 | 20247218 | 20247319 | 20 | 6 | [illegible] |
| SE | 104253 | ENSG00000128487 | SPECC1 | chr17 | + | 20245925 | 20246574 | 20232199 | 20232405 | 20247218 | 20247319 | 7 | 0 | [illegible] |
| SE | 104257 | ENSG00000128487 | SPECC1 | chr17 | + | 20306022 | 20306082 | 20296960 | 20297077 | 20313975 | 20314138 | 5 | 12 | [illegible] |
| SE | 104264 | ENSG00000132305 | IMMT | chr2 | - | 86151296 | 86151520 | 86147701 | 86147833 | 86159535 | 86159671 | 58 | 50 | [illegible] |
| SE | 104267 | ENSG00000132305 | IMMT | chr2 | - | 86158591 | 86158721 | 86151296 | 86151520 | 86159535 | 86159671 | 251 | 156 | [illegible] |
| SE | 104271 | ENSG00000132305 | IMMT | chr2 | - | 86166507 | 86166644 | 86161951 | 86162079 | 86170748 | 86170801 | 203 | 133 | [illegible] |
| SE | 104276 | ENSG00000132305 | IMMT | chr2 | - | 86173649 | 86173761 | 86171207 | 86171312 | 86179432 | 86179622 | 211 | 119 | [illegible] |
| SE | 104277 | ENSG00000132305 | IMMT | chr2 | - | 86179432 | 86179622 | 86173507 | 86173761 | 86181298 | 86181372 | 265 | 139 | [illegible] |
| SE | 104281 | ENSG00000110107 | PRPF19 | chr11 | - | 60899148 | 60899304 | 60898861 | 60898931 | 60900581 | 60900691 | 229 | 134 | [illegible] |
| SE | 104282 | ENSG00000110107 | PRPF19 | chr11 | - | 60903458 | 60903535 | 60902739 | 60902881 | 60903711 | 60903861 | 184 | 80 | [illegible] |
| SE | 104283 | ENSG00000110107 | PRPF19 | chr11 | - | 60905364 | 60905452 | 60903711 | 60903861 | 60906363 | 60906541 | 0 | 0 | [illegible] |
| SE | 104290 | ENSG00000135655 | USP15 | chr12 | + | 62294178 | 62294306 | 62260337 | 62260503 | 62302789 | 62302920 | 1410 | 951 | [illegible] |
| SE | 104291 | ENSG00000135655 | USP15 | chr12 | + | 62312285 | 62312309 | 62302789 | 62302920 | 62314789 | 62314916 | 5 | 2 | [illegible] |
| SE | 104292 | ENSG00000135655 | USP15 | chr12 | + | 62312285 | 62312309 | 62305594 | 62306058 | 62314789 | 62314916 | 0 | 1 | [illegible] |
| SE | 104293 | ENSG00000135655 | USP15 | chr12 | + | 62314789 | 62314916 | 62302789 | 62302920 | 62321463 | 62321489 | 1935 | 1470 | [illegible] |
| SE | 104294 | ENSG00000135655 | USP15 | chr12 | + | 62321463 | 62321609 | 62314789 | 62314916 | 62325871 | 62325933 | 1926 | 1445 | [illegible] |
| SE | 104296 | ENSG00000135655 | USP15 | chr12 | + | 62325871 | 62325933 | 62321460 | 62321609 | 62355330 | 62355475 | 399 | 259 | [illegible] |
| SE | 104297 | ENSG00000135655 | USP15 | chr12 | + | 62325871 | 62325933 | 62321460 | 62321609 | 62374389 | 62374535 | 324 | 205 | [illegible] |
| SE | 104299 | ENSG00000135655 | USP15 | chr12 | + | 62349220 | 62349307 | 62321463 | 62321609 | 62355330 | 62355475 | 259 | 216 | [illegible] |
| SE | 104301 | ENSG00000135655 | USP15 | chr12 | + | 62355330 | 62355475 | 62325871 | 62325933 | 62374389 | 62374535 | 198 | 137 | [illegible] |
| SE | 104303 | ENSG00000135655 | USP15 | chr12 | + | 62374389 | 62374535 | 62325871 | 62325933 | 62381489 | 62381633 | 71 | 57 | [illegible] |
| SE | 104305 | ENSG00000135655 | USP15 | chr12 | + | 62374389 | 62374535 | 62355330 | 62355475 | 62383839 | 62384302 | 109 | 70 | [illegible] |
| SE | 104306 | ENSG00000135655 | USP15 | chr12 | + | 62381489 | 62381663 | 62355330 | 62355475 | 62383839 | 62383998 | 1117 | 935 | [illegible] |
| SE | 104307 | ENSG00000135655 | USP15 | chr12 | + | 62381489 | 62381663 | 62374389 | 62374535 | 62383839 | 62383998 | 777 | 661 | [illegible] |
| SE | 104308 | ENSG00000135655 | USP15 | chr12 | + | 62384077 | 62384302 | 62383839 | 62383998 | 62389430 | 62389514 | 701 | 461 | [illegible] |
| SE | 104309 | ENSG00000135655 | USP15 | chr12 | + | 62389430 | 62389514 | 62383839 | 62384302 | 62389604 | 62389699 | 1114 | 798 | [illegible] |
| SE | 104310 | ENSG00000135655 | USP15 | chr12 | + | 62389604 | 62389699 | 62389430 | 62389514 | 62389796 | 62389873 | 1251 | 908 | [illegible] |
| SE | 104311 | ENSG00000135655 | USP15 | chr12 | + | 62391156 | 62391429 | 62390863 | 62390979 | 62391815 | 62391886 | 825 | 653 | [illegible] |
| SE | 104316 | ENSG00000122884 | P4HA1 | chr10 | - | 73016845 | 73016899 | 73014223 | 73014289 | 73030270 | 73030370 | 18 | 12 | [illegible] |
| SE | 104317 | ENSG00000122884 | P4HA1 | chr10 | - | 73030270 | 73030370 | 73016845 | 73016899 | 73044980 | 73045051 | 38 | 26 | [illegible] |
| SE | 104319 | ENSG00000122884 | P4HA1 | chr10 | - | 73043886 | 73043957 | 73030266 | 73030370 | 73046924 | 73047101 | 61 | 27 | [illegible] |
| SE | 104321 | ENSG00000122884 | P4HA1 | chr10 | - | 73044980 | 73045051 | 73043886 | 73043957 | 73046924 | 73047101 | 20 | 24 | [illegible] |
| SE | 104323 | ENSG00000122884 | P4HA1 | chr10 | - | 73068845 | 73068983 | 73053350 | 73053590 | 73072028 | 73072180 | 249 | 179 | [illegible] |
| SE | 104324 | ENSG00000122884 | P4HA1 | chr10 | - | 73073730 | 73073827 | 73072028 | 73072180 | 73074807 | 73074915 | 204 | 153 | [illegible] |
| SE | 104325 | ENSG00000243660 | ZNF487 | chr10 | + | 43476106 | 43476202 | 43475720 | 43475847 | 43481428 | 43481678 | 5 | 6 | [illegible] |
| SE | 104329 | ENSG00000155256 | ZFYVE27 | chr10 | + | 97743093 | 97743164 | 97738476 | 97738649 | 97748268 | 97748364 | 0 | 0 | [illegible] |
| SE | 104331 | ENSG00000155256 | ZFYVE27 | chr10 | + | 97743093 | 97743164 | 97738476 | 97738674 | 97748268 | 97748364 | 5 | 1 | [illegible] |
| SE | 104332 | ENSG00000155256 | ZFYVE27 | chr10 | + | 97744728 | 97744915 | 97738476 | 97738649 | 97748268 | 97748364 | 12 | 8 | [illegible] |
| SE | 104333 | ENSG00000155256 | ZFYVE27 | chr10 | + | 97744728 | 97744915 | 97738476 | 97738649 | 97749473 | 97749586 | 0 | 0 | [illegible] |
| SE | 104334 | ENSG00000155256 | ZFYVE27 | chr10 | + | 97744728 | 97744915 | 97738476 | 97738674 | 97748268 | 97748364 | 13 | 11 | [illegible] |
| SE | 104335 | ENSG00000155256 | ZFYVE27 | chr10 | + | 97744728 | 97744915 | 97738476 | 97738674 | 97749473 | 97749586 | 1 | 3 | [illegible] |
| SE | 104336 | ENSG00000155256 | ZFYVE27 | chr10 | + | 97744728 | 97744915 | 97743093 | 97743164 | 97748268 | 97748364 | 30 | 13 | [illegible] |
| SE | 104338 | ENSG00000155256 | ZFYVE27 | chr10 | + | 97748268 | 97748364 | 97738476 | 97738674 | 97749473 | 97749586 | 16 | 10 | [illegible] |
| SE | 104340 | ENSG00000155256 | ZFYVE27 | chr10 | + | 97749473 | 97749586 | 97748268 | 97748364 | 97750330 | 97750470 | 51 | 32 | [illegible] |
| SE | 104341 | ENSG00000155256 | ZFYVE27 | chr10 | + | 97752856 | 97752877 | 97751375 | 97751462 | 97753037 | 97753182 | 1 | 0 | [illegible] |
| SE | 104342 | ENSG00000260729 | NA | chr15 | - | 72319612 | 72320702 | 72315844 | 72315927 | 72345445 | 72345550 | 1 | 3 | [illegible] |
| SE | 104348 | ENSG00000150054 | MPP7 | chr10 | - | 28059515 | 28059743 | 28058494 | 28058603 | 28069771 | 28069852 | 57 | 42 | [illegible] |
| SE | 104349 | ENSG00000150054 | MPP7 | chr10 | - | 28059649 | 28059743 | 28058494 | 28058603 | 28069771 | 28069852 | 154 | 114 | [illegible] |
| SE | 104350 | ENSG00000150054 | MPP7 | chr10 | - | 28089670 | 28089841 | 28069771 | 28069852 | 28119650 | 28119715 | 182 | 198 | [illegible] |
| SE | 104351 | ENSG00000150054 | MPP7 | chr10 | - | 28120193 | 28120390 | 28119650 | 28119715 | 28120593 | 28120668 | 224 | 240 | [illegible] |
| SE | 104352 | ENSG00000150054 | MPP7 | chr10 | - | 28147482 | 28147563 | 28131559 | 28131691 | 28149981 | 28150059 | 171 | 128 | [illegible] |
| SE | 104353 | ENSG00000150054 | MPP7 | chr10 | - | 28149981 | 28150059 | 28131559 | 28131691 | 28202152 | 28202271 | 37 | 18 | [illegible] |
| SE | 104355 | ENSG00000150054 | MPP7 | chr10 | - | 28254773 | 28254876 | 28238567 | 28238735 | 28280099 | 28280140 | 0 | 0 | [illegible] |
| SE | 104372 | ENSG00000095637 | SORBS1 | chr10 | - | 95375972 | 95376056 | 95367649 | 95367699 | 95381684 | 95381799 | 1 | 0 | [illegible] |
| SE | 104382 | ENSG00000095637 | SORBS1 | chr10 | - | 95422189 | 95422216 | 95421960 | 95422073 | 95432447 | 95432576 | 0 | 0 | [illegible] |
| SE | 104384 | ENSG00000169813 | HNRNPF | chr10 | - | 43394629 | 43394688 | 43385616 | 43387936 | 43396455 | 43396590 | 525 | 184 | [illegible] |
| SE | 104385 | ENSG00000169813 | HNRNPF | chr10 | - | 43394629 | 43394688 | 43385616 | 43387936 | 43409089 | 43409166 | 119 | 47 | [illegible] |
| SE | 104386 | ENSG00000169813 | HNRNPF | chr10 | - | 43396455 | 43396590 | 43385616 | 43387936 | 43407716 | 43407819 | 25 | 27 | [illegible] |
| SE | 104387 | ENSG00000169813 | HNRNPF | chr10 | - | 43396455 | 43396590 | 43385616 | 43387936 | 43409089 | 43409166 | 15 | 6 | [illegible] |
| SE | 104388 | ENSG00000169813 | HNRNPF | chr10 | - | 43396455 | 43396590 | 43385616 | 43387936 | 43409130 | 43409186 | 55 | 9 | [illegible] |
| SE | 104389 | ENSG00000169813 | HNRNPF | chr10 | - | 43396455 | 43396590 | 43394629 | 43394688 | 43409089 | 43409166 | 404 | 140 | [illegible] |
| SE | 104391 | ENSG00000204179 | PTPN20 | chr10 | + | 46932376 | 46932533 | 46911427 | 46911501 | 46943917 | 46944015 | 0 | 0 | [illegible] |
| SE | 104393 | ENSG00000204179 | PTPN20 | chr10 | + | 46932376 | 46932533 | 46911780 | 46911842 | 46943917 | 46944015 | 0 | 0 | [illegible] |
| SE | 104394 | ENSG00000204179 | PTPN20 | chr10 | + | 46940581 | 46940717 | 46911780 | 46911842 | 46943917 | 46944015 | 1 | 0 | [illegible] |
| SE | 104395 | ENSG00000204179 | PTPN20 | chr10 | + | 46940581 | 46940717 | 46932416 | 46932533 | 46943917 | 46944015 | 1 | 0 | [illegible] |
| SE | 104396 | ENSG00000204179 | PTPN20 | chr10 | + | 46940622 | 46940717 | 46911437 | 46911501 | 46943917 | 46943992 | 1 | 0 | [illegible] |
| SE | 104397 | ENSG00000204179 | PTPN20 | chr10 | + | 46940622 | 46940717 | 46911780 | 46911842 | 46943917 | 46943992 | 1 | 0 | [illegible] |
| SE | 104398 | ENSG00000204179 | PTPN20 | chr10 | + | 46940622 | 46940717 | 46932376 | 46932533 | 46943917 | 46943992 | 1 | 0 | [illegible] |
| SE | 104400 | ENSG00000204179 | PTPN20 | chr10 | + | 46947840 | 46947955 | 46946562 | 46946675 | 46964985 | 46965137 | 2 | 0 | [illegible] |
| SE | 104402 | ENSG00000204179 | PTPN20 | chr10 | + | 46964985 | 46965137 | 46946562 | 46946675 | 46999911 | 46999974 | 0 | 2 | [illegible] |
| SE | 104404 | ENSG00000204179 | PTPN20 | chr10 | + | 46967956 | 46968047 | 46964985 | 46965137 | 46999911 | 46999950 | 0 | 0 | [illegible] |
| SE | 104405 | ENSG00000204179 | PTPN20 | chr10 | + | 46984229 | 46984564 | 46964985 | 46965137 | 46987339 | 46987555 | 0 | 0 | [illegible] |
| SE | 104409 | ENSG00000204179 | PTPN20 | chr10 | + | 46987339 | 46987555 | 46984229 | 46984564 | 46999911 | 46999974 | 0 | 0 | [illegible] |
| SE | 104414 | ENSG00000184047 | DIABLO | chr12 | - | 122218265 | 122218397 | 122216758 | 122216869 | 122225964 | 122226568 | 138 | 59 | [illegible] |
| SE | 104415 | ENSG00000184047 | DIABLO | chr12 | - | 122220529 | 122220627 | 122218265 | 122218397 | 122224511 | 122224644 | 0 | 0 | [illegible] |
| SE | 104416 | ENSG00000184047 | DIABLO | chr12 | - | 122220529 | 122220627 | 122218265 | 122218397 | 122224644 | 122226062 | 0 | 0 | [illegible] |
| SE | 104417 | ENSG00000184047 | DIABLO | chr12 | - | 122224511 | 122224644 | 122216758 | 122216869 | 122225964 | 122226009 | 39 | 7 | [illegible] |
| SE | 104418 | ENSG00000184047 | DIABLO | chr12 | - | 122224511 | 122224644 | 122218265 | 122218397 | 122225964 | 122226009 | 105 | 39 | [illegible] |
| SE | 104419 | ENSG00000184047 | DIABLO | chr12 | - | 122224511 | 122224644 | 122218265 | 122218397 | 122227355 | 122227456 | 70 | 32 | [illegible] |
| SE | 104420 | ENSG00000184047 | DIABLO | chr12 | - | 122224644 | 122224748 | 122216758 | 122216869 | 122225964 | 122226039 | 3 | 2 | [illegible] |
| SE | 104423 | ENSG00000142920 | AZIN2 | chr1 | + | 33081600 | 33081812 | 33081387 | 33081508 | 33082177 | 33082354 | 0 | 0 | [illegible] |
| SE | 104430 | ENSG00000142920 | AZIN2 | chr1 | + | 33083953 | 33084127 | 33082177 | 33082354 | 33092049 | 33092210 | 3 | 3 | [illegible] |
| SE | 104432 | ENSG00000142920 | AZIN2 | chr1 | + | 33101832 | 33101902 | 33098066 | 33098179 | 33117901 | 33118116 | 0 | 0 | [illegible] |
| SE | 104434 | ENSG00000255639 | NA | chr12 | + | 4685258 | 4685325 | 4682204 | 4682300 | 4726531 | 4726829 | 35 | 21 | [illegible] |
| SE | 104440 | ENSG00000149100 | EIF3M | chr11 | + | 32588593 | 32588732 | 32586933 | 32587144 | 32589546 | 32589641 | 591 | 416 | [illegible] |
| SE | 104442 | ENSG00000149100 | EIF3M | chr11 | + | 32589546 | 32589641 | 32583831 | 32583929 | 32593865 | 32593949 | 528 | 364 | [illegible] |
| SE | 104444 | ENSG00000149100 | EIF3M | chr11 | + | 32594913 | 32595013 | 32593768 | 32593949 | 32600688 | 32600832 | 749 | 436 | [illegible] |
| SE | 104445 | ENSG00000149100 | EIF3M | chr11 | + | 32595965 | 32596047 | 32593768 | 32593949 | 32600688 | 32600793 | 775 | 540 | [illegible] |
| SE | 104446 | ENSG00000149100 | EIF3M | chr11 | + | 32595965 | 32596047 | 32593865 | 32595013 | 32600688 | 32600793 | 1732 | 1107 | [illegible] |
| SE | 104447 | ENSG00000149100 | EIF3M | chr11 | + | 32601761 | 32601822 | 32600688 | 32600832 | 32602278 | 32602302 | 1294 | 751 | [illegible] |
| SE | 104449 | ENSG00000205846 | CLEC6A | chr12 | + | 8465483 | 8465629 | 8459596 | 8459698 | 8476124 | 8476240 | 28 | 20 | [illegible] |
| SE | 104450 | ENSG00000205846 | CLEC6A | chr12 | + | 8476124 | 8476240 | 8465483 | 8465629 | 8477319 | 8478330 | 7 | 14 | [illegible] |
| SE | 104451 | ENSG00000066651 | TRMT11 | chr6 | + | 125993756 | 125993822 | 125986478 | 125986622 | 125995966 | 125996040 | 52 | 32 | [illegible] |
| SE | 104452 | ENSG00000066651 | TRMT11 | chr6 | + | 125993792 | 125993822 | 125986550 | 125986622 | 125995966 | 125996040 | 17 | 9 | [illegible] |
| SE | 104454 | ENSG00000066651 | TRMT11 | chr6 | + | 125999456 | 125999613 | 125998549 | 125998684 | 126008391 | 126008472 | 110 | 102 | [illegible] |
| SE | 104456 | ENSG00000066651 | TRMT11 | chr6 | + | 126008656 | 126008745 | 126008391 | 126008472 | 126010924 | 126011417 | 6 | 7 | [illegible] |
| SE | 104457 | ENSG00000066651 | TRMT11 | chr6 | + | 126008656 | 126008745 | 126008391 | 126008472 | 126011252 | 126011274 | 10 | 6 | [illegible] |
| SE | 104458 | ENSG00000066651 | TRMT11 | chr6 | + | 126009334 | 126009434 | 126008656 | 126008745 | 126011252 | 126011417 | 0 | 2 | [illegible] |
| SE | 104459 | ENSG00000066651 | TRMT11 | chr6 | + | 126040794 | 126040845 | 126038704 | 126038925 | 126174824 | 126174911 | 0 | 1 | [illegible] |
| SE | 104461 | ENSG00000066651 | TRMT11 | chr6 | + | 126115344 | 126115493 | 126114704 | 126114739 | 126115742 | 126115855 | 1 | 3 | [illegible] |
| SE | 104464 | ENSG00000228063 | LYPLAL1-DT | chr1 | - | 218991270 | 218991354 | 218983022 | 218983283 | 219086494 | 219086660 | 0 | 0 | [illegible] |
| SE | 104467 | ENSG00000228063 | LYPLAL1-DT | chr1 | - | 219149938 | 219150031 | 219086494 | 219086660 | 219173469 | 219173517 | 0 | 5 | [illegible] |
| SE | 104468 | ENSG00000228063 | LYPLAL1-DT | chr1 | - | 219149938 | 219150031 | 219100625 | 219100754 | 219156119 | 219156199 | 0 | 30 | [illegible] |
| SE | 104469 | ENSG00000228063 | LYPLAL1-DT | chr1 | - | 219149938 | 219150031 | 219100625 | 219100754 | 219173469 | 219173517 | 0 | 0 | [illegible] |
| SE | 104470 | ENSG00000228063 | LYPLAL1-DT | chr1 | - | 219156119 | 219156199 | 219086494 | 219086660 | 219173469 | 219173493 | 12 | 23 | [illegible] |
| SE | 104471 | ENSG00000228063 | LYPLAL1-DT | chr1 | - | 219156119 | 219156199 | 219100625 | 219100754 | 219173469 | 219173493 | 5 | 21 | [illegible] |
| SE | 104473 | ENSG00000226711 | FAM66C | chr12 | + | 8187731 | 8188225 | 8180208 | 8180752 | 8188471 | 8188645 | 0 | 0 | [illegible] |
| SE | 104474 | ENSG00000226711 | FAM66C | chr12 | + | 8193519 | 8193655 | 8188471 | 8188645 | 8194202 | 8194481 | 8 | 0 | [illegible] |
| SE | 104475 | ENSG00000226711 | FAM66C | chr12 | + | 8193519 | 8193655 | 8188484 | 8188535 | 8194202 | 8194481 | 6 | 0 | [illegible] |
| SE | 104476 | ENSG00000226711 | FAM66C | chr12 | + | 8194728 | 8194796 | 8194202 | 8194481 | 8195276 | 8195412 | 7 | 1 | [illegible] |
| SE | 104477 | ENSG00000226711 | FAM66C | chr12 | + | 8195276 | 8195412 | 8194728 | 8194796 | 8196256 | 8196518 | 1 | 0 | [illegible] |
| SE | 104480 | ENSG00000013561 | RNF14 | chr5 | + | 141970703 | 141970877 | 141969094 | 141969167 | 141973582 | 141973634 | 85 | 54 | [illegible] |
| SE | 104485 | ENSG00000013561 | RNF14 | chr5 | + | 141978302 | 141978830 | 141973582 | 141973742 | 141983379 | 141983552 | 19 | 11 | [illegible] |
| SE | 104490 | ENSG00000013561 | RNF14 | chr5 | + | 141980122 | 141980351 | 141978302 | 141978830 | 141983379 | 141983552 | 116 | 68 | [illegible] |
| SE | 104491 | ENSG00000249790 | NA | chr12 | + | 8789999 | 8790263 | 8788273 | 8788786 | 8794304 | 8794336 | 0 | 0 | [illegible] |
| SE | 104492 | ENSG00000103966 | EHD4 | chr15 | - | 41919209 | 41919622 | 41909698 | 41909863 | 41943066 | 41943164 | 131 | 48 | [illegible] |
| SE | 104493 | ENSG00000103966 | EHD4 | chr15 | - | 41943066 | 41943164 | 41919209 | 41919622 | 41953763 | 41953940 | 142 | 70 | [illegible] |
| SE | 104495 | ENSG00000151490 | PTPRO | chr12 | + | 15551550 | 15551671 | 15549093 | 15549226 | 15557454 | 15557523 | 23 | 27 | [illegible] |
| SE | 104496 | ENSG00000151490 | PTPRO | chr12 | + | 15557454 | 15557523 | 15551550 | 15551671 | 15565592 | 15565628 | 16 | 14 | [illegible] |
| SE | 104497 | ENSG00000151490 | PTPRO | chr12 | + | 15560192 | 15560276 | 15557454 | 15557523 | 15565592 | 15565628 | 8 | 11 | [illegible] |
| SE | 104500 | ENSG00000151490 | PTPRO | chr12 | + | 15570088 | 15570135 | 15569416 | 15569498 | 15578852 | 15578943 | 1 | 0 | [illegible] |
| SE | 104502 | ENSG00000120647 | CCDC77 | chr12 | + | 395123 | 395221 | 389361 | 389527 | 405510 | 405564 | 0 | 0 | [illegible] |
| SE | 104503 | ENSG00000120647 | CCDC77 | chr12 | + | 395123 | 395221 | 389418 | 389486 | 405510 | 405564 | 0 | 0 | [illegible] |
| SE | 104504 | ENSG00000120647 | CCDC77 | chr12 | + | 395123 | 395221 | 389418 | 389486 | 409367 | 409421 | 0 | 0 | [illegible] |
| SE | 104505 | ENSG00000120647 | CCDC77 | chr12 | + | 405510 | 405564 | 389349 | 389486 | 409367 | 409421 | 4 | 0 | [illegible] |
| SE | 104506 | ENSG00000120647 | CCDC77 | chr12 | + | 405510 | 405564 | 389349 | 389486 | 411746 | 411978 | 4 | 0 | [illegible] |
| SE | 104510 | ENSG00000120647 | CCDC77 | chr12 | + | 428768 | 428865 | 418493 | 418636 | 430663 | 430736 | 27 | 31 | [illegible] |
| SE | 104511 | ENSG00000120647 | CCDC77 | chr12 | + | 430663 | 430736 | 428768 | 428865 | 431865 | 431954 | 23 | 25 | [illegible] |
| SE | 104513 | ENSG00000175806 | MSRA | chr8 | + | 10245103 | 10245223 | 10207832 | 10207901 | 10301533 | 10301638 | 56 | 47 | [illegible] |

| | | | | | | | | | | | | | | | | | | | | | | | | | | | | | | | | | | | | | | | | | | | | | | |
|---|---|---|---|---|---|---|---|---|---|---|---|---|---|---|---|---|---|---|---|---|---|---|---|---|---|---|---|---|---|---|---|---|---|---|---|---|---|---|---|---|---|---|---|---|---|---|
| SE | 104514 | ENSG00000175806 | MSRA | chr8 | + | 10301533 | 10301638 | 10245103 | 10245223 | 10319882 | 10319891 | 51 | 51 | 85 | 56 | 79 | 1 | 0 | 0 | 0 | 0 | 104 | 85 | 83 | 185 | 231 | 0 | 0 | 0 | 0 | 0 | 254 | 149 | 1 | 1 | 0.968 | 1 | 1 | 1 | 1 | 1 | 1 | 1 | 1 | 1 | -0.006 |
| SE | 104518 | ENSG00000079739 | PGM1 | chr1 | + | 63651668 | 63651852 | 63648516 | 63648652 | 63654331 | 63654466 | 219 | 96 | 118 | 140 | 144 | 0 | 0 | 0 | 4 | 0 | 216 | 111 | 118 | 256 | 281 | 0 | 0 | 0 | 0 | 0 | 298 | 149 | 1 | 1 | 1 | 1 | 1 | 0.946 | 1 | 1 | 1 | 1 | 1 | 1 | -0.011 |
| SE | 104521 | ENSG00000171792 | RHNO1 | chr12 | + | 2885161 | 2885534 | 2877198 | 2877282 | 2887910 | 2888057 | 11 | 2 | 27 | 12 | 8 | 1 | 2 | 0 | 0 | 0 | 0 | 19 | 20 | 28 | 47 | 0 | 0 | 0 | 0 | 0 | 298 | 149 | 1 | 1 | 0.846 | 0.333 | 1 | 1 | 1 | NA | 1 | 1 | 1 | 1 | -0.164 |
| SE | 104522 | ENSG00000171792 | RHNO1 | chr12 | + | 2885262 | 2885534 | 2877198 | 2877282 | 2887910 | 2888137 | 11 | 2 | 27 | 12 | 8 | 1 | 2 | 0 | 0 | 0 | 0 | 19 | 20 | 28 | 47 | 0 | 0 | 0 | 0 | 0 | 298 | 149 | 1 | 1 | 0.846 | 0.333 | 1 | 1 | 1 | NA | 1 | 1 | 1 | 1 | -0.164 |
| SE | 104523 | ENSG00000171792 | RHNO1 | chr12 | + | 2885282 | 2885492 | 2877198 | 2877282 | 2887910 | 2889271 | 13 | 2 | 13 | 5 | 2 | 1 | 2 | 0 | 0 | 0 | 14 | 9 | 15 | 10 | 0 | 0 | 0 | 0 | 0 | 0 | 298 | 149 | 1 | 1 | 0.867 | 0.333 | 1 | 1 | 1 | 1 | 1 | 1 | 1 | NA | -0.16 |
| SE | 104524 | ENSG00000171792 | RHNO1 | chr12 | + | 2885282 | 2885534 | 2877222 | 2877282 | 2887910 | 2887934 | 24 | 4 | 37 | 17 | 10 | 1 | 2 | 0 | 0 | 0 | 14 | 26 | 32 | 38 | 47 | 0 | 0 | 0 | 0 | 0 | 298 | 149 | 1 | 1 | 0.923 | 0.5 | 1 | 1 | 1 | 1 | 1 | 1 | 1 | 1 | -0.115 |
| SE | 104525 | ENSG00000171792 | RHNO1 | chr12 | + | 2886977 | 2887111 | 2885282 | 2885492 | 2887910 | 2888007 | 0 | 0 | 0 | 0 | 0 | 0 | 0 | 3 | 0 | 0 | 0 | 0 | 1 | 0 | 0 | 0 | 2 | 3 | 0 | 0 | 283 | 149 | 1 | 1 | NA | NA | 0 | NA | NA | NA | 0 | 0.149 | NA | NA | -0.074 |
| SE | 104528 | ENSG00000214776 | NA | chr12 | + | 9545095 | 9545241 | 9544237 | 9544292 | 9545597 | 9545747 | 0 | 0 | 0 | 3 | 0 | 0 | 0 | 0 | 0 | 0 | 0 | 0 | 0 | 0 | 0 | 0 | 0 | 4 | 0 | 0 | 295 | 149 | 1 | 1 | NA | NA | NA | 1 | NA | NA | NA | 0 | NA | NA | 1 |
| SE | 104529 | ENSG00000214776 | NA | chr12 | + | 9545597 | 9545747 | 9544237 | 9544292 | 9551452 | 9551623 | 0 | 1 | 0 | 3 | 2 | 0 | 0 | 1 | 0 | 0 | 0 | 0 | 6 | 0 | 0 | 0 | 0 | 0 | 0 | 0 | 298 | 149 | 1 | 1 | NA | 1 | 0 | 1 | 1 | NA | NA | 1 | NA | NA | -0.25 |
| SE | 104533 | ENSG00000145354 | CISD2 | chr4 | + | 102882770 | 102882873 | 102869053 | 102869187 | 102885215 | 102885430 | 0 | 0 | 0 | 4 | 0 | 63 | 19 | 55 | 34 | 32 | 0 | 0 | 0 | 0 | 0 | 62 | 77 | 28 | 42 | 35 | 252 | 149 | 1 | 1 | 0 | 0 | 0 | 0.065 | 0 | 0 | 0 | 0 | 0 | 0 | 0.013 |
| SE | 104534 | ENSG00000145354 | CISD2 | chr4 | + | 102882791 | 102882873 | 102868973 | 102869187 | 102885215 | 102885430 | 0 | 0 | 0 | 0 | 0 | 63 | 19 | 55 | 34 | 32 | 0 | 2 | 0 | 0 | 0 | 62 | 77 | 28 | 42 | 35 | 231 | 149 | 1 | 1 | 0 | 0 | 0 | 0 | 0 | 0 | 0.016 | 0 | 0 | 0 | -0.003 |
| SE | 104535 | ENSG00000145354 | CISD2 | chr4 | + | 102885215 | 102885430 | 102868991 | 102869187 | 102887340 | 102887359 | 175 | 84 | 155 | 145 | 204 | 2 | 0 | 0 | 0 | 0 | 212 | 211 | 95 | 197 | 137 | 0 | 0 | 0 | 0 | 0 | 298 | 149 | 1 | 1 | 0.978 | 1 | 1 | 1 | 1 | 1 | 1 | 1 | 1 | 1 | -0.004 |
| SE | 104536 | ENSG00000228705 | LINC00659 | chr20 | - | 62775314 | 62775479 | 62774120 | 62774569 | 62776805 | 62776949 | 0 | 3 | 4 | 2 | 7 | 1 | 0 | 0 | 0 | 0 | 0 | 0 | 2 | 6 | 9 | 0 | 0 | 0 | 0 | 0 | 298 | 149 | 1 | 1 | 0 | 1 | 1 | 1 | 1 | NA | NA | 1 | 1 | 1 | -0.2 |
| SE | 104537 | ENSG00000228705 | LINC00659 | chr20 | - | 62775314 | 62775520 | 62774120 | 62774569 | 62776805 | 62776921 | 0 | 3 | 8 | 3 | 7 | 1 | 0 | 0 | 0 | 0 | 0 | 0 | 2 | 12 | 9 | 0 | 0 | 0 | 0 | 0 | 298 | 149 | 1 | 1 | 0 | 1 | 1 | 1 | 1 | NA | NA | 1 | 1 | 1 | -0.2 |
| SE | 104539 | ENSG00000075303 | SLC25A40 | chr7 | - | 87841931 | 87842041 | 87841632 | 87841714 | 87843753 | 87843864 | 1 | 0 | 2 | 1 | 2 | 49 | 51 | 51 | 38 | 131 | 0 | 2 | 3 | 0 | 0 | 40 | 29 | 25 | 42 | 22 | 259 | 149 | 1 | 1 | 0.012 | 0 | 0.022 | 0.015 | 0.009 | 0 | 0.038 | 0.065 | 0 | 0 | -0.009 |
| SE | 104541 | ENSG00000075303 | SLC25A40 | chr7 | - | 87849880 | 87849948 | 87847852 | 87847977 | 87854203 | 87854310 | 108 | 85 | 91 | 118 | 171 | 1 | 0 | 0 | 0 | 1 | 52 | 74 | 45 | 106 | 79 | 0 | 0 | 2 | 0 | 0 | 217 | 149 | 1 | 1 | 0.987 | 1 | 1 | 1 | 0.992 | 1 | 1 | 0.939 | 1 | 1 | 0.008 |
| SE | 104542 | ENSG00000075303 | SLC25A40 | chr7 | - | 87856291 | 87856351 | 87854203 | 87854310 | 87858630 | 87858751 | 128 | 86 | 104 | 95 | 139 | 0 | 0 | 0 | 2 | 0 | 128 | 90 | 44 | 126 | 59 | 0 | 0 | 0 | 0 | 0 | 209 | 149 | 1 | 1 | 1 | 1 | 1 | 0.971 | 1 | 1 | 1 | 1 | 1 | 1 | -0.006 |
| SE | 104543 | ENSG00000075303 | SLC25A40 | chr7 | - | 87856291 | 87856389 | 87854203 | 87854310 | 87858630 | 87858751 | 59 | 41 | 41 | 44 | 42 | 0 | 0 | 0 | 2 | 0 | 51 | 35 | 26 | 77 | 21 | 0 | 0 | 0 | 0 | 0 | 247 | 149 | 1 | 1 | 1 | 1 | 1 | 0.93 | 1 | 1 | 1 | 1 | 1 | 1 | -0.014 |
| SE | 104545 | ENSG00000150045 | KLRF1 | chr12 | + | 9832315 | 9832414 | 9827480 | 9827629 | 9833302 | 9833452 | 1984 | 665 | 591 | 702 | 1023 | 1 | 0 | 0 | 0 | 0 | 52 | 125 | 169 | 9 | 0 | 0 | 0 | 0 | 0 | 0 | 248 | 149 | 1 | 1 | 0.999 | 1 | 1 | 1 | 1 | 1 | 1 | 1 | 1 | NA | 0 |
| SE | 104546 | ENSG00000150045 | KLRF1 | chr12 | + | 9833302 | 9833452 | 9832315 | 9832414 | 9841811 | 9841951 | 2020 | 707 | 584 | 726 | 965 | 3 | 3 | 1 | 2 | 9 | 59 | 167 | 178 | 18 | 12 | 0 | 1 | 1 | 0 | 0 | 298 | 149 | 1 | 1 | 0.997 | 0.992 | 0.997 | 0.995 | 0.982 | 1 | 0.988 | 0.989 | 1 | 1 | -0.003 |
| SE | 104547 | ENSG00000150045 | KLRF1 | chr12 | + | 9833302 | 9833452 | 9832315 | 9832414 | 9842320 | 9842433 | 1199 | 394 | 348 | 407 | 560 | 14 | 4 | 6 | 5 | 0 | 39 | 85 | 98 | 14 | 0 | 0 | 0 | 0 | 0 | 0 | 298 | 149 | 1 | 1 | 0.977 | 0.98 | 0.967 | 0.976 | 1 | 1 | 1 | 1 | 1 | NA | -0.02 |
| SE | 104548 | ENSG00000150045 | KLRF1 | chr12 | + | 9833302 | 9833452 | 9832315 | 9832414 | 9844417 | 9844657 | 1153 | 383 | 331 | 398 | 572 | 2 | 1 | 0 | 4 | 0 | 39 | 91 | 90 | 7 | 4 | 0 | 0 | 1 | 0 | 0 | 298 | 149 | 1 | 1 | 0.997 | 0.995 | 1 | 0.98 | 1 | 1 | 1 | 0.978 | 1 | 1 | -0.001 |
| SE | 104550 | ENSG00000150045 | KLRF1 | chr12 | + | 9841811 | 9841951 | 9832315 | 9832414 | 9844417 | 9845005 | 5 | 3 | 3 | 5 | 10 | 2 | 1 | 0 | 4 | 0 | 0 | 1 | 1 | 0 | 0 | 0 | 0 | 1 | 0 | 0 | 289 | 149 | 1 | 1 | 0.563 | 0.607 | 1 | 0.392 | 1 | NA | 1 | 0.34 | NA | NA | 0.042 |
| SE | 104553 | ENSG00000150045 | KLRF1 | chr12 | + | 9842301 | 9842433 | 9833302 | 9833452 | 9844417 | 9844657 | 382 | 132 | 125 | 160 | 190 | 30 | 9 | 3 | 12 | 17 | 0 | 29 | 14 | 10 | 6 | 0 | 10 | 0 | 0 | 4 | 281 | 149 | 1 | 1 | 0.871 | 0.886 | 0.957 | 0.876 | 0.856 | NA | 0.606 | 1 | 1 | 0.443 | 0.127 |
| SE | 104554 | ENSG00000150045 | KLRF1 | chr12 | + | 9842301 | 9842433 | 9841813 | 9841951 | 9844417 | 9844657 | 408 | 146 | 124 | 168 | 204 | 3 | 1 | 2 | 3 | 1 | 0 | 32 | 24 | 10 | 6 | 0 | 0 | 0 | 0 | 0 | 281 | 149 | 1 | 1 | 0.986 | 0.987 | 0.97 | 0.967 | 0.991 | NA | 1 | 1 | 1 | 1 | -0.02 |
| SE | 104555 | ENSG00000150045 | KLRF1 | chr12 | + | 9842320 | 9842433 | 9832315 | 9832414 | 9844417 | 9845002 | 353 | 118 | 124 | 151 | 187 | 2 | 1 | 0 | 4 | 0 | 0 | 29 | 14 | 10 | 6 | 0 | 0 | 1 | 0 | 0 | 262 | 149 | 1 | 1 | 0.99 | 0.985 | 1 | 0.955 | 1 | NA | 1 | 0.888 | 1 | 1 | 0.014 |
| SE | 104556 | ENSG00000150045 | KLRF1 | chr12 | + | 9842320 | 9842433 | 9833302 | 9833452 | 9844417 | 9845002 | 415 | 134 | 138 | 167 | 192 | 30 | 9 | 3 | 12 | 17 | 0 | 33 | 22 | 17 | 6 | 0 | 10 | 0 | 0 | 4 | 262 | 149 | 1 | 1 | 0.887 | 0.894 | 0.963 | 0.888 | 0.865 | NA | 0.652 | 1 | 1 | 0.46 | 0.121 |
| SE | 104557 | ENSG00000150045 | KLRF1 | chr12 | + | 9842320 | 9842433 | 9841811 | 9841951 | 9844417 | 9845002 | 1478 | 487 | 455 | 588 | 609 | 3 | 1 | 2 | 3 | 1 | 18 | 158 | 93 | 41 | 13 | 0 | 0 | 0 | 0 | 0 | 262 | 149 | 1 | 1 | 0.996 | 0.996 | 0.992 | 0.991 | 0.997 | 1 | 1 | 1 | 1 | 1 | -0.006 |
| SE | 104562 | ENSG00000241484 | ARHGAP8 | chr22 | + | 44809005 | 44809098 | 44808306 | 44808438 | 44814671 | 44814758 | 0 | 0 | 0 | 0 | 0 | 7 | 0 | 1 | 2 | 0 | 0 | 0 | 3 | 0 | 0 | 0 | 32 | 38 | 4 | 0 | 242 | 149 | 1 | 1 | 0 | NA | 0 | 0 | NA | NA | 0 | 0.046 | 0 | NA | -0.015 |
| SE | 104564 | ENSG00000256039 | LINC02446 | chr12 | + | 10557335 | 10557677 | 10553361 | 10553516 | 10572237 | 10572688 | 131 | 89 | 88 | 67 | 208 | 28 | 16 | 4 | 3 | 23 | 0 | 107 | 95 | 22 | 3 | 0 | 3 | 11 | 0 | 6 | 298 | 149 | 1 | 1 | 0.701 | 0.736 | 0.917 | 0.918 | 0.819 | NA | 0.947 | 0.812 | 1 | 0.2 | 0.078 |
| SE | 104566 | ENSG00000284697 | NA | chr12 | + | 8125560 | 8125677 | 8120907 | 8121106 | 8129263 | 8129362 | 152 | 117 | 94 | 113 | 103 | 2 | 0 | 0 | 0 | 0 | 172 | 72 | 65 | 113 | 65 | 0 | 0 | 0 | 0 | 0 | 266 | 149 | 1 | 1 | 0.977 | 1 | 1 | 1 | 1 | 1 | 1 | 1 | 1 | 1 | -0.005 |
| SE | 104568 | ENSG00000153495 | TEX29 | chr13 | + | 111328182 | 111328293 | 111320856 | 111320948 | 111339862 | 111339932 | 0 | 0 | 2 | 2 | 0 | 0 | 0 | 0 | 0 | 0 | 0 | 0 | 4 | 0 | 0 | 0 | 0 | 0 | 2 | 0 | 260 | 149 | 1 | 1 | NA | NA | 1 | 1 | NA | NA | NA | 1 | 0 | NA | 0.5 |
| SE | 104569 | ENSG00000153495 | TEX29 | chr13 | + | 111339293 | 111339438 | 111328182 | 111328293 | 111339862 | 111339932 | 0 | 1 | 0 | 0 | 0 | 0 | 0 | 2 | 1 | 0 | 0 | 0 | 0 | 0 | 0 | 0 | 0 | 2 | 0 | 0 | 294 | 149 | 1 | 1 | NA | 1 | 0 | 0 | NA | NA | NA | 0 | NA | NA | 0.333 |
| SE | 104573 | ENSG00000214212 | C19orf38 | chr19 | + | 10850258 | 10850567 | 10848414 | 10848539 | 10856264 | 10856357 | 196 | 61 | 84 | 99 | 57 | 0 | 1 | 0 | 0 | 0 | 261 | 233 | 89 | 212 | 395 | 0 | 1 | 0 | 6 | 0 | 298 | 149 | 1 | 1 | 1 | 0.968 | 1 | 1 | 1 | 1 | 0.991 | 1 | 0.946 | 1 | 0.006 |
| SE | 104574 | ENSG00000214212 | C19orf38 | chr19 | + | 10858315 | 10858343 | 10856264 | 10856357 | 10859914 | 10859958 | 112 | 68 | 54 | 66 | 43 | 0 | 0 | 0 | 0 | 0 | 219 | 176 | 97 | 172 | 285 | 0 | 0 | 0 | 0 | 4 | 177 | 149 | 1 | 1 | 1 | 1 | 1 | 1 | 1 | 1 | 1 | 1 | 1 | 0.984 | 0.003 |
| SE | 104575 | ENSG00000214212 | C19orf38 | chr19 | + | 10858315 | 10858343 | 10856264 | 10856357 | 10863169 | 10863207 | 16 | 9 | 11 | 6 | 19 | 0 | 0 | 0 | 0 | 0 | 13 | 30 | 15 | 23 | 31 | 0 | 1 | 0 | 0 | 0 | 177 | 149 | 1 | 1 | 1 | 1 | 1 | 1 | 1 | 1 | 0.962 | 1 | 1 | 1 | 0.008 |
| SE | 104576 | ENSG00000214212 | C19orf38 | chr19 | + | 10859914 | 10859958 | 10856264 | 10856357 | 10863169 | 10863207 | 7 | 11 | 10 | 7 | 10 | 0 | 0 | 0 | 0 | 0 | 53 | 24 | 9 | 19 | 20 | 0 | 1 | 0 | 0 | 0 | 193 | 149 | 1 | 1 | 1 | 1 | 1 | 1 | 1 | 1 | 0.949 | 1 | 1 | 1 | 0.01 |
| SE | 104578 | ENSG00000214212 | C19orf38 | chr19 | + | 10863169 | 10863207 | 10859914 | 10859958 | 10869217 | 10869470 | 73 | 59 | 49 | 52 | 44 | 0 | 1 | 0 | 0 | 0 | 241 | 152 | 80 | 156 | 169 | 0 | 0 | 0 | 0 | 0 | 187 | 149 | 1 | 1 | 1 | 0.979 | 1 | 1 | 1 | 1 | 1 | 1 | 1 | 1 | -0.004 |
| SE | 104583 | ENSG00000111796 | KLRB1 | chr12 | - | 9598045 | 9598161 | 9594550 | 9595421 | 9598498 | 9598653 | 1473 | 346 | 864 | 763 | 1517 | 0 | 1 | 0 | 0 | 0 | 1 | 116 | 145 | 45 | 22 | 0 | 0 | 0 | 0 | 0 | 265 | 149 | 1 | 1 | 1 | 0.995 | 1 | 1 | 1 | 1 | 1 | 1 | 1 | 1 | -0.001 |
| SE | 104584 | ENSG00000111796 | KLRB1 | chr12 | - | 9601500 | 9601599 | 9599766 | 9599841 | 9607754 | 9607916 | 5644 | 1838 | 3812 | 3516 | 2958 | 2 | 0 | 0 | 0 | 6 | 90 | 976 | 1073 | 246 | 184 | 0 | 0 | 1 | 0 | 0 | 248 | 149 | 1 | 1 | 0.999 | 1 | 1 | 1 | 0.997 | 1 | 1 | 0.998 | 1 | 1 | 0 |
| SE | 104598 | ENSG00000166527 | CLEC4D | chr12 | + | 8518163 | 8518274 | 8515235 | 8515328 | 8521123 | 8522366 | 50 | 39 | 21 | 32 | 55 | 0 | 1 | 0 | 0 | 0 | 37 | 56 | 59 | 121 | 15 | 0 | 0 | 0 | 0 | 0 | 260 | 149 | 1 | 1 | 1 | 0.957 | 1 | 1 | 1 | 1 | 1 | 1 | 1 | 1 | -0.009 |
| SE | 104599 | ENSG00000166527 | CLEC4D | chr12 | + | 8519008 | 8519160 | 8518163 | 8518274 | 8520225 | 8520273 | 164 | 104 | 85 | 113 | 127 | 0 | 0 | 0 | 0 | 0 | 68 | 209 | 181 | 249 | 38 | 0 | 0 | 1 | 0 | 0 | 298 | 149 | 1 | 1 | 1 | 1 | 1 | 1 | 1 | 1 | 1 | 0.989 | 1 | 1 | 0.002 |
| SE | 104600 | ENSG00000166527 | CLEC4D | chr12 | + | 8520225 | 8520341 | 8518163 | 8518274 | 8521123 | 8522366 | 57 | 38 | 28 | 41 | 45 | 0 | 0 | 0 | 0 | 0 | 45 | 60 | 72 | 58 | 9 | 0 | 0 | 4 | 0 | 0 | 265 | 149 | 1 | 1 | 1 | 1 | 1 | 1 | 1 | 1 | 1 | 0.91 | 1 | 1 | 0.018 |
| SE | 104601 | ENSG00000066336 | SPI1 | chr11 | - | 47359852 | 47360040 | 47358180 | 47359006 | 47375632 | 47375729 | 321 | 239 | 193 | 225 | 276 | 0 | 0 | 1 | 1 | 0 | 1473 | 585 | 499 | 1441 | 1217 | 0 | 0 | 0 | 6 | 0 | 298 | 149 | 1 | 1 | 1 | 1 | 0.99 | 0.991 | 1 | 1 | 1 | 1 | 0.992 | 1 | -0.002 |
| SE | 104602 | ENSG00000066336 | SPI1 | chr11 | - | 47375632 | 47375729 | 47359852 | 47360040 | 47378308 | 47378487 | 201 | 119 | 105 | 107 | 196 | 0 | 0 | 0 | 0 | 0 | 798 | 301 | 242 | 852 | 666 | 0 | 0 | 1 | 0 | 0 | 246 | 149 | 1 | 1 | 1 | 1 | 1 | 1 | 1 | 1 | 1 | 0.993 | 1 | 1 | 0.001 |
| SE | 104603 | ENSG00000066336 | SPI1 | chr11 | - | 47375632 | 47375732 | 47359852 | 47360040 | 47378308 | 47378391 | 145 | 98 | 68 | 83 | 134 | 0 | 0 | 0 | 0 | 0 | 630 | 251 | 191 | 791 | 503 | 0 | 0 | 1 | 0 | 0 | 249 | 149 | 1 | 1 | 1 | 1 | 1 | 1 | 1 | 1 | 1 | 0.991 | 1 | 1 | 0.002 |
| SE | 104609 | ENSG00000172243 | CLEC7A | chr12 | - | 10125296 | 10125448 | 10123244 | 10123363 | 10126570 | 10126708 | 366 | 527 | 270 | 276 | 654 | 0 | 0 | 0 | 0 | 1 | 520 | 361 | 945 | 568 | 359 | 0 | 0 | 0 | 1 | 0 | 298 | 149 | 1 | 1 | 1 | 1 | 1 | 1 | 0.997 | 1 | 1 | 1 | 0.996 | 1 | 0 |
| SE | 104610 | ENSG00000172243 | CLEC7A | chr12 | - | 10125296 | 10125448 | 10123244 | 10123363 | 10127746 | 10127845 | 390 | 502 | 285 | 264 | 601 | 0 | 0 | 0 | 0 | 0 | 656 | 313 | 662 | 496 | 402 | 0 | 0 | 1 | 2 | 0 | 298 | 149 | 1 | 1 | 1 | 1 | 1 | 1 | 1 | 1 | 1 | 0.997 | 0.992 | 1 | 0.002 |
| SE | 104612 | ENSG00000172243 | CLEC7A | chr12 | - | 10126570 | 10126708 | 10123244 | 10123363 | 10127746 | 10127845 | 44 | 34 | 45 | 39 | 96 | 0 | 0 | 0 | 0 | 0 | 96 | 55 | 168 | 74 | 80 | 0 | 0 | 1 | 2 | 0 | 287 | 149 | 1 | 1 | 1 | 1 | 1 | 1 | 1 | 1 | 1 | 0.989 | 0.951 | 1 | 0.012 |
| SE | 104615 | ENSG00000172243 | CLEC7A | chr12 | - | 10126570 | 10127845 | 10125296 | 10125448 | 10129979 | 10130257 | 330 | 348 | 225 | 228 | 373 | 0 | 2 | 0 | 1 | 1 | 248 | 209 | 699 | 429 | 202 | 0 | 0 | 0 | 1 | 0 | 298 | 149 | 1 | 1 | 1 | 0.989 | 1 | 0.991 | 0.995 | 1 | 1 | 1 | 0.995 | 1 | -0.004 |
| SE | 104618 | ENSG00000172243 | CLEC7A | chr12 | - | 10127746 | 10127845 | 10124745 | 10125448 | 10129979 | 10130082 | 294 | 275 | 200 | 183 | 279 | 0 | 2 | 0 | 1 | 1 | 396 | 136 | 359 | 345 | 237 | 0 | 0 | 0 | 1 | 0 | 248 | 149 | 1 | 1 | 1 | 0.988 | 1 | 0.991 | 0.994 | 1 | 1 | 1 | 0.995 | 1 | -0.004 |
| SE | 104619 | ENSG00000172243 | CLEC7A | chr12 | - | 10127746 | 10127845 | 10126570 | 10126708 | 10130202 | 10130258 | 42 | 28 | 40 | 28 | 89 | 0 | 0 | 0 | 0 | 0 | 98 | 44 | 149 | 64 | 78 | 0 | 0 | 1 | 0 | 0 | 248 | 149 | 1 | 1 | 1 | 1 | 1 | 1 | 1 | 1 | 1 | 0.989 | 1 | 1 | 0.002 |
| SE | 104620 | ENSG00000172243 | CLEC7A | chr12 | - | 10129979 | 10130097 | 10127445 | 10127845 | 10130202 | 10130257 | 244 | 213 | 147 | 167 | 217 | 0 | 0 | 0 | 0 | 0 | 237 | 99 | 329 | 258 | 167 | 5 | 0 | 0 | 0 | 0 | 267 | 149 | 1 | 1 | 1 | 1 | 1 | 1 | 1 | 0.964 | 1 | 1 | 1 | 1 | 0.007 |
| SE | 104623 | ENSG00000172322 | CLEC12A | chr12 | + | 9980581 | 9980733 | 9978965 | 9979524 | 9982019 | 9982129 | 1203 | 1027 | 699 | 1359 | 1872 | 0 | 1 | 0 | 0 | 0 | 918 | 354 | 756 | 1419 | 247 | 0 | 0 | 0 | 3 | 0 | 298 | 149 | 1 | 1 | 1 | 0.998 | 1 | 1 | 1 | 1 | 1 | 1 | 0.996 | 1 | 0 |
| SE | 104636 | ENSG00000084444 | FAM234B | chr12 | + | 13067154 | 13067296 | 13066639 | 13066787 | 13068303 | 13068447 | 23 | 23 | 25 | 10 | 49 | 0 | 0 | 0 | 1 | 0 | 45 | 18 | 21 | 14 | 67 | 0 | 0 | 0 | 0 | 0 | 291 | 149 | 1 | 1 | 1 | 1 | 1 | 0.837 | 1 | 1 | 1 | 1 | 1 | 1 | -0.033 |
| SE | 104641 | ENSG00000079974 | RABL2B | chr22 | - | 50769904 | 50770016 | 50769424 | 50769549 | 50775771 | 50775851 | 42 | 14 | 26 | 34 | 62 | 1 | 0 | 0 | 0 | 0 | 6 | 22 | 24 | 24 | 49 | 0 | 0 | 0 | 0 | 0 | 261 | 149 | 1 | 1 | 0.96 | 1 | 1 | 1 | 1 | 1 | 1 | 1 | 1 | 1 | -0.008 |
| SE | 104645 | ENSG00000079974 | RABL2B | chr22 | - | 50782890 | 50783045 | 50782187 | 50782351 | 50783500 | 50783642 | 1 | 0 | 0 | 0 | 0 | 0 | 0 | 0 | 0 | 0 | 0 | 0 | 3 | 0 | 0 | 0 | 0 | 2 | 0 | 0 | 298 | 149 | 1 | 1 | 1 | NA | NA | NA | NA | NA | NA | 0.429 | NA | NA | 0.571 |
| SE | 104646 | ENSG00000079974 | RABL2B | chr22 | - | 50783038 | 50783286 | 50782187 | 50782351 | 50783500 | 50783636 | 3 | 4 | 6 | 8 | 12 | 0 | 0 | 0 | 0 | 0 | 20 | 13 | 4 | 9 | 15 | 0 | 0 | 2 | 0 | 0 | 298 | 149 | 1 | 1 | 1 | 1 | 1 | 1 | 1 | 1 | 1 | 0.5 | 1 | 1 | 0.1 |
| SE | 104649 | ENSG00000110925 | CSRNP2 | chr12 | - | 51081427 | 51081496 | 51076568 | 51076647 | 51082520 | 51082694 | 0 | 0 | 0 | 1 | 0 | 0 | 2 | 4 | 0 | 0 | 0 | 0 | 0 | 0 | 0 | 0 | 0 | 0 | 5 | 0 | 218 | 149 | 1 | 1 | NA | 0 | 0 | 1 | NA | NA | NA | NA | 0 | NA | 0.333 |
| SE | 104650 | ENSG00000110925 | CSRNP2 | chr12 | - | 51082520 | 51082694 | 51076410 | 51076647 | 51083338 | 51083583 | 0 | 2 | 5 | 2 | 0 | 0 | 0 | 1 | 1 | 0 | 0 | 0 | 0 | 5 | 0 | 0 | 0 | 0 | 0 | 0 | 298 | 149 | 1 | 1 | NA | 1 | 0.714 | 0.5 | NA | NA | NA | NA | 1 | NA | -0.262 |
| SE | 104651 | ENSG00000060982 | BCAT1 | chr12 | - | 24878529 | 24878649 | 24849785 | 24849949 | 24881300 | 24881398 | 44 | 40 | 31 | 35 | 75 | 0 | 0 | 0 | 1 | 0 | 12 | 158 | 57 | 22 | 23 | 0 | 0 | 0 | 0 | 0 | 269 | 149 | 1 | 1 | 1 | 1 | 1 | 0.951 | 1 | 1 | 1 | 1 | 1 | 1 | -0.01 |
| SE | 104652 | ENSG00000060982 | BCAT1 | chr12 | - | 24881300 | 24881411 | 24878529 | 24878649 | 24894274 | 24894475 | 33 | 28 | 26 | 30 | 58 | 0 | 0 | 1 | 0 | 0 | 12 | 161 | 55 | 18 | 18 | 0 | 3 | 1 | 0 | 0 | 260 | 149 | 1 | 1 | 1 | 1 | 0.937 | 1 | 1 | 1 | 0.969 | 0.969 | 1 | 1 | 0 |
| SE | 104654 | ENSG00000144579 | CTDSP1 | chr2 | + | 218399376 | 218399474 | 218398255 | 218398332 | 218401563 | 218401712 | 0 | 0 | 0 | 0 | 0 | 0 | 0 | 0 | 1 | 0 | 9 | 0 | 0 | 0 | 0 | 0 | 0 | 0 | 0 | 0 | 247 | 149 | 1 | 1 | NA | NA | NA | 0 | NA | 1 | NA | NA | NA | NA | -1 |
| SE | 104655 | ENSG00000144579 | CTDSP1 | chr2 | + | 218402110 | 218402215 | 218401563 | 218401712 | 218403034 | 218403127 | 47 | 16 | 30 | 33 | 2 | 0 | 0 | 0 | 0 | 0 | 32 | 37 | 22 | 7 | 59 | 0 | 0 | 0 | 1 | 0 | 254 | 149 | 1 | 1 | 1 | 1 | 1 | 1 | 1 | 1 | 1 | 1 | 0.804 | 1 | 0.039 |
| SE | 104656 | ENSG00000144579 | CTDSP1 | chr2 | + | 218402113 | 218402215 | 218401166 | 218401712 | 218403034 | 218403127 | 80 | 32 | 55 | 58 | 9 | 0 | 0 | 0 | 0 | 0 | 58 | 64 | 22 | 28 | 115 | 0 | 0 | 0 | 1 | 0 | 251 | 149 | 1 | 1 | 1 | 1 | 1 | 1 | 1 | 1 | 1 | 1 | 0.943 | 1 | 0.011 |
| SE | 104657 | ENSG00000144579 | CTDSP1 | chr2 | + | 218402348 | 218402405 | 218402110 | 218402215 | 218403034 | 218403127 | 412 | 150 | 308 | 291 | 150 | 0 | 0 | 3 | 0 | 0 | 201 | 311 | 145 | 283 | 506 | 0 | 0 | 5 | 0 | 0 | 206 | 149 | 1 | 1 | 1 | 1 | 0.987 | 1 | 1 | 1 | 1 | 0.954 | 1 | 1 | 0.007 |
| SE | 104658 | ENSG00000144579 | CTDSP1 | chr2 | + | 218402348 | 218402405 | 218402110 | 218402215 | 218403231 | 218403400 | 72 | 37 | 50 | 43 | 12 | 0 | 0 | 0 | 0 | 0 | 44 | 99 | 32 | 44 | 112 | 0 | 2 | 0 | 0 | 0 | 206 | 149 | 1 | 1 | 1 | 1 | 1 | 1 | 1 | 1 | 0.973 | 1 | 1 | 1 | 0.005 |
| SE | 104659 | ENSG00000144579 | CTDSP1 | chr2 | + | 218403010 | 218403127 | 218402348 | 218402405 | 218403231 | 218403417 | 218 | 97 | 154 | 167 | 72 | 0 | 0 | 0 | 0 | 0 | 153 | 169 | 69 | 133 | 333 | 0 | 3 | 0 | 0 | 0 | 266 | 149 | 1 | 1 | 1 | 1 | 1 | 1 | 1 | 1 | 0.969 | 1 | 1 | 1 | 0.006 |
| SE | 104660 | ENSG00000144579 | CTDSP1 | chr2 | + | 218403034 | 218403127 | 218402110 | 218402215 | 218403231 | 218403400 | 217 | 93 | 157 | 163 | 72 | 0 | 0 | 0 | 0 | 0 | 153 | 169 | 72 | 133 | 333 | 0 | 2 | 0 | 0 | 0 | 242 | 149 | 1 | 1 | 1 | 1 | 1 | 1 | 1 | 1 | 0.981 | 1 | 1 | 1 | 0.004 |
| SE | 104661 | ENSG00000144579 | CTDSP1 | chr2 | + | 218403034 | 218403127 | 218402348 | 218402405 | 218403231 | 218403400 | 556 | 206 | 412 | 411 | 210 | 0 | 0 | 0 | 0 | 0 | 310 | 381 | 180 | 372 | 727 | 0 | 3 | 0 | 0 | 0 | 242 | 149 | 1 | 1 | 1 | 1 | 1 | 1 | 1 | 1 | 0.987 | 1 | 1 | 1 | 0.003 |
| SE | 104662 | ENSG00000100354 | TNRC6B | chr22 | + | 40117054 | 40117128 | 40044816 | 40044998 | 40156114 | 40156182 | 65 | 34 | 76 | 58 | 23 | 2 | 0 | 0 | 0 | 0 | 78 | 53 | 122 | 92 | 37 | 0 | 0 | 0 | 0 | 0 | 223 | 149 | 1 | 1 | 0.956 | 1 | 1 | 1 | 1 | 1 | 1 | 1 | 1 | 1 | -0.009 |
| SE | 104664 | ENSG00000100354 | TNRC6B | chr22 | + | 40156114 | 40156182 | 40125771 | 40125862 | 40246014 | 40246102 | 118 | 91 | 122 | 143 | 59 | 0 | 0 | 0 | 0 | 0 | 140 | 104 | 193 | 142 | 70 | 0 | 0 | 1 | 0 | 0 | 217 | 149 | 1 | 1 | 1 | 1 | 1 | 1 | 1 | 1 | 1 | 0.993 | 1 | 1 | 0.001 |
| SE | 104665 | ENSG00000100354 | TNRC6B | chr22 | + | 40246014 | 40246102 | 40156114 | 40156182 | 40251178 | 40251200 | 112 | 75 | 96 | 122 | 95 | 0 | 4 | 0 | 0 | 0 | 156 | 79 | 161 | 108 | 46 | 0 | 0 | 0 | 0 | 0 | 237 | 149 | 1 | 1 | 1 | 0.922 | 1 | 1 | 1 | 1 | 1 | 1 | 1 | 1 | -0.016 |
| SE | 104666 | ENSG00000100354 | TNRC6B | chr22 | + | 40246014 | 40246102 | 40156114 | 40156182 | 40261831 | 40262173 | 70 | 59 | 66 | 77 | 49 | 0 | 1 | 0 | 3 | 0 | 120 | 51 | 116 | 84 | 20 | 0 | 0 | 0 | 0 | 0 | 237 | 149 | 1 | 1 | 1 | 0.974 | 1 | 0.942 | 1 | 1 | 1 | 1 | 1 | 1 | -0.017 |
| SE | 104667 | ENSG00000100354 | TNRC6B | chr22 | + | 40251178 | 40251200 | 40156114 | 40156182 | 40261831 | 40262173 | 21 | 8 | 12 | 17 | 16 | 0 | 1 | 0 | 3 | 0 | 0 | 17 | 10 | 32 | 17 | 0 | 0 | 0 | 0 | 0 | 171 | 149 | 1 | 1 | 1 | 0.875 | 1 | 0.832 | 1 | NA | 1 | 1 | 1 | 1 | -0.059 |
| SE | 104668 | ENSG00000100354 | TNRC6B | chr22 | + | 40251178 | 40251200 | 40246014 | 40246102 | 40261831 | 40262173 | 102 | 32 | 54 | 91 | 83 | 0 | 0 | 1 | 0 | 0 | 55 | 51 | 58 | 68 | 47 | 0 | 0 | 0 | 0 | 0 | 171 | 149 | 1 | 1 | 1 | 1 | 0.979 | 1 | 1 | 1 | 1 | 1 | 1 | 1 | -0.004 |
| SE | 104669 | ENSG00000100354 | TNRC6B | chr22 | + | 40261831 | 40262173 | 40251178 | 40251200 | 40264687 | 40267036 | 173 | 116 | 172 | 215 | 139 | 2 | 2 | 0 | 0 | 0 | 154 | 171 | 201 | 150 | 68 | 0 | 0 | 0 | 0 | 0 | 298 | 149 | 1 | 1 | 0.977 | 0.967 | 1 | 1 | 1 | 1 | 1 | 1 | 1 | 1 | -0.011 |
| SE | 104670 | ENSG00000100354 | TNRC6B | chr22 | + | 40261831 | 40262173 | 40251178 | 40251200 | 40270121 | 40270280 | 81 | 30 | 51 | 71 | 74 | 0 | 0 | 0 | 0 | 0 | 49 | 48 | 58 | 64 | 49 | 4 | 0 | 0 | 0 | 0 | 298 | 149 | 1 | 1 | 1 | 1 | 1 | 1 | 1 | 0.86 | 1 | 1 | 1 | 1 | 0.028 |
| SE | 104671 | ENSG00000100354 | TNRC6B | chr22 | + | 40261831 | 40262173 | 40251178 | 40251200 | 40273424 | 40273600 | 81 | 30 | 51 | 71 | 74 | 0 | 0 | 0 | 0 | 0 | 49 | 51 | 58 | 65 | 49 | 0 | 0 | 1 | 0 | 0 | 298 | 149 | 1 | 1 | 1 | 1 | 1 | 1 | 1 | 1 | 1 | 0.967 | 1 | 1 | 0.007 |
| SE | 104673 | ENSG00000100354 | TNRC6B | chr22 | + | 40264687 | 40267036 | 40251178 | 40251200 | 40273424 | 40273600 | 124 | 115 | 110 | 150 | 131 | 0 | 0 | 0 | 0 | 0 | 111 | 147 | 140 | 153 | 50 | 0 | 0 | 1 | 0 | 0 | 298 | 149 | 1 | 1 | 1 | 1 | 1 | 1 | 1 | 1 | 1 | 0.986 | 1 | 1 | 0.003 |
| SE | 104674 | ENSG00000100354 | TNRC6B | chr22 | + | 40264687 | 40267036 | 40251178 | 40251200 | 40277076 | 40277151 | 5 | 2 | 0 | 0 | 4 | 1 | 0 | 0 | 0 | 0 | 0 | 0 | 0 | 0 | 3 | 0 | 0 | 0 | 0 | 0 | 298 | 149 | 1 | 1 | 0.714 | 1 | NA | NA | 1 | NA | NA | NA | NA | 1 | -0.095 |
| SE | 104676 | ENSG00000100354 | TNRC6B | chr22 | + | 40264687 | 40267036 | 40261831 | 40262173 | 40273424 | 40273600 | 214 | 199 | 231 | 294 | 196 | 0 | 0 | 0 | 0 | 0 | 216 | 270 | 283 | 239 | 69 | 0 | 3 | 0 | 1 | 0 | 298 | 149 | 1 | 1 | 1 | 1 | 1 | 1 | 1 | 1 | 0.978 | 1 | 0.992 | 1 | 0.006 |
| SE | 104677 | ENSG00000100354 | TNRC6B | chr22 | + | 40270121 | 40270280 | 40251178 | 40251200 | 40273424 | 40273600 | 30 | 31 | 65 | 54 | 37 | 0 | 0 | 0 | 0 | 0 | 48 | 66 | 67 | 54 | 46 | 0 | 0 | 1 | 0 | 0 | 298 | 149 | 1 | 1 | 1 | 1 | 1 | 1 | 1 | 1 | 1 | 0.971 | 1 | 1 | 0.006 |
| SE | 104678 | ENSG00000100354 | TNRC6B | chr22 | + | 40270121 | 40270280 | 40261831 | 40262173 | 40273424 | 40273600 | 30 | 31 | 65 | 54 | 37 | 0 | 0 | 0 | 0 | 0 | 44 | 66 | 67 | 54 | 46 | 0 | 3 | 0 | 1 | 0 | 298 | 149 | 1 | 1 | 1 | 1 | 1 | 1 | 1 | 1 | 0.917 | 1 | 0.964 | 1 | 0.024 |
| SE | 104680 | ENSG00000100354 | TNRC6B | chr22 | + | 40273424 | 40273600 | 40251178 | 40251200 | 40277076 | 40277151 | 161 | 86 | 144 | 168 | 149 | 1 | 0 | 0 | 0 | 0 | 86 | 181 | 157 | 126 | 78 | 0 | 0 | 0 | 0 | 0 | 298 | 149 | 1 | 1 | 0.988 | 1 | 1 | 1 | 1 | 1 | 1 | 1 | 1 | 1 | -0.002 |
| SE | 104681 | ENSG00000100354 | TNRC6B | chr22 | + | 40273424 | 40273600 | 40264687 | 40267036 | 40277076 | 40277151 | 283 | 199 | 254 | 318 | 280 | 3 | 0 | 0 | 0 | 4 | 197 | 328 | 296 | 279 | 128 | 0 | 0 | 0 | 0 | 3 | 298 | 149 | 1 | 1 | 0.979 | 1 | 1 | 1 | 0.972 | 1 | 1 | 1 | 1 | 0.955 | -0.001 |
| SE | 104682 | ENSG00000100354 | TNRC6B | chr22 | + | 40277076 | 40277151 | 40273424 | 40273600 | 40277998 | 40278044 | 372 | 222 | 348 | 373 | 393 | 0 | 0 | 0 | 0 | 0 | 214 | 438 | 333 | 361 | 218 | 0 | 0 | 0 | 7 | 0 | 224 | 149 | 1 | 1 | 1 | 1 | 1 | 1 | 1 | 1 | 1 | 1 | 0.972 | 1 | 0.006 |
| SE | 104683 | ENSG00000100354 | TNRC6B | chr22 | + | 40277998 | 40278044 | 40273424 | 40273600 | 40279994 | 40280143 | 63 | 43 | 63 | 68 | 105 | 0 | 0 | 1 | 0 | 0 | 15 | 75 | 48 | 48 | 35 | 0 | 0 | 0 | 0 | 0 | 195 | 149 | 1 | 1 | 1 | 1 | 0.98 | 1 | 1 | 1 | 1 | 1 | 1 | 1 | -0.004 |
| SE | 104686 | ENSG00000100354 | TNRC6B | chr22 | + | 40308511 | 40308649 | 40301149 | 40301333 | 40310816 | 40310993 | 123 | 68 | 148 | 188 | 158 | 0 | 0 | 1 | 0 | 0 | 45 | 154 | 114 | 137 | 42 | 0 | 0 | 0 | 0 | 0 | 287 | 149 | 1 | 1 | 1 | 1 | 0.987 | 1 | 1 | 1 | 1 | 1 | 1 | 1 | -0.003 |
| SE | 104691 | ENSG00000211455 | STK38L | chr12 | + | 27297029 | 27297158 | 27244292 | 27244332 | 27297709 | 27297854 | 0 | 1 | 0 | 6 | 2 | 13 | 9 | 8 | 11 | 11 | 4 | 4 | 0 | 0 | 0 | 0 | 0 | 10 | 0 | 2 | 278 | 149 | 1 | 1 | 0 | 0.056 | 0 | 0.226 | 0.089 | 1 | 1 | 0 | NA | 0 | -0.426 |
| SE | 104692 | ENSG00000211455 | STK38L | chr12 | + | 27300553 | 27300601 | 27297709 | 27297854 | 27302136 | 27302188 | 0 | 2 | 0 | 5 | 0 | 34 | 24 | 21 | 26 | 21 | 0 | 4 | 7 | 0 | 0 | 0 | 13 | 22 | 1 | 0 | 197 | 149 | 1 | 1 | 0 | 0.059 | 0 | 0.127 | 0 | NA | 0.189 | 0.194 | 0 | NA | -0.09 |
| SE | 104693 | ENSG00000211455 | STK38L | chr12 | + | 27308338 | 27308461 | 27302135 | 27302188 | 27309113 | 27309197 | 92 | 126 | 72 | 96 | 137 | 2 | 9 | 2 | 2 | 1 | 22 | 44 | 79 | 27 | 0 | 0 | 0 | 0 | 0 | 0 | 272 | 149 | 1 | 1 | 0.962 | 0.885 | 0.952 | 0.963 | 0.987 | 1 | 1 | 1 | 1 | NA | -0.05 |
| SE | 104697 | ENSG00000211455 | STK38L | chr12 | + | 27319327 | 27319423 | 27317581 | 27318019 | 27322142 | 27322234 | 185 | 143 | 100 | 143 | 322 | 0 | 0 | 0 | 2 | 0 | 19 | 53 | 54 | 35 | 0 | 0 | 0 | 0 | 0 | 0 | 245 | 149 | 1 | 1 | 1 | 1 | 1 | 0.978 | 1 | 1 | 1 | 1 | 1 | NA | -0.004 |
| SE | 104698 | ENSG00000211455 | STK38L | chr12 | + | 27322142 | 27322234 | 27319327 | 27319423 | 27322327 | 27322683 | 170 | 128 | 95 | 144 | 266 | 1 | 4 | 1 | 4 | 4 | 8 | 54 | 56 | 26 | 0 | 0 | 0 | 0 | 0 | 0 | 241 | 149 | 1 | 1 | 0.991 | 0.952 | 0.983 | 0.957 | 0.976 | 1 | 1 | 1 | 1 | NA | -0.028 |
| SE | 104699 | ENSG00000110888 | CAPRIN2 | chr12 | - | 30711738 | 30711782 | 30711565 | 30711629 | 30713784 | 30713885 | 2 | 1 | 2 | 2 | 5 | 30 | 22 | 24 | 26 | 23 | 7 | 6 | 5 | 0 | 0 | 0 | 18 | 10 | 14 | 0 | 193 | 149 | 1 | 1 | 0.049 | 0.034 | 0.06 | 0.056 | 0.144 | 1 | 0.205 | 0.279 | 0 | NA | -0.302 |
| SE | 104700 | ENSG00000110888 | CAPRIN2 | chr12 | - | 30719078 | 30719225 | 30716289 | 30716676 | 30720810 | 30720915 | 64 | 66 | 65 | 65 | 117 | 1 | 0 | 1 | 1 | 6 | 12 | 54 | 23 | 11 | 0 | 0 | 14 | 2 | 0 | 0 | 296 | 149 | 1 | 1 | 0.97 | 1 | 0.97 | 0.97 | 0.908 | 1 | 0.66 | 0.853 | 1 | NA | 0.085 |
| SE | 104701 | ENSG00000110888 | CAPRIN2 | chr12 | - | 30719078 | 30719225 | 30716289 | 30716676 | 30723258 | 30723314 | 45 | 37 | 47 | 44 | 66 | 10 | 9 | 6 | 3 | 30 | 31 | 15 | 12 | 21 | 0 | 6 | 7 | 1 | 12 | 0 | 296 | 149 | 1 | 1 | 0.694 | 0.674 | 0.798 | 0.881 | 0.525 | 0.722 | 0.519 | 0.858 | 0.468 | NA | 0.073 |
| SE | 104702 | ENSG00000110888 | CAPRIN2 | chr12 | - | 30719078 | 30720915 | 30714958 | 30716676 | 30723258 | 30723314 | 43 | 35 | 45 | 58 | 71 | 10 | 9 | 6 | 3 | 30 | 12 | 14 | 15 | 11 | 0 | 6 | 7 | 1 | 12 | 0 | 298 | 149 | 1 | 1 | 0.683 | 0.66 | 0.789 | 0.906 | 0.542 | 0.5 | 0.5 | 0.882 | 0.314 | NA | 0.167 |
| SE | 104703 | ENSG00000110888 | CAPRIN2 | chr12 | - | 30720810 | 30720915 | 30716507 | 30716676 | 30723258 | 30723314 | 11 | 8 | 10 | 26 | 23 | 10 | 9 | 6 | 3 | 30 | 0 | 16 | 9 | 1 | 0 | 6 | 7 | 1 | 12 | 0 | 254 | 149 | 1 | 1 | 0.392 | 0.343 | 0.494 | 0.836 | 0.31 | 0 | 0.573 | 0.841 | 0.047 | NA | 0.11 |
| SE | 104704 | ENSG00000110888 | CAPRIN2 | chr12 | - | 30720810 | 30720915 | 30719078 | 30719225 | 30723258 | 30723314 | 39 | 47 | 38 | 56 | 82 | 12 | 10 | 11 | 11 | 14 | 0 | 44 | 23 | 2 | 0 | 19 | 3 | 5 | 11 | 0 | 254 | 149 | 1 | 1 | 0.656 | 0.734 | 0.67 | 0.749 | 0.775 | 0 | 0.896 | 0.73 | 0.096 | NA | 0.286 |
| SE | 104705 | ENSG00000110888 | CAPRIN2 | chr12 | - | 30725965 | 30726088 | 30724369 | 30724451 | 30728647 | 30728778 | 70 | 83 | 68 | 80 | 220 | 0 | 0 | 0 | 0 | 4 | 24 | 33 | 39 | 34 | 0 | 0 | 0 | 0 | 0 | 0 | 272 | 149 | 1 | 1 | 1 | 1 | 1 | 1 | 0.968 | 1 | 1 | 1 | 1 | NA | -0.006 |
| SE | 104706 | ENSG00000110888 | CAPRIN2 | chr12 | - | 30730238 | 30730282 | 30728378 | 30729325 | 30731342 | 30731448 | 97 | 72 | 80 | 121 | 172 | 1 | 0 | 0 | 0 | 0 | 34 | 90 | 46 | 37 | 39 | 0 | 0 | 0 | 0 | 0 | 193 | 149 | 1 | 1 | 0.987 | 1 | 1 | 1 | 1 | 1 | 1 | 1 | 1 | 1 | -0.003 |
| SE | 104709 | ENSG00000110888 | CAPRIN2 | chr12 | - | 30741015 | 30741106 | 30734967 | 30735206 | 30751070 | 30751133 | 34 | 29 | 34 | 38 | 39 | 1 | 0 | 0 | 0 | 0 | 48 | 44 | 26 | 10 | 20 | 0 | 0 | 0 | 0 | 0 | 240 | 149 | 1 | 1 | 0.955 | 1 | 1 | 1 | 1 | 1 | 1 | 1 | 1 | 1 | -0.009 |
| SE | 104710 | ENSG00000110888 | CAPRIN2 | chr12 | - | 30741019 | 30741106 | 30734954 | 30735206 | 30751070 | 30751124 | 188 | 156 | 144 | 161 | 119 | 1 | 0 | 0 | 0 | 0 | 146 | 175 | 109 | 81 | 119 | 0 | 0 | 0 | 0 | 0 | 236 | 149 | 1 | 1 | 0.992 | 1 | 1 | 1 | 1 | 1 | 1 | 1 | 1 | 1 | -0.002 |
| SE | 104712 | ENSG00000110888 | CAPRIN2 | chr12 | - | 30753343 | 30753559 | 30751070 | 30751133 | 30754688 | 30754860 | 35 | 31 | 33 | 48 | 54 | 1 | 0 | 0 | 1 | 0 | 57 | 50 | 32 | 13 | 24 | 0 | 0 | 0 | 0 | 0 | 298 | 149 | 1 | 1 | 0.946 | 1 | 1 | 0.96 | 1 | 1 | 1 | 1 | 1 | 1 | -0.019 |
| SE | 104713 | ENSG00000110888 | CAPRIN2 | chr12 | - | 30753343 | 30753559 | 30751070 | 30751133 | 30754750 | 30754888 | 36 | 30 | 37 | 52 | 54 | 0 | 0 | 2 | 0 | 0 | 57 | 50 | 32 | 13 | 24 | 0 | 0 | 0 | 0 | 1 | 298 | 149 | 1 | 1 | 1 | 1 | 0.902 | 1 | 1 | 1 | 1 | 1 | 1 | 0.923 | -0.004 |
| SE | 104717 | ENSG00000005189 | REXO5 | chr16 | + | 20806951 | 20807091 | 20806495 | 20806705 | 20813189 | 20813302 | 7 | 1 | 2 | 1 | 2 | 0 | 0 | 1 | 0 | 0 | 16 | 13 | 2 | 0 | 0 | 0 | 0 | 0 | 0 | 0 | 289 | 149 | 1 | 1 | 1 | 1 | 0.508 | 1 | 1 | 1 | 1 | 1 | NA | NA | -0.098 |
| SE | 104719 | ENSG00000005189 | REXO5 | chr16 | + | 20813189 | 20813302 | 20806551 | 20806721 | 20821761 | 20821902 | 0 | 0 | 0 | 0 | 0 | 1 | 0 | 0 | 0 | 0 | 0 | 1 | 0 | 0 | 0 | 0 | 0 | 0 | 0 | 0 | 262 | 149 | 1 | 1 | 0 | NA | NA | NA | NA | NA | 1 | NA | NA | NA | -1 |
| SE | 104721 | ENSG00000005189 | REXO5 | chr16 | + | 20824438 | 20824527 | 20821761 | 20821902 | 20825832 | 20825871 | 2 | 1 | 3 | 1 | 5 | 1 | 0 | 0 | 0 | 2 | 33 | 0 | 1 | 0 | 6 | 0 | 0 | 0 | 3 | 0 | 238 | 149 | 1 | 1 | 0.556 | 1 | 1 | 1 | 0.61 | 1 | NA | 1 | 0 | 1 | 0.083 |
| SE | 104722 | ENSG00000005189 | REXO5 | chr16 | + | 20828434 | 20828537 | 20827352 | 20827447 | 20832155 | 20832199 | 6 | 2 | 1 | 6 | 7 | 0 | 0 | 0 | 0 | 0 | 11 | 13 | 2 | 14 | 13 | 0 | 0 | 2 | 0 | 0 | 252 | 149 | 1 | 1 | 1 | 1 | 1 | 1 | 1 | 1 | 1 | 0.372 | 1 | 1 | 0.126 |
| SE | 104723 | ENSG00000005189 | REXO5 | chr16 | + | 20839754 | 20839859 | 20833002 | 20833123 | 20840330 | 20840468 | 2 | 5 | 0 | 6 | 4 | 1 | 0 | 0 | 0 | 0 | 28 | 4 | 1 | 4 | 6 | 0 | 0 | 0 | 6 | 0 | 254 | 149 | 1 | 1 | 0.54 | 1 | NA | 1 | 1 | 1 | 1 | 1 | 0.281 | 1 | 0.029 |
| SE | 104724 | ENSG00000005189 | REXO5 | chr16 | + | 20843933 | 20844026 | 20839852 | 20840468 | 20844628 | 20844662 | 6 | 2 | 1 | 3 | 0 | 0 | 0 | 0 | 4 | 0 | 9 | 0 | 0 | 4 | 6 | 0 | 1 | 1 | 5 | 0 | 242 | 149 | 1 | 1 | 1 | 1 | 1 | 0.316 | NA | 1 | 0 | 0 | 0.33 | 1 | 0.363 |
| SE | 104728 | ENSG00000226210 | WASH8P | chr12 | - | 15084 | 15153 | 14521 | 14944 | 15912 | 16065 | 19 | 44 | 66 | 41 | 1 | 0 | 0 | 0 | 0 | 0 | 19 | 42 | 12 | 47 | 88 | 0 | 0 | 0 | 4 | 0 | 218 | 149 | 1 | 1 | 1 | 1 | 1 | 1 | 1 | 1 | 1 | 1 | 0.889 | 1 | 0.022 |
| SE | 104729 | ENSG00000226210 | WASH8P | chr12 | - | 17347 | 17483 | 16968 | 17170 | 17722 | 17859 | 30 | 63 | 118 | 97 | 47 | 0 | 0 | 0 | 0 | 2 | 85 | 61 | 28 | 31 | 88 | 0 | 2 | 0 | 2 | 0 | 285 | 149 | 1 | 1 | 1 | 1 | 1 | 1 | 0.925 | 1 | 0.941 | 1 | 0.89 | 1 | 0.019 |
| SE | 104733 | ENSG00000226210 | WASH8P | chr12 | - | 18372 | 18471 | 18036 | 18183 | 26800 | 26954 | 30 | 51 | 107 | 62 | 14 | 1 | 0 | 4 | 0 | 0 | 60 | 60 | 23 | 26 | 103 | 0 | 2 | 2 | 0 | 0 | 248 | 149 | 1 | 1 | 0.947 | 1 | 0.941 | 1 | 1 | 1 | 0.947 | 0.874 | 1 | 1 | 0.013 |
| SE | 104734 | ENSG00000122694 | GLIPR2 | chr9 | + | 36146111 | 36146215 | 36144275 | 36145041 | 36147785 | 36147894 | 2 | 0 | 1 | 1 | 1 | 2 | 2 | 0 | 9 | 1 | 0 | 0 | 0 | 0 | 0 | 16 | 0 | 12 | 4 | 15 | 253 | 149 | 1 | 1 | 0.371 | 0 | 1 | 0.061 | 0.371 | 0 | NA | 0 | 0 | 0 | 0.361 |
| SE | 104735 | ENSG00000122694 | GLIPR2 | chr9 | + | 36147785 | 36147894 | 36136535 | 36136791 | 36148546 | 36148650 | 381 | 227 | 240 | 328 | 205 | 0 | 0 | 0 | 0 | 0 | 467 | 441 | 206 | 437 | 786 | 7 | 0 | 0 | 0 | 0 | 258 | 149 | 1 | 1 | 1 | 1 | 1 | 1 | 1 | 0.975 | 1 | 1 | 1 | 1 | 0.005 |
| SE | 104736 | ENSG00000122694 | GLIPR2 | chr9 | + | 36147785 | 36147894 | 36136535 | 36136791 | 36162361 | 36163913 | 31 | 15 | 17 | 21 | 8 | 0 | 1 | 1 | 0 | 1 | 45 | 41 | 19 | 33 | 24 | 4 | 0 | 0 | 0 | 0 | 258 | 149 | 1 | 1 | 1 | 0.897 | 0.908 | 1 | 0.822 | 0.867 | 1 | 1 | 1 | 1 | -0.048 |
| SE | 104738 | ENSG00000122694 | GLIPR2 | chr9 | + | 36148546 | 36148650 | 36136732 | 36136791 | 36162361 | 36162910 | 3 | 0 | 0 | 1 | 0 | 0 | 1 | 1 | 0 | 1 | 9 | 10 | 3 | 1 | 0 | 4 | 0 | 0 | 0 | 0 | 253 | 149 | 1 | 1 | 1 | 0 | 0 | 1 | 0 | 0.57 | 1 | 1 | 1 | NA | -0.492 |
| SE | 104743 | ENSG00000136783 | NIPSNAP3A | chr9 | + | 104750955 | 104751166 | 104747682 | 104747852 | 104752905 | 104753064 | 300 | 163 | 142 | 204 | 322 | 0 | 0 | 0 | 0 | 0 | 155 | 137 | 112 | 137 | 175 | 0 | 0 | 1 | 0 | 0 | 298 | 149 | 1 | 1 | 1 | 1 | 1 | 1 | 1 | 1 | 1 | 0.982 | 1 | 1 | 0.004 |
| SE | 104746 | ENSG00000164845 | FAM86FP | chr12 | - | 8235252 | 8235518 | 8234341 | 8234425 | 8235600 | 8235734 | 5 | 1 | 0 | 0 | 0 | 0 | 5 | 2 | 0 | 0 | 0 | 5 | 0 | 0 | 0 | 0 | 8 | 0 | 0 | 0 | 298 | 149 | 1 | 1 | 1 | 0.091 | 0 | NA | NA | NA | 0.238 | NA | NA | NA | 0.126 |
| SE | 104747 | ENSG00000164845 | FAM86FP | chr12 | - | 8235600 | 8235734 | 8234341 | 8234425 | 8238666 | 8238749 | 4 | 6 | 2 | 2 | 0 | 0 | 0 | 0 | 2 | 0 | 0 | 11 | 0 | 0 | 0 | 0 | 0 | 0 | 0 | 0 | 283 | 149 | 1 | 1 | 1 | 1 | 1 | 0.345 | NA | NA | 1 | NA | NA | NA | -0.164 |
| SE | 104748 | ENSG00000164845 | FAM86FP | chr12 | - | 8236970 | 8237072 | 8235600 | 8235734 | 8238666 | 8238749 | 1 | 0 | 0 | 0 | 1 | 4 | 1 | 0 | 2 | 0 | 0 | 0 | 0 | 0 | 0 | 0 | 3 | 0 | 0 | 0 | 251 | 149 | 1 | 1 | 0.129 | 0 | NA | 0 | 1 | NA | 0 | NA | NA | NA | 0.282 |
| SE | 104755 | ENSG00000011258 | MBTD1 | chr17 | - | 51193427 | 51193510 | 51191583 | 51193016 | 51201591 | 51201696 | 26 | 43 | 31 | 53 | 79 | 0 | 0 | 0 | 1 | 0 | 18 | 59 | 74 | 31 | 8 | 0 | 0 | 1 | 0 | 0 | 232 | 149 | 1 | 1 | 1 | 1 | 1 | 0.971 | 1 | 1 | 1 | 0.979 | 1 | 1 | -0.002 |
| SE | 104756 | ENSG00000011258 | MBTD1 | chr17 | - | 51195213 | 51195361 | 51191583 | 51193016 | 51201591 | 51201696 | 55 | 47 | 60 | 76 | 115 | 0 | 0 | 0 | 1 | 0 | 8 | 61 | 59 | 43 | 15 | 0 | 0 | 1 | 0 | 0 | 297 | 149 | 1 | 1 | 1 | 1 | 1 | 0.974 | 1 | 1 | 1 | 0.967 | 1 | 1 | 0.001 |
| SE | 104759 | ENSG00000011258 | MBTD1 | chr17 | - | 51202700 | 51202935 | 51201591 | 51201696 | 51203139 | 51203228 | 96 | 73 | 59 | 101 | 146 | 4 | 0 | 1 | 0 | 0 | 38 | 114 | 93 | 66 | 27 | 0 | 0 | 0 | 0 | 0 | 298 | 149 | 1 | 1 | 0.923 | 1 | 0.967 | 1 | 1 | 1 | 1 | 1 | 1 | 1 | -0.022 |
| SE | 104760 | ENSG00000011258 | MBTD1 | chr17 | - | 51202700 | 51202935 | 51201591 | 51201696 | 51203790 | 51203925 | 9 | 8 | 6 | 8 | 14 | 1 | 0 | 0 | 0 | 0 | 7 | 19 | 14 | 18 | 0 | 0 | 0 | 0 | 0 | 0 | 298 | 149 | 1 | 1 | 0.818 | 1 | 1 | 1 | 1 | 1 | 1 | 1 | 1 | NA | -0.036 |
| SE | 104761 | ENSG00000011258 | MBTD1 | chr17 | - | 51203139 | 51203228 | 51201591 | 51201696 | 51203790 | 51203925 | 33 | 30 | 33 | 38 | 62 | 1 | 0 | 0 | 0 | 0 | 0 | 19 | 27 | 16 | 0 | 0 | 0 | 0 | 0 | 0 | 238 | 149 | 1 | 1 | 0.954 | 1 | 1 | 1 | 1 | NA | 1 | 1 | 1 | NA | -0.009 |
| SE | 104762 | ENSG00000011258 | MBTD1 | chr17 | - | 51203139 | 51203228 | 51202700 | 51202935 | 51203790 | 51203925 | 116 | 95 | 86 | 131 | 194 | 0 | 0 | 1 | 0 | 0 | 31 | 114 | 106 | 64 | 27 | 0 | 0 | 0 | 0 | 0 | 238 | 149 | 1 | 1 | 1 | 1 | 0.982 | 1 | 1 | 1 | 1 | 1 | 1 | 1 | -0.004 |
| SE | 104763 | ENSG00000011258 | MBTD1 | chr17 | - | 51206887 | 51207005 | 51203790 | 51203925 | 51209313 | 51209404 | 28 | 25 | 36 | 29 | 60 | 2 | 0 | 0 | 0 | 0 | 1 | 32 | 32 | 21 | 16 | 0 | 1 | 2 | 0 | 0 | 267 | 149 | 1 | 1 | 0.887 | 1 | 1 | 1 | 1 | 1 | 0.947 | 0.899 | 1 | 1 | 0.008 |
| SE | 104765 | ENSG00000011258 | MBTD1 | chr17 | - | 51209313 | 51209430 | 51203790 | 51203925 | 51217333 | 51217416 | 7 | 3 | 2 | 3 | 4 | 0 | 3 | 1 | 0 | 0 | 0 | 6 | 9 | 1 | 0 | 0 | 4 | 2 | 2 | 0 | 266 | 149 | 1 | 1 | 1 | 0.359 | 0.528 | 1 | 1 | NA | 0.457 | 0.716 | 0.219 | NA | 0.313 |
| SE | 104769 | ENSG00000151491 | EPS8 | chr12 | - | 15623157 | 15623287 | 15620133 | 15621430 | 15624226 | 15624407 | 21 | 15 | 16 | 20 | 37 | 0 | 0 | 0 | 0 | 0 | 9 | 5 | 13 | 31 | 7 | 0 | 0 | 0 | 4 | 0 | 279 | 149 | 1 | 1 | 1 | 1 | 1 | 1 | 1 | 1 | 1 | 1 | 0.805 | 1 | 0.039 |
| SE | 104781 | ENSG00000130119 | GNL3L | chrX | + | 54540134 | 54540238 | 54539039 | 54539101 | 54541272 | 54541389 | 95 | 58 | 73 | 54 | 87 | 0 | 0 | 0 | 0 | 1 | 176 | 88 | 47 | 105 | 154 | 0 | 0 | 0 | 4 | 0 | 253 | 149 | 1 | 1 | 1 | 1 | 1 | 1 | 0.981 | 1 | 1 | 1 | 0.939 | 1 | 0.008 |
| SE | 104782 | ENSG00000130119 | GNL3L | chrX | + | 54540134 | 54540242 | 54539039 | 54539101 | 54541272 | 54541389 | 208 | 157 | 175 | 158 | 173 | 0 | 0 | 0 | 0 | 1 | 525 | 211 | 141 | 312 | 392 | 0 | 0 | 0 | 4 | 0 | 257 | 149 | 1 | 1 | 1 | 1 | 1 | 1 | 0.99 | 1 | 1 | 1 | 0.978 | 1 | 0.002 |
| SE | 104783 | ENSG00000130119 | GNL3L | chrX | + | 54541272 | 54541389 | 54540134 | 54540242 | 54542954 | 54543038 | 215 | 178 | 209 | 203 | 158 | 0 | 0 | 0 | 0 | 0 | 445 | 192 | 157 | 315 | 353 | 0 | 0 | 0 | 0 | 4 | 266 | 149 | 1 | 1 | 1 | 1 | 1 | 1 | 1 | 1 | 1 | 1 | 1 | 0.98 | 0.004 |
| SE | 104784 | ENSG00000130119 | GNL3L | chrX | + | 54544005 | 54544120 | 54543206 | 54543342 | 54544222 | 54544322 | 0 | 0 | 0 | 0 | 0 | 85 | 46 | 72 | 85 | 70 | 0 | 0 | 3 | 0 | 0 | 122 | 65 | 48 | 66 | 98 | 264 | 149 | 1 | 1 | 0 | 0 | 0 | 0 | 0 | 0 | 0 | 0.034 | 0 | 0 | -0.007 |
| SE | 104786 | ENSG00000130119 | GNL3L | chrX | + | 54550365 | 54550616 | 54548228 | 54548373 | 54550962 | 54551050 | 1 | 0 | 0 | 1 | 6 | 60 | 24 | 33 | 49 | 56 | 0 | 0 | 0 | 0 | 0 | 44 | 48 | 38 | 54 | 122 | 298 | 149 | 1 | 1 | 0.008 | 0 | 0 | 0.01 | 0.051 | 0 | 0 | 0 | 0 | 0 | 0.014 |
| SE | 104787 | ENSG00000130119 | GNL3L | chrX | + | 54551831 | 54551974 | 54551567 | 54551742 | 54552291 | 54552428 | 120 | 73 | 144 | 177 | 139 | 0 | 0 | 0 | 0 | 0 | 246 | 134 | 103 | 131 | 365 | 0 | 0 | 0 | 3 | 0 | 292 | 149 | 1 | 1 | 1 | 1 | 1 | 1 | 1 | 1 | 1 | 1 | 0.957 | 1 | 0.009 |
| SE | 104790 | ENSG00000228784 | LINC00954 | chr2 | + | 19870401 | 19870546 | 19869495 | 19869613 | 19872392 | 19872596 | 13 | 2 | 2 | 6 | 0 | 1 | 0 | 5 | 1 | 0 | 15 | 0 | 0 | 5 | 0 | 0 | 0 | 1 | 12 | 0 | 294 | 149 | 1 | 1 | 0.868 | 1 | 0.169 | 0.753 | NA | 1 | NA | 0 | 0.174 | NA | 0.306 |
| SE | 104791 | ENSG00000228784 | LINC00954 | chr2 | + | 19870424 | 19870546 | 19869495 | 19869613 | 19872392 | 19872530 | 17 | 4 | 6 | 5 | 0 | 1 | 0 | 5 | 1 | 0 | 11 | 1 | 0 | 5 | 0 | 0 | 0 | 1 | 12 | 0 | 271 | 149 | 1 | 1 | 0.903 | 1 | 0.398 | 0.733 | NA | 1 | 1 | 0 | 0.186 | NA | 0.212 |
| SE | 104793 | ENSG00000228784 | LINC00954 | chr2 | + | 19881705 | 19881886 | 19881566 | 19881618 | 19882214 | 19882260 | 1 | 0 | 2 | 2 | 0 | 0 | 0 | 2 | 2 | 3 | 0 | 0 | 0 | 0 | 0 | 0 | 0 | 0 | 6 | 0 | 298 | 149 | 1 | 1 | 1 | NA | 0.333 | 0.333 | 0 | NA | NA | NA | 0 | NA | 0.416 |
| SE | 104794 | ENSG00000228784 | LINC00954 | chr2 | + | 19881705 | 19881886 | 19881566 | 19881618 | 19882614 | 19882735 | 1 | 0 | 1 | 2 | 0 | 1 | 0 | 0 | 2 | 2 | 0 | 0 | 1 | 1 | 0 | 0 | 0 | 1 | 0 | 0 | 298 | 149 | 1 | 1 | 0.333 | NA | 1 | 0.333 | 0 | NA | NA | 0.333 | 1 | NA | -0.25 |
| SE | 104795 | ENSG00000228784 | LINC00954 | chr2 | + | 19882614 | 19882735 | 19881566 | 19881618 | 19883557 | 19883648 | 3 | 0 | 7 | 11 | 9 | 0 | 0 | 0 | 0 | 0 | 0 | 4 | 4 | 1 | 0 | 0 | 0 | 1 | 0 | 0 | 270 | 149 | 1 | 1 | 1 | NA | 1 | 1 | 1 | NA | 1 | 0.688 | 1 | NA | 0.104 |
| SE | 104796 | ENSG00000067560 | RHOA | chr3 | - | 49360927 | 49361066 | 49359138 | 49360382 | 49362495 | 49362626 | 12 | 3 | 4 | 18 | 9 | 1983 | 1460 | 1368 | 1848 | 2203 | 18 | 7 | 6 | 16 | 13 | 3723 | 2538 | 1701 | 3047 | 6660 | 288 | 149 | 1 | 1 | 0.003 | 0.001 | 0.002 | 0.005 | 0.002 | 0.002 | 0.001 | 0.002 | 0.003 | 0.001 | 0.001 |
| SE | 104797 | ENSG00000067560 | RHOA | chr3 | - | 49362495 | 49362626 | 49359138 | 49360382 | 49368427 | 49368542 | 3421 | 2549 | 2347 | 3152 | 3517 | 7 | 4 | 4 | 3 | 2 | 6965 | 4249 | 3040 | 5418 | 11691 | 4 | 5 | 5 | 12 | 20 | 280 | 149 | 1 | 1 | 0.996 | 0.997 | 0.997 | 0.998 | 0.999 | 0.999 | 0.998 | 0.997 | 0.996 | 0.997 | 0 |
| SE | 104798 | ENSG00000067560 | RHOA | chr3 | - | 49362495 | 49362626 | 49359138 | 49360382 | 49375433 | 49375591 | 1818 | 1325 | 1231 | 1704 | 1996 | 7 | 0 | 5 | 3 | 3 | 3445 | 2293 | 1495 | 2740 | 5917 | 10 | 1 | 6 | 4 | 15 | 280 | 149 | 1 | 1 | 0.993 | 1 | 0.992 | 0.997 | 0.997 | 0.995 | 0.999 | 0.993 | 0.997 | 0.995 | 0 |
| SE | 104799 | ENSG00000067560 | RHOA | chr3 | - | 49368427 | 49368548 | 49359138 | 49360382 | 49375433 | 49375591 | 1935 | 1082 | 1062 | 1530 | 1382 | 7 | 0 | 5 | 3 | 3 | 3929 | 2304 | 1566 | 2609 | 6405 | 10 | 1 | 6 | 4 | 15 | 270 | 149 | 1 | 1 | 0.993 | 1 | 0.992 | 0.996 | 0.996 | 0.995 | 0.999 | 0.993 | 0.997 | 0.996 | -0.001 |
| SE | 104800 | ENSG00000067560 | RHOA | chr3 | - | 49368427 | 49368548 | 49359139 | 49362626 | 49375433 | 49375591 | 3667 | 2442 | 2245 | 3095 | 2980 | 3 | 2 | 5 | 12 | 0 | 7927 | 4468 | 3256 | 5519 | 12872 | 44 | 13 | 1 | 0 | 8 | 270 | 149 | 1 | 1 | 0.999 | 0.999 | 0.996 | 0.993 | 1 | 0.99 | 0.995 | 0.999 | 1 | 0.999 | 0.001 |
| SE | 104801 | ENSG00000067560 | RHOA | chr3 | - | 49406711 | 49406806 | 49375433 | 49375591 | 49411819 | 49411968 | 1 | 0 | 1 | 1 | 0 | 1288 | 688 | 786 | 1068 | 265 | 0 | 2 | 2 | 1 | 9 | 2848 | 1540 | 947 | 1779 | 4416 | 244 | 149 | 1 | 1 | 0 | 0 | 0.001 | 0.001 | 0 | 0 | 0.001 | 0.001 | 0 | 0.001 | 0 |
| SE | 104809 | ENSG00000136404 | TM6SF1 | chr15 | + | 83112796 | 83112900 | 83107571 | 83107772 | 83119577 | 83119681 | 20 | 1 | 12 | 17 | 2 | 1 | 0 | 0 | 0 | 0 | 22 | 11 | 8 | 12 | 22 | 0 | 0 | 0 | 0 | 0 | 253 | 149 | 1 | 1 | 0.922 | 1 | 1 | 1 | 1 | 1 | 1 | 1 | 1 | 1 | -0.016 |
| SE | 104810 | ENSG00000136404 | TM6SF1 | chr15 | + | 83115844 | 83115942 | 83112796 | 83112900 | 83119577 | 83119681 | 53 | 36 | 34 | 84 | 20 | 0 | 0 | 0 | 0 | 0 | 40 | 48 | 49 | 125 | 142 | 0 | 0 | 0 | 1 | 3 | 247 | 149 | 1 | 1 | 1 | 1 | 1 | 1 | 1 | 1 | 1 | 1 | 0.987 | 0.966 | 0.009 |
| SE | 104813 | ENSG00000136404 | TM6SF1 | chr15 | + | 83121920 | 83122003 | 83119577 | 83119734 | 83124671 | 83124776 | 1 | 7 | 3 | 6 | 7 | 0 | 1 | 0 | 0 | 0 | 10 | 5 | 1 | 0 | 4 | 0 | 0 | 0 | 0 | 0 | 232 | 149 | 1 | 1 | 1 | 0.818 | 1 | 1 | 1 | 1 | 1 | 1 | NA | 1 | -0.036 |
| SE | 104820 | ENSG00000087502 | ERGIC2 | chr12 | - | 29356381 | 29356477 | 29350012 | 29350068 | 29357622 | 29357668 | 300 | 207 | 279 | 225 | 441 | 1 | 0 | 0 | 3 | 0 | 6 | 93 | 63 | 17 | 8 | 0 | 0 | 0 | 0 | 0 | 245 | 149 | 1 | 1 | 0.995 | 1 | 1 | 0.979 | 1 | 1 | 1 | 1 | 1 | 1 | -0.005 |
| SE | 104823 | ENSG00000205356 | TECPR1 | chr7 | - | 98217691 | 98217811 | 98214623 | 98217503 | 98217935 | 98218042 | 104 | 59 | 86 | 83 | 92 | 0 | 0 | 1 | 0 | 0 | 30 | 61 | 34 | 20 | 73 | 0 | 0 | 0 | 0 | 4 | 269 | 149 | 1 | 1 | 1 | 1 | 0.979 | 1 | 1 | 1 | 1 | 1 | 1 | 0.91 | 0.014 |
| SE | 104825 | ENSG00000205356 | TECPR1 | chr7 | - | 98230960 | 98231118 | 98229038 | 98229166 | 98231223 | 98231373 | 80 | 39 | 80 | 54 | 35 | 2 | 0 | 0 | 0 | 0 | 97 | 73 | 50 | 19 | 62 | 0 | 0 | 2 | 2 | 0 | 298 | 149 | 1 | 1 | 0.952 | 1 | 1 | 1 | 1 | 1 | 1 | 0.926 | 0.826 | 1 | 0.04 |
| SE | 104826 | ENSG00000205356 | TECPR1 | chr7 | - | 98233420 | 98233911 | 98232826 | 98232972 | 98236775 | 98236921 | 67 | 42 | 124 | 88 | 52 | 0 | 0 | 0 | 0 | 0 | 89 | 106 | 42 | 43 | 128 | 0 | 0 | 1 | 1 | 0 | 298 | 149 | 1 | 1 | 1 | 1 | 1 | 1 | 1 | 1 | 1 | 0.955 | 0.956 | 1 | 0.018 |
| SE | 104829 | ENSG00000205356 | TECPR1 | chr7 | - | 98236775 | 98236921 | 98232826 | 98232972 | 98240850 | 98240951 | 2 | 0 | 0 | 1 | 5 | 0 | 0 | 0 | 2 | 2 | 0 | 0 | 1 | 1 | 0 | 4 | 0 | 0 | 2 | 0 | 295 | 149 | 1 | 1 | 1 | NA | NA | 0.202 | 0.558 | 0 | NA | 1 | 0.202 | NA | 0.186 |
| SE | 104830 | ENSG00000205356 | TECPR1 | chr7 | - | 98236775 | 98236921 | 98233420 | 98233911 | 98238508 | 98238610 | 95 | 73 | 175 | 128 | 142 | 0 | 0 | 0 | 0 | 0 | 137 | 161 | 89 | 69 | 197 | 0 | 3 | 0 | 0 | 0 | 295 | 149 | 1 | 1 | 1 | 1 | 1 | 1 | 1 | 1 | 0.964 | 1 | 1 | 1 | 0.007 |
| SE | 104831 | ENSG00000205356 | TECPR1 | chr7 | - | 98238508 | 98238610 | 98232826 | 98232972 | 98240850 | 98240951 | 53 | 38 | 67 | 50 | 63 | 0 | 0 | 0 | 2 | 2 | 66 | 86 | 58 | 32 | 98 | 4 | 0 | 0 | 2 | 0 | 251 | 149 | 1 | 1 | 1 | 1 | 1 | 0.937 | 0.949 | 0.907 | 1 | 1 | 0.905 | 1 | 0.015 |
| SE | 104834 | ENSG00000205356 | TECPR1 | chr7 | - | 98244570 | 98244693 | 98243466 | 98243592 | 98251393 | 98251548 | 103 | 73 | 122 | 102 | 110 | 0 | 0 | 0 | 0 | 0 | 136 | 96 | 69 | 64 | 223 | 0 | 0 | 0 | 0 | 2 | 272 | 149 | 1 | 1 | 1 | 1 | 1 | 1 | 1 | 1 | 1 | 1 | 1 | 0.984 | 0.003 |
| SE | 104835 | ENSG00000205356 | TECPR1 | chr7 | - | 98244884 | 98245067 | 98244570 | 98244693 | 98251393 | 98251548 | 51 | 38 | 38 | 44 | 72 | 0 | 2 | 0 | 0 | 0 | 82 | 45 | 19 | 24 | 94 | 0 | 0 | 0 | 2 | 0 | 298 | 149 | 1 | 1 | 1 | 0.905 | 1 | 1 | 1 | 1 | 1 | 1 | 0.857 | 1 | 0.01 |
| SE | 104836 | ENSG00000205356 | TECPR1 | chr7 | - | 98245921 | 98246165 | 98244884 | 98245067 | 98251393 | 98251548 | 94 | 44 | 92 | 77 | 89 | 0 | 0 | 0 | 0 | 1 | 147 | 70 | 77 | 64 | 143 | 0 | 0 | 0 | 0 | 0 | 298 | 149 | 1 | 1 | 1 | 1 | 1 | 1 | 0.978 | 1 | 1 | 1 | 1 | 1 | -0.004 |
| SE | 104838 | ENSG00000205356 | TECPR1 | chr7 | - | 98251393 | 98251548 | 98245921 | 98246165 | 98252125 | 98252206 | 50 | 25 | 71 | 47 | 45 | 3 | 0 | 0 | 0 | 0 | 35 | 45 | 39 | 27 | 57 | 0 | 0 | 0 | 0 | 0 | 298 | 149 | 1 | 1 | 0.893 | 1 | 1 | 1 | 1 | 1 | 1 | 1 | 1 | 1 | -0.021 |
| SE | 104839 | ENSG00000205356 | TECPR1 | chr7 | - | 98251418 | 98251548 | 98245921 | 98246165 | 98252125 | 98252218 | 28 | 23 | 52 | 30 | 8 | 3 | 0 | 0 | 0 | 0 | 32 | 32 | 9 | 19 | 44 | 0 | 0 | 0 | 0 | 0 | 279 | 149 | 1 | 1 | 0.833 | 1 | 1 | 1 | 1 | 1 | 1 | 1 | 1 | 1 | -0.033 |
| SE | 104841 | ENSG00000011105 | TSPAN9 | chr12 | + | 3083652 | 3083719 | 3077376 | 3077453 | 3278420 | 3278612 | 1 | 0 | 1 | 0 | 7 | 1 | 0 | 1 | 0 | 0 | 0 | 0 | 0 | 2 | 4 | 0 | 0 | 0 | 0 | 3 | 216 | 149 | 1 | 1 | 0.408 | NA | 0.408 | NA | 1 | NA | NA | NA | 1 | 0.479 | -0.134 |
| SE | 104842 | ENSG00000011105 | TSPAN9 | chr12 | + | 3201176 | 3201256 | 3077354 | 3077453 | 3278420 | 3278612 | 1 | 14 | 2 | 5 | 17 | 1 | 0 | 1 | 0 | 0 | 26 | 2 | 4 | 3 | 93 | 0 | 0 | 0 | 0 | 3 | 229 | 149 | 1 | 1 | 0.394 | 1 | 0.565 | 1 | 1 | 1 | 1 | 1 | 1 | 0.953 | -0.199 |
| SE | 104846 | ENSG00000257176 | NA | chr12 | - | 29307849 | 29308044 | 29293065 | 29293540 | 29309770 | 29309903 | 8 | 1 | 3 | 7 | 3 | 0 | 8 | 0 | 0 | 0 | 0 | 0 | 3 | 5 | 7 | 0 | 0 | 0 | 0 | 0 | 298 | 149 | 1 | 1 | 1 | 0.059 | 1 | 1 | 1 | NA | NA | 1 | 1 | 1 | -0.188 |
| SE | 104847 | ENSG00000139044 | B4GALNT3 | chr12 | + | 556546 | 556866 | 552466 | 553983 | 557607 | 557761 | 11 | 3 | 1 | 0 | 0 | 1 | 0 | 0 | 0 | 0 | 0 | 0 | 0 | 11 | 0 | 0 | 0 | 0 | 0 | 0 | 298 | 149 | 1 | 1 | 0.846 | 1 | 1 | NA | NA | NA | NA | NA | 1 | NA | -0.051 |
| SE | 104848 | ENSG00000151743 | AMN1 | chr12 | - | 31708917 | 31709010 | 31701998 | 31702007 | 31709091 | 31709167 | 0 | 0 | 0 | 0 | 0 | 0 | 1 | 0 | 0 | 3 | 0 | 0 | 0 | 0 | 9 | 0 | 3 | 1 | 0 | 0 | 242 | 149 | 1 | 1 | NA | 0 | NA | NA | 0 | NA | 0 | 0 | NA | 1 | -0.333 |
| SE | 104850 | ENSG00000151743 | AMN1 | chr12 | - | 31709292 | 31709425 | 31701862 | 31702007 | 31714881 | 31714951 | 28 | 29 | 27 | 25 | 99 | 2 | 0 | 1 | 1 | 0 | 7 | 32 | 14 | 15 | 10 | 0 | 3 | 4 | 0 | 0 | 282 | 149 | 1 | 1 | 0.881 | 1 | 0.934 | 0.93 | 1 | 1 | 0.849 | 0.649 | 1 | 1 | 0.049 |
| SE | 104851 | ENSG00000151743 | AMN1 | chr12 | - | 31709292 | 31709425 | 31701862 | 31702007 | 31728970 | 31729021 | 25 | 20 | 33 | 18 | 71 | 1 | 0 | 0 | 1 | 2 | 8 | 45 | 8 | 8 | 13 | 0 | 6 | 0 | 1 | 0 | 282 | 149 | 1 | 1 | 0.93 | 1 | 1 | 0.905 | 0.949 | 1 | 0.798 | 1 | 0.809 | 1 | 0.035 |
| SE | 104852 | ENSG00000151743 | AMN1 | chr12 | - | 31714440 | 31714631 | 31709292 | 31709425 | 31714881 | 31714951 | 0 | 0 | 0 | 0 | 0 | 7 | 13 | 4 | 9 | 35 | 0 | 1 | 0 | 0 | 0 | 3 | 1 | 8 | 8 | 0 | 298 | 149 | 1 | 1 | 0 | 0 | 0 | 0 | 0 | 0 | 0.333 | 0 | 0 | NA | -0.083 |
| SE | 104853 | ENSG00000151743 | AMN1 | chr12 | - | 31714881 | 31714951 | 31701862 | 31702007 | 31728970 | 31729019 | 2 | 0 | 1 | 1 | 0 | 1 | 0 | 0 | 1 | 2 | 0 | 3 | 2 | 0 | 0 | 0 | 6 | 0 | 1 | 0 | 219 | 149 | 1 | 1 | 0.576 | NA | 1 | 0.405 | 0 | NA | 0.254 | 1 | 0 | NA | 0.077 |
| SE | 104855 | ENSG00000151743 | AMN1 | chr12 | - | 31719217 | 31719356 | 31714881 | 31714951 | 31728970 | 31729010 | 2 | 8 | 1 | 9 | 20 | 0 | 0 | 0 | 0 | 0 | 6 | 1 | 5 | 0 | 0 | 0 | 0 | 0 | 2 | 0 | 288 | 149 | 1 | 1 | 1 | 1 | 1 | 1 | 1 | 1 | 1 | 1 | 0 | NA | 0.25 |
| SE | 104857 | ENSG00000109685 | NSD2 | chr4 | + | 1893729 | 1893909 | 1871433 | 1871542 | 1899220 | 1899395 | 0 | 0 | 0 | 0 | 1 | 0 | 1 | 0 | 0 | 0 | 18 | 0 | 0 | 0 | 0 | 0 | 0 | 0 | 0 | 0 | 298 | 149 | 1 | 1 | NA | 0 | NA | NA | 1 | 1 | NA | NA | NA | NA | -0.5 |
| SE | 104859 | ENSG00000109685 | NSD2 | chr4 | + | 1893729 | 1893909 | 1892165 | 1892263 | 1900625 | 1900766 | 3 | 0 | 0 | 1 | 6 | 3 | 4 | 0 | 1 | 0 | 40 | 3 | 1 | 4 | 8 | 0 | 0 | 0 | 4 | 0 | 298 | 149 | 1 | 1 | 0.333 | 0 | NA | 0.333 | 1 | 1 | 1 | 1 | 0.333 | 1 | -0.45 |
| SE | 104861 | ENSG00000109685 | NSD2 | chr4 | + | 1893729 | 1893909 | 1892781 | 1892891 | 1900625 | 1900766 | 3 | 0 | 0 | 1 | 6 | 0 | 0 | 1 | 0 | 0 | 40 | 3 | 1 | 4 | 8 | 0 | 0 | 0 | 0 | 0 | 298 | 149 | 1 | 1 | 1 | NA | 0 | 1 | 1 | 1 | 1 | 1 | 1 | 1 | -0.25 |
| SE | 104862 | ENSG00000109685 | NSD2 | chr4 | + | 1899220 | 1899395 | 1871460 | 1871542 | 1900625 | 1901251 | 1 | 1 | 0 | 0 | 0 | 22 | 1 | 13 | 9 | 0 | 0 | 0 | 1 | 0 | 0 | 10 | 24 | 5 | 15 | 3 | 298 | 149 | 1 | 1 | 0.022 | 0.333 | 0 | 0 | NA | 0 | 0 | 0.091 | 0 | 0 | 0.071 |
| SE | 104863 | ENSG00000109685 | NSD2 | chr4 | + | 1899220 | 1899395 | 1892781 | 1892891 | 1900625 | 1901251 | 1 | 0 | 0 | 0 | 0 | 0 | 0 | 1 | 0 | 0 | 0 | 0 | 1 | 0 | 0 | 0 | 0 | 0 | 0 | 0 | 298 | 149 | 1 | 1 | 1 | NA | 0 | NA | NA | NA | NA | 1 | NA | NA | -0.5 |
| SE | 104864 | ENSG00000109685 | NSD2 | chr4 | + | 1899220 | 1899395 | 1893729 | 1893909 | 1900625 | 1901251 | 1 | 0 | 0 | 0 | 0 | 3 | 0 | 0 | 1 | 6 | 0 | 0 | 1 | 0 | 0 | 40 | 3 | 1 | 4 | 8 | 298 | 149 | 1 | 1 | 0.143 | NA | NA | 0 | 0 | 0 | 0 | 0.333 | 0 | 0 | -0.019 |
| SE | 104868 | ENSG00000109685 | NSD2 | chr4 | + | 1916870 | 1917037 | 1904215 | 1904378 | 1918140 | 1918389 | 168 | 139 | 183 | 170 | 204 | 1 | 1 | 1 | 0 | 0 | 476 | 195 | 177 | 403 | 312 | 0 | 3 | 0 | 0 | 2 | 298 | 149 | 1 | 1 | 0.988 | 0.986 | 0.989 | 1 | 1 | 1 | 0.97 | 1 | 1 | 0.987 | 0.001 |
| SE | 104869 | ENSG00000109685 | NSD2 | chr4 | + | 1916870 | 1917037 | 1912090 | 1912194 | 1918140 | 1918389 | 107 | 87 | 117 | 137 | 126 | 0 | 0 | 2 | 0 | 0 | 349 | 151 | 120 | 241 | 222 | 0 | 0 | 0 | 0 | 0 | 298 | 149 | 1 | 1 | 1 | 1 | 0.967 | 1 | 1 | 1 | 1 | 1 | 1 | 1 | -0.007 |
| SE | 104870 | ENSG00000109685 | NSD2 | chr4 | + | 1918140 | 1918623 | 1904215 | 1904378 | 1930625 | 1930770 | 58 | 38 | 63 | 66 | 58 | 0 | 0 | 0 | 0 | 0 | 115 | 78 | 111 | 201 | 76 | 0 | 0 | 0 | 1 | 0 | 298 | 149 | 1 | 1 | 1 | 1 | 1 | 1 | 1 | 1 | 1 | 1 | 0.99 | 1 | 0.002 |
| SE | 104871 | ENSG00000109685 | NSD2 | chr4 | + | 1935143 | 1935262 | 1930625 | 1930770 | 1951071 | 1951203 | 78 | 46 | 85 | 70 | 73 | 0 | 0 | 0 | 0 | 0 | 150 | 100 | 134 | 177 | 108 | 0 | 0 | 2 | 0 | 0 | 268 | 149 | 1 | 1 | 1 | 1 | 1 | 1 | 1 | 1 | 1 | 0.974 | 1 | 1 | 0.005 |
| SE | 104873 | ENSG00000109685 | NSD2 | chr4 | + | 1939653 | 1939778 | 1935143 | 1935262 | 1942338 | 1942432 | 10 | 2 | 6 | 5 | 12 | 0 | 0 | 0 | 1 | 0 | 15 | 1 | 8 | 28 | 5 | 0 | 0 | 0 | 0 | 0 | 274 | 149 | 1 | 1 | 1 | 1 | 1 | 0.731 | 1 | 1 | 1 | 1 | 1 | 1 | -0.054 |
| SE | 104876 | ENSG00000109685 | NSD2 | chr4 | + | 1942338 | 1942432 | 1939653 | 1939778 | 1951071 | 1951203 | 3 | 0 | 1 | 1 | 0 | 54 | 42 | 52 | 55 | 41 | 0 | 0 | 0 | 3 | 0 | 96 | 54 | 50 | 84 | 54 | 243 | 149 | 1 | 1 | 0.033 | 0 | 0.012 | 0.011 | 0 | 0 | 0 | 0 | 0.021 | 0 | 0.007 |
| SE | 104877 | ENSG00000109685 | NSD2 | chr4 | + | 1942338 | 1942565 | 1935143 | 1935262 | 1951071 | 1951203 | 2 | 0 | 0 | 3 | 0 | 1 | 6 | 3 | 6 | 2 | 0 | 0 | 0 | 2 | 5 | 33 | 0 | 4 | 19 | 1 | 298 | 149 | 1 | 1 | 0.5 | 0 | 0 | 0.2 | 0 | 0 | NA | 0 | 0.05 | 0.714 | -0.051 |
| SE | 104879 | ENSG00000109685 | NSD2 | chr4 | + | 1951998 | 1952231 | 1951071 | 1951203 | 1953323 | 1953446 | 21 | 9 | 17 | 16 | 7 | 0 | 0 | 0 | 0 | 1 | 10 | 6 | 26 | 21 | 16 | 0 | 0 | 0 | 2 | 0 | 298 | 149 | 1 | 1 | 1 | 1 | 1 | 1 | 0.778 | 1 | 1 | 1 | 0.84 | 1 | -0.012 |
| SE | 104880 | ENSG00000109685 | NSD2 | chr4 | + | 1952107 | 1952231 | 1951071 | 1951203 | 1953323 | 1953524 | 55 | 35 | 42 | 55 | 39 | 0 | 0 | 0 | 0 | 1 | 46 | 31 | 60 | 86 | 51 | 0 | 0 | 0 | 2 | 0 | 273 | 149 | 1 | 1 | 1 | 1 | 1 | 1 | 0.955 | 1 | 1 | 1 | 0.959 | 1 | -0.001 |
| SE | 104881 | ENSG00000109685 | NSD2 | chr4 | + | 1952751 | 1953524 | 1952104 | 1952231 | 1955160 | 1955340 | 20 | 9 | 24 | 15 | 13 | 0 | 1 | 0 | 0 | 0 | 20 | 17 | 10 | 32 | 6 | 0 | 0 | 0 | 0 | 0 | 298 | 149 | 1 | 1 | 1 | 0.818 | 1 | 1 | 1 | 1 | 1 | 1 | 1 | 1 | -0.036 |
| SE | 104882 | ENSG00000109685 | NSD2 | chr4 | + | 1953323 | 1953524 | 1952104 | 1952231 | 1955160 | 1955221 | 43 | 19 | 42 | 30 | 18 | 0 | 1 | 0 | 0 | 0 | 24 | 24 | 35 | 56 | 17 | 0 | 0 | 0 | 0 | 0 | 298 | 149 | 1 | 1 | 1 | 0.905 | 1 | 1 | 1 | 1 | 1 | 1 | 1 | 1 | -0.019 |
| SE | 104885 | ENSG00000109685 | NSD2 | chr4 | + | 1961034 | 1961151 | 1959295 | 1959740 | 1974862 | 1975004 | 114 | 65 | 128 | 118 | 94 | 0 | 0 | 0 | 0 | 0 | 412 | 179 | 178 | 294 | 239 | 0 | 0 | 0 | 1 | 6 | 266 | 149 | 1 | 1 | 1 | 1 | 1 | 1 | 1 | 1 | 1 | 1 | 0.994 | 0.957 | 0.01 |
| SE | 104888 | ENSG00000142166 | IFNAR1 | chr21 | + | 33340998 | 33341174 | 33335523 | 33335647 | 33343534 | 33343676 | 130 | 98 | 138 | 170 | 244 | 1 | 0 | 0 | 0 | 0 | 78 | 85 | 125 | 132 | 4 | 0 | 0 | 0 | 0 | 0 | 298 | 149 | 1 | 1 | 0.985 | 1 | 1 | 1 | 1 | 1 | 1 | 1 | 1 | 1 | -0.003 |
| SE | 104890 | ENSG00000142166 | IFNAR1 | chr21 | + | 33343267 | 33343422 | 33335523 | 33335647 | 33343534 | 33343604 | 421 | 282 | 370 | 322 | 818 | 1 | 0 | 0 | 0 | 0 | 252 | 297 | 413 | 335 | 57 | 0 | 0 | 0 | 0 | 0 | 298 | 149 | 1 | 1 | 0.995 | 1 | 1 | 1 | 1 | 1 | 1 | 1 | 1 | 1 | -0.001 |
| SE | 104894 | ENSG00000142166 | IFNAR1 | chr21 | + | 33343534 | 33343676 | 33343267 | 33343422 | 33345245 | 33345358 | 775 | 484 | 623 | 568 | 1384 | 0 | 0 | 0 | 0 | 0 | 417 | 504 | 646 | 580 | 74 | 0 | 0 | 1 | 0 | 0 | 291 | 149 | 1 | 1 | 1 | 1 | 1 | 1 | 1 | 1 | 1 | 0.997 | 1 | 1 | 0.001 |
| SE | 104895 | ENSG00000142166 | IFNAR1 | chr21 | + | 33345245 | 33345360 | 33343534 | 33343676 | 33348549 | 33349077 | 333 | 184 | 260 | 246 | 574 | 0 | 0 | 0 | 0 | 3 | 146 | 216 | 266 | 260 | 25 | 0 | 0 | 6 | 0 | 0 | 264 | 149 | 1 | 1 | 1 | 1 | 1 | 1 | 0.991 | 1 | 1 | 0.962 | 1 | 1 | 0.006 |
| SE | 104896 | ENSG00000142166 | IFNAR1 | chr21 | + | 33345245 | 33345360 | 33343534 | 33343676 | 33349090 | 33349166 | 549 | 334 | 382 | 403 | 895 | 0 | 0 | 0 | 1 | 2 | 288 | 308 | 286 | 369 | 34 | 0 | 0 | 0 | 0 | 0 | 264 | 149 | 1 | 1 | 1 | 1 | 1 | 0.996 | 0.996 | 1 | 1 | 1 | 1 | 1 | -0.002 |
| SE | 104897 | ENSG00000142166 | IFNAR1 | chr21 | + | 33345245 | 33345360 | 33343534 | 33343676 | 33352757 | 33352908 | 324 | 181 | 243 | 230 | 546 | 0 | 0 | 0 | 0 | 0 | 122 | 206 | 224 | 251 | 14 | 0 | 0 | 0 | 1 | 0 | 264 | 149 | 1 | 1 | 1 | 1 | 1 | 1 | 1 | 1 | 1 | 1 | 0.993 | 1 | 0.001 |
| SE | 104898 | ENSG00000142166 | IFNAR1 | chr21 | + | 33349090 | 33349290 | 33343534 | 33343676 | 33352757 | 33352908 | 0 | 1 | 0 | 1 | 2 | 0 | 0 | 0 | 0 | 0 | 0 | 0 | 0 | 0 | 0 | 0 | 0 | 0 | 1 | 0 | 298 | 149 | 1 | 1 | NA | 1 | NA | 1 | 1 | NA | NA | NA | 0 | NA | 1 |
| SE | 104899 | ENSG00000142166 | IFNAR1 | chr21 | + | 33349090 | 33349290 | 33345245 | 33345360 | 33352757 | 33352908 | 227 | 155 | 139 | 176 | 357 | 0 | 0 | 0 | 0 | 0 | 166 | 102 | 67 | 118 | 20 | 0 | 0 | 5 | 0 | 0 | 298 | 149 | 1 | 1 | 1 | 1 | 1 | 1 | 1 | 1 | 1 | 0.87 | 1 | 1 | 0.026 |
| SE | 104900 | ENSG00000142166 | IFNAR1 | chr21 | + | 33349388 | 33349543 | 33349090 | 33349290 | 33352757 | 33352908 | 568 | 406 | 443 | 580 | 1146 | 0 | 1 | 0 | 0 | 0 | 308 | 392 | 288 | 431 | 57 | 0 | 0 | 0 | 0 | 0 | 298 | 149 | 1 | 1 | 1 | 0.995 | 1 | 1 | 1 | 1 | 1 | 1 | 1 | 1 | -0.001 |
| SE | 104901 | ENSG00000142166 | IFNAR1 | chr21 | + | 33353637 | 33353783 | 33352757 | 33352908 | 33355315 | 33355703 | 216 | 161 | 217 | 250 | 329 | 0 | 0 | 0 | 0 | 1 | 159 | 166 | 149 | 170 | 43 | 0 | 0 | 0 | 0 | 0 | 295 | 149 | 1 | 1 | 1 | 1 | 1 | 1 | 0.994 | 1 | 1 | 1 | 1 | 1 | -0.001 |
| SE | 104904 | ENSG00000170456 | DENND5B | chr12 | - | 31399653 | 31399772 | 31398174 | 31398362 | 31402497 | 31402643 | 20 | 11 | 16 | 12 | 19 | 0 | 0 | 3 | 0 | 0 | 11 | 22 | 41 | 19 | 0 | 0 | 0 | 0 | 3 | 0 | 268 | 149 | 1 | 1 | 1 | 1 | 0.748 | 1 | 1 | 1 | 1 | 1 | 0.779 | NA | 0.005 |
| SE | 104907 | ENSG00000170456 | DENND5B | chr12 | - | 31433154 | 31433248 | 31426292 | 31426424 | 31447537 | 31447769 | 8 | 7 | 10 | 5 | 14 | 0 | 0 | 0 | 0 | 0 | 10 | 5 | 6 | 3 | 0 | 0 | 2 | 0 | 0 | 0 | 243 | 149 | 1 | 1 | 1 | 1 | 1 | 1 | 1 | 1 | 0.605 | 1 | 1 | NA | 0.099 |
| SE | 104908 | ENSG00000170456 | DENND5B | chr12 | - | 31442774 | 31442925 | 31432374 | 31433248 | 31447537 | 31447769 | 66 | 47 | 135 | 69 | 115 | 0 | 2 | 0 | 0 | 1 | 187 | 116 | 198 | 118 | 21 | 0 | 4 | 0 | 0 | 0 | 298 | 149 | 1 | 1 | 1 | 0.922 | 1 | 1 | 0.983 | 1 | 0.935 | 1 | 1 | 1 | -0.006 |
| SE | 104909 | ENSG00000170456 | DENND5B | chr12 | - | 31442774 | 31442925 | 31432374 | 31433248 | 31451939 | 31452476 | 44 | 34 | 83 | 46 | 77 | 2 | 0 | 1 | 0 | 0 | 111 | 85 | 132 | 82 | 15 | 0 | 0 | 0 | 0 | 0 | 298 | 149 | 1 | 1 | 0.917 | 1 | 0.976 | 1 | 1 | 1 | 1 | 1 | 1 | 1 | -0.021 |
| SE | 104910 | ENSG00000170456 | DENND5B | chr12 | - | 31447537 | 31447769 | 31432374 | 31433248 | 31451939 | 31452476 | 9 | 20 | 38 | 11 | 23 | 2 | 0 | 1 | 0 | 0 | 5 | 62 | 44 | 26 | 12 | 0 | 0 | 0 | 0 | 0 | 298 | 149 | 1 | 1 | 0.692 | 1 | 0.95 | 1 | 1 | 1 | 1 | 1 | 1 | 1 | -0.072 |
| SE | 104912 | ENSG00000170456 | DENND5B | chr12 | - | 31451939 | 31452476 | 31447537 | 31447769 | 31460193 | 31460381 | 25 | 32 | 79 | 35 | 48 | 0 | 0 | 1 | 0 | 0 | 29 | 117 | 60 | 32 | 55 | 0 | 0 | 0 | 2 | 0 | 298 | 149 | 1 | 1 | 1 | 1 | 0.975 | 1 | 1 | 1 | 1 | 1 | 0.889 | 1 | 0.017 |
| SE | 104913 | ENSG00000170456 | DENND5B | chr12 | - | 31460193 | 31460381 | 31451701 | 31452476 | 31479588 | 31480255 | 30 | 18 | 60 | 37 | 32 | 0 | 0 | 1 | 0 | 0 | 41 | 85 | 54 | 15 | 53 | 0 | 0 | 1 | 0 | 0 | 298 | 149 | 1 | 1 | 1 | 1 | 0.968 | 1 | 1 | 1 | 1 | 0.964 | 1 | 1 | 0.001 |

| | | | | | | | | | | | | | | | | | | | | | | | | | | | | | | | | | | | | | | | | | | | | | | |
|---|---|---|---|---|---|---|---|---|---|---|---|---|---|---|---|---|---|---|---|---|---|---|---|---|---|---|---|---|---|---|---|---|---|---|---|---|---|---|---|---|---|---|---|---|---|---|
| SE | 104914 | ENSG00000170456 | DENND5B | chr12 | - | 31479588 | 31480255 | 31460193 | 31460381 | 31495809 | 31495902 | 21 | 6 | 37 | 23 | 18 | 0 | 0 | 0 | 0 | 0 | 42 | 47 | 80 | 27 | 19 | 0 | 0 | 1 | 0 | 0 | 298 | 149 | 1 | 1 | 1 | 1 | 1 | 1 | 1 | 1 | 1 | 0.976 | 1 | 1 | 0.005 |
| SE | 104915 | ENSG00000170456 | DENND5B | chr12 | - | 31499605 | 31499671 | 31495809 | 31495919 | 31540919 | 31541031 | 1 | 0 | 5 | 2 | 2 | 0 | 0 | 0 | 0 | 1 | 0 | 3 | 9 | 5 | 7 | 0 | 0 | 0 | 0 | 0 | 215 | 149 | 1 | 1 | 1 | NA | 1 | 1 | 0.581 | NA | 1 | 1 | 1 | 1 | -0.105 |
| SE | 104916 | ENSG00000170456 | DENND5B | chr12 | - | 31499605 | 31499671 | 31495809 | 31495919 | 31590705 | 31591012 | 1 | 0 | 6 | 5 | 2 | 2 | 0 | 1 | 2 | 0 | 0 | 10 | 20 | 5 | 7 | 0 | 3 | 13 | 0 | 2 | 215 | 149 | 1 | 1 | 0.257 | NA | 0.806 | 0.634 | 1 | NA | 0.698 | 0.516 | 1 | 0.708 | -0.056 |
| SE | 104917 | ENSG00000170456 | DENND5B | chr12 | - | 31540919 | 31541031 | 31495809 | 31495919 | 31590705 | 31590999 | 0 | 0 | 0 | 0 | 1 | 2 | 0 | 1 | 2 | 0 | 0 | 0 | 0 | 0 | 0 | 0 | 3 | 13 | 0 | 2 | 261 | 149 | 1 | 1 | 0 | NA | 0 | 0 | 1 | NA | 0 | 0 | NA | 0 | 0.25 |
| SE | 104918 | ENSG00000170456 | DENND5B | chr12 | - | 31540919 | 31541031 | 31499605 | 31499671 | 31590705 | 31590999 | 0 | 0 | 0 | 0 | 0 | 0 | 0 | 1 | 3 | 0 | 0 | 0 | 1 | 0 | 0 | 0 | 7 | 12 | 0 | 0 | 261 | 149 | 1 | 1 | NA | NA | 0 | 0 | NA | NA | 0 | 0.045 | NA | NA | -0.022 |
| SE | 104921 | ENSG00000064102 | INTS13 | chr12 | - | 26913973 | 26914128 | 26913456 | 26913687 | 26914407 | 26914578 | 51 | 46 | 79 | 73 | 108 | 1 | 0 | 0 | 0 | 0 | 28 | 34 | 53 | 38 | 12 | 0 | 0 | 0 | 0 | 0 | 298 | 149 | 1 | 1 | 0.962 | 1 | 1 | 1 | 1 | 1 | 1 | 1 | 1 | 1 | -0.008 |
| SE | 104922 | ENSG00000064102 | INTS13 | chr12 | - | 26913973 | 26914128 | 26913456 | 26913687 | 26916001 | 26916180 | 28 | 29 | 51 | 38 | 83 | 0 | 0 | 1 | 0 | 0 | 10 | 17 | 32 | 34 | 0 | 0 | 0 | 0 | 0 | 0 | 298 | 149 | 1 | 1 | 1 | 1 | 0.962 | 1 | 1 | 1 | 1 | 1 | 1 | NA | -0.008 |
| SE | 104923 | ENSG00000064102 | INTS13 | chr12 | - | 26913973 | 26914128 | 26913456 | 26913687 | 26917351 | 26917441 | 28 | 23 | 47 | 37 | 75 | 0 | 0 | 0 | 0 | 0 | 10 | 13 | 27 | 36 | 0 | 0 | 4 | 0 | 0 | 0 | 298 | 149 | 1 | 1 | 1 | 1 | 1 | 1 | 1 | 1 | 0.619 | 1 | 1 | NA | 0.095 |
| SE | 104925 | ENSG00000064102 | INTS13 | chr12 | - | 26914407 | 26914532 | 26914045 | 26914128 | 26917351 | 26917441 | 23 | 20 | 27 | 40 | 39 | 0 | 0 | 0 | 0 | 0 | 18 | 18 | 30 | 3 | 12 | 0 | 0 | 1 | 2 | 0 | 274 | 149 | 1 | 1 | 1 | 1 | 1 | 1 | 1 | 1 | 1 | 0.942 | 0.449 | 1 | 0.122 |
| SE | 104926 | ENSG00000064102 | INTS13 | chr12 | - | 26914407 | 26914578 | 26913456 | 26913687 | 26916001 | 26916180 | 49 | 24 | 21 | 34 | 59 | 0 | 0 | 1 | 0 | 0 | 0 | 19 | 6 | 5 | 0 | 0 | 0 | 0 | 0 | 0 | 298 | 149 | 1 | 1 | 1 | 1 | 0.913 | 1 | 1 | NA | 1 | 1 | 1 | NA | -0.017 |
| SE | 104927 | ENSG00000064102 | INTS13 | chr12 | - | 26914407 | 26914578 | 26913456 | 26913687 | 26917351 | 26917441 | 1 | 0 | 0 | 0 | 4 | 0 | 0 | 0 | 0 | 0 | 0 | 0 | 0 | 0 | 0 | 0 | 4 | 0 | 0 | 0 | 298 | 149 | 1 | 1 | 1 | NA | NA | NA | 1 | NA | 0 | NA | NA | NA | 1 |
| SE | 104929 | ENSG00000064102 | INTS13 | chr12 | - | 26914407 | 26914578 | 26913973 | 26914128 | 26917351 | 26917441 | 23 | 23 | 32 | 36 | 36 | 0 | 0 | 0 | 0 | 0 | 18 | 18 | 27 | 4 | 12 | 0 | 0 | 1 | 2 | 0 | 298 | 149 | 1 | 1 | 1 | 1 | 1 | 1 | 1 | 1 | 1 | 0.931 | 0.5 | 1 | 0.114 |
| SE | 104930 | ENSG00000064102 | INTS13 | chr12 | - | 26916001 | 26916180 | 26913456 | 26913687 | 26917351 | 26917441 | 111 | 86 | 81 | 86 | 167 | 0 | 0 | 0 | 0 | 0 | 66 | 50 | 84 | 27 | 6 | 0 | 4 | 0 | 0 | 0 | 298 | 149 | 1 | 1 | 1 | 1 | 1 | 1 | 1 | 1 | 0.862 | 1 | 1 | 1 | 0.028 |
| SE | 104931 | ENSG00000064102 | INTS13 | chr12 | - | 26916001 | 26916180 | 26913973 | 26914128 | 26917351 | 26917441 | 111 | 92 | 84 | 87 | 175 | 0 | 0 | 0 | 0 | 0 | 66 | 54 | 90 | 27 | 6 | 0 | 0 | 1 | 2 | 0 | 298 | 149 | 1 | 1 | 1 | 1 | 1 | 1 | 1 | 1 | 1 | 0.978 | 0.871 | 1 | 0.03 |
| SE | 104933 | ENSG00000064102 | INTS13 | chr12 | - | 26916001 | 26916180 | 26914407 | 26914578 | 26917351 | 26917441 | 159 | 110 | 101 | 120 | 226 | 0 | 0 | 0 | 0 | 4 | 66 | 69 | 90 | 32 | 6 | 0 | 0 | 0 | 0 | 0 | 298 | 149 | 1 | 1 | 1 | 1 | 1 | 1 | 0.966 | 1 | 1 | 1 | 1 | 1 | -0.007 |
| SE | 104934 | ENSG00000064102 | INTS13 | chr12 | - | 26917643 | 26917733 | 26917351 | 26917441 | 26922615 | 26922700 | 103 | 76 | 76 | 88 | 198 | 0 | 0 | 1 | 0 | 4 | 7 | 49 | 65 | 11 | 0 | 0 | 0 | 2 | 0 | 0 | 239 | 149 | 1 | 1 | 1 | 1 | 0.979 | 1 | 0.969 | 1 | 1 | 0.953 | 1 | NA | 0.001 |
| SE | 104935 | ENSG00000064102 | INTS13 | chr12 | - | 26924354 | 26924483 | 26922615 | 26922700 | 26925760 | 26925851 | 87 | 45 | 70 | 82 | 163 | 0 | 0 | 0 | 0 | 0 | 31 | 43 | 30 | 16 | 5 | 0 | 0 | 1 | 0 | 0 | 278 | 149 | 1 | 1 | 1 | 1 | 1 | 1 | 1 | 1 | 1 | 0.941 | 1 | 1 | 0.012 |
| SE | 104936 | ENSG00000064102 | INTS13 | chr12 | - | 26934555 | 26934630 | 26928702 | 26928905 | 26936578 | 26936814 | 76 | 44 | 74 | 60 | 75 | 0 | 0 | 0 | 0 | 0 | 2 | 37 | 26 | 13 | 0 | 0 | 1 | 0 | 0 | 0 | 224 | 149 | 1 | 1 | 1 | 1 | 1 | 1 | 1 | 1 | 0.961 | 1 | 1 | NA | 0.01 |
| SE | 104937 | ENSG00000064102 | INTS13 | chr12 | - | 26936578 | 26936814 | 26928702 | 26928905 | 26937795 | 26937981 | 35 | 15 | 20 | 15 | 15 | 1 | 0 | 0 | 0 | 0 | 11 | 24 | 9 | 7 | 0 | 0 | 0 | 0 | 0 | 0 | 298 | 149 | 1 | 1 | 0.946 | 1 | 1 | 1 | 1 | 1 | 1 | 1 | 1 | NA | -0.011 |
| SE | 104941 | ENSG00000257261 | NA | chr12 | + | 46455011 | 46455141 | 46387746 | 46387972 | 46470152 | 46470603 | 0 | 0 | 0 | 2 | 0 | 3 | 5 | 2 | 1 | 2 | 0 | 0 | 0 | 0 | 21 | 0 | 2 | 0 | 5 | 28 | 279 | 149 | 1 | 1 | 0 | 0 | 0 | 0.516 | 0 | NA | 0 | NA | 0 | 0.286 | 0.008 |
| SE | 104952 | ENSG00000101290 | CDS2 | chr20 | + | 5173522 | 5173659 | 5128149 | 5128750 | 5175182 | 5175279 | 90 | 41 | 50 | 80 | 80 | 0 | 0 | 0 | 0 | 0 | 83 | 70 | 109 | 80 | 220 | 0 | 0 | 1 | 0 | 0 | 286 | 149 | 1 | 1 | 1 | 1 | 1 | 1 | 1 | 1 | 1 | 0.983 | 1 | 1 | 0.003 |
| SE | 104953 | ENSG00000101290 | CDS2 | chr20 | + | 5175182 | 5175279 | 5126878 | 5127149 | 5182386 | 5182445 | 1 | 0 | 0 | 2 | 0 | 1 | 0 | 0 | 0 | 0 | 0 | 0 | 0 | 1 | 0 | 0 | 0 | 0 | 0 | 0 | 246 | 149 | 1 | 1 | 0.377 | NA | NA | 1 | NA | NA | NA | NA | 1 | NA | -0.311 |
| SE | 104954 | ENSG00000101290 | CDS2 | chr20 | + | 5176647 | 5176745 | 5175182 | 5175279 | 5182386 | 5182445 | 92 | 44 | 70 | 82 | 90 | 1 | 0 | 0 | 2 | 0 | 82 | 96 | 89 | 74 | 167 | 0 | 0 | 0 | 2 | 0 | 247 | 149 | 1 | 1 | 0.982 | 1 | 1 | 0.961 | 1 | 1 | 1 | 1 | 0.957 | 1 | -0.003 |
| SE | 104955 | ENSG00000101290 | CDS2 | chr20 | + | 5178816 | 5178956 | 5126878 | 5127149 | 5182386 | 5182445 | 118 | 48 | 104 | 88 | 353 | 1 | 0 | 0 | 0 | 0 | 32 | 58 | 73 | 96 | 151 | 0 | 0 | 0 | 0 | 0 | 289 | 149 | 1 | 1 | 0.984 | 1 | 1 | 1 | 1 | 1 | 1 | 1 | 1 | 1 | -0.003 |
| SE | 104956 | ENSG00000101290 | CDS2 | chr20 | + | 5178816 | 5178956 | 5176101 | 5176745 | 5182386 | 5182445 | 210 | 98 | 197 | 179 | 567 | 0 | 0 | 0 | 0 | 0 | 96 | 189 | 155 | 161 | 379 | 0 | 0 | 0 | 0 | 5 | 289 | 149 | 1 | 1 | 1 | 1 | 1 | 1 | 1 | 1 | 1 | 1 | 1 | 0.975 | 0.005 |
| SE | 104957 | ENSG00000101290 | CDS2 | chr20 | + | 5184857 | 5184945 | 5183060 | 5183143 | 5186686 | 5186774 | 122 | 51 | 136 | 117 | 307 | 0 | 0 | 0 | 0 | 1 | 126 | 123 | 109 | 124 | 231 | 0 | 2 | 0 | 0 | 0 | 237 | 149 | 1 | 1 | 1 | 1 | 1 | 1 | 0.995 | 1 | 0.975 | 1 | 1 | 1 | 0.004 |
| SE | 104958 | ENSG00000101290 | CDS2 | chr20 | + | 5185757 | 5185826 | 5184857 | 5184945 | 5186686 | 5186774 | 172 | 83 | 184 | 176 | 499 | 0 | 0 | 0 | 0 | 0 | 131 | 213 | 144 | 147 | 387 | 0 | 1 | 1 | 5 | 0 | 218 | 149 | 1 | 1 | 1 | 1 | 1 | 1 | 1 | 1 | 0.993 | 0.99 | 0.953 | 1 | 0.013 |
| SE | 104959 | ENSG00000101290 | CDS2 | chr20 | + | 5185757 | 5185826 | 5184857 | 5184945 | 5189066 | 5189186 | 23 | 19 | 50 | 39 | 137 | 0 | 0 | 0 | 0 | 0 | 7 | 37 | 38 | 37 | 96 | 0 | 0 | 1 | 0 | 0 | 218 | 149 | 1 | 1 | 1 | 1 | 1 | 1 | 1 | 1 | 1 | 0.963 | 1 | 1 | 0.007 |
| SE | 104960 | ENSG00000101290 | CDS2 | chr20 | + | 5186686 | 5186839 | 5184857 | 5184945 | 5189066 | 5189186 | 205 | 91 | 163 | 164 | 227 | 0 | 0 | 0 | 0 | 0 | 142 | 257 | 171 | 216 | 592 | 0 | 0 | 1 | 0 | 0 | 298 | 149 | 1 | 1 | 1 | 1 | 1 | 1 | 1 | 1 | 1 | 0.988 | 1 | 1 | 0.002 |
| SE | 104962 | ENSG00000101290 | CDS2 | chr20 | + | 5189066 | 5189186 | 5186686 | 5186839 | 5189734 | 5189838 | 393 | 211 | 296 | 325 | 458 | 0 | 0 | 0 | 0 | 0 | 471 | 512 | 331 | 434 | 1230 | 0 | 0 | 0 | 0 | 6 | 269 | 149 | 1 | 1 | 1 | 1 | 1 | 1 | 1 | 1 | 1 | 1 | 1 | 0.991 | 0.002 |
| SE | 104963 | ENSG00000123200 | ZC3H13 | chr13 | - | 45963841 | 45964042 | 45959482 | 45959646 | 45965279 | 45965429 | 323 | 204 | 244 | 293 | 372 | 0 | 0 | 0 | 1 | 1 | 0 | 125 | 78 | 2 | 0 | 0 | 0 | 0 | 0 | 0 | 298 | 149 | 1 | 1 | 1 | 1 | 1 | 0.993 | 0.995 | NA | 1 | 1 | 1 | NA | -0.002 |
| SE | 104964 | ENSG00000123200 | ZC3H13 | chr13 | - | 46003138 | 46003336 | 45988786 | 45989097 | 46010347 | 46010505 | 158 | 146 | 106 | 150 | 256 | 0 | 0 | 0 | 0 | 1 | 49 | 63 | 96 | 11 | 80 | 0 | 0 | 0 | 0 | 0 | 298 | 149 | 1 | 1 | 1 | 1 | 1 | 1 | 0.992 | 1 | 1 | 1 | 1 | 1 | -0.002 |
| SE | 104965 | ENSG00000123200 | ZC3H13 | chr13 | - | 46020448 | 46020557 | 46011414 | 46011554 | 46042163 | 46042275 | 391 | 316 | 284 | 332 | 592 | 3 | 3 | 5 | 0 | 0 | 179 | 190 | 252 | 82 | 110 | 0 | 2 | 0 | 0 | 0 | 258 | 149 | 1 | 1 | 0.987 | 0.984 | 0.97 | 1 | 1 | 1 | 0.982 | 1 | 1 | 1 | -0.008 |
| SE | 104966 | ENSG00000123200 | ZC3H13 | chr13 | - | 46020448 | 46020557 | 46011414 | 46011554 | 46052403 | 46052746 | 255 | 193 | 188 | 189 | 440 | 0 | 0 | 0 | 0 | 0 | 116 | 106 | 130 | 45 | 79 | 0 | 0 | 3 | 0 | 0 | 258 | 149 | 1 | 1 | 1 | 1 | 1 | 1 | 1 | 1 | 1 | 0.962 | 1 | 1 | 0.008 |
| SE | 104967 | ENSG00000123200 | ZC3H13 | chr13 | - | 46020448 | 46020557 | 46011414 | 46011554 | 46052697 | 46052759 | 255 | 193 | 187 | 191 | 440 | 0 | 0 | 0 | 0 | 0 | 116 | 106 | 130 | 45 | 79 | 0 | 2 | 0 | 0 | 0 | 258 | 149 | 1 | 1 | 1 | 1 | 1 | 1 | 1 | 1 | 0.968 | 1 | 1 | 1 | 0.006 |
| SE | 104968 | ENSG00000123200 | ZC3H13 | chr13 | - | 46042163 | 46042275 | 46011414 | 46011554 | 46052403 | 46052746 | 1 | 3 | 6 | 0 | 0 | 0 | 0 | 0 | 0 | 0 | 0 | 2 | 0 | 0 | 0 | 0 | 0 | 3 | 0 | 0 | 261 | 149 | 1 | 1 | 1 | 1 | 1 | NA | NA | NA | 1 | 0 | NA | NA | 0.5 |
| SE | 104969 | ENSG00000123200 | ZC3H13 | chr13 | - | 46042163 | 46042275 | 46020448 | 46020557 | 46044954 | 46045064 | 268 | 217 | 210 | 277 | 271 | 0 | 0 | 0 | 3 | 0 | 109 | 164 | 203 | 68 | 95 | 0 | 0 | 2 | 0 | 0 | 261 | 149 | 1 | 1 | 1 | 1 | 1 | 0.981 | 1 | 1 | 1 | 0.983 | 1 | 1 | 0 |
| SE | 104970 | ENSG00000123200 | ZC3H13 | chr13 | - | 46042163 | 46042275 | 46020448 | 46020557 | 46052403 | 46052746 | 84 | 78 | 56 | 111 | 122 | 0 | 0 | 1 | 0 | 0 | 45 | 42 | 70 | 28 | 20 | 0 | 0 | 0 | 0 | 0 | 261 | 149 | 1 | 1 | 1 | 1 | 0.97 | 1 | 1 | 1 | 1 | 1 | 1 | 1 | -0.006 |
| SE | 104971 | ENSG00000123200 | ZC3H13 | chr13 | - | 46051931 | 46051989 | 46020448 | 46020557 | 46052403 | 46052685 | 6 | 0 | 0 | 5 | 0 | 0 | 0 | 1 | 1 | 0 | 0 | 3 | 0 | 0 | 0 | 0 | 0 | 0 | 0 | 0 | 207 | 149 | 1 | 1 | 1 | NA | 0 | 1 | NA | NA | 1 | NA | NA | NA | -0.333 |
| SE | 104973 | ENSG00000123200 | ZC3H13 | chr13 | - | 46051931 | 46051989 | 46045390 | 46045513 | 46052403 | 46052685 | 12 | 4 | 5 | 14 | 3 | 4 | 2 | 5 | 4 | 7 | 0 | 0 | 0 | 0 | 0 | 0 | 5 | 3 | 0 | 0 | 207 | 149 | 1 | 1 | 0.683 | 0.59 | 0.419 | 0.716 | 0.236 | NA | 0 | 0 | NA | NA | 0.529 |
| SE | 104974 | ENSG00000123200 | ZC3H13 | chr13 | - | 46051931 | 46051989 | 46045390 | 46045513 | 46052697 | 46052759 | 4 | 4 | 5 | 5 | 3 | 7 | 3 | 5 | 3 | 6 | 0 | 0 | 0 | 0 | 0 | 0 | 1 | 2 | 0 | 0 | 207 | 149 | 1 | 1 | 0.291 | 0.49 | 0.419 | 0.545 | 0.265 | NA | 0 | 0 | NA | NA | 0.402 |
| SE | 104975 | ENSG00000123200 | ZC3H13 | chr13 | - | 46051931 | 46051989 | 46045390 | 46045516 | 46052403 | 46052685 | 16 | 13 | 5 | 18 | 4 | 28 | 14 | 28 | 23 | 28 | 0 | 0 | 0 | 0 | 0 | 0 | 10 | 22 | 6 | 2 | 207 | 149 | 1 | 1 | 0.291 | 0.401 | 0.114 | 0.36 | 0.093 | NA | 0 | 0 | 0 | 0 | 0.252 |
| SE | 104978 | ENSG00000111371 | SLC38A1 | chr12 | - | 46197719 | 46197817 | 46183062 | 46189071 | 46197918 | 46198060 | 1019 | 823 | 1118 | 1134 | 2403 | 0 | 0 | 0 | 0 | 0 | 225 | 1012 | 1294 | 450 | 54 | 0 | 0 | 1 | 0 | 0 | 247 | 149 | 1 | 1 | 1 | 1 | 1 | 1 | 1 | 1 | 1 | 0.999 | 1 | 1 | 0 |
| SE | 104980 | ENSG00000111371 | SLC38A1 | chr12 | - | 46201097 | 46201198 | 46198624 | 46198743 | 46204300 | 46204417 | 261 | 167 | 345 | 316 | 546 | 2 | 0 | 0 | 1 | 1 | 38 | 270 | 344 | 101 | 21 | 0 | 0 | 0 | 0 | 0 | 250 | 149 | 1 | 1 | 0.987 | 1 | 1 | 0.995 | 0.997 | 1 | 1 | 1 | 1 | 1 | -0.004 |
| SE | 104981 | ENSG00000111371 | SLC38A1 | chr12 | - | 46203009 | 46203089 | 46198624 | 46198743 | 46204300 | 46204417 | 414 | 259 | 465 | 379 | 582 | 2 | 0 | 0 | 1 | 1 | 34 | 251 | 335 | 137 | 5 | 0 | 0 | 0 | 0 | 0 | 229 | 149 | 1 | 1 | 0.993 | 1 | 1 | 0.996 | 0.997 | 1 | 1 | 1 | 1 | 1 | -0.003 |
| SE | 104982 | ENSG00000111371 | SLC38A1 | chr12 | - | 46203009 | 46203089 | 46201097 | 46201198 | 46204300 | 46204417 | 769 | 462 | 906 | 754 | 1137 | 1 | 0 | 0 | 0 | 0 | 27 | 534 | 641 | 232 | 14 | 0 | 0 | 0 | 0 | 0 | 229 | 149 | 1 | 1 | 0.998 | 1 | 1 | 1 | 1 | 1 | 1 | 1 | 1 | 1 | 0 |
| SE | 104983 | ENSG00000111371 | SLC38A1 | chr12 | - | 46207154 | 46207236 | 46206079 | 46206162 | 46207528 | 46207621 | 524 | 348 | 645 | 496 | 970 | 0 | 0 | 0 | 2 | 0 | 82 | 334 | 476 | 124 | 17 | 0 | 0 | 0 | 0 | 0 | 231 | 149 | 1 | 1 | 1 | 1 | 1 | 0.994 | 1 | 1 | 1 | 1 | 1 | 1 | -0.001 |
| SE | 104984 | ENSG00000111371 | SLC38A1 | chr12 | - | 46209053 | 46209127 | 46207528 | 46207621 | 46229152 | 46229268 | 1155 | 834 | 1290 | 1217 | 1346 | 3 | 0 | 1 | 2 | 0 | 292 | 1277 | 1427 | 524 | 68 | 0 | 5 | 1 | 0 | 0 | 223 | 149 | 1 | 1 | 0.996 | 1 | 0.999 | 0.998 | 1 | 1 | 0.994 | 0.999 | 1 | 1 | 0 |
| SE | 104985 | ENSG00000111371 | SLC38A1 | chr12 | - | 46239678 | 46239893 | 46229563 | 46229639 | 46243199 | 46243314 | 1710 | 1198 | 1842 | 1803 | 2512 | 0 | 0 | 2 | 0 | 4 | 457 | 1398 | 1574 | 629 | 356 | 0 | 1 | 2 | 0 | 0 | 298 | 149 | 1 | 1 | 1 | 1 | 0.998 | 1 | 0.997 | 1 | 0.999 | 0.997 | 1 | 1 | 0 |
| SE | 104986 | ENSG00000111371 | SLC38A1 | chr12 | - | 46239678 | 46239893 | 46229563 | 46229639 | 46268525 | 46268995 | 1033 | 703 | 1135 | 1163 | 1671 | 4 | 2 | 6 | 3 | 0 | 282 | 881 | 1129 | 415 | 207 | 0 | 0 | 0 | 2 | 0 | 298 | 149 | 1 | 1 | 0.992 | 0.994 | 0.99 | 0.995 | 1 | 1 | 1 | 1 | 0.99 | 1 | -0.004 |
| SE | 104987 | ENSG00000111371 | SLC38A1 | chr12 | - | 46239678 | 46239893 | 46229563 | 46229639 | 46268864 | 46269130 | 1033 | 702 | 1136 | 1162 | 1681 | 5 | 0 | 6 | 9 | 0 | 282 | 881 | 1127 | 415 | 207 | 0 | 0 | 1 | 0 | 0 | 298 | 149 | 1 | 1 | 0.99 | 1 | 0.99 | 0.985 | 1 | 1 | 1 | 0.998 | 1 | 1 | -0.007 |
| SE | 104988 | ENSG00000111371 | SLC38A1 | chr12 | - | 46243199 | 46243314 | 46229563 | 46229639 | 46268525 | 46269043 | 75 | 63 | 45 | 55 | 33 | 4 | 2 | 6 | 3 | 0 | 6 | 63 | 43 | 18 | 0 | 0 | 0 | 0 | 2 | 0 | 264 | 149 | 1 | 1 | 0.914 | 0.947 | 0.809 | 0.912 | 1 | 1 | 1 | 1 | 0.836 | NA | -0.043 |
| SE | 104990 | ENSG00000111371 | SLC38A1 | chr12 | - | 46243199 | 46243314 | 46239678 | 46239893 | 46268525 | 46269043 | 717 | 511 | 707 | 666 | 795 | 2 | 1 | 2 | 2 | 0 | 180 | 540 | 458 | 220 | 149 | 0 | 0 | 2 | 0 | 0 | 264 | 149 | 1 | 1 | 0.995 | 0.997 | 0.995 | 0.995 | 1 | 1 | 1 | 0.992 | 1 | 1 | -0.002 |
| SE | 104991 | ENSG00000111371 | SLC38A1 | chr12 | - | 46243199 | 46243314 | 46239678 | 46239893 | 46268864 | 46269119 | 827 | 623 | 904 | 800 | 890 | 2 | 0 | 3 | 1 | 10 | 182 | 632 | 539 | 251 | 159 | 0 | 0 | 0 | 0 | 0 | 264 | 149 | 1 | 1 | 0.996 | 1 | 0.994 | 0.998 | 0.98 | 1 | 1 | 1 | 1 | 1 | -0.006 |
| SE | 104994 | ENSG00000178449 | COX14 | chr12 | + | 50112831 | 50112903 | 50112081 | 50112301 | 50120035 | 50120451 | 0 | 1 | 0 | 0 | 0 | 62 | 13 | 36 | 51 | 26 | 0 | 0 | 0 | 0 | 0 | 45 | 58 | 35 | 44 | 95 | 221 | 149 | 1 | 1 | 0 | 0.049 | 0 | 0 | 0 | 0 | 0 | 0 | 0 | 0 | 0.01 |
| SE | 104996 | ENSG00000129315 | CCNT1 | chr12 | - | 48695758 | 48695829 | 48688457 | 48694436 | 48695998 | 48696162 | 88 | 52 | 80 | 108 | 55 | 0 | 0 | 2 | 0 | 0 | 32 | 73 | 92 | 64 | 14 | 0 | 0 | 0 | 0 | 2 | 220 | 149 | 1 | 1 | 1 | 1 | 0.964 | 1 | 1 | 1 | 1 | 1 | 1 | 0.826 | 0.028 |
| SE | 104997 | ENSG00000129315 | CCNT1 | chr12 | - | 48695758 | 48695829 | 48688457 | 48694436 | 48698137 | 48698183 | 98 | 50 | 75 | 143 | 77 | 0 | 0 | 0 | 0 | 0 | 46 | 125 | 189 | 115 | 0 | 0 | 1 | 7 | 0 | 0 | 220 | 149 | 1 | 1 | 1 | 1 | 1 | 1 | 1 | 1 | 0.988 | 0.948 | 1 | NA | 0.016 |
| SE | 105001 | ENSG00000129315 | CCNT1 | chr12 | - | 48699777 | 48699840 | 48698137 | 48698183 | 48705767 | 48705896 | 47 | 33 | 33 | 63 | 51 | 0 | 0 | 1 | 0 | 0 | 19 | 31 | 102 | 45 | 0 | 0 | 0 | 0 | 0 | 0 | 212 | 149 | 1 | 1 | 1 | 1 | 0.959 | 1 | 1 | 1 | 1 | 1 | 1 | NA | -0.008 |
| SE | 105002 | ENSG00000129315 | CCNT1 | chr12 | - | 48701012 | 48701073 | 48698137 | 48698183 | 48705767 | 48705896 | 7 | 8 | 27 | 15 | 5 | 0 | 0 | 1 | 0 | 0 | 14 | 25 | 23 | 10 | 12 | 0 | 0 | 0 | 0 | 0 | 210 | 149 | 1 | 1 | 1 | 1 | 0.95 | 1 | 1 | 1 | 1 | 1 | 1 | 1 | -0.01 |
| SE | 105003 | ENSG00000129315 | CCNT1 | chr12 | - | 48701012 | 48701073 | 48699777 | 48699840 | 48705767 | 48705896 | 156 | 101 | 149 | 176 | 192 | 0 | 0 | 0 | 0 | 1 | 30 | 186 | 238 | 75 | 40 | 0 | 0 | 1 | 0 | 0 | 210 | 149 | 1 | 1 | 1 | 1 | 1 | 1 | 0.993 | 1 | 1 | 0.994 | 1 | 1 | 0 |
| SE | 105004 | ENSG00000129315 | CCNT1 | chr12 | - | 48705767 | 48705896 | 48701012 | 48701073 | 48714442 | 48714524 | 197 | 105 | 177 | 206 | 201 | 1 | 0 | 1 | 2 | 1 | 64 | 206 | 278 | 79 | 43 | 0 | 0 | 6 | 0 | 0 | 278 | 149 | 1 | 1 | 0.991 | 1 | 0.99 | 0.982 | 0.991 | 1 | 1 | 0.961 | 1 | 1 | -0.001 |
| SE | 105015 | ENSG00000284770 | TBCE | chr1 | + | 235435744 | 235436608 | 235433350 | 235434280 | 235437321 | 235437474 | 47 | 39 | 48 | 72 | 89 | 0 | 0 | 1 | 0 | 1 | 76 | 61 | 55 | 32 | 42 | 0 | 0 | 0 | 0 | 0 | 298 | 149 | 1 | 1 | 1 | 1 | 0.96 | 1 | 0.978 | 1 | 1 | 1 | 1 | 1 | -0.012 |
| SE | 105018 | ENSG00000188223 | NA | chr19 | + | 35746418 | 35746523 | 35745834 | 35745991 | 35752175 | 35752251 | 67 | 7 | 54 | 43 | 41 | 0 | 0 | 0 | 1 | 0 | 46 | 38 | 30 | 48 | 62 | 0 | 0 | 0 | 0 | 0 | 254 | 149 | 1 | 1 | 1 | 1 | 1 | 0.962 | 1 | 1 | 1 | 1 | 1 | 1 | -0.008 |
| SE | 105044 | ENSG00000133704 | IPO8 | chr12 | - | 30631894 | 30632011 | 30628987 | 30630957 | 30634082 | 30634286 | 218 | 162 | 176 | 182 | 280 | 0 | 0 | 2 | 2 | 0 | 77 | 188 | 71 | 106 | 1 | 0 | 0 | 0 | 0 | 0 | 266 | 149 | 1 | 1 | 1 | 1 | 0.98 | 0.981 | 1 | 1 | 1 | 1 | 1 | 1 | -0.008 |
| SE | 105045 | ENSG00000133704 | IPO8 | chr12 | - | 30652191 | 30652289 | 30649136 | 30649232 | 30656683 | 30656750 | 145 | 121 | 139 | 173 | 184 | 0 | 0 | 0 | 0 | 0 | 57 | 145 | 69 | 100 | 21 | 0 | 3 | 0 | 0 | 0 | 247 | 149 | 1 | 1 | 1 | 1 | 1 | 1 | 1 | 1 | 0.967 | 1 | 1 | 1 | 0.007 |
| SE | 105046 | ENSG00000133704 | IPO8 | chr12 | - | 30652966 | 30653092 | 30652191 | 30652289 | 30656683 | 30656750 | 257 | 169 | 255 | 218 | 282 | 0 | 0 | 0 | 1 | 0 | 36 | 182 | 83 | 89 | 49 | 0 | 0 | 0 | 0 | 0 | 275 | 149 | 1 | 1 | 1 | 1 | 1 | 0.992 | 1 | 1 | 1 | 1 | 1 | 1 | -0.002 |
| SE | 105048 | ENSG00000133704 | IPO8 | chr12 | - | 30670961 | 30671096 | 30669182 | 30669282 | 30673989 | 30674074 | 163 | 132 | 138 | 207 | 284 | 0 | 1 | 0 | 0 | 0 | 12 | 174 | 64 | 107 | 31 | 0 | 0 | 0 | 0 | 0 | 284 | 149 | 1 | 1 | 1 | 0.986 | 1 | 1 | 1 | 1 | 1 | 1 | 1 | 1 | -0.003 |
| SE | 105049 | ENSG00000133704 | IPO8 | chr12 | - | 30670961 | 30671096 | 30669182 | 30669282 | 30676497 | 30676587 | 85 | 65 | 58 | 94 | 127 | 0 | 0 | 0 | 0 | 0 | 0 | 70 | 25 | 56 | 22 | 0 | 0 | 0 | 0 | 1 | 284 | 149 | 1 | 1 | 1 | 1 | 1 | 1 | 1 | NA | 1 | 1 | 1 | 0.92 | 0.02 |
| SE | 105050 | ENSG00000133704 | IPO8 | chr12 | - | 30676497 | 30676587 | 30674658 | 30674753 | 30680481 | 30680638 | 194 | 109 | 164 | 189 | 204 | 0 | 0 | 0 | 1 | 0 | 0 | 182 | 80 | 83 | 0 | 0 | 0 | 0 | 0 | 0 | 239 | 149 | 1 | 1 | 1 | 1 | 1 | 0.992 | 1 | NA | 1 | 1 | 1 | NA | -0.002 |
| SE | 105051 | ENSG00000133704 | IPO8 | chr12 | - | 30681658 | 30681817 | 30680371 | 30680638 | 30684300 | 30684457 | 189 | 143 | 184 | 193 | 339 | 0 | 0 | 1 | 0 | 0 | 14 | 194 | 94 | 87 | 6 | 0 | 0 | 3 | 0 | 0 | 298 | 149 | 1 | 1 | 1 | 1 | 0.989 | 1 | 1 | 1 | 1 | 0.94 | 1 | 1 | 0.01 |
| SE | 105052 | ENSG00000133704 | IPO8 | chr12 | - | 30684300 | 30684457 | 30681658 | 30681817 | 30690495 | 30690548 | 238 | 174 | 247 | 254 | 386 | 1 | 0 | 0 | 0 | 0 | 32 | 225 | 139 | 164 | 17 | 0 | 0 | 0 | 0 | 0 | 298 | 149 | 1 | 1 | 0.992 | 1 | 1 | 1 | 1 | 1 | 1 | 1 | 1 | 1 | -0.002 |
| SE | 105059 | ENSG00000119801 | YPEL5 | chr2 | + | 30148244 | 30148541 | 30147956 | 30148091 | 30155849 | 30155910 | 2 | 0 | 3 | 9 | 10 | 0 | 0 | 0 | 0 | 0 | 13 | 14 | 13 | 7 | 0 | 4 | 0 | 0 | 0 | 0 | 298 | 149 | 1 | 1 | 1 | NA | 1 | 1 | 1 | 0.619 | 1 | 1 | 1 | NA | 0.095 |
| SE | 105066 | ENSG00000119801 | YPEL5 | chr2 | + | 30156627 | 30156792 | 30147006 | 30147062 | 30158618 | 30158747 | 846 | 696 | 647 | 1005 | 713 | 2 | 1 | 0 | 2 | 1 | 778 | 1136 | 1053 | 847 | 652 | 0 | 3 | 5 | 0 | 0 | 298 | 149 | 1 | 1 | 0.995 | 0.997 | 1 | 0.996 | 0.997 | 1 | 0.995 | 0.991 | 1 | 1 | 0 |
| SE | 105067 | ENSG00000119801 | YPEL5 | chr2 | + | 30156627 | 30156792 | 30148244 | 30148541 | 30158618 | 30158747 | 573 | 463 | 430 | 678 | 662 | 0 | 0 | 0 | 0 | 2 | 551 | 704 | 683 | 554 | 366 | 0 | 0 | 0 | 0 | 0 | 298 | 149 | 1 | 1 | 1 | 1 | 1 | 1 | 0.994 | 1 | 1 | 1 | 1 | 1 | -0.001 |
| SE | 105087 | ENSG00000136026 | CKAP4 | chr12 | - | 106254077 | 106254219 | 106240321 | 106240349 | 106276414 | 106276549 | 2 | 0 | 1 | 0 | 0 | 0 | 1 | 0 | 0 | 0 | 2 | 0 | 0 | 10 | 1 | 0 | 0 | 0 | 0 | 0 | 291 | 149 | 1 | 1 | 1 | 0 | 1 | NA | NA | 1 | NA | NA | 1 | 1 | -0.333 |
| SE | 105089 | ENSG00000172687 | ZNF738 | chr19 | + | 21375237 | 21375364 | 21361765 | 21361858 | 21375868 | 21375964 | 53 | 22 | 33 | 49 | 45 | 0 | 0 | 1 | 0 | 0 | 13 | 34 | 38 | 31 | 34 | 0 | 0 | 0 | 0 | 0 | 276 | 149 | 1 | 1 | 1 | 1 | 0.947 | 1 | 1 | 1 | 1 | 1 | 1 | 1 | -0.011 |
| SE | 105090 | ENSG00000172687 | ZNF738 | chr19 | + | 21375868 | 21375964 | 21361765 | 21361858 | 21382865 | 21388582 | 5 | 5 | 6 | 11 | 6 | 0 | 1 | 0 | 0 | 0 | 3 | 3 | 0 | 0 | 0 | 0 | 0 | 0 | 0 | 0 | 245 | 149 | 1 | 1 | 1 | 0.753 | 1 | 1 | 1 | 1 | 1 | NA | NA | NA | -0.049 |
| SE | 105092 | ENSG00000112297 | CRYBG1 | chr6 | + | 106525132 | 106525180 | 106519130 | 106521453 | 106525267 | 106525386 | 651 | 555 | 502 | 566 | 1217 | 2 | 0 | 0 | 0 | 1 | 237 | 666 | 1282 | 336 | 61 | 0 | 2 | 0 | 0 | 0 | 197 | 149 | 1 | 1 | 0.996 | 1 | 1 | 1 | 0.999 | 1 | 0.996 | 1 | 1 | 1 | 0 |
| SE | 105094 | ENSG00000112297 | CRYBG1 | chr6 | + | 106539402 | 106539529 | 106530175 | 106530315 | 106541585 | 106541621 | 723 | 651 | 571 | 647 | 780 | 0 | 0 | 0 | 0 | 0 | 483 | 692 | 1174 | 660 | 102 | 0 | 0 | 0 | 1 | 0 | 276 | 149 | 1 | 1 | 1 | 1 | 1 | 1 | 1 | 1 | 1 | 1 | 0.997 | 1 | 0.001 |
| SE | 105095 | ENSG00000112297 | CRYBG1 | chr6 | + | 106543439 | 106543597 | 106541585 | 106541621 | 106544570 | 106544697 | 697 | 611 | 581 | 647 | 762 | 4 | 1 | 0 | 3 | 0 | 463 | 779 | 1293 | 598 | 88 | 0 | 2 | 10 | 0 | 0 | 298 | 149 | 1 | 1 | 0.989 | 0.997 | 1 | 0.991 | 1 | 1 | 0.995 | 0.985 | 1 | 1 | -0.001 |
| SE | 105096 | ENSG00000112297 | CRYBG1 | chr6 | + | 106544787 | 106544933 | 106544570 | 106544697 | 106551851 | 106551978 | 692 | 526 | 501 | 550 | 886 | 0 | 0 | 0 | 0 | 0 | 180 | 581 | 993 | 459 | 6 | 0 | 3 | 3 | 7 | 0 | 295 | 149 | 1 | 1 | 1 | 1 | 1 | 1 | 1 | 1 | 0.99 | 0.994 | 0.971 | 1 | 0.009 |
| SE | 105097 | ENSG00000112297 | CRYBG1 | chr6 | + | 106551851 | 106551978 | 106544787 | 106544933 | 106552183 | 106552216 | 534 | 410 | 370 | 472 | 872 | 2 | 0 | 0 | 0 | 0 | 112 | 433 | 808 | 346 | 4 | 0 | 0 | 0 | 0 | 0 | 276 | 149 | 1 | 1 | 0.993 | 1 | 1 | 1 | 1 | 1 | 1 | 1 | 1 | 1 | -0.001 |
| SE | 105098 | ENSG00000112297 | CRYBG1 | chr6 | + | 106552183 | 106552216 | 106551851 | 106551978 | 106553454 | 106553567 | 316 | 214 | 223 | 287 | 619 | 0 | 0 | 0 | 2 | 0 | 45 | 241 | 403 | 160 | 0 | 0 | 0 | 0 | 0 | 0 | 182 | 149 | 1 | 1 | 1 | 1 | 1 | 0.992 | 1 | 1 | 1 | 1 | 1 | NA | -0.002 |
| SE | 105099 | ENSG00000112297 | CRYBG1 | chr6 | + | 106555767 | 106555897 | 106553454 | 106553567 | 106558485 | 106558625 | 501 | 331 | 389 | 426 | 839 | 0 | 0 | 0 | 4 | 0 | 70 | 343 | 341 | 214 | 0 | 0 | 0 | 7 | 0 | 0 | 279 | 149 | 1 | 1 | 1 | 1 | 1 | 0.983 | 1 | 1 | 1 | 0.963 | 1 | NA | 0.006 |
| SE | 105100 | ENSG00000112297 | CRYBG1 | chr6 | + | 106555767 | 106555897 | 106553454 | 106553567 | 106560802 | 106560926 | 197 | 129 | 160 | 160 | 305 | 0 | 0 | 0 | 0 | 0 | 17 | 119 | 158 | 106 | 0 | 0 | 5 | 0 | 0 | 0 | 279 | 149 | 1 | 1 | 1 | 1 | 1 | 1 | 1 | 1 | 0.927 | 1 | 1 | NA | 0.018 |
| SE | 105101 | ENSG00000112297 | CRYBG1 | chr6 | + | 106558485 | 106558625 | 106553454 | 106553567 | 106560802 | 106560926 | 404 | 316 | 262 | 323 | 444 | 0 | 0 | 0 | 0 | 0 | 114 | 331 | 455 | 228 | 16 | 0 | 5 | 0 | 0 | 0 | 289 | 149 | 1 | 1 | 1 | 1 | 1 | 1 | 1 | 1 | 0.972 | 1 | 1 | 1 | 0.006 |
| SE | 105102 | ENSG00000112297 | CRYBG1 | chr6 | + | 106558485 | 106558625 | 106555767 | 106555897 | 106560802 | 106560926 | 709 | 523 | 498 | 589 | 979 | 0 | 0 | 5 | 0 | 0 | 167 | 557 | 636 | 343 | 16 | 0 | 0 | 5 | 4 | 0 | 289 | 149 | 1 | 1 | 1 | 1 | 0.981 | 1 | 1 | 1 | 1 | 0.985 | 0.978 | 1 | 0.004 |
| SE | 105103 | ENSG00000112297 | CRYBG1 | chr6 | + | 106560802 | 106560926 | 106558485 | 106558625 | 106561341 | 106561500 | 835 | 642 | 541 | 732 | 955 | 1 | 0 | 2 | 2 | 1 | 265 | 796 | 988 | 507 | 36 | 0 | 0 | 6 | 0 | 0 | 273 | 149 | 1 | 1 | 0.998 | 1 | 0.993 | 0.995 | 0.998 | 1 | 1 | 0.989 | 1 | 1 | -0.001 |
| SE | 105105 | ENSG00000112297 | CRYBG1 | chr6 | + | 106563763 | 106563926 | 106561341 | 106561500 | 106568471 | 106572017 | 491 | 491 | 410 | 495 | 640 | 1 | 0 | 3 | 0 | 0 | 182 | 479 | 832 | 488 | 105 | 0 | 0 | 0 | 0 | 0 | 298 | 149 | 1 | 1 | 0.996 | 1 | 0.986 | 1 | 1 | 1 | 1 | 1 | 1 | 1 | -0.004 |
| SE | 105106 | ENSG00000182199 | SHMT2 | chr12 | + | 57230802 | 57231000 | 57229572 | 57229811 | 57231712 | 57231913 | 112 | 49 | 81 | 100 | 37 | 0 | 0 | 1 | 0 | 0 | 178 | 142 | 85 | 104 | 181 | 0 | 0 | 0 | 0 | 0 | 298 | 149 | 1 | 1 | 1 | 1 | 0.976 | 1 | 1 | 1 | 1 | 1 | 1 | 1 | -0.005 |
| SE | 105114 | ENSG00000278311 | GGNBP2 | chr17 | + | 36581343 | 36581538 | 36579244 | 36579419 | 36585299 | 36585450 | 713 | 737 | 630 | 637 | 1079 | 0 | 0 | 1 | 3 | 0 | 335 | 521 | 1025 | 456 | 85 | 0 | 0 | 8 | 2 | 0 | 298 | 149 | 1 | 1 | 1 | 1 | 0.997 | 0.991 | 1 | 1 | 1 | 0.985 | 0.991 | 1 | 0.002 |
| SE | 105115 | ENSG00000278311 | GGNBP2 | chr17 | + | 36585839 | 36585965 | 36585299 | 36585450 | 36586049 | 36586153 | 836 | 840 | 825 | 797 | 1190 | 0 | 0 | 1 | 0 | 4 | 403 | 722 | 1250 | 534 | 59 | 0 | 0 | 0 | 8 | 0 | 275 | 149 | 1 | 1 | 1 | 1 | 0.998 | 1 | 0.994 | 1 | 1 | 1 | 0.973 | 1 | 0.004 |
| SE | 105116 | ENSG00000139117 | CPNE8 | chr12 | - | 38839915 | 38839955 | 38829378 | 38829455 | 38848558 | 38848662 | 100 | 120 | 58 | 75 | 69 | 0 | 0 | 0 | 1 | 0 | 43 | 81 | 65 | 32 | 25 | 0 | 0 | 0 | 0 | 0 | 189 | 149 | 1 | 1 | 1 | 1 | 1 | 0.983 | 1 | 1 | 1 | 1 | 1 | 1 | -0.003 |
| SE | 105119 | ENSG00000161970 | RPL26 | chr17 | - | 8379795 | 8379936 | 8377515 | 8377692 | 8382142 | 8382309 | 38452 | 23782 | 37978 | 27183 | 63988 | 21 | 3 | 12 | 16 | 18 | 42308 | 24391 | 25170 | 24195 | 37394 | 31 | 25 | 15 | 3 | 34 | 290 | 149 | 1 | 1 | 0.999 | 1 | 0.999 | 0.999 | 0.999 | 0.999 | 0.998 | 0.999 | 1 | 0.998 | 0 |
| SE | 105120 | ENSG00000161970 | RPL26 | chr17 | - | 8380586 | 8380703 | 8379795 | 8379936 | 8382142 | 8382315 | 9 | 4 | 8 | 8 | 12 | 22486 | 13871 | 21153 | 15813 | 33243 | 0 | 38 | 5 | 24 | 1 | 22301 | 12948 | 13039 | 12691 | 20071 | 266 | 149 | 1 | 1 | 0 | 0 | 0 | 0 | 0 | 0 | 0.002 | 0 | 0.001 | 0 | -0.001 |
| SE | 105121 | ENSG00000161970 | RPL26 | chr17 | - | 8381997 | 8382090 | 8379795 | 8379936 | 8383156 | 8383192 | 2 | 3 | 1 | 5 | 0 | 0 | 0 | 0 | 1 | 0 | 0 | 0 | 0 | 5 | 0 | 0 | 0 | 0 | 0 | 0 | 242 | 149 | 1 | 1 | 1 | 1 | 1 | 0.755 | NA | NA | NA | NA | 1 | NA | -0.061 |
| SE | 105122 | ENSG00000161970 | RPL26 | chr17 | - | 8382142 | 8382315 | 8379144 | 8379936 | 8383156 | 8383193 | 36479 | 21274 | 34999 | 26335 | 42418 | 0 | 0 | 0 | 1 | 0 | 31321 | 20417 | 19403 | 18431 | 29396 | 0 | 0 | 0 | 0 | 0 | 298 | 149 | 1 | 1 | 1 | 1 | 1 | 1 | 1 | 1 | 1 | 1 | 1 | 1 | 0 |
| SE | 105123 | ENSG00000161970 | RPL26 | chr17 | - | 8382142 | 8382338 | 8379795 | 8379936 | 8383156 | 8383213 | 22487 | 13872 | 21150 | 15812 | 33243 | 0 | 0 | 0 | 1 | 0 | 22301 | 12948 | 13040 | 12691 | 20074 | 0 | 0 | 0 | 0 | 0 | 298 | 149 | 1 | 1 | 1 | 1 | 1 | 1 | 1 | 1 | 1 | 1 | 1 | 1 | 0 |
| SE | 105126 | ENSG00000243335 | KCTD7 | chr7 | + | 66639127 | 66639228 | 66638855 | 66638929 | 66643534 | 66643611 | 0 | 0 | 2 | 1 | 2 | 1 | 0 | 1 | 0 | 0 | 0 | 2 | 0 | 0 | 0 | 0 | 0 | 0 | 0 | 0 | 250 | 149 | 1 | 1 | 0 | NA | 0.544 | 1 | 1 | NA | 1 | NA | NA | NA | -0.364 |
| SE | 105132 | ENSG00000120253 | NUP43 | chr6 | - | 149736470 | 149736622 | 149731370 | 149731735 | 149742389 | 149742570 | 97 | 53 | 107 | 50 | 201 | 0 | 0 | 1 | 0 | 0 | 34 | 38 | 26 | 32 | 34 | 0 | 0 | 0 | 0 | 0 | 298 | 149 | 1 | 1 | 1 | 1 | 0.982 | 1 | 1 | 1 | 1 | 1 | 1 | 1 | -0.004 |
| SE | 105134 | ENSG00000120253 | NUP43 | chr6 | - | 149742389 | 149742570 | 149738642 | 149738778 | 149743637 | 149743715 | 228 | 234 | 253 | 216 | 415 | 0 | 4 | 0 | 0 | 3 | 140 | 231 | 88 | 96 | 90 | 0 | 0 | 0 | 0 | 0 | 298 | 149 | 1 | 1 | 1 | 0.967 | 1 | 1 | 0.986 | 1 | 1 | 1 | 1 | 1 | -0.009 |
| SE | 105135 | ENSG00000120253 | NUP43 | chr6 | - | 149743637 | 149743715 | 149742389 | 149742570 | 149745939 | 149746062 | 227 | 253 | 232 | 238 | 386 | 1 | 0 | 0 | 0 | 0 | 133 | 203 | 72 | 94 | 70 | 0 | 0 | 0 | 0 | 0 | 227 | 149 | 1 | 1 | 0.993 | 1 | 1 | 1 | 1 | 1 | 1 | 1 | 1 | 1 | -0.001 |
| SE | 105136 | ENSG00000161328 | LRRC56 | chr11 | + | 551650 | 551827 | 551130 | 551302 | 551902 | 551967 | 0 | 0 | 2 | 3 | 0 | 0 | 0 | 1 | 0 | 0 | 0 | 11 | 0 | 0 | 0 | 0 | 0 | 0 | 0 | 0 | 298 | 149 | 1 | 1 | NA | NA | 0.5 | 1 | NA | NA | 1 | NA | NA | NA | -0.25 |
| SE | 105140 | ENSG00000247137 | NA | chr11 | - | 83192427 | 83192541 | 83191062 | 83191168 | 83192735 | 83192792 | 0 | 1 | 1 | 0 | 0 | 5 | 0 | 0 | 1 | 2 | 0 | 4 | 0 | 0 | 7 | 0 | 10 | 3 | 0 | 8 | 263 | 149 | 1 | 1 | 0 | 1 | 1 | 0 | 0 | NA | 0.185 | 0 | NA | 0.331 | 0.228 |
| SE | 105143 | ENSG00000242516 | LINC00960 | chr3 | + | 75675967 | 75676128 | 75675763 | 75675880 | 75678832 | 75679314 | 0 | 0 | 0 | 2 | 4 | 0 | 0 | 0 | 0 | 0 | 4 | 0 | 0 | 0 | 0 | 0 | 0 | 0 | 0 | 8 | 298 | 149 | 1 | 1 | NA | NA | NA | 1 | 1 | 1 | NA | NA | NA | 0 | 0.5 |
| SE | 105146 | ENSG00000135407 | AVIL | chr12 | - | 57807330 | 57807727 | 57806359 | 57806539 | 57808193 | 57808294 | 4 | 5 | 3 | 12 | 1 | 0 | 1 | 0 | 0 | 0 | 0 | 1 | 2 | 7 | 0 | 0 | 0 | 0 | 0 | 0 | 298 | 149 | 1 | 1 | 1 | 0.714 | 1 | 1 | 1 | NA | 1 | 1 | 1 | NA | -0.057 |
| SE | 105149 | ENSG00000170946 | DNAJC24 | chr11 | + | 31374102 | 31374205 | 31370715 | 31370859 | 31426286 | 31427978 | 0 | 0 | 1 | 0 | 2 | 4 | 1 | 4 | 4 | 6 | 0 | 0 | 0 | 0 | 0 | 5 | 0 | 4 | 1 | 0 | 252 | 149 | 1 | 1 | 0 | 0 | 0.129 | 0 | 0.165 | 0 | NA | 0 | 0 | NA | 0.059 |
| SE | 105150 | ENSG00000170946 | DNAJC24 | chr11 | + | 31396282 | 31396394 | 31370715 | 31370859 | 31414810 | 31414906 | 0 | 1 | 1 | 0 | 0 | 57 | 38 | 61 | 54 | 25 | 0 | 0 | 0 | 0 | 0 | 0 | 10 | 15 | 10 | 0 | 261 | 149 | 1 | 1 | 0 | 0.015 | 0.009 | 0 | 0 | NA | 0 | 0 | 0 | NA | 0.005 |
| SE | 105151 | ENSG00000170946 | DNAJC24 | chr11 | + | 31407455 | 31407543 | 31370715 | 31370859 | 31408109 | 31408175 | 1 | 4 | 0 | 1 | 0 | 2 | 0 | 0 | 0 | 1 | 0 | 4 | 0 | 0 | 0 | 0 | 2 | 0 | 1 | 0 | 237 | 149 | 1 | 1 | 0.239 | 1 | NA | 1 | 0 | NA | 0.557 | NA | 0 | NA | 0.281 |
| SE | 105152 | ENSG00000170946 | DNAJC24 | chr11 | + | 31408109 | 31408330 | 31370806 | 31370859 | 31414810 | 31414949 | 5 | 0 | 2 | 5 | 10 | 57 | 38 | 61 | 54 | 25 | 0 | 6 | 0 | 1 | 0 | 0 | 10 | 15 | 10 | 0 | 298 | 149 | 1 | 1 | 0.042 | 0 | 0.016 | 0.044 | 0.167 | NA | 0.231 | 0 | 0.048 | NA | -0.039 |
| SE | 105153 | ENSG00000170946 | DNAJC24 | chr11 | + | 31414810 | 31414949 | 31370715 | 31370859 | 31421971 | 31422132 | 56 | 37 | 63 | 55 | 25 | 0 | 0 | 0 | 0 | 2 | 0 | 10 | 15 | 10 | 0 | 0 | 0 | 0 | 0 | 0 | 288 | 149 | 1 | 1 | 1 | 1 | 1 | 1 | 0.866 | NA | 1 | 1 | 1 | NA | -0.027 |
| SE | 105154 | ENSG00000170946 | DNAJC24 | chr11 | + | 31414810 | 31414949 | 31370715 | 31370859 | 31426286 | 31426355 | 89 | 69 | 100 | 97 | 75 | 4 | 1 | 4 | 4 | 6 | 0 | 31 | 21 | 15 | 0 | 5 | 0 | 4 | 1 | 0 | 288 | 149 | 1 | 1 | 0.92 | 0.973 | 0.928 | 0.926 | 0.866 | 0 | 1 | 0.731 | 0.886 | NA | 0.268 |
| SE | 105155 | ENSG00000170946 | DNAJC24 | chr11 | + | 31421971 | 31422132 | 31370715 | 31370859 | 31426286 | 31427978 | 7 | 6 | 4 | 9 | 6 | 4 | 1 | 4 | 4 | 6 | 0 | 0 | 3 | 0 | 0 | 5 | 0 | 4 | 1 | 0 | 298 | 149 | 1 | 1 | 0.467 | 0.75 | 0.333 | 0.529 | 0.333 | 0 | NA | 0.273 | 0 | NA | 0.391 |
| SE | 105156 | ENSG00000170946 | DNAJC24 | chr11 | + | 31421971 | 31422132 | 31414810 | 31414949 | 31426286 | 31427978 | 7 | 8 | 6 | 10 | 4 | 33 | 34 | 39 | 43 | 50 | 0 | 0 | 3 | 0 | 0 | 0 | 21 | 6 | 5 | 0 | 298 | 149 | 1 | 1 | 0.096 | 0.105 | 0.071 | 0.104 | 0.038 | NA | 0 | 0.2 | 0 | NA | 0.016 |
| SE | 105160 | ENSG00000174718 | RESF1 | chr12 | + | 31970188 | 31970356 | 31959414 | 31959491 | 31980877 | 31982979 | 1263 | 1061 | 1083 | 1144 | 3053 | 0 | 4 | 1 | 3 | 2 | 148 | 422 | 313 | 190 | 128 | 0 | 0 | 1 | 0 | 0 | 298 | 149 | 1 | 1 | 1 | 0.993 | 0.998 | 0.995 | 0.999 | 1 | 1 | 0.994 | 1 | 1 | -0.002 |
| SE | 105163 | ENSG00000174718 | RESF1 | chr12 | + | 31970188 | 31970356 | 31960776 | 31960871 | 31992377 | 31992941 | 1039 | 779 | 826 | 828 | 1752 | 2 | 1 | 0 | 1 | 0 | 70 | 264 | 199 | 105 | 80 | 0 | 0 | 0 | 0 | 0 | 298 | 149 | 1 | 1 | 0.996 | 0.997 | 1 | 0.998 | 1 | 1 | 1 | 1 | 1 | 1 | -0.002 |
| SE | 105165 | ENSG00000174718 | RESF1 | chr12 | + | 31980877 | 31985957 | 31970188 | 31970356 | 31992377 | 31993107 | 1252 | 1059 | 1082 | 1140 | 3060 | 7 | 0 | 6 | 4 | 13 | 148 | 425 | 306 | 187 | 117 | 1 | 0 | 5 | 1 | 0 | 298 | 149 | 1 | 1 | 0.989 | 1 | 0.989 | 0.993 | 0.992 | 0.987 | 1 | 0.968 | 0.989 | 1 | 0.004 |
| SE | 105166 | ENSG00000174718 | RESF1 | chr12 | + | 31987238 | 31987322 | 31960776 | 31960871 | 31992377 | 31992941 | 460 | 363 | 406 | 445 | 750 | 2 | 1 | 0 | 1 | 0 | 18 | 97 | 56 | 12 | 5 | 0 | 0 | 0 | 0 | 0 | 233 | 149 | 1 | 1 | 0.993 | 0.996 | 1 | 0.996 | 1 | 1 | 1 | 1 | 1 | 1 | -0.003 |
| SE | 105174 | ENSG00000117151 | CTBS | chr1 | - | 84563734 | 84563832 | 84563256 | 84563418 | 84565840 | 84566012 | 213 | 208 | 228 | 143 | 318 | 0 | 0 | 3 | 0 | 0 | 136 | 191 | 189 | 138 | 46 | 0 | 0 | 0 | 0 | 0 | 247 | 149 | 1 | 1 | 1 | 1 | 0.979 | 1 | 1 | 1 | 1 | 1 | 1 | 1 | -0.004 |
| SE | 105176 | ENSG00000117151 | CTBS | chr1 | - | 84570581 | 84570720 | 84565840 | 84566012 | 84574238 | 84574440 | 84 | 62 | 96 | 127 | 155 | 0 | 1 | 1 | 0 | 0 | 111 | 98 | 120 | 69 | 41 | 0 | 0 | 0 | 0 | 0 | 288 | 149 | 1 | 1 | 1 | 0.97 | 0.98 | 1 | 1 | 1 | 1 | 1 | 1 | 1 | -0.01 |
| SE | 105181 | ENSG00000177990 | DPY19L2 | chr12 | - | 63590998 | 63591127 | 63582405 | 63582525 | 63594086 | 63594133 | 0 | 4 | 5 | 6 | 0 | 0 | 0 | 1 | 0 | 0 | 3 | 1 | 2 | 3 | 28 | 0 | 0 | 0 | 0 | 0 | 278 | 149 | 1 | 1 | NA | 1 | 0.728 | 1 | NA | 1 | 1 | 1 | 1 | 1 | -0.091 |
| SE | 105183 | ENSG00000177990 | DPY19L2 | chr12 | - | 63594086 | 63594133 | 63582405 | 63582525 | 63595965 | 63596037 | 0 | 1 | 3 | 1 | 9 | 0 | 0 | 0 | 0 | 0 | 0 | 0 | 0 | 0 | 4 | 0 | 0 | 0 | 2 | 0 | 196 | 149 | 1 | 1 | NA | 1 | 1 | 1 | 1 | NA | NA | NA | 0 | 1 | 0.5 |
| SE | 105184 | ENSG00000177990 | DPY19L2 | chr12 | - | 63608615 | 63608675 | 63600305 | 63600386 | 63617303 | 63617390 | 7 | 10 | 4 | 15 | 15 | 0 | 0 | 1 | 0 | 0 | 8 | 1 | 1 | 5 | 22 | 0 | 0 | 0 | 0 | 0 | 209 | 149 | 1 | 1 | 1 | 1 | 0.74 | 1 | 1 | 1 | 1 | 1 | 1 | 1 | -0.052 |
| SE | 105185 | ENSG00000177990 | DPY19L2 | chr12 | - | 63618150 | 63618228 | 63617303 | 63617390 | 63621237 | 63621337 | 9 | 22 | 15 | 16 | 5 | 0 | 0 | 0 | 0 | 0 | 32 | 11 | 2 | 7 | 44 | 0 | 1 | 0 | 0 | 0 | 227 | 149 | 1 | 1 | 1 | 1 | 1 | 1 | 1 | 1 | 0.878 | 1 | 1 | 1 | 0.024 |
| SE | 105190 | ENSG00000139624 | CERS5 | chr12 | - | 50132961 | 50133064 | 50129791 | 50130694 | 50134545 | 50134702 | 0 | 0 | 1 | 0 | 0 | 16 | 16 | 16 | 27 | 20 | 0 | 1 | 0 | 0 | 0 | 24 | 31 | 19 | 15 | 51 | 252 | 149 | 1 | 1 | 0 | 0 | 0.036 | 0 | 0 | 0 | 0.019 | 0 | 0 | 0 | 0.003 |
| SE | 105196 | ENSG00000139624 | CERS5 | chr12 | - | 50135940 | 50136069 | 50135439 | 50135838 | 50137727 | 50137793 | 181 | 106 | 136 | 158 | 196 | 0 | 0 | 0 | 0 | 0 | 221 | 214 | 111 | 189 | 190 | 0 | 0 | 6 | 0 | 0 | 278 | 149 | 1 | 1 | 1 | 1 | 1 | 1 | 1 | 1 | 1 | 0.908 | 1 | 1 | 0.018 |
| SE | 105197 | ENSG00000139624 | CERS5 | chr12 | - | 50137727 | 50137820 | 50135940 | 50136069 | 50138566 | 50138617 | 120 | 60 | 106 | 103 | 129 | 1 | 0 | 0 | 2 | 0 | 131 | 124 | 78 | 131 | 153 | 0 | 0 | 0 | 0 | 0 | 242 | 149 | 1 | 1 | 0.987 | 1 | 1 | 0.969 | 1 | 1 | 1 | 1 | 1 | 1 | -0.009 |
| SE | 105198 | ENSG00000139624 | CERS5 | chr12 | - | 50138566 | 50138617 | 50135940 | 50136069 | 50142052 | 50142110 | 35 | 11 | 29 | 34 | 35 | 0 | 1 | 2 | 0 | 1 | 49 | 36 | 26 | 33 | 16 | 0 | 0 | 5 | 3 | 11 | 200 | 149 | 1 | 1 | 1 | 0.891 | 0.915 | 1 | 0.963 | 1 | 1 | 0.795 | 0.891 | 0.52 | 0.113 |
| SE | 105203 | ENSG00000139624 | CERS5 | chr12 | - | 50148471 | 50148588 | 50143951 | 50144057 | 50167100 | 50167359 | 0 | 0 | 0 | 0 | 0 | 33 | 18 | 25 | 17 | 18 | 0 | 0 | 0 | 4 | 0 | 21 | 35 | 28 | 37 | 18 | 266 | 149 | 1 | 1 | 0 | 0 | 0 | 0 | 0 | 0 | 0 | 0 | 0.057 | 0 | -0.011 |
| SE | 105206 | ENSG00000139624 | CERS5 | chr12 | - | 50165821 | 50165943 | 50143951 | 50144057 | 50167100 | 50167326 | 0 | 0 | 2 | 0 | 0 | 33 | 18 | 25 | 17 | 18 | 0 | 0 | 0 | 0 | 0 | 21 | 35 | 28 | 37 | 18 | 271 | 149 | 1 | 1 | 0 | 0 | 0.042 | 0 | 0 | 0 | 0 | 0 | 0 | 0 | 0.008 |
| SE | 105210 | ENSG00000118596 | SLC16A7 | chr12 | + | 59704771 | 59705018 | 59655151 | 59655250 | 59771218 | 59771362 | 102 | 68 | 78 | 102 | 122 | 1 | 0 | 0 | 0 | 2 | 20 | 71 | 48 | 16 | 13 | 0 | 0 | 0 | 0 | 0 | 298 | 149 | 1 | 1 | 0.981 | 1 | 1 | 1 | 0.968 | 1 | 1 | 1 | 1 | 1 | -0.01 |
| SE | 105211 | ENSG00000118596 | SLC16A7 | chr12 | + | 59704791 | 59705018 | 59655151 | 59655250 | 59771218 | 59771362 | 17 | 16 | 16 | 34 | 13 | 1 | 0 | 0 | 0 | 2 | 0 | 19 | 11 | 6 | 0 | 0 | 0 | 0 | 0 | 0 | 298 | 149 | 1 | 1 | 0.895 | 1 | 1 | 1 | 0.765 | NA | 1 | 1 | 1 | NA | -0.068 |
| SE | 105214 | ENSG00000118596 | SLC16A7 | chr12 | + | 59735929 | 59735988 | 59704771 | 59705018 | 59761124 | 59761146 | 0 | 0 | 0 | 0 | 3 | 2 | 1 | 1 | 1 | 0 | 0 | 0 | 0 | 1 | 0 | 0 | 0 | 0 | 0 | 0 | 208 | 149 | 1 | 1 | 0 | 0 | 0 | 0 | 1 | NA | NA | NA | 1 | NA | -0.8 |
| SE | 105215 | ENSG00000118596 | SLC16A7 | chr12 | + | 59735929 | 59735988 | 59704771 | 59705018 | 59771218 | 59771362 | 0 | 0 | 0 | 0 | 3 | 17 | 16 | 16 | 34 | 13 | 0 | 0 | 0 | 1 | 0 | 0 | 19 | 11 | 6 | 0 | 208 | 149 | 1 | 1 | 0 | 0 | 0 | 0 | 0.142 | NA | 0 | 0 | 0.107 | NA | -0.007 |
| SE | 105216 | ENSG00000118596 | SLC16A7 | chr12 | + | 59761124 | 59761202 | 59704771 | 59705018 | 59771218 | 59771362 | 2 | 0 | 2 | 2 | 0 | 17 | 16 | 16 | 34 | 13 | 0 | 3 | 0 | 0 | 0 | 0 | 19 | 11 | 6 | 0 | 227 | 149 | 1 | 1 | 0.072 | 0 | 0.076 | 0.037 | 0 | NA | 0.094 | 0 | 0 | NA | 0.006 |
| SE | 105217 | ENSG00000118596 | SLC16A7 | chr12 | + | 59761124 | 59761202 | 59735929 | 59735988 | 59771218 | 59771362 | 0 | 0 | 1 | 1 | 0 | 1 | 0 | 0 | 0 | 0 | 0 | 3 | 0 | 0 | 0 | 0 | 0 | 0 | 0 | 0 | 227 | 149 | 1 | 1 | 0 | NA | 1 | 1 | NA | NA | 1 | NA | NA | NA | -0.333 |
| SE | 105218 | ENSG00000118596 | SLC16A7 | chr12 | + | 59768125 | 59768323 | 59761124 | 59761202 | 59771218 | 59771362 | 0 | 1 | 1 | 0 | 0 | 0 | 0 | 3 | 2 | 0 | 0 | 0 | 0 | 0 | 0 | 0 | 3 | 0 | 0 | 0 | 298 | 149 | 1 | 1 | NA | 1 | 0.143 | 0 | NA | NA | 0 | NA | NA | NA | 0.381 |
| SE | 105224 | ENSG00000102967 | DHODH | chr16 | + | 72021123 | 72021311 | 72017023 | 72017106 | 72023164 | 72023318 | 12 | 5 | 6 | 11 | 15 | 0 | 0 | 1 | 0 | 0 | 12 | 22 | 7 | 8 | 0 | 0 | 0 | 0 | 2 | 0 | 298 | 149 | 1 | 1 | 1 | 1 | 0.75 | 1 | 1 | 1 | 1 | 1 | 0.667 | NA | 0.033 |
| SE | 105225 | ENSG00000102967 | DHODH | chr16 | + | 72022361 | 72022475 | 72017023 | 72017106 | 72023164 | 72023318 | 5 | 8 | 17 | 8 | 8 | 0 | 0 | 1 | 0 | 0 | 24 | 3 | 3 | 8 | 18 | 0 | 0 | 0 | 2 | 0 | 263 | 149 | 1 | 1 | 1 | 1 | 0.906 | 1 | 1 | 1 | 1 | 1 | 0.694 | 1 | 0.042 |
| SE | 105226 | ENSG00000102967 | DHODH | chr16 | + | 72022361 | 72022475 | 72021123 | 72021311 | 72023164 | 72023318 | 15 | 6 | 28 | 24 | 14 | 0 | 0 | 0 | 3 | 0 | 11 | 20 | 6 | 15 | 43 | 0 | 0 | 2 | 0 | 0 | 263 | 149 | 1 | 1 | 1 | 1 | 1 | 0.819 | 1 | 1 | 1 | 0.63 | 1 | 1 | 0.038 |
| SE | 105229 | ENSG00000076067 | RBMS2 | chr12 | + | 56527864 | 56527963 | 56521994 | 56522089 | 56562416 | 56562583 | 0 | 0 | 0 | 0 | 0 | 9 | 8 | 6 | 4 | 14 | 0 | 0 | 0 | 3 | 0 | 30 | 1 | 13 | 14 | 24 | 248 | 149 | 1 | 1 | 0 | 0 | 0 | 0 | 0 | 0 | 0 | 0 | 0.114 | 0 | -0.023 |
| SE | 105235 | ENSG00000076067 | RBMS2 | chr12 | + | 56581183 | 56581263 | 56571697 | 56571855 | 56581398 | 56581508 | 45 | 40 | 44 | 38 | 34 | 0 | 0 | 0 | 4 | 0 | 131 | 25 | 21 | 102 | 173 | 0 | 0 | 0 | 1 | 0 | 229 | 149 | 1 | 1 | 1 | 1 | 1 | 0.861 | 1 | 1 | 1 | 1 | 0.985 | 1 | -0.025 |
| SE | 105237 | ENSG00000076067 | RBMS2 | chr12 | + | 56588931 | 56589018 | 56588293 | 56588374 | 56589139 | 56589503 | 20 | 10 | 3 | 22 | 19 | 3 | 0 | 0 | 0 | 2 | 38 | 0 | 11 | 12 | 58 | 0 | 0 | 0 | 6 | 0 | 236 | 149 | 1 | 1 | 0.808 | 1 | 1 | 1 | 0.857 | 1 | NA | 1 | 0.558 | 1 | 0.044 |
| SE | 105238 | ENSG00000082898 | XPO1 | chr2 | - | 61481184 | 61481281 | 61476031 | 61478966 | 61482379 | 61482426 | 284 | 194 | 233 | 242 | 539 | 1 | 0 | 0 | 0 | 0 | 32 | 120 | 108 | 59 | 18 | 0 | 0 | 0 | 0 | 0 | 246 | 149 | 1 | 1 | 0.994 | 1 | 1 | 1 | 1 | 1 | 1 | 1 | 1 | 1 | -0.001 |
| SE | 105239 | ENSG00000082898 | XPO1 | chr2 | - | 61485767 | 61485962 | 61483936 | 61484105 | 61488164 | 61488213 | 943 | 745 | 879 | 941 | 1752 | 5 | 2 | 3 | 1 | 3 | 174 | 400 | 396 | 308 | 131 | 0 | 0 | 6 | 0 | 0 | 298 | 149 | 1 | 1 | 0.99 | 0.995 | 0.993 | 0.998 | 0.997 | 1 | 1 | 0.971 | 1 | 1 | 0 |
| SE | 105240 | ENSG00000082898 | XPO1 | chr2 | - | 61485767 | 61485962 | 61483936 | 61484105 | 61492034 | 61492196 | 550 | 452 | 508 | 520 | 839 | 0 | 0 | 0 | 0 | 0 | 88 | 273 | 231 | 183 | 76 | 0 | 0 | 3 | 0 | 0 | 298 | 149 | 1 | 1 | 1 | 1 | 1 | 1 | 1 | 1 | 1 | 0.975 | 1 | 1 | 0.005 |
| SE | 105241 | ENSG00000082898 | XPO1 | chr2 | - | 61488164 | 61488271 | 61483936 | 61484105 | 61492034 | 61492196 | 6 | 0 | 2 | 1 | 3 | 0 | 0 | 0 | 0 | 0 | 0 | 0 | 2 | 0 | 0 | 0 | 0 | 3 | 0 | 0 | 256 | 149 | 1 | 1 | 1 | NA | 1 | 1 | 1 | NA | NA | 0.28 | NA | NA | 0.72 |
| SE | 105242 | ENSG00000082898 | XPO1 | chr2 | - | 61488164 | 61488271 | 61483936 | 61485962 | 61490641 | 61490776 | 331 | 248 | 310 | 367 | 770 | 0 | 1 | 0 | 0 | 0 | 69 | 98 | 145 | 111 | 46 | 0 | 0 | 0 | 0 | 0 | 256 | 149 | 1 | 1 | 1 | 0.993 | 1 | 1 | 1 | 1 | 1 | 1 | 1 | 1 | -0.001 |
| SE | 105243 | ENSG00000082898 | XPO1 | chr2 | - | 61488164 | 61488271 | 61483936 | 61485962 | 61492034 | 61492196 | 332 | 247 | 309 | 361 | 770 | 0 | 0 | 0 | 0 | 3 | 69 | 98 | 143 | 111 | 46 | 0 | 0 | 0 | 0 | 0 | 256 | 149 | 1 | 1 | 1 | 1 | 1 | 1 | 0.993 | 1 | 1 | 1 | 1 | 1 | -0.001 |
| SE | 105244 | ENSG00000082898 | XPO1 | chr2 | - | 61488587 | 61488771 | 61488164 | 61488271 | 61490641 | 61490776 | 463 | 284 | 413 | 513 | 936 | 0 | 2 | 2 | 6 | 0 | 74 | 155 | 158 | 99 | 38 | 0 | 0 | 2 | 0 | 0 | 298 | 149 | 1 | 1 | 1 | 0.986 | 0.99 | 0.977 | 1 | 1 | 1 | 0.975 | 1 | 1 | -0.004 |
| SE | 105245 | ENSG00000082898 | XPO1 | chr2 | - | 61488587 | 61488771 | 61488164 | 61488271 | 61492034 | 61492198 | 192 | 128 | 173 | 211 | 420 | 1 | 0 | 0 | 0 | 0 | 34 | 72 | 74 | 52 | 30 | 0 | 0 | 0 | 0 | 0 | 298 | 149 | 1 | 1 | 0.99 | 1 | 1 | 1 | 1 | 1 | 1 | 1 | 1 | 1 | -0.002 |
| SE | 105247 | ENSG00000082898 | XPO1 | chr2 | - | 61490641 | 61490776 | 61485767 | 61485962 | 61492034 | 61492196 | 599 | 406 | 525 | 570 | 926 | 0 | 0 | 0 | 0 | 3 | 56 | 251 | 226 | 161 | 26 | 0 | 0 | 0 | 0 | 0 | 284 | 149 | 1 | 1 | 1 | 1 | 1 | 1 | 0.994 | 1 | 1 | 1 | 1 | 1 | -0.001 |
| SE | 105248 | ENSG00000082898 | XPO1 | chr2 | - | 61490641 | 61490776 | 61488164 | 61488271 | 61492034 | 61492196 | 599 | 407 | 527 | 576 | 926 | 1 | 0 | 0 | 0 | 0 | 56 | 251 | 228 | 161 | 26 | 0 | 0 | 0 | 0 | 0 | 284 | 149 | 1 | 1 | 0.997 | 1 | 1 | 1 | 1 | 1 | 1 | 1 | 1 | 1 | -0.001 |
| SE | 105250 | ENSG00000082898 | XPO1 | chr2 | - | 61492034 | 61492196 | 61490641 | 61490776 | 61492324 | 61492481 | 623 | 411 | 544 | 587 | 951 | 0 | 0 | 3 | 0 | 0 | 56 | 262 | 226 | 163 | 26 | 0 | 0 | 0 | 0 | 0 | 298 | 149 | 1 | 1 | 1 | 1 | 0.989 | 1 | 1 | 1 | 1 | 1 | 1 | 1 | -0.002 |
| SE | 105251 | ENSG00000082898 | XPO1 | chr2 | - | 61492034 | 61492198 | 61490641 | 61490776 | 61492324 | 61492481 | 1109 | 777 | 981 | 1084 | 1932 | 0 | 0 | 3 | 0 | 0 | 114 | 449 | 419 | 294 | 99 | 0 | 0 | 0 | 0 | 0 | 298 | 149 | 1 | 1 | 1 | 1 | 0.994 | 1 | 1 | 1 | 1 | 1 | 1 | 1 | -0.001 |
| SE | 105252 | ENSG00000082898 | XPO1 | chr2 | - | 61492914 | 61493053 | 61492566 | 61492748 | 61493893 | 61494091 | 1333 | 1069 | 1091 | 1252 | 1929 | 0 | 1 | 1 | 2 | 2 | 287 | 837 | 837 | 618 | 127 | 0 | 7 | 4 | 2 | 0 | 288 | 149 | 1 | 1 | 1 | 0.998 | 0.998 | 0.997 | 0.998 | 1 | 0.984 | 0.991 | 0.994 | 1 | 0.004 |
| SE | 105253 | ENSG00000082898 | XPO1 | chr2 | - | 61493893 | 61494091 | 61492914 | 61493053 | 61495454 | 61495613 | 1033 | 912 | 863 | 1038 | 1542 | 13 | 3 | 1 | 3 | 1 | 223 | 757 | 754 | 562 | 119 | 0 | 10 | 1 | 0 | 0 | 298 | 149 | 1 | 1 | 0.975 | 0.993 | 0.998 | 0.994 | 0.999 | 1 | 0.974 | 0.997 | 1 | 1 | -0.002 |
| SE | 105254 | ENSG00000082898 | XPO1 | chr2 | - | 61493893 | 61495613 | 61492914 | 61493053 | 61496878 | 61498913 | 907 | 815 | 795 | 951 | 1331 | 0 | 0 | 0 | 1 | 0 | 252 | 663 | 617 | 462 | 152 | 0 | 0 | 0 | 0 | 0 | 298 | 149 | 1 | 1 | 1 | 1 | 1 | 0.998 | 1 | 1 | 1 | 1 | 1 | 1 | 0 |
| SE | 105255 | ENSG00000082898 | XPO1 | chr2 | - | 61495454 | 61495613 | 61492914 | 61493053 | 61496878 | 61497007 | 194 | 177 | 179 | 210 | 391 | 0 | 0 | 0 | 1 | 0 | 51 | 131 | 71 | 50 | 61 | 0 | 0 | 0 | 0 | 0 | 298 | 149 | 1 | 1 | 1 | 1 | 1 | 0.991 | 1 | 1 | 1 | 1 | 1 | 1 | -0.002 |
| SE | 105256 | ENSG00000082898 | XPO1 | chr2 | - | 61498672 | 61498913 | 61496878 | 61497007 | 61499712 | 61500402 | 336 | 200 | 227 | 262 | 592 | 0 | 0 | 2 | 0 | 0 | 80 | 163 | 121 | 125 | 55 | 0 | 0 | 0 | 0 | 0 | 298 | 149 | 1 | 1 | 1 | 1 | 0.983 | 1 | 1 | 1 | 1 | 1 | 1 | 1 | -0.003 |
| SE | 105257 | ENSG00000082898 | XPO1 | chr2 | - | 61499712 | 61499894 | 61498864 | 61498913 | 61522610 | 61522683 | 412 | 241 | 295 | 336 | 595 | 3 | 3 | 0 | 1 | 3 | 79 | 188 | 136 | 132 | 63 | 0 | 0 | 4 | 0 | 0 | 298 | 149 | 1 | 1 | 0.986 | 0.976 | 1 | 0.994 | 0.99 | 1 | 1 | 0.944 | 1 | 1 | 0 |
| SE | 105260 | ENSG00000082898 | XPO1 | chr2 | - | 61514061 | 61514200 | 61502248 | 61502310 | 61520390 | 61520475 | 0 | 0 | 2 | 1 | 4 | 1 | 3 | 0 | 4 | 4 | 0 | 0 | 0 | 3 | 0 | 0 | 0 | 1 | 0 | 0 | 288 | 149 | 1 | 1 | 0 | 0 | 1 | 0.115 | 0.341 | NA | NA | 0 | 1 | NA | -0.209 |
| SE | 105261 | ENSG00000082898 | XPO1 | chr2 | - | 61520390 | 61520475 | 61502248 | 61502310 | 61522610 | 61522683 | 2 | 4 | 2 | 5 | 5 | 248 | 178 | 254 | 256 | 267 | 0 | 0 | 1 | 0 | 0 | 13 | 112 | 85 | 74 | 59 | 234 | 149 | 1 | 1 | 0.005 | 0.014 | 0.005 | 0.012 | 0.012 | 0 | 0 | 0.007 | 0 | 0 | 0.008 |
| SE | 105267 | ENSG00000082898 | XPO1 | chr2 | - | 61533771 | 61533903 | 61525491 | 61526521 | 61537561 | 61537587 | 960 | 697 | 752 | 917 | 783 | 2 | 1 | 0 | 1 | 0 | 618 | 637 | 476 | 493 | 446 | 0 | 0 | 0 | 0 | 0 | 281 | 149 | 1 | 1 | 0.996 | 0.997 | 1 | 0.998 | 1 | 1 | 1 | 1 | 1 | 1 | -0.002 |
| SE | 105269 | ENSG00000082898 | XPO1 | chr2 | - | 61534557 | 61534654 | 61533771 | 61533903 | 61537561 | 61537668 | 4 | 5 | 10 | 2 | 4 | 334 | 259 | 307 | 321 | 240 | 0 | 0 | 0 | 0 | 2 | 55 | 197 | 184 | 153 | 33 | 246 | 149 | 1 | 1 | 0.007 | 0.012 | 0.019 | 0.004 | 0.01 | 0 | 0 | 0 | 0 | 0.035 | 0.003 |
| SE | 105271 | ENSG00000082898 | XPO1 | chr2 | - | 61537561 | 61537587 | 61533771 | 61533903 | 61538491 | 61538626 | 59 | 50 | 70 | 64 | 43 | 4 | 0 | 2 | 3 | 0 | 0 | 30 | 23 | 38 | 5 | 0 | 0 | 0 | 0 | 0 | 175 | 149 | 1 | 1 | 0.926 | 1 | 0.968 | 0.948 | 1 | NA | 1 | 1 | 1 | 1 | -0.032 |
| SE | 105272 | ENSG00000082898 | XPO1 | chr2 | - | 61537561 | 61537668 | 61533771 | 61533903 | 61538491 | 61538516 | 256 | 205 | 255 | 247 | 188 | 4 | 0 | 2 | 3 | 0 | 55 | 139 | 138 | 110 | 24 | 0 | 0 | 0 | 0 | 0 | 256 | 149 | 1 | 1 | 0.974 | 1 | 0.987 | 0.98 | 1 | 1 | 1 | 1 | 1 | 1 | -0.012 |
| SE | 105275 | ENSG00000161638 | ITGA5 | chr12 | - | 54401584 | 54401665 | 54401372 | 54401478 | 54401775 | 54401855 | 428 | 223 | 325 | 374 | 245 | 0 | 0 | 0 | 0 | 0 | 965 | 585 | 287 | 447 | 1003 | 0 | 4 | 0 | 0 | 9 | 230 | 149 | 1 | 1 | 1 | 1 | 1 | 1 | 1 | 1 | 0.99 | 1 | 1 | 0.986 | 0.005 |
| SE | 105276 | ENSG00000161638 | ITGA5 | chr12 | - | 54402000 | 54402093 | 54401775 | 54401855 | 54402982 | 54403050 | 144 | 87 | 109 | 125 | 81 | 0 | 0 | 0 | 1 | 0 | 255 | 179 | 84 | 174 | 227 | 0 | 8 | 0 | 0 | 0 | 242 | 149 | 1 | 1 | 1 | 1 | 1 | 0.987 | 1 | 1 | 0.932 | 1 | 1 | 1 | 0.011 |
| SE | 105278 | ENSG00000161638 | ITGA5 | chr12 | - | 54402982 | 54403050 | 54402179 | 54402330 | 54403186 | 54403324 | 199 | 84 | 173 | 173 | 209 | 0 | 0 | 0 | 2 | 0 | 329 | 208 | 88 | 143 | 251 | 0 | 0 | 1 | 7 | 0 | 217 | 149 | 1 | 1 | 1 | 1 | 1 | 0.983 | 1 | 1 | 1 | 0.984 | 0.933 | 1 | 0.013 |
| SE | 105281 | ENSG00000161638 | ITGA5 | chr12 | - | 54411833 | 54411964 | 54409484 | 54409597 | 54418980 | 54419127 | 92 | 20 | 49 | 44 | 25 | 0 | 1 | 1 | 2 | 0 | 78 | 39 | 14 | 13 | 61 | 0 | 0 | 3 | 8 | 0 | 280 | 149 | 1 | 1 | 1 | 0.914 | 0.963 | 0.921 | 1 | 1 | 1 | 0.713 | 0.464 | 1 | 0.124 |
| SE | 105282 | ENSG00000241370 | RPP21 | chr6 | + | 30345297 | 30345398 | 30345130 | 30345228 | 30345490 | 30345573 | 36 | 24 | 22 | 29 | 10 | 0 | 0 | 0 | 0 | 0 | 28 | 42 | 12 | 17 | 30 | 0 | 0 | 0 | 0 | 2 | 250 | 149 | 1 | 1 | 1 | 1 | 1 | 1 | 1 | 1 | 1 | 1 | 1 | 0.899 | 0.02 |
| SE | 105284 | ENSG00000241370 | RPP21 | chr6 | + | 30345490 | 30345929 | 30345297 | 30345398 | 30346431 | 30346493 | 4 | 7 | 6 | 6 | 2 | 0 | 0 | 0 | 0 | 0 | 19 | 6 | 3 | 7 | 9 | 2 | 0 | 0 | 0 | 0 | 298 | 149 | 1 | 1 | 1 | 1 | 1 | 1 | 1 | 0.826 | 1 | 1 | 1 | 1 | 0.035 |
| SE | 105286 | ENSG00000105221 | AKT2 | chr19 | - | 40235262 | 40235350 | 40234497 | 40235147 | 40235889 | 40235958 | 328 | 122 | 267 | 222 | 293 | 0 | 0 | 0 | 0 | 0 | 369 | 355 | 195 | 271 | 508 | 0 | 0 | 1 | 0 | 0 | 237 | 149 | 1 | 1 | 1 | 1 | 1 | 1 | 1 | 1 | 1 | 0.992 | 1 | 1 | 0.002 |
| SE | 105287 | ENSG00000105221 | AKT2 | chr19 | - | 40235262 | 40235467 | 40235044 | 40235147 | 40235889 | 40236031 | 193 | 70 | 172 | 141 | 204 | 0 | 0 | 0 | 0 | 0 | 233 | 271 | 130 | 182 | 378 | 0 | 0 | 1 | 0 | 0 | 298 | 149 | 1 | 1 | 1 | 1 | 1 | 1 | 1 | 1 | 1 | 0.985 | 1 | 1 | 0.003 |
| SE | 105288 | ENSG00000105221 | AKT2 | chr19 | - | 40236237 | 40236385 | 40236031 | 40236104 | 40237968 | 40238091 | 106 | 47 | 131 | 116 | 120 | 2 | 0 | 3 | 0 | 0 | 146 | 141 | 61 | 77 | 212 | 0 | 0 | 0 | 3 | 0 | 297 | 149 | 1 | 1 | 0.964 | 1 | 0.956 | 1 | 1 | 1 | 1 | 1 | 0.928 | 1 | -0.002 |
| SE | 105289 | ENSG00000105221 | AKT2 | chr19 | - | 40236256 | 40236385 | 40235889 | 40236104 | 40237968 | 40238091 | 177 | 96 | 224 | 195 | 218 | 2 | 0 | 3 | 0 | 0 | 244 | 261 | 147 | 159 | 335 | 0 | 0 | 0 | 3 | 0 | 278 | 149 | 1 | 1 | 0.979 | 1 | 0.976 | 1 | 1 | 1 | 1 | 1 | 0.966 | 1 | -0.002 |
| SE | 105290 | ENSG00000105221 | AKT2 | chr19 | - | 40236256 | 40236385 | 40236023 | 40236142 | 40237968 | 40238091 | 100 | 46 | 131 | 115 | 120 | 0 | 0 | 3 | 0 | 0 | 146 | 141 | 61 | 77 | 206 | 0 | 0 | 0 | 0 | 0 | 278 | 149 | 1 | 1 | 1 | 1 | 0.959 | 1 | 1 | 1 | 1 | 1 | 1 | 1 | -0.008 |
| SE | 105293 | ENSG00000105221 | AKT2 | chr19 | - | 40240044 | 40240110 | 40238904 | 40238973 | 40241937 | 40242069 | 321 | 223 | 338 | 351 | 292 | 0 | 0 | 7 | 1 | 1 | 400 | 480 | 208 | 370 | 533 | 0 | 0 | 0 | 0 | 0 | 215 | 149 | 1 | 1 | 1 | 1 | 0.971 | 0.996 | 0.995 | 1 | 1 | 1 | 1 | 1 | -0.008 |
| SE | 105300 | ENSG00000105221 | AKT2 | chr19 | - | 40268583 | 40268685 | 40265221 | 40265351 | 40269422 | 40269540 | 2 | 0 | 2 | 0 | 1 | 1 | 0 | 2 | 0 | 0 | 0 | 0 | 0 | 0 | 0 | 0 | 0 | 4 | 0 | 14 | 251 | 149 | 1 | 1 | 0.543 | NA | 0.372 | NA | 1 | NA | NA | 0 | NA | 0 | 0.638 |
| SE | 105301 | ENSG00000105221 | AKT2 | chr19 | - | 40269422 | 40269540 | 40265221 | 40265351 | 40270497 | 40270769 | 1 | 0 | 1 | 0 | 0 | 0 | 3 | 0 | 0 | 0 | 0 | 0 | 4 | 0 | 5 | 0 | 0 | 0 | 0 | 0 | 267 | 149 | 1 | 1 | 1 | 0 | 1 | NA | NA | NA | NA | 1 | NA | 1 | -0.333 |
| SE | 105308 | ENSG00000127325 | BEST3 | chr12 | - | 69676915 | 69677068 | 69672884 | 69672965 | 69677179 | 69677257 | 0 | 0 | 0 | 0 | 0 | 0 | 3 | 0 | 2 | 0 | 0 | 0 | 0 | 0 | 8 | 4 | 0 | 0 | 15 | 85 | 298 | 149 | 1 | 1 | NA | 0 | NA | 0 | NA | 0 | NA | NA | 0 | 0.045 | -0.015 |
| SE | 105314 | ENSG00000240505 | TNFRSF13B | chr17 | - | 16940325 | 16940511 | 16939080 | 16939797 | 16948737 | 16948983 | 12 | 3 | 23 | 19 | 21 | 0 | 0 | 0 | 2 | 0 | 17 | 29 | 14 | 6 | 8 | 0 | 0 | 3 | 0 | 0 | 298 | 149 | 1 | 1 | 1 | 1 | 1 | 0.826 | 1 | 1 | 1 | 0.7 | 1 | 1 | 0.025 |
| SE | 105319 | ENSG00000123268 | ATF1 | chr12 | + | 50780139 | 50780238 | 50764077 | 50764307 | 50795908 | 50796009 | 101 | 53 | 78 | 80 | 68 | 0 | 0 | 1 | 0 | 0 | 9 | 51 | 24 | 32 | 26 | 0 | 0 | 0 | 0 | 0 | 248 | 149 | 1 | 1 | 1 | 1 | 0.979 | 1 | 1 | 1 | 1 | 1 | 1 | 1 | -0.004 |
| SE | 105320 | ENSG00000123268 | ATF1 | chr12 | + | 50788207 | 50788325 | 50780139 | 50780238 | 50795908 | 50796009 | 2 | 6 | 3 | 5 | 0 | 71 | 44 | 59 | 68 | 64 | 0 | 0 | 0 | 0 | 0 | 0 | 42 | 22 | 21 | 24 | 267 | 149 | 1 | 1 | 0.015 | 0.071 | 0.028 | 0.039 | 0 | NA | 0 | 0 | 0 | 0 | 0.031 |
| SE | 105321 | ENSG00000123268 | ATF1 | chr12 | + | 50795908 | 50796009 | 50780139 | 50780238 | 50809455 | 50809475 | 113 | 67 | 104 | 113 | 156 | 0 | 0 | 2 | 0 | 0 | 19 | 79 | 42 | 45 | 24 | 0 | 0 | 0 | 0 | 0 | 250 | 149 | 1 | 1 | 1 | 1 | 0.969 | 1 | 1 | 1 | 1 | 1 | 1 | 1 | -0.006 |
| SE | 105324 | ENSG00000123268 | ATF1 | chr12 | + | 50809455 | 50809589 | 50795908 | 50796009 | 50814009 | 50814192 | 188 | 137 | 186 | 205 | 419 | 1 | 0 | 0 | 0 | 0 | 30 | 117 | 73 | 87 | 45 | 0 | 0 | 0 | 0 | 0 | 283 | 149 | 1 | 1 | 0.99 | 1 | 1 | 1 | 1 | 1 | 1 | 1 | 1 | 1 | -0.002 |
| SE | 105337 | ENSG00000172819 | RARG | chr12 | - | 53215645 | 53215794 | 53215292 | 53215434 | 53219949 | 53220259 | 30 | 14 | 58 | 41 | 8 | 2 | 0 | 0 | 0 | 0 | 17 | 12 | 7 | 26 | 47 | 0 | 0 | 0 | 0 | 0 | 298 | 149 | 1 | 1 | 0.882 | 1 | 1 | 1 | 1 | 1 | 1 | 1 | 1 | 1 | -0.024 |
| SE | 105338 | ENSG00000172819 | RARG | chr12 | - | 53215645 | 53215794 | 53215292 | 53215434 | 53227361 | 53227687 | 27 | 13 | 46 | 44 | 8 | 0 | 0 | 0 | 0 | 0 | 17 | 11 | 4 | 17 | 44 | 0 | 3 | 0 | 0 | 0 | 298 | 149 | 1 | 1 | 1 | 1 | 1 | 1 | 1 | 1 | 0.647 | 1 | 1 | 1 | 0.071 |
| SE | 105341 | ENSG00000172819 | RARG | chr12 | - | 53227361 | 53227687 | 53215645 | 53215794 | 53231168 | 53231235 | 10 | 1 | 9 | 4 | 1 | 2 | 0 | 1 | 0 | 0 | 0 | 2 | 1 | 1 | 9 | 0 | 0 | 0 | 0 | 18 | 298 | 149 | 1 | 1 | 0.714 | 1 | 0.818 | 1 | 1 | NA | 1 | 1 | 1 | 0.2 | 0.106 |
| SE | 105342 | ENSG00000172819 | RARG | chr12 | - | 53229841 | 53230053 | 53227634 | 53227687 | 53231168 | 53231232 | 1 | 0 | 3 | 0 | 3 | 7 | 1 | 2 | 3 | 1 | 0 | 0 | 2 | 0 | 0 | 0 | 0 | 0 | 1 | 1 | 298 | 149 | 1 | 1 | 0.067 | 0 | 0.429 | 0 | 0.6 | NA | NA | 1 | 0 | 0 | -0.114 |
| SE | 105343 | ENSG00000172819 | RARG | chr12 | - | 53229910 | 53230053 | 53227557 | 53227687 | 53231168 | 53231232 | 1 | 0 | 4 | 0 | 3 | 7 | 1 | 2 | 3 | 1 | 0 | 0 | 3 | 0 | 0 | 0 | 0 | 0 | 1 | 1 | 292 | 149 | 1 | 1 | 0.068 | 0 | 0.505 | 0 | 0.605 | NA | NA | 1 | 0 | 0 | -0.098 |
| SE | 105344 | ENSG00000172819 | RARG | chr12 | - | 53229910 | 53230074 | 53227607 | 53227687 | 53231168 | 53231235 | 0 | 0 | 1 | 0 | 0 | 7 | 1 | 2 | 3 | 1 | 0 | 0 | 1 | 0 | 0 | 0 | 0 | 0 | 1 | 1 | 298 | 149 | 1 | 1 | 0 | 0 | 0.2 | 0 | 0 | NA | NA | 1 | 0 | 0 | -0.293 |
| SE | 105347 | ENSG00000090382 | LYZ | chr12 | + | 69350107 | 69350272 | 69348380 | 69348544 | 69352219 | 69352298 | 36123 | 27746 | 23363 | 24912 | 37020 | 35 | 28 | 10 | 20 | 42 | 61073 | 14216 | 16126 | 48185 | 37366 | 47 | 17 | 13 | 44 | 5 | 298 | 149 | 1 | 1 | 0.998 | 0.998 | 0.999 | 0.998 | 0.998 | 0.998 | 0.998 | 0.998 | 0.998 | 1 | 0 |
| SE | 105348 | ENSG00000090382 | LYZ | chr12 | + | 69352219 | 69352298 | 69350107 | 69350272 | 69353152 | 69354234 | 29736 | 22636 | 18903 | 20441 | 29095 | 89 | 58 | 43 | 82 | 85 | 51319 | 11210 | 12949 | 39569 | 32668 | 162 | 17 | 41 | 118 | 69 | 228 | 149 | 1 | 1 | 0.995 | 0.996 | 0.997 | 0.994 | 0.996 | 0.995 | 0.998 | 0.995 | 0.995 | 0.997 | 0 |
| SE | 105355 | ENSG00000211829 | TRDC | chr14 | + | 22463773 | 22463839 | 22462931 | 22463210 | 22464203 | 22464323 | 1415 | 499 | 802 | 569 | 3503 | 0 | 1 | 0 | 0 | 0 | 27 | 196 | 168 | 129 | 72 | 0 | 0 | 0 | 0 | 0 | 215 | 149 | 1 | 1 | 1 | 0.997 | 1 | 1 | 1 | 1 | 1 | 1 | 1 | 1 | -0.001 |
| SE | 105356 | ENSG00000175782 | SLC35E3 | chr12 | + | 68759156 | 68759239 | 68752031 | 68752190 | 68764703 | 68767158 | 69 | 38 | 71 | 55 | 57 | 0 | 0 | 0 | 0 | 0 | 95 | 66 | 33 | 34 | 95 | 0 | 0 | 2 | 0 | 0 | 232 | 149 | 1 | 1 | 1 | 1 | 1 | 1 | 1 | 1 | 1 | 0.914 | 1 | 1 | 0.017 |
| SE | 105361 | ENSG00000122432 | SPATA1 | chr1 | + | 84522389 | 84522507 | 84520584 | 84520691 | 84525695 | 84525749 | 4 | 7 | 0 | 0 | 4 | 0 | 0 | 0 | 1 | 0 | 0 | 11 | 0 | 2 | 34 | 0 | 0 | 0 | 0 | 0 | 267 | 149 | 1 | 1 | 1 | 1 | NA | 0 | 1 | NA | 1 | NA | 1 | 1 | -0.25 |
| SE | 105362 | ENSG00000122432 | SPATA1 | chr1 | + | 84525695 | 84525749 | 84522389 | 84522507 | 84532859 | 84532974 | 1 | 2 | 0 | 0 | 1 | 0 | 1 | 0 | 0 | 0 | 0 | 7 | 0 | 1 | 10 | 0 | 0 | 0 | 0 | 0 | 203 | 149 | 1 | 1 | 1 | 0.595 | NA | NA | 1 | NA | 1 | NA | 1 | 1 | -0.135 |
| SE | 105363 | ENSG00000122432 | SPATA1 | chr1 | + | 84525844 | 84526073 | 84525695 | 84525749 | 84532859 | 84532974 | 4 | 0 | 0 | 0 | 9 | 0 | 0 | 0 | 0 | 0 | 0 | 0 | 0 | 3 | 35 | 0 | 2 | 0 | 1 | 0 | 298 | 149 | 1 | 1 | 1 | NA | NA | NA | 1 | NA | 0 | NA | 0.6 | 1 | 0.467 |
| SE | 105364 | ENSG00000122432 | SPATA1 | chr1 | + | 84545633 | 84545759 | 84544201 | 84544304 | 84548785 | 84548964 | 6 | 9 | 0 | 2 | 7 | 0 | 0 | 0 | 0 | 2 | 0 | 0 | 8 | 19 | 10 | 0 | 0 | 0 | 0 | 0 | 275 | 149 | 1 | 1 | 1 | 1 | NA | 1 | 0.655 | NA | NA | 1 | 1 | 1 | -0.086 |
| SE | 105370 | ENSG00000110719 | TCIRG1 | chr11 | + | 68047881 | 68047972 | 68047646 | 68047804 | 68048878 | 68048927 | 383 | 167 | 266 | 276 | 277 | 0 | 0 | 0 | 1 | 0 | 482 | 556 | 267 | 408 | 482 | 0 | 0 | 0 | 0 | 0 | 240 | 149 | 1 | 1 | 1 | 1 | 1 | 0.994 | 1 | 1 | 1 | 1 | 1 | 1 | -0.001 |
| SE | 105371 | ENSG00000110719 | TCIRG1 | chr11 | + | 68049080 | 68049294 | 68048878 | 68048997 | 68049662 | 68049788 | 382 | 191 | 248 | 317 | 332 | 0 | 0 | 0 | 3 | 0 | 522 | 653 | 299 | 476 | 585 | 0 | 0 | 2 | 0 | 0 | 298 | 149 | 1 | 1 | 1 | 1 | 1 | 0.981 | 1 | 1 | 1 | 0.987 | 1 | 1 | -0.001 |
| SE | 105372 | ENSG00000110719 | TCIRG1 | chr11 | + | 68050136 | 68050254 | 68049961 | 68050066 | 68050486 | 68050664 | 201 | 76 | 135 | 115 | 75 | 0 | 0 | 0 | 0 | 0 | 168 | 259 | 90 | 130 | 310 | 0 | 4 | 2 | 0 | 0 | 267 | 149 | 1 | 1 | 1 | 1 | 1 | 1 | 1 | 1 | 0.973 | 0.962 | 1 | 1 | 0.013 |
| SE | 105375 | ENSG00000114302 | PRKAR2A | chr3 | - | 48752175 | 48752317 | 48746585 | 48751718 | 48765003 | 48765078 | 77 | 60 | 64 | 77 | 136 | 0 | 0 | 0 | 0 | 0 | 122 | 65 | 54 | 114 | 109 | 0 | 5 | 0 | 0 | 0 | 291 | 149 | 1 | 1 | 1 | 1 | 1 | 1 | 1 | 1 | 0.869 | 1 | 1 | 1 | 0.026 |
| SE | 105378 | ENSG00000114302 | PRKAR2A | chr3 | - | 48765003 | 48765078 | 48756378 | 48756444 | 48772954 | 48773108 | 60 | 46 | 43 | 47 | 68 | 0 | 0 | 0 | 0 | 3 | 106 | 61 | 69 | 83 | 99 | 0 | 0 | 0 | 0 | 0 | 224 | 149 | 1 | 1 | 1 | 1 | 1 | 1 | 0.938 | 1 | 1 | 1 | 1 | 1 | -0.012 |
| SE | 105379 | ENSG00000114302 | PRKAR2A | chr3 | - | 48765247 | 48765349 | 48765003 | 48765078 | 48772954 | 48773108 | 390 | 328 | 302 | 364 | 488 | 0 | 2 | 0 | 2 | 0 | 1010 | 329 | 444 | 512 | 754 | 0 | 0 | 1 | 3 | 0 | 251 | 149 | 1 | 1 | 1 | 0.99 | 1 | 0.991 | 1 | 1 | 1 | 0.996 | 0.99 | 1 | -0.001 |
| SE | 105380 | ENSG00000114302 | PRKAR2A | chr3 | - | 48772954 | 48773108 | 48765247 | 48765349 | 48782985 | 48783092 | 453 | 333 | 332 | 400 | 641 | 4 | 1 | 2 | 3 | 0 | 1097 | 352 | 544 | 586 | 786 | 2 | 0 | 0 | 7 | 6 | 298 | 149 | 1 | 1 | 0.983 | 0.994 | 0.988 | 0.985 | 1 | 0.996 | 1 | 1 | 0.977 | 0.985 | -0.002 |
| SE | 105382 | ENSG00000114302 | PRKAR2A | chr3 | - | 48772954 | 48773108 | 48765247 | 48765349 | 48793996 | 48794013 | 300 | 255 | 244 | 288 | 395 | 0 | 0 | 0 | 0 | 0 | 835 | 261 | 361 | 410 | 609 | 0 | 0 | 0 | 4 | 0 | 298 | 149 | 1 | 1 | 1 | 1 | 1 | 1 | 1 | 1 | 1 | 1 | 0.981 | 1 | 0.004 |

| | | | | | | | | | | | | | | | | |
|---|---|---|---|---|---|---|---|---|---|---|---|---|---|---|---|---|
| SE | 105384 | ENSG00000114302 | PRKAR2A | chr3 | - | 48782985 | 48783092 | 48765247 | 48765349 | 48793996 | 48794013 | 5 | 0 | 2 | 7 | 1 |
| SE | 105386 | ENSG00000114302 | PRKAR2A | chr3 | - | 48782985 | 48783092 | 48772954 | 48773108 | 48793996 | 48794013 | 111 | 60 | 63 | 83 | 137 |
| SE | 105387 | ENSG00000114302 | PRKAR2A | chr3 | - | 48790543 | 48790627 | 48765247 | 48765349 | 48793996 | 48794013 | 60 | 51 | 59 | 61 | 73 |
| SE | 105388 | ENSG00000114302 | PRKAR2A | chr3 | - | 48790543 | 48790627 | 48772954 | 48773108 | 48793996 | 48794013 | 60 | 54 | 61 | 66 | 75 |
| SE | 105389 | ENSG00000114302 | PRKAR2A | chr3 | - | 48790543 | 48790627 | 48782985 | 48783092 | 48793996 | 48794013 | 260 | 181 | 180 | 208 | 363 |
| SE | 105390 | ENSG00000114302 | PRKAR2A | chr3 | - | 48793996 | 48794049 | 48790543 | 48790627 | 48807648 | 48807684 | 156 | 105 | 107 | 127 | 232 |
| SE | 105391 | ENSG00000114302 | PRKAR2A | chr3 | - | 48807648 | 48807684 | 48793996 | 48794049 | 48847334 | 48847804 | 83 | 57 | 73 | 75 | 96 |
| SE | 105403 | ENSG00000132141 | CCT6B | chr17 | - | 34942795 | 34942906 | 34942483 | 34942643 | 34951949 | 34952053 | 6 | 10 | 21 | 2 | 10 |
| SE | 105405 | ENSG00000132141 | CCT6B | chr17 | - | 34959586 | 34959650 | 34954425 | 34954599 | 34980898 | 34981078 | 2 | 0 | 0 | 0 | 0 |
| SE | 105406 | ENSG00000050438 | SLC4A8 | chr12 | + | 51407802 | 51407897 | 51391347 | 51391488 | 51440707 | 51440789 | 0 | 0 | 1 | 1 | 0 |
| SE | 105409 | ENSG00000050438 | SLC4A8 | chr12 | + | 51471286 | 51471532 | 51470391 | 51470525 | 51474341 | 51474447 | 7 | 11 | 3 | 8 | 7 |
| SE | 105410 | ENSG00000050438 | SLC4A8 | chr12 | + | 51489699 | 51489951 | 51488698 | 51488860 | 51493703 | 51493772 | 19 | 10 | 15 | 6 | 6 |
| SE | 105412 | ENSG00000040531 | CTNS | chr17 | + | 3637106 | 3637316 | 3636725 | 3636831 | 3640187 | 3640267 | 34 | 24 | 20 | 52 | 27 |
| SE | 105420 | ENSG00000170486 | KRT72 | chr12 | - | 52599270 | 52599302 | 52598953 | 52599112 | 52601026 | 52601409 | 0 | 0 | 0 | 3 | 0 |
| SE | 105421 | ENSG00000170486 | KRT72 | chr12 | - | 52599270 | 52599409 | 52598897 | 52599112 | 52601026 | 52601432 | 1 | 0 | 0 | 4 | 0 |
| SE | 105422 | ENSG00000011523 | CEP68 | chr2 | + | 65069398 | 65069801 | 65056415 | 65056528 | 65071453 | 65072569 | 36 | 2 | 13 | 19 | 16 |
| SE | 105424 | ENSG00000011523 | CEP68 | chr2 | + | 65071453 | 65072569 | 65069398 | 65069801 | 65074281 | 65074404 | 19 | 2 | 13 | 10 | 16 |
| SE | 105428 | ENSG00000011523 | CEP68 | chr2 | + | 65082535 | 65082709 | 65074281 | 65074404 | 65083638 | 65087004 | 25 | 29 | 64 | 39 | 60 |
| SE | 105429 | ENSG00000145901 | TNIP1 | chr5 | - | 151032286 | 151032383 | 151029944 | 151030747 | 151033607 | 151033799 | 224 | 109 | 161 | 202 | 101 |
| SE | 105430 | ENSG00000145901 | TNIP1 | chr5 | - | 151032286 | 151032383 | 151029944 | 151030747 | 151035001 | 151035067 | 153 | 72 | 112 | 129 | 66 |
| SE | 105431 | ENSG00000145901 | TNIP1 | chr5 | - | 151032286 | 151032400 | 151029946 | 151030747 | 151033607 | 151033799 | 154 | 74 | 114 | 118 | 71 |
| SE | 105432 | ENSG00000145901 | TNIP1 | chr5 | - | 151032286 | 151032400 | 151029946 | 151030747 | 151035001 | 151035067 | 149 | 71 | 104 | 117 | 65 |
| SE | 105434 | ENSG00000145901 | TNIP1 | chr5 | - | 151033607 | 151033799 | 151029944 | 151030747 | 151035001 | 151035067 | 206 | 136 | 163 | 193 | 395 |
| SE | 105436 | ENSG00000145901 | TNIP1 | chr5 | - | 151033607 | 151033799 | 151032286 | 151032400 | 151035001 | 151035067 | 211 | 139 | 174 | 194 | 401 |
| SE | 105438 | ENSG00000145901 | TNIP1 | chr5 | - | 151056765 | 151056957 | 151052164 | 151052259 | 151060317 | 151060395 | 479 | 201 | 330 | 280 | 215 |
| SE | 105439 | ENSG00000145901 | TNIP1 | chr5 | - | 151060317 | 151060395 | 151056765 | 151056957 | 151062126 | 151062212 | 266 | 142 | 193 | 200 | 129 |
| SE | 105440 | ENSG00000145901 | TNIP1 | chr5 | - | 151064959 | 151065131 | 151063612 | 151063747 | 151080323 | 151080359 | 197 | 129 | 158 | 185 | 194 |
| SE | 105444 | ENSG00000145901 | TNIP1 | chr5 | - | 151080323 | 151080359 | 151063612 | 151063747 | 151080879 | 151081034 | 1 | 1 | 0 | 0 | 0 |
| SE | 105453 | ENSG00000179889 | PDXDC1 | chr16 | + | 15004186 | 15004333 | 14998339 | 14998405 | 15008778 | 15008840 | 8 | 5 | 6 | 1 | 18 |
| SE | 105454 | ENSG00000179889 | PDXDC1 | chr16 | + | 15006393 | 15006583 | 14998339 | 14998405 | 15008778 | 15008840 | 104 | 124 | 197 | 134 | 151 |
| SE | 105455 | ENSG00000179889 | PDXDC1 | chr16 | + | 15006393 | 15006583 | 15004186 | 15004333 | 15008778 | 15008840 | 187 | 222 | 355 | 264 | 277 |
| SE | 105457 | ENSG00000179889 | PDXDC1 | chr16 | + | 15017119 | 15017168 | 15016128 | 15016213 | 15017320 | 15017387 | 63 | 33 | 96 | 90 | 91 |
| SE | 105458 | ENSG00000179889 | PDXDC1 | chr16 | + | 15022703 | 15022754 | 15018839 | 15018965 | 15026642 | 15026706 | 134 | 171 | 291 | 269 | 133 |
| SE | 105461 | ENSG00000135720 | DYNC1LI2 | chr16 | - | 66724661 | 66724748 | 66723600 | 66723822 | 66725827 | 66725944 | 5 | 1 | 2 | 2 | 3 |
| SE | 105463 | ENSG00000135720 | DYNC1LI2 | chr16 | - | 66727687 | 66727805 | 66725827 | 66725944 | 66728200 | 66728219 | 190 | 186 | 185 | 200 | 251 |
| SE | 105466 | ENSG00000135720 | DYNC1LI2 | chr16 | - | 66728200 | 66728242 | 66727687 | 66727805 | 66729039 | 66729099 | 164 | 181 | 161 | 189 | 254 |
| SE | 105467 | ENSG00000135720 | DYNC1LI2 | chr16 | - | 66729039 | 66729099 | 66725827 | 66725944 | 66730111 | 66730219 | 38 | 29 | 27 | 35 | 47 |
| SE | 105468 | ENSG00000135720 | DYNC1LI2 | chr16 | - | 66742437 | 66742554 | 66736101 | 66736244 | 66749196 | 66749313 | 83 | 68 | 70 | 62 | 71 |
| SE | 105469 | ENSG00000135720 | DYNC1LI2 | chr16 | - | 66742437 | 66742668 | 66736074 | 66736244 | 66749196 | 66749313 | 174 | 167 | 181 | 160 | 141 |
| SE | 105470 | ENSG00000135720 | DYNC1LI2 | chr16 | - | 66749196 | 66749313 | 66742437 | 66742668 | 66751272 | 66751303 | 150 | 125 | 162 | 114 | 89 |
| SE | 105471 | ENSG00000183281 | PLGLB1 | chr2 | - | 87013004 | 87013119 | 87011129 | 87011206 | 87016301 | 87016408 | 8 | 7 | 15 | 13 | 1 |
| SE | 105474 | ENSG00000111145 | ELK3 | chr12 | + | 96246939 | 96247734 | 96223564 | 96223773 | 96259730 | 96259853 | 227 | 127 | 175 | 209 | 224 |
| SE | 105475 | ENSG00000111145 | ELK3 | chr12 | + | 96247602 | 96247734 | 96247130 | 96247283 | 96259730 | 96259853 | 87 | 14 | 50 | 63 | 35 |
| SE | 105476 | ENSG00000111145 | ELK3 | chr12 | + | 96259730 | 96259853 | 96246939 | 96247734 | 96267081 | 96267357 | 152 | 50 | 110 | 105 | 75 |
| SE | 105477 | ENSG00000152256 | PDK1 | chr2 | + | 172562219 | 172562291 | 172558707 | 172558849 | 172564502 | 172564687 | 54 | 66 | 83 | 90 | 149 |
| SE | 105478 | ENSG00000152256 | PDK1 | chr2 | + | 172562740 | 172562800 | 172562219 | 172562291 | 172564051 | 172564130 | 5 | 13 | 14 | 8 | 8 |
| SE | 105481 | ENSG00000152256 | PDK1 | chr2 | + | 172564051 | 172564130 | 172562219 | 172562291 | 172564502 | 172564687 | 1 | 0 | 0 | 0 | 0 |
| SE | 105482 | ENSG00000152256 | PDK1 | chr2 | + | 172564051 | 172564130 | 172562219 | 172562291 | 172564977 | 172565073 | 0 | 1 | 0 | 0 | 0 |
| SE | 105483 | ENSG00000152256 | PDK1 | chr2 | + | 172564051 | 172564130 | 172562740 | 172562800 | 172564502 | 172564687 | 1 | 3 | 4 | 1 | 0 |
| SE | 105488 | ENSG00000152256 | PDK1 | chr2 | + | 172564977 | 172565073 | 172562219 | 172562291 | 172570725 | 172570812 | 0 | 7 | 0 | 7 | 6 |
| SE | 105489 | ENSG00000152256 | PDK1 | chr2 | + | 172566855 | 172566933 | 172564977 | 172565073 | 172568740 | 172568817 | 81 | 113 | 149 | 126 | 364 |
| SE | 105491 | ENSG00000152256 | PDK1 | chr2 | + | 172568740 | 172568817 | 172564977 | 172565073 | 172570725 | 172570812 | 19 | 27 | 29 | 23 | 28 |
| SE | 105493 | ENSG00000152256 | PDK1 | chr2 | + | 172586277 | 172586388 | 172570584 | 172570824 | 172592934 | 172593048 | 88 | 92 | 162 | 112 | 201 |
| SE | 105502 | ENSG00000174206 | C12orf66 | chr12 | - | 64215677 | 64215963 | 64191079 | 64194658 | 64222002 | 64222265 | 54 | 33 | 76 | 32 | 134 |
| SE | 105504 | ENSG00000111647 | UHRF1BP1L | chr12 | - | 100039595 | 100039789 | 100037071 | 100037755 | 100047529 | 100047638 | 127 | 138 | 123 | 172 | 160 |
| SE | 105505 | ENSG00000111647 | UHRF1BP1L | chr12 | - | 100059849 | 100060043 | 100058028 | 100059533 | 100072671 | 100072813 | 161 | 130 | 103 | 113 | 364 |
| SE | 105507 | ENSG00000111647 | UHRF1BP1L | chr12 | - | 100095666 | 100095837 | 100088914 | 100089095 | 100097364 | 100097505 | 233 | 220 | 159 | 232 | 429 |
| SE | 105509 | ENSG00000111647 | UHRF1BP1L | chr12 | - | 100108385 | 100108548 | 100103912 | 100103951 | 100142591 | 100142847 | 176 | 156 | 170 | 196 | 240 |
| SE | 105511 | ENSG00000174840 | PDE12 | chr3 | + | 57559309 | 57559388 | 57556273 | 57557687 | 57559561 | 57566848 | 179 | 92 | 157 | 164 | 104 |
| SE | 105512 | ENSG00000153179 | RASSF3 | chr12 | + | 64684786 | 64684894 | 64610494 | 64610743 | 64691469 | 64691579 | 73 | 35 | 48 | 60 | 31 |
| SE | 105515 | ENSG00000084110 | HAL | chr12 | - | 95976428 | 95976498 | 95972661 | 95974372 | 95976597 | 95976706 | 20 | 24 | 24 | 58 | 21 |
| SE | 105516 | ENSG00000084110 | HAL | chr12 | - | 95977943 | 95978078 | 95976597 | 95976706 | 95980555 | 95980668 | 25 | 17 | 20 | 38 | 25 |
| SE | 105517 | ENSG00000084110 | HAL | chr12 | - | 95987066 | 95987214 | 95986064 | 95986160 | 95988192 | 95988240 | 33 | 15 | 30 | 45 | 52 |
| SE | 105524 | ENSG00000188342 | GTF2F2 | chr13 | + | 45149769 | 45149788 | 45136732 | 45136806 | 45151686 | 45151831 | 105 | 74 | 58 | 83 | 66 |
| SE | 105525 | ENSG00000188342 | GTF2F2 | chr13 | + | 45151686 | 45151831 | 45149769 | 45149788 | 45207423 | 45207505 | 136 | 116 | 90 | 123 | 114 |
| SE | 105527 | ENSG00000188342 | GTF2F2 | chr13 | + | 45252870 | 45252970 | 45206354 | 45207505 | 45267232 | 45267333 | 46 | 44 | 48 | 63 | 56 |
| SE | 105529 | ENSG00000196653 | ZNF502 | chr3 | + | 44714590 | 44714688 | 44712659 | 44712740 | 44720202 | 44720316 | 5 | 3 | 4 | 5 | 1 |
| SE | 105530 | ENSG00000224429 | LINC00539 | chr13 | - | 21311041 | 21311118 | 21303169 | 21303820 | 21330573 | 21330658 | 2 | 1 | 2 | 0 | 0 |
| SE | 105532 | ENSG00000224429 | LINC00539 | chr13 | - | 21312145 | 21312360 | 21311041 | 21311118 | 21312451 | 21312555 | 1 | 0 | 5 | 4 | 0 |
| SE | 105533 | ENSG00000224429 | LINC00539 | chr13 | - | 21312145 | 21312365 | 21311041 | 21311118 | 21312451 | 21312555 | 8 | 1 | 12 | 5 | 4 |
| SE | 105534 | ENSG00000224429 | LINC00539 | chr13 | - | 21312150 | 21312365 | 21311041 | 21311118 | 21312451 | 21312555 | 7 | 1 | 7 | 1 | 4 |
| SE | 105535 | ENSG00000224429 | LINC00539 | chr13 | - | 21312278 | 21312365 | 21311041 | 21311118 | 21312451 | 21312555 | 8 | 1 | 4 | 5 | 5 |
| SE | 105536 | ENSG00000224429 | LINC00539 | chr13 | - | 21312451 | 21312555 | 21311041 | 21311118 | 21330573 | 21330658 | 7 | 1 | 3 | 2 | 2 |
| SE | 105537 | ENSG00000224429 | LINC00539 | chr13 | - | 21312638 | 21312761 | 21312451 | 21312555 | 21330573 | 21330658 | 2 | 1 | 0 | 0 | 0 |
| SE | 105538 | ENSG00000258102 | MAP1LC3B2 | chr12 | + | 116551114 | 116551172 | 116550370 | 116550774 | 116554294 | 116554381 | 0 | 2 | 0 | 0 | 0 |
| SE | 105547 | ENSG00000012822 | CALCOCO1 | chr12 | - | 53715199 | 53715325 | 53714597 | 53714693 | 53716259 | 53716415 | 258 | 192 | 214 | 359 | 230 |
| SE | 105549 | ENSG00000012822 | CALCOCO1 | chr12 | - | 53715792 | 53716047 | 53715199 | 53715325 | 53719738 | 53719829 | 247 | 141 | 195 | 310 | 216 |
| SE | 105550 | ENSG00000012822 | CALCOCO1 | chr12 | - | 53716259 | 53716415 | 53715199 | 53715325 | 53719738 | 53719778 | 187 | 113 | 179 | 259 | 201 |
| SE | 105551 | ENSG00000012822 | CALCOCO1 | chr12 | - | 53716259 | 53716415 | 53715674 | 53716047 | 53719738 | 53719778 | 370 | 234 | 348 | 492 | 447 |
| SE | 105552 | ENSG00000012822 | CALCOCO1 | chr12 | - | 53719738 | 53719829 | 53716259 | 53716415 | 53721466 | 53721615 | 269 | 164 | 271 | 356 | 340 |
| SE | 105554 | ENSG00000012822 | CALCOCO1 | chr12 | - | 53725689 | 53725759 | 53725086 | 53725266 | 53726199 | 53726340 | 1 | 0 | 2 | 0 | 7 |

| | | | | | | | | | | | | | |
|---|---|---|---|---|---|---|---|---|---|---|---|---|---|
| SE | 105804 | ENSG00000102189 | EEA1 | chr12 | - | 92905485 | 92905596 | 92891628 | 92891721 | 92929042 | 92929331 | 0 | 1 |
| SE | 105806 | ENSG00000111785 | RIC8B | chr12 | + | 106783996 | 106784044 | 106774639 | 106774829 | 106814695 | 106814870 | 27 | 19 |
| SE | 105809 | ENSG00000111785 | RIC8B | chr12 | + | 106825725 | 106825820 | 106814695 | 106815304 | 106842588 | 106842817 | 65 | 69 |
| SE | 105810 | ENSG00000111785 | RIC8B | chr12 | + | 106825725 | 106825820 | 106814695 | 106815304 | 106851449 | 106851594 | 18 | 17 |
| SE | 105813 | ENSG00000111785 | RIC8B | chr12 | + | 106843851 | 106843947 | 106825725 | 106825820 | 106860267 | 106860316 | 4 | 9 |
| SE | 105816 | ENSG00000111785 | RIC8B | chr12 | + | 106868300 | 106868423 | 106860267 | 106860412 | 106870822 | 106870942 | 1 | 0 |
| SE | 105817 | ENSG00000111785 | RIC8B | chr12 | + | 106870822 | 106870942 | 106860267 | 106860412 | 106885903 | 106889316 | 17 | 9 |
| SE | 105818 | ENSG00000111785 | RIC8B | chr12 | + | 106870822 | 106870942 | 106868300 | 106868423 | 106878986 | 106879125 | 2 | 3 |
| SE | 105819 | ENSG00000111785 | RIC8B | chr12 | + | 106873031 | 106873174 | 106870822 | 106870942 | 106874480 | 106874567 | 1 | 0 |
| SE | 105820 | ENSG00000111785 | RIC8B | chr12 | + | 106873031 | 106873174 | 106870822 | 106870942 | 106885903 | 106886360 | 0 | 0 |
| SE | 105821 | ENSG00000111785 | RIC8B | chr12 | + | 106874480 | 106874567 | 106870822 | 106870942 | 106878986 | 106879125 | 3 | 1 |
| SE | 105822 | ENSG00000111785 | RIC8B | chr12 | + | 106874480 | 106874567 | 106870822 | 106870942 | 106879774 | 106879947 | 3 | 1 |
| SE | 105823 | ENSG00000111785 | RIC8B | chr12 | + | 106874480 | 106874567 | 106870822 | 106870942 | 106885903 | 106887284 | 3 | 1 |
| SE | 105825 | ENSG00000111785 | RIC8B | chr12 | + | 106878986 | 106879125 | 106870822 | 106870942 | 106879774 | 106879947 | 6 | 12 |
| SE | 105826 | ENSG00000111785 | RIC8B | chr12 | + | 106878986 | 106879125 | 106870822 | 106870942 | 106885903 | 106887282 | 2 | 5 |
| SE | 105827 | ENSG00000111785 | RIC8B | chr12 | + | 106878986 | 106879125 | 106874480 | 106874567 | 106879774 | 106879947 | 4 | 8 |
| SE | 105829 | ENSG00000111785 | RIC8B | chr12 | + | 106879774 | 106879947 | 106870822 | 106870942 | 106885903 | 106887282 | 5 | 3 |
| SE | 105830 | ENSG00000111785 | RIC8B | chr12 | + | 106879774 | 106879947 | 106874480 | 106874567 | 106885903 | 106887282 | 5 | 3 |
| SE | 105831 | ENSG00000111785 | RIC8B | chr12 | + | 106879774 | 106879947 | 106878986 | 106879125 | 106885903 | 106887282 | 10 | 11 |
| SE | 105834 | ENSG00000257489 | NA | chr12 | - | 100173656 | 100173684 | 100173311 | 100173343 | 100177042 | 100177120 | 3 | 5 |
| SE | 105835 | ENSG00000257489 | NA | chr12 | - | 100180552 | 100180716 | 100177042 | 100177120 | 100195581 | 100195673 | 23 | 27 |
| SE | 105836 | ENSG00000257489 | NA | chr12 | - | 100180552 | 100180716 | 100177042 | 100177120 | 100198843 | 100198972 | 16 | 23 |
| SE | 105838 | ENSG00000257489 | NA | chr12 | - | 100195581 | 100195673 | 100180552 | 100180716 | 100198843 | 100198972 | 5 | 1 |
| SE | 105839 | ENSG00000257489 | NA | chr12 | - | 100195581 | 100195673 | 100180552 | 100180716 | 100199519 | 100199783 | 5 | 1 |
| SE | 105840 | ENSG00000257489 | NA | chr12 | - | 100198843 | 100198972 | 100180552 | 100180716 | 100199519 | 100199783 | 4 | 0 |
| SE | 105841 | ENSG00000257489 | NA | chr12 | - | 100198843 | 100198972 | 100195581 | 100195673 | 100199519 | 100199783 | 4 | 0 |
| SE | 105842 | ENSG00000111727 | HCFC2 | chr12 | + | 104079444 | 104079653 | 104067946 | 104068107 | 104080745 | 104080830 | 170 | 159 |
| SE | 105843 | ENSG00000111727 | HCFC2 | chr12 | + | 104079444 | 104079653 | 104067946 | 104068107 | 104082499 | 104082606 | 73 | 69 |
| SE | 105844 | ENSG00000111727 | HCFC2 | chr12 | + | 104080745 | 104080830 | 104067946 | 104068107 | 104082499 | 104082606 | 33 | 9 |
| SE | 105845 | ENSG00000111727 | HCFC2 | chr12 | + | 104080745 | 104080830 | 104079444 | 104079653 | 104082499 | 104082606 | 124 | 97 |
| SE | 105846 | ENSG00000111727 | HCFC2 | chr12 | + | 104086846 | 104087014 | 104082712 | 104082901 | 104087985 | 104088038 | 125 | 73 |
| SE | 105848 | ENSG00000111727 | HCFC2 | chr12 | + | 104097610 | 104097691 | 104096359 | 104096433 | 104098342 | 104098480 | 18 | 5 |
| SE | 105849 | ENSG00000111727 | HCFC2 | chr12 | + | 104098342 | 104098480 | 104096359 | 104096433 | 104101967 | 104102153 | 84 | 35 |
| SE | 105850 | ENSG00000111727 | HCFC2 | chr12 | + | 104101967 | 104102153 | 104098123 | 104098480 | 104102958 | 104103301 | 88 | 57 |
| SE | 105852 | ENSG00000188596 | CFAP54 | chr12 | + | 96554675 | 96554802 | 96554181 | 96554310 | 96564467 | 96564554 | 0 | 0 |
| SE | 105855 | ENSG00000090006 | LTBP4 | chr19 | + | 40611151 | 40611394 | 40610212 | 40610657 | 40611858 | 40611922 | 17 | 7 |
| SE | 105856 | ENSG00000090006 | LTBP4 | chr19 | + | 40612072 | 40612192 | 40611858 | 40611984 | 40613064 | 40613180 | 45 | 6 |
| SE | 105857 | ENSG00000090006 | LTBP4 | chr19 | + | 40615337 | 40615377 | 40614141 | 40614446 | 40616888 | 40617020 | 2 | 0 |
| SE | 105858 | ENSG00000090006 | LTBP4 | chr19 | + | 40615337 | 40615377 | 40614141 | 40614446 | 40619346 | 40619493 | 0 | 0 |
| SE | 105861 | ENSG00000090006 | LTBP4 | chr19 | + | 40616888 | 40617020 | 40615337 | 40615377 | 40619346 | 40619382 | 5 | 7 |
| SE | 105866 | ENSG00000090006 | LTBP4 | chr19 | + | 40622949 | 40623021 | 40619346 | 40619493 | 40623603 | 40623732 | 15 | 0 |
| SE | 105867 | ENSG00000090006 | LTBP4 | chr19 | + | 40622949 | 40623021 | 40622400 | 40622667 | 40623603 | 40623732 | 40 | 5 |
| SE | 105899 | ENSG00000137955 | RABGGTB | chr1 | + | 75786699 | 75786797 | 75786227 | 75786274 | 75787496 | 75787604 | 0 | 0 |
| SE | 105900 | ENSG00000137955 | RABGGTB | chr1 | + | 75787496 | 75787604 | 75786227 | 75786274 | 75791284 | 75791337 | 2 | 2 |
| SE | 105901 | ENSG00000137955 | RABGGTB | chr1 | + | 75787496 | 75790517 | 75786196 | 75786274 | 75791284 | 75791337 | 78 | 74 |
| SE | 105906 | ENSG00000137955 | RABGGTB | chr1 | + | 75789951 | 75790057 | 75789158 | 75789356 | 75791284 | 75791337 | 156 | 102 |
| SE | 105907 | ENSG00000137955 | RABGGTB | chr1 | + | 75792180 | 75792306 | 75791460 | 75791571 | 75794083 | 75794233 | 310 | 254 |
| SE | 105910 | ENSG00000111144 | LTA4H | chr12 | - | 96009093 | 96009148 | 96006313 | 96006409 | 96013187 | 96013252 | 1276 | 823 |
| SE | 105912 | ENSG00000111144 | LTA4H | chr12 | - | 96013749 | 96013853 | 96009093 | 96013258 | 96014854 | 96014999 | 1157 | 712 |
| SE | 105913 | ENSG00000111144 | LTA4H | chr12 | - | 96019167 | 96019240 | 96018762 | 96018903 | 96022146 | 96022251 | 600 | 422 |
| SE | 105915 | ENSG00000111144 | LTA4H | chr12 | - | 96021084 | 96021137 | 96019167 | 96019240 | 96022146 | 96022251 | 1083 | 676 |
| SE | 105916 | ENSG00000111144 | LTA4H | chr12 | - | 96022146 | 96022251 | 96021084 | 96021137 | 96024478 | 96024547 | 1524 | 1087 |
| SE | 105917 | ENSG00000111144 | LTA4H | chr12 | - | 96024478 | 96024547 | 96022146 | 96022251 | 96027443 | 96027503 | 1185 | 875 |
| SE | 105918 | ENSG00000111144 | LTA4H | chr12 | - | 96027443 | 96027503 | 96024478 | 96024547 | 96029054 | 96029185 | 668 | 511 |
| SE | 105919 | ENSG00000111144 | LTA4H | chr12 | - | 96027443 | 96027564 | 96024478 | 96024547 | 96029054 | 96029185 | 1587 | 1204 |
| SE | 105927 | ENSG00000110801 | PSMD9 | chr12 | + | 121899633 | 121899845 | 121894738 | 121894841 | 121903005 | 121903107 | 194 | 144 |
| SE | 105928 | ENSG00000110801 | PSMD9 | chr12 | + | 121899633 | 121899889 | 121893816 | 121894841 | 121903005 | 121903107 | 124 | 91 |
| SE | 105933 | ENSG00000127314 | RAP1B | chr12 | + | 68624670 | 68624873 | 68610898 | 68611052 | 68650399 | 68650468 | 0 | 0 |
| SE | 105935 | ENSG00000127314 | RAP1B | chr12 | + | 68624670 | 68624873 | 68610907 | 68611043 | 68650399 | 68650468 | 1 | 0 |
| SE | 105938 | ENSG00000127314 | RAP1B | chr12 | + | 68624674 | 68624873 | 68610907 | 68611052 | 68650399 | 68650440 | 0 | 0 |
| SE | 105940 | ENSG00000127314 | RAP1B | chr12 | + | 68624674 | 68624873 | 68610924 | 68611043 | 68650399 | 68650440 | 0 | 0 |
| SE | 105945 | ENSG00000127314 | RAP1B | chr12 | + | 68648669 | 68648781 | 68610932 | 68611043 | 68650399 | 68650468 | 1361 | 1282 |
| SE | 105946 | ENSG00000127314 | RAP1B | chr12 | + | 68648698 | 68648781 | 68610854 | 68611043 | 68650399 | 68650440 | 1984 | 1737 |
| SE | 105948 | ENSG00000127314 | RAP1B | chr12 | + | 68648698 | 68648781 | 68610857 | 68611052 | 68650399 | 68650440 | 1591 | 1474 |
| SE | 105949 | ENSG00000127314 | RAP1B | chr12 | + | 68648698 | 68648781 | 68610857 | 68611052 | 68654111 | 68654252 | 60 | 53 |
| SE | 105950 | ENSG00000127314 | RAP1B | chr12 | + | 68648698 | 68648781 | 68624670 | 68624873 | 68650399 | 68650440 | 1382 | 1305 |
| SE | 105951 | ENSG00000127314 | RAP1B | chr12 | + | 68648705 | 68648781 | 68610907 | 68611043 | 68650399 | 68650468 | 1307 | 1210 |
| SE | 105952 | ENSG00000127314 | RAP1B | chr12 | + | 68648705 | 68648781 | 68610924 | 68611052 | 68650399 | 68650468 | 1303 | 1210 |
| SE | 105953 | ENSG00000127314 | RAP1B | chr12 | + | 68650399 | 68650468 | 68610857 | 68611052 | 68654111 | 68654252 | 1 | 0 |
| SE | 105955 | ENSG00000127314 | RAP1B | chr12 | + | 68650399 | 68650468 | 68648669 | 68648781 | 68654111 | 68654252 | 644 | 546 |
| SE | 105956 | ENSG00000127314 | RAP1B | chr12 | + | 68651994 | 68652051 | 68650243 | 68650468 | 68654111 | 68654182 | 2474 | 2412 |
| SE | 105957 | ENSG00000127314 | RAP1B | chr12 | + | 68654111 | 68654252 | 68651994 | 68652051 | 68656305 | 68656348 | 2801 | 3000 |
| SE | 105959 | ENSG00000127314 | RAP1B | chr12 | + | 68656305 | 68656449 | 68654111 | 68654252 | 68657100 | 68657139 | 2287 | 2351 |
| SE | 105960 | ENSG00000127314 | RAP1B | chr12 | + | 68657100 | 68657217 | 68656305 | 68656449 | 68659279 | 68659401 | 2885 | 2403 |
| SE | 105962 | ENSG00000102225 | CDK16 | chrX | + | 47222260 | 47222380 | 47218412 | 47218664 | 47223551 | 47223759 | 0 | 1 |
| SE | 105967 | ENSG00000102225 | CDK16 | chrX | + | 47224384 | 47224518 | 47222852 | 47223759 | 47224617 | 47224731 | 147 | 62 |
| SE | 105968 | ENSG00000102225 | CDK16 | chrX | + | 47224833 | 47224890 | 47224617 | 47224743 | 47224987 | 47225036 | 93 | 77 |
| SE | 105971 | ENSG00000102225 | CDK16 | chrX | + | 47226811 | 47226909 | 47225964 | 47226703 | 47226993 | 47227099 | 83 | 40 |
| SE | 105972 | ENSG00000102225 | CDK16 | chrX | + | 47227378 | 47227469 | 47226993 | 47227223 | 47228566 | 47228644 | 95 | 41 |
| SE | 105975 | ENSG00000136014 | USP44 | chr12 | - | 95528806 | 95529002 | 95524679 | 95524788 | 95532828 | 95534326 | 19 | 3 |
| SE | 105976 | ENSG00000136014 | USP44 | chr12 | - | 95535372 | 95535426 | 95533763 | 95534326 | 95548770 | 95548819 | 0 | 0 |
| SE | 105978 | ENSG00000088992 | TESC | chr12 | - | 117041946 | 117041994 | 117038922 | 117039210 | 117046558 | 117046666 | 77 | 44 |
| SE | 105979 | ENSG00000088992 | TESC | chr12 | - | 117046558 | 117046666 | 117041946 | 117041994 | 117049018 | 117049158 | 44 | 18 |
| SE | 105980 | ENSG00000088992 | TESC | chr12 | - | 117046771 | 117046838 | 117046558 | 117046666 | 117049018 | 117049158 | 27 | 6 |
| SE | 105981 | ENSG00000088992 | TESC | chr12 | - | 117046776 | 117046838 | 117046479 | 117046666 | 117049018 | 117049158 | 60 | 25 |
| SE | 105983 | ENSG00000088992 | TESC | chr12 | - | 117049018 | 117049158 | 117046771 | 117046838 | 117056805 | 117056886 | 92 | 34 |
| SE | 105984 | ENSG00000088992 | TESC | chr12 | - | 117056805 | 117056886 | 117048797 | 117049158 | 117075270 | 117075340 | 51 | 22 |
| SE | 105985 | ENSG00000088992 | TESC | chr12 | - | 117083968 | 117084090 | 117075270 | 117075340 | 117099224 | 117099431 | 0 | 0 |
| SE | 105986 | ENSG00000122965 | RBM19 | chr12 | - | 113844667 | 113844788 | 113822056 | 113823321 | 113858790 | 113858896 | 90 | 57 |
| SE | 105988 | ENSG00000122965 | RBM19 | chr12 | - | 113948832 | 113949036 | 113947333 | 113947464 | 113950082 | 113950154 | 201 | 118 |
| SE | 105992 | ENSG00000007392 | LUC7L | chr16 | - | 190540 | 190570 | 188968 | 190135 | 192926 | 193015 | 131 | 65 |
| SE | 105993 | ENSG00000007392 | LUC7L | chr16 | - | 190661 | 190802 | 190540 | 190570 | 192926 | 193015 | 3 | 0 |
| SE | 105994 | ENSG00000007392 | LUC7L | chr16 | - | 192926 | 193015 | 190540 | 190570 | 199061 | 199238 | 222 | 176 |
| SE | 105995 | ENSG00000007392 | LUC7L | chr16 | - | 192926 | 193015 | 190540 | 190570 | 206003 | 206087 | 56 | 27 |
| SE | 105996 | ENSG00000007392 | LUC7L | chr16 | - | 199061 | 199238 | 190540 | 190570 | 206003 | 206147 | 170 | 134 |
| SE | 105998 | ENSG00000007392 | LUC7L | chr16 | - | 206003 | 206147 | 199061 | 199238 | 208077 | 208188 | 285 | 220 |
| SE | 106000 | ENSG00000007392 | LUC7L | chr16 | - | 220648 | 220747 | 208077 | 208188 | 227241 | 227336 | 254 | 271 |
| SE | 106003 | ENSG00000007392 | LUC7L | chr16 | - | 227241 | 227336 | 220648 | 220747 | 229278 | 229449 | 144 | 160 |
| SE | 106005 | ENSG00000007392 | LUC7L | chr16 | - | 227241 | 228402 | 220648 | 220747 | 229278 | 229451 | 194 | 203 |
| SE | 106007 | ENSG00000139344 | AMDHD1 | chr12 | + | 95954910 | 95954975 | 95952716 | 95952823 | 95956684 | 95956696 | 5 | 3 |
| SE | 106008 | ENSG00000139405 | RITA1 | chr12 | + | 113186184 | 113186316 | 113185720 | 113186021 | 113186682 | 113187048 | 32 | 17 |
| SE | 106009 | ENSG00000139405 | RITA1 | chr12 | + | 113186192 | 113186316 | 113185525 | 113186021 | 113186682 | 113187048 | 25 | 13 |
| SE | 106013 | ENSG00000163312 | HELQ | chr4 | - | 83418092 | 83418206 | 83416730 | 83416865 | 83421562 | 83421736 | 91 | 83 |
| SE | 106014 | ENSG00000163312 | HELQ | chr4 | - | 83421562 | 83421736 | 83418092 | 83418206 | 83425993 | 83426092 | 70 | 74 |
| SE | 106016 | ENSG00000163312 | HELQ | chr4 | - | 83436857 | 83437097 | 83432125 | 83432267 | 83439862 | 83440008 | 50 | 57 |
| SE | 106018 | ENSG00000163312 | HELQ | chr4 | - | 83446013 | 83446086 | 83443516 | 83443614 | 83446834 | 83447035 | 54 | 47 |
| SE | 106023 | ENSG00000111361 | EIF2B1 | chr12 | - | 123624786 | 123624862 | 123622635 | 123622761 | 123626424 | 123626493 | 334 | 238 |
| SE | 106024 | ENSG00000111361 | EIF2B1 | chr12 | - | 123624786 | 123624862 | 123622635 | 123622761 | 123630168 | 123630285 | 142 | 88 |
| SE | 106025 | ENSG00000111361 | EIF2B1 | chr12 | - | 123626424 | 123626493 | 123624786 | 123624862 | 123627043 | 123627156 | 234 | 188 |
| SE | 106028 | ENSG00000177707 | NECTIN3 | chr3 | + | 111126183 | 111126335 | 111122120 | 111122238 | 111133634 | 111137560 | 3 | 20 |
| SE | 106043 | ENSG00000204482 | LST1 | chr6 | + | 31587948 | 31587966 | 31587315 | 31587318 | 31588562 | 31588757 | 0 | 0 |
| SE | 106045 | ENSG00000204482 | LST1 | chr6 | + | 31587948 | 31587966 | 31587640 | 31587733 | 31588562 | 31588757 | 0 | 0 |
| SE | 106057 | ENSG00000130177 | CDC16 | chr13 | + | 114265149 | 114265240 | 114262878 | 114263014 | 114269562 | 114269620 | 62 | 42 |
| SE | 106059 | ENSG00000130177 | CDC16 | chr13 | + | 114269562 | 114269620 | 114262878 | 114263014 | 114272183 | 114272481 | 1 | 0 |
| SE | 106061 | ENSG00000123384 | LRP1 | chr12 | + | 57156776 | 57156920 | 57156093 | 57156283 | 57158401 | 57158638 | 396 | 249 |
| SE | 106062 | ENSG00000123384 | LRP1 | chr12 | + | 57168184 | 57168235 | 57167444 | 57167524 | 57169139 | 57169300 | 2 | 0 |
| SE | 106063 | ENSG00000123384 | LRP1 | chr12 | + | 57180666 | 57180807 | 57180329 | 57180479 | 57181156 | 57181291 | 222 | 120 |
| SE | 106064 | ENSG00000123384 | LRP1 | chr12 | + | 57191319 | 57191512 | 57190804 | 57191009 | 57192844 | 57192970 | 184 | 95 |
| SE | 106065 | ENSG00000123384 | LRP1 | chr12 | + | 57194353 | 57194503 | 57193898 | 57194012 | 57194576 | 57194699 | 308 | 114 |
| SE | 106066 | ENSG00000123384 | LRP1 | chr12 | + | 57198155 | 57198343 | 57197544 | 57197664 | 57198464 | 57198670 | 303 | 124 |
| SE | 106071 | ENSG00000196498 | NCOR2 | chr12 | - | 124330844 | 124330898 | 124327408 | 124327633 | 124332318 | 124332345 | 131 | 67 |
| SE | 106073 | ENSG00000196498 | NCOR2 | chr12 | - | 124340601 | 124340694 | 124340293 | 124340443 | 124341822 | 124342055 | 3 | 3 |
| SE | 106076 | ENSG00000196498 | NCOR2 | chr12 | - | 124350586 | 124350737 | 124348173 | 124348314 | 124354092 | 124354196 | 110 | 51 |
| SE | 106080 | ENSG00000196498 | NCOR2 | chr12 | - | 124363678 | 124363799 | 124362125 | 124362297 | 124378236 | 124378384 | 4 | 0 |
| SE | 106081 | ENSG00000196498 | NCOR2 | chr12 | - | 124372021 | 124372610 | 124348173 | 124348314 | 124385744 | 124385887 | 0 | 0 |
| SE | 106083 | ENSG00000196498 | NCOR2 | chr12 | - | 124372021 | 124372610 | 124362125 | 124362297 | 124378236 | 124378384 | 35 | 36 |
| SE | 106086 | ENSG00000196498 | NCOR2 | chr12 | - | 124378236 | 124378384 | 124372021 | 124372610 | 124385744 | 124385887 | 304 | 206 |
| SE | 106091 | ENSG00000196498 | NCOR2 | chr12 | - | 124385744 | 124385887 | 124378236 | 124378384 | 124419956 | 124420055 | 254 | 167 |
| SE | 106092 | ENSG00000196498 | NCOR2 | chr12 | - | 124426621 | 124426800 | 124422500 | 124422555 | 124437929 | 124437996 | 93 | 49 |
| SE | 106096 | ENSG00000196498 | NCOR2 | chr12 | - | 124437929 | 124437996 | 124422500 | 124422555 | 124449814 | 124449867 | 55 | 20 |
| SE | 106097 | ENSG00000196498 | NCOR2 | chr12 | - | 124437929 | 124437996 | 124430609 | 124430787 | 124466172 | 124466286 | 37 | 26 |
| SE | 106099 | ENSG00000196498 | NCOR2 | chr12 | - | 124449814 | 124449867 | 124437929 | 124437996 | 124466172 | 124466286 | 106 | 56 |
| SE | 106100 | ENSG00000196498 | NCOR2 | chr12 | - | 124457105 | 124457162 | 124437929 | 124437996 | 124466172 | 124466286 | 45 | 24 |
| SE | 106103 | ENSG00000108515 | ENO3 | chr17 | + | 4951827 | 4951914 | 4950189 | 4950630 | 4955074 | 4955093 | 0 | 0 |
| SE | 106105 | ENSG00000108515 | ENO3 | chr17 | + | 4956572 | 4956681 | 4955799 | 4956143 | 4956830 | 4956889 | 59 | 12 |
| SE | 106111 | ENSG00000089157 | RPLP0 | chr12 | - | 120200045 | 120200150 | 120199309 | 120199485 | 120200718 | 120200852 | 0 | 1 |
| SE | 106112 | ENSG00000089157 | RPLP0 | chr12 | - | 120200045 | 120200150 | 120199309 | 120199485 | 120200729 | 120200831 | 0 | 1 |
| SE | 106114 | ENSG00000089157 | RPLP0 | chr12 | - | 120200729 | 120200831 | 120199117 | 120199485 | 120201082 | 120201097 | 5766 | 2739 |
| SE | 106115 | ENSG00000089157 | RPLP0 | chr12 | - | 120200729 | 120200891 | 120199309 | 120199485 | 120201082 | 120201211 | 3560 | 1621 |
| SE | 106117 | ENSG00000187778 | MCRS1 | chr12 | - | 49559213 | 49559301 | 49558842 | 49558970 | 49559728 | 49559821 | 58 | 18 |
| SE | 106118 | ENSG00000187778 | MCRS1 | chr12 | - | 49559452 | 49559535 | 49559213 | 49559301 | 49559728 | 49559780 | 131 | 41 |
| SE | 106119 | ENSG00000187778 | MCRS1 | chr12 | - | 49563437 | 49563546 | 49563000 | 49563139 | 49564480 | 49564590 | 83 | 64 |
| SE | 106121 | ENSG00000187778 | MCRS1 | chr12 | - | 49565528 | 49565667 | 49564736 | 49564895 | 49566076 | 49566215 | 140 | 75 |
| SE | 106122 | ENSG00000187778 | MCRS1 | chr12 | - | 49565528 | 49565667 | 49564736 | 49564895 | 49568047 | 49568131 | 39 | 34 |
| SE | 106123 | ENSG00000187778 | MCRS1 | chr12 | - | 49565528 | 49565679 | 49564762 | 49564895 | 49566076 | 49566215 | 38 | 34 |
| SE | 106124 | ENSG00000187778 | MCRS1 | chr12 | - | 49566076 | 49566215 | 49564736 | 49564895 | 49568047 | 49568100 | 8 | 1 |
| SE | 106126 | ENSG00000187778 | MCRS1 | chr12 | - | 49566076 | 49566841 | 49564736 | 49564895 | 49568047 | 49568128 | 51 | 24 |
| SE | 106131 | ENSG00000136021 | SCYL2 | chr12 | + | 100313421 | 100313538 | 100312431 | 100312653 | 100314488 | 100314614 | 257 | 215 |
| SE | 106132 | ENSG00000136021 | SCYL2 | chr12 | + | 100317680 | 100317692 | 100315557 | 100315734 | 100317802 | 100317925 | 2 | 0 |
| SE | 106140 | ENSG00000089022 | MAPKAPK5 | chr12 | + | 111865249 | 111865323 | 111842227 | 111842769 | 111866155 | 111866231 | 139 | 87 |
| SE | 106141 | ENSG00000089022 | MAPKAPK5 | chr12 | + | 111865249 | 111865323 | 111842227 | 111842769 | 111867571 | 111867669 | 4 | 3 |
| SE | 106142 | ENSG00000089022 | MAPKAPK5 | chr12 | + | 111866155 | 111866231 | 111842227 | 111842769 | 111867571 | 111867669 | 138 | 68 |
| SE | 106147 | ENSG00000089022 | MAPKAPK5 | chr12 | + | 111867571 | 111867669 | 111866155 | 111866231 | 111870270 | 111870307 | 98 | 63 |
| SE | 106148 | ENSG00000089022 | MAPKAPK5 | chr12 | + | 111867571 | 111867669 | 111866155 | 111866231 | 111871084 | 111871180 | 92 | 69 |
| SE | 106149 | ENSG00000089022 | MAPKAPK5 | chr12 | + | 111868752 | 111868861 | 111866155 | 111866231 | 111870270 | 111870360 | 60 | 27 |
| SE | 106150 | ENSG00000089022 | MAPKAPK5 | chr12 | + | 111868752 | 111868861 | 111866155 | 111866231 | 111871084 | 111871180 | 14 | 8 |
| SE | 106153 | ENSG00000089022 | MAPKAPK5 | chr12 | + | 111870270 | 111870360 | 111866155 | 111866231 | 111871084 | 111871180 | 51 | 18 |
| SE | 106157 | ENSG00000089022 | MAPKAPK5 | chr12 | + | 111888884 | 111889000 | 111888487 | 111888618 | 111890039 | 111890144 | 196 | 112 |
| SE | 106158 | ENSG00000089022 | MAPKAPK5 | chr12 | + | 111888884 | 111889006 | 111887577 | 111888618 | 111890039 | 111890141 | 126 | 74 |
| SE | 106159 | ENSG00000120805 | ARL1 | chr12 | - | 101400290 | 101400381 | 101396539 | 101396577 | 101401061 | 101401173 | 0 | 0 |
| SE | 106161 | ENSG00000120805 | ARL1 | chr12 | - | 101401061 | 101401173 | 101396142 | 101396577 | 101405843 | 101405981 | 134 | 99 |
| SE | 106162 | ENSG00000120805 | ARL1 | chr12 | - | 101401061 | 101401173 | 101396142 | 101396577 | 101407641 | 101407746 | 126 | 99 |
| SE | 106163 | ENSG00000120805 | ARL1 | chr12 | - | 101402864 | 101402946 | 101396142 | 101396577 | 101405843 | 101405981 | 112 | 79 |
| SE | 106164 | ENSG00000120805 | ARL1 | chr12 | - | 101402864 | 101402946 | 101396142 | 101396577 | 101407641 | 101407746 | 1 | 0 |
| SE | 106168 | ENSG00000120805 | ARL1 | chr12 | - | 101403141 | 101403255 | 101402864 | 101402946 | 101407641 | 101407762 | 5 | 2 |
| SE | 106171 | ENSG00000120805 | ARL1 | chr12 | - | 101405843 | 101405981 | 101402864 | 101402946 | 101407641 | 101407746 | 300 | 195 |
| SE | 106172 | ENSG00000120805 | ARL1 | chr12 | - | 101405843 | 101405981 | 101403141 | 101403255 | 101407641 | 101407746 | 94 | 48 |
| SE | 106173 | ENSG00000168916 | ZNF608 | chr5 | - | 124638781 | 124638823 | 124636912 | 124637906 | 124639132 | 124639214 | 2 | 0 |
| SE | 106174 | ENSG00000168916 | ZNF608 | chr5 | - | 124641251 | 124641405 | 124639132 | 124639214 | 124643510 | 124643683 | 2 | 0 |
| SE | 106176 | ENSG00000168916 | ZNF608 | chr5 | - | 124646678 | 124649133 | 124644243 | 124644661 | 124649609 | 124649697 | 4 | 6 |
| SE | 106177 | ENSG00000168916 | ZNF608 | chr5 | - | 124701013 | 124701269 | 124649609 | 124649697 | 124748532 | 124748807 | 7 | 3 |
| SE | 106178 | ENSG00000168916 | ZNF608 | chr5 | - | 124744083 | 124745172 | 124701013 | 124701269 | 124746194 | 124746639 | 6 | 1 |
| SE | 106179 | ENSG00000168916 | ZNF608 | chr5 | - | 124744083 | 124745172 | 124701013 | 124701269 | 124748532 | 124748807 | 8 | 5 |
| SE | 106180 | ENSG00000089094 | KDM2B | chr12 | - | 121442991 | 121443030 | 121442156 | 121442836 | 121443679 | 121443793 | 110 | 112 |
| SE | 106182 | ENSG00000089094 | KDM2B | chr12 | - | 121453119 | 121453344 | 121444449 | 121445418 | 121494578 | 121494665 | 184 | 97 |
| SE | 106183 | ENSG00000089094 | KDM2B | chr12 | - | 121494578 | 121494665 | 121453119 | 121453344 | 121509566 | 121509706 | 120 | 57 |
| SE | 106185 | ENSG00000089094 | KDM2B | chr12 | - | 121520984 | 121521100 | 121513275 | 121513402 | 121532805 | 121532959 | 108 | 61 |
| SE | 106189 | ENSG00000135090 | TAOK3 | chr12 | - | 118160145 | 118160358 | 118152226 | 118152409 | 118161787 | 118162027 | 416 | 371 |
| SE | 106190 | ENSG00000135090 | TAOK3 | chr12 | - | 118161787 | 118162027 | 118152226 | 118152409 | 118172456 | 118172660 | 188 | 109 |
| SE | 106191 | ENSG00000135090 | TAOK3 | chr12 | - | 118172456 | 118172660 | 118152226 | 118152409 | 118177200 | 118177329 | 90 | 53 |
| SE | 106193 | ENSG00000135090 | TAOK3 | chr12 | - | 118189806 | 118189941 | 118181370 | 118181607 | 118199050 | 118199257 | 355 | 215 |
| SE | 106197 | ENSG00000135090 | TAOK3 | chr12 | - | 118243414 | 118243516 | 118239226 | 118239272 | 118244893 | 118244965 | 363 | 269 |
| SE | 106198 | ENSG00000135090 | TAOK3 | chr12 | - | 118244893 | 118244965 | 118243414 | 118243516 | 118255447 | 118255655 | 276 | 198 |
| SE | 106199 | ENSG00000135090 | TAOK3 | chr12 | - | 118255447 | 118255655 | 118244893 | 118244965 | 118266654 | 118266759 | 375 | 311 |
| SE | 106200 | ENSG00000135090 | TAOK3 | chr12 | - | 118261496 | 118261673 | 118244913 | 118244965 | 118266654 | 118266759 | 48 | 48 |
| SE | 106202 | ENSG00000135090 | TAOK3 | chr12 | - | 118266654 | 118266759 | 118255447 | 118255655 | 118295239 | 118295381 | 82 | 70 |
| SE | 106203 | ENSG00000135090 | TAOK3 | chr12 | - | 118266654 | 118266759 | 118255447 | 118255655 | 118359520 | 118359676 | 111 | 111 |
| SE | 106204 | ENSG00000135090 | TAOK3 | chr12 | - | 118283604 | 118283702 | 118266654 | 118266759 | 118359505 | 118359720 | 0 | 0 |
| SE | 106205 | ENSG00000135090 | TAOK3 | chr12 | - | 118283604 | 118283702 | 118266654 | 118266759 | 118372647 | 118372889 | 0 | 0 |
| SE | 106206 | ENSG00000135090 | TAOK3 | chr12 | - | 118295239 | 118295381 | 118266654 | 118266759 | 118296941 | 118297163 | 6 | 6 |
| SE | 106208 | ENSG00000135090 | TAOK3 | chr12 | - | 118296941 | 118297163 | 118266654 | 118266759 | 118358918 | 118359105 | 1 | 0 |
| SE | 106210 | ENSG00000135090 | TAOK3 | chr12 | - | 118296941 | 118297163 | 118266654 | 118266759 | 118359520 | 118359720 | 2 | 1 |
| SE | 106215 | ENSG00000123444 | KBTBD4 | chr11 | - | 47575592 | 47575699 | 47572196 | 47573790 | 47577410 | 47578024 | 32 | 18 |
| SE | 106216 | ENSG00000257433 | NA | chr12 | + | 47717491 | 47717636 | 47706084 | 47706163 | 47719146 | 47719272 | 0 | 0 |
| SE | 106217 | ENSG00000257433 | NA | chr12 | + | 47719599 | 47719699 | 47719146 | 47719272 | 47728886 | 47729203 | 3 | 0 |
| SE | 106218 | ENSG00000257433 | NA | chr12 | + | 47719599 | 47719699 | 47719146 | 47719272 | 47738914 | 47739011 | 0 | 0 |
| SE | 106220 | ENSG00000070214 | SLC44A1 | chr9 | + | 105309723 | 105309866 | 105299219 | 105299309 | 105335562 | 105335587 | 76 | 68 |
| SE | 106222 | ENSG00000070214 | SLC44A1 | chr9 | + | 105365482 | 105365639 | 105364554 | 105364720 | 105366345 | 105366429 | 108 | 93 |
| SE | 106224 | ENSG00000070214 | SLC44A1 | chr9 | + | 105385421 | 105385502 | 105383122 | 105383359 | 105389903 | 105397346 | 16 | 18 |
| SE | 106226 | ENSG00000083937 | CHMP2B | chr3 | + | 87240322 | 87240453 | 87227308 | 87227556 | 87240698 | 87240790 | 2 | 1 |
| SE | 106227 | ENSG00000083937 | CHMP2B | chr3 | + | 87240326 | 87240453 | 87227311 | 87227556 | 87240698 | 87240790 | 2 | 1 |
| SE | 106231 | ENSG00000089154 | GCN1 | chr12 | - | 120138694 | 120138856 | 120138322 | 120138415 | 120140858 | 120141023 | 145 | 61 |
| SE | 106232 | ENSG00000089154 | GCN1 | chr12 | - | 120142506 | 120142722 | 120140858 | 120141023 | 120142823 | 120142941 | 163 | 86 |
| SE | 106234 | ENSG00000089154 | GCN1 | chr12 | - | 120147051 | 120147272 | 120145261 | 120145330 | 120148166 | 120148346 | 128 | 60 |
| SE | 106236 | ENSG00000089154 | GCN1 | chr12 | - | 120151144 | 120151391 | 120149921 | 120150043 | 120153212 | 120153407 | 154 | 64 |
| SE | 106245 | ENSG00000089154 | GCN1 | chr12 | - | 120184111 | 120184243 | 120183568 | 120183677 | 120184823 | 120184887 | 100 | 65 |
| SE | 106246 | ENSG00000089154 | GCN1 | chr12 | - | 120184823 | 120184887 | 120178851 | 120178950 | 120190297 | 120190400 | 12 | 6 |
| SE | 106247 | ENSG00000272636 | DOC2B | chr17 | - | 164129 | 164204 | 162080 | 162190 | 172536 | 172616 | 0 | 0 |
| SE | 106248 | ENSG00000272636 | DOC2B | chr17 | - | 172536 | 172616 | 164129 | 164204 | 181106 | 181170 | 0 | 0 |

| | | | | | | | | | | | | | | | | | | | | | | | | | | | | | | | | | | | | | | | | | | | | | | |
|---|---|---|---|---|---|---|---|---|---|---|---|---|---|---|---|---|---|---|---|---|---|---|---|---|---|---|---|---|---|---|---|---|---|---|---|---|---|---|---|---|---|---|---|---|---|---|
| SE | 106261 | ENSG00000116198 | CEP104 | chr1 | - | 3823173 | 3823241 | 3816148 | 3816370 | 3823423 | 3823562 | 44 | 48 | 42 | 45 | 30 | 0 | 0 | 0 | 0 | 0 | 36 | 43 | 30 | 55 | 54 | 0 | 3 | 1 | 0 | 0 | 217 | 149 | 1 | 1 | 1 | 1 | 1 | 1 | 1 | 1 | 0.908 | 0.954 | 1 | 1 | 0.028 |
| SE | 106264 | ENSG00000116198 | CEP104 | chr1 | - | 3829790 | 3829997 | 3829132 | 3829373 | 3831045 | 3831222 | 237 | 179 | 198 | 172 | 303 | 0 | 3 | 2 | 0 | 0 | 353 | 202 | 133 | 323 | 339 | 0 | 1 | 0 | 0 | 0 | 298 | 149 | 1 | 1 | 1 | 0.968 | 0.98 | 1 | 1 | 1 | 0.99 | 1 | 1 | 1 | -0.008 |
| SE | 106265 | ENSG00000116198 | CEP104 | chr1 | - | 3831045 | 3831222 | 3829790 | 3829997 | 3833861 | 3834035 | 293 | 224 | 252 | 232 | 435 | 3 | 0 | 0 | 0 | 0 | 406 | 287 | 149 | 356 | 504 | 0 | 0 | 0 | 0 | 0 | 298 | 149 | 1 | 1 | 0.98 | 1 | 1 | 1 | 1 | 1 | 1 | 1 | 1 | 1 | -0.004 |
| SE | 106267 | ENSG00000116198 | CEP104 | chr1 | - | 3841081 | 3841169 | 3839607 | 3839776 | 3844906 | 3844983 | 0 | 0 | 0 | 0 | 3 | 93 | 78 | 96 | 124 | 175 | 0 | 0 | 0 | 0 | 0 | 226 | 200 | 117 | 141 | 144 | 237 | 149 | 1 | 1 | 0 | 0 | 0 | 0 | 0.011 | 0 | 0 | 0 | 0 | 0 | 0.002 |
| SE | 106268 | ENSG00000116198 | CEP104 | chr1 | - | 3848607 | 3848781 | 3846576 | 3847613 | 3852294 | 3852421 | 175 | 136 | 123 | 178 | 156 | 0 | 1 | 1 | 0 | 0 | 246 | 116 | 88 | 129 | 142 | 0 | 0 | 0 | 0 | 0 | 298 | 149 | 1 | 1 | 1 | 0.986 | 0.984 | 1 | 1 | 1 | 1 | 1 | 1 | 1 | -0.006 |
| SE | 106270 | ENSG00000116198 | CEP104 | chr1 | - | 3855873 | 3856001 | 3852294 | 3852421 | 3856888 | 3857211 | 1 | 0 | 0 | 2 | 0 | 11 | 9 | 21 | 9 | 4 | 0 | 0 | 0 | 0 | 0 | 2 | 6 | 6 | 6 | 8 | 277 | 149 | 1 | 1 | 0.047 | 0 | 0 | 0.107 | 0 | 0 | 0 | 0 | 0 | 0 | 0.031 |
| SE | 106272 | ENSG00000251301 | LINC02384 | chr12 | - | 68335654 | 68335780 | 68332887 | 68333664 | 68442063 | 68442216 | 0 | 0 | 0 | 0 | 0 | 4 | 9 | 3 | 4 | 9 | 0 | 0 | 6 | 0 | 0 | 0 | 24 | 1 | 2 | 3 | 275 | 149 | 1 | 1 | 0 | 0 | 0 | 0 | 0 | NA | 0 | 0.765 | 0 | 0 | -0.191 |
| SE | 106276 | ENSG00000251301 | LINC02384 | chr12 | - | 68433283 | 68433371 | 68380462 | 68383602 | 68451366 | 68451385 | 12 | 2 | 0 | 3 | 10 | 4 | 0 | 0 | 2 | 0 | 14 | 4 | 0 | 4 | 26 | 0 | 0 | 0 | 0 | 0 | 237 | 149 | 1 | 1 | 0.654 | 1 | NA | 0.485 | 1 | 1 | 1 | NA | 1 | 1 | -0.215 |
| SE | 106290 | ENSG00000197057 | DTHD1 | chr4 | + | 36283975 | 36284591 | 36281615 | 36282029 | 36290372 | 36290703 | 178 | 52 | 45 | 70 | 44 | 10 | 4 | 9 | 4 | 0 | 10 | 42 | 37 | 8 | 0 | 0 | 0 | 2 | 0 | 0 | 298 | 149 | 1 | 1 | 0.899 | 0.867 | 0.714 | 0.897 | 1 | 1 | 1 | 0.902 | 1 | NA | -0.1 |
| SE | 106291 | ENSG00000197057 | DTHD1 | chr4 | + | 36290372 | 36290703 | 36281621 | 36282029 | 36293525 | 36293705 | 135 | 45 | 36 | 47 | 94 | 0 | 1 | 0 | 0 | 0 | 0 | 33 | 25 | 0 | 10 | 0 | 0 | 0 | 0 | 0 | 298 | 149 | 1 | 1 | 1 | 0.957 | 1 | 1 | 1 | NA | 1 | 1 | NA | 1 | -0.009 |
| SE | 106292 | ENSG00000197057 | DTHD1 | chr4 | + | 36306190 | 36306352 | 36294794 | 36295039 | 36308203 | 36308493 | 192 | 52 | 66 | 56 | 81 | 18 | 2 | 6 | 4 | 2 | 0 | 65 | 10 | 6 | 19 | 0 | 4 | 1 | 0 | 0 | 298 | 149 | 1 | 1 | 0.842 | 0.929 | 0.846 | 0.875 | 0.953 | NA | 0.89 | 0.833 | 1 | 1 | -0.042 |
| SE | 106293 | ENSG00000197057 | DTHD1 | chr4 | + | 36308203 | 36308493 | 36294794 | 36295039 | 36339111 | 36339169 | 18 | 9 | 6 | 4 | 2 | 0 | 0 | 3 | 0 | 0 | 0 | 9 | 2 | 0 | 0 | 0 | 0 | 0 | 0 | 0 | 298 | 149 | 1 | 1 | 1 | 1 | 0.5 | 1 | 1 | NA | 1 | 1 | NA | NA | -0.1 |
| SE | 106296 | ENSG00000162482 | AKR7A3 | chr1 | - | 19285017 | 19285114 | 19284685 | 19284785 | 19285887 | 19285992 | 0 | 0 | 2 | 2 | 3 | 0 | 0 | 0 | 0 | 0 | 0 | 5 | 0 | 8 | 0 | 0 | 0 | 0 | 2 | 14 | 246 | 149 | 1 | 1 | NA | NA | 1 | 1 | 1 | NA | 1 | NA | 0.708 | 0 | 0.431 |
| SE | 106297 | ENSG00000111837 | MAK | chr6 | - | 10770110 | 10770230 | 10762722 | 10764606 | 10775327 | 10775459 | 3 | 4 | 0 | 2 | 8 | 2 | 2 | 1 | 2 | 9 | 0 | 0 | 15 | 0 | 0 | 0 | 0 | 1 | 0 | 0 | 269 | 149 | 1 | 1 | 0.454 | 0.526 | 0 | 0.356 | 0.33 | NA | NA | 0.893 | NA | NA | -0.56 |
| SE | 106298 | ENSG00000111837 | MAK | chr6 | - | 10773033 | 10773108 | 10770110 | 10770230 | 10775327 | 10775459 | 0 | 0 | 0 | 0 | 0 | 0 | 1 | 0 | 0 | 2 | 0 | 0 | 2 | 0 | 0 | 0 | 0 | 6 | 0 | 0 | 224 | 149 | 1 | 1 | NA | 0 | NA | NA | 0 | NA | NA | 0.181 | NA | NA | -0.181 |
| SE | 106301 | ENSG00000111837 | MAK | chr6 | - | 10808809 | 10808942 | 10803719 | 10803891 | 10813643 | 10813723 | 15 | 20 | 2 | 7 | 23 | 1 | 0 | 0 | 1 | 0 | 8 | 2 | 48 | 19 | 0 | 0 | 0 | 2 | 0 | 0 | 282 | 149 | 1 | 1 | 0.888 | 1 | 1 | 0.787 | 1 | 1 | 1 | 0.927 | 1 | NA | -0.047 |
| SE | 106302 | ENSG00000111837 | MAK | chr6 | - | 10808809 | 10808942 | 10803719 | 10803891 | 10817849 | 10817971 | 4 | 13 | 1 | 6 | 14 | 0 | 0 | 0 | 1 | 0 | 4 | 0 | 45 | 13 | 0 | 0 | 0 | 0 | 0 | 0 | 282 | 149 | 1 | 1 | 1 | 1 | 1 | 0.76 | 1 | 1 | NA | 1 | 1 | NA | -0.048 |
| SE | 106303 | ENSG00000111837 | MAK | chr6 | - | 10813643 | 10813723 | 10803719 | 10803891 | 10817849 | 10817971 | 7 | 0 | 0 | 3 | 9 | 0 | 0 | 0 | 1 | 0 | 0 | 0 | 26 | 0 | 0 | 0 | 0 | 0 | 0 | 0 | 229 | 149 | 1 | 1 | 1 | NA | NA | 0.661 | 1 | NA | NA | 1 | NA | NA | -0.113 |
| SE | 106304 | ENSG00000111837 | MAK | chr6 | - | 10813643 | 10813723 | 10808809 | 10808942 | 10817849 | 10817971 | 17 | 10 | 1 | 4 | 18 | 0 | 3 | 0 | 1 | 0 | 4 | 2 | 29 | 3 | 0 | 0 | 0 | 2 | 0 | 0 | 229 | 149 | 1 | 1 | 1 | 0.684 | 1 | 0.722 | 1 | 1 | 1 | 0.904 | 1 | NA | -0.095 |
| SE | 106305 | ENSG00000111837 | MAK | chr6 | - | 10830547 | 10830877 | 10818885 | 10818940 | 10838502 | 10838531 | 6 | 0 | 4 | 8 | 10 | 3 | 0 | 0 | 1 | 0 | 7 | 0 | 12 | 3 | 0 | 0 | 0 | 0 | 0 | 0 | 298 | 149 | 1 | 1 | 0.5 | NA | 1 | 0.8 | 1 | 1 | NA | 1 | 1 | NA | -0.175 |
| SE | 106312 | ENSG00000145439 | CBR4 | chr4 | - | 169008951 | 169008999 | 169006801 | 169006891 | 169009947 | 169010250 | 1 | 0 | 0 | 0 | 0 | 0 | 0 | 7 | 2 | 0 | 0 | 0 | 0 | 7 | 0 | 0 | 0 | 0 | 0 | 3 | 197 | 149 | 1 | 1 | 1 | NA | 0 | 0 | NA | NA | NA | NA | 1 | 0 | -0.167 |
| SE | 106316 | ENSG00000111450 | STX2 | chr12 | - | 130801414 | 130801488 | 130801152 | 130801290 | 130806981 | 130807090 | 87 | 66 | 89 | 102 | 93 | 1 | 0 | 0 | 0 | 0 | 162 | 61 | 70 | 70 | 62 | 0 | 0 | 0 | 0 | 0 | 223 | 149 | 1 | 1 | 0.983 | 1 | 1 | 1 | 1 | 1 | 1 | 1 | 1 | 1 | -0.003 |
| SE | 106317 | ENSG00000111450 | STX2 | chr12 | - | 130806981 | 130807090 | 130801152 | 130801290 | 130812956 | 130813031 | 1 | 0 | 0 | 0 | 0 | 0 | 0 | 0 | 0 | 0 | 0 | 0 | 1 | 0 | 0 | 0 | 0 | 0 | 0 | 10 | 258 | 149 | 1 | 1 | 1 | NA | NA | NA | NA | NA | NA | 1 | NA | 0 | 0.5 |
| SE | 106318 | ENSG00000111450 | STX2 | chr12 | - | 130806981 | 130807090 | 130801414 | 130801488 | 130808630 | 130808704 | 94 | 98 | 94 | 107 | 133 | 0 | 0 | 0 | 2 | 1 | 231 | 93 | 115 | 128 | 139 | 0 | 4 | 0 | 0 | 0 | 258 | 149 | 1 | 1 | 1 | 1 | 1 | 0.969 | 0.987 | 1 | 0.931 | 1 | 1 | 1 | 0.005 |
| SE | 106319 | ENSG00000111450 | STX2 | chr12 | - | 130808630 | 130808704 | 130806981 | 130807090 | 130812956 | 130813031 | 126 | 124 | 119 | 138 | 179 | 0 | 0 | 0 | 0 | 0 | 255 | 111 | 142 | 182 | 197 | 0 | 0 | 1 | 0 | 0 | 223 | 149 | 1 | 1 | 1 | 1 | 1 | 1 | 1 | 1 | 1 | 0.99 | 1 | 1 | 0.002 |
| SE | 106320 | ENSG00000111450 | STX2 | chr12 | - | 130812309 | 130812443 | 130808630 | 130808704 | 130812956 | 130813031 | 0 | 0 | 1 | 0 | 1 | 99 | 90 | 98 | 110 | 145 | 0 | 0 | 0 | 0 | 0 | 164 | 88 | 128 | 133 | 188 | 283 | 149 | 1 | 1 | 0 | 0 | 0.005 | 0 | 0.004 | 0 | 0 | 0 | 0 | 0 | 0.002 |
| SE | 106321 | ENSG00000229117 | RPL41 | chr12 | + | 56117188 | 56117211 | 56116632 | 56116799 | 56117481 | 56117710 | 1838 | 1022 | 1786 | 1613 | 1965 | 3 | 1 | 6 | 3 | 2 | 1247 | 1097 | 945 | 1082 | 1724 | 11 | 7 | 2 | 8 | 22 | 172 | 149 | 1 | 1 | 0.998 | 0.999 | 0.996 | 0.998 | 0.999 | 0.99 | 0.993 | 0.998 | 0.992 | 0.985 | 0.006 |
| SE | 106322 | ENSG00000125458 | NT5C | chr17 | - | 75130752 | 75130867 | 75130224 | 75130642 | 75131044 | 75131105 | 83 | 36 | 63 | 61 | 62 | 0 | 1 | 0 | 0 | 0 | 90 | 89 | 51 | 36 | 145 | 0 | 0 | 1 | 0 | 0 | 264 | 149 | 1 | 1 | 1 | 0.953 | 1 | 1 | 1 | 1 | 1 | 0.966 | 1 | 1 | -0.003 |
| SE | 106334 | ENSG00000111231 | GPN3 | chr12 | - | 110457509 | 110457634 | 110455814 | 110455930 | 110459694 | 110459862 | 39 | 44 | 70 | 49 | 75 | 1 | 0 | 2 | 1 | 0 | 33 | 2 | 5 | 24 | 2 | 0 | 0 | 0 | 0 | 0 | 274 | 149 | 1 | 1 | 0.955 | 1 | 0.95 | 0.964 | 1 | 1 | 1 | 1 | 1 | 1 | -0.026 |
| SE | 106335 | ENSG00000111231 | GPN3 | chr12 | - | 110468155 | 110468375 | 110465185 | 110465214 | 110468471 | 110468719 | 7 | 5 | 14 | 16 | 0 | 0 | 0 | 0 | 2 | 0 | 0 | 13 | 8 | 6 | 0 | 0 | 0 | 0 | 0 | 0 | 298 | 149 | 1 | 1 | 1 | 1 | 1 | 0.8 | NA | NA | 1 | 1 | 1 | NA | -0.05 |
| SE | 106338 | ENSG00000198563 | DDX39B | chr6 | - | 31538762 | 31538939 | 31536608 | 31536683 | 31539146 | 31539274 | 304 | 176 | 280 | 314 | 195 | 0 | 0 | 0 | 0 | 0 | 458 | 377 | 282 | 387 | 440 | 0 | 3 | 0 | 0 | 0 | 298 | 149 | 1 | 1 | 1 | 1 | 1 | 1 | 1 | 1 | 0.984 | 1 | 1 | 1 | 0.003 |
| SE | 106340 | ENSG00000198563 | DDX39B | chr6 | - | 31541152 | 31541327 | 31540321 | 31540664 | 31541949 | 31541986 | 2 | 1 | 5 | 2 | 0 | 19 | 21 | 34 | 22 | 0 | 0 | 4 | 7 | 1 | 7 | 14 | 13 | 1 | 36 | 34 | 298 | 149 | 1 | 1 | 0.05 | 0.023 | 0.068 | 0.043 | NA | 0 | 0.133 | 0.778 | 0.014 | 0.093 | -0.158 |
| SE | 106341 | ENSG00000076513 | ANKRD13A | chr12 | + | 110013124 | 110013249 | 110012004 | 110012137 | 110016387 | 110016433 | 389 | 216 | 302 | 334 | 528 | 1 | 0 | 0 | 0 | 0 | 720 | 364 | 289 | 416 | 427 | 0 | 0 | 0 | 0 | 1 | 274 | 149 | 1 | 1 | 0.995 | 1 | 1 | 1 | 1 | 1 | 1 | 1 | 1 | 0.996 | 0 |
| SE | 106346 | ENSG00000076513 | ANKRD13A | chr12 | + | 110019138 | 110019328 | 110013124 | 110013249 | 110024045 | 110024112 | 700 | 512 | 682 | 828 | 1433 | 0 | 0 | 0 | 0 | 0 | 2149 | 960 | 917 | 1044 | 1691 | 0 | 0 | 0 | 6 | 0 | 298 | 149 | 1 | 1 | 1 | 1 | 1 | 1 | 1 | 1 | 1 | 1 | 0.989 | 1 | 0.002 |
| SE | 106348 | ENSG00000076513 | ANKRD13A | chr12 | + | 110021107 | 110021301 | 110019138 | 110019328 | 110025741 | 110025823 | 6 | 4 | 3 | 10 | 4 | 7 | 4 | 2 | 4 | 15 | 4 | 4 | 16 | 31 | 0 | 25 | 23 | 16 | 16 | 0 | 298 | 149 | 1 | 1 | 0.3 | 0.333 | 0.429 | 0.556 | 0.118 | 0.074 | 0.08 | 0.333 | 0.492 | NA | 0.102 |
| SE | 106350 | ENSG00000076513 | ANKRD13A | chr12 | + | 110024045 | 110024112 | 110021107 | 110021301 | 110025741 | 110025823 | 337 | 285 | 300 | 340 | 545 | 0 | 0 | 0 | 0 | 0 | 879 | 380 | 366 | 472 | 679 | 1 | 0 | 4 | 0 | 0 | 216 | 149 | 1 | 1 | 1 | 1 | 1 | 1 | 1 | 0.998 | 1 | 0.984 | 1 | 1 | 0.004 |
| SE | 106351 | ENSG00000076513 | ANKRD13A | chr12 | + | 110025741 | 110025823 | 110023618 | 110024112 | 110027704 | 110027766 | 908 | 680 | 839 | 976 | 1673 | 0 | 1 | 0 | 0 | 0 | 2584 | 1093 | 1120 | 1304 | 2000 | 0 | 0 | 0 | 0 | 0 | 231 | 149 | 1 | 1 | 1 | 0.998 | 1 | 1 | 1 | 1 | 1 | 1 | 1 | 1 | 0 |
| SE | 106352 | ENSG00000076513 | ANKRD13A | chr12 | + | 110025741 | 110025823 | 110023618 | 110024112 | 110028511 | 110028642 | 459 | 385 | 459 | 562 | 870 | 0 | 0 | 0 | 0 | 0 | 1519 | 634 | 656 | 751 | 1137 | 0 | 0 | 0 | 6 | 0 | 231 | 149 | 1 | 1 | 1 | 1 | 1 | 1 | 1 | 1 | 1 | 1 | 0.988 | 1 | 0.002 |
| SE | 106353 | ENSG00000076513 | ANKRD13A | chr12 | + | 110027704 | 110027766 | 110023618 | 110024112 | 110028511 | 110028642 | 225 | 213 | 207 | 234 | 244 | 0 | 0 | 0 | 0 | 0 | 563 | 284 | 261 | 322 | 461 | 0 | 0 | 0 | 6 | 0 | 211 | 149 | 1 | 1 | 1 | 1 | 1 | 1 | 1 | 1 | 1 | 1 | 0.974 | 1 | 0.005 |
| SE | 106357 | ENSG00000076513 | ANKRD13A | chr12 | + | 110030644 | 110030758 | 110028946 | 110029635 | 110036260 | 110036328 | 184 | 156 | 224 | 236 | 303 | 0 | 1 | 0 | 2 | 0 | 453 | 273 | 278 | 349 | 426 | 0 | 0 | 0 | 0 | 0 | 263 | 149 | 1 | 1 | 1 | 0.989 | 1 | 0.985 | 1 | 1 | 1 | 1 | 1 | 1 | -0.005 |
| SE | 106358 | ENSG00000076513 | ANKRD13A | chr12 | + | 110033796 | 110033957 | 110028946 | 110029635 | 110036260 | 110036328 | 407 | 286 | 325 | 411 | 641 | 0 | 1 | 0 | 2 | 0 | 707 | 437 | 407 | 458 | 738 | 0 | 0 | 0 | 0 | 0 | 298 | 149 | 1 | 1 | 1 | 0.993 | 1 | 0.99 | 1 | 1 | 1 | 1 | 1 | 1 | -0.003 |
| SE | 106361 | ENSG00000132341 | RAN | chr12 | + | 130872583 | 130872617 | 130872075 | 130872126 | 130872835 | 130872920 | 0 | 0 | 0 | 0 | 3 | 7 | 3 | 4 | 5 | 7 | 0 | 0 | 0 | 0 | 4 | 0 | 2 | 1 | 3 | 5 | 183 | 149 | 1 | 1 | 0 | 0 | 0 | 0 | 0.259 | NA | 0 | 0 | 0 | 0.394 | -0.047 |
| SE | 106363 | ENSG00000132341 | RAN | chr12 | + | 130874033 | 130874084 | 130873002 | 130873128 | 130874545 | 130874733 | 0 | 0 | 1 | 0 | 1 | 544 | 351 | 604 | 622 | 1315 | 0 | 0 | 0 | 0 | 0 | 613 | 534 | 369 | 516 | 697 | 200 | 149 | 1 | 1 | 0 | 0 | 0.001 | 0 | 0.001 | 0 | 0 | 0 | 0 | 0 | 0 |
| SE | 106364 | ENSG00000112053 | SLC26A8 | chr6 | - | 35960842 | 35960912 | 35959713 | 35959806 | 35960992 | 35961099 | 0 | 4 | 0 | 1 | 4 | 0 | 0 | 0 | 0 | 0 | 0 | 6 | 2 | 0 | 0 | 0 | 0 | 0 | 4 | 0 | 219 | 149 | 1 | 1 | NA | 1 | NA | 1 | 1 | NA | 1 | 1 | 0 | NA | 0.333 |
| SE | 106365 | ENSG00000112053 | SLC26A8 | chr6 | - | 35977203 | 35977351 | 35975374 | 35975488 | 35982120 | 35982203 | 3 | 2 | 0 | 0 | 9 | 0 | 0 | 0 | 0 | 0 | 0 | 10 | 12 | 6 | 0 | 0 | 0 | 0 | 0 | 4 | 297 | 149 | 1 | 1 | 1 | 1 | NA | NA | 1 | NA | 1 | 1 | 1 | 0 | 0.25 |
| SE | 106366 | ENSG00000112053 | SLC26A8 | chr6 | - | 35977203 | 35977351 | 35975374 | 35975488 | 35991658 | 35991808 | 6 | 1 | 0 | 0 | 3 | 0 | 0 | 1 | 0 | 0 | 0 | 4 | 5 | 2 | 0 | 0 | 0 | 0 | 0 | 0 | 297 | 149 | 1 | 1 | 1 | 1 | 0 | NA | 1 | NA | 1 | 1 | 1 | NA | -0.25 |
| SE | 106367 | ENSG00000112053 | SLC26A8 | chr6 | - | 35982120 | 35982203 | 35975374 | 35975488 | 35991658 | 35991808 | 0 | 0 | 0 | 0 | 0 | 0 | 0 | 1 | 0 | 0 | 0 | 5 | 3 | 0 | 8 | 0 | 0 | 0 | 0 | 0 | 232 | 149 | 1 | 1 | NA | NA | 0 | NA | NA | NA | 1 | 1 | NA | 1 | -1 |
| SE | 106368 | ENSG00000112053 | SLC26A8 | chr6 | - | 35982120 | 35982203 | 35977203 | 35977351 | 35991658 | 35991808 | 0 | 1 | 0 | 0 | 5 | 3 | 0 | 0 | 0 | 1 | 0 | 11 | 6 | 6 | 4 | 0 | 0 | 0 | 2 | 0 | 232 | 149 | 1 | 1 | 0 | 1 | NA | NA | 0.763 | NA | 1 | 1 | 0.658 | 1 | -0.327 |
| SE | 106369 | ENSG00000112053 | SLC26A8 | chr6 | - | 35991658 | 35991808 | 35982120 | 35982203 | 35992509 | 35992674 | 0 | 1 | 1 | 1 | 6 | 0 | 0 | 0 | 0 | 2 | 0 | 16 | 12 | 15 | 4 | 0 | 0 | 4 | 0 | 0 | 298 | 149 | 1 | 1 | NA | 1 | 1 | 1 | 0.6 | NA | 1 | 0.6 | 1 | 1 | 0 |
| SE | 106371 | ENSG00000112053 | SLC26A8 | chr6 | - | 36012232 | 36012319 | 35999991 | 36000108 | 36019519 | 36019709 | 1 | 1 | 0 | 0 | 0 | 0 | 0 | 0 | 0 | 0 | 0 | 1 | 1 | 0 | 0 | 0 | 0 | 0 | 11 | 0 | 236 | 149 | 1 | 1 | 1 | 1 | NA | NA | NA | NA | 1 | 1 | 0 | NA | 0.333 |
| SE | 106372 | ENSG00000112053 | SLC26A8 | chr6 | - | 36012232 | 36012372 | 35999991 | 36000108 | 36019519 | 36019709 | 2 | 1 | 0 | 1 | 0 | 0 | 0 | 0 | 0 | 0 | 0 | 7 | 2 | 0 | 7 | 0 | 0 | 0 | 11 | 0 | 289 | 149 | 1 | 1 | 1 | 1 | NA | 1 | NA | NA | 1 | 1 | 0 | 1 | 0.25 |
| SE | 106373 | ENSG00000092531 | SNAP23 | chr15 | + | 42511832 | 42511903 | 42495305 | 42495713 | 42512954 | 42512996 | 395 | 440 | 283 | 466 | 626 | 0 | 1 | 0 | 2 | 0 | 688 | 365 | 435 | 556 | 1093 | 12 | 0 | 0 | 0 | 0 | 220 | 149 | 1 | 1 | 1 | 0.997 | 1 | 0.994 | 1 | 0.975 | 1 | 1 | 1 | 1 | 0.003 |
| SE | 106374 | ENSG00000092531 | SNAP23 | chr15 | + | 42512954 | 42512996 | 42511832 | 42511903 | 42513173 | 42513209 | 92 | 83 | 82 | 108 | 79 | 0 | 0 | 0 | 0 | 0 | 64 | 78 | 106 | 79 | 172 | 0 | 0 | 1 | 0 | 0 | 191 | 149 | 1 | 1 | 1 | 1 | 1 | 1 | 1 | 1 | 1 | 0.988 | 1 | 1 | 0.002 |
| SE | 106378 | ENSG00000092531 | SNAP23 | chr15 | + | 42515236 | 42515354 | 42512954 | 42513447 | 42531412 | 42531827 | 240 | 236 | 164 | 271 | 458 | 0 | 0 | 0 | 0 | 0 | 519 | 270 | 242 | 424 | 743 | 0 | 0 | 3 | 0 | 0 | 267 | 149 | 1 | 1 | 1 | 1 | 1 | 1 | 1 | 1 | 1 | 0.978 | 1 | 1 | 0.004 |
| SE | 106379 | ENSG00000092531 | SNAP23 | chr15 | + | 42515236 | 42515354 | 42512954 | 42513447 | 42545135 | 42545347 | 240 | 237 | 164 | 271 | 458 | 0 | 0 | 0 | 0 | 0 | 519 | 270 | 243 | 424 | 743 | 0 | 0 | 0 | 3 | 0 | 267 | 149 | 1 | 1 | 1 | 1 | 1 | 1 | 1 | 1 | 1 | 1 | 0.987 | 1 | 0.003 |
| SE | 106380 | ENSG00000092531 | SNAP23 | chr15 | + | 42521613 | 42521708 | 42515236 | 42515354 | 42528261 | 42528343 | 5 | 3 | 6 | 11 | 2 | 550 | 697 | 411 | 708 | 954 | 1 | 3 | 12 | 11 | 0 | 1211 | 516 | 501 | 665 | 1716 | 244 | 149 | 1 | 1 | 0.006 | 0.003 | 0.009 | 0.009 | 0.001 | 0.001 | 0.004 | 0.014 | 0.01 | 0 | 0 |
| SE | 106384 | ENSG00000092531 | SNAP23 | chr15 | + | 42529674 | 42529819 | 42515236 | 42515354 | 42545135 | 42545345 | 4 | 2 | 1 | 4 | 13 | 0 | 1 | 0 | 0 | 0 | 0 | 18 | 3 | 0 | 0 | 0 | 0 | 1 | 0 | 0 | 294 | 149 | 1 | 1 | 1 | 0.503 | 1 | 1 | 1 | NA | 1 | 0.603 | NA | NA | 0.099 |
| SE | 106385 | ENSG00000092531 | SNAP23 | chr15 | + | 42529674 | 42529819 | 42527942 | 42528420 | 42531412 | 42531444 | 809 | 836 | 608 | 1078 | 1431 | 3 | 1 | 0 | 0 | 4 | 1597 | 676 | 683 | 961 | 2490 | 0 | 0 | 0 | 0 | 0 | 294 | 149 | 1 | 1 | 0.993 | 0.998 | 1 | 1 | 0.995 | 1 | 1 | 1 | 1 | 1 | -0.003 |
| SE | 106399 | ENSG00000089041 | P2RX7 | chr12 | + | 121160901 | 121160974 | 121156078 | 121156147 | 121162270 | 121162430 | 21 | 22 | 20 | 16 | 45 | 1 | 0 | 0 | 0 | 0 | 124 | 38 | 17 | 21 | 20 | 0 | 0 | 0 | 0 | 0 | 222 | 149 | 1 | 1 | 0.934 | 1 | 1 | 1 | 1 | 1 | 1 | 1 | 1 | 1 | -0.013 |
| SE | 106406 | ENSG00000089041 | P2RX7 | chr12 | + | 121166057 | 121166187 | 121165356 | 121165437 | 121175387 | 121175478 | 43 | 32 | 19 | 32 | 53 | 0 | 0 | 0 | 1 | 0 | 109 | 56 | 49 | 46 | 18 | 0 | 2 | 0 | 0 | 0 | 279 | 149 | 1 | 1 | 1 | 1 | 1 | 0.945 | 1 | 1 | 0.937 | 1 | 1 | 1 | 0.002 |
| SE | 106411 | ENSG00000135297 | MTO1 | chr6 | + | 73466208 | 73466408 | 73461577 | 73462071 | 73466488 | 73466606 | 185 | 142 | 193 | 210 | 185 | 0 | 0 | 0 | 2 | 0 | 203 | 177 | 126 | 140 | 108 | 0 | 0 | 3 | 0 | 0 | 298 | 149 | 1 | 1 | 1 | 1 | 1 | 0.981 | 1 | 1 | 1 | 0.955 | 1 | 1 | 0.005 |
| SE | 106412 | ENSG00000135297 | MTO1 | chr6 | + | 73466213 | 73466408 | 73461736 | 73462071 | 73466488 | 73466606 | 132 | 116 | 141 | 133 | 152 | 0 | 0 | 0 | 2 | 0 | 143 | 129 | 93 | 93 | 87 | 0 | 0 | 3 | 0 | 0 | 298 | 149 | 1 | 1 | 1 | 1 | 1 | 0.971 | 1 | 1 | 1 | 0.939 | 1 | 1 | 0.006 |
| SE | 106417 | ENSG00000135297 | MTO1 | chr6 | + | 73473364 | 73473654 | 73466208 | 73466606 | 73480674 | 73480805 | 143 | 115 | 164 | 161 | 156 | 0 | 0 | 2 | 0 | 0 | 134 | 169 | 93 | 124 | 71 | 0 | 0 | 0 | 0 | 0 | 298 | 149 | 1 | 1 | 1 | 1 | 0.976 | 1 | 1 | 1 | 1 | 1 | 1 | 1 | -0.005 |
| SE | 106418 | ENSG00000135297 | MTO1 | chr6 | + | 73473364 | 73473654 | 73471459 | 73471557 | 73479731 | 73479844 | 104 | 88 | 80 | 77 | 148 | 0 | 0 | 3 | 0 | 0 | 105 | 78 | 48 | 51 | 63 | 0 | 0 | 0 | 0 | 0 | 298 | 149 | 1 | 1 | 1 | 1 | 0.93 | 1 | 1 | 1 | 1 | 1 | 1 | 1 | -0.014 |
| SE | 106419 | ENSG00000135297 | MTO1 | chr6 | + | 73479731 | 73479844 | 73473364 | 73473654 | 73479935 | 73480126 | 175 | 135 | 134 | 145 | 209 | 0 | 0 | 0 | 1 | 0 | 146 | 125 | 114 | 77 | 98 | 0 | 0 | 0 | 0 | 0 | 262 | 149 | 1 | 1 | 1 | 1 | 1 | 0.988 | 1 | 1 | 1 | 1 | 1 | 1 | -0.002 |
| SE | 106420 | ENSG00000135297 | MTO1 | chr6 | + | 73479731 | 73480126 | 73473364 | 73473654 | 73480674 | 73480805 | 200 | 168 | 169 | 164 | 242 | 0 | 0 | 0 | 0 | 0 | 171 | 154 | 119 | 98 | 102 | 0 | 0 | 0 | 0 | 0 | 298 | 149 | 1 | 1 | 1 | 1 | 0.988 | 1 | 1 | 1 | 1 | 1 | 1 | 1 | -0.002 |
| SE | 106421 | ENSG00000135297 | MTO1 | chr6 | + | 73479935 | 73480126 | 73473364 | 73473654 | 73480674 | 73480805 | 128 | 99 | 111 | 107 | 147 | 0 | 0 | 1 | 0 | 0 | 90 | 98 | 82 | 74 | 59 | 0 | 0 | 0 | 0 | 0 | 298 | 149 | 1 | 1 | 1 | 1 | 0.982 | 1 | 1 | 1 | 1 | 1 | 1 | 1 | -0.004 |
| SE | 106428 | ENSG00000135297 | MTO1 | chr6 | + | 73481000 | 73481116 | 73480674 | 73480805 | 73482448 | 73482620 | 1 | 0 | 0 | 5 | 5 | 0 | 3 | 3 | 3 | 6 | 0 | 0 | 8 | 0 | 0 | 8 | 0 | 3 | 0 | 0 | 265 | 149 | 1 | 1 | 1 | 0 | 0 | 0.484 | 0.319 | 0 | NA | 0.6 | NA | NA | 0.061 |
| SE | 106436 | ENSG00000135297 | MTO1 | chr6 | + | 73482448 | 73482620 | 73482028 | 73482244 | 73492233 | 73492352 | 129 | 90 | 138 | 146 | 236 | 3 | 0 | 1 | 0 | 0 | 92 | 113 | 75 | 37 | 29 | 0 | 0 | 0 | 0 | 3 | 298 | 149 | 1 | 1 | 0.956 | 1 | 0.986 | 1 | 1 | 1 | 1 | 1 | 1 | 0.829 | 0.023 |
| SE | 106437 | ENSG00000135297 | MTO1 | chr6 | + | 73482448 | 73482620 | 73482028 | 73482244 | 73497735 | 73497896 | 41 | 25 | 35 | 43 | 66 | 0 | 0 | 1 | 0 | 0 | 66 | 29 | 31 | 12 | 9 | 0 | 0 | 0 | 0 | 0 | 298 | 149 | 1 | 1 | 1 | 1 | 0.946 | 1 | 1 | 1 | 1 | 1 | 1 | 1 | -0.011 |
| SE | 106440 | ENSG00000135297 | MTO1 | chr6 | + | 73492233 | 73492352 | 73482028 | 73482244 | 73497735 | 73497896 | 56 | 55 | 70 | 66 | 128 | 0 | 0 | 1 | 0 | 0 | 38 | 36 | 43 | 41 | 27 | 0 | 0 | 0 | 0 | 0 | 268 | 149 | 1 | 1 | 1 | 1 | 0.975 | 1 | 1 | 1 | 1 | 1 | 1 | 1 | -0.005 |
| SE | 106441 | ENSG00000135297 | MTO1 | chr6 | + | 73492233 | 73492352 | 73482448 | 73482620 | 73497735 | 73497896 | 138 | 120 | 171 | 167 | 298 | 0 | 0 | 0 | 0 | 0 | 64 | 120 | 90 | 66 | 44 | 0 | 0 | 3 | 0 | 0 | 268 | 149 | 1 | 1 | 1 | 1 | 1 | 1 | 1 | 1 | 1 | 0.943 | 1 | 1 | 0.011 |
| SE | 106443 | ENSG00000135297 | MTO1 | chr6 | + | 73498910 | 73499057 | 73492233 | 73492352 | 73500573 | 73501131 | 1 | 0 | 0 | 0 | 0 | 5 | 1 | 0 | 2 | 7 | 0 | 0 | 0 | 0 | 0 | 0 | 12 | 5 | 4 | 0 | 296 | 149 | 1 | 1 | 0.091 | 0 | NA | 0 | 0 | NA | 0 | 0 | 0 | NA | 0.023 |
| SE | 106444 | ENSG00000135127 | BICDL1 | chr12 | + | 120071621 | 120071801 | 120064636 | 120064879 | 120080886 | 120081017 | 67 | 38 | 87 | 75 | 47 | 1 | 0 | 0 | 2 | 0 | 61 | 55 | 46 | 81 | 166 | 0 | 0 | 0 | 0 | 0 | 298 | 149 | 1 | 1 | 0.971 | | 1 | 0.949 | | 1 | 1 | 1 | 1 | 1 | -0.016 |
| SE | 106445 | ENSG00000135127 | BICDL1 | chr12 | + | 120072510 | 120072729 | 120071621 | 120071801 | 120074442 | 120074586 | 47 | 19 | 29 | 48 | 28 | 0 | 0 | 0 | 2 | 0 | 28 | 46 | 17 | 31 | 37 | 0 | 0 | 0 | 0 | 0 | 298 | 149 | 1 | 1 | 1 | | 1 | 0.923 | | 1 | 1 | 1 | 1 | 1 | -0.015 |
| SE | 106449 | ENSG00000135127 | BICDL1 | chr12 | + | 120090910 | 120091019 | 120089950 | 120090071 | 120092999 | 120093456 | 0 | 0 | 0 | 1 | 0 | 65 | 47 | 52 | 58 | 37 | 0 | 4 | 0 | 0 | 0 | 33 | 48 | 27 | 20 | 54 | 258 | 149 | 1 | 1 | 0 | 0 | 0 | 0.01 | 0 | 0 | 0.046 | 0 | 0 | 0 | -0.007 |
| SE | 106452 | ENSG00000198270 | TMEM116 | chr12 | - | 111932585 | 111932659 | 111931281 | 111931827 | 111933885 | 111934030 | 39 | 37 | 52 | 62 | 96 | 0 | 2 | 0 | 0 | 0 | 5 | 15 | 29 | 12 | 25 | 0 | 0 | 0 | 0 | 0 | 223 | 149 | 1 | 1 | 1 | 0.925 | 1 | 1 | 1 | 1 | 1 | 1 | 1 | 1 | -0.015 |
| SE | 106453 | ENSG00000198270 | TMEM116 | chr12 | - | 111932585 | 111932731 | 111931379 | 111931827 | 111933885 | 111934030 | 13 | 20 | 38 | 37 | 48 | 0 | 2 | 0 | 0 | 0 | 8 | 18 | 7 | 11 | 0 | 0 | 0 | 0 | 0 | 0 | 295 | 149 | 1 | 1 | 1 | 0.835 | 1 | 1 | 1 | 1 | 1 | 1 | 1 | NA | -0.033 |
| SE | 106454 | ENSG00000198270 | TMEM116 | chr12 | - | 111933885 | 111934030 | 111932585 | 111932659 | 111936691 | 111936830 | 64 | 46 | 52 | 67 | 92 | 0 | 0 | 0 | 0 | 2 | 5 | 7 | 35 | 13 | 47 | 0 | 0 | 0 | 0 | 0 | 294 | 149 | 1 | 1 | 1 | 1 | 1 | 1 | 0.959 | 1 | 1 | 1 | 1 | 1 | -0.008 |
| SE | 106455 | ENSG00000198270 | TMEM116 | chr12 | - | 111933885 | 111934030 | 111932585 | 111932659 | 111937159 | 111937243 | 52 | 34 | 50 | 46 | 73 | 0 | 1 | 2 | 0 | 0 | 5 | 13 | 33 | 26 | 36 | 0 | 0 | 5 | 0 | 0 | 294 | 149 | 1 | 1 | 1 | 0.945 | 0.927 | 1 | 1 | 1 | 1 | 0.77 | 1 | 1 | 0.02 |
| SE | 106462 | ENSG00000198270 | TMEM116 | chr12 | - | 111936691 | 111936830 | 111933885 | 111934030 | 111938160 | 111938210 | 25 | 22 | 19 | 30 | 34 | 0 | 0 | 1 | 0 | 0 | 0 | 1 | 8 | 5 | 18 | 0 | 0 | 0 | 0 | 0 | 288 | 149 | 1 | 1 | 1 | 1 | 0.908 | 1 | 1 | NA | 1 | 1 | 1 | 1 | -0.018 |
| SE | 106466 | ENSG00000198270 | TMEM116 | chr12 | - | 111937159 | 111937243 | 111933885 | 111934030 | 111938160 | 111938210 | 39 | 32 | 40 | 30 | 78 | 0 | 0 | 1 | 0 | 0 | 8 | 26 | 19 | 43 | 37 | 0 | 0 | 0 | 0 | 0 | 233 | 149 | 1 | 1 | 1 | 1 | 0.962 | 1 | 1 | 1 | 1 | 1 | 1 | 1 | -0.008 |
| SE | 106467 | ENSG00000198270 | TMEM116 | chr12 | - | 111937159 | 111937243 | 111933885 | 111936830 | 111938160 | 111938210 | 80 | 62 | 70 | 62 | 190 | 0 | 0 | 1 | 0 | 1 | 16 | 50 | 32 | 55 | 72 | 0 | 0 | 0 | 0 | 0 | 233 | 149 | 1 | 1 | 1 | 1 | 0.978 | 1 | 0.992 | 1 | 1 | 1 | 1 | 1 | -0.006 |
| SE | 106468 | ENSG00000198270 | TMEM116 | chr12 | - | 111938160 | 111938210 | 111933885 | 111934030 | 111943264 | 111943369 | 17 | 13 | 21 | 15 | 39 | 0 | 0 | 0 | 1 | 0 | 10 | 8 | 27 | 11 | 9 | 0 | 0 | 0 | 0 | 0 | 199 | 149 | 1 | 1 | 1 | 1 | 1 | 0.918 | 1 | 1 | 1 | 1 | 1 | 1 | -0.016 |
| SE | 106470 | ENSG00000198270 | TMEM116 | chr12 | - | 111943264 | 111943369 | 111938160 | 111938210 | 111978735 | 111978787 | 52 | 31 | 39 | 34 | 82 | 0 | 0 | 1 | 0 | 0 | 8 | 28 | 21 | 34 | 26 | 0 | 0 | 0 | 0 | 0 | 254 | 149 | 1 | 1 | 1 | 1 | 0.958 | 1 | 1 | 1 | 1 | 1 | 1 | 1 | -0.008 |
| SE | 106472 | ENSG00000198270 | TMEM116 | chr12 | - | 111943264 | 111943369 | 111938160 | 111938210 | 112003799 | 112003863 | 46 | 27 | 34 | 30 | 80 | 0 | 1 | 0 | 1 | 0 | 8 | 19 | 16 | 31 | 26 | 0 | 0 | 0 | 0 | 0 | 254 | 149 | 1 | 1 | 1 | 0.941 | 1 | 0.946 | 1 | 1 | 1 | 1 | 1 | 1 | -0.023 |
| SE | 106474 | ENSG00000198270 | TMEM116 | chr12 | - | 111965594 | 111965759 | 111938183 | 111938210 | 111991757 | 111991889 | 2 | 2 | 4 | 2 | 2 | 2 | 1 | 1 | 0 | 8 | 0 | 2 | 0 | 0 | 0 | 0 | 0 | 0 | 0 | 0 | 298 | 149 | 1 | 1 | 0.333 | 0.5 | 0.667 | 1 | 0.111 | NA | 1 | NA | NA | NA | -0.478 |
| SE | 106475 | ENSG00000198270 | TMEM116 | chr12 | - | 111965594 | 111965759 | 111943264 | 111943369 | 111978735 | 111978787 | 6 | 3 | 7 | 10 | 4 | 6 | 4 | 6 | 4 | 2 | 0 | 5 | 1 | 0 | 9 | 0 | 9 | 5 | 3 | 0 | 298 | 149 | 1 | 1 | 0.333 | 0.273 | 0.368 | 0.556 | 0.5 | NA | 0.217 | 0.091 | 0 | 1 | 0.079 |
| SE | 106477 | ENSG00000198270 | TMEM116 | chr12 | - | 111978735 | 111978787 | 111938160 | 111938210 | 111991757 | 111991889 | 0 | 0 | 3 | 3 | 3 | 2 | 1 | 1 | 0 | 8 | 0 | 5 | 0 | 0 | 0 | 0 | 0 | 0 | 0 | 0 | 201 | 149 | 1 | 1 | 0 | 0 | 0.69 | 1 | 0.218 | NA | 1 | NA | NA | NA | -0.618 |
| SE | 106479 | ENSG00000198270 | TMEM116 | chr12 | - | 111978735 | 111978787 | 111965594 | 111965759 | 111991757 | 111991889 | 1 | 0 | 4 | 5 | 3 | 2 | 2 | 4 | 2 | 2 | 0 | 5 | 0 | 0 | 0 | 0 | 2 | 0 | 0 | 0 | 201 | 149 | 1 | 1 | 0.27 | 0 | 0.426 | 0.65 | 0.527 | NA | 0.65 | NA | NA | NA | -0.275 |
| SE | 106480 | ENSG00000198270 | TMEM116 | chr12 | - | 111991757 | 111991889 | 111938046 | 111938210 | 112003799 | 112003863 | 24 | 1 | 7 | 1 | 23 | 0 | 1 | 0 | 1 | 0 | 0 | 6 | 4 | 0 | 0 | 0 | 0 | 0 | 0 | 0 | 281 | 149 | 1 | 1 | 1 | 0.347 | 1 | 0.347 | 1 | NA | 1 | 1 | NA | NA | -0.261 |
| SE | 106481 | ENSG00000198270 | TMEM116 | chr12 | - | 111991757 | 111991889 | 111943260 | 111943369 | 112003799 | 112003863 | 43 | 18 | 25 | 13 | 45 | 0 | 0 | 1 | 0 | 0 | 19 | 10 | 20 | 15 | 11 | 0 | 0 | 0 | 0 | 0 | 281 | 149 | 1 | 1 | 1 | 1 | 0.93 | 1 | 1 | 1 | 1 | 1 | 1 | 1 | -0.014 |
| SE | 106482 | ENSG00000198270 | TMEM116 | chr12 | - | 112003799 | 112003863 | 111991743 | 111991889 | 112005256 | 112005303 | 28 | 0 | 9 | 1 | 20 | 6 | 6 | 10 | 8 | 16 | 0 | 13 | 5 | 0 | 0 | 0 | 0 | 1 | 6 | 0 | 213 | 149 | 1 | 1 | 0.766 | 0 | 0.386 | 0.08 | 0.466 | NA | 1 | 0.778 | 0 | NA | -0.253 |
| SE | 106486 | ENSG00000176915 | ANKLE2 | chr12 | - | 132735405 | 132735512 | 132734384 | 132734575 | 132741418 | 132741485 | 40 | 32 | 42 | 17 | 36 | 0 | 0 | 0 | 0 | 0 | 30 | 38 | 27 | 22 | 33 | 0 | 0 | 0 | 0 | 2 | 256 | 149 | 1 | 1 | 1 | 1 | 1 | 1 | 1 | 1 | 1 | 1 | 1 | 0.906 | 0.019 |
| SE | 106490 | ENSG00000176915 | ANKLE2 | chr12 | - | 132750642 | 132750849 | 132748137 | 132748331 | 132754674 | 132755133 | 262 | 143 | 232 | 217 | 311 | 0 | 3 | 1 | 0 | 2 | 431 | 262 | 259 | 285 | 331 | 0 | 0 | 0 | 0 | 0 | 298 | 149 | 1 | 1 | 1 | 0.96 | 0.991 | 1 | 0.987 | 1 | 1 | 1 | 1 | 1 | -0.012 |
| SE | 106491 | ENSG00000185482 | STAC3 | chr12 | - | 57244314 | 57244366 | 57244087 | 57244225 | 57244915 | 57244965 | 0 | 2 | 4 | 2 | 16 | 0 | 0 | 0 | 0 | 0 | 8 | 9 | 4 | 5 | 0 | 5 | 0 | 2 | 0 | 0 | 201 | 149 | 1 | 1 | NA | 1 | 1 | 1 | 1 | 0.543 | 1 | 0.597 | 1 | NA | 0.215 |
| SE | 106496 | ENSG00000185482 | STAC3 | chr12 | - | 57248125 | 57248198 | 57245144 | 57245211 | 57248705 | 57248803 | 51 | 29 | 27 | 29 | 80 | 1 | 0 | 0 | 0 | 0 | 189 | 50 | 39 | 153 | 93 | 0 | 0 | 1 | 2 | 0 | 222 | 149 | 1 | 1 | 0.972 | 1 | 1 | 1 | 1 | 1 | 1 | 0.963 | 0.981 | 1 | 0.006 |
| SE | 106498 | ENSG00000185482 | STAC3 | chr12 | - | 57249040 | 57249308 | 57248705 | 57248803 | 57250992 | 57251184 | 96 | 61 | 45 | 70 | 116 | 1 | 0 | 0 | 0 | 0 | 370 | 165 | 83 | 323 | 236 | 0 | 0 | 2 | 0 | 0 | 298 | 149 | 1 | 1 | 0.98 | 1 | 1 | 1 | 1 | 1 | 1 | 0.954 | 1 | 1 | 0.005 |
| SE | 106499 | ENSG00000185482 | STAC3 | chr12 | - | 57249570 | 57249637 | 57249018 | 57249308 | 57250992 | 57251186 | 4 | 0 | 0 | 0 | 2 | 0 | 1 | 0 | 0 | 0 | 13 | 0 | 1 | 15 | 6 | 4 | 0 | 0 | 6 | 0 | 216 | 149 | 1 | 1 | 1 | 0 | NA | NA | 1 | 0.692 | NA | 1 | 0.633 | 1 | -0.165 |
| SE | 106500 | ENSG00000185482 | STAC3 | chr12 | - | 57249570 | 57249730 | 57249040 | 57249308 | 57250992 | 57251188 | 2 | 0 | 0 | 0 | 0 | 0 | 1 | 0 | 0 | 0 | 13 | 0 | 0 | 12 | 0 | 4 | 0 | 0 | 6 | 0 | 298 | 149 | 1 | 1 | 1 | 0 | NA | NA | NA | 0.619 | NA | NA | 0.5 | NA | -0.059 |
| SE | 106501 | ENSG00000139725 | RHOF | chr12 | - | 121799846 | 121799956 | 121793150 | 121793239 | 121801571 | 121802784 | 4 | 2 | 8 | 7 | 10 | 0 | 0 | 0 | 0 | 0 | 16 | 0 | 3 | 7 | 22 | 0 | 0 | 3 | 0 | 0 | 259 | 149 | 1 | 1 | 1 | 1 | 1 | 1 | 1 | 1 | NA | 0.365 | 1 | 1 | 0.159 |
| SE | 106506 | ENSG00000111666 | CHPT1 | chr12 | + | 101720122 | 101720254 | 101716727 | 101716812 | 101723167 | 101723254 | 177 | 67 | 105 | 138 | 87 | 0 | 0 | 0 | 0 | 0 | 287 | 207 | 159 | 223 | 165 | 0 | 0 | 1 | 0 | 0 | 281 | 149 | 1 | 1 | 1 | 1 | 1 | 1 | 1 | 1 | 1 | 0.988 | 1 | 1 | 0.002 |
| SE | 106507 | ENSG00000111666 | CHPT1 | chr12 | + | 101726293 | 101726404 | 101723721 | 101723847 | 101728900 | 101729051 | 114 | 59 | 71 | 83 | 177 | 0 | 3 | 0 | 0 | 0 | 202 | 124 | 127 | 158 | 97 | 0 | 0 | 0 | 0 | 0 | 260 | 149 | 1 | 1 | 1 | 0.919 | 1 | 1 | 1 | 1 | 1 | 1 | 1 | 1 | -0.016 |
| SE | 106508 | ENSG00000111666 | CHPT1 | chr12 | + | 101742396 | 101742506 | 101730520 | 101730636 | 101743714 | 101744010 | 2 | 0 | 0 | 0 | 0 | 0 | 0 | 1 | 2 | 0 | 0 | 6 | 0 | 3 | 0 | 0 | 0 | 0 | 0 | 0 | 259 | 149 | 1 | 1 | 1 | NA | 0 | 0 | NA | NA | 1 | NA | 1 | NA | -0.667 |
| SE | 106516 | ENSG00000184967 | NOC4L | chr12 | + | 132151737 | 132151820 | 132151483 | 132151644 | 132152083 | 132152197 | 10 | 5 | 6 | 6 | 21 | 0 | 0 | 0 | 1 | 0 | 15 | 18 | 7 | 23 | 8 | 0 | 0 | 0 | 0 | 0 | 232 | 149 | 1 | 1 | 1 | 1 | 1 | 0.794 | 1 | 1 | 1 | 1 | 1 | 1 | -0.041 |
| SE | 106518 | ENSG00000272369 | NA | chr12 | + | 46571255 | 46571412 | 46537501 | 46537631 | 46652389 | 46652550 | 16 | 7 | 11 | 4 | 11 | 1 | 0 | 0 | 0 | 7 | 4 | 9 | 7 | 0 | 7 | 0 | 0 | 0 | 0 | 0 | 298 | 149 | 1 | 1 | 0.889 | 1 | 1 | 1 | 0.44 | 1 | 1 | 1 | NA | 1 | -0.134 |
| SE | 106524 | ENSG00000186496 | ZNF396 | chr18 | - | 35374469 | 35374823 | 35374173 | 35374364 | 35377277 | 35377310 | 0 | 0 | 1 | 0 | 0 | 3 | 2 | 0 | 1 | 0 | 0 | 0 | 4 | 0 | 0 | 0 | 0 | 0 | 0 | 0 | 298 | 149 | 1 | 1 | 0 | 0 | 1 | 0 | NA | NA | NA | 1 | NA | NA | -0.75 |
| SE | 106534 | ENSG00000176834 | VSIG10 | chr12 | - | 118082126 | 118082429 | 118079345 | 118079606 | 118095532 | 118095814 | 3 | 0 | 1 | 1 | 0 | 0 | 0 | 0 | 0 | 1 | 0 | 3 | 0 | 20 | 6 | 0 | 0 | 0 | 0 | 0 | 298 | 149 | 1 | 1 | 1 | NA | 1 | 1 | 0 | NA | 1 | NA | 1 | 1 | -0.25 |
| SE | 106535 | ENSG00000176834 | VSIG10 | chr12 | - | 118102988 | 118103170 | 118095532 | 118095814 | 118103592 | 118103862 | 0 | 0 | 1 | 0 | 1 | 2 | 0 | 0 | 0 | 0 | 0 | 0 | 0 | 3 | 0 | 0 | 0 | 0 | 0 | 0 | 298 | 149 | 1 | 1 | 0 | NA | 1 | NA | 1 | NA | NA | NA | 1 | NA | -0.333 |
| SE | 106537 | ENSG00000176834 | VSIG10 | chr12 | - | 118135638 | 118135742 | 118126480 | 118126670 | 118135856 | 118135948 | 0 | 1 | 7 | 0 | 0 | 2 | 3 | 0 | 0 | 1 | 0 | 0 | 0 | 0 | 0 | 0 | 0 | 0 | 5 | 0 | 253 | 149 | 1 | 1 | 0 | 0.164 | 1 | NA | 0 | NA | NA | NA | 0 | NA | 0.291 |
| SE | 106539 | ENSG00000183955 | KMT5A | chr12 | + | 123389432 | 123389554 | 123383772 | 123384208 | 123390629 | 123390786 | 2 | 3 | 4 | 10 | 2 | 0 | 0 | 0 | 0 | 1 | 0 | 16 | 10 | 10 | 18 | 0 | 7 | 0 | 0 | 0 | 271 | 149 | 1 | 1 | 1 | 1 | 1 | 1 | 0.524 | NA | 0.557 | 1 | 1 | 1 | 0.016 |
| SE | 106541 | ENSG00000183955 | KMT5A | chr12 | + | 123404883 | 123405074 | 123403572 | 123403632 | 123407492 | 123407568 | 222 | 121 | 255 | 241 | 250 | 1 | 0 | 0 | 0 | 0 | 387 | 214 | 183 | 474 | 355 | 0 | 0 | 1 | 5 | 0 | 298 | 149 | 1 | 1 | 0.991 | 1 | 1 | 1 | 1 | 1 | 1 | 0.989 | 0.979 | 1 | 0.005 |
| SE | 106542 | ENSG00000111696 | NT5DC3 | chr12 | - | 103785570 | 103785747 | 103785334 | 103785475 | 103787440 | 103787527 | 3 | 0 | 0 | 0 | 0 | 12 | 19 | 4 | 4 | 11 | 0 | 0 | 5 | 4 | 0 | 6 | 8 | 11 | 17 | 0 | 298 | 149 | 1 | 1 | 0.111 | 0 | 0 | 0 | 0 | 0 | 0 | 0.185 | 0.105 | NA | -0.05 |
| SE | 106543 | ENSG00000111696 | NT5DC3 | chr12 | - | 103787440 | 103787527 | 103785334 | 103785475 | 103788837 | 103788919 | 18 | 23 | 6 | 11 | 14 | 0 | 2 | 0 | 0 | 0 | 13 | 10 | 14 | 21 | 0 | 0 | 0 | 0 | 0 | 0 | 236 | 149 | 1 | 1 | 1 | 0.879 | 1 | 1 | 1 | 1 | 1 | 1 | 1 | NA | -0.024 |
| SE | 106544 | ENSG00000111696 | NT5DC3 | chr12 | - | 103788837 | 103788919 | 103787440 | 103787527 | 103793163 | 103793265 | 13 | 17 | 11 | 9 | 16 | 1 | 0 | 0 | 0 | 0 | 12 | 6 | 15 | 11 | 0 | 0 | 0 | 0 | 0 | 0 | 231 | 149 | 1 | 1 | 0.893 | 1 | 1 | 1 | 1 | 1 | 1 | 1 | 1 | NA | -0.021 |
| SE | 106548 | ENSG00000129562 | DAD1 | chr14 | - | 22575058 | 22575233 | 22564906 | 22565137 | 22588946 | 22589224 | 426 | 203 | 360 | 381 | 393 | 0 | 2 | 1 | 1 | 0 | 440 | 337 | 198 | 415 | 823 | 0 | 6 | 1 | 2 | 0 | 298 | 149 | 1 | 1 | 1 | 0.981 | 0.994 | 0.995 | 1 | 1 | 0.966 | 0.99 | 0.99 | 1 | 0.005 |
| SE | 106551 | ENSG00000134532 | SOX5 | chr12 | - | 23665443 | 23665564 | 23640811 | 23640897 | 23734683 | 23734752 | 0 | 6 | 2 | 1 | 10 | 0 | 1 | 0 | 3 | 0 | 0 | 3 | 2 | 0 | 0 | 0 | 0 | 0 | 0 | 0 | 270 | 149 | 1 | 1 | NA | 0.768 | 1 | 0.155 | 1 | NA | 1 | 1 | NA | NA | -0.269 |
| SE | 106552 | ENSG00000134532 | SOX5 | chr12 | - | 23665443 | 23665567 | 23640811 | 23640897 | 23734683 | 23734715 | 0 | 2 | 0 | 0 | 6 | 0 | 1 | 0 | 3 | 0 | 0 | 3 | 0 | 0 | 0 | 0 | 0 | 0 | 0 | 0 | 273 | 149 | 1 | 1 | NA | 0.522 | NA | 0 | 1 | NA | 1 | NA | NA | NA | -0.493 |
| SE | 106555 | ENSG00000134532 | SOX5 | chr12 | - | 24277215 | 24277312 | 24213342 | 24213417 | 24368562 | 24368639 | 10 | 20 | 12 | 11 | 7 | 0 | 0 | 0 | 3 | 0 | 0 | 9 | 3 | 1 | 0 | 0 | 0 | 0 | 0 | 0 | 246 | 149 | 1 | 1 | 1 | 1 | 1 | 0.69 | 1 | NA | 1 | 1 | 1 | NA | -0.062 |
| SE | 106557 | ENSG00000134532 | SOX5 | chr12 | - | 24407484 | 24407566 | 24367934 | 24368639 | 24562337 | 24562382 | 2 | 7 | 0 | 8 | 4 | 0 | 0 | 0 | 3 | 0 | 0 | 0 | 1 | 0 | 0 | 0 | 8 | 0 | 0 | 0 | 231 | 149 | 1 | 1 | 1 | 1 | NA | 1 | 1 | NA | 0 | 1 | NA | NA | 0.5 |
| SE | 106560 | ENSG00000111790 | FGFR1OP2 | chr12 | + | 26953548 | 26953706 | 26938480 | 26938710 | 26954144 | 26954293 | 0 | 3 | 0 | 0 | 0 | 126 | 77 | 116 | 121 | 141 | 0 | 0 | 0 | 0 | 0 | 72 | 65 | 68 | 19 | 39 | 298 | 149 | 1 | 1 | 0 | 0.019 | 0 | 0 | 0 | 0 | 0 | 0 | 0 | 0 | 0.004 |
| SE | 106561 | ENSG00000111790 | FGFR1OP2 | chr12 | + | 26954144 | 26954293 | 26938469 | 26938710 | 26956542 | 26956587 | 504 | 386 | 439 | 498 | 571 | 4 | 0 | 0 | 1 | 3 | 298 | 292 | 191 | 174 | 180 | 0 | 0 | 0 | 0 | 0 | 298 | 149 | 1 | 1 | 0.984 | 1 | 1 | 0.996 | 0.99 | 1 | 1 | 1 | 1 | 1 | -0.006 |
| SE | 106572 | ENSG00000103356 | EARS2 | chr16 | - | 23546381 | 23546434 | 23544540 | 23544703 | 23552148 | 23552304 | 1 | 0 | 0 | 0 | 0 | 14 | 1 | 9 | 8 | 2 | 0 | 0 | 0 | 0 | 0 | 39 | 16 | 11 | 2 | 10 | 202 | 149 | 1 | 1 | 0.05 | 0 | 0 | 0 | 0 | 0 | 0 | 0 | 0 | 0 | 0.01 |
| SE | 106573 | ENSG00000103356 | EARS2 | chr16 | - | 23553006 | 23553123 | 23552148 | 23552304 | 23557204 | 23557364 | 1 | 0 | 0 | 0 | 0 | 8 | 3 | 3 | 2 | 2 | 0 | 0 | 0 | 0 | 0 | 7 | 2 | 4 | 0 | 22 | 266 | 149 | 1 | 1 | 0.065 | 0 | 0 | 0 | 0 | 0 | 0 | 0 | NA | 0 | 0.013 |
| SE | 106574 | ENSG00000103356 | EARS2 | chr16 | - | 23553043 | 23553123 | 23552148 | 23552304 | 23556889 | 23557177 | 0 | 0 | 0 | 4 | 0 | 0 | 0 | 2 | 2 | 0 | 0 | 0 | 2 | 5 | 0 | 0 | 1 | 0 | 0 | 0 | 229 | 149 | 1 | 1 | NA | NA | 0 | 0.565 | NA | NA | NA | 0.565 | 1 | NA | -0.5 |
| SE | 106577 | ENSG00000166012 | TAF1D | chr11 | - | 93730676 | 93730712 | 93730193 | 93730447 | 93731149 | 93731216 | 182 | 169 | 167 | 151 | 450 | 0 | 1 | 0 | 4 | 4 | 10 | 51 | 64 | 18 | 28 | 0 | 5 | 0 | 0 | 0 | 185 | 149 | 1 | 1 | 1 | 0.993 | 1 | 0.968 | 0.989 | 1 | 0.891 | 1 | 1 | 1 | 0.012 |
| SE | 106580 | ENSG00000166012 | TAF1D | chr11 | - | 93733086 | 93733121 | 93732243 | 93732291 | 93734624 | 93734660 | 21 | 17 | 14 | 17 | 79 | 1 | 0 | 0 | 1 | 4 | 10 | 15 | 17 | 1 | 0 | 0 | 0 | 0 | 0 | 0 | 184 | 149 | 1 | 1 | 0.944 | 1 | 1 | 0.932 | 0.941 | 1 | 1 | 1 | 1 | NA | -0.037 |
| SE | 106581 | ENSG00000166012 | TAF1D | chr11 | - | 93733349 | 93733397 | 93732799 | 93733121 | 93734624 | 93734660 | 419 | 363 | 390 | 362 | 871 | 0 | 0 | 0 | 0 | 4 | 22 | 187 | 172 | 55 | 11 | 0 | 0 | 0 | 0 | 0 | 197 | 149 | 1 | 1 | 1 | 1 | 1 | 1 | 0.994 | 1 | 1 | 1 | 1 | 1 | -0.001 |
| SE | 106582 | ENSG00000166012 | TAF1D | chr11 | - | 93733349 | 93733419 | 93733086 | 93733121 | 93734624 | 93734660 | 39 | 37 | 45 | 47 | 60 | 0 | 0 | 0 | 0 | 4 | 17 | 10 | 19 | 3 | 26 | 0 | 0 | 0 | 0 | 0 | 219 | 149 | 1 | 1 | 1 | 1 | 1 | 1 | 0.911 | 1 | 1 | 1 | 1 | 1 | -0.018 |
| SE | 106583 | ENSG00000166012 | TAF1D | chr11 | - | 93733349 | 93733505 | 93733086 | 93733121 | 93734624 | 93734660 | 44 | 29 | 40 | 34 | 58 | 0 | 0 | 0 | 0 | 4 | 23 | 9 | 22 | 8 | 17 | 0 | 0 | 0 | 0 | 0 | 298 | 149 | 1 | 1 | 1 | 1 | 1 | 1 | 0.879 | 1 | 1 | 1 | 1 | 1 | -0.024 |
| SE | 106584 | ENSG00000166012 | TAF1D | chr11 | - | 93734624 | 93734660 | 93730676 | 93730712 | 93736140 | 93736239 | 70 | 60 | 87 | 71 | 148 | 1 | 5 | 3 | 4 | 0 | 7 | 30 | 28 | 7 | 0 | 0 | 0 | 3 | 0 | 0 | 185 | 149 | 1 | 1 | 0.983 | 0.906 | 0.959 | 0.935 | 1 | 1 | 1 | 0.883 | 1 | NA | -0.014 |
| SE | 106585 | ENSG00000166012 | TAF1D | chr11 | - | 93734624 | 93734660 | 93730676 | 93730712 | 93737063 | 93737239 | 18 | 6 | 12 | 8 | 42 | 0 | 0 | 0 | 2 | 0 | 0 | 8 | 25 | 0 | 0 | 0 | 0 | 0 | 0 | 0 | 185 | 149 | 1 | 1 | 1 | 1 | 1 | 0.763 | 1 | NA | 1 | 1 | NA | NA | -0.047 |
| SE | 106586 | ENSG00000166012 | TAF1D | chr11 | - | 93734624 | 93734660 | 93732243 | 93732291 | 93736140 | 93736239 | 57 | 51 | 69 | 65 | 121 | 1 | 0 | 0 | 0 | 0 | 7 | 17 | 24 | 4 | 0 | 0 | 0 | 0 | 0 | 0 | 185 | 149 | 1 | 1 | 0.979 | 1 | 1 | 1 | 1 | 1 | 1 | 1 | 1 | NA | -0.004 |
| SE | 106587 | ENSG00000166012 | TAF1D | chr11 | - | 93734624 | 93734660 | 93732243 | 93732291 | 93737063 | 93737239 | 14 | 5 | 9 | 7 | 43 | 1 | 0 | 0 | 0 | 0 | 0 | 6 | 14 | 0 | 0 | 0 | 0 | 0 | 0 | 0 | 185 | 149 | 1 | 1 | 0.919 | 1 | 1 | 1 | 1 | NA | 1 | 1 | NA | NA | -0.016 |
| SE | 106588 | ENSG00000166012 | TAF1D | chr11 | - | 93734624 | 93734660 | 93732976 | 93733397 | 93734962 | 93736175 | 254 | 211 | 220 | 266 | 431 | 0 | 0 | 3 | 0 | 0 | 13 | 123 | 98 | 36 | 9 | 0 | 0 | 0 | 0 | 0 | 185 | 149 | 1 | 1 | 1 | 1 | 0.983 | 1 | 1 | 1 | 1 | 1 | 1 | 1 | -0.003 |
| SE | 106589 | ENSG00000166012 | TAF1D | chr11 | - | 93734624 | 93734660 | 93732976 | 93733397 | 93736140 | 93736239 | 441 | 414 | 424 | 416 | 835 | 0 | 0 | 1 | 0 | 1 | 20 | 173 | 175 | 47 | 11 | 0 | 0 | 3 | 0 | 0 | 185 | 149 | 1 | 1 | 1 | 1 | 0.997 | 1 | 0.999 | 1 | 1 | 0.979 | 1 | 1 | 0.003 |
| SE | 106590 | ENSG00000166012 | TAF1D | chr11 | - | 93734624 | 93734660 | 93732976 | 93733397 | 93737063 | 93737239 | 298 | 252 | 255 | 290 | 525 | 0 | 0 | 0 | 0 | 2 | 19 | 140 | 129 | 41 | 9 | 0 | 0 | 0 | 0 | 0 | 185 | 149 | 1 | 1 | 1 | 1 | 1 | 1 | 0.995 | 1 | 1 | 1 | 1 | 1 | -0.001 |
| SE | 106591 | ENSG00000166012 | TAF1D | chr11 | - | 93736140 | 93736304 | 93730676 | 93730712 | 93737063 | 93737239 | 1 | 7 | 6 | 7 | 6 | 0 | 0 | 0 | 2 | 0 | 6 | 1 | 10 | 0 | 0 | 0 | 0 | 0 | 0 | 0 | 298 | 149 | 1 | 1 | 1 | 1 | 1 | 0.636 | 1 | 1 | 1 | 1 | NA | NA | -0.073 |
| SE | 106592 | ENSG00000166012 | TAF1D | chr11 | - | 93736140 | 93736304 | 93732243 | 93732291 | 93737063 | 93737239 | 1 | 2 | 3 | 3 | 6 | 1 | 0 | 0 | 0 | 0 | 6 | 1 | 7 | 0 | 0 | 0 | 0 | 0 | 0 | 0 | 298 | 149 | 1 | 1 | 0.333 | 1 | 1 | 1 | 1 | 1 | 1 | 1 | NA | NA | -0.133 |
| SE | 106593 | ENSG00000166012 | TAF1D | chr11 | - | 93736140 | 93736304 | 93733304 | 93733397 | 93737063 | 93737239 | 0 | 2 | 4 | 3 | 7 | 0 | 0 | 0 | 0 | 2 | 6 | 1 | 10 | 0 | 0 | 0 | 0 | 0 | 0 | 0 | 298 | 149 | 1 | 1 | NA | 1 | 1 | 1 | 0.636 | 1 | 1 | 1 | NA | NA | -0.091 |
| SE | 106596 | ENSG00000166012 | TAF1D | chr11 | - | 93737063 | 93737239 | 93734624 | 93734660 | 93738108 | 93738499 | 264 | 198 | 255 | 222 | 438 | 0 | 0 | 2 | 0 | 2 | 11 | 110 | 132 | 51 | 15 | 0 | 0 | 0 | 0 | 0 | 298 | 149 | 1 | 1 | 1 | 1 | 0.985 | 1 | 0.991 | 1 | 1 | 1 | 1 | 1 | -0.005 |
| SE | 106597 | ENSG00000166012 | TAF1D | chr11 | - | 93738108 | 93738440 | 93737088 | 93737239 | 93739236 | 93739331 | 217 | 161 | 221 | 195 | 348 | 1 | 0 | 2 | 0 | 5 | 5 | 89 | 93 | 46 | 2 | 0 | 0 | 0 | 0 | 0 | 298 | 149 | 1 | 1 | 0.991 | 1 | 0.982 | 1 | 0.972 | 1 | 1 | 1 | 1 | 1 | -0.011 |
| SE | 106598 | ENSG00000166012 | TAF1D | chr11 | - | 93738108 | 93738499 | 93734624 | 93734660 | 93739236 | 93739331 | 233 | 185 | 197 | 202 | 445 | 0 | 0 | 1 | 0 | 1 | 37 | 81 | 78 | 38 | 9 | 0 | 0 | 0 | 0 | 0 | 298 | 149 | 1 | 1 | 1 | 1 | 0.99 | 1 | 0.996 | 1 | 1 | 1 | 1 | 1 | -0.003 |
| SE | 106599 | ENSG00000166012 | TAF1D | chr11 | - | 93738108 | 93738499 | 93737063 | 93737239 | 93739236 | 93739331 | 445 | 341 | 411 | 394 | 778 | 1 | 0 | 2 | 0 | 5 | 42 | 168 | 168 | 84 | 11 | 0 | 0 | 0 | 0 | 0 | 298 | 149 | 1 | 1 | 0.996 | 1 | 0.99 | 1 | 0.987 | 1 | 1 | 1 | 1 | 1 | -0.005 |
| SE | 106606 | ENSG00000112200 | ZNF451 | chr6 | + | 57090832 | 57090894 | 57090210 | 57090274 | 57099060 | 57099141 | 38 | 43 | 36 | 25 | 91 | 7 | 1 | 2 | 2 | 5 | 21 | 29 | 29 | 18 | 3 | 0 | 1 | 2 | 0 | 0 | 211 | 149 | 1 | 1 | 0.793 | 0.968 | 0.927 | 0.898 | 0.928 | 1 | 0.953 | 0.911 | 1 | 1 | -0.07 |
| SE | 106608 | ENSG00000112200 | ZNF451 | chr6 | + | 57099060 | 57099141 | 57090782 | 57090894 | 57124733 | 57124859 | 89 | 108 | 79 | 93 | 138 | 0 | 0 | 2 | 0 | 0 | 16 | 52 | 54 | 25 | 5 | 0 | 0 | 0 | 0 | 0 | 230 | 149 | 1 | 1 | 1 | 1 | 0.962 | 1 | 1 | 1 | 1 | 1 | 1 | 1 | -0.008 |
| SE | 106610 | ENSG00000112200 | ZNF451 | chr6 | + | 57124733 | 57124859 | 57099060 | 57099141 | 57133041 | 57133154 | 57 | 79 | 49 | 54 | 104 | 0 | 0 | 0 | 2 | 0 | 6 | 26 | 36 | 14 | 5 | 0 | 0 | 0 | 0 | 0 | 275 | 149 | 1 | 1 | 1 | 1 | 1 | 0.936 | 1 | 1 | 1 | 1 | 1 | 1 | -0.013 |
| SE | 106612 | ENSG00000112200 | ZNF451 | chr6 | + | 57133041 | 57133192 | 57128728 | 57128840 | 57134743 | 57134870 | 241 | 168 | 172 | 241 | 400 | 0 | 0 | 0 | 0 | 2 | 47 | 101 | 85 | 54 | 45 | 0 | 0 | 0 | 0 | 0 | 298 | 149 | 1 | 1 | 1 | 1 | 1 | 1 | 0.99 | 1 | 1 | 1 | 1 | 1 | -0.002 |
| SE | 106613 | ENSG00000112200 | ZNF451 | chr6 | + | 57134743 | 57134870 | 57133041 | 57133192 | 57141301 | 57141368 | 166 | 132 | 129 | 165 | 411 | 1 | 0 | 0 | 0 | 0 | 23 | 86 | 65 | 30 | 19 | 0 | 0 | 1 | 0 | 0 | 276 | 149 | 1 | 1 | 0.989 | 1 | 1 | 1 | 1 | 1 | 1 | 0.972 | 1 | 1 | 0.003 |
| SE | 106614 | ENSG00000112200 | ZNF451 | chr6 | + | 57141301 | 57141455 | 57134743 | 57134870 | 57141947 | 57142095 | 216 | 156 | 191 | 199 | 584 | 0 | 0 | 0 | 2 | 0 | 14 | 150 | 72 | 32 | 41 | 0 | 0 | 0 | 0 | 0 | 298 | 149 | 1 | 1 | 1 | 1 | 1 | 0.98 | 1 | 1 | 1 | 1 | 1 | 1 | -0.004 |
| SE | 106615 | ENSG00000112200 | ZNF451 | chr6 | + | 57147089 | 57148693 | 57141947 | 57142095 | 57150718 | 57150862 | 229 | 160 | 215 | 241 | 385 | 0 | 2 | 0 | 0 | 0 | 0 | 146 | 101 | 44 | 5 | 0 | 0 | 0 | 0 | 0 | 298 | 149 | 1 | 1 | 1 | 0.976 | 1 | 1 | 1 | NA | 1 | 1 | 1 | 1 | -0.005 |
| SE | 106617 | ENSG00000112200 | ZNF451 | chr6 | + | 57150718 | 57150862 | 57147089 | 57148693 | 57152220 | 57152351 | 166 | 131 | 161 | 192 | 217 | 2 | 0 | 6 | 2 | 8 | 0 | 91 | 97 | 9 | 5 | 0 | 0 | 5 | 0 | 0 | 293 | 149 | 1 | 1 | 0.977 | 1 | 0.932 | 0.98 | 0.932 | NA | 1 | 0.908 | 1 | 1 | -0.013 |
| SE | 106618 | ENSG00000112200 | ZNF451 | chr6 | + | 57152220 | 57152351 | 57147089 | 57148693 | 57161083 | 57161152 | 12 | 11 | 12 | 14 | 11 | 0 | 0 | 0 | 2 | 0 | 0 | 4 | 8 | 0 | 0 | 0 | 0 | 0 | 0 | 0 | 280 | 149 | 1 | 1 | 1 | 1 | 1 | 0.788 | 1 | NA | 1 | 1 | NA | NA | -0.042 |
| SE | 106621 | ENSG00000196081 | ZNF724 | chr19 | - | 23232166 | 23232293 | 23230859 | 23231361 | 23250239 | 23250390 | 3 | 2 | 8 | 16 | 1 | 0 | 0 | 1 | 0 | 0 | 0 | 0 | 0 | 1 | 0 | 0 | 0 | 0 | 0 | 0 | 276 | 149 | 1 | 1 | 1 | 1 | 0.812 | 1 | 1 | NA | NA | NA | 1 | NA | -0.038 |
| SE | 106625 | ENSG00000106133 | NSUN5P2 | chr7 | - | 72952151 | 72952295 | 72950057 | 72950196 | 72953510 | 72953677 | 26 | 15 | 11 | 6 | 15 | 0 | 0 | 2 | 0 | 0 | 9 | 5 | 4 | 26 | 46 | 0 | 0 | 0 | 0 | 0 | 293 | 149 | 1 | 1 | 1 | 1 | 0.737 | 1 | 1 | 1 | 1 | 1 | 1 | 1 | -0.053 |
| SE | 106626 | ENSG00000106133 | NSUN5P2 | chr7 | - | 72953325 | 72953430 | 72950057 | 72950196 | 72953510 | 72953677 | 14 | 12 | 14 | 24 | 10 | 0 | 0 | 2 | 0 | 0 | 13 | 31 | 12 | 14 | 20 | 0 | 0 | 0 | 0 | 0 | 254 | 149 | 1 | 1 | 1 | 1 | 0.804 | 1 | 1 | 1 | 1 | 1 | 1 | 1 | -0.039 |
| SE | 106627 | ENSG00000106133 | NSUN5P2 | chr7 | - | 72953325 | 72953430 | 72952151 | 72952295 | 72953510 | 72953677 | 47 | 42 | 46 | 30 | 40 | 0 | 0 | 0 | 0 | 1 | 17 | 57 | 18 | 67 | 91 | 0 | 0 | 0 | 0 | 0 | 254 | 149 | 1 | 1 | 1 | 1 | 1 | 1 | 0.959 | 1 | 1 | 1 | 1 | 1 | -0.008 |
| SE | 106628 | ENSG00000106133 | NSUN5P2 | chr7 | - | 72953510 | 72953677 | 72950057 | 72950196 | 72954358 | 72954479 | 0 | 0 | 2 | 0 | 0 | 5 | 0 | 1 | 0 | 0 | 0 | 0 | 0 | 0 | 0 | 0 | 1 | 0 | 0 | 0 | 298 | 149 | 1 | 1 | 0 | NA | 0.5 | NA | NA | NA | 0 | NA | NA | NA | 0.25 |
| SE | 106629 | ENSG00000106133 | NSUN5P2 | chr7 | - | 72954358 | 72954479 | 72950057 | 72950196 | 72954624 | 72954717 | 9 | 0 | 2 | 0 | 0 | 6 | 4 | 5 | 0 | 0 | 0 | 3 | 0 | 0 | 0 | 0 | 5 | 1 | 2 | 0 | 270 | 149 | 1 | 1 | 0.453 | 0 | 0.181 | NA | NA | NA | 0.249 | 0 | 0 | NA | 0.128 |
| SE | 106633 | ENSG00000135968 | GCC2 | chr2 | + | 108452398 | 108452466 | 108451027 | 108451112 | 108468979 | 108469084 | 205 | 256 | 248 | 322 | 275 | 0 | 0 | 1 | 0 | 0 | 38 | 66 | 111 | 30 | 28 | 0 | 0 | 0 | 0 | 0 | 217 | 149 | 1 | 1 | 1 | 1 | 0.994 | 1 | 1 | 1 | 1 | 1 | 1 | 1 | -0.001 |
| SE | 106634 | ENSG00000135968 | GCC2 | chr2 | + | 108462587 | 108462685 | 108452398 | 108452466 | 108468979 | 108469084 | 4 | 2 | 2 | 0 | 1 | 162 | 232 | 208 | 271 | 248 | 0 | 0 | 0 | 0 | 0 | 32 | 46 | 101 | 33 | 24 | 247 | 149 | 1 | 1 | 0.015 | 0.005 | 0.006 | 0 | 0.002 | 0 | 0 | 0 | 0 | 0 | 0.006 |
| SE | 106636 | ENSG00000135968 | GCC2 | chr2 | + | 108472826 | 108472899 | 108469650 | 108472116 | 108475534 | 108475635 | 183 | 230 | 175 | 207 | 354 | 1 | 1 | 0 | 0 | 0 | 2 | 30 | 50 | 1 | 2 | 0 | 0 | 0 | 0 | 0 | 222 | 149 | 1 | 1 | 0.992 | 0.994 | 1 | 1 | 1 | 1 | 1 | 1 | 1 | 1 | -0.003 |
| SE | 106637 | ENSG00000135968 | GCC2 | chr2 | + | 108481696 | 108481816 | 108475751 | 108475850 | 108482286 | 108482451 | 250 | 206 | 234 | 332 | 467 | 0 | 0 | 0 | 1 | 0 | 2 | 35 | 41 | 8 | 1 | 0 | 0 | 0 | 0 | 0 | 269 | 149 | 1 | 1 | 1 | 1 | 1 | 0.995 | 1 | 1 | 1 | 1 | 1 | 1 | -0.001 |
| SE | 106638 | ENSG00000135968 | GCC2 | chr2 | + | 108481709 | 108481816 | 108475751 | 108475850 | 108482286 | 108482451 | 111 | 104 | 111 | 150 | 200 | 0 | 0 | 0 | 1 | 0 | 2 | 17 | 22 | 4 | 0 | 0 | 0 | 0 | 0 | 0 | 256 | 149 | 1 | 1 | 1 | 1 | 1 | 0.989 | 1 | 1 | 1 | 1 | 1 | NA | -0.002 |
| SE | 106639 | ENSG00000135968 | GCC2 | chr2 | + | 108482286 | 108482451 | 108481696 | 108481816 | 108483061 | 108483156 | 278 | 245 | 288 | 364 | 642 | 0 | 0 | 0 | 0 | 4 | 4 | 22 | 55 | 4 | 3 | 0 | 0 | 0 | 0 | 0 | 298 | 149 | 1 | 1 | 1 | 1 | 1 | 1 | 0.988 | 1 | 1 | 1 | 1 | 1 | -0.002 |
| SE | 106640 | ENSG00000135968 | GCC2 | chr2 | + | 108483061 | 108483166 | 108482286 | 108482451 | 108484148 | 108484311 | 172 | 188 | 216 | 251 | 527 | 5 | 0 | 2 | 5 | 3 | 7 | 20 | 49 | 0 | 16 | 0 | 0 | 0 | 0 | 0 | 254 | 149 | 1 | 1 | 0.953 | 1 | 0.984 | 0.967 | 0.99 | 1 | 1 | 1 | NA | 1 | -0.021 |
| SE | 106641 | ENSG00000135968 | GCC2 | chr2 | + | 108486434 | 108486648 | 108485830 | 108485908 | 108487698 | 108487820 | 147 | 138 | 129 | 147 | 234 | 0 | 1 | 2 | 0 | 3 | 0 | 12 | 48 | 2 | 10 | 0 | 0 | 0 | 0 | 0 | 298 | 149 | 1 | 1 | 1 | 0.986 | 0.97 | 1 | 0.975 | NA | 1 | 1 | 1 | 1 | -0.014 |
| SE | 106642 | ENSG00000135968 | GCC2 | chr2 | + | 108486510 | 108486648 | 108485830 | 108485908 | 108487698 | 108487820 | 281 | 255 | 288 | 338 | 529 | 0 | 1 | 2 | 0 | 3 | 0 | 22 | 82 | 8 | 10 | 0 | 0 | 0 | 0 | 0 | 287 | 149 | 1 | 1 | 1 | 0.993 | 0.987 | 1 | 0.989 | NA | 1 | 1 | 1 | 1 | -0.006 |
| SE | 106643 | ENSG00000135968 | GCC2 | chr2 | + | 108487698 | 108487820 | 108486434 | 108486648 | 108489837 | 108490014 | 217 | 193 | 192 | 229 | 362 | 5 | 5 | 6 | 11 | 5 | 0 | 19 | 64 | 2 | 10 | 0 | 4 | 0 | 0 | 0 | 271 | 149 | 1 | 1 | 0.96 | 0.955 | 0.946 | 0.92 | 0.975 | NA | 0.723 | 1 | 1 | 1 | 0.02 |
| SE | 106645 | ENSG00000139178 | C1RL | chr12 | - | 7099685 | 7099760 | 7094553 | 7097163 | 7101897 | 7102087 | 2 | 1 | 3 | 5 | 0 | 1 | 1 | 0 | 0 | 0 | 8 | 5 | 4 | 10 | 13 | 0 | 0 | 0 | 0 | 0 | 224 | 149 | 1 | 1 | 0.571 | 0.399 | 1 | 1 | NA | 1 | 1 | 1 | 1 | 1 | -0.258 |
| SE | 106649 | ENSG00000149256 | TENM4 | chr11 | - | 78712481 | 78712714 | 78708360 | 78708515 | 78722667 | 78722917 | 2 | 0 | 0 | 3 | 3 | 0 | 0 | 0 | 1 | 0 | 0 | 0 | 0 | 0 | 5 | 0 | 0 | 0 | 0 | 5 | 298 | 149 | 1 | 1 | 1 | NA | NA | 0.6 | 1 | NA | NA | NA | NA | 0.333 | 0.534 |
| SE | 106654 | ENSG00000139350 | NEDD1 | chr12 | + | 96907603 | 96907624 | 96907229 | 96907300 | 96909751 | 96909895 | 3 | 0 | 1 | 0 | 0 | 6 | 9 | 7 | 1 | 0 | 0 | 1 | 0 | 0 | 0 | 0 | 8 | 6 | 1 | 4 | 170 | 149 | 1 | 1 | 0.305 | 0 | 0.111 | 0 | NA | NA | 0.099 | 0 | 0 | 0 | 0.079 |
| SE | 106656 | ENSG00000139350 | NEDD1 | chr12 | + | 96907603 | 96907624 | 96907229 | 96907300 | 96919984 | 96920118 | 0 | 0 | 0 | 0 | 0 | 0 | 2 | 0 | 0 | 1 | 0 | 0 | 0 | 5 | 0 | 0 | 0 | 0 | 0 | 0 | 170 | 149 | 1 | 1 | NA | 0 | NA | NA | 0 | NA | NA | NA | 1 | NA | -1 |
| SE | 106658 | ENSG00000139350 | NEDD1 | chr12 | + | 96907603 | 96907856 | 96907237 | 96907300 | 96909751 | 96909895 | 28 | 21 | 13 | 35 | 36 | 6 | 9 | 7 | 1 | 0 | 0 | 8 | 1 | 3 | 0 | 0 | 8 | 6 | 1 | 4 | 298 | 149 | 1 | 1 | 0.7 | 0.538 | 0.481 | 0.946 | 1 | NA | 0.333 | 0.077 | 0.6 | 0 | 0.48 |
| SE | 106661 | ENSG00000139350 | NEDD1 | chr12 | + | 96907701 | 96907856 | 96907603 | 96907624 | 96909751 | 96909895 | 23 | 20 | 12 | 27 | 30 | 3 | 1 | 3 | 1 | 0 | 0 | 8 | 1 | 3 | 0 | 2 | 1 | 1 | 0 | 0 | 298 | 149 | 1 | 1 | 0.793 | 0.909 | 0.667 | 0.931 | 1 | 0 | 0.8 | 0.333 | 1 | NA | 0.327 |
| SE | 106662 | ENSG00000139350 | NEDD1 | chr12 | + | 96917620 | 96917737 | 96907223 | 96907300 | 96919984 | 96920118 | 59 | 34 | 75 | 64 | 73 | 0 | 2 | 0 | 0 | 1 | 7 | 37 | 20 | 4 | 25 | 0 | 0 | 0 | 0 | 0 | 266 | 149 | 1 | 1 | 1 | 0.905 | 1 | 1 | 0.976 | 1 | 1 | 1 | 1 | 1 | -0.024 |
| SE | 106665 | ENSG00000139350 | NEDD1 | chr12 | + | 96917620 | 96917737 | 96912575 | 96912817 | 96919984 | 96920118 | 102 | 77 | 112 | 99 | 96 | 0 | 0 | 0 | 2 | 0 | 7 | 56 | 38 | 9 | 25 | 0 | 0 | 0 | 0 | 0 | 266 | 149 | 1 | 1 | 1 | 1 | 1 | 0.965 | 1 | 1 | 1 | 1 | 1 | 1 | -0.007 |
| SE | 106666 | ENSG00000139350 | NEDD1 | chr12 | + | 96924832 | 96924942 | 96919984 | 96920125 | 96934975 | 96935205 | 12 | 5 | 13 | 8 | 20 | 52 | 28 | 74 | 56 | 51 | 0 | 1 | 2 | 2 | 12 | 0 | 22 | 18 | 6 | 0 | 259 | 149 | 1 | 1 | 0.117 | 0.093 | 0.092 | 0.076 | 0.184 | NA | 0.025 | 0.06 | 0.161 | 1 | -0.199 |
| SE | 106667 | ENSG00000139350 | NEDD1 | chr12 | + | 96924832 | 96924942 | 96919984 | 96920125 | 96936610 | 96936718 | 12 | 5 | 5 | 5 | 18 | 2 | 3 | 0 | 3 | 2 | 0 | 1 | 2 | 2 | 12 | 0 | 0 | 0 | 0 | 0 | 259 | 149 | 1 | 1 | 0.775 | 0.489 | 1 | 0.489 | 0.838 | NA | 1 | 1 | 1 | 1 | -0.282 |
| SE | 106668 | ENSG00000139350 | NEDD1 | chr12 | + | 96934975 | 96935205 | 96919984 | 96920125 | 96936610 | 96936638 | 132 | 94 | 154 | 123 | 147 | 2 | 3 | 0 | 3 | 2 | 0 | 34 | 52 | 10 | 9 | 0 | 0 | 0 | 0 | 0 | 298 | 149 | 1 | 1 | 0.971 | 0.94 | 1 | 0.953 | 0.974 | NA | 1 | 1 | 1 | 1 | -0.032 |
| SE | 106670 | ENSG00000139350 | NEDD1 | chr12 | + | 96936610 | 96936812 | 96934975 | 96935205 | 96937197 | 96937393 | 188 | 150 | 178 | 172 | 307 | 0 | 0 | 0 | 2 | 0 | 13 | 42 | 96 | 9 | 17 | 0 | 0 | 0 | 0 | 0 | 298 | 149 | 1 | 1 | 1 | 1 | 1 | 0.977 | 1 | 1 | 1 | 1 | 1 | 1 | -0.005 |
| SE | 106673 | ENSG00000120662 | MTRF1 | chr13 | - | 41252948 | 41253030 | 41252281 | 41252752 | 41254528 | 41254620 | 294 | 179 | 234 | 238 | 581 | 0 | 0 | 1 | 0 | 0 | 175 | 146 | 149 | 177 | 133 | 0 | 0 | 0 | 0 | 0 | 231 | 149 | 1 | 1 | 1 | 1 | 0.993 | 1 | 1 | 1 | 1 | 1 | 1 | 1 | -0.001 |
| SE | 106676 | ENSG00000120662 | MTRF1 | chr13 | - | 41261690 | 41261825 | 41260492 | 41260915 | 41262214 | 41262331 | 20 | 20 | 28 | 35 | 67 | 3 | 0 | 0 | 0 | 0 | 4 | 38 | 9 | 16 | 3 | 0 | 0 | 0 | 0 | 0 | 284 | 149 | 1 | 1 | 0.778 | 1 | 1 | 1 | 1 | 1 | 1 | 1 | 1 | 1 | -0.044 |
| SE | 106677 | ENSG00000120662 | MTRF1 | chr13 | - | 41262047 | 41262331 | 41260492 | 41260915 | 41263484 | 41263569 | 3 | 2 | 5 | 3 | 6 | 13 | 2 | 12 | 11 | 25 | 8 | 5 | 0 | 0 | 0 | 0 | 0 | 0 | 0 | 0 | 298 | 149 | 1 | 1 | 0.103 | 0.333 | 0.172 | 0.12 | 0.107 | 1 | 1 | NA | NA | NA | -0.833 |
| SE | 106679 | ENSG00000120662 | MTRF1 | chr13 | - | 41262214 | 41262331 | 41261432 | 41261825 | 41263272 | 41263490 | 4 | 7 | 3 | 7 | 16 | 0 | 0 | 0 | 0 | 0 | 0 | 13 | 1 | 7 | 7 | 0 | 17 | 0 | 0 | 0 | 266 | 149 | 1 | 1 | 1 | 1 | 1 | 1 | 1 | NA | 0.3 | 1 | 1 | 1 | 0.175 |
| SE | 106680 | ENSG00000120662 | MTRF1 | chr13 | - | 41262214 | 41262331 | 41261432 | 41261825 | 41263274 | 41263484 | 4 | 7 | 3 | 8 | 16 | 0 | 2 | 0 | 0 | 0 | 0 | 13 | 1 | 7 | 7 | 0 | 0 | 0 | 0 | 0 | 266 | 149 | 1 | 1 | 1 | 0.662 | 1 | 1 | 1 | NA | 1 | 1 | 1 | 1 | -0.068 |
| SE | 106681 | ENSG00000120662 | MTRF1 | chr13 | - | 41262410 | 41262481 | 41260492 | 41260915 | 41263272 | 41263490 | 2 | 1 | 5 | 0 | 6 | 0 | 1 | 0 | 0 | 0 | 0 | 0 | 1 | 0 | 6 | 0 | 0 | 0 | 0 | 0 | 220 | 149 | 1 | 1 | 1 | 0.404 | 1 | NA | 1 | NA | NA | 1 | NA | 1 | -0.149 |
| SE | 106682 | ENSG00000120662 | MTRF1 | chr13 | - | 41262410 | 41262481 | 41260492 | 41260915 | 41263484 | 41263538 | 4 | 3 | 3 | 0 | 7 | 13 | 2 | 12 | 11 | 25 | 0 | 2 | 0 | 0 | 6 | 0 | 0 | 0 | 0 | 0 | 220 | 149 | 1 | 1 | 0.172 | 0.504 | 0.145 | 0 | 0.159 | NA | 1 | NA | NA | 1 | -0.804 |
| SE | 106683 | ENSG00000120662 | MTRF1 | chr13 | - | 41262410 | 41262481 | 41261332 | 41261825 | 41263272 | 41263490 | 4 | 11 | 2 | 5 | 10 | 0 | 0 | 0 | 0 | 0 | 0 | 3 | 2 | 5 | 0 | 0 | 17 | 0 | 0 | 0 | 220 | 149 | 1 | 1 | 1 | 1 | 1 | 1 | 1 | NA | 0.107 | 1 | 1 | NA | 0.298 |
| SE | 106684 | ENSG00000120662 | MTRF1 | chr13 | - | 41262410 | 41262481 | 41261332 | 41261825 | 41263274 | 41263484 | 5 | 11 | 0 | 7 | 16 | 0 | 2 | 0 | 0 | 0 | 4 | 7 | 1 | 11 | 0 | 0 | 0 | 0 | 0 | 0 | 220 | 149 | 1 | 1 | 1 | 0.788 | NA | 1 | 1 | 1 | 1 | 1 | 1 | NA | -0.053 |
| SE | 106685 | ENSG00000120662 | MTRF1 | chr13 | - | 41262410 | 41262481 | 41261332 | 41261825 | 41263484 | 41263538 | 10 | 12 | 2 | 6 | 18 | 0 | 0 | 1 | 0 | 0 | 0 | 3 | 3 | 5 | 0 | 0 | 0 | 0 | 0 | 0 | 220 | 149 | 1 | 1 | 1 | 1 | 0.575 | 1 | 1 | NA | 1 | 1 | 1 | NA | -0.085 |
| SE | 106688 | ENSG00000174405 | LIG4 | chr13 | - | 108214537 | 108214610 | 108207441 | 108211296 | 108215483 | 108215523 | 19 | 2 | 15 | 13 | 3 | 0 | 0 | 1 | 4 | 0 | 0 | 6 | 2 | 0 | 0 | 0 | 0 | 0 | 0 | 0 | 222 | 149 | 1 | 1 | 1 | 1 | 0.91 | 0.686 | 1 | NA | 1 | 1 | NA | NA | -0.081 |
| SE | 106689 | ENSG00000174405 | LIG4 | chr13 | - | 108214537 | 108214610 | 108207441 | 108211296 | 108218161 | 108218368 | 12 | 1 | 14 | 10 | 1 | 6 | 1 | 1 | 11 | 5 | 0 | 6 | 2 | 0 | 0 | 0 | 1 | 0 | 0 | 0 | 222 | 149 | 1 | 1 | 0.573 | 0.402 | 0.904 | 0.379 | 0.118 | NA | 0.801 | 1 | NA | NA | -0.425 |
| SE | 106691 | ENSG00000174405 | LIG4 | chr13 | - | 108214537 | 108214807 | 108207438 | 108211296 | 108215483 | 108215535 | 13 | 1 | 16 | 12 | 3 | 0 | 0 | 1 | 4 | 0 | 0 | 8 | 2 | 0 | 8 | 0 | 0 | 0 | 0 | 0 | 298 | 149 | 1 | 1 | 1 | 1 | 0.889 | 0.6 | 1 | NA | 1 | 1 | NA | 1 | -0.102 |
| SE | 106692 | ENSG00000141376 | BCAS3 | chr17 | + | 60679452 | 60679540 | 60677847 | 60677914 | 60683981 | 60684036 | 154 | 86 | 80 | 74 | 101 | 1 | 0 | 0 | 0 | 0 | 108 | 107 | 86 | 36 | 114 | 0 | 0 | 0 | 0 | 0 | 237 | 149 | 1 | 1 | 0.99 | 1 | 1 | 1 | 1 | 1 | 1 | 1 | 1 | 1 | -0.002 |
| SE | 106696 | ENSG00000141376 | BCAS3 | chr17 | + | 60688931 | 60689046 | 60688019 | 60688195 | 60689685 | 60689761 | 8 | 4 | 1 | 2 | 12 | 0 | 1 | 0 | 2 | 7 | 7 | 0 | 2 | 1 | 4 | 0 | 0 | 4 | 0 | 0 | 264 | 149 | 1 | 1 | 1 | 0.693 | 1 | 0.361 | 0.492 | 1 | NA | 0.22 | 1 | 1 | -0.096 |
| SE | 106697 | ENSG00000141376 | BCAS3 | chr17 | + | 60709434 | 60709552 | 60709218 | 60709325 | 60747197 | 60747222 | 0 | 0 | 0 | 0 | 0 | 77 | 37 | 57 | 54 | 97 | 0 | 0 | 1 | 0 | 6 | 47 | 32 | 48 | 25 | 101 | 267 | 149 | 1 | 1 | 0 | 0 | 0 | 0 | 0 | 0 | 0 | 0.011 | 0 | 0.032 | -0.009 |
| SE | 106702 | ENSG00000141376 | BCAS3 | chr17 | + | 61037888 | 61038049 | 61034665 | 61034790 | 61040791 | 61040892 | 46 | 47 | 42 | 58 | 93 | 0 | 0 | 0 | 0 | 2 | 43 | 50 | 61 | 57 | 177 | 0 | 0 | 0 | 0 | 0 | 298 | 149 | 1 | 1 | 1 | 1 | 1 | 1 | 0.959 | 1 | 1 | 1 | 1 | 1 | -0.008 |
| SE | 106703 | ENSG00000141376 | BCAS3 | chr17 | + | 61037888 | 61038054 | 61034665 | 61034790 | 61040791 | 61040892 | 98 | 89 | 102 | 105 | 155 | 0 | 0 | 0 | 0 | 2 | 84 | 110 | 105 | 82 | 225 | 0 | 0 | 0 | 0 | 0 | 298 | 149 | 1 | 1 | 1 | 1 | 1 | 1 | 0.975 | 1 | 1 | 1 | 1 | 1 | -0.005 |
| SE | 106704 | ENSG00000141376 | BCAS3 | chr17 | + | 61084466 | 61084564 | 61078332 | 61078529 | 61368326 | 61368342 | 83 | 60 | 74 | 97 | 96 | 0 | 0 | 0 | 0 | 0 | 82 | 90 | 57 | 79 | 119 | 0 | 1 | 0 | 0 | 0 | 247 | 149 | 1 | 1 | 1 | 1 | 1 | 1 | 1 | 1 | 0.982 | 1 | 1 | 1 | 0.004 |
| SE | 106706 | ENSG00000141376 | BCAS3 | chr17 | + | 61353587 | 61354502 | 61245215 | 61245371 | 61368326 | 61368494 | 2 | 0 | 0 | 1 | 0 | 0 | 0 | 0 | 0 | 0 | 3 | 7 | 3 | 0 | 0 | 0 | 0 | 0 | 3 | 0 | 298 | 149 | 1 | 1 | 1 | NA | NA | 1 | NA | 1 | 1 | 1 | 0 | NA | 0.25 |
| SE | 106708 | ENSG00000141376 | BCAS3 | chr17 | + | 61354361 | 61354502 | 61245259 | 61245371 | 61368326 | 61368494 | 2 | 0 | 0 | 1 | 0 | 0 | 0 | 0 | 0 | 0 | 3 | 7 | 3 | 0 | 0 | 0 | 0 | 0 | 3 | 0 | 290 | 149 | 1 | 1 | 1 | NA | NA | 1 | NA | 1 | 1 | 1 | 0 | NA | 0.25 |
| SE | 106709 | ENSG00000141376 | BCAS3 | chr17 | + | 61356848 | 61356986 | 61354361 | 61354502 | 61368326 | 61368410 | 0 | 0 | 4 | 2 | 3 | 2 | 0 | 0 | 1 | 0 | 0 | 0 | 0 | 0 | 0 | 3 | 7 | 3 | 0 | 0 | 287 | 149 | 1 | 1 | 0 | NA | 1 | 0.509 | 1 | 0 | 0 | 0 | NA | NA | 0.627 |
| SE | 106710 | ENSG00000141376 | BCAS3 | chr17 | + | 61368326 | 61368494 | 61084466 | 61084564 | 61391976 | 61392125 | 79 | 56 | 68 | 66 | 74 | 0 | 0 | 1 | 0 | 0 | 69 | 75 | 35 | 45 | 90 | 0 | 0 | 3 | 0 | 0 | 298 | 149 | 1 | 1 | 1 | 1 | 0.971 | 1 | 1 | 1 | 1 | 0.854 | 1 | 1 | 0.023 |
| SE | 106711 | ENSG00000141376 | BCAS3 | chr17 | + | 61378950 | 61379140 | 61368326 | 61368494 | 61380505 | 61380571 | 0 | 0 | 0 | 0 | 0 | 4 | 1 | 3 | 2 | 10 | 0 | 1 | 0 | 0 | 0 | 12 | 14 | 3 | 2 | 11 | 298 | 149 | 1 | 1 | 0 | 0 | 0 | 0 | 0 | 0 | 0.034 | 0 | 0 | 0 | -0.007 |
| SE | 106717 | ENSG00000274349 | ZNF658 | chr9 | + | 66903360 | 66903576 | 66900737 | 66900836 | 66908237 | 66908364 | 2 | 1 | 5 | 1 | 1 | 0 | 0 | 0 | 1 | 0 | 0 | 0 | 1 | 0 | 0 | 0 | 0 | 0 | 0 | 0 | 298 | 149 | 1 | 1 | 1 | 1 | 1 | 0.333 | 1 | NA | NA | 1 | NA | NA | -0.133 |
| SE | 106718 | ENSG00000274349 | ZNF658 | chr9 | + | 66903517 | 66903576 | 66900724 | 66900836 | 66908237 | 66908364 | 0 | 1 | 5 | 0 | 1 | 0 | 0 | 0 | 1 | 0 | 0 | 0 | 1 | 0 | 0 | 0 | 0 | 0 | 0 | 0 | 208 | 149 | 1 | 1 | NA | 1 | 1 | 0 | 1 | NA | NA | 1 | NA | NA | -0.25 |
| SE | 106721 | ENSG00000136111 | TBC1D4 | chr13 | - | 75288933 | 75289110 | 75283502 | 75287025 | 75292101 | 75292271 | 214 | 267 | 440 | 360 | 586 | 0 | 0 | 0 | 1 | 0 | 79 | 46 | 90 | 94 | 27 | 0 | 0 | 0 | 0 | 0 | 298 | 149 | 1 | 1 | 1 | 1 | 1 | 0.994 | 1 | 1 | 1 | 1 | 1 | 1 | -0.001 |
| SE | 106722 | ENSG00000136111 | TBC1D4 | chr13 | - | 75294853 | 75295013 | 75292101 | 75292271 | 75299329 | 75299574 | 232 | 187 | 393 | 237 | 348 | 1 | 0 | 0 | 1 | 2 | 35 | 29 | 52 | 25 | 50 | 0 | 0 | 0 | 0 | 0 | 298 | 149 | 1 | 1 | 0.991 | 1 | 1 | 0.992 | 0.989 | 1 | 1 | 1 | 1 | 1 | -0.006 |

| | | | | | | | | | | | | | |
|---|---|---|---|---|---|---|---|---|---|---|---|---|---|
| SE | 106724 | ENSG00000136111 | TBC1D4 | chr13 | - | 75309941 | 75310151 | 75306312 | 75306471 | 75326196 | 75326423 | 150 | 152 |
| SE | 106725 | ENSG00000136111 | TBC1D4 | chr13 | - | 75312737 | 75312898 | 75309941 | 75310151 | 75326196 | 75326423 | 127 | 100 |
| SE | 106727 | ENSG00000136111 | TBC1D4 | chr13 | - | 75324236 | 75324401 | 75312737 | 75312898 | 75326196 | 75326250 | 0 | 0 |
| SE | 106728 | ENSG00000136111 | TBC1D4 | chr13 | - | 75327751 | 75327826 | 75326196 | 75326423 | 75336920 | 75337040 | 21 | 14 |
| SE | 106729 | ENSG00000136111 | TBC1D4 | chr13 | - | 75341124 | 75341235 | 75336920 | 75337040 | 75341495 | 75341531 | 15 | 24 |
| SE | 106731 | ENSG00000145781 | COMMD10 | chr5 | + | 116134067 | 116134178 | 116092544 | 116092700 | 116291516 | 116291576 | 231 | 142 |
| SE | 106732 | ENSG00000145781 | COMMD10 | chr5 | + | 116298246 | 116298392 | 116292450 | 116292560 | 116303529 | 116303683 | 44 | 48 |
| SE | 106734 | ENSG00000145781 | COMMD10 | chr5 | + | 116382277 | 116382454 | 116303529 | 116303683 | 116399481 | 116399622 | 0 | 1 |
| SE | 106735 | ENSG00000145781 | COMMD10 | chr5 | + | 116399481 | 116399622 | 116303529 | 116303683 | 116412642 | 116412762 | 37 | 14 |
| SE | 106736 | ENSG00000105810 | CDK6 | chr7 | - | 92623035 | 92623086 | 92618071 | 92618207 | 92671425 | 92671535 | 57 | 41 |
| SE | 106737 | ENSG00000105810 | CDK6 | chr7 | - | 92671425 | 92671535 | 92623035 | 92623086 | 92725625 | 92725793 | 96 | 74 |
| SE | 106740 | ENSG00000215483 | LINC00598 | chr13 | - | 40451063 | 40451203 | 40350108 | 40350302 | 40460281 | 40460406 | 0 | 3 |
| SE | 106741 | ENSG00000215483 | LINC00598 | chr13 | - | 40457513 | 40458668 | 40450933 | 40451203 | 40460281 | 40460406 | 3 | 2 |
| SE | 106742 | ENSG00000215483 | LINC00598 | chr13 | - | 40466506 | 40466677 | 40460281 | 40460406 | 40469995 | 40470162 | 4 | 2 |
| SE | 106743 | ENSG00000215483 | LINC00598 | chr13 | - | 40466506 | 40466677 | 40460281 | 40460406 | 40473993 | 40474095 | 4 | 2 |
| SE | 106744 | ENSG00000215483 | LINC00598 | chr13 | - | 40469995 | 40470162 | 40460281 | 40460406 | 40473993 | 40474095 | 0 | 1 |
| SE | 106745 | ENSG00000215483 | LINC00598 | chr13 | - | 40469995 | 40470162 | 40466506 | 40466677 | 40473993 | 40474095 | 0 | 1 |
| SE | 106746 | ENSG00000215483 | LINC00598 | chr13 | - | 40473993 | 40474095 | 40460281 | 40460406 | 40480235 | 40480468 | 0 | 0 |
| SE | 106749 | ENSG00000184162 | NR2C2AP | chr19 | - | 19202469 | 19202575 | 19202057 | 19202398 | 19202790 | 19202881 | 20 | 12 |
| SE | 106752 | ENSG00000078674 | PCM1 | chr8 | + | 17935588 | 17935706 | 17934705 | 17934858 | 17937133 | 17937321 | 185 | 192 |
| SE | 106753 | ENSG00000078674 | PCM1 | chr8 | + | 17938739 | 17939009 | 17937133 | 17937379 | 17939690 | 17939861 | 640 | 585 |
| SE | 106754 | ENSG00000078674 | PCM1 | chr8 | + | 17939690 | 17939861 | 17937133 | 17937379 | 17947185 | 17947363 | 361 | 310 |
| SE | 106756 | ENSG00000078674 | PCM1 | chr8 | + | 17947185 | 17947363 | 17939690 | 17939861 | 17950614 | 17950724 | 663 | 596 |
| SE | 106757 | ENSG00000078674 | PCM1 | chr8 | + | 17950614 | 17950724 | 17947185 | 17947363 | 17952969 | 17953035 | 437 | 391 |
| SE | 106758 | ENSG00000078674 | PCM1 | chr8 | + | 17957263 | 17957421 | 17956603 | 17956777 | 17957539 | 17957775 | 788 | 952 |

[illegible]

| | | | | | | | | | | | | | | | | | | | | | | | | | | | | | | | | | | | | | | | | | | | | | | |
|---|---|---|---|---|---|---|---|---|---|---|---|---|---|---|---|---|---|---|---|---|---|---|---|---|---|---|---|---|---|---|---|---|---|---|---|---|---|---|---|---|---|---|---|---|---|---|
| SE | 107136 | ENSG00000175582 | RAB6A | chr11 | - | 73679653 | 73679720 | 73675637 | 73677962 | 73707419 | 73707513 | 542 | 393 | 395 | 526 | 844 | 1 | 0 | 0 | 0 | 5 | 469 | 335 | 310 | 419 | 286 | 0 | 1 | 0 | 3 | 0 | 216 | 149 | 1 | 1 | 0.997 | 1 | 1 | 1 | 0.991 | 1 | 0.996 | 1 | 0.99 | 1 | 0 |
| SE | 107137 | ENSG00000175582 | RAB6A | chr11 | - | 73707419 | 73707513 | 73679653 | 73679720 | 73716250 | 73716362 | 1135 | 811 | 808 | 1138 | 1483 | 0 | 0 | 0 | 1 | 0 | 1805 | 995 | 937 | 1420 | 1465 | 0 | 0 | 0 | 0 | 0 | 243 | 149 | 1 | 1 | 1 | 1 | 1 | 0.999 | 1 | 1 | 1 | 1 | 1 | 1 | 0 |
| SE | 107138 | ENSG00000175582 | RAB6A | chr11 | - | 73716250 | 73716362 | 73679653 | 73679720 | 73718784 | 73718890 | 203 | 165 | 150 | 272 | 368 | 0 | 1 | 0 | 0 | 0 | 281 | 180 | 115 | 260 | 285 | 0 | 0 | 0 | 0 | 0 | 261 | 149 | 1 | 1 | 1 | 0.989 | 1 | 1 | 1 | 1 | 1 | 1 | 1 | 1 | -0.002 |
| SE | 107139 | ENSG00000175582 | RAB6A | chr11 | - | 73716250 | 73716362 | 73679653 | 73679720 | 73720845 | 73720899 | 1 | 2 | 1 | 2 | 0 | 1 | 1 | 0 | 1 | 0 | 0 | 0 | 3 | 3 | 0 | 0 | 0 | 0 | 0 | 8 | 261 | 149 | 1 | 1 | 0.363 | 0.533 | 1 | 0.533 | NA | NA | NA | 1 | 1 | 0 | -0.059 |
| SE | 107141 | ENSG00000175582 | RAB6A | chr11 | - | 73718612 | 73718718 | 73716250 | 73716362 | 73720845 | 73720899 | 989 | 681 | 681 | 940 | 1138 | 3 | 4 | 1 | 2 | 0 | 1922 | 958 | 1052 | 1303 | 1336 | 0 | 0 | 4 | 6 | 0 | 255 | 149 | 1 | 1 | 0.995 | 0.99 | 0.997 | 0.996 | 1 | 1 | 1 | 0.994 | 0.992 | 1 | -0.002 |
| SE | 107142 | ENSG00000175582 | RAB6A | chr11 | - | 73718784 | 73718890 | 73679653 | 73679720 | 73720845 | 73720899 | 178 | 127 | 107 | 160 | 217 | 1 | 1 | 0 | 1 | 0 | 210 | 139 | 114 | 187 | 213 | 0 | 0 | 0 | 0 | 8 | 255 | 149 | 1 | 1 | 0.99 | 0.987 | 1 | 0.989 | 1 | 1 | 1 | 1 | 1 | 0.94 | 0.005 |
| SE | 107143 | ENSG00000175582 | RAB6A | chr11 | - | 73718784 | 73718890 | 73716250 | 73716362 | 73720845 | 73720899 | 648 | 481 | 435 | 698 | 841 | 3 | 4 | 1 | 2 | 0 | 870 | 531 | 385 | 807 | 950 | 0 | 0 | 4 | 6 | 0 | 255 | 149 | 1 | 1 | 0.992 | 0.986 | 0.996 | 0.995 | 1 | 1 | 1 | 0.983 | 0.987 | 1 | 0 |
| SE | 107151 | ENSG00000061794 | MRPS35 | chr12 | + | 27714779 | 27714820 | 27710831 | 27710955 | 27716290 | 27716458 | 122 | 77 | 120 | 119 | 144 | 1 | 1 | 0 | 2 | 0 | 53 | 101 | 70 | 42 | 39 | 0 | 0 | 0 | 0 | 0 | 190 | 149 | 1 | 1 | 0.99 | 0.984 | 1 | 0.979 | 1 | 1 | 1 | 1 | 1 | 1 | -0.009 |
| SE | 107154 | ENSG00000061794 | MRPS35 | chr12 | + | 27737538 | 27737608 | 27724046 | 27724186 | 27755180 | 27756294 | 110 | 64 | 81 | 82 | 235 | 0 | 0 | 0 | 0 | 0 | 62 | 94 | 51 | 104 | 8 | 0 | 1 | 0 | 0 | 0 | 219 | 149 | 1 | 1 | 1 | 1 | 1 | 1 | 1 | 1 | 0.985 | 1 | 1 | 1 | 0.003 |
| SE | 107158 | ENSG00000110799 | VWF | chr12 | - | 6075334 | 6075551 | 6073618 | 6073741 | 6095459 | 6095584 | 1 | 0 | 0 | 1 | 0 | 0 | 0 | 0 | 0 | 0 | 0 | 0 | 0 | 0 | 53 | 0 | 0 | 0 | 0 | 8 | 298 | 149 | 1 | 1 | 1 | NA | NA | 1 | NA | NA | NA | NA | NA | 0.768 | 0.232 |
| SE | 107159 | ENSG00000110799 | VWF | chr12 | - | 6122724 | 6122966 | 6121173 | 6121338 | 6123141 | 6123196 | 1 | 0 | 0 | 0 | 0 | 2 | 1 | 0 | 4 | 0 | 0 | 0 | 0 | 0 | 0 | 2 | 0 | 0 | 0 | 33 | 298 | 149 | 1 | 1 | 0.2 | 0 | NA | 0 | NA | 0 | NA | NA | NA | 0 | 0.067 |
| SE | 107162 | ENSG00000136003 | ISCU | chr12 | + | 108567657 | 108567743 | 108567189 | 108567268 | 108568830 | 108569099 | 1 | 0 | 2 | 3 | 1 | 194 | 120 | 216 | 224 | 526 | 0 | 1 | 0 | 0 | 0 | 227 | 250 | 238 | 173 | 288 | 235 | 149 | 1 | 1 | 0.003 | 0 | 0.006 | 0.008 | 0.001 | 0 | 0.003 | 0 | 0 | 0 | 0.003 |
| SE | 107163 | ENSG00000136003 | ISCU | chr12 | + | 108567657 | 108567757 | 108567189 | 108567268 | 108568830 | 108569368 | 1 | 0 | 2 | 3 | 0 | 194 | 120 | 216 | 224 | 526 | 0 | 1 | 2 | 0 | 0 | 227 | 250 | 238 | 173 | 288 | 249 | 149 | 1 | 1 | 0.003 | 0 | 0.006 | 0.008 | 0 | 0 | 0.002 | 0.005 | 0 | 0 | 0.002 |
| SE | 107164 | ENSG00000136003 | ISCU | chr12 | + | 108567919 | 108568016 | 108567189 | 108567268 | 108568830 | 108569092 | 0 | 0 | 0 | 0 | 0 | 194 | 120 | 216 | 224 | 526 | 0 | 0 | 3 | 0 | 0 | 227 | 250 | 238 | 173 | 288 | 246 | 149 | 1 | 1 | 0 | 0 | 0 | 0 | 0 | 0 | 0 | 0.008 | 0 | 0 | -0.002 |
| SE | 107173 | ENSG00000063046 | EIF4B | chr12 | + | 53016472 | 53016610 | 53006430 | 53006496 | 53018797 | 53019006 | 3447 | 1975 | 3592 | 3185 | 5753 | 1 | 0 | 0 | 3 | 0 | 1451 | 2003 | 2086 | 1184 | 919 | 0 | 0 | 0 | 0 | 0 | 287 | 149 | 1 | 1 | 0.999 | 1 | 1 | 0.998 | 1 | 1 | 1 | 1 | 1 | 1 | -0.001 |
| SE | 107174 | ENSG00000063046 | EIF4B | chr12 | + | 53019909 | 53020026 | 53018797 | 53019006 | 53021805 | 53021835 | 4632 | 2850 | 4928 | 4361 | 5683 | 1 | 3 | 3 | 1 | 0 | 3040 | 2381 | 2768 | 1827 | 1454 | 0 | 0 | 0 | 0 | 11 | 266 | 149 | 1 | 1 | 1 | 0.998 | 0.999 | 1 | 1 | 1 | 1 | 1 | 1 | 0.987 | 0.002 |
| SE | 107175 | ENSG00000063046 | EIF4B | chr12 | + | 53021805 | 53021860 | 53018797 | 53019006 | 53027781 | 53027919 | 0 | 4 | 4 | 6 | 3 | 0 | 0 | 0 | 0 | 0 | 0 | 5 | 0 | 0 | 5 | 0 | 0 | 1 | 0 | 0 | 204 | 149 | 1 | 1 | NA | 1 | 1 | 1 | 1 | NA | 1 | 0 | NA | 1 | 0.333 |
| SE | 107176 | ENSG00000063046 | EIF4B | chr12 | + | 53022253 | 53022366 | 53021805 | 53021860 | 53022492 | 53022627 | 0 | 4 | 7 | 2 | 2 | 3201 | 1943 | 3271 | 2964 | 4541 | 0 | 4 | 14 | 0 | 0 | 2088 | 1625 | 2002 | 1344 | 900 | 262 | 149 | 1 | 1 | 0 | 0.001 | 0.001 | 0 | 0 | 0 | 0.001 | 0.004 | 0 | 0 | -0.001 |
| SE | 107177 | ENSG00000063046 | EIF4B | chr12 | + | 53022492 | 53022627 | 53021715 | 53021860 | 53027781 | 53027856 | 5580 | 3628 | 5864 | 5414 | 8309 | 6 | 7 | 10 | 11 | 22 | 4113 | 3006 | 3748 | 2345 | 1733 | 0 | 6 | 3 | 0 | 10 | 284 | 149 | 1 | 1 | 0.998 | 0.996 | 0.997 | 0.996 | 0.995 | 1 | 0.996 | 0.998 | 1 | 0.989 | 0 |
| SE | 107180 | ENSG00000198715 | GLMP | chr1 | - | 156290976 | 156291106 | 156290095 | 156290360 | 156291402 | 156291550 | 7 | 0 | 0 | 0 | 0 | 0 | 0 | 0 | 1 | 0 | 0 | 0 | 2 | 0 | 0 | 0 | 0 | 0 | 0 | 0 | 279 | 149 | 1 | 1 | 1 | NA | NA | 0 | NA | NA | NA | 1 | NA | NA | -0.5 |
| [illegible] | | | | | | | | | | | | | | | | | | | | | | | | | | | | | | | | | | | | | | | | | | | | | | |

| | | | | | | | | | | | | | | | | | | | | | | | | | | | | | | | | | | | | | | | | | | | | | | |
|---|---|---|---|---|---|---|---|---|---|---|---|---|---|---|---|---|---|---|---|---|---|---|---|---|---|---|---|---|---|---|---|---|---|---|---|---|---|---|---|---|---|---|---|---|---|---|
| SE | 107641 | ENSG00000116396 | KCNC4 | chr1 | + | 110232220 | 110232597 | 110225974 | 110226178 | 110232910 | 110234044 | 0 | 0 | 3 | 1 | 0 | 0 | 0 | 0 | 0 | 0 | 0 | 0 | 0 | 0 | 0 | 0 | 0 | 0 | 0 | 7 | 298 | 149 | 1 | 1 | NA | NA | 1 | 1 | NA | NA | NA | NA | NA | 0 | 1 |
| SE | 107642 | ENSG00000139193 | CD27 | chr12 | + | 6450172 | 6450352 | 6445351 | 6445555 | 6450540 | 6450630 | 121 | 72 | 207 | 148 | 199 | 1 | 0 | 0 | 1 | 0 | 61 | 143 | 71 | 34 | 47 | 0 | 1 | 0 | 3 | 0 | 298 | 149 | 1 | 1 | 0.984 | 1 | 1 | 0.987 | 1 | 1 | 0.986 | 1 | 0.85 | 1 | 0.027 |
| SE | 107649 | ENSG00000198040 | ZNF84 | chr12 | + | 133038874 | 133039012 | 133037508 | 133037545 | 133041277 | 133041482 | 1 | 9 | 5 | 6 | 11 | 3 | 8 | 4 | 6 | 0 | 10 | 12 | 2 | 0 | 0 | 0 | 4 | 4 | 0 | 3 | 287 | 149 | 1 | 1 | 0.148 | 0.369 | 0.394 | 0.342 | 1 | 1 | 0.609 | 0.206 | NA | 0 | -0.003 |
| SE | 107653 | ENSG00000139718 | SETD1B | chr12 | + | 121817369 | 121817704 | 121817032 | 121817294 | 121817798 | 121817904 | 360 | 288 | 361 | 406 | 363 | 0 | 0 | 0 | 0 | 0 | 953 | 575 | 484 | 610 | 933 | 0 | 0 | 0 | 0 | 4 | 298 | 149 | 1 | 1 | 1 | 1 | 1 | 1 | 1 | 1 | 1 | 1 | 1 | 0.991 | 0.002 |
| SE | 107654 | ENSG00000139718 | SETD1B | chr12 | + | 121819403 | 121819895 | 121817798 | 121817904 | 121822489 | 121823749 | 384 | 316 | 414 | 409 | 513 | 0 | 2 | 0 | 1 | 0 | 1087 | 607 | 550 | 690 | 1042 | 0 | 0 | 0 | 0 | 12 | 298 | 149 | 1 | 1 | 1 | 0.988 | 1 | 0.995 | 1 | 1 | 1 | 1 | 1 | 0.977 | 0.001 |
| SE | 107664 | ENSG00000111247 | RAD51AP1 | chr12 | + | 4545218 | 4545312 | 4543762 | 4543904 | 4545781 | 4545832 | 0 | 0 | 0 | 1 | 0 | 0 | 0 | 0 | 2 | 2 | 9 | 0 | 0 | 0 | 0 | 0 | 0 | 3 | 0 | 0 | 243 | 149 | 1 | 1 | NA | NA | NA | 0.235 | 0 | 1 | NA | 0 | NA | NA | -0.383 |
| SE | 107665 | ENSG00000111247 | RAD51AP1 | chr12 | + | 4545218 | 4545312 | 4543762 | 4543904 | 4546308 | 4546418 | 0 | 0 | 0 | 1 | 0 | 2 | 6 | 4 | 1 | 3 | 9 | 0 | 0 | 0 | 0 | 0 | 0 | 9 | 3 | 0 | 243 | 149 | 1 | 1 | 0 | 0 | 0 | 0.38 | 0 | 1 | NA | 0 | 0 | NA | -0.257 |
| SE | 107668 | ENSG00000111247 | RAD51AP1 | chr12 | + | 4545781 | 4545832 | 4543762 | 4543904 | 4546308 | 4546418 | 1 | 1 | 0 | 3 | 2 | 2 | 6 | 4 | 1 | 3 | 0 | 0 | 7 | 0 | 0 | 0 | 0 | 9 | 3 | 0 | 200 | 149 | 1 | 1 | 0.271 | 0.11 | 0 | 0.691 | 0.332 | NA | NA | 0.367 | 0 | NA | 0.097 |
| SE | 107669 | ENSG00000111247 | RAD51AP1 | chr12 | + | 4545781 | 4545832 | 4543762 | 4543911 | 4546308 | 4546418 | 2 | 1 | 0 | 1 | 0 | 2 | 0 | 0 | 0 | 0 | 0 | 0 | 4 | 0 | 0 | 0 | 0 | 0 | 0 | 0 | 200 | 149 | 1 | 1 | 0.427 | 1 | NA | 1 | NA | NA | NA | 1 | NA | NA | -0.191 |
| SE | 107673 | ENSG00000111247 | RAD51AP1 | chr12 | + | 4556352 | 4556502 | 4552982 | 4553147 | 4558856 | 4560018 | 6 | 3 | 4 | 14 | 9 | 1 | 0 | 0 | 0 | 4 | 0 | 2 | 2 | 10 | 0 | 0 | 0 | 0 | 0 | 0 | 298 | 149 | 1 | 1 | 0.75 | 1 | 1 | 1 | 0.529 | NA | 1 | 1 | 1 | NA | -0.144 |
| SE | 107674 | ENSG00000111247 | RAD51AP1 | chr12 | + | 4556352 | 4556550 | 4552982 | 4553147 | 4558856 | 4559019 | 6 | 3 | 4 | 13 | 9 | 1 | 0 | 0 | 0 | 4 | 0 | 0 | 2 | 4 | 0 | 0 | 0 | 0 | 0 | 0 | 298 | 149 | 1 | 1 | 0.75 | 1 | 1 | 1 | 0.529 | NA | NA | 1 | 1 | NA | -0.144 |
| SE | 107676 | ENSG00000198015 | MRPL42 | chr12 | + | 93470485 | 93470590 | 93469191 | 93469355 | 93476953 | 93477017 | 3 | 1 | 3 | 4 | 0 | 163 | 105 | 102 | 107 | 265 | 0 | 0 | 0 | 0 | 0 | 11 | 59 | 67 | 38 | 15 | 254 | 149 | 1 | 1 | 0.011 | 0.006 | 0.017 | 0.021 | 0 | 0 | 0 | 0 | 0 | 0 | 0.011 |
| SE | 107679 | ENSG00000198015 | MRPL42 | chr12 | + | 93501175 | 93501311 | 93487496 | 93487660 | 93502705 | 93502864 | 87 | 56 | 96 | 87 | 100 | 0 | 0 | 0 | 0 | 4 | 53 | 59 | 53 | 36 | 0 | 0 | 0 | 0 | 0 | 0 | 285 | 149 | 1 | 1 | 1 | 1 | 1 | 1 | 0.929 | 1 | 1 | 1 | 1 | NA | -0.014 |
| SE | 107681 | ENSG00000128585 | MKLN1 | chr7 | + | 131375423 | 131375493 | 131327875 | 131327997 | 131387119 | 131387262 | 87 | 81 | 74 | 60 | 57 | 0 | 1 | 0 | 0 | 0 | 47 | 85 | 90 | 31 | 62 | 0 | 0 | 0 | 0 | 0 | 219 | 149 | 1 | 1 | 1 | 0.982 | 1 | 1 | 1 | 1 | 1 | 1 | 1 | 1 | -0.004 |
| SE | 107683 | ENSG00000128585 | MKLN1 | chr7 | + | 131387119 | 131387262 | 131375423 | 131375493 | 131388883 | 131388972 | 383 | 237 | 284 | 295 | 432 | 0 | 1 | 0 | 1 | 0 | 181 | 304 | 239 | 180 | 255 | 7 | 0 | 1 | 0 | 0 | 292 | 149 | 1 | 1 | 1 | 1 | 1 | 0.993 | 1 | 0.93 | 1 | 0.992 | 1 | 1 | 0.014 |
| SE | 107684 | ENSG00000128585 | MKLN1 | chr7 | + | 131388883 | 131388972 | 131387119 | 131387262 | 131397266 | 131397376 | 377 | 239 | 266 | 323 | 588 | 0 | 2 | 0 | 1 | 0 | 218 | 276 | 188 | 177 | 245 | 0 | 0 | 0 | 0 | 0 | 238 | 149 | 1 | 1 | 1 | 0.987 | 1 | 0.995 | 1 | 1 | 1 | 1 | 1 | 1 | -0.004 |
| SE | 107686 | ENSG00000128585 | MKLN1 | chr7 | + | 131443480 | 131443702 | 131437784 | 131437997 | 131445773 | 131445903 | 260 | 270 | 191 | 289 | 266 | 3 | 0 | 0 | 1 | 0 | 45 | 205 | 234 | 115 | 15 | 0 | 1 | 5 | 0 | 0 | 298 | 149 | 1 | 1 | 0.977 | 1 | 1 | 0.993 | 1 | 1 | 0.99 | 0.959 | 1 | 1 | 0.004 |
| SE | 107690 | ENSG00000128585 | MKLN1 | chr7 | + | 131464280 | 131464408 | 131463324 | 131463364 | 131466275 | 131466415 | 127 | 126 | 104 | 135 | 230 | 0 | 0 | 0 | 1 | 0 | 14 | 85 | 49 | 33 | 30 | 0 | 0 | 1 | 0 | 0 | 277 | 149 | 1 | 1 | 1 | 1 | 1 | 0.986 | 1 | 1 | 1 | 0.963 | 1 | 1 | 0.005 |
| SE | 107691 | ENSG00000128585 | MKLN1 | chr7 | + | 131464293 | 131464408 | 131463216 | 131463364 | 131466275 | 131466415 | 206 | 203 | 174 | 221 | 340 | 0 | 0 | 0 | 1 | 0 | 16 | 127 | 87 | 42 | 31 | 0 | 0 | 1 | 0 | 0 | 264 | 149 | 1 | 1 | 1 | 1 | 1 | 0.992 | 1 | 1 | 1 | 0.98 | 1 | 1 | 0.002 |
| SE | 107692 | ENSG00000128585 | MKLN1 | chr7 | + | 131470841 | 131470944 | 131466275 | 131466415 | 131478622 | 131478677 | 202 | 131 | 157 | 175 | 366 | 0 | 0 | 0 | 0 | 0 | 2 | 119 | 120 | 31 | 23 | 0 | 0 | 1 | 0 | 0 | 252 | 149 | 1 | 1 | 1 | 1 | 1 | 1 | 1 | 1 | 1 | 0.986 | 1 | 1 | 0.003 |
| SE | 107693 | ENSG00000128585 | MKLN1 | chr7 | + | 131478622 | 131478677 | 131470841 | 131470944 | 131487606 | 131487844 | 105 | 53 | 93 | 99 | 218 | 0 | 1 | 0 | 1 | 0 | 0 | 38 | 70 | 12 | 6 | 0 | 0 | 1 | 0 | 0 | 204 | 149 | 1 | 1 | 1 | 0.975 | 1 | 0.986 | 1 | NA | 1 | 0.981 | 1 | 1 | -0.003 |
| SE | 107694 | ENSG00000129317 | PUS7L | chr12 | - | 43731704 | 43731758 | 43718991 | 43730702 | 43736380 | 43736661 | 91 | 58 | 66 | 51 | 143 | 0 | 0 | 0 | 1 | 0 | 0 | 38 | 25 | 1 | 0 | 0 | 0 | 0 | 0 | 0 | 203 | 149 | 1 | 1 | 1 | 1 | 1 | 0.974 | 1 | NA | 1 | 1 | 1 | NA | -0.005 |
| SE | 107695 | ENSG00000129317 | PUS7L | chr12 | - | 43736380 | 43736661 | 43731704 | 43731758 | 43742456 | 43742555 | 88 | 65 | 68 | 55 | 136 | 1 | 1 | 6 | 0 | 1 | 0 | 33 | 23 | 1 | 0 | 0 | 0 | 0 | 0 | 0 | 298 | 149 | 1 | 1 | 0.978 | 0.97 | 0.85 | 1 | 0.986 | NA | 1 | 1 | 1 | NA | -0.043 |
| SE | 107696 | ENSG00000129317 | PUS7L | chr12 | - | 43738309 | 43738391 | 43736380 | 43736661 | 43742456 | 43742555 | 50 | 72 | 83 | 90 | 210 | 2 | 2 | 1 | 2 | 0 | 0 | 67 | 33 | 8 | 0 | 0 | 0 | 0 | 0 | 0 | 231 | 149 | 1 | 1 | 0.942 | 0.959 | 0.982 | 0.967 | 1 | NA | 1 | 1 | 1 | NA | -0.03 |
| SE | 107697 | ENSG00000129317 | PUS7L | chr12 | - | 43742456 | 43742555 | 43731704 | 43731758 | 43746045 | 43746238 | 36 | 66 | 60 | 71 | 71 | 2 | 0 | 0 | 0 | 0 | 0 | 40 | 15 | 1 | 4 | 0 | 0 | 0 | 0 | 0 | 248 | 149 | 1 | 1 | 0.915 | 1 | 1 | 1 | 1 | NA | 1 | 1 | 1 | 1 | -0.017 |
| SE | 107698 | ENSG00000129317 | PUS7L | chr12 | - | 43742456 | 43742555 | 43738309 | 43738391 | 43746045 | 43746238 | 60 | 125 | 109 | 133 | 228 | 23 | 0 | 0 | 2 | 1 | 0 | 94 | 28 | 9 | 4 | 0 | 0 | 0 | 0 | 0 | 248 | 149 | 1 | 1 | 0.61 | 1 | 1 | 0.976 | 0.993 | NA | 1 | 1 | 1 | 1 | -0.084 |
| SE | 107699 | ENSG00000129317 | PUS7L | chr12 | - | 43748449 | 43748609 | 43746045 | 43746238 | 43754335 | 43754442 | 147 | 91 | 102 | 117 | 128 | 1 | 1 | 0 | 0 | 1 | 10 | 42 | 28 | 1 | 1 | 0 | 0 | 0 | 0 | 0 | 298 | 149 | 1 | 1 | 0.987 | 0.978 | 1 | 1 | 0.985 | 1 | 1 | 1 | 1 | 1 | -0.01 |
| SE | 107700 | ENSG00000129317 | PUS7L | chr12 | - | 43754335 | 43755261 | 43748449 | 43748609 | 43758729 | 43758767 | 88 | 63 | 79 | 80 | 105 | 2 | 3 | 1 | 0 | 0 | 10 | 36 | 15 | 0 | 0 | 0 | 0 | 0 | 0 | 0 | 298 | 149 | 1 | 1 | 0.957 | 0.913 | 0.975 | 1 | 1 | 1 | 1 | 1 | NA | NA | -0.031 |
| SE | 107701 | ENSG00000129317 | PUS7L | chr12 | - | 43755216 | 43755261 | 43748449 | 43748609 | 43758729 | 43758759 | 1 | 1 | 0 | 4 | 4 | 2 | 3 | 1 | 0 | 0 | 0 | 4 | 0 | 0 | 0 | 0 | 0 | 0 | 0 | 0 | 194 | 149 | 1 | 1 | 0.277 | 0.204 | 0 | 1 | 1 | NA | 1 | NA | NA | NA | -0.504 |
| SE | 107710 | ENSG00000177575 | CD163 | chr12 | - | 7479859 | 7479913 | 7470810 | 7471397 | 7481160 | 7481256 | 136 | 97 | 70 | 170 | 180 | 0 | 0 | 0 | 0 | 0 | 395 | 854 | 529 | 293 | 184 | 0 | 9 | 0 | 0 | 0 | 203 | 149 | 1 | 1 | 1 | 1 | 1 | 1 | 1 | 1 | 0.986 | 1 | 1 | 1 | 0.003 |
| SE | 107712 | ENSG00000177575 | CD163 | chr12 | - | 7481160 | 7481256 | 7479859 | 7479913 | 7482642 | 7482762 | 249 | 163 | 111 | 293 | 244 | 0 | 0 | 0 | 2 | 0 | 844 | 1672 | 1049 | 572 | 355 | 0 | 8 | 4 | 3 | 2 | 245 | 149 | 1 | 1 | 1 | 1 | 1 | 0.989 | 1 | 1 | 0.992 | 0.994 | 0.991 | 0.991 | 0.004 |
| SE | 107713 | ENSG00000177575 | CD163 | chr12 | - | 7481160 | 7481256 | 7479859 | 7479913 | 7482965 | 7483004 | 107 | 81 | 52 | 125 | 105 | 0 | 1 | 0 | 0 | 0 | 287 | 690 | 389 | 200 | 138 | 0 | 0 | 0 | 0 | 0 | 245 | 149 | 1 | 1 | 1 | 0.98 | 1 | 1 | 1 | 1 | 1 | 1 | 1 | 1 | -0.004 |
| SE | 107716 | ENSG00000177575 | CD163 | chr12 | - | 7482642 | 7482762 | 7479859 | 7479913 | 7482965 | 7483004 | 137 | 81 | 54 | 150 | 144 | 0 | 1 | 0 | 0 | 0 | 341 | 820 | 500 | 350 | 170 | 0 | 0 | 0 | 0 | 0 | 269 | 149 | 1 | 1 | 1 | 0.978 | 1 | 1 | 1 | 1 | 1 | 1 | 1 | 1 | -0.004 |
| SE | 107717 | ENSG00000177575 | CD163 | chr12 | - | 7482642 | 7482762 | 7479859 | 7479996 | 7482965 | 7483004 | 142 | 84 | 56 | 157 | 145 | 0 | 0 | 0 | 2 | 0 | 362 | 837 | 496 | 378 | 168 | 6 | 3 | 0 | 0 | 0 | 269 | 149 | 1 | 1 | 1 | 1 | 1 | 0.978 | 1 | 0.971 | 0.994 | 1 | 1 | 1 | 0.003 |
| SE | 107719 | ENSG00000177575 | CD163 | chr12 | - | 7482642 | 7482762 | 7479859 | 7481256 | 7485095 | 7485416 | 138 | 67 | 44 | 158 | 123 | 0 | 0 | 0 | 0 | 0 | 519 | 873 | 599 | 343 | 202 | 0 | 0 | 2 | 0 | 0 | 269 | 149 | 1 | 1 | 1 | 1 | 1 | 1 | 1 | 1 | 1 | 0.994 | 1 | 1 | 0.001 |
| SE | 107720 | ENSG00000177575 | CD163 | chr12 | - | 7482965 | 7483004 | 7479859 | 7481256 | 7485095 | 7485416 | 0 | 3 | 2 | 1 | 0 | 0 | 0 | 0 | 0 | 0 | 4 | 35 | 26 | 25 | 0 | 0 | 0 | 2 | 0 | 0 | 188 | 149 | 1 | 1 | NA | 1 | 1 | 1 | NA | 1 | 1 | 0.912 | 1 | NA | 0.022 |
| SE | 107721 | ENSG00000177575 | CD163 | chr12 | - | 7482965 | 7483004 | 7482642 | 7482762 | 7485095 | 7485416 | 18 | 30 | 17 | 25 | 32 | 0 | 0 | 0 | 0 | 0 | 71 | 230 | 153 | 94 | 30 | 0 | 3 | 0 | 0 | 0 | 188 | 149 | 1 | 1 | 1 | 1 | 1 | 1 | 1 | 1 | 0.984 | 1 | 1 | 1 | 0.003 |
| SE | 107723 | ENSG00000177575 | CD163 | chr12 | - | 7486893 | 7486986 | 7486498 | 7486813 | 7487358 | 7487673 | 313 | 194 | 136 | 358 | 170 | 0 | 3 | 0 | 0 | 0 | 1418 | 2294 | 1579 | 833 | 608 | 0 | 2 | 2 | 7 | 0 | 242 | 149 | 1 | 1 | 1 | 0.975 | 1 | 1 | 1 | 1 | 0.999 | 0.998 | 0.987 | 1 | -0.002 |
| SE | 107724 | ENSG00000177575 | CD163 | chr12 | - | 7495080 | 7495401 | 7487358 | 7488087 | 7496812 | 7497133 | 750 | 421 | 301 | 641 | 609 | 1 | 1 | 0 | 0 | 0 | 2318 | 4453 | 2547 | 1657 | 769 | 6 | 17 | 15 | 0 | 0 | 298 | 149 | 1 | 1 | 0.997 | 0.995 | 1 | 1 | 1 | 0.995 | 0.992 | 0.988 | 1 | 1 | 0.003 |
| SE | 107725 | ENSG00000177575 | CD163 | chr12 | - | 7496812 | 7497133 | 7495080 | 7495401 | 7498867 | 7499188 | 1040 | 700 | 467 | 1105 | 1222 | 4 | 0 | 0 | 1 | 0 | 5589 | 9045 | 5594 | 3751 | 1920 | 0 | 1 | 10 | 0 | 0 | 298 | 149 | 1 | 1 | 0.992 | 1 | 1 | 0.998 | 1 | 1 | 1 | 0.996 | 1 | 1 | -0.001 |
| SE | 107726 | ENSG00000177575 | CD163 | chr12 | - | 7501138 | 7501462 | 7498867 | 7499188 | 7502477 | 7502564 | 794 | 512 | 338 | 827 | 586 | 0 | 0 | 0 | 0 | 0 | 2898 | 5366 | 4289 | 2225 | 1138 | 0 | 4 | 0 | 0 | 0 | 298 | 149 | 1 | 1 | 1 | 1 | 1 | 1 | 1 | 1 | 0.999 | 1 | 1 | 1 | 0 |
| SE | 107727 | ENSG00000177575 | CD163 | chr12 | - | 7501138 | 7501462 | 7498867 | 7499188 | 7503644 | 7503776 | 293 | 164 | 142 | 276 | 150 | 0 | 0 | 0 | 1 | 0 | 1125 | 2000 | 1576 | 649 | 438 | 0 | 8 | 7 | 0 | 0 | 298 | 149 | 1 | 1 | 1 | 1 | 1 | 0.993 | 1 | 1 | 0.992 | 0.991 | 1 | 1 | 0.002 |
| SE | 107728 | ENSG00000177575 | CD163 | chr12 | - | 7502477 | 7502564 | 7498867 | 7499188 | 7503644 | 7503776 | 443 | 313 | 166 | 422 | 390 | 0 | 0 | 0 | 1 | 0 | 1687 | 2685 | 2558 | 1192 | 522 | 0 | 8 | 7 | 0 | 0 | 236 | 149 | 1 | 1 | 1 | 1 | 1 | 0.996 | 1 | 1 | 0.995 | 0.996 | 1 | 1 | 0.001 |
| SE | 107729 | ENSG00000177575 | CD163 | chr12 | - | 7502477 | 7502564 | 7501138 | 7501462 | 7503644 | 7503776 | 948 | 658 | 362 | 974 | 825 | 4 | 0 | 0 | 2 | 0 | 3460 | 6044 | 5272 | 2768 | 1222 | 0 | 4 | 16 | 0 | 0 | 236 | 149 | 1 | 1 | 0.993 | 1 | 1 | 0.997 | 1 | 1 | 0.999 | 0.995 | 1 | 1 | -0.001 |
| SE | 107730 | ENSG00000196876 | SCN8A | chr12 | + | 51688757 | 51688849 | 51687090 | 51687219 | 51699569 | 51699791 | 1 | 0 | 0 | 0 | 3 | 0 | 0 | 1 | 3 | 0 | 0 | 0 | 0 | 4 | 0 | 0 | 0 | 0 | 6 | 4 | 241 | 149 | 1 | 1 | 1 | NA | 0 | 0 | 1 | NA | NA | NA | 0.292 | 0 | 0.354 |
| SE | 107731 | ENSG00000196876 | SCN8A | chr12 | + | 51688757 | 51690019 | 51687090 | 51687219 | 51699569 | 51699791 | 1 | 0 | 0 | 0 | 3 | 0 | 0 | 1 | 3 | 0 | 0 | 0 | 0 | 4 | 0 | 0 | 0 | 0 | 6 | 4 | 298 | 149 | 1 | 1 | 1 | NA | 0 | 0 | 1 | NA | NA | NA | 0.25 | 0 | 0.375 |
| SE | 107733 | ENSG00000196876 | SCN8A | chr12 | + | 51751354 | 51751593 | 51745900 | 51746035 | 51762502 | 51762676 | 2 | 1 | 1 | 0 | 0 | 0 | 0 | 0 | 0 | 0 | 10 | 0 | 2 | 24 | 0 | 0 | 0 | 0 | 1 | 0 | 298 | 149 | 1 | 1 | 1 | 1 | 1 | NA | NA | 1 | NA | 1 | 0.923 | NA | 0.026 |
| SE | 107735 | ENSG00000196876 | SCN8A | chr12 | + | 51780201 | 51780271 | 51774188 | 51774362 | 51786541 | 51786826 | 0 | 0 | 0 | 1 | 0 | 0 | 0 | 0 | 0 | 0 | 0 | 8 | 0 | 4 | 0 | 0 | 0 | 1 | 0 | 0 | 219 | 149 | 1 | 1 | NA | NA | NA | 1 | NA | NA | 1 | 0 | 1 | NA | 0.333 |
| SE | 107737 | ENSG00000135677 | GNS | chr12 | - | 64723005 | 64723113 | 64721594 | 64721705 | 64728955 | 64729057 | 612 | 476 | 357 | 461 | 597 | 0 | 1 | 2 | 1 | 3 | 1296 | 779 | 443 | 746 | 964 | 0 | 2 | 0 | 3 | 0 | 257 | 149 | 1 | 1 | 1 | 0.996 | 0.99 | 0.996 | 0.991 | 1 | 0.996 | 1 | 0.993 | 1 | -0.003 |
| SE | 107738 | ENSG00000135677 | GNS | chr12 | - | 64728955 | 64729057 | 64723005 | 64723113 | 64737003 | 64737107 | 795 | 550 | 453 | 598 | 825 | 1 | 0 | 0 | 0 | 2 | 1526 | 948 | 544 | 923 | 1198 | 0 | 9 | 4 | 0 | 0 | 251 | 149 | 1 | 1 | 0.998 | 1 | 1 | 1 | 0.996 | 1 | 0.984 | 0.988 | 1 | 1 | 0.004 |
| SE | 107739 | ENSG00000135677 | GNS | chr12 | - | 64737003 | 64737107 | 64728955 | 64729057 | 64739380 | 64739499 | 785 | 523 | 418 | 582 | 830 | 0 | 0 | 0 | 0 | 0 | 1370 | 831 | 497 | 884 | 1114 | 0 | 2 | 0 | 0 | 0 | 253 | 149 | 1 | 1 | 1 | 1 | 1 | 1 | 1 | 1 | 0.996 | 1 | 1 | 1 | 0.001 |
| SE | 107740 | ENSG00000135677 | GNS | chr12 | - | 64737003 | 64737107 | 64728955 | 64729057 | 64743140 | 64743308 | 362 | 252 | 227 | 272 | 406 | 1 | 1 | 0 | 0 | 0 | 685 | 427 | 268 | 386 | 542 | 0 | 0 | 1 | 0 | 0 | 253 | 149 | 1 | 1 | 0.995 | 0.993 | 1 | 1 | 1 | 1 | 1 | 0.994 | 1 | 1 | -0.001 |
| SE | 107741 | ENSG00000135677 | GNS | chr12 | - | 64739380 | 64739499 | 64728955 | 64729057 | 64743140 | 64743308 | 0 | 0 | 0 | 0 | 1 | 1 | 1 | 0 | 0 | 0 | 0 | 2 | 0 | 0 | 0 | 0 | 0 | 1 | 0 | 0 | 268 | 149 | 1 | 1 | 0 | 0 | NA | NA | 1 | NA | 1 | 0 | NA | NA | -0.167 |
| SE | 107742 | ENSG00000135677 | GNS | chr12 | - | 64739380 | 64739499 | 64737003 | 64737107 | 64740605 | 64740688 | 1071 | 643 | 553 | 781 | 855 | 0 | 1 | 1 | 1 | 0 | 1881 | 1146 | 759 | 1286 | 1496 | 0 | 3 | 0 | 10 | 0 | 268 | 149 | 1 | 1 | 1 | 0.997 | 0.997 | 0.998 | 1 | 1 | 0.995 | 1 | 0.986 | 1 | 0.002 |
| SE | 107743 | ENSG00000135677 | GNS | chr12 | - | 64739380 | 64739499 | 64737003 | 64737107 | 64743140 | 64743308 | 387 | 223 | 174 | 275 | 355 | 1 | 2 | 3 | 0 | 0 | 582 | 376 | 186 | 435 | 489 | 0 | 3 | 0 | 2 | 0 | 268 | 149 | 1 | 1 | 0.995 | 0.984 | 0.97 | 1 | 1 | 1 | 0.986 | 1 | 0.992 | 1 | -0.006 |
| SE | 107744 | ENSG00000135677 | GNS | chr12 | - | 64740605 | 64740688 | 64737003 | 64737107 | 64743140 | 64743308 | 189 | 140 | 105 | 154 | 176 | 1 | 2 | 3 | 0 | 0 | 308 | 219 | 108 | 296 | 326 | 0 | 3 | 0 | 2 | 0 | 232 | 149 | 1 | 1 | 0.992 | 0.978 | 0.957 | 1 | 1 | 1 | 0.979 | 1 | 0.99 | 1 | -0.008 |
| SE | 107745 | ENSG00000135677 | GNS | chr12 | - | 64740605 | 64740688 | 64739380 | 64739499 | 64743140 | 64743308 | 871 | 557 | 478 | 652 | 677 | 0 | 0 | 0 | 0 | 1 | 1606 | 980 | 673 | 1125 | 1333 | 0 | 0 | 0 | 0 | 0 | 232 | 149 | 1 | 1 | 1 | 1 | 1 | 1 | 0.998 | 1 | 1 | 1 | 1 | 1 | 0 |
| SE | 107746 | ENSG00000135677 | GNS | chr12 | - | 64752697 | 64752757 | 64747711 | 64747918 | 64759084 | 64759406 | 247 | 190 | 158 | 152 | 167 | 4 | 0 | 3 | 0 | 0 | 397 | 300 | 177 | 342 | 371 | 0 | 0 | 0 | 0 | 0 | 209 | 149 | 1 | 1 | 0.978 | 1 | 0.974 | 1 | 1 | 1 | 1 | 1 | 1 | 1 | -0.01 |
| SE | 107756 | ENSG00000174106 | LEMD3 | chr12 | + | 65218551 | 65218619 | 65215976 | 65216043 | 65238501 | 65238581 | 311 | 181 | 296 | 267 | 362 | 0 | 0 | 0 | 0 | 0 | 133 | 193 | 243 | 164 | 84 | 0 | 1 | 0 | 0 | 0 | 217 | 149 | 1 | 1 | 1 | 1 | 1 | 1 | 1 | 1 | 0.993 | 1 | 1 | 1 | 0.001 |
| SE | 107757 | ENSG00000174106 | LEMD3 | chr12 | + | 65238668 | 65238814 | 65238501 | 65238581 | 65239928 | 65240030 | 450 | 250 | 422 | 438 | 495 | 1 | 1 | 1 | 0 | 0 | 110 | 261 | 317 | 272 | 88 | 0 | 0 | 0 | 0 | 0 | 295 | 149 | 1 | 1 | 0.996 | 0.992 | 0.995 | 1 | 1 | 1 | 1 | 1 | 1 | 1 | -0.003 |
| SE | 107758 | ENSG00000174106 | LEMD3 | chr12 | + | 65239928 | 65240030 | 65238668 | 65238814 | 65240135 | 65240238 | 310 | 147 | 275 | 319 | 444 | 0 | 0 | 0 | 1 | 0 | 69 | 154 | 184 | 170 | 50 | 0 | 0 | 0 | 0 | 0 | 251 | 149 | 1 | 1 | 1 | 1 | 1 | 0.995 | 1 | 1 | 1 | 1 | 1 | 1 | -0.001 |
| SE | 107759 | ENSG00000174106 | LEMD3 | chr12 | + | 65245668 | 65245774 | 65243387 | 65243469 | 65245860 | 65245939 | 279 | 161 | 250 | 230 | 675 | 0 | 0 | 0 | 2 | 0 | 96 | 138 | 156 | 94 | 0 | 0 | 0 | 0 | 0 | 0 | 255 | 149 | 1 | 1 | 1 | 1 | 1 | 0.985 | 1 | 1 | 1 | 1 | 1 | NA | -0.003 |
| SE | 107760 | ENSG00000174106 | LEMD3 | chr12 | + | 65245668 | 65245774 | 65243387 | 65243469 | 65246161 | 65248355 | 153 | 79 | 149 | 133 | 319 | 1 | 0 | 0 | 2 | 5 | 60 | 87 | 88 | 56 | 0 | 0 | 6 | 0 | 0 | 0 | 255 | 149 | 1 | 1 | 0.989 | 1 | 1 | 0.975 | 0.974 | 1 | 0.894 | 1 | 1 | NA | 0.014 |
| SE | 107761 | ENSG00000174106 | LEMD3 | chr12 | + | 65245668 | 65245939 | 65243448 | 65243469 | 65246161 | 65246281 | 259 | 152 | 247 | 233 | 627 | 1 | 0 | 0 | 2 | 5 | 78 | 137 | 146 | 81 | 0 | 0 | 6 | 0 | 0 | 0 | 298 | 149 | 1 | 1 | 0.992 | 1 | 1 | 0.983 | 0.984 | 1 | 0.919 | 1 | 1 | NA | 0.012 |
| SE | 107762 | ENSG00000174106 | LEMD3 | chr12 | + | 65245860 | 65245939 | 65243387 | 65243469 | 65246161 | 65246539 | 97 | 58 | 82 | 77 | 289 | 1 | 0 | 0 | 2 | 5 | 14 | 34 | 47 | 16 | 0 | 0 | 6 | 0 | 0 | 0 | 228 | 149 | 1 | 1 | 0.984 | 1 | 1 | 0.962 | 0.974 | 1 | 0.787 | 1 | 1 | NA | 0.037 |
| SE | 107763 | ENSG00000174106 | LEMD3 | chr12 | + | 65245860 | 65245939 | 65245299 | 65245774 | 65246161 | 65246539 | 229 | 139 | 183 | 170 | 643 | 8 | 0 | 0 | 0 | 4 | 50 | 84 | 117 | 54 | 0 | 0 | 2 | 2 | 0 | 0 | 228 | 149 | 1 | 1 | 0.949 | 1 | 1 | 1 | 0.991 | 1 | 0.965 | 0.975 | 1 | NA | 0.003 |
| SE | 107764 | ENSG00000261609 | GAN | chr16 | + | 81351582 | 81351697 | 81314961 | 81315280 | 81354404 | 81354755 | 22 | 9 | 29 | 17 | 38 | 0 | 0 | 0 | 0 | 6 | 1 | 14 | 13 | 6 | 0 | 0 | 0 | 0 | 0 | 0 | 264 | 149 | 1 | 1 | 1 | 1 | 1 | 1 | 0.781 | 1 | 1 | 1 | 1 | NA | -0.044 |
| SE | 107767 | ENSG00000073060 | SCARB1 | chr12 | - | 124786356 | 124786503 | 124782682 | 124782811 | 124787405 | 124787457 | 15 | 9 | 20 | 14 | 24 | 0 | 0 | 0 | 1 | 0 | 27 | 10 | 1 | 18 | 57 | 0 | 0 | 0 | 0 | 0 | 296 | 149 | 1 | 1 | 1 | 1 | 1 | 0.876 | 1 | 1 | 1 | 1 | 1 | 1 | -0.025 |
| SE | 107768 | ENSG00000073060 | SCARB1 | chr12 | - | 124786440 | 124786503 | 124782682 | 124782811 | 124787405 | 124787457 | 2 | 1 | 9 | 2 | 7 | 0 | 0 | 0 | 1 | 0 | 0 | 1 | 1 | 12 | 29 | 0 | 0 | 0 | 0 | 0 | 212 | 149 | 1 | 1 | 1 | 1 | 1 | 0.584 | 1 | NA | 1 | 1 | 1 | 1 | -0.083 |
| SE | 107770 | ENSG00000073060 | SCARB1 | chr12 | - | 124795194 | 124795268 | 124787405 | 124787457 | 124807760 | 124807927 | 9 | 0 | 4 | 6 | 4 | 0 | 0 | 0 | 1 | 0 | 24 | 1 | 0 | 10 | 5 | 0 | 0 | 0 | 0 | 0 | 223 | 149 | 1 | 1 | 1 | NA | 1 | 0.8 | 1 | 1 | 1 | NA | 1 | 1 | -0.05 |
| SE | 107771 | ENSG00000073060 | SCARB1 | chr12 | - | 124800123 | 124800242 | 124795194 | 124795268 | 124807760 | 124807927 | 23 | 13 | 30 | 15 | 9 | 0 | 0 | 0 | 2 | 0 | 33 | 17 | 6 | 26 | 61 | 0 | 0 | 0 | 0 | 0 | 268 | 149 | 1 | 1 | 1 | 1 | 1 | 0.807 | 1 | 1 | 1 | 1 | 1 | 1 | -0.039 |
| SE | 107772 | ENSG00000073060 | SCARB1 | chr12 | - | 124807760 | 124807927 | 124800123 | 124800242 | 124811869 | 124811965 | 10 | 10 | 15 | 12 | 9 | 0 | 0 | 0 | 0 | 0 | 33 | 16 | 6 | 19 | 36 | 0 | 0 | 0 | 1 | 0 | 298 | 149 | 1 | 1 | 1 | 1 | 1 | 1 | 1 | 1 | 1 | 1 | 0.905 | 1 | 0.019 |
| SE | 107773 | ENSG00000073060 | SCARB1 | chr12 | - | 124810173 | 124810265 | 124807760 | 124807927 | 124811869 | 124811965 | 8 | 6 | 13 | 11 | 15 | 0 | 0 | 0 | 0 | 1 | 27 | 19 | 5 | 24 | 12 | 0 | 0 | 0 | 0 | 0 | 241 | 149 | 1 | 1 | 1 | 1 | 1 | 1 | 0.903 | 1 | 1 | 1 | 1 | 1 | -0.019 |
| SE | 107774 | ENSG00000073060 | SCARB1 | chr12 | - | 124810173 | 124810289 | 124805764 | 124807927 | 124811869 | 124811965 | 27 | 17 | 42 | 34 | 47 | 0 | 0 | 0 | 0 | 1 | 90 | 37 | 10 | 59 | 58 | 0 | 0 | 0 | 0 | 0 | 265 | 149 | 1 | 1 | 1 | 1 | 1 | 1 | 0.964 | 1 | 1 | 1 | 1 | 1 | -0.007 |
| SE | 107778 | ENSG00000102780 | DGKH | chr13 | + | 42153749 | 42153915 | 42129551 | 42129632 | 42155290 | 42155395 | 4 | 7 | 0 | 0 | 0 | 7 | 5 | 9 | 13 | 15 | 1 | 5 | 0 | 0 | 27 | 0 | 13 | 14 | 0 | 3 | 298 | 149 | 1 | 1 | 0.222 | 0.412 | 0 | 0 | 0 | 1 | 0.161 | 0 | NA | 0.818 | -0.368 |
| SE | 107779 | ENSG00000102780 | DGKH | chr13 | + | 42159265 | 42159372 | 42155666 | 42155799 | 42160010 | 42160136 | 14 | 18 | 18 | 12 | 15 | 0 | 1 | 2 | 0 | 0 | 0 | 6 | 22 | 5 | 8 | 0 | 0 | 0 | 0 | 43 | 256 | 149 | 1 | 1 | 1 | 0.913 | 0.84 | 1 | 1 | NA | 1 | 1 | 1 | 0.098 | 0.176 |
| SE | 107785 | ENSG00000102780 | DGKH | chr13 | + | 42221161 | 42221394 | 42219685 | 42219794 | 42229098 | 42229250 | 0 | 1 | 0 | 1 | 0 | 10 | 9 | 9 | 10 | 10 | 0 | 0 | 0 | 0 | 0 | 24 | 8 | 10 | 8 | 6 | 298 | 149 | 1 | 1 | 0 | 0.053 | 0 | 0.048 | 0 | 0 | 0 | 0 | 0 | 0 | 0.02 |
| SE | 107786 | ENSG00000102780 | DGKH | chr13 | + | 42221263 | 42221394 | 42219685 | 42219794 | 42229098 | 42229250 | 0 | 1 | 0 | 1 | 0 | 10 | 9 | 9 | 10 | 10 | 0 | 0 | 0 | 0 | 0 | 24 | 8 | 10 | 8 | 6 | 280 | 149 | 1 | 1 | 0 | 0.056 | 0 | 0.051 | 0 | 0 | 0 | 0 | 0 | 0 | 0.021 |
| SE | 107788 | ENSG00000050426 | LETMD1 | chr12 | + | 51049033 | 51049185 | 51048300 | 51048478 | 51055834 | 51055848 | 41 | 17 | 44 | 35 | 28 | 1 | 2 | 0 | 0 | 0 | 16 | 40 | 26 | 34 | 32 | 0 | 0 | 0 | 4 | 0 | 298 | 149 | 1 | 1 | 0.953 | 0.81 | 1 | 1 | 1 | 1 | 1 | 1 | 0.81 | 1 | -0.009 |
| SE | 107789 | ENSG00000050426 | LETMD1 | chr12 | + | 51049033 | 51049185 | 51048300 | 51048478 | 51056349 | 51056502 | 69 | 28 | 66 | 79 | 85 | 2 | 0 | 3 | 0 | 0 | 29 | 52 | 45 | 40 | 37 | 0 | 0 | 0 | 0 | 0 | 298 | 149 | 1 | 1 | 0.945 | 1 | 0.917 | 1 | 1 | 1 | 1 | 1 | 1 | 1 | -0.028 |
| SE | 107794 | ENSG00000050426 | LETMD1 | chr12 | + | 51052091 | 51052207 | 51048342 | 51049185 | 51056143 | 51056237 | 36 | 17 | 44 | 29 | 94 | 0 | 0 | 0 | 3 | 0 | 13 | 35 | 23 | 18 | 51 | 0 | 0 | 3 | 0 | 0 | 265 | 149 | 1 | 1 | 1 | 1 | 1 | 0.845 | 1 | 1 | 1 | 0.812 | 1 | 1 | 0.007 |
| SE | 107798 | ENSG00000050426 | LETMD1 | chr12 | + | 51053777 | 51053860 | 51049033 | 51049185 | 51056143 | 51056245 | 5 | 1 | 1 | 9 | 11 | 0 | 0 | 0 | 3 | 0 | 0 | 17 | 10 | 10 | 0 | 0 | 0 | 3 | 0 | 0 | 232 | 149 | 1 | 1 | 1 | 1 | 1 | 0.658 | 1 | NA | 1 | 0.682 | 1 | NA | 0.038 |
| SE | 107800 | ENSG00000050426 | LETMD1 | chr12 | + | 51053777 | 51053860 | 51052087 | 51052207 | 51055834 | 51055848 | 102 | 52 | 104 | 61 | 218 | 0 | 0 | 0 | 1 | 4 | 36 | 46 | 67 | 38 | 56 | 0 | 0 | 0 | 0 | 1 | 232 | 149 | 1 | 1 | 1 | 1 | 1 | 0.975 | 0.972 | 1 | 1 | 1 | 1 | 0.973 | -0.005 |
| SE | 107801 | ENSG00000050426 | LETMD1 | chr12 | + | 51053777 | 51053860 | 51052087 | 51052207 | 51056143 | 51056245 | 49 | 27 | 47 | 22 | 86 | 0 | 0 | 3 | 0 | 2 | 14 | 19 | 29 | 8 | 17 | 0 | 0 | 0 | 0 | 0 | 232 | 149 | 1 | 1 | 1 | 1 | 0.91 | 1 | 0.965 | 1 | 1 | 1 | 1 | 1 | -0.025 |
| SE | 107804 | ENSG00000050426 | LETMD1 | chr12 | + | 51053811 | 51053860 | 51049033 | 51049185 | 51055986 | 51056021 | 5 | 4 | 3 | 1 | 4 | 0 | 0 | 0 | 0 | 0 | 18 | 7 | 2 | 0 | 0 | 0 | 1 | 1 | 0 | 4 | 198 | 149 | 1 | 1 | 1 | 1 | 1 | 1 | 1 | 1 | 0.84 | 0.601 | NA | 0 | 0.39 |
| SE | 107806 | ENSG00000050426 | LETMD1 | chr12 | + | 51053811 | 51053860 | 51052091 | 51052207 | 51055834 | 51056021 | 31 | 15 | 34 | 18 | 74 | 0 | 0 | 0 | 1 | 4 | 0 | 15 | 32 | 28 | 14 | 0 | 0 | 0 | 0 | 1 | 198 | 149 | 1 | 1 | 1 | 1 | 1 | 0.931 | 0.933 | NA | 1 | 1 | 1 | 0.913 | -0.005 |
| SE | 107807 | ENSG00000050426 | LETMD1 | chr12 | + | 51053811 | 51053860 | 51052091 | 51052207 | 51056349 | 51056502 | 3 | 0 | 0 | 3 | 1 | 6 | 0 | 3 | 9 | 17 | 0 | 0 | 2 | 12 | 0 | 0 | 0 | 0 | 0 | 13 | 198 | 149 | 1 | 1 | 0.273 | NA | 0 | 0.201 | 0.042 | NA | NA | 1 | 1 | 0 | -0.538 |
| SE | 107808 | ENSG00000050426 | LETMD1 | chr12 | + | 51055834 | 51056021 | 51049033 | 51049185 | 51056143 | 51056221 | 55 | 30 | 51 | 46 | 84 | 0 | 0 | 0 | 3 | 0 | 10 | 46 | 46 | 36 | 75 | 0 | 0 | 3 | 0 | 0 | 298 | 149 | 1 | 1 | 1 | 1 | 1 | 0.885 | 1 | 1 | 1 | 0.885 | 1 | 1 | 0 |
| SE | 107809 | ENSG00000050426 | LETMD1 | chr12 | + | 51055834 | 51056021 | 51052091 | 51052207 | 51056143 | 51056221 | 51 | 28 | 41 | 47 | 85 | 0 | 0 | 3 | 0 | 2 | 10 | 42 | 46 | 25 | 64 | 0 | 0 | 0 | 0 | 0 | 298 | 149 | 1 | 1 | 1 | 1 | 0.872 | 1 | 0.955 | 1 | 1 | 1 | 1 | 1 | -0.035 |
| SE | 107810 | ENSG00000050426 | LETMD1 | chr12 | + | 51055834 | 51056021 | 51053777 | 51053860 | 51056143 | 51056221 | 117 | 59 | 112 | 89 | 219 | 1 | 0 | 0 | 0 | 0 | 47 | 94 | 90 | 59 | 114 | 0 | 0 | 1 | 0 | 0 | 298 | 149 | 1 | 1 | 0.983 | 1 | 1 | 1 | 1 | 1 | 1 | 0.978 | 1 | 1 | 0.001 |
| SE | 107811 | ENSG00000050426 | LETMD1 | chr12 | + | 51055986 | 51056021 | 51049033 | 51049185 | 51056143 | 51056245 | 9 | 11 | 20 | 18 | 25 | 0 | 0 | 0 | 3 | 0 | 3 | 7 | 13 | 2 | 28 | 0 | 0 | 3 | 0 | 0 | 184 | 149 | 1 | 1 | 1 | 1 | 1 | 0.829 | 1 | 1 | 1 | 0.778 | 1 | 1 | 0.01 |
| SE | 107812 | ENSG00000050426 | LETMD1 | chr12 | + | 51055986 | 51056021 | 51053811 | 51053860 | 51056143 | 51056245 | 9 | 11 | 20 | 18 | 25 | 1 | 0 | 0 | 0 | 0 | 3 | 6 | 12 | 2 | 24 | 0 | 0 | 1 | 0 | 0 | 184 | 149 | 1 | 1 | 0.879 | 1 | 1 | 1 | 1 | 1 | 1 | 0.907 | 1 | 1 | -0.006 |
| SE | 107817 | ENSG00000050426 | LETMD1 | chr12 | + | 51056143 | 51056245 | 51055834 | 51056021 | 51056349 | 51056428 | 101 | 54 | 89 | 84 | 142 | 0 | 0 | 0 | 0 | 0 | 59 | 120 | 88 | 45 | 97 | 0 | 0 | 0 | 0 | 4 | 251 | 149 | 1 | 1 | 1 | 1 | 1 | 1 | 1 | 1 | 1 | 1 | 1 | 0.935 | 0.013 |
| SE | 107819 | ENSG00000050426 | LETMD1 | chr12 | + | 51056149 | 51056245 | 51055986 | 51056021 | 51056349 | 51056502 | 48 | 28 | 45 | 36 | 60 | 0 | 0 | 0 | 0 | 0 | 41 | 73 | 46 | 18 | 28 | 0 | 0 | 0 | 0 | 4 | 245 | 149 | 1 | 1 | 1 | 1 | 1 | 1 | 1 | 1 | 1 | 1 | 1 | 0.81 | 0.038 |
| SE | 107827 | ENSG00000135457 | TFCP2 | chr12 | - | 51103669 | 51103763 | 51101934 | 51102025 | 51106524 | 51106613 | 56 | 54 | 53 | 65 | 74 | 0 | 0 | 0 | 3 | 0 | 74 | 61 | 17 | 67 | 30 | 0 | 0 | 0 | 0 | 0 | 243 | 149 | 1 | 1 | 1 | 1 | 1 | 0.93 | 1 | 1 | 1 | 1 | 1 | 1 | -0.014 |
| SE | 107830 | ENSG00000135457 | TFCP2 | chr12 | - | 51117670 | 51117747 | 51116314 | 51116420 | 51118620 | 51118772 | 208 | 122 | 146 | 184 | 159 | 0 | 0 | 0 | 2 | 0 | 160 | 123 | 83 | 184 | 47 | 0 | 0 | 0 | 0 | 0 | 226 | 149 | 1 | 1 | 1 | 1 | 1 | 0.984 | 1 | 1 | 1 | 1 | 1 | 1 | -0.003 |
| SE | 107831 | ENSG00000135457 | TFCP2 | chr12 | - | 51118620 | 51118772 | 51117670 | 51117747 | 51172300 | 51172434 | 203 | 98 | 165 | 183 | 186 | 0 | 2 | 1 | 0 | 3 | 125 | 137 | 67 | 118 | 36 | 0 | 0 | 0 | 0 | 0 | 298 | 149 | 1 | 1 | 1 | 0.961 | 0.988 | 1 | 0.969 | 1 | 1 | 1 | 1 | 1 | -0.016 |
| SE | 107832 | ENSG00000125629 | INSIG2 | chr2 | + | 118096418 | 118096426 | 118088473 | 118088541 | 118103196 | 118103321 | 5 | 0 | 3 | 5 | 3 | 0 | 0 | 4 | 3 | 0 | 0 | 8 | 6 | 0 | 0 | 0 | 0 | 0 | 0 | 0 | 157 | 149 | 1 | 1 | 1 | NA | 0.416 | 0.613 | 1 | NA | 1 | 1 | NA | NA | -0.243 |
| SE | 107833 | ENSG00000125629 | INSIG2 | chr2 | + | 118096418 | 118096800 | 118088470 | 118088541 | 118103196 | 118103321 | 45 | 31 | 34 | 51 | 56 | 0 | 0 | 4 | 3 | 0 | 24 | 35 | 22 | 26 | 0 | 0 | 0 | 0 | 0 | 0 | 298 | 149 | 1 | 1 | 1 | 1 | 0.81 | 0.895 | 1 | 1 | 1 | | 1 | NA | -0.059 |
| SE | 107834 | ENSG00000125629 | INSIG2 | chr2 | + | 118107089 | 118107189 | 118106658 | 118106903 | 118108280 | 118108304 | 46 | 36 | 41 | 57 | 77 | 1 | 1 | 0 | 0 | 0 | 0 | 29 | 29 | 20 | 4 | 0 | 0 | 2 | 3 | 0 | 249 | 149 | 1 | 1 | 0.965 | 0.956 | 1 | 1 | 1 | NA | 1 | 0.897 | 0.8 | 1 | 0.06 |
| SE | 107837 | ENSG00000160124 | MIX23 | chr3 | - | 122365335 | 122365466 | 122362967 | 122363027 | 122368175 | 122368322 | 9 | 11 | 9 | 16 | 11 | 2 | 1 | 9 | 4 | 1 | 0 | 1 | 4 | 10 | 3 | 0 | 0 | 0 | 0 | 0 | 280 | 149 | 1 | 1 | 0.705 | 0.854 | 0.347 | 0.68 | 0.854 | NA | 1 | 1 | | 1 | -0.312 |
| SE | 107838 | ENSG00000160124 | MIX23 | chr3 | - | 122365564 | 122365633 | 122365335 | 122365466 | 122366452 | 122366762 | 0 | 2 | 1 | 1 | 0 | 0 | 1 | 0 | 1 | 3 | 0 | 0 | 0 | 0 | 0 | 0 | 0 | 0 | 4 | 0 | 218 | 149 | 1 | 1 | NA | 0.578 | 1 | 0.406 | 0 | NA | NA | NA | 0 | NA | 0.496 |
| SE | 107839 | ENSG00000160124 | MIX23 | chr3 | - | 122372982 | 122373022 | 122371674 | 122371800 | 122383173 | 122383231 | 0 | 0 | 0 | 0 | 2 | 19 | 3 | 5 | 4 | 11 | 0 | 0 | 0 | 0 | 0 | 0 | 6 | 2 | 2 | 0 | 189 | 149 | 1 | 1 | 0 | 0 | 0 | 0 | 0.125 | NA | 0 | 0 | 0 | NA | 0.025 |
| SE | 107840 | ENSG00000061936 | SFSWAP | chr12 | + | 131719453 | 131719539 | 131711099 | 131714953 | 131725404 | 131725609 | 131 | 102 | 154 | 155 | 241 | 0 | 0 | 1 | 1 | 0 | 226 | 169 | 231 | 276 | 221 | 0 | 0 | 0 | 0 | 0 | 235 | 149 | 1 | 1 | 1 | 1 | 0.99 | 0.99 | 1 | 1 | 1 | 1 | 1 | 1 | -0.004 |
| SE | 107844 | ENSG00000169429 | CXCL8 | chr4 | + | 73741541 | 73741677 | 73740540 | 73740722 | 73741948 | 73742032 | 563 | 264 | 439 | 1255 | 1161 | 0 | 0 | 0 | 1 | 0 | 247 | 196 | 1454 | 654 | 52 | 0 | 0 | 0 | 0 | 0 | 285 | 149 | 1 | 1 | 1 | 1 | 1 | 0.998 | 1 | 1 | 1 | 1 | 1 | 1 | 0 |
| SE | 107845 | ENSG00000225342 | LRRK2-DT | chr12 | - | 40190252 | 40190394 | 40189082 | 40189205 | 40223658 | 40223803 | 0 | 0 | 0 | 1 | 0 | 0 | 0 | 0 | 0 | 0 | 0 | 0 | 0 | 0 | 0 | 0 | 0 | 2 | 0 | 0 | 291 | 149 | 1 | 1 | NA | NA | NA | 1 | NA | NA | NA | 0 | NA | NA | 1 |
| SE | 107846 | ENSG00000166598 | HSP90B1 | chr12 | + | 103931710 | 103931748 | 103931520 | 103931623 | 103932276 | 103932418 | 7 | 1 | 2 | 7 | 2 | 498 | 400 | 617 | 485 | 709 | 0 | 0 | 6 | 0 | 0 | 71 | 347 | 879 | 174 | 22 | 187 | 149 | 1 | 1 | 0.011 | 0.002 | 0.003 | 0.011 | 0.002 | 0 | 0 | 0.005 | 0 | 0 | 0.005 |
| SE | 107847 | ENSG00000166598 | HSP90B1 | chr12 | + | 103932276 | 103932418 | 103931520 | 103931623 | 103932825 | 103932929 | 1791 | 1306 | 2094 | 1821 | 3068 | 3 | 0 | 1 | 6 | 5 | 230 | 1146 | 2716 | 457 | 100 | 2 | 0 | 2 | 4 | 2 | 291 | 149 | 1 | 1 | 0.997 | 1 | 0.999 | 0.994 | 0.997 | 0.983 | 1 | 0.999 | 0.983 | 0.962 | 0.012 |
| SE | 107848 | ENSG00000166598 | HSP90B1 | chr12 | + | 103932278 | 103932418 | 103931520 | 103931623 | 103932825 | 103932942 | 1299 | 917 | 1486 | 1340 | 2373 | 3 | 0 | 1 | 6 | 5 | 159 | 803 | 1843 | 285 | 78 | 2 | 0 | 2 | 4 | 2 | 289 | 149 | 1 | 1 | 0.996 | 1 | 0.999 | 0.991 | 0.996 | 0.976 | 1 | 0.998 | 0.973 | 0.953 | 0.016 |
| SE | 107849 | ENSG00000166598 | HSP90B1 | chr12 | + | 103932280 | 103932418 | 103931520 | 103931623 | 103932825 | 103932942 | 1285 | 901 | 1459 | 1334 | 2316 | 3 | 0 | 1 | 6 | 5 | 159 | 789 | 1816 | 281 | 78 | 2 | 0 | 2 | 4 | 2 | 287 | 149 | 1 | 1 | 0.996 | 1 | 0.999 | 0.991 | 0.996 | 0.976 | 1 | 0.998 | 0.973 | 0.953 | 0.016 |
| SE | 107850 | ENSG00000166598 | HSP90B1 | chr12 | + | 103932825 | 103932942 | 103931520 | 103932418 | 103933955 | 103934287 | 2365 | 1540 | 2745 | 2412 | 4437 | 1 | 0 | 0 | 1 | 0 | 210 | 1271 | 3073 | 465 | 132 | 0 | 3 | 0 | 3 | 0 | 266 | 149 | 1 | 1 | 0.999 | 1 | 1 | 0.999 | 1 | 1 | 0.996 | 1 | 0.989 | 1 | 0.003 |
| SE | 107851 | ENSG00000166598 | HSP90B1 | chr12 | + | 103937694 | 103937806 | 103933955 | 103934287 | 103938339 | 103938459 | 1333 | 930 | 1598 | 1145 | 1660 | 0 | 3 | 2 | 2 | 1 | 128 | 710 | 1983 | 318 | 42 | 0 | 0 | 7 | 0 | 0 | 261 | 149 | 1 | 1 | 1 | 0.994 | 0.998 | 0.997 | 0.999 | 1 | 1 | 0.994 | 1 | 1 | -0.001 |
| SE | 107853 | ENSG00000166598 | HSP90B1 | chr12 | + | 103941831 | 103941897 | 103941628 | 103941706 | 103942526 | 103942796 | 1441 | 854 | 1731 | 1413 | 1416 | 0 | 0 | 0 | 2 | 0 | 118 | 823 | 1742 | 246 | 42 | 0 | 2 | 0 | 0 | 0 | 215 | 149 | 1 | 1 | 1 | 1 | 1 | 0.998 | 1 | 1 | 0.997 | 1 | 1 | 1 | 0 |
| SE | 107854 | ENSG00000166598 | HSP90B1 | chr12 | + | 103944199 | 103944229 | 103943737 | 103943874 | 103946617 | 103946696 | 2 | 0 | 0 | 2 | 0 | 1023 | 711 | 1346 | 1026 | 1107 | 0 | 0 | 0 | 0 | 0 | 99 | 587 | 1371 | 228 | 61 | 179 | 149 | 1 | 1 | 0.002 | 0 | 0 | 0.002 | 0 | 0 | 0 | 0 | 0 | 0 | 0.001 |
| SE | 107868 | ENSG00000188906 | LRRK2 | chr12 | + | 40227890 | 40227916 | 40225554 | 40225640 | 40232273 | 40232383 | 2 | 0 | 1 | 1 | 0 | 269 | 209 | 129 | 180 | 180 | 0 | 0 | 1 | 7 | 0 | 123 | 129 | 169 | 60 | 21 | 175 | 149 | 1 | 1 | 0.006 | 0 | 0.007 | 0.005 | 0 | 0 | 0 | 0.005 | 0.09 | 0 | -0.015 |
| SE | 107869 | ENSG00000188906 | LRRK2 | chr12 | + | 40232273 | 40232383 | 40225554 | 40225640 | 40235625 | 40235714 | 480 | 399 | 231 | 309 | 308 | 0 | 0 | 1 | 0 | 0 | 181 | 225 | 282 | 96 | 56 | 0 | 0 | 0 | 0 | 0 | 259 | 149 | 1 | 1 | 1 | 1 | 0.993 | 1 | 1 | 1 | 1 | 1 | 1 | 1 | -0.001 |
| SE | 107870 | ENSG00000188906 | LRRK2 | chr12 | + | 40237968 | 40238103 | 40235625 | 40235714 | 40240482 | 40240617 | 553 | 504 | 264 | 304 | 586 | 0 | 2 | 0 | 0 | 0 | 132 | 282 | 298 | 99 | 60 | 0 | 0 | 0 | 1 | 0 | 284 | 149 | 1 | 1 | 1 | 0.992 | 1 | 1 | 1 | 1 | 1 | 1 | 0.981 | 1 | 0.002 |
| SE | 107871 | ENSG00000188906 | LRRK2 | chr12 | + | 40237968 | 40238103 | 40235625 | 40235714 | 40243549 | 40243681 | 257 | 231 | 114 | 139 | 238 | 0 | 0 | 1 | 0 | 0 | 56 | 125 | 131 | 55 | 46 | 0 | 0 | 0 | 0 | 0 | 284 | 149 | 1 | 1 | 1 | 1 | 0.984 | 1 | 1 | 1 | 1 | 1 | 1 | 1 | -0.003 |
| SE | 107872 | ENSG00000188906 | LRRK2 | chr12 | + | 40238046 | 40238103 | 40235625 | 40235714 | 40240482 | 40240617 | 68 | 49 | 33 | 22 | 83 | 0 | 2 | 0 | 0 | 0 | 5 | 17 | 33 | 12 | 2 | 0 | 0 | 0 | 1 | 0 | 206 | 149 | 1 | 1 | 1 | 0.947 | 1 | 1 | 1 | 1 | 1 | 1 | 0.897 | 1 | 0.01 |
| SE | 107873 | ENSG00000188906 | LRRK2 | chr12 | + | 40240482 | 40240617 | 40235625 | 40235714 | 40243549 | 40243681 | 154 | 155 | 77 | 112 | 228 | 0 | 0 | 1 | 0 | 0 | 38 | 122 | 95 | 38 | 11 | 0 | 0 | 0 | 0 | 0 | 284 | 149 | 1 | 1 | 1 | 1 | 0.976 | 1 | 1 | 1 | 1 | 1 | 1 | 1 | -0.005 |
| SE | 107876 | ENSG00000188906 | LRRK2 | chr12 | + | 40249825 | 40249945 | 40243549 | 40243681 | 40251231 | 40251374 | 375 | 465 | 194 | 298 | 661 | 0 | 0 | 0 | 4 | 0 | 106 | 337 | 303 | 115 | 53 | 0 | 0 | 0 | 0 | 0 | 269 | 149 | 1 | 1 | 1 | 1 | 1 | 0.976 | 1 | 1 | 1 | 1 | 1 | 1 | -0.005 |
| SE | 107877 | ENSG00000188906 | LRRK2 | chr12 | + | 40251464 | 40251544 | 40251231 | 40251374 | 40252909 | 40253016 | 204 | 231 | 126 | 165 | 390 | 0 | 0 | 0 | 0 | 0 | 29 | 143 | 116 | 45 | 2 | 0 | 0 | 1 | 0 | 0 | 229 | 149 | 1 | 1 | 1 | 1 | 1 | 1 | 1 | 1 | 1 | 0.987 | 1 | 1 | 0.003 |
| SE | 107878 | ENSG00000188906 | LRRK2 | chr12 | + | 40252004 | 40252125 | 40251464 | 40251544 | 40252909 | 40253016 | 8 | 4 | 1 | 3 | 0 | 127 | 165 | 99 | 120 | 280 | 8 | 1 | 2 | 0 | 0 | 29 | 86 | 87 | 36 | 0 | 270 | 149 | 1 | 1 | 0.034 | 0.013 | 0.006 | 0.014 | 0 | 0.132 | 0.006 | 0.013 | 0 | NA | -0.024 |
| SE | 107879 | ENSG00000188906 | LRRK2 | chr12 | + | 40252909 | 40253016 | 40251464 | 40251544 | 40257247 | 40257377 | 198 | 252 | 137 | 190 | 379 | 0 | 0 | 0 | 0 | 0 | 48 | 128 | 157 | 35 | 0 | 0 | 0 | 0 | 3 | 0 | 256 | 149 | 1 | 1 | 1 | 1 | 1 | 1 | 1 | 1 | 1 | 1 | 0.872 | NA | 0.032 |
| SE | 107880 | ENSG00000188906 | LRRK2 | chr12 | + | 40257247 | 40257377 | 40252909 | 40253016 | 40259479 | 40259604 | 203 | 231 | 124 | 177 | 337 | 0 | 0 | 0 | 1 | 0 | 59 | 174 | 154 | 29 | 4 | 0 | 0 | 0 | 0 | 0 | 279 | 149 | 1 | 1 | 1 | 1 | 1 | 0.99 | 1 | 1 | 1 | 1 | 1 | 1 | -0.002 |
| SE | 107881 | ENSG00000188906 | LRRK2 | chr12 | + | 40257247 | 40257377 | 40252909 | 40253016 | 40263788 | 40263901 | 111 | 118 | 54 | 89 | 201 | 0 | 0 | 4 | 0 | 0 | 59 | 68 | 90 | 9 | 0 | 0 | 5 | 0 | 5 | 0 | 279 | 149 | 1 | 1 | 1 | 1 | 0.878 | 1 | 1 | 1 | 0.879 | 1 | 0.49 | NA | 0.133 |
| SE | 107884 | ENSG00000188906 | LRRK2 | chr12 | + | 40263788 | 40263901 | 40259479 | 40259604 | 40274582 | 40274727 | 126 | 147 | 71 | 81 | 237 | 0 | 0 | 0 | 0 | 2 | 24 | 70 | 90 | 42 | 4 | 0 | 2 | 0 | 0 | 0 | 262 | 149 | 1 | 1 | 1 | 1 | 1 | 1 | 0.985 | 1 | 0.952 | 1 | 1 | 1 | 0.007 |
| SE | 107885 | ENSG00000188906 | LRRK2 | chr12 | + | 40278090 | 40278261 | 40277887 | 40278016 | 40283874 | 40283952 | 322 | 324 | 148 | 209 | 538 | 0 | 0 | 0 | 0 | 0 | 54 | 235 | 186 | 47 | 44 | 0 | 0 | 1 | 0 | 0 | 298 | 149 | 1 | 1 | 1 | 1 | 1 | 1 | 1 | 1 | 1 | 0.989 | 1 | 1 | 0.002 |
| SE | 107886 | ENSG00000188906 | LRRK2 | chr12 | + | 40283874 | 40284133 | 40278090 | 40278261 | 40287350 | 40287539 | 457 | 483 | 239 | 364 | 636 | 0 | 0 | 0 | 0 | 2 | 37 | 342 | 242 | 75 | 36 | 0 | 0 | 0 | 0 | 0 | 298 | 149 | 1 | 1 | 1 | 1 | 1 | 1 | 0.994 | 1 | 1 | 1 | 1 | 1 | -0.001 |
| SE | 107887 | ENSG00000188906 | LRRK2 | chr12 | + | 40295426 | 40295644 | 40294844 | 40294914 | 40298242 | 40298493 | 385 | 345 | 207 | 259 | 655 | 2 | 4 | 0 | 4 | 1 | 0 | 225 | 171 | 62 | 18 | 0 | 2 | 0 | 0 | 0 | 298 | 149 | 1 | 1 | 0.99 | 0.977 | 1 | 0.97 | 0.997 | NA | 0.983 | 1 | 1 | 1 | -0.009 |
| SE | 107888 | ENSG00000188906 | LRRK2 | chr12 | + | 40298242 | 40298493 | 40295426 | 40295644 | 40299108 | 40299257 | 451 | 428 | 209 | 313 | 940 | 2 | 3 | 0 | 0 | 0 | 14 | 288 | 246 | 77 | 45 | 0 | 0 | 2 | 1 | 0 | 298 | 149 | 1 | 1 | 0.991 | 0.986 | 1 | 1 | 1 | 1 | 1 | 0.984 | 0.975 | 1 | 0.004 |
| SE | 107889 | ENSG00000188906 | LRRK2 | chr12 | + | 40299108 | 40299257 | 40295426 | 40295644 | 40302788 | 40302882 | 210 | 183 | 89 | 150 | 401 | 0 | 0 | 0 | 0 | 4 | 0 | 117 | 85 | 59 | 20 | 0 | 0 | 0 | 0 | 0 | 298 | 149 | 1 | 1 | 1 | 1 | 1 | 1 | 0.98 | NA | 1 | 1 | 1 | 1 | -0.004 |
| SE | 107890 | ENSG00000188906 | LRRK2 | chr12 | + | 40300812 | 40301064 | 40299108 | 40299257 | 40302788 | 40302882 | 0 | 1 | 1 | 1 | 10 | 208 | 180 | 89 | 150 | 401 | 0 | 5 | 0 | 0 | 0 | 0 | 117 | 83 | 58 | 20 | 298 | 149 | 1 | 1 | 0 | 0.003 | 0.006 | 0.003 | 0.012 | NA | 0.021 | 0 | 0 | 0 | 0 |
| SE | 107891 | ENSG00000188906 | LRRK2 | chr12 | + | 40303947 | 40304134 | 40302788 | 40302882 | 40308466 | 40308696 | 204 | 133 | 72 | 111 | 238 | 0 | 0 | 0 | 2 | 0 | 3 | 77 | 77 | 18 | 16 | 0 | 4 | 0 | 0 | 0 | 298 | 149 | 1 | 1 | 1 | 1 | 1 | 0.965 | 1 | 1 | 0.906 | 1 | 1 | 1 | 0.012 |
| SE | 107893 | ENSG00000188906 | LRRK2 | chr12 | + | 40310430 | 40310649 | 40309105 | 40309233 | 40313971 | 40314173 | 389 | 431 | 220 | 280 | 587 | 0 | 0 | 0 | 1 | 0 | 22 | 278 | 210 | 27 | 36 | 0 | 0 | 0 | 0 | 0 | 298 | 149 | 1 | 1 | 1 | 1 | 1 | 0.993 | 1 | 1 | 1 | 1 | 1 | 1 | -0.001 |
| SE | 107894 | ENSG00000188906 | LRRK2 | chr12 | + | 40313971 | 40314173 | 40309105 | 40309233 | 40315211 | 40315300 | 262 | 305 | 174 | 191 | 458 | 0 | 0 | 0 | 1 | 0 | 33 | 150 | 173 | 31 | 5 | 0 | 0 | 0 | 0 | 0 | 298 | 149 | 1 | 1 | 1 | 1 | 1 | 0.99 | 1 | 1 | 1 | 1 | 1 | 1 | -0.002 |
| SE | 107897 | ENSG00000188906 | LRRK2 | chr12 | + | 40316277 | 40316403 | 40315211 | 40315300 | 40321033 | 40321188 | 5 | 3 | 0 | 2 | 2 | 5 | 8 | 1 | 5 | 16 | 0 | 3 | 4 | 0 | 0 | 0 | 2 | 6 | 0 | 0 | 275 | 149 | 1 | 1 | 0.351 | 0.169 | 0 | 0.178 | 0.063 | NA | 0.448 | 0.265 | NA | NA | -0.204 |
| SE | 107899 | ENSG00000188906 | LRRK2 | chr12 | + | 40319987 | 40320175 | 40316277 | 40316403 | 40321033 | 40321188 | 164 | 185 | 110 | 109 | 248 | 2 | 1 | 0 | 0 | 0 | 13 | 80 | 69 | 20 | 1 | 0 | 0 | 0 | 0 | 0 | 298 | 149 | 1 | 1 | 0.976 | 0.989 | 1 | 1 | 1 | 1 | 1 | 1 | 1 | 1 | -0.007 |
| SE | 107900 | ENSG00000188906 | LRRK2 | chr12 | + | 40322034 | 40322181 | 40321033 | 40321188 | 40322318 | 40322510 | 535 | 627 | 277 | 380 | 754 | 0 | 0 | 0 | 0 | 2 | 122 | 383 | 390 | 64 | 0 | 0 | 0 | 0 | 0 | 0 | 296 | 149 | 1 | 1 | 1 | 1 | 1 | 1 | 0.995 | 1 | 1 | 1 | 1 | NA | -0.001 |
| SE | 107901 | ENSG00000188906 | LRRK2 | chr12 | + | 40322034 | 40322181 | 40321033 | 40321188 | 40323159 | 40323306 | 208 | 232 | 100 | 148 | 309 | 0 | 0 | 0 | 0 | 3 | 71 | 136 | 138 | 28 | 0 | 0 | 0 | 0 | 0 | 0 | 296 | 149 | 1 | 1 | 1 | 1 | 1 | 1 | 0.981 | 1 | 1 | 1 | 1 | NA | -0.004 |
| SE | 107902 | ENSG00000188906 | LRRK2 | chr12 | + | 40322318 | 40322510 | 40321033 | 40321188 | 40323159 | 40323306 | 137 | 142 | 100 | 102 | 303 | 0 | 0 | 0 | 0 | 3 | 8 | 69 | 41 | 7 | 8 | 0 | 0 | 0 | 0 | 0 | 298 | 149 | 1 | 1 | 1 | 1 | 1 | 1 | 0.981 | 1 | 1 | 1 | 1 | 1 | -0.004 |
| SE | 107903 | ENSG00000188906 | LRRK2 | chr12 | + | 40322318 | 40322510 | 40322034 | 40322181 | 40323159 | 40323306 | 465 | 537 | 277 | 334 | 746 | 1 | 0 | 0 | 0 | 0 | 59 | 316 | 293 | 43 | 8 | 0 | 0 | 0 | 0 | 0 | 298 | 149 | 1 | 1 | 0.996 | 1 | 1 | 1 | 1 | 1 | 1 | 1 | 1 | 1 | -0.001 |
| SE | 107904 | ENSG00000188906 | LRRK2 | chr12 | + | 40340293 | 40340454 | 40334966 | 40335157 | 40346752 | 40346923 | 410 | 358 | 222 | 307 | 663 | 1 | 2 | 0 | 3 | 1 | 47 | 342 | 294 | 61 | 51 | 0 | 2 | 0 | 0 | 0 | 298 | 149 | 1 | 1 | 0.995 | 0.989 | 1 | 0.981 | 0.997 | 1 | 0.988 | 1 | 1 | 1 | -0.005 |
| SE | 107905 | ENSG00000188906 | LRRK2 | chr12 | + | 40348408 | 40348509 | 40346752 | 40346923 | 40351538 | 40351733 | 396 | 406 | 196 | 281 | 563 | 3 | 3 | 0 | 4 | 1 | 83 | 270 | 212 | 14 | 9 | 0 | 0 | 0 | 0 | 0 | 250 | 149 | 1 | 1 | 0.987 | 0.988 | 1 | 0.977 | 0.997 | 1 | 1 | 1 | 1 | 1 | -0.01 |
| SE | 107906 | ENSG00000188906 | LRRK2 | chr12 | + | 40351538 | 40351733 | 40348408 | 40348509 | 40354298 | 40354492 | 386 | 356 | 175 | 274 | 529 | 0 | 0 | 0 | 2 | 0 | 66 | 251 | 201 | 20 | 5 | 0 | 0 | 0 | 0 | 0 | 298 | 149 | 1 | 1 | 1 | 1 | 1 | 0.986 | 1 | 1 | 1 | 1 | 1 | 1 | -0.003 |
| SE | 107907 | ENSG00000188906 | LRRK2 | chr12 | + | 40354298 | 40354492 | 40351538 | 40351733 | 40356114 | 40356187 | 295 | 260 | 140 | 196 | 441 | 1 | 0 | 1 | 4 | 1 | 32 | 193 | 161 | 14 | 27 | 0 | 1 | 0 | 0 | 0 | 298 | 149 | 1 | 1 | 0.993 | 1 | 0.986 | 0.961 | 0.995 | 1 | 0.99 | 1 | 1 | 1 | -0.011 |
| SE | 107908 | ENSG00000188906 | LRRK2 | chr12 | + | 40356114 | 40356187 | 40354298 | 40354492 | 40359259 | 40359444 | 269 | 281 | 147 | 197 | 410 | 0 | 1 | 0 | 0 | 0 | 28 | 183 | 169 | 13 | 45 | 0 | 0 | 0 | 0 | 0 | 222 | 149 | 1 | 1 | 1 | 0.995 | 1 | 1 | 1 | 1 | 1 | 1 | 1 | 1 | -0.001 |
| SE | 107909 | ENSG00000188906 | LRRK2 | chr12 | + | 40367005 | 40367059 | 40364841 | 40365050 | 40367643 | 40368879 | 92 | 65 | 42 | 41 | 181 | 0 | 1 | 0 | 4 | 6 | 0 | 37 | 46 | 15 | 0 | 0 | 1 | 3 | 0 | 0 | 203 | 149 | 1 | 1 | 1 | 0.979 | 1 | 0.883 | 0.957 | NA | 0.964 | 0.918 | 1 | NA | 0.003 |
| SE | 107910 | ENSG00000188906 | LRRK2 | chr12 | + | 40367005 | 40367077 | 40364841 | 40365050 | 40367643 | 40368330 | 228 | 240 | 145 | 125 | 443 | 0 | 1 | 0 | 4 | 6 | 19 | 112 | 131 | 29 | 10 | 0 | 1 | 3 | 0 | 0 | 221 | 149 | 1 | 1 | 1 | 0.994 | 1 | 0.955 | 0.98 | 1 | 0.987 | 0.967 | 1 | 1 | -0.005 |
| SE | 107911 | ENSG00000188906 | LRRK2 | chr12 | + | 40367009 | 40367077 | 40364841 | 40365050 | 40367643 | 40368330 | 61 | 71 | 49 | 46 | 128 | 0 | 1 | 0 | 4 | 6 | 1 | 45 | 36 | 7 | 0 | 0 | 1 | 3 | 0 | 0 | 217 | 149 | 1 | 1 | 1 | 0.98 | 1 | 0.888 | 0.936 | 1 | 0.969 | 0.892 | 1 | NA | -0.004 |
| SE | 107923 | ENSG00000111335 | OAS2 | chr12 | + | 112978671 | 112978785 | 112978444 | 112978477 | 112987037 | 112987308 | 162 | 112 | 121 | 193 | 161 | 0 | 0 | 0 | 0 | 0 | 471 | 308 | 97 | 442 | 104 | 0 | 0 | 0 | 2 | 0 | 263 | 149 | 1 | 1 | 1 | 1 | 1 | 1 | 1 | 1 | 1 | 1 | 0.992 | 1 | 0.002 |
| SE | 107926 | ENSG00000111335 | OAS2 | chr12 | + | 112987037 | 112987308 | 112978394 | 112978785 | 112995295 | 112995474 | 409 | 300 | 309 | 473 | 480 | 0 | 1 | 2 | 0 | 0 | 1234 | 734 | 213 | 1067 | 298 | 0 | 4 | 2 | 3 | 6 | 298 | 149 | 1 | 1 | 1 | 0.993 | 0.987 | 1 | 1 | 1 | 0.989 | 0.982 | 0.994 | 0.961 | 0.011 |
| SE | 107929 | ENSG00000111335 | OAS2 | chr12 | + | 112998265 | 112998410 | 112997519 | 112997755 | 113002931 | 113003102 | 692 | 704 | 581 | 891 | 1114 | 0 | 0 | 2 | 0 | 0 | 3172 | 1274 | 338 | 2093 | 516 | 0 | 0 | 0 | 0 | 0 | 294 | 149 | 1 | 1 | 1 | 1 | 0.993 | 1 | 1 | 1 | 1 | 1 | 1 | 1 | -0.001 |
| SE | 107930 | ENSG00000111335 | OAS2 | chr12 | + | 113002931 | 113003102 | 112997519 | 112998410 | 113004933 | 113005222 | 768 | 745 | 644 | 949 | 1092 | 1 | 0 | 0 | 0 | 0 | 3232 | 1278 | 393 | 2040 | 478 | 0 | 0 | 0 | 0 | 0 | 298 | 149 | 1 | 1 | 0.997 | 1 | 1 | 1 | 1 | 1 | 1 | 1 | 1 | 1 | -0.001 |
| SE | 107931 | ENSG00000111335 | OAS2 | chr12 | + | 113004474 | 113004673 | 113002931 | 113003102 | 113004933 | 113005222 | 7 | 8 | 1 | 4 | 2 | 291 | 268 | 257 | 358 | 370 | 5 | 2 | 2 | 8 | 0 | 1124 | 510 | 165 | 676 | 157 | 298 | 149 | 1 | 1 | 0.012 | 0.015 | 0.002 | 0.006 | 0.003 | 0.002 | 0.002 | 0.006 | 0.006 | 0 | 0.004 |
| SE | 107932 | ENSG00000111335 | OAS2 | chr12 | + | 113004474 | 113005222 | 113002931 | 113003102 | 113006412 | 113006600 | 205 | 180 | 189 | 260 | 198 | 0 | 0 | 0 | 1 | 1 | 597 | 383 | 93 | 544 | 178 | 0 | 0 | 1 | 0 | 0 | 298 | 149 | 1 | 1 | 1 | 1 | 1 | 0.992 | 0.99 | 1 | 1 | 0.979 | 1 | 1 | 0.001 |
| SE | 107933 | ENSG00000111335 | OAS2 | chr12 | + | 113004933 | 113005222 | 113002931 | 113003102 | 113006412 | 113006600 | 489 | 442 | 445 | 616 | 566 | 0 | 0 | 0 | 1 | 1 | 1716 | 891 | 256 | 1212 | 335 | 0 | 0 | 1 | 0 | 0 | 298 | 149 | 1 | 1 | 1 | 1 | 1 | 0.997 | 0.996 | 1 | 1 | 0.992 | 1 | 1 | 0 |
| SE | 107934 | ENSG00000111335 | OAS2 | chr12 | + | 113006412 | 113006600 | 113003304 | 113005222 | 113007704 | 113007943 | 359 | 342 | 325 | 480 | 396 | 0 | 0 | 0 | 1 | 0 | 1321 | 626 | 186 | 1093 | 368 | 0 | 0 | 0 | 0 | 0 | 298 | 149 | 1 | 1 | 1 | 1 | 1 | 0.996 | 1 | 1 | 1 | 1 | 1 | 1 | -0.001 |
| SE | 107935 | ENSG00000111335 | OAS2 | chr12 | + | 113006417 | 113006600 | 113004933 | 113005222 | 113007704 | 113007943 | 161 | 170 | 138 | 222 | 200 | 0 | 0 | 0 | 1 | 0 | 729 | 245 | 95 | 557 | 190 | 0 | 0 | 0 | 0 | 0 | 298 | 149 | 1 | 1 | 1 | 1 | 1 | 0.991 | 1 | 1 | 1 | 1 | 1 | 1 | -0.002 |
| SE | 107936 | ENSG00000111335 | OAS2 | chr12 | + | 113007704 | 113007943 | 113006412 | 113006600 | 113009086 | 113009240 | 404 | 429 | 352 | 543 | 563 | 1 | 0 | 0 | 0 | 0 | 1939 | 740 | 243 | 1485 | 488 | 0 | 0 | 0 | 3 | 0 | 298 | 149 | 1 | 1 | 0.995 | 1 | 1 | 1 | 1 | 1 | 1 | 1 | 0.996 | 1 | 0 |
| SE | 107937 | ENSG00000135114 | OASL | chr12 | - | 121023989 | 121024137 | 121019110 | 121021058 | 121031441 | 121031617 | 38 | 22 | 21 | 69 | 63 | 0 | 0 | 0 | 1 | 0 | 125 | 176 | 60 | 265 | 46 | 0 | 0 | 2 | 0 | 0 | 297 | 149 | 1 | 1 | 1 | 1 | 1 | 0.972 | 1 | 1 | 1 | 0.938 | 1 | 1 | 0.007 |
| SE | 107947 | ENSG00000177119 | ANO6 | chr12 | + | 45270461 | 45270524 | 45228167 | 45228273 | 45331294 | 45331423 | 3 | 0 | 0 | 4 | 0 | 1 | 2 | 0 | 0 | 3 | 18 | 4 | 3 | 2 | 0 | 0 | 1 | 0 | 0 | 0 | 212 | 149 | 1 | 1 | 0.678 | 0 | NA | 1 | 0 | 1 | 0.738 | 1 | 1 | NA | -0.515 |
| SE | 107949 | ENSG00000177119 | ANO6 | chr12 | + | 45302013 | 45302093 | 45216019 | 45216391 | 45347021 | 45347087 | 56 | 52 | 51 | 77 | 47 | 1 | 0 | 0 | 2 | 0 | 102 | 87 | 80 | 83 | 48 | 0 | 0 | 0 | 0 | 0 | 229 | 149 | 1 | 1 | 0.973 | 1 | | 0.962 | 1 | 1 | 1 | 1 | 1 | 1 | -0.013 |
| SE | 107950 | ENSG00000177119 | ANO6 | chr12 | + | 45302013 | 45302093 | 45228187 | 45228273 | 45331294 | 45331423 | 117 | 148 | 78 | 183 | 290 | 1 | 2 | 0 | 0 | 3 | 199 | 171 | 143 | 191 | 115 | 0 | 1 | 0 | 0 | 0 | 229 | 149 | 1 | 1 | 0.987 | 0.98 | 1 | 1 | 0.984 | 1 | 0.991 | 1 | 1 | 1 | -0.008 |
| SE | 107952 | ENSG00000177119 | ANO6 | chr12 | + | 45302013 | 45302093 | 45292682 | 45292934 | 45331294 | 45331423 | 116 | 146 | 77 | 178 | 298 | 0 | 1 | 0 | 0 | 0 | 206 | 161 | 137 | 187 | 115 | 0 | 0 | 0 | 0 | 0 | 229 | 149 | 1 | 1 | 1 | 0.99 | 1 | 1 | 1 | 1 | 1 | 1 | 1 | 1 | -0.002 |
| SE | 107953 | ENSG00000177119 | ANO6 | chr12 | + | 45331294 | 45331423 | 45215986 | 45216391 | 45347021 | 45347087 | 301 | 341 | 140 | 451 | 729 | 1 | 0 | 0 | 2 | 0 | 490 | 319 | 262 | 466 | 506 | 0 | 0 | 0 | 0 | 0 | 278 | 149 | 1 | 1 | 0.994 | 1 | 1 | 0.992 | 1 | 1 | 1 | 1 | 1 | 1 | -0.003 |
| SE | 107954 | ENSG00000177119 | ANO6 | chr12 | + | 45331294 | 45331423 | 45302013 | 45302093 | 45347021 | 45347087 | 465 | 524 | 263 | 690 | 1059 | 0 | 0 | 0 | 0 | 0 | 743 | 516 | 440 | 727 | 659 | 0 | 1 | 0 | 0 | 0 | 278 | 149 | 1 | 1 | 1 | 1 | 1 | 1 | 1 | 1 | 0.996 | 1 | 1 | 1 | 0.001 |
| SE | 107955 | ENSG00000177119 | ANO6 | chr12 | + | 45348517 | 45348631 | 45348027 | 45348315 | 45350658 | 45350774 | 140 | 131 | 60 | 186 | 209 | 0 | 0 | 0 | 0 | 0 | 118 | 168 | 87 | 140 | 227 | 0 | 2 | 0 | 0 | 0 | 263 | 149 | 1 | 1 | 1 | 1 | 1 | 1 | 1 | 1 | 0.979 | 1 | 1 | 1 | 0.004 |
| SE | 107956 | ENSG00000177119 | ANO6 | chr12 | + | 45350658 | 45350774 | 45348517 | 45348631 | 45357289 | 45357424 | 218 | 186 | 93 | 277 | 199 | 1 | 0 | 0 | 0 | 0 | 208 | 262 | 145 | 264 | 266 | 0 | 0 | 0 | 0 | 0 | 265 | 149 | 1 | 1 | 0.992 | 1 | 1 | 1 | 1 | 1 | 1 | 1 | 1 | 1 | -0.002 |
| SE | 107957 | ENSG00000177119 | ANO6 | chr12 | + | 45378052 | 45378113 | 45367687 | 45367793 | 45401794 | 45402020 | 115 | 132 | 74 | 139 | 212 | 0 | 0 | 0 | 0 | 0 | 105 | 123 | 106 | 203 | 88 | 6 | 0 | 0 | 0 | 0 | 210 | 149 | 1 | 1 | 1 | 1 | 1 | 1 | 1 | 0.925 | 1 | 1 | 1 | 1 | 0.015 |
| SE | 107958 | ENSG00000177119 | ANO6 | chr12 | + | 45383194 | 45383355 | 45378052 | 45378113 | 45388160 | 45388303 | 0 | 0 | 1 | 4 | 0 | 282 | 317 | 162 | 468 | 474 | 4 | 0 | 0 | 0 | 0 | 370 | 359 | 276 | 475 | 387 | 298 | 149 | 1 | 1 | 0 | 0 | 0.003 | 0.004 | 0 | 0.005 | 0 | 0 | 0 | 0 | 0 |
| SE | 107959 | ENSG00000177119 | ANO6 | chr12 | + | 45388160 | 45388303 | 45378052 | 45378113 | 45390420 | 45390498 | 489 | 522 | 267 | 741 | 822 | 1 | 0 | 0 | 0 | 0 | 633 | 668 | 485 | 704 | 637 | 0 | 0 | 0 | 2 | 0 | 292 | 149 | 1 | 1 | 0.996 | 1 | 1 | 1 | 1 | 1 | 1 | 1 | 0.994 | 1 | 0 |
| SE | 107960 | ENSG00000177119 | ANO6 | chr12 | + | 45388160 | 45388303 | 45378052 | 45378113 | 45401794 | 45402020 | 281 | 311 | 161 | 453 | 431 | 0 | 0 | 0 | 0 | 4 | 358 | 355 | 270 | 463 | 385 | 0 | 0 | 1 | 0 | 0 | 292 | 149 | 1 | 1 | 1 | 1 | 1 | 1 | 0.982 | 1 | 1 | 0.993 | 1 | 1 | -0.002 |
| SE | 107961 | ENSG00000177119 | ANO6 | chr12 | + | 45390420 | 45390498 | 45378052 | 45378113 | 45401794 | 45402020 | 138 | 154 | 75 | 211 | 286 | 0 | 0 | 0 | 0 | 4 | 195 | 172 | 172 | 183 | 164 | 0 | 0 | 1 | 0 | 0 | 227 | 149 | 1 | 1 | 1 | 1 | 1 | 1 | 0.979 | 1 | 1 | 0.991 | 1 | 1 | -0.002 |
| SE | 107962 | ENSG00000177119 | ANO6 | chr12 | + | 45390420 | 45390498 | 45388160 | 45388303 | 45401794 | 45402020 | 352 | 367 | 181 | 503 | 678 | 0 | 0 | 0 | 0 | 0 | 470 | 489 | 389 | 435 | 420 | 0 | 0 | 1 | 0 | 0 | 227 | 149 | 1 | 1 | 1 | 1 | 1 | 1 | 1 | 1 | 1 | 0.996 | 1 | 1 | 0.001 |
| SE | 107963 | ENSG00000177119 | ANO6 | chr12 | + | 45401794 | 45402020 | 45390420 | 45390498 | 45403071 | 45403241 | 430 | 467 | 193 | 646 | 714 | 4 | 0 | 0 | 0 | 0 | 630 | 622 | 568 | 650 | 543 | 0 | 0 | 0 | 0 | 0 | 298 | 149 | 1 | 1 | 0.982 | 1 | 1 | 1 | 1 | 1 | 1 | 1 | 1 | 1 | -0.004 |
| SE | 107964 | ENSG00000177119 | ANO6 | chr12 | + | 45409356 | 45409487 | 45403438 | 45403536 | 45416698 | 45416904 | 523 | 596 | 333 | 799 | 922 | 0 | 0 | 0 | 1 | 0 | 548 | 670 | 535 | 754 | 486 | 0 | 0 | 0 | 0 | 0 | 280 | 149 | 1 | 1 | 1 | 1 | 1 | 0.998 | 1 | 1 | 1 | 1 | 1 | 1 | 0 |
| SE | 107965 | ENSG00000177119 | ANO6 | chr12 | + | 45416698 | 45416904 | 45409356 | 45409487 | 45421070 | 45421217 | 525 | 525 | 360 | 848 | 988 | 0 | 4 | 1 | 0 | 8 | 557 | 665 | 555 | 854 | 610 | 0 | 0 | 2 | 0 | 0 | 298 | 149 | 1 | 1 | 1 | 0.985 | 0.994 | 1 | 0.984 | 1 | 1 | 0.993 | 1 | 1 | -0.006 |
| SE | 107966 | ENSG00000177119 | ANO6 | chr12 | + | 45421070 | 45421273 | 45416698 | 45416904 | 45422956 | 45423062 | 481 | 469 | 315 | 762 | 846 | 0 | 0 | 1 | 0 | 4 | 429 | 625 | 533 | 722 | 456 | 0 | 6 | 2 | 0 | 0 | 298 | 149 | 1 | 1 | 1 | 1 | 0.994 | 1 | 0.991 | 1 | 0.981 | 0.993 | 1 | 1 | 0.002 |
| SE | 107968 | ENSG00000168778 | TCTN2 | chr12 | + | 123679188 | 123679289 | 123673614 | 123673810 | 123686835 | 123687035 | 14 | 14 | 16 | 11 | 21 | 0 | 0 | 0 | 0 | 0 | 0 | 2 | 11 | 2 | 14 | 0 | 3 | 0 | 0 | 0 | 250 | 149 | 1 | 1 | 1 | 1 | 1 | 1 | 1 | NA | 0.284 | 1 | 1 | 1 | 0.179 |
| SE | 107969 | ENSG00000168778 | TCTN2 | chr12 | + | 123688050 | 123688177 | 123686835 | 123687035 | 123690532 | 123690674 | 31 | 7 | 13 | 27 | 18 | 2 | 2 | 0 | 0 | 0 | 0 | 27 | 17 | 13 | 28 | 0 | 0 | 4 | 5 | 0 | 276 | 149 | 1 | 1 | 0.893 | 0.654 | 1 | 1 | 1 | NA | 1 | 0.696 | 0.584 | 1 | 0.089 |
| SE | 107970 | ENSG00000168778 | TCTN2 | chr12 | + | 123694841 | 123694976 | 123692657 | 123692723 | 123695219 | 123695297 | 16 | 11 | 34 | 21 | 39 | 0 | 0 | 0 | 0 | 8 | 0 | 13 | 12 | 3 | 0 | 0 | 8 | 0 | 0 | 0 | 284 | 149 | 1 | 1 | 1 | 1 | 1 | 1 | 0.719 | NA | 0.46 | 1 | 1 | NA | 0.124 |
| SE | 107971 | ENSG00000168778 | TCTN2 | chr12 | + | 123695219 | 123695297 | 123694841 | 123694976 | 123696414 | 123696495 | 10 | 4 | 16 | 10 | 46 | 0 | 0 | 1 | 0 | 2 | 0 | 9 | 11 | 1 | 0 | 0 | 0 | 0 | 0 | 0 | 227 | 149 | 1 | 1 | 1 | 1 | 0.913 | 1 | 0.938 | NA | 1 | 1 | 1 | NA | -0.03 |
| SE | 107975 | ENSG00000111229 | ARPC3 | chr12 | - | 110440311 | 110440388 | 110437083 | 110437152 | 110445451 | 110445551 | 3616 | 2446 | 2703 | 3059 | 4427 | 8 | 0 | 5 | 3 | 6 | 4021 | 3410 | 2320 | 3654 | 3420 | 13 | 3 | 5 | 10 | 5 | 226 | 149 | 1 | 1 | 0.997 | 1 | 0.997 | 0.999 | 0.998 | 0.995 | 0.999 | 0.997 | 0.996 | 0.998 | 0.001 |
| SE | 107980 | ENSG00000204852 | TCTN1 | chr12 | + | 110619835 | 110619956 | 110614026 | 110614402 | 110626361 | 110626382 | 0 | 1 | 0 | 0 | 5 | 0 | 0 | 0 | 0 | 0 | 0 | 0 | 0 | 0 | 0 | 4 | 0 | 2 | 0 | 0 | 270 | 149 | 1 | 1 | NA | 1 | NA | NA | 1 | 0 | NA | 0 | NA | NA | 1 |
| SE | 107981 | ENSG00000204852 | TCTN1 | chr12 | + | 110619835 | 110619956 | 110614106 | 110614463 | 110626361 | 110626382 | 0 | 1 | 0 | 0 | 5 | 0 | 0 | 0 | 0 | 0 | 0 | 0 | 0 | 0 | 0 | 4 | 0 | 2 | 0 | 0 | 270 | 149 | 1 | 1 | NA | 1 | NA | NA | 1 | NA | NA | 0 | NA | NA | 1 |
| SE | 107983 | ENSG00000204852 | TCTN1 | chr12 | + | 110627929 | 110628224 | 110626361 | 110626492 | 110628766 | 110628918 | 1 | 0 | 1 | 0 | 4 | 7 | 8 | 8 | 4 | 6 | 0 | 0 | 0 | 4 | 0 | 10 | 0 | 0 | 3 | 14 | 298 | 149 | 1 | 1 | 0.067 | 0 | 0.059 | 0 | 0.25 | 0 | NA | NA | 0.4 | 0 | -0.058 |
| SE | 107984 | ENSG00000204852 | TCTN1 | chr12 | + | 110628051 | 110628224 | 110626361 | 110626492 | 110628766 | 110628918 | 1 | 0 | 4 | 0 | 6 | 7 | 8 | 8 | 4 | 6 | 0 | 0 | 0 | 8 | 0 | 10 | 0 | 0 | 3 | 14 | 298 | 149 | 1 | 1 | 0.067 | 0 | 0.2 | 0 | 0.333 | 0 | NA | NA | 0.571 | 0 | -0.07 |
| SE | 107985 | ENSG00000204852 | TCTN1 | chr12 | + | 110628051 | 110628224 | 110626361 | 110626492 | 110628796 | 110630216 | 0 | 0 | 4 | 1 | 5 | 1 | 0 | 1 | 0 | 7 | 0 | 0 | 1 | 5 | 0 | 0 | 2 | 5 | 0 | 0 | 298 | 149 | 1 | 1 | 0 | NA | 0.667 | 1 | 0.263 | NA | 0 | 0.091 | 1 | NA | 0.119 |
| SE | 107986 | ENSG00000204852 | TCTN1 | chr12 | + | 110634395 | 110634463 | 110632471 | 110632559 | 110634669 | 110634779 | 0 | 0 | 0 | 1 | 0 | 16 | 17 | 10 | 16 | 43 | 0 | 0 | 0 | 0 | 0 | 10 | 9 | 4 | 8 | 27 | 217 | 149 | 1 | 1 | 0 | 0 | 0 | 0.041 | 0 | 0 | 0 | 0 | 0 | 0 | 0.008 |
| SE | 107987 | ENSG00000204852 | TCTN1 | chr12 | + | 110634395 | 110634463 | 110632471 | 110632559 | 110636480 | 110636501 | 1 | 1 | 0 | 2 | 0 | 4 | 1 | 4 | 3 | 3 | 0 | 0 | 0 | 0 | 0 | 0 | 3 | 0 | 1 | 0 | 217 | 149 | 1 | 1 | 0.147 | 0.407 | 0 | 0.314 | 0 | NA | 0 | NA | 0 | NA | 0.174 |
| SE | 107990 | ENSG00000204852 | TCTN1 | chr12 | + | 110634669 | 110634779 | 110634395 | 110634463 | 110636480 | 110636501 | 9 | 7 | 9 | 2 | 28 | 1 | 1 | 0 | 1 | 0 | 0 | 1 | 3 | 5 | 0 | 0 | 0 | 0 | 0 | 0 | 259 | 149 | 1 | 1 | 0.838 | 0.801 | 1 | 0.535 | 1 | NA | 1 | 1 | 1 | NA | -0.165 |
| SE | 107991 | ENSG00000204852 | TCTN1 | chr12 | + | 110635950 | 110636244 | 110634669 | 110634779 | 110636480 | 110636501 | 0 | 0 | 3 | 1 | 0 | 14 | 16 | 14 | 7 | 41 | 0 | 0 | 0 | 0 | 0 | 0 | 10 | 4 | 8 | 8 | 298 | 149 | 1 | 1 | 0 | 0 | 0.097 | 0.067 | 0 | NA | 0 | 0 | 0 | 0 | 0.033 |
| SE | 107993 | ENSG00000204852 | TCTN1 | chr12 | + | 110641023 | 110641149 | 110640382 | 110640517 | 110641541 | 110641586 | 16 | 15 | 15 | 9 | 33 | 0 | 0 | 0 | 2 | 0 | 12 | 12 | 7 | 11 | 8 | 0 | 0 | 0 | 0 | 0 | 275 | 149 | 1 | 1 | 1 | 1 | 1 | 0.709 | 1 | 1 | 1 | 1 | 1 | 1 | -0.058 |
| SE | 107995 | ENSG00000204852 | TCTN1 | chr12 | + | 110641023 | 110641149 | 110640382 | 110640517 | 110647210 | 110647336 | 6 | 11 | 9 | 6 | 16 | 2 | 0 | 1 | 0 | 0 | 6 | 3 | 11 | 7 | 5 | 0 | 2 | 0 | 0 | 0 | 275 | 149 | 1 | 1 | 0.619 | 1 | 0.83 | 1 | 1 | 1 | 0.448 | 1 | 1 | 1 | 0 |
| SE | 107996 | ENSG00000204852 | TCTN1 | chr12 | + | 110641541 | 110642389 | 110640382 | 110640517 | 110647210 | 110647336 | 0 | 1 | 0 | 1 | 0 | 2 | 0 | 1 | 0 | 0 | 0 | 0 | 0 | 0 | 0 | 0 | 2 | 0 | 0 | 0 | 298 | 149 | 1 | 1 | 0 | 1 | 0 | 1 | NA | NA | 0 | NA | NA | NA | 0.5 |
| SE | 107997 | ENSG00000204852 | TCTN1 | chr12 | + | 110641541 | 110642389 | 110641023 | 110641149 | 110644991 | 110645129 | 7 | 4 | 7 | 0 | 14 | 1 | 0 | 0 | 0 | 0 | 0 | 7 | 1 | 4 | 0 | 0 | 0 | 0 | 0 | 0 | 298 | 149 | 1 | 1 | 0.778 | 1 | 1 | NA | 1 | NA | 1 | 1 | 1 | NA | -0.055 |
| SE | 107998 | ENSG00000204852 | TCTN1 | chr12 | + | 110641541 | 110642389 | 110641023 | 110641149 | 110647210 | 110647336 | 7 | 5 | 7 | 0 | 14 | 1 | 0 | 4 | 0 | 0 | 0 | 7 | 1 | 4 | 0 | 0 | 0 | 5 | 0 | 0 | 298 | 149 | 1 | 1 | 0.778 | 1 | 0.467 | NA | 1 | NA | 1 | 0.091 | 1 | NA | 0.114 |
| SE | 107999 | ENSG00000204852 | TCTN1 | chr12 | + | 110644966 | 110645129 | 110640382 | 110640517 | 110647195 | 110647336 | 0 | 0 | 0 | 0 | 2 | 0 | 0 | 2 | 0 | 0 | 0 | 0 | 3 | 10 | 0 | 0 | 0 | 0 | 0 | 0 | 298 | 149 | 1 | 1 | NA | NA | 0 | NA | 1 | NA | NA | NA | 1 | 1 | -0.5 |
| SE | 108001 | ENSG00000204852 | TCTN1 | chr12 | + | 110644966 | 110645129 | 110641023 | 110641149 | 110647210 | 110647336 | 14 | 13 | 13 | 3 | 4 | 1 | 0 | 4 | 0 | 0 | 6 | 0 | 4 | 11 | 19 | 0 | 0 | 5 | 0 | 0 | 298 | 149 | 1 | 1 | 0.875 | 1 | 0.619 | 1 | 1 | 1 | NA | 0.286 | 1 | 1 | 0.077 |
| SE | 108002 | ENSG00000204852 | TCTN1 | chr12 | + | 110644966 | 110645129 | 110642248 | 110642389 | 110647210 | 110647336 | 15 | 18 | 16 | 5 | 18 | 0 | 1 | 0 | 0 | 0 | 6 | 7 | 6 | 22 | 32 | 0 | 0 | 0 | 0 | 0 | 298 | 149 | 1 | 1 | 1 | 0.9 | 1 | 1 | 1 | 1 | 1 | 1 | 1 | 1 | -0.02 |
| SE | 108003 | ENSG00000204852 | TCTN1 | chr12 | + | 110644966 | 110645143 | 110640382 | 110640517 | 110647210 | 110647262 | 0 | 0 | 0 | 1 | 2 | 2 | 0 | 1 | 0 | 0 | 0 | 0 | 0 | 0 | 7 | 0 | 2 | 0 | 0 | 0 | 298 | 149 | 1 | 1 | 0 | NA | 0 | 1 | 1 | NA | 0 | NA | NA | 1 | 0 |
| SE | 108004 | ENSG00000204852 | TCTN1 | chr12 | + | 110644991 | 110645129 | 110641023 | 110641149 | 110647210 | 110647336 | 15 | 12 | 11 | 3 | 4 | 1 | 0 | 4 | 0 | 0 | 6 | 0 | 4 | 11 | 16 | 0 | 0 | 5 | 0 | 0 | 287 | 149 | 1 | 1 | 0.886 | 1 | 0.588 | 1 | 1 | 1 | NA | 0.293 | 1 | 1 | 0.072 |
| SE | 108005 | ENSG00000204852 | TCTN1 | chr12 | + | 110644991 | 110645129 | 110641541 | 110642389 | 110647210 | 110647336 | 14 | 12 | 11 | 3 | 4 | 0 | 1 | 0 | 0 | 0 | 6 | 0 | 4 | 11 | 16 | 0 | 0 | 0 | 0 | 0 | 287 | 149 | 1 | 1 | 1 | 0.862 | 1 | 1 | 1 | 1 | NA | 1 | 1 | 1 | -0.028 |
| SE | 108009 | ENSG00000135451 | TROAP | chr12 | + | 49323844 | 49324037 | 49323373 | 49323752 | 49324211 | 49324296 | 2 | 0 | 11 | 6 | 0 | 0 | 0 | 0 | 0 | 0 | 5 | 19 | 6 | 17 | 1 | 0 | 1 | 2 | 2 | 0 | 298 | 149 | 1 | 1 | 1 | NA | 1 | 1 | NA | 1 | 0.905 | 0.6 | 0.81 | 1 | 0.137 |
| SE | 108014 | ENSG00000135451 | TROAP | chr12 | + | 49326075 | 49326158 | 49325746 | 49325884 | 49326667 | 49326720 | 0 | 0 | 2 | 1 | 0 | 1 | 0 | 0 | 0 | 0 | 0 | 4 | 1 | 5 | 8 | 0 | 0 | 0 | 0 | 0 | 232 | 149 | 1 | 1 | 0 | NA | 1 | 1 | NA | NA | 1 | 1 | 1 | 1 | -0.333 |
| SE | 108018 | ENSG00000111540 | RAB5B | chr12 | + | 55979371 | 55979474 | 55974083 | 55974139 | 55986868 | 55987123 | 7 | 4 | 1 | 3 | 0 | 2 | 2 | 2 | 1 | 2 | 0 | 1 | 5 | 8 | 1 | 0 | 3 | 1 | 0 | 2 | 252 | 149 | 1 | 1 | 0.674 | 0.542 | 0.228 | 0.639 | 0 | NA | 0.165 | 0.747 | 1 | 0.228 | -0.118 |
| SE | 108019 | ENSG00000111540 | RAB5B | chr12 | + | 55980888 | 55980922 | 55979371 | 55979474 | 55986868 | 55987123 | 0 | 0 | 0 | 0 | 1 | 3 | 4 | 0 | 2 | 0 | 0 | 4 | 2 | 4 | 2 | 0 | 0 | 5 | 4 | 1 | 183 | 149 | 1 | 1 | 0 | 0 | NA | 0 | 1 | NA | 1 | 0.246 | 0.449 | 0.62 | -0.329 |
| SE | 108020 | ENSG00000111540 | RAB5B | chr12 | + | 55985599 | 55985723 | 55979371 | 55979474 | 55986868 | 55987092 | 1 | 0 | 0 | 0 | 0 | 3 | 4 | 0 | 2 | 0 | 0 | 1 | 0 | 0 | 0 | 0 | 0 | 5 | 4 | 1 | 273 | 149 | 1 | 1 | 0.154 | 0 | NA | 0 | NA | NA | 1 | 0 | 0 | 0 | -0.199 |

| | | | | | | | | | | | | | | | | | | | | | | | | | | | | | | | | | | | | | | | | | | | | | | | | |
|---|---|---|---|---|---|---|---|---|---|---|---|---|---|---|---|---|---|---|---|---|---|---|---|---|---|---|---|---|---|---|---|---|---|---|---|---|---|---|---|---|---|---|---|---|---|---|---|---|
| SE | 108021 | ENSG00000111540 | RAB5B | chr12 | + | 55989946 | 55990098 | 55986868 | 55987123 | 55990681 | 55990711 | 672 | 371 | 614 | 735 | 580 | 2 | 0 | 0 | 0 | 0 | 744 | 607 | 364 | 652 | 1250 | 0 | 0 | 0 | 5 | 0 | 5 | 0 | 298 | 149 | 1 | 1 | 0.994 | 1 | 1 | 1 | 1 | 1 | 1 | 1 | 0.985 | 1 | 0.002 |
| SE | 108024 | ENSG00000123411 | IKZF4 | chr12 | + | 56019317 | 56019415 | 56018125 | 56018177 | 56021430 | 56021580 | 0 | 5 | 4 | 0 | 0 | 2 | 0 | 0 | 0 | 1 | 32 | 0 | 5 | 7 | 0 | 0 | 0 | 0 | 0 | 0 | 0 | 0 | 247 | 149 | 1 | 1 | 0 | 1 | 1 | NA | 0 | 1 | NA | 1 | 1 | NA | -0.5 |
| SE | 108025 | ENSG00000123411 | IKZF4 | chr12 | + | 56019317 | 56019415 | 56018125 | 56018177 | 56023670 | 56023764 | 0 | 5 | 2 | 0 | 0 | 0 | 2 | 2 | 0 | 0 | 34 | 0 | 5 | 7 | 0 | 8 | 0 | 0 | 0 | 0 | 0 | 0 | 247 | 149 | 1 | 1 | NA | 0.601 | 0.376 | NA | NA | 0.719 | NA | 1 | 1 | NA | -0.418 |
| SE | 108026 | ENSG00000123411 | IKZF4 | chr12 | + | 56019320 | 56019415 | 56018125 | 56018177 | 56021430 | 56021523 | 3 | 0 | 5 | 0 | 0 | 2 | 0 | 0 | 0 | 1 | 0 | 2 | 8 | 0 | 0 | 0 | 0 | 0 | 0 | 0 | 0 | 0 | 244 | 149 | 1 | 1 | 0.478 | NA | 1 | NA | 0 | NA | 1 | 1 | NA | NA | -0.507 |
| SE | 108027 | ENSG00000123411 | IKZF4 | chr12 | + | 56021430 | 56021580 | 56018125 | 56018177 | 56023670 | 56023764 | 5 | 1 | 1 | 0 | 2 | 0 | 2 | 2 | 0 | 0 | 0 | 0 | 0 | 0 | 0 | 8 | 0 | 0 | 0 | 0 | 0 | 0 | 298 | 149 | 1 | 1 | 1 | 0.2 | 0.2 | NA | 1 | 0 | NA | NA | NA | NA | 0.6 |
| SE | 108028 | ENSG00000123411 | IKZF4 | chr12 | + | 56021430 | 56021580 | 56019317 | 56019415 | 56023670 | 56023764 | 3 | 1 | 3 | 0 | 1 | 0 | 0 | 0 | 0 | 0 | 0 | 0 | 0 | 0 | 0 | 2 | 0 | 0 | 0 | 0 | 0 | 0 | 298 | 149 | 1 | 1 | 1 | 1 | 1 | NA | 1 | 0 | NA | NA | NA | NA | 1 |
| SE | 108029 | ENSG00000123411 | IKZF4 | chr12 | + | 56023670 | 56023764 | 56018125 | 56018177 | 56025053 | 56025158 | 2 | 4 | 6 | 0 | 0 | 2 | 1 | 0 | 2 | 4 | 34 | 0 | 0 | 0 | 0 | 8 | 4 | 0 | 0 | 0 | 0 | 0 | 243 | 149 | 1 | 1 | 0.38 | 0.71 | 1 | 0 | 0 | 0.723 | 0 | NA | NA | NA | 0.056 |
| SE | 108031 | ENSG00000123411 | IKZF4 | chr12 | + | 56023670 | 56023764 | 56021042 | 56021580 | 56025053 | 56025158 | 4 | 3 | 5 | 0 | 1 | 0 | 0 | 0 | 0 | 0 | 26 | 0 | 0 | 0 | 0 | 0 | 0 | 0 | 2 | 0 | 2 | 6 | 243 | 149 | 1 | 1 | 1 | 1 | 1 | NA | 1 | 1 | NA | NA | 0 | 0 | 0.667 |
| SE | 108034 | ENSG00000139613 | SMARCC2 | chr12 | - | 56169775 | 56169911 | 56168059 | 56168194 | 56171270 | 56171432 | 1 | 0 | 1 | 2 | 2 | 0 | 0 | 0 | 0 | 0 | 20 | 4 | 4 | 40 | 1 | 0 | 0 | 1 | 0 | 1 | 0 | 0 | 285 | 149 | 1 | 1 | 1 | NA | 1 | 1 | 1 | 1 | 1 | 0.677 | 1 | 1 | 0.065 |
| SE | 108038 | ENSG00000139613 | SMARCC2 | chr12 | - | 56172584 | 56172704 | 56172238 | 56172490 | 56173695 | 56173849 | 241 | 165 | 256 | 252 | 254 | 1 | 0 | 0 | 0 | 0 | 417 | 322 | 253 | 306 | 515 | 0 | 0 | 0 | 4 | 0 | 4 | 0 | 269 | 149 | 1 | 1 | 0.993 | 1 | 1 | 1 | 1 | 1 | 1 | 1 | 0.977 | 1 | 0.003 |
| SE | 108040 | ENSG00000139613 | SMARCC2 | chr12 | - | 56174650 | 56174764 | 56173695 | 56173849 | 56178021 | 56178093 | 427 | 245 | 434 | 464 | 631 | 0 | 0 | 0 | 2 | 0 | 506 | 447 | 278 | 316 | 835 | 0 | 0 | 0 | 0 | 0 | 0 | 0 | 263 | 149 | 1 | 1 | 1 | 1 | 1 | 0.992 | 1 | 1 | 1 | 1 | 1 | 1 | -0.002 |
| SE | 108041 | ENSG00000139613 | SMARCC2 | chr12 | - | 56178021 | 56178093 | 56174530 | 56174764 | 56178403 | 56178534 | 386 | 224 | 398 | 420 | 573 | 0 | 0 | 2 | 0 | 0 | 473 | 356 | 253 | 292 | 770 | 2 | 0 | 0 | 2 | 0 | 2 | 0 | 221 | 149 | 1 | 1 | 1 | 1 | 0.993 | 1 | 1 | 0.994 | 1 | 1 | 0.99 | 1 | 0.002 |
| SE | 108044 | ENSG00000139613 | SMARCC2 | chr12 | - | 56184174 | 56184244 | 56182003 | 56182079 | 56184843 | 56184936 | 190 | 107 | 184 | 208 | 226 | 0 | 0 | 1 | 0 | 0 | 260 | 218 | 120 | 209 | 297 | 0 | 0 | 0 | 0 | 0 | 0 | 0 | 219 | 149 | 1 | 1 | 1 | 1 | 0.992 | 1 | 1 | 1 | 1 | 1 | 1 | 1 | -0.002 |
| SE | 108046 | ENSG00000139613 | SMARCC2 | chr12 | - | 56184843 | 56184936 | 56182003 | 56182079 | 56186154 | 56186240 | 0 | 1 | 1 | 0 | 0 | 0 | 0 | 0 | 2 | 0 | 0 | 7 | 1 | 1 | 8 | 7 | 1 | 0 | 0 | 0 | 0 | 0 | 242 | 149 | 1 | 1 | NA | 1 | 1 | 0 | NA | 0 | 0.812 | 1 | 1 | 1 | -0.096 |
| SE | 108047 | ENSG00000139613 | SMARCC2 | chr12 | - | 56184843 | 56184936 | 56184174 | 56184244 | 56186154 | 56186240 | 209 | 170 | 177 | 251 | 426 | 0 | 0 | 0 | 2 | 0 | 420 | 209 | 163 | 208 | 335 | 0 | 0 | 0 | 0 | 0 | 0 | 0 | 242 | 149 | 1 | 1 | 1 | 1 | 1 | 0.987 | 1 | 1 | 1 | 1 | 1 | 1 | -0.003 |
| SE | 108050 | ENSG00000139613 | SMARCC2 | chr12 | - | 56187186 | 56187306 | 56185789 | 56186240 | 56189350 | 56189391 | 277 | 205 | 303 | 340 | 210 | 0 | 0 | 0 | 2 | 0 | 427 | 338 | 195 | 280 | 386 | 0 | 2 | 0 | 0 | 0 | 0 | 0 | 269 | 149 | 1 | 1 | 1 | 1 | 1 | 0.989 | 1 | 1 | 0.989 | 1 | 1 | 1 | 0 |
| SE | 108051 | ENSG00000103550 | KNOP1 | chr16 | - | 19710508 | 19710586 | 19701936 | 19707221 | 19714117 | 19714577 | 23 | 12 | 43 | 25 | 30 | 0 | 0 | 0 | 0 | 0 | 36 | 12 | 15 | 5 | 23 | 0 | 0 | 1 | 0 | 1 | 0 | 0 | 227 | 149 | 1 | 1 | 1 | 1 | 1 | 1 | 1 | 1 | 1 | 0.908 | 1 | 1 | 0.018 |
| SE | 108053 | ENSG00000139579 | NABP2 | chr12 | + | 56223583 | 56223712 | 56222014 | 56222199 | 56224833 | 56224935 | 1 | 0 | 1 | 0 | 0 | 1 | 0 | 1 | 0 | 0 | 0 | 4 | 4 | 0 | 14 | 0 | 0 | 1 | 0 | 1 | 0 | 6 | 278 | 149 | 1 | 1 | 0.349 | NA | 0.349 | NA | NA | NA | 1 | 0.682 | NA | 0.556 | -0.397 |
| SE | 108054 | ENSG00000139579 | NABP2 | chr12 | + | 56225623 | 56225695 | 56225372 | 56225511 | 56226178 | 56226260 | 87 | 33 | 65 | 88 | 149 | 0 | 0 | 1 | 0 | 0 | 154 | 98 | 79 | 71 | 177 | 0 | 0 | 1 | 0 | 0 | 0 | 0 | 221 | 149 | 1 | 1 | 1 | 1 | 0.978 | 1 | 1 | 1 | 1 | 1 | 1 | 1 | -0.004 |
| SE | 108056 | ENSG00000257727 | CNPY2 | chr12 | - | 56311210 | 56311453 | 56310957 | 56311054 | 56314850 | 56314968 | 129 | 64 | 131 | 116 | 119 | 0 | 0 | 0 | 1 | 0 | 205 | 84 | 80 | 163 | 119 | 0 | 0 | 0 | 0 | 0 | 0 | 0 | 298 | 149 | 1 | 1 | 1 | 1 | 1 | 0.983 | 1 | 1 | 1 | 1 | 1 | 1 | -0.003 |
| [illegible] | | | | | | | | | | | | | | | | | | | | | | | | | | | | | | | | | | | | | | | | | | | | | | | | |

| | | | | | | | | | | | | | | |
|---|---|---|---|---|---|---|---|---|---|---|---|---|---|---|
| SE | 108423 | ENSG00000197111 | PCBP2 | chr12 | + | 53471637 | 53471807 | 53467722 | 53468832 | 53479405 | 53479614 | 2921 | 1455 | [illegible] |
| SE | 108426 | ENSG00000067167 | TRAM1 | chr8 | - | 70583163 | 70583324 | 70573217 | 70575005 | 70586894 | 70586999 | 1733 | 1215 | [illegible] |
| SE | 108427 | ENSG00000067167 | TRAM1 | chr8 | - | 70583649 | 70583793 | 70583163 | 70583324 | 70586894 | 70586999 | 3053 | 2428 | [illegible] |
| SE | 108428 | ENSG00000067167 | TRAM1 | chr8 | - | 70586894 | 70586999 | 70573217 | 70575005 | 70587105 | 70587176 | 672 | 441 | [illegible] |
| SE | 108429 | ENSG00000067167 | TRAM1 | chr8 | - | 70586894 | 70586999 | 70583649 | 70583793 | 70587105 | 70587176 | 2049 | 1432 | [illegible] |
| SE | 108430 | ENSG00000067167 | TRAM1 | chr8 | - | 70587105 | 70587176 | 70583649 | 70583793 | 70594505 | 70594590 | 404 | 229 | [illegible] |
| SE | 108432 | ENSG00000067167 | TRAM1 | chr8 | - | 70594505 | 70594590 | 70587105 | 70587176 | 70596262 | 70596321 | 1396 | 821 | [illegible] |
| SE | 108433 | ENSG00000067167 | TRAM1 | chr8 | - | 70594505 | 70594590 | 70587105 | 70587176 | 70597894 | 70598011 | 657 | 440 | [illegible] |
| SE | 108436 | ENSG00000067167 | TRAM1 | chr8 | - | 70596262 | 70596321 | 70594500 | 70594590 | 70600018 | 70600082 | 307 | 145 | [illegible] |
| SE | 108437 | ENSG00000067167 | TRAM1 | chr8 | - | 70597890 | 70598011 | 70596262 | 70596321 | 70598133 | 70598255 | 307 | 153 | [illegible] |
| SE | 108438 | ENSG00000067167 | TRAM1 | chr8 | - | 70597894 | 70598011 | 70594500 | 70594590 | 70600018 | 70600082 | 12 | 10 | [illegible] |
| SE | 108439 | ENSG00000067167 | TRAM1 | chr8 | - | 70597894 | 70598011 | 70596262 | 70596321 | 70598133 | 70598255 | 938 | 504 | [illegible] |
| SE | 108440 | ENSG00000067167 | TRAM1 | chr8 | - | 70597894 | 70598011 | 70596262 | 70596321 | 70600018 | 70600082 | 487 | 276 | [illegible] |
| SE | 108443 | ENSG00000067167 | TRAM1 | chr8 | - | 70600018 | 70600082 | 70598133 | 70598255 | 70608076 | 70608136 | 300 | 137 | [illegible] |
| SE | 108447 | ENSG00000092978 | GPATCH2 | chr1 | - | 217491679 | 217491750 | 217449248 | 217449337 | 217498355 | 217498395 | 16 | 11 | [illegible] |
| SE | 108448 | ENSG00000092978 | GPATCH2 | chr1 | - | 217491679 | 217491750 | 217449248 | 217449337 | 217514821 | 217514889 | 4 | 3 | [illegible] |
| SE | 108449 | ENSG00000092978 | GPATCH2 | chr1 | - | 217495014 | 217495075 | 217491505 | 217491750 | 217498355 | 217498395 | 2 | 0 | [illegible] |
| SE | 108450 | ENSG00000092978 | GPATCH2 | chr1 | - | 217495014 | 217495075 | 217492500 | 217492693 | 217498355 | 217498395 | 1 | 8 | [illegible] |
| SE | 108451 | ENSG00000092978 | GPATCH2 | chr1 | - | 217498355 | 217498395 | 217449248 | 217449337 | 217514821 | 217514889 | 4 | 10 | [illegible] |
| SE | 108452 | ENSG00000092978 | GPATCH2 | chr1 | - | 217498355 | 217498395 | 217491679 | 217491750 | 217514821 | 217514889 | 13 | 18 | [illegible] |
| SE | 108454 | ENSG00000092978 | GPATCH2 | chr1 | - | 217614140 | 217614202 | 217610320 | 217610400 | 217619782 | 217620499 | 34 | 33 | [illegible] |
| SE | 108457 | ENSG00000244649 | LINC02086 | chr17 | + | 48683238 | 48683417 | 48652870 | 48652974 | 48687496 | 48687710 | 3 | 0 | [illegible] |
| SE | 108458 | ENSG00000244649 | LINC02086 | chr17 | + | 48683238 | 48683417 | 48652870 | 48652974 | 48700611 | 48700696 | 3 | 0 | [illegible] |
| SE | 108461 | ENSG00000196199 | MPHOSPH8 | chr13 | + | 19642114 | 19642270 | 19633658 | 19633961 | 19646442 | 19647291 | 203 | 228 | [illegible] |
| SE | 108462 | ENSG00000196199 | MPHOSPH8 | chr13 | + | 19670839 | 19670964 | 19670235 | 19670363 | 19671205 | 19671289 | 8 | 10 | [illegible] |
| SE | 108463 | ENSG00000196199 | MPHOSPH8 | chr13 | + | 19670839 | 19671289 | 19670339 | 19670363 | 19671833 | 19671942 | 72 | 66 | [illegible] |
| SE | 108464 | ENSG00000196199 | MPHOSPH8 | chr13 | + | 19671205 | 19671289 | 19670235 | 19670363 | 19671833 | 19672072 | 103 | 83 | [illegible] |
| SE | 108465 | ENSG00000146281 | PM20D2 | chr6 | + | 89153042 | 89153185 | 89149264 | 89149413 | 89158324 | 89158460 | 13 | 7 | [illegible] |
| SE | 108466 | ENSG00000146281 | PM20D2 | chr6 | + | 89154747 | 89154902 | 89153042 | 89153185 | 89158324 | 89158460 | 34 | 51 | [illegible] |
| [illegible] | | | | | | | | | | | | | | |

| | | | | | | | | | | | | | | |
|---|---|---|---|---|---|---|---|---|---|---|---|---|---|---|
| SE | 108922 | ENSG00000196683 | TOMM7 | chr7 | - | 22817411 | 22817548 | 22812972 | 22813185 | 22817999 | 22818048 | 1 | 1 | [illegible] |
| SE | 108923 | ENSG00000196683 | TOMM7 | chr7 | - | 22817999 | 22818048 | 22812970 | 22813185 | 22822153 | 22822344 | 81 | 46 | [illegible] |
| SE | 108925 | ENSG00000102471 | NDFIP2 | chr13 | + | 79534391 | 79534554 | 79533322 | 79533456 | 79539681 | 79539775 | 0 | 0 | [illegible] |
| SE | 108928 | ENSG00000102471 | NDFIP2 | chr13 | + | 79551016 | 79551122 | 79548327 | 79548394 | 79552515 | 79552578 | 14 | 23 | [illegible] |
| SE | 108934 | ENSG00000197558 | SSPOP | chr7 | + | 149812265 | 149812436 | 149811664 | 149811717 | 149812560 | 149812635 | 0 | 0 | [illegible] |
| SE | 108944 | ENSG00000136141 | LRCH1 | chr13 | + | 46681740 | 46681846 | 46669029 | 46669156 | 46692535 | 46692577 | 23 | 19 | [illegible] |
| SE | 108945 | ENSG00000136141 | LRCH1 | chr13 | + | 46685904 | 46686041 | 46681740 | 46681846 | 46687851 | 46687979 | 101 | 73 | [illegible] |
| SE | 108946 | ENSG00000136141 | LRCH1 | chr13 | + | 46687851 | 46687979 | 46681740 | 46681846 | 46692535 | 46692577 | 0 | 0 | [illegible] |
| SE | 108948 | ENSG00000136141 | LRCH1 | chr13 | + | 46699335 | 46699403 | 46694892 | 46695017 | 46701120 | 46701207 | 96 | 72 | [illegible] |
| SE | 108952 | ENSG00000139496 | NUP58 | chr13 | + | 25303101 | 25303155 | 25301624 | 25301880 | 25307805 | 25307948 | 0 | 2 | [illegible] |
| SE | 108955 | ENSG00000139496 | NUP58 | chr13 | + | 25312882 | 25313032 | 25309246 | 25309282 | 25313613 | 25313751 | 200 | 150 | [illegible] |
| SE | 108956 | ENSG00000139496 | NUP58 | chr13 | + | 25313613 | 25313751 | 25309246 | 25309282 | 25315356 | 25315467 | 104 | 68 | [illegible] |
| SE | 108961 | ENSG00000102543 | CDADC1 | chr13 | + | 49259345 | 49259523 | 49255838 | 49255913 | 49267489 | 49268059 | 47 | 39 | [illegible] |
| SE | 108963 | ENSG00000102543 | CDADC1 | chr13 | + | 49259485 | 49259523 | 49255838 | 49255913 | 49267489 | 49267785 | 7 | 9 | [illegible] |
| SE | 108970 | ENSG00000111877 | MCM9 | chr6 | - | 118931419 | 118931738 | 118923810 | 118924127 | 118934890 | 118935114 | 50 | 26 | [illegible] |
| SE | 108973 | ENSG00000180370 | PAK2 | chr3 | + | 196801926 | 196802027 | 196782625 | 196782833 | 196803016 | 196803164 | 199 | 196 | [illegible] |
| SE | 108974 | ENSG00000180370 | PAK2 | chr3 | + | 196810589 | 196810653 | 196807781 | 196807914 | 196812218 | 196812267 | 611 | 395 | [illegible] |
| SE | 108975 | ENSG00000180370 | PAK2 | chr3 | + | 196814450 | 196814568 | 196812738 | 196812851 | 196818056 | 196818156 | 921 | 595 | [illegible] |
| SE | 108976 | ENSG00000180370 | PAK2 | chr3 | + | 196818056 | 196818156 | 196814450 | 196814568 | 196820370 | 196820567 | 598 | 366 | [illegible] |
| SE | 108978 | ENSG00000180370 | PAK2 | chr3 | + | 196827195 | 196827333 | 196820370 | 196820567 | 196828318 | 196832647 | 687 | 351 | [illegible] |
| SE | 108980 | ENSG00000133103 | COG6 | chr13 | + | 39656116 | 39656275 | 39655650 | 39655879 | 39659363 | 39659503 | 4 | 0 | [illegible] |
| SE | 108981 | ENSG00000133103 | COG6 | chr13 | + | 39656825 | 39656890 | 39655723 | 39655879 | 39659363 | 39659467 | 11 | 0 | [illegible] |
| SE | 108982 | ENSG00000133103 | COG6 | chr13 | + | 39656825 | 39656890 | 39656116 | 39656275 | 39659363 | 39659467 | 5 | 0 | [illegible] |
| SE | 108984 | ENSG00000133103 | COG6 | chr13 | + | 39694633 | 39694725 | 39689759 | 39689824 | 39699500 | 39699618 | 126 | 52 | [illegible] |
| SE | 108985 | ENSG00000133103 | COG6 | chr13 | + | 39699500 | 39699618 | 39694633 | 39694725 | 39719235 | 39719367 | 125 | 31 | [illegible] |
| SE | 108986 | ENSG00000102738 | MRPS31 | chr13 | - | 40749137 | 40749281 | 40729127 | 40729601 | 40756872 | 40757013 | 83 | 61 | [illegible] |
| SE | 108987 | ENSG00000102738 | MRPS31 | chr13 | - | 40754018 | 40754092 | 40749137 | 40749281 | 40756872 | 40757013 | 161 | 125 | [illegible] |
| SE | 108988 | ENSG00000102738 | MRPS31 | chr13 | - | 40758947 | 40759106 | 40756872 | 40757013 | 40766745 | 40767033 | 179 | 141 | [illegible] |
| SE | 108989 | ENSG00000102738 | MRPS31 | chr13 | - | 40766745 | 40767033 | 40758947 | 40759106 | 40770984 | 40771190 | 178 | 111 | [illegible] |
| SE | 108990 | ENSG00000160961 | ZNF333 | chr19 | + | 14695009 | 14695133 | 14693450 | 14693494 | 14695565 | 14695661 | 27 | 15 | [illegible] |
| SE | 108992 | ENSG00000160961 | ZNF333 | chr19 | + | 14706685 | 14706773 | 14705053 | 14705170 | 14715381 | 14715470 | 31 | 29 | [illegible] |
| SE | 109002 | ENSG00000105352 | CEACAM4 | chr19 | - | 41619669 | 41619711 | 41618970 | 41619395 | 41620210 | 41620242 | 18 | 14 | [illegible] |
| SE | 109003 | ENSG00000105352 | CEACAM4 | chr19 | - | 41620210 | 41620242 | 41619544 | 41619711 | 41620574 | 41620621 | 19 | 12 | [illegible] |
| SE | 109008 | ENSG00000256771 | ZNF253 | chr19 | + | 19878480 | 19878607 | 19865881 | 19865999 | 19880050 | 19880146 | 24 | 13 | [illegible] |
| SE | 109009 | ENSG00000256771 | ZNF253 | chr19 | + | 19880050 | 19880146 | 19865885 | 19865999 | 19891473 | 19891917 | 10 | 11 | [illegible] |
| SE | 109010 | ENSG00000186056 | MATN1-AS1 | chr1 | + | 30720976 | 30721174 | 30719173 | 30719232 | 30721287 | 30721722 | 2 | 0 | [illegible] |
| SE | 109011 | ENSG00000137822 | TUBGCP4 | chr15 | + | 43383302 | 43383504 | 43380083 | 43380163 | 43385790 | 43385956 | 36 | 52 | [illegible] |
| SE | 109012 | ENSG00000137822 | TUBGCP4 | chr15 | + | 43385367 | 43385508 | 43383302 | 43383504 | 43385790 | 43385916 | 0 | 2 | [illegible] |
| SE | 109015 | ENSG00000137822 | TUBGCP4 | chr15 | + | 43401715 | 43401850 | 43400043 | 43400221 | 43403682 | 43403799 | 16 | 38 | [illegible] |
| SE | 109025 | ENSG00000125457 | MIF4GD | chr17 | - | 75267745 | 75267901 | 75266511 | 75267630 | 75268082 | 75268192 | 83 | 62 | [illegible] |
| SE | 109028 | ENSG00000125457 | MIF4GD | chr17 | - | 75268082 | 75268192 | 75267745 | 75267901 | 75269346 | 75269448 | 42 | 32 | [illegible] |
| SE | 109029 | ENSG00000125457 | MIF4GD | chr17 | - | 75268082 | 75268192 | 75267745 | 75267901 | 75270113 | 75270227 | 68 | 46 | [illegible] |
| SE | 109030 | ENSG00000125457 | MIF4GD | chr17 | - | 75269346 | 75269448 | 75267745 | 75267901 | 75270113 | 75270245 | 4 | 0 | [illegible] |
| SE | 109032 | ENSG00000125457 | MIF4GD | chr17 | - | 75269346 | 75269469 | 75267745 | 75267901 | 75270113 | 75270245 | 9 | 5 | [illegible] |
| SE | 109034 | ENSG00000102595 | UGGT2 | chr13 | - | 95887272 | 95887393 | 95884490 | 95884680 | 95887891 | 95887971 | 7 | 5 | [illegible] |
| SE | 109035 | ENSG00000102595 | UGGT2 | chr13 | - | 95900806 | 95900938 | 95895179 | 95895304 | 95902853 | 95903060 | 14 | 10 | [illegible] |
| SE | 109036 | ENSG00000102595 | UGGT2 | chr13 | - | 95983803 | 95983864 | 95972579 | 95972671 | 95986332 | 95986432 | 14 | 7 | [illegible] |
| SE | 109037 | ENSG00000102595 | UGGT2 | chr13 | - | 95986332 | 95986432 | 95972579 | 95972671 | 95996062 | 95996135 | 2 | 2 | [illegible] |
| SE | 109038 | ENSG00000102595 | UGGT2 | chr13 | - | 95989520 | 95989598 | 95986332 | 95986432 | 95989972 | 95990073 | 1 | 0 | [illegible] |
| SE | 109039 | ENSG00000102595 | UGGT2 | chr13 | - | 95989972 | 95990073 | 95986332 | 95986432 | 95996062 | 95996135 | 50 | 58 | [illegible] |
| SE | 109040 | ENSG00000083844 | ZNF264 | chr19 | + | 57191698 | 57191946 | 57191500 | 57191527 | 57193874 | 57193958 | 27 | 7 | [illegible] |
| SE | 109041 | ENSG00000083844 | ZNF264 | chr19 | + | 57193874 | 57194001 | 57191499 | 57191946 | 57205396 | 57205492 | 67 | 24 | [illegible] |
| SE | 109043 | ENSG00000173391 | OLR1 | chr12 | - | 10160346 | 10160462 | 10158300 | 10160021 | 10160785 | 10160925 | 2 | 0 | [illegible] |
| SE | 109044 | ENSG00000173391 | OLR1 | chr12 | - | 10160346 | 10160462 | 10158300 | 10160021 | 10166711 | 10166957 | 1 | 0 | [illegible] |
| SE | 109045 | ENSG00000173391 | OLR1 | chr12 | - | 10160785 | 10160925 | 10159617 | 10160021 | 10166711 | 10166949 | 0 | 0 | [illegible] |
| SE | 109049 | ENSG00000139668 | WDFY2 | chr13 | + | 51660595 | 51660663 | 51584461 | 51584824 | 51675169 | 51675243 | 98 | 98 | [illegible] |
| SE | 109051 | ENSG00000139668 | WDFY2 | chr13 | + | 51703595 | 51703650 | 51675169 | 51675243 | 51719197 | 51719348 | 145 | 136 | [illegible] |
| SE | 109052 | ENSG00000139668 | WDFY2 | chr13 | + | 51703595 | 51703650 | 51675169 | 51675243 | 51739048 | 51739175 | 36 | 34 | [illegible] |
| SE | 109060 | ENSG00000164649 | CDCA7L | chr7 | - | 21906288 | 21906456 | 21905505 | 21905631 | 21906567 | 21906639 | 112 | 77 | [illegible] |
| SE | 109066 | ENSG00000198176 | TFDP1 | chr13 | + | 113610995 | 113611062 | 113585773 | 113585849 | 113623179 | 113623286 | 57 | 39 | [illegible] |
| SE | 109071 | ENSG00000198176 | TFDP1 | chr13 | + | 113631622 | 113631905 | 113610995 | 113611062 | 113633119 | 113633285 | 0 | 0 | [illegible] |
| SE | 109077 | ENSG00000144567 | RETREG2 | chr2 | + | 219180669 | 219180754 | 219180109 | 219180245 | 219181368 | 219181463 | 34 | 19 | [illegible] |
| SE | 109079 | ENSG00000144567 | RETREG2 | chr2 | + | 219181061 | 219181205 | 219180669 | 219180754 | 219181368 | 219181463 | 330 | 186 | [illegible] |
| SE | 109080 | ENSG00000144567 | RETREG2 | chr2 | + | 219181061 | 219181205 | 219180669 | 219180754 | 219181639 | 219181663 | 127 | 63 | [illegible] |
| SE | 109082 | ENSG00000144567 | RETREG2 | chr2 | + | 219181061 | 219181463 | 219180667 | 219180754 | 219181639 | 219181693 | 302 | 169 | [illegible] |
| SE | 109084 | ENSG00000144567 | RETREG2 | chr2 | + | 219181368 | 219181463 | 219180669 | 219180754 | 219181639 | 219181663 | 175 | 106 | [illegible] |
| SE | 109086 | ENSG00000144567 | RETREG2 | chr2 | + | 219181639 | 219181775 | 219180669 | 219180754 | 219182012 | 219182120 | 245 | 163 | [illegible] |
| SE | 109089 | ENSG00000285534 | NA | chr13 | - | 109571073 | 109571158 | 109560400 | 109560490 | 109730942 | 109731700 | 0 | 1 | [illegible] |
| SE | 109090 | ENSG00000013364 | MVP | chr16 | + | 29825941 | 29826092 | 29820472 | 29820510 | 29830473 | 29830674 | 0 | 0 | [illegible] |
| SE | 109091 | ENSG00000013364 | MVP | chr16 | + | 29825941 | 29826092 | 29820472 | 29820510 | 29830514 | 29830675 | 2 | 1 | [illegible] |
| SE | 109092 | ENSG00000013364 | MVP | chr16 | + | 29830877 | 29831073 | 29830473 | 29830674 | 29833732 | 29833856 | 331 | 133 | [illegible] |
| SE | 109093 | ENSG00000013364 | MVP | chr16 | + | 29835703 | 29835798 | 29833934 | 29834066 | 29836721 | 29836958 | 530 | 334 | [illegible] |
| SE | 109094 | ENSG00000013364 | MVP | chr16 | + | 29835703 | 29835798 | 29834185 | 29834437 | 29836721 | 29836958 | 296 | 171 | [illegible] |
| SE | 109095 | ENSG00000013364 | MVP | chr16 | + | 29836721 | 29836958 | 29835703 | 29835798 | 29840177 | 29840267 | 480 | 277 | [illegible] |
| SE | 109096 | ENSG00000013364 | MVP | chr16 | + | 29846157 | 29846284 | 29845862 | 29845979 | 29847196 | 29847385 | 149 | 69 | [illegible] |
| SE | 109097 | ENSG00000196312 | MFSD14C | chr9 | - | 96902916 | 96903108 | 96897916 | 96898127 | 96918642 | 96918826 | 13 | 8 | [illegible] |
| SE | 109102 | ENSG00000196312 | MFSD14C | chr9 | - | 96972852 | 96972948 | 96958834 | 96958897 | 97013258 | 97013366 | 35 | 28 | [illegible] |
| SE | 109104 | ENSG00000184497 | TMEM255B | chr13 | + | 113768886 | 113768969 | 113766163 | 113766257 | 113769097 | 113769160 | 1 | 0 | [illegible] |
| SE | 109105 | ENSG00000184497 | TMEM255B | chr13 | + | 113795147 | 113795237 | 113769097 | 113769160 | 113799338 | 113799419 | 1 | 0 | [illegible] |
| SE | 109114 | ENSG00000089486 | CDIP1 | chr16 | - | 4514045 | 4514144 | 4513693 | 4513851 | 4514574 | 4514664 | 34 | 18 | [illegible] |
| SE | 109116 | ENSG00000166529 | ZSCAN21 | chr7 | + | 100051477 | 100051610 | 100049793 | 100049841 | 100056910 | 100057405 | 9 | 4 | [illegible] |
| SE | 109117 | ENSG00000166529 | ZSCAN21 | chr7 | + | 100063787 | 100063979 | 100057697 | 100057890 | 100064082 | 100065038 | 23 | 13 | [illegible] |
| SE | 109119 | ENSG00000006625 | GGCT | chr7 | - | 30497740 | 30497896 | 30496783 | 30497235 | 30500535 | 30500681 | 0 | 0 | [illegible] |
| SE | 109125 | ENSG00000126217 | MCF2L | chr13 | + | 113024643 | 113024758 | 113014762 | 113014846 | 113045270 | 113045361 | 5 | 8 | [illegible] |
| SE | 109130 | ENSG00000126217 | MCF2L | chr13 | + | 113091120 | 113091195 | 113089609 | 113089728 | 113094513 | 113094635 | 0 | 0 | [illegible] |
| SE | 109132 | ENSG00000185989 | RASA3 | chr13 | - | 113981674 | 113981858 | 113977782 | 113979422 | 113992484 | 113992588 | 440 | 170 | [illegible] |
| SE | 109133 | ENSG00000185989 | RASA3 | chr13 | - | 114007532 | 114007606 | 114000825 | 114000932 | 114009386 | 114009464 | 753 | 425 | [illegible] |
| SE | 109135 | ENSG00000185989 | RASA3 | chr13 | - | 114018762 | 114018919 | 114018103 | 114018252 | 114021403 | 114021508 | 955 | 530 | [illegible] |
| SE | 109136 | ENSG00000185989 | RASA3 | chr13 | - | 114024278 | 114024355 | 114021403 | 114021508 | 114040999 | 114041094 | 247 | 157 | [illegible] |
| SE | 109137 | ENSG00000185989 | RASA3 | chr13 | - | 114024278 | 114024355 | 114021403 | 114021508 | 114052051 | 114052155 | 246 | 157 | [illegible] |
| SE | 109138 | ENSG00000185989 | RASA3 | chr13 | - | 114029810 | 114029887 | 114027842 | 114027927 | 114040999 | 114041094 | 572 | 228 | [illegible] |
| SE | 109139 | ENSG00000185989 | RASA3 | chr13 | - | 114029810 | 114029887 | 114027842 | 114027927 | 114052051 | 114052155 | 212 | 85 | [illegible] |
| SE | 109140 | ENSG00000185989 | RASA3 | chr13 | - | 114040999 | 114041094 | 114000825 | 114000932 | 114073719 | 114073837 | 2 | 2 | [illegible] |
| SE | 109141 | ENSG00000185989 | RASA3 | chr13 | - | 114040999 | 114041094 | 114021403 | 114021508 | 114052051 | 114052155 | 490 | 254 | [illegible] |
| SE | 109142 | ENSG00000185989 | RASA3 | chr13 | - | 114040999 | 114041094 | 114024278 | 114024355 | 114052051 | 114052155 | 491 | 254 | [illegible] |
| SE | 109143 | ENSG00000185989 | RASA3 | chr13 | - | 114040999 | 114041094 | 114024278 | 114024355 | 114073719 | 114073837 | 3 | 1 | [illegible] |
| SE | 109144 | ENSG00000185989 | RASA3 | chr13 | - | 114040999 | 114041094 | 114027846 | 114027927 | 114052051 | 114052155 | 490 | 254 | [illegible] |
| SE | 109146 | ENSG00000185989 | RASA3 | chr13 | - | 114040999 | 114041094 | 114029810 | 114029887 | 114073719 | 114073837 | 226 | 79 | [illegible] |
| SE | 109147 | ENSG00000185989 | RASA3 | chr13 | - | 114052051 | 114052155 | 114024278 | 114024355 | 114073719 | 114073837 | 228 | 124 | [illegible] |
| SE | 109148 | ENSG00000185989 | RASA3 | chr13 | - | 114052051 | 114052155 | 114029810 | 114029887 | 114073719 | 114073837 | 241 | 129 | [illegible] |
| SE | 109150 | ENSG00000185989 | RASA3 | chr13 | - | 114052051 | 114052155 | 114040999 | 114041094 | 114073719 | 114073837 | 849 | 438 | [illegible] |
| SE | 109151 | ENSG00000185989 | RASA3 | chr13 | - | 114073719 | 114073837 | 114029810 | 114029887 | 114132434 | 114132623 | 78 | 40 | [illegible] |
| SE | 109152 | ENSG00000185989 | RASA3 | chr13 | - | 114073719 | 114073837 | 114052051 | 114052155 | 114132434 | 114132623 | 505 | 275 | [illegible] |
| SE | 109154 | ENSG00000197442 | MAP3K5 | chr6 | - | 136558799 | 136558876 | 136557045 | 136557818 | 136561532 | 136561645 | 421 | 334 | [illegible] |
| SE | 109156 | ENSG00000197442 | MAP3K5 | chr6 | - | 136601021 | 136601042 | 136592436 | 136592614 | 136605208 | 136605366 | 5 | 6 | [illegible] |
| SE | 109157 | ENSG00000197442 | MAP3K5 | chr6 | - | 136601801 | 136601979 | 136592436 | 136592614 | 136605208 | 136605366 | 107 | 76 | [illegible] |
| SE | 109158 | ENSG00000197442 | MAP3K5 | chr6 | - | 136601801 | 136601979 | 136601021 | 136601042 | 136605208 | 136605366 | 196 | 129 | [illegible] |
| SE | 109159 | ENSG00000197442 | MAP3K5 | chr6 | - | 136611281 | 136611387 | 136605208 | 136605366 | 136613119 | 136613256 | 266 | 176 | [illegible] |
| SE | 109161 | ENSG00000197442 | MAP3K5 | chr6 | - | 136637324 | 136637406 | 136622847 | 136622981 | 136656306 | 136656460 | 87 | 62 | [illegible] |
| SE | 109162 | ENSG00000197442 | MAP3K5 | chr6 | - | 136639542 | 136639638 | 136637324 | 136637406 | 136642519 | 136642569 | 416 | 326 | [illegible] |
| SE | 109163 | ENSG00000197442 | MAP3K5 | chr6 | - | 136639542 | 136639638 | 136637324 | 136637406 | 136650983 | 136651091 | 135 | 69 | [illegible] |
| SE | 109165 | ENSG00000197442 | MAP3K5 | chr6 | - | 136642519 | 136642569 | 136637324 | 136637406 | 136650983 | 136651091 | 30 | 14 | [illegible] |
| SE | 109166 | ENSG00000197442 | MAP3K5 | chr6 | - | 136642519 | 136642569 | 136637324 | 136637406 | 136656306 | 136656460 | 3 | 4 | [illegible] |
| SE | 109168 | ENSG00000197442 | MAP3K5 | chr6 | - | 136642519 | 136642569 | 136639542 | 136639638 | 136656306 | 136656460 | 136 | 129 | [illegible] |
| SE | 109170 | ENSG00000197442 | MAP3K5 | chr6 | - | 136650983 | 136651091 | 136639542 | 136639638 | 136656306 | 136656460 | 82 | 74 | [illegible] |
| SE | 109172 | ENSG00000197442 | MAP3K5 | chr6 | - | 136669282 | 136669395 | 136659218 | 136659378 | 136694139 | 136694310 | 214 | 141 | [illegible] |
| SE | 109173 | ENSG00000197442 | MAP3K5 | chr6 | - | 136705109 | 136705133 | 136698488 | 136698682 | 136720449 | 136720589 | 160 | 121 | [illegible] |
| SE | 109177 | ENSG00000124496 | TRERF1 | chr6 | - | 42232680 | 42232892 | 42224930 | 42228669 | 42236204 | 42236411 | 63 | 42 | [illegible] |
| SE | 109179 | ENSG00000124496 | TRERF1 | chr6 | - | 42232680 | 42232928 | 42228148 | 42228669 | 42236204 | 42236411 | 46 | 33 | [illegible] |
| SE | 109183 | ENSG00000124496 | TRERF1 | chr6 | - | 42236204 | 42236411 | 42232680 | 42232928 | 42243247 | 42243361 | 45 | 46 | [illegible] |
| SE | 109184 | ENSG00000124496 | TRERF1 | chr6 | - | 42254850 | 42254926 | 42246455 | 42246544 | 42256727 | 42256831 | 62 | 37 | [illegible] |
| SE | 109185 | ENSG00000124496 | TRERF1 | chr6 | - | 42259338 | 42259723 | 42258134 | 42258201 | 42264703 | 42264854 | 59 | 18 | [illegible] |
| SE | 109190 | ENSG00000124496 | TRERF1 | chr6 | - | 42362946 | 42363079 | 42300637 | 42300749 | 42451176 | 42451261 | 71 | 40 | [illegible] |
| SE | 109191 | ENSG00000124496 | TRERF1 | chr6 | - | 42362996 | 42363079 | 42300637 | 42300749 | 42451176 | 42451261 | 55 | 57 | [illegible] |
| SE | 109207 | ENSG00000085662 | AKR1B1 | chr7 | - | 134448996 | 134449119 | 134448386 | 134448493 | 134449719 | 134449797 | 226 | 94 | [illegible] |
| SE | 109208 | ENSG00000085662 | AKR1B1 | chr7 | - | 134449719 | 134449797 | 134448996 | 134449119 | 134450785 | 134450902 | 160 | 60 | [illegible] |
| SE | 109211 | ENSG00000189057 | FAM111B | chr11 | + | 59109539 | 59109706 | 59108667 | 59108712 | 59124178 | 59127412 | 3 | 5 | [illegible] |
| SE | 109212 | ENSG00000173473 | SMARCC1 | chr3 | - | 47590660 | 47590837 | 47585268 | 47588306 | 47610065 | 47610327 | 108 | 33 | [illegible] |
| SE | 109213 | ENSG00000173473 | SMARCC1 | chr3 | - | 47635189 | 47635344 | 47622206 | 47622341 | 47636021 | 47636136 | 322 | 189 | [illegible] |
| SE | 109214 | ENSG00000173473 | SMARCC1 | chr3 | - | 47636021 | 47636136 | 47622206 | 47622341 | 47638724 | 47638780 | 131 | 118 | [illegible] |
| SE | 109215 | ENSG00000173473 | SMARCC1 | chr3 | - | 47638724 | 47638780 | 47636021 | 47636136 | 47661293 | 47661455 | 161 | 126 | [illegible] |
| SE | 109216 | ENSG00000173473 | SMARCC1 | chr3 | - | 47661293 | 47661455 | 47638724 | 47638780 | 47662333 | 47662592 | 272 | 200 | [illegible] |
| SE | 109218 | ENSG00000173473 | SMARCC1 | chr3 | - | 47670657 | 47670717 | 47662333 | 47662592 | 47675474 | 47675588 | 147 | 94 | [illegible] |
| SE | 109220 | ENSG00000173473 | SMARCC1 | chr3 | - | 47675474 | 47675588 | 47670349 | 47670717 | 47678197 | 47678311 | 89 | 58 | [illegible] |
| SE | 109221 | ENSG00000173473 | SMARCC1 | chr3 | - | 47676628 | 47676782 | 47670349 | 47670717 | 47678197 | 47678311 | 210 | 160 | [illegible] |
| SE | 109223 | ENSG00000173473 | SMARCC1 | chr3 | - | 47678197 | 47678311 | 47676628 | 47676782 | 47680436 | 47680508 | 449 | 356 | [illegible] |
| SE | 109224 | ENSG00000173473 | SMARCC1 | chr3 | - | 47680436 | 47680508 | 47678195 | 47678311 | 47686048 | 47686170 | 483 | 401 | [illegible] |
| SE | 109225 | ENSG00000173473 | SMARCC1 | chr3 | - | 47689386 | 47689424 | 47686048 | 47686170 | 47693240 | 47693300 | 219 | 149 | [illegible] |
| SE | 109226 | ENSG00000173473 | SMARCC1 | chr3 | - | 47689386 | 47689424 | 47686048 | 47686170 | 47701277 | 47701402 | 37 | 20 | [illegible] |
| SE | 109227 | ENSG00000173473 | SMARCC1 | chr3 | - | 47693240 | 47693300 | 47686048 | 47686170 | 47701277 | 47701402 | 29 | 28 | [illegible] |
| SE | 109228 | ENSG00000173473 | SMARCC1 | chr3 | - | 47693240 | 47693300 | 47689386 | 47689424 | 47701277 | 47701402 | 205 | 167 | [illegible] |
| SE | 109230 | ENSG00000173473 | SMARCC1 | chr3 | - | 47720665 | 47720735 | 47714414 | 47714490 | 47729024 | 47729094 | 346 | 278 | [illegible] |
| SE | 109231 | ENSG00000173473 | SMARCC1 | chr3 | - | 47720665 | 47720735 | 47714414 | 47714490 | 47736033 | 47736126 | 201 | 172 | [illegible] |
| SE | 109232 | ENSG00000173473 | SMARCC1 | chr3 | - | 47729024 | 47729094 | 47714414 | 47714490 | 47736033 | 47736126 | 57 | 43 | [illegible] |
| SE | 109234 | ENSG00000173473 | SMARCC1 | chr3 | - | 47736033 | 47736126 | 47729024 | 47729094 | 47738028 | 47738110 | 297 | 179 | [illegible] |
| SE | 109235 | ENSG00000173473 | SMARCC1 | chr3 | - | 47738028 | 47738110 | 47736033 | 47736126 | 47745907 | 47745993 | 254 | 134 | [illegible] |
| SE | 109236 | ENSG00000173473 | SMARCC1 | chr3 | - | 47745907 | 47745993 | 47738028 | 47738110 | 47772816 | 47772936 | 196 | 106 | [illegible] |
| SE | 109237 | ENSG00000173473 | SMARCC1 | chr3 | - | 47772816 | 47772936 | 47745907 | 47745993 | 47781602 | 47781881 | 138 | 75 | [illegible] |
| SE | 109241 | ENSG00000275066 | SYNRG | chr17 | - | 37535978 | 37536127 | 37520537 | 37520648 | 37538323 | 37538420 | 235 | 206 | [illegible] |
| SE | 109243 | ENSG00000275066 | SYNRG | chr17 | - | 37538323 | 37538420 | 37520537 | 37520648 | 37539191 | 37539245 | 142 | 74 | [illegible] |
| SE | 109244 | ENSG00000275066 | SYNRG | chr17 | - | 37538323 | 37538420 | 37535978 | 37536127 | 37539191 | 37539245 | 306 | 214 | [illegible] |
| SE | 109245 | ENSG00000275066 | SYNRG | chr17 | - | 37539191 | 37539245 | 37535978 | 37536127 | 37540379 | 37540480 | 95 | 74 | [illegible] |
| SE | 109247 | ENSG00000275066 | SYNRG | chr17 | - | 37553114 | 37554059 | 37541971 | 37542430 | 37561194 | 37561257 | 434 | 259 | [illegible] |
| SE | 109248 | ENSG00000275066 | SYNRG | chr17 | - | 37553114 | 37554059 | 37541971 | 37542565 | 37561194 | 37561257 | 599 | 382 | [illegible] |
| SE | 109249 | ENSG00000275066 | SYNRG | chr17 | - | 37557139 | 37557203 | 37553986 | 37554059 | 37561194 | 37561257 | 2 | 0 | [illegible] |
| SE | 109251 | ENSG00000275066 | SYNRG | chr17 | - | 37570636 | 37570885 | 37566848 | 37568924 | 37576340 | 37576418 | 133 | 82 | [illegible] |
| SE | 109252 | ENSG00000275066 | SYNRG | chr17 | - | 37571790 | 37571987 | 37568790 | 37568924 | 37576340 | 37576418 | 343 | 218 | [illegible] |
| SE | 109253 | ENSG00000275066 | SYNRG | chr17 | - | 37571790 | 37571987 | 37570636 | 37570885 | 37576340 | 37576418 | 532 | 334 | [illegible] |
| SE | 109254 | ENSG00000275066 | SYNRG | chr17 | - | 37576340 | 37576418 | 37571790 | 37571987 | 37584647 | 37584759 | 286 | 181 | [illegible] |
| SE | 109259 | ENSG00000275066 | SYNRG | chr17 | - | 37586418 | 37586549 | 37585324 | 37585430 | 37596222 | 37596341 | 339 | 185 | [illegible] |
| SE | 109261 | ENSG00000055147 | FAM114A2 | chr5 | - | 153994591 | 153994743 | 153992687 | 153993110 | 153994918 | 153994972 | 0 | 0 | [illegible] |
| SE | 109262 | ENSG00000055147 | FAM114A2 | chr5 | - | 154011240 | 154011320 | 154002846 | 154002969 | 154026398 | 154026522 | 90 | 54 | [illegible] |
| SE | 109263 | ENSG00000055147 | FAM114A2 | chr5 | - | 154029488 | 154029580 | 154028148 | 154028283 | 154033790 | 154033877 | 98 | 77 | [illegible] |
| SE | 109264 | ENSG00000055147 | FAM114A2 | chr5 | - | 154036241 | 154036385 | 154034743 | 154034967 | 154036721 | 154037310 | 0 | 0 | [illegible] |
| SE | 109265 | ENSG00000055147 | FAM114A2 | chr5 | - | 154036241 | 154036385 | 154034743 | 154034967 | 154038292 | 154038434 | 0 | 0 | [illegible] |
| SE | 109267 | ENSG00000055147 | FAM114A2 | chr5 | - | 154038292 | 154038434 | 154034743 | 154034967 | 154038856 | 154038880 | 31 | 4 | [illegible] |
| SE | 109268 | ENSG00000055147 | FAM114A2 | chr5 | - | 154038292 | 154038434 | 154034875 | 154034946 | 154038856 | 154038880 | 8 | 3 | [illegible] |
| SE | 109269 | ENSG00000055147 | FAM114A2 | chr5 | - | 154038387 | 154038434 | 154034743 | 154034967 | 154038856 | 154038921 | 2 | 0 | [illegible] |
| SE | 109270 | ENSG00000099821 | POLRMT | chr19 | - | 619585 | 619765 | 619209 | 619296 | 619957 | 620080 | 12 | 7 | [illegible] |
| SE | 109275 | ENSG00000198553 | KCNRG | chr13 | + | 50018822 | 50018920 | 50015253 | 50016071 | 50020213 | 50020920 | 0 | 0 | [illegible] |
| SE | 109276 | ENSG00000085832 | EPS15 | chr1 | - | 51361170 | 51361355 | 51354262 | 51356846 | 51363865 | 51364028 | 727 | 443 | [illegible] |
| SE | 109278 | ENSG00000085832 | EPS15 | chr1 | - | 51397110 | 51397211 | 51394211 | 51394447 | 51399031 | 51399165 | 2 | 1 | [illegible] |
| SE | 109280 | ENSG00000085832 | EPS15 | chr1 | - | 51400917 | 51400953 | 51394211 | 51394447 | 51402434 | 51402525 | 41 | 21 | [illegible] |
| SE | 109281 | ENSG00000085832 | EPS15 | chr1 | - | 51402434 | 51402525 | 51400917 | 51400953 | 51403418 | 51403451 | 325 | 187 | [illegible] |
| SE | 109283 | ENSG00000085832 | EPS15 | chr1 | - | 51405904 | 51406108 | 51403418 | 51403532 | 51409534 | 51409696 | 190 | 141 | [illegible] |
| SE | 109284 | ENSG00000085832 | EPS15 | chr1 | - | 51408134 | 51408332 | 51403418 | 51403532 | 51409534 | 51409696 | 157 | 113 | [illegible] |
| SE | 109285 | ENSG00000085832 | EPS15 | chr1 | - | 51408134 | 51408332 | 51405904 | 51406108 | 51409534 | 51409696 | 664 | 394 | [illegible] |
| SE | 109286 | ENSG00000085832 | EPS15 | chr1 | - | 51421785 | 51421858 | 51409534 | 51409696 | 51440346 | 51440432 | 1232 | 912 | [illegible] |
| SE | 109288 | ENSG00000085832 | EPS15 | chr1 | - | 51421785 | 51421858 | 51409534 | 51409696 | 51461090 | 51461150 | 144 | 111 | [illegible] |
| SE | 109289 | ENSG00000085832 | EPS15 | chr1 | - | 51440346 | 51440432 | 51409534 | 51409696 | 51444888 | 51445045 | 483 | 359 | [illegible] |
| SE | 109290 | ENSG00000085832 | EPS15 | chr1 | - | 51440346 | 51440432 | 51421785 | 51421858 | 51444888 | 51445045 | 1571 | 1169 | [illegible] |
| SE | 109291 | ENSG00000085832 | EPS15 | chr1 | - | 51444888 | 51445045 | 51409534 | 51409696 | 51461090 | 51461150 | 1 | 1 | [illegible] |
| SE | 109292 | ENSG00000085832 | EPS15 | chr1 | - | 51444888 | 51445045 | 51421785 | 51421858 | 51461090 | 51461150 | 4 | 7 | [illegible] |
| SE | 109293 | ENSG00000085832 | EPS15 | chr1 | - | 51446959 | 51447105 | 51444888 | 51445045 | 51461090 | 51461150 | 642 | 453 | [illegible] |
| SE | 109294 | ENSG00000085832 | EPS15 | chr1 | - | 51448045 | 51448135 | 51446959 | 51447105 | 51459040 | 51459148 | 114 | 104 | [illegible] |
| SE | 109298 | ENSG00000085832 | EPS15 | chr1 | - | 51459040 | 51459148 | 51458395 | 51458733 | 51461090 | 51461150 | 6 | 5 | [illegible] |
| SE | 109299 | ENSG00000085832 | EPS15 | chr1 | - | 51463672 | 51463798 | 51460943 | 51461150 | 51465260 | 51465326 | 526 | 266 | [illegible] |
| SE | 109300 | ENSG00000085832 | EPS15 | chr1 | - | 51465260 | 51465326 | 51463672 | 51463798 | 51468472 | 51468568 | 322 | 155 | [illegible] |
| SE | 109301 | ENSG00000085832 | EPS15 | chr1 | - | 51468472 | 51468568 | 51465260 | 51465326 | 51471689 | 51471737 | 441 | 231 | [illegible] |
| SE | 109303 | ENSG00000188549 | CCDC9B | chr15 | - | 40336573 | 40336664 | 40335783 | 40335826 | 40336759 | 40336813 | 0 | 3 | [illegible] |
| SE | 109310 | ENSG00000139597 | N4BP2L1 | chr13 | - | 32411527 | 32411592 | 32407644 | 32407772 | 32415958 | 32416029 | 0 | 0 | [illegible] |
| SE | 109311 | ENSG00000139597 | N4BP2L1 | chr13 | - | 32415958 | 32416029 | 32407644 | 32407772 | 32427903 | 32428130 | 0 | 2 | [illegible] |
| SE | 109312 | ENSG00000139597 | N4BP2L1 | chr13 | - | 32416550 | 32416703 | 32415958 | 32416029 | 32427903 | 32428311 | 7 | 10 | [illegible] |
| SE | 109313 | ENSG00000139597 | N4BP2L1 | chr13 | - | 32420370 | 32420576 | 32416498 | 32416703 | 32427903 | 32428130 | 4 | 1 | [illegible] |
| SE | 109316 | ENSG00000123104 | ITPR2 | chr12 | - | 26411319 | 26411412 | 26400127 | 26400258 | 26415302 | 26415498 | 101 | 64 | [illegible] |
| SE | 109317 | ENSG00000123104 | ITPR2 | chr12 | - | 26419048 | 26419213 | 26415302 | 26415498 | 26427912 | 26427937 | 103 | 45 | [illegible] |
| SE | 109318 | ENSG00000123104 | ITPR2 | chr12 | - | 26427912 | 26428088 | 26415302 | 26415498 | 26436220 | 26436346 | 68 | 49 | [illegible] |
| SE | 109320 | ENSG00000123104 | ITPR2 | chr12 | - | 26475295 | 26475418 | 26443542 | 26443650 | 26481130 | 26481241 | 75 | 97 | [illegible] |
| SE | 109321 | ENSG00000123104 | ITPR2 | chr12 | - | 26476911 | 26477007 | 26443542 | 26443650 | 26481130 | 26481241 | 45 | 27 | [illegible] |
| SE | 109322 | ENSG00000123104 | ITPR2 | chr12 | - | 26476911 | 26477007 | 26475295 | 26475418 | 26481130 | 26481241 | 126 | 98 | [illegible] |
| SE | 109324 | ENSG00000123104 | ITPR2 | chr12 | - | 26487067 | 26487251 | 26486103 | 26486360 | 26494152 | 26494233 | 247 | 359 | [illegible] |
| SE | 109328 | ENSG00000123104 | ITPR2 | chr12 | - | 26659112 | 26659285 | 26658010 | 26658130 | 26663684 | 26663846 | 155 | 218 | [illegible] |
| SE | 109329 | ENSG00000123104 | ITPR2 | chr12 | - | 26681873 | 26682034 | 26665909 | 26666051 | 26682573 | 26682673 | 351 | 405 | [illegible] |
| SE | 109331 | ENSG00000123104 | ITPR2 | chr12 | - | 26695605 | 26695650 | 26686480 | 26686632 | 26711172 | 26711268 | 165 | 202 | [illegible] |
| SE | 109332 | ENSG00000123104 | ITPR2 | chr12 | - | 26711172 | 26711268 | 26695601 | 26695650 | 26715298 | 26715445 | 209 | 285 | [illegible] |
| SE | 109333 | ENSG00000123104 | ITPR2 | chr12 | - | 26715298 | 26715445 | 26711172 | 26711268 | 26715751 | 26715835 | 196 | 272 | [illegible] |
| SE | 109334 | ENSG00000123104 | ITPR2 | chr12 | - | 26724655 | 26724742 | 26722396 | 26722555 | 26725649 | 26725765 | 284 | 265 | [illegible] |
| SE | 109335 | ENSG00000123104 | ITPR2 | chr12 | - | 26790156 | 26790227 | 26725649 | 26725765 | 26832689 | 26832781 | 113 | 100 | [illegible] |
| SE | 109337 | ENSG00000102710 | SUPT20H | chr13 | - | 37017244 | 37017328 | 37012191 | 37012297 | 37019341 | 37019397 | 49 | 34 | [illegible] |

| | | | | | | | | | | | | | | | | | | | | | | | | | | | | | | | | | | | | | | | | | | | | | |
|---|---|---|---|---|---|---|---|---|---|---|---|---|---|---|---|---|---|---|---|---|---|---|---|---|---|---|---|---|---|---|---|---|---|---|---|---|---|---|---|---|---|---|---|---|---|
| SE | 109340 | ENSG00000102710 | SUPT20H | chr13 | - | 37024034 | 37024193 | 37022010 | 37022080 | 37024339 | 37024442 | 188 | 130 | 209 | 189 | 412 | 0 | 1 | 0 | 0 | 0 | 101 | 198 | 153 | 109 | 27 | 0 | 0 | 0 | 0 | 0 | 298 | 149 | 1 | 1 | 1 | 0.985 | 1 | 1 | 1 | 1 | 1 | 1 | 1 | 1 | -0.003 |
| SE | 109341 | ENSG00000102710 | SUPT20H | chr13 | - | 37024339 | 37024442 | 37023548 | 37024193 | 37025319 | 37025437 | 211 | 133 | 210 | 187 | 513 | 0 | 1 | 0 | 0 | 1 | 100 | 132 | 149 | 82 | 20 | 0 | 0 | 0 | 0 | 0 | 252 | 149 | 1 | 1 | 1 | 0.987 | 1 | 1 | 0.997 | 1 | 1 | 1 | 1 | 1 | -0.003 |
| SE | 109343 | ENSG00000102710 | SUPT20H | chr13 | - | 37028147 | 37028305 | 37026789 | 37026816 | 37029764 | 37029836 | 344 | 233 | 354 | 352 | 589 | 0 | 0 | 0 | 0 | 0 | 101 | 304 | 175 | 181 | 13 | 0 | 1 | 0 | 0 | 0 | 298 | 149 | 1 | 1 | 1 | 1 | 1 | 1 | 1 | 1 | 0.993 | 1 | 1 | 1 | 0.001 |
| SE | 109344 | ENSG00000102710 | SUPT20H | chr13 | - | 37029764 | 37029836 | 37028147 | 37028305 | 37031566 | 37031623 | 158 | 104 | 162 | 212 | 240 | 1 | 0 | 6 | 0 | 2 | 55 | 185 | 111 | 91 | 19 | 0 | 0 | 0 | 0 | 0 | 221 | 149 | 1 | 1 | 0.991 | 1 | 0.948 | 1 | 0.988 | 1 | 1 | 1 | 1 | 1 | -0.015 |
| SE | 109345 | ENSG00000102710 | SUPT20H | chr13 | - | 37031566 | 37031623 | 37029764 | 37029836 | 37031738 | 37031895 | 107 | 76 | 104 | 149 | 121 | 4 | 0 | 0 | 0 | 0 | 57 | 135 | 80 | 73 | 19 | 0 | 0 | 0 | 0 | 0 | 206 | 149 | 1 | 1 | 0.951 | 1 | 1 | 1 | 1 | 1 | 1 | 1 | 1 | 1 | -0.01 |
| SE | 109347 | ENSG00000102710 | SUPT20H | chr13 | - | 37045246 | 37045373 | 37044077 | 37044181 | 37047345 | 37047408 | 157 | 100 | 146 | 187 | 376 | 0 | 0 | 0 | 2 | 2 | 63 | 75 | 71 | 73 | 23 | 0 | 0 | 0 | 0 | 0 | 276 | 149 | 1 | 1 | 1 | 1 | 1 | 0.981 | 0.99 | 1 | 1 | 1 | 1 | 1 | -0.006 |
| SE | 109349 | ENSG00000102710 | SUPT20H | chr13 | - | 37045246 | 37045376 | 37044077 | 37044181 | 37047345 | 37047408 | 167 | 95 | 141 | 176 | 364 | 0 | 0 | 0 | 2 | 2 | 63 | 76 | 76 | 71 | 25 | 0 | 0 | 0 | 0 | 0 | 279 | 149 | 1 | 1 | 1 | 1 | 1 | 0.979 | 0.99 | 1 | 1 | 1 | 1 | 1 | -0.006 |
| SE | 109354 | ENSG00000102710 | SUPT20H | chr13 | - | 37047877 | 37047936 | 37047534 | 37047601 | 37048563 | 37048599 | 307 | 217 | 202 | 259 | 341 | 0 | 0 | 0 | 1 | 0 | 103 | 225 | 122 | 171 | 103 | 0 | 0 | 0 | 0 | 0 | 208 | 149 | 1 | 1 | 1 | 1 | 1 | 0.995 | 1 | 1 | 1 | 1 | 1 | 1 | -0.001 |
| SE | 109355 | ENSG00000102710 | SUPT20H | chr13 | - | 37047877 | 37047936 | 37047534 | 37047601 | 37051487 | 37051583 | 143 | 143 | 114 | 154 | 175 | 1 | 0 | 0 | 0 | 0 | 52 | 76 | 67 | 78 | 21 | 0 | 0 | 0 | 0 | 0 | 208 | 149 | 1 | 1 | 0.99 | 1 | 1 | 1 | 1 | 1 | 1 | 1 | 1 | 1 | -0.002 |
| SE | 109356 | ENSG00000102710 | SUPT20H | chr13 | - | 37048563 | 37048599 | 37047534 | 37047601 | 37051487 | 37051559 | 14 | 9 | 10 | 13 | 7 | 1 | 0 | 0 | 0 | 0 | 7 | 10 | 3 | 13 | 5 | 0 | 0 | 0 | 0 | 0 | 185 | 149 | 1 | 1 | 0.919 | 1 | 1 | 1 | 1 | 1 | 1 | 1 | 1 | 1 | -0.016 |
| SE | 109359 | ENSG00000186184 | POLR1D | chr13 | + | 27621336 | 27621437 | 27620742 | 27621140 | 27648378 | 27648444 | 7 | 0 | 3 | 4 | 2 | 0 | 0 | 0 | 0 | 0 | 0 | 0 | 4 | 4 | 10 | 0 | 0 | 4 | 0 | 0 | 250 | 149 | 1 | 1 | 1 | NA | 1 | 1 | 1 | NA | NA | 0.373 | 1 | 1 | 0.209 |
| SE | 109360 | ENSG00000186184 | POLR1D | chr13 | + | 27646193 | 27646218 | 27621892 | 27622009 | 27648378 | 27648453 | 5 | 1 | 0 | 0 | 0 | 127 | 50 | 97 | 67 | 129 | 0 | 0 | 0 | 0 | 0 | 117 | 78 | 83 | 109 | 144 | 174 | 149 | 1 | 1 | 0.033 | 0.017 | 0 | 0 | 0 | 0 | 0 | 0 | 0 | 0 | 0.01 |
| SE | 109361 | ENSG00000186184 | POLR1D | chr13 | + | 27648378 | 27648449 | 27621874 | 27622009 | 27665685 | 27667410 | 28 | 8 | 25 | 25 | 3 | 2 | 0 | 1 | 0 | 0 | 30 | 27 | 15 | 54 | 56 | 0 | 0 | 2 | 0 | 0 | 220 | 149 | 1 | 1 | 0.905 | 1 | 0.944 | 1 | 1 | 1 | 1 | 0.836 | 1 | 1 | 0.003 |
| SE | 109362 | ENSG00000186184 | POLR1D | chr13 | + | 27648378 | 27648453 | 27621778 | 27622009 | 27665685 | 27667022 | 210 | 87 | 140 | 131 | 234 | 2 | 0 | 1 | 0 | 0 | 232 | 128 | 139 | 183 | 255 | 0 | 0 | 2 | 0 | 0 | 224 | 149 | 1 | 1 | 0.986 | 1 | 0.989 | 1 | 1 | 1 | 1 | 0.979 | 1 | 1 | -0.001 |
| SE | 109380 | ENSG00000166226 | CCT2 | chr12 | + | 69585750 | 69586009 | 69585459 | 69585524 | 69586269 | 69586344 | 1 | 0 | 0 | 3 | 0 | 53 | 40 | 82 | 69 | 51 | 0 | 4 | 1 | 0 | 0 | 51 | 51 | 58 | 43 | 34 | 298 | 149 | 1 | 1 | 0.009 | 0 | 0 | 0.021 | 0 | 0 | 0.038 | 0.009 | 0 | 0 | -0.003 |
| SE | 109381 | ENSG00000166226 | CCT2 | chr12 | + | 69586269 | 69586344 | 69585425 | 69585524 | 69586752 | 69586818 | 112 | 79 | 158 | 114 | 110 | 0 | 0 | 1 | 0 | 0 | 81 | 119 | 119 | 75 | 55 | 0 | 0 | 0 | 0 | 0 | 224 | 149 | 1 | 1 | 1 | 1 | 0.991 | 1 | 1 | 1 | 1 | 1 | 1 | 1 | -0.002 |
| SE | 109382 | ENSG00000166226 | CCT2 | chr12 | + | 69589484 | 69589687 | 69588149 | 69588262 | 69592058 | 69592159 | 594 | 370 | 573 | 525 | 554 | 0 | 0 | 1 | 1 | 0 | 517 | 321 | 402 | 380 | 352 | 0 | 0 | 0 | 0 | 0 | 298 | 149 | 1 | 1 | 1 | 1 | 0.997 | 0.996 | 1 | 1 | 1 | 1 | 1 | 1 | -0.001 |
| SE | 109385 | ENSG00000166226 | CCT2 | chr12 | + | 69598321 | 69598421 | 69597967 | 69598071 | 69599862 | 69600004 | 266 | 177 | 248 | 268 | 314 | 1 | 0 | 0 | 0 | 0 | 241 | 219 | 188 | 202 | 152 | 0 | 0 | 0 | 0 | 0 | 249 | 149 | 1 | 1 | 0.994 | 1 | 1 | 1 | 1 | 1 | 1 | 1 | 1 | 1 | -0.001 |
| SE | 109389 | ENSG00000167186 | COQ7 | chr16 | + | 19075720 | 19075860 | 19071927 | 19072106 | 19077305 | 19077374 | 18 | 7 | 4 | 14 | 12 | 0 | 1 | 0 | 2 | 0 | 26 | 28 | 8 | 3 | 12 | 0 | 0 | 0 | 0 | 0 | 289 | 149 | 1 | 1 | 1 | 0.783 | 1 | 0.783 | 1 | 1 | 1 | 1 | 1 | 1 | -0.087 |
| SE | 109390 | ENSG00000167186 | COQ7 | chr16 | + | 19077305 | 19077374 | 19075720 | 19075860 | 19078080 | 19078159 | 19 | 8 | 4 | 17 | 12 | 0 | 1 | 0 | 0 | 0 | 26 | 26 | 8 | 3 | 12 | 0 | 0 | 0 | 0 | 0 | 218 | 149 | 1 | 1 | 1 | 0.845 | 1 | 1 | 1 | 1 | 1 | 1 | 1 | 1 | -0.031 |
| SE | 109391 | ENSG00000134900 | TPP2 | chr13 | + | 102614100 | 102614196 | 102604792 | 102604921 | 102616395 | 102616500 | 422 | 391 | 530 | 476 | 790 | 2 | 0 | 0 | 0 | 0 | 253 | 416 | 515 | 293 | 282 | 0 | 0 | 6 | 0 | 0 | 245 | 149 | 1 | 1 | 0.992 | 1 | 1 | 1 | 1 | 1 | 1 | 0.981 | 1 | 1 | 0.002 |
| SE | 109393 | ENSG00000134900 | TPP2 | chr13 | + | 102622876 | 102623040 | 102618721 | 102618846 | 102627011 | 102627166 | 844 | 658 | 759 | 656 | 813 | 1 | 2 | 1 | 0 | 1 | 329 | 668 | 833 | 367 | 222 | 0 | 0 | 0 | 0 | 0 | 298 | 149 | 1 | 1 | 0.998 | 0.994 | 0.997 | 1 | 0.998 | 1 | 1 | 1 | 1 | 1 | -0.003 |
| SE | 109394 | ENSG00000134900 | TPP2 | chr13 | + | 102627011 | 102627166 | 102622876 | 102623040 | 102633949 | 102634098 | 414 | 322 | 379 | 303 | 517 | 0 | 0 | 0 | 2 | 0 | 133 | 320 | 384 | 175 | 135 | 0 | 0 | 0 | 0 | 0 | 298 | 149 | 1 | 1 | 1 | 1 | 1 | 0.987 | 1 | 1 | 1 | 1 | 1 | 1 | -0.003 |
| SE | 109395 | ENSG00000134900 | TPP2 | chr13 | + | 102627847 | 102627924 | 102627011 | 102627166 | 102629481 | 102629609 | 406 | 256 | 370 | 376 | 480 | 0 | 0 | 0 | 0 | 0 | 94 | 276 | 268 | 178 | 66 | 0 | 2 | 0 | 0 | 0 | 226 | 149 | 1 | 1 | 1 | 1 | 1 | 1 | 1 | 1 | 0.989 | 1 | 1 | 1 | 0.002 |
| SE | 109396 | ENSG00000134900 | TPP2 | chr13 | + | 102627847 | 102627924 | 102627011 | 102627166 | 102633949 | 102634098 | 150 | 78 | 108 | 131 | 123 | 0 | 1 | 0 | 0 | 0 | 27 | 109 | 107 | 70 | 26 | 0 | 0 | 5 | 0 | 0 | 226 | 149 | 1 | 1 | 1 | 0.981 | 1 | 1 | 1 | 1 | 1 | 0.934 | 1 | 1 | 0.009 |
| SE | 109397 | ENSG00000134900 | TPP2 | chr13 | + | 102629481 | 102629609 | 102627011 | 102627166 | 102633949 | 102634098 | 1 | 0 | 2 | 2 | 0 | 0 | 1 | 0 | 0 | 0 | 0 | 2 | 2 | 2 | 0 | 0 | 0 | 5 | 0 | 0 | 277 | 149 | 1 | 1 | 1 | 0 | 1 | 1 | NA | NA | 1 | 0.177 | 1 | NA | 0.024 |
| SE | 109403 | ENSG00000134900 | TPP2 | chr13 | + | 102638238 | 102638315 | 102637081 | 102637239 | 102640269 | 102640376 | 182 | 144 | 204 | 173 | 264 | 0 | 0 | 2 | 1 | 4 | 49 | 147 | 74 | 51 | 47 | 0 | 0 | 0 | 1 | 0 | 226 | 149 | 1 | 1 | 1 | 1 | 0.985 | 0.991 | 0.978 | 1 | 1 | 1 | 0.971 | 1 | -0.003 |
| SE | 109404 | ENSG00000134900 | TPP2 | chr13 | + | 102643221 | 102643376 | 102640269 | 102640376 | 102644556 | 102644673 | 285 | 188 | 272 | 302 | 316 | 0 | 0 | 0 | 1 | 0 | 70 | 138 | 131 | 63 | 31 | 0 | 0 | 0 | 0 | 0 | 298 | 149 | 1 | 1 | 1 | 1 | 1 | 0.993 | 1 | 1 | 1 | 1 | 1 | 1 | -0.001 |
| SE | 109405 | ENSG00000134900 | TPP2 | chr13 | + | 102644556 | 102644673 | 102643221 | 102643376 | 102644908 | 102645009 | 369 | 267 | 348 | 396 | 432 | 0 | 0 | 0 | 0 | 0 | 54 | 186 | 216 | 89 | 35 | 0 | 2 | 0 | 0 | 0 | 266 | 149 | 1 | 1 | 1 | 1 | 1 | 1 | 1 | 1 | 0.981 | 1 | 1 | 1 | 0.004 |
| SE | 109406 | ENSG00000134900 | TPP2 | chr13 | + | 102644908 | 102645009 | 102644556 | 102644673 | 102646293 | 102646390 | 266 | 195 | 268 | 305 | 341 | 0 | 0 | 0 | 0 | 0 | 35 | 167 | 168 | 71 | 29 | 0 | 0 | 1 | 0 | 0 | 250 | 149 | 1 | 1 | 1 | 1 | 1 | 1 | 1 | 1 | 1 | 0.99 | 1 | 1 | 0.002 |
| SE | 109409 | ENSG00000134900 | TPP2 | chr13 | + | 102647206 | 102647344 | 102644908 | 102646390 | 102649407 | 102649486 | 201 | 180 | 234 | 222 | 327 | 0 | 0 | 0 | 1 | 0 | 67 | 106 | 123 | 60 | 6 | 0 | 0 | 0 | 0 | 0 | 287 | 149 | 1 | 1 | 1 | 1 | 1 | 0.991 | 1 | 1 | 1 | 1 | 1 | 1 | -0.002 |
| SE | 109410 | ENSG00000134900 | TPP2 | chr13 | + | 102648906 | 102649151 | 102647206 | 102647344 | 102649407 | 102649486 | 682 | 484 | 650 | 634 | 1267 | 0 | 0 | 0 | 0 | 0 | 136 | 376 | 364 | 180 | 19 | 0 | 0 | 3 | 3 | 0 | 298 | 149 | 1 | 1 | 1 | 1 | 1 | 1 | 1 | 1 | 1 | 0.984 | 0.968 | 1 | 0.01 |
| SE | 109411 | ENSG00000134900 | TPP2 | chr13 | + | 102649407 | 102649486 | 102648906 | 102649151 | 102657055 | 102657207 | 327 | 218 | 295 | 301 | 504 | 0 | 0 | 1 | 2 | 0 | 65 | 214 | 174 | 106 | 9 | 0 | 0 | 0 | 2 | 0 | 228 | 149 | 1 | 1 | 1 | 1 | 0.995 | 0.99 | 1 | 1 | 1 | 1 | 0.972 | 1 | 0.003 |
| SE | 109415 | ENSG00000134900 | TPP2 | chr13 | + | 102651358 | 102651397 | 102649407 | 102649486 | 102663647 | 102663744 | 7 | 9 | 5 | 10 | 9 | 0 | 0 | 1 | 0 | 0 | 8 | 7 | 17 | 3 | 0 | 0 | 0 | 0 | 0 | 0 | 188 | 149 | 1 | 1 | 1 | 1 | 0.798 | 1 | 1 | 1 | 1 | 1 | 1 | NA | -0.04 |
| SE | 109416 | ENSG00000134900 | TPP2 | chr13 | + | 102657055 | 102657207 | 102649407 | 102649486 | 102663647 | 102663744 | 425 | 320 | 378 | 360 | 674 | 0 | 0 | 1 | 0 | 0 | 89 | 257 | 192 | 111 | 45 | 0 | 0 | 0 | 0 | 0 | 298 | 149 | 1 | 1 | 1 | 1 | 0.995 | 1 | 1 | 1 | 1 | 1 | 1 | 1 | -0.001 |
| SE | 109417 | ENSG00000134900 | TPP2 | chr13 | + | 102657055 | 102657207 | 102651338 | 102651397 | 102663647 | 102663744 | 287 | 237 | 284 | 255 | 404 | 0 | 0 | 0 | 0 | 1 | 47 | 169 | 158 | 81 | 43 | 0 | 0 | 0 | 0 | 0 | 298 | 149 | 1 | 1 | 1 | 1 | 1 | 1 | 0.995 | 1 | 1 | 1 | 1 | 1 | -0.001 |
| SE | 109418 | ENSG00000134900 | TPP2 | chr13 | + | 102663647 | 102663744 | 102656941 | 102657207 | 102664794 | 102664925 | 420 | 321 | 392 | 365 | 481 | 0 | 0 | 1 | 0 | 0 | 59 | 222 | 189 | 106 | 66 | 0 | 0 | 0 | 0 | 0 | 246 | 149 | 1 | 1 | 1 | 1 | 0.996 | 1 | 1 | 1 | 1 | 1 | 1 | 1 | -0.001 |
| SE | 109419 | ENSG00000134900 | TPP2 | chr13 | + | 102667757 | 102667883 | 102664794 | 102664925 | 102674282 | 102674411 | 13 | 8 | 22 | 8 | 11 | 228 | 170 | 260 | 253 | 283 | 0 | 5 | 6 | 12 | 0 | 0 | 196 | 174 | 82 | 22 | 275 | 149 | 1 | 1 | 0.03 | 0.025 | 0.044 | 0.017 | 0.021 | NA | 0.014 | 0.018 | 0.073 | 0 | 0.001 |
| SE | 109420 | ENSG00000134900 | TPP2 | chr13 | + | 102667757 | 102667883 | 102664794 | 102664936 | 102674282 | 102674411 | 14 | 7 | 10 | 5 | 3 | 35 | 30 | 38 | 36 | 50 | 0 | 3 | 3 | 8 | 0 | 7 | 12 | 22 | 0 | 0 | 275 | 149 | 1 | 1 | 0.178 | 0.112 | 0.125 | 0.07 | 0.031 | 0 | 0.119 | 0.069 | 1 | NA | -0.194 |
| SE | 109421 | ENSG00000134900 | TPP2 | chr13 | + | 102676295 | 102676415 | 102673835 | 102674490 | 102678226 | 102678232 | 436 | 342 | 436 | 348 | 611 | 1 | 0 | 0 | 0 | 0 | 52 | 295 | 213 | 124 | 27 | 0 | 0 | 0 | 0 | 0 | 269 | 149 | 1 | 1 | 0.996 | 1 | 1 | 1 | 1 | 1 | 1 | 1 | 1 | 1 | -0.001 |
| SE | 109422 | ENSG00000134901 | POGLUT2 | chr13 | - | 102786231 | 102786339 | 102784280 | 102784512 | 102787833 | 102787923 | 1 | 0 | 6 | 6 | 4 | 0 | 0 | 1 | 0 | 0 | 0 | 3 | 1 | 0 | 0 | 0 | 0 | 0 | 0 | 0 | 257 | 149 | 1 | 1 | 1 | NA | 0.777 | 1 | 1 | NA | 1 | 1 | NA | NA | -0.056 |
| SE | 109434 | ENSG00000270181 | BIVM-ERCC5 | chr13 | + | 102853756 | 102853872 | 102852117 | 102852293 | 102854287 | 102854374 | 330 | 258 | 348 | 338 | 454 | 0 | 0 | 0 | 0 | 1 | 72 | 231 | 210 | 174 | 39 | 0 | 0 | 0 | 6 | 0 | 265 | 149 | 1 | 1 | 1 | 1 | 1 | 1 | 0.996 | 1 | 1 | 1 | 0.942 | 1 | 0.011 |
| SE | 109435 | ENSG00000270181 | BIVM-ERCC5 | chr13 | + | 102856051 | 102856112 | 102854287 | 102854374 | 102858274 | 102858418 | 517 | 428 | 556 | 478 | 674 | 3 | 1 | 0 | 0 | 0 | 86 | 410 | 452 | 251 | 51 | 0 | 0 | 10 | 0 | 0 | 210 | 149 | 1 | 1 | 0.992 | 0.997 | 1 | 1 | 1 | 1 | 1 | 0.97 | 1 | 1 | 0.004 |
| SE | 109436 | ENSG00000270181 | BIVM-ERCC5 | chr13 | + | 102858274 | 102858418 | 102856051 | 102856112 | 102861506 | 102861714 | 899 | 801 | 982 | 889 | 993 | 0 | 0 | 0 | 0 | 0 | 380 | 760 | 1013 | 560 | 120 | 0 | 0 | 0 | 2 | 0 | 293 | 149 | 1 | 1 | 1 | 1 | 1 | 1 | 1 | 1 | 1 | 1 | 0.993 | 1 | 0.001 |
| SE | 109437 | ENSG00000270181 | BIVM-ERCC5 | chr13 | + | 102865666 | 102865911 | 102862029 | 102863103 | 102866261 | 102866381 | 389 | 300 | 368 | 380 | 503 | 0 | 0 | 0 | 0 | 1 | 132 | 363 | 404 | 215 | 37 | 0 | 0 | 1 | 0 | 0 | 298 | 149 | 1 | 1 | 1 | 1 | 1 | 1 | 0.996 | 1 | 1 | 0.995 | 1 | 1 | 0 |
| SE | 109438 | ENSG00000121644 | DESI2 | chr1 | + | 244686596 | 244686669 | 244653102 | 244653355 | 244689248 | 244689342 | 65 | 39 | 38 | 49 | 76 | 2 | 0 | 0 | 0 | 0 | 17 | 36 | 43 | 27 | 5 | 0 | 0 | 0 | 0 | 0 | 222 | 149 | 1 | 1 | 0.956 | 1 | 1 | 1 | 1 | 1 | 1 | 1 | 1 | 1 | -0.009 |
| SE | 109441 | ENSG00000136146 | MED4 | chr13 | - | 48077152 | 48077311 | 48053322 | 48053491 | 48079843 | 48079975 | 559 | 460 | 484 | 518 | 675 | 1 | 0 | 0 | 3 | 1 | 182 | 352 | 402 | 258 | 87 | 0 | 0 | 3 | 0 | 0 | 298 | 149 | 1 | 1 | 0.996 | 1 | 1 | 0.989 | 0.997 | 1 | 1 | 0.985 | 1 | 1 | -0.001 |
| SE | 109442 | ENSG00000136146 | MED4 | chr13 | - | 48079843 | 48079975 | 48075723 | 48077311 | 48081644 | 48081731 | 942 | 744 | 782 | 838 | 1085 | 1 | 0 | 0 | 0 | 0 | 294 | 610 | 633 | 392 | 171 | 0 | 0 | 0 | 0 | 0 | 281 | 149 | 1 | 1 | 0.998 | 1 | 1 | 1 | 1 | 1 | 1 | 1 | 1 | 1 | 0 |
| SE | 109443 | ENSG00000136146 | MED4 | chr13 | - | 48081644 | 48081731 | 48079843 | 48079975 | 48086281 | 48086452 | 290 | 231 | 235 | 236 | 341 | 0 | 2 | 0 | 0 | 0 | 113 | 196 | 197 | 123 | 75 | 0 | 0 | 0 | 0 | 0 | 236 | 149 | 1 | 1 | 1 | 0.986 | 1 | 1 | 1 | 1 | 1 | 1 | 1 | 1 | -0.003 |
| SE | 109445 | ENSG00000136146 | MED4 | chr13 | - | 48090351 | 48090418 | 48086281 | 48086452 | 48094953 | 48095104 | 252 | 244 | 198 | 199 | 242 | 0 | 2 | 0 | 0 | 0 | 120 | 148 | 225 | 160 | 80 | 0 | 0 | 3 | 0 | 0 | 216 | 149 | 1 | 1 | 1 | 0.988 | 1 | 1 | 1 | 1 | 1 | 0.981 | 1 | 1 | 0.001 |
| SE | 109446 | ENSG00000136146 | MED4 | chr13 | - | 48093547 | 48093687 | 48090351 | 48090418 | 48094953 | 48095068 | 2 | 3 | 0 | 5 | 0 | 191 | 159 | 118 | 132 | 168 | 0 | 0 | 5 | 0 | 0 | 67 | 83 | 128 | 67 | 42 | 289 | 149 | 1 | 1 | 0.005 | 0.01 | 0 | 0.019 | 0 | 0 | 0 | 0.02 | 0 | 0 | 0.003 |
| SE | 109447 | ENSG00000136146 | MED4 | chr13 | - | 48093547 | 48093687 | 48090351 | 48090418 | 48095059 | 48095131 | 0 | 3 | 0 | 4 | 0 | 30 | 21 | 25 | 8 | 12 | 0 | 0 | 5 | 0 | 0 | 5 | 8 | 15 | 31 | 0 | 289 | 149 | 1 | 1 | 0 | 0.069 | 0 | 0.205 | 0 | 0 | 0 | 0.147 | 0 | NA | 0.018 |
| SE | 109448 | ENSG00000139687 | RB1 | chr13 | + | 48307279 | 48307406 | 48303743 | 48304049 | 48342598 | 48342714 | 146 | 166 | 85 | 125 | 153 | 1 | 1 | 0 | 0 | 0 | 96 | 115 | 54 | 70 | 29 | 0 | 0 | 3 | 0 | 0 | 276 | 149 | 1 | 1 | 0.987 | 0.989 | 1 | 1 | 1 | 1 | 1 | 0.907 | 1 | 1 | 0.014 |
| SE | 109449 | ENSG00000139687 | RB1 | chr13 | + | 48342598 | 48342714 | 48307279 | 48307406 | 48345079 | 48345199 | 433 | 446 | 275 | 408 | 501 | 0 | 0 | 0 | 3 | 0 | 258 | 324 | 238 | 245 | 67 | 0 | 0 | 0 | 0 | 0 | 265 | 149 | 1 | 1 | 1 | 1 | 1 | 0.987 | 1 | 1 | 1 | 1 | 1 | 1 | -0.003 |
| SE | 109450 | ENSG00000139687 | RB1 | chr13 | + | 48347824 | 48347863 | 48345079 | 48345199 | 48348955 | 48349023 | 408 | 417 | 290 | 370 | 649 | 0 | 0 | 0 | 0 | 0 | 272 | 309 | 210 | 184 | 71 | 0 | 0 | 2 | 0 | 0 | 188 | 149 | 1 | 1 | 1 | 1 | 1 | 1 | 1 | 1 | 1 | 0.988 | 1 | 1 | 0.002 |
| SE | 109452 | ENSG00000139687 | RB1 | chr13 | + | 48364893 | 48364971 | 48362814 | 48362957 | 48367493 | 48367603 | 272 | 276 | 190 | 303 | 834 | 2 | 0 | 0 | 0 | 0 | 63 | 192 | 143 | 140 | 27 | 0 | 0 | 0 | 0 | 0 | 227 | 149 | 1 | 1 | 0.989 | 1 | 1 | 1 | 1 | 1 | 1 | 1 | 1 | 1 | -0.002 |
| SE | 109453 | ENSG00000139687 | RB1 | chr13 | + | 48376917 | 48377034 | 48373404 | 48373492 | 48380052 | 48380084 | 169 | 157 | 110 | 183 | 331 | 0 | 0 | 0 | 1 | 0 | 45 | 70 | 64 | 62 | 28 | 0 | 0 | 2 | 0 | 0 | 266 | 149 | 1 | 1 | 1 | 1 | 1 | 0.99 | 1 | 1 | 1 | 0.947 | 1 | 1 | 0.009 |
| SE | 109454 | ENSG00000139687 | RB1 | chr13 | + | 48379593 | 48379650 | 48376917 | 48377034 | 48380052 | 48380084 | 317 | 230 | 177 | 322 | 475 | 1 | 0 | 0 | 0 | 4 | 57 | 155 | 133 | 125 | 64 | 0 | 0 | 0 | 5 | 0 | 206 | 149 | 1 | 1 | 0.996 | 1 | 1 | 1 | 0.988 | 1 | 1 | 1 | 0.948 | 1 | 0.007 |
| SE | 109456 | ENSG00000139687 | RB1 | chr13 | + | 48464997 | 48465111 | 48459687 | 48459833 | 48465204 | 48465368 | 67 | 50 | 36 | 39 | 163 | 2 | 0 | 0 | 0 | 0 | 18 | 32 | 9 | 20 | 11 | 0 | 0 | 0 | 0 | 0 | 263 | 149 | 1 | 1 | 0.95 | 1 | 1 | 1 | 1 | 1 | 1 | 1 | 1 | 1 | -0.01 |
| SE | 109457 | ENSG00000153395 | LPCAT1 | chr5 | - | 1470825 | 1470924 | 1466748 | 1466890 | 1477403 | 1477486 | 203 | 44 | 102 | 88 | 80 | 0 | 0 | 0 | 0 | 0 | 86 | 90 | 29 | 102 | 86 | 0 | 0 | 3 | 0 | 0 | 248 | 149 | 1 | 1 | 1 | 1 | 1 | 1 | 1 | 1 | 1 | 0.853 | 1 | 1 | 0.029 |
| SE | 109458 | ENSG00000153395 | LPCAT1 | chr5 | - | 1473956 | 1474110 | 1470825 | 1470924 | 1474559 | 1474685 | 735 | 373 | 532 | 666 | 439 | 0 | 0 | 0 | 0 | 0 | 881 | 694 | 314 | 647 | 755 | 0 | 0 | 3 | 1 | 0 | 298 | 149 | 1 | 1 | 1 | 1 | 1 | 1 | 1 | 1 | 1 | 0.981 | 0.997 | 1 | 0.004 |
| SE | 109459 | ENSG00000153395 | LPCAT1 | chr5 | - | 1473956 | 1474110 | 1470825 | 1470924 | 1477403 | 1477486 | 449 | 204 | 286 | 331 | 293 | 0 | 0 | 2 | 0 | 0 | 438 | 277 | 130 | 341 | 377 | 0 | 0 | 0 | 0 | 0 | 298 | 149 | 1 | 1 | 1 | 1 | 0.986 | 1 | 1 | 1 | 1 | 1 | 1 | 1 | -0.003 |
| SE | 109460 | ENSG00000153395 | LPCAT1 | chr5 | - | 1474559 | 1474685 | 1470825 | 1470924 | 1477403 | 1477486 | 187 | 135 | 150 | 214 | 113 | 0 | 0 | 2 | 0 | 0 | 168 | 290 | 92 | 211 | 228 | 0 | 0 | 0 | 0 | 0 | 275 | 149 | 1 | 1 | 1 | 1 | 0.976 | 1 | 1 | 1 | 1 | 1 | 1 | 1 | -0.005 |
| SE | 109461 | ENSG00000153395 | LPCAT1 | chr5 | - | 1474559 | 1474685 | 1473956 | 1474110 | 1477403 | 1477486 | 472 | 305 | 396 | 549 | 259 | 0 | 1 | 0 | 0 | 0 | 611 | 707 | 273 | 517 | 595 | 0 | 0 | 0 | 1 | 0 | 275 | 149 | 1 | 1 | 1 | 0.994 | 1 | 1 | 1 | 1 | 1 | 1 | 0.996 | 1 | 0 |
| SE | 109462 | ENSG00000153395 | LPCAT1 | chr5 | - | 1479620 | 1479675 | 1477403 | 1477486 | 1480941 | 1480976 | 346 | 207 | 227 | 328 | 273 | 1 | 0 | 0 | 0 | 0 | 258 | 300 | 86 | 213 | 281 | 0 | 1 | 0 | 0 | 0 | 204 | 149 | 1 | 1 | 0.996 | 1 | 1 | 1 | 1 | 1 | 0.995 | 1 | 1 | 1 | 0 |
| SE | 109463 | ENSG00000153395 | LPCAT1 | chr5 | - | 1494699 | 1494914 | 1489745 | 1489858 | 1501460 | 1501603 | 306 | 83 | 157 | 242 | 106 | 1 | 0 | 0 | 0 | 0 | 124 | 201 | 81 | 124 | 163 | 0 | 0 | 1 | 0 | 0 | 298 | 149 | 1 | 1 | 0.994 | 1 | 1 | 1 | 1 | 1 | 1 | 0.976 | 1 | 1 | 0.004 |
| SE | 109465 | ENSG00000139684 | ESD | chr13 | - | 46779934 | 46780033 | 46775385 | 46777623 | 46781495 | 46781615 | 435 | 267 | 350 | 416 | 893 | 0 | 0 | 0 | 1 | 5 | 162 | 212 | 215 | 207 | 72 | 0 | 0 | 2 | 0 | 0 | 248 | 149 | 1 | 1 | 1 | 1 | 1 | 0.996 | 0.991 | 1 | 1 | 0.985 | 1 | 1 | 0 |
| SE | 109466 | ENSG00000139684 | ESD | chr13 | - | 46784251 | 46784350 | 46782666 | 46782791 | 46787020 | 46787109 | 316 | 142 | 301 | 196 | 698 | 0 | 0 | 0 | 0 | 1 | 129 | 118 | 103 | 152 | 44 | 0 | 0 | 0 | 0 | 0 | 248 | 149 | 1 | 1 | 1 | 1 | 1 | 1 | 0.998 | 1 | 1 | 1 | 1 | 1 | 0 |
| SE | 109467 | ENSG00000139684 | ESD | chr13 | - | 46785648 | 46785719 | 46784251 | 46784350 | 46787020 | 46787109 | 4 | 0 | 0 | 5 | 2 | 168 | 75 | 174 | 98 | 339 | 0 | 0 | 0 | 4 | 0 | 81 | 58 | 50 | 72 | 14 | 220 | 149 | 1 | 1 | 0.016 | 0 | 0 | 0.033 | 0.004 | 0 | 0 | 0 | 0.036 | 0 | 0.003 |
| SE | 109468 | ENSG00000139684 | ESD | chr13 | - | 46790596 | 46790661 | 46787020 | 46787109 | 46791345 | 46791420 | 4 | 0 | 1 | 5 | 1 | 123 | 78 | 157 | 111 | 180 | 0 | 2 | 1 | 4 | 0 | 99 | 60 | 43 | 51 | 9 | 214 | 149 | 1 | 1 | 0.022 | 0 | 0.004 | 0.03 | 0.004 | 0 | 0.023 | 0.016 | 0 | 0 | 0.004 |
| SE | 109471 | ENSG00000133104 | SPART | chr13 | - | 36308418 | 36308444 | 36304450 | 36304632 | 36312144 | 36312235 | 0 | 0 | 0 | 0 | 0 | 230 | 163 | 173 | 221 | 342 | 0 | 0 | 1 | 4 | 0 | 31 | 133 | 163 | 160 | 5 | 175 | 149 | 1 | 1 | 0 | 0 | 0 | 0 | 0 | 0 | 0 | 0.005 | 0.021 | 0 | -0.005 |
| SE | 109472 | ENSG00000133104 | SPART | chr13 | - | 36313967 | 36314082 | 36312318 | 36312477 | 36314226 | 36314441 | 9 | 11 | 2 | 5 | 4 | 62 | 33 | 62 | 50 | 64 | 0 | 0 | 4 | 0 | 0 | 2 | 33 | 29 | 30 | 0 | 264 | 149 | 1 | 1 | 0.076 | 0.158 | 0.018 | 0.053 | 0.034 | 0 | 0 | 0.072 | 0 | NA | 0.05 |
| SE | 109473 | ENSG00000133104 | SPART | chr13 | - | 36314221 | 36314421 | 36312318 | 36312477 | 36326574 | 36326623 | 198 | 148 | 137 | 196 | 321 | 0 | 2 | 0 | 2 | 3 | 18 | 105 | 86 | 171 | 18 | 0 | 0 | 1 | 0 | 0 | 298 | 149 | 1 | 1 | 1 | 0.974 | 1 | 0.98 | 0.982 | 1 | 1 | 0.977 | 1 | 1 | -0.008 |
| SE | 109474 | ENSG00000133104 | SPART | chr13 | - | 36314226 | 36314421 | 36312318 | 36312477 | 36326574 | 36326698 | 260 | 181 | 199 | 246 | 385 | 0 | 2 | 0 | 2 | 3 | 20 | 138 | 115 | 201 | 18 | 0 | 0 | 1 | 0 | 0 | 298 | 149 | 1 | 1 | 1 | 0.978 | 1 | 0.984 | 0.985 | 1 | 1 | 0.983 | 1 | 1 | -0.007 |
| SE | 109476 | ENSG00000133104 | SPART | chr13 | - | 36336286 | 36336358 | 36335020 | 36335832 | 36346313 | 36346493 | 1 | 6 | 4 | 4 | 1 | 0 | 1 | 1 | 1 | 1 | 0 | 3 | 10 | 2 | 3 | 0 | 0 | 4 | 1 | 0 | 221 | 149 | 1 | 1 | 1 | 0.802 | 0.729 | 0.729 | 0.403 | NA | 1 | 0.628 | 0.574 | 1 | -0.068 |
| SE | 109477 | ENSG00000133104 | SPART | chr13 | - | 36336286 | 36336358 | 36335020 | 36335832 | 36346581 | 36346797 | 1 | 12 | 4 | 4 | 1 | 1 | 3 | 4 | 3 | 2 | 0 | 4 | 10 | 4 | 3 | 0 | 1 | 0 | 0 | 0 | 221 | 149 | 1 | 1 | 0.403 | 0.729 | 0.403 | 0.473 | 0.252 | NA | 0.729 | 1 | 1 | 1 | -0.48 |
| SE | 109479 | ENSG00000102572 | STK24 | chr13 | - | 98457167 | 98457304 | 98445184 | 98453209 | 98460371 | 98460440 | 427 | 286 | 379 | 436 | 594 | 2 | 3 | 0 | 3 | 1 | 825 | 466 | 326 | 392 | 1376 | 0 | 2 | 0 | 0 | 0 | 286 | 149 | 1 | 1 | 0.991 | 0.98 | 1 | 0.987 | 0.997 | 1 | 0.992 | 1 | 1 | 1 | -0.007 |
| SE | 109480 | ENSG00000102572 | STK24 | chr13 | - | 98460371 | 98460440 | 98455420 | 98457304 | 98461773 | 98461897 | 388 | 244 | 339 | 394 | 543 | 4 | 1 | 2 | 5 | 0 | 682 | 384 | 303 | 338 | 1064 | 0 | 0 | 0 | 0 | 6 | 218 | 149 | 1 | 1 | 0.985 | 0.994 | 0.991 | 0.982 | 1 | 1 | 1 | 1 | 1 | 0.992 | -0.008 |
| SE | 109481 | ENSG00000102572 | STK24 | chr13 | - | 98482264 | 98482321 | 98475249 | 98475358 | 98519242 | 98519267 | 833 | 542 | 673 | 827 | 1039 | 2 | 0 | 0 | 3 | 0 | 1427 | 831 | 671 | 951 | 2865 | 0 | 0 | 0 | 0 | 7 | 206 | 149 | 1 | 1 | 0.997 | 1 | 1 | 0.995 | 1 | 1 | 1 | 1 | 1 | 0.997 | -0.001 |
| SE | 109482 | ENSG00000102572 | STK24 | chr13 | - | 98508778 | 98508967 | 98505027 | 98505053 | 98519242 | 98519473 | 0 | 0 | 0 | 3 | 0 | 10 | 2 | 7 | 13 | 24 | 0 | 0 | 0 | 0 | 0 | 21 | 11 | 13 | 28 | 36 | 298 | 149 | 1 | 1 | 0 | 0 | 0 | 0.103 | 0 | 0 | 0 | 0 | 0 | 0 | 0.021 |
| SE | 109483 | ENSG00000132965 | ALOX5AP | chr13 | + | 30752051 | 30752122 | 30744059 | 30744159 | 30755943 | 30756025 | 480 | 212 | 301 | 412 | 476 | 0 | 0 | 1 | 4 | 0 | 508 | 972 | 1077 | 1447 | 752 | 7 | 2 | 0 | 4 | 0 | 220 | 149 | 1 | 1 | 1 | 1 | 0.995 | 0.986 | 1 | 0.98 | 0.997 | 1 | 0.996 | 1 | 0.002 |
| SE | 109484 | ENSG00000132965 | ALOX5AP | chr13 | + | 30752051 | 30752122 | 30744059 | 30744159 | 30763943 | 30764425 | 92 | 45 | 63 | 73 | 99 | 0 | 0 | 0 | 0 | 0 | 69 | 189 | 157 | 243 | 117 | 0 | 0 | 0 | 2 | 0 | 220 | 149 | 1 | 1 | 1 | 1 | 1 | 1 | 1 | 1 | 1 | 1 | 0.988 | 1 | 0.002 |
| SE | 109485 | ENSG00000132965 | ALOX5AP | chr13 | + | 30755943 | 30756025 | 30744059 | 30744159 | 30763943 | 30764425 | 234 | 73 | 173 | 225 | 227 | 0 | 0 | 0 | 0 | 0 | 234 | 446 | 498 | 532 | 279 | 0 | 0 | 0 | 2 | 0 | 231 | 149 | 1 | 1 | 1 | 1 | 1 | 1 | 1 | 1 | 1 | 1 | 0.994 | 1 | 0.001 |
| SE | 109486 | ENSG00000132965 | ALOX5AP | chr13 | + | 30755943 | 30756025 | 30752051 | 30752122 | 30763943 | 30764425 | 621 | 243 | 414 | 560 | 606 | 0 | 0 | 0 | 5 | 0 | 666 | 1232 | 1438 | 1731 | 914 | 0 | 0 | 7 | 0 | 0 | 231 | 149 | 1 | 1 | 1 | 1 | 1 | 0.986 | 1 | 1 | 1 | 0.993 | 1 | 1 | -0.001 |
| SE | 109487 | ENSG00000148362 | PAXX | chr9 | + | 136992639 | 136992700 | 136992421 | 136992562 | 136992924 | 136992974 | 18 | 7 | 11 | 24 | 3 | 1 | 0 | 0 | 0 | 0 | 6 | 14 | 6 | 0 | 30 | 0 | 0 | 2 | 0 | 0 | 210 | 149 | 1 | 1 | 0.927 | 1 | 1 | 1 | 1 | 1 | 1 | 0.68 | NA | 1 | 0.065 |
| SE | 109488 | ENSG00000148362 | PAXX | chr9 | + | 136992924 | 136992974 | 136992639 | 136992700 | 136993052 | 136993243 | 12 | 5 | 9 | 16 | 7 | 0 | 0 | 0 | 0 | 0 | 0 | 7 | 9 | 0 | 16 | 9 | 5 | 0 | 0 | 0 | 199 | 149 | 1 | 1 | 1 | 1 | 1 | 1 | 1 | 0 | 0.512 | 1 | NA | 1 | 0.372 |
| SE | 109489 | ENSG00000148362 | PAXX | chr9 | + | 136993342 | 136993411 | 136992639 | 136993243 | 136993579 | 136993664 | 109 | 37 | 36 | 53 | 29 | 0 | 0 | 0 | 1 | 0 | 36 | 83 | 27 | 27 | 112 | 0 | 0 | 0 | 0 | 0 | 218 | 149 | 1 | 1 | 1 | 1 | 1 | 0.973 | 1 | 1 | 1 | 1 | 1 | 1 | -0.005 |
| SE | 109491 | ENSG00000005893 | LAMP2 | chrX | - | 120456650 | 120456769 | 120455356 | 120455570 | 120469105 | 120469296 | 353 | 244 | 274 | 334 | 259 | 0 | 0 | 1 | 0 | 0 | 379 | 288 | 383 | 523 | 274 | 0 | 0 | 0 | 0 | 0 | 268 | 149 | 1 | 1 | 1 | 1 | 0.993 | 1 | 1 | 1 | 1 | 1 | 1 | 1 | -0.001 |
| SE | 109495 | ENSG00000143155 | TIPRL | chr1 | + | 168191368 | 168191500 | 168184778 | 168184878 | 168196546 | 168196642 | 778 | 611 | 593 | 698 | 1147 | 4 | 1 | 0 | 0 | 0 | 718 | 572 | 668 | 741 | 568 | 0 | 0 | 3 | 0 | 0 | 281 | 149 | 1 | 1 | 0.99 | 0.997 | 1 | 1 | 1 | 1 | 1 | 0.992 | 1 | 1 | -0.001 |
| SE | 109496 | ENSG00000139793 | MBNL2 | chr13 | + | 97334275 | 97334440 | 97275631 | 97276409 | 97343015 | 97343216 | 382 | 321 | 353 | 359 | 830 | 0 | 1 | 0 | 0 | 0 | 79 | 263 | 168 | 98 | 28 | 0 | 1 | 0 | 0 | 0 | 298 | 149 | 1 | 1 | 1 | 0.994 | 1 | 1 | 1 | 1 | 0.992 | 1 | 1 | 1 | 0 |
| SE | 109497 | ENSG00000139793 | MBNL2 | chr13 | + | 97343015 | 97343216 | 97275631 | 97276409 | 97357481 | 97357635 | 2 | 2 | 0 | 0 | 0 | 0 | 2 | 0 | 0 | 0 | 0 | 2 | 1 | 0 | 0 | 0 | 0 | 0 | 0 | 0 | 298 | 149 | 1 | 1 | 1 | 0.333 | NA | NA | NA | NA | 1 | 1 | NA | NA | -0.334 |
| SE | 109498 | ENSG00000139793 | MBNL2 | chr13 | + | 97346803 | 97347067 | 97343015 | 97343216 | 97357481 | 97357635 | 121 | 73 | 112 | 123 | 111 | 2 | 1 | 0 | 0 | 0 | 0 | 87 | 62 | 25 | 11 | 0 | 1 | 1 | 0 | 0 | 298 | 149 | 1 | 1 | 0.968 | 0.973 | 1 | 1 | 1 | NA | 0.978 | 0.969 | 1 | 1 | 0.001 |
| SE | 109499 | ENSG00000139793 | MBNL2 | chr13 | + | 97356795 | 97356849 | 97346803 | 97347067 | 97357481 | 97357635 | 0 | 1 | 2 | 0 | 0 | 53 | 29 | 44 | 65 | 40 | 0 | 0 | 0 | 0 | 0 | 0 | 27 | 21 | 4 | 0 | 203 | 149 | 1 | 1 | 0 | 0.025 | 0.032 | 0 | 0 | NA | 0 | 0 | 0 | NA | 0.011 |
| SE | 109500 | ENSG00000139793 | MBNL2 | chr13 | + | 97365135 | 97365171 | 97357481 | 97357635 | 97366458 | 97366553 | 0 | 0 | 0 | 0 | 4 | 4 | 7 | 2 | 7 | 1 | 0 | 0 | 0 | 0 | 0 | 1 | 0 | 7 | 0 | 0 | 185 | 149 | 1 | 1 | 0 | 0 | 0 | 0 | 0.763 | 0 | NA | 0 | NA | NA | 0.153 |
| SE | 109501 | ENSG00000139793 | MBNL2 | chr13 | + | 97365135 | 97365171 | 97357481 | 97357635 | 97391321 | 97394119 | 0 | 0 | 0 | 0 | 5 | 25 | 14 | 7 | 21 | 24 | 0 | 0 | 0 | 0 | 0 | 0 | 5 | 0 | 0 | 0 | 185 | 149 | 1 | 1 | 0 | 0 | 0 | 0 | 0.144 | NA | 0 | NA | NA | NA | 0.029 |
| SE | 109502 | ENSG00000139793 | MBNL2 | chr13 | + | 97366458 | 97366553 | 97357481 | 97357635 | 97391321 | 97391599 | 8 | 7 | 7 | 11 | 1 | 25 | 14 | 7 | 21 | 24 | 1 | 0 | 9 | 0 | 0 | 0 | 5 | 0 | 0 | 0 | 244 | 149 | 1 | 1 | 0.163 | 0.234 | 0.379 | 0.242 | 0.025 | 1 | 0 | 1 | NA | NA | -0.458 |
| SE | 109503 | ENSG00000139793 | MBNL2 | chr13 | + | 97366458 | 97366553 | 97365135 | 97365171 | 97391321 | 97391599 | 4 | 0 | 5 | 4 | 0 | 0 | 0 | 0 | 0 | 1 | 0 | 0 | 2 | 0 | 0 | 0 | 0 | 0 | 0 | 0 | 244 | 149 | 1 | 1 | 1 | NA | 1 | 1 | 0 | NA | NA | 1 | NA | NA | -0.25 |
| SE | 109509 | ENSG00000163519 | TRAT1 | chr3 | + | 108830669 | 108830780 | 108822769 | 108822934 | 108838933 | 108838967 | 636 | 518 | 819 | 809 | 2008 | 0 | 3 | 1 | 2 | 0 | 8 | 270 | 186 | 24 | 0 | 0 | 0 | 8 | 0 | 0 | 260 | 149 | 1 | 1 | 1 | 0.99 | 0.998 | 0.996 | 1 | 1 | 1 | 0.93 | 1 | NA | 0.014 |
| SE | 109510 | ENSG00000163519 | TRAT1 | chr3 | + | 108830669 | 108830780 | 108822769 | 108822934 | 108847067 | 108847338 | 302 | 260 | 367 | 339 | 950 | 1 | 0 | 0 | 0 | 0 | 0 | 136 | 83 | 14 | 0 | 0 | 0 | 0 | 0 | 0 | 260 | 149 | 1 | 1 | 0.994 | 1 | 1 | 1 | 1 | NA | 1 | 1 | 1 | NA | -0.001 |
| SE | 109511 | ENSG00000163519 | TRAT1 | chr3 | + | 108838933 | 108838967 | 108822771 | 108822934 | 108847067 | 108847129 | 117 | 82 | 168 | 155 | 517 | 1 | 0 | 0 | 0 | 0 | 9 | 34 | 39 | 0 | 0 | 0 | 0 | 0 | 0 | 0 | 183 | 149 | 1 | 1 | 0.99 | 1 | 1 | 1 | 1 | 1 | 1 | 1 | NA | NA | -0.002 |
| SE | 109512 | ENSG00000163519 | TRAT1 | chr3 | + | 108838933 | 108838967 | 108830669 | 108830780 | 108847067 | 108847129 | 447 | 329 | 611 | 623 | 1545 | 1 | 0 | 4 | 4 | 0 | 17 | 157 | 133 | 15 | 0 | 0 | 0 | 0 | 0 | 0 | 183 | 149 | 1 | 1 | 0.997 | 1 | 0.992 | 0.992 | 1 | 1 | 1 | 1 | 1 | NA | -0.004 |
| SE | 109513 | ENSG00000163519 | TRAT1 | chr3 | + | 108847067 | 108847129 | 108838933 | 108838967 | 108849165 | 108849254 | 469 | 343 | 598 | 592 | 1470 | 4 | 0 | 1 | 3 | 0 | 9 | 160 | 152 | 24 | 0 | 0 | 2 | 0 | 0 | 0 | 211 | 149 | 1 | 1 | 0.988 | 1 | 0.998 | 0.993 | 1 | 1 | 0.983 | 1 | 1 | NA | 0 |
| SE | 109515 | ENSG00000119917 | IFIT3 | chr10 | + | 89328548 | 89328629 | 89327996 | 89328078 | 89338660 | 89340968 | 0 | 0 | 0 | 0 | 4 | 50 | 154 | 59 | 203 | 105 | 0 | 0 | 0 | 0 | 0 | 49 | 93 | 122 | 389 | 0 | 230 | 149 | 1 | 1 | 0 | 0 | 0 | 0 | 0.024 | 0 | 0 | 0 | 0 | NA | 0.005 |
| SE | 109516 | ENSG00000119917 | IFIT3 | chr10 | + | 89337420 | 89337542 | 89332511 | 89332627 | 89338660 | 89340968 | 1 | 0 | 1 | 2 | 0 | 16 | 40 | 21 | 49 | 23 | 0 | 0 | 2 | 8 | 0 | 64 | 28 | 48 | 145 | 0 | 271 | 149 | 1 | 1 | 0.033 | 0 | 0.026 | 0.022 | 0 | 0 | 0 | 0.022 | 0.029 | NA | 0.003 |
| SE | 109517 | ENSG00000119917 | IFIT3 | chr10 | + | 89337424 | 89337542 | 89327981 | 89328078 | 89338660 | 89340968 | 1 | 0 | 1 | 4 | 0 | 50 | 154 | 59 | 203 | 105 | 0 | 0 | 2 | 8 | 0 | 49 | 93 | 122 | 389 | 0 | 267 | 149 | 1 | 1 | 0.011 | 0 | 0.009 | 0.011 | 0 | 0 | 0 | 0.009 | 0.011 | NA | 0.001 |
| SE | 109518 | ENSG00000119917 | IFIT3 | chr10 | + | 89337424 | 89337542 | 89332488 | 89332627 | 89338660 | 89340968 | 1 | 0 | 1 | 2 | 0 | 16 | 40 | 21 | 49 | 23 | 0 | 0 | 2 | 8 | 0 | 64 | 28 | 48 | 145 | 0 | 267 | 149 | 1 | 1 | 0.034 | 0 | 0.026 | 0.022 | 0 | 0 | 0 | 0.023 | 0.03 | NA | 0.003 |
| SE | 109522 | ENSG00000286724 | NA | chrX | + | 19051703 | 19052145 | 19050834 | 19050981 | 19054931 | 19055716 | 0 | 0 | 0 | 1 | 0 | 1 | 0 | 2 | 0 | 0 | 0 | 0 | 0 | 0 | 0 | 0 | 5 | 1 | 0 | 0 | 298 | 149 | 1 | 1 | 0 | NA | 0 | 1 | NA | NA | 0 | 0 | NA | NA | 0.333 |
| SE | 109523 | ENSG00000010319 | SEMA3G | chr3 | - | 52440376 | 52440521 | 52439866 | 52440098 | 52440753 | 52440823 | 1 | 0 | 4 | 4 | 0 | 0 | 0 | 0 | 0 | 0 | 0 | 5 | 8 | 0 | 0 | 0 | 0 | 3 | 5 | 6 | 294 | 149 | 1 | 1 | 1 | NA | 1 | 1 | NA | NA | 1 | 0.575 | 0 | 0 | 0.606 |
| SE | 109525 | ENSG00000151849 | CENPJ | chr13 | - | 24886234 | 24886385 | 24885605 | 24885670 | 24889315 | 24889355 | 2 | 1 | 2 | 0 | 4 | 6 | 2 | 4 | 5 | 6 | 0 | 0 | 0 | 0 | 0 | 0 | 3 | 3 | 0 | 0 | 298 | 149 | 1 | 1 | 0.143 | 0.2 | 0.2 | 0 | 0.25 | NA | 0 | 0 | NA | NA | 0.159 |
| SE | 109526 | ENSG00000151849 | CENPJ | chr13 | - | 24889315 | 24889400 | 24885605 | 24885670 | 24892642 | 24892867 | 15 | 7 | 10 | 11 | 11 | 0 | 1 | 0 | 0 | 0 | 0 | 3 | 4 | 0 | 0 | 0 | 0 | 0 | 0 | 0 | 234 | 149 | 1 | 1 | 1 | 0.817 | 1 | 1 | 1 | NA | 1 | 1 | NA | NA | -0.037 |
| SE | 109527 | ENSG00000151849 | CENPJ | chr13 | - | 24909789 | 24910086 | 24908017 | 24908126 | 24911945 | 24912069 | 70 | 50 | 25 | 75 | 75 | 1 | 0 | 0 | 6 | 0 | 9 | 89 | 49 | 28 | 0 | 0 | 0 | 0 | 0 | 0 | 298 | 149 | 1 | 1 | 0.972 | 1 | 1 | 0.862 | 1 | 1 | 1 | 1 | 1 | NA | -0.033 |
| SE | 109528 | ENSG00000151849 | CENPJ | chr13 | - | 24912581 | 24913091 | 24911945 | 24912069 | 24922760 | 24922810 | 50 | 27 | 31 | 43 | 37 | 0 | 0 | 0 | 0 | 0 | 16 | 31 | 29 | 15 | 0 | 0 | 0 | 2 | 0 | 0 | 298 | 149 | 1 | 1 | 1 | 1 | 1 | 1 | 1 | 1 | 1 | 0.879 | 1 | NA | 0.03 |
| SE | 109532 | ENSG00000127022 | CANX | chr5 | + | 179699574 | 179699770 | 179698955 | 179699102 | 179705678 | 179705852 | 3 | 0 | 6 | 1 | 0 | 449 | 231 | 375 | 425 | 112 | 5 | 2 | 9 | 3 | 0 | 200 | 317 | 303 | 401 | 101 | 298 | 149 | 1 | 1 | 0.003 | 0 | 0.008 | 0.001 | 0 | 0.012 | 0.003 | 0.015 | 0.004 | 0 | -0.004 |
| SE | 109533 | ENSG00000127022 | CANX | chr5 | + | 179700383 | 179700533 | 179699118 | 179699770 | 179705678 | 179705852 | 0 | 7 | 0 | 0 | 0 | 3 | 0 | 5 | 1 | 0 | 0 | 0 | 0 | 0 | 0 | 5 | 2 | 9 | 3 | 0 | 298 | 149 | 1 | 1 | 0 | 1 | 0 | 0 | NA | 0 | 0 | 0 | 0 | NA | 0.25 |
| SE | 109535 | ENSG00000127022 | CANX | chr5 | + | 179702492 | 179702630 | 179699008 | 179699102 | 179705678 | 179705852 | 2 | 0 | 0 | 0 | 0 | 449 | 231 | 375 | 425 | 112 | 0 | 0 | 2 | 2 | 0 | 200 | 317 | 303 | 401 | 101 | 287 | 149 | 1 | 1 | 0.002 | 0 | 0 | 0 | 0 | 0 | 0 | 0.003 | 0.003 | 0 | -0.001 |
| SE | 109549 | ENSG00000127022 | CANX | chr5 | + | 179707131 | 179707190 | 179706257 | 179706331 | 179708238 | 179708380 | 2234 | 1195 | 1720 | 1861 | 3006 | 0 | 2 | 1 | 1 | 0 | 896 | 1407 | 1626 | 1759 | 682 | 0 | 3 | 2 | 1 | 0 | 208 | 149 | 1 | 1 | 1 | 0.998 | 0.999 | 0.999 | 1 | 1 | 0.997 | 0.998 | 0.999 | 1 | 0 |
| SE | 109550 | ENSG00000127022 | CANX | chr5 | + | 179708238 | 179708380 | 179698933 | 179699102 | 179709872 | 179710065 | 22 | 21 | 17 | 36 | 59 | 0 | 0 | 2 | 0 | 0 | 42 | 48 | 75 | 43 | 24 | 0 | 0 | 0 | 0 | 0 | 291 | 149 | 1 | 1 | 1 | 1 | 0.813 | 1 | 1 | 1 | 1 | 1 | 1 | 1 | -0.037 |
| SE | 109552 | ENSG00000127022 | CANX | chr5 | + | 179708977 | 179709059 | 179708238 | 179708380 | 179709918 | 179710065 | 914 | 505 | 758 | 795 | 1917 | 0 | 0 | 0 | 0 | 7 | 300 | 491 | 585 | 638 | 256 | 0 | 3 | 4 | 0 | 0 | 231 | 149 | 1 | 1 | 1 | 1 | 1 | 1 | 0.994 | 1 | 0.991 | 0.99 | 1 | 1 | 0.003 |
| SE | 109553 | ENSG00000127022 | CANX | chr5 | + | 179709872 | 179710065 | 179708238 | 179708380 | 179716104 | 179716294 | 2156 | 1482 | 1900 | 1864 | 3408 | 0 | 1 | 0 | 0 | 0 | 1386 | 1638 | 1762 | 1984 | 493 | 0 | 0 | 0 | 0 | 0 | 298 | 149 | 1 | 1 | 1 | 0.999 | 1 | 1 | 1 | 1 | 1 | 1 | 1 | 1 | 0 |
| SE | 109554 | ENSG00000127022 | CANX | chr5 | + | 179709918 | 179710065 | 179708238 | 179708380 | 179716104 | 179716294 | 2134 | 1461 | 1883 | 1828 | 3356 | 0 | 1 | 0 | 0 | 0 | 1344 | 1593 | 1689 | 1939 | 469 | 0 | 0 | 0 | 0 | 0 | 296 | 149 | 1 | 1 | 1 | 0.999 | 1 | 1 | 1 | 1 | 1 | 1 | 1 | 1 | 0 |
| SE | 109555 | ENSG00000127022 | CANX | chr5 | + | 179715898 | 179716000 | 179709872 | 179710065 | 179716104 | 179716294 | 7 | 5 | 6 | 10 | 4 | 2134 | 1461 | 1883 | 1828 | 3349 | 0 | 10 | 5 | 0 | 0 | 1344 | 1591 | 1685 | 1939 | 469 | 251 | 149 | 1 | 1 | 0.002 | 0.002 | 0.002 | 0.003 | 0.001 | 0 | 0.004 | 0.002 | 0 | 0 | 0.001 |
| SE | 109556 | ENSG00000127022 | CANX | chr5 | + | 179715936 | 179716000 | 179709872 | 179710065 | 179716104 | 179716294 | 4 | 1 | 2 | 5 | 3 | 2134 | 1461 | 1883 | 1828 | 3349 | 0 | 0 | 2 | 0 | 2 | 1344 | 1591 | 1685 | 1939 | 469 | 213 | 149 | 1 | 1 | 0.001 | 0 | 0.001 | 0.002 | 0.001 | 0 | 0 | 0.001 | 0 | 0.003 | 0 |
| SE | 109562 | ENSG00000127022 | CANX | chr5 | + | 179724656 | 179724783 | 179722803 | 179723779 | 179726679 | 179726759 | 2333 | 1780 | 1897 | 1885 | 2598 | 0 | 0 | 1 | 0 | 0 | 1308 | 1811 | 2457 | 2991 | 430 | 0 | 0 | 2 | 0 | 0 | 276 | 149 | 1 | 1 | 1 | 1 | 0.999 | 1 | 1 | 1 | 1 | 0.998 | 1 | 1 | 0 |
| SE | 109563 | ENSG00000127022 | CANX | chr5 | + | 179724662 | 179724783 | 179723659 | 179723779 | 179726679 | 179726759 | 1013 | 753 | 815 | 795 | 1235 | 0 | 0 | 1 | 0 | 0 | 516 | 780 | 1052 | 1309 | 140 | 0 | 0 | 2 | 0 | 0 | 270 | 149 | 1 | 1 | 1 | 1 | 0.998 | 1 | 1 | 1 | 1 | 0.997 | 1 | 1 | 0 |
| SE | 109564 | ENSG00000127022 | CANX | chr5 | + | 179726659 | 179726759 | 179724656 | 179724783 | 179728590 | 179731453 | 608 | 365 | 486 | 433 | 809 | 0 | 0 | 0 | 0 | 0 | 101 | 280 | 331 | 365 | 46 | 0 | 0 | 0 | 1 | 0 | 249 | 149 | 1 | 1 | 1 | 1 | 1 | 1 | 1 | 1 | 1 | 1 | 0.995 | 1 | 0.001 |
| SE | 109565 | ENSG00000127022 | CANX | chr5 | + | 179726679 | 179726759 | 179724656 | 179724783 | 179728590 | 179728682 | 1655 | 1144 | 1353 | 1285 | 2170 | 0 | 0 | 0 | 0 | 0 | 629 | 1103 | 1385 | 1726 | 206 | 0 | 0 | 0 | 1 | 0 | 229 | 149 | 1 | 1 | 1 | 1 | 1 | 1 | 1 | 1 | 1 | 1 | 0.999 | 1 | 0 |
| SE | 109566 | ENSG00000127022 | CANX | chr5 | + | 179726682 | 179726759 | 179724656 | 179724783 | 179728590 | 179731453 | 608 | 368 | 479 | 433 | 792 | 0 | 0 | 0 | 0 | 0 | 101 | 296 | 339 | 367 | 46 | 0 | 0 | 0 | 1 | 0 | 226 | 149 | 1 | 1 | 1 | 1 | 1 | 1 | 1 | 1 | 1 | 1 | 0.996 | 1 | 0.001 |
| SE | 109568 | ENSG00000051128 | HOMER3 | chr19 | - | 18940320 | 18940404 | 18938968 | 18939049 | 18941050 | 18941122 | 0 | 0 | 0 | 0 | 0 | 5 | 3 | 3 | 0 | 0 | 0 | 1 | 0 | 0 | 9 | 0 | 1 | 0 | 0 | 2 | 233 | 149 | 1 | 1 | 0 | 0 | 0 | NA | NA | NA | 0.39 | NA | NA | 0.742 | -0.566 |
| SE | 109569 | ENSG00000133027 | PEMT | chr17 | - | 17506226 | 17506301 | 17505562 | 17505848 | 17509433 | 17509545 | 30 | 4 | 5 | 12 | 0 | 1 | 0 | 0 | 1 | 0 | 16 | 9 | 18 | 10 | 46 | 0 | 0 | 0 | 0 | 0 | 224 | 149 | 1 | 1 | 0.952 | 1 | 1 | 0.889 | NA | 1 | 1 | 1 | 1 | 1 | -0.04 |
| SE | 109570 | ENSG00000133027 | PEMT | chr17 | - | 17507168 | 17507199 | 17506226 | 17506301 | 17509433 | 17509545 | 1 | 0 | 0 | 0 | 0 | 26 | 4 | 4 | 10 | 0 | 0 | 0 | 0 | 0 | 0 | 0 | 4 | 11 | 8 | 40 | 180 | 149 | 1 | 1 | 0.031 | 0 | 0 | 0 | NA | NA | 0 | 0 | 0 | 0 | 0.008 |
| SE | 109572 | ENSG00000133027 | PEMT | chr17 | - | 17509433 | 17509545 | 17506226 | 17506301 | 17576919 | 17577027 | 16 | 4 | 3 | 6 | 0 | 0 | 0 | 0 | 0 | 0 | 0 | 0 | 11 | 6 | 36 | 0 | 0 | 0 | 0 | 1 | 261 | 149 | 1 | 1 | 1 | 1 | 1 | 1 | NA | NA | NA | 1 | 1 | 0.954 | 0.015 |
| SE | 109573 | ENSG00000133027 | PEMT | chr17 | - | 17512508 | 17512654 | 17509433 | 17509545 | 17522279 | 17522395 | 26 | 2 | 9 | 17 | 12 | 1 | 0 | 0 | 0 | 0 | 14 | 15 | 3 | 15 | 44 | 0 | 0 | 0 | 0 | 0 | 295 | 149 | 1 | 1 | 0.929 | 1 | 1 | 1 | 1 | 1 | 1 | 1 | 1 | 1 | -0.014 |
| SE | 109578 | ENSG00000102804 | TSC22D1 | chr13 | - | 44435792 | 44436095 | 44434708 | 44434883 | 44436503 | 44436596 | 138 | 178 | 49 | 193 | 249 | 0 | 0 | 0 | 2 | 0 | 473 | 477 | 100 | 289 | 1041 | 0 | 0 | 0 | 0 | 0 | 298 | 149 | 1 | 1 | 1 | 1 | 1 | 0.98 | 1 | 1 | 1 | 1 | 1 | 1 | -0.004 |
| SE | 109579 | ENSG00000102804 | TSC22D1 | chr13 | - | 44436043 | 44436095 | 44432142 | 44434883 | 44436503 | 44436845 | 453 | 692 | 192 | 744 | 1246 | 0 | 0 | 0 | 2 | 0 | 1838 | 1716 | 418 | 1178 | 4076 | 0 | 0 | 0 | 0 | 0 | 201 | 149 | 1 | 1 | 1 | 1 | 1 | 0.996 | 1 | 1 | 1 | 1 | 1 | 1 | -0.001 |
| SE | 109580 | ENSG00000102804 | TSC22D1 | chr13 | - | 44539766 | 44539923 | 44538325 | 44538888 | 44573162 | 44573345 | 4 | 6 | 6 | 0 | 8 | 0 | 0 | 1 | 0 | 0 | 0 | 2 | 3 | 0 | 0 | 0 | 0 | 0 | 0 | 0 | 298 | 149 | 1 | 1 | 1 | 1 | 0.75 | NA | 1 | NA | 1 | 1 | NA | NA | -0.062 |
| SE | 109581 | ENSG00000102804 | TSC22D1 | chr13 | - | 44539766 | 44539936 | 44538325 | 44538888 | 44573162 | 44573522 | 4 | 6 | 10 | 8 | 10 | 0 | 0 | 1 | 0 | 0 | 16 | 10 | 6 | 0 | 0 | 0 | 0 | 0 | 0 | 0 | 298 | 149 | 1 | 1 | 1 | 1 | 0.833 | 1 | 1 | 1 | 1 | 1 | NA | NA | -0.033 |
| SE | 109588 | ENSG00000236778 | INTS6-AS1 | chr13 | + | 51467508 | 51467684 | 51454163 | 51454329 | 51468267 | 51468367 | 1 | 0 | 0 | 0 | 0 | 0 | 0 | 0 | 1 | 0 | 0 | 0 | 0 | 0 | 0 | 0 | 0 | 3 | 0 | 0 | 298 | 149 | 1 | 1 | 1 | NA | NA | 0 | NA | NA | NA | 0 | NA | NA | 0.5 |
| SE | 109589 | ENSG00000236778 | INTS6-AS1 | chr13 | + | 51467537 | 51467684 | 51454149 | 51454329 | 51468267 | 51468367 | 1 | 0 | 0 | 0 | 0 | 0 | 0 | 0 | 1 | 0 | 0 | 0 | 0 | 0 | 0 | 0 | 0 | 3 | 0 | 0 | 296 | 149 | 1 | 1 | 1 | NA | NA | 0 | NA | NA | NA | 0 | NA | NA | 0.5 |
| SE | 109590 | ENSG00000236778 | INTS6-AS1 | chr13 | + | 51467595 | 51467684 | 51454002 | 51454329 | 51468267 | 51468367 | 1 | 0 | 0 | 0 | 0 | 0 | 0 | 0 | 1 | 0 | 0 | 0 | 0 | 0 | 0 | 0 | 0 | 3 | 0 | 0 | 238 | 149 | 1 | 1 | 1 | NA | NA | 0 | NA | NA | NA | 0 | NA | NA | 0.5 |
| SE | 109595 | ENSG00000236778 | INTS6-AS1 | chr13 | + | 51506542 | 51506638 | 51468267 | 51468367 | 51549709 | 51549875 | 1 | 0 | 0 | 0 | 0 | 0 | 0 | 0 | 0 | 0 | 0 | 0 | 0 | 0 | 0 | 0 | 6 | 0 | 0 | 0 | 245 | 149 | 1 | 1 | 1 | NA | NA | NA | NA | NA | 0 | NA | NA | NA | 1 |
| SE | 109602 | ENSG00000285444 | NA | chr13 | - | 51799533 | 51799667 | 51791164 | 51791257 | 51813170 | 51813339 | 15 | 11 | 15 | 21 | 37 | 0 | 0 | 2 | 0 | 0 | 33 | 5 | 9 | 12 | 25 | 0 | 0 | 0 | 0 | 0 | 283 | 149 | 1 | 1 | 1 | 1 | 0.798 | 1 | 1 | 1 | 1 | 1 | 1 | 1 | -0.04 |
| SE | 109603 | ENSG00000285444 | NA | chr13 | - | 51799533 | 51799667 | 51791164 | 51791257 | 51838472 | 51838553 | 17 | 12 | 15 | 20 | 35 | 0 | 0 | 0 | 0 | 0 | 33 | 5 | 22 | 12 | 25 | 0 | 0 | 1 | 0 | 0 | 283 | 149 | 1 | 1 | 1 | 1 | 1 | 1 | 1 | 1 | 1 | 0.921 | 1 | 1 | 0.016 |
| SE | 109604 | ENSG00000285444 | NA | chr13 | - | 51813170 | 51813339 | 51791164 | 51791257 | 51838472 | 51838553 | 0 | 0 | 2 | 0 | 0 | 0 | 0 | 0 | 0 | 0 | 0 | 0 | 0 | 0 | 0 | 0 | 0 | 1 | 0 | 0 | 298 | 149 | 1 | 1 | NA | NA | 1 | NA | NA | NA | NA | 0 | NA | NA | 1 |
| SE | 109605 | ENSG00000285444 | NA | chr13 | - | 51813170 | 51813339 | 51799533 | 51799667 | 51838472 | 51838553 | 0 | 0 | 0 | 1 | 2 | 2 | 1 | 0 | 0 | 0 | 0 | 0 | 2 | 0 | 0 | 0 | 0 | 15 | 0 | 0 | 298 | 149 | 1 | 1 | 0 | 0 | NA | 1 | 1 | NA | NA | 0.062 | NA | NA | 0.438 |
| SE | 109606 | ENSG00000285444 | NA | chr13 | - | 51838472 | 51838553 | 51799533 | 51799667 | 51845015 | 51845159 | 4 | 1 | 0 | 0 | 0 | 0 | 0 | 0 | 0 | 0 | 0 | 3 | 28 | 0 | 0 | 0 | 0 | 2 | 0 | 0 | 230 | 149 | 1 | 1 | 1 | 1 | NA | NA | NA | NA | 1 | 0.901 | NA | NA | 0.049 |
| SE | 109608 | ENSG00000005810 | MYCBP2 | chr13 | - | 77058217 | 77058406 | 77056985 | 77057093 | 77059522 | 77059626 | 230 | 162 | 237 | 282 | 382 | 0 | 0 | 0 | 0 | 0 | 46 | 149 | 139 | 32 | 20 | 0 | 2 | 0 | 0 | 0 | 298 | 149 | 1 | 1 | 1 | 1 | 1 | 1 | 1 | 1 | 0.974 | 1 | 1 | 1 | 0.005 |
| SE | 109609 | ENSG00000005810 | MYCBP2 | chr13 | - | 77059522 | 77059626 | 77058217 | 77058406 | 77061168 | 77061301 | 187 | 158 | 208 | 212 | 294 | 0 | 0 | 1 | 0 | 0 | 91 | 147 | 138 | 29 | 23 | 0 | 0 | 0 | 0 | 0 | 253 | 149 | 1 | 1 | 1 | 1 | 0.992 | 1 | 1 | 1 | 1 | 1 | 1 | 1 | -0.002 |
| SE | 109610 | ENSG00000005810 | MYCBP2 | chr13 | - | 77061168 | 77061301 | 77059522 | 77059626 | 77061661 | 77061790 | 306 | 319 | 329 | 349 | 627 | 0 | 0 | 0 | 3 | 0 | 88 | 304 | 260 | 65 | 25 | 0 | 0 | 0 | 0 | 0 | 282 | 149 | 1 | 1 | 1 | 1 | 1 | 0.984 | 1 | 1 | 1 | 1 | 1 | 1 | -0.003 |
| SE | 109611 | ENSG00000005810 | MYCBP2 | chr13 | - | 77061661 | 77061790 | 77061168 | 77061301 | 77062595 | 77062697 | 401 | 433 | 436 | 500 | 736 | 0 | 0 | 2 | 0 | 0 | 59 | 433 | 425 | 129 | 33 | 0 | 1 | 0 | 0 | 0 | 278 | 149 | 1 | 1 | 1 | 1 | 0.992 | 1 | 1 | 1 | 0.996 | 1 | 1 | 1 | -0.001 |
| SE | 109612 | ENSG00000005810 | MYCBP2 | chr13 | - | 77062595 | 77062697 | 77061661 | 77061790 | 77064614 | 77064734 | 369 | 370 | 394 | 467 | 635 | 0 | 0 | 0 | 0 | 0 | 102 | 335 | 428 | 127 | 22 | 0 | 0 | 0 | 3 | 0 | 251 | 149 | 1 | 1 | 1 | 1 | 1 | 1 | 1 | 1 | 1 | 1 | 0.962 | 1 | 0.008 |
| SE | 109613 | ENSG00000005810 | MYCBP2 | chr13 | - | 77076750 | 77076849 | 77070630 | 77070711 | 77077147 | 77077387 | 216 | 161 | 199 | 213 | 446 | 0 | 0 | 0 | 0 | 0 | 37 | 135 | 197 | 59 | 22 | 0 | 0 | 3 | 0 | 0 | 248 | 149 | 1 | 1 | 1 | 1 | 1 | 1 | 1 | 1 | 1 | 0.975 | 1 | 1 | 0.005 |
| SE | 109614 | ENSG00000005810 | MYCBP2 | chr13 | - | 77076759 | 77076849 | 77070630 | 77070711 | 77077147 | 77077387 | 213 | 179 | 207 | 234 | 436 | 0 | 0 | 0 | 0 | 0 | 24 | 120 | 203 | 51 | 28 | 0 | 0 | 3 | 0 | 0 | 239 | 149 | 1 | 1 | 1 | 1 | 1 | 1 | 1 | 1 | 1 | 0.977 | 1 | 1 | 0.005 |
| SE | 109617 | ENSG00000005810 | MYCBP2 | chr13 | - | 77090105 | 77090263 | 77087483 | 77087633 | 77093164 | 77093332 | 287 | 217 | 251 | 235 | 630 | 4 | 1 | 1 | 0 | 0 | 0 | 172 | 135 | 83 | 2 | 0 | 0 | 0 | 0 | 18 | 298 | 149 | 1 | 1 | 0.973 | 0.991 | 0.992 | 1 | 1 | NA | 1 | 1 | 1 | 0.053 | 0.228 |
| SE | 109618 | ENSG00000005810 | MYCBP2 | chr13 | - | 77090105 | 77090263 | 77088831 | 77089031 | 77093164 | 77093332 | 531 | 433 | 471 | 468 | 1167 | 0 | 0 | 0 | 0 | 2 | 16 | 362 | 297 | 121 | 13 | 0 | 0 | 0 | 0 | 0 | 298 | 149 | 1 | 1 | 1 | 1 | 1 | 1 | 0.997 | 1 | 1 | 1 | 1 | 1 | -0.001 |
| SE | 109619 | ENSG00000005810 | MYCBP2 | chr13 | - | 77095357 | 77095602 | 77093164 | 77093332 | 77096311 | 77096481 | 387 | 297 | 319 | 340 | 569 | 0 | 0 | 0 | 0 | 0 | 43 | 225 | 208 | 99 | 21 | 0 | 0 | 1 | 0 | 0 | 298 | 149 | 1 | 1 | 1 | 1 | 1 | 1 | 1 | 1 | 1 | 0.99 | 1 | 1 | 0.002 |
| SE | 109620 | ENSG00000005810 | MYCBP2 | chr13 | - | 77095530 | 77095602 | 77093285 | 77093332 | 77096311 | 77096481 | 116 | 73 | 101 | 101 | 167 | 0 | 0 | 0 | 0 | 0 | 6 | 45 | 47 | 36 | 0 | 0 | 0 | 1 | 0 | 0 | 221 | 149 | 1 | 1 | 1 | 1 | 1 | 1 | 1 | 1 | 1 | 0.969 | 1 | NA | 0.008 |
| SE | 109621 | ENSG00000005810 | MYCBP2 | chr13 | - | 77096311 | 77096481 | 77093164 | 77093332 | 77121372 | 77121495 | 1 | 0 | 0 | 0 | 0 | 0 | 0 | 0 | 0 | 0 | 0 | 0 | 1 | 3 | 0 | 0 | 0 | 0 | 1 | 0 | 298 | 149 | 1 | 1 | 1 | NA | NA | NA | NA | NA | NA | 1 | 0.6 | NA | 0.2 |
| SE | 109622 | ENSG00000005810 | MYCBP2 | chr13 | - | 77097369 | 77099013 | 77096311 | 77096481 | 77121372 | 77121495 | 503 | 404 | 399 | 491 | 664 | 1 | 0 | 0 | 0 | 0 | 58 | 266 | 257 | 130 | 13 | 0 | 0 | 0 | 3 | 0 | 298 | 149 | 1 | 1 | 0.996 | 1 | 1 | 1 | 1 | 1 | 1 | 1 | 0.956 | 1 | 0.008 |
| SE | 109623 | ENSG00000005810 | MYCBP2 | chr13 | - | 77103208 | 77103310 | 77097369 | 77099013 | 77118339 | 77118519 | 0 | 2 | 0 | 2 | 0 | 48 | 37 | 26 | 53 | 72 | 0 | 2 | 1 | 0 | 0 | 8 | 54 | 23 | 3 | 31 | 251 | 149 | 1 | 1 | 0 | 0.031 | 0 | 0.022 | 0 | 0 | 0.022 | 0.025 | 0 | 0 | 0.001 |
| SE | 109624 | ENSG00000005810 | MYCBP2 | chr13 | - | 77103208 | 77103310 | 77097369 | 77099013 | 77121372 | 77121495 | 0 | 7 | 0 | 2 | 0 | 242 | 191 | 186 | 239 | 344 | 0 | 1 | 1 | 0 | 0 | 27 | 121 | 159 | 75 | 0 | 251 | 149 | 1 | 1 | 0 | 0.021 | 0 | 0.005 | 0 | 0 | 0.005 | 0.004 | 0 | NA | 0.003 |
| SE | 109627 | ENSG00000005810 | MYCBP2 | chr13 | - | 77118339 | 77118519 | 77103208 | 77103310 | 77121372 | 77121495 | 8 | 3 | 0 | 10 | 5 | 0 | 5 | 0 | 0 | 0 | 0 | 3 | 1 | 0 | 0 | 0 | 0 | 0 | 0 | 0 | 298 | 149 | 1 | 1 | 1 | 0.231 | NA | 1 | 1 | NA | 1 | 1 | NA | NA | -0.192 |
| SE | 109630 | ENSG00000005810 | MYCBP2 | chr13 | - | 77140046 | 77140163 | 77139195 | 77139336 | 77140845 | 77140943 | 249 | 229 | 258 | 236 | 326 | 0 | 0 | 0 | 0 | 0 | 55 | 217 | 149 | 51 | 20 | 0 | 1 | 0 | 1 | 0 | 266 | 149 | 1 | 1 | 1 | 1 | 1 | 1 | 1 | 1 | 0.992 | 1 | 0.966 | 1 | 0.008 |
| SE | 109631 | ENSG00000005810 | MYCBP2 | chr13 | - | 77140845 | 77140943 | 77140046 | 77140163 | 77144444 | 77144560 | 302 | 228 | 298 | 261 | 345 | 0 | 1 | 1 | 0 | 1 | 81 | 274 | 162 | 50 | 21 | 0 | 0 | 2 | 0 | 0 | 247 | 149 | 1 | 1 | 1 | 0.993 | 0.994 | 1 | 0.995 | 1 | 1 | 0.98 | 1 | 1 | 0 |
| SE | 109632 | ENSG00000005810 | MYCBP2 | chr13 | - | 77156057 | 77156202 | 77150733 | 77150949 | 77157936 | 77158109 | 465 | 443 | 514 | 604 | 894 | 0 | 0 | 1 | 0 | 0 | 75 | 343 | 303 | 123 | 73 | 0 | 0 | 1 | 0 | 0 | 294 | 149 | 1 | 1 | 1 | 1 | 0.996 | 1 | 1 | 1 | 1 | 0.994 | 1 | 1 | 0 |
| SE | 109633 | ENSG00000005810 | MYCBP2 | chr13 | - | 77157936 | 77158109 | 77156057 | 77156202 | 77161905 | 77161955 | 403 | 402 | 413 | 509 | 852 | 4 | 0 | 0 | 0 | 2 | 54 | 275 | 270 | 139 | 83 | 0 | 0 | 0 | 0 | 0 | 298 | 149 | 1 | 1 | 0.981 | 1 | 1 | 1 | 0.995 | 1 | 1 | 1 | 1 | 1 | -0.005 |
| SE | 109634 | ENSG00000005810 | MYCBP2 | chr13 | - | 77161905 | 77161955 | 77157936 | 77158109 | 77164453 | 77164541 | 193 | 169 | 163 | 197 | 339 | 0 | 0 | 1 | 0 | 0 | 15 | 88 | 116 | 87 | 21 | 0 | 0 | 0 | 0 | 0 | 199 | 149 | 1 | 1 | 1 | 1 | 0.992 | 1 | 1 | 1 | 1 | 1 | 1 | 1 | -0.002 |
| SE | 109638 | ENSG00000005810 | MYCBP2 | chr13 | - | 77165272 | 77165391 | 77164453 | 77164541 | 77166328 | 77166554 | 504 | 499 | 428 | 524 | 773 | 1 | 0 | 0 | 1 | 0 | 177 | 499 | 374 | 176 | 68 | 0 | 0 | 0 | 4 | 0 | 268 | 149 | 1 | 1 | 0.996 | 1 | 1 | 0.997 | 1 | 1 | 1 | 1 | 0.961 | 1 | 0.006 |
| SE | 109639 | ENSG00000005810 | MYCBP2 | chr13 | - | 77169613 | 77169714 | 77168427 | 77168646 | 77171491 | 77171634 | 427 | 350 | 370 | 375 | 685 | 4 | 2 | 0 | 2 | 0 | 108 | 310 | 268 | 136 | 29 | 0 | 0 | 0 | 0 | 0 | 250 | 149 | 1 | 1 | 0.985 | 0.991 | 1 | 0.991 | 1 | 1 | 1 | 1 | 1 | 1 | -0.007 |
| SE | 109640 | ENSG00000005810 | MYCBP2 | chr13 | - | 77174310 | 77174489 | 77171491 | 77171634 | 77176496 | 77176628 | 491 | 518 | 463 | 462 | 889 | 0 | 0 | 0 | 1 | 0 | 189 | 333 | 339 | 154 | 34 | 0 | 0 | 0 | 0 | 0 | 298 | 149 | 1 | 1 | 1 | 1 | 1 | 0.996 | 1 | 1 | 1 | 1 | 1 | 1 | -0.001 |
| SE | 109641 | ENSG00000005810 | MYCBP2 | chr13 | - | 77177747 | 77177954 | 77176496 | 77176628 | 77180126 | 77180318 | 992 | 948 | 832 | 1014 | 1412 | 2 | 2 | 0 | 0 | 0 | 509 | 798 | 666 | 365 | 135 | 0 | 0 | 0 | 2 | 0 | 298 | 149 | 1 | 1 | 0.996 | 0.996 | 1 | 1 | 1 | 1 | 1 | 1 | 0.989 | 1 | 0.001 |
| SE | 109642 | ENSG00000005810 | MYCBP2 | chr13 | - | 77181700 | 77181922 | 77180126 | 77180318 | 77185102 | 77185377 | 519 | 462 | 493 | 504 | 683 | 0 | 0 | 0 | 0 | 0 | 160 | 348 | 294 | 128 | 73 | 0 | 0 | 1 | 0 | 0 | 298 | 149 | 1 | 1 | 1 | 1 | 1 | 1 | 1 | 1 | 1 | 0.993 | 1 | 1 | 0.001 |
| SE | 109644 | ENSG00000005810 | MYCBP2 | chr13 | - | 77211955 | 77212160 | 77211166 | 77211320 | 77217839 | 77217957 | 396 | 292 | 371 | 378 | 402 | 0 | 0 | 0 | 1 | 0 | 125 | 252 | 289 | 145 | 125 | 0 | 0 | 0 | 0 | 0 | 298 | 149 | 1 | 1 | 1 | 1 | 1 | 0.995 | 1 | 1 | 1 | 1 | 1 | 1 | -0.001 |
| SE | 109645 | ENSG00000005810 | MYCBP2 | chr13 | - | 77217839 | 77217957 | 77211955 | 77212160 | 77224450 | 77224532 | 397 | 298 | 377 | 450 | 396 | 0 | 7 | 1 | 1 | 2 | 93 | 258 | 235 | 111 | 127 | 0 | 3 | 0 | 5 | 0 | 267 | 149 | 1 | 1 | 1 | 0.96 | 0.995 | 0.996 | 0.991 | 1 | 0.98 | 1 | 0.925 | 1 | 0.007 |
| SE | 109647 | ENSG00000005810 | MYCBP2 | chr13 | - | 77251150 | 77251355 | 77243805 | 77243951 | 77257670 | 77257829 | 668 | 836 | 721 | 851 | 851 | 0 | 0 | 2 | 0 | 1 | 324 | 626 | 598 | 271 | 94 | 0 | 0 | 0 | 0 | 0 | 298 | 149 | 1 | 1 | 1 | 1 | 0.994 | 1 | 0.998 | 1 | 1 | 1 | 1 | 1 | -0.002 |
| SE | 109648 | ENSG00000005810 | MYCBP2 | chr13 | - | 77263650 | 77263789 | 77262052 | 77262129 | 77263928 | 77264002 | 623 | 677 | 530 | 591 | 759 | 0 | 0 | 1 | 0 | 0 | 311 | 429 | 374 | 301 | 65 | 0 | 0 | 0 | 0 | 0 | 288 | 149 | 1 | 1 | 1 | 1 | 0.996 | 1 | 1 | 1 | 1 | 1 | 1 | 1 | -0.001 |
| SE | 109649 | ENSG00000005810 | MYCBP2 | chr13 | - | 77270295 | 77270538 | 77269991 | 77270063 | 77273471 | 77273668 | 567 | 453 | 571 | 645 | 758 | 1 | 0 | 0 | 0 | 2 | 179 | 389 | 268 | 177 | 158 | 0 | 0 | 0 | 0 | 0 | 298 | 149 | 1 | 1 | 0.996 | 1 | 1 | 1 | 0.995 | 1 | 1 | 1 | 1 | 1 | -0.002 |
| SE | 109651 | ENSG00000005810 | MYCBP2 | chr13 | - | 77278757 | 77278911 | 77273471 | 77273668 | 77288160 | 77288376 | 408 | 285 | 355 | 445 | 380 | 1 | 0 | 2 | 1 | 3 | 137 | 186 | 218 | 132 | 134 | 0 | 0 | 0 | 1 | 0 | 298 | 149 | 1 | 1 | 0.995 | 1 | 0.989 | 0.996 | 0.984 | 1 | 1 | 1 | 0.985 | 1 | -0.004 |
| SE | 109652 | ENSG00000005810 | MYCBP2 | chr13 | - | 77288160 | 77288376 | 77273471 | 77273668 | 77296598 | 77296674 | 205 | 196 | 168 | 189 | 190 | 0 | 0 | 0 | 0 | 2 | 53 | 118 | 103 | 51 | 48 | 0 | 0 | 0 | 0 | 0 | 298 | 149 | 1 | 1 | 1 | 1 | 1 | 1 | 0.979 | 1 | 1 | 1 | 1 | 1 | -0.004 |
| SE | 109653 | ENSG00000005810 | MYCBP2 | chr13 | - | 77296598 | 77296674 | 77288160 | 77288376 | 77326473 | 77327094 | 214 | 217 | 177 | 203 | 181 | 0 | 0 | 2 | 0 | 0 | 64 | 132 | 115 | 55 | 61 | 0 | 0 | 0 | 0 | 0 | 225 | 149 | 1 | 1 | 1 | 1 | 0.983 | 1 | 1 | 1 | 1 | 1 | 1 | 1 | -0.003 |
| SE | 109654 | ENSG00000139182 | CLSTN3 | chr12 | + | 7135803 | 7135953 | 7135326 | 7135535 | 7136205 | 7136391 | 95 | 41 | 78 | 55 | 66 | 0 | 0 | 0 | 0 | 0 | 67 | 201 | 85 | 147 | 162 | 0 | 4 | 0 | 0 | 0 | 298 | 149 | 1 | 1 | 1 | 1 | 1 | 1 | 1 | 1 | 1 | 0.914 | 1 | 1 | 0.017 |
| SE | 109657 | ENSG00000139182 | CLSTN3 | chr12 | + | 7142085 | 7142139 | 7141241 | 7141404 | 7142868 | 7143026 | 92 | 24 | 43 | 62 | 128 | 1 | 0 | 0 | 0 | 0 | 80 | 151 | 61 | 107 | 119 | 0 | 3 | 0 | 0 | 0 | 203 | 149 | 1 | 1 | 0.985 | 1 | 1 | 1 | 1 | 1 | 0.974 | 1 | 1 | 1 | 0.002 |
| SE | 109658 | ENSG00000139182 | CLSTN3 | chr12 | + | 7142868 | 7143026 | 7142085 | 7142139 | 7148971 | 7149198 | 29 | 3 | 14 | 13 | 46 | 0 | 0 | 0 | 0 | 0 | 18 | 29 | 19 | 24 | 28 | 0 | 2 | 0 | 0 | 0 | 298 | 149 | 1 | 1 | 1 | 1 | 1 | 1 | 1 | 1 | 0.879 | 1 | 1 | 1 | 0.024 |
| SE | 109659 | ENSG00000139182 | CLSTN3 | chr12 | + | 7143162 | 7143311 | 7142868 | 7143026 | 7148971 | 7149198 | 29 | 15 | 23 | 16 | 4 | 0 | 0 | 0 | 0 | 0 | 47 | 58 | 33 | 26 | 87 | 0 | 0 | 0 | 0 | 3 | 298 | 149 | 1 | 1 | 1 | 1 | 1 | 1 | 1 | 1 | 1 | 1 | 1 | 0.935 | 0.013 |
| SE | 109661 | ENSG00000154122 | ANKH | chr5 | - | 14749171 | 14749306 | 14745869 | 14745962 | 14751068 | 14751239 | 140 | 51 | 153 | 161 | 135 | 0 | 0 | 0 | 0 | 0 | 2 | 55 | 33 | 32 | 84 | 0 | 0 | 0 | 3 | 0 | 284 | 149 | 1 | 1 | 1 | 1 | 1 | 1 | 1 | 1 | 1 | 1 | 0.848 | 1 | 0.03 |
| SE | 109663 | ENSG00000198331 | HYLS1 | chr11 | + | 125891422 | 125891472 | 125887667 | 125887765 | 125899343 | 125900617 | 4 | 3 | 6 | 5 | 0 | 0 | 0 | 2 | 0 | 0 | 0 | 0 | 4 | 3 | 0 | 0 | 0 | 0 | 0 | 0 | 199 | 149 | 1 | 1 | 1 | 1 | 0.692 | 1 | NA | NA | NA | 1 | 1 | NA | -0.077 |
| SE | 109664 | ENSG00000198331 | HYLS1 | chr11 | + | 125899103 | 125899158 | 125891422 | 125891472 | 125899343 | 125900644 | 0 | 0 | 0 | 0 | 0 | 4 | 3 | 7 | 5 | 0 | 0 | 0 | 2 | 0 | 0 | 0 | 0 | 4 | 3 | 0 | 204 | 149 | 1 | 1 | 0 | 0 | 0 | 0 | NA | NA | NA | 0.268 | 0 | NA | -0.134 |
| SE | 109665 | ENSG00000136114 | THSD1 | chr13 | - | 52386027 | 52386186 | 52377166 | 52378789 | 52397231 | 52398194 | 1 | 0 | 2 | 0 | 0 | 1 | 0 | 2 | 0 | 3 | 0 | 0 | 0 | 4 | 0 | 8 | 3 | 0 | 0 | 0 | 298 | 149 | 1 | 1 | 0.333 | NA | 0.333 | NA | 0 | 0 | 0 | NA | 1 | NA | -0.111 |

**U12_SE_filtered**

| Splice_type | ID | Gene_ID | Gene_symbol | Chromosome | Strand | LongExonStart_0base | LongExonEnd | ShortES | ShortEE | FlankingES | FlankingEE |
|---|---|---|---|---|---|---|---|---|---|---|---|
| SE | 7693 | ENSG00000170004 | CHD3 | chr17 | + | 7902936 | 7903061 | 7902183 | 7902727 | 7903271 | 7903503 |
| SE | 15465 | ENSG00000140199 | SLC12A6 | chr15 | - | 34257641 | 34257788 | 34256228 | 34256283 | 34258812 | 34258944 |
| SE | 30535 | ENSG00000132182 | NUP210 | chr3 | - | 13335453 | 13335612 | 13332292 | 13332384 | 13336786 | 13336918 |
| SE | 35452 | ENSG00000111642 | CHD4 | chr12 | - | 6582136 | 6582281 | 6581648 | 6581814 | 6582614 | 6582748 |
| SE | 35456 | ENSG00000111642 | CHD4 | chr12 | - | 6588297 | 6588422 | 6587660 | 6587949 | 6591465 | 6591583 |
| SE | 47672 | ENSG00000163479 | SSR2 | chr1 | - | 156019084 | 156019266 | 156018131 | 156018368 | 156020012 | 156020167 |
| SE | 65650 | ENSG00000143924 | EML4 | chr2 | + | 42301240 | 42301392 | 42295380 | 42295516 | 42303103 | 42303229 |
| SE | 98435 | ENSG00000099783 | HNRNPM | chr19 | + | 8473663 | 8473708 | 8471325 | 8471427 | 8474166 | 8474244 |
| SE | 100288 | ENSG00000167986 | DDB1 | chr11 | - | 61324237 | 61324387 | 61323978 | 61324137 | 61325610 | 61325708 |
| SE | 102579 | ENSG00000120438 | TCP1 | chr6 | - | 159780934 | 159781110 | 159780442 | 159780566 | 159783940 | 159784067 |
| SE | 109055 | ENSG00000139668 | WDFY2 | chr13 | + | 51739048 | 51739175 | 51727677 | 51727790 | 51751309 | 51751415 |
| SE | 96611 | ENSG00000030066 | NUP160 | chr11 | - | 47812881 | 47813047 | 47812063 | 47812429 | 47813315 | 47813415 |
| SE | 22551 | ENSG00000167881 | SRP68 | chr17 | - | 76050427 | 76050526 | 76047905 | 76047970 | 76057402 | 76057543 |
| SE | 16961 | ENSG00000172575 | RASGRP1 | chr15 | - | 38503271 | 38503376 | 38502311 | 38502421 | 38505839 | 38505920 |
| SE | 35458 | ENSG00000111642 | CHD4 | chr12 | - | 6600925 | 6601053 | 6600533 | 6600669 | 6601288 | 6601530 |
| SE | 8273 | ENSG00000161654 | LSM12 | chr17 | - | 44036861 | 44037102 | 44036142 | 44036300 | 44037411 | 44037538 |
| SE | 98439 | ENSG00000099783 | HNRNPM | chr19 | + | 8483157 | 8483211 | 8474166 | 8474244 | 8485602 | 8485648 |
| [illegible] | [illegible] | [illegible] | [illegible] | [illegible] | [illegible] | [illegible] | [illegible] | [illegible] | [illegible] | [illegible] | [illegible] |

| ID | Inclusion_counts_HC SRR13214146 | SRR13214147 | SRR13214148 | SRR13214149 | SRR13214150 | Skipping_counts_HC SRR13214146 | SRR13214147 | SRR13214148 | SRR13214149 | SRR13214150 | Inclusion counts_SLE SRR13214141 | SRR13214242 | SRR13214143 | SRR13214144 | SRR13214145 |
|---|---|---|---|---|---|---|---|---|---|---|---|---|---|---|---|
| 7693 | 447 | 286 | 590 | 611 | 417 | 4 | 3 | 3 | 10 | 0 | 686 | 429 | 349 | 465 | 804 |
| 15465 | 131 | 109 | 202 | 158 | 237 | 4 | 6 | 5 | 7 | 14 | 120 | 71 | 75 | 80 | 99 |
| 30535 | 460 | 214 | 397 | 368 | 282 | 2 | 0 | 1 | 3 | 2 | 337 | 479 | 251 | 232 | 500 |
| 35452 | 370 | 295 | 475 | 553 | 431 | 1 | 4 | 1 | 9 | 0 | 860 | 522 | 540 | 637 | 777 |
| 35456 | 495 | 323 | 504 | 485 | 409 | 1 | 4 | 1 | 11 | 12 | 888 | 705 | 439 | 624 | 813 |
| 47672 | 3 | 1 | 2 | 7 | 14 | 757 | 301 | 762 | 630 | 820 | 33 | 23 | 40 | 24 | 37 |
| 65650 | 792 | 844 | 998 | 986 | 1068 | 4 | 1 | 1 | 4 | 2 | 255 | 733 | 870 | 514 | 93 |
| 98435 | 976 | 712 | 836 | 1030 | 1010 | 20 | 15 | 18 | 27 | 25 | 1985 | 1058 | 1077 | 1400 | 1109 |
| 100288 | 4 | 4 | 4 | 9 | 2 | 187 | 86 | 160 | 147 | 179 | 49 | 48 | 51 | 31 | 15 |
| 102579 | 894 | 774 | 773 | 825 | 973 | 6 | 5 | 3 | 22 | 21 | 628 | 492 | 741 | 679 | 311 |
| 109055 | 76 | 48 | 60 | 75 | 73 | 0 | 0 | 3 | 0 | 2 | 91 | 88 | 70 | 105 | 137 |
| 96611 | 239 | 133 | 192 | 231 | 276 | 0 | 0 | 0 | 1 | 2 | 53 | 97 | 139 | 130 | 16 |
| 22551 | 270 | 219 | 282 | 279 | 358 | 0 | 2 | 3 | 4 | 3 | 259 | 234 | 254 | 219 | 290 |
| 16961 | 439 | 277 | 482 | 444 | 645 | 21 | 12 | 32 | 30 | 38 | 9 | 240 | 139 | 77 | 42 |
| 35458 | 455 | 262 | 382 | 402 | 372 | 0 | 0 | 7 | 3 | 3 | 495 | 431 | 410 | 468 | 713 |
| 8273 | 0 | 1 | 2 | 3 | 3 | 228 | 145 | 208 | 219 | 340 | 34 | 24 | 9 | 17 | 16 |
| 98439 | 146 | 136 | 152 | 156 | 178 | 20 | 8 | 14 | 33 | 17 | 323 | 166 | 251 | 237 | 256 |
| [illegible] | [illegible] | [illegible] | [illegible] | [illegible] | [illegible] | [illegible] | [illegible] | [illegible] | [illegible] | [illegible] | [illegible] | [illegible] | [illegible] | [illegible] | [illegible] |

| ID | Skipping_counts_SLE SRR13214141 | SRR13214242 | SRR13214143 | SRR13214144 | SRR13214145 | IncFormLen | SkipFormLen | p value | FDR | Inclusion_level(normalized) SRR13214146 | SRR13214147 | SRR13214148 | SRR13214149 | SRR13214150 | SRR13214141 | SRR13214242 | SRR13214143 | SRR13214144 | SRR13214145 | Inclevel_diff |
|---|---|---|---|---|---|---|---|---|---|---|---|---|---|---|---|---|---|---|---|---|
| 7693 | 38 | 31 | 20 | 45 | 20 | 274 | 149 | 0 | 0 | 0.984 | 0.981 | 0.991 | 0.971 | 1 | 0.908 | 0.883 | 0.905 | 0.849 | 0.956 | 0.085 |
| 15465 | 35 | 15 | 32 | 25 | 18 | 296 | 149 | 0 | 0 | 0.943 | 0.901 | 0.953 | 0.919 | 0.895 | 0.633 | 0.704 | 0.541 | 0.617 | 0.735 | 0.276 |
| 30535 | 22 | 10 | 8 | 29 | 17 | 298 | 149 | 0 | 0 | 0.991 | 1 | 0.995 | 0.984 | 0.986 | 0.885 | 0.96 | 0.94 | 0.8 | 0.936 | 0.087 |
| 35452 | 103 | 36 | 41 | 63 | 1 | 294 | 149 | 0 | 0 | 0.995 | 0.974 | 0.996 | 0.969 | 1 | 0.809 | 0.88 | 0.87 | 0.837 | 0.997 | 0.108 |
| 35456 | 60 | 71 | 37 | 89 | 15 | 274 | 149 | 0 | 0 | 0.996 | 0.978 | 0.996 | 0.96 | 0.949 | 0.889 | 0.844 | 0.866 | 0.792 | 0.967 | 0.104 |
| 47672 | 463 | 591 | 434 | 454 | 680 | 298 | 149 | 0 | 0 | 0.002 | 0.002 | 0.001 | 0.006 | 0.008 | 0.034 | 0.019 | 0.044 | 0.026 | 0.026 | -0.026 |
| 65650 | 0 | 19 | 35 | 20 | 5 | 298 | 149 | 0 | 0 | 0.99 | 0.998 | 0.998 | 0.992 | 0.996 | 1 | 0.951 | 0.926 | 0.928 | 0.903 | 0.053 |
| 98435 | 134 | 58 | 80 | 130 | 70 | 194 | 149 | 0 | 0 | 0.974 | 0.973 | 0.973 | 0.967 | 0.969 | 0.919 | 0.933 | 0.912 | 0.892 | 0.924 | 0.055 |
| 100288 | 322 | 172 | 98 | 129 | 222 | 298 | 149 | 0 | 0 | 0.011 | 0.023 | 0.012 | 0.03 | 0.006 | 0.071 | 0.122 | 0.206 | 0.107 | 0.033 | -0.091 |
| 102579 | 17 | 55 | 24 | 121 | 12 | 298 | 149 | 0 | 0 | 0.987 | 0.987 | 0.992 | 0.949 | 0.959 | 0.949 | 0.817 | 0.939 | 0.737 | 0.928 | 0.101 |
| 109055 | 33 | 16 | 25 | 24 | 11 | 276 | 149 | 0 | 0 | 1 | 1 | 0.915 | 1 | 0.952 | 0.598 | 0.748 | 0.602 | 0.703 | 0.871 | 0.269 |
| 96611 | 22 | 12 | 5 | 15 | 0 | 298 | 149 | 2.22045E-16 | 9.9915E-14 | 1 | 1 | 1 | 0.991 | 0.986 | 0.546 | 0.802 | 0.933 | 0.812 | 1 | 0.177 |
| 22551 | 35 | 6 | 14 | 10 | 17 | 248 | 149 | 7.77156E-16 | 3.13651E-13 | 1 | 0.985 | 0.983 | 0.977 | 0.986 | 0.816 | 0.959 | 0.916 | 0.929 | 0.911 | 0.08 |
| 16961 | 8 | 47 | 73 | 16 | 31 | 254 | 149 | 8.02691E-14 | 2.32769E-11 | 0.925 | 0.931 | 0.898 | 0.897 | 0.909 | 0.398 | 0.75 | 0.528 | 0.738 | 0.443 | 0.341 |
| 35458 | 31 | 11 | 10 | 36 | 12 | 277 | 149 | 8.59313E-14 | 2.4555E-11 | 1 | 1 | 0.967 | 0.986 | 0.985 | 0.896 | 0.955 | 0.957 | 0.875 | 0.97 | 0.057 |
| 8273 | 352 | 260 | 202 | 237 | 330 | 298 | 149 | 1.76525E-13 | 4.80206E-11 | 0 | 0.003 | 0.005 | 0.007 | 0.004 | 0.046 | 0.044 | 0.022 | 0.035 | 0.024 | -0.03 |
| 98439 | 5 | 16 | 2 | 5 | 5 | 203 | 149 | 1.75449E-12 | 3.89148E-10 | 0.843 | 0.926 | 0.889 | 0.776 | 0.885 | 0.979 | 0.884 | 0.989 | 0.972 | 0.974 | -0.096 |
| [illegible] | [illegible] | [illegible] | [illegible] | [illegible] | [illegible] | [illegible] | [illegible] | [illegible] | [illegible] | [illegible] | [illegible] | [illegible] | [illegible] | [illegible] | [illegible] | [illegible] | [illegible] | [illegible] | [illegible] | [illegible] |

| | | | | | | | | | | | | | | | | | | | | | | | | | | | | | | | | | | | | | | | | | | | | | | |
|---|---|---|---|---|---|---|---|---|---|---|---|---|---|---|---|---|---|---|---|---|---|---|---|---|---|---|---|---|---|---|---|---|---|---|---|---|---|---|---|---|---|---|---|---|---|---|
| SE | 55696 | ENSG00000135749 | PCNX2 | chr1 | - | 233160282 | 233160433 | 233139713 | 233139855 | 233161270 | 233161363 | 60 | 35 | 106 | 63 | 79 | 2 | 0 | 6 | 0 | 5 | 29 | 39 | 47 | 30 | 17 | 0 | 0 | 0 | 0 | 0 | 298 | 149 | 0.00066348 | 0.010984398 | 0.938 | 1 | 0.898 | 1 | 0.888 | 1 | 1 | 1 | 1 | 1 | -0.055 |
| SE | 1567 | ENSG00000198089 | SFI1 | chr22 | + | 31546860 | 31546971 | 31508254 | 31508376 | 31556941 | 31557059 | 16 | 19 | 24 | 18 | 12 | 0 | 0 | 0 | 0 | 0 | 53 | 10 | 14 | 23 | 7 | 11 | 0 | 0 | 0 | 0 | 260 | 149 | 0.000715219 | 0.011695654 | 1 | 1 | 1 | 1 | 1 | 0.734 | 1 | 1 | 1 | 1 | 0.053 |
| SE | 16689 | ENSG00000137841 | PLCB2 | chr15 | - | 40290024 | 40290082 | 40289065 | 40289358 | 40290576 | 40290672 | 656 | 487 | 502 | 713 | 652 | 35 | 29 | 12 | 29 | 19 | 1576 | 897 | 681 | 1447 | 1701 | 37 | 30 | 24 | 39 | 27 | 207 | 149 | 0.000732055 | 0.011928616 | 0.931 | 0.924 | 0.968 | 0.947 | 0.961 | 0.968 | 0.956 | 0.953 | 0.964 | 0.978 | -0.018 |
| SE | 105518 | ENSG00000084110 | HAL | chr12 | - | 95990392 | 95990532 | 95988192 | 95988240 | 95992679 | 95992805 | 38 | 20 | 29 | 48 | 49 | 1 | 0 | 0 | 0 | 4 | 64 | 61 | 95 | 93 | 94 | 0 | 3 | 9 | 13 | 0 | 289 | 149 | 0.00075769 | 0.012277337 | 0.951 | 1 | 1 | 1 | 0.863 | 1 | 0.913 | 0.845 | 0.787 | 1 | 0.054 |
| SE | 13612 | ENSG00000038358 | EDC4 | chr16 | + | 67875737 | 67875837 | 67873051 | 67873343 | 67875944 | 67876101 | 1 | 2 | 3 | 0 | 0 | 25 | 9 | 16 | 24 | 4 | 0 | 0 | 9 | 4 | 9 | 14 | 27 | 14 | 15 | 26 | 249 | 149 | 0.000784141 | 0.012628392 | 0.023 | 0.117 | 0.101 | 0 | 0 | 0 | 0 | 0.278 | 0.138 | 0.172 | -0.069 |
| SE | 69003 | ENSG00000107521 | HPS1 | chr10 | - | 98431130 | 98431291 | 98430570 | 98430670 | 98433939 | 98434091 | 28 | 13 | 35 | 20 | 23 | 18 | 8 | 7 | 8 | 9 | 44 | 37 | 29 | 42 | 77 | 0 | 7 | 1 | 6 | 27 | 298 | 149 | 0.000789927 | 0.012686788 | 0.437 | 0.448 | 0.714 | 0.556 | 0.561 | 1 | 0.725 | 0.935 | 0.778 | 0.588 | -0.262 |
| SE | 80842 | ENSG00000164885 | CDK5 | chr7 | - | 151055752 | 151055848 | 151055531 | 151055606 | 151056579 | 151056636 | 46 | 28 | 38 | 54 | 69 | 2 | 4 | 2 | 0 | 4 | 127 | 117 | 33 | 104 | 74 | 25 | 4 | 4 | 10 | 15 | 245 | 149 | 0.000792299 | 0.012719667 | 0.933 | 0.81 | 0.92 | 1 | 0.913 | 0.755 | 0.947 | 0.834 | 0.863 | 0.75 | 0.085 |
| SE | 45891 | ENSG00000145740 | SLC30A5 | chr5 | + | 69128003 | 69128132 | 69123198 | 69123425 | 69129446 | 69131065 | 256 | 227 | 241 | 237 | 329 | 6 | 1 | 6 | 5 | 6 | 78 | 103 | 148 | 97 | 69 | 0 | 0 | 0 | 1 | 0 | 278 | 149 | 0.000824598 | 0.013138526 | 0.958 | 0.992 | 0.956 | 0.962 | 0.967 | 1 | 1 | 1 | 0.981 | 1 | -0.029 |
| SE | 86265 | ENSG00000134516 | DOCK2 | chr5 | + | 169761518 | 169761625 | 169747395 | 169747504 | 169803057 | 169803206 | 676 | 317 | 391 | 619 | 1027 | 8 | 3 | 5 | 13 | 6 | 1563 | 841 | 876 | 709 | 546 | 3 | 18 | 31 | 33 | 10 | 256 | 149 | 0.000829343 | 0.013203382 | 0.98 | 0.984 | 0.979 | 0.965 | 0.99 | 0.997 | 0.965 | 0.943 | 0.926 | 0.969 | 0.02 |
| SE | 14341 | ENSG00000007545 | CRAMP1 | chr16 | + | 1667334 | 1667400 | 1666421 | 1666600 | 1667961 | 1668193 | 31 | 3 | 27 | 15 | 8 | 2 | 11 | 0 | 3 | 0 | 15 | 38 | 12 | 8 | 21 | 0 | 0 | 0 | 3 | 0 | 215 | 149 | 0.000891876 | 0.014047546 | 0.915 | 0.159 | 1 | 0.776 | 1 | 1 | 1 | 1 | 0.649 | 1 | -0.16 |
| SE | 1747 | ENSG00000100325 | ASCC2 | chr22 | - | 29832244 | 29832342 | 29825621 | 29825780 | 29834488 | 29834550 | 18 | 6 | 17 | 9 | 12 | 4 | 0 | 4 | 2 | 0 | 10 | 9 | 8 | 9 | 24 | 0 | 1 | 2 | 0 | 0 | 247 | 149 | 0.000930672 | 0.014591088 | 0.731 | 1 | 0.719 | 0.731 | 1 | 1 | 0.844 | 0.707 | 1 | 1 | -0.074 |
| SE | 57889 | ENSG00000162604 | TM2D1 | chr1 | - | 61723712 | 61723772 | 61709328 | 61709437 | 61724919 | 61725067 | 64 | 38 | 42 | 43 | 36 | 1 | 0 | 1 | 0 | 0 | 94 | 38 | 47 | 37 | 20 | 10 | 2 | 1 | 0 | 0 | 209 | 149 | 0.000939988 | 0.014713568 | 0.979 | 1 | 0.968 | 1 | 1 | 0.87 | 0.931 | 0.971 | 1 | 1 | 0.035 |
| SE | 69319 | ENSG00000112367 | FIG4 | chr6 | + | 109760249 | 109760383 | 109743109 | 109743772 | 109762090 | 109762140 | 145 | 109 | 84 | 123 | 197 | 4 | 3 | 3 | 0 | 2 | 157 | 131 | 50 | 110 | 31 | 22 | 0 | 2 | 5 | 0 | 283 | 149 | 0.000941667 | 0.01472325 | 0.95 | 0.95 | 0.936 | 1 | 0.981 | 0.79 | 1 | 0.929 | 0.921 | 1 | 0.035 |
| SE | 50705 | ENSG00000113318 | MSH3 | chr5 | + | 80744505 | 80744615 | 80728850 | 80728965 | 80761545 | 80761610 | 38 | 24 | 37 | 36 | 115 | 0 | 0 | 5 | 3 | 4 | 13 | 57 | 44 | 35 | 15 | 0 | 0 | 0 | 0 | 0 | 259 | 149 | 0.000999976 | 0.015457868 | 1 | 1 | 0.81 | 0.873 | 0.943 | 1 | 1 | 1 | 1 | 1 | -0.075 |
| SE | 73941 | ENSG00000144134 | RABL2A | chr2 | + | 113635050 | 113635130 | 113634152 | 113634232 | 113640893 | 113640938 | 28 | 4 | 9 | 11 | 24 | 0 | 0 | 2 | 2 | 0 | 21 | 40 | 9 | 19 | 34 | 0 | 6 | 0 | 0 | 10 | 229 | 149 | 0.001008471 | 0.015573289 | 1 | 1 | 0.745 | 0.782 | 1 | 1 | 0.813 | 1 | 1 | 0.689 | 0.005 |
| SE | 82950 | ENSG00000075142 | SRI | chr7 | - | 88214794 | 88214936 | 88210881 | 88210925 | 88217121 | 88217191 | 10 | 2 | 4 | 3 | 7 | 145 | 86 | 125 | 134 | 181 | 23 | 6 | 8 | 14 | 0 | 157 | 136 | 83 | 116 | 112 | 291 | 149 | 0.001008636 | 0.015573289 | 0.034 | 0.012 | 0.016 | 0.011 | 0.019 | 0.07 | 0.022 | 0.047 | 0.058 | 0 | -0.021 |
| SE | 41972 | ENSG00000120008 | WDR11 | chr10 | + | 120890715 | 120890887 | 120889894 | 120890009 | 120900028 | 120900137 | 207 | 170 | 160 | 168 | 349 | 4 | 1 | 9 | 7 | 4 | 81 | 140 | 63 | 102 | 30 | 0 | 1 | 2 | 1 | 0 | 298 | 149 | 0.001069546 | 0.016346116 | 0.963 | 0.988 | 0.899 | 0.923 | 0.978 | 1 | 0.986 | 0.94 | 0.981 | 1 | -0.031 |
| SE | 93923 | ENSG00000184465 | WDR27 | chr6 | - | 169689126 | 169689308 | 169688816 | 169689012 | 169701550 | 169701930 | 23 | 16 | 22 | 32 | 39 | 6 | 0 | 2 | 4 | 3 | 23 | 38 | 13 | 21 | 11 | 0 | 3 | 0 | 2 | 0 | 298 | 149 | 0.001115131 | 0.016914041 | 0.657 | 1 | 0.846 | 0.8 | 0.867 | 1 | 0.864 | 1 | 0.84 | 1 | -0.107 |
| SE | 33411 | ENSG00000096063 | SRPK1 | chr6 | - | 35869009 | 35869110 | 35857260 | 35857368 | 35869481 | 35869901 | 141 | 91 | 110 | 113 | 151 | 1 | 0 | 2 | 1 | 0 | 19 | 163 | 197 | 235 | 14 | 0 | 2 | 7 | 9 | 0 | 250 | 149 | 0.00112966 | 0.017094673 | 0.988 | 1 | 0.97 | 0.985 | 1 | 1 | 0.98 | 0.944 | 0.94 | 1 | 0.016 |
| SE | 85067 | ENSG00000147459 | DOCK5 | chr8 | + | 25368570 | 25368687 | 25366869 | 25366970 | 25369555 | 25369641 | 1 | 4 | 2 | 10 | 11 | 4 | 1 | 1 | 0 | 5 | 0 | 0 | 15 | 14 | 1 | 20 | 5 | 15 | 24 | 3 | 266 | 149 | 0.001145262 | 0.017290673 | 0.123 | 0.691 | 0.528 | 1 | 0.552 | 0 | 0 | 0.359 | 0.246 | 0.157 | 0.426 |
| SE | 96616 | ENSG00000030066 | NUP160 | chr11 | - | 47839842 | 47840065 | 47837544 | 47837623 | 47840377 | 47840588 | 127 | 78 | 118 | 99 | 125 | 2 | 2 | 0 | 0 | 3 | 29 | 115 | 36 | 59 | 16 | 0 | 5 | 8 | 2 | 0 | 298 | 149 | 0.001153851 | 0.017380137 | 0.969 | 0.951 | 1 | 1 | 0.954 | 1 | 0.92 | 0.692 | 0.937 | 1 | 0.065 |
| SE | 1553 | ENSG00000198089 | SFI1 | chr22 | + | 31497247 | 31497348 | 31496557 | 31496637 | 31508258 | 31508376 | 0 | 1 | 4 | 0 | 0 | 16 | 13 | 24 | 12 | 6 | 12 | 0 | 3 | 5 | 0 | 23 | 16 | 26 | 15 | 32 | 250 | 149 | 0.001185545 | 0.017791527 | 0 | 0.044 | 0.09 | 0 | 0 | 0.237 | 0 | 0.064 | 0.166 | 0 | -0.067 |
| SE | 29421 | ENSG00000169762 | TAPT1 | chr4 | - | 16202461 | 16202580 | 16191360 | 16191523 | 16213767 | 16213898 | 43 | 35 | 30 | 22 | 85 | 5 | 1 | 4 | 4 | 4 | 16 | 73 | 20 | 17 | 12 | 7 | 8 | 5 | 9 | 3 | 268 | 149 | 0.001185935 | 0.017791527 | 0.827 | 0.951 | 0.807 | 0.754 | 0.922 | 0.56 | 0.835 | 0.69 | 0.512 | 0.69 | 0.195 |
| SE | 12269 | ENSG00000103194 | USP10 | chr16 | + | 84704739 | 84704860 | 84699994 | 84700111 | 84733434 | 84733503 | 0 | 0 | 0 | 0 | 0 | 32 | 31 | 24 | 40 | 18 | 0 | 2 | 0 | 2 | 10 | 17 | 46 | 6 | 20 | 29 | 270 | 149 | 0.001203775 | 0.018028076 | 0 | 0 | 0 | 0 | 0 | 0 | 0.023 | 0 | 0.052 | 0.16 | -0.047 |
| SE | 42793 | ENSG00000139842 | CUL4A | chr13 | + | 113239432 | 113239551 | 113236822 | 113236890 | 113242967 | 113243160 | 475 | 302 | 339 | 376 | 434 | 1 | 3 | 1 | 2 | 1 | 506 | 372 | 218 | 446 | 211 | 0 | 0 | 5 | 23 | 0 | 268 | 149 | 0.00122346 | 0.018287904 | 0.996 | 0.982 | 0.995 | 0.991 | 0.996 | 1 | 1 | 0.96 | 0.915 | 1 | 0.017 |
| SE | 59366 | ENSG00000162927 | PUS10 | chr2 | - | 61010655 | 61010888 | 61008780 | 61009015 | 61011764 | 61011905 | 8 | 5 | 8 | 9 | 17 | 20 | 16 | 7 | 11 | 20 | 17 | 3 | 9 | 2 | 3 | 0 | 2 | 8 | 1 | 0 | 298 | 149 | 0.001228871 | 0.018365062 | 0.167 | 0.135 | 0.364 | 0.29 | 0.298 | 1 | 0.429 | 0.36 | 0.5 | 1 | -0.407 |
| SE | 22799 | ENSG00000131504 | DIAPH1 | chr5 | - | 141529602 | 141529697 | 141529171 | 141529273 | 141534334 | 141534433 | 623 | 382 | 397 | 483 | 988 | 4 | 4 | 1 | 1 | 7 | 890 | 475 | 337 | 452 | 1156 | 6 | 8 | 5 | 19 | 8 | 244 | 149 | 0.001291581 | 0.019138089 | 0.99 | 0.983 | 0.996 | 0.997 | 0.989 | 0.989 | 0.973 | 0.976 | 0.936 | 0.989 | 0.018 |
| SE | 19023 | ENSG00000131966 | ACTR10 | chr14 | + | 58213630 | 58213698 | 58208998 | 58209107 | 58215204 | 58215284 | 33 | 23 | 29 | 19 | 54 | 12 | 14 | 4 | 13 | 51 | 3 | 23 | 11 | 12 | 20 | 6 | 31 | 21 | 23 | 13 | 217 | 149 | 0.001324975 | 0.019573629 | 0.654 | 0.53 | 0.833 | 0.501 | 0.421 | 0.256 | 0.338 | 0.265 | 0.264 | 0.514 | 0.26 |
| SE | 21236 | ENSG00000136986 | DERL1 | chr8 | - | 123030604 | 123030716 | 123024985 | 123025050 | 123041969 | 123042122 | 274 | 122 | 188 | 198 | 138 | 1 | 2 | 4 | 0 | 2 | 188 | 175 | 201 | 144 | 226 | 0 | 12 | 7 | 6 | 0 | 261 | 149 | 0.001347741 | 0.019864979 | 0.994 | 0.972 | 0.964 | 1 | 0.975 | 1 | 0.893 | 0.943 | 0.932 | 1 | 0.027 |
| SE | 38017 | ENSG00000197386 | HTT | chr4 | + | 3199731 | 3199939 | 3187650 | 3189093 | 3204006 | 3204148 | 219 | 133 | 156 | 236 | 353 | 2 | 2 | 0 | 0 | 4 | 387 | 187 | 182 | 244 | 546 | 10 | 1 | 2 | 5 | 11 | 298 | 149 | 0.001379972 | 0.020275159 | 0.982 | 0.971 | 1 | 1 | 0.978 | 0.951 | 0.989 | 0.978 | 0.961 | 0.961 | 0.018 |
| SE | 78634 | ENSG00000168014 | C2CD3 | chr11 | - | 74057405 | 74057544 | 74054606 | 74054671 | 74074252 | 74074505 | 23 | 17 | 52 | 31 | 21 | 2 | 0 | 0 | 3 | 5 | 17 | 59 | 19 | 39 | 37 | 0 | 0 | 1 | 0 | 0 | 288 | 149 | 0.001399374 | 0.020510178 | 0.856 | 1 | 1 | 0.842 | 0.685 | 1 | 1 | 0.908 | 1 | 1 | -0.105 |
| SE | 88037 | ENSG00000156171 | DRAM2 | chr1 | - | 111124741 | 111124881 | 111119876 | 111119959 | 111126226 | 111126294 | 18 | 8 | 6 | 20 | 38 | 0 | 0 | 3 | 3 | 0 | 23 | 16 | 3 | 5 | 4 | 0 | 3 | 2 | 4 | 5 | 289 | 149 | 0.001430859 | 0.020905121 | 1 | 1 | 0.508 | 0.775 | 1 | 1 | 0.733 | 0.436 | 0.392 | 0.292 | 0.286 |
| SE | 107652 | ENSG00000139718 | SETD1B | chr12 | + | 121808207 | 121808320 | 121805834 | 121806105 | 121809602 | 121810835 | 73 | 60 | 85 | 101 | 91 | 0 | 0 | 1 | 0 | 0 | 188 | 150 | 64 | 101 | 184 | 9 | 1 | 2 | 5 | 0 | 262 | 149 | 0.001442204 | 0.021035551 | 1 | 1 | 0.98 | 1 | 1 | 0.922 | 0.988 | 0.948 | 0.92 | 1 | 0.04 |
| SE | 55707 | ENSG00000135749 | PCNX2 | chr1 | - | 233227225 | 233227371 | 233218030 | 233218184 | 233236844 | 233236980 | 124 | 102 | 146 | 162 | 277 | 7 | 7 | 11 | 5 | 6 | 89 | 93 | 137 | 107 | 99 | 1 | 14 | 18 | 5 | 18 | 295 | 149 | 0.001475607 | 0.021430924 | 0.899 | 0.88 | 0.87 | 0.942 | 0.959 | 0.978 | 0.77 | 0.794 | 0.915 | 0.735 | 0.072 |
| SE | 30542 | ENSG00000132182 | NUP210 | chr3 | - | 13373717 | 13373873 | 13365945 | 13366091 | 13375503 | 13375641 | 155 | 57 | 138 | 126 | 107 | 0 | 0 | 0 | 0 | 0 | 109 | 106 | 70 | 50 | 149 | 0 | 3 | 0 | 4 | 7 | 298 | 149 | 0.001493631 | 0.02163251 | 1 | 1 | 1 | 1 | 1 | 1 | 0.946 | 1 | 0.862 | 0.914 | 0.056 |
| SE | 56082 | ENSG00000153207 | AHCTF1 | chr1 | - | 246842693 | 246842776 | 246839097 | 246840998 | 246843794 | 246843928 | 94 | 95 | 114 | 114 | 106 | 4 | 0 | 0 | 3 | 1 | 39 | 49 | 68 | 4 | 11 | 0 | 0 | 16 | 0 | 0 | 232 | 149 | 0.001510447 | 0.021815519 | 0.938 | 1 | 1 | 0.961 | 0.986 | 1 | 1 | 0.732 | 1 | 1 | 0.031 |
| SE | 55477 | ENSG00000143258 | USP21 | chr1 | + | 161160365 | 161160506 | 161159449 | 161159678 | 161160619 | 161161240 | 18 | 15 | 11 | 16 | 19 | 1 | 0 | 2 | 2 | 0 | 11 | 43 | 3 | 24 | 39 | 0 | 16 | 8 | 6 | 0 | 290 | 149 | 0.001542093 | 0.022227183 | 0.902 | 1 | 0.739 | 0.804 | 1 | 1 | 0.58 | 0.162 | 0.673 | 1 | 0.206 |
| SE | 57173 | ENSG00000129084 | PSMA1 | chr11 | - | 14519904 | 14520211 | 14518996 | 14519041 | 14520296 | 14520445 | 17 | 13 | 17 | 10 | 8 | 241 | 119 | 181 | 226 | 47 | 14 | 12 | 18 | 16 | 39 | 160 | 115 | 125 | 170 | 35 | 298 | 149 | 0.001628964 | 0.023290972 | 0.034 | 0.052 | 0.045 | 0.022 | 0.078 | 0.042 | 0.05 | 0.067 | 0.045 | 0.358 | -0.066 |
| SE | 97616 | ENSG00000118363 | SPCS2 | chr11 | + | 74969639 | 74969699 | 74965033 | 74965117 | 74976856 | 74977348 | 75 | 56 | 86 | 80 | 89 | 0 | 0 | 0 | 2 | 0 | 59 | 32 | 87 | 47 | 10 | 0 | 14 | 1 | 0 | 0 | 209 | 149 | 0.0016603 | 0.023656934 | 1 | 1 | 1 | 0.966 | 1 | 1 | 0.62 | 0.984 | 1 | 1 | 0.072 |
| SE | 108853 | ENSG00000117226 | GBP3 | chr1 | - | 89013184 | 89013427 | 89011746 | 89012027 | 89014082 | 89014279 | 139 | 153 | 133 | 103 | 156 | 0 | 0 | 6 | 11 | 0 | 6 | 126 | 75 | 53 | 42 | 0 | 0 | 2 | 0 | 0 | 298 | 149 | 0.001662306 | 0.023672593 | 1 | 1 | 0.917 | 0.824 | 1 | 1 | 1 | 0.949 | 1 | 1 | -0.042 |
| SE | 99109 | ENSG00000110274 | CEP164 | chr11 | + | 117408889 | 117409004 | 117407924 | 117408032 | 117409617 | 117409965 | 30 | 33 | 60 | 47 | 110 | 0 | 0 | 0 | 0 | 3 | 76 | 41 | 27 | 37 | 109 | 14 | 0 | 0 | 0 | 0 | 264 | 149 | 0.001668565 | 0.023740136 | 1 | 1 | 1 | 1 | 0.954 | 0.754 | 1 | 1 | 1 | 1 | 0.04 |
| SE | 605 | ENSG00000099949 | LZTR1 | chr22 | + | 20987503 | 20987583 | 20985840 | 20985897 | 20988009 | 20988113 | 89 | 34 | 64 | 69 | 35 | 4 | 2 | 6 | 5 | 1 | 52 | 87 | 32 | 62 | 121 | 11 | 2 | 12 | 18 | 9 | 229 | 149 | 0.001759995 | 0.024855801 | 0.935 | 0.917 | 0.874 | 0.9 | 0.958 | 0.755 | 0.966 | 0.634 | 0.691 | 0.897 | 0.128 |
| SE | 8837 | ENSG00000108587 | GOSR1 | chr17 | + | 30495425 | 30495582 | 30492678 | 30492753 | 30510879 | 30510909 | 9 | 1 | 0 | 4 | 8 | 63 | 54 | 67 | 76 | 78 | 0 | 0 | 0 | 0 | 0 | 79 | 75 | 55 | 41 | 46 | 298 | 149 | 0.001760395 | 0.024856968 | 0.067 | 0.009 | 0 | 0.026 | 0.049 | 0 | 0 | 0 | 0 | 0 | 0.03 |
| SE | 102593 | ENSG00000175470 | PPP2R2D | chr10 | + | 131940581 | 131940694 | 131939775 | 131940196 | 131944094 | 131944145 | 107 | 84 | 91 | 102 | 177 | 0 | 0 | 0 | 0 | 0 | 160 | 82 | 80 | 103 | 248 | 16 | 0 | 1 | 0 | 0 | 262 | 149 | 0.001809989 | 0.025410598 | 1 | 1 | 1 | 1 | 1 | 0.85 | 1 | 0.978 | 1 | 1 | 0.034 |
| SE | 61973 | ENSG00000205423 | CNEP1R1 | chr16 | + | 50025650 | 50025701 | 50024409 | 50025340 | 50026395 | 50026467 | 0 | 10 | 3 | 2 | 2 | 8 | 1 | 4 | 5 | 0 | 23 | 13 | 7 | 8 | 2 | 0 | 2 | 6 | 0 | 0 | 200 | 149 | 0.001924127 | 0.026796766 | 0 | 0.882 | 0.358 | 0.23 | 1 | 1 | 0.829 | 0.465 | 1 | 1 | -0.365 |
| SE | 28215 | ENSG00000204084 | INPP5B | chr1 | - | 37869924 | 37870014 | 37868500 | 37868614 | 37872929 | 37873028 | 1 | 3 | 1 | 1 | 6 | 72 | 29 | 65 | 47 | 62 | 8 | 8 | 2 | 9 | 0 | 36 | 91 | 41 | 27 | 36 | 239 | 149 | 0.002045551 | 0.028221758 | 0.009 | 0.061 | 0.01 | 0.013 | 0.057 | 0.122 | 0.052 | 0.03 | 0.172 | 0 | -0.045 |
| SE | 92578 | ENSG00000137054 | POLR1E | chr9 | + | 37492281 | 37492375 | 37489314 | 37489400 | 37492656 | 37492715 | 1 | 3 | 0 | 1 | 5 | 100 | 58 | 106 | 95 | 163 | 12 | 0 | 4 | 0 | 4 | 67 | 62 | 59 | 58 | 63 | 243 | 149 | 0.002192619 | 0.029840663 | 0.006 | 0.031 | 0 | 0.006 | 0.018 | 0.099 | 0 | 0.04 | 0 | 0.037 | -0.023 |
| SE | 11028 | ENSG00000072849 | DERL2 | chr17 | - | 5480386 | 5480543 | 5480053 | 5480144 | 5481100 | 5481142 | 124 | 65 | 113 | 107 | 195 | 2 | 7 | 1 | 0 | 3 | 44 | 94 | 86 | 80 | 57 | 0 | 0 | 14 | 16 | 0 | 298 | 149 | 0.00226971 | 0.030713659 | 0.969 | 0.823 | 0.983 | 1 | 0.97 | 1 | 1 | 0.754 | 0.714 | 1 | 0.055 |
| SE | 82044 | ENSG00000104133 | SPG11 | chr15 | - | 44610839 | 44610985 | 44608443 | 44608605 | 44613429 | 44613536 | 384 | 303 | 329 | 379 | 754 | 4 | 5 | 3 | 2 | 0 | 148 | 405 | 375 | 304 | 92 | 0 | 5 | 5 | 4 | 10 | 295 | 149 | 0.002408709 | 0.032287254 | 0.98 | 0.968 | 0.982 | 0.99 | 1 | 1 | 0.976 | 0.974 | 0.975 | 0.823 | 0.034 |
| SE | 50707 | ENSG00000113318 | MSH3 | chr5 | + | 80761545 | 80761678 | 80744505 | 80744615 | 80767932 | 80768120 | 139 | 74 | 99 | 110 | 305 | 3 | 0 | 1 | 6 | 1 | 39 | 124 | 96 | 54 | 27 | 1 | 12 | 5 | 0 | 0 | 282 | 149 | 0.002412674 | 0.03232422 | 0.961 | 1 | 0.981 | 0.906 | 0.994 | 0.954 | 0.845 | 0.91 | 1 | 1 | 0.027 |
| SE | 15470 | ENSG00000140199 | SLC12A6 | chr15 | - | 34275344 | 34275389 | 34260925 | 34261020 | 34336409 | 34336601 | 258 | 203 | 227 | 259 | 277 | 16 | 11 | 9 | 10 | 9 | 522 | 205 | 430 | 454 | 309 | 32 | 24 | 15 | 20 | 50 | 194 | 149 | 0.002416795 | 0.032368351 | 0.925 | 0.934 | 0.951 | 0.952 | 0.959 | 0.926 | 0.868 | 0.957 | 0.946 | 0.826 | 0.04 |
| SE | 107364 | ENSG00000033030 | ZCCHC8 | chr12 | - | 122482634 | 122482695 | 122481944 | 122482087 | 122489385 | 122489463 | 75 | 60 | 64 | 95 | 109 | 3 | 3 | 3 | 1 | 7 | 50 | 57 | 54 | 72 | 0 | 0 | 5 | 8 | 25 | 11 | 210 | 149 | 0.002506732 | 0.033344734 | 0.947 | 0.934 | 0.938 | 0.985 | 0.917 | 1 | 0.89 | 0.827 | 0.671 | 0 | 0.267 |
| SE | 53582 | ENSG00000188529 | SRSF10 | chr1 | - | 23971572 | 23971626 | 23964803 | 23967723 | 23971849 | 23972012 | 163 | 144 | 131 | 186 | 165 | 8 | 5 | 7 | 11 | 5 | 8 | 114 | 113 | 49 | 21 | 0 | 0 | 3 | 4 | 0 | 203 | 149 | 0.002585977 | 0.03422442 | 0.937 | 0.955 | 0.932 | 0.925 | 0.96 | 1 | 1 | 0.965 | 0.9 | 1 | -0.031 |
| SE | 65683 | ENSG00000186141 | POLR3C | chr1 | + | 145826605 | 145826709 | 145826453 | 145826492 | 145826819 | 145827005 | 72 | 47 | 84 | 101 | 156 | 0 | 0 | 0 | 0 | 5 | 86 | 110 | 71 | 94 | 62 | 6 | 0 | 3 | 6 | 0 | 253 | 149 | 0.002677127 | 0.035145593 | 1 | 1 | 1 | 1 | 0.948 | 0.894 | 1 | 0.933 | 0.902 | 1 | 0.044 |
| SE | 86271 | ENSG00000134516 | DOCK2 | chr5 | + | 170008687 | 170008746 | 170008496 | 170008597 | 170018959 | 170019108 | 887 | 545 | 742 | 757 | 816 | 3 | 2 | 2 | 3 | 0 | 1749 | 1051 | 957 | 1251 | 745 | 12 | 7 | 7 | 29 | 15 | 208 | 149 | 0.002728527 | 0.035700661 | 0.995 | 0.995 | 0.996 | 0.994 | 1 | 0.991 | 0.991 | 0.99 | 0.969 | 0.973 | 0.013 |
| SE | 70234 | ENSG00000160695 | VPS11 | chr11 | + | 119071595 | 119071843 | 119070233 | 119070397 | 119073799 | 119073951 | 55 | 27 | 78 | 50 | 84 | 1 | 3 | 1 | 6 | 5 | 58 | 46 | 47 | 53 | 65 | 28 | 1 | 5 | 5 | 11 | 298 | 149 | 0.002733736 | 0.035746873 | 0.965 | 0.818 | 0.975 | 0.806 | 0.894 | 0.509 | 0.958 | 0.825 | 0.841 | 0.747 | 0.116 |
| SE | 79025 | ENSG00000160218 | TRAPPC10 | chr21 | + | 44074323 | 44074470 | 44063537 | 44063785 | 44075038 | 44075153 | 243 | 121 | 187 | 225 | 173 | 3 | 1 | 2 | 1 | 0 | 245 | 254 | 182 | 207 | 214 | 0 | 5 | 4 | 13 | 0 | 296 | 149 | 0.002791041 | 0.036348531 | 0.976 | 0.984 | 0.979 | 0.991 | 1 | 1 | 0.962 | 0.958 | 0.889 | 1 | 0.024 |
| SE | 89483 | ENSG00000064313 | TAF2 | chr8 | - | 119793365 | 119793451 | 119791323 | 119791459 | 119795531 | 119795631 | 95 | 66 | 97 | 114 | 116 | 4 | 2 | 12 | 2 | 5 | 10 | 87 | 46 | 30 | 53 | 0 | 4 | 26 | 2 | 16 | 235 | 149 | 0.002796333 | 0.036385287 | 0.938 | 0.954 | 0.837 | 0.973 | 0.936 | 1 | 0.932 | 0.529 | 0.905 | 0.677 | 0.119 |
| SE | 70235 | ENSG00000160695 | VPS11 | chr11 | + | 119073197 | 119073399 | 119070233 | 119070397 | 119073799 | 119073951 | 61 | 73 | 58 | 72 | 128 | 1 | 3 | 1 | 6 | 5 | 119 | 83 | 60 | 100 | 108 | 28 | 1 | 5 | 5 | 11 | 298 | 149 | 0.00281463 | 0.036558438 | 0.968 | 0.924 | 0.967 | 0.857 | 0.928 | 0.68 | 0.976 | 0.857 | 0.909 | 0.831 | 0.078 |
| SE | 57888 | ENSG00000162604 | TM2D1 | chr1 | - | 61723712 | 61723772 | 61709328 | 61709437 | 61724901 | 61725144 | 65 | 32 | 50 | 45 | 37 | 3 | 2 | 1 | 6 | 0 | 94 | 38 | 46 | 36 | 19 | 0 | 1 | 0 | 2 | 0 | 209 | 149 | 0.002874997 | 0.037232853 | 0.939 | 0.919 | 0.973 | 0.842 | 1 | 1 | 0.964 | 1 | 0.928 | 1 | -0.044 |
| SE | 1571 | ENSG00000198089 | SFI1 | chr22 | + | 31546860 | 31546971 | 31531057 | 31531129 | 31550253 | 31550348 | 289 | 241 | 325 | 294 | 298 | 1 | 0 | 0 | 2 | 1 | 284 | 343 | 244 | 293 | 507 | 0 | 1 | 1 | 10 | 14 | 260 | 149 | 0.002914908 | 0.037698316 | 0.994 | 1 | 1 | 0.988 | 0.994 | 1 | 0.995 | 0.993 | 0.944 | 0.954 | 0.018 |
| SE | 107988 | ENSG00000204852 | TCTN1 | chr12 | + | 110634669 | 110634779 | 110632471 | 110632559 | 110636480 | 110636501 | 25 | 24 | 19 | 17 | 71 | 4 | 1 | 4 | 3 | 3 | 10 | 10 | 7 | 13 | 27 | 0 | 3 | 0 | 1 | 0 | 259 | 149 | 0.002928803 | 0.037840518 | 0.782 | 0.932 | 0.732 | 0.765 | 0.932 | 1 | 0.657 | 1 | 0.882 | 1 | -0.079 |
| SE | 100118 | ENSG00000159063 | ALG8 | chr11 | - | 78102873 | 78102948 | 78100931 | 78101195 | 78103979 | 78104052 | 8 | 3 | 27 | 4 | 22 | 19 | 14 | 19 | 25 | 40 | 8 | 9 | 1 | 2 | 0 | 59 | 12 | 9 | 23 | 14 | 224 | 149 | 0.002963495 | 0.038187918 | 0.219 | 0.125 | 0.486 | 0.096 | 0.268 | 0.083 | 0.333 | 0.069 | 0.055 | 0 | 0.131 |
| SE | 28221 | ENSG00000204084 | INPP5B | chr1 | - | 37885637 | 37885825 | 37882806 | 37882918 | 37886887 | 37887004 | 168 | 99 | 155 | 126 | 214 | 1 | 0 | 4 | 0 | 2 | 220 | 120 | 82 | 192 | 189 | 0 | 22 | 1 | 0 | 0 | 298 | 149 | 0.003251634 | 0.041369831 | 0.988 | 1 | 0.951 | 1 | 0.982 | 1 | 0.732 | 0.976 | 1 | 1 | 0.043 |
| SE | 61910 | ENSG00000124067 | SLC12A4 | chr16 | - | 67950653 | 67951060 | 67950318 | 67950493 | 67951139 | 67951304 | 7 | 2 | 3 | 9 | 4 | 2 | 0 | 1 | 0 | 0 | 5 | 6 | 3 | 11 | 8 | 7 | 8 | 0 | 0 | 0 | 298 | 149 | 0.003364944 | 0.042555592 | 0.636 | 1 | 0.6 | 1 | 1 | 0.263 | 0.273 | 1 | 1 | 1 | 0.14 |
| SE | 81071 | ENSG00000157764 | BRAF | chr7 | - | 140753274 | 140754233 | 140749286 | 140749418 | 140776911 | 140777088 | 152 | 115 | 142 | 171 | 222 | 5 | 9 | 5 | 5 | 8 | 149 | 106 | 173 | 58 | 58 | 0 | 7 | 2 | 1 | 0 | 298 | 149 | 0.003467415 | 0.043555935 | 0.938 | 0.865 | 0.934 | 0.945 | 0.933 | 1 | 0.883 | 0.977 | 0.967 | 1 | -0.042 |
| SE | 99110 | ENSG00000110274 | CEP164 | chr11 | + | 117408889 | 117409028 | 117407924 | 117408032 | 117409617 | 117409965 | 46 | 43 | 61 | 52 | 101 | 0 | 0 | 0 | 0 | 3 | 123 | 47 | 41 | 57 | 150 | 14 | 0 | 0 | 0 | 0 | 288 | 149 | 0.003500548 | 0.043852625 | 1 | 1 | 1 | 1 | 0.946 | 0.82 | 1 | 1 | 1 | 1 | 0.025 |
| SE | 30540 | ENSG00000132182 | NUP210 | chr3 | - | 13371833 | 13372032 | 13365945 | 13366091 | 13373717 | 13373873 | 591 | 209 | 434 | 441 | 516 | 0 | 0 | 0 | 0 | 0 | 335 | 554 | 217 | 249 | 503 | 0 | 10 | 1 | 5 | 9 | 298 | 149 | 0.003857221 | 0.047530192 | 1 | 1 | 1 | 1 | 1 | 1 | 0.965 | 0.991 | 0.961 | 0.965 | 0.024 |
| SE | 16947 | ENSG00000172575 | RASGRP1 | chr15 | - | 38494381 | 38494767 | 38488102 | 38490688 | 38500102 | 38500139 | 190 | 134 | 178 | 153 | 300 | 3 | 2 | 1 | 0 | 0 | 47 | 120 | 111 | 25 | 26 | 0 | 11 | 1 | 0 | 0 | 298 | 149 | 0.003859553 | 0.047536186 | 0.969 | 0.971 | 0.989 | 1 | 1 | 1 | 0.845 | 0.982 | 1 | 1 | 0.02 |
| SE | 58929 | ENSG00000135336 | ORC3 | chr6 | + | 87663002 | 87663144 | 87657920 | 87658018 | 87664742 | 87664859 | 127 | 87 | 142 | 109 | 325 | 0 | 4 | 6 | 4 | 6 | 1 | 48 | 26 | 8 | 4 | 0 | 1 | 0 | 1 | 0 | 291 | 149 | 0.003977079 | 0.048723106 | 1 | 0.918 | 0.924 | 0.933 | 0.965 | 1 | 0.961 | 1 | 0.804 | 1 | -0.005 |
| SE | 22561 | ENSG00000167881 | SRP68 | chr17 | - | 76070377 | 76070444 | 76067216 | 76067330 | 76072344 | 76072497 | 45 | 19 | 34 | 35 | 48 | 0 | 1 | 2 | 0 | 0 | 21 | 38 | 43 | 47 | 31 | 0 | 0 | 1 | 3 | 7 | 216 | 149 | 0.004121182 | 0.050229151 | 1 | 0.929 | 0.921 | 1 | 1 | 1 | 1 | 0.967 | 0.915 | 0.753 | 0.043 |
| SE | 66973 | ENSG00000162645 | GBP2 | chr1 | - | 89108022 | 89108291 | 89106131 | 89107052 | 89109676 | 89109870 | 331 | 227 | 284 | 288 | 483 | 7 | 6 | 6 | 6 | 25 | 314 | 171 | 162 | 260 | 161 | 7 | 0 | 6 | 4 | 0 | 298 | 149 | 0.004189353 | 0.050948982 | 0.959 | 0.95 | 0.959 | 0.96 | 0.906 | 0.957 | 1 | 0.931 | 0.97 | 1 | -0.025 |
| SE | 74954 | ENSG00000105393 | BABAM1 | chr19 | + | 17271596 | 17271655 | 17268684 | 17269091 | 17273903 | 17273980 | 269 | 143 | 208 | 169 | 269 | 6 | 0 | 0 | 4 | 4 | 365 | 214 | 179 | 194 | 293 | 9 | 6 | 9 | 11 | 0 | 208 | 149 | 0.004201389 | 0.051055711 | 0.97 | 1 | 1 | 0.968 | 0.98 | 0.967 | 0.962 | 0.934 | 0.927 | 1 | 0.026 |
| SE | 66496 | ENSG00000132676 | DAP3 | chr1 | + | 155721516 | 155721618 | 155709772 | 155709824 | 155725381 | 155725411 | 117 | 78 | 103 | 138 | 109 | 5 | 1 | 5 | 4 | 0 | 243 | 112 | 106 | 73 | 182 | 0 | 2 | 1 | 2 | 0 | 251 | 149 | 0.004208175 | 0.051106443 | 0.933 | 0.979 | 0.924 | 0.953 | 1 | 1 | 0.971 | 0.984 | 0.956 | 1 | -0.024 |
| SE | 4405 | ENSG00000130475 | FCHO1 | chr19 | + | 17781230 | 17781343 | 17778440 | 17778884 | 17781451 | 17781539 | 21 | 3 | 19 | 15 | 11 | 0 | 0 | 0 | 0 | 0 | 8 | 16 | 13 | 5 | 38 | 0 | 0 | 2 | 0 | 7 | 262 | 149 | 0.004261156 | 0.051612411 | 1 | 1 | 1 | 1 | 1 | 1 | 1 | 0.787 | 1 | 0.755 | 0.092 |
| SE | 100091 | ENSG00000149262 | INTS4 | chr11 | - | 77938650 | 77938825 | 77928341 | 77928547 | 77941179 | 77941251 | 207 | 134 | 179 | 200 | 233 | 1 | 0 | 0 | 0 | 6 | 222 | 204 | 153 | 206 | 150 | 0 | 4 | 8 | 5 | 2 | 298 | 149 | 0.004330057 | 0.052278507 | 0.99 | 1 | 1 | 1 | 0.951 | 1 | 0.962 | 0.905 | 0.954 | 0.974 | 0.029 |
| SE | 42662 | ENSG00000088854 | C20orf194 | chr20 | - | 3304439 | 3304543 | 3295493 | 3296949 | 3315032 | 3315120 | 9 | 12 | 14 | 8 | 15 | 1 | 0 | 0 | 0 | 4 | 13 | 13 | 8 | 15 | 15 | 9 | 4 | 0 | 2 | 0 | 253 | 149 | 0.004525127 | 0.054276178 | 0.841 | 1 | 1 | 1 | 0.688 | 0.46 | 0.657 | 1 | 0.815 | 1 | 0.119 |
| SE | 53581 | ENSG00000188529 | SRSF10 | chr1 | - | 23971572 | 23971623 | 23966444 | 23967723 | 23971849 | 23972012 | 47 | 33 | 60 | 61 | 50 | 8 | 5 | 7 | 11 | 5 | 8 | 17 | 31 | 33 | 3 | 0 | 0 | 3 | 4 | 0 | 200 | 149 | 0.004608212 | 0.055042649 | 0.814 | 0.831 | 0.865 | 0.805 | 0.882 | 1 | 1 | 0.885 | 0.86 | 1 | -0.11 |
| SE | 45108 | ENSG00000088035 | ALG6 | chr1 | + | 63411681 | 63411806 | 63411145 | 63411331 | 63411925 | 63412061 | 0 | 5 | 3 | 5 | 12 | 17 | 3 | 23 | 19 | 38 | 11 | 5 | 8 | 12 | 0 | 6 | 5 | 7 | 2 | 10 | 274 | 149 | 0.005055832 | 0.059206017 | 0 | 0.475 | 0.066 | 0.125 | 0.147 | 0.499 | 0.352 | 0.383 | 0.765 | 0 | -0.237 |
| SE | 100097 | ENSG00000149262 | INTS4 | chr11 | - | 77955941 | 77956062 | 77941179 | 77941251 | 77958745 | 77958834 | 266 | 227 | 255 | 325 | 261 | 3 | 0 | 0 | 0 | 0 | 337 | 394 | 270 | 272 | 238 | 0 | 2 | 7 | 0 | 14 | 270 | 149 | 0.005176166 | 0.060339705 | 0.98 | 1 | 1 | 1 | 1 | 1 | 0.991 | 0.955 | 1 | 0.904 | 0.026 |
| SE | 108801 | ENSG00000105983 | LMBR1 | chr7 | - | 156779654 | 156779732 | 156763668 | 156763795 | 156796388 | 156796492 | 10 | 6 | 7 | 10 | 21 | 100 | 43 | 87 | 69 | 135 | 16 | 23 | 2 | 7 | 47 | 33 | 90 | 57 | 62 | 34 | 227 | 149 | 0.005274517 | 0.061244789 | 0.062 | 0.084 | 0.05 | 0.087 | 0.093 | 0.241 | 0.144 | 0.023 | 0.069 | 0.476 | -0.115 |
| SE | 38021 | ENSG00000197386 | HTT | chr4 | + | 3223991 | 3224131 | 3223405 | 3223560 | 3225660 | 3225743 | 128 | 70 | 124 | 109 | 68 | 0 | 0 | 0 | 0 | 0 | 204 | 115 | 67 | 151 | 231 | 0 | 0 | 0 | 6 | 10 | 289 | 149 | 0.005358502 | 0.062008461 | 1 | 1 | 1 | 1 | 1 | 1 | 1 | 1 | 0.928 | 0.923 | 0.03 |
| SE | 74955 | ENSG00000105393 | BABAM1 | chr19 | + | 17273903 | 17274024 | 17271596 | 17271655 | 17276494 | 17276555 | 157 | 82 | 114 | 99 | 192 | 6 | 2 | 1 | 3 | 5 | 167 | 116 | 98 | 134 | 188 | 17 | 0 | 1 | 5 | 18 | 270 | 149 | 0.005405382 | 0.062394194 | 0.935 | 0.958 | 0.984 | 0.948 | 0.955 | 0.844 | 1 | 0.982 | 0.937 | 0.852 | 0.033 |
| SE | 65365 | ENSG00000153214 | TMEM87B | chr2 | + | 112081334 | 112081518 | 112081056 | 112081118 | 112086004 | 112086104 | 327 | 304 | 217 | 412 | 418 | 6 | 11 | 7 | 18 | 16 | 285 | 239 | 135 | 182 | 177 | 2 | 6 | 11 | 16 | 41 | 298 | 149 | 0.005441235 | 0.062743293 | 0.965 | 0.933 | 0.939 | 0.92 | 0.929 | 0.986 | 0.952 | 0.86 | 0.85 | 0.683 | 0.071 |
| SE | 68468 | ENSG00000143799 | PARP1 | chr1 | - | 226392916 | 226392980 | 226392257 | 226392314 | 226402213 | 226402379 | 5 | 0 | 1 | 4 | 1 | 237 | 139 | 178 | 264 | 129 | 9 | 14 | 2 | 7 | 2 | 263 | 199 | 153 | 164 | 285 | 213 | 149 | 0.005481335 | 0.063089652 | 0.015 | 0 | 0.004 | 0.01 | 0.005 | 0.023 | 0.047 | 0.009 | 0.029 | 0.005 | -0.016 |
| SE | 82046 | ENSG00000104133 | SPG11 | chr15 | - | 44610917 | 44610985 | 44608443 | 44608605 | 44613429 | 44613536 | 91 | 99 | 89 | 91 | 162 | 4 | 5 | 3 | 2 | 0 | 25 | 144 | 70 | 74 | 18 | 0 | 5 | 5 | 4 | 10 | 217 | 149 | 0.005482553 | 0.063089652 | 0.94 | 0.931 | 0.953 | 0.969 | 1 | 1 | 0.952 | 0.906 | 0.927 | 0.553 | 0.091 |
| SE | 24103 | ENSG00000143889 | HNRNPLL | chr2 | - | 38591529 | 38591648 | 38585643 | 38585881 | 38602437 | 38602600 | 106 | 65 | 91 | 72 | 68 | 2 | 0 | 2 | 2 | 0 | 59 | 79 | 69 | 110 | 68 | 0 | 3 | 1 | 3 | 9 | 268 | 149 | 0.005500768 | 0.063234201 | 0.967 | 1 | 0.962 | 0.952 | 1 | 1 | 0.936 | 0.975 | 0.953 | 0.808 | 0.042 |
| SE | 58915 | ENSG00000135336 | ORC3 | chr6 | + | 87594526 | 87594613 | 87594352 | 87594407 | 87601783 | 87601881 | 22 | 24 | 34 | 16 | 38 | 77 | 47 | 97 | 88 | 94 | 0 | 11 | 18 | 4 | 0 | 25 | 67 | 40 | 18 | 18 | 236 | 149 | 0.005601993 | 0.06419989 | 0.153 | 0.244 | 0.181 | 0.103 | 0.203 | 0 | 0.094 | 0.221 | 0.123 | 0 | 0.089 |
| SE | 39532 | ENSG00000111707 | SUDS3 | chr12 | + | 118404055 | 118404473 | 118403500 | 118403517 | 118411072 | 118411147 | 4 | 0 | 3 | 5 | 0 | 81 | 57 | 126 | 92 | 124 | 0 | 6 | 5 | 21 | 0 | 165 | 115 | 84 | 119 | 93 | 298 | 149 | 0.005727137 | 0.065394771 | 0.024 | 0 | 0.012 | 0.026 | 0 | 0 | 0.025 | 0.029 | 0.081 | 0 | -0.015 |
| SE | 6053 | ENSG00000132792 | CTNNBL1 | chr20 | + | 37694926 | 37695033 | 37694031 | 37694152 | 37732878 | 37733067 | 2 | 0 | 4 | 8 | 3 | 26 | 47 | 46 | 72 | 39 | 0 | 5 | 0 | 5 | 0 | 134 | 48 | 57 | 85 | 106 | 256 | 149 | 0.005995699 | 0.06797557 | 0.043 | 0 | 0.048 | 0.061 | 0.043 | 0 | 0.057 | 0 | 0.033 | 0 | 0.021 |
| SE | 100701 | ENSG00000109917 | ZPR1 | chr11 | - | 116784377 | 116784498 | 116783529 | 116783619 | 116784858 | 116784921 | 78 | 45 | 71 | 91 | 50 | 0 | 0 | 0 | 0 | 0 | 70 | 94 | 65 | 110 | 110 | 6 | 1 | 1 | 5 | 0 | 270 | 149 | 0.006126525 | 0.069168334 | 1 | 1 | 1 | 1 | 1 | 0.866 | 0.981 | 0.973 | 0.924 | 1 | 0.051 |
| SE | 68825 | ENSG00000082258 | CCNT2 | chr2 | + | 134945786 | 134945982 | 134942611 | 134942674 | 134946100 | 134946214 | 6 | 3 | 12 | 3 | 15 | 29 | 27 | 26 | 27 | 40 | 0 | 9 | 0 | 1 | 0 | 7 | 32 | 29 | 7 | 2 | 298 | 149 | 0.006146531 | 0.069354195 | 0.094 | 0.053 | 0.188 | 0.053 | 0.158 | 0 | 0.123 | 0 | 0.067 | 0 | 0.071 |
| SE | 12388 | ENSG00000188559 | RALGAPA2 | chr20 | - | 20396694 | 20396734 | 20389529 | 20393253 | 20412026 | 20412113 | 23 | 17 | 23 | 20 | 41 | 5 | 6 | 0 | 4 | 9 | 18 | 24 | 19 | 45 | 39 | 14 | 15 | 17 | 10 | 8 | 189 | 149 | 0.006242917 | 0.070219143 | 0.784 | 0.691 | 1 | 0.798 | 0.782 | 0.503 | 0.558 | 0.468 | 0.78 | 0.794 | 0.19 |
| SE | 31556 | ENSG00000163913 | IFT122 | chr3 | + | 129458598 | 129458677 | 129451913 | 129451998 | 129461227 | 129461304 | 37 | 21 | 23 | 31 | 17 | 0 | 0 | 1 | 0 | 0 | 106 | 36 | 23 | 67 | 65 | 6 | 6 | 0 | 0 | 0 | 228 | 149 | 0.006307707 | 0.070785182 | 1 | 1 | 0.938 | 1 | 1 | 0.92 | 0.797 | 1 | 1 | 1 | 0.044 |
| SE | 73313 | ENSG00000136699 | SMPD4 | chr2 | - | 130173513 | 130173656 | 130172786 | 130172895 | 130174913 | 130175000 | 30 | 12 | 24 | 40 | 17 | 0 | 0 | 0 | 0 | 0 | 55 | 18 | 24 | 27 | 11 | 0 | 0 | 1 | 2 | 6 | 292 | 149 | 0.006377022 | 0.071378886 | 1 | 1 | 1 | 1 | 1 | 1 | 1 | 0.925 | 0.873 | 0.483 | 0.144 |
| SE | 79996 | ENSG00000084070 | SMAP2 | chr1 | + | 40362299 | 40362436 | 40344849 | 40344910 | 40406735 | 40406869 | 126 | 92 | 34 | 81 | 60 | 7 | 2 | 3 | 4 | 0 | 268 | 126 | 30 | 159 | 181 | 0 | 9 | 1 | 0 | 0 | 286 | 149 | 0.006532858 | 0.072893681 | 0.904 | 0.96 | 0.855 | 0.913 | 1 | 1 | 0.879 | 0.94 | 1 | 1 | -0.037 |
| SE | 55693 | ENSG00000135749 | PCNX2 | chr1 | - | 233112844 | 233112966 | 233057231 | 233057290 | 233135012 | 233135190 | 12 | 11 | 29 | 32 | 46 | 1 | 0 | 0 | 0 | 1 | 21 | 5 | 12 | 11 | 7 | 14 | 0 | 0 | 0 | 0 | 271 | 149 | 0.006776595 | 0.075099826 | 0.868 | 1 | 1 | 1 | 0.962 | 0.452 | 1 | 1 | 1 | 1 | 0.076 |
| SE | 52283 | ENSG00000007402 | CACNA2D2 | chr3 | - | 50384195 | 50384337 | 50380994 | 50381126 | 50387567 | 50387612 | 106 | 43 | 35 | 84 | 34 | 0 | 0 | 0 | 0 | 0 | 16 | 44 | 29 | 29 | 56 | 0 | 10 | 0 | 0 | 0 | 291 | 149 | 0.006866599 | 0.075968238 | 1 | 1 | 1 | 1 | 1 | 1 | 0.693 | 1 | 1 | 1 | 0.061 |
| SE | 42661 | ENSG00000088854 | C20orf194 | chr20 | - | 3298028 | 3298175 | 3295493 | 3296949 | 3315032 | 3315120 | 19 | 17 | 8 | 14 | 9 | 1 | 0 | 0 | 0 | 4 | 25 | 16 | 7 | 21 | 15 | 9 | 4 | 0 | 2 | 0 | 296 | 149 | 0.006956847 | 0.076836403 | 0.905 | 1 | 1 | 1 | 0.531 | 0.583 | 0.668 | 1 | 0.841 | 1 | 0.069 |
| SE | 59868 | ENSG00000103168 | TAF1C | chr16 | - | 84184850 | 84184899 | 84183696 | 84183778 | 84186900 | 84187049 | 18 | 9 | 16 | 10 | 10 | 3 | 1 | 4 | 5 | 0 | 23 | 21 | 25 | 24 | 38 | 0 | 3 | 1 | 4 | 0 | 198 | 149 | 0.007116142 | 0.078330591 | 0.819 | 0.871 | 0.751 | 0.601 | 1 | 1 | 0.84 | 0.95 | 0.819 | 1 | -0.113 |
| SE | 31565 | ENSG00000163913 | IFT122 | chr3 | + | 129492140 | 129492194 | 129488256 | 129488397 | 129495445 | 129495607 | 7 | 9 | 17 | 12 | 10 | 1 | 0 | 1 | 1 | 0 | 31 | 24 | 18 | 15 | 37 | 0 | 0 | 10 | 2 | 0 | 203 | 149 | 0.007137094 | 0.07850603 | 0.837 | 1 | 0.926 | 0.898 | 1 | 1 | 1 | 0.569 | 0.846 | 1 | 0.049 |
| SE | 101644 | ENSG00000198258 | UBL5 | chr19 | + | 9828591 | 9828675 | 9828326 | 9828389 | 9828836 | 9828874 | 103 | 56 | 91 | 99 | 132 | 0 | 0 | 0 | 0 | 0 | 79 | 77 | 46 | 59 | 181 | 14 | 0 | 0 | 0 | 0 | 233 | 149 | 0.007717896 | 0.083860984 | 1 | 1 | 1 | 1 | 1 | 0.783 | 1 | 1 | 1 | 1 | 0.043 |
| SE | 42656 | ENSG00000088854 | C20orf194 | chr20 | - | 3270426 | 3270562 | 3264437 | 3264524 | 3278911 | 3278949 | 18 | 9 | 15 | 20 | 9 | 0 | 0 | 1 | 0 | 3 | 23 | 6 | 2 | 5 | 19 | 7 | 0 | 3 | 1 | 0 | 285 | 149 | 0.007832394 | 0.084866466 | 1 | 1 | 0.887 | 1 | 0.611 | 0.632 | 1 | 0.258 | 0.723 | 1 | 0.177 |
| SE | 103092 | ENSG00000108055 | SMC3 | chr10 | + | 110581922 | 110582098 | 110580903 | 110581021 | 110582561 | 110582642 | 312 | 332 | 283 | 396 | 439 | 4 | 3 | 0 | 6 | 0 | 63 | 240 | 257 | 85 | 20 | 0 | 12 | 3 | 1 | 0 | 298 | 149 | 0.00787261 | 0.085290425 | 0.975 | 0.982 | 1 | 0.971 | 1 | 1 | 0.909 | 0.977 | 0.977 | 1 | 0.013 |
| SE | 85074 | ENSG00000147459 | DOCK5 | chr8 | + | 25407982 | 25408154 | 25403557 | 25403724 | 25408801 | 25408940 | 129 | 93 | 57 | 166 | 113 | 1 | 0 | 0 | 0 | 0 | 155 | 184 | 214 | 145 | 49 | 0 | 9 | 2 | 4 | 0 | 298 | 149 | 0.007940763 | 0.085921772 | 0.985 | 1 | 1 | 1 | 1 | 1 | 0.911 | 0.982 | 0.948 | 1 | 0.029 |
| SE | 27150 | ENSG00000109534 | GAR1 | chr4 | + | 109816152 | 109816378 | 109815634 | 109815804 | 109817935 | 109818090 | 82 | 101 | 80 | 74 | 73 | 0 | 0 | 0 | 0 | 0 | 176 | 77 | 98 | 93 | 77 | 0 | 1 | 3 | 9 | 0 | 298 | 149 | 0.008000291 | 0.086422572 | 1 | 1 | 1 | 1 | 1 | 1 | 0.975 | 0.942 | 0.838 | 1 | 0.049 |
| SE | 99092 | ENSG00000110274 | CEP164 | chr11 | + | 117362403 | 117362538 | 117361834 | 117361993 | 117371079 | 117371466 | 46 | 30 | 41 | 64 | 51 | 0 | 0 | 0 | 0 | 0 | 124 | 59 | 45 | 57 | 185 | 11 | 0 | 0 | 0 | 2 | 284 | 149 | 0.008058867 | 0.086899474 | 1 | 1 | 1 | 1 | 1 | 0.855 | 1 | 1 | 1 | 0.98 | 0.033 |
| SE | 31564 | ENSG00000163913 | IFT122 | chr3 | + | 129466889 | 129467066 | 129464634 | 129464781 | 129469341 | 129469417 | 1 | 0 | 4 | 4 | 0 | 11 | 3 | 6 | 10 | 1 | 0 | 0 | 0 | 2 | 0 | 13 | 2 | 11 | 10 | 6 | 298 | 149 | 0.008130795 | 0.087566542 | 0.043 | 0 | 0.25 | 0.167 | 0 | 0 | 0 | 0 | 0.091 | 0 | 0.074 |
| SE | 69926 | ENSG00000169016 | E2F6 | chr2 | - | 11455341 | 11455474 | 11453581 | 11453798 | 11457178 | 11457233 | 17 | 3 | 4 | 6 | 15 | 19 | 26 | 8 | 16 | 6 | 0 | 1 | 2 | 10 | 6 | 35 | 18 | 14 | 23 | 5 | 282 | 149 | 0.008229695 | 0.088400635 | 0.321 | 0.057 | 0.209 | 0.165 | 0.569 | 0 | 0.029 | 0.07 | 0.187 | 0.388 | 0.129 |
| SE | 54677 | ENSG00000070010 | UFD1 | chr22 | - | 19475067 | 19475100 | 19471686 | 19471808 | 19479082 | 19479172 | 46 | 22 | 29 | 45 | 45 | 7 | 5 | 17 | 11 | 2 | 28 | 51 | 26 | 33 | 50 | 15 | 24 | 4 | 14 | 43 | 182 | 149 | 0.00832705 | 0.089274915 | 0.843 | 0.783 | 0.583 | 0.77 | 0.949 | 0.604 | 0.635 | 0.842 | 0.659 | 0.488 | 0.14 |
| SE | 88160 | ENSG00000130227 | XPO7 | chr8 | + | 21990810 | 21990919 | 21990343 | 21990407 | 21991867 | 21991974 | 173 | 139 | 130 | 192 | 199 | 10 | 8 | 0 | 15 | 9 | 194 | 125 | 94 | 189 | 332 | 18 | 7 | 24 | 22 | 9 | 258 | 149 | 0.008881644 | 0.094227263 | 0.909 | 0.909 | 1 | 0.881 | 0.927 | 0.862 | 0.912 | 0.693 | 0.832 | 0.955 | 0.074 |
| SE | 9893 | ENSG00000185813 | PCYT2 | chr17 | - | 81907203 | 81907257 | 81906759 | 81906898 | 81907553 | 81907598 | 9 | 5 | 11 | 9 | 12 | 15 | 4 | 23 | 13 | 10 | 16 | 20 | 4 | 13 | 29 | 5 | 13 | 5 | 4 | 23 | 203 | 149 | 0.009187689 | 0.096922573 | 0.306 | 0.478 | 0.26 | 0.337 | 0.468 | 0.701 | 0.53 | 0.37 | 0.705 | 0.481 | -0.188 |
| SE | 13613 | ENSG00000038358 | EDC4 | chr16 | + | 67876487 | 67876599 | 67875944 | 67876101 | 67876872 | 67876972 | 98 | 65 | 110 | 79 | 74 | 0 | 0 | 0 | 0 | 0 | 131 | 135 | 69 | 85 | 102 | 14 | 0 | 0 | 0 | 0 | 261 | 149 | 0.009577081 | 0.100193365 | 1 | 1 | 1 | 1 | 1 | 0.842 | 1 | 1 | 1 | 1 | 0.032 |
| SE | 107368 | ENSG00000033030 | ZCCHC8 | chr12 | - | 122483278 | 122483563 | 122482634 | 122482695 | 122489385 | 122489463 | 94 | 66 | 76 | 113 | 186 | 6 | 12 | 11 | 13 | 23 | 69 | 38 | 56 | 43 | 10 | 16 | 24 | 14 | 20 | 0 | 298 | 149 | 0.00970328 | 0.101256561 | 0.887 | 0.733 | 0.776 | 0.813 | 0.802 | 0.683 | 0.442 | 0.667 | 0.518 | 1 | 0.14 |
| SE | 74717 | ENSG00000112282 | MED23 | chr6 | - | 131602217 | 131602381 | 131600037 | 131600162 | 131603029 | 131603204 | 136 | 100 | 134 | 173 | 146 | 0 | 0 | 3 | 0 | 0 | 4 | 79 | 91 | 24 | 1 | 0 | 0 | 10 | 0 | 0 | 298 | 149 | 0.009812851 | 0.102141312 | 1 | 1 | 0.957 | 1 | 1 | 1 | 1 | 0.82 | 1 | 1 | 0.027 |
| SE | 99364 | ENSG00000156599 | ZDHHC5 | chr11 | + | 57682421 | 57682543 | 57672020 | 57673194 | 57688507 | 57688665 | 260 | 112 | 202 | 233 | 136 | 12 | 6 | 8 | 9 | 12 | 228 | 217 | 151 | 224 | 267 | 24 | 10 | 16 | 19 | 18 | 271 | 149 | 0.010093219 | 0.10467002 | 0.923 | 0.911 | 0.933 | 0.934 | 0.862 | 0.839 | 0.923 | 0.838 | 0.866 | 0.891 | 0.041 |
| SE | 13000 | ENSG00000122390 | NAA60 | chr16 | + | 3476221 | 3476337 | 3448470 | 3448540 | 3479470 | 3479591 | 155 | 91 | 139 | 168 | 109 | 0 | 0 | 3 | 0 | 0 | 212 | 225 | 174 | 280 | 418 | 13 | 6 | 0 | 0 | 2 | 265 | 149 | 0.010183826 | 0.105344544 | 1 | 1 | 0.963 | 1 | 1 | 0.902 | 0.955 | 1 | 1 | 0.992 | 0.023 |
| SE | 41220 | ENSG00000157181 | ODR4 | chr1 | + | 186385987 | 186386083 | 186383021 | 186383156 | 186388441 | 186388548 | 76 | 65 | 89 | 75 | 153 | 6 | 3 | 2 | 4 | 5 | 48 | 52 | 42 | 37 | 13 | 0 | 9 | 6 | 9 | 0 | 245 | 149 | 0.010230788 | 0.105746502 | 0.885 | 0.929 | 0.964 | 0.919 | 0.949 | 1 | 0.778 | 0.81 | 0.714 | 1 | 0.069 |
| SE | 79036 | ENSG00000160218 | TRAPPC10 | chr21 | + | 44087698 | 44087928 | 44084121 | 44084263 | 44089832 | 44089933 | 224 | 112 | 155 | 166 | 200 | 0 | 0 | 0 | 1 | 0 | 132 | 182 | 110 | 129 | 76 | 11 | 2 | 1 | 0 | 0 | 298 | 149 | 0.010322346 | 0.106552196 | 1 | 1 | 1 | 0.988 | 1 | 0.857 | 0.978 | 0.982 | 1 | 1 | 0.034 |
| SE | 16949 | ENSG00000172575 | RASGRP1 | chr15 | - | 38498793 | 38498946 | 38490553 | 38490688 | 38500102 | 38500139 | 278 | 183 | 308 | 341 | 311 | 3 | 2 | 1 | 0 | 0 | 30 | 174 | 104 | 59 | 32 | 0 | 11 | 1 | 0 | 0 | 298 | 149 | 0.010612386 | 0.108914322 | 0.979 | 0.979 | 0.994 | 1 | 1 | 1 | 0.888 | 0.981 | 1 | 1 | 0.017 |
| SE | 103326 | ENSG00000148688 | RPP30 | chr10 | + | 90895453 | 90895483 | 90894772 | 90894891 | 90895879 | 90895917 | 54 | 16 | 73 | 34 | 51 | 6 | 12 | 19 | 23 | 27 | 37 | 31 | 17 | 24 | 30 | 29 | 37 | 16 | 28 | 10 | 179 | 149 | 0.010769912 | 0.110126813 | 0.882 | 0.526 | 0.762 | 0.552 | 0.611 | 0.515 | 0.411 | 0.469 | 0.416 | 0.714 | 0.162 |
| SE | 55090 | ENSG00000086200 | IPO11 | chr5 | + | 62601763 | 62601848 | 62591576 | 62591672 | 62627153 | 62627569 | 68 | 62 | 62 | 107 | 130 | 1 | 3 | 1 | 1 | 1 | 61 | 143 | 54 | 73 | 392 | 0 | 4 | 3 | 10 | 6 | 234 | 149 | 0.011446722 | 0.115927115 | 0.977 | 0.929 | 0.975 | 0.986 | 0.988 | 1 | 0.958 | 0.92 | 0.823 | 0.977 | 0.035 |
| SE | 63665 | ENSG00000144451 | SPAG16 | chr2 | + | 213310058 | 213310177 | 213297261 | 213297357 | 213317218 | 213317356 | 67 | 50 | 114 | 74 | 250 | 0 | 0 | 0 | 0 | 14 | 18 | 32 | 19 | 13 | 48 | 0 | 0 | 0 | 0 | 0 | 268 | 149 | 0.011643244 | 0.117535246 | 1 | 1 | 1 | 1 | 0.908 | 1 | 1 | 1 | 1 | 1 | -0.018 |
| SE | 55476 | ENSG00000143258 | USP21 | chr1 | + | 161160328 | 161160506 | 161159520 | 161159678 | 161160619 | 161160992 | 9 | 13 | 6 | 9 | 13 | 1 | 0 | 2 | 2 | 0 | 11 | 38 | 1 | 17 | 67 | 0 | 16 | 8 | 6 | 0 | 298 | 149 | 0.011715509 | 0.11816018 | 0.818 | 1 | 0.6 | 0.692 | 1 | 1 | 0.543 | 0.059 | 0.586 | 1 | 0.184 |
| SE | 76687 | ENSG00000223501 | VPS52 | chr6 | - | 33269998 | 33270051 | 33269743 | 33269819 | 33270198 | 33270283 | 54 | 30 | 46 | 62 | 59 | 0 | 3 | 6 | 6 | 5 | 103 | 134 | 73 | 80 | 171 | 12 | 8 | 32 | 16 | 14 | 202 | 149 | 0.012155386 | 0.121734214 | 1 | 0.881 | 0.85 | 0.884 | 0.897 | 0.864 | 0.925 | 0.627 | 0.787 | 0.9 | 0.082 |
| SE | 96390 | ENSG00000197563 | PIGN | chr18 | - | 62123506 | 62123544 | 62114560 | 62114639 | 62138242 | 62138298 | 2 | 2 | 1 | 0 | 1 | 26 | 19 | 27 | 25 | 65 | 8 | 1 | 2 | 0 | 1 | 0 | 44 | 4 | 20 | 5 | 187 | 149 | 0.012682346 | 0.12612434 | 0.058 | 0.077 | 0.029 | 0 | 0.012 | 1 | 0.018 | 0.285 | 0 | 0.137 | -0.253 |
| SE | 86862 | ENSG00000090316 | MAEA | chr4 | + | 1322380 | 1322503 | 1315396 | 1315600 | 1327626 | 1327703 | 159 | 49 | 98 | 129 | 93 | 4 | 0 | 3 | 0 | 0 | 78 | 105 | 73 | 80 | 116 | 6 | 4 | 3 | 2 | 0 | 272 | 149 | 0.012720294 | 0.126373319 | 0.956 | 1 | 0.947 | 1 | 1 | 0.877 | 0.935 | 0.93 | 0.956 | 1 | 0.041 |
| SE | 29708 | ENSG00000182621 | PLCB1 | chr20 | + | 8739260 | 8739360 | 8737027 | 8737192 | 8740343 | 8740448 | 75 | 62 | 46 | 54 | 74 | 3 | 0 | 0 | 1 | 0 | 31 | 41 | 49 | 13 | 10 | 0 | 10 | 0 | 0 | 0 | 249 | 149 | 0.013465477 | 0.132415593 | 0.937 | 1 | 1 | 0.97 | 1 | 1 | 0.71 | 1 | 1 | 1 | 0.039 |
| SE | 88654 | ENSG00000187735 | TCEA1 | chr8 | - | 54002891 | 54002940 | 53999944 | 54000050 | 54010429 | 54010492 | 6 | 7 | 2 | 6 | 8 | 637 | 473 | 501 | 587 | 626 | 0 | 22 | 9 | 4 | 8 | 179 | 509 | 625 | 358 | 90 | 198 | 149 | 0.013840771 | 0.135425768 | 0.007 | 0.011 | 0.003 | 0.008 | 0.01 | 0 | 0.032 | 0.011 | 0.008 | 0.063 | -0.015 |
| SE | 38018 | ENSG00000197386 | HTT | chr4 | + | 3202976 | 3203066 | 3199731 | 3199939 | 3204006 | 3204148 | 35 | 22 | 21 | 43 | 37 | 73 | 33 | 43 | 80 | 52 | 23 | 57 | 33 | 15 | 44 | 108 | 56 | 73 | 82 | 205 | 239 | 149 | 0.014026713 | 0.136878495 | 0.23 | 0.294 | 0.233 | 0.251 | 0.307 | 0.117 | 0.388 | 0.22 | 0.102 | 0.118 | 0.074 |
| SE | 100278 | ENSG00000167986 | DDB1 | chr11 | - | 61310294 | 61310418 | 61309795 | 61309960 | 61311783 | 61311895 | 557 | 266 | 417 | 472 | 612 | 0 | 5 | 2 | 0 | 2 | 453 | 479 | 352 | 353 | 673 | 10 | 3 | 3 | 9 | 3 | 273 | 149 | 0.014161075 | 0.137984081 | 1 | 0.967 | 0.991 | 1 | 0.994 | 0.961 | 0.989 | 0.985 | 0.955 | 0.992 | 0.014 |
| SE | 50572 | ENSG00000139636 | LMBR1L | chr12 | - | 49104447 | 49104551 | 49103686 | 49103813 | 49105923 | 49105957 | 16 | 10 | 13 | 16 | 9 | 0 | 0 | 0 | 0 | 0 | 10 | 17 | 6 | 17 | 9 | 0 | 0 | 1 | 0 | 8 | 253 | 149 | 0.014243747 | 0.138641027 | 1 | 1 | 1 | 1 | 1 | 1 | 1 | 0.779 | 1 | 0.399 | 0.164 |
| SE | 100088 | ENSG00000149262 | INTS4 | chr11 | - | 77918030 | 77918264 | 77907716 | 77907810 | 77918820 | 77918978 | 2 | 1 | 0 | 3 | 0 | 34 | 20 | 41 | 61 | 30 | 2 | 4 | 6 | 3 | 0 | 31 | 47 | 25 | 41 | 32 | 298 | 149 | 0.01453945 | 0.14090615 | 0.029 | 0.024 | 0 | 0.024 | 0 | 0.031 | 0.041 | 0.107 | 0.035 | 0 | -0.027 |
| SE | 32332 | ENSG00000068745 | IP6K2 | chr3 | - | 48694458 | 48694525 | 48693452 | 48694240 | 48695089 | 48695421 | 19 | 8 | 0 | 10 | 9 | 4 | 6 | 2 | 12 | 13 | 0 | 6 | 0 | 16 | 0 | 10 | 19 | 3 | 0 | 10 | 216 | 149 | 0.014717002 | 0.142186866 | 0.766 | 0.479 | 0 | 0.365 | 0.323 | 0 | 0.179 | 0 | 1 | 0 | 0.151 |
| SE | 55710 | ENSG00000135749 | PCNX2 | chr1 | - | 233262044 | 233262165 | 233261284 | 233261321 | 233262957 | 233263163 | 40 | 33 | 39 | 30 | 73 | 9 | 17 | 10 | 12 | 13 | 38 | 44 | 30 | 31 | 41 | 0 | 14 | 9 | 0 | 6 | 270 | 149 | 0.014837853 | 0.143124862 | 0.71 | 0.517 | 0.683 | 0.58 | 0.756 | 1 | 0.634 | 0.648 | 1 | 0.79 | -0.165 |
| SE | 16951 | ENSG00000172575 | RASGRP1 | chr15 | - | 38498793 | 38498946 | 38494381 | 38494767 | 38500102 | 38500139 | 376 | 228 | 420 | 439 | 445 | 17 | 7 | 21 | 11 | 12 | 30 | 218 | 156 | 85 | 44 | 10 | 7 | 18 | 6 | 0 | 298 | 149 | 0.014903392 | 0.143562499 | 0.917 | 0.942 | 0.909 | 0.952 | 0.949 | 0.6 | 0.94 | 0.812 | 0.876 | 1 | 0.088 |
| SE | 1740 | ENSG00000100325 | ASCC2 | chr22 | - | 29827476 | 29827600 | 29825621 | 29825780 | 29832244 | 29832342 | 3 | 5 | 1 | 0 | 0 | 15 | 7 | 18 | 9 | 7 | 0 | 14 | 17 | 0 | 17 | 27 | 10 | 7 | 11 | 50 | 273 | 149 | 0.015020111 | 0.144509041 | 0.098 | 0.28 | 0.029 | 0 | 0 | 0 | 0.433 | 0.57 | 0 | 0.157 | -0.151 |
| SE | 54635 | ENSG00000140992 | PDPK1 | chr16 | + | 2538618 | 2538771 | 2538020 | 2538136 | 2557702 | 2557963 | 8 | 0 | 0 | 0 | 1 | 10 | 3 | 5 | 12 | 0 | 6 | 0 | 0 | 5 | 14 | 16 | 12 | 13 | 6 | 10 | 298 | 149 | 0.015114066 | 0.145305855 | 0.286 | 0 | 0 | 0 | 1 | 0.158 | 0 | 0 | 0.294 | 0.412 | 0.084 |
| SE | 50580 | ENSG00000139636 | LMBR1L | chr12 | - | 49106510 | 49106746 | 49105923 | 49105957 | 49106960 | 49107011 | 4 | 3 | 13 | 3 | 10 | 25 | 33 | 37 | 32 | 66 | 16 | 16 | 2 | 8 | 11 | 26 | 35 | 14 | 17 | 83 | 298 | 149 | 0.015146805 | 0.145566989 | 0.074 | 0.043 | 0.149 | 0.045 | 0.07 | 0.235 | 0.186 | 0.067 | 0.19 | 0.062 | -0.072 |
| SE | 50574 | ENSG00000139636 | LMBR1L | chr12 | - | 49104745 | 49104885 | 49103700 | 49103813 | 49105923 | 49105957 | 23 | 32 | 47 | 25 | 67 | 0 | 0 | 0 | 0 | 0 | 36 | 21 | 14 | 6 | 57 | 0 | 0 | 1 | 0 | 8 | 289 | 149 | 0.015400172 | 0.147657586 | 1 | 1 | 1 | 1 | 1 | 1 | 1 | 0.878 | 1 | 0.786 | 0.067 |
| SE | 42206 | ENSG00000132155 | RAF1 | chr3 | - | 12599112 | 12599225 | 12591707 | 12591792 | 12599690 | 12599808 | 5 | 2 | 0 | 4 | 11 | 138 | 87 | 130 | 124 | 185 | 16 | 5 | 5 | 13 | 14 | 201 | 122 | 169 | 238 | 207 | 262 | 149 | 0.015465516 | 0.148211513 | 0.02 | 0.013 | 0 | 0.018 | 0.033 | 0.043 | 0.023 | 0.017 | 0.03 | 0.037 | -0.013 |
| SE | 88400 | ENSG00000145041 | DCAF1 | chr3 | - | 51466805 | 51466876 | 51463113 | 51463227 | 51470928 | 51471005 | 64 | 51 | 61 | 55 | 51 | 3 | 0 | 5 | 3 | 8 | 60 | 74 | 67 | 64 | 33 | 4 | 0 | 1 | 4 | 0 | 220 | 149 | 0.015530311 | 0.148650516 | 0.935 | 1 | 0.892 | 0.925 | 0.812 | 0.91 | 1 | 0.978 | 0.916 | 1 | -0.048 |
| SE | 29419 | ENSG00000169762 | TAPT1 | chr4 | - | 16196657 | 16196723 | 16191360 | 16191523 | 16202461 | 16202582 | 5 | 6 | 3 | 9 | 1 | 28 | 21 | 14 | 17 | 56 | 0 | 0 | 1 | 6 | 0 | 4 | 42 | 8 | 8 | 12 | 215 | 149 | 0.015671509 | 0.149682432 | 0.11 | 0.165 | 0.129 | 0.268 | 0.012 | 0 | 0 | 0.08 | 0.342 | 0 | 0.052 |
| SE | 5039 | ENSG00000173418 | NAA20 | chr20 | + | 20032507 | 20032653 | 20026783 | 20026919 | 20033101 | 20033629 | 231 | 144 | 204 | 275 | 340 | 3 | 2 | 6 | 11 | 27 | 317 | 160 | 103 | 220 | 248 | 22 | 2 | 16 | 8 | 24 | 295 | 149 | 0.016036272 | 0.152431221 | 0.975 | 0.973 | 0.945 | 0.927 | 0.864 | 0.879 | 0.976 | 0.765 | 0.933 | 0.839 | 0.058 |
| SE | 30157 | ENSG00000177054 | ZDHHC13 | chr11 | + | 19166301 | 19166385 | 19165051 | 19165145 | 19170410 | 19170568 | 44 | 23 | 32 | 42 | 45 | 1 | 2 | 1 | 6 | 0 | 4 | 26 | 22 | 14 | 9 | 0 | 3 | 0 | 0 | 0 | 233 | 149 | 0.016752261 | 0.15778119 | 0.966 | 0.88 | 0.953 | 0.817 | 1 | 1 | 0.847 | 1 | 1 | 1 | -0.046 |
| SE | 66190 | ENSG00000157483 | MYO1E | chr15 | - | 59163156 | 59163303 | 59161072 | 59161230 | 59171896 | 59172042 | 76 | 41 | 41 | 69 | 113 | 0 | 0 | 0 | 0 | 6 | 122 | 24 | 45 | 51 | 23 | 11 | 0 | 0 | 0 | 0 | 296 | 149 | 0.017392172 | 0.162906746 | 1 | 1 | 1 | 1 | 0.905 | 0.848 | 1 | 1 | 1 | 1 | 0.011 |
| SE | 29806 | ENSG00000141452 | RMC1 | chr18 | + | 23520201 | 23520313 | 23519068 | 23519174 | 23524129 | 23524158 | 159 | 96 | 101 | 152 | 178 | 0 | 0 | 0 | 0 | 15 | 268 | 140 | 127 | 139 | 315 | 0 | 1 | 0 | 0 | 0 | 261 | 149 | 0.017412529 | 0.163058408 | 1 | 1 | 1 | 1 | 0.871 | 1 | 0.988 | 1 | 1 | 1 | -0.023 |
| SE | 84347 | ENSG00000159256 | MORC3 | chr21 | + | 36368987 | 36369876 | 36364092 | 36364259 | 36372373 | 36372531 | 424 | 352 | 308 | 359 | 656 | 0 | 5 | 2 | 3 | 15 | 64 | 152 | 166 | 45 | 82 | 4 | 0 | 7 | 6 | 0 | 298 | 149 | 0.017560682 | 0.164230784 | 1 | 0.972 | 0.987 | 0.984 | 0.956 | 0.889 | 1 | 0.922 | 0.789 | 1 | 0.06 |
| SE | 92134 | ENSG00000108666 | C17orf75 | chr17 | - | 32333416 | 32333520 | 32328440 | 32331978 | 32334468 | 32334605 | 69 | 68 | 88 | 81 | 90 | 0 | 0 | 0 | 0 | 1 | 63 | 25 | 49 | 28 | 10 | 0 | 5 | 0 | 0 | 4 | 253 | 149 | 0.017888457 | 0.166816891 | 1 | 1 | 1 | 1 | 0.981 | 1 | 0.746 | 1 | 1 | 0.596 | 0.128 |
| SE | 57162 | ENSG00000011021 | CLCN6 | chr1 | + | 11807130 | 11807190 | 11806095 | 11806349 | 11815845 | 11815911 | 28 | 22 | 31 | 30 | 50 | 0 | 0 | 0 | 0 | 0 | 62 | 32 | 26 | 48 | 138 | 0 | 0 | 0 | 0 | 12 | 209 | 149 | 0.017964146 | 0.167440166 | 1 | 1 | 1 | 1 | 1 | 1 | 1 | 1 | 1 | 0.891 | 0.022 |
| SE | 38010 | ENSG00000197386 | HTT | chr4 | + | 3154292 | 3154419 | 3148004 | 3148207 | 3157071 | 3157199 | 152 | 96 | 136 | 149 | 183 | 2 | 0 | 0 | 0 | 2 | 303 | 188 | 113 | 178 | 316 | 10 | 0 | 2 | 0 | 6 | 276 | 149 | 0.018065057 | 0.168127115 | 0.976 | 1 | 1 | 1 | 0.98 | 0.942 | 1 | 0.968 | 1 | 0.966 | 0.016 |
| SE | 107503 | ENSG00000123349 | PFDN5 | chr12 | + | 53296429 | 53296551 | 53295524 | 53295639 | 53298044 | 53298150 | 3 | 1 | 4 | 2 | 0 | 21 | 6 | 15 | 20 | 7 | 12 | 3 | 0 | 0 | 9 | 14 | 11 | 12 | 8 | 9 | 271 | 149 | 0.018240864 | 0.169417159 | 0.073 | 0.084 | 0.128 | 0.052 | 0 | 0.32 | 0.13 | 0 | 0 | 0.355 | -0.094 |
| SE | 104998 | ENSG00000129315 | CCNT1 | chr12 | - | 48695998 | 48696162 | 48688457 | 48694436 | 48698137 | 48698183 | 55 | 25 | 40 | 44 | 55 | 0 | 0 | 0 | 0 | 0 | 13 | 14 | 30 | 6 | 2 | 0 | 1 | 7 | 0 | 0 | 298 | 149 | 0.018763236 | 0.173404901 | 1 | 1 | 1 | 1 | 1 | 1 | 0.875 | 0.682 | 1 | 1 | 0.089 |
| SE | 13617 | ENSG00000038358 | EDC4 | chr16 | + | 67882957 | 67883006 | 67882779 | 67882865 | 67883567 | 67883731 | 26 | 17 | 21 | 11 | 7 | 0 | 0 | 0 | 0 | 0 | 2 | 19 | 12 | 25 | 16 | 0 | 4 | 2 | 2 | 0 | 198 | 149 | 0.01894764 | 0.174758505 | 1 | 1 | 1 | 1 | 1 | 1 | 0.781 | 0.819 | 0.904 | 1 | 0.099 |
| SE | 12087 | ENSG00000205250 | E2F4 | chr16 | + | 67193421 | 67193515 | 67192760 | 67192870 | 67194397 | 67194459 | 38 | 24 | 30 | 49 | 48 | 0 | 3 | 0 | 1 | 0 | 50 | 53 | 10 | 89 | 70 | 0 | 10 | 1 | 1 | 0 | 243 | 149 | 0.019050651 | 0.175543102 | 1 | 0.831 | 1 | 0.968 | 1 | 1 | 0.765 | 0.86 | 0.982 | 1 | 0.038 |
| SE | 87766 | ENSG00000196459 | TRAPPC2 | chrX | - | 13720427 | 13720497 | 13719870 | 13719982 | 13734524 | 13734599 | 6 | 1 | 0 | 4 | 2 | 39 | 25 | 23 | 30 | 13 | 16 | 11 | 1 | 0 | 6 | 36 | 19 | 11 | 37 | 59 | 219 | 149 | 0.019607727 | 0.179640373 | 0.095 | 0.026 | 0 | 0.083 | 0.095 | 0.232 | 0.283 | 0.058 | 0 | 0.065 | -0.068 |
| SE | 58939 | ENSG00000131778 | CHD1L | chr1 | + | 147254869 | 147254976 | 147242672 | 147242830 | 147255812 | 147255927 | 50 | 30 | 30 | 30 | 67 | 1 | 0 | 0 | 0 | 0 | 38 | 21 | 8 | 31 | 40 | 0 | 0 | 0 | 3 | 6 | 256 | 149 | 0.019750358 | 0.180672283 | 0.967 | 1 | 1 | 1 | 1 | 1 | 1 | 1 | 0.857 | 0.795 | 0.063 |
| SE | 28224 | ENSG00000204084 | INPP5B | chr1 | - | 37942082 | 37942264 | 37940762 | 37940798 | 37943639 | 37943669 | 0 | 4 | 6 | 1 | 1 | 6 | 1 | 4 | 1 | 7 | 0 | 0 | 4 | 0 | 0 | 13 | 20 | 0 | 2 | 19 | 298 | 149 | 0.02018855 | 0.183928635 | 0 | 0.667 | 0.429 | 0.333 | 0.067 | 0 | 0 | 1 | 0 | 0 | 0.099 |
| SE | 89484 | ENSG00000064313 | TAF2 | chr8 | - | 119796989 | 119797103 | 119795531 | 119795631 | 119797661 | 119797846 | 164 | 105 | 160 | 206 | 198 | 3 | 12 | 2 | 2 | 9 | 36 | 141 | 128 | 70 | 95 | 0 | 4 | 1 | 4 | 0 | 263 | 149 | 0.020692301 | 0.187536102 | 0.969 | 0.832 | 0.978 | 0.983 | 0.926 | 1 | 0.952 | 0.986 | 0.908 | 1 | -0.032 |
| SE | 105693 | ENSG00000028203 | VEZT | chr12 | + | 95256549 | 95256622 | 95251939 | 95252071 | 95257149 | 95257181 | 55 | 52 | 62 | 60 | 138 | 13 | 15 | 17 | 23 | 29 | 17 | 45 | 43 | 21 | 27 | 0 | 3 | 9 | 13 | 0 | 222 | 149 | 0.020859603 | 0.18880712 | 0.74 | 0.699 | 0.71 | 0.636 | 0.762 | 1 | 0.91 | 0.762 | 0.52 | 1 | -0.129 |
| SE | 66491 | ENSG00000132676 | DAP3 | chr1 | + | 155709032 | 155709125 | 155689094 | 155689174 | 155709772 | 155709824 | 7 | 12 | 8 | 4 | 5 | 168 | 107 | 189 | 159 | 59 | 27 | 11 | 2 | 8 | 13 | 211 | 116 | 107 | 84 | 183 | 242 | 149 | 0.020970087 | 0.189570941 | 0.025 | 0.065 | 0.025 | 0.015 | 0.05 | 0.073 | 0.055 | 0.011 | 0.055 | 0.042 | -0.011 |
| SE | 17329 | ENSG00000092529 | CAPN3 | chr15 | + | 42389952 | 42390096 | 42388927 | 42389096 | 42392638 | 42392722 | 34 | 21 | 25 | 24 | 12 | 9 | 13 | 3 | 3 | 10 | 56 | 63 | 35 | 132 | 60 | 0 | 12 | 5 | 43 | 2 | 293 | 149 | 0.021087742 | 0.190414696 | 0.658 | 0.451 | 0.809 | 0.803 | 0.379 | 1 | 0.728 | 0.781 | 0.61 | 0.938 | -0.191 |
| SE | 107501 | ENSG00000123349 | PFDN5 | chr12 | + | 53296429 | 53296467 | 53295838 | 53295941 | 53297849 | 53297924 | 0 | 1 | 8 | 0 | 0 | 3 | 13 | 11 | 3 | 11 | 0 | 2 | 0 | 0 | 0 | 34 | 5 | 5 | 4 | 20 | 187 | 149 | 0.02129409 | 0.191990102 | 0 | 0.058 | 0.367 | 0 | 0 | 0 | 0.242 | 0 | 0 | 0 | 0.037 |
| SE | 73317 | ENSG00000136699 | SMPD4 | chr2 | - | 130174913 | 130175000 | 130172786 | 130172895 | 130176553 | 130176637 | 23 | 10 | 23 | 31 | 19 | 0 | 0 | 0 | 0 | 0 | 26 | 27 | 10 | 18 | 37 | 8 | 1 | 0 | 0 | 0 | 236 | 149 | 0.021922364 | 0.196613222 | 1 | 1 | 1 | 1 | 1 | 0.672 | 0.945 | 1 | 1 | 1 | 0.077 |
| SE | 37124 | ENSG00000144848 | ATG3 | chr3 | - | 112553279 | 112553329 | 112550191 | 112550262 | 112558375 | 112558417 | 245 | 211 | 242 | 262 | 314 | 1 | 6 | 2 | 8 | 1 | 468 | 212 | 232 | 311 | 213 | 1 | 2 | 12 | 8 | 13 | 199 | 149 | 0.021984929 | 0.19699382 | 0.995 | 0.963 | 0.989 | 0.961 | 0.996 | 0.997 | 0.988 | 0.935 | 0.967 | 0.925 | 0.018 |
| SE | 87765 | ENSG00000196459 | TRAPPC2 | chrX | - | 13720427 | 13720497 | 13719870 | 13719982 | 13734043 | 13734185 | 8 | 1 | 5 | 8 | 5 | 69 | 24 | 51 | 54 | 44 | 43 | 18 | 0 | 7 | 17 | 17 | 140 | 51 | 29 | 52 | 219 | 149 | 0.022008663 | 0.1971388 | 0.073 | 0.028 | 0.063 | 0.092 | 0.072 | 0.632 | 0.08 | 0 | 0.141 | 0.182 | -0.141 |
| SE | 64457 | ENSG00000065150 | IPO5 | chr13 | + | 97990125 | 97990222 | 97989061 | 97989164 | 97990432 | 97990502 | 228 | 100 | 220 | 215 | 357 | 0 | 0 | 3 | 0 | 2 | 84 | 104 | 92 | 111 | 15 | 0 | 9 | 1 | 0 | 2 | 246 | 149 | 0.022726319 | 0.202248226 | 1 | 1 | 0.978 | 1 | 0.991 | 1 | 0.875 | 0.982 | 1 | 0.82 | 0.058 |
| SE | 15781 | ENSG00000167077 | MEI1 | chr22 | + | 41795410 | 41795542 | 41793828 | 41794477 | 41795734 | 41795773 | 134 | 43 | 126 | 85 | 79 | 5 | 8 | 9 | 9 | 11 | 35 | 91 | 42 | 47 | 74 | 13 | 10 | 12 | 9 | 5 | 281 | 149 | 0.02275209 | 0.202406424 | 0.934 | 0.74 | 0.881 | 0.834 | 0.792 | 0.588 | 0.828 | 0.65 | 0.735 | 0.887 | 0.099 |
| SE | 69657 | ENSG00000143390 | RFX5 | chr1 | - | 151346481 | 151346606 | 151346204 | 151346333 | 151347210 | 151347251 | 96 | 73 | 89 | 60 | 131 | 0 | 0 | 0 | 0 | 0 | 116 | 89 | 66 | 76 | 108 | 4 | 0 | 0 | 0 | 8 | 274 | 149 | 0.023137679 | 0.205232549 | 1 | 1 | 1 | 1 | 1 | 0.94 | 1 | 1 | 1 | 0.88 | 0.036 |
| SE | 30152 | ENSG00000177054 | ZDHHC13 | chr11 | + | 19142977 | 19143123 | 19117167 | 19117276 | 19146180 | 19146303 | 50 | 24 | 51 | 35 | 39 | 2 | 0 | 2 | 4 | 0 | 10 | 41 | 8 | 28 | 12 | 0 | 4 | 0 | 13 | 0 | 295 | 149 | 0.023313533 | 0.206393487 | 0.927 | 1 | 0.928 | 0.815 | 1 | 1 | 0.838 | 1 | 0.521 | 1 | 0.062 |
| SE | 41065 | ENSG00000160741 | CRTC2 | chr1 | - | 153953265 | 153953369 | 153952804 | 153952834 | 153953537 | 153953606 | 208 | 105 | 119 | 174 | 175 | 6 | 0 | 1 | 0 | 0 | 319 | 250 | 179 | 171 | 364 | 14 | 1 | 2 | 2 | 3 | 253 | 149 | 0.023538916 | 0.207920012 | 0.953 | 1 | 0.986 | 1 | 1 | 0.931 | 0.993 | 0.981 | 0.981 | 0.986 | 0.013 |
| SE | 79021 | ENSG00000160218 | TRAPPC10 | chr21 | + | 44063052 | 44063168 | 44059102 | 44059214 | 44063537 | 44063785 | 2 | 8 | 8 | 15 | 2 | 80 | 35 | 68 | 58 | 66 | 16 | 14 | 5 | 27 | 0 | 28 | 77 | 41 | 74 | 77 | 265 | 149 | 0.023617686 | 0.208474676 | 0.014 | 0.114 | 0.062 | 0.127 | 0.017 | 0.243 | 0.093 | 0.064 | 0.17 | 0 | -0.047 |
| SE | 29797 | ENSG00000141452 | RMC1 | chr18 | + | 23506969 | 23507054 | 23504370 | 23504447 | 23515855 | 23515996 | 32 | 31 | 46 | 49 | 14 | 1 | 0 | 0 | 0 | 3 | 58 | 51 | 24 | 60 | 27 | 0 | 1 | 0 | 0 | 12 | 234 | 149 | 0.023876559 | 0.21014378 | 0.953 | 1 | 1 | 1 | 0.748 | 1 | 0.97 | 1 | 1 | 0.589 | 0.028 |
| SE | 102590 | ENSG00000175470 | PPP2R2D | chr10 | + | 131934893 | 131935006 | 131934457 | 131934555 | 131940030 | 131940196 | 30 | 34 | 10 | 14 | 11 | 69 | 31 | 78 | 56 | 75 | 24 | 18 | 41 | 25 | 214 | 92 | 53 | 41 | 61 | 88 | 262 | 149 | 0.024016757 | 0.211140357 | 0.198 | 0.384 | 0.068 | 0.124 | 0.077 | 0.129 | 0.162 | 0.363 | 0.189 | 0.58 | -0.114 |
| SE | 12403 | ENSG00000188559 | RALGAPA2 | chr20 | - | 20637362 | 20637501 | 20635350 | 20635617 | 20639784 | 20639900 | 126 | 134 | 163 | 193 | 196 | 6 | 4 | 5 | 2 | 4 | 113 | 193 | 163 | 270 | 69 | 0 | 3 | 48 | 16 | 0 | 288 | 149 | 0.024358079 | 0.213565525 | 0.916 | 0.945 | 0.944 | 0.98 | 0.962 | 1 | 0.971 | 0.637 | 0.897 | 1 | 0.048 |
| SE | 78270 | ENSG00000173715 | C11orf80 | chr11 | + | 66822563 | 66822674 | 66816052 | 66816199 | 66828265 | 66828364 | 20 | 12 | 33 | 29 | 59 | 0 | 4 | 4 | 2 | 12 | 31 | 51 | 28 | 40 | 24 | 0 | 1 | 0 | 10 | 0 | 260 | 149 | 0.02466684 | 0.215885913 | 1 | 0.632 | 0.825 | 0.893 | 0.738 | 1 | 0.967 | 1 | 0.696 | 1 | -0.115 |
| SE | 58955 | ENSG00000131778 | CHD1L | chr1 | + | 147272381 | 147272475 | 147272170 | 147272281 | 147275353 | 147275468 | 1 | 0 | 0 | 0 | 0 | 49 | 28 | 53 | 41 | 46 | 16 | 0 | 0 | 0 | 0 | 61 | 52 | 23 | 34 | 121 | 243 | 149 | 0.024701332 | 0.216139413 | 0.012 | 0 | 0 | 0 | 0 | 0.139 | 0 | 0 | 0 | 0 | -0.025 |
| SE | 42669 | ENSG00000088854 | C20orf194 | chr20 | - | 3359515 | 3359593 | 3348524 | 3348623 | 3374047 | 3374154 | 34 | 54 | 47 | 46 | 37 | 0 | 4 | 0 | 0 | 0 | 147 | 37 | 24 | 85 | 41 | 0 | 0 | 4 | 8 | 0 | 227 | 149 | 0.024808977 | 0.216911631 | 1 | 0.899 | 1 | 1 | 1 | 1 | 1 | 0.798 | 0.875 | 1 | 0.045 |
| SE | 53351 | ENSG00000131778 | CHD1L | chr1 | + | 147284350 | 147284499 | 147276103 | 147276257 | 147285323 | 147285487 | 160 | 198 | 170 | 212 | 411 | 2 | 5 | 2 | 5 | 1 | 493 | 285 | 232 | 301 | 872 | 24 | 10 | 4 | 5 | 3 | 298 | 149 | 0.025511547 | 0.221550894 | 0.976 | 0.952 | 0.977 | 0.955 | 0.995 | 0.911 | 0.934 | 0.967 | 0.968 | 0.993 | 0.016 |
| SE | 47783 | ENSG00000154358 | OBSCN | chr1 | + | 228288624 | 228288888 | 228288060 | 228288324 | 228291897 | 228292161 | 47 | 45 | 61 | 54 | 43 | 0 | 2 | 0 | 0 | 0 | 17 | 8 | 18 | 49 | 30 | 0 | 0 | 0 | 0 | 9 | 298 | 149 | 0.025558293 | 0.221901982 | 1 | 0.918 | 1 | 1 | 1 | 1 | 1 | 1 | 1 | 0.625 | 0.059 |
| SE | 29800 | ENSG00000141452 | RMC1 | chr18 | + | 23507984 | 23508041 | 23506969 | 23507054 | 23509192 | 23509279 | 192 | 132 | 156 | 176 | 223 | 1 | 0 | 0 | 2 | 0 | 176 | 201 | 145 | 190 | 252 | 3 | 0 | 6 | 3 | 7 | 206 | 149 | 0.026251737 | 0.226765891 | 0.993 | 1 | 1 | 0.985 | 1 | 0.977 | 1 | 0.946 | 0.979 | 0.963 | 0.023 |
| SE | 98873 | ENSG00000175294 | CATSPER1 | chr11 | - | 66020563 | 66020627 | 66020316 | 66020389 | 66020810 | 66020954 | 22 | 5 | 2 | 2 | 2 | 8 | 10 | 3 | 8 | 0 | 11 | 8 | 9 | 2 | 31 | 6 | 5 | 1 | 0 | 5 | 213 | 149 | 0.026868393 | 0.230895366 | 0.658 | 0.259 | 0.318 | 0.149 | 1 | 0.562 | 0.528 | 0.863 | 1 | 0.813 | -0.276 |
| SE | 12389 | ENSG00000188559 | RALGAPA2 | chr20 | - | 20524405 | 20524543 | 20520916 | 20521100 | 20524829 | 20524898 | 151 | 79 | 110 | 107 | 150 | 1 | 1 | 0 | 1 | 0 | 109 | 126 | 114 | 203 | 78 | 0 | 5 | 7 | 1 | 0 | 287 | 149 | 0.02709016 | 0.23236757 | 0.987 | 0.976 | 1 | 0.982 | 1 | 1 | 0.929 | 0.894 | 0.991 | 1 | 0.026 |
| SE | 92580 | ENSG00000137054 | POLR1E | chr9 | + | 37500839 | 37500921 | 37498090 | 37498224 | 37501712 | 37501844 | 177 | 78 | 183 | 169 | 205 | 2 | 3 | 8 | 9 | 6 | 95 | 71 | 82 | 53 | 133 | 0 | 5 | 15 | 11 | 0 | 231 | 149 | 0.027517958 | 0.235212448 | 0.983 | 0.944 | 0.937 | 0.924 | 0.957 | 1 | 0.902 | 0.779 | 0.757 | 1 | 0.061 |
| SE | 38567 | ENSG00000198925 | ATG9A | chr2 | - | 219227921 | 219228079 | 219227769 | 219227813 | 219228433 | 219228485 | 28 | 13 | 26 | 16 | 14 | 0 | 0 | 1 | 0 | 0 | 16 | 13 | 5 | 8 | 15 | 1 | 2 | 4 | 0 | 0 | 298 | 149 | 0.027607558 | 0.235823843 | 1 | 1 | 0.929 | 1 | 1 | 0.889 | 0.765 | 0.385 | 1 | 1 | 0.178 |
| SE | 83154 | ENSG00000055130 | CUL1 | chr7 | + | 148760332 | 148760496 | 148759547 | 148759638 | 148766560 | 148766723 | 428 | 388 | 344 | 407 | 539 | 4 | 4 | 5 | 7 | 11 | 362 | 300 | 263 | 603 | 31 | 4 | 18 | 3 | 15 | 0 | 298 | 149 | 0.027626684 | 0.23593574 | 0.982 | 0.98 | 0.972 | 0.967 | 0.961 | 0.978 | 0.893 | 0.978 | 0.953 | 1 | 0.012 |
| SE | 22135 | ENSG00000175727 | MLXIP | chr12 | + | 122137468 | 122137590 | 122135488 | 122135666 | 122138193 | 122138295 | 78 | 35 | 95 | 119 | 65 | 5 | 1 | 2 | 3 | 3 | 22 | 94 | 22 | 67 | 325 | 1 | 0 | 2 | 15 | 18 | 271 | 149 | 0.028140073 | 0.239640546 | 0.896 | 0.951 | 0.963 | 0.956 | 0.923 | 0.924 | 1 | 0.858 | 0.711 | 0.908 | 0.058 |
| SE | 82048 | ENSG00000104133 | SPG11 | chr15 | - | 44613429 | 44613536 | 44610839 | 44610985 | 44615362 | 44615566 | 355 | 291 | 301 | 325 | 602 | 9 | 17 | 4 | 16 | 8 | 84 | 350 | 241 | 247 | 86 | 8 | 0 | 17 | 26 | 0 | 256 | 149 | 0.02843038 | 0.241299191 | 0.958 | 0.909 | 0.978 | 0.922 | 0.978 | 0.859 | 1 | 0.892 | 0.847 | 1 | 0.029 |
| SE | 103884 | ENSG00000119977 | TCTN3 | chr10 | - | 95682650 | 95682804 | 95680471 | 95680609 | 95683100 | 95683180 | 69 | 49 | 77 | 69 | 102 | 2 | 3 | 3 | 0 | 3 | 49 | 56 | 18 | 24 | 25 | 0 | 0 | 3 | 0 | 0 | 298 | 149 | 0.030001857 | 0.252149571 | 0.945 | 0.891 | 0.928 | 1 | 0.944 | 1 | 1 | 0.75 | 1 | 1 | -0.008 |
| SE | 1586 | ENSG00000198089 | SFI1 | chr22 | + | 31580271 | 31580364 | 31578381 | 31578452 | 31583874 | 31583877 | 72 | 72 | 121 | 96 | 134 | 11 | 2 | 6 | 5 | 7 | 67 | 95 | 63 | 84 | 153 | 13 | 13 | 13 | 0 | 12 | 242 | 149 | 0.030418483 | 0.255021473 | 0.801 | 0.957 | 0.925 | 0.922 | 0.922 | 0.76 | 0.818 | 0.749 | 1 | 0.887 | 0.063 |
| SE | 78269 | ENSG00000173715 | C11orf80 | chr11 | + | 66821642 | 66821764 | 66815833 | 66816199 | 66828265 | 66828364 | 18 | 8 | 25 | 18 | 55 | 0 | 4 | 4 | 2 | 12 | 36 | 13 | 13 | 46 | 33 | 0 | 1 | 0 | 10 | 0 | 271 | 149 | 0.030484758 | 0.255331045 | 1 | 0.524 | 0.775 | 0.832 | 0.716 | 1 | 0.877 | 1 | 0.717 | 1 | -0.149 |
| SE | 86202 | ENSG00000171311 | EXOSC1 | chr10 | - | 97445452 | 97445606 | 97443236 | 97443311 | 97445731 | 97445847 | 0 | 0 | 2 | 0 | 0 | 56 | 32 | 49 | 51 | 14 | 14 | 0 | 0 | 0 | 0 | 39 | 43 | 31 | 41 | 21 | 298 | 149 | 0.030518753 | 0.25553377 | 0 | 0 | 0.02 | 0 | 0 | 0.152 | 0 | 0 | 0 | 0 | -0.026 |
| SE | 52023 | ENSG00000135250 | SRPK2 | chr7 | - | 105126247 | 105126340 | 105125781 | 105125874 | 105126992 | 105127062 | 54 | 28 | 37 | 61 | 47 | 32 | 10 | 11 | 18 | 16 | 32 | 40 | 35 | 12 | 5 | 10 | 4 | 3 | 7 | 0 | 242 | 149 | 0.031128933 | 0.25965203 | 0.51 | 0.633 | 0.674 | 0.676 | 0.644 | 0.663 | 0.86 | 0.878 | 0.513 | 1 | -0.155 |
| SE | 96392 | ENSG00000197563 | PIGN | chr18 | - | 62140419 | 62140479 | 62138982 | 62139075 | 62143305 | 62143346 | 53 | 22 | 48 | 40 | 105 | 2 | 2 | 0 | 2 | 6 | 40 | 69 | 26 | 35 | 37 | 0 | 0 | 0 | 5 | 0 | 209 | 149 | 0.031419694 | 0.261678154 | 0.95 | 0.887 | 1 | 0.934 | 0.926 | 1 | 1 | 1 | 0.833 | 1 | -0.027 |
| SE | 108798 | ENSG00000105983 | LMBR1 | chr7 | - | 156762133 | 156762198 | 156756392 | 156756465 | 156763107 | 156763176 | 144 | 113 | 136 | 131 | 202 | 0 | 2 | 0 | 0 | 0 | 68 | 118 | 146 | 88 | 46 | 0 | 0 | 7 | 3 | 3 | 214 | 149 | 0.03217281 | 0.266928292 | 1 | 0.975 | 1 | 1 | 1 | 1 | 1 | 0.936 | 0.953 | 0.914 | 0.034 |
| SE | 29420 | ENSG00000169762 | TAPT1 | chr4 | - | 16196657 | 16196723 | 16191360 | 16191523 | 16213767 | 16213898 | 3 | 4 | 1 | 0 | 0 | 5 | 1 | 4 | 4 | 4 | 0 | 0 | 1 | 3 | 0 | 7 | 8 | 5 | 9 | 3 | 215 | 149 | 0.032308399 | 0.267854554 | 0.294 | 0.735 | 0.148 | 0 | 0 | 0 | 0 | 0.122 | 0.188 | 0 | 0.173 |
| SE | 26443 | ENSG00000083312 | TNPO1 | chr5 | + | 72855773 | 72855923 | 72851204 | 72851319 | 72861807 | 72861914 | 361 | 257 | 255 | 309 | 408 | 0 | 0 | 0 | 3 | 0 | 130 | 227 | 141 | 84 | 64 | 10 | 2 | 1 | 0 | 0 | 298 | 149 | 0.032484918 | 0.268964959 | 1 | 1 | 1 | 0.981 | 1 | 0.867 | 0.983 | 0.986 | 1 | 1 | 0.029 |
| SE | 41969 | ENSG00000120008 | WDR11 | chr10 | + | 120873838 | 120873923 | 120871169 | 120871346 | 120883779 | 120883805 | 69 | 59 | 48 | 66 | 128 | 0 | 0 | 0 | 0 | 0 | 1 | 35 | 15 | 17 | 2 | 12 | 0 | 0 | 0 | 0 | 234 | 149 | 0.033227279 | 0.2741662 | 1 | 1 | 1 | 1 | 1 | 0.05 | 1 | 1 | 1 | 1 | 0.19 |
| SE | 82004 | ENSG00000104133 | SPG11 | chr15 | - | 44569397 | 44569505 | 44567423 | 44567592 | 44572682 | 44572820 | 102 | 57 | 81 | 94 | 104 | 2 | 0 | 0 | 2 | 11 | 46 | 63 | 110 | 82 | 62 | 0 | 0 | 1 | 6 | 0 | 257 | 149 | 0.033363396 | 0.274912702 | 0.967 | 1 | 1 | 0.965 | 0.846 | 1 | 1 | 0.985 | 0.888 | 1 | -0.019 |
| SE | 84866 | ENSG00000164074 | ABHD18 | chr4 | + | 128008919 | 128008994 | 127984338 | 127984423 | 128009106 | 128009191 | 46 | 46 | 38 | 42 | 64 | 0 | 0 | 0 | 0 | 0 | 20 | 30 | 49 | 39 | 43 | 10 | 0 | 0 | 0 | 0 | 228 | 149 | 0.033437684 | 0.275335004 | 1 | 1 | 1 | 1 | 1 | 0.567 | 1 | 1 | 1 | 1 | 0.087 |
| SE | 16965 | ENSG00000172575 | RASGRP1 | chr15 | - | 38511603 | 38511720 | 38507725 | 38508001 | 38512782 | 38512956 | 449 | 240 | 469 | 544 | 555 | 4 | 2 | 10 | 17 | 6 | 95 | 203 | 274 | 116 | 66 | 0 | 7 | 1 | 1 | 0 | 266 | 149 | 0.034093338 | 0.279691115 | 0.984 | 0.985 | 0.963 | 0.947 | 0.981 | 1 | 0.942 | 0.994 | 0.985 | 1 | -0.012 |
| SE | 91181 | ENSG00000164941 | INTS8 | chr8 | + | 94866157 | 94866191 | 94859510 | 94859632 | 94867139 | 94867196 | 34 | 39 | 28 | 43 | 55 | 1 | 5 | 2 | 2 | 0 | 6 | 4 | 18 | 10 | 1 | 0 | 0 | 0 | 0 | 0 | 183 | 149 | 0.035740723 | 0.289985042 | 0.965 | 0.864 | 0.919 | 0.946 | 1 | 1 | 1 | 1 | 1 | 1 | -0.061 |
| SE | 87610 | ENSG00000102078 | SLC25A14 | chrX | + | 130346543 | 130346691 | 130345181 | 130345275 | 130349250 | 130349345 | 30 | 31 | 26 | 44 | 25 | 1 | 0 | 3 | 1 | 3 | 40 | 26 | 34 | 29 | 4 | 0 | 0 | 0 | 1 | 0 | 297 | 149 | 0.035926296 | 0.29118895 | 0.938 | 1 | 0.813 | 0.957 | 0.807 | 1 | 1 | 1 | 0.936 | 1 | -0.084 |
| SE | 96888 | ENSG00000100731 | PCNX1 | chr14 | + | 70995740 | 70995843 | 70976941 | 70978648 | 71009633 | 71009724 | 8 | 3 | 7 | 6 | 5 | 82 | 47 | 69 | 69 | 92 | 14 | 6 | 10 | 8 | 0 | 7 | 112 | 90 | 17 | 28 | 252 | 149 | 0.036346314 | 0.293954242 | 0.055 | 0.036 | 0.057 | 0.049 | 0.031 | 0.542 | 0.031 | 0.062 | 0.218 | 0 | -0.125 |
| SE | 29422 | ENSG00000169762 | TAPT1 | chr4 | - | 16202461 | 16202580 | 16191360 | 16191523 | 16226258 | 16226471 | 21 | 17 | 10 | 13 | 38 | 3 | 0 | 0 | 2 | 5 | 4 | 36 | 7 | 3 | 12 | 0 | 1 | 0 | 3 | 0 | 268 | 149 | 0.037323743 | 0.300030777 | 0.796 | 1 | 1 | 0.783 | 0.809 | 1 | 0.952 | 1 | 0.357 | 1 | 0.016 |
| SE | 94160 | ENSG00000198000 | NOL8 | chr9 | - | 92311145 | 92311259 | 92310552 | 92310675 | 92314266 | 92316138 | 136 | 166 | 145 | 171 | 161 | 9 | 6 | 8 | 4 | 20 | 5 | 86 | 79 | 16 | 25 | 0 | 5 | 4 | 0 | 0 | 263 | 149 | 0.039609198 | 0.314623291 | 0.895 | 0.94 | 0.911 | 0.96 | 0.82 | 1 | 0.907 | 0.918 | 1 | 1 | -0.06 |
| SE | 52067 | ENSG00000043514 | TRIT1 | chr1 | - | 39847547 | 39847660 | 39847219 | 39847297 | 39847985 | 39848091 | 81 | 60 | 56 | 70 | 101 | 1 | 2 | 4 | 3 | 1 | 61 | 70 | 54 | 51 | 60 | 0 | 0 | 0 | 4 | 0 | 262 | 149 | 0.040107036 | 0.317708715 | 0.979 | 0.945 | 0.888 | 0.93 | 0.983 | 1 | 1 | 1 | 0.879 | 1 | -0.031 |
| SE | 46333 | ENSG00000095564 | BTAF1 | chr10 | + | 91966636 | 91966757 | 91962537 | 91962678 | 91980453 | 91980558 | 150 | 116 | 165 | 171 | 130 | 3 | 0 | 1 | 1 | 0 | 25 | 59 | 120 | 37 | 14 | 0 | 0 | 5 | 0 | 5 | 270 | 149 | 0.04069636 | 0.32145307 | 0.965 | 1 | 0.989 | 0.99 | 1 | 1 | 1 | 0.93 | 1 | 0.607 | 0.081 |
| SE | 33416 | ENSG00000096063 | SRPK1 | chr6 | - | 35904878 | 35904974 | 35890894 | 35891041 | 35920467 | 35920528 | 0 | 0 | 2 | 2 | 1 | 16 | 9 | 7 | 12 | 3 | 0 | 4 | 1 | 7 | 0 | 13 | 30 | 8 | 27 | 4 | 245 | 149 | 0.040943837 | 0.323093999 | 0 | 0 | 0.148 | 0.092 | 0.169 | 0 | 0.075 | 0.071 | 0.136 | 0 | 0.025 |

| | | | | | | | | | | | | | | | | | | | | | | | | | | | | | | | | | | | | | | | | | | | | | | |
|---|---|---|---|---|---|---|---|---|---|---|---|---|---|---|---|---|---|---|---|---|---|---|---|---|---|---|---|---|---|---|---|---|---|---|---|---|---|---|---|---|---|---|---|---|---|---|
| SE | 26435 | ENSG00000158290 | CUL4B | chrX | - | 120544480 | 120544643 | 120544113 | 120544203 | 120545443 | 120545517 | 378 | 264 | 368 | 391 | 548 | 9 | 6 | 2 | 5 | 0 | 92 | 188 | 174 | 89 | 59 | 0 | 0 | 1 | 0 | 0 | 298 | 149 | 0.041708143 | 0.32760642 | 0.955 | 0.957 | 0.989 | 0.975 | 1 | 1 | 1 | 0.989 | 1 | 1 | -0.023 |
| SE | 57885 | ENSG00000162604 | TM2D1 | chr1 | - | 61686791 | 61686921 | 61683451 | 61683546 | 61694696 | 61694770 | 15 | 4 | 20 | 24 | 11 | 76 | 40 | 30 | 49 | 118 | 25 | 11 | 12 | 0 | 21 | 21 | 43 | 27 | 12 | 9 | 279 | 149 | 0.041855754 | 0.328568087 | 0.095 | 0.051 | 0.263 | 0.207 | 0.047 | 0.389 | 0.12 | 0.192 | 0 | 0.555 | -0.119 |
| SE | 76683 | ENSG00000223501 | VPS52 | chr6 | - | 33267681 | 33267739 | 33266556 | 33266712 | 33267864 | 33267997 | 27 | 11 | 21 | 19 | 36 | 2 | 1 | 4 | 0 | 2 | 6 | 24 | 17 | 20 | 43 | 0 | 3 | 7 | 1 | 7 | 207 | 149 | 0.042422928 | 0.332254234 | 0.907 | 0.888 | 0.791 | 1 | 0.928 | 1 | 0.852 | 0.636 | 0.935 | 0.816 | 0.055 |
| SE | 57884 | ENSG00000162604 | TM2D1 | chr1 | - | 61686791 | 61686895 | 61683496 | 61683546 | 61694696 | 61694770 | 10 | 3 | 15 | 16 | 12 | 76 | 40 | 30 | 49 | 118 | 17 | 6 | 10 | 0 | 21 | 21 | 43 | 27 | 12 | 9 | 253 | 149 | 0.043460148 | 0.338683758 | 0.072 | 0.042 | 0.227 | 0.161 | 0.057 | 0.323 | 0.076 | 0.179 | 0 | 0.579 | -0.12 |
| SE | 65655 | ENSG00000143924 | EML4 | chr2 | + | 42317426 | 42317524 | 42304483 | 42304551 | 42325466 | 42325554 | 95 | 62 | 122 | 104 | 92 | 3 | 1 | 8 | 0 | 0 | 22 | 57 | 94 | 94 | 8 | 0 | 0 | 0 | 0 | 0 | 247 | 149 | 0.043478787 | 0.338761581 | 0.95 | 0.974 | 0.902 | | 1 | 1 | 1 | 1 | 0 | 1 | -0.035 |
| SE | 108805 | ENSG00000105983 | LMBR1 | chr7 | - | 156834477 | 156834595 | 156833752 | 156833792 | 156836812 | 156836885 | 12 | 6 | 3 | 7 | 21 | 88 | 64 | 60 | 63 | 49 | 0 | 5 | 11 | 3 | 8 | 36 | 75 | 74 | 82 | 28 | 267 | 149 | 0.043659978 | 0.339782063 | 0.071 | 0.05 | 0.027 | 0.058 | 0.193 | 0 | 0.036 | 0.077 | 0.02 | 0.138 | 0.026 |
| SE | 77125 | ENSG00000164654 | MIOS | chr7 | + | 7589404 | 7589563 | 7588497 | 7588563 | 7596256 | 7596461 | 98 | 76 | 89 | 95 | 122 | 2 | 0 | 6 | 1 | 5 | 52 | 74 | 40 | 44 | 5 | 0 | 0 | 5 | 0 | 0 | 298 | 149 | 0.043863065 | 0.341009092 | 0.961 | 1 | 0.881 | 0.979 | 0.924 | 1 | 1 | 0.8 | 1 | 1 | -0.011 |
| SE | 11029 | ENSG00000072849 | DERL2 | chr17 | - | 5481295 | 5481389 | 5480386 | 5480582 | 5482808 | 5482879 | 193 | 120 | 170 | 122 | 250 | 13 | 9 | 1 | 7 | 26 | 110 | 131 | 106 | 156 | 123 | 0 | 14 | 7 | 2 | 3 | 243 | 149 | 0.044970382 | 0.347959387 | 0.901 | 0.891 | 0.99 | 0.914 | 0.855 | 1 | 0.852 | 0.903 | 0.98 | 0.962 | -0.029 |
| SE | 108790 | ENSG00000105983 | LMBR1 | chr7 | - | 156680442 | 156680574 | 156679076 | 156679306 | 156681828 | 156682093 | 40 | 23 | 41 | 27 | 40 | 0 | 0 | 0 | 0 | 9 | 11 | 32 | 49 | 9 | 6 | 0 | 0 | 0 | 0 | 0 | 281 | 149 | 0.045531178 | 0.351395675 | 1 | 1 | 1 | 1 | 0.702 | 1 | 1 | 1 | 1 | 1 | -0.06 |
| SE | 7694 | ENSG00000170004 | CHD3 | chr17 | + | 7905620 | 7905706 | 7905099 | 7905165 | 7905855 | 7905989 | 303 | 216 | 444 | 419 | 397 | 0 | 0 | 0 | 4 | 0 | 397 | 307 | 197 | 354 | 603 | 5 | 4 | 0 | 3 | 13 | 235 | 149 | 0.045586264 | 0.351716802 | 1 | 1 | 1 | 0.985 | 1 | 0.981 | 0.98 | 1 | 0.987 | 0.967 | 0.014 |
| SE | 89477 | ENSG00000064313 | TAF2 | chr8 | - | 119778018 | 119778129 | 119762414 | 119762608 | 119781052 | 119781193 | 125 | 93 | 88 | 104 | 124 | 4 | 0 | 2 | 1 | 5 | 21 | 73 | 48 | 12 | 19 | 0 | 0 | 2 | 0 | 0 | 260 | 149 | 0.045766975 | 0.35272872 | 0.947 | 1 | 0.962 | 0.983 | 0.934 | 1 | 1 | 0.932 | 1 | 1 | -0.021 |
| SE | 47548 | ENSG00000144357 | UBR3 | chr2 | + | 170013840 | 170013972 | 170008839 | 170008940 | 170015280 | 170015366 | 0 | 2 | 1 | 6 | 5 | 108 | 108 | 75 | 108 | 239 | 0 | 9 | 4 | 0 | 0 | 6 | 79 | 65 | 18 | 4 | 281 | 149 | 0.045999589 | 0.353978487 | 0 | 0.01 | 0.007 | 0.029 | 0.011 | 0 | 0.057 | 0.032 | 0 | 0 | -0.006 |
| SE | 86864 | ENSG00000090316 | MAEA | chr4 | + | 1322380 | 1322503 | 1315396 | 1315600 | 1339073 | 1339177 | 58 | 19 | 43 | 61 | 22 | 1 | 0 | 0 | 0 | 0 | 35 | 31 | 39 | 43 | 35 | 0 | 1 | 0 | 0 | 8 | 272 | 149 | 0.046084633 | 0.354500733 | 0.969 | 1 | 1 | 1 | 1 | 1 | 0.944 | 1 | 1 | 0.706 | 0.064 |
| SE | 1588 | ENSG00000198089 | SFI1 | chr22 | + | 31602211 | 31602293 | 31589446 | 31589577 | 31604868 | 31604945 | 16 | 25 | 36 | 42 | 28 | 0 | 0 | 0 | 0 | 0 | 0 | 65 | 19 | 23 | 32 | 8 | 0 | 0 | 0 | 0 | 231 | 149 | 0.047872211 | 0.365714482 | 1 | 1 | 1 | 1 | 1 | 0 | 1 | 1 | 1 | 1 | 0.2 |
| SE | 55650 | ENSG00000163463 | KRTCAP2 | chr1 | - | 155172350 | 155172489 | 155169790 | 155169857 | 155172564 | 155172628 | 6 | 4 | 4 | 1 | 1 | 162 | 54 | 121 | 141 | 86 | 0 | 6 | 5 | 3 | 17 | 97 | 162 | 87 | 122 | 173 | 288 | 149 | 0.049406046 | 0.374798547 | 0.019 | 0.037 | 0.017 | 0.004 | 0.006 | 0 | 0.019 | 0.029 | 0.013 | 0.048 | -0.005 |
| SE | 108804 | ENSG00000105983 | LMBR1 | chr7 | - | 156834477 | 156834575 | 156833752 | 156833792 | 156836812 | 156836885 | 13 | 11 | 5 | 13 | 16 | 88 | 64 | 60 | 63 | 49 | 1 | 10 | 12 | 3 | 8 | 36 | 75 | 74 | 82 | 28 | 247 | 149 | 0.049882253 | 0.377739805 | 0.082 | 0.094 | 0.048 | 0.111 | 0.165 | 0.016 | 0.074 | 0.089 | 0.022 | 0.147 | 0.03 |
| SE | 22805 | ENSG00000131504 | DIAPH1 | chr5 | - | 141588223 | 141588250 | 141587041 | 141587197 | 141618797 | 141618999 | 38 | 18 | 29 | 39 | 34 | 140 | 78 | 99 | 120 | 150 | 49 | 35 | 7 | 19 | 43 | 140 | 111 | 81 | 119 | 350 | 176 | 149 | 0.050206388 | 0.379560589 | 0.187 | 0.163 | 0.199 | 0.216 | 0.161 | 0.229 | 0.211 | 0.068 | 0.119 | 0.094 | 0.041 |
| SE | 99087 | ENSG00000110274 | CEP164 | chr11 | + | 117335604 | 117335680 | 117327928 | 117328208 | 117344165 | 117344277 | 11 | 28 | 11 | 10 | 3 | 2 | 0 | 3 | 1 | 1 | 7 | 8 | 8 | 13 | 10 | 0 | 0 | 0 | 1 | 0 | 225 | 149 | 0.051094512 | 0.384589112 | 0.785 | 1 | 0.708 | 0.869 | 0.665 | 1 | 1 | 1 | 0.896 | 1 | -0.174 |
| SE | 107492 | ENSG00000123349 | PFDN5 | chr12 | + | 53295838 | 53295941 | 53295290 | 53295639 | 53296453 | 53296551 | 669 | 413 | 531 | 553 | 335 | 4 | 1 | 9 | 16 | 0 | 701 | 451 | 363 | 357 | 826 | 8 | 3 | 2 | 1 | 4 | 252 | 149 | 0.051174933 | 0.385083376 | 0.99 | 0.996 | 0.972 | 0.953 | 1 | 0.981 | 0.989 | 0.991 | 0.995 | 0.992 | -0.007 |
| SE | 66503 | ENSG00000132676 | DAP3 | chr1 | + | 155725991 | 155726019 | 155725381 | 155725411 | 155727607 | 155727738 | 19 | 11 | 9 | 12 | 23 | 1 | 0 | 3 | 4 | 0 | 22 | 21 | 26 | 12 | 6 | 0 | 0 | 0 | 1 | 0 | 177 | 149 | 0.051963169 | 0.389703855 | 0.941 | 1 | 0.716 | 0.716 | 1 | 1 | 1 | 1 | 0.91 | 1 | -0.107 |
| SE | 63630 | ENSG00000198520 | ARMH1 | chr1 | + | 44724761 | 44724839 | 44724319 | 44724392 | 44725032 | 44725217 | 4 | 3 | 7 | 8 | 1 | 4 | 0 | 1 | 4 | 4 | 21 | 0 | 0 | 13 | 0 | 0 | 1 | 2 | 12 | 10 | 227 | 149 | 0.053365872 | 0.398240025 | 0.396 | 1 | 0.821 | 0.568 | 0.141 | 1 | 0 | 0 | 0.416 | 0 | 0.302 |
| SE | 89246 | ENSG00000137574 | TGS1 | chr8 | + | 55786237 | 55787060 | 55785718 | 55785891 | 55790181 | 55790299 | 153 | 114 | 144 | 200 | 452 | 0 | 1 | 0 | 1 | 9 | 42 | 109 | 182 | 129 | 15 | 4 | 0 | 2 | 0 | 6 | 298 | 149 | 0.053891443 | 0.401289059 | 1 | 0.983 | 1 | 0.99 | 0.962 | 0.84 | 1 | 0.978 | 1 | 0.556 | 0.112 |
| SE | 32329 | ENSG00000068745 | IP6K2 | chr3 | - | 48693269 | 48693338 | 48693058 | 48693179 | 48695089 | 48695421 | 7 | 6 | 0 | 1 | 10 | 142 | 77 | 92 | 123 | 136 | 0 | 0 | 1 | 16 | 0 | 226 | 110 | 161 | 121 | 132 | 218 | 149 | 0.054648067 | 0.405758655 | 0.033 | 0.051 | 0 | 0.006 | 0.048 | 0 | 0 | 0.004 | 0.083 | 0 | 0.01 |
| SE | 93384 | ENSG00000065923 | SLC9A7 | chrX | - | 46682335 | 46682535 | 46679677 | 46679755 | 46758704 | 46759055 | 26 | 26 | 17 | 11 | 20 | 0 | 0 | 0 | 0 | 0 | 21 | 50 | 16 | 21 | 20 | 0 | 3 | 5 | 0 | 0 | 298 | 149 | 0.055230067 | 0.409110056 | 1 | 1 | 1 | 1 | 1 | 1 | 0.893 | 0.615 | 1 | 1 | 0.098 |
| SE | 63643 | ENSG00000112855 | HARS2 | chr5 | + | 140693490 | 140693508 | 140691457 | 140691756 | 140693590 | 140693665 | 0 | 2 | 1 | 0 | 2 | 28 | 14 | 31 | 22 | 10 | 2 | 0 | 8 | 0 | 0 | 5 | 9 | 19 | 19 | 36 | 167 | 149 | 0.055385953 | 0.410148357 | 0 | 0.113 | 0.028 | 0 | 0.151 | 0.263 | 0 | 0.273 | 0 | 0 | -0.049 |
| SE | 46324 | ENSG00000095564 | BTAF1 | chr10 | + | 91953736 | 91953873 | 91951402 | 91951566 | 91956527 | 91956657 | 470 | 486 | 509 | 568 | 448 | 7 | 3 | 1 | 1 | 1 | 221 | 413 | 403 | 202 | 94 | 0 | 8 | 10 | 0 | 0 | 286 | 149 | 0.056283911 | 0.41557951 | 0.972 | 0.988 | 0.996 | 0.997 | 0.996 | 1 | 0.964 | 0.955 | 1 | 1 | 0.006 |
| SE | 59089 | ENSG00000180182 | MED14 | chrX | - | 40666719 | 40666851 | 40664306 | 40664489 | 40671860 | 40671972 | 54 | 46 | 43 | 64 | 69 | 3 | 4 | 1 | 3 | 11 | 32 | 85 | 54 | 46 | 21 | 0 | 10 | 1 | 0 | 0 | 281 | 149 | 0.056539559 | 0.41703451 | 0.905 | 0.859 | 0.958 | 0.919 | 0.769 | 1 | 0.818 | 0.966 | 1 | 1 | -0.075 |
| SE | 30368 | ENSG00000145348 | TBCK | chr4 | - | 106164391 | 106164461 | 106116318 | 106116378 | 106171094 | 106171240 | 14 | 4 | 5 | 5 | 12 | 84 | 53 | 48 | 57 | 90 | 0 | 7 | 8 | 0 | 0 | 17 | 66 | 24 | 54 | 32 | 219 | 149 | 0.056850972 | 0.418857977 | 0.102 | 0.049 | 0.066 | 0.056 | 0.083 | 0 | 0.067 | 0.185 | 0 | 0 | 0.021 |
| SE | 19506 | ENSG00000100479 | POLE2 | chr14 | - | 49654783 | 49654838 | 49654154 | 49654214 | 49655004 | 49655094 | 5 | 4 | 8 | 11 | 5 | 4 | 2 | 1 | 0 | 18 | 26 | 8 | 6 | 12 | 5 | 4 | 0 | 0 | 7 | 0 | 204 | 149 | 0.057008595 | 0.419861249 | 0.477 | 0.594 | 0.854 | 1 | 0.169 | 0.826 | 1 | 1 | 0.556 | 1 | -0.258 |
| SE | 63651 | ENSG00000112855 | HARS2 | chr5 | + | 140696102 | 140696201 | 140695737 | 140695845 | 140696520 | 140696614 | 81 | 44 | 88 | 83 | 86 | 1 | 1 | 3 | 2 | 11 | 67 | 55 | 41 | 33 | 35 | 0 | 0 | 2 | 5 | 0 | 248 | 149 | 0.057256149 | 0.42132777 | 0.98 | 0.964 | 0.946 | 0.961 | 0.824 | 1 | 1 | 0.925 | 0.799 | 1 | -0.01 |
| SE | 82050 | ENSG00000104133 | SPG11 | chr15 | - | 44620124 | 44620403 | 44615362 | 44615566 | 44621758 | 44621934 | 310 | 279 | 245 | 272 | 452 | 7 | 2 | 2 | 2 | 7 | 63 | 319 | 282 | 219 | 49 | 0 | 1 | 3 | 3 | 0 | 298 | 149 | 0.057331793 | 0.421725861 | 0.957 | 0.986 | 0.984 | 0.986 | 0.97 | 1 | 0.994 | 0.979 | 0.973 | 1 | -0.013 |
| SE | 65648 | ENSG00000143924 | EML4 | chr2 | + | 42264918 | 42264951 | 42264705 | 42264731 | 42280849 | 42280973 | 5 | 6 | 9 | 6 | 1 | 198 | 176 | 198 | 168 | 189 | 1 | 1 | 13 | 9 | 7 | 56 | 126 | 184 | 148 | 93 | 182 | 149 | 0.057726667 | 0.423921882 | 0.02 | 0.027 | 0.036 | 0.028 | 0.004 | 0.014 | 0.006 | 0.055 | 0.047 | 0.058 | -0.013 |
| SE | 79040 | ENSG00000160218 | TRAPPC10 | chr21 | + | 44102777 | 44102972 | 44096579 | 44098534 | 44103292 | 44104733 | 67 | 21 | 48 | 52 | 59 | 0 | 0 | 0 | 1 | 0 | 23 | 75 | 47 | 40 | 11 | 0 | 0 | 0 | 0 | 8 | 298 | 149 | 0.057789844 | 0.424099129 | 1 | 1 | 1 | 0.963 | 1 | 1 | 1 | 1 | 1 | 0.407 | 0.111 |
| SE | 31566 | ENSG00000163913 | IFT122 | chr3 | + | 129495445 | 129495607 | 129492140 | 129492194 | 129499901 | 129500068 | 21 | 9 | 22 | 16 | 19 | 0 | 0 | 0 | 0 | 8 | 65 | 32 | 19 | 27 | 34 | 0 | 0 | 0 | 0 | 0 | 298 | 149 | 0.058130408 | 0.426238849 | 1 | 1 | 1 | 1 | 0.543 | 1 | 1 | 1 | 1 | 1 | -0.091 |
| SE | 95853 | ENSG00000107263 | RAPGEF1 | chr9 | - | 131603960 | 131604053 | 131602060 | 131602149 | 131621795 | 131621838 | 21 | 1 | 0 | 0 | 9 | 179 | 120 | 187 | 160 | 102 | 3 | 5 | 19 | 2 | 0 | 326 | 300 | 194 | 135 | 277 | 242 | 149 | 0.058284594 | 0.427209376 | 0.067 | 0.005 | 0 | 0 | 0.052 | 0.006 | 0.01 | 0.057 | 0.009 | 0 | 0.008 |
| SE | 30285 | ENSG00000114480 | GBE1 | chr3 | - | 81646391 | 81646504 | 81642570 | 81642990 | 81648855 | 81648991 | 25 | 17 | 19 | 15 | 21 | 2 | 0 | 1 | 5 | 0 | 17 | 4 | 10 | 8 | 18 | 0 | 2 | 1 | 0 | 0 | 262 | 149 | 0.058919016 | 0.430810913 | 0.877 | 1 | 0.915 | 0.63 | 1 | 1 | 0.532 | 0.85 | 1 | 1 | 0.008 |
| SE | 86242 | ENSG00000134516 | DOCK2 | chr5 | + | 169669287 | 169669328 | 169654402 | 169654486 | 169671077 | 169671174 | 61 | 20 | 42 | 28 | 6 | 0 | 0 | 0 | 1 | 7 | 14 | 63 | 21 | 61 | 45 | 0 | 0 | 6 | 9 | 4 | 190 | 149 | 0.059177301 | 0.432376446 | 1 | 1 | 1 | 0.956 | 0.402 | 1 | 1 | 0.733 | 0.842 | 0.898 | -0.023 |
| SE | 73310 | ENSG00000136699 | SMPD4 | chr2 | - | 130173278 | 130173354 | 130172786 | 130172895 | 130174913 | 130175000 | 30 | 11 | 26 | 21 | 65 | 0 | 0 | 0 | 0 | 0 | 43 | 40 | 39 | 20 | 56 | 0 | 0 | 1 | 2 | 6 | 225 | 149 | 0.059380688 | 0.433660141 | 1 | 1 | 1 | 1 | 1 | 1 | 1 | 0.963 | 0.869 | 0.861 | 0.061 |
| SE | 65654 | ENSG00000143924 | EML4 | chr2 | + | 42315961 | 42316050 | 42304483 | 42304551 | 42325466 | 42325554 | 80 | 71 | 104 | 103 | 131 | 3 | 1 | 8 | 0 | 0 | 27 | 40 | 84 | 53 | 12 | 0 | 0 | 0 | 0 | 0 | 238 | 149 | 0.059488933 | 0.434248132 | 0.943 | 0.978 | 0.891 | 1 | 1 | 1 | 1 | 1 | 1 | 1 | -0.038 |
| SE | 1603 | ENSG00000198089 | SFI1 | chr22 | + | 31604868 | 31604945 | 31589446 | 31589577 | 31606327 | 31606430 | 8 | 8 | 11 | 8 | 3 | 0 | 4 | 3 | 0 | 0 | 22 | 8 | 4 | 2 | 11 | 0 | 0 | 0 | 0 | 0 | 226 | 149 | 0.059861728 | 0.436409832 | 1 | 0.569 | 0.707 | 1 | 1 | 1 | 1 | 1 | 1 | 1 | -0.145 |
| SE | 7390 | ENSG00000101557 | USP14 | chr18 | + | 178932 | 179037 | 166786 | 166819 | 180235 | 180339 | 154 | 74 | 117 | 103 | 143 | 6 | 3 | 5 | 5 | 0 | 58 | 36 | 135 | 94 | 62 | 13 | 0 | 4 | 8 | 0 | 254 | 149 | 0.060216641 | 0.438578799 | 0.938 | 0.935 | 0.932 | 0.924 | 1 | 0.724 | 1 | 0.952 | 0.873 | 1 | 0.036 |
| SE | 58304 | ENSG00000117262 | GPR89A | chr1 | + | 145646203 | 145646272 | 145644085 | 145645743 | 145647174 | 145647267 | 5 | 2 | 4 | 1 | 3 | 2 | 0 | 0 | 2 | 0 | 14 | 0 | 6 | 0 | 16 | 0 | 7 | 0 | 2 | 0 | 218 | 149 | 0.06070082 | 0.441489214 | 0.631 | 1 | 1 | 0.255 | 1 | 1 | 0 | 1 | 0 | 1 | 0.177 |
| SE | 100255 | ENSG00000166471 | TMEM41B | chr11 | - | 9286454 | 9286593 | 9280653 | 9283593 | 9287701 | 9287806 | 125 | 100 | 181 | 131 | 214 | 0 | 1 | 2 | 4 | 4 | 87 | 121 | 139 | 93 | 4 | 8 | 1 | 0 | 6 | 0 | 288 | 149 | 0.060719622 | 0.441584948 | 1 | 0.981 | 0.979 | 0.944 | 0.965 | 0.849 | 0.984 | 1 | 0.889 | 1 | 0.029 |
| SE | 19132 | ENSG00000090061 | CCNK | chr14 | + | 99491945 | 99492320 | 99481424 | 99481479 | 99492625 | 99492874 | 9 | 11 | 6 | 6 | 11 | 75 | 40 | 85 | 74 | 49 | 7 | 31 | 28 | 15 | 10 | 124 | 95 | 82 | 121 | 56 | 298 | 149 | 0.060960595 | 0.443049356 | 0.057 | 0.121 | 0.034 | 0.039 | 0.101 | 0.027 | 0.14 | 0.146 | 0.058 | 0.082 | -0.02 |
| SE | 30541 | ENSG00000132182 | NUP210 | chr3 | - | 13371833 | 13372032 | 13365945 | 13366091 | 13375503 | 13375641 | 277 | 109 | 204 | 216 | 251 | 0 | 0 | 0 | 0 | 0 | 179 | 261 | 93 | 123 | 212 | 0 | 3 | 0 | 4 | 7 | 298 | 149 | 0.061282877 | 0.444566303 | 1 | 1 | 1 | 1 | 1 | 1 | 0.978 | 1 | 0.939 | 0.938 | 0.029 |
| SE | 77119 | ENSG00000164654 | MIOS | chr7 | + | 7583117 | 7583372 | 7574097 | 7574196 | 7585635 | 7585805 | 196 | 134 | 175 | 190 | 282 | 0 | 0 | 1 | 1 | 2 | 88 | 131 | 138 | 134 | 8 | 3 | 0 | 1 | 7 | 0 | 298 | 149 | 0.062587231 | 0.451352107 | 1 | 1 | 0.989 | 0.99 | 0.986 | 0.936 | 1 | 0.986 | 0.905 | 1 | 0.028 |
| SE | 49544 | ENSG00000120158 | RCL1 | chr9 | + | 4826857 | 4827033 | 4823547 | 4823619 | 4833153 | 4833228 | 125 | 44 | 152 | 126 | 116 | 0 | 0 | 0 | 5 | 10 | 251 | 212 | 120 | 185 | 122 | 9 | 0 | 0 | 2 | 0 | 298 | 149 | 0.062672764 | 0.451760881 | 1 | 1 | 1 | 0.926 | 0.853 | 0.933 | 1 | 1 | 0.979 | 1 | -0.027 |
| SE | 31037 | ENSG00000172795 | DCP2 | chr5 | + | 113007937 | 113008042 | 113003941 | 113004077 | 113010755 | 113010807 | 406 | 300 | 322 | 429 | 412 | 5 | 0 | 2 | 16 | 7 | 125 | 277 | 209 | 165 | 114 | 7 | 0 | 8 | 7 | 4 | 254 | 149 | 0.062832903 | 0.452665159 | 0.979 | 1 | 0.99 | 0.94 | 0.972 | 0.913 | 1 | 0.939 | 0.933 | 0.944 | 0.03 |
| SE | 88034 | ENSG00000156171 | DRAM2 | chr1 | - | 111120515 | 111120693 | 111119876 | 111119959 | 111126226 | 111126294 | 292 | 195 | 264 | 343 | 520 | 0 | 0 | 3 | 3 | 0 | 217 | 233 | 182 | 163 | 82 | 0 | 3 | 2 | 4 | 5 | 298 | 149 | 0.063006173 | 0.453537865 | 1 | 1 | 0.978 | 0.983 | 1 | 1 | 0.975 | 0.978 | 0.953 | 0.891 | 0.033 |
| SE | 84867 | ENSG00000164074 | ABHD18 | chr4 | + | 128017357 | 128017501 | 128011674 | 128011700 | 128020079 | 128020169 | 84 | 50 | 83 | 106 | 48 | 2 | 2 | 1 | 3 | 1 | 115 | 98 | 90 | 77 | 96 | 0 | 0 | 4 | 0 | 0 | 293 | 149 | 0.063947148 | 0.458582698 | 0.955 | 0.927 | 0.977 | 0.947 | 0.961 | 1 | 1 | 0.92 | 1 | 1 | -0.031 |
| SE | 45883 | ENSG00000145740 | SLC30A5 | chr5 | + | 69117238 | 69117396 | 69115925 | 69116214 | 69118498 | 69118628 | 75 | 35 | 80 | 81 | 131 | 0 | 0 | 0 | 0 | 1 | 36 | 35 | 8 | 35 | 16 | 0 | 3 | 5 | 0 | 0 | 298 | 149 | 0.064007568 | 0.458862588 | 1 | 0.814 | 1 | 1 | 0.985 | 1 | 0.854 | 0.444 | 1 | 1 | 0.1 |
| SE | 69320 | ENSG00000112367 | FIG4 | chr6 | + | 109763936 | 109763982 | 109762090 | 109762207 | 109765012 | 109765161 | 115 | 69 | 46 | 97 | 137 | 0 | 4 | 0 | 0 | 0 | 81 | 97 | 36 | 70 | 37 | 8 | 3 | 0 | 0 | 0 | 195 | 149 | 0.064275395 | 0.460241679 | 1 | 1 | 1 | 1 | 1 | 0.886 | 0.961 | 1 | 1 | 1 | 0.031 |
| SE | 96901 | ENSG00000100731 | PCNX1 | chr14 | + | 71011491 | 71011549 | 71009633 | 71009724 | 71012999 | 71013202 | 79 | 38 | 72 | 63 | 121 | 0 | 1 | 0 | 2 | 3 | 7 | 92 | 71 | 24 | 15 | 14 | 0 | 0 | 0 | 0 | 207 | 149 | 0.064323503 | 0.460396102 | 1 | 0.965 | 1 | 0.958 | 0.967 | 0.265 | 1 | 1 | 1 | 1 | 0.125 |
| SE | 97614 | ENSG00000118363 | SPCS2 | chr11 | + | 74969610 | 74969699 | 74965033 | 74965117 | 74976856 | 74977156 | 212 | 127 | 259 | 195 | 284 | 0 | 0 | 0 | 2 | 0 | 206 | 166 | 230 | 149 | 48 | 0 | 14 | 1 | 0 | 0 | 238 | 149 | 0.064879095 | 0.46352529 | 1 | 1 | 1 | 0.984 | 1 | 1 | 0.881 | 0.993 | 1 | 1 | 0.022 |
| SE | 1560 | ENSG00000198089 | SFI1 | chr22 | + | 31528689 | 31528863 | 31508254 | 31508376 | 31556941 | 31557059 | 105 | 96 | 101 | 90 | 100 | 0 | 0 | 0 | 0 | 0 | 52 | 75 | 86 | 97 | 213 | 11 | 0 | 0 | 0 | 0 | 298 | 149 | 0.064954307 | 0.463851006 | 1 | 1 | 1 | 1 | 1 | 0.703 | 1 | 1 | 1 | 1 | 0.059 |
| SE | 103095 | ENSG00000108055 | SMC3 | chr10 | + | 110582561 | 110582642 | 110581922 | 110582098 | 110583383 | 110583548 | 366 | 424 | 369 | 465 | 428 | 4 | 6 | 4 | 7 | 1 | 34 | 241 | 266 | 118 | 18 | 0 | 6 | 9 | 2 | 0 | 230 | 149 | 0.06622695 | 0.470985986 | 0.983 | 0.979 | 0.984 | 0.977 | 0.996 | 1 | 0.963 | 0.95 | 0.975 | 1 | 0.006 |
| SE | 32349 | ENSG00000068745 | IP6K2 | chr3 | - | 48715314 | 48715527 | 48693520 | 48695421 | 48717156 | 48717188 | 4 | 9 | 8 | 8 | 3 | 33 | 7 | 7 | 17 | 18 | 11 | 7 | 1 | 6 | 13 | 32 | 4 | 1 | 1 | 12 | 298 | 149 | 0.066924268 | 0.475016092 | 0.057 | 0.391 | 0.364 | 0.19 | 0.077 | 0.147 | 0.467 | 0.333 | 0.75 | 0.351 | -0.194 |
| SE | 59869 | ENSG00000103168 | TAF1C | chr16 | - | 84184850 | 84184989 | 84183696 | 84183778 | 84186900 | 84187045 | 42 | 10 | 30 | 7 | 14 | 3 | 1 | 4 | 5 | 0 | 19 | 47 | 13 | 25 | 16 | 0 | 3 | 1 | 4 | 0 | 288 | 149 | 0.067494011 | 0.478322869 | 0.879 | 0.838 | 0.795 | 0.42 | 1 | 1 | 0.89 | 0.871 | 0.764 | 1 | -0.119 |
| SE | 96896 | ENSG00000100731 | PCNX1 | chr14 | + | 71009633 | 71009724 | 70988566 | 70988699 | 71011491 | 71011549 | 140 | 83 | 121 | 121 | 150 | 10 | 1 | 0 | 2 | 0 | 27 | 140 | 109 | 42 | 43 | 0 | 0 | 2 | 0 | 0 | 240 | 149 | 0.067530714 | 0.478496363 | 0.897 | 0.981 | 1 | 0.974 | 1 | 1 | 1 | 0.971 | 1 | 1 | -0.024 |
| SE | 49685 | ENSG00000256269 | HMBS | chr11 | + | 119092403 | 119092523 | 119092124 | 119092163 | 119092757 | 119092811 | 23 | 4 | 13 | 17 | 10 | 2 | 0 | 2 | 0 | 1 | 15 | 25 | 6 | 13 | 10 | 0 | 22 | 0 | 1 | 0 | 269 | 149 | 0.068345994 | 0.482692076 | 0.864 | 1 | 0.783 | 1 | 0.847 | 1 | 0.386 | 1 | 0.878 | 1 | 0.046 |
| SE | 51125 | ENSG00000233355 | CHRM3-AS2 | chr1 | - | 239707614 | 239707760 | 239706424 | 239706496 | 239714176 | 239714264 | 38 | 57 | 109 | 45 | 85 | 1 | 2 | 8 | 8 | 12 | 8 | 17 | 19 | 6 | 3 | 0 | 0 | 0 | 0 | 15 | 295 | 149 | 0.068798081 | 0.485193165 | 0.95 | 0.935 | 0.873 | 0.74 | 0.782 | 1 | 1 | 1 | 1 | 0.092 | 0.038 |
| SE | 47766 | ENSG00000154358 | OBSCN | chr1 | + | 228219323 | 228219599 | 228217012 | 228217288 | 228243182 | 228243458 | 15 | 3 | 32 | 11 | 6 | 2 | 4 | 2 | 0 | 0 | 13 | 14 | 0 | 4 | 2 | 21 | 5 | 2 | 0 | 0 | 298 | 149 | 0.069101227 | 0.486972335 | 0.789 | 0.273 | 0.889 | 1 | 1 | 0.236 | 0.583 | 0 | 1 | 1 | 0.226 |
| SE | 99154 | ENSG00000103544 | VPS35L | chr16 | + | 19616113 | 19616191 | 19610321 | 19610415 | 19626176 | 19626202 | 71 | 45 | 38 | 62 | 64 | 3 | 0 | 1 | 5 | 2 | 44 | 83 | 16 | 43 | 40 | 0 | 0 | 4 | 1 | 0 | 227 | 149 | 0.069112555 | 0.486972335 | 0.94 | 1 | 0.961 | 0.891 | 0.955 | 1 | 1 | 0.724 | 0.966 | 1 | 0.011 |
| SE | 103937 | ENSG00000171862 | PTEN | chr10 | + | 87952117 | 87952259 | 87933012 | 87933251 | 87957852 | 87958019 | 1686 | 1372 | 1241 | 1758 | 3367 | 35 | 20 | 22 | 30 | 42 | 617 | 1549 | 2112 | 1069 | 260 | 29 | 21 | 49 | 19 | 0 | 291 | 149 | 0.06961336 | 0.490012389 | 0.961 | 0.972 | 0.967 | 0.968 | 0.976 | 0.916 | 0.974 | 0.957 | 0.966 | 1 | 0.006 |
| SE | 34620 | ENSG00000198700 | IPO9 | chr1 | + | 201848392 | 201848594 | 201847551 | 201847638 | 201852103 | 201852192 | 141 | 69 | 95 | 107 | 99 | 1 | 0 | 0 | 0 | 0 | 154 | 152 | 92 | 99 | 136 | 0 | 4 | 5 | 3 | 0 | 298 | 149 | 0.069694798 | 0.490280673 | 0.986 | 1 | 1 | 1 | 1 | 1 | 0.95 | 0.902 | 0.943 | 1 | 0.038 |
| SE | 52941 | ENSG00000144744 | UBA3 | chr3 | - | 69057255 | 69057309 | 69056778 | 69056815 | 69061813 | 69061927 | 262 | 253 | 192 | 236 | 475 | 2 | 0 | 4 | 9 | 6 | 5 | 166 | 87 | 68 | 11 | 0 | 7 | 7 | 0 | 0 | 203 | 149 | 0.070361849 | 0.493908136 | 0.99 | 1 | 0.972 | 0.951 | 0.983 | 1 | 0.946 | 0.901 | 1 | 1 | 0.01 |
| SE | 45879 | ENSG00000145740 | SLC30A5 | chr5 | + | 69100806 | 69100929 | 69093948 | 69094338 | 69103061 | 69103128 | 101 | 64 | 107 | 88 | 86 | 24 | 10 | 13 | 21 | 16 | 43 | 56 | 28 | 88 | 75 | 20 | 6 | 31 | 14 | 26 | 272 | 149 | 0.070612815 | 0.495485217 | 0.697 | 0.778 | 0.818 | 0.697 | 0.746 | 0.541 | 0.836 | 0.331 | 0.775 | 0.612 | 0.128 |
| SE | 67145 | ENSG00000089060 | SLC8B1 | chr12 | - | 113320330 | 113320498 | 113318963 | 113319071 | 113320580 | 113320686 | 29 | 9 | 26 | 24 | 21 | 0 | 0 | 0 | 0 | 8 | 39 | 3 | 9 | 10 | 20 | 0 | 0 | 0 | 0 | 0 | 298 | 149 | 0.071757343 | 0.50181021 | 1 | 1 | 1 | 1 | 0.568 | 1 | 1 | 1 | 1 | 1 | -0.086 |
| SE | 99099 | ENSG00000110274 | CEP164 | chr11 | + | 117382795 | 117382942 | 117381700 | 117381868 | 117390776 | 117390908 | 4 | 2 | 3 | 5 | 3 | 0 | 1 | 0 | 4 | 0 | 18 | 4 | 8 | 6 | 25 | 10 | 1 | 1 | 0 | 0 | 296 | 149 | 0.071855898 | 0.502234365 | 1 | 0.502 | 1 | 0.386 | 1 | 0.475 | 0.668 | 0.801 | 1 | 1 | -0.011 |
| SE | 51123 | ENSG00000233355 | CHRM3-AS2 | chr1 | - | 239707533 | 239707760 | 239706321 | 239706496 | 239714176 | 239714264 | 47 | 53 | 113 | 42 | 89 | 1 | 2 | 8 | 8 | 12 | 8 | 17 | 19 | 6 | 3 | 0 | 0 | 0 | 0 | 15 | 298 | 149 | 0.073068828 | 0.508848425 | 0.959 | 0.93 | 0.876 | 0.724 | 0.788 | 1 | 1 | 1 | 1 | 0.091 | 0.037 |
| SE | 45659 | ENSG00000163608 | NEPRO | chr3 | - | 113009497 | 113009584 | 113008170 | 113008430 | 113010642 | 113010704 | 3 | 0 | 0 | 1 | 0 | 120 | 101 | 104 | 74 | 223 | 9 | 0 | 0 | 0 | 0 | 3 | 87 | 121 | 35 | 9 | 236 | 149 | 0.073838171 | 0.5130589 | 0.016 | 0 | 0 | 0.008 | 0 | 0.654 | 0 | 0 | 0 | 0 | -0.126 |
| SE | 6743 | ENSG00000150477 | KIAA1328 | chr18 | + | 36844207 | 36844302 | 36835233 | 36835376 | 36885556 | 36885672 | 66 | 102 | 78 | 119 | 94 | 6 | 2 | 5 | 7 | 3 | 39 | 58 | 123 | 71 | 20 | 0 | 0 | 2 | 8 | 0 | 244 | 149 | 0.073882597 | 0.513284104 | 0.87 | 0.969 | 0.905 | 0.912 | 0.95 | 1 | 1 | 0.974 | 0.844 | 1 | -0.042 |
| SE | 79039 | ENSG00000160218 | TRAPPC10 | chr21 | + | 44098356 | 44098534 | 44094062 | 44094233 | 44102777 | 44102972 | 230 | 91 | 195 | 218 | 254 | 0 | 1 | 0 | 0 | 0 | 76 | 217 | 160 | 115 | 82 | 0 | 0 | 10 | 2 | 0 | 298 | 149 | 0.076229338 | 0.525623269 | 1 | 0.978 | 1 | 1 | 1 | 1 | 1 | 0.889 | 0.966 | 1 | 0.025 |
| SE | 42211 | ENSG00000132155 | RAF1 | chr3 | - | 12611949 | 12612062 | 12609232 | 12609335 | 12618514 | 12618747 | 272 | 151 | 239 | 259 | 301 | 7 | 3 | 5 | 6 | 6 | 202 | 178 | 336 | 241 | 227 | 4 | 9 | 1 | 2 | 0 | 262 | 149 | 0.077540668 | 0.533069112 | 0.957 | 0.966 | 0.965 | 0.961 | 0.966 | 0.966 | 0.918 | 0.995 | 0.986 | 1 | -0.01 |
| SE | 31467 | ENSG00000178950 | GAK | chr4 | - | 877614 | 877809 | 877089 | 877207 | 881906 | 882040 | 187 | 89 | 167 | 173 | 92 | 0 | 2 | 0 | 0 | 0 | 256 | 231 | 160 | 232 | 323 | 0 | 0 | 2 | 13 | 1 | 298 | 149 | 0.078404023 | 0.537588352 | 1 | 0.957 | 1 | 1 | 1 | 1 | 1 | 0.976 | 0.899 | 0.994 | 0.018 |
| SE | 17333 | ENSG00000092529 | CAPN3 | chr15 | + | 42408210 | 42408324 | 42403740 | 42403777 | 42409302 | 42409380 | 22 | 27 | 18 | 19 | 17 | 8 | 0 | 7 | 7 | 2 | 24 | 56 | 19 | 120 | 28 | 6 | 11 | 0 | 9 | 0 | 263 | 149 | 0.078860064 | 0.539461269 | 0.609 | 1 | 0.593 | 0.606 | 0.828 | 0.694 | 0.743 | 1 | 0.883 | 1 | -0.137 |
| SE | 26307 | ENSG00000105875 | WDR91 | chr7 | - | 135188432 | 135188545 | 135186971 | 135187169 | 135189343 | 135189452 | 26 | 17 | 35 | 44 | 16 | 5 | 0 | 2 | 2 | 0 | 25 | 44 | 45 | 28 | 33 | 8 | 0 | 14 | 3 | 0 | 262 | 149 | 0.079843625 | 0.544692558 | 0.747 | 1 | 0.909 | 0.926 | 1 | 0.64 | 1 | 0.646 | 0.841 | 1 | 0.091 |
| SE | 64423 | ENSG00000143919 | CAMKMT | chr2 | + | 44706286 | 44706341 | 44704282 | 44704343 | 44707398 | 44707462 | 23 | 14 | 33 | 28 | 49 | 3 | 0 | 4 | 3 | 0 | 8 | 20 | 14 | 16 | 31 | 0 | 8 | 0 | 10 | 0 | 204 | 149 | 0.080368178 | 0.54743578 | 0.848 | 1 | 0.858 | 0.872 | 1 | 1 | 0.646 | 1 | 0.539 | 1 | 0.079 |
| SE | 84864 | ENSG00000164074 | ABHD18 | chr4 | + | 128001715 | 128001760 | 127984338 | 127984423 | 128008919 | 128008998 | 3 | 2 | 3 | 3 | 3 | 15 | 25 | 19 | 16 | 26 | 3 | 3 | 15 | 0 | 2 | 8 | 15 | 25 | 18 | 8 | 194 | 149 | 0.081641171 | 0.554443327 | 0.133 | 0.058 | 0.108 | 0.126 | 0.081 | 0.224 | 0.133 | 0.315 | 0 | 0.161 | -0.065 |
| SE | 107498 | ENSG00000123349 | PFDN5 | chr12 | + | 53296218 | 53296275 | 53295838 | 53295941 | 53297849 | 53297924 | 626 | 367 | 549 | 516 | 649 | 3 | 13 | 11 | 3 | 11 | 848 | 569 | 331 | 482 | 930 | 34 | 5 | 5 | 4 | 20 | 206 | 149 | 0.083176236 | 0.562137813 | 0.993 | 0.953 | 0.973 | 0.992 | 0.977 | 0.947 | 0.988 | 0.98 | 0.989 | 0.971 | 0.003 |
| SE | 75565 | ENSG00000221838 | AP4M1 | chr7 | + | 100106240 | 100106291 | 100105958 | 100106003 | 100106402 | 100106514 | 9 | 10 | 10 | 12 | 4 | 0 | 0 | 0 | 0 | 0 | 20 | 16 | 4 | 6 | 6 | 0 | 0 | 0 | 0 | 7 | 200 | 149 | 0.083369782 | 0.56315421 | 1 | 1 | 1 | 1 | 1 | 1 | 1 | 1 | 1 | 0.39 | 0.122 |
| SE | 100159 | ENSG00000159063 | ALG8 | chr11 | - | 78138709 | 78138767 | 78127357 | 78127436 | 78139493 | 78139630 | 0 | 1 | 1 | 0 | 4 | 13 | 9 | 3 | 12 | 9 | 5 | 4 | 3 | 0 | 0 | 0 | 5 | 15 | 4 | 9 | 207 | 149 | 0.084958932 | 0.571775714 | 0 | 0.074 | 0.194 | 0 | 0.242 | 1 | 0.365 | 0.126 | 0 | 0 | -0.196 |
| SE | 16953 | ENSG00000172575 | RASGRP1 | chr15 | - | 38498793 | 38499184 | 38494451 | 38494767 | 38500102 | 38500139 | 104 | 51 | 116 | 115 | 146 | 17 | 7 | 21 | 11 | 12 | 0 | 44 | 54 | 26 | 12 | 10 | 7 | 18 | 6 | 0 | 298 | 149 | 0.085030222 | 0.572050717 | 0.754 | 0.785 | 0.734 | 0.839 | 0.859 | 0 | 0.759 | 0.6 | 0.684 | 1 | 0.186 |
| SE | 11027 | ENSG00000072849 | DERL2 | chr17 | - | 5480386 | 5480459 | 5480053 | 5480144 | 5481100 | 5481142 | 38 | 27 | 37 | 35 | 92 | 2 | 7 | 1 | 0 | 3 | 22 | 38 | 42 | 27 | 15 | 0 | 0 | 14 | 16 | 0 | 222 | 149 | 0.085046453 | 0.572070531 | 0.927 | 0.721 | 0.961 | 1 | 0.954 | 1 | 1 | 0.668 | 0.531 | 1 | 0.073 |
| SE | 79677 | ENSG00000186591 | UBE2H | chr7 | - | 129948958 | 129949102 | 129880894 | 129880971 | 129952502 | 129952722 | 0 | 0 | 0 | 0 | 0 | 46 | 28 | 58 | 38 | 63 | 18 | 0 | 0 | 0 | 0 | 37 | 116 | 102 | 68 | 120 | 293 | 149 | 0.08700765 | 0.582850012 | 0 | 0 | 0 | 0 | 0 | 0.198 | 0 | 0 | 0 | 0 | -0.04 |
| SE | 50581 | ENSG00000139636 | LMBR1L | chr12 | - | 49106544 | 49106746 | 49105923 | 49105957 | 49106960 | 49107045 | 5 | 3 | 20 | 3 | 10 | 25 | 33 | 37 | 32 | 66 | 16 | 16 | 2 | 9 | 11 | 26 | 35 | 14 | 17 | 83 | 298 | 149 | 0.088067643 | 0.588236565 | 0.091 | 0.043 | 0.213 | 0.045 | 0.07 | 0.235 | 0.186 | 0.067 | 0.209 | 0.062 | -0.059 |
| SE | 47801 | ENSG00000154358 | OBSCN | chr1 | + | 228336199 | 228336270 | 228335790 | 228335851 | 228337003 | 228337126 | 18 | 8 | 16 | 8 | 23 | 1 | 7 | 7 | 1 | 3 | 0 | 1 | 7 | 10 | 14 | 5 | 0 | 0 | 2 | 0 | 220 | 149 | 0.088805008 | 0.592204831 | 0.924 | 0.436 | 0.608 | 0.844 | 0.839 | 0 | 1 | 1 | 0.772 | 1 | -0.024 |
| SE | 47768 | ENSG00000154358 | OBSCN | chr1 | + | 228224458 | 228224734 | 228217012 | 228217288 | 228243182 | 228243458 | 6 | 6 | 22 | 18 | 6 | 2 | 4 | 2 | 0 | 0 | 21 | 0 | 7 | 9 | 1 | 21 | 5 | 2 | 0 | 0 | 298 | 149 | 0.089446646 | 0.595266858 | 0.6 | 0.429 | 0.846 | 1 | 1 | 0.333 | 1 | 0.636 | 1 | 1 | 0.181 |
| SE | 6088 | ENSG00000105576 | TNPO2 | chr19 | - | 12706009 | 12706109 | 12705681 | 12705768 | 12706195 | 12706282 | 5 | 1 | 1 | 3 | 3 | 13 | 15 | 16 | 33 | 10 | 11 | 0 | 0 | 9 | 11 | 28 | 12 | 8 | 12 | 64 | 249 | 149 | 0.090858059 | 0.603428841 | 0.187 | 0.038 | 0.036 | 0.052 | 0.152 | 0.19 | 0 | 0 | 0.31 | 0.093 | -0.026 |
| SE | 7686 | ENSG00000170004 | CHD3 | chr17 | + | 7895338 | 7895542 | 7894916 | 7895150 | 7897082 | 7897294 | 471 | 334 | 577 | 458 | 523 | 17 | 6 | 15 | 12 | 8 | 438 | 334 | 280 | 487 | 777 | 22 | 9 | 14 | 25 | 6 | 298 | 149 | 0.092162697 | 0.610281677 | 0.933 | 0.965 | 0.951 | 0.95 | 0.97 | 0.909 | 0.949 | 0.909 | 0.907 | 0.985 | 0.022 |
| SE | 58956 | ENSG00000131778 | CHD1L | chr1 | + | 147272446 | 147272630 | 147272170 | 147272281 | 147275353 | 147275468 | 4 | 1 | 0 | 2 | 2 | 49 | 28 | 53 | 41 | 46 | 0 | 8 | 3 | 8 | 0 | 61 | 52 | 23 | 34 | 121 | 298 | 149 | 0.09417598 | 0.620882668 | 0.039 | 0.018 | 0 | 0.024 | 0.021 | 0 | 0.071 | 0.061 | 0.105 | 0 | -0.027 |
| SE | 34039 | ENSG00000159147 | DONSON | chr21 | - | 33582164 | 33582246 | 33581950 | 33582055 | 33584589 | 33584768 | 3 | 4 | 8 | 9 | 10 | 5 | 0 | 0 | 1 | 0 | 12 | 2 | 7 | 15 | 10 | 0 | 0 | 0 | 0 | 0 | 231 | 149 | 0.095044027 | 0.625499467 | 0.279 | 1 | 1 | 0.853 | 1 | 1 | 1 | 1 | 1 | 1 | -0.174 |
| SE | 6594 | ENSG00000134444 | RELCH | chr18 | + | 62264728 | 62264852 | 62263988 | 62264145 | 62266700 | 62266749 | 172 | 132 | 143 | 187 | 173 | 9 | 5 | 9 | 14 | 3 | 71 | 111 | 114 | 76 | 94 | 0 | 6 | 2 | 8 | 0 | 273 | 149 | 0.095096428 | 0.625633975 | 0.913 | 0.935 | 0.897 | 0.879 | 0.969 | 1 | 0.91 | 0.969 | 0.838 | 1 | -0.025 |
| SE | 88399 | ENSG00000145041 | DCAF1 | chr3 | - | 51466802 | 51466876 | 51463113 | 51463227 | 51470928 | 51471005 | 268 | 204 | 268 | 258 | 246 | 3 | 0 | 5 | 3 | 8 | 480 | 324 | 309 | 400 | 271 | 4 | 0 | 1 | 4 | 0 | 223 | 149 | 0.095665455 | 0.628532553 | 0.984 | 1 | 0.973 | 0.983 | 0.954 | 0.988 | 1 | 0.995 | 0.985 | 1 | -0.015 |
| SE | 45884 | ENSG00000145740 | SLC30A5 | chr5 | + | 69117238 | 69117396 | 69116393 | 69116602 | 69118498 | 69118628 | 165 | 88 | 147 | 185 | 229 | 3 | 0 | 5 | 2 | 0 | 52 | 66 | 34 | 61 | 34 | 0 | 1 | 10 | 0 | 0 | 298 | 149 | 0.097322011 | 0.636811192 | 0.965 | 1 | 0.936 | 0.979 | 1 | 1 | 0.971 | 0.63 | 1 | 1 | 0.056 |
| SE | 34626 | ENSG00000198700 | IPO9 | chr1 | + | 201874831 | 201874936 | 201874249 | 201874372 | 201875151 | 201875228 | 122 | 73 | 145 | 103 | 124 | 0 | 0 | 0 | 1 | 1 | 254 | 153 | 72 | 128 | 177 | 14 | 0 | 0 | 0 | 0 | 254 | 149 | 0.098346201 | 0.642211358 | 1 | 1 | 1 | 0.984 | 0.986 | 0.914 | 1 | 1 | 1 | 1 | 0.011 |
| SE | 99102 | ENSG00000110274 | CEP164 | chr11 | + | 117390776 | 117390908 | 117384596 | 117387412 | 117392225 | 117392303 | 23 | 27 | 21 | 35 | 18 | 0 | 0 | 1 | 1 | 0 | 89 | 34 | 61 | 74 | 164 | 0 | 0 | 2 | 0 | 8 | 281 | 149 | 0.098927899 | 0.645148992 | 1 | 1 | 0.918 | 0.949 | 1 | 1 | 1 | 0.942 | 1 | 0.916 | 0.002 |
| SE | 99155 | ENSG00000103544 | VPS35L | chr16 | + | 19616685 | 19616808 | 19616113 | 19616191 | 19626176 | 19626202 | 80 | 75 | 90 | 112 | 155 | 1 | 1 | 1 | 0 | 0 | 183 | 85 | 72 | 155 | 253 | 0 | 0 | 0 | 13 | 0 | 272 | 149 | 0.099248146 | 0.646860308 | 0.978 | 0.976 | 0.98 | 1 | 1 | 1 | 1 | 1 | 0.867 | 1 | 0.013 |
| SE | 24280 | ENSG00000104671 | DCTN6 | chr8 | + | 30165715 | 30165901 | 30164110 | 30164175 | 30175084 | 30175190 | 0 | 1 | 1 | 4 | 1 | 63 | 44 | 53 | 59 | 70 | 0 | 0 | 8 | 7 | 0 | 16 | 76 | 115 | 62 | 4 | 298 | 149 | 0.099758859 | 0.649540096 | 0 | 0.011 | 0.009 | 0.033 | 0.007 | 0 | 0 | 0.034 | 0.053 | 0 | -0.005 |
| SE | 21428 | ENSG00000164180 | TMEM161B | chr5 | - | 88221673 | 88221760 | 88220562 | 88220719 | 88225768 | 88225866 | 29 | 6 | 21 | 15 | 25 | 33 | 30 | 51 | 49 | 70 | 8 | 18 | 3 | 2 | 12 | 2 | 18 | 11 | 11 | 4 | 236 | 149 | 0.103901932 | 0.671157965 | 0.357 | 0.112 | 0.206 | 0.162 | 0.184 | 0.716 | 0.387 | 0.147 | 0.103 | 0.654 | -0.197 |
| SE | 107494 | ENSG00000123349 | PFDN5 | chr12 | + | 53295838 | 53295941 | 53295290 | 53295639 | 53298044 | 53298150 | 669 | 407 | 530 | 553 | 332 | 21 | 6 | 15 | 20 | 7 | 701 | 454 | 363 | 357 | 826 | 14 | 11 | 12 | 8 | 9 | 252 | 149 | 0.10473753 | 0.674718585 | 0.95 | 0.976 | 0.954 | 0.942 | 0.966 | 0.967 | 0.961 | 0.947 | 0.963 | 0.982 | -0.006 |
| SE | 34623 | ENSG00000198700 | IPO9 | chr1 | + | 201863447 | 201863607 | 201858854 | 201858994 | 201866732 | 201866959 | 92 | 52 | 83 | 79 | 68 | 0 | 2 | 3 | 3 | 3 | 145 | 80 | 41 | 77 | 128 | 0 | 2 | 5 | 0 | 0 | 298 | 149 | 0.104840076 | 0.675176999 | 1 | 0.929 | 0.933 | 0.929 | 0.919 | 1 | 0.952 | 0.804 | 1 | 1 | -0.009 |
| SE | 102563 | ENSG00000171307 | ZDHHC16 | chr10 | + | 97453798 | 97453846 | 97453529 | 97453663 | 97454713 | 97454799 | 39 | 22 | 24 | 31 | 34 | 1 | 0 | 3 | 4 | 16 | 62 | 28 | 17 | 35 | 50 | 11 | 0 | 0 | 0 | 4 | 197 | 149 | 0.106587194 | 0.684161195 | 0.967 | 1 | 0.858 | 0.854 | 0.616 | 0.81 | 1 | 1 | 1 | 0.904 | -0.084 |
| SE | 77118 | ENSG00000164654 | MIOS | chr7 | + | 7582629 | 7582819 | 7574097 | 7574196 | 7583117 | 7583372 | 17 | 14 | 15 | 24 | 24 | 119 | 95 | 111 | 121 | 218 | 6 | 17 | 32 | 9 | 10 | 50 | 73 | 123 | 97 | 8 | 298 | 149 | 0.107723314 | 0.689753401 | 0.067 | 0.069 | 0.063 | 0.09 | 0.052 | 0.057 | 0.104 | 0.115 | 0.044 | 0.385 | -0.073 |
| SE | 24099 | ENSG00000143889 | HNRNPLL | chr2 | - | 38577919 | 38578002 | 38577460 | 38577532 | 38581912 | 38581985 | 25 | 27 | 18 | 28 | 62 | 62 | 37 | 66 | 88 | 130 | 36 | 41 | 51 | 36 | 36 | 45 | 96 | 55 | 107 | 66 | 232 | 149 | 0.107962652 | 0.691059824 | 0.206 | 0.319 | 0.149 | 0.17 | 0.234 | 0.339 | 0.215 | 0.373 | 0.178 | 0.259 | -0.057 |
| SE | 19131 | ENSG00000090061 | CCNK | chr14 | + | 99491945 | 99492320 | 99481411 | 99481614 | 99492625 | 99492874 | 9 | 9 | 6 | 6 | 11 | 14 | 9 | 9 | 17 | 16 | 7 | 25 | 28 | 16 | 10 | 21 | 22 | 9 | 27 | 3 | 298 | 149 | 0.107997987 | 0.691229492 | 0.243 | 0.333 | 0.25 | 0.15 | 0.256 | 0.143 | 0.362 | 0.609 | 0.229 | 0.625 | -0.147 |
| SE | 39955 | ENSG00000164253 | WDR41 | chr5 | - | 77453801 | 77453928 | 77451323 | 77451353 | 77459061 | 77459124 | 112 | 37 | 46 | 67 | 132 | 0 | 0 | 0 | 0 | 0 | 68 | 37 | 34 | 37 | 27 | 0 | 0 | 2 | 6 | 0 | 276 | 149 | 0.10825636 | 0.692543508 | 1 | 1 | 1 | 1 | 1 | 1 | 1 | 0.902 | 0.769 | 1 | 0.066 |
| SE | 88447 | ENSG00000221914 | PPP2R2A | chr8 | + | 26350598 | 26350681 | 26339902 | 26339973 | 26354467 | 26354637 | 2 | 6 | 0 | 2 | 15 | 12 | 3 | 10 | 1 | 7 | 6 | 0 | 8 | 0 | 0 | 12 | 7 | 2 | 6 | 8 | 232 | 149 | 0.108305758 | 0.692802911 | 0.097 | 0.562 | 0 | 0.562 | 0.579 | 0.243 | 0 | 0.72 | 0 | 0 | 0.167 |
| SE | 51121 | ENSG00000233355 | CHRM3-AS2 | chr1 | - | 239706321 | 239706496 | 239703891 | 239705715 | 239714176 | 239714264 | 45 | 44 | 92 | 74 | 68 | 2 | 0 | 0 | 5 | 3 | 18 | 10 | 20 | 8 | 31 | 0 | 0 | 4 | 0 | 0 | 298 | 149 | 0.109788731 | 0.70005851 | 0.918 | 1 | 1 | 0.881 | 0.919 | 1 | 1 | 0.714 | 1 | 1 | 0.001 |
| SE | 47778 | ENSG00000154358 | OBSCN | chr1 | + | 228271924 | 228272221 | 228264115 | 228264391 | 228273271 | 228273547 | 20 | 21 | 31 | 20 | 19 | 0 | 2 | 2 | 1 | 2 | 4 | 5 | 13 | 11 | 17 | 0 | 0 | 0 | 0 | 0 | 298 | 149 | 0.110863577 | 0.705384717 | 1 | 0.84 | 0.886 | 0.909 | 0.826 | 1 | 1 | 1 | 1 | 1 | -0.108 |
| SE | 100294 | ENSG00000167986 | DDB1 | chr11 | - | 61329957 | 61330074 | 61326778 | 61326893 | 61331542 | 61331691 | 189 | 73 | 143 | 108 | 259 | 2 | 0 | 2 | 0 | 3 | 122 | 133 | 79 | 149 | 152 | 0 | 4 | 1 | 0 | 7 | 266 | 149 | 0.111359737 | 0.707655383 | 0.981 | 1 | 0.976 | 1 | 0.98 | 1 | 0.949 | 0.978 | 1 | 0.924 | 0.017 |
| SE | 82008 | ENSG00000104133 | SPG11 | chr15 | - | 44570524 | 44570658 | 44567423 | 44567592 | 44572682 | 44572820 | 107 | 61 | 75 | 115 | 113 | 2 | 0 | 0 | 2 | 11 | 21 | 99 | 63 | 35 | 14 | 0 | 0 | 1 | 6 | 0 | 283 | 149 | 0.112779339 | 0.714588585 | 0.966 | 1 | 1 | 0.968 | 0.844 | 1 | 1 | 0.971 | 0.754 | 1 | 0.011 |
| SE | 105683 | ENSG00000028203 | VEZT | chr12 | + | 95242163 | 95242264 | 95217779 | 95217886 | 95251939 | 95252071 | 3 | 0 | 0 | 4 | 2 | 49 | 32 | 32 | 52 | 39 | 13 | 5 | 1 | 0 | 0 | 2 | 35 | 39 | 27 | 8 | 250 | 149 | 0.112950315 | 0.715244459 | 0.035 | 0 | 0 | 0.044 | 0.03 | 0.795 | 0.078 | 0.015 | 0 | 0 | -0.156 |
| SE | 29795 | ENSG00000141452 | RMC1 | chr18 | + | 23506969 | 23507054 | 23504370 | 23504447 | 23507984 | 23508041 | 163 | 126 | 149 | 171 | 150 | 1 | 1 | 3 | 3 | 6 | 186 | 201 | 126 | 183 | 208 | 0 | 4 | 0 | 1 | 0 | 234 | 149 | 0.113351492 | 0.716702631 | 0.99 | 0.988 | 0.969 | 0.973 | 0.941 | 1 | 0.97 | 1 | 0.991 | 1 | -0.02 |
| SE | 14219 | ENSG00000167965 | MLST8 | chr16 | + | 2207034 | 2207110 | 2206357 | 2206659 | 2207192 | 2207250 | 33 | 12 | 27 | 18 | 24 | 1 | 0 | 0 | 0 | 0 | 66 | 36 | 17 | 5 | 54 | 0 | 0 | 0 | 6 | 0 | 225 | 149 | 0.113358142 | 0.716702631 | 0.956 | 1 | 1 | 1 | 1 | 1 | 1 | 1 | 0.356 | 1 | 0.12 |
| SE | 49676 | ENSG00000256269 | HMBS | chr11 | + | 119088254 | 119088308 | 119084865 | 119085066 | 119088634 | 119088707 | 19 | 14 | 19 | 26 | 13 | 0 | 0 | 0 | 3 | 0 | 41 | 40 | 13 | 7 | 15 | 3 | 0 | 0 | 5 | 0 | 203 | 149 | 0.114935909 | 0.724303457 | 1 | 1 | 1 | 0.864 | 1 | 0.909 | 1 | 1 | 0.507 | 1 | 0.09 |
| SE | 62918 | ENSG00000185420 | SMYD3 | chr1 | - | 245814341 | 245814415 | 245764040 | 245764149 | 245858495 | 245858670 | 0 | 5 | 2 | 3 | 0 | 32 | 31 | 18 | 26 | 37 | 0 | 0 | 0 | 0 | 0 | 46 | 10 | 10 | 11 | 42 | 223 | 149 | 0.115507363 | 0.726815436 | 0 | 0.097 | 0.069 | 0.072 | 0 | 0 | 0 | 0 | 0 | 0 | 0.048 |
| SE | 93902 | ENSG00000184465 | WDR27 | chr6 | - | 169662303 | 169662424 | 169660662 | 169660766 | 169672254 | 169672396 | 9 | 1 | 16 | 13 | 14 | 0 | 0 | 0 | 0 | 0 | 2 | 1 | 9 | 14 | 0 | 0 | 0 | 0 | 0 | 4 | 270 | 149 | 0.11588384 | 0.728247942 | 1 | 1 | 1 | 1 | 1 | 1 | 1 | 1 | 1 | 0 | 0.2 |
| SE | 107994 | ENSG00000204852 | TCTN1 | chr12 | + | 110641023 | 110641149 | 110640382 | 110640517 | 110644966 | 110645057 | 5 | 12 | 7 | 6 | 16 | 0 | 0 | 0 | 0 | 2 | 6 | 3 | 6 | 7 | 8 | 0 | 0 | 0 | 0 | 7 | 275 | 149 | 0.115941863 | 0.72854635 | 1 | 1 | 1 | 1 | 0.813 | 1 | 1 | 1 | 1 | 0.382 | 0.086 |
| SE | 96403 | ENSG00000197563 | PIGN | chr18 | - | 62162252 | 62162329 | 62161132 | 62161385 | 62163530 | 62163656 | 11 | 15 | 22 | 32 | 16 | 17 | 8 | 4 | 9 | 15 | 9 | 17 | 8 | 6 | 22 | 4 | 5 | 1 | 8 | 0 | 226 | 149 | 0.115949934 | 0.72854635 | 0.299 | 0.553 | 0.784 | 0.701 | 0.413 | 0.597 | 0.692 | 0.841 | 0.331 | 1 | -0.142 |
| SE | 83158 | ENSG00000055130 | CUL1 | chr7 | + | 148767618 | 148767749 | 148766560 | 148766723 | 148783782 | 148783890 | 441 | 326 | 357 | 410 | 495 | 0 | 0 | 0 | 0 | 1 | 283 | 295 | 308 | 414 | 49 | 0 | 14 | 2 | 1 | 0 | 280 | 149 | 0.115962805 | 0.728568753 | 1 | 1 | 1 | 1 | 0.996 | 1 | 0.918 | 0.988 | 0.995 | 1 | 0.019 |
| SE | 54641 | ENSG00000140992 | PDPK1 | chr16 | + | 2597122 | 2597275 | 2586675 | 2586893 | 2597650 | 2597928 | 155 | 96 | 137 | 134 | 195 | 0 | 0 | 0 | 1 | 0 | 192 | 159 | 86 | 136 | 157 | 0 | 3 | 6 | 3 | 0 | 298 | 149 | 0.121151028 | 0.754739434 | 1 | 1 | 1 | 0.985 | 1 | 1 | 0.964 | 0.878 | 0.958 | 1 | 0.037 |
| SE | 51124 | ENSG00000233355 | CHRM3-AS2 | chr1 | - | 239707614 | 239707760 | 239703891 | 239705715 | 239714176 | 239714264 | 38 | 52 | 103 | 41 | 78 | 2 | 0 | 0 | 5 | 3 | 8 | 18 | 19 | 6 | 3 | 0 | 0 | 4 | 0 | 0 | 295 | 149 | 0.12170125 | 0.757329601 | 0.906 | 1 | 1 | 0.806 | 0.929 | 1 | 1 | 0.706 | 1 | 1 | -0.013 |
| SE | 51122 | ENSG00000233355 | CHRM3-AS2 | chr1 | - | 239707533 | 239707760 | 239703891 | 239705715 | 239714176 | 239714264 | 38 | 52 | 103 | 40 | 82 | 2 | 0 | 0 | 5 | 3 | 8 | 17 | 19 | 6 | 3 | 0 | 0 | 4 | 0 | 0 | 298 | 149 | 0.123214509 | 0.764740639 | 0.905 | 1 | 1 | 0.8 | 0.932 | 1 | 1 | 0.704 | 1 | 1 | -0.013 |
| SE | 66494 | ENSG00000132676 | DAP3 | chr1 | + | 155717005 | 155717128 | 155709772 | 155709824 | 155725381 | 155725490 | 235 | 167 | 281 | 239 | 136 | 5 | 1 | 5 | 4 | 0 | 225 | 185 | 142 | 136 | 186 | 0 | 2 | 1 | 2 | 0 | 272 | 149 | 0.123893322 | 0.768101316 | 0.963 | 0.989 | 0.969 | 0.97 | 1 | 1 | 0.981 | 0.987 | 0.974 | 1 | -0.01 |
| SE | 96884 | ENSG00000100731 | PCNX1 | chr14 | + | 70988566 | 70988699 | 70976941 | 70978648 | 70995740 | 70995843 | 54 | 26 | 51 | 52 | 41 | 9 | 7 | 7 | 8 | 10 | 23 | 32 | 35 | 34 | 16 | 14 | 9 | 16 | 12 | 0 | 282 | 149 | 0.128878366 | 0.792487911 | 0.76 | 0.662 | 0.794 | 0.774 | 0.684 | 0.465 | 0.653 | 0.536 | 0.6 | 1 | 0.084 |
| SE | 37933 | ENSG00000148843 | PDCD11 | chr10 | + | 103399646 | 103399830 | 103398415 | 103398528 | 103400396 | 103400528 | 13 | 2 | 15 | 8 | 10 | 42 | 28 | 16 | 43 | 18 | 21 | 0 | 6 | 18 | 4 | 77 | 31 | 11 | 32 | 86 | 298 | 149 | 0.128879831 | 0.792487911 | 0.134 | 0.034 | 0.319 | 0.085 | 0.217 | 0.12 | 0 | 0.214 | 0.22 | 0.023 | 0.042 |
| SE | 31032 | ENSG00000172795 | DCP2 | chr5 | + | 112981041 | 112981171 | 112976812 | 112976986 | 112983973 | 112984651 | 0 | 2 | 4 | 5 | 4 | 9 | 2 | 6 | 13 | 3 | 3 | 10 | 4 | 6 | 0 | 42 | 23 | 20 | 15 | 28 | 279 | 149 | 0.129252425 | 0.794336857 | 0 | 0.348 | 0.263 | 0.17 | 0.416 | 0.037 | 0.188 | 0.097 | 0.176 | 0 | 0.14 |
| SE | 45882 | ENSG00000145740 | SLC30A5 | chr5 | + | 69116393 | 69116602 | 69115925 | 69116214 | 69118498 | 69118628 | 68 | 26 | 60 | 60 | 58 | 0 | 4 | 0 | 0 | 1 | 41 | 28 | 47 | 37 | 33 | 0 | 3 | 5 | 0 | 0 | 298 | 149 | 0.130467333 | 0.799935094 | 1 | 0.765 | 1 | 1 | 0.967 | 1 | 0.824 | 0.825 | 1 | 1 | 0.017 |
| SE | 66928 | ENSG00000024862 | CCDC28A | chr6 | + | 138785226 | 138785381 | 138779821 | 138779985 | 138788365 | 138788388 | 172 | 111 | 156 | 130 | 197 | 0 | 0 | 0 | 0 | 0 | 121 | 95 | 204 | 95 | 86 | 8 | 3 | 0 | 0 | 0 | 298 | 149 | 0.131448583 | 0.804872002 | 1 | 1 | 1 | 1 | 1 | 0.883 | 0.941 | 1 | 1 | 1 | 0.035 |
| SE | 102597 | ENSG00000175470 | PPP2R2D | chr10 | + | 131943967 | 131943997 | 131940581 | 131940694 | 131944094 | 131944145 | 19 | 12 | 15 | 14 | 18 | 9 | 6 | 6 | 8 | 14 | 7 | 19 | 12 | 23 | 28 | 0 | 1 | 9 | 11 | 13 | 179 | 149 | 0.133526939 | 0.815040545 | 0.637 | 0.625 | 0.675 | 0.593 | 0.517 | 1 | 0.941 | 0.526 | 0.635 | 0.642 | -0.139 |
| SE | 19510 | ENSG00000100479 | POLE2 | chr14 | - | 49669523 | 49669598 | 49666329 | 49666413 | 49674122 | 49674216 | 35 | 39 | 35 | 38 | 82 | 0 | 0 | 0 | 6 | 0 | 139 | 71 | 24 | 81 | 90 | 8 | 0 | 0 | 0 | 5 | 224 | 149 | 0.133901764 | 0.816699583 | 1 | 1 | 1 | 0.808 | 1 | 0.92 | 1 | 1 | 1 | 0.923 | -0.007 |
| SE | 66490 | ENSG00000132676 | DAP3 | chr1 | + | 155709032 | 155709125 | 155689090 | 155689243 | 155709772 | 155709824 | 5 | 9 | 2 | 1 | 2 | 65 | 35 | 84 | 71 | 35 | 17 | 4 | 1 | 0 | 1 | 45 | 52 | 28 | 48 | 26 | 242 | 149 | 0.134228218 | 0.818117124 | 0.045 | 0.137 | 0.014 | 0.009 | 0.034 | 0.189 | 0.045 | 0.022 | 0 | 0.023 | -0.008 |
| SE | 93252 | ENSG00000136937 | NCBP1 | chr9 | + | 97662041 | 97662144 | 97660945 | 97661068 | 97662953 | 97663047 | 142 | 97 | 109 | 119 | 248 | 0 | 0 | 1 | 0 | 1 | 44 | 49 | 53 | 30 | 15 | 0 | 4 | 4 | 0 | 0 | 252 | 149 | 0.134904545 | 0.82134419 | 1 | 1 | 0.985 | 1 | 0.993 | 1 | 0.879 | 0.887 | 1 | 1 | 0.042 |
| SE | 91166 | ENSG00000164941 | INTS8 | chr8 | + | 94838493 | 94838618 | 94831991 | 94832174 | 94841490 | 94841591 | 100 | 69 | 80 | 67 | 103 | 0 | 0 | 3 | 8 | 6 | 72 | 80 | 57 | 45 | 22 | 8 | 0 | 0 | 0 | 0 | 274 | 149 | 0.135286262 | 0.822836416 | 1 | 1 | 0.935 | 0.82 | 0.903 | 0.83 | 1 | 1 | 1 | 1 | -0.034 |
| SE | 42659 | ENSG00000088854 | C20orf194 | chr20 | - | 3296860 | 3296949 | 3294527 | 3294629 | 3304439 | 3304543 | 15 | 24 | 16 | 19 | 22 | 0 | 0 | 0 | 0 | 1 | 57 | 33 | 12 | 39 | 47 | 0 | 0 | 7 | 0 | 0 | 238 | 149 | 0.136768644 | 0.829339975 | 1 | 1 | 1 | 1 | 0.932 | 1 | 1 | 0.518 | 1 | 1 | 0.083 |
| SE | 1572 | ENSG00000198089 | SFI1 | chr22 | + | 31550253 | 31550348 | 31508254 | 31508376 | 31556941 | 31557059 | 72 | 48 | 113 | 70 | 105 | 0 | 0 | 0 | 0 | 0 | 133 | 82 | 61 | 84 | 108 | 11 | 0 | 0 | 0 | 0 | 244 | 149 | 0.137147482 | 0.831035993 | 1 | 1 | 1 | 1 | 1 | 0.881 | 1 | 1 | 1 | 1 | 0.024 |
| SE | 59871 | ENSG00000103168 | TAF1C | chr16 | - | 84184850 | 84185060 | 84183696 | 84183778 | 84186900 | 84186981 | 45 | 12 | 18 | 20 | 10 | 3 | 1 | 4 | 5 | 0 | 19 | 51 | 17 | 24 | 16 | 0 | 3 | 1 | 4 | 0 | 298 | 149 | 0.137384407 | 0.832364741 | 0.882 | 0.857 | 0.692 | 0.667 | 1 | 1 | 0.895 | 0.895 | 0.75 | 1 | -0.088 |
| SE | 107496 | ENSG00000123349 | PFDN5 | chr12 | + | 53295838 | 53297924 | 53295555 | 53295639 | 53298044 | 53298150 | 1463 | 818 | 1261 | 1166 | 1018 | 21 | 6 | 15 | 20 | 7 | 1659 | 985 | 806 | 889 | 1674 | 14 | 11 | 12 | 8 | 9 | 298 | 149 | 0.141153091 | 0.850267213 | 0.972 | 0.986 | 0.977 | 0.967 | 0.986 | 0.983 | 0.978 | 0.971 | 0.982 | 0.989 | -0.003 |
| SE | 91185 | ENSG00000164941 | INTS8 | chr8 | + | 94868691 | 94868813 | 94867275 | 94867337 | 94871883 | 94872002 | 5 | 4 | 2 | 3 | 0 | 238 | 180 | 175 | 200 | 192 | 6 | 0 | 2 | 0 | 0 | 0 | 86 | 97 | 61 | 11 | 271 | 149 | 0.141346545 | 0.851166674 | 0.011 | 0.012 | 0.006 | 0.008 | 0 | 1 | 0 | 0.011 | 0 | 0 | -0.195 |
| SE | 78260 | ENSG00000173715 | C11orf80 | chr11 | + | 66762078 | 66762450 | 66759042 | 66759096 | 66788159 | 66788269 | 7 | 2 | 2 | 15 | 3 | 5 | 7 | 8 | 19 | 10 | 7 | 5 | 8 | 19 | 0 | 0 | 2 | 11 | 6 | 12 | 298 | 149 | 0.14178067 | 0.853321498 | 0.412 | 0.125 | 0.111 | 0.283 | 0.13 | 1 | 0.556 | 0.267 | 0.613 | 0 | -0.275 |
| SE | 58951 | ENSG00000131778 | CHD1L | chr1 | + | 147270931 | 147271195 | 147268781 | 147268878 | 147272170 | 147272281 | 12 | 5 | 18 | 15 | 17 | 7 | 7 | 7 | 16 | 11 | 2 | 11 | 4 | 13 | 19 | 7 | 8 | 9 | 18 | 12 | 298 | 149 | 0.142173057 | 0.85463199 | 0.462 | 0.263 | 0.562 | 0.319 | 0.436 | 0.125 | 0.407 | 0.182 | 0.265 | 0.442 | 0.124 |
| SE | 100290 | ENSG00000167986 | DDB1 | chr11 | - | 61326778 | 61326893 | 61323660 | 61324137 | 61329362 | 61329584 | 113 | 87 | 116 | 157 | 111 | 0 | 0 | 0 | 0 | 0 | 211 | 150 | 120 | 109 | 195 | 0 | 9 | 0 | 0 | 4 | 264 | 149 | 0.143402886 | 0.86070313 | 1 | 1 | 1 | 1 | 1 | 1 | 0.904 | 1 | 1 | 0.965 | 0.026 |
| SE | 86465 | ENSG00000215021 | PHB2 | chr12 | - | 6967170 | 6967779 | 6966423 | 6966500 | 6967891 | 6968021 | 433 | 273 | 472 | 428 | 667 | 0 | 2 | 1 | 0 | 1 | 627 | 288 | 314 | 404 | 606 | 16 | 0 | 0 | 6 | 0 | 298 | 149 | 0.143614815 | 0.861644871 | 1 | 0.986 | 0.996 | 1 | 0.997 | 0.951 | 1 | 1 | 0.971 | 1 | 0.011 |
| SE | 4413 | ENSG00000105662 | CRTC1 | chr19 | + | 18765403 | 18765528 | 18760007 | 18760228 | 18775640 | 18775821 | 10 | 14 | 6 | 3 | 7 | 0 | 0 | 0 | 0 | 0 | 8 | 2 | 4 | 5 | 20 | 0 | 0 | 2 | 0 | 4 | 274 | 149 | 0.143946259 | 0.863302667 | 1 | 1 | 1 | 1 | 1 | 1 | 1 | 0.521 | 1 | 0.731 | 0.15 |
| SE | 45885 | ENSG00000145740 | SLC30A5 | chr5 | + | 69118498 | 69118628 | 69116393 | 69116602 | 69121693 | 69121895 | 44 | 29 | 62 | 52 | 81 | 0 | 0 | 0 | 0 | 0 | 31 | 10 | 23 | 19 | 15 | 0 | 0 | 0 | 0 | 7 | 279 | 149 | 0.145025287 | 0.86824437 | 1 | 1 | 1 | 1 | 1 | 1 | 1 | 1 | 1 | 0.534 | 0.093 |
| SE | 21424 | ENSG00000164180 | TMEM161B | chr5 | - | 88205800 | 88205954 | 88202961 | 88203075 | 88206438 | 88206499 | 86 | 37 | 72 | 66 | 173 | 0 | 0 | 3 | 0 | 8 | 11 | 27 | 12 | 7 | 3 | 0 | 0 | 0 | 3 | 0 | 298 | 149 | 0.145068157 | 0.868368229 | 1 | 1 | 0.923 | 1 | 0.915 | 1 | 1 | 1 | 0.538 | 1 | 0.06 |
| SE | 6592 | ENSG00000134444 | RELCH | chr18 | + | 62252663 | 62252754 | 62244763 | 62244876 | 62255406 | 62255478 | 105 | 83 | 105 | 115 | 149 | 5 | 0 | 3 | 4 | 0 | 67 | 51 | 50 | 26 | 32 | 0 | 3 | 2 | 7 | 0 | 240 | 149 | 0.146518909 | 0.874978607 | 0.929 | 1 | 0.956 | 0.947 | 1 | 1 | 0.913 | 0.939 | 0.698 | 1 | 0.056 |
| SE | 79675 | ENSG00000186591 | UBE2H | chr7 | - | 129858901 | 129858941 | 129839206 | 129839335 | 129879567 | 129879642 | 34 | 31 | 42 | 37 | 108 | 3 | 3 | 1 | 0 | 7 | 73 | 82 | 59 | 49 | 104 | 0 | 0 | 9 | 0 | 2 | 189 | 149 | 0.148165972 | 0.882324532 | 0.899 | 0.891 | 0.971 | 1 | 0.924 | 1 | 1 | 0.838 | 1 | 0.976 | -0.026 |
| SE | 107511 | ENSG00000123349 | PFDN5 | chr12 | + | 53297849 | 53297924 | 53295524 | 53295639 | 53298044 | 53298150 | 916 | 481 | 863 | 714 | 745 | 21 | 6 | 15 | 20 | 7 | 1102 | 587 | 534 | 583 | 1073 | 14 | 11 | 12 | 8 | 9 | 224 | 149 | 0.148820961 | 0.885147825 | 0.967 | 0.982 | 0.975 | 0.96 | 0.986 | 0.981 | 0.973 | 0.967 | 0.98 | 0.988 | -0.004 |
| SE | 97605 | ENSG00000179832 | MROH1 | chr8 | + | 144191714 | 144191855 | 144190783 | 144190935 | 144192308 | 144192401 | 15 | 14 | 12 | 15 | 7 | 0 | 0 | 0 | 0 | 0 | 65 | 17 | 14 | 25 | 34 | 0 | 4 | 0 | 3 | 0 | 290 | 149 | 0.149172267 | 0.886631126 | 1 | 1 | 1 | 1 | 1 | 1 | 0.686 | 1 | 0.811 | 1 | 0.101 |
| SE | 33415 | ENSG00000096063 | SRPK1 | chr6 | - | 35904878 | 35904974 | 35890894 | 35891013 | 35920467 | 35920528 | 9 | 6 | 12 | 16 | 3 | 134 | 91 | 138 | 125 | 112 | 8 | 12 | 38 | 19 | 0 | 67 | 196 | 282 | 211 | 10 | 245 | 149 | 0.149454382 | 0.887970883 | 0.039 | 0.039 | 0.05 | 0.072 | 0.016 | 0.068 | 0.036 | 0.076 | 0.052 | 0 | -0.003 |
| SE | 37698 | ENSG00000163507 | CIP2A | chr3 | - | 108582976 | 108583083 | 108582107 | 108582202 | 108585064 | 108585212 | 6 | 5 | 22 | 8 | 5 | 1 | 0 | 0 | 0 | 0 | 8 | 13 | 10 | 5 | 4 | 0 | 2 | 0 | 4 | 0 | 256 | 149 | 0.149928193 | 0.889773195 | 0.777 | 1 | 1 | 1 | 1 | 1 | 0.791 | 1 | 0.421 | 1 | 0.113 |
| SE | 102414 | ENSG00000138346 | DNA2 | chr10 | - | 68465666 | 68465812 | 68459103 | 68459235 | 68468122 | 68468306 | 74 | 51 | 19 | 67 | 89 | 0 | 0 | 0 | 0 | 9 | 105 | 83 | 63 | 108 | 33 | 0 | 0 | 0 | 0 | 0 | 295 | 149 | 0.150023647 | 0.890137274 | 1 | 1 | 1 | 1 | 0.833 | 1 | 1 | 1 | 1 | 1 | -0.033 |
| SE | 107366 | ENSG00000033030 | ZCCHC8 | chr12 | - | 122483278 | 122483344 | 122482634 | 122482695 | 122489385 | 122489463 | 36 | 12 | 23 | 45 | 75 | 6 | 12 | 11 | 13 | 23 | 23 | 18 | 27 | 19 | 4 | 16 | 24 | 14 | 20 | 0 | 215 | 149 | 0.150177038 | 0.890844864 | 0.806 | 0.409 | 0.592 | 0.706 | 0.693 | 0.499 | 0.342 | 0.572 | 0.397 | 1 | 0.079 |
| SE | 34625 | ENSG00000198700 | IPO9 | chr1 | + | 201872827 | 201872961 | 201871160 | 201871327 | 201874249 | 201874372 | 84 | 45 | 83 | 92 | 65 | 0 | 0 | 1 | 0 | 7 | 138 | 94 | 81 | 91 | 129 | 0 | 0 | 3 | 5 | 8 | 283 | 149 | 0.150228766 | 0.891084199 | 1 | 1 | 0.978 | 1 | 0.83 | 1 | 1 | 0.934 | 0.906 | 0.895 | 0.015 |
| SE | 21423 | ENSG00000164180 | TMEM161B | chr5 | - | 88205753 | 88205954 | 88197668 | 88203075 | 88206438 | 88206499 | 87 | 37 | 72 | 69 | 181 | 0 | 0 | 3 | 0 | 8 | 11 | 27 | 12 | 7 | 2 | 0 | 0 | 0 | 3 | 0 | 298 | 149 | 0.154239795 | 0.90847815 | 1 | 1 | 0.923 | 1 | 0.919 | 1 | 1 | 1 | 0.538 | 1 | 0.061 |
| SE | 106663 | ENSG00000139350 | NEDD1 | chr12 | + | 96917620 | 96917737 | 96907603 | 96907624 | 96919984 | 96920118 | 61 | 34 | 74 | 64 | 73 | 0 | 0 | 0 | 0 | 0 | 7 | 37 | 20 | 4 | 25 | 0 | 0 | 0 | 5 | 0 | 266 | 149 | 0.154472359 | 0.90950277 | 1 | 1 | 1 | 1 | 1 | 1 | 1 | 1 | 0.309 | 1 | 0.138 |
| SE | 42664 | ENSG00000088854 | C20orf194 | chr20 | - | 3316722 | 3316793 | 3315734 | 3315785 | 3322216 | 3322262 | 11 | 11 | 5 | 11 | 6 | 1 | 0 | 0 | 0 | 0 | 17 | 4 | 10 | 8 | 13 | 0 | 2 | 4 | 0 | 0 | 220 | 149 | 0.155400819 | 0.913938444 | 0.882 | 1 | 1 | 1 | 1 | 1 | 0.575 | 0.629 | 1 | 1 | 0.136 |
| SE | 99086 | ENSG00000110274 | CEP164 | chr11 | + | 117335604 | 117335680 | 117327928 | 117328208 | 117338565 | 117338668 | 33 | 21 | 29 | 41 | 15 | 14 | 22 | 16 | 20 | 7 | 48 | 35 | 21 | 28 | 34 | 27 | 59 | 22 | 13 | 24 | 225 | 149 | 0.15568705 | 0.91493457 | 0.61 | 0.387 | 0.546 | 0.576 | 0.587 | 0.541 | 0.282 | 0.387 | 0.588 | 0.484 | 0.085 |
| SE | 82032 | ENSG00000104133 | SPG11 | chr15 | - | 44589251 | 44589414 | 44585635 | 44585850 | 44592330 | 44592438 | 314 | 227 | 189 | 195 | 322 | 18 | 21 | 8 | 7 | 19 | 100 | 198 | 121 | 185 | 25 | 2 | 18 | 6 | 0 | 3 | 298 | 149 | 0.157743657 | 0.924315024 | 0.897 | 0.844 | 0.922 | 0.933 | 0.894 | 0.962 | 0.846 | 0.91 | 1 | 0.806 | -0.007 |
| SE | 30855 | ENSG00000109133 | TMEM33 | chr4 | + | 41939195 | 41939383 | 41938601 | 41938696 | 41943746 | 41943814 | 348 | 185 | 234 | 250 | 271 | 5 | 4 | 2 | 0 | 9 | 149 | 251 | 217 | 171 | 125 | 0 | 5 | 15 | 0 | 3 | 298 | 149 | 0.157820734 | 0.924432645 | 0.972 | 0.959 | 0.983 | 1 | 0.938 | 1 | 0.962 | 0.879 | 1 | 0.954 | 0.011 |
| SE | 68824 | ENSG00000082258 | CCNT2 | chr2 | + | 134942611 | 134942674 | 134939001 | 134939062 | 134946100 | 134946146 | 59 | 40 | 66 | 75 | 110 | 5 | 2 | 6 | 0 | 1 | 7 | 55 | 55 | 23 | 10 | 0 | 1 | 1 | 4 | 0 | 212 | 149 | 0.159506258 | 0.932549429 | 0.892 | 0.934 | 0.885 | 1 | 0.987 | 1 | 0.975 | 0.975 | 0.802 | 1 | -0.011 |
| SE | 17486 | ENSG00000185024 | BRF1 | chr14 | - | 105241264 | 105241414 | 105228819 | 105228913 | 105252506 | 105252579 | 71 | 46 | 38 | 42 | 46 | 3 | 1 | 1 | 1 | 0 | 41 | 46 | 9 | 19 | 67 | 0 | 24 | 0 | 0 | 0 | 298 | 149 | 0.159887093 | 0.933939111 | 0.922 | 0.958 | 0.95 | 0.955 | 1 | 1 | 0.489 | 1 | 1 | 1 | 0.059 |
| SE | 99100 | ENSG00000110274 | CEP164 | chr11 | + | 117387412 | 117387412 | 117381700 | 117381868 | 117390776 | 117390908 | 33 | 33 | 29 | 49 | 26 | 0 | 1 | 0 | 4 | 0 | 137 | 48 | 68 | 81 | 183 | 10 | 1 | 1 | 0 | 0 | 298 | 149 | 0.160563783 | 0.936634028 | 1 | 0.943 | 1 | 0.86 | 1 | 0.873 | 0.96 | 0.971 | 1 | 1 | 0 |
| SE | 109053 | ENSG00000139668 | WDFY2 | chr13 | + | 51719197 | 51719348 | 51703595 | 51703650 | 51727677 | 51727790 | 215 | 180 | 166 | 210 | 266 | 0 | 3 | 0 | 0 | 2 | 326 | 268 | 249 | 293 | 221 | 0 | 5 | 0 | 11 | 0 | 298 | 149 | 0.162310779 | 0.944292226 | 1 | 0.968 | 1 | 1 | 0.985 | 1 | 0.964 | 1 | 0.93 | 1 | 0.012 |
| SE | 106783 | ENSG00000151062 | CACNA2D4 | chr12 | - | 1801573 | 1801644 | 1801042 | 1801118 | 1810277 | 1810340 | 4 | 3 | 12 | 12 | 15 | 8 | 0 | 2 | 3 | 3 | 6 | 0 | 5 | 10 | 4 | 8 | 2 | 5 | 4 | 0 | 220 | 149 | 0.163296868 | 0.94883644 | 0.253 | 1 | 0.803 | 0.73 | 0.772 | 0.337 | 0 | 0.404 | 0.629 | 1 | 0.238 |
| SE | 52022 | ENSG00000135250 | SRPK2 | chr7 | - | 105126247 | 105126340 | 105116373 | 105118022 | 105126992 | 105127062 | 101 | 63 | 82 | 94 | 81 | 0 | 2 | 1 | 2 | 5 | 24 | 70 | 73 | 35 | 5 | 0 | 4 | 0 | 0 | 0 | 242 | 149 | 0.163297883 | 0.94883644 | 1 | 0.951 | 0.981 | 0.967 | 0.909 | 1 | 0.915 | 1 | 1 | 1 | -0.021 |
| SE | 107144 | ENSG00000175582 | RAB6A | chr11 | - | 73718784 | 73718890 | 73718612 | 73718718 | 73720845 | 73720899 | 179 | 126 | 107 | 160 | 217 | 349 | 232 | 272 | 346 | 436 | 210 | 139 | 114 | 187 | 221 | 557 | 332 | 365 | 410 | 377 | 255 | 149 | 0.163511432 | 0.949724856 | 0.231 | 0.241 | 0.187 | 0.213 | 0.225 | 0.181 | 0.197 | 0.154 | 0.21 | 0.255 | 0.02 |
| SE | 63076 | ENSG00000132953 | XPO4 | chr13 | - | 20799164 | 20799339 | 20796763 | 20797057 | 20800155 | 20800325 | 134 | 112 | 104 | 118 | 179 | 0 | 0 | 2 | 0 | 0 | 9 | 117 | 164 | 37 | 21 | 0 | 8 | 1 | 0 | 0 | 298 | 149 | 0.163698907 | 0.95060221 | 1 | 1 | 0.963 | 1 | 1 | 1 | 0.88 | 0.988 | 1 | 1 | 0.019 |
| SE | 4393 | ENSG00000130475 | FCHO1 | chr19 | + | 17754591 | 17754758 | 17754316 | 17754361 | 17755117 | 17755191 | 42 | 11 | 29 | 36 | 2 | 1 | 1 | 8 | 5 | 0 | 30 | 36 | 4 | 20 | 106 | 0 | 3 | 7 | 10 | 23 | 298 | 149 | 0.163946761 | 0.951593586 | 0.955 | 0.846 | 0.644 | 0.783 | 1 | 1 | 0.857 | 0.222 | 0.5 | 0.697 | 0.19 |
| SE | 99373 | ENSG00000156599 | ZDHHC5 | chr11 | + | 57696760 | 57696873 | 57695919 | 57696043 | 57698558 | 57698835 | 195 | 100 | 171 | 118 | 236 | 20 | 2 | 7 | 6 | 23 | 236 | 198 | 106 | 169 | 259 | 23 | 17 | 17 | 16 | 12 | 262 | 149 | 0.165230196 | 0.957417504 | 0.847 | 0.966 | 0.933 | 0.918 | 0.854 | 0.854 | 0.869 | 0.78 | 0.857 | 0.925 | 0.047 |
| SE | 12928 | ENSG00000197818 | SLC9A8 | chr20 | + | 49839540 | 49839599 | 49823060 | 49823141 | 49845035 | 49845119 | 64 | 47 | 64 | 57 | 47 | 1 | 0 | 3 | 0 | 0 | 122 | 125 | 140 | 182 | 98 | 4 | 0 | 1 | 3 | 5 | 208 | 149 | 0.166636517 | 0.963510095 | 0.979 | 1 | 0.939 | 1 | 1 | 0.956 | 1 | 0.99 | 0.978 | 0.934 | 0.012 |
| SE | 62921 | ENSG00000185420 | SMYD3 | chr1 | - | 246186742 | 246186942 | 245929869 | 245929937 | 246327200 | 246327337 | 1 | 0 | 8 | 1 | 2 | 25 | 17 | 17 | 21 | 18 | 6 | 2 | 9 | 0 | 2 | 5 | 18 | 23 | 7 | 46 | 298 | 149 | 0.168054533 | 0.969638744 | 0.02 | 0 | 0.19 | 0.023 | 0.053 | 0.375 | 0.053 | 0.164 | 0 | 0.021 | -0.065 |
| SE | 46332 | ENSG00000095564 | BTAF1 | chr10 | + | 91964076 | 91964201 | 91962537 | 91962678 | 91980453 | 91980558 | 290 | 237 | 273 | 309 | 403 | 3 | 0 | 1 | 1 | 0 | 19 | 130 | 216 | 102 | 27 | 0 | 0 | 5 | 0 | 5 | 274 | 149 | 0.170319359 | 0.979602183 | 0.981 | 1 | 0.993 | 0.994 | 1 | 1 | 1 | 0.959 | 1 | 0.746 | 0.053 |
| SE | 81065 | ENSG00000157764 | BRAF | chr7 | - | 140734616 | 140734770 | 140719326 | 140726516 | 140739811 | 140739946 | 49 | 55 | 75 | 98 | 236 | 34 | 24 | 33 | 19 | 58 | 37 | 41 | 64 | 25 | 7 | 39 | 30 | 47 | 25 | 0 | 298 | 149 | 0.173531888 | 0.993983161 | 0.419 | 0.534 | 0.532 | 0.721 | 0.67 | 0.322 | 0.406 | 0.405 | 0.333 | 1 | 0.082 |
| SE | 10342 | ENSG00000198496 | NBR2 | chr17 | + | 43132955 | 43133137 | 43131207 | 43131270 | 43138656 | 43138922 | 6 | 2 | 3 | 4 | 8 | 11 | 12 | 14 | 12 | 15 | 0 | 4 | 1 | 6 | 12 | 34 | 18 | 10 | 16 | 30 | 298 | 149 | 0.173993714 | 0.995834197 | 0.214 | 0.077 | 0.097 | 0.143 | 0.211 | 0 | 0.1 | 0.048 | 0.158 | 0.167 | 0.054 |
| SE | 100138 | ENSG00000159063 | ALG8 | chr11 | - | 78112649 | 78112770 | 78109441 | 78109581 | 78113885 | 78113989 | 101 | 61 | 67 | 59 | 113 | 0 | 0 | 0 | 0 | 0 | 92 | 48 | 42 | 48 | 52 | 0 | 0 | 0 | 9 | 0 | 270 | 149 | 0.177399363 | 1 | 1 | 1 | 1 | 1 | 1 | 1 | 1 | 1 | 0.746 | 1 | 0.051 |
| SE | 15789 | ENSG00000103978 | TMEM87A | chr15 | - | 42272062 | 42272123 | 42267946 | 42268032 | 42273254 | 42273433 | 214 | 158 | 170 | 195 | 222 | 5 | 6 | 8 | 13 | 2 | 280 | 279 | 254 | 242 | 64 | 0 | 13 | 27 | 11 | 2 | 210 | 149 | 0.179431045 | 1 | 0.968 | 0.949 | 0.938 | 0.914 | 0.987 | 1 | 0.938 | 0.87 | 0.94 | 0.958 | 0.01 |
| SE | 84863 | ENSG00000164074 | ABHD18 | chr4 | + | 127989720 | 127989821 | 127984338 | 127984423 | 128008919 | 128008957 | 143 | 90 | 132 | 128 | 137 | 15 | 25 | 19 | 16 | 26 | 148 | 105 | 111 | 107 | 126 | 8 | 15 | 25 | 18 | 8 | 250 | 149 | 0.180938457 | 1 | 0.85 | 0.682 | 0.805 | 0.827 | 0.758 | 0.917 | 0.807 | 0.726 | 0.78 | 0.904 | -0.042 |
| SE | 56431 | ENSG00000050748 | MAPK9 | chr5 | - | 180279795 | 180280036 | 180269279 | 180269409 | 180280439 | 180280581 | 42 | 12 | 30 | 36 | 27 | 60 | 38 | 35 | 58 | 76 | 19 | 17 | 34 | 18 | 47 | 53 | 22 | 42 | 33 | 24 | 298 | 149 | 0.182202714 | 1 | 0.259 | 0.136 | 0.3 | 0.237 | 0.151 | 0.152 | 0.279 | 0.288 | 0.214 | 0.495 | -0.069 |
| SE | 101961 | ENSG00000197771 | MCMBP | chr10 | - | 119835539 | 119835704 | 119832011 | 119832100 | 119836895 | 119837029 | 367 | 198 | 262 | 299 | 572 | 8 | 4 | 2 | 1 | 12 | 392 | 303 | 200 | 277 | 265 | 0 | 5 | 4 | 7 | 0 | 298 | 149 | 0.182435788 | 1 | 0.958 | 0.961 | 0.985 | 0.993 | 0.96 | 1 | 0.968 | 0.962 | 0.952 | 1 | -0.005 |
| SE | 98436 | ENSG00000099783 | HNRNPM | chr19 | + | 8474166 | 8474244 | 8471325 | 8471427 | 8483157 | 8483211 | 53 | 36 | 38 | 64 | 66 | 1 | 1 | 0 | 4 | 7 | 161 | 70 | 97 | 133 | 89 | 32 | 0 | 2 | 9 | 0 | 227 | 149 | 0.184578971 | 1 | 0.972 | 0.959 | 1 | 0.913 | 0.861 | 0.768 | 1 | 0.97 | 0.907 | 1 | 0.012 |
| SE | 14218 | ENSG00000167965 | MLST8 | chr16 | + | 2206742 | 2206929 | 2206496 | 2206659 | 2207034 | 2207110 | 2 | 0 | 0 | 1 | 6 | 26 | 11 | 23 | 14 | 17 | 0 | 0 | 1 | 0 | 0 | 48 | 26 | 18 | 5 | 34 | 298 | 149 | 0.186791779 | 1 | 0.037 | 0 | 0 | 0.034 | 0.15 | 0 | 0 | 0.027 | 0 | 0 | 0.039 |
| SE | 70237 | ENSG00000160695 | VPS11 | chr11 | + | 119078564 | 119078704 | 119078172 | 119078334 | 119078794 | 119078997 | 152 | 80 | 162 | 179 | 136 | 0 | 0 | 0 | 0 | 0 | 260 | 168 | 142 | 166 | 172 | 10 | 0 | 0 | 3 | 0 | 289 | 149 | 0.18689352 | 1 | 1 | 1 | 1 | 1 | 1 | 0.931 | 1 | 1 | 0.966 | 1 | 0.021 |
| SE | 34627 | ENSG00000198700 | IPO9 | chr1 | + | 201875151 | 201875228 | 201874831 | 201874936 | 201875943 | 201884291 | 108 | 66 | 134 | 91 | 118 | 0 | 0 | 1 | 0 | 0 | 166 | 125 | 66 | 99 | 134 | 0 | 6 | 2 | 0 | 4 | 226 | 149 | 0.187111881 | 1 | 1 | 1 | 0.989 | 1 | 1 | 1 | 0.932 | 0.956 | 1 | 0.957 | 0.029 |
| SE | 16695 | ENSG00000137841 | PLCB2 | chr15 | - | 40296521 | 40296634 | 40296295 | 40296392 | 40296757 | 40296908 | 197 | 174 | 178 | 190 | 351 | 1 | 0 | 0 | 0 | 0 | 271 | 206 | 123 | 273 | 358 | 0 | 0 | 1 | 0 | 15 | 262 | 149 | 0.188991329 | 1 | 0.991 | 1 | 1 | 1 | 1 | 1 | 1 | 0.986 | 1 | 0.931 | 0.015 |
| SE | 66651 | ENSG00000084623 | EIF3I | chr1 | + | 32226170 | 32226320 | 32222439 | 32222630 | 32228498 | 32228609 | 1 | 2 | 2 | 5 | 7 | 0 | 1 | 0 | 0 | 1 | 6 | 11 | 15 | 14 | 12 | 0 | 1 | 0 | 1 | 4 | 298 | 149 | 0.190710872 | 1 | 1 | 0.5 | 1 | 1 | 0.778 | 1 | 0.846 | 1 | 0.875 | 0.6 | -0.009 |
| SE | 47787 | ENSG00000154358 | OBSCN | chr1 | + | 228299258 | 228299522 | 228298453 | 228298717 | 228299881 | 228300145 | 11 | 10 | 17 | 19 | 17 | 6 | 6 | 27 | 21 | 9 | 14 | 26 | 7 | 17 | 0 | 0 | 0 | 8 | 7 | 16 | 298 | 149 | 0.190831407 | 1 | 0.478 | 0.455 | 0.239 | 0.311 | 0.486 | 1 | 1 | 0.304 | 0.548 | 0 | -0.177 |
| SE | 96890 | ENSG00000100731 | PCNX1 | chr14 | + | 70995740 | 70995925 | 70976941 | 70978648 | 71009633 | 71009724 | 43 | 25 | 42 | 36 | 43 | 82 | 47 | 69 | 69 | 92 | 14 | 37 | 39 | 34 | 20 | 7 | 112 | 90 | 17 | 28 | 298 | 149 | 0.191805043 | 1 | 0.208 | 0.21 | 0.233 | 0.207 | 0.189 | 0.5 | 0.142 | 0.178 | 0.5 | 0.263 | -0.107 |
| SE | 104999 | ENSG00000129315 | CCNT1 | chr12 | - | 48695998 | 48696162 | 48695758 | 48695829 | 48698137 | 48698183 | 102 | 60 | 75 | 88 | 77 | 57 | 35 | 32 | 79 | 44 | 23 | 39 | 73 | 14 | 14 | 24 | 81 | 144 | 59 | 0 | 298 | 149 | 0.192124267 | 1 | 0.472 | 0.462 | 0.54 | 0.358 | 0.467 | 0.324 | 0.194 | 0.202 | 0.106 | 1 | 0.095 |
| SE | 60210 | ENSG00000154451 | GBP5 | chr1 | - | 89262682 | 89262785 | 89262219 | 89262401 | 89263735 | 89263948 | 824 | 660 | 767 | 762 | 970 | 4 | 4 | 5 | 10 | 15 | 637 | 677 | 381 | 522 | 183 | 3 | 5 | 9 | 0 | 0 | 252 | 149 | 0.193160561 | 1 | 0.992 | 0.99 | 0.989 | 0.978 | 0.975 | 0.992 | 0.988 | 0.962 | 1 | 1 | -0.004 |
| SE | 30343 | ENSG00000108591 | DRG2 | chr17 | + | 18091917 | 18092074 | 18087964 | 18088087 | 18093812 | 18093973 | 0 | 0 | 0 | 0 | 0 | 43 | 24 | 33 | 39 | 33 | 0 | 0 | 11 | 0 | 0 | 44 | 79 | 16 | 43 | 54 | 298 | 149 | 0.19332295 | 1 | 0 | 0 | 0 | 0 | 0 | 0 | 0 | 0.256 | 0 | 0 | -0.051 |
| SE | 77023 | ENSG00000146433 | TMEM181 | chr6 | + | 158607243 | 158607343 | 158605266 | 158605347 | 158608332 | 158608463 | 127 | 64 | 113 | 135 | 179 | 1 | 6 | 3 | 4 | 18 | 41 | 90 | 101 | 36 | 92 | 0 | 11 | 13 | 6 | 0 | 249 | 149 | 0.193379353 | 1 | 0.987 | 0.865 | 0.958 | 0.953 | 0.856 | 1 | 0.83 | 0.823 | 0.782 | 1 | 0.037 |
| SE | 42794 | ENSG00000139842 | CUL4A | chr13 | + | 113245955 | 113246063 | 113245151 | 113245237 | 113253081 | 113253195 | 358 | 213 | 307 | 358 | 376 | 8 | 8 | 5 | 3 | 3 | 227 | 345 | 141 | 300 | 104 | 9 | 4 | 6 | 9 | 0 | 257 | 149 | 0.194155871 | 1 | 0.963 | 0.939 | 0.973 | 0.986 | 0.986 | 0.936 | 0.98 | 0.932 | 0.951 | 1 | 0.01 |
| SE | 85061 | ENSG00000147459 | DOCK5 | chr8 | + | 25340876 | 25340988 | 25336238 | 25336373 | 25341738 | 25341809 | 104 | 96 | 57 | 99 | 97 | 0 | 0 | 0 | 0 | 0 | 135 | 57 | 112 | 101 | 49 | 8 | 1 | 1 | 0 | 0 | 261 | 149 | 0.194600579 | 1 | 1 | 1 | 1 | 1 | 1 | 0.906 | 0.97 | 0.985 | 1 | 1 | 0.028 |
| SE | 31466 | ENSG00000178950 | GAK | chr4 | - | 868538 | 868685 | 866955 | 867432 | 870710 | 870904 | 269 | 125 | 189 | 271 | 144 | 17 | 1 | 0 | 0 | 5 | 270 | 241 | 129 | 248 | 539 | 0 | 3 | 12 | 10 | 18 | 296 | 149 | 0.194988283 | 1 | 0.888 | 0.984 | 1 | 1 | 0.935 | 1 | 0.976 | 0.844 | 0.926 | 0.938 | 0.025 |
| SE | 69653 | ENSG00000143390 | RFX5 | chr1 | - | 151345927 | 151345957 | 151344961 | 151345188 | 151346204 | 151346333 | 29 | 6 | 15 | 18 | 9 | 0 | 0 | 7 | 1 | 2 | 6 | 18 | 18 | 5 | 15 | 8 | 0 | 3 | 7 | 0 | 179 | 149 | 0.195155717 | 1 | 1 | 1 | 0.641 | 0.937 | 0.789 | 0.384 | 1 | 0.833 | 0.373 | 1 | 0.155 |
| SE | 29706 | ENSG00000182621 | PLCB1 | chr20 | + | 8646101 | 8646181 | 8628287 | 8628431 | 8647899 | 8647908 | 64 | 28 | 47 | 44 | 46 | 0 | 0 | 0 | 0 | 0 | 8 | 12 | 21 | 11 | 6 | 0 | 0 | 0 | 6 | 0 | 229 | 149 | 0.19653423 | 1 | 1 | 1 | 1 | 1 | 1 | 1 | 1 | 1 | 0.544 | 1 | 0.091 |
| SE | 99206 | ENSG00000074266 | EED | chr11 | + | 86252147 | 86252240 | 86250295 | 86250448 | 86255221 | 86255287 | 129 | 82 | 97 | 130 | 299 | 5 | 1 | 4 | 3 | 0 | 70 | 117 | 71 | 57 | 108 | 0 | 0 | 5 | 1 | 0 | 242 | 149 | 0.196827746 | 1 | 0.941 | 0.981 | 0.937 | 0.964 | 1 | 1 | 1 | 0.897 | 0.972 | 1 | -0.009 |
| SE | 19455 | ENSG00000125746 | EML2 | chr19 | - | 45614825 | 45615036 | 45614604 | 45614700 | 45615801 | 45615889 | 0 | 0 | 0 | 1 | 0 | 28 | 19 | 25 | 35 | 16 | 0 | 12 | 0 | 0 | 0 | 86 | 35 | 26 | 26 | 33 | 298 | 149 | 0.197105796 | 1 | 0 | 0 | 0 | 0.014 | 0 | 0 | 0.146 | 0 | 0 | 0 | -0.026 |
| SE | 84012 | ENSG00000111581 | NUP107 | chr12 | + | 68690630 | 68690746 | 68689532 | 68689619 | 68696850 | 68696922 | 53 | 43 | 32 | 55 | 95 | 0 | 0 | 0 | 0 | 0 | 15 | 24 | 14 | 14 | 6 | 0 | 0 | 0 | 6 | 0 | 265 | 149 | 0.197921293 | 1 | 1 | 1 | 1 | 1 | 1 | 1 | 1 | 1 | 0.567 | 1 | 0.087 |
| SE | 1584 | ENSG00000198089 | SFI1 | chr22 | + | 31561289 | 31561392 | 31550253 | 31550348 | 31573057 | 31573214 | 29 | 25 | 51 | 28 | 38 | 0 | 0 | 0 | 0 | 0 | 67 | 32 | 21 | 42 | 80 | 0 | 0 | 0 | 8 | 0 | 252 | 149 | 0.201059833 | 1 | 1 | 1 | 1 | 1 | 1 | 1 | 1 | 1 | 0.756 | 1 | 0.049 |
| SE | 85064 | ENSG00000147459 | DOCK5 | chr8 | + | 25364625 | 25364704 | 25363046 | 25363141 | 25366869 | 25366970 | 187 | 149 | 116 | 189 | 178 | 1 | 0 | 0 | 1 | 0 | 160 | 143 | 247 | 205 | 86 | 7 | 3 | 0 | 4 | 0 | 228 | 149 | 0.202120614 | 1 | 0.992 | 1 | 1 | 0.992 | 1 | 0.937 | 0.969 | 1 | 0.971 | 1 | 0.021 |
| SE | 94014 | ENSG00000099381 | SETD1A | chr16 | + | 30966883 | 30967060 | 30965600 | 30966386 | 30967500 | 30967588 | 136 | 83 | 133 | 155 | 187 | 0 | 0 | 0 | 0 | 0 | 162 | 194 | 168 | 158 | 307 | 0 | 0 | 2 | 11 | 0 | 298 | 149 | 0.203591003 | 1 | 1 | 1 | 1 | 1 | 1 | 1 | 1 | 0.977 | 0.878 | 1 | 0.029 |
| SE | 65652 | ENSG00000143924 | EML4 | chr2 | + | 42304483 | 42304551 | 42303314 | 42303446 | 42315961 | 42316050 | 237 | 204 | 291 | 316 | 346 | 2 | 1 | 1 | 4 | 2 | 73 | 189 | 268 | 122 | 29 | 4 | 2 | 7 | 0 | 0 | 217 | 149 | 0.205319989 | 1 | 0.988 | 0.993 | 0.995 | 0.982 | 0.992 | 0.926 | 0.985 | 0.963 | 1 | 1 | 0.015 |
| SE | 102599 | ENSG00000175470 | PPP2R2D | chr10 | + | 131947529 | 131947791 | 131945294 | 131945459 | 131955683 | 131956242 | 220 | 134 | 152 | 184 | 157 | 0 | 0 | 1 | 2 | 0 | 196 | 151 | 104 | 119 | 240 | 0 | 0 | 0 | 0 | 12 | 298 | 149 | 0.206155859 | 1 | 1 | 1 | 0.987 | 0.979 | 1 | 1 | 1 | 1 | 1 | 0.909 | 0.011 |
| SE | 64456 | ENSG00000065150 | IPO5 | chr13 | + | 97987989 | 97988101 | 97985420 | 97985613 | 97989061 | 97989099 | 5 | 2 | 6 | 3 | 2 | 188 | 131 | 233 | 216 | 333 | 0 | 0 | 4 | 14 | 0 | 77 | 130 | 111 | 117 | 49 | 261 | 149 | 0.206513732 | 1 | 0.015 | 0.009 | 0.014 | 0.008 | 0.003 | 0 | 0 | 0.02 | 0.064 | 0 | -0.007 |
| SE | 99363 | ENSG00000156599 | ZDHHC5 | chr11 | + | 57682421 | 57682543 | 57667746 | 57668187 | 57688507 | 57688665 | 107 | 44 | 92 | 107 | 65 | 0 | 0 | 0 | 0 | 0 | 107 | 100 | 71 | 72 | 109 | 0 | 4 | 0 | 2 | 4 | 271 | 149 | 0.20696848 | 1 | 1 | 1 | 1 | 1 | 1 | 1 | 0.932 | 1 | 0.952 | 0.937 | 0.036 |

| | | | | | | | | | | | | | | | | | | | | | | | | | | | | | | | | | | | | | | | | | | | | | | |
|---|---|---|---|---|---|---|---|---|---|---|---|---|---|---|---|---|---|---|---|---|---|---|---|---|---|---|---|---|---|---|---|---|---|---|---|---|---|---|---|---|---|---|---|---|---|---|
| SE | 88445 | ENSG00000221914 | PPP2R2A | chr8 | + | 26339902 | 26339973 | 26338889 | 26338987 | 26354467 | 26354633 | 13 | 5 | 16 | 2 | 11 | 163 | 135 | 155 | 134 | 185 | 12 | 8 | 11 | 6 | 8 | 83 | 118 | 202 | 89 | 57 | 220 | 149 | 0.20755474 | 1 | 0.051 | 0.024 | 0.065 | 0.01 | 0.039 | 0.089 | 0.044 | 0.036 | 0.044 | 0.087 | -0.022 |
| SE | 16962 | ENSG00000172575 | RASGRP1 | chr15 | - | 38505839 | 38505920 | 38502311 | 38502421 | 38507725 | 38508001 | 252 | 162 | 253 | 293 | 342 | 1 | 9 | 5 | 7 | 9 | 37 | 176 | 206 | 91 | 49 | 17 | 4 | 4 | 0 | 0 | 230 | 149 | 0.216735256 | 1 | 0.994 | 0.921 | 0.97 | 0.964 | 0.961 | 0.585 | 0.966 | 0.971 | 1 | 1 | 0.058 |
| SE | 55709 | ENSG00000135749 | PCNX2 | chr1 | - | 233250738 | 233250832 | 233236844 | 233236980 | 233252353 | 233252499 | 130 | 94 | 117 | 106 | 327 | 0 | 1 | 5 | 2 | 5 | 121 | 101 | 135 | 131 | 88 | 0 | 0 | 4 | 0 | 0 | 243 | 149 | 0.218239597 | 1 | 1 | 0.983 | 0.935 | 0.97 | 0.976 | 1 | 1 | 0.954 | 1 | 1 | -0.018 |
| SE | 94924 | ENSG00000148655 | LRMDA | chr10 | + | 76346429 | 76346553 | 76324400 | 76324485 | 76557208 | 76557306 | 2 | 0 | 4 | 1 | 4 | 3 | 14 | 5 | 7 | 19 | 0 | 0 | 6 | 14 | 16 | 35 | 7 | 13 | 6 | 41 | 273 | 149 | 0.218774957 | 1 | 0.267 | 0 | 0.304 | 0.072 | 0.103 | 0 | 0 | 0.201 | 0.56 | 0.176 | -0.038 |
| SE | 12991 | ENSG00000122390 | NAA60 | chr16 | + | 3447481 | 3447620 | 3443693 | 3443837 | 3448470 | 3448540 | 5 | 6 | 3 | 0 | 3 | 8 | 1 | 7 | 11 | 10 | 15 | 8 | 5 | 11 | 18 | 23 | 18 | 2 | 16 | 21 | 288 | 149 | 0.218933188 | 1 | 0.244 | 0.756 | 0.181 | 0 | 0.134 | 0.252 | 0.187 | 0.564 | 0.262 | 0.307 | -0.051 |
| SE | 84773 | ENSG00000125676 | THOC2 | chrX | - | 123623643 | 123623709 | 123623104 | 123623283 | 123623786 | 123623971 | 15 | 19 | 13 | 25 | 33 | 82 | 90 | 98 | 138 | 214 | 0 | 23 | 10 | 0 | 0 | 1 | 53 | 34 | 1 | 6 | 215 | 149 | 0.219708444 | 1 | 0.113 | 0.128 | 0.084 | 0.112 | 0.097 | 0 | 0.231 | 0.169 | 0 | 0 | 0.027 |
| SE | 55786 | ENSG00000012174 | MBTPS2 | chrX | + | 21869497 | 21869678 | 21853375 | 21853503 | 21878041 | 21878136 | 23 | 11 | 33 | 18 | 54 | 1 | 0 | 0 | 0 | 0 | 22 | 13 | 14 | 13 | 4 | 0 | 0 | 6 | 0 | 0 | 298 | 149 | 0.220217863 | 1 | 0.92 | 1 | 1 | 1 | 1 | 1 | 1 | 0.538 | 1 | 1 | 0.076 |
| SE | 99108 | ENSG00000110274 | CEP164 | chr11 | + | 117408525 | 117408755 | 117407924 | 117408032 | 117408889 | 117409028 | 4 | 0 | 4 | 1 | 6 | 26 | 31 | 49 | 31 | 84 | 4 | 3 | 4 | 0 | 0 | 72 | 36 | 24 | 30 | 98 | 298 | 149 | 0.222973108 | 1 | 0.071 | 0 | 0.039 | 0.016 | 0.034 | 0.027 | 0.04 | 0.077 | 0 | 0 | 0.003 |
| SE | 86258 | ENSG00000134516 | DOCK2 | chr5 | + | 169708168 | 169708267 | 169702302 | 169702427 | 169711934 | 169712007 | 1126 | 819 | 902 | 1130 | 1334 | 1 | 8 | 0 | 5 | 40 | 1771 | 1425 | 1240 | 1337 | 870 | 21 | 15 | 0 | 9 | 6 | 248 | 149 | 0.223110077 | 1 | 0.999 | 0.984 | 1 | 0.993 | 0.952 | 0.981 | 0.983 | 1 | 0.989 | 0.989 | -0.003 |
| SE | 19457 | ENSG00000125746 | EML2 | chr19 | - | 45617629 | 45617697 | 45616764 | 45616853 | 45619059 | 45619090 | 69 | 27 | 35 | 48 | 14 | 4 | 3 | 1 | 0 | 0 | 40 | 43 | 22 | 22 | 62 | 0 | 0 | 0 | 3 | 0 | 217 | 149 | 0.224347249 | 1 | 0.922 | 0.861 | 0.96 | 1 | 1 | 1 | 1 | 1 | 0.834 | 1 | -0.018 |
| SE | 76085 | ENSG00000112697 | TMEM30A | chr6 | - | 75267640 | 75267748 | 75265230 | 75265338 | 75284401 | 75284763 | 506 | 278 | 423 | 439 | 578 | 8 | 4 | 1 | 4 | 6 | 56 | 292 | 376 | 135 | 112 | 0 | 0 | 2 | 0 | 0 | 257 | 149 | 0.224350475 | 1 | 0.973 | 0.976 | 0.996 | 0.985 | 0.982 | 1 | 1 | 0.991 | 1 | 1 | -0.016 |
| SE | 64421 | ENSG00000143919 | CAMKMT | chr2 | + | 44549554 | 44549728 | 44390240 | 44390305 | 44704282 | 44704343 | 1 | 5 | 13 | 5 | 2 | 15 | 12 | 28 | 28 | 36 | 0 | 6 | 4 | 9 | 4 | 6 | 17 | 11 | 2 | 15 | 298 | 149 | 0.22497154 | 1 | 0.032 | 0.172 | 0.188 | 0.082 | 0.027 | 0 | 0.15 | 0.154 | 0.692 | 0.118 | -0.123 |
| SE | 38564 | ENSG00000198925 | ATG9A | chr2 | - | 219227921 | 219228053 | 219227769 | 219227813 | 219228433 | 219228482 | 40 | 19 | 38 | 22 | 16 | 0 | 0 | 1 | 0 | 0 | 16 | 24 | 9 | 15 | 47 | 1 | 2 | 4 | 0 | 0 | 281 | 149 | 0.22732695 | 1 | 1 | 1 | 0.953 | 1 | 1 | 0.895 | 0.864 | 0.544 | 1 | 1 | 0.13 |
| SE | 29803 | ENSG00000141452 | RMC1 | chr18 | + | 23509192 | 23509279 | 23504370 | 23504447 | 23515855 | 23515886 | 131 | 102 | 85 | 102 | 143 | 1 | 0 | 0 | 0 | 3 | 182 | 140 | 95 | 151 | 211 | 0 | 1 | 0 | 0 | 12 | 236 | 149 | 0.227992972 | 1 | 0.988 | 1 | 1 | 1 | 0.968 | 1 | 0.989 | 1 | 1 | 0.917 | 0.01 |
| SE | 8066 | ENSG00000214087 | ARL16 | chr17 | - | 81682490 | 81682675 | 81682033 | 81682149 | 81683012 | 81683126 | 27 | 11 | 29 | 17 | 18 | 30 | 8 | 8 | 6 | 34 | 10 | 44 | 12 | 11 | 48 | 34 | 18 | 8 | 43 | 32 | 298 | 149 | 0.229446432 | 1 | 0.31 | 0.407 | 0.644 | 0.586 | 0.209 | 0.128 | 0.55 | 0.429 | 0.113 | 0.429 | 0.101 |
| SE | 76679 | ENSG00000223501 | VPS52 | chr6 | - | 33251517 | 33251636 | 33250271 | 33250987 | 33251859 | 33251971 | 82 | 46 | 86 | 57 | 111 | 0 | 0 | 0 | 6 | 0 | 116 | 65 | 70 | 82 | 90 | 10 | 0 | 0 | 0 | 0 | 268 | 149 | 0.229890326 | 1 | 1 | 1 | 1 | 0.841 | 1 | 0.866 | 1 | 1 | 1 | 1 | -0.005 |
| SE | 21430 | ENSG00000164180 | TMEM161B | chr5 | - | 88225768 | 88225866 | 88220562 | 88220719 | 88228444 | 88228528 | 48 | 45 | 70 | 65 | 79 | 4 | 2 | 6 | 9 | 15 | 21 | 36 | 14 | 13 | 11 | 0 | 3 | 0 | 0 | 8 | 247 | 149 | 0.231183679 | 1 | 0.879 | 0.931 | 0.876 | 0.813 | 0.761 | 1 | 0.879 | 1 | 1 | 0.453 | -0.014 |
| SE | 58950 | ENSG00000131778 | CHD1L | chr1 | + | 147270931 | 147271113 | 147268781 | 147268878 | 147272170 | 147272281 | 16 | 5 | 25 | 15 | 18 | 7 | 7 | 7 | 16 | 11 | 2 | 22 | 4 | 13 | 23 | 7 | 8 | 9 | 18 | 12 | 298 | 149 | 0.233391153 | 1 | 0.533 | 0.263 | 0.641 | 0.319 | 0.45 | 0.125 | 0.579 | 0.182 | 0.265 | 0.489 | 0.113 |
| SE | 8029 | ENSG00000073350 | LLGL2 | chr17 | + | 75574452 | 75574495 | 75574110 | 75574260 | 75574609 | 75574668 | 34 | 14 | 15 | 21 | 24 | 3 | 1 | 0 | 8 | 0 | 4 | 31 | 8 | 17 | 22 | 0 | 13 | 0 | 8 | 0 | 192 | 149 | 0.2352208 | 1 | 0.898 | 0.916 | 1 | 0.671 | 1 | 1 | 0.649 | 1 | 0.623 | 1 | 0.043 |
| SE | 69004 | ENSG00000107521 | HPS1 | chr10 | - | 98431130 | 98431291 | 98430570 | 98430670 | 98433982 | 98434091 | 68 | 28 | 57 | 77 | 46 | 3 | 1 | 1 | 5 | 0 | 80 | 90 | 100 | 65 | 239 | 0 | 3 | 0 | 0 | 7 | 298 | 149 | 0.23545253 | 1 | 0.919 | 0.933 | 0.966 | 0.885 | 1 | 1 | 0.938 | 1 | 1 | 0.945 | -0.036 |
| SE | 51295 | ENSG00000174780 | SRP72 | chr4 | + | 56478591 | 56478649 | 56476670 | 56476702 | 56483138 | 56483270 | 51 | 45 | 39 | 60 | 120 | 4 | 0 | 1 | 0 | 6 | 29 | 28 | 40 | 28 | 4 | 0 | 0 | 6 | 0 | 8 | 207 | 149 | 0.236221306 | 1 | 0.902 | 1 | 0.966 | 1 | 0.935 | 1 | 1 | 0.828 | 1 | 0.265 | 0.142 |
| SE | 101744 | ENSG00000198056 | PRIM1 | chr12 | - | 56740011 | 56740128 | 56739293 | 56739363 | 56741434 | 56741576 | 6 | 3 | 9 | 4 | 3 | 97 | 69 | 54 | 88 | 120 | 0 | 3 | 2 | 15 | 0 | 38 | 42 | 33 | 23 | 45 | 266 | 149 | 0.237417035 | 1 | 0.033 | 0.024 | 0.085 | 0.025 | 0.014 | 0 | 0.038 | 0.033 | 0.268 | 0 | -0.032 |
| SE | 73304 | ENSG00000136699 | SMPD4 | chr2 | - | 130161185 | 130161272 | 130157250 | 130157396 | 130164373 | 130164445 | 12 | 10 | 22 | 10 | 25 | 26 | 9 | 25 | 31 | 55 | 17 | 20 | 12 | 19 | 11 | 62 | 41 | 26 | 19 | 58 | 236 | 149 | 0.238301795 | 1 | 0.226 | 0.412 | 0.357 | 0.169 | 0.223 | 0.148 | 0.235 | 0.226 | 0.387 | 0.107 | 0.057 |
| SE | 56430 | ENSG00000050748 | MAPK9 | chr5 | - | 180247662 | 180247728 | 180246544 | 180247510 | 180248972 | 180249138 | 1 | 5 | 5 | 1 | 4 | 99 | 46 | 90 | 88 | 157 | 0 | 7 | 6 | 8 | 0 | 90 | 109 | 73 | 79 | 77 | 215 | 149 | 0.239265734 | 1 | 0.007 | 0.07 | 0.037 | 0.008 | 0.017 | 0 | 0.043 | 0.054 | 0.066 | 0 | -0.005 |
| SE | 108198 | ENSG00000186908 | ZDHHC17 | chr12 | + | 76809042 | 76809120 | 76805316 | 76805439 | 76809712 | 76809750 | 215 | 168 | 168 | 189 | 244 | 3 | 1 | 0 | 2 | 9 | 58 | 84 | 106 | 55 | 23 | 0 | 0 | 0 | 0 | 0 | 227 | 149 | 0.239357581 | 1 | 0.979 | 0.991 | 1 | 0.984 | 0.947 | 1 | 1 | 1 | 1 | 1 | -0.02 |
| SE | 12281 | ENSG00000103194 | USP10 | chr16 | + | 84759890 | 84759946 | 84744632 | 84745673 | 84760171 | 84760236 | 13 | 13 | 30 | 19 | 28 | 1 | 2 | 2 | 0 | 1 | 12 | 24 | 28 | 27 | 40 | 0 | 0 | 0 | 8 | 3 | 205 | 149 | 0.240547281 | 1 | 0.904 | 0.825 | 0.916 | 1 | 0.953 | 1 | 1 | 1 | 0.71 | 0.906 | -0.004 |
| SE | 16203 | ENSG00000140598 | EFL1 | chr15 | - | 82238306 | 82238521 | 82230847 | 82230971 | 82240417 | 82240555 | 130 | 87 | 91 | 129 | 105 | 0 | 2 | 1 | 1 | 6 | 55 | 106 | 50 | 43 | 20 | 0 | 1 | 2 | 8 | 0 | 298 | 149 | 0.241224316 | 1 | 1 | 0.956 | 0.978 | 0.985 | 0.897 | 1 | 0.981 | 0.926 | 0.729 | 1 | 0.036 |
| SE | 19456 | ENSG00000125746 | EML2 | chr19 | - | 45616460 | 45616558 | 45615801 | 45615889 | 45616764 | 45616853 | 47 | 28 | 36 | 35 | 16 | 3 | 1 | 4 | 0 | 3 | 45 | 34 | 16 | 18 | 75 | 22 | 3 | 1 | 0 | 0 | 247 | 149 | 0.241507819 | 1 | 0.904 | 0.944 | 0.844 | 1 | 0.763 | 0.552 | 0.872 | 0.906 | 1 | 1 | 0.025 |
| SE | 12277 | ENSG00000103194 | USP10 | chr16 | + | 84740308 | 84740369 | 84733137 | 84733503 | 84758715 | 84758807 | 15 | 21 | 17 | 27 | 14 | 0 | 0 | 0 | 0 | 1 | 33 | 45 | 14 | 15 | 16 | 0 | 4 | 0 | 0 | 3 | 210 | 149 | 0.244108156 | 1 | 1 | 1 | 1 | 1 | 0.909 | 1 | 0.889 | 1 | 1 | 0.791 | 0.046 |
| SE | 65203 | ENSG00000010244 | ZNF207 | chr17 | + | 32366664 | 32366757 | 32365329 | 32365487 | 32367771 | 32368014 | 334 | 248 | 378 | 351 | 514 | 323 | 277 | 286 | 363 | 452 | 507 | 510 | 539 | 602 | 140 | 342 | 262 | 298 | 303 | 343 | 242 | 149 | 0.244482313 | 1 | 0.389 | 0.355 | 0.449 | 0.373 | 0.412 | 0.477 | 0.545 | 0.527 | 0.55 | 0.201 | -0.064 |
| SE | 52020 | ENSG00000135250 | SRPK2 | chr7 | - | 105125781 | 105125874 | 105117387 | 105118022 | 105126247 | 105126340 | 22 | 8 | 18 | 27 | 21 | 58 | 38 | 52 | 49 | 45 | 24 | 21 | 15 | 2 | 5 | 0 | 44 | 49 | 23 | 5 | 242 | 149 | 0.246165111 | 1 | 0.189 | 0.115 | 0.176 | 0.253 | 0.223 | 1 | 0.227 | 0.159 | 0.051 | 0.381 | -0.172 |
| SE | 86244 | ENSG00000134516 | DOCK2 | chr5 | + | 169670541 | 169670597 | 169654402 | 169654486 | 169671077 | 169671174 | 252 | 182 | 159 | 173 | 207 | 0 | 0 | 0 | 1 | 7 | 315 | 247 | 294 | 299 | 181 | 0 | 0 | 6 | 9 | 4 | 205 | 149 | 0.246541475 | 1 | 1 | 1 | 1 | 0.992 | 0.956 | 1 | 1 | 0.973 | 0.96 | 0.97 | 0.009 |
| SE | 78631 | ENSG00000168014 | C2CD3 | chr11 | - | 74054606 | 74054671 | 74049336 | 74049542 | 74057405 | 74057544 | 43 | 11 | 40 | 33 | 44 | 1 | 0 | 2 | 0 | 0 | 26 | 52 | 17 | 37 | 44 | 0 | 3 | 0 | 4 | 0 | 214 | 149 | 0.250451281 | 1 | 0.968 | 1 | 0.933 | 1 | 1 | 1 | 0.923 | 1 | 0.866 | 1 | 0.022 |
| SE | 84868 | ENSG00000164074 | ABHD18 | chr4 | + | 128017362 | 128017501 | 128011672 | 128011700 | 128020079 | 128020169 | 111 | 75 | 113 | 148 | 65 | 2 | 2 | 1 | 3 | 1 | 146 | 126 | 117 | 101 | 126 | 0 | 0 | 4 | 0 | 0 | 288 | 149 | 0.251038016 | 1 | 0.966 | 0.951 | 0.983 | 0.962 | 0.971 | 1 | 1 | 0.938 | 1 | 1 | -0.021 |
| SE | 1564 | ENSG00000198089 | SFI1 | chr22 | + | 31531057 | 31531129 | 31528689 | 31528863 | 31546860 | 31546971 | 139 | 116 | 118 | 109 | 131 | 52 | 44 | 35 | 66 | 40 | 89 | 141 | 92 | 92 | 152 | 46 | 35 | 57 | 60 | 65 | 221 | 149 | 0.254056447 | 1 | 0.643 | 0.64 | 0.694 | 0.527 | 0.688 | 0.566 | 0.731 | 0.521 | 0.508 | 0.612 | 0.051 |

[illegible]

| | | | | | | | | | | | | | | | | | | | | | | | | | | | | | | | | | | | | | | | | | | | | | | |
|---|---|---|---|---|---|---|---|---|---|---|---|---|---|---|---|---|---|---|---|---|---|---|---|---|---|---|---|---|---|---|---|---|---|---|---|---|---|---|---|---|---|---|---|---|---|---|
| SE | 52033 | ENSG00000135250 | SRPK2 | chr7 | - | 105297420 | 105297533 | 105203627 | 105203785 | 105388647 | 105388702 | 0 | 4 | 1 | 1 | 0 | 17 | 17 | 9 | 13 | 10 | 0 | 0 | 0 | 0 | 0 | 15 | 15 | 13 | 9 | 17 | 262 | 149 | 0.548879634 | 1 | 0 | 0.118 | 0.059 | 0.042 | 0 | 0 | 0 | 0 | 0 | 0 | 0.044 |
| SE | 107655 | ENSG00000139718 | SETD1B | chr12 | + | 121827518 | 121827650 | 121825199 | 121825366 | 121827734 | 121827854 | 79 | 44 | 125 | 88 | 55 | 0 | 0 | 1 | 0 | 0 | 74 | 141 | 86 | 80 | 162 | 0 | 0 | 0 | 0 | 9 | 281 | 149 | 0.553032083 | 1 | 1 | 1 | 0.985 | 1 | 1 | 1 | 1 | 1 | 1 | 0.905 | 0.016 |
| SE | 1582 | ENSG00000198089 | SFI1 | chr22 | + | 31556941 | 31557059 | 31550253 | 31550348 | 31573057 | 31573214 | 67 | 47 | 74 | 69 | 77 | 0 | 0 | 0 | 0 | 0 | 60 | 73 | 75 | 75 | 143 | 0 | 0 | 0 | 8 | 0 | 267 | 149 | 0.553059294 | 1 | 1 | 1 | 1 | 1 | 1 | 1 | 1 | 1 | 0.84 | 1 | 0.032 |
| SE | 107507 | ENSG00000123349 | PFDN5 | chr12 | + | 53296453 | 53296551 | 53295541 | 53295639 | 53297849 | 53297924 | 18 | 7 | 25 | 23 | 11 | 208 | 128 | 218 | 168 | 123 | 14 | 12 | 8 | 23 | 26 | 212 | 136 | 153 | 132 | 306 | 247 | 149 | 0.553172426 | 1 | 0.05 | 0.032 | 0.065 | 0.076 | 0.051 | 0.038 | 0.051 | 0.031 | 0.095 | 0.049 | 0.002 |
| SE | 1579 | ENSG00000198089 | SFI1 | chr22 | + | 31556941 | 31557059 | 31546860 | 31546971 | 31561289 | 31561333 | 134 | 83 | 109 | 102 | 103 | 0 | 0 | 0 | 0 | 0 | 114 | 88 | 59 | 82 | 173 | 0 | 0 | 0 | 0 | 9 | 267 | 149 | 0.553278735 | 1 | 1 | 1 | 1 | 1 | 1 | 1 | 1 | 1 | 1 | 0.915 | 0.017 |
| SE | 79032 | ENSG00000160218 | TRAPPC10 | chr21 | + | 44084121 | 44084263 | 44082787 | 44083302 | 44089832 | 44089933 | 117 | 46 | 104 | 92 | 101 | 0 | 0 | 0 | 1 | 0 | 80 | 91 | 92 | 63 | 68 | 0 | 1 | 2 | 3 | 0 | 291 | 149 | 0.554877123 | 1 | 1 | 1 | 1 | 0.979 | 1 | 1 | 0.979 | 0.959 | 0.915 | 1 | 0.025 |
| SE | 97606 | ENSG00000179832 | MROH1 | chr8 | + | 144242601 | 144242628 | 144242368 | 144242515 | 144243493 | 144243616 | 34 | 37 | 39 | 17 | 35 | 33 | 20 | 35 | 19 | 36 | 10 | 52 | 31 | 53 | 48 | 52 | 11 | 30 | 43 | 84 | 176 | 149 | 0.55515376 | 1 | 0.466 | 0.61 | 0.485 | 0.431 | 0.451 | 0.14 | 0.8 | 0.467 | 0.511 | 0.326 | 0.04 |
| SE | 91179 | ENSG00000164941 | INTS8 | chr8 | + | 94865505 | 94865690 | 94859510 | 94859632 | 94867139 | 94867196 | 171 | 108 | 112 | 137 | 160 | 1 | 5 | 2 | 2 | 0 | 62 | 97 | 60 | 22 | 7 | 0 | 0 | 0 | 0 | 0 | 298 | 149 | 0.557333351 | 1 | 0.988 | 0.915 | 0.966 | 0.972 | 1 | 1 | 1 | 1 | 1 | 1 | -0.032 |
| SE | 89482 | ENSG00000064313 | TAF2 | chr8 | - | 119793365 | 119793451 | 119789591 | 119789746 | 119795531 | 119795631 | 53 | 45 | 45 | 72 | 65 | 0 | 0 | 0 | 0 | 0 | 4 | 52 | 33 | 21 | 34 | 0 | 0 | 6 | 0 | 0 | 235 | 149 | 0.558866413 | 1 | 1 | 1 | 1 | 1 | 1 | 1 | 1 | 0.777 | 1 | 1 | 0.045 |
| SE | 4780 | ENSG00000064932 | SBNO2 | chr19 | - | 1112188 | 1112301 | 1111995 | 1112067 | 1112401 | 1112537 | 62 | 22 | 31 | 41 | 1 | 0 | 0 | 0 | 0 | 0 | 16 | 68 | 38 | 44 | 58 | 0 | 0 | 2 | 0 | 5 | 262 | 149 | 0.559024012 | 1 | 1 | 1 | 1 | 1 | 1 | 1 | 1 | 0.915 | 1 | 0.868 | 0.043 |
| SE | 10339 | ENSG00000198496 | NBR2 | chr17 | + | 43131207 | 43131270 | 43125609 | 43125770 | 43138656 | 43138922 | 18 | 15 | 25 | 22 | 22 | 0 | 4 | 4 | 4 | 5 | 41 | 38 | 18 | 22 | 33 | 0 | 5 | 9 | 0 | 6 | 212 | 149 | 0.562206857 | 1 | 1 | 0.725 | 0.815 | 0.794 | 0.756 | 1 | 0.842 | 0.584 | 1 | 0.794 | -0.026 |
| SE | 73946 | ENSG00000144134 | RABL2A | chr2 | + | 113640893 | 113641005 | 113635050 | 113635130 | 113641352 | 113641450 | 15 | 3 | 7 | 10 | 18 | 3 | 2 | 4 | 0 | 1 | 16 | 23 | 6 | 11 | 40 | 0 | 0 | 0 | 2 | 19 | 261 | 149 | 0.562733202 | 1 | 0.741 | 0.461 | 0.5 | 1 | 0.911 | 1 | 1 | 1 | 0.758 | 0.546 | -0.138 |
| SE | 100706 | ENSG00000187147 | RNF220 | chr1 | + | 44635544 | 44635666 | 44632234 | 44632385 | 44636029 | 44636152 | 209 | 137 | 156 | 176 | 172 | 9 | 5 | 6 | 9 | 2 | 269 | 205 | 175 | 160 | 280 | 15 | 5 | 5 | 6 | 2 | 271 | 149 | 0.563136492 | 1 | 0.927 | 0.938 | 0.935 | 0.915 | 0.979 | 0.908 | 0.958 | 0.951 | 0.936 | 0.987 | -0.009 |
| SE | 53430 | ENSG00000111358 | GTF2H3 | chr12 | + | 123654923 | 123654998 | 123652706 | 123652735 | 123659515 | 123659584 | 20 | 18 | 25 | 29 | 43 | 0 | 0 | 1 | 0 | 0 | 10 | 9 | 12 | 7 | 14 | 4 | 0 | 1 | 0 | 0 | 224 | 149 | 0.56455486 | 1 | 1 | 1 | 0.943 | 1 | 1 | 0.624 | 1 | 0.889 | 1 | 1 | 0.086 |
| SE | 96907 | ENSG00000100731 | PCNX1 | chr14 | + | 71051882 | 71052012 | 71047806 | 71047984 | 71055503 | 71055562 | 125 | 80 | 90 | 127 | 165 | 5 | 0 | 1 | 3 | 10 | 28 | 111 | 138 | 63 | 59 | 0 | 4 | 2 | 6 | 0 | 279 | 149 | 0.567501949 | 1 | 0.93 | 1 | 0.98 | 0.958 | 0.898 | 1 | 0.937 | 0.974 | 0.849 | 1 | 0.001 |
| SE | 4398 | ENSG00000130475 | FCHO1 | chr19 | + | 17772456 | 17772555 | 17770791 | 17770896 | 17774238 | 17774241 | 114 | 85 | 110 | 84 | 78 | 0 | 0 | 2 | 3 | 2 | 142 | 148 | 73 | 100 | 178 | 0 | 1 | 0 | 0 | 0 | 248 | 149 | 0.568806014 | 1 | 1 | 1 | 0.971 | 0.944 | 0.959 | 1 | 0.989 | 1 | 1 | 1 | -0.023 |
| SE | 105465 | ENSG00000135720 | DYNC1LI2 | chr16 | - | 66728200 | 66728242 | 66725827 | 66725944 | 66729039 | 66729099 | 29 | 34 | 26 | 19 | 59 | 0 | 0 | 0 | 0 | 0 | 18 | 44 | 47 | 26 | 12 | 4 | 0 | 0 | 2 | 0 | 191 | 149 | 0.573838813 | 1 | 1 | 1 | 1 | 1 | 1 | 0.778 | 1 | 1 | 0.91 | 1 | 0.062 |
| SE | 79948 | ENSG00000186480 | INSIG1 | chr7 | + | 155302746 | 155302846 | 155298258 | 155298697 | 155308240 | 155308757 | 83 | 33 | 38 | 53 | 50 | 1 | 1 | 1 | 5 | 0 | 46 | 39 | 30 | 47 | 60 | 0 | 0 | 18 | 0 | 0 | 249 | 149 | 0.574444869 | 1 | 0.98 | 0.952 | 0.958 | 0.864 | 1 | 1 | 1 | 0.499 | 1 | 1 | 0.051 |
| SE | 95855 | ENSG00000107263 | RAPGEF1 | chr9 | - | 131627912 | 131628096 | 131625921 | 131626422 | 131629101 | 131629254 | 247 | 88 | 148 | 183 | 177 | 1 | 0 | 0 | 8 | 0 | 236 | 242 | 196 | 188 | 413 | 0 | 2 | 1 | 0 | 0 | 298 | 149 | 0.575280041 | 1 | 0.992 | 1 | 1 | 0.92 | 1 | 1 | 0.984 | 0.99 | 1 | 1 | -0.012 |
| SE | 88036 | ENSG00000156171 | DRAM2 | chr1 | - | 111122615 | 111122643 | 111120515 | 111120693 | 111126226 | 111126294 | 16 | 8 | 10 | 14 | 26 | 7 | 11 | 15 | 11 | 29 | 10 | 2 | 5 | 5 | 5 | 4 | 14 | 16 | 13 | 0 | 177 | 149 | 0.5756803 | 1 | 0.658 | 0.38 | 0.359 | 0.517 | 0.43 | 0.678 | 0.107 | 0.208 | 0.245 | 1 | 0.021 |
| SE | 96894 | ENSG00000100731 | PCNX1 | chr14 | + | 71009633 | 71009724 | 70976941 | 70978648 | 71011491 | 71011549 | 220 | 123 | 189 | 187 | 239 | 0 | 0 | 2 | 1 | 1 | 34 | 250 | 190 | 59 | 71 | 0 | 10 | 0 | 0 | 0 | 240 | 149 | 0.575688514 | 1 | 1 | 1 | 0.983 | 0.991 | 0.993 | 1 | 0.939 | 1 | 1 | 1 | 0.006 |
| SE | 52567 | ENSG00000111445 | RFC5 | chr12 | + | 118017689 | 118017852 | 118016728 | 118016892 | 118019071 | 118019136 | 2 | 1 | 0 | 4 | 2 | 13 | 12 | 15 | 23 | 8 | 9 | 0 | 3 | 1 | 0 | 20 | 25 | 10 | 18 | 31 | 298 | 149 | 0.576143452 | 1 | 0.071 | 0.04 | 0 | 0.08 | 0.111 | 0.184 | 0 | 0.13 | 0.027 | 0 | -0.008 |
| SE | 86872 | ENSG00000090316 | MAEA | chr4 | + | 1332756 | 1332865 | 1327626 | 1327703 | 1339073 | 1339177 | 105 | 22 | 57 | 59 | 67 | 0 | 0 | 0 | 2 | 0 | 49 | 67 | 26 | 39 | 81 | 0 | 2 | 1 | 0 | 3 | 258 | 149 | 0.577435636 | 1 | 1 | 1 | 1 | 0.945 | 1 | 1 | 0.951 | 0.938 | 1 | 0.94 | 0.023 |
| SE | 35721 | ENSG00000130787 | HIP1R | chr12 | + | 122855550 | 122855612 | 122855264 | 122855405 | 122855830 | 122855903 | 86 | 23 | 86 | 85 | 26 | 0 | 0 | 2 | 0 | 0 | 14 | 110 | 49 | 51 | 164 | 0 | 0 | 0 | 0 | 7 | 211 | 149 | 0.577593519 | 1 | 1 | 1 | 0.968 | 1 | 1 | 1 | 1 | 1 | 1 | 0.943 | 0.005 |
| SE | 12935 | ENSG00000197818 | SLC9A8 | chr20 | + | 49874704 | 49874821 | 49864738 | 49864844 | 49877980 | 49878063 | 87 | 62 | 90 | 74 | 121 | 3 | 0 | 0 | 0 | 0 | 108 | 123 | 200 | 167 | 161 | 0 | 5 | 2 | 2 | 0 | 266 | 149 | 0.578531036 | 1 | 0.942 | 1 | 1 | 1 | 1 | 1 | 0.932 | 0.982 | 0.979 | 1 | 0.01 |
| SE | 108802 | ENSG00000105983 | LMBR1 | chr7 | - | 156826604 | 156826744 | 156796388 | 156796492 | 156833752 | 156833792 | 167 | 101 | 121 | 117 | 197 | 1 | 2 | 0 | 3 | 2 | 78 | 162 | 106 | 129 | 111 | 0 | 1 | 0 | 0 | 0 | 289 | 149 | 0.579439159 | 1 | 0.989 | 0.963 | 1 | 0.953 | 0.981 | 1 | 0.988 | 1 | 1 | 1 | -0.02 |
| SE | 107512 | ENSG00000123349 | PFDN5 | chr12 | + | 53297849 | 53297924 | 53295524 | 53295639 | 53298048 | 53298150 | 61 | 42 | 71 | 54 | 41 | 0 | 0 | 0 | 0 | 0 | 34 | 23 | 34 | 52 | 78 | 0 | 0 | 0 | 0 | 6 | 224 | 149 | 0.579760196 | 1 | 1 | 1 | 1 | 1 | 1 | 1 | 1 | 1 | 1 | 0.896 | 0.021 |
| SE | 96407 | ENSG00000197563 | PIGN | chr18 | - | 62163530 | 62163656 | 62161132 | 62161385 | 62186843 | 62186989 | 28 | 16 | 9 | 16 | 27 | 12 | 7 | 3 | 6 | 3 | 4 | 12 | 7 | 13 | 0 | 0 | 2 | 4 | 4 | 6 | 275 | 149 | 0.579897712 | 1 | 0.558 | 0.553 | 0.619 | 0.591 | 0.83 | 1 | 0.765 | 0.487 | 0.638 | 0 | 0.052 |
| SE | 107490 | ENSG00000123349 | PFDN5 | chr12 | + | 53295838 | 53295941 | 53295290 | 53295639 | 53296243 | 53296275 | 1673 | 937 | 1395 | 1374 | 1303 | 7 | 7 | 9 | 16 | 16 | 1833 | 1073 | 827 | 1053 | 1936 | 12 | 4 | 19 | 17 | 9 | 252 | 149 | 0.581738313 | 1 | 0.993 | 0.988 | 0.989 | 0.981 | 0.98 | 0.989 | 0.994 | 0.963 | 0.973 | 0.992 | 0.004 |
| SE | 19696 | ENSG00000119599 | DCAF4 | chr14 | + | 72935022 | 72935186 | 72926500 | 72926543 | 72937970 | 72938070 | 0 | 1 | 1 | 6 | 2 | 21 | 11 | 28 | 18 | 25 | 0 | 1 | 7 | 0 | 5 | 28 | 16 | 24 | 11 | 29 | 298 | 149 | 0.582787964 | 1 | 0 | 0.043 | 0.018 | 0.143 | 0.038 | 0 | 0.03 | 0.127 | 0 | 0.079 | 0.001 |
| SE | 78648 | ENSG00000168014 | C2CD3 | chr11 | - | 74133424 | 74133557 | 74131867 | 74132972 | 74138719 | 74138967 | 108 | 76 | 90 | 77 | 138 | 0 | 0 | 0 | 0 | 0 | 24 | 129 | 92 | 58 | 34 | 0 | 0 | 0 | 7 | 0 | 282 | 149 | 0.586486661 | 1 | 1 | 1 | 1 | 1 | 1 | 1 | 1 | 1 | 0.814 | 1 | 0.037 |
| SE | 26450 | ENSG00000083312 | TNPO1 | chr5 | + | 72891809 | 72891896 | 72889785 | 72889957 | 72893138 | 72893246 | 265 | 125 | 181 | 264 | 428 | 1 | 0 | 0 | 1 | 3 | 32 | 140 | 103 | 53 | 35 | 4 | 2 | 2 | 0 | 0 | 236 | 149 | 0.589784817 | 1 | 0.994 | 1 | 1 | 0.994 | 0.989 | 0.835 | 0.978 | 0.97 | 1 | 1 | 0.039 |
| SE | 5031 | ENSG00000101343 | CRNKL1 | chr20 | - | 20040685 | 20040766 | 20039608 | 20039848 | 20041565 | 20041625 | 208 | 181 | 204 | 186 | 258 | 2 | 0 | 1 | 4 | 0 | 19 | 114 | 145 | 70 | 9 | 0 | 5 | 0 | 3 | 0 | 230 | 149 | 0.591095868 | 1 | 0.985 | 1 | 0.992 | 0.968 | 1 | 1 | 0.937 | 1 | 0.938 | 1 | 0.014 |
| SE | 102581 | ENSG00000120438 | TCP1 | chr6 | - | 159785899 | 159785997 | 159783940 | 159784067 | 159787742 | 159787871 | 128 | 86 | 143 | 137 | 139 | 2 | 0 | 1 | 0 | 3 | 159 | 104 | 98 | 152 | 45 | 0 | 6 | 2 | 0 | 0 | 247 | 149 | 0.591357941 | 1 | 0.975 | 1 | 0.989 | 1 | 0.965 | 1 | 0.913 | 0.967 | 1 | 1 | 0.01 |
| SE | 15778 | ENSG00000167077 | MEI1 | chr22 | + | 41784338 | 41784420 | 41781398 | 41781845 | 41793828 | 41793910 | 18 | 7 | 13 | 15 | 14 | 11 | 4 | 15 | 9 | 8 | 15 | 27 | 8 | 9 | 20 | 4 | 10 | 8 | 20 | 32 | 231 | 149 | 0.591596645 | 1 | 0.513 | 0.53 | 0.359 | 0.518 | 0.53 | 0.708 | 0.635 | 0.392 | 0.225 | 0.287 | 0.041 |

[illegible]

| | | | | | | | | | | | | | | | |
|---|---|---|---|---|---|---|---|---|---|---|---|---|---|---|---|
| SE | 84022 | ENSG00000111581 | NUP107 | chr12 | + | 68719577 | 68719654 | 68719340 | 68719431 | 68721117 | 68721177 | 89 | 65 | 70 | [illegible] |
| SE | 103093 | ENSG00000108055 | SMC3 | chr10 | + | 110581922 | 110582098 | 110580903 | 110581021 | 110583383 | 110583548 | 65 | 90 | 59 | [illegible] |
| SE | 84453 | ENSG00000137522 | RNF121 | chr11 | + | 71957226 | 71957264 | 71929017 | 71929124 | 71960749 | 71960891 | 19 | 7 | 21 | [illegible] |
| SE | 1726 | ENSG00000100325 | ASCC2 | chr22 | - | 29800990 | 29801110 | 29793359 | 29793676 | 29804637 | 29804824 | 68 | 35 | 41 | [illegible] |
| SE | 63851 | ENSG00000148690 | FRA10AC1 | chr10 | - | 93692645 | 93692729 | 93692008 | 93692093 | 93694860 | 93694937 | 90 | 145 | 102 | [illegible] |
| SE | 29807 | ENSG00000141452 | RMC1 | chr18 | + | 23526636 | 23526765 | 23524409 | 23524482 | 23527794 | 23527901 | 177 | 122 | 124 | [illegible] |
| SE | 17650 | ENSG00000100949 | RABGGTA | chr14 | - | 24270836 | 24270998 | 24270333 | 24270458 | 24271112 | 24271169 | 35 | 5 | 39 | [illegible] |
| SE | 88032 | ENSG00000156171 | DRAM2 | chr1 | - | 111119614 | 111119681 | 111118864 | 111118897 | 111119876 | 111119959 | 3 | 4 | 2 | [illegible] |
| SE | 70233 | ENSG00000160695 | VPS11 | chr11 | + | 119071595 | 119071843 | 119070233 | 119070397 | 119073197 | 119073399 | 112 | 65 | 115 | [illegible] |
| SE | 96397 | ENSG00000197563 | PIGN | chr18 | - | 62157128 | 62157227 | 62148213 | 62148253 | 62157686 | 62157808 | 41 | 50 | 33 | [illegible] |
| SE | 39952 | ENSG00000164253 | WDR41 | chr5 | - | 77449759 | 77449870 | 77440812 | 77440997 | 77451290 | 77451353 | 248 | 108 | 131 | [illegible] |
| SE | 82057 | ENSG00000104133 | SPG11 | chr15 | - | 44629232 | 44629388 | 44627635 | 44628844 | 44633504 | 44633637 | 588 | 351 | 426 | [illegible] |
| SE | 12393 | ENSG00000188559 | RALGAPA2 | chr20 | - | 20536655 | 20536784 | 20531686 | 20531795 | 20546703 | 20546832 | 65 | 49 | 57 | [illegible] |
| SE | 101968 | ENSG00000197771 | MCMBP | chr10 | - | 119842471 | 119842595 | 119840842 | 119840960 | 119843253 | 119843426 | 371 | 192 | 210 | [illegible] |
| SE | 99105 | ENSG00000110274 | CEP164 | chr11 | + | 117394919 | 117395003 | 117394349 | 117394493 | 117395122 | 117395191 | 38 | 19 | 37 | [illegible] |
| SE | 73303 | ENSG00000136699 | SMPD4 | chr2 | - | 130161090 | 130161272 | 130157250 | 130157396 | 130164373 | 130164445 | 10 | 4 | 12 | [illegible] |
| SE | 87767 | ENSG00000196459 | TRAPPC2 | chrX | - | 13734043 | 13734185 | 13719870 | 13719982 | 13734524 | 13734620 | 92 | 34 | 63 | [illegible] |
| SE | 57166 | ENSG00000011021 | CLCN6 | chr1 | + | 11828457 | 11828624 | 11828105 | 11828219 | 11829195 | 11829322 | 17 | 6 | 2 | [illegible] |
| SE | 19460 | ENSG00000125746 | EML2 | chr19 | - | 45621482 | 45621637 | 45620811 | 45621332 | 45624718 | 45624782 | 92 | 29 | 48 | [illegible] |
| SE | 65686 | ENSG00000186141 | POLR3C | chr1 | + | 145836493 | 145836574 | 145833489 | 145833582 | 145839889 | 145839991 | 27 | 19 | 34 | [illegible] |
| SE | 54749 | ENSG00000083720 | OXCT1 | chr5 | - | 41803068 | 41803163 | 41801021 | 41801070 | 41807330 | 41807438 | 46 | 29 | 34 | [illegible] |
| SE | 74725 | ENSG00000112282 | MED23 | chr6 | - | 131624864 | 131624989 | 131623350 | 131623462 | 131627395 | 131627483 | 600 | 575 | 463 | [illegible] |
| SE | 108854 | ENSG00000117226 | GBP3 | chr1 | - | 89020125 | 89020211 | 89015286 | 89015414 | 89020531 | 89020743 | 8 | 15 | 19 | [illegible] |
| SE | 98003 | ENSG00000205758 | CRYZL1 | chr21 | - | 33631485 | 33631557 | 33624682 | 33624760 | 33641680 | 33641720 | 150 | 110 | 116 | [illegible] |
| SE | 6600 | ENSG00000134444 | RELCH | chr18 | + | 62280359 | 62280442 | 62275373 | 62275473 | 62280645 | 62280709 | 19 | 22 | 37 | [illegible] |
| SE | 84013 | ENSG00000111581 | NUP107 | chr12 | + | 68691967 | 68692112 | 68690467 | 68690746 | 68696818 | 68696910 | 180 | 98 | 126 | [illegible] |
| SE | 21433 | ENSG00000164180 | TMEM161B | chr5 | - | 88240812 | 88240916 | 88228440 | 88228528 | 88268720 | 88268799 | 80 | 61 | 121 | [illegible] |
| SE | 96994 | ENSG00000153339 | TRAPPC8 | chr18 | - | 31832083 | 31832173 | 31829196 | 31830989 | 31839311 | 31839457 | 265 | 185 | 186 | [illegible] |
| SE | 31001 | ENSG00000113504 | SLC12A7 | chr5 | - | 1088305 | 1088360 | 1086902 | 1087033 | 1093532 | 1093575 | 23 | 8 | 10 | [illegible] |
| SE | 102580 | ENSG00000120438 | TCP1 | chr6 | - | 159783940 | 159784067 | 159780934 | 159781110 | 159785899 | 159785997 | 577 | 480 | 468 | [illegible] |
| SE | 69622 | ENSG00000156711 | MAPK13 | chr6 | + | 36136018 | 36136048 | 36135752 | 36135861 | 36136483 | 36136515 | 89 | 28 | 91 | [illegible] |
| SE | 4399 | ENSG00000130475 | FCHO1 | chr19 | + | 17772644 | 17772741 | 17772456 | 17772555 | 17774238 | 17774241 | 198 | 168 | 214 | [illegible] |
| SE | 95860 | ENSG00000107263 | RAPGEF1 | chr9 | - | 131638634 | 131638791 | 131629101 | 131629254 | 131643238 | 131643391 | 518 | 247 | 373 | [illegible] |
| SE | 34969 | ENSG00000117133 | RPF1 | chr1 | + | 84495881 | 84496063 | 84495372 | 84495455 | 84496243 | 84496370 | 832 | 545 | 632 | [illegible] |
| SE | 107003 | ENSG00000133858 | ZFC3H1 | chr12 | - | 71642435 | 71642559 | 71638417 | 71638515 | 71644094 | 71644318 | 539 | 448 | 532 | [illegible] |
| SE | 90773 | ENSG00000174744 | BRMS1 | chr11 | - | 66340966 | 66341046 | 66340773 | 66340870 | 66341205 | 66341333 | 211 | 123 | 164 | [illegible] |
| SE | 22133 | ENSG00000175727 | MLXIP | chr12 | + | 122132291 | 122132383 | 122130843 | 122130933 | 122133347 | 122133987 | 64 | 48 | 99 | [illegible] |
| SE | 30347 | ENSG00000108591 | DRG2 | chr17 | + | 18099632 | 18099715 | 18099016 | 18099077 | 18100362 | 18100435 | 26 | 12 | 25 | [illegible] |
| SE | 3911 | ENSG00000072958 | AP1M1 | chr19 | + | 16234198 | 16234274 | 16233492 | 16233618 | 16234412 | 16234784 | 212 | 80 | 169 | [illegible] |
| SE | 54680 | ENSG00000070010 | UFD1 | chr22 | - | 19475469 | 19475602 | 19475067 | 19475100 | 19479082 | 19479172 | 95 | 54 | 76 | [illegible] |
| SE | 57839 | ENSG00000079246 | XRCC5 | chr2 | + | 216110466 | 216110551 | 216109350 | 216109457 | 216113015 | 216113129 | 1 | 0 | 0 | [illegible] |
| SE | 99153 | ENSG00000103544 | VPS35L | chr16 | + | 19611727 | 19611798 | 19610321 | 19610415 | 19616113 | 19616191 | 0 | 1 | 2 | [illegible] |
| SE | 42207 | ENSG00000132155 | RAF1 | chr3 | - | 12600151 | 12600279 | 12599690 | 12599808 | 12600387 | 12600415 | 412 | 303 | 437 | [illegible] |
| SE | 104232 | ENSG00000139620 | KANSL2 | chr12 | - | 48679035 | 48679238 | 48671798 | 48671962 | 48679654 | 48679833 | 85 | 62 | 93 | [illegible] |
| SE | 99377 | ENSG00000156599 | ZDHHC5 | chr11 | + | 57698558 | 57699418 | 57696760 | 57696873 | 57699865 | 57700363 | 182 | 109 | 184 | [illegible] |
| SE | 65299 | ENSG00000149499 | EML3 | chr11 | - | 62607665 | 62607821 | 62606957 | 62607099 | 62608200 | 62608296 | 244 | 156 | 232 | [illegible] |
| SE | 100102 | ENSG00000149262 | INTS4 | chr11 | - | 77987603 | 77987698 | 77981458 | 77981576 | 77991107 | 77991299 | 3 | 3 | 1 | [illegible] |
| SE | 47784 | ENSG00000154358 | OBSCN | chr1 | + | 228292522 | 228292786 | 228291897 | 228292161 | 228293352 | 228293616 | 24 | 30 | 79 | [illegible] |
| SE | 2404 | ENSG00000126247 | CAPNS1 | chr19 | + | 36146246 | 36146312 | 36145975 | 36146054 | 36149577 | 36149636 | 1083 | 514 | 570 | [illegible] |
| SE | 31560 | ENSG00000163913 | IFT122 | chr3 | + | 129461227 | 129461304 | 129451913 | 129451998 | 129463559 | 129463626 | 14 | 6 | 7 | [illegible] |
| SE | 99094 | ENSG00000110274 | CEP164 | chr11 | + | 117373750 | 117373831 | 117371079 | 117371466 | 117375707 | 117375791 | 79 | 46 | 53 | [illegible] |
| SE | 86245 | ENSG00000134516 | DOCK2 | chr5 | + | 169670541 | 169670597 | 169669287 | 169669328 | 169671077 | 169671174 | 480 | 310 | 293 | [illegible] |
| SE | 79034 | ENSG00000160218 | TRAPPC10 | chr21 | + | 44086801 | 44086960 | 44084121 | 44084263 | 44087698 | 44087928 | 499 | 219 | 367 | [illegible] |
| SE | 88452 | ENSG00000221914 | PPP2R2A | chr8 | + | 26360168 | 26360281 | 26353810 | 26354633 | 26366314 | 26366406 | 212 | 218 | 188 | [illegible] |
| SE | 86280 | ENSG00000134516 | DOCK2 | chr5 | + | 170075946 | 170076084 | 170067509 | 170067686 | 170077709 | 170077837 | 355 | 157 | 193 | [illegible] |
| SE | 7697 | ENSG00000170004 | CHD3 | chr17 | + | 7907600 | 7907702 | 7907352 | 7907488 | 7907890 | 7908019 | 141 | 136 | 178 | [illegible] |
| SE | 98001 | ENSG00000205758 | CRYZL1 | chr21 | - | 33624682 | 33624760 | 33621995 | 33622068 | 33631485 | 33631527 | 193 | 163 | 172 | [illegible] |
| SE | 22560 | ENSG00000167881 | SRP68 | chr17 | - | 76067216 | 76067330 | 76063975 | 76064171 | 76070377 | 76070436 | 256 | 207 | 273 | [illegible] |
| SE | 14279 | ENSG00000103249 | CLCN7 | chr16 | - | 1452754 | 1452893 | 1451622 | 1451716 | 1453833 | 1453894 | 310 | 217 | 267 | [illegible] |
| SE | 109050 | ENSG00000139668 | WDFY2 | chr13 | + | 51675169 | 51675243 | 51660595 | 51660663 | 51703595 | 51703650 | 149 | 145 | 142 | [illegible] |
| SE | 98047 | ENSG00000139697 | SBNO1 | chr12 | - | 123348028 | 123348133 | 123345257 | 123345570 | 123350309 | 123350441 | 158 | 128 | 176 | [illegible] |
| SE | 6056 | ENSG00000132792 | CTNNBL1 | chr20 | + | 37767952 | 37768044 | 37757558 | 37757656 | 37777344 | 37777417 | 98 | 51 | 91 | [illegible] |
| SE | 95864 | ENSG00000107263 | RAPGEF1 | chr9 | - | 131643238 | 131643417 | 131638634 | 131638791 | 131650128 | 131650242 | 727 | 349 | 490 | [illegible] |
| SE | 79450 | ENSG00000121680 | PEX16 | chr11 | - | 45910897 | 45910962 | 45909662 | 45910312 | 45913818 | 45913938 | 30 | 21 | 35 | [illegible] |
| SE | 89407 | ENSG00000133740 | E2F5 | chr8 | + | 85209141 | 85209409 | 85207424 | 85207489 | 85212156 | 85212204 | 87 | 46 | 71 | [illegible] |
| SE | 38001 | ENSG00000197386 | HTT | chr4 | + | 3099273 | 3099394 | 3086938 | 3087022 | 3105356 | 3105436 | 137 | 85 | 89 | [illegible] |
| SE | 54673 | ENSG00000070010 | UFD1 | chr22 | - | 19471686 | 19471808 | 19467402 | 19468003 | 19479082 | 19479144 | 191 | 103 | 207 | [illegible] |
| SE | 100101 | ENSG00000149262 | INTS4 | chr11 | - | 77987528 | 77987698 | 77981458 | 77981576 | 77991107 | 77991299 | 5 | 7 | 13 | [illegible] |
| SE | 7385 | ENSG00000101557 | USP14 | chr18 | + | 162982 | 163174 | 158556 | 158714 | 163307 | 163431 | 0 | 0 | 4 | [illegible] |
| SE | 87612 | ENSG00000102078 | SLC25A14 | chrX | + | 130350645 | 130350731 | 130349247 | 130349345 | 130358639 | 130358735 | 21 | 22 | 18 | [illegible] |
| SE | 38782 | ENSG00000162601 | MYSM1 | chr1 | - | 58689037 | 58689116 | 58685152 | 58685251 | 58690225 | 58690249 | 139 | 102 | 118 | [illegible] |
| SE | 107495 | ENSG00000123349 | PFDN5 | chr12 | + | 53295838 | 53296275 | 53295478 | 53295639 | 53297849 | 53297924 | 1395 | 849 | 1210 | [illegible] |
| SE | 79016 | ENSG00000160218 | TRAPPC10 | chr21 | + | 44052279 | 44052476 | 44037791 | 44037927 | 44055697 | 44055893 | 510 | 235 | 286 | [illegible] |
| SE | 30857 | ENSG00000109133 | TMEM33 | chr4 | + | 41939720 | 41939828 | 41939254 | 41939383 | 41943746 | 41943814 | 6 | 4 | 3 | [illegible] |
| SE | 104230 | ENSG00000139620 | KANSL2 | chr12 | - | 48662559 | 48662654 | 48660467 | 48660619 | 48667692 | 48667789 | 33 | 11 | 20 | [illegible] |
| SE | 108201 | ENSG00000186908 | ZDHHC17 | chr12 | + | 76821057 | 76821167 | 76815856 | 76816019 | 76822405 | 76822531 | 40 | 36 | 32 | [illegible] |
| SE | 22076 | ENSG00000174032 | SLC25A30 | chr13 | - | 45408926 | 45409074 | 45405882 | 45405977 | 45418299 | 45418380 | 94 | 52 | 79 | [illegible] |
| SE | 91163 | ENSG00000164941 | INTS8 | chr8 | + | 94836523 | 94836631 | 94831991 | 94832174 | 94838493 | 94838618 | 437 | 375 | 341 | [illegible] |
| SE | 91164 | ENSG00000164941 | INTS8 | chr8 | + | 94836523 | 94836631 | 94831991 | 94832174 | 94841490 | 94841591 | 432 | 372 | 332 | [illegible] |
| SE | 99374 | ENSG00000156599 | ZDHHC5 | chr11 | + | 57696760 | 57696873 | 57695919 | 57696043 | 57699865 | 57700363 | 106 | 48 | 96 | [illegible] |
| SE | 63083 | ENSG00000132953 | XPO4 | chr13 | - | 20862968 | 20863123 | 20862716 | 20862858 | 20868595 | 20868701 | 59 | 18 | 63 | [illegible] |
| SE | 64868 | ENSG00000170310 | STX8 | chr17 | - | 9505037 | 9505162 | 9491828 | 9491921 | 9545171 | 9545282 | 329 | 217 | 239 | [illegible] |
| SE | 24628 | ENSG00000157212 | PAXIP1 | chr7 | - | 154975695 | 154976331 | 154968402 | 154969126 | 154983218 | 154983332 | 108 | 75 | 83 | [illegible] |
| SE | 65354 | ENSG00000153214 | TMEM87B | chr2 | + | 112064161 | 112064253 | 112059976 | 112060037 | 112066935 | 112067067 | 134 | 105 | 119 | [illegible] |
| SE | 54748 | ENSG00000083720 | OXCT1 | chr5 | - | 41794676 | 41794749 | 41794002 | 41794078 | 41801021 | 41801070 | 49 | 34 | 47 | [illegible] |
| SE | 88405 | ENSG00000145041 | DCAF1 | chr3 | - | 51496733 | 51496780 | 51483718 | 51483836 | 51499872 | 51500015 | 31 | 24 | 42 | [illegible] |
| SE | 105695 | ENSG00000028203 | VEZT | chr12 | + | 95256549 | 95256622 | 95251939 | 95252071 | 95262905 | 95263081 | 44 | 41 | 28 | [illegible] |
| SE | 69654 | ENSG00000143390 | RFX5 | chr1 | - | 151345927 | 151345961 | 151344988 | 151345188 | 151346204 | 151346333 | 313 | 198 | 273 | [illegible] |
| SE | 42208 | ENSG00000132155 | RAF1 | chr3 | - | 12603477 | 12603537 | 12600387 | 12600415 | 12604135 | 12604289 | 3 | 0 | 0 | [illegible] |
| SE | 19133 | ENSG00000090061 | CCNK | chr14 | + | 99501355 | 99501413 | 99500765 | 99500871 | 99502206 | 99502312 | 484 | 304 | 401 | [illegible] |
| SE | 7698 | ENSG00000170004 | CHD3 | chr17 | + | 7907600 | 7907702 | 7907352 | 7907488 | 7907893 | 7908019 | 189 | 190 | 220 | [illegible] |
| SE | 99093 | ENSG00000110274 | CEP164 | chr11 | + | 117363428 | 117363506 | 117362403 | 117362538 | 117371079 | 117371466 | 26 | 8 | 36 | [illegible] |
| SE | 29418 | ENSG00000169762 | TAPT1 | chr4 | - | 16190046 | 16190270 | 16188231 | 16188355 | 16191360 | 16191523 | 6 | 0 | 1 | [illegible] |
| SE | 606 | ENSG00000099949 | LZTR1 | chr22 | + | 20988788 | 20988872 | 20988009 | 20988118 | 20989624 | 20989675 | 109 | 58 | 90 | [illegible] |
| SE | 806 | ENSG00000100147 | CCDC134 | chr22 | + | 41808874 | 41808993 | 41800678 | 41800766 | 41809878 | 41810000 | 31 | 13 | 24 | [illegible] |
| SE | 807 | ENSG00000100147 | CCDC134 | chr22 | + | 41810206 | 41810291 | 41809878 | 41810000 | 41813263 | 41813445 | 61 | 13 | 50 | [illegible] |
| SE | 808 | ENSG00000100147 | CCDC134 | chr22 | + | 41810206 | 41810291 | 41809878 | 41810000 | 41825697 | 41832164 | 19 | 4 | 11 | [illegible] |
| SE | 810 | ENSG00000100147 | CCDC134 | chr22 | + | 41813263 | 41813445 | 41810206 | 41810291 | 41813750 | 41813822 | 83 | 26 | 80 | [illegible] |
| SE | 811 | ENSG00000100147 | CCDC134 | chr22 | + | 41813263 | 41813445 | 41810206 | 41810291 | 41825697 | 41832164 | 43 | 9 | 40 | [illegible] |
| SE | 812 | ENSG00000100147 | CCDC134 | chr22 | + | 41813750 | 41813822 | 41810206 | 41810291 | 41825697 | 41832164 | 7 | 7 | 5 | [illegible] |
| SE | 813 | ENSG00000100147 | CCDC134 | chr22 | + | 41813750 | 41813822 | 41813263 | 41813445 | 41825697 | 41832164 | 48 | 24 | 45 | [illegible] |
| SE | 855 | ENSG00000185386 | MAPK11 | chr22 | - | 50267568 | 50267627 | 50267370 | 50267482 | 50267819 | 50267949 | 0 | 0 | 2 | [illegible] |
| SE | 858 | ENSG00000188130 | MAPK12 | chr22 | - | 50248365 | 50248472 | 50248224 | 50248292 | 50249102 | 50249454 | 5 | 4 | 5 | [illegible] |
| SE | 859 | ENSG00000188130 | MAPK12 | chr22 | - | 50248661 | 50248751 | 50248224 | 50248292 | 50249102 | 50249454 | 0 | 0 | 0 | [illegible] |
| SE | 860 | ENSG00000188130 | MAPK12 | chr22 | - | 50248661 | 50248751 | 50248365 | 50248472 | 50248830 | 50248890 | 20 | 1 | 15 | [illegible] |
| SE | 862 | ENSG00000188130 | MAPK12 | chr22 | - | 50248830 | 50248890 | 50248365 | 50248472 | 50249102 | 50249454 | 8 | 2 | 8 | [illegible] |
| SE | 864 | ENSG00000188130 | MAPK12 | chr22 | - | 50249102 | 50249454 | 50248365 | 50248472 | 50249634 | 50249703 | 20 | 13 | 11 | [illegible] |
| SE | 888 | ENSG00000093009 | CDC45 | chr22 | + | 19480952 | 19481045 | 19479737 | 19480217 | 19482689 | 19482766 | 4 | 1 | 11 | [illegible] |
| SE | 890 | ENSG00000093009 | CDC45 | chr22 | + | 19494473 | 19494602 | 19494326 | 19494382 | 19495980 | 19496029 | 0 | 0 | 0 | [illegible] |
| SE | 891 | ENSG00000093009 | CDC45 | chr22 | + | 19494506 | 19494602 | 19494326 | 19494382 | 19495980 | 19495986 | 0 | 0 | 0 | [illegible] |
| SE | 892 | ENSG00000093009 | CDC45 | chr22 | + | 19499100 | 19499151 | 19497385 | 19497447 | 19505361 | 19505370 | 0 | 0 | 13 | [illegible] |
| SE | 893 | ENSG00000093009 | CDC45 | chr22 | + | 19514748 | 19514887 | 19508529 | 19508691 | 19514964 | 19515048 | 1 | 2 | 5 | [illegible] |
| SE | 894 | ENSG00000093009 | CDC45 | chr22 | + | 19514748 | 19514887 | 19508529 | 19508691 | 19516526 | 19516645 | 1 | 2 | 4 | [illegible] |
| SE | 895 | ENSG00000093009 | CDC45 | chr22 | + | 19514964 | 19515048 | 19508529 | 19508691 | 19516526 | 19516645 | 0 | 2 | 3 | [illegible] |
| SE | 896 | ENSG00000093009 | CDC45 | chr22 | + | 19514964 | 19515048 | 19514748 | 19514887 | 19516526 | 19516645 | 0 | 2 | 4 | [illegible] |
| SE | 960 | ENSG00000100413 | POLR3H | chr22 | - | 41532603 | 41532745 | 41532093 | 41532157 | 41540698 | 41540795 | 35 | 7 | 29 | [illegible] |
| SE | 961 | ENSG00000100413 | POLR3H | chr22 | - | 41532658 | 41532745 | 41532093 | 41532157 | 41540698 | 41540795 | 72 | 46 | 67 | [illegible] |
| SE | 963 | ENSG00000100413 | POLR3H | chr22 | - | 41540698 | 41540795 | 41532603 | 41532745 | 41543990 | 41544121 | 93 | 37 | 82 | [illegible] |
| SE | 1089 | ENSG00000100030 | MAPK1 | chr22 | - | 21772872 | 21772982 | 21769039 | 21769320 | 21788256 | 21788388 | 649 | 284 | 458 | [illegible] |
| SE | 1091 | ENSG00000100030 | MAPK1 | chr22 | - | 21788256 | 21788388 | 21772872 | 21772982 | 21799011 | 21799128 | 423 | 230 | 336 | [illegible] |
| SE | 1092 | ENSG00000100030 | MAPK1 | chr22 | - | 21788693 | 21788808 | 21772872 | 21772982 | 21799011 | 21799128 | 426 | 176 | 300 | [illegible] |
| SE | 1094 | ENSG00000100030 | MAPK1 | chr22 | - | 21799011 | 21799128 | 21788693 | 21788808 | 21805849 | 21806039 | 806 | 366 | 567 | [illegible] |
| SE | 1557 | ENSG00000198089 | SFI1 | chr22 | + | 31528689 | 31528863 | 31508254 | 31508376 | 31531057 | 31531129 | 204 | 184 | 175 | [illegible] |
| SE | 1563 | ENSG00000198089 | SFI1 | chr22 | + | 31531057 | 31531129 | 31508254 | 31508376 | 31550253 | 31550348 | 1 | 0 | 0 | [illegible] |
| SE | 1565 | ENSG00000198089 | SFI1 | chr22 | + | 31531057 | 31531129 | 31528689 | 31528863 | 31550253 | 31550348 | 27 | 22 | 21 | [illegible] |
| SE | 1566 | ENSG00000198089 | SFI1 | chr22 | + | 31546860 | 31546971 | 31508254 | 31508376 | 31550253 | 31550348 | 198 | 163 | 256 | [illegible] |
| SE | 1569 | ENSG00000198089 | SFI1 | chr22 | + | 31546860 | 31546971 | 31528689 | 31528863 | 31556941 | 31557059 | 46 | 34 | 25 | [illegible] |
| SE | 1570 | ENSG00000198089 | SFI1 | chr22 | + | 31546860 | 31546971 | 31528689 | 31528863 | 31573057 | 31573214 | 46 | 35 | 25 | [illegible] |
| SE | 1573 | ENSG00000198089 | SFI1 | chr22 | + | 31550253 | 31550348 | 31528689 | 31528863 | 31556941 | 31557059 | 73 | 49 | 111 | [illegible] |
| SE | 1574 | ENSG00000198089 | SFI1 | chr22 | + | 31550253 | 31550348 | 31528689 | 31528863 | 31573057 | 31573214 | 1 | 1 | 0 | [illegible] |
| SE | 1575 | ENSG00000198089 | SFI1 | chr22 | + | 31550253 | 31550348 | 31546860 | 31546971 | 31556941 | 31557059 | 267 | 217 | 378 | [illegible] |
| SE | 1576 | ENSG00000198089 | SFI1 | chr22 | + | 31550253 | 31550348 | 31546860 | 31546971 | 31561289 | 31561333 | 173 | 141 | 238 | [illegible] |
| SE | 1577 | ENSG00000198089 | SFI1 | chr22 | + | 31550253 | 31550348 | 31546860 | 31546971 | 31573057 | 31573214 | 173 | 140 | 235 | [illegible] |
| SE | 1578 | ENSG00000198089 | SFI1 | chr22 | + | 31556941 | 31557059 | 31528689 | 31528863 | 31573057 | 31573214 | 7 | 0 | 5 | [illegible] |
| SE | 1580 | ENSG00000198089 | SFI1 | chr22 | + | 31556941 | 31557059 | 31546860 | 31546971 | 31573057 | 31573214 | 6 | 2 | 5 | [illegible] |
| SE | 1581 | ENSG00000198089 | SFI1 | chr22 | + | 31556941 | 31557059 | 31550253 | 31550348 | 31561289 | 31561333 | 228 | 157 | 243 | [illegible] |
| SE | 1583 | ENSG00000198089 | SFI1 | chr22 | + | 31561289 | 31561392 | 31546860 | 31546971 | 31573057 | 31573214 | 29 | 24 | 48 | [illegible] |
| SE | 1587 | ENSG00000198089 | SFI1 | chr22 | + | 31602211 | 31602293 | 31589446 | 31589577 | 31604308 | 31604404 | 16 | 25 | 36 | [illegible] |
| SE | 1589 | ENSG00000198089 | SFI1 | chr22 | + | 31602211 | 31602293 | 31589446 | 31589577 | 31606327 | 31606430 | 17 | 27 | 40 | [illegible] |
| SE | 1590 | ENSG00000198089 | SFI1 | chr22 | + | 31602211 | 31602293 | 31589446 | 31589577 | 31607936 | 31608033 | 16 | 25 | 36 | [illegible] |
| SE | 1591 | ENSG00000198089 | SFI1 | chr22 | + | 31602606 | 31602785 | 31602211 | 31602293 | 31604308 | 31604404 | 88 | 93 | 100 | [illegible] |
| SE | 1592 | ENSG00000198089 | SFI1 | chr22 | + | 31602606 | 31602785 | 31602211 | 31602293 | 31607936 | 31608033 | 84 | 87 | 91 | [illegible] |
| SE | 1593 | ENSG00000198089 | SFI1 | chr22 | + | 31603743 | 31603819 | 31589446 | 31589577 | 31604308 | 31604404 | 18 | 6 | 20 | [illegible] |
| SE | 1597 | ENSG00000198089 | SFI1 | chr22 | + | 31604308 | 31604404 | 31589446 | 31589577 | 31604868 | 31604945 | 9 | 4 | 12 | [illegible] |
| SE | 1598 | ENSG00000198089 | SFI1 | chr22 | + | 31604308 | 31604404 | 31589446 | 31589577 | 31606327 | 31606430 | 0 | 0 | 2 | [illegible] |
| SE | 1599 | ENSG00000198089 | SFI1 | chr22 | + | 31604308 | 31604404 | 31602211 | 31602293 | 31604868 | 31604945 | 9 | 4 | 12 | [illegible] |
| SE | 1600 | ENSG00000198089 | SFI1 | chr22 | + | 31604308 | 31604404 | 31602211 | 31602293 | 31606327 | 31606430 | 0 | 0 | 2 | [illegible] |
| SE | 1602 | ENSG00000198089 | SFI1 | chr22 | + | 31604308 | 31604404 | 31603743 | 31603819 | 31606327 | 31606430 | 29 | 22 | 39 | [illegible] |
| SE | 1605 | ENSG00000198089 | SFI1 | chr22 | + | 31604868 | 31604945 | 31603743 | 31603819 | 31606327 | 31606430 | 8 | 8 | 11 | [illegible] |
| SE | 1607 | ENSG00000198089 | SFI1 | chr22 | + | 31604868 | 31606430 | 31589446 | 31589577 | 31607936 | 31608033 | 17 | 13 | 26 | [illegible] |
| SE | 1608 | ENSG00000198089 | SFI1 | chr22 | + | 31604868 | 31606430 | 31602211 | 31602293 | 31607936 | 31608033 | 17 | 13 | 26 | [illegible] |
| SE | 1609 | ENSG00000198089 | SFI1 | chr22 | + | 31606327 | 31606430 | 31589446 | 31589577 | 31607936 | 31608033 | 17 | 16 | 29 | [illegible] |
| SE | 1610 | ENSG00000198089 | SFI1 | chr22 | + | 31606327 | 31606430 | 31602211 | 31602293 | 31607936 | 31608033 | 18 | 15 | 29 | [illegible] |
| SE | 1611 | ENSG00000198089 | SFI1 | chr22 | + | 31614788 | 31614860 | 31613601 | 31613855 | 31615047 | 31615279 | 38 | 5 | 45 | [illegible] |
| SE | 1725 | ENSG00000100325 | ASCC2 | chr22 | - | 29800990 | 29801110 | 29793359 | 29793676 | 29801993 | 29802208 | 91 | 54 | 55 | [illegible] |
| SE | 1728 | ENSG00000100325 | ASCC2 | chr22 | - | 29801993 | 29802208 | 29800990 | 29801110 | 29804637 | 29804824 | 95 | 58 | 68 | [illegible] |
| SE | 1729 | ENSG00000100325 | ASCC2 | chr22 | - | 29802114 | 29802208 | 29800990 | 29801110 | 29804637 | 29804830 | 52 | 25 | 38 | [illegible] |
| SE | 1730 | ENSG00000100325 | ASCC2 | chr22 | - | 29808110 | 29808185 | 29806796 | 29806904 | 29813429 | 29813542 | 116 | 46 | 107 | [illegible] |
| SE | 1731 | ENSG00000100325 | ASCC2 | chr22 | - | 29815171 | 29815289 | 29814656 | 29814767 | 29816005 | 29816073 | 4 | 0 | 0 | [illegible] |
| SE | 1733 | ENSG00000100325 | ASCC2 | chr22 | - | 29822334 | 29822464 | 29816005 | 29816073 | 29825086 | 29825257 | 206 | 90 | 177 | [illegible] |
| SE | 1736 | ENSG00000100325 | ASCC2 | chr22 | - | 29825086 | 29825257 | 29822141 | 29822464 | 29832244 | 29832342 | 109 | 49 | 105 | [illegible] |
| SE | 1737 | ENSG00000100325 | ASCC2 | chr22 | - | 29825086 | 29825257 | 29822141 | 29822464 | 29838177 | 29838274 | 108 | 49 | 102 | [illegible] |
| SE | 1739 | ENSG00000100325 | ASCC2 | chr22 | - | 29825621 | 29825780 | 29825086 | 29825257 | 29834441 | 29834550 | 56 | 30 | 64 | [illegible] |
| SE | 1741 | ENSG00000100325 | ASCC2 | chr22 | - | 29827476 | 29827600 | 29825621 | 29825780 | 29834488 | 29834550 | 2 | 1 | 0 | [illegible] |
| SE | 1743 | ENSG00000100325 | ASCC2 | chr22 | - | 29832244 | 29832342 | 29825086 | 29825257 | 29834441 | 29834550 | 5 | 0 | 4 | [illegible] |
| SE | 1748 | ENSG00000100325 | ASCC2 | chr22 | - | 29832244 | 29832342 | 29827476 | 29827600 | 29834488 | 29834550 | 11 | 5 | 4 | [illegible] |
| SE | 1749 | ENSG00000100325 | ASCC2 | chr22 | - | 29833540 | 29833614 | 29825621 | 29825780 | 29834488 | 29834550 | 1 | 0 | 1 | [illegible] |
| SE | 1750 | ENSG00000100325 | ASCC2 | chr22 | - | 29833540 | 29833614 | 29832244 | 29832342 | 29834488 | 29834550 | 2 | 1 | 1 | [illegible] |
| SE | 1751 | ENSG00000100325 | ASCC2 | chr22 | - | 29833540 | 29833614 | 29832244 | 29832342 | 29838177 | 29838228 | 0 | 1 | 0 | [illegible] |
| SE | 1753 | ENSG00000100325 | ASCC2 | chr22 | - | 29834441 | 29834550 | 29832244 | 29832342 | 29838177 | 29838274 | 5 | 0 | 0 | [illegible] |
| SE | 1754 | ENSG00000100325 | ASCC2 | chr22 | - | 29834488 | 29834550 | 29832244 | 29832342 | 29838177 | 29838261 | 11 | 3 | 6 | [illegible] |
| SE | 2274 | ENSG00000104824 | HNRNPL | chr19 | - | 38838893 | 38839015 | 38838396 | 38838598 | 38840487 | 38840559 | 366 | 187 | 347 | [illegible] |
| SE | 2275 | ENSG00000104824 | HNRNPL | chr19 | - | 38840095 | 38840376 | 38838396 | 38838598 | 38840487 | 38840559 | 270 | 162 | 235 | [illegible] |
| SE | 2276 | ENSG00000104824 | HNRNPL | chr19 | - | 38840095 | 38840376 | 38838893 | 38839015 | 38840487 | 38840559 | 471 | 261 | 458 | [illegible] |
| SE | 2278 | ENSG00000104824 | HNRNPL | chr19 | - | 38845852 | 38846090 | 38843370 | 38843914 | 38847315 | 38847434 | 382 | 188 | 312 | [illegible] |
| SE | 2402 | ENSG00000126247 | CAPNS1 | chr19 | + | 36142651 | 36142741 | 36142299 | 36142333 | 36142908 | 36142966 | 2229 | 930 | 1367 | [illegible] |
| SE | 2403 | ENSG00000126247 | CAPNS1 | chr19 | + | 36146195 | 36146312 | 36145975 | 36146054 | 36149577 | 36149580 | 2674 | 1207 | 1476 | [illegible] |
| SE | 2527 | ENSG00000178982 | EIF3K | chr19 | + | 38620336 | 38620435 | 38619185 | 38619327 | 38624076 | 38624197 | 2516 | 1056 | 2210 | [illegible] |
| SE | 2696 | ENSG00000005007 | UPF1 | chr19 | + | 18851049 | 18851129 | 18850687 | 18850868 | 18852134 | 18852182 | 0 | 1 | 0 | [illegible] |
| SE | 2697 | ENSG00000005007 | UPF1 | chr19 | + | 18860825 | 18860982 | 18859382 | 18860438 | 18862009 | 18862152 | 237 | 99 | 241 | [illegible] |
| SE | 2698 | ENSG00000005007 | UPF1 | chr19 | + | 18863437 | 18863612 | 18862009 | 18862152 | 18865288 | 18865450 | 101 | 37 | 92 | [illegible] |
| SE | 2699 | ENSG00000005007 | UPF1 | chr19 | + | 18864169 | 18864251 | 18862932 | 18863612 | 18865288 | 18865450 | 78 | 34 | 89 | [illegible] |
| SE | 3203 | ENSG00000013275 | PSMC4 | chr19 | + | 39972145 | 39972244 | 39971164 | 39971238 | 39972368 | 39972555 | 116 | 37 | 81 | [illegible] |
| SE | 3204 | ENSG00000013275 | PSMC4 | chr19 | + | 39972145 | 39972244 | 39971164 | 39971238 | 39972461 | 39972555 | 19 | 12 | 23 | [illegible] |
| SE | 3205 | ENSG00000013275 | PSMC4 | chr19 | + | 39980069 | 39980146 | 39979816 | 39979984 | 39980285 | 39980454 | 182 | 92 | 147 | [illegible] |
| SE | 3240 | ENSG00000180061 | TMEM150B | chr19 | - | 55322647 | 55322743 | 55320968 | 55321093 | 55325271 | 55325305 | 1 | 10 | 0 | [illegible] |
| SE | 3906 | ENSG00000072958 | AP1M1 | chr19 | + | 16202946 | 16203101 | 16198010 | 16198068 | 16203458 | 16203615 | 1 | 0 | 2 | [illegible] |
| SE | 3908 | ENSG00000072958 | AP1M1 | chr19 | + | 16202988 | 16203101 | 16197970 | 16198068 | 16203458 | 16203615 | 0 | 0 | 0 | [illegible] |
| SE | 3910 | ENSG00000072958 | AP1M1 | chr19 | + | 16228769 | 16228928 | 16228136 | 16228208 | 16233492 | 16233618 | 258 | 139 | 211 | [illegible] |
| SE | 4380 | ENSG00000064545 | TMEM161A | chr19 | - | 19134783 | 19134887 | 19132879 | 19133210 | 19138425 | 19138447 | 3 | 0 | 1 | [illegible] |
| SE | 4381 | ENSG00000064545 | TMEM161A | chr19 | - | 19134783 | 19134916 | 19133129 | 19133210 | 19138425 | 19138467 | 2 | 0 | 1 | [illegible] |
| SE | 4392 | ENSG00000130475 | FCHO1 | chr19 | + | 17754591 | 17754621 | 17754316 | 17754361 | 17755117 | 17755191 | 5 | 2 | 13 | [illegible] |
| SE | 4394 | ENSG00000130475 | FCHO1 | chr19 | + | 17755075 | 17755191 | 17747717 | 17747814 | 17762761 | 17762853 | 26 | 16 | 39 | [illegible] |
| SE | 4395 | ENSG00000130475 | FCHO1 | chr19 | + | 17755117 | 17755191 | 17754316 | 17754361 | 17762761 | 17762805 | 25 | 17 | 45 | [illegible] |
| SE | 4401 | ENSG00000130475 | FCHO1 | chr19 | + | 17776246 | 17776271 | 17775982 | 17776161 | 17776634 | 17776686 | 60 | 30 | 43 | [illegible] |
| SE | 4402 | ENSG00000130475 | FCHO1 | chr19 | + | 17776634 | 17776686 | 17776156 | 17776271 | 17778136 | 17778228 | 67 | 23 | 43 | [illegible] |
| SE | 4404 | ENSG00000130475 | FCHO1 | chr19 | + | 17778608 | 17778884 | 17778136 | 17778228 | 17781230 | 17781343 | 40 | 8 | 27 | [illegible] |
| SE | 4406 | ENSG00000130475 | FCHO1 | chr19 | + | 17784724 | 17784924 | 17784102 | 17784235 | 17786573 | 17786629 | 91 | 46 | 68 | [illegible] |
| SE | 4412 | ENSG00000105662 | CRTC1 | chr19 | + | 18744107 | 18744155 | 18742909 | 18743026 | 18745822 | 18745960 | 0 | 1 | 1 | [illegible] |
| SE | 4414 | ENSG00000105662 | CRTC1 | chr19 | + | 18768484 | 18768793 | 18765403 | 18765528 | 18775640 | 18775821 | 3 | 11 | 4 | [illegible] |
| SE | 4415 | ENSG00000105662 | CRTC1 | chr19 | + | 18771441 | 18771546 | 18768484 | 18768793 | 18775640 | 18775821 | 0 | 2 | 1 | [illegible] |
| SE | 4416 | ENSG00000105662 | CRTC1 | chr19 | + | 18774899 | 18774986 | 18765403 | 18765528 | 18775640 | 18775821 | 9 | 2 | 3 | [illegible] |
| SE | 4417 | ENSG00000105662 | CRTC1 | chr19 | + | 18774899 | 18774986 | 18771441 | 18771546 | 18775640 | 18775821 | 19 | 4 | 4 | [illegible] |
| SE | 4781 | ENSG00000064932 | SBNO2 | chr19 | - | 1122893 | 1123045 | 1122657 | 1122791 | 1123533 | 1123639 | 110 | 37 | 72 | [illegible] |
| SE | 4782 | ENSG00000064932 | SBNO2 | chr19 | - | 1123533 | 1123639 | 1122893 | 1123045 | 1123941 | 1124022 | 110 | 36 | 77 | [illegible] |
| SE | 5026 | ENSG00000101343 | CRNKL1 | chr20 | - | 20037322 | 20037534 | 20036005 | 20036362 | 20038348 | 20038452 | 134 | 77 | 96 | [illegible] |
| SE | 5027 | ENSG00000101343 | CRNKL1 | chr20 | - | 20037322 | 20037571 | 20034367 | 20036362 | 20038348 | 20038450 | 207 | 132 | 184 | [illegible] |
| SE | 5030 | ENSG00000101343 | CRNKL1 | chr20 | - | 20040685 | 20040766 | 20038348 | 20038450 | 20041565 | 20041625 | 67 | 64 | 70 | [illegible] |
| SE | 5032 | ENSG00000101343 | CRNKL1 | chr20 | - | 20043491 | 20043662 | 20042324 | 20042516 | 20045307 | 20045486 | 235 | 126 | 322 | [illegible] |

| | | | | | | | | | | | | | | | |
|---|---|---|---|---|---|---|---|---|---|---|---|---|---|---|---|
| SE | 5033 | ENSG00000173418 | NAA20 | chr20 | + | 20018361 | 20018517 | 20017309 | 20017449 | 20022455 | 20022480 | 1 | 0 | 1 | [illegible] |
| SE | 5035 | ENSG00000173418 | NAA20 | chr20 | + | 20025668 | 20025767 | 20017358 | 20017449 | 20026783 | 20026919 | 116 | 82 | 103 | [illegible] |
| SE | 5036 | ENSG00000173418 | NAA20 | chr20 | + | 20025668 | 20025767 | 20017358 | 20017449 | 20032507 | 20032653 | 0 | 0 | 0 | [illegible] |
| SE | 5037 | ENSG00000173418 | NAA20 | chr20 | + | 20026783 | 20026919 | 20017309 | 20017449 | 20032507 | 20032653 | 88 | 60 | 70 | [illegible] |
| SE | 5038 | ENSG00000173418 | NAA20 | chr20 | + | 20026783 | 20026919 | 20025668 | 20025767 | 20032507 | 20032653 | 274 | 220 | 252 | [illegible] |
| SE | 6054 | ENSG00000132792 | CTNNBL1 | chr20 | + | 37727301 | 37727402 | 37693954 | 37694152 | 37732878 | 37733067 | 1 | 0 | 0 | [illegible] |
| SE | 6055 | ENSG00000132792 | CTNNBL1 | chr20 | + | 37765196 | 37765290 | 37757558 | 37757656 | 37767952 | 37768044 | 189 | 166 | 185 | [illegible] |
| SE | 6057 | ENSG00000132792 | CTNNBL1 | chr20 | + | 37832247 | 37832345 | 37802866 | 37803048 | 37840101 | 37840199 | 2 | 0 | 0 | [illegible] |
| SE | 6083 | ENSG00000105576 | TNPO2 | chr19 | - | 12702071 | 12702177 | 12701751 | 12701851 | 12702822 | 12702918 | 17 | 12 | 15 | [illegible] |
| SE | 6084 | ENSG00000105576 | TNPO2 | chr19 | - | 12703713 | 12703801 | 12703427 | 12703526 | 12705239 | 12705398 | 46 | 33 | 43 | [illegible] |
| SE | 6085 | ENSG00000105576 | TNPO2 | chr19 | - | 12705491 | 12705599 | 12705239 | 12705398 | 12705681 | 12705768 | 53 | 46 | 52 | [illegible] |
| SE | 6086 | ENSG00000105576 | TNPO2 | chr19 | - | 12705681 | 12705768 | 12705491 | 12705599 | 12706195 | 12706367 | 29 | 32 | 33 | [illegible] |
| SE | 6087 | ENSG00000105576 | TNPO2 | chr19 | - | 12705681 | 12705830 | 12705491 | 12705599 | 12706195 | 12706367 | 23 | 24 | 20 | [illegible] |
| SE | 6089 | ENSG00000105576 | TNPO2 | chr19 | - | 12711295 | 12711461 | 12710620 | 12710773 | 12711552 | 12711613 | 97 | 59 | 92 | [illegible] |
| SE | 6090 | ENSG00000105576 | TNPO2 | chr19 | - | 12711552 | 12711613 | 12710620 | 12710773 | 12714820 | 12714939 | 13 | 17 | 23 | [illegible] |
| SE | 6554 | ENSG00000134265 | NAPG | chr18 | + | 10534465 | 10534496 | 10533535 | 10533553 | 10539761 | 10539871 | 138 | 61 | 70 | [illegible] |
| SE | 6586 | ENSG00000134444 | RELCH | chr18 | + | 62221036 | 62221108 | 62211152 | 62211242 | 62221218 | 62221274 | 127 | 84 | 91 | [illegible] |
| SE | 6587 | ENSG00000134444 | RELCH | chr18 | + | 62221218 | 62221274 | 62211152 | 62211242 | 62221383 | 62221497 | 86 | 54 | 64 | [illegible] |
| SE | 6588 | ENSG00000134444 | RELCH | chr18 | + | 62221383 | 62221497 | 62211152 | 62211242 | 62227288 | 62227450 | 189 | 147 | 146 | [illegible] |
| SE | 6589 | ENSG00000134444 | RELCH | chr18 | + | 62227288 | 62227492 | 62221383 | 62221497 | 62228304 | 62228598 | 221 | 169 | 176 | [illegible] |
| SE | 6591 | ENSG00000134444 | RELCH | chr18 | + | 62232331 | 62232427 | 62231193 | 62231269 | 62244763 | 62244876 | 212 | 187 | 239 | [illegible] |
| SE | 6596 | ENSG00000134444 | RELCH | chr18 | + | 62273979 | 62274086 | 62268868 | 62268948 | 62275373 | 62275473 | 131 | 118 | 123 | [illegible] |
| SE | 6597 | ENSG00000134444 | RELCH | chr18 | + | 62279773 | 62279856 | 62275373 | 62275473 | 62280359 | 62280442 | 5 | 7 | 8 | [illegible] |
| SE | 6598 | ENSG00000134444 | RELCH | chr18 | + | 62279773 | 62279856 | 62275373 | 62275473 | 62280645 | 62280709 | 5 | 10 | 5 | [illegible] |
| SE | 6599 | ENSG00000134444 | RELCH | chr18 | + | 62279773 | 62279856 | 62277595 | 62277655 | 62280359 | 62280442 | 0 | 3 | 3 | [illegible] |
| SE | 6601 | ENSG00000134444 | RELCH | chr18 | + | 62280359 | 62280442 | 62279773 | 62279856 | 62280645 | 62280709 | 9 | 14 | 28 | [illegible] |
| SE | 6602 | ENSG00000134444 | RELCH | chr18 | + | 62282305 | 62282442 | 62280645 | 62280709 | 62287350 | 62287467 | 68 | 81 | 86 | [illegible] |
| SE | 6603 | ENSG00000134444 | RELCH | chr18 | + | 62282305 | 62282444 | 62280645 | 62280709 | 62287350 | 62287467 | 140 | 159 | 161 | [illegible] |
| SE | 6605 | ENSG00000134444 | RELCH | chr18 | + | 62284238 | 62284361 | 62282315 | 62282444 | 62287350 | 62287467 | 3 | 4 | 7 | [illegible] |
| SE | 6606 | ENSG00000134444 | RELCH | chr18 | + | 62298789 | 62298860 | 62291542 | 62291631 | 62305413 | 62307112 | 304 | 325 | 260 | [illegible] |
| SE | 6746 | ENSG00000150477 | KIAA1328 | chr18 | + | 37172972 | 37173081 | 37160199 | 37160381 | 37222016 | 37222117 | 22 | 14 | 18 | [illegible] |
| SE | 7386 | ENSG00000101557 | USP14 | chr18 | + | 166786 | 166819 | 163307 | 163453 | 178932 | 179037 | 74 | 60 | 55 | [illegible] |
| SE | 7391 | ENSG00000101557 | USP14 | chr18 | + | 180235 | 180339 | 166786 | 166819 | 196636 | 196767 | 4 | 0 | 3 | [illegible] |
| SE | 7684 | ENSG00000170004 | CHD3 | chr17 | + | 7893804 | 7893935 | 7893285 | 7893569 | 7894126 | 7894265 | 183 | 137 | 278 | [illegible] |
| SE | 7687 | ENSG00000170004 | CHD3 | chr17 | + | 7897970 | 7898102 | 7897082 | 7897294 | 7898495 | 7898595 | 415 | 358 | 656 | [illegible] |
| SE | 7688 | ENSG00000170004 | CHD3 | chr17 | + | 7898495 | 7898595 | 7897970 | 7898102 | 7899010 | 7899202 | 308 | 236 | 509 | [illegible] |
| SE | 7689 | ENSG00000170004 | CHD3 | chr17 | + | 7899895 | 7900033 | 7899342 | 7899543 | 7900289 | 7900411 | 544 | 449 | 767 | [illegible] |
| SE | 7690 | ENSG00000170004 | CHD3 | chr17 | + | 7899901 | 7900033 | 7899342 | 7899543 | 7900289 | 7900411 | 249 | 203 | 311 | [illegible] |
| SE | 7691 | ENSG00000170004 | CHD3 | chr17 | + | 7900289 | 7900411 | 7899342 | 7899543 | 7900557 | 7900731 | 202 | 127 | 255 | [illegible] |
| SE | 7692 | ENSG00000170004 | CHD3 | chr17 | + | 7901243 | 7901375 | 7900851 | 7900993 | 7902609 | 7902727 | 314 | 224 | 509 | [illegible] |
| SE | 7695 | ENSG00000170004 | CHD3 | chr17 | + | 7906868 | 7907031 | 7906552 | 7906697 | 7907125 | 7907247 | 547 | 400 | 663 | [illegible] |
| SE | 7696 | ENSG00000170004 | CHD3 | chr17 | + | 7907352 | 7907488 | 7907125 | 7907247 | 7907893 | 7908019 | 186 | 173 | 251 | [illegible] |
| SE | 7699 | ENSG00000170004 | CHD3 | chr17 | + | 7907890 | 7908019 | 7907352 | 7907488 | 7909142 | 7909338 | 0 | 1 | 3 | [illegible] |
| SE | 7700 | ENSG00000170004 | CHD3 | chr17 | + | 7907893 | 7908019 | 7907352 | 7907488 | 7909142 | 7909338 | 50 | 49 | 80 | [illegible] |
| SE | 7701 | ENSG00000170004 | CHD3 | chr17 | + | 7910427 | 7910591 | 7909142 | 7909338 | 7910846 | 7910973 | 130 | 95 | 135 | [illegible] |
| SE | 7702 | ENSG00000170004 | CHD3 | chr17 | + | 7910438 | 7910591 | 7909142 | 7909338 | 7910846 | 7910895 | 52 | 38 | 59 | [illegible] |
| SE | 7703 | ENSG00000170004 | CHD3 | chr17 | + | 7910846 | 7910973 | 7909500 | 7910591 | 7911463 | 7911750 | 139 | 103 | 148 | [illegible] |
| SE | 8023 | ENSG00000073350 | LLGL2 | chr17 | + | 75533641 | 75533705 | 75525778 | 75525825 | 75543396 | 75543501 | 0 | 0 | 0 | [illegible] |
| SE | 8024 | ENSG00000073350 | LLGL2 | chr17 | + | 75538530 | 75538636 | 75525788 | 75525825 | 75543396 | 75543501 | 7 | 0 | 3 | [illegible] |
| SE | 8025 | ENSG00000073350 | LLGL2 | chr17 | + | 75538530 | 75538636 | 75533641 | 75533705 | 75543396 | 75543501 | 6 | 3 | 4 | [illegible] |
| SE | 8026 | ENSG00000073350 | LLGL2 | chr17 | + | 75564352 | 75564507 | 75563751 | 75563806 | 75568475 | 75568693 | 40 | 15 | 20 | [illegible] |
| SE | 8030 | ENSG00000073350 | LLGL2 | chr17 | + | 75574452 | 75574668 | 75574212 | 75574260 | 75574870 | 75575121 | 10 | 3 | 6 | [illegible] |
| SE | 8031 | ENSG00000073350 | LLGL2 | chr17 | + | 75574609 | 75574668 | 75574110 | 75574260 | 75574870 | 75574921 | 3 | 1 | 1 | [illegible] |
| SE | 8032 | ENSG00000073350 | LLGL2 | chr17 | + | 75574609 | 75574668 | 75574110 | 75574260 | 75574873 | 75574906 | 3 | 0 | 0 | [illegible] |
| SE | 8067 | ENSG00000214087 | ARL16 | chr17 | - | 81683012 | 81683126 | 81682033 | 81682149 | 81683535 | 81683594 | 37 | 11 | 14 | [illegible] |
| SE | 8274 | ENSG00000161654 | LSM12 | chr17 | - | 44037411 | 44037538 | 44034327 | 44036300 | 44040146 | 44040256 | 495 | 326 | 451 | [illegible] |
| SE | 8833 | ENSG00000108587 | GOSR1 | chr17 | + | 30481142 | 30481257 | 30477407 | 30477464 | 30484213 | 30484301 | 101 | 91 | 84 | [illegible] |
| SE | 8834 | ENSG00000108587 | GOSR1 | chr17 | + | 30481142 | 30481263 | 30477361 | 30477464 | 30484213 | 30484301 | 70 | 56 | 61 | [illegible] |
| SE | 8866 | ENSG00000180891 | CUEDC1 | chr17 | - | 57871285 | 57871369 | 57869121 | 57869193 | 57872662 | 57872670 | 26 | 24 | 16 | [illegible] |
| SE | 9260 | ENSG00000179029 | TMEM107 | chr17 | - | 8175958 | 8176026 | 8174002 | 8174272 | 8176199 | 8176316 | 3 | 1 | 1 | [illegible] |
| SE | 9890 | ENSG00000185813 | PCYT2 | chr17 | - | 81905381 | 81905447 | 81905065 | 81905154 | 81905669 | 81905735 | 21 | 6 | 16 | [illegible] |
| SE | 9891 | ENSG00000185813 | PCYT2 | chr17 | - | 81905669 | 81905735 | 81905381 | 81905447 | 81906099 | 81906177 | 25 | 5 | 20 | [illegible] |
| SE | 9894 | ENSG00000185813 | PCYT2 | chr17 | - | 81907553 | 81907598 | 81906759 | 81906898 | 81907772 | 81907794 | 32 | 13 | 36 | [illegible] |
| SE | 9895 | ENSG00000185813 | PCYT2 | chr17 | - | 81908875 | 81909037 | 81908567 | 81908634 | 81909513 | 81909602 | 53 | 29 | 43 | [illegible] |
| SE | 9896 | ENSG00000185813 | PCYT2 | chr17 | - | 81908875 | 81909321 | 81908567 | 81908634 | 81909513 | 81909602 | 46 | 18 | 34 | [illegible] |
| SE | 9897 | ENSG00000185813 | PCYT2 | chr17 | - | 81909161 | 81909321 | 81908875 | 81909037 | 81909513 | 81909602 | 1 | 0 | 0 | [illegible] |
| SE | 10340 | ENSG00000198496 | NBR2 | chr17 | + | 43132598 | 43132723 | 43131207 | 43131270 | 43132955 | 43133137 | 5 | 0 | 2 | [illegible] |
| SE | 10341 | ENSG00000198496 | NBR2 | chr17 | + | 43132955 | 43133137 | 43131207 | 43131270 | 43138155 | 43138360 | 4 | 1 | 1 | [illegible] |
| SE | 10343 | ENSG00000198496 | NBR2 | chr17 | + | 43138155 | 43138360 | 43131207 | 43131270 | 43138656 | 43138922 | 0 | 2 | 0 | [illegible] |
| SE | 10746 | ENSG00000131899 | LLGL1 | chr17 | + | 18241450 | 18241715 | 18240577 | 18240873 | 18241884 | 18241999 | 41 | 10 | 24 | [illegible] |
| SE | 10762 | ENSG00000118046 | STK11 | chr19 | + | 1220372 | 1220505 | 1219323 | 1219413 | 1220580 | 1220617 | 173 | 105 | 181 | [illegible] |
| SE | 10763 | ENSG00000118046 | STK11 | chr19 | + | 1223637 | 1226663 | 1223147 | 1223172 | 1227592 | 1228095 | 54 | 25 | 35 | [illegible] |
| SE | 10764 | ENSG00000118046 | STK11 | chr19 | + | 1226453 | 1226663 | 1222984 | 1223172 | 1227592 | 1228431 | 167 | 63 | 99 | [illegible] |
| SE | 11030 | ENSG00000072849 | DERL2 | chr17 | - | 5481295 | 5481389 | 5481100 | 5481142 | 5482808 | 5482879 | 78 | 53 | 78 | [illegible] |
| SE | 11031 | ENSG00000072849 | DERL2 | chr17 | - | 5485150 | 5485216 | 5480053 | 5480144 | 5486068 | 5486148 | 16 | 5 | 17 | [illegible] |
| SE | 11032 | ENSG00000072849 | DERL2 | chr17 | - | 5485150 | 5485216 | 5482269 | 5482882 | 5486068 | 5486148 | 96 | 65 | 98 | [illegible] |
| SE | 12089 | ENSG00000205250 | E2F4 | chr16 | + | 67194397 | 67194459 | 67192760 | 67192870 | 67194685 | 67194819 | 39 | 22 | 42 | [illegible] |
| SE | 12090 | ENSG00000205250 | E2F4 | chr16 | + | 67194397 | 67194459 | 67193421 | 67193515 | 67194685 | 67194819 | 201 | 95 | 168 | [illegible] |
| SE | 12271 | ENSG00000103194 | USP10 | chr16 | + | 84716194 | 84716518 | 84704739 | 84704860 | 84733434 | 84733503 | 0 | 3 | 0 | [illegible] |
| SE | 12272 | ENSG00000103194 | USP10 | chr16 | + | 84732467 | 84732589 | 84700007 | 84700111 | 84733052 | 84733105 | 0 | 0 | 0 | [illegible] |
| SE | 12273 | ENSG00000103194 | USP10 | chr16 | + | 84732467 | 84732589 | 84700007 | 84700111 | 84733434 | 84733503 | 0 | 0 | 0 | [illegible] |
| SE | 12276 | ENSG00000103194 | USP10 | chr16 | + | 84740308 | 84740369 | 84733137 | 84733503 | 84744632 | 84744780 | 84 | 115 | 82 | [illegible] |
| SE | 12279 | ENSG00000103194 | USP10 | chr16 | + | 84744632 | 84745673 | 84733434 | 84733503 | 84759890 | 84759946 | 4 | 2 | 10 | [illegible] |
| SE | 12282 | ENSG00000103194 | USP10 | chr16 | + | 84759890 | 84759946 | 84759277 | 84759472 | 84760171 | 84760236 | 101 | 73 | 118 | [illegible] |
| SE | 12284 | ENSG00000103194 | USP10 | chr16 | + | 84772540 | 84772685 | 84768192 | 84768358 | 84778894 | 84778987 | 202 | 155 | 174 | [illegible] |
| SE | 12285 | ENSG00000103194 | USP10 | chr16 | + | 84775159 | 84775225 | 84768264 | 84768358 | 84778894 | 84779015 | 42 | 29 | 46 | [illegible] |
| SE | 12286 | ENSG00000103194 | USP10 | chr16 | + | 84775159 | 84775225 | 84772540 | 84772685 | 84778894 | 84779015 | 270 | 245 | 286 | [illegible] |
| SE | 12390 | ENSG00000188559 | RALGAPA2 | chr20 | - | 20531686 | 20531795 | 20526251 | 20526362 | 20535744 | 20535803 | 87 | 51 | 106 | [illegible] |
| SE | 12391 | ENSG00000188559 | RALGAPA2 | chr20 | - | 20535744 | 20535803 | 20531686 | 20531795 | 20536655 | 20536784 | 68 | 37 | 86 | [illegible] |
| SE | 12394 | ENSG00000188559 | RALGAPA2 | chr20 | - | 20536655 | 20536784 | 20531686 | 20531795 | 20571457 | 20571613 | 2 | 0 | 2 | [illegible] |
| SE | 12395 | ENSG00000188559 | RALGAPA2 | chr20 | - | 20536655 | 20536784 | 20535744 | 20535803 | 20546703 | 20546832 | 119 | 71 | 107 | [illegible] |
| SE | 12396 | ENSG00000188559 | RALGAPA2 | chr20 | - | 20546703 | 20546832 | 20531686 | 20531795 | 20571457 | 20571613 | 61 | 52 | 86 | [illegible] |
| SE | 12400 | ENSG00000188559 | RALGAPA2 | chr20 | - | 20619276 | 20619414 | 20616042 | 20616191 | 20620462 | 20620630 | 93 | 114 | 122 | [illegible] |
| SE | 12401 | ENSG00000188559 | RALGAPA2 | chr20 | - | 20620462 | 20620630 | 20616042 | 20616191 | 20629362 | 20629590 | 92 | 102 | 126 | [illegible] |
| SE | 12402 | ENSG00000188559 | RALGAPA2 | chr20 | - | 20629362 | 20629590 | 20620462 | 20620630 | 20635417 | 20635617 | 200 | 225 | 215 | [illegible] |
| SE | 12404 | ENSG00000188559 | RALGAPA2 | chr20 | - | 20653529 | 20653587 | 20643505 | 20643549 | 20676235 | 20676288 | 69 | 71 | 74 | [illegible] |
| SE | 12405 | ENSG00000188559 | RALGAPA2 | chr20 | - | 20653529 | 20653587 | 20643505 | 20643549 | 20712374 | 20712644 | 23 | 30 | 34 | [illegible] |
| SE | 12408 | ENSG00000188559 | RALGAPA2 | chr20 | - | 20676235 | 20676288 | 20653529 | 20653587 | 20712374 | 20712644 | 14 | 18 | 9 | [illegible] |
| SE | 12409 | ENSG00000188559 | RALGAPA2 | chr20 | - | 20680690 | 20680801 | 20653529 | 20653587 | 20712374 | 20712644 | 11 | 10 | 12 | [illegible] |
| SE | 12410 | ENSG00000188559 | RALGAPA2 | chr20 | - | 20680690 | 20680801 | 20676235 | 20676288 | 20712374 | 20712644 | 63 | 47 | 47 | [illegible] |
| SE | 12445 | ENSG00000103043 | VAC14 | chr16 | - | 70698636 | 70698811 | 70696889 | 70697257 | 70731494 | 70731627 | 143 | 67 | 100 | [illegible] |
| SE | 12446 | ENSG00000103043 | VAC14 | chr16 | - | 70784775 | 70784838 | 70784112 | 70784220 | 70785701 | 70785869 | 79 | 38 | 58 | [illegible] |
| SE | 12926 | ENSG00000197818 | SLC9A8 | chr20 | + | 49815007 | 49815189 | 49812712 | 49812948 | 49823060 | 49823141 | 84 | 58 | 80 | [illegible] |
| SE | 12927 | ENSG00000197818 | SLC9A8 | chr20 | + | 49823060 | 49823141 | 49815007 | 49815189 | 49839540 | 49839599 | 72 | 55 | 65 | [illegible] |
| SE | 12929 | ENSG00000197818 | SLC9A8 | chr20 | + | 49845035 | 49845119 | 49839540 | 49839599 | 49849578 | 49849680 | 49 | 43 | 53 | [illegible] |
| SE | 12930 | ENSG00000197818 | SLC9A8 | chr20 | + | 49845035 | 49845119 | 49839540 | 49839599 | 49855437 | 49855581 | 16 | 12 | 16 | [illegible] |
| SE | 12931 | ENSG00000197818 | SLC9A8 | chr20 | + | 49849578 | 49849680 | 49839540 | 49839599 | 49855437 | 49855581 | 6 | 7 | 9 | [illegible] |
| SE | 12936 | ENSG00000197818 | SLC9A8 | chr20 | + | 49877980 | 49878063 | 49874704 | 49874821 | 49880923 | 49881035 | 44 | 40 | 54 | [illegible] |
| SE | 12990 | ENSG00000122390 | NAA60 | chr16 | + | 3447481 | 3447620 | 3443687 | 3443893 | 3448470 | 3448540 | 5 | 4 | 1 | [illegible] |
| SE | 12992 | ENSG00000122390 | NAA60 | chr16 | + | 3448470 | 3448540 | 3443677 | 3443827 | 3476221 | 3476337 | 13 | 22 | 29 | [illegible] |
| SE | 12997 | ENSG00000122390 | NAA60 | chr16 | + | 3467676 | 3468245 | 3458075 | 3458181 | 3476221 | 3476299 | 0 | 1 | 0 | [illegible] |
| SE | 13002 | ENSG00000122390 | NAA60 | chr16 | + | 3476221 | 3476337 | 3457984 | 3458181 | 3479470 | 3479591 | 183 | 96 | 140 | [illegible] |
| SE | 13003 | ENSG00000122390 | NAA60 | chr16 | + | 3476221 | 3476337 | 3457984 | 3458181 | 3482501 | 3482598 | 31 | 20 | 20 | [illegible] |
| SE | 13004 | ENSG00000122390 | NAA60 | chr16 | + | 3479470 | 3479600 | 3448470 | 3448540 | 3482501 | 3482540 | 103 | 44 | 72 | [illegible] |
| SE | 13005 | ENSG00000122390 | NAA60 | chr16 | + | 3479470 | 3479600 | 3457992 | 3458181 | 3482501 | 3482540 | 104 | 44 | 70 | [illegible] |
| SE | 13006 | ENSG00000122390 | NAA60 | chr16 | + | 3479470 | 3479600 | 3476221 | 3476337 | 3482501 | 3482540 | 258 | 128 | 191 | [illegible] |
| SE | 13007 | ENSG00000122390 | NAA60 | chr16 | + | 3482501 | 3482598 | 3479470 | 3479600 | 3483362 | 3483385 | 183 | 90 | 122 | [illegible] |
| SE | 13008 | ENSG00000122390 | NAA60 | chr16 | + | 3482501 | 3482598 | 3479470 | 3479600 | 3484690 | 3485083 | 51 | 28 | 40 | [illegible] |
| SE | 13009 | ENSG00000122390 | NAA60 | chr16 | + | 3484690 | 3485083 | 3482501 | 3482598 | 3485466 | 3485578 | 3 | 0 | 3 | [illegible] |
| SE | 13010 | ENSG00000122390 | NAA60 | chr16 | + | 3484761 | 3485061 | 3482501 | 3482598 | 3485466 | 3485604 | 12 | 4 | 7 | [illegible] |
| SE | 13011 | ENSG00000122390 | NAA60 | chr16 | + | 3484761 | 3485083 | 3482501 | 3482598 | 3485466 | 3485786 | 3 | 0 | 4 | [illegible] |
| SE | 13283 | ENSG00000051523 | CYBA | chr16 | - | 88646519 | 88646603 | 88646131 | 88646197 | 88646754 | 88646838 | 4 | 0 | 2 | [illegible] |
| SE | 13284 | ENSG00000051523 | CYBA | chr16 | - | 88646754 | 88646838 | 88645086 | 88646197 | 88647100 | 88647175 | 1380 | 446 | 688 | [illegible] |
| SE | 13614 | ENSG00000038358 | EDC4 | chr16 | + | 67881496 | 67881533 | 67881264 | 67881417 | 67881667 | 67881845 | 109 | 40 | 79 | [illegible] |
| SE | 13615 | ENSG00000038358 | EDC4 | chr16 | + | 67881667 | 67882109 | 67881496 | 67881533 | 67882211 | 67882327 | 125 | 53 | 103 | [illegible] |
| SE | 13619 | ENSG00000038358 | EDC4 | chr16 | + | 67883114 | 67883177 | 67882957 | 67883006 | 67883567 | 67883731 | 12 | 3 | 7 | [illegible] |
| SE | 13890 | ENSG00000069329 | VPS35 | chr16 | - | 46662982 | 46663162 | 46662242 | 46662482 | 46668929 | 46669052 | 381 | 286 | 281 | [illegible] |
| SE | 13893 | ENSG00000069329 | VPS35 | chr16 | - | 46674563 | 46674660 | 46674313 | 46674462 | 46676582 | 46676692 | 619 | 416 | 510 | [illegible] |
| SE | 13894 | ENSG00000069329 | VPS35 | chr16 | - | 46677314 | 46677398 | 46676504 | 46676692 | 46678942 | 46679043 | 479 | 322 | 397 | [illegible] |
| SE | 13895 | ENSG00000069329 | VPS35 | chr16 | - | 46678942 | 46679156 | 46677314 | 46677398 | 46680670 | 46680853 | 870 | 778 | 703 | [illegible] |
| SE | 13896 | ENSG00000069329 | VPS35 | chr16 | - | 46682477 | 46682666 | 46682078 | 46682175 | 46683507 | 46683606 | 0 | 0 | 0 | [illegible] |
| SE | 14220 | ENSG00000167965 | MLST8 | chr16 | + | 2207856 | 2208024 | 2207196 | 2207345 | 2208209 | 2208328 | 0 | 0 | 2 | [illegible] |
| SE | 14221 | ENSG00000167965 | MLST8 | chr16 | + | 2207856 | 2208024 | 2207196 | 2207345 | 2208449 | 2209412 | 0 | 2 | 0 | [illegible] |
| SE | 14280 | ENSG00000103249 | CLCN7 | chr16 | - | 1457693 | 1457756 | 1457253 | 1457337 | 1459106 | 1459187 | 119 | 73 | 77 | [illegible] |
| SE | 14281 | ENSG00000103249 | CLCN7 | chr16 | - | 1465266 | 1465338 | 1461602 | 1461674 | 1474833 | 1475025 | 49 | 35 | 54 | [illegible] |
| SE | 14340 | ENSG00000007545 | CRAMP1 | chr16 | + | 1666072 | 1666177 | 1665056 | 1665138 | 1666421 | 1666513 | 38 | 8 | 12 | [illegible] |
| SE | 14498 | ENSG00000181827 | RFX7 | chr15 | - | 56101358 | 56101566 | 56098080 | 56098376 | 56102168 | 56102253 | 87 | 97 | 89 | [illegible] |
| SE | 14499 | ENSG00000181827 | RFX7 | chr15 | - | 56160715 | 56160747 | 56144400 | 56144483 | 56179058 | 56179133 | 0 | 0 | 0 | [illegible] |
| SE | 14501 | ENSG00000181827 | RFX7 | chr15 | - | 56179269 | 56179303 | 56144400 | 56144483 | 56243124 | 56243287 | 49 | 27 | 37 | [illegible] |
| SE | 14827 | ENSG00000069943 | PIGB | chr15 | + | 55324779 | 55324834 | 55319272 | 55319413 | 55327530 | 55327635 | 2 | 3 | 1 | [illegible] |
| SE | 14830 | ENSG00000069943 | PIGB | chr15 | + | 55327530 | 55327635 | 55321269 | 55321390 | 55329723 | 55329854 | 30 | 23 | 30 | [illegible] |
| SE | 14831 | ENSG00000069943 | PIGB | chr15 | + | 55329723 | 55329854 | 55321269 | 55321390 | 55333866 | 55334007 | 35 | 26 | 38 | [illegible] |
| SE | 14832 | ENSG00000069943 | PIGB | chr15 | + | 55333866 | 55334007 | 55329723 | 55329854 | 55339266 | 55339289 | 56 | 42 | 55 | [illegible] |
| SE | 15273 | ENSG00000166262 | FAM227B | chr15 | - | 49576740 | 49576845 | 49574737 | 49575109 | 49577628 | 49577664 | 9 | 2 | 3 | [illegible] |
| SE | 15274 | ENSG00000166262 | FAM227B | chr15 | - | 49577171 | 49577305 | 49576493 | 49576845 | 49577628 | 49577664 | 4 | 0 | 3 | [illegible] |
| SE | 15275 | ENSG00000166262 | FAM227B | chr15 | - | 49577628 | 49577664 | 49574737 | 49575109 | 49588015 | 49588083 | 6 | 8 | 20 | [illegible] |
| SE | 15276 | ENSG00000166262 | FAM227B | chr15 | - | 49577628 | 49577664 | 49577171 | 49577305 | 49588015 | 49588083 | 6 | 8 | 22 | [illegible] |
| SE | 15277 | ENSG00000166262 | FAM227B | chr15 | - | 49588015 | 49588083 | 49577628 | 49577664 | 49611214 | 49611268 | 7 | 4 | 4 | [illegible] |
| SE | 15278 | ENSG00000166262 | FAM227B | chr15 | - | 49589775 | 49590007 | 49588015 | 49588083 | 49606155 | 49606218 | 31 | 23 | 40 | [illegible] |
| SE | 15279 | ENSG00000166262 | FAM227B | chr15 | - | 49589775 | 49590007 | 49588015 | 49588083 | 49611214 | 49611268 | 35 | 25 | 45 | [illegible] |
| SE | 15280 | ENSG00000166262 | FAM227B | chr15 | - | 49606155 | 49606218 | 49588015 | 49588083 | 49611214 | 49611268 | 2 | 1 | 0 | [illegible] |
| SE | 15281 | ENSG00000166262 | FAM227B | chr15 | - | 49606155 | 49606218 | 49589775 | 49590007 | 49611214 | 49611268 | 5 | 3 | 7 | [illegible] |
| SE | 15282 | ENSG00000166262 | FAM227B | chr15 | - | 49620133 | 49620190 | 49615120 | 49615243 | 49620630 | 49620747 | 0 | 0 | 1 | [illegible] |
| SE | 15283 | ENSG00000166262 | FAM227B | chr15 | - | 49620133 | 49620190 | 49615120 | 49615243 | 49620699 | 49620818 | 0 | 0 | 2 | [illegible] |
| SE | 15458 | ENSG00000140199 | SLC12A6 | chr15 | - | 34238964 | 34239160 | 34238231 | 34238331 | 34240660 | 34240829 | 152 | 111 | 203 | [illegible] |
| SE | 15459 | ENSG00000140199 | SLC12A6 | chr15 | - | 34238964 | 34239160 | 34238231 | 34238331 | 34241232 | 34241337 | 9 | 0 | 2 | [illegible] |
| SE | 15461 | ENSG00000140199 | SLC12A6 | chr15 | - | 34240660 | 34240829 | 34238231 | 34238331 | 34241232 | 34241337 | 97 | 52 | 78 | [illegible] |
| SE | 15464 | ENSG00000140199 | SLC12A6 | chr15 | - | 34245284 | 34245403 | 34243973 | 34244072 | 34245692 | 34245867 | 216 | 137 | 227 | [illegible] |
| SE | 15466 | ENSG00000140199 | SLC12A6 | chr15 | - | 34260925 | 34261020 | 34255261 | 34255392 | 34275344 | 34275389 | 176 | 157 | 231 | [illegible] |
| SE | 15467 | ENSG00000140199 | SLC12A6 | chr15 | - | 34260925 | 34261020 | 34258812 | 34258944 | 34275344 | 34275389 | 388 | 380 | 535 | [illegible] |
| SE | 15468 | ENSG00000140199 | SLC12A6 | chr15 | - | 34260925 | 34261020 | 34258812 | 34258944 | 34336409 | 34336844 | 191 | 202 | 258 | [illegible] |
| SE | 15469 | ENSG00000140199 | SLC12A6 | chr15 | - | 34275344 | 34275389 | 34258812 | 34258944 | 34336409 | 34336601 | 80 | 55 | 73 | [illegible] |
| SE | 15772 | ENSG00000167077 | MEI1 | chr22 | + | 41753948 | 41754046 | 41752459 | 41752651 | 41758364 | 41758533 | 0 | 1 | 8 | [illegible] |
| SE | 15773 | ENSG00000167077 | MEI1 | chr22 | + | 41758364 | 41758533 | 41753948 | 41754046 | 41763173 | 41763321 | 0 | 2 | 5 | [illegible] |
| SE | 15774 | ENSG00000167077 | MEI1 | chr22 | + | 41758364 | 41758547 | 41753948 | 41754046 | 41763173 | 41763321 | 0 | 1 | 4 | [illegible] |
| SE | 15776 | ENSG00000167077 | MEI1 | chr22 | + | 41778707 | 41778812 | 41776048 | 41776267 | 41781283 | 41781394 | 0 | 0 | 1 | [illegible] |
| SE | 15777 | ENSG00000167077 | MEI1 | chr22 | + | 41778707 | 41778877 | 41776046 | 41776267 | 41781283 | 41781363 | 0 | 0 | 0 | [illegible] |
| SE | 15779 | ENSG00000167077 | MEI1 | chr22 | + | 41784607 | 41784783 | 41784338 | 41784420 | 41793828 | 41793910 | 27 | 3 | 33 | [illegible] |
| SE | 15780 | ENSG00000167077 | MEI1 | chr22 | + | 41794370 | 41794477 | 41793828 | 41793910 | 41795734 | 41795847 | 14 | 7 | 13 | [illegible] |
| SE | 15782 | ENSG00000103978 | TMEM87A | chr15 | - | 42226805 | 42226909 | 42220061 | 42220135 | 42227710 | 42227769 | 291 | 193 | 244 | [illegible] |
| SE | 15783 | ENSG00000103978 | TMEM87A | chr15 | - | 42233212 | 42233306 | 42231191 | 42231260 | 42236319 | 42236419 | 116 | 73 | 103 | [illegible] |
| SE | 15784 | ENSG00000103978 | TMEM87A | chr15 | - | 42233212 | 42233306 | 42231191 | 42231260 | 42237431 | 42237475 | 39 | 21 | 47 | [illegible] |
| SE | 15785 | ENSG00000103978 | TMEM87A | chr15 | - | 42236319 | 42236419 | 42231191 | 42231260 | 42237431 | 42237475 | 44 | 17 | 37 | [illegible] |
| SE | 15786 | ENSG00000103978 | TMEM87A | chr15 | - | 42236319 | 42236419 | 42232878 | 42233306 | 42237431 | 42237475 | 123 | 72 | 90 | [illegible] |
| SE | 15787 | ENSG00000103978 | TMEM87A | chr15 | - | 42239669 | 42239731 | 42237431 | 42237615 | 42244049 | 42244167 | 101 | 31 | 59 | [illegible] |
| SE | 15836 | ENSG00000104177 | MYEF2 | chr15 | - | 48149031 | 48149083 | 48134631 | 48143071 | 48149162 | 48149326 | 13 | 25 | 20 | [illegible] |
| SE | 15837 | ENSG00000104177 | MYEF2 | chr15 | - | 48151099 | 48151171 | 48149162 | 48149371 | 48151472 | 48151571 | 12 | 8 | 12 | [illegible] |
| SE | 15838 | ENSG00000104177 | MYEF2 | chr15 | - | 48151873 | 48151942 | 48151472 | 48151571 | 48152233 | 48152280 | 14 | 9 | 11 | [illegible] |
| SE | 15839 | ENSG00000104177 | MYEF2 | chr15 | - | 48152233 | 48152284 | 48151873 | 48151942 | 48153791 | 48153893 | 13 | 5 | 12 | [illegible] |
| SE | 15840 | ENSG00000104177 | MYEF2 | chr15 | - | 48167348 | 48167401 | 48166120 | 48166128 | 48168630 | 48168839 | 29 | 27 | 22 | [illegible] |
| SE | 15898 | ENSG00000140577 | CRTC3 | chr15 | + | 90540038 | 90540137 | 90529922 | 90530203 | 90593635 | 90593755 | 39 | 20 | 45 | [illegible] |
| SE | 15899 | ENSG00000140577 | CRTC3 | chr15 | + | 90604384 | 90604447 | 90602323 | 90602385 | 90614452 | 90614488 | 23 | 22 | 24 | [illegible] |
| SE | 15902 | ENSG00000140577 | CRTC3 | chr15 | + | 90617882 | 90617968 | 90614452 | 90614488 | 90619740 | 90619790 | 189 | 107 | 189 | [illegible] |
| SE | 15903 | ENSG00000140577 | CRTC3 | chr15 | + | 90619740 | 90619790 | 90617882 | 90617968 | 90625775 | 90625786 | 223 | 99 | 179 | [illegible] |
| SE | 15904 | ENSG00000140577 | CRTC3 | chr15 | + | 90629233 | 90629532 | 90625775 | 90625993 | 90638445 | 90638646 | 70 | 64 | 113 | [illegible] |
| SE | 15915 | ENSG00000103852 | TTC23 | chr15 | - | 99161739 | 99161867 | 99155363 | 99156297 | 99175049 | 99175120 | 2 | 2 | 9 | [illegible] |
| SE | 15922 | ENSG00000103852 | TTC23 | chr15 | - | 99245388 | 99245510 | 99228532 | 99228732 | 99249621 | 99249799 | 1 | 0 | 0 | [illegible] |
| SE | 15924 | ENSG00000103852 | TTC23 | chr15 | - | 99245388 | 99245510 | 99241364 | 99241559 | 99249416 | 99249518 | 4 | 0 | 0 | [illegible] |
| SE | 16197 | ENSG00000140598 | EFL1 | chr15 | - | 82138657 | 82138842 | 82130232 | 82130561 | 82157712 | 82157860 | 67 | 57 | 43 | [illegible] |
| SE | 16199 | ENSG00000140598 | EFL1 | chr15 | - | 82151464 | 82152423 | 82138657 | 82138842 | 82163852 | 82163984 | 75 | 57 | 72 | [illegible] |
| SE | 16200 | ENSG00000140598 | EFL1 | chr15 | - | 82157712 | 82157860 | 82138657 | 82138842 | 82163852 | 82163984 | 35 | 32 | 29 | [illegible] |
| SE | 16201 | ENSG00000140598 | EFL1 | chr15 | - | 82157712 | 82157860 | 82151464 | 82152423 | 82163852 | 82163984 | 99 | 78 | 58 | [illegible] |
| SE | 16202 | ENSG00000140598 | EFL1 | chr15 | - | 82225164 | 82225264 | 82220077 | 82220229 | 82227449 | 82227572 | 121 | 80 | 78 | [illegible] |
| SE | 16204 | ENSG00000140598 | EFL1 | chr15 | - | 82240417 | 82240555 | 82238306 | 82238521 | 82241269 | 82241403 | 198 | 111 | 141 | [illegible] |
| SE | 16206 | ENSG00000140598 | EFL1 | chr15 | - | 82259087 | 82259155 | 82241269 | 82241403 | 82261687 | 82261797 | 53 | 33 | 40 | [illegible] |
| SE | 16659 | ENSG00000174444 | RPL4 | chr15 | - | 66500292 | 66500376 | 66500055 | 66500176 | 66500948 | 66501105 | 12187 | 7612 | 17237 | [illegible] |
| SE | 16660 | ENSG00000174444 | RPL4 | chr15 | - | 66501787 | 66501912 | 66500948 | 66501504 | 66502611 | 66502750 | 15659 | 8907 | 21174 | [illegible] |
| SE | 16661 | ENSG00000174444 | RPL4 | chr15 | - | 66502611 | 66502871 | 66501787 | 66501912 | 66503357 | 66503529 | 6093 | 3376 | 7861 | [illegible] |
| SE | 16662 | ENSG00000174444 | RPL4 | chr15 | - | 66502611 | 66503164 | 66501787 | 66501912 | 66503357 | 66503529 | 7749 | 4123 | 9667 | [illegible] |
| SE | 16693 | ENSG00000137841 | PLCB2 | chr15 | - | 40292338 | 40292443 | 40291873 | 40292158 | 40292925 | 40293025 | 338 | 184 | 212 | [illegible] |
| SE | 16694 | ENSG00000137841 | PLCB2 | chr15 | - | 40296521 | 40296634 | 40296295 | 40296392 | 40296745 | 40296908 | 350 | 230 | 272 | [illegible] |
| SE | 16696 | ENSG00000137841 | PLCB2 | chr15 | - | 40296745 | 40296908 | 40296295 | 40296392 | 40297520 | 40297605 | 201 | 110 | 107 | [illegible] |
| SE | 16697 | ENSG00000137841 | PLCB2 | chr15 | - | 40296757 | 40296908 | 40296295 | 40296392 | 40297520 | 40297605 | 194 | 110 | 107 | [illegible] |
| SE | 16698 | ENSG00000137841 | PLCB2 | chr15 | - | 40298797 | 40298964 | 40298561 | 40298708 | 40299127 | 40299228 | 918 | 604 | 713 | [illegible] |
| SE | 16699 | ENSG00000137841 | PLCB2 | chr15 | - | 40299127 | 40299228 | 40298797 | 40298964 | 40301956 | 40302032 | 544 | 326 | 427 | [illegible] |
| SE | 16700 | ENSG00000137841 | PLCB2 | chr15 | - | 40302135 | 40302189 | 40301956 | 40302032 | 40302269 | 40302349 | 347 | 150 | 199 | [illegible] |
| SE | 16701 | ENSG00000137841 | PLCB2 | chr15 | - | 40302468 | 40302609 | 40302269 | 40302349 | 40303287 | 40303356 | 326 | 145 | 219 | [illegible] |
| SE | 16948 | ENSG00000172575 | RASGRP1 | chr15 | - | 38494381 | 38494767 | 38488102 | 38490688 | 38502311 | 38502421 | 173 | 127 | 157 | [illegible] |
| SE | 16950 | ENSG00000172575 | RASGRP1 | chr15 | - | 38498793 | 38498946 | 38490553 | 38490688 | 38502311 | 38502421 | 6 | 7 | 4 | [illegible] |

| | | | | | | | | | | | | | | | |
|---|---|---|---|---|---|---|---|---|---|---|---|---|---|---|---|
| SE | 16952 | ENSG00000172575 | RASGRP1 | chr15 | - | 38498793 | 38498946 | 38494381 | 38494767 | 38502311 | 38502421 | 104 | 52 | 116 | 115 | 146 |
| SE | 16954 | ENSG00000172575 | RASGRP1 | chr15 | - | 38500102 | 38500139 | 38490553 | 38490688 | 38502311 | 38502421 | 3 | 2 | 5 | 0 | 20 |
| SE | 16955 | ENSG00000172575 | RASGRP1 | chr15 | - | 38500102 | 38500139 | 38494381 | 38494767 | 38502311 | 38502421 | 4 | 4 | 15 | 3 | 21 |
| SE | 16956 | ENSG00000172575 | RASGRP1 | chr15 | - | 38500102 | 38500139 | 38498793 | 38498946 | 38502311 | 38502421 | 10 | 18 | 35 | 26 | 73 |
| SE | 16959 | ENSG00000172575 | RASGRP1 | chr15 | - | 38502311 | 38502421 | 38500102 | 38500139 | 38507725 | 38508001 | 8 | 18 | 15 | 20 | 23 |
| SE | 16960 | ENSG00000172575 | RASGRP1 | chr15 | - | 38502311 | 38502421 | 38500102 | 38500139 | 38511603 | 38511705 | 7 | 10 | 11 | 16 | 14 |
| SE | 16963 | ENSG00000172575 | RASGRP1 | chr15 | - | 38507725 | 38508001 | 38500102 | 38500139 | 38511603 | 38511705 | 261 | 152 | 298 | 337 | 357 |
| SE | 16964 | ENSG00000172575 | RASGRP1 | chr15 | - | 38507725 | 38508001 | 38502311 | 38502421 | 38511603 | 38511705 | 262 | 161 | 303 | 342 | 366 |
| SE | 16966 | ENSG00000172575 | RASGRP1 | chr15 | - | 38518291 | 38518423 | 38516196 | 38516350 | 38519308 | 38519364 | 432 | 296 | 478 | 514 | 512 |
| SE | 16967 | ENSG00000172575 | RASGRP1 | chr15 | - | 38518291 | 38518423 | 38516196 | 38516350 | 38559820 | 38560005 | 157 | 76 | 135 | 189 | 224 |
| SE | 17482 | ENSG00000185024 | BRF1 | chr14 | - | 105226250 | 105226290 | 105226068 | 105226161 | 105226633 | 105226760 | 58 | 21 | 62 | 32 | 42 |
| SE | 17483 | ENSG00000185024 | BRF1 | chr14 | - | 105228819 | 105228913 | 105226633 | 105226760 | 105247070 | 105247193 | 12 | 6 | 18 | 12 | 16 |
| SE | 17595 | ENSG00000174373 | RALGAPA1 | chr14 | - | 35542026 | 35542029 | 35538356 | 35539690 | 35549109 | 35549234 | 1 | 0 | 0 | 0 | 0 |
| SE | 17597 | ENSG00000174373 | RALGAPA1 | chr14 | - | 35595633 | 35595789 | 35572559 | 35572718 | 35605585 | 35605709 | 140 | 119 | 145 | 151 | 244 |
| SE | 17599 | ENSG00000174373 | RALGAPA1 | chr14 | - | 35634573 | 35634757 | 35627089 | 35627951 | 35635463 | 35635598 | 140 | 86 | 155 | 154 | 216 |
| SE | 17600 | ENSG00000174373 | RALGAPA1 | chr14 | - | 35635463 | 35635598 | 35634573 | 35634757 | 35651804 | 35651873 | 111 | 60 | 111 | 108 | 166 |
| SE | 17601 | ENSG00000174373 | RALGAPA1 | chr14 | - | 35659137 | 35659196 | 35655806 | 35655915 | 35664641 | 35664767 | 99 | 78 | 102 | 120 | 190 |
| SE | 17602 | ENSG00000174373 | RALGAPA1 | chr14 | - | 35659137 | 35659196 | 35655806 | 35655915 | 35671388 | 35671517 | 46 | 34 | 55 | 53 | 115 |
| SE | 17603 | ENSG00000174373 | RALGAPA1 | chr14 | - | 35664641 | 35664767 | 35655806 | 35655915 | 35671388 | 35671517 | 38 | 34 | 36 | 46 | 97 |
| SE | 17604 | ENSG00000174373 | RALGAPA1 | chr14 | - | 35664641 | 35664767 | 35659137 | 35659196 | 35671388 | 35671517 | 91 | 78 | 84 | 112 | 173 |
| SE | 17605 | ENSG00000174373 | RALGAPA1 | chr14 | - | 35677949 | 35678102 | 35674515 | 35674709 | 35683808 | 35683985 | 123 | 67 | 122 | 104 | 138 |
| SE | 17606 | ENSG00000174373 | RALGAPA1 | chr14 | - | 35688458 | 35688540 | 35686541 | 35686623 | 35689835 | 35690003 | 2 | 1 | 4 | 0 | 7 |
| SE | 17607 | ENSG00000174373 | RALGAPA1 | chr14 | - | 35700161 | 35700302 | 35688458 | 35690003 | 35721687 | 35721849 | 0 | 0 | 1 | 0 | 0 |
| SE | 17608 | ENSG00000174373 | RALGAPA1 | chr14 | - | 35728361 | 35728510 | 35725023 | 35725153 | 35738512 | 35738650 | 291 | 256 | 280 | 283 | 485 |
| SE | 17610 | ENSG00000174373 | RALGAPA1 | chr14 | - | 35756792 | 35756908 | 35752023 | 35752162 | 35760828 | 35760857 | 187 | 120 | 150 | 129 | 317 |
| SE | 17611 | ENSG00000174373 | RALGAPA1 | chr14 | - | 35760828 | 35761006 | 35756792 | 35756908 | 35762709 | 35762753 | 151 | 119 | 161 | 148 | 324 |
| SE | 17612 | ENSG00000174373 | RALGAPA1 | chr14 | - | 35762709 | 35762753 | 35760828 | 35761006 | 35770941 | 35770999 | 120 | 96 | 141 | 144 | 249 |
| SE | 17613 | ENSG00000174373 | RALGAPA1 | chr14 | - | 35770941 | 35770999 | 35762709 | 35762753 | 35775005 | 35775055 | 105 | 70 | 131 | 141 | 223 |
| SE | 17648 | ENSG00000100949 | RABGGTA | chr14 | - | 24269079 | 24269163 | 24268826 | 24268993 | 24269490 | 24269694 | 44 | 17 | 25 | 18 | 18 |
| SE | 17649 | ENSG00000100949 | RABGGTA | chr14 | - | 24270836 | 24270947 | 24270333 | 24270458 | 24271112 | 24271169 | 54 | 8 | 46 | 42 | 12 |
| SE | 17656 | ENSG00000100916 | BRMS1L | chr14 | + | 35829794 | 35829907 | 35826440 | 35826658 | 35831409 | 35831500 | 7 | 19 | 4 | 19 | 0 |
| SE | 17657 | ENSG00000100916 | BRMS1L | chr14 | + | 35831409 | 35831500 | 35829794 | 35829907 | 35832977 | 35833105 | 20 | 28 | 25 | 32 | 28 |
| SE | 17658 | ENSG00000100916 | BRMS1L | chr14 | + | 35832977 | 35833105 | 35831409 | 35831500 | 35862589 | 35862686 | 20 | 35 | 29 | 33 | 29 |
| SE | 17659 | ENSG00000100916 | BRMS1L | chr14 | + | 35834843 | 35834923 | 35832977 | 35833105 | 35858576 | 35858681 | 18 | 14 | 8 | 15 | 7 |
| SE | 17660 | ENSG00000100916 | BRMS1L | chr14 | + | 35834843 | 35834923 | 35832977 | 35833105 | 35862589 | 35862686 | 49 | 50 | 44 | 54 | 49 |
| SE | 18419 | ENSG00000196497 | IPO4 | chr14 | - | 24182791 | 24182842 | 24182277 | 24182403 | 24182975 | 24183161 | 30 | 8 | 24 | 50 | 28 |
| SE | 18421 | ENSG00000196497 | IPO4 | chr14 | - | 24187084 | 24187150 | 24186877 | 24186979 | 24187399 | 24187579 | 27 | 18 | 39 | 37 | 40 |
| SE | 18655 | ENSG00000099904 | ZDHHC8 | chr22 | + | 20139193 | 20139315 | 20131803 | 20132051 | 20139477 | 20139635 | 24 | 6 | 30 | 30 | 12 |
| SE | 18757 | ENSG00000197045 | GMFB | chr14 | - | 54482121 | 54482202 | 54481394 | 54481458 | 54483670 | 54483767 | 28 | 12 | 18 | 39 | 27 |
| SE | 18758 | ENSG00000197045 | GMFB | chr14 | - | 54482152 | 54482202 | 54481325 | 54481458 | 54483670 | 54483767 | 85 | 68 | 70 | 89 | 143 |
| SE | 18759 | ENSG00000197045 | GMFB | chr14 | - | 54483932 | 54484060 | 54483670 | 54483767 | 54488924 | 54489026 | 1 | 3 | 2 | 1 | 4 |
| SE | 19019 | ENSG00000131966 | ACTR10 | chr14 | + | 58202854 | 58202927 | 58200079 | 58200294 | 58207935 | 58208018 | 62 | 46 | 48 | 77 | 112 |
| SE | 19021 | ENSG00000131966 | ACTR10 | chr14 | + | 58207935 | 58208018 | 58202854 | 58202927 | 58208998 | 58209107 | 98 | 71 | 60 | 112 | 259 |
| SE | 19022 | ENSG00000131966 | ACTR10 | chr14 | + | 58211291 | 58211399 | 58208998 | 58209107 | 58213630 | 58213698 | 141 | 79 | 119 | 119 | 195 |
| SE | 19024 | ENSG00000131966 | ACTR10 | chr14 | + | 58215204 | 58215284 | 58208998 | 58209107 | 58219693 | 58219729 | 120 | 99 | 98 | 123 | 270 |
| SE | 19025 | ENSG00000131966 | ACTR10 | chr14 | + | 58219693 | 58219729 | 58208998 | 58209107 | 58230398 | 58230480 | 0 | 0 | 0 | 0 | 0 |
| SE | 19028 | ENSG00000131966 | ACTR10 | chr14 | + | 58223621 | 58223701 | 58219669 | 58219729 | 58223782 | 58223827 | 169 | 116 | 152 | 163 | 312 |
| SE | 19029 | ENSG00000131966 | ACTR10 | chr14 | + | 58223782 | 58223856 | 58219669 | 58219729 | 58230398 | 58230480 | 192 | 116 | 112 | 162 | 252 |
| SE | 19031 | ENSG00000131966 | ACTR10 | chr14 | + | 58223782 | 58223856 | 58223598 | 58223701 | 58230398 | 58230480 | 299 | 189 | 210 | 257 | 449 |
| SE | 19033 | ENSG00000131966 | ACTR10 | chr14 | + | 58230398 | 58230480 | 58223492 | 58223856 | 58232065 | 58232108 | 342 | 205 | 237 | 318 | 540 |
| SE | 19035 | ENSG00000131966 | ACTR10 | chr14 | + | 58231079 | 58231153 | 58230398 | 58230480 | 58232065 | 58232267 | 0 | 0 | 0 | 0 | 0 |
| SE | 19036 | ENSG00000131966 | ACTR10 | chr14 | + | 58232065 | 58232267 | 58223782 | 58223856 | 58234369 | 58234734 | 216 | 121 | 176 | 213 | 306 |
| SE | 19461 | ENSG00000125746 | EML2 | chr19 | - | 45624718 | 45624818 | 45621206 | 45621332 | 45629950 | 45630046 | 1 | 1 | 5 | 7 | 5 |
| SE | 19462 | ENSG00000125746 | EML2 | chr19 | - | 45624718 | 45624818 | 45621482 | 45621637 | 45626704 | 45626752 | 90 | 43 | 75 | 100 | 52 |
| SE | 19464 | ENSG00000125746 | EML2 | chr19 | - | 45628697 | 45628795 | 45626704 | 45626839 | 45629950 | 45630046 | 2 | 0 | 2 | 3 | 0 |
| SE | 19465 | ENSG00000125746 | EML2 | chr19 | - | 45629950 | 45630046 | 45626704 | 45626839 | 45632860 | 45632971 | 98 | 42 | 58 | 107 | 39 |
| SE | 19466 | ENSG00000125746 | EML2 | chr19 | - | 45632860 | 45633139 | 45629963 | 45630046 | 45634321 | 45634496 | 48 | 20 | 36 | 70 | 34 |
| SE | 19507 | ENSG00000100479 | POLE2 | chr14 | - | 49655670 | 49655843 | 49655004 | 49655094 | 49663314 | 49663387 | 10 | 12 | 10 | 9 | 30 |
| SE | 19508 | ENSG00000100479 | POLE2 | chr14 | - | 49663314 | 49663387 | 49655670 | 49655843 | 49664625 | 49664663 | 13 | 18 | 19 | 17 | 24 |
| SE | 19509 | ENSG00000100479 | POLE2 | chr14 | - | 49666329 | 49666413 | 49665093 | 49665163 | 49669523 | 49669598 | 23 | 35 | 25 | 27 | 92 |
| SE | 19511 | ENSG00000100479 | POLE2 | chr14 | - | 49674122 | 49674216 | 49669523 | 49669598 | 49674349 | 49674427 | 25 | 29 | 27 | 34 | 59 |
| SE | 20746 | ENSG00000175048 | ZDHHC14 | chr6 | + | 157592987 | 157593146 | 157542584 | 157542745 | 157628348 | 157628374 | 26 | 32 | 28 | 44 | 45 |
| SE | 20747 | ENSG00000175048 | ZDHHC14 | chr6 | + | 157628348 | 157628374 | 157592987 | 157593146 | 157632833 | 157632882 | 2 | 3 | 1 | 7 | 2 |
| SE | 20749 | ENSG00000175048 | ZDHHC14 | chr6 | + | 157628447 | 157628486 | 157593008 | 157593146 | 157632833 | 157632882 | 8 | 5 | 5 | 8 | 13 |
| SE | 21233 | ENSG00000136986 | DERL1 | chr8 | - | 123019194 | 123019305 | 123013169 | 123015585 | 123021446 | 123021499 | 526 | 347 | 339 | 412 | 561 |
| SE | 21234 | ENSG00000136986 | DERL1 | chr8 | - | 123021446 | 123021499 | 123019194 | 123019245 | 123022683 | 123022779 | 130 | 72 | 108 | 106 | 78 |
| SE | 21235 | ENSG00000136986 | DERL1 | chr8 | - | 123024985 | 123025050 | 123021446 | 123021499 | 123041969 | 123042122 | 0 | 0 | 1 | 0 | 0 |
| SE | 21422 | ENSG00000164180 | TMEM161B | chr5 | - | 88197668 | 88197765 | 88195046 | 88196488 | 88198975 | 88199069 | 60 | 25 | 47 | 34 | 48 |
| SE | 21426 | ENSG00000164180 | TMEM161B | chr5 | - | 88221300 | 88221471 | 88220614 | 88220719 | 88225768 | 88225866 | 0 | 0 | 2 | 1 | 0 |
| SE | 21427 | ENSG00000164180 | TMEM161B | chr5 | - | 88221300 | 88221471 | 88220614 | 88220719 | 88228444 | 88228528 | 1 | 0 | 1 | 0 | 0 |
| SE | 21429 | ENSG00000164180 | TMEM161B | chr5 | - | 88221673 | 88221760 | 88220562 | 88220719 | 88228444 | 88228528 | 11 | 3 | 1 | 1 | 6 |
| SE | 21431 | ENSG00000164180 | TMEM161B | chr5 | - | 88225768 | 88225866 | 88221300 | 88221471 | 88228444 | 88228528 | 16 | 15 | 23 | 18 | 11 |
| SE | 21432 | ENSG00000164180 | TMEM161B | chr5 | - | 88225768 | 88225866 | 88221673 | 88221760 | 88228444 | 88228528 | 34 | 21 | 42 | 29 | 24 |
| SE | 21434 | ENSG00000164180 | TMEM161B | chr5 | - | 88240812 | 88240916 | 88228440 | 88228528 | 88268787 | 88269070 | 50 | 31 | 58 | 50 | 41 |
| SE | 22069 | ENSG00000174032 | SLC25A30 | chr13 | - | 45397257 | 45397338 | 45393315 | 45396015 | 45398939 | 45399078 | 39 | 33 | 36 | 30 | 42 |
| SE | 22070 | ENSG00000174032 | SLC25A30 | chr13 | - | 45401082 | 45401207 | 45393315 | 45399078 | 45402274 | 45402357 | 64 | 31 | 43 | 69 | 69 |
| SE | 22072 | ENSG00000174032 | SLC25A30 | chr13 | - | 45404326 | 45404412 | 45402274 | 45402370 | 45408926 | 45409074 | 49 | 49 | 52 | 57 | 54 |
| SE | 22073 | ENSG00000174032 | SLC25A30 | chr13 | - | 45405882 | 45405977 | 45402274 | 45402370 | 45408926 | 45409074 | 72 | 38 | 58 | 70 | 80 |
| SE | 22074 | ENSG00000174032 | SLC25A30 | chr13 | - | 45405882 | 45405977 | 45404326 | 45404412 | 45408926 | 45409074 | 156 | 78 | 118 | 153 | 224 |
| SE | 22078 | ENSG00000174032 | SLC25A30 | chr13 | - | 45411361 | 45411480 | 45408926 | 45409074 | 45418299 | 45418373 | 67 | 32 | 35 | 38 | 25 |
| SE | 22134 | ENSG00000175727 | MLXIP | chr12 | + | 122135488 | 122135666 | 122134320 | 122135345 | 122138193 | 122138295 | 78 | 43 | 93 | 97 | 90 |
| SE | 22549 | ENSG00000167881 | SRP68 | chr17 | - | 76040902 | 76040978 | 76040418 | 76040474 | 76043828 | 76043958 | 227 | 164 | 256 | 197 | 432 |
| SE | 22554 | ENSG00000167881 | SRP68 | chr17 | - | 76061109 | 76061219 | 76057402 | 76057543 | 76061491 | 76061574 | 179 | 143 | 170 | 194 | 279 |
| SE | 22555 | ENSG00000167881 | SRP68 | chr17 | - | 76061109 | 76061219 | 76057402 | 76057543 | 76063975 | 76064171 | 2 | 0 | 2 | 6 | 6 |
| SE | 22556 | ENSG00000167881 | SRP68 | chr17 | - | 76061109 | 76061219 | 76060307 | 76060390 | 76061491 | 76061574 | 482 | 353 | 448 | 491 | 613 |
| SE | 22557 | ENSG00000167881 | SRP68 | chr17 | - | 76061491 | 76061574 | 76057402 | 76057543 | 76063975 | 76064171 | 105 | 81 | 147 | 158 | 164 |
| SE | 22793 | ENSG00000131504 | DIAPH1 | chr5 | - | 141518943 | 141518996 | 141515020 | 141517008 | 141524142 | 141524229 | 1 | 0 | 1 | 0 | 0 |
| SE | 22794 | ENSG00000131504 | DIAPH1 | chr5 | - | 141524142 | 141524229 | 141515020 | 141517008 | 141526037 | 141526173 | 1099 | 580 | 753 | 826 | 1461 |
| SE | 22795 | ENSG00000131504 | DIAPH1 | chr5 | - | 141526037 | 141526173 | 141524142 | 141524229 | 141526296 | 141526459 | 2354 | 1606 | 1458 | 2107 | 3013 |
| SE | 22796 | ENSG00000131504 | DIAPH1 | chr5 | - | 141529171 | 141529273 | 141528701 | 141528941 | 141529602 | 141529697 | 985 | 585 | 614 | 775 | 1480 |
| SE | 22797 | ENSG00000131504 | DIAPH1 | chr5 | - | 141529171 | 141529273 | 141528701 | 141528941 | 141534334 | 141534433 | 501 | 312 | 308 | 400 | 743 |
| SE | 22801 | ENSG00000131504 | DIAPH1 | chr5 | - | 141571427 | 141571436 | 141534334 | 141534433 | 141571925 | 141572040 | 13 | 5 | 8 | 10 | 31 |
| SE | 22802 | ENSG00000131504 | DIAPH1 | chr5 | - | 141580743 | 141580883 | 141579087 | 141579196 | 141582311 | 141582375 | 1849 | 1268 | 1275 | 1687 | 2299 |
| SE | 22803 | ENSG00000131504 | DIAPH1 | chr5 | - | 141581914 | 141582056 | 141580743 | 141580883 | 141582311 | 141582375 | 6 | 0 | 4 | 0 | 3 |
| SE | 22804 | ENSG00000131504 | DIAPH1 | chr5 | - | 141584123 | 141584225 | 141583484 | 141583615 | 141587041 | 141587176 | 836 | 433 | 524 | 656 | 915 |
| SE | 23007 | ENSG00000206077 | ZDHHC11B | chr5 | - | 716800 | 716865 | 710354 | 712282 | 730433 | 730468 | 2 | 3 | 14 | 13 | 14 |
| SE | 23009 | ENSG00000206077 | ZDHHC11B | chr5 | - | 749251 | 750296 | 748403 | 748559 | 751132 | 751257 | 0 | 0 | 1 | 0 | 2 |
| SE | 23010 | ENSG00000206077 | ZDHHC11B | chr5 | - | 769954 | 770117 | 768834 | 768930 | 784667 | 784729 | 0 | 0 | 0 | 0 | 0 |
| SE | 23057 | ENSG00000111641 | NOP2 | chr12 | - | 6558015 | 6558137 | 6556870 | 6557642 | 6560097 | 6560326 | 0 | 0 | 0 | 2 | 0 |
| SE | 23059 | ENSG00000111641 | NOP2 | chr12 | - | 6564149 | 6564248 | 6563890 | 6563946 | 6566528 | 6566617 | 0 | 0 | 2 | 0 | 6 |
| SE | 23062 | ENSG00000111641 | NOP2 | chr12 | - | 6566100 | 6566336 | 6564149 | 6564248 | 6566528 | 6566559 | 97 | 69 | 83 | 81 | 82 |
| SE | 23224 | ENSG00000183763 | TRAIP | chr3 | - | 49842452 | 49842547 | 49841825 | 49841939 | 49843800 | 49843928 | 2 | 0 | 3 | 3 | 5 |
| SE | 23226 | ENSG00000183763 | TRAIP | chr3 | - | 49848136 | 49848200 | 49847524 | 49847608 | 49856355 | 49856564 | 1 | 0 | 1 | 0 | 1 |
| SE | 23227 | ENSG00000183763 | TRAIP | chr3 | - | 49848142 | 49848200 | 49847524 | 49847608 | 49856355 | 49856474 | 2 | 2 | 5 | 2 | 1 |
| SE | 24097 | ENSG00000143889 | HNRNPLL | chr2 | - | 38569803 | 38569925 | 38569132 | 38569334 | 38573209 | 38573427 | 227 | 138 | 144 | 192 | 227 |
| SE | 24098 | ENSG00000143889 | HNRNPLL | chr2 | - | 38577460 | 38577532 | 38573209 | 38573427 | 38581912 | 38581980 | 119 | 80 | 96 | 137 | 219 |
| SE | 24102 | ENSG00000143889 | HNRNPLL | chr2 | - | 38585643 | 38585881 | 38583840 | 38583926 | 38602437 | 38602600 | 139 | 80 | 81 | 126 | 153 |
| SE | 24282 | ENSG00000104671 | DCTN6 | chr8 | + | 30175084 | 30175190 | 30164110 | 30164175 | 30177125 | 30177214 | 100 | 89 | 80 | 110 | 120 |
| SE | 24283 | ENSG00000104671 | DCTN6 | chr8 | + | 30175084 | 30175190 | 30165715 | 30165901 | 30177125 | 30177214 | 37 | 46 | 28 | 55 | 51 |
| SE | 24627 | ENSG00000157212 | PAXIP1 | chr7 | - | 154957223 | 154957294 | 154955528 | 154955631 | 154959889 | 154959933 | 45 | 41 | 67 | 80 | 143 |
| SE | 25740 | ENSG00000157107 | FCHO2 | chr5 | + | 72990711 | 72990864 | 72990477 | 72990619 | 73006444 | 73006549 | 212 | 187 | 110 | 217 | 141 |
| SE | 25741 | ENSG00000157107 | FCHO2 | chr5 | + | 73006444 | 73006549 | 72990711 | 72990864 | 73015625 | 73015724 | 116 | 83 | 68 | 90 | 65 |
| SE | 25742 | ENSG00000157107 | FCHO2 | chr5 | + | 73015625 | 73015724 | 73006444 | 73006549 | 73017211 | 73017282 | 135 | 94 | 86 | 87 | 103 |
| SE | 25743 | ENSG00000157107 | FCHO2 | chr5 | + | 73063840 | 73063944 | 73058420 | 73058524 | 73068649 | 73068779 | 112 | 66 | 34 | 66 | 171 |
| SE | 26050 | ENSG00000198689 | SLC9A6 | chrX | + | 135998481 | 135998558 | 135998107 | 135998185 | 135998855 | 135998883 | 75 | 56 | 64 | 51 | 92 |
| SE | 26051 | ENSG00000198689 | SLC9A6 | chrX | + | 136000577 | 136000651 | 135998855 | 135998968 | 136002107 | 136002213 | 2 | 0 | 1 | 0 | 2 |
| SE | 26052 | ENSG00000198689 | SLC9A6 | chrX | + | 136002107 | 136002213 | 135998855 | 135998968 | 136010441 | 136010578 | 42 | 30 | 31 | 43 | 40 |
| SE | 26053 | ENSG00000198689 | SLC9A6 | chrX | + | 136013348 | 136013437 | 136012948 | 136013054 | 136016644 | 136016758 | 58 | 31 | 59 | 56 | 62 |
| SE | 26054 | ENSG00000198689 | SLC9A6 | chrX | + | 136028885 | 136028975 | 136024329 | 136024483 | 136030131 | 136030162 | 0 | 0 | 0 | 0 | 0 |
| SE | 26140 | ENSG00000172244 | C5orf34 | chr5 | - | 43502371 | 43502495 | 43494509 | 43494601 | 43503664 | 43503760 | 8 | 5 | 5 | 4 | 24 |
| SE | 26141 | ENSG00000172244 | C5orf34 | chr5 | - | 43508576 | 43508666 | 43505549 | 43506394 | 43509144 | 43509375 | 9 | 6 | 9 | 13 | 7 |
| SE | 26142 | ENSG00000172244 | C5orf34 | chr5 | - | 43508576 | 43508666 | 43505549 | 43506394 | 43514805 | 43515078 | 7 | 5 | 6 | 8 | 5 |
| SE | 26144 | ENSG00000172244 | C5orf34 | chr5 | - | 43509144 | 43509375 | 43508576 | 43508666 | 43514805 | 43515148 | 5 | 1 | 5 | 6 | 2 |
| SE | 26429 | ENSG00000158290 | CUL4B | chrX | - | 120530101 | 120530254 | 120505919 | 120526856 | 120532421 | 120532594 | 564 | 475 | 466 | 557 | 926 |
| SE | 26430 | ENSG00000158290 | CUL4B | chrX | - | 120532421 | 120532594 | 120529975 | 120530254 | 120534480 | 120534586 | 554 | 483 | 463 | 551 | 1089 |
| SE | 26432 | ENSG00000158290 | CUL4B | chrX | - | 120538347 | 120538409 | 120538123 | 120538209 | 120538659 | 120538770 | 1 | 0 | 2 | 2 | 0 |
| SE | 26433 | ENSG00000158290 | CUL4B | chrX | - | 120539267 | 120539372 | 120538123 | 120538770 | 120540369 | 120540562 | 334 | 245 | 271 | 331 | 491 |
| SE | 26438 | ENSG00000158290 | CUL4B | chrX | - | 120545443 | 120545517 | 120544480 | 120544643 | 120546546 | 120546616 | 279 | 199 | 256 | 263 | 279 |
| SE | 26439 | ENSG00000158290 | CUL4B | chrX | - | 120554907 | 120554953 | 120547135 | 120547239 | 120557923 | 120558039 | 2 | 4 | 0 | 1 | 0 |
| SE | 26440 | ENSG00000158290 | CUL4B | chrX | - | 120558248 | 120558292 | 120557923 | 120558039 | 120560082 | 120560776 | 1 | 0 | 0 | 0 | 0 |
| SE | 26444 | ENSG00000083312 | TNPO1 | chr5 | + | 72855773 | 72855923 | 72851204 | 72851319 | 72872638 | 72872720 | 200 | 135 | 131 | 150 | 186 |
| SE | 26445 | ENSG00000083312 | TNPO1 | chr5 | + | 72861807 | 72861914 | 72855773 | 72855923 | 72865595 | 72865693 | 255 | 162 | 192 | 246 | 338 |
| SE | 26446 | ENSG00000083312 | TNPO1 | chr5 | + | 72865595 | 72865729 | 72855773 | 72855923 | 72872638 | 72872720 | 123 | 72 | 114 | 116 | 167 |
| SE | 26447 | ENSG00000083312 | TNPO1 | chr5 | + | 72872638 | 72872720 | 72865595 | 72865729 | 72875614 | 72875737 | 211 | 125 | 185 | 180 | 243 |
| SE | 26448 | ENSG00000083312 | TNPO1 | chr5 | + | 72875614 | 72875737 | 72872638 | 72872720 | 72877227 | 72877346 | 269 | 157 | 246 | 285 | 351 |
| SE | 26451 | ENSG00000083312 | TNPO1 | chr5 | + | 72893138 | 72893246 | 72891809 | 72891896 | 72893376 | 72893535 | 402 | 205 | 302 | 384 | 591 |
| SE | 26452 | ENSG00000083312 | TNPO1 | chr5 | + | 72893615 | 72893703 | 72893376 | 72893535 | 72896457 | 72896556 | 266 | 181 | 226 | 268 | 416 |
| SE | 26453 | ENSG00000083312 | TNPO1 | chr5 | + | 72896457 | 72896556 | 72893615 | 72893703 | 72897055 | 72897151 | 364 | 294 | 345 | 386 | 482 |
| SE | 26454 | ENSG00000083312 | TNPO1 | chr5 | + | 72896457 | 72896556 | 72893615 | 72893703 | 72900005 | 72900081 | 172 | 126 | 151 | 174 | 220 |
| SE | 26455 | ENSG00000083312 | TNPO1 | chr5 | + | 72896457 | 72896556 | 72893615 | 72893703 | 72900973 | 72901073 | 167 | 126 | 150 | 171 | 216 |
| SE | 26456 | ENSG00000083312 | TNPO1 | chr5 | + | 72897055 | 72897151 | 72893615 | 72893703 | 72900005 | 72900081 | 81 | 60 | 60 | 73 | 66 |
| SE | 26457 | ENSG00000083312 | TNPO1 | chr5 | + | 72897055 | 72897151 | 72893615 | 72893703 | 72900973 | 72901073 | 3 | 2 | 0 | 4 | 3 |
| SE | 26458 | ENSG00000083312 | TNPO1 | chr5 | + | 72897055 | 72897151 | 72895458 | 72896556 | 72900005 | 72900081 | 280 | 227 | 259 | 287 | 333 |
| SE | 26459 | ENSG00000083312 | TNPO1 | chr5 | + | 72897055 | 72897151 | 72895458 | 72896556 | 72900973 | 72901073 | 169 | 136 | 156 | 173 | 252 |
| SE | 26460 | ENSG00000083312 | TNPO1 | chr5 | + | 72900005 | 72900081 | 72893615 | 72893703 | 72900973 | 72901073 | 17 | 7 | 9 | 14 | 3 |
| SE | 26461 | ENSG00000083312 | TNPO1 | chr5 | + | 72900005 | 72900081 | 72896457 | 72896556 | 72900973 | 72901073 | 23 | 7 | 13 | 21 | 9 |
| SE | 26462 | ENSG00000083312 | TNPO1 | chr5 | + | 72900005 | 72900081 | 72897055 | 72897151 | 72900973 | 72901073 | 125 | 101 | 109 | 128 | 87 |
| SE | 26463 | ENSG00000083312 | TNPO1 | chr5 | + | 72900150 | 72900224 | 72897055 | 72897151 | 72900973 | 72901073 | 0 | 0 | 0 | 0 | 0 |
| SE | 26559 | ENSG00000032742 | IFT88 | chr13 | + | 20567669 | 20567997 | 20567165 | 20567256 | 20574379 | 20574475 | 9 | 10 | 4 | 9 | 7 |
| SE | 26560 | ENSG00000032742 | IFT88 | chr13 | + | 20589810 | 20589867 | 20582956 | 20583019 | 20590966 | 20591020 | 51 | 36 | 26 | 40 | 39 |
| SE | 26561 | ENSG00000032742 | IFT88 | chr13 | + | 20592334 | 20592404 | 20591617 | 20591681 | 20596149 | 20596240 | 68 | 63 | 37 | 46 | 70 |
| SE | 26562 | ENSG00000032742 | IFT88 | chr13 | + | 20598650 | 20598753 | 20597013 | 20597119 | 20599450 | 20599565 | 110 | 68 | 58 | 54 | 162 |
| SE | 26563 | ENSG00000032742 | IFT88 | chr13 | + | 20599450 | 20599565 | 20597013 | 20597119 | 20601704 | 20601748 | 45 | 33 | 27 | 21 | 65 |
| SE | 26564 | ENSG00000032742 | IFT88 | chr13 | + | 20614356 | 20614435 | 20605055 | 20605105 | 20615792 | 20615879 | 1 | 0 | 2 | 0 | 0 |
| SE | 26565 | ENSG00000032742 | IFT88 | chr13 | + | 20625749 | 20625849 | 20615790 | 20615879 | 20631015 | 20631102 | 25 | 33 | 26 | 30 | 52 |
| SE | 26566 | ENSG00000032742 | IFT88 | chr13 | + | 20638331 | 20638518 | 20631015 | 20631102 | 20641289 | 20641398 | 38 | 27 | 19 | 34 | 90 |
| SE | 26567 | ENSG00000032742 | IFT88 | chr13 | + | 20644842 | 20644958 | 20643454 | 20643605 | 20653875 | 20653928 | 44 | 32 | 31 | 46 | 97 |
| SE | 26568 | ENSG00000032742 | IFT88 | chr13 | + | 20663262 | 20663386 | 20656364 | 20656430 | 20663497 | 20663604 | 0 | 4 | 5 | 0 | 7 |
| SE | 26569 | ENSG00000032742 | IFT88 | chr13 | + | 20690704 | 20690815 | 20670972 | 20671039 | 20691053 | 20691217 | 10 | 3 | 5 | 8 | 9 |
| SE | 26618 | ENSG00000188818 | ZDHHC11 | chr5 | - | 823388 | 823809 | 821860 | 821895 | 825163 | 825251 | 4 | 1 | 1 | 2 | 0 |
| SE | 26619 | ENSG00000188818 | ZDHHC11 | chr5 | - | 823388 | 824093 | 821860 | 821895 | 825163 | 825251 | 4 | 1 | 1 | 2 | 0 |
| SE | 26976 | ENSG00000187091 | PLCD1 | chr3 | - | 38008457 | 38008636 | 38008234 | 38008367 | 38009041 | 38009158 | 42 | 6 | 27 | 31 | 31 |
| SE | 26977 | ENSG00000187091 | PLCD1 | chr3 | - | 38010360 | 38010562 | 38009903 | 38010275 | 38011213 | 38011445 | 19 | 4 | 20 | 7 | 11 |
| SE | 27149 | ENSG00000109534 | GAR1 | chr4 | + | 109816152 | 109816378 | 109815509 | 109815545 | 109817935 | 109818090 | 70 | 85 | 61 | 61 | 59 |
| SE | 28211 | ENSG00000204084 | INPP5B | chr1 | - | 37864311 | 37864423 | 37860696 | 37862430 | 37865760 | 37865888 | 154 | 87 | 149 | 109 | 234 |
| SE | 28212 | ENSG00000204084 | INPP5B | chr1 | - | 37864311 | 37864423 | 37860696 | 37862430 | 37868500 | 37868614 | 95 | 56 | 79 | 69 | 124 |
| SE | 28213 | ENSG00000204084 | INPP5B | chr1 | - | 37868500 | 37868614 | 37860696 | 37862430 | 37872929 | 37873165 | 49 | 16 | 41 | 35 | 51 |
| SE | 28214 | ENSG00000204084 | INPP5B | chr1 | - | 37868500 | 37868614 | 37866458 | 37866543 | 37872929 | 37873165 | 120 | 53 | 108 | 89 | 152 |
| SE | 28216 | ENSG00000204084 | INPP5B | chr1 | - | 37870669 | 37870720 | 37869924 | 37870014 | 37872929 | 37873028 | 0 | 0 | 1 | 1 | 4 |
| SE | 28217 | ENSG00000204084 | INPP5B | chr1 | - | 37873992 | 37874155 | 37872929 | 37873165 | 37875605 | 37875716 | 99 | 80 | 107 | 85 | 152 |
| SE | 28218 | ENSG00000204084 | INPP5B | chr1 | - | 37882806 | 37882918 | 37880084 | 37880194 | 37885637 | 37885825 | 121 | 92 | 112 | 91 | 185 |
| SE | 28219 | ENSG00000204084 | INPP5B | chr1 | - | 37882806 | 37882918 | 37880084 | 37880194 | 37886887 | 37887004 | 65 | 46 | 56 | 50 | 108 |
| SE | 28220 | ENSG00000204084 | INPP5B | chr1 | - | 37885637 | 37885825 | 37880084 | 37880194 | 37886887 | 37887004 | 112 | 53 | 95 | 85 | 135 |
| SE | 28222 | ENSG00000204084 | INPP5B | chr1 | - | 37931672 | 37932053 | 37891357 | 37891454 | 37940687 | 37940798 | 58 | 19 | 38 | 48 | 29 |
| SE | 28223 | ENSG00000204084 | INPP5B | chr1 | - | 37931912 | 37932053 | 37891357 | 37891454 | 37940687 | 37940798 | 23 | 8 | 14 | 14 | 45 |
| SE | 28316 | ENSG00000088538 | DOCK3 | chr3 | + | 51089242 | 51089284 | 51075355 | 51075440 | 51090229 | 51090384 | 4 | 5 | 3 | 5 | 11 |
| SE | 28317 | ENSG00000088538 | DOCK3 | chr3 | + | 51146548 | 51146630 | 51090229 | 51090384 | 51159243 | 51159304 | 6 | 3 | 5 | 9 | 1 |
| SE | 28318 | ENSG00000088538 | DOCK3 | chr3 | + | 51227282 | 51227445 | 51225648 | 51225773 | 51227981 | 51228088 | 9 | 2 | 14 | 2 | 11 |
| SE | 28319 | ENSG00000088538 | DOCK3 | chr3 | + | 51270814 | 51271007 | 51260155 | 51260326 | 51275078 | 51275206 | 10 | 2 | 4 | 2 | 9 |
| SE | 29569 | ENSG00000146350 | TBC1D32 | chr6 | - | 121090852 | 121091041 | 121079493 | 121080890 | 121106022 | 121106163 | 11 | 16 | 14 | 16 | 20 |
| SE | 29570 | ENSG00000146350 | TBC1D32 | chr6 | - | 121129868 | 121129924 | 121126377 | 121126461 | 121131626 | 121131752 | 0 | 0 | 5 | 1 | 1 |
| SE | 29571 | ENSG00000146350 | TBC1D32 | chr6 | - | 121139773 | 121139836 | 121131626 | 121131752 | 121160009 | 121160103 | 4 | 5 | 4 | 1 | 15 |
| SE | 29572 | ENSG00000146350 | TBC1D32 | chr6 | - | 121170404 | 121170527 | 121160947 | 121161056 | 121205074 | 121205163 | 24 | 29 | 9 | 16 | 51 |
| SE | 29573 | ENSG00000146350 | TBC1D32 | chr6 | - | 121170404 | 121170527 | 121160947 | 121161056 | 121223235 | 121223352 | 7 | 14 | 4 | 6 | 31 |
| SE | 29575 | ENSG00000146350 | TBC1D32 | chr6 | - | 121205074 | 121205163 | 121160947 | 121161056 | 121223235 | 121223352 | 11 | 5 | 9 | 6 | 4 |
| SE | 29576 | ENSG00000146350 | TBC1D32 | chr6 | - | 121205074 | 121205163 | 121170404 | 121170527 | 121223235 | 121223352 | 29 | 20 | 14 | 17 | 28 |
| SE | 29577 | ENSG00000146350 | TBC1D32 | chr6 | - | 121224362 | 121224451 | 121223235 | 121223352 | 121239069 | 121239188 | 0 | 0 | 0 | 1 | 0 |
| SE | 29578 | ENSG00000146350 | TBC1D32 | chr6 | - | 121241464 | 121241552 | 121239069 | 121239188 | 121242200 | 121242339 | 14 | 16 | 15 | 20 | 15 |
| SE | 29579 | ENSG00000146350 | TBC1D32 | chr6 | - | 121242200 | 121242339 | 121239069 | 121239188 | 121255327 | 121255410 | 20 | 13 | 16 | 15 | 37 |
| SE | 29580 | ENSG00000146350 | TBC1D32 | chr6 | - | 121242200 | 121242339 | 121239069 | 121239188 | 121279120 | 121279245 | 0 | 1 | 4 | 2 | 0 |
| SE | 29581 | ENSG00000146350 | TBC1D32 | chr6 | - | 121255327 | 121255410 | 121239069 | 121239188 | 121279120 | 121279245 | 9 | 3 | 3 | 3 | 7 |
| SE | 29582 | ENSG00000146350 | TBC1D32 | chr6 | - | 121255327 | 121255410 | 121242200 | 121242339 | 121279120 | 121279245 | 13 | 10 | 7 | 2 | 10 |
| SE | 29583 | ENSG00000146350 | TBC1D32 | chr6 | - | 121256083 | 121256285 | 121255327 | 121255410 | 121279120 | 121279245 | 33 | 37 | 20 | 28 | 58 |
| SE | 29584 | ENSG00000146350 | TBC1D32 | chr6 | - | 121307975 | 121308101 | 121304754 | 121304833 | 121310778 | 121310847 | 46 | 35 | 30 | 60 | 89 |
| SE | 29585 | ENSG00000146350 | TBC1D32 | chr6 | - | 121310778 | 121310847 | 121304754 | 121304833 | 121317494 | 121317672 | 8 | 13 | 9 | 10 | 12 |
| SE | 29586 | ENSG00000146350 | TBC1D32 | chr6 | - | 121317494 | 121317672 | 121310778 | 121310847 | 121321632 | 121321794 | 60 | 68 | 62 | 55 | 101 |
| SE | 29587 | ENSG00000146350 | TBC1D32 | chr6 | - | 121317494 | 121317690 | 121310778 | 121310847 | 121321632 | 121321794 | 25 | 24 | 19 | 30 | 46 |
| SE | 29705 | ENSG00000182621 | PLCB1 | chr20 | + | 8141870 | 8141948 | 8132648 | 8132750 | 8150293 | 8150371 | 2 | 0 | 0 | 0 | 3 |
| SE | 29707 | ENSG00000182621 | PLCB1 | chr20 | + | 8657183 | 8657284 | 8649373 | 8649449 | 8658537 | 8658555 | 57 | 30 | 46 | 63 | 46 |
| SE | 29709 | ENSG00000182621 | PLCB1 | chr20 | + | 8757045 | 8757178 | 8741463 | 8741573 | 8760406 | 8760460 | 58 | 26 | 42 | 50 | 77 |
| SE | 29710 | ENSG00000182621 | PLCB1 | chr20 | + | 8760406 | 8760460 | 8757045 | 8757178 | 8765138 | 8765358 | 26 | 28 | 23 | 27 | 24 |
| SE | 29711 | ENSG00000182621 | PLCB1 | chr20 | + | 8765138 | 8765358 | 8760406 | 8760460 | 8774538 | 8774719 | 65 | 57 | 48 | 48 | 42 |
| SE | 29712 | ENSG00000182621 | PLCB1 | chr20 | + | 8789517 | 8789575 | 8788632 | 8788722 | 8790174 | 8790261 | 54 | 60 | 41 | 56 | 74 |
| SE | 29713 | ENSG00000182621 | PLCB1 | chr20 | + | 8802055 | 8802173 | 8790174 | 8790261 | 8881621 | 8883199 | 23 | 32 | 28 | 35 | 53 |
| SE | 29715 | ENSG00000182621 | PLCB1 | chr20 | + | 8962485 | 8963010 | 8962307 | 8962336 | 8965095 | 8965151 | 1 | 0 | 0 | 1 | 0 |
| SE | 29716 | ENSG00000182621 | PLCB1 | chr20 | + | 8962892 | 8963010 | 8962195 | 8962336 | 8965095 | 8965435 | 1 | 0 | 0 | 1 | 0 |
| SE | 29794 | ENSG00000141452 | RMC1 | chr18 | + | 23504370 | 23504447 | 23503469 | 23503720 | 23515855 | 23515886 | 12 | 5 | 6 | 8 | 5 |
| SE | 29799 | ENSG00000141452 | RMC1 | chr18 | + | 23507984 | 23508041 | 23504370 | 23504447 | 23515855 | 23515886 | 0 | 0 | 0 | 3 | 2 |
| SE | 29801 | ENSG00000141452 | RMC1 | chr18 | + | 23507984 | 23508041 | 23506969 | 23507054 | 23515855 | 23515886 | 42 | 33 | 38 | 50 | 46 |

| | | | | | | | | | | | | | | | | | | | | | | | | | | | | | | | | | | | | | | | | | | | | | | |
|---|---|---|---|---|---|---|---|---|---|---|---|---|---|---|---|---|---|---|---|---|---|---|---|---|---|---|---|---|---|---|---|---|---|---|---|---|---|---|---|---|---|---|---|---|---|---|
| SE | 29802 | ENSG00000141452 | RMC1 | chr18 | + | 23508654 | 23508776 | 23507984 | 23508041 | 23509192 | 23509279 | 0 | 0 | 0 | 0 | 1 | 153 | 103 | 131 | 133 | 200 | 0 | 0 | 0 | 0 | 0 | 157 | 187 | 104 | 155 | 210 | 271 | 149 | 1 | 1 | 0 | 0 | 0 | 0 | 0.003 | 0 | 0 | 0 | 0 | 0 | 0.001 |
| SE | 29805 | ENSG00000141452 | RMC1 | chr18 | + | 23509192 | 23509279 | 23507984 | 23508041 | 23515855 | 23515886 | 280 | 199 | 210 | 230 | 333 | 0 | 0 | 0 | 2 | 2 | 326 | 327 | 198 | 301 | 414 | 0 | 3 | 0 | 0 | 0 | 236 | 149 | 1 | 1 | 1 | 1 | 1 | 0.986 | 0.991 | 1 | 0.986 | 1 | 1 | 1 | -0.002 |
| SE | 29809 | ENSG00000141452 | RMC1 | chr18 | + | 23529634 | 23529712 | 23529178 | 23529298 | 23530027 | 23530132 | 80 | 50 | 83 | 89 | 75 | 0 | 0 | 0 | 0 | 0 | 80 | 48 | 52 | 51 | 35 | 0 | 0 | 0 | 1 | 0 | 227 | 149 | 1 | 1 | 1 | 1 | 1 | 1 | 1 | 1 | 1 | 1 | 0.971 | 1 | 0.006 |
| SE | 30153 | ENSG00000177054 | ZDHHC13 | chr11 | + | 19152157 | 19152320 | 19150726 | 19150791 | 19152558 | 19152625 | 101 | 83 | 122 | 86 | 148 | 1 | 0 | 0 | 0 | 0 | 27 | 40 | 46 | 91 | 15 | 0 | 0 | 0 | 0 | 0 | 298 | 149 | 1 | 1 | 0.981 | 1 | 1 | 1 | 1 | 1 | 1 | 1 | 1 | 1 | -0.004 |
| SE | 30154 | ENSG00000177054 | ZDHHC13 | chr11 | + | 19152157 | 19152320 | 19150726 | 19150791 | 19155795 | 19155929 | 50 | 43 | 57 | 49 | 80 | 0 | 0 | 0 | 0 | 0 | 7 | 28 | 16 | 38 | 4 | 0 | 0 | 0 | 1 | 0 | 298 | 149 | 1 | 1 | 1 | 1 | 1 | 1 | 1 | 1 | 1 | 1 | 0.95 | 1 | 0.01 |
| SE | 30155 | ENSG00000177054 | ZDHHC13 | chr11 | + | 19152558 | 19152684 | 19150726 | 19150791 | 19155795 | 19155929 | 34 | 12 | 23 | 17 | 22 | 0 | 0 | 0 | 0 | 0 | 0 | 30 | 7 | 29 | 7 | 0 | 0 | 0 | 1 | 0 | 275 | 149 | 1 | 1 | 1 | 1 | 1 | 1 | 1 | NA | 1 | 1 | 0.94 | 1 | 0.015 |
| SE | 30156 | ENSG00000177054 | ZDHHC13 | chr11 | + | 19152558 | 19152684 | 19152157 | 19152320 | 19155795 | 19155929 | 84 | 52 | 87 | 54 | 90 | 2 | 0 | 0 | 0 | 0 | 20 | 46 | 37 | 82 | 18 | 0 | 4 | 0 | 0 | 0 | 275 | 149 | 1 | 1 | 0.958 | 1 | 1 | 1 | 1 | 1 | 0.862 | 1 | 1 | 1 | 0.019 |
| SE | 30286 | ENSG00000114480 | GBE1 | chr3 | - | 81648855 | 81648991 | 81642570 | 81642990 | 81649795 | 81649878 | 46 | 43 | 55 | 33 | 44 | 0 | 0 | 0 | 1 | 3 | 27 | 45 | 42 | 51 | 29 | 0 | 3 | 0 | 0 | 0 | 285 | 149 | 1 | 1 | 1 | 1 | 1 | 0.945 | 0.885 | 1 | 0.887 | 1 | 1 | 1 | -0.011 |
| SE | 30348 | ENSG00000108591 | DRG2 | chr17 | + | 18099632 | 18099723 | 18099016 | 18099077 | 18100362 | 18100435 | 57 | 18 | 47 | 66 | 62 | 3 | 0 | 0 | 1 | 0 | 86 | 119 | 23 | 56 | 100 | 0 | 0 | 0 | 0 | 0 | 240 | 149 | 1 | 1 | 0.922 | 1 | 0.967 | 0.976 | 1 | 1 | 1 | 1 | 1 | 1 | -0.027 |
| SE | 30350 | ENSG00000108591 | DRG2 | chr17 | + | 18106432 | 18106470 | 18104622 | 18104681 | 18107153 | 18107959 | 34 | 13 | 16 | 20 | 7 | 2 | 0 | 0 | 1 | 0 | 37 | 23 | 10 | 11 | 17 | 0 | 0 | 0 | 0 | 0 | 187 | 149 | 1 | 1 | 0.931 | 1 | 1 | 0.941 | 1 | 1 | 1 | 1 | 1 | 1 | -0.026 |
| SE | 30351 | ENSG00000108591 | DRG2 | chr17 | + | 18106432 | 18106486 | 18104559 | 18104681 | 18107153 | 18107600 | 98 | 41 | 48 | 63 | 60 | 2 | 0 | 0 | 1 | 0 | 136 | 109 | 33 | 58 | 109 | 0 | 0 | 0 | 0 | 0 | 203 | 149 | 1 | 1 | 0.973 | 1 | 1 | 0.979 | 1 | 1 | 1 | 1 | 1 | 1 | -0.01 |
| SE | 30369 | ENSG00000145348 | TBCK | chr4 | - | 106194717 | 106194754 | 106193608 | 106193770 | 106212749 | 106212835 | 98 | 48 | 74 | 94 | 145 | 0 | 2 | 0 | 4 | 0 | 5 | 30 | 24 | 37 | 10 | 0 | 0 | 0 | 0 | 0 | 186 | 149 | 1 | 1 | 1 | 0.951 | 1 | 0.95 | 1 | 1 | 1 | 1 | 1 | 1 | -0.02 |
| SE | 30370 | ENSG00000145348 | TBCK | chr4 | - | 106233587 | 106233650 | 106232937 | 106233064 | 106235268 | 106235367 | 67 | 53 | 51 | 57 | 180 | 0 | 1 | 0 | 0 | 0 | 11 | 22 | 28 | 18 | 19 | 0 | 0 | 0 | 0 | 0 | 212 | 149 | 1 | 1 | 1 | 0.974 | 1 | 1 | 1 | 1 | 1 | 1 | 1 | 1 | -0.005 |
| SE | 30372 | ENSG00000145348 | TBCK | chr4 | - | 106247138 | 106247287 | 106244625 | 106244764 | 106248244 | 106248306 | 124 | 94 | 67 | 97 | 191 | 0 | 0 | 0 | 1 | 2 | 19 | 82 | 45 | 36 | 34 | 0 | 0 | 0 | 2 | 0 | 298 | 149 | 1 | 1 | 1 | 1 | 1 | 0.98 | 0.979 | 1 | 1 | 1 | 0.9 | 1 | 0.012 |
| SE | 30373 | ENSG00000145348 | TBCK | chr4 | - | 106247138 | 106247287 | 106244625 | 106244764 | 106248920 | 106248982 | 90 | 58 | 49 | 69 | 134 | 0 | 0 | 0 | 0 | 5 | 7 | 45 | 27 | 13 | 33 | 0 | 0 | 0 | 0 | 0 | 298 | 149 | 1 | 1 | 1 | 1 | 1 | 1 | 0.931 | 1 | 1 | 1 | 1 | 1 | -0.014 |
| SE | 30374 | ENSG00000145348 | TBCK | chr4 | - | 106247138 | 106247287 | 106244625 | 106244764 | 106250417 | 106250470 | 90 | 57 | 49 | 67 | 136 | 0 | 0 | 0 | 0 | 4 | 7 | 45 | 27 | 13 | 33 | 0 | 0 | 0 | 0 | 0 | 298 | 149 | 1 | 1 | 1 | 1 | 1 | 1 | 0.944 | 1 | 1 | 1 | 1 | 1 | -0.011 |
| SE | 30375 | ENSG00000145348 | TBCK | chr4 | - | 106248244 | 106248306 | 106244625 | 106244764 | 106248920 | 106248982 | 22 | 15 | 9 | 18 | 30 | 0 | 0 | 0 | 0 | 5 | 0 | 14 | 8 | 9 | 0 | 0 | 0 | 0 | 0 | 0 | 211 | 149 | 1 | 1 | 1 | 1 | 1 | 1 | 0.809 | NA | 1 | 1 | 1 | NA | -0.038 |
| SE | 30376 | ENSG00000145348 | TBCK | chr4 | - | 106248244 | 106248306 | 106247138 | 106247287 | 106248920 | 106248982 | 53 | 51 | 27 | 47 | 86 | 0 | 1 | 0 | 2 | 0 | 12 | 51 | 26 | 30 | 1 | 0 | 0 | 0 | 0 | 0 | 211 | 149 | 1 | 1 | 1 | 0.973 | 1 | 0.943 | 1 | 1 | 1 | 1 | 1 | 1 | -0.017 |
| SE | 30377 | ENSG00000145348 | TBCK | chr4 | - | 106248920 | 106248982 | 106244625 | 106244764 | 106250417 | 106250478 | 14 | 13 | 5 | 12 | 25 | 0 | 0 | 0 | 0 | 4 | 6 | 6 | 8 | 6 | 0 | 0 | 0 | 0 | 0 | 0 | 211 | 149 | 1 | 1 | 1 | 1 | 1 | 1 | 0.815 | 1 | 1 | 1 | 1 | NA | -0.037 |
| SE | 30378 | ENSG00000145348 | TBCK | chr4 | - | 106248920 | 106248982 | 106247138 | 106247287 | 106250417 | 106250478 | 14 | 14 | 5 | 14 | 20 | 0 | 0 | 0 | 0 | 2 | 6 | 6 | 8 | 6 | 0 | 0 | 0 | 0 | 0 | 0 | 211 | 149 | 1 | 1 | 1 | 1 | 1 | 1 | 0.876 | 1 | 1 | 1 | 1 | NA | -0.025 |
| SE | 30380 | ENSG00000145348 | TBCK | chr4 | - | 106254946 | 106255047 | 106251865 | 106252007 | 106260436 | 106260510 | 32 | 32 | 50 | 28 | 76 | 26 | 12 | 18 | 35 | 36 | 0 | 32 | 31 | 2 | 9 | 0 | 15 | 12 | 13 | 0 | 250 | 149 | 1 | 1 | 0.423 | 0.614 | 0.623 | 0.323 | 0.557 | NA | 0.56 | 0.606 | 0.084 | 1 | -0.054 |
| SE | 30381 | ENSG00000145348 | TBCK | chr4 | - | 106260436 | 106260510 | 106251865 | 106252007 | 106295093 | 106295166 | 12 | 7 | 5 | 9 | 12 | 0 | 0 | 4 | 1 | 3 | 0 | 1 | 8 | 10 | 0 | 0 | 0 | 0 | 0 | 1 | 223 | 149 | 1 | 1 | 1 | 1 | 0.455 | 0.857 | 0.728 | NA | 1 | 1 | 1 | 0 | 0.058 |
| SE | 30382 | ENSG00000145348 | TBCK | chr4 | - | 106262097 | 106262212 | 106260436 | 106260510 | 106295093 | 106295166 | 116 | 93 | 118 | 121 | 176 | 0 | 0 | 0 | 0 | 1 | 30 | 125 | 70 | 47 | 35 | 0 | 0 | 0 | 0 | 0 | 264 | 149 | 1 | 1 | 1 | 1 | 1 | 1 | 0.99 | 1 | 1 | 1 | 1 | 1 | -0.002 |
| SE | 30383 | ENSG00000145348 | TBCK | chr4 | - | 106309926 | 106309967 | 106308767 | 106308989 | 106315930 | 106316235 | 1 | 0 | 2 | 4 | 8 | 16 | 29 | 36 | 34 | 21 | 0 | 0 | 0 | 0 | 0 | 0 | 22 | 4 | 7 | 0 | 190 | 149 | 1 | 1 | 0.047 | 0 | 0.042 | 0.084 | 0.23 | NA | 0 | 0 | 0 | NA | 0.081 |
| SE | 30536 | ENSG00000132182 | NUP210 | chr3 | - | 13351878 | 13351980 | 13343174 | 13343303 | 13352079 | 13352184 | 242 | 80 | 181 | 165 | 69 | 2 | 0 | 0 | 0 | 0 | 121 | 270 | 99 | 123 | 206 | 0 | 0 | 0 | 0 | 0 | 251 | 149 | 1 | 1 | 0.986 | 1 | 1 | 1 | 1 | 1 | 1 | 1 | 1 | 1 | -0.003 |
| SE | 30537 | ENSG00000132182 | NUP210 | chr3 | - | 13365945 | 13366091 | 13360269 | 13360491 | 13371833 | 13372032 | 427 | 148 | 303 | 332 | 342 | 0 | 1 | 0 | 0 | 0 | 300 | 403 | 177 | 207 | 378 | 0 | 0 | 0 | 0 | 0 | 295 | 149 | 1 | 1 | 1 | 0.987 | 1 | 1 | 1 | 1 | 1 | 1 | 1 | 1 | -0.003 |
| SE | 30538 | ENSG00000132182 | NUP210 | chr3 | - | 13365945 | 13366091 | 13360269 | 13360491 | 13373717 | 13373873 | 150 | 41 | 99 | 116 | 91 | 1 | 0 | 0 | 0 | 0 | 121 | 157 | 86 | 93 | 175 | 0 | 0 | 0 | 0 | 0 | 295 | 149 | 1 | 1 | 0.987 | 1 | 1 | 1 | 1 | 1 | 1 | 1 | 1 | 1 | -0.003 |
| SE | 30539 | ENSG00000132182 | NUP210 | chr3 | - | 13371833 | 13372032 | 13360269 | 13360491 | 13373717 | 13373873 | 314 | 103 | 230 | 225 | 265 | 1 | 0 | 0 | 0 | 0 | 156 | 293 | 125 | 130 | 291 | 0 | 0 | 0 | 0 | 0 | 298 | 149 | 1 | 1 | 0.994 | 1 | 1 | 1 | 1 | 1 | 1 | 1 | 1 | 1 | -0.001 |
| SE | 30543 | ENSG00000132182 | NUP210 | chr3 | - | 13373717 | 13373873 | 13371833 | 13372032 | 13375503 | 13375641 | 469 | 159 | 368 | 351 | 372 | 0 | 2 | 0 | 0 | 0 | 265 | 389 | 194 | 175 | 431 | 0 | 0 | 1 | 4 | 0 | 298 | 149 | 1 | 1 | 1 | 0.975 | 1 | 1 | 1 | 1 | 1 | 0.99 | 0.956 | 1 | 0.006 |
| SE | 30544 | ENSG00000132182 | NUP210 | chr3 | - | 13386274 | 13386407 | 13379562 | 13379721 | 13388302 | 13388453 | 510 | 300 | 527 | 483 | 557 | 2 | 0 | 2 | 0 | 0 | 439 | 675 | 330 | 334 | 643 | 0 | 0 | 0 | 4 | 0 | 282 | 149 | 1 | 1 | 0.993 | 1 | 0.993 | 1 | 1 | 1 | 1 | 1 | 0.978 | 1 | 0.002 |
| SE | 30545 | ENSG00000132182 | NUP210 | chr3 | - | 13391210 | 13391307 | 13388302 | 13388453 | 13397356 | 13397488 | 286 | 92 | 233 | 242 | 165 | 0 | 0 | 0 | 1 | 0 | 173 | 277 | 176 | 290 | 341 | 0 | 0 | 0 | 0 | 0 | 246 | 149 | 1 | 1 | 1 | 1 | 1 | 0.993 | 1 | 1 | 1 | 1 | 1 | 1 | -0.001 |
| SE | 30853 | ENSG00000109133 | TMEM33 | chr4 | + | 41938601 | 41938696 | 41935128 | 41935529 | 41943746 | 41943814 | 22 | 20 | 22 | 16 | 24 | 0 | 0 | 0 | 0 | 0 | 25 | 23 | 41 | 7 | 18 | 0 | 0 | 3 | 0 | 0 | 244 | 149 | 1 | 1 | 1 | 1 | 1 | 1 | 1 | 1 | 1 | 0.893 | 1 | 1 | 0.021 |
| SE | 30854 | ENSG00000109133 | TMEM33 | chr4 | + | 41938601 | 41938696 | 41935128 | 41935529 | 41944792 | 41944926 | 18 | 22 | 20 | 21 | 19 | 1 | 0 | 0 | 0 | 0 | 25 | 18 | 27 | 7 | 15 | 0 | 0 | 0 | 0 | 0 | 244 | 149 | 1 | 1 | 0.917 | 1 | 1 | 1 | 1 | 1 | 1 | 1 | 1 | 1 | -0.017 |
| SE | 30856 | ENSG00000109133 | TMEM33 | chr4 | + | 41939720 | 41939828 | 41938601 | 41938696 | 41943746 | 41943814 | 6 | 4 | 3 | 5 | 3 | 5 | 4 | 2 | 0 | 9 | 0 | 9 | 1 | 0 | 0 | 0 | 5 | 15 | 0 | 3 | 257 | 149 | 1 | 1 | 0.41 | 0.367 | 0.465 | 1 | 0.162 | NA | 0.511 | 0.037 | NA | 0 | 0.298 |
| SE | 30858 | ENSG00000109133 | TMEM33 | chr4 | + | 41943746 | 41943814 | 41935128 | 41935529 | 41944792 | 41944926 | 97 | 53 | 46 | 74 | 99 | 1 | 0 | 0 | 0 | 0 | 10 | 62 | 58 | 60 | 13 | 0 | 0 | 0 | 0 | 0 | 217 | 149 | 1 | 1 | 0.985 | 1 | 1 | 1 | 1 | 1 | 1 | 1 | 1 | 1 | -0.003 |
| SE | 30998 | ENSG00000113504 | SLC12A7 | chr5 | - | 1056581 | 1056596 | 1053348 | 1053482 | 1057470 | 1057649 | 2 | 0 | 2 | 2 | 0 | 101 | 38 | 18 | 74 | 29 | 0 | 0 | 0 | 0 | 1 | 72 | 52 | 39 | 31 | 54 | 164 | 149 | 1 | 1 | 0.018 | 0 | 0.092 | 0.024 | 0 | 0 | 0 | 0 | 0 | 0.017 | 0.023 |
| SE | 30999 | ENSG00000113504 | SLC12A7 | chr5 | - | 1085231 | 1085473 | 1065282 | 1065478 | 1093532 | 1093575 | 1 | 0 | 0 | 0 | 0 | 0 | 0 | 0 | 2 | 0 | 0 | 2 | 0 | 0 | 0 | 0 | 0 | 1 | 0 | 0 | 298 | 149 | 1 | 1 | 1 | NA | NA | 0 | NA | NA | 1 | 0 | NA | NA | 0 |
| SE | 31000 | ENSG00000113504 | SLC12A7 | chr5 | - | 1086902 | 1087033 | 1085231 | 1085473 | 1093532 | 1093575 | 34 | 16 | 9 | 15 | 18 | 0 | 0 | 0 | 0 | 0 | 28 | 16 | 13 | 12 | 26 | 0 | 2 | 0 | 0 | 0 | 280 | 149 | 1 | 1 | 1 | 1 | 1 | 1 | 1 | 1 | 0.81 | 1 | 1 | 1 | 0.038 |
| SE | 31002 | ENSG00000113504 | SLC12A7 | chr5 | - | 1088981 | 1089128 | 1088305 | 1088360 | 1093532 | 1093575 | 118 | 62 | 24 | 67 | 73 | 0 | 0 | 0 | 0 | 0 | 77 | 93 | 40 | 43 | 127 | 0 | 1 | 0 | 0 | 0 | 296 | 149 | 1 | 1 | 1 | 1 | 1 | 1 | 1 | 1 | 0.979 | 1 | 1 | 1 | 0.004 |
| SE | 31097 | ENSG00000099949 | LZTR1 | chr22 | + | 20987503 | 20987583 | 20983026 | 20983089 | 20988009 | 20988113 | 33 | 16 | 25 | 23 | 6 | 1 | 0 | 0 | 0 | 0 | 24 | 30 | 11 | 23 | 33 | 0 | 1 | 0 | 0 | 0 | 229 | 149 | 1 | 1 | 0.955 | 1 | 1 | 1 | 1 | 1 | 1 | 1 | 1 | 1 | -0.009 |
| SE | 31100 | ENSG00000099949 | LZTR1 | chr22 | + | 20995367 | 20995484 | 20994869 | 20995026 | 20995762 | 20995872 | 3 | 0 | 2 | 3 | 2 | 0 | 0 | 0 | 1 | 0 | 4 | 0 | 0 | 0 | 0 | 0 | 0 | 0 | 0 | 3 | 266 | 149 | 1 | 1 | 1 | NA | 1 | 0.627 | 1 | 1 | NA | NA | NA | 0 | 0.407 |
| SE | 31135 | ENSG00000134825 | TMEM258 | chr11 | - | 61789784 | 61789990 | 61789129 | 61789235 | 61790374 | 61790602 | 184 | 88 | 197 | 179 | 91 | 0 | 0 | 1 | 1 | 0 | 248 | 217 | 86 | 179 | 166 | 2 | 0 | 0 | 0 | 0 | 298 | 149 | 1 | 1 | 1 | 1 | 0.99 | 0.989 | 1 | 0.984 | 1 | 1 | 1 | 1 | -0.001 |
| SE | 31461 | ENSG00000178950 | GAK | chr4 | - | 851749 | 851974 | 850935 | 851084 | 859605 | 859722 | 162 | 86 | 107 | 146 | 53 | 1 | 0 | 2 | 0 | 0 | 221 | 167 | 127 | 138 | 197 | 0 | 0 | 0 | 3 | 0 | 298 | 149 | 1 | 1 | 0.988 | 1 | 0.964 | 1 | 1 | 1 | 1 | 1 | 0.958 | 1 | -0.001 |
| SE | 31462 | ENSG00000178950 | GAK | chr4 | - | 859361 | 859493 | 850935 | 851084 | 859605 | 859722 | 0 | 0 | 1 | 1 | 0 | 1 | 0 | 0 | 0 | 0 | 0 | 0 | 0 | 8 | 0 | 0 | 0 | 0 | 3 | 0 | 281 | 149 | 1 | 1 | 0 | NA | 0.21 | 1 | NA | NA | NA | NA | 0.586 | NA | -0.183 |
| SE | 31463 | ENSG00000178950 | GAK | chr4 | - | 859361 | 859493 | 851749 | 851974 | 859605 | 859722 | 4 | 0 | 1 | 3 | 0 | 44 | 19 | 36 | 34 | 13 | 0 | 4 | 0 | 0 | 3 | 68 | 25 | 33 | 36 | 35 | 281 | 149 | 1 | 1 | 0.046 | 0 | 0.015 | 0.045 | 0 | 0 | 0.078 | 0 | 0 | 0.043 | -0.003 |
| SE | 31468 | ENSG00000178950 | GAK | chr4 | - | 888846 | 888970 | 884036 | 884086 | 890531 | 890622 | 169 | 58 | 118 | 138 | 71 | 0 | 0 | 0 | 0 | 0 | 149 | 133 | 89 | 105 | 248 | 0 | 0 | 2 | 0 | 0 | 273 | 149 | 1 | 1 | 1 | 1 | 1 | 1 | 1 | 1 | 1 | 0.96 | 1 | 1 | 0.008 |
| SE | 31469 | ENSG00000178950 | GAK | chr4 | - | 896459 | 896549 | 893873 | 894009 | 898032 | 898158 | 393 | 219 | 355 | 339 | 520 | 2 | 0 | 0 | 0 | 0 | 471 | 481 | 225 | 390 | 640 | 10 | 0 | 0 | 0 | 0 | 239 | 149 | 1 | 1 | 0.992 | 1 | 1 | 1 | 1 | 0.967 | 1 | 1 | 1 | 1 | 0.005 |
| SE | 31470 | ENSG00000178950 | GAK | chr4 | - | 898032 | 898158 | 893942 | 894009 | 904636 | 904779 | 126 | 74 | 106 | 116 | 120 | 0 | 0 | 0 | 0 | 0 | 201 | 136 | 73 | 140 | 217 | 0 | 0 | 0 | 5 | 0 | 275 | 149 | 1 | 1 | 1 | 1 | 1 | 1 | 1 | 1 | 1 | 1 | 0.938 | 1 | 0.012 |
| SE | 31474 | ENSG00000178950 | GAK | chr4 | - | 912734 | 912794 | 912029 | 912174 | 913606 | 913668 | 34 | 11 | 21 | 23 | 13 | 0 | 0 | 1 | 0 | 0 | 39 | 25 | 20 | 33 | 17 | 0 | 0 | 0 | 0 | 0 | 209 | 149 | 1 | 1 | 1 | 1 | 0.937 | 1 | 1 | 1 | 1 | 1 | 1 | 1 | -0.013 |
| SE | 31476 | ENSG00000178950 | GAK | chr4 | - | 913606 | 913668 | 904636 | 904779 | 932042 | 932220 | 68 | 27 | 44 | 41 | 15 | 2 | 0 | 1 | 0 | 0 | 76 | 47 | 26 | 47 | 71 | 0 | 1 | 0 | 0 | 0 | 211 | 149 | 1 | 1 | 0.96 | 1 | 0.969 | 1 | 1 | 1 | 0.971 | 1 | 1 | 1 | -0.008 |
| SE | 31554 | ENSG00000163913 | IFT122 | chr3 | + | 129456113 | 129456264 | 129451913 | 129451998 | 129458598 | 129458677 | 2 | 0 | 1 | 0 | 2 | 26 | 19 | 15 | 16 | 15 | 0 | 0 | 0 | 0 | 0 | 50 | 32 | 19 | 40 | 74 | 298 | 149 | 1 | 1 | 0.037 | 0 | 0.032 | 0 | 0.062 | 0 | 0 | 0 | 0 | 0 | 0.026 |
| SE | 31555 | ENSG00000163913 | IFT122 | chr3 | + | 129457921 | 129458015 | 129451913 | 129451998 | 129458598 | 129458677 | 0 | 0 | 0 | 0 | 0 | 26 | 19 | 15 | 16 | 15 | 0 | 3 | 0 | 0 | 2 | 50 | 32 | 19 | 40 | 74 | 243 | 149 | 1 | 1 | 0 | 0 | 0 | 0 | 0 | 0 | 0.054 | 0 | 0 | 0.016 | -0.014 |
| SE | 31559 | ENSG00000163913 | IFT122 | chr3 | + | 129460853 | 129461006 | 129458598 | 129458677 | 129461227 | 129461304 | 0 | 0 | 0 | 0 | 0 | 25 | 15 | 17 | 26 | 9 | 0 | 2 | 2 | 0 | 0 | 100 | 24 | 14 | 41 | 55 | 298 | 149 | 1 | 1 | 0 | 0 | 0 | 0 | 0 | 0 | 0.04 | 0.067 | 0 | 0 | -0.021 |
| SE | 31567 | ENSG00000163913 | IFT122 | chr3 | + | 129499901 | 129500068 | 129495445 | 129495607 | 129502710 | 129502764 | 23 | 7 | 21 | 19 | 23 | 0 | 0 | 0 | 0 | 0 | 78 | 22 | 17 | 29 | 35 | 0 | 0 | 0 | 2 | 0 | 298 | 149 | 1 | 1 | 1 | 1 | 1 | 1 | 1 | 1 | 1 | 1 | 0.879 | 1 | 0.024 |
| SE | 31568 | ENSG00000163913 | IFT122 | chr3 | + | 129504318 | 129504421 | 129502710 | 129502882 | 129506408 | 129506549 | 4 | 5 | 6 | 9 | 4 | 0 | 0 | 0 | 2 | 0 | 0 | 3 | 8 | 19 | 2 | 0 | 2 | 0 | 2 | 0 | 252 | 149 | 1 | 1 | 1 | 1 | 1 | 0.727 | 1 | NA | 0.47 | 1 | 0.849 | 1 | 0.116 |
| SE | 32333 | ENSG00000068745 | IP6K2 | chr3 | - | 48694693 | 48694824 | 48693487 | 48694240 | 48695089 | 48695418 | 0 | 0 | 4 | 0 | 0 | 4 | 6 | 2 | 12 | 13 | 0 | 4 | 0 | 0 | 0 | 10 | 19 | 3 | 0 | 10 | 280 | 149 | 1 | 1 | 0 | 0 | 0.516 | 0 | 0 | 0 | 0.101 | 0 | NA | 0 | 0.078 |
| SE | 32334 | ENSG00000068745 | IP6K2 | chr3 | - | 48694701 | 48694824 | 48694174 | 48694240 | 48695089 | 48695418 | 0 | 0 | 4 | 0 | 0 | 4 | 6 | 2 | 12 | 13 | 0 | 4 | 0 | 0 | 0 | 10 | 19 | 3 | 0 | 10 | 272 | 149 | 1 | 1 | 0 | 0 | 0.523 | 0 | 0 | 0 | 0.103 | 0 | NA | 0 | 0.079 |
| SE | 32335 | ENSG00000068745 | IP6K2 | chr3 | - | 48708119 | 48708237 | 48695089 | 48695418 | 48711366 | 48711459 | 5 | 0 | 0 | 0 | 0 | 0 | 0 | 1 | 0 | 0 | 0 | 2 | 0 | 0 | 0 | 0 | 0 | 0 | 0 | 8 | 267 | 149 | 1 | 1 | 1 | NA | 0 | NA | NA | NA | 1 | NA | NA | 0 | 0 |
| SE | 32336 | ENSG00000068745 | IP6K2 | chr3 | - | 48708119 | 48708237 | 48695089 | 48695418 | 48715314 | 48715527 | 3 | 0 | 0 | 0 | 0 | 5 | 1 | 5 | 4 | 5 | 0 | 4 | 0 | 0 | 0 | 0 | 5 | 2 | 4 | 10 | 267 | 149 | 1 | 1 | 0.251 | 0 | 0 | 0 | 0 | NA | 0.309 | 0 | 0 | 0 | -0.027 |
| SE | 32337 | ENSG00000068745 | IP6K2 | chr3 | - | 48708119 | 48708237 | 48695089 | 48695418 | 48717156 | 48717210 | 4 | 0 | 2 | 0 | 0 | 5 | 6 | 11 | 5 | 8 | 0 | 2 | 0 | 0 | 0 | 0 | 16 | 1 | 11 | 11 | 267 | 149 | 1 | 1 | 0.309 | 0 | 0.092 | 0 | 0 | NA | 0.065 | 0 | 0 | 0 | 0.064 |
| SE | 32338 | ENSG00000068745 | IP6K2 | chr3 | - | 48708119 | 48708237 | 48695089 | 48695421 | 48711366 | 48711459 | 2 | 0 | 1 | 1 | 0 | 0 | 0 | 0 | 0 | 0 | 0 | 0 | 0 | 0 | 0 | 0 | 5 | 0 | 0 | 0 | 267 | 149 | 1 | 1 | 1 | NA | 1 | 1 | NA | NA | 0 | NA | NA | NA | 1 |
| SE | 32339 | ENSG00000068745 | IP6K2 | chr3 | - | 48708119 | 48708237 | 48695089 | 48695421 | 48715314 | 48715527 | 0 | 0 | 1 | 1 | 0 | 5 | 3 | 0 | 2 | 0 | 0 | 2 | 0 | 0 | 0 | 11 | 2 | 0 | 0 | 0 | 267 | 149 | 1 | 1 | NA | 0 | 1 | 0.218 | NA | 0 | 0.358 | NA | NA | NA | 0.227 |
| SE | 32340 | ENSG00000068745 | IP6K2 | chr3 | - | 48708119 | 48708237 | 48695089 | 48695421 | 48717156 | 48717210 | 1 | 0 | 3 | 1 | 0 | 33 | 7 | 7 | 17 | 18 | 0 | 0 | 0 | 0 | 0 | 32 | 4 | 1 | 1 | 12 | 267 | 149 | 1 | 1 | 0.017 | 0 | 0.193 | 0.032 | 0 | 0 | 0 | 0 | 0 | 0 | 0.048 |
| SE | 32341 | ENSG00000068745 | IP6K2 | chr3 | - | 48711366 | 48711459 | 48695089 | 48695418 | 48715314 | 48715527 | 0 | 0 | 1 | 0 | 0 | 5 | 1 | 5 | 4 | 5 | 0 | 5 | 0 | 2 | 0 | 0 | 5 | 2 | 4 | 10 | 242 | 149 | 1 | 1 | 0 | 0 | 0.11 | 0 | 0 | NA | 0.381 | 0 | 0.235 | 0 | -0.132 |
| SE | 32343 | ENSG00000068745 | IP6K2 | chr3 | - | 48711366 | 48711459 | 48695089 | 48695418 | 48717156 | 48717188 | 0 | 0 | 1 | 0 | 0 | 5 | 6 | 11 | 5 | 8 | 0 | 0 | 0 | 0 | 0 | 0 | 16 | 1 | 11 | 11 | 242 | 149 | 1 | 1 | 0 | 0 | 0.053 | 0 | 0 | NA | 0 | 0 | 0 | 0 | 0.011 |
| SE | 32344 | ENSG00000068745 | IP6K2 | chr3 | - | 48711366 | 48711459 | 48695089 | 48695421 | 48715314 | 48715527 | 0 | 0 | 0 | 0 | 0 | 5 | 3 | 0 | 2 | 0 | 0 | 5 | 0 | 2 | 0 | 11 | 2 | 0 | 0 | 0 | 242 | 149 | 1 | 1 | NA | 0 | NA | 0 | NA | 0 | 0.606 | NA | 1 | NA | -0.535 |
| SE | 32345 | ENSG00000068745 | IP6K2 | chr3 | - | 48711366 | 48711459 | 48695089 | 48695421 | 48715388 | 48715529 | 0 | 0 | 1 | 0 | 0 | 1 | 0 | 3 | 0 | 1 | 0 | 0 | 0 | 0 | 0 | 0 | 4 | 0 | 1 | 4 | 242 | 149 | 1 | 1 | 0 | NA | 0.17 | NA | 0 | NA | 0 | NA | 0 | 0 | 0.057 |
| SE | 32347 | ENSG00000068745 | IP6K2 | chr3 | - | 48711366 | 48711459 | 48708119 | 48708237 | 48715314 | 48715527 | 1 | 0 | 0 | 0 | 0 | 0 | 0 | 0 | 0 | 0 | 0 | 5 | 0 | 2 | 0 | 0 | 2 | 0 | 0 | 0 | 242 | 149 | 1 | 1 | 1 | NA | NA | NA | NA | NA | 0.606 | NA | 1 | NA | 0.197 |
| SE | 32350 | ENSG00000068745 | IP6K2 | chr3 | - | 48715314 | 48715527 | 48695089 | 48695418 | 48717156 | 48717188 | 9 | 7 | 13 | 10 | 8 | 5 | 6 | 11 | 5 | 8 | 0 | 10 | 3 | 10 | 23 | 0 | 16 | 1 | 11 | 11 | 298 | 149 | 1 | 1 | 0.474 | 0.368 | 0.371 | 0.5 | 0.333 | NA | 0.238 | 0.6 | 0.312 | 0.511 | -0.006 |
| SE | 32351 | ENSG00000068745 | IP6K2 | chr3 | - | 48715314 | 48715527 | 48708119 | 48708237 | 48717470 | 48717188 | 4 | 6 | 8 | 6 | 3 | 1 | 0 | 2 | 0 | 0 | 0 | 7 | 1 | 6 | 13 | 0 | 1 | 0 | 0 | 0 | 298 | 149 | 1 | 1 | 0.667 | 1 | 0.667 | 1 | 1 | NA | 1 | 1 | 1 | 1 | -0.133 |
| SE | 32352 | ENSG00000068745 | IP6K2 | chr3 | - | 48715314 | 48715527 | 48711366 | 48711459 | 48717156 | 48717188 | 4 | 6 | 8 | 6 | 3 | 1 | 0 | 0 | 0 | 0 | 0 | 10 | 1 | 8 | 13 | 0 | 0 | 0 | 0 | 0 | 298 | 149 | 1 | 1 | 0.667 | 1 | 1 | 1 | 1 | NA | 1 | 1 | 1 | 1 | -0.067 |
| SE | 32353 | ENSG00000068745 | IP6K2 | chr3 | - | 48715388 | 48715527 | 48694050 | 48695418 | 48717156 | 48717192 | 5 | 7 | 8 | 7 | 3 | 5 | 6 | 11 | 5 | 8 | 0 | 5 | 1 | 6 | 13 | 0 | 16 | 1 | 11 | 11 | 288 | 149 | 1 | 1 | 0.341 | 0.376 | 0.273 | 0.42 | 0.162 | NA | 0.139 | 0.341 | 0.22 | 0.379 | 0.045 |
| SE | 32355 | ENSG00000068745 | IP6K2 | chr3 | - | 48716431 | 48716477 | 48715314 | 48715527 | 48717156 | 48717215 | 1 | 0 | 3 | 0 | 0 | 4 | 6 | 8 | 6 | 3 | 0 | 0 | 0 | 0 | 0 | 0 | 5 | 1 | 6 | 13 | 195 | 149 | 1 | 1 | 0.16 | 0 | 0.223 | 0 | 0 | NA | 0 | 0 | 0 | 0 | 0.077 |
| SE | 32550 | ENSG00000211456 | SACM1L | chr3 | + | 45690347 | 45690387 | 45689240 | 45689497 | 45703437 | 45703535 | 15 | 12 | 19 | 8 | 5 | 55 | 24 | 52 | 37 | 16 | 4 | 1 | 18 | 2 | 0 | 7 | 23 | 12 | 6 | 0 | 189 | 149 | 1 | 1 | 0.177 | 0.283 | 0.224 | 0.146 | 0.198 | 0.311 | 0.033 | 0.542 | 0.208 | NA | -0.068 |
| SE | 32552 | ENSG00000211456 | SACM1L | chr3 | + | 45731300 | 45731380 | 45723474 | 45723543 | 45732052 | 45732151 | 415 | 304 | 304 | 382 | 282 | 2 | 2 | 1 | 8 | 0 | 51 | 146 | 153 | 90 | 12 | 0 | 6 | 0 | 0 | 0 | 229 | 149 | 1 | 1 | 0.993 | 0.99 | 0.995 | 0.969 | 1 | 1 | 0.941 | 1 | 1 | 1 | 0.001 |
| SE | 32553 | ENSG00000211456 | SACM1L | chr3 | + | 45732052 | 45732151 | 45723474 | 45723543 | 45735234 | 45735373 | 127 | 85 | 98 | 135 | 131 | 0 | 0 | 0 | 0 | 3 | 16 | 30 | 29 | 21 | 4 | 0 | 0 | 0 | 0 | 0 | 248 | 149 | 1 | 1 | 1 | 1 | 1 | 1 | 0.963 | 1 | 1 | 1 | 1 | 1 | -0.007 |
| SE | 32556 | ENSG00000211456 | SACM1L | chr3 | + | 45739586 | 45739644 | 45738780 | 45738873 | 45743532 | 45743735 | 73 | 54 | 55 | 81 | 60 | 1 | 0 | 3 | 0 | 6 | 0 | 35 | 7 | 20 | 5 | 0 | 1 | 0 | 0 | 0 | 207 | 149 | 1 | 1 | 0.981 | 1 | 0.93 | 1 | 0.878 | NA | 0.962 | 1 | 1 | 1 | -0.033 |
| SE | 32842 | ENSG00000184470 | TXNRD2 | chr22 | - | 19919542 | 19919599 | 19918859 | 19919004 | 19931029 | 19931095 | 44 | 19 | 19 | 28 | 9 | 0 | 0 | 1 | 0 | 0 | 34 | 22 | 18 | 21 | 90 | 0 | 1 | 0 | 0 | 0 | 206 | 149 | 1 | 1 | 1 | 1 | 0.932 | 1 | 1 | 1 | 1 | 1 | 1 | 1 | -0.014 |
| SE | 33041 | ENSG00000154917 | RAB6B | chr3 | - | 133864583 | 133864642 | 133841609 | 133841663 | 133895396 | 133895595 | 28 | 24 | 4 | 36 | 40 | 0 | 0 | 0 | 1 | 0 | 187 | 52 | 26 | 36 | 422 | 0 | 0 | 0 | 0 | 0 | 208 | 149 | 1 | 1 | 1 | 1 | 1 | 0.963 | 1 | 1 | 1 | 1 | 1 | 1 | -0.007 |
| SE | 33410 | ENSG00000096063 | SRPK1 | chr6 | - | 35857260 | 35857368 | 35842534 | 35842604 | 35869009 | 35869110 | 185 | 102 | 123 | 151 | 181 | 0 | 0 | 0 | 0 | 2 | 72 | 188 | 235 | 237 | 20 | 0 | 0 | 0 | 0 | 0 | 257 | 149 | 1 | 1 | 1 | 1 | 1 | 1 | 0.981 | 1 | 1 | 1 | 1 | 1 | -0.004 |
| SE | 33412 | ENSG00000096063 | SRPK1 | chr6 | - | 35869009 | 35869138 | 35856933 | 35857368 | 35869481 | 35869551 | 81 | 52 | 54 | 51 | 82 | 1 | 0 | 2 | 1 | 0 | 13 | 55 | 100 | 106 | 14 | 0 | 2 | 7 | 9 | 0 | 278 | 149 | 1 | 1 | 0.977 | 1 | 0.935 | 0.965 | 1 | 1 | 0.936 | 0.884 | 0.863 | 1 | 0.039 |
| SE | 33414 | ENSG00000096063 | SRPK1 | chr6 | - | 35872562 | 35872728 | 35870933 | 35870959 | 35874232 | 35874339 | 111 | 139 | 138 | 157 | 239 | 0 | 1 | 0 | 0 | 0 | 48 | 159 | 350 | 230 | 10 | 0 | 0 | 0 | 0 | 0 | 298 | 149 | 1 | 1 | 1 | 0.986 | 1 | 1 | 1 | 1 | 1 | 1 | 1 | 1 | -0.003 |
| SE | 33494 | ENSG00000164068 | RNF123 | chr3 | + | 49691424 | 49691509 | 49691129 | 49691247 | 49697142 | 49697204 | 29 | 28 | 34 | 42 | 24 | 0 | 0 | 0 | 0 | 0 | 86 | 85 | 25 | 41 | 79 | 0 | 0 | 0 | 0 | 0 | 234 | 149 | 1 | 1 | 1 | 1 | 1 | 0.964 | 1 | 1 | 1 | 1 | 1 | 1 | -0.007 |
| SE | 33495 | ENSG00000164068 | RNF123 | chr3 | + | 49697044 | 49697222 | 49691424 | 49691509 | 49697362 | 49697412 | 32 | 17 | 20 | 30 | 11 | 0 | 0 | 0 | 0 | 0 | 13 | 22 | 13 | 38 | 43 | 0 | 0 | 2 | 0 | 0 | 298 | 149 | 1 | 1 | 1 | 1 | 1 | 1 | 1 | 1 | 1 | 0.765 | 1 | 1 | 0.047 |
| SE | 33496 | ENSG00000164068 | RNF123 | chr3 | + | 49697142 | 49697222 | 49691424 | 49691509 | 49697362 | 49697457 | 54 | 34 | 41 | 62 | 21 | 0 | 0 | 0 | 0 | 0 | 70 | 89 | 20 | 62 | 88 | 0 | 0 | 2 | 0 | 0 | 229 | 149 | 1 | 1 | 1 | 1 | 1 | 1 | 1 | 1 | 1 | 0.867 | 1 | 1 | 0.027 |
| SE | 33497 | ENSG00000164068 | RNF123 | chr3 | + | 49698754 | 49698822 | 49698439 | 49698526 | 49698979 | 49699105 | 37 | 10 | 26 | 22 | 10 | 1 | 0 | 3 | 0 | 0 | 9 | 37 | 13 | 23 | 34 | 0 | 0 | 0 | 0 | 0 | 217 | 149 | 1 | 1 | 0.962 | 1 | 1 | 0.834 | 1 | 1 | 1 | 1 | 1 | 1 | -0.041 |
| SE | 33498 | ENSG00000164068 | RNF123 | chr3 | + | 49713737 | 49713825 | 49712478 | 49713587 | 49713909 | 49713997 | 26 | 12 | 19 | 25 | 9 | 0 | 0 | 0 | 0 | 0 | 11 | 17 | 21 | 9 | 74 | 0 | 0 | 2 | 0 | 0 | 237 | 149 | 1 | 1 | 1 | 1 | 1 | 1 | 1 | 1 | 1 | 0.868 | 1 | 1 | 0.026 |
| SE | 33828 | ENSG00000041802 | LSG1 | chr3 | - | 194670008 | 194670135 | 194666451 | 194666572 | 194672063 | 194672162 | 177 | 113 | 132 | 118 | 182 | 0 | 0 | 4 | 0 | 0 | 224 | 131 | 137 | 195 | 77 | 0 | 0 | 0 | 3 | 0 | 276 | 149 | 1 | 1 | 1 | 0.938 | 1 | 1 | 1 | 1 | 1 | 1 | 0.972 | 1 | -0.007 |
| SE | 34621 | ENSG00000198700 | IPO9 | chr1 | + | 201855782 | 201855934 | 201855123 | 201855182 | 201857095 | 201857194 | 163 | 99 | 114 | 156 | 327 | 1 | 1 | 0 | 0 | 1 | 79 | 105 | 55 | 181 | 157 | 0 | 2 | 1 | 0 | 0 | 298 | 149 | 1 | 1 | 0.988 | 0.98 | 1 | 1 | 0.994 | 1 | 0.963 | 0.965 | 1 | 1 | 0.007 |
| SE | 34622 | ENSG00000198700 | IPO9 | chr1 | + | 201857095 | 201857194 | 201855782 | 201855934 | 201858446 | 201858553 | 200 | 89 | 132 | 158 | 315 | 0 | 0 | 0 | 0 | 1 | 105 | 128 | 72 | 150 | 158 | 0 | 0 | 0 | 0 | 0 | 248 | 149 | 1 | 1 | 1 | 1 | 1 | 1 | 0.995 | 1 | 1 | 1 | 1 | 1 | -0.001 |
| SE | 34624 | ENSG00000198700 | IPO9 | chr1 | + | 201871160 | 201871327 | 201870582 | 201870858 | 201872827 | 201872961 | 94 | 42 | 91 | 107 | 47 | 0 | 1 | 0 | 0 | 0 | 76 | 55 | 53 | 75 | 99 | 0 | 0 | 0 | 2 | 0 | 298 | 149 | 1 | 1 | 1 | 0.955 | 1 | 1 | 1 | 1 | 1 | 1 | 1 | 0.961 | -0.001 |
| SE | 34970 | ENSG00000117133 | RPF1 | chr1 | + | 84496243 | 84496370 | 84495881 | 84496063 | 84497428 | 84498352 | 727 | 480 | 571 | 561 | 886 | 0 | 0 | 0 | 0 | 0 | 244 | 541 | 591 | 389 | 53 | 6 | 0 | 0 | 0 | 0 | 276 | 149 | 1 | 1 | 1 | 1 | 1 | 1 | 1 | 0.956 | 1 | 1 | 1 | 1 | 0.009 |
| SE | 35448 | ENSG00000111642 | CHD4 | chr12 | - | 6578845 | 6578917 | 6577784 | 6578546 | 6581043 | 6581173 | 356 | 276 | 328 | 441 | 462 | 0 | 0 | 0 | 0 | 0 | 1090 | 520 | 442 | 698 | 876 | 0 | 0 | 0 | 5 | 0 | 221 | 149 | 1 | 1 | 1 | 1 | 1 | 1 | 1 | 1 | 1 | 1 | 0.989 | 1 | 0.002 |
| SE | 35449 | ENSG00000111642 | CHD4 | chr12 | - | 6580581 | 6580679 | 6578845 | 6578917 | 6581043 | 6581173 | 0 | 5 | 3 | 1 | 3 | 320 | 254 | 303 | 402 | 418 | 0 | 1 | 4 | 3 | 0 | 1002 | 488 | 397 | 640 | 830 | 247 | 149 | 1 | 1 | 0 | 0.012 | 0.006 | 0.001 | 0.004 | 0 | 0.001 | 0.006 | 0.003 | 0 | 0.003 |
| SE | 35450 | ENSG00000111642 | CHD4 | chr12 | - | 6580581 | 6580699 | 6578845 | 6578917 | 6581043 | 6581173 | 0 | 1 | 1 | 1 | 0 | 320 | 254 | 303 | 402 | 418 | 0 | 1 | 0 | 3 | 0 | 1002 | 488 | 397 | 640 | 830 | 267 | 149 | 1 | 1 | 0 | 0.002 | 0.002 | 0.001 | 0 | 0 | 0.001 | 0 | 0.003 | 0 | 0 |
| SE | 35451 | ENSG00000111642 | CHD4 | chr12 | - | 6581648 | 6581814 | 6581290 | 6581388 | 6582136 | 6582281 | 671 | 522 | 773 | 881 | 734 | 0 | 0 | 0 | 0 | 0 | 1753 | 1041 | 897 | 1357 | 1556 | 0 | 7 | 0 | 2 | 0 | 298 | 149 | 1 | 1 | 1 | 1 | 1 | 1 | 1 | 1 | 0.987 | 1 | 0.997 | 1 | 0.003 |
| SE | 35457 | ENSG00000111642 | CHD4 | chr12 | - | 6590072 | 6590213 | 6588297 | 6588422 | 6591465 | 6591583 | 1 | 0 | 0 | 0 | 6 | 310 | 184 | 316 | 300 | 241 | 0 | 0 | 0 | 0 | 0 | 451 | 389 | 231 | 280 | 404 | 290 | 149 | 1 | 1 | 0.002 | 0 | 0 | 0 | 0.013 | 0 | 0 | 0 | 0 | 0 | 0.003 |
| SE | 35722 | ENSG00000130787 | HIP1R | chr12 | + | 122855979 | 122856163 | 122855830 | 122855903 | 122856255 | 122856344 | 160 | 83 | 141 | 130 | 51 | 0 | 0 | 0 | 0 | 0 | 146 | 242 | 76 | 87 | 231 | 0 | 0 | 3 | 0 | 0 | 298 | 149 | 1 | 1 | 1 | 1 | 1 | 1 | 1 | 1 | 1 | 0.927 | 1 | 1 | 0.015 |
| SE | 35724 | ENSG00000130787 | HIP1R | chr12 | + | 122861307 | 122861514 | 122861130 | 122861192 | 122861705 | 122862084 | 105 | 49 | 121 | 108 | 91 | 0 | 0 | 0 | 1 | 0 | 151 | 237 | 77 | 50 | 192 | 0 | 0 | 0 | 0 | 0 | 298 | 149 | 1 | 1 | 1 | 1 | 1 | 0.982 | 1 | 1 | 1 | 1 | 1 | 1 | -0.004 |
| SE | 35992 | ENSG00000145428 | RNF175 | chr4 | - | 153728206 | 153728361 | 153723350 | 153723458 | 153745668 | 153745727 | 0 | 4 | 1 | 5 | 0 | 1 | 0 | 0 | 0 | 0 | 3 | 0 | 0 | 0 | 0 | 0 | 3 | 0 | 0 | 0 | 298 | 149 | 1 | 1 | 0 | 1 | 1 | 1 | NA | 1 | 0 | NA | NA | NA | 0.25 |
| SE | 35994 | ENSG00000145428 | RNF175 | chr4 | - | 153728206 | 153728361 | 153723350 | 153723458 | 153751437 | 153751475 | 0 | 4 | 1 | 5 | 0 | 1 | 0 | 0 | 0 | 0 | 3 | 0 | 0 | 0 | 0 | 0 | 0 | 0 | 0 | 0 | 298 | 149 | 1 | 1 | 0 | 1 | 1 | 1 | NA | 1 | NA | NA | NA | NA | -0.25 |
| SE | 35997 | ENSG00000145428 | RNF175 | chr4 | - | 153748644 | 153748786 | 153723350 | 153723458 | 153751437 | 153751475 | 0 | 0 | 0 | 1 | 0 | 1 | 0 | 0 | 0 | 0 | 0 | 3 | 0 | 0 | 0 | 0 | 0 | 0 | 0 | 0 | 291 | 149 | 1 | 1 | 0 | NA | NA | 1 | NA | NA | 1 | NA | NA | NA | -0.5 |
| SE | 36661 | ENSG00000180329 | CCDC43 | chr17 | - | 44680584 | 44680643 | 44677439 | 44679043 | 44682002 | 44682138 | 114 | 101 | 82 | 122 | 189 | 6 | 11 | 9 | 11 | 15 | 232 | 88 | 171 | 161 | 0 | 0 | 3 | 13 | 6 | 0 | 208 | 149 | 1 | 1 | 0.932 | 0.868 | 0.867 | 0.888 | 0.9 | 1 | 0.955 | 0.904 | 0.951 | NA | -0.061 |
| SE | 36662 | ENSG00000180329 | CCDC43 | chr17 | - | 44680584 | 44680652 | 44677482 | 44679043 | 44682002 | 44682138 | 61 | 37 | 29 | 68 | 95 | 6 | 11 | 9 | 11 | 15 | 109 | 42 | 83 | 85 | 0 | 0 | 3 | 13 | 6 | 0 | 217 | 149 | 1 | 1 | 0.875 | 0.698 | 0.689 | 0.809 | 0.813 | 1 | 0.906 | 0.814 | 0.907 | NA | -0.13 |
| SE | 36663 | ENSG00000180329 | CCDC43 | chr17 | - | 44683871 | 44683959 | 44681730 | 44682138 | 44689549 | 44689725 | 93 | 80 | 78 | 67 | 98 | 0 | 0 | 2 | 0 | 0 | 130 | 113 | 98 | 103 | 20 | 0 | 0 | 0 | 2 | 0 | 237 | 149 | 1 | 1 | 1 | 1 | 0.961 | 1 | 1 | 1 | 1 | 1 | 0.97 | 1 | -0.002 |
| SE | 37121 | ENSG00000144848 | ATG3 | chr3 | - | 112550191 | 112550262 | 112548532 | 112548640 | 112553279 | 112553329 | 453 | 413 | 453 | 470 | 689 | 0 | 1 | 0 | 5 | 0 | 932 | 331 | 396 | 623 | 434 | 0 | 0 | 0 | 4 | 0 | 220 | 149 | 1 | 1 | 1 | 0.996 | 1 | 0.985 | 1 | 1 | 1 | 1 | 0.991 | 1 | -0.002 |
| SE | 37122 | ENSG00000144848 | ATG3 | chr3 | - | 112550191 | 112550262 | 112548532 | 112548640 | 112558375 | 112558417 | 202 | 211 | 215 | 222 | 377 | 0 | 0 | 0 | 0 | 0 | 432 | 108 | 197 | 321 | 225 | 0 | 2 | 0 | 0 | 0 | 220 | 149 | 1 | 1 | 1 | 1 | 1 | 1 | 1 | 1 | 0.973 | 1 | 1 | 1 | 0.005 |
| SE | 37123 | ENSG00000144848 | ATG3 | chr3 | - | 112553279 | 112553329 | 112548532 | 112548640 | 112558375 | 112558417 | 10 | 22 | 14 | 26 | 19 | 0 | 0 | 0 | 0 | 0 | 27 | 25 | 43 | 18 | 2 | 0 | 2 | 0 | 0 | 0 | 199 | 149 | 1 | 1 | 1 | 1 | 1 | 1 | 1 | 1 | 0.903 | 1 | 1 | 1 | 0.019 |
| SE | 37237 | ENSG00000189283 | FHIT | chr3 | - | 59886219 | 59886287 | 59749701 | 59752321 | 59922345 | 59922414 | 0 | 0 | 1 | 0 | 0 | 20 | 6 | 27 | 36 | 90 | 0 | 0 | 0 | 0 | 0 | 12 | 3 | 24 | 8 | 7 | 217 | 149 | 1 | 1 | 0 | 0 | 0.025 | 0 | 0 | 0 | 0 | 0 | 0 | 0 | 0.005 |
| SE | 37238 | ENSG00000189283 | FHIT | chr3 | - | 59922345 | 59922414 | 59751406 | 59752321 | 60080790 | 60080840 | 7 | 2 | 14 | 18 | 31 | 0 | 0 | 0 | 0 | 3 | 6 | 3 | 11 | 3 | 0 | 0 | 0 | 0 | 0 | 0 | 218 | 149 | 1 | 1 | 1 | 1 | 1 | 1 | 0.876 | 1 | 1 | 1 | 1 | NA | -0.025 |
| SE | 37239 | ENSG00000189283 | FHIT | chr3 | - | 60011370 | 60011400 | 59922345 | 59922414 | 60014006 | 60014075 | 19 | 14 | 31 | 15 | 37 | 0 | 0 | 0 | 0 | 0 | 6 | 4 | 36 | 19 | 15 | 0 | 0 | 0 | 3 | 0 | 179 | 149 | 1 | 1 | 1 | 1 | 1 | 1 | 1 | 1 | 1 | 1 | 0.841 | 1 | 0.032 |
| SE | 37240 | ENSG00000189283 | FHIT | chr3 | - | 60011370 | 60011400 | 59922345 | 59922414 | 60080790 | 60080840 | 3 | 1 | 4 | 3 | 12 | 3 | 0 | 0 | 6 | 1 | 6 | 0 | 6 | 5 | 0 | 0 | 0 | 2 | 0 | 0 | 179 | 149 | 1 | 1 | 0.454 | 1 | 1 | 0.294 | 0.909 | 1 | NA | 0.714 | 1 | NA | -0.173 |
| SE | 37242 | ENSG00000189283 | FHIT | chr3 | - | 60014006 | 60014075 | 59922345 | 59922414 | 60080790 | 60080840 | 0 | 0 | 0 | 0 | 3 | 3 | 0 | 0 | 6 | 1 | 0 | 0 | 0 | 0 | 0 | 0 | 0 | 2 | 0 | 0 | 218 | 149 | 1 | 1 | 0 | NA | NA | 0 | 0.672 | NA | NA | 0 | NA | NA | 0.224 |
| SE | 37243 | ENSG00000189283 | FHIT | chr3 | - | 60014006 | 60014075 | 60011370 | 60011400 | 60080790 | 60080840 | 8 | 8 | 12 | 6 | 14 | 1 | 0 | 0 | 0 | 0 | 0 | 0 | 13 | 5 | 7 | 0 | 0 | 0 | 0 | 0 | 218 | 149 | 1 | 1 | 0.845 | 1 | 1 | 1 | 1 | NA | NA | 1 | 1 | 1 | -0.031 |
| SE | 37244 | ENSG00000189283 | FHIT | chr3 | - | 60077369 | 60077526 | 59922345 | 59922414 | 60080790 | 60080840 | 4 | 0 | 2 | 1 | 1 | 3 | 0 | 0 | 6 | 1 | 0 | 0 | 0 | 2 | 0 | 0 | 0 | 2 | 0 | 0 | 298 | 149 | 1 | 1 | 0.4 | NA | 1 | 0.077 | 0.333 | NA | NA | 0 | 1 | NA | -0.048 |
| SE | 37245 | ENSG00000189283 | FHIT | chr3 | - | 60077369 | 60077526 | 60014006 | 60014075 | 60080790 | 60080840 | 4 | 0 | 2 | 1 | 1 | 0 | 0 | 0 | 0 | 3 | 0 | 0 | 0 | 2 | 0 | 0 | 0 | 0 | 0 | 0 | 298 | 149 | 1 | 1 | 1 | NA | 1 | 1 | 0.143 | NA | NA | NA | 1 | NA | -0.214 |
| SE | 37246 | ENSG00000189283 | FHIT | chr3 | - | 60765523 | 60765563 | 60536859 | 60536979 | 60821918 | 60822011 | 0 | 0 | 0 | 1 | 0 | 5 | 7 | 4 | 3 | 10 | 0 | 0 | 0 | 0 | 0 | 12 | 10 | 1 | 8 | 15 | 189 | 149 | 1 | 1 | 0 | 0 | 0 | 0.208 | 0 | 0 | 0 | 0 | 0 | 0 | 0.042 |
| SE | 37247 | ENSG00000189283 | FHIT | chr3 | - | 61166972 | 61167042 | 61042046 | 61042099 | 61200616 | 61200665 | 1 | 2 | 0 | 0 | 0 | 3 | 11 | 2 | 4 | 5 | 0 | 0 | 0 | 0 | 0 | 21 | 15 | 12 | 0 | 4 | 219 | 149 | 1 | 1 | 0.185 | 0.11 | 0 | 0 | 0 | 0 | 0 | 0 | NA | 0 | 0.059 |
| SE | 37426 | ENSG00000114850 | SSR3 | chr3 | - | 156548865 | 156549003 | 156544307 | 156544439 | 156553654 | 156553781 | 186 | 93 | 207 | 157 | 254 | 1 | 0 | 1 | 0 | 0 | 182 | 122 | 180 | 108 | 59 | 0 | 0 | 0 | 0 | 0 | 287 | 149 | 1 | 1 | 0.99 | 1 | 0.991 | 1 | 1 | 1 | 1 | 1 | 1 | 1 | -0.004 |
| SE | 37427 | ENSG00000114850 | SSR3 | chr3 | - | 156548904 | 156549003 | 156543953 | 156544439 | 156553654 | 156553781 | 551 | 383 | 669 | 499 | 1043 | 1 | 0 | 1 | 0 | 0 | 573 | 320 | 628 | 347 | 228 | 0 | 0 | 0 | 0 | 0 | 248 | 149 | 1 | 1 | 0.997 | 1 | 0.998 | 1 | 1 | 1 | 1 | 1 | 1 | 1 | -0.001 |
| SE | 37428 | ENSG00000114850 | SSR3 | chr3 | - | 156553654 | 156553781 | 156548865 | 156549003 | 156554956 | 156555117 | 359 | 191 | 419 | 322 | 411 | 0 | 0 | 0 | 0 | 0 | 314 | 236 | 342 | 274 | 90 | 0 | 0 | 0 | 7 | 0 | 276 | 149 | 1 | 1 | 1 | 1 | 1 | 1 | 1 | 1 | 1 | 1 | 0.955 | 1 | 0.009 |
| SE | 37691 | ENSG00000163507 | CIP2A | chr3 | - | 108553647 | 108553730 | 108552233 | 108552373 | 108554375 | 108554489 | 24 | 30 | 51 | 32 | 57 | 0 | 0 | 0 | 0 | 3 | 0 | 22 | 14 | 7 | 0 | 0 | 0 | 0 | 0 | 0 | 232 | 149 | 1 | 1 | 1 | 1 | 1 | 1 | 0.924 | NA | 1 | 1 | 1 | NA | -0.015 |
| SE | 37692 | ENSG00000163507 | CIP2A | chr3 | - | 108557217 | 108557414 | 108554375 | 108554489 | 108559756 | 108559867 | 28 | 34 | 54 | 34 | 61 | 0 | 0 | 3 | 0 | 0 | 9 | 21 | 31 | 8 | 0 | 0 | 0 | 0 | 0 | 0 | 298 | 149 | 1 | 1 | 1 | 1 | 0.9 | 1 | 1 | 1 | 1 | 1 | 1 | NA | -0.02 |
| SE | 37693 | ENSG00000163507 | CIP2A | chr3 | - | 108563121 | 108563244 | 108560734 | 108560841 | 108565354 | 108565454 | 8 | 7 | 18 | 16 | 36 | 0 | 0 | 0 | 1 | 0 | 0 | 0 | 1 | 7 | 0 | 0 | 0 | 0 | 0 | 0 | 272 | 149 | 1 | 1 | 1 | 1 | 1 | 0.898 | 1 | NA | NA | 1 | 1 | NA | -0.02 |
| SE | 37694 | ENSG00000163507 | CIP2A | chr3 | - | 108563125 | 108563244 | 108560648 | 108560841 | 108565354 | 108565454 | 18 | 14 | 40 | 29 | 72 | 0 | 0 | 0 | 1 | 0 | 0 | 12 | 2 | 11 | 2 | 0 | 0 | 0 | 0 | 0 | 268 | 149 | 1 | 1 | 1 | 1 | 1 | 0.942 | 1 | NA | 1 | 1 | 1 | 1 | -0.012 |
| SE | 37695 | ENSG00000163507 | CIP2A | chr3 | - | 108582107 | 108582138 | 108581414 | 108581511 | 108582976 | 108583083 | 3 | 6 | 5 | 1 | 0 | 0 | 0 | 1 | 1 | 0 | 0 | 2 | 3 | 3 | 4 | 0 | 0 | 0 | 0 | 0 | 180 | 149 | 1 | 1 | 1 | 1 | 0.805 | 0.453 | NA | NA | 1 | 1 | 1 | 1 | -0.185 |
| SE | 37696 | ENSG00000163507 | CIP2A | chr3 | - | 108582107 | 108582202 | 108581414 | 108581511 | 108582976 | 108583083 | 18 | 16 | 32 | 12 | 9 | 0 | 0 | 0 | 1 | 0 | 9 | 22 | 5 | 13 | 15 | 0 | 0 | 0 | 0 | 0 | 244 | 149 | 1 | 1 | 1 | 1 | 0.951 | 0.88 | 1 | 1 | 1 | 1 | 1 | 1 | -0.034 |
| SE | 37697 | ENSG00000163507 | CIP2A | chr3 | - | 108582976 | 108583083 | 108582107 | 108582138 | 108585064 | 108585212 | 0 | 0 | 13 | 4 | 2 | 0 | 1 | 0 | 0 | 0 | 0 | 2 | 0 | 0 | 0 | 0 | 0 | 0 | 1 | 0 | 256 | 149 | 1 | 1 | NA | 0 | 1 | 1 | 1 | NA | NA | NA | 0 | NA | 0.75 |
| SE | 37699 | ENSG00000163507 | CIP2A | chr3 | - | 108585665 | 108585749 | 108585064 | 108585212 | 108589273 | 108589456 | 1 | 0 | 0 | 0 | 0 | 2 | 1 | 2 | 5 | 2 | 0 | 0 | 0 | 0 | 0 | 0 | 14 | 1 | 4 | 0 | 233 | 149 | 1 | 1 | 0.242 | 0 | 0 | 0 | 0 | NA | 0 | 0 | 0 | NA | 0.048 |
| SE | 37858 | ENSG00000114573 | ATP6V1A | chr3 | + | 113786231 | 113786383 | 113778740 | 113778835 | 113788712 | 113788875 | 73 | 67 | 64 | 75 | 110 | 1 | 0 | 0 | 0 | 0 | 142 | 78 | 57 | 87 | 72 | 0 | 0 | 0 | 0 | 0 | 298 | 149 | 1 | 1 | 0.973 | 1 | 1 | 1 | 1 | 1 | 1 | 1 | 1 | 1 | -0.005 |
| SE | 37859 | ENSG00000114573 | ATP6V1A | chr3 | + | 113794871 | 113794994 | 113789731 | 113789840 | 113795089 | 113795204 | 92 | 67 | 88 | 96 | 99 | 0 | 0 | 0 | 0 | 0 | 73 | 113 | 75 | 91 | 20 | 0 | 0 | 3 | 0 | 0 | 272 | 149 | 1 | 1 | 1 | 1 | 1 | 1 | 1 | 1 | 1 | 0.932 | 1 | 1 | 0.014 |
| SE | 37934 | ENSG00000148843 | PDCD11 | chr10 | + | 103432128 | 103432234 | 103427328 | 103427391 | 103433947 | 103434037 | 81 | 54 | 92 | 63 | 69 | 1 | 0 | 0 | 0 | 0 | 60 | 42 | 31 | 43 | 93 | 0 | 0 | 2 | 3 | 0 | 255 | 149 | 1 | 1 | 0.979 | 1 | 1 | 1 | 1 | 1 | 1 | 0.901 | 0.893 | 1 | 0.037 |
| SE | 37936 | ENSG00000148843 | PDCD11 | chr10 | + | 103438014 | 103438071 | 103434797 | 103434975 | 103439745 | 103439868 | 22 | 8 | 27 | 15 | 8 | 0 | 0 | 0 | 0 | 2 | 14 | 4 | 3 | 20 | 37 | 0 | 0 | 0 | 0 | 0 | 206 | 149 | 1 | 1 | 1 | 1 | 1 | 1 | 0.743 | 1 | 1 | 1 | 1 | 1 | -0.051 |
| SE | 37937 | ENSG00000148843 | PDCD11 | chr10 | + | 103438685 | 103438808 | 103438014 | 103438071 | 103439745 | 103439868 | 112 | 64 | 111 | 105 | 125 | 1 | 0 | 0 | 0 | 0 | 193 | 120 | 80 | 78 | 173 | 0 | 0 | 0 | 0 | 0 | 272 | 149 | 1 | 1 | 0.984 | 1 | 1 | 1 | 1 | 1 | 1 | 1 | 1 | 1 | -0.003 |
| SE | 37997 | ENSG00000197386 | HTT | chr4 | + | 3062475 | 3062692 | 3041421 | 3042245 | 3071235 | 3071396 | 2 | 0 | 0 | 0 | 0 | 1 | 0 | 0 | 0 | 0 | 0 | 5 | 0 | 0 | 0 | 0 | 0 | 1 | 0 | 0 | 298 | 149 | 1 | 1 | 0.5 | NA | NA | NA | NA | NA | 1 | 0 | NA | NA | 0 |
| SE | 37998 | ENSG00000197386 | HTT | chr4 | + | 3071235 | 3071396 | 3041421 | 3042245 | 3086938 | 3087022 | 6 | 0 | 0 | 0 | 0 | 3 | 0 | 0 | 0 | 0 | 0 | 0 | 1 | 0 | 0 | 0 | 0 | 0 | 0 | 0 | 298 | 149 | 1 | 1 | 0.5 | NA | NA | NA | NA | NA | NA | 1 | NA | NA | -0.5 |
| SE | 37999 | ENSG00000197386 | HTT | chr4 | + | 3086938 | 3087022 | 3074680 | 3075088 | 3099273 | 3099394 | 145 | 99 | 104 | 146 | 208 | 0 | 0 | 0 | 0 | 0 | 273 | 149 | 113 | 177 | 344 | 0 | 0 | 0 | 0 | 2 | 233 | 149 | 1 | 1 | 1 | 1 | 1 | 1 | 1 | 1 | 1 | 1 | 1 | 0.991 | 0.002 |
| SE | 38000 | ENSG00000197386 | HTT | chr4 | + | 3099273 | 3099394 | 3086938 | 3087022 | 3103823 | 3103883 | 275 | 193 | 179 | 226 | 344 | 0 | 2 | 1 | 0 | 0 | 479 | 241 | 246 | 330 | 507 | 0 | 0 | 0 | 0 | 5 | 270 | 149 | 1 | 1 | 1 | 0.982 | 0.99 | 1 | 1 | 1 | 1 | 1 | 1 | 0.982 | -0.002 |
| SE | 38002 | ENSG00000197386 | HTT | chr4 | + | 3103823 | 3103883 | 3086938 | 3087022 | 3105356 | 3105436 | 129 | 79 | 90 | 113 | 145 | 0 | 0 | 0 | 0 | 0 | 233 | 171 | 119 | 103 | 358 | 8 | 0 | 0 | 0 | 0 | 209 | 149 | 1 | 1 | 1 | 1 | 1 | 1 | 1 | 0.954 | 1 | 1 | 1 | 1 | 0.009 |
| SE | 38003 | ENSG00000197386 | HTT | chr4 | + | 3103823 | 3103883 | 3099273 | 3099394 | 3105356 | 3105436 | 267 | 185 | 179 | 216 | 319 | 0 | 0 | 0 | 0 | 1 | 472 | 310 | 275 | 284 | 594 | 0 | 2 | 4 | 3 | 2 | 209 | 149 | 1 | 1 | 1 | 1 | 1 | 1 | 0.996 | 1 | 0.991 | 0.98 | 0.985 | 0.995 | 0.009 |
| SE | 38004 | ENSG00000197386 | HTT | chr4 | + | 3115303 | 3115445 | 3107284 | 3107423 | 3116084 | 3116263 | 178 | 103 | 128 | 157 | 183 | 2 | 0 | 0 | 1 | 0 | 330 | 202 | 120 | 211 | 447 | 0 | 0 | 0 | 7 | 0 | 291 | 149 | 1 | 1 | 0.979 | 1 | 1 | 0.988 | 1 | 1 | 1 | 1 | 0.939 | 1 | 0.006 |
| SE | 38005 | ENSG00000197386 | HTT | chr4 | + | 3122888 | 3122936 | 3121227 | 3121432 | 3125548 | 3125629 | 107 | 60 | 80 | 108 | 102 | 0 | 0 | 1 | 0 | 0 | 161 | 158 | 97 | 141 | 361 | 0 | 0 | 3 | 0 | 0 | 197 | 149 | 1 | 1 | 1 | 1 | 0.984 | 1 | 1 | 1 | 1 | 0.961 | 1 | 1 | 0.005 |
| SE | 38008 | ENSG00000197386 | HTT | chr4 | + | 3135903 | 3135967 | 3134400 | 3134540 | 3136225 | 3136326 | 156 | 114 | 107 | 100 | 160 | 0 | 0 | 0 | 1 | 0 | 263 | 179 | 131 | 187 | 362 | 0 | 0 | 0 | 0 | 0 | 213 | 149 | 1 | 1 | 1 | 1 | 1 | 0.986 | 1 | 1 | 1 | 1 | 1 | 1 | -0.003 |
| SE | 38009 | ENSG00000197386 | HTT | chr4 | + | 3142765 | 3142886 | 3140509 | 3140656 | 3145151 | 3145228 | 182 | 152 | 182 | 194 | 266 | 0 | 0 | 0 | 0 | 0 | 381 | 241 | 151 | 283 | 392 | 0 | 1 | 0 | 0 | 0 | 270 | 149 | 1 | 1 | 1 | 1 | 1 | 1 | 1 | 1 | 0.993 | 1 | 1 | 1 | 0.001 |
| SE | 38011 | ENSG00000197386 | HTT | chr4 | + | 3154292 | 3154419 | 3148004 | 3148207 | 3160281 | 3160392 | 49 | 21 | 37 | 37 | 56 | 0 | 0 | 4 | 0 | 0 | 118 | 66 | 17 | 59 | 89 | 0 | 0 | 0 | 0 | 0 | 276 | 149 | 1 | 1 | 1 | 1 | 0.833 | 1 | 1 | 1 | 1 | 1 | 1 | 1 | -0.033 |
| SE | 38012 | ENSG00000197386 | HTT | chr4 | + | 3157071 | 3157199 | 3148004 | 3148207 | 3160281 | 3160392 | 131 | 94 | 94 | 119 | 210 | 0 | 0 | 4 | 0 | 0 | 231 | 145 | 112 | 126 | 405 | 0 | 0 | 0 | 0 | 0 | 277 | 149 | 1 | 1 | 1 | 1 | 0.927 | 1 | 1 | 1 | 1 | 1 | 1 | 1 | -0.015 |
| SE | 38013 | ENSG00000197386 | HTT | chr4 | + | 3157071 | 3157199 | 3154292 | 3154419 | 3160281 | 3160392 | 228 | 168 | 192 | 230 | 335 | 0 | 0 | 0 | 0 | 0 | 403 | 262 | 204 | 240 | 621 | 9 | 0 | 0 | 0 | 0 | 277 | 149 | 1 | 1 | 1 | 1 | 1 | 1 | 1 | 0.96 | 1 | 1 | 1 | 1 | 0.008 |
| SE | 38014 | ENSG00000197386 | HTT | chr4 | + | 3160281 | 3160392 | 3157071 | 3157199 | 3172319 | 3172397 | 306 | 256 | 261 | 295 | 637 | 0 | 0 | 0 | 0 | 0 | 568 | 408 | 324 | 340 | 1091 | 0 | 5 | 0 | 0 | 0 | 260 | 149 | 1 | 1 | 1 | 1 | 1 | 1 | 1 | 1 | 0.979 | 1 | 1 | 1 | 0.004 |
| SE | 38015 | ENSG00000197386 | HTT | chr4 | + | 3174945 | 3175107 | 3174720 | 3174799 | 3177331 | 3177387 | 237 | 200 | 217 | 255 | 201 | 0 | 0 | 0 | 0 | 0 | 287 | 245 | 187 | 278 | 502 | 0 | 0 | 0 | 5 | 0 | 298 | 149 | 1 | 1 | 1 | 1 | 1 | 1 | 1 | 1 | 1 | 1 | 0.965 | 1 | 0.007 |
| SE | 38016 | ENSG00000197386 | HTT | chr4 | + | 3182353 | 3182470 | 3180514 | 3180651 | 3186596 | 3186719 | 135 | 74 | 101 | 119 | 219 | 0 | 0 | 1 | 0 | 0 | 188 | 136 | 94 | 130 | 242 | 0 | 1 | 0 | 0 | 0 | 266 | 149 | 1 | 1 | 1 | 1 | 0.983 | 1 | 1 | 1 | 0.987 | 1 | 1 | 1 | -0.001 |
| SE | 38019 | ENSG00000197386 | HTT | chr4 | + | 3207280 | 3207357 | 3206806 | 3206983 | 3208772 | 3208911 | 126 | 59 | 89 | 137 | 130 | 1 | 0 | 0 | 0 | 0 | 155 | 139 | 90 | 174 | 339 | 0 | 1 | 0 | 5 | 0 | 226 | 149 | 1 | 1 | 0.988 | 1 | 1 | 1 | 1 | 1 | 1 | 1 | 0.958 | 1 | 0.006 |
| SE | 38020 | ENSG00000197386 | HTT | chr4 | + | 3217764 | 3217952 | 3215109 | 3215211 | 3220181 | 3220308 | 137 | 58 | 96 | 142 | 162 | 0 | 0 | 0 | 0 | 0 | 242 | 164 | 97 | 96 | 289 | 0 | 0 | 2 | 5 | 0 | 298 | 149 | 1 | 1 | 1 | 1 | 1 | 1 | 1 | 1 | 1 | 0.96 | 0.906 | 1 | 0.027 |
| SE | 38023 | ENSG00000197386 | HTT | chr4 | + | 3236148 | 3236254 | 3235564 | 3235778 | 3238446 | 3238609 | 177 | 104 | 139 | 201 | 168 | 0 | 0 | 0 | 0 | 0 | 144 | 250 | 91 | 109 | 501 | 5 | 0 | 2 | 0 | 0 | 255 | 149 | 1 | 1 | 1 | 1 | 1 | 1 | 1 | 0.944 | 1 | 0.964 | 1 | 1 | 0.018 |
| SE | 38563 | ENSG00000198925 | ATG9A | chr2 | - | 219222049 | 219222167 | 219221079 | 219221302 | 219222271 | 219222450 | 42 | 16 | 24 | 28 | 8 | 0 | 0 | 0 | 0 | 0 | 4 | 16 | 17 | 25 | 68 | 0 | 0 | 2 | 0 | 0 | 267 | 149 | 1 | 1 | 1 | 1 | 1 | 1 | 1 | 1 | 1 | 0.826 | 1 | 1 | 0.035 |
| SE | 38565 | ENSG00000198925 | ATG9A | chr2 | - | 219227921 | 219228053 | 219227769 | 219227813 | 219229009 | 219229342 | 28 | 14 | 27 | 19 | 14 | 0 | 0 | 0 | 2 | 0 | 16 | 8 | 5 | 8 | 15 | 0 | 0 | 0 | 0 | 0 | 281 | 149 | 1 | 1 | 1 | 1 | 1 | 0.834 | 1 | 1 | 1 | 1 | 1 | 1 | -0.033 |
| SE | 38570 | ENSG00000198925 | ATG9A | chr2 | - | 219228433 | 219228482 | 219227921 | 219228053 | 219229009 | 219229325 | 7 | 3 | 5 | 3 | 1 | 0 | 1 | 1 | 4 | 0 | 0 | 8 | 0 | 1 | 13 | 0 | 0 | 0 | 0 | 0 | 198 | 149 | 1 | 1 | 1 | 0.693 | 0.79 | 0.361 | 1 | NA | 1 | NA | 1 | 1 | -0.231 |
| SE | 38571 | ENSG00000198925 | ATG9A | chr2 | - | 219228433 | 219228482 | 219227921 | 219228053 | 219229534 | 219229639 | 7 | 3 | 6 | 3 | 1 | 5 | 4 | 0 | 2 | 0 | 0 | 10 | 0 | 3 | 13 | 0 | 1 | 3 | 0 | 0 | 198 | 149 | 1 | 1 | 0.513 | 0.361 | 1 | 0.53 | 1 | NA | 0.883 | 0 | 1 | 1 | -0.04 |
| SE | 38572 | ENSG00000198925 | ATG9A | chr2 | - | 219228433 | 219228485 | 219227769 | 219227813 | 219229009 | 219229174 | 0 | 3 | 0 | 1 | 0 | 0 | 0 | 0 | 2 | 0 | 1 | 2 | 2 | 0 | 3 | 0 | 0 | 0 | 0 | 0 | 201 | 149 | 1 | 1 | NA | 1 | NA | 0.27 | NA | 1 | 1 | 1 | NA | 1 | -0.365 |
| SE | 38574 | ENSG00000198925 | ATG9A | chr2 | - | 219228433 | 219228485 | 219227921 | 219228053 | 219229009 | 219229174 | 9 | 8 | 7 | 3 | 2 | 0 | 1 | 1 | 4 | 0 | 0 | 14 | 2 | 1 | 24 | 0 | 0 | 0 | 0 | 0 | 201 | 149 | 1 | 1 | 1 | 0.856 | 0.838 | 0.357 | 1 | NA | 1 | 1 | 1 | 1 | -0.19 |
| SE | 38575 | ENSG00000198925 | ATG9A | chr2 | - | 219228433 | 219228485 | 219227921 | 219228053 | 219229534 | 219229594 | 9 | 4 | 8 | 2 | 1 | 5 | 4 | 0 | 2 | 0 | 0 | 8 | 4 | 2 | 22 | 0 | 1 | 3 | 0 | 0 | 201 | 149 | 1 | 1 | 0.572 | 0.426 | 1 | 0.426 | 1 | NA | 0.856 | 0.497 | 1 | 1 | -0.153 |
| SE | 38576 | ENSG00000198925 | ATG9A | chr2 | - | 219228433 | 219228485 | 219227921 | 219228079 | 219229534 | 219229594 | 0 | 0 | 0 | 0 | 0 | 0 | 0 | 0 | 1 | 0 | 0 | 5 | 0 | 0 | 0 | 0 | 0 | 0 | 0 | 0 | 201 | 149 | 1 | 1 | NA | NA | NA | 0 | NA | NA | 1 | NA | NA | NA | -1 |
| SE | 38578 | ENSG00000198925 | ATG9A | chr2 | - | 219229009 | 219229174 | 219228433 | 219228485 | 219229534 | 219229618 | 3 | 8 | 4 | 3 | 1 | 2 | 1 | 3 | 1 | 0 | 0 | 8 | 4 | 0 | 11 | 0 | 0 | 8 | 1 | 9 | 298 | 149 | 1 | 1 | 0.429 | 0.8 | 0.4 | 0.6 | 1 | NA | 1 | 0.2 | 0 | 0.379 | 0.251 |
| SE | 38768 | ENSG00000181804 | SLC9A9 | chr3 | - | 143382059 | 143382114 | 143363483 | 143363563 | 143467036 | 143467190 | 45 | 40 | 26 | 39 | 11 | 0 | 0 | 1 | 0 | 0 | 29 | 36 | 39 | 54 | 18 | 0 | 0 | 0 | 0 | 0 | 204 | 149 | 1 | 1 | 1 | 1 | 0.95 | 1 | 1 | 1 | 1 | 1 | 1 | 1 | -0.01 |
| SE | 38769 | ENSG00000181804 | SLC9A9 | chr3 | - | 143652254 | 143652361 | 143552361 | 143552450 | 143693191 | 143693307 | 25 | 10 | 16 | 34 | 47 | 1 | 0 | 0 | 0 | 0 | 12 | 24 | 24 | 29 | 21 | 0 | 1 | 0 | 0 | 0 | 255 | 149 | 1 | 1 | 0.936 | 1 | 1 | 1 | 1 | 1 | 0.933 | 1 | 1 | 1 | 0.001 |
| SE | 38770 | ENSG00000181804 | SLC9A9 | chr3 | - | 143652254 | 143652360 | 143578584 | 143578723 | 143693191 | 143693307 | 55 | 52 | 50 | 86 | 107 | 0 | 0 | 0 | 0 | 0 | 67 | 54 | 31 | 57 | 56 | 0 | 1 | 0 | 0 | 0 | 255 | 149 | 1 | 1 | 1 | 1 | 1 | 1 | 1 | 1 | 0.969 | 1 | 1 | 1 | 0.006 |
| SE | 38771 | ENSG00000181804 | SLC9A9 | chr3 | - | 143693191 | 143693307 | 143652254 | 143652360 | 143795000 | 143795077 | 67 | 40 | 45 | 87 | 116 | 0 | 0 | 2 | 0 | 0 | 59 | 28 | 27 | 55 | 46 | 0 | 0 | 0 | 0 | 0 | 265 | 149 | 1 | 1 | 1 | 0.918 | 1 | 1 | 1 | 1 | 1 | 1 | 1 | 1 | -0.016 |
| SE | 38772 | ENSG00000181804 | SLC9A9 | chr3 | - | 143796825 | 143796903 | 143795000 | 143795077 | 143832018 | 143832221 | 158 | 129 | 101 | 181 | 190 | 0 | 0 | 0 | 0 | 0 | 103 | 59 | 42 | 155 | 140 | 0 | 0 | 0 | 2 | 0 | 227 | 149 | 1 | 1 | 1 | 1 | 1 | 1 | 1 | 1 | 1 | 1 | 0.981 | 1 | 0.004 |
| SE | 38773 | ENSG00000181804 | SLC9A9 | chr3 | - | 143832018 | 143832221 | 143796825 | 143796903 | 143848147 | 143848458 | 276 | 240 | 153 | 274 | 280 | 2 | 1 | 1 | 0 | 1 | 148 | 149 | 99 | 247 | 166 | 0 | 0 | 0 | 0 | 4 | 298 | 149 | 1 | 1 | 0.986 | 0.992 | 0.987 | 1 | 0.993 | 1 | 1 | 1 | 1 | 0.954 | 0.001 |
| SE | 38775 | ENSG00000162601 | MYSM1 | chr1 | - | 58663449 | 58663522 | 58661405 | 58661511 | 58665498 | 58665631 | 3 | 9 | 9 | 23 | 5 | 84 | 87 | 91 | 82 | 85 | 0 | 9 | 6 | 5 | 0 | 4 | 20 | 34 | 13 | 0 | 222 | 149 | 1 | 1 | 0.023 | 0.065 | 0.062 | 0.158 | 0.038 | 0 | 0.232 | 0.106 | 0.205 | NA | -0.067 |
| SE | 38776 | ENSG00000162601 | MYSM1 | chr1 | - | 58665498 | 58665631 | 58661405 | 58661511 | 58667037 | 58667226 | 166 | 137 | 158 | 140 | 184 | 0 | 0 | 0 | 0 | 0 | 24 | 41 | 56 | 19 | 5 | 0 | 0 | 2 | 0 | 0 | 282 | 149 | 1 | 1 | 1 | 1 | 1 | 1 | 1 | 1 | 1 | 0.937 | 1 | 1 | 0.013 |
| SE | 38777 | ENSG00000162601 | MYSM1 | chr1 | - | 58667037 | 58667226 | 58661405 | 58661511 | 58667846 | 58667921 | 69 | 65 | 68 | 46 | 148 | 2 | 0 | 0 | 3 | 0 | 5 | 21 | 34 | 2 | 5 | 0 | 0 | 0 | 0 | 0 | 298 | 149 | 1 | 1 | 0.945 | 1 | 1 | 0.885 | 1 | 1 | 1 | 1 | 1 | 1 | -0.034 |
| SE | 38778 | ENSG00000162601 | MYSM1 | chr1 | - | 58668429 | 58668566 | 58667846 | 58667921 | 58668983 | 58669038 | 39 | 31 | 24 | 49 | 47 | 0 | 0 | 1 | 0 | 3 | 0 | 6 | 8 | 3 | 0 | 0 | 0 | 0 | 0 | 0 | 286 | 149 | 1 | 1 | 1 | 1 | 0.926 | 1 | 0.891 | NA | 1 | 1 | 1 | NA | -0.037 |
| SE | 38779 | ENSG00000162601 | MYSM1 | chr1 | - | 58668631 | 58668682 | 58667846 | 58667921 | 58668983 | 58669038 | 91 | 79 | 94 | 87 | 159 | 0 | 0 | 1 | 0 | 3 | 0 | 52 | 40 | 7 | 0 | 0 | 0 | 0 | 0 | 0 | 200 | 149 | 1 | 1 | 1 | 1 | 0.986 | 1 | 0.975 | NA | 1 | 1 | 1 | NA | -0.008 |
| SE | 38783 | ENSG00000162601 | MYSM1 | chr1 | - | 58689293 | 58689367 | 58689037 | 58689116 | 58690225 | 58690249 | 6 | 11 | 11 | 11 | 22 | 77 | 63 | 71 | 58 | 77 | 0 | 5 | 6 | 2 | 11 | 0 | 36 | 53 | 0 | 0 | 223 | 149 | 1 | 1 | 0.049 | 0.104 | 0.094 | 0.112 | 0.16 | NA | 0.085 | 0.07 | 1 | 1 | -0.435 |
| SE | 39106 | ENSG00000117228 | GBP1 | chr1 | - | 89055112 | 89056228 | 89054681 | 89054875 | 89056853 | 89057134 | 412 | 521 | 432 | 444 | 500 | 0 | 0 | 0 | 3 | 0 | 493 | 362 | 272 | 774 | 16 | 0 | 2 | 0 | 0 | 0 | 298 | 149 | 1 | 1 | 1 | 1 | 1 | 0.987 | 1 | 1 | 0.989 | 1 | 1 | 1 | 0 |
| SE | 39507 | ENSG00000141179 | PCTP | chr17 | + | 55771105 | 55771201 | 55767334 | 55767452 | 55773655 | 55773895 | 23 | 26 | 9 | 27 | 13 | 0 | 0 | 0 | 0 | 0 | 32 | 29 | 22 | 69 | 49 | 0 | 0 | 2 | 0 | 0 | 245 | 149 | 1 | 1 | 1 | 1 | 1 | 1 | 1 | 1 | 1 | 0.87 | 1 | 1 | 0.026 |
| SE | 39650 | ENSG00000127946 | HIP1 | chr7 | - | 75546938 | 75547032 | 75545087 | 75545188 | 75547754 | 75547813 | 17 | 35 | 22 | 33 | 18 | 0 | 0 | 0 | 0 | 0 | 67 | 5 | 32 | 89 | 62 | 0 | 0 | 1 | 4 | 0 | 243 | 149 | 1 | 1 | 1 | 1 | 1 | 1 | 1 | 1 | 1 | 0.952 | 0.932 | 1 | 0.023 |
| SE | 39652 | ENSG00000127946 | HIP1 | chr7 | - | 75559731 | 75559915 | 75558166 | 75558255 | 75561328 | 75561401 | 49 | 67 | 41 | 44 | 34 | 2 | 0 | 1 | 0 | 1 | 230 | 62 | 67 | 127 | 166 | 6 | 0 | 5 | 0 | 0 | 298 | 149 | 1 | 1 | 0.925 | 1 | 0.953 | 1 | 0.944 | 0.95 | 1 | 0.87 | 1 | 1 | 0 |
| SE | 39653 | ENSG00000127946 | HIP1 | chr7 | - | 75573760 | 75573901 | 75568198 | 75568256 | 75581236 | 75581298 | 79 | 68 | 60 | 47 | 70 | 0 | 0 | 3 | 0 | 0 | 206 | 37 | 43 | 166 | 66 | 3 | 0 | 0 | 0 | 0 | 290 | 149 | 1 | 1 | 1 | 1 | 0.911 | 1 | 1 | 0.972 | 1 | 1 | 1 | 1 | -0.012 |
| SE | 39930 | ENSG00000206530 | CFAP44 | chr3 | - | 113401583 | 113401739 | 113401239 | 113401287 | 113403851 | 113404016 | 22 | 54 | 46 | 46 | 25 | 1 | 1 | 0 | 0 | 2 | 0 | 0 | 4 | 4 | 15 | 0 | 0 | 0 | 0 | 0 | 298 | 149 | 1 | 1 | 0.917 | 0.964 | 1 | 1 | 0.862 | NA | NA | 1 | 1 | 1 | -0.051 |
| SE | 39946 | ENSG00000164253 | WDR41 | chr5 | - | 77426943 | 77427034 | 77425969 | 77426191 | 77432892 | 77432955 | 2 | 0 | 6 | 0 | 0 | 2 | 1 | 3 | 0 | 4 | 0 | 2 | 0 | 0 | 0 | 0 | 0 | 0 | 6 | 0 | 240 | 149 | 1 | 1 | 0.383 | 0 | 0.554 | NA | 0 | NA | 1 | NA | 0 | NA | -0.266 |
| SE | 39948 | ENSG00000164253 | WDR41 | chr5 | - | 77437335 | 77437424 | 77436260 | 77436394 | 77440812 | 77440997 | 61 | 27 | 18 | 42 | 35 | 1 | 2 | 0 | 1 | 0 | 25 | 28 | 8 | 28 | 5 | 0 | 3 | 3 | 1 | 0 | 238 | 149 | 1 | 1 | 0.974 | 0.894 | 1 | 0.963 | 1 | 1 | 0.854 | 0.625 | 0.946 | 1 | 0.081 |
| SE | 39949 | ENSG00000164253 | WDR41 | chr5 | - | 77438239 | 77438361 | 77436359 | 77436394 | 77440812 | 77440997 | 94 | 43 | 60 | 56 | 105 | 1 | 2 | 0 | 1 | 0 | 17 | 23 | 29 | 56 | 28 | 0 | 3 | 3 | 1 | 0 | 271 | 149 | 1 | 1 | 0.981 | 0.922 | 1 | 0.969 | 1 | 1 | 0.808 | 0.842 | 0.969 | 1 | 0.051 |
| SE | 39950 | ENSG00000164253 | WDR41 | chr5 | - | 77438239 | 77438361 | 77437335 | 77437424 | 77440812 | 77440997 | 193 | 85 | 108 | 111 | 200 | 0 | 0 | 0 | 1 | 0 | 42 | 69 | 46 | 93 | 66 | 0 | 0 | 1 | 0 | 0 | 271 | 149 | 1 | 1 | 1 | 1 | 1 | 0.984 | 1 | 1 | 1 | 1 | 1 | 1 | -0.003 |
| SE | 39951 | ENSG00000164253 | WDR41 | chr5 | - | 77449759 | 77449860 | 77440812 | 77440997 | 77453816 | 77453928 | 86 | 46 | 62 | 64 | 135 | 2 | 0 | 1 | 0 | 1 | 21 | 41 | 23 | 74 | 0 | 0 | 0 | 1 | 0 | 0 | 250 | 149 | 1 | 1 | 0.962 | 1 | 0.974 | 1 | 0.988 | 1 | 1 | 0.932 | 1 | NA | 0.002 |
| SE | 39953 | ENSG00000164253 | WDR41 | chr5 | - | 77451290 | 77451353 | 77440812 | 77440997 | 77453816 | 77453910 | 39 | 20 | 22 | 34 | 76 | 2 | 0 | 1 | 0 | 1 | 34 | 43 | 14 | 23 | 1 | 0 | 0 | 1 | 0 | 0 | 212 | 149 | 1 | 1 | 0.932 | 1 | 0.939 | 1 | 0.982 | 1 | 1 | 0.908 | 1 | 1 | -0.011 |
| SE | 39959 | ENSG00000164253 | WDR41 | chr5 | - | 77464760 | 77464809 | 77463092 | 77463226 | 77489456 | 77489557 | 95 | 54 | 57 | 56 | 68 | 2 | 0 | 0 | 0 | 0 | 54 | 39 | 28 | 54 | 27 | 0 | 0 | 0 | 0 | 0 | 198 | 149 | 1 | 1 | 0.973 | 1 | 1 | 1 | 1 | 1 | 1 | 1 | 1 | 1 | -0.005 |
| SE | 39961 | ENSG00000164253 | WDR41 | chr5 | - | 77480080 | 77480189 | 77464760 | 77464809 | 77489456 | 77489572 | 6 | 0 | 0 | 0 | 0 | 78 | 48 | 41 | 46 | 52 | 0 | 4 | 0 | 0 | 0 | 41 | 20 | 26 | 41 | 22 | 258 | 149 | 1 | 1 | 0.043 | 0 | 0 | 0 | 0 | 0 | 0.104 | 0 | 0 | 0 | -0.012 |
| SE | 39962 | ENSG00000164253 | WDR41 | chr5 | - | 77480080 | 77480189 | 77464760 | 77464809 | 77492169 | 77492220 | 3 | 0 | 0 | 1 | 0 | 0 | 1 | 0 | 0 | 0 | 0 | 4 | 0 | 0 | 0 | 0 | 0 | 0 | 0 | 0 | 258 | 149 | 1 | 1 | 1 | 0 | NA | 1 | NA | NA | 1 | NA | NA | NA | -0.333 |
| SE | 39963 | ENSG00000164253 | WDR41 | chr5 | - | 77489456 | 77489557 | 77464760 | 77464809 | 77492169 | 77492487 | 66 | 41 | 30 | 34 | 48 | 0 | 1 | 0 | 0 | 0 | 29 | 19 | 21 | 38 | 22 | 0 | 0 | 0 | 0 | 0 | 250 | 149 | 1 | 1 | 1 | 0.961 | 1 | 1 | 1 | 1 | 1 | 1 | 1 | 1 | -0.008 |
| SE | 39965 | ENSG00000164253 | WDR41 | chr5 | - | 77489456 | 77489572 | 77464760 | 77464809 | 77492169 | 77492195 | 109 | 51 | 66 | 60 | 57 | 0 | 1 | 0 | 0 | 0 | 55 | 21 | 33 | 46 | 30 | 0 | 0 | 0 | 0 | 0 | 265 | 149 | 1 | 1 | 1 | 0.966 | 1 | 1 | 1 | 1 | 1 | 1 | 1 | 1 | -0.007 |
| SE | 39966 | ENSG00000164253 | WDR41 | chr5 | - | 77489456 | 77489572 | 77480080 | 77480189 | 77492169 | 77492195 | 31 | 3 | 25 | 15 | 5 | 0 | 0 | 0 | 1 | 0 | 14 | 1 | 7 | 5 | 8 | 0 | 0 | 0 | 0 | 0 | 265 | 149 | 1 | 1 | 1 | 1 | 1 | 0.894 | 1 | 1 | 1 | 1 | 1 | 1 | -0.021 |
| SE | 40930 | ENSG00000204219 | TCEA3 | chr1 | - | 23419076 | 23419139 | 23417903 | 23418009 | 23424564 | 23424618 | 11 | 3 | 15 | 13 | 3 | 0 | 0 | 0 | 2 | 1 | 0 | 0 | 1 | 4 | 10 | 0 | 1 | 0 | 0 | 0 | 212 | 149 | 1 | 1 | 1 | 1 | 1 | 0.82 | 0.678 | NA | 0 | 1 | 1 | 1 | 0.15 |
| SE | 40978 | ENSG00000114485 | PPP5C | chr19 | + | 46387092 | 46387235 | 46384376 | 46384909 | 46387365 | 46387453 | 271 | 158 | 239 | 273 | 318 | 0 | 0 | 0 | 1 | 0 | 350 | 309 | 132 | 194 | 334 | 0 | 0 | 0 | 0 | 0 | 292 | 149 | 1 | 1 | 1 | 1 | 1 | 0.993 | 1 | 1 | 1 | 1 | 1 | 1 | -0.001 |
| SE | 41060 | ENSG00000160741 | CRTC2 | chr1 | - | 153949114 | 153949384 | 153948457 | 153948644 | 153951259 | 153951604 | 72 | 27 | 68 | 80 | 57 | 0 | 0 | 0 | 0 | 0 | 146 | 114 | 102 | 119 | 299 | 0 | 0 | 0 | 2 | 5 | 298 | 149 | 1 | 1 | 1 | 1 | 1 | 1 | 1 | 1 | 1 | 1 | 0.967 | 0.968 | 0.013 |
| SE | 41061 | ENSG00000160741 | CRTC2 | chr1 | - | 153949114 | 153949384 | 153948457 | 153948644 | 153952017 | 153952262 | 66 | 26 | 62 | 71 | 53 | 0 | 0 | 0 | 0 | 0 | 105 | 103 | 79 | 106 | 221 | 0 | 0 | 0 | 2 | 0 | 298 | 149 | 1 | 1 | 1 | 1 | 1 | 1 | 1 | 1 | 1 | 1 | 0.964 | 1 | 0.007 |
| SE | 41062 | ENSG00000160741 | CRTC2 | chr1 | - | 153951259 | 153951666 | 153948457 | 153948644 | 153952017 | 153952262 | 12 | 8 | 4 | 19 | 5 | 0 | 0 | 0 | 0 | 0 | 89 | 24 | 27 | 44 | 61 | 0 | 0 | 0 | 2 | 0 | 298 | 149 | 1 | 1 | 1 | 1 | 1 | 1 | 1 | 1 | 1 | 1 | 0.917 | 1 | 0.017 |
| SE | 41064 | ENSG00000160741 | CRTC2 | chr1 | - | 153952017 | 153952262 | 153951259 | 153951666 | 153952396 | 153952446 | 141 | 55 | 106 | 133 | 170 | 2 | 0 | 1 | 0 | 2 | 338 | 217 | 194 | 232 | 321 | 0 | 0 | 0 | 0 | 0 | 298 | 149 | 1 | 1 | 0.972 | 1 | 0.981 | 1 | 0.977 | 1 | 1 | 1 | 1 | 1 | -0.014 |
| SE | 41066 | ENSG00000160741 | CRTC2 | chr1 | - | 153953537 | 153953606 | 153953265 | 153953369 | 153954254 | 153954316 | 157 | 90 | 95 | 150 | 113 | 0 | 0 | 0 | 1 | 0 | 210 | 222 | 147 | 134 | 277 | 0 | 0 | 0 | 0 | 0 | 218 | 149 | 1 | 1 | 1 | 1 | 1 | 0.99 | 1 | 1 | 1 | 1 | 1 | 1 | -0.002 |
| SE | 41219 | ENSG00000157181 | ODR4 | chr1 | + | 186379766 | 186379884 | 186375837 | 186375974 | 186383021 | 186383156 | 109 | 122 | 119 | 136 | 168 | 0 | 0 | 1 | 0 | 0 | 25 | 46 | 96 | 43 | 38 | 0 | 0 | 0 | 0 | 0 | 267 | 149 | 1 | 1 | 1 | 1 | 0.985 | 1 | 1 | 1 | 1 | 1 | 1 | 1 | -0.003 |
| SE | 41221 | ENSG00000157181 | ODR4 | chr1 | + | 186385987 | 186386083 | 186383021 | 186383156 | 186390710 | 186390851 | 28 | 25 | 41 | 39 | 53 | 1 | 0 | 0 | 0 | 1 | 48 | 8 | 22 | 26 | 8 | 0 | 0 | 4 | 0 | 0 | 245 | 149 | 1 | 1 | 0.945 | 1 | 1 | 1 | 0.97 | 1 | 1 | 1 | 0.798 | 1 | 0.023 |
| SE | 41222 | ENSG00000157181 | ODR4 | chr1 | + | 186393946 | 186394015 | 186391695 | 186391791 | 186398312 | 186398441 | 71 | 64 | 66 | 111 | 133 | 0 | 0 | 4 | 0 | 0 | 7 | 27 | 62 | 35 | 0 | 0 | 0 | 0 | 0 | 0 | 218 | 149 | 1 | 1 | 1 | 1 | 1 | 0.95 | 1 | 1 | 1 | 1 | 1 | NA | -0.01 |
| SE | 41264 | ENSG00000164506 | STXBP5 | chr6 | + | 147291093 | 147291172 | 147278080 | 147278204 | 147310083 | 147310238 | 180 | 120 | 163 | 174 | 271 | 0 | 0 | 0 | 4 | 0 | 134 | 95 | 156 | 182 | 88 | 0 | 0 | 0 | 0 | 0 | 228 | 149 | 1 | 1 | 1 | 1 | 1 | 1 | 0.978 | 1 | 1 | 1 | 1 | 1 | -0.004 |
| SE | 41266 | ENSG00000164506 | STXBP5 | chr6 | + | 147324958 | 147325084 | 147316228 | 147316407 | 147327124 | 147327276 | 179 | 167 | 176 | 155 | 171 | 0 | 0 | 0 | 0 | 0 | 30 | 110 | 135 | 89 | 31 | 0 | 0 | 5 | 0 | 0 | 275 | 149 | 1 | 1 | 1 | 1 | 1 | 1 | 1 | 1 | 1 | 0.936 | 1 | 1 | 0.013 |
| SE | 41267 | ENSG00000164506 | STXBP5 | chr6 | + | 147324958 | 147325084 | 147316228 | 147316407 | 147334156 | 147334222 | 80 | 69 | 72 | 58 | 72 | 0 | 0 | 1 | 0 | 0 | 21 | 51 | 45 | 34 | 17 | 0 | 0 | 0 | 0 | 0 | 275 | 149 | 1 | 1 | 1 | 1 | 0.975 | 1 | 1 | 1 | 1 | 1 | 1 | 1 | -0.005 |
| SE | 41268 | ENSG00000164506 | STXBP5 | chr6 | + | 147327124 | 147327276 | 147316228 | 147316407 | 147334156 | 147334222 | 85 | 62 | 65 | 81 | 93 | 0 | 0 | 1 | 0 | 0 | 5 | 34 | 60 | 45 | 17 | 0 | 0 | 0 | 0 | 0 | 298 | 149 | 1 | 1 | 1 | 1 | 0.97 | 1 | 1 | 1 | 1 | 1 | 1 | 1 | -0.006 |
| SE | 41269 | ENSG00000164506 | STXBP5 | chr6 | + | 147327124 | 147327276 | 147324958 | 147325084 | 147334156 | 147334222 | 184 | 160 | 169 | 179 | 192 | 0 | 0 | 1 | 0 | 0 | 14 | 93 | 145 | 101 | 31 | 0 | 0 | 0 | 1 | 0 | 298 | 149 | 1 | 1 | 1 | 1 | 1 | 0.989 | 1 | 1 | 1 | 1 | 0.981 | 1 | 0.002 |
| SE | 41271 | ENSG00000164506 | STXBP5 | chr6 | + | 147339178 | 147339238 | 147334156 | 147334222 | 147353322 | 147353373 | 12 | 16 | 16 | 14 | 17 | 67 | 33 | 43 | 63 | 71 | 0 | 4 | 10 | 6 | 0 | 0 | 26 | 33 | 14 | 9 | 209 | 149 | 1 | 1 | 0.113 | 0.257 | 0.21 | 0.137 | 0.146 | NA | 0.099 | 0.178 | 0.234 | 0 | 0.045 |
| SE | 41272 | ENSG00000164506 | STXBP5 | chr6 | + | 147339336 | 147339384 | 147339178 | 147339238 | 147353322 | 147353373 | 15 | 9 | 13 | 10 | 24 | 5 | 10 | 8 | 11 | 3 | 0 | 17 | 14 | 18 | 0 | 0 | 2 | 6 | 3 | 0 | 197 | 149 | 1 | 1 | 0.694 | 0.405 | 0.551 | 0.407 | 0.858 | NA | 0.865 | 0.638 | 0.819 | NA | -0.191 |

| | | | | | | | | | | | | | | | |
|---|---|---|---|---|---|---|---|---|---|---|---|---|---|---|---|
| SE | 41274 | ENSG00000164506 | STXBP5 | chr6 | + | 147364000 | 147364166 | 147363334 | 147363704 | 147373730 | 147373842 | 252 | 260 | 208 | [illegible] |
| SE | 41275 | ENSG00000164506 | STXBP5 | chr6 | + | 147373730 | 147373842 | 147364000 | 147364166 | 147382777 | 147382998 | 251 | 195 | 217 | [illegible] |
| SE | 41538 | ENSG00000071127 | WDR1 | chr4 | - | 10075688 | 10077186 | 10075148 | 10075484 | 10077303 | 10077448 | 2 | 1 | 2 | [illegible] |
| SE | 41539 | ENSG00000071127 | WDR1 | chr4 | - | 10087706 | 10087935 | 10084442 | 10084530 | 10088292 | 10088373 | 830 | 654 | 643 | [illegible] |
| SE | 41540 | ENSG00000071127 | WDR1 | chr4 | - | 10087706 | 10087940 | 10084423 | 10084530 | 10088292 | 10088373 | 1878 | 1290 | 1368 | [illegible] |
| SE | 41541 | ENSG00000071127 | WDR1 | chr4 | - | 10088292 | 10088373 | 10087705 | 10087940 | 10088663 | 10088741 | 1436 | 1000 | 1078 | [illegible] |
| SE | 41542 | ENSG00000071127 | WDR1 | chr4 | - | 10088663 | 10088741 | 10088286 | 10088373 | 10097710 | 10097891 | 1068 | 802 | 780 | [illegible] |
| SE | 41544 | ENSG00000071127 | WDR1 | chr4 | - | 10103895 | 10103986 | 10098989 | 10099139 | 10116112 | 10116234 | 1760 | 1555 | 1440 | [illegible] |
| SE | 41545 | ENSG00000071127 | WDR1 | chr4 | - | 10108217 | 10108395 | 10103895 | 10103986 | 10116112 | 10116234 | 5 | 0 | 6 | [illegible] |
| SE | 41967 | ENSG00000120008 | WDR11 | chr10 | + | 120866568 | 120866764 | 120865629 | 120865744 | 120867065 | 120867169 | 301 | 187 | 209 | [illegible] |
| SE | 41968 | ENSG00000120008 | WDR11 | chr10 | + | 120867065 | 120867169 | 120866568 | 120866764 | 120871169 | 120871346 | 244 | 175 | 187 | [illegible] |
| SE | 41970 | ENSG00000120008 | WDR11 | chr10 | + | 120878352 | 120878459 | 120873838 | 120873923 | 120883779 | 120883805 | 62 | 40 | 42 | [illegible] |
| SE | 41971 | ENSG00000120008 | WDR11 | chr10 | + | 120880825 | 120880901 | 120878352 | 120878459 | 120883779 | 120883805 | 193 | 123 | 145 | [illegible] |
| SE | 41973 | ENSG00000120008 | WDR11 | chr10 | + | 120900028 | 120900137 | 120890715 | 120890887 | 120901035 | 120901098 | 186 | 143 | 137 | [illegible] |
| SE | 41974 | ENSG00000120008 | WDR11 | chr10 | + | 120904645 | 120904811 | 120904046 | 120904142 | 120905318 | 120905416 | 216 | 170 | 141 | [illegible] |
| SE | 41975 | ENSG00000120008 | WDR11 | chr10 | + | 120904649 | 120904811 | 120904046 | 120904142 | 120905318 | 120905416 | 108 | 92 | 74 | [illegible] |
| SE | 41976 | ENSG00000120008 | WDR11 | chr10 | + | 120905875 | 120906021 | 120905318 | 120905416 | 120906775 | 120906855 | 183 | 126 | 122 | [illegible] |
| SE | 42203 | ENSG00000132155 | RAF1 | chr3 | - | 12584846 | 12584981 | 12583600 | 12584657 | 12585121 | 12585253 | 295 | 184 | 259 | [illegible] |
| SE | 42204 | ENSG00000132155 | RAF1 | chr3 | - | 12590797 | 12590974 | 12586846 | 12587637 | 12591707 | 12591792 | 253 | 165 | 284 | [illegible] |
| SE | 42205 | ENSG00000132155 | RAF1 | chr3 | - | 12591707 | 12591792 | 12586846 | 12587637 | 12599690 | 12599755 | 108 | 55 | 93 | [illegible] |
| [illegible] | | | | | | | | | | | | | | | |

| | | | | | | | | | | | | | | | | | | | | | | | | |
|---|---|---|---|---|---|---|---|---|---|---|---|---|---|---|---|---|---|---|---|---|---|---|---|---|
| SE | 52035 | ENSG00000135250 | SRPK2 | chr7 | - | 105297420 | 105297533 | 105268805 | 105268869 | 105388647 | 105388702 | 1 | 2 | 1 | [illegible] | [illegible] | [illegible] | [illegible] | [illegible] | [illegible] | [illegible] | [illegible] | [illegible] | [illegible] |
| SE | 52037 | ENSG00000135250 | SRPK2 | chr7 | - | 105306435 | 105306571 | 105268805 | 105268869 | 105388647 | 105388702 | 1 | 4 | 3 | [illegible] | [illegible] | [illegible] | [illegible] | [illegible] | [illegible] | [illegible] | [illegible] | [illegible] | [illegible] |
| SE | 52038 | ENSG00000135250 | SRPK2 | chr7 | - | 105306435 | 105306571 | 105297420 | 105297533 | 105388647 | 105388702 | 1 | 4 | 3 | [illegible] | [illegible] | [illegible] | [illegible] | [illegible] | [illegible] | [illegible] | [illegible] | [illegible] | [illegible] |
| SE | 52064 | ENSG00000043514 | TRIT1 | chr1 | - | 39844530 | 39844640 | 39844100 | 39844218 | 39847219 | 39847297 | 69 | 43 | 70 | 73 | 134 | 0 | 0 | 0 | 0 | 0 | 48 | 54 | 61 |
| SE | 52065 | ENSG00000043514 | TRIT1 | chr1 | - | 39847219 | 39847297 | 39844530 | 39844640 | 39847547 | 39847660 | 51 | 35 | 62 | 56 | 103 | 0 | 0 | 0 | 2 | 0 | 29 | 55 | 55 |
| SE | 52068 | ENSG00000043514 | TRIT1 | chr1 | - | 39847985 | 39848091 | 39847547 | 39847660 | 39850118 | 39850261 | 46 | 34 | 29 | 35 | 44 | 0 | 0 | 0 | 0 | 0 | 26 | 27 | 19 |
| SE | 52070 | ENSG00000043514 | TRIT1 | chr1 | - | 39847985 | 39848097 | 39847547 | 39847660 | 39850118 | 39850261 | 82 | 47 | 44 | 49 | 67 | 0 | 0 | 0 | 0 | 0 | 57 | 51 | 25 |
| SE | 52071 | ENSG00000043514 | TRIT1 | chr1 | - | 39850118 | 39850261 | 39847985 | 39848097 | 39853969 | 39854068 | 32 | 15 | 16 | 17 | 30 | 0 | 0 | 0 | 2 | 0 | 33 | 23 | 7 |
| SE | 52072 | ENSG00000043514 | TRIT1 | chr1 | - | 39850118 | 39850261 | 39847985 | 39848097 | 39883317 | 39883511 | 29 | 13 | 16 | 14 | 22 | 0 | 1 | 0 | 0 | 0 | 31 | 23 | 6 |
| SE | 52073 | ENSG00000043514 | TRIT1 | chr1 | - | 39850122 | 39850261 | 39847985 | 39848097 | 39853969 | 39854068 | 8 | 11 | 11 | 7 | 17 | 0 | 0 | 0 | 2 | 0 | 8 | 0 | 15 |
| SE | 52075 | ENSG00000043514 | TRIT1 | chr1 | - | 39852730 | 39852876 | 39850118 | 39850261 | 39883317 | 39883491 | 26 | 18 | 26 | 18 | 15 | 0 | 0 | 0 | 1 | 0 | 7 | 7 | 14 |
| SE | 52077 | ENSG00000043514 | TRIT1 | chr1 | - | 39876061 | 39876132 | 39857276 | 39857417 | 39883317 | 39883475 | 0 | 0 | 0 | 0 | 0 | 12 | 6 | 7 | 2 | 6 | 0 | 0 | 6 |
| SE | 52279 | ENSG00000007402 | CACNA2D2 | chr3 | - | 50370319 | 50370380 | 50368137 | 50368235 | 50375643 | 50375705 | 34 | 11 | 15 | 25 | 23 | 0 | 0 | 1 | 2 | 0 | 19 | 17 | 8 |
| SE | 52281 | ENSG00000007402 | CACNA2D2 | chr3 | - | 50374736 | 50374813 | 50370319 | 50370380 | 50375643 | 50375705 | 80 | 36 | 32 | 58 | 77 | 0 | 0 | 0 | 0 | 0 | 35 | 45 | 37 |
| SE | 52282 | ENSG00000007402 | CACNA2D2 | chr3 | - | 50379967 | 50380018 | 50379724 | 50379824 | 50380747 | 50380805 | 42 | 15 | 8 | 19 | 10 | 0 | 0 | 0 | 1 | 0 | 0 | 2 | 16 |
| SE | 52284 | ENSG00000007402 | CACNA2D2 | chr3 | - | 50387567 | 50387612 | 50380994 | 50381126 | 50394108 | 50394168 | 22 | 4 | 6 | 12 | 0 | 0 | 0 | 0 | 0 | 0 | 0 | 20 | 2 |
| SE | 52564 | ENSG00000111445 | RFC5 | chr12 | + | 118017689 | 118017815 | 118016700 | 118016892 | 118017994 | 118018053 | 4 | 1 | 0 | 3 | 2 | 1 | 0 | 7 | 1 | 0 | 9 | 0 | 2 |
| SE | 52566 | ENSG00000111445 | RFC5 | chr12 | + | 118017689 | 118017852 | 118016728 | 118016892 | 118017994 | 118018053 | 2 | 1 | 0 | 2 | 2 | 1 | 0 | 7 | 1 | 0 | 9 | 0 | 3 |
| SE | 52570 | ENSG00000111445 | RFC5 | chr12 | + | 118017994 | 118018053 | 118017689 | 118017815 | 118019071 | 118019136 | 5 | 1 | 3 | 2 | 5 | 2 | 1 | 1 | 2 | 0 | 8 | 3 | 1 |
| SE | 52571 | ENSG00000111445 | RFC5 | chr12 | + | 118017994 | 118018053 | 118017689 | 118017852 | 118019071 | 118019136 | 3 | 1 | 3 | 1 | 5 | 0 | 0 | 0 | 2 | 0 | 8 | 3 | 1 |
| SE | 52572 | ENSG00000111445 | RFC5 | chr12 | + | 118022285 | 118022359 | 118020905 | 118020985 | 118024850 | 118024981 | 23 | 27 | 38 | 14 | 60 | 0 | 0 | 0 | 0 | 0 | 55 | 42 | 30 |
| SE | 52939 | ENSG00000144744 | UBA3 | chr3 | - | 69056611 | 69056693 | 69056182 | 69056283 | 69056778 | 69056815 | 453 | 467 | 357 | 423 | 936 | 0 | 0 | 0 | 1 | 0 | 28 | 333 | 165 |
| SE | 52940 | ENSG00000144744 | UBA3 | chr3 | - | 69056778 | 69056815 | 69056611 | 69056693 | 69057255 | 69057309 | 248 | 241 | 176 | 228 | 452 | 0 | 0 | 0 | 1 | 0 | 5 | 161 | 90 |
| SE | 52942 | ENSG00000144744 | UBA3 | chr3 | - | 69061813 | 69061927 | 69057255 | 69057309 | 69062981 | 69063137 | 116 | 96 | 88 | 110 | 220 | 0 | 0 | 1 | 0 | 0 | 0 | 69 | 46 |
| SE | 52943 | ENSG00000144744 | UBA3 | chr3 | - | 69062076 | 69062179 | 69061582 | 69061927 | 69062981 | 69063137 | 428 | 327 | 298 | 373 | 661 | 1 | 0 | 0 | 0 | 1 | 90 | 288 | 271 |
| SE | 52944 | ENSG00000144744 | UBA3 | chr3 | - | 69062981 | 69063137 | 69062076 | 69062179 | 69063438 | 69063503 | 488 | 313 | 336 | 359 | 631 | 0 | 0 | 0 | 0 | 0 | 100 | 303 | 232 |
| SE | 52945 | ENSG00000144744 | UBA3 | chr3 | - | 69071534 | 69071617 | 69063438 | 69063503 | 69075429 | 69075510 | 82 | 66 | 77 | 94 | 92 | 1 | 0 | 0 | 0 | 1 | 6 | 25 | 19 |
| SE | 52946 | ENSG00000144744 | UBA3 | chr3 | - | 69071534 | 69071617 | 69063438 | 69063503 | 69077797 | 69077918 | 1 | 2 | 0 | 1 | 0 | 0 | 3 | 1 | 0 | 1 | 0 | 2 | 0 |
| SE | 52948 | ENSG00000144744 | UBA3 | chr3 | - | 69071534 | 69071617 | 69067927 | 69068008 | 69077797 | 69077918 | 78 | 38 | 46 | 50 | 114 | 1 | 0 | 0 | 0 | 4 | 0 | 29 | 19 |
| SE | 52949 | ENSG00000144744 | UBA3 | chr3 | - | 69075429 | 69075510 | 69063438 | 69063503 | 69077797 | 69077918 | 46 | 34 | 41 | 54 | 45 | 0 | 3 | 1 | 0 | 1 | 12 | 22 | 21 |
| SE | 52950 | ENSG00000144744 | UBA3 | chr3 | - | 69075429 | 69075510 | 69067927 | 69068008 | 69077797 | 69077918 | 47 | 36 | 45 | 57 | 51 | 1 | 0 | 0 | 0 | 4 | 25 | 24 | 24 |
| SE | 52951 | ENSG00000144744 | UBA3 | chr3 | - | 69075429 | 69075510 | 69071092 | 69071617 | 69077797 | 69077918 | 166 | 134 | 155 | 193 | 188 | 0 | 1 | 0 | 2 | 0 | 18 | 69 | 55 |
| SE | 53192 | ENSG00000149527 | PLCH2 | chr1 | + | 2487597 | 2487718 | 2487172 | 2487376 | 2489206 | 2489378 | 32 | 12 | 16 | 43 | 40 | 0 | 0 | 0 | 0 | 3 | 9 | 24 | 7 |
| SE | 53279 | ENSG00000164151 | ICE1 | chr5 | + | 5454551 | 5454638 | 5447840 | 5447897 | 5457331 | 5457741 | 186 | 161 | 195 | 305 | 348 | 0 | 0 | 0 | 0 | 0 | 1 | 162 | 114 |
| SE | 53341 | ENSG00000131778 | CHD1L | chr1 | + | 147259836 | 147259918 | 147256530 | 147256562 | 147264421 | 147264532 | 91 | 45 | 66 | 65 | 125 | 0 | 0 | 0 | 2 | 0 | 98 | 65 | 86 |
| SE | 53343 | ENSG00000131778 | CHD1L | chr1 | + | 147264421 | 147264584 | 147259836 | 147259918 | 147265931 | 147266087 | 136 | 96 | 92 | 112 | 168 | 1 | 0 | 1 | 2 | 2 | 148 | 128 | 134 |
| SE | 53426 | ENSG00000111358 | GTF2H3 | chr12 | + | 123639263 | 123639343 | 123633738 | 123633872 | 123645454 | 123645561 | 44 | 31 | 51 | 31 | 106 | 1 | 0 | 0 | 0 | 0 | 26 | 23 | 37 |
| SE | 53427 | ENSG00000111358 | GTF2H3 | chr12 | + | 123639293 | 123639343 | 123633828 | 123633872 | 123645454 | 123645561 | 17 | 9 | 12 | 9 | 22 | 1 | 0 | 0 | 0 | 0 | 8 | 2 | 5 |
| SE | 53428 | ENSG00000111358 | GTF2H3 | chr12 | + | 123639956 | 123640001 | 123639263 | 123639343 | 123645454 | 123645561 | 0 | 0 | 2 | 0 | 0 | 44 | 31 | 52 | 31 | 106 | 0 | 0 | 0 |
| SE | 53429 | ENSG00000111358 | GTF2H3 | chr12 | + | 123647962 | 123648126 | 123645454 | 123645561 | 123650993 | 123651056 | 93 | 83 | 108 | 83 | 219 | 5 | 0 | 0 | 0 | 0 | 50 | 58 | 59 |
| SE | 53431 | ENSG00000111358 | GTF2H3 | chr12 | + | 123655770 | 123655824 | 123654923 | 123654998 | 123659515 | 123659584 | 65 | 36 | 72 | 84 | 174 | 0 | 0 | 1 | 2 | 0 | 16 | 47 | 32 |
| SE | 53899 | ENSG00000114805 | PLCH1 | chr3 | - | 155504554 | 155504626 | 155500702 | 155500794 | 155514722 | 155514884 | 0 | 6 | 1 | 7 | 1 | 0 | 0 | 1 | 0 | 0 | 8 | 0 | 6 |
| SE | 54335 | ENSG00000176095 | IP6K1 | chr3 | - | 49738211 | 49738422 | 49732790 | 49732972 | 49747817 | 49748168 | 161 | 68 | 119 | 115 | 94 | 0 | 0 | 0 | 1 | 0 | 118 | 100 | 121 |
| SE | 54636 | ENSG00000140992 | PDPK1 | chr16 | + | 2566355 | 2566453 | 2561479 | 2561522 | 2577424 | 2577500 | 0 | 0 | 3 | 3 | 5 | 2 | 1 | 3 | 2 | 0 | 0 | 4 | 2 |
| SE | 54637 | ENSG00000140992 | PDPK1 | chr16 | + | 2577424 | 2577500 | 2561479 | 2561522 | 2581294 | 2581363 | 4 | 5 | 6 | 7 | 0 | 0 | 0 | 0 | 0 | 0 | 0 | 9 | 5 |
| SE | 54638 | ENSG00000140992 | PDPK1 | chr16 | + | 2583411 | 2583585 | 2581606 | 2581703 | 2586675 | 2586893 | 120 | 38 | 91 | 106 | 60 | 0 | 0 | 1 | 0 | 0 | 108 | 79 | 63 |
| SE | 54639 | ENSG00000140992 | PDPK1 | chr16 | + | 2595792 | 2595850 | 2586675 | 2586893 | 2597122 | 2597275 | 144 | 84 | 135 | 165 | 101 | 0 | 0 | 0 | 0 | 0 | 146 | 102 | 85 |
| SE | 54642 | ENSG00000140992 | PDPK1 | chr16 | + | 2597122 | 2597275 | 2595792 | 2595850 | 2597650 | 2597928 | 250 | 160 | 226 | 231 | 279 | 0 | 0 | 0 | 0 | 0 | 305 | 229 | 136 |
| SE | 54669 | ENSG00000070010 | UFD1 | chr22 | - | 19455679 | 19455768 | 19450620 | 19454830 | 19456586 | 19456634 | 145 | 115 | 158 | 175 | 218 | 0 | 0 | 0 | 0 | 0 | 341 | 230 | 279 |
| SE | 54670 | ENSG00000070010 | UFD1 | chr22 | - | 19458070 | 19458139 | 19456852 | 19456918 | 19465201 | 19465274 | 151 | 118 | 131 | 197 | 229 | 0 | 0 | 0 | 0 | 0 | 333 | 227 | 245 |
| SE | 54671 | ENSG00000070010 | UFD1 | chr22 | - | 19471686 | 19471808 | 19467402 | 19468003 | 19475067 | 19475100 | 300 | 174 | 285 | 294 | 432 | 0 | 0 | 0 | 0 | 0 | 477 | 388 | 349 |
| SE | 54672 | ENSG00000070010 | UFD1 | chr22 | - | 19471686 | 19471808 | 19467402 | 19468003 | 19475469 | 19475602 | 184 | 98 | 190 | 180 | 281 | 0 | 0 | 0 | 0 | 0 | 365 | 243 | 211 |
| SE | 54674 | ENSG00000070010 | UFD1 | chr22 | - | 19475067 | 19475100 | 19467402 | 19468003 | 19475469 | 19475602 | 6 | 0 | 3 | 1 | 3 | 0 | 0 | 0 | 0 | 0 | 4 | 14 | 8 |
| SE | 54675 | ENSG00000070010 | UFD1 | chr22 | - | 19475067 | 19475100 | 19467402 | 19468003 | 19479082 | 19479172 | 0 | 0 | 0 | 0 | 0 | 0 | 0 | 2 | 1 | 4 | 0 | 0 | 1 |
| SE | 54676 | ENSG00000070010 | UFD1 | chr22 | - | 19475067 | 19475100 | 19471686 | 19471808 | 19475469 | 19475602 | 103 | 69 | 71 | 100 | 140 | 0 | 0 | 0 | 0 | 0 | 104 | 147 | 118 |
| SE | 54744 | ENSG00000083720 | OXCT1 | chr5 | - | 41749526 | 41749607 | 41739389 | 41739491 | 41762110 | 41762200 | 66 | 50 | 86 | 75 | 48 | 2 | 0 | 0 | 0 | 0 | 0 | 40 | 49 |
| SE | 54745 | ENSG00000083720 | OXCT1 | chr5 | - | 41749526 | 41749607 | 41739389 | 41739491 | 41794002 | 41794078 | 26 | 14 | 27 | 24 | 16 | 0 | 0 | 0 | 0 | 0 | 0 | 10 | 17 |
| SE | 54746 | ENSG00000083720 | OXCT1 | chr5 | - | 41762110 | 41762200 | 41739389 | 41739491 | 41794002 | 41794078 | 25 | 12 | 35 | 39 | 24 | 0 | 0 | 0 | 0 | 0 | 7 | 41 | 50 |
| SE | 54747 | ENSG00000083720 | OXCT1 | chr5 | - | 41762110 | 41762200 | 41749526 | 41749607 | 41794002 | 41794078 | 66 | 52 | 99 | 96 | 58 | 0 | 3 | 0 | 0 | 2 | 7 | 71 | 82 |
| SE | 54750 | ENSG00000083720 | OXCT1 | chr5 | - | 41803068 | 41803163 | 41801021 | 41801070 | 41840450 | 41840511 | 43 | 29 | 34 | 39 | 33 | 2 | 0 | 0 | 1 | 0 | 12 | 42 | 33 |
| SE | 54751 | ENSG00000083720 | OXCT1 | chr5 | - | 41805566 | 41805681 | 41803068 | 41803163 | 41807330 | 41807438 | 124 | 79 | 126 | 101 | 160 | 4 | 0 | 0 | 0 | 0 | 34 | 80 | 86 |
| SE | 54753 | ENSG00000083720 | OXCT1 | chr5 | - | 41807330 | 41807438 | 41801021 | 41801070 | 41840450 | 41840511 | 61 | 27 | 79 | 54 | 69 | 2 | 0 | 0 | 1 | 0 | 46 | 52 | 59 |

[illegible]

| | | | | | | | | | | | | | | | | | | | | | | | | | | | | | | | | | | | | | | | | | | | | | | |
|---|---|---|---|---|---|---|---|---|---|---|---|---|---|---|---|---|---|---|---|---|---|---|---|---|---|---|---|---|---|---|---|---|---|---|---|---|---|---|---|---|---|---|---|---|---|---|
| SE | 60179 | ENSG00000162852 | CNST | chr1 | + | 246641748 | 246641770 | 246633923 | 246634587 | 246647138 | 246648037 | 4 | 22 | 8 | 15 | 33 | 3 | 1 | 0 | 3 | 1 | 6 | 7 | 6 | 18 | 0 | 0 | 16 | 16 | 14 | 0 | 171 | 149 | 1 | 1 | 0.537 | 0.95 | 1 | 0.813 | 0.966 | 1 | 0.276 | 0.246 | 0.528 | NA | 0.341 |
| SE | 60180 | ENSG00000162852 | CNST | chr1 | + | 246641940 | 246642037 | 246641748 | 246641770 | 246647138 | 246648037 | 309 | 414 | 252 | 411 | 652 | 0 | 0 | 1 | 0 | 11 | 249 | 582 | 358 | 371 | 128 | 0 | 1 | 0 | 5 | 0 | 246 | 149 | 1 | 1 | 1 | 1 | 0.993 | 1 | 0.973 | 1 | 0.997 | 1 | 0.978 | 1 | -0.002 |
| SE | 60211 | ENSG00000154451 | GBP5 | chr1 | - | 89268728 | 89268856 | 89267416 | 89267526 | 89269365 | 89269574 | 960 | 886 | 1154 | 862 | 972 | 2 | 0 | 2 | 0 | 1 | 823 | 755 | 533 | 739 | 203 | 0 | 8 | 4 | 0 | 0 | 277 | 149 | 1 | 1 | 0.996 | 1 | 0.997 | 1 | 0.998 | 1 | 0.981 | 0.986 | 1 | 1 | 0.005 |
| SE | 60212 | ENSG00000154451 | GBP5 | chr1 | - | 89269365 | 89269574 | 89268728 | 89268856 | 89270754 | 89270863 | 973 | 941 | 1160 | 886 | 1015 | 0 | 0 | 2 | 0 | 0 | 1016 | 852 | 618 | 804 | 274 | 0 | 0 | 0 | 2 | 0 | 298 | 149 | 1 | 1 | 1 | 1 | 0.997 | 1 | 1 | 1 | 1 | 1 | 0.995 | 1 | 0 |
| SE | 60352 | ENSG00000172766 | NAA16 | chr13 | + | 41316845 | 41316930 | 41311266 | 41311582 | 41318805 | 41318910 | 159 | 128 | 210 | 161 | 284 | 0 | 0 | 0 | 0 | 0 | 34 | 111 | 139 | 48 | 44 | 0 | 3 | 1 | 0 | 0 | 234 | 149 | 1 | 1 | 1 | 1 | 1 | 1 | 1 | 1 | 0.959 | 0.989 | 1 | 1 | 0.01 |
| SE | 60353 | ENSG00000172766 | NAA16 | chr13 | + | 41323055 | 41323190 | 41320666 | 41320824 | 41325697 | 41325851 | 423 | 396 | 476 | 490 | 464 | 1 | 0 | 0 | 0 | 0 | 182 | 307 | 419 | 265 | 219 | 0 | 2 | 1 | 0 | 0 | 284 | 149 | 1 | 1 | 0.996 | 1 | 1 | 1 | 1 | 1 | 0.988 | 0.995 | 1 | 1 | 0.003 |
| SE | 60354 | ENSG00000172766 | NAA16 | chr13 | + | 41331273 | 41331369 | 41328723 | 41328843 | 41336649 | 41336687 | 107 | 119 | 112 | 135 | 172 | 0 | 0 | 0 | 2 | 4 | 48 | 127 | 84 | 62 | 69 | 0 | 0 | 0 | 0 | 0 | 245 | 149 | 1 | 1 | 1 | 1 | 1 | 0.976 | 0.963 | 1 | 1 | 1 | 1 | 1 | -0.012 |
| SE | 60357 | ENSG00000172766 | NAA16 | chr13 | + | 41355143 | 41355216 | 41331273 | 41331369 | 41358303 | 41358473 | 16 | 20 | 16 | 23 | 62 | 5 | 8 | 5 | 2 | 7 | 3 | 7 | 28 | 1 | 0 | 6 | 0 | 12 | 1 | 0 | 222 | 149 | 1 | 1 | 0.682 | 0.627 | 0.682 | 0.885 | 0.856 | 0.251 | 1 | 0.61 | 0.402 | NA | 0.181 |
| SE | 60358 | ENSG00000172766 | NAA16 | chr13 | + | 41355143 | 41355216 | 41336649 | 41336756 | 41358303 | 41358473 | 49 | 59 | 60 | 61 | 147 | 6 | 4 | 3 | 17 | 7 | 0 | 26 | 16 | 4 | 0 | 5 | 6 | 6 | 0 | 0 | 222 | 149 | 1 | 1 | 0.846 | 0.908 | 0.931 | 0.707 | 0.934 | 0 | 0.744 | 0.642 | 1 | NA | 0.269 |
| SE | 60360 | ENSG00000172766 | NAA16 | chr13 | + | 41369089 | 41369283 | 41367438 | 41367652 | 41372202 | 41372311 | 77 | 77 | 110 | 109 | 219 | 0 | 1 | 0 | 1 | 3 | 8 | 75 | 41 | 22 | 19 | 0 | 0 | 0 | 0 | 0 | 298 | 149 | 1 | 1 | 1 | 0.975 | 1 | 0.982 | 0.973 | 1 | 1 | 1 | 1 | 1 | -0.014 |
| SE | 60361 | ENSG00000172766 | NAA16 | chr13 | + | 41372202 | 41372311 | 41369089 | 41369283 | 41372731 | 41372830 | 69 | 33 | 78 | 68 | 143 | 0 | 0 | 0 | 0 | 3 | 28 | 51 | 34 | 18 | 36 | 0 | 0 | 1 | 0 | 0 | 258 | 149 | 1 | 1 | 1 | 1 | 1 | 1 | 0.965 | 1 | 1 | 0.952 | 1 | 1 | 0.003 |
| SE | 60362 | ENSG00000172766 | NAA16 | chr13 | + | 41373636 | 41373780 | 41372731 | 41372830 | 41374741 | 41374839 | 62 | 32 | 77 | 62 | 119 | 2 | 0 | 0 | 0 | 2 | 8 | 35 | 35 | 10 | 9 | 0 | 0 | 0 | 0 | 0 | 293 | 149 | 1 | 1 | 0.94 | 1 | 1 | 1 | 0.968 | 1 | 1 | 1 | 1 | 1 | -0.018 |
| SE | 60363 | ENSG00000172766 | NAA16 | chr13 | + | 41373767 | 41373880 | 41372814 | 41372830 | 41374741 | 41374839 | 4 | 1 | 0 | 5 | 7 | 2 | 0 | 0 | 0 | 2 | 0 | 1 | 3 | 0 | 0 | 0 | 0 | 0 | 0 | 0 | 262 | 149 | 1 | 1 | 0.532 | 1 | NA | 1 | 0.666 | NA | 1 | 1 | NA | NA | -0.201 |
| SE | 60364 | ENSG00000172766 | NAA16 | chr13 | + | 41374741 | 41374839 | 41373767 | 41373880 | 41375404 | 41376182 | 36 | 19 | 22 | 36 | 58 | 1 | 0 | 0 | 0 | 0 | 0 | 8 | 16 | 5 | 8 | 0 | 0 | 0 | 0 | 0 | 247 | 149 | 1 | 1 | 0.956 | 1 | 1 | 1 | 1 | NA | 1 | 1 | 1 | 1 | -0.009 |
| SE | 60746 | ENSG00000198216 | CACNA1E | chr1 | + | 181580594 | 181580776 | 181579071 | 181579224 | 181651337 | 181651441 | 0 | 0 | 0 | 0 | 2 | 0 | 0 | 0 | 1 | 0 | 0 | 0 | 4 | 2 | 0 | 0 | 0 | 0 | 0 | 0 | 298 | 149 | 1 | 1 | NA | NA | NA | 0 | 1 | NA | NA | 1 | 1 | NA | -0.5 |
| SE | 60914 | ENSG00000130713 | EXOSC2 | chr9 | + | 130703052 | 130703181 | 130700841 | 130702310 | 130703693 | 130703774 | 239 | 124 | 221 | 215 | 173 | 0 | 0 | 0 | 1 | 0 | 282 | 230 | 192 | 268 | 372 | 0 | 0 | 4 | 0 | 0 | 278 | 149 | 1 | 1 | 1 | 1 | 1 | 1 | 1 | 1 | 1 | 0.963 | 1 | 1 | 0.007 |
| SE | 61790 | ENSG00000174371 | EXO1 | chr1 | + | 241849108 | 241849216 | 241848950 | 241849027 | 241850408 | 241850586 | 1 | 2 | 3 | 0 | 0 | 1 | 0 | 0 | 1 | 2 | 0 | 2 | 3 | 5 | 0 | 0 | 0 | 6 | 1 | 0 | 257 | 149 | 1 | 1 | 0.367 | 1 | 1 | 0 | 0 | NA | 1 | 0.225 | 0.744 | NA | -0.183 |
| SE | 61791 | ENSG00000174371 | EXO1 | chr1 | + | 241849111 | 241849216 | 241848730 | 241849027 | 241850408 | 241850586 | 0 | 0 | 2 | 0 | 0 | 1 | 0 | 0 | 1 | 2 | 0 | 2 | 0 | 4 | 0 | 0 | 0 | 6 | 1 | 0 | 254 | 149 | 1 | 1 | 0 | NA | 1 | 0 | 0 | NA | 1 | 0 | 0.701 | NA | -0.317 |
| SE | 61792 | ENSG00000174371 | EXO1 | chr1 | + | 241852291 | 241852411 | 241850408 | 241850586 | 241853357 | 241853481 | 6 | 0 | 22 | 4 | 4 | 0 | 0 | 0 | 1 | 2 | 0 | 22 | 20 | 10 | 0 | 11 | 0 | 3 | 2 | 0 | 269 | 149 | 1 | 1 | 1 | NA | 1 | 0.689 | 0.526 | 0 | 1 | 0.787 | 0.735 | NA | 0.173 |
| SE | 61908 | ENSG00000124067 | SLC12A4 | chr16 | - | 67946437 | 67946633 | 67946170 | 67946340 | 67946936 | 67947105 | 41 | 13 | 33 | 31 | 10 | 0 | 0 | 0 | 0 | 0 | 28 | 43 | 16 | 30 | 80 | 0 | 0 | 2 | 0 | 0 | 298 | 149 | 1 | 1 | 1 | 1 | 1 | 1 | 1 | 1 | 1 | 0.8 | 1 | 1 | 0.04 |
| SE | 61909 | ENSG00000124067 | SLC12A4 | chr16 | - | 67947330 | 67947435 | 67946936 | 67947105 | 67947668 | 67947788 | 28 | 7 | 27 | 22 | 15 | 0 | 0 | 1 | 3 | 0 | 44 | 37 | 16 | 31 | 75 | 0 | 0 | 0 | 2 | 0 | 254 | 149 | 1 | 1 | 1 | 1 | 0.941 | 0.811 | 1 | 1 | 1 | 1 | 0.901 | 1 | -0.03 |
| SE | 61974 | ENSG00000205423 | CNEP1R1 | chr16 | + | 50029724 | 50029798 | 50026159 | 50026467 | 50033396 | 50033506 | 50 | 32 | 40 | 37 | 45 | 2 | 0 | 0 | 0 | 0 | 33 | 28 | 40 | 34 | 22 | 0 | 0 | 0 | 0 | 0 | 223 | 149 | 1 | 1 | 0.944 | 1 | 1 | 1 | 1 | 1 | 1 | 1 | 1 | 1 | -0.011 |
| SE | 62098 | ENSG00000092529 | CAPN3 | chr15 | + | 42405925 | 42405943 | 42403740 | 42403777 | 42408210 | 42408324 | 0 | 0 | 0 | 1 | 2 | 12 | 17 | 3 | 7 | 9 | 0 | 0 | 3 | 1 | 0 | 12 | 20 | 10 | 44 | 19 | 167 | 149 | 1 | 1 | 0 | 0 | 0 | 0.113 | 0.165 | 0 | 0 | 0.211 | 0.02 | 0 | 0.009 |
| SE | 62099 | ENSG00000092529 | CAPN3 | chr15 | + | 42405925 | 42405943 | 42403740 | 42403777 | 42409302 | 42409380 | 0 | 0 | 0 | 1 | 2 | 8 | 0 | 7 | 7 | 2 | 0 | 0 | 3 | 0 | 0 | 6 | 11 | 0 | 9 | 0 | 167 | 149 | 1 | 1 | 0 | NA | 0 | 0 | 0.472 | 0 | 0 | 1 | 0 | NA | -0.132 |
| SE | 62734 | ENSG00000101333 | PLCB4 | chr20 | + | 9373046 | 9373104 | 9372302 | 9372403 | 9380053 | 9380162 | 0 | 0 | 0 | 1 | 0 | 0 | 2 | 0 | 0 | 1 | 0 | 2 | 2 | 11 | 0 | 0 | 0 | 2 | 6 | 0 | 207 | 149 | 1 | 1 | NA | 0 | NA | 1 | 0 | NA | 1 | 0.419 | 0.569 | NA | -0.329 |
| SE | 62920 | ENSG00000185420 | SMYD3 | chr1 | - | 246186742 | 246186942 | 245929869 | 245929937 | 246193674 | 246193727 | 1 | 0 | 0 | 4 | 2 | 0 | 0 | 0 | 0 | 0 | 6 | 4 | 0 | 0 | 2 | 0 | 0 | 0 | 4 | 0 | 298 | 149 | 1 | 1 | 1 | NA | NA | 1 | 1 | 1 | 1 | NA | 0 | 1 | 0.25 |
| SE | 62922 | ENSG00000185420 | SMYD3 | chr1 | - | 246335366 | 246335474 | 246330479 | 246330537 | 246355030 | 246355094 | 57 | 31 | 42 | 49 | 51 | 1 | 0 | 0 | 0 | 0 | 86 | 35 | 39 | 22 | 70 | 0 | 0 | 0 | 0 | 0 | 257 | 149 | 1 | 1 | 0.971 | 1 | 1 | 1 | 1 | 1 | 1 | 1 | 1 | 1 | -0.006 |
| SE | 62923 | ENSG00000185420 | SMYD3 | chr1 | - | 246355030 | 246355094 | 246335366 | 246335474 | 246507053 | 246507144 | 26 | 5 | 17 | 20 | 20 | 2 | 0 | 2 | 0 | 0 | 29 | 18 | 16 | 8 | 21 | 0 | 0 | 0 | 0 | 0 | 213 | 149 | 1 | 1 | 0.901 | 1 | 0.856 | 1 | 1 | 1 | 1 | 1 | 1 | 1 | -0.049 |
| SE | 63071 | ENSG00000132953 | XPO4 | chr13 | - | 20786964 | 20787057 | 20777328 | 20783919 | 20790461 | 20790580 | 46 | 29 | 38 | 38 | 60 | 3 | 2 | 2 | 4 | 4 | 5 | 38 | 24 | 2 | 0 | 0 | 0 | 8 | 0 | 0 | 242 | 149 | 1 | 1 | 0.904 | 0.899 | 0.921 | 0.854 | 0.902 | 1 | 1 | 0.649 | 1 | NA | -0.016 |
| SE | 63072 | ENSG00000132953 | XPO4 | chr13 | - | 20787480 | 20787598 | 20786964 | 20787057 | 20790461 | 20790580 | 37 | 41 | 46 | 43 | 85 | 3 | 3 | 2 | 1 | 1 | 0 | 43 | 23 | 8 | 0 | 0 | 0 | 0 | 0 | 0 | 267 | 149 | 1 | 1 | 0.873 | 0.884 | 0.928 | 0.96 | 0.979 | NA | 1 | 1 | 1 | NA | -0.075 |
| SE | 63073 | ENSG00000132953 | XPO4 | chr13 | - | 20788485 | 20788616 | 20787480 | 20787598 | 20790461 | 20790580 | 107 | 82 | 94 | 111 | 332 | 0 | 3 | 0 | 0 | 0 | 4 | 105 | 72 | 21 | 20 | 0 | 1 | 1 | 0 | 0 | 280 | 149 | 1 | 1 | 1 | 0.936 | 1 | 1 | 1 | 1 | 0.982 | 0.975 | 1 | 1 | -0.004 |
| SE | 63074 | ENSG00000132953 | XPO4 | chr13 | - | 20790461 | 20790580 | 20777328 | 20783919 | 20796075 | 20796256 | 82 | 64 | 54 | 59 | 123 | 0 | 0 | 1 | 0 | 0 | 0 | 28 | 50 | 7 | 7 | 0 | 0 | 0 | 0 | 0 | 268 | 149 | 1 | 1 | 1 | 1 | 0.968 | 1 | 1 | NA | 1 | 1 | 1 | 1 | -0.006 |
| SE | 63075 | ENSG00000132953 | XPO4 | chr13 | - | 20790461 | 20790580 | 20788485 | 20788616 | 20796075 | 20796256 | 130 | 100 | 93 | 105 | 289 | 1 | 0 | 0 | 0 | 0 | 0 | 67 | 84 | 14 | 18 | 0 | 0 | 0 | 0 | 0 | 268 | 149 | 1 | 1 | 0.986 | 1 | 1 | 1 | 1 | NA | 1 | 1 | 1 | 1 | -0.003 |
| SE | 63077 | ENSG00000132953 | XPO4 | chr13 | - | 20800155 | 20800325 | 20796763 | 20797057 | 20807456 | 20807634 | 1 | 5 | 2 | 0 | 0 | 0 | 0 | 0 | 0 | 0 | 0 | 8 | 1 | 0 | 0 | 0 | 0 | 3 | 0 | 0 | 298 | 149 | 1 | 1 | 1 | 1 | 1 | NA | NA | NA | 1 | 0.143 | NA | NA | 0.428 |
| SE | 63078 | ENSG00000132953 | XPO4 | chr13 | - | 20800830 | 20800990 | 20800155 | 20800325 | 20807456 | 20807634 | 128 | 88 | 133 | 100 | 190 | 1 | 5 | 0 | 0 | 0 | 9 | 110 | 142 | 25 | 0 | 0 | 0 | 0 | 0 | 0 | 298 | 149 | 1 | 1 | 0.985 | 0.898 | 1 | 1 | 1 | 1 | 1 | 1 | 1 | NA | -0.023 |
| SE | 63079 | ENSG00000132953 | XPO4 | chr13 | - | 20807456 | 20807634 | 20796763 | 20797057 | 20808435 | 20808581 | 56 | 44 | 65 | 36 | 74 | 0 | 0 | 0 | 0 | 0 | 10 | 38 | 30 | 8 | 0 | 0 | 0 | 0 | 2 | 0 | 298 | 149 | 1 | 1 | 1 | 1 | 1 | 1 | 1 | 1 | 1 | 1 | 0.667 | NA | 0.083 |
| SE | 63080 | ENSG00000132953 | XPO4 | chr13 | - | 20807456 | 20807634 | 20800830 | 20800990 | 20808435 | 20808581 | 116 | 77 | 121 | 77 | 142 | 1 | 5 | 0 | 1 | 2 | 10 | 105 | 102 | 20 | 0 | 0 | 0 | 1 | 0 | 0 | 298 | 149 | 1 | 1 | 0.983 | 0.885 | 1 | 0.975 | 0.973 | 1 | 1 | 0.981 | 1 | NA | -0.032 |
| SE | 63081 | ENSG00000132953 | XPO4 | chr13 | - | 20843769 | 20843886 | 20842894 | 20843048 | 20855626 | 20855765 | 88 | 82 | 86 | 72 | 175 | 0 | 4 | 0 | 0 | 0 | 16 | 62 | 58 | 32 | 7 | 0 | 0 | 0 | 0 | 0 | 266 | 149 | 1 | 1 | 1 | 0.92 | 1 | 1 | 1 | 1 | 1 | 1 | 1 | 1 | -0.016 |
| SE | 63084 | ENSG00000132953 | XPO4 | chr13 | - | 20868595 | 20868701 | 20862968 | 20863123 | 20902669 | 20902810 | 32 | 21 | 44 | 32 | 22 | 1 | 0 | 0 | 0 | 0 | 12 | 32 | 41 | 11 | 0 | 0 | 0 | 0 | 0 | 0 | 255 | 149 | 1 | 1 | 0.949 | 1 | 1 | 1 | 1 | 1 | 1 | 1 | 1 | NA | -0.01 |
| SE | 63125 | ENSG00000164134 | NAA15 | chr4 | + | 139334173 | 139334258 | 139301504 | 139301831 | 139336847 | 139336952 | 143 | 127 | 140 | 159 | 183 | 0 | 0 | 0 | 0 | 0 | 26 | 115 | 133 | 76 | 27 | 0 | 0 | 0 | 1 | 0 | 234 | 149 | 1 | 1 | 1 | 1 | 1 | 1 | 1 | 1 | 1 | 1 | 0.98 | 1 | 0.004 |
| SE | 63126 | ENSG00000164134 | NAA15 | chr4 | + | 139342825 | 139342960 | 139340911 | 139341069 | 139344185 | 139344339 | 289 | 228 | 253 | 320 | 510 | 0 | 0 | 0 | 0 | 0 | 54 | 166 | 201 | 78 | 29 | 0 | 0 | 0 | 1 | 0 | 284 | 149 | 1 | 1 | 1 | 1 | 1 | 1 | 1 | 1 | 1 | 1 | 0.976 | 1 | 0.005 |
| SE | 63127 | ENSG00000164134 | NAA15 | chr4 | + | 139344185 | 139344339 | 139340911 | 139341069 | 139349461 | 139349581 | 140 | 150 | 136 | 134 | 241 | 0 | 0 | 0 | 0 | 1 | 20 | 76 | 99 | 34 | 5 | 0 | 0 | 0 | 0 | 0 | 298 | 149 | 1 | 1 | 1 | 1 | 1 | 1 | 0.992 | 1 | 1 | 1 | 1 | 1 | -0.002 |
| SE | 63128 | ENSG00000164134 | NAA15 | chr4 | + | 139354025 | 139354098 | 139351504 | 139351611 | 139357385 | 139357555 | 168 | 86 | 143 | 162 | 235 | 0 | 3 | 0 | 2 | 6 | 0 | 63 | 97 | 8 | 17 | 0 | 0 | 1 | 0 | 0 | 222 | 149 | 1 | 1 | 1 | 0.951 | 1 | 0.982 | 0.963 | NA | 1 | 0.985 | 1 | 1 | -0.017 |
| SE | 63130 | ENSG00000164134 | NAA15 | chr4 | + | 139357385 | 139357555 | 139354025 | 139354098 | 139359742 | 139359857 | 294 | 193 | 241 | 274 | 450 | 0 | 0 | 1 | 0 | 0 | 0 | 145 | 153 | 30 | 35 | 0 | 0 | 0 | 0 | 0 | 298 | 149 | 1 | 1 | 1 | 1 | 0.992 | 1 | 1 | NA | 1 | 1 | 1 | 1 | -0.002 |
| SE | 63131 | ENSG00000164134 | NAA15 | chr4 | + | 139370210 | 139370404 | 139361723 | 139361937 | 139376364 | 139376473 | 180 | 163 | 145 | 187 | 204 | 0 | 0 | 1 | 1 | 0 | 0 | 112 | 190 | 15 | 5 | 0 | 0 | 3 | 0 | 0 | 298 | 149 | 1 | 1 | 1 | 1 | 0.986 | 0.989 | 1 | NA | 1 | 0.969 | 1 | 1 | 0.003 |
| SE | 63132 | ENSG00000164134 | NAA15 | chr4 | + | 139376364 | 139376473 | 139370210 | 139370404 | 139378755 | 139378854 | 89 | 58 | 79 | 79 | 127 | 0 | 0 | 0 | 0 | 2 | 0 | 24 | 52 | 3 | 5 | 0 | 0 | 0 | 0 | 0 | 258 | 149 | 1 | 1 | 1 | 1 | 1 | 1 | 0.973 | NA | 1 | 1 | 1 | 1 | -0.005 |
| SE | 63133 | ENSG00000164134 | NAA15 | chr4 | + | 139378755 | 139378854 | 139376364 | 139376473 | 139384831 | 139384978 | 191 | 144 | 156 | 175 | 274 | 0 | 0 | 0 | 1 | 0 | 0 | 65 | 101 | 10 | 0 | 0 | 0 | 0 | 0 | 0 | 248 | 149 | 1 | 1 | 1 | 1 | 1 | 0.991 | 1 | NA | 1 | 1 | 1 | NA | -0.002 |
| SE | 63134 | ENSG00000164134 | NAA15 | chr4 | + | 139384831 | 139384978 | 139378755 | 139378854 | 139386132 | 139386230 | 351 | 272 | 291 | 328 | 559 | 3 | 0 | 0 | 0 | 0 | 0 | 144 | 200 | 45 | 0 | 0 | 0 | 0 | 0 | 0 | 296 | 149 | 1 | 1 | 0.983 | 1 | 1 | 1 | 1 | NA | 1 | 1 | 1 | NA | -0.003 |
| SE | 63135 | ENSG00000164134 | NAA15 | chr4 | + | 139384834 | 139384978 | 139378755 | 139378854 | 139386132 | 139386230 | 229 | 167 | 194 | 189 | 364 | 3 | 0 | 0 | 0 | 0 | 0 | 89 | 129 | 39 | 0 | 0 | 0 | 0 | 0 | 0 | 293 | 149 | 1 | 1 | 0.975 | 1 | 1 | 1 | 1 | NA | 1 | 1 | 1 | NA | -0.005 |
| SE | 63136 | ENSG00000164134 | NAA15 | chr4 | + | 139386132 | 139386230 | 139384831 | 139384978 | 139387883 | 139391368 | 290 | 219 | 224 | 230 | 500 | 2 | 0 | 0 | 0 | 0 | 18 | 123 | 175 | 41 | 0 | 0 | 0 | 0 | 0 | 0 | 247 | 149 | 1 | 1 | 0.989 | 1 | 1 | 1 | 1 | 1 | 1 | 1 | 1 | NA | -0.002 |
| SE | 63137 | ENSG00000164134 | NAA15 | chr4 | + | 139393590 | 139393722 | 139386167 | 139386230 | 139404397 | 139404490 | 2 | 1 | 3 | 2 | 0 | 0 | 0 | 0 | 2 | 0 | 0 | 1 | 0 | 0 | 0 | 0 | 0 | 0 | 0 | 0 | 281 | 149 | 1 | 1 | 1 | 1 | 1 | 0.347 | NA | NA | 1 | NA | NA | NA | -0.163 |
| SE | 63618 | ENSG00000198520 | ARMH1 | chr1 | + | 44684392 | 44684656 | 44674712 | 44674873 | 44689675 | 44689903 | 0 | 2 | 0 | 0 | 0 | 0 | 0 | 0 | 0 | 0 | 0 | 0 | 0 | 0 | 0 | 0 | 0 | 0 | 0 | 11 | 298 | 149 | 1 | 1 | NA | 1 | NA | NA | NA | NA | NA | NA | NA | 0 | 1 |
| SE | 63627 | ENSG00000198520 | ARMH1 | chr1 | + | 44723802 | 44723906 | 44704088 | 44704173 | 44724121 | 44724244 | 17 | 9 | 26 | 22 | 22 | 0 | 0 | 0 | 0 | 0 | 28 | 12 | 9 | 62 | 84 | 0 | 3 | 0 | 0 | 0 | 253 | 149 | 1 | 1 | 1 | 1 | 1 | 1 | 1 | 1 | 0.702 | 1 | 1 | 1 | 0.06 |
| SE | 63628 | ENSG00000198520 | ARMH1 | chr1 | + | 44724319 | 44724392 | 44724121 | 44724244 | 44724538 | 44724668 | 14 | 8 | 25 | 10 | 23 | 1 | 0 | 0 | 0 | 0 | 10 | 15 | 6 | 33 | 38 | 0 | 0 | 0 | 0 | 0 | 222 | 149 | 1 | 1 | 0.904 | 1 | 1 | 1 | 1 | 1 | 1 | 1 | 1 | 1 | -0.019 |
| SE | 63635 | ENSG00000112855 | HARS2 | chr5 | + | 140691606 | 140691756 | 140691429 | 140691515 | 140693590 | 140693665 | 29 | 15 | 35 | 29 | 13 | 0 | 0 | 0 | 1 | 0 | 7 | 9 | 21 | 19 | 36 | 0 | 0 | 0 | 0 | 0 | 298 | 149 | 1 | 1 | 1 | 1 | 1 | 0.935 | 1 | 1 | 1 | 1 | 1 | 1 | -0.013 |
| SE | 63639 | ENSG00000112855 | HARS2 | chr5 | + | 140691794 | 140692017 | 140691454 | 140691515 | 140693590 | 140693665 | 0 | 0 | 2 | 3 | 9 | 0 | 0 | 0 | 1 | 0 | 0 | 4 | 0 | 0 | 0 | 0 | 0 | 0 | 0 | 0 | 298 | 149 | 1 | 1 | NA | NA | 1 | 0.6 | 1 | NA | 1 | NA | NA | NA | -0.133 |
| SE | 63642 | ENSG00000112855 | HARS2 | chr5 | + | 140693490 | 140693508 | 140691457 | 140691515 | 140693590 | 140693665 | 0 | 0 | 0 | 0 | 2 | 0 | 0 | 0 | 1 | 0 | 0 | 0 | 1 | 0 | 0 | 0 | 0 | 0 | 0 | 0 | 167 | 149 | 1 | 1 | NA | NA | NA | 0 | 1 | NA | NA | 1 | NA | NA | -0.5 |
| SE | 63645 | ENSG00000112855 | HARS2 | chr5 | + | 140693490 | 140693665 | 140691791 | 140692017 | 140693934 | 140694054 | 41 | 24 | 22 | 37 | 50 | 0 | 0 | 0 | 2 | 0 | 32 | 26 | 12 | 23 | 52 | 0 | 0 | 0 | 0 | 0 | 298 | 149 | 1 | 1 | 1 | 1 | 1 | 0.902 | 1 | 1 | 1 | 1 | 1 | 1 | -0.02 |
| SE | 63647 | ENSG00000112855 | HARS2 | chr5 | + | 140693590 | 140693665 | 140691562 | 140692017 | 140693934 | 140693987 | 38 | 18 | 19 | 34 | 40 | 0 | 0 | 0 | 2 | 0 | 27 | 17 | 11 | 23 | 52 | 0 | 0 | 0 | 0 | 0 | 224 | 149 | 1 | 1 | 1 | 1 | 1 | 0.919 | 1 | 1 | 1 | 1 | 1 | 1 | -0.016 |
| SE | 63663 | ENSG00000144451 | SPAG16 | chr2 | + | 213297021 | 213297139 | 213296039 | 213296110 | 213297261 | 213297357 | 2 | 0 | 0 | 2 | 2 | 17 | 17 | 21 | 16 | 24 | 0 | 0 | 2 | 0 | 0 | 0 | 8 | 0 | 7 | 0 | 267 | 149 | 1 | 1 | 0.062 | 0 | 0 | 0.065 | 0.044 | NA | 0 | 1 | 0 | NA | -0.299 |
| SE | 63664 | ENSG00000144451 | SPAG16 | chr2 | + | 213302512 | 213302582 | 213297261 | 213297357 | 213310058 | 213310177 | 2 | 0 | 2 | 0 | 4 | 27 | 26 | 56 | 35 | 104 | 0 | 0 | 0 | 0 | 0 | 0 | 18 | 4 | 7 | 21 | 219 | 149 | 1 | 1 | 0.048 | 0 | 0.024 | 0 | 0.026 | NA | 0 | 0 | 0 | 0 | 0.02 |
| SE | 63667 | ENSG00000144451 | SPAG16 | chr2 | + | 213340162 | 213340270 | 213317218 | 213317356 | 213364075 | 213364145 | 10 | 9 | 11 | 10 | 18 | 0 | 0 | 0 | 0 | 0 | 5 | 14 | 5 | 8 | 16 | 0 | 0 | 2 | 0 | 0 | 257 | 149 | 1 | 1 | 1 | 1 | 1 | 1 | 1 | 1 | 1 | 0.592 | 1 | 1 | 0.082 |
| SE | 63668 | ENSG00000144451 | SPAG16 | chr2 | + | 213350527 | 213350645 | 213340162 | 213340270 | 213364075 | 213364145 | 23 | 9 | 3 | 8 | 37 | 1 | 2 | 1 | 0 | 2 | 40 | 16 | 2 | 0 | 0 | 0 | 0 | 0 | 1 | 0 | 267 | 149 | 1 | 1 | 0.928 | 0.715 | 0.626 | 1 | 0.912 | 1 | 1 | 1 | 0 | NA | 0.086 |
| SE | 63669 | ENSG00000144451 | SPAG16 | chr2 | + | 213929959 | 213930145 | 213862484 | 213862628 | 214013950 | 214014077 | 1 | 0 | 0 | 0 | 0 | 0 | 0 | 0 | 0 | 0 | 0 | 0 | 0 | 0 | 0 | 0 | 1 | 0 | 0 | 0 | 298 | 149 | 1 | 1 | 1 | NA | NA | NA | NA | NA | 0 | NA | NA | NA | 1 |
| SE | 63694 | ENSG00000112242 | E2F3 | chr6 | + | 20488112 | 20488248 | 20481205 | 20481425 | 20490167 | 20493714 | 58 | 54 | 49 | 62 | 112 | 0 | 0 | 0 | 0 | 0 | 80 | 97 | 63 | 88 | 22 | 1 | 0 | 0 | 1 | 0 | 285 | 149 | 1 | 1 | 1 | 1 | 1 | 1 | 1 | 0.977 | 1 | 1 | 0.979 | 1 | 0.009 |
| SE | 63776 | ENSG00000133131 | MORC4 | chrX | - | 106999676 | 106999749 | 106993229 | 106993362 | 106999867 | 107000078 | 3 | 0 | 7 | 2 | 2 | 0 | 2 | 0 | 0 | 0 | 0 | 5 | 0 | 0 | 0 | 0 | 0 | 0 | 0 | 0 | 222 | 149 | 1 | 1 | 1 | 0 | 1 | 1 | 1 | NA | 1 | NA | NA | NA | -0.2 |
| SE | 63846 | ENSG00000148690 | FRA10AC1 | chr10 | - | 93681479 | 93681598 | 93676652 | 93676691 | 93684055 | 93684098 | 127 | 120 | 105 | 94 | 152 | 2 | 2 | 0 | 0 | 0 | 62 | 62 | 55 | 43 | 13 | 0 | 0 | 0 | 0 | 0 | 268 | 149 | 1 | 1 | 0.972 | 0.971 | 1 | 1 | 1 | 1 | 1 | 1 | 1 | 1 | -0.011 |
| SE | 63847 | ENSG00000148690 | FRA10AC1 | chr10 | - | 93684055 | 93684098 | 93681479 | 93681598 | 93685245 | 93685359 | 83 | 103 | 67 | 71 | 131 | 0 | 6 | 0 | 1 | 0 | 30 | 69 | 52 | 49 | 12 | 0 | 0 | 4 | 1 | 0 | 192 | 149 | 1 | 1 | 1 | 0.93 | 1 | 0.982 | 1 | 1 | 0.93 | 0.976 | 1 | 1 | 0.001 |
| SE | 63849 | ENSG00000148690 | FRA10AC1 | chr10 | - | 93692008 | 93692093 | 93687403 | 93687449 | 93694860 | 93694937 | 53 | 55 | 45 | 33 | 91 | 0 | 0 | 0 | 0 | 0 | 1 | 25 | 25 | 18 | 2 | 0 | 0 | 3 | 0 | 0 | 234 | 149 | 1 | 1 | 1 | 1 | 1 | 1 | 1 | 1 | 0.841 | 1 | 1 | 1 | 0.032 |
| SE | 63850 | ENSG00000148690 | FRA10AC1 | chr10 | - | 93692645 | 93692729 | 93687403 | 93687449 | 93694860 | 93694937 | 37 | 60 | 43 | 68 | 76 | 0 | 0 | 0 | 0 | 0 | 14 | 13 | 33 | 27 | 0 | 0 | 0 | 3 | 0 | 0 | 233 | 149 | 1 | 1 | 1 | 1 | 1 | 1 | 1 | 1 | 0.735 | 1 | 1 | NA | 0.066 |
| SE | 63852 | ENSG00000148690 | FRA10AC1 | chr10 | - | 93698135 | 93698181 | 93694860 | 93694937 | 93698300 | 93698396 | 78 | 119 | 93 | 116 | 142 | 0 | 1 | 1 | 1 | 0 | 41 | 38 | 77 | 70 | 6 | 0 | 0 | 0 | 0 | 0 | 195 | 149 | 1 | 1 | 1 | 0.989 | 0.986 | 0.989 | 1 | 1 | 1 | 1 | 1 | 1 | -0.007 |
| SE | 64422 | ENSG00000143919 | CAMKMT | chr2 | + | 44704282 | 44704343 | 44549554 | 44549728 | 44707398 | 44707462 | 3 | 2 | 2 | 3 | 0 | 0 | 0 | 0 | 0 | 0 | 0 | 12 | 4 | 6 | 0 | 0 | 0 | 3 | 0 | 0 | 210 | 149 | 1 | 1 | 1 | 1 | 1 | 1 | NA | NA | 1 | 0.486 | 1 | NA | 0.171 |
| SE | 64425 | ENSG00000143919 | CAMKMT | chr2 | + | 44715286 | 44715353 | 44704282 | 44704343 | 44743621 | 44743696 | 12 | 4 | 5 | 10 | 15 | 0 | 5 | 1 | 0 | 5 | 2 | 4 | 8 | 0 | 0 | 0 | 4 | 0 | 0 | 3 | 216 | 149 | 1 | 1 | 1 | 0.356 | 0.775 | 1 | 0.674 | 1 | 0.408 | 1 | NA | 0 | 0.159 |
| SE | 64426 | ENSG00000143919 | CAMKMT | chr2 | + | 44723728 | 44723816 | 44715286 | 44715353 | 44743621 | 44743657 | 2 | 1 | 0 | 0 | 0 | 23 | 9 | 16 | 14 | 53 | 0 | 0 | 0 | 0 | 0 | 17 | 20 | 19 | 14 | 5 | 237 | 149 | 1 | 1 | 0.052 | 0.065 | 0 | 0 | 0 | 0 | 0 | 0 | 0 | 0 | 0.023 |
| SE | 64427 | ENSG00000143919 | CAMKMT | chr2 | + | 44743621 | 44743696 | 44704282 | 44704343 | 44754054 | 44754118 | 4 | 16 | 9 | 9 | 18 | 0 | 0 | 0 | 0 | 0 | 10 | 27 | 6 | 7 | 7 | 2 | 0 | 0 | 0 | 0 | 224 | 149 | 1 | 1 | 1 | 1 | 1 | 1 | 1 | 0.769 | 1 | 1 | 1 | 1 | 0.046 |
| SE | 64453 | ENSG00000065150 | IPO5 | chr13 | + | 97981241 | 97981400 | 97976692 | 97976786 | 97982502 | 97982569 | 2 | 0 | 0 | 0 | 0 | 98 | 74 | 132 | 102 | 58 | 0 | 0 | 1 | 0 | 0 | 43 | 73 | 63 | 42 | 17 | 298 | 149 | 1 | 1 | 0.01 | 0 | 0 | 0 | 0 | 0 | 0 | 0.008 | 0 | 0 | 0 |
| SE | 64458 | ENSG00000065150 | IPO5 | chr13 | + | 98007361 | 98007587 | 98006129 | 98006348 | 98008058 | 98008142 | 0 | 1 | 0 | 6 | 1 | 108 | 112 | 112 | 105 | 145 | 0 | 0 | 0 | 0 | 0 | 34 | 67 | 109 | 57 | 13 | 298 | 149 | 1 | 1 | 0 | 0.004 | 0 | 0.028 | 0.003 | 0 | 0 | 0 | 0 | 0 | 0.007 |
| SE | 64459 | ENSG00000065150 | IPO5 | chr13 | + | 98016728 | 98016851 | 98015725 | 98015781 | 98018484 | 98018704 | 427 | 308 | 444 | 404 | 969 | 0 | 0 | 0 | 0 | 4 | 124 | 295 | 257 | 210 | 52 | 0 | 0 | 0 | 0 | 0 | 272 | 149 | 1 | 1 | 1 | 1 | 1 | 1 | 0.993 | 1 | 1 | 1 | 1 | 1 | -0.001 |
| SE | 64541 | ENSG00000147533 | GOLGA7 | chr8 | + | 41490832 | 41490965 | 41490481 | 41490617 | 41497508 | 41497661 | 153 | 82 | 176 | 166 | 143 | 0 | 0 | 2 | 0 | 0 | 63 | 173 | 137 | 182 | 69 | 0 | 0 | 0 | 0 | 0 | 282 | 149 | 1 | 1 | 1 | 1 | 0.979 | 1 | 1 | 1 | 1 | 1 | 1 | 1 | -0.004 |
| SE | 64542 | ENSG00000147533 | GOLGA7 | chr8 | + | 41490869 | 41490965 | 41490561 | 41490617 | 41497508 | 41497661 | 124 | 61 | 144 | 125 | 124 | 0 | 0 | 2 | 0 | 0 | 45 | 145 | 108 | 161 | 52 | 0 | 0 | 0 | 0 | 0 | 245 | 149 | 1 | 1 | 1 | 1 | 0.978 | 1 | 1 | 1 | 1 | 1 | 1 | 1 | -0.004 |
| SE | 64851 | ENSG00000206530 | CFAP44 | chr3 | - | 113293951 | 113294038 | 113290852 | 113291748 | 113294686 | 113294821 | 5 | 11 | 8 | 7 | 7 | 2 | 4 | 0 | 3 | 3 | 0 | 0 | 2 | 0 | 0 | 0 | 0 | 0 | 2 | 10 | 236 | 149 | 1 | 1 | 0.612 | 0.635 | 1 | 0.596 | 0.596 | NA | NA | 1 | 0 | 0 | 0.354 |
| SE | 64852 | ENSG00000206530 | CFAP44 | chr3 | - | 113296724 | 113296885 | 113294686 | 113294821 | 113303915 | 113304117 | 9 | 14 | 17 | 23 | 22 | 0 | 0 | 2 | 0 | 0 | 20 | 2 | 0 | 9 | 4 | 0 | 0 | 0 | 0 | 0 | 298 | 149 | 1 | 1 | 1 | 1 | 0.81 | 1 | 1 | 1 | 1 | NA | 1 | 1 | -0.038 |
| SE | 64853 | ENSG00000206530 | CFAP44 | chr3 | - | 113296724 | 113296902 | 113294686 | 113294821 | 113303915 | 113304067 | 6 | 2 | 4 | 10 | 17 | 0 | 0 | 2 | 0 | 0 | 0 | 0 | 0 | 2 | 4 | 0 | 0 | 0 | 0 | 0 | 298 | 149 | 1 | 1 | 1 | 1 | 0.5 | 1 | 1 | NA | NA | NA | 1 | 1 | -0.1 |
| SE | 64854 | ENSG00000206530 | CFAP44 | chr3 | - | 113327615 | 113327819 | 113326444 | 113326640 | 113330167 | 113330668 | 28 | 51 | 35 | 31 | 36 | 0 | 0 | 0 | 0 | 2 | 0 | 11 | 5 | 0 | 12 | 0 | 0 | 0 | 0 | 0 | 298 | 149 | 1 | 1 | 1 | 1 | 1 | 1 | 0.9 | NA | 1 | 1 | NA | 1 | -0.02 |
| SE | 64855 | ENSG00000206530 | CFAP44 | chr3 | - | 113333405 | 113333583 | 113330167 | 113330668 | 113341743 | 113341918 | 7 | 9 | 9 | 7 | 19 | 1 | 0 | 1 | 3 | 12 | 0 | 3 | 10 | 1 | 0 | 0 | 0 | 0 | 0 | 0 | 298 | 149 | 1 | 1 | 0.778 | 1 | 0.818 | 0.538 | 0.442 | NA | 1 | 1 | 1 | NA | -0.285 |
| SE | 64856 | ENSG00000206530 | CFAP44 | chr3 | - | 113333405 | 113333583 | 113330167 | 113330668 | 113358744 | 113358875 | 3 | 7 | 7 | 6 | 8 | 0 | 0 | 0 | 1 | 6 | 0 | 3 | 9 | 1 | 0 | 0 | 0 | 0 | 0 | 0 | 298 | 149 | 1 | 1 | 1 | 1 | 1 | 0.75 | 0.4 | NA | 1 | 1 | 1 | NA | -0.17 |
| SE | 64858 | ENSG00000206530 | CFAP44 | chr3 | - | 113341743 | 113341918 | 113333405 | 113333583 | 113358744 | 113358875 | 4 | 4 | 3 | 3 | 11 | 0 | 1 | 0 | 1 | 0 | 0 | 0 | 1 | 0 | 0 | 0 | 0 | 0 | 0 | 0 | 298 | 149 | 1 | 1 | 1 | 0.667 | 1 | 0.6 | 1 | NA | NA | 1 | NA | NA | -0.147 |
| SE | 64859 | ENSG00000206530 | CFAP44 | chr3 | - | 113344515 | 113344712 | 113341743 | 113341918 | 113358744 | 113358875 | 13 | 23 | 38 | 31 | 32 | 0 | 1 | 1 | 1 | 0 | 0 | 6 | 4 | 1 | 6 | 0 | 0 | 0 | 0 | 0 | 298 | 149 | 1 | 1 | 1 | 0.92 | 0.95 | 0.939 | 1 | NA | 1 | 1 | 1 | 1 | -0.038 |
| SE | 64861 | ENSG00000206530 | CFAP44 | chr3 | - | 113373410 | 113373556 | 113366038 | 113366309 | 113379305 | 113379551 | 0 | 0 | 0 | 0 | 0 | 3 | 25 | 17 | 8 | 10 | 0 | 0 | 1 | 7 | 0 | 0 | 1 | 1 | 0 | 0 | 295 | 149 | 1 | 1 | 0 | 0 | 0 | 0 | 0 | NA | 0 | 0.336 | 1 | NA | -0.445 |
| SE | 64862 | ENSG00000206530 | CFAP44 | chr3 | - | 113379305 | 113379551 | 113363476 | 113363532 | 113380898 | 113381060 | 36 | 33 | 30 | 30 | 63 | 0 | 1 | 0 | 0 | 0 | 7 | 15 | 3 | 22 | 3 | 0 | 0 | 0 | 0 | 0 | 298 | 149 | 1 | 1 | 1 | 0.943 | 1 | 1 | 1 | 1 | 1 | 1 | 1 | 1 | -0.011 |
| SE | 64867 | ENSG00000170310 | STX8 | chr17 | - | 9378551 | 9378653 | 9250470 | 9250645 | 9491828 | 9491921 | 163 | 163 | 123 | 159 | 447 | 0 | 0 | 2 | 0 | 0 | 278 | 129 | 83 | 178 | 208 | 0 | 0 | 0 | 0 | 2 | 251 | 149 | 1 | 1 | 1 | 1 | 0.973 | 1 | 1 | 1 | 1 | 1 | 1 | 0.984 | -0.002 |
| SE | 64870 | ENSG00000170310 | STX8 | chr17 | - | 9568370 | 9568470 | 9557433 | 9557528 | 9575791 | 9575811 | 311 | 191 | 193 | 265 | 230 | 2 | 0 | 1 | 0 | 0 | 364 | 289 | 223 | 310 | 330 | 0 | 0 | 2 | 0 | 0 | 249 | 149 | 1 | 1 | 0.989 | 1 | 0.991 | 1 | 1 | 1 | 1 | 0.985 | 1 | 1 | -0.001 |
| SE | 65137 | ENSG00000152689 | RASGRP3 | chr2 | + | 33493053 | 33493172 | 33447819 | 33447943 | 33511709 | 33511842 | 0 | 0 | 0 | 3 | 0 | 0 | 0 | 0 | 1 | 0 | 0 | 0 | 0 | 0 | 0 | 0 | 1 | 0 | 0 | 0 | 268 | 149 | 1 | 1 | NA | NA | NA | 0.625 | NA | NA | 0 | NA | NA | NA | 0.625 |
| SE | 65139 | ENSG00000152689 | RASGRP3 | chr2 | + | 33511709 | 33511842 | 33476596 | 33476707 | 33515009 | 33515206 | 23 | 30 | 40 | 31 | 58 | 0 | 0 | 2 | 1 | 2 | 4 | 35 | 23 | 27 | 12 | 0 | 1 | 1 | 0 | 0 | 282 | 149 | 1 | 1 | 1 | 1 | 0.914 | 0.942 | 0.939 | 1 | 0.949 | 0.924 | 1 | 1 | -0.016 |
| SE | 65140 | ENSG00000152689 | RASGRP3 | chr2 | + | 33511811 | 33511842 | 33476648 | 33476707 | 33515009 | 33515206 | 7 | 10 | 17 | 12 | 21 | 0 | 0 | 2 | 1 | 2 | 2 | 7 | 14 | 7 | 0 | 0 | 1 | 1 | 0 | 0 | 180 | 149 | 1 | 1 | 1 | 1 | 0.876 | 0.909 | 0.897 | 1 | 0.853 | 0.921 | 1 | NA | -0.007 |
| SE | 65141 | ENSG00000152689 | RASGRP3 | chr2 | + | 33534322 | 33534400 | 33527136 | 33527412 | 33539093 | 33539210 | 24 | 25 | 47 | 46 | 71 | 0 | 0 | 0 | 0 | 0 | 33 | 67 | 27 | 50 | 28 | 0 | 0 | 0 | 1 | 0 | 227 | 149 | 1 | 1 | 1 | 1 | 1 | 1 | 1 | 1 | 1 | 1 | 0.97 | 1 | 0.006 |
| SE | 65142 | ENSG00000152689 | RASGRP3 | chr2 | + | 33543511 | 33543627 | 33539650 | 33540108 | 33549603 | 33549751 | 11 | 18 | 15 | 17 | 30 | 0 | 0 | 0 | 0 | 0 | 48 | 36 | 10 | 21 | 15 | 4 | 0 | 0 | 0 | 0 | 265 | 149 | 1 | 1 | 1 | 1 | 1 | 1 | 1 | 0.871 | 1 | 1 | 1 | 1 | 0.026 |
| SE | 65143 | ENSG00000152689 | RASGRP3 | chr2 | + | 33555530 | 33555567 | 33549603 | 33549751 | 33558210 | 33558336 | 28 | 25 | 43 | 42 | 33 | 0 | 0 | 0 | 1 | 0 | 47 | 54 | 37 | 21 | 17 | 0 | 0 | 0 | 0 | 0 | 186 | 149 | 1 | 1 | 1 | 1 | 1 | 0.971 | 1 | 1 | 1 | 1 | 1 | 1 | -0.006 |
| SE | 65144 | ENSG00000152689 | RASGRP3 | chr2 | + | 33558671 | 33559030 | 33558210 | 33558336 | 33562728 | 33564605 | 33 | 26 | 40 | 37 | 37 | 0 | 0 | 0 | 1 | 0 | 30 | 106 | 51 | 25 | 13 | 0 | 0 | 0 | 0 | 0 | 298 | 149 | 1 | 1 | 1 | 1 | 1 | 0.949 | 1 | 1 | 1 | 1 | 1 | 1 | -0.01 |
| SE | 65198 | ENSG00000010244 | ZNF207 | chr17 | + | 32360891 | 32360967 | 32360597 | 32360765 | 32361467 | 32361515 | 667 | 393 | 569 | 651 | 1455 | 0 | 1 | 0 | 1 | 0 | 521 | 643 | 599 | 490 | 422 | 0 | 0 | 4 | 0 | 0 | 225 | 149 | 1 | 1 | 1 | 0.996 | 1 | 0.998 | 1 | 1 | 1 | 0.99 | 1 | 1 | 0.001 |
| SE | 65199 | ENSG00000010244 | ZNF207 | chr17 | + | 32360891 | 32360967 | 32360597 | 32360765 | 32362913 | 32362984 | 249 | 146 | 196 | 217 | 491 | 0 | 0 | 0 | 1 | 0 | 155 | 181 | 190 | 134 | 153 | 0 | 0 | 0 | 0 | 0 | 225 | 149 | 1 | 1 | 1 | 1 | 1 | 0.993 | 1 | 1 | 1 | 1 | 1 | 1 | -0.001 |
| SE | 65200 | ENSG00000010244 | ZNF207 | chr17 | + | 32361467 | 32361515 | 32360597 | 32360765 | 32362913 | 32362984 | 56 | 73 | 89 | 108 | 165 | 0 | 0 | 0 | 1 | 0 | 106 | 75 | 71 | 89 | 73 | 0 | 0 | 0 | 0 | 0 | 197 | 149 | 1 | 1 | 1 | 1 | 1 | 0.988 | 1 | 1 | 1 | 1 | 1 | 1 | -0.002 |
| SE | 65202 | ENSG00000010244 | ZNF207 | chr17 | + | 32365329 | 32365487 | 32362913 | 32362984 | 32367771 | 32368014 | 761 | 628 | 744 | 864 | 1568 | 0 | 0 | 0 | 0 | 0 | 869 | 806 | 847 | 785 | 769 | 0 | 2 | 0 | 0 | 0 | 298 | 149 | 1 | 1 | 1 | 1 | 1 | 1 | 1 | 1 | 0.995 | 1 | 1 | 1 | 0.001 |
| SE | 65295 | ENSG00000149499 | EML3 | chr11 | - | 62604112 | 62604201 | 62603943 | 62604041 | 62605112 | 62605180 | 103 | 44 | 102 | 93 | 85 | 1 | 0 | 0 | 0 | 0 | 116 | 98 | 104 | 90 | 135 | 0 | 0 | 2 | 0 | 0 | 238 | 149 | 1 | 1 | 0.985 | 1 | 1 | 1 | 1 | 1 | 1 | 0.97 | 1 | 1 | 0.003 |
| SE | 65296 | ENSG00000149499 | EML3 | chr11 | - | 62605641 | 62605773 | 62605112 | 62605180 | 62605854 | 62605980 | 72 | 16 | 58 | 58 | 68 | 0 | 0 | 0 | 0 | 3 | 27 | 64 | 37 | 57 | 114 | 0 | 0 | 0 | 0 | 0 | 281 | 149 | 1 | 1 | 1 | 1 | 1 | 1 | 0.923 | 1 | 1 | 1 | 1 | 1 | -0.015 |
| SE | 65355 | ENSG00000153214 | TMEM87B | chr2 | + | 112066935 | 112067067 | 112059976 | 112060037 | 112074911 | 112074962 | 206 | 135 | 149 | 179 | 315 | 0 | 0 | 0 | 0 | 0 | 116 | 171 | 79 | 95 | 75 | 0 | 0 | 0 | 0 | 4 | 281 | 149 | 1 | 1 | 1 | 1 | 1 | 1 | 1 | 1 | 1 | 1 | 1 | 0.909 | 0.018 |
| SE | 65356 | ENSG00000153214 | TMEM87B | chr2 | + | 112066935 | 112067067 | 112059976 | 112060037 | 112081056 | 112081118 | 2 | 1 | 5 | 6 | 1 | 0 | 0 | 0 | 1 | 0 | 0 | 8 | 4 | 1 | 0 | 0 | 0 | 0 | 0 | 0 | 281 | 149 | 1 | 1 | 1 | 1 | 1 | 0.761 | 1 | NA | 1 | 1 | 1 | NA | -0.048 |
| SE | 65357 | ENSG00000153214 | TMEM87B | chr2 | + | 112074911 | 112074962 | 112059976 | 112060037 | 112081056 | 112081118 | 1 | 0 | 0 | 0 | 0 | 0 | 0 | 0 | 1 | 0 | 0 | 0 | 1 | 0 | 4 | 0 | 0 | 0 | 0 | 0 | 200 | 149 | 1 | 1 | 1 | NA | NA | 0 | NA | NA | NA | 1 | NA | 1 | -0.5 |
| SE | 65359 | ENSG00000153214 | TMEM87B | chr2 | + | 112074911 | 112074962 | 112066935 | 112067067 | 112081056 | 112081118 | 50 | 32 | 23 | 60 | 69 | 2 | 0 | 1 | 3 | 0 | 11 | 30 | 30 | 45 | 0 | 0 | 0 | 2 | 1 | 0 | 200 | 149 | 1 | 1 | 0.949 | 1 | 0.945 | 0.937 | 1 | 1 | 1 | 0.918 | 0.971 | NA | -0.006 |
| SE | 65360 | ENSG00000153214 | TMEM87B | chr2 | + | 112077191 | 112077282 | 112066935 | 112067067 | 112081056 | 112081118 | 83 | 83 | 60 | 98 | 154 | 2 | 0 | 1 | 3 | 0 | 96 | 37 | 27 | 41 | 51 | 0 | 0 | 2 | 1 | 0 | 240 | 149 | 1 | 1 | 0.963 | 1 | 0.974 | 0.953 | 1 | 1 | 1 | 0.893 | 0.962 | 1 | 0.007 |
| SE | 65361 | ENSG00000153214 | TMEM87B | chr2 | + | 112077191 | 112077282 | 112066935 | 112067067 | 112086004 | 112086104 | 1 | 3 | 1 | 3 | 2 | 0 | 0 | 0 | 0 | 0 | 7 | 2 | 1 | 4 | 0 | 0 | 0 | 1 | 0 | 0 | 240 | 149 | 1 | 1 | 1 | 1 | 1 | 1 | 1 | 1 | 1 | 0.383 | 1 | NA | 0.154 |
| SE | 65362 | ENSG00000153214 | TMEM87B | chr2 | + | 112077191 | 112077282 | 112074911 | 112074962 | 112081056 | 112081118 | 278 | 203 | 224 | 262 | 463 | 2 | 2 | 0 | 0 | 0 | 190 | 210 | 84 | 104 | 149 | 0 | 1 | 1 | 0 | 0 | 240 | 149 | 1 | 1 | 0.989 | 0.984 | 1 | 1 | 1 | 1 | 0.992 | 0.981 | 1 | 1 | 0 |
| SE | 65363 | ENSG00000153214 | TMEM87B | chr2 | + | 112081056 | 112081118 | 112066935 | 112067067 | 112086004 | 112086104 | 4 | 6 | 4 | 11 | 8 | 0 | 0 | 0 | 0 | 0 | 0 | 4 | 6 | 11 | 27 | 0 | 0 | 1 | 0 | 0 | 211 | 149 | 1 | 1 | 1 | 1 | 1 | 1 | 1 | NA | 1 | 0.809 | 1 | 1 | 0.048 |
| SE | 65364 | ENSG00000153214 | TMEM87B | chr2 | + | 112081056 | 112081118 | 112077191 | 112077282 | 112086004 | 112086104 | 44 | 36 | 56 | 64 | 82 | 0 | 2 | 0 | 0 | 0 | 41 | 48 | 19 | 32 | 72 | 0 | 0 | 0 | 0 | 0 | 211 | 149 | 1 | 1 | 1 | 0.927 | 1 | 1 | 1 | 1 | 1 | 1 | 1 | 1 | -0.015 |
| SE | 65366 | ENSG00000153214 | TMEM87B | chr2 | + | 112097232 | 112097291 | 112097043 | 112097152 | 112098594 | 112098698 | 207 | 163 | 156 | 198 | 172 | 0 | 0 | 0 | 0 | 0 | 110 | 202 | 127 | 126 | 47 | 0 | 0 | 0 | 0 | 5 | 208 | 149 | 1 | 1 | 1 | 1 | 1 | 1 | 1 | 1 | 1 | 1 | 1 | 0.871 | 0.026 |
| SE | 65367 | ENSG00000153214 | TMEM87B | chr2 | + | 112107787 | 112107840 | 112106001 | 112106075 | 112116083 | 112119273 | 34 | 14 | 29 | 30 | 90 | 0 | 0 | 1 | 0 | 0 | 13 | 18 | 14 | 15 | 0 | 0 | 0 | 0 | 0 | 0 | 202 | 149 | 1 | 1 | 1 | 1 | 0.955 | 1 | 1 | 1 | 1 | 1 | 1 | NA | -0.009 |
| SE | 65368 | ENSG00000153214 | TMEM87B | chr2 | + | 112112898 | 112112929 | 112107787 | 112107840 | 112116083 | 112119273 | 141 | 152 | 148 | 138 | 318 | 0 | 0 | 0 | 0 | 1 | 36 | 98 | 106 | 116 | 68 | 3 | 0 | 0 | 0 | 0 | 180 | 149 | 1 | 1 | 1 | 1 | 1 | 1 | 0.996 | 0.909 | 1 | 1 | 1 | 1 | 0.017 |
| SE | 65645 | ENSG00000143924 | EML4 | chr2 | + | 42207928 | 42208082 | 42169390 | 42169636 | 42245504 | 42245688 | 0 | 0 | 0 | 1 | 1 | 96 | 49 | 74 | 91 | 57 | 0 | 1 | 1 | 0 | 0 | 10 | 56 | 71 | 51 | 29 | 298 | 149 | 1 | 1 | 0 | 0 | 0 | 0.005 | 0.009 | 0 | 0.009 | 0.007 | 0 | 0 | 0 |
| SE | 65646 | ENSG00000143924 | EML4 | chr2 | + | 42256500 | 42256630 | 42245504 | 42245687 | 42261120 | 42261294 | 330 | 316 | 444 | 384 | 955 | 0 | 0 | 0 | 0 | 0 | 158 | 274 | 368 | 274 | 50 | 0 | 0 | 1 | 0 | 0 | 279 | 149 | 1 | 1 | 1 | 1 | 1 | 1 | 1 | 1 | 1 | 0.995 | 1 | 1 | 0.001 |
| SE | 65647 | ENSG00000143924 | EML4 | chr2 | + | 42261120 | 42261294 | 42256500 | 42256630 | 42263177 | 42263306 | 329 | 303 | 435 | 390 | 1089 | 0 | 2 | 1 | 0 | 0 | 157 | 275 | 406 | 251 | 48 | 0 | 0 | 0 | 0 | 0 | 298 | 149 | 1 | 1 | 1 | 0.987 | 0.995 | 1 | 1 | 1 | 1 | 1 | 1 | 1 | -0.004 |
| SE | 65649 | ENSG00000143924 | EML4 | chr2 | + | 42282822 | 42282972 | 42280849 | 42280973 | 42284633 | 42284703 | 489 | 432 | 510 | 481 | 581 | 0 | 0 | 3 | 0 | 0 | 65 | 292 | 332 | 267 | 200 | 0 | 0 | 0 | 0 | 5 | 298 | 149 | 1 | 1 | 1 | 1 | 0.988 | 1 | 1 | 1 | 1 | 1 | 1 | 0.952 | 0.007 |
| SE | 65651 | ENSG00000143924 | EML4 | chr2 | + | 42303314 | 42303446 | 42303103 | 42303229 | 42304483 | 42304551 | 483 | 438 | 590 | 541 | 664 | 0 | 0 | 1 | 0 | 0 | 271 | 364 | 597 | 290 | 66 | 0 | 0 | 0 | 0 | 0 | 281 | 149 | 1 | 1 | 1 | 1 | 0.997 | 1 | 1 | 1 | 1 | 1 | 1 | 1 | -0.001 |
| SE | 65653 | ENSG00000143924 | EML4 | chr2 | + | 42315961 | 42316050 | 42304483 | 42304551 | 42317426 | 42317524 | 156 | 127 | 195 | 216 | 273 | 0 | 0 | 0 | 0 | 0 | 57 | 114 | 167 | 98 | 37 | 0 | 0 | 2 | 0 | 0 | 238 | 149 | 1 | 1 | 1 | 1 | 1 | 1 | 1 | 1 | 1 | 0.981 | 1 | 1 | 0.004 |
| SE | 65656 | ENSG00000143924 | EML4 | chr2 | + | 42317426 | 42317524 | 42315961 | 42316050 | 42325466 | 42325554 | 174 | 122 | 216 | 218 | 238 | 1 | 1 | 0 | 0 | 0 | 52 | 140 | 181 | 139 | 33 | 0 | 0 | 0 | 0 | 0 | 247 | 149 | 1 | 1 | 0.991 | 0.987 | 1 | 1 | 1 | 1 | 1 | 1 | 1 | 1 | -0.004 |
| SE | 65657 | ENSG00000143924 | EML4 | chr2 | + | 42328885 | 42329016 | 42326153 | 42326252 | 42329733 | 42330744 | 441 | 302 | 548 | 480 | 1000 | 0 | 0 | 1 | 0 | 0 | 113 | 251 | 326 | 196 | 49 | 0 | 0 | 0 | 0 | 0 | 280 | 149 | 1 | 1 | 1 | 1 | 0.997 | 1 | 1 | 1 | 1 | 1 | 1 | 1 | -0.001 |
| SE | 65684 | ENSG00000186141 | POLR3C | chr1 | + | 145836493 | 145836574 | 145833489 | 145833582 | 145836814 | 145836854 | 93 | 84 | 106 | 113 | 159 | 2 | 0 | 1 | 0 | 0 | 132 | 168 | 89 | 88 | 108 | 0 | 0 | 0 | 0 | 0 | 230 | 149 | 1 | 1 | 0.968 | 1 | 0.986 | 1 | 1 | 1 | 1 | 1 | 1 | 1 | -0.009 |
| SE | 65687 | ENSG00000186141 | POLR3C | chr1 | + | 145838055 | 145838206 | 145833489 | 145833582 | 145839889 | 145839991 | 107 | 79 | 83 | 107 | 149 | 0 | 0 | 0 | 1 | 1 | 231 | 106 | 135 | 101 | 125 | 6 | 0 | 0 | 0 | 0 | 298 | 149 | 1 | 1 | 1 | 1 | 1 | 0.982 | 0.987 | 0.951 | 1 | 1 | 1 | 1 | 0.004 |
| SE | 65689 | ENSG00000186141 | POLR3C | chr1 | + | 145838055 | 145838206 | 145836493 | 145836574 | 145842338 | 145842493 | 7 | 0 | 6 | 6 | 4 | 0 | 0 | 0 | 1 | 0 | 5 | 7 | 5 | 2 | 9 | 0 | 0 | 0 | 0 | 0 | 298 | 149 | 1 | 1 | 1 | NA | 1 | 0.75 | 1 | 1 | 1 | 1 | 1 | 1 | -0.062 |
| SE | 65690 | ENSG00000186141 | POLR3C | chr1 | + | 145839889 | 145839991 | 145836493 | 145836574 | 145842338 | 145844402 | 2 | 0 | 2 | 4 | 2 | 0 | 0 | 0 | 1 | 0 | 0 | 5 | 5 | 0 | 14 | 0 | 0 | 0 | 0 | 0 | 251 | 149 | 1 | 1 | 1 | NA | 1 | 0.704 | 1 | NA | 1 | 1 | NA | 1 | -0.074 |
| SE | 65691 | ENSG00000186141 | POLR3C | chr1 | + | 145839889 | 145839991 | 145838055 | 145838206 | 145842338 | 145844402 | 89 | 77 | 71 | 97 | 134 | 0 | 0 | 1 | 2 | 2 | 208 | 97 | 121 | 100 | 125 | 0 | 0 | 0 | 0 | 0 | 251 | 149 | 1 | 1 | 1 | 1 | 0.977 | 0.966 | 0.975 | 1 | 1 | 1 | 1 | 1 | -0.016 |
| SE | 65692 | ENSG00000186141 | POLR3C | chr1 | + | 145840115 | 145840165 | 145839889 | 145839991 | 145842338 | 145844402 | 17 | 5 | 14 | 12 | 21 | 0 | 0 | 0 | 2 | 0 | 10 | 9 | 13 | 3 | 0 | 0 | 0 | 0 | 0 | 0 | 199 | 149 | 1 | 1 | 1 | 1 | 1 | 0.818 | 1 | 1 | 1 | 1 | 1 | NA | -0.036 |
| SE | 65693 | ENSG00000186141 | POLR3C | chr1 | + | 145840921 | 145841071 | 145840115 | 145840165 | 145842338 | 145844402 | 202 | 106 | 165 | 187 | 277 | 0 | 0 | 0 | 1 | 0 | 245 | 152 | 171 | 135 | 145 | 0 | 0 | 0 | 2 | 0 | 298 | 149 | 1 | 1 | 1 | 1 | 1 | 0.989 | 1 | 1 | 1 | 1 | 0.971 | 1 | 0.004 |
| SE | 65837 | ENSG00000088205 | DDX18 | chr2 | + | 117819648 | 117819792 | 117817443 | 117817728 | 117821160 | 117821296 | 770 | 682 | 702 | 701 | 1181 | 0 | 1 | 0 | 0 | 0 | 56 | 509 | 510 | 207 | 15 | 0 | 1 | 0 | 0 | 0 | 293 | 149 | 1 | 1 | 1 | 0.997 | 1 | 1 | 1 | 1 | 0.996 | 1 | 1 | 1 | 0 |
| SE | 65965 | ENSG00000135931 | ARMC9 | chr2 | + | 231206197 | 231206289 | 231198545 | 231198698 | 231208126 | 231208252 | 1 | 1 | 1 | 0 | 3 | 0 | 1 | 0 | 0 | 0 | 3 | 0 | 4 | 0 | 0 | 0 | 0 | 0 | 0 | 0 | 241 | 149 | 1 | 1 | 1 | 0.382 | 1 | NA | 1 | 1 | NA | 1 | NA | NA | -0.154 |
| SE | 65966 | ENSG00000135931 | ARMC9 | chr2 | + | 231206199 | 231206289 | 231198630 | 231198698 | 231208126 | 231208252 | 1 | 1 | 1 | 0 | 3 | 0 | 1 | 0 | 0 | 0 | 3 | 0 | 2 | 0 | 0 | 0 | 0 | 0 | 0 | 0 | 239 | 149 | 1 | 1 | 1 | 0.384 | 1 | NA | 1 | 1 | NA | 1 | NA | NA | -0.154 |
| SE | 65984 | ENSG00000168890 | TMEM150A | chr2 | - | 85599524 | 85599702 | 85598546 | 85599317 | 85599890 | 85600018 | 12 | 2 | 6 | 17 | 2 | 1 | 0 | 0 | 0 | 0 | 11 | 3 | 3 | 6 | 4 | 0 | 0 | 0 | 0 | 0 | 298 | 149 | 1 | 1 | 0.857 | 1 | 1 | 1 | 1 | 1 | 1 | 1 | 1 | 1 | -0.029 |
| SE | 65985 | ENSG00000168890 | TMEM150A | chr2 | - | 85600344 | 85600412 | 85599890 | 85600018 | 85601020 | 85601107 | 8 | 1 | 2 | 10 | 2 | 1 | 0 | 2 | 4 | 0 | 5 | 0 | 0 | 0 | 0 | 0 | 1 | 0 | 0 | 0 | 217 | 149 | 1 | 1 | 0.846 | 1 | 0.407 | 0.632 | 1 | 1 | 0 | NA | NA | NA | 0.277 |
| SE | 65986 | ENSG00000168890 | TMEM150A | chr2 | - | 85600499 | 85600759 | 85600344 | 85600412 | 85601020 | 85601107 | 2 | 1 | 1 | 1 | 0 | 6 | 1 | 1 | 8 | 0 | 3 | 0 | 4 | 0 | 0 | 5 | 0 | 0 | 0 | 0 | 298 | 149 | 1 | 1 | 0.143 | 0.333 | 0.333 | 0.059 | NA | 0.231 | NA | 1 | NA | NA | -0.399 |
| SE | 65987 | ENSG00000168890 | TMEM150A | chr2 | - | 85601305 | 85601482 | 85601020 | 85601107 | 85601883 | 85602064 | 4 | 0 | 6 | 7 | 3 | 1 | 0 | 0 | 0 | 0 | 6 | 1 | 3 | 0 | 0 | 0 | 0 | 0 | 0 | 0 | 298 | 149 | 1 | 1 | 0.667 | NA | 1 | 1 | 1 | 1 | 1 | 1 | NA | NA | -0.083 |
| SE | 65988 | ENSG00000168890 | TMEM150A | chr2 | - | 85601434 | 85601482 | 85601020 | 85601107 | 85601883 | 85602064 | 4 | 3 | 10 | 9 | 1 | 1 | 0 | 0 | 0 | 0 | 7 | 0 | 5 | 0 | 10 | 0 | 0 | 0 | 0 | 0 | 197 | 149 | 1 | 1 | 0.752 | 1 | 1 | 1 | 1 | 1 | NA | 1 | NA | 1 | -0.05 |
| SE | 66189 | ENSG00000157483 | MYO1E | chr15 | - | 59138197 | 59138367 | 59132433 | 59137456 | 59153589 | 59153791 | 52 | 37 | 36 | 47 | 42 | 0 | 0 | 0 | 0 | 0 | 23 | 5 | 20 | 28 | 35 | 0 | 0 | 0 | 1 | 0 | 298 | 149 | 1 | 1 | 1 | 1 | 1 | 1 | 1 | 1 | 1 | 1 | 0.933 | 1 | 0.013 |
| SE | 66191 | ENSG00000157483 | MYO1E | chr15 | - | 59171896 | 59172042 | 59163156 | 59163303 | 59173745 | 59173915 | 68 | 28 | 41 | 51 | 86 | 0 | 0 | 0 | 0 | 0 | 55 | 16 | 24 | 51 | 43 | 0 | 0 | 2 | 2 | 0 | 295 | 149 | 1 | 1 | 1 | 1 | 1 | 1 | 1 | 1 | 1 | 0.858 | 0.928 | 1 | 0.043 |
| SE | 66192 | ENSG00000157483 | MYO1E | chr15 | - | 59178392 | 59178537 | 59174125 | 59174240 | 59188117 | 59188216 | 99 | 88 | 71 | 98 | 79 | 0 | 0 | 0 | 0 | 0 | 348 | 71 | 98 | 169 | 255 | 0 | 0 | 1 | 3 | 0 | 294 | 149 | 1 | 1 | 1 | 1 | 1 | 1 | 1 | 1 | 1 | 0.98 | 0.966 | 1 | 0.011 |
| SE | 66193 | ENSG00000157483 | MYO1E | chr15 | - | 59188117 | 59188216 | 59178392 | 59178537 | 59195460 | 59195567 | 75 | 70 | 59 | 64 | 70 | 0 | 0 | 0 | 0 | 3 | 309 | 58 | 54 | 137 | 159 | 0 | 0 | 3 | 1 | 0 | 248 | 149 | 1 | 1 | 1 | 1 | 1 | 1 | 0.933 | 1 | 1 | 0.915 | 0.988 | 1 | 0.006 |
| SE | 66194 | ENSG00000157483 | MYO1E | chr15 | - | 59217890 | 59218087 | 59214639 | 59214720 | 59223058 | 59223191 | 28 | 39 | 26 | 27 | 32 | 0 | 2 | 0 | 0 | 0 | 61 | 10 | 22 | 51 | 87 | 0 | 0 | 0 | 0 | 0 | 298 | 149 | 1 | 1 | 1 | 0.907 | 1 | 1 | 1 | 1 | 1 | 1 | 1 | 1 | -0.019 |
| SE | 66195 | ENSG00000157483 | MYO1E | chr15 | - | 59236584 | 59236672 | 59231701 | 59231791 | 59256283 | 59256378 | 70 | 63 | 40 | 75 | 100 | 1 | 0 | 0 | 0 | 0 | 104 | 6 | 42 | 93 | 55 | 0 | 0 | 0 | 0 | 5 | 237 | 149 | 1 | 1 | 0.978 | 1 | 1 | 1 | 1 | 1 | 1 | 1 | 1 | 0.874 | 0.021 |
| SE | 66196 | ENSG00000157483 | MYO1E | chr15 | - | 59256283 | 59256378 | 59236584 | 59236672 | 59261419 | 59261509 | 72 | 82 | 42 | 82 | 91 | 0 | 0 | 0 | 0 | 0 | 119 | 21 | 50 | 104 | 48 | 0 | 0 | 0 | 3 | 0 | 244 | 149 | 1 | 1 | 1 | 1 | 1 | 1 | 1 | 1 | 1 | 1 | 0.955 | 1 | 0.009 |
| SE | 66479 | ENSG00000155561 | NUP205 | chr7 | + | 135571104 | 135571247 | 135557916 | 135557972 | 135576269 | 135576414 | 42 | 32 | 37 | 23 | 29 | 1 | 0 | 0 | 0 | 0 | 46 | 40 | 34 | 33 | 0 | 0 | 0 | 0 | 0 | 0 | 292 | 149 | 1 | 1 | 0.955 | 1 | 1 | 1 | 1 | 1 | 1 | 1 | 1 | NA | -0.009 |
| SE | 66480 | ENSG00000155561 | NUP205 | chr7 | + | 135573653 | 135573825 | 135571104 | 135571247 | 135576269 | 135576414 | 325 | 249 | 329 | 337 | 262 | 4 | 0 | 0 | 0 | 0 | 197 | 306 | 275 | 198 | 38 | 0 | 0 | 0 | 0 | 0 | 298 | 149 | 1 | 1 | 0.976 | 1 | 1 | 1 | 1 | 1 | 1 | 1 | 1 | 1 | -0.005 |
| SE | 66481 | ENSG00000155561 | NUP205 | chr7 | + | 135597997 | 135598207 | 135597238 | 135597418 | 135600869 | 135600969 | 93 | 74 | 84 | 86 | 107 | 0 | 0 | 2 | 0 | 0 | 26 | 29 | 36 | 56 | 6 | 0 | 0 | 2 | 0 | 0 | 298 | 149 | 1 | 1 | 1 | 1 | 0.955 | 1 | 1 | 1 | 1 | 0.9 | 1 | 1 | 0.011 |
| SE | 66482 | ENSG00000155561 | NUP205 | chr7 | + | 135598796 | 135598896 | 135597739 | 135598207 | 135600869 | 135600969 | 0 | 0 | 1 | 2 | 1 | 27 | 21 | 16 | 32 | 27 | 0 | 0 | 0 | 0 | 0 | 10 | 8 | 13 | 9 | 0 | 249 | 149 | 1 | 1 | 0 | 0 | 0.036 | 0.036 | 0.022 | 0 | 0 | 0 | 0 | NA | 0.019 |
| SE | 66483 | ENSG00000155561 | NUP205 | chr7 | + | 135602804 | 135602994 | 135601369 | 135601507 | 135604339 | 135604460 | 107 | 58 | 145 | 111 | 130 | 0 | 0 | 0 | 1 | 0 | 28 | 40 | 54 | 38 | 0 | 0 | 2 | 1 | 0 | 0 | 298 | 149 | 1 | 1 | 1 | 1 | 1 | 0.982 | 1 | 1 | 0.909 | 0.964 | 1 | NA | 0.028 |
| SE | 66484 | ENSG00000155561 | NUP205 | chr7 | + | 135604339 | 135604460 | 135601369 | 135601507 | 135606144 | 135606210 | 109 | 54 | 109 | 88 | 105 | 0 | 0 | 0 | 0 | 0 | 8 | 52 | 78 | 59 | 16 | 0 | 0 | 0 | 2 | 0 | 270 | 149 | 1 | 1 | 1 | 1 | 1 | 1 | 1 | 1 | 1 | 1 | 0.942 | 1 | 0.012 |
| SE | 66486 | ENSG00000155561 | NUP205 | chr7 | + | 135615915 | 135616065 | 135614158 | 135614273 | 135616654 | 135616701 | 223 | 164 | 234 | 207 | 248 | 0 | 0 | 0 | 0 | 0 | 138 | 123 | 153 | 139 | 22 | 0 | 2 | 0 | 0 | 0 | 298 | 149 | 1 | 1 | 1 | 1 | 1 | 1 | 1 | 1 | 0.969 | 1 | 1 | 1 | 0.006 |
| SE | 66487 | ENSG00000155561 | NUP205 | chr7 | + | 135630343 | 135630470 | 135627972 | 135628111 | 135635580 | 135635657 | 44 | 32 | 102 | 86 | 52 | 0 | 0 | 0 | 2 | 0 | 56 | 80 | 35 | 43 | 19 | 0 | 0 | 2 | 0 | 0 | 276 | 149 | 1 | 1 | 1 | 1 | 1 | 0.959 | 1 | 1 | 1 | 0.904 | 1 | 1 | 0.011 |
| SE | 66489 | ENSG00000132676 | DAP3 | chr1 | + | 155689094 | 155689174 | 155687966 | 155688192 | 155709772 | 155709824 | 149 | 96 | 155 | 153 | 43 | 0 | 0 | 2 | 0 | 0 | 196 | 96 | 98 | 76 | 170 | 0 | 0 | 0 | 0 | 0 | 229 | 149 | 1 | 1 | 1 | 1 | 0.981 | 1 | 1 | 1 | 1 | 1 | 1 | 1 | -0.004 |
| SE | 66492 | ENSG00000132676 | DAP3 | chr1 | + | 155709772 | 155709824 | 155689090 | 155689174 | 155717005 | 155717128 | 238 | 165 | 280 | 229 | 139 | 1 | 0 | 0 | 0 | 0 | 241 | 187 | 150 | 132 | 218 | 0 | 0 | 0 | 0 | 0 | 201 | 149 | 1 | 1 | 0.994 | 1 | 1 | 1 | 1 | 1 | 1 | 1 | 1 | 1 | -0.001 |
| SE | 66495 | ENSG00000132676 | DAP3 | chr1 | + | 155717005 | 155717128 | 155709772 | 155709824 | 155727607 | 155727738 | 226 | 160 | 264 | 225 | 136 | 0 | 1 | 0 | 2 | 0 | 225 | 175 | 137 | 135 | 187 | 0 | 0 | 0 | 0 | 0 | 272 | 149 | 1 | 1 | 1 | 0.989 | 1 | 0.984 | 1 | 1 | 1 | 1 | 1 | 1 | -0.005 |
| SE | 66497 | ENSG00000132676 | DAP3 | chr1 | + | 155721516 | 155721618 | 155709772 | 155709824 | 155731355 | 155731411 | 8 | 7 | 7 | 13 | 1 | 0 | 0 | 2 | 0 | 0 | 38 | 1 | 7 | 3 | 29 | 0 | 0 | 0 | 0 | 0 | 251 | 149 | 1 | 1 | 1 | 1 | 0.675 | 1 | 1 | 1 | 1 | 1 | 1 | 1 | -0.065 |
| SE | 66499 | ENSG00000132676 | DAP3 | chr1 | + | 155725381 | 155725411 | 155709772 | 155709824 | 155727607 | 155727738 | 0 | 0 | 0 | 4 | 0 | 0 | 1 | 0 | 2 | 0 | 0 | 0 | 1 | 2 | 0 | 0 | 0 | 0 | 0 | 0 | 179 | 149 | 1 | 1 | NA | 0 | NA | 0.625 | NA | NA | NA | 1 | 1 | NA | -0.688 |
| SE | 66501 | ENSG00000132676 | DAP3 | chr1 | + | 155725381 | 155725411 | 155721516 | 155721618 | 155736945 | 155737063 | 19 | 9 | 23 | 23 | 24 | 0 | 0 | 1 | 0 | 0 | 45 | 11 | 17 | 18 | 8 | 0 | 0 | 0 | 0 | 0 | 179 | 149 | 1 | 1 | 1 | 1 | 0.95 | 1 | 1 | 1 | 1 | 1 | 1 | 1 | -0.01 |
| SE | 66504 | ENSG00000132676 | DAP3 | chr1 | + | 155727607 | 155727738 | 155725926 | 155726019 | 155729041 | 155729064 | 498 | 480 | 514 | 490 | 784 | 0 | 0 | 1 | 0 | 1 | 875 | 433 | 495 | 487 | 857 | 0 | 5 | 0 | 0 | 0 | 280 | 149 | 1 | 1 | 1 | 1 | 0.996 | 1 | 0.998 | 1 | 0.979 | 1 | 1 | 1 | 0.003 |
| SE | 66506 | ENSG00000132676 | DAP3 | chr1 | + | 155729207 | 155729366 | 155721516 | 155721618 | 155731355 | 155731411 | 192 | 104 | 268 | 194 | 319 | 0 | 0 | 0 | 2 | 0 | 229 | 163 | 138 | 132 | 268 | 0 | 0 | 0 | 0 | 0 | 298 | 149 | 1 | 1 | 1 | 1 | 1 | 0.98 | 1 | 1 | 1 | 1 | 1 | 1 | -0.004 |
| SE | 66507 | ENSG00000132676 | DAP3 | chr1 | + | 155729207 | 155729366 | 155721516 | 155721618 | 155736945 | 155737063 | 0 | 0 | 0 | 1 | 0 | 0 | 0 | 1 | 0 | 0 | 0 | 3 | 1 | 0 | 2 | 0 | 0 | 0 | 0 | 0 | 298 | 149 | 1 | 1 | NA | NA | 0 | 1 | NA | NA | 1 | 1 | NA | 1 | -0.5 |
| SE | 66508 | ENSG00000132676 | DAP3 | chr1 | + | 155731355 | 155731415 | 155721516 | 155721618 | 155736945 | 155737063 | 0 | 0 | 0 | 1 | 3 | 0 | 0 | 1 | 0 | 0 | 0 | 0 | 0 | 1 | 0 | 0 | 0 | 0 | 0 | 0 | 209 | 149 | 1 | 1 | NA | NA | 0 | 1 | 1 | NA | NA | NA | 1 | NA | -0.333 |
| SE | 66509 | ENSG00000132676 | DAP3 | chr1 | + | 155731355 | 155731415 | 155729207 | 155729366 | 155736945 | 155737063 | 58 | 44 | 86 | 70 | 115 | 0 | 0 | 0 | 1 | 0 | 82 | 33 | 40 | 53 | 91 | 0 | 3 | 1 | 0 | 0 | 209 | 149 | 1 | 1 | 1 | 1 | 1 | 0.98 | 1 | 1 | 0.887 | 0.966 | 1 | 1 | 0.025 |
| SE | 66510 | ENSG00000132676 | DAP3 | chr1 | + | 155731943 | 155732033 | 155731355 | 155731415 | 155736945 | 155737063 | 370 | 198 | 424 | 355 | 445 | 0 | 0 | 1 | 2 | 9 | 376 | 300 | 262 | 242 | 379 | 0 | 2 | 0 | 1 | 0 | 239 | 149 | 1 | 1 | 1 | 1 | 0.996 | 0.991 | 0.969 | 1 | 0.989 | 1 | 0.993 | 1 | -0.005 |
| SE | 66511 | ENSG00000132676 | DAP3 | chr1 | + | 155736607 | 155736694 | 155731943 | 155732033 | 155736945 | 155737063 | 1 | 0 | 1 | 1 | 0 | 236 | 138 | 241 | 230 | 242 | 0 | 1 | 0 | 2 | 7 | 229 | 167 | 165 | 162 | 218 | 236 | 149 | 1 | 1 | 0.003 | 0 | 0.003 | 0.003 | 0 | 0 | 0.004 | 0 | 0.008 | 0.02 | -0.005 |
| SE | 66512 | ENSG00000132676 | DAP3 | chr1 | + | 155736945 | 155737063 | 155731943 | 155732033 | 155738156 | 155738426 | 406 | 236 | 419 | 400 | 379 | 0 | 0 | 2 | 1 | 0 | 323 | 323 | 274 | 272 | 399 | 0 | 0 | 0 | 8 | 0 | 267 | 149 | 1 | 1 | 1 | 1 | 0.992 | 0.996 | 1 | 1 | 1 | 1 | 0.95 | 1 | 0.008 |
| SE | 66647 | ENSG00000084623 | EIF3I | chr1 | + | 32226104 | 32226320 | 32224409 | 32224475 | 32226402 | 32226530 | 277 | 137 | 260 | 256 | 250 | 0 | 0 | 1 | 2 | 0 | 373 | 288 | 216 | 217 | 547 | 0 | 0 | 1 | 0 | 6 | 298 | 149 | 1 | 1 | 1 | 1 | 0.992 | 0.985 | 1 | 1 | 1 | 0.991 | 1 | 0.979 | 0.001 |
| SE | 66648 | ENSG00000084623 | EIF3I | chr1 | + | 32226148 | 32226320 | 32224409 | 32224475 | 32226402 | 32226530 | 278 | 137 | 261 | 256 | 250 | 0 | 0 | 1 | 2 | 0 | 373 | 288 | 216 | 217 | 547 | 0 | 0 | 1 | 0 | 6 | 298 | 149 | 1 | 1 | 1 | 1 | 0.992 | 0.985 | 1 | 1 | 1 | 0.991 | 1 | 0.979 | 0.001 |
| SE | 66649 | ENSG00000084623 | EIF3I | chr1 | + | 32226170 | 32226283 | 32224409 | 32224475 | 32226402 | 32226530 | 151 | 78 | 166 | 152 | 72 | 0 | 0 | 1 | 2 | 0 | 177 | 167 | 107 | 100 | 272 | 0 | 0 | 1 | 0 | 6 | 262 | 149 | 1 | 1 | 1 | 1 | 0.99 | 0.977 | 1 | 1 | 1 | 0.984 | 1 | 0.963 | 0.004 |
| SE | 66652 | ENSG00000084623 | EIF3I | chr1 | + | 32226170 | 32226320 | 32224033 | 32224475 | 32226402 | 32226530 | 445 | 233 | 449 | 428 | 351 | 0 | 0 | 1 | 2 | 0 | 562 | 479 | 335 | 341 | 867 | 0 | 0 | 1 | 0 | 6 | 298 | 149 | 1 | 1 | 1 | 1 | 0.996 | 0.991 | 1 | 1 | 1 | 0.994 | 1 | 0.986 | 0.001 |
| SE | 66786 | ENSG00000169432 | SCN9A | chr2 | - | 166243489 | 166243529 | 166242574 | 166242656 | 166251764 | 166251885 | 0 | 1 | 0 | 0 | 0 | 8 | 2 | 0 | 5 | 12 | 0 | 0 | 0 | 0 | 0 | 0 | 0 | 0 | 15 | 0 | 189 | 149 | 1 | 1 | 0 | 0.283 | NA | 0 | 0 | NA | NA | NA | 0 | NA | 0.071 |
| SE | 66789 | ENSG00000169432 | SCN9A | chr2 | - | 166293230 | 166293372 | 166288436 | 166288643 | 166294598 | 166294662 | 10 | 10 | 7 | 24 | 17 | 0 | 0 | 0 | 2 | 0 | 0 | 1 | 9 | 4 | 20 | 0 | 0 | 0 | 1 | 0 | 291 | 149 | 1 | 1 | 1 | 1 | 1 | 0.86 | 1 | NA | 1 | 1 | 0.672 | 1 | 0.054 |
| SE | 66790 | ENSG00000169432 | SCN9A | chr2 | - | 166294598 | 166294662 | 166288436 | 166288643 | 166303089 | 166303302 | 1 | 0 | 1 | 2 | 1 | 1 | 0 | 0 | 0 | 0 | 0 | 0 | 7 | 1 | 0 | 0 | 0 | 0 | 0 | 0 | 213 | 149 | 1 | 1 | 0.412 | NA | 1 | 1 | 1 | NA | NA | 1 | 1 | NA | -0.147 |
| SE | 66792 | ENSG00000169432 | SCN9A | chr2 | - | 166304237 | 166304329 | 166302062 | 166303302 | 166305791 | 166305920 | 8 | 5 | 6 | 18 | 8 | 0 | 0 | 0 | 0 | 1 | 13 | 2 | 2 | 16 | 0 | 0 | 0 | 0 | 1 | 0 | 241 | 149 | 1 | 1 | 1 | 1 | 1 | 1 | 0.832 | 1 | 1 | 1 | 0.908 | NA | -0.011 |
| SE | 66974 | ENSG00000162645 | GBP2 | chr1 | - | 89114015 | 89114296 | 89112471 | 89112684 | 89116991 | 89117234 | 1050 | 841 | 951 | 989 | 960 | 0 | 0 | 1 | 0 | 0 | 1662 | 607 | 689 | 1016 | 520 | 0 | 0 | 0 | 0 | 0 | 298 | 149 | 1 | 1 | 1 | 1 | 0.998 | 1 | 1 | 1 | 1 | 1 | 1 | 1 | 0 |
| SE | 66975 | ENSG00000162645 | GBP2 | chr1 | - | 89116991 | 89117234 | 89114015 | 89114296 | 89117576 | 89117773 | 864 | 540 | 763 | 772 | 704 | 0 | 0 | 2 | 5 | 0 | 882 | 396 | 512 | 487 | 319 | 0 | 0 | 4 | 8 | 0 | 298 | 149 | 1 | 1 | 1 | 1 | 0.995 | 0.987 | 1 | 1 | 1 | 0.985 | 0.968 | 1 | 0.006 |
| SE | 67066 | ENSG00000188177 | ZC3H6 | chr2 | + | 112324963 | 112325197 | 112324151 | 112324663 | 112331004 | 112340059 | 55 | 45 | 72 | 54 | 53 | 2 | 1 | 0 | 0 | 0 | 0 | 45 | 30 | 21 | 0 | 0 | 0 | 0 | 0 | 0 | 298 | 149 | 1 | 1 | 0.932 | 0.957 | 1 | 1 | 1 | NA | 1 | 1 | 1 | NA | -0.022 |
| SE | 67147 | ENSG00000089060 | SLC8B1 | chr12 | - | 113334442 | 113334835 | 113332722 | 113332960 | 113334992 | 113335119 | 148 | 107 | 176 | 163 | 143 | 1 | 0 | 0 | 1 | 0 | 238 | 186 | 89 | 242 | 288 | 0 | 0 | 3 | 0 | 0 | 298 | 149 | 1 | 1 | 0.987 | 1 | 1 | 0.988 | 1 | 1 | 1 | 0.937 | 1 | 1 | 0.008 |
| SE | 67148 | ENSG00000089060 | SLC8B1 | chr12 | - | 113334442 | 113334838 | 113332722 | 113332960 | 113334992 | 113335030 | 148 | 107 | 176 | 163 | 143 | 1 | 0 | 0 | 1 | 0 | 238 | 186 | 89 | 242 | 288 | 0 | 0 | 3 | 0 | 0 | 298 | 149 | 1 | 1 | 0.987 | 1 | 1 | 0.988 | 1 | 1 | 1 | 0.937 | 1 | 1 | 0.008 |
| SE | 67299 | ENSG00000204764 | RANBP17 | chr5 | + | 170916464 | 170916584 | 170914166 | 170914240 | 170918712 | 170918859 | 13 | 6 | 1 | 8 | 8 | 0 | 0 | 0 | 0 | 0 | 0 | 13 | 0 | 9 | 11 | 0 | 0 | 0 | 1 | 0 | 269 | 149 | 1 | 1 | 1 | 1 | 1 | 1 | 1 | NA | 1 | NA | 0.833 | 1 | 0.056 |
| SE | 67302 | ENSG00000204764 | RANBP17 | chr5 | + | 171028871 | 171028989 | 170968241 | 170968377 | 171170129 | 171170203 | 0 | 0 | 0 | 3 | 0 | 0 | 0 | 0 | 1 | 0 | 0 | 4 | 0 | 0 | 3 | 0 | 0 | 0 | 0 | 0 | 267 | 149 | 1 | 1 | NA | NA | NA | 0.626 | NA | NA | 1 | NA | NA | 1 | -0.374 |
| SE | 67418 | ENSG00000204160 | ZDHHC18 | chr1 | + | 26850300 | 26850438 | 26848607 | 26848757 | 26850557 | 26850606 | 138 | 52 | 107 | 154 | 173 | 0 | 0 | 0 | 0 | 2 | 49 | 162 | 156 | 113 | 153 | 0 | 0 | 0 | 0 | 0 | 287 | 149 | 1 | 1 | 1 | 1 | 1 | 1 | 0.978 | 1 | 1 | 1 | 1 | 1 | -0.004 |
| SE | 67419 | ENSG00000204160 | ZDHHC18 | chr1 | + | 26856184 | 26856262 | 26853725 | 26853825 | 26856847 | 26857602 | 0 | 0 | 0 | 0 | 2 | 0 | 0 | 0 | 2 | 0 | 0 | 2 | 0 | 0 | 0 | 0 | 1 | 0 | 0 | 0 | 227 | 149 | 1 | 1 | NA | NA | NA | 0 | 1 | NA | 0.568 | NA | NA | NA | -0.068 |
| SE | 67967 | ENSG00000166762 | CATSPER2 | chr15 | - | 43631429 | 43631538 | 43630570 | 43630732 | 43632198 | 43632363 | 0 | 0 | 0 | 3 | 0 | 6 | 8 | 3 | 3 | 7 | 0 | 0 | 0 | 0 | 0 | 0 | 0 | 1 | 2 | 0 | 258 | 149 | 1 | 1 | 0 | 0 | 0 | 0.366 | 0 | NA | NA | 0 | 0 | NA | 0.073 |
| SE | 67968 | ENSG00000166762 | CATSPER2 | chr15 | - | 43632716 | 43632928 | 43632198 | 43632363 | 43635359 | 43635416 | 0 | 3 | 0 | 3 | 14 | 2 | 0 | 0 | 0 | 0 | 0 | 0 | 1 | 0 | 0 | 0 | 0 | 0 | 0 | 0 | 298 | 149 | 1 | 1 | 0 | 1 | NA | 1 | 1 | NA | NA | 1 | NA | NA | -0.25 |
| SE | 67969 | ENSG00000166762 | CATSPER2 | chr15 | - | 43632716 | 43632934 | 43632198 | 43632363 | 43635359 | 43635416 | 0 | 3 | 0 | 3 | 14 | 2 | 0 | 0 | 0 | 0 | 0 | 0 | 1 | 0 | 0 | 0 | 0 | 0 | 0 | 0 | 298 | 149 | 1 | 1 | 0 | 1 | NA | 1 | 1 | NA | NA | 1 | NA | NA | -0.25 |
| SE | 67971 | ENSG00000166762 | CATSPER2 | chr15 | - | 43635726 | 43635826 | 43635359 | 43635416 | 43636040 | 43636219 | 0 | 0 | 1 | 0 | 0 | 4 | 2 | 0 | 2 | 1 | 0 | 0 | 10 | 0 | 0 | 0 | 0 | 1 | 0 | 0 | 249 | 149 | 1 | 1 | 0 | 0 | 1 | 0 | 0 | NA | NA | 0.857 | NA | NA | -0.657 |
| SE | 67974 | ENSG00000166762 | CATSPER2 | chr15 | - | 43647916 | 43648063 | 43647293 | 43647467 | 43664417 | 43664543 | 0 | 0 | 3 | 0 | 0 | 0 | 0 | 1 | 0 | 0 | 14 | 0 | 0 | 0 | 0 | 0 | 0 | 0 | 0 | 0 | 296 | 149 | 1 | 1 | NA | NA | 0.602 | NA | NA | 1 | NA | NA | NA | NA | -0.398 |
| SE | 67976 | ENSG00000166762 | CATSPER2 | chr15 | - | 43649726 | 43649879 | 43647916 | 43648063 | 43664417 | 43664543 | 0 | 2 | 0 | 1 | 0 | 0 | 0 | 1 | 0 | 0 | 0 | 0 | 0 | 1 | 0 | 0 | 0 | 0 | 0 | 0 | 298 | 149 | 1 | 1 | NA | 1 | 0 | 1 | NA | NA | NA | 1 | NA | NA | -0.333 |
| SE | 67977 | ENSG00000166762 | CATSPER2 | chr15 | - | 43649726 | 43649879 | 43647916 | 43648066 | 43668027 | 43668118 | 0 | 3 | 2 | 1 | 0 | 0 | 0 | 1 | 0 | 0 | 0 | 0 | 0 | 0 | 5 | 0 | 2 | 0 | 0 | 0 | 298 | 149 | 1 | 1 | NA | 1 | 1 | 1 | NA | NA | 0 | NA | NA | 1 | 0.5 |
| SE | 67978 | ENSG00000166762 | CATSPER2 | chr15 | - | 43664417 | 43664543 | 43649726 | 43649879 | 43668027 | 43668118 | 0 | 2 | 3 | 1 | 0 | 0 | 0 | 3 | 0 | 0 | 0 | 0 | 7 | 1 | 0 | 0 | 0 | 0 | 0 | 5 | 275 | 149 | 1 | 1 | NA | 0.265 | 1 | 1 | NA | NA | NA | 1 | 1 | 0 | 0.088 |
| SE | 68009 | ENSG00000214517 | PPME1 | chr11 | + | 74222311 | 74222369 | 74204352 | 74204445 | 74225204 | 74225256 | 45 | 13 | 23 | 26 | 42 | 0 | 0 | 0 | 4 | 0 | 15 | 7 | 16 | 32 | 13 | 0 | 0 | 0 | 0 | 0 | 207 | 149 | 1 | 1 | 1 | 1 | 1 | 0.824 | 1 | 1 | 1 | 1 | 1 | 1 | -0.035 |
| SE | 68210 | ENSG00000107643 | MAPK8 | chr10 | + | 48308094 | 48308288 | 48306654 | 48306821 | 48325896 | 48325965 | 4 | 6 | 4 | 1 | 5 | 4 | 3 | 3 | 0 | 1 | 0 | 0 | 0 | 0 | 6 | 0 | 1 | 0 | 0 | 0 | 298 | 149 | 1 | 1 | 0.333 | 0.5 | 0.4 | 1 | 0.714 | NA | 0 | NA | NA | 1 | 0.089 |
| SE | 68211 | ENSG00000107643 | MAPK8 | chr10 | + | 48325896 | 48325965 | 48306654 | 48306821 | 48401611 | 48401782 | 10 | 9 | 16 | 4 | 18 | 7 | 5 | 2 | 12 | 1 | 0 | 7 | 4 | 0 | 9 | 0 | 3 | 0 | 3 | 18 | 218 | 149 | 1 | 1 | 0.494 | 0.552 | 0.845 | 0.186 | 0.925 | NA | 0.615 | 1 | 0 | 0.255 | 0.133 |
| SE | 68212 | ENSG00000107643 | MAPK8 | chr10 | + | 48325896 | 48325965 | 48306654 | 48306821 | 48404851 | 48404981 | 2 | 0 | 0 | 0 | 1 | 1 | 0 | 0 | 0 | 0 | 0 | 1 | 1 | 0 | 0 | 0 | 0 | 0 | 0 | 0 | 218 | 149 | 1 | 1 | 0.578 | NA | NA | NA | 1 | NA | 1 | 1 | NA | NA | -0.211 |
| SE | 68213 | ENSG00000107643 | MAPK8 | chr10 | + | 48325896 | 48325965 | 48308094 | 48308288 | 48401611 | 48401782 | 10 | 12 | 17 | 4 | 20 | 6 | 1 | 4 | 2 | 9 | 0 | 6 | 4 | 0 | 9 | 0 | 2 | 4 | 6 | 0 | 218 | 149 | 1 | 1 | 0.533 | 0.891 | 0.744 | 0.578 | 0.603 | NA | 0.672 | 0.406 | 0 | 1 | 0.15 |
| SE | 68215 | ENSG00000107643 | MAPK8 | chr10 | + | 48401611 | 48401782 | 48306638 | 48306821 | 48404851 | 48404981 | 112 | 64 | 106 | 84 | 153 | 1 | 0 | 0 | 0 | 0 | 40 | 59 | 34 | 43 | 60 | 0 | 0 | 0 | 0 | 0 | 298 | 149 | 1 | 1 | 0.982 | 1 | 1 | 1 | 1 | 1 | 1 | 1 | 1 | 1 | -0.004 |
| SE | 68216 | ENSG00000107643 | MAPK8 | chr10 | + | 48401611 | 48401782 | 48325896 | 48325965 | 48404851 | 48404981 | 113 | 70 | 121 | 76 | 170 | 0 | 0 | 0 | 0 | 0 | 40 | 62 | 38 | 40 | 51 | 0 | 0 | 1 | 0 | 0 | 298 | 149 | 1 | 1 | 1 | 1 | 1 | 1 | 1 | 1 | 1 | 0.95 | 1 | 1 | 0.01 |
| SE | 68218 | ENSG00000107643 | MAPK8 | chr10 | + | 48420154 | 48420320 | 48410029 | 48410168 | 48425178 | 48425219 | 83 | 45 | 63 | 54 | 110 | 0 | 0 | 0 | 1 | 0 | 21 | 55 | 58 | 28 | 1 | 0 | 0 | 0 | 0 | 0 | 298 | 149 | 1 | 1 | 1 | 1 | 1 | 0.964 | 1 | 1 | 1 | 1 | 1 | 1 | -0.007 |
| SE | 68219 | ENSG00000107643 | MAPK8 | chr10 | + | 48426379 | 48426504 | 48425887 | 48426070 | 48427079 | 48427143 | 140 | 115 | 134 | 164 | 161 | 0 | 1 | 0 | 0 | 0 | 96 | 94 | 129 | 80 | 56 | 0 | 0 | 0 | 0 | 0 | 274 | 149 | 1 | 1 | 1 | 0.984 | 1 | 1 | 1 | 1 | 1 | 1 | 1 | 1 | -0.003 |
| SE | 68220 | ENSG00000107643 | MAPK8 | chr10 | + | 48431192 | 48431270 | 48425188 | 48427143 | 48434878 | 48434898 | 83 | 66 | 70 | 94 | 80 | 0 | 1 | 0 | 0 | 0 | 36 | 78 | 93 | 63 | 31 | 0 | 0 | 0 | 0 | 0 | 227 | 149 | 1 | 1 | 1 | 0.977 | 1 | 1 | 1 | 1 | 1 | 1 | 1 | 1 | -0.005 |
| SE | 68221 | ENSG00000107643 | MAPK8 | chr10 | + | 48431192 | 48431270 | 48425188 | 48427143 | 48434883 | 48439357 | 77 | 61 | 68 | 86 | 66 | 0 | 0 | 0 | 3 | 0 | 36 | 76 | 93 | 54 | 31 | 0 | 0 | 0 | 0 | 0 | 227 | 149 | 1 | 1 | 1 | 1 | 1 | 1 | 0.935 | 1 | 1 | 1 | 1 | 1 | -0.013 |
| SE | 68462 | ENSG00000143799 | PARP1 | chr1 | - | 226371015 | 226371106 | 226370433 | 226370517 | 226374225 | 226376067 | 0 | 0 | 1 | 0 | 2 | 274 | 159 | 234 | 265 | 436 | 0 | 9 | 0 | 2 | 0 | 172 | 233 | 126 | 121 | 106 | 240 | 149 | 1 | 1 | 0 | 0 | 0.003 | 0 | 0.003 | 0 | 0.023 | 0 | 0.01 | 0 | -0.005 |
| SE | 68464 | ENSG00000143799 | PARP1 | chr1 | - | 226383035 | 226383183 | 226379921 | 226381208 | 226385503 | 226385680 | 528 | 314 | 498 | 517 | 460 | 0 | 0 | 1 | 0 | 0 | 424 | 576 | 372 | 345 | 560 | 0 | 0 | 1 | 0 | 0 | 297 | 149 | 1 | 1 | 1 | 1 | 0.996 | 1 | 1 | 1 | 1 | 0.995 | 1 | 1 | 0 |
| SE | 68465 | ENSG00000143799 | PARP1 | chr1 | - | 226390409 | 226390624 | 226388655 | 226388755 | 226392198 | 226392314 | 336 | 189 | 252 | 320 | 215 | 0 | 0 | 0 | 0 | 0 | 403 | 234 | 191 | 195 | 269 | 7 | 1 | 0 | 1 | 0 | 298 | 149 | 1 | 1 | 1 | 1 | 1 | 1 | 1 | 0.966 | 0.992 | 1 | 0.99 | 1 | 0.01 |
| SE | 68466 | ENSG00000143799 | PARP1 | chr1 | - | 226392198 | 226392314 | 226388655 | 226388755 | 226402213 | 226402379 | 200 | 121 | 144 | 210 | 96 | 0 | 0 | 0 | 0 | 0 | 227 | 153 | 137 | 147 | 251 | 0 | 0 | 3 | 0 | 0 | 265 | 149 | 1 | 1 | 1 | 1 | 1 | 1 | 1 | 1 | 1 | 0.963 | 1 | 1 | 0.007 |

| | | | | | | | | | | | | | | | |
|---|---|---|---|---|---|---|---|---|---|---|---|---|---|---|---|
| SE | 68652 | ENSG00000184465 | WDR27 | chr6 | - | 169613558 | 169613656 | 169602218 | 169602321 | 169632946 | 169633068 | 10 | 3 | 0 | [illegible] |
| SE | 68823 | ENSG00000082258 | CCNT2 | chr2 | + | 134936840 | 134936969 | 134919809 | 134919891 | 134939001 | 134939062 | 163 | 130 | 160 | [illegible] |
| SE | 68827 | ENSG00000082258 | CCNT2 | chr2 | + | 134946100 | 134946146 | 134942611 | 134942674 | 134947735 | 134947899 | 24 | 31 | 26 | [illegible] |
| SE | 68828 | ENSG00000082258 | CCNT2 | chr2 | + | 134947735 | 134947899 | 134942611 | 134942674 | 134952640 | 134952711 | 129 | 94 | 100 | [illegible] |
| SE | 68829 | ENSG00000082258 | CCNT2 | chr2 | + | 134952640 | 134952711 | 134947729 | 134947899 | 134953229 | 134953268 | 181 | 134 | 149 | [illegible] |
| SE | 68976 | ENSG00000009954 | BAZ1B | chr7 | - | 73450846 | 73450994 | 73449541 | 73449689 | 73459535 | 73459718 | 125 | 132 | 88 | [illegible] |
| SE | 68977 | ENSG00000009954 | BAZ1B | chr7 | - | 73459535 | 73459718 | 73450846 | 73450994 | 73462921 | 73463099 | 275 | 180 | 217 | [illegible] |
| SE | 68978 | ENSG00000009954 | BAZ1B | chr7 | - | 73469516 | 73469650 | 73466295 | 73466401 | 73470344 | 73470483 | 382 | 305 | 313 | [illegible] |
| SE | 68980 | ENSG00000009954 | BAZ1B | chr7 | - | 73489193 | 73489391 | 73476867 | 73478569 | 73492799 | 73492921 | 656 | 545 | 708 | [illegible] |
| SE | 69000 | ENSG00000107521 | HPS1 | chr10 | - | 98424312 | 98424374 | 98423752 | 98423887 | 98425540 | 98425720 | 91 | 55 | 87 | [illegible] |
| SE | 69305 | ENSG00000112367 | FIG4 | chr6 | + | 109715077 | 109715176 | 109691215 | 109691501 | 109727108 | 109727265 | 28 | 12 | 31 | [illegible] |
| SE | 69307 | ENSG00000112367 | FIG4 | chr6 | + | 109716444 | 109716568 | 109715077 | 109715176 | 109727108 | 109727149 | 232 | 181 | 140 | [illegible] |
| SE | 69308 | ENSG00000112367 | FIG4 | chr6 | + | 109716444 | 109716568 | 109715077 | 109715176 | 109732636 | 109732687 | 125 | 91 | 75 | [illegible] |
| SE | 69310 | ENSG00000112367 | FIG4 | chr6 | + | 109727108 | 109727265 | 109715077 | 109715176 | 109732636 | 109732687 | 112 | 71 | 56 | [illegible] |
| SE | 69313 | ENSG00000112367 | FIG4 | chr6 | + | 109730379 | 109730506 | 109727108 | 109727265 | 109732636 | 109732687 | 3 | 0 | 0 | [illegible] |
| SE | 69314 | ENSG00000112367 | FIG4 | chr6 | + | 109738324 | 109738453 | 109735131 | 109735298 | 109741443 | 109741544 | 156 | 84 | 116 | [illegible] |
| SE | 69316 | ENSG00000112367 | FIG4 | chr6 | + | 109743109 | 109743272 | 109735149 | 109735298 | 109743674 | 109743772 | 76 | 53 | 45 | [illegible] |
| SE | 69317 | ENSG00000112367 | FIG4 | chr6 | + | 109743674 | 109743772 | 109743109 | 109743272 | 109762090 | 109762207 | 41 | 28 | 37 | [illegible] |
| SE | 69322 | ENSG00000112367 | FIG4 | chr6 | + | 109785670 | 109785769 | 109784969 | 109785028 | 109786301 | 109786449 | 3 | 1 | 2 | [illegible] |
| SE | 69325 | ENSG00000112367 | FIG4 | chr6 | + | 109796764 | 109796851 | 109789593 | 109792664 | 109825087 | 109825199 | 186 | 104 | 123 | [illegible] |
| SE | 69620 | ENSG00000156711 | MAPK13 | chr6 | + | 36131270 | 36131400 | 36130483 | 36130701 | 36135752 | 36135832 | 9 | 5 | 19 | [illegible] |
| SE | 69621 | ENSG00000156711 | MAPK13 | chr6 | + | 36132620 | 36132679 | 36131270 | 36131400 | 36135752 | 36135832 | 82 | 24 | 80 | [illegible] |
| SE | 69650 | ENSG00000143390 | RFX5 | chr1 | - | 151343680 | 151343882 | 151343341 | 151343442 | 151344196 | 151344278 | 168 | 102 | 159 | [illegible] |
| SE | 69651 | ENSG00000143390 | RFX5 | chr1 | - | 151344727 | 151344847 | 151344332 | 151344536 | 151345105 | 151345177 | 191 | 82 | 160 | [illegible] |
| SE | 69655 | ENSG00000143390 | RFX5 | chr1 | - | 151345927 | 151345961 | 151344988 | 151345188 | 151346214 | 151346333 | 28 | 11 | 22 | [illegible] |
| SE | 69737 | ENSG00000149782 | PLCB3 | chr11 | + | 64255233 | 64255313 | 64254897 | 64255038 | 64255395 | 64255449 | 17 | 26 | 26 | [illegible] |
| SE | 69738 | ENSG00000149782 | PLCB3 | chr11 | + | 64258472 | 64258713 | 64256617 | 64256764 | 64258884 | 64258969 | 48 | 21 | 15 | [illegible] |
| SE | 69740 | ENSG00000149782 | PLCB3 | chr11 | + | 64264020 | 64264112 | 64263690 | 64263795 | 64264950 | 64265006 | 19 | 9 | 18 | [illegible] |
| SE | 69741 | ENSG00000149782 | PLCB3 | chr11 | + | 64266494 | 64266552 | 64266314 | 64266404 | 64267184 | 64267271 | 28 | 27 | 23 | [illegible] |
| [illegible] | | | | | | | | | | | | | | | |

| | | | | | | | | | | | | | | | |
|---|---|---|---|---|---|---|---|---|---|---|---|---|---|---|---|
| SE | 81066 | ENSG00000157764 | BRAF | chr7 | - | 140739811 | 140739946 | 140725144 | 140726516 | 140749286 | 140749418 | 96 | 74 | 86 | [illegible] |
| SE | 81067 | ENSG00000157764 | BRAF | chr7 | - | 140747365 | 140747447 | 140739811 | 140739946 | 140748931 | 140749052 | 5 | 0 | 2 | [illegible] |
| SE | 81069 | ENSG00000157764 | BRAF | chr7 | - | 140748931 | 140749052 | 140739811 | 140739946 | 140749286 | 140749418 | 1 | 1 | 0 | [illegible] |
| SE | 81070 | ENSG00000157764 | BRAF | chr7 | - | 140748931 | 140749052 | 140747365 | 140747447 | 140749286 | 140749418 | 0 | 1 | 0 | [illegible] |
| SE | 81072 | ENSG00000157764 | BRAF | chr7 | - | 140777990 | 140778075 | 140776911 | 140777088 | 140781575 | 140781677 | 260 | 170 | 200 | [illegible] |
| SE | 81075 | ENSG00000157764 | BRAF | chr7 | - | 140800361 | 140800462 | 140794307 | 140794467 | 140801411 | 140801560 | 99 | 68 | 91 | [illegible] |
| SE | 81076 | ENSG00000157764 | BRAF | chr7 | - | 140800361 | 140800481 | 140794307 | 140794467 | 140801411 | 140801560 | 286 | 218 | 272 | [illegible] |
| SE | 81077 | ENSG00000157764 | BRAF | chr7 | - | 140808236 | 140808316 | 140807959 | 140808062 | 140808882 | 140808995 | 21 | 7 | 20 | [illegible] |
| SE | 81079 | ENSG00000157764 | BRAF | chr7 | - | 140808882 | 140808995 | 140807959 | 140808062 | 140834608 | 140834872 | 35 | 34 | 43 | [illegible] |
| SE | 81080 | ENSG00000157764 | BRAF | chr7 | - | 140808891 | 140808995 | 140807959 | 140808062 | 140834608 | 140834703 | 113 | 130 | 120 | [illegible] |
| SE | 81081 | ENSG00000157764 | BRAF | chr7 | - | 140850110 | 140850212 | 140833971 | 140834872 | 140924565 | 140924703 | 88 | 57 | 76 | [illegible] |
| SE | 81467 | ENSG00000106477 | CEP41 | chr7 | - | 130426628 | 130426759 | 130416918 | 130416966 | 130427954 | 130428018 | 0 | 1 | 0 | [illegible] |
| SE | 81468 | ENSG00000106477 | CEP41 | chr7 | - | 130426628 | 130426759 | 130421852 | 130422007 | 130427954 | 130428018 | 0 | 0 | 0 | [illegible] |
| SE | 81469 | ENSG00000106477 | CEP41 | chr7 | - | 130426645 | 130426759 | 130416918 | 130416966 | 130427954 | 130428018 | 0 | 0 | 0 | [illegible] |
| SE | 81470 | ENSG00000106477 | CEP41 | chr7 | - | 130426645 | 130426759 | 130421852 | 130422007 | 130427954 | 130428018 | 0 | 0 | 0 | [illegible] |
| SE | 82001 | ENSG00000104133 | SPG11 | chr15 | - | 44567423 | 44567592 | 44566216 | 44566305 | 44569397 | 44569505 | 225 | 128 | 159 | [illegible] |
| SE | 82002 | ENSG00000104133 | SPG11 | chr15 | - | 44569397 | 44569505 | 44566216 | 44566305 | 44572682 | 44572820 | 0 | 1 | 3 | [illegible] |
| SE | 82005 | ENSG00000104133 | SPG11 | chr15 | - | 44569397 | 44569505 | 44567423 | 44567592 | 44574901 | 44575041 | 102 | 57 | 81 | [illegible] |
| SE | 82006 | ENSG00000104133 | SPG11 | chr15 | - | 44570524 | 44570652 | 44569397 | 44569505 | 44572682 | 44572824 | 74 | 48 | 61 | [illegible] |
| SE | 82007 | ENSG00000104133 | SPG11 | chr15 | - | 44570524 | 44570652 | 44569397 | 44569505 | 44574901 | 44575041 | 73 | 48 | 59 | [illegible] |
| SE | 82009 | ENSG00000104133 | SPG11 | chr15 | - | 44570524 | 44570658 | 44567423 | 44567592 | 44573546 | 44573745 | 4 | 3 | 1 | [illegible] |
| SE | 82010 | ENSG00000104133 | SPG11 | chr15 | - | 44570524 | 44570658 | 44567423 | 44567592 | 44574901 | 44575041 | 1 | 3 | 1 | [illegible] |
| SE | 82011 | ENSG00000104133 | SPG11 | chr15 | - | 44570524 | 44570658 | 44567423 | 44567592 | 44583813 | 44584558 | 1 | 3 | 1 | [illegible] |
| SE | 82012 | ENSG00000104133 | SPG11 | chr15 | - | 44570524 | 44570658 | 44569397 | 44569505 | 44572682 | 44572820 | 183 | 107 | 135 | [illegible] |
| SE | 82013 | ENSG00000104133 | SPG11 | chr15 | - | 44570524 | 44570658 | 44569397 | 44569505 | 44574901 | 44575041 | 77 | 49 | 59 | [illegible] |
| SE | 82014 | ENSG00000104133 | SPG11 | chr15 | - | 44572682 | 44572820 | 44567337 | 44567592 | 44573546 | 44573683 | 78 | 40 | 42 | [illegible] |
| SE | 82015 | ENSG00000104133 | SPG11 | chr15 | - | 44572682 | 44572820 | 44567337 | 44567592 | 44574901 | 44575041 | 2 | 0 | 0 | [illegible] |
| SE | 82016 | ENSG00000104133 | SPG11 | chr15 | - | 44572682 | 44572820 | 44567337 | 44567592 | 44583813 | 44584558 | 2 | 0 | 0 | [illegible] |
| SE | 82017 | ENSG00000104133 | SPG11 | chr15 | - | 44572682 | 44572820 | 44569397 | 44569505 | 44574901 | 44575041 | 0 | 0 | 1 | [illegible] |
| SE | 82018 | ENSG00000104133 | SPG11 | chr15 | - | 44572682 | 44572820 | 44570524 | 44570658 | 44573546 | 44573683 | 179 | 93 | 115 | [illegible] |
| SE | 82019 | ENSG00000104133 | SPG11 | chr15 | - | 44572682 | 44572820 | 44570524 | 44570658 | 44574901 | 44575041 | 101 | 44 | 66 | [illegible] |
| SE | 82020 | ENSG00000104133 | SPG11 | chr15 | - | 44572682 | 44572820 | 44570524 | 44570658 | 44583813 | 44584558 | 101 | 44 | 66 | [illegible] |
| SE | 82021 | ENSG00000104133 | SPG11 | chr15 | - | 44573546 | 44573745 | 44567423 | 44567592 | 44574901 | 44575041 | 71 | 50 | 59 | [illegible] |
| SE | 82022 | ENSG00000104133 | SPG11 | chr15 | - | 44573546 | 44573745 | 44567423 | 44567592 | 44583813 | 44584558 | 1 | 0 | 0 | [illegible] |
| SE | 82023 | ENSG00000104133 | SPG11 | chr15 | - | 44573546 | 44573745 | 44570524 | 44570658 | 44574901 | 44575041 | 73 | 50 | 59 | [illegible] |
| SE | 82024 | ENSG00000104133 | SPG11 | chr15 | - | 44573546 | 44573745 | 44570524 | 44570658 | 44583813 | 44584558 | 3 | 0 | 0 | [illegible] |
| SE | 82025 | ENSG00000104133 | SPG11 | chr15 | - | 44573546 | 44573745 | 44572682 | 44572820 | 44574901 | 44575041 | 149 | 100 | 108 | [illegible] |
| SE | 82027 | ENSG00000104133 | SPG11 | chr15 | - | 44574901 | 44575041 | 44567423 | 44567592 | 44583813 | 44583935 | 44 | 17 | 28 | [illegible] |
| SE | 82028 | ENSG00000104133 | SPG11 | chr15 | - | 44574901 | 44575041 | 44570524 | 44570658 | 44583813 | 44583935 | 44 | 17 | 28 | [illegible] |
| SE | 82030 | ENSG00000104133 | SPG11 | chr15 | - | 44574901 | 44575041 | 44573118 | 44573745 | 44583813 | 44583935 | 113 | 67 | 87 | [illegible] |
| SE | 82033 | ENSG00000104133 | SPG11 | chr15 | - | 44595258 | 44595459 | 44592330 | 44592438 | 44596059 | 44596355 | 186 | 94 | 126 | [illegible] |
| SE | 82034 | ENSG00000104133 | SPG11 | chr15 | - | 44596783 | 44596943 | 44596059 | 44596355 | 44598264 | 44598373 | 233 | 193 | 209 | [illegible] |
| SE | 82036 | ENSG00000104133 | SPG11 | chr15 | - | 44600466 | 44600498 | 44598264 | 44598373 | 44608443 | 44608605 | 0 | 1 | 4 | [illegible] |
| SE | 82037 | ENSG00000104133 | SPG11 | chr15 | - | 44600466 | 44600632 | 44597384 | 44598373 | 44608443 | 44608605 | 2 | 13 | 7 | [illegible] |
| SE | 82038 | ENSG00000104133 | SPG11 | chr15 | - | 44601738 | 44601824 | 44600466 | 44600632 | 44606024 | 44606091 | 0 | 2 | 3 | [illegible] |
| SE | 82039 | ENSG00000104133 | SPG11 | chr15 | - | 44604104 | 44604197 | 44600466 | 44600632 | 44606024 | 44606091 | 6 | 4 | 9 | [illegible] |
| SE | 82040 | ENSG00000104133 | SPG11 | chr15 | - | 44604104 | 44604197 | 44600466 | 44600632 | 44608443 | 44608605 | 5 | 4 | 6 | [illegible] |
| SE | 82041 | ENSG00000104133 | SPG11 | chr15 | - | 44604802 | 44604927 | 44600466 | 44600632 | 44606024 | 44606091 | 4 | 0 | 2 | [illegible] |
| SE | 82042 | ENSG00000104133 | SPG11 | chr15 | - | 44606024 | 44606091 | 44600259 | 44600632 | 44608443 | 44608605 | 81 | 51 | 53 | [illegible] |
| SE | 82043 | ENSG00000104133 | SPG11 | chr15 | - | 44606024 | 44606091 | 44604104 | 44604197 | 44608443 | 44608605 | 14 | 11 | 6 | [illegible] |
| SE | 82045 | ENSG00000104133 | SPG11 | chr15 | - | 44610839 | 44610985 | 44608443 | 44608605 | 44615362 | 44615566 | 196 | 154 | 150 | [illegible] |
| SE | 82047 | ENSG00000104133 | SPG11 | chr15 | - | 44613429 | 44613536 | 44608443 | 44608605 | 44615362 | 44615566 | 161 | 130 | 120 | [illegible] |
| SE | 82049 | ENSG00000104133 | SPG11 | chr15 | - | 44615362 | 44615566 | 44613429 | 44613536 | 44620189 | 44620403 | 405 | 285 | 312 | [illegible] |
| SE | 82052 | ENSG00000104133 | SPG11 | chr15 | - | 44621758 | 44621934 | 44620124 | 44620403 | 44622219 | 44622347 | 572 | 600 | 445 | [illegible] |
| SE | 82055 | ENSG00000104133 | SPG11 | chr15 | - | 44622219 | 44622347 | 44621758 | 44621934 | 44622727 | 44622799 | 458 | 473 | 336 | [illegible] |
| SE | 82056 | ENSG00000104133 | SPG11 | chr15 | - | 44622727 | 44622799 | 44622219 | 44622347 | 44626330 | 44626507 | 338 | 280 | 265 | [illegible] |
| SE | 82058 | ENSG00000104133 | SPG11 | chr15 | - | 44633504 | 44633637 | 44629232 | 44629388 | 44648865 | 44649011 | 630 | 356 | 452 | [illegible] |
| SE | 82059 | ENSG00000104133 | SPG11 | chr15 | - | 44648865 | 44649011 | 44633469 | 44633637 | 44651490 | 44651644 | 466 | 298 | 341 | [illegible] |
| SE | 82060 | ENSG00000104133 | SPG11 | chr15 | - | 44657094 | 44657263 | 44652128 | 44652266 | 44660431 | 44660616 | 301 | 209 | 186 | [illegible] |
| SE | 82113 | ENSG00000136048 | DRAM1 | chr12 | + | 101914173 | 101914232 | 101901290 | 101901433 | 101921215 | 101921542 | 1 | 0 | 1 | [illegible] |
| SE | 82114 | ENSG00000136048 | DRAM1 | chr12 | + | 101920108 | 101920201 | 101901290 | 101901433 | 101921215 | 101921542 | 52 | 59 | 30 | [illegible] |
| SE | 82153 | ENSG00000146828 | SLC12A9 | chr7 | + | 100854155 | 100854378 | 100852712 | 100852835 | 100858834 | 100858942 | 3 | 1 | 2 | [illegible] |
| SE | 82155 | ENSG00000146828 | SLC12A9 | chr7 | + | 100856867 | 100857176 | 100852732 | 100852835 | 100858834 | 100858942 | 18 | 11 | 14 | [illegible] |
| SE | 82156 | ENSG00000146828 | SLC12A9 | chr7 | + | 100856867 | 100857176 | 100854155 | 100854378 | 100858834 | 100858942 | 18 | 12 | 16 | [illegible] |
| SE | 82157 | ENSG00000146828 | SLC12A9 | chr7 | + | 100856867 | 100857176 | 100855705 | 100855837 | 100858834 | 100858942 | 26 | 12 | 16 | [illegible] |
| SE | 82159 | ENSG00000146828 | SLC12A9 | chr7 | + | 100861137 | 100861262 | 100860149 | 100860232 | 100861391 | 100861584 | 52 | 19 | 40 | [illegible] |
| SE | 82356 | ENSG00000114503 | NCBP2 | chr3 | - | 196937509 | 196937631 | 196935433 | 196937082 | 196939392 | 196939432 | 70 | 90 | 121 | [illegible] |
| SE | 82357 | ENSG00000114503 | NCBP2 | chr3 | - | 196941486 | 196941602 | 196939250 | 196939432 | 196942369 | 196942436 | 1 | 0 | 1 | [illegible] |
| SE | 82566 | ENSG00000104915 | STX10 | chr19 | - | 13145287 | 13145395 | 13144763 | 13144870 | 13149028 | 13149091 | 195 | 109 | 166 | [illegible] |
| SE | 82948 | ENSG00000075142 | SRI | chr7 | - | 88209982 | 88210130 | 88208859 | 88209452 | 88210881 | 88210925 | 501 | 371 | 382 | [illegible] |
| SE | 82949 | ENSG00000075142 | SRI | chr7 | - | 88210881 | 88210925 | 88208859 | 88209452 | 88217121 | 88217191 | 26 | 12 | 29 | [illegible] |
| SE | 83151 | ENSG00000055130 | CUL1 | chr7 | + | 148759303 | 148759354 | 148756967 | 148757150 | 148759547 | 148759638 | 216 | 154 | 188 | [illegible] |
| SE | 83153 | ENSG00000055130 | CUL1 | chr7 | + | 148759547 | 148759638 | 148759303 | 148759354 | 148760332 | 148760484 | 322 | 249 | 293 | [illegible] |
| SE | 83161 | ENSG00000055130 | CUL1 | chr7 | + | 148792725 | 148792818 | 148790309 | 148790441 | 148797811 | 148797859 | 183 | 144 | 179 | [illegible] |
| SE | 84014 | ENSG00000111581 | NUP107 | chr12 | + | 68691967 | 68692112 | 68690467 | 68690746 | 68696850 | 68696922 | 79 | 55 | 49 | [illegible] |
| SE | 84015 | ENSG00000111581 | NUP107 | chr12 | + | 68696818 | 68696922 | 68691967 | 68692112 | 68700725 | 68700853 | 145 | 79 | 103 | [illegible] |
| SE | 84016 | ENSG00000111581 | NUP107 | chr12 | + | 68696850 | 68696922 | 68691967 | 68692112 | 68700725 | 68700814 | 35 | 34 | 23 | [illegible] |
| SE | 84017 | ENSG00000111581 | NUP107 | chr12 | + | 68700725 | 68700853 | 68696818 | 68696922 | 68702735 | 68702784 | 175 | 118 | 140 | [illegible] |
| SE | 84018 | ENSG00000111581 | NUP107 | chr12 | + | 68700725 | 68700853 | 68696818 | 68696922 | 68709237 | 68709309 | 94 | 52 | 66 | [illegible] |
| SE | 84019 | ENSG00000111581 | NUP107 | chr12 | + | 68702735 | 68702784 | 68696818 | 68696922 | 68709237 | 68709265 | 33 | 18 | 29 | [illegible] |
| SE | 84020 | ENSG00000111581 | NUP107 | chr12 | + | 68702735 | 68702784 | 68700725 | 68700853 | 68709237 | 68709265 | 118 | 86 | 105 | [illegible] |
| SE | 84021 | ENSG00000111581 | NUP107 | chr12 | + | 68709237 | 68709309 | 68702735 | 68702784 | 68710004 | 68710093 | 138 | 91 | 129 | [illegible] |
| SE | 84023 | ENSG00000111581 | NUP107 | chr12 | + | 68721117 | 68721177 | 68719577 | 68719654 | 68721840 | 68721986 | 75 | 50 | 55 | [illegible] |
| SE | 84024 | ENSG00000111581 | NUP107 | chr12 | + | 68725726 | 68725796 | 68722103 | 68722152 | 68726498 | 68726617 | 175 | 141 | 134 | [illegible] |
| SE | 84025 | ENSG00000111581 | NUP107 | chr12 | + | 68731109 | 68731260 | 68727350 | 68727389 | 68731606 | 68731719 | 255 | 250 | 263 | [illegible] |
| SE | 84026 | ENSG00000111581 | NUP107 | chr12 | + | 68731606 | 68731719 | 68727350 | 68727389 | 68732636 | 68732739 | 65 | 65 | 38 | [illegible] |
| SE | 84027 | ENSG00000111581 | NUP107 | chr12 | + | 68735230 | 68735344 | 68734707 | 68734833 | 68741812 | 68741980 | 164 | 137 | 96 | [illegible] |
| SE | 84235 | ENSG00000112305 | SMAP1 | chr6 | + | 70798656 | 70798737 | 70773349 | 70773425 | 70836940 | 70837028 | 58 | 52 | 57 | [illegible] |
| SE | 84236 | ENSG00000112305 | SMAP1 | chr6 | + | 70856858 | 70857030 | 70852539 | 70852664 | 70860199 | 70861166 | 34 | 33 | 51 | [illegible] |
| SE | 84238 | ENSG00000112305 | SMAP1 | chr6 | + | 70859328 | 70859416 | 70857921 | 70858229 | 70860199 | 70861166 | 1 | 0 | 1 | [illegible] |
| SE | 84300 | ENSG00000147202 | DIAPH2 | chrX | + | 96735757 | 96735790 | 96684662 | 96685190 | 96738585 | 96738762 | 12 | 26 | 15 | [illegible] |
| SE | 84301 | ENSG00000147202 | DIAPH2 | chrX | + | 96758153 | 96758258 | 96738585 | 96738762 | 96763099 | 96763120 | 203 | 204 | 127 | [illegible] |
| SE | 84303 | ENSG00000147202 | DIAPH2 | chrX | + | 96763099 | 96763120 | 96738585 | 96738762 | 96881578 | 96881718 | 4 | 7 | 5 | [illegible] |
| SE | 84306 | ENSG00000147202 | DIAPH2 | chrX | + | 96948934 | 96949039 | 96945526 | 96945591 | 96957827 | 96958148 | 300 | 364 | 180 | [illegible] |
| SE | 84340 | ENSG00000159256 | MORC3 | chr21 | + | 36333645 | 36333718 | 36320188 | 36320303 | 36336873 | 36337006 | 127 | 96 | 126 | [illegible] |
| SE | 84341 | ENSG00000159256 | MORC3 | chr21 | + | 36341398 | 36341546 | 36338773 | 36338921 | 36344578 | 36344707 | 2092 | 1508 | 1346 | [illegible] |
| SE | 84342 | ENSG00000159256 | MORC3 | chr21 | + | 36349310 | 36349408 | 36344911 | 36345031 | 36356619 | 36356724 | 365 | 317 | 286 | [illegible] |
| SE | 84343 | ENSG00000159256 | MORC3 | chr21 | + | 36349310 | 36349408 | 36344911 | 36345031 | 36359954 | 36360049 | 148 | 118 | 110 | [illegible] |
| SE | 84344 | ENSG00000159256 | MORC3 | chr21 | + | 36356619 | 36356724 | 36344911 | 36345031 | 36359954 | 36360049 | 98 | 130 | 113 | [illegible] |
| SE | 84345 | ENSG00000159256 | MORC3 | chr21 | + | 36356619 | 36356724 | 36349310 | 36349408 | 36359954 | 36360049 | 312 | 318 | 281 | [illegible] |
| SE | 84346 | ENSG00000159256 | MORC3 | chr21 | + | 36362182 | 36362228 | 36360183 | 36360258 | 36364092 | 36364259 | 264 | 212 | 212 | [illegible] |
| SE | 84454 | ENSG00000137522 | RNF121 | chr11 | + | 71962151 | 71962289 | 71960749 | 71960891 | 71982760 | 71982915 | 2 | 0 | 5 | [illegible] |
| SE | 84456 | ENSG00000137522 | RNF121 | chr11 | + | 71978085 | 71978235 | 71960749 | 71960891 | 71987003 | 71987028 | 0 | 0 | 5 | [illegible] |
| SE | 84457 | ENSG00000137522 | RNF121 | chr11 | + | 71982760 | 71982915 | 71960749 | 71960891 | 71987003 | 71987028 | 35 | 17 | 48 | [illegible] |
| SE | 84458 | ENSG00000137522 | RNF121 | chr11 | + | 71982760 | 71982915 | 71960749 | 71960891 | 71990596 | 71990717 | 12 | 11 | 21 | [illegible] |
| SE | 84459 | ENSG00000137522 | RNF121 | chr11 | + | 71982760 | 71982915 | 71978085 | 71978235 | 71987003 | 71987028 | 32 | 7 | 28 | [illegible] |
| SE | 84460 | ENSG00000137522 | RNF121 | chr11 | + | 71982806 | 71982915 | 71960749 | 71960891 | 71987003 | 71987111 | 20 | 5 | 30 | [illegible] |
| SE | 84461 | ENSG00000137522 | RNF121 | chr11 | + | 71982806 | 71982915 | 71960749 | 71960891 | 71990596 | 71990663 | 2 | 0 | 3 | [illegible] |
| SE | 84464 | ENSG00000137522 | RNF121 | chr11 | + | 71990596 | 71990717 | 71960749 | 71960891 | 71994718 | 71994777 | 40 | 13 | 27 | [illegible] |
| SE | 84465 | ENSG00000137522 | RNF121 | chr11 | + | 71995449 | 71995551 | 71994718 | 71994852 | 71996194 | 71996275 | 70 | 34 | 39 | [illegible] |
| SE | 84774 | ENSG00000125676 | THOC2 | chrX | - | 123625911 | 123626069 | 123624540 | 123624669 | 123626520 | 123626636 | 361 | 269 | 310 | [illegible] |
| SE | 84775 | ENSG00000125676 | THOC2 | chrX | - | 123632860 | 123633040 | 123631687 | 123631852 | 123633952 | 123634070 | 278 | 164 | 204 | [illegible] |
| SE | 84776 | ENSG00000125676 | THOC2 | chrX | - | 123668158 | 123668314 | 123667105 | 123667278 | 123671668 | 123671761 | 304 | 327 | 239 | [illegible] |
| SE | 84777 | ENSG00000125676 | THOC2 | chrX | - | 123686547 | 123686714 | 123671668 | 123671761 | 123696020 | 123696154 | 422 | 389 | 367 | [illegible] |
| SE | 84778 | ENSG00000125676 | THOC2 | chrX | - | 123696020 | 123696154 | 123671668 | 123671761 | 123696720 | 123696835 | 205 | 133 | 150 | [illegible] |
| SE | 84779 | ENSG00000125676 | THOC2 | chrX | - | 123703453 | 123703505 | 123697680 | 123697751 | 123706857 | 123706949 | 193 | 170 | 150 | [illegible] |
| SE | 84861 | ENSG00000164074 | ABHD18 | chr4 | + | 127982938 | 127983047 | 127965305 | 127965606 | 127984338 | 127984423 | 88 | 78 | 112 | [illegible] |
| SE | 84869 | ENSG00000164074 | ABHD18 | chr4 | + | 128020079 | 128020169 | 128011672 | 128011700 | 128028474 | 128028853 | 3 | 1 | 2 | [illegible] |
| SE | 84871 | ENSG00000164074 | ABHD18 | chr4 | + | 128028474 | 128028853 | 128021136 | 128021238 | 128030509 | 128030672 | 154 | 103 | 115 | [illegible] |
| SE | 84872 | ENSG00000164074 | ABHD18 | chr4 | + | 128030509 | 128030672 | 128028474 | 128028853 | 128035761 | 128036286 | 182 | 142 | 154 | [illegible] |
| SE | 85053 | ENSG00000147459 | DOCK5 | chr8 | + | 25268844 | 25268885 | 25243673 | 25243757 | 25275385 | 25275441 | 74 | 56 | 30 | [illegible] |
| SE | 85054 | ENSG00000147459 | DOCK5 | chr8 | + | 25268844 | 25268885 | 25243673 | 25243757 | 25292023 | 25292172 | 12 | 5 | 4 | [illegible] |
| SE | 85056 | ENSG00000147459 | DOCK5 | chr8 | + | 25275385 | 25275441 | 25268844 | 25268885 | 25292023 | 25292172 | 42 | 33 | 19 | [illegible] |
| SE | 85057 | ENSG00000147459 | DOCK5 | chr8 | + | 25278568 | 25278665 | 25275385 | 25275441 | 25292023 | 25292172 | 83 | 72 | 30 | [illegible] |
| SE | 85058 | ENSG00000147459 | DOCK5 | chr8 | + | 25296512 | 25296648 | 25292023 | 25292172 | 25298943 | 25299101 | 166 | 101 | 97 | [illegible] |
| SE | 85059 | ENSG00000147459 | DOCK5 | chr8 | + | 25300575 | 25300657 | 25298943 | 25299101 | 25302324 | 25302454 | 100 | 73 | 63 | [illegible] |
| SE | 85060 | ENSG00000147459 | DOCK5 | chr8 | + | 25317006 | 25317131 | 25310406 | 25310532 | 25319577 | 25319676 | 287 | 251 | 119 | [illegible] |
| SE | 85062 | ENSG00000147459 | DOCK5 | chr8 | + | 25341738 | 25341809 | 25336238 | 25336373 | 25342400 | 25342507 | 6 | 15 | 4 | [illegible] |
| SE | 85063 | ENSG00000147459 | DOCK5 | chr8 | + | 25342400 | 25342507 | 25336238 | 25336373 | 25345474 | 25345611 | 33 | 27 | 19 | [illegible] |
| SE | 85068 | ENSG00000147459 | DOCK5 | chr8 | + | 25368570 | 25368687 | 25368191 | 25368250 | 25369555 | 25369641 | 69 | 35 | 38 | [illegible] |
| SE | 85070 | ENSG00000147459 | DOCK5 | chr8 | + | 25368570 | 25368725 | 25368191 | 25368250 | 25369555 | 25369641 | 240 | 176 | 113 | [illegible] |
| SE | 85071 | ENSG00000147459 | DOCK5 | chr8 | + | 25374563 | 25374654 | 25373617 | 25373658 | 25377304 | 25377424 | 88 | 60 | 52 | [illegible] |
| SE | 85073 | ENSG00000147459 | DOCK5 | chr8 | + | 25395542 | 25395719 | 25392795 | 25392882 | 25399910 | 25399994 | 123 | 84 | 76 | [illegible] |
| SE | 85075 | ENSG00000147459 | DOCK5 | chr8 | + | 25408801 | 25408940 | 25403557 | 25403724 | 25410098 | 25410202 | 21 | 13 | 3 | [illegible] |
| SE | 85175 | ENSG00000162669 | HFM1 | chr1 | - | 91316390 | 91316476 | 91316100 | 91316184 | 91319077 | 91319209 | 1 | 1 | 0 | [illegible] |
| SE | 85180 | ENSG00000162669 | HFM1 | chr1 | - | 91350737 | 91350871 | 91347428 | 91347476 | 91352505 | 91352651 | 0 | 2 | 0 | [illegible] |
| SE | 85181 | ENSG00000162669 | HFM1 | chr1 | - | 91351548 | 91351643 | 91350737 | 91350871 | 91352505 | 91352651 | 1 | 4 | 0 | [illegible] |
| SE | 85426 | ENSG00000122126 | OCRL | chrX | + | 129557860 | 129557950 | 129557324 | 129557435 | 129558632 | 129558753 | 37 | 23 | 33 | [illegible] |
| SE | 85427 | ENSG00000122126 | OCRL | chrX | + | 129560549 | 129560651 | 129558839 | 129559001 | 129561178 | 129561293 | 42 | 29 | 14 | [illegible] |
| SE | 85430 | ENSG00000122126 | OCRL | chrX | + | 129584343 | 129584367 | 129576316 | 129576552 | 129587001 | 129587118 | 4 | 0 | 1 | [illegible] |
| SE | 85431 | ENSG00000122126 | OCRL | chrX | + | 129589236 | 129589364 | 129588885 | 129589013 | 129589844 | 129589956 | 2 | 2 | 0 | [illegible] |
| SE | 85529 | ENSG00000100802 | C14orf93 | chr14 | - | 22987902 | 22988015 | 22985893 | 22987634 | 22989741 | 22989845 | 68 | 52 | 56 | [illegible] |
| SE | 85530 | ENSG00000100802 | C14orf93 | chr14 | - | 22989741 | 22989845 | 22987902 | 22988015 | 22990065 | 22990127 | 68 | 63 | 55 | [illegible] |
| SE | 85531 | ENSG00000100802 | C14orf93 | chr14 | - | 22999956 | 23000117 | 22998772 | 22999156 | 23000958 | 23001060 | 4 | 0 | 0 | [illegible] |
| SE | 85532 | ENSG00000100802 | C14orf93 | chr14 | - | 22999956 | 23000117 | 22998772 | 22999156 | 23010016 | 23010166 | 4 | 4 | 0 | [illegible] |
| SE | 85533 | ENSG00000100802 | C14orf93 | chr14 | - | 22999956 | 23000117 | 22998772 | 22999156 | 23010100 | 23010143 | 4 | 0 | 0 | [illegible] |
| SE | 85534 | ENSG00000100802 | C14orf93 | chr14 | - | 22999956 | 23000117 | 22999044 | 22999362 | 23000958 | 23001060 | 0 | 4 | 2 | [illegible] |
| SE | 85535 | ENSG00000100802 | C14orf93 | chr14 | - | 22999956 | 23000117 | 22999044 | 22999362 | 23010016 | 23010166 | 0 | 8 | 2 | [illegible] |
| SE | 85536 | ENSG00000100802 | C14orf93 | chr14 | - | 22999956 | 23000117 | 22999044 | 22999362 | 23010100 | 23010143 | 0 | 4 | 2 | [illegible] |
| SE | 85537 | ENSG00000100802 | C14orf93 | chr14 | - | 22999956 | 23000117 | 22999120 | 22999402 | 23000958 | 23001060 | 1 | 0 | 0 | [illegible] |
| SE | 85538 | ENSG00000100802 | C14orf93 | chr14 | - | 22999956 | 23000117 | 22999120 | 22999402 | 23010100 | 23010143 | 1 | 0 | 0 | [illegible] |
| SE | 85539 | ENSG00000100802 | C14orf93 | chr14 | - | 23000958 | 23001031 | 22998704 | 22999156 | 23010100 | 23010143 | 3 | 1 | 3 | [illegible] |
| SE | 85540 | ENSG00000100802 | C14orf93 | chr14 | - | 23000958 | 23001037 | 22998426 | 22999156 | 23010016 | 23010143 | 3 | 1 | 3 | [illegible] |
| SE | 85541 | ENSG00000100802 | C14orf93 | chr14 | - | 23000958 | 23001037 | 22998426 | 22999156 | 23010100 | 23010128 | 4 | 2 | 5 | [illegible] |
| SE | 85542 | ENSG00000100802 | C14orf93 | chr14 | - | 23000958 | 23001037 | 22999044 | 22999362 | 23010016 | 23010143 | 2 | 0 | 0 | [illegible] |
| SE | 85543 | ENSG00000100802 | C14orf93 | chr14 | - | 23000958 | 23001037 | 22999044 | 22999362 | 23010100 | 23010128 | 2 | 0 | 1 | [illegible] |
| SE | 85546 | ENSG00000100802 | C14orf93 | chr14 | - | 23005064 | 23005130 | 23000958 | 23001037 | 23010016 | 23010143 | 2 | 3 | 0 | [illegible] |
| SE | 85548 | ENSG00000100532 | CGRRF1 | chr14 | + | 54522453 | 54522593 | 54509811 | 54510063 | 54530048 | 54530226 | 92 | 38 | 43 | [illegible] |
| SE | 85973 | ENSG00000144161 | ZC3H8 | chr2 | - | 112240367 | 112240492 | 112238314 | 112238528 | 112250190 | 112250272 | 2 | 0 | 2 | [illegible] |
| SE | 86197 | ENSG00000171311 | EXOSC1 | chr10 | - | 97438662 | 97438703 | 97437699 | 97437750 | 97441170 | 97441259 | 32 | 24 | 30 | [illegible] |
| SE | 86198 | ENSG00000171311 | EXOSC1 | chr10 | - | 97438669 | 97438703 | 97437699 | 97437750 | 97441170 | 97441259 | 116 | 80 | 126 | [illegible] |
| SE | 86200 | ENSG00000171311 | EXOSC1 | chr10 | - | 97441170 | 97441259 | 97438662 | 97438703 | 97445731 | 97445847 | 79 | 44 | 75 | [illegible] |
| SE | 86228 | ENSG00000177042 | TMEM80 | chr11 | + | 700614 | 700707 | 698868 | 700235 | 702944 | 703146 | 122 | 45 | 58 | [illegible] |
| SE | 86241 | ENSG00000134516 | DOCK2 | chr5 | + | 169669287 | 169669328 | 169654402 | 169654486 | 169670541 | 169670597 | 277 | 140 | 174 | [illegible] |
| SE | 86243 | ENSG00000134516 | DOCK2 | chr5 | + | 169669287 | 169669328 | 169654402 | 169654486 | 169674296 | 169674445 | 62 | 21 | 41 | [illegible] |
| SE | 86246 | ENSG00000134516 | DOCK2 | chr5 | + | 169671077 | 169671174 | 169654402 | 169654486 | 169674296 | 169674445 | 295 | 230 | 214 | [illegible] |
| SE | 86247 | ENSG00000134516 | DOCK2 | chr5 | + | 169671077 | 169671174 | 169654402 | 169654486 | 169681743 | 169681879 | 0 | 3 | 0 | [illegible] |
| SE | 86248 | ENSG00000134516 | DOCK2 | chr5 | + | 169671077 | 169671174 | 169669287 | 169669328 | 169674296 | 169674445 | 296 | 230 | 215 | [illegible] |
| SE | 86249 | ENSG00000134516 | DOCK2 | chr5 | + | 169674296 | 169674445 | 169654402 | 169654486 | 169681743 | 169681879 | 601 | 565 | 514 | [illegible] |
| SE | 86250 | ENSG00000134516 | DOCK2 | chr5 | + | 169674296 | 169674445 | 169671077 | 169671174 | 169681743 | 169681879 | 1150 | 961 | 873 | [illegible] |
| SE | 86252 | ENSG00000134516 | DOCK2 | chr5 | + | 169684195 | 169684350 | 169681743 | 169681879 | 169695802 | 169695938 | 767 | 536 | 649 | [illegible] |
| SE | 86253 | ENSG00000134516 | DOCK2 | chr5 | + | 169689251 | 169689333 | 169681743 | 169681879 | 169695802 | 169695938 | 364 | 244 | 291 | [illegible] |
| SE | 86254 | ENSG00000134516 | DOCK2 | chr5 | + | 169689251 | 169689333 | 169684195 | 169684350 | 169695802 | 169695938 | 947 | 648 | 826 | [illegible] |
| SE | 86255 | ENSG00000134516 | DOCK2 | chr5 | + | 169695802 | 169695938 | 169689251 | 169689333 | 169698373 | 169698449 | 1690 | 1177 | 1319 | [illegible] |
| SE | 86256 | ENSG00000134516 | DOCK2 | chr5 | + | 169699381 | 169699458 | 169698373 | 169698449 | 169700013 | 169700039 | 1617 | 1108 | 1047 | [illegible] |
| SE | 86257 | ENSG00000134516 | DOCK2 | chr5 | + | 169702302 | 169702427 | 169700013 | 169700139 | 169708168 | 169708267 | 1335 | 894 | 927 | [illegible] |
| SE | 86260 | ENSG00000134516 | DOCK2 | chr5 | + | 169714027 | 169714211 | 169712119 | 169712223 | 169714359 | 169714403 | 2050 | 1487 | 1512 | [illegible] |
| SE | 86261 | ENSG00000134516 | DOCK2 | chr5 | + | 169714359 | 169714457 | 169714027 | 169714211 | 169716212 | 169716302 | 1103 | 890 | 764 | [illegible] |
| SE | 86262 | ENSG00000134516 | DOCK2 | chr5 | + | 169717383 | 169717484 | 169716212 | 169716302 | 169718656 | 169718791 | 1036 | 849 | 676 | [illegible] |
| SE | 86263 | ENSG00000134516 | DOCK2 | chr5 | + | 169759704 | 169759775 | 169747395 | 169747504 | 169761518 | 169761625 | 973 | 606 | 592 | [illegible] |
| SE | 86267 | ENSG00000134516 | DOCK2 | chr5 | + | 169840756 | 169840852 | 169803057 | 169803206 | 169983067 | 169983166 | 565 | 345 | 496 | [illegible] |
| SE | 86268 | ENSG00000134516 | DOCK2 | chr5 | + | 169983067 | 169983166 | 169840756 | 169840852 | 169985827 | 169985922 | 569 | 334 | 467 | [illegible] |
| SE | 86269 | ENSG00000134516 | DOCK2 | chr5 | + | 169997973 | 169998024 | 169996085 | 169996164 | 170008496 | 170008597 | 0 | 0 | 0 | [illegible] |
| SE | 86272 | ENSG00000134516 | DOCK2 | chr5 | + | 170034398 | 170034555 | 170027862 | 170027948 | 170036514 | 170036555 | 837 | 551 | 657 | [illegible] |
| SE | 86273 | ENSG00000134516 | DOCK2 | chr5 | + | 170034398 | 170034555 | 170027862 | 170027948 | 170041054 | 170041145 | 551 | 411 | 483 | [illegible] |
| SE | 86274 | ENSG00000134516 | DOCK2 | chr5 | + | 170036514 | 170036555 | 170027862 | 170027948 | 170041054 | 170041145 | 95 | 80 | 76 | [illegible] |
| SE | 86275 | ENSG00000134516 | DOCK2 | chr5 | + | 170036514 | 170036555 | 170034398 | 170034555 | 170041054 | 170041145 | 375 | 222 | 238 | [illegible] |
| SE | 86276 | ENSG00000134516 | DOCK2 | chr5 | + | 170045815 | 170045905 | 170042012 | 170042132 | 170047509 | 170047614 | 460 | 252 | 269 | [illegible] |
| SE | 86277 | ENSG00000134516 | DOCK2 | chr5 | + | 170067509 | 170067686 | 170056437 | 170057666 | 170069136 | 170069220 | 652 | 330 | 486 | [illegible] |
| SE | 86278 | ENSG00000134516 | DOCK2 | chr5 | + | 170069136 | 170069220 | 170067509 | 170067686 | 170075946 | 170076084 | 345 | 169 | 235 | [illegible] |
| SE | 86281 | ENSG00000134516 | DOCK2 | chr5 | + | 170075946 | 170076084 | 170069136 | 170069220 | 170077709 | 170077837 | 626 | 280 | 372 | [illegible] |
| SE | 86282 | ENSG00000134516 | DOCK2 | chr5 | + | 170077709 | 170077837 | 170069136 | 170069220 | 170078974 | 170079146 | 225 | 112 | 207 | [illegible] |
| SE | 86283 | ENSG00000134516 | DOCK2 | chr5 | + | 170080162 | 170080283 | 170078974 | 170079146 | 170081841 | 170081984 | 435 | 193 | 336 | [illegible] |
| SE | 86397 | ENSG00000102001 | CACNA1F | chrX | - | 49217897 | 49218005 | 49217754 | 49217807 | 49218454 | 49218542 | 1 | 0 | 3 | [illegible] |
| SE | 86398 | ENSG00000102001 | CACNA1F | chrX | - | 49224760 | 49224986 | 49222928 | 49223136 | 49225908 | 49226069 | 2 | 0 | 1 | [illegible] |
| SE | 86399 | ENSG00000102001 | CACNA1F | chrX | - | 49225908 | 49226069 | 49222928 | 49223136 | 49226216 | 49226243 | 0 | 0 | 2 | [illegible] |
| SE | 86464 | ENSG00000215021 | PHB2 | chr12 | - | 6967170 | 6967335 | 6966423 | 6966500 | 6967895 | 6967962 | 193 | 117 | 198 | [illegible] |
| SE | 86855 | ENSG00000090316 | MAEA | chr4 | + | 1297955 | 1298115 | 1289890 | 1289982 | 1311978 | 1312161 | 0 | 0 | 0 | [illegible] |
| SE | 86857 | ENSG00000090316 | MAEA | chr4 | + | 1315396 | 1315600 | 1311978 | 1312161 | 1332756 | 1332865 | 42 | 24 | 50 | [illegible] |
| SE | 86858 | ENSG00000090316 | MAEA | chr4 | + | 1315396 | 1315600 | 1311978 | 1312238 | 1322380 | 1322414 | 61 | 25 | 53 | [illegible] |
| SE | 86859 | ENSG00000090316 | MAEA | chr4 | + | 1315396 | 1315600 | 1311978 | 1312238 | 1332756 | 1332865 | 4 | 5 | 15 | [illegible] |
| SE | 86860 | ENSG00000090316 | MAEA | chr4 | + | 1322380 | 1322503 | 1311978 | 1312161 | 1332756 | 1332865 | 1 | 6 | 3 | [illegible] |
| SE | 86861 | ENSG00000090316 | MAEA | chr4 | + | 1322380 | 1322503 | 1311978 | 1312238 | 1332756 | 1332865 | 2 | 6 | 3 | [illegible] |
| SE | 86867 | ENSG00000090316 | MAEA | chr4 | + | 1327626 | 1327703 | 1315396 | 1315600 | 1339073 | 1339177 | 2 | 0 | 3 | [illegible] |
| SE | 86870 | ENSG00000090316 | MAEA | chr4 | + | 1332756 | 1332865 | 1315396 | 1315600 | 1339073 | 1339177 | 2 | 0 | 9 | [illegible] |
| SE | 86871 | ENSG00000090316 | MAEA | chr4 | + | 1332756 | 1332865 | 1322380 | 1322503 | 1339073 | 1339177 | 1 | 6 | 3 | [illegible] |
| SE | 86873 | ENSG00000090316 | MAEA | chr4 | + | 1336860 | 1338617 | 1332560 | 1332865 | 1339073 | 1340147 | 251 | 106 | 161 | [illegible] |
| SE | 86915 | ENSG00000188706 | ZDHHC9 | chrX | - | 129841778 | 129841956 | 129828980 | 129829141 | 129843695 | 129843909 | 1 | 5 | 7 | [illegible] |
| SE | 86994 | ENSG00000232119 | MCTS1 | chrX | + | 120608224 | 120608463 | 120606078 | 120606176 | 120612182 | 120612387 | 109 | 54 | 82 | [illegible] |

| | | | | | | | | | | | | | | |
|---|---|---|---|---|---|---|---|---|---|---|---|---|---|---|
| SE | 86995 | ENSG00000232119 | MCTS1 | chrX | + | 120611010 | 120611078 | 120608224 | 120608463 | 120612182 | 120612312 | 59 | 41 | [illegible] |
| SE | 87129 | ENSG00000104613 | INTS10 | chr8 | + | 19818193 | 19818342 | 19817436 | 19817666 | 19819572 | 19819676 | 29 | 23 | [illegible] |
| SE | 87130 | ENSG00000104613 | INTS10 | chr8 | + | 19818274 | 19818342 | 19817415 | 19817666 | 19819572 | 19819676 | 60 | 27 | [illegible] |
| SE | 87131 | ENSG00000104613 | INTS10 | chr8 | + | 19818274 | 19818473 | 19817390 | 19817666 | 19819572 | 19819653 | 7 | 7 | [illegible] |
| SE | 87133 | ENSG00000104613 | INTS10 | chr8 | + | 19820378 | 19820518 | 19819572 | 19819676 | 19822438 | 19822520 | 315 | 169 | [illegible] |
| SE | 87136 | ENSG00000104613 | INTS10 | chr8 | + | 19826425 | 19826559 | 19824663 | 19824972 | 19830405 | 19830526 | 259 | 154 | [illegible] |
| SE | 87137 | ENSG00000104613 | INTS10 | chr8 | + | 19830405 | 19830559 | 19826425 | 19826559 | 19832027 | 19832110 | 236 | 178 | [illegible] |
| SE | 87138 | ENSG00000104613 | INTS10 | chr8 | + | 19832027 | 19832110 | 19830405 | 19830559 | 19833168 | 19833321 | 284 | 211 | [illegible] |
| SE | 87139 | ENSG00000104613 | INTS10 | chr8 | + | 19832027 | 19832110 | 19830405 | 19830559 | 19837051 | 19837160 | 89 | 74 | [illegible] |
| SE | 87140 | ENSG00000104613 | INTS10 | chr8 | + | 19833168 | 19833321 | 19830405 | 19830559 | 19837051 | 19837160 | 145 | 107 | [illegible] |
| SE | 87141 | ENSG00000104613 | INTS10 | chr8 | + | 19833168 | 19833321 | 19832027 | 19832110 | 19837051 | 19837160 | 345 | 248 | [illegible] |
| SE | 87142 | ENSG00000104613 | INTS10 | chr8 | + | 19837051 | 19837160 | 19832027 | 19832110 | 19842847 | 19842927 | 73 | 41 | [illegible] |
| SE | 87143 | ENSG00000104613 | INTS10 | chr8 | + | 19841826 | 19841955 | 19837080 | 19837160 | 19842847 | 19842927 | 1 | 3 | [illegible] |
| SE | 87144 | ENSG00000104613 | INTS10 | chr8 | + | 19842847 | 19842927 | 19837051 | 19837160 | 19844075 | 19844237 | 179 | 107 | [illegible] |
| SE | 87145 | ENSG00000104613 | INTS10 | chr8 | + | 19844075 | 19844238 | 19837051 | 19837160 | 19845703 | 19845797 | 178 | 66 | [illegible] |
| SE | 87146 | ENSG00000104613 | INTS10 | chr8 | + | 19849161 | 19849239 | 19845703 | 19845797 | 19851648 | 19852061 | 0 | 0 | [illegible] |
| SE | 87459 | ENSG00000102172 | SMS | chrX | + | 21972506 | 21972571 | 21967195 | 21967316 | 21977060 | 21977236 | 308 | 208 | [illegible] |
| SE | 87606 | ENSG00000102078 | SLC25A14 | chrX | + | 130340106 | 130340344 | 130339887 | 130339976 | 130345181 | 130345275 | 0 | 0 | [illegible] |
| SE | 87607 | ENSG00000102078 | SLC25A14 | chrX | + | 130340106 | 130340353 | 130339912 | 130339976 | 130345181 | 130345275 | 0 | 0 | [illegible] |
| SE | 87608 | ENSG00000102078 | SLC25A14 | chrX | + | 130345181 | 130345275 | 130339887 | 130339976 | 130346543 | 130346691 | 7 | 9 | [illegible] |
| SE | 87609 | ENSG00000102078 | SLC25A14 | chrX | + | 130346543 | 130346691 | 130339887 | 130339976 | 130349250 | 130349345 | 21 | 22 | [illegible] |
| SE | 87768 | ENSG00000196459 | TRAPPC2 | chrX | - | 13734043 | 13734185 | 13720427 | 13720497 | 13734524 | 13734620 | 27 | 11 | [illegible] |
| SE | 87770 | ENSG00000102158 | MAGT1 | chrX | - | 77830804 | 77830895 | 77825746 | 77829235 | 77841245 | 77841320 | 226 | 165 | [illegible] |
| SE | 87771 | ENSG00000102158 | MAGT1 | chrX | - | 77855500 | 77855590 | 77853900 | 77853964 | 77856732 | 77856873 | 172 | 112 | [illegible] |
| SE | 88033 | ENSG00000156171 | DRAM2 | chr1 | - | 111120515 | 111120693 | 111119876 | 111119959 | 111124741 | 111124881 | 498 | 326 | [illegible] |
| SE | 88038 | ENSG00000156171 | DRAM2 | chr1 | - | 111124741 | 111124881 | 111119876 | 111119959 | 111131423 | 111131568 | 1 | 2 | [illegible] |
| SE | 88042 | ENSG00000156171 | DRAM2 | chr1 | - | 111126226 | 111126294 | 111124741 | 111124881 | 111131423 | 111131568 | 57 | 40 | [illegible] |
| SE | 88044 | ENSG00000156171 | DRAM2 | chr1 | - | 111139500 | 111139666 | 111137522 | 111137586 | 111140037 | 111140090 | 90 | 76 | [illegible] |
| SE | 88045 | ENSG00000156171 | DRAM2 | chr1 | - | 111139587 | 111139711 | 111137522 | 111137586 | 111140037 | 111140062 | 5 | 4 | [illegible] |
| SE | 88155 | ENSG00000130227 | XPO7 | chr8 | + | 21969482 | 21969576 | 21966856 | 21967003 | 21970143 | 21970310 | 152 | 111 | [illegible] |
| SE | 88158 | ENSG00000130227 | XPO7 | chr8 | + | 21977769 | 21977843 | 21976355 | 21976521 | 21980083 | 21980203 | 202 | 141 | [illegible] |
| SE | 88159 | ENSG00000130227 | XPO7 | chr8 | + | 21987140 | 21987276 | 21985585 | 21985691 | 21987783 | 21987857 | 220 | 161 | [illegible] |
| SE | 88390 | ENSG00000145041 | DCAF1 | chr3 | - | 51403142 | 51403395 | 51397841 | 51398827 | 51412378 | 51412480 | 203 | 191 | [illegible] |
| SE | 88391 | ENSG00000145041 | DCAF1 | chr3 | - | 51412378 | 51412480 | 51403142 | 51403395 | 51412992 | 51413066 | 236 | 269 | [illegible] |
| SE | 88392 | ENSG00000145041 | DCAF1 | chr3 | - | 51427371 | 51427541 | 51422306 | 51422431 | 51429260 | 51429470 | 88 | 50 | [illegible] |
| SE | 88393 | ENSG00000145041 | DCAF1 | chr3 | - | 51429260 | 51429470 | 51422306 | 51422431 | 51430032 | 51430212 | 44 | 25 | [illegible] |
| SE | 88396 | ENSG00000145041 | DCAF1 | chr3 | - | 51441384 | 51441897 | 51440969 | 51441071 | 51463113 | 51463227 | 49 | 42 | [illegible] |
| SE | 88397 | ENSG00000145041 | DCAF1 | chr3 | - | 51443765 | 51443903 | 51440969 | 51441071 | 51463113 | 51463227 | 136 | 93 | [illegible] |
| SE | 88398 | ENSG00000145041 | DCAF1 | chr3 | - | 51443765 | 51443903 | 51441384 | 51441897 | 51463113 | 51463227 | 274 | 185 | [illegible] |
| SE | 88401 | ENSG00000145041 | DCAF1 | chr3 | - | 51470928 | 51471005 | 51466802 | 51466876 | 51483718 | 51483836 | 277 | 180 | [illegible] |
| SE | 88402 | ENSG00000145041 | DCAF1 | chr3 | - | 51483718 | 51483836 | 51470928 | 51471005 | 51496733 | 51496780 | 144 | 102 | [illegible] |
| SE | 88403 | ENSG00000145041 | DCAF1 | chr3 | - | 51483718 | 51483836 | 51470928 | 51471005 | 51499872 | 51499994 | 121 | 82 | [illegible] |
| SE | 88404 | ENSG00000145041 | DCAF1 | chr3 | - | 51496733 | 51496780 | 51470928 | 51471005 | 51499872 | 51500015 | 2 | 0 | [illegible] |
| SE | 88438 | ENSG00000221914 | PPP2R2A | chr8 | + | 26293665 | 26293740 | 26291507 | 26291826 | 26338889 | 26338987 | 108 | 107 | [illegible] |
| SE | 88439 | ENSG00000221914 | PPP2R2A | chr8 | + | 26293665 | 26293860 | 26291527 | 26291826 | 26338889 | 26338987 | 25 | 17 | [illegible] |
| SE | 88441 | ENSG00000221914 | PPP2R2A | chr8 | + | 26333485 | 26333570 | 26293665 | 26293860 | 26338889 | 26338987 | 12 | 3 | [illegible] |
| SE | 88442 | ENSG00000221914 | PPP2R2A | chr8 | + | 26338889 | 26338987 | 26293571 | 26293740 | 26354467 | 26354550 | 201 | 179 | [illegible] |
| SE | 88443 | ENSG00000221914 | PPP2R2A | chr8 | + | 26338889 | 26338987 | 26293665 | 26293860 | 26354467 | 26354550 | 117 | 91 | [illegible] |
| SE | 88444 | ENSG00000221914 | PPP2R2A | chr8 | + | 26339902 | 26339973 | 26338889 | 26338987 | 26350598 | 26350681 | 2 | 2 | [illegible] |
| SE | 88448 | ENSG00000221914 | PPP2R2A | chr8 | + | 26354467 | 26354633 | 26338889 | 26338987 | 26360168 | 26360182 | 450 | 395 | [illegible] |
| SE | 88449 | ENSG00000221914 | PPP2R2A | chr8 | + | 26358906 | 26359032 | 26354467 | 26354633 | 26360168 | 26360281 | 6 | 2 | [illegible] |
| SE | 88450 | ENSG00000221914 | PPP2R2A | chr8 | + | 26358911 | 26359032 | 26354467 | 26354633 | 26360168 | 26360281 | 3 | 2 | [illegible] |
| SE | 88451 | ENSG00000221914 | PPP2R2A | chr8 | + | 26360168 | 26360281 | 26353810 | 26354633 | 26360973 | 26361001 | 432 | 357 | [illegible] |
| SE | 88646 | ENSG00000187735 | TCEA1 | chr8 | - | 53970391 | 53970463 | 53966551 | 53968112 | 53984362 | 53984517 | 60 | 23 | [illegible] |
| SE | 88647 | ENSG00000187735 | TCEA1 | chr8 | - | 53978522 | 53978697 | 53970391 | 53970463 | 53979024 | 53979171 | 0 | 0 | [illegible] |
| SE | 88648 | ENSG00000187735 | TCEA1 | chr8 | - | 53979024 | 53979171 | 53970391 | 53970463 | 53984362 | 53984517 | 893 | 632 | [illegible] |
| SE | 88651 | ENSG00000187735 | TCEA1 | chr8 | - | 53993667 | 53993755 | 53988113 | 53988259 | 53999944 | 54000050 | 1035 | 859 | [illegible] |
| SE | 88652 | ENSG00000187735 | TCEA1 | chr8 | - | 53999944 | 54000050 | 53993375 | 53993755 | 54010429 | 54010492 | 951 | 740 | [illegible] |
| SE | 88653 | ENSG00000187735 | TCEA1 | chr8 | - | 53999944 | 54000050 | 53993375 | 53993755 | 54022062 | 54022448 | 323 | 270 | [illegible] |
| SE | 88655 | ENSG00000187735 | TCEA1 | chr8 | - | 54002891 | 54003086 | 54000009 | 54000050 | 54010429 | 54010492 | 3 | 2 | [illegible] |
| SE | 88656 | ENSG00000187735 | TCEA1 | chr8 | - | 54002891 | 54003092 | 53999944 | 54000050 | 54010429 | 54010492 | 3 | 2 | [illegible] |
| SE | 88657 | ENSG00000187735 | TCEA1 | chr8 | - | 54010429 | 54010492 | 53993375 | 53993755 | 54022062 | 54022125 | 25 | 9 | [illegible] |
| SE | 88659 | ENSG00000187735 | TCEA1 | chr8 | - | 54021525 | 54021586 | 54010429 | 54010492 | 54022062 | 54022310 | 1 | 3 | [illegible] |
| SE | 88882 | ENSG00000167302 | TEPSIN | chr17 | - | 81232314 | 81232518 | 81231846 | 81232021 | 81233431 | 81233503 | 42 | 12 | [illegible] |
| SE | 88883 | ENSG00000167302 | TEPSIN | chr17 | - | 81237550 | 81237660 | 81237386 | 81237459 | 81238513 | 81238873 | 6 | 4 | [illegible] |
| SE | 89247 | ENSG00000137574 | TGS1 | chr8 | + | 55795977 | 55796152 | 55792697 | 55792784 | 55798913 | 55798998 | 139 | 85 | [illegible] |
| SE | 89248 | ENSG00000137574 | TGS1 | chr8 | + | 55798913 | 55799220 | 55795977 | 55796152 | 55802456 | 55802606 | 118 | 97 | [illegible] |
| SE | 89249 | ENSG00000137574 | TGS1 | chr8 | + | 55813054 | 55813118 | 55804892 | 55805036 | 55824580 | 55825448 | 45 | 37 | [illegible] |
| SE | 89472 | ENSG00000064313 | TAF2 | chr8 | - | 119744880 | 119745036 | 119744287 | 119744393 | 119746704 | 119746934 | 6 | 4 | [illegible] |
| SE | 89473 | ENSG00000064313 | TAF2 | chr8 | - | 119744880 | 119745036 | 119744287 | 119744393 | 119756005 | 119756115 | 2 | 2 | [illegible] |
| SE | 89474 | ENSG00000064313 | TAF2 | chr8 | - | 119746704 | 119746934 | 119744287 | 119744393 | 119756005 | 119756115 | 94 | 79 | [illegible] |
| SE | 89475 | ENSG00000064313 | TAF2 | chr8 | - | 119746704 | 119746934 | 119744880 | 119745036 | 119756005 | 119756115 | 56 | 47 | [illegible] |
| SE | 89478 | ENSG00000064313 | TAF2 | chr8 | - | 119783380 | 119783633 | 119781052 | 119781193 | 119785200 | 119785266 | 156 | 136 | [illegible] |
| SE | 89479 | ENSG00000064313 | TAF2 | chr8 | - | 119788337 | 119788447 | 119785200 | 119785266 | 119788789 | 119788904 | 135 | 121 | [illegible] |
| SE | 89480 | ENSG00000064313 | TAF2 | chr8 | - | 119791323 | 119791459 | 119789591 | 119789746 | 119793365 | 119793451 | 206 | 149 | [illegible] |
| SE | 89481 | ENSG00000064313 | TAF2 | chr8 | - | 119791323 | 119791459 | 119789591 | 119789746 | 119795531 | 119795631 | 149 | 126 | [illegible] |
| SE | 89485 | ENSG00000064313 | TAF2 | chr8 | - | 119801793 | 119802025 | 119797661 | 119797846 | 119803877 | 119804019 | 186 | 69 | [illegible] |
| SE | 89486 | ENSG00000064313 | TAF2 | chr8 | - | 119803877 | 119804019 | 119801793 | 119802025 | 119806282 | 119806401 | 120 | 62 | [illegible] |
| SE | 89600 | ENSG00000172167 | MTBP | chr8 | + | 120470819 | 120470937 | 120463686 | 120463761 | 120488158 | 120488332 | 14 | 7 | [illegible] |
| SE | 89601 | ENSG00000172167 | MTBP | chr8 | + | 120509933 | 120510029 | 120506705 | 120506861 | 120515924 | 120516191 | 36 | 4 | [illegible] |
| SE | 90635 | ENSG00000014216 | CAPN1 | chr11 | + | 65183127 | 65183197 | 65182461 | 65182968 | 65183473 | 65183503 | 251 | 118 | [illegible] |
| SE | 90636 | ENSG00000014216 | CAPN1 | chr11 | + | 65186169 | 65186338 | 65185916 | 65186050 | 65187214 | 65187298 | 324 | 126 | [illegible] |
| SE | 90637 | ENSG00000014216 | CAPN1 | chr11 | + | 65187954 | 65188040 | 65187214 | 65187298 | 65188413 | 65188488 | 304 | 125 | [illegible] |
| SE | 90768 | ENSG00000174744 | BRMS1 | chr11 | - | 66338242 | 66338282 | 66337332 | 66337807 | 66338720 | 66338785 | 23 | 9 | [illegible] |
| SE | 90770 | ENSG00000174744 | BRMS1 | chr11 | - | 66338242 | 66338785 | 66337368 | 66337889 | 66340773 | 66340870 | 4 | 5 | [illegible] |
| SE | 90771 | ENSG00000174744 | BRMS1 | chr11 | - | 66338720 | 66338785 | 66337338 | 66337889 | 66340773 | 66340870 | 0 | 0 | [illegible] |
| SE | 90772 | ENSG00000174744 | BRMS1 | chr11 | - | 66340120 | 66340213 | 66338242 | 66338785 | 66340773 | 66340870 | 86 | 18 | [illegible] |
| SE | 91156 | ENSG00000164941 | INTS8 | chr8 | + | 94824892 | 94825067 | 94823286 | 94823561 | 94827262 | 94827353 | 295 | 182 | [illegible] |
| SE | 91159 | ENSG00000164941 | INTS8 | chr8 | + | 94827262 | 94827403 | 94824892 | 94825067 | 94827721 | 94827793 | 486 | 304 | [illegible] |
| SE | 91160 | ENSG00000164941 | INTS8 | chr8 | + | 94827721 | 94827793 | 94824892 | 94825067 | 94828974 | 94829026 | 120 | 121 | [illegible] |
| SE | 91161 | ENSG00000164941 | INTS8 | chr8 | + | 94828974 | 94829026 | 94827721 | 94827793 | 94831991 | 94832001 | 375 | 361 | [illegible] |
| SE | 91162 | ENSG00000164941 | INTS8 | chr8 | + | 94831991 | 94832174 | 94828974 | 94829026 | 94836523 | 94836631 | 761 | 702 | [illegible] |
| SE | 91165 | ENSG00000164941 | INTS8 | chr8 | + | 94838462 | 94838618 | 94836523 | 94836631 | 94841490 | 94841591 | 141 | 91 | [illegible] |
| SE | 91167 | ENSG00000164941 | INTS8 | chr8 | + | 94838493 | 94838618 | 94836523 | 94836631 | 94841490 | 94841591 | 107 | 72 | [illegible] |
| SE | 91168 | ENSG00000164941 | INTS8 | chr8 | + | 94841490 | 94841591 | 94838462 | 94838618 | 94842346 | 94842488 | 276 | 206 | [illegible] |
| SE | 91169 | ENSG00000164941 | INTS8 | chr8 | + | 94841490 | 94841591 | 94838462 | 94838618 | 94849461 | 94849532 | 76 | 48 | [illegible] |
| SE | 91170 | ENSG00000164941 | INTS8 | chr8 | + | 94842346 | 94842488 | 94838462 | 94838618 | 94849461 | 94849532 | 312 | 236 | [illegible] |
| SE | 91172 | ENSG00000164941 | INTS8 | chr8 | + | 94849915 | 94850091 | 94849461 | 94849532 | 94851552 | 94851686 | 511 | 371 | [illegible] |
| SE | 91173 | ENSG00000164941 | INTS8 | chr8 | + | 94849915 | 94850091 | 94849461 | 94849532 | 94853804 | 94853915 | 389 | 301 | [illegible] |
| SE | 91174 | ENSG00000164941 | INTS8 | chr8 | + | 94851552 | 94851686 | 94849461 | 94849532 | 94853804 | 94853915 | 122 | 96 | [illegible] |
| SE | 91175 | ENSG00000164941 | INTS8 | chr8 | + | 94851552 | 94851686 | 94849915 | 94850091 | 94853804 | 94853915 | 240 | 166 | [illegible] |
| SE | 91177 | ENSG00000164941 | INTS8 | chr8 | + | 94859510 | 94859632 | 94856776 | 94856978 | 94865505 | 94865690 | 278 | 181 | [illegible] |
| SE | 91178 | ENSG00000164941 | INTS8 | chr8 | + | 94865505 | 94865690 | 94859510 | 94859632 | 94866157 | 94866191 | 348 | 197 | [illegible] |
| SE | 91180 | ENSG00000164941 | INTS8 | chr8 | + | 94865505 | 94865700 | 94859510 | 94859632 | 94866157 | 94866191 | 174 | 105 | [illegible] |
| SE | 91182 | ENSG00000164941 | INTS8 | chr8 | + | 94866157 | 94866191 | 94865505 | 94865690 | 94867139 | 94867196 | 210 | 130 | [illegible] |
| SE | 91183 | ENSG00000164941 | INTS8 | chr8 | + | 94866311 | 94866351 | 94866157 | 94866191 | 94867139 | 94867196 | 0 | 0 | [illegible] |
| SE | 91186 | ENSG00000164941 | INTS8 | chr8 | + | 94871883 | 94872002 | 94867139 | 94867337 | 94873373 | 94873477 | 476 | 333 | [illegible] |
| SE | 91188 | ENSG00000164941 | INTS8 | chr8 | + | 94876073 | 94876147 | 94873373 | 94873477 | 94876220 | 94876285 | 180 | 141 | [illegible] |
| SE | 91280 | ENSG00000155265 | GOLGA7B | chr10 | + | 97859457 | 97859583 | 97849842 | 97850315 | 97863929 | 97864082 | 11 | 0 | [illegible] |
| SE | 91342 | ENSG00000167632 | TRAPPC9 | chr8 | - | 140252776 | 140252929 | 140221458 | 140221583 | 140275657 | 140275821 | 230 | 153 | [illegible] |
| SE | 91343 | ENSG00000167632 | TRAPPC9 | chr8 | - | 140283888 | 140284021 | 140275657 | 140275821 | 140287607 | 140287734 | 96 | 27 | [illegible] |
| SE | 91344 | ENSG00000167632 | TRAPPC9 | chr8 | - | 140287607 | 140287734 | 140283888 | 140284021 | 140290992 | 140291078 | 98 | 52 | [illegible] |
| SE | 91346 | ENSG00000167632 | TRAPPC9 | chr8 | - | 140300468 | 140300614 | 140290992 | 140291078 | 140311247 | 140311374 | 162 | 107 | [illegible] |
| SE | 91347 | ENSG00000167632 | TRAPPC9 | chr8 | - | 140311247 | 140311374 | 140300468 | 140300614 | 140360049 | 140360193 | 160 | 75 | [illegible] |
| SE | 91348 | ENSG00000167632 | TRAPPC9 | chr8 | - | 140360049 | 140360193 | 140311247 | 140311374 | 140370963 | 140371180 | 79 | 34 | [illegible] |
| SE | 91349 | ENSG00000167632 | TRAPPC9 | chr8 | - | 140370963 | 140371180 | 140360049 | 140360193 | 140397619 | 140397745 | 110 | 63 | [illegible] |
| SE | 91351 | ENSG00000167632 | TRAPPC9 | chr8 | - | 140426614 | 140426641 | 140405576 | 140405698 | 140439051 | 140439197 | 12 | 15 | [illegible] |
| SE | 91352 | ENSG00000167632 | TRAPPC9 | chr8 | - | 140435111 | 140435240 | 140405576 | 140405698 | 140439051 | 140439197 | 86 | 40 | [illegible] |
| SE | 91353 | ENSG00000167632 | TRAPPC9 | chr8 | - | 140435111 | 140435240 | 140426614 | 140426641 | 140439051 | 140439197 | 138 | 68 | [illegible] |
| SE | 91354 | ENSG00000167632 | TRAPPC9 | chr8 | - | 140439051 | 140439197 | 140435111 | 140435240 | 140450789 | 140450930 | 124 | 67 | [illegible] |
| SE | 92133 | ENSG00000108666 | C17orf75 | chr17 | - | 32333416 | 32333520 | 32324564 | 32324774 | 32334468 | 32334605 | 37 | 27 | [illegible] |
| SE | 92444 | ENSG00000136950 | ARPC5L | chr9 | + | 124874974 | 124875151 | 124872588 | 124873764 | 124876877 | 124877401 | 382 | 254 | [illegible] |
| SE | 92579 | ENSG00000137054 | POLR1E | chr9 | + | 37498090 | 37498224 | 37495889 | 37495986 | 37500839 | 37500921 | 143 | 59 | [illegible] |
| SE | 93244 | ENSG00000136937 | NCBP1 | chr9 | + | 97643203 | 97643360 | 97641561 | 97641662 | 97645116 | 97645224 | 157 | 126 | [illegible] |
| SE | 93245 | ENSG00000136937 | NCBP1 | chr9 | + | 97643203 | 97643360 | 97641561 | 97641662 | 97647491 | 97647561 | 49 | 40 | [illegible] |
| SE | 93246 | ENSG00000136937 | NCBP1 | chr9 | + | 97645116 | 97645224 | 97641561 | 97641662 | 97647491 | 97647561 | 6 | 4 | [illegible] |
| SE | 93248 | ENSG00000136937 | NCBP1 | chr9 | + | 97645116 | 97645224 | 97643203 | 97643360 | 97647491 | 97647561 | 98 | 70 | [illegible] |
| SE | 93249 | ENSG00000136937 | NCBP1 | chr9 | + | 97645610 | 97645732 | 97643203 | 97643360 | 97647491 | 97647561 | 64 | 58 | [illegible] |
| SE | 93253 | ENSG00000136937 | NCBP1 | chr9 | + | 97662953 | 97663047 | 97662041 | 97662144 | 97664339 | 97664443 | 196 | 171 | [illegible] |
| SE | 93892 | ENSG00000184465 | WDR27 | chr6 | - | 169572418 | 169572540 | 169558320 | 169559927 | 169582835 | 169582934 | 0 | 0 | [illegible] |
| SE | 93893 | ENSG00000184465 | WDR27 | chr6 | - | 169636370 | 169636504 | 169634427 | 169634525 | 169638538 | 169638660 | 0 | 2 | [illegible] |
| SE | 93894 | ENSG00000184465 | WDR27 | chr6 | - | 169638538 | 169638660 | 169634427 | 169634525 | 169643696 | 169643786 | 7 | 7 | [illegible] |
| SE | 93895 | ENSG00000184465 | WDR27 | chr6 | - | 169638538 | 169638660 | 169634427 | 169634525 | 169658275 | 169658358 | 2 | 2 | [illegible] |
| SE | 93896 | ENSG00000184465 | WDR27 | chr6 | - | 169643696 | 169643786 | 169634427 | 169634525 | 169658275 | 169658358 | 1 | 2 | [illegible] |
| SE | 93897 | ENSG00000184465 | WDR27 | chr6 | - | 169643696 | 169643786 | 169638538 | 169638660 | 169647772 | 169647870 | 7 | 12 | [illegible] |
| SE | 93898 | ENSG00000184465 | WDR27 | chr6 | - | 169643696 | 169643786 | 169638538 | 169638660 | 169658275 | 169658358 | 4 | 1 | [illegible] |
| SE | 93899 | ENSG00000184465 | WDR27 | chr6 | - | 169651929 | 169652008 | 169649197 | 169649275 | 169658275 | 169658358 | 4 | 4 | [illegible] |
| SE | 93900 | ENSG00000184465 | WDR27 | chr6 | - | 169659085 | 169659207 | 169658275 | 169658358 | 169659450 | 169659518 | 8 | 11 | [illegible] |
| SE | 93901 | ENSG00000184465 | WDR27 | chr6 | - | 169659450 | 169659518 | 169659085 | 169659207 | 169660662 | 169660692 | 11 | 6 | [illegible] |
| SE | 93903 | ENSG00000184465 | WDR27 | chr6 | - | 169663562 | 169663649 | 169662303 | 169662424 | 169664165 | 169664286 | 0 | 4 | [illegible] |
| SE | 93904 | ENSG00000184465 | WDR27 | chr6 | - | 169663562 | 169663706 | 169662303 | 169662424 | 169664165 | 169664632 | 0 | 2 | [illegible] |
| SE | 93905 | ENSG00000184465 | WDR27 | chr6 | - | 169663562 | 169663878 | 169662303 | 169662424 | 169664165 | 169664286 | 0 | 6 | [illegible] |
| SE | 93906 | ENSG00000184465 | WDR27 | chr6 | - | 169664165 | 169664286 | 169662303 | 169662424 | 169672254 | 169672396 | 9 | 2 | [illegible] |
| SE | 93907 | ENSG00000184465 | WDR27 | chr6 | - | 169665485 | 169665556 | 169664165 | 169664286 | 169667135 | 169667187 | 1 | 3 | [illegible] |
| SE | 93911 | ENSG00000184465 | WDR27 | chr6 | - | 169667135 | 169667187 | 169664285 | 169665556 | 169672254 | 169672396 | 2 | 0 | [illegible] |
| SE | 93914 | ENSG00000184465 | WDR27 | chr6 | - | 169667135 | 169668776 | 169665134 | 169665556 | 169672254 | 169672396 | 2 | 0 | [illegible] |
| SE | 93916 | ENSG00000184465 | WDR27 | chr6 | - | 169667981 | 169668185 | 169667135 | 169667187 | 169670568 | 169670693 | 1 | 4 | [illegible] |
| SE | 93918 | ENSG00000184465 | WDR27 | chr6 | - | 169670568 | 169670693 | 169665485 | 169665556 | 169672254 | 169672396 | 5 | 6 | [illegible] |
| SE | 93919 | ENSG00000184465 | WDR27 | chr6 | - | 169670568 | 169670693 | 169666917 | 169668776 | 169672254 | 169672396 | 8 | 7 | [illegible] |
| SE | 93920 | ENSG00000184465 | WDR27 | chr6 | - | 169670568 | 169670693 | 169667135 | 169667187 | 169672254 | 169672396 | 5 | 6 | [illegible] |
| SE | 93924 | ENSG00000184465 | WDR27 | chr6 | - | 169694462 | 169695065 | 169689126 | 169689308 | 169701550 | 169701854 | 3 | 2 | [illegible] |
| SE | 94012 | ENSG00000099381 | SETD1A | chr16 | + | 30963432 | 30963554 | 30961266 | 30961537 | 30964093 | 30964323 | 108 | 54 | [illegible] |
| SE | 94013 | ENSG00000099381 | SETD1A | chr16 | + | 30964093 | 30964323 | 30963432 | 30963554 | 30964611 | 30965461 | 76 | 25 | [illegible] |
| SE | 94016 | ENSG00000099381 | SETD1A | chr16 | + | 30967500 | 30967588 | 30966883 | 30967060 | 30969304 | 30969462 | 147 | 92 | [illegible] |
| SE | 94017 | ENSG00000099381 | SETD1A | chr16 | + | 30969304 | 30969462 | 30967500 | 30967588 | 30969601 | 30969689 | 194 | 115 | [illegible] |
| SE | 94161 | ENSG00000198000 | NOL8 | chr9 | - | 92319942 | 92320137 | 92319220 | 92319356 | 92321667 | 92321746 | 18 | 5 | [illegible] |
| SE | 94162 | ENSG00000198000 | NOL8 | chr9 | - | 92319942 | 92320379 | 92319220 | 92319356 | 92321667 | 92321746 | 19 | 7 | [illegible] |
| SE | 94163 | ENSG00000198000 | NOL8 | chr9 | - | 92321667 | 92321746 | 92319987 | 92320382 | 92323440 | 92323503 | 36 | 27 | [illegible] |
| SE | 94164 | ENSG00000198000 | NOL8 | chr9 | - | 92323235 | 92323351 | 92321667 | 92321746 | 92323440 | 92323503 | 29 | 22 | [illegible] |
| SE | 94165 | ENSG00000198000 | NOL8 | chr9 | - | 92323440 | 92323503 | 92321667 | 92321746 | 92324022 | 92324080 | 93 | 86 | [illegible] |
| SE | 94445 | ENSG00000099958 | DERL3 | chr22 | - | 23838351 | 23838445 | 23837658 | 23837854 | 23838563 | 23838637 | 1 | 2 | [illegible] |
| SE | 94923 | ENSG00000148655 | LRMDA | chr10 | + | 76058665 | 76058783 | 76047163 | 76047303 | 76324400 | 76324485 | 29 | 30 | [illegible] |
| SE | 94926 | ENSG00000148655 | LRMDA | chr10 | + | 76535797 | 76535987 | 76324400 | 76324485 | 76557208 | 76557366 | 0 | 1 | [illegible] |
| SE | 95110 | ENSG00000096872 | IFT74 | chr9 | + | 26992382 | 26992677 | 26990133 | 26990195 | 27009019 | 27009158 | 0 | 3 | [illegible] |
| SE | 95111 | ENSG00000096872 | IFT74 | chr9 | + | 27009019 | 27009158 | 26990133 | 26990195 | 27011905 | 27011968 | 45 | 68 | [illegible] |
| SE | 95850 | ENSG00000107263 | RAPGEF1 | chr9 | - | 131598198 | 131598310 | 131588800 | 131588986 | 131621795 | 131621838 | 1 | 9 | [illegible] |
| SE | 95852 | ENSG00000107263 | RAPGEF1 | chr9 | - | 131603960 | 131604053 | 131602060 | 131602149 | 131604930 | 131605188 | 5 | 0 | [illegible] |
| SE | 95854 | ENSG00000107263 | RAPGEF1 | chr9 | - | 131625921 | 131626422 | 131621795 | 131621998 | 131643238 | 131643417 | 251 | 184 | [illegible] |
| SE | 95856 | ENSG00000107263 | RAPGEF1 | chr9 | - | 131628548 | 131628672 | 131627912 | 131628096 | 131629101 | 131629254 | 462 | 167 | [illegible] |
| SE | 95857 | ENSG00000107263 | RAPGEF1 | chr9 | - | 131629101 | 131629254 | 131625921 | 131626422 | 131643238 | 131643391 | 1 | 0 | [illegible] |
| SE | 95861 | ENSG00000107263 | RAPGEF1 | chr9 | - | 131638634 | 131638791 | 131630235 | 131630324 | 131643238 | 131643391 | 989 | 474 | [illegible] |
| SE | 95862 | ENSG00000107263 | RAPGEF1 | chr9 | - | 131643238 | 131643417 | 131625921 | 131626422 | 131650128 | 131650242 | 210 | 102 | [illegible] |
| SE | 95863 | ENSG00000107263 | RAPGEF1 | chr9 | - | 131643238 | 131643417 | 131625921 | 131626422 | 131739769 | 131739944 | 2 | 0 | [illegible] |
| SE | 95865 | ENSG00000107263 | RAPGEF1 | chr9 | - | 131643238 | 131643417 | 131638634 | 131638791 | 131650809 | 131650949 | 623 | 313 | [illegible] |
| SE | 95868 | ENSG00000107263 | RAPGEF1 | chr9 | - | 131650809 | 131650949 | 131643238 | 131643417 | 131739769 | 131739944 | 135 | 91 | [illegible] |
| SE | 96382 | ENSG00000197563 | PIGN | chr18 | - | 62070389 | 62070506 | 62052548 | 62052957 | 62072672 | 62072725 | 1 | 0 | [illegible] |
| SE | 96384 | ENSG00000197563 | PIGN | chr18 | - | 62072320 | 62072365 | 62070389 | 62070506 | 62072672 | 62072725 | 0 | 1 | [illegible] |
| SE | 96385 | ENSG00000197563 | PIGN | chr18 | - | 62090475 | 62090578 | 62088390 | 62088842 | 62095847 | 62095950 | 142 | 53 | [illegible] |
| SE | 96388 | ENSG00000197563 | PIGN | chr18 | - | 62107467 | 62107586 | 62106985 | 62107085 | 62109833 | 62110150 | 1 | 0 | [illegible] |
| SE | 96391 | ENSG00000197563 | PIGN | chr18 | - | 62123506 | 62123544 | 62118546 | 62118657 | 62138242 | 62138298 | 0 | 3 | [illegible] |
| SE | 96394 | ENSG00000197563 | PIGN | chr18 | - | 62154544 | 62154651 | 62148213 | 62148253 | 62157128 | 62157227 | 48 | 59 | [illegible] |
| SE | 96395 | ENSG00000197563 | PIGN | chr18 | - | 62154544 | 62154651 | 62148213 | 62148338 | 62157128 | 62157227 | 154 | 122 | [illegible] |
| SE | 96396 | ENSG00000197563 | PIGN | chr18 | - | 62154544 | 62154651 | 62148213 | 62148338 | 62161132 | 62161337 | 84 | 56 | [illegible] |
| SE | 96398 | ENSG00000197563 | PIGN | chr18 | - | 62157128 | 62157227 | 62148213 | 62148338 | 62157686 | 62157808 | 41 | 50 | [illegible] |
| SE | 96399 | ENSG00000197563 | PIGN | chr18 | - | 62157128 | 62157227 | 62148213 | 62148338 | 62161132 | 62161226 | 0 | 0 | [illegible] |
| SE | 96401 | ENSG00000197563 | PIGN | chr18 | - | 62157686 | 62157808 | 62148213 | 62148338 | 62161132 | 62161226 | 23 | 28 | [illegible] |
| SE | 96402 | ENSG00000197563 | PIGN | chr18 | - | 62157686 | 62157808 | 62157128 | 62157227 | 62161132 | 62161226 | 80 | 109 | [illegible] |
| SE | 96406 | ENSG00000197563 | PIGN | chr18 | - | 62163530 | 62163656 | 62161132 | 62161226 | 62186843 | 62186989 | 11 | 8 | [illegible] |
| SE | 96607 | ENSG00000030066 | NUP160 | chr11 | - | 47780342 | 47780447 | 47778086 | 47779194 | 47783072 | 47783198 | 181 | 130 | [illegible] |
| SE | 96608 | ENSG00000030066 | NUP160 | chr11 | - | 47788181 | 47788305 | 47786452 | 47786554 | 47788500 | 47788611 | 66 | 34 | [illegible] |
| SE | 96610 | ENSG00000030066 | NUP160 | chr11 | - | 47807069 | 47807140 | 47806152 | 47806312 | 47808395 | 47808529 | 53 | 40 | [illegible] |
| SE | 96612 | ENSG00000030066 | NUP160 | chr11 | - | 47818055 | 47818124 | 47815945 | 47816029 | 47819373 | 47819458 | 129 | 92 | [illegible] |
| SE | 96613 | ENSG00000030066 | NUP160 | chr11 | - | 47819373 | 47819458 | 47818055 | 47818124 | 47821723 | 47821821 | 118 | 86 | [illegible] |
| SE | 96614 | ENSG00000030066 | NUP160 | chr11 | - | 47821723 | 47821821 | 47819373 | 47819458 | 47822086 | 47822164 | 141 | 93 | [illegible] |
| SE | 96617 | ENSG00000030066 | NUP160 | chr11 | - | 47840377 | 47840588 | 47837544 | 47837623 | 47847847 | 47847959 | 78 | 72 | [illegible] |
| SE | 96618 | ENSG00000030066 | NUP160 | chr11 | - | 47840377 | 47840588 | 47839842 | 47840065 | 47847847 | 47847959 | 154 | 123 | [illegible] |
| SE | 96619 | ENSG00000030066 | NUP160 | chr11 | - | 47847847 | 47847959 | 47837544 | 47837623 | 47848218 | 47848313 | 28 | 13 | [illegible] |
| SE | 96637 | ENSG00000135372 | NAT10 | chr11 | + | 34108210 | 34108333 | 34105628 | 34105792 | 34108741 | 34108833 | 163 | 86 | [illegible] |
| SE | 96639 | ENSG00000135372 | NAT10 | chr11 | + | 34122458 | 34122592 | 34118395 | 34118503 | 34127462 | 34127599 | 71 | 33 | [illegible] |
| SE | 96640 | ENSG00000135372 | NAT10 | chr11 | + | 34130812 | 34130937 | 34127462 | 34127599 | 34131380 | 34131531 | 149 | 95 | [illegible] |
| SE | 96641 | ENSG00000135372 | NAT10 | chr11 | + | 34139384 | 34139495 | 34139190 | 34139287 | 34140399 | 34140572 | 139 | 77 | [illegible] |
| SE | 96882 | ENSG00000100731 | PCNX1 | chr14 | + | 70962225 | 70962331 | 70946881 | 70947123 | 70976941 | 70978648 | 229 | 161 | [illegible] |
| SE | 96883 | ENSG00000100731 | PCNX1 | chr14 | + | 70969858 | 70969999 | 70969020 | 70969110 | 70976941 | 70977151 | 0 | 4 | [illegible] |
| SE | 96886 | ENSG00000100731 | PCNX1 | chr14 | + | 70988566 | 70988699 | 70976941 | 70978648 | 71011491 | 71011549 | 48 | 20 | [illegible] |
| SE | 96891 | ENSG00000100731 | PCNX1 | chr14 | + | 70995740 | 70995925 | 70976941 | 70978648 | 71012999 | 71013202 | 9 | 7 | [illegible] |
| SE | 96893 | ENSG00000100731 | PCNX1 | chr14 | + | 70995740 | 70995925 | 70988566 | 70988699 | 71012999 | 71013202 | 15 | 7 | [illegible] |
| SE | 96897 | ENSG00000100731 | PCNX1 | chr14 | + | 71009633 | 71009724 | 70988566 | 70988699 | 71012999 | 71013202 | 0 | 1 | [illegible] |
| SE | 96898 | ENSG00000100731 | PCNX1 | chr14 | + | 71009633 | 71009724 | 70995740 | 70995925 | 71012999 | 71013202 | 20 | 8 | [illegible] |
| SE | 96899 | ENSG00000100731 | PCNX1 | chr14 | + | 71011491 | 71011549 | 70976941 | 70978648 | 71012999 | 71013202 | 9 | 5 | [illegible] |

| | | | | | | | | | | | | | | | | | | | | | | | | | | | | | | | | | | | | | | | | | | | | | | |
|---|---|---|---|---|---|---|---|---|---|---|---|---|---|---|---|---|---|---|---|---|---|---|---|---|---|---|---|---|---|---|---|---|---|---|---|---|---|---|---|---|---|---|---|---|---|---|
| SE | 96900 | ENSG00000100731 | PCNX1 | chr14 | + | 71011491 | 71011549 | 70988566 | 70988699 | 71012999 | 71013202 | 10 | 5 | 11 | 1 | 12 | 1 | 0 | 0 | 0 | 0 | 0 | 11 | 5 | 0 | 0 | 0 | 0 | 0 | 0 | 0 | 207 | 149 | 1 | 1 | 0.878 | 1 | 1 | 1 | 1 | NA | 1 | 1 | NA | NA | -0.024 |
| SE | 96902 | ENSG00000100731 | PCNX1 | chr14 | + | 71019008 | 71019162 | 71012984 | 71013202 | 71023199 | 71023232 | 229 | 143 | 233 | 253 | 267 | 0 | 0 | 0 | 1 | 0 | 122 | 245 | 239 | 64 | 100 | 0 | 0 | 0 | 0 | 0 | 298 | 149 | 1 | 1 | 1 | 1 | 1 | 0.992 | 1 | 1 | 1 | 1 | 1 | 1 | -0.002 |
| SE | 96903 | ENSG00000100731 | PCNX1 | chr14 | + | 71026116 | 71026288 | 71023199 | 71023232 | 71026771 | 71026882 | 213 | 121 | 194 | 205 | 298 | 3 | 0 | 0 | 0 | 0 | 31 | 191 | 200 | 94 | 36 | 0 | 0 | 0 | 0 | 0 | 298 | 149 | 1 | 1 | 0.973 | 1 | 1 | 1 | 1 | 1 | 1 | 1 | 1 | 1 | -0.005 |
| SE | 96904 | ENSG00000100731 | PCNX1 | chr14 | + | 71036064 | 71036157 | 71033930 | 71034036 | 71045132 | 71045283 | 107 | 64 | 133 | 115 | 243 | 1 | 0 | 0 | 2 | 3 | 20 | 105 | 88 | 30 | 19 | 0 | 0 | 0 | 0 | 0 | 242 | 149 | 1 | 1 | 0.985 | 1 | 1 | 0.973 | 0.98 | 1 | 1 | 1 | 1 | 1 | -0.012 |
| SE | 96908 | ENSG00000100731 | PCNX1 | chr14 | + | 71051882 | 71052012 | 71047806 | 71047984 | 71073544 | 71073759 | 0 | 0 | 0 | 2 | 5 | 0 | 1 | 0 | 0 | 0 | 8 | 0 | 5 | 0 | 0 | 0 | 0 | 0 | 0 | 0 | 279 | 149 | 1 | 1 | NA | 0 | NA | 1 | 1 | 1 | NA | 1 | NA | NA | -0.333 |
| SE | 96909 | ENSG00000100731 | PCNX1 | chr14 | + | 71053296 | 71053412 | 71051882 | 71052012 | 71055503 | 71055562 | 1 | 1 | 0 | 0 | 0 | 138 | 89 | 111 | 147 | 204 | 0 | 0 | 1 | 0 | 0 | 28 | 130 | 167 | 80 | 66 | 265 | 149 | 1 | 1 | 0.004 | 0.006 | 0 | 0 | 0 | 0 | 0 | 0.003 | 0 | 0 | 0.001 |
| SE | 96910 | ENSG00000100731 | PCNX1 | chr14 | + | 71053296 | 71053412 | 71051882 | 71052012 | 71073544 | 71073759 | 1 | 0 | 0 | 0 | 0 | 0 | 0 | 0 | 0 | 0 | 0 | 0 | 1 | 0 | 0 | 0 | 0 | 1 | 0 | 0 | 265 | 149 | 1 | 1 | 1 | NA | NA | NA | NA | NA | NA | 0.36 | NA | NA | 0.64 |
| SE | 96911 | ENSG00000100731 | PCNX1 | chr14 | + | 71055503 | 71055562 | 71047806 | 71047984 | 71073544 | 71073798 | 6 | 0 | 1 | 0 | 4 | 0 | 1 | 0 | 0 | 0 | 0 | 4 | 2 | 0 | 0 | 0 | 0 | 0 | 0 | 0 | 208 | 149 | 1 | 1 | 1 | 0 | 1 | NA | 1 | NA | 1 | 1 | NA | NA | -0.25 |
| SE | 96912 | ENSG00000100731 | PCNX1 | chr14 | + | 71055503 | 71055562 | 71051882 | 71052012 | 71073544 | 71073798 | 67 | 45 | 56 | 83 | 128 | 0 | 0 | 0 | 0 | 0 | 28 | 54 | 90 | 54 | 41 | 0 | 0 | 1 | 0 | 0 | 208 | 149 | 1 | 1 | 1 | 1 | 1 | 1 | 1 | 1 | 1 | 0.985 | 1 | 1 | 0.003 |
| SE | 96915 | ENSG00000100731 | PCNX1 | chr14 | + | 71101989 | 71102220 | 71089191 | 71089342 | 71103394 | 71103669 | 221 | 141 | 222 | 205 | 412 | 0 | 0 | 0 | 0 | 0 | 133 | 229 | 270 | 116 | 94 | 0 | 7 | 0 | 0 | 0 | 298 | 149 | 1 | 1 | 1 | 1 | 1 | 1 | 1 | 1 | 0.942 | 1 | 1 | 1 | 0.012 |
| SE | 96916 | ENSG00000100731 | PCNX1 | chr14 | + | 71103394 | 71103669 | 71101989 | 71102220 | 71105234 | 71105440 | 310 | 131 | 219 | 280 | 441 | 0 | 0 | 0 | 0 | 3 | 89 | 225 | 324 | 127 | 107 | 0 | 0 | 0 | 0 | 0 | 298 | 149 | 1 | 1 | 1 | 1 | 1 | 1 | 0.987 | 1 | 1 | 1 | 1 | 1 | -0.003 |
| SE | 96917 | ENSG00000100731 | PCNX1 | chr14 | + | 71108603 | 71109046 | 71105234 | 71105440 | 71109451 | 71109592 | 464 | 359 | 436 | 492 | 364 | 0 | 0 | 0 | 0 | 0 | 188 | 559 | 667 | 278 | 91 | 0 | 0 | 4 | 2 | 0 | 298 | 149 | 1 | 1 | 1 | 1 | 1 | 1 | 1 | 1 | 1 | 0.988 | 0.986 | 1 | 0.005 |
| SE | 96995 | ENSG00000153339 | TRAPPC8 | chr18 | - | 31832083 | 31832173 | 31829196 | 31830989 | 31846715 | 31846817 | 77 | 54 | 62 | 58 | 59 | 1 | 5 | 0 | 0 | 4 | 0 | 29 | 19 | 3 | 0 | 0 | 0 | 1 | 2 | 0 | 239 | 149 | 1 | 1 | 0.98 | 0.871 | 1 | 1 | 0.902 | NA | 1 | 0.922 | 0.483 | NA | 0.149 |
| SE | 96997 | ENSG00000153339 | TRAPPC8 | chr18 | - | 31839311 | 31839457 | 31832083 | 31832173 | 31846715 | 31846817 | 432 | 345 | 326 | 376 | 513 | 2 | 1 | 0 | 0 | 2 | 32 | 201 | 175 | 77 | 57 | 0 | 0 | 1 | 0 | 0 | 295 | 149 | 1 | 1 | 0.991 | 0.994 | 1 | 1 | 0.992 | 1 | 1 | 0.989 | 1 | 1 | -0.002 |
| SE | 96998 | ENSG00000153339 | TRAPPC8 | chr18 | - | 31849565 | 31849739 | 31846715 | 31846817 | 31852445 | 31852504 | 537 | 418 | 393 | 465 | 579 | 0 | 1 | 0 | 1 | 2 | 62 | 228 | 196 | 113 | 50 | 0 | 0 | 0 | 0 | 0 | 298 | 149 | 1 | 1 | 1 | 0.995 | 1 | 0.996 | 0.993 | 1 | 1 | 1 | 1 | 1 | -0.003 |
| SE | 96999 | ENSG00000153339 | TRAPPC8 | chr18 | - | 31853848 | 31853945 | 31852594 | 31852663 | 31855659 | 31855807 | 308 | 223 | 182 | 281 | 389 | 0 | 0 | 1 | 0 | 0 | 7 | 85 | 130 | 27 | 0 | 0 | 0 | 0 | 0 | 0 | 246 | 149 | 1 | 1 | 1 | 1 | 0.991 | 1 | 1 | 1 | 1 | 1 | 1 | NA | -0.002 |
| SE | 97000 | ENSG00000153339 | TRAPPC8 | chr18 | - | 31855659 | 31855807 | 31853848 | 31853945 | 31857539 | 31857720 | 319 | 263 | 203 | 276 | 406 | 0 | 1 | 2 | 0 | 2 | 11 | 135 | 154 | 27 | 0 | 0 | 0 | 0 | 0 | 0 | 297 | 149 | 1 | 1 | 1 | 0.992 | 0.981 | 1 | 0.99 | 1 | 1 | 1 | 1 | NA | -0.007 |
| SE | 97001 | ENSG00000153339 | TRAPPC8 | chr18 | - | 31857539 | 31857720 | 31855659 | 31855807 | 31864626 | 31864781 | 141 | 108 | 85 | 106 | 136 | 0 | 2 | 0 | 1 | 0 | 8 | 85 | 67 | 14 | 0 | 0 | 0 | 2 | 0 | 0 | 298 | 149 | 1 | 1 | 1 | 0.964 | 1 | 0.981 | 1 | 1 | 1 | 0.944 | 1 | NA | 0.003 |
| SE | 97002 | ENSG00000153339 | TRAPPC8 | chr18 | - | 31857539 | 31857982 | 31855659 | 31855807 | 31864626 | 31864781 | 204 | 146 | 124 | 142 | 236 | 0 | 2 | 0 | 1 | 0 | 4 | 108 | 75 | 17 | 6 | 0 | 0 | 2 | 0 | 0 | 298 | 149 | 1 | 1 | 1 | 0.973 | 1 | 0.986 | 1 | 1 | 1 | 0.949 | 1 | 1 | 0.002 |
| SE | 97003 | ENSG00000153339 | TRAPPC8 | chr18 | - | 31866848 | 31866975 | 31864626 | 31864781 | 31867401 | 31867476 | 223 | 149 | 155 | 174 | 421 | 1 | 0 | 1 | 0 | 2 | 2 | 52 | 58 | 24 | 3 | 0 | 4 | 0 | 0 | 0 | 276 | 149 | 1 | 1 | 0.992 | 1 | 0.988 | 1 | 0.991 | 1 | 0.875 | 1 | 1 | 1 | 0.019 |
| SE | 97004 | ENSG00000153339 | TRAPPC8 | chr18 | - | 31874479 | 31874704 | 31873429 | 31873538 | 31890734 | 31890866 | 281 | 234 | 281 | 261 | 382 | 0 | 2 | 0 | 0 | 0 | 63 | 168 | 75 | 54 | 16 | 0 | 0 | 0 | 0 | 0 | 298 | 149 | 1 | 1 | 1 | 0.983 | 1 | 1 | 1 | 1 | 1 | 1 | 1 | 1 | -0.003 |
| SE | 97005 | ENSG00000153339 | TRAPPC8 | chr18 | - | 31890734 | 31890866 | 31873569 | 31874704 | 31897785 | 31897891 | 126 | 126 | 150 | 142 | 144 | 0 | 2 | 0 | 0 | 0 | 40 | 84 | 52 | 19 | 21 | 0 | 0 | 0 | 0 | 0 | 281 | 149 | 1 | 1 | 1 | 0.971 | 1 | 1 | 1 | 1 | 1 | 1 | 1 | 1 | -0.006 |
| SE | 97007 | ENSG00000153339 | TRAPPC8 | chr18 | - | 31913368 | 31913522 | 31909666 | 31909760 | 31931328 | 31931523 | 215 | 138 | 201 | 198 | 290 | 2 | 1 | 0 | 1 | 7 | 50 | 144 | 98 | 43 | 45 | 2 | 2 | 1 | 0 | 0 | 298 | 149 | 1 | 1 | 0.982 | 0.986 | 1 | 0.99 | 0.954 | 0.926 | 0.973 | 0.98 | 1 | 1 | 0.007 |
| SE | 97221 | ENSG00000213465 | ARL2 | chr11 | + | 65020418 | 65020499 | 65018570 | 65018733 | 65021720 | 65021888 | 36 | 12 | 39 | 23 | 39 | 1 | 0 | 0 | 0 | 0 | 17 | 32 | 12 | 14 | 63 | 0 | 0 | 0 | 0 | 0 | 230 | 149 | 1 | 1 | 0.959 | 1 | 1 | 1 | 1 | 1 | 1 | 1 | 1 | 1 | -0.008 |
| SE | 97607 | ENSG00000179832 | MROH1 | chr8 | + | 144243862 | 144243942 | 144243493 | 144243616 | 144244221 | 144244336 | 72 | 79 | 106 | 58 | 85 | 2 | 0 | 1 | 0 | 0 | 84 | 85 | 51 | 123 | 163 | 0 | 0 | 0 | 1 | 0 | 229 | 149 | 1 | 1 | 0.959 | 1 | 0.986 | 1 | 1 | 1 | 1 | 1 | 0.988 | 1 | -0.009 |
| SE | 97609 | ENSG00000179832 | MROH1 | chr8 | + | 144260185 | 144260374 | 144259910 | 144260057 | 144260676 | 144260832 | 12 | 6 | 25 | 24 | 6 | 0 | 0 | 1 | 1 | 0 | 18 | 17 | 6 | 0 | 28 | 0 | 5 | 0 | 0 | 0 | 298 | 149 | 1 | 1 | 1 | 1 | 0.926 | 0.923 | 1 | 1 | 0.63 | 1 | NA | 1 | 0.062 |
| SE | 97610 | ENSG00000179832 | MROH1 | chr8 | + | 144261283 | 144261349 | 144261113 | 144261216 | 144261654 | 144261809 | 9 | 6 | 7 | 9 | 2 | 0 | 0 | 0 | 0 | 2 | 8 | 19 | 2 | 3 | 0 | 0 | 0 | 0 | 0 | 0 | 215 | 149 | 1 | 1 | 1 | 1 | 1 | 1 | 0.409 | 1 | 1 | 1 | 1 | NA | -0.118 |
| SE | 97611 | ENSG00000118363 | SPCS2 | chr11 | + | 74963594 | 74963687 | 74949264 | 74949399 | 74965033 | 74965117 | 0 | 0 | 1 | 4 | 0 | 152 | 72 | 179 | 152 | 135 | 0 | 0 | 0 | 0 | 0 | 113 | 93 | 129 | 118 | 64 | 242 | 149 | 1 | 1 | 0 | 0 | 0.003 | 0.016 | 0 | 0 | 0 | 0 | 0 | 0 | 0.004 |
| SE | 97612 | ENSG00000118363 | SPCS2 | chr11 | + | 74965033 | 74965117 | 74949246 | 74949399 | 74969639 | 74969699 | 68 | 30 | 78 | 51 | 42 | 2 | 0 | 2 | 0 | 0 | 21 | 22 | 61 | 46 | 21 | 0 | 0 | 0 | 0 | 0 | 233 | 149 | 1 | 1 | 0.956 | 1 | 0.961 | 1 | 1 | 1 | 1 | 1 | 1 | 1 | -0.017 |
| SE | 97613 | ENSG00000118363 | SPCS2 | chr11 | + | 74965033 | 74965117 | 74949246 | 74949399 | 74976856 | 74977087 | 60 | 25 | 67 | 48 | 39 | 1 | 0 | 0 | 0 | 0 | 13 | 30 | 37 | 37 | 21 | 0 | 0 | 0 | 0 | 0 | 233 | 149 | 1 | 1 | 0.975 | 1 | 1 | 1 | 1 | 1 | 1 | 1 | 1 | 1 | -0.005 |
| SE | 97615 | ENSG00000118363 | SPCS2 | chr11 | + | 74969639 | 74969699 | 74949263 | 74949399 | 74976856 | 74977348 | 69 | 51 | 78 | 75 | 86 | 1 | 0 | 0 | 0 | 0 | 51 | 26 | 62 | 38 | 10 | 0 | 0 | 0 | 0 | 0 | 209 | 149 | 1 | 1 | 0.98 | 1 | 1 | 1 | 1 | 1 | 1 | 1 | 1 | 1 | -0.004 |
| SE | 97894 | ENSG00000145555 | MYO10 | chr5 | - | 16670525 | 16670978 | 16668276 | 16668468 | 16671421 | 16671542 | 7 | 0 | 2 | 3 | 3 | 0 | 0 | 0 | 1 | 0 | 0 | 4 | 0 | 0 | 0 | 0 | 0 | 0 | 0 | 0 | 298 | 149 | 1 | 1 | 1 | NA | 1 | 0.6 | 1 | NA | 1 | NA | NA | NA | -0.1 |
| SE | 97995 | ENSG00000205758 | CRYZL1 | chr21 | - | 33591161 | 33591371 | 33589467 | 33589921 | 33595355 | 33595614 | 18 | 17 | 26 | 26 | 16 | 0 | 0 | 0 | 0 | 0 | 0 | 25 | 1 | 36 | 3 | 0 | 6 | 0 | 0 | 0 | 298 | 149 | 1 | 1 | 1 | 1 | 1 | 1 | 1 | NA | 0.676 | 1 | 1 | 1 | 0.081 |
| SE | 97996 | ENSG00000205758 | CRYZL1 | chr21 | - | 33597279 | 33597401 | 33595130 | 33595528 | 33599149 | 33599248 | 116 | 101 | 112 | 85 | 101 | 2 | 1 | 4 | 0 | 0 | 46 | 90 | 60 | 98 | 0 | 0 | 0 | 0 | 2 | 0 | 271 | 149 | 1 | 1 | 0.97 | 0.982 | 0.939 | 1 | 1 | 1 | 1 | 1 | 0.964 | NA | -0.013 |
| SE | 97998 | ENSG00000205758 | CRYZL1 | chr21 | - | 33597279 | 33597401 | 33596061 | 33596213 | 33599149 | 33599248 | 116 | 103 | 114 | 89 | 105 | 0 | 0 | 0 | 2 | 0 | 50 | 111 | 73 | 98 | 6 | 0 | 0 | 0 | 0 | 0 | 271 | 149 | 1 | 1 | 1 | 1 | 1 | 0.961 | 1 | 1 | 1 | 1 | 1 | 1 | -0.008 |
| SE | 97999 | ENSG00000205758 | CRYZL1 | chr21 | - | 33613537 | 33613606 | 33603155 | 33603537 | 33621995 | 33622068 | 18 | 14 | 24 | 29 | 46 | 0 | 0 | 0 | 0 | 0 | 11 | 10 | 7 | 12 | 0 | 0 | 0 | 1 | 0 | 0 | 218 | 149 | 1 | 1 | 1 | 1 | 1 | 1 | 1 | 1 | 1 | 0.827 | 1 | NA | 0.043 |
| SE | 98000 | ENSG00000205758 | CRYZL1 | chr21 | - | 33616705 | 33616750 | 33613537 | 33613606 | 33621995 | 33622042 | 164 | 165 | 163 | 165 | 194 | 0 | 0 | 0 | 1 | 0 | 146 | 125 | 111 | 131 | 67 | 0 | 0 | 0 | 0 | 0 | 194 | 149 | 1 | 1 | 1 | 1 | 1 | 0.992 | 1 | 1 | 1 | 1 | 1 | 1 | -0.002 |
| SE | 98004 | ENSG00000205758 | CRYZL1 | chr21 | - | 33641178 | 33641305 | 33631485 | 33631557 | 33641680 | 33641721 | 0 | 0 | 0 | 4 | 0 | 67 | 47 | 54 | 45 | 28 | 0 | 0 | 0 | 2 | 0 | 21 | 58 | 21 | 39 | 12 | 276 | 149 | 1 | 1 | 0 | 0 | 0 | 0.046 | 0 | 0 | 0 | 0 | 0.027 | 0 | 0.004 |
| SE | 98038 | ENSG00000139697 | SBNO1 | chr12 | - | 123304581 | 123304719 | 123302823 | 123302900 | 123309309 | 123309402 | 257 | 180 | 185 | 261 | 309 | 1 | 0 | 0 | 0 | 1 | 48 | 154 | 185 | 56 | 0 | 0 | 0 | 0 | 0 | 0 | 287 | 149 | 1 | 1 | 0.993 | 1 | 1 | 1 | 0.994 | 1 | 1 | 1 | 1 | NA | -0.003 |
| SE | 98039 | ENSG00000139697 | SBNO1 | chr12 | - | 123309309 | 123309402 | 123304581 | 123304719 | 123309488 | 123309583 | 188 | 138 | 148 | 219 | 298 | 0 | 2 | 0 | 0 | 0 | 28 | 144 | 151 | 55 | 0 | 0 | 0 | 0 | 0 | 0 | 242 | 149 | 1 | 1 | 1 | 0.977 | 1 | 1 | 1 | 1 | 1 | 1 | 1 | NA | -0.005 |
| SE | 98040 | ENSG00000139697 | SBNO1 | chr12 | - | 123311054 | 123311129 | 123309709 | 123309856 | 123313619 | 123313719 | 258 | 225 | 266 | 256 | 446 | 1 | 0 | 2 | 0 | 7 | 33 | 233 | 215 | 104 | 0 | 0 | 0 | 0 | 0 | 0 | 224 | 149 | 1 | 1 | 0.994 | 1 | 0.989 | 1 | 0.977 | 1 | 1 | 1 | 1 | NA | -0.008 |
| SE | 98041 | ENSG00000139697 | SBNO1 | chr12 | - | 123313619 | 123313719 | 123311054 | 123311129 | 123315372 | 123315444 | 253 | 214 | 252 | 245 | 394 | 0 | 0 | 0 | 3 | 3 | 36 | 200 | 177 | 101 | 0 | 0 | 1 | 0 | 0 | 0 | 249 | 149 | 1 | 1 | 1 | 1 | 1 | 0.98 | 0.987 | 1 | 0.992 | 1 | 1 | NA | -0.005 |
| SE | 98042 | ENSG00000139697 | SBNO1 | chr12 | - | 123317220 | 123317356 | 123315547 | 123315660 | 123319899 | 123320031 | 544 | 531 | 475 | 672 | 478 | 9 | 3 | 0 | 0 | 0 | 165 | 531 | 830 | 415 | 42 | 4 | 3 | 1 | 4 | 0 | 285 | 149 | 1 | 1 | 0.969 | 0.989 | 1 | 1 | 1 | 0.956 | 0.989 | 0.998 | 0.982 | 1 | 0.007 |
| SE | 98043 | ENSG00000139697 | SBNO1 | chr12 | - | 123334052 | 123334213 | 123331241 | 123331375 | 123336394 | 123336491 | 399 | 349 | 362 | 343 | 482 | 0 | 0 | 0 | 1 | 0 | 221 | 367 | 542 | 280 | 73 | 0 | 0 | 0 | 0 | 0 | 298 | 149 | 1 | 1 | 1 | 1 | 1 | 0.994 | 1 | 1 | 1 | 1 | 1 | 1 | -0.001 |
| SE | 98044 | ENSG00000139697 | SBNO1 | chr12 | - | 123345257 | 123345567 | 123340987 | 123341088 | 123348028 | 123348133 | 428 | 351 | 333 | 496 | 454 | 5 | 0 | 2 | 0 | 0 | 364 | 517 | 405 | 314 | 129 | 0 | 0 | 0 | 1 | 0 | 298 | 149 | 1 | 1 | 0.977 | 1 | 0.988 | 1 | 1 | 1 | 1 | 1 | 0.994 | 1 | -0.006 |
| SE | 98045 | ENSG00000139697 | SBNO1 | chr12 | - | 123345257 | 123345570 | 123340987 | 123341088 | 123348028 | 123348133 | 413 | 363 | 339 | 462 | 490 | 5 | 0 | 2 | 0 | 0 | 337 | 511 | 373 | 263 | 113 | 0 | 0 | 0 | 1 | 0 | 298 | 149 | 1 | 1 | 0.976 | 1 | 0.988 | 1 | 1 | 1 | 1 | 1 | 0.992 | 1 | -0.006 |
| SE | 98046 | ENSG00000139697 | SBNO1 | chr12 | - | 123348028 | 123348133 | 123345257 | 123345567 | 123350309 | 123350441 | 173 | 116 | 167 | 194 | 274 | 2 | 4 | 0 | 0 | 0 | 104 | 119 | 149 | 160 | 74 | 0 | 0 | 1 | 0 | 0 | 254 | 149 | 1 | 1 | 0.981 | 0.944 | 1 | 1 | 1 | 1 | 1 | 0.989 | 1 | 1 | -0.013 |
| SE | 98048 | ENSG00000139697 | SBNO1 | chr12 | - | 123350309 | 123350441 | 123348028 | 123348133 | 123364700 | 123364843 | 162 | 105 | 155 | 176 | 232 | 1 | 3 | 0 | 0 | 0 | 79 | 107 | 115 | 119 | 59 | 0 | 0 | 0 | 1 | 0 | 281 | 149 | 1 | 1 | 0.988 | 0.949 | 1 | 1 | 1 | 1 | 1 | 1 | 0.984 | 1 | -0.009 |
| SE | 98050 | ENSG00000110318 | CEP126 | chr11 | + | 101955791 | 101956943 | 101948030 | 101948142 | 101958167 | 101958366 | 12 | 12 | 12 | 12 | 14 | 7 | 1 | 7 | 13 | 4 | 0 | 2 | 0 | 0 | 0 | 0 | 3 | 0 | 0 | 0 | 298 | 149 | 1 | 1 | 0.462 | 0.857 | 0.462 | 0.316 | 0.636 | NA | 0.25 | NA | NA | NA | 0.297 |
| SE | 98051 | ENSG00000110318 | CEP126 | chr11 | + | 101956261 | 101956943 | 101948030 | 101948142 | 101958167 | 101958366 | 13 | 12 | 11 | 13 | 10 | 7 | 1 | 7 | 13 | 4 | 0 | 4 | 0 | 0 | 0 | 0 | 3 | 0 | 0 | 0 | 298 | 149 | 1 | 1 | 0.481 | 0.857 | 0.44 | 0.333 | 0.556 | NA | 0.4 | NA | NA | NA | 0.133 |
| SE | 98052 | ENSG00000110318 | CEP126 | chr11 | + | 101956867 | 101956943 | 101948030 | 101948142 | 101958167 | 101958366 | 6 | 4 | 2 | 7 | 4 | 7 | 1 | 7 | 13 | 4 | 0 | 2 | 0 | 0 | 0 | 0 | 3 | 0 | 0 | 0 | 225 | 149 | 1 | 1 | 0.362 | 0.726 | 0.159 | 0.263 | 0.398 | NA | 0.306 | NA | NA | NA | 0.076 |
| SE | 98053 | ENSG00000110318 | CEP126 | chr11 | + | 101958167 | 101958366 | 101948030 | 101948142 | 101978346 | 101978459 | 8 | 1 | 9 | 13 | 4 | 0 | 0 | 2 | 0 | 0 | 0 | 3 | 0 | 0 | 0 | 0 | 0 | 0 | 0 | 0 | 298 | 149 | 1 | 1 | 1 | 1 | 0.692 | 1 | 1 | NA | 1 | NA | NA | NA | -0.062 |
| SE | 98054 | ENSG00000110318 | CEP126 | chr11 | + | 101961740 | 101963880 | 101958167 | 101958366 | 101978346 | 101978459 | 48 | 33 | 24 | 28 | 29 | 1 | 0 | 2 | 0 | 0 | 0 | 5 | 0 | 3 | 0 | 0 | 0 | 0 | 0 | 0 | 298 | 149 | 1 | 1 | 0.96 | 1 | 0.857 | 1 | 1 | NA | 1 | NA | 1 | NA | -0.037 |
| SE | 98055 | ENSG00000110318 | CEP126 | chr11 | + | 101978346 | 101978459 | 101961740 | 101963880 | 101981888 | 101981964 | 37 | 33 | 18 | 19 | 21 | 5 | 0 | 0 | 6 | 10 | 0 | 0 | 0 | 6 | 0 | 0 | 0 | 0 | 0 | 0 | 262 | 149 | 1 | 1 | 0.808 | 1 | 1 | 0.643 | 0.544 | NA | NA | NA | 1 | NA | -0.201 |
| SE | 98093 | ENSG00000135249 | RINT1 | chr7 | + | 105535571 | 105535691 | 105532823 | 105532869 | 105536564 | 105536749 | 5 | 0 | 3 | 1 | 1 | 36 | 13 | 17 | 27 | 18 | 0 | 1 | 4 | 0 | 0 | 15 | 14 | 25 | 28 | 24 | 269 | 149 | 1 | 1 | 0.071 | 0 | 0.089 | 0.02 | 0.03 | 0 | 0.038 | 0.081 | 0 | 0 | 0.018 |
| SE | 98094 | ENSG00000135249 | RINT1 | chr7 | + | 105546909 | 105547083 | 105542407 | 105542649 | 105547183 | 105547321 | 133 | 95 | 128 | 95 | 287 | 0 | 0 | 0 | 0 | 0 | 114 | 127 | 151 | 173 | 68 | 0 | 0 | 0 | 1 | 0 | 298 | 149 | 1 | 1 | 1 | 1 | 1 | 1 | 1 | 1 | 1 | 1 | 0.989 | 1 | 0.002 |
| SE | 98095 | ENSG00000135249 | RINT1 | chr7 | + | 105550260 | 105550486 | 105550054 | 105550165 | 105551569 | 105551707 | 78 | 61 | 74 | 65 | 123 | 0 | 0 | 1 | 0 | 0 | 66 | 65 | 97 | 97 | 35 | 0 | 0 | 0 | 0 | 0 | 298 | 149 | 1 | 1 | 1 | 1 | 0.974 | 1 | 1 | 1 | 1 | 1 | 1 | 1 | -0.005 |
| SE | 98096 | ENSG00000135249 | RINT1 | chr7 | + | 105555027 | 105555227 | 105551569 | 105551707 | 105563732 | 105563947 | 110 | 86 | 128 | 94 | 207 | 3 | 1 | 0 | 0 | 1 | 115 | 107 | 91 | 146 | 56 | 0 | 0 | 0 | 0 | 0 | 298 | 149 | 1 | 1 | 0.948 | 0.977 | 1 | 1 | 0.99 | 1 | 1 | 1 | 1 | 1 | -0.017 |
| SE | 98097 | ENSG00000135249 | RINT1 | chr7 | + | 105563732 | 105563947 | 105555027 | 105555227 | 105565276 | 105565457 | 96 | 77 | 90 | 93 | 134 | 1 | 0 | 0 | 1 | 0 | 88 | 90 | 104 | 197 | 45 | 0 | 0 | 0 | 4 | 0 | 298 | 149 | 1 | 1 | 0.98 | 1 | 1 | 0.979 | 1 | 1 | 1 | 1 | 0.961 | 1 | 0 |
| SE | 98281 | ENSG00000068831 | RASGRP2 | chr11 | - | 64727295 | 64727360 | 64726910 | 64727131 | 64728862 | 64729042 | 127 | 68 | 145 | 172 | 43 | 1 | 0 | 0 | 1 | 0 | 123 | 105 | 113 | 92 | 256 | 0 | 0 | 0 | 0 | 0 | 214 | 149 | 1 | 1 | 1 | 1 | 1 | 0.992 | 1 | 1 | 1 | 1 | 1 | 1 | -0.002 |
| SE | 98282 | ENSG00000068831 | RASGRP2 | chr11 | - | 64727295 | 64727363 | 64726910 | 64727131 | 64728862 | 64729042 | 11 | 3 | 12 | 15 | 3 | 0 | 0 | 0 | 1 | 0 | 5 | 12 | 6 | 14 | 23 | 0 | 0 | 0 | 0 | 0 | 217 | 149 | 1 | 1 | 1 | 1 | 1 | 0.912 | 1 | 1 | 1 | 1 | 1 | 1 | -0.018 |
| SE | 98283 | ENSG00000068831 | RASGRP2 | chr11 | - | 64729761 | 64729798 | 64728862 | 64729042 | 64730052 | 64730194 | 413 | 212 | 463 | 458 | 407 | 1 | 0 | 0 | 0 | 0 | 180 | 260 | 268 | 230 | 577 | 0 | 0 | 0 | 0 | 0 | 186 | 149 | 1 | 1 | 0.997 | 1 | 1 | 1 | 1 | 1 | 1 | 1 | 1 | 1 | -0.001 |
| SE | 98287 | ENSG00000068831 | RASGRP2 | chr11 | - | 64743816 | 64743868 | 64742793 | 64742937 | 64744002 | 64744118 | 2 | 1 | 2 | 2 | 0 | 35 | 10 | 37 | 20 | 20 | 0 | 6 | 0 | 5 | 10 | 0 | 17 | 29 | 1 | 43 | 201 | 149 | 1 | 1 | 0.041 | 0.069 | 0.039 | 0.069 | 0 | NA | 0.207 | 0 | 0.788 | 0.147 | -0.242 |
| SE | 98288 | ENSG00000068831 | RASGRP2 | chr11 | - | 64743816 | 64743868 | 64742793 | 64742937 | 64744741 | 64744867 | 0 | 0 | 1 | 2 | 0 | 6 | 3 | 6 | 6 | 1 | 0 | 0 | 0 | 0 | 0 | 0 | 15 | 0 | 6 | 0 | 201 | 149 | 1 | 1 | 0 | 0 | 0.11 | 0.198 | 0 | NA | 0 | NA | 0 | NA | 0.062 |
| SE | 98348 | ENSG00000131379 | C3orf20 | chr3 | + | 14726900 | 14727024 | 14721652 | 14721784 | 14728438 | 14728688 | 0 | 0 | 0 | 1 | 0 | 3 | 1 | 0 | 0 | 0 | 0 | 6 | 0 | 2 | 0 | 0 | 0 | 0 | 1 | 0 | 273 | 149 | 1 | 1 | 0 | 0 | NA | 1 | NA | NA | 1 | NA | 0.522 | NA | -0.428 |
| SE | 98431 | ENSG00000099783 | HNRNPM | chr19 | + | 8465323 | 8465362 | 8463592 | 8463686 | 8465479 | 8465515 | 363 | 231 | 277 | 327 | 437 | 0 | 0 | 1 | 0 | 3 | 414 | 253 | 218 | 248 | 241 | 0 | 1 | 2 | 0 | 0 | 188 | 149 | 1 | 1 | 1 | 1 | 0.995 | 1 | 0.991 | 1 | 0.995 | 0.989 | 1 | 1 | 0 |
| SE | 98432 | ENSG00000099783 | HNRNPM | chr19 | + | 8465479 | 8465515 | 8465323 | 8465362 | 8466234 | 8466388 | 419 | 260 | 320 | 374 | 526 | 0 | 0 | 0 | 1 | 0 | 569 | 304 | 298 | 362 | 270 | 2 | 0 | 3 | 0 | 3 | 185 | 149 | 1 | 1 | 1 | 1 | 1 | 0.997 | 1 | 0.996 | 1 | 0.988 | 1 | 0.986 | 0.005 |
| SE | 98433 | ENSG00000099783 | HNRNPM | chr19 | + | 8471325 | 8471427 | 8468773 | 8468834 | 8473663 | 8473708 | 1174 | 825 | 953 | 1238 | 1180 | 0 | 0 | 0 | 0 | 0 | 2259 | 1238 | 1172 | 1724 | 1259 | 0 | 6 | 2 | 0 | 0 | 251 | 149 | 1 | 1 | 1 | 1 | 1 | 1 | 1 | 1 | 0.992 | 0.997 | 1 | 1 | 0.002 |
| SE | 98437 | ENSG00000099783 | HNRNPM | chr19 | + | 8474166 | 8474244 | 8471325 | 8471427 | 8485602 | 8486273 | 28 | 14 | 25 | 36 | 23 | 1 | 0 | 0 | 0 | 0 | 73 | 39 | 49 | 73 | 40 | 0 | 0 | 0 | 0 | 0 | 227 | 149 | 1 | 1 | 0.948 | 1 | 1 | 1 | 1 | 1 | 1 | 1 | 1 | 1 | -0.01 |
| SE | 98438 | ENSG00000099783 | HNRNPM | chr19 | + | 8483157 | 8483211 | 8471325 | 8471427 | 8485602 | 8485648 | 18 | 20 | 28 | 31 | 32 | 1 | 0 | 0 | 0 | 0 | 71 | 28 | 30 | 23 | 18 | 0 | 0 | 0 | 0 | 0 | 203 | 149 | 1 | 1 | 0.93 | 1 | 1 | 1 | 1 | 1 | 1 | 1 | 1 | 1 | -0.014 |
| SE | 98530 | ENSG00000130935 | NOL11 | chr17 | + | 67719110 | 67719167 | 67717930 | 67718088 | 67719673 | 67719787 | 0 | 4 | 0 | 0 | 0 | 97 | 104 | 109 | 108 | 91 | 0 | 0 | 0 | 0 | 0 | 10 | 99 | 85 | 48 | 19 | 206 | 149 | 1 | 1 | 0 | 0.027 | 0 | 0 | 0 | 0 | 0 | 0 | 0 | 0 | 0.005 |
| SE | 98531 | ENSG00000130935 | NOL11 | chr17 | + | 67720346 | 67720444 | 67719905 | 67719962 | 67721377 | 67721440 | 1 | 2 | 0 | 0 | 0 | 185 | 132 | 195 | 160 | 203 | 5 | 1 | 0 | 0 | 0 | 78 | 135 | 105 | 51 | 31 | 247 | 149 | 1 | 1 | 0.003 | 0.009 | 0 | 0 | 0 | 0.037 | 0.004 | 0 | 0 | 0 | -0.006 |
| SE | 98532 | ENSG00000130935 | NOL11 | chr17 | + | 67722579 | 67722637 | 67721377 | 67721526 | 67724048 | 67724193 | 267 | 151 | 238 | 200 | 522 | 0 | 1 | 0 | 4 | 0 | 35 | 102 | 112 | 39 | 10 | 0 | 0 | 0 | 0 | 0 | 207 | 149 | 1 | 1 | 1 | 0.991 | 1 | 0.973 | 1 | 1 | 1 | 1 | 1 | 1 | -0.007 |
| SE | 98533 | ENSG00000130935 | NOL11 | chr17 | + | 67726459 | 67726648 | 67724048 | 67724193 | 67734362 | 67734439 | 343 | 217 | 350 | 292 | 720 | 0 | 0 | 0 | 0 | 1 | 30 | 157 | 192 | 60 | 28 | 0 | 0 | 0 | 0 | 0 | 298 | 149 | 1 | 1 | 1 | 1 | 1 | 1 | 0.997 | 1 | 1 | 1 | 1 | 1 | -0.001 |
| SE | 98534 | ENSG00000130935 | NOL11 | chr17 | + | 67734362 | 67734439 | 67726459 | 67726648 | 67735899 | 67736023 | 121 | 73 | 136 | 94 | 313 | 2 | 0 | 0 | 0 | 1 | 18 | 63 | 79 | 25 | 40 | 0 | 0 | 0 | 0 | 0 | 226 | 149 | 1 | 1 | 0.976 | 1 | 1 | 1 | 0.995 | 1 | 1 | 1 | 1 | 1 | -0.006 |
| SE | 98536 | ENSG00000130935 | NOL11 | chr17 | + | 67735899 | 67736023 | 67734362 | 67734439 | 67737070 | 67737145 | 66 | 55 | 69 | 69 | 153 | 0 | 0 | 0 | 0 | 0 | 66 | 79 | 46 | 32 | 40 | 0 | 3 | 0 | 0 | 0 | 273 | 149 | 1 | 1 | 1 | 1 | 1 | 1 | 1 | 1 | 0.935 | 1 | 1 | 1 | 0.013 |
| SE | 98538 | ENSG00000130935 | NOL11 | chr17 | + | 67737070 | 67737145 | 67736579 | 67736754 | 67737507 | 67737689 | 378 | 211 | 347 | 307 | 522 | 1 | 0 | 1 | 0 | 0 | 35 | 145 | 192 | 82 | 11 | 0 | 0 | 0 | 0 | 0 | 224 | 149 | 1 | 1 | 0.996 | 1 | 0.996 | 1 | 1 | 1 | 1 | 1 | 1 | 1 | -0.002 |
| SE | 98869 | ENSG00000175294 | CATSPER1 | chr11 | - | 66020139 | 66020200 | 66018787 | 66018902 | 66020810 | 66020954 | 9 | 6 | 4 | 1 | 7 | 0 | 0 | 0 | 1 | 0 | 21 | 14 | 10 | 5 | 6 | 0 | 0 | 0 | 0 | 0 | 210 | 149 | 1 | 1 | 1 | 1 | 1 | 0.415 | 1 | 1 | 1 | 1 | 1 | 1 | -0.117 |
| SE | 98870 | ENSG00000175294 | CATSPER1 | chr11 | - | 66020316 | 66020389 | 66020139 | 66020200 | 66020563 | 66020627 | 33 | 7 | 10 | 13 | 3 | 0 | 0 | 0 | 1 | 0 | 5 | 19 | 16 | 10 | 24 | 0 | 0 | 0 | 0 | 0 | 222 | 149 | 1 | 1 | 1 | 1 | 1 | 0.897 | 1 | 1 | 1 | 1 | 1 | 1 | -0.021 |
| SE | 98871 | ENSG00000175294 | CATSPER1 | chr11 | - | 66020316 | 66020389 | 66020139 | 66020200 | 66020810 | 66020954 | 26 | 13 | 10 | 16 | 1 | 0 | 0 | 0 | 0 | 0 | 9 | 17 | 12 | 8 | 7 | 0 | 0 | 0 | 3 | 0 | 222 | 149 | 1 | 1 | 1 | 1 | 1 | 1 | 1 | 1 | 1 | 1 | 0.642 | 1 | 0.072 |
| SE | 98872 | ENSG00000175294 | CATSPER1 | chr11 | - | 66020563 | 66020627 | 66020139 | 66020200 | 66020810 | 66020954 | 5 | 1 | 0 | 1 | 0 | 0 | 0 | 0 | 0 | 0 | 9 | 0 | 4 | 0 | 9 | 0 | 0 | 0 | 3 | 0 | 213 | 149 | 1 | 1 | 1 | 1 | NA | 1 | NA | 1 | NA | 1 | 0 | 1 | 0.25 |
| SE | 99084 | ENSG00000110274 | CEP164 | chr11 | + | 117335604 | 117335680 | 117327853 | 117327904 | 117338565 | 117338668 | 18 | 7 | 19 | 25 | 11 | 0 | 0 | 0 | 1 | 0 | 40 | 29 | 19 | 11 | 20 | 0 | 0 | 1 | 1 | 0 | 225 | 149 | 1 | 1 | 1 | 1 | 1 | 0.943 | 1 | 1 | 1 | 0.926 | 0.879 | 1 | 0.028 |
| SE | 99095 | ENSG00000110274 | CEP164 | chr11 | + | 117373750 | 117373831 | 117371079 | 117371466 | 117380613 | 117380705 | 30 | 26 | 20 | 24 | 54 | 0 | 0 | 0 | 2 | 0 | 52 | 42 | 28 | 23 | 50 | 0 | 0 | 0 | 0 | 0 | 230 | 149 | 1 | 1 | 1 | 1 | 1 | 0.886 | 1 | 1 | 1 | 1 | 1 | 1 | -0.023 |
| SE | 99096 | ENSG00000110274 | CEP164 | chr11 | + | 117375707 | 117375791 | 117371079 | 117371466 | 117380613 | 117380705 | 20 | 7 | 13 | 18 | 3 | 0 | 0 | 0 | 2 | 0 | 12 | 12 | 9 | 8 | 24 | 0 | 0 | 0 | 0 | 0 | 233 | 149 | 1 | 1 | 1 | 1 | 1 | 0.852 | 1 | 1 | 1 | 1 | 1 | 1 | -0.03 |
| SE | 99097 | ENSG00000110274 | CEP164 | chr11 | + | 117375707 | 117375791 | 117373750 | 117373831 | 117380613 | 117380705 | 67 | 26 | 43 | 61 | 40 | 0 | 0 | 0 | 0 | 0 | 62 | 57 | 60 | 42 | 89 | 3 | 0 | 0 | 0 | 0 | 233 | 149 | 1 | 1 | 1 | 1 | 1 | 1 | 1 | 0.93 | 1 | 1 | 1 | 1 | 0.014 |
| SE | 99101 | ENSG00000110274 | CEP164 | chr11 | + | 117387202 | 117387412 | 117382795 | 117382942 | 117390776 | 117390908 | 40 | 35 | 25 | 52 | 33 | 0 | 0 | 0 | 1 | 0 | 171 | 50 | 85 | 85 | 222 | 0 | 0 | 1 | 0 | 0 | 298 | 149 | 1 | 1 | 1 | 1 | 1 | 0.963 | 1 | 1 | 1 | 0.977 | 1 | 1 | -0.003 |
| SE | 99103 | ENSG00000110274 | CEP164 | chr11 | + | 117390998 | 117391215 | 117390776 | 117390908 | 117392225 | 117392303 | 88 | 98 | 91 | 118 | 84 | 0 | 3 | 0 | 0 | 0 | 303 | 174 | 162 | 175 | 314 | 0 | 0 | 1 | 4 | 0 | 298 | 149 | 1 | 1 | 1 | 0.942 | 1 | 1 | 1 | 1 | 1 | 0.988 | 0.956 | 1 | 0 |
| SE | 99106 | ENSG00000110274 | CEP164 | chr11 | + | 117395122 | 117395191 | 117394919 | 117395003 | 117395546 | 117395722 | 30 | 14 | 29 | 31 | 23 | 0 | 0 | 0 | 0 | 0 | 70 | 42 | 20 | 23 | 65 | 0 | 0 | 0 | 1 | 0 | 218 | 149 | 1 | 1 | 1 | 1 | 1 | 1 | 1 | 1 | 1 | 1 | 0.94 | 1 | 0.012 |
| SE | 99144 | ENSG00000103544 | VPS35L | chr16 | + | 19569243 | 19569591 | 19564850 | 19564950 | 19573118 | 19573163 | 87 | 30 | 55 | 48 | 66 | 1 | 0 | 0 | 0 | 0 | 100 | 73 | 53 | 44 | 58 | 0 | 0 | 0 | 0 | 0 | 298 | 149 | 1 | 1 | 0.978 | 1 | 1 | 1 | 1 | 1 | 1 | 1 | 1 | 1 | -0.004 |
| SE | 99145 | ENSG00000103544 | VPS35L | chr16 | + | 19569423 | 19569591 | 19564850 | 19564950 | 19573118 | 19573241 | 158 | 55 | 105 | 82 | 171 | 1 | 0 | 0 | 0 | 0 | 137 | 148 | 81 | 88 | 133 | 0 | 0 | 0 | 0 | 0 | 298 | 149 | 1 | 1 | 0.988 | 1 | 1 | 1 | 1 | 1 | 1 | 1 | 1 | 1 | -0.002 |
| SE | 99146 | ENSG00000103544 | VPS35L | chr16 | + | 19573118 | 19573241 | 19569423 | 19569591 | 19575097 | 19575122 | 156 | 87 | 105 | 145 | 152 | 0 | 0 | 0 | 0 | 0 | 209 | 163 | 133 | 142 | 212 | 0 | 0 | 2 | 0 | 0 | 272 | 149 | 1 | 1 | 1 | 1 | 1 | 1 | 1 | 1 | 1 | 0.973 | 1 | 1 | 0.005 |
| SE | 99147 | ENSG00000103544 | VPS35L | chr16 | + | 19573123 | 19573241 | 19569423 | 19569591 | 19575097 | 19575122 | 69 | 61 | 53 | 98 | 87 | 0 | 0 | 0 | 0 | 0 | 109 | 97 | 90 | 94 | 154 | 0 | 0 | 2 | 0 | 0 | 267 | 149 | 1 | 1 | 1 | 1 | 1 | 1 | 1 | 1 | 1 | 0.962 | 1 | 1 | 0.008 |
| SE | 99148 | ENSG00000103544 | VPS35L | chr16 | + | 19575097 | 19575122 | 19573118 | 19573241 | 19579051 | 19579128 | 139 | 92 | 87 | 148 | 146 | 0 | 1 | 0 | 0 | 0 | 164 | 133 | 103 | 154 | 196 | 0 | 1 | 0 | 0 | 0 | 174 | 149 | 1 | 1 | 1 | 0.987 | 1 | 1 | 1 | 1 | 0.991 | 1 | 1 | 1 | -0.001 |
| SE | 99150 | ENSG00000103544 | VPS35L | chr16 | + | 19579051 | 19579128 | 19575097 | 19575122 | 19581524 | 19581608 | 160 | 102 | 109 | 180 | 183 | 0 | 0 | 0 | 1 | 0 | 203 | 178 | 123 | 172 | 240 | 0 | 0 | 0 | 0 | 0 | 226 | 149 | 1 | 1 | 1 | 1 | 1 | 0.992 | 1 | 1 | 1 | 1 | 1 | 1 | -0.002 |
| SE | 99151 | ENSG00000103544 | VPS35L | chr16 | + | 19591789 | 19591874 | 19581524 | 19581653 | 19601663 | 19601723 | 249 | 173 | 218 | 247 | 256 | 0 | 0 | 0 | 0 | 0 | 270 | 265 | 142 | 295 | 317 | 3 | 0 | 0 | 0 | 0 | 234 | 149 | 1 | 1 | 1 | 1 | 1 | 1 | 1 | 0.983 | 1 | 1 | 1 | 1 | 0.003 |
| SE | 99152 | ENSG00000103544 | VPS35L | chr16 | + | 19601663 | 19601723 | 19591789 | 19591874 | 19608177 | 19608274 | 149 | 107 | 147 | 161 | 160 | 0 | 1 | 0 | 0 | 2 | 171 | 181 | 83 | 172 | 194 | 0 | 0 | 0 | 0 | 0 | 209 | 149 | 1 | 1 | 1 | 0.987 | 1 | 1 | 0.983 | 1 | 1 | 1 | 1 | 1 | -0.006 |
| SE | 99156 | ENSG00000103544 | VPS35L | chr16 | + | 19626176 | 19626223 | 19616685 | 19616808 | 19627693 | 19627802 | 108 | 82 | 103 | 121 | 170 | 1 | 1 | 0 | 0 | 0 | 202 | 100 | 66 | 160 | 204 | 0 | 0 | 0 | 0 | 0 | 196 | 149 | 1 | 1 | 0.988 | 0.984 | 1 | 1 | 1 | 1 | 1 | 1 | 1 | 1 | -0.006 |
| SE | 99157 | ENSG00000103544 | VPS35L | chr16 | + | 19640014 | 19640100 | 19637593 | 19637656 | 19642395 | 19642476 | 157 | 104 | 126 | 154 | 225 | 0 | 0 | 0 | 1 | 0 | 295 | 156 | 110 | 141 | 271 | 0 | 0 | 0 | 0 | 5 | 235 | 149 | 1 | 1 | 1 | 1 | 1 | 0.99 | 0.993 | 1 | 1 | 1 | 1 | 0.972 | 0.002 |
| SE | 99158 | ENSG00000103544 | VPS35L | chr16 | + | 19642395 | 19642476 | 19640014 | 19640100 | 19644885 | 19644949 | 195 | 123 | 172 | 183 | 261 | 2 | 0 | 2 | 0 | 1 | 408 | 171 | 139 | 191 | 351 | 3 | 0 | 0 | 0 | 0 | 230 | 149 | 1 | 1 | 0.984 | 1 | 0.982 | 1 | 1 | 0.989 | 1 | 1 | 1 | 1 | -0.005 |
| SE | 99159 | ENSG00000103544 | VPS35L | chr16 | + | 19644885 | 19644949 | 19642395 | 19642476 | 19647783 | 19647882 | 174 | 137 | 178 | 155 | 240 | 3 | 0 | 0 | 2 | 0 | 486 | 171 | 133 | 194 | 343 | 0 | 0 | 0 | 4 | 0 | 213 | 149 | 1 | 1 | 0.976 | 1 | 1 | 0.982 | 1 | 1 | 1 | 1 | 0.971 | 1 | -0.003 |
| SE | 99160 | ENSG00000103544 | VPS35L | chr16 | + | 19647783 | 19647882 | 19644885 | 19644949 | 19651975 | 19652090 | 77 | 38 | 69 | 61 | 80 | 0 | 0 | 0 | 0 | 0 | 141 | 67 | 38 | 74 | 118 | 0 | 0 | 0 | 4 | 0 | 248 | 149 | 1 | 1 | 1 | 1 | 1 | 1 | 1 | 1 | 1 | 1 | 0.917 | 1 | 0.017 |
| SE | 99162 | ENSG00000103544 | VPS35L | chr16 | + | 19691352 | 19691471 | 19682224 | 19682390 | 19699501 | 19699648 | 93 | 80 | 104 | 100 | 161 | 0 | 0 | 0 | 0 | 0 | 115 | 107 | 61 | 132 | 195 | 0 | 0 | 0 | 0 | 2 | 268 | 149 | 1 | 1 | 1 | 1 | 1 | 1 | 1 | 1 | 1 | 1 | 1 | 0.982 | 0.004 |
| SE | 99163 | ENSG00000103544 | VPS35L | chr16 | + | 19699501 | 19699648 | 19691352 | 19691471 | 19700377 | 19700687 | 90 | 65 | 81 | 86 | 150 | 1 | 1 | 0 | 0 | 1 | 135 | 64 | 57 | 109 | 114 | 0 | 0 | 0 | 4 | 0 | 296 | 149 | 1 | 1 | 0.978 | 0.97 | 1 | 1 | 0.987 | 1 | 1 | 1 | 0.932 | 1 | 0.001 |
| SE | 99205 | ENSG00000074266 | EED | chr11 | + | 86250295 | 86250448 | 86244752 | 86245343 | 86255221 | 86255287 | 43 | 24 | 31 | 33 | 41 | 0 | 3 | 1 | 0 | 0 | 0 | 9 | 19 | 19 | 9 | 0 | 0 | 0 | 0 | 0 | 298 | 149 | 1 | 1 | 1 | 0.8 | 0.939 | 1 | 1 | NA | 1 | 1 | 1 | 1 | -0.052 |
| SE | 99207 | ENSG00000074266 | EED | chr11 | + | 86257514 | 86257596 | 86256386 | 86256512 | 86264171 | 86264263 | 80 | 36 | 93 | 103 | 116 | 5 | 2 | 3 | 0 | 0 | 4 | 48 | 29 | 27 | 0 | 0 | 6 | 5 | 0 | 0 | 231 | 149 | 1 | 1 | 0.912 | 0.921 | 0.952 | 1 | 1 | 1 | 0.838 | 0.789 | 1 | NA | 0.05 |
| SE | 99209 | ENSG00000074266 | EED | chr11 | + | 86264171 | 86264263 | 86257514 | 86257596 | 86276979 | 86277138 | 61 | 23 | 65 | 67 | 60 | 0 | 1 | 0 | 0 | 1 | 0 | 41 | 20 | 12 | 0 | 0 | 0 | 0 | 0 | 0 | 241 | 149 | 1 | 1 | 1 | 0.934 | 1 | 1 | 0.974 | NA | 1 | 1 | 1 | NA | -0.018 |
| SE | 99210 | ENSG00000074266 | EED | chr11 | + | 86266082 | 86266216 | 86263944 | 86264263 | 86276979 | 86277138 | 55 | 26 | 57 | 68 | 71 | 1 | 0 | 0 | 0 | 0 | 5 | 13 | 47 | 24 | 0 | 0 | 0 | 3 | 6 | 0 | 283 | 149 | 1 | 1 | 0.967 | 1 | 1 | 1 | 1 | 1 | 1 | 0.892 | 0.678 | NA | 0.101 |
| SE | 99211 | ENSG00000074266 | EED | chr11 | + | 86268455 | 86268561 | 86263944 | 86266216 | 86276979 | 86277138 | 62 | 34 | 99 | 87 | 124 | 12 | 4 | 8 | 10 | 13 | 26 | 35 | 47 | 27 | 0 | 0 | 4 | 10 | 6 | 0 | 255 | 149 | 1 | 1 | 0.751 | 0.832 | 0.879 | 0.836 | 0.848 | 1 | 0.836 | 0.733 | 0.724 | NA | 0.006 |
| SE | 99212 | ENSG00000074266 | EED | chr11 | + | 86270082 | 86270157 | 86267894 | 86268561 | 86276979 | 86277138 | 24 | 5 | 9 | 22 | 15 | 34 | 19 | 53 | 40 | 55 | 0 | 0 | 8 | 21 | 0 | 11 | 23 | 24 | 4 | 0 | 224 | 149 | 1 | 1 | 0.32 | 0.149 | 0.101 | 0.268 | 0.154 | 0 | 0 | 0.181 | 0.777 | NA | -0.041 |
| SE | 99362 | ENSG00000156599 | ZDHHC5 | chr11 | + | 57672020 | 57673194 | 57668113 | 57668187 | 57682421 | 57682543 | 226 | 93 | 159 | 173 | 106 | 0 | 0 | 0 | 0 | 0 | 166 | 166 | 138 | 180 | 216 | 0 | 0 | 5 | 0 | 0 | 298 | 149 | 1 | 1 | 1 | 1 | 1 | 1 | 1 | 1 | 1 | 0.932 | 1 | 1 | 0.014 |
| SE | 99365 | ENSG00000156599 | ZDHHC5 | chr11 | + | 57682421 | 57682543 | 57681478 | 57681698 | 57688507 | 57688665 | 107 | 44 | 93 | 109 | 65 | 0 | 0 | 0 | 0 | 1 | 107 | 100 | 66 | 72 | 109 | 0 | 0 | 0 | 0 | 0 | 271 | 149 | 1 | 1 | 1 | 1 | 1 | 1 | 0.973 | 1 | 1 | 1 | 1 | 1 | -0.005 |
| SE | 99366 | ENSG00000156599 | ZDHHC5 | chr11 | + | 57688507 | 57688665 | 57682421 | 57682543 | 57690030 | 57690203 | 310 | 111 | 274 | 262 | 232 | 0 | 1 | 1 | 0 | 0 | 311 | 287 | 183 | 226 | 279 | 0 | 0 | 0 | 0 | 0 | 298 | 149 | 1 | 1 | 1 | 0.982 | 0.993 | 1 | 1 | 1 | 1 | 1 | 1 | 1 | -0.005 |
| SE | 99368 | ENSG00000156599 | ZDHHC5 | chr11 | + | 57695919 | 57696043 | 57693782 | 57693915 | 57696760 | 57696873 | 254 | 129 | 233 | 177 | 359 | 2 | 0 | 0 | 2 | 0 | 456 | 263 | 164 | 252 | 391 | 0 | 0 | 0 | 0 | 0 | 273 | 149 | 1 | 1 | 0.986 | 1 | 1 | 0.98 | 1 | 1 | 1 | 1 | 1 | 1 | -0.007 |
| SE | 99369 | ENSG00000156599 | ZDHHC5 | chr11 | + | 57695919 | 57696043 | 57693782 | 57693915 | 57698558 | 57698835 | 143 | 59 | 120 | 95 | 214 | 0 | 1 | 0 | 2 | 0 | 269 | 159 | 115 | 133 | 215 | 0 | 1 | 0 | 0 | 0 | 273 | 149 | 1 | 1 | 1 | 0.97 | 1 | 0.963 | 1 | 1 | 0.989 | 1 | 1 | 1 | -0.011 |
| SE | 99370 | ENSG00000156599 | ZDHHC5 | chr11 | + | 57695919 | 57696043 | 57693782 | 57693915 | 57699865 | 57700363 | 123 | 57 | 113 | 89 | 191 | 0 | 0 | 2 | 0 | 0 | 246 | 148 | 98 | 119 | 203 | 0 | 0 | 0 | 0 | 0 | 273 | 149 | 1 | 1 | 1 | 1 | 0.969 | 1 | 1 | 1 | 1 | 1 | 1 | 1 | -0.006 |
| SE | 99371 | ENSG00000156599 | ZDHHC5 | chr11 | + | 57696760 | 57696873 | 57693782 | 57693915 | 57698558 | 57698835 | 67 | 25 | 51 | 34 | 68 | 0 | 1 | 0 | 2 | 0 | 22 | 77 | 37 | 39 | 70 | 0 | 1 | 0 | 0 | 0 | 262 | 149 | 1 | 1 | 1 | 0.934 | 1 | 0.906 | 1 | 1 | 0.978 | 1 | 1 | 1 | -0.028 |
| SE | 99372 | ENSG00000156599 | ZDHHC5 | chr11 | + | 57696760 | 57696873 | 57693782 | 57693915 | 57699865 | 57700363 | 4 | 0 | 0 | 2 | 0 | 0 | 0 | 2 | 0 | 0 | 0 | 0 | 0 | 5 | 0 | 0 | 0 | 0 | 0 | 0 | 262 | 149 | 1 | 1 | 1 | NA | 0 | 1 | NA | NA | NA | NA | 1 | NA | -0.333 |
| SE | 99375 | ENSG00000156599 | ZDHHC5 | chr11 | + | 57698558 | 57699418 | 57693782 | 57693915 | 57699865 | 57700363 | 91 | 58 | 109 | 76 | 138 | 0 | 0 | 2 | 0 | 0 | 84 | 87 | 88 | 79 | 228 | 0 | 0 | 0 | 0 | 0 | 298 | 149 | 1 | 1 | 1 | 1 | 0.965 | 1 | 1 | 1 | 1 | 1 | 1 | 1 | -0.007 |
| SE | 99376 | ENSG00000156599 | ZDHHC5 | chr11 | + | 57698558 | 57699418 | 57695919 | 57696043 | 57699865 | 57700363 | 111 | 59 | 116 | 80 | 161 | 0 | 0 | 0 | 0 | 0 | 107 | 103 | 105 | 95 | 240 | 0 | 6 | 0 | 0 | 0 | 298 | 149 | 1 | 1 | 1 | 1 | 1 | 1 | 1 | 1 | 0.896 | 1 | 1 | 1 | 0.021 |
| SE | 100084 | ENSG00000149262 | INTS4 | chr11 | - | 77894289 | 77894349 | 77891680 | 77891840 | 77901420 | 77901551 | 92 | 84 | 84 | 99 | 118 | 0 | 0 | 0 | 1 | 3 | 89 | 84 | 57 | 58 | 122 | 0 | 0 | 4 | 0 | 0 | 209 | 149 | 1 | 1 | 1 | 1 | 1 | 0.986 | 0.966 | 1 | 1 | 0.91 | 1 | 1 | 0.008 |
| SE | 100085 | ENSG00000149262 | INTS4 | chr11 | - | 77903539 | 77903620 | 77901356 | 77901551 | 77907716 | 77907810 | 112 | 80 | 151 | 166 | 110 | 1 | 0 | 1 | 0 | 5 | 182 | 117 | 102 | 118 | 113 | 0 | 0 | 0 | 0 | 0 | 230 | 149 | 1 | 1 | 0.986 | 1 | 0.99 | 1 | 0.934 | 1 | 1 | 1 | 1 | 1 | -0.018 |
| SE | 100086 | ENSG00000149262 | INTS4 | chr11 | - | 77910969 | 77911152 | 77907716 | 77907810 | 77918030 | 77918264 | 8 | 3 | 11 | 10 | 17 | 2 | 1 | 0 | 3 | 0 | 29 | 4 | 1 | 6 | 0 | 2 | 4 | 4 | 3 | 0 | 298 | 149 | 1 | 1 | 0.667 | 0.6 | 1 | 0.625 | 1 | 0.879 | 0.333 | 0.111 | 0.5 | NA | 0.323 |
| SE | 100089 | ENSG00000149262 | INTS4 | chr11 | - | 77918030 | 77918264 | 77910969 | 77911152 | 77918820 | 77918978 | 1 | 0 | 1 | 1 | 0 | 11 | 7 | 5 | 1 | 4 | 0 | 0 | 2 | 0 | 0 | 0 | 2 | 10 | 4 | 8 | 298 | 149 | 1 | 1 | 0.043 | 0 | 0.091 | 0.333 | 0 | NA | 0 | 0.091 | 0 | 0 | 0.071 |
| SE | 100090 | ENSG00000149262 | INTS4 | chr11 | - | 77928341 | 77928547 | 77924441 | 77924892 | 77938650 | 77938771 | 119 | 65 | 110 | 92 | 254 | 0 | 0 | 0 | 1 | 0 | 122 | 123 | 100 | 94 | 60 | 0 | 0 | 0 | 0 | 0 | 298 | 149 | 1 | 1 | 1 | 1 | 1 | 0.979 | 1 | 1 | 1 | 1 | 1 | 1 | -0.004 |
| SE | 100092 | ENSG00000149262 | INTS4 | chr11 | - | 77938650 | 77938825 | 77928341 | 77928547 | 77955941 | 77956062 | 58 | 30 | 41 | 40 | 104 | 0 | 1 | 0 | 0 | 0 | 45 | 35 | 51 | 45 | 34 | 0 | 2 | 0 | 0 | 0 | 298 | 149 | 1 | 1 | 1 | 0.938 | 1 | 1 | 1 | 1 | 0.897 | 1 | 1 | 1 | 0.008 |
| SE | 100093 | ENSG00000149262 | INTS4 | chr11 | - | 77941179 | 77941251 | 77928341 | 77928547 | 77955941 | 77956062 | 76 | 61 | 72 | 86 | 37 | 0 | 1 | 0 | 0 | 0 | 57 | 89 | 85 | 71 | 41 | 0 | 2 | 0 | 0 | 0 | 221 | 149 | 1 | 1 | 1 | 0.976 | 1 | 1 | 1 | 1 | 0.968 | 1 | 1 | 1 | 0.002 |
| SE | 100094 | ENSG00000149262 | INTS4 | chr11 | - | 77941179 | 77941251 | 77928341 | 77928547 | 77958745 | 77958834 | 3 | 0 | 0 | 0 | 0 | 0 | 0 | 0 | 0 | 0 | 0 | 4 | 5 | 3 | 7 | 0 | 0 | 0 | 4 | 0 | 221 | 149 | 1 | 1 | 1 | NA | NA | NA | NA | NA | 1 | 1 | 0.336 | 1 | 0.166 |
| SE | 100095 | ENSG00000149262 | INTS4 | chr11 | - | 77941179 | 77941251 | 77938650 | 77938825 | 77955941 | 77956062 | 225 | 165 | 212 | 247 | 161 | 1 | 1 | 2 | 1 | 1 | 224 | 249 | 179 | 227 | 155 | 0 | 0 | 0 | 0 | 0 | 221 | 149 | 1 | 1 | 0.993 | 0.991 | 0.986 | 0.994 | 0.991 | 1 | 1 | 1 | 1 | 1 | -0.009 |
| SE | 100096 | ENSG00000149262 | INTS4 | chr11 | - | 77955941 | 77956062 | 77928341 | 77928547 | 77958745 | 77958834 | 108 | 97 | 113 | 148 | 123 | 0 | 0 | 0 | 0 | 0 | 171 | 202 | 125 | 115 | 125 | 0 | 0 | 0 | 4 | 0 | 270 | 149 | 1 | 1 | 1 | 1 | 1 | 1 | 1 | 1 | 1 | 1 | 0.941 | 1 | 0.012 |
| SE | 100098 | ENSG00000149262 | INTS4 | chr11 | - | 77958745 | 77958834 | 77955941 | 77956062 | 77960340 | 77960391 | 192 | 165 | 208 | 261 | 173 | 0 | 0 | 0 | 0 | 0 | 286 | 309 | 179 | 217 | 187 | 0 | 3 | 0 | 3 | 0 | 238 | 149 | 1 | 1 | 1 | 1 | 1 | 1 | 1 | 1 | 0.985 | 1 | 0.978 | 1 | 0.007 |
| SE | 100099 | ENSG00000149262 | INTS4 | chr11 | - | 77960340 | 77960391 | 77955941 | 77956062 | 77960952 | 77961138 | 24 | 8 | 20 | 30 | 33 | 0 | 0 | 0 | 0 | 0 | 19 | 32 | 21 | 43 | 17 | 0 | 0 | 0 | 1 | 0 | 200 | 149 | 1 | 1 | 1 | 1 | 1 | 1 | 1 | 1 | 1 | 1 | 0.97 | 1 | 0.006 |
| SE | 100100 | ENSG00000149262 | INTS4 | chr11 | - | 77978995 | 77979102 | 77960952 | 77961138 | 77981458 | 77981576 | 314 | 181 | 264 | 261 | 362 | 3 | 0 | 2 | 2 | 0 | 325 | 292 | 223 | 261 | 261 | 0 | 0 | 0 | 0 | 0 | 256 | 149 | 1 | 1 | 0.984 | 1 | 0.987 | 0.987 | 1 | 1 | 1 | 1 | 1 | 1 | -0.008 |
| SE | 100120 | ENSG00000159063 | ALG8 | chr11 | - | 78103979 | 78104052 | 78100935 | 78101195 | 78104355 | 78104453 | 31 | 23 | 34 | 39 | 54 | 0 | 0 | 0 | 0 | 5 | 59 | 15 | 12 | 35 | 16 | 0 | 0 | 0 | 0 | 0 | 222 | 149 | 1 | 1 | 1 | 1 | 1 | 1 | 0.879 | 1 | 1 | 1 | 1 | 1 | -0.024 |
| SE | 100124 | ENSG00000159063 | ALG8 | chr11 | - | 78106628 | 78106714 | 78104355 | 78104453 | 78106806 | 78106946 | 1 | 0 | 0 | 0 | 0 | 50 | 27 | 67 | 64 | 41 | 0 | 0 | 0 | 0 | 0 | 44 | 31 | 20 | 21 | 17 | 235 | 149 | 1 | 1 | 0.013 | 0 | 0 | 0 | 0 | 0 | 0 | 0 | 0 | 0 | 0.003 |
| SE | 100125 | ENSG00000159063 | ALG8 | chr11 | - | 78106806 | 78106912 | 78104355 | 78104453 | 78109441 | 78109581 | 46 | 25 | 57 | 52 | 42 | 1 | 0 | 0 | 1 | 2 | 43 | 27 | 18 | 10 | 11 | 0 | 0 | 0 | 0 | 0 | 255 | 149 | 1 | 1 | 0.964 | 1 | 1 | 0.968 | 0.925 | 1 | 1 | 1 | 1 | 1 | -0.029 |
| SE | 100126 | ENSG00000159063 | ALG8 | chr11 | - | 78106806 | 78106912 | 78104355 | 78104453 | 78112649 | 78112770 | 45 | 23 | 55 | 49 | 39 | 1 | 0 | 0 | 0 | 0 | 39 | 27 | 14 | 10 | 11 | 0 | 4 | 0 | 0 | 0 | 255 | 149 | 1 | 1 | 0.963 | 1 | 1 | 1 | 1 | 1 | 0.798 | 1 | 1 | 1 | 0.033 |
| SE | 100127 | ENSG00000159063 | ALG8 | chr11 | - | 78106806 | 78106946 | 78104355 | 78104453 | 78109441 | 78109581 | 79 | 40 | 117 | 86 | 75 | 1 | 0 | 0 | 1 | 2 | 67 | 55 | 27 | 42 | 18 | 0 | 0 | 0 | 0 | 0 | 289 | 149 | 1 | 1 | 0.976 | 1 | 1 | 0.978 | 0.951 | 1 | 1 | 1 | 1 | 1 | -0.019 |
| SE | 100128 | ENSG00000159063 | ALG8 | chr11 | - | 78106806 | 78106946 | 78104355 | 78104453 | 78112649 | 78112770 | 48 | 27 | 61 | 63 | 41 | 1 | 0 | 0 | 0 | 0 | 44 | 31 | 15 | 21 | 17 | 0 | 4 | 0 | 0 | 0 | 289 | 149 | 1 | 1 | 0.961 | 1 | 1 | 1 | 1 | 1 | 0.8 | 1 | 1 | 1 | 0.032 |
| SE | 100129 | ENSG00000159063 | ALG8 | chr11 | - | 78106806 | 78106946 | 78104355 | 78104453 | 78113885 | 78113989 | 48 | 27 | 61 | 63 | 41 | 0 | 0 | 0 | 0 | 2 | 44 | 31 | 15 | 21 | 17 | 0 | 0 | 0 | 0 | 0 | 289 | 149 | 1 | 1 | 1 | 1 | 1 | 1 | 0.914 | 1 | 1 | 1 | 1 | 1 | -0.017 |
| SE | 100131 | ENSG00000159063 | ALG8 | chr11 | - | 78109441 | 78109581 | 78104355 | 78104453 | 78112649 | 78112770 | 46 | 28 | 35 | 28 | 61 | 1 | 0 | 0 | 0 | 0 | 59 | 31 | 24 | 29 | 14 | 0 | 4 | 0 | 0 | 0 | 289 | 149 | 1 | 1 | 0.96 | 1 | 1 | 1 | 1 | 1 | 0.8 | 1 | 1 | 1 | 0.032 |
| SE | 100132 | ENSG00000159063 | ALG8 | chr11 | - | 78109441 | 78109581 | 78104355 | 78104453 | 78113885 | 78113989 | 1 | 0 | 0 | 1 | 2 | 0 | 0 | 0 | 0 | 2 | 0 | 0 | 0 | 9 | 0 | 0 | 0 | 0 | 0 | 0 | 289 | 149 | 1 | 1 | 1 | NA | NA | 1 | 0.34 | NA | NA | NA | 1 | NA | -0.22 |
| SE | 100136 | ENSG00000159063 | ALG8 | chr11 | - | 78112649 | 78112770 | 78104355 | 78104453 | 78113885 | 78113989 | 57 | 33 | 31 | 31 | 52 | 0 | 0 | 0 | 0 | 2 | 29 | 19 | 16 | 19 | 38 | 0 | 0 | 0 | 0 | 0 | 270 | 149 | 1 | 1 | 1 | 1 | 1 | 1 | 0.935 | 1 | 1 | 1 | 1 | 1 | -0.013 |
| SE | 100140 | ENSG00000159063 | ALG8 | chr11 | - | 78113885 | 78113989 | 78109441 | 78109581 | 78114265 | 78114392 | 66 | 55 | 75 | 59 | 60 | 0 | 0 | 0 | 0 | 0 | 53 | 31 | 27 | 62 | 16 | 0 | 4 | 0 | 1 | 0 | 253 | 149 | 1 | 1 | 1 | 1 | 1 | 1 | 1 | 1 | 0.82 | 1 | 0.973 | 1 | 0.041 |
| SE | 100141 | ENSG00000159063 | ALG8 | chr11 | - | 78113885 | 78113989 | 78112383 | 78112770 | 78114265 | 78114392 | 124 | 95 | 107 | 93 | 118 | 0 | 0 | 0 | 4 | 0 | 82 | 52 | 44 | 83 | 54 | 0 | 0 | 0 | 0 | 0 | 253 | 149 | 1 | 1 | 1 | 1 | 1 | 0.932 | 1 | 1 | 1 | 1 | 1 | 1 | -0.014 |
| SE | 100144 | ENSG00000159063 | ALG8 | chr11 | - | 78124020 | 78124214 | 78121064 | 78121174 | 78127357 | 78127436 | 115 | 68 | 68 | 99 | 132 | 0 | 2 | 0 | 0 | 0 | 53 | 62 | 63 | 56 | 57 | 0 | 0 | 0 | 3 | 0 | 298 | 149 | 1 | 1 | 1 | 0.944 | 1 | 1 | 1 | 1 | 1 | 1 | 0.903 | 1 | 0.008 |
| SE | 100145 | ENSG00000159063 | ALG8 | chr11 | - | 78124024 | 78124214 | 78120094 | 78121174 | 78127357 | 78127436 | 46 | 26 | 29 | 46 | 77 | 0 | 2 | 0 | 0 | 0 | 25 | 34 | 34 | 23 | 35 | 0 | 0 | 0 | 3 | 0 | 298 | 149 | 1 | 1 | 1 | 0.867 | 1 | 1 | 1 | 1 | 1 | 1 | 0.793 | 1 | 0.015 |
| SE | 100147 | ENSG00000159063 | ALG8 | chr11 | - | 78127357 | 78127436 | 78124020 | 78124214 | 78138709 | 78138767 | 43 | 22 | 25 | 32 | 65 | 0 | 0 | 0 | 0 | 2 | 25 | 25 | 25 | 20 | 26 | 0 | 0 | 3 | 0 | 0 | 228 | 149 | 1 | 1 | 1 | 1 | 1 | 1 | 0.955 | 1 | 1 | 0.845 | 1 | 1 | 0.022 |
| SE | 100151 | ENSG00000159063 | ALG8 | chr11 | - | 78134027 | 78135347 | 78127357 | 78127436 | 78139493 | 78139611 | 0 | 0 | 0 | 0 | 3 | 13 | 9 | 3 | 12 | 9 | 0 | 0 | 0 | 0 | 0 | 0 | 5 | 15 | 4 | 9 | 298 | 149 | 1 | 1 | 0 | 0 | 0 | 0 | 0.143 | NA | 0 | 0 | 0 | 0 | 0.029 |
| SE | 100152 | ENSG00000159063 | ALG8 | chr11 | - | 78135229 | 78135347 | 78127357 | 78127436 | 78138709 | 78139512 | 0 | 6 | 0 | 0 | 0 | 2 | 1 | 2 | 2 | 8 | 0 | 0 | 0 | 0 | 0 | 5 | 4 | 3 | 0 | 0 | 267 | 149 | 1 | 1 | 0 | 0.77 | 0 | 0 | 0 | 0 | 0 | 0 | NA | NA | 0.154 |
| SE | 100154 | ENSG00000159063 | ALG8 | chr11 | - | 78137241 | 78137645 | 78127357 | 78127436 | 78138709 | 78139626 | 2 | 2 | 0 | 1 | 1 | 2 | 1 | 2 | 2 | 8 | 0 | 11 | 0 | 4 | 0 | 5 | 4 | 3 | 0 | 0 | 298 | 149 | 1 | 1 | 0.333 | 0.5 | 0 | 0.2 | 0.059 | 0 | 0.579 | 0 | 1 | NA | -0.176 |
| SE | 100155 | ENSG00000159063 | ALG8 | chr11 | - | 78137241 | 78137645 | 78127357 | 78127436 | 78139493 | 78139611 | 1 | 0 | 0 | 0 | 3 | 13 | 9 | 3 | 12 | 9 | 0 | 0 | 0 | 0 | 0 | 0 | 5 | 15 | 4 | 9 | 298 | 149 | 1 | 1 | 0.037 | 0 | 0 | 0 | 0.143 | NA | 0 | 0 | 0 | 0 | 0.036 |
| SE | 100156 | ENSG00000159063 | ALG8 | chr11 | - | 78137241 | 78137645 | 78135288 | 78135347 | 78138709 | 78139626 | 1 | 2 | 0 | 1 | 1 | 0 | 6 | 0 | 0 | 0 | 0 | 11 | 0 | 4 | 0 | 0 | 0 | 0 | 0 | 0 | 298 | 149 | 1 | 1 | 1 | 0.143 | NA | 1 | 1 | NA | 1 | NA | 1 | NA | -0.214 |
| SE | 100254 | ENSG00000166471 | TMEM41B | chr11 | - | 9283370 | 9283593 | 9280653 | 9281041 | 9286454 | 9286593 | 82 | 79 | 109 | 93 | 112 | 0 | 0 | 0 | 0 | 0 | 63 | 120 | 127 | 102 | 4 | 0 | 0 | 0 | 2 | 0 | 298 | 149 | 1 | 1 | 1 | 1 | 1 | 1 | 1 | 1 | 1 | 1 | 0.962 | 1 | 0.008 |
| SE | 100256 | ENSG00000166471 | TMEM41B | chr11 | - | 9299583 | 9299701 | 9295095 | 9295387 | 9314320 | 9314567 | 38 | 26 | 59 | 45 | 77 | 0 | 0 | 0 | 1 | 0 | 0 | 31 | 23 | 23 | 20 | 0 | 0 | 0 | 0 | 0 | 267 | 149 | 1 | 1 | 1 | 1 | 1 | 0.962 | 1 | NA | 1 | 1 | 1 | 1 | -0.008 |
| SE | 100271 | ENSG00000167986 | DDB1 | chr11 | - | 61302581 | 61302751 | 61301711 | 61302359 | 61303045 | 61303155 | 462 | 253 | 431 | 402 | 611 | 0 | 0 | 0 | 2 | 0 | 580 | 361 | 201 | 236 | 446 | 8 | 1 | 0 | 0 | 2 | 298 | 149 | 1 | 1 | 1 | 1 | 1 | 0.99 | 1 | 0.973 | 0.994 | 1 | 1 | 0.991 | 0.006 |
| SE | 100272 | ENSG00000167986 | DDB1 | chr11 | - | 61308982 | 61309077 | 61303750 | 61304035 | 61310294 | 61310418 | 104 | 41 | 103 | 90 | 208 | 0 | 0 | 0 | 0 | 0 | 114 | 97 | 50 | 61 | 156 | 0 | 0 | 1 | 0 | 0 | 244 | 149 | 1 | 1 | 1 | 1 | 1 | 1 | 1 | 1 | 1 | 1 | 0.974 | 1 | 0.005 |
| SE | 100273 | ENSG00000167986 | DDB1 | chr11 | - | 61309795 | 61309861 | 61308982 | 61309077 | 61310294 | 61310418 | 106 | 63 | 91 | 100 | 90 | 0 | 0 | 0 | 1 | 0 | 92 | 89 | 57 | 101 | 99 | 0 | 0 | 0 | 0 | 0 | 215 | 149 | 1 | 1 | 1 | 1 | 1 | 0.986 | 1 | 1 | 1 | 1 | 1 | 1 | -0.003 |
| SE | 100274 | ENSG00000167986 | DDB1 | chr11 | - | 61309795 | 61309960 | 61308982 | 61309077 | 61310294 | 61310418 | 604 | 309 | 473 | 524 | 557 | 0 | 0 | 0 | 1 | 0 | 484 | 490 | 351 | 402 | 649 | 0 | 0 | 0 | 0 | 0 | 298 | 149 | 1 | 1 | 1 | 1 | 1 | 0.996 | 1 | 1 | 1 | 1 | 1 | 1 | -0.001 |
| SE | 100275 | ENSG00000167986 | DDB1 | chr11 | - | 61309795 | 61309960 | 61308982 | 61309077 | 61311783 | 61311895 | 308 | 164 | 236 | 285 | 276 | 0 | 0 | 0 | 3 | 0 | 281 | 251 | 172 | 234 | 375 | 0 | 1 | 0 | 0 | 0 | 298 | 149 | 1 | 1 | 1 | 1 | 1 | 0.979 | 1 | 1 | 0.992 | 1 | 1 | 1 | -0.003 |
| SE | 100276 | ENSG00000167986 | DDB1 | chr11 | - | 61309795 | 61309979 | 61308982 | 61309077 | 61310294 | 61310418 | 308 | 159 | 234 | 285 | 274 | 0 | 0 | 0 | 1 | 0 | 271 | 248 | 169 | 225 | 372 | 0 | 0 | 0 | 0 | 0 | 298 | 149 | 1 | 1 | 1 | 1 | 1 | 0.993 | 1 | 1 | 1 | 1 | 1 | 1 | -0.001 |
| SE | 100277 | ENSG00000167986 | DDB1 | chr11 | - | 61310294 | 61310418 | 61308982 | 61309077 | 61311783 | 61311895 | 262 | 116 | 181 | 237 | 329 | 0 | 0 | 0 | 3 | 0 | 240 | 237 | 170 | 183 | 396 | 0 | 1 | 0 | 0 | 0 | 273 | 149 | 1 | 1 | 1 | 1 | 1 | 0.977 | 1 | 1 | 0.992 | 1 | 1 | 1 | -0.003 |
| SE | 100279 | ENSG00000167986 | DDB1 | chr11 | - | 61313861 | 61313969 | 61313498 | 61313706 | 61314046 | 61314202 | 345 | 160 | 307 | 275 | 313 | 0 | 0 | 0 | 0 | 0 | 253 | 327 | 150 | 261 | 417 | 0 | 0 | 2 | 0 | 0 | 257 | 149 | 1 | 1 | 1 | 1 | 1 | 1 | 1 | 1 | 1 | 0.978 | 1 | 1 | 0.004 |
| SE | 100280 | ENSG00000167986 | DDB1 | chr11 | - | 61315331 | 61315523 | 61314452 | 61314486 | 61316284 | 61316393 | 0 | 0 | 0 | 1 | 0 | 269 | 137 | 240 | 221 | 509 | 0 | 0 | 0 | 0 | 0 | 173 | 203 | 128 | 146 | 306 | 298 | 149 | 1 | 1 | 0 | 0 | 0 | 0.002 | 0 | 0 | 0 | 0 | 0 | 0 | 0 |
| SE | 100282 | ENSG00000167986 | DDB1 | chr11 | - | 61316284 | 61316393 | 61314046 | 61314486 | 61321594 | 61321697 | 202 | 99 | 169 | 150 | 293 | 0 | 0 | 2 | 0 | 0 | 104 | 136 | 79 | 103 | 203 | 0 | 0 | 1 | 0 | 0 | 258 | 149 | 1 | 1 | 1 | 1 | 0.98 | 1 | 1 | 1 | 1 | 0.979 | 1 | 1 | 0 |
| SE | 100283 | ENSG00000167986 | DDB1 | chr11 | - | 61316491 | 61316567 | 61314883 | 61316393 | 61321594 | 61321697 | 384 | 208 | 336 | 346 | 502 | 0 | 1 | 0 | 0 | 1 | 384 | 312 | 253 | 283 | 483 | 0 | 0 | 0 | 0 | 0 | 225 | 149 | 1 | 1 | 1 | 0.993 | 1 | 1 | 0.997 | 1 | 1 | 1 | 1 | 1 | -0.002 |
| SE | 100284 | ENSG00000167986 | DDB1 | chr11 | - | 61321594 | 61321697 | 61316491 | 61316567 | 61322295 | 61322412 | 452 | 249 | 357 | 394 | 492 | 0 | 0 | 0 | 1 | 0 | 385 | 381 | 260 | 336 | 501 | 0 | 0 | 0 | 1 | 0 | 252 | 149 | 1 | 1 | 1 | 1 | 1 | 0.996 | 1 | 1 | 1 | 1 | 0.995 | 1 | 0 |
| SE | 100285 | ENSG00000167986 | DDB1 | chr11 | - | 61322295 | 61322412 | 61316491 | 61316567 | 61323978 | 61324137 | 0 | 0 | 0 | 1 | 0 | 1 | 0 | 0 | 0 | 0 | 18 | 0 | 9 | 5 | 4 | 0 | 0 | 1 | 0 | 0 | 266 | 149 | 1 | 1 | 0 | NA | NA | 1 | NA | 1 | NA | 0.834 | 1 | 1 | -0.459 |
| SE | 100286 | ENSG00000167986 | DDB1 | chr11 | - | 61323010 | 61323094 | 61314046 | 61314486 | 61323978 | 61324137 | 178 | 124 | 161 | 147 | 157 | 0 | 0 | 2 | 2 | 0 | 155 | 103 | 100 | 110 | 183 | 0 | 0 | 0 | 0 | 0 | 233 | 149 | 1 | 1 | 1 | 1 | 0.981 | 0.979 | 1 | 1 | 1 | 1 | 1 | 1 | -0.008 |
| SE | 100289 | ENSG00000167986 | DDB1 | chr11 | - | 61325610 | 61325708 | 61323660 | 61324137 | 61326778 | 61326893 | 257 | 125 | 215 | 207 | 292 | 0 | 0 | 1 | 2 | 0 | 456 | 281 | 148 | 197 | 399 | 0 | 0 | 0 | 0 | 0 | 247 | 149 | 1 | 1 | 1 | 1 | 0.992 | 0.984 | 1 | 1 | 1 | 1 | 1 | 1 | -0.005 |
| SE | 100291 | ENSG00000167986 | DDB1 | chr11 | - | 61326778 | 61326893 | 61323660 | 61324137 | 61331542 | 61331691 | 2 | 0 | 2 | 2 | 3 | 0 | 0 | 0 | 0 | 0 | 0 | 4 | 1 | 0 | 7 | 0 | 6 | 0 | 0 | 0 | 264 | 149 | 1 | 1 | 1 | NA | 1 | 1 | 1 | NA | 0.273 | 1 | NA | 1 | 0.242 |
| SE | 100295 | ENSG00000167986 | DDB1 | chr11 | - | 61331542 | 61331691 | 61326778 | 61326893 | 61332907 | 61333008 | 81 | 31 | 72 | 74 | 45 | 0 | 0 | 0 | 0 | 0 | 52 | 98 | 50 | 52 | 124 | 0 | 0 | 0 | 2 | 0 | 298 | 149 | 1 | 1 | 1 | 1 | 1 | 1 | 1 | 1 | 1 | 1 | 0.929 | 1 | 0.014 |
| SE | 100700 | ENSG00000109917 | ZPR1 | chr11 | - | 116782918 | 116783029 | 116782157 | 116782244 | 116783529 | 116783619 | 316 | 229 | 286 | 359 | 230 | 0 | 0 | 0 | 1 | 0 | 466 | 400 | 329 | 384 | 339 | 0 | 0 | 0 | 0 | 0 | 260 | 149 | 1 | 1 | 1 | 1 | 1 | 0.995 | 1 | 1 | 1 | 1 | 1 | 1 | -0.001 |
| SE | 100703 | ENSG00000109917 | ZPR1 | chr11 | - | 116787481 | 116787602 | 116786968 | 116787059 | 116787819 | 116787898 | 72 | 44 | 74 | 93 | 95 | 1 | 2 | 0 | 0 | 0 | 114 | 72 | 85 | 114 | 109 | 0 | 0 | 0 | 0 | 2 | 270 | 149 | 1 | 1 | 0.975 | 0.924 | 1 | 1 | 1 | 1 | 1 | 1 | 1 | 0.968 | -0.014 |
| SE | 100704 | ENSG00000109917 | ZPR1 | chr11 | - | 116787481 | 116787643 | 116786968 | 116787059 | 116787819 | 116787989 | 122 | 68 | 110 | 161 | 138 | 1 | 2 | 0 | 0 | 0 | 182 | 113 | 134 | 181 | 136 | 0 | 0 | 0 | 0 | 2 | 298 | 149 | 1 | 1 | 0.984 | 0.944 | 1 | 1 | 1 | 1 | 1 | 1 | 1 | 0.971 | -0.009 |
| SE | 100707 | ENSG00000187147 | RNF220 | chr1 | + | 44649882 | 44649957 | 44649660 | 44649769 | 44650703 | 44650808 | 392 | 156 | 363 | 355 | 439 | 1 | 2 | 0 | 1 | 1 | 371 | 338 | 251 | 163 | 484 | 0 | 0 | 1 | 3 | 0 | 224 | 149 | 1 | 1 | 0.996 | 0.981 | 1 | 0.996 | 0.997 | 1 | 1 | 0.994 | 0.973 | 1 | 0.001 |
| SE | 101368 | ENSG00000107537 | PHYH | chr10 | - | 13298186 | 13298245 | 13295227 | 13295606 | 13299419 | 13299549 | 9 | 9 | 4 | 4 | 14 | 1 | 0 | 0 | 0 | 0 | 13 | 0 | 10 | 12 | 0 | 0 | 0 | 0 | 0 | 0 | 208 | 149 | 1 | 1 | 0.866 | 1 | 1 | 1 | 1 | 1 | NA | 1 | 1 | NA | -0.027 |
| SE | 101609 | ENSG00000143702 | CEP170 | chr1 | - | 243126377 | 243126494 | 243125917 | 243126134 | 243126612 | 243126698 | 0 | 0 | 0 | 0 | 0 | 1 | 0 | 2 | 2 | 7 | 0 | 2 | 0 | 0 | 0 | 10 | 1 | 0 | 0 | 0 | 266 | 149 | 1 | 1 | 0 | NA | 0 | 0 | 0 | 0 | 0.528 | NA | NA | NA | -0.264 |
| SE | 101610 | ENSG00000143702 | CEP170 | chr1 | - | 243142285 | 243142463 | 243139936 | 243140107 | 243156220 | 243156347 | 46 | 32 | 32 | 46 | 53 | 0 | 2 | 1 | 4 | 1 | 0 | 17 | 19 | 9 | 0 | 0 | 5 | 1 | 0 | 0 | 298 | 149 | 1 | 1 | 1 | 0.889 | 0.941 | 0.852 | 0.964 | NA | 0.63 | 0.905 | 1 | NA | 0.084 |
| SE | 101611 | ENSG00000143702 | CEP170 | chr1 | - | 243142285 | 243142463 | 243139936 | 243140107 | 243164283 | 243166116 | 4 | 3 | 5 | 5 | 9 | 0 | 0 | 3 | 0 | 0 | 0 | 0 | 2 | 0 | 0 | 0 | 0 | 0 | 0 | 0 | 298 | 149 | 1 | 1 | 1 | 1 | 0.455 | 1 | 1 | NA | NA | 1 | NA | NA | -0.109 |
| SE | 101612 | ENSG00000143702 | CEP170 | chr1 | - | 243142315 | 243142463 | 243139936 | 243140107 | 243156220 | 243156455 | 56 | 42 | 33 | 55 | 66 | 0 | 2 | 1 | 4 | 1 | 0 | 20 | 24 | 9 | 0 | 0 | 5 | 1 | 0 | 0 | 297 | 149 | 1 | 1 | 1 | 0.913 | 0.943 | 0.873 | 0.971 | NA | 0.667 | 0.923 | 1 | NA | 0.077 |
| SE | 101613 | ENSG00000143702 | CEP170 | chr1 | - | 243142315 | 243142463 | 243139936 | 243140107 | 243164283 | 243166116 | 14 | 13 | 6 | 14 | 22 | 0 | 0 | 3 | 0 | 0 | 0 | 3 | 7 | 0 | 0 | 0 | 0 | 0 | 0 | 0 | 297 | 149 | 1 | 1 | 1 | 1 | 0.501 | 1 | 1 | NA | 1 | 1 | NA | NA | -0.1 |
| SE | 101614 | ENSG00000143702 | CEP170 | chr1 | - | 243142366 | 243142463 | 243139936 | 243140107 | 243156220 | 243156347 | 39 | 18 | 20 | 47 | 35 | 0 | 2 | 1 | 4 | 1 | 0 | 6 | 15 | 12 | 0 | 0 | 5 | 1 | 0 | 0 | 246 | 149 | 1 | 1 | 1 | 0.845 | 0.924 | 0.877 | 0.955 | NA | 0.421 | 0.901 | 1 | NA | 0.146 |
| SE | 101615 | ENSG00000143702 | CEP170 | chr1 | - | 243142366 | 243142463 | 243139936 | 243140107 | 243164283 | 243164494 | 13 | 2 | 3 | 10 | 6 | 0 | 0 | 3 | 0 | 0 | 0 | 0 | 4 | 3 | 0 | 0 | 0 | 0 | 0 | 0 | 246 | 149 | 1 | 1 | 1 | 1 | 0.377 | 1 | 1 | NA | NA | 1 | 1 | NA | -0.125 |
| SE | 101616 | ENSG00000143702 | CEP170 | chr1 | - | 243156220 | 243156347 | 243139936 | 243140107 | 243164283 | 243164494 | 10 | 9 | 10 | 2 | 14 | 0 | 0 | 3 | 0 | 0 | 0 | 5 | 1 | 0 | 0 | 0 | 0 | 0 | 0 | 0 | 276 | 149 | 1 | 1 | 1 | 1 | 0.643 | 1 | 1 | NA | 1 | 1 | NA | NA | -0.071 |
| SE | 101617 | ENSG00000143702 | CEP170 | chr1 | - | 243156220 | 243156347 | 243142285 | 243142463 | 243164283 | 243164494 | 50 | 36 | 38 | 45 | 55 | 0 | 0 | 0 | 1 | 0 | 0 | 17 | 16 | 9 | 0 | 0 | 0 | 0 | 0 | 0 | 276 | 149 | 1 | 1 | 1 | 1 | 1 | 0.96 | 1 | NA | 1 | 1 | 1 | NA | -0.008 |
| SE | 101618 | ENSG00000143702 | CEP170 | chr1 | - | 243156220 | 243156455 | 243139936 | 243140107 | 243164283 | 243166116 | 58 | 71 | 45 | 74 | 153 | 0 | 0 | 3 | 0 | 0 | 8 | 35 | 36 | 0 | 7 | 0 | 0 | 0 | 0 | 0 | 298 | 149 | 1 | 1 | 1 | 1 | 0.882 | 1 | 1 | 1 | 1 | 1 | NA | 1 | -0.024 |
| SE | 101619 | ENSG00000143702 | CEP170 | chr1 | - | 243156220 | 243156455 | 243142285 | 243142463 | 243164283 | 243166116 | 100 | 98 | 76 | 117 | 198 | 0 | 0 | 0 | 1 | 0 | 8 | 47 | 53 | 9 | 7 | 0 | 0 | 0 | 0 | 0 | 298 | 149 | 1 | 1 | 1 | 1 | 1 | 0.983 | 1 | 1 | 1 | 1 | 1 | 1 | -0.003 |
| SE | 101620 | ENSG00000143702 | CEP170 | chr1 | - | 243176701 | 243176816 | 243172696 | 243172846 | 243185778 | 243186072 | 0 | 3 | 0 | 0 | 0 | 64 | 65 | 58 | 47 | 109 | 0 | 0 | 0 | 0 | 0 | 0 | 24 | 21 | 2 | 8 | 264 | 149 | 1 | 1 | 0 | 0.025 | 0 | 0 | 0 | NA | 0 | 0 | 0 | 0 | 0.005 |
| SE | 101621 | ENSG00000143702 | CEP170 | chr1 | - | 243185778 | 243186072 | 243172696 | 243172846 | 243186258 | 243186280 | 135 | 113 | 108 | 104 | 191 | 24 | 24 | 17 | 18 | 32 | 0 | 66 | 40 | 13 | 30 | 0 | 15 | 8 | 0 | 0 | 298 | 149 | 1 | 1 | 0.738 | 0.702 | 0.761 | 0.743 | 0.749 | NA | 0.688 | 0.714 | 1 | 1 | -0.112 |
| SE | 101743 | ENSG00000198056 | PRIM1 | chr12 | - | 56739293 | 56739363 | 56738433 | 56738525 | 56741434 | 56741576 | 110 | 79 | 68 | 110 | 129 | 0 | 0 | 0 | 0 | 0 | 53 | 55 | 39 | 41 | 45 | 0 | 0 | 0 | 3 | 0 | 219 | 149 | 1 | 1 | 1 | 1 | 1 | 1 | 1 | 1 | 1 | 1 | 0.903 | 1 | 0.019 |
| SE | 101962 | ENSG00000197771 | MCMBP | chr10 | - | 119836895 | 119837029 | 119835539 | 119835704 | 119838534 | 119838700 | 495 | 307 | 356 | 363 | 730 | 4 | 0 | 0 | 2 | 0 | 583 | 295 | 306 | 477 | 351 | 0 | 0 | 0 | 0 | 0 | 283 | 149 | 1 | 1 | 0.985 | 1 | 1 | 0.99 | 1 | 1 | 1 | 1 | 1 | 1 | -0.005 |
| SE | 101963 | ENSG00000197771 | MCMBP | chr10 | - | 119838534 | 119838700 | 119835539 | 119835704 | 119840842 | 119840960 | 450 | 280 | 299 | 403 | 908 | 1 | 0 | 0 | 0 | 0 | 483 | 303 | 293 | 411 | 297 | 0 | 0 | 3 | 0 | 0 | 298 | 149 | 1 | 1 | 0.996 | 1 | 1 | 1 | 1 | 1 | 1 | 0.98 | 1 | 1 | 0.003 |
| SE | 101964 | ENSG00000197771 | MCMBP | chr10 | - | 119840842 | 119840960 | 119838534 | 119838700 | 119842471 | 119842595 | 667 | 392 | 425 | 599 | 1373 | 0 | 0 | 1 | 0 | 0 | 644 | 449 | 414 | 588 | 382 | 0 | 0 | 0 | 0 | 0 | 267 | 149 | 1 | 1 | 1 | 1 | 0.996 | 1 | 1 | 1 | 1 | 1 | 1 | 1 | -0.001 |
| SE | 101965 | ENSG00000197771 | MCMBP | chr10 | - | 119840842 | 119840960 | 119838534 | 119838700 | 119843253 | 119843426 | 418 | 256 | 286 | 378 | 843 | 0 | 0 | 2 | 0 | 1 | 452 | 285 | 270 | 381 | 261 | 0 | 0 | 1 | 0 | 0 | 267 | 149 | 1 | 1 | 1 | 1 | 0.988 | 1 | 0.998 | 1 | 1 | 0.993 | 1 | 1 | -0.001 |
| SE | 101966 | ENSG00000197771 | MCMBP | chr10 | - | 119842471 | 119842595 | 119838534 | 119838700 | 119843253 | 119843426 | 122 | 57 | 76 | 123 | 195 | 0 | 0 | 2 | 0 | 1 | 32 | 95 | 77 | 66 | 20 | 0 | 0 | 1 | 0 | 0 | 273 | 149 | 1 | 1 | 1 | 1 | 0.954 | 1 | 0.991 | 1 | 1 | 0.977 | 1 | 1 | -0.006 |
| SE | 102319 | ENSG00000075945 | KIFAP3 | chr1 | - | 169984581 | 169984690 | 169983269 | 169983382 | 169992154 | 169992255 | 278 | 98 | 138 | 157 | 136 | 0 | 0 | 0 | 0 | 0 | 99 | 97 | 130 | 21 | 274 | 0 | 0 | 0 | 2 | 0 | 258 | 149 | 1 | 1 | 1 | 1 | 1 | 1 | 1 | 1 | 1 | 1 | 0.858 | 1 | 0.028 |
| SE | 102321 | ENSG00000075945 | KIFAP3 | chr1 | - | 170041676 | 170041840 | 170039232 | 170039288 | 170046711 | 170046866 | 1 | 3 | 0 | 1 | 3 | 97 | 85 | 89 | 71 | 173 | 2 | 4 | 0 | 0 | 0 | 26 | 88 | 53 | 28 | 61 | 298 | 149 | 1 | 1 | 0.005 | 0.017 | 0 | 0.007 | 0.009 | 0.037 | 0.022 | 0 | 0 | 0 | -0.004 |
| SE | 102322 | ENSG00000075945 | KIFAP3 | chr1 | - | 170046711 | 170046866 | 170039232 | 170039288 | 170055304 | 170055436 | 187 | 155 | 157 | 138 | 376 | 0 | 0 | 0 | 0 | 0 | 59 | 176 | 85 | 49 | 140 | 0 | 1 | 0 | 0 | 0 | 298 | 149 | 1 | 1 | 1 | 1 | 1 | 1 | 1 | 1 | 0.989 | 1 | 1 | 1 | 0.002 |
| SE | 102323 | ENSG00000075945 | KIFAP3 | chr1 | - | 170055304 | 170055436 | 170046711 | 170046866 | 170074435 | 170074695 | 115 | 84 | 89 | 75 | 220 | 0 | 2 | 0 | 2 | 0 | 47 | 100 | 48 | 40 | 79 | 0 | 0 | 0 | 0 | 0 | 281 | 149 | 1 | 1 | 1 | 0.957 | 1 | 0.952 | 1 | 1 | 1 | 1 | 1 | 1 | -0.018 |
| SE | 102408 | ENSG00000138346 | DNA2 | chr10 | - | 68419802 | 68419892 | 68419033 | 68419213 | 68422224 | 68422429 | 13 | 5 | 7 | 22 | 17 | 2 | 0 | 0 | 0 | 0 | 0 | 1 | 7 | 4 | 0 | 0 | 0 | 0 | 4 | 0 | 239 | 149 | 1 | 1 | 0.802 | 1 | 1 | 1 | 1 | NA | 1 | 1 | 0.384 | NA | 0.166 |
| SE | 102410 | ENSG00000138346 | DNA2 | chr10 | - | 68437010 | 68437241 | 68432393 | 68432510 | 68442916 | 68443111 | 14 | 19 | 15 | 23 | 29 | 0 | 0 | 0 | 3 | 0 | 19 | 10 | 17 | 18 | 5 | 0 | 0 | 0 | 0 | 0 | 298 | 149 | 1 | 1 | 1 | 1 | 1 | 0.793 | 1 | 1 | 1 | 1 | 1 | 1 | -0.041 |
| SE | 102411 | ENSG00000138346 | DNA2 | chr10 | - | 68442916 | 68443111 | 68437010 | 68437241 | 68450027 | 68450247 | 12 | 17 | 15 | 13 | 16 | 0 | 0 | 0 | 0 | 1 | 19 | 4 | 11 | 17 | 5 | 0 | 0 | 3 | 0 | 0 | 298 | 149 | 1 | 1 | 1 | 1 | 1 | 1 | 0.889 | 1 | 1 | 0.647 | 1 | 1 | 0.048 |
| SE | 102412 | ENSG00000138346 | DNA2 | chr10 | - | 68444920 | 68445083 | 68442916 | 68443111 | 68450027 | 68450247 | 11 | 9 | 19 | 15 | 27 | 0 | 1 | 1 | 0 | 0 | 0 | 9 | 9 | 12 | 0 | 0 | 0 | 0 | 0 | 0 | 298 | 149 | 1 | 1 | 1 | 0.818 | 0.905 | 1 | 1 | NA | 1 | 1 | 1 | NA | -0.055 |
| SE | 102415 | ENSG00000138346 | DNA2 | chr10 | - | 68468122 | 68468306 | 68465666 | 68465812 | 68469980 | 68470163 | 59 | 42 | 31 | 62 | 97 | 0 | 2 | 0 | 0 | 0 | 135 | 96 | 59 | 102 | 95 | 0 | 0 | 0 | 0 | 0 | 298 | 149 | 1 | 1 | 1 | 0.913 | 1 | 1 | 1 | 1 | 1 | 1 | 1 | 1 | -0.017 |
| SE | 102416 | ENSG00000138346 | DNA2 | chr10 | - | 68470488 | 68470612 | 68469980 | 68470163 | 68471790 | 68471886 | 0 | 0 | 0 | 0 | 3 | 2 | 1 | 2 | 1 | 4 | 0 | 0 | 0 | 0 | 0 | 0 | 4 | 3 | 2 | 9 | 273 | 149 | 1 | 1 | 0 | 0 | 0 | 0 | 0.29 | NA | 0 | 0 | 0 | 0 | 0.058 |
| SE | 102560 | ENSG00000171307 | ZDHHC16 | chr10 | + | 97450356 | 97450536 | 97446174 | 97446353 | 97451670 | 97451918 | 0 | 6 | 4 | 0 | 0 | 4 | 4 | 0 | 3 | 3 | 0 | 0 | 0 | 4 | 0 | 0 | 0 | 0 | 0 | 0 | 298 | 149 | 1 | 1 | 0 | 0.429 | 1 | 0 | 0 | NA | NA | NA | 1 | NA | -0.714 |
| SE | 102561 | ENSG00000171307 | ZDHHC16 | chr10 | + | 97451670 | 97451918 | 97446169 | 97446353 | 97452414 | 97452503 | 4 | 4 | 1 | 3 | 3 | 0 | 0 | 0 | 0 | 0 | 0 | 0 | 0 | 0 | 0 | 0 | 0 | 0 | 0 | 8 | 298 | 149 | 1 | 1 | 1 | 1 | 1 | 1 | 1 | NA | NA | NA | NA | 0 | 1 |
| SE | 102562 | ENSG00000171307 | ZDHHC16 | chr10 | + | 97452089 | 97452284 | 97451670 | 97451918 | 97452414 | 97452465 | 50 | 17 | 28 | 43 | 25 | 0 | 0 | 1 | 0 | 0 | 59 | 52 | 27 | 31 | 45 | 0 | 0 | 0 | 0 | 0 | 298 | 149 | 1 | 1 | 1 | 1 | 0.933 | 1 | 1 | 1 | 1 | 1 | 1 | 1 | -0.013 |
| SE | 102564 | ENSG00000171307 | ZDHHC16 | chr10 | + | 97455638 | 97455783 | 97454713 | 97454799 | 97455973 | 97456044 | 47 | 16 | 22 | 32 | 36 | 0 | 0 | 1 | 0 | 0 | 34 | 30 | 13 | 18 | 34 | 0 | 2 | 0 | 0 | 0 | 294 | 149 | 1 | 1 | 1 | 1 | 0.918 | 1 | 1 | 1 | 0.884 | 1 | 1 | 1 | 0.007 |

| | | | | | | | | | | | | | | | | | | | | | | | | | | | | | | | | | | | | | | | | | | | | | | |
|---|---|---|---|---|---|---|---|---|---|---|---|---|---|---|---|---|---|---|---|---|---|---|---|---|---|---|---|---|---|---|---|---|---|---|---|---|---|---|---|---|---|---|---|---|---|---|
| SE | 102565 | ENSG00000171307 | ZDHHC16 | chr10 | + | 97455659 | 97455783 | 97454713 | 97454799 | 97455973 | 97456044 | 83 | 33 | 50 | 59 | 71 | 0 | 0 | 1 | 0 | 0 | 83 | 55 | 20 | 31 | 52 | 0 | 2 | 0 | 0 | 0 | 273 | 149 | 1 | 1 | 1 | 1 | 0.965 | 1 | 1 | 1 | 0.938 | 1 | 1 | 1 | 0.005 |
| SE | 102566 | ENSG00000171307 | ZDHHC16 | chr10 | + | 97455713 | 97455783 | 97454713 | 97454799 | 97455973 | 97456044 | 39 | 17 | 16 | 22 | 24 | 0 | 0 | 1 | 0 | 0 | 10 | 30 | 4 | 6 | 30 | 0 | 2 | 0 | 0 | 0 | 219 | 149 | 1 | 1 | 1 | 1 | 0.916 | 1 | 1 | 1 | 0.911 | 1 | 1 | 1 | 0.001 |
| SE | 102567 | ENSG00000171307 | ZDHHC16 | chr10 | + | 97455973 | 97456044 | 97455601 | 97455783 | 97456776 | 97456924 | 87 | 25 | 46 | 71 | 70 | 0 | 0 | 4 | 0 | 0 | 69 | 75 | 18 | 40 | 76 | 0 | 0 | 0 | 0 | 0 | 220 | 149 | 1 | 1 | 1 | 0.809 | 1 | 1 | 1 | 1 | 1 | 1 | 1 | 1 | -0.038 |
| SE | 102578 | ENSG00000120438 | TCP1 | chr6 | - | 159779894 | 159780087 | 159779626 | 159779790 | 159780442 | 159780566 | 922 | 698 | 1047 | 1113 | 913 | 0 | 0 | 1 | 0 | 0 | 758 | 764 | 1033 | 856 | 320 | 0 | 0 | 0 | 11 | 0 | 298 | 149 | 1 | 1 | 1 | 1 | 0.998 | 1 | 1 | 1 | 1 | 1 | 0.975 | 1 | 0.005 |
| SE | 102586 | ENSG00000120438 | TCP1 | chr6 | - | 159788057 | 159788143 | 159787742 | 159787871 | 159789404 | 159789572 | 183 | 107 | 156 | 159 | 210 | 0 | 0 | 0 | 0 | 0 | 82 | 67 | 90 | 104 | 13 | 0 | 0 | 3 | 0 | 0 | 235 | 149 | 1 | 1 | 1 | 1 | 1 | 1 | 1 | 1 | 1 | 0.95 | 1 | 1 | 0.01 |
| SE | 102587 | ENSG00000120438 | TCP1 | chr6 | - | 159788609 | 159788696 | 159788057 | 159788143 | 159789404 | 159789672 | 0 | 1 | 4 | 1 | 0 | 76 | 54 | 66 | 70 | 70 | 0 | 0 | 2 | 3 | 0 | 19 | 31 | 60 | 44 | 0 | 236 | 149 | 1 | 1 | 0 | 0.012 | 0.037 | 0.009 | 0 | 0 | 0 | 0.021 | 0.041 | NA | -0.004 |
| SE | 102591 | ENSG00000175470 | PPP2R2D | chr10 | + | 131940030 | 131940196 | 131901237 | 131901330 | 131955683 | 131956555 | 0 | 3 | 4 | 3 | 7 | 0 | 0 | 0 | 1 | 0 | 10 | 0 | 3 | 3 | 1 | 0 | 0 | 0 | 0 | 3 | 298 | 149 | 1 | 1 | NA | 1 | 1 | 0.6 | 1 | 1 | NA | 1 | 1 | 0.143 | 0.114 |
| SE | 102592 | ENSG00000175470 | PPP2R2D | chr10 | + | 131940581 | 131940694 | 131939775 | 131940196 | 131943967 | 131943997 | 188 | 133 | 149 | 152 | 302 | 0 | 0 | 0 | 2 | 0 | 226 | 162 | 110 | 178 | 318 | 0 | 0 | 0 | 0 | 0 | 262 | 149 | 1 | 1 | 1 | 1 | 1 | 0.977 | 1 | 1 | 1 | 1 | 1 | 1 | -0.005 |
| SE | 102595 | ENSG00000175470 | PPP2R2D | chr10 | + | 131940581 | 131940841 | 131940158 | 131940196 | 131943967 | 131944145 | 103 | 82 | 90 | 98 | 167 | 0 | 0 | 0 | 2 | 0 | 176 | 84 | 73 | 92 | 238 | 0 | 0 | 0 | 0 | 0 | 298 | 149 | 1 | 1 | 1 | 1 | 1 | 0.961 | 1 | 1 | 1 | 1 | 1 | 1 | -0.008 |
| SE | 102596 | ENSG00000175470 | PPP2R2D | chr10 | + | 131943967 | 131943997 | 131940162 | 131940196 | 131944094 | 131944145 | 0 | 0 | 0 | 1 | 0 | 0 | 0 | 0 | 0 | 0 | 0 | 0 | 0 | 0 | 0 | 16 | 0 | 1 | 0 | 0 | 179 | 149 | 1 | 1 | NA | NA | NA | 1 | NA | 0 | NA | 0 | NA | NA | 1 |
| SE | 103091 | ENSG00000108055 | SMC3 | chr10 | + | 110573706 | 110573745 | 110568937 | 110569013 | 110575335 | 110575403 | 62 | 42 | 58 | 76 | 129 | 0 | 0 | 0 | 0 | 6 | 24 | 43 | 14 | 12 | 0 | 0 | 0 | 0 | 0 | 0 | 188 | 149 | 1 | 1 | 1 | 1 | 1 | 1 | 0.945 | 1 | 1 | 1 | 1 | NA | -0.011 |
| SE | 103094 | ENSG00000108055 | SMC3 | chr10 | + | 110582561 | 110582642 | 110580903 | 110581021 | 110583383 | 110583548 | 125 | 182 | 145 | 174 | 160 | 0 | 0 | 0 | 1 | 3 | 0 | 78 | 81 | 54 | 0 | 0 | 0 | 5 | 0 | 0 | 230 | 149 | 1 | 1 | 1 | 1 | 1 | 0.991 | 0.972 | NA | 1 | 0.913 | 1 | NA | 0.022 |
| SE | 103096 | ENSG00000108055 | SMC3 | chr10 | + | 110590411 | 110590572 | 110589891 | 110589991 | 110590990 | 110591132 | 294 | 308 | 300 | 293 | 397 | 0 | 0 | 1 | 0 | 0 | 40 | 63 | 107 | 39 | 9 | 0 | 0 | 0 | 0 | 0 | 298 | 149 | 1 | 1 | 1 | 1 | 0.993 | 1 | 1 | 1 | 1 | 1 | 1 | 1 | -0.001 |
| SE | 103097 | ENSG00000108055 | SMC3 | chr10 | + | 110590990 | 110591132 | 110590411 | 110590572 | 110593072 | 110593223 | 354 | 310 | 341 | 331 | 421 | 1 | 2 | 2 | 0 | 0 | 21 | 110 | 114 | 49 | 3 | 0 | 3 | 0 | 0 | 0 | 291 | 149 | 1 | 1 | 0.995 | 0.988 | 0.989 | 1 | 1 | 1 | 0.949 | 1 | 1 | 1 | 0.005 |
| SE | 103098 | ENSG00000108055 | SMC3 | chr10 | + | 110593072 | 110593223 | 110590990 | 110591132 | 110596397 | 110596550 | 372 | 258 | 365 | 398 | 303 | 0 | 0 | 0 | 2 | 0 | 20 | 179 | 89 | 31 | 5 | 0 | 0 | 0 | 0 | 0 | 298 | 149 | 1 | 1 | 1 | 1 | 1 | 0.99 | 1 | 1 | 1 | 1 | 1 | 1 | -0.002 |
| SE | 103099 | ENSG00000108055 | SMC3 | chr10 | + | 110600438 | 110600546 | 110599653 | 110599812 | 110601021 | 110601130 | 199 | 162 | 149 | 211 | 295 | 0 | 0 | 0 | 0 | 0 | 15 | 47 | 48 | 35 | 0 | 0 | 0 | 0 | 2 | 0 | 257 | 149 | 1 | 1 | 1 | 1 | 1 | 1 | 1 | 1 | 1 | 1 | 0.91 | NA | 0.022 |
| SE | 103100 | ENSG00000108055 | SMC3 | chr10 | + | 110602473 | 110602665 | 110601965 | 110602178 | 110602824 | 110603002 | 408 | 442 | 299 | 376 | 630 | 4 | 2 | 1 | 2 | 2 | 0 | 114 | 126 | 25 | 7 | 0 | 0 | 2 | 0 | 0 | 298 | 149 | 1 | 1 | 0.981 | 0.991 | 0.993 | 0.989 | 0.994 | NA | 1 | 0.969 | 1 | 1 | -0.003 |
| SE | 103324 | ENSG00000148688 | RPP30 | chr10 | + | 90874868 | 90874924 | 90871951 | 90872068 | 90875557 | 90875614 | 128 | 50 | 72 | 91 | 101 | 1 | 0 | 0 | 0 | 0 | 44 | 76 | 61 | 76 | 72 | 0 | 0 | 0 | 0 | 0 | 205 | 149 | 1 | 1 | 0.989 | 1 | 1 | 1 | 1 | 1 | 1 | 1 | 1 | 1 | -0.002 |
| SE | 103327 | ENSG00000148688 | RPP30 | chr10 | + | 90902242 | 90902344 | 90900569 | 90900668 | 90903191 | 90903262 | 3 | 9 | 17 | 13 | 26 | 0 | 0 | 1 | 0 | 0 | 20 | 4 | 11 | 3 | 10 | 0 | 0 | 0 | 0 | 0 | 251 | 149 | 1 | 1 | 1 | 1 | 0.91 | 1 | 1 | 1 | 1 | 1 | 1 | 1 | -0.018 |
| SE | 103328 | ENSG00000148688 | RPP30 | chr10 | + | 90903191 | 90903276 | 90900569 | 90900668 | 90905301 | 90906065 | 6 | 4 | 7 | 7 | 19 | 0 | 1 | 0 | 0 | 0 | 0 | 4 | 0 | 5 | 15 | 0 | 0 | 0 | 0 | 0 | 234 | 149 | 1 | 1 | 1 | 0.718 | 1 | 1 | 1 | NA | 1 | NA | 1 | 1 | -0.056 |
| SE | 103329 | ENSG00000150760 | DOCK1 | chr10 | + | 127447393 | 127447545 | 127444125 | 127444279 | 127451331 | 127452516 | 2 | 1 | 0 | 0 | 0 | 0 | 0 | 0 | 2 | 0 | 0 | 4 | 2 | 0 | 0 | 0 | 0 | 0 | 0 | 0 | 298 | 149 | 1 | 1 | 1 | 1 | NA | 0 | NA | NA | 1 | 1 | NA | NA | -0.333 |
| SE | 103339 | ENSG00000108094 | CUL2 | chr10 | - | 35011847 | 35011964 | 35008550 | 35010442 | 35013698 | 35013800 | 121 | 64 | 77 | 82 | 154 | 0 | 2 | 0 | 0 | 7 | 16 | 70 | 40 | 31 | 0 | 0 | 4 | 0 | 0 | 0 | 266 | 149 | 1 | 1 | 1 | 0.947 | 1 | 1 | 0.925 | 1 | 0.907 | 1 | 1 | NA | -0.002 |
| SE | 103340 | ENSG00000108094 | CUL2 | chr10 | - | 35013698 | 35013800 | 35011847 | 35011964 | 35016191 | 35016394 | 167 | 116 | 116 | 149 | 311 | 3 | 0 | 0 | 0 | 0 | 19 | 108 | 69 | 45 | 17 | 0 | 0 | 0 | 0 | 0 | 251 | 149 | 1 | 1 | 0.971 | 1 | 1 | 1 | 1 | 1 | 1 | 1 | 1 | 1 | -0.006 |
| SE | 103341 | ENSG00000108094 | CUL2 | chr10 | - | 35029487 | 35029640 | 35028809 | 35028887 | 35031299 | 35031386 | 266 | 237 | 279 | 348 | 495 | 1 | 1 | 0 | 0 | 0 | 49 | 191 | 230 | 84 | 28 | 0 | 0 | 0 | 0 | 0 | 298 | 149 | 1 | 1 | 0.993 | 0.992 | 1 | 1 | 1 | 1 | 1 | 1 | 1 | 1 | -0.003 |
| SE | 103342 | ENSG00000108094 | CUL2 | chr10 | - | 35038919 | 35039082 | 35035171 | 35035296 | 35044565 | 35044676 | 216 | 134 | 161 | 209 | 400 | 0 | 2 | 0 | 0 | 0 | 29 | 91 | 128 | 27 | 23 | 0 | 0 | 0 | 0 | 0 | 298 | 149 | 1 | 1 | 1 | 0.971 | 1 | 1 | 1 | 1 | 1 | 1 | 1 | 1 | -0.006 |
| SE | 103343 | ENSG00000108094 | CUL2 | chr10 | - | 35044771 | 35044868 | 35044565 | 35044676 | 35049682 | 35049765 | 187 | 144 | 153 | 159 | 313 | 1 | 0 | 0 | 1 | 0 | 42 | 137 | 153 | 72 | 17 | 0 | 0 | 0 | 0 | 0 | 246 | 149 | 1 | 1 | 0.991 | 1 | 1 | 0.99 | 1 | 1 | 1 | 1 | 1 | 1 | -0.004 |
| SE | 103344 | ENSG00000108094 | CUL2 | chr10 | - | 35049682 | 35049765 | 35044771 | 35044868 | 35054433 | 35054539 | 133 | 123 | 106 | 98 | 229 | 0 | 3 | 0 | 1 | 0 | 26 | 110 | 116 | 42 | 12 | 0 | 0 | 0 | 0 | 0 | 232 | 149 | 1 | 1 | 1 | 0.963 | 1 | 0.984 | 1 | 1 | 1 | 1 | 1 | 1 | -0.011 |
| SE | 103345 | ENSG00000108094 | CUL2 | chr10 | - | 35049682 | 35049765 | 35044771 | 35044868 | 35060873 | 35060924 | 56 | 38 | 45 | 32 | 94 | 0 | 1 | 0 | 0 | 0 | 15 | 44 | 33 | 21 | 12 | 0 | 0 | 0 | 0 | 0 | 232 | 149 | 1 | 1 | 1 | 0.961 | 1 | 1 | 1 | 1 | 1 | 1 | 1 | 1 | -0.008 |
| SE | 103346 | ENSG00000108094 | CUL2 | chr10 | - | 35054433 | 35054539 | 35044771 | 35044868 | 35060873 | 35060924 | 156 | 132 | 124 | 134 | 144 | 0 | 1 | 0 | 0 | 0 | 65 | 96 | 166 | 67 | 14 | 0 | 0 | 0 | 0 | 0 | 255 | 149 | 1 | 1 | 1 | 0.987 | 1 | 1 | 1 | 1 | 1 | 1 | 1 | 1 | -0.003 |
| SE | 103349 | ENSG00000108094 | CUL2 | chr10 | - | 35089895 | 35090001 | 35071198 | 35071339 | 35090178 | 35090237 | 9 | 6 | 13 | 2 | 10 | 14 | 7 | 7 | 8 | 5 | 2 | 6 | 21 | 0 | 0 | 0 | 9 | 3 | 3 | 0 | 255 | 149 | 1 | 1 | 0.273 | 0.334 | 0.52 | 0.127 | 0.539 | 1 | 0.28 | 0.804 | 0 | NA | -0.162 |
| SE | 103880 | ENSG00000119977 | TCTN3 | chr10 | - | 95669046 | 95669157 | 95663416 | 95664300 | 95680471 | 95680607 | 2 | 1 | 3 | 0 | 6 | 22 | 10 | 32 | 24 | 30 | 0 | 0 | 3 | 0 | 12 | 0 | 11 | 26 | 19 | 10 | 260 | 149 | 1 | 1 | 0.05 | 0.054 | 0.051 | 0 | 0.103 | NA | 0 | 0.062 | 0 | 0.407 | -0.066 |
| SE | 103881 | ENSG00000119977 | TCTN3 | chr10 | - | 95680471 | 95680609 | 95659822 | 95664300 | 95682650 | 95682804 | 46 | 23 | 59 | 44 | 56 | 1 | 0 | 0 | 3 | 0 | 0 | 22 | 38 | 22 | 14 | 0 | 0 | 0 | 0 | 0 | 287 | 149 | 1 | 1 | 0.96 | 1 | 1 | 0.884 | 1 | NA | 1 | 1 | 1 | 1 | -0.031 |
| SE | 103882 | ENSG00000119977 | TCTN3 | chr10 | - | 95680471 | 95680609 | 95659822 | 95664300 | 95683100 | 95683180 | 24 | 12 | 34 | 17 | 31 | 0 | 0 | 0 | 2 | 0 | 0 | 11 | 29 | 15 | 10 | 0 | 0 | 0 | 0 | 0 | 287 | 149 | 1 | 1 | 1 | 1 | 1 | 0.815 | 1 | NA | 1 | 1 | 1 | 1 | -0.037 |
| SE | 103883 | ENSG00000119977 | TCTN3 | chr10 | - | 95682650 | 95682804 | 95663400 | 95664300 | 95683100 | 95683180 | 46 | 35 | 49 | 45 | 74 | 0 | 0 | 0 | 2 | 0 | 49 | 45 | 6 | 17 | 21 | 0 | 0 | 0 | 0 | 0 | 298 | 149 | 1 | 1 | 1 | 1 | 1 | 0.918 | 1 | 1 | 1 | 1 | 1 | 1 | -0.016 |
| SE | 103889 | ENSG00000119977 | TCTN3 | chr10 | - | 95686494 | 95686530 | 95684498 | 95684624 | 95687043 | 95687159 | 11 | 7 | 13 | 14 | 40 | 0 | 0 | 1 | 0 | 0 | 13 | 6 | 12 | 2 | 14 | 0 | 0 | 0 | 0 | 0 | 185 | 149 | 1 | 1 | 1 | 1 | 0.913 | 1 | 1 | 1 | 1 | 1 | 1 | 1 | -0.017 |
| SE | 103890 | ENSG00000119977 | TCTN3 | chr10 | - | 95687043 | 95687159 | 95684532 | 95684624 | 95692919 | 95693038 | 0 | 0 | 1 | 0 | 0 | 0 | 0 | 0 | 0 | 0 | 0 | 0 | 0 | 0 | 0 | 0 | 1 | 0 | 0 | 0 | 265 | 149 | 1 | 1 | NA | NA | 1 | NA | NA | NA | 0 | NA | NA | NA | 1 |
| SE | 103895 | ENSG00000119977 | TCTN3 | chr10 | - | 95693352 | 95693476 | 95692863 | 95693038 | 95693643 | 95693899 | 46 | 25 | 63 | 54 | 50 | 0 | 0 | 0 | 1 | 0 | 54 | 51 | 69 | 23 | 52 | 0 | 0 | 3 | 0 | 0 | 273 | 149 | 1 | 1 | 1 | 1 | 1 | 0.967 | 1 | 1 | 1 | 0.926 | 1 | 1 | 0.008 |
| SE | 103936 | ENSG00000171862 | PTEN | chr10 | + | 87925512 | 87925557 | 87894024 | 87894109 | 87931045 | 87931089 | 773 | 530 | 464 | 602 | 1270 | 0 | 3 | 2 | 1 | 8 | 338 | 452 | 628 | 333 | 173 | 0 | 0 | 0 | 3 | 0 | 194 | 149 | 1 | 1 | 1 | 0.993 | 0.994 | 0.998 | 0.992 | 1 | 1 | 1 | 0.988 | 1 | -0.002 |
| SE | 103938 | ENSG00000171862 | PTEN | chr10 | + | 87952117 | 87952259 | 87933012 | 87933251 | 87960893 | 87961118 | 556 | 355 | 367 | 574 | 754 | 1 | 0 | 0 | 0 | 0 | 138 | 443 | 526 | 284 | 90 | 0 | 0 | 0 | 0 | 0 | 291 | 149 | 1 | 1 | 0.996 | 1 | 1 | 1 | 1 | 1 | 1 | 1 | 1 | 1 | -0.001 |
| SE | 103939 | ENSG00000171862 | PTEN | chr10 | + | 87957852 | 87958019 | 87933012 | 87933251 | 87960893 | 87961118 | 1684 | 1398 | 1225 | 1774 | 3131 | 1 | 0 | 0 | 0 | 0 | 691 | 1701 | 2273 | 1002 | 266 | 0 | 0 | 0 | 0 | 0 | 298 | 149 | 1 | 1 | 0.999 | 1 | 1 | 1 | 1 | 1 | 1 | 1 | 1 | 1 | 0 |
| SE | 103940 | ENSG00000171862 | PTEN | chr10 | + | 87957852 | 87958019 | 87952117 | 87952259 | 87960893 | 87961118 | 2790 | 2400 | 2084 | 2934 | 5745 | 0 | 0 | 0 | 1 | 0 | 1146 | 2798 | 3819 | 1775 | 436 | 0 | 0 | 0 | 0 | 0 | 298 | 149 | 1 | 1 | 1 | 1 | 1 | 0.999 | 1 | 1 | 1 | 1 | 1 | 1 | 0 |
| SE | 104079 | ENSG00000139734 | DIAPH3 | chr13 | - | 60112009 | 60112186 | 60093627 | 60093732 | 60163586 | 60163766 | 0 | 0 | 2 | 7 | 0 | 0 | 0 | 1 | 0 | 0 | 10 | 0 | 3 | 10 | 0 | 0 | 0 | 0 | 0 | 0 | 298 | 149 | 1 | 1 | NA | NA | 0.5 | 1 | NA | 1 | NA | 1 | 1 | NA | -0.25 |
| SE | 104080 | ENSG00000139734 | DIAPH3 | chr13 | - | 60132956 | 60132989 | 60112009 | 60112186 | 60163586 | 60163766 | 0 | 0 | 1 | 1 | 0 | 0 | 0 | 0 | 1 | 0 | 0 | 0 | 1 | 0 | 0 | 0 | 0 | 0 | 0 | 0 | 182 | 149 | 1 | 1 | NA | NA | 1 | 0.45 | NA | NA | NA | 1 | NA | NA | -0.275 |
| SE | 104227 | ENSG00000139620 | KANSL2 | chr12 | - | 48654940 | 48655060 | 48653210 | 48654175 | 48660365 | 48660619 | 94 | 60 | 85 | 109 | 97 | 0 | 0 | 0 | 1 | 0 | 93 | 84 | 120 | 130 | 160 | 0 | 0 | 0 | 1 | 0 | 269 | 149 | 1 | 1 | 1 | 1 | 1 | 0.984 | 1 | 1 | 1 | 1 | 0.986 | 1 | 0 |
| SE | 104229 | ENSG00000139620 | KANSL2 | chr12 | - | 48661194 | 48661243 | 48660365 | 48660619 | 48662559 | 48662592 | 16 | 11 | 25 | 23 | 22 | 0 | 1 | 2 | 1 | 0 | 19 | 21 | 13 | 4 | 30 | 0 | 0 | 0 | 1 | 0 | 198 | 149 | 1 | 1 | 1 | 0.892 | 0.904 | 0.945 | 1 | 1 | 1 | 1 | 0.751 | 1 | -0.002 |
| SE | 104233 | ENSG00000139620 | KANSL2 | chr12 | - | 48679654 | 48679833 | 48679035 | 48679150 | 48681381 | 48681416 | 116 | 73 | 94 | 122 | 119 | 0 | 0 | 0 | 0 | 0 | 161 | 126 | 106 | 100 | 132 | 0 | 0 | 0 | 1 | 0 | 298 | 149 | 1 | 1 | 1 | 1 | 1 | 1 | 1 | 1 | 1 | 1 | 0.98 | 1 | 0.004 |
| SE | 104234 | ENSG00000139620 | KANSL2 | chr12 | - | 48681381 | 48681641 | 48679654 | 48679833 | 48682186 | 48682211 | 65 | 40 | 67 | 80 | 66 | 1 | 0 | 1 | 0 | 0 | 112 | 76 | 43 | 49 | 86 | 0 | 0 | 0 | 2 | 0 | 298 | 149 | 1 | 1 | 0.97 | 1 | 0.971 | 1 | 1 | 1 | 1 | 1 | 0.925 | 1 | 0.003 |
| SE | 104235 | ENSG00000139620 | KANSL2 | chr12 | - | 48681570 | 48681641 | 48679654 | 48679833 | 48682186 | 48682238 | 2 | 2 | 0 | 0 | 0 | 1 | 0 | 1 | 0 | 0 | 0 | 1 | 0 | 0 | 0 | 0 | 0 | 0 | 2 | 0 | 220 | 149 | 1 | 1 | 0.575 | 1 | 0 | NA | NA | NA | 1 | NA | 0 | NA | 0.025 |
| SE | 104236 | ENSG00000139620 | KANSL2 | chr12 | - | 48681857 | 48682065 | 48681381 | 48681641 | 48682186 | 48682238 | 0 | 0 | 0 | 0 | 0 | 15 | 6 | 16 | 15 | 2 | 0 | 2 | 2 | 0 | 0 | 22 | 16 | 6 | 5 | 5 | 298 | 149 | 1 | 1 | 0 | 0 | 0 | 0 | 0 | 0 | 0.059 | 0.143 | 0 | 0 | -0.04 |
| SE | 104641 | ENSG00000079974 | RABL2B | chr22 | - | 50769904 | 50770016 | 50769424 | 50769549 | 50775771 | 50775851 | 42 | 14 | 26 | 34 | 62 | 1 | 0 | 0 | 0 | 0 | 6 | 22 | 24 | 24 | 49 | 0 | 0 | 0 | 0 | 0 | 261 | 149 | 1 | 1 | 0.96 | 1 | 1 | 1 | 1 | 1 | 1 | 1 | 1 | 1 | -0.008 |
| SE | 104645 | ENSG00000079974 | RABL2B | chr22 | - | 50782890 | 50783045 | 50782187 | 50782351 | 50783500 | 50783642 | 1 | 0 | 0 | 0 | 0 | 0 | 0 | 0 | 0 | 0 | 0 | 0 | 3 | 0 | 0 | 0 | 0 | 2 | 0 | 0 | 298 | 149 | 1 | 1 | 1 | NA | NA | NA | NA | NA | NA | 0.429 | NA | NA | 0.571 |
| SE | 104646 | ENSG00000079974 | RABL2B | chr22 | - | 50783038 | 50783286 | 50782187 | 50782351 | 50783500 | 50783636 | 3 | 4 | 6 | 8 | 12 | 0 | 0 | 0 | 0 | 0 | 20 | 13 | 4 | 9 | 15 | 0 | 0 | 2 | 0 | 0 | 298 | 149 | 1 | 1 | 1 | 1 | 1 | 1 | 1 | 1 | 1 | 0.5 | 1 | 1 | 0.1 |
| SE | 104996 | ENSG00000129315 | CCNT1 | chr12 | - | 48695758 | 48695829 | 48688457 | 48694436 | 48695998 | 48696162 | 88 | 52 | 80 | 108 | 55 | 0 | 0 | 2 | 0 | 0 | 32 | 73 | 92 | 64 | 14 | 0 | 0 | 0 | 0 | 2 | 220 | 149 | 1 | 1 | 1 | 1 | 0.964 | 1 | 1 | 1 | 1 | 1 | 1 | 0.826 | 0.028 |
| SE | 104997 | ENSG00000129315 | CCNT1 | chr12 | - | 48695758 | 48695829 | 48688457 | 48694436 | 48698137 | 48698183 | 98 | 50 | 75 | 143 | 77 | 0 | 0 | 0 | 0 | 0 | 46 | 125 | 189 | 115 | 0 | 0 | 1 | 7 | 0 | 0 | 220 | 149 | 1 | 1 | 1 | 1 | 1 | 1 | 1 | 1 | 0.988 | 0.948 | 1 | NA | 0.016 |
| SE | 105001 | ENSG00000129315 | CCNT1 | chr12 | - | 48699777 | 48699840 | 48698137 | 48698183 | 48705767 | 48705896 | 47 | 33 | 33 | 63 | 51 | 0 | 0 | 1 | 0 | 0 | 19 | 31 | 102 | 45 | 0 | 0 | 0 | 0 | 0 | 0 | 212 | 149 | 1 | 1 | 1 | 1 | 0.959 | 1 | 1 | 1 | 1 | 1 | 1 | NA | -0.008 |
| SE | 105002 | ENSG00000129315 | CCNT1 | chr12 | - | 48701012 | 48701073 | 48698137 | 48698183 | 48705767 | 48705896 | 7 | 8 | 27 | 15 | 5 | 0 | 0 | 1 | 0 | 0 | 14 | 25 | 23 | 10 | 12 | 0 | 0 | 0 | 0 | 0 | 210 | 149 | 1 | 1 | 1 | 1 | 0.95 | 1 | 1 | 1 | 1 | 1 | 1 | 1 | -0.01 |
| SE | 105003 | ENSG00000129315 | CCNT1 | chr12 | - | 48701012 | 48701073 | 48699777 | 48699840 | 48705767 | 48705896 | 156 | 101 | 149 | 176 | 192 | 0 | 0 | 0 | 0 | 1 | 30 | 186 | 238 | 75 | 40 | 0 | 0 | 1 | 0 | 0 | 210 | 149 | 1 | 1 | 1 | 1 | 1 | 1 | 0.993 | 1 | 1 | 0.994 | 1 | 1 | 0 |
| SE | 105004 | ENSG00000129315 | CCNT1 | chr12 | - | 48705767 | 48705896 | 48701012 | 48701073 | 48714442 | 48714524 | 197 | 105 | 177 | 206 | 201 | 1 | 0 | 1 | 2 | 1 | 64 | 206 | 278 | 79 | 43 | 0 | 0 | 6 | 0 | 0 | 278 | 149 | 1 | 1 | 0.991 | 1 | 0.99 | 0.982 | 0.991 | 1 | 1 | 0.961 | 1 | 1 | -0.001 |
| SE | 105356 | ENSG00000175782 | SLC35E3 | chr12 | + | 68759156 | 68759239 | 68752031 | 68752190 | 68764703 | 68767158 | 69 | 38 | 71 | 55 | 57 | 0 | 0 | 0 | 0 | 0 | 95 | 66 | 33 | 34 | 95 | 0 | 0 | 2 | 0 | 0 | 232 | 149 | 1 | 1 | 1 | 1 | 1 | 1 | 1 | 1 | 1 | 0.914 | 1 | 1 | 0.017 |
| SE | 105461 | ENSG00000135720 | DYNC1LI2 | chr16 | - | 66724661 | 66724748 | 66723600 | 66723822 | 66725827 | 66725944 | 5 | 1 | 2 | 2 | 3 | 85 | 70 | 69 | 62 | 34 | 0 | 0 | 0 | 3 | 3 | 0 | 49 | 93 | 53 | 27 | 236 | 149 | 1 | 1 | 0.036 | 0.009 | 0.018 | 0.02 | 0.053 | NA | 0 | 0 | 0.035 | 0.066 | 0.002 |
| SE | 105463 | ENSG00000135720 | DYNC1LI2 | chr16 | - | 66727687 | 66727805 | 66725827 | 66725944 | 66728200 | 66728219 | 190 | 186 | 185 | 200 | 251 | 0 | 0 | 0 | 0 | 0 | 148 | 255 | 300 | 272 | 86 | 0 | 0 | 0 | 2 | 0 | 267 | 149 | 1 | 1 | 1 | 1 | 1 | 1 | 1 | 1 | 1 | 1 | 0.987 | 1 | 0.003 |
| SE | 105466 | ENSG00000135720 | DYNC1LI2 | chr16 | - | 66728200 | 66728242 | 66727687 | 66727805 | 66729039 | 66729099 | 164 | 181 | 161 | 189 | 254 | 0 | 2 | 0 | 0 | 1 | 148 | 239 | 291 | 237 | 87 | 0 | 0 | 2 | 1 | 7 | 191 | 149 | 1 | 1 | 1 | 0.986 | 1 | 1 | 0.995 | 1 | 1 | 0.991 | 0.995 | 0.907 | 0.018 |
| SE | 105467 | ENSG00000135720 | DYNC1LI2 | chr16 | - | 66729039 | 66729099 | 66725827 | 66725944 | 66730111 | 66730219 | 38 | 29 | 27 | 35 | 47 | 0 | 2 | 1 | 0 | 0 | 16 | 52 | 70 | 39 | 6 | 0 | 0 | 0 | 1 | 0 | 209 | 149 | 1 | 1 | 1 | 0.912 | 0.951 | 1 | 1 | 1 | 1 | 1 | 1 | 1 | -0.027 |
| SE | 105468 | ENSG00000135720 | DYNC1LI2 | chr16 | - | 66742437 | 66742554 | 66736101 | 66736244 | 66749196 | 66749313 | 83 | 68 | 70 | 62 | 71 | 0 | 0 | 0 | 1 | 0 | 126 | 96 | 101 | 72 | 44 | 0 | 0 | 2 | 1 | 0 | 266 | 149 | 1 | 1 | 1 | 1 | 1 | 0.972 | 1 | 1 | 1 | 0.966 | 0.976 | 1 | 0.006 |
| SE | 105469 | ENSG00000135720 | DYNC1LI2 | chr16 | - | 66742437 | 66742668 | 66736074 | 66736244 | 66749196 | 66749313 | 174 | 167 | 181 | 160 | 141 | 0 | 0 | 0 | 1 | 0 | 239 | 210 | 231 | 194 | 111 | 0 | 0 | 2 | 1 | 0 | 298 | 149 | 1 | 1 | 1 | 1 | 1 | 0.988 | 1 | 1 | 1 | 0.983 | 0.99 | 1 | 0.003 |
| SE | 105470 | ENSG00000135720 | DYNC1LI2 | chr16 | - | 66749196 | 66749313 | 66742437 | 66742668 | 66751272 | 66751303 | 150 | 125 | 162 | 114 | 89 | 0 | 0 | 0 | 0 | 0 | 113 | 138 | 148 | 139 | 80 | 0 | 2 | 3 | 0 | 0 | 266 | 149 | 1 | 1 | 1 | 1 | 1 | 1 | 1 | 1 | 0.975 | 0.965 | 1 | 1 | 0.012 |
| SE | 105515 | ENSG00000084110 | HAL | chr12 | - | 95976428 | 95976498 | 95972661 | 95974372 | 95976597 | 95976706 | 20 | 24 | 24 | 58 | 21 | 0 | 0 | 0 | 0 | 0 | 22 | 54 | 60 | 49 | 48 | 0 | 0 | 0 | 2 | 0 | 219 | 149 | 1 | 1 | 1 | 1 | 1 | 1 | 1 | 1 | 1 | 1 | 0.943 | 1 | 0.011 |
| SE | 105516 | ENSG00000084110 | HAL | chr12 | - | 95977943 | 95978078 | 95976597 | 95976706 | 95980555 | 95980668 | 25 | 17 | 20 | 38 | 25 | 0 | 0 | 0 | 0 | 0 | 51 | 68 | 42 | 37 | 48 | 0 | 0 | 0 | 3 | 0 | 284 | 149 | 1 | 1 | 1 | 1 | 1 | 1 | 1 | 1 | 1 | 1 | 0.866 | 1 | 0.027 |
| SE | 105517 | ENSG00000084110 | HAL | chr12 | - | 95987066 | 95987214 | 95986064 | 95986160 | 95988192 | 95988240 | 33 | 15 | 30 | 45 | 52 | 3 | 0 | 2 | 0 | 0 | 85 | 60 | 144 | 106 | 119 | 0 | 0 | 0 | 0 | 0 | 297 | 149 | 1 | 1 | 0.847 | 1 | 0.883 | 1 | 1 | 1 | 1 | 1 | 1 | 1 | -0.054 |
| SE | 105679 | ENSG00000028203 | VEZT | chr12 | + | 95224179 | 95224236 | 95217809 | 95217886 | 95242163 | 95242314 | 3 | 1 | 0 | 2 | 0 | 1 | 0 | 0 | 2 | 2 | 0 | 0 | 0 | 0 | 0 | 7 | 0 | 0 | 0 | 0 | 206 | 149 | 1 | 1 | 0.685 | 1 | NA | 0.42 | 0 | 0 | NA | NA | NA | NA | 0.526 |
| SE | 105680 | ENSG00000028203 | VEZT | chr12 | + | 95224179 | 95224236 | 95217809 | 95217886 | 95243940 | 95244009 | 4 | 3 | 0 | 2 | 0 | 0 | 1 | 0 | 0 | 0 | 3 | 0 | 0 | 0 | 0 | 0 | 0 | 0 | 1 | 0 | 206 | 149 | 1 | 1 | 1 | 0.685 | NA | 1 | NA | 1 | NA | NA | 0 | NA | 0.395 |
| SE | 105682 | ENSG00000028203 | VEZT | chr12 | + | 95242163 | 95242264 | 95217779 | 95217886 | 95243940 | 95244009 | 1 | 0 | 0 | 0 | 0 | 0 | 1 | 0 | 0 | 0 | 0 | 0 | 2 | 0 | 0 | 0 | 0 | 0 | 1 | 0 | 250 | 149 | 1 | 1 | 1 | 0 | NA | NA | NA | NA | NA | 1 | 0 | NA | 0 |
| SE | 105684 | ENSG00000028203 | VEZT | chr12 | + | 95242163 | 95242264 | 95218064 | 95218277 | 95243940 | 95244009 | 0 | 0 | 0 | 0 | 0 | 0 | 1 | 0 | 0 | 0 | 0 | 0 | 2 | 0 | 0 | 0 | 0 | 0 | 0 | 0 | 250 | 149 | 1 | 1 | NA | 0 | NA | NA | NA | NA | NA | 1 | NA | NA | -1 |
| SE | 105686 | ENSG00000028203 | VEZT | chr12 | + | 95242163 | 95242314 | 95217745 | 95217886 | 95243940 | 95244009 | 0 | 0 | 0 | 0 | 0 | 0 | 1 | 0 | 0 | 0 | 7 | 0 | 0 | 0 | 0 | 0 | 0 | 0 | 1 | 0 | 298 | 149 | 1 | 1 | NA | 0 | NA | NA | NA | 1 | NA | NA | 0 | NA | -0.5 |
| SE | 105687 | ENSG00000028203 | VEZT | chr12 | + | 95242163 | 95242314 | 95224179 | 95224236 | 95243940 | 95244009 | 1 | 0 | 0 | 0 | 0 | 2 | 2 | 0 | 0 | 0 | 7 | 0 | 0 | 0 | 0 | 3 | 0 | 0 | 0 | 0 | 298 | 149 | 1 | 1 | 0.2 | 0 | NA | NA | NA | 0 | NA | NA | NA | NA | 0.1 |
| SE | 105691 | ENSG00000028203 | VEZT | chr12 | + | 95243940 | 95244009 | 95224179 | 95224236 | 95251939 | 95252071 | 3 | 4 | 0 | 0 | 0 | 3 | 2 | 7 | 3 | 2 | 7 | 0 | 0 | 1 | 0 | 0 | 0 | 1 | 0 | 0 | 218 | 149 | 1 | 1 | 0.406 | 0.578 | 0 | 0 | 0 | 1 | NA | 0 | 1 | NA | -0.47 |
| SE | 105692 | ENSG00000028203 | VEZT | chr12 | + | 95243940 | 95244009 | 95242163 | 95242264 | 95251939 | 95252071 | 2 | 2 | 0 | 0 | 0 | 3 | 0 | 0 | 4 | 2 | 4 | 0 | 2 | 1 | 0 | 13 | 5 | 1 | 0 | 0 | 218 | 149 | 1 | 1 | 0.313 | 1 | NA | 0 | 0 | 0.174 | 0 | 0.578 | 1 | NA | -0.11 |
| SE | 105696 | ENSG00000028203 | VEZT | chr12 | + | 95256549 | 95256622 | 95251939 | 95252071 | 95266356 | 95266633 | 32 | 26 | 25 | 35 | 53 | 1 | 0 | 0 | 0 | 0 | 17 | 39 | 30 | 17 | 8 | 0 | 0 | 0 | 0 | 0 | 222 | 149 | 1 | 1 | 0.956 | 1 | 1 | 1 | 1 | 1 | 1 | 1 | 1 | 1 | -0.009 |
| SE | 105697 | ENSG00000028203 | VEZT | chr12 | + | 95257149 | 95257239 | 95251939 | 95252071 | 95258244 | 95258282 | 13 | 12 | 13 | 20 | 43 | 1 | 0 | 0 | 0 | 2 | 0 | 3 | 5 | 11 | 0 | 0 | 0 | 0 | 0 | 0 | 239 | 149 | 1 | 1 | 0.89 | 1 | 1 | 1 | 0.931 | NA | 1 | 1 | 1 | NA | -0.036 |
| SE | 105698 | ENSG00000028203 | VEZT | chr12 | + | 95257149 | 95257239 | 95251939 | 95252071 | 95262905 | 95263059 | 73 | 68 | 50 | 67 | 106 | 9 | 0 | 3 | 4 | 4 | 0 | 36 | 36 | 30 | 6 | 0 | 15 | 2 | 0 | 0 | 239 | 149 | 1 | 1 | 0.835 | 1 | 0.912 | 0.913 | 0.943 | NA | 0.599 | 0.918 | 1 | 1 | 0.041 |
| SE | 105699 | ENSG00000028203 | VEZT | chr12 | + | 95257149 | 95257239 | 95251939 | 95252071 | 95266356 | 95266384 | 11 | 11 | 14 | 21 | 29 | 1 | 0 | 0 | 0 | 0 | 0 | 3 | 4 | 11 | 0 | 0 | 0 | 0 | 0 | 0 | 239 | 149 | 1 | 1 | 0.873 | 1 | 1 | 1 | 1 | NA | 1 | 1 | 1 | NA | -0.025 |
| SE | 105700 | ENSG00000028203 | VEZT | chr12 | + | 95257149 | 95257239 | 95256549 | 95256622 | 95262905 | 95263059 | 81 | 78 | 69 | 68 | 157 | 12 | 15 | 5 | 11 | 49 | 0 | 39 | 42 | 21 | 25 | 0 | 10 | 22 | 19 | 0 | 239 | 149 | 1 | 1 | 0.808 | 0.764 | 0.896 | 0.794 | 0.666 | NA | 0.709 | 0.543 | 0.408 | 1 | 0.121 |
| SE | 105701 | ENSG00000028203 | VEZT | chr12 | + | 95257149 | 95257239 | 95256549 | 95256622 | 95266356 | 95266384 | 17 | 17 | 17 | 12 | 47 | 0 | 0 | 2 | 1 | 0 | 0 | 6 | 10 | 3 | 19 | 0 | 0 | 3 | 0 | 0 | 239 | 149 | 1 | 1 | 1 | 1 | 0.841 | 0.882 | 1 | NA | 1 | 0.675 | 1 | 1 | 0.026 |
| SE | 105702 | ENSG00000028203 | VEZT | chr12 | + | 95257153 | 95257239 | 95251939 | 95252071 | 95262905 | 95263081 | 60 | 53 | 33 | 44 | 77 | 9 | 0 | 3 | 4 | 4 | 0 | 33 | 27 | 17 | 6 | 0 | 15 | 2 | 0 | 0 | 235 | 149 | 1 | 1 | 0.809 | 1 | 0.875 | 0.875 | 0.924 | NA | 0.582 | 0.895 | 1 | 1 | 0.027 |
| SE | 105704 | ENSG00000028203 | VEZT | chr12 | + | 95257153 | 95257239 | 95256549 | 95256622 | 95262905 | 95263081 | 62 | 53 | 33 | 44 | 81 | 12 | 15 | 5 | 11 | 49 | 0 | 33 | 28 | 17 | 6 | 0 | 10 | 22 | 19 | 0 | 235 | 149 | 1 | 1 | 0.766 | 0.691 | 0.807 | 0.717 | 0.512 | NA | 0.677 | 0.447 | 0.362 | 1 | 0.077 |
| SE | 105705 | ENSG00000028203 | VEZT | chr12 | + | 95257153 | 95257239 | 95256549 | 95256622 | 95266356 | 95266632 | 3 | 0 | 2 | 1 | 3 | 0 | 0 | 2 | 1 | 0 | 0 | 0 | 0 | 0 | 0 | 0 | 0 | 3 | 0 | 0 | 235 | 149 | 1 | 1 | 1 | NA | 0.388 | 0.388 | 1 | NA | NA | 0 | NA | NA | 0.694 |
| SE | 105706 | ENSG00000028203 | VEZT | chr12 | + | 95258244 | 95258282 | 95251939 | 95252071 | 95262905 | 95263081 | 4 | 0 | 3 | 1 | 6 | 9 | 0 | 3 | 4 | 4 | 0 | 0 | 0 | 0 | 0 | 0 | 15 | 2 | 0 | 0 | 187 | 149 | 1 | 1 | 0.262 | NA | 0.443 | 0.166 | 0.544 | NA | 0 | 0 | NA | NA | 0.354 |
| SE | 105707 | ENSG00000028203 | VEZT | chr12 | + | 95258244 | 95258282 | 95257149 | 95257239 | 95262905 | 95263081 | 8 | 2 | 3 | 4 | 15 | 68 | 65 | 60 | 60 | 117 | 0 | 0 | 1 | 0 | 0 | 0 | 33 | 38 | 20 | 6 | 187 | 149 | 1 | 1 | 0.086 | 0.024 | 0.038 | 0.05 | 0.093 | NA | 0 | 0.021 | 0 | 0 | 0.053 |
| SE | 105708 | ENSG00000028203 | VEZT | chr12 | + | 95258244 | 95258287 | 95251939 | 95252071 | 95262905 | 95263074 | 4 | 3 | 0 | 0 | 6 | 9 | 0 | 3 | 4 | 4 | 0 | 0 | 0 | 0 | 0 | 0 | 15 | 2 | 0 | 0 | 192 | 149 | 1 | 1 | 0.256 | 1 | 0 | 0 | 0.538 | NA | 0 | 0 | NA | NA | 0.359 |
| SE | 105709 | ENSG00000028203 | VEZT | chr12 | + | 95258244 | 95258287 | 95257149 | 95257239 | 95262905 | 95263074 | 4 | 4 | 1 | 2 | 18 | 68 | 65 | 60 | 60 | 117 | 0 | 0 | 0 | 4 | 0 | 0 | 33 | 38 | 20 | 6 | 192 | 149 | 1 | 1 | 0.044 | 0.046 | 0.013 | 0.025 | 0.107 | NA | 0 | 0 | 0.134 | 0 | 0.013 |
| SE | 105710 | ENSG00000028203 | VEZT | chr12 | + | 95262905 | 95263081 | 95251939 | 95252071 | 95266356 | 95266384 | 64 | 67 | 48 | 73 | 88 | 1 | 0 | 0 | 0 | 0 | 17 | 72 | 39 | 29 | 0 | 0 | 0 | 0 | 0 | 0 | 298 | 149 | 1 | 1 | 0.97 | 1 | 1 | 1 | 1 | 1 | 1 | 1 | 1 | NA | -0.006 |
| SE | 105711 | ENSG00000028203 | VEZT | chr12 | + | 95262905 | 95263081 | 95256549 | 95256622 | 95266356 | 95266384 | 67 | 82 | 50 | 80 | 133 | 0 | 0 | 2 | 1 | 0 | 17 | 67 | 59 | 48 | 0 | 0 | 0 | 3 | 0 | 0 | 298 | 149 | 1 | 1 | 1 | 1 | 0.926 | 0.976 | 1 | 1 | 1 | 0.908 | 1 | NA | 0.003 |
| SE | 105712 | ENSG00000028203 | VEZT | chr12 | + | 95262905 | 95263081 | 95257149 | 95257239 | 95266356 | 95266384 | 123 | 132 | 105 | 129 | 201 | 1 | 0 | 2 | 1 | 2 | 17 | 90 | 75 | 49 | 6 | 0 | 0 | 0 | 0 | 0 | 298 | 149 | 1 | 1 | 0.984 | 1 | 0.963 | 0.985 | 0.98 | 1 | 1 | 1 | 1 | 1 | -0.018 |
| SE | 105714 | ENSG00000028203 | VEZT | chr12 | + | 95282312 | 95282359 | 95266356 | 95266632 | 95287663 | 95287857 | 2 | 0 | 0 | 0 | 0 | 0 | 1 | 0 | 0 | 0 | 0 | 0 | 6 | 0 | 0 | 0 | 0 | 0 | 0 | 0 | 196 | 149 | 1 | 1 | 1 | 0 | NA | NA | NA | NA | NA | 1 | NA | NA | -0.5 |
| SE | 105715 | ENSG00000028203 | VEZT | chr12 | + | 95282312 | 95282359 | 95274741 | 95274889 | 95287663 | 95287857 | 35 | 27 | 18 | 27 | 49 | 2 | 0 | 1 | 0 | 0 | 0 | 35 | 35 | 10 | 0 | 0 | 8 | 0 | 0 | 0 | 196 | 149 | 1 | 1 | 0.93 | 1 | 0.932 | 1 | 1 | NA | 0.769 | 1 | 1 | NA | 0.049 |
| SE | 105716 | ENSG00000028203 | VEZT | chr12 | + | 95282312 | 95282644 | 95266356 | 95266632 | 95287663 | 95287719 | 156 | 124 | 147 | 127 | 202 | 0 | 1 | 0 | 0 | 0 | 28 | 93 | 91 | 27 | 5 | 0 | 0 | 0 | 0 | 0 | 298 | 149 | 1 | 1 | 1 | 0.984 | 1 | 1 | 1 | 1 | 1 | 1 | 1 | 1 | -0.003 |
| SE | 105718 | ENSG00000028203 | VEZT | chr12 | + | 95287663 | 95287857 | 95282312 | 95282644 | 95294271 | 95294372 | 254 | 220 | 243 | 233 | 390 | 1 | 2 | 3 | 0 | 3 | 53 | 161 | 160 | 42 | 11 | 0 | 0 | 0 | 0 | 0 | 298 | 149 | 1 | 1 | 0.992 | 0.982 | 0.976 | 1 | 0.985 | 1 | 1 | 1 | 1 | 1 | -0.013 |
| SE | 106654 | ENSG00000139350 | NEDD1 | chr12 | + | 96907603 | 96907624 | 96907229 | 96907300 | 96909751 | 96909895 | 3 | 0 | 1 | 0 | 0 | 6 | 9 | 7 | 1 | 0 | 0 | 1 | 0 | 0 | 0 | 0 | 8 | 6 | 1 | 4 | 170 | 149 | 1 | 1 | 0.305 | 0 | 0.111 | 0 | NA | NA | 0.099 | 0 | 0 | 0 | 0.079 |
| SE | 106656 | ENSG00000139350 | NEDD1 | chr12 | + | 96907603 | 96907624 | 96907229 | 96907300 | 96919984 | 96920118 | 0 | 0 | 0 | 0 | 0 | 0 | 2 | 0 | 0 | 1 | 0 | 0 | 0 | 5 | 0 | 0 | 0 | 0 | 0 | 0 | 170 | 149 | 1 | 1 | NA | 0 | NA | NA | 0 | NA | NA | NA | 1 | NA | -1 |
| SE | 106658 | ENSG00000139350 | NEDD1 | chr12 | + | 96907603 | 96907856 | 96907237 | 96907300 | 96909751 | 96909895 | 28 | 21 | 13 | 35 | 36 | 6 | 9 | 7 | 1 | 0 | 0 | 8 | 1 | 3 | 0 | 0 | 8 | 6 | 1 | 4 | 298 | 149 | 1 | 1 | 0.7 | 0.538 | 0.481 | 0.946 | 1 | NA | 0.333 | 0.077 | 0.6 | 0 | 0.48 |
| SE | 106661 | ENSG00000139350 | NEDD1 | chr12 | + | 96907701 | 96907856 | 96907603 | 96907624 | 96909751 | 96909895 | 23 | 20 | 12 | 27 | 30 | 3 | 1 | 3 | 1 | 0 | 0 | 8 | 1 | 3 | 0 | 2 | 1 | 1 | 0 | 0 | 298 | 149 | 1 | 1 | 0.793 | 0.909 | 0.667 | 0.931 | 1 | 0 | 0.8 | 0.333 | 1 | NA | 0.327 |
| SE | 106662 | ENSG00000139350 | NEDD1 | chr12 | + | 96917620 | 96917737 | 96907223 | 96907300 | 96919984 | 96920118 | 59 | 34 | 75 | 64 | 73 | 0 | 2 | 0 | 0 | 1 | 7 | 37 | 20 | 4 | 25 | 0 | 0 | 0 | 0 | 0 | 266 | 149 | 1 | 1 | 1 | 0.905 | 1 | 1 | 0.976 | 1 | 1 | 1 | 1 | 1 | -0.024 |
| SE | 106665 | ENSG00000139350 | NEDD1 | chr12 | + | 96917620 | 96917737 | 96912575 | 96912817 | 96919984 | 96920118 | 102 | 77 | 112 | 99 | 96 | 0 | 0 | 0 | 2 | 0 | 7 | 56 | 38 | 9 | 25 | 0 | 0 | 0 | 0 | 0 | 266 | 149 | 1 | 1 | 1 | 1 | 1 | 0.965 | 1 | 1 | 1 | 1 | 1 | 1 | -0.007 |
| SE | 106666 | ENSG00000139350 | NEDD1 | chr12 | + | 96924832 | 96924942 | 96919984 | 96920125 | 96934975 | 96935205 | 12 | 5 | 13 | 8 | 20 | 52 | 28 | 74 | 56 | 51 | 0 | 1 | 2 | 2 | 12 | 0 | 22 | 18 | 6 | 0 | 259 | 149 | 1 | 1 | 0.117 | 0.093 | 0.092 | 0.076 | 0.184 | NA | 0.025 | 0.06 | 0.161 | 1 | -0.199 |
| SE | 106667 | ENSG00000139350 | NEDD1 | chr12 | + | 96924832 | 96924942 | 96919984 | 96920125 | 96936610 | 96936718 | 12 | 5 | 5 | 5 | 18 | 2 | 3 | 0 | 3 | 2 | 0 | 1 | 2 | 2 | 12 | 0 | 0 | 0 | 0 | 0 | 259 | 149 | 1 | 1 | 0.775 | 0.489 | 1 | 0.489 | 0.838 | NA | 1 | 1 | 1 | 1 | -0.282 |
| SE | 106668 | ENSG00000139350 | NEDD1 | chr12 | + | 96934975 | 96935205 | 96919984 | 96920125 | 96936610 | 96936638 | 132 | 94 | 154 | 123 | 147 | 2 | 3 | 0 | 3 | 2 | 0 | 34 | 52 | 10 | 9 | 0 | 0 | 0 | 0 | 0 | 298 | 149 | 1 | 1 | 0.971 | 0.94 | 1 | 0.953 | 0.974 | NA | 1 | 1 | 1 | 1 | -0.032 |
| SE | 106670 | ENSG00000139350 | NEDD1 | chr12 | + | 96936610 | 96936812 | 96934975 | 96935205 | 96937197 | 96937393 | 188 | 150 | 178 | 172 | 307 | 0 | 0 | 0 | 2 | 0 | 13 | 42 | 96 | 9 | 17 | 0 | 0 | 0 | 0 | 0 | 298 | 149 | 1 | 1 | 1 | 1 | 1 | 0.977 | 1 | 1 | 1 | 1 | 1 | 1 | -0.005 |
| SE | 106749 | ENSG00000184162 | NR2C2AP | chr19 | - | 19202469 | 19202575 | 19202057 | 19202398 | 19202790 | 19202881 | 20 | 12 | 11 | 30 | 57 | 5 | 1 | 7 | 6 | 6 | 0 | 32 | 7 | 6 | 29 | 0 | 33 | 6 | 7 | 19 | 255 | 149 | 1 | 1 | 0.7 | 0.875 | 0.479 | 0.745 | 0.847 | NA | 0.362 | 0.405 | 0.334 | 0.471 | 0.336 |
| SE | 106784 | ENSG00000151062 | CACNA2D4 | chr12 | - | 1810542 | 1810587 | 1810277 | 1810340 | 1811661 | 1811720 | 4 | 0 | 4 | 7 | 12 | 5 | 0 | 3 | 1 | 0 | 14 | 1 | 0 | 2 | 4 | 0 | 2 | 2 | 0 | 0 | 194 | 149 | 1 | 1 | 0.381 | NA | 0.506 | 0.843 | 1 | 1 | 0.277 | 0 | 1 | 1 | 0.027 |
| SE | 106790 | ENSG00000151062 | CACNA2D4 | chr12 | - | 1887008 | 1887069 | 1886222 | 1886373 | 1907439 | 1907571 | 0 | 0 | 2 | 4 | 2 | 0 | 0 | 0 | 1 | 0 | 0 | 0 | 1 | 0 | 0 | 0 | 0 | 0 | 0 | 0 | 210 | 149 | 1 | 1 | NA | NA | 1 | 0.739 | 1 | NA | NA | 1 | NA | NA | -0.087 |
| SE | 106860 | ENSG00000135452 | TSPAN31 | chr12 | + | 57745744 | 57745873 | 57745022 | 57745217 | 57746588 | 57748205 | 7 | 7 | 15 | 6 | 8 | 2 | 0 | 0 | 0 | 2 | 19 | 2 | 0 | 23 | 21 | 0 | 0 | 0 | 0 | 6 | 278 | 149 | 1 | 1 | 0.652 | 1 | 1 | 1 | 0.682 | 1 | 1 | NA | 1 | 0.652 | -0.046 |
| SE | 106861 | ENSG00000135452 | TSPAN31 | chr12 | + | 57746349 | 57746448 | 57746175 | 57746256 | 57746588 | 57746720 | 1 | 0 | 1 | 2 | 0 | 62 | 36 | 35 | 31 | 43 | 0 | 0 | 0 | 0 | 0 | 53 | 25 | 14 | 43 | 78 | 248 | 149 | 1 | 1 | 0.01 | 0 | 0.017 | 0.037 | 0 | 0 | 0 | 0 | 0 | 0 | 0.013 |
| SE | 106994 | ENSG00000133858 | ZFC3H1 | chr12 | - | 71611057 | 71611097 | 71610694 | 71610757 | 71611785 | 71611887 | 83 | 53 | 73 | 67 | 149 | 0 | 1 | 0 | 1 | 3 | 0 | 53 | 14 | 1 | 0 | 0 | 0 | 1 | 0 | 0 | 189 | 149 | 1 | 1 | 1 | 0.977 | 1 | 0.981 | 0.975 | NA | 1 | 0.917 | 1 | NA | 0.014 |
| SE | 106995 | ENSG00000133858 | ZFC3H1 | chr12 | - | 71614534 | 71614700 | 71613334 | 71613435 | 71615205 | 71615316 | 109 | 88 | 86 | 86 | 156 | 0 | 0 | 0 | 2 | 0 | 16 | 67 | 19 | 11 | 2 | 0 | 0 | 0 | 0 | 0 | 298 | 149 | 1 | 1 | 1 | 1 | 1 | 0.956 | 1 | 1 | 1 | 1 | 1 | 1 | -0.009 |
| SE | 106997 | ENSG00000133858 | ZFC3H1 | chr12 | - | 71619314 | 71619409 | 71615070 | 71615316 | 71619925 | 71619965 | 378 | 334 | 337 | 409 | 311 | 1 | 0 | 1 | 3 | 3 | 8 | 187 | 172 | 33 | 28 | 0 | 0 | 0 | 2 | 0 | 244 | 149 | 1 | 1 | 0.996 | 1 | 0.995 | 0.988 | 0.984 | 1 | 1 | 1 | 0.91 | 1 | 0.011 |
| SE | 106998 | ENSG00000133858 | ZFC3H1 | chr12 | - | 71619925 | 71620124 | 71619314 | 71619409 | 71620209 | 71620315 | 531 | 430 | 467 | 584 | 359 | 0 | 0 | 0 | 0 | 1 | 13 | 257 | 235 | 44 | 20 | 0 | 0 | 0 | 0 | 0 | 298 | 149 | 1 | 1 | 1 | 1 | 1 | 1 | 0.994 | 1 | 1 | 1 | 1 | 1 | -0.001 |
| SE | 106999 | ENSG00000133858 | ZFC3H1 | chr12 | - | 71627750 | 71627934 | 71626259 | 71626446 | 71628917 | 71629037 | 333 | 356 | 332 | 377 | 594 | 0 | 2 | 2 | 0 | 3 | 79 | 164 | 228 | 51 | 56 | 0 | 0 | 1 | 0 | 0 | 298 | 149 | 1 | 1 | 1 | 0.989 | 0.988 | 1 | 0.99 | 1 | 1 | 0.991 | 1 | 1 | -0.005 |
| SE | 107000 | ENSG00000133858 | ZFC3H1 | chr12 | - | 71631877 | 71631887 | 71630822 | 71630954 | 71631971 | 71632514 | 362 | 258 | 339 | 310 | 416 | 0 | 2 | 2 | 0 | 1 | 18 | 128 | 93 | 16 | 10 | 0 | 2 | 0 | 2 | 0 | 259 | 149 | 1 | 1 | 1 | 0.987 | 0.99 | 1 | 0.996 | 1 | 0.974 | 1 | 0.822 | 1 | 0.035 |
| SE | 107001 | ENSG00000133858 | ZFC3H1 | chr12 | - | 71632885 | 71633017 | 71631971 | 71632514 | 71633263 | 71633438 | 429 | 433 | 374 | 416 | 622 | 0 | 0 | 1 | 0 | 4 | 44 | 257 | 195 | 35 | 51 | 0 | 1 | 0 | 0 | 0 | 281 | 149 | 1 | 1 | 1 | 1 | 1 | 0.995 | 0.988 | 1 | 0.993 | 1 | 1 | 1 | -0.002 |
| SE | 107002 | ENSG00000133858 | ZFC3H1 | chr12 | - | 71636849 | 71637059 | 71636489 | 71636654 | 71638417 | 71638515 | 313 | 328 | 418 | 446 | 1023 | 0 | 0 | 0 | 1 | 0 | 87 | 233 | 214 | 56 | 88 | 0 | 0 | 0 | 0 | 0 | 298 | 149 | 1 | 1 | 1 | 1 | 1 | 0.996 | 1 | 1 | 1 | 1 | 1 | 1 | -0.001 |
| SE | 107004 | ENSG00000133858 | ZFC3H1 | chr12 | - | 71644876 | 71645075 | 71644094 | 71644318 | 71647748 | 71647813 | 345 | 472 | 391 | 393 | 608 | 1 | 0 | 2 | 1 | 1 | 126 | 433 | 310 | 264 | 57 | 0 | 0 | 0 | 3 | 0 | 298 | 149 | 1 | 1 | 0.994 | 1 | 0.99 | 0.995 | 0.997 | 1 | 1 | 1 | 0.978 | 1 | 0 |
| SE | 107005 | ENSG00000133858 | ZFC3H1 | chr12 | - | 71647748 | 71647813 | 71644876 | 71645075 | 71656884 | 71657301 | 301 | 357 | 309 | 350 | 729 | 0 | 2 | 0 | 0 | 0 | 79 | 327 | 234 | 185 | 27 | 0 | 0 | 0 | 0 | 0 | 214 | 149 | 1 | 1 | 1 | 0.992 | 1 | 1 | 1 | 1 | 1 | 1 | 1 | 1 | -0.002 |
| SE | 107006 | ENSG00000133858 | ZFC3H1 | chr12 | - | 71656884 | 71657301 | 71647748 | 71647813 | 71660288 | 71660427 | 213 | 219 | 221 | 254 | 554 | 0 | 0 | 1 | 1 | 2 | 48 | 228 | 155 | 127 | 15 | 0 | 0 | 0 | 0 | 0 | 298 | 149 | 1 | 1 | 1 | 1 | 0.991 | 0.992 | 0.993 | 1 | 1 | 1 | 1 | 1 | -0.005 |
| SE | 107136 | ENSG00000175582 | RAB6A | chr11 | - | 73679653 | 73679720 | 73675637 | 73677962 | 73707419 | 73707513 | 542 | 393 | 395 | 526 | 844 | 1 | 0 | 0 | 0 | 5 | 469 | 335 | 310 | 419 | 286 | 0 | 1 | 0 | 3 | 0 | 216 | 149 | 1 | 1 | 0.997 | 1 | 1 | 1 | 0.991 | 1 | 0.996 | 1 | 0.99 | 1 | 0 |
| SE | 107137 | ENSG00000175582 | RAB6A | chr11 | - | 73707419 | 73707513 | 73679653 | 73679720 | 73716250 | 73716362 | 1135 | 811 | 808 | 1138 | 1483 | 0 | 0 | 0 | 1 | 0 | 1805 | 995 | 937 | 1420 | 1465 | 0 | 0 | 0 | 0 | 0 | 243 | 149 | 1 | 1 | 1 | 1 | 1 | 0.999 | 1 | 1 | 1 | 1 | 1 | 1 | 0 |
| SE | 107138 | ENSG00000175582 | RAB6A | chr11 | - | 73716250 | 73716362 | 73679653 | 73679720 | 73718784 | 73718890 | 203 | 165 | 150 | 272 | 368 | 0 | 1 | 0 | 0 | 0 | 281 | 180 | 115 | 260 | 285 | 0 | 0 | 0 | 0 | 0 | 261 | 149 | 1 | 1 | 1 | 0.989 | 1 | 1 | 1 | 1 | 1 | 1 | 1 | 1 | -0.002 |
| SE | 107139 | ENSG00000175582 | RAB6A | chr11 | - | 73716250 | 73716362 | 73679653 | 73679720 | 73720845 | 73720899 | 1 | 2 | 1 | 2 | 0 | 1 | 1 | 0 | 1 | 0 | 0 | 0 | 3 | 3 | 0 | 0 | 0 | 0 | 0 | 8 | 261 | 149 | 1 | 1 | 0.363 | 0.533 | 1 | 0.533 | NA | NA | NA | | 1 | 0 | -0.059 |
| SE | 107141 | ENSG00000175582 | RAB6A | chr11 | - | 73718612 | 73718718 | 73716250 | 73716362 | 73720845 | 73720899 | 989 | 681 | 681 | 940 | 1138 | 3 | 4 | 1 | 2 | 0 | 1922 | 958 | 1052 | 1303 | 1336 | 0 | 0 | 4 | 6 | 0 | 255 | 149 | 1 | 1 | 0.995 | 0.99 | 0.997 | 0.996 | 1 | 1 | 1 | 0.994 | 0.992 | 1 | -0.002 |
| SE | 107142 | ENSG00000175582 | RAB6A | chr11 | - | 73718784 | 73718890 | 73679653 | 73679720 | 73720845 | 73720899 | 178 | 127 | 107 | 160 | 217 | 1 | 1 | 0 | 1 | 0 | 210 | 139 | 114 | 187 | 213 | 0 | 0 | 0 | 0 | 8 | 255 | 149 | 1 | 1 | 0.99 | 0.987 | 1 | 0.989 | 1 | 1 | 1 | 1 | 1 | 0.94 | 0.005 |
| SE | 107143 | ENSG00000175582 | RAB6A | chr11 | - | 73718784 | 73718890 | 73716250 | 73716362 | 73720845 | 73720899 | 648 | 481 | 435 | 698 | 841 | 3 | 4 | 1 | 2 | 0 | 870 | 531 | 385 | 807 | 950 | 0 | 0 | 4 | 6 | 0 | 255 | 149 | 1 | 1 | 0.992 | 0.986 | 0.996 | 0.995 | 1 | 1 | 1 | 0.983 | 0.987 | 1 | 0 |
| SE | 107362 | ENSG00000033030 | ZCCHC8 | chr12 | - | 122478205 | 122478292 | 122471598 | 122474275 | 122480189 | 122480311 | 52 | 41 | 57 | 63 | 123 | 0 | 0 | 0 | 1 | 0 | 24 | 57 | 40 | 50 | 12 | 0 | 0 | 0 | 0 | 0 | 236 | 149 | 1 | 1 | 1 | 1 | 1 | 0.975 | 1 | 1 | 1 | 1 | 1 | 1 | -0.005 |
| SE | 107363 | ENSG00000033030 | ZCCHC8 | chr12 | - | 122482634 | 122482695 | 122481944 | 122482087 | 122483278 | 122483344 | 120 | 69 | 91 | 142 | 180 | 2 | 0 | 0 | 0 | 0 | 65 | 65 | 75 | 80 | 9 | 0 | 0 | 0 | 0 | 0 | 210 | 149 | 1 | 1 | 0.977 | 1 | 1 | 1 | 1 | 1 | 1 | 1 | 1 | 1 | -0.005 |
| SE | 107365 | ENSG00000033030 | ZCCHC8 | chr12 | - | 122483278 | 122483344 | 122481944 | 122482087 | 122489385 | 122489463 | 2 | 0 | 0 | 0 | 2 | 3 | 3 | 3 | 1 | 7 | 0 | 0 | 0 | 0 | 0 | 0 | 5 | 8 | 25 | 11 | 215 | 149 | 1 | 1 | 0.316 | 0 | 0 | 0 | 0.165 | NA | 0 | 0 | 0 | 0 | 0.096 |
| SE | 107369 | ENSG00000033030 | ZCCHC8 | chr12 | - | 122483459 | 122483563 | 122483278 | 122483344 | 122489385 | 122489463 | 107 | 80 | 88 | 107 | 207 | 0 | 0 | 0 | 0 | 2 | 59 | 61 | 47 | 34 | 32 | 0 | 0 | 0 | 0 | 0 | 253 | 149 | 1 | 1 | 1 | 1 | 1 | 1 | 0.984 | 1 | 1 | 1 | 1 | 1 | -0.003 |
| SE | 107370 | ENSG00000033030 | ZCCHC8 | chr12 | - | 122490461 | 122490567 | 122489385 | 122489463 | 122492714 | 122492789 | 214 | 170 | 160 | 209 | 314 | 1 | 3 | 0 | 1 | 0 | 171 | 176 | 193 | 210 | 51 | 0 | 2 | 0 | 0 | 0 | 255 | 149 | 1 | 1 | 0.992 | 0.971 | 1 | 0.992 | 1 | 1 | 0.981 | 1 | 1 | 1 | -0.005 |
| SE | 107371 | ENSG00000033030 | ZCCHC8 | chr12 | - | 122490934 | 122491053 | 122490461 | 122490567 | 122492714 | 122492789 | 3 | 4 | 1 | 0 | 7 | 167 | 143 | 114 | 162 | 227 | 0 | 2 | 2 | 5 | 2 | 139 | 141 | 147 | 154 | 36 | 268 | 149 | 1 | 1 | 0.01 | 0.015 | 0.005 | 0 | 0.017 | 0 | 0.008 | 0.008 | 0.018 | 0.03 | -0.003 |
| SE | 107372 | ENSG00000033030 | ZCCHC8 | chr12 | - | 122492714 | 122492789 | 122490461 | 122490567 | 122498826 | 122498867 | 180 | 146 | 127 | 171 | 227 | 0 | 2 | 0 | 1 | 0 | 135 | 163 | 167 | 178 | 39 | 0 | 0 | 0 | 0 | 0 | 224 | 149 | 1 | 1 | 1 | 0.98 | 1 | 0.991 | 1 | 1 | 1 | 1 | 1 | 1 | -0.006 |
| SE | 107505 | ENSG00000123349 | PFDN5 | chr12 | + | 53296429 | 53296551 | 53295838 | 53295941 | 53298044 | 53298150 | 0 | 0 | 3 | 1 | 0 | 0 | 0 | 0 | 0 | 0 | 0 | 0 | 0 | 3 | 0 | 0 | 4 | 0 | 0 | 0 | 271 | 149 | 1 | 1 | NA | NA | 1 | 1 | NA | NA | 0 | NA | 1 | NA | 0.5 |
| SE | 107506 | ENSG00000123349 | PFDN5 | chr12 | + | 53296429 | 53297924 | 53295838 | 53295941 | 53298044 | 53298150 | 726 | 362 | 663 | 564 | 647 | 0 | 0 | 0 | 0 | 0 | 900 | 464 | 393 | 479 | 767 | 0 | 4 | 0 | 0 | 0 | 298 | 149 | 1 | 1 | 1 | 1 | 1 | 1 | 1 | 1 | 0.983 | 1 | 1 | 1 | 0.003 |
| SE | 107510 | ENSG00000123349 | PFDN5 | chr12 | + | 53296453 | 53296551 | 53295838 | 53295941 | 53298044 | 53298150 | 0 | 3 | 0 | 0 | 2 | 0 | 0 | 0 | 0 | 0 | 0 | 0 | 0 | 0 | 0 | 0 | 4 | 0 | 0 | 0 | 247 | 149 | 1 | 1 | NA | 1 | NA | NA | 1 | NA | 0 | NA | NA | NA | 1 |
| SE | 107513 | ENSG00000123349 | PFDN5 | chr12 | + | 53297849 | 53297924 | 53295838 | 53295941 | 53298044 | 53298150 | 714 | 367 | 656 | 550 | 634 | 0 | 0 | 0 | 0 | 0 | 924 | 456 | 386 | 455 | 787 | 0 | 4 | 0 | 0 | 0 | 224 | 149 | 1 | 1 | 1 | 1 | 1 | 1 | 1 | 1 | 0.987 | 1 | 1 | 1 | 0.003 |
| SE | 107514 | ENSG00000123349 | PFDN5 | chr12 | + | 53297849 | 53297924 | 53295890 | 53296551 | 53298044 | 53298150 | 726 | 367 | 666 | 562 | 635 | 0 | 0 | 0 | 0 | 0 | 904 | 464 | 387 | 465 | 793 | 0 | 3 | 0 | 0 | 0 | 224 | 149 | 1 | 1 | 1 | 1 | 1 | 1 | 1 | 1 | 0.99 | 1 | 1 | 1 | 0.002 |
| SE | 107653 | ENSG00000139718 | SETD1B | chr12 | + | 121817369 | 121817704 | 121817032 | 121817294 | 121817798 | 121817904 | 360 | 288 | 361 | 406 | 363 | 0 | 0 | 0 | 0 | 0 | 953 | 575 | 484 | 610 | 933 | 0 | 0 | 0 | 0 | 4 | 298 | 149 | 1 | 1 | 1 | 1 | 1 | 1 | 1 | 1 | 1 | 1 | 1 | 0.991 | 0.002 |
| SE | 107654 | ENSG00000139718 | SETD1B | chr12 | + | 121819403 | 121819895 | 121817798 | 121817904 | 121822489 | 121823749 | 384 | 316 | 414 | 409 | 513 | 0 | 2 | 0 | 1 | 0 | 1087 | 607 | 550 | 690 | 1042 | 0 | 0 | 0 | 0 | 12 | 298 | 149 | 1 | 1 | 1 | 0.988 | 1 | 0.995 | 1 | 1 | 1 | 1 | 1 | 0.977 | 0.001 |
| SE | 107730 | ENSG00000196876 | SCN8A | chr12 | + | 51688757 | 51688849 | 51687090 | 51687219 | 51699569 | 51699791 | 1 | 0 | 0 | 0 | 3 | 0 | 0 | 1 | 3 | 0 | 0 | 0 | 0 | 4 | 0 | 0 | 0 | 0 | 6 | 4 | 241 | 149 | 1 | 1 | 1 | NA | 0 | 0 | 1 | NA | NA | NA | 0.292 | 0 | 0.354 |
| SE | 107731 | ENSG00000196876 | SCN8A | chr12 | + | 51688757 | 51690019 | 51687090 | 51687219 | 51699569 | 51699791 | 1 | 0 | 0 | 0 | 3 | 0 | 0 | 1 | 3 | 0 | 0 | 0 | 0 | 4 | 0 | 0 | 0 | 0 | 6 | 4 | 298 | 149 | 1 | 1 | 1 | NA | 0 | 0 | 1 | NA | NA | NA | 0.25 | 0 | 0.375 |
| SE | 107733 | ENSG00000196876 | SCN8A | chr12 | + | 51751354 | 51751593 | 51745900 | 51746035 | 51762502 | 51762676 | 2 | 1 | 1 | 0 | 0 | 0 | 0 | 0 | 0 | 0 | 10 | 0 | 2 | 24 | 0 | 0 | 0 | 0 | 1 | 0 | 298 | 149 | 1 | 1 | 1 | 1 | 1 | NA | NA | 1 | NA | 1 | 0.923 | NA | 0.026 |
| SE | 107735 | ENSG00000196876 | SCN8A | chr12 | + | 51780201 | 51780271 | 51774188 | 51774362 | 51786541 | 51786826 | 0 | 0 | 0 | 1 | 0 | 0 | 0 | 0 | 0 | 0 | 0 | 8 | 0 | 4 | 0 | 0 | 0 | 1 | 0 | 0 | 219 | 149 | 1 | 1 | NA | NA | NA | 1 | NA | | 1 | 0 | 1 | NA | 0.333 |
| SE | 107832 | ENSG00000125629 | INSIG2 | chr2 | + | 118096418 | 118096426 | 118088473 | 118088541 | 118103196 | 118103321 | 5 | 0 | 3 | 5 | 3 | 0 | 0 | 4 | 3 | 0 | 0 | 8 | 6 | 0 | 0 | 0 | 0 | 0 | 0 | 0 | 157 | 149 | 1 | 1 | 1 | NA | 0.416 | 0.613 | 1 | NA | 1 | 1 | NA | NA | -0.243 |
| SE | 107833 | ENSG00000125629 | INSIG2 | chr2 | + | 118096418 | 118096800 | 118088470 | 118088541 | 118103196 | 118103321 | 45 | 31 | 34 | 51 | 56 | 0 | 0 | 4 | 3 | 0 | 24 | 35 | 22 | 26 | 0 | 0 | 0 | 0 | 0 | 0 | 298 | 149 | 1 | 1 | 1 | 1 | 0.81 | 0.895 | 1 | 1 | 1 | 1 | 1 | NA | -0.059 |
| SE | 107834 | ENSG00000125629 | INSIG2 | chr2 | + | 118107089 | 118107189 | 118106658 | 118106903 | 118108280 | 118108304 | 46 | 36 | 41 | 57 | 77 | 1 | 1 | 0 | 0 | 0 | 0 | 29 | 29 | 20 | 4 | 0 | 0 | 2 | 3 | 0 | 249 | 149 | 1 | 1 | 0.965 | 0.956 | 1 | 1 | 1 | NA | 1 | 0.897 | 0.8 | 1 | 0.06 |
| SE | 107980 | ENSG00000204852 | TCTN1 | chr12 | + | 110619835 | 110619956 | 110614026 | 110614402 | 110626361 | 110626382 | 0 | 1 | 0 | 0 | 5 | 0 | 0 | 0 | 0 | 0 | 0 | 0 | 0 | 0 | 0 | 4 | 0 | 2 | 0 | 0 | 270 | 149 | 1 | 1 | NA | 1 | NA | NA | 1 | 0 | NA | 0 | NA | NA | 1 |
| SE | 107981 | ENSG00000204852 | TCTN1 | chr12 | + | 110619835 | 110619956 | 110614106 | 110614463 | 110626361 | 110626382 | 0 | 1 | 0 | 0 | 5 | 0 | 0 | 0 | 0 | 0 | 0 | 0 | 0 | 0 | 0 | 0 | 0 | 2 | 0 | 0 | 270 | 149 | 1 | 1 | NA | 1 | NA | NA | 1 | NA | NA | 0 | NA | NA | 1 |
| SE | 107983 | ENSG00000204852 | TCTN1 | chr12 | + | 110627929 | 110628224 | 110626361 | 110626492 | 110628766 | 110628918 | 1 | 0 | 1 | 0 | 4 | 7 | 8 | 8 | 4 | 6 | 0 | 0 | 0 | 4 | 0 | 10 | 0 | 0 | 3 | 14 | 298 | 149 | 1 | 1 | 0.067 | 0 | 0.059 | 0 | 0.25 | 0 | NA | NA | 0.4 | 0 | -0.058 |
| SE | 107984 | ENSG00000204852 | TCTN1 | chr12 | + | 110628051 | 110628224 | 110626361 | 110626492 | 110628766 | 110628918 | 1 | 0 | 4 | 0 | 6 | 7 | 8 | 8 | 4 | 6 | 0 | 0 | 0 | 8 | 0 | 10 | 0 | 0 | 3 | 14 | 298 | 149 | 1 | 1 | 0.067 | 0 | 0.2 | 0 | 0.333 | 0 | NA | NA | 0.571 | 0 | -0.07 |
| SE | 107985 | ENSG00000204852 | TCTN1 | chr12 | + | 110628051 | 110628224 | 110626361 | 110626492 | 110628796 | 110630216 | 0 | 0 | 4 | 1 | 5 | 1 | 0 | 1 | 0 | 7 | 0 | 0 | 1 | 5 | 0 | 0 | 2 | 5 | 0 | 0 | 298 | 149 | 1 | 1 | 0 | NA | 0.667 | 1 | 0.263 | NA | 0 | 0.091 | 1 | NA | 0.119 |
| SE | 107986 | ENSG00000204852 | TCTN1 | chr12 | + | 110634395 | 110634463 | 110632471 | 110632559 | 110636669 | 110634779 | 0 | 0 | 0 | 1 | 0 | 16 | 17 | 10 | 16 | 43 | 0 | 0 | 0 | 0 | 0 | 10 | 9 | 4 | 8 | 27 | 217 | 149 | 1 | 1 | 0 | 0 | 0 | 0.041 | 0 | 0 | 0 | 0 | 0 | 0 | 0.008 |
| SE | 107987 | ENSG00000204852 | TCTN1 | chr12 | + | 110634395 | 110634463 | 110632471 | 110632559 | 110636480 | 110636501 | 1 | 1 | 0 | 2 | 0 | 4 | 1 | 4 | 3 | 3 | 0 | 0 | 0 | 0 | 0 | 0 | 3 | 0 | 1 | 0 | 217 | 149 | 1 | 1 | 0.147 | 0.407 | 0 | 0.314 | 0 | NA | 0 | NA | 0 | NA | 0.174 |
| SE | 107990 | ENSG00000204852 | TCTN1 | chr12 | + | 110634669 | 110634779 | 110634395 | 110634463 | 110636480 | 110636501 | 9 | 7 | 9 | 2 | 28 | 1 | 1 | 0 | 1 | 0 | 0 | 1 | 3 | 5 | 0 | 0 | 0 | 0 | 0 | 0 | 259 | 149 | 1 | 1 | 0.838 | 0.801 | 1 | 0.535 | 1 | NA | 1 | 1 | 1 | NA | -0.165 |
| SE | 107991 | ENSG00000204852 | TCTN1 | chr12 | + | 110635950 | 110636244 | 110634669 | 110634779 | 110636480 | 110636501 | 0 | 0 | 3 | 1 | 0 | 14 | 16 | 14 | 7 | 41 | 0 | 0 | 0 | 0 | 0 | 0 | 10 | 4 | 8 | 8 | 298 | 149 | 1 | 1 | 0 | 0 | 0.097 | 0.067 | 0 | NA | 0 | 0 | 0 | 0 | 0.033 |
| SE | 107993 | ENSG00000204852 | TCTN1 | chr12 | + | 110641023 | 110641149 | 110640382 | 110640517 | 110641541 | 110641586 | 16 | 15 | 15 | 9 | 33 | 0 | 0 | 0 | 2 | 0 | 12 | 12 | 7 | 11 | 8 | 0 | 0 | 0 | 0 | 0 | 275 | 149 | 1 | 1 | 1 | 1 | 1 | 0.709 | 1 | 1 | 1 | 1 | 1 | 1 | -0.058 |
| SE | 107995 | ENSG00000204852 | TCTN1 | chr12 | + | 110641023 | 110641149 | 110640382 | 110640517 | 110647210 | 110647336 | 6 | 11 | 9 | 6 | 16 | 2 | 0 | 1 | 0 | 0 | 6 | 3 | 11 | 7 | 5 | 0 | 2 | 0 | 0 | 0 | 275 | 149 | 1 | 1 | 0.619 | 1 | 0.83 | 1 | 1 | 1 | 0.448 | 1 | 1 | 1 | 0 |
| SE | 107996 | ENSG00000204852 | TCTN1 | chr12 | + | 110641541 | 110642389 | 110640382 | 110640517 | 110647210 | 110647336 | 0 | 1 | 0 | 1 | 0 | 2 | 0 | 1 | 0 | 0 | 0 | 0 | 0 | 0 | 0 | 0 | 2 | 0 | 0 | 0 | 298 | 149 | 1 | 1 | 0 | 1 | 0 | 1 | NA | NA | 0 | NA | NA | NA | 0.5 |
| SE | 107997 | ENSG00000204852 | TCTN1 | chr12 | + | 110641541 | 110642389 | 110641023 | 110641149 | 110644991 | 110645129 | 7 | 4 | 7 | 0 | 14 | 1 | 0 | 0 | 0 | 0 | 0 | 7 | 1 | 4 | 0 | 0 | 0 | 0 | 0 | 0 | 298 | 149 | 1 | 1 | 0.778 | 1 | 1 | NA | 1 | NA | 1 | 1 | 1 | NA | -0.055 |
| SE | 107998 | ENSG00000204852 | TCTN1 | chr12 | + | 110641541 | 110642389 | 110641023 | 110641149 | 110647210 | 110647336 | 7 | 5 | 7 | 0 | 14 | 1 | 0 | 4 | 0 | 0 | 0 | 7 | 1 | 4 | 0 | 0 | 0 | 5 | 0 | 0 | 298 | 149 | 1 | 1 | 0.778 | 1 | 0.467 | NA | 1 | NA | 1 | 0.091 | 1 | NA | 0.114 |
| SE | 107999 | ENSG00000204852 | TCTN1 | chr12 | + | 110644966 | 110645129 | 110640382 | 110640517 | 110647195 | 110647336 | 0 | 0 | 0 | 0 | 2 | 0 | 0 | 2 | 0 | 0 | 0 | 0 | 0 | 3 | 10 | 0 | 0 | 0 | 0 | 0 | 298 | 149 | 1 | 1 | NA | NA | 0 | NA | 1 | NA | NA | NA | 1 | 1 | -0.5 |
| SE | 108001 | ENSG00000204852 | TCTN1 | chr12 | + | 110644966 | 110645129 | 110641023 | 110641149 | 110647210 | 110647336 | 14 | 13 | 13 | 3 | 4 | 1 | 0 | 4 | 0 | 0 | 6 | 0 | 4 | 11 | 19 | 0 | 0 | 5 | 0 | 0 | 298 | 149 | 1 | 1 | 0.875 | 1 | 0.619 | 1 | 1 | 1 | NA | 0.286 | 1 | 1 | 0.077 |
| SE | 108002 | ENSG00000204852 | TCTN1 | chr12 | + | 110644966 | 110645129 | 110642248 | 110642389 | 110647210 | 110647336 | 15 | 18 | 16 | 5 | 18 | 0 | 1 | 0 | 0 | 0 | 6 | 7 | 6 | 22 | 32 | 0 | 0 | 0 | 0 | 0 | 298 | 149 | 1 | 1 | 1 | 0.9 | 1 | 1 | 1 | 1 | 1 | 1 | 1 | 1 | -0.02 |
| SE | 108003 | ENSG00000204852 | TCTN1 | chr12 | + | 110644966 | 110645143 | 110640382 | 110640517 | 110647210 | 110647262 | 0 | 0 | 0 | 1 | 2 | 2 | 0 | 1 | 0 | 0 | 0 | 0 | 0 | 0 | 7 | 0 | 2 | 0 | 0 | 0 | 298 | 149 | 1 | 1 | 0 | NA | 0 | 1 | 1 | NA | 0 | NA | NA | 1 | 0 |
| SE | 108004 | ENSG00000204852 | TCTN1 | chr12 | + | 110644991 | 110645129 | 110641023 | 110641149 | 110647210 | 110647336 | 15 | 12 | 11 | 3 | 4 | 1 | 0 | 4 | 0 | 0 | 6 | 0 | 4 | 11 | 16 | 0 | 0 | 5 | 0 | 0 | 287 | 149 | 1 | 1 | 0.886 | 1 | 0.588 | 1 | 1 | 1 | NA | 0.293 | 1 | 1 | 0.072 |
| SE | 108005 | ENSG00000204852 | TCTN1 | chr12 | + | 110644991 | 110645129 | 110641541 | 110642389 | 110647210 | 110647336 | 14 | 12 | 11 | 3 | 4 | 0 | 1 | 0 | 0 | 0 | 6 | 0 | 4 | 11 | 16 | 0 | 0 | 0 | 0 | 0 | 287 | 149 | 1 | 1 | 1 | 0.862 | 1 | 1 | 1 | 1 | NA | 1 | 1 | 1 | -0.028 |
| SE | 108196 | ENSG00000186908 | ZDHHC17 | chr12 | + | 76781628 | 76781683 | 76764252 | 76764329 | 76797433 | 76797537 | 2 | 0 | 0 | 1 | 1 | 48 | 25 | 24 | 45 | 39 | 0 | 0 | 0 | 0 | 0 | 22 | 17 | 17 | 35 | 15 | 204 | 149 | 1 | 1 | 0.03 | 0 | 0 | 0.016 | 0.018 | 0 | 0 | 0 | 0 | 0 | 0.013 |
| SE | 108200 | ENSG00000186908 | ZDHHC17 | chr12 | + | 76815856 | 76816019 | 76815145 | 76815210 | 76821057 | 76821167 | 124 | 102 | 89 | 125 | 98 | 1 | 0 | 0 | 0 | 0 | 29 | 39 | 52 | 27 | 11 | 0 | 0 | 0 | 0 | 0 | 298 | 149 | 1 | 1 | 0.984 | 1 | 1 | 1 | 1 | 1 | 1 | 1 | 1 | 1 | -0.003 |
| SE | 108202 | ENSG00000186908 | ZDHHC17 | chr12 | + | 76822405 | 76822531 | 76815856 | 76816019 | 76826907 | 76826914 | 107 | 114 | 91 | 147 | 186 | 0 | 1 | 0 | 0 | 0 | 49 | 44 | 72 | 23 | 16 | 0 | 0 | 0 | 0 | 0 | 275 | 149 | 1 | 1 | 1 | 0.984 | 1 | 1 | 1 | 1 | 1 | 1 | 1 | 1 | -0.003 |
| SE | 108203 | ENSG00000186908 | ZDHHC17 | chr12 | + | 76826907 | 76827050 | 76822405 | 76822531 | 76828389 | 76828490 | 132 | 111 | 109 | 120 | 200 | 0 | 1 | 0 | 0 | 0 | 5 | 49 | 57 | 21 | 4 | 0 | 0 | 2 | 0 | 0 | 292 | 149 | 1 | 1 | 1 | 0.983 | 1 | 1 | 1 | 1 | 1 | 0.936 | 1 | 1 | 0.009 |
| SE | 108204 | ENSG00000186908 | ZDHHC17 | chr12 | + | 76840964 | 76841106 | 76840390 | 76840483 | 76842024 | 76842106 | 5 | 9 | 4 | 5 | 2 | 2 | 3 | 6 | 4 | 1 | 0 | 0 | 1 | 0 | 0 | 0 | 8 | 1 | 3 | 0 | 291 | 149 | 1 | 1 | 0.561 | 0.606 | 0.254 | 0.39 | 0.506 | NA | 0 | 0.339 | 0 | NA | 0.35 |
| SE | 108205 | ENSG00000186908 | ZDHHC17 | chr12 | + | 76842918 | 76842981 | 76841981 | 76842106 | 76845708 | 76845802 | 74 | 66 | 76 | 92 | 117 | 0 | 1 | 0 | 0 | 0 | 2 | 39 | 14 | 22 | 11 | 0 | 0 | 0 | 0 | 0 | 212 | 149 | 1 | 1 | 1 | 0.979 | 1 | 1 | 1 | 1 | 1 | 1 | 1 | 1 | -0.004 |
| SE | 108206 | ENSG00000186908 | ZDHHC17 | chr12 | + | 76848232 | 76848390 | 76846290 | 76846679 | 76849375 | 76849466 | 280 | 228 | 228 | 258 | 334 | 0 | 0 | 0 | 3 | 0 | 43 | 135 | 93 | 35 | 7 | 0 | 0 | 0 | 0 | 0 | 298 | 149 | 1 | 1 | 1 | 1 | 1 | 0.977 | 1 | 1 | 1 | 1 | 1 | 1 | -0.005 |
| SE | 108207 | ENSG00000186908 | ZDHHC17 | chr12 | + | 76849375 | 76849470 | 76848232 | 76848390 | 76850846 | 76851392 | 211 | 164 | 182 | 180 | 245 | 0 | 2 | 1 | 1 | 0 | 26 | 86 | 74 | 21 | 15 | 0 | 0 | 0 | 0 | 0 | 244 | 149 | 1 | 1 | 1 | 0.98 | 0.991 | 0.991 | 1 | 1 | 1 | 1 | 1 | 1 | -0.008 |
| SE | 108792 | ENSG00000105983 | LMBR1 | chr7 | - | 156724111 | 156724178 | 156688029 | 156688191 | 156734176 | 156734257 | 47 | 35 | 47 | 36 | 56 | 0 | 0 | 1 | 0 | 0 | 53 | 41 | 38 | 26 | 16 | 0 | 0 | 0 | 0 | 0 | 216 | 149 | 1 | 1 | 1 | 1 | 0.97 | 1 | 1 | 1 | 1 | 1 | 1 | 1 | -0.006 |
| SE | 108793 | ENSG00000105983 | LMBR1 | chr7 | - | 156727929 | 156728007 | 156724111 | 156724178 | 156734176 | 156734257 | 0 | 2 | 1 | 1 | 0 | 6 | 5 | 20 | 13 | 17 | 0 | 0 | 0 | 0 | 0 | 14 | 8 | 16 | 3 | 7 | 227 | 149 | 1 | 1 | 0 | 0.208 | 0.032 | 0.048 | 0 | 0 | 0 | 0 | 0 | 0 | 0.058 |
| SE | 108796 | ENSG00000105983 | LMBR1 | chr7 | - | 156734176 | 156734257 | 156724111 | 156724178 | 156756392 | 156756465 | 53 | 46 | 61 | 65 | 71 | 1 | 1 | 0 | 0 | 0 | 29 | 24 | 64 | 41 | 24 | 0 | 2 | 0 | 0 | 0 | 230 | 149 | 1 | 1 | 0.972 | 0.968 | 1 | 1 | 1 | 1 | 0.886 | 1 | 1 | 1 | 0.011 |
| SE | 108797 | ENSG00000105983 | LMBR1 | chr7 | - | 156736492 | 156736615 | 156734176 | 156734257 | 156756392 | 156756465 | 0 | 0 | 1 | 1 | 1 | 93 | 74 | 90 | 116 | 100 | 0 | 0 | 6 | 0 | 0 | 59 | 74 | 116 | 77 | 43 | 272 | 149 | 1 | 1 | 0 | 0 | 0.006 | 0.005 | 0.005 | 0 | 0 | 0.028 | 0 | 0 | -0.002 |
| SE | 108799 | ENSG00000105983 | LMBR1 | chr7 | - | 156763107 | 156763176 | 156762133 | 156762198 | 156763668 | 156763795 | 117 | 83 | 120 | 102 | 167 | 0 | 0 | 0 | 0 | 1 | 61 | 121 | 137 | 80 | 42 | 0 | 0 | 0 | 0 | 0 | 218 | 149 | 1 | 1 | 1 | 1 | 1 | 1 | 0.991 | 1 | 1 | 1 | 1 | 1 | -0.002 |
| SE | 108806 | ENSG00000105983 | LMBR1 | chr7 | - | 156836808 | 156836885 | 156833752 | 156833792 | 156892927 | 156893216 | 1 | 1 | 7 | 5 | 0 | 0 | 0 | 1 | 0 | 0 | 0 | 3 | 11 | 1 | 2 | 0 | 0 | 0 | 0 | 0 | 226 | 149 | 1 | 1 | 1 | 1 | 0.822 | 1 | NA | NA | 1 | 1 | 1 | 1 | -0.044 |
| SE | 108807 | ENSG00000105983 | LMBR1 | chr7 | - | 156836812 | 156836885 | 156833477 | 156833792 | 156892927 | 156892987 | 84 | 62 | 64 | 67 | 49 | 0 | 0 | 1 | 0 | 0 | 36 | 78 | 85 | 82 | 30 | 0 | 0 | 0 | 0 | 0 | 222 | 149 | 1 | 1 | 1 | 1 | 0.977 | 1 | 1 | 1 | 1 | 1 | 1 | 1 | -0.005 |
| SE | 108850 | ENSG00000117226 | GBP3 | chr1 | - | 89010903 | 89011116 | 89009391 | 89009494 | 89013184 | 89013427 | 72 | 114 | 89 | 125 | 127 | 30 | 0 | 0 | 0 | 0 | 0 | 46 | 22 | 18 | 0 | 0 | 0 | 0 | 22 | 0 | 298 | 149 | 1 | 1 | 0.545 | 1 | 1 | 1 | 1 | NA | 1 | 1 | 0.29 | NA | 0.146 |
| SE | 108851 | ENSG00000117226 | GBP3 | chr1 | - | 89011746 | 89012027 | 89008946 | 89009140 | 89014082 | 89014279 | 0 | 0 | 6 | 15 | 0 | 1 | 1 | 0 | 0 | 0 | 0 | 0 | 4 | 0 | 0 | 0 | 0 | 11 | 0 | 0 | 298 | 149 | 1 | 1 | 0 | 0 | 1 | 1 | NA | NA | NA | 0.154 | NA | NA | 0.346 |
| SE | 108852 | ENSG00000117226 | GBP3 | chr1 | - | 89011746 | 89012027 | 89010903 | 89011116 | 89013184 | 89013427 | 91 | 154 | 107 | 108 | 88 | 0 | 2 | 2 | 0 | 2 | 0 | 87 | 62 | 31 | 26 | 0 | 5 | 0 | 0 | 0 | 298 | 149 | 1 | 1 | 1 | 0.975 | 0.964 | 1 | 0.957 | NA | 0.897 | 1 | 1 | 1 | 0.005 |
| SE | 108884 | ENSG00000145014 | TMEM44 | chr3 | - | 194615568 | 194615670 | 194610915 | 194611020 | 194623223 | 194623310 | 5 | 0 | 0 | 0 | 1 | 0 | 0 | 0 | 1 | 1 | 0 | 4 | 4 | 0 | 0 | 0 | 0 | 0 | 0 | 0 | 251 | 149 | 1 | 1 | 1 | NA | NA | 0 | 0.372 | NA | 1 | 1 | NA | NA | -0.543 |
| SE | 109049 | ENSG00000139668 | WDFY2 | chr13 | + | 51660595 | 51660663 | 51584461 | 51584824 | 51675169 | 51675243 | 98 | 98 | 96 | 111 | 107 | 0 | 0 | 0 | 1 | 0 | 92 | 123 | 138 | 97 | 51 | 0 | 5 | 0 | 0 | 0 | 217 | 149 | 1 | 1 | 1 | 1 | 1 | 0.987 | 1 | 1 | 0.944 | 1 | 1 | 1 | 0.009 |
| SE | 109051 | ENSG00000139668 | WDFY2 | chr13 | + | 51703595 | 51703650 | 51675169 | 51675243 | 51719197 | 51719348 | 145 | 136 | 115 | 131 | 202 | 0 | 0 | 0 | 0 | 0 | 202 | 206 | 180 | 214 | 127 | 0 | 0 | 1 | 0 | 0 | 204 | 149 | 1 | 1 | 1 | 1 | 1 | 1 | 1 | 1 | 1 | 0.992 | 1 | 1 | 0.002 |
| SE | 109052 | ENSG00000139668 | WDFY2 | chr13 | + | 51703595 | 51703650 | 51675169 | 51675243 | 51739048 | 51739175 | 36 | 34 | 31 | 28 | 63 | 0 | 1 | 0 | 0 | 0 | 21 | 44 | 39 | 59 | 25 | 0 | 0 | 0 | 0 | 0 | 204 | 149 | 1 | 1 | 1 | 0.961 | 1 | 1 | 1 | 1 | 1 | 1 | 1 | 1 | -0.008 |

RI

| | | | | | | | | | | | | Inclusion_counts_HC | | | | | Skipping_counts_HC | | | | | Inclusion counts_SLE | | | | | Skipping_counts_SLE | | | | | | | | | Inclusion_level(normalized) | | | | | | | | | | |
|---|---|---|---|---|---|---|---|---|---|---|---|---|---|---|---|---|---|---|---|---|---|---|---|---|---|---|---|---|---|---|---|---|---|---|---|---|---|---|---|---|---|---|---|---|---|---|
| Splice_type | ID | Gene_ID | Gene_symbol | Chromosome | Strand | LongExonStart_0base | LongExonEnd | ShortES | ShortEE | FlankingES | FlankingEE | SRR13214146 | SRR13214147 | SRR13214148 | SRR13214149 | SRR13214150 | SRR13214146 | SRR13214147 | SRR13214148 | SRR13214149 | SRR13214150 | SRR13214141 | SRR13214242 | SRR13214143 | SRR13214144 | SRR13214145 | SRR13214141 | SRR13214242 | SRR13214143 | SRR13214144 | SRR13214145 | IncFormLen | SkipFormLen | p value | FDR | SRR13214146 | SRR13214147 | SRR13214148 | SRR13214149 | SRR13214150 | SRR13214141 | SRR13214242 | SRR13214143 | SRR13214144 | SRR13214145 | Inclevel_diff |
| RI | 7539 | ENSG00000130714 | POMT1 | chr9 | + | 131509902 | 131510415 | 131509902 | 131509996 | 131510259 | 131510415 | 13 | 5 | 18 | 18 | 5 | 8 | 1 | 12 | 6 | 4 | 47 | 40 | 19 | 17 | 23 | 0 | 0 | 0 | 0 | 0 | 298 | 149 | 0 | 0 | 0.448 | 0.714 | 0.429 | 0.6 | 0.385 | 1 | 1 | 1 | 1 | 1 | -0.485 |
| RI | 1430 | ENSG00000197170 | PSMD12 | chr17 | - | 67357302 | 67357578 | 67357302 | 67357431 | 67357518 | 67357578 | 61 | 77 | 43 | 69 | 29 | 128 | 74 | 109 | 128 | 89 | 239 | 185 | 227 | 286 | 36 | 45 | 73 | 110 | 39 | 17 | 236 | 149 | 0 | 0 | 0.231 | 0.396 | 0.199 | 0.254 | 0.171 | 0.77 | 0.615 | 0.566 | 0.822 | 0.572 | -0.419 |
| RI | 2447 | ENSG00000125089 | SH3TC1 | chr4 | + | 8235432 | 8237670 | 8235432 | 8235555 | 8236277 | 8237670 | 16 | 15 | 5 | 14 | 9 | 46 | 22 | 23 | 30 | 17 | 91 | 56 | 25 | 51 | 100 | 29 | 24 | 10 | 20 | 33 | 298 | 149 | 0 | 0 | 0.148 | 0.254 | 0.098 | 0.189 | 0.209 | 0.611 | 0.538 | 0.556 | 0.56 | 0.602 | -0.394 |
| RI | 8365 | ENSG00000166012 | TAF1D | chr11 | - | 93730676 | 93731216 | 93730676 | 93730712 | 93731149 | 93731216 | 354 | 209 | 266 | 302 | 1156 | 150 | 155 | 141 | 134 | 301 | 384 | 441 | 254 | 226 | 725 | 10 | 40 | 58 | 17 | 28 | 298 | 149 | 0 | 0 | 0.541 | 0.403 | 0.485 | 0.53 | 0.658 | 0.95 | 0.846 | 0.686 | 0.869 | 0.928 | -0.332 |
| RI | 6534 | ENSG00000106263 | EIF3B | chr7 | + | 2366316 | 2366591 | 2366316 | 2366448 | 2366524 | 2366591 | 28 | 22 | 19 | 33 | 4 | 100 | 71 | 125 | 123 | 126 | 236 | 110 | 68 | 182 | 156 | 157 | 113 | 100 | 116 | 137 | 225 | 149 | 0 | 0 | 0.156 | 0.17 | 0.091 | 0.151 | 0.021 | 0.499 | 0.392 | 0.31 | 0.51 | 0.43 | -0.31 |
| RI | 4891 | ENSG00000070814 | TCOF1 | chr5 | + | 150375720 | 150376330 | 150375720 | 150375909 | 150376081 | 150376330 | 16 | 5 | 24 | 21 | 10 | 55 | 31 | 53 | 68 | 22 | 63 | 49 | 55 | 76 | 134 | 25 | 37 | 45 | 43 | 90 | 298 | 149 | 0 | 0 | 0.127 | 0.075 | 0.185 | 0.134 | 0.185 | 0.558 | 0.398 | 0.379 | 0.469 | 0.427 | -0.305 |
| RI | 3116 | ENSG00000161204 | ABCF3 | chr3 | + | 184187396 | 184187761 | 184187396 | 184187443 | 184187663 | 184187761 | 62 | 47 | 43 | 42 | 48 | 104 | 50 | 58 | 85 | 61 | 221 | 179 | 105 | 140 | 195 | 89 | 53 | 52 | 66 | 61 | 298 | 149 | 0 | 0 | 0.23 | 0.32 | 0.27 | 0.198 | 0.282 | 0.554 | 0.628 | 0.502 | 0.515 | 0.615 | -0.303 |
| RI | 4630 | ENSG00000003756 | RBM5 | chr3 | + | 50117073 | 50117379 | 50117073 | 50117171 | 50117249 | 50117379 | 58 | 65 | 53 | 84 | 64 | 171 | 130 | 178 | 148 | 257 | 170 | 174 | 175 | 271 | 120 | 155 | 141 | 138 | 127 | 90 | 227 | 149 | 0 | 0 | 0.182 | 0.247 | 0.163 | 0.271 | 0.14 | 0.419 | 0.448 | 0.454 | 0.583 | 0.467 | -0.274 |
| RI | 2880 | ENSG00000178035 | IMPDH2 | chr3 | - | 49024342 | 49024578 | 49024342 | 49024404 | 49024494 | 49024578 | 41 | 31 | 35 | 33 | 18 | 83 | 57 | 133 | 107 | 130 | 97 | 62 | 65 | 64 | 103 | 62 | 50 | 52 | 48 | 105 | 239 | 149 | 0 | 0 | 0.235 | 0.253 | 0.141 | 0.161 | 0.079 | 0.494 | 0.436 | 0.438 | 0.454 | 0.379 | -0.266 |
| RI | 6924 | ENSG00000152332 | UHMK1 | chr1 | + | 162499954 | 162501104 | 162499954 | 162500247 | 162500912 | 162501104 | 69 | 33 | 78 | 59 | 51 | 191 | 115 | 150 | 170 | 258 | 67 | 133 | 146 | 110 | 35 | 38 | 122 | 93 | 83 | 31 | 298 | 149 | 0 | 0 | 0.153 | 0.125 | 0.206 | 0.148 | 0.09 | 0.469 | 0.353 | 0.44 | 0.399 | 0.361 | -0.26 |
| RI | 7904 | ENSG00000054654 | SYNE2 | chr14 | + | 64022750 | 64024459 | 64022750 | 64022863 | 64024256 | 64024459 | 51 | 42 | 27 | 16 | 16 | 498 | 305 | 291 | 345 | 304 | 14 | 51 | 83 | 25 | 12 | 17 | 127 | 93 | 14 | 28 | 298 | 149 | 0 | 0 | 0.049 | 0.064 | 0.044 | 0.023 | 0.026 | 0.292 | 0.167 | 0.309 | 0.472 | 0.176 | -0.242 |
| RI | 3908 | ENSG00000126749 | EMG1 | chr12 | + | 6974551 | 6975148 | 6974551 | 6974693 | 6975089 | 6975148 | 20 | 14 | 26 | 22 | 43 | 126 | 43 | 140 | 106 | 190 | 46 | 38 | 45 | 42 | 44 | 49 | 70 | 39 | 40 | 41 | 298 | 149 | 0 | 0 | 0.074 | 0.14 | 0.085 | 0.094 | 0.102 | 0.319 | 0.213 | 0.366 | 0.344 | 0.349 | -0.219 |
| RI | 3093 | ENSG00000111642 | CHD4 | chr12 | - | 6578028 | 6578546 | 6578028 | 6578137 | 6578408 | 6578546 | 52 | 59 | 39 | 58 | 56 | 384 | 226 | 344 | 359 | 525 | 383 | 139 | 146 | 347 | 175 | 361 | 239 | 246 | 215 | 405 | 298 | 149 | 0 | 0 | 0.063 | 0.115 | 0.054 | 0.075 | 0.051 | 0.347 | 0.225 | 0.229 | 0.447 | 0.178 | -0.214 |
| RI | 1066 | ENSG00000168259 | DNAJC7 | chr17 | - | 41987818 | 41988896 | 41987818 | 41987910 | 41988731 | 41988896 | 99 | 78 | 99 | 139 | 78 | 300 | 272 | 286 | 257 | 556 | 750 | 192 | 251 | 448 | 273 | 633 | 210 | 402 | 477 | 182 | 298 | 149 | 0 | 0 | 0.142 | 0.125 | 0.148 | 0.213 | 0.066 | 0.372 | 0.314 | 0.238 | 0.32 | 0.429 | -0.196 |
| RI | 5341 | ENSG00000084623 | EIF3I | chr1 | + | 32228726 | 32229208 | 32228726 | 32228816 | 32229134 | 32229208 | 21 | 20 | 21 | 30 | 12 | 136 | 82 | 153 | 117 | 130 | 103 | 69 | 44 | 94 | 110 | 112 | 86 | 84 | 107 | 255 | 298 | 149 | 0 | 0 | 0.072 | 0.109 | 0.064 | 0.114 | 0.044 | 0.315 | 0.286 | 0.208 | 0.305 | 0.177 | -0.178 |
| RI | 2166 | ENSG00000065357 | DGKA | chr12 | + | 55940081 | 55940722 | 55940081 | 55940433 | 55940623 | 55940722 | 48 | 35 | 93 | 81 | 27 | 163 | 90 | 255 | 242 | 216 | 53 | 84 | 91 | 135 | 122 | 77 | 114 | 103 | 111 | 129 | 298 | 149 | 0 | 0 | 0.128 | 0.163 | 0.154 | 0.143 | 0.059 | 0.256 | 0.269 | 0.306 | 0.378 | 0.321 | -0.177 |
| RI | 256 | ENSG00000104774 | MAN2B1 | chr19 | - | 12655693 | 12656687 | 12655693 | 12655879 | 12656570 | 12656687 | 55 | 39 | 45 | 68 | 34 | 251 | 157 | 173 | 182 | 194 | 323 | 105 | 69 | 160 | 346 | 328 | 224 | 122 | 174 | 391 | 298 | 149 | 0 | 0 | 0.099 | 0.11 | 0.115 | 0.157 | 0.081 | 0.33 | 0.19 | 0.22 | 0.315 | 0.307 | -0.16 |
| RI | 7531 | ENSG00000180104 | EXOC3 | chr5 | + | 464289 | 465272 | 464289 | 464412 | 465110 | 465272 | 10 | 13 | 13 | 7 | 25 | 111 | 81 | 79 | 109 | 204 | 52 | 54 | 28 | 52 | 64 | 72 | 118 | 55 | 112 | 219 | 298 | 149 | 0 | 0 | 0.043 | 0.074 | 0.076 | 0.031 | 0.058 | 0.265 | 0.186 | 0.203 | 0.188 | 0.127 | -0.137 |
| RI | 5665 | ENSG00000165280 | VCP | chr9 | - | 35062002 | 35062350 | 35062002 | 35062138 | 35062216 | 35062350 | 46 | 39 | 59 | 64 | 28 | 386 | 236 | 313 | 362 | 267 | 361 | 161 | 82 | 297 | 256 | 632 | 412 | 338 | 441 | 672 | 227 | 149 | 0 | 0 | 0.073 | 0.098 | 0.11 | 0.104 | 0.064 | 0.273 | 0.204 | 0.137 | 0.307 | 0.2 | -0.134 |
| RI | 192 | ENSG00000099875 | MKNK2 | chr19 | - | 2042606 | 2042870 | 2042606 | 2042662 | 2042765 | 2042870 | 37 | 15 | 24 | 39 | 22 | 248 | 109 | 217 | 261 | 200 | 23 | 98 | 87 | 75 | 114 | 92 | 248 | 145 | 122 | 320 | 252 | 149 | 0 | 0 | 0.081 | 0.075 | 0.061 | 0.081 | 0.061 | 0.129 | 0.189 | 0.262 | 0.267 | 0.174 | -0.132 |
| RI | 1259 | ENSG00000005844 | ITGAL | chr16 | + | 30496096 | 30496566 | 30496096 | 30496294 | 30496435 | 30496566 | 158 | 98 | 100 | 103 | 67 | 594 | 262 | 305 | 418 | 415 | 324 | 214 | 153 | 193 | 364 | 387 | 525 | 224 | 269 | 571 | 290 | 149 | 0 | 0 | 0.12 | 0.161 | 0.144 | 0.112 | 0.077 | 0.301 | 0.173 | 0.26 | 0.269 | 0.247 | -0.127 |
| RI | 5336 | ENSG00000084623 | EIF3I | chr1 | + | 32224033 | 32224475 | 32224033 | 32224121 | 32224409 | 32224475 | 26 | 17 | 22 | 48 | 13 | 198 | 124 | 225 | 216 | 151 | 99 | 74 | 61 | 137 | 122 | 245 | 213 | 141 | 163 | 377 | 298 | 149 | 0 | 0 | 0.062 | 0.064 | 0.047 | 0.1 | 0.041 | 0.168 | 0.148 | 0.178 | 0.296 | 0.139 | -0.123 |
| RI | 5113 | ENSG00000181163 | NPM1 | chr5 | + | 171400152 | 171400925 | 171400152 | 171400210 | 171400838 | 171400925 | 37 | 39 | 51 | 72 | 74 | 1044 | 698 | 1317 | 1051 | 2497 | 116 | 76 | 53 | 74 | 30 | 130 | 525 | 589 | 399 | 54 | 298 | 149 | 0 | 0 | 0.017 | 0.027 | 0.019 | 0.033 | 0.015 | 0.309 | 0.067 | 0.043 | 0.085 | 0.217 | -0.122 |
| RI | 6002 | ENSG00000197081 | IGF2R | chr6 | + | 160088032 | 160089253 | 160088032 | 160088147 | 160089106 | 160089253 | 38 | 36 | 36 | 41 | 6 | 347 | 131 | 174 | 261 | 205 | 150 | 78 | 106 | 125 | 90 | 248 | 267 | 279 | 173 | 274 | 298 | 149 | 0 | 0 | 0.052 | 0.121 | 0.094 | 0.073 | 0.014 | 0.232 | 0.127 | 0.16 | 0.265 | 0.141 | -0.114 |
| RI | 8500 | ENSG00000139613 | SMARCC2 | chr12 | - | 56178403 | 56178847 | 56178403 | 56178534 | 56178809 | 56178847 | 71 | 43 | 80 | 78 | 72 | 316 | 203 | 282 | 319 | 557 | 166 | 155 | 97 | 148 | 136 | 371 | 267 | 129 | 279 | 348 | 298 | 149 | 0 | 0 | 0.101 | 0.096 | 0.124 | 0.109 | 0.061 | 0.183 | 0.225 | 0.273 | 0.21 | 0.163 | -0.113 |
| RI | 4058 | ENSG00000198821 | CD247 | chr1 | - | 167434019 | 167435434 | 167434019 | 167434076 | 167435398 | 167435434 | 126 | 51 | 87 | 111 | 70 | 1363 | 454 | 706 | 774 | 978 | 112 | 40 | 164 | 101 | 92 | 222 | 380 | 380 | 160 | 308 | 298 | 149 | 0 | 0 | 0.044 | 0.053 | 0.058 | 0.067 | 0.035 | 0.201 | 0.05 | 0.177 | 0.24 | 0.13 | -0.108 |
| RI | 5690 | ENSG00000160710 | ADAR | chr1 | - | 154588124 | 154588673 | 154588124 | 154588258 | 154588550 | 154588673 | 51 | 40 | 53 | 65 | 58 | 428 | 303 | 363 | 433 | 680 | 307 | 118 | 97 | 290 | 81 | 600 | 378 | 282 | 522 | 312 | 298 | 149 | 0 | 0 | 0.056 | 0.062 | 0.068 | 0.07 | 0.041 | 0.204 | 0.135 | 0.147 | 0.217 | 0.115 | -0.104 |
| RI | 125 | ENSG00000100242 | SUN2 | chr22 | - | 38750897 | 38751373 | 38750897 | 38751035 | 38751209 | 38751373 | 89 | 45 | 68 | 83 | 20 | 587 | 257 | 486 | 473 | 165 | 100 | 97 | 91 | 89 | 134 | 186 | 395 | 190 | 174 | 401 | 298 | 149 | 0 | 0 | 0.07 | 0.081 | 0.065 | 0.081 | 0.057 | 0.212 | 0.109 | 0.193 | 0.204 | 0.143 | -0.101 |
| RI | 1992 | ENSG00000175061 | SNHG29 | chr17 | + | 16439326 | 16440253 | 16439326 | 16439414 | 16440184 | 16440253 | 65 | 46 | 51 | 61 | 44 | 617 | 324 | 818 | 518 | 513 | 116 | 47 | 76 | 128 | 114 | 272 | 214 | 229 | 323 | 462 | 298 | 149 | 0 | 0 | 0.05 | 0.066 | 0.03 | 0.056 | 0.041 | 0.176 | 0.099 | 0.142 | 0.165 | 0.11 | -0.09 |
| RI | 6745 | ENSG00000285171 | NA | chrX | - | 71110896 | 71111571 | 71110896 | 71111050 | 71111424 | 71111571 | 88 | 60 | 135 | 145 | 123 | 891 | 365 | 538 | 638 | 961 | 199 | 163 | 201 | 162 | 177 | 406 | 474 | 419 | 535 | 660 | 298 | 149 | 0 | 0 | 0.047 | 0.076 | 0.111 | 0.102 | 0.06 | 0.197 | 0.147 | 0.193 | 0.131 | 0.118 | -0.078 |
| RI | 2169 | ENSG00000065357 | DGKA | chr12 | + | 55941997 | 55942263 | 55941997 | 55942083 | 55942173 | 55942263 | 18 | 23 | 36 | 20 | 32 | 365 | 427 | 645 | 519 | 537 | 46 | 77 | 48 | 77 | 70 | 437 | 331 | 353 | 423 | 463 | 239 | 149 | 0 | 0 | 0.03 | 0.032 | 0.034 | 0.023 | 0.036 | 0.062 | 0.127 | 0.078 | 0.102 | 0.086 | -0.06 |
| RI | 3097 | ENSG00000111642 | CHD4 | chr12 | - | 6587383 | 6587949 | 6587383 | 6587559 | 6587711 | 6587949 | 21 | 27 | 30 | 51 | 17 | 359 | 260 | 345 | 423 | 371 | 148 | 167 | 92 | 192 | 237 | 1058 | 572 | 515 | 874 | 977 | 298 | 149 | 0 | 0 | 0.028 | 0.049 | 0.042 | 0.057 | 0.022 | 0.065 | 0.127 | 0.082 | 0.099 | 0.108 | -0.057 |
| RI | 5499 | ENSG00000081059 | TCF7 | chr5 | + | 134142720 | 134143100 | 134142720 | 134142883 | 134142992 | 134143100 | 19 | 20 | 62 | 44 | 5 | 493 | 436 | 1108 | 711 | 999 | 10 | 44 | 11 | 36 | 59 | 63 | 245 | 267 | 191 | 337 | 258 | 149 | 0 | 0 | 0.022 | 0.026 | 0.031 | 0.035 | 0.003 | 0.084 | 0.094 | 0.023 | 0.098 | 0.092 | -0.055 |
| RI | 5240 | ENSG00000131143 | COX4I1 | chr16 | + | 85804936 | 85805864 | 85804936 | 85805104 | 85805732 | 85805864 | 51 | 47 | 83 | 87 | 54 | 841 | 392 | 804 | 758 | 778 | 263 | 125 | 192 | 147 | 195 | 1035 | 772 | 485 | 831 | 1555 | 298 | 149 | 0 | 0 | 0.029 | 0.057 | 0.049 | 0.054 | 0.034 | 0.113 | 0.075 | 0.165 | 0.081 | 0.059 | -0.054 |
| RI | 6850 | ENSG00000215021 | PHB2 | chr12 | - | 6967170 | 6967779 | 6967170 | 6967248 | 6967675 | 6967779 | 14 | 18 | 19 | 35 | 64 | 593 | 281 | 561 | 447 | 725 | 51 | 64 | 62 | 62 | 68 | 534 | 279 | 280 | 305 | 534 | 298 | 149 | 0 | 0 | 0.012 | 0.031 | 0.017 | 0.038 | 0.042 | 0.046 | 0.103 | 0.1 | 0.092 | 0.06 | -0.052 |
| RI | 2308 | ENSG00000204628 | RACK1 | chr5 | - | 181241491 | 181242345 | 181241491 | 181241639 | 181242173 | 181242345 | 88 | 98 | 140 | 154 | 102 | 2027 | 923 | 2060 | 1639 | 1595 | 385 | 120 | 149 | 252 | 308 | 1502 | 1264 | 952 | 942 | 2166 | 298 | 149 | 0 | 0 | 0.021 | 0.05 | 0.033 | 0.045 | 0.031 | 0.114 | 0.045 | 0.073 | 0.118 | 0.066 | -0.047 |
| RI | 2304 | ENSG00000204628 | RACK1 | chr5 | - | 181236928 | 181237719 | 181236928 | 181237042 | 181237608 | 181237719 | 66 | 39 | 153 | 54 | 106 | 1572 | 555 | 1691 | 745 | 979 | 146 | 131 | 66 | 114 | 308 | 544 | 1254 | 760 | 617 | 1386 | 298 | 149 | 0 | 0 | 0.021 | 0.034 | 0.043 | 0.035 | 0.051 | 0.118 | 0.05 | 0.042 | 0.085 | 0.1 | -0.042 |
| RI | 3959 | ENSG00000147403 | RPL10 | chrX | + | 154399337 | 154399594 | 154399337 | 154399396 | 154399486 | 154399594 | 272 | 144 | 213 | 292 | 310 | 9609 | 4215 | 11461 | 8678 | 12617 | 426 | 319 | 174 | 408 | 611 | 3925 | 4724 | 3946 | 2927 | 7551 | 239 | 149 | 0 | 0 | 0.017 | 0.021 | 0.011 | 0.021 | 0.015 | 0.063 | 0.04 | 0.027 | 0.08 | 0.048 | -0.035 |
| RI | 8453 | ENSG00000123349 | PFDN5 | chr12 | + | 53295838 | 53297924 | 53295838 | 53296275 | 53297849 | 53297924 | 56 | 52 | 86 | 116 | 142 | 1544 | 881 | 1393 | 1315 | 1587 | 347 | 138 | 115 | 204 | 194 | 2006 | 1172 | 805 | 1133 | 1967 | 298 | 149 | 0 | 0 | 0.018 | 0.029 | 0.03 | 0.042 | 0.043 | 0.08 | 0.056 | 0.067 | 0.083 | 0.047 | -0.034 |
| RI | 373 | ENSG00000105193 | RPS16 | chr19 | - | 39433664 | 39435708 | 39433664 | 39433761 | 39435606 | 39435708 | 167 | 73 | 161 | 239 | 111 | 3550 | 1216 | 4274 | 3254 | 2693 | 175 | 184 | 165 | 197 | 338 | 1368 | 2273 | 1544 | 1050 | 3360 | 298 | 149 | 0 | 0 | 0.023 | 0.029 | 0.018 | 0.035 | 0.02 | 0.06 | 0.039 | 0.051 | 0.086 | 0.048 | -0.032 |
| RI | 5109 | ENSG00000181163 | NPM1 | chr5 | + | 171391304 | 171391799 | 171391304 | 171391424 | 171391705 | 171391799 | 171 | 178 | 230 | 232 | 200 | 4105 | 3548 | 5449 | 4941 | 8235 | 181 | 287 | 279 | 314 | 114 | 1944 | 3624 | 4980 | 3284 | 503 | 298 | 149 | 0 | 0 | 0.02 | 0.024 | 0.021 | 0.023 | 0.012 | 0.044 | 0.038 | 0.027 | 0.046 | 0.102 | -0.031 |
| RI | 3863 | ENSG00000108654 | DDX5 | chr17 | - | 64500548 | 64501143 | 64500548 | 64500773 | 64501056 | 64501143 | 2939 | 1762 | 2823 | 3111 | 2639 | 126 | 80 | 88 | 99 | 101 | 403 | 2364 | 2518 | 1519 | 167 | 9 | 40 | 12 | 9 | 10 | 298 | 149 | 0 | 0 | 0.921 | 0.917 | 0.941 | 0.94 | 0.929 | 0.957 | 0.967 | 0.991 | 0.988 | 0.893 | -0.03 |
| RI | 489 | ENSG00000142347 | MYO1F | chr19 | - | 8553138 | 8553437 | 8553138 | 8553228 | 8553349 | 8553437 | 98 | 43 | 52 | 84 | 45 | 2055 | 1058 | 1314 | 1592 | 1249 | 415 | 135 | 187 | 363 | 511 | 3812 | 2937 | 1655 | 2910 | 4780 | 270 | 149 | 0 | 0 | 0.026 | 0.022 | 0.021 | 0.028 | 0.019 | 0.057 | 0.025 | 0.059 | 0.064 | 0.056 | -0.029 |
| RI | 2472 | ENSG00000070756 | PABPC1 | chr8 | - | 100712651 | 100713181 | 100712651 | 100712789 | 100713086 | 100713181 | 303 | 515 | 301 | 555 | 430 | 10942 | 8709 | 11477 | 10639 | 17216 | 674 | 906 | 1009 | 1273 | 235 | 13212 | 6355 | 8286 | 7930 | 13649 | 298 | 149 | 0 | 0 | 0.014 | 0.029 | 0.013 | 0.025 | 0.012 | 0.025 | 0.067 | 0.057 | 0.074 | 0.009 | -0.028 |
| RI | 5108 | ENSG00000181163 | NPM1 | chr5 | + | 171390050 | 171391424 | 171390050 | 171390130 | 171391304 | 171391424 | 81 | 71 | 126 | 119 | 139 | 3892 | 3080 | 5046 | 4516 | 5974 | 151 | 156 | 133 | 171 | 73 | 1614 | 3298 | 4362 | 2833 | 414 | 298 | 149 | 0 | 0 | 0.01 | 0.011 | 0.012 | 0.013 | 0.011 | 0.045 | 0.023 | 0.015 | 0.029 | 0.081 | -0.027 |
| RI | 4612 | ENSG00000019582 | CD74 | chr5 | - | 150405084 | 150406321 | 150405084 | 150405180 | 150406258 | 150406321 | 46 | 51 | 38 | 43 | 47 | 3501 | 1525 | 2441 | 3131 | 1791 | 385 | 153 | 63 | 230 | 557 | 5579 | 3547 | 1674 | 3067 | 8990 | 298 | 149 | 0 | 0 | 0.007 | 0.016 | 0.008 | 0.007 | 0.013 | 0.033 | 0.021 | 0.018 | 0.036 | 0.03 | -0.017 |
| RI | 4046 | ENSG00000142534 | RPS11 | chr19 | + | 49497193 | 49498046 | 49497193 | 49497595 | 49497916 | 49498046 | 244 | 209 | 310 | 379 | 227 | 8196 | 4842 | 8502 | 7585 | 8692 | 893 | 492 | 365 | 605 | 861 | 11074 | 7952 | 6609 | 6879 | 13936 | 298 | 149 | 0 | 0 | 0.015 | 0.021 | 0.018 | 0.024 | 0.013 | 0.039 | 0.03 | 0.027 | 0.042 | 0.03 | -0.015 |
| RI | 3960 | ENSG00000147403 | RPL10 | chrX | + | 154399802 | 154400626 | 154399802 | 154399941 | 154400463 | 154400626 | 156 | 66 | 126 | 192 | 140 | 7308 | 3453 | 8995 | 7146 | 8875 | 196 | 150 | 119 | 163 | 334 | 4248 | 4604 | 4101 | 2575 | 7342 | 298 | 149 | 0 | 0 | 0.011 | 0.009 | 0.007 | 0.013 | 0.008 | 0.023 | 0.016 | 0.014 | 0.031 | 0.022 | -0.012 |
| RI | 3786 | ENSG00000026025 | VIM | chr10 | + | 17235168 | 17236379 | 17235168 | 17235389 | 17235845 | 17236379 | 395 | 287 | 217 | 269 | 728 | 5414 | 3441 | 4286 | 4861 | 10142 | 275 | 167 | 149 | 425 | 66 | 13433 | 7773 | 5486 | 9019 | 13008 | 298 | 149 | 0 | 0 | 0.035 | 0.04 | 0.025 | 0.027 | 0.035 | 0.01 | 0.011 | 0.013 | 0.023 | 0.003 | 0.02 |
| RI | 3788 | ENSG00000026025 | VIM | chr10 | + | 17235845 | 17236379 | 17235845 | 17235889 | 17236293 | 17236379 | 723 | 703 | 451 | 548 | 1355 | 4625 | 2859 | 3631 | 4144 | 7983 | 309 | 411 | 323 | 729 | 83 | 11320 | 6639 | 4497 | 7663 | 11033 | 298 | 149 | 0 | 0 | 0.072 | 0.109 | 0.058 | 0.062 | 0.078 | 0.013 | 0.03 | 0.035 | 0.045 | 0.004 | 0.05 |
| RI | 5081 | ENSG00000110934 | BIN2 | chr12 | - | 51302019 | 51302780 | 51302019 | 51302115 | 51302685 | 51302780 | 259 | 237 | 115 | 218 | 204 | 1459 | 1099 | 567 | 1753 | 1965 | 68 | 90 | 141 | 144 | 120 | 3315 | 2003 | 1695 | 2370 | 3839 | 298 | 149 | 0 | 0 | 0.082 | 0.097 | 0.092 | 0.059 | 0.049 | 0.01 | 0.022 | 0.04 | 0.029 | 0.015 | 0.053 |
| RI | 3622 | ENSG00000133226 | SRRM1 | chr1 | + | 24646666 | 24649029 | 24646666 | 24646789 | 24648858 | 24649029 | 155 | 179 | 143 | 147 | 171 | 589 | 450 | 486 | 645 | 808 | 9 | 132 | 92 | 65 | 4 | 1160 | 606 | 695 | 876 | 651 | 298 | 149 | 0 | 0 | 0.116 | 0.166 | 0.128 | 0.102 | 0.096 | 0.004 | 0.098 | 0.062 | 0.036 | 0.003 | 0.081 |
| RI | 7770 | ENSG00000160593 | JAML | chr11 | - | 118196734 | 118198091 | 118196734 | 118196821 | 118197997 | 118198091 | 139 | 139 | 168 | 172 | 223 | 530 | 335 | 349 | 396 | 637 | 51 | 50 | 76 | 98 | 19 | 1043 | 321 | 173 | 416 | 626 | 298 | 149 | 0 | 0 | 0.116 | 0.172 | 0.194 | 0.178 | 0.149 | 0.024 | 0.072 | 0.18 | 0.105 | 0.015 | 0.083 |
| RI | 5885 | ENSG00000137959 | IFI44L | chr1 | + | 78627905 | 78628999 | 78627905 | 78628393 | 78628950 | 78628999 | 36 | 127 | 28 | 128 | 29 | 176 | 583 | 146 | 316 | 276 | 18 | 65 | 30 | 39 | 0 | 737 | 1220 | 543 | 1390 | 24 | 298 | 149 | 0 | 0 | 0.093 | 0.098 | 0.088 | 0.168 | 0.05 | 0.012 | 0.026 | 0.027 | 0.014 | 0 | 0.084 |
| RI | 5487 | ENSG00000198830 | HMGN2 | chr1 | + | 26473482 | 26473732 | 26473482 | 26473527 | 26473702 | 26473732 | 118 | 58 | 128 | 95 | 143 | 346 | 189 | 344 | 342 | 266 | 63 | 61 | 84 | 134 | 71 | 795 | 383 | 308 | 1063 | 734 | 298 | 149 | 0 | 0 | 0.146 | 0.133 | 0.157 | 0.122 | 0.212 | 0.038 | 0.074 | 0.12 | 0.059 | 0.046 | 0.087 |
| RI | 7064 | ENSG00000205581 | HMGN1 | chr21 | - | 39345145 | 39348339 | 39345145 | 39345274 | 39348291 | 39348339 | 102 | 95 | 109 | 120 | 85 | 261 | 222 | 299 | 339 | 241 | 32 | 33 | 105 | 41 | 7 | 419 | 230 | 252 | 469 | 342 | 298 | 149 | 0 | 0 | 0.163 | 0.176 | 0.154 | 0.15 | 0.15 | 0.037 | 0.067 | 0.172 | 0.042 | 0.01 | 0.093 |
| RI | 4637 | ENSG00000115271 | GCA | chr2 | + | 162359043 | 162359552 | 162359043 | 162359157 | 162359493 | 162359552 | 149 | 125 | 67 | 89 | 76 | 396 | 283 | 259 | 282 | 439 | 0 | 21 | 33 | 8 | 4 | 94 | 387 | 260 | 359 | 65 | 298 | 149 | 0 | 0 | 0.158 | 0.181 | 0.115 | 0.136 | 0.08 | 0 | 0.026 | 0.06 | 0.011 | 0.03 | 0.109 |
| RI | 5489 | ENSG00000198830 | HMGN2 | chr1 | + | 26473482 | 26474135 | 26473482 | 26473732 | 26474084 | 26474135 | 217 | 120 | 212 | 173 | 265 | 475 | 274 | 443 | 478 | 355 | 171 | 101 | 122 | 191 | 128 | 1159 | 494 | 367 | 1466 | 891 | 298 | 149 | 0 | 0 | 0.186 | 0.18 | 0.193 | 0.153 | 0.272 | 0.069 | 0.093 | 0.143 | 0.061 | 0.067 | 0.11 |
| RI | 5513 | ENSG00000112367 | FIG4 | chr6 | + | 109786301 | 109789677 | 109786301 | 109786449 | 109789593 | 109789677 | 19 | 15 | 14 | 24 | 13 | 74 | 62 | 41 | 65 | 151 | 0 | 1 | 0 | 0 | 0 | 76 | 49 | 33 | 69 | 17 | 298 | 149 | 0 | 0 | 0.114 | 0.108 | 0.146 | 0.156 | 0.041 | 0 | 0.01 | 0 | 0 | 0 | 0.111 |
| RI | 2643 | ENSG00000203875 | SNHG5 | chr6 | - | 85677423 | 85678032 | 85677423 | 85677875 | 85677953 | 85678032 | 75 | 81 | 107 | 102 | 98 | 244 | 602 | 318 | 241 | 452 | 5 | 43 | 60 | 28 | 5 | 202 | 384 | 497 | 599 | 968 | 227 | 149 | 0 | 0 | 0.168 | 0.081 | 0.181 | 0.217 | 0.125 | 0.016 | 0.068 | 0.073 | 0.03 | 0.003 | 0.116 |
| RI | 4610 | ENSG00000060138 | YBX3 | chr12 | - | 10709907 | 10713333 | 10709907 | 10710114 | 10713210 | 10713333 | 65 | 53 | 44 | 42 | 79 | 172 | 71 | 85 | 107 | 137 | 11 | 90 | 37 | 13 | 2 | 353 | 201 | 103 | 215 | 805 | 298 | 149 | 0 | 0 | 0.159 | 0.272 | 0.206 | 0.164 | 0.224 | 0.015 | 0.183 | 0.152 | 0.029 | 0.001 | 0.129 |
| RI | 8229 | ENSG00000213809 | KLRK1 | chr12 | - | 10379446 | 10379792 | 10379446 | 10379482 | 10379699 | 10379792 | 611 | 436 | 395 | 591 | 736 | 1023 | 503 | 562 | 782 | 914 | 34 | 42 | 96 | 19 | 19 | 108 | 564 | 348 | 221 | 36 | 298 | 149 | 0 | 0 | 0.23 | 0.302 | 0.26 | 0.274 | 0.287 | 0.136 | 0.036 | 0.121 | 0.041 | 0.209 | 0.162 |
| RI | 6521 | ENSG00000105835 | NAMPT | chr7 | - | 106269153 | 106272658 | 106269153 | 106269312 | 106272529 | 106272658 | 285 | 514 | 146 | 464 | 539 | 623 | 538 | 504 | 785 | 1646 | 3 | 135 | 298 | 18 | 1 | 275 | 539 | 3464 | 445 | 48 | 298 | 149 | 0 | 0 | 0.186 | 0.323 | 0.127 | 0.228 | 0.141 | 0.005 | 0.111 | 0.041 | 0.02 | 0.01 | 0.164 |
| RI | 8221 | ENSG00000172322 | CLEC12A | chr12 | + | 9978965 | 9979524 | 9978965 | 9979064 | 9979335 | 9979524 | 403 | 467 | 254 | 408 | 659 | 420 | 327 | 254 | 492 | 934 | 64 | 45 | 127 | 105 | 48 | 347 | 143 | 333 | 608 | 65 | 298 | 149 | 0 | 0 | 0.324 | 0.417 | 0.333 | 0.293 | 0.261 | 0.084 | 0.136 | 0.16 | 0.079 | 0.27 | 0.18 |
| RI | 8578 | ENSG00000134884 | ARGLU1 | chr13 | - | 106557047 | 106559657 | 106557047 | 106557131 | 106559431 | 106559657 | 1165 | 950 | 1138 | 1007 | 1721 | 42 | 38 | 35 | 59 | 107 | 235 | 424 | 504 | 223 | 119 | 185 | 68 | 79 | 118 | 57 | 298 | 149 | 0 | 0 | 0.933 | 0.926 | 0.942 | 0.895 | 0.889 | 0.388 | 0.757 | 0.761 | 0.486 | 0.511 | 0.336 |
| RI | 7769 | ENSG00000160593 | JAML | chr11 | - | 118196734 | 118198091 | 118196734 | 118196821 | 118197967 | 118198091 | 144 | 152 | 169 | 178 | 205 | 68 | 52 | 44 | 52 | 96 | 47 | 67 | 65 | 85 | 27 | 166 | 72 | 81 | 167 | 77 | 298 | 149 | 0 | 0 | 0.514 | 0.594 | 0.658 | 0.631 | 0.516 | 0.124 | 0.318 | 0.286 | 0.203 | 0.149 | 0.367 |
| RI | 2840 | ENSG00000013441 | CLK1 | chr2 | - | 200856890 | 200857884 | 200856890 | 200856985 | 200857717 | 200857884 | 593 | 710 | 475 | 729 | 351 | 449 | 307 | 488 | 513 | 1069 | 997 | 1361 | 1439 | 1630 | 180 | 53 | 235 | 210 | 113 | 59 | 298 | 149 | 1.11E-16 | 9.37E-15 | 0.398 | 0.536 | 0.327 | 0.415 | 0.141 | 0.904 | 0.743 | 0.774 | 0.878 | 0.604 | -0.417 |
| RI | 2882 | ENSG00000178035 | IMPDH2 | chr3 | - | 49024494 | 49025040 | 49024494 | 49024802 | 49024895 | 49025040 | 30 | 33 | 47 | 58 | 47 | 150 | 114 | 228 | 191 | 196 | 120 | 58 | 81 | 41 | 133 | 73 | 80 | 69 | 72 | 142 | 242 | 149 | 1.11E-16 | 9.37E-15 | 0.11 | 0.151 | 0.113 | 0.158 | 0.129 | 0.503 | 0.309 | 0.42 | 0.26 | 0.366 | -0.239 |
| RI | 7353 | ENSG00000168003 | SLC3A2 | chr11 | + | 62881892 | 62882999 | 62881892 | 62882066 | 62882907 | 62882999 | 18 | 20 | 21 | 26 | 16 | 125 | 60 | 81 | 122 | 180 | 68 | 48 | 72 | 89 | 134 | 151 | 114 | 78 | 132 | 167 | 298 | 149 | 1.11E-16 | 9.37E-15 | 0.067 | 0.143 | 0.115 | 0.096 | 0.043 | 0.184 | 0.174 | 0.316 | 0.252 | 0.286 | -0.15 |
| RI | 7184 | ENSG00000110077 | MS4A6A | chr11 | - | 60179490 | 60181741 | 60179490 | 60179965 | 60181580 | 60181741 | 120 | 116 | 75 | 77 | 65 | 1081 | 710 | 599 | 809 | 1490 | 84 | 113 | 91 | 46 | 3 | 3302 | 1078 | 716 | 1412 | 1772 | 298 | 149 | 1.11E-16 | 9.37E-15 | 0.053 | 0.076 | 0.059 | 0.045 | 0.021 | 0.013 | 0.05 | 0.06 | 0.016 | 0.001 | 0.023 |
| RI | 2444 | ENSG00000197728 | RPS26 | chr12 | + | 56042424 | 56043493 | 56042424 | 56042602 | 56043362 | 56043493 | 130 | 82 | 141 | 159 | 248 | 799 | 2368 | 4045 | 3476 | 1179 | 177 | 123 | 99 | 171 | 151 | 3301 | 571 | 513 | 699 | 907 | 298 | 149 | 3.33E-16 | 2.77E-14 | 0.075 | 0.017 | 0.017 | 0.022 | 0.095 | 0.026 | 0.097 | 0.088 | 0.109 | 0.077 | -0.034 |
| RI | 2474 | ENSG00000070756 | PABPC1 | chr8 | - | 100717772 | 100718280 | 100717772 | 100717888 | 100718182 | 100718280 | 2524 | 1779 | 2515 | 2500 | 1890 | 25 | 11 | 45 | 18 | 16 | 2026 | 1577 | 2142 | 1891 | 2332 | 77 | 48 | 35 | 46 | 72 | 298 | 149 | 4.44E-16 | 3.59E-14 | 0.981 | 0.988 | 0.965 | 0.986 | 0.983 | 0.929 | 0.943 | 0.968 | 0.954 | 0.942 | 0.033 |
| RI | 6082 | ENSG00000167996 | FTH1 | chr11 | - | 61964986 | 61965515 | 61964986 | 61965105 | 61965368 | 61965515 | 666 | 861 | 879 | 755 | 789 | 9 | 4 | 6 | 14 | 7 | 1769 | 754 | 760 | 1521 | 2267 | 91 | 25 | 38 | 37 | 66 | 298 | 149 | 4.44E-16 | 3.59E-14 | 0.974 | 0.991 | 0.987 | 0.964 | 0.983 | 0.907 | 0.938 | 0.909 | 0.954 | 0.945 | 0.049 |
| RI | 6244 | ENSG00000075975 | MKRN2 | chr3 | + | 12581807 | 12582342 | 12581807 | 12581952 | 12582115 | 12582342 | 23 | 13 | 18 | 25 | 10 | 30 | 53 | 54 | 49 | 58 | 76 | 56 | 34 | 102 | 36 | 23 | 26 | 13 | 20 | 30 | 298 | 149 | 1.33E-15 | 1.05E-13 | 0.277 | 0.109 | 0.143 | 0.203 | 0.079 | 0.623 | 0.519 | 0.567 | 0.718 | 0.375 | -0.398 |
| RI | 6388 | ENSG00000197694 | SPTAN1 | chr9 | + | 128598962 | 128600115 | 128598962 | 128598986 | 128600079 | 128600115 | 61 | 35 | 63 | 76 | 50 | 210 | 135 | 236 | 207 | 291 | 60 | 109 | 97 | 93 | 136 | 109 | 169 | 121 | 163 | 230 | 298 | 149 | 1.33E-15 | 1.05E-13 | 0.127 | 0.115 | 0.118 | 0.155 | 0.079 | 0.216 | 0.244 | 0.286 | 0.222 | 0.228 | -0.12 |
| RI | 2167 | ENSG00000065357 | DGKA | chr12 | + | 55940896 | 55941325 | 55940896 | 55940980 | 55941251 | 55941325 | 39 | 30 | 78 | 38 | 24 | 187 | 150 | 313 | 290 | 373 | 88 | 37 | 54 | 96 | 88 | 205 | 151 | 154 | 164 | 140 | 298 | 149 | 1.78E-15 | 1.35E-13 | 0.094 | 0.091 | 0.111 | 0.061 | 0.031 | 0.177 | 0.109 | 0.149 | 0.226 | 0.239 | -0.102 |
| RI | 3132 | ENSG00000109861 | CTSC | chr11 | - | 88296132 | 88300645 | 88296132 | 88296264 | 88300529 | 88300645 | 108 | 64 | 39 | 84 | 78 | 858 | 469 | 467 | 590 | 1375 | 20 | 46 | 56 | 29 | 10 | 1272 | 608 | 404 | 833 | 932 | 298 | 149 | 1.78E-15 | 1.35E-13 | 0.059 | 0.064 | 0.04 | 0.066 | 0.028 | 0.008 | 0.036 | 0.065 | 0.017 | 0.005 | 0.025 |
| RI | 2641 | ENSG00000203875 | SNHG5 | chr6 | - | 85677078 | 85677875 | 85677078 | 85677492 | 85677794 | 85677875 | 137 | 106 | 173 | 215 | 250 | 202 | 470 | 289 | 229 | 465 | 57 | 102 | 72 | 68 | 46 | 169 | 312 | 365 | 417 | 788 | 298 | 149 | 1.78E-15 | 1.35E-13 | 0.253 | 0.101 | 0.23 | 0.319 | 0.212 | 0.144 | 0.14 | 0.09 | 0.075 | 0.028 | 0.128 |
| RI | 8153 | ENSG00000107862 | GBF1 | chr10 | + | 102379852 | 102380362 | 102379852 | 102379954 | 102380248 | 102380362 | 21 | 6 | 22 | 10 | 0 | 53 | 18 | 23 | 23 | 25 | 73 | 50 | 34 | 88 | 157 | 9 | 19 | 11 | 29 | 70 | 298 | 149 | 2.00E-15 | 1.48E-13 | 0.165 | 0.143 | 0.324 | 0.179 | 0 | 0.802 | 0.568 | 0.607 | 0.603 | 0.529 | -0.46 |
| RI | 5222 | ENSG00000125875 | TBC1D20 | chr20 | - | 439107 | 442043 | 439107 | 439295 | 440247 | 442043 | 17 | 4 | 11 | 18 | 5 | 129 | 66 | 112 | 161 | 139 | 67 | 29 | 25 | 55 | 74 | 129 | 175 | 59 | 104 | 292 | 298 | 149 | 2.00E-15 | 1.48E-13 | 0.062 | 0.029 | 0.047 | 0.053 | 0.018 | 0.206 | 0.077 | 0.175 | 0.209 | 0.112 | -0.114 |
| RI | 1364 | ENSG00000149923 | PPP4C | chr16 | + | 30082745 | 30083567 | 30082745 | 30082847 | 30083369 | 30083567 | 12 | 5 | 34 | 20 | 14 | 102 | 54 | 79 | 116 | 106 | 107 | 87 | 45 | 76 | 90 | 155 | 143 | 84 | 110 | 191 | 298 | 149 | 2.11E-15 | 1.54E-13 | 0.056 | 0.044 | 0.177 | 0.079 | 0.062 | 0.257 | 0.233 | 0.211 | 0.257 | 0.191 | -0.146 |
| RI | 161 | ENSG00000099331 | MYO9B | chr19 | + | 17202841 | 17203258 | 17202841 | 17202883 | 17203146 | 17203258 | 63 | 29 | 61 | 50 | 23 | 271 | 119 | 207 | 280 | 281 | 186 | 87 | 66 | 108 | 214 | 213 | 248 | 173 | 198 | 389 | 298 | 149 | 2.22E-15 | 1.60E-13 | 0.104 | 0.109 | 0.128 | 0.082 | 0.039 | 0.304 | 0.149 | 0.16 | 0.214 | 0.216 | -0.116 |
| RI | 5225 | ENSG00000125875 | TBC1D20 | chr20 | - | 440247 | 441689 | 440247 | 440389 | 441587 | 441689 | 25 | 11 | 15 | 19 | 20 | 111 | 66 | 96 | 149 | 136 | 105 | 60 | 40 | 46 | 58 | 125 | 108 | 55 | 113 | 161 | 298 | 149 | 2.55E-15 | 1.82E-13 | 0.101 | 0.077 | 0.072 | 0.06 | 0.068 | 0.296 | 0.217 | 0.267 | 0.169 | 0.153 | -0.145 |
| RI | 2648 | ENSG00000203875 | SNHG5 | chr6 | - | 85677423 | 85678188 | 85677423 | 85678032 | 85678135 | 85678188 | 69 | 87 | 104 | 99 | 125 | 198 | 345 | 252 | 183 | 303 | 31 | 62 | 54 | 45 | 20 | 92 | 270 | 339 | 333 | 535 | 252 | 149 | 2.89E-15 | 2.03E-13 | 0.171 | 0.13 | 0.196 | 0.242 | 0.196 | 0.166 | 0.12 | 0.086 | 0.074 | 0.022 | 0.093 |
| RI | 990 | ENSG00000141030 | COPS3 | chr17 | - | 17270757 | 17271008 | 17270757 | 17270807 | 17270895 | 17271008 | 2 | 1 | 6 | 7 | 2 | 181 | 58 | 131 | 100 | 135 | 61 | 20 | 4 | 30 | 26 | 149 | 186 | 150 | 153 | 114 | 237 | 149 | 3.00E-15 | 2.08E-13 | 0.007 | 0.011 | 0.028 | 0.042 | 0.009 | 0.205 | 0.063 | 0.016 | 0.11 | 0.125 | -0.084 |
| RI | 4043 | ENSG00000142534 | RPS11 | chr19 | + | 49497193 | 49497595 | 49497193 | 49497325 | 49497519 | 49497595 | 360 | 239 | 430 | 473 | 350 | 8493 | 5453 | 8276 | 7903 | 7616 | 1071 | 613 | 626 | 889 | 1228 | 12991 | 8645 | 7392 | 8089 | 15804 | 298 | 149 | 3.33E-15 | 2.29E-13 | 0.021 | 0.021 | 0.025 | 0.029 | 0.022 | 0.04 | 0.034 | 0.041 | 0.052 | 0.037 | -0.017 |
| RI | 5237 | ENSG00000148303 | RPL7A | chr9 | + | 133350239 | 133350727 | 133350239 | 133350319 | 133350596 | 133350727 | 200 | 107 | 202 | 213 | 307 | 8060 | 4485 | 9184 | 7580 | 8492 | 420 | 282 | 255 | 302 | 220 | 7311 | 5421 | 4870 | 4274 | 9108 | 298 | 149 | 6.00E-15 | 4.07E-13 | 0.012 | 0.012 | 0.011 | 0.014 | 0.018 | 0.028 | 0.025 | 0.026 | 0.034 | 0.012 | -0.012 |
| RI | 8573 | ENSG00000144567 | RETREG2 | chr2 | + | 219181061 | 219181463 | 219181061 | 219181205 | 219181368 | 219181463 | 28 | 25 | 23 | 30 | 17 | 203 | 123 | 191 | 178 | 90 | 70 | 42 | 64 | 106 | 204 | 184 | 188 | 128 | 213 | 331 | 298 | 149 | 8.22E-15 | 5.39E-13 | 0.065 | 0.092 | 0.057 | 0.078 | 0.086 | 0.16 | 0.1 | 0.2 | 0.199 | 0.236 | -0.103 |
| RI | 806 | ENSG00000185624 | P4HB | chr17 | - | 81845143 | 81845742 | 81845143 | 81845230 | 81845560 | 81845742 | 46 | 28 | 41 | 63 | 26 | 1352 | 705 | 1012 | 1085 | 1281 | 241 | 109 | 106 | 182 | 225 | 2766 | 1697 | 1290 | 1999 | 2849 | 298 | 149 | 8.22E-15 | 5.39E-13 | 0.017 | 0.019 | 0.02 | 0.028 | 0.01 | 0.042 | 0.031 | 0.039 | 0.044 | 0.038 | -0.02 |
| RI | 3743 | ENSG00000136450 | SRSF1 | chr17 | - | 58004922 | 58005973 | 58004922 | 58005041 | 58005800 | 58005973 | 1731 | 1310 | 1509 | 1547 | 1805 | 55 | 35 | 46 | 63 | 32 | 1212 | 940 | 1082 | 1153 | 292 | 40 | 67 | 106 | 95 | 45 | 298 | 149 | 8.22E-15 | 5.39E-13 | 0.94 | 0.949 | 0.943 | 0.925 | 0.966 | 0.938 | 0.875 | 0.836 | 0.859 | 0.764 | 0.09 |
| RI | 2351 | ENSG00000011243 | AKAP8L | chr19 | - | 15397713 | 15398766 | 15397713 | 15397855 | 15398629 | 15398766 | 38 | 17 | 23 | 31 | 19 | 19 | 12 | 14 | 8 | 8 | 38 | 71 | 59 | 111 | 89 | 3 | 0 | 0 | 10 | 6 | 298 | 149 | 8.55E-15 | 5.55E-13 | 0.5 | 0.415 | 0.451 | 0.66 | 0.543 | 0.864 | 1 | 1 | 0.847 | 0.881 | -0.405 |
| RI | 7446 | ENSG00000165688 | PMPCA | chr9 | + | 136421831 | 136423760 | 136421831 | 136421976 | 136423094 | 136423760 | 8 | 4 | 11 | 7 | 4 | 41 | 21 | 39 | 59 | 32 | 22 | 42 | 18 | 13 | 38 | 26 | 24 | 24 | 16 | 26 | 298 | 149 | 8.77E-15 | 5.63E-13 | 0.089 | 0.087 | 0.124 | 0.056 | 0.059 | 0.297 | 0.467 | 0.273 | 0.289 | 0.422 | -0.267 |
| RI | 7775 | ENSG00000087053 | MTMR2 | chr11 | - | 95862271 | 95865676 | 95862271 | 95862366 | 95865600 | 95865676 | 11 | 10 | 9 | 11 | 8 | 39 | 32 | 28 | 51 | 34 | 0 | 1 | 0 | 0 | 0 | 39 | 51 | 29 | 33 | 29 | 298 | 149 | 9.44E-15 | 5.99E-13 | 0.124 | 0.135 | 0.138 | 0.097 | 0.105 | 0 | 0.01 | 0 | 0 | 0 | 0.118 |
| RI | 4971 | ENSG00000231500 | RPS18 | chr6 | + | 33275964 | 33276267 | 33275964 | 33276066 | 33276175 | 33276267 | 532 | 329 | 567 | 475 | 286 | 4335 | 2137 | 4860 | 3744 | 8319 | 249 | 359 | 310 | 271 | 594 | 1785 | 1973 | 1796 | 1588 | 2854 | 258 | 149 | 1.63E-14 | 1.03E-12 | 0.066 | 0.082 | 0.063 | 0.068 | 0.019 | 0.075 | 0.095 | 0.091 | 0.09 | 0.107 | -0.032 |
| RI | 3687 | ENSG00000099622 | CIRBP | chr19 | + | 1271980 | 1273172 | 1271980 | 1272051 | 1272426 | 1273172 | 189 | 122 | 198 | 165 | 84 | 431 | 215 | 411 | 361 | 418 | 48 | 131 | 82 | 58 | 114 | 428 | 598 | 399 | 268 | 598 | 298 | 149 | 1.89E-14 | 1.17E-12 | 0.18 | 0.221 | 0.194 | 0.186 | 0.091 | 0.053 | 0.099 | 0.093 | 0.098 | 0.087 | 0.088 |
| RI | 6638 | ENSG00000167778 | SPRYD3 | chr12 | - | 53074648 | 53075219 | 53074648 | 53074784 | 53075094 | 53075219 | 25 | 18 | 20 | 38 | 24 | 123 | 58 | 129 | 131 | 104 | 89 | 41 | 37 | 49 | 55 | 119 | 64 | 47 | 70 | 112 | 298 | 149 | 2.18E-14 | 1.34E-12 | 0.092 | 0.134 | 0.072 | 0.127 | 0.103 | 0.272 | 0.243 | 0.282 | 0.259 | 0.197 | -0.145 |
| RI | 3787 | ENSG00000026025 | VIM | chr10 | + | 17235168 | 17236379 | 17235168 | 17235389 | 17236293 | 17236379 | 545 | 508 | 285 | 345 | 664 | 16 | 1 | 4 | 17 | 31 | 297 | 297 | 222 | 533 | 32 | 50 | 65 | 8 | 109 | 13 | 298 | 149 | 3.42E-14 | 2.08E-12 | 0.945 | 0.996 | 0.973 | 0.91 | 0.915 | 0.748 | 0.696 | 0.933 | 0.71 | 0.552 | 0.22 |
| RI | 5716 | ENSG00000115053 | NCL | chr2 | - | 231458265 | 231459125 | 231458265 | 231458389 | 231459000 | 231459125 | 216 | 194 | 207 | 307 | 273 | 1841 | 1584 | 1802 | 1962 | 2837 | 53 | 200 | 109 | 86 | 5 | 1511 | 1478 | 1675 | 1874 | 325 | 298 | 149 | 4.35E-14 | 2.62E-12 | 0.055 | 0.058 | 0.054 | 0.073 | 0.046 | 0.017 | 0.063 | 0.032 | 0.022 | 0.008 | 0.029 |
| RI | 6876 | ENSG00000147140 | NONO | chrX | + | 71296564 | 71297047 | 71296564 | 71296660 | 71296850 | 71297047 | 48 | 23 | 38 | 40 | 53 | 245 | 251 | 304 | 232 | 354 | 252 | 129 | 37 | 158 | 136 | 349 | 312 | 228 | 293 | 455 | 298 | 149 | 4.61E-14 | 2.75E-12 | 0.089 | 0.044 | 0.059 | 0.079 | 0.07 | 0.265 | 0.171 | 0.075 | 0.212 | 0.13 | -0.102 |
| RI | 4611 | ENSG00000019582 | CD74 | chr5 | - | 150403120 | 150404767 | 150403120 | 150403312 | 150404679 | 150404767 | 197 | 187 | 145 | 244 | 226 | 490 | 271 | 284 | 350 | 408 | 998 | 514 | 165 | 550 | 1273 | 785 | 441 | 192 | 334 | 1231 | 298 | 149 | 4.65E-14 | 2.75E-12 | 0.167 | 0.257 | 0.203 | 0.258 | 0.217 | 0.389 | 0.368 | 0.301 | 0.452 | 0.341 | -0.15 |
| RI | 5967 | ENSG00000114423 | CBLB | chr3 | - | 105680991 | 105681818 | 105680991 | 105681610 | 105681723 | 105681818 | 45 | 38 | 25 | 19 | 25 | 210 | 138 | 156 | 157 | 323 | 0 | 5 | 3 | 7 | 0 | 115 | 76 | 184 | 74 | 40 | 262 | 149 | 8.27E-14 | 4.84E-12 | 0.109 | 0.135 | 0.084 | 0.064 | 0.042 | 0 | 0.036 | 0.009 | 0.051 | 0 | 0.068 |
| RI | 7528 | ENSG00000108424 | KPNB1 | chr17 | + | 47678307 | 47680134 | 47678307 | 47678413 | 47680019 | 47680134 | 61 | 63 | 38 | 92 | 20 | 528 | 378 | 498 | 531 | 647 | 101 | 125 | 119 | 213 | 6 | 438 | 417 | 424 | 348 | 105 | 298 | 149 | 8.48E-14 | 4.91E-12 | 0.055 | 0.077 | 0.037 | 0.08 | 0.015 | 0.103 | 0.13 | 0.123 | 0.234 | 0.028 | -0.071 |
| RI | 6833 | ENSG00000147168 | IL2RG | chrX | - | 71110155 | 71110688 | 71110155 | 71110295 | 71110503 | 71110688 | 110 | 76 | 119 | 132 | 185 | 906 | 343 | 606 | 731 | 958 | 156 | 150 | 174 | 170 | 184 | 392 | 686 | 284 | 399 | 587 | 298 | 149 | 9.14E-14 | 5.24E-12 | 0.057 | 0.1 | 0.089 | 0.083 | 0.088 | 0.166 | 0.099 | 0.235 | 0.176 | 0.135 | -0.079 |
| RI | 4345 | ENSG00000087191 | PSMC5 | chr17 | + | 63828137 | 63829563 | 63828137 | 63828209 | 63829493 | 63829563 | 29 | 26 | 27 | 43 | 19 | 305 | 154 | 272 | 321 | 183 | 57 | 118 | 47 | 91 | 63 | 302 | 199 | 190 | 205 | 273 | 298 | 149 | 1.46E-13 | 8.29E-12 | 0.045 | 0.078 | 0.047 | 0.063 | 0.049 | 0.086 | 0.229 | 0.11 | 0.182 | 0.103 | -0.086 |
| RI | 8348 | ENSG00000198563 | DDX39B | chr6 | - | 31532779 | 31535485 | 31532779 | 31532911 | 31535366 | 31535485 | 316 | 240 | 282 | 357 | 474 | 291 | 214 | 241 | 274 | 345 | 735 | 439 | 384 | 621 | 718 | 332 | 226 | 167 | 237 | 362 | 298 | 149 | 1.68E-13 | 9.45E-12 | 0.352 | 0.359 | 0.369 | 0.394 | 0.407 | 0.525 | 0.493 | 0.535 | 0.567 | 0.498 | -0.147 |
| RI | 8127 | ENSG00000176986 | SEC24C | chr10 | + | 73769346 | 73769733 | 73769346 | 73769485 | 73769614 | 73769733 | 2 | 4 | 7 | 11 | 0 | 124 | 50 | 120 | 123 | 160 | 47 | 27 | 15 | 27 | 14 | 71 | 121 | 87 | 156 | 166 | 278 | 149 | 1.73E-13 | 9.65E-12 | 0.009 | 0.041 | 0.03 | 0.046 | 0 | 0.262 | 0.107 | 0.085 | 0.085 | 0.043 | -0.091 |
| RI | 2559 | ENSG00000145907 | G3BP1 | chr5 | + | 151794158 | 151795575 | 151794158 | 151794249 | 151795478 | 151795575 | 47 | 41 | 41 | 32 | 32 | 395 | 249 | 390 | 407 | 466 | 52 | 63 | 102 | 28 | 10 | 167 | 233 | 247 | 211 | 70 | 298 | 149 | 2.14E-13 | 1.18E-11 | 0.056 | 0.076 | 0.05 | 0.038 | 0.033 | 0.135 | 0.119 | 0.171 | 0.062 | 0.067 | -0.06 |
| RI | 4155 | ENSG00000134909 | ARHGAP32 | chr11 | - | 128972452 | 128975002 | 128972452 | 128973432 | 128974123 | 128975002 | 6 | 4 | 4 | 6 | 18 | 11 | 18 | 9 | 13 | 21 | 0 | 0 | 0 | 3 | 0 | 28 | 15 | 3 | 21 | 46 | 298 | 149 | 2.30E-13 | 1.25E-11 | 0.214 | 0.1 | 0.182 | 0.188 | 0.3 | 0 | 0 | 0 | 0.067 | 0 | 0.183 |
| RI | 3878 | ENSG00000162434 | JAK1 | chr1 | - | 64845512 | 64846736 | 64845512 | 64845640 | 64846648 | 64846736 | 41 | 37 | 61 | 43 | 29 | 941 | 492 | 761 | 980 | 1496 | 13 | 50 | 117 | 26 | 26 | 434 | 458 | 531 | 403 | 114 | 298 | 149 | 2.30E-13 | 1.25E-11 | 0.021 | 0.036 | 0.039 | 0.021 | 0.01 | 0.015 | 0.052 | 0.099 | 0.031 | 0.102 | -0.034 |
| RI | 5163 | ENSG00000198755 | RPL10A | chr6 | + | 35468946 | 35469529 | 35468946 | 35469027 | 35469380 | 35469529 | 132 | 56 | 167 | 145 | 125 | 2053 | 806 | 2614 | 2166 | 2237 | 124 | 121 | 119 | 103 | 287 | 1166 | 1405 | 1141 | 808 | 1614 | 298 | 149 | 2.51E-13 | 1.33E-11 | 0.031 | 0.034 | 0.031 | 0.032 | 0.027 | 0.05 | 0.041 | 0.05 | 0.06 | 0.082 | -0.026 |
| RI | 4736 | ENSG00000140416 | TPM1 | chr15 | + | 63061197 | 63062277 | 63061197 | 63061273 | 63062214 | 63062277 | 8 | 11 | 12 | 9 | 8 | 46 | 33 | 18 | 29 | 38 | 7 | 6 | 6 | 12 | 3 | 100 | 64 | 40 | 95 | 586 | 298 | 149 | 2.51E-13 | 1.33E-11 | 0.08 | 0.143 | 0.25 | 0.134 | 0.095 | 0.034 | 0.045 | 0.07 | 0.059 | 0.003 | 0.098 |
| RI | 3007 | ENSG00000114867 | EIF4G1 | chr3 | + | 184331265 | 184331606 | 184331265 | 184331364 | 184331471 | 184331606 | 17 | 27 | 18 | 24 | 12 | 209 | 103 | 177 | 195 | 111 | 121 | 61 | 38 | 88 | 88 | 307 | 199 | 143 | 167 | 378 | 256 | 149 | 2.53E-13 | 1.33E-11 | 0.045 | 0.132 | 0.056 | 0.067 | 0.059 | 0.187 | 0.151 | 0.134 | 0.235 | 0.119 | -0.093 |
| RI | 4147 | ENSG00000156482 | RPL30 | chr8 | - | 98044942 | 98045399 | 98044942 | 98045088 | 98045346 | 98045399 | 257 | 135 | 259 | 226 | 228 | 3756 | 1656 | 4425 | 3203 | 2299 | 278 | 286 | 189 | 268 | 132 | 1795 | 1934 | 1533 | 1312 | 2299 | 298 | 149 | 3.30E-13 | 1.73E-11 | 0.033 | 0.039 | 0.028 | 0.034 | 0.047 | 0.072 | 0.069 | 0.058 | 0.093 | 0.028 | -0.028 |
| RI | 5670 | ENSG00000115234 | SNX17 | chr2 | + | 27374084 | 27374433 | 27374084 | 27374175 | 27374345 | 27374433 | 10 | 13 | 10 | 11 | 13 | 219 | 118 | 158 | 165 | 276 | 52 | 28 | 21 | 24 | 44 | 265 | 177 | 81 | 78 | 275 | 298 | 149 | 6.09E-13 | 3.16E-11 | 0.022 | 0.052 | 0.031 | 0.032 | 0.023 | 0.089 | 0.073 | 0.115 | 0.133 | 0.074 | -0.065 |
| RI | 5221 | ENSG00000125875 | TBC1D20 | chr20 | - | 437439 | 438841 | 437439 | 437598 | 438549 | 438841 | 130 | 63 | 73 | 112 | 111 | 7 | 8 | 12 | 15 | 22 | 170 | 116 | 74 | 130 | 267 | 0 | 6 | 2 | 0 | 7 | 298 | 149 | 7.02E-13 | 3.61E-11 | 0.903 | 0.797 | 0.753 | 0.789 | 0.716 | 1 | 0.906 | 0.949 | 1 | 0.95 | -0.169 |
| RI | 4634 | ENSG00000159377 | PSMB4 | chr1 | + | 151401238 | 151401630 | 151401238 | 151401355 | 151401540 | 151401630 | 203 | 110 | 169 | 250 | 73 | 799 | 366 | 693 | 852 | 528 | 377 | 354 | 207 | 424 | 455 | 1127 | 825 | 493 | 606 | 1034 | 298 | 149 | 1.00E-12 | 5.10E-11 | 0.113 | 0.131 | 0.109 | 0.128 | 0.065 | 0.143 | 0.177 | 0.174 | 0.259 | 0.18 | -0.077 |
| RI | 5410 | ENSG00000137726 | FXYD6 | chr11 | - | 117841989 | 117842261 | 117841989 | 117842025 | 117842106 | 117842261 | 7 | 1 | 2 | 1 | 2 | 0 | 0 | 0 | 0 | 0 | 23 | 15 | 3 | 5 | 12 | 3 | 22 | 2 | 5 | 6 | 230 | 149 | 1.12E-12 | 5.67E-11 | 1 | 1 | 1 | 1 | 1 | 0.832 | 0.306 | 0.493 | 0.393 | 0.564 | 0.482 |
| RI | 6873 | ENSG00000147140 | NONO | chrX | + | 71290628 | 71291972 | 71290628 | 71290791 | 71291778 | 71291972 | 71 | 56 | 75 | 98 | 61 | 216 | 137 | 244 | 246 | 259 | 304 | 139 | 116 | 196 | 124 | 268 | 180 | 152 | 180 | 276 | 298 | 149 | 1.13E-12 | 5.68E-11 | 0.141 | 0.17 | 0.133 | 0.166 | 0.105 | 0.362 | 0.279 | 0.276 | 0.353 | 0.183 | -0.148 |
| RI | 6322 | ENSG00000139641 | ESYT1 | chr12 | + | 56131039 | 56131316 | 56131039 | 56131113 | 56131243 | 56131316 | 8 | 3 | 15 | 5 | 9 | 240 | 139 | 242 | 217 | 263 | 10 | 7 | 18 | 21 | 29 | 111 | 129 | 81 | 65 | 169 | 279 | 149 | 1.20E-12 | 5.95E-11 | 0.017 | 0.011 | 0.032 | 0.012 | 0.018 | 0.046 | 0.028 | 0.106 | 0.147 | 0.084 | -0.064 |
| RI | 3095 | ENSG00000111642 | CHD4 | chr12 | - | 6583026 | 6583378 | 6583026 | 6583113 | 6583197 | 6583378 | 25 | 15 | 21 | 17 | 23 | 323 | 203 | 328 | 349 | 402 | 57 | 48 | 42 | 76 | 75 | 407 | 290 | 237 | 413 | 561 | 233 | 149 | 1.42E-12 | 6.98E-11 | 0.047 | 0.045 | 0.039 | 0.03 | 0.035 | 0.082 | 0.096 | 0.102 | 0.105 | 0.079 | -0.054 |
| RI | 5300 | ENSG00000197037 | ZSCAN25 | chr7 | + | 99619560 | 99621574 | 99619560 | 99619993 | 99621372 | 99621574 | 45 | 29 | 22 | 31 | 11 | 16 | 28 | 32 | 20 | 13 | 104 | 74 | 80 | 93 | 93 | 0 | 19 | 13 | 8 | 7 | 298 | 149 | 1.45E-12 | 7.09E-11 | 0.584 | 0.341 | 0.256 | 0.437 | 0.297 | 1 | 0.661 | 0.755 | 0.853 | 0.869 | -0.445 |
| RI | 5662 | ENSG00000165280 | VCP | chr9 | - | 35057375 | 35059219 | 35057375 | 35057530 | 35059063 | 35059219 | 73 | 45 | 70 | 84 | 64 | 315 | 169 | 292 | 264 | 286 | 284 | 170 | 89 | 238 | 217 | 309 | 343 | 211 | 256 | 484 | 298 | 149 | 1.84E-12 | 8.93E-11 | 0.104 | 0.117 | 0.107 | 0.137 | 0.101 | 0.315 | 0.199 | 0.174 | 0.317 | 0.183 | -0.124 |
| RI | 2203 | ENSG00000086189 | DIMT1 | chr5 | - | 62390875 | 62392234 | 62390875 | 62390982 | 62392170 | 62392234 | 63 | 86 | 76 | 99 | 155 | 155 | 133 | 161 | 168 | 245 | 0 | 26 | 66 | 8 | 4 | 200 | 129 | 60 | 146 | 183 | 298 | 149 | 2.18E-12 | 1.05E-10 | 0.169 | 0.244 | 0.191 | 0.228 | 0.24 | 0 | 0.092 | 0.355 | 0.027 | 0.011 | 0.117 |
| RI | 1055 | ENSG00000073584 | SMARCE1 | chr17 | - | 40630713 | 40632367 | 40630713 | 40631693 | 40632194 | 40632367 | 49 | 46 | 37 | 52 | 41 | 259 | 226 | 261 | 254 | 430 | 0 | 18 | 42 | 10 | 0 | 188 | 304 | 351 | 277 | 59 | 298 | 149 | 3.15E-12 | 1.50E-10 | 0.086 | 0.092 | 0.066 | 0.093 | 0.046 | 0 | 0.029 | 0.056 | 0.018 | 0 | 0.056 |
| RI | 495 | ENSG00000104904 | OAZ1 | chr19 | + | 2271384 | 2271954 | 2271384 | 2271530 | 2271781 | 2271954 | 68 | 52 | 66 | 127 | 54 | 2552 | 1334 | 1723 | 2574 | 1994 | 296 | 127 | 245 | 264 | 436 | 4225 | 3286 | 1715 | 2859 | 12529 | 298 | 149 | 4.42E-12 | 2.09E-10 | 0.013 | 0.019 | 0.019 | 0.024 | 0.013 | 0.034 | 0.019 | 0.067 | 0.044 | 0.017 | -0.019 |
| RI | 3506 | ENSG00000124614 | RPS10 | chr6 | - | 34417453 | 34418424 | 34417453 | 34417547 | 34418368 | 34418424 | 19 | 1 | 25 | 19 | 19 | 1325 | 637 | 1509 | 1244 | 1960 | 73 | 38 | 31 | 43 | 80 | 1442 | 1232 | 985 | 821 | 2126 | 298 | 149 | 4.84E-12 | 2.27E-10 | 0.007 | 0.001 | 0.008 | 0.008 | 0.005 | 0.025 | 0.015 | 0.015 | 0.026 | 0.018 | -0.014 |
| RI | 3889 | ENSG00000213593 | TMX2 | chr11 | + | 57738663 | 57739039 | 57738663 | 57738770 | 57738973 | 57739039 | 61 | 65 | 66 | 99 | 68 | 156 | 108 | 157 | 165 | 203 | 238 | 134 | 127 | 227 | 142 | 209 | 165 | 110 | 206 | 193 | 298 | 149 | 5.30E-12 | 2.47E-10 | 0.164 | 0.231 | 0.174 | 0.231 | 0.143 | 0.363 | 0.289 | 0.366 | 0.355 | 0.269 | -0.14 |
| RI | 2340 | ENSG00000126456 | IRF3 | chr19 | - | 49664673 | 49665857 | 49664673 | 49664846 | 49665630 | 49665857 | 172 | 163 | 191 | 194 | 167 | 16 | 7 | 21 | 17 | 11 | 382 | 162 | 193 | 303 | 484 | 0 | 10 | 7 | 2 | 7 | 298 | 149 | 5.61E-12 | 2.59E-10 | 0.843 | 0.921 | 0.82 | 0.851 | 0.884 | 1 | 0.89 | 0.932 | 0.987 | 0.972 | -0.092 |
| RI | 1746 | ENSG00000197102 | DYNC1H1 | chr14 | + | 102040597 | 102041734 | 102040597 | 102040673 | 102041573 | 102041734 | 9 | 18 | 2 | 2 | 3 | 108 | 59 | 93 | 81 | 91 | 20 | 25 | 12 | 10 | 34 | 17 | 69 | 42 | 50 | 46 | 298 | 149 | 5.73E-12 | 2.62E-10 | 0.04 | 0.132 | 0.011 | 0.012 | 0.016 | 0.37 | 0.153 | 0.125 | 0.091 | 0.27 | -0.16 |
| RI | 2931 | ENSG00000105373 | NOP53 | chr19 | + | 47755755 | 47756604 | 47755755 | 47755822 | 47756527 | 47756604 | 6 | 2 | 9 | 17 | 8 | 276 | 67 | 273 | 240 | 59 | 4 | 28 | 12 | 9 | 21 | 54 | 107 | 57 | 49 | 106 | 298 | 149 | 5.76E-12 | 2.62E-10 | 0.011 | 0.015 | 0.016 | 0.034 | 0.063 | 0.036 | 0.116 | 0.095 | 0.084 | 0.09 | -0.056 |
| RI | 6083 | ENSG00000167996 | FTH1 | chr11 | - | 61964986 | 61965515 | 61964986 | 61965112 | 61965368 | 61965515 | 640 | 842 | 845 | 727 | 730 | 8743 | 6766 | 4530 | 9038 | 7552 | 1698 | 737 | 728 | 1496 | 2157 | 30859 | 9177 | 8276 | 16052 | 56532 | 298 | 149 | 6.85E-12 | 3.09E-10 | 0.035 | 0.059 | 0.085 | 0.039 | 0.046 | 0.027 | 0.039 | 0.042 | 0.045 | 0.019 | 0.018 |
| RI | 5450 | ENSG00000143799 | PARP1 | chr1 | - | 226365953 | 226367608 | 226365953 | 226366052 | 226367479 | 226367608 | 18 | 8 | 21 | 11 | 9 | 358 | 206 | 282 | 377 | 468 | 13 | 34 | 5 | 42 | 39 | 232 | 223 | 178 | 165 | 280 | 298 | 149 | 7.04E-12 | 3.15E-10 | 0.025 | 0.019 | 0.036 | 0.014 | 0.01 | 0.027 | 0.071 | 0.014 | 0.113 | 0.065 | -0.037 |
| RI | 3763 | ENSG00000115524 | SF3B1 | chr2 | - | 197418508 | 197420542 | 197418508 | 197418588 | 197418895 | 197420542 | 978 | 777 | 1057 | 895 | 966 | 46 | 80 | 69 | 67 | 94 | 82 | 302 | 263 | 115 | 86 | 0 | 8 | 0 | 0 | 7 | 298 | 149 | 7.11E-12 | 3.16E-10 | 0.914 | 0.829 | 0.885 | 0.87 | 0.837 | 1 | 0.95 | 1 | 1 | 0.86 | -0.095 |
| RI | 6878 | ENSG00000147140 | NONO | chrX | + | 71296850 | 71298508 | 71296850 | 71297047 | 71297376 | 71298508 | 55 | 62 | 59 | 74 | 32 | 234 | 181 | 218 | 200 | 173 | 353 | 125 | 118 | 166 | 147 | 526 | 252 | 211 | 223 | 364 | 298 | 149 | 7.50E-12 | 3.31E-10 | 0.105 | 0.146 | 0.119 | 0.156 | 0.085 | 0.251 | 0.199 | 0.219 | 0.271 | 0.168 | -0.099 |
| RI | 6535 | ENSG00000106263 | EIF3B | chr7 | + | 2378688 | 2379242 | 2378688 | 2378766 | 2379133 | 2379242 | 7 | 10 | 3 | 8 | 5 | 208 | 136 | 205 | 172 | 166 | 55 | 29 | 7 | 47 | 44 | 367 | 233 | 169 | 249 | 439 | 298 | 149 | 8.42E-12 | 3.69E-10 | 0.017 | 0.035 | 0.007 | 0.023 | 0.015 | 0.07 | 0.059 | 0.02 | 0.086 | 0.048 | -0.037 |
| RI | 5699 | ENSG00000079785 | DDX1 | chr2 | + | 15620207 | 15621116 | 15620207 | 15620396 | 15621064 | 15621116 | 2 | 9 | 6 | 8 | 10 | 241 | 168 | 209 | 232 | 365 | 22 | 14 | 18 | 6 | 0 | 41 | 168 | 117 | 49 | 14 | 298 | 149 | 8.76E-12 | 3.81E-10 | 0.004 | 0.026 | 0.014 | 0.017 | 0.014 | 0.212 | 0.04 | 0.071 | 0.058 | 0 | -0.061 |
| RI | 1625 | ENSG00000175826 | CTDNEP1 | chr17 | - | 7243590 | 7244245 | 7243590 | 7243951 | 7244188 | 7244245 | 306 | 112 | 255 | 252 | 149 | 1 | 2 | 2 | 1 | 1 | 220 | 271 | 157 | 310 | 384 | 1 | 12 | 8 | 5 | 49 | 298 | 149 | 9.53E-12 | 4.11E-10 | 0.994 | 0.966 | 0.985 | 0.992 | 0.987 | 0.991 | 0.919 | 0.908 | 0.969 | 0.797 | 0.068 |
| RI | 5288 | ENSG00000136718 | IMP4 | chr2 | + | 130345566 | 130345933 | 130345566 | 130345699 | 130345778 | 130345933 | 2 | 0 | 0 | 1 | 2 | 104 | 49 | 130 | 96 | 113 | 12 | 2 | 6 | 10 | 15 | 62 | 83 | 44 | 83 | 117 | 228 | 149 | 9.85E-12 | 4.21E-10 | 0.012 | 0 | 0 | 0.007 | 0.011 | 0.112 | 0.016 | 0.082 | 0.073 | 0.077 | -0.066 |
| RI | 1744 | ENSG00000197102 | DYNC1H1 | chr14 | + | 102015855 | 102016489 | 102015855 | 102016086 | 102016348 | 102016489 | 5 | 10 | 6 | 13 | 13 | 77 | 31 | 78 | 71 | 32 | 52 | 44 | 46 | 47 | 51 | 104 | 90 | 57 | 90 | 83 | 298 | 149 | 1.43E-11 | 6.09E-10 | 0.031 | 0.139 | 0.037 | 0.084 | 0.169 | 0.2 | 0.196 | 0.287 | 0.207 | 0.235 | -0.133 |
| RI | 3657 | ENSG00000186470 | BTN3A2 | chr6 | + | 26373275 | 26373413 | 26373275 | 26373296 | 26373386 | 26373413 | 33 | 25 | 36 | 26 | 49 | 359 | 220 | 249 | 165 | 490 | 26 | 22 | 11 | 6 | 0 | 287 | 300 | 163 | 141 | 800 | 239 | 149 | 2.01E-11 | 8.48E-10 | 0.054 | 0.066 | 0.083 | 0.089 | 0.059 | 0.053 | 0.044 | 0.04 | 0.026 | 0 | 0.038 |
| RI | 6518 | ENSG00000204257 | HLA-DMA | chr6 | - | 32949610 | 32950803 | 32949610 | 32949889 | 32950518 | 32950803 | 99 | 90 | 115 | 104 | 112 | 286 | 158 | 230 | 316 | 439 | 452 | 227 | 111 | 214 | 636 | 753 | 214 | 140 | 269 | 993 | 298 | 149 | 2.19E-11 | 9.18E-10 | 0.148 | 0.222 | 0.2 | 0.141 | 0.113 | 0.231 | 0.347 | 0.284 | 0.285 | 0.243 | -0.113 |
| RI | 2889 | ENSG00000178035 | IMPDH2 | chr3 | - | 49027709 | 49028322 | 49027709 | 49027916 | 49028247 | 49028322 | 62 | 32 | 53 | 60 | 46 | 100 | 66 | 112 | 82 | 106 | 178 | 102 | 96 | 69 | 128 | 95 | 60 | 34 | 66 | 103 | 298 | 149 | 2.26E-11 | 9.40E-10 | 0.237 | 0.195 | 0.191 | 0.268 | 0.178 | 0.484 | 0.459 | 0.585 | 0.343 | 0.383 | -0.237 |
| RI | 6899 | ENSG00000068308 | OTUD5 | chrX | - | 48934463 | 48934867 | 48934463 | 48934612 | 48934710 | 48934867 | 41 | 41 | 49 | 63 | 35 | 201 | 135 | 184 | 196 | 185 | 242 | 93 | 122 | 140 | 202 | 489 | 216 | 208 | 216 | 465 | 247 | 149 | 2.91E-11 | 1.20E-09 | 0.11 | 0.155 | 0.138 | 0.162 | 0.102 | 0.23 | 0.206 | 0.261 | 0.281 | 0.208 | -0.104 |
| RI | 8276 | ENSG00000161638 | ITGA5 | chr12 | - | 54403186 | 54403779 | 54403186 | 54403324 | 54403624 | 54403779 | 8 | 8 | 5 | 20 | 4 | 170 | 61 | 134 | 136 | 146 | 19 | 30 | 58 | 28 | 35 | 169 | 138 | 69 | 103 | 162 | 298 | 149 | 2.94E-11 | 1.21E-09 | 0.023 | 0.062 | 0.018 | 0.068 | 0.014 | 0.053 | 0.098 | 0.296 | 0.12 | 0.097 | -0.096 |
| RI | 8181 | ENSG00000151923 | TIAL1 | chr10 | - | 119576610 | 119577203 | 119576610 | 119576750 | 119577079 | 119577203 | 239 | 181 | 194 | 218 | 309 | 379 | 256 | 342 | 407 | 597 | 48 | 139 | 227 | 39 | 11 | 344 | 331 | 314 | 292 | 341 | 298 | 149 | 3.15E-11 | 1.28E-09 | 0.24 | 0.261 | 0.221 | 0.211 | 0.206 | 0.065 | 0.174 | 0.265 | 0.063 | 0.016 | 0.111 |
| RI | 493 | ENSG00000160633 | SAFB | chr19 | + | 5667352 | 5667886 | 5667352 | 5667450 | 5667819 | 5667886 | 12 | 8 | 12 | 9 | 5 | 85 | 55 | 71 | 75 | 31 | 45 | 26 | 47 | 69 | 23 | 90 | 70 | 59 | 55 | 70 | 298 | 149 | 3.21E-11 | 1.30E-09 | 0.066 | 0.068 | 0.078 | 0.057 | 0.075 | 0.2 | 0.157 | 0.285 | 0.385 | 0.141 | -0.165 |
| RI | 4881 | ENSG00000143418 | CERS2 | chr1 | - | 150967391 | 150967714 | 150967391 | 150967484 | 150967663 | 150967714 | 14 | 18 | 30 | 67 | 26 | 329 | 191 | 224 | 282 | 456 | 103 | 64 | 48 | 141 | 124 | 349 | 322 | 254 | 230 | 829 | 298 | 149 | 4.18E-11 | 1.68E-09 | 0.021 | 0.045 | 0.063 | 0.106 | 0.028 | 0.129 | 0.09 | 0.086 | 0.235 | 0.07 | -0.069 |
| RI | 5099 | ENSG00000131748 | STARD3 | chr17 | + | 39657972 | 39658522 | 39657972 | 39658026 | 39658404 | 39658522 | 31 | 19 | 27 | 47 | 20 | 65 | 22 | 62 | 44 | 84 | 89 | 89 | 81 | 118 | 82 | 39 | 59 | 51 | 58 | 49 | 298 | 149 | 4.63E-11 | 1.85E-09 | 0.193 | 0.302 | 0.179 | 0.348 | 0.106 | 0.533 | 0.43 | 0.443 | 0.504 | 0.456 | -0.248 |
| RI | 8405 | ENSG00000002016 | RAD52 | chr12 | - | 926795 | 927263 | 926795 | 926946 | 927144 | 927263 | 19 | 7 | 24 | 16 | 24 | 8 | 2 | 2 | 9 | 3 | 35 | 33 | 6 | 63 | 31 | 0 | 0 | 1 | 1 | 0 | 298 | 149 | 5.26E-11 | 2.09E-09 | 0.543 | 0.636 | 0.857 | 0.471 | 0.8 | 1 | 1 | 0.75 | 0.969 | 1 | -0.282 |
| RI | 3884 | ENSG00000158517 | NCF1 | chr7 | + | 74782938 | 74783632 | 74782938 | 74783061 | 74783524 | 74783632 | 76 | 50 | 60 | 47 | 24 | 450 | 227 | 247 | 238 | 125 | 301 | 148 | 109 | 162 | 273 | 565 | 496 | 167 | 441 | 825 | 298 | 149 | 5.96E-11 | 2.35E-09 | 0.078 | 0.099 | 0.108 | 0.09 | 0.088 | 0.21 | 0.13 | 0.246 | 0.155 | 0.142 | -0.084 |
| RI | 5449 | ENSG00000143799 | PARP1 | chr1 | - | 226363098 | 226365154 | 226363098 | 226364070 | 226365001 | 226365154 | 9 | 4 | 9 | 4 | 3 | 132 | 49 | 120 | 100 | 124 | 29 | 15 | 13 | 12 | 27 | 100 | 64 | 65 | 90 | 130 | 298 | 149 | 6.18E-11 | 2.42E-09 | 0.033 | 0.039 | 0.036 | 0.02 | 0.012 | 0.127 | 0.105 | 0.091 | 0.062 | 0.094 | -0.068 |
| RI | 4509 | ENSG00000168610 | STAT3 | chr17 | - | 42316788 | 42317224 | 42316788 | 42316901 | 42317181 | 42317224 | 14 | 15 | 13 | 19 | 15 | 291 | 156 | 193 | 297 | 327 | 68 | 14 | 62 | 75 | 60 | 392 | 350 | 301 | 355 | 470 | 298 | 149 | 8.50E-11 | 3.30E-09 | 0.023 | 0.046 | 0.033 | 0.031 | 0.022 | 0.08 | 0.02 | 0.093 | 0.096 | 0.06 | -0.039 |
| RI | 5549 | ENSG00000108883 | EFTUD2 | chr17 | - | 44859904 | 44860543 | 44859904 | 44860045 | 44860431 | 44860543 | 23 | 16 | 16 | 23 | 23 | 150 | 85 | 91 | 155 | 235 | 113 | 52 | 19 | 68 | 42 | 159 | 125 | 62 | 96 | 249 | 298 | 149 | 8.67E-11 | 3.35E-09 | 0.071 | 0.086 | 0.081 | 0.069 | 0.047 | 0.262 | 0.172 | 0.133 | 0.262 | 0.078 | -0.111 |
| RI | 2907 | ENSG00000198218 | QRICH1 | chr3 | - | 49032621 | 49033228 | 49032621 | 49032773 | 49033119 | 49033228 | 57 | 55 | 91 | 94 | 55 | 170 | 130 | 217 | 151 | 266 | 180 | 170 | 124 | 144 | 121 | 189 | 168 | 178 | 145 | 208 | 298 | 149 | 9.36E-11 | 3.59E-09 | 0.144 | 0.175 | 0.173 | 0.237 | 0.094 | 0.323 | 0.336 | 0.258 | 0.332 | 0.225 | -0.13 |
| RI | 2619 | ENSG00000173020 | GRK2 | chr11 | + | 67281821 | 67282542 | 67281821 | 67281952 | 67282270 | 67282542 | 57 | 46 | 54 | 77 | 33 | 168 | 62 | 193 | 134 | 100 | 109 | 134 | 113 | 142 | 242 | 149 | 142 | 77 | 99 | 238 | 298 | 149 | 1.03E-10 | 3.92E-09 | 0.145 | 0.271 | 0.123 | 0.223 | 0.142 | 0.268 | 0.321 | 0.423 | 0.418 | 0.337 | -0.173 |
| RI | 7063 | ENSG00000205581 | HMGN1 | chr21 | - | 39345829 | 39347292 | 39345829 | 39345958 | 39347246 | 39347292 | 348 | 399 | 383 | 375 | 372 | 1 | 2 | 6 | 2 | 8 | 41 | 169 | 236 | 94 | 53 | 0 | 7 | 8 | 21 | 11 | 298 | 149 | 1.06E-10 | 4.03E-09 | 0.994 | 0.99 | 0.97 | 0.989 | 0.959 | 1 | 0.923 | 0.937 | 0.691 | 0.707 | 0.129 |
| RI | 1136 | ENSG00000142512 | SIGLEC10 | chr19 | - | 51415897 | 51416357 | 51415897 | 51416167 | 51416309 | 51416357 | 20 | 10 | 2 | 8 | 16 | 32 | 8 | 10 | 25 | 6 | 1 | 0 | 0 | 6 | 0 | 14 | 13 | 8 | 50 | 38 | 291 | 149 | 1.33E-10 | 5.02E-09 | 0.242 | 0.39 | 0.093 | 0.141 | 0.577 | 0.035 | 0 | 0 | 0.058 | 0 | 0.27 |
| RI | 7748 | ENSG00000077458 | FAM76B | chr11 | - | 95788498 | 95789782 | 95788498 | 95788563 | 95789391 | 95789782 | 58 | 66 | 57 | 102 | 77 | 47 | 33 | 43 | 50 | 27 | 82 | 40 | 64 | 57 | 21 | 4 | 13 | 4 | 3 | 0 | 298 | 149 | 1.42E-10 | 5.32E-09 | 0.382 | 0.5 | 0.399 | 0.505 | 0.588 | 0.911 | 0.606 | 0.889 | 0.905 | 1 | -0.387 |
| RI | 3352 | ENSG00000158864 | NDUFS2 | chr1 | + | 161212350 | 161213475 | 161212350 | 161212480 | 161213379 | 161213475 | 5 | 7 | 6 | 11 | 5 | 126 | 53 | 120 | 133 | 116 | 26 | 17 | 20 | 9 | 56 | 104 | 160 | 85 | 90 | 129 | 298 | 149 | 1.46E-10 | 5.44E-09 | 0.019 | 0.062 | 0.024 | 0.04 | 0.021 | 0.111 | 0.05 | 0.105 | 0.048 | 0.178 | -0.065 |
| RI | 4005 | ENSG00000197530 | MIB2 | chr1 | + | 1628272 | 1628722 | 1628272 | 1628399 | 1628488 | 1628722 | 14 | 6 | 11 | 22 | 8 | 44 | 16 | 13 | 37 | 21 | 1 | 0 | 11 | 0 | 3 | 31 | 28 | 28 | 48 | 82 | 238 | 149 | 1.49E-10 | 5.50E-09 | 0.166 | 0.19 | 0.346 | 0.271 | 0.193 | 0.02 | 0 | 0.197 | 0 | 0.022 | 0.185 |
| RI | 338 | ENSG00000013275 | PSMC4 | chr19 | + | 39980285 | 39980717 | 39980285 | 39980454 | 39980661 | 39980717 | 26 | 7 | 30 | 23 | 24 | 149 | 88 | 115 | 114 | 228 | 91 | 28 | 34 | 56 | 68 | 186 | 149 | 88 | 97 | 161 | 298 | 149 | 1.51E-10 | 5.53E-09 | 0.08 | 0.038 | 0.115 | 0.092 | 0.05 | 0.197 | 0.086 | 0.162 | 0.224 | 0.174 | -0.094 |
| RI | 5314 | ENSG00000117632 | STMN1 | chr1 | - | 25901490 | 25903813 | 25901490 | 25901682 | 25903640 | 25903813 | 12 | 11 | 17 | 25 | 14 | 121 | 133 | 217 | 230 | 277 | 0 | 9 | 4 | 15 | 0 | 356 | 206 | 263 | 439 | 107 | 298 | 149 | 1.55E-10 | 5.66E-09 | 0.047 | 0.04 | 0.038 | 0.052 | 0.025 | 0 | 0.021 | 0.008 | 0.017 | 0 | 0.031 |
| RI | 2786 | ENSG00000181789 | COPG1 | chr3 | + | 129252622 | 129252955 | 129252622 | 129252694 | 129252875 | 129252955 | 14 | 8 | 7 | 21 | 8 | 85 | 44 | 73 | 107 | 76 | 83 | 49 | 21 | 37 | 60 | 101 | 93 | 92 | 120 | 95 | 298 | 149 | 1.59E-10 | 5.76E-09 | 0.076 | 0.083 | 0.046 | 0.089 | 0.05 | 0.291 | 0.209 | 0.102 | 0.134 | 0.24 | -0.126 |
| RI | 7292 | ENSG00000137509 | PRCP | chr11 | - | 82850323 | 82853278 | 82850323 | 82850505 | 82853176 | 82853278 | 32 | 24 | 10 | 10 | 41 | 186 | 114 | 153 | 149 | 299 | 5 | 6 | 15 | 6 | 0 | 272 | 97 | 60 | 219 | 246 | 298 | 149 | 1.70E-10 | 6.12E-09 | 0.079 | 0.095 | 0.032 | 0.032 | 0.064 | 0.009 | 0.03 | 0.111 | 0.014 | 0 | 0.028 |
| RI | 3870 | ENSG00000108654 | DDX5 | chr17 | - | 64502161 | 64502549 | 64502161 | 64502223 | 64502438 | 64502549 | 1109 | 910 | 1093 | 1307 | 902 | 2926 | 2360 | 2849 | 3523 | 4477 | 533 | 1603 | 1397 | 1011 | 117 | 1331 | 1990 | 2591 | 1705 | 253 | 298 | 149 | 1.93E-10 | 6.93E-09 | 0.159 | 0.162 | 0.161 | 0.156 | 0.092 | 0.167 | 0.287 | 0.212 | 0.229 | 0.188 | -0.071 |
| RI | 8315 | ENSG00000007392 | LUC7L | chr16 | - | 188968 | 190135 | 188968 | 189339 | 189967 | 190135 | 41 | 20 | 32 | 45 | 14 | 120 | 95 | 92 | 157 | 78 | 60 | 44 | 39 | 66 | 95 | 80 | 80 | 83 | 74 | 81 | 298 | 149 | 2.04E-10 | 7.27E-09 | 0.146 | 0.095 | 0.148 | 0.125 | 0.082 | 0.273 | 0.216 | 0.19 | 0.308 | 0.37 | -0.152 |
| RI | 6294 | ENSG00000145191 | EIF2B5 | chr3 | + | 184140079 | 184140730 | 184140079 | 184140157 | 184140417 | 184140730 | 40 | 41 | 33 | 33 | 29 | 55 | 32 | 69 | 63 | 97 | 166 | 77 | 111 | 100 | 116 | 85 | 47 | 31 | 62 | 60 | 298 | 149 | 2.10E-10 | 7.43E-09 | 0.267 | 0.39 | 0.193 | 0.208 | 0.13 | 0.494 | 0.45 | 0.642 | 0.446 | 0.492 | -0.267 |
| RI | 5737 | ENSG00000186635 | ARAP1 | chr11 | - | 72687438 | 72687738 | 72687438 | 72687502 | 72687687 | 72687738 | 57 | 39 | 40 | 39 | 21 | 400 | 162 | 175 | 196 | 270 | 120 | 96 | 71 | 75 | 135 | 421 | 166 | 89 | 203 | 554 | 298 | 149 | 2.58E-10 | 9.06E-09 | 0.067 | 0.107 | 0.103 | 0.09 | 0.037 | 0.125 | 0.224 | 0.285 | 0.156 | 0.109 | -0.099 |
| RI | 810 | ENSG00000185624 | P4HB | chr17 | - | 81845143 | 81845991 | 81845143 | 81845742 | 81845870 | 81845991 | 16 | 10 | 7 | 25 | 24 | 1242 | 744 | 1021 | 1002 | 1415 | 130 | 35 | 32 | 142 | 63 | 2889 | 1624 | 1248 | 1918 | 3143 | 277 | 149 | 2.94E-10 | 1.03E-08 | 0.007 | 0.007 | 0.004 | 0.013 | 0.009 | 0.024 | 0.011 | 0.014 | 0.038 | 0.011 | -0.012 |
| RI | 1053 | ENSG00000073584 | SMARCE1 | chr17 | - | 40630713 | 40631693 | 40630713 | 40630924 | 40631591 | 40631693 | 101 | 107 | 83 | 97 | 83 | 271 | 321 | 323 | 341 | 452 | 7 | 65 | 148 | 23 | 0 | 257 | 365 | 462 | 409 | 68 | 298 | 149 | 2.99E-10 | 1.04E-08 | 0.157 | 0.143 | 0.114 | 0.125 | 0.084 | 0.013 | 0.082 | 0.138 | 0.027 | 0 | 0.073 |
| RI | 1525 | ENSG00000213614 | HEXA | chr15 | - | 72353067 | 72355624 | 72353067 | 72353178 | 72355558 | 72355624 | 9 | 9 | 12 | 16 | 11 | 12 | 12 | 5 | 8 | 16 | 39 | 40 | 36 | 54 | 12 | 0 | 11 | 3 | 4 | 0 | 298 | 149 | 3.00E-10 | 1.04E-08 | 0.273 | 0.273 | 0.545 | 0.5 | 0.256 | 1 | 0.645 | 0.857 | 0.871 | 1 | -0.505 |
| RI | 2881 | ENSG00000178035 | IMPDH2 | chr3 | - | 49024494 | 49024802 | 49024494 | 49024578 | 49024658 | 49024802 | 35 | 26 | 39 | 24 | 24 | 186 | 136 | 283 | 251 | 278 | 29 | 20 | 44 | 34 | 91 | 127 | 82 | 99 | 145 | 171 | 229 | 149 | 3.23E-10 | 1.11E-08 | 0.109 | 0.111 | 0.082 | 0.059 | 0.053 | 0.129 | 0.137 | 0.224 | 0.132 | 0.257 | -0.093 |
| RI | 2275 | ENSG00000104731 | KLHDC4 | chr16 | - | 87755193 | 87756477 | 87755193 | 87755292 | 87756398 | 87756477 | 26 | 17 | 24 | 21 | 10 | 67 | 41 | 60 | 69 | 74 | 43 | 43 | 61 | 74 | 88 | 67 | 56 | 40 | 77 | 48 | 298 | 149 | 3.27E-10 | 1.12E-08 | 0.163 | 0.172 | 0.167 | 0.132 | 0.063 | 0.243 | 0.277 | 0.433 | 0.325 | 0.478 | -0.212 |
| RI | 2056 | ENSG00000133706 | LARS1 | chr5 | - | 146130017 | 146131109 | 146130017 | 146130158 | 146131018 | 146131109 | 21 | 37 | 19 | 27 | 41 | 194 | 177 | 188 | 197 | 444 | 0 | 3 | 9 | 0 | 0 | 26 | 117 | 158 | 100 | 7 | 298 | 149 | 3.34E-10 | 1.13E-08 | 0.051 | 0.095 | 0.048 | 0.064 | 0.044 | 0 | 0.013 | 0.028 | 0 | 0 | 0.052 |
| RI | 6052 | ENSG00000135486 | HNRNPA1 | chr12 | + | 54282393 | 54282874 | 54282393 | 54282665 | 54282799 | 54282874 | 287 | 273 | 333 | 350 | 240 | 2803 | 2712 | 3192 | 3019 | 3029 | 379 | 532 | 269 | 696 | 233 | 3271 | 2331 | 4028 | 3061 | 867 | 283 | 149 | 3.68E-10 | 1.24E-08 | 0.051 | 0.05 | 0.052 | 0.058 | 0.04 | 0.057 | 0.107 | 0.034 | 0.107 | 0.124 | -0.036 |
| RI | 5664 | ENSG00000165280 | VCP | chr9 | - | 35061014 | 35061689 | 35061014 | 35061179 | 35061576 | 35061689 | 39 | 28 | 30 | 37 | 23 | 342 | 174 | 299 | 319 | 237 | 58 | 102 | 79 | 75 | 123 | 458 | 389 | 251 | 302 | 470 | 298 | 149 | 3.76E-10 | 1.26E-08 | 0.054 | 0.074 | 0.048 | 0.055 | 0.046 | 0.06 | 0.116 | 0.136 | 0.11 | 0.116 | -0.052 |
| RI | 5176 | ENSG00000104907 | TRMT1 | chr19 | - | 13115996 | 13116431 | 13115996 | 13116052 | 13116145 | 13116431 | 7 | 8 | 11 | 14 | 9 | 57 | 59 | 63 | 72 | 84 | 70 | 17 | 28 | 18 | 28 | 87 | 51 | 54 | 50 | 72 | 242 | 149 | 4.05E-10 | 1.35E-08 | 0.07 | 0.077 | 0.097 | 0.107 | 0.062 | 0.331 | 0.17 | 0.242 | 0.181 | 0.193 | -0.141 |
| RI | 8240 | ENSG00000067560 | RHOA | chr3 | - | 49359139 | 49362626 | 49359139 | 49360382 | 49362495 | 49362626 | 174 | 170 | 120 | 182 | 134 | 1983 | 1460 | 1368 | 1848 | 2203 | 726 | 318 | 232 | 390 | 899 | 3723 | 2538 | 1701 | 3047 | 6660 | 298 | 149 | 4.23E-10 | 1.41E-08 | 0.042 | 0.055 | 0.042 | 0.047 | 0.03 | 0.089 | 0.059 | 0.064 | 0.06 | 0.063 | -0.024 |
| RI | 4892 | ENSG00000070814 | TCOF1 | chr5 | + | 150391543 | 150392176 | 150391543 | 150391657 | 150391956 | 150392176 | 35 | 31 | 38 | 40 | 27 | 57 | 10 | 27 | 39 | 23 | 284 | 141 | 59 | 159 | 287 | 12 | 36 | 21 | 14 | 43 | 298 | 149 | 4.47E-10 | 1.47E-08 | 0.235 | 0.608 | 0.413 | 0.339 | 0.37 | 0.922 | 0.662 | 0.584 | 0.85 | 0.769 | -0.364 |
| RI | 7311 | ENSG00000147162 | OGT | chrX | + | 71544566 | 71548023 | 71544566 | 71544635 | 71547906 | 71548023 | 1226 | 992 | 1266 | 1226 | 679 | 106 | 42 | 71 | 81 | 47 | 709 | 1221 | 1264 | 1229 | 418 | 50 | 48 | 15 | 33 | 0 | 298 | 149 | 4.76E-10 | 1.56E-08 | 0.853 | 0.922 | 0.899 | 0.883 | 0.878 | 0.876 | 0.927 | 0.977 | 0.949 | 1 | -0.059 |
| RI | 6957 | ENSG00000215301 | DDX3X | chrX | + | 41345179 | 41345548 | 41345179 | 41345324 | 41345403 | 41345548 | 161 | 124 | 123 | 181 | 142 | 1099 | 750 | 1044 | 1238 | 1610 | 21 | 144 | 137 | 181 | 26 | 131 | 459 | 762 | 304 | 89 | 228 | 149 | 4.90E-10 | 1.60E-08 | 0.087 | 0.098 | 0.071 | 0.087 | 0.054 | 0.095 | 0.17 | 0.105 | 0.28 | 0.16 | -0.083 |
| RI | 8128 | ENSG00000176986 | SEC24C | chr10 | + | 73770279 | 73770798 | 73770279 | 73770471 | 73770708 | 73770798 | 118 | 89 | 89 | 91 | 105 | 203 | 112 | 135 | 194 | 190 | 278 | 214 | 205 | 292 | 273 | 188 | 190 | 100 | 146 | 317 | 298 | 149 | 6.27E-10 | 2.04E-08 | 0.225 | 0.284 | 0.248 | 0.19 | 0.216 | 0.425 | 0.36 | 0.506 | 0.5 | 0.301 | -0.186 |
| RI | 5909 | ENSG00000165792 | METTL17 | chr14 | + | 20993117 | 20994063 | 20993117 | 20993191 | 20993968 | 20994063 | 70 | 53 | 89 | 86 | 55 | 55 | 29 | 60 | 59 | 54 | 202 | 71 | 124 | 109 | 209 | 6 | 21 | 31 | 17 | 70 | 298 | 149 | 6.54E-10 | 2.11E-08 | 0.389 | 0.477 | 0.426 | 0.422 | 0.337 | 0.944 | 0.628 | 0.667 | 0.762 | 0.599 | -0.31 |
| RI | 5364 | ENSG00000123136 | DDX39A | chr19 | - | 14409302 | 14409645 | 14409302 | 14409447 | 14409535 | 14409645 | 55 | 37 | 54 | 96 | 32 | 166 | 106 | 140 | 175 | 63 | 178 | 100 | 97 | 159 | 181 | 159 | 118 | 109 | 146 | 296 | 237 | 149 | 6.78E-10 | 2.18E-08 | 0.172 | 0.18 | 0.195 | 0.256 | 0.242 | 0.413 | 0.348 | 0.359 | 0.406 | 0.278 | -0.152 |
| RI | 8485 | ENSG00000166598 | HSP90B1 | chr12 | + | 103942526 | 103943319 | 103942526 | 103942796 | 103943073 | 103943319 | 35 | 27 | 35 | 38 | 26 | 547 | 402 | 707 | 609 | 631 | 6 | 38 | 102 | 25 | 13 | 106 | 258 | 930 | 127 | 46 | 298 | 149 | 6.81E-10 | 2.18E-08 | 0.031 | 0.032 | 0.024 | 0.03 | 0.02 | 0.028 | 0.069 | 0.052 | 0.09 | 0.124 | -0.045 |
| RI | 5728 | ENSG00000139531 | SUOX | chr12 | + | 56002211 | 56002720 | 56002211 | 56002271 | 56002542 | 56002720 | 0 | 1 | 3 | 0 | 0 | 14 | 8 | 5 | 3 | 4 | 13 | 8 | 8 | 2 | 18 | 12 | 2 | 5 | 5 | 4 | 298 | 149 | 7.35E-10 | 2.33E-08 | 0 | 0.059 | 0.231 | 0 | 0 | 0.351 | 0.667 | 0.444 | 0.167 | 0.692 | -0.406 |
| RI | 778 | ENSG00000182173 | TSEN54 | chr17 | + | 75521408 | 75522333 | 75521408 | 75521510 | 75521704 | 75522333 | 20 | 5 | 23 | 12 | 15 | 76 | 37 | 66 | 86 | 50 | 33 | 50 | 45 | 31 | 30 | 22 | 75 | 32 | 36 | 67 | 298 | 149 | 7.40E-10 | 2.34E-08 | 0.116 | 0.063 | 0.148 | 0.065 | 0.13 | 0.429 | 0.25 | 0.413 | 0.301 | 0.183 | -0.211 |
| RI | 2891 | ENSG00000178035 | IMPDH2 | chr3 | - | 49028247 | 49028532 | 49028247 | 49028322 | 49028430 | 49028532 | 43 | 13 | 40 | 55 | 37 | 63 | 44 | 73 | 82 | 56 | 140 | 100 | 114 | 72 | 93 | 101 | 53 | 43 | 37 | 86 | 257 | 149 | 7.61E-10 | 2.39E-08 | 0.284 | 0.146 | 0.241 | 0.28 | 0.277 | 0.446 | 0.522 | 0.606 | 0.53 | 0.385 | -0.252 |
| RI | 6678 | ENSG00000102226 | USP11 | chrX | + | 47247071 | 47247419 | 47247071 | 47247221 | 47247303 | 47247419 | 17 | 7 | 14 | 15 | 6 | 70 | 33 | 67 | 74 | 63 | 6 | 45 | 39 | 53 | 35 | 29 | 39 | 22 | 27 | 28 | 231 | 149 | 8.47E-10 | 2.65E-08 | 0.135 | 0.12 | 0.119 | 0.116 | 0.058 | 0.118 | 0.427 | 0.533 | 0.559 | 0.446 | -0.307 |
| RI | 3121 | ENSG00000161204 | ABCF3 | chr3 | + | 184192600 | 184192896 | 184192600 | 184192689 | 184192804 | 184192896 | 20 | 12 | 22 | 16 | 25 | 33 | 16 | 39 | 36 | 31 | 67 | 41 | 39 | 40 | 49 | 33 | 23 | 24 | 15 | 29 | 264 | 149 | 8.57E-10 | 2.66E-08 | 0.255 | 0.297 | 0.241 | 0.201 | 0.313 | 0.534 | 0.502 | 0.478 | 0.601 | 0.488 | -0.259 |
| RI | 2170 | ENSG00000065357 | DGKA | chr12 | + | 55951622 | 55952099 | 55951622 | 55951783 | 55952034 | 55952099 | 25 | 52 | 55 | 50 | 26 | 331 | 341 | 622 | 447 | 886 | 75 | 27 | 42 | 45 | 54 | 274 | 308 | 177 | 316 | 358 | 298 | 149 | 9.06E-10 | 2.80E-08 | 0.036 | 0.071 | 0.042 | 0.053 | 0.014 | 0.12 | 0.042 | 0.106 | 0.066 | 0.07 | -0.038 |
| RI | 3702 | ENSG00000058673 | ZC3H11A | chr1 | + | 203801574 | 203803016 | 203801574 | 203801628 | 203802956 | 203803016 | 408 | 288 | 360 | 423 | 559 | 35 | 12 | 20 | 26 | 37 | 124 | 380 | 449 | 266 | 52 | 7 | 4 | 7 | 5 | 1 | 298 | 149 | 9.31E-10 | 2.86E-08 | 0.854 | 0.923 | 0.9 | 0.891 | 0.883 | 0.899 | 0.979 | 0.97 | 0.964 | 0.963 | -0.065 |
| RI | 4116 | ENSG00000116337 | AMPD2 | chr1 | + | 109628082 | 109628806 | 109628082 | 109628495 | 109628642 | 109628806 | 54 | 34 | 49 | 58 | 28 | 134 | 79 | 77 | 98 | 122 | 228 | 72 | 62 | 110 | 198 | 222 | 89 | 64 | 54 | 142 | 296 | 149 | 9.63E-10 | 2.95E-08 | 0.169 | 0.178 | 0.243 | 0.23 | 0.104 | 0.341 | 0.289 | 0.328 | 0.506 | 0.412 | -0.19 |
| RI | 4218 | ENSG00000104231 | ZFAND1 | chr8 | - | 81713917 | 81714895 | 81713917 | 81714039 | 81714803 | 81714895 | 44 | 52 | 55 | 36 | 64 | 264 | 303 | 268 | 297 | 497 | 2 | 8 | 14 | 0 | 2 | 84 | 162 | 99 | 90 | 67 | 298 | 149 | 1.03E-09 | 3.13E-08 | 0.077 | 0.079 | 0.093 | 0.057 | 0.06 | 0.012 | 0.024 | 0.066 | 0 | 0.015 | 0.05 |
| RI | 5596 | ENSG00000095059 | DHPS | chr19 | - | 12679456 | 12679719 | 12679456 | 12679543 | 12679622 | 12679719 | 32 | 43 | 55 | 63 | 21 | 197 | 103 | 170 | 149 | 221 | 123 | 127 | 45 | 104 | 148 | 216 | 195 | 86 | 158 | 283 | 228 | 149 | 1.17E-09 | 3.53E-08 | 0.096 | 0.214 | 0.175 | 0.216 | 0.058 | 0.271 | 0.299 | 0.255 | 0.301 | 0.255 | -0.124 |
| RI | 6243 | ENSG00000075975 | MKRN2 | chr3 | + | 12574791 | 12576741 | 12574791 | 12575006 | 12576630 | 12576741 | 2 | 2 | 4 | 7 | 0 | 117 | 89 | 111 | 115 | 144 | 29 | 23 | 6 | 5 | 0 | 164 | 99 | 79 | 85 | 77 | 298 | 149 | 1.19E-09 | 3.58E-08 | 0.008 | 0.011 | 0.018 | 0.03 | 0 | 0.081 | 0.104 | 0.037 | 0.029 | 0 | -0.037 |
| RI | 4512 | ENSG00000168610 | STAT3 | chr17 | - | 42329409 | 42329647 | 42329409 | 42329457 | 42329553 | 42329647 | 13 | 16 | 10 | 21 | 6 | 384 | 222 | 315 | 381 | 597 | 69 | 51 | 56 | 49 | 34 | 749 | 463 | 466 | 627 | 845 | 245 | 149 | 1.26E-09 | 3.78E-08 | 0.02 | 0.042 | 0.019 | 0.032 | 0.006 | 0.053 | 0.063 | 0.068 | 0.045 | 0.024 | -0.027 |
| RI | 1977 | ENSG00000006744 | ELAC2 | chr17 | - | 13002273 | 13002579 | 13002273 | 13002359 | 13002440 | 13002579 | 10 | 8 | 17 | 32 | 5 | 103 | 61 | 83 | 121 | 128 | 42 | 59 | 24 | 32 | 84 | 146 | 104 | 78 | 111 | 161 | 230 | 149 | 1.27E-09 | 3.80E-08 | 0.059 | 0.078 | 0.117 | 0.146 | 0.025 | 0.157 | 0.269 | 0.166 | 0.157 | 0.253 | -0.115 |
| RI | 5764 | ENSG00000118495 | PLAGL1 | chr6 | - | 143964786 | 143966198 | 143964786 | 143964818 | 143966157 | 143966198 | 26 | 31 | 20 | 29 | 41 | 16 | 18 | 5 | 8 | 22 | 0 | 22 | 8 | 27 | 0 | 27 | 43 | 19 | 37 | 4 | 298 | 149 | 1.34E-09 | 3.99E-08 | 0.448 | 0.463 | 0.667 | 0.644 | 0.482 | 0 | 0.204 | 0.174 | 0.267 | 0 | 0.412 |
| RI | 8358 | ENSG00000135297 | MTO1 | chr6 | + | 73479731 | 73480126 | 73479731 | 73479844 | 73479935 | 73480126 | 14 | 13 | 10 | 9 | 8 | 105 | 66 | 79 | 87 | 118 | 69 | 32 | 20 | 16 | 105 | 65 | 71 | 77 | 53 | 55 | 240 | 149 | 1.36E-09 | 4.02E-08 | 0.076 | 0.109 | 0.073 | 0.06 | 0.04 | 0.397 | 0.219 | 0.139 | 0.158 | 0.542 | -0.219 |
| RI | 1679 | ENSG00000174444 | RPL4 | chr15 | - | 66502611 | 66503164 | 66502611 | 66502750 | 66503057 | 66503164 | 88 | 56 | 98 | 85 | 111 | 1518 | 660 | 1745 | 1247 | 1546 | 106 | 96 | 81 | 141 | 83 | 765 | 811 | 915 | 663 | 1305 | 298 | 149 | 1.65E-09 | 4.84E-08 | 0.028 | 0.041 | 0.027 | 0.033 | 0.035 | 0.065 | 0.056 | 0.042 | 0.096 | 0.031 | -0.025 |
| RI | 2209 | ENSG00000160201 | U2AF1 | chr21 | - | 43092955 | 43094563 | 43092955 | 43093249 | 43094470 | 43094563 | 20 | 24 | 12 | 8 | 5 | 0 | 0 | 0 | 0 | 0 | 56 | 27 | 44 | 23 | 13 | 9 | 0 | 13 | 13 | 0 | 298 | 149 | 1.71E-09 | 4.99E-08 | 1 | 1 | 1 | 1 | 1 | 0.757 | 1 | 0.629 | 0.469 | 1 | 0.229 |
| RI | 2164 | ENSG00000065357 | DGKA | chr12 | + | 55939414 | 55940433 | 55939414 | 55940170 | 55940313 | 55940433 | 114 | 115 | 234 | 155 | 241 | 112 | 94 | 193 | 165 | 164 | 136 | 150 | 172 | 186 | 111 | 44 | 86 | 63 | 43 | 44 | 292 | 149 | 1.76E-09 | 5.14E-08 | 0.342 | 0.384 | 0.382 | 0.324 | 0.429 | 0.612 | 0.471 | 0.582 | 0.688 | 0.563 | -0.211 |
| RI | 748 | ENSG00000109118 | PHF12 | chr17 | - | 28923908 | 28927063 | 28923908 | 28924302 | 28926990 | 28927063 | 40 | 32 | 30 | 28 | 31 | 249 | 113 | 197 | 242 | 177 | 109 | 53 | 95 | 66 | 74 | 239 | 220 | 258 | 160 | 301 | 298 | 149 | 1.84E-09 | 5.33E-08 | 0.074 | 0.124 | 0.071 | 0.055 | 0.081 | 0.186 | 0.108 | 0.155 | 0.171 | 0.109 | -0.065 |
| RI | 1963 | ENSG00000112659 | CUL9 | chr6 | + | 43196068 | 43196862 | 43196068 | 43196265 | 43196644 | 43196862 | 19 | 5 | 13 | 15 | 18 | 31 | 30 | 38 | 43 | 77 | 23 | 27 | 20 | 57 | 61 | 37 | 18 | 17 | 26 | 49 | 298 | 149 | 1.86E-09 | 5.37E-08 | 0.235 | 0.077 | 0.146 | 0.149 | 0.105 | 0.237 | 0.429 | 0.37 | 0.523 | 0.384 | -0.246 |
| RI | 6072 | ENSG00000204348 | DXO | chr6 | - | 31970605 | 31971147 | 31970605 | 31970752 | 31970911 | 31971147 | 30 | 25 | 32 | 15 | 20 | 2 | 2 | 14 | 4 | 4 | 14 | 24 | 26 | 24 | 30 | 0 | 0 | 1 | 1 | 0 | 298 | 149 | 2.19E-09 | 6.28E-08 | 0.882 | 0.862 | 0.533 | 0.652 | 0.714 | 1 | 1 | 0.929 | 0.923 | 1 | -0.242 |

| | | | | | | | | | | | | | | | |
|---|---|---|---|---|---|---|---|---|---|---|---|---|---|---|---|
| RI | 3203 | ENSG00000143862 | ARL8A | chr1 | - | 202135458 | 202135800 | 202135458 | 202135526 | 202135706 | 202135800 | 25 | 11 | 35 | [illegible] |
| RI | 5318 | ENSG00000170445 | HARS1 | chr5 | - | 140675016 | 140676896 | 140675016 | 140675133 | 140676653 | 140676896 | 12 | 11 | 8 | [illegible] |
| RI | 6660 | ENSG00000203965 | EFCAB7 | chr1 | + | 63568309 | 63571128 | 63568309 | 63568519 | 63571020 | 63571128 | 23 | 21 | 7 | [illegible] |
| RI | 3342 | ENSG00000158864 | NDUFS2 | chr1 | + | 161209482 | 161209931 | 161209482 | 161209595 | 161209856 | 161209931 | 70 | 73 | 63 | [illegible] |
| RI | 8180 | ENSG00000151651 | ADAM8 | chr10 | - | 133273253 | 133273838 | 133273253 | 133273443 | 133273761 | 133273838 | 4 | 4 | 10 | [illegible] |
| RI | 1445 | ENSG00000068903 | SIRT2 | chr19 | - | 38889688 | 38889961 | 38889688 | 38889745 | 38889854 | 38889961 | 2 | 0 | 1 | [illegible] |
| RI | 4608 | ENSG00000060138 | YBX3 | chr12 | - | 10701959 | 10704148 | 10701959 | 10702134 | 10704050 | 10704148 | 11 | 21 | 12 | [illegible] |
| RI | 3456 | ENSG00000144591 | GMPPA | chr2 | + | 219504082 | 219505761 | 219504082 | 219505362 | 219505457 | 219505761 | 17 | 6 | 9 | [illegible] |
| RI | 8032 | ENSG00000090621 | PABPC4 | chr1 | - | 39567750 | 39568939 | 39567750 | 39567846 | 39568801 | 39568939 | 113 | 97 | 96 | [illegible] |
| RI | 5124 | ENSG00000109332 | UBE2D3 | chr4 | - | 102801453 | 102802638 | 102801453 | 102801559 | 102802560 | 102802638 | 291 | 218 | 265 | [illegible] |
| RI | 8017 | ENSG00000198258 | UBL5 | chr19 | + | 9828326 | 9828675 | 9828326 | 9828393 | 9828591 | 9828675 | 29 | 8 | 31 | [illegible] |
| RI | 4618 | ENSG00000117616 | RSRP1 | chr1 | - | 25243549 | 25245301 | 25243549 | 25243633 | 25245149 | 25245301 | 656 | 598 | 544 | [illegible] |
| RI | 8197 | ENSG00000023041 | ZDHHC6 | chr10 | - | 112432375 | 112434464 | 112432375 | 112432521 | 112434296 | 112434464 | 142 | 123 | 163 | [illegible] |
| RI | 4050 | ENSG00000132463 | GRSF1 | chr4 | - | 70825295 | 70826245 | 70825295 | 70825431 | 70826123 | 70826245 | 38 | 24 | 31 | [illegible] |
| RI | 3564 | ENSG00000175203 | DCTN2 | chr12 | - | 57532983 | 57533303 | 57532983 | 57533022 | 57533237 | 57533303 | 19 | 16 | 8 | [illegible] |
| RI | 6448 | ENSG00000123505 | AMD1 | chr6 | + | 110892298 | 110892827 | 110892298 | 110892443 | 110892734 | 110892827 | 74 | 58 | 45 | [illegible] |
| RI | 3182 | ENSG00000149480 | MTA2 | chr11 | - | 62597315 | 62597712 | 62597315 | 62597415 | 62597609 | 62597712 | 30 | 14 | 28 | [illegible] |
| RI | 4208 | ENSG00000215788 | TNFRSF25 | chr1 | - | 6465439 | 6466099 | 6465439 | 6465560 | 6466068 | 6466099 | 11 | 1 | 11 | [illegible] |
| RI | 6536 | ENSG00000106263 | EIF3B | chr7 | + | 2379393 | 2380740 | 2379393 | 2379511 | 2380203 | 2380740 | 35 | 20 | 43 | [illegible] |
| RI | 2205 | ENSG00000086189 | DIMT1 | chr5 | - | 62390875 | 62392990 | 62390875 | 62392234 | 62392925 | 62392990 | 22 | 14 | 26 | [illegible] |
| RI | 5313 | ENSG00000168438 | CDC40 | chr6 | + | 110210706 | 110212272 | 110210706 | 110210803 | 110212132 | 110212272 | 100 | 92 | 95 | [illegible] |
| RI | 6939 | ENSG00000147133 | TAF1 | chrX | + | 71423116 | 71424066 | 71423116 | 71423239 | 71423973 | 71424066 | 85 | 82 | 68 | [illegible] |
| RI | 6090 | ENSG00000112739 | PRPF4B | chr6 | + | 4056320 | 4057187 | 4056320 | 4056435 | 4057035 | 4057187 | 88 | 93 | 104 | [illegible] |
| RI | 1814 | ENSG00000196497 | IPO4 | chr14 | - | 24183455 | 24183667 | 24183455 | 24183513 | 24183589 | 24183667 | 9 | 6 | 14 | [illegible] |
| RI | 4711 | ENSG00000232956 | SNHG15 | chr7 | - | 44983821 | 44986097 | 44983821 | 44984046 | 44986020 | 44986097 | 24 | 18 | 37 | [illegible] |
| RI | 5372 | ENSG00000003509 | NDUFAF7 | chr2 | + | 37241577 | 37242693 | 37241577 | 37241791 | 37242634 | 37242693 | 68 | 92 | 58 | [illegible] |
| RI | 6293 | ENSG00000145191 | EIF2B5 | chr3 | + | 184137619 | 184138075 | 184137619 | 184137805 | 184137897 | 184138075 | 48 | 14 | 64 | [illegible] |
| RI | 5821 | ENSG00000124356 | STAMBP | chr2 | + | 73828975 | 73831059 | 73828975 | 73828996 | 73830844 | 73831059 | 11 | 11 | 15 | [illegible] |
| RI | 2105 | ENSG00000137070 | IL11RA | chr9 | + | 34655217 | 34655665 | 34655217 | 34655317 | 34655604 | 34655665 | 28 | 18 | 49 | [illegible] |
| [illegible] | | | | | | | | | | | | | | | |

| | | | | | | | | | | | | | | | | | | | | | | | | | | | | | | | | | | | | | | | | | | | | | | |
|---|---|---|---|---|---|---|---|---|---|---|---|---|---|---|---|---|---|---|---|---|---|---|---|---|---|---|---|---|---|---|---|---|---|---|---|---|---|---|---|---|---|---|---|---|---|---|
| RI | 909 | ENSG00000076382 | SPAG5 | chr17 | - | 28579750 | 28580120 | 28579750 | 28579837 | 28580008 | 28580120 | 2 | 0 | 0 | 5 | 0 | 11 | 9 | 22 | 6 | 4 | 16 | 8 | 5 | 39 | 18 | 15 | 18 | 11 | 18 | 0 | 298 | 149 | 1.04E-06 | 1.48E-05 | 0.083 | 0 | 0 | 0.294 | 0 | 0.348 | 0.182 | 0.185 | 0.52 | 1 | -0.372 |
| RI | 1126 | ENSG00000167840 | ZNF232 | chr17 | - | 5108925 | 5109868 | 5108925 | 5109052 | 5109393 | 5109868 | 7 | 13 | 12 | 17 | 29 | 5 | 0 | 1 | 2 | 5 | 17 | 22 | 18 | 26 | 3 | 0 | 0 | 2 | 0 | 0 | 298 | 149 | 1.04E-06 | 1.48E-05 | 0.412 | 1 | 0.857 | 0.81 | 0.744 | 1 | 1 | 0.818 | 1 | 1 | -0.199 |
| RI | 349 | ENSG00000221946 | FXYD7 | chr19 | + | 35151253 | 35151482 | 35151253 | 35151328 | 35151439 | 35151482 | 2 | 3 | 5 | 1 | 0 | 6 | 8 | 9 | 0 | 11 | 7 | 16 | 10 | 5 | 4 | 0 | 1 | 10 | 0 | 0 | 260 | 149 | 1.08E-06 | 1.53E-05 | 0.16 | 0.177 | 0.241 | 1 | 0 | 1 | 0.902 | 0.364 | 1 | 1 | -0.538 |
| RI | 6883 | ENSG00000147140 | NONO | chrX | + | 71297376 | 71298508 | 71297376 | 71297938 | 71298468 | 71298508 | 86 | 69 | 113 | 133 | 63 | 248 | 135 | 239 | 259 | 216 | 287 | 191 | 224 | 223 | 132 | 363 | 244 | 188 | 212 | 333 | 298 | 149 | 1.13E-06 | 1.59E-05 | 0.148 | 0.204 | 0.191 | 0.204 | 0.127 | 0.283 | 0.281 | 0.373 | 0.345 | 0.165 | -0.115 |
| RI | 2883 | ENSG00000178035 | IMPDH2 | chr3 | - | 49025125 | 49026419 | 49025125 | 49025269 | 49026323 | 49026419 | 62 | 34 | 74 | 63 | 45 | 57 | 51 | 84 | 77 | 53 | 94 | 119 | 54 | 82 | 72 | 17 | 38 | 33 | 25 | 59 | 298 | 149 | 1.13E-06 | 1.59E-05 | 0.352 | 0.25 | 0.306 | 0.29 | 0.298 | 0.734 | 0.61 | 0.45 | 0.621 | 0.379 | -0.26 |
| RI | 5195 | ENSG00000115355 | CCDC88A | chr2 | - | 55295596 | 55296056 | 55295596 | 55295727 | 55295949 | 55296056 | 236 | 285 | 175 | 216 | 307 | 4 | 10 | 8 | 12 | 5 | 43 | 72 | 68 | 35 | 18 | 0 | 0 | 0 | 0 | 0 | 298 | 149 | 1.15E-06 | 1.62E-05 | 0.967 | 0.934 | 0.916 | 0.9 | 0.968 | 1 | 1 | 1 | 1 | 1 | -0.063 |
| RI | 2562 | ENSG00000145907 | G3BP1 | chr5 | + | 151795478 | 151797428 | 151795478 | 151795575 | 151797226 | 151797428 | 42 | 25 | 28 | 55 | 31 | 408 | 258 | 365 | 376 | 440 | 23 | 47 | 66 | 44 | 12 | 181 | 262 | 289 | 172 | 126 | 298 | 149 | 1.23E-06 | 1.72E-05 | 0.049 | 0.046 | 0.037 | 0.068 | 0.034 | 0.06 | 0.082 | 0.102 | 0.113 | 0.045 | -0.034 |
| RI | 5776 | ENSG00000110700 | RPS13 | chr11 | - | 17077428 | 17077667 | 17077428 | 17077477 | 17077618 | 17077667 | 34 | 25 | 28 | 44 | 29 | 1620 | 737 | 1769 | 1437 | 674 | 54 | 39 | 25 | 28 | 130 | 1324 | 1077 | 795 | 831 | 1614 | 290 | 149 | 1.33E-06 | 1.85E-05 | 0.011 | 0.017 | 0.008 | 0.015 | 0.022 | 0.021 | 0.018 | 0.016 | 0.017 | 0.04 | -0.008 |
| RI | 8175 | ENSG00000119977 | TCTN3 | chr10 | - | 95683100 | 95683629 | 95683100 | 95683180 | 95683521 | 95683629 | 17 | 12 | 24 | 12 | 35 | 1 | 2 | 4 | 1 | 4 | 22 | 17 | 3 | 8 | 10 | 0 | 0 | 0 | 0 | 0 | 298 | 149 | 1.39E-06 | 1.93E-05 | 0.895 | 0.75 | 0.75 | 0.857 | 0.814 | 1 | 1 | 1 | 1 | 1 | -0.187 |
| RI | 6948 | ENSG00000215301 | DDX3X | chrX | + | 41337407 | 41339083 | 41337407 | 41337465 | 41339035 | 41339083 | 126 | 123 | 117 | 162 | 127 | 1127 | 591 | 871 | 1034 | 437 | 81 | 127 | 140 | 234 | 20 | 177 | 530 | 614 | 413 | 178 | 298 | 149 | 1.41E-06 | 1.96E-05 | 0.053 | 0.094 | 0.063 | 0.073 | 0.127 | 0.186 | 0.107 | 0.102 | 0.221 | 0.053 | -0.052 |
| RI | 550 | ENSG00000125826 | RBCK1 | chr20 | + | 428490 | 429094 | 428490 | 428589 | 428950 | 429094 | 11 | 7 | 19 | 19 | 14 | 40 | 23 | 66 | 37 | 57 | 49 | 34 | 21 | 38 | 66 | 79 | 36 | 25 | 53 | 105 | 298 | 149 | 1.42E-06 | 1.97E-05 | 0.121 | 0.132 | 0.126 | 0.204 | 0.109 | 0.237 | 0.321 | 0.296 | 0.264 | 0.239 | -0.133 |
| RI | 7736 | ENSG00000175166 | PSMD2 | chr3 | + | 184303319 | 184303749 | 184303319 | 184303466 | 184303642 | 184303749 | 15 | 12 | 14 | 33 | 5 | 85 | 38 | 59 | 60 | 82 | 80 | 38 | 59 | 51 | 61 | 109 | 68 | 25 | 61 | 153 | 298 | 149 | 1.42E-06 | 1.97E-05 | 0.081 | 0.136 | 0.106 | 0.216 | 0.03 | 0.268 | 0.218 | 0.541 | 0.295 | 0.166 | -0.184 |
| RI | 420 | ENSG00000104856 | RELB | chr19 | + | 45034243 | 45034528 | 45034243 | 45034312 | 45034450 | 45034528 | 2 | 1 | 2 | 4 | 6 | 38 | 12 | 22 | 19 | 34 | 32 | 4 | 10 | 81 | 44 | 45 | 56 | 39 | 45 | 58 | 287 | 149 | 1.45E-06 | 2.00E-05 | 0.027 | 0.041 | 0.045 | 0.099 | 0.084 | 0.27 | 0.036 | 0.117 | 0.483 | 0.283 | -0.179 |
| RI | 3460 | ENSG00000144591 | GMPPA | chr2 | + | 219504082 | 219506072 | 219504082 | 219505761 | 219505979 | 219506072 | 0 | 5 | 10 | 2 | 2 | 21 | 11 | 13 | 19 | 9 | 0 | 0 | 0 | 3 | 0 | 15 | 29 | 15 | 21 | 33 | 298 | 149 | 1.52E-06 | 2.09E-05 | 0 | 0.185 | 0.278 | 0.05 | 0.1 | 0 | 0 | 0 | 0.067 | 0 | 0.109 |
| RI | 4547 | ENSG00000117410 | ATP6V0B | chr1 | + | 43976301 | 43976659 | 43976301 | 43976379 | 43976589 | 43976659 | 38 | 18 | 33 | 40 | 52 | 80 | 32 | 58 | 56 | 47 | 136 | 86 | 53 | 82 | 265 | 101 | 77 | 39 | 66 | 115 | 298 | 149 | 1.56E-06 | 2.15E-05 | 0.192 | 0.22 | 0.221 | 0.263 | 0.356 | 0.402 | 0.358 | 0.405 | 0.383 | 0.535 | -0.166 |
| RI | 1217 | ENSG00000090581 | GNPTG | chr16 | + | 1352101 | 1352306 | 1352101 | 1352159 | 1352238 | 1352306 | 3 | 0 | 1 | 4 | 2 | 17 | 1 | 17 | 18 | 8 | 12 | 8 | 28 | 3 | 32 | 5 | 9 | 5 | 29 | 9 | 228 | 149 | 1.61E-06 | 2.20E-05 | 0.103 | 0 | 0.037 | 0.127 | 0.14 | 0.611 | 0.367 | 0.785 | 0.063 | 0.699 | -0.424 |
| RI | 6264 | ENSG00000004059 | ARF5 | chr7 | + | 127589484 | 127590137 | 127589484 | 127589594 | 127590065 | 127590137 | 65 | 53 | 78 | 97 | 104 | 394 | 178 | 236 | 326 | 581 | 225 | 123 | 140 | 117 | 226 | 606 | 384 | 250 | 323 | 715 | 298 | 149 | 1.62E-06 | 2.21E-05 | 0.076 | 0.13 | 0.142 | 0.13 | 0.082 | 0.157 | 0.138 | 0.219 | 0.153 | 0.136 | -0.049 |
| RI | 6818 | ENSG00000090520 | DNAJB11 | chr3 | + | 186577667 | 186582077 | 186577667 | 186581513 | 186581994 | 186582077 | 42 | 41 | 37 | 49 | 83 | 64 | 41 | 75 | 71 | 44 | 0 | 45 | 73 | 16 | 2 | 66 | 76 | 118 | 78 | 37 | 298 | 149 | 1.63E-06 | 2.22E-05 | 0.247 | 0.333 | 0.198 | 0.257 | 0.485 | 0 | 0.228 | 0.236 | 0.093 | 0.026 | 0.187 |
| RI | 4240 | ENSG00000197121 | PGAP1 | chr2 | - | 196847002 | 196848037 | 196847002 | 196847200 | 196847946 | 196848037 | 3 | 10 | 7 | 13 | 4 | 1 | 5 | 9 | 4 | 6 | 13 | 0 | 7 | 20 | 3 | 0 | 2 | 0 | 0 | 0 | 298 | 149 | 1.63E-06 | 2.22E-05 | 0.6 | 0.5 | 0.28 | 0.619 | 0.25 | 1 | 0 | 1 | 1 | 1 | -0.35 |
| RI | 6844 | ENSG00000104921 | FCER2 | chr19 | - | 7698355 | 7698854 | 7698355 | 7698409 | 7698740 | 7698854 | 8 | 5 | 7 | 4 | 6 | 115 | 42 | 81 | 64 | 84 | 7 | 11 | 2 | 9 | 75 | 50 | 57 | 17 | 40 | 319 | 298 | 149 | 1.73E-06 | 2.35E-05 | 0.034 | 0.056 | 0.041 | 0.03 | 0.034 | 0.065 | 0.088 | 0.056 | 0.101 | 0.105 | -0.044 |
| RI | 6304 | ENSG00000170632 | ARMC10 | chr7 | + | 103083681 | 103084062 | 103083681 | 103083830 | 103084029 | 103084062 | 33 | 38 | 23 | 9 | 33 | 4 | 3 | 6 | 3 | 4 | 0 | 20 | 18 | 34 | 0 | 15 | 6 | 13 | 11 | 18 | 298 | 149 | 1.74E-06 | 2.35E-05 | 0.805 | 0.864 | 0.657 | 0.6 | 0.805 | 0 | 0.625 | 0.409 | 0.607 | 0 | 0.418 |
| RI | 4694 | ENSG00000273136 | NBPF26 | chr1 | + | 120805417 | 120805765 | 120805417 | 120805458 | 120805555 | 120805765 | 23 | 16 | 2 | 5 | 6 | 49 | 49 | 21 | 39 | 21 | 0 | 11 | 7 | 0 | 10 | 36 | 51 | 96 | 135 | 52 | 246 | 149 | 1.84E-06 | 2.49E-05 | 0.221 | 0.165 | 0.055 | 0.072 | 0.148 | 0 | 0.116 | 0.042 | 0 | 0.104 | 0.08 |
| RI | 5685 | ENSG00000077380 | DYNC1I2 | chr2 | + | 171725918 | 171726293 | 171725918 | 171726081 | 171726193 | 171726293 | 79 | 38 | 51 | 29 | 57 | 268 | 144 | 183 | 192 | 396 | 1 | 16 | 22 | 8 | 0 | 88 | 118 | 90 | 135 | 36 | 261 | 149 | 1.90E-06 | 2.56E-05 | 0.144 | 0.131 | 0.137 | 0.079 | 0.076 | 0.006 | 0.072 | 0.122 | 0.033 | 0 | 0.067 |
| RI | 5787 | ENSG00000119535 | CSF3R | chr1 | - | 36466043 | 36467311 | 36466043 | 36466827 | 36467229 | 36467311 | 11 | 8 | 6 | 13 | 10 | 225 | 103 | 114 | 224 | 157 | 72 | 42 | 69 | 34 | 32 | 573 | 235 | 321 | 425 | 523 | 298 | 149 | 1.94E-06 | 2.61E-05 | 0.024 | 0.037 | 0.026 | 0.028 | 0.031 | 0.059 | 0.082 | 0.097 | 0.038 | 0.03 | -0.032 |
| RI | 3588 | ENSG00000197343 | ZNF655 | chr7 | + | 99561887 | 99562494 | 99561887 | 99561973 | 99562367 | 99562494 | 167 | 77 | 114 | 193 | 97 | 104 | 52 | 72 | 74 | 95 | 195 | 237 | 249 | 195 | 253 | 77 | 95 | 47 | 48 | 57 | 298 | 149 | 1.99E-06 | 2.67E-05 | 0.445 | 0.425 | 0.442 | 0.566 | 0.338 | 0.559 | 0.555 | 0.726 | 0.67 | 0.689 | -0.197 |
| RI | 5897 | ENSG00000188846 | RPL14 | chr3 | + | 40461406 | 40461661 | 40461406 | 40461506 | 40461607 | 40461661 | 57 | 56 | 71 | 52 | 54 | 1885 | 966 | 2466 | 2075 | 2153 | 127 | 60 | 35 | 71 | 34 | 1548 | 1061 | 1200 | 1029 | 1418 | 250 | 149 | 2.02E-06 | 2.71E-05 | 0.018 | 0.033 | 0.017 | 0.015 | 0.015 | 0.047 | 0.033 | 0.017 | 0.039 | 0.014 | -0.01 |
| RI | 5681 | ENSG00000169247 | SH3TC2 | chr5 | - | 149006880 | 149009001 | 149006880 | 149007077 | 149008850 | 149009001 | 0 | 1 | 0 | 0 | 0 | 5 | 7 | 2 | 7 | 7 | 18 | 1 | 4 | 4 | 4 | 44 | 19 | 7 | 6 | 49 | 298 | 149 | 2.12E-06 | 2.83E-05 | 0 | 0.067 | 0 | 0 | 0 | 0.17 | 0.026 | 0.222 | 0.25 | 0.039 | -0.128 |
| RI | 889 | ENSG00000141295 | SCRN2 | chr17 | - | 47837691 | 47838450 | 47837691 | 47838002 | 47838269 | 47838450 | 35 | 15 | 34 | 50 | 21 | 6 | 9 | 18 | 7 | 4 | 31 | 40 | 11 | 24 | 33 | 0 | 1 | 5 | 0 | 0 | 298 | 149 | 2.13E-06 | 2.84E-05 | 0.745 | 0.455 | 0.486 | 0.781 | 0.724 | 1 | 0.952 | 0.524 | 1 | 1 | -0.257 |
| RI | 65 | ENSG00000182944 | EWSR1 | chr22 | + | 29298732 | 29299851 | 29298732 | 29299280 | 29299598 | 29299851 | 354 | 215 | 282 | 334 | 221 | 9 | 7 | 12 | 5 | 2 | 840 | 590 | 539 | 814 | 396 | 54 | 29 | 19 | 29 | 52 | 298 | 149 | 2.18E-06 | 2.90E-05 | 0.952 | 0.939 | 0.922 | 0.971 | 0.982 | 0.886 | 0.91 | 0.934 | 0.933 | 0.792 | 0.062 |
| [illegible] | | | | | | | | | | | | | | | | | | | | | | | | | | | | | | | | | | | | | | | | | | | | | | |

| | | | | | | | | | | | | | | |
|---|---|---|---|---|---|---|---|---|---|---|---|---|---|---|
| RI | 3470 | ENSG00000143793 | C1orf35 | chr1 | - | 228102079 | 228102382 | 228102079 | 228102165 | 228102309 | 228102382 | 3 | 0 | 9 |
| RI | 8330 | ENSG00000089094 | KDM2B | chr12 | - | 121444449 | 121445418 | 121444449 | 121444536 | 121445274 | 121445418 | 8 | 2 | 9 |
| RI | 4885 | ENSG00000115875 | SRSF7 | chr2 | - | 38748578 | 38749705 | 38748578 | 38748653 | 38749528 | 38749705 | 670 | 419 | 538 |
| RI | 7999 | ENSG00000150433 | TMEM218 | chr11 | - | 125102131 | 125102809 | 125102131 | 125102351 | 125102733 | 125102809 | 29 | 22 | 16 |
| RI | 3019 | ENSG00000169714 | CNBP | chr3 | - | 129171078 | 129171538 | 129171078 | 129171277 | 129171445 | 129171538 | 130 | 141 | 166 |
| RI | 2298 | ENSG00000170144 | HNRNPA3 | chr2 | + | 177216675 | 177216940 | 177216675 | 177216747 | 177216907 | 177216940 | 219 | 309 | 246 |
| RI | 8268 | ENSG00000082898 | XPO1 | chr2 | - | 61483936 | 61485962 | 61483936 | 61484105 | 61485767 | 61485962 | 30 | 47 | 38 |
| RI | 3589 | ENSG00000163818 | LZTFL1 | chr3 | - | 45834237 | 45835784 | 45834237 | 45834298 | 45835589 | 45835784 | 3 | 1 | 2 |
| RI | 3644 | ENSG00000239900 | ADSL | chr22 | + | 40360401 | 40361342 | 40360401 | 40360492 | 40361272 | 40361342 | 10 | 3 | 12 |
| RI | 6584 | ENSG00000196305 | IARS1 | chr9 | - | 92285721 | 92287910 | 92285721 | 92285839 | 92287790 | 92287910 | 20 | 18 | 26 |
| RI | 3813 | ENSG00000138074 | SLC5A6 | chr2 | - | 27202812 | 27203345 | 27202812 | 27202880 | 27203232 | 27203345 | 16 | 4 | 8 |
| RI | 5356 | ENSG00000167393 | PPP2R3B | chrX | - | 341306 | 341931 | 341306 | 341396 | 341882 | 341931 | 10 | 5 | 12 |
| RI | 249 | ENSG00000130816 | DNMT1 | chr19 | - | 10139675 | 10140328 | 10139675 | 10139817 | 10140045 | 10140328 | 3 | 2 | 22 |
| RI | 7115 | ENSG00000166946 | CCNDBP1 | chr15 | + | 43185819 | 43186233 | 43185819 | 43185879 | 43186153 | 43186233 | 50 | 13 | 61 |
| RI | 3809 | ENSG00000135469 | COQ10A | chr12 | + | 56269058 | 56269561 | 56269058 | 56269251 | 56269459 | 56269561 | 23 | 10 | 20 |
| RI | 5672 | ENSG00000123983 | ACSL3 | chr2 | + | 222916318 | 222918155 | 222916318 | 222916496 | 222918045 | 222918155 | 49 | 42 | 36 |
| RI | 8344 | ENSG00000116198 | CEP104 | chr1 | - | 3837291 | 3839119 | 3837291 | 3837519 | 3838963 | 3839119 | 11 | 6 | 8 |
| RI | 7278 | ENSG00000147789 | ZNF7 | chr8 | + | 144829042 | 144829604 | 144829042 | 144829090 | 144829444 | 144829604 | 14 | 4 | 10 |
| RI | 6434 | ENSG00000198556 | ZNF789 | chr7 | + | 99478273 | 99482326 | 99478273 | 99478356 | 99479660 | 99482326 | 21 | 19 | 11 |
| RI | 4030 | ENSG00000143543 | JTB | chr1 | - | 153976692 | 153977013 | 153976692 | 153976775 | 153976975 | 153977013 | 87 | 40 | 89 |
| RI | 701 | ENSG00000154889 | MPPE1 | chr18 | - | 11885675 | 11887025 | 11885675 | 11885816 | 11886916 | 11887025 | 70 | 36 | 66 |
| RI | 2852 | ENSG00000114857 | NKTR | chr3 | + | 42627199 | 42630575 | 42627199 | 42627227 | 42630545 | 42630575 | 166 | 160 | 117 |
| RI | 4379 | ENSG00000172531 | PPP1CA | chr11 | - | 67398721 | 67399163 | 67398721 | 67398856 | 67398939 | 67399163 | 20 | 2 | 12 |
| RI | 5064 | ENSG00000151092 | NGLY1 | chr3 | - | 25736308 | 25737455 | 25736308 | 25736400 | 25737333 | 25737455 | 87 | 66 | 59 |
| RI | 6758 | ENSG00000104870 | FCGRT | chr19 | + | 49513881 | 49514486 | 49513881 | 49514133 | 49514210 | 49514486 | 7 | 11 | 13 |
| RI | 5875 | ENSG00000111011 | RSRC2 | chr12 | - | 122518838 | 122521428 | 122518838 | 122519029 | 122521384 | 122521428 | 222 | 214 | 204 |
| RI | 953 | ENSG00000129657 | SEC14L1 | chr17 | + | 77213917 | 77216802 | 77213917 | 77214019 | 77216548 | 77216802 | 207 | 113 | 133 |
| RI | 6024 | ENSG00000124523 | SIRT5 | chr6 | + | 13599031 | 13600949 | 13599031 | 13599155 | 13600833 | 13600949 | 6 | 11 | 6 |
| RI | 3794 | ENSG00000167065 | DUSP18 | chr22 | - | 30661444 | 30664080 | 30661444 | 30663728 | 30663838 | 30664080 | 42 | 36 | 59 |
| RI | 6853 | ENSG00000215021 | PHB2 | chr12 | - | 6967170 | 6968021 | 6967170 | 6967779 | 6967891 | 6968021 | 17 | 14 | 39 |
| RI | 3919 | ENSG00000155463 | OXA1L | chr14 | + | 22770192 | 22770625 | 22770192 | 22770278 | 22770460 | 22770625 | 183 | 106 | 140 |
| RI | 3920 | ENSG00000055609 | KMT2C | chr7 | - | 152144712 | 152145295 | 152144712 | 152144881 | 152145152 | 152145295 | 36 | 32 | 21 |
| RI | 2300 | ENSG00000172785 | CBWD1 | chr9 | - | 121088 | 122090 | 121088 | 121573 | 121960 | 122090 | 17 | 52 | 10 |
| RI | 1411 | ENSG00000051523 | CYBA | chr16 | - | 88646115 | 88646838 | 88646115 | 88646197 | 88646754 | 88646838 | 35 | 23 | 47 |
| RI | 4577 | ENSG00000156675 | RAB11FIP1 | chr8 | - | 37870419 | 37873278 | 37870419 | 37870528 | 37871277 | 37873278 | 23 | 22 | 27 |
| RI | 5789 | ENSG00000119535 | CSF3R | chr1 | - | 36472516 | 36473622 | 36472516 | 36472686 | 36473434 | 36473622 | 44 | 29 | 40 |
| RI | 3612 | ENSG00000116353 | MECR | chr1 | - | 29195940 | 29196258 | 29195940 | 29196013 | 29196197 | 29196258 | 7 | 3 | 5 |
| RI | 2750 | ENSG00000103174 | NAGPA | chr16 | - | 5027134 | 5027379 | 5027134 | 5027198 | 5027277 | 5027379 | 7 | 2 | 2 |
| RI | 3117 | ENSG00000161204 | ABCF3 | chr3 | + | 184187663 | 184187983 | 184187663 | 184187761 | 184187860 | 184187983 | 20 | 32 | 26 |
| RI | 288 | ENSG00000142546 | NOSIP | chr19 | - | 49556548 | 49556993 | 49556548 | 49556657 | 49556874 | 49556993 | 76 | 24 | 49 |
| RI | 6727 | ENSG00000148229 | POLE3 | chr9 | - | 113409609 | 113410140 | 113409609 | 113409728 | 113410054 | 113410140 | 19 | 9 | 15 |
| RI | 4124 | ENSG00000171793 | CTPS1 | chr1 | + | 41008658 | 41008890 | 41008658 | 41008714 | 41008793 | 41008890 | 33 | 18 | 16 |
| RI | 6648 | ENSG00000178761 | FAM219B | chr15 | - | 74899991 | 74902063 | 74899991 | 74900137 | 74901949 | 74902063 | 63 | 66 | 71 |
| RI | 8413 | ENSG00000089692 | LAG3 | chr12 | + | 6777790 | 6778455 | 6777790 | 6777921 | 6778243 | 6778455 | 2 | 3 | 0 |
| RI | 2640 | ENSG00000203875 | SNHG5 | chr6 | - | 85677423 | 85677875 | 85677423 | 85677492 | 85677790 | 85677875 | 138 | 106 | 176 |
| RI | 7821 | ENSG00000110321 | EIF4G2 | chr11 | - | 10799551 | 10800348 | 10799551 | 10799756 | 10800089 | 10800348 | 186 | 147 | 157 |
| RI | 3631 | ENSG00000125637 | PSD4 | chr2 | + | 113185364 | 113186255 | 113185364 | 113185440 | 113185876 | 113186255 | 85 | 89 | 82 |
| RI | 7093 | ENSG00000143321 | HDGF | chr1 | - | 156745007 | 156745373 | 156745007 | 156745146 | 156745296 | 156745373 | 63 | 27 | 70 |
| RI | 8026 | ENSG00000090621 | PABPC4 | chr1 | - | 39562322 | 39564542 | 39562322 | 39563922 | 39564422 | 39564542 | 17 | 27 | 30 |
| RI | 4926 | ENSG00000117155 | SSX2IP | chr1 | - | 84662197 | 84662530 | 84662197 | 84662375 | 84662454 | 84662530 | 17 | 18 | 6 |
| RI | 267 | ENSG00000142541 | RPL13A | chr19 | + | 49490778 | 49491099 | 49490778 | 49490864 | 49491039 | 49491099 | 209 | 130 | 264 |
| RI | 2885 | ENSG00000178035 | IMPDH2 | chr3 | - | 49026518 | 49027047 | 49026518 | 49026609 | 49026686 | 49027047 | 37 | 23 | 36 |
| RI | 836 | ENSG00000108788 | MLX | chr17 | + | 42569203 | 42569606 | 42569203 | 42569303 | 42569506 | 42569606 | 97 | 50 | 54 |
| RI | 6062 | ENSG00000243649 | CFB | chr6 | + | 31950285 | 31950772 | 31950285 | 31950403 | 31950618 | 31950772 | 8 | 8 | 11 |
| RI | 8162 | ENSG00000150093 | ITGB1 | chr10 | - | 32932514 | 32933583 | 32932514 | 32932600 | 32933350 | 32933583 | 102 | 86 | 51 |
| RI | 5926 | ENSG00000114388 | NPRL2 | chr3 | - | 50348526 | 50348782 | 50348526 | 50348563 | 50348684 | 50348782 | 8 | 3 | 8 |
| RI | 2930 | ENSG00000188313 | PLSCR1 | chr3 | - | 146521543 | 146522053 | 146521543 | 146521705 | 146521832 | 146522053 | 106 | 138 | 58 |
| RI | 437 | ENSG00000187994 | RINL | chr19 | - | 38876330 | 38876781 | 38876330 | 38876474 | 38876692 | 38876781 | 25 | 3 | 18 |
| RI | 1574 | ENSG00000242028 | HYPK | chr15 | + | 43801517 | 43804427 | 43801517 | 43801569 | 43801710 | 43804427 | 24 | 10 | 20 |
| RI | 5274 | ENSG00000116809 | ZBTB17 | chr1 | - | 15943598 | 15943895 | 15943598 | 15943715 | 15943807 | 15943895 | 17 | 2 | 8 |
| RI | 2536 | ENSG00000049656 | CLPTM1L | chr5 | - | 1324762 | 1325816 | 1324762 | 1324813 | 1325750 | 1325816 | 20 | 15 | 18 |
| RI | 4677 | ENSG00000173846 | PLK3 | chr1 | + | 44802954 | 44803391 | 44802954 | 44803153 | 44803267 | 44803391 | 22 | 12 | 33 |
| RI | 1178 | ENSG00000125124 | BBS2 | chr16 | - | 56501352 | 56506036 | 56501352 | 56502808 | 56505949 | 56506036 | 22 | 23 | 23 |
| RI | 124 | ENSG00000100242 | SUN2 | chr22 | - | 38740266 | 38741050 | 38740266 | 38740432 | 38741006 | 38741050 | 68 | 28 | 75 |
| RI | 5614 | ENSG00000105325 | FZR1 | chr19 | + | 3526119 | 3526386 | 3526119 | 3526183 | 3526258 | 3526386 | 15 | 3 | 10 |
| RI | 8188 | ENSG00000099246 | RAB18 | chr10 | + | 27526827 | 27532579 | 27526827 | 27526889 | 27532506 | 27532579 | 117 | 126 | 123 |
| RI | 1913 | ENSG00000089737 | DDX24 | chr14 | - | 94079024 | 94079747 | 94079024 | 94079069 | 94079234 | 94079747 | 653 | 380 | 669 |
| RI | 6006 | ENSG00000103363 | ELOB | chr16 | - | 2771413 | 2772102 | 2771413 | 2771604 | 2771993 | 2772102 | 83 | 31 | 303 |
| RI | 6049 | ENSG00000135486 | HNRNPA1 | chr12 | + | 54281385 | 54281941 | 54281385 | 54281502 | 54281794 | 54281941 | 336 | 233 | 399 |
| RI | 1924 | ENSG00000125746 | EML2 | chr19 | - | 45638504 | 45638873 | 45638504 | 45638634 | 45638844 | 45638873 | 11 | 2 | 19 |
| RI | 4153 | ENSG00000126012 | KDM5C | chrX | - | 53198489 | 53198888 | 53198489 | 53198637 | 53198763 | 53198888 | 79 | 52 | 69 |
| RI | 2676 | ENSG00000070423 | RNF126 | chr19 | - | 648881 | 649748 | 648881 | 648975 | 649678 | 649748 | 18 | 7 | 12 |
| RI | 5918 | ENSG00000089818 | NECAP1 | chr12 | + | 8090194 | 8093055 | 8090194 | 8090299 | 8092871 | 8093055 | 203 | 155 | 182 |
| RI | 1994 | ENSG00000175061 | SNHG29 | chr17 | + | 16441073 | 16442018 | 16441073 | 16441130 | 16441356 | 16442018 | 35 | 16 | 28 |
| RI | 3772 | ENSG00000163346 | PBXIP1 | chr1 | - | 154951470 | 154951921 | 154951470 | 154951535 | 154951794 | 154951921 | 32 | 26 | 47 |
| RI | 5453 | ENSG00000143799 | PARP1 | chr1 | - | 226385503 | 226386442 | 226385503 | 226385680 | 226386325 | 226386442 | 5 | 3 | 10 |
| RI | 2499 | ENSG00000162413 | KLHL21 | chr1 | - | 6590723 | 6595557 | 6590723 | 6593658 | 6595484 | 6595557 | 25 | 13 | 21 |
| RI | 3160 | ENSG00000107404 | DVL1 | chr1 | - | 1339286 | 1339649 | 1339286 | 1339439 | 1339581 | 1339649 | 4 | 2 | 2 |
| RI | 7412 | ENSG00000107331 | ABCA2 | chr9 | - | 137010192 | 137010737 | 137010192 | 137010371 | 137010619 | 137010737 | 18 | 2 | 8 |
| RI | 7202 | ENSG00000104763 | ASAH1 | chr8 | - | 18064456 | 18067298 | 18064456 | 18064531 | 18067219 | 18067298 | 294 | 155 | 94 |
| RI | 901 | ENSG00000076604 | TRAF4 | chr17 | + | 28747842 | 28748178 | 28747842 | 28747947 | 28748016 | 28748178 | 7 | 1 | 10 |
| RI | 3020 | ENSG00000169714 | CNBP | chr3 | - | 129171078 | 129171538 | 129171078 | 129171280 | 129171445 | 129171538 | 118 | 143 | 158 |
| RI | 2419 | ENSG00000214655 | ZSWIM8 | chr10 | + | 73800639 | 73801195 | 73800639 | 73800759 | 73801016 | 73801195 | 13 | 6 | 7 |
| RI | 1870 | ENSG00000105122 | RASAL3 | chr19 | - | 15451623 | 15452815 | 15451623 | 15451938 | 15452044 | 15452815 | 8 | 0 | 5 |
| RI | 6777 | ENSG00000072778 | ACADVL | chr17 | + | 7220121 | 7220676 | 7220121 | 7220197 | 7220603 | 7220676 | 21 | 8 | 10 |
| RI | 4815 | ENSG00000167671 | UBXN6 | chr19 | - | 4446499 | 4446920 | 4446499 | 4446719 | 4446835 | 4446920 | 13 | 14 | 19 |
| RI | 4738 | ENSG00000164828 | SUN1 | chr7 | + | 852608 | 852952 | 852608 | 852667 | 852812 | 852952 | 58 | 29 | 56 |
| RI | 6198 | ENSG00000120314 | WDR55 | chr5 | + | 140668890 | 140669248 | 140668890 | 140668990 | 140669078 | 140669248 | 21 | 10 | 35 |
| RI | 7431 | ENSG00000070061 | ELP1 | chr9 | - | 108916203 | 108919349 | 108916203 | 108916297 | 108917546 | 108919349 | 7 | 4 | 4 |
| RI | 5639 | ENSG00000163516 | ANKZF1 | chr2 | + | 219234132 | 219235312 | 219234132 | 219234288 | 219234825 | 219235312 | 273 | 203 | 249 |
| RI | 217 | ENSG00000160953 | PWWP3A | chr19 | + | 1357008 | 1358464 | 1357008 | 1357094 | 1358393 | 1358464 | 13 | 17 | 23 |
| RI | 2001 | ENSG00000196655 | TRAPPC4 | chr11 | + | 119018765 | 119019317 | 119018765 | 119018970 | 119019142 | 119019317 | 4 | 3 | 11 |
| RI | 4040 | ENSG00000138095 | LRPPRC | chr2 | - | 43976142 | 43977052 | 43976142 | 43976229 | 43976993 | 43977052 | 58 | 83 | 58 |
| RI | 3344 | ENSG00000158864 | NDUFS2 | chr1 | + | 161209482 | 161210188 | 161209482 | 161209931 | 161210110 | 161210188 | 37 | 16 | 35 |
| RI | 6363 | ENSG00000012124 | CD22 | chr19 | + | 35332018 | 35332924 | 35332018 | 35332074 | 35332552 | 35332924 | 39 | 46 | 38 |
| RI | 5400 | ENSG00000126768 | TIMM17B | chrX | - | 48893899 | 48894225 | 48893899 | 48894010 | 48894096 | 48894225 | 14 | 4 | 16 |
| RI | 1983 | ENSG00000143379 | SETDB1 | chr1 | + | 150962973 | 150963741 | 150962973 | 150963139 | 150963529 | 150963741 | 23 | 18 | 28 |
| RI | 5122 | ENSG00000044574 | HSPA5 | chr9 | - | 125239420 | 125240309 | 125239420 | 125239533 | 125240171 | 125240309 | 155 | 108 | 175 |
| RI | 2796 | ENSG00000163703 | CRELD1 | chr3 | + | 9933829 | 9934612 | 9933829 | 9933920 | 9934419 | 9934612 | 35 | 16 | 29 |
| RI | 5747 | ENSG00000163214 | DHX57 | chr2 | - | 38854053 | 38855252 | 38854053 | 38854178 | 38855056 | 38855252 | 26 | 12 | 11 |
| RI | 3294 | ENSG00000165801 | ARHGEF40 | chr14 | + | 21080872 | 21082119 | 21080872 | 21081016 | 21081508 | 21082119 | 2 | 2 | 8 |
| RI | 1838 | ENSG00000099797 | TECR | chr19 | + | 14564785 | 14564992 | 14564785 | 14564858 | 14564948 | 14564992 | 5 | 1 | 5 |
| RI | 8370 | ENSG00000139190 | VAMP1 | chr12 | - | 6462236 | 6464941 | 6462236 | 6462842 | 6464889 | 6464941 | 190 | 141 | 275 |
| RI | 2252 | ENSG00000161960 | EIF4A1 | chr17 | + | 7574259 | 7574678 | 7574259 | 7574308 | 7574545 | 7574678 | 89 | 90 | 84 |
| RI | 3924 | ENSG00000055609 | KMT2C | chr7 | - | 152154266 | 152156057 | 152154266 | 152154445 | 152155909 | 152156057 | 33 | 30 | 18 |
| RI | 6862 | ENSG00000109111 | SUPT6H | chr17 | + | 28700345 | 28701128 | 28700345 | 28700512 | 28700940 | 28701128 | 6 | 4 | 7 |
| RI | 4291 | ENSG00000155363 | MOV10 | chr1 | + | 112698286 | 112698789 | 112698286 | 112698478 | 112698714 | 112698789 | 12 | 3 | 15 |
| RI | 3855 | ENSG00000175390 | EIF3F | chr11 | + | 7994425 | 7995118 | 7994425 | 7994517 | 7994981 | 7995118 | 41 | 34 | 36 |
| RI | 5097 | ENSG00000166986 | MARS1 | chr12 | + | 57516244 | 57516655 | 57516244 | 57516337 | 57516434 | 57516655 | 18 | 18 | 16 |
| RI | 2272 | ENSG00000103496 | STX4 | chr16 | + | 31034203 | 31034536 | 31034203 | 31034325 | 31034461 | 31034536 | 46 | 19 | 47 |
| RI | 4701 | ENSG00000089127 | OAS1 | chr12 | + | 112911050 | 112917700 | 112911050 | 112911235 | 112916508 | 112917700 | 25 | 18 | 3 |
| RI | 4800 | ENSG00000137101 | CD72 | chr9 | - | 35615942 | 35616689 | 35615942 | 35616386 | 35616599 | 35616689 | 19 | 8 | 10 |
| RI | 6537 | ENSG00000106263 | EIF3B | chr7 | + | 2379393 | 2380740 | 2379393 | 2379511 | 2380278 | 2380740 | 85 | 37 | 76 |
| RI | 6900 | ENSG00000163682 | RPL9 | chr4 | - | 39458193 | 39458440 | 39458193 | 39458309 | 39458393 | 39458440 | 74 | 52 | 78 |
| RI | 2392 | ENSG00000181929 | PRKAG1 | chr12 | - | 49004506 | 49005018 | 49004506 | 49004633 | 49004963 | 49005018 | 8 | 6 | 7 |
| RI | 600 | ENSG00000215440 | NPEPL1 | chr20 | + | 58713419 | 58714093 | 58713419 | 58713543 | 58713916 | 58714093 | 18 | 6 | 13 |
| RI | 3368 | ENSG00000186517 | ARHGAP30 | chr1 | - | 161052285 | 161052543 | 161052285 | 161052363 | 161052439 | 161052543 | 32 | 24 | 41 |
| RI | 7578 | ENSG00000182208 | MOB2 | chr11 | - | 1480392 | 1480885 | 1480392 | 1480486 | 1480724 | 1480885 | 21 | 4 | 26 |
| RI | 4037 | ENSG00000138095 | LRPPRC | chr2 | - | 43957384 | 43960634 | 43957384 | 43957451 | 43960540 | 43960634 | 53 | 46 | 54 |
| RI | 3187 | ENSG00000168028 | RPSA | chr3 | + | 39411895 | 39412434 | 39411895 | 39412061 | 39412273 | 39412434 | 56 | 45 | 66 |
| RI | 4475 | ENSG00000141456 | PELP1 | chr17 | - | 4675273 | 4675884 | 4675273 | 4675362 | 4675796 | 4675884 | 4 | 6 | 8 |
| RI | 5211 | ENSG00000152291 | TGOLN2 | chr2 | - | 85326507 | 85327685 | 85326507 | 85326804 | 85326978 | 85327685 | 528 | 303 | 401 |
| RI | 5235 | ENSG00000138434 | ITPRID2 | chr2 | + | 181915215 | 181918675 | 181915215 | 181916427 | 181918597 | 181918675 | 56 | 63 | 47 |
| RI | 4662 | ENSG00000001497 | LAS1L | chrX | - | 65523559 | 65524614 | 65523559 | 65523707 | 65524055 | 65524614 | 7 | 5 | 7 |
| RI | 3379 | ENSG00000182866 | LCK | chr1 | + | 32279670 | 32279994 | 32279670 | 32279747 | 32279840 | 32279994 | 31 | 9 | 55 |
| RI | 2184 | ENSG00000113240 | CLK4 | chr5 | - | 178617343 | 178618778 | 178617343 | 178617434 | 178618555 | 178618778 | 57 | 62 | 72 |
| RI | 5633 | ENSG00000178719 | GRINA | chr8 | + | 143991691 | 143992373 | 143991691 | 143991804 | 143992244 | 143992373 | 13 | 3 | 10 |
| RI | 1928 | ENSG00000029364 | SLC39A9 | chr14 | + | 69458362 | 69462381 | 69458362 | 69458729 | 69461696 | 69462381 | 202 | 118 | 162 |
| RI | 1058 | ENSG00000073584 | SMARCE1 | chr17 | - | 40631591 | 40636102 | 40631591 | 40632367 | 40635930 | 40636102 | 28 | 25 | 32 |
| RI | 6465 | ENSG00000113013 | HSPA9 | chr5 | - | 138556773 | 138557496 | 138556773 | 138556866 | 138557401 | 138557496 | 113 | 110 | 108 |
| RI | 8198 | ENSG00000023041 | ZDHHC6 | chr10 | - | 112433239 | 112434464 | 112433239 | 112433281 | 112434296 | 112434464 | 169 | 156 | 192 |
| RI | 6442 | ENSG00000110497 | AMBRA1 | chr11 | - | 46541944 | 46543398 | 46541944 | 46542981 | 46543251 | 46543398 | 17 | 9 | 14 |
| RI | 1947 | ENSG00000103811 | CTSH | chr15 | - | 78931450 | 78935082 | 78931450 | 78932458 | 78934977 | 78935082 | 19 | 40 | 33 |
| RI | 3524 | ENSG00000126107 | HECTD3 | chr1 | - | 45005793 | 45006116 | 45005793 | 45005883 | 45005996 | 45006116 | 18 | 10 | 28 |
| RI | 5023 | ENSG00000112651 | MRPL2 | chr6 | - | 43055896 | 43056196 | 43055896 | 43056007 | 43056080 | 43056196 | 4 | 12 | 12 |
| RI | 5301 | ENSG00000197037 | ZSCAN25 | chr7 | + | 99619560 | 99621574 | 99619560 | 99619997 | 99621372 | 99621574 | 43 | 28 | 18 |
| RI | 4858 | ENSG00000149532 | CPSF7 | chr11 | - | 61420469 | 61421608 | 61420469 | 61420573 | 61421389 | 61421608 | 15 | 7 | 18 |
| RI | 7813 | ENSG00000255717 | SNHG1 | chr11 | - | 62855423 | 62855885 | 62855423 | 62855466 | 62855864 | 62855885 | 110 | 51 | 164 |
| RI | 371 | ENSG00000105193 | RPS16 | chr19 | - | 39433222 | 39433569 | 39433222 | 39433418 | 39433521 | 39433569 | 44 | 12 | 59 |
| RI | 5067 | ENSG00000167595 | PROSER3 | chr19 | + | 35766767 | 35768065 | 35766767 | 35766955 | 35767071 | 35768065 | 2 | 1 | 1 |
| RI | 7492 | ENSG00000213930 | GALT | chr9 | + | 34647088 | 34647567 | 34647088 | 34647258 | 34647540 | 34647567 | 64 | 43 | 65 |
| RI | 877 | ENSG00000136448 | NMT1 | chr17 | + | 45103708 | 45104996 | 45103708 | 45103876 | 45104858 | 45104996 | 10 | 11 | 12 |
| RI | 1033 | ENSG00000182087 | TMEM259 | chr19 | - | 1012065 | 1012573 | 1012065 | 1012188 | 1012462 | 1012573 | 66 | 29 | 81 |
| RI | 3459 | ENSG00000144591 | GMPPA | chr2 | + | 219505457 | 219505761 | 219505457 | 219505555 | 219505714 | 219505761 | 22 | 5 | 16 |
| RI | 6716 | ENSG00000094631 | HDAC6 | chrX | + | 48805630 | 48806464 | 48805630 | 48805671 | 48806367 | 48806464 | 20 | 4 | 7 |
| RI | 2911 | ENSG00000163930 | BAP1 | chr3 | - | 52402778 | 52403298 | 52402778 | 52402871 | 52403137 | 52403298 | 52 | 22 | 31 |
| RI | 3670 | ENSG00000090863 | GLG1 | chr16 | - | 74462487 | 74463479 | 74462487 | 74462630 | 74463355 | 74463479 | 56 | 35 | 43 |
| RI | 3137 | ENSG00000164463 | CREBRF | chr5 | + | 173108623 | 173110711 | 173108623 | 173108818 | 173110521 | 173110711 | 27 | 43 | 30 |
| RI | 6897 | ENSG00000166707 | ZCCHC18 | chrX | + | 104113433 | 104115842 | 104113433 | 104113573 | 104115251 | 104115842 | 11 | 10 | 15 |
| RI | 4636 | ENSG00000197323 | TRIM33 | chr1 | - | 114397607 | 114397990 | 114397607 | 114397860 | 114397939 | 114397990 | 28 | 32 | 20 |
| RI | 3115 | ENSG00000161204 | ABCF3 | chr3 | + | 184186506 | 184186875 | 184186506 | 184186654 | 184186795 | 184186875 | 25 | 16 | 22 |
| RI | 2633 | ENSG00000100650 | SRSF5 | chr14 | + | 69768797 | 69770540 | 69768797 | 69769251 | 69770466 | 69770540 | 2274 | 2548 | 2168 |
| RI | 3297 | ENSG00000198925 | ATG9A | chr2 | - | 219227769 | 219228053 | 219227769 | 219227813 | 219227921 | 219228053 | 12 | 7 | 14 |
| RI | 3472 | ENSG00000177954 | RPS27 | chr1 | + | 153990762 | 153991034 | 153990762 | 153990802 | 153990913 | 153991034 | 141 | 115 | 237 |
| RI | 6041 | ENSG00000137312 | FLOT1 | chr6 | - | 30741189 | 30741704 | 30741189 | 30741333 | 30741613 | 30741704 | 54 | 48 | 51 |
| RI | 6053 | ENSG00000135486 | HNRNPA1 | chr12 | + | 54283078 | 54283234 | 54283078 | 54283093 | 54283207 | 54283234 | 548 | 483 | 572 |
| RI | 5164 | ENSG00000198755 | RPL10A | chr6 | + | 35468946 | 35469529 | 35468946 | 35469285 | 35469380 | 35469529 | 71 | 30 | 102 |
| RI | 4361 | ENSG00000158941 | CCAR2 | chr8 | + | 22619149 | 22620468 | 22619149 | 22619355 | 22619637 | 22620468 | 13 | 25 | 33 |
| RI | 3635 | ENSG00000114861 | FOXP1 | chr3 | - | 70972010 | 70972676 | 70972010 | 70972180 | 70972554 | 70972676 | 21 | 40 | 38 |
| RI | 1616 | ENSG00000284946 | NA | chr15 | - | 90980883 | 90981669 | 90980883 | 90981033 | 90981498 | 90981669 | 10 | 13 | 10 |
| RI | 5111 | ENSG00000181163 | NPM1 | chr5 | + | 171392709 | 171392978 | 171392709 | 171392816 | 171392913 | 171392978 | 86 | 93 | 118 |
| RI | 4625 | ENSG00000130775 | THEMIS2 | chr1 | + | 27881970 | 27883043 | 27881970 | 27882077 | 27882512 | 27883043 | 398 | 258 | 232 |
| RI | 481 | ENSG00000123146 | ADGRE5 | chr19 | + | 14397732 | 14397935 | 14397732 | 14397804 | 14397888 | 14397935 | 9 | 27 | 5 |
| RI | 936 | ENSG00000132475 | H3-3B | chr17 | - | 75778809 | 75779184 | 75778809 | 75778963 | 75779046 | 75779184 | 79 | 30 | 115 |
| RI | 6711 | ENSG00000137801 | THBS1 | chr15 | + | 39593396 | 39594196 | 39593396 | 39593668 | 39594098 | 39594196 | 21 | 3 | 16 |
| RI | 2615 | ENSG00000214174 | AMZ2P1 | chr17 | - | 64972753 | 64973517 | 64972753 | 64972927 | 64973241 | 64973517 | 17 | 14 | 18 |
| RI | 2324 | ENSG00000185627 | PSMD13 | chr11 | + | 244040 | 244469 | 244040 | 244216 | 244425 | 244469 | 34 | 46 | 17 |
| RI | 3726 | ENSG00000142208 | AKT1 | chr14 | - | 104772877 | 104773379 | 104772877 | 104773092 | 104773250 | 104773379 | 10 | 0 | 1 |
| RI | 2253 | ENSG00000161960 | EIF4A1 | chr17 | + | 7574259 | 7574678 | 7574259 | 7574308 | 7574547 | 7574678 | 90 | 88 | 86 |
| RI | 1666 | ENSG00000140474 | ULK3 | chr15 | - | 74840233 | 74840641 | 74840233 | 74840316 | 74840497 | 74840641 | 46 | 17 | 86 |
| RI | 4004 | ENSG00000197530 | MIB2 | chr1 | + | 1628018 | 1628399 | 1628018 | 1628179 | 1628272 | 1628399 | 57 | 10 | 18 |
| RI | 6139 | ENSG00000118515 | SGK1 | chr6 | - | 134173273 | 134173566 | 134173273 | 134173357 | 134173461 | 134173566 | 82 | 50 | 95 |
| RI | 8481 | ENSG00000166598 | HSP90B1 | chr12 | + | 103931520 | 103932418 | 103931520 | 103931623 | 103932278 | 103932418 | 100 | 62 | 127 |
| RI | 931 | ENSG00000180891 | CUEDC1 | chr17 | - | 57866473 | 57867415 | 57866473 | 57866544 | 57867356 | 57867415 | 9 | 15 | 9 |
| RI | 1119 | ENSG00000177731 | FLII | chr17 | - | 18245554 | 18246062 | 18245554 | 18245660 | 18245837 | 18246062 | 171 | 85 | 133 |
| RI | 3967 | ENSG00000163479 | SSR2 | chr1 | - | 156011809 | 156012565 | 156011809 | 156011887 | 156012384 | 156012565 | 22 | 6 | 8 |
| RI | 417 | ENSG00000105369 | CD79A | chr19 | + | 41878989 | 41879653 | 41878989 | 41879175 | 41879534 | 41879653 | 125 | 35 | 99 |
| RI | 8425 | ENSG00000135124 | P2RX4 | chr12 | + | 121222093 | 121223043 | 121222093 | 121222166 | 121222946 | 121223043 | 25 | 20 | 17 |
| RI | 4071 | ENSG00000178498 | DTX3 | chr12 | + | 57605988 | 57606252 | 57605988 | 57606029 | 57606189 | 57606252 | 13 | 14 | 18 |
| RI | 3249 | ENSG00000064961 | HMG20B | chr19 | + | 3573691 | 3574586 | 3573691 | 3573800 | 3574369 | 3574586 | 5 | 12 | 13 |
| RI | 4140 | ENSG00000154222 | CC2D1B | chr1 | - | 52357780 | 52358461 | 52357780 | 52357916 | 52358330 | 52358461 | 28 | 23 | 30 |
| RI | 2449 | ENSG00000125089 | SH3TC1 | chr4 | + | 8236277 | 8237670 | 8236277 | 8236428 | 8237473 | 8237670 | 7 | 9 | 6 |
| RI | 4236 | ENSG00000197785 | ATAD3A | chr1 | + | 1517310 | 1517775 | 1517310 | 1517412 | 1517715 | 1517775 | 1 | 0 | 0 |
| RI | 5609 | ENSG00000144199 | FAHD2B | chr2 | - | 97085698 | 97086198 | 97085698 | 97085861 | 97086138 | 97086198 | 6 | 3 | 3 |
| RI | 3651 | ENSG00000177728 | TMEM94 | chr17 | + | 75498443 | 75498722 | 75498443 | 75498538 | 75498628 | 75498722 | 0 | 1 | 3 |
| RI | 7756 | ENSG00000174851 | YIF1A | chr11 | - | 66284579 | 66284966 | 66284579 | 66284783 | 66284872 | 66284966 | 4 | 0 | 4 |
| RI | 1908 | ENSG00000182979 | MTA1 | chr14 | + | 105464462 | 105464863 | 105464462 | 105464567 | 105464673 | 105464863 | 14 | 14 | 4 |
| RI | 3279 | ENSG00000163002 | NUP35 | chr2 | + | 183159487 | 183161680 | 183159487 | 183159652 | 183161053 | 183161680 | 0 | 2 | 3 |
| RI | 5187 | ENSG00000135945 | REV1 | chr2 | - | 99402243 | 99402800 | 99402243 | 99402346 | 99402643 | 99402800 | 76 | 86 | 91 |
| RI | 2917 | ENSG00000221978 | CCNL2 | chr1 | - | 1390765 | 1392803 | 1390765 | 1390865 | 1392678 | 1392803 | 250 | 192 | 336 |
| RI | 5671 | ENSG00000115234 | SNX17 | chr2 | + | 27374688 | 27375153 | 27374688 | 27374758 | 27375060 | 27375153 | 25 | 19 | 32 |
| RI | 2119 | ENSG00000172262 | ZNF131 | chr5 | + | 43161248 | 43161931 | 43161248 | 43161613 | 43161708 | 43161931 | 144 | 91 | 166 |
| RI | 1988 | ENSG00000175061 | SNHG29 | chr17 | + | 16439326 | 16439703 | 16439326 | 16439414 | 16439659 | 16439703 | 184 | 94 | 232 |
| RI | 3295 | ENSG00000165801 | ARHGEF40 | chr14 | + | 21087000 | 21087463 | 21087000 | 21087105 | 21087319 | 21087463 | 44 | 25 | 35 |
| RI | 3172 | ENSG00000233452 | STXBP5-AS1 | chr6 | - | 146850824 | 146851331 | 146850824 | 146851023 | 146851117 | 146851331 | 9 | 2 | 2 |
| RI | 5603 | ENSG00000102125 | TAZ | chrX | + | 154413206 | 154413567 | 154413206 | 154413358 | 154413481 | 154413567 | 8 | 3 | 2 |
| RI | 1004 | ENSG00000100360 | IFT27 | chr22 | - | 36758210 | 36764036 | 36758210 | 36758409 | 36763918 | 36764036 | 34 | 35 | 32 |
| RI | 2095 | ENSG00000141279 | NPEPPS | chr17 | + | 47585499 | 47586398 | 47585499 | 47585700 | 47586364 | 47586398 | 25 | 26 | 36 |
| RI | 386 | ENSG00000129347 | KRI1 | chr19 | - | 10564928 | 10565790 | 10564928 | 10565034 | 10565716 | 10565790 | 3 | 3 | 2 |
| RI | 5917 | ENSG00000089818 | NECAP1 | chr12 | + | 8090194 | 8092784 | 8090194 | 8091850 | 8092675 | 8092784 | 215 | 163 | 167 |
| RI | 1995 | ENSG00000175061 | SNHG29 | chr17 | + | 16441073 | 16441502 | 16441073 | 16441130 | 16441367 | 16441502 | 70 | 28 | 69 |
| RI | 844 | ENSG00000173757 | STAT5B | chr17 | - | 42210170 | 42210497 | 42210170 | 42210301 | 42210402 | 42210497 | 6 | 10 | 19 |
| RI | 7991 | ENSG00000175224 | ATG13 | chr11 | + | 46668498 | 46668910 | 46668498 | 46668576 | 46668793 | 46668910 | 19 | 6 | 10 |
| RI | 5089 | ENSG00000166986 | MARS1 | chr12 | + | 57489018 | 57489345 | 57489018 | 57489109 | 57489266 | 57489345 | 10 | 10 | 20 |
| RI | 5758 | ENSG00000074582 | BCS1L | chr2 | + | 218662196 | 218662679 | 218662196 | 218662260 | 218662509 | 218662679 | 5 | 13 | 2 |
| RI | 8501 | ENSG00000139613 | SMARCC2 | chr12 | - | 56180976 | 56181597 | 56180976 | 56181101 | 56181481 | 56181597 | 63 | 42 | 55 |
| RI | 369 | ENSG00000205744 | DENND1C | chr19 | - | 6476856 | 6477127 | 6476856 | 6476967 | 6477073 | 6477127 | 8 | 1 | 10 |
| RI | 7699 | ENSG00000168067 | MAP4K2 | chr11 | - | 64791908 | 64792422 | 64791908 | 64792086 | 64792171 | 64792422 | 3 | 1 | 2 |
| RI | 3147 | ENSG00000169715 | MT1E | chr16 | + | 56626465 | 56627112 | 56626465 | 56626531 | 56626879 | 56627112 | 1 | 0 | 2 |

| | | | | | | | | | | | | | | | | | | | | | | | | | | | | | | | | | | | | | | | | | | | | | | |
|---|---|---|---|---|---|---|---|---|---|---|---|---|---|---|---|---|---|---|---|---|---|---|---|---|---|---|---|---|---|---|---|---|---|---|---|---|---|---|---|---|---|---|---|---|---|---|
| RI | 7018 | ENSG00000182196 | ARL6IP4 | chr12 | + | 122982468 | 122982908 | 122982468 | 122982538 | 122982619 | 122982908 | 57 | 72 | 120 | 118 | 76 | 190 | 69 | 142 | 126 | 68 | 113 | 31 | 47 | 35 | 140 | 202 | 132 | 110 | 202 | 217 | 230 | 149 | 0.000265604 | 0.001881201 | 0.163 | 0.403 | 0.354 | 0.378 | 0.42 | 0.266 | 0.132 | 0.217 | 0.101 | 0.295 | 0.141 |
| RI | 1219 | ENSG00000090581 | GNPTG | chr16 | + | 1362451 | 1362742 | 1362451 | 1362534 | 1362610 | 1362742 | 1 | 1 | 7 | 12 | 5 | 49 | 21 | 37 | 38 | 10 | 0 | 3 | 6 | 3 | 3 | 157 | 32 | 30 | 32 | 95 | 225 | 149 | 0.000266499 | 0.001885279 | 0.013 | 0.031 | 0.111 | 0.173 | 0.249 | 0 | 0.058 | 0.117 | 0.058 | 0.02 | 0.065 |
| RI | 5470 | ENSG00000223960 | CHROMR | chr2 | + | 178426365 | 178430889 | 178426365 | 178426505 | 178430694 | 178430889 | 65 | 28 | 34 | 41 | 38 | 11 | 10 | 5 | 9 | 11 | 48 | 35 | 47 | 32 | 11 | 0 | 8 | 5 | 0 | 0 | 298 | 149 | 0.000268116 | 0.00189445 | 0.747 | 0.583 | 0.773 | 0.695 | 0.633 | 1 | 0.686 | 0.825 | 1 | 1 | -0.216 |
| RI | 112 | ENSG00000231711 | LINC00899 | chr22 | - | 46042573 | 46043699 | 46042573 | 46042640 | 46043562 | 46043699 | 0 | 0 | 1 | 1 | 1 | 3 | 1 | 0 | 6 | 3 | 12 | 2 | 1 | 5 | 0 | 1 | 2 | 2 | 2 | 6 | 298 | 149 | 0.000271484 | 0.001915957 | 0 | 0 | 1 | 0.077 | 0.143 | 0.857 | 0.333 | 0.2 | 0.556 | 0 | -0.145 |
| RI | 1598 | ENSG00000102901 | CENPT | chr16 | - | 67832227 | 67832545 | 67832227 | 67832315 | 67832454 | 67832545 | 7 | 6 | 11 | 18 | 5 | 39 | 32 | 30 | 58 | 58 | 71 | 26 | 20 | 8 | 34 | 14 | 37 | 22 | 25 | 120 | 288 | 149 | 0.000274125 | 0.001932289 | 0.085 | 0.088 | 0.159 | 0.138 | 0.043 | 0.724 | 0.267 | 0.32 | 0.142 | 0.128 | -0.214 |
| RI | 2241 | ENSG00000104886 | PLEKHJ1 | chr19 | - | 2235761 | 2235990 | 2235761 | 2235828 | 2235922 | 2235990 | 0 | 0 | 0 | 5 | 0 | 93 | 26 | 53 | 52 | 27 | 17 | 2 | 1 | 0 | 0 | 59 | 38 | 19 | 59 | 37 | 243 | 149 | 0.000274937 | 0.0019357 | 0 | 0 | 0 | 0.056 | 0 | 0.15 | 0.031 | 0.031 | 0 | 0 | -0.031 |
| RI | 2211 | ENSG00000166888 | STAT6 | chr12 | - | 57105467 | 57106339 | 57105467 | 57105599 | 57106190 | 57106339 | 47 | 24 | 39 | 54 | 37 | 319 | 209 | 286 | 309 | 219 | 64 | 19 | 26 | 25 | 75 | 682 | 370 | 197 | 296 | 645 | 298 | 149 | 0.000275483 | 0.001937234 | 0.069 | 0.054 | 0.064 | 0.08 | 0.078 | 0.045 | 0.025 | 0.062 | 0.041 | 0.055 | 0.023 |
| RI | 5026 | ENSG00000115525 | ST3GAL5 | chr2 | - | 85847860 | 85848204 | 85847860 | 85847984 | 85848115 | 85848204 | 229 | 251 | 239 | 330 | 356 | 3 | 5 | 2 | 1 | 0 | 753 | 343 | 213 | 307 | 257 | 14 | 3 | 4 | 10 | 8 | 280 | 149 | 0.000277068 | 0.001943789 | 0.976 | 0.964 | 0.985 | 0.994 | 1 | 0.966 | 0.984 | 0.966 | 0.942 | 0.945 | 0.023 |
| RI | 6121 | ENSG00000112511 | PHF1 | chr6 | + | 33414724 | 33415144 | 33414724 | 33414829 | 33414954 | 33415144 | 37 | 19 | 36 | 61 | 39 | 72 | 53 | 67 | 53 | 100 | 45 | 54 | 50 | 59 | 78 | 52 | 87 | 31 | 34 | 36 | 274 | 149 | 0.000277073 | 0.001943789 | 0.218 | 0.163 | 0.226 | 0.385 | 0.175 | 0.32 | 0.252 | 0.467 | 0.486 | 0.541 | -0.18 |
| RI | 4568 | ENSG00000122678 | POLM | chr7 | - | 44073782 | 44074530 | 44073782 | 44074025 | 44074397 | 44074530 | 16 | 5 | 26 | 15 | 18 | 4 | 3 | 6 | 1 | 9 | 19 | 39 | 16 | 24 | 39 | 0 | 3 | 5 | 0 | 0 | 298 | 149 | 0.000277906 | 0.00194732 | 0.667 | 0.455 | 0.684 | 0.882 | 0.5 | 1 | 0.867 | 0.615 | 1 | 1 | -0.259 |
| RI | 2356 | ENSG00000011243 | AKAP8L | chr19 | - | 15397713 | 15400358 | 15397713 | 15397855 | 15400294 | 15400358 | 51 | 18 | 35 | 51 | 11 | 0 | 0 | 0 | 1 | 0 | 59 | 49 | 69 | 112 | 112 | 25 | 1 | 0 | 1 | 0 | 298 | 149 | 0.000288206 | 0.002017103 | 1 | 1 | 1 | 0.962 | 1 | 0.541 | 0.961 | 1 | 0.982 | 1 | 0.096 |
| RI | 341 | ENSG00000115286 | NDUFS7 | chr19 | + | 1388524 | 1388938 | 1388524 | 1388593 | 1388832 | 1388938 | 0 | 0 | 1 | 8 | 0 | 44 | 12 | 23 | 20 | 11 | 0 | 0 | 5 | 10 | 21 | 28 | 17 | 9 | 29 | 18 | 298 | 149 | 0.000291025 | 0.002034418 | 0 | 0 | 0.021 | 0.167 | 0 | 0 | 0 | 0.217 | 0.147 | 0.368 | -0.109 |
| RI | 3021 | ENSG00000169714 | CNBP | chr3 | - | 129171078 | 129171538 | 129171078 | 129171283 | 129171445 | 129171538 | 116 | 143 | 158 | 186 | 125 | 11 | 11 | 18 | 11 | 22 | 113 | 85 | 163 | 120 | 73 | 44 | 14 | 15 | 39 | 17 | 298 | 149 | 0.000291913 | 0.002038213 | 0.841 | 0.867 | 0.814 | 0.894 | 0.74 | 0.562 | 0.752 | 0.845 | 0.606 | 0.682 | 0.142 |
| RI | 6310 | ENSG00000133606 | MKRN1 | chr7 | - | 140455094 | 140455900 | 140455094 | 140455233 | 140455789 | 140455900 | 66 | 91 | 45 | 116 | 65 | 489 | 451 | 435 | 627 | 996 | 397 | 123 | 99 | 235 | 162 | 1603 | 580 | 666 | 1021 | 1696 | 298 | 149 | 0.000295847 | 0.002062519 | 0.063 | 0.092 | 0.049 | 0.085 | 0.032 | 0.11 | 0.096 | 0.069 | 0.103 | 0.046 | -0.021 |
| RI | 3478 | ENSG00000160741 | CRTC2 | chr1 | - | 153954872 | 153955166 | 153954872 | 153954989 | 153955064 | 153955166 | 39 | 21 | 45 | 29 | 47 | 41 | 33 | 36 | 36 | 28 | 105 | 106 | 81 | 82 | 158 | 56 | 83 | 40 | 16 | 43 | 224 | 149 | 0.000296593 | 0.002062519 | 0.388 | 0.297 | 0.454 | 0.349 | 0.528 | 0.555 | 0.459 | 0.574 | 0.773 | 0.71 | -0.211 |
| RI | 7962 | ENSG00000133895 | MEN1 | chr11 | - | 64807551 | 64808099 | 64807551 | 64807680 | 64807890 | 64808099 | 10 | 4 | 12 | 5 | 11 | 23 | 10 | 25 | 28 | 12 | 30 | 24 | 12 | 26 | 7 | 0 | 29 | 4 | 7 | 16 | 298 | 149 | 0.000296766 | 0.002062519 | 0.179 | 0.167 | 0.194 | 0.082 | 0.314 | 1 | 0.293 | 0.6 | 0.65 | 0.179 | -0.357 |
| RI | 971 | ENSG00000178971 | CTC1 | chr17 | - | 8229890 | 8230468 | 8229890 | 8229968 | 8230293 | 8230468 | 81 | 44 | 71 | 68 | 57 | 58 | 42 | 54 | 67 | 74 | 71 | 63 | 83 | 82 | 139 | 41 | 33 | 20 | 56 | 63 | 298 | 149 | 0.00029679 | 0.002062519 | 0.411 | 0.344 | 0.397 | 0.337 | 0.278 | 0.464 | 0.488 | 0.675 | 0.423 | 0.525 | -0.162 |
| RI | 5436 | ENSG00000197858 | GPAA1 | chr8 | + | 144085288 | 144085743 | 144085288 | 144085479 | 144085572 | 144085743 | 7 | 0 | 4 | 2 | 5 | 51 | 8 | 42 | 45 | 40 | 1 | 10 | 8 | 8 | 8 | 9 | 56 | 17 | 11 | 27 | 242 | 149 | 0.000306338 | 0.002126367 | 0.078 | 0 | 0.055 | 0.027 | 0.071 | 0.064 | 0.099 | 0.225 | 0.309 | 0.154 | -0.124 |
| RI | 1990 | ENSG00000175061 | SNHG29 | chr17 | + | 16439037 | 16440253 | 16439037 | 16439060 | 16440184 | 16440253 | 53 | 43 | 45 | 63 | 42 | 8 | 5 | 8 | 7 | 0 | 73 | 42 | 69 | 105 | 126 | 0 | 3 | 0 | 1 | 10 | 298 | 149 | 0.000310769 | 0.002154591 | 0.768 | 0.811 | 0.738 | 0.818 | 1 | 1 | 0.875 | 1 | 0.981 | 0.863 | -0.117 |
| RI | 6010 | ENSG00000112146 | FBXO9 | chr6 | + | 53092733 | 53093561 | 53092733 | 53092824 | 53093465 | 53093561 | 139 | 117 | 158 | 102 | 151 | 92 | 56 | 98 | 90 | 195 | 38 | 113 | 101 | 141 | 2 | 160 | 94 | 52 | 166 | 163 | 298 | 149 | 0.00031909 | 0.002209689 | 0.43 | 0.511 | 0.446 | 0.362 | 0.279 | 0.106 | 0.375 | 0.493 | 0.298 | 0.006 | 0.15 |
| RI | 5981 | ENSG00000204256 | BRD2 | chr6 | + | 32976030 | 32976464 | 32976030 | 32976169 | 32976249 | 32976464 | 82 | 129 | 83 | 193 | 154 | 526 | 317 | 444 | 455 | 1256 | 634 | 232 | 123 | 293 | 254 | 1602 | 573 | 500 | 731 | 963 | 229 | 149 | 0.000319892 | 0.002212646 | 0.092 | 0.209 | 0.108 | 0.216 | 0.074 | 0.205 | 0.209 | 0.138 | 0.207 | 0.146 | -0.041 |
| RI | 105 | ENSG00000100359 | SGSM3 | chr22 | + | 40408932 | 40409372 | 40408932 | 40409018 | 40409249 | 40409372 | 35 | 27 | 44 | 46 | 27 | 14 | 8 | 8 | 14 | 15 | 181 | 58 | 35 | 35 | 83 | 0 | 15 | 0 | 4 | 28 | 298 | 149 | 0.000323951 | 0.00223563 | 0.556 | 0.628 | 0.733 | 0.622 | 0.474 | 1 | 0.659 | 1 | 0.814 | 0.597 | -0.211 |
| RI | 2485 | ENSG00000077232 | DNAJC10 | chr2 | + | 182728575 | 182728994 | 182728575 | 182728658 | 182728862 | 182728994 | 22 | 39 | 39 | 12 | 39 | 207 | 190 | 131 | 165 | 397 | 8 | 15 | 19 | 20 | 12 | 43 | 78 | 76 | 40 | 20 | 298 | 149 | 0.000323972 | 0.00223563 | 0.05 | 0.093 | 0.13 | 0.035 | 0.047 | 0.085 | 0.088 | 0.111 | 0.2 | 0.231 | -0.072 |
| RI | 7772 | ENSG00000087053 | MTMR2 | chr11 | - | 95857551 | 95858632 | 95857551 | 95857635 | 95858530 | 95858632 | 8 | 13 | 14 | 8 | 9 | 97 | 62 | 44 | 67 | 95 | 0 | 10 | 8 | 2 | 0 | 79 | 91 | 59 | 29 | 75 | 298 | 149 | 0.000325922 | 0.002246465 | 0.04 | 0.095 | 0.137 | 0.056 | 0.045 | 0 | 0.052 | 0.063 | 0.033 | 0 | 0.045 |
| RI | 6246 | ENSG00000171777 | RASGRP4 | chr19 | - | 38411114 | 38411381 | 38411114 | 38411249 | 38411344 | 38411381 | 2 | 12 | 7 | 7 | 0 | 44 | 25 | 33 | 33 | 23 | 56 | 22 | 8 | 39 | 55 | 62 | 56 | 46 | 117 | 140 | 244 | 149 | 0.000331383 | 0.002281446 | 0.027 | 0.227 | 0.115 | 0.115 | 0 | 0.355 | 0.193 | 0.096 | 0.169 | 0.193 | -0.104 |
| RI | 283 | ENSG00000126458 | RRAS | chr19 | - | 49636823 | 49637130 | 49636823 | 49636926 | 49637042 | 49637130 | 7 | 0 | 4 | 13 | 2 | 33 | 20 | 8 | 33 | 13 | 0 | 7 | 0 | 3 | 7 | 58 | 67 | 7 | 36 | 92 | 265 | 149 | 0.000332225 | 0.002283899 | 0.107 | 0 | 0.219 | 0.181 | 0.08 | 0 | 0.055 | 0 | 0.045 | 0.041 | 0.089 |
| RI | 5079 | ENSG00000112855 | HARS2 | chr5 | + | 140691452 | 140691756 | 140691452 | 140691515 | 140691606 | 140691756 | 39 | 18 | 39 | 28 | 22 | 1 | 1 | 4 | 7 | 3 | 8 | 11 | 26 | 15 | 56 | 2 | 0 | 2 | 0 | 0 | 240 | 149 | 0.000332513 | 0.002283899 | 0.96 | 0.918 | 0.858 | 0.713 | 0.82 | 0.713 | 1 | 0.89 | 1 | 1 | -0.067 |
| RI | 968 | ENSG00000178921 | PFAS | chr17 | + | 8256266 | 8256648 | 8256266 | 8256407 | 8256523 | 8256648 | 2 | 1 | 2 | 0 | 2 | 23 | 1 | 22 | 16 | 27 | 0 | 11 | 1 | 3 | 26 | 36 | 5 | 8 | 7 | 0 | 265 | 149 | 0.00033402 | 0.002291588 | 0.047 | 0.36 | 0.049 | 0 | 0.04 | 0 | 0.553 | 0.066 | 0.194 | 1 | -0.263 |
| RI | 5319 | ENSG00000170445 | HARS1 | chr5 | - | 140677654 | 140679127 | 140677654 | 140677753 | 140677907 | 140679127 | 45 | 24 | 32 | 33 | 42 | 92 | 38 | 100 | 67 | 123 | 77 | 42 | 38 | 65 | 45 | 27 | 81 | 44 | 54 | 73 | 298 | 149 | 0.000335212 | 0.002297098 | 0.197 | 0.24 | 0.138 | 0.198 | 0.146 | 0.588 | 0.206 | 0.302 | 0.376 | 0.236 | -0.158 |

| | | | | | | | | | | | | | | | | | | | | | | | | | | | | | | | | | | | | | | | | | | | | | | |
|---|---|---|---|---|---|---|---|---|---|---|---|---|---|---|---|---|---|---|---|---|---|---|---|---|---|---|---|---|---|---|---|---|---|---|---|---|---|---|---|---|---|---|---|---|---|---|
| RI | 5780 | ENSG00000204843 | DCTN1 | chr2 | - | 74366243 | 74366620 | 74366243 | 74366375 | 74366458 | 74366620 | 27 | 19 | 35 | 25 | 32 | 184 | 122 | 175 | 193 | 130 | 33 | 24 | 38 | 48 | 36 | 496 | 291 | 191 | 249 | 453 | 232 | 149 | 0.001478909 | 0.008375759 | 0.086 | 0.091 | 0.114 | 0.077 | 0.137 | 0.041 | 0.05 | 0.113 | 0.11 | 0.049 | 0.028 |
| RI | 705 | ENSG00000073605 | GSDMB | chr17 | - | 39906110 | 39906987 | 39906110 | 39906271 | 39906960 | 39906987 | 39 | 30 | 51 | 39 | 19 | 43 | 26 | 29 | 57 | 69 | 10 | 34 | 17 | 20 | 17 | 39 | 63 | 28 | 62 | 58 | 298 | 149 | 0.001482649 | 0.008388897 | 0.312 | 0.366 | 0.468 | 0.255 | 0.121 | 0.114 | 0.212 | 0.233 | 0.139 | 0.128 | 0.139 |
| RI | 7267 | ENSG00000104518 | GSDMD | chr8 | + | 143559331 | 143559969 | 143559331 | 143559552 | 143559776 | 143559969 | 0 | 5 | 6 | 4 | 1 | 83 | 19 | 55 | 55 | 23 | 9 | 5 | 1 | 7 | 32 | 62 | 67 | 25 | 102 | 88 | 298 | 149 | 0.001493814 | 0.00844398 | 0 | 0.116 | 0.052 | 0.035 | 0.021 | 0.068 | 0.036 | 0.02 | 0.033 | 0.154 | -0.017 |
| RI | 2833 | ENSG00000114745 | GORASP1 | chr3 | - | 39100304 | 39100877 | 39100304 | 39100503 | 39100746 | 39100877 | 12 | 8 | 18 | 8 | 10 | 10 | 3 | 14 | 18 | 6 | 39 | 40 | 14 | 24 | 77 | 11 | 25 | 2 | 7 | 16 | 298 | 149 | 0.001507191 | 0.008511453 | 0.375 | 0.571 | 0.391 | 0.182 | 0.455 | 0.639 | 0.444 | 0.778 | 0.632 | 0.706 | -0.245 |
| RI | 7979 | ENSG00000187531 | SIRT7 | chr17 | - | 81913773 | 81914167 | 81913773 | 81913880 | 81914086 | 81914167 | 5 | 0 | 4 | 4 | 0 | 63 | 45 | 73 | 76 | 30 | 8 | 14 | 2 | 4 | 21 | 96 | 77 | 92 | 82 | 155 | 298 | 149 | 0.00154021 | 0.008689609 | 0.038 | 0 | 0.027 | 0.026 | 0 | 0.04 | 0.083 | 0.011 | 0.024 | 0.063 | -0.026 |
| RI | 874 | ENSG00000125450 | NUP85 | chr17 | + | 75234636 | 75235201 | 75234636 | 75234788 | 75235099 | 75235201 | 21 | 27 | 31 | 35 | 41 | 14 | 15 | 22 | 28 | 10 | 107 | 49 | 87 | 75 | 78 | 14 | 5 | 9 | 17 | 59 | 298 | 149 | 0.001549372 | 0.00873296 | 0.429 | 0.474 | 0.413 | 0.385 | 0.672 | 0.793 | 0.831 | 0.829 | 0.688 | 0.398 | -0.233 |
| RI | 7442 | ENSG00000233016 | SNHG7 | chr9 | - | 136726310 | 136727086 | 136726310 | 136726416 | 136727045 | 136727086 | 21 | 13 | 22 | 17 | 14 | 44 | 19 | 45 | 46 | 13 | 46 | 33 | 22 | 33 | 48 | 14 | 41 | 18 | 22 | 69 | 298 | 149 | 0.001551135 | 0.008734559 | 0.193 | 0.255 | 0.196 | 0.156 | 0.35 | 0.622 | 0.287 | 0.379 | 0.429 | 0.258 | -0.165 |
| RI | 1792 | ENSG00000092108 | SCFD1 | chr14 | + | 30700187 | 30702375 | 30700187 | 30700258 | 30702295 | 30702375 | 48 | 49 | 32 | 36 | 27 | 71 | 30 | 69 | 95 | 137 | 19 | 12 | 25 | 7 | 13 | 4 | 18 | 12 | 7 | 0 | 298 | 149 | 0.001588029 | 0.008929609 | 0.253 | 0.45 | 0.188 | 0.159 | 0.09 | 0.704 | 0.25 | 0.51 | 0.333 | 1 | -0.331 |
| RI | 3316 | ENSG00000137818 | RPLP1 | chr15 | + | 69453646 | 69455287 | 69453646 | 69453721 | 69455169 | 69455287 | 107 | 65 | 150 | 114 | 152 | 4970 | 2358 | 5262 | 4646 | 5971 | 133 | 117 | 86 | 128 | 137 | 3463 | 3361 | 2701 | 2770 | 6076 | 298 | 149 | 0.001588796 | 0.008929609 | 0.011 | 0.014 | 0.014 | 0.012 | 0.013 | 0.019 | 0.017 | 0.016 | 0.023 | 0.011 | -0.004 |
| RI | 637 | ENSG00000196839 | ADA | chr20 | - | 44622828 | 44623078 | 44622828 | 44622930 | 44623006 | 44623078 | 20 | 17 | 12 | 24 | 21 | 35 | 15 | 18 | 26 | 32 | 31 | 18 | 21 | 15 | 41 | 13 | 29 | 15 | 3 | 3 | 225 | 149 | 0.001591015 | 0.008933581 | 0.275 | 0.429 | 0.306 | 0.379 | 0.303 | 0.612 | 0.291 | 0.481 | 0.768 | 0.901 | -0.272 |
| RI | 646 | ENSG00000141644 | MBD1 | chr18 | - | 50276661 | 50276998 | 50276661 | 50276744 | 50276831 | 50276998 | 25 | 20 | 18 | 23 | 14 | 61 | 36 | 49 | 46 | 35 | 48 | 36 | 34 | 45 | 49 | 76 | 28 | 41 | 52 | 89 | 236 | 149 | 0.001593604 | 0.008939621 | 0.206 | 0.26 | 0.188 | 0.24 | 0.202 | 0.285 | 0.448 | 0.344 | 0.353 | 0.258 | -0.118 |
| RI | 8455 | ENSG00000123349 | PFDN5 | chr12 | + | 53296429 | 53297924 | 53296429 | 53296551 | 53297849 | 53297924 | 52 | 40 | 62 | 82 | 101 | 15 | 13 | 21 | 15 | 12 | 254 | 116 | 102 | 163 | 121 | 14 | 13 | 6 | 24 | 26 | 298 | 149 | 0.00162206 | 0.009084791 | 0.634 | 0.606 | 0.596 | 0.732 | 0.808 | 0.901 | 0.817 | 0.895 | 0.773 | 0.699 | -0.142 |
| RI | 3355 | ENSG00000213024 | NUP62 | chr19 | - | 49908104 | 49909884 | 49908104 | 49908745 | 49908973 | 49909884 | 85 | 26 | 54 | 55 | 18 | 2 | 0 | 4 | 4 | 1 | 84 | 57 | 34 | 66 | 132 | 0 | 0 | 1 | 0 | 0 | 298 | 149 | 0.001622559 | 0.009084791 | 0.955 | 1 | 0.871 | 0.873 | 0.9 | 1 | 1 | 0.944 | 1 | 1 | -0.069 |
| RI | 330 | ENSG00000076662 | ICAM3 | chr19 | - | 10335065 | 10335976 | 10335065 | 10335353 | 10335670 | 10335976 | 24 | 21 | 27 | 26 | 14 | 236 | 95 | 159 | 222 | 94 | 35 | 18 | 18 | 8 | 23 | 253 | 193 | 148 | 204 | 415 | 298 | 149 | 0.001631085 | 0.00912388 | 0.048 | 0.1 | 0.078 | 0.055 | 0.069 | 0.065 | 0.045 | 0.057 | 0.019 | 0.027 | 0.027 |
| RI | 4387 | ENSG00000145020 | AMT | chr3 | - | 49418970 | 49420342 | 49418970 | 49419151 | 49419709 | 49420342 | 13 | 7 | 20 | 11 | 7 | 0 | 0 | 0 | 0 | 0 | 22 | 23 | 16 | 10 | 14 | 0 | 0 | 4 | 5 | 0 | 298 | 149 | 0.001635824 | 0.009141732 | 1 | 1 | 1 | 1 | 1 | 1 | 1 | 0.667 | 0.5 | 1 | 0.167 |
| RI | 6600 | ENSG00000188878 | FBF1 | chr17 | - | 75914121 | 75914932 | 75914121 | 75914298 | 75914746 | 75914932 | 0 | 0 | 0 | 0 | 1 | 5 | 7 | 2 | 7 | 1 | 9 | 0 | 0 | 9 | 0 | 0 | 16 | 4 | 0 | 4 | 298 | 149 | 0.001654763 | 0.009236575 | 0 | 0 | 0 | 0 | 0.333 | 1 | 0 | 0 | 1 | 0 | -0.333 |
| RI | 2997 | ENSG00000060971 | ACAA1 | chr3 | - | 38125825 | 38126700 | 38125825 | 38125881 | 38126509 | 38126700 | 32 | 20 | 41 | 39 | 20 | 62 | 24 | 43 | 34 | 17 | 40 | 43 | 39 | 59 | 90 | 114 | 72 | 68 | 118 | 166 | 298 | 149 | 0.001655922 | 0.009236575 | 0.205 | 0.294 | 0.323 | 0.364 | 0.37 | 0.149 | 0.23 | 0.223 | 0.2 | 0.213 | 0.108 |
| RI | 5803 | ENSG00000136699 | SMPD4 | chr2 | - | 130156034 | 130156675 | 130156034 | 130156135 | 130156584 | 130156675 | 15 | 5 | 22 | 12 | 10 | 39 | 9 | 27 | 30 | 29 | 49 | 23 | 8 | 40 | 28 | 28 | 36 | 18 | 21 | 25 | 298 | 149 | 0.001685799 | 0.009394352 | 0.161 | 0.217 | 0.289 | 0.167 | 0.147 | 0.467 | 0.242 | 0.182 | 0.488 | 0.359 | -0.151 |
| RI | 8304 | ENSG00000125651 | GTF2F1 | chr19 | - | 6381358 | 6381615 | 6381358 | 6381478 | 6381553 | 6381615 | 10 | 13 | 6 | 8 | 0 | 63 | 61 | 67 | 35 | 51 | 58 | 5 | 7 | 47 | 59 | 170 | 117 | 61 | 121 | 137 | 224 | 149 | 0.001697543 | 0.009450881 | 0.096 | 0.124 | 0.056 | 0.132 | 0 | 0.185 | 0.028 | 0.071 | 0.205 | 0.223 | -0.061 |
| RI | 2139 | ENSG00000140853 | NLRC5 | chr16 | + | 57028071 | 57028385 | 57028071 | 57028155 | 57028301 | 57028385 | 75 | 52 | 89 | 94 | 67 | 62 | 36 | 79 | 80 | 63 | 183 | 179 | 127 | 98 | 310 | 99 | 79 | 51 | 92 | 151 | 295 | 149 | 0.001722048 | 0.009578284 | 0.379 | 0.422 | 0.363 | 0.372 | 0.349 | 0.483 | 0.534 | 0.557 | 0.35 | 0.509 | -0.11 |
| RI | 7827 | ENSG00000149743 | TRPT1 | chr11 | - | 64224099 | 64224341 | 64224099 | 64224210 | 64224284 | 64224341 | 3 | 4 | 7 | 3 | 4 | 17 | 7 | 16 | 7 | 26 | 36 | 8 | 11 | 5 | 32 | 53 | 11 | 21 | 11 | 10 | 223 | 149 | 0.001730117 | 0.009614114 | 0.105 | 0.276 | 0.226 | 0.223 | 0.093 | 0.312 | 0.327 | 0.259 | 0.233 | 0.681 | -0.178 |
| RI | 1406 | ENSG00000174177 | CTU2 | chr16 | + | 88714382 | 88714737 | 88714382 | 88714486 | 88714586 | 88714737 | 2 | 3 | 6 | 2 | 3 | 7 | 3 | 15 | 8 | 2 | 0 | 0 | 0 | 8 | 0 | 29 | 10 | 1 | 2 | 19 | 249 | 149 | 0.00173385 | 0.009625799 | 0.146 | 0.374 | 0.193 | 0.13 | 0.473 | 0 | 0 | 0 | 0.705 | 0 | 0.122 |
| RI | 7534 | ENSG00000130714 | POMT1 | chr9 | + | 131506402 | 131509022 | 131506402 | 131506453 | 131507367 | 131509022 | 17 | 21 | 30 | 14 | 9 | 16 | 3 | 15 | 12 | 18 | 45 | 30 | 40 | 25 | 50 | 0 | 24 | 9 | 0 | 13 | 298 | 149 | 0.001753945 | 0.009728218 | 0.347 | 0.778 | 0.5 | 0.368 | 0.2 | 1 | 0.385 | 0.69 | 1 | 0.658 | -0.308 |
| RI | 3891 | ENSG00000139645 | ANKRD52 | chr12 | - | 56237806 | 56243392 | 56237806 | 56237968 | 56243237 | 56243392 | 43 | 13 | 46 | 38 | 6 | 0 | 0 | 0 | 0 | 0 | 39 | 44 | 22 | 37 | 52 | 14 | 0 | 0 | 0 | 0 | 298 | 149 | 0.001759908 | 0.009752135 | 1 | 1 | 1 | 1 | 1 | 0.582 | 1 | 1 | 1 | 1 | 0.084 |
| RI | 6941 | ENSG00000147133 | TAF1 | chrX | + | 71423116 | 71424066 | 71423116 | 71423239 | 71424016 | 71424066 | 103 | 101 | 78 | 67 | 88 | 4 | 2 | 2 | 0 | 2 | 136 | 63 | 115 | 111 | 22 | 0 | 4 | 12 | 9 | 2 | 298 | 149 | 0.001777756 | 0.009841801 | 0.928 | 0.962 | 0.951 | 1 | 0.957 | 1 | 0.887 | 0.827 | 0.86 | 0.846 | 0.076 |
| RI | 8038 | ENSG00000138326 | RPS24 | chr10 | + | 78037438 | 78040225 | 78037438 | 78037441 | 78040203 | 78040225 | 73 | 51 | 63 | 60 | 189 | 9 | 3 | 6 | 2 | 9 | 5 | 26 | 37 | 25 | 14 | 8 | 7 | 0 | 5 | 11 | 298 | 149 | 0.001808283 | 0.010001429 | 0.802 | 0.895 | 0.84 | 0.938 | 0.913 | 0.238 | 0.65 | 1 | 0.714 | 0.389 | 0.279 |
| RI | 8040 | ENSG00000176155 | CCDC57 | chr17 | - | 82193755 | 82194139 | 82193755 | 82193830 | 82193981 | 82194139 | 7 | 16 | 15 | 21 | 4 | 3 | 14 | 12 | 23 | 21 | 51 | 27 | 15 | 43 | 12 | 14 | 12 | 4 | 16 | 11 | 298 | 149 | 0.00182191 | 0.010067374 | 0.538 | 0.364 | 0.385 | 0.313 | 0.087 | 0.646 | 0.529 | 0.652 | 0.573 | 0.353 | -0.213 |
| RI | 7521 | ENSG00000115459 | ELMOD3 | chr2 | + | 85355075 | 85357252 | 85355075 | 85355170 | 85356966 | 85357252 | 32 | 25 | 15 | 29 | 24 | 18 | 1 | 8 | 9 | 12 | 37 | 20 | 23 | 35 | 37 | 2 | 3 | 3 | 9 | 0 | 298 | 149 | 0.001841321 | 0.010165123 | 0.471 | 0.926 | 0.484 | 0.617 | 0.5 | 0.902 | 0.769 | 0.793 | 0.66 | 1 | -0.225 |
| RI | 4177 | ENSG00000134574 | DDB2 | chr11 | + | 47234756 | 47235412 | 47234756 | 47234934 | 47235269 | 47235412 | 37 | 33 | 38 | 21 | 32 | 41 | 19 | 45 | 23 | 38 | 63 | 101 | 48 | 73 | 49 | 20 | 65 | 25 | 26 | 45 | 298 | 149 | 0.001846636 | 0.01018495 | 0.311 | 0.465 | 0.297 | 0.313 | 0.296 | 0.612 | 0.437 | 0.49 | 0.584 | 0.353 | -0.159 |
| RI | 6589 | ENSG00000234741 | GAS5 | chr1 | - | 173864256 | 173864704 | 173864256 | 173864304 | 173864674 | 173864704 | 300 | 274 | 340 | 320 | 354 | 9 | 5 | 26 | 22 | 40 | 170 | 158 | 245 | 212 | 51 | 4 | 1 | 8 | 15 | 0 | 298 | 149 | 0.001857966 | 0.010237875 | 0.943 | 0.965 | 0.867 | 0.879 | 0.816 | 0.955 | 0.988 | 0.939 | 0.876 | 1 | -0.058 |
| RI | 7167 | ENSG00000213782 | DDX47 | chr12 | + | 12823202 | 12824016 | 12823202 | 12823319 | 12823869 | 12824016 | 131 | 79 | 171 | 180 | 208 | 139 | 94 | 133 | 128 | 360 | 69 | 215 | 128 | 124 | 34 | 90 | 86 | 62 | 55 | 19 | 298 | 149 | 0.001928143 | 0.010614669 | 0.32 | 0.296 | 0.391 | 0.413 | 0.224 | 0.277 | 0.556 | 0.508 | 0.53 | 0.472 | -0.14 |
| RI | 1513 | ENSG00000175265 | GOLGA8A | chr15 | - | 34391122 | 34391426 | 34391122 | 34391265 | 34391352 | 34391426 | 24 | 22 | 18 | 19 | 21 | 11 | 17 | 25 | 15 | 23 | 15 | 6 | 16 | 3 | 3 | 0 | 0 | 7 | 0 | 5 | 236 | 149 | 0.001947142 | 0.010709282 | 0.579 | 0.45 | 0.313 | 0.444 | 0.366 | 1 | 1 | 0.591 | 1 | 0.275 | -0.343 |
| RI | 4646 | ENSG00000147274 | RBMX | chrX | - | 136879016 | 136879453 | 136879016 | 136879123 | 136879318 | 136879453 | 35 | 29 | 48 | 22 | 34 | 363 | 295 | 326 | 403 | 359 | 46 | 69 | 45 | 57 | 44 | 480 | 416 | 377 | 299 | 308 | 298 | 149 | 0.001949117 | 0.010710169 | 0.046 | 0.047 | 0.069 | 0.027 | 0.045 | 0.046 | 0.077 | 0.056 | 0.087 | 0.067 | -0.02 |
| RI | 4455 | ENSG00000117139 | KDM5B | chr1 | - | 202730908 | 202731939 | 202730908 | 202731063 | 202731827 | 202731939 | 12 | 3 | 16 | 12 | 7 | 37 | 33 | 30 | 27 | 55 | 0 | 10 | 7 | 6 | 0 | 16 | 41 | 49 | 67 | 15 | 298 | 149 | 0.001974089 | 0.01083731 | 0.14 | 0.043 | 0.211 | 0.182 | 0.06 | 0 | 0.109 | 0.067 | 0.043 | 0 | 0.083 |
| RI | 5691 | ENSG00000160710 | ADAR | chr1 | - | 154589756 | 154590409 | 154589756 | 154589928 | 154590183 | 154590409 | 23 | 17 | 13 | 19 | 31 | 304 | 197 | 284 | 348 | 243 | 60 | 47 | 37 | 86 | 10 | 572 | 362 | 254 | 504 | 414 | 298 | 149 | 0.001977271 | 0.0108447 | 0.036 | 0.041 | 0.022 | 0.027 | 0.06 | 0.05 | 0.061 | 0.068 | 0.079 | 0.012 | -0.017 |
| RI | 1841 | ENSG00000099797 | TECR | chr19 | + | 14565065 | 14565290 | 14565065 | 14565123 | 14565201 | 14565290 | 7 | 3 | 11 | 9 | 10 | 59 | 27 | 79 | 60 | 40 | 0 | 2 | 1 | 6 | 11 | 26 | 69 | 63 | 57 | 84 | 227 | 149 | 0.001989139 | 0.01088471 | 0.072 | 0.068 | 0.084 | 0.09 | 0.141 | 0 | 0.019 | 0.01 | 0.065 | 0.079 | 0.056 |
| RI | 3770 | ENSG00000072210 | ALDH3A2 | chr17 | + | 19663332 | 19665047 | 19663332 | 19663499 | 19664947 | 19665047 | 2 | 6 | 7 | 2 | 7 | 95 | 51 | 64 | 85 | 130 | 4 | 2 | 11 | 30 | 3 | 98 | 51 | 39 | 98 | 111 | 298 | 149 | 0.001989772 | 0.01088471 | 0.01 | 0.056 | 0.052 | 0.012 | 0.026 | 0.02 | 0.019 | 0.124 | 0.133 | 0.013 | -0.031 |
| RI | 2418 | ENSG00000214655 | ZSWIM8 | chr10 | + | 73794146 | 73794639 | 73794146 | 73794330 | 73794555 | 73794639 | 36 | 35 | 21 | 30 | 23 | 39 | 44 | 37 | 41 | 40 | 65 | 44 | 36 | 74 | 79 | 55 | 59 | 25 | 51 | 60 | 298 | 149 | 0.001991917 | 0.01088471 | 0.316 | 0.285 | 0.221 | 0.268 | 0.223 | 0.371 | 0.272 | 0.419 | 0.42 | 0.397 | -0.113 |
| RI | 3222 | ENSG00000184014 | DENND5A | chr11 | - | 9141939 | 9143485 | 9141939 | 9142108 | 9143402 | 9143485 | 17 | 30 | 17 | 14 | 14 | 0 | 1 | 0 | 0 | 4 | 42 | 20 | 24 | 31 | 27 | 15 | 0 | 1 | 11 | 0 | 298 | 149 | 0.001991937 | 0.01088471 | 1 | 0.938 | 1 | 1 | 0.636 | 0.583 | 1 | 0.923 | 0.585 | 1 | 0.097 |
| RI | 1518 | ENSG00000130703 | OSBPL2 | chr20 | + | 62281057 | 62281879 | 62281057 | 62281165 | 62281789 | 62281879 | 20 | 9 | 12 | 17 | 14 | 58 | 50 | 56 | 69 | 62 | 21 | 33 | 30 | 37 | 47 | 80 | 73 | 64 | 106 | 97 | 298 | 149 | 0.001995295 | 0.010892983 | 0.147 | 0.083 | 0.097 | 0.11 | 0.101 | 0.116 | 0.184 | 0.19 | 0.149 | 0.195 | -0.059 |
| RI | 5412 | ENSG00000018280 | SLC11A1 | chr2 | + | 218382959 | 218385266 | 218382959 | 218383102 | 218385146 | 218385266 | 59 | 62 | 35 | 45 | 26 | 0 | 0 | 0 | 1 | 0 | 78 | 91 | 152 | 104 | 227 | 12 | 0 | 0 | 4 | 0 | 298 | 149 | 0.001998641 | 0.010901176 | 1 | 1 | 1 | 0.957 | 1 | 0.765 | 1 | 1 | 0.929 | 1 | 0.053 |
| RI | 7892 | ENSG00000204356 | NELFE | chr6 | - | 31954554 | 31955096 | 31954554 | 31954892 | 31955058 | 31955096 | 20 | 13 | 12 | 25 | 5 | 69 | 32 | 63 | 57 | 76 | 54 | 33 | 34 | 28 | 63 | 108 | 94 | 70 | 114 | 109 | 298 | 149 | 0.002002328 | 0.010911208 | 0.127 | 0.169 | 0.087 | 0.18 | 0.032 | 0.2 | 0.149 | 0.195 | 0.109 | 0.224 | -0.056 |
| RI | 7791 | ENSG00000060749 | QSER1 | chr11 | + | 32931742 | 32935435 | 32931742 | 32932374 | 32933091 | 32935435 | 175 | 156 | 135 | 207 | 346 | 0 | 0 | 0 | 2 | 0 | 44 | 110 | 146 | 81 | 33 | 7 | 0 | 3 | 3 | 0 | 298 | 149 | 0.002008571 | 0.010932846 | 1 | 1 | 1 | 0.981 | 1 | 0.759 | 1 | 0.961 | 0.931 | 1 | 0.066 |
| RI | 2187 | ENSG00000165819 | METTL3 | chr14 | - | 21503172 | 21503829 | 21503172 | 21503577 | 21503663 | 21503829 | 20 | 16 | 17 | 20 | 13 | 65 | 37 | 50 | 69 | 67 | 12 | 22 | 47 | 43 | 27 | 67 | 62 | 22 | 23 | 24 | 235 | 149 | 0.00201 | 0.010932846 | 0.163 | 0.215 | 0.177 | 0.155 | 0.11 | 0.102 | 0.184 | 0.575 | 0.542 | 0.416 | -0.2 |
| RI | 1523 | ENSG00000213614 | HEXA | chr15 | - | 72347685 | 72348134 | 72347685 | 72347758 | 72348047 | 72348134 | 10 | 4 | 3 | 11 | 4 | 194 | 134 | 165 | 150 | 215 | 16 | 5 | 28 | 20 | 30 | 473 | 189 | 200 | 281 | 370 | 298 | 149 | 0.002015167 | 0.010950866 | 0.025 | 0.015 | 0.009 | 0.035 | 0.009 | 0.017 | 0.013 | 0.065 | 0.034 | 0.039 | -0.015 |
| RI | 2096 | ENSG00000141279 | NPEPPS | chr17 | + | 47585499 | 47586398 | 47585499 | 47585745 | 47586364 | 47586398 | 19 | 23 | 27 | 30 | 59 | 0 | 0 | 0 | 1 | 0 | 4 | 41 | 50 | 53 | 12 | 0 | 1 | 0 | 2 | 8 | 298 | 149 | 0.002039195 | 0.011071254 | 1 | 1 | 1 | 0.938 | 1 | 1 | 0.953 | 1 | 0.93 | 0.429 | 0.125 |
| RI | 7920 | ENSG00000179532 | DNHD1 | chr11 | + | 6570008 | 6572027 | 6570008 | 6571423 | 6571635 | 6572027 | 1 | 6 | 8 | 4 | 2 | 1 | 1 | 3 | 6 | 4 | 0 | 7 | 13 | 12 | 7 | 3 | 0 | 1 | 0 | 1 | 298 | 149 | 0.002054666 | 0.011145005 | 0.333 | 0.75 | 0.571 | 0.25 | 0.2 | 0 | 1 | 0.867 | 1 | 0.778 | -0.308 |
| RI | 8487 | ENSG00000166598 | HSP90B1 | chr12 | + | 103943073 | 103943874 | 103943073 | 103943573 | 103943737 | 103943874 | 178 | 136 | 167 | 170 | 194 | 9 | 9 | 3 | 9 | 10 | 16 | 35 | 133 | 26 | 21 | 0 | 2 | 4 | 0 | 0 | 298 | 149 | 0.002063286 | 0.011181493 | 0.908 | 0.883 | 0.965 | 0.904 | 0.907 | 1 | 0.897 | 0.943 | 1 | 1 | -0.055 |
| RI | 8266 | ENSG00000211451 | GNRHR2 | chr1 | + | 145919343 | 145919900 | 145919343 | 145919419 | 145919679 | 145919900 | 1 | 5 | 8 | 4 | 3 | 2 | 0 | 1 | 3 | 9 | 6 | 0 | 1 | 8 | 9 | 0 | 8 | 0 | 0 | 0 | 298 | 149 | 0.00207144 | 0.011213159 | 0.2 | 1 | 0.8 | 0.4 | 0.143 | 1 | 0 | 1 | 1 | 1 | -0.291 |
| RI | 7381 | ENSG00000167272 | POP5 | chr12 | - | 120579513 | 120579923 | 120579513 | 120579597 | 120579773 | 120579923 | 15 | 21 | 24 | 46 | 9 | 63 | 24 | 47 | 62 | 94 | 28 | 12 | 15 | 37 | 39 | 37 | 24 | 15 | 55 | 26 | 298 | 149 | 0.002072925 | 0.011213159 | 0.106 | 0.304 | 0.203 | 0.271 | 0.046 | 0.275 | 0.2 | 0.333 | 0.252 | 0.429 | -0.112 |
| RI | 4643 | ENSG00000227057 | WDR46 | chr6 | - | 33288147 | 33288470 | 33288147 | 33288235 | 33288357 | 33288470 | 7 | 16 | 16 | 14 | 2 | 27 | 34 | 33 | 20 | 58 | 22 | 18 | 14 | 9 | 36 | 23 | 28 | 16 | 18 | 63 | 271 | 149 | 0.002096797 | 0.01133191 | 0.125 | 0.206 | 0.21 | 0.278 | 0.019 | 0.345 | 0.261 | 0.325 | 0.216 | 0.239 | -0.11 |
| RI | 8029 | ENSG00000090621 | PABPC4 | chr1 | - | 39562322 | 39564542 | 39562322 | 39563922 | 39564470 | 39564542 | 76 | 55 | 86 | 107 | 61 | 38 | 14 | 42 | 27 | 20 | 89 | 113 | 85 | 118 | 73 | 9 | 14 | 6 | 17 | 39 | 298 | 149 | 0.002100754 | 0.01134292 | 0.5 | 0.663 | 0.506 | 0.665 | 0.604 | 0.832 | 0.801 | 0.876 | 0.776 | 0.483 | -0.166 |
| RI | 8350 | ENSG00000132341 | RAN | chr12 | + | 130872583 | 130872920 | 130872583 | 130872629 | 130872835 | 130872920 | 42 | 24 | 55 | 60 | 38 | 338 | 217 | 323 | 361 | 253 | 36 | 20 | 35 | 47 | 58 | 464 | 357 | 260 | 387 | 489 | 298 | 149 | 0.002107138 | 0.011367 | 0.058 | 0.052 | 0.078 | 0.077 | 0.07 | 0.037 | 0.027 | 0.063 | 0.057 | 0.056 | 0.019 |
| RI | 4224 | ENSG00000132781 | MUTYH | chr1 | - | 45331660 | 45332086 | 45331660 | 45331849 | 45332022 | 45332086 | 15 | 14 | 13 | 16 | 9 | 14 | 1 | 4 | 5 | 5 | 22 | 0 | 17 | 20 | 30 | 0 | 9 | 0 | 0 | 1 | 298 | 149 | 0.002120322 | 0.011427687 | 0.349 | 0.875 | 0.619 | 0.615 | 0.474 | 1 | 0 | 1 | 1 | 0.938 | -0.201 |
| RI | 572 | ENSG00000185513 | L3MBTL1 | chr20 | + | 43536096 | 43536458 | 43536096 | 43536294 | 43536408 | 43536458 | 0 | 0 | 0 | 2 | 0 | 7 | 1 | 3 | 8 | 1 | 8 | 0 | 0 | 3 | 0 | 7 | 2 | 5 | 4 | 12 | 263 | 149 | 0.002144402 | 0.011546932 | 0 | 0 | 0 | 0.124 | 0 | 0.393 | 0 | 0 | 0.298 | 0 | -0.113 |
| RI | 3669 | ENSG00000135597 | REPS1 | chr6 | - | 138926400 | 138930098 | 138926400 | 138926481 | 138929976 | 138930098 | 52 | 88 | 54 | 58 | 42 | 43 | 41 | 48 | 57 | 98 | 6 | 32 | 49 | 3 | 3 | 53 | 42 | 29 | 13 | 27 | 298 | 149 | 0.002152512 | 0.011580045 | 0.377 | 0.518 | 0.36 | 0.337 | 0.176 | 0.054 | 0.276 | 0.458 | 0.103 | 0.053 | 0.165 |
| RI | 2212 | ENSG00000171530 | TBCA | chr5 | - | 77691240 | 77693352 | 77691240 | 77691498 | 77693265 | 77693352 | 39 | 14 | 32 | 22 | 15 | 872 | 511 | 780 | 805 | 1051 | 0 | 1 | 2 | 3 | 0 | 1272 | 806 | 608 | 871 | 815 | 298 | 149 | 0.002166015 | 0.01163176 | 0.022 | 0.014 | 0.02 | 0.013 | 0.007 | 0 | 0.001 | 0.002 | 0.002 | 0 | 0.014 |
| RI | 5375 | ENSG00000163050 | COQ8A | chr1 | + | 226983760 | 226984235 | 226983760 | 226983854 | 226984093 | 226984235 | 3 | 5 | 7 | 5 | 4 | 36 | 20 | 23 | 49 | 45 | 8 | 4 | 2 | 5 | 51 | 23 | 37 | 27 | 12 | 46 | 298 | 149 | 0.002166063 | 0.01163176 | 0.04 | 0.111 | 0.132 | 0.049 | 0.043 | 0.148 | 0.051 | 0.036 | 0.172 | 0.357 | -0.078 |
| RI | 6498 | ENSG00000128604 | IRF5 | chr7 | + | 128947022 | 128947535 | 128947022 | 128947056 | 128947229 | 128947535 | 35 | 23 | 21 | 8 | 10 | 39 | 9 | 18 | 14 | 23 | 80 | 36 | 29 | 45 | 115 | 51 | 29 | 5 | 16 | 29 | 298 | 149 | 0.002181726 | 0.011705226 | 0.31 | 0.561 | 0.368 | 0.222 | 0.179 | 0.44 | 0.383 | 0.744 | 0.584 | 0.665 | -0.235 |
| RI | 2665 | ENSG00000154743 | TSEN2 | chr3 | + | 12529761 | 12531659 | 12529761 | 12529873 | 12531569 | 12531659 | 4 | 3 | 20 | 6 | 35 | 23 | 16 | 32 | 32 | 38 | 0 | 3 | 3 | 6 | 0 | 8 | 13 | 27 | 4 | 37 | 298 | 149 | 0.002231944 | 0.011963788 | 0.08 | 0.086 | 0.238 | 0.086 | 0.315 | 0 | 0.103 | 0.053 | 0.429 | 0 | 0.044 |
| RI | 8217 | ENSG00000069493 | CLEC2D | chr12 | + | 9692827 | 9693948 | 9692827 | 9692931 | 9693824 | 9693948 | 281 | 264 | 233 | 264 | 226 | 3 | 0 | 1 | 5 | 20 | 12 | 58 | 78 | 20 | 37 | 0 | 1 | 2 | 0 | 0 | 298 | 149 | 0.002241108 | 0.012002019 | 0.979 | 1 | 0.991 | 0.964 | 0.85 | 1 | 0.967 | 0.951 | 1 | 1 | -0.027 |
| RI | 784 | ENSG00000182173 | TSEN54 | chr17 | + | 75521704 | 75523335 | 75521704 | 75522521 | 75523274 | 75523335 | 43 | 32 | 37 | 54 | 47 | 15 | 1 | 1 | 16 | 2 | 14 | 59 | 89 | 60 | 85 | 0 | 12 | 1 | 0 | 0 | 298 | 149 | 0.002246891 | 0.012022085 | 0.589 | 0.941 | 0.949 | 0.628 | 0.922 | 1 | 0.711 | 0.978 | 1 | 1 | -0.132 |
| RI | 1662 | ENSG00000185088 | RPS27L | chr15 | - | 63156412 | 63157477 | 63156412 | 63156521 | 63157399 | 63157477 | 36 | 24 | 32 | 26 | 21 | 96 | 38 | 85 | 59 | 59 | 14 | 0 | 21 | 15 | 18 | 109 | 55 | 44 | 89 | 25 | 298 | 149 | 0.002268621 | 0.012127373 | 0.158 | 0.24 | 0.158 | 0.181 | 0.151 | 0.06 | 0 | 0.193 | 0.078 | 0.265 | 0.058 |
| RI | 6849 | ENSG00000102103 | PQBP1 | chrX | + | 48902232 | 48903084 | 48902232 | 48902795 | 48902927 | 48903084 | 11 | 5 | 21 | 8 | 16 | 104 | 59 | 107 | 94 | 95 | 0 | 46 | 23 | 20 | 17 | 145 | 75 | 56 | 36 | 99 | 281 | 149 | 0.00227712 | 0.012161799 | 0.053 | 0.043 | 0.094 | 0.043 | 0.082 | 0 | 0.245 | 0.179 | 0.228 | 0.083 | -0.084 |
| RI | 1358 | ENSG00000160014 | CALM3 | chr19 | + | 46605826 | 46608340 | 46605826 | 46605857 | 46608196 | 46608340 | 41 | 16 | 29 | 50 | 26 | 578 | 415 | 433 | 678 | 297 | 194 | 94 | 67 | 106 | 143 | 1403 | 739 | 564 | 724 | 4542 | 298 | 149 | 0.002279419 | 0.012162123 | 0.034 | 0.019 | 0.032 | 0.036 | 0.042 | 0.065 | 0.06 | 0.056 | 0.068 | 0.015 | -0.02 |
| RI | 5930 | ENSG00000114388 | NPRL2 | chr3 | - | 50349385 | 50349828 | 50349385 | 50349494 | 50349664 | 50349828 | 44 | 26 | 37 | 69 | 28 | 38 | 16 | 39 | 27 | 35 | 63 | 82 | 48 | 48 | 42 | 18 | 25 | 2 | 21 | 51 | 298 | 149 | 0.002281299 | 0.012162123 | 0.367 | 0.448 | 0.322 | 0.561 | 0.286 | 0.636 | 0.621 | 0.923 | 0.5 | 0.46 | -0.231 |
| RI | 6031 | ENSG00000196730 | DAPK1 | chr9 | + | 87639649 | 87639815 | 87639649 | 87639698 | 87639788 | 87639815 | 22 | 33 | 12 | 12 | 11 | 104 | 120 | 54 | 138 | 222 | 117 | 21 | 25 | 43 | 19 | 277 | 104 | 101 | 149 | 249 | 239 | 149 | 0.002286852 | 0.012180736 | 0.117 | 0.146 | 0.122 | 0.051 | 0.03 | 0.208 | 0.112 | 0.134 | 0.152 | 0.045 | -0.037 |
| RI | 8054 | ENSG00000052749 | RRP12 | chr10 | - | 97358536 | 97359010 | 97358536 | 97358619 | 97358942 | 97359010 | 6 | 1 | 8 | 3 | 0 | 71 | 45 | 28 | 48 | 46 | 18 | 10 | 7 | 33 | 13 | 129 | 107 | 91 | 120 | 110 | 298 | 149 | 0.002293919 | 0.012207368 | 0.041 | 0.011 | 0.125 | 0.03 | 0 | 0.065 | 0.045 | 0.037 | 0.121 | 0.056 | -0.023 |
| RI | 8424 | ENSG00000136003 | ISCU | chr12 | + | 108564059 | 108564392 | 108564059 | 108564155 | 108564278 | 108564392 | 196 | 161 | 241 | 283 | 354 | 7 | 2 | 6 | 10 | 10 | 500 | 312 | 432 | 444 | 305 | 39 | 16 | 18 | 15 | 11 | 272 | 149 | 0.002297779 | 0.012216903 | 0.939 | 0.978 | 0.957 | 0.939 | 0.951 | 0.875 | 0.914 | 0.929 | 0.942 | 0.938 | 0.033 |
| RI | 5994 | ENSG00000156508 | EEF1A1 | chr6 | - | 73517514 | 73518264 | 73517514 | 73517934 | 73518029 | 73518264 | 432 | 281 | 497 | 403 | 1352 | 25700 | 14994 | 30394 | 24309 | 44917 | 376 | 318 | 238 | 361 | 278 | 14786 | 11614 | 11488 | 9559 | 18309 | 244 | 149 | 0.002327654 | 0.012364615 | 0.01 | 0.011 | 0.01 | 0.01 | 0.018 | 0.015 | 0.016 | 0.012 | 0.023 | 0.009 | -0.003 |
| RI | 3882 | ENSG00000117593 | DARS2 | chr1 | + | 173826686 | 173828399 | 173826686 | 173826786 | 173828332 | 173828399 | 2 | 12 | 3 | 1 | 2 | 30 | 18 | 30 | 37 | 39 | 4 | 5 | 18 | 14 | 0 | 41 | 14 | 24 | 24 | 5 | 298 | 149 | 0.002352006 | 0.01248275 | 0.032 | 0.25 | 0.048 | 0.013 | 0.025 | 0.047 | 0.152 | 0.273 | 0.226 | 0 | -0.066 |
| RI | 1847 | ENSG00000100722 | ZC3H14 | chr14 | + | 88609266 | 88609803 | 88609266 | 88609403 | 88609714 | 88609803 | 23 | 34 | 31 | 35 | 84 | 60 | 55 | 50 | 70 | 152 | 27 | 46 | 18 | 9 | 23 | 19 | 11 | 12 | 30 | 0 | 298 | 149 | 0.002360232 | 0.012515164 | 0.161 | 0.236 | 0.237 | 0.2 | 0.216 | 0.415 | 0.676 | 0.429 | 0.13 | 1 | -0.32 |
| RI | 5908 | ENSG00000165792 | METTL17 | chr14 | + | 20990229 | 20990598 | 20990229 | 20990383 | 20990463 | 20990598 | 25 | 15 | 28 | 37 | 5 | 46 | 35 | 42 | 59 | 29 | 42 | 41 | 43 | 65 | 64 | 64 | 48 | 28 | 54 | 89 | 229 | 149 | 0.002373051 | 0.01257185 | 0.261 | 0.218 | 0.303 | 0.29 | 0.101 | 0.299 | 0.357 | 0.5 | 0.439 | 0.319 | -0.148 |
| RI | 6332 | ENSG00000136286 | MYO1G | chr7 | - | 44962969 | 44964162 | 44962969 | 44963124 | 44964048 | 44964162 | 46 | 22 | 15 | 27 | 6 | 198 | 75 | 77 | 124 | 66 | 24 | 59 | 24 | 41 | 60 | 103 | 217 | 65 | 130 | 124 | 298 | 149 | 0.00237811 | 0.012587363 | 0.104 | 0.128 | 0.089 | 0.098 | 0.043 | 0.104 | 0.12 | 0.156 | 0.136 | 0.195 | -0.05 |
| RI | 7263 | ENSG00000104365 | IKBKB | chr8 | + | 42322053 | 42322494 | 42322053 | 42322153 | 42322346 | 42322494 | 45 | 38 | 51 | 27 | 24 | 64 | 46 | 55 | 50 | 60 | 65 | 39 | 93 | 61 | 58 | 63 | 45 | 30 | 37 | 69 | 298 | 149 | 0.002380758 | 0.012590094 | 0.26 | 0.292 | 0.317 | 0.213 | 0.167 | 0.34 | 0.302 | 0.608 | 0.452 | 0.296 | -0.15 |
| RI | 5950 | ENSG00000197756 | RPL37A | chr2 | + | 216498824 | 216499398 | 216498824 | 216498877 | 216499269 | 216499398 | 109 | 27 | 99 | 97 | 106 | 1215 | 435 | 1365 | 1117 | 770 | 113 | 96 | 44 | 50 | 85 | 662 | 924 | 459 | 413 | 735 | 298 | 149 | 0.002403442 | 0.012698687 | 0.043 | 0.03 | 0.035 | 0.042 | 0.064 | 0.079 | 0.049 | 0.046 | 0.057 | 0.055 | -0.014 |
| RI | 7536 | ENSG00000130714 | POMT1 | chr9 | + | 131507367 | 131509022 | 131507367 | 131507514 | 131508910 | 131509022 | 14 | 8 | 24 | 8 | 3 | 5 | 4 | 11 | 3 | 6 | 26 | 17 | 7 | 19 | 30 | 0 | 11 | 4 | 0 | 1 | 298 | 149 | 0.002420646 | 0.012778153 | 0.583 | 0.5 | 0.522 | 0.571 | 0.2 | 1 | 0.436 | 0.467 | 1 | 0.938 | -0.293 |
| RI | 3641 | ENSG00000239900 | ADSL | chr22 | + | 40349831 | 40353117 | 40349831 | 40350035 | 40353072 | 40353117 | 15 | 14 | 19 | 23 | 26 | 91 | 54 | 90 | 57 | 121 | 23 | 19 | 32 | 48 | 26 | 104 | 47 | 28 | 67 | 105 | 298 | 149 | 0.002440189 | 0.012869819 | 0.076 | 0.115 | 0.095 | 0.168 | 0.097 | 0.1 | 0.168 | 0.364 | 0.264 | 0.11 | -0.091 |
| RI | 7324 | ENSG00000137207 | YIPF3 | chr6 | - | 43513068 | 43513451 | 43513068 | 43513200 | 43513358 | 43513451 | 29 | 12 | 17 | 25 | 6 | 94 | 41 | 65 | 97 | 124 | 35 | 25 | 22 | 35 | 68 | 97 | 139 | 48 | 91 | 167 | 298 | 149 | 0.002454417 | 0.012933311 | 0.134 | 0.128 | 0.116 | 0.114 | 0.024 | 0.153 | 0.083 | 0.186 | 0.161 | 0.169 | -0.047 |
| RI | 5567 | ENSG00000115282 | TTC31 | chr2 | + | 74490243 | 74491184 | 74490243 | 74490473 | 74491127 | 74491184 | 54 | 45 | 54 | 40 | 78 | 1 | 0 | 3 | 5 | 0 | 51 | 56 | 76 | 51 | 133 | 0 | 4 | 10 | 6 | 10 | 298 | 149 | 0.002492533 | 0.013122451 | 0.964 | 1 | 0.9 | 0.8 | 1 | 1 | 0.875 | 0.792 | 0.81 | 0.869 | 0.064 |
| RI | 6394 | ENSG00000099901 | RANBP1 | chr22 | + | 20126302 | 20127181 | 20126302 | 20126368 | 20126951 | 20127181 | 22 | 9 | 24 | 17 | 19 | 128 | 77 | 130 | 110 | 113 | 23 | 7 | 8 | 23 | 6 | 177 | 99 | 131 | 163 | 133 | 298 | 149 | 0.002505347 | 0.013171831 | 0.079 | 0.055 | 0.085 | 0.072 | 0.078 | 0.061 | 0.034 | 0.03 | 0.066 | 0.022 | 0.031 |
| RI | 8284 | ENSG00000110719 | TCIRG1 | chr11 | + | 68042945 | 68043497 | 68042945 | 68043031 | 68043370 | 68043497 | 96 | 24 | 43 | 46 | 12 | 16 | 1 | 4 | 5 | 5 | 36 | 110 | 75 | 78 | 112 | 32 | 52 | 8 | 23 | 47 | 298 | 149 | 0.002506372 | 0.013171831 | 0.75 | 0.923 | 0.843 | 0.821 | 0.545 | 0.36 | 0.514 | 0.824 | 0.629 | 0.544 | 0.202 |
| RI | 4635 | ENSG00000129084 | PSMA1 | chr11 | - | 14513816 | 14514491 | 14513816 | 14513887 | 14514402 | 14514491 | 233 | 168 | 143 | 205 | 415 | 356 | 185 | 288 | 292 | 314 | 48 | 133 | 175 | 114 | 34 | 258 | 233 | 230 | 216 | 52 | 298 | 149 | 0.002509661 | 0.013177393 | 0.247 | 0.312 | 0.199 | 0.26 | 0.398 | 0.085 | 0.222 | 0.276 | 0.209 | 0.246 | 0.076 |
| RI | 123 | ENSG00000100242 | SUN2 | chr22 | - | 38740266 | 38741050 | 38740266 | 38740432 | 38740888 | 38741050 | 70 | 27 | 85 | 106 | 22 | 5 | 1 | 3 | 1 | 1 | 78 | 70 | 67 | 96 | 98 | 0 | 0 | 0 | 0 | 0 | 298 | 149 | 0.002517402 | 0.013206298 | 0.875 | 0.931 | 0.934 | 0.981 | 0.917 | 1 | 1 | 1 | 1 | 1 | -0.072 |
| RI | 5648 | ENSG00000161202 | DVL3 | chr3 | + | 184166445 | 184166673 | 184166445 | 184166522 | 184166605 | 184166673 | 15 | 3 | 14 | 22 | 16 | 96 | 47 | 70 | 81 | 118 | 48 | 12 | 14 | 37 | 55 | 132 | 133 | 60 | 81 | 154 | 232 | 149 | 0.002523058 | 0.013224229 | 0.091 | 0.039 | 0.114 | 0.149 | 0.08 | 0.189 | 0.055 | 0.13 | 0.227 | 0.187 | -0.063 |
| RI | 3650 | ENSG00000177728 | TMEM94 | chr17 | + | 75492928 | 75493593 | 75492928 | 75493102 | 75493490 | 75493593 | 12 | 3 | 12 | 8 | 12 | 17 | 9 | 7 | 8 | 13 | 34 | 22 | 8 | 19 | 34 | 10 | 13 | 2 | 14 | 14 | 298 | 149 | 0.002527949 | 0.013238113 | 0.261 | 0.143 | 0.462 | 0.333 | 0.316 | 0.63 | 0.458 | 0.667 | 0.404 | 0.548 | -0.238 |
| RI | 1418 | ENSG00000039523 | RIPOR1 | chr16 | + | 67545948 | 67546231 | 67545948 | 67546032 | 67546140 | 67546231 | 5 | 2 | 11 | 11 | 5 | 11 | 15 | 22 | 18 | 15 | 20 | 10 | 16 | 18 | 42 | 1 | 38 | 15 | 17 | 12 | 257 | 149 | 0.002545706 | 0.013319294 | 0.209 | 0.072 | 0.225 | 0.262 | 0.162 | 0.921 | 0.132 | 0.382 | 0.38 | 0.67 | -0.311 |
| RI | 4647 | ENSG00000147274 | RBMX | chrX | - | 136879016 | 136879453 | 136879016 | 136879158 | 136879318 | 136879453 | 32 | 28 | 44 | 19 | 33 | 7 | 9 | 1 | 7 | 1 | 32 | 77 | 44 | 48 | 58 | 0 | 0 | 8 | 4 | 0 | 298 | 149 | 0.002549397 | 0.013326805 | 0.696 | 0.609 | 0.957 | 0.576 | 0.943 | 1 | 1 | 0.733 | 0.857 | 1 | -0.162 |
| RI | 3566 | ENSG00000265241 | RBM8A | chr1 | - | 145926481 | 145927077 | 145926481 | 145926618 | 145926808 | 145927077 | 95 | 100 | 76 | 134 | 91 | 543 | 364 | 448 | 613 | 534 | 299 | 142 | 99 | 269 | 90 | 803 | 542 | 654 | 839 | 467 | 298 | 149 | 0.002558221 | 0.013361107 | 0.08 | 0.121 | 0.078 | 0.099 | 0.079 | 0.157 | 0.116 | 0.07 | 0.138 | 0.088 | -0.022 |
| RI | 2790 | ENSG00000206573 | THUMPD3-AS1 | chr3 | - | 9390289 | 9391826 | 9390289 | 9390497 | 9391426 | 9391826 | 75 | 71 | 33 | 69 | 115 | 2 | 4 | 1 | 5 | 0 | 19 | 62 | 49 | 42 | 10 | 10 | 6 | 1 | 0 | 12 | 298 | 149 | 0.002563784 | 0.01337833 | 0.949 | 0.899 | 0.943 | 0.873 | 1 | 0.487 | 0.838 | 0.961 | 1 | 0.294 | 0.217 |
| RI | 6588 | ENSG00000234741 | GAS5 | chr1 | - | 173864256 | 173864506 | 173864256 | 173864304 | 173864483 | 173864506 | 149 | 120 | 179 | 144 | 345 | 494 | 279 | 634 | 432 | 1019 | 19 | 52 | 71 | 83 | 0 | 148 | 190 | 214 | 150 | 198 | 298 | 149 | 0.002571553 | 0.013407028 | 0.131 | 0.177 | 0.124 | 0.143 | 0.145 | 0.06 | 0.12 | 0.142 | 0.217 | 0 | 0.036 |
| RI | 4809 | ENSG00000111801 | BTN3A3 | chr6 | + | 26443956 | 26445985 | 26443956 | 26444304 | 26445703 | 26445985 | 104 | 67 | 79 | 115 | 51 | 107 | 46 | 79 | 85 | 137 | 24 | 60 | 29 | 27 | 36 | 52 | 96 | 39 | 37 | 115 | 298 | 149 | 0.00257874 | 0.013422911 | 0.327 | 0.421 | 0.333 | 0.404 | 0.157 | 0.188 | 0.238 | 0.271 | 0.267 | 0.135 | 0.109 |
| RI | 3789 | ENSG00000157978 | LDLRAP1 | chr1 | + | 25563660 | 25565207 | 25563660 | 25563791 | 25565172 | 25565207 | 20 | 10 | 34 | 22 | 19 | 84 | 52 | 119 | 115 | 79 | 8 | 5 | 16 | 3 | 7 | 32 | 47 | 29 | 57 | 195 | 298 | 149 | 0.002579144 | 0.013422911 | 0.106 | 0.088 | 0.125 | 0.087 | 0.107 | 0.111 | 0.051 | 0.216 | 0.026 | 0.018 | 0.018 |
| RI | 674 | ENSG00000130813 | SHFL | chr19 | + | 10089110 | 10089378 | 10089110 | 10089208 | 10089306 | 10089378 | 26 | 17 | 27 | 44 | 37 | 13 | 6 | 25 | 7 | 8 | 118 | 43 | 57 | 125 | 61 | 10 | 17 | 9 | 11 | 7 | 247 | 149 | 0.002599874 | 0.013518887 | 0.547 | 0.631 | 0.394 | 0.791 | 0.736 | 0.877 | 0.604 | 0.793 | 0.873 | 0.84 | -0.178 |
| RI | 3455 | ENSG00000144591 | GMPPA | chr2 | + | 219504082 | 219505761 | 219504082 | 219504213 | 219505457 | 219505761 | 25 | 9 | 13 | 19 | 3 | 3 | 0 | 2 | 0 | 1 | 0 | 7 | 17 | 23 | 11 | 11 | 0 | 2 | 4 | 5 | 298 | 149 | 0.002615511 | 0.013588235 | 0.806 | 1 | 0.765 | 1 | 0.6 | 0 | 1 | 0.81 | 0.742 | 0.524 | 0.219 |
| RI | 6541 | ENSG00000197448 | GSTK1 | chr7 | + | 143264547 | 143265092 | 143264547 | 143264676 | 143264991 | 143265092 | 99 | 88 | 114 | 117 | 99 | 739 | 373 | 535 | 643 | 621 | 213 | 151 | 141 | 180 | 195 | 934 | 704 | 478 | 540 | 1251 | 298 | 149 | 0.002647932 | 0.01374458 | 0.063 | 0.106 | 0.096 | 0.083 | 0.074 | 0.102 | 0.097 | 0.129 | 0.143 | 0.072 | -0.024 |
| RI | 5928 | ENSG00000114388 | NPRL2 | chr3 | - | 50348684 | 50349010 | 50348684 | 50348782 | 50348873 | 50349010 | 23 | 1 | 6 | 13 | 8 | 43 | 21 | 40 | 48 | 38 | 18 | 19 | 16 | 37 | 20 | 58 | 41 | 23 | 46 | 51 | 240 | 149 | 0.002684214 | 0.013920676 | 0.249 | 0.029 | 0.085 | 0.144 | 0.116 | 0.162 | 0.223 | 0.302 | 0.333 | 0.196 | -0.119 |
| RI | 5365 | ENSG00000123136 | DDX39A | chr19 | - | 14409741 | 14410334 | 14409741 | 14409873 | 14410215 | 14410334 | 83 | 51 | 57 | 73 | 39 | 56 | 25 | 51 | 54 | 58 | 149 | 134 | 85 | 214 | 157 | 45 | 64 | 51 | 63 | 105 | 298 | 149 | 0.002688034 | 0.013928263 | 0.426 | 0.505 | 0.358 | 0.403 | 0.252 | 0.623 | 0.511 | 0.455 | 0.629 | 0.428 | -0.14 |
| RI | 5622 | ENSG00000119912 | IDE | chr10 | - | 92507574 | 92508205 | 92507574 | 92507666 | 92508112 | 92508205 | 23 | 13 | 18 | 16 | 32 | 84 | 59 | 99 | 82 | 64 | 15 | 13 | 28 | 9 | 14 | 0 | 68 | 46 | 23 | 0 | 298 | 149 | 0.002730563 | 0.014136227 | 0.12 | 0.099 | 0.083 | 0.089 | 0.2 | 1 | 0.087 | 0.233 | 0.164 | 1 | -0.379 |
| RI | 1234 | ENSG00000162066 | AMDHD2 | chr16 | + | 2528451 | 2528718 | 2528451 | 2528559 | 2528649 | 2528718 | 14 | 2 | 6 | 6 | 0 | 7 | 1 | 8 | 2 | 1 | 28 | 19 | 4 | 28 | 35 | 0 | 4 | 1 | 3 | 8 | 239 | 149 | 0.002776656 | 0.014362264 | 0.555 | 0.555 | 0.319 | 0.652 | 0 | 1 | 0.748 | 0.714 | 0.853 | 0.732 | -0.393 |
| RI | 2618 | ENSG00000173020 | GRK2 | chr11 | + | 67280731 | 67281184 | 67280731 | 67280783 | 67281092 | 67281184 | 122 | 42 | 105 | 115 | 96 | 718 | 374 | 621 | 666 | 664 | 126 | 111 | 81 | 194 | 235 | 1016 | 655 | 397 | 704 | 861 | 298 | 149 | 0.00278441 | 0.014389773 | 0.078 | 0.053 | 0.078 | 0.079 | 0.067 | 0.058 | 0.078 | 0.093 | 0.121 | 0.12 | -0.023 |
| RI | 4721 | ENSG00000129968 | ABHD17A | chr19 | - | 1877507 | 1880115 | 1877507 | 1877687 | 1879920 | 1880115 | 31 | 14 | 32 | 13 | 19 | 186 | 29 | 105 | 153 | 63 | 10 | 19 | 9 | 19 | 1 | 52 | 149 | 33 | 78 | 191 | 298 | 149 | 0.002814285 | 0.01453145 | 0.077 | 0.194 | 0.132 | 0.041 | 0.131 | 0.088 | 0.06 | 0.12 | 0.109 | 0.003 | 0.039 |
| RI | 1268 | ENSG00000168701 | TMEM208 | chr16 | + | 67228494 | 67228881 | 67228494 | 67228657 | 67228796 | 67228881 | 17 | 2 | 19 | 30 | 5 | 0 | 0 | 4 | 0 | 0 | 48 | 28 | 23 | 46 | 35 | 0 | 5 | 6 | 0 | 0 | 288 | 149 | 0.002847116 | 0.014688136 | 1 | 1 | 0.711 | 1 | 1 | 1 | 0.743 | 0.665 | 1 | 1 | 0.061 |
| RI | 6835 | ENSG00000184634 | MED12 | chrX | + | 71137725 | 71137943 | 71137725 | 71137780 | 71137855 | 71137943 | 59 | 33 | 50 | 53 | 30 | 2 | 7 | 4 | 0 | 0 | 57 | 56 | 40 | 92 | 69 | 0 | 5 | 0 | 0 | 0 | 224 | 149 | 0.002850981 | 0.01469524 | 0.952 | 0.758 | 0.893 | 1 | 1 | 1 | 0.882 | 1 | 1 | 1 | -0.056 |
| RI | 1661 | ENSG00000185088 | RPS27L | chr15 | - | 63153852 | 63155731 | 63153852 | 63154060 | 63155620 | 63155731 | 52 | 32 | 38 | 31 | 25 | 236 | 211 | 209 | 158 | 116 | 17 | 11 | 43 | 20 | 13 | 175 | 204 | 124 | 186 | 155 | 298 | 149 | 0.002860053 | 0.014729149 | 0.099 | 0.07 | 0.083 | 0.089 | 0.097 | 0.046 | 0.026 | 0.148 | 0.051 | 0.04 | 0.025 |
| RI | 6352 | ENSG00000198198 | SZT2 | chr1 | + | 43447848 | 43448484 | 43447848 | 43447971 | 43448078 | 43448484 | 3 | 4 | 3 | 3 | 0 | 38 | 8 | 33 | 21 | 7 | 0 | 2 | 2 | 0 | 0 | 41 | 33 | 14 | 15 | 27 | 256 | 149 | 0.002863769 | 0.014735437 | 0.044 | 0.225 | 0.05 | 0.077 | 0 | 0 | 0.034 | 0.077 | 0 | 0 | 0.057 |
| RI | 7031 | ENSG00000147475 | ERLIN2 | chr8 | + | 37744570 | 37746550 | 37744570 | 37744696 | 37745568 | 37746550 | 5 | 31 | 15 | 14 | 32 | 11 | 13 | 15 | 7 | 17 | 0 | 6 | 13 | 13 | 0 | 24 | 6 | 4 | 21 | 15 | 298 | 149 | 0.002899052 | 0.014904005 | 0.185 | 0.544 | 0.333 | 0.5 | 0.485 | 0 | 0.333 | 0.619 | 0.236 | 0 | 0.172 |
| RI | 5175 | ENSG00000104907 | TRMT1 | chr19 | - | 13105266 | 13105606 | 13105266 | 13105396 | 13105486 | 13105606 | 40 | 25 | 41 | 45 | 30 | 27 | 6 | 25 | 29 | 16 | 86 | 29 | 53 | 61 | 126 | 12 | 23 | 4 | 4 | 46 | 239 | 149 | 0.002904874 | 0.014920947 | 0.48 | 0.722 | 0.506 | 0.492 | 0.539 | 0.817 | 0.44 | 0.892 | 0.905 | 0.631 | -0.189 |
| RI | 4624 | ENSG00000269713 | NBPF9 | chr1 | - | 149081964 | 149082311 | 149081964 | 149082174 | 149082271 | 149082311 | 4 | 1 | 0 | 1 | 12 | 20 | 22 | 51 | 31 | 50 | 18 | 11 | 17 | 1 | 0 | 77 | 26 | 32 | 29 | 15 | 246 | 149 | 0.002921103 | 0.014991275 | 0.108 | 0.027 | 0 | 0.019 | 0.127 | 0.124 | 0.204 | 0.243 | 0.02 | 0 | -0.062 |
| RI | 8193 | ENSG00000130640 | TUBGCP2 | chr10 | - | 133293038 | 133293769 | 133293038 | 133293238 | 133293561 | 133293769 | 18 | 6 | 6 | 7 | 5 | 94 | 46 | 61 | 50 | 19 | 42 | 16 | 9 | 9 | 59 | 89 | 84 | 72 | 57 | 170 | 298 | 149 | 0.002931639 | 0.015032284 | 0.087 | 0.061 | 0.047 | 0.065 | 0.116 | 0.191 | 0.087 | 0.059 | 0.073 | 0.148 | -0.036 |
| RI | 1738 | ENSG00000172590 | MRPL52 | chr14 | + | 22829886 | 22830110 | 22829886 | 22829940 | 22830052 | 22830110 | 0 | 2 | 0 | 2 | 0 | 1 | 2 | 1 | 5 | 3 | 2 | 3 | 9 | 5 | 0 | 16 | 8 | 4 | 7 | 15 | 261 | 149 | 0.002945878 | 0.015083402 | 0 | 0.363 | 0 | 0.186 | 0 | 0.067 | 0.176 | 0.562 | 0.29 | 0 | -0.109 |
| RI | 4524 | ENSG00000204469 | PRRC2A | chr6 | + | 31635581 | 31636049 | 31635581 | 31635749 | 31635966 | 31636049 | 2 | 5 | 6 | 13 | 6 | 339 | 145 | 228 | 269 | 281 | 15 | 6 | 17 | 11 | 19 | 388 | 272 | 226 | 215 | 351 | 298 | 149 | 0.002946715 | 0.015083402 | 0.003 | 0.017 | 0.013 | 0.024 | 0.011 | 0.019 | 0.011 | 0.036 | 0.025 | 0.026 | -0.01 |
| RI | 2780 | ENSG00000178950 | GAK | chr4 | - | 850935 | 851974 | 850935 | 851084 | 851749 | 851974 | 26 | 25 | 51 | 29 | 21 | 118 | 67 | 71 | 112 | 40 | 104 | 49 | 53 | 82 | 106 | 153 | 142 | 94 | 102 | 162 | 298 | 149 | 0.002954819 | 0.01511179 | 0.099 | 0.157 | 0.264 | 0.115 | 0.208 | 0.254 | 0.147 | 0.22 | 0.287 | 0.247 | -0.062 |
| RI | 5615 | ENSG00000243970 | PPIEL | chr1 | - | 39547578 | 39548951 | 39547578 | 39547634 | 39548861 | 39548951 | 28 | 17 | 20 | 20 | 21 | 3 | 0 | 4 | 2 | 0 | 72 | 38 | 16 | 25 | 34 | 0 | 0 | 0 | 1 | 0 | 298 | 149 | 0.002966615 | 0.015158992 | 0.824 | 1 | 0.714 | 0.833 | 1 | 1 | 1 | 1 | 0.926 | 1 | -0.111 |
| RI | 7928 | ENSG00000166340 | TPP1 | chr11 | - | 6616974 | 6617776 | 6616974 | 6617428 | 6617625 | 6617776 | 4 | 0 | 0 | 5 | 0 | 278 | 125 | 169 | 222 | 243 | 29 | 19 | 6 | 0 | 10 | 278 | 300 | 147 | 283 | 871 | 298 | 149 | 0.002990647 | 0.015268585 | 0.007 | 0 | 0 | 0.011 | 0 | 0.05 | 0.031 | 0.02 | 0 | 0.006 | -0.018 |
| RI | 1560 | ENSG00000137876 | RSL24D1 | chr15 | - | 55183314 | 55185425 | 55183314 | 55183400 | 55185361 | 55185425 | 47 | 43 | 60 | 50 | 155 | 282 | 159 | 284 | 281 | 553 | 3 | 26 | 34 | 4 | 2 | 43 | 149 | 142 | 95 | 27 | 298 | 149 | 0.003020345 | 0.015406889 | 0.077 | 0.119 | 0.096 | 0.082 | 0.123 | 0.034 | 0.08 | 0.107 | 0.021 | 0.036 | 0.044 |
| RI | 7692 | ENSG00000168071 | CCDC88B | chr11 | + | 64351159 | 64351616 | 64351159 | 64351255 | 64351475 | 64351616 | 26 | 14 | 18 | 12 | 4 | 30 | 17 | 19 | 25 | 1 | 48 | 26 | 22 | 32 | 76 | 7 | 18 | 4 | 25 | 31 | 298 | 149 | 0.003043883 | 0.015513558 | 0.302 | 0.292 | 0.321 | 0.194 | 0.667 | 0.774 | 0.419 | 0.733 | 0.39 | 0.551 | -0.218 |
| RI | 6837 | ENSG00000147224 | PRPS1 | chrX | + | 107649939 | 107651026 | 107649939 | 107650522 | 107650577 | 107651026 | 23 | 14 | 21 | 26 | 13 | 0 | 1 | 0 | 0 | 0 | 16 | 11 | 8 | 12 | 1 | 0 | 0 | 4 | 0 | 5 | 204 | 149 | 0.003052988 | 0.015546552 | 1 | 0.911 | 1 | 1 | 1 | 1 | 1 | 0.594 | 1 | 0.127 | 0.238 |
| RI | 271 | ENSG00000011304 | PTBP1 | chr19 | + | 804291 | 804702 | 804291 | 804438 | 804531 | 804702 | 14 | 4 | 19 | 18 | 3 | 110 | 61 | 103 | 143 | 79 | 2 | 13 | 12 | 3 | 7 | 91 | 144 | 80 | 126 | 170 | 242 | 149 | 0.003105002 | 0.015773857 | 0.073 | 0.039 | 0.102 | 0.072 | 0.023 | 0.013 | 0.053 | 0.085 | 0.014 | 0.025 | 0.024 |
| RI | 6894 | ENSG00000102393 | GLA | chrX | - | 101400665 | 101401809 | 101400665 | 101400757 | 101401631 | 101401809 | 20 | 33 | 19 | 38 | 43 | 51 | 58 | 35 | 52 | 66 | 55 | 77 | 52 | 138 | 138 | 51 | 106 | 25 | 89 | 243 | 298 | 149 | 0.003105846 | 0.015773857 | 0.164 | 0.221 | 0.213 | 0.268 | 0.246 | 0.35 | 0.266 | 0.51 | 0.437 | 0.221 | -0.134 |
| RI | 6464 | ENSG00000113013 | HSPA9 | chr5 | - | 138553755 | 138556592 | 138553755 | 138556114 | 138556451 | 138556592 | 36 | 30 | 28 | 52 | 19 | 406 | 407 | 378 | 514 | 571 | 57 | 80 | 27 | 85 | 0 | 519 | 431 | 653 | 673 | 80 | 298 | 149 | 0.003107321 | 0.015773857 | 0.042 | 0.036 | 0.036 | 0.048 | 0.016 | 0.052 | 0.085 | 0.02 | 0.059 | 0 | -0.008 |
| RI | 8371 | ENSG00000139190 | VAMP1 | chr12 | - | 6462236 | 6464941 | 6462236 | 6464486 | 6464889 | 6464941 | 172 | 123 | 160 | 218 | 188 | 0 | 0 | 1 | 0 | 10 | 382 | 156 | 123 | 232 | 190 | 0 | 3 | 6 | 0 | 23 | 298 | 149 | 0.003108307 | 0.015773857 | 1 | 1 | 0.988 | 1 | 0.904 | 1 | 0.963 | 0.911 | 1 | 0.805 | 0.043 |
| RI | 3729 | ENSG00000179335 | CLK3 | chr15 | + | 74622119 | 74622560 | 74622119 | 74622216 | 74622493 | 74622560 | 127 | 49 | 84 | 124 | 123 | 76 | 28 | 64 | 76 | 117 | 306 | 111 | 104 | 277 | 323 | 69 | 91 | 65 | 32 | 99 | 298 | 149 | 0.003123791 | 0.015838827 | 0.455 | 0.467 | 0.396 | 0.449 | 0.345 | 0.689 | 0.379 | 0.444 | 0.812 | 0.62 | -0.166 |
| RI | 7227 | ENSG00000121022 | COPS5 | chr8 | - | 67043081 | 67045960 | 67043081 | 67043317 | 67045811 | 67045960 | 6 | 10 | 7 | 15 | 13 | 48 | 24 | 40 | 54 | 158 | 9 | 8 | 19 | 2 | 0 | 8 | 18 | 6 | 9 | 3 | 298 | 149 | 0.003141007 | 0.015912462 | 0.059 | 0.172 | 0.08 | 0.122 | 0.04 | 0.36 | 0.182 | 0.613 | 0.1 | 0 | -0.156 |
| RI | 1325 | ENSG00000214078 | CPNE1 | chr20 | - | 35626043 | 35626803 | 35626043 | 35626381 | 35626562 | 35626803 | 23 | 20 | 49 | 61 | 29 | 1 | 0 | 0 | 1 | 0 | 16 | 29 | 13 | 23 | 54 | 0 | 0 | 2 | 8 | 0 | 298 | 149 | 0.003172527 | 0.016058372 | 0.92 | 1 | 1 | 0.968 | 1 | 1 | 1 | 0.765 | 0.59 | 1 | 0.107 |
| RI | 2871 | ENSG00000075711 | DLG1 | chr3 | - | 197065275 | 197065809 | 197065275 | 197065448 | 197065707 | 197065809 | 100 | 90 | 110 | 103 | 168 | 178 | 129 | 136 | 144 | 407 | 2 | 37 | 56 | 32 | 0 | 23 | 119 | 89 | 49 | 28 | 298 | 149 | 0.003179985 | 0.01608234 | 0.219 | 0.259 | 0.288 | 0.263 | 0.171 | 0.042 | 0.135 | 0.239 | 0.246 | 0 | 0.108 |
| RI | 3563 | ENSG00000175203 | DCTN2 | chr12 | - | 57532014 | 57532643 | 57532014 | 57532106 | 57532571 | 57532643 | 4 | 6 | 6 | 7 | 8 | 0 | 0 | 0 | 2 | 0 | 52 | 21 | 34 | 14 | 8 | 3 | 8 | 0 | 0 | 0 | 298 | 149 | 0.003184547 | 0.016091633 | 1 | 1 | 1 | 0.636 | 1 | 0.897 | 0.568 | 1 | 1 | 1 | 0.034 |
| RI | 1984 | ENSG00000175061 | SNHG29 | chr17 | + | 16439036 | 16439414 | 16439036 | 16439060 | 16439326 | 16439414 | 78 | 37 | 68 | 68 | 55 | 194 | 92 | 239 | 135 | 134 | 50 | 41 | 56 | 77 | 55 | 85 | 89 | 73 | 82 | 151 | 298 | 149 | 0.00321522 | 0.01623274 | 0.167 | 0.167 | 0.125 | 0.201 | 0.17 | 0.227 | 0.187 | 0.277 | 0.32 | 0.154 | -0.067 |
| RI | 227 | ENSG00000214046 | SMIM7 | chr19 | - | 16646147 | 16647261 | 16646147 | 16646800 | 16647095 | 16647261 | 217 | 130 | 166 | 169 | 290 | 15 | 1 | 14 | 11 | 9 | 383 | 189 | 238 | 301 | 481 | 8 | 13 | 0 | 7 | 13 | 298 | 149 | 0.003220697 | 0.016246506 | 0.879 | 0.985 | 0.856 | 0.885 | 0.942 | 0.96 | 0.879 | 1 | 0.956 | 0.949 | -0.039 |
| RI | 1766 | ENSG00000100906 | NFKBIA | chr14 | - | 35402393 | 35402859 | 35402393 | 35402663 | 35402770 | 35402859 | 223 | 141 | 330 | 211 | 291 | 730 | 434 | 1004 | 1006 | 742 | 513 | 414 | 316 | 440 | 559 | 2611 | 1299 | 1346 | 2362 | 1738 | 256 | 149 | 0.003229943 | 0.016279245 | 0.151 | 0.159 | 0.161 | 0.109 | 0.186 | 0.103 | 0.156 | 0.12 | 0.098 | 0.158 | 0.026 |
| RI | 2430 | ENSG00000159199 | ATP5MC1 | chr17 | + | 48893408 | 48895334 | 48893408 | 48893456 | 48895155 | 48895334 | 6 | 12 | 13 | 2 | 11 | 2 | 1 | 4 | 0 | 2 | 28 | 9 | 6 | 10 | 23 | 0 | 1 | 3 | 0 | 0 | 298 | 149 | 0.0032344 | 0.01628781 | 0.6 | 0.857 | 0.619 | 1 | 0.733 | 1 | 0.818 | 0.5 | 1 | 1 | -0.102 |
| RI | 7401 | ENSG00000147684 | NDUFB9 | chr8 | + | 124543086 | 124547113 | 124543086 | 124543279 | 124546999 | 124547113 | 38 | 5 | 23 | 26 | 11 | 231 | 122 | 200 | 181 | 148 | 71 | 28 | 66 | 59 | 10 | 283 | 234 | 179 | 265 | 357 | 298 | 149 | 0.003253408 | 0.016369573 | 0.076 | 0.02 | 0.054 | 0.067 | 0.036 | 0.111 | 0.056 | 0.156 | 0.1 | 0.014 | -0.037 |
| RI | 4014 | ENSG00000069424 | KCNAB2 | chr1 | + | 6094399 | 6095443 | 6094399 | 6094485 | 6095322 | 6095443 | 30 | 17 | 22 | 26 | 2 | 120 | 53 | 75 | 102 | 65 | 13 | 45 | 49 | 32 | 117 | 74 | 146 | 55 | 102 | 248 | 298 | 149 | 0.003262434 | 0.01640102 | 0.111 | 0.138 | 0.128 | 0.113 | 0.015 | 0.081 | 0.134 | 0.308 | 0.136 | 0.191 | -0.069 |
| RI | 7518 | ENSG00000165699 | TSC1 | chr9 | - | 132910570 | 132911568 | 132910570 | 132910689 | 132911452 | 132911568 | 64 | 46 | 58 | 74 | 110 | 0 | 0 | 0 | 1 | 0 | 44 | 65 | 98 | 97 | 50 | 0 | 6 | 1 | 5 | 0 | 298 | 149 | 0.003291205 | 0.016531588 | 1 | 1 | 1 | 0.974 | 1 | 1 | 0.844 | 0.98 | 0.907 | 1 | 0.049 |
| RI | 6270 | ENSG00000183735 | TBK1 | chr12 | + | 64485454 | 64486017 | 64485454 | 64485513 | 64485925 | 64486017 | 16 | 21 | 10 | 19 | 11 | 177 | 113 | 111 | 152 | 65 | 0 | 45 | 33 | 3 | 0 | 17 | 132 | 82 | 15 | 33 | 298 | 149 | 0.003315398 | 0.016625926 | 0.043 | 0.085 | 0.043 | 0.059 | 0.078 | 0 | 0.146 | 0.168 | 0.091 | 1 | -0.019 |
| RI | 1368 | ENSG00000169592 | INO80E | chr16 | + | 29996243 | 29996617 | 29996243 | 29996391 | 29996546 | 29996617 | 26 | 12 | 23 | 37 | 16 | 148 | 55 | 117 | 100 | 116 | 82 | 23 | 46 | 12 | 43 | 50 | 124 | 53 | 83 | 123 | 298 | 149 | 0.003315616 | 0.016625926 | 0.081 | 0.098 | 0.089 | 0.156 | 0.065 | 0.451 | 0.085 | 0.303 | 0.067 | 0.149 | -0.113 |
| RI | 1197 | ENSG00000033011 | ALG1 | chr16 | + | 5082558 | 5083757 | 5082558 | 5082673 | 5083681 | 5083757 | 3 | 3 | 2 | 3 | 8 | 25 | 18 | 17 | 29 | 14 | 0 | 0 | 5 | 1 | 0 | 48 | 12 | 12 | 7 | 1 | 298 | 149 | 0.003318681 | 0.016627185 | 0.057 | 0.077 | 0.056 | 0.049 | 0.222 | 0 | 0 | 0.172 | 0.067 | 0 | 0.044 |
| RI | 8043 | ENSG00000166987 | MBD6 | chr12 | + | 57524028 | 57524416 | 57524028 | 57524092 | 57524279 | 57524416 | 12 | 11 | 11 | 14 | 4 | 23 | 12 | 39 | 18 | 13 | 39 | 32 | 21 | 35 | 82 | 52 | 49 | 20 | 29 | 18 | 298 | 149 | 0.003334582 | 0.016677925 | 0.207 | 0.314 | 0.124 | 0.28 | 0.133 | 0.273 | 0.246 | 0.344 | 0.376 | 0.695 | -0.175 |
| RI | 3458 | ENSG00000144591 | GMPPA | chr2 | + | 219505457 | 219505761 | 219505457 | 219505555 | 219505694 | 219505761 | 20 | 5 | 16 | 13 | 16 | 0 | 0 | 1 | 0 | 0 | 8 | 10 | 12 | 7 | 8 | 0 | 4 | 0 | 0 | 4 | 288 | 149 | 0.003335015 | 0.016677925 | 1 | 1 | 0.892 | 1 | 1 | 1 | 0.564 | 1 | 1 | 0.509 | 0.164 |
| RI | 819 | ENSG00000184009 | ACTG1 | chr17 | - | 81511187 | 81512142 | 81511187 | 81511578 | 81511926 | 81512142 | 977 | 358 | 773 | 660 | 620 | 40 | 23 | 53 | 43 | 22 | 657 | 691 | 463 | 706 | 2317 | 45 | 47 | 41 | 58 | 166 | 298 | 149 | 0.003337279 | 0.016677925 | 0.924 | 0.886 | 0.879 | 0.885 | 0.934 | 0.88 | 0.88 | 0.85 | 0.859 | 0.875 | 0.033 |
| RI | 8322 | ENSG00000123384 | LRP1 | chr12 | + | 57208037 | 57208817 | 57208037 | 57208216 | 57208710 | 57208817 | 6 | 2 | 4 | 1 | 0 | 139 | 64 | 77 | 92 | 65 | 10 | 14 | 12 | 8 | 13 | 324 | 139 | 81 | 141 | 337 | 298 | 149 | 0.0033796 | 0.016875145 | 0.021 | 0.015 | 0.025 | 0.005 | 0 | 0.015 | 0.048 | 0.069 | 0.028 | 0.019 | -0.023 |
| RI | 1978 | ENSG00000006744 | ELAC2 | chr17 | - | 13002273 | 13002579 | 13002273 | 13002359 | 13002445 | 13002579 | 13 | 8 | 17 | 32 | 5 | 1 | 2 | 2 | 0 | 0 | 42 | 59 | 24 | 32 | 84 | 11 | 5 | 6 | 4 | 0 | 235 | 149 | 0.003394236 | 0.016933914 | 0.892 | 0.717 | 0.843 | 1 | 1 | 0.708 | 0.882 | 0.717 | 0.835 | 1 | 0.062 |
| RI | 6675 | ENSG00000164062 | APEH | chr3 | + | 49676055 | 49676515 | 49676055 | 49676219 | 49676377 | 49676515 | 42 | 14 | 51 | 58 | 10 | 366 | 189 | 233 | 279 | 275 | 72 | 58 | 35 | 61 | 118 | 521 | 320 | 172 | 280 | 631 | 298 | 149 | 0.00346163 | 0.017255566 | 0.054 | 0.036 | 0.099 | 0.094 | 0.018 | 0.065 | 0.083 | 0.092 | 0.098 | 0.086 | -0.025 |
| RI | 2696 | ENSG00000178678 | GPAT3 | chr4 | + | 83536107 | 83536763 | 83536107 | 83536206 | 83536503 | 83536763 | 20 | 11 | 16 | 10 | 22 | 5 | 0 | 0 | 0 | 0 | 20 | 17 | 8 | 14 | 24 | 6 | 1 | 0 | 3 | 0 | 298 | 149 | 0.003472453 | 0.017294925 | 0.667 | 1 | 1 | 1 | 1 | 0.625 | 0.895 | 1 | 0.7 | 1 | 0.089 |
| RI | 253 | ENSG00000130816 | DNMT1 | chr19 | - | 10160017 | 10160418 | 10160017 | 10160063 | 10160383 | 10160418 | 21 | 16 | 33 | 14 | 10 | 18 | 9 | 24 | 19 | 46 | 93 | 23 | 29 | 57 | 77 | 60 | 12 | 14 | 43 | 28 | 298 | 149 | 0.003493515 | 0.017385169 | 0.368 | 0.471 | 0.407 | 0.269 | 0.098 | 0.437 | 0.489 | 0.509 | 0.399 | 0.579 | -0.16 |
| RI | 1902 | ENSG00000090061 | CCNK | chr14 | + | 99502718 | 99503644 | 99502718 | 99502984 | 99503610 | 99503644 | 29 | 14 | 19 | 29 | 32 | 18 | 17 | 14 | 21 | 26 | 12 | 62 | 39 | 55 | 33 | 61 | 46 | 37 | 58 | 136 | 298 | 149 | 0.003498498 | 0.017395311 | 0.446 | 0.292 | 0.404 | 0.408 | 0.381 | 0.09 | 0.403 | 0.345 | 0.322 | 0.108 | 0.133 |
| RI | 8480 | ENSG00000166598 | HSP90B1 | chr12 | + | 103931520 | 103932418 | 103931520 | 103931748 | 103932276 | 103932418 | 82 | 39 | 126 | 115 | 190 | 7 | 1 | 2 | 7 | 2 | 81 | 88 | 94 | 90 | 43 | 0 | 0 | 6 | 0 | 0 | 298 | 149 | 0.003515951 | 0.017467384 | 0.854 | 0.951 | 0.969 | 0.891 | 0.979 | 1 | 1 | 0.887 | 1 | 1 | -0.049 |
| RI | 7028 | ENSG00000111364 | DDX55 | chr12 | + | 123607431 | 123607662 | 123607431 | 123607523 | 123607599 | 123607662 | 21 | 16 | 33 | 29 | 34 | 41 | 33 | 38 | 45 | 37 | 100 | 21 | 24 | 109 | 46 | 46 | 37 | 19 | 53 | 43 | 225 | 149 | 0.003524488 | 0.017495086 | 0.253 | 0.243 | 0.365 | 0.299 | 0.378 | 0.59 | 0.273 | 0.455 | 0.577 | 0.415 | -0.154 |
| RI | 5642 | ENSG00000163516 | ANKZF1 | chr2 | + | 219236009 | 219236651 | 219236009 | 219236095 | 219236321 | 219236651 | 6 | 6 | 18 | 7 | 4 | 60 | 29 | 75 | 74 | 44 | 15 | 8 | 42 | 16 | 10 | 49 | 46 | 57 | 55 | 118 | 298 | 149 | 0.003583069 | 0.017770939 | 0.048 | 0.094 | 0.107 | 0.045 | 0.043 | 0.133 | 0.08 | 0.269 | 0.127 | 0.041 | -0.063 |
| RI | 1292 | ENSG00000197912 | SPG7 | chr16 | + | 89529476 | 89530808 | 89529476 | 89529579 | 89530682 | 89530808 | 16 | 6 | 17 | 9 | 2 | 23 | 21 | 38 | 19 | 17 | 21 | 13 | 22 | 37 | 45 | 48 | 33 | 11 | 30 | 20 | 298 | 149 | 0.003606904 | 0.017874144 | 0.258 | 0.125 | 0.183 | 0.191 | 0.056 | 0.179 | 0.165 | 0.5 | 0.381 | 0.529 | -0.188 |
| RI | 4054 | ENSG00000011295 | TTC19 | chr17 | + | 15999823 | 16000245 | 15999823 | 16000032 | 16000117 | 16000245 | 1 | 1 | 1 | 2 | 0 | 9 | 6 | 4 | 3 | 2 | 11 | 5 | 0 | 4 | 0 | 9 | 7 | 9 | 4 | 13 | 234 | 149 | 0.003634813 | 0.017981365 | 0.066 | 0.096 | 0.137 | 0.298 | 0 | 0.438 | 0.313 | 0 | 0.389 | 0 | -0.109 |
| RI | 7987 | ENSG00000103510 | KAT8 | chr16 | + | 31130455 | 31130900 | 31130455 | 31130606 | 31130745 | 31130900 | 19 | 18 | 16 | 24 | 13 | 71 | 31 | 58 | 86 | 89 | 64 | 51 | 27 | 23 | 10 | 70 | 95 | 32 | 45 | 90 | 288 | 149 | 0.003637234 | 0.017981365 | 0.122 | 0.231 | 0.125 | 0.126 | 0.07 | 0.321 | 0.217 | 0.304 | 0.209 | 0.054 | -0.086 |
| RI | 5334 | ENSG00000064703 | DDX20 | chr1 | + | 111760705 | 111763007 | 111760705 | 111762782 | 111762905 | 111763007 | 19 | 13 | 32 | 15 | 30 | 40 | 60 | 64 | 38 | 112 | 21 | 32 | 39 | 14 | 7 | 33 | 44 | 52 | 19 | 27 | 272 | 149 | 0.003637673 | 0.017981365 | 0.206 | 0.106 | 0.215 | 0.178 | 0.128 | 0.258 | 0.285 | 0.291 | 0.288 | 0.124 | -0.083 |
| RI | 8471 | ENSG00000177575 | CD163 | chr12 | - | 7479859 | 7481256 | 7479859 | 7479913 | 7481160 | 7481256 | 43 | 31 | 16 | 31 | 43 | 122 | 91 | 63 | 160 | 132 | 83 | 130 | 133 | 63 | 7 | 362 | 818 | 490 | 254 | 171 | 298 | 149 | 0.003645041 | 0.018002724 | 0.15 | 0.146 | 0.113 | 0.088 | 0.14 | 0.103 | 0.074 | 0.119 | 0.11 | 0.02 | 0.042 |
| RI | 1987 | ENSG00000175061 | SNHG29 | chr17 | + | 16439326 | 16439703 | 16439326 | 16439414 | 16439580 | 16439703 | 132 | 69 | 149 | 143 | 107 | 18 | 13 | 36 | 18 | 11 | 210 | 182 | 154 | 248 | 166 | 11 | 18 | 19 | 25 | 16 | 298 | 149 | 0.003670902 | 0.018115303 | 0.786 | 0.726 | 0.674 | 0.799 | 0.829 | 0.905 | 0.835 | 0.802 | 0.832 | 0.838 | -0.08 |
| RI | 8119 | ENSG00000119969 | HELLS | chr10 | + | 94592228 | 94592514 | 94592228 | 94592312 | 94592394 | 94592514 | 0 | 0 | 1 | 1 | 5 | 17 | 23 | 16 | 38 | 56 | 0 | 11 | 3 | 4 | 0 | 14 | 33 | 58 | 16 | 9 | 231 | 149 | 0.003676146 | 0.018126037 | 0 | 0 | 0.039 | 0.017 | 0.054 | 0 | 0.177 | 0.032 | 0.139 | 0 | -0.048 |
| RI | 401 | ENSG00000090372 | STRN4 | chr19 | - | 46721985 | 46722340 | 46721985 | 46722072 | 46722241 | 46722340 | 10 | 15 | 8 | 14 | 14 | 78 | 48 | 33 | 72 | 39 | 24 | 22 | 16 | 38 | 52 | 64 | 68 | 53 | 69 | 158 | 298 | 149 | 0.003680519 | 0.018131018 | 0.06 | 0.135 | 0.108 | 0.089 | 0.152 | 0.158 | 0.139 | 0.131 | 0.216 | 0.141 | -0.048 |
| RI | 7701 | ENSG00000168067 | MAP4K2 | chr11 | - | 64797274 | 64797534 | 64797274 | 64797380 | 64797500 | 64797534 | 24 | 17 | 23 | 37 | 23 | 66 | 41 | 55 | 59 | 84 | 8 | 43 | 17 | 16 | 31 | 82 | 100 | 30 | 64 | 262 | 269 | 149 | 0.003683295 | 0.018131018 | 0.168 | 0.187 | 0.188 | 0.258 | 0.132 | 0.051 | 0.192 | 0.239 | 0.122 | 0.062 | 0.053 |
| RI | 2160 | ENSG00000065357 | DGKA | chr12 | + | 55938914 | 55939305 | 55938914 | 55938989 | 55939185 | 55939305 | 56 | 44 | 73 | 55 | 39 | 145 | 133 | 276 | 245 | 241 | 47 | 70 | 38 | 61 | 79 | 172 | 124 | 166 | 178 | 142 | 298 | 149 | 0.00370301 | 0.01821289 | 0.162 | 0.142 | 0.117 | 0.101 | 0.075 | 0.12 | 0.22 | 0.103 | 0.146 | 0.218 | -0.042 |
| RI | 669 | ENSG00000099956 | SMARCB1 | chr22 | + | 23791755 | 23793688 | 23791755 | 23791867 | 23793558 | 23793688 | 56 | 38 | 55 | 44 | 55 | 83 | 67 | 82 | 75 | 147 | 167 | 77 | 29 | 89 | 126 | 150 | 162 | 39 | 103 | 122 | 298 | 149 | 0.003717813 | 0.018270482 | 0.252 | 0.221 | 0.251 | 0.227 | 0.158 | 0.358 | 0.192 | 0.271 | 0.302 | 0.341 | -0.071 |
| RI | 3247 | ENSG00000064961 | HMG20B | chr19 | + | 3572978 | 3573347 | 3572978 | 3572994 | 3573291 | 3573347 | 6 | 6 | 3 | 0 | 1 | 8 | 8 | 9 | 13 | 1 | 0 | 4 | 0 | 1 | 5 | 34 | 16 | 11 | 4 | 30 | 298 | 149 | 0.003728067 | 0.018291369 | 0.273 | 0.273 | 0.143 | 0 | 0.333 | 0 | 0.111 | 0 | 0.111 | 0.077 | 0.145 |
| RI | 4471 | ENSG00000072135 | PTPN18 | chr2 | + | 130370556 | 130370782 | 130370556 | 130370623 | 130370704 | 130370782 | 5 | 14 | 10 | 30 | 3 | 45 | 42 | 37 | 57 | 35 | 9 | 2 | 8 | 26 | 25 | 154 | 94 | 31 | 96 | 159 | 230 | 149 | 0.003728256 | 0.018291369 | 0.067 | 0.178 | 0.149 | 0.254 | 0.053 | 0.036 | 0.014 | 0.143 | 0.149 | 0.092 | 0.053 |
| RI | 1008 | ENSG00000173821 | RNF213 | chr17 | + | 80388611 | 80389367 | 80388611 | 80388689 | 80389172 | 80389367 | 49 | 30 | 42 | 43 | 72 | 337 | 162 | 256 | 335 | 401 | 159 | 71 | 60 | 87 | 31 | 601 | 341 | 226 | 403 | 268 | 298 | 149 | 0.003763762 | 0.018443705 | 0.068 | 0.085 | 0.076 | 0.06 | 0.082 | 0.117 | 0.094 | 0.117 | 0.097 | 0.055 | -0.022 |
| RI | 5569 | ENSG00000115282 | TTC31 | chr2 | + | 74492003 | 74494559 | 74492003 | 74492055 | 74492138 | 74494559 | 11 | 13 | 2 | 6 | 13 | 56 | 42 | 30 | 51 | 91 | 6 | 5 | 5 | 8 | 6 | 85 | 66 | 63 | 57 | 106 | 232 | 149 | 0.003765551 | 0.018443705 | 0.112 | 0.166 | 0.041 | 0.07 | 0.084 | 0.043 | 0.046 | 0.048 | 0.083 | 0.035 | 0.044 |
| RI | 7515 | ENSG00000165699 | TSC1 | chr9 | - | 132897183 | 132897610 | 132897183 | 132897345 | 132897422 | 132897610 | 5 | 8 | 3 | 7 | 5 | 45 | 31 | 42 | 51 | 64 | 8 | 0 | 4 | 0 | 0 | 38 | 50 | 52 | 70 | 28 | 226 | 149 | 0.003782804 | 0.018512859 | 0.068 | 0.145 | 0.045 | 0.083 | 0.049 | 0.122 | 0 | 0.048 | 0 | 0 | 0.044 |
| RI | 7178 | ENSG00000100023 | PPIL2 | chr22 | + | 21694754 | 21695070 | 21694754 | 21694817 | 21694936 | 21695070 | 13 | 5 | 11 | 10 | 2 | 96 | 60 | 64 | 69 | 52 | 14 | 10 | 14 | 16 | 26 | 115 | 59 | 20 | 55 | 123 | 268 | 149 | 0.003797958 | 0.018571639 | 0.07 | 0.044 | 0.087 | 0.075 | 0.021 | 0.063 | 0.086 | 0.28 | 0.139 | 0.105 | -0.075 |
| RI | 1758 | ENSG00000101384 | JAG1 | chr20 | - | 10648548 | 10649107 | 10648548 | 10648722 | 10649060 | 10649107 | 1 | 1 | 3 | 0 | 0 | 10 | 6 | 4 | 6 | 6 | 0 | 0 | 3 | 7 | 19 | 17 | 1 | 13 | 28 | 6 | 298 | 149 | 0.003807532 | 0.018603052 | 0.048 | 0.077 | 0.273 | 0 | 0 | 0 | 0 | 0.103 | 0.111 | 0.613 | -0.086 |
| RI | 2471 | ENSG00000105755 | ETHE1 | chr19 | - | 43526200 | 43526659 | 43526200 | 43526349 | 43526514 | 43526659 | 0 | 0 | 5 | 0 | 0 | 26 | 2 | 7 | 14 | 5 | 0 | 5 | 2 | 9 | 0 | 31 | 21 | 3 | 17 | 11 | 298 | 149 | 0.003832177 | 0.018707992 | 0 | 0 | 0.263 | 0 | 0 | 0 | 0.106 | 0.25 | 0.209 | 0 | -0.06 |
| RI | 8158 | ENSG00000107862 | GBF1 | chr10 | + | 102379852 | 102380686 | 102379852 | 102380362 | 102380505 | 102380686 | 8 | 2 | 6 | 12 | 0 | 42 | 24 | 34 | 28 | 23 | 24 | 7 | 8 | 34 | 15 | 30 | 55 | 29 | 31 | 70 | 292 | 149 | 0.003842139 | 0.018741138 | 0.089 | 0.041 | 0.083 | 0.179 | 0 | 0.29 | 0.061 | 0.123 | 0.359 | 0.099 | -0.108 |
| RI | 1433 | ENSG00000169189 | NSMCE1 | chr16 | - | 27225725 | 27226836 | 27225725 | 27225846 | 27226719 | 27226836 | 12 | 10 | 10 | 9 | 17 | 67 | 32 | 62 | 57 | 142 | 1 | 7 | 1 | 0 | 15 | 57 | 72 | 19 | 47 | 86 | 298 | 149 | 0.003890149 | 0.018959661 | 0.082 | 0.135 | 0.075 | 0.073 | 0.056 | 0.009 | 0.046 | 0.026 | 0 | 0.08 | 0.052 |
| RI | 7037 | ENSG00000100918 | REC8 | chr14 | + | 24172898 | 24173198 | 24172898 | 24173041 | 24173125 | 24173198 | 8 | 3 | 12 | 15 | 7 | 10 | 5 | 14 | 18 | 22 | 101 | 25 | 14 | 46 | 13 | 30 | 7 | 29 | 18 | 13 | 233 | 149 | 0.003900756 | 0.018995686 | 0.338 | 0.277 | 0.354 | 0.348 | 0.169 | 0.683 | 0.695 | 0.236 | 0.62 | 0.39 | -0.228 |
| RI | 6944 | ENSG00000172732 | MUS81 | chr11 | + | 65864719 | 65865323 | 65864719 | 65865145 | 65865219 | 65865323 | 19 | 10 | 20 | 18 | 7 | 77 | 37 | 49 | 57 | 42 | 47 | 23 | 13 | 20 | 36 | 52 | 47 | 41 | 49 | 87 | 223 | 149 | 0.003953142 | 0.019234933 | 0.142 | 0.153 | 0.214 | 0.174 | 0.1 | 0.377 | 0.246 | 0.175 | 0.214 | 0.217 | -0.089 |
| RI | 622 | ENSG00000062598 | ELMO2 | chr20 | - | 46368890 | 46370525 | 46368890 | 46368968 | 46370442 | 46370525 | 16 | 18 | 16 | 23 | 29 | 140 | 66 | 63 | 104 | 268 | 42 | 16 | 50 | 41 | 24 | 94 | 120 | 108 | 168 | 180 | 298 | 149 | 0.003980862 | 0.019353871 | 0.054 | 0.12 | 0.113 | 0.1 | 0.051 | 0.183 | 0.062 | 0.188 | 0.109 | 0.062 | -0.033 |
| RI | 200 | ENSG00000239382 | ALKBH6 | chr19 | - | 36013020 | 36013422 | 36013020 | 36013089 | 36013343 | 36013422 | 19 | 10 | 17 | 26 | 8 | 29 | 21 | 25 | 26 | 47 | 36 | 39 | 32 | 20 | 34 | 51 | 40 | 17 | 27 | 14 | 298 | 149 | 0.004008781 | 0.019468997 | 0.247 | 0.192 | 0.254 | 0.333 | 0.078 | 0.261 | 0.328 | 0.485 | 0.27 | 0.548 | -0.158 |
| RI | 4426 | ENSG00000197989 | SNHG12 | chr1 | - | 28579532 | 28579981 | 28579532 | 28579587 | 28579911 | 28579981 | 61 | 31 | 59 | 60 | 49 | 34 | 19 | 30 | 35 | 74 | 57 | 31 | 90 | 52 | 108 | 9 | 28 | 52 | 18 | 0 | 298 | 149 | 0.004011134 | 0.019468997 | 0.473 | 0.449 | 0.496 | 0.462 | 0.249 | 0.76 | 0.356 | 0.464 | 0.591 | 1 | -0.208 |
| RI | 2229 | ENSG00000113263 | ITK | chr5 | + | 157240061 | 157241720 | 157240061 | 157240195 | 157241645 | 157241720 | 184 | 186 | 262 | 174 | 240 | 267 | 153 | 301 | 233 | 496 | 148 | 113 | 409 | 143 | 94 | 105 | 177 | 306 | 101 | 26 | 298 | 149 | 0.004077024 | 0.01977264 | 0.256 | 0.378 | 0.303 | 0.272 | 0.195 | 0.413 | 0.242 | 0.401 | 0.414 | 0.644 | -0.142 |
| RI | 7927 | ENSG00000166340 | TPP1 | chr11 | - | 6616974 | 6617776 | 6616974 | 6617153 | 6617625 | 6617776 | 16 | 14 | 11 | 28 | 10 | 3 | 0 | 3 | 0 | 3 | 25 | 19 | 14 | 18 | 44 | 3 | 0 | 1 | 0 | 0 | 298 | 149 | 0.004103314 | 0.019883736 | 0.727 | 1 | 0.647 | 1 | 0.625 | 0.806 | 1 | 0.875 | 1 | 1 | -0.136 |
| RI | 7988 | ENSG00000103510 | KAT8 | chr16 | + | 31130745 | 31131359 | 31130745 | 31130900 | 31131194 | 31131359 | 7 | 8 | 16 | 15 | 16 | 69 | 24 | 56 | 40 | 42 | 6 | 27 | 7 | 12 | 55 | 12 | 57 | 38 | 25 | 63 | 298 | 149 | 0.004182368 | 0.020250203 | 0.048 | 0.143 | 0.125 | 0.158 | 0.16 | 0.2 | 0.191 | 0.084 | 0.194 | 0.304 | -0.068 |
| RI | 290 | ENSG00000142546 | NOSIP | chr19 | - | 49557089 | 49558978 | 49557089 | 49557249 | 49558896 | 49558978 | 42 | 22 | 82 | 21 | 32 | 238 | 126 | 286 | 219 | 181 | 15 | 10 | 8 | 14 | 13 | 108 | 121 | 82 | 123 | 64 | 298 | 149 | 0.004189291 | 0.020267111 | 0.081 | 0.08 | 0.125 | 0.046 | 0.081 | 0.065 | 0.04 | 0.047 | 0.054 | 0.092 | 0.023 |
| RI | 3225 | ENSG00000184014 | DENND5A | chr11 | - | 9141939 | 9144278 | 9141939 | 9142108 | 9144096 | 9144278 | 15 | 27 | 13 | 7 | 13 | 1 | 4 | 2 | 1 | 0 | 36 | 20 | 34 | 24 | 16 | 7 | 3 | 6 | 16 | 4 | 298 | 149 | 0.004226546 | 0.020430612 | 0.882 | 0.771 | 0.765 | 0.778 | 1 | 0.72 | 0.769 | 0.739 | 0.429 | 0.667 | 0.174 |
| RI | 7765 | ENSG00000172932 | ANKRD13D | chr11 | + | 67290077 | 67290446 | 67290077 | 67290213 | 67290321 | 67290446 | 23 | 19 | 14 | 24 | 18 | 80 | 40 | 89 | 94 | 36 | 54 | 21 | 38 | 43 | 40 | 129 | 49 | 30 | 69 | 118 | 257 | 149 | 0.004258709 | 0.020541971 | 0.143 | 0.216 | 0.084 | 0.129 | 0.225 | 0.195 | 0.199 | 0.423 | 0.265 | 0.164 | -0.09 |
| RI | 3118 | ENSG00000161204 | ABCF3 | chr3 | + | 184189087 | 184189277 | 184189087 | 184189144 | 184189254 | 184189277 | 7 | 5 | 7 | 10 | 18 | 51 | 13 | 43 | 27 | 35 | 84 | 14 | 33 | 26 | 8 | 60 | 41 | 54 | 39 | 41 | 259 | 149 | 0.004259544 | 0.020541971 | 0.073 | 0.181 | 0.086 | 0.176 | 0.228 | 0.446 | 0.164 | 0.26 | 0.277 | 0.101 | -0.101 |
| RI | 6898 | ENSG00000068308 | OTUD5 | chrX | - | 48934463 | 48934867 | 48934463 | 48934612 | 48934695 | 48934867 | 39 | 35 | 49 | 58 | 31 | 5 | 4 | 7 | 18 | 15 | 220 | 85 | 114 | 130 | 186 | 14 | 3 | 13 | 13 | 31 | 232 | 149 | 0.004260016 | 0.020541971 | 0.834 | 0.849 | 0.818 | 0.674 | 0.57 | 0.91 | 0.948 | 0.849 | 0.865 | 0.794 | -0.124 |
| RI | 1366 | ENSG00000149923 | PPP4C | chr16 | + | 30083393 | 30083781 | 30083393 | 30083567 | 30083654 | 30083781 | 5 | 5 | 12 | 26 | 5 | 142 | 60 | 112 | 112 | 75 | 51 | 18 | 5 | 33 | 29 | 154 | 124 | 104 | 116 | 268 | 236 | 149 | 0.004297894 | 0.020707716 | 0.022 | 0.05 | 0.063 | 0.128 | 0.04 | 0.173 | 0.084 | 0.029 | 0.152 | 0.064 | -0.04 |
| RI | 64 | ENSG00000182944 | EWSR1 | chr22 | + | 29298732 | 29299851 | 29298732 | 29298895 | 29299598 | 29299851 | 70 | 61 | 68 | 94 | 54 | 0 | 0 | 0 | 2 | 0 | 370 | 235 | 269 | 445 | 141 | 17 | 0 | 5 | 0 | 2 | 298 | 149 | 0.004315546 | 0.020775818 | 1 | 1 | 1 | 0.959 | 1 | 0.916 | 1 | 0.964 | 1 | 0.972 | 0.021 |
| RI | 1985 | ENSG00000175061 | SNHG29 | chr17 | + | 16439326 | 16439703 | 16439326 | 16439414 | 16439527 | 16439703 | 78 | 42 | 92 | 80 | 74 | 79 | 23 | 54 | 56 | 52 | 113 | 140 | 100 | 140 | 74 | 52 | 46 | 64 | 29 | 43 | 262 | 149 | 0.004324124 | 0.020800166 | 0.36 | 0.509 | 0.492 | 0.448 | 0.447 | 0.553 | 0.634 | 0.471 | 0.733 | 0.495 | -0.126 |
| RI | 7479 | ENSG00000148334 | PTGES2 | chr9 | - | 128122361 | 128123134 | 128122361 | 128122479 | 128122933 | 128123134 | 16 | 14 | 16 | 23 | 8 | 27 | 13 | 29 | 21 | 15 | 28 | 22 | 16 | 27 | 84 | 6 | 27 | 6 | 25 | 45 | 298 | 149 | 0.004343078 | 0.020859234 | 0.229 | 0.35 | 0.216 | 0.354 | 0.211 | 0.7 | 0.289 | 0.571 | 0.351 | 0.483 | -0.207 |
| RI | 3293 | ENSG00000165801 | ARHGEF40 | chr14 | + | 21075331 | 21075765 | 21075331 | 21075499 | 21075644 | 21075765 | 6 | 8 | 5 | 2 | 0 | 25 | 7 | 10 | 19 | 14 | 26 | 11 | 7 | 31 | 53 | 73 | 21 | 18 | 24 | 57 | 294 | 149 | 0.004343734 | 0.020859234 | 0.108 | 0.367 | 0.202 | 0.051 | 0 | 0.153 | 0.21 | 0.165 | 0.396 | 0.32 | -0.103 |
| RI | 7529 | ENSG00000137145 | DENND4C | chr9 | + | 19336269 | 19336832 | 19336269 | 19336414 | 19336685 | 19336832 | 92 | 54 | 59 | 71 | 29 | 235 | 202 | 161 | 206 | 260 | 16 | 65 | 85 | 26 | 16 | 19 | 170 | 142 | 75 | 5 | 298 | 149 | 0.004346998 | 0.020859234 | 0.164 | 0.118 | 0.155 | 0.147 | 0.053 | 0.296 | 0.16 | 0.23 | 0.148 | 0.615 | -0.162 |
| RI | 4015 | ENSG00000069424 | KCNAB2 | chr1 | + | 6094399 | 6095443 | 6094399 | 6094962 | 6095322 | 6095443 | 24 | 8 | 23 | 15 | 9 | 2 | 0 | 1 | 0 | 0 | 6 | 27 | 22 | 19 | 39 | 10 | 0 | 4 | 0 | 0 | 298 | 149 | 0.00439607 | 0.021077584 | 0.857 | 1 | 0.92 | 1 | 1 | 0.231 | 1 | 0.733 | 1 | 1 | 0.163 |
| RI | 228 | ENSG00000196684 | HSH2D | chr19 | + | 16152551 | 16153208 | 16152551 | 16152641 | 16153031 | 16153208 | 83 | 52 | 76 | 104 | 28 | 15 | 3 | 1 | 11 | 0 | 237 | 96 | 110 | 227 | 119 | 11 | 5 | 0 | 1 | 7 | 298 | 149 | 0.004433688 | 0.021240708 | 0.735 | 0.897 | 0.974 | 0.825 | 1 | 0.915 | 0.906 | 1 | 0.991 | 0.895 | -0.055 |
| RI | 3069 | ENSG00000076108 | BAZ2A | chr12 | - | 56615013 | 56615607 | 56615013 | 56615128 | 56615218 | 56615607 | 257 | 131 | 244 | 246 | 209 | 2 | 7 | 1 | 10 | 11 | 620 | 334 | 287 | 397 | 370 | 0 | 6 | 2 | 12 | 5 | 239 | 149 | 0.004438894 | 0.021245953 | 0.988 | 0.921 | 0.993 | 0.939 | 0.922 | 1 | 0.972 | 0.989 | 0.954 | 0.979 | -0.026 |
| RI | 4275 | ENSG00000042429 | MED17 | chr11 | + | 93809716 | 93812378 | 93809716 | 93809876 | 93811852 | 93812378 | 30 | 33 | 29 | 27 | 25 | 69 | 41 | 30 | 57 | 29 | 76 | 41 | 37 | 58 | 0 | 24 | 21 | 28 | 11 | 1 | 298 | 149 | 0.004441976 | 0.021245953 | 0.179 | 0.287 | 0.326 | 0.191 | 0.301 | 0.613 | 0.494 | 0.398 | 0.725 | 0 | -0.189 |
| RI | 8588 | ENSG00000148362 | PAXX | chr9 | + | 136992639 | 136992974 | 136992639 | 136992700 | 136992924 | 136992974 | 37 | 6 | 11 | 28 | 1 | 5 | 3 | 6 | 14 | 3 | 10 | 35 | 5 | 3 | 24 | 0 | 5 | 0 | 0 | 6 | 298 | 149 | 0.004478879 | 0.02140513 | 0.787 | 0.5 | 0.478 | 0.5 | 0.143 | 1 | 0.778 | 1 | 1 | 0.667 | -0.407 |
| RI | 2884 | ENSG00000178035 | IMPDH2 | chr3 | - | 49025125 | 49026609 | 49025125 | 49026419 | 49026518 | 49026609 | 33 | 29 | 39 | 26 | 58 | 186 | 84 | 235 | 175 | 254 | 7 | 35 | 37 | 36 | 47 | 91 | 92 | 70 | 83 | 166 | 248 | 149 | 0.004572962 | 0.021837095 | 0.096 | 0.172 | 0.091 | 0.082 | 0.121 | 0.044 | 0.186 | 0.241 | 0.207 | 0.145 | -0.052 |
| RI | 5005 | ENSG00000162650 | ATXN7L2 | chr1 | + | 109488846 | 109490128 | 109488846 | 109489100 | 109489929 | 109490128 | 7 | 2 | 6 | 16 | 8 | 11 | 3 | 9 | 11 | 7 | 6 | 16 | 1 | 2 | 1 | 17 | 39 | 7 | 4 | 3 | 298 | 149 | 0.004588533 | 0.021893751 | 0.241 | 0.25 | 0.25 | 0.421 | 0.364 | 0.15 | 0.17 | 0.067 | 0.2 | 0.143 | 0.159 |
| RI | 6666 | ENSG00000068354 | TBC1D25 | chrX | + | 48558896 | 48559346 | 48558896 | 48559024 | 48559157 | 48559346 | 3 | 2 | 0 | 5 | 7 | 42 | 20 | 35 | 30 | 60 | 0 | 0 | 5 | 2 | 1 | 19 | 65 | 44 | 41 | 98 | 282 | 149 | 0.004695445 | 0.022376479 | 0.036 | 0.05 | 0 | 0.081 | 0.058 | 0 | 0 | 0.057 | 0.025 | 0.005 | 0.028 |
| RI | 5190 | ENSG00000204642 | HLA-F | chr6 | + | 29726010 | 29726526 | 29726010 | 29726043 | 29726402 | 29726526 | 456 | 256 | 353 | 382 | 569 | 78 | 29 | 46 | 42 | 66 | 442 | 467 | 449 | 384 | 544 | 28 | 48 | 42 | 31 | 65 | 298 | 149 | 0.004700382 | 0.022376479 | 0.745 | 0.815 | 0.793 | 0.82 | 0.812 | 0.888 | 0.829 | 0.842 | 0.861 | 0.807 | -0.048 |
| RI | 6127 | ENSG00000168036 | CTNNB1 | chr3 | + | 41239133 | 41239955 | 41239133 | 41239354 | 41239812 | 41239955 | 252 | 150 | 255 | 285 | 218 | 0 | 0 | 3 | 0 | 0 | 82 | 227 | 236 | 193 | 55 | 0 | 9 | 1 | 3 | 3 | 298 | 149 | 0.004701068 | 0.022376479 | 1 | 1 | 0.977 | 1 | 1 | 1 | 0.927 | 0.992 | 0.97 | 0.902 | 0.037 |
| RI | 5892 | ENSG00000112787 | FBRSL1 | chr12 | + | 132574318 | 132574564 | 132574318 | 132574348 | 132574492 | 132574564 | 0 | 0 | 3 | 0 | 0 | 48 | 20 | 30 | 41 | 40 | 0 | 0 | 3 | 5 | 10 | 11 | 31 | 45 | 33 | 125 | 293 | 149 | 0.004723355 | 0.022464459 | 0 | 0 | 0.048 | 0 | 0 | 0 | 0 | 0.033 | 0.072 | 0.039 | -0.019 |
| RI | 82 | ENSG00000100417 | PMM1 | chr22 | - | 41583958 | 41584372 | 41583958 | 41584058 | 41584280 | 41584372 | 21 | 1 | 9 | 13 | 10 | 14 | 11 | 38 | 29 | 28 | 18 | 32 | 8 | 13 | 42 | 34 | 17 | 14 | 7 | 33 | 298 | 149 | 0.004777913 | 0.022705659 | 0.429 | 0.043 | 0.106 | 0.183 | 0.152 | 0.209 | 0.485 | 0.222 | 0.481 | 0.389 | -0.175 |
| RI | 4988 | ENSG00000080947 | CROCCP3 | chr1 | - | 16476929 | 16478437 | 16476929 | 16477074 | 16478254 | 16478437 | 9 | 5 | 2 | 1 | 4 | 4 | 5 | 2 | 7 | 1 | 12 | 6 | 0 | 8 | 7 | 0 | 4 | 3 | 0 | 0 | 298 | 149 | 0.004788155 | 0.022736038 | 0.529 | 0.333 | 0.333 | 0.067 | 0.667 | 1 | 0.429 | 0 | 1 | 1 | -0.3 |
| RI | 8407 | ENSG00000002016 | RAD52 | chr12 | - | 929763 | 930144 | 929763 | 929886 | 930050 | 930144 | 20 | 9 | 8 | 14 | 2 | 6 | 6 | 13 | 19 | 4 | 31 | 24 | 16 | 23 | 21 | 20 | 3 | 3 | 11 | 4 | 298 | 149 | 0.004801795 | 0.022782493 | 0.625 | 0.429 | 0.235 | 0.269 | 0.2 | 0.437 | 0.8 | 0.727 | 0.511 | 0.724 | -0.288 |
| RI | 374 | ENSG00000105193 | RPS16 | chr19 | - | 39433664 | 39435708 | 39433664 | 39433761 | 39435657 | 39435708 | 516 | 214 | 599 | 570 | 235 | 88 | 39 | 135 | 88 | 52 | 311 | 448 | 314 | 328 | 656 | 64 | 49 | 51 | 33 | 80 | 298 | 149 | 0.004812066 | 0.022812902 | 0.746 | 0.733 | 0.689 | 0.764 | 0.693 | 0.708 | 0.821 | 0.755 | 0.832 | 0.804 | -0.059 |
| RI | 334 | ENSG00000161847 | RAVER1 | chr19 | - | 10320654 | 10320951 | 10320654 | 10320782 | 10320857 | 10320951 | 7 | 3 | 6 | 2 | 1 | 2 | 0 | 1 | 0 | 0 | 0 | 7 | 0 | 9 | 20 | 9 | 0 | 3 | 0 | 0 | 224 | 149 | 0.004832328 | 0.022890584 | 0.7 | 1 | 0.8 | 1 | 1 | 0 | 1 | 0 | 1 | 1 | 0.3 |
| RI | 5980 | ENSG00000204256 | BRD2 | chr6 | + | 32975383 | 32976464 | 32975383 | 32975521 | 32976030 | 32976464 | 49 | 58 | 90 | 113 | 107 | 931 | 620 | 756 | 796 | 1563 | 280 | 121 | 53 | 190 | 156 | 1828 | 1073 | 1061 | 1293 | 1294 | 298 | 149 | 0.004893966 | 0.02316399 | 0.026 | 0.045 | 0.056 | 0.066 | 0.033 | 0.071 | 0.053 | 0.024 | 0.068 | 0.057 | -0.009 |
| RI | 3759 | ENSG00000161547 | SRSF2 | chr17 | - | 76734114 | 76736464 | 76734114 | 76735158 | 76736153 | 76736464 | 464 | 297 | 423 | 536 | 396 | 154 | 78 | 102 | 113 | 66 | 291 | 405 | 629 | 387 | 153 | 214 | 145 | 104 | 169 | 186 | 298 | 149 | 0.004908635 | 0.023214816 | 0.601 | 0.656 | 0.675 | 0.703 | 0.75 | 0.405 | 0.583 | 0.751 | 0.534 | 0.291 | 0.164 |
| RI | 6972 | ENSG00000147050 | KDM6A | chrX | + | 45062646 | 45063817 | 45062646 | 45062748 | 45063421 | 45063817 | 88 | 90 | 70 | 66 | 41 | 81 | 67 | 106 | 101 | 111 | 21 | 37 | 86 | 31 | 0 | 57 | 77 | 114 | 115 | 10 | 298 | 149 | 0.004941802 | 0.02335298 | 0.352 | 0.402 | 0.248 | 0.246 | 0.156 | 0.156 | 0.194 | 0.274 | 0.119 | 0 | 0.132 |
| RI | 4799 | ENSG00000137101 | CD72 | chr9 | - | 35615942 | 35616689 | 35615942 | 35616278 | 35616599 | 35616689 | 19 | 14 | 18 | 35 | 22 | 9 | 5 | 6 | 9 | 6 | 13 | 111 | 28 | 53 | 59 | 8 | 7 | 0 | 1 | 14 | 298 | 149 | 0.00497006 | 0.02346774 | 0.514 | 0.583 | 0.6 | 0.66 | 0.647 | 0.448 | 0.888 | 1 | 0.964 | 0.678 | -0.195 |

| | | | | | | | | | | | | | | |
|---|---|---|---|---|---|---|---|---|---|---|---|---|---|---|
| RI | 4495 | ENSG00000072110 | ACTN1 | chr14 | - | 68909317 | 68910042 | 68909317 | 68909396 | 68909954 | 68910042 | 9 | 13 | 19 |
| RI | 5606 | ENSG00000102125 | TAZ | chrX | + | 154419542 | 154420094 | 154419542 | 154419746 | 154420031 | 154420094 | 42 | 16 | 58 |
| RI | 4380 | ENSG00000172531 | PPP1CA | chr11 | - | 67398939 | 67399665 | 67398939 | 67399163 | 67399560 | 67399665 | 23 | 17 | 23 |
| RI | 5754 | ENSG00000169045 | HNRNPH1 | chr5 | - | 179623036 | 179624669 | 179623036 | 179623164 | 179623635 | 179624669 | 72 | 50 | 52 |
| RI | 5793 | ENSG00000115307 | AUP1 | chr2 | - | 74526940 | 74527363 | 74526940 | 74527059 | 74527247 | 74527363 | 51 | 37 | 45 |
| RI | 1753 | ENSG00000197102 | DYNC1H1 | chr14 | + | 102050070 | 102050773 | 102050070 | 102050198 | 102050434 | 102050773 | 17 | 12 | 17 |
| RI | 1627 | ENSG00000137845 | ADAM10 | chr15 | - | 58610296 | 58611991 | 58610296 | 58610517 | 58611807 | 58611991 | 109 | 106 | 69 |
| RI | 5962 | ENSG00000168000 | BSCL2 | chr11 | - | 62690786 | 62691141 | 62690786 | 62690867 | 62691074 | 62691141 | 6 | 5 | 6 |
| RI | 1919 | ENSG00000092203 | TOX4 | chr14 | + | 21487450 | 21488850 | 21487450 | 21487693 | 21488589 | 21488850 | 26 | 22 | 26 |
| RI | 6258 | ENSG00000196465 | MYL6B | chr12 | + | 56154812 | 56155198 | 56154812 | 56154840 | 56155054 | 56155198 | 5 | 0 | 4 |
| RI | 3921 | ENSG00000055609 | KMT2C | chr7 | - | 152144712 | 152145295 | 152144712 | 152144881 | 152145164 | 152145295 | 37 | 32 | 23 |
| RI | 8568 | ENSG00000185896 | LAMP1 | chr13 | + | 113320344 | 113321470 | 113320344 | 113320470 | 113321403 | 113321470 | 72 | 69 | 55 |
| RI | 2236 | ENSG00000113758 | DBN1 | chr5 | - | 177467480 | 177467817 | 177467480 | 177467627 | 177467742 | 177467817 | 2 | 0 | 2 |
| RI | 3554 | ENSG00000127054 | INTS11 | chr1 | - | 1312592 | 1313124 | 1312592 | 1312949 | 1313034 | 1313124 | 7 | 0 | 7 |
| RI | 6622 | ENSG00000146707 | POMZP3 | chr7 | - | 76618182 | 76618615 | 76618182 | 76618300 | 76618496 | 76618615 | 3 | 0 | 3 |
| RI | 5435 | ENSG00000197858 | GPAA1 | chr8 | + | 144083388 | 144083861 | 144083388 | 144083500 | 144083817 | 144083861 | 55 | 23 | 53 |
| RI | 5427 | ENSG00000189046 | ALKBH2 | chr12 | - | 109092506 | 109093472 | 109092506 | 109092937 | 109093246 | 109093472 | 15 | 11 | 11 |
| RI | 1867 | ENSG00000100823 | APEX1 | chr14 | + | 20455577 | 20456101 | 20455577 | 20455773 | 20455913 | 20456101 | 41 | 34 | 63 |
| RI | 427 | ENSG00000269858 | EGLN2 | chr19 | + | 40807137 | 40807551 | 40807137 | 40807274 | 40807483 | 40807551 | 123 | 63 | 111 |
| RI | 5485 | ENSG00000144895 | EIF2A | chr3 | + | 150567901 | 150568046 | 150567901 | 150567929 | 150568016 | 150568046 | 352 | 311 | 495 |
| RI | 5782 | ENSG00000204843 | DCTN1 | chr2 | - | 74367970 | 74368880 | 74367970 | 74368131 | 74368727 | 74368880 | 21 | 16 | 23 |
| RI | 6768 | ENSG00000177646 | ACAD9 | chr3 | + | 128909343 | 128910813 | 128909343 | 128910149 | 128910740 | 128910813 | 6 | 0 | 3 |
| RI | 3229 | ENSG00000184014 | DENND5A | chr11 | - | 9144096 | 9147151 | 9144096 | 9145113 | 9147029 | 9147151 | 7 | 11 | 9 |
| RI | 342 | ENSG00000115286 | NDUFS7 | chr19 | + | 1388832 | 1391050 | 1388832 | 1388938 | 1390870 | 1391050 | 5 | 0 | 1 |
| RI | 8075 | ENSG00000175197 | DDIT3 | chr12 | - | 57517268 | 57517753 | 57517268 | 57517595 | 57517705 | 57517753 | 51 | 31 | 73 |
| RI | 4927 | ENSG00000117155 | SSX2IP | chr1 | - | 84669680 | 84670815 | 84669680 | 84669893 | 84670645 | 84670815 | 13 | 7 | 14 |
| RI | 2572 | ENSG00000104972 | LILRB1 | chr19 | + | 54635556 | 54636652 | 54635556 | 54635609 | 54636493 | 54636652 | 23 | 19 | 8 |
| RI | 5270 | ENSG00000145476 | CYP4V2 | chr4 | + | 186201156 | 186205302 | 186201156 | 186201342 | 186205199 | 186205302 | 10 | 23 | 7 |
| RI | 1829 | ENSG00000100897 | DCAF11 | chr14 | + | 24115381 | 24115726 | 24115381 | 24115392 | 24115542 | 24115726 | 181 | 104 | 143 |
| RI | 895 | ENSG00000108963 | DPH1 | chr17 | + | 2035969 | 2036686 | 2035969 | 2036091 | 2036428 | 2036686 | 10 | 11 | 16 |
| RI | 6962 | ENSG00000215301 | DDX3X | chrX | + | 41347311 | 41349886 | 41347311 | 41347451 | 41347639 | 41349886 | 381 | 339 | 351 |
| RI | 6472 | ENSG00000106588 | PSMA2 | chr7 | - | 42917777 | 42921931 | 42917777 | 42917835 | 42921857 | 42921931 | 42 | 27 | 33 |
| RI | 4213 | ENSG00000124256 | ZBP1 | chr20 | - | 57615511 | 57616468 | 57615511 | 57615580 | 57616243 | 57616468 | 39 | 14 | 20 |
| RI | 3290 | ENSG00000116871 | MAP7D1 | chr1 | + | 36176198 | 36176581 | 36176198 | 36176321 | 36176417 | 36176581 | 77 | 24 | 39 |
| RI | 8269 | ENSG00000082898 | XPO1 | chr2 | - | 61492566 | 61494091 | 61492566 | 61492748 | 61493893 | 61494091 | 66 | 104 | 105 |
| RI | 7800 | ENSG00000255717 | SNHG1 | chr11 | - | 62851987 | 62852681 | 62851987 | 62852436 | 62852655 | 62852681 | 600 | 401 | 846 |
| RI | 938 | ENSG00000161526 | SAP30BP | chr17 | + | 75705610 | 75706092 | 75705610 | 75705693 | 75706007 | 75706092 | 28 | 28 | 27 |
| RI | 6797 | ENSG00000005302 | MSL3 | chrX | + | 11759633 | 11759875 | 11759633 | 11759712 | 11759792 | 11759875 | 166 | 125 | 128 |
| RI | 7753 | ENSG00000159352 | PSMD4 | chr1 | + | 151266003 | 151266439 | 151266003 | 151266112 | 151266307 | 151266439 | 3 | 1 | 3 |
| RI | 6999 | ENSG00000182774 | RPS17 | chr15 | - | 82539980 | 82540457 | 82539980 | 82540132 | 82540425 | 82540457 | 105 | 35 | 70 |
| RI | 7657 | ENSG00000149823 | VPS51 | chr11 | + | 65109704 | 65110603 | 65109704 | 65109923 | 65110481 | 65110603 | 27 | 12 | 34 |
| RI | 4411 | ENSG00000143575 | HAX1 | chr1 | + | 154273335 | 154273961 | 154273335 | 154273598 | 154273773 | 154273961 | 18 | 9 | 37 |
| RI | 4687 | ENSG00000175634 | RPS6KB2 | chr11 | + | 67429526 | 67432657 | 67429526 | 67429595 | 67432599 | 67432657 | 28 | 14 | 28 |
| RI | 4959 | ENSG00000096080 | MRPS18A | chr6 | - | 43675201 | 43675617 | 43675201 | 43675271 | 43675493 | 43675617 | 26 | 20 | 17 |
| RI | 6418 | ENSG00000164896 | FASTK | chr7 | - | 151077100 | 151077401 | 151077100 | 151077236 | 151077309 | 151077401 | 43 | 14 | 52 |
| RI | 8435 | ENSG00000137343 | ATAT1 | chr6 | + | 30645894 | 30646109 | 30645894 | 30645974 | 30646066 | 30646109 | 13 | 7 | 18 |
| RI | 6160 | ENSG00000168944 | CEP120 | chr5 | - | 123389923 | 123391337 | 123389923 | 123390140 | 123391109 | 123391337 | 57 | 43 | 30 |
| RI | 2995 | ENSG00000114742 | WDR48 | chr3 | + | 39093873 | 39096664 | 39093873 | 39094066 | 39094647 | 39096664 | 18 | 16 | 26 |
| RI | 1167 | ENSG00000125124 | BBS2 | chr16 | - | 56500853 | 56501497 | 56500853 | 56501025 | 56501352 | 56501497 | 10 | 9 | 34 |
| RI | 4678 | ENSG00000173846 | PLK3 | chr1 | + | 44803599 | 44804024 | 44803599 | 44803691 | 44803930 | 44804024 | 28 | 29 | 34 |
| RI | 5692 | ENSG00000160710 | ADAR | chr1 | - | 154589756 | 154590409 | 154589756 | 154589928 | 154590261 | 154590409 | 138 | 90 | 101 |
| RI | 4255 | ENSG00000036549 | ZZZ3 | chr1 | - | 77579526 | 77581069 | 77579526 | 77579628 | 77580997 | 77581069 | 18 | 21 | 19 |
| RI | 633 | ENSG00000196756 | SNHG17 | chr20 | - | 38425934 | 38426503 | 38425934 | 38426048 | 38426418 | 38426503 | 19 | 13 | 6 |
| RI | 976 | ENSG00000179094 | PER1 | chr17 | - | 8146896 | 8147381 | 8146896 | 8147002 | 8147249 | 8147381 | 66 | 33 | 43 |
| RI | 2213 | ENSG00000196670 | ZFP62 | chr5 | - | 180847610 | 180851493 | 180847610 | 180848082 | 180851380 | 180851493 | 71 | 58 | 86 |
| RI | 8519 | ENSG00000075415 | SLC25A3 | chr12 | + | 98593974 | 98595848 | 98593974 | 98594135 | 98595726 | 98595848 | 105 | 69 | 113 |
| RI | 5945 | ENSG00000094880 | CDC23 | chr5 | - | 138191861 | 138192388 | 138191861 | 138191937 | 138192268 | 138192388 | 12 | 12 | 8 |
| RI | 5657 | ENSG00000121741 | ZMYM2 | chr13 | + | 19993062 | 19993919 | 19993062 | 19993564 | 19993825 | 19993919 | 473 | 499 | 522 |
| RI | 2877 | ENSG00000183323 | CCDC125 | chr5 | - | 69320236 | 69320580 | 69320236 | 69320345 | 69320464 | 69320580 | 74 | 55 | 74 |
| RI | 672 | ENSG00000099956 | SMARCB1 | chr22 | + | 23800943 | 23803422 | 23800943 | 23801135 | 23803294 | 23803422 | 10 | 6 | 14 |
| RI | 1454 | ENSG00000090905 | TNRC6A | chr16 | + | 24805013 | 24805733 | 24805013 | 24805151 | 24805604 | 24805733 | 17 | 19 | 14 |
| RI | 4246 | ENSG00000172432 | GTPBP2 | chr6 | - | 43624887 | 43625560 | 43624887 | 43625062 | 43625362 | 43625560 | 76 | 46 | 81 |
| RI | 4272 | ENSG00000042429 | MED17 | chr11 | + | 93793727 | 93794035 | 93793727 | 93793864 | 93793950 | 93794035 | 63 | 46 | 46 |
| RI | 3429 | ENSG00000008128 | CDK11A | chr1 | - | 1704222 | 1704655 | 1704222 | 1704344 | 1704549 | 1704655 | 24 | 6 | 20 |
| RI | 3417 | ENSG00000170581 | STAT2 | chr12 | - | 56354777 | 56355351 | 56354777 | 56354863 | 56355275 | 56355351 | 73 | 86 | 72 |
| RI | 5218 | ENSG00000204344 | STK19 | chr6 | + | 31979413 | 31980548 | 31979413 | 31979553 | 31980450 | 31980548 | 17 | 24 | 15 |
| RI | 6575 | ENSG00000265681 | RPL17 | chr18 | - | 49491531 | 49491833 | 49491531 | 49491584 | 49491735 | 49491833 | 1164 | 663 | 1069 |
| RI | 362 | ENSG00000125755 | SYMPK | chr19 | - | 45827836 | 45829205 | 45827836 | 45827918 | 45828969 | 45829205 | 17 | 10 | 25 |
| RI | 1344 | ENSG00000168872 | DDX19A | chr16 | + | 70370222 | 70371563 | 70370222 | 70370385 | 70371371 | 70371563 | 13 | 20 | 10 |
| RI | 636 | ENSG00000196839 | ADA | chr20 | - | 44620298 | 44621147 | 44620298 | 44620401 | 44621017 | 44621147 | 14 | 4 | 4 |
| RI | 7131 | ENSG00000104783 | KCNN4 | chr19 | - | 43766532 | 43767707 | 43766532 | 43767089 | 43767539 | 43767707 | 0 | 4 | 1 |
| RI | 1586 | ENSG00000140521 | POLG | chr15 | - | 89317375 | 89318749 | 89317375 | 89317536 | 89317938 | 89318749 | 87 | 78 | 62 |
| RI | 8176 | ENSG00000119977 | TCTN3 | chr10 | - | 95683100 | 95683629 | 95683100 | 95683195 | 95683521 | 95683629 | 5 | 5 | 14 |
| RI | 8048 | ENSG00000196531 | NACA | chr12 | - | 56712439 | 56712908 | 56712439 | 56712552 | 56712785 | 56712908 | 70 | 53 | 61 |
| RI | 1332 | ENSG00000161981 | SNRNP25 | chr16 | + | 55776 | 56613 | 55776 | 55882 | 56538 | 56613 | 7 | 2 | 2 |
| RI | 2846 | ENSG00000196378 | ZNF34 | chr8 | - | 144778037 | 144778525 | 144778037 | 144778164 | 144778438 | 144778525 | 0 | 2 | 0 |
| RI | 1739 | ENSG00000087111 | PIGS | chr17 | - | 28570851 | 28571188 | 28570851 | 28570963 | 28571048 | 28571188 | 1 | 2 | 10 |
| RI | 5973 | ENSG00000204410 | MSH5 | chr6 | + | 31743903 | 31744299 | 31743903 | 31744025 | 31744189 | 31744299 | 10 | 15 | 4 |
| RI | 738 | ENSG00000004975 | DVL2 | chr17 | - | 7230284 | 7230797 | 7230284 | 7230430 | 7230727 | 7230797 | 8 | 1 | 8 |
| RI | 1355 | ENSG00000213658 | LAT | chr16 | + | 28986539 | 28986714 | 28986539 | 28986569 | 28986656 | 28986714 | 66 | 37 | 57 |
| RI | 1993 | ENSG00000175061 | SNHG29 | chr17 | + | 16439326 | 16440253 | 16439326 | 16439703 | 16440184 | 16440253 | 129 | 108 | 202 |
| RI | 8469 | ENSG00000089693 | MLF2 | chr12 | - | 6750176 | 6750766 | 6750176 | 6750305 | 6750712 | 6750766 | 17 | 19 | 32 |
| RI | 6556 | ENSG00000127914 | AKAP9 | chr7 | + | 92098108 | 92099869 | 92098108 | 92098214 | 92099686 | 92099869 | 10 | 21 | 5 |
| RI | 7696 | ENSG00000160226 | CFAP410 | chr21 | - | 44330462 | 44332014 | 44330462 | 44330916 | 44331842 | 44332014 | 20 | 13 | 38 |
| RI | 7546 | ENSG00000130714 | POMT1 | chr9 | + | 131522931 | 131523799 | 131522931 | 131523170 | 131523346 | 131523799 | 74 | 53 | 109 |
| RI | 4126 | ENSG00000241878 | PISD | chr22 | - | 31620995 | 31621472 | 31620995 | 31621142 | 31621333 | 31621472 | 10 | 10 | 11 |
| RI | 3073 | ENSG00000007923 | DNAJC11 | chr1 | - | 6637446 | 6638364 | 6637446 | 6637504 | 6638082 | 6638364 | 10 | 4 | 5 |
| RI | 136 | ENSG00000100325 | ASCC2 | chr22 | - | 29793359 | 29793676 | 29793359 | 29793490 | 29793576 | 29793676 | 9 | 9 | 9 |
| RI | 4125 | ENSG00000215305 | VPS16 | chr20 | + | 2863948 | 2864287 | 2863948 | 2864083 | 2864178 | 2864287 | 23 | 15 | 27 |
| RI | 870 | ENSG00000125447 | GGA3 | chr17 | - | 75239788 | 75241057 | 75239788 | 75240108 | 75240341 | 75241057 | 3 | 1 | 7 |
| RI | 4990 | ENSG00000128699 | ORMDL1 | chr2 | - | 189782421 | 189783123 | 189782421 | 189782602 | 189783013 | 189783123 | 261 | 232 | 330 |
| RI | 6080 | ENSG00000225791 | TRAM2-AS1 | chr6 | + | 52578049 | 52578594 | 52578049 | 52578393 | 52578501 | 52578594 | 9 | 9 | 14 |
| RI | 7573 | ENSG00000170909 | OSCAR | chr19 | - | 54094669 | 54096153 | 54094669 | 54095357 | 54095871 | 54096153 | 26 | 16 | 9 |
| RI | 5171 | ENSG00000108387 | SEPTIN4 | chr17 | - | 58521735 | 58522101 | 58521735 | 58521853 | 58521966 | 58522101 | 1 | 0 | 0 |
| RI | 578 | ENSG00000088970 | KIZ | chr20 | + | 21244244 | 21246622 | 21244244 | 21244288 | 21246478 | 21246622 | 28 | 23 | 20 |
| RI | 8169 | ENSG00000107937 | GTPBP4 | chr10 | + | 996105 | 997308 | 996105 | 996242 | 997207 | 997308 | 32 | 34 | 31 |
| RI | 6185 | ENSG00000214021 | TTLL3 | chr3 | + | 9834680 | 9835404 | 9834680 | 9834907 | 9835093 | 9835404 | 33 | 17 | 38 |
| RI | 119 | ENSG00000100201 | DDX17 | chr22 | - | 38487878 | 38492115 | 38487878 | 38488115 | 38492055 | 38492115 | 1446 | 1384 | 1781 |
| RI | 4923 | ENSG00000138398 | PPIG | chr2 | + | 169603641 | 169604102 | 169603641 | 169603694 | 169604025 | 169604102 | 33 | 50 | 26 |
| RI | 2140 | ENSG00000140853 | NLRC5 | chr16 | + | 57028071 | 57028385 | 57028071 | 57028203 | 57028301 | 57028385 | 69 | 48 | 77 |
| RI | 3126 | ENSG00000130787 | HIP1R | chr12 | + | 122860046 | 122860210 | 122860046 | 122860077 | 122860147 | 122860210 | 8 | 1 | 8 |
| RI | 2762 | ENSG00000185000 | DGAT1 | chr8 | - | 144318260 | 144318566 | 144318260 | 144318362 | 144318460 | 144318566 | 8 | 0 | 9 |
| RI | 5523 | ENSG00000035720 | STAP1 | chr4 | + | 67606295 | 67607337 | 67606295 | 67606389 | 67607193 | 67607337 | 139 | 92 | 110 |
| RI | 5259 | ENSG00000224078 | SNHG14 | chr15 | + | 25103858 | 25106294 | 25103858 | 25103970 | 25106163 | 25106294 | 1 | 7 | 4 |
| RI | 6132 | ENSG00000135317 | SNX14 | chr6 | - | 85507967 | 85513895 | 85507967 | 85508313 | 85513799 | 85513895 | 53 | 43 | 36 |
| RI | 6506 | ENSG00000122566 | HNRNPA2B1 | chr7 | - | 26190992 | 26192577 | 26190992 | 26191128 | 26192494 | 26192577 | 550 | 594 | 639 |
| RI | 7272 | ENSG00000104529 | EEF1D | chr8 | - | 143581053 | 143581328 | 143581053 | 143581154 | 143581285 | 143581328 | 697 | 306 | 659 |
| RI | 1249 | ENSG00000188603 | CLN3 | chr16 | - | 28482104 | 28482545 | 28482104 | 28482382 | 28482476 | 28482545 | 16 | 12 | 18 |
| RI | 7583 | ENSG00000132017 | DCAF15 | chr19 | + | 13960286 | 13960580 | 13960286 | 13960391 | 13960464 | 13960580 | 11 | 4 | 6 |
| RI | 6302 | ENSG00000145191 | EIF2B5 | chr3 | + | 184143051 | 184144707 | 184143051 | 184144224 | 184144596 | 184144707 | 5 | 3 | 4 |
| RI | 6330 | ENSG00000168255 | POLR2J3 | chr7 | - | 102566996 | 102568184 | 102566996 | 102567083 | 102568009 | 102568184 | 11 | 6 | 17 |
| RI | 4534 | ENSG00000143258 | USP21 | chr1 | + | 161164533 | 161164942 | 161164533 | 161164612 | 161164834 | 161164942 | 35 | 19 | 40 |
| RI | 1927 | ENSG00000119723 | COQ6 | chr14 | + | 73959414 | 73961375 | 73959414 | 73959522 | 73961172 | 73961375 | 8 | 2 | 1 |
| RI | 3300 | ENSG00000167085 | PHB | chr17 | - | 49406758 | 49409158 | 49406758 | 49406854 | 49409041 | 49409158 | 5 | 6 | 8 |
| RI | 3728 | ENSG00000179335 | CLK3 | chr15 | + | 74620008 | 74622216 | 74620008 | 74620225 | 74622119 | 74622216 | 112 | 30 | 61 |
| RI | 2405 | ENSG00000105875 | WDR91 | chr7 | - | 135186971 | 135189452 | 135186971 | 135188545 | 135189343 | 135189452 | 14 | 7 | 7 |
| RI | 925 | ENSG00000011143 | MKS1 | chr17 | - | 58214740 | 58216243 | 58214740 | 58214838 | 58216087 | 58216243 | 3 | 5 | 2 |
| RI | 4139 | ENSG00000154222 | CC2D1B | chr1 | - | 52357780 | 52358461 | 52357780 | 52357898 | 52358330 | 52358461 | 28 | 23 | 30 |
| RI | 5540 | ENSG00000144635 | DYNC1LI1 | chr3 | - | 32570345 | 32570858 | 32570345 | 32570419 | 32570624 | 32570858 | 2 | 3 | 2 |
| RI | 2638 | ENSG00000156876 | SASS6 | chr1 | - | 100107627 | 100108004 | 100107627 | 100107717 | 100107809 | 100108004 | 20 | 18 | 12 |
| RI | 5561 | ENSG00000120697 | ALG5 | chr13 | - | 36993603 | 36995596 | 36993603 | 36993672 | 36994988 | 36995596 | 40 | 10 | 38 |
| RI | 3958 | ENSG00000147403 | RPL10 | chrX | + | 154398376 | 154398542 | 154398376 | 154398394 | 154398496 | 154398542 | 315 | 125 | 334 |
| RI | 2280 | ENSG00000185404 | SP140L | chr2 | + | 230400126 | 230401443 | 230400126 | 230400242 | 230401365 | 230401443 | 43 | 44 | 48 |
| RI | 3586 | ENSG00000197343 | ZNF655 | chr7 | + | 99561887 | 99562494 | 99561887 | 99561973 | 99562343 | 99562494 | 135 | 67 | 107 |
| RI | 6544 | ENSG00000106144 | CASP2 | chr7 | + | 143300203 | 143303933 | 143300203 | 143300294 | 143303783 | 143303933 | 9 | 8 | 14 |
| RI | 2921 | ENSG00000247596 | TWF2 | chr3 | - | 52229919 | 52230995 | 52229919 | 52230070 | 52230869 | 52230995 | 2 | 4 | 11 |
| RI | 5638 | ENSG00000163516 | ANKZF1 | chr2 | + | 219233078 | 219233433 | 219233078 | 219233191 | 219233285 | 219233433 | 23 | 40 | 48 |
| RI | 1516 | ENSG00000103653 | CSK | chr15 | + | 74800680 | 74800922 | 74800680 | 74800746 | 74800822 | 74800922 | 32 | 36 | 36 |
| RI | 7170 | ENSG00000014216 | CAPN1 | chr11 | + | 65209848 | 65210096 | 65209848 | 65209917 | 65210017 | 65210096 | 5 | 7 | 4 |
| RI | 7828 | ENSG00000149743 | TRPT1 | chr11 | - | 64224284 | 64224717 | 64224284 | 64224341 | 64224542 | 64224717 | 5 | 4 | 1 |
| RI | 5050 | ENSG00000242485 | MRPL20 | chr1 | - | 1405808 | 1407019 | 1405808 | 1405886 | 1406908 | 1407019 | 71 | 64 | 86 |
| RI | 8179 | ENSG00000151651 | ADAM8 | chr10 | - | 133273253 | 133273838 | 133273253 | 133273430 | 133273761 | 133273838 | 29 | 17 | 13 |
| RI | 93 | ENSG00000100099 | HPS4 | chr22 | - | 26466225 | 26468623 | 26466225 | 26466262 | 26468550 | 26468623 | 11 | 18 | 23 |
| RI | 5229 | ENSG00000125875 | TBC1D20 | chr20 | - | 440247 | 442043 | 440247 | 441689 | 441856 | 442043 | 19 | 17 | 20 |
| RI | 4969 | ENSG00000186652 | PRG2 | chr11 | - | 57388576 | 57389317 | 57388576 | 57388708 | 57389009 | 57389317 | 2 | 3 | 5 |
| RI | 8402 | ENSG00000002016 | RAD52 | chr12 | - | 916343 | 916820 | 916343 | 916483 | 916638 | 916820 | 12 | 2 | 7 |
| RI | 2820 | ENSG00000242086 | MUC20-OT1 | chr3 | + | 195708153 | 195708435 | 195708153 | 195708268 | 195708428 | 195708435 | 17 | 10 | 27 |
| RI | 1388 | ENSG00000131652 | THOC6 | chr16 | + | 3025923 | 3026166 | 3025923 | 3025988 | 3026062 | 3026166 | 4 | 1 | 7 |
| RI | 8271 | ENSG00000082898 | XPO1 | chr2 | - | 61496878 | 61498913 | 61496878 | 61497007 | 61498672 | 61498913 | 54 | 68 | 35 |
| RI | 4702 | ENSG00000089127 | OAS1 | chr12 | + | 112916508 | 112917700 | 112916508 | 112916738 | 112917546 | 112917700 | 24 | 22 | 11 |
| RI | 126 | ENSG00000100055 | CYTH4 | chr22 | + | 37311455 | 37312174 | 37311455 | 37311527 | 37312019 | 37312174 | 53 | 44 | 58 |
| RI | 5877 | ENSG00000111679 | PTPN6 | chr12 | + | 6951608 | 6952177 | 6951608 | 6951731 | 6951911 | 6952177 | 51 | 25 | 39 |
| RI | 2490 | ENSG00000166913 | YWHAB | chr20 | + | 44904967 | 44906096 | 44904967 | 44905131 | 44906000 | 44906096 | 255 | 192 | 245 |
| RI | 7750 | ENSG00000175467 | SART1 | chr11 | + | 65966075 | 65966556 | 65966075 | 65966218 | 65966349 | 65966556 | 16 | 4 | 6 |
| RI | 1750 | ENSG00000197102 | DYNC1H1 | chr14 | + | 102049439 | 102049882 | 102049439 | 102049582 | 102049713 | 102049882 | 13 | 6 | 12 |
| RI | 7636 | ENSG00000162300 | ZFPL1 | chr11 | + | 65084241 | 65084800 | 65084241 | 65084378 | 65084690 | 65084800 | 35 | 17 | 23 |
| RI | 2468 | ENSG00000137802 | MAPKBP1 | chr15 | + | 41819245 | 41819650 | 41819245 | 41819379 | 41819594 | 41819650 | 9 | 2 | 3 |
| RI | 1603 | ENSG00000140199 | SLC12A6 | chr15 | - | 34336409 | 34338057 | 34336409 | 34336686 | 34337960 | 34338057 | 113 | 114 | 138 |
| RI | 770 | ENSG00000129197 | RPAIN | chr17 | + | 5425970 | 5426299 | 5425970 | 5426082 | 5426235 | 5426299 | 64 | 30 | 96 |
| RI | 6934 | ENSG00000046651 | OFD1 | chrX | + | 13753367 | 13755242 | 13753367 | 13753441 | 13755150 | 13755242 | 60 | 153 | 101 |
| RI | 5704 | ENSG00000079785 | DDX1 | chr2 | + | 15627053 | 15629697 | 15627053 | 15628839 | 15629601 | 15629697 | 22 | 15 | 9 |
| RI | 1823 | ENSG00000100911 | PSME2 | chr14 | - | 24144399 | 24145273 | 24144399 | 24145155 | 24145242 | 24145273 | 16 | 43 | 31 |
| RI | 7017 | ENSG00000182196 | ARL6IP4 | chr12 | + | 122981956 | 122982538 | 122981956 | 122982074 | 122982468 | 122982538 | 32 | 24 | 38 |
| RI | 7102 | ENSG00000094914 | AAAS | chr12 | - | 53314297 | 53315782 | 53314297 | 53315426 | 53315726 | 53315782 | 6 | 13 | 14 |
| RI | 6291 | ENSG00000145191 | EIF2B5 | chr3 | + | 184136611 | 184137805 | 184136611 | 184136736 | 184137619 | 184137805 | 56 | 39 | 47 |
| RI | 5548 | ENSG00000108883 | EFTUD2 | chr17 | - | 44853516 | 44854356 | 44853516 | 44853635 | 44854268 | 44854356 | 36 | 19 | 31 |
| RI | 1067 | ENSG00000168961 | LGALS9 | chr17 | + | 27647029 | 27647432 | 27647029 | 27647118 | 27647269 | 27647432 | 14 | 15 | 10 |
| RI | 7656 | ENSG00000149823 | VPS51 | chr11 | + | 65107580 | 65108022 | 65107580 | 65107727 | 65107802 | 65108022 | 21 | 5 | 16 |
| RI | 8311 | ENSG00000102225 | CDK16 | chrX | + | 47225964 | 47226703 | 47225964 | 47226027 | 47226582 | 47226703 | 29 | 18 | 48 |
| RI | 947 | ENSG00000132522 | GPS2 | chr17 | - | 7313028 | 7313291 | 7313028 | 7313124 | 7313211 | 7313291 | 39 | 20 | 30 |
| RI | 8448 | ENSG00000123349 | PFDN5 | chr12 | + | 53295838 | 53296275 | 53295838 | 53295941 | 53296218 | 53296275 | 212 | 165 | 266 |
| RI | 3665 | ENSG00000167962 | ZNF598 | chr16 | - | 1999495 | 2000481 | 1999495 | 2000256 | 2000390 | 2000481 | 7 | 7 | 5 |
| RI | 464 | ENSG00000105726 | ATP13A1 | chr19 | - | 19649740 | 19651797 | 19649740 | 19649940 | 19651688 | 19651797 | 3 | 2 | 1 |
| RI | 6778 | ENSG00000072778 | ACADVL | chr17 | + | 7220463 | 7220676 | 7220463 | 7220529 | 7220603 | 7220676 | 35 | 7 | 10 |
| RI | 1763 | ENSG00000100949 | RABGGTA | chr14 | - | 24270836 | 24271169 | 24270836 | 24270947 | 24271112 | 24271169 | 7 | 3 | 10 |
| RI | 7340 | ENSG00000158850 | B4GALT3 | chr1 | - | 161176433 | 161177106 | 161176433 | 161176579 | 161176911 | 161177106 | 19 | 18 | 31 |
| RI | 7491 | ENSG00000213930 | GALT | chr9 | + | 34647088 | 34647567 | 34647088 | 34647258 | 34647491 | 34647567 | 46 | 18 | 32 |
| RI | 8256 | ENSG00000133316 | WDR74 | chr11 | - | 62833617 | 62833937 | 62833617 | 62833674 | 62833791 | 62833937 | 7 | 11 | 21 |
| RI | 4227 | ENSG00000132781 | MUTYH | chr1 | - | 45332762 | 45333324 | 45332762 | 45332834 | 45332917 | 45333324 | 6 | 1 | 10 |
| RI | 6333 | ENSG00000136286 | MYO1G | chr7 | - | 44966072 | 44966838 | 44966072 | 44966280 | 44966671 | 44966838 | 116 | 43 | 75 |
| RI | 2803 | ENSG00000163703 | CRELD1 | chr3 | + | 9943076 | 9943515 | 9943076 | 9943172 | 9943380 | 9943515 | 6 | 6 | 9 |
| RI | 3425 | ENSG00000164088 | PPM1M | chr3 | + | 52248152 | 52248453 | 52248152 | 52248237 | 52248334 | 52248453 | 43 | 18 | 27 |
| RI | 1172 | ENSG00000125124 | BBS2 | chr16 | - | 56498436 | 56502456 | 56498436 | 56501497 | 56502316 | 56502456 | 101 | 100 | 106 |
| RI | 3056 | ENSG00000198034 | RPS4X | chrX | - | 72273800 | 72275150 | 72273800 | 72273972 | 72275052 | 72275150 | 712 | 475 | 913 |
| RI | 196 | ENSG00000105618 | PRPF31 | chr19 | + | 54129056 | 54129370 | 54129056 | 54129185 | 54129271 | 54129370 | 11 | 4 | 6 |
| RI | 264 | ENSG00000142541 | RPL13A | chr19 | + | 49490231 | 49490576 | 49490231 | 49490297 | 49490479 | 49490576 | 225 | 143 | 255 |
| RI | 7481 | ENSG00000148334 | PTGES2 | chr9 | - | 128124491 | 128125441 | 128124491 | 128124550 | 128125243 | 128125441 | 1 | 0 | 5 |
| RI | 2148 | ENSG00000105698 | USF2 | chr19 | + | 35278697 | 35279074 | 35278697 | 35278792 | 35278969 | 35279074 | 207 | 136 | 194 |
| RI | 1196 | ENSG00000033011 | ALG1 | chr16 | + | 5078756 | 5079807 | 5078756 | 5078878 | 5079747 | 5079807 | 2 | 7 | 6 |
| RI | 7683 | ENSG00000149273 | RPS3 | chr11 | + | 75401639 | 75402446 | 75401639 | 75401705 | 75402379 | 75402446 | 4728 | 2273 | 4742 |
| RI | 8108 | ENSG00000103024 | NME3 | chr16 | - | 1771069 | 1771410 | 1771069 | 1771171 | 1771279 | 1771410 | 10 | 2 | 1 |
| RI | 2044 | ENSG00000126777 | KTN1 | chr14 | + | 55671565 | 55671877 | 55671565 | 55671655 | 55671784 | 55671877 | 79 | 97 | 77 |
| RI | 4528 | ENSG00000130717 | UCK1 | chr9 | - | 131528943 | 131529270 | 131528943 | 131529038 | 131529127 | 131529270 | 5 | 5 | 12 |
| RI | 7810 | ENSG00000255717 | SNHG1 | chr11 | - | 62854518 | 62855216 | 62854518 | 62854551 | 62855132 | 62855216 | 132 | 95 | 242 |
| RI | 5422 | ENSG00000141522 | ARHGDIA | chr17 | - | 81869329 | 81869625 | 81869329 | 81869406 | 81869541 | 81869625 | 44 | 28 | 63 |
| RI | 3658 | ENSG00000198590 | C3orf35 | chr3 | + | 37417018 | 37418101 | 37417018 | 37417142 | 37417252 | 37418101 | 8 | 14 | 16 |
| RI | 4292 | ENSG00000155363 | MOV10 | chr1 | + | 112698286 | 112699810 | 112698286 | 112698789 | 112699684 | 112699810 | 27 | 11 | 12 |
| RI | 5161 | ENSG00000198755 | RPL10A | chr6 | + | 35468419 | 35468873 | 35468419 | 35468439 | 35468798 | 35468873 | 73 | 27 | 83 |
| RI | 7552 | ENSG00000137073 | UBAP2 | chr9 | - | 33922965 | 33923478 | 33922965 | 33923293 | 33923378 | 33923478 | 11 | 5 | 7 |
| RI | 5082 | ENSG00000162882 | HAAO | chr2 | - | 42770142 | 42770582 | 42770142 | 42770186 | 42770492 | 42770582 | 3 | 5 | 6 |
| RI | 6502 | ENSG00000122566 | HNRNPA2B1 | chr7 | - | 26189926 | 26192338 | 26189926 | 26190460 | 26190992 | 26192338 | 507 | 407 | 504 |
| RI | 3099 | ENSG00000161217 | PCYT1A | chr3 | - | 196247366 | 196248323 | 196247366 | 196247518 | 196248206 | 196248323 | 38 | 26 | 41 |
| RI | 3052 | ENSG00000156709 | AIFM1 | chrX | - | 130136044 | 130136731 | 130136044 | 130136185 | 130136642 | 130136731 | 3 | 8 | 0 |
| RI | 2388 | ENSG00000005156 | LIG3 | chr17 | + | 34998603 | 34999449 | 34998603 | 34998727 | 34999306 | 34999449 | 29 | 15 | 13 |
| RI | 1414 | ENSG00000158805 | ZNF276 | chr16 | + | 89723136 | 89723709 | 89723136 | 89723183 | 89723259 | 89723709 | 58 | 25 | 58 |
| RI | 4594 | ENSG00000131165 | CHMP1A | chr16 | - | 89646526 | 89647331 | 89646526 | 89646714 | 89647202 | 89647331 | 21 | 19 | 30 |
| RI | 7579 | ENSG00000123360 | PDE1B | chr12 | + | 54570240 | 54572741 | 54570240 | 54570357 | 54572600 | 54572741 | 4 | 5 | 5 |
| RI | 6151 | ENSG00000106009 | BRAT1 | chr7 | - | 2543203 | 2543962 | 2543203 | 2543323 | 2543589 | 2543962 | 12 | 12 | 21 |
| RI | 3250 | ENSG00000064961 | HMG20B | chr19 | + | 3573691 | 3574586 | 3573691 | 3573800 | 3574382 | 3574586 | 7 | 15 | 14 |
| RI | 4788 | ENSG00000058453 | CROCC | chr1 | + | 16953301 | 16954357 | 16953301 | 16953481 | 16954222 | 16954357 | 2 | 0 | 0 |
| RI | 5588 | ENSG00000164548 | TRA2A | chr7 | - | 23521706 | 23522432 | 23521706 | 23521840 | 23522131 | 23522432 | 453 | 499 | 528 |
| RI | 3080 | ENSG00000103064 | SLC7A6 | chr16 | + | 68297233 | 68301819 | 68297233 | 68297476 | 68298490 | 68301819 | 100 | 103 | 170 |
| RI | 4794 | ENSG00000160310 | PRMT2 | chr21 | + | 46660832 | 46661936 | 46660832 | 46660962 | 46661799 | 46661936 | 106 | 43 | 74 |
| RI | 1933 | ENSG00000103811 | CTSH | chr15 | - | 78921771 | 78923118 | 78921771 | 78922205 | 78922992 | 78923118 | 44 | 34 | 57 |
| RI | 4667 | ENSG00000001497 | LAS1L | chrX | - | 65528259 | 65529258 | 65528259 | 65528369 | 65529211 | 65529258 | 42 | 18 | 31 |
| RI | 297 | ENSG00000063177 | RPL18 | chr19 | - | 48615330 | 48615946 | 48615330 | 48615447 | 48615876 | 48615946 | 9 | 8 | 11 |
| RI | 878 | ENSG00000185359 | HGS | chr17 | + | 81694814 | 81695067 | 81694814 | 81694853 | 81694923 | 81695067 | 32 | 14 | 21 |
| RI | 3304 | ENSG00000228794 | LINC01128 | chr1 | + | 851926 | 852766 | 851926 | 852110 | 852670 | 852766 | 14 | 1 | 17 |
| RI | 772 | ENSG00000129197 | RPAIN | chr17 | + | 5425970 | 5428211 | 5425970 | 5426299 | 5428070 | 5428211 | 41 | 35 | 55 |

| | | | | | | | | | | | | | | | |
|---|---|---|---|---|---|---|---|---|---|---|---|---|---|---|---|
| RI | 7607 | ENSG00000266173 | STRADA | chr17 | - | 63710490 | 63710836 | 63710490 | 63710614 | 63710727 | 63710836 | 31 | 28 | 34 | [illegible] |
| RI | 7553 | ENSG00000137073 | UBAP2 | chr9 | - | 33996222 | 33998864 | 33996222 | 33996333 | 33998786 | 33998864 | 10 | 5 | 16 | [illegible] |
| RI | 1599 | ENSG00000102901 | CENPT | chr16 | - | 67832227 | 67832545 | 67832227 | 67832350 | 67832454 | 67832545 | 6 | 3 | 7 | [illegible] |
| RI | 555 | ENSG00000226648 | PLCG1-AS1 | chr20 | - | 41100657 | 41101289 | 41100657 | 41100792 | 41101171 | 41101289 | 5 | 8 | 7 | [illegible] |
| RI | 3145 | ENSG00000174243 | DDX23 | chr12 | - | 48836568 | 48837037 | 48836568 | 48836794 | 48836893 | 48837037 | 21 | 7 | 23 | [illegible] |
| RI | 7490 | ENSG00000213930 | GALT | chr9 | + | 34647088 | 34647567 | 34647088 | 34647258 | 34647474 | 34647567 | 40 | 14 | 32 | [illegible] |
| RI | 892 | ENSG00000006025 | OSBPL7 | chr17 | - | 47816106 | 47816482 | 47816106 | 47816202 | 47816387 | 47816482 | 8 | 11 | 0 | [illegible] |
| RI | 8362 | ENSG00000198270 | TMEM116 | chr12 | - | 111933885 | 111936830 | 111933885 | 111934030 | 111936691 | 111936830 | 20 | 22 | 34 | [illegible] |
| RI | 6325 | ENSG00000134905 | CARS2 | chr13 | - | 110642314 | 110644182 | 110642314 | 110642521 | 110644064 | 110644182 | 28 | 29 | 16 | [illegible] |
| RI | 1014 | ENSG00000169738 | DCXR | chr17 | - | 82036190 | 82036448 | 82036190 | 82036308 | 82036383 | 82036448 | 10 | 11 | 8 | [illegible] |
| RI | 2266 | ENSG00000124357 | NAGK | chr2 | + | 71068649 | 71070586 | 71068649 | 71068712 | 71070501 | 71070586 | 73 | 88 | 39 | [illegible] |
| RI | 4181 | ENSG00000087095 | NLK | chr17 | + | 28194581 | 28196381 | 28194581 | 28194708 | 28196179 | 28196381 | 61 | 77 | 76 | [illegible] |
| RI | 3196 | ENSG00000075826 | SEC31B | chr10 | - | 100505360 | 100506201 | 100505360 | 100505495 | 100506039 | 100506201 | 104 | 104 | 145 | [illegible] |
| RI | 159 | ENSG00000179909 | ZNF154 | chr19 | - | 57697366 | 57701145 | 57697366 | 57697841 | 57700987 | 57701145 | 40 | 50 | 21 | [illegible] |
| RI | 3022 | ENSG00000102349 | KLF8 | chrX | + | 56265179 | 56269489 | 56265179 | 56265744 | 56269377 | 56269489 | 5 | 4 | 0 | [illegible] |
| RI | 8459 | ENSG00000123349 | PFDN5 | chr12 | + | 53297849 | 53298150 | 53297849 | 53297924 | 53298048 | 53298150 | 112 | 79 | 101 | [illegible] |
| RI | 7258 | ENSG00000133794 | ARNTL | chr11 | + | 13378336 | 13381254 | 13378336 | 13378445 | 13381160 | 13381254 | 51 | 29 | 44 | [illegible] |
| RI | 3553 | ENSG00000127054 | INTS11 | chr1 | - | 1312592 | 1312949 | 1312592 | 1312700 | 1312786 | 1312949 | 10 | 5 | 13 | [illegible] |
| RI | 7367 | ENSG00000140988 | RPS2 | chr16 | - | 1962496 | 1963256 | 1962496 | 1962656 | 1963148 | 1963256 | 210 | 78 | 309 | [illegible] |
| RI | 934 | ENSG00000092929 | UNC13D | chr17 | - | 75827224 | 75828086 | 75827224 | 75827445 | 75827956 | 75828086 | 30 | 20 | 21 | [illegible] |
| RI | 5996 | ENSG00000156508 | EEF1A1 | chr6 | - | 73518029 | 73518610 | 73518029 | 73518264 | 73518353 | 73518610 | 524 | 290 | 643 | [illegible] |
| RI | 6893 | ENSG00000102393 | GLA | chrX | - | 101398369 | 101401809 | 101398369 | 101398946 | 101401631 | 101401809 | 12 | 30 | 27 | [illegible] |
| RI | 3120 | ENSG00000161204 | ABCF3 | chr3 | + | 184189387 | 184189757 | 184189387 | 184189443 | 184189556 | 184189757 | 3 | 10 | 16 | [illegible] |
| RI | 2273 | ENSG00000103496 | STX4 | chr16 | + | 31039540 | 31039818 | 31039540 | 31039651 | 31039722 | 31039818 | 19 | 5 | 26 | [illegible] |
| RI | 7262 | ENSG00000104365 | IKBKB | chr8 | + | 42319269 | 42320844 | 42319269 | 42319421 | 42320734 | 42320844 | 97 | 50 | 41 | [illegible] |
| RI | 1477 | ENSG00000156858 | PRR14 | chr16 | + | 30651595 | 30651964 | 30651595 | 30651668 | 30651795 | 30651964 | 7 | 2 | 3 | [illegible] |
| RI | 7746 | ENSG00000149091 | DGKZ | chr11 | + | 46379202 | 46379568 | 46379202 | 46379236 | 46379453 | 46379568 | 32 | 29 | 19 | [illegible] |
| RI | 7795 | ENSG00000110079 | MS4A4A | chr11 | + | 60302558 | 60306201 | 60302558 | 60302717 | 60306099 | 60306201 | 2 | 8 | 4 | [illegible] |
| RI | 4230 | ENSG00000132781 | MUTYH | chr1 | - | 45332762 | 45333324 | 45332762 | 45333170 | 45333284 | 45333324 | 2 | 2 | 10 | [illegible] |
| RI | 5684 | ENSG00000169247 | SH3TC2 | chr5 | - | 149026859 | 149028718 | 149026859 | 149028554 | 149028676 | 149028718 | 0 | 0 | 3 | [illegible] |
| RI | 2609 | ENSG00000067082 | KLF6 | chr10 | - | 3775995 | 3779590 | 3775995 | 3776403 | 3779346 | 3779590 | 804 | 604 | 753 | [illegible] |
| RI | 969 | ENSG00000178971 | CTC1 | chr17 | - | 8229141 | 8229446 | 8229141 | 8229206 | 8229301 | 8229446 | 61 | 53 | 50 | [illegible] |
| RI | 355 | ENSG00000076928 | ARHGEF1 | chr19 | + | 41892024 | 41892373 | 41892024 | 41892123 | 41892330 | 41892373 | 24 | 25 | 38 | [illegible] |
| RI | 7239 | ENSG00000104408 | EIF3E | chr8 | - | 108202982 | 108203503 | 108202982 | 108203117 | 108203400 | 108203503 | 35 | 56 | 30 | [illegible] |
| RI | 8303 | ENSG00000125651 | GTF2F1 | chr19 | - | 6380903 | 6381195 | 6380903 | 6381042 | 6381121 | 6381195 | 5 | 5 | 0 | [illegible] |
| RI | 5337 | ENSG00000084623 | EIF3I | chr1 | + | 32224409 | 32226530 | 32224409 | 32224475 | 32226170 | 32226530 | 43 | 30 | 29 | [illegible] |
| RI | 6947 | ENSG00000215301 | DDX3X | chrX | + | 41337407 | 41339083 | 41337407 | 41337465 | 41337690 | 41339083 | 83 | 64 | 108 | [illegible] |
| RI | 5460 | ENSG00000184840 | TMED9 | chr5 | + | 177593649 | 177594285 | 177593649 | 177593775 | 177594138 | 177594285 | 17 | 8 | 11 | [illegible] |
| RI | 7519 | ENSG00000165699 | TSC1 | chr9 | - | 132910570 | 132911568 | 132910570 | 132910692 | 132911452 | 132911568 | 64 | 47 | 58 | [illegible] |
| RI | 6344 | ENSG00000107175 | CREB3 | chr9 | + | 35732995 | 35733282 | 35732995 | 35733143 | 35733214 | 35733282 | 33 | 12 | 24 | [illegible] |
| RI | 3523 | ENSG00000270629 | NBPF14 | chr1 | - | 148595542 | 148595890 | 148595542 | 148595752 | 148595854 | 148595890 | 10 | 1 | 10 | [illegible] |
| RI | 2583 | ENSG00000182093 | GET1 | chr21 | + | 39390697 | 39391476 | 39390697 | 39390863 | 39391226 | 39391476 | 11 | 14 | 10 | [illegible] |
| RI | 7613 | ENSG00000062822 | POLD1 | chr19 | + | 50415723 | 50416528 | 50415723 | 50415826 | 50416395 | 50416528 | 7 | 0 | 2 | [illegible] |
| RI | 7177 | ENSG00000100023 | PPIL2 | chr22 | + | 21694592 | 21694817 | 21694592 | 21694665 | 21694754 | 21694817 | 36 | 13 | 12 | [illegible] |
| RI | 6790 | ENSG00000126903 | SLC10A3 | chrX | - | 154487305 | 154489082 | 154487305 | 154488496 | 154488583 | 154489082 | 35 | 7 | 18 | [illegible] |
| RI | 8552 | ENSG00000105967 | TFEC | chr7 | - | 115939948 | 115942040 | 115939948 | 115940444 | 115941892 | 115942040 | 189 | 234 | 109 | [illegible] |
| RI | 4239 | ENSG00000166484 | MAPK7 | chr17 | + | 19380607 | 19382466 | 19380607 | 19381686 | 19381780 | 19382466 | 46 | 36 | 45 | [illegible] |
| RI | 6802 | ENSG00000005302 | MSL3 | chrX | + | 11763779 | 11768682 | 11763779 | 11765729 | 11768572 | 11768682 | 82 | 78 | 87 | [illegible] |
| RI | 4392 | ENSG00000145020 | AMT | chr3 | - | 49418970 | 49421572 | 49418970 | 49420342 | 49421491 | 49421572 | 24 | 15 | 23 | [illegible] |
| RI | 6573 | ENSG00000186088 | GSAP | chr7 | - | 77329332 | 77330367 | 77329332 | 77329391 | 77330181 | 77330367 | 432 | 457 | 325 | [illegible] |
| RI | 2323 | ENSG00000185627 | PSMD13 | chr11 | + | 244040 | 244216 | 244040 | 244075 | 244160 | 244216 | 10 | 20 | 17 | [illegible] |
| RI | 4690 | ENSG00000175634 | RPS6KB2 | chr11 | + | 67434581 | 67435401 | 67434581 | 67434694 | 67434988 | 67435401 | 15 | 2 | 22 | [illegible] |
| RI | 7981 | ENSG00000187531 | SIRT7 | chr17 | - | 81914293 | 81915512 | 81914293 | 81914702 | 81915439 | 81915512 | 9 | 7 | 3 | [illegible] |
| RI | 8379 | ENSG00000123329 | ARHGAP9 | chr12 | - | 57477155 | 57477680 | 57477155 | 57477269 | 57477458 | 57477680 | 25 | 37 | 47 | [illegible] |
| RI | 601 | ENSG00000171703 | TCEA2 | chr20 | + | 64069764 | 64070414 | 64069764 | 64069821 | 64070259 | 64070414 | 3 | 10 | 6 | [illegible] |
| RI | 7339 | ENSG00000158850 | B4GALT3 | chr1 | - | 161173604 | 161174049 | 161173604 | 161173727 | 161173858 | 161174049 | 32 | 35 | 46 | [illegible] |
| RI | 3226 | ENSG00000184014 | DENND5A | chr11 | - | 9141939 | 9144278 | 9141939 | 9143485 | 9144096 | 9144278 | 6 | 13 | 9 | [illegible] |
| RI | 6488 | ENSG00000087077 | TRIP6 | chr7 | + | 100870369 | 100870743 | 100870369 | 100870463 | 100870573 | 100870743 | 0 | 2 | 3 | [illegible] |
| RI | 3037 | ENSG00000206503 | HLA-A | chr6 | + | 29944499 | 29945091 | 29944499 | 29944616 | 29945062 | 29945091 | 452 | 189 | 1024 | [illegible] |
| RI | 6719 | ENSG00000094631 | HDAC6 | chrX | + | 48814961 | 48815490 | 48814961 | 48815051 | 48815383 | 48815490 | 25 | 12 | 27 | [illegible] |
| RI | 5388 | ENSG00000106636 | YKT6 | chr7 | + | 44208132 | 44211124 | 44208132 | 44208198 | 44211022 | 44211124 | 40 | 25 | 14 | [illegible] |
| RI | 8528 | ENSG00000197111 | PCBP2 | chr12 | + | 53455346 | 53455493 | 53455346 | 53455370 | 53455460 | 53455493 | 207 | 167 | 182 | [illegible] |
| RI | 7640 | ENSG00000113163 | CERT1 | chr5 | - | 75400204 | 75403058 | 75400204 | 75400297 | 75402971 | 75403058 | 12 | 8 | 4 | [illegible] |
| RI | 7907 | ENSG00000102317 | RBM3 | chrX | + | 48575560 | 48576083 | 48575560 | 48575667 | 48575814 | 48576083 | 54 | 41 | 55 | [illegible] |
| RI | 7480 | ENSG00000148334 | PTGES2 | chr9 | - | 128122361 | 128123134 | 128122361 | 128122646 | 128122933 | 128123134 | 21 | 15 | 17 | [illegible] |
| RI | 7601 | ENSG00000198561 | CTNND1 | chr11 | + | 57811398 | 57814373 | 57811398 | 57811486 | 57814310 | 57814373 | 2 | 7 | 3 | [illegible] |
| RI | 5710 | ENSG00000115053 | NCL | chr2 | - | 231456630 | 231457800 | 231456630 | 231456764 | 231457000 | 231457800 | 53 | 81 | 62 | [illegible] |
| RI | 5851 | ENSG00000108465 | CDK5RAP3 | chr17 | + | 47973518 | 47974031 | 47973518 | 47973650 | 47973930 | 47974031 | 28 | 21 | 29 | [illegible] |
| RI | 872 | ENSG00000125447 | GGA3 | chr17 | - | 75241399 | 75241696 | 75241399 | 75241516 | 75241614 | 75241696 | 45 | 42 | 28 | [illegible] |
| RI | 5117 | ENSG00000136463 | TACO1 | chr17 | + | 63606312 | 63607464 | 63606312 | 63606440 | 63607286 | 63607464 | 5 | 5 | 4 | [illegible] |
| RI | 1642 | ENSG00000140400 | MAN2C1 | chr15 | - | 75359058 | 75359425 | 75359058 | 75359153 | 75359327 | 75359425 | 44 | 47 | 45 | [illegible] |
| RI | 1734 | ENSG00000185024 | BRF1 | chr14 | - | 105219150 | 105220130 | 105219150 | 105219232 | 105220068 | 105220130 | 13 | 5 | 3 | [illegible] |
| RI | 4609 | ENSG00000060138 | YBX3 | chr12 | - | 10701959 | 10704148 | 10701959 | 10702134 | 10704059 | 10704148 | 24 | 20 | 17 | [illegible] |
| RI | 1322 | ENSG00000103061 | SLC7A6OS | chr16 | - | 68310334 | 68310946 | 68310334 | 68310613 | 68310734 | 68310946 | 6 | 5 | 5 | [illegible] |
| RI | 4664 | ENSG00000001497 | LAS1L | chrX | - | 65524055 | 65524614 | 65524055 | 65524262 | 65524563 | 65524614 | 32 | 15 | 25 | [illegible] |
| RI | 3400 | ENSG00000095321 | CRAT | chr9 | - | 129100510 | 129102057 | 129100510 | 129100689 | 129101882 | 129102057 | 3 | 1 | 4 | [illegible] |
| RI | 5431 | ENSG00000214517 | PPME1 | chr11 | + | 74250953 | 74251715 | 74250953 | 74251018 | 74251647 | 74251715 | 2 | 3 | 4 | [illegible] |
| RI | 2063 | ENSG00000079616 | KIF22 | chr16 | + | 29798592 | 29799184 | 29798592 | 29798747 | 29798974 | 29799184 | 32 | 19 | 29 | [illegible] |
| RI | 6230 | ENSG00000164054 | SHISA5 | chr3 | - | 48467875 | 48469573 | 48467875 | 48468217 | 48469360 | 48469573 | 717 | 322 | 393 | [illegible] |
| RI | 235 | ENSG00000129353 | SLC44A2 | chr19 | + | 10634755 | 10635073 | 10634755 | 10634887 | 10634973 | 10635073 | 64 | 36 | 84 | [illegible] |
| RI | 2344 | ENSG00000184584 | STING1 | chr5 | - | 139481158 | 139481704 | 139481158 | 139481342 | 139481477 | 139481704 | 16 | 4 | 27 | [illegible] |
| RI | 8436 | ENSG00000137343 | ATAT1 | chr6 | + | 30646066 | 30646823 | 30646066 | 30646109 | 30646468 | 30646823 | 11 | 12 | 10 | [illegible] |
| RI | 6122 | ENSG00000112511 | PHF1 | chr6 | + | 33414724 | 33415144 | 33414724 | 33414829 | 33414965 | 33415144 | 42 | 19 | 42 | [illegible] |
| RI | 604 | ENSG00000198276 | UCKL1 | chr20 | - | 63940949 | 63941208 | 63940949 | 63941043 | 63941109 | 63941208 | 10 | 3 | 16 | [illegible] |
| RI | 5823 | ENSG00000124356 | STAMBP | chr2 | + | 73844812 | 73845262 | 73844812 | 73844888 | 73845166 | 73845262 | 9 | 7 | 10 | [illegible] |
| RI | 6021 | ENSG00000167967 | E4F1 | chr16 | + | 2229569 | 2232364 | 2229569 | 2229675 | 2232170 | 2232364 | 9 | 5 | 8 | [illegible] |
| RI | 29 | ENSG00000160294 | MCM3AP | chr21 | - | 46246306 | 46246886 | 46246306 | 46246404 | 46246627 | 46246886 | 25 | 17 | 33 | [illegible] |
| RI | 5509 | ENSG00000112367 | FIG4 | chr6 | + | 109735149 | 109738453 | 109735149 | 109735298 | 109738324 | 109738453 | 54 | 31 | 29 | [illegible] |
| RI | 3482 | ENSG00000092841 | MYL6 | chr12 | + | 56158683 | 56159730 | 56158683 | 56158711 | 56159586 | 56159730 | 116 | 84 | 100 | [illegible] |
| RI | 1117 | ENSG00000177731 | FLII | chr17 | - | 18245353 | 18245660 | 18245353 | 18245419 | 18245554 | 18245660 | 15 | 6 | 18 | [illegible] |
| RI | 6623 | ENSG00000122545 | SEPTIN7 | chr7 | + | 35898247 | 35903215 | 35898247 | 35898383 | 35903075 | 35903215 | 83 | 63 | 65 | [illegible] |
| RI | 1748 | ENSG00000197102 | DYNC1H1 | chr14 | + | 102042383 | 102042748 | 102042383 | 102042507 | 102042634 | 102042748 | 3 | 2 | 2 | [illegible] |
| RI | 4507 | ENSG00000168610 | STAT3 | chr17 | - | 42316788 | 42317224 | 42316788 | 42316851 | 42317181 | 42317224 | 112 | 81 | 119 | [illegible] |
| RI | 6736 | ENSG00000131871 | SELENOS | chr15 | - | 101274591 | 101275361 | 101274591 | 101274681 | 101275254 | 101275361 | 52 | 39 | 25 | [illegible] |
| RI | 6462 | ENSG00000160856 | FCRL3 | chr1 | - | 157697139 | 157697919 | 157697139 | 157697424 | 157697658 | 157697919 | 119 | 70 | 46 | [illegible] |
| RI | 5784 | ENSG00000239665 | NA | chr10 | + | 13654352 | 13655482 | 13654352 | 13654427 | 13655377 | 13655482 | 8 | 7 | 7 | [illegible] |
| RI | 7467 | ENSG00000135047 | CTSL | chr9 | + | 87726212 | 87727729 | 87726212 | 87726398 | 87727593 | 87727729 | 1 | 4 | 2 | [illegible] |
| RI | 7749 | ENSG00000175550 | DRAP1 | chr11 | + | 65919779 | 65920041 | 65919779 | 65919852 | 65919947 | 65920041 | 32 | 12 | 17 | [illegible] |
| RI | 2630 | ENSG00000100650 | SRSF5 | chr14 | + | 69768797 | 69769251 | 69768797 | 69768896 | 69769181 | 69769251 | 830 | 690 | 724 | [illegible] |
| RI | 529 | ENSG00000064601 | CTSA | chr20 | + | 45892272 | 45892484 | 45892272 | 45892323 | 45892397 | 45892484 | 8 | 6 | 6 | [illegible] |
| RI | 6551 | ENSG00000103145 | HCFC1R1 | chr16 | - | 3023232 | 3023968 | 3023232 | 3023530 | 3023846 | 3023968 | 2 | 1 | 1 | [illegible] |
| RI | 5092 | ENSG00000166986 | MARS1 | chr12 | + | 57512007 | 57512353 | 57512007 | 57512103 | 57512235 | 57512353 | 18 | 22 | 32 | [illegible] |
| RI | 378 | ENSG00000130755 | GMFG | chr19 | - | 39328358 | 39329073 | 39328358 | 39328548 | 39328999 | 39329073 | 119 | 62 | 121 | [illegible] |
| RI | 2379 | ENSG00000247774 | PCED1B-AS1 | chr12 | - | 47216164 | 47216434 | 47216164 | 47216264 | 47216383 | 47216434 | 64 | 38 | 64 | [illegible] |
| RI | 2948 | ENSG00000164068 | RNF123 | chr3 | + | 49705533 | 49706065 | 49705533 | 49705679 | 49705981 | 49706065 | 8 | 3 | 4 | [illegible] |
| RI | 1250 | ENSG00000188603 | CLN3 | chr16 | - | 28482326 | 28482672 | 28482326 | 28482382 | 28482607 | 28482672 | 18 | 12 | 18 | [illegible] |
| RI | 2986 | ENSG00000110203 | FOLR3 | chr11 | + | 72138960 | 72139149 | 72138960 | 72139110 | 72139112 | 72139149 | 0 | 31 | 0 | [illegible] |
| RI | 109 | ENSG00000100241 | SBF1 | chr22 | - | 50447321 | 50447609 | 50447321 | 50447453 | 50447521 | 50447609 | 3 | 0 | 0 | [illegible] |
| RI | 6586 | ENSG00000184863 | RBM33 | chr7 | + | 155738059 | 155740026 | 155738059 | 155738403 | 155739714 | 155740026 | 8 | 6 | 7 | [illegible] |
| RI | 2964 | ENSG00000155975 | VPS37A | chr8 | + | 17280238 | 17280443 | 17280238 | 17280297 | 17280374 | 17280443 | 15 | 25 | 24 | [illegible] |
| RI | 547 | ENSG00000022277 | RTF2 | chr20 | + | 56516934 | 56517201 | 56516934 | 56516989 | 56517105 | 56517201 | 14 | 14 | 21 | [illegible] |
| RI | 4073 | ENSG00000142684 | ZNF593 | chr1 | + | 26170417 | 26170873 | 26170417 | 26170480 | 26170574 | 26170873 | 2 | 5 | 0 | [illegible] |
| RI | 274 | ENSG00000090674 | MCOLN1 | chr19 | + | 7527519 | 7527960 | 7527519 | 7527628 | 7527863 | 7527960 | 6 | 9 | 8 | [illegible] |
| RI | 5384 | ENSG00000114786 | ABHD14A-ACY1 | chr3 | + | 51987555 | 51988603 | 51987555 | 51987624 | 51988523 | 51988603 | 0 | 0 | 3 | [illegible] |
| RI | 129 | ENSG00000100365 | NCF4 | chr22 | + | 36875652 | 36876094 | 36875652 | 36875783 | 36876028 | 36876094 | 21 | 21 | 27 | [illegible] |
| RI | 2963 | ENSG00000010295 | IFFO1 | chr12 | - | 6548064 | 6548545 | 6548064 | 6548160 | 6548424 | 6548545 | 46 | 9 | 23 | [illegible] |
| RI | 1822 | ENSG00000100911 | PSME2 | chr14 | - | 24144399 | 24145273 | 24144399 | 24144468 | 24145242 | 24145273 | 98 | 82 | 72 | [illegible] |
| RI | 4367 | ENSG00000144381 | HSPD1 | chr2 | - | 197494656 | 197495376 | 197494656 | 197494752 | 197495293 | 197495376 | 71 | 65 | 77 | [illegible] |
| RI | 716 | ENSG00000169733 | RFNG | chr17 | - | 82049676 | 82050006 | 82049676 | 82049842 | 82049917 | 82050006 | 4 | 2 | 9 | [illegible] |
| RI | 6925 | ENSG00000102312 | PORCN | chrX | + | 48511891 | 48512507 | 48511891 | 48511935 | 48512325 | 48512507 | 4 | 2 | 0 | [illegible] |
| RI | 1160 | ENSG00000125124 | BBS2 | chr16 | - | 56496966 | 56497880 | 56496966 | 56497079 | 56497742 | 56497880 | 22 | 26 | 23 | [illegible] |
| RI | 3679 | ENSG00000160789 | LMNA | chr1 | + | 156137653 | 156138757 | 156137653 | 156137743 | 156138487 | 156138757 | 2 | 3 | 6 | [illegible] |
| RI | 6858 | ENSG00000071889 | FAM3A | chrX | - | 154515759 | 154516242 | 154515759 | 154515812 | 154515909 | 154516242 | 25 | 11 | 22 | [illegible] |
| RI | 6011 | ENSG00000204420 | MPIG6B | chr6 | + | 31724586 | 31724844 | 31724586 | 31724627 | 31724745 | 31724844 | 2 | 1 | 1 | [illegible] |
| RI | 1719 | ENSG00000165795 | NDRG2 | chr14 | - | 21017986 | 21018814 | 21017986 | 21018038 | 21018203 | 21018814 | 1 | 5 | 0 | [illegible] |
| RI | 5396 | ENSG00000115159 | GPD2 | chr2 | + | 156578888 | 156579164 | 156578888 | 156579001 | 156579085 | 156579164 | 3 | 6 | 4 | [illegible] |
| RI | 4391 | ENSG00000145020 | AMT | chr3 | - | 49419709 | 49420342 | 49419709 | 49419788 | 49420210 | 49420342 | 23 | 11 | 23 | [illegible] |
| RI | 3291 | ENSG00000180098 | TRNAU1AP | chr1 | + | 28560632 | 28561398 | 28560632 | 28560732 | 28561345 | 28561398 | 28 | 24 | 23 | [illegible] |
| RI | 4714 | ENSG00000172500 | FIBP | chr11 | - | 65884569 | 65884998 | 65884569 | 65884656 | 65884934 | 65884998 | 14 | 6 | 15 | [illegible] |
| RI | 1285 | ENSG00000167526 | RPL13 | chr16 | + | 89561226 | 89561751 | 89561226 | 89561368 | 89561577 | 89561751 | 52 | 50 | 106 | [illegible] |
| RI | 1422 | ENSG00000167508 | MVD | chr16 | - | 88657435 | 88658029 | 88657435 | 88657582 | 88657914 | 88658029 | 7 | 0 | 2 | [illegible] |
| RI | 5939 | ENSG00000134440 | NARS1 | chr18 | - | 57602354 | 57602943 | 57602354 | 57602486 | 57602811 | 57602943 | 122 | 99 | 98 | [illegible] |
| RI | 1840 | ENSG00000099797 | TECR | chr19 | + | 14564785 | 14565123 | 14564785 | 14564992 | 14565065 | 14565123 | 10 | 1 | 6 | [illegible] |
| RI | 6523 | ENSG00000105835 | NAMPT | chr7 | - | 106274945 | 106277179 | 106274945 | 106275049 | 106277022 | 106277179 | 256 | 411 | 135 | [illegible] |
| RI | 641 | ENSG00000088826 | SMOX | chr20 | + | 4182088 | 4182848 | 4182088 | 4182323 | 4182482 | 4182848 | 22 | 37 | 9 | [illegible] |
| RI | 618 | ENSG00000080189 | SLC35C2 | chr20 | - | 46349527 | 46350890 | 46349527 | 46350524 | 46350760 | 46350890 | 42 | 32 | 37 | [illegible] |
| RI | 3626 | ENSG00000143376 | SNX27 | chr1 | + | 151693423 | 151694484 | 151693423 | 151693483 | 151693592 | 151694484 | 25 | 21 | 13 | [illegible] |
| RI | 593 | ENSG00000160410 | SHKBP1 | chr19 | + | 40577157 | 40577441 | 40577157 | 40577284 | 40577395 | 40577441 | 5 | 3 | 13 | [illegible] |
| RI | 7430 | ENSG00000070061 | ELP1 | chr9 | - | 108906302 | 108908404 | 108906302 | 108906485 | 108908304 | 108908404 | 17 | 13 | 39 | [illegible] |
| RI | 399 | ENSG00000130244 | FAM98C | chr19 | + | 38405518 | 38407077 | 38405518 | 38405635 | 38406909 | 38407077 | 26 | 11 | 20 | [illegible] |
| RI | 276 | ENSG00000076944 | STXBP2 | chr19 | + | 7647161 | 7647511 | 7647161 | 7647247 | 7647353 | 7647511 | 8 | 17 | 12 | [illegible] |
| RI | 4791 | ENSG00000115042 | FAHD2A | chr2 | + | 95410526 | 95411026 | 95410526 | 95410586 | 95410863 | 95411026 | 34 | 8 | 20 | [illegible] |
| RI | 5600 | ENSG00000102125 | TAZ | chrX | + | 154413206 | 154413567 | 154413206 | 154413252 | 154413481 | 154413567 | 8 | 3 | 7 | [illegible] |
| RI | 5796 | ENSG00000115307 | AUP1 | chr2 | - | 74529131 | 74529499 | 74529131 | 74529282 | 74529361 | 74529499 | 21 | 7 | 11 | [illegible] |
| RI | 3548 | ENSG00000168389 | MFSD2A | chr1 | + | 39967627 | 39967916 | 39967627 | 39967711 | 39967803 | 39967916 | 0 | 4 | 4 | [illegible] |
| RI | 8493 | ENSG00000111229 | ARPC3 | chr12 | - | 110436109 | 110436683 | 110436109 | 110436204 | 110436556 | 110436683 | 104 | 95 | 96 | [illegible] |
| RI | 2540 | ENSG00000198707 | CEP290 | chr12 | - | 88102837 | 88106905 | 88102837 | 88103011 | 88106674 | 88106905 | 35 | 57 | 16 | [illegible] |
| RI | 6744 | ENSG00000285171 | NA | chrX | - | 71105999 | 71106426 | 71105999 | 71106137 | 71106358 | 71106426 | 50 | 18 | 55 | [illegible] |
| RI | 6916 | ENSG00000248323 | LUCAT1 | chr5 | - | 91310626 | 91313229 | 91310626 | 91312413 | 91313077 | 91313229 | 28 | 22 | 19 | [illegible] |
| RI | 2377 | ENSG00000128694 | OSGEPL1 | chr2 | - | 189753915 | 189754345 | 189753915 | 189754064 | 189754140 | 189754345 | 11 | 8 | 20 | [illegible] |
| RI | 7237 | ENSG00000104408 | EIF3E | chr8 | - | 108201742 | 108203117 | 108201742 | 108201923 | 108202982 | 108203117 | 50 | 30 | 36 | [illegible] |
| RI | 910 | ENSG00000109103 | UNC119 | chr17 | - | 28546707 | 28547849 | 28546707 | 28547409 | 28547676 | 28547849 | 103 | 36 | 60 | [illegible] |
| RI | 7524 | ENSG00000148356 | LRSAM1 | chr9 | + | 127451514 | 127452084 | 127451514 | 127451669 | 127451928 | 127452084 | 22 | 2 | 10 | [illegible] |
| RI | 1237 | ENSG00000167978 | SRRM2 | chr16 | + | 2757471 | 2757945 | 2757471 | 2757579 | 2757780 | 2757945 | 23 | 21 | 39 | [illegible] |
| RI | 8343 | ENSG00000116198 | CEP104 | chr1 | - | 3837291 | 3839119 | 3837291 | 3837345 | 3838963 | 3839119 | 91 | 51 | 89 | [illegible] |
| RI | 2989 | ENSG00000109065 | NAT9 | chr17 | - | 74771959 | 74772277 | 74771959 | 74772054 | 74772217 | 74772277 | 23 | 13 | 28 | [illegible] |
| RI | 2256 | ENSG00000161960 | EIF4A1 | chr17 | + | 7577568 | 7578244 | 7577568 | 7577706 | 7577826 | 7578244 | 42 | 46 | 81 | [illegible] |
| RI | 7782 | ENSG00000149136 | SSRP1 | chr11 | - | 57328296 | 57330138 | 57328296 | 57328426 | 57330092 | 57330138 | 5 | 9 | 20 | [illegible] |
| RI | 6180 | ENSG00000214021 | TTLL3 | chr3 | + | 9817644 | 9818920 | 9817644 | 9817759 | 9818554 | 9818920 | 67 | 57 | 61 | [illegible] |
| RI | 5483 | ENSG00000196295 | GARS1-DT | chr7 | - | 30576563 | 30578091 | 30576563 | 30576710 | 30577648 | 30578091 | 28 | 37 | 29 | [illegible] |
| RI | 8247 | ENSG00000284770 | TBCE | chr1 | + | 235448348 | 235448951 | 235448348 | 235448440 | 235448669 | 235448951 | 24 | 31 | 29 | [illegible] |
| RI | 679 | ENSG00000133313 | CNDP2 | chr18 | + | 74516227 | 74519096 | 74516227 | 74516392 | 74518948 | 74519096 | 13 | 13 | 6 | [illegible] |
| RI | 7482 | ENSG00000148334 | PTGES2 | chr9 | - | 128125243 | 128125441 | 128125243 | 128125283 | 128125375 | 128125441 | 50 | 23 | 59 | [illegible] |
| RI | 6142 | ENSG00000288612 | NA | chr6 | + | 3026752 | 3027426 | 3026752 | 3026871 | 3026944 | 3027426 | 8 | 7 | 5 | [illegible] |
| RI | 5936 | ENSG00000179222 | MAGED1 | chrX | + | 51901552 | 51902354 | 51901552 | 51901938 | 51902145 | 51902354 | 3 | 20 | 20 | [illegible] |
| RI | 4424 | ENSG00000173575 | CHD2 | chr15 | + | 92946037 | 92946340 | 92946037 | 92946082 | 92946247 | 92946340 | 439 | 536 | 449 | [illegible] |
| RI | 1474 | ENSG00000140983 | RHOT2 | chr16 | + | 672253 | 672566 | 672253 | 672384 | 672488 | 672566 | 17 | 2 | 14 | [illegible] |
| RI | 8321 | ENSG00000123384 | LRP1 | chr12 | + | 57204626 | 57205249 | 57204626 | 57204749 | 57205108 | 57205249 | 31 | 17 | 10 | [illegible] |
| RI | 4388 | ENSG00000145020 | AMT | chr3 | - | 49418970 | 49420342 | 49418970 | 49419405 | 49419709 | 49420342 | 17 | 14 | 14 | [illegible] |
| RI | 7847 | ENSG00000174483 | BBS1 | chr11 | + | 66531655 | 66532219 | 66531655 | 66531742 | 66531950 | 66532219 | 1 | 0 | 2 | [illegible] |
| RI | 1941 | ENSG00000103811 | CTSH | chr15 | - | 78931450 | 78932458 | 78931450 | 78932038 | 78932371 | 78932458 | 7 | 8 | 3 | [illegible] |
| RI | 6619 | ENSG00000130305 | NSUN5 | chr7 | - | 73307582 | 73308553 | 73307582 | 73307757 | 73308430 | 73308553 | 22 | 17 | 29 | [illegible] |
| RI | 1980 | ENSG00000164080 | RAD54L2 | chr3 | + | 51630271 | 51630931 | 51630271 | 51630388 | 51630704 | 51630931 | 13 | 5 | 12 | [illegible] |
| RI | 970 | ENSG00000178971 | CTC1 | chr17 | - | 8229890 | 8230468 | 8229890 | 8229922 | 8230293 | 8230468 | 86 | 51 | 81 | [illegible] |
| RI | 5128 | ENSG00000115464 | USP34 | chr2 | - | 61206759 | 61208977 | 61206759 | 61206886 | 61208898 | 61208977 | 52 | 43 | 38 | [illegible] |
| RI | 6283 | ENSG00000026950 | BTN3A1 | chr6 | + | 26409893 | 26410032 | 26409893 | 26409914 | 26410005 | 26410032 | 52 | 28 | 79 | [illegible] |
| RI | 3331 | ENSG00000117228 | GBP1 | chr1 | - | 89055112 | 89056228 | 89055112 | 89055215 | 89056015 | 89056228 | 51 | 48 | 34 | [illegible] |
| RI | 7295 | ENSG00000128567 | PODXL | chr7 | - | 131508950 | 131509585 | 131508950 | 131509028 | 131509364 | 131509585 | 4 | 0 | 3 | [illegible] |
| RI | 1527 | ENSG00000099341 | PSMD8 | chr19 | + | 38376159 | 38376454 | 38376159 | 38376232 | 38376351 | 38376454 | 10 | 7 | 13 | [illegible] |
| RI | 6203 | ENSG00000223705 | NSUN5P1 | chr7 | + | 75415591 | 75416191 | 75415591 | 75415770 | 75415980 | 75416191 | 45 | 28 | 72 | [illegible] |
| RI | 4336 | ENSG00000160055 | TMEM234 | chr1 | - | 32215135 | 32215795 | 32215135 | 32215216 | 32215672 | 32215795 | 7 | 12 | 17 | [illegible] |
| RI | 7767 | ENSG00000172932 | ANKRD13D | chr11 | + | 67299057 | 67299611 | 67299057 | 67299124 | 67299529 | 67299611 | 72 | 38 | 44 | [illegible] |
| RI | 7758 | ENSG00000101158 | NELFCD | chr20 | + | 58988913 | 58989640 | 58988913 | 58989021 | 58989487 | 58989640 | 52 | 35 | 44 | [illegible] |
| RI | 2613 | ENSG00000127419 | TMEM175 | chr4 | + | 948115 | 948615 | 948115 | 948154 | 948410 | 948615 | 23 | 3 | 16 | [illegible] |
| RI | 3547 | ENSG00000136059 | VILL | chr3 | + | 38006180 | 38006700 | 38006180 | 38006252 | 38006448 | 38006700 | 9 | 4 | 8 | [illegible] |
| RI | 3867 | ENSG00000108654 | DDX5 | chr17 | - | 64500548 | 64502069 | 64500548 | 64500882 | 64502009 | 64502069 | 2224 | 1398 | 2103 | [illegible] |
| RI | 5482 | ENSG00000137309 | HMGA1 | chr6 | + | 34236872 | 34237317 | 34236872 | 34236963 | 34237203 | 34237317 | 1 | 0 | 1 | [illegible] |
| RI | 1668 | ENSG00000140474 | ULK3 | chr15 | - | 74842074 | 74842420 | 74842074 | 74842195 | 74842279 | 74842420 | 12 | 7 | 9 | [illegible] |
| RI | 8489 | ENSG00000166598 | HSP90B1 | chr12 | + | 103946617 | 103946941 | 103946617 | 103946696 | 103946785 | 103946941 | 43 | 39 | 55 | [illegible] |
| RI | 6834 | ENSG00000177042 | TMEM80 | chr11 | + | 698868 | 700235 | 698868 | 698888 | 700141 | 700235 | 3 | 2 | 12 | [illegible] |
| RI | 1648 | ENSG00000196547 | MAN2A2 | chr15 | + | 90912042 | 90912664 | 90912042 | 90912279 | 90912516 | 90912664 | 9 | 9 | 7 | [illegible] |
| RI | 405 | ENSG00000105677 | TMEM147 | chr19 | + | 35546671 | 35547240 | 35546671 | 35547029 | 35547118 | 35547240 | 18 | 6 | 10 | [illegible] |
| RI | 1975 | ENSG00000006744 | ELAC2 | chr17 | - | 12994763 | 12995062 | 12994763 | 12994884 | 12994962 | 12995062 | 11 | 5 | 15 | [illegible] |
| RI | 4752 | ENSG00000147813 | NAPRT | chr8 | - | 143574784 | 143575093 | 143574784 | 143574900 | 143574985 | 143575093 | 4 | 3 | 3 | [illegible] |
| RI | 2783 | ENSG00000161013 | MGAT4B | chr5 | - | 179799505 | 179800068 | 179799505 | 179799636 | 179799953 | 179800068 | 38 | 18 | 20 | [illegible] |
| RI | 5752 | ENSG00000169045 | HNRNPH1 | chr5 | - | 179617798 | 179618060 | 179617798 | 179617932 | 179617988 | 179618060 | 424 | 508 | 415 | [illegible] |
| RI | 6361 | ENSG00000122515 | ZMIZ2 | chr7 | + | 44761448 | 44761905 | 44761448 | 44761593 | 44761694 | 44761905 | 11 | 7 | 16 | [illegible] |
| RI | 3738 | ENSG00000114626 | ABTB1 | chr3 | + | 127676535 | 127677286 | 127676535 | 127676581 | 127677167 | 127677286 | 27 | 30 | 26 | [illegible] |
| RI | 8025 | ENSG00000090621 | PABPC4 | chr1 | - | 39563613 | 39563922 | 39563613 | 39563741 | 39563835 | 39563922 | 18 | 18 | 34 | [illegible] |
| RI | 3369 | ENSG00000186517 | ARHGAP30 | chr1 | - | 161052625 | 161053385 | 161052625 | 161052797 | 161053257 | 161053385 | 99 | 69 | 67 | [illegible] |
| RI | 6026 | ENSG00000204264 | PSMB8 | chr6 | - | 32842671 | 32843089 | 32842671 | 32842783 | 32842941 | 32843089 | 63 | 54 | 93 | [illegible] |
| RI | 1967 | ENSG00000112640 | PPP2R5D | chr6 | + | 43008366 | 43009227 | 43008366 | 43008475 | 43009056 | 43009227 | 18 | 18 | 17 | [illegible] |
| RI | 2938 | ENSG00000156976 | EIF4A2 | chr3 | + | 186784563 | 186786051 | 186784563 | 186784696 | 186785882 | 186786051 | 454 | 323 | 496 | [illegible] |
| RI | 4456 | ENSG00000117139 | KDM5B | chr1 | - | 202731827 | 202733886 | 202731827 | 202731939 | 202733400 | 202733886 | 7 | 6 | 7 | [illegible] |
| RI | 2949 | ENSG00000164068 | RNF123 | chr3 | + | 49712478 | 49713587 | 49712478 | 49712656 | 49713512 | 49713587 | 12 | 2 | 7 | [illegible] |
| RI | 1077 | ENSG00000126581 | BECN1 | chr17 | - | 42818786 | 42819609 | 42818786 | 42818877 | 42819547 | 42819609 | 70 | 48 | 66 | [illegible] |
| RI | 7845 | ENSG00000174483 | BBS1 | chr11 | + | 66515539 | 66515731 | 66515539 | 66515586 | 66515692 | 66515731 | 4 | 2 | 7 | [illegible] |
| RI | 7503 | ENSG00000148288 | GBGT1 | chr9 | - | 133152952 | 133155312 | 133152952 | 133154242 | 133155177 | 133155312 | 12 | 8 | 9 | [illegible] |
| RI | 2998 | ENSG00000060971 | ACAA1 | chr3 | - | 38126161 | 38126700 | 38126161 | 38126341 | 38126509 | 38126700 | 47 | 19 | 42 | [illegible] |

| | | | | | | | | | | | | | | | | | | | | | | | | | | | | | | | | | | | | | | | | | | | | | | |
|---|---|---|---|---|---|---|---|---|---|---|---|---|---|---|---|---|---|---|---|---|---|---|---|---|---|---|---|---|---|---|---|---|---|---|---|---|---|---|---|---|---|---|---|---|---|---|
| RI | 1165 | ENSG00000125124 | BBS2 | chr16 | - | 56498436 | 56501025 | 56498436 | 56499907 | 56500853 | 56501025 | 32 | 23 | 25 | 33 | 5 | 35 | 33 | 67 | 59 | 86 | 20 | 43 | 36 | 21 | 0 | 14 | 25 | 34 | 12 | 30 | 298 | 149 | 0.031745003 | 0.112286067 | 0.314 | 0.258 | 0.157 | 0.219 | 0.028 | 0.417 | 0.462 | 0.346 | 0.467 | 0 | -0.143 |
| RI | 7379 | ENSG00000198000 | NOL8 | chr9 | - | 92298256 | 92298954 | 92298256 | 92298336 | 92298883 | 92298954 | 73 | 68 | 85 | 57 | 96 | 145 | 153 | 115 | 127 | 341 | 10 | 27 | 70 | 11 | 8 | 0 | 34 | 75 | 6 | 23 | 298 | 149 | 0.031818556 | 0.112478881 | 0.201 | 0.182 | 0.27 | 0.183 | 0.123 | 1 | 0.284 | 0.318 | 0.478 | 0.148 | -0.254 |
| RI | 1175 | ENSG00000125124 | BBS2 | chr16 | - | 56501944 | 56502456 | 56501944 | 56502079 | 56502316 | 56502456 | 118 | 134 | 141 | 137 | 60 | 3 | 1 | 7 | 2 | 0 | 32 | 139 | 174 | 75 | 10 | 0 | 1 | 4 | 0 | 0 | 298 | 149 | 0.031980965 | 0.112967071 | 0.952 | 0.985 | 0.91 | 0.972 | 1 | 1 | 0.986 | 0.956 | 1 | 1 | -0.025 |
| RI | 3468 | ENSG00000117394 | SLC2A1 | chr1 | - | 42929209 | 42929780 | 42929209 | 42929314 | 42929592 | 42929780 | 15 | 13 | 10 | 3 | 8 | 37 | 9 | 39 | 22 | 29 | 15 | 18 | 5 | 17 | 75 | 36 | 41 | 10 | 23 | 23 | 298 | 149 | 0.031994906 | 0.112967071 | 0.169 | 0.419 | 0.114 | 0.064 | 0.121 | 0.172 | 0.18 | 0.2 | 0.27 | 0.62 | -0.111 |
| RI | 4810 | ENSG00000111801 | BTN3A3 | chr6 | + | 26448247 | 26448754 | 26448247 | 26448448 | 26448727 | 26448754 | 64 | 103 | 82 | 68 | 91 | 20 | 18 | 13 | 22 | 15 | 55 | 62 | 67 | 37 | 25 | 14 | 29 | 9 | 19 | 37 | 298 | 149 | 0.032046952 | 0.113083241 | 0.615 | 0.741 | 0.759 | 0.607 | 0.752 | 0.663 | 0.517 | 0.788 | 0.493 | 0.253 | 0.152 |
| RI | 8166 | ENSG00000168209 | DDIT4 | chr10 | + | 72273923 | 72274421 | 72273923 | 72274060 | 72274156 | 72274421 | 38 | 2 | 17 | 10 | 7 | 92 | 24 | 48 | 63 | 3 | 0 | 11 | 8 | 6 | 20 | 13 | 78 | 141 | 18 | 69 | 245 | 149 | 0.032169856 | 0.113449158 | 0.201 | 0.048 | 0.177 | 0.088 | 0.587 | 0 | 0.079 | 0.033 | 0.169 | 0.15 | 0.134 |
| RI | 4113 | ENSG00000116337 | AMPD2 | chr1 | + | 109628082 | 109628495 | 109628082 | 109628277 | 109628363 | 109628495 | 48 | 17 | 46 | 38 | 20 | 79 | 56 | 52 | 67 | 67 | 124 | 27 | 52 | 65 | 161 | 144 | 78 | 52 | 74 | 195 | 235 | 149 | 0.032242794 | 0.113638535 | 0.278 | 0.161 | 0.359 | 0.264 | 0.159 | 0.353 | 0.18 | 0.388 | 0.358 | 0.344 | -0.08 |
| RI | 2456 | ENSG00000138760 | SCARB2 | chr4 | - | 76179516 | 76181101 | 76179516 | 76179705 | 76180953 | 76181101 | 21 | 6 | 21 | 31 | 18 | 156 | 120 | 116 | 172 | 217 | 4 | 20 | 22 | 18 | 6 | 231 | 139 | 140 | 224 | 87 | 298 | 149 | 0.032349018 | 0.113944931 | 0.063 | 0.024 | 0.083 | 0.083 | 0.04 | 0.009 | 0.067 | 0.073 | 0.039 | 0.033 | 0.014 |
| RI | 7008 | ENSG00000198373 | WWP2 | chr16 | + | 69931508 | 69931890 | 69931508 | 69931580 | 69931801 | 69931890 | 9 | 6 | 9 | 4 | 1 | 81 | 38 | 83 | 61 | 132 | 8 | 27 | 4 | 10 | 16 | 168 | 92 | 52 | 89 | 167 | 298 | 149 | 0.032520847 | 0.114481909 | 0.053 | 0.073 | 0.051 | 0.032 | 0.004 | 0.023 | 0.128 | 0.037 | 0.053 | 0.046 | -0.015 |
| RI | 1883 | ENSG00000213983 | AP1G2 | chr14 | - | 23562503 | 23563638 | 23562503 | 23562593 | 23563379 | 23563638 | 68 | 28 | 38 | 44 | 12 | 64 | 31 | 55 | 37 | 32 | 67 | 73 | 38 | 61 | 48 | 22 | 71 | 21 | 28 | 84 | 298 | 149 | 0.032609802 | 0.114726682 | 0.347 | 0.311 | 0.257 | 0.373 | 0.158 | 0.604 | 0.34 | 0.475 | 0.521 | 0.222 | -0.143 |
| RI | 4951 | ENSG00000171503 | ETFDH | chr4 | + | 158680466 | 158682424 | 158680466 | 158680607 | 158682194 | 158682424 | 16 | 35 | 30 | 28 | 51 | 9 | 24 | 11 | 17 | 16 | 22 | 39 | 85 | 32 | 6 | 19 | 0 | 8 | 26 | 0 | 298 | 149 | 0.032836619 | 0.115455899 | 0.471 | 0.422 | 0.577 | 0.452 | 0.614 | 0.367 | 1 | 0.842 | 0.381 | 1 | -0.211 |
| RI | 1328 | ENSG00000214078 | CPNE1 | chr20 | - | 35632162 | 35632385 | 35632162 | 35632234 | 35632310 | 35632385 | 34 | 36 | 38 | 29 | 14 | 206 | 90 | 225 | 154 | 207 | 30 | 22 | 10 | 28 | 16 | 134 | 182 | 128 | 147 | 271 | 225 | 149 | 0.033011733 | 0.116002562 | 0.099 | 0.209 | 0.101 | 0.111 | 0.043 | 0.129 | 0.074 | 0.049 | 0.112 | 0.038 | 0.032 |
| RI | 1382 | ENSG00000103423 | DNAJA3 | chr16 | + | 4442267 | 4443164 | 4442267 | 4442420 | 4443016 | 4443164 | 11 | 15 | 17 | 19 | 18 | 42 | 14 | 40 | 19 | 46 | 50 | 18 | 24 | 5 | 24 | 26 | 17 | 15 | 11 | 53 | 298 | 149 | 0.033179515 | 0.116522826 | 0.116 | 0.349 | 0.175 | 0.333 | 0.164 | 0.49 | 0.346 | 0.444 | 0.185 | 0.185 | -0.103 |
| RI | 1534 | ENSG00000172345 | STARD5 | chr15 | - | 81322407 | 81324141 | 81322407 | 81322540 | 81323801 | 81324141 | 28 | 17 | 19 | 21 | 27 | 0 | 0 | 3 | 5 | 0 | 57 | 33 | 61 | 21 | 32 | 0 | 0 | 0 | 0 | 0 | 298 | 149 | 0.033561425 | 0.117794022 | 1 | 1 | 0.76 | 0.677 | 1 | 1 | 1 | 1 | 1 | 1 | -0.113 |
| RI | 3849 | ENSG00000109323 | MANBA | chr4 | - | 102664684 | 102669049 | 102664684 | 102664852 | 102668962 | 102669049 | 48 | 61 | 23 | 36 | 24 | 180 | 155 | 101 | 133 | 170 | 21 | 44 | 63 | 27 | 9 | 349 | 153 | 106 | 217 | 95 | 298 | 149 | 0.03384686 | 0.118725298 | 0.118 | 0.164 | 0.102 | 0.119 | 0.066 | 0.029 | 0.126 | 0.229 | 0.059 | 0.045 | 0.016 |
| RI | 8417 | ENSG00000182796 | TMEM198B | chr12 | + | 55834850 | 55836246 | 55834850 | 55835615 | 55836125 | 55836246 | 26 | 5 | 14 | 23 | 18 | 0 | 3 | 0 | 0 | 0 | 43 | 38 | 21 | 23 | 62 | 25 | 0 | 0 | 0 | 0 | 298 | 149 | 0.034049434 | 0.119364991 | 1 | 0.455 | 1 | 1 | 1 | 0.462 | 1 | 1 | 1 | 1 | -0.001 |
| RI | 7934 | ENSG00000137504 | CREBZF | chr11 | - | 85660334 | 85661749 | 85660334 | 85660597 | 85661636 | 85661749 | 477 | 396 | 402 | 400 | 376 | 13 | 7 | 13 | 22 | 33 | 88 | 153 | 194 | 44 | 48 | 0 | 16 | 9 | 9 | 7 | 298 | 149 | 0.03412508 | 0.11955922 | 0.948 | 0.966 | 0.939 | 0.901 | 0.851 | 1 | 0.827 | 0.915 | 0.71 | 0.774 | 0.076 |
| RI | 7176 | ENSG00000100023 | PPIL2 | chr22 | + | 21688072 | 21688849 | 21688072 | 21688106 | 21688731 | 21688849 | 5 | 1 | 9 | 11 | 0 | 108 | 79 | 91 | 107 | 36 | 2 | 6 | 0 | 2 | 33 | 92 | 62 | 55 | 76 | 70 | 298 | 149 | 0.034315113 | 0.120153749 | 0.023 | 0.006 | 0.047 | 0.049 | 0 | 0.011 | 0.046 | 0 | 0.013 | 0.191 | -0.027 |
| RI | 2582 | ENSG00000010256 | UQCRC1 | chr3 | - | 48600679 | 48601118 | 48600679 | 48600840 | 48600974 | 48601118 | 16 | 3 | 34 | 17 | 9 | 90 | 47 | 63 | 87 | 29 | 49 | 15 | 15 | 33 | 40 | 125 | 77 | 39 | 104 | 80 | 283 | 149 | 0.034512595 | 0.120773637 | 0.086 | 0.033 | 0.221 | 0.093 | 0.14 | 0.171 | 0.093 | 0.168 | 0.143 | 0.208 | -0.042 |
| RI | 5406 | ENSG00000004455 | AK2 | chr1 | - | 33012073 | 33013402 | 33012073 | 33012122 | 33013206 | 33013402 | 53 | 54 | 63 | 44 | 119 | 0 | 0 | 0 | 1 | 0 | 22 | 36 | 38 | 74 | 48 | 0 | 0 | 0 | 2 | 7 | 298 | 149 | 0.034565932 | 0.120888668 | 1 | 1 | 1 | 0.957 | 1 | 1 | 1 | 1 | 0.949 | 0.774 | 0.047 |
| RI | 7109 | ENSG00000103266 | STUB1 | chr16 | + | 681792 | 682076 | 681792 | 681903 | 682019 | 682076 | 121 | 76 | 114 | 105 | 79 | 2 | 2 | 1 | 1 | 0 | 133 | 140 | 103 | 123 | 282 | 14 | 0 | 0 | 3 | 0 | 265 | 149 | 0.034730529 | 0.121392446 | 0.971 | 0.955 | 0.985 | 0.983 | 1 | 0.842 | 1 | 1 | 0.958 | 1 | 0.019 |
| RI | 7195 | ENSG00000174744 | BRMS1 | chr11 | - | 66340773 | 66341333 | 66340773 | 66340870 | 66341205 | 66341333 | 12 | 7 | 17 | 16 | 12 | 0 | 0 | 0 | 0 | 0 | 11 | 26 | 4 | 15 | 34 | 10 | 0 | 0 | 0 | 0 | 298 | 149 | 0.034763137 | 0.121424469 | 1 | 1 | 1 | 1 | 1 | 0.355 | 1 | 1 | 1 | 1 | 0.129 |
| RI | 7133 | ENSG00000104783 | KCNN4 | chr19 | - | 43768962 | 43769560 | 43768962 | 43769032 | 43769441 | 43769560 | 2 | 1 | 1 | 9 | 0 | 47 | 18 | 31 | 51 | 46 | 2 | 1 | 5 | 5 | 6 | 43 | 15 | 17 | 37 | 18 | 298 | 149 | 0.034793919 | 0.121424469 | 0.021 | 0.027 | 0.016 | 0.081 | 0 | 0.023 | 0.032 | 0.128 | 0.063 | 0.143 | -0.049 |
| RI | 223 | ENSG00000104814 | MAP4K1 | chr19 | - | 38595484 | 38595729 | 38595484 | 38595555 | 38595639 | 38595729 | 16 | 1 | 11 | 12 | 5 | 46 | 15 | 36 | 23 | 31 | 23 | 49 | 3 | 5 | 37 | 13 | 35 | 26 | 25 | 58 | 233 | 149 | 0.034801359 | 0.121424469 | 0.182 | 0.041 | 0.163 | 0.25 | 0.093 | 0.531 | 0.472 | 0.069 | 0.113 | 0.29 | -0.149 |
| RI | 3820 | ENSG00000107099 | DOCK8 | chr9 | + | 368017 | 370300 | 368017 | 368135 | 370229 | 370300 | 107 | 141 | 140 | 133 | 116 | 727 | 622 | 631 | 816 | 2308 | 148 | 164 | 169 | 164 | 60 | 1074 | 965 | 582 | 825 | 727 | 298 | 149 | 0.035122581 | 0.122472897 | 0.069 | 0.102 | 0.1 | 0.075 | 0.025 | 0.064 | 0.078 | 0.127 | 0.09 | 0.04 | -0.006 |
| RI | 1613 | ENSG00000167077 | MEI1 | chr22 | + | 41793828 | 41794477 | 41793828 | 41793910 | 41794370 | 41794477 | 41 | 13 | 33 | 39 | 37 | 13 | 2 | 18 | 13 | 6 | 42 | 39 | 48 | 33 | 62 | 0 | 18 | 4 | 2 | 18 | 298 | 149 | 0.035201404 | 0.122675334 | 0.612 | 0.765 | 0.478 | 0.6 | 0.755 | 1 | 0.52 | 0.857 | 0.892 | 0.633 | -0.138 |
| RI | 7650 | ENSG00000149761 | NUDT22 | chr11 | + | 64226289 | 64227132 | 64226289 | 64226427 | 64226634 | 64227132 | 22 | 3 | 5 | 21 | 5 | 21 | 11 | 7 | 14 | 2 | 16 | 55 | 18 | 8 | 29 | 30 | 0 | 1 | 10 | 23 | 298 | 149 | 0.035254635 | 0.122788402 | 0.344 | 0.12 | 0.263 | 0.429 | 0.556 | 0.211 | 1 | 0.9 | 0.286 | 0.387 | -0.214 |
| RI | 873 | ENSG00000125447 | GGA3 | chr17 | - | 75241614 | 75242473 | 75241614 | 75241696 | 75242335 | 75242473 | 42 | 29 | 36 | 40 | 14 | 139 | 83 | 118 | 142 | 151 | 86 | 44 | 51 | 52 | 90 | 293 | 172 | 79 | 195 | 240 | 298 | 149 | 0.03534239 | 0.123021506 | 0.131 | 0.149 | 0.132 | 0.123 | 0.044 | 0.128 | 0.113 | 0.244 | 0.118 | 0.158 | -0.036 |
| RI | 4375 | ENSG00000066654 | THUMPD1 | chr16 | - | 20733663 | 20737286 | 20733663 | 20736358 | 20736859 | 20737286 | 221 | 154 | 176 | 221 | 243 | 28 | 15 | 32 | 21 | 33 | 42 | 192 | 178 | 64 | 8 | 0 | 21 | 21 | 3 | 0 | 298 | 149 | 0.035364742 | 0.123026813 | 0.798 | 0.837 | 0.733 | 0.84 | 0.786 | 1 | 0.821 | 0.809 | 0.914 | 1 | -0.11 |
| RI | 7937 | ENSG00000187147 | RNF220 | chr1 | + | 44649882 | 44651723 | 44649882 | 44649957 | 44650703 | 44651723 | 10 | 5 | 21 | 13 | 7 | 231 | 106 | 214 | 212 | 209 | 14 | 8 | 12 | 21 | 29 | 255 | 183 | 143 | 103 | 321 | 298 | 149 | 0.035477119 | 0.123345112 | 0.021 | 0.023 | 0.047 | 0.03 | 0.016 | 0.027 | 0.021 | 0.04 | 0.093 | 0.043 | -0.017 |
| RI | 894 | ENSG00000136490 | LIMD2 | chr17 | - | 63698798 | 63699069 | 63698798 | 63698938 | 63699027 | 63699069 | 68 | 27 | 73 | 70 | 25 | 184 | 56 | 179 | 135 | 46 | 10 | 51 | 41 | 34 | 98 | 134 | 154 | 86 | 82 | 233 | 238 | 149 | 0.035572199 | 0.123602928 | 0.188 | 0.232 | 0.203 | 0.245 | 0.254 | 0.045 | 0.172 | 0.23 | 0.206 | 0.208 | 0.052 |
| RI | 1864 | ENSG00000100823 | APEX1 | chr14 | + | 20455225 | 20455703 | 20455225 | 20455389 | 20455577 | 20455703 | 128 | 78 | 108 | 95 | 65 | 5 | 4 | 6 | 0 | 1 | 113 | 45 | 75 | 89 | 220 | 0 | 8 | 0 | 18 | 10 | 298 | 149 | 0.035649431 | 0.123735537 | 0.928 | 0.907 | 0.9 | 1 | 0.97 | 1 | 0.738 | 1 | 0.712 | 0.917 | 0.068 |
| RI | 4085 | ENSG00000138496 | PARP9 | chr3 | - | 122536167 | 122537073 | 122536167 | 122536342 | 122536933 | 122537073 | 40 | 47 | 32 | 50 | 48 | 170 | 223 | 150 | 226 | 357 | 103 | 68 | 90 | 134 | 0 | 396 | 296 | 188 | 623 | 23 | 298 | 149 | 0.035656854 | 0.123735537 | 0.105 | 0.095 | 0.096 | 0.1 | 0.063 | 0.115 | 0.103 | 0.193 | 0.097 | 0 | -0.01 |
| RI | 852 | ENSG00000141503 | MINK1 | chr17 | + | 4890221 | 4890735 | 4890221 | 4890308 | 4890516 | 4890735 | 10 | 14 | 16 | 11 | 13 | 2 | 3 | 3 | 1 | 0 | 14 | 19 | 20 | 0 | 21 | 5 | 0 | 2 | 7 | 25 | 298 | 149 | 0.035673205 | 0.123735537 | 0.714 | 0.7 | 0.727 | 0.846 | 1 | 0.583 | 1 | 0.833 | 0 | 0.296 | 0.255 |
| RI | 3868 | ENSG00000108654 | DDX5 | chr17 | - | 64500548 | 64502069 | 64500548 | 64501143 | 64502009 | 64502069 | 1685 | 1095 | 1660 | 1833 | 1607 | 10 | 12 | 4 | 14 | 21 | 389 | 1323 | 1463 | 809 | 241 | 4 | 7 | 4 | 0 | 0 | 298 | 149 | 0.035774465 | 0.124013946 | 0.988 | 0.979 | 0.995 | 0.985 | 0.975 | 0.98 | 0.99 | 0.995 | 1 | 1 | -0.009 |
| RI | 3684 | ENSG00000099622 | CIRBP | chr19 | + | 1270927 | 1271467 | 1270927 | 1271036 | 1271328 | 1271467 | 62 | 33 | 63 | 75 | 63 | 1 | 0 | 7 | 2 | 4 | 88 | 100 | 74 | 84 | 125 | 0 | 1 | 5 | 0 | 3 | 298 | 149 | 0.036012874 | 0.12471701 | 0.969 | 1 | 0.818 | 0.949 | 0.887 | 1 | 0.98 | 0.881 | 1 | 0.954 | -0.038 |
| RI | 3575 | ENSG00000141526 | SLC16A3 | chr17 | + | 82235982 | 82236231 | 82235982 | 82236050 | 82236128 | 82236231 | 31 | 13 | 47 | 23 | 9 | 0 | 0 | 0 | 0 | 0 | 21 | 21 | 19 | 23 | 28 | 0 | 9 | 0 | 0 | 0 | 227 | 149 | 0.036019506 | 0.12471701 | 1 | 1 | 1 | 1 | 1 | 1 | 0.605 | 1 | 1 | 1 | 0.079 |
| RI | 6467 | ENSG00000176454 | LPCAT4 | chr15 | - | 34359588 | 34360209 | 34359588 | 34359745 | 34360110 | 34360209 | 41 | 21 | 21 | 27 | 51 | 20 | 14 | 26 | 32 | 27 | 80 | 32 | 36 | 27 | 80 | 21 | 21 | 14 | 24 | 23 | 298 | 149 | 0.036082663 | 0.124862501 | 0.506 | 0.429 | 0.288 | 0.297 | 0.486 | 0.656 | 0.432 | 0.562 | 0.36 | 0.635 | -0.128 |
| RI | 3951 | ENSG00000156873 | PHKG2 | chr16 | + | 30751105 | 30751603 | 30751105 | 30751281 | 30751548 | 30751603 | 29 | 17 | 18 | 24 | 3 | 33 | 12 | 24 | 36 | 20 | 54 | 43 | 18 | 48 | 78 | 15 | 41 | 24 | 31 | 92 | 298 | 149 | 0.036195479 | 0.125179563 | 0.305 | 0.415 | 0.273 | 0.25 | 0.07 | 0.643 | 0.344 | 0.273 | 0.436 | 0.298 | -0.136 |
| RI | 5395 | ENSG00000068654 | POLR1A | chr2 | - | 86081600 | 86083168 | 86081600 | 86081706 | 86083081 | 86083168 | 5 | 6 | 5 | 4 | 3 | 39 | 18 | 29 | 32 | 33 | 9 | 13 | 3 | 3 | 9 | 56 | 26 | 11 | 16 | 27 | 298 | 149 | 0.036270462 | 0.125365487 | 0.06 | 0.143 | 0.079 | 0.059 | 0.043 | 0.074 | 0.2 | 0.12 | 0.086 | 0.143 | -0.048 |
| RI | 1009 | ENSG00000214176 | PLEKHM1P1 | chr17 | - | 64821766 | 64822393 | 64821766 | 64822052 | 64822253 | 64822393 | 18 | 8 | 6 | 14 | 4 | 102 | 47 | 96 | 127 | 60 | 39 | 26 | 15 | 8 | 56 | 278 | 90 | 100 | 240 | 186 | 298 | 149 | 0.036303674 | 0.125406902 | 0.081 | 0.078 | 0.03 | 0.052 | 0.032 | 0.066 | 0.126 | 0.07 | 0.016 | 0.131 | -0.027 |
| RI | 1571 | ENSG00000168970 | JMJD7-PLA2G4B | chr15 | + | 41842192 | 41842591 | 41842192 | 41842276 | 41842553 | 41842591 | 8 | 7 | 21 | 11 | 2 | 9 | 8 | 25 | 8 | 3 | 0 | 9 | 8 | 13 | 24 | 12 | 11 | 7 | 34 | 53 | 298 | 149 | 0.036655226 | 0.126547294 | 0.308 | 0.304 | 0.296 | 0.407 | 0.25 | 0 | 0.29 | 0.364 | 0.16 | 0.185 | 0.113 |
| RI | 501 | ENSG00000099817 | POLR2E | chr19 | - | 1093903 | 1095375 | 1093903 | 1094078 | 1095258 | 1095375 | 1 | 1 | 7 | 3 | 1 | 128 | 84 | 87 | 83 | 47 | 17 | 6 | 1 | 9 | 1 | 138 | 134 | 59 | 60 | 303 | 298 | 149 | 0.036707299 | 0.12660734 | 0.004 | 0.006 | 0.039 | 0.018 | 0.011 | 0.058 | 0.022 | 0.008 | 0.07 | 0.002 | -0.016 |
| RI | 191 | ENSG00000105699 | LSR | chr19 | + | 35266678 | 35266967 | 35266678 | 35266738 | 35266838 | 35266967 | 17 | 2 | 22 | 8 | 11 | 3 | 2 | 2 | 2 | 4 | 10 | 6 | 18 | 26 | 20 | 0 | 0 | 0 | 0 | 11 | 249 | 149 | 0.036715486 | 0.12660734 | 0.772 | 0.374 | 0.868 | 0.705 | 0.622 | 1 | 1 | 1 | 1 | 0.521 | -0.236 |
| RI | 2158 | ENSG00000111641 | NOP2 | chr12 | - | 6560097 | 6560569 | 6560097 | 6560326 | 6560446 | 6560569 | 30 | 17 | 41 | 33 | 23 | 24 | 29 | 39 | 53 | 70 | 33 | 39 | 26 | 38 | 45 | 30 | 28 | 27 | 37 | 51 | 269 | 149 | 0.036805899 | 0.126845068 | 0.409 | 0.245 | 0.368 | 0.256 | 0.154 | 0.379 | 0.436 | 0.348 | 0.363 | 0.328 | -0.084 |
| RI | 4099 | ENSG00000256269 | HMBS | chr11 | + | 119089682 | 119090067 | 119089682 | 119089760 | 119089989 | 119090067 | 0 | 1 | 2 | 5 | 2 | 28 | 9 | 20 | 13 | 26 | 14 | 0 | 7 | 0 | 0 | 26 | 15 | 8 | 13 | 14 | 298 | 149 | 0.037024372 | 0.127507838 | 0 | 0.053 | 0.048 | 0.161 | 0.037 | 0.212 | 0 | 0.304 | 0 | 0 | -0.043 |
| RI | 1013 | ENSG00000108592 | FTSJ3 | chr17 | - | 63826257 | 63826672 | 63826257 | 63826304 | 63826566 | 63826672 | 5 | 8 | 4 | 5 | 5 | 36 | 18 | 35 | 31 | 27 | 6 | 8 | 14 | 9 | 6 | 54 | 26 | 6 | 12 | 34 | 298 | 149 | 0.037041383 | 0.127507838 | 0.065 | 0.182 | 0.054 | 0.075 | 0.085 | 0.053 | 0.133 | 0.538 | 0.273 | 0.081 | -0.123 |
| RI | 4328 | ENSG00000115548 | KDM3A | chr2 | + | 86477876 | 86478265 | 86477876 | 86478029 | 86478169 | 86478265 | 18 | 22 | 21 | 9 | 24 | 285 | 185 | 257 | 317 | 393 | 14 | 25 | 25 | 6 | 15 | 111 | 203 | 384 | 186 | 38 | 289 | 149 | 0.037086195 | 0.127587743 | 0.032 | 0.058 | 0.04 | 0.014 | 0.031 | 0.061 | 0.06 | 0.032 | 0.016 | 0.169 | -0.033 |
| RI | 3241 | ENSG00000100258 | LMF2 | chr22 | - | 50504339 | 50504727 | 50504339 | 50504451 | 50504558 | 50504727 | 9 | 1 | 11 | 13 | 0 | 24 | 9 | 19 | 19 | 13 | 20 | 10 | 5 | 20 | 19 | 12 | 21 | 3 | 32 | 44 | 256 | 149 | 0.037143959 | 0.127712089 | 0.179 | 0.061 | 0.252 | 0.285 | 0 | 0.492 | 0.217 | 0.492 | 0.267 | 0.201 | -0.178 |
| RI | 7407 | ENSG00000165802 | NSMF | chr9 | - | 137452735 | 137453820 | 137452735 | 137453180 | 137453730 | 137453820 | 5 | 3 | 2 | 7 | 2 | 18 | 7 | 18 | 14 | 6 | 2 | 26 | 7 | 18 | 20 | 27 | 16 | 22 | 7 | 42 | 298 | 149 | 0.037248978 | 0.127938642 | 0.122 | 0.176 | 0.053 | 0.2 | 0.143 | 0.036 | 0.448 | 0.137 | 0.562 | 0.192 | -0.136 |
| [illegible] | | | | | | | | | | | | | | | | | | | | | | | | | | | | | | | | | | | | | | | | | | | | | | |

| | | | | | | | | | | | | | | |
|---|---|---|---|---|---|---|---|---|---|---|---|---|---|---|
| RI | 8092 | ENSG00000177239 | MAN1B1 | chr9 | + | 137101483 | 137106315 | 137101483 | 137101672 | 137106124 | 137106315 | 17 | 12 | 10 |
| RI | 357 | ENSG00000105518 | TMEM205 | chr19 | - | 11345519 | 11346268 | 11345519 | 11345628 | 11346207 | 11346268 | 38 | 20 | 25 |
| RI | 4337 | ENSG00000160055 | TMEM234 | chr1 | - | 32216199 | 32216947 | 32216199 | 32216517 | 32216888 | 32216947 | 44 | 27 | 49 |
| RI | 6005 | ENSG00000204386 | NEU1 | chr6 | - | 31860438 | 31862191 | 31860438 | 31861450 | 31861998 | 31862191 | 7 | 6 | 5 |
| RI | 1017 | ENSG00000169738 | DCXR | chr17 | - | 82036190 | 82036643 | 82036190 | 82036448 | 82036543 | 82036643 | 13 | 9 | 10 |
| RI | 4708 | ENSG00000232956 | SNHG15 | chr7 | - | 44983821 | 44984426 | 44983821 | 44984046 | 44984356 | 44984426 | 23 | 24 | 30 |
| RI | 1716 | ENSG00000100429 | HDAC10 | chr22 | - | 50246678 | 50246966 | 50246678 | 50246735 | 50246874 | 50246966 | 10 | 2 | 4 |
| RI | 955 | ENSG00000129667 | RHBDF2 | chr17 | - | 76472685 | 76473105 | 76472685 | 76472839 | 76473004 | 76473105 | 17 | 5 | 18 |
| RI | 1634 | ENSG00000140319 | SRP14 | chr15 | - | 40038281 | 40038948 | 40038281 | 40038394 | 40038875 | 40038948 | 34 | 26 | 36 |
| RI | 2414 | ENSG00000158290 | CUL4B | chrX | - | 120538123 | 120538770 | 120538123 | 120538209 | 120538659 | 120538770 | 96 | 114 | 108 |
| RI | 2874 | ENSG00000146192 | FGD2 | chr6 | + | 37022238 | 37025938 | 37022238 | 37022370 | 37025791 | 37025938 | 8 | 7 | 6 |
| RI | 1030 | ENSG00000182087 | TMEM259 | chr19 | - | 1011366 | 1011612 | 1011366 | 1011499 | 1011579 | 1011612 | 7 | 0 | 2 |
| RI | 7373 | ENSG00000100296 | THOC5 | chr22 | - | 29544459 | 29544603 | 29544459 | 29544508 | 29544579 | 29544603 | 137 | 130 | 129 |
| RI | 2677 | ENSG00000177054 | ZDHHC13 | chr11 | + | 19166301 | 19170568 | 19166301 | 19166385 | 19170410 | 19170568 | 11 | 18 | 10 |
| RI | 5136 | ENSG00000267796 | LIN37 | chr19 | + | 35753086 | 35754157 | 35753086 | 35753253 | 35754016 | 35754157 | 16 | 12 | 18 |
| RI | 2417 | ENSG00000214655 | ZSWIM8 | chr10 | + | 73794146 | 73794639 | 73794146 | 73794330 | 73794540 | 73794639 | 20 | 26 | 9 |
| RI | 7253 | ENSG00000147687 | TATDN1 | chr8 | - | 124493832 | 124495542 | 124493832 | 124493959 | 124495471 | 124495542 | 19 | 28 | 10 |
| RI | 24 | ENSG00000141959 | PFKL | chr21 | + | 44313912 | 44316339 | 44313912 | 44314021 | 44316243 | 44316339 | 8 | 2 | 9 |
| RI | 811 | ENSG00000185624 | P4HB | chr17 | - | 81846428 | 81847068 | 81846428 | 81846629 | 81846946 | 81847068 | 21 | 8 | 12 |
| RI | 260 | ENSG00000142541 | RPL13A | chr19 | + | 49489849 | 49490297 | 49489849 | 49490090 | 49490231 | 49490297 | 75 | 55 | 79 |
| RI | 3526 | ENSG00000150991 | UBC | chr12 | - | 124911645 | 124913774 | 124911645 | 124912180 | 124913320 | 124913774 | 1027 | 603 | 811 |
| RI | 1262 | ENSG00000132603 | NIP7 | chr16 | + | 69340005 | 69340332 | 69340005 | 69340092 | 69340193 | 69340332 | 2 | 8 | 1 |
| RI | 1476 | ENSG00000140990 | NDUFB10 | chr16 | + | 1961496 | 1961975 | 1961496 | 1961636 | 1961796 | 1961975 | 25 | 16 | 30 |
| RI | 571 | ENSG00000185513 | L3MBTL1 | chr20 | + | 43534283 | 43534942 | 43534283 | 43534394 | 43534827 | 43534942 | 2 | 1 | 1 |
| RI | 1938 | ENSG00000103811 | CTSH | chr15 | - | 78931450 | 78932038 | 78931450 | 78931506 | 78931900 | 78932038 | 15 | 13 | 11 |
| RI | 4190 | ENSG00000173039 | RELA | chr11 | - | 65661936 | 65662205 | 65661936 | 65662088 | 65662178 | 65662205 | 17 | 0 | 10 |
| RI | 7693 | ENSG00000168071 | CCDC88B | chr11 | + | 64353053 | 64353496 | 64353053 | 64353240 | 64353350 | 64353496 | 4 | 12 | 2 |
| RI | 6838 | ENSG00000035681 | NSMAF | chr8 | - | 58585651 | 58586000 | 58585651 | 58585761 | 58585897 | 58586000 | 46 | 22 | 35 |
| RI | 4143 | ENSG00000132300 | PTCD3 | chr2 | + | 86130647 | 86131106 | 86130647 | 86130737 | 86131077 | 86131106 | 48 | 19 | 30 |
| RI | 2996 | ENSG00000060971 | ACAA1 | chr3 | - | 38125825 | 38126700 | 38125825 | 38125881 | 38126161 | 38126700 | 27 | 11 | 31 |
| RI | 6993 | ENSG00000072506 | HSD17B10 | chrX | - | 53431260 | 53431906 | 53431260 | 53431594 | 53431797 | 53431906 | 11 | 15 | 14 |
| RI | 1884 | ENSG00000213983 | AP1G2 | chr14 | - | 23563379 | 23563638 | 23563379 | 23563502 | 23563583 | 23563638 | 77 | 42 | 36 |
| RI | 7535 | ENSG00000130714 | POMT1 | chr9 | + | 131506402 | 131509022 | 131506402 | 131506453 | 131508910 | 131509022 | 22 | 25 | 42 |
| RI | 4762 | ENSG00000214941 | ZSWIM7 | chr17 | - | 15976559 | 15978163 | 15976559 | 15977092 | 15978043 | 15978163 | 49 | 33 | 42 |
| RI | 6742 | ENSG00000285171 | NA | chrX | - | 71105999 | 71106426 | 71105999 | 71106137 | 71106339 | 71106426 | 49 | 17 | 48 |
| RI | 6059 | ENSG00000132424 | PNISR | chr6 | - | 99414571 | 99416428 | 99414571 | 99414690 | 99416348 | 99416428 | 228 | 201 | 164 |
| RI | 1140 | ENSG00000008517 | IL32 | chr16 | + | 3068989 | 3069551 | 3068989 | 3069109 | 3069280 | 3069551 | 1398 | 294 | 1347 |
| RI | 6590 | ENSG00000234741 | GAS5 | chr1 | - | 173864483 | 173864704 | 173864483 | 173864506 | 173864674 | 173864704 | 263 | 253 | 320 |
| RI | 66 | ENSG00000161179 | YDJC | chr22 | - | 21629009 | 21629407 | 21629009 | 21629187 | 21629307 | 21629407 | 26 | 5 | 17 |
| RI | 3485 | ENSG00000092841 | MYL6 | chr12 | + | 56159586 | 56160148 | 56159586 | 56159730 | 56159974 | 56160148 | 63 | 40 | 52 |
| RI | 7365 | ENSG00000140988 | RPS2 | chr16 | - | 1962496 | 1962909 | 1962496 | 1962656 | 1962735 | 1962909 | 210 | 77 | 318 |
| RI | 4133 | ENSG00000162298 | SYVN1 | chr11 | - | 65132247 | 65132780 | 65132247 | 65132351 | 65132731 | 65132780 | 30 | 5 | 27 |
| RI | 4428 | ENSG00000197989 | SNHG12 | chr1 | - | 28579532 | 28580646 | 28579532 | 28579981 | 28580559 | 28580646 | 72 | 44 | 76 |
| RI | 3339 | ENSG00000144559 | TAMM41 | chr3 | - | 11807832 | 11809682 | 11807832 | 11807895 | 11809516 | 11809682 | 14 | 17 | 31 |
| RI | 7152 | ENSG00000172269 | DPAGT1 | chr11 | - | 119101017 | 119101860 | 119101017 | 119101138 | 119101494 | 119101860 | 9 | 3 | 17 |
| RI | 2858 | ENSG00000169696 | ASPSCR1 | chr17 | + | 82016799 | 82017113 | 82016799 | 82016869 | 82016940 | 82017113 | 2 | 1 | 4 |
| RI | 4286 | ENSG00000071859 | FAM50A | chrX | + | 154449680 | 154449931 | 154449680 | 154449735 | 154449882 | 154449931 | 39 | 30 | 41 |
| RI | 4141 | ENSG00000154222 | CC2D1B | chr1 | - | 52360423 | 52361132 | 52360423 | 52360549 | 52360973 | 52361132 | 38 | 19 | 33 |
| RI | 5702 | ENSG00000079785 | DDX1 | chr2 | + | 15627053 | 15628839 | 15627053 | 15628710 | 15628796 | 15628839 | 13 | 8 | 18 |
| RI | 3963 | ENSG00000110619 | CARS1 | chr11 | - | 3001969 | 3002600 | 3001969 | 3002053 | 3002540 | 3002600 | 30 | 50 | 44 |
| RI | 3499 | ENSG00000013573 | DDX11 | chr12 | + | 31103576 | 31104791 | 31103576 | 31103731 | 31103806 | 31104791 | 0 | 1 | 1 |
| RI | 1001 | ENSG00000083290 | ULK2 | chr17 | - | 19770829 | 19776407 | 19770829 | 19771067 | 19776308 | 19776407 | 21 | 15 | 18 |
| RI | 3843 | ENSG00000117335 | CD46 | chr1 | + | 207767012 | 207770362 | 207767012 | 207767195 | 207770320 | 207770362 | 275 | 272 | 301 |
| RI | 7205 | ENSG00000185122 | HSF1 | chr8 | + | 144313845 | 144314054 | 144313845 | 144313911 | 144313984 | 144314054 | 6 | 5 | 10 |
| RI | 1915 | ENSG00000129460 | NGDN | chr14 | + | 23475170 | 23476152 | 23475170 | 23475308 | 23476028 | 23476152 | 54 | 47 | 81 |
| RI | 1065 | ENSG00000168259 | DNAJC7 | chr17 | - | 41987818 | 41988896 | 41987818 | 41987905 | 41988731 | 41988896 | 167 | 156 | 182 |
| RI | 3645 | ENSG00000239900 | ADSL | chr22 | + | 40361272 | 40363071 | 40361272 | 40361342 | 40361487 | 40363071 | 16 | 12 | 26 |
| RI | 6882 | ENSG00000147140 | NONO | chrX | + | 71297376 | 71298508 | 71297376 | 71297461 | 71298468 | 71298508 | 91 | 72 | 110 |
| RI | 73 | ENSG00000128159 | TUBGCP6 | chr22 | - | 50217917 | 50218402 | 50217917 | 50218117 | 50218188 | 50218402 | 7 | 7 | 7 |
| RI | 7013 | ENSG00000182196 | ARL6IP4 | chr12 | + | 122981128 | 122981879 | 122981128 | 122981266 | 122981570 | 122981879 | 24 | 13 | 36 |
| RI | 8273 | ENSG00000082898 | XPO1 | chr2 | - | 61498672 | 61498913 | 61498672 | 61498792 | 61498864 | 61498913 | 53 | 42 | 56 |
| RI | 99 | ENSG00000100031 | GGT1 | chr22 | + | 24628080 | 24628388 | 24628080 | 24628193 | 24628274 | 24628388 | 0 | 0 | 2 |
| RI | 6473 | ENSG00000106588 | PSMA2 | chr7 | - | 42921857 | 42923406 | 42921857 | 42921931 | 42923324 | 42923406 | 26 | 20 | 20 |
| RI | 4908 | ENSG00000143303 | RRNAD1 | chr1 | + | 156731990 | 156732473 | 156731990 | 156732115 | 156732280 | 156732473 | 41 | 24 | 44 |
| RI | 4105 | ENSG00000101444 | AHCY | chr20 | - | 34290331 | 34290638 | 34290331 | 34290449 | 34290550 | 34290638 | 3 | 8 | 6 |
| RI | 3123 | ENSG00000100151 | PICK1 | chr22 | + | 38073772 | 38074451 | 38073772 | 38073823 | 38074306 | 38074451 | 14 | 10 | 14 |
| RI | 4876 | ENSG00000154451 | GBP5 | chr1 | - | 89262219 | 89263948 | 89262219 | 89262401 | 89263735 | 89263948 | 71 | 89 | 125 |
| RI | 1831 | ENSG00000100889 | PCK2 | chr14 | + | 24098202 | 24098678 | 24098202 | 24098387 | 24098474 | 24098678 | 4 | 2 | 5 |
| RI | 568 | ENSG00000185513 | L3MBTL1 | chr20 | + | 43530283 | 43530889 | 43530283 | 43530419 | 43530797 | 43530889 | 4 | 6 | 5 |
| RI | 8244 | ENSG00000284770 | TBCE | chr1 | + | 235435744 | 235436608 | 235435744 | 235435840 | 235436385 | 235436608 | 24 | 15 | 17 |
| RI | 1472 | ENSG00000140983 | RHOT2 | chr16 | + | 670891 | 671203 | 670891 | 671000 | 671114 | 671203 | 91 | 38 | 82 |
| RI | 1878 | ENSG00000182809 | CRIP2 | chr14 | + | 105479124 | 105479493 | 105479124 | 105479219 | 105479435 | 105479493 | 7 | 3 | 11 |
| RI | 2359 | ENSG00000011243 | AKAP8L | chr19 | - | 15397713 | 15400864 | 15397713 | 15397855 | 15400793 | 15400864 | 33 | 15 | 19 |
| RI | 678 | ENSG00000104884 | ERCC2 | chr19 | - | 45368929 | 45369147 | 45368929 | 45368992 | 45369069 | 45369147 | 0 | 2 | 0 |
| RI | 2334 | ENSG00000112759 | SLC29A1 | chr6 | + | 44229906 | 44230481 | 44229906 | 44230046 | 44230346 | 44230481 | 1 | 0 | 4 |
| RI | 6350 | ENSG00000198198 | SZT2 | chr1 | + | 43432988 | 43434485 | 43432988 | 43433190 | 43434385 | 43434485 | 14 | 18 | 29 |
| RI | 1890 | ENSG00000213983 | AP1G2 | chr14 | - | 23565815 | 23566160 | 23565815 | 23565892 | 23566063 | 23566160 | 136 | 68 | 124 |
| RI | 1132 | ENSG00000072849 | DERL2 | chr17 | - | 5481100 | 5481389 | 5481100 | 5481142 | 5481295 | 5481389 | 74 | 63 | 91 |
| RI | 3654 | ENSG00000136878 | USP20 | chr9 | + | 129880112 | 129881828 | 129880112 | 129880289 | 129880466 | 129881828 | 18 | 17 | 42 |
| RI | 5350 | ENSG00000129292 | PHF20L1 | chr8 | + | 132814636 | 132817545 | 132814636 | 132814889 | 132817338 | 132817545 | 137 | 133 | 110 |
| RI | 3428 | ENSG00000008128 | CDK11A | chr1 | - | 1704038 | 1704344 | 1704038 | 1704146 | 1704222 | 1704344 | 1 | 2 | 1 |
| RI | 6412 | ENSG00000160813 | PPP1R35 | chr7 | - | 100435630 | 100436064 | 100435630 | 100435767 | 100435847 | 100436064 | 30 | 20 | 35 |
| RI | 5517 | ENSG00000171720 | HDAC3 | chr5 | - | 141628113 | 141628639 | 141628113 | 141628187 | 141628558 | 141628639 | 11 | 5 | 10 |
| RI | 6903 | ENSG00000090316 | MAEA | chr4 | + | 1336860 | 1338617 | 1336860 | 1336994 | 1338425 | 1338617 | 20 | 12 | 16 |
| RI | 1821 | ENSG00000100911 | PSME2 | chr14 | - | 24144399 | 24145273 | 24144399 | 24144468 | 24145226 | 24145273 | 95 | 79 | 71 |
| RI | 2182 | ENSG00000113240 | CLK4 | chr5 | - | 178616881 | 178618778 | 178616881 | 178616948 | 178617343 | 178618778 | 162 | 109 | 157 |
| RI | 6974 | ENSG00000147050 | KDM6A | chrX | + | 45069578 | 45076826 | 45069578 | 45070357 | 45076696 | 45076826 | 45 | 38 | 31 |
| RI | 520 | ENSG00000101246 | ARFRP1 | chr20 | - | 63698642 | 63700702 | 63698642 | 63700530 | 63700601 | 63700702 | 16 | 9 | 15 |
| RI | 4430 | ENSG00000197989 | SNHG12 | chr1 | - | 28580559 | 28581229 | 28580559 | 28580646 | 28581110 | 28581229 | 85 | 71 | 89 |
| RI | 7841 | ENSG00000162032 | SPSB3 | chr16 | - | 1777746 | 1778612 | 1777746 | 1777872 | 1777945 | 1778612 | 5 | 1 | 2 |
| RI | 1517 | ENSG00000103653 | CSK | chr15 | + | 74801694 | 74802083 | 74801694 | 74801890 | 74801996 | 74802083 | 36 | 18 | 29 |
| RI | 1351 | ENSG00000089280 | FUS | chr16 | + | 31184938 | 31188357 | 31184938 | 31185179 | 31188324 | 31188357 | 167 | 168 | 215 |
| RI | 3036 | ENSG00000206503 | HLA-A | chr6 | + | 29944499 | 29945091 | 29944499 | 29944634 | 29945058 | 29945091 | 385 | 184 | 928 |
| RI | 5595 | ENSG00000168214 | RBPJ | chr4 | + | 26429897 | 26430432 | 26429897 | 26430053 | 26430418 | 26430432 | 111 | 122 | 121 |
| RI | 523 | ENSG00000026036 | RTEL1-TNFRSF6B | chr20 | + | 63691741 | 63693003 | 63691741 | 63691837 | 63692804 | 63693003 | 3 | 4 | 6 |
| RI | 7887 | ENSG00000132275 | RRP8 | chr11 | - | 6600925 | 6601602 | 6600925 | 6601055 | 6601148 | 6601602 | 4 | 4 | 21 |
| RI | 3139 | ENSG00000114416 | FXR1 | chr3 | + | 180962882 | 180963090 | 180962882 | 180962940 | 180963027 | 180963090 | 186 | 187 | 218 |
| RI | 6181 | ENSG00000214021 | TTLL3 | chr3 | + | 9817644 | 9818920 | 9817644 | 9817759 | 9818821 | 9818920 | 129 | 93 | 109 |
| RI | 7754 | ENSG00000159352 | PSMD4 | chr1 | + | 151266003 | 151266439 | 151266003 | 151266121 | 151266307 | 151266439 | 2 | 1 | 3 |
| RI | 2936 | ENSG00000010278 | CD9 | chr12 | + | 6232631 | 6233486 | 6232631 | 6232729 | 6233411 | 6233486 | 0 | 3 | 0 |
| RI | 7323 | ENSG00000137207 | YIPF3 | chr6 | - | 43513068 | 43513451 | 43513068 | 43513098 | 43513358 | 43513451 | 71 | 25 | 48 |
| RI | 7780 | ENSG00000110274 | CEP164 | chr11 | + | 117394919 | 117395191 | 117394919 | 117395003 | 117395122 | 117395191 | 4 | 0 | 5 |
| RI | 8063 | ENSG00000148824 | MTG1 | chr10 | + | 133401528 | 133402245 | 133401528 | 133401590 | 133402148 | 133402245 | 54 | 18 | 41 |
| RI | 5121 | ENSG00000044574 | HSPA5 | chr9 | - | 125238940 | 125239533 | 125238940 | 125239331 | 125239420 | 125239533 | 62 | 31 | 82 |
| RI | 4454 | ENSG00000117139 | KDM5B | chr1 | - | 202729706 | 202731063 | 202729706 | 202730027 | 202730908 | 202731063 | 7 | 4 | 4 |
| RI | 5374 | ENSG00000163050 | COQ8A | chr1 | + | 226982026 | 226982763 | 226982026 | 226982149 | 226982677 | 226982763 | 33 | 17 | 41 |
| RI | 7848 | ENSG00000213619 | NDUFS3 | chr11 | + | 47579268 | 47580622 | 47579268 | 47579334 | 47580524 | 47580622 | 8 | 1 | 1 |
| RI | 6593 | ENSG00000234741 | GAS5 | chr1 | - | 173865228 | 173865547 | 173865228 | 173865282 | 173865470 | 173865547 | 173 | 100 | 178 |
| RI | 244 | ENSG00000104969 | SGTA | chr19 | - | 2757692 | 2759294 | 2757692 | 2757782 | 2759256 | 2759294 | 10 | 13 | 9 |
| RI | 3706 | ENSG00000070761 | CFAP20 | chr16 | - | 58114809 | 58115457 | 58114809 | 58114920 | 58115268 | 58115457 | 50 | 20 | 51 |
| RI | 7162 | ENSG00000184575 | XPOT | chr12 | + | 64425037 | 64425457 | 64425037 | 64425138 | 64425337 | 64425457 | 80 | 59 | 59 |
| RI | 3682 | ENSG00000099622 | CIRBP | chr19 | + | 1270927 | 1271467 | 1270927 | 1271036 | 1271139 | 1271467 | 35 | 18 | 54 |
| RI | 183 | ENSG00000105607 | GCDH | chr19 | + | 12891486 | 12891974 | 12891486 | 12891522 | 12891830 | 12891974 | 11 | 11 | 10 |
| RI | 6483 | ENSG00000172354 | GNB2 | chr7 | + | 100677497 | 100677818 | 100677497 | 100677660 | 100677751 | 100677818 | 10 | 2 | 8 |
| RI | 7550 | ENSG00000137073 | UBAP2 | chr9 | - | 33922965 | 33923293 | 33922965 | 33923033 | 33923185 | 33923293 | 15 | 9 | 4 |
| RI | 6917 | ENSG00000248323 | LUCAT1 | chr5 | - | 91310626 | 91313229 | 91310626 | 91312495 | 91313077 | 91313229 | 31 | 18 | 24 |
| RI | 833 | ENSG00000100038 | TOP3B | chr22 | - | 21962428 | 21962893 | 21962428 | 21962602 | 21962746 | 21962893 | 7 | 5 | 23 |
| RI | 3649 | ENSG00000177728 | TMEM94 | chr17 | + | 75485870 | 75486426 | 75485870 | 75485998 | 75486289 | 75486426 | 2 | 4 | 6 |
| RI | 4134 | ENSG00000162298 | SYVN1 | chr11 | - | 65132921 | 65133320 | 65132921 | 65133074 | 65133159 | 65133320 | 12 | 9 | 10 |
| RI | 1435 | ENSG00000077235 | GTF3C1 | chr16 | - | 27463540 | 27464836 | 27463540 | 27463592 | 27464319 | 27464836 | 19 | 4 | 14 |
| RI | 807 | ENSG00000185624 | P4HB | chr17 | - | 81845143 | 81845742 | 81845143 | 81845303 | 81845560 | 81845742 | 46 | 24 | 33 |
| RI | 989 | ENSG00000131469 | RPL27 | chr17 | + | 42998418 | 43000102 | 42998418 | 42998471 | 42999932 | 43000102 | 50 | 12 | 50 |
| RI | 5929 | ENSG00000114388 | NPRL2 | chr3 | - | 50349385 | 50349828 | 50349385 | 50349490 | 50349664 | 50349828 | 44 | 26 | 40 |
| RI | 1402 | ENSG00000130304 | SLC27A1 | chr19 | + | 17486562 | 17487335 | 17486562 | 17486957 | 17487173 | 17487335 | 0 | 2 | 4 |
| RI | 957 | ENSG00000129667 | RHBDF2 | chr17 | - | 76487711 | 76488132 | 76487711 | 76487851 | 76487973 | 76488132 | 98 | 56 | 34 |
| RI | 656 | ENSG00000176014 | TUBB6 | chr18 | + | 12308240 | 12308795 | 12308240 | 12308349 | 12308686 | 12308795 | 6 | 2 | 0 |
| RI | 6202 | ENSG00000223705 | NSUN5P1 | chr7 | + | 75412788 | 75415023 | 75412788 | 75412932 | 75414884 | 75415023 | 117 | 69 | 168 |
| RI | 6868 | ENSG00000196924 | FLNA | chrX | - | 154350030 | 154351696 | 154350030 | 154350207 | 154350908 | 154351696 | 99 | 57 | 77 |
| RI | 2814 | ENSG00000100003 | SEC14L2 | chr22 | + | 30415758 | 30416087 | 30415758 | 30415865 | 30415947 | 30416087 | 10 | 6 | 20 |
| RI | 1334 | ENSG00000090861 | AARS1 | chr16 | - | 70253267 | 70253800 | 70253267 | 70253381 | 70253713 | 70253800 | 7 | 1 | 11 |
| RI | 6131 | ENSG00000135317 | SNX14 | chr6 | - | 85507967 | 85513895 | 85507967 | 85508059 | 85513799 | 85513895 | 73 | 57 | 56 |
| RI | 5011 | ENSG00000197043 | ANXA6 | chr5 | - | 151138677 | 151139447 | 151138677 | 151138791 | 151139352 | 151139447 | 51 | 31 | 37 |
| RI | 3667 | ENSG00000167962 | ZNF598 | chr16 | - | 2002205 | 2002732 | 2002205 | 2002423 | 2002520 | 2002732 | 11 | 0 | 0 |
| RI | 6315 | ENSG00000147894 | C9orf72 | chr9 | - | 27558490 | 27560299 | 27558490 | 27558607 | 27560226 | 27560299 | 387 | 310 | 195 |
| RI | 7958 | ENSG00000103479 | RBL2 | chr16 | + | 53469915 | 53470663 | 53469915 | 53470185 | 53470382 | 53470663 | 278 | 388 | 342 |
| RI | 4127 | ENSG00000241878 | PISD | chr22 | - | 31620995 | 31621885 | 31620995 | 31621142 | 31621648 | 31621885 | 21 | 12 | 21 |
| RI | 1960 | ENSG00000213780 | GTF2H4 | chr6 | + | 30913109 | 30913387 | 30913109 | 30913157 | 30913308 | 30913387 | 14 | 7 | 15 |
| RI | 3188 | ENSG00000004534 | RBM6 | chr3 | + | 50061461 | 50062108 | 50061461 | 50061547 | 50061961 | 50062108 | 128 | 165 | 120 |
| RI | 2829 | ENSG00000113971 | NPHP3 | chr3 | - | 132684553 | 132686387 | 132684553 | 132684794 | 132686259 | 132686387 | 22 | 8 | 26 |
| RI | 1623 | ENSG00000182253 | SYNM | chr15 | + | 99129366 | 99135593 | 99129366 | 99131813 | 99132749 | 99135593 | 6 | 12 | 12 |
| RI | 630 | ENSG00000196756 | SNHG17 | chr20 | - | 38421005 | 38422241 | 38421005 | 38421098 | 38422091 | 38422241 | 24 | 11 | 22 |
| RI | 8520 | ENSG00000121073 | SLC35B1 | chr17 | - | 49705123 | 49706309 | 49705123 | 49705281 | 49705865 | 49706309 | 17 | 10 | 17 |
| RI | 7468 | ENSG00000135047 | CTSL | chr9 | + | 87728026 | 87728396 | 87728026 | 87728149 | 87728249 | 87728396 | 0 | 6 | 0 |
| RI | 2835 | ENSG00000114745 | GORASP1 | chr3 | - | 39100746 | 39101102 | 39100746 | 39100877 | 39101015 | 39101102 | 22 | 5 | 12 |
| RI | 6223 | ENSG00000127980 | PEX1 | chr7 | - | 92501506 | 92502079 | 92501506 | 92501673 | 92501889 | 92502079 | 30 | 34 | 23 |
| RI | 6114 | ENSG00000237441 | RGL2 | chr6 | - | 33293794 | 33294066 | 33293794 | 33293916 | 33294033 | 33294066 | 16 | 11 | 5 |
| RI | 1582 | ENSG00000114062 | UBE3A | chr15 | - | 25333727 | 25340228 | 25333727 | 25339257 | 25340084 | 25340228 | 72 | 97 | 78 |
| RI | 7279 | ENSG00000147789 | ZNF7 | chr8 | + | 144829042 | 144829604 | 144829042 | 144829090 | 144829477 | 144829604 | 14 | 5 | 11 |
| RI | 2574 | ENSG00000155229 | MMS19 | chr10 | - | 97461495 | 97461896 | 97461495 | 97461622 | 97461827 | 97461896 | 26 | 28 | 38 |
| RI | 7006 | ENSG00000120699 | EXOSC8 | chr13 | + | 37002933 | 37004561 | 37002933 | 37003007 | 37004515 | 37004561 | 77 | 46 | 48 |
| RI | 8433 | ENSG00000033030 | ZCCHC8 | chr12 | - | 122483278 | 122483563 | 122483278 | 122483344 | 122483459 | 122483563 | 26 | 52 | 48 |
| RI | 3678 | ENSG00000160789 | LMNA | chr1 | + | 156136920 | 156137232 | 156136920 | 156137028 | 156137112 | 156137232 | 0 | 2 | 1 |
| RI | 1121 | ENSG00000177731 | FLII | chr17 | - | 18253543 | 18254182 | 18253543 | 18253719 | 18254078 | 18254182 | 52 | 27 | 34 |
| RI | 2942 | ENSG00000173540 | GMPPB | chr3 | - | 49722595 | 49723114 | 49722595 | 49722754 | 49722971 | 49723114 | 24 | 8 | 31 |
| RI | 6092 | ENSG00000112739 | PRPF4B | chr6 | + | 4060412 | 4061147 | 4060412 | 4060686 | 4061014 | 4061147 | 455 | 318 | 369 |
| RI | 1981 | ENSG00000164080 | RAD54L2 | chr3 | + | 51638143 | 51639670 | 51638143 | 51638321 | 51639418 | 51639670 | 36 | 18 | 15 |
| RI | 4551 | ENSG00000131788 | PIAS3 | chr1 | - | 145848521 | 145850269 | 145848521 | 145849712 | 145850231 | 145850269 | 18 | 13 | 18 |
| RI | 2185 | ENSG00000165819 | METTL3 | chr14 | - | 21499024 | 21499600 | 21499024 | 21499137 | 21499491 | 21499600 | 237 | 234 | 338 |
| RI | 6752 | ENSG00000167716 | WDR81 | chr17 | + | 1735571 | 1736218 | 1735571 | 1735717 | 1736038 | 1736218 | 6 | 0 | 9 |
| RI | 5694 | ENSG00000160710 | ADAR | chr1 | - | 154601040 | 154602626 | 154601040 | 154601888 | 154602035 | 154602626 | 873 | 681 | 843 |
| RI | 8192 | ENSG00000130640 | TUBGCP2 | chr10 | - | 133285084 | 133285628 | 133285084 | 133285213 | 133285455 | 133285628 | 26 | 12 | 21 |
| RI | 3737 | ENSG00000114626 | ABTB1 | chr3 | + | 127676535 | 127677286 | 127676535 | 127676581 | 127676966 | 127677286 | 27 | 26 | 36 |
| RI | 3330 | ENSG00000117228 | GBP1 | chr1 | - | 89054681 | 89055215 | 89054681 | 89054875 | 89055112 | 89055215 | 26 | 45 | 14 |
| RI | 7193 | ENSG00000174744 | BRMS1 | chr11 | - | 66338242 | 66338785 | 66338242 | 66338282 | 66338720 | 66338785 | 4 | 2 | 12 |
| RI | 2352 | ENSG00000011243 | AKAP8L | chr19 | - | 15397713 | 15399410 | 15397713 | 15397855 | 15399301 | 15399410 | 58 | 39 | 45 |
| RI | 384 | ENSG00000129347 | KRI1 | chr19 | - | 10559358 | 10559708 | 10559358 | 10559529 | 10559612 | 10559708 | 12 | 3 | 23 |
| RI | 3474 | ENSG00000064687 | ABCA7 | chr19 | + | 1056906 | 1057429 | 1056906 | 1057084 | 1057313 | 1057429 | 5 | 10 | 9 |
| RI | 4335 | ENSG00000123416 | TUBA1B | chr12 | - | 49129499 | 49131395 | 49129499 | 49129722 | 49131297 | 49131395 | 14 | 2 | 12 |
| RI | 3872 | ENSG00000108654 | DDX5 | chr17 | - | 64502161 | 64503098 | 64502161 | 64502549 | 64502925 | 64503098 | 1232 | 1079 | 1232 |
| RI | 4837 | ENSG00000175054 | ATR | chr3 | - | 142457603 | 142459111 | 142457603 | 142457755 | 142458957 | 142459111 | 16 | 27 | 12 |
| RI | 8398 | ENSG00000125967 | NECAB3 | chr20 | - | 33659884 | 33660395 | 33659884 | 33660003 | 33660258 | 33660395 | 16 | 2 | 1 |
| RI | 6366 | ENSG00000012124 | CD22 | chr19 | + | 35332018 | 35332924 | 35332018 | 35332074 | 35332652 | 35332924 | 74 | 65 | 83 |
| RI | 3689 | ENSG00000099622 | CIRBP | chr19 | + | 1271980 | 1274810 | 1271980 | 1274440 | 1274648 | 1274810 | 44 | 28 | 35 |
| RI | 204 | ENSG00000167645 | YIF1B | chr19 | - | 38307427 | 38307752 | 38307427 | 38307521 | 38307596 | 38307752 | 5 | 0 | 2 |
| RI | 5992 | ENSG00000004139 | SARM1 | chr17 | + | 28388173 | 28388539 | 28388173 | 28388276 | 28388349 | 28388539 | 6 | 3 | 8 |
| RI | 8428 | ENSG00000127328 | RAB3IP | chr12 | + | 69812777 | 69813033 | 69812777 | 69812877 | 69812963 | 69813033 | 15 | 14 | 40 |
| RI | 5934 | ENSG00000114388 | NPRL2 | chr3 | - | 50349664 | 50350022 | 50349664 | 50349833 | 50349930 | 50350022 | 16 | 10 | 15 |
| RI | 3758 | ENSG00000071894 | CPSF1 | chr8 | - | 144400165 | 144400493 | 144400165 | 144400276 | 144400353 | 144400493 | 3 | 5 | 2 |
| RI | 6526 | ENSG00000105835 | NAMPT | chr7 | - | 106274945 | 106277179 | 106274945 | 106275049 | 106277083 | 106277179 | 240 | 384 | 160 |
| RI | 6808 | ENSG00000149600 | COMMD7 | chr20 | - | 32702702 | 32704071 | 32702702 | 32703458 | 32704022 | 32704071 | 120 | 96 | 61 |
| RI | 945 | ENSG00000132522 | GPS2 | chr17 | - | 7312660 | 7313124 | 7312660 | 7312839 | 7313028 | 7313124 | 22 | 27 | 29 |
| RI | 5866 | ENSG00000120896 | SORBS3 | chr8 | + | 22566660 | 22566868 | 22566660 | 22566713 | 22566821 | 22566868 | 8 | 15 | 13 |
| RI | 4686 | ENSG00000211584 | SLC48A1 | chr12 | + | 47780144 | 47782751 | 47780144 | 47780289 | 47782217 | 47782751 | 9 | 9 | 17 |
| RI | 1837 | ENSG00000099797 | TECR | chr19 | + | 14563981 | 14564287 | 14563981 | 14564097 | 14564181 | 14564287 | 21 | 8 | 24 |
| RI | 6720 | ENSG00000094631 | HDAC6 | chrX | + | 48818040 | 48818412 | 48818040 | 48818109 | 48818219 | 48818412 | 5 | 1 | 1 |
| RI | 6715 | ENSG00000107185 | RGP1 | chr9 | + | 35750657 | 35750989 | 35750657 | 35750741 | 35750839 | 35750989 | 13 | 5 | 10 |
| RI | 824 | ENSG00000275700 | AATF | chr17 | + | 36989246 | 36990857 | 36989246 | 36989411 | 36990773 | 36990857 | 10 | 0 | 7 |
| RI | 4921 | ENSG00000130713 | EXOSC2 | chr9 | + | 130697581 | 130698251 | 130697581 | 130697627 | 130698161 | 130698251 | 31 | 15 | 34 |
| RI | 2400 | ENSG00000108039 | XPNPEP1 | chr10 | - | 109875527 | 109877867 | 109875527 | 109875599 | 109877789 | 109877867 | 59 | 36 | 58 |
| RI | 4735 | ENSG00000140416 | TPM1 | chr15 | + | 63060868 | 63061273 | 63060868 | 63060939 | 63061197 | 63061273 | 16 | 12 | 17 |
| RI | 949 | ENSG00000132522 | GPS2 | chr17 | - | 7313567 | 7314159 | 7313567 | 7313988 | 7314079 | 7314159 | 42 | 16 | 42 |
| RI | 4427 | ENSG00000197989 | SNHG12 | chr1 | - | 28579532 | 28579981 | 28579532 | 28579587 | 28579941 | 28579981 | 73 | 37 | 75 |
| RI | 448 | ENSG00000081665 | ZNF506 | chr19 | - | 19791561 | 19792484 | 19791561 | 19791731 | 19792410 | 19792484 | 19 | 13 | 37 |
| RI | 797 | ENSG00000131462 | TUBG1 | chr17 | + | 42610422 | 42612143 | 42610422 | 42610590 | 42612074 | 42612143 | 8 | 3 | 8 |
| RI | 5802 | ENSG00000136699 | SMPD4 | chr2 | - | 130154276 | 130155259 | 130154276 | 130154482 | 130155095 | 130155259 | 10 | 2 | 14 |
| RI | 4649 | ENSG00000226822 | LINC02785 | chr1 | + | 108071483 | 108072697 | 108071483 | 108071637 | 108072548 | 108072697 | 13 | 25 | 2 |
| RI | 7500 | ENSG00000213930 | GALT | chr9 | + | 34648333 | 34648894 | 34648333 | 34648456 | 34648761 | 34648894 | 34 | 30 | 51 |
| RI | 6767 | ENSG00000177646 | ACAD9 | chr3 | + | 128909343 | 128910149 | 128909343 | 128909421 | 128910020 | 128910149 | 28 | 12 | 39 |
| RI | 7038 | ENSG00000100918 | REC8 | chr14 | + | 24178605 | 24178916 | 24178605 | 24178672 | 24178776 | 24178916 | 7 | 2 | 3 |
| RI | 466 | ENSG00000105726 | ATP13A1 | chr19 | - | 19656659 | 19656916 | 19656659 | 19656766 | 19656846 | 19656916 | 28 | 8 | 15 |
| RI | 6240 | ENSG00000135218 | CD36 | chr7 | + | 80669952 | 80671164 | 80669952 | 80670022 | 80670976 | 80671164 | 108 | 66 | 42 |
| RI | 5415 | ENSG00000018280 | SLC11A1 | chr2 | + | 218387564 | 218388385 | 218387564 | 218387632 | 218387799 | 218388385 | 15 | 8 | 7 |
| RI | 5731 | ENSG00000172936 | MYD88 | chr3 | + | 38140387 | 38140848 | 38140387 | 38140568 | 38140756 | 38140848 | 72 | 51 | 36 |
| RI | 7135 | ENSG00000140525 | FANCI | chr15 | + | 89281764 | 89283250 | 89281764 | 89281835 | 89283135 | 89283250 | 7 | 3 | 20 |
| RI | 7041 | ENSG00000147471 | PLPBP | chr8 | + | 37765525 | 37765746 | 37765525 | 37765633 | 37765710 | 37765746 | 10 | 6 | 18 |
| RI | 7098 | ENSG00000094914 | AAAS | chr12 | - | 53307844 | 53308133 | 53307844 | 53307929 | 53308051 | 53308133 | 17 | 6 | 22 |
| RI | 4978 | ENSG00000196975 | ANXA4 | chr2 | + | 69807905 | 69810673 | 69807905 | 69807996 | 69810593 | 69810673 | 8 | 9 | 7 |
| RI | 305 | ENSG00000161265 | U2AF1L4 | chr19 | - | 35742528 | 35743904 | 35742528 | 35742803 | 35743808 | 35743904 | 32 | 13 | 19 |
| RI | 2007 | ENSG00000135596 | MICAL1 | chr6 | - | 109446135 | 109446772 | 109446135 | 109446357 | 109446695 | 109446772 | 54 | 24 | 74 |

| | | | | | | | | | | | | | | | | | | | | | | | | | | | | | | | | | | | | | | | | | | | | | | |
|---|---|---|---|---|---|---|---|---|---|---|---|---|---|---|---|---|---|---|---|---|---|---|---|---|---|---|---|---|---|---|---|---|---|---|---|---|---|---|---|---|---|---|---|---|---|---|
| RI | 280 | ENSG00000183207 | RUVBL2 | chr19 | + | 49009972 | 49010611 | 49009972 | 49010066 | 49010487 | 49010611 | 4 | 2 | 3 | 5 | 7 | 48 | 35 | 41 | 56 | 21 | 0 | 6 | 4 | 3 | 13 | 155 | 64 | 73 | 76 | 118 | 298 | 149 | 0.088776448 | 0.251136674 | 0.04 | 0.028 | 0.035 | 0.043 | 0.143 | 0 | 0.045 | 0.027 | 0.019 | 0.052 | 0.029 |
| RI | 928 | ENSG00000181038 | METTL23 | chr17 | + | 76732977 | 76733377 | 76732977 | 76733215 | 76733292 | 76733377 | 54 | 26 | 67 | 73 | 78 | 177 | 75 | 144 | 132 | 249 | 68 | 68 | 105 | 55 | 13 | 69 | 108 | 105 | 88 | 136 | 226 | 149 | 0.088814036 | 0.251136674 | 0.167 | 0.186 | 0.235 | 0.267 | 0.171 | 0.394 | 0.293 | 0.397 | 0.292 | 0.059 | -0.082 |
| RI | 2751 | ENSG00000099949 | LZTR1 | chr22 | + | 20994103 | 20994727 | 20994103 | 20994269 | 20994557 | 20994727 | 13 | 9 | 15 | 23 | 5 | 13 | 7 | 10 | 5 | 0 | 0 | 11 | 0 | 3 | 19 | 4 | 5 | 2 | 0 | 21 | 298 | 149 | 0.089210509 | 0.252130266 | 0.333 | 0.391 | 0.429 | 0.697 | 1 | 0 | 0.524 | 0 | 1 | 0.311 | 0.203 |
| RI | 978 | ENSG00000170037 | CNTROB | chr17 | + | 7948486 | 7949137 | 7948486 | 7948619 | 7949084 | 7949137 | 4 | 3 | 5 | 13 | 12 | 27 | 22 | 30 | 64 | 30 | 20 | 6 | 7 | 18 | 0 | 40 | 36 | 11 | 32 | 21 | 298 | 149 | 0.089250785 | 0.252130266 | 0.069 | 0.064 | 0.077 | 0.092 | 0.167 | 0.2 | 0.077 | 0.241 | 0.22 | 0 | -0.054 |
| RI | 1793 | ENSG00000092108 | SCFD1 | chr14 | + | 30705822 | 30708065 | 30705822 | 30705885 | 30707989 | 30708065 | 75 | 77 | 53 | 57 | 49 | 92 | 71 | 73 | 108 | 104 | 1 | 27 | 19 | 25 | 3 | 15 | 41 | 27 | 20 | 34 | 298 | 149 | 0.089559717 | 0.252882051 | 0.29 | 0.352 | 0.266 | 0.209 | 0.191 | 0.032 | 0.248 | 0.26 | 0.385 | 0.042 | 0.068 |
| RI | 2149 | ENSG00000133112 | TPT1 | chr13 | - | 45340711 | 45341158 | 45340711 | 45340820 | 45341041 | 45341158 | 8 | 18 | 7 | 24 | 13 | 0 | 4 | 23 | 19 | 16 | 30 | 28 | 21 | 10 | 6 | 63 | 37 | 38 | 4 | 18 | 298 | 149 | 0.090341372 | 0.254967263 | 1 | 0.692 | 0.132 | 0.387 | 0.289 | 0.192 | 0.275 | 0.216 | 0.556 | 0.143 | 0.224 |
| RI | 1807 | ENSG00000100888 | CHD8 | chr14 | - | 21428963 | 21431858 | 21428963 | 21429335 | 21430800 | 21431858 | 26 | 27 | 27 | 28 | 60 | 36 | 22 | 36 | 44 | 42 | 49 | 81 | 92 | 50 | 4 | 43 | 26 | 22 | 22 | 19 | 298 | 149 | 0.09070024 | 0.255857842 | 0.265 | 0.38 | 0.273 | 0.241 | 0.417 | 0.363 | 0.609 | 0.676 | 0.532 | 0.095 | -0.14 |
| RI | 712 | ENSG00000169718 | DUS1L | chr17 | - | 82058340 | 82058818 | 82058340 | 82058416 | 82058780 | 82058818 | 74 | 42 | 86 | 67 | 31 | 375 | 181 | 306 | 296 | 536 | 39 | 67 | 59 | 71 | 85 | 320 | 314 | 229 | 245 | 366 | 298 | 149 | 0.091320688 | 0.257485108 | 0.09 | 0.104 | 0.123 | 0.102 | 0.028 | 0.057 | 0.096 | 0.114 | 0.127 | 0.104 | -0.01 |
| RI | 8126 | ENSG00000148688 | RPP30 | chr10 | + | 90894774 | 90895483 | 90894774 | 90894891 | 90895453 | 90895483 | 54 | 54 | 53 | 65 | 53 | 48 | 9 | 61 | 26 | 47 | 28 | 33 | 76 | 16 | 0 | 24 | 29 | 17 | 24 | 30 | 298 | 149 | 0.091383723 | 0.25753991 | 0.36 | 0.75 | 0.303 | 0.556 | 0.361 | 0.368 | 0.363 | 0.691 | 0.25 | 0 | 0.132 |
| RI | 4600 | ENSG00000185946 | RNPC3 | chr1 | + | 103525698 | 103526262 | 103525698 | 103525930 | 103526066 | 103526262 | 114 | 65 | 92 | 104 | 86 | 5 | 5 | 5 | 2 | 1 | 101 | 63 | 70 | 36 | 15 | 0 | 5 | 0 | 7 | 16 | 285 | 149 | 0.09173208 | 0.258335395 | 0.923 | 0.872 | 0.906 | 0.965 | 0.978 | 1 | 0.868 | 1 | 0.729 | 0.329 | 0.144 |
| RI | 2394 | ENSG00000181929 | PRKAG1 | chr12 | - | 49005305 | 49005543 | 49005305 | 49005391 | 49005461 | 49005543 | 19 | 11 | 10 | 16 | 25 | 0 | 2 | 3 | 0 | 2 | 1 | 2 | 39 | 15 | 21 | 0 | 0 | 0 | 0 | 0 | 219 | 149 | 0.091753455 | 0.258335395 | 1 | 0.789 | 0.694 | 1 | 0.895 | 1 | 1 | 1 | 1 | 1 | -0.124 |
| RI | 4835 | ENSG00000103168 | TAF1C | chr16 | - | 84181745 | 84182058 | 84181745 | 84181863 | 84181941 | 84182058 | 13 | 10 | 23 | 27 | 5 | 5 | 8 | 13 | 16 | 0 | 61 | 21 | 30 | 52 | 76 | 1 | 15 | 8 | 14 | 16 | 227 | 149 | 0.092622295 | 0.260657407 | 0.631 | 0.451 | 0.537 | 0.526 | 1 | 0.976 | 0.479 | 0.711 | 0.709 | 0.757 | -0.097 |
| RI | 1148 | ENSG00000168488 | ATXN2L | chr16 | + | 28835932 | 28837232 | 28835932 | 28836176 | 28836750 | 28837232 | 28 | 22 | 16 | 37 | 2 | 4 | 0 | 0 | 0 | 2 | 45 | 34 | 15 | 21 | 35 | 8 | 0 | 3 | 0 | 0 | 298 | 149 | 0.093102942 | 0.261885274 | 0.778 | 1 | 1 | 1 | 0.333 | 0.738 | 1 | 0.714 | 1 | 1 | -0.068 |
| RI | 6365 | ENSG00000012124 | CD22 | chr19 | + | 35332018 | 35332924 | 35332018 | 35332074 | 35332593 | 35332924 | 51 | 56 | 55 | 98 | 140 | 2 | 0 | 0 | 0 | 8 | 203 | 242 | 35 | 157 | 202 | 8 | 1 | 0 | 0 | 20 | 298 | 149 | 0.093504386 | 0.262889295 | 0.927 | 1 | 1 | 1 | 0.897 | 0.927 | 0.992 | 1 | 1 | 0.835 | 0.014 |
| RI | 8089 | ENSG00000177239 | MAN1B1 | chr9 | + | 137099695 | 137101153 | 137099695 | 137099881 | 137101004 | 137101153 | 15 | 2 | 14 | 10 | 26 | 124 | 100 | 126 | 140 | 81 | 14 | 20 | 10 | 14 | 81 | 181 | 238 | 84 | 115 | 221 | 298 | 149 | 0.093626607 | 0.26310769 | 0.057 | 0.01 | 0.053 | 0.034 | 0.138 | 0.037 | 0.04 | 0.056 | 0.057 | 0.155 | -0.011 |
| RI | 717 | ENSG00000169733 | RFNG | chr17 | - | 82049676 | 82050555 | 82049676 | 82050006 | 82050401 | 82050555 | 13 | 5 | 22 | 12 | 16 | 12 | 4 | 11 | 21 | 27 | 21 | 18 | 4 | 12 | 9 | 37 | 20 | 3 | 14 | 53 | 298 | 149 | 0.0939069 | 0.263706436 | 0.351 | 0.385 | 0.5 | 0.222 | 0.229 | 0.221 | 0.31 | 0.4 | 0.3 | 0.078 | 0.076 |
| RI | 2477 | ENSG00000070756 | PABPC1 | chr8 | - | 100721390 | 100721886 | 100721390 | 100721619 | 100721733 | 100721886 | 1035 | 458 | 790 | 889 | 739 | 9 | 1 | 18 | 2 | 9 | 1204 | 465 | 618 | 887 | 2178 | 11 | 4 | 13 | 2 | 0 | 263 | 149 | 0.093928956 | 0.263706436 | 0.985 | 0.996 | 0.961 | 0.996 | 0.979 | 0.984 | 0.985 | 0.964 | 0.996 | 1 | -0.002 |
| RI | 5546 | ENSG00000109171 | SLAIN2 | chr4 | + | 48382567 | 48382927 | 48382567 | 48382639 | 48382727 | 48382927 | 559 | 372 | 403 | 638 | 1298 | 10 | 5 | 4 | 15 | 17 | 731 | 519 | 428 | 485 | 582 | 8 | 8 | 3 | 9 | 2 | 237 | 149 | 0.094162484 | 0.26397622 | 0.972 | 0.979 | 0.984 | 0.964 | 0.98 | 0.983 | 0.976 | 0.989 | 0.971 | 0.995 | -0.007 |
| RI | 8318 | ENSG00000204482 | LST1 | chr6 | + | 31587199 | 31588909 | 31587199 | 31587318 | 31588562 | 31588909 | 273 | 183 | 115 | 242 | 333 | 181 | 84 | 59 | 117 | 172 | 594 | 162 | 149 | 276 | 557 | 361 | 89 | 75 | 179 | 397 | 298 | 149 | 0.094172802 | 0.26397622 | 0.43 | 0.521 | 0.494 | 0.508 | 0.492 | 0.451 | 0.476 | 0.498 | 0.435 | 0.412 | 0.035 |
| RI | 3784 | ENSG00000166526 | ZNF3 | chr7 | - | 100075134 | 100075630 | 100075134 | 100075261 | 100075541 | 100075630 | 13 | 6 | 17 | 25 | 4 | 40 | 19 | 29 | 40 | 22 | 24 | 9 | 22 | 10 | 20 | 20 | 23 | 39 | 30 | 19 | 298 | 149 | 0.094196951 | 0.26397622 | 0.14 | 0.136 | 0.227 | 0.238 | 0.083 | 0.375 | 0.164 | 0.22 | 0.143 | 0.345 | -0.085 |
| RI | 5145 | ENSG00000138035 | PNPT1 | chr2 | - | 55644636 | 55645432 | 55644636 | 55644720 | 55645348 | 55645432 | 10 | 15 | 11 | 7 | 2 | 83 | 99 | 100 | 124 | 129 | 15 | 14 | 13 | 8 | 2 | 38 | 73 | 76 | 118 | 0 | 298 | 149 | 0.094203804 | 0.26397622 | 0.057 | 0.07 | 0.052 | 0.027 | 0.008 | 0.165 | 0.088 | 0.079 | 0.033 | 1 | -0.23 |
| RI | 254 | ENSG00000104889 | RNASEH2A | chr19 | + | 12807205 | 12807506 | 12807205 | 12807329 | 12807418 | 12807506 | 0 | 0 | 2 | 5 | 0 | 20 | 8 | 11 | 23 | 12 | 15 | 0 | 0 | 0 | 8 | 0 | 15 | 16 | 6 | 10 | 238 | 149 | 0.094345492 | 0.264247901 | 0 | 0 | 0.102 | 0.12 | 0 | 1 | 0 | 0 | 0 | 0.334 | -0.222 |
| RI | 7015 | ENSG00000182196 | ARL6IP4 | chr12 | + | 122981128 | 122981879 | 122981128 | 122981266 | 122981594 | 122981879 | 30 | 17 | 39 | 39 | 15 | 79 | 39 | 62 | 64 | 78 | 67 | 30 | 32 | 25 | 89 | 81 | 95 | 63 | 66 | 73 | 298 | 149 | 0.094511721 | 0.264588027 | 0.16 | 0.179 | 0.239 | 0.234 | 0.088 | 0.293 | 0.136 | 0.203 | 0.159 | 0.379 | -0.054 |
| RI | 7331 | ENSG00000056586 | RC3H2 | chr9 | - | 122865348 | 122865657 | 122865348 | 122865541 | 122865623 | 122865657 | 200 | 160 | 176 | 191 | 351 | 3 | 1 | 2 | 2 | 9 | 51 | 187 | 147 | 22 | 79 | 0 | 1 | 4 | 2 | 9 | 231 | 149 | 0.09465979 | 0.264877027 | 0.977 | 0.99 | 0.983 | 0.984 | 0.962 | 1 | 0.992 | 0.96 | 0.876 | 0.85 | 0.044 |
| RI | 4818 | ENSG00000149231 | CCDC82 | chr11 | - | 96383961 | 96384761 | 96383961 | 96384293 | 96384362 | 96384761 | 204 | 263 | 169 | 171 | 372 | 99 | 139 | 61 | 70 | 178 | 29 | 143 | 55 | 32 | 20 | 22 | 33 | 14 | 5 | 9 | 218 | 149 | 0.094944917 | 0.265549064 | 0.585 | 0.564 | 0.654 | 0.625 | 0.588 | 0.474 | 0.748 | 0.729 | 0.814 | 0.603 | -0.07 |
| RI | 7185 | ENSG00000168404 | MLKL | chr16 | - | 74674959 | 74675404 | 74674959 | 74675100 | 74675323 | 74675404 | 16 | 8 | 13 | 10 | 5 | 12 | 10 | 19 | 19 | 9 | 85 | 16 | 26 | 49 | 11 | 46 | 23 | 30 | 20 | 18 | 298 | 149 | 0.09515104 | 0.265999617 | 0.4 | 0.286 | 0.255 | 0.208 | 0.217 | 0.48 | 0.258 | 0.302 | 0.551 | 0.234 | -0.092 |
| RI | 4429 | ENSG00000197989 | SNHG12 | chr1 | - | 28580559 | 28581229 | 28580559 | 28580646 | 28581093 | 28581229 | 83 | 63 | 85 | 85 | 43 | 11 | 3 | 5 | 8 | 2 | 137 | 55 | 109 | 70 | 86 | 34 | 0 | 18 | 6 | 9 | 298 | 149 | 0.095589249 | 0.267098247 | 0.79 | 0.913 | 0.895 | 0.842 | 0.915 | 0.668 | 1 | 0.752 | 0.854 | 0.827 | 0.051 |
| RI | 2914 | ENSG00000163930 | BAP1 | chr3 | - | 52406252 | 52406907 | 52406252 | 52406376 | 52406828 | 52406907 | 12 | 7 | 11 | 24 | 7 | 66 | 24 | 39 | 60 | 21 | 24 | 28 | 16 | 17 | 20 | 66 | 61 | 28 | 53 | 61 | 298 | 149 | 0.095900632 | 0.267841624 | 0.083 | 0.127 | 0.124 | 0.167 | 0.143 | 0.154 | 0.187 | 0.222 | 0.138 | 0.141 | -0.04 |
| RI | 1176 | ENSG00000125124 | BBS2 | chr16 | - | 56498436 | 56502808 | 56498436 | 56502456 | 56502672 | 56502808 | 83 | 97 | 80 | 84 | 66 | 81 | 45 | 54 | 68 | 83 | 114 | 79 | 102 | 42 | 22 | 32 | 27 | 18 | 42 | 33 | 298 | 149 | 0.096360063 | 0.268997587 | 0.339 | 0.519 | 0.426 | 0.382 | 0.284 | 0.64 | 0.594 | 0.739 | 0.333 | 0.25 | -0.121 |
| [illegible] | | | | | | | | | | | | | | | | | | | | | | | | | | | | | | | | | | | | | | | | | | | | | | |

| | | | | | | | | | | | | | | | |
|---|---|---|---|---|---|---|---|---|---|---|---|---|---|---|---|
| RI | 4186 | ENSG00000173039 | RELA | chr11 | - | 65653600 | 65655762 | 65653600 | 65655000 | 65655687 | 65655762 | 30 | 32 | 37 | [illegible] |
| RI | 4872 | ENSG00000163534 | FCRL1 | chr1 | - | 157801914 | 157802664 | 157801914 | 157802162 | 157802376 | 157802664 | 61 | 38 | 67 | [illegible] |
| RI | 8326 | ENSG00000187778 | MCRS1 | chr12 | - | 49566076 | 49566841 | 49566076 | 49566215 | 49566721 | 49566841 | 47 | 25 | 58 | [illegible] |
| RI | 6771 | ENSG00000147955 | SIGMAR1 | chr9 | - | 34636996 | 34637420 | 34636996 | 34637089 | 34637219 | 34637420 | 3 | 0 | 2 | [illegible] |
| RI | 6942 | ENSG00000172732 | MUS81 | chr11 | + | 65864719 | 65865145 | 65864719 | 65864815 | 65865016 | 65865145 | 37 | 10 | 21 | [illegible] |
| RI | 5028 | ENSG00000138646 | HERC5 | chr4 | + | 88504231 | 88504597 | 88504231 | 88504415 | 88504494 | 88504597 | 11 | 9 | 8 | [illegible] |
| RI | 5398 | ENSG00000115317 | HTRA2 | chr2 | + | 74531338 | 74531702 | 74531338 | 74531371 | 74531596 | 74531702 | 18 | 4 | 17 | [illegible] |
| RI | 3420 | ENSG00000184432 | COPB2 | chr3 | - | 139373665 | 139374588 | 139373665 | 139373808 | 139374488 | 139374588 | 155 | 99 | 110 | [illegible] |
| RI | 2299 | ENSG00000170144 | HNRNPA3 | chr2 | + | 177219246 | 177221976 | 177219246 | 177219314 | 177219407 | 177221976 | 295 | 415 | 267 | [illegible] |
| RI | 3666 | ENSG00000167962 | ZNF598 | chr16 | - | 1999495 | 2000481 | 1999495 | 2000256 | 2000408 | 2000481 | 11 | 7 | 7 | [illegible] |
| RI | 8440 | ENSG00000168010 | ATG16L2 | chr11 | + | 72828358 | 72828776 | 72828358 | 72828508 | 72828728 | 72828776 | 33 | 8 | 16 | [illegible] |
| RI | 6798 | ENSG00000005302 | MSL3 | chrX | + | 11760836 | 11762252 | 11760836 | 11760937 | 11762129 | 11762252 | 124 | 90 | 136 | [illegible] |
| RI | 4284 | ENSG00000197355 | UAP1L1 | chr9 | + | 137081997 | 137084538 | 137081997 | 137082064 | 137082636 | 137084538 | 2 | 3 | 0 | [illegible] |
| RI | 1866 | ENSG00000100823 | APEX1 | chr14 | + | 20455577 | 20456101 | 20455577 | 20455703 | 20455913 | 20456101 | 35 | 27 | 59 | [illegible] |
| RI | 4759 | ENSG00000103365 | GGA2 | chr16 | - | 23493359 | 23494378 | 23493359 | 23493587 | 23494302 | 23494378 | 35 | 40 | 34 | [illegible] |
| RI | 1811 | ENSG00000213920 | MDP1 | chr14 | - | 24215551 | 24215797 | 24215551 | 24215662 | 24215736 | 24215797 | 19 | 7 | 11 | [illegible] |
| RI | 2411 | ENSG00000139625 | MAP3K12 | chr12 | - | 53484256 | 53485214 | 53484256 | 53484365 | 53485055 | 53485214 | 22 | 25 | 28 | [illegible] |
| RI | 1457 | ENSG00000140694 | PARN | chr16 | - | 14627105 | 14628251 | 14627105 | 14627187 | 14628171 | 14628251 | 13 | 6 | 17 | [illegible] |
| RI | 4462 | ENSG00000151694 | ADAM17 | chr2 | - | 9521202 | 9523338 | 9521202 | 9521316 | 9523248 | 9523338 | 34 | 59 | 21 | [illegible] |
| RI | 5473 | ENSG00000223960 | CHROMR | chr2 | + | 178426365 | 178430889 | 178426365 | 178426505 | 178430769 | 178430889 | 60 | 38 | 36 | [illegible] |
| RI | 8567 | ENSG00000137822 | TUBGCP4 | chr15 | + | 43401715 | 43403799 | 43401715 | 43401850 | 43403682 | 43403799 | 26 | 43 | 26 | [illegible] |
| RI | 7859 | ENSG00000142102 | PGGHG | chr11 | + | 290677 | 292095 | 290677 | 291113 | 291975 | 292095 | 30 | 19 | 32 | [illegible] |
| RI | 5077 | ENSG00000198520 | ARMH1 | chr1 | + | 44724319 | 44724668 | 44724319 | 44724392 | 44724538 | 44724668 | 24 | 26 | 22 | [illegible] |
| RI | 4182 | ENSG00000142784 | WDTC1 | chr1 | + | 27303620 | 27305193 | 27303620 | 27303795 | 27305000 | 27305193 | 18 | 17 | 21 | [illegible] |
| RI | 5651 | ENSG00000142733 | MAP3K6 | chr1 | - | 27356589 | 27357114 | 27356589 | 27356749 | 27357008 | 27357114 | 13 | 8 | 5 | [illegible] |
| RI | 4950 | ENSG00000171503 | ETFDH | chr4 | + | 158680466 | 158681614 | 158680466 | 158680607 | 158681570 | 158681614 | 20 | 19 | 18 | [illegible] |
| RI | 4135 | ENSG00000157873 | TNFRSF14 | chr1 | + | 2558342 | 2559978 | 2558342 | 2558468 | 2559822 | 2559978 | 28 | 13 | 24 | [illegible] |
| RI | 313 | ENSG00000115268 | RPS15 | chr19 | + | 1438806 | 1440253 | 1438806 | 1438892 | 1440025 | 1440253 | 80 | 24 | 76 | [illegible] |
| RI | 2240 | ENSG00000104886 | PLEKHJ1 | chr19 | - | 2233153 | 2234061 | 2233153 | 2233905 | 2233997 | 2234061 | 9 | 6 | 12 | [illegible] |
| RI | 2863 | ENSG00000168291 | PDHB | chr3 | - | 58431876 | 58433684 | 58431876 | 58431984 | 58433630 | 58433684 | 23 | 25 | 35 | [illegible] |
| RI | 3053 | ENSG00000145431 | PDGFC | chr4 | - | 156767772 | 156767990 | 156767772 | 156767876 | 156767961 | 156767990 | 72 | 93 | 28 | [illegible] |
| RI | 7869 | ENSG00000166333 | ILK | chr11 | + | 6608693 | 6608967 | 6608693 | 6608790 | 6608883 | 6608967 | 31 | 25 | 40 | [illegible] |
| RI | 1290 | ENSG00000185324 | CDK10 | chr16 | + | 89693273 | 89694232 | 89693273 | 89693467 | 89694172 | 89694232 | 14 | 7 | 10 | [illegible] |
| RI | 6235 | ENSG00000234545 | FAM133B | chr7 | - | 92577654 | 92578182 | 92577654 | 92577717 | 92578149 | 92578182 | 73 | 68 | 57 | [illegible] |
| RI | 3587 | ENSG00000197343 | ZNF655 | chr7 | + | 99561887 | 99562494 | 99561887 | 99561973 | 99562348 | 99562494 | 137 | 66 | 104 | [illegible] |
| RI | 7881 | ENSG00000149483 | TMEM138 | chr11 | + | 61364249 | 61364518 | 61364249 | 61364315 | 61364432 | 61364518 | 165 | 57 | 91 | [illegible] |
| RI | 3451 | ENSG00000144591 | GMPPA | chr2 | + | 219504082 | 219505362 | 219504082 | 219504213 | 219505227 | 219505362 | 15 | 9 | 9 | [illegible] |
| RI | 3311 | ENSG00000117054 | ACADM | chr1 | + | 75732852 | 75733628 | 75732852 | 75732922 | 75733527 | 75733628 | 34 | 22 | 17 | [illegible] |
| RI | 4363 | ENSG00000124574 | ABCC10 | chr6 | + | 43442969 | 43444010 | 43442969 | 43443159 | 43443932 | 43444010 | 34 | 26 | 37 | [illegible] |
| RI | 5774 | ENSG00000163938 | GNL3 | chr3 | + | 52690591 | 52691071 | 52690591 | 52690704 | 52690944 | 52691071 | 88 | 65 | 114 | [illegible] |
| RI | 3322 | ENSG00000157216 | SSBP3 | chr1 | - | 54242163 | 54243299 | 54242163 | 54242212 | 54243234 | 54243299 | 13 | 7 | 15 | [illegible] |
| RI | 61 | ENSG00000182944 | EWSR1 | chr22 | + | 29288605 | 29291599 | 29288605 | 29288786 | 29291561 | 29291599 | 152 | 107 | 130 | [illegible] |
| RI | 2011 | ENSG00000135506 | OS9 | chr12 | + | 57695961 | 57696373 | 57695961 | 57695976 | 57696274 | 57696373 | 131 | 135 | 118 | [illegible] |
| RI | 1767 | ENSG00000119682 | AREL1 | chr14 | - | 74669946 | 74670871 | 74669946 | 74670126 | 74670761 | 74670871 | 37 | 28 | 43 | [illegible] |
| RI | 5811 | ENSG00000177463 | NR2C2 | chr3 | + | 15037999 | 15039227 | 15037999 | 15038137 | 15039121 | 15039227 | 55 | 21 | 47 | [illegible] |
| RI | 364 | ENSG00000125755 | SYMPK | chr19 | - | 45854174 | 45854507 | 45854174 | 45854240 | 45854390 | 45854507 | 14 | 3 | 4 | [illegible] |
| RI | 3219 | ENSG00000015479 | MATR3 | chr5 | + | 139314674 | 139316188 | 139314674 | 139314736 | 139316075 | 139316188 | 166 | 166 | 112 | [illegible] |
| RI | 8047 | ENSG00000156650 | KAT6B | chr10 | + | 74981786 | 74985241 | 74981786 | 74981928 | 74985079 | 74985241 | 36 | 30 | 19 | [illegible] |
| RI | 4587 | ENSG00000106462 | EZH2 | chr7 | - | 148816683 | 148818117 | 148816683 | 148817391 | 148817876 | 148818117 | 18 | 14 | 14 | [illegible] |
| RI | 1280 | ENSG00000141002 | TCF25 | chr16 | + | 89904117 | 89905096 | 89904117 | 89904205 | 89904937 | 89905096 | 7 | 7 | 11 | [illegible] |
| RI | 1248 | ENSG00000188603 | CLN3 | chr16 | - | 28482104 | 28482382 | 28482104 | 28482198 | 28482326 | 28482382 | 31 | 14 | 17 | [illegible] |
| RI | 1609 | ENSG00000166411 | IDH3A | chr15 | + | 78156907 | 78157631 | 78156907 | 78157162 | 78157547 | 78157631 | 48 | 36 | 35 | [illegible] |
| RI | 2779 | ENSG00000159496 | RGL4 | chr22 | + | 23698554 | 23698978 | 23698554 | 23698645 | 23698843 | 23698978 | 24 | 15 | 39 | [illegible] |
| RI | 4720 | ENSG00000129968 | ABHD17A | chr19 | - | 1877507 | 1880115 | 1877507 | 1877687 | 1878663 | 1880115 | 48 | 18 | 39 | [illegible] |
| RI | 5857 | ENSG00000108465 | CDK5RAP3 | chr17 | + | 47975513 | 47976013 | 47975513 | 47975653 | 47975868 | 47976013 | 302 | 201 | 280 | [illegible] |
| RI | 3299 | ENSG00000238227 | TMEM250 | chr9 | - | 136114588 | 136117024 | 136114588 | 136114950 | 136115455 | 136117024 | 274 | 110 | 176 | [illegible] |
| RI | 972 | ENSG00000178971 | CTC1 | chr17 | - | 8230293 | 8230468 | 8230293 | 8230348 | 8230446 | 8230468 | 245 | 176 | 227 | [illegible] |
| RI | 3533 | ENSG00000183258 | DDX41 | chr5 | - | 177513352 | 177513847 | 177513352 | 177513484 | 177513684 | 177513847 | 8 | 6 | 7 | [illegible] |
| RI | 1932 | ENSG00000119688 | ABCD4 | chr14 | - | 74296332 | 74298069 | 74296332 | 74296449 | 74297929 | 74298069 | 19 | 10 | 16 | [illegible] |
| RI | 5141 | ENSG00000143387 | CTSK | chr1 | - | 150799167 | 150799709 | 150799167 | 150799273 | 150799543 | 150799709 | 8 | 12 | 8 | [illegible] |
| RI | 4467 | ENSG00000177409 | SAMD9L | chr7 | - | 93145384 | 93146040 | 93145384 | 93145532 | 93145908 | 93146040 | 289 | 315 | 194 | [illegible] |
| RI | 7042 | ENSG00000087088 | BAX | chr19 | + | 48955547 | 48955833 | 48955547 | 48955599 | 48955686 | 48955833 | 26 | 12 | 10 | [illegible] |
| RI | 3086 | ENSG00000164073 | MFSD8 | chr4 | - | 127917888 | 127920836 | 127917888 | 127919338 | 127920456 | 127920836 | 56 | 42 | 71 | [illegible] |
| RI | 5536 | ENSG00000198901 | PRC1 | chr15 | - | 90981498 | 90981981 | 90981498 | 90981669 | 90981747 | 90981981 | 14 | 2 | 18 | [illegible] |
| RI | 5123 | ENSG00000044574 | HSPA5 | chr9 | - | 125239420 | 125240907 | 125239420 | 125240309 | 125240675 | 125240907 | 66 | 50 | 43 | [illegible] |
| RI | 6508 | ENSG00000122566 | HNRNPA2B1 | chr7 | - | 26190992 | 26192577 | 26190992 | 26192338 | 26192494 | 26192577 | 1088 | 1468 | 1162 | [illegible] |
| RI | 6459 | ENSG00000160856 | FCRL3 | chr1 | - | 157676480 | 157678856 | 157676480 | 157676768 | 157678594 | 157678856 | 272 | 81 | 99 | [illegible] |
| RI | 5353 | ENSG00000066923 | STAG3 | chr7 | + | 100202171 | 100202590 | 100202171 | 100202340 | 100202453 | 100202590 | 38 | 14 | 17 | [illegible] |
| RI | 2922 | ENSG00000247596 | TWF2 | chr3 | - | 52229919 | 52231231 | 52229919 | 52230070 | 52231126 | 52231231 | 8 | 7 | 9 | [illegible] |
| RI | 2420 | ENSG00000214655 | ZSWIM8 | chr10 | + | 73800639 | 73801195 | 73800639 | 73800759 | 73801084 | 73801195 | 32 | 14 | 22 | [illegible] |
| RI | 6199 | ENSG00000141258 | SGSM2 | chr17 | + | 2362837 | 2363134 | 2362837 | 2362905 | 2362988 | 2363134 | 16 | 13 | 23 | [illegible] |
| RI | 7557 | ENSG00000106397 | PLOD3 | chr7 | - | 101211845 | 101212374 | 101211845 | 101211950 | 101212252 | 101212374 | 8 | 2 | 0 | [illegible] |
| RI | 2100 | ENSG00000141279 | NPEPPS | chr17 | + | 47619277 | 47619784 | 47619277 | 47619322 | 47619736 | 47619784 | 30 | 37 | 28 | [illegible] |
| RI | 5327 | ENSG00000064703 | DDX20 | chr1 | + | 111760705 | 111761125 | 111760705 | 111760848 | 111760986 | 111761125 | 32 | 21 | 63 | [illegible] |
| RI | 2859 | ENSG00000168291 | PDHB | chr3 | - | 58429707 | 58430238 | 58429707 | 58429799 | 58430127 | 58430238 | 45 | 14 | 30 | [illegible] |
| RI | 5469 | ENSG00000223960 | CHROMR | chr2 | + | 178426365 | 178430889 | 178426365 | 178426505 | 178430677 | 178430889 | 66 | 26 | 32 | [illegible] |
| RI | 2793 | ENSG00000204463 | BAG6 | chr6 | - | 31640644 | 31640938 | 31640644 | 31640704 | 31640791 | 31640938 | 14 | 9 | 12 | [illegible] |
| RI | 4413 | ENSG00000143575 | HAX1 | chr1 | + | 154274949 | 154275260 | 154274949 | 154275001 | 154275153 | 154275260 | 42 | 31 | 44 | [illegible] |
| RI | 2821 | ENSG00000242086 | MUC20-OT1 | chr3 | + | 195708133 | 195708807 | 195708133 | 195708268 | 195708746 | 195708807 | 16 | 4 | 22 | [illegible] |
| RI | 2401 | ENSG00000048140 | TSPAN17 | chr5 | + | 176652742 | 176655020 | 176652742 | 176652913 | 176654894 | 176655020 | 10 | 8 | 12 | [illegible] |
| RI | 5636 | ENSG00000115649 | CNPPD1 | chr2 | - | 219174777 | 219175108 | 219174777 | 219174906 | 219174987 | 219175108 | 16 | 4 | 16 | [illegible] |
| RI | 1231 | ENSG00000198736 | MSRB1 | chr16 | - | 1940777 | 1941405 | 1940777 | 1940892 | 1941256 | 1941405 | 5 | 6 | 2 | [illegible] |
| RI | 5862 | ENSG00000164609 | SLU7 | chr5 | - | 160418595 | 160419081 | 160418595 | 160418685 | 160419022 | 160419081 | 87 | 48 | 99 | [illegible] |
| RI | 143 | ENSG00000099917 | MED15 | chr22 | + | 20554935 | 20555148 | 20554935 | 20554995 | 20555076 | 20555148 | 126 | 30 | 137 | [illegible] |
| RI | 6799 | ENSG00000005302 | MSL3 | chrX | + | 11763779 | 11765729 | 11763779 | 11763938 | 11765466 | 11765729 | 206 | 149 | 211 | [illegible] |
| RI | 4128 | ENSG00000241878 | PISD | chr22 | - | 31620995 | 31621885 | 31620995 | 31621472 | 31621648 | 31621885 | 18 | 8 | 26 | [illegible] |
| RI | 2080 | ENSG00000120318 | ARAP3 | chr5 | - | 141661683 | 141662255 | 141661683 | 141661789 | 141662042 | 141662255 | 0 | 4 | 0 | [illegible] |
| RI | 670 | ENSG00000099956 | SMARCB1 | chr22 | + | 23791755 | 23793688 | 23791755 | 23791894 | 23793558 | 23793688 | 49 | 41 | 52 | [illegible] |
| RI | 522 | ENSG00000026036 | RTEL1-TNFRSF6B | chr20 | + | 63689749 | 63690210 | 63689749 | 63689865 | 63690086 | 63690210 | 7 | 4 | 8 | [illegible] |
| RI | 2913 | ENSG00000163930 | BAP1 | chr3 | - | 52406252 | 52406907 | 52406252 | 52406322 | 52406828 | 52406907 | 47 | 16 | 35 | [illegible] |
| RI | 4939 | ENSG00000129933 | MAU2 | chr19 | + | 19344848 | 19345369 | 19344848 | 19344926 | 19345303 | 19345369 | 22 | 10 | 22 | [illegible] |
| RI | 5465 | ENSG00000237765 | FAM200B | chr4 | + | 15686235 | 15687148 | 15686235 | 15686462 | 15687000 | 15687148 | 87 | 69 | 98 | [illegible] |
| RI | 7906 | ENSG00000084207 | GSTP1 | chr11 | + | 67586103 | 67586653 | 67586103 | 67586211 | 67586388 | 67586653 | 14 | 7 | 15 | [illegible] |
| RI | 3569 | ENSG00000265241 | RBM8A | chr1 | - | 145926808 | 145927077 | 145926808 | 145926886 | 145927017 | 145927077 | 48 | 43 | 31 | [illegible] |
| RI | 62 | ENSG00000182944 | EWSR1 | chr22 | + | 29288605 | 29291599 | 29288605 | 29288786 | 29291564 | 29291599 | 156 | 112 | 132 | [illegible] |
| RI | 4764 | ENSG00000159658 | EFCAB14 | chr1 | - | 46684490 | 46686870 | 46684490 | 46684602 | 46686783 | 46686870 | 57 | 33 | 28 | [illegible] |
| RI | 2008 | ENSG00000135596 | MICAL1 | chr6 | - | 109446135 | 109446772 | 109446135 | 109446412 | 109446695 | 109446772 | 11 | 6 | 35 | [illegible] |
| RI | 2031 | ENSG00000135423 | GLS2 | chr12 | - | 56475623 | 56475977 | 56475623 | 56475682 | 56475944 | 56475977 | 21 | 12 | 8 | [illegible] |
| RI | 2571 | ENSG00000174652 | ZNF266 | chr19 | - | 9418761 | 9419299 | 9418761 | 9418895 | 9419050 | 9419299 | 160 | 85 | 89 | [illegible] |
| RI | 8227 | ENSG00000213809 | KLRK1 | chr12 | - | 10378553 | 10379792 | 10378553 | 10378705 | 10379446 | 10379792 | 403 | 387 | 274 | [illegible] |
| RI | 5417 | ENSG00000039650 | PNKP | chr19 | - | 49861448 | 49861881 | 49861448 | 49861510 | 49861607 | 49861881 | 10 | 2 | 5 | [illegible] |
| RI | 4414 | ENSG00000143575 | HAX1 | chr1 | + | 154275153 | 154275483 | 154275153 | 154275260 | 154275392 | 154275483 | 29 | 25 | 38 | [illegible] |
| RI | 108 | ENSG00000100395 | L3MBTL2 | chr22 | + | 41227088 | 41227869 | 41227088 | 41227323 | 41227803 | 41227869 | 5 | 4 | 9 | [illegible] |
| RI | 4330 | ENSG00000115548 | KDM3A | chr2 | + | 86482457 | 86484158 | 86482457 | 86482694 | 86483986 | 86484158 | 69 | 61 | 58 | [illegible] |
| RI | 4946 | ENSG00000167578 | RAB4B | chr19 | + | 40780018 | 40780499 | 40780018 | 40780099 | 40780384 | 40780499 | 25 | 8 | 10 | [illegible] |
| RI | 5355 | ENSG00000162645 | GBP2 | chr1 | - | 89106131 | 89108291 | 89106131 | 89107052 | 89108022 | 89108291 | 128 | 139 | 90 | [illegible] |
| RI | 2894 | ENSG00000172053 | QARS1 | chr3 | - | 49097991 | 49098480 | 49097991 | 49098117 | 49098191 | 49098480 | 18 | 13 | 28 | [illegible] |
| RI | 3254 | ENSG00000137074 | APTX | chr9 | - | 32973375 | 32974561 | 32973375 | 32973652 | 32974457 | 32974561 | 12 | 7 | 7 | [illegible] |
| RI | 3065 | ENSG00000159433 | STARD9 | chr15 | + | 42662793 | 42663917 | 42662793 | 42662891 | 42663819 | 42663917 | 6 | 13 | 8 | [illegible] |
| RI | 2851 | ENSG00000114857 | NKTR | chr3 | + | 42620015 | 42621516 | 42620015 | 42620043 | 42621428 | 42621516 | 337 | 321 | 305 | [illegible] |
| RI | 2770 | ENSG00000159788 | RGS12 | chr4 | + | 3414071 | 3414844 | 3414071 | 3414241 | 3414751 | 3414844 | 0 | 0 | 0 | [illegible] |
| RI | 643 | ENSG00000069399 | BCL3 | chr19 | + | 44758245 | 44758841 | 44758245 | 44758413 | 44758723 | 44758841 | 6 | 3 | 17 | [illegible] |
| RI | 7302 | ENSG00000100353 | EIF3D | chr22 | - | 36525663 | 36526131 | 36525663 | 36525709 | 36525998 | 36526131 | 36 | 19 | 51 | [illegible] |
| RI | 2357 | ENSG00000011243 | AKAP8L | chr19 | - | 15397713 | 15400358 | 15397713 | 15399410 | 15400294 | 15400358 | 53 | 19 | 54 | [illegible] |
| RI | 2823 | ENSG00000242086 | MUC20-OT1 | chr3 | + | 195708133 | 195708807 | 195708133 | 195708435 | 195708746 | 195708807 | 11 | 5 | 12 | [illegible] |
| RI | 5511 | ENSG00000112367 | FIG4 | chr6 | + | 109743109 | 109743772 | 109743109 | 109743272 | 109743674 | 109743772 | 38 | 10 | 7 | [illegible] |
| RI | 3245 | ENSG00000161904 | LEMD2 | chr6 | - | 33778241 | 33780179 | 33778241 | 33778387 | 33780099 | 33780179 | 51 | 42 | 31 | [illegible] |
| RI | 4648 | ENSG00000226822 | LINC02785 | chr1 | + | 108071483 | 108072697 | 108071483 | 108071637 | 108072528 | 108072697 | 13 | 25 | 2 | [illegible] |
| RI | 4879 | ENSG00000182400 | TRAPPC6B | chr14 | - | 39158284 | 39159550 | 39158284 | 39158402 | 39159482 | 39159550 | 38 | 36 | 27 | [illegible] |
| RI | 7328 | ENSG00000110455 | ACCS | chr11 | + | 44077844 | 44078784 | 44077844 | 44077922 | 44078683 | 44078784 | 98 | 23 | 55 | [illegible] |
| RI | 8594 | ENSG00000127022 | CANX | chr5 | + | 179720403 | 179723019 | 179720403 | 179720560 | 179722803 | 179723019 | 29 | 6 | 26 | [illegible] |
| RI | 6371 | ENSG00000077454 | LRCH4 | chr7 | - | 100576693 | 100577005 | 100576693 | 100576777 | 100576901 | 100577005 | 55 | 50 | 48 | [illegible] |
| RI | 7725 | ENSG00000130935 | NOL11 | chr17 | + | 67737507 | 67737972 | 67737507 | 67737692 | 67737846 | 67737972 | 62 | 50 | 59 | [illegible] |
| RI | 7362 | ENSG00000089053 | ANAPC5 | chr12 | - | 121327095 | 121328497 | 121327095 | 121327231 | 121328315 | 121328497 | 38 | 16 | 24 | [illegible] |
| RI | 881 | ENSG00000185359 | HGS | chr17 | + | 81695785 | 81696529 | 81695785 | 81695999 | 81696356 | 81696529 | 14 | 11 | 2 | [illegible] |
| RI | 2118 | ENSG00000172262 | ZNF131 | chr5 | + | 43161248 | 43161931 | 43161248 | 43161606 | 43161708 | 43161931 | 154 | 106 | 184 | [illegible] |
| RI | 1272 | ENSG00000205250 | E2F4 | chr16 | + | 67192760 | 67193170 | 67192760 | 67192870 | 67193008 | 67193170 | 70 | 37 | 31 | [illegible] |
| RI | 1412 | ENSG00000051523 | CYBA | chr16 | - | 88646754 | 88647175 | 88646754 | 88646838 | 88647100 | 88647175 | 55 | 29 | 49 | [illegible] |
| RI | 7144 | ENSG00000172269 | DPAGT1 | chr11 | - | 119097141 | 119098043 | 119097141 | 119097551 | 119097854 | 119098043 | 6 | 0 | 4 | [illegible] |
| RI | 7111 | ENSG00000147536 | GINS4 | chr8 | + | 41541808 | 41545030 | 41541808 | 41541899 | 41541990 | 41545030 | 10 | 1 | 1 | [illegible] |
| RI | 6779 | ENSG00000072778 | ACADVL | chr17 | + | 7220765 | 7221058 | 7220765 | 7220830 | 7220923 | 7221058 | 21 | 6 | 27 | [illegible] |
| RI | 3494 | ENSG00000013573 | DDX11 | chr12 | + | 31089043 | 31090094 | 31089043 | 31089151 | 31089402 | 31090094 | 1 | 1 | 7 | [illegible] |
| RI | 7884 | ENSG00000167770 | OTUB1 | chr11 | + | 63996529 | 63996941 | 63996529 | 63996648 | 63996856 | 63996941 | 68 | 19 | 73 | [illegible] |
| RI | 5901 | ENSG00000159618 | ADGRG5 | chr16 | + | 57565033 | 57566751 | 57565033 | 57565150 | 57566598 | 57566751 | 8 | 18 | 3 | [illegible] |
| RI | 7016 | ENSG00000182196 | ARL6IP4 | chr12 | + | 122981128 | 122981879 | 122981128 | 122981299 | 122981594 | 122981879 | 31 | 18 | 44 | [illegible] |
| RI | 7216 | ENSG00000160948 | VPS28 | chr8 | - | 144425682 | 144426208 | 144425682 | 144425772 | 144426025 | 144426208 | 27 | 13 | 24 | [illegible] |
| RI | 6985 | ENSG00000102119 | EMD | chrX | + | 154379941 | 154380367 | 154379941 | 154380019 | 154380233 | 154380367 | 39 | 39 | 41 | [illegible] |
| RI | 5432 | ENSG00000172890 | NADSYN1 | chr11 | + | 71484311 | 71485648 | 71484311 | 71484447 | 71485541 | 71485648 | 12 | 16 | 12 | [illegible] |
| RI | 1282 | ENSG00000140854 | KATNB1 | chr16 | + | 57755789 | 57756066 | 57755789 | 57755917 | 57755991 | 57756066 | 6 | 0 | 3 | [illegible] |
| RI | 2621 | ENSG00000173020 | GRK2 | chr11 | + | 67283706 | 67283949 | 67283706 | 67283773 | 67283853 | 67283949 | 27 | 10 | 36 | [illegible] |
| RI | 645 | ENSG00000141644 | MBD1 | chr18 | - | 50268845 | 50271540 | 50268845 | 50269818 | 50271468 | 50271540 | 189 | 141 | 181 | [illegible] |
| RI | 521 | ENSG00000026036 | RTEL1-TNFRSF6B | chr20 | + | 63688527 | 63689132 | 63688527 | 63688605 | 63689054 | 63689132 | 6 | 9 | 7 | [illegible] |
| RI | 296 | ENSG00000130159 | ECSIT | chr19 | - | 11507701 | 11508048 | 11507701 | 11507850 | 11507990 | 11508048 | 20 | 6 | 7 | [illegible] |
| RI | 5910 | ENSG00000165792 | METTL17 | chr14 | + | 20994793 | 20995233 | 20994793 | 20994901 | 20995164 | 20995233 | 16 | 45 | 94 | [illegible] |
| RI | 3755 | ENSG00000071894 | CPSF1 | chr8 | - | 144393882 | 144394160 | 144393882 | 144394038 | 144394112 | 144394160 | 14 | 3 | 12 | [illegible] |
| RI | 6033 | ENSG00000100316 | RPL3 | chr22 | - | 39312883 | 39313310 | 39312883 | 39312984 | 39313190 | 39313310 | 84 | 62 | 103 | [illegible] |
| RI | 6705 | ENSG00000130734 | ATG4D | chr19 | + | 10551896 | 10552121 | 10551896 | 10551975 | 10552044 | 10552121 | 15 | 6 | 9 | [illegible] |
| RI | 92 | ENSG00000100099 | HPS4 | chr22 | - | 26465454 | 26466262 | 26465454 | 26465551 | 26466225 | 26466262 | 10 | 8 | 25 | [illegible] |
| RI | 6236 | ENSG00000088038 | CNOT3 | chr19 | + | 54143659 | 54144134 | 54143659 | 54143749 | 54144005 | 54144134 | 8 | 9 | 18 | [illegible] |
| RI | 741 | ENSG00000070413 | DGCR2 | chr22 | - | 19041057 | 19041959 | 19041057 | 19041294 | 19041806 | 19041959 | 17 | 8 | 18 | [illegible] |
| RI | 3854 | ENSG00000175390 | EIF3F | chr11 | + | 7992886 | 7994517 | 7992886 | 7993024 | 7994425 | 7994517 | 35 | 12 | 24 | [illegible] |
| RI | 3720 | ENSG00000138002 | IFT172 | chr2 | - | 27457840 | 27458223 | 27457840 | 27457976 | 27458125 | 27458223 | 12 | 7 | 6 | [illegible] |
| RI | 1852 | ENSG00000100664 | EIF5 | chr14 | + | 103334236 | 103334597 | 103334236 | 103334260 | 103334388 | 103334597 | 22 | 8 | 39 | [illegible] |
| RI | 4845 | ENSG00000159363 | ATP13A2 | chr1 | - | 16986462 | 16986956 | 16986462 | 16986632 | 16986804 | 16986956 | 0 | 2 | 3 | [illegible] |
| RI | 7260 | ENSG00000156738 | MS4A1 | chr11 | + | 60462184 | 60462533 | 60462184 | 60462298 | 60462417 | 60462533 | 706 | 500 | 618 | [illegible] |
| RI | 3402 | ENSG00000095321 | CRAT | chr9 | - | 129102399 | 129103066 | 129102399 | 129102565 | 129103012 | 129103066 | 3 | 0 | 1 | [illegible] |
| RI | 804 | ENSG00000185624 | P4HB | chr17 | - | 81843168 | 81845742 | 81843168 | 81844092 | 81845143 | 81845742 | 11 | 3 | 14 | [illegible] |
| RI | 721 | ENSG00000161955 | TNFSF13 | chr17 | + | 7560723 | 7561601 | 7560723 | 7560817 | 7560998 | 7561601 | 490 | 289 | 287 | [illegible] |
| RI | 2587 | ENSG00000180979 | LRRC57 | chr15 | - | 42547260 | 42548244 | 42547260 | 42547529 | 42548105 | 42548244 | 49 | 46 | 31 | [illegible] |
| RI | 7425 | ENSG00000070061 | ELP1 | chr9 | - | 108892985 | 108894066 | 108892985 | 108893083 | 108893942 | 108894066 | 7 | 2 | 8 | [illegible] |
| RI | 6659 | ENSG00000102316 | MAGED2 | chrX | + | 54815247 | 54816007 | 54815247 | 54815690 | 54815880 | 54816007 | 36 | 10 | 23 | [illegible] |
| RI | 111 | ENSG00000100241 | SBF1 | chr22 | - | 50467531 | 50467923 | 50467531 | 50467693 | 50467785 | 50467923 | 5 | 0 | 5 | [illegible] |
| RI | 1326 | ENSG00000214078 | CPNE1 | chr20 | - | 35626043 | 35626803 | 35626043 | 35626381 | 35626566 | 35626803 | 23 | 20 | 48 | [illegible] |
| RI | 5695 | ENSG00000160710 | ADAR | chr1 | - | 154601040 | 154602626 | 154601040 | 154602248 | 154602384 | 154602626 | 836 | 523 | 789 | [illegible] |
| RI | 5443 | ENSG00000170092 | SPDYE5 | chr7 | + | 75501361 | 75501755 | 75501361 | 75501476 | 75501551 | 75501755 | 10 | 10 | 8 | [illegible] |
| RI | 2306 | ENSG00000204628 | RACK1 | chr5 | - | 181241491 | 181242345 | 181241491 | 181241639 | 181242169 | 181242345 | 58 | 83 | 113 | [illegible] |
| RI | 5779 | ENSG00000204843 | DCTN1 | chr2 | - | 74365074 | 74365657 | 74365074 | 74365241 | 74365514 | 74365657 | 19 | 21 | 8 | [illegible] |
| RI | 4570 | ENSG00000122678 | POLM | chr7 | - | 44074130 | 44074530 | 44074130 | 44074233 | 44074397 | 44074530 | 22 | 14 | 31 | [illegible] |
| RI | 3484 | ENSG00000092841 | MYL6 | chr12 | + | 56159586 | 56160148 | 56159586 | 56159622 | 56159974 | 56160148 | 460 | 352 | 335 | [illegible] |
| RI | 482 | ENSG00000105401 | CDC37 | chr19 | - | 10395227 | 10395543 | 10395227 | 10395343 | 10395434 | 10395543 | 142 | 156 | 205 | [illegible] |
| RI | 7670 | ENSG00000129255 | MPDU1 | chr17 | + | 7586691 | 7587271 | 7586691 | 7586838 | 7587160 | 7587271 | 19 | 15 | 42 | [illegible] |
| RI | 3383 | ENSG00000117385 | P3H1 | chr1 | - | 42754868 | 42755217 | 42754868 | 42754990 | 42755164 | 42755217 | 88 | 95 | 121 | [illegible] |
| RI | 3211 | ENSG00000168615 | ADAM9 | chr8 | + | 39018852 | 39021714 | 39018852 | 39018918 | 39021642 | 39021714 | 11 | 8 | 10 | [illegible] |
| RI | 7634 | ENSG00000204577 | LILRB3 | chr19 | - | 54222746 | 54223087 | 54222746 | 54222782 | 54222942 | 54223087 | 23 | 30 | 8 | [illegible] |
| RI | 468 | ENSG00000130299 | GTPBP3 | chr19 | + | 17338953 | 17339266 | 17338953 | 17339026 | 17339122 | 17339266 | 0 | 3 | 7 | [illegible] |
| RI | 8437 | ENSG00000168010 | ATG16L2 | chr11 | + | 72822477 | 72824842 | 72822477 | 72822543 | 72822847 | 72824842 | 45 | 31 | 27 | [illegible] |
| RI | 1487 | ENSG00000168096 | ANKS3 | chr16 | - | 4701433 | 4702242 | 4701433 | 4701543 | 4702101 | 4702242 | 1 | 8 | 11 | [illegible] |
| RI | 1948 | ENSG00000100714 | MTHFD1 | chr14 | + | 64440125 | 64441453 | 64440125 | 64440266 | 64441384 | 64441453 | 26 | 22 | 22 | [illegible] |
| RI | 1075 | ENSG00000126581 | BECN1 | chr17 | - | 42818543 | 42819609 | 42818543 | 42818680 | 42818786 | 42819609 | 35 | 30 | 45 | [illegible] |
| RI | 1808 | ENSG00000100888 | CHD8 | chr14 | - | 21428963 | 21431858 | 21428963 | 21429335 | 21431810 | 21431858 | 58 | 31 | 55 | [illegible] |
| RI | 1937 | ENSG00000103811 | CTSH | chr15 | - | 78931450 | 78932038 | 78931450 | 78931506 | 78931868 | 78932038 | 13 | 16 | 11 | [illegible] |
| RI | 4081 | ENSG00000116691 | MIIP | chr1 | + | 12029222 | 12029894 | 12029222 | 12029281 | 12029764 | 12029894 | 25 | 12 | 11 | [illegible] |
| RI | 2013 | ENSG00000135506 | OS9 | chr12 | + | 57715759 | 57716193 | 57715759 | 57715970 | 57716088 | 57716193 | 12 | 17 | 20 | [illegible] |
| RI | 2688 | ENSG00000108591 | DRG2 | chr17 | + | 18099016 | 18100435 | 18099016 | 18099077 | 18099632 | 18100435 | 19 | 14 | 24 | [illegible] |
| RI | 2224 | ENSG00000254996 | ANKHD1-EIF4EBP3 | chr5 | + | 140538918 | 140542281 | 140538918 | 140539083 | 140542171 | 140542281 | 21 | 36 | 31 | [illegible] |
| RI | 6222 | ENSG00000127980 | PEX1 | chr7 | - | 92492952 | 92494396 | 92492952 | 92493129 | 92494292 | 92494396 | 68 | 45 | 37 | [illegible] |
| RI | 6113 | ENSG00000237441 | RGL2 | chr6 | - | 33293412 | 33293699 | 33293412 | 33293524 | 33293603 | 33293699 | 36 | 16 | 12 | [illegible] |
| RI | 1488 | ENSG00000172366 | MCRIP2 | chr16 | + | 647416 | 647884 | 647416 | 647544 | 647782 | 647884 | 9 | 1 | 11 | [illegible] |
| RI | 6674 | ENSG00000164062 | APEH | chr3 | + | 49675182 | 49675787 | 49675182 | 49675309 | 49675693 | 49675787 | 8 | 6 | 16 | [illegible] |
| RI | 8133 | ENSG00000107959 | PITRM1 | chr10 | - | 3156929 | 3157531 | 3156929 | 3157064 | 3157434 | 3157531 | 13 | 13 | 21 | [illegible] |
| RI | 5137 | ENSG00000267796 | LIN37 | chr19 | + | 35754245 | 35754519 | 35754245 | 35754319 | 35754392 | 35754519 | 5 | 5 | 1 | [illegible] |
| RI | 799 | ENSG00000131462 | TUBG1 | chr17 | + | 42610422 | 42612506 | 42610422 | 42612143 | 42612426 | 42612506 | 6 | 1 | 2 | [illegible] |
| RI | 6272 | ENSG00000183735 | TBK1 | chr12 | + | 64495482 | 64496406 | 64495482 | 64495604 | 64495698 | 64496406 | 5 | 6 | 2 | [illegible] |
| RI | 4179 | ENSG00000133275 | CSNK1G2 | chr19 | + | 1978858 | 1979248 | 1978858 | 1979093 | 1979162 | 1979248 | 89 | 19 | 67 | [illegible] |
| RI | 7384 | ENSG00000137098 | SPAG8 | chr9 | - | 35810438 | 35810682 | 35810438 | 35810553 | 35810636 | 35810682 | 5 | 2 | 3 | [illegible] |
| RI | 6809 | ENSG00000244754 | N4BP2L2 | chr13 | - | 32538617 | 32538792 | 32538617 | 32538684 | 32538777 | 32538792 | 350 | 316 | 308 | [illegible] |
| RI | 1935 | ENSG00000103811 | CTSH | chr15 | - | 78925333 | 78927781 | 78925333 | 78925440 | 78927712 | 78927781 | 24 | 19 | 25 | [illegible] |
| RI | 6545 | ENSG00000171863 | RPS7 | chr2 | + | 3575324 | 3575684 | 3575324 | 3575448 | 3575591 | 3575684 | 120 | 50 | 132 | [illegible] |
| RI | 4996 | ENSG00000196642 | RABL6 | chr9 | + | 136831720 | 136832370 | 136831720 | 136831861 | 136832261 | 136832370 | 4 | 2 | 3 | [illegible] |
| RI | 8145 | ENSG00000107862 | GBF1 | chr10 | + | 102362474 | 102363396 | 102362474 | 102362671 | 102363262 | 102363396 | 21 | 19 | 17 | [illegible] |
| RI | 1561 | ENSG00000182718 | ANXA2 | chr15 | - | 60351192 | 60351819 | 60351192 | 60351251 | 60351723 | 60351819 | 28 | 34 | 13 | [illegible] |
| RI | 815 | ENSG00000184009 | ACTG1 | chr17 | - | 81510926 | 81511626 | 81510926 | 81511108 | 81511187 | 81511626 | 90 | 40 | 72 | [illegible] |
| RI | 4645 | ENSG00000147274 | RBMX | chrX | - | 136872267 | 136875168 | 136872267 | 136872345 | 136875085 | 136875168 | 246 | 248 | 247 | [illegible] |
| RI | 8441 | ENSG00000157538 | VPS26C | chr21 | - | 37223419 | 37227806 | 37223419 | 37225626 | 37227653 | 37227806 | 15 | 6 | 26 | [illegible] |
| RI | 6497 | ENSG00000123066 | MED13L | chr12 | - | 115980749 | 115982603 | 115980749 | 115980938 | 115982383 | 115982603 | 63 | 49 | 59 | [illegible] |
| RI | 682 | ENSG00000263753 | LINC00667 | chr18 | + | 5238080 | 5241889 | 5238080 | 5239196 | 5241698 | 5241889 | 111 | 65 | 89 | [illegible] |
| RI | 6542 | ENSG00000197448 | GSTK1 | chr7 | + | 143264991 | 143265296 | 143264991 | 143265092 | 143265260 | 143265296 | 286 | 182 | 280 | [illegible] |
| RI | 8281 | ENSG00000241370 | RPP21 | chr6 | + | 30345297 | 30346557 | 30345297 | 30345573 | 30346431 | 30346557 | 8 | 5 | 12 | [illegible] |
| RI | 4317 | ENSG00000173209 | AHSA2P | chr2 | + | 61185501 | 61186497 | 61185501 | 61185618 | 61186445 | 61186497 | 347 | 493 | 460 | [illegible] |

| | | | | | | | | | | | | | | | |
|---|---|---|---|---|---|---|---|---|---|---|---|---|---|---|---|
| RI | 2404 | ENSG00000105875 | WDR91 | chr7 | - | 135186971 | 135189452 | 135186971 | 135187169 | 135189343 | 135189452 | 12 | 10 | 7 | [illegible] |
| RI | 5289 | ENSG00000136718 | IMP4 | chr2 | + | 130345778 | 130346112 | 130345778 | 130345933 | 130346017 | 130346112 | 19 | 6 | 10 | [illegible] |
| RI | 1537 | ENSG00000140263 | SORD | chr15 | + | 45040407 | 45043421 | 45040407 | 45040441 | 45043256 | 45043421 | 1 | 0 | 1 | [illegible] |
| RI | 4904 | ENSG00000147099 | HDAC8 | chrX | - | 72567581 | 72568030 | 72567581 | 72567796 | 72567888 | 72568030 | 12 | 6 | 4 | [illegible] |
| RI | 1182 | ENSG00000140650 | PMM2 | chr16 | + | 8849848 | 8851050 | 8849848 | 8849993 | 8850932 | 8851050 | 12 | 11 | 14 | [illegible] |
| RI | 3221 | ENSG00000184014 | DENND5A | chr11 | - | 9141939 | 9143485 | 9141939 | 9142108 | 9142721 | 9143485 | 18 | 25 | 14 | [illegible] |
| RI | 449 | ENSG00000089639 | GMIP | chr19 | - | 19635414 | 19635721 | 19635414 | 19635569 | 19635643 | 19635721 | 7 | 3 | 5 | [illegible] |
| RI | 7099 | ENSG00000094914 | AAAS | chr12 | - | 53308724 | 53309020 | 53308724 | 53308815 | 53308959 | 53309020 | 1 | 1 | 2 | [illegible] |
| RI | 3543 | ENSG00000247828 | TMEM161B-AS1 | chr5 | + | 88269467 | 88282106 | 88269467 | 88270585 | 88282041 | 88282106 | 52 | 62 | 61 | [illegible] |
| RI | 583 | ENSG00000149636 | DSN1 | chr20 | - | 36770872 | 36771473 | 36770872 | 36771193 | 36771424 | 36771473 | 19 | 7 | 11 | [illegible] |
| RI | 6661 | ENSG00000140543 | DET1 | chr15 | - | 88530622 | 88531715 | 88530622 | 88531473 | 88531612 | 88531715 | 28 | 45 | 53 | [illegible] |
| RI | 7622 | ENSG00000110025 | SNX15 | chr11 | + | 65035519 | 65038829 | 65035519 | 65035663 | 65038571 | 65038829 | 11 | 9 | 13 | [illegible] |
| RI | 5628 | ENSG00000131781 | FMO5 | chr1 | - | 147224894 | 147225339 | 147224894 | 147225066 | 147225286 | 147225339 | 15 | 7 | 12 | [illegible] |
| RI | 6860 | ENSG00000085224 | ATRX | chrX | - | 77681519 | 77684593 | 77681519 | 77683451 | 77683538 | 77684593 | 444 | 345 | 321 | [illegible] |
| RI | 1635 | ENSG00000140553 | UNC45A | chr15 | + | 90935945 | 90936460 | 90935945 | 90935982 | 90936136 | 90936460 | 3 | 2 | 4 | [illegible] |
| RI | 2614 | ENSG00000127419 | TMEM175 | chr4 | + | 948115 | 948615 | 948115 | 948154 | 948510 | 948615 | 42 | 14 | 37 | [illegible] |
| RI | 6902 | ENSG00000090316 | MAEA | chr4 | + | 1336860 | 1338617 | 1336860 | 1336994 | 1338421 | 1338617 | 15 | 9 | 15 | [illegible] |
| RI | 4896 | ENSG00000116984 | MTR | chr1 | + | 236886291 | 236889336 | 236886291 | 236886367 | 236889180 | 236889336 | 0 | 7 | 7 | [illegible] |
| RI | 3643 | ENSG00000239900 | ADSL | chr22 | + | 40360401 | 40361342 | 40360401 | 40360492 | 40361087 | 40361342 | 8 | 8 | 8 | [illegible] |
| RI | 3692 | ENSG00000018510 | AGPS | chr2 | + | 177440964 | 177442486 | 177440964 | 177441036 | 177442406 | 177442486 | 15 | 26 | 33 | [illegible] |
| RI | 5419 | ENSG00000039650 | PNKP | chr19 | - | 49861607 | 49861881 | 49861607 | 49861695 | 49861771 | 49861881 | 11 | 4 | 7 | [illegible] |
| RI | 6032 | ENSG00000234745 | HLA-B | chr6 | - | 31356166 | 31356957 | 31356166 | 31356442 | 31356687 | 31356957 | 142 | 53 | 51 | [illegible] |
| RI | 7893 | ENSG00000204356 | NELFE | chr6 | - | 31956692 | 31957010 | 31956692 | 31956838 | 31956940 | 31957010 | 6 | 2 | 12 | [illegible] |
| RI | 7214 | ENSG00000160948 | VPS28 | chr8 | - | 144424945 | 144425772 | 144424945 | 144425000 | 144425682 | 144425772 | 118 | 43 | 100 | [illegible] |
| RI | 1741 | ENSG00000139998 | RAB15 | chr14 | - | 64950324 | 64951151 | 64950324 | 64950414 | 64950942 | 64951151 | 20 | 7 | 13 | [illegible] |
| RI | 5942 | ENSG00000112282 | MED23 | chr6 | - | 131592387 | 131593171 | 131592387 | 131592460 | 131593005 | 131593171 | 38 | 34 | 44 | [illegible] |
| RI | 186 | ENSG00000104824 | HNRNPL | chr19 | - | 38840095 | 38840559 | 38840095 | 38840376 | 38840487 | 38840559 | 66 | 64 | 72 | [illegible] |
| RI | 4107 | ENSG00000077549 | CAPZB | chr1 | - | 19344357 | 19345252 | 19344357 | 19344434 | 19345186 | 19345252 | 125 | 41 | 82 | [illegible] |
| RI | 8307 | ENSG00000137955 | RABGGTB | chr1 | + | 75789951 | 75790274 | 75789951 | 75790057 | 75790151 | 75790274 | 247 | 244 | 198 | [illegible] |
| RI | 7556 | ENSG00000067064 | IDI1 | chr10 | - | 1043300 | 1044171 | 1043300 | 1043393 | 1043998 | 1044171 | 110 | 80 | 69 | [illegible] |
| RI | 7241 | ENSG00000008988 | RPS20 | chr8 | - | 56073694 | 56074159 | 56073694 | 56073768 | 56074059 | 56074159 | 248 | 208 | 208 | [illegible] |
| RI | 3893 | ENSG00000028310 | BRD9 | chr5 | - | 868606 | 869119 | 868606 | 868804 | 869037 | 869119 | 0 | 10 | 3 | [illegible] |
| RI | 5599 | ENSG00000102125 | TAZ | chrX | + | 154412085 | 154413567 | 154412085 | 154412214 | 154413206 | 154413567 | 9 | 1 | 4 | [illegible] |
| RI | 5057 | ENSG00000106105 | GARS1 | chr7 | + | 30603495 | 30609730 | 30603495 | 30603572 | 30609584 | 30609730 | 35 | 36 | 32 | [illegible] |
| RI | 8328 | ENSG00000089094 | KDM2B | chr12 | - | 121439856 | 121440977 | 121439856 | 121440075 | 121440815 | 121440977 | 15 | 8 | 21 | [illegible] |
| RI | 4867 | ENSG00000135924 | DNAJB2 | chr2 | + | 219280577 | 219281771 | 219280577 | 219280687 | 219281717 | 219281771 | 11 | 7 | 13 | [illegible] |
| RI | 465 | ENSG00000105726 | ATP13A1 | chr19 | - | 19655315 | 19655654 | 19655315 | 19655453 | 19655527 | 19655654 | 3 | 3 | 5 | [illegible] |
| RI | 2955 | ENSG00000134072 | CAMK1 | chr3 | - | 9757346 | 9757846 | 9757346 | 9757621 | 9757728 | 9757846 | 11 | 4 | 23 | [illegible] |
| RI | 3686 | ENSG00000099622 | CIRBP | chr19 | + | 1271139 | 1271632 | 1271139 | 1271467 | 1271550 | 1271632 | 27 | 9 | 16 | [illegible] |
| RI | 3839 | ENSG00000169994 | MYO7B | chr2 | + | 127634175 | 127634683 | 127634175 | 127634289 | 127634595 | 127634683 | 1 | 5 | 2 | [illegible] |
| RI | 7142 | ENSG00000172269 | DPAGT1 | chr11 | - | 119097141 | 119097551 | 119097141 | 119097297 | 119097463 | 119097551 | 24 | 12 | 4 | [illegible] |
| RI | 863 | ENSG00000213339 | QTRT1 | chr19 | + | 10712160 | 10712628 | 10712160 | 10712299 | 10712552 | 10712628 | 16 | 18 | 25 | [illegible] |
| RI | 7978 | ENSG00000187531 | SIRT7 | chr17 | - | 81911938 | 81913880 | 81911938 | 81912614 | 81913773 | 81913880 | 2 | 1 | 6 | [illegible] |
| RI | 6411 | ENSG00000147535 | PLPP5 | chr8 | - | 38267896 | 38268990 | 38267896 | 38267960 | 38268881 | 38268990 | 20 | 24 | 43 | [illegible] |
| RI | 4287 | ENSG00000169228 | RAB24 | chr5 | - | 177301924 | 177302496 | 177301924 | 177301987 | 177302127 | 177302496 | 19 | 15 | 13 | [illegible] |
| RI | 7436 | ENSG00000204394 | VARS1 | chr6 | - | 31781690 | 31781952 | 31781690 | 31781761 | 31781846 | 31781952 | 1 | 0 | 3 | [illegible] |
| RI | 1478 | ENSG00000156858 | PRR14 | chr16 | + | 30652720 | 30653103 | 30652720 | 30652842 | 30652913 | 30653103 | 4 | 6 | 12 | [illegible] |
| RI | 3030 | ENSG00000116350 | SRSF4 | chr1 | - | 29159373 | 29160517 | 29159373 | 29159486 | 29160374 | 29160517 | 38 | 50 | 32 | [illegible] |
| RI | 2979 | ENSG00000009830 | POMT2 | chr14 | - | 77283796 | 77285041 | 77283796 | 77283873 | 77284949 | 77285041 | 7 | 7 | 8 | [illegible] |
| RI | 3237 | ENSG00000095906 | NUBP2 | chr16 | + | 1786536 | 1787831 | 1786536 | 1786955 | 1787676 | 1787831 | 4 | 3 | 10 | [illegible] |
| RI | 3618 | ENSG00000168256 | NKIRAS2 | chr17 | + | 42022398 | 42022640 | 42022398 | 42022472 | 42022564 | 42022640 | 103 | 60 | 59 | [illegible] |
| RI | 7255 | ENSG00000092094 | OSGEP | chr14 | - | 20447614 | 20448171 | 20447614 | 20447690 | 20448105 | 20448171 | 40 | 33 | 73 | [illegible] |
| RI | 1254 | ENSG00000188603 | CLN3 | chr16 | - | 28491481 | 28491835 | 28491481 | 28491560 | 28491713 | 28491835 | 9 | 5 | 14 | [illegible] |
| RI | 8511 | ENSG00000101160 | CTSZ | chr20 | - | 58996638 | 58997753 | 58996638 | 58996801 | 58997602 | 58997753 | 25 | 26 | 5 | [illegible] |
| RI | 1448 | ENSG00000180096 | SEPTIN1 | chr16 | - | 30382092 | 30382365 | 30382092 | 30382179 | 30382274 | 30382365 | 8 | 11 | 14 | [illegible] |
| RI | 4632 | ENSG00000173276 | ZBTB21 | chr21 | - | 41986830 | 41994108 | 41986830 | 41991889 | 41992492 | 41994108 | 61 | 35 | 65 | [illegible] |
| RI | 1399 | ENSG00000182224 | CYB5D1 | chr17 | + | 7858402 | 7858824 | 7858402 | 7858479 | 7858605 | 7858824 | 11 | 0 | 3 | [illegible] |
| RI | 3698 | ENSG00000010292 | NCAPD2 | chr12 | + | 6514264 | 6514587 | 6514264 | 6514392 | 6514463 | 6514587 | 6 | 2 | 10 | [illegible] |
| RI | 668 | ENSG00000173482 | PTPRM | chr18 | + | 8376461 | 8378414 | 8376461 | 8376597 | 8378264 | 8378414 | 11 | 5 | 3 | [illegible] |
| RI | 5547 | ENSG00000283761 | NA | chr1 | + | 100078450 | 100080659 | 100078450 | 100078548 | 100080495 | 100080659 | 34 | 33 | 25 | [illegible] |
| RI | 8160 | ENSG00000150093 | ITGB1 | chr10 | - | 32920244 | 32922346 | 32920244 | 32920385 | 32922256 | 32922346 | 166 | 152 | 101 | [illegible] |
| RI | 7487 | ENSG00000165283 | STOML2 | chr9 | - | 35100597 | 35101011 | 35100597 | 35100726 | 35100931 | 35101011 | 7 | 2 | 13 | [illegible] |
| RI | 2489 | ENSG00000166913 | YWHAB | chr20 | + | 44901530 | 44904116 | 44901530 | 44901833 | 44903992 | 44904116 | 125 | 83 | 111 | [illegible] |
| RI | 4093 | ENSG00000076685 | NT5C2 | chr10 | - | 103097290 | 103098984 | 103097290 | 103097374 | 103098930 | 103098984 | 89 | 106 | 53 | [illegible] |
| RI | 1241 | ENSG00000260565 | ERVK13-1 | chr16 | - | 2669410 | 2672562 | 2669410 | 2669561 | 2671344 | 2672562 | 60 | 39 | 44 | [illegible] |
| RI | 1522 | ENSG00000213614 | HEXA | chr15 | - | 72347685 | 72348134 | 72347685 | 72347758 | 72348030 | 72348134 | 4 | 3 | 2 | [illegible] |
| RI | 5474 | ENSG00000223960 | CHROMR | chr2 | + | 178426365 | 178430889 | 178426365 | 178426661 | 178430769 | 178430889 | 43 | 37 | 24 | [illegible] |
| RI | 1795 | ENSG00000075413 | MARK3 | chr14 | + | 103490951 | 103492034 | 103490951 | 103491107 | 103491776 | 103492034 | 38 | 89 | 48 | [illegible] |
| RI | 6364 | ENSG00000012124 | CD22 | chr19 | + | 35332018 | 35332924 | 35332018 | 35332074 | 35332577 | 35332924 | 44 | 53 | 51 | [illegible] |
| RI | 2940 | ENSG00000165782 | PIP4P1 | chr14 | - | 20457857 | 20459296 | 20457857 | 20458702 | 20459205 | 20459296 | 55 | 28 | 71 | [illegible] |
| RI | 8174 | ENSG00000119977 | TCTN3 | chr10 | - | 95682650 | 95683629 | 95682650 | 95682804 | 95683100 | 95683629 | 5 | 17 | 9 | [illegible] |
| RI | 6490 | ENSG00000146872 | TLK2 | chr17 | + | 62578476 | 62580192 | 62578476 | 62578574 | 62580110 | 62580192 | 32 | 47 | 42 | [illegible] |
| RI | 6335 | ENSG00000136286 | MYO1G | chr7 | - | 44972114 | 44975227 | 44972114 | 44972225 | 44975173 | 44975227 | 76 | 62 | 60 | [illegible] |
| RI | 409 | ENSG00000090013 | BLVRB | chr19 | - | 40458173 | 40458545 | 40458173 | 40458244 | 40458380 | 40458545 | 15 | 13 | 9 | [illegible] |
| RI | 507 | ENSG00000087460 | GNAS | chr20 | + | 58903671 | 58909216 | 58903671 | 58905480 | 58909161 | 58909216 | 41 | 22 | 62 | [illegible] |
| RI | 1462 | ENSG00000140983 | RHOT2 | chr16 | + | 668352 | 668569 | 668352 | 668411 | 668487 | 668569 | 7 | 2 | 7 | [illegible] |
| RI | 7898 | ENSG00000172375 | C2CD2L | chr11 | + | 119111520 | 119112392 | 119111520 | 119111629 | 119112327 | 119112392 | 19 | 10 | 7 | [illegible] |
| RI | 2865 | ENSG00000157601 | MX1 | chr21 | + | 41441715 | 41443866 | 41441715 | 41441914 | 41443787 | 41443866 | 25 | 49 | 30 | [illegible] |
| RI | 7116 | ENSG00000166946 | CCNDBP1 | chr15 | + | 43194080 | 43194461 | 43194080 | 43194137 | 43194414 | 43194461 | 47 | 30 | 30 | [illegible] |
| RI | 8510 | ENSG00000101160 | CTSZ | chr20 | - | 58995184 | 58997753 | 58995184 | 58995759 | 58997602 | 58997753 | 20 | 21 | 19 | [illegible] |
| RI | 4490 | ENSG00000104894 | CD37 | chr19 | + | 49339329 | 49340606 | 49339329 | 49339413 | 49340250 | 49340606 | 69 | 43 | 84 | [illegible] |
| RI | 1415 | ENSG00000158805 | ZNF276 | chr16 | + | 89723136 | 89723709 | 89723136 | 89723183 | 89723278 | 89723709 | 72 | 29 | 67 | [illegible] |
| RI | 7080 | ENSG00000110697 | PITPNM1 | chr11 | - | 67496177 | 67497436 | 67496177 | 67496345 | 67497230 | 67497436 | 10 | 6 | 3 | [illegible] |
| RI | 5985 | ENSG00000196262 | PPIA | chr7 | + | 44799391 | 44799874 | 44799391 | 44799480 | 44799701 | 44799874 | 307 | 196 | 312 | [illegible] |
| RI | 7604 | ENSG00000152556 | PFKM | chr12 | + | 48134942 | 48135383 | 48134942 | 48135038 | 48135290 | 48135383 | 7 | 7 | 13 | [illegible] |
| RI | 2163 | ENSG00000065357 | DGKA | chr12 | + | 55939414 | 55940433 | 55939414 | 55939529 | 55940313 | 55940433 | 122 | 113 | 221 | [illegible] |
| RI | 5562 | ENSG00000120697 | ALG5 | chr13 | - | 36993603 | 36995596 | 36993603 | 36993672 | 36995424 | 36995596 | 48 | 23 | 35 | [illegible] |
| RI | 6670 | ENSG00000109736 | MFSD10 | chr4 | - | 2932599 | 2933209 | 2932599 | 2932758 | 2933105 | 2933209 | 8 | 15 | 11 | [illegible] |
| RI | 5490 | ENSG00000198830 | HMGN2 | chr1 | + | 26473702 | 26474135 | 26473702 | 26473817 | 26474084 | 26474135 | 303 | 153 | 325 | [illegible] |
| RI | 5854 | ENSG00000108465 | CDK5RAP3 | chr17 | + | 47974399 | 47975337 | 47974399 | 47974448 | 47975158 | 47975337 | 366 | 302 | 337 | [illegible] |
| RI | 4650 | ENSG00000101361 | NOP56 | chr20 | + | 2657080 | 2658393 | 2657080 | 2657218 | 2657928 | 2658393 | 22 | 23 | 20 | [illegible] |
| RI | 2899 | ENSG00000172053 | QARS1 | chr3 | - | 49098352 | 49098692 | 49098352 | 49098480 | 49098599 | 49098692 | 24 | 7 | 22 | [illegible] |
| RI | 6653 | ENSG00000182220 | ATP6AP2 | chrX | + | 40599591 | 40600881 | 40599591 | 40599741 | 40600761 | 40600881 | 27 | 9 | 20 | [illegible] |
| RI | 1554 | ENSG00000182511 | FES | chr15 | + | 90885036 | 90885585 | 90885036 | 90885258 | 90885411 | 90885585 | 6 | 5 | 13 | [illegible] |
| RI | 7398 | ENSG00000197070 | ARRDC1 | chr9 | + | 137613614 | 137614214 | 137613614 | 137613769 | 137614139 | 137614214 | 127 | 42 | 73 | [illegible] |
| RI | 26 | ENSG00000141959 | PFKL | chr21 | + | 44319350 | 44321875 | 44319350 | 44320147 | 44321728 | 44321875 | 3 | 1 | 8 | [illegible] |
| RI | 6813 | ENSG00000169359 | SLC33A1 | chr3 | - | 155833467 | 155834041 | 155833467 | 155833585 | 155833856 | 155834041 | 72 | 43 | 68 | [illegible] |
| RI | 4144 | ENSG00000132300 | PTCD3 | chr2 | + | 86133177 | 86133436 | 86133177 | 86133256 | 86133345 | 86133436 | 59 | 49 | 56 | [illegible] |
| RI | 6124 | ENSG00000168036 | CTNNB1 | chr3 | + | 41233340 | 41233867 | 41233340 | 41233444 | 41233528 | 41233867 | 33 | 41 | 47 | [illegible] |
| RI | 6338 | ENSG00000106617 | PRKAG2 | chr7 | - | 151565719 | 151566699 | 151565719 | 151565885 | 151566613 | 151566699 | 34 | 23 | 19 | [illegible] |
| RI | 7493 | ENSG00000213930 | GALT | chr9 | + | 34647491 | 34647705 | 34647491 | 34647567 | 34647656 | 34647705 | 30 | 16 | 35 | [illegible] |
| RI | 7118 | ENSG00000226091 | LINC00937 | chr12 | - | 8390269 | 8390969 | 8390269 | 8390511 | 8390901 | 8390969 | 4 | 5 | 6 | [illegible] |
| RI | 6128 | ENSG00000168036 | CTNNB1 | chr3 | + | 41239133 | 41239904 | 41239133 | 41239354 | 41239818 | 41239904 | 232 | 140 | 237 | [illegible] |
| RI | 4149 | ENSG00000178467 | P4HTM | chr3 | + | 49004860 | 49005867 | 49004860 | 49005046 | 49005776 | 49005867 | 28 | 4 | 36 | [illegible] |
| RI | 7368 | ENSG00000140988 | RPS2 | chr16 | - | 1962496 | 1963256 | 1962496 | 1962909 | 1963148 | 1963256 | 408 | 188 | 433 | [illegible] |
| RI | 1576 | ENSG00000159459 | UBR1 | chr15 | - | 43025373 | 43026663 | 43025373 | 43025429 | 43026560 | 43026663 | 34 | 36 | 15 | [illegible] |
| RI | 6486 | ENSG00000065000 | AP3D1 | chr19 | - | 2112859 | 2113413 | 2112859 | 2112931 | 2113335 | 2113413 | 14 | 9 | 13 | [illegible] |
| RI | 5935 | ENSG00000179222 | MAGED1 | chrX | + | 51897794 | 51898193 | 51897794 | 51897886 | 51898113 | 51898193 | 0 | 0 | 4 | [illegible] |
| RI | 3976 | ENSG00000164405 | UQCRQ | chr5 | + | 132866868 | 132867687 | 132866868 | 132867035 | 132867487 | 132867687 | 137 | 64 | 97 | [illegible] |
| RI | 5632 | ENSG00000178719 | GRINA | chr8 | + | 143991691 | 143992373 | 143991691 | 143991804 | 143991877 | 143992373 | 18 | 4 | 7 | [illegible] |
| RI | 8411 | ENSG00000159403 | C1R | chr12 | - | 7091451 | 7092441 | 7091451 | 7091680 | 7092386 | 7092441 | 4 | 8 | 10 | [illegible] |
| RI | 5194 | ENSG00000115355 | CCDC88A | chr2 | - | 55290654 | 55291775 | 55290654 | 55291164 | 55291675 | 55291775 | 23 | 22 | 15 | [illegible] |
| RI | 3026 | ENSG00000178149 | DALRD3 | chr3 | - | 49015972 | 49016340 | 49015972 | 49016086 | 49016157 | 49016340 | 14 | 13 | 15 | [illegible] |
| RI | 753 | ENSG00000005961 | ITGA2B | chr17 | - | 44384310 | 44384585 | 44384310 | 44384354 | 44384537 | 44384585 | 6 | 11 | 0 | [illegible] |
| RI | 2834 | ENSG00000114745 | GORASP1 | chr3 | - | 39100304 | 39100877 | 39100304 | 39100503 | 39100787 | 39100877 | 23 | 9 | 23 | [illegible] |
| RI | 3296 | ENSG00000165801 | ARHGEF40 | chr14 | + | 21088829 | 21089545 | 21088829 | 21088876 | 21089013 | 21089545 | 16 | 15 | 6 | [illegible] |
| RI | 4847 | ENSG00000142327 | RNPEPL1 | chr2 | + | 240575029 | 240575610 | 240575029 | 240575142 | 240575501 | 240575610 | 3 | 2 | 15 | [illegible] |
| RI | 4288 | ENSG00000169228 | RAB24 | chr5 | - | 177302127 | 177302496 | 177302127 | 177302178 | 177302396 | 177302496 | 108 | 82 | 65 | [illegible] |
| RI | 6686 | ENSG00000174080 | CTSF | chr11 | - | 66564087 | 66564648 | 66564087 | 66564146 | 66564557 | 66564648 | 6 | 5 | 8 | [illegible] |
| RI | 2952 | ENSG00000164068 | RNF123 | chr3 | + | 49720799 | 49721105 | 49720799 | 49720894 | 49721019 | 49721105 | 32 | 14 | 49 | [illegible] |
| RI | 2924 | ENSG00000243989 | ACY1 | chr3 | + | 51985851 | 51986331 | 51985851 | 51985946 | 51986254 | 51986331 | 1 | 0 | 2 | [illegible] |
| RI | 383 | ENSG00000129347 | KRI1 | chr19 | - | 10557551 | 10557895 | 10557551 | 10557682 | 10557768 | 10557895 | 33 | 14 | 40 | [illegible] |
| RI | 233 | ENSG00000129351 | ILF3 | chr19 | + | 10683916 | 10685765 | 10683916 | 10683970 | 10684415 | 10685765 | 30 | 7 | 35 | [illegible] |
| RI | 6099 | ENSG00000246067 | RAB30-DT | chr11 | + | 83072622 | 83073604 | 83072622 | 83072804 | 83073302 | 83073604 | 41 | 18 | 27 | [illegible] |
| RI | 2441 | ENSG00000083838 | ZNF446 | chr19 | + | 58479642 | 58480019 | 58479642 | 58479727 | 58479929 | 58480019 | 9 | 8 | 10 | [illegible] |
| RI | 8401 | ENSG00000125967 | NECAB3 | chr20 | - | 33669374 | 33669712 | 33669374 | 33669472 | 33669686 | 33669712 | 6 | 7 | 5 | [illegible] |
| RI | 8262 | ENSG00000182199 | SHMT2 | chr12 | + | 57234002 | 57234726 | 57234002 | 57234110 | 57234233 | 57234726 | 4 | 2 | 2 | [illegible] |
| RI | 3605 | ENSG00000104381 | GDAP1 | chr8 | + | 74361883 | 74363053 | 74361883 | 74361978 | 74362938 | 74363053 | 7 | 7 | 2 | [illegible] |
| RI | 796 | ENSG00000141252 | VPS53 | chr17 | - | 708140 | 710613 | 708140 | 708787 | 710532 | 710613 | 22 | 17 | 22 | [illegible] |
| RI | 948 | ENSG00000132522 | GPS2 | chr17 | - | 7313567 | 7313988 | 7313567 | 7313721 | 7313905 | 7313988 | 98 | 62 | 70 | [illegible] |
| RI | 6015 | ENSG00000130340 | SNX9 | chr6 | + | 157937433 | 157938747 | 157937433 | 157937523 | 157938632 | 157938747 | 17 | 31 | 30 | [illegible] |
| RI | 2984 | ENSG00000134107 | BHLHE40 | chr3 | + | 4979961 | 4980408 | 4979961 | 4980031 | 4980300 | 4980408 | 70 | 29 | 69 | [illegible] |
| RI | 903 | ENSG00000007202 | KIAA0100 | chr17 | - | 28639574 | 28640025 | 28639574 | 28639642 | 28639900 | 28640025 | 31 | 39 | 44 | [illegible] |
| RI | 3727 | ENSG00000142208 | AKT1 | chr14 | - | 104773454 | 104773980 | 104773454 | 104773580 | 104773911 | 104773980 | 23 | 1 | 10 | [illegible] |
| RI | 5463 | ENSG00000239779 | WBP1 | chr2 | + | 74459437 | 74459745 | 74459437 | 74459562 | 74459642 | 74459745 | 22 | 10 | 50 | [illegible] |
| RI | 8312 | ENSG00000102225 | CDK16 | chrX | + | 47226993 | 47227223 | 47226993 | 47227099 | 47227159 | 47227223 | 8 | 12 | 13 | [illegible] |
| RI | 5786 | ENSG00000119535 | CSF3R | chr1 | - | 36466042 | 36466827 | 36466042 | 36466200 | 36466620 | 36466827 | 228 | 119 | 200 | [illegible] |
| RI | 1336 | ENSG00000090861 | AARS1 | chr16 | - | 70255727 | 70258217 | 70255727 | 70255836 | 70258032 | 70258217 | 5 | 2 | 10 | [illegible] |
| RI | 6950 | ENSG00000215301 | DDX3X | chrX | + | 41337407 | 41339083 | 41337407 | 41337465 | 41339038 | 41339083 | 127 | 119 | 115 | [illegible] |
| RI | 8583 | ENSG00000188549 | CCDC9B | chr15 | - | 40338747 | 40339619 | 40338747 | 40338903 | 40339511 | 40339619 | 2 | 0 | 2 | [illegible] |
| RI | 6288 | ENSG00000169660 | HEXD | chr17 | + | 82440996 | 82441266 | 82440996 | 82441075 | 82441164 | 82441266 | 16 | 7 | 23 | [illegible] |
| RI | 6718 | ENSG00000094631 | HDAC6 | chrX | + | 48814674 | 48814893 | 48814674 | 48814740 | 48814833 | 48814893 | 17 | 15 | 21 | [illegible] |
| RI | 5058 | ENSG00000106105 | GARS1 | chr7 | + | 30621392 | 30622462 | 30621392 | 30621500 | 30622316 | 30622462 | 22 | 18 | 20 | [illegible] |
| RI | 6370 | ENSG00000010318 | PHF7 | chr3 | + | 52421647 | 52422881 | 52421647 | 52421754 | 52422759 | 52422881 | 6 | 18 | 1 | [illegible] |
| RI | 7045 | ENSG00000221914 | PPP2R2A | chr8 | + | 26362683 | 26363890 | 26362683 | 26362848 | 26363720 | 26363890 | 27 | 17 | 18 | [illegible] |
| RI | 1213 | ENSG00000086504 | MRPL28 | chr16 | - | 368500 | 370225 | 368500 | 369220 | 369930 | 370225 | 3 | 1 | 5 | [illegible] |
| RI | 8456 | ENSG00000123349 | PFDN5 | chr12 | + | 53297849 | 53298150 | 53297849 | 53297924 | 53298018 | 53298150 | 18 | 26 | 28 | [illegible] |
| RI | 7801 | ENSG00000255717 | SNHG1 | chr11 | - | 62852810 | 62853115 | 62852810 | 62852839 | 62853074 | 62853115 | 375 | 253 | 462 | [illegible] |
| RI | 5879 | ENSG00000111679 | PTPN6 | chr12 | + | 6951608 | 6952177 | 6951608 | 6951731 | 6952016 | 6952177 | 250 | 137 | 205 | [illegible] |
| RI | 5299 | ENSG00000143207 | COP1 | chr1 | - | 175988287 | 175989479 | 175988287 | 175988412 | 175989361 | 175989479 | 35 | 56 | 26 | [illegible] |
| RI | 2767 | ENSG00000087266 | SH3BP2 | chr4 | + | 2832989 | 2834289 | 2832989 | 2833049 | 2833696 | 2834289 | 25 | 22 | 26 | [illegible] |
| RI | 5403 | ENSG00000114735 | HEMK1 | chr3 | + | 50579843 | 50580229 | 50579843 | 50579939 | 50580115 | 50580229 | 16 | 0 | 13 | [illegible] |
| RI | 1370 | ENSG00000149929 | HIRIP3 | chr16 | - | 29995102 | 29995463 | 29995102 | 29995217 | 29995342 | 29995463 | 0 | 2 | 4 | [illegible] |
| RI | 6065 | ENSG00000204351 | SKIV2L | chr6 | + | 31962713 | 31963091 | 31962713 | 31962798 | 31962984 | 31963091 | 16 | 8 | 3 | [illegible] |
| RI | 3358 | ENSG00000213024 | NUP62 | chr19 | - | 49927693 | 49929504 | 49927693 | 49927815 | 49929416 | 49929504 | 29 | 21 | 42 | [illegible] |
| RI | 1048 | ENSG00000006282 | SPATA20 | chr17 | + | 50549984 | 50550312 | 50549984 | 50550115 | 50550207 | 50550312 | 12 | 1 | 10 | [illegible] |
| RI | 7169 | ENSG00000213782 | DDX47 | chr12 | + | 12823202 | 12824677 | 12823202 | 12824016 | 12824539 | 12824677 | 40 | 55 | 97 | [illegible] |
| RI | 6852 | ENSG00000215021 | PHB2 | chr12 | - | 6967170 | 6968021 | 6967170 | 6967335 | 6967891 | 6968021 | 30 | 19 | 46 | [illegible] |
| RI | 4169 | ENSG00000120889 | TNFRSF10B | chr8 | - | 23028330 | 23028602 | 23028330 | 23028441 | 23028528 | 23028602 | 135 | 115 | 88 | [illegible] |
| RI | 8323 | ENSG00000196498 | NCOR2 | chr12 | - | 124334423 | 124335280 | 124334423 | 124334617 | 124335134 | 124335280 | 0 | 0 | 1 | [illegible] |
| RI | 2492 | ENSG00000167995 | BEST1 | chr11 | + | 61958145 | 61959578 | 61958145 | 61958298 | 61959497 | 61959578 | 16 | 9 | 8 | [illegible] |
| RI | 6514 | ENSG00000122566 | HNRNPA2B1 | chr7 | - | 26196806 | 26197461 | 26196806 | 26197017 | 26197314 | 26197461 | 425 | 389 | 445 | [illegible] |
| RI | 2841 | ENSG00000013441 | CLK1 | chr2 | - | 200859679 | 200861466 | 200859679 | 200859746 | 200861237 | 200861466 | 1040 | 1074 | 939 | [illegible] |
| RI | 5150 | ENSG00000177000 | MTHFR | chr1 | - | 11802880 | 11803526 | 11802880 | 11803129 | 11803404 | 11803526 | 31 | 18 | 29 | [illegible] |
| RI | 8600 | ENSG00000134899 | ERCC5 | chr13 | + | 102872197 | 102873343 | 102872197 | 102872398 | 102873258 | 102873343 | 21 | 38 | 16 | [illegible] |
| RI | 1341 | ENSG00000157353 | FCSK | chr16 | + | 70466881 | 70467966 | 70466881 | 70467471 | 70467885 | 70467966 | 5 | 3 | 3 | [illegible] |
| RI | 5993 | ENSG00000004139 | SARM1 | chr17 | + | 28388173 | 28388539 | 28388173 | 28388276 | 28388359 | 28388539 | 8 | 5 | 8 | [illegible] |
| RI | 7101 | ENSG00000094914 | AAAS | chr12 | - | 53315093 | 53315426 | 53315093 | 53315140 | 53315334 | 53315426 | 21 | 9 | 11 | [illegible] |
| RI | 3450 | ENSG00000144591 | GMPPA | chr2 | + | 219501475 | 219502037 | 219501475 | 219501579 | 219501911 | 219502037 | 34 | 17 | 37 | [illegible] |
| RI | 6568 | ENSG00000164610 | RP9 | chr7 | - | 33097270 | 33099436 | 33097270 | 33097362 | 33099306 | 33099436 | 21 | 30 | 25 | [illegible] |
| RI | 7461 | ENSG00000266714 | MYO15B | chr17 | + | 75590906 | 75591246 | 75590906 | 75591016 | 75591171 | 75591246 | 42 | 29 | 28 | [illegible] |
| RI | 7707 | ENSG00000132274 | TRIM22 | chr11 | + | 5697247 | 5698545 | 5697247 | 5697343 | 5698314 | 5698545 | 362 | 491 | 431 | [illegible] |
| RI | 1581 | ENSG00000157734 | SNX22 | chr15 | + | 64152242 | 64152742 | 64152242 | 64152326 | 64152637 | 64152742 | 21 | 5 | 8 | [illegible] |
| RI | 1114 | ENSG00000205560 | CPT1B | chr22 | - | 50571157 | 50571539 | 50571157 | 50571292 | 50571374 | 50571539 | 3 | 3 | 2 | [illegible] |
| RI | 1649 | ENSG00000196547 | MAN2A2 | chr15 | + | 90912042 | 90912664 | 90912042 | 90912279 | 90912541 | 90912664 | 11 | 11 | 8 | [illegible] |
| RI | 8246 | ENSG00000284770 | TBCE | chr1 | + | 235436385 | 235436608 | 235436385 | 235436450 | 235436543 | 235436608 | 32 | 10 | 13 | [illegible] |
| RI | 7409 | ENSG00000159069 | FBXW5 | chr9 | - | 136942045 | 136942695 | 136942045 | 136942466 | 136942546 | 136942695 | 20 | 4 | 20 | [illegible] |
| RI | 3102 | ENSG00000175110 | MRPS22 | chr3 | + | 139350178 | 139351060 | 139350178 | 139350322 | 139351030 | 139351060 | 63 | 37 | 74 | [illegible] |
| RI | 3571 | ENSG00000120008 | WDR11 | chr10 | + | 120905875 | 120906855 | 120905875 | 120906021 | 120906775 | 120906855 | 60 | 62 | 64 | [illegible] |
| RI | 2651 | ENSG00000203875 | SNHG5 | chr6 | - | 85678135 | 85678733 | 85678135 | 85678188 | 85678693 | 85678733 | 121 | 143 | 203 | [illegible] |
| RI | 2669 | ENSG00000143624 | INTS3 | chr1 | + | 153773192 | 153774808 | 153773192 | 153773331 | 153774517 | 153774808 | 88 | 45 | 60 | [illegible] |
| RI | 8006 | ENSG00000158545 | ZC3H18 | chr16 | + | 88622196 | 88623344 | 88622196 | 88622388 | 88623218 | 88623344 | 9 | 0 | 13 | [illegible] |
| RI | 1494 | ENSG00000008710 | PKD1 | chr16 | - | 2091780 | 2092188 | 2091780 | 2091906 | 2092046 | 2092188 | 5 | 0 | 2 | [illegible] |
| RI | 2766 | ENSG00000087266 | SH3BP2 | chr4 | + | 2831922 | 2832412 | 2831922 | 2831978 | 2832330 | 2832412 | 29 | 25 | 26 | [illegible] |
| RI | 4088 | ENSG00000136068 | FLNB | chr3 | + | 58130740 | 58132931 | 58130740 | 58130908 | 58132807 | 58132931 | 15 | 15 | 9 | [illegible] |
| RI | 3357 | ENSG00000213024 | NUP62 | chr19 | - | 49927693 | 49929504 | 49927693 | 49927848 | 49929325 | 49929504 | 22 | 18 | 37 | [illegible] |
| RI | 5564 | ENSG00000120697 | ALG5 | chr13 | - | 36993603 | 36995596 | 36993603 | 36993672 | 36995426 | 36995596 | 49 | 23 | 39 | [illegible] |
| RI | 8200 | ENSG00000108528 | SLC25A11 | chr17 | - | 4938153 | 4938429 | 4938153 | 4938253 | 4938338 | 4938429 | 4 | 0 | 6 | [illegible] |
| RI | 5658 | ENSG00000123143 | PKN1 | chr19 | + | 14467564 | 14467793 | 14467564 | 14467641 | 14467717 | 14467793 | 9 | 3 | 12 | [illegible] |
| RI | 7471 | ENSG00000135047 | CTSL | chr9 | + | 87728584 | 87729735 | 87728584 | 87728809 | 87729572 | 87729735 | 2 | 2 | 0 | [illegible] |
| RI | 4408 | ENSG00000160050 | CCDC28B | chr1 | + | 32204597 | 32205387 | 32204597 | 32204620 | 32205193 | 32205387 | 32 | 13 | 21 | [illegible] |
| RI | 8566 | ENSG00000160961 | ZNF333 | chr19 | + | 14716993 | 14717733 | 14716993 | 14717089 | 14717656 | 14717733 | 11 | 11 | 10 | [illegible] |
| RI | 2680 | ENSG00000005700 | IBTK | chr6 | - | 82191072 | 82191879 | 82191072 | 82191216 | 82191786 | 82191879 | 41 | 23 | 37 | [illegible] |
| RI | 827 | ENSG00000141349 | G6PC3 | chr17 | + | 44074968 | 44075451 | 44074968 | 44075087 | 44075309 | 44075451 | 1 | 0 | 2 | [illegible] |
| RI | 4048 | ENSG00000116670 | MAD2L2 | chr1 | - | 11680352 | 11680613 | 11680352 | 11680471 | 11680561 | 11680613 | 1 | 2 | 0 | [illegible] |
| RI | 3332 | ENSG00000117228 | GBP1 | chr1 | - | 89057991 | 89059043 | 89057991 | 89058234 | 89058840 | 89059043 | 24 | 66 | 37 | [illegible] |
| RI | 793 | ENSG00000141252 | VPS53 | chr17 | - | 511531 | 512625 | 511531 | 511743 | 512371 | 512625 | 16 | 14 | 13 | [illegible] |
| RI | 6014 | ENSG00000130340 | SNX9 | chr6 | + | 157906127 | 157909790 | 157906127 | 157906212 | 157909664 | 157909790 | 26 | 37 | 50 | [illegible] |
| RI | 6701 | ENSG00000130734 | ATG4D | chr19 | + | 10546838 | 10547253 | 10546838 | 10547071 | 10547188 | 10547253 | 17 | 8 | 16 | [illegible] |
| RI | 8143 | ENSG00000107862 | GBF1 | chr10 | + | 102362474 | 102363396 | 102362474 | 102362628 | 102363255 | 102363396 | 45 | 45 | 33 | [illegible] |
| RI | 7595 | ENSG00000143549 | TPM3 | chr1 | - | 154170648 | 154172978 | 154170648 | 154170711 | 154172907 | 154172978 | 175 | 174 | 139 | [illegible] |
| RI | 2426 | ENSG00000032742 | IFT88 | chr13 | + | 20601704 | 20601933 | 20601704 | 20601748 | 20601850 | 20601933 | 52 | 56 | 46 | [illegible] |
| RI | 8555 | ENSG00000117226 | GBP3 | chr1 | - | 89013184 | 89014279 | 89013184 | 89013427 | 89014082 | 89014279 | 59 | 25 | 46 | [illegible] |
| RI | 2060 | ENSG00000133706 | LARS1 | chr5 | - | 146140203 | 146143084 | 146140203 | 146140261 | 146142871 | 146143084 | 48 | 55 | 53 | [illegible] |
| RI | 4585 | ENSG00000106462 | EZH2 | chr7 | - | 148816683 | 148817391 | 148816683 | 148816778 | 148817221 | 148817391 | 11 | 11 | 21 | [illegible] |
| RI | 3171 | ENSG00000233452 | STXBP5-AS1 | chr6 | - | 146844318 | 146844712 | 146844318 | 146844522 | 146844637 | 146844712 | 11 | 7 | 3 | [illegible] |
| RI | 4565 | ENSG00000132510 | KDM6B | chr17 | + | 7853209 | 7854796 | 7853209 | 7853380 | 7853497 | 7854796 | 15 | 11 | 11 | [illegible] |
| RI | 1069 | ENSG00000197381 | ADARB1 | chr21 | + | 45222017 | 45226560 | 45222017 | 45222107 | 45225546 | 45226560 | 66 | 52 | 59 | [illegible] |
| RI | 6905 | ENSG00000214026 | MRPL23 | chr11 | + | 1956255 | 1956609 | 1956255 | 1956448 | 1956541 | 1956609 | 26 | 5 | 13 | [illegible] |

| | | | | | | | | | | | | | | | | | | | | | | | | | | | | | | |
|---|---|---|---|---|---|---|---|---|---|---|---|---|---|---|---|---|---|---|---|---|---|---|---|---|---|---|---|---|---|---|
| RI | 2668 | ENSG00000143624 | INTS3 | chr1 | + | 153772339 | 153772711 | 153772339 | 153772440 | 153772638 | 153772711 | 7 | 13 | 10 | 17 | 8 | 36 | 8 | 15 | 19 | 17 | 15 | 12 | 17 | 14 | 26 | 36 | 37 | 37 | 24 | 59 |
| RI | 3695 | ENSG00000145916 | RMND5B | chr5 | + | 178142582 | 178142992 | 178142582 | 178142728 | 178142851 | 178142992 | 9 | 7 | 8 | 16 | 10 | 24 | 18 | 31 | 18 | 4 | 0 | 6 | 6 | 22 | 7 | 31 | 7 | 7 | 19 | 42 |
| RI | 5761 | ENSG00000157456 | CCNB2 | chr15 | + | 59107321 | 59107670 | 59107321 | 59107450 | 59107556 | 59107670 | 1 | 0 | 2 | 0 | 0 | 3 | 1 | 10 | 2 | 4 | 1 | 0 | 0 | 13 | 0 | 0 | 14 | 22 | 38 | 8 |
| RI | 3785 | ENSG00000142676 | RPL11 | chr1 | + | 23694659 | 23695908 | 23694659 | 23694791 | 23695797 | 23695908 | 123 | 76 | 141 | 126 | 104 | 8051 | 5759 | 8203 | 7479 | 11598 | 136 | 135 | 125 | 196 | 101 | 11980 | 5067 | 7115 | 7297 | 10394 |
| RI | 860 | ENSG00000132382 | MYBBP1A | chr17 | - | 4553809 | 4554093 | 4553809 | 4553917 | 4554018 | 4554093 | 11 | 1 | 0 | 1 | 3 | 28 | 12 | 40 | 21 | 39 | 0 | 8 | 8 | 3 | 4 | 43 | 35 | 3 | 34 | 34 |
| RI | 7452 | ENSG00000167106 | FAM102A | chr9 | - | 127940880 | 127944684 | 127940880 | 127943237 | 127944614 | 127944684 | 6 | 20 | 31 | 13 | 8 | 198 | 84 | 210 | 144 | 119 | 0 | 2 | 7 | 1 | 46 | 12 | 139 | 61 | 22 | 68 |
| RI | 3061 | ENSG00000169727 | GPS1 | chr17 | + | 82056628 | 82057470 | 82056628 | 82056766 | 82056839 | 82057470 | 4 | 1 | 5 | 6 | 6 | 74 | 36 | 76 | 86 | 60 | 0 | 9 | 3 | 2 | 5 | 131 | 84 | 45 | 50 | 98 |
| RI | 7033 | ENSG00000136279 | DBNL | chr7 | + | 44058431 | 44058983 | 44058431 | 44058480 | 44058901 | 44058983 | 56 | 43 | 73 | 45 | 45 | 266 | 207 | 307 | 318 | 366 | 53 | 73 | 82 | 62 | 101 | 618 | 403 | 230 | 335 | 908 |
| RI | 1641 | ENSG00000140400 | MAN2C1 | chr15 | - | 75358461 | 75358808 | 75358461 | 75358618 | 75358703 | 75358808 | 49 | 2 | 67 | 43 | 8 | 43 | 14 | 10 | 20 | 22 | 106 | 38 | 68 | 1 | 0 | 11 | 37 | 17 | 32 | 72 |
| RI | 5749 | ENSG00000169045 | HNRNPH1 | chr5 | - | 179616125 | 179617110 | 179616125 | 179616218 | 179617020 | 179617110 | 600 | 706 | 618 | 615 | 497 | 128 | 111 | 62 | 109 | 80 | 132 | 746 | 981 | 311 | 57 | 66 | 40 | 38 | 57 | 10 |
| RI | 7274 | ENSG00000104522 | GFUS | chr8 | - | 143613523 | 143613817 | 143613523 | 143613603 | 143613750 | 143613817 | 1 | 2 | 3 | 2 | 0 | 50 | 19 | 45 | 50 | 17 | 0 | 1 | 7 | 4 | 6 | 86 | 78 | 20 | 51 | 107 |
| RI | 498 | ENSG00000104897 | SF3A2 | chr19 | + | 2246752 | 2247022 | 2246752 | 2246802 | 2246881 | 2247022 | 20 | 11 | 7 | 28 | 9 | 141 | 55 | 139 | 127 | 120 | 9 | 16 | 13 | 18 | 16 | 168 | 137 | 107 | 81 | 198 |
| RI | 3223 | ENSG00000184014 | DENND5A | chr11 | - | 9142721 | 9143485 | 9142721 | 9142845 | 9143402 | 9143485 | 9 | 25 | 18 | 16 | 19 | 146 | 97 | 134 | 129 | 146 | 49 | 21 | 36 | 29 | 22 | 297 | 168 | 121 | 216 | 223 |
| RI | 5196 | ENSG00000115355 | CCDC88A | chr2 | - | 55299838 | 55301277 | 55299838 | 55299919 | 55301205 | 55301277 | 10 | 21 | 21 | 17 | 10 | 110 | 113 | 85 | 117 | 141 | 0 | 7 | 14 | 1 | 7 | 33 | 34 | 12 | 11 | 0 |
| RI | 567 | ENSG00000185513 | L3MBTL1 | chr20 | + | 43528656 | 43529368 | 43528656 | 43528745 | 43529263 | 43529368 | 6 | 3 | 4 | 12 | 3 | 0 | 1 | 4 | 1 | 2 | 5 | 7 | 10 | 9 | 12 | 0 | 0 | 5 | 3 | 0 |
| RI | 4072 | ENSG00000010270 | STARD3NL | chr7 | + | 38215027 | 38217305 | 38215027 | 38215105 | 38217187 | 38217305 | 19 | 18 | 9 | 26 | 14 | 17 | 13 | 12 | 10 | 30 | 15 | 9 | 13 | 13 | 2 | 13 | 4 | 7 | 14 | 53 |
| RI | 1278 | ENSG00000103051 | COG4 | chr16 | - | 70508405 | 70509388 | 70508405 | 70508464 | 70509230 | 70509388 | 30 | 27 | 45 | 39 | 26 | 60 | 26 | 34 | 39 | 49 | 76 | 21 | 36 | 41 | 80 | 64 | 57 | 20 | 32 | 72 |
| RI | 5283 | ENSG00000155366 | RHOC | chr1 | - | 112702562 | 112703119 | 112702562 | 112702693 | 112702998 | 112703119 | 13 | 1 | 5 | 13 | 2 | 427 | 219 | 145 | 281 | 131 | 49 | 5 | 6 | 2 | 5 | 443 | 215 | 232 | 149 | 1006 |
| RI | 3740 | ENSG00000114626 | ABTB1 | chr3 | + | 127677464 | 127677843 | 127677464 | 127677563 | 127677675 | 127677843 | 19 | 7 | 42 | 19 | 9 | 45 | 4 | 34 | 29 | 24 | 17 | 32 | 25 | 13 | 60 | 11 | 46 | 18 | 27 | 58 |
| RI | 1163 | ENSG00000125124 | BBS2 | chr16 | - | 56498436 | 56499907 | 56498436 | 56498693 | 56499777 | 56499907 | 38 | 32 | 22 | 32 | 18 | 9 | 5 | 5 | 4 | 9 | 3 | 14 | 35 | 3 | 14 | 0 | 2 | 5 | 5 | 0 |
| RI | 5563 | ENSG00000120697 | ALG5 | chr13 | - | 36994988 | 36995596 | 36994988 | 36995035 | 36995424 | 36995596 | 29 | 25 | 20 | 19 | 28 | 96 | 54 | 73 | 65 | 102 | 4 | 29 | 31 | 11 | 2 | 35 | 32 | 49 | 35 | 13 |
| RI | 2962 | ENSG00000010295 | IFFO1 | chr12 | - | 6548064 | 6548545 | 6548064 | 6548157 | 6548424 | 6548545 | 47 | 9 | 24 | 17 | 15 | 34 | 29 | 37 | 57 | 23 | 33 | 4 | 20 | 20 | 47 | 98 | 29 | 28 | 50 | 61 |
| RI | 4145 | ENSG00000025708 | TYMP | chr22 | - | 50526000 | 50526476 | 50526000 | 50526141 | 50526245 | 50526476 | 5 | 5 | 0 | 6 | 0 | 37 | 26 | 15 | 33 | 1 | 24 | 0 | 1 | 19 | 0 | 89 | 43 | 8 | 34 | 41 |
| RI | 4826 | ENSG00000185163 | DDX51 | chr12 | - | 132141497 | 132141956 | 132141497 | 132141606 | 132141849 | 132141956 | 18 | 9 | 8 | 25 | 6 | 13 | 10 | 11 | 12 | 10 | 16 | 21 | 17 | 37 | 2 | 8 | 28 | 16 | 9 | 0 |
| RI | 2875 | ENSG00000146192 | FGD2 | chr6 | + | 37027428 | 37029069 | 37027428 | 37027575 | 37027947 | 37029069 | 13 | 6 | 6 | 5 | 8 | 76 | 49 | 40 | 77 | 20 | 9 | 27 | 7 | 8 | 40 | 151 | 109 | 18 | 42 | 209 |
| RI | 4986 | ENSG00000014914 | MTMR11 | chr1 | - | 149930364 | 149930965 | 149930364 | 149930547 | 149930791 | 149930965 | 17 | 21 | 7 | 8 | 7 | 66 | 47 | 32 | 55 | 37 | 28 | 21 | 7 | 19 | 24 | 259 | 68 | 25 | 68 | 103 |
| RI | 6182 | ENSG00000214021 | TTLL3 | chr3 | + | 9817644 | 9820741 | 9817644 | 9817759 | 9820545 | 9820741 | 107 | 85 | 85 | 100 | 68 | 2 | 3 | 0 | 2 | 5 | 188 | 121 | 153 | 184 | 232 | 0 | 5 | 2 | 6 | 0 |
| RI | 8382 | ENSG00000135452 | TSPAN31 | chr12 | + | 57746175 | 57746720 | 57746175 | 57746256 | 57746588 | 57746720 | 20 | 20 | 23 | 28 | 20 | 62 | 36 | 35 | 31 | 43 | 56 | 14 | 38 | 22 | 20 | 53 | 25 | 14 | 43 | 78 |
| RI | 6863 | ENSG00000126759 | CFP | chrX | - | 47628101 | 47629674 | 47628101 | 47628277 | 47629523 | 47629674 | 6 | 3 | 2 | 3 | 6 | 200 | 81 | 98 | 114 | 51 | 0 | 7 | 0 | 10 | 24 | 196 | 149 | 81 | 133 | 348 |
| RI | 7698 | ENSG00000068831 | RASGRP2 | chr11 | - | 64735541 | 64735980 | 64735541 | 64735664 | 64735902 | 64735980 | 7 | 5 | 6 | 7 | 3 | 174 | 73 | 192 | 171 | 101 | 0 | 6 | 7 | 6 | 0 | 179 | 122 | 71 | 78 | 184 |
| RI | 8556 | ENSG00000121310 | ECHDC2 | chr1 | - | 52897436 | 52899224 | 52897436 | 52897484 | 52899173 | 52899224 | 7 | 7 | 18 | 11 | 3 | 36 | 10 | 14 | 20 | 14 | 2 | 11 | 10 | 26 | 11 | 30 | 6 | 4 | 22 | 24 |
| RI | 2073 | ENSG00000137221 | TJAP1 | chr6 | + | 43500743 | 43501687 | 43500743 | 43500772 | 43501525 | 43501687 | 6 | 3 | 16 | 3 | 1 | 50 | 23 | 46 | 33 | 54 | 5 | 6 | 11 | 1 | 13 | 43 | 37 | 32 | 39 | 45 |
| RI | 7020 | ENSG00000142046 | TMEM91 | chr19 | + | 41382771 | 41384083 | 41382771 | 41382921 | 41383714 | 41384083 | 27 | 25 | 35 | 35 | 18 | 5 | 0 | 2 | 9 | 2 | 82 | 44 | 58 | 61 | 172 | 11 | 2 | 1 | 2 | 15 |
| RI | 7797 | ENSG00000162231 | NXF1 | chr11 | - | 62798538 | 62800486 | 62798538 | 62798575 | 62800376 | 62800486 | 179 | 148 | 173 | 253 | 292 | 89 | 52 | 79 | 110 | 229 | 465 | 259 | 194 | 361 | 521 | 184 | 162 | 78 | 140 | 211 |
| RI | 2850 | ENSG00000114857 | NKTR | chr3 | + | 42619663 | 42621516 | 42619663 | 42619708 | 42621428 | 42621516 | 425 | 467 | 363 | 408 | 340 | 17 | 13 | 6 | 13 | 19 | 57 | 171 | 203 | 41 | 15 | 7 | 3 | 5 | 0 | 0 |

| | | | | | | | | | | | | | | | | | | | | | | | | | | | | | | | | | | | | | | | | | | | | | | |
|---|---|---|---|---|---|---|---|---|---|---|---|---|---|---|---|---|---|---|---|---|---|---|---|---|---|---|---|---|---|---|---|---|---|---|---|---|---|---|---|---|---|---|---|---|---|---|
| RI | 5952 | ENSG00000197756 | RPL37A | chr2 | + | 216499948 | 216501467 | 216499948 | 216500031 | 216501340 | 216501467 | 136 | 83 | 151 | 129 | 161 | 3800 | 1721 | 4187 | 3369 | 5599 | 62 | 80 | 91 | 108 | 63 | 2146 | 2450 | 1609 | 1634 | 2542 | 298 | 149 | 0.398563144 | 0.805168431 | 0.018 | 0.024 | 0.018 | 0.019 | 0.014 | 0.014 | 0.016 | 0.028 | 0.032 | 0.012 | -0.002 |
| RI | 1790 | ENSG00000092108 | SCFD1 | chr14 | + | 30673256 | 30673997 | 30673256 | 30673347 | 30673930 | 30673997 | 24 | 44 | 33 | 31 | 36 | 4 | 5 | 4 | 2 | 10 | 1 | 17 | 10 | 12 | 3 | 0 | 0 | 3 | 0 | 5 | 298 | 149 | 0.398754524 | 0.805279649 | 0.75 | 0.815 | 0.805 | 0.886 | 0.643 | 1 | 1 | 0.625 | 1 | 0.231 | 0.009 |
| RI | 7812 | ENSG00000255717 | SNHG1 | chr11 | - | 62855132 | 62855885 | 62855132 | 62855216 | 62855423 | 62855885 | 162 | 69 | 233 | 255 | 147 | 81 | 41 | 88 | 74 | 106 | 142 | 199 | 180 | 147 | 39 | 63 | 60 | 116 | 38 | 17 | 298 | 149 | 0.39963212 | 0.806473304 | 0.5 | 0.457 | 0.57 | 0.633 | 0.409 | 0.53 | 0.624 | 0.437 | 0.659 | 0.534 | -0.043 |
| RI | 5688 | ENSG00000138385 | SSB | chr2 | + | 169808476 | 169808902 | 169808476 | 169808553 | 169808859 | 169808902 | 48 | 41 | 62 | 83 | 66 | 237 | 236 | 192 | 211 | 294 | 26 | 29 | 34 | 40 | 21 | 40 | 140 | 219 | 158 | 26 | 298 | 149 | 0.399636825 | 0.806473304 | 0.092 | 0.08 | 0.139 | 0.164 | 0.101 | 0.245 | 0.094 | 0.072 | 0.112 | 0.288 | -0.047 |
| RI | 4251 | ENSG00000011009 | LYPLA2 | chr1 | + | 23794223 | 23794600 | 23794223 | 23794325 | 23794426 | 23794600 | 20 | 6 | 18 | 28 | 29 | 33 | 13 | 14 | 15 | 36 | 12 | 53 | 15 | 16 | 37 | 38 | 30 | 9 | 6 | 31 | 250 | 149 | 0.399866394 | 0.806473304 | 0.265 | 0.216 | 0.434 | 0.527 | 0.324 | 0.158 | 0.513 | 0.498 | 0.614 | 0.416 | -0.087 |
| RI | 8014 | ENSG00000143702 | CEP170 | chr1 | - | 243185778 | 243186422 | 243185778 | 243186072 | 243186258 | 243186422 | 170 | 182 | 126 | 148 | 163 | 71 | 48 | 50 | 57 | 82 | 26 | 118 | 141 | 31 | 11 | 0 | 42 | 19 | 11 | 22 | 298 | 149 | 0.399891706 | 0.806473304 | 0.545 | 0.655 | 0.558 | 0.565 | 0.498 | 1 | 0.584 | 0.788 | 0.585 | 0.2 | -0.067 |
| RI | 3527 | ENSG00000084072 | PPIE | chr1 | + | 39741894 | 39743297 | 39741894 | 39741921 | 39743215 | 39743297 | 31 | 21 | 28 | 29 | 28 | 139 | 88 | 72 | 119 | 235 | 38 | 20 | 26 | 23 | 40 | 138 | 80 | 92 | 130 | 163 | 298 | 149 | 0.400161385 | 0.806513255 | 0.1 | 0.107 | 0.163 | 0.109 | 0.056 | 0.121 | 0.111 | 0.124 | 0.081 | 0.109 | -0.002 |
| RI | 2484 | ENSG00000077232 | DNAJC10 | chr2 | + | 182717015 | 182718290 | 182717015 | 182717072 | 182718156 | 182718290 | 352 | 262 | 259 | 333 | 560 | 7 | 4 | 4 | 3 | 5 | 145 | 254 | 219 | 72 | 84 | 5 | 4 | 6 | 1 | 0 | 298 | 149 | 0.400277773 | 0.806513255 | 0.962 | 0.97 | 0.97 | 0.982 | 0.982 | 0.935 | 0.969 | 0.948 | 0.973 | 1 | 0.008 |
| RI | 6336 | ENSG00000136286 | MYO1G | chr7 | - | 44972114 | 44975227 | 44972114 | 44972274 | 44975173 | 44975227 | 86 | 67 | 64 | 70 | 72 | 16 | 4 | 5 | 8 | 12 | 199 | 138 | 125 | 105 | 204 | 32 | 21 | 6 | 14 | 43 | 298 | 149 | 0.400321121 | 0.806513255 | 0.729 | 0.893 | 0.865 | 0.814 | 0.75 | 0.757 | 0.767 | 0.912 | 0.789 | 0.703 | 0.025 |
| RI | 3647 | ENSG00000239900 | ADSL | chr22 | + | 40361487 | 40363071 | 40361487 | 40361666 | 40362980 | 40363071 | 22 | 18 | 19 | 21 | 21 | 2 | 2 | 1 | 3 | 0 | 15 | 52 | 46 | 95 | 40 | 0 | 3 | 5 | 1 | 0 | 298 | 149 | 0.401620128 | 0.808854448 | 0.846 | 0.818 | 0.905 | 0.778 | 1 | 1 | 0.897 | 0.821 | 0.979 | 1 | -0.07 |
| RI | 4556 | ENSG00000135749 | PCNX2 | chr1 | - | 233217898 | 233218184 | 233217898 | 233217931 | 233218030 | 233218184 | 19 | 4 | 19 | 7 | 23 | 79 | 44 | 71 | 85 | 156 | 13 | 3 | 6 | 2 | 10 | 24 | 46 | 73 | 56 | 45 | 248 | 149 | 0.402579019 | 0.810509293 | 0.126 | 0.052 | 0.139 | 0.047 | 0.081 | 0.246 | 0.038 | 0.047 | 0.021 | 0.118 | -0.005 |
| RI | 197 | ENSG00000105258 | POLR2I | chr19 | - | 36114338 | 36114713 | 36114338 | 36114412 | 36114658 | 36114713 | 11 | 6 | 2 | 10 | 13 | 22 | 6 | 31 | 26 | 13 | 0 | 13 | 4 | 6 | 11 | 13 | 35 | 21 | 10 | 18 | 298 | 149 | 0.403015962 | 0.810930954 | 0.2 | 0.333 | 0.031 | 0.161 | 0.333 | 0 | 0.157 | 0.087 | 0.231 | 0.234 | 0.07 |
| RI | 956 | ENSG00000129667 | RHBDF2 | chr17 | - | 76478805 | 76479277 | 76478805 | 76479009 | 76479081 | 76479277 | 2 | 4 | 4 | 0 | 0 | 61 | 26 | 25 | 32 | 13 | 0 | 3 | 0 | 4 | 4 | 52 | 78 | 29 | 30 | 142 | 221 | 149 | 0.403063024 | 0.810930954 | 0.022 | 0.094 | 0.097 | 0 | 0 | 0 | 0.025 | 0 | 0.082 | 0.019 | 0.017 |
| RI | 1473 | ENSG00000140983 | RHOT2 | chr16 | + | 671859 | 672181 | 671859 | 672002 | 672083 | 672181 | 1 | 0 | 5 | 7 | 0 | 17 | 8 | 13 | 14 | 3 | 8 | 5 | 2 | 0 | 34 | 0 | 24 | 29 | 15 | 47 | 230 | 149 | 0.403913545 | 0.812022449 | 0.037 | 0 | 0.199 | 0.245 | 0 | 1 | 0.119 | 0.043 | 0 | 0.319 | -0.2 |
| RI | 510 | ENSG00000087460 | GNAS | chr20 | + | 58905382 | 58909579 | 58905382 | 58909423 | 58909520 | 58909579 | 26 | 22 | 24 | 31 | 23 | 1830 | 1183 | 1706 | 2476 | 2134 | 38 | 52 | 33 | 41 | 1 | 2704 | 2487 | 1412 | 1776 | 9318 | 246 | 149 | 0.403999562 | 0.812022449 | 0.009 | 0.011 | 0.008 | 0.008 | 0.006 | 0.008 | 0.013 | 0.014 | 0.014 | 0 | -0.001 |
| RI | 1865 | ENSG00000100823 | APEX1 | chr14 | + | 20455225 | 20455703 | 20455225 | 20455394 | 20455577 | 20455703 | 85 | 64 | 83 | 80 | 54 | 167 | 72 | 147 | 131 | 78 | 101 | 38 | 64 | 70 | 164 | 167 | 84 | 65 | 103 | 200 | 298 | 149 | 0.404017941 | 0.812022449 | 0.203 | 0.308 | 0.22 | 0.234 | 0.257 | 0.232 | 0.184 | 0.33 | 0.254 | 0.291 | -0.014 |
| RI | 2249 | ENSG00000241106 | HLA-DOB | chr6 | - | 32813722 | 32814601 | 32813722 | 32813833 | 32814319 | 32814601 | 39 | 18 | 11 | 34 | 19 | 15 | 5 | 15 | 12 | 27 | 27 | 38 | 13 | 14 | 36 | 25 | 24 | 5 | 7 | 60 | 298 | 149 | 0.404256973 | 0.81222651 | 0.565 | 0.643 | 0.268 | 0.586 | 0.26 | 0.351 | 0.442 | 0.565 | 0.5 | 0.231 | 0.047 |
| RI | 259 | ENSG00000142541 | RPL13A | chr19 | + | 49489849 | 49490297 | 49489849 | 49489922 | 49490231 | 49490297 | 81 | 55 | 99 | 89 | 70 | 12021 | 4703 | 13109 | 9474 | 13052 | 169 | 101 | 72 | 78 | 221 | 5818 | 7716 | 5279 | 4104 | 9979 | 298 | 149 | 0.404470219 | 0.812378641 | 0.003 | 0.006 | 0.004 | 0.005 | 0.003 | 0.014 | 0.007 | 0.007 | 0.009 | 0.011 | -0.005 |
| RI | 1432 | ENSG00000140848 | CPNE2 | chr16 | + | 57146084 | 57147964 | 57146084 | 57146321 | 57147550 | 57147964 | 3 | 2 | 3 | 0 | 0 | 36 | 10 | 27 | 18 | 6 | 0 | 0 | 3 | 7 | 8 | 47 | 25 | 20 | 42 | 85 | 298 | 149 | 0.405902073 | 0.814977412 | 0.04 | 0.091 | 0.053 | 0 | 0 | 0 | 0 | 0.07 | 0.077 | 0.045 | -0.002 |
| RI | 394 | ENSG00000167613 | LAIR1 | chr19 | - | 54356229 | 54356398 | 54356229 | 54356267 | 54356355 | 54356398 | 12 | 13 | 10 | 13 | 8 | 205 | 101 | 147 | 120 | 114 | 17 | 6 | 12 | 12 | 17 | 186 | 107 | 72 | 109 | 192 | 237 | 149 | 0.410327368 | 0.823582658 | 0.035 | 0.075 | 0.041 | 0.064 | 0.042 | 0.054 | 0.034 | 0.095 | 0.065 | 0.053 | -0.009 |
| RI | 2189 | ENSG00000145604 | SKP2 | chr5 | + | 36152108 | 36153042 | 36152108 | 36152270 | 36152770 | 36153042 | 9 | 0 | 4 | 5 | 1 | 16 | 4 | 11 | 17 | 3 | 9 | 4 | 4 | 0 | 0 | 0 | 2 | 3 | 5 | 5 | 298 | 149 | 0.41084171 | 0.824334912 | 0.22 | 0 | 0.154 | 0.128 | 0.143 | 1 | 0.5 | 0.4 | 0 | 0 | -0.251 |
| RI | 1658 | ENSG00000140365 | COMMD4 | chr15 | + | 75338061 | 75338420 | 75338061 | 75338133 | 75338354 | 75338420 | 9 | 7 | 11 | 17 | 2 | 45 | 10 | 27 | 22 | 41 | 9 | 8 | 7 | 7 | 17 | 30 | 30 | 42 | 36 | 53 | 298 | 149 | 0.411160205 | 0.824693831 | 0.091 | 0.259 | 0.169 | 0.279 | 0.024 | 0.13 | 0.118 | 0.077 | 0.089 | 0.138 | 0.054 |
| RI | 2667 | ENSG00000143624 | INTS3 | chr1 | + | 153746956 | 153748755 | 153746956 | 153747070 | 153748688 | 153748755 | 52 | 50 | 72 | 64 | 43 | 0 | 0 | 2 | 0 | 1 | 97 | 51 | 49 | 82 | 95 | 0 | 0 | 0 | 8 | 0 | 298 | 149 | 0.41160822 | 0.825312206 | 1 | 1 | 0.947 | 1 | 0.956 | 1 | 1 | 1 | 0.837 | 1 | 0.013 |
| RI | 6762 | ENSG00000119760 | SUPT7L | chr2 | - | 27660983 | 27662281 | 27660983 | 27661388 | 27662178 | 27662281 | 122 | 118 | 118 | 188 | 346 | 48 | 37 | 44 | 60 | 62 | 155 | 222 | 253 | 130 | 65 | 69 | 31 | 52 | 72 | 62 | 298 | 149 | 0.411812594 | 0.825441802 | 0.56 | 0.615 | 0.573 | 0.61 | 0.736 | 0.529 | 0.782 | 0.709 | 0.474 | 0.344 | 0.051 |
| RI | 617 | ENSG00000124224 | PPP4R1L | chr20 | - | 58245782 | 58247506 | 58245782 | 58246252 | 58247321 | 58247506 | 13 | 8 | 7 | 23 | 10 | 1 | 4 | 4 | 2 | 0 | 1 | 10 | 27 | 32 | 5 | 0 | 2 | 8 | 18 | 0 | 298 | 149 | 0.412611092 | 0.826682391 | 0.867 | 0.5 | 0.467 | 0.852 | 1 | 1 | 0.714 | 0.628 | 0.471 | 1 | -0.025 |
| RI | 1566 | ENSG00000166783 | MARF1 | chr16 | - | 15634756 | 15636342 | 15634756 | 15634931 | 15635655 | 15636342 | 50 | 22 | 51 | 36 | 74 | 138 | 149 | 175 | 179 | 217 | 103 | 49 | 56 | 52 | 11 | 259 | 183 | 212 | 194 | 50 | 298 | 149 | 0.412711422 | 0.826682391 | 0.153 | 0.069 | 0.127 | 0.091 | 0.146 | 0.166 | 0.118 | 0.117 | 0.118 | 0.099 | -0.006 |
| RI | 1644 | ENSG00000177082 | WDR73 | chr15 | - | 84645470 | 84646348 | 84645470 | 84645836 | 84645972 | 84646348 | 48 | 33 | 31 | 52 | 22 | 1 | 0 | 5 | 3 | 0 | 44 | 34 | 44 | 43 | 55 | 0 | 4 | 3 | 2 | 4 | 285 | 149 | 0.412971165 | 0.826702917 | 0.962 | 1 | 0.764 | 0.901 | 1 | 1 | 0.816 | 0.885 | 0.918 | 0.878 | 0.026 |
| RI | 7588 | ENSG00000159314 | ARHGAP27 | chr17 | - | 45404600 | 45405106 | 45404600 | 45404681 | 45405027 | 45405106 | 144 | 71 | 87 | 122 | 59 | 12 | 2 | 0 | 6 | 4 | 126 | 139 | 93 | 116 | 139 | 0 | 6 | 4 | 11 | 2 | 298 | 149 | 0.413001576 | 0.826702917 | 0.857 | 0.947 | 1 | 0.91 | 0.881 | 1 | 0.921 | 0.921 | 0.841 | 0.972 | -0.012 |
| RI | 791 | ENSG00000214087 | ARL16 | chr17 | - | 81682033 | 81683126 | 81682033 | 81682675 | 81683012 | 81683126 | 15 | 10 | 5 | 12 | 4 | 17 | 9 | 24 | 12 | 10 | 0 | 13 | 34 | 12 | 2 | 5 | 35 | 12 | 11 | 45 | 298 | 149 | 0.413684312 | 0.82764407 | 0.306 | 0.357 | 0.094 | 0.333 | 0.167 | 0 | 0.157 | 0.586 | 0.353 | 0.022 | 0.028 |
| RI | 68 | ENSG00000161179 | YDJC | chr22 | - | 21629009 | 21629761 | 21629009 | 21629407 | 21629601 | 21629761 | 43 | 20 | 36 | 44 | 14 | 2 | 0 | 2 | 1 | 1 | 6 | 40 | 39 | 29 | 45 | 0 | 0 | 1 | 0 | 0 | 298 | 149 | 0.413832768 | 0.82764407 | 0.915 | 1 | 0.9 | 0.957 | 0.875 | 1 | 1 | 0.951 | 1 | 1 | -0.061 |
| RI | 1345 | ENSG00000090857 | PDPR | chr16 | + | 70128783 | 70129122 | 70128783 | 70128865 | 70128958 | 70129122 | 14 | 4 | 0 | 17 | 3 | 22 | 7 | 30 | 33 | 23 | 8 | 5 | 3 | 0 | 106 | 17 | 17 | 23 | 7 | 62 | 242 | 149 | 0.413892091 | 0.82764407 | 0.282 | 0.26 | 0 | 0.241 | 0.074 | 0.225 | 0.153 | 0.074 | 0 | 0.513 | -0.022 |
| RI | 5166 | ENSG00000167895 | TMC8 | chr17 | + | 78137234 | 78137814 | 78137234 | 78137358 | 78137716 | 78137814 | 72 | 55 | 54 | 77 | 54 | 136 | 90 | 131 | 235 | 61 | 28 | 42 | 60 | 55 | 64 | 86 | 183 | 82 | 72 | 158 | 298 | 149 | 0.414419558 | 0.827721197 | 0.209 | 0.234 | 0.171 | 0.141 | 0.307 | 0.14 | 0.103 | 0.268 | 0.276 | 0.168 | 0.021 |
| RI | 5238 | ENSG00000138073 | PREB | chr2 | - | 27131671 | 27132082 | 27131671 | 27131831 | 27132009 | 27132082 | 18 | 5 | 5 | 11 | 12 | 17 | 11 | 17 | 13 | 6 | 13 | 0 | 4 | 2 | 23 | 0 | 17 | 8 | 12 | 36 | 298 | 149 | 0.414480255 | 0.827721197 | 0.346 | 0.185 | 0.128 | 0.297 | 0.5 | 1 | 0 | 0.2 | 0.077 | 0.242 | -0.013 |
| RI | 3110 | ENSG00000054967 | RELT | chr11 | + | 73394476 | 73395285 | 73394476 | 73394734 | 73395086 | 73395285 | 23 | 2 | 13 | 13 | 6 | 29 | 28 | 34 | 50 | 12 | 38 | 25 | 10 | 22 | 24 | 104 | 87 | 71 | 64 | 90 | 298 | 149 | 0.414525243 | 0.827721197 | 0.284 | 0.034 | 0.16 | 0.115 | 0.2 | 0.154 | 0.126 | 0.066 | 0.147 | 0.118 | 0.036 |
| RI | 7410 | ENSG00000159069 | FBXW5 | chr9 | - | 136942546 | 136942943 | 136942546 | 136942695 | 136942768 | 136942943 | 36 | 20 | 37 | 34 | 14 | 51 | 25 | 44 | 46 | 22 | 28 | 28 | 23 | 18 | 51 | 21 | 81 | 21 | 32 | 117 | 222 | 149 | 0.414705186 | 0.827721197 | 0.321 | 0.349 | 0.361 | 0.332 | 0.299 | 0.472 | 0.188 | 0.424 | 0.274 | 0.226 | 0.016 |
| RI | 473 | ENSG00000105135 | ILVBL | chr19 | - | 15116407 | 15118015 | 15116407 | 15116522 | 15117880 | 15118015 | 1 | 8 | 3 | 17 | 3 | 41 | 17 | 29 | 10 | 2 | 0 | 3 | 5 | 5 | 7 | 13 | 28 | 3 | 23 | 18 | 298 | 149 | 0.414768074 | 0.827721197 | 0.012 | 0.19 | 0.049 | 0.459 | 0.429 | 0 | 0.051 | 0.455 | 0.098 | 0.163 | 0.074 |
| RI | 1647 | ENSG00000177082 | WDR73 | chr15 | - | 84647889 | 84648625 | 84647889 | 84647954 | 84648536 | 84648625 | 42 | 37 | 49 | 40 | 48 | 24 | 24 | 42 | 29 | 29 | 33 | 22 | 46 | 54 | 33 | 27 | 31 | 42 | 21 | 29 | 298 | 149 | 0.414817752 | 0.827721197 | 0.467 | 0.435 | 0.368 | 0.408 | 0.453 | 0.379 | 0.262 | 0.354 | 0.562 | 0.363 | 0.042 |
| RI | 943 | ENSG00000215041 | NEURL4 | chr17 | - | 7318268 | 7318674 | 7318268 | 7318356 | 7318494 | 7318674 | 4 | 3 | 2 | 7 | 0 | 12 | 5 | 11 | 11 | 5 | 0 | 10 | 0 | 0 | 9 | 6 | 31 | 11 | 12 | 8 | 287 | 149 | 0.414911539 | 0.827721197 | 0.148 | 0.238 | 0.086 | 0.248 | 0 | 0 | 0.143 | 0 | 0 | 0.369 | 0.042 |
| RI | 4998 | ENSG00000110446 | SLC15A3 | chr11 | - | 60941121 | 60942145 | 60941121 | 60941290 | 60942034 | 60942145 | 18 | 11 | 19 | 26 | 11 | 34 | 19 | 28 | 19 | 5 | 39 | 5 | 14 | 35 | 53 | 47 | 72 | 31 | 25 | 33 | 298 | 149 | 0.41624554 | 0.830102094 | 0.209 | 0.224 | 0.253 | 0.406 | 0.524 | 0.293 | 0.034 | 0.184 | 0.412 | 0.445 | 0.05 |
| RI | 6359 | ENSG00000075624 | ACTB | chr7 | - | 5528280 | 5529400 | 5528280 | 5528719 | 5529160 | 5529400 | 152 | 90 | 148 | 138 | 176 | 5712 | 2377 | 4193 | 4574 | 4615 | 299 | 196 | 149 | 170 | 564 | 6280 | 5672 | 2822 | 5355 | 24163 | 298 | 149 | 0.416557889 | 0.830444634 | 0.013 | 0.019 | 0.017 | 0.015 | 0.019 | 0.023 | 0.017 | 0.026 | 0.016 | 0.012 | -0.002 |
| RI | 368 | ENSG00000205744 | DENND1C | chr19 | - | 6475483 | 6475751 | 6475483 | 6475585 | 6475705 | 6475751 | 7 | 3 | 9 | 5 | 11 | 58 | 49 | 72 | 70 | 8 | 31 | 0 | 14 | 15 | 12 | 58 | 88 | 56 | 87 | 82 | 269 | 149 | 0.416875005 | 0.830796442 | 0.063 | 0.033 | 0.065 | 0.038 | 0.432 | 0.228 | 0 | 0.122 | 0.087 | 0.075 | 0.024 |
| RI | 1211 | ENSG00000086504 | MRPL28 | chr16 | - | 368500 | 369220 | 368500 | 368635 | 369067 | 369220 | 27 | 6 | 14 | 22 | 32 | 107 | 46 | 74 | 85 | 82 | 35 | 14 | 21 | 36 | 78 | 152 | 178 | 130 | 158 | 196 | 298 | 149 | 0.418146825 | 0.833050015 | 0.112 | 0.061 | 0.086 | 0.115 | 0.163 | 0.103 | 0.038 | 0.075 | 0.102 | 0.166 | 0.011 |
| RI | 2147 | ENSG00000105698 | USF2 | chr19 | + | 35278697 | 35279074 | 35278697 | 35278792 | 35278945 | 35279074 | 48 | 30 | 63 | 33 | 117 | 566 | 469 | 416 | 580 | 706 | 102 | 59 | 52 | 96 | 54 | 722 | 604 | 344 | 457 | 1269 | 298 | 149 | 0.419109898 | 0.834687177 | 0.041 | 0.031 | 0.07 | 0.028 | 0.077 | 0.066 | 0.047 | 0.07 | 0.095 | 0.021 | -0.01 |
| RI | 8163 | ENSG00000150093 | ITGB1 | chr10 | - | 32932514 | 32933583 | 32932514 | 32932600 | 32933481 | 32933583 | 100 | 85 | 40 | 63 | 77 | 5 | 9 | 7 | 3 | 2 | 16 | 53 | 44 | 12 | 0 | 0 | 13 | 5 | 0 | 4 | 298 | 149 | 0.41944346 | 0.834960951 | 0.909 | 0.825 | 0.741 | 0.913 | 0.951 | 1 | 0.671 | 0.815 | 1 | 0 | 0.171 |
| RI | 5153 | ENSG00000264538 | SUZ12P1 | chr17 | + | 30768833 | 30769064 | 30768833 | 30768867 | 30769000 | 30769064 | 173 | 155 | 203 | 163 | 321 | 5 | 13 | 8 | 14 | 21 | 158 | 231 | 121 | 267 | 266 | 45 | 11 | 1 | 1 | 60 | 282 | 149 | 0.419664249 | 0.834960951 | 0.948 | 0.863 | 0.931 | 0.86 | 0.89 | 0.65 | 0.917 | 0.985 | 0.993 | 0.701 | 0.049 |
| RI | 851 | ENSG00000141503 | MINK1 | chr17 | + | 4889646 | 4890735 | 4889646 | 4889763 | 4890516 | 4890735 | 6 | 8 | 8 | 13 | 12 | 34 | 5 | 18 | 43 | 10 | 13 | 21 | 17 | 2 | 37 | 18 | 28 | 12 | 14 | 77 | 298 | 149 | 0.419671418 | 0.834960951 | 0.081 | 0.444 | 0.182 | 0.131 | 0.375 | 0.265 | 0.273 | 0.415 | 0.067 | 0.194 | 0 |
| RI | 2960 | ENSG00000009790 | TRAF3IP3 | chr1 | + | 209775348 | 209775736 | 209775348 | 209775489 | 209775598 | 209775736 | 134 | 89 | 138 | 106 | 45 | 576 | 474 | 594 | 584 | 565 | 43 | 108 | 76 | 69 | 31 | 378 | 498 | 278 | 307 | 496 | 258 | 149 | 0.420372998 | 0.836075184 | 0.118 | 0.098 | 0.118 | 0.095 | 0.044 | 0.062 | 0.111 | 0.136 | 0.115 | 0.035 | 0.003 |
| RI | 5641 | ENSG00000163516 | ANKZF1 | chr2 | + | 219234132 | 219235875 | 219234132 | 219235312 | 219235707 | 219235875 | 145 | 124 | 203 | 172 | 104 | 4 | 4 | 13 | 5 | 8 | 225 | 300 | 222 | 214 | 248 | 10 | 4 | 5 | 9 | 15 | 298 | 149 | 0.420730039 | 0.836503649 | 0.948 | 0.939 | 0.886 | 0.945 | 0.867 | 0.918 | 0.974 | 0.957 | 0.922 | 0.892 | -0.016 |
| RI | 7007 | ENSG00000168066 | SF1 | chr11 | - | 64764605 | 64766155 | 64764605 | 64765518 | 64765950 | 64766155 | 158 | 86 | 186 | 134 | 108 | 0 | 2 | 5 | 3 | 0 | 305 | 285 | 159 | 211 | 207 | 10 | 0 | 0 | 0 | 0 | 298 | 149 | 0.421064316 | 0.836886579 | 1 | 0.956 | 0.949 | 0.957 | 1 | 0.938 | 1 | 1 | 1 | 1 | -0.015 |
| RI | 1959 | ENSG00000213780 | GTF2H4 | chr6 | + | 30911683 | 30912146 | 30911683 | 30911767 | 30912013 | 30912146 | 16 | 15 | 21 | 18 | 32 | 17 | 15 | 14 | 18 | 27 | 9 | 17 | 7 | 3 | 10 | 32 | 9 | 5 | 5 | 6 | 298 | 149 | 0.422427977 | 0.839063137 | 0.32 | 0.333 | 0.429 | 0.333 | 0.372 | 0.123 | 0.486 | 0.412 | 0.231 | 0.455 | 0.016 |
| RI | 5159 | ENSG00000166801 | FAM111A | chr11 | + | 59143496 | 59145856 | 59143496 | 59143695 | 59145826 | 59145856 | 373 | 453 | 262 | 473 | 447 | 23 | 32 | 33 | 36 | 91 | 98 | 278 | 249 | 74 | 40 | 0 | 25 | 26 | 25 | 0 | 298 | 149 | 0.422459128 | 0.839063137 | 0.89 | 0.876 | 0.799 | 0.868 | 0.711 | 1 | 0.848 | 0.827 | 0.597 | 1 | -0.026 |
| RI | 4329 | ENSG00000115548 | KDM3A | chr2 | + | 86478607 | 86480362 | 86478607 | 86478735 | 86480166 | 86480362 | 64 | 50 | 76 | 85 | 60 | 40 | 25 | 33 | 39 | 81 | 82 | 76 | 81 | 43 | 4 | 4 | 41 | 82 | 24 | 5 | 298 | 149 | 0.422585548 | 0.839063137 | 0.444 | 0.5 | 0.535 | 0.521 | 0.27 | 0.911 | 0.481 | 0.331 | 0.473 | 0.286 | -0.042 |
| RI | 3721 | ENSG00000138002 | IFT172 | chr2 | - | 27458778 | 27459522 | 27458778 | 27458868 | 27459377 | 27459522 | 7 | 3 | 2 | 4 | 3 | 8 | 10 | 7 | 9 | 3 | 17 | 0 | 1 | 4 | 5 | 0 | 4 | 3 | 2 | 13 | 298 | 149 | 0.423717764 | 0.841028505 | 0.304 | 0.13 | 0.125 | 0.182 | 0.333 | 1 | 0 | 0.143 | 0.5 | 0.161 | -0.146 |
| RI | 4975 | ENSG00000163382 | NAXE | chr1 | + | 156592556 | 156593555 | 156592556 | 156592670 | 156593407 | 156593555 | 28 | 8 | 6 | 22 | 16 | 65 | 51 | 65 | 72 | 62 | 8 | 38 | 18 | 8 | 19 | 42 | 31 | 74 | 71 | 66 | 298 | 149 | 0.424101687 | 0.841099024 | 0.177 | 0.073 | 0.044 | 0.133 | 0.114 | 0.087 | 0.38 | 0.108 | 0.053 | 0.126 | -0.043 |
| RI | 6192 | ENSG00000172638 | EFEMP2 | chr11 | - | 65866440 | 65867079 | 65866440 | 65866712 | 65866844 | 65867079 | 32 | 15 | 26 | 31 | 15 | 3 | 0 | 6 | 2 | 0 | 37 | 32 | 28 | 61 | 62 | 0 | 0 | 1 | 5 | 6 | 281 | 149 | 0.424154357 | 0.841099024 | 0.85 | 1 | 0.697 | 0.892 | 1 | 1 | 1 | 0.937 | 0.866 | 0.846 | -0.042 |
| RI | 2817 | ENSG00000126970 | ZC4H2 | chrX | - | 64919041 | 64919204 | 64919041 | 64919063 | 64919192 | 64919204 | 37 | 21 | 36 | 34 | 53 | 0 | 0 | 0 | 0 | 5 | 61 | 27 | 13 | 18 | 75 | 0 | 0 | 0 | 0 | 7 | 278 | 149 | 0.424180463 | 0.841099024 | 1 | 1 | 1 | 1 | 0.85 | 1 | 1 | 1 | 1 | 0.852 | 0 |
| RI | 5421 | ENSG00000123374 | CDK2 | chr12 | + | 55966830 | 55967934 | 55966830 | 55967124 | 55967856 | 55967934 | 13 | 16 | 30 | 20 | 7 | 22 | 8 | 28 | 37 | 67 | 18 | 13 | 12 | 40 | 44 | 33 | 22 | 22 | 17 | 77 | 298 | 149 | 0.424356514 | 0.841165748 | 0.228 | 0.5 | 0.349 | 0.213 | 0.05 | 0.214 | 0.228 | 0.214 | 0.541 | 0.222 | -0.016 |
| RI | 7844 | ENSG00000254986 | DPP3 | chr11 | + | 66495205 | 66495489 | 66495205 | 66495268 | 66495364 | 66495489 | 0 | 4 | 2 | 2 | 1 | 19 | 5 | 17 | 15 | 8 | 0 | 4 | 4 | 12 | 0 | 36 | 39 | 12 | 23 | 26 | 245 | 149 | 0.425560936 | 0.843270194 | 0 | 0.327 | 0.067 | 0.075 | 0.071 | 0 | 0.059 | 0.169 | 0.241 | 0 | 0.014 |
| RI | 3152 | ENSG00000146830 | GIGYF1 | chr7 | - | 100684019 | 100684337 | 100684019 | 100684157 | 100684236 | 100684337 | 7 | 4 | 3 | 7 | 2 | 63 | 38 | 56 | 44 | 16 | 0 | 0 | 1 | 22 | 0 | 20 | 47 | 18 | 18 | 99 | 228 | 149 | 0.426120557 | 0.844095953 | 0.068 | 0.064 | 0.034 | 0.094 | 0.076 | 0 | 0 | 0.035 | 0.444 | 0 | -0.029 |
| RI | 4846 | ENSG00000142327 | RNPEPL1 | chr2 | + | 240573774 | 240574628 | 240573774 | 240573891 | 240574514 | 240574628 | 8 | 7 | 8 | 15 | 2 | 0 | 7 | 1 | 0 | 0 | 45 | 3 | 8 | 10 | 7 | 0 | 0 | 0 | 0 | 10 | 298 | 149 | 0.426390653 | 0.844266368 | 1 | 0.333 | 0.8 | 1 | 1 | 1 | 1 | 1 | 1 | 0.259 | -0.025 |
| RI | 5527 | ENSG00000168818 | STX18 | chr4 | - | 4418967 | 4420944 | 4418967 | 4420129 | 4420863 | 4420944 | 23 | 2 | 8 | 12 | 9 | 46 | 44 | 70 | 57 | 72 | 9 | 18 | 31 | 20 | 0 | 73 | 80 | 71 | 63 | 58 | 298 | 149 | 0.42649244 | 0.844266368 | 0.2 | 0.022 | 0.054 | 0.095 | 0.059 | 0.058 | 0.101 | 0.179 | 0.137 | 0 | -0.009 |
| RI | 6495 | ENSG00000123066 | MED13L | chr12 | - | 115975170 | 115975738 | 115975170 | 115975313 | 115975514 | 115975738 | 74 | 126 | 53 | 80 | 59 | 121 | 71 | 103 | 78 | 99 | 58 | 167 | 157 | 133 | 61 | 179 | 106 | 128 | 149 | 38 | 298 | 149 | 0.427077892 | 0.844927898 | 0.234 | 0.47 | 0.205 | 0.339 | 0.23 | 0.139 | 0.441 | 0.38 | 0.309 | 0.445 | -0.047 |
| RI | 1815 | ENSG00000196497 | IPO4 | chr14 | - | 24188215 | 24188423 | 24188215 | 24188257 | 24188343 | 24188423 | 3 | 2 | 2 | 7 | 7 | 3 | 3 | 4 | 5 | 6 | 0 | 13 | 5 | 19 | 13 | 1 | 7 | 5 | 8 | 12 | 235 | 149 | 0.427112697 | 0.844927898 | 0.388 | 0.297 | 0.241 | 0.47 | 0.425 | 0 | 0.541 | 0.388 | 0.601 | 0.407 | -0.023 |
| RI | 7783 | ENSG00000149136 | SSRP1 | chr11 | - | 57330290 | 57330927 | 57330290 | 57330429 | 57330854 | 57330927 | 10 | 11 | 12 | 17 | 8 | 153 | 124 | 145 | 145 | 173 | 2 | 6 | 10 | 0 | 44 | 250 | 168 | 131 | 166 | 197 | 298 | 149 | 0.428477228 | 0.847185482 | 0.032 | 0.042 | 0.04 | 0.055 | 0.023 | 0.004 | 0.018 | 0.037 | 0 | 0.1 | 0.007 |
| RI | 224 | ENSG00000104814 | MAP4K1 | chr19 | - | 38607863 | 38608033 | 38607863 | 38607911 | 38607989 | 38608033 | 11 | 12 | 17 | 25 | 15 | 79 | 40 | 66 | 77 | 109 | 6 | 28 | 13 | 7 | 17 | 52 | 104 | 69 | 38 | 115 | 227 | 149 | 0.428634888 | 0.847185482 | 0.084 | 0.165 | 0.145 | 0.176 | 0.083 | 0.07 | 0.15 | 0.11 | 0.108 | 0.088 | 0.025 |
| RI | 8473 | ENSG00000177575 | CD163 | chr12 | - | 7487358 | 7488087 | 7487358 | 7487673 | 7487772 | 7488087 | 0 | 9 | 5 | 2 | 0 | 126 | 64 | 56 | 123 | 122 | 0 | 0 | 10 | 0 | 0 | 310 | 838 | 376 | 223 | 160 | 248 | 149 | 0.428684172 | 0.847185482 | 0 | 0.078 | 0.051 | 0.01 | 0 | 0 | 0 | 0.016 | 0 | 0 | 0.025 |
| RI | 734 | ENSG00000270647 | TAF15 | chr17 | + | 35820023 | 35820437 | 35820023 | 35820075 | 35820331 | 35820437 | 39 | 24 | 14 | 27 | 29 | 0 | 1 | 1 | 1 | 4 | 0 | 10 | 70 | 8 | 14 | 5 | 4 | 0 | 0 | 0 | 298 | 149 | 0.429828787 | 0.849163425 | 1 | 0.923 | 0.875 | 0.931 | 0.784 | 0 | 0.556 | 1 | 1 | 1 | 0.191 |
| RI | 306 | ENSG00000161265 | U2AF1L4 | chr19 | - | 35744069 | 35744421 | 35744069 | 35744145 | 35744322 | 35744421 | 77 | 55 | 61 | 94 | 76 | 54 | 30 | 62 | 31 | 99 | 233 | 115 | 67 | 55 | 72 | 65 | 65 | 64 | 42 | 49 | 298 | 149 | 0.431274331 | 0.851734362 | 0.416 | 0.478 | 0.33 | 0.603 | 0.277 | 0.642 | 0.469 | 0.344 | 0.396 | 0.424 | -0.034 |
| RI | 5673 | ENSG00000123983 | ACSL3 | chr2 | + | 222916318 | 222918155 | 222916318 | 222916496 | 222918069 | 222918155 | 53 | 46 | 44 | 55 | 21 | 1 | 0 | 3 | 4 | 17 | 24 | 63 | 81 | 33 | 3 | 6 | 11 | 1 | 7 | 0 | 298 | 149 | 0.431431605 | 0.851760191 | 0.964 | 1 | 0.88 | 0.873 | 0.382 | 0.667 | 0.741 | 0.976 | 0.702 | 1 | 0.003 |
| RI | 8292 | ENSG00000012822 | CALCOCO1 | chr12 | - | 53713700 | 53714241 | 53713700 | 53713900 | 53714132 | 53714241 | 58 | 29 | 39 | 60 | 37 | 54 | 48 | 24 | 42 | 43 | 91 | 72 | 69 | 69 | 69 | 97 | 95 | 47 | 50 | 161 | 298 | 149 | 0.431711484 | 0.852001874 | 0.349 | 0.232 | 0.448 | 0.417 | 0.301 | 0.319 | 0.275 | 0.423 | 0.408 | 0.176 | 0.029 |
| RI | 7429 | ENSG00000070061 | ELP1 | chr9 | - | 108897147 | 108898581 | 108897147 | 108897285 | 108898501 | 108898581 | 5 | 8 | 3 | 5 | 2 | 85 | 52 | 108 | 85 | 182 | 0 | 3 | 5 | 0 | 0 | 30 | 21 | 49 | 40 | 16 | 298 | 149 | 0.431967248 | 0.852001874 | 0.029 | 0.071 | 0.014 | 0.029 | 0.005 | 0 | 0.067 | 0.049 | 0 | 0 | 0.006 |
| RI | 835 | ENSG00000108788 | MLX | chr17 | + | 42568469 | 42568943 | 42568469 | 42568559 | 42568836 | 42568943 | 46 | 14 | 54 | 62 | 22 | 14 | 9 | 13 | 15 | 19 | 92 | 41 | 53 | 43 | 164 | 70 | 12 | 4 | 14 | 39 | 298 | 149 | 0.43198673 | 0.852001874 | 0.622 | 0.437 | 0.675 | 0.674 | 0.367 | 0.397 | 0.631 | 0.869 | 0.606 | 0.678 | -0.081 |
| RI | 4655 | ENSG00000001497 | LAS1L | chrX | - | 65517986 | 65522285 | 65517986 | 65518465 | 65520701 | 65522285 | 19 | 15 | 15 | 25 | 27 | 2 | 0 | 1 | 3 | 0 | 86 | 44 | 31 | 33 | 42 | 0 | 0 | 0 | 0 | 0 | 298 | 149 | 0.43388652 | 0.855264919 | 0.826 | 1 | 0.882 | 0.806 | 1 | 1 | 1 | 1 | 1 | 1 | -0.097 |
| RI | 7463 | ENSG00000266714 | MYO15B | chr17 | + | 75621344 | 75621570 | 75621344 | 75621408 | 75621500 | 75621570 | 1 | 1 | 3 | 2 | 3 | 21 | 17 | 17 | 21 | 5 | 0 | 3 | 3 | 7 | 0 | 42 | 42 | 28 | 58 | 39 | 241 | 149 | 0.433930754 | 0.855264919 | 0.029 | 0.035 | 0.098 | 0.056 | 0.271 | 0 | 0.042 | 0.062 | 0.069 | 0 | 0.063 |
| RI | 5332 | ENSG00000064703 | DDX20 | chr1 | + | 111761225 | 111762782 | 111761225 | 111761284 | 111762676 | 111762782 | 19 | 30 | 28 | 44 | 59 | 0 | 0 | 0 | 0 | 0 | 55 | 13 | 34 | 22 | 2 | 0 | 0 | 0 | 0 | 3 | 298 | 149 | 0.434335198 | 0.855776523 | 1 | 1 | 1 | 1 | 1 | 1 | 1 | 1 | 1 | 0.25 | 0.15 |
| RI | 538 | ENSG00000064601 | CTSA | chr20 | + | 45894993 | 45898109 | 45894993 | 45895138 | 45896964 | 45898109 | 9 | 3 | 14 | 9 | 2 | 0 | 0 | 0 | 0 | 3 | 23 | 40 | 15 | 53 | 30 | 0 | 0 | 0 | 1 | 0 | 298 | 149 | 0.434604483 | 0.856021567 | 1 | 1 | 1 | 1 | 0.25 | 1 | 1 | 1 | 0.964 | 1 | -0.143 |
| RI | 2171 | ENSG00000065357 | DGKA | chr12 | + | 55951622 | 55952431 | 55951622 | 55951783 | 55952340 | 55952431 | 26 | 57 | 57 | 47 | 28 | 9 | 8 | 21 | 17 | 46 | 96 | 44 | 37 | 51 | 45 | 12 | 12 | 42 | 24 | 5 | 298 | 149 | 0.435535239 | 0.857568887 | 0.591 | 0.781 | 0.576 | 0.58 | 0.233 | 0.8 | 0.647 | 0.306 | 0.515 | 0.818 | -0.065 |
| RI | 2947 | ENSG00000164068 | RNF123 | chr3 | + | 49704649 | 49705186 | 49704649 | 49704756 | 49705081 | 49705186 | 17 | 6 | 25 | 17 | 6 | 4 | 1 | 0 | 0 | 0 | 10 | 23 | 3 | 12 | 11 | 0 | 0 | 0 | 7 | 0 | 298 | 149 | 0.43656097 | 0.859302115 | 0.68 | 0.75 | 1 | 1 | 1 | 1 | 1 | 1 | 0.462 | 1 | -0.006 |
| RI | 6557 | ENSG00000127914 | AKAP9 | chr7 | + | 92099686 | 92101056 | 92099686 | 92099869 | 92100855 | 92101056 | 4 | 18 | 11 | 10 | 8 | 37 | 32 | 46 | 51 | 123 | 1 | 3 | 6 | 0 | 0 | 31 | 7 | 2 | 4 | 9 | 298 | 149 | 0.436729733 | 0.859347945 | 0.051 | 0.22 | 0.107 | 0.089 | 0.031 | 0.016 | 0.176 | 0.6 | 0 | 0 | -0.059 |
| RI | 8316 | ENSG00000007392 | LUC7L | chr16 | - | 227241 | 228402 | 227241 | 227336 | 228332 | 228402 | 420 | 347 | 387 | 451 | 507 | 6 | 19 | 10 | 21 | 10 | 553 | 462 | 322 | 412 | 578 | 18 | 16 | 14 | 14 | 21 | 298 | 149 | 0.436960927 | 0.859516549 | 0.972 | 0.901 | 0.951 | 0.915 | 0.962 | 0.939 | 0.935 | 0.92 | 0.936 | 0.932 | 0.008 |
| RI | 1031 | ENSG00000182087 | TMEM259 | chr19 | - | 1011579 | 1011798 | 1011579 | 1011612 | 1011740 | 1011798 | 50 | 10 | 34 | 43 | 39 | 21 | 5 | 13 | 49 | 18 | 37 | 59 | 31 | 31 | 59 | 0 | 14 | 24 | 24 | 79 | 277 | 149 | 0.437504013 | 0.860239468 | 0.562 | 0.518 | 0.585 | 0.321 | 0.538 | 1 | 0.694 | 0.41 | 0.41 | 0.287 | -0.055 |
| RI | 2068 | ENSG00000125459 | MSTO1 | chr1 | + | 155611215 | 155611603 | 155611215 | 155611355 | 155611548 | 155611603 | 7 | 13 | 22 | 29 | 10 | 0 | 1 | 1 | 1 | 5 | 0 | 19 | 16 | 2 | 15 | 16 | 0 | 0 | 0 | 0 | 298 | 149 | 0.437619706 | 0.860239468 | 1 | 0.867 | 0.917 | 0.935 | 0.5 | 0 | 1 | 1 | 1 | 1 | 0.044 |
| RI | 1835 | ENSG00000099797 | TECR | chr19 | + | 14563657 | 14563903 | 14563657 | 14563702 | 14563799 | 14563903 | 18 | 5 | 13 | 15 | 6 | 136 | 65 | 170 | 130 | 107 | 9 | 3 | 29 | 9 | 31 | 91 | 154 | 124 | 106 | 282 | 246 | 149 | 0.438388694 | 0.861464409 | 0.074 | 0.045 | 0.044 | 0.065 | 0.033 | 0.057 | 0.012 | 0.124 | 0.049 | 0.062 | -0.009 |
| RI | 4479 | ENSG00000156983 | BRPF1 | chr3 | + | 9745572 | 9745930 | 9745572 | 9745709 | 9745811 | 9745930 | 35 | 22 | 23 | 25 | 34 | 79 | 45 | 53 | 76 | 84 | 38 | 11 | 31 | 59 | 149 | 88 | 86 | 51 | 126 | 152 | 251 | 149 | 0.43905085 | 0.862285574 | 0.208 | 0.225 | 0.205 | 0.163 | 0.194 | 0.204 | 0.071 | 0.265 | 0.218 | 0.368 | -0.026 |
| RI | 2523 | ENSG00000106003 | LFNG | chr7 | + | 2525218 | 2525567 | 2525218 | 2525318 | 2525413 | 2525567 | 54 | 16 | 45 | 37 | 14 | 82 | 39 | 40 | 56 | 36 | 5 | 13 | 11 | 27 | 35 | 25 | 17 | 5 | 17 | 48 | 244 | 149 | 0.439098528 | 0.862285574 | 0.287 | 0.2 | 0.407 | 0.287 | 0.192 | 0.109 | 0.318 | 0.573 | 0.492 | 0.308 | -0.085 |
| RI | 7271 | ENSG00000104529 | EEF1D | chr8 | - | 143581053 | 143581328 | 143581053 | 143581154 | 143581228 | 143581328 | 100 | 35 | 83 | 80 | 69 | 1722 | 902 | 1608 | 1470 | 1879 | 53 | 61 | 69 | 83 | 84 | 1715 | 1317 | 1019 | 929 | 1875 | 223 | 149 | 0.440613938 | 0.864973923 | 0.037 | 0.025 | 0.033 | 0.035 | 0.024 | 0.02 | 0.03 | 0.043 | 0.056 | 0.029 | -0.005 |
| RI | 1887 | ENSG00000213983 | AP1G2 | chr14 | - | 23563715 | 23564221 | 23563715 | 23563856 | 23564045 | 23564221 | 82 | 65 | 71 | 116 | 42 | 58 | 34 | 39 | 69 | 36 | 85 | 67 | 62 | 72 | 87 | 47 | 53 | 37 | 33 | 129 | 298 | 149 | 0.441785673 | 0.866986037 | 0.414 | 0.489 | 0.477 | 0.457 | 0.368 | 0.475 | 0.387 | 0.456 | 0.522 | 0.252 | 0.023 |
| RI | 3158 | ENSG00000100462 | PRMT5 | chr14 | - | 22928125 | 22928615 | 22928125 | 22928211 | 22928496 | 22928615 | 6 | 7 | 25 | 13 | 30 | 41 | 28 | 55 | 53 | 83 | 19 | 18 | 10 | 7 | 14 | 33 | 61 | 45 | 48 | 74 | 298 | 149 | 0.442501361 | 0.868102138 | 0.068 | 0.111 | 0.185 | 0.109 | 0.153 | 0.224 | 0.129 | 0.1 | 0.068 | 0.086 | 0.004 |
| RI | 4055 | ENSG00000011295 | TTC19 | chr17 | + | 16026539 | 16028786 | 16026539 | 16026702 | 16027373 | 16028786 | 70 | 113 | 73 | 63 | 80 | 116 | 49 | 98 | 74 | 106 | 23 | 103 | 142 | 72 | 50 | 44 | 81 | 69 | 65 | 59 | 298 | 149 | 0.443349551 | 0.869477356 | 0.232 | 0.536 | 0.271 | 0.299 | 0.274 | 0.207 | 0.389 | 0.507 | 0.356 | 0.298 | -0.029 |
| RI | 6296 | ENSG00000145191 | EIF2B5 | chr3 | + | 184143051 | 184143565 | 184143051 | 184143142 | 184143430 | 184143565 | 56 | 76 | 105 | 90 | 136 | 1 | 3 | 1 | 1 | 4 | 171 | 107 | 106 | 110 | 131 | 12 | 0 | 3 | 2 | 0 | 298 | 149 | 0.443637149 | 0.869664017 | 0.966 | 0.927 | 0.981 | 0.978 | 0.944 | 0.877 | 1 | 0.946 | 0.965 | 1 | 0.002 |
| RI | 5530 | ENSG00000157379 | DHRS1 | chr14 | - | 24292183 | 24292784 | 24292183 | 24292330 | 24292651 | 24292784 | 37 | 11 | 23 | 25 | 3 | 19 | 26 | 12 | 15 | 6 | 7 | 33 | 38 | 31 | 24 | 20 | 7 | 12 | 13 | 31 | 298 | 149 | 0.443739182 | 0.869664017 | 0.493 | 0.175 | 0.489 | 0.455 | 0.2 | 0.149 | 0.702 | 0.613 | 0.544 | 0.279 | -0.095 |
| RI | 5170 | ENSG00000108387 | SEPTIN4 | chr17 | - | 58520742 | 58521160 | 58520742 | 58520842 | 58520997 | 58521160 | 3 | 2 | 0 | 1 | 0 | 7 | 2 | 2 | 8 | 10 | 0 | 0 | 0 | 0 | 0 | 26 | 1 | 7 | 15 | 80 | 298 | 149 | 0.444647477 | 0.871002493 | 0.176 | 0.333 | 0 | 0.059 | 0 | 0 | 0 | 0 | 0 | 0 | 0.114 |
| RI | 7781 | ENSG00000149136 | SSRP1 | chr11 | - | 57326702 | 57327514 | 57326702 | 57326889 | 57327425 | 57327514 | 4 | 2 | 0 | 13 | 0 | 42 | 29 | 57 | 63 | 101 | 10 | 8 | 11 | 3 | 1 | 136 | 78 | 74 | 69 | 79 | 298 | 149 | 0.444717034 | 0.871002493 | 0.045 | 0.033 | 0 | 0.094 | 0 | 0.035 | 0.049 | 0.069 | 0.021 | 0.006 | -0.002 |
| RI | 2733 | ENSG00000109133 | TMEM33 | chr4 | + | 41935128 | 41935529 | 41935128 | 41935262 | 41935466 | 41935529 | 134 | 94 | 122 | 100 | 37 | 4 | 3 | 2 | 5 | 11 | 86 | 76 | 69 | 72 | 101 | 0 | 11 | 5 | 2 | 0 | 298 | 149 | 0.445151105 | 0.871563665 | 0.944 | 0.94 | 0.968 | 0.909 | 0.627 | 1 | 0.776 | 0.873 | 0.947 | 1 | -0.042 |
| RI | 4011 | ENSG00000111602 | TIMELESS | chr12 | - | 56420962 | 56421809 | 56420962 | 56421134 | 56421726 | 56421809 | 9 | 4 | 15 | 5 | 6 | 3 | 0 | 0 | 3 | 0 | 20 | 2 | 1 | 19 | 15 | 0 | 0 | 0 | 3 | 6 | 298 | 149 | 0.446013069 | 0.872734918 | 0.6 | 1 | 1 | 0.455 | 1 | 1 | 1 | 1 | 0.76 | 0.556 | -0.052 |
| RI | 6998 | ENSG00000182774 | RPS17 | chr15 | - | 82538305 | 82538985 | 82538305 | 82538371 | 82538879 | 82538985 | 187 | 79 | 217 | 152 | 157 | 10220 | 5418 | 10595 | 10044 | 8709 | 304 | 119 | 106 | 98 | 237 | 10998 | 6893 | 6833 | 5901 | 10977 | 298 | 149 | 0.446138132 | 0.872734918 | 0.009 | 0.007 | 0.01 | 0.008 | 0.009 | 0.014 | 0.009 | 0.008 | 0.008 | 0.011 | -0.001 |
| RI | 1307 | ENSG00000197912 | SPG7 | chr16 | + | 89554485 | 89557747 | 89554485 | 89554563 | 89556886 | 89557747 | 0 | 1 | 2 | 19 | 3 | 42 | 16 | 34 | 48 | 26 | 13 | 8 | 10 | 4 | 38 | 117 | 101 | 66 | 84 | 91 | 298 | 149 | 0.44619256 | 0.872734918 | 0 | 0.03 | 0.029 | 0.165 | 0.055 | 0.053 | 0.038 | 0.07 | 0.023 | 0.173 | -0.016 |
| RI | 4854 | ENSG00000096093 | EFHC1 | chr6 | + | 52452687 | 52454287 | 52452687 | 52452837 | 52454094 | 52454287 | 73 | 57 | 25 | 36 | 69 | 2 | 2 | 2 | 0 | 12 | 15 | 11 | 23 | 1 | 1 | 0 | 3 | 0 | 1 | 5 | 298 | 149 | 0.44657735 | 0.873198414 | 0.948 | 0.934 | 0.862 | 1 | 0.742 | 1 | 0.647 | 1 | 0.333 | 0.091 | 0.283 |
| RI | 8521 | ENSG00000121073 | SLC35B1 | chr17 | - | 49705865 | 49706309 | 49705865 | 49705896 | 49706203 | 49706309 | 22 | 2 | 31 | 27 | 30 | 43 | 21 | 55 | 40 | 49 | 46 | 5 | 34 | 44 | 73 | 56 | 49 | 69 | 52 | 65 | 298 | 149 | 0.44690566 | 0.873551202 | 0.204 | 0.045 | 0.22 | 0.252 | 0.234 | 0.291 | 0.049 | 0.198 | 0.297 | 0.36 | -0.048 |
| RI | 1377 | ENSG00000125148 | MT2A | chr16 | + | 56608583 | 56609057 | 56608583 | 56608683 | 56608991 | 56609057 | 1 | 2 | 10 | 12 | 2 | 58 | 38 | 61 | 83 | 11 | 0 | 30 | 9 | 4 | 17 | 150 | 73 | 89 | 88 | 111 | 298 | 149 | 0.447235542 | 0.873818053 | 0.009 | 0.026 | 0.076 | 0.067 | 0.083 | 0 | 0.17 | 0.048 | 0.022 | 0.071 | -0.01 |
| RI | 4422 | ENSG00000111653 | ING4 | chr12 | - | 6652661 | 6653050 | 6652661 | 6652767 | 6652944 | 6653050 | 15 | 12 | 12 | 13 | 31 | 31 | 23 | 21 | 41 | 92 | 8 | 11 | 25 | 9 | 6 | 43 | 24 | 18 | 25 | 77 | 298 | 149 | 0.447338038 | 0.873818053 | 0.195 | 0.207 | 0.222 | 0.137 | 0.144 | 0.085 | 0.186 | 0.41 | 0.153 | 0.037 | 0.007 |
| RI | 134 | ENSG00000235954 | TTC28-AS1 | chr22 | + | 27997410 | 27998838 | 27997410 | 27998152 | 27998603 | 27998838 | 14 | 9 | 16 | 18 | 11 | 1 | 2 | 1 | 0 | 1 | 7 | 13 | 0 | 15 | 19 | 0 | 0 | 4 | 0 | 0 | 298 | 149 | 0.448245371 | 0.875300961 | 0.875 | 0.692 | 0.889 | 1 | 0.846 | 1 | 1 | 0 | 1 | 1 | 0.06 |
| RI | 1417 | ENSG00000039523 | RIPOR1 | chr16 | + | 67540457 | 67540704 | 67540457 | 67540501 | 67540578 | 67540704 | 25 | 17 | 28 | 14 | 21 | 87 | 47 | 63 | 75 | 21 | 40 | 14 | 12 | 47 | 62 | 99 | 75 | 14 | 56 | 157 | 226 | 149 | 0.449864137 | 0.877800686 | 0.159 | 0.193 | 0.227 | 0.11 | 0.397 | 0.21 | 0.11 | 0.361 | 0.356 | 0.207 | -0.032 |
| RI | 3559 | ENSG00000171163 | ZNF692 | chr1 | - | 248855557 | 248855946 | 248855557 | 248855635 | 248855724 | 248855946 | 14 | 6 | 15 | 15 | 6 | 12 | 5 | 8 | 16 | 14 | 15 | 7 | 15 | 41 | 48 | 64 | 19 | 2 | 3 | 37 | 238 | 149 | 0.449955162 | 0.877800686 | 0.422 | 0.429 | 0.54 | 0.37 | 0.212 | 0.128 | 0.187 | 0.824 | 0.895 | 0.448 | -0.102 |
| RI | 5080 | ENSG00000112855 | HARS2 | chr5 | + | 140693490 | 140693665 | 140693490 | 140693508 | 140693590 | 140693665 | 26 | 29 | 26 | 29 | 39 | 0 | 3 | 0 | 0 | 2 | 76 | 46 | 79 | 79 | 71 | 2 | 0 | 1 | 4 | 0 | 231 | 149 | 0.449971301 | 0.877800686 | 1 | 0.862 | 1 | 1 | 0.926 | 0.961 | 1 | 0.981 | 0.927 | 1 | -0.016 |
| RI | 7273 | ENSG00000104522 | GFUS | chr8 | - | 143612617 | 143613295 | 143612617 | 143612965 | 143613195 | 143613295 | 4 | 0 | 0 | 4 | 2 | 56 | 17 | 31 | 31 | 22 | 6 | 0 | 1 | 5 | 5 | 37 | 53 | 14 | 35 | 107 | 298 | 149 | 0.450422518 | 0.878390827 | 0.034 | 0 | 0 | 0.061 | 0.043 | 0.075 | 0 | 0.034 | 0.067 | 0.023 | -0.012 |
| RI | 2752 | ENSG00000099949 | LZTR1 | chr22 | + | 20994869 | 20995872 | 20994869 | 20995026 | 20995745 | 20995872 | 10 | 6 | 11 | 10 | 10 | 62 | 20 | 32 | 47 | 10 | 9 | 4 | 3 | 7 | 21 | 66 | 20 | 14 | 35 | 48 | 298 | 149 | 0.450664114 | 0.878487297 | 0.075 | 0.13 | 0.147 | 0.096 | 0.333 | 0.064 | 0.091 | 0.097 | 0.091 | 0.179 | 0.052 |
| RI | 3996 | ENSG00000198912 | C1orf174 | chr1 | - | 3890568 | 3892996 | 3890568 | 3891057 | 3892882 | 3892996 | 57 | 22 | 44 | 64 | 55 | 21 | 12 | 11 | 23 | 0 | 45 | 51 | 73 | 87 | 106 | 39 | 5 | 8 | 51 | 47 | 298 | 149 | 0.450769426 | 0.878487297 | 0.576 | 0.478 | 0.667 | 0.582 | 1 | 0.366 | 0.836 | 0.82 | 0.46 | 0.53 | 0.058 |
| RI | 4395 | ENSG00000249915 | PDCD6 | chr5 | + | 271620 | 271821 | 271620 | 271690 | 271775 | 271821 | 71 | 39 | 32 | 42 | 8 | 0 | 0 | 0 | 0 | 0 | 14 | 57 | 36 | 48 | 94 | 5 | 0 | 0 | 0 | 0 | 234 | 149 | 0.451130695 | 0.878896768 | 1 | 1 | 1 | 1 | 1 | 0.641 | 1 | 1 | 1 | 1 | 0.072 |
| RI | 3722 | ENSG00000138002 | IFT172 | chr2 | - | 27476640 | 27477320 | 27476640 | 27476726 | 27477216 | 27477320 | 2 | 4 | 0 | 3 | 1 | 10 | 6 | 13 | 11 | 8 | 0 | 5 | 0 | 1 | 2 | 9 | 7 | 6 | 11 | 17 | 298 | 149 | 0.451277111 | 0.878896768 | 0.091 | 0.25 | 0 | 0.12 | 0.059 | 0 | 0.263 | 0 | 0.043 | 0.056 | 0.032 |
| RI | 4870 | ENSG00000162852 | CNST | chr1 | + | 246633923 | 246634587 | 246633923 | 246634010 | 246634472 | 246634587 | 17 | 29 | 23 | 26 | 34 | 168 | 216 | 157 | 237 | 305 | 0 | 14 | 37 | 12 | 25 | 123 | 216 | 172 | 185 | 115 | 298 | 149 | 0.451700861 | 0.879432097 | 0.048 | 0.063 | 0.068 | 0.052 | 0.053 | 0 | 0.031 | 0.097 | 0.031 | 0.098 | 0.005 |
| RI | 3017 | ENSG00000204427 | ABHD16A | chr6 | - | 31688040 | 31688305 | 31688040 | 31688103 | 31688248 | 31688305 | 28 | 16 | 33 | 30 | 16 | 65 | 40 | 55 | 66 | 20 | 29 | 43 | 32 | 37 | 78 | 101 | 69 | 57 | 54 | 268 | 294 | 149 | 0.452600705 | 0.880718918 | 0.179 | 0.169 | 0.233 | 0.187 | 0.288 | 0.127 | 0.24 | 0.221 | 0.258 | 0.129 | 0.016 |
| RI | 8478 | ENSG00000166598 | HSP90B1 | chr12 | + | 103931520 | 103931748 | 103931520 | 103931623 | 103931710 | 103931748 | 115 | 59 | 134 | 147 | 203 | 7 | 0 | 1 | 2 | 1 | 122 | 135 | 122 | 109 | 27 | 0 | 0 | 5 | 0 | 0 | 236 | 149 | 0.452660003 | 0.880718918 | 0.912 | 1 | 0.988 | 0.979 | 0.992 | 1 | 1 | 0.939 | 1 | 1 | -0.014 |
| RI | 5404 | ENSG00000198805 | PNP | chr14 | + | 20474471 | 20474948 | 20474471 | 20474575 | 20474772 | 20474948 | 38 | 46 | 32 | 61 | 28 | 190 | 74 | 125 | 163 | 101 | 38 | 25 | 93 | 97 | 78 | 152 | 97 | 68 | 195 | 491 | 298 | 149 | 0.453019002 | 0.88112718 | 0.091 | 0.237 | 0.113 | 0.158 | 0.122 | 0.111 | 0.114 | 0.406 | 0.199 | 0.074 | -0.037 |
| RI | 7066 | ENSG00000140632 | GLYR1 | chr16 | - | 4803202 | 4805310 | 4803202 | 4804148 | 4804521 | 4805310 | 142 | 103 | 123 | 143 | 87 | 11 | 12 | 15 | 15 | 12 | 174 | 145 | 109 | 208 | 293 | 11 | 21 | 33 | 22 | 27 | 298 | 149 | 0.453439476 | 0.881473334 | 0.866 | 0.811 | 0.804 | 0.827 | 0.784 | 0.888 | 0.775 | 0.623 | 0.825 | 0.844 | 0.027 |
| RI | 6564 | ENSG00000181513 | ACBD4 | chr17 | + | 45136499 | 45136776 | 45136499 | 45136620 | 45136691 | 45136776 | 9 | 2 | 15 | 13 | 0 | 21 | 15 | 31 | 25 | 41 | 20 | 3 | 0 | 6 | 15 | 29 | 8 | 6 | 26 | 44 | 220 | 149 | 0.453495423 | 0.881473334 | 0.225 | 0.083 | 0.247 | 0.26 | 0 | 0.318 | 0.203 | 0 | 0.135 | 0.188 | -0.006 |
| RI | 4013 | ENSG00000069424 | KCNAB2 | chr1 | + | 6087466 | 6089051 | 6087466 | 6087511 | 6089007 | 6089051 | 18 | 11 | 21 | 19 | 6 | 273 | 143 | 210 | 203 | 200 | 38 | 27 | 14 | 7 | 20 | 294 | 413 | 196 | 344 | 452 | 298 | 149 | 0.453792593 | 0.881760804 | 0.032 | 0.037 | 0.048 | 0.045 | 0.015 | 0.061 | 0.032 | 0.034 | 0.01 | 0.022 | 0.004 |
| RI | 3088 | ENSG00000164073 | MFSD8 | chr4 | - | 127920556 | 127921771 | 127920556 | 127920836 | 127921523 | 127921771 | 65 | 45 | 57 | 50 | 57 | 10 | 1 | 9 | 11 | 17 | 14 | 34 | 27 | 20 | 24 | 0 | 10 | 3 | 7 | 0 | 298 | 149 | 0.454643993 | 0.883080145 | 0.765 | 0.957 | 0.76 | 0.694 | 0.626 | 1 | 0.63 | 0.818 | 0.588 | 1 | -0.047 |
| RI | 1470 | ENSG00000140983 | RHOT2 | chr16 | + | 670122 | 670357 | 670122 | 670175 | 670265 | 670357 | 57 | 41 | 28 | 39 | 39 | 32 | 5 | 25 | 13 | 9 | 122 | 46 | 34 | 44 | 96 | 49 | 41 | 23 | 17 | 22 | 239 | 149 | 0.454770577 | 0.883080145 | 0.526 | 0.836 | 0.411 | 0.652 | 0.73 | 0.608 | 0.412 | 0.48 | 0.617 | 0.731 | 0.061 |
| RI | 7902 | ENSG00000247708 | STX18-AS1 | chr4 | + | 4648976 | 4649542 | 4648976 | 4649109 | 4649372 | 4649542 | 6 | 3 | 4 | 3 | 1 | 8 | 3 | 4 | 4 | 1 | 12 | 11 | 8 | 3 | 0 | 0 | 6 | 2 | 7 | 10 | 298 | 149 | 0.45539051 | 0.883993343 | 0.273 | 0.333 | 0.333 | 0.273 | 0.333 | 1 | 0.478 | 0.667 | 0.176 | 0 | -0.155 |
| RI | 6353 | ENSG00000198198 | SZT2 | chr1 | + | 43448611 | 43450171 | 43448611 | 43448728 | 43450102 | 43450171 | 11 | 8 | 23 | 17 | 10 | 16 | 14 | 19 | 20 | 17 | 33 | 37 | 18 | 30 | 25 | 30 | 12 | 36 | 44 | 33 | 298 | 149 | 0.456190413 | 0.885239389 | 0.256 | 0.222 | 0.377 | 0.298 | 0.227 | 0.355 | 0.607 | 0.2 | 0.254 | 0.275 | -0.062 |
| RI | 6493 | ENSG00000104823 | ECH1 | chr19 | - | 38815856 | 38816355 | 38815856 | 38816007 | 38816283 | 38816355 | 8 | 4 | 8 | 8 | 2 | 199 | 68 | 147 | 145 | 96 | 0 | 11 | 12 | 7 | 11 | 122 | 225 | 98 | 117 | 205 | 298 | 149 | 0.456377711 | 0.885239389 | 0.02 | 0.029 | 0.026 | 0.027 | 0.01 | 0 | 0.024 | 0.058 | 0.029 | 0.026 | -0.005 |
| RI | 5234 | ENSG00000186141 | POLR3C | chr1 | + | 145826453 | 145826709 | 145826453 | 145826492 | 145826605 | 145826709 | 117 | 95 | 126 | 146 | 165 | 14 | 12 | 15 | 19 | 28 | 104 | 160 | 91 | 166 | 150 | 13 | 20 | 4 | 30 | 14 | 262 | 149 | 0.456482001 | 0.885239389 | 0.826 | 0.818 | 0.827 | 0.814 | 0.77 | 0.82 | 0.82 | 0.928 | 0.759 | 0.859 | -0.026 |
| RI | 3133 | ENSG00000109861 | CTSC | chr11 | - | 88328111 | 88335082 | 88328111 | 88328196 | 88334936 | 88335082 | 114 | 60 | 62 | 70 | 50 | 198 | 105 | 126 | 147 | 147 | 61 | 95 | 41 | 51 | 39 | 153 | 145 | 81 | 78 | 199 | 298 | 149 | 0.45702855 | 0.886008417 | 0.224 | 0.222 | 0.197 | 0.192 | 0.145 | 0.166 | 0.247 | 0.202 | 0.246 | 0.089 | 0.006 |
| RI | 8414 | ENSG00000182796 | TMEM198B | chr12 | + | 55830840 | 55831349 | 55830840 | 55831083 | 55831256 | 55831349 | 14 | 6 | 6 | 19 | 9 | 0 | 0 | 2 | 3 | 7 | 38 | 19 | 22 | 8 | 12 | 0 | 7 | 6 | 0 | 0 | 298 | 149 | 0.45799381 | 0.887429338 | 1 | 1 | 0.6 | 0.76 | 0.391 | 1 | 0.576 | 0.647 | 1 | 1 | -0.094 |
| RI | 4041 | ENSG00000106610 | STAG3L4 | chr7 | + | 67308576 | 67309139 | 67308576 | 67308652 | 67308934 | 67309139 | 52 | 21 | 52 | 39 | 58 | 2 | 0 | 1 | 0 | 2 | 90 | 74 | 68 | 52 | 155 | 0 | 0 | 0 | 0 | 0 | 298 | 149 | 0.458061969 | 0.887429338 | 0.929 | 1 | 0.963 | 1 | 0.935 | 1 | 1 | 1 | 1 | 1 | -0.035 |
| RI | 5589 | ENSG00000164548 | TRA2A | chr7 | - | 23521706 | 23522432 | 23521706 | 23521840 | 23522353 | 23522432 | 424 | 434 | 438 | 594 | 556 | 3 | 4 | 4 | 2 | 11 | 325 | 638 | 847 | 720 | 77 | 5 | 1 | 6 | 2 | 6 | 298 | 149 | 0.458433741 | 0.887858396 | 0.986 | 0.982 | 0.982 | 0.993 | 0.962 | 0.97 | 0.997 | 0.986 | 0.994 | 0.865 | 0.019 |
| RI | 6444 | ENSG00000115677 | HDLBP | chr2 | - | 241235120 | 241235594 | 241235120 | 241235255 | 241235489 | 241235594 | 8 | 8 | 22 | 23 | 9 | 276 | 207 | 328 | 297 | 357 | 25 | 12 | 13 | 8 | 32 | 566 | 322 | 375 | 369 | 556 | 298 | 149 | 0.458825358 | 0.888325596 | 0.014 | 0.019 | 0.032 | 0.037 | 0.012 | 0.022 | 0.018 | 0.017 | 0.011 | 0.028 | 0.004 |
| RI | 2502 | ENSG00000126522 | ASL | chr7 | + | 66086743 | 66088921 | 66086743 | 66087791 | 66088806 | 66088921 | 5 | 1 | 3 | 6 | 6 | 31 | 10 | 32 | 24 | 20 | 2 | 17 | 3 | 9 | 8 | 36 | 43 | 20 | 38 | 47 | 298 | 149 | 0.460140224 | 0.890579392 | 0.075 | 0.048 | 0.045 | 0.111 | 0.13 | 0.027 | 0.165 | 0.07 | 0.106 | 0.078 | -0.007 |
| RI | 1142 | ENSG00000168488 | ATXN2L | chr16 | + | 28835277 | 28835758 | 28835277 | 28835399 | 28835548 | 28835758 | 10 | 5 | 6 | 3 | 0 | 11 | 18 | 5 | 14 | 4 | 0 | 6 | 1 | 7 | 11 | 61 | 4 | 14 | 25 | 4 | 298 | 149 | 0.460572762 | 0.891124567 | 0.312 | 0.122 | 0.375 | 0.097 | 0 | 0 | 0.429 | 0.034 | 0.123 | 0.579 | -0.052 |
| RI | 2040 | ENSG00000206560 | ANKRD28 | chr3 | - | 15686001 | 15686309 | 15686001 | 15686119 | 15686221 | 15686309 | 10 | 19 | 14 | 8 | 15 | 165 | 180 | 130 | 239 | 169 | 3 | 12 | 27 | 3 | 12 | 70 | 189 | 177 | 162 | 37 | 251 | 149 | 0.462158248 | 0.893899401 | 0.035 | 0.059 | 0.06 | 0.019 | 0.05 | 0.025 | 0.036 | 0.083 | 0.011 | 0.161 | -0.019 |
| RI | 5224 | ENSG00000125875 | TBC1D20 | chr20 | - | 440247 | 441689 | 440247 | 440317 | 441587 | 441689 | 70 | 24 | 39 | 64 | 48 | 2 | 0 | 0 | 5 | 6 | 163 | 98 | 41 | 56 | 149 | 5 | 4 | 1 | 6 | 0 | 298 | 149 | 0.462569097 | 0.894401197 | 0.946 | 1 | 1 | 0.865 | 0.8 | 0.942 | 0.925 | 0.953 | 0.824 | 1 | -0.007 |
| RI | 1520 | ENSG00000213614 | HEXA | chr15 | - | 72345445 | 72346325 | 72345445 | 72346128 | 72346234 | 72346325 | 33 | 13 | 33 | 26 | 18 | 0 | 0 | 0 | 1 | 0 | 62 | 14 | 31 | 68 | 33 | 0 | 4 | 2 | 0 | 0 | 255 | 149 | 0.464218314 | 0.897296329 | 1 | 1 | 1 | 0.938 | 1 | 1 | 0.672 | 0.901 | 1 | 1 | 0.073 |
| RI | 7863 | ENSG00000185101 | ANO9 | chr11 | - | 419581 | 420615 | 419581 | 419729 | 420462 | 420615 | 41 | 11 | 17 | 19 | 13 | 7 | 3 | 9 | 3 | 2 | 15 | 16 | 2 | 19 | 32 | 0 | 1 | 0 | 8 | 14 | 298 | 149 | 0.464625086 | 0.897600468 | 0.745 | 0.647 | 0.486 | 0.76 | 0.765 | 1 | 0.889 | 1 | 0.543 | 0.533 | -0.112 |
| RI | 5946 | ENSG00000184154 | LRTOMT | chr11 | + | 72094947 | 72096895 | 72094947 | 72095096 | 72095378 | 72096895 | 25 | 26 | 15 | 25 | 20 | 0 | 2 | 1 | 8 | 3 | 32 | 31 | 59 | 11 | 8 | 0 | 3 | 5 | 0 | 4 | 298 | 149 | 0.464679572 | 0.897600468 | 1 | 0.867 | 0.882 | 0.61 | 0.769 | 1 | 0.838 | 0.855 | 1 | 0.5 | -0.013 |
| RI | 1600 | ENSG00000102901 | CENPT | chr16 | - | 67832227 | 67832545 | 67832227 | 67832354 | 67832454 | 67832545 | 6 | 3 | 7 | 16 | 5 | 2 | 0 | 1 | 1 | 7 | 71 | 21 | 12 | 8 | 19 | 0 | 0 | 4 | 7 | 3 | 249 | 149 | 0.465162818 | 0.898240198 | 0.642 | 1 | 0.807 | 0.905 | 0.299 | 1 | 1 | 0.642 | 0.406 | 0.791 | -0.037 |
| RI | 7786 | ENSG00000115414 | FN1 | chr2 | - | 215379129 | 215382325 | 215379129 | 215379317 | 215382211 | 215382325 | 2 | 3 | 4 | 5 | 7 | 0 | 45 | 3 | 7 | 6 | 0 | 7 | 3 | 9 | 22 | 27 | 8 | 21 | 29 | 14 | 298 | 149 | 0.465318422 | 0.898247033 | 1 | 0.032 | 0.4 | 0.263 | 0.368 | 0 | 0.304 | 0.067 | 0.134 | 0.44 | 0.224 |
| RI | 4963 | ENSG00000138757 | G3BP2 | chr4 | - | 75653982 | 75655246 | 75653982 | 75654081 | 75655065 | 75655246 | 64 | 61 | 53 | 57 | 141 | 38 | 31 | 37 | 30 | 68 | 48 | 79 | 60 | 53 | 9 | 26 | 38 | 31 | 28 | 21 | 298 | 149 | 0.465766626 | 0.898733315 | 0.457 | 0.496 | 0.417 | 0.487 | 0.509 | 0.48 | 0.51 | 0.492 | 0.486 | 0.176 | 0.044 |
| RI | 7462 | ENSG00000266714 | MYO15B | chr17 | + | 75620466 | 75621176 | 75620466 | 75620636 | 75621030 | 75621176 | 15 | 14 | 11 | 12 | 6 | 13 | 5 | 3 | 10 | 0 | 21 | 12 | 8 | 38 | 14 | 0 | 0 | 6 | 20 | 18 | 298 | 149 | 0.46587836 | 0.898733315 | 0.366 | 0.583 | 0.647 | 0.375 | 1 | 1 | 1 | 0.4 | 0.487 | 0.28 | -0.039 |
| RI | 7441 | ENSG00000233016 | SNHG7 | chr9 | - | 136724657 | 136725110 | 136724657 | 136724939 | 136725054 | 136725110 | 162 | 51 | 143 | 152 | 158 | 5 | 8 | 15 | 6 | 18 | 57 | 191 | 59 | 54 | 203 | 6 | 28 | 1 | 2 | 20 | 264 | 149 | 0.466026772 | 0.898733315 | 0.948 | 0.783 | 0.843 | 0.935 | 0.832 | 0.843 | 0.794 | 0.971 | 0.938 | 0.851 | -0.011 |
| RI | 2343 | ENSG00000184584 | STING1 | chr5 | - | 139480789 | 139481704 | 139480789 | 139480898 | 139481158 | 139481704 | 13 | 11 | 19 | 12 | 2 | 86 | 37 | 86 | 64 | 35 | 28 | 12 | 10 | 13 | 17 | 142 | 93 | 62 | 71 | 154 | 298 | 149 | 0.466628637 | 0.899307906 | 0.07 | 0.129 | 0.099 | 0.086 | 0.028 | 0.09 | 0.061 | 0.075 | 0.084 | 0.052 | 0.01 |
| RI | 7857 | ENSG00000172543 | CTSW | chr11 | + | 65883214 | 65883741 | 65883214 | 65883424 | 65883507 | 65883741 | 178 | 25 | 75 | 82 | 69 | 1045 | 256 | 434 | 563 | 281 | 66 | 101 | 48 | 31 | 94 | 211 | 349 | 383 | 247 | 756 | 232 | 149 | 0.466629208 | 0.899307906 | 0.099 | 0.059 | 0.1 | 0.086 | 0.136 | 0.167 | 0.157 | 0.074 | 0.075 | 0.074 | -0.013 |
| RI | 3194 | ENSG00000075826 | SEC31B | chr10 | - | 100488857 | 100489398 | 100488857 | 100488974 | 100489251 | 100489398 | 29 | 18 | 28 | 16 | 26 | 0 | 2 | 2 | 3 | 4 | 19 | 8 | 26 | 41 | 33 | 0 | 3 | 0 | 2 | 4 | 298 | 149 | 0.46695233 | 0.899637121 | 1 | 0.818 | 0.875 | 0.727 | 0.765 | 1 | 0.571 | 1 | 0.911 | 0.805 | -0.02 |
| RI | 5810 | ENSG00000115942 | ORC2 | chr2 | - | 200913294 | 200913992 | 200913294 | 200913413 | 200913930 | 200913992 | 68 | 34 | 51 | 51 | 19 | 26 | 30 | 41 | 37 | 37 | 14 | 50 | 44 | 24 | 1 | 13 | 32 | 17 | 47 | 8 | 298 | 149 | 0.467864218 | 0.901100077 | 0.567 | 0.362 | 0.383 | 0.408 | 0.204 | 0.35 | 0.439 | 0.564 | 0.203 | 0.059 | 0.062 |
| RI | 2195 | ENSG00000168268 | NT5DC2 | chr3 | - | 52527616 | 52527931 | 52527616 | 52527718 | 52527828 | 52527931 | 0 | 1 | 0 | 3 | 0 | 3 | 0 | 8 | 7 | 2 | 0 | 3 | 2 | 0 | 0 | 19 | 17 | 13 | 4 | 12 | 259 | 149 | 0.468043725 | 0.901151983 | 0 | 1 | 0 | 0.198 | 0 | 0 | 0.092 | 0.081 | 0 | 0 | 0.205 |
| RI | 8329 | ENSG00000089094 | KDM2B | chr12 | - | 121442156 | 121443793 | 121442156 | 121442836 | 121443679 | 121443793 | 9 | 6 | 11 | 8 | 17 | 0 | 0 | 3 | 2 | 0 | 14 | 14 | 10 | 21 | 89 | 0 | 0 | 0 | 0 | 0 | 298 | 149 | 0.468206001 | 0.901170691 | 1 | 1 | 0.647 | 0.667 | 1 | 1 | 1 | 1 | 1 | 1 | -0.137 |
| RI | 5308 | ENSG00000186318 | BACE1 | chr11 | - | 117293053 | 117294008 | 117293053 | 117293188 | 117293870 | 117294008 | 8 | 23 | 17 | 12 | 13 | 3 | 4 | 2 | 1 | 1 | 9 | 24 | 9 | 33 | 4 | 0 | 5 | 0 | 5 | 22 | 298 | 149 | 0.46884385 | 0.902104437 | 0.571 | 0.742 | 0.81 | 0.857 | 0.867 | 1 | 0.706 | 1 | 0.767 | 0.083 | 0.058 |
| RI | 7146 | ENSG00000172269 | DPAGT1 | chr11 | - | 119097141 | 119098487 | 119097141 | 119098043 | 119098402 | 119098487 | 7 | 3 | 4 | 9 | 19 | 27 | 9 | 16 | 23 | 21 | 18 | 3 | 10 | 14 | 5 | 8 | 22 | 20 | 14 | 24 | 298 | 149 | 0.469064016 | 0.902234172 | 0.115 | 0.143 | 0.111 | 0.164 | 0.311 | 0.529 | 0.064 | 0.2 | 0.333 | 0.094 | -0.075 |
| RI | 8095 | ENSG00000177239 | MAN1B1 | chr9 | + | 137106124 | 137106809 | 137106124 | 137106315 | 137106688 | 137106809 | 3 | 6 | 6 | 7 | 4 | 23 | 9 | 24 | 9 | 16 | 9 | 2 | 8 | 0 | 12 | 13 | 25 | 5 | 31 | 70 | 298 | 149 | 0.469512167 | 0.902802204 | 0.061 | 0.25 | 0.111 | 0.28 | 0.111 | 0.257 | 0.038 | 0.444 | 0 | 0.079 | -0.001 |
| RI | 5214 | ENSG00000149499 | EML3 | chr11 | - | 62603943 | 62604201 | 62603943 | 62604041 | 62604112 | 62604201 | 14 | 6 | 5 | 26 | 8 | 89 | 41 | 94 | 83 | 85 | 15 | 14 | 11 | 21 | 17 | 104 | 83 | 75 | 78 | 120 | 220 | 149 | 0.469934252 | 0.903319762 | 0.096 | 0.09 | 0.035 | 0.175 | 0.06 | 0.089 | 0.103 | 0.09 | 0.154 | 0.088 | -0.014 |
| RI | 2868 | ENSG00000184470 | TXNRD2 | chr22 | - | 19875521 | 19877234 | 19875521 | 19875807 | 19877039 | 19877234 | 0 | 1 | 3 | 9 | 6 | 15 | 1 | 4 | 10 | 1 | 18 | 2 | 2 | 3 | 22 | 5 | 10 | 8 | 2 | 22 | 298 | 149 | 0.4704967 | 0.904106704 | 0 | 0.333 | 0.273 | 0.31 | 0.75 | 0.643 | 0.091 | 0.111 | 0.429 | 0.333 | 0.012 |
| RI | 8416 | ENSG00000182796 | TMEM198B | chr12 | + | 55832958 | 55833541 | 55832958 | 55833038 | 55833446 | 55833541 | 15 | 1 | 7 | 7 | 7 | 0 | 0 | 0 | 1 | 0 | 8 | 17 | 8 | 19 | 19 | 0 | 0 | 0 | 5 | 0 | 298 | 149 | 0.471658516 | 0.906044506 | 1 | 1 | 1 | 0.778 | 1 | 1 | 1 | 1 | 0.655 | 1 | 0.025 |
| RI | 7714 | ENSG00000177697 | CD151 | chr11 | + | 836250 | 836843 | 836250 | 836442 | 836768 | 836843 | 13 | 4 | 15 | 12 | 17 | 27 | 9 | 23 | 40 | 25 | 17 | 19 | 19 | 21 | 9 | 42 | 35 | 25 | 15 | 195 | 298 | 149 | 0.472061522 | 0.906523865 | 0.194 | 0.182 | 0.246 | 0.13 | 0.254 | 0.168 | 0.213 | 0.275 | 0.412 | 0.023 | -0.017 |
| RI | 790 | ENSG00000214087 | ARL16 | chr17 | - | 81682033 | 81683126 | 81682033 | 81682149 | 81683012 | 81683126 | 32 | 26 | 36 | 45 | 17 | 30 | 8 | 8 | 6 | 34 | 76 | 91 | 54 | 81 | 57 | 34 | 18 | 8 | 43 | 32 | 298 | 149 | 0.473156612 | 0.908331526 | 0.348 | 0.619 | 0.692 | 0.789 | 0.2 | 0.528 | 0.717 | 0.771 | 0.485 | 0.471 | -0.065 |
| RI | 4860 | ENSG00000160326 | SLC2A6 | chr9 | - | 133478253 | 133479096 | 133478253 | 133478416 | 133478967 | 133479096 | 8 | 5 | 12 | 6 | 6 | 7 | 1 | 4 | 6 | 4 | 8 | 1 | 10 | 6 | 17 | 1 | 11 | 6 | 3 | 17 | 298 | 149 | 0.473362273 | 0.908347908 | 0.364 | 0.714 | 0.6 | 0.333 | 0.429 | 0.8 | 0.043 | 0.455 | 0.5 | 0.333 | 0.062 |
| RI | 1046 | ENSG00000006282 | SPATA20 | chr17 | + | 50548282 | 50548618 | 50548282 | 50548453 | 50548553 | 50548618 | 12 | 2 | 7 | 4 | 11 | 62 | 20 | 57 | 48 | 55 | 15 | 16 | 11 | 8 | 16 | 68 | 100 | 59 | 70 | 103 | 249 | 149 | 0.473472695 | 0.908347908 | 0.104 | 0.056 | 0.068 | 0.047 | 0.107 | 0.117 | 0.087 | 0.1 | 0.064 | 0.085 | -0.014 |
| RI | 4616 | ENSG00000117616 | RSRP1 | chr1 | - | 25242248 | 25242705 | 25242248 | 25242426 | 25242538 | 25242705 | 744 | 562 | 665 | 714 | 938 | 15 | 26 | 20 | 22 | 21 | 334 | 192 | 362 | 288 | 275 | 7 | 3 | 9 | 9 | 11 | 261 | 149 | 0.473815986 | 0.908711373 | 0.966 | 0.925 | 0.95 | 0.949 | 0.962 | 0.965 | 0.973 | 0.958 | 0.948 | 0.935 | -0.005 |
| RI | 6938 | ENSG00000046651 | OFD1 | chrX | + | 13767126 | 13768224 | 13767126 | 13767284 | 13768053 | 13768224 | 27 | 40 | 33 | 24 | 46 | 46 | 40 | 45 | 19 | 41 | 8 | 34 | 29 | 1 | 10 | 10 | 10 | 27 | 13 | 0 | 298 | 149 | 0.474609457 | 0.909937703 | 0.227 | 0.333 | 0.268 | 0.387 | 0.359 | 0.286 | 0.63 | 0.349 | 0.037 | 1 | -0.146 |
| RI | 4359 | ENSG00000158941 | CCAR2 | chr8 | + | 22618616 | 22619015 | 22618616 | 22618728 | 22618826 | 22619015 | 6 | 5 | 19 | 16 | 1 | 39 | 15 | 60 | 34 | 28 | 2 | 12 | 29 | 47 | 17 | 71 | 81 | 21 | 41 | 90 | 247 | 149 | 0.475118138 | 0.910617404 | 0.085 | 0.167 | 0.16 | 0.221 | 0.021 | 0.017 | 0.082 | 0.454 | 0.409 | 0.102 | -0.082 |
| RI | 8367 | ENSG00000106133 | NSUN5P2 | chr7 | - | 72948891 | 72949102 | 72948891 | 72948980 | 72949052 | 72949102 | 25 | 7 | 27 | 33 | 8 | 0 | 0 | 0 | 0 | 0 | 18 | 11 | 14 | 29 | 19 | 0 | 0 | 0 | 6 | 0 | 221 | 149 | 0.475347738 | 0.910761948 | 1 | 1 | 1 | 1 | 1 | 1 | 1 | 1 | 0.765 | 1 | 0.047 |
| RI | 7543 | ENSG00000130714 | POMT1 | chr9 | + | 131511336 | 131512136 | 131511336 | 131511467 | 131512040 | 131512136 | 6 | 3 | 7 | 4 | 0 | 19 | 9 | 20 | 12 | 10 | 0 | 17 | 3 | 5 | 39 | 10 | 36 | 27 | 28 | 9 | 298 | 149 | 0.476658753 | 0.91297771 | 0.136 | 0.143 | 0.149 | 0.143 | 0 | 0 | 0.191 | 0.053 | 0.082 | 0.684 | -0.088 |
| RI | 644 | ENSG00000069399 | BCL3 | chr19 | + | 44758245 | 44758841 | 44758245 | 44758473 | 44758723 | 44758841 | 7 | 3 | 9 | 15 | 6 | 0 | 0 | 0 | 0 | 0 | 47 | 9 | 13 | 50 | 33 | 0 | 4 | 0 | 0 | 0 | 298 | 149 | 0.477385563 | 0.914073427 | 1 | 1 | 1 | 1 | 1 | 1 | 0.529 | 1 | 1 | 1 | 0.094 |
| RI | 2153 | ENSG00000106123 | EPHB6 | chr7 | + | 142862755 | 142863327 | 142862755 | 142862833 | 142863130 | 142863327 | 1 | 0 | 6 | 7 | 10 | 0 | 2 | 2 | 1 | 0 | 2 | 1 | 3 | 1 | 13 | 0 | 0 | 0 | 3 | 0 | 298 | 149 | 0.477924114 | 0.914808083 | 1 | 0 | 0.6 | 0.778 | 1 | 1 | 1 | 1 | 0.143 | 1 | -0.153 |
| RI | 2749 | ENSG00000145247 | OCIAD2 | chr4 | - | 48892771 | 48894053 | 48892771 | 48892889 | 48894005 | 48894053 | 4 | 4 | 10 | 10 | 7 | 35 | 24 | 67 | 55 | 96 | 0 | 10 | 8 | 1 | 0 | 19 | 33 | 41 | 30 | 43 | 298 | 149 | 0.478203058 | 0.915045502 | 0.054 | 0.077 | 0.069 | 0.083 | 0.035 | 0 | 0.132 | 0.089 | 0.016 | 0 | 0.016 |
| RI | 5002 | ENSG00000092529 | CAPN3 | chr15 | + | 42410427 | 42410666 | 42410427 | 42410496 | 42410587 | 42410666 | 2 | 5 | 5 | 3 | 2 | 24 | 11 | 20 | 40 | 45 | 21 | 9 | 3 | 8 | 0 | 25 | 64 | 38 | 166 | 19 | 240 | 149 | 0.479537699 | 0.917302198 | 0.049 | 0.22 | 0.134 | 0.044 | 0.027 | 0.343 | 0.08 | 0.047 | 0.029 | 0 | -0.005 |
| RI | 3660 | ENSG00000204252 | HLA-DOA | chr6 | - | 33007079 | 33007592 | 33007079 | 33007215 | 33007310 | 33007592 | 1 | 3 | 2 | 9 | 0 | 18 | 12 | 20 | 19 | 18 | 1 | 4 | 3 | 5 | 11 | 29 | 11 | 3 | 17 | 69 | 244 | 149 | 0.480394188 | 0.918643078 | 0.033 | 0.132 | 0.058 | 0.224 | 0 | 0.021 | 0.182 | 0.379 | 0.152 | 0.089 | -0.075 |
| RI | 5022 | ENSG00000049167 | ERCC8 | chr5 | - | 60887439 | 60891086 | 60887439 | 60887520 | 60890888 | 60891086 | 7 | 18 | 12 | 12 | 10 | 23 | 4 | 17 | 23 | 39 | 0 | 13 | 6 | 3 | 5 | 10 | 9 | 2 | 3 | 1 | 298 | 149 | 0.480936538 | 0.919382567 | 0.132 | 0.692 | 0.261 | 0.207 | 0.114 | 0 | 0.419 | 0.6 | 0.333 | 0.714 | -0.132 |
| RI | 3931 | ENSG00000095787 | WAC | chr10 | + | 28614566 | 28616362 | 28614566 | 28614685 | 28616175 | 28616362 | 49 | 46 | 35 | 42 | 36 | 114 | 66 | 93 | 132 | 120 | 0 | 45 | 67 | 12 | 0 | 26 | 68 | 120 | 43 | 7 | 298 | 149 | 0.481933417 | 0.920990195 | 0.177 | 0.258 | 0.158 | 0.137 | 0.13 | 0 | 0.249 | 0.218 | 0.122 | 0 | 0.054 |
| RI | 6073 | ENSG00000204348 | DXO | chr6 | - | 31970605 | 31971147 | 31970605 | 31970825 | 31970911 | 31971147 | 16 | 22 | 28 | 10 | 18 | 6 | 0 | 6 | 1 | 7 | 8 | 18 | 28 | 22 | 3 | 10 | 3 | 2 | 1 | 6 | 235 | 149 | 0.482837645 | 0.922201069 | 0.628 | 1 | 0.747 | 0.864 | 0.62 | 0.337 | 0.792 | 0.899 | 0.933 | 0.241 | 0.131 |
| RI | 3363 | ENSG00000103507 | BCKDK | chr16 | + | 31111868 | 31112567 | 31111868 | 31112027 | 31112120 | 31112567 | 23 | 5 | 16 | 6 | 8 | 31 | 4 | 13 | 15 | 20 | 50 | 15 | 14 | 34 | 50 | 46 | 46 | 10 | 28 | 72 | 242 | 149 | 0.48287928 | 0.922201069 | 0.314 | 0.435 | 0.431 | 0.198 | 0.198 | 0.401 | 0.167 | 0.463 | 0.428 | 0.3 | -0.037 |
| RI | 3157 | ENSG00000100462 | PRMT5 | chr14 | - | 22920528 | 22921056 | 22920528 | 22920731 | 22920987 | 22921056 | 50 | 18 | 33 | 48 | 111 | 0 | 1 | 0 | 1 | 4 | 58 | 40 | 30 | 22 | 15 | 0 | 0 | 0 | 0 | 0 | 298 | 149 | 0.483974459 | 0.9239939 | 1 | 0.9 | 1 | 0.96 | 0.933 | 1 | 1 | 1 | 1 | 1 | -0.041 |
| RI | 1715 | ENSG00000100429 | HDAC10 | chr22 | - | 50246297 | 50246735 | 50246297 | 50246376 | 50246678 | 50246735 | 8 | 0 | 12 | 6 | 1 | 8 | 6 | 17 | 9 | 8 | 1 | 14 | 5 | 6 | 7 | 14 | 10 | 9 | 8 | 7 | 298 | 149 | 0.484196954 | 0.924120002 | 0.333 | 0 | 0.261 | 0.25 | 0.059 | 0.034 | 0.412 | 0.217 | 0.273 | 0.333 | -0.073 |
| RI | 206 | ENSG00000167642 | SPINT2 | chr19 | + | 38290536 | 38292468 | 38290536 | 38290575 | 38291839 | 38292468 | 2 | 1 | 4 | 5 | 2 | 65 | 43 | 63 | 103 | 54 | 4 | 0 | 0 | 0 | 6 | 76 | 65 | 25 | 74 | 417 | 298 | 149 | 0.484503008 | 0.924405448 | 0.015 | 0.011 | 0.031 | 0.024 | 0.018 | 0.026 | 0 | 0 | 0 | 0.007 | 0.013 |
| RI | 7793 | ENSG00000086848 | ALG9 | chr11 | - | 111782194 | 111786520 | 111782194 | 111786094 | 111786401 | 111786520 | 19 | 15 | 29 | 17 | 26 | 4 | 4 | 2 | 0 | 1 | 10 | 35 | 24 | 26 | 2 | 0 | 4 | 3 | 0 | 18 | 298 | 149 | 0.484777762 | 0.924446923 | 0.704 | 0.652 | 0.879 | 1 | 0.929 | 1 | 0.814 | 0.8 | 1 | 0.053 | 0.099 |
| RI | 4108 | ENSG00000116337 | AMPD2 | chr1 | + | 109626725 | 109627316 | 109626725 | 109626912 | 109627174 | 109627316 | 24 | 15 | 17 | 10 | 21 | 66 | 20 | 21 | 53 | 16 | 27 | 9 | 23 | 27 | 20 | 62 | 55 | 25 | 37 | 63 | 298 | 149 | 0.48486845 | 0.924446923 | 0.154 | 0.273 | 0.288 | 0.086 | 0.396 | 0.179 | 0.076 | 0.315 | 0.267 | 0.137 | 0.045 |
| RI | 1464 | ENSG00000140983 | RHOT2 | chr16 | + | 668352 | 668699 | 668352 | 668569 | 668655 | 668699 | 7 | 3 | 10 | 7 | 2 | 79 | 41 | 55 | 70 | 29 | 0 | 11 | 12 | 9 | 13 | 51 | 67 | 58 | 61 | 137 | 235 | 149 | 0.484994246 | 0.924446923 | 0.053 | 0.044 | 0.103 | 0.06 | 0.042 | 0 | 0.094 | 0.116 | 0.086 | 0.057 | -0.01 |
| RI | 7189 | ENSG00000214706 | IFRD2 | chr3 | - | 50289928 | 50290294 | 50289928 | 50290086 | 50290169 | 50290294 | 3 | 0 | 6 | 4 | 1 | 8 | 7 | 13 | 13 | 17 | 5 | 0 | 4 | 3 | 0 | 0 | 9 | 5 | 1 | 17 | 232 | 149 | 0.487836838 | 0.929565226 | 0.194 | 0 | 0.229 | 0.165 | 0.036 | 1 | 0 | 0.339 | 0.658 | 0 | -0.275 |
| RI | 209 | ENSG00000132024 | CC2D1A | chr19 | + | 13926977 | 13928030 | 13926977 | 13927265 | 13927892 | 13928030 | 4 | 9 | 7 | 7 | 2 | 14 | 14 | 24 | 17 | 6 | 7 | 7 | 3 | 22 | 36 | 16 | 34 | 17 | 4 | 99 | 298 | 149 | 0.488262112 | 0.930075555 | 0.125 | 0.243 | 0.127 | 0.171 | 0.143 | 0.179 | 0.093 | 0.081 | 0.733 | 0.154 | -0.086 |
| RI | 4814 | ENSG00000167671 | UBXN6 | chr19 | - | 4446048 | 4446413 | 4446048 | 4446197 | 4446282 | 4446413 | 7 | 5 | 5 | 3 | 1 | 29 | 25 | 19 | 27 | 8 | 0 | 1 | 0 | 10 | 15 | 21 | 30 | 5 | 12 | 108 | 234 | 149 | 0.488671161 | 0.930554657 | 0.133 | 0.113 | 0.144 | 0.066 | 0.074 | 0 | 0.021 | 0 | 0.347 | 0.081 | 0.016 |
| RI | 642 | ENSG00000283633 | NA | chr22 | + | 16614076 | 16615111 | 16614076 | 16614178 | 16614698 | 16615111 | 85 | 128 | 10 | 207 | 96 | 9 | 32 | 6 | 35 | 17 | 1305 | 117 | 181 | 172 | 1991 | 213 | 57 | 27 | 27 | 668 | 298 | 149 | 0.488831075 | 0.930559187 | 0.825 | 0.667 | 0.455 | 0.747 | 0.738 | 0.754 | 0.506 | 0.77 | 0.761 | 0.598 | 0.009 |
| RI | 2900 | ENSG00000172053 | QARS1 | chr3 | - | 49098599 | 49098989 | 49098599 | 49098692 | 49098884 | 49098989 | 30 | 4 | 21 | 22 | 23 | 105 | 67 | 122 | 97 | 87 | 9 | 34 | 18 | 31 | 50 | 139 | 100 | 97 | 70 | 209 | 298 | 149 | 0.489069716 | 0.930713535 | 0.125 | 0.029 | 0.079 | 0.102 | 0.117 | 0.031 | 0.145 | 0.085 | 0.181 | 0.107 | -0.019 |
| RI | 6851 | ENSG00000215021 | PHB2 | chr12 | - | 6967170 | 6968021 | 6967170 | 6967238 | 6967891 | 6968021 | 55 | 49 | 81 | 85 | 116 | 14 | 6 | 2 | 5 | 13 | 166 | 93 | 101 | 113 | 114 | 31 | 14 | 22 | 11 | 0 | 298 | 149 | 0.489343042 | 0.930933768 | 0.663 | 0.803 | 0.953 | 0.895 | 0.817 | 0.728 | 0.769 | 0.697 | 0.837 | 1 | 0.02 |
| RI | 5927 | ENSG00000114388 | NPRL2 | chr3 | - | 50348522 | 50348782 | 50348522 | 50348563 | 50348710 | 50348782 | 13 | 6 | 15 | 17 | 16 | 5 | 1 | 2 | 0 | 3 | 21 | 16 | 27 | 29 | 34 | 0 | 4 | 3 | 8 | 11 | 296 | 149 | 0.489726813 | 0.931363903 | 0.567 | 0.751 | 0.791 | 1 | 0.729 | 1 | 0.668 | 0.819 | 0.646 | 0.609 | 0.019 |
| RI | 1111 | ENSG00000118046 | STK11 | chr19 | + | 1220372 | 1220717 | 1220372 | 1220505 | 1220580 | 1220717 | 15 | 4 | 17 | 9 | 16 | 64 | 32 | 70 | 51 | 48 | 28 | 11 | 3 | 1 | 10 | 19 | 68 | 14 | 15 | 42 | 224 | 149 | 0.490133114 | 0.931836597 | 0.135 | 0.077 | 0.139 | 0.105 | 0.181 | 0.495 | 0.097 | 0.125 | 0.042 | 0.137 | -0.052 |
| RI | 3560 | ENSG00000089327 | FXYD5 | chr19 | + | 35166141 | 35166325 | 35166141 | 35166171 | 35166250 | 35166325 | 40 | 28 | 46 | 33 | 31 | 1134 | 589 | 991 | 873 | 738 | 86 | 60 | 55 | 38 | 89 | 1695 | 1118 | 819 | 1044 | 3026 | 228 | 149 | 0.490550052 | 0.9323292 | 0.023 | 0.03 | 0.029 | 0.024 | 0.027 | 0.032 | 0.034 | 0.042 | 0.023 | 0.019 | -0.003 |
| RI | 4695 | ENSG00000273136 | NBPF26 | chr1 | + | 120805555 | 120805765 | 120805555 | 120805710 | 120805712 | 120805765 | 83 | 88 | 34 | 25 | 39 | 0 | 0 | 8 | 0 | 0 | 69 | 56 | 121 | 125 | 64 | 0 | 0 | 0 | 0 | 0 | 151 | 149 | 0.491661804 | 0.934141613 | 1 | 1 | 0.807 | 1 | 1 | 1 | 1 | 1 | 1 | 1 | -0.039 |
| RI | 3124 | ENSG00000185507 | IRF7 | chr11 | - | 613784 | 614037 | 613784 | 613865 | 613950 | 614037 | 11 | 7 | 3 | 16 | 16 | 21 | 10 | 11 | 21 | 2 | 4 | 12 | 7 | 39 | 24 | 8 | 23 | 6 | 13 | 15 | 234 | 149 | 0.492335513 | 0.934826885 | 0.25 | 0.308 | 0.148 | 0.327 | 0.836 | 0.241 | 0.249 | 0.426 | 0.656 | 0.505 | -0.042 |
| RI | 3251 | ENSG00000064961 | HMG20B | chr19 | + | 3576260 | 3576625 | 3576260 | 3576307 | 3576552 | 3576625 | 5 | 3 | 6 | 1 | 19 | 248 | 228 | 191 | 286 | 240 | 21 | 1 | 3 | 3 | 19 | 483 | 282 | 164 | 255 | 1201 | 298 | 149 | 0.492338994 | 0.934826885 | 0.01 | 0.007 | 0.015 | 0.002 | 0.038 | 0.021 | 0.002 | 0.009 | 0.006 | 0.008 | 0.005 |
| RI | 4984 | ENSG00000124067 | SLC12A4 | chr16 | - | 67950961 | 67951304 | 67950961 | 67951060 | 67951139 | 67951304 | 5 | 8 | 8 | 11 | 1 | 9 | 4 | 5 | 5 | 2 | 0 | 9 | 4 | 6 | 26 | 5 | 7 | 7 | 11 | 4 | 228 | 149 | 0.492531237 | 0.934891394 | 0.266 | 0.567 | 0.511 | 0.59 | 0.246 | 0 | 0.457 | 0.272 | 0.263 | 0.809 | 0.076 |
| RI | 3930 | ENSG00000095787 | WAC | chr10 | + | 28614566 | 28616362 | 28614566 | 28614685 | 28616172 | 28616362 | 44 | 47 | 31 | 40 | 31 | 317 | 192 | 316 | 281 | 418 | 0 | 39 | 68 | 12 | 0 | 41 | 242 | 225 | 183 | 31 | 298 | 149 | 0.494664866 | 0.938639693 | 0.065 | 0.109 | 0.047 | 0.066 | 0.036 | 0 | 0.075 | 0.131 | 0.032 | 0 | 0.017 |
| RI | 2856 | ENSG00000113851 | CRBN | chr3 | - | 3153423 | 3156281 | 3153423 | 3153488 | 3156218 | 3156281 | 86 | 64 | 66 | 54 | 94 | 9 | 4 | 7 | 5 | 6 | 28 | 16 | 42 | 10 | 6 | 19 | 0 | 1 | 0 | 0 | 298 | 149 | 0.495213538 | 0.939379052 | 0.827 | 0.889 | 0.825 | 0.844 | 0.887 | 0.424 | 1 | 0.955 | 1 | 1 | -0.021 |
| RI | 3771 | ENSG00000134419 | RPS15A | chr16 | - | 18782967 | 18784823 | 18782967 | 18783102 | 18784737 | 18784823 | 43 | 34 | 77 | 61 | 28 | 5149 | 2875 | 6322 | 5046 | 7458 | 117 | 25 | 32 | 59 | 59 | 4104 | 3393 | 3473 | 2544 | 5579 | 298 | 149 | 0.495851482 | 0.940287224 | 0.004 | 0.006 | 0.006 | 0.006 | 0.002 | 0.014 | 0.004 | 0.005 | 0.011 | 0.005 | -0.003 |
| RI | 1553 | ENSG00000166794 | PPIB | chr15 | - | 64156724 | 64160197 | 64156724 | 64156909 | 64160103 | 64160197 | 91 | 31 | 76 | 91 | 112 | 635 | 309 | 719 | 551 | 884 | 141 | 67 | 94 | 70 | 124 | 674 | 824 | 723 | 500 | 869 | 298 | 149 | 0.496504545 | 0.941099129 | 0.067 | 0.048 | 0.05 | 0.076 | 0.06 | 0.095 | 0.039 | 0.061 | 0.065 | 0.067 | -0.005 |
| RI | 188 | ENSG00000142002 | DPP9 | chr19 | - | 4684662 | 4685771 | 4684662 | 4684809 | 4685625 | 4685771 | 13 | 3 | 4 | 3 | 9 | 35 | 10 | 11 | 23 | 4 | 16 | 21 | 8 | 1 | 28 | 47 | 46 | 15 | 14 | 23 | 298 | 149 | 0.496598271 | 0.941099129 | 0.157 | 0.13 | 0.154 | 0.061 | 0.529 | 0.145 | 0.186 | 0.211 | 0.034 | 0.378 | 0.015 |
| RI | 3885 | ENSG00000158517 | NCF1 | chr7 | + | 74787983 | 74788704 | 74787983 | 74788088 | 74788558 | 74788704 | 4 | 4 | 7 | 4 | 0 | 240 | 130 | 132 | 99 | 63 | 4 | 7 | 4 | 0 | 13 | 148 | 192 | 71 | 150 | 280 | 298 | 149 | 0.499041551 | 0.945426056 | 0.008 | 0.015 | 0.026 | 0.02 | 0 | 0.013 | 0.018 | 0.027 | 0 | 0.023 | -0.002 |
| RI | 7713 | ENSG00000177666 | PNPLA2 | chr11 | + | 823693 | 824130 | 823693 | 823855 | 823997 | 824130 | 15 | 11 | 12 | 11 | 4 | 73 | 27 | 65 | 42 | 30 | 14 | 15 | 21 | 30 | 26 | 78 | 104 | 44 | 95 | 111 | 291 | 149 | 0.500431937 | 0.947756156 | 0.095 | 0.173 | 0.086 | 0.118 | 0.064 | 0.084 | 0.069 | 0.196 | 0.139 | 0.107 | -0.012 |
| RI | 8366 | ENSG00000166012 | TAF1D | chr11 | - | 93736693 | 93737239 | 93736693 | 93736751 | 93737063 | 93737239 | 274 | 326 | 238 | 308 | 372 | 288 | 234 | 305 | 275 | 599 | 32 | 73 | 193 | 23 | 17 | 31 | 119 | 136 | 26 | 5 | 298 | 149 | 0.501860878 | 0.950157758 | 0.322 | 0.411 | 0.281 | 0.359 | 0.237 | 0.34 | 0.235 | 0.415 | 0.307 | 0.63 | -0.063 |
| RI | 5814 | ENSG00000054148 | PHPT1 | chr9 | + | 136850012 | 136850560 | 136850012 | 136850137 | 136850502 | 136850560 | 21 | 4 | 15 | 17 | 2 | 1 | 1 | 2 | 0 | 1 | 7 | 12 | 3 | 7 | 36 | 0 | 0 | 0 | 5 | 0 | 298 | 149 | 0.502317578 | 0.950717697 | 0.913 | 0.667 | 0.789 | 1 | 0.5 | 1 | 1 | 1 | 0.412 | 1 | -0.109 |
| RI | 7882 | ENSG00000118894 | EEF2KMT | chr16 | - | 5090083 | 5090565 | 5090083 | 5090349 | 5090431 | 5090565 | 2 | 1 | 3 | 0 | 0 | 2 | 0 | 1 | 5 | 3 | 0 | 0 | 10 | 0 | 8 | 7 | 8 | 2 | 6 | 0 | 231 | 149 | 0.50251316 | 0.950731967 | 0.392 | 1 | 0.659 | 0 | 0 | 0 | 0 | 0.763 | 0 | 1 | 0.058 |
| RI | 8150 | ENSG00000107862 | GBF1 | chr10 | + | 102379520 | 102379954 | 102379520 | 102379729 | 102379852 | 102379954 | 5 | 4 | 7 | 4 | 2 | 3 | 0 | 4 | 0 | 0 | 57 | 24 | 20 | 38 | 27 | 15 | 2 | 13 | 12 | 0 | 272 | 149 | 0.502647018 | 0.950731967 | 0.477 | 1 | 0.489 | 1 | 1 | 0.675 | 0.868 | 0.457 | 0.634 | 1 | 0.066 |
| RI | 277 | ENSG00000099860 | GADD45B | chr19 | + | 2476355 | 2476630 | 2476355 | 2476431 | 2476528 | 2476630 | 14 | 5 | 14 | 7 | 4 | 5 | 2 | 1 | 1 | 0 | 10 | 9 | 24 | 15 | 54 | 9 | 1 | 6 | 8 | 0 | 246 | 149 | 0.502844147 | 0.950800377 | 0.629 | 0.602 | 0.895 | 0.809 | 1 | 0.402 | 0.845 | 0.708 | 0.532 | 1 | 0.09 |
| RI | 5414 | ENSG00000018280 | SLC11A1 | chr2 | + | 218386634 | 218387230 | 218386634 | 218386741 | 218387159 | 218387230 | 64 | 75 | 54 | 85 | 92 | 77 | 35 | 14 | 51 | 14 | 161 | 184 | 247 | 176 | 402 | 71 | 154 | 114 | 106 | 158 | 298 | 149 | 0.503336319 | 0.951426444 | 0.294 | 0.517 | 0.659 | 0.455 | 0.767 | 0.531 | 0.374 | 0.52 | 0.454 | 0.56 | 0.051 |
| RI | 4952 | ENSG00000171503 | ETFDH | chr4 | + | 158680466 | 158682424 | 158680466 | 158681614 | 158682194 | 158682424 | 16 | 26 | 19 | 17 | 56 | 11 | 7 | 8 | 3 | 1 | 14 | 34 | 81 | 37 | 13 | 0 | 14 | 14 | 10 | 0 | 298 | 149 | 0.503665601 | 0.951744307 | 0.421 | 0.65 | 0.543 | 0.739 | 0.966 | 1 | 0.548 | 0.743 | 0.649 | 1 | -0.124 |
| RI | 8057 | ENSG00000156671 | SAMD8 | chr10 | + | 75176065 | 75182123 | 75176065 | 75176216 | 75176387 | 75182123 | 34 | 25 | 23 | 7 | 33 | 46 | 37 | 80 | 51 | 69 | 10 | 33 | 57 | 48 | 0 | 22 | 38 | 74 | 56 | 54 | 298 | 149 | 0.505449247 | 0.954488133 | 0.27 | 0.253 | 0.126 | 0.064 | 0.193 | 0.185 | 0.303 | 0.278 | 0.3 | 0 | -0.032 |
| RI | 3035 | ENSG00000206503 | HLA-A | chr6 | + | 29944499 | 29945091 | 29944499 | 29944616 | 29945058 | 29945091 | 457 | 190 | 963 | 391 | 653 | 7613 | 2121 | 5050 | 5130 | 2449 | 1823 | 963 | 1640 | 1842 | 1379 | 13612 | 7888 | 6627 | 13599 | 25534 | 298 | 149 | 0.505480735 | 0.954488133 | 0.029 | 0.043 | 0.087 | 0.037 | 0.118 | 0.063 | 0.058 | 0.11 | 0.063 | 0.026 | -0.001 |
| RI | 4086 | ENSG00000107551 | RASSF4 | chr10 | + | 44970164 | 44971848 | 44970164 | 44970264 | 44971772 | 44971848 | 202 | 93 | 77 | 82 | 52 | 92 | 43 | 43 | 60 | 47 | 554 | 81 | 79 | 113 | 225 | 149 | 64 | 22 | 75 | 183 | 298 | 149 | 0.505602399 | 0.954488133 | 0.523 | 0.52 | 0.472 | 0.406 | 0.356 | 0.65 | 0.388 | 0.642 | 0.43 | 0.381 | -0.043 |
| RI | 6420 | ENSG00000164896 | FASTK | chr7 | - | 151076912 | 151078092 | 151076912 | 151077780 | 151077878 | 151078092 | 10 | 7 | 20 | 15 | 9 | 77 | 9 | 43 | 51 | 58 | 8 | 10 | 23 | 5 | 39 | 17 | 116 | 31 | 54 | 47 | 247 | 149 | 0.507105725 | 0.957020294 | 0.073 | 0.319 | 0.219 | 0.151 | 0.086 | 0.221 | 0.049 | 0.309 | 0.053 | 0.334 | -0.024 |
| RI | 1393 | ENSG00000123815 | COQ8B | chr19 | - | 40703540 | 40703855 | 40703540 | 40703622 | 40703714 | 40703855 | 5 | 3 | 6 | 2 | 1 | 15 | 1 | 15 | 13 | 0 | 5 | 0 | 0 | 2 | 8 | 0 | 5 | 2 | 3 | 0 | 241 | 149 | 0.507438575 | 0.957075177 | 0.171 | 0.65 | 0.198 | 0.087 | 1 | 1 | 0 | 0 | 0.292 | 1 | -0.037 |
| RI | 4712 | ENSG00000232956 | SNHG15 | chr7 | - | 44983821 | 44986097 | 44983821 | 44984426 | 44986020 | 44986097 | 12 | 12 | 30 | 22 | 33 | 14 | 7 | 20 | 13 | 9 | 43 | 39 | 21 | 20 | 75 | 15 | 5 | 21 | 33 | 17 | 298 | 149 | 0.507566229 | 0.957075177 | 0.3 | 0.462 | 0.429 | 0.458 | 0.647 | 0.589 | 0.796 | 0.333 | 0.233 | 0.688 | -0.069 |

| | | | | | | | | | | | | | | | | | | | | | | | | | | | | | | | | | | | | | | | | | | | | | | |
|---|---|---|---|---|---|---|---|---|---|---|---|---|---|---|---|---|---|---|---|---|---|---|---|---|---|---|---|---|---|---|---|---|---|---|---|---|---|---|---|---|---|---|---|---|---|---|
| RI | 376 | ENSG00000006712 | PAF1 | chr19 | - | 39390068 | 39390289 | 39390068 | 39390161 | 39390259 | 39390289 | 7 | 1 | 17 | 16 | 31 | 37 | 20 | 42 | 38 | 10 | 14 | 26 | 6 | 21 | 55 | 31 | 37 | 36 | 51 | 54 | 247 | 149 | 0.507620879 | 0.957075177 | 0.102 | 0.029 | 0.196 | 0.203 | 0.652 | 0.214 | 0.298 | 0.091 | 0.199 | 0.381 | 0 |
| RI | 7124 | ENSG00000198851 | CD3E | chr11 | + | 118312152 | 118313874 | 118312152 | 118312170 | 118313706 | 118313874 | 66 | 42 | 55 | 50 | 155 | 7 | 3 | 15 | 15 | 33 | 36 | 52 | 40 | 25 | 35 | 11 | 5 | 13 | 4 | 0 | 298 | 149 | 0.507877032 | 0.957252593 | 0.825 | 0.875 | 0.647 | 0.625 | 0.701 | 0.621 | 0.839 | 0.606 | 0.758 | 1 | -0.03 |
| RI | 3491 | ENSG00000143578 | CREB3L4 | chr1 | + | 153972743 | 153973078 | 153972743 | 153972836 | 153972971 | 153973078 | 4 | 9 | 11 | 1 | 2 | 5 | 2 | 2 | 2 | 0 | 8 | 0 | 4 | 4 | 3 | 0 | 10 | 0 | 5 | 0 | 284 | 149 | 0.509935989 | 0.960826758 | 0.296 | 0.702 | 0.743 | 0.208 | 1 | 1 | 0 | 1 | 0.296 | 1 | -0.069 |
| RI | 4910 | ENSG00000160818 | GPATCH4 | chr1 | - | 156596080 | 156596478 | 156596080 | 156596131 | 156596386 | 156596478 | 25 | 15 | 32 | 22 | 66 | 17 | 5 | 15 | 19 | 16 | 43 | 33 | 18 | 25 | 49 | 69 | 18 | 21 | 17 | 4 | 298 | 149 | 0.511357071 | 0.963001564 | 0.424 | 0.6 | 0.516 | 0.367 | 0.673 | 0.238 | 0.478 | 0.3 | 0.424 | 0.86 | 0.056 |
| RI | 4277 | ENSG00000165487 | MICU2 | chr13 | - | 21495160 | 21496160 | 21495160 | 21495318 | 21496051 | 21496160 | 18 | 28 | 22 | 37 | 30 | 88 | 117 | 91 | 102 | 131 | 0 | 16 | 32 | 5 | 9 | 8 | 50 | 72 | 36 | 6 | 298 | 149 | 0.511540832 | 0.963001564 | 0.093 | 0.107 | 0.108 | 0.154 | 0.103 | 0 | 0.138 | 0.182 | 0.065 | 0.429 | -0.05 |
| RI | 7276 | ENSG00000161016 | RPL8 | chr8 | - | 144791276 | 144791495 | 144791276 | 144791307 | 144791370 | 144791495 | 5393 | 2283 | 4919 | 4415 | 3576 | 158 | 77 | 115 | 130 | 52 | 4555 | 4504 | 3228 | 2588 | 6536 | 117 | 111 | 91 | 61 | 131 | 212 | 149 | 0.511579297 | 0.963001564 | 0.96 | 0.954 | 0.968 | 0.96 | 0.98 | 0.965 | 0.966 | 0.961 | 0.968 | 0.972 | -0.002 |
| RI | 927 | ENSG00000011143 | MKS1 | chr17 | - | 58214258 | 58216736 | 58214258 | 58216243 | 58216665 | 58216736 | 2 | 0 | 1 | 1 | 4 | 32 | 7 | 29 | 23 | 32 | 0 | 0 | 0 | 0 | 2 | 12 | 22 | 14 | 2 | 12 | 298 | 149 | 0.511910488 | 0.963039622 | 0.03 | 0 | 0.017 | 0.021 | 0.059 | 0 | 0 | 0 | 0 | 0.077 | 0.01 |
| RI | 4264 | ENSG00000198837 | DENND4B | chr1 | - | 153930538 | 153931064 | 153930538 | 153930603 | 153930691 | 153931064 | 9 | 10 | 17 | 15 | 0 | 85 | 19 | 62 | 52 | 22 | 19 | 16 | 8 | 9 | 15 | 71 | 68 | 24 | 30 | 80 | 237 | 149 | 0.511948292 | 0.963039622 | 0.062 | 0.249 | 0.147 | 0.154 | 0 | 0.144 | 0.129 | 0.173 | 0.159 | 0.105 | -0.02 |
| RI | 6373 | ENSG00000168137 | SETD5 | chr3 | + | 9474448 | 9475156 | 9474448 | 9474582 | 9475067 | 9475156 | 29 | 18 | 24 | 33 | 33 | 129 | 63 | 105 | 105 | 100 | 46 | 25 | 32 | 28 | 20 | 108 | 117 | 103 | 87 | 85 | 298 | 149 | 0.512088616 | 0.963039622 | 0.101 | 0.125 | 0.103 | 0.136 | 0.142 | 0.176 | 0.097 | 0.134 | 0.139 | 0.105 | -0.009 |
| RI | 719 | ENSG00000169733 | RFNG | chr17 | - | 82050661 | 82051342 | 82050661 | 82050764 | 82051293 | 82051342 | 17 | 1 | 3 | 6 | 0 | 14 | 1 | 6 | 10 | 4 | 4 | 11 | 1 | 4 | 12 | 6 | 13 | 5 | 3 | 10 | 298 | 149 | 0.513332011 | 0.965070715 | 0.378 | 0.333 | 0.2 | 0.231 | 0 | 0.25 | 0.297 | 0.091 | 0.4 | 0.375 | -0.054 |
| RI | 787 | ENSG00000196704 | AMZ2 | chr17 | + | 68250187 | 68250967 | 68250187 | 68250470 | 68250793 | 68250967 | 22 | 10 | 31 | 38 | 19 | 61 | 36 | 57 | 58 | 89 | 22 | 30 | 38 | 30 | 8 | 63 | 63 | 62 | 74 | 105 | 298 | 149 | 0.513533467 | 0.96514228 | 0.153 | 0.122 | 0.214 | 0.247 | 0.096 | 0.149 | 0.192 | 0.235 | 0.169 | 0.037 | 0.01 |
| RI | 5290 | ENSG00000136718 | IMP4 | chr2 | + | 130346017 | 130346249 | 130346017 | 130346112 | 130346200 | 130346249 | 11 | 7 | 13 | 16 | 9 | 32 | 26 | 25 | 23 | 37 | 13 | 8 | 9 | 37 | 36 | 40 | 38 | 26 | 40 | 71 | 237 | 149 | 0.513925786 | 0.965408027 | 0.178 | 0.145 | 0.246 | 0.304 | 0.133 | 0.17 | 0.117 | 0.179 | 0.368 | 0.242 | -0.014 |
| RI | 5777 | ENSG00000272752 | STAG3L5P-PVRIG2P-PILRB | chr7 | + | 100352563 | 100352914 | 100352563 | 100352667 | 100352760 | 100352914 | 119 | 45 | 125 | 83 | 64 | 1 | 1 | 1 | 1 | 3 | 40 | 96 | 46 | 48 | 97 | 0 | 0 | 0 | 0 | 0 | 242 | 149 | 0.514060002 | 0.965408027 | 0.987 | 0.965 | 0.987 | 0.981 | 0.929 | 1 | 1 | 1 | 1 | 1 | -0.03 |
| RI | 5494 | ENSG00000147454 | SLC25A37 | chr8 | + | 23566107 | 23568378 | 23566107 | 23566336 | 23568321 | 23568378 | 92 | 104 | 119 | 209 | 167 | 7 | 7 | 9 | 8 | 15 | 365 | 235 | 918 | 555 | 257 | 12 | 24 | 23 | 32 | 27 | 298 | 149 | 0.51421543 | 0.965408027 | 0.868 | 0.881 | 0.869 | 0.929 | 0.848 | 0.938 | 0.83 | 0.952 | 0.897 | 0.826 | -0.01 |
| RI | 3515 | ENSG00000130826 | DKC1 | chrX | + | 154774601 | 154775273 | 154774601 | 154774705 | 154775194 | 154775273 | 8 | 31 | 13 | 40 | 33 | 42 | 34 | 43 | 43 | 63 | 42 | 36 | 23 | 63 | 8 | 52 | 43 | 38 | 36 | 89 | 298 | 149 | 0.514328604 | 0.965408027 | 0.087 | 0.313 | 0.131 | 0.317 | 0.208 | 0.288 | 0.295 | 0.232 | 0.467 | 0.043 | -0.054 |
| RI | 5965 | ENSG00000168000 | BSCL2 | chr11 | - | 62690786 | 62692473 | 62690786 | 62691421 | 62692375 | 62692473 | 1 | 1 | 2 | 4 | 0 | 43 | 18 | 27 | 37 | 39 | 0 | 0 | 3 | 1 | 0 | 73 | 55 | 11 | 37 | 159 | 298 | 149 | 0.514826955 | 0.966036475 | 0.011 | 0.027 | 0.036 | 0.051 | 0 | 0 | 0 | 0.12 | 0.013 | 0 | -0.002 |
| RI | 675 | ENSG00000130813 | SHFL | chr19 | + | 10089306 | 10089695 | 10089306 | 10089378 | 10089656 | 10089695 | 30 | 25 | 31 | 24 | 50 | 15 | 4 | 20 | 7 | 5 | 74 | 48 | 41 | 81 | 71 | 37 | 20 | 18 | 24 | 14 | 298 | 149 | 0.515126907 | 0.966141103 | 0.5 | 0.758 | 0.437 | 0.632 | 0.833 | 0.5 | 0.545 | 0.532 | 0.628 | 0.717 | 0.048 |
| RI | 7374 | ENSG00000148346 | LCN2 | chr9 | + | 128153099 | 128153453 | 128153099 | 128153126 | 128153310 | 128153453 | 0 | 5 | 6 | 5 | 11 | 59 | 53 | 29 | 132 | 71 | 24 | 27 | 7 | 357 | 135 | 306 | 339 | 106 | 4707 | 1676 | 298 | 149 | 0.515365771 | 0.966141103 | 0 | 0.045 | 0.094 | 0.019 | 0.072 | 0.038 | 0.038 | 0.032 | 0.037 | 0.039 | 0.009 |
| RI | 2866 | ENSG00000157601 | MX1 | chr21 | + | 41445999 | 41449295 | 41445999 | 41446141 | 41449136 | 41449295 | 59 | 116 | 72 | 134 | 59 | 156 | 312 | 146 | 359 | 362 | 344 | 147 | 158 | 492 | 47 | 1343 | 526 | 173 | 996 | 247 | 298 | 149 | 0.515396393 | 0.966141103 | 0.159 | 0.157 | 0.198 | 0.157 | 0.075 | 0.114 | 0.123 | 0.313 | 0.198 | 0.087 | -0.018 |
| RI | 5812 | ENSG00000198721 | ECI2 | chr6 | - | 4117307 | 4119275 | 4117307 | 4117451 | 4119185 | 4119275 | 7 | 12 | 28 | 23 | 10 | 28 | 44 | 47 | 30 | 50 | 10 | 11 | 27 | 16 | 0 | 43 | 26 | 18 | 18 | 10 | 298 | 149 | 0.515536949 | 0.966141103 | 0.111 | 0.12 | 0.23 | 0.277 | 0.091 | 0.104 | 0.175 | 0.429 | 0.308 | 0 | -0.037 |
| RI | 292 | ENSG00000104872 | PIH1D1 | chr19 | - | 49447826 | 49448010 | 49447826 | 49447908 | 49448000 | 49448010 | 26 | 23 | 21 | 35 | 19 | 107 | 90 | 101 | 110 | 110 | 27 | 59 | 20 | 19 | 48 | 174 | 172 | 95 | 136 | 222 | 241 | 149 | 0.515769344 | 0.966270066 | 0.131 | 0.136 | 0.114 | 0.164 | 0.096 | 0.088 | 0.175 | 0.115 | 0.08 | 0.118 | 0.013 |
| RI | 8601 | ENSG00000134899 | ERCC5 | chr13 | + | 102872197 | 102873343 | 102872197 | 102872398 | 102873260 | 102873343 | 21 | 39 | 16 | 40 | 56 | 7 | 5 | 6 | 5 | 4 | 20 | 31 | 41 | 4 | 3 | 0 | 3 | 5 | 7 | 0 | 298 | 149 | 0.516131201 | 0.966641409 | 0.6 | 0.796 | 0.571 | 0.8 | 0.875 | 1 | 0.838 | 0.804 | 0.222 | 1 | -0.044 |
| RI | 96 | ENSG00000100031 | GGT1 | chr22 | + | 24610249 | 24610531 | 24610249 | 24610336 | 24610413 | 24610531 | 13 | 4 | 16 | 23 | 33 | 4 | 0 | 0 | 1 | 0 | 33 | 27 | 8 | 17 | 39 | 8 | 0 | 0 | 0 | 0 | 226 | 149 | 0.516494112 | 0.967014491 | 0.682 | 1 | 1 | 0.938 | 1 | 0.731 | 1 | 1 | 1 | 1 | -0.022 |
| RI | 7150 | ENSG00000172269 | DPAGT1 | chr11 | - | 119100629 | 119101138 | 119100629 | 119100843 | 119101017 | 119101138 | 12 | 13 | 11 | 28 | 3 | 9 | 8 | 1 | 16 | 6 | 16 | 14 | 1 | 15 | 14 | 16 | 7 | 4 | 5 | 15 | 298 | 149 | 0.516662548 | 0.967023344 | 0.4 | 0.448 | 0.846 | 0.467 | 0.2 | 0.333 | 0.5 | 0.111 | 0.6 | 0.318 | 0.1 |
| RI | 7651 | ENSG00000142186 | SCYL1 | chr11 | + | 65538269 | 65538704 | 65538269 | 65538324 | 65538441 | 65538704 | 11 | 3 | 6 | 12 | 2 | 104 | 40 | 84 | 74 | 46 | 7 | 24 | 14 | 15 | 11 | 132 | 114 | 101 | 87 | 257 | 266 | 149 | 0.517023612 | 0.967392612 | 0.056 | 0.04 | 0.038 | 0.083 | 0.024 | 0.029 | 0.105 | 0.072 | 0.088 | 0.023 | -0.015 |
| RI | 7697 | ENSG00000144354 | CDCA7 | chr2 | + | 173363817 | 173364989 | 173363817 | 173363895 | 173364794 | 173364989 | 6 | 6 | 6 | 10 | 3 | 7 | 4 | 2 | 11 | 9 | 0 | 12 | 20 | 18 | 5 | 19 | 11 | 5 | 19 | 15 | 298 | 149 | 0.518169467 | 0.969172494 | 0.3 | 0.429 | 0.6 | 0.312 | 0.143 | 0 | 0.353 | 0.667 | 0.321 | 0.143 | 0.06 |
| RI | 8360 | ENSG00000135297 | MTO1 | chr6 | + | 73479935 | 73480367 | 73479935 | 73480126 | 73480292 | 73480367 | 26 | 34 | 44 | 32 | 42 | 0 | 0 | 0 | 1 | 0 | 56 | 38 | 22 | 32 | 109 | 0 | 0 | 0 | 6 | 0 | 298 | 149 | 0.518462736 | 0.969172494 | 1 | 1 | 1 | 0.941 | 1 | 1 | 1 | 1 | 0.727 | 1 | 0.043 |
| RI | 3676 | ENSG00000160789 | LMNA | chr1 | + | 156136213 | 156137028 | 156136213 | 156136436 | 156136920 | 156137028 | 3 | 3 | 1 | 9 | 1 | 31 | 27 | 15 | 50 | 19 | 10 | 0 | 2 | 3 | 22 | 42 | 19 | 25 | 37 | 181 | 298 | 149 | 0.518467087 | 0.969172494 | 0.046 | 0.053 | 0.032 | 0.083 | 0.026 | 0.106 | 0 | 0.038 | 0.039 | 0.057 | 0 |
| RI | 4488 | ENSG00000104894 | CD37 | chr19 | + | 49335713 | 49337033 | 49335713 | 49335782 | 49336908 | 49337033 | 65 | 57 | 45 | 60 | 29 | 68 | 39 | 76 | 54 | 60 | 147 | 80 | 47 | 98 | 234 | 135 | 119 | 41 | 73 | 296 | 298 | 149 | 0.519122934 | 0.97009148 | 0.323 | 0.422 | 0.228 | 0.357 | 0.195 | 0.353 | 0.252 | 0.364 | 0.402 | 0.283 | -0.026 |
| RI | 2260 | ENSG00000143889 | HNRNPLL | chr2 | - | 38602437 | 38603036 | 38602437 | 38602600 | 38602856 | 38603036 | 34 | 16 | 14 | 31 | 4 | 1 | 1 | 0 | 0 | 1 | 18 | 19 | 9 | 28 | 12 | 0 | 2 | 3 | 0 | 0 | 298 | 149 | 0.519780301 | 0.971012726 | 0.944 | 0.889 | 1 | 1 | 0.667 | 1 | 0.826 | 0.6 | 1 | 1 | 0.015 |
| RI | 5765 | ENSG00000197223 | C1D | chr2 | - | 68045987 | 68046410 | 68045987 | 68046043 | 68046343 | 68046410 | 37 | 43 | 28 | 38 | 47 | 340 | 161 | 234 | 269 | 308 | 25 | 44 | 42 | 26 | 0 | 168 | 171 | 224 | 164 | 83 | 298 | 149 | 0.520088788 | 0.971043659 | 0.052 | 0.118 | 0.056 | 0.066 | 0.071 | 0.069 | 0.114 | 0.086 | 0.073 | 0 | 0.004 |
| RI | 4447 | ENSG00000142765 | SYTL1 | chr1 | + | 27349397 | 27349765 | 27349397 | 27349498 | 27349651 | 27349765 | 24 | 4 | 14 | 16 | 3 | 67 | 19 | 51 | 42 | 32 | 0 | 20 | 2 | 6 | 10 | 18 | 38 | 11 | 30 | 45 | 298 | 149 | 0.520125637 | 0.971043659 | 0.152 | 0.095 | 0.121 | 0.16 | 0.045 | 0 | 0.208 | 0.083 | 0.091 | 0.1 | 0.018 |
| RI | 8211 | ENSG00000149100 | EIF3M | chr11 | + | 32593865 | 32595013 | 32593865 | 32593949 | 32594913 | 32595013 | 294 | 248 | 253 | 305 | 275 | 917 | 566 | 701 | 688 | 732 | 174 | 253 | 333 | 178 | 97 | 662 | 482 | 518 | 494 | 582 | 298 | 149 | 0.520596115 | 0.971614929 | 0.138 | 0.18 | 0.153 | 0.181 | 0.158 | 0.116 | 0.208 | 0.243 | 0.153 | 0.077 | 0.003 |
| RI | 4198 | ENSG00000082512 | TRAF5 | chr1 | + | 211356366 | 211360779 | 211356366 | 211356468 | 211360701 | 211360779 | 40 | 20 | 31 | 52 | 35 | 2 | 0 | 0 | 0 | 2 | 21 | 42 | 22 | 55 | 24 | 0 | 0 | 6 | 0 | 0 | 298 | 149 | 0.521151633 | 0.971930252 | 0.909 | 1 | 1 | 1 | 0.897 | 1 | 1 | 0.647 | 1 | 1 | 0.032 |
| RI | 4421 | ENSG00000111653 | ING4 | chr12 | - | 6652661 | 6653050 | 6652661 | 6652770 | 6652935 | 6653050 | 14 | 9 | 12 | 7 | 14 | 64 | 31 | 47 | 51 | 102 | 2 | 7 | 23 | 12 | 2 | 43 | 58 | 28 | 62 | 83 | 298 | 149 | 0.521372539 | 0.971930252 | 0.099 | 0.127 | 0.113 | 0.064 | 0.064 | 0.023 | 0.057 | 0.291 | 0.088 | 0.012 | -0.001 |
| RI | 150 | ENSG00000093072 | ADA2 | chr22 | - | 17189941 | 17191810 | 17189941 | 17190032 | 17191682 | 17191810 | 67 | 53 | 51 | 57 | 32 | 774 | 427 | 532 | 573 | 1028 | 197 | 80 | 25 | 88 | 150 | 2070 | 896 | 499 | 1012 | 1715 | 298 | 149 | 0.521401049 | 0.971930252 | 0.041 | 0.058 | 0.046 | 0.047 | 0.015 | 0.045 | 0.043 | 0.024 | 0.042 | 0.042 | 0.002 |
| RI | 6669 | ENSG00000109736 | MFSD10 | chr4 | - | 2931824 | 2932153 | 2931824 | 2931936 | 2932044 | 2932153 | 10 | 5 | 12 | 8 | 0 | 6 | 0 | 3 | 1 | 5 | 6 | 8 | 4 | 0 | 6 | 0 | 18 | 0 | 5 | 6 | 257 | 149 | 0.521455025 | 0.971930252 | 0.491 | 1 | 0.699 | 0.823 | 0 | 1 | 0.205 | 1 | 0 | 0.367 | 0.088 |
| RI | 8161 | ENSG00000150093 | ITGB1 | chr10 | - | 32920244 | 32922346 | 32920244 | 32920397 | 32922256 | 32922346 | 166 | 148 | 100 | 161 | 59 | 3 | 2 | 1 | 1 | 1 | 43 | 82 | 190 | 22 | 58 | 0 | 6 | 3 | 0 | 0 | 298 | 149 | 0.52158776 | 0.971930252 | 0.965 | 0.974 | 0.98 | 0.988 | 0.967 | 1 | 0.872 | 0.969 | 1 | 1 | 0.007 |
| RI | 4256 | ENSG00000143801 | PSEN2 | chr1 | + | 226888828 | 226890133 | 226888828 | 226889049 | 226890034 | 226890133 | 0 | 0 | 2 | 2 | 2 | 4 | 5 | 5 | 6 | 1 | 0 | 7 | 3 | 0 | 0 | 28 | 7 | 3 | 2 | 8 | 298 | 149 | 0.522038516 | 0.972463423 | 0 | 0 | 0.167 | 0.143 | 0.5 | 0 | 0.333 | 0.333 | 0 | 0 | 0.029 |
| RI | 1224 | ENSG00000103042 | SLC38A7 | chr16 | - | 58675939 | 58676346 | 58675939 | 58676054 | 58676288 | 58676346 | 5 | 1 | 5 | 4 | 9 | 6 | 5 | 0 | 3 | 1 | 19 | 6 | 2 | 8 | 4 | 5 | 1 | 0 | 3 | 6 | 298 | 149 | 0.522355286 | 0.972601013 | 0.294 | 0.091 | 1 | 0.4 | 0.818 | 0.655 | 0.75 | 1 | 0.571 | 0.25 | -0.125 |
| RI | 4750 | ENSG00000159348 | CYB5R1 | chr1 | - | 202965886 | 202966600 | 202965886 | 202965993 | 202966527 | 202966600 | 22 | 5 | 19 | 15 | 19 | 75 | 14 | 31 | 39 | 47 | 13 | 64 | 17 | 19 | 41 | 51 | 60 | 35 | 30 | 202 | 298 | 149 | 0.522441682 | 0.972601013 | 0.128 | 0.152 | 0.235 | 0.161 | 0.168 | 0.113 | 0.348 | 0.195 | 0.241 | 0.092 | -0.029 |
| RI | 6782 | ENSG00000072778 | ACADVL | chr17 | + | 7223812 | 7224069 | 7223812 | 7223875 | 7223967 | 7224069 | 32 | 28 | 42 | 31 | 11 | 321 | 165 | 216 | 308 | 341 | 40 | 100 | 28 | 66 | 16 | 505 | 498 | 285 | 428 | 871 | 241 | 149 | 0.522933347 | 0.972980141 | 0.058 | 0.095 | 0.107 | 0.059 | 0.02 | 0.047 | 0.11 | 0.057 | 0.087 | 0.011 | 0.005 |
| RI | 3213 | ENSG00000168615 | ADAM9 | chr8 | + | 39090046 | 39091346 | 39090046 | 39090188 | 39091258 | 39091346 | 12 | 14 | 11 | 7 | 18 | 50 | 57 | 34 | 61 | 139 | 0 | 10 | 41 | 13 | 0 | 8 | 87 | 65 | 66 | 15 | 298 | 149 | 0.522974767 | 0.972980141 | 0.107 | 0.109 | 0.139 | 0.054 | 0.061 | 0 | 0.054 | 0.24 | 0.09 | 0 | 0.017 |
| RI | 626 | ENSG00000174365 | SNHG11 | chr20 | + | 38447465 | 38450447 | 38447465 | 38448730 | 38450335 | 38450447 | 9 | 8 | 3 | 2 | 4 | 7 | 3 | 5 | 8 | 0 | 12 | 4 | 5 | 10 | 12 | 10 | 19 | 10 | 7 | 7 | 298 | 149 | 0.523554209 | 0.973381545 | 0.391 | 0.571 | 0.231 | 0.111 | 1 | 0.375 | 0.095 | 0.2 | 0.417 | 0.462 | 0.151 |
| RI | 7039 | ENSG00000100918 | REC8 | chr14 | + | 24179799 | 24180254 | 24179799 | 24179906 | 24180009 | 24180254 | 8 | 10 | 2 | 23 | 16 | 7 | 2 | 13 | 9 | 2 | 33 | 45 | 8 | 54 | 15 | 5 | 18 | 10 | 19 | 7 | 252 | 149 | 0.523609446 | 0.973381545 | 0.403 | 0.747 | 0.083 | 0.602 | 0.825 | 0.796 | 0.596 | 0.321 | 0.627 | 0.559 | -0.048 |
| RI | 7760 | ENSG00000101158 | NELFCD | chr20 | + | 58993623 | 58994239 | 58993623 | 58993675 | 58994109 | 58994239 | 62 | 56 | 64 | 61 | 47 | 4 | 3 | 7 | 7 | 2 | 116 | 103 | 50 | 84 | 67 | 0 | 13 | 3 | 3 | 7 | 298 | 149 | 0.523684874 | 0.973381545 | 0.886 | 0.903 | 0.821 | 0.813 | 0.922 | 1 | 0.798 | 0.893 | 0.933 | 0.827 | -0.021 |
| RI | 5464 | ENSG00000237765 | FAM200B | chr4 | + | 15686235 | 15687148 | 15686235 | 15686462 | 15686931 | 15687148 | 91 | 67 | 113 | 103 | 217 | 1 | 0 | 1 | 0 | 0 | 130 | 38 | 65 | 42 | 75 | 0 | 6 | 0 | 0 | 0 | 298 | 149 | 0.523947935 | 0.973564157 | 0.978 | 1 | 0.983 | 1 | 1 | 1 | 0.76 | 1 | 1 | 1 | 0.04 |
| RI | 438 | ENSG00000187994 | RINL | chr19 | - | 38876330 | 38876781 | 38876330 | 38876490 | 38876692 | 38876781 | 26 | 3 | 19 | 16 | 6 | 21 | 11 | 6 | 24 | 5 | 33 | 21 | 33 | 18 | 10 | 22 | 35 | 0 | 18 | 35 | 298 | 149 | 0.524166401 | 0.973581175 | 0.382 | 0.12 | 0.613 | 0.25 | 0.375 | 0.429 | 0.231 | 1 | 0.333 | 0.125 | -0.076 |
| RI | 2697 | ENSG00000103502 | CDIPT | chr16 | - | 29860580 | 29861259 | 29860580 | 29860662 | 29861105 | 29861259 | 39 | 19 | 20 | 30 | 15 | 48 | 18 | 30 | 46 | 52 | 6 | 42 | 28 | 13 | 51 | 55 | 46 | 22 | 55 | 57 | 298 | 149 | 0.524286731 | 0.973581175 | 0.289 | 0.345 | 0.25 | 0.246 | 0.126 | 0.052 | 0.313 | 0.389 | 0.106 | 0.309 | 0.017 |
| RI | 2950 | ENSG00000164068 | RNF123 | chr3 | + | 49712478 | 49713587 | 49712478 | 49712892 | 49713512 | 49713587 | 3 | 1 | 4 | 1 | 1 | 3 | 0 | 1 | 0 | 3 | 6 | 11 | 4 | 2 | 12 | 0 | 2 | 2 | 0 | 9 | 298 | 149 | 0.524921023 | 0.974452698 | 0.333 | 1 | 0.667 | 1 | 0.143 | 1 | 0.733 | 0.5 | 1 | 0.4 | -0.098 |
| RI | 6540 | ENSG00000127824 | TUBA4A | chr2 | - | 219251193 | 219251713 | 219251193 | 219251323 | 219251564 | 219251713 | 48 | 36 | 39 | 50 | 40 | 401 | 233 | 295 | 500 | 283 | 48 | 109 | 86 | 22 | 161 | 434 | 643 | 425 | 337 | 2145 | 298 | 149 | 0.52588645 | 0.975821011 | 0.056 | 0.072 | 0.062 | 0.048 | 0.066 | 0.052 | 0.078 | 0.092 | 0.032 | 0.036 | 0.003 |
| RI | 6118 | ENSG00000044090 | CUL7 | chr6 | - | 43046510 | 43047107 | 43046510 | 43046601 | 43046879 | 43047107 | 17 | 13 | 24 | 15 | 15 | 10 | 2 | 2 | 2 | 5 | 13 | 33 | 12 | 19 | 70 | 10 | 3 | 7 | 18 | 3 | 298 | 149 | 0.525988505 | 0.975821011 | 0.459 | 0.765 | 0.857 | 0.789 | 0.6 | 0.394 | 0.846 | 0.462 | 0.345 | 0.921 | 0.1 |
| RI | 8452 | ENSG00000123349 | PFDN5 | chr12 | + | 53295838 | 53297924 | 53295838 | 53295941 | 53297849 | 53297924 | 192 | 157 | 217 | 240 | 285 | 3 | 13 | 11 | 3 | 11 | 433 | 302 | 224 | 293 | 324 | 34 | 5 | 5 | 4 | 20 | 298 | 149 | 0.526469632 | 0.976204343 | 0.97 | 0.858 | 0.908 | 0.976 | 0.928 | 0.864 | 0.968 | 0.957 | 0.973 | 0.89 | -0.002 |
| RI | 8141 | ENSG00000107862 | GBF1 | chr10 | + | 102358505 | 102359435 | 102358505 | 102358729 | 102359269 | 102359435 | 10 | 5 | 9 | 14 | 12 | 112 | 48 | 103 | 61 | 59 | 6 | 32 | 21 | 14 | 17 | 200 | 72 | 73 | 142 | 164 | 298 | 149 | 0.526525654 | 0.976204343 | 0.043 | 0.05 | 0.042 | 0.103 | 0.092 | 0.015 | 0.182 | 0.126 | 0.047 | 0.049 | -0.018 |
| RI | 2753 | ENSG00000099949 | LZTR1 | chr22 | + | 20995962 | 20996801 | 20995962 | 20996112 | 20996695 | 20996801 | 10 | 3 | 9 | 12 | 13 | 41 | 5 | 23 | 18 | 6 | 8 | 12 | 2 | 6 | 24 | 44 | 8 | 6 | 19 | 42 | 298 | 149 | 0.52729852 | 0.977330518 | 0.109 | 0.231 | 0.164 | 0.25 | 0.52 | 0.083 | 0.429 | 0.143 | 0.136 | 0.222 | 0.052 |
| RI | 5616 | ENSG00000120833 | SOCS2 | chr12 | - | 93572684 | 93576206 | 93572684 | 93573036 | 93574721 | 93576206 | 9 | 1 | 9 | 8 | 15 | 36 | 13 | 17 | 38 | 11 | 14 | 0 | 1 | 10 | 0 | 19 | 12 | 3 | 2 | 21 | 298 | 149 | 0.52856632 | 0.979373041 | 0.111 | 0.037 | 0.209 | 0.095 | 0.405 | 0.269 | 0 | 0.143 | 0.714 | 0 | -0.054 |
| RI | 74 | ENSG00000128159 | TUBGCP6 | chr22 | - | 50219287 | 50219791 | 50219287 | 50219456 | 50219643 | 50219791 | 8 | 7 | 20 | 7 | 0 | 4 | 5 | 0 | 6 | 1 | 0 | 10 | 7 | 11 | 30 | 26 | 16 | 13 | 0 | 1 | 298 | 149 | 0.52966147 | 0.981094483 | 0.5 | 0.412 | 1 | 0.368 | 0 | 0 | 0.238 | 0.212 | 1 | 0.938 | -0.022 |
| RI | 7956 | ENSG00000149311 | ATM | chr11 | + | 108331443 | 108333968 | 108331443 | 108331557 | 108333885 | 108333968 | 96 | 119 | 117 | 132 | 78 | 4 | 1 | 2 | 0 | 0 | 14 | 67 | 32 | 12 | 7 | 0 | 0 | 0 | 0 | 0 | 298 | 149 | 0.530846033 | 0.982980413 | 0.923 | 0.983 | 0.967 | 1 | 1 | 1 | 1 | 1 | 1 | 1 | -0.025 |
| RI | 7231 | ENSG00000164941 | INTS8 | chr8 | + | 94867139 | 94867337 | 94867139 | 94867196 | 94867275 | 94867337 | 13 | 26 | 16 | 28 | 30 | 213 | 145 | 138 | 116 | 199 | 12 | 12 | 10 | 2 | 0 | 8 | 69 | 59 | 32 | 11 | 228 | 149 | 0.531060799 | 0.983069928 | 0.038 | 0.105 | 0.07 | 0.136 | 0.09 | 0.495 | 0.102 | 0.1 | 0.039 | 0 | -0.059 |
| RI | 4234 | ENSG00000165458 | INPPL1 | chr11 | + | 72233440 | 72233744 | 72233440 | 72233522 | 72233654 | 72233744 | 41 | 25 | 22 | 22 | 24 | 123 | 82 | 84 | 109 | 79 | 150 | 46 | 58 | 28 | 73 | 324 | 199 | 83 | 149 | 412 | 281 | 149 | 0.531339054 | 0.983276877 | 0.15 | 0.139 | 0.122 | 0.097 | 0.139 | 0.197 | 0.109 | 0.27 | 0.091 | 0.086 | -0.021 |
| RI | 2094 | ENSG00000141279 | NPEPPS | chr17 | + | 47579389 | 47582849 | 47579389 | 47579511 | 47582741 | 47582849 | 39 | 36 | 21 | 35 | 36 | 202 | 132 | 143 | 192 | 377 | 69 | 50 | 22 | 68 | 21 | 270 | 260 | 185 | 218 | 366 | 298 | 149 | 0.532228554 | 0.984614491 | 0.088 | 0.12 | 0.068 | 0.084 | 0.046 | 0.113 | 0.088 | 0.056 | 0.135 | 0.028 | -0.003 |
| RI | 5989 | ENSG00000171453 | POLR1C | chr6 | + | 43519705 | 43520427 | 43519705 | 43520185 | 43520274 | 43520427 | 13 | 5 | 21 | 20 | 17 | 36 | 31 | 34 | 47 | 73 | 11 | 8 | 12 | 19 | 0 | 41 | 31 | 21 | 36 | 29 | 238 | 149 | 0.534321332 | 0.988176616 | 0.184 | 0.092 | 0.279 | 0.21 | 0.127 | 0.144 | 0.139 | 0.263 | 0.248 | 0 | 0.02 |
| RI | 6210 | ENSG00000187796 | CARD9 | chr9 | - | 136364482 | 136366845 | 136364482 | 136365217 | 136366799 | 136366845 | 7 | 4 | 4 | 4 | 5 | 14 | 6 | 12 | 14 | 1 | 13 | 3 | 0 | 15 | 69 | 6 | 24 | 7 | 18 | 15 | 298 | 149 | 0.534891919 | 0.988922243 | 0.2 | 0.25 | 0.143 | 0.125 | 0.714 | 0.52 | 0.059 | 0 | 0.294 | 0.697 | -0.028 |
| RI | 1109 | ENSG00000154760 | SLFN13 | chr17 | - | 35444614 | 35445693 | 35444614 | 35444701 | 35445372 | 35445693 | 125 | 71 | 112 | 135 | 79 | 4 | 1 | 2 | 1 | 0 | 54 | 57 | 160 | 91 | 28 | 0 | 0 | 0 | 0 | 0 | 298 | 149 | 0.535749122 | 0.990012658 | 0.94 | 0.973 | 0.966 | 0.985 | 1 | 1 | 1 | 1 | 1 | 1 | -0.027 |
| RI | 1204 | ENSG00000166747 | AP1G1 | chr16 | - | 71808504 | 71808813 | 71808504 | 71808677 | 71808762 | 71808813 | 6 | 3 | 12 | 11 | 6 | 3 | 4 | 1 | 1 | 1 | 35 | 18 | 10 | 18 | 28 | 6 | 6 | 1 | 2 | 2 | 234 | 149 | 0.535816907 | 0.990012658 | 0.56 | 0.323 | 0.884 | 0.875 | 0.793 | 0.788 | 0.656 | 0.864 | 0.851 | 0.899 | -0.125 |
| RI | 339 | ENSG00000160007 | ARHGAP35 | chr19 | + | 46987988 | 46989675 | 46987988 | 46988066 | 46989543 | 46989675 | 32 | 19 | 26 | 31 | 26 | 173 | 122 | 168 | 162 | 128 | 40 | 32 | 29 | 28 | 46 | 164 | 164 | 148 | 174 | 260 | 298 | 149 | 0.53824442 | 0.994186927 | 0.085 | 0.072 | 0.072 | 0.087 | 0.092 | 0.109 | 0.089 | 0.089 | 0.074 | 0.081 | -0.007 |
| RI | 5533 | ENSG00000184922 | FMNL1 | chr17 | + | 45246209 | 45246604 | 45246209 | 45246330 | 45246504 | 45246604 | 53 | 40 | 50 | 63 | 45 | 12 | 17 | 8 | 21 | 19 | 136 | 64 | 88 | 89 | 145 | 49 | 27 | 30 | 28 | 43 | 298 | 149 | 0.538731993 | 0.994776456 | 0.688 | 0.541 | 0.758 | 0.6 | 0.542 | 0.581 | 0.542 | 0.595 | 0.614 | 0.628 | 0.034 |
| RI | 1387 | ENSG00000131652 | THOC6 | chr16 | + | 3025707 | 3026288 | 3025707 | 3025823 | 3025923 | 3026288 | 3 | 1 | 11 | 1 | 0 | 13 | 6 | 12 | 13 | 23 | 5 | 5 | 2 | 2 | 5 | 17 | 23 | 7 | 7 | 11 | 249 | 149 | 0.538905341 | 0.994785578 | 0.121 | 0.091 | 0.354 | 0.044 | 0 | 0.15 | 0.115 | 0.146 | 0.146 | 0.214 | -0.032 |
| RI | 7110 | ENSG00000103266 | STUB1 | chr16 | + | 681792 | 682076 | 681792 | 681880 | 682058 | 682076 | 167 | 91 | 133 | 157 | 149 | 1 | 0 | 7 | 0 | 2 | 173 | 221 | 135 | 142 | 364 | 0 | 0 | 2 | 0 | 0 | 298 | 149 | 0.539306011 | 0.995214184 | 0.988 | 1 | 0.905 | 1 | 0.974 | 1 | 1 | 0.971 | 1 | 1 | -0.021 |
| RI | 7594 | ENSG00000025434 | NR1H3 | chr11 | + | 47261546 | 47262018 | 47261546 | 47261726 | 47261918 | 47262018 | 5 | 2 | 1 | 2 | 1 | 17 | 2 | 3 | 12 | 4 | 6 | 0 | 2 | 0 | 3 | 16 | 19 | 7 | 1 | 14 | 298 | 149 | 0.540093703 | 0.995877073 | 0.128 | 0.333 | 0.143 | 0.077 | 0.111 | 0.158 | 0 | 0.125 | 0 | 0.097 | 0.082 |
| RI | 7858 | ENSG00000142102 | PGGHG | chr11 | + | 290677 | 291113 | 290677 | 290781 | 291001 | 291113 | 75 | 13 | 60 | 78 | 26 | 5 | 0 | 0 | 2 | 0 | 16 | 26 | 71 | 41 | 86 | 0 | 0 | 3 | 0 | 0 | 298 | 149 | 0.540122299 | 0.995877073 | 0.882 | 1 | 1 | 0.951 | 1 | 1 | 1 | 0.922 | 1 | 1 | -0.018 |
| RI | 6953 | ENSG00000215301 | DDX3X | chrX | + | 41343215 | 41344128 | 41343215 | 41343351 | 41343736 | 41344128 | 229 | 247 | 249 | 309 | 457 | 384 | 300 | 351 | 483 | 598 | 24 | 229 | 215 | 191 | 6 | 67 | 202 | 330 | 217 | 30 | 298 | 149 | 0.540171008 | 0.995877073 | 0.23 | 0.292 | 0.262 | 0.242 | 0.276 | 0.152 | 0.362 | 0.246 | 0.306 | 0.091 | 0.029 |
| RI | 2735 | ENSG00000164934 | DCAF13 | chr8 | + | 103427096 | 103430689 | 103427096 | 103427252 | 103430611 | 103430689 | 30 | 41 | 32 | 40 | 23 | 114 | 73 | 111 | 100 | 162 | 0 | 25 | 40 | 9 | 0 | 20 | 52 | 36 | 23 | 7 | 298 | 149 | 0.540628317 | 0.996159162 | 0.116 | 0.219 | 0.126 | 0.167 | 0.066 | 0 | 0.194 | 0.357 | 0.164 | 0 | -0.004 |
| RI | 8112 | ENSG00000205542 | TMSB4X | chrX | + | 12975109 | 12976361 | 12975109 | 12975168 | 12976245 | 12976361 | 248 | 187 | 243 | 221 | 251 | 7292 | 4872 | 6142 | 9498 | 4500 | 405 | 234 | 270 | 535 | 500 | 11443 | 7415 | 4031 | 7936 | 30016 | 298 | 149 | 0.540661296 | 0.996159162 | 0.017 | 0.019 | 0.019 | 0.012 | 0.027 | 0.017 | 0.016 | 0.032 | 0.033 | 0.008 | -0.002 |
| RI | 1803 | ENSG00000100938 | GMPR2 | chr14 | + | 24238245 | 24239239 | 24238245 | 24238405 | 24238588 | 24239239 | 92 | 67 | 81 | 75 | 127 | 122 | 67 | 104 | 109 | 190 | 61 | 126 | 116 | 129 | 68 | 139 | 119 | 61 | 70 | 222 | 298 | 149 | 0.541156195 | 0.996760102 | 0.274 | 0.333 | 0.28 | 0.256 | 0.25 | 0.18 | 0.346 | 0.487 | 0.48 | 0.133 | -0.047 |
| RI | 2920 | ENSG00000247596 | TWF2 | chr3 | - | 52229660 | 52230995 | 52229660 | 52229782 | 52229919 | 52230995 | 11 | 4 | 8 | 15 | 6 | 110 | 53 | 95 | 84 | 69 | 0 | 4 | 9 | 9 | 28 | 136 | 94 | 86 | 69 | 163 | 286 | 149 | 0.542423133 | 0.998782247 | 0.05 | 0.038 | 0.042 | 0.085 | 0.043 | 0 | 0.022 | 0.052 | 0.064 | 0.082 | 0.008 |
| RI | 234 | ENSG00000129351 | ILF3 | chr19 | + | 10683916 | 10685765 | 10683916 | 10683970 | 10684858 | 10685765 | 140 | 89 | 116 | 153 | 187 | 1 | 1 | 0 | 4 | 2 | 201 | 224 | 150 | 180 | 153 | 0 | 1 | 1 | 0 | 0 | 298 | 149 | 0.544370095 | 1 | 0.986 | 0.978 | 1 | 0.95 | 0.979 | 1 | 0.991 | 0.987 | 1 | 1 | -0.017 |
| RI | 4719 | ENSG00000141458 | NPC1 | chr18 | - | 23533354 | 23534559 | 23533354 | 23533517 | 23534445 | 23534559 | 62 | 31 | 30 | 44 | 25 | 76 | 19 | 40 | 32 | 18 | 25 | 52 | 79 | 40 | 0 | 17 | 18 | 18 | 21 | 20 | 298 | 149 | 0.544653544 | 1 | 0.29 | 0.449 | 0.273 | 0.407 | 0.41 | 0.424 | 0.591 | 0.687 | 0.488 | 0 | -0.072 |
| RI | 4964 | ENSG00000131876 | SNRPA1 | chr15 | - | 101285725 | 101287010 | 101285725 | 101285801 | 101286907 | 101287010 | 439 | 476 | 456 | 580 | 180 | 12 | 17 | 21 | 24 | 40 | 560 | 623 | 893 | 1012 | 137 | 15 | 32 | 60 | 39 | 12 | 298 | 149 | 0.545638212 | 1 | 0.948 | 0.933 | 0.916 | 0.924 | 0.692 | 0.949 | 0.907 | 0.882 | 0.928 | 0.851 | -0.021 |
| RI | 2439 | ENSG00000160179 | ABCG1 | chr21 | + | 42287888 | 42288312 | 42287888 | 42288037 | 42288210 | 42288312 | 4 | 2 | 16 | 17 | 12 | 5 | 3 | 15 | 2 | 0 | 15 | 8 | 31 | 19 | 0 | 3 | 8 | 1 | 0 | 5 | 298 | 149 | 0.545757608 | 1 | 0.286 | 0.25 | 0.348 | 0.81 | 1 | 0.714 | 0.333 | 0.939 | 1 | 0 | -0.058 |
| RI | 5739 | ENSG00000186635 | ARAP1 | chr11 | - | 72696548 | 72697195 | 72696548 | 72696654 | 72696982 | 72697195 | 12 | 2 | 4 | 12 | 12 | 478 | 232 | 209 | 288 | 107 | 12 | 9 | 10 | 15 | 19 | 509 | 284 | 226 | 547 | 902 | 298 | 149 | 0.546079688 | 1 | 0.012 | 0.004 | 0.009 | 0.02 | 0.053 | 0.012 | 0.016 | 0.022 | 0.014 | 0.01 | 0.005 |
| RI | 6587 | ENSG00000234741 | GAS5 | chr1 | - | 173863900 | 173864304 | 173863900 | 173864075 | 173864256 | 173864304 | 96 | 66 | 92 | 120 | 157 | 419 | 188 | 513 | 309 | 628 | 13 | 54 | 41 | 35 | 23 | 88 | 112 | 149 | 77 | 134 | 298 | 149 | 0.546275952 | 1 | 0.103 | 0.149 | 0.082 | 0.163 | 0.111 | 0.069 | 0.194 | 0.121 | 0.185 | 0.079 | -0.008 |
| RI | 4562 | ENSG00000186364 | NUDT17 | chr1 | + | 145846432 | 145846690 | 145846432 | 145846458 | 145846597 | 145846690 | 11 | 8 | 9 | 23 | 1 | 0 | 0 | 0 | 0 | 0 | 2 | 15 | 27 | 23 | 23 | 0 | 0 | 0 | 5 | 0 | 288 | 149 | 0.546441729 | 1 | 1 | 1 | 1 | 1 | 1 | 1 | 1 | 1 | 0.704 | 1 | 0.059 |
| RI | 6485 | ENSG00000065000 | AP3D1 | chr19 | - | 2109085 | 2109958 | 2109085 | 2109207 | 2109872 | 2109958 | 18 | 7 | 20 | 7 | 15 | 42 | 21 | 61 | 71 | 26 | 18 | 39 | 6 | 20 | 45 | 70 | 71 | 23 | 54 | 164 | 298 | 149 | 0.54726468 | 1 | 0.176 | 0.143 | 0.141 | 0.047 | 0.224 | 0.114 | 0.215 | 0.115 | 0.156 | 0.121 | 0.002 |
| RI | 7437 | ENSG00000204394 | VARS1 | chr6 | - | 31782529 | 31782845 | 31782529 | 31782633 | 31782720 | 31782845 | 9 | 6 | 8 | 12 | 4 | 24 | 15 | 20 | 24 | 10 | 0 | 28 | 8 | 23 | 23 | 43 | 30 | 17 | 7 | 44 | 236 | 149 | 0.548133715 | 1 | 0.191 | 0.202 | 0.202 | 0.24 | 0.202 | 0 | 0.371 | 0.229 | 0.675 | 0.248 | -0.097 |
| RI | 2010 | ENSG00000198399 | ITSN2 | chr2 | - | 24202863 | 24204418 | 24202863 | 24203783 | 24204244 | 24204418 | 8 | 4 | 3 | 3 | 0 | 56 | 40 | 34 | 58 | 55 | 7 | 4 | 0 | 0 | 4 | 18 | 34 | 17 | 56 | 15 | 298 | 149 | 0.548406746 | 1 | 0.067 | 0.048 | 0.042 | 0.025 | 0 | 0.163 | 0.056 | 0 | 0 | 0.118 | -0.031 |
| RI | 4450 | ENSG00000090097 | PCBP4 | chr3 | - | 51959894 | 51960320 | 51959894 | 51960023 | 51960188 | 51960320 | 5 | 0 | 5 | 11 | 0 | 24 | 6 | 16 | 15 | 22 | 8 | 10 | 1 | 0 | 7 | 15 | 23 | 8 | 10 | 31 | 298 | 149 | 0.54849541 | 1 | 0.094 | 0 | 0.135 | 0.268 | 0 | 0.211 | 0.179 | 0.059 | 0 | 0.101 | -0.011 |
| RI | 530 | ENSG00000064601 | CTSA | chr20 | + | 45892724 | 45893311 | 45892724 | 45892880 | 45893219 | 45893311 | 31 | 32 | 38 | 44 | 30 | 326 | 212 | 179 | 369 | 574 | 135 | 58 | 55 | 66 | 333 | 743 | 769 | 267 | 605 | 3887 | 298 | 149 | 0.549035026 | 1 | 0.045 | 0.07 | 0.096 | 0.056 | 0.025 | 0.083 | 0.036 | 0.093 | 0.052 | 0.041 | -0.003 |
| RI | 4094 | ENSG00000081760 | AACS | chr12 | + | 125124704 | 125125024 | 125124704 | 125124769 | 125124901 | 125125024 | 0 | 6 | 5 | 8 | 2 | 3 | 4 | 7 | 1 | 17 | 0 | 9 | 6 | 6 | 12 | 45 | 2 | 5 | 3 | 5 | 281 | 149 | 0.549150149 | 1 | 0 | 0.443 | 0.275 | 0.809 | 0.059 | 0 | 0.705 | 0.389 | 0.515 | 0.56 | -0.117 |
| RI | 2516 | ENSG00000156253 | RWDD2B | chr21 | - | 29008239 | 29008621 | 29008239 | 29008307 | 29008394 | 29008621 | 11 | 6 | 6 | 15 | 25 | 8 | 3 | 11 | 4 | 24 | 4 | 10 | 6 | 0 | 3 | 0 | 5 | 8 | 3 | 0 | 236 | 149 | 0.549862341 | 1 | 0.465 | 0.558 | 0.256 | 0.703 | 0.397 | 1 | 0.558 | 0.321 | 0 | 1 | -0.1 |
| RI | 3704 | ENSG00000196628 | TCF4 | chr18 | - | 55228846 | 55232671 | 55228846 | 55229076 | 55232508 | 55232671 | 29 | 50 | 30 | 51 | 45 | 31 | 34 | 39 | 61 | 60 | 33 | 50 | 47 | 13 | 9 | 11 | 42 | 41 | 23 | 16 | 298 | 149 | 0.550306777 | 1 | 0.319 | 0.424 | 0.278 | 0.295 | 0.273 | 0.6 | 0.373 | 0.364 | 0.22 | 0.22 | -0.038 |
| RI | 2905 | ENSG00000172053 | QARS1 | chr3 | - | 49103344 | 49103706 | 49103344 | 49103431 | 49103630 | 49103706 | 84 | 67 | 100 | 114 | 71 | 3 | 2 | 2 | 1 | 4 | 172 | 120 | 160 | 163 | 308 | 12 | 7 | 0 | 0 | 0 | 298 | 149 | 0.550435933 | 1 | 0.933 | 0.944 | 0.962 | 0.983 | 0.899 | 0.878 | 0.896 | 1 | 1 | 1 | -0.011 |
| RI | 6984 | ENSG00000140678 | ITGAX | chr16 | + | 31360309 | 31361213 | 31360309 | 31360463 | 31361062 | 31361213 | 169 | 122 | 95 | 85 | 109 | 147 | 160 | 111 | 186 | 311 | 301 | 131 | 119 | 70 | 290 | 462 | 262 | 146 | 183 | 428 | 298 | 149 | 0.550675565 | 1 | 0.365 | 0.276 | 0.3 | 0.186 | 0.149 | 0.246 | 0.2 | 0.29 | 0.161 | 0.253 | 0.025 |
| RI | 5326 | ENSG00000064703 | DDX20 | chr1 | + | 111760705 | 111761284 | 111760705 | 111760848 | 111760973 | 111761284 | 29 | 21 | 53 | 57 | 67 | 0 | 0 | 0 | 1 | 6 | 13 | 24 | 41 | 45 | 4 | 0 | 0 | 0 | 0 | 0 | 274 | 149 | 0.550988131 | 1 | 1 | 1 | 1 | 0.969 | 0.859 | 1 | 1 | 1 | 1 | 1 | -0.034 |
| RI | 1519 | ENSG00000213614 | HEXA | chr15 | - | 72345445 | 72346325 | 72345445 | 72345550 | 72346234 | 72346325 | 54 | 23 | 48 | 45 | 21 | 118 | 58 | 100 | 93 | 122 | 71 | 46 | 56 | 94 | 65 | 251 | 146 | 109 | 119 | 183 | 298 | 149 | 0.551537605 | 1 | 0.186 | 0.165 | 0.194 | 0.195 | 0.079 | 0.124 | 0.136 | 0.204 | 0.283 | 0.151 | -0.016 |
| RI | 3568 | ENSG00000265241 | RBM8A | chr1 | - | 145926808 | 145927077 | 145926808 | 145926883 | 145927015 | 145927077 | 54 | 49 | 36 | 76 | 47 | 65 | 69 | 79 | 86 | 141 | 114 | 60 | 57 | 105 | 24 | 171 | 127 | 107 | 138 | 85 | 283 | 149 | 0.551634728 | 1 | 0.304 | 0.272 | 0.193 | 0.318 | 0.149 | 0.26 | 0.199 | 0.219 | 0.286 | 0.129 | 0.029 |
| RI | 1146 | ENSG00000168488 | ATXN2L | chr16 | + | 28835932 | 28837237 | 28835932 | 28836485 | 28836727 | 28837237 | 30 | 24 | 26 | 36 | 4 | 1 | 0 | 2 | 1 | 3 | 48 | 34 | 23 | 29 | 35 | 0 | 7 | 2 | 0 | 4 | 298 | 149 | 0.551877749 | 1 | 0.938 | 1 | 0.867 | 0.947 | 0.4 | 1 | 0.708 | 0.852 | 1 | 0.814 | -0.044 |
| RI | 2110 | ENSG00000159753 | CARMIL2 | chr16 | + | 67645539 | 67645777 | 67645539 | 67645631 | 67645723 | 67645777 | 4 | 3 | 7 | 3 | 8 | 2 | 0 | 5 | 3 | 0 | 12 | 6 | 0 | 12 | 10 | 0 | 3 | 3 | 0 | 2 | 241 | 149 | 0.551930025 | 1 | 0.553 | 1 | 0.464 | 0.382 | 1 | 1 | 0.553 | 0 | 1 | 0.756 | 0.018 |
| RI | 5501 | ENSG00000105374 | NKG7 | chr19 | - | 51371939 | 51372202 | 51371939 | 51372074 | 51372160 | 51372202 | 268 | 48 | 87 | 113 | 49 | 1232 | 171 | 366 | 468 | 132 | 9 | 200 | 111 | 163 | 182 | 284 | 909 | 362 | 394 | 676 | 235 | 149 | 0.552864919 | 1 | 0.121 | 0.151 | 0.131 | 0.133 | 0.191 | 0.02 | 0.122 | 0.163 | 0.208 | 0.146 | 0.014 |
| RI | 207 | ENSG00000132024 | CC2D1A | chr19 | + | 13926977 | 13927265 | 13926977 | 13927077 | 13927174 | 13927265 | 6 | 13 | 6 | 11 | 11 | 26 | 18 | 30 | 28 | 22 | 0 | 10 | 10 | 12 | 43 | 19 | 58 | 24 | 43 | 59 | 246 | 149 | 0.555160308 | 1 | 0.123 | 0.304 | 0.108 | 0.192 | 0.232 | 0 | 0.095 | 0.202 | 0.145 | 0.306 | 0.042 |
| RI | 4222 | ENSG00000132002 | DNAJB1 | chr19 | - | 14514781 | 14517046 | 14514781 | 14516170 | 14516465 | 14517046 | 86 | 45 | 99 | 93 | 112 | 678 | 469 | 662 | 861 | 1269 | 46 | 101 | 108 | 88 | 131 | 683 | 1131 | 816 | 802 | 1026 | 298 | 149 | 0.555172391 | 1 | 0.06 | 0.046 | 0.07 | 0.051 | 0.042 | 0.033 | 0.043 | 0.062 | 0.052 | 0.06 | 0.004 |
| RI | 380 | ENSG00000105483 | CARD8 | chr19 | - | 48232452 | 48234543 | 48232452 | 48232493 | 48234402 | 48234543 | 293 | 301 | 348 | 368 | 276 | 167 | 112 | 83 | 192 | 211 | 153 | 211 | 201 | 185 | 50 | 67 | 57 | 22 | 76 | 77 | 298 | 149 | 0.555204003 | 1 | 0.467 | 0.573 | 0.677 | 0.489 | 0.395 | 0.533 | 0.649 | 0.82 | 0.549 | 0.245 | -0.039 |
| RI | 8450 | ENSG00000123349 | PFDN5 | chr12 | + | 53295838 | 53296467 | 53295838 | 53295941 | 53296429 | 53296467 | 146 | 118 | 157 | 161 | 186 | 0 | 1 | 7 | 1 | 0 | 195 | 197 | 138 | 147 | 203 | 0 | 2 | 0 | 3 | 0 | 298 | 149 | 0.555750848 | 1 | 1 | 0.983 | 0.918 | 0.988 | 1 | 1 | 0.98 | 1 | 0.961 | 1 | -0.01 |
| RI | 6419 | ENSG00000164896 | FASTK | chr7 | - | 151077309 | 151077780 | 151077309 | 151077401 | 151077620 | 151077780 | 25 | 4 | 26 | 26 | 7 | 43 | 14 | 29 | 44 | 21 | 13 | 11 | 11 | 10 | 45 | 5 | 67 | 16 | 27 | 24 | 298 | 149 | 0.556154825 | 1 | 0.225 | 0.125 | 0.31 | 0.228 | 0.143 | 0.565 | 0.076 | 0.256 | 0.156 | 0.484 | -0.101 |
| RI | 8195 | ENSG00000023041 | ZDHHC6 | chr10 | - | 112432375 | 112434464 | 112432375 | 112432521 | 112433239 | 112434464 | 52 | 74 | 41 | 73 | 54 | 244 | 282 | 223 | 205 | 469 | 36 | 91 | 118 | 35 | 0 | 351 | 250 | 252 | 277 | 92 | 298 | 149 | 0.556208031 | 1 | 0.096 | 0.116 | 0.084 | 0.151 | 0.054 | 0.049 | 0.154 | 0.19 | 0.059 | 0 | 0.01 |
| RI | 7585 | ENSG00000159314 | ARHGAP27 | chr17 | - | 45396206 | 45396585 | 45396206 | 45396284 | 45396486 | 45396585 | 32 | 19 | 28 | 33 | 8 | 58 | 14 | 43 | 45 | 20 | 9 | 32 | 11 | 30 | 12 | 43 | 27 | 15 | 20 | 59 | 298 | 149 | 0.556699001 | 1 | 0.216 | 0.404 | 0.246 | 0.268 | 0.167 | 0.095 | 0.372 | 0.268 | 0.429 | 0.092 | 0.009 |
| RI | 5899 | ENSG00000168237 | GLYCTK | chr3 | + | 52290959 | 52291922 | 52290959 | 52291111 | 52291746 | 52291922 | 41 | 9 | 36 | 26 | 16 | 6 | 0 | 5 | 3 | 0 | 5 | 55 | 34 | 24 | 80 | 9 | 4 | 0 | 1 | 1 | 298 | 149 | 0.556713548 | 1 | 0.774 | 1 | 0.783 | 0.812 | 1 | 0.217 | 0.873 | 1 | 0.923 | 0.976 | 0.076 |
| RI | 5461 | ENSG00000138442 | WDR12 | chr2 | - | 202884197 | 202884535 | 202884197 | 202884303 | 202884394 | 202884535 | 2 | 4 | 5 | 8 | 0 | 29 | 20 | 35 | 16 | 57 | 0 | 9 | 0 | 0 | 0 | 8 | 12 | 11 | 20 | 28 | 240 | 149 | 0.556975272 | 1 | 0.041 | 0.11 | 0.081 | 0.237 | 0 | 0 | 0.318 | 0 | 0 | 0 | 0.03 |
| RI | 245 | ENSG00000104969 | SGTA | chr19 | - | 2767135 | 2767686 | 2767135 | 2767220 | 2767579 | 2767686 | 3 | 0 | 2 | 5 | 3 | 28 | 19 | 50 | 33 | 15 | 0 | 9 | 1 | 2 | 4 | 17 | 22 | 34 | 33 | 25 | 298 | 149 | 0.557539792 | 1 | 0.051 | 0 | 0.02 | 0.07 | 0.091 | 0 | 0.17 | 0.014 | 0.029 | 0.074 | -0.011 |
| RI | 7171 | ENSG00000120885 | CLU | chr8 | - | 27598459 | 27600009 | 27598459 | 27598635 | 27599779 | 27600009 | 10 | 8 | 13 | 28 | 8 | 265 | 274 | 63 | 697 | 295 | 74 | 49 | 18 | 18 | 189 | 1170 | 1395 | 509 | 558 | 6838 | 298 | 149 | 0.558251275 | 1 | 0.019 | 0.014 | 0.094 | 0.02 | 0.013 | 0.031 | 0.017 | 0.017 | 0.016 | 0.014 | 0.013 |
| RI | 5604 | ENSG00000102125 | TAZ | chrX | + | 154419542 | 154419746 | 154419542 | 154419623 | 154419704 | 154419746 | 18 | 10 | 28 | 30 | 28 | 12 | 3 | 23 | 16 | 24 | 13 | 45 | 19 | 26 | 34 | 2 | 27 | 2 | 36 | 37 | 230 | 149 | 0.558376795 | 1 | 0.493 | 0.683 | 0.441 | 0.548 | 0.43 | 0.808 | 0.519 | 0.86 | 0.319 | 0.373 | -0.057 |
| RI | 1732 | ENSG00000126822 | PLEKHG3 | chr14 | + | 64736852 | 64737375 | 64736852 | 64736891 | 64737355 | 64737375 | 23 | 8 | 3 | 11 | 1 | 207 | 70 | 102 | 149 | 71 | 21 | 8 | 7 | 6 | 11 | 177 | 72 | 112 | 115 | 141 | 298 | 149 | 0.558700645 | 1 | 0.053 | 0.054 | 0.014 | 0.036 | 0.007 | 0.056 | 0.053 | 0.03 | 0.025 | 0.038 | -0.008 |
| RI | 2237 | ENSG00000115661 | STK16 | chr2 | + | 219245895 | 219246876 | 219245895 | 219246085 | 219246656 | 219246876 | 13 | 11 | 7 | 16 | 11 | 34 | 6 | 16 | 27 | 14 | 8 | 12 | 13 | 13 | 19 | 14 | 39 | 26 | 20 | 25 | 298 | 149 | 0.559200033 | 1 | 0.16 | 0.478 | 0.179 | 0.229 | 0.282 | 0.222 | 0.133 | 0.2 | 0.245 | 0.275 | 0.051 |
| RI | 375 | ENSG00000006712 | PAF1 | chr19 | - | 39389479 | 39389761 | 39389479 | 39389546 | 39389639 | 39389761 | 34 | 9 | 15 | 39 | 14 | 69 | 32 | 58 | 90 | 79 | 92 | 65 | 21 | 8 | 68 | 153 | 132 | 78 | 94 | 161 | 242 | 149 | 0.559664411 | 1 | 0.233 | 0.148 | 0.137 | 0.211 | 0.098 | 0.27 | 0.233 | 0.142 | 0.05 | 0.206 | -0.015 |
| RI | 6676 | ENSG00000188917 | TRMT2B | chrX | - | 101051250 | 101051733 | 101051250 | 101051366 | 101051643 | 101051733 | 37 | 45 | 32 | 46 | 31 | 3 | 1 | 0 | 3 | 0 | 68 | 60 | 44 | 43 | 70 | 0 | 8 | 0 | 0 | 0 | 298 | 149 | 0.559749872 | 1 | 0.86 | 0.957 | 1 | 0.885 | 1 | 1 | 0.789 | 1 | 1 | 1 | -0.017 |
| RI | 2593 | ENSG00000145495 | MARCHF6 | chr5 | + | 10400783 | 10402139 | 10400783 | 10400842 | 10402058 | 10402139 | 206 | 212 | 214 | 213 | 272 | 99 | 53 | 74 | 83 | 143 | 81 | 244 | 313 | 151 | 39 | 93 | 70 | 29 | 57 | 89 | 298 | 149 | 0.560029831 | 1 | 0.51 | 0.667 | 0.591 | 0.562 | 0.487 | 0.303 | 0.635 | 0.844 | 0.57 | 0.18 | 0.057 |
| RI | 1112 | ENSG00000118046 | STK11 | chr19 | + | 1221212 | 1222006 | 1221212 | 1221340 | 1221948 | 1222006 | 1 | 4 | 10 | 4 | 0 | 118 | 53 | 79 | 95 | 109 | 11 | 12 | 2 | 3 | 0 | 148 | 108 | 35 | 98 | 212 | 298 | 149 | 0.56003204 | 1 | 0.004 | 0.036 | 0.06 | 0.021 | 0 | 0.036 | 0.053 | 0.028 | 0.015 | 0 | -0.002 |
| RI | 8072 | ENSG00000175197 | DDIT3 | chr12 | - | 57517268 | 57517753 | 57517268 | 57517424 | 57517705 | 57517753 | 63 | 33 | 65 | 59 | 88 | 4 | 0 | 0 | 1 | 1 | 140 | 186 | 160 | 139 | 200 | 0 | 0 | 9 | 0 | 0 | 298 | 149 | 0.561249209 | 1 | 0.887 | 1 | 1 | 0.967 | 0.978 | 1 | 1 | 0.899 | 1 | 1 | -0.013 |
| RI | 8383 | ENSG00000135452 | TSPAN31 | chr12 | + | 57746175 | 57746720 | 57746175 | 57746256 | 57746592 | 57746720 | 31 | 24 | 32 | 33 | 24 | 4 | 1 | 2 | 6 | 0 | 58 | 17 | 47 | 27 | 37 | 10 | 0 | 0 | 4 | 0 | 298 | 149 | 0.561255074 | 1 | 0.795 | 0.923 | 0.889 | 0.733 | 1 | 0.744 | 1 | 1 | 0.771 | 1 | -0.035 |
| RI | 2032 | ENSG00000148218 | ALAD | chr9 | - | 113390592 | 113390933 | 113390592 | 113390676 | 113390797 | 113390933 | 4 | 20 | 15 | 19 | 4 | 33 | 5 | 24 | 14 | 13 | 5 | 19 | 5 | 16 | 5 | 17 | 13 | 0 | 7 | 29 | 270 | 149 | 0.561847466 | 1 | 0.063 | 0.688 | 0.256 | 0.428 | 0.145 | 0.14 | 0.446 | 1 | 0.558 | 0.087 | -0.13 |
| RI | 2507 | ENSG00000131591 | C1orf159 | chr1 | - | 1084352 | 1084506 | 1084352 | 1084383 | 1084480 | 1084506 | 0 | 0 | 0 | 8 | 0 | 4 | 6 | 7 | 6 | 2 | 0 | 2 | 2 | 0 | 2 | 15 | 0 | 2 | 2 | 5 | 246 | 149 | 0.562274427 | 1 | 0 | 0 | 0 | 0.447 | 0 | 0 | 1 | 0.377 | 0 | 0.195 | -0.225 |
| RI | 1400 | ENSG00000123154 | WDR83 | chr19 | + | 12669754 | 12670080 | 12669754 | 12669893 | 12669976 | 12670080 | 13 | 3 | 16 | 18 | 7 | 19 | 2 | 7 | 19 | 5 | 41 | 16 | 9 | 8 | 44 | 50 | 25 | 5 | 10 | 4 | 232 | 149 | 0.562317878 | 1 | 0.305 | 0.491 | 0.595 | 0.378 | 0.473 | 0.345 | 0.291 | 0.536 | 0.339 | 0.876 | -0.029 |
| RI | 4673 | ENSG00000182578 | CSF1R | chr5 | - | 150061490 | 150061849 | 150061490 | 150061595 | 150061722 | 150061849 | 1 | 10 | 1 | 2 | 4 | 144 | 102 | 70 | 188 | 56 | 5 | 0 | 0 | 0 | 12 | 308 | 71 | 31 | 164 | 368 | 276 | 149 | 0.563209338 | 1 | 0.004 | 0.05 | 0.008 | 0.006 | 0.037 | 0.009 | 0 | 0 | 0 | 0.017 | 0.016 |
| RI | 3447 | ENSG00000144591 | GMPPA | chr2 | + | 219500120 | 219502037 | 219500120 | 219500218 | 219501475 | 219502037 | 3 | 1 | 6 | 12 | 6 | 52 | 28 | 59 | 64 | 68 | 20 | 4 | 7 | 8 | 2 | 65 | 84 | 54 | 43 | 124 | 298 | 149 | 0.564306681 | 1 | 0.028 | 0.018 | 0.048 | 0.086 | 0.042 | 0.133 | 0.023 | 0.061 | 0.085 | 0.008 | -0.018 |
| RI | 3465 | ENSG00000123473 | STIL | chr1 | - | 47280240 | 47281209 | 47280240 | 47280363 | 47280945 | 47281209 | 28 | 20 | 23 | 33 | 56 | 0 | 0 | 1 | 0 | 0 | 98 | 59 | 46 | 106 | 54 | 0 | 0 | 1 | 5 | 0 | 298 | 149 | 0.564347519 | 1 | 1 | 1 | 0.92 | 1 | 1 | 1 | 1 | 0.958 | 0.914 | 1 | 0.01 |
| RI | 4704 | ENSG00000089127 | OAS1 | chr12 | + | 112917546 | 112919903 | 112917546 | 112917700 | 112919486 | 112919903 | 160 | 166 | 152 | 199 | 261 | 26 | 7 | 14 | 26 | 11 | 608 | 298 | 114 | 545 | 228 | 29 | 33 | 20 | 24 | 21 | 298 | 149 | 0.564598588 | 1 | 0.755 | 0.922 | 0.844 | 0.793 | 0.922 | 0.913 | 0.819 | 0.74 | 0.919 | 0.844 | 0 |
| RI | 5721 | ENSG00000130414 | NDUFA10 | chr2 | - | 240011616 | 240014860 | 240011616 | 240011696 | 240014738 | 240014860 | 89 | 68 | 126 | 111 | 111 | 60 | 23 | 97 | 69 | 99 | 183 | 117 | 110 | 174 | 61 | 114 | 70 | 45 | 66 | 71 | 298 | 149 | 0.565220359 | 1 | 0.426 | 0.596 | 0.394 | 0.446 | 0.359 | 0.445 | 0.455 | 0.55 | 0.569 | 0.3 | -0.02 |
| RI | 2503 | ENSG00000126522 | ASL | chr7 | + | 66089090 | 66089335 | 66089090 | 66089175 | 66089275 | 66089335 | 4 | 6 | 1 | 1 | 0 | 28 | 11 | 18 | 30 | 13 | 5 | 10 | 0 | 5 | 0 | 44 | 69 | 34 | 41 | 106 | 249 | 149 | 0.565276256 | 1 | 0.079 | 0.246 | 0.032 | 0.02 | 0 | 0.064 | 0.08 | 0 | 0.068 | 0 | 0.033 |
| RI | 2990 | ENSG00000171109 | MFN1 | chr3 | + | 179378340 | 179378814 | 179378340 | 179378443 | 179378584 | 179378814 | 29 | 18 | 20 | 28 | 26 | 74 | 40 | 38 | 60 | 135 | 7 | 16 | 29 | 2 | 0 | 7 | 21 | 36 | 22 | 8 | 290 | 149 | 0.56624126 | 1 | 0.168 | 0.188 | 0.213 | 0.193 | 0.09 | 0.339 | 0.281 | 0.293 | 0.045 | 0 | -0.021 |
| RI | 8602 | ENSG00000071564 | TCF3 | chr19 | - | 1609289 | 1611849 | 1609289 | 1611369 | 1611703 | 1611849 | 107 | 33 | 134 | 138 | 121 | 20 | 21 | 10 | 17 | 27 | 132 | 124 | 79 | 66 | 188 | 39 | 29 | 30 | 17 | 14 | 298 | 149 | 0.566513337 | 1 | 0.728 | 0.44 | 0.87 | 0.802 | 0.691 | 0.629 | 0.681 | 0.568 | 0.66 | 0.87 | 0.025 |
| RI | 2064 | ENSG00000079616 | KIF22 | chr16 | + | 29799263 | 29799781 | 29799263 | 29799494 | 29799627 | 29799781 | 22 | 7 | 21 | 19 | 5 | 52 | 19 | 47 | 45 | 50 | 18 | 8 | 15 | 24 | 38 | 75 | 41 | 71 | 27 | 47 | 282 | 149 | 0.566578652 | 1 | 0.183 | 0.163 | 0.191 | 0.182 | 0.05 | 0.113 | 0.093 | 0.1 | 0.32 | 0.299 | -0.031 |
| RI | 5103 | ENSG00000160753 | RUSC1 | chr1 | + | 155325101 | 155325490 | 155325101 | 155325178 | 155325315 | 155325490 | 6 | 0 | 7 | 8 | 0 | 16 | 4 | 16 | 11 | 17 | 12 | 0 | 0 | 8 | 21 | 3 | 12 | 11 | 30 | 16 | 286 | 149 | 0.567561448 | 1 | 0.163 | 0 | 0.186 | 0.275 | 0 | 0.676 | 0 | 0 | 0.122 | 0.406 | -0.116 |
| RI | 4836 | ENSG00000103168 | TAF1C | chr16 | - | 84183075 | 84183328 | 84183075 | 84183149 | 84183243 | 84183328 | 7 | 1 | 12 | 17 | 6 | 28 | 15 | 23 | 30 | 11 | 13 | 27 | 7 | 1 | 9 | 15 | 29 | 32 | 12 | 23 | 243 | 149 | 0.567599424 | 1 | 0.133 | 0.039 | 0.242 | 0.258 | 0.251 | 0.347 | 0.363 | 0.118 | 0.049 | 0.194 | -0.03 |
| RI | 6956 | ENSG00000215301 | DDX3X | chrX | + | 41343736 | 41344128 | 41343736 | 41343822 | 41344029 | 41344128 | 231 | 326 | 274 | 402 | 372 | 422 | 358 | 368 | 544 | 623 | 47 | 268 | 214 | 271 | 0 | 106 | 243 | 280 | 208 | 34 | 298 | 149 | 0.567969465 | 1 | 0.215 | 0.313 | 0.271 | 0.27 | 0.23 | 0.181 | 0.355 | 0.276 | 0.394 | 0 | 0.019 |
| RI | 6346 | ENSG00000288558 | DUS4L-BCAP29 | chr7 | + | 107563970 | 107564676 | 107563970 | 107564209 | 107564587 | 107564676 | 18 | 15 | 26 | 14 | 16 | 1 | 0 | 1 | 0 | 0 | 11 | 20 | 13 | 12 | 12 | 0 | 0 | 0 | 0 | 4 | 298 | 149 | 0.568004519 | 1 | 0.9 | 1 | 0.929 | 1 | 1 | 1 | 1 | 1 | 1 | 0.6 | 0.046 |
| RI | 4642 | ENSG00000227057 | WDR46 | chr6 | - | 33280427 | 33280987 | 33280427 | 33280522 | 33280673 | 33280987 | 39 | 14 | 37 | 18 | 23 | 29 | 11 | 11 | 23 | 8 | 0 | 20 | 34 | 29 | 51 | 13 | 8 | 7 | 18 | 45 | 298 | 149 | 0.568063343 | 1 | 0.402 | 0.389 | 0.627 | 0.281 | 0.59 | 0 | 0.556 | 0.708 | 0.446 | 0.362 | 0.043 |
| RI | 4138 | ENSG00000154222 | CC2D1B | chr1 | - | 52355397 | 52355666 | 52355397 | 52355449 | 52355607 | 52355666 | 17 | 11 | 12 | 16 | 7 | 82 | 44 | 60 | 76 | 72 | 49 | 10 | 9 | 2 | 10 | 58 | 62 | 59 | 41 | 84 | 298 | 149 | 0.568310729 | 1 | 0.094 | 0.111 | 0.091 | 0.095 | 0.046 | 0.297 | 0.075 | 0.071 | 0.024 | 0.056 | -0.017 |
| RI | 6671 | ENSG00000169598 | DFFB | chr1 | + | 3883506 | 3885429 | 3883506 | 3883800 | 3885037 | 3885429 | 103 | 71 | 63 | 61 | 52 | 6 | 4 | 4 | 0 | 1 | 145 | 97 | 58 | 117 | 96 | 4 | 8 | 0 | 4 | 0 | 298 | 149 | 0.568451324 | 1 | 0.896 | 0.899 | 0.887 | 1 | 0.963 | 0.948 | 0.858 | 1 | 0.936 | 1 | -0.019 |
| RI | 7975 | ENSG00000174775 | HRAS | chr11 | - | 532241 | 532755 | 532241 | 532522 | 532630 | 532755 | 11 | 0 | 16 | 7 | 1 | 30 | 20 | 36 | 31 | 20 | 15 | 6 | 3 | 4 | 14 | 47 | 46 | 35 | 29 | 56 | 257 | 149 | 0.568594349 | 1 | 0.175 | 0 | 0.205 | 0.116 | 0.028 | 0.156 | 0.07 | 0.047 | 0.074 | 0.127 | 0.01 |
| RI | 7811 | ENSG00000255717 | SNHG1 | chr11 | - | 62854518 | 62855216 | 62854518 | 62854938 | 62855132 | 62855216 | 147 | 77 | 220 | 244 | 177 | 143 | 72 | 167 | 141 | 168 | 116 | 158 | 169 | 167 | 25 | 98 | 108 | 165 | 45 | 34 | 298 | 149 | 0.568897448 | 1 | 0.339 | 0.348 | 0.397 | 0.464 | 0.345 | 0.372 | 0.422 | 0.339 | 0.65 | 0.269 | -0.032 |
| RI | 5550 | ENSG00000135775 | COG2 | chr1 | + | 230669355 | 230671640 | 230669355 | 230669535 | 230671515 | 230671640 | 23 | 25 | 16 | 21 | 25 | 58 | 35 | 60 | 53 | 91 | 16 | 9 | 24 | 5 | 9 | 54 | 53 | 25 | 39 | 16 | 298 | 149 | 0.569694129 | 1 | 0.165 | 0.263 | 0.118 | 0.165 | 0.121 | 0.129 | 0.078 | 0.324 | 0.06 | 0.22 | 0.004 |
| RI | 6098 | ENSG00000155906 | RMND1 | chr6 | - | 151421244 | 151423631 | 151421244 | 151422605 | 151423524 | 151423631 | 15 | 15 | 18 | 22 | 15 | 49 | 37 | 30 | 44 | 58 | 0 | 14 | 13 | 5 | 4 | 17 | 39 | 28 | 31 | 0 | 298 | 149 | 0.569708087 | 1 | 0.133 | 0.169 | 0.231 | 0.2 | 0.115 | 0 | 0.152 | 0.188 | 0.075 | 1 | -0.113 |
| RI | 893 | ENSG00000136490 | LIMD2 | chr17 | - | 63698798 | 63699069 | 63698798 | 63698818 | 63699027 | 63699069 | 225 | 67 | 213 | 173 | 54 | 5 | 0 | 1 | 3 | 0 | 94 | 194 | 126 | 92 | 244 | 0 | 0 | 0 | 0 | 0 | 298 | 149 | 0.569991759 | 1 | 0.957 | 1 | 0.991 | 0.966 | 1 | 1 | 1 | 1 | 1 | 1 | -0.017 |
| RI | 3373 | ENSG00000106330 | MOSPD3 | chr7 | + | 100612729 | 100613269 | 100612729 | 100612996 | 100613193 | 100613269 | 0 | 0 | 8 | 1 | 0 | 15 | 1 | 14 | 24 | 5 | 1 | 11 | 0 | 0 | 0 | 19 | 31 | 11 | 16 | 34 | 298 | 149 | 0.570499034 | 1 | 0 | 0 | 0.222 | 0.02 | 0 | 0.026 | 0.151 | 0 | 0 | 0 | 0.013 |
| RI | 4878 | ENSG00000182400 | TRAPPC6B | chr14 | - | 39158284 | 39159550 | 39158284 | 39158402 | 39159449 | 39159550 | 38 | 35 | 28 | 49 | 23 | 39 | 27 | 29 | 22 | 38 | 47 | 20 | 34 | 22 | 0 | 10 | 28 | 15 | 18 | 43 | 298 | 149 | 0.570787743 | 1 | 0.328 | 0.393 | 0.326 | 0.527 | 0.232 | 0.701 | 0.263 | 0.531 | 0.379 | 0 | -0.014 |
| RI | 3167 | ENSG00000138459 | SLC35A5 | chr3 | + | 112580545 | 112581326 | 112580545 | 112580706 | 112581178 | 112581326 | 127 | 102 | 147 | 149 | 188 | 1 | 0 | 0 | 0 | 7 | 24 | 126 | 67 | 41 | 38 | 0 | 0 | 0 | 0 | 0 | 298 | 149 | 0.571743679 | 1 | 0.984 | 1 | 1 | 1 | 0.931 | 1 | 1 | 1 | 1 | 1 | -0.017 |
| RI | 614 | ENSG00000101194 | SLC17A9 | chr20 | + | 62966702 | 62968597 | 62966702 | 62966732 | 62967336 | 62968597 | 3 | 0 | 2 | 2 | 0 | 3 | 2 | 8 | 10 | 7 | 0 | 7 | 0 | 0 | 0 | 8 | 21 | 6 | 5 | 12 | 298 | 149 | 0.57326682 | 1 | 0.333 | 0 | 0.111 | 0.091 | 0 | 0 | 0.143 | 0 | 0 | 0 | 0.078 |
| RI | 309 | ENSG00000161265 | U2AF1L4 | chr19 | - | 35744322 | 35745187 | 35744322 | 35744738 | 35745124 | 35745187 | 50 | 10 | 32 | 33 | 42 | 9 | 1 | 2 | 7 | 4 | 108 | 50 | 19 | 57 | 29 | 13 | 7 | 7 | 0 | 13 | 298 | 149 | 0.574393315 | 1 | 0.735 | 0.833 | 0.889 | 0.702 | 0.84 | 0.806 | 0.781 | 0.576 | 1 | 0.527 | 0.062 |
| RI | 7610 | ENSG00000126934 | MAP2K2 | chr19 | - | 4094452 | 4095449 | 4094452 | 4094498 | 4095387 | 4095449 | 18 | 10 | 24 | 8 | 15 | 250 | 67 | 155 | 171 | 271 | 10 | 3 | 11 | 20 | 19 | 163 | 211 | 86 | 118 | 271 | 298 | 149 | 0.574939582 | 1 | 0.035 | 0.069 | 0.072 | 0.023 | 0.027 | 0.03 | 0.007 | 0.06 | 0.078 | 0.034 | 0.003 |
| RI | 8569 | ENSG00000125457 | MIF4GD | chr17 | - | 75267745 | 75268192 | 75267745 | 75267901 | 75268082 | 75268192 | 7 | 4 | 8 | 19 | 6 | 44 | 31 | 38 | 36 | 40 | 9 | 11 | 14 | 11 | 10 | 16 | 68 | 19 | 40 | 89 | 298 | 149 | 0.575429089 | 1 | 0.074 | 0.061 | 0.095 | 0.209 | 0.07 | 0.22 | 0.075 | 0.269 | 0.121 | 0.053 | -0.046 |
| RI | 6360 | ENSG00000075624 | ACTB | chr7 | - | 5529160 | 5529663 | 5529160 | 5529400 | 5529534 | 5529663 | 188 | 86 | 175 | 158 | 195 | 13622 | 6280 | 8649 | 10509 | 7774 | 212 | 244 | 96 | 180 | 802 | 17451 | 15301 | 6579 | 13364 | 65078 | 283 | 149 | 0.575539983 | 1 | 0.007 | 0.007 | 0.011 | 0.008 | 0.013 | 0.006 | 0.008 | 0.008 | 0.007 | 0.006 | 0.002 |
| RI | 831 | ENSG00000100038 | TOP3B | chr22 | - | 21957026 | 21958693 | 21957026 | 21957595 | 21958491 | 21958693 | 4 | 5 | 13 | 10 | 17 | 16 | 7 | 14 | 11 | 1 | 11 | 9 | 14 | 0 | 32 | 8 | 11 | 5 | 3 | 7 | 298 | 149 | 0.575714755 | 1 | 0.111 | 0.263 | 0.317 | 0.312 | 0.895 | 0.407 | 0.29 | 0.583 | 0 | 0.696 | -0.016 |
| RI | 7580 | ENSG00000117751 | PPP1R8 | chr1 | + | 27850092 | 27851676 | 27850092 | 27850414 | 27851539 | 27851676 | 20 | 19 | 21 | 12 | 22 | 0 | 0 | 1 | 0 | 0 | 32 | 17 | 7 | 15 | 15 | 5 | 0 | 0 | 0 | 0 | 298 | 149 | 0.575769496 | 1 | 1 | 1 | 0.913 | 1 | 1 | 0.762 | 1 | 1 | 1 | 1 | 0.03 |
| RI | 4522 | ENSG00000065457 | ADAT1 | chr16 | - | 75622402 | 75623281 | 75622402 | 75622464 | 75623237 | 75623281 | 44 | 31 | 45 | 66 | 49 | 14 | 13 | 8 | 23 | 9 | 73 | 74 | 26 | 27 | 54 | 7 | 16 | 8 | 11 | 18 | 298 | 149 | 0.577268822 | 1 | 0.611 | 0.544 | 0.738 | 0.589 | 0.731 | 0.839 | 0.698 | 0.619 | 0.551 | 0.6 | -0.019 |
| RI | 3614 | ENSG00000126088 | UROD | chr1 | + | 45013135 | 45013354 | 45013135 | 45013215 | 45013291 | 45013354 | 26 | 17 | 27 | 38 | 32 | 73 | 53 | 82 | 104 | 75 | 26 | 36 | 35 | 18 | 36 | 108 | 94 | 44 | 68 | 179 | 225 | 149 | 0.5778987 | 1 | 0.191 | 0.175 | 0.179 | 0.195 | 0.22 | 0.138 | 0.202 | 0.345 | 0.149 | 0.118 | 0.002 |
| RI | 8461 | ENSG00000139168 | ZCRB1 | chr12 | - | 42322417 | 42324104 | 42322417 | 42322446 | 42324018 | 42324104 | 16 | 12 | 16 | 18 | 16 | 200 | 103 | 148 | 146 | 110 | 1 | 19 | 29 | 8 | 4 | 79 | 161 | 189 | 119 | 44 | 298 | 149 | 0.578188529 | 1 | 0.038 | 0.055 | 0.051 | 0.058 | 0.068 | 0.006 | 0.056 | 0.071 | 0.033 | 0.043 | 0.012 |
| RI | 8502 | ENSG00000139579 | NABP2 | chr12 | + | 56226178 | 56226419 | 56226178 | 56226260 | 56226355 | 56226419 | 2 | 3 | 2 | 4 | 6 | 70 | 27 | 44 | 57 | 143 | 4 | 8 | 3 | 2 | 1 | 106 | 70 | 53 | 84 | 154 | 244 | 149 | 0.578720197 | 1 | 0.017 | 0.064 | 0.027 | 0.041 | 0.025 | 0.023 | 0.065 | 0.033 | 0.014 | 0.004 | 0.007 |
| RI | 3988 | ENSG00000197077 | KIAA1671 | chr22 | + | 25174239 | 25177522 | 25174239 | 25174489 | 25177347 | 25177522 | 4 | 1 | 4 | 3 | 3 | 10 | 3 | 4 | 11 | 9 | 0 | 5 | 0 | 5 | 15 | 7 | 9 | 7 | 14 | 32 | 298 | 149 | 0.580392157 | 1 | 0.167 | 0.143 | 0.333 | 0.12 | 0.143 | 0 | 0.217 | 0 | 0.152 | 0.19 | 0.069 |
| RI | 3430 | ENSG00000008128 | CDK11A | chr1 | - | 1722587 | 1722831 | 1722587 | 1722634 | 1722707 | 1722831 | 18 | 43 | 30 | 72 | 32 | 0 | 4 | 8 | 8 | 14 | 127 | 60 | 17 | 112 | 79 | 8 | 20 | 0 | 21 | 6 | 222 | 149 | 0.582611373 | 1 | 1 | 0.878 | 0.716 | 0.858 | 0.605 | 0.914 | 0.668 | 1 | 0.782 | 0.898 | -0.041 |
| RI | 7275 | ENSG00000104522 | GFUS | chr8 | - | 143614319 | 143614697 | 143614319 | 143614453 | 143614623 | 143614697 | 23 | 16 | 25 | 12 | 14 | 42 | 14 | 46 | 53 | 55 | 40 | 23 | 11 | 16 | 24 | 45 | 89 | 25 | 44 | 102 | 298 | 149 | 0.582782214 | 1 | 0.215 | 0.364 | 0.214 | 0.102 | 0.113 | 0.308 | 0.114 | 0.18 | 0.154 | 0.105 | 0.029 |
| RI | 7228 | ENSG00000180900 | SCRIB | chr8 | - | 143792484 | 143792867 | 143792484 | 143792635 | 143792707 | 143792867 | 3 | 0 | 0 | 3 | 0 | 17 | 7 | 13 | 20 | 4 | 10 | 0 | 0 | 0 | 0 | 5 | 6 | 23 | 5 | 5 | 221 | 149 | 0.583405262 | 1 | 0.106 | 0 | 0 | 0.092 | 0 | 0.574 | 0 | 0 | 0 | 0 | -0.075 |
| RI | 2434 | ENSG00000215845 | TSTD1 | chr1 | - | 161037630 | 161038075 | 161037630 | 161037826 | 161037912 | 161038075 | 46 | 19 | 20 | 44 | 14 | 70 | 20 | 58 | 85 | 37 | 9 | 30 | 28 | 21 | 9 | 28 | 51 | 24 | 32 | 76 | 235 | 149 | 0.5836165 | 1 | 0.294 | 0.376 | 0.179 | 0.247 | 0.193 | 0.169 | 0.272 | 0.425 | 0.294 | 0.07 | 0.012 |
| RI | 4365 | ENSG00000144381 | HSPD1 | chr2 | - | 197494156 | 197495376 | 197494156 | 197494250 | 197494656 | 197495376 | 56 | 48 | 62 | 52 | 45 | 184 | 125 | 185 | 199 | 234 | 0 | 26 | 63 | 22 | 24 | 39 | 100 | 118 | 91 | 1 | 298 | 149 | 0.584829919 | 1 | 0.132 | 0.161 | 0.144 | 0.116 | 0.088 | 0 | 0.115 | 0.211 | 0.108 | 0.923 | -0.143 |
| RI | 496 | ENSG00000196365 | LONP1 | chr19 | - | 5696248 | 5696757 | 5696248 | 5696371 | 5696669 | 5696757 | 9 | 5 | 13 | 11 | 10 | 10 | 12 | 9 | 14 | 14 | 0 | 15 | 1 | 31 | 23 | 7 | 4 | 9 | 10 | 5 | 298 | 149 | 0.585520899 | 1 | 0.31 | 0.172 | 0.419 | 0.282 | 0.263 | 0 | 0.652 | 0.053 | 0.608 | 0.697 | -0.113 |
| RI | 1357 | ENSG00000102879 | CORO1A | chr16 | + | 30186815 | 30187223 | 30186815 | 30186945 | 30187038 | 30187223 | 124 | 91 | 105 | 151 | 87 | 1411 | 492 | 1062 | 1107 | 422 | 168 | 197 | 135 | 153 | 272 | 1488 | 1474 | 697 | 1398 | 1802 | 242 | 149 | 0.586964428 | 1 | 0.051 | 0.102 | 0.057 | 0.077 | 0.113 | 0.065 | 0.076 | 0.107 | 0.063 | 0.085 | 0.001 |
| RI | 4342 | ENSG00000149476 | TKFC | chr11 | + | 61342460 | 61342658 | 61342460 | 61342495 | 61342573 | 61342658 | 1 | 3 | 4 | 3 | 4 | 9 | 9 | 15 | 8 | 4 | 8 | 2 | 5 | 7 | 21 | 36 | 17 | 8 | 19 | 44 | 227 | 149 | 0.58699364 | 1 | 0.068 | 0.18 | 0.149 | 0.198 | 0.396 | 0.127 | 0.072 | 0.291 | 0.195 | 0.239 | 0.013 |
| RI | 951 | ENSG00000132507 | EIF5A | chr17 | + | 7311349 | 7311651 | 7311349 | 7311481 | 7311577 | 7311651 | 118 | 84 | 127 | 103 | 83 | 1018 | 573 | 942 | 796 | 882 | 236 | 121 | 75 | 130 | 171 | 1487 | 1000 | 817 | 1127 | 1781 | 245 | 149 | 0.587743712 | 1 | 0.066 | 0.082 | 0.076 | 0.073 | 0.054 | 0.088 | 0.069 | 0.053 | 0.066 | 0.055 | 0.004 |
| RI | 1284 | ENSG00000167526 | RPL13 | chr16 | + | 89560939 | 89561368 | 89560939 | 89561063 | 89561226 | 89561368 | 4 | 18 | 23 | 13 | 7 | 1238 | 404 | 840 | 853 | 318 | 1 | 14 | 1 | 1 | 7 | 503 | 785 | 452 | 397 | 956 | 298 | 149 | 0.587982985 | 1 | 0.002 | 0.022 | 0.014 | 0.008 | 0.011 | 0.001 | 0.009 | 0.001 | 0.001 | 0.004 | 0.008 |
| RI | 4591 | ENSG00000165637 | VDAC2 | chr10 | + | 75219062 | 75219356 | 75219062 | 75219215 | 75219303 | 75219356 | 47 | 71 | 32 | 56 | 50 | 433 | 231 | 346 | 406 | 557 | 90 | 78 | 45 | 110 | 95 | 667 | 494 | 348 | 591 | 909 | 237 | 149 | 0.589006947 | 1 | 0.064 | 0.162 | 0.055 | 0.08 | 0.053 | 0.078 | 0.09 | 0.075 | 0.105 | 0.062 | 0.001 |
| RI | 6461 | ENSG00000160856 | FCRL3 | chr1 | - | 157697139 | 157697919 | 157697139 | 157697329 | 157697658 | 157697919 | 130 | 62 | 65 | 75 | 74 | 4 | 0 | 0 | 5 | 12 | 54 | 118 | 55 | 46 | 38 | 7 | 3 | 4 | 0 | 0 | 298 | 149 | 0.589586137 | 1 | 0.942 | 1 | 1 | 0.882 | 0.755 | 0.794 | 0.952 | 0.873 | 1 | 1 | -0.008 |
| RI | 8444 | ENSG00000100599 | RIN3 | chr14 | + | 92684986 | 92688049 | 92684986 | 92685150 | 92687685 | 92688049 | 30 | 8 | 17 | 29 | 13 | 2 | 3 | 2 | 0 | 3 | 35 | 17 | 16 | 33 | 25 | 0 | 1 | 0 | 2 | 8 | 298 | 149 | 0.590200875 | 1 | 0.882 | 0.571 | 0.81 | 1 | 0.684 | 1 | 0.895 | 1 | 0.892 | 0.61 | -0.09 |
| RI | 4855 | ENSG00000149532 | CPSF7 | chr11 | - | 61415665 | 61416519 | 61415665 | 61415784 | 61416104 | 61416519 | 35 | 48 | 36 | 45 | 25 | 84 | 49 | 67 | 85 | 38 | 121 | 59 | 67 | 70 | 59 | 147 | 97 | 46 | 169 | 145 | 298 | 149 | 0.591929687 | 1 | 0.172 | 0.329 | 0.212 | 0.209 | 0.248 | 0.292 | 0.233 | 0.421 | 0.172 | 0.169 | -0.023 |
| RI | 1214 | ENSG00000167522 | ANKRD11 | chr16 | - | 89286038 | 89288670 | 89286038 | 89286186 | 89288527 | 89288670 | 43 | 19 | 23 | 39 | 25 | 30 | 23 | 20 | 30 | 4 | 78 | 38 | 55 | 71 | 72 | 8 | 53 | 40 | 65 | 29 | 298 | 149 | 0.593336142 | 1 | 0.417 | 0.292 | 0.365 | 0.394 | 0.758 | 0.83 | 0.264 | 0.407 | 0.353 | 0.554 | -0.036 |
| RI | 8173 | ENSG00000152778 | IFIT5 | chr10 | + | 89417204 | 89420997 | 89417204 | 89417551 | 89418063 | 89420997 | 213 | 225 | 273 | 343 | 418 | 0 | 0 | 1 | 1 | 3 | 67 | 318 | 214 | 145 | 18 | 0 | 9 | 0 | 0 | 0 | 298 | 149 | 0.593499096 | 1 | 1 | 1 | 0.993 | 0.994 | 0.986 | 1 | 0.946 | 1 | 1 | 1 | 0.005 |
| RI | 423 | ENSG00000249115 | HAUS5 | chr19 | + | 35613867 | 35614059 | 35613867 | 35613900 | 35614034 | 35614059 | 2 | 0 | 6 | 3 | 0 | 13 | 1 | 5 | 2 | 4 | 0 | 5 | 2 | 4 | 0 | 4 | 20 | 2 | 6 | 9 | 283 | 149 | 0.5947346 | 1 | 0.075 | 0 | 0.387 | 0.441 | 0 | 0 | 0.116 | 0.345 | 0.26 | 0 | 0.036 |
| RI | 488 | ENSG00000142347 | MYO1F | chr19 | - | 8526452 | 8526935 | 8526452 | 8526601 | 8526788 | 8526935 | 30 | 2 | 3 | 7 | 11 | 97 | 42 | 59 | 60 | 46 | 24 | 11 | 8 | 28 | 12 | 101 | 111 | 40 | 82 | 127 | 298 | 149 | 0.597184892 | 1 | 0.134 | 0.023 | 0.025 | 0.055 | 0.107 | 0.106 | 0.047 | 0.091 | 0.146 | 0.045 | -0.018 |
| RI | 1854 | ENSG00000166887 | VPS39 | chr15 | - | 42164357 | 42165816 | 42164357 | 42164486 | 42165717 | 42165816 | 33 | 29 | 30 | 42 | 52 | 0 | 0 | 1 | 2 | 2 | 74 | 76 | 65 | 95 | 69 | 0 | 2 | 4 | 0 | 2 | 298 | 149 | 0.597531429 | 1 | 1 | 1 | 0.938 | 0.913 | 0.929 | 1 | 0.95 | 0.89 | 1 | 0.945 | -0.001 |
| RI | 1252 | ENSG00000188603 | CLN3 | chr16 | - | 28486577 | 28487541 | 28486577 | 28486650 | 28487455 | 28487541 | 1 | 8 | 10 | 3 | 0 | 43 | 20 | 14 | 17 | 21 | 10 | 3 | 6 | 8 | 5 | 47 | 27 | 18 | 22 | 32 | 298 | 149 | 0.59780198 | 1 | 0.011 | 0.167 | 0.263 | 0.081 | 0 | 0.096 | 0.053 | 0.143 | 0.154 | 0.072 | 0.001 |
| RI | 5915 | ENSG00000089818 | NECAP1 | chr12 | + | 8090194 | 8092784 | 8090194 | 8090299 | 8092675 | 8092784 | 239 | 175 | 188 | 287 | 253 | 13 | 5 | 9 | 13 | 38 | 515 | 449 | 619 | 692 | 125 | 28 | 39 | 40 | 37 | 18 | 298 | 149 | 0.598368483 | 1 | 0.902 | 0.946 | 0.913 | 0.917 | 0.769 | 0.902 | 0.852 | 0.886 | 0.903 | 0.776 | 0.026 |
| RI | 7660 | ENSG00000110711 | AIP | chr11 | + | 67489266 | 67490214 | 67489266 | 67489455 | 67490037 | 67490214 | 19 | 6 | 14 | 21 | 7 | 190 | 87 | 166 | 170 | 91 | 30 | 8 | 16 | 12 | 6 | 101 | 191 | 123 | 84 | 207 | 298 | 149 | 0.598806104 | 1 | 0.048 | 0.033 | 0.04 | 0.058 | 0.037 | 0.129 | 0.021 | 0.061 | 0.067 | 0.014 | -0.015 |
| RI | 4271 | ENSG00000114841 | DNAH1 | chr3 | + | 52388417 | 52388937 | 52388417 | 52388609 | 52388805 | 52388937 | 19 | 2 | 17 | 22 | 9 | 13 | 4 | 10 | 11 | 12 | 16 | 24 | 27 | 0 | 12 | 6 | 18 | 5 | 8 | 33 | 298 | 149 | 0.598964916 | 1 | 0.422 | 0.2 | 0.459 | 0.5 | 0.273 | 0.571 | 0.4 | 0.73 | 0 | 0.154 | 0 |
| RI | 3995 | ENSG00000198912 | C1orf174 | chr1 | - | 3890568 | 3892996 | 3890568 | 3891057 | 3892740 | 3892996 | 49 | 16 | 31 | 46 | 47 | 0 | 2 | 3 | 0 | 0 | 24 | 41 | 49 | 61 | 49 | 0 | 0 | 0 | 0 | 7 | 298 | 149 | 0.599777856 | 1 | 1 | 0.8 | 0.838 | 1 | 1 | 1 | 1 | 1 | 1 | 0.778 | -0.028 |
| RI | 8605 | ENSG00000071564 | TCF3 | chr19 | - | 1619110 | 1619474 | 1619110 | 1619234 | 1619315 | 1619474 | 4 | 1 | 6 | 3 | 1 | 45 | 16 | 46 | 52 | 18 | 5 | 2 | 1 | 0 | 5 | 18 | 52 | 26 | 43 | 52 | 230 | 149 | 0.600433265 | 1 | 0.054 | 0.039 | 0.078 | 0.036 | 0.035 | 0.153 | 0.024 | 0.024 | 0 | 0.059 | -0.004 |
| RI | 7496 | ENSG00000213930 | GALT | chr9 | + | 34647491 | 34647961 | 34647491 | 34647705 | 34647831 | 34647961 | 53 | 31 | 42 | 41 | 23 | 43 | 23 | 52 | 38 | 45 | 30 | 58 | 27 | 36 | 83 | 54 | 41 | 31 | 43 | 39 | 275 | 149 | 0.600918471 | 1 | 0.4 | 0.422 | 0.304 | 0.369 | 0.217 | 0.231 | 0.434 | 0.321 | 0.312 | 0.536 | -0.024 |
| RI | 6058 | ENSG00000132424 | PNISR | chr6 | - | 99414571 | 99416428 | 99414571 | 99414686 | 99416348 | 99416428 | 254 | 225 | 193 | 239 | 288 | 69 | 75 | 106 | 82 | 148 | 96 | 316 | 292 | 142 | 25 | 43 | 41 | 35 | 53 | 42 | 298 | 149 | 0.601933875 | 1 | 0.648 | 0.6 | 0.477 | 0.593 | 0.493 | 0.527 | 0.794 | 0.807 | 0.573 | 0.229 | -0.024 |
| RI | 7211 | ENSG00000160948 | VPS28 | chr8 | - | 144424214 | 144424819 | 144424214 | 144424268 | 144424717 | 144424819 | 34 | 10 | 29 | 34 | 17 | 232 | 110 | 165 | 165 | 158 | 25 | 20 | 31 | 30 | 40 | 102 | 254 | 115 | 155 | 309 | 298 | 149 | 0.603973386 | 1 | 0.068 | 0.043 | 0.081 | 0.093 | 0.051 | 0.109 | 0.038 | 0.119 | 0.088 | 0.061 | -0.016 |
| RI | 8557 | ENSG00000121310 | ECHDC2 | chr1 | - | 52905033 | 52906611 | 52905033 | 52905090 | 52906518 | 52906611 | 15 | 10 | 13 | 22 | 15 | 14 | 9 | 18 | 19 | 15 | 46 | 5 | 7 | 6 | 6 | 6 | 7 | 13 | 8 | 36 | 298 | 149 | 0.604128358 | 1 | 0.349 | 0.357 | 0.265 | 0.367 | 0.333 | 0.793 | 0.263 | 0.212 | 0.273 | 0.077 | 0.011 |
| RI | 7454 | ENSG00000188283 | ZNF383 | chr19 | + | 37235548 | 37236074 | 37235548 | 37235758 | 37235978 | 37236074 | 29 | 24 | 39 | 34 | 30 | 0 | 0 | 0 | 1 | 2 | 42 | 18 | 42 | 30 | 16 | 0 | 0 | 6 | 0 | 0 | 298 | 149 | 0.605660324 | 1 | 1 | 1 | 1 | 0.944 | 0.882 | 1 | 1 | 0.778 | 1 | 1 | 0.01 |
| RI | 6067 | ENSG00000204351 | SKIV2L | chr6 | + | 31969279 | 31969751 | 31969279 | 31969420 | 31969514 | 31969751 | 27 | 25 | 42 | 41 | 34 | 9 | 10 | 13 | 6 | 4 | 14 | 35 | 6 | 34 | 67 | 3 | 16 | 5 | 2 | 24 | 243 | 149 | 0.605894768 | 1 | 0.648 | 0.605 | 0.665 | 0.807 | 0.839 | 0.741 | 0.573 | 0.424 | 0.912 | 0.631 | 0.057 |
| RI | 5746 | ENSG00000125503 | PPP1R12C | chr19 | - | 55093015 | 55093233 | 55093015 | 55093076 | 55093152 | 55093233 | 36 | 25 | 39 | 20 | 16 | 87 | 25 | 62 | 47 | 24 | 57 | 51 | 15 | 45 | 70 | 74 | 79 | 62 | 71 | 71 | 225 | 149 | 0.606295041 | 1 | 0.215 | 0.398 | 0.294 | 0.22 | 0.306 | 0.338 | 0.299 | 0.138 | 0.296 | 0.395 | -0.007 |
| RI | 794 | ENSG00000141252 | VPS53 | chr17 | - | 551871 | 553462 | 551871 | 551950 | 553379 | 553462 | 35 | 23 | 21 | 21 | 16 | 71 | 37 | 64 | 57 | 156 | 25 | 7 | 40 | 22 | 27 | 164 | 95 | 31 | 74 | 84 | 298 | 149 | 0.608157367 | 1 | 0.198 | 0.237 | 0.141 | 0.156 | 0.049 | 0.071 | 0.036 | 0.392 | 0.129 | 0.138 | 0.003 |
| RI | 76 | ENSG00000128159 | TUBGCP6 | chr22 | - | 50219956 | 50221874 | 50219956 | 50220015 | 50220250 | 50221874 | 32 | 4 | 26 | 28 | 23 | 17 | 0 | 12 | 18 | 0 | 37 | 12 | 12 | 18 | 82 | 29 | 6 | 0 | 6 | 24 | 298 | 149 | 0.60829237 | 1 | 0.485 | 1 | 0.52 | 0.437 | 1 | 0.389 | 0.5 | 1 | 0.6 | 0.631 | 0.064 |
| RI | 3319 | ENSG00000198818 | SFT2D1 | chr6 | - | 166322856 | 166324595 | 166322856 | 166322886 | 166324536 | 166324595 | 74 | 89 | 51 | 82 | 79 | 88 | 67 | 84 | 65 | 144 | 64 | 53 | 189 | 104 | 9 | 70 | 46 | 39 | 85 | 68 | 298 | 149 | 0.608310215 | 1 | 0.296 | 0.399 | 0.233 | 0.387 | 0.215 | 0.314 | 0.366 | 0.708 | 0.38 | 0.062 | -0.06 |
| RI | 5278 | ENSG00000059804 | SLC2A3 | chr12 | - | 7929683 | 7930642 | 7929683 | 7929871 | 7930479 | 7930642 | 45 | 62 | 38 | 57 | 38 | 149 | 150 | 168 | 281 | 154 | 312 | 123 | 163 | 282 | 138 | 519 | 695 | 695 | 629 | 935 | 298 | 149 | 0.608375319 | 1 | 0.131 | 0.171 | 0.102 | 0.092 | 0.11 | 0.231 | 0.081 | 0.105 | 0.183 | 0.069 | -0.013 |
| RI | 6028 | ENSG00000010704 | HFE | chr6 | + | 26090840 | 26091589 | 26090840 | 26091104 | 26091313 | 26091589 | 10 | 0 | 5 | 0 | 2 | 25 | 13 | 12 | 13 | 34 | 0 | 5 | 1 | 8 | 9 | 16 | 7 | 0 | 15 | 48 | 298 | 149 | 0.608814006 | 1 | 0.167 | 0 | 0.172 | 0 | 0.029 | 0 | 0.263 | 1 | 0.211 | 0.086 | -0.238 |
| RI | 1168 | ENSG00000125124 | BBS2 | chr16 | - | 56498436 | 56502456 | 56498436 | 56499907 | 56501352 | 56502456 | 31 | 23 | 28 | 28 | 15 | 6 | 1 | 4 | 5 | 16 | 22 | 52 | 33 | 17 | 8 | 0 | 11 | 9 | 0 | 10 | 298 | 149 | 0.613467039 | 1 | 0.721 | 0.92 | 0.778 | 0.737 | 0.319 | 1 | 0.703 | 0.647 | 1 | 0.286 | -0.032 |
| RI | 2156 | ENSG00000183474 | GTF2H2C | chr5 | + | 69578711 | 69579108 | 69578711 | 69578802 | 69579020 | 69579108 | 4 | 2 | 3 | 5 | 4 | 52 | 32 | 60 | 64 | 86 | 0 | 2 | 12 | 0 | 0 | 79 | 73 | 28 | 50 | 43 | 298 | 149 | 0.613913958 | 1 | 0.037 | 0.03 | 0.024 | 0.038 | 0.023 | 0 | 0.014 | 0.176 | 0 | 0 | -0.008 |
| RI | 7160 | ENSG00000182307 | C8orf33 | chr8 | + | 145052466 | 145052897 | 145052466 | 145052497 | 145052585 | 145052897 | 6 | 2 | 5 | 6 | 0 | 0 | 0 | 1 | 0 | 3 | 7 | 10 | 6 | 0 | 9 | 0 | 3 | 0 | 3 | 0 | 237 | 149 | 0.615061775 | 1 | 1 | 1 | 0.759 | 1 | 0 | 1 | 0.677 | 1 | 0 | 1 | 0.016 |
| RI | 5581 | ENSG00000132323 | ILKAP | chr2 | - | 238194274 | 238194870 | 238194274 | 238194331 | 238194804 | 238194870 | 100 | 62 | 75 | 74 | 97 | 20 | 5 | 27 | 13 | 16 | 196 | 138 | 120 | 234 | 71 | 7 | 33 | 16 | 18 | 62 | 298 | 149 | 0.615528043 | 1 | 0.714 | 0.861 | 0.581 | 0.74 | 0.752 | 0.933 | 0.676 | 0.789 | 0.867 | 0.364 | 0.004 |
| RI | 4110 | ENSG00000116337 | AMPD2 | chr1 | + | 109626725 | 109627518 | 109626725 | 109627316 | 109627428 | 109627518 | 8 | 5 | 9 | 8 | 0 | 56 | 20 | 20 | 21 | 13 | 3 | 3 | 12 | 18 | 16 | 42 | 68 | 47 | 25 | 106 | 261 | 149 | 0.617184763 | 1 | 0.075 | 0.125 | 0.204 | 0.179 | 0 | 0.039 | 0.025 | 0.127 | 0.291 | 0.079 | 0.004 |
| RI | 6509 | ENSG00000122566 | HNRNPA2B1 | chr7 | - | 26190992 | 26193694 | 26190992 | 26192577 | 26193250 | 26193694 | 335 | 481 | 338 | 474 | 224 | 746 | 670 | 566 | 781 | 703 | 119 | 431 | 625 | 198 | 38 | 214 | 484 | 777 | 726 | 86 | 298 | 149 | 0.617547919 | 1 | 0.183 | 0.264 | 0.23 | 0.233 | 0.137 | 0.218 | 0.308 | 0.287 | 0.12 | 0.181 | -0.013 |
| RI | 5700 | ENSG00000079785 | DDX1 | chr2 | + | 15627053 | 15628710 | 15627053 | 15627145 | 15628444 | 15628710 | 14 | 24 | 23 | 23 | 20 | 54 | 37 | 58 | 56 | 82 | 7 | 17 | 15 | 4 | 0 | 0 | 28 | 26 | 17 | 5 | 298 | 149 | 0.618800872 | 1 | 0.115 | 0.245 | 0.165 | 0.17 | 0.109 | 1 | 0.233 | 0.224 | 0.105 | 0 | -0.152 |
| RI | 7065 | ENSG00000205581 | HMGN1 | chr21 | - | 39348421 | 39348577 | 39348421 | 39348451 | 39348544 | 39348577 | 6 | 5 | 7 | 9 | 18 | 208 | 136 | 225 | 265 | 91 | 0 | 18 | 4 | 2 | 28 | 299 | 167 | 173 | 306 | 276 | 242 | 149 | 0.619750041 | 1 | 0.017 | 0.022 | 0.019 | 0.02 | 0.109 | 0 | 0.062 | 0.014 | 0.004 | 0.059 | 0.01 |
| RI | 786 | ENSG00000196704 | AMZ2 | chr17 | + | 68248042 | 68248705 | 68248042 | 68248086 | 68248639 | 68248705 | 178 | 100 | 102 | 147 | 193 | 1 | 1 | 0 | 1 | 0 | 81 | 230 | 128 | 151 | 307 | 0 | 0 | 0 | 0 | 9 | 298 | 149 | 0.620116763 | 1 | 0.989 | 0.98 | 1 | 0.987 | 1 | 1 | 1 | 1 | 1 | 0.945 | 0.002 |
| RI | 834 | ENSG00000267278 | MAP3K14-AS1 | chr17 | + | 45267112 | 45267490 | 45267112 | 45267328 | 45267415 | 45267490 | 34 | 9 | 3 | 19 | 26 | 2 | 4 | 1 | 4 | 6 | 80 | 21 | 14 | 3 | 67 | 5 | 5 | 2 | 0 | 17 | 236 | 149 | 0.621200992 | 1 | 0.915 | 0.587 | 0.654 | 0.75 | 0.732 | 0.91 | 0.726 | 0.815 | 1 | 0.713 | -0.105 |
| RI | 7475 | ENSG00000135047 | CTSL | chr9 | + | 87730380 | 87731320 | 87730380 | 87730498 | 87731007 | 87731320 | 3 | 9 | 1 | 4 | 0 | 73 | 101 | 31 | 142 | 117 | 0 | 0 | 2 | 8 | 15 | 322 | 144 | 78 | 218 | 213 | 298 | 149 | 0.621512664 | 1 | 0.02 | 0.043 | 0.016 | 0.014 | 0 | 0 | 0 | 0.013 | 0.018 | 0.034 | 0.006 |
| RI | 5781 | ENSG00000204843 | DCTN1 | chr2 | - | 74366458 | 74366932 | 74366458 | 74366620 | 74366782 | 74366932 | 30 | 10 | 39 | 36 | 22 | 99 | 71 | 96 | 89 | 88 | 40 | 14 | 29 | 66 | 41 | 104 | 153 | 50 | 83 | 196 | 298 | 149 | 0.62175634 | 1 | 0.132 | 0.066 | 0.169 | 0.168 | 0.111 | 0.161 | 0.044 | 0.225 | 0.284 | 0.095 | -0.033 |
| RI | 2705 | ENSG00000163161 | ERCC3 | chr2 | - | 127289336 | 127289824 | 127289336 | 127289519 | 127289688 | 127289824 | 60 | 40 | 52 | 71 | 58 | 3 | 0 | 1 | 4 | 0 | 248 | 90 | 75 | 85 | 68 | 0 | 1 | 0 | 12 | 0 | 298 | 149 | 0.622948071 | 1 | 0.909 | 1 | 0.963 | 0.899 | 1 | 1 | 0.978 | 1 | 0.78 | 1 | 0.003 |
| RI | 7494 | ENSG00000213930 | GALT | chr9 | + | 34647088 | 34647961 | 34647088 | 34647258 | 34647831 | 34647961 | 33 | 15 | 31 | 26 | 22 | 3 | 0 | 4 | 0 | 0 | 37 | 58 | 16 | 33 | 37 | 10 | 1 | 0 | 0 | 0 | 298 | 149 | 0.623147486 | 1 | 0.846 | 1 | 0.795 | 1 | 1 | 0.649 | 0.967 | 1 | 1 | 1 | 0.005 |

| | | | | | | | | | | | | | |
|---|---|---|---|---|---|---|---|---|---|---|---|---|---|
| RI | 418 | ENSG00000105369 | CD79A | chr19 | + | 41878989 | 41879653 | 41878989 | 41879289 | 41879534 | 41879653 | 7 | 1 |
| RI | 4083 | ENSG00000084754 | HADHA | chr2 | - | 26190654 | 26191628 | 26190654 | 26191395 | 26191482 | 26191628 | 28 | 16 |
| RI | 1998 | ENSG00000226067 | LINC00623 | chr1 | + | 120952438 | 120953460 | 120952438 | 120952625 | 120953222 | 120953460 | 183 | 67 |
| RI | 5858 | ENSG00000108465 | CDK5RAP3 | chr17 | + | 47981436 | 47981768 | 47981436 | 47981459 | 47981586 | 47981768 | 257 | 186 |
| RI | 8342 | ENSG00000116198 | CEP104 | chr1 | - | 3836494 | 3837519 | 3836494 | 3836692 | 3837291 | 3837519 | 14 | 15 |
| RI | 7862 | ENSG00000185101 | ANO9 | chr11 | - | 418719 | 418989 | 418719 | 418813 | 418887 | 418989 | 18 | 7 |
| RI | 3228 | ENSG00000184014 | DENND5A | chr11 | - | 9144096 | 9145113 | 9144096 | 9144278 | 9144994 | 9145113 | 13 | 15 |
| RI | 6278 | ENSG00000183735 | TBK1 | chr12 | + | 64497647 | 64498039 | 64497647 | 64497754 | 64497967 | 64498039 | 11 | 6 |
| RI | 6784 | ENSG00000072778 | ACADVL | chr17 | + | 7224320 | 7224552 | 7224320 | 7224393 | 7224483 | 7224552 | 8 | 10 |
| RI | 7210 | ENSG00000160948 | VPS28 | chr8 | - | 144423616 | 144424132 | 144423616 | 144423922 | 144424040 | 144424132 | 16 | 6 |
| RI | 661 | ENSG00000124155 | PIGT | chr20 | + | 45416516 | 45418979 | 45416516 | 45416694 | 45418851 | 45418979 | 44 | 15 |
| RI | 4968 | ENSG00000061273 | HDAC7 | chr12 | - | 47791870 | 47792669 | 47791870 | 47792004 | 47792537 | 47792669 | 23 | 9 |
| RI | 5095 | ENSG00000166986 | MARS1 | chr12 | + | 57515919 | 57516337 | 57515919 | 57515991 | 57516244 | 57516337 | 87 | 73 |
| RI | 1873 | ENSG00000105122 | RASAL3 | chr19 | - | 15457291 | 15458426 | 15457291 | 15457834 | 15458327 | 15458426 | 41 | 15 |
| RI | 6677 | ENSG00000102226 | USP11 | chrX | + | 47239350 | 47239907 | 47239350 | 47239481 | 47239789 | 47239907 | 14 | 9 |
| RI | 8359 | ENSG00000135297 | MTO1 | chr6 | + | 73479731 | 73480126 | 73479731 | 73479844 | 73480038 | 73480126 | 82 | 49 |
| RI | 6100 | ENSG00000153406 | NMRAL1 | chr16 | - | 4474092 | 4474792 | 4474092 | 4474166 | 4474553 | 4474792 | 22 | 5 |
| RI | 2724 | ENSG00000100412 | ACO2 | chr22 | + | 41517596 | 41518572 | 41517596 | 41517631 | 41518480 | 41518572 | 33 | 12 |
| RI | 2173 | ENSG00000065357 | DGKA | chr12 | + | 55953349 | 55954021 | 55953349 | 55953410 | 55953684 | 55954021 | 25 | 28 |
| RI | 4418 | ENSG00000111653 | ING4 | chr12 | - | 6652270 | 6652770 | 6652270 | 6652309 | 6652661 | 6652770 | 98 | 35 |
| RI | 3498 | ENSG00000013573 | DDX11 | chr12 | + | 31089402 | 31090094 | 31089402 | 31089490 | 31089890 | 31090094 | 6 | 2 |
| RI | 6574 | ENSG00000186088 | GSAP | chr7 | - | 77329332 | 77330367 | 77329332 | 77329391 | 77330238 | 77330367 | 349 | 376 |
| RI | 3045 | ENSG00000187514 | PTMA | chr2 | + | 231711347 | 231711983 | 231711347 | 231711419 | 231711886 | 231711983 | 111 | 93 |
| RI | 2897 | ENSG00000172053 | QARS1 | chr3 | - | 49098191 | 49098480 | 49098191 | 49098258 | 49098352 | 49098480 | 12 | 15 |
| RI | 858 | ENSG00000132382 | MYBBP1A | chr17 | - | 4538901 | 4539967 | 4538901 | 4538975 | 4539438 | 4539967 | 68 | 27 |
| RI | 2612 | ENSG00000127419 | TMEM175 | chr4 | + | 947708 | 948154 | 947708 | 947892 | 948115 | 948154 | 4 | 2 |
| RI | 6776 | ENSG00000072778 | ACADVL | chr17 | + | 7220121 | 7220676 | 7220121 | 7220197 | 7220463 | 7220676 | 21 | 5 |
| RI | 4696 | ENSG00000160688 | FLAD1 | chr1 | + | 154990158 | 154990528 | 154990158 | 154990257 | 154990338 | 154990528 | 33 | 14 |
| RI | 5832 | ENSG00000119707 | RBM25 | chr14 | + | 73110830 | 73111802 | 73110830 | 73111155 | 73111527 | 73111802 | 70 | 82 |

| | | | | | | | | | | | | | | | |
|---|---|---|---|---|---|---|---|---|---|---|---|---|---|---|---|
| RI | 2890 | ENSG00000178035 | IMPDH2 | chr3 | - | 49027709 | 49028532 | 49027709 | 49027916 | 49028430 | 49028532 | 72 | 37 | 63 | [illegible] |
| RI | 5411 | ENSG00000137726 | FXYD6 | chr11 | - | 117841989 | 117842261 | 117841989 | 117842028 | 117842106 | 117842261 | 7 | 0 | 2 | [illegible] |
| RI | 3103 | ENSG00000172830 | SSH3 | chr11 | + | 67306837 | 67307113 | 67306837 | 67306962 | 67307041 | 67307113 | 4 | 8 | 1 | [illegible] |
| RI | 7586 | ENSG00000159314 | ARHGAP27 | chr17 | - | 45403618 | 45404096 | 45403618 | 45403709 | 45404028 | 45404096 | 11 | 4 | 9 | [illegible] |
| RI | 2953 | ENSG00000145029 | NICN1 | chr3 | - | 49424948 | 49425438 | 49424948 | 49425053 | 49425366 | 49425438 | 20 | 8 | 5 | [illegible] |
| RI | 63 | ENSG00000182944 | EWSR1 | chr22 | + | 29298732 | 29299280 | 29298732 | 29298895 | 29299194 | 29299280 | 98 | 83 | 88 | [illegible] |
| RI | 7747 | ENSG00000149091 | DGKZ | chr11 | + | 46379202 | 46379568 | 46379202 | 46379236 | 46379468 | 46379568 | 31 | 25 | 17 | [illegible] |
| RI | 2004 | ENSG00000184436 | THAP7 | chr22 | - | 21000646 | 21001411 | 21000646 | 21000787 | 21001255 | 21001411 | 61 | 27 | 85 | [illegible] |
| RI | 3916 | ENSG00000167257 | RNF214 | chr11 | + | 117238600 | 117239860 | 117238600 | 117239111 | 117239800 | 117239860 | 9 | 0 | 1 | [illegible] |
| RI | 1020 | ENSG00000169738 | DCXR | chr17 | - | 82036858 | 82037547 | 82036858 | 82037013 | 82037449 | 82037547 | 6 | 5 | 4 | [illegible] |
| RI | 3320 | ENSG00000160298 | C21orf58 | chr21 | - | 46301129 | 46302154 | 46301129 | 46301713 | 46301996 | 46302154 | 11 | 3 | 4 | [illegible] |
| RI | 602 | ENSG00000171703 | TCEA2 | chr20 | + | 64069764 | 64070635 | 64069764 | 64070414 | 64070488 | 64070635 | 5 | 6 | 7 | [illegible] |
| RI | 1471 | ENSG00000140983 | RHOT2 | chr16 | + | 670891 | 671203 | 670891 | 671000 | 671082 | 671203 | 81 | 29 | 68 | [illegible] |
| RI | 2234 | ENSG00000104805 | NUCB1 | chr19 | + | 48919029 | 48919286 | 48919029 | 48919122 | 48919193 | 48919286 | 39 | 26 | 44 | [illegible] |
| RI | 3012 | ENSG00000075785 | RAB7A | chr3 | + | 128806371 | 128807671 | 128806371 | 128806590 | 128807542 | 128807671 | 27 | 6 | 12 | [illegible] |
| RI | 5605 | ENSG00000102125 | TAZ | chrX | + | 154419542 | 154420094 | 154419542 | 154419623 | 154420031 | 154420094 | 33 | 10 | 43 | [illegible] |
| RI | 3962 | ENSG00000213672 | NCKIPSD | chr3 | - | 48678876 | 48679183 | 48678876 | 48678969 | 48679054 | 48679183 | 4 | 1 | 3 | [illegible] |
| RI | 4477 | ENSG00000156983 | BRPF1 | chr3 | + | 9742024 | 9742541 | 9742024 | 9742153 | 9742351 | 9742541 | 28 | 21 | 15 | [illegible] |
| RI | 8046 | ENSG00000156650 | KAT6B | chr10 | + | 74975398 | 74977437 | 74975398 | 74976330 | 74977315 | 74977437 | 28 | 29 | 17 | [illegible] |
| RI | 651 | ENSG00000049759 | NEDD4L | chr18 | + | 58383245 | 58385586 | 58383245 | 58383319 | 58385525 | 58385586 | 1 | 1 | 0 | [illegible] |
| RI | 6913 | ENSG00000197114 | ZGPAT | chr20 | + | 63733586 | 63734824 | 63733586 | 63733739 | 63734704 | 63734824 | 28 | 16 | 50 | [illegible] |
| RI | 7905 | ENSG00000110011 | DNAJC4 | chr11 | + | 64233893 | 64234281 | 64233893 | 64233980 | 64234072 | 64234281 | 40 | 20 | 32 | [illegible] |
| RI | 1727 | ENSG00000079134 | THOC1 | chr18 | - | 224923 | 225403 | 224923 | 224994 | 225088 | 225403 | 8 | 33 | 20 | [illegible] |
| RI | 3638 | ENSG00000076258 | FMO4 | chr1 | + | 171341412 | 171342084 | 171341412 | 171341493 | 171341944 | 171342084 | 22 | 11 | 18 | [illegible] |
| RI | 4857 | ENSG00000149532 | CPSF7 | chr11 | - | 61420469 | 61421608 | 61420469 | 61420530 | 61421389 | 61421608 | 65 | 30 | 61 | [illegible] |
| RI | 1335 | ENSG00000090861 | AARS1 | chr16 | - | 70253918 | 70254734 | 70253918 | 70254038 | 70254620 | 70254734 | 4 | 5 | 6 | [illegible] |
| RI | 2428 | ENSG00000164889 | SLC4A2 | chr7 | + | 151074038 | 151074841 | 151074038 | 151074099 | 151074674 | 151074841 | 14 | 14 | 15 | [illegible] |
| RI | 5143 | ENSG00000143387 | CTSK | chr1 | - | 150804020 | 150806016 | 150804020 | 150804239 | 150805860 | 150806016 | 8 | 13 | 5 | [illegible] |
| RI | 4658 | ENSG00000001497 | LAS1L | chrX | - | 65520701 | 65523707 | 65520701 | 65522285 | 65523559 | 65523707 | 5 | 15 | 11 | [illegible] |
| RI | 1836 | ENSG00000099797 | TECR | chr19 | + | 14563657 | 14563903 | 14563657 | 14563702 | 14563834 | 14563903 | 82 | 22 | 102 | [illegible] |
| RI | 3409 | ENSG00000114544 | SLC41A3 | chr3 | - | 126006356 | 126007225 | 126006356 | 126006564 | 126007113 | 126007225 | 26 | 14 | 28 | [illegible] |
| RI | 7413 | ENSG00000107331 | ABCA2 | chr9 | - | 137010972 | 137011309 | 137010972 | 137011105 | 137011185 | 137011309 | 1 | 0 | 2 | [illegible] |
| RI | 7277 | ENSG00000147789 | ZNF7 | chr8 | + | 144829042 | 144829604 | 144829042 | 144829090 | 144829426 | 144829604 | 13 | 4 | 10 | [illegible] |
| RI | 1725 | ENSG00000258461 | NA | chr15 | + | 42402123 | 42403002 | 42402123 | 42402135 | 42402793 | 42403002 | 19 | 10 | 8 | [illegible] |
| RI | 4419 | ENSG00000111653 | ING4 | chr12 | - | 6652270 | 6652770 | 6652270 | 6652418 | 6652661 | 6652770 | 21 | 8 | 8 | [illegible] |
| RI | 6910 | ENSG00000197114 | ZGPAT | chr20 | + | 63733586 | 63734824 | 63733586 | 63733739 | 63734644 | 63734824 | 27 | 19 | 51 | [illegible] |
| RI | 6824 | ENSG00000115275 | MOGS | chr2 | - | 74463189 | 74463386 | 74463189 | 74463219 | 74463379 | 74463386 | 146 | 56 | 113 | [illegible] |
| RI | 4057 | ENSG00000198821 | CD247 | chr1 | - | 167430639 | 167431746 | 167430639 | 167431013 | 167431576 | 167431746 | 1251 | 353 | 690 | [illegible] |
| RI | 605 | ENSG00000198276 | UCKL1 | chr20 | - | 63945804 | 63946267 | 63945804 | 63945975 | 63946160 | 63946267 | 7 | 2 | 11 | [illegible] |
| RI | 4281 | ENSG00000153187 | HNRNPU | chr1 | - | 244859274 | 244862381 | 244859274 | 244859374 | 244860334 | 244862381 | 178 | 161 | 181 | [illegible] |
| RI | 8206 | ENSG00000135655 | USP15 | chr12 | + | 62383839 | 62384302 | 62383839 | 62383998 | 62384077 | 62384302 | 104 | 100 | 56 | [illegible] |
| RI | 2493 | ENSG00000167995 | BEST1 | chr11 | + | 61958145 | 61959578 | 61958145 | 61958501 | 61959497 | 61959578 | 13 | 7 | 8 | [illegible] |
| RI | 3255 | ENSG00000137074 | APTX | chr9 | - | 32987543 | 32988129 | 32987543 | 32987842 | 32988078 | 32988129 | 19 | 45 | 27 | [illegible] |
| RI | 6734 | ENSG00000147400 | CETN2 | chrX | - | 152828536 | 152829277 | 152828536 | 152828674 | 152829148 | 152829277 | 6 | 0 | 4 | [illegible] |
| RI | 7865 | ENSG00000185101 | ANO9 | chr11 | - | 431693 | 431906 | 431693 | 431767 | 431847 | 431906 | 6 | 9 | 21 | [illegible] |
| RI | 6173 | ENSG00000164620 | RELL2 | chr5 | + | 141636949 | 141638409 | 141636949 | 141637741 | 141638169 | 141638409 | 13 | 7 | 29 | [illegible] |
| RI | 1389 | ENSG00000131652 | THOC6 | chr16 | + | 3025923 | 3026288 | 3025923 | 3025988 | 3026250 | 3026288 | 10 | 6 | 13 | [illegible] |
| RI | 5601 | ENSG00000102125 | TAZ | chrX | + | 154413206 | 154413567 | 154413206 | 154413339 | 154413481 | 154413567 | 8 | 3 | 2 | [illegible] |
| RI | 3497 | ENSG00000013573 | DDX11 | chr12 | + | 31089402 | 31090094 | 31089402 | 31089490 | 31089885 | 31090094 | 6 | 2 | 4 | [illegible] |
| RI | 2067 | ENSG00000125459 | MSTO1 | chr1 | + | 155611215 | 155611603 | 155611215 | 155611291 | 155611548 | 155611603 | 7 | 14 | 20 | [illegible] |
| RI | 3398 | ENSG00000095321 | CRAT | chr9 | - | 129097249 | 129098148 | 129097249 | 129097312 | 129098012 | 129098148 | 0 | 0 | 7 | [illegible] |
| RI | 4539 | ENSG00000082014 | SMARCD3 | chr7 | - | 151242136 | 151242603 | 151242136 | 151242232 | 151242480 | 151242603 | 5 | 5 | 2 | [illegible] |
| RI | 3992 | ENSG00000158863 | FAM160B2 | chr8 | + | 22101707 | 22102315 | 22101707 | 22101851 | 22102174 | 22102315 | 24 | 26 | 44 | [illegible] |
| RI | 275 | ENSG00000076944 | STXBP2 | chr19 | + | 7642428 | 7642823 | 7642428 | 7642536 | 7642765 | 7642823 | 12 | 4 | 4 | [illegible] |
| RI | 239 | ENSG00000172009 | THOP1 | chr19 | + | 2807441 | 2808444 | 2807441 | 2807808 | 2808242 | 2808444 | 0 | 2 | 1 | [illegible] |
| RI | 7878 | ENSG00000167986 | DDB1 | chr11 | - | 61314046 | 61314486 | 61314046 | 61314202 | 61314307 | 61314486 | 19 | 7 | 22 | [illegible] |
| RI | 5532 | ENSG00000184922 | FMNL1 | chr17 | + | 45243790 | 45244244 | 45243790 | 45244025 | 45244175 | 45244244 | 5 | 5 | 13 | [illegible] |
| RI | 2172 | ENSG00000065357 | DGKA | chr12 | + | 55952340 | 55952932 | 55952340 | 55952431 | 55952733 | 55952932 | 11 | 22 | 34 | [illegible] |
| RI | 2693 | ENSG00000108591 | DRG2 | chr17 | + | 18099632 | 18100435 | 18099632 | 18099723 | 18100362 | 18100435 | 15 | 11 | 18 | [illegible] |
| RI | 1686 | ENSG00000137841 | PLCB2 | chr15 | - | 40291264 | 40291924 | 40291264 | 40291487 | 40291848 | 40291924 | 81 | 43 | 70 | [illegible] |
| RI | 4811 | ENSG00000158828 | PINK1 | chr1 | + | 20648504 | 20649231 | 20648504 | 20648632 | 20648994 | 20649231 | 43 | 9 | 36 | [illegible] |
| RI | 5114 | ENSG00000181163 | NPM1 | chr5 | + | 171405301 | 171407774 | 171405301 | 171405403 | 171406396 | 171407774 | 58 | 58 | 65 | [illegible] |
| RI | 6312 | ENSG00000001631 | KRIT1 | chr7 | - | 92235402 | 92236542 | 92235402 | 92235646 | 92236412 | 92236542 | 32 | 51 | 22 | [illegible] |
| RI | 4059 | ENSG00000159079 | CFAP298 | chr21 | - | 32601666 | 32603292 | 32601666 | 32601973 | 32603160 | 32603292 | 22 | 30 | 23 | [illegible] |
| RI | 7268 | ENSG00000104518 | GSDMD | chr8 | + | 143561001 | 143561423 | 143561001 | 143561104 | 143561369 | 143561423 | 10 | 2 | 5 | [illegible] |
| RI | 5193 | ENSG00000159840 | ZYX | chr7 | + | 143382247 | 143382685 | 143382247 | 143382447 | 143382592 | 143382685 | 27 | 18 | 23 | [illegible] |
| RI | 1171 | ENSG00000125124 | BBS2 | chr16 | - | 56498436 | 56502456 | 56498436 | 56499907 | 56502316 | 56502456 | 110 | 112 | 99 | [illegible] |
| RI | 5828 | ENSG00000124356 | STAMBP | chr2 | + | 73847386 | 73849487 | 73847386 | 73847572 | 73849362 | 73849487 | 86 | 57 | 63 | [illegible] |
| RI | 1510 | ENSG00000103197 | TSC2 | chr16 | + | 2088047 | 2088325 | 2088047 | 2088139 | 2088226 | 2088325 | 1 | 0 | 2 | [illegible] |
| RI | 475 | ENSG00000167460 | TPM4 | chr19 | + | 16088026 | 16089120 | 16088026 | 16088097 | 16089044 | 16089120 | 31 | 36 | 28 | [illegible] |
| RI | 3556 | ENSG00000127054 | INTS11 | chr1 | - | 1320455 | 1321093 | 1320455 | 1320529 | 1320995 | 1321093 | 10 | 2 | 10 | [illegible] |
| RI | 270 | ENSG00000011304 | PTBP1 | chr19 | + | 804035 | 804438 | 804035 | 804208 | 804291 | 804438 | 75 | 32 | 61 | [illegible] |
| RI | 3765 | ENSG00000115524 | SF3B1 | chr2 | - | 197418795 | 197420542 | 197418795 | 197418951 | 197420427 | 197420542 | 1224 | 934 | 1328 | [illegible] |
| RI | 4974 | ENSG00000163382 | NAXE | chr1 | + | 156592100 | 156592475 | 156592100 | 156592209 | 156592364 | 156592475 | 3 | 1 | 4 | [illegible] |
| RI | 5420 | ENSG00000039650 | PNKP | chr19 | - | 49864178 | 49864403 | 49864178 | 49864236 | 49864323 | 49864403 | 48 | 14 | 42 | [illegible] |
| RI | 6881 | ENSG00000147140 | NONO | chrX | + | 71297376 | 71297938 | 71297376 | 71297461 | 71297839 | 71297938 | 60 | 44 | 43 | [illegible] |
| RI | 5835 | ENSG00000114770 | ABCC5 | chr3 | - | 183983752 | 183987917 | 183983752 | 183984894 | 183987769 | 183987917 | 27 | 23 | 15 | [illegible] |
| RI | 6920 | ENSG00000248323 | LUCAT1 | chr5 | - | 91314079 | 91314386 | 91314079 | 91314244 | 91314331 | 91314386 | 22 | 12 | 24 | [illegible] |
| RI | 6070 | ENSG00000204348 | DXO | chr6 | - | 31970342 | 31970825 | 31970342 | 31970478 | 31970605 | 31970825 | 32 | 23 | 35 | [illegible] |
| RI | 7581 | ENSG00000125611 | CHCHD5 | chr2 | + | 112585973 | 112586365 | 112585973 | 112586114 | 112586199 | 112586365 | 3 | 1 | 5 | [illegible] |
| RI | 2700 | ENSG00000055955 | ITIH4 | chr3 | - | 52819753 | 52819990 | 52819753 | 52819792 | 52819939 | 52819990 | 54 | 20 | 44 | [illegible] |
| RI | 214 | ENSG00000105127 | AKAP8 | chr19 | - | 15372217 | 15373340 | 15372217 | 15372347 | 15372850 | 15373340 | 16 | 20 | 18 | [illegible] |
| RI | 436 | ENSG00000187994 | RINL | chr19 | - | 38871077 | 38871870 | 38871077 | 38871227 | 38871797 | 38871870 | 12 | 10 | 10 | [illegible] |
| RI | 7802 | ENSG00000255717 | SNHG1 | chr11 | - | 62852810 | 62853115 | 62852810 | 62852993 | 62853074 | 62853115 | 286 | 192 | 341 | [illegible] |
| RI | 5178 | ENSG00000104907 | TRMT1 | chr19 | - | 13115996 | 13116431 | 13115996 | 13116052 | 13116186 | 13116431 | 22 | 19 | 25 | [illegible] |
| RI | 1371 | ENSG00000177455 | CD19 | chr16 | + | 28932345 | 28933114 | 28932345 | 28932612 | 28932910 | 28933114 | 6 | 7 | 8 | [illegible] |
| RI | 2541 | ENSG00000138796 | HADH | chr4 | + | 108023473 | 108027760 | 108023473 | 108023563 | 108027687 | 108027760 | 24 | 21 | 36 | [illegible] |
| RI | 4766 | ENSG00000047410 | TPR | chr1 | - | 186341251 | 186343473 | 186341251 | 186341389 | 186343325 | 186343473 | 35 | 45 | 39 | [illegible] |
| RI | 2297 | ENSG00000170144 | HNRNPA3 | chr2 | + | 177215977 | 177216940 | 177215977 | 177216771 | 177216859 | 177216940 | 100 | 199 | 120 | [illegible] |
| RI | 2946 | ENSG00000164068 | RNF123 | chr3 | + | 49704649 | 49705186 | 49704649 | 49704756 | 49704983 | 49705186 | 9 | 2 | 9 | [illegible] |
| RI | 7809 | ENSG00000255717 | SNHG1 | chr11 | - | 62854518 | 62854938 | 62854518 | 62854551 | 62854887 | 62854938 | 98 | 80 | 193 | [illegible] |
| RI | 6066 | ENSG00000204351 | SKIV2L | chr6 | + | 31968622 | 31969089 | 31968622 | 31968796 | 31968870 | 31969089 | 1 | 3 | 3 | [illegible] |
| RI | 3756 | ENSG00000071894 | CPSF1 | chr8 | - | 144394233 | 144394555 | 144394233 | 144394300 | 144394378 | 144394555 | 9 | 2 | 13 | [illegible] |
| RI | 4084 | ENSG00000120733 | KDM3B | chr5 | + | 138386021 | 138386621 | 138386021 | 138386190 | 138386288 | 138386621 | 296 | 185 | 273 | [illegible] |
| RI | 6713 | ENSG00000233093 | LINC00892 | chrX | + | 136642426 | 136640773 | 136640840 | 136642102 | 136642426 | [illegible] | 16 | 12 | 16 | [illegible] |
| RI | 5344 | ENSG00000084623 | EIF3I | chr1 | + | 32228726 | 32231019 | 32228726 | 32229208 | 32230926 | 32231019 | 8 | 7 | 5 | [illegible] |
| RI | 6886 | ENSG00000182484 | WASH6P | chrX | + | 156021998 | 156022459 | 156021998 | 156022145 | 156022322 | 156022459 | 9 | 0 | 2 | [illegible] |
| RI | 7434 | ENSG00000070061 | ELP1 | chr9 | - | 108930996 | 108934124 | 108930996 | 108931201 | 108933863 | 108934124 | 12 | 19 | 14 | [illegible] |
| RI | 5217 | ENSG00000164172 | MOCS2 | chr5 | - | 53100410 | 53101509 | 53100410 | 53100534 | 53101358 | 53101509 | 7 | 6 | 9 | [illegible] |
| RI | 5806 | ENSG00000145736 | GTF2H2 | chr5 | - | 71048761 | 71049158 | 71048761 | 71048849 | 71049067 | 71049158 | 0 | 7 | 0 | [illegible] |
| RI | 7334 | ENSG00000127084 | FGD3 | chr9 | + | 93032768 | 93034681 | 93032768 | 93032873 | 93034543 | 93034681 | 37 | 20 | 28 | [illegible] |
| RI | 7470 | ENSG00000135047 | CTSL | chr9 | + | 87728026 | 87728809 | 87728026 | 87728396 | 87728584 | 87728809 | 2 | 1 | 4 | [illegible] |
| RI | 2644 | ENSG00000203875 | SNHG5 | chr6 | - | 85677794 | 85678032 | 85677794 | 85677868 | 85677953 | 85678032 | 81 | 81 | 108 | [illegible] |
| RI | 6804 | ENSG00000111203 | ITFG2 | chr12 | + | 2817222 | 2817950 | 2817222 | 2817318 | 2817908 | 2817950 | 4 | 2 | 4 | [illegible] |
| RI | 4120 | ENSG00000116337 | AMPD2 | chr1 | + | 109629326 | 109629916 | 109629326 | 109629490 | 109629795 | 109629916 | 15 | 5 | 8 | [illegible] |
| RI | 2807 | ENSG00000163703 | CRELD1 | chr3 | + | 9943076 | 9944095 | 9943076 | 9943515 | 9943674 | 9944095 | 25 | 13 | 14 | [illegible] |
| RI | 817 | ENSG00000184009 | ACTG1 | chr17 | - | 81511187 | 81512142 | 81511187 | 81511626 | 81511902 | 81512142 | 22 | 8 | 26 | [illegible] |
| RI | 8146 | ENSG00000107862 | GBF1 | chr10 | + | 102375358 | 102375584 | 102375358 | 102375479 | 102375527 | 102375584 | 50 | 32 | 50 | [illegible] |
| RI | 6887 | ENSG00000182484 | WASH6P | chrX | + | 156022313 | 156022834 | 156022313 | 156022459 | 156022698 | 156022834 | 25 | 14 | 19 | [illegible] |
| RI | 2794 | ENSG00000204463 | BAG6 | chr6 | - | 31641319 | 31641592 | 31641319 | 31641422 | 31641538 | 31641592 | 35 | 16 | 23 | [illegible] |
| RI | 8492 | ENSG00000111335 | OAS2 | chr12 | + | 113009086 | 113011723 | 113009086 | 113009240 | 113010373 | 113011723 | 351 | 397 | 341 | [illegible] |
| RI | 1047 | ENSG00000006282 | SPATA20 | chr17 | + | 50548809 | 50549186 | 50548809 | 50548964 | 50549042 | 50549186 | 1 | 2 | 2 | [illegible] |
| RI | 1966 | ENSG00000112640 | PPP2R5D | chr6 | + | 43007934 | 43008260 | 43007934 | 43008065 | 43008200 | 43008260 | 16 | 11 | 9 | [illegible] |
| RI | 3257 | ENSG00000137074 | APTX | chr9 | - | 32987543 | 32988129 | 32987543 | 32987731 | 32988082 | 32988129 | 31 | 52 | 30 | [illegible] |
| RI | 2928 | ENSG00000125247 | TMTC4 | chr13 | - | 100634804 | 100635195 | 100634804 | 100634936 | 100635023 | 100635195 | 9 | 0 | 5 | [illegible] |
| RI | 1664 | ENSG00000214331 | PDPR2P | chr16 | - | 74352039 | 74352378 | 74352039 | 74352203 | 74352296 | 74352378 | 9 | 13 | 18 | [illegible] |
| RI | 8024 | ENSG00000090621 | PABPC4 | chr1 | - | 39562322 | 39563922 | 39562322 | 39562416 | 39563613 | 39563922 | 27 | 12 | 24 | [illegible] |
| RI | 2577 | ENSG00000105202 | FBL | chr19 | - | 39834457 | 39834813 | 39834457 | 39834562 | 39834667 | 39834813 | 6 | 2 | 8 | [illegible] |
| RI | 2473 | ENSG00000070756 | PABPC1 | chr8 | - | 100717772 | 100718280 | 100717772 | 100717888 | 100718086 | 100718280 | 632 | 745 | 693 | [illegible] |
| RI | 1588 | ENSG00000140521 | POLG | chr15 | - | 89317375 | 89318749 | 89317375 | 89317536 | 89318540 | 89318749 | 40 | 26 | 37 | [illegible] |
| RI | 5162 | ENSG00000198755 | RPL10A | chr6 | + | 35468798 | 35469027 | 35468798 | 35468873 | 35468946 | 35469027 | 12 | 1 | 11 | [illegible] |
| RI | 2122 | ENSG00000177595 | PIDD1 | chr11 | - | 800761 | 801120 | 800761 | 800912 | 800984 | 801120 | 3 | 2 | 3 | [illegible] |
| RI | 1372 | ENSG00000177455 | CD19 | chr16 | + | 28937454 | 28937680 | 28937454 | 28937523 | 28937623 | 28937680 | 5 | 2 | 0 | [illegible] |
| RI | 170 | ENSG00000104936 | DMPK | chr19 | - | 45778492 | 45779359 | 45778492 | 45778641 | 45779263 | 45779359 | 9 | 3 | 6 | [illegible] |
| RI | 6879 | ENSG00000147140 | NONO | chrX | + | 71297376 | 71298508 | 71297376 | 71297461 | 71297587 | 71298508 | 46 | 42 | 33 | [illegible] |
| RI | 6631 | ENSG00000196998 | WDR45 | chrX | - | 49076644 | 49077911 | 49076644 | 49076750 | 49077836 | 49077911 | 42 | 14 | 31 | [illegible] |
| RI | 6814 | ENSG00000102054 | RBBP7 | chrX | - | 16845827 | 16849301 | 16845827 | 16845938 | 16849243 | 16849301 | 36 | 28 | 22 | [illegible] |
| RI | 8546 | ENSG00000168883 | USP39 | chr2 | + | 85648760 | 85649280 | 85648760 | 85648853 | 85649016 | 85649280 | 123 | 65 | 96 | [illegible] |
| RI | 8030 | ENSG00000090621 | PABPC4 | chr1 | - | 39563613 | 39564542 | 39563613 | 39563741 | 39564470 | 39564542 | 74 | 48 | 88 | [illegible] |
| RI | 6237 | ENSG00000088038 | CNOT3 | chr19 | + | 54153714 | 54155680 | 54153714 | 54153840 | 54155308 | 54155680 | 31 | 14 | 31 | [illegible] |
| RI | 6594 | ENSG00000234741 | GAS5 | chr1 | - | 173865228 | 173865547 | 173865228 | 173865282 | 173865509 | 173865547 | 200 | 129 | 222 | [illegible] |
| RI | 6503 | ENSG00000122566 | HNRNPA2B1 | chr7 | - | 26190992 | 26191650 | 26190992 | 26191128 | 26191421 | 26191650 | 554 | 511 | 553 | [illegible] |
| RI | 385 | ENSG00000129347 | KRI1 | chr19 | - | 10561666 | 10561845 | 10561666 | 10561716 | 10561790 | 10561845 | 1 | 0 | 2 | [illegible] |
| RI | 7998 | ENSG00000150433 | TMEM218 | chr11 | - | 125102131 | 125102809 | 125102131 | 125102317 | 125102733 | 125102809 | 24 | 18 | 16 | [illegible] |
| RI | 676 | ENSG00000130813 | SHFL | chr19 | + | 10091249 | 10091630 | 10091249 | 10091353 | 10091475 | 10091630 | 69 | 21 | 62 | [illegible] |
| RI | 743 | ENSG00000161939 | RNASEK-C17orf49 | chr17 | + | 7014144 | 7015156 | 7014144 | 7014878 | 7015061 | 7015156 | 4 | 1 | 9 | [illegible] |
| RI | 7648 | ENSG00000076984 | MAP2K7 | chr19 | + | 7910062 | 7910373 | 7910062 | 7910129 | 7910259 | 7910373 | 3 | 6 | 6 | [illegible] |
| RI | 6016 | ENSG00000130340 | SNX9 | chr6 | + | 157938632 | 157940974 | 157938632 | 157938747 | 157940882 | 157940974 | 9 | 21 | 32 | [illegible] |
| RI | 7523 | ENSG00000115459 | ELMOD3 | chr2 | + | 85390137 | 85391748 | 85390137 | 85390265 | 85390759 | 85391748 | 9 | 18 | 16 | [illegible] |
| RI | 8535 | ENSG00000185189 | NRBP2 | chr8 | - | 143839508 | 143839825 | 143839508 | 143839549 | 143839735 | 143839825 | 10 | 12 | 15 | [illegible] |
| RI | 6148 | ENSG00000106009 | BRAT1 | chr7 | - | 2537880 | 2539351 | 2537880 | 2538764 | 2539178 | 2539351 | 34 | 6 | 17 | [illegible] |
| RI | 4249 | ENSG00000132604 | TERF2 | chr16 | - | 69368375 | 69368595 | 69368375 | 69368482 | 69368586 | 69368595 | 8 | 8 | 12 | [illegible] |
| RI | 4894 | ENSG00000115211 | EIF2B4 | chr2 | - | 27369413 | 27369919 | 27369413 | 27369549 | 27369875 | 27369919 | 15 | 12 | 21 | [illegible] |
| RI | 8537 | ENSG00000132286 | TIMM10B | chr11 | + | 6481500 | 6481852 | 6481500 | 6481555 | 6481756 | 6481852 | 16 | 7 | 16 | [illegible] |
| RI | 4243 | ENSG00000143621 | ILF2 | chr1 | - | 153663018 | 153663276 | 153663018 | 153663133 | 153663214 | 153663276 | 41 | 31 | 73 | [illegible] |
| RI | 5407 | ENSG00000049768 | FOXP3 | chrX | - | 49251333 | 49251765 | 49251333 | 49251483 | 49251663 | 49251765 | 2 | 4 | 3 | [illegible] |
| RI | 2014 | ENSG00000135506 | OS9 | chr12 | + | 57715759 | 57716193 | 57715759 | 57715970 | 57716091 | 57716193 | 12 | 17 | 22 | [illegible] |
| RI | 3059 | ENSG00000169727 | GPS1 | chr17 | + | 82053867 | 82054810 | 82053867 | 82054049 | 82054509 | 82054810 | 3 | 2 | 5 | [illegible] |
| RI | 4396 | ENSG00000249915 | PDCD6 | chr5 | + | 304176 | 306754 | 304176 | 304221 | 306601 | 306754 | 9 | 7 | 9 | [illegible] |
| RI | 8165 | ENSG00000120616 | EPC1 | chr10 | - | 32291162 | 32292644 | 32291162 | 32291322 | 32292495 | 32292644 | 81 | 88 | 44 | [illegible] |
| RI | 1965 | ENSG00000121851 | POLR3GL | chr1 | + | 145977777 | 145978096 | 145977777 | 145977851 | 145977982 | 145978096 | 12 | 15 | 28 | [illegible] |
| RI | 1506 | ENSG00000103197 | TSC2 | chr16 | + | 2053341 | 2054440 | 2053341 | 2053452 | 2054295 | 2054440 | 5 | 7 | 4 | [illegible] |
| RI | 6458 | ENSG00000181090 | EHMT1 | chr9 | + | 137817438 | 137818138 | 137817438 | 137817525 | 137818059 | 137818138 | 3 | 1 | 1 | [illegible] |
| RI | 897 | ENSG00000108963 | DPH1 | chr17 | + | 2040217 | 2040605 | 2040217 | 2040374 | 2040504 | 2040605 | 7 | 9 | 10 | [illegible] |
| RI | 2255 | ENSG00000161960 | EIF4A1 | chr17 | + | 7576523 | 7577165 | 7576523 | 7576692 | 7577055 | 7577165 | 216 | 148 | 208 | [illegible] |
| RI | 3672 | ENSG00000092010 | PSME1 | chr14 | + | 24138048 | 24138263 | 24138048 | 24138117 | 24138195 | 24138263 | 122 | 62 | 118 | [illegible] |
| RI | 6565 | ENSG00000161048 | NAPEPLD | chr7 | - | 103099775 | 103103554 | 103099775 | 103103122 | 103103423 | 103103554 | 91 | 47 | 40 | [illegible] |
| RI | 346 | ENSG00000108107 | RPL28 | chr19 | + | 55386569 | 55388048 | 55386569 | 55386674 | 55387929 | 55388048 | 410 | 216 | 333 | [illegible] |
| RI | 1242 | ENSG00000260565 | ERVK13-1 | chr16 | - | 2671023 | 2672570 | 2671023 | 2671231 | 2671344 | 2672570 | 67 | 47 | 56 | [illegible] |
| RI | 1976 | ENSG00000006744 | ELAC2 | chr17 | - | 12995939 | 12996685 | 12995939 | 12995978 | 12996546 | 12996685 | 5 | 0 | 8 | [illegible] |
| RI | 3048 | ENSG00000156709 | AIFM1 | chrX | - | 130129969 | 130131799 | 130129969 | 130130102 | 130131674 | 130131799 | 37 | 14 | 37 | [illegible] |
| RI | 6596 | ENSG00000234741 | GAS5 | chr1 | - | 173866527 | 173866796 | 173866527 | 173866567 | 173866760 | 173866796 | 106 | 67 | 132 | [illegible] |
| RI | 4106 | ENSG00000154832 | CXXC1 | chr18 | - | 50285748 | 50286257 | 50285748 | 50285928 | 50286021 | 50286257 | 6 | 1 | 10 | [illegible] |
| RI | 2567 | ENSG00000145907 | G3BP1 | chr5 | + | 151803884 | 151812785 | 151803884 | 151803996 | 151804405 | 151812785 | 397 | 174 | 337 | [illegible] |
| RI | 662 | ENSG00000124155 | PIGT | chr20 | + | 45418771 | 45419395 | 45418771 | 45418979 | 45419294 | 45419395 | 29 | 13 | 35 | [illegible] |
| RI | 677 | ENSG00000130813 | SHFL | chr19 | + | 10091249 | 10091630 | 10091249 | 10091353 | 10091583 | 10091630 | 103 | 41 | 91 | [illegible] |
| RI | 424 | ENSG00000249115 | HAUS5 | chr19 | + | 35619612 | 35620123 | 35619612 | 35619758 | 35620011 | 35620123 | 6 | 6 | 5 | [illegible] |
| RI | 1338 | ENSG00000090861 | AARS1 | chr16 | - | 70258032 | 70259186 | 70258032 | 70258217 | 70258979 | 70259186 | 4 | 2 | 8 | [illegible] |
| RI | 6785 | ENSG00000147130 | ZMYM3 | chrX | - | 71247344 | 71247906 | 71247344 | 71247510 | 71247733 | 71247906 | 2 | 0 | 2 | [illegible] |
| RI | 8100 | ENSG00000177239 | MAN1B1 | chr9 | + | 137107530 | 137109180 | 137107530 | 137107662 | 137108387 | 137109180 | 2 | 4 | 3 | [illegible] |
| RI | 1215 | ENSG00000167522 | ANKRD11 | chr16 | - | 89290624 | 89291183 | 89290624 | 89290828 | 89291012 | 89291183 | 10 | 2 | 7 | [illegible] |
| RI | 5852 | ENSG00000108465 | CDK5RAP3 | chr17 | + | 47973518 | 47974031 | 47973518 | 47973693 | 47973930 | 47974031 | 28 | 20 | 28 | [illegible] |
| RI | 1239 | ENSG00000260565 | ERVK13-1 | chr16 | - | 2660361 | 2663407 | 2660361 | 2661381 | 2663083 | 2663407 | 141 | 119 | 90 | [illegible] |
| RI | 6211 | ENSG00000187796 | CARD9 | chr9 | - | 136367215 | 136367976 | 136367215 | 136367257 | 136367636 | 136367976 | 4 | 0 | 0 | [illegible] |
| RI | 2681 | ENSG00000178104 | PDE4DIP | chr1 | + | 148929196 | 148931923 | 148929196 | 148929273 | 148931799 | 148931923 | 3 | 3 | 1 | [illegible] |
| RI | 6576 | ENSG00000265681 | RPL17 | chr18 | - | 49491531 | 49491833 | 49491531 | 49491584 | 49491745 | 49491833 | 1146 | 667 | 1035 | [illegible] |
| RI | 4765 | ENSG00000159658 | EFCAB14 | chr1 | - | 46686783 | 46689691 | 46686783 | 46686870 | 46689586 | 46689691 | 19 | 12 | 16 | [illegible] |
| RI | 286 | ENSG00000142546 | NOSIP | chr19 | - | 49556316 | 49556993 | 49556316 | 49556425 | 49556548 | 49556993 | 10 | 0 | 0 | [illegible] |
| RI | 3621 | ENSG00000182473 | EXOC7 | chr17 | - | 76087653 | 76088562 | 76087653 | 76087720 | 76088463 | 76088562 | 24 | 10 | 21 | [illegible] |
| RI | 2264 | ENSG00000143774 | GUK1 | chr1 | + | 228147405 | 228147699 | 228147405 | 228147544 | 228147614 | 228147699 | 7 | 7 | 12 | [illegible] |
| RI | 4763 | ENSG00000214941 | ZSWIM7 | chr17 | - | 15977726 | 15978163 | 15977726 | 15977913 | 15978043 | 15978163 | 15 | 12 | 32 | [illegible] |
| RI | 5836 | ENSG00000144224 | UBXN4 | chr2 | + | 135753538 | 135754277 | 135753538 | 135753567 | 135754158 | 135754277 | 193 | 222 | 176 | [illegible] |
| RI | 7672 | ENSG00000137500 | CCDC90B | chr11 | - | 83273792 | 83273992 | 83273792 | 83273864 | 83273950 | 83273992 | 6 | 2 | 5 | [illegible] |
| RI | 1479 | ENSG00000167965 | MLST8 | chr16 | + | 2206357 | 2206659 | 2206357 | 2206409 | 2206478 | 2206659 | 5 | 7 | 2 | [illegible] |
| RI | 6951 | ENSG00000215301 | DDX3X | chrX | + | 41341483 | 41342653 | 41341483 | 41341616 | 41342494 | 41342653 | 109 | 117 | 120 | [illegible] |
| RI | 4982 | ENSG00000124067 | SLC12A4 | chr16 | - | 67945086 | 67945553 | 67945086 | 67945220 | 67945368 | 67945553 | 11 | 9 | 8 | [illegible] |
| RI | 8454 | ENSG00000123349 | PFDN5 | chr12 | + | 53296429 | 53297924 | 53296429 | 53296467 | 53297849 | 53297924 | 54 | 44 | 73 | [illegible] |
| RI | 7289 | ENSG00000115091 | ACTR3 | chr2 | + | 113951719 | 113955706 | 113951719 | 113951845 | 113955622 | 113955706 | 208 | 238 | 142 | [illegible] |
| RI | 1011 | ENSG00000108592 | FTSJ3 | chr17 | - | 63820838 | 63821115 | 63820838 | 63820938 | 63821029 | 63821115 | 10 | 21 | 15 | [illegible] |
| RI | 5434 | ENSG00000197858 | GPAA1 | chr8 | + | 144083388 | 144083861 | 144083388 | 144083500 | 144083713 | 144083861 | 18 | 12 | 22 | [illegible] |
| RI | 6513 | ENSG00000122566 | HNRNPA2B1 | chr7 | - | 26196556 | 26197017 | 26196556 | 26196658 | 26196806 | 26197017 | 602 | 759 | 580 | [illegible] |
| RI | 5620 | ENSG00000119912 | IDE | chr10 | - | 92506441 | 92508205 | 92506441 | 92506522 | 92507574 | 92508205 | 7 | 3 | 9 | [illegible] |
| RI | 5528 | ENSG00000168818 | STX18 | chr4 | - | 4457190 | 4457500 | 4457190 | 4457257 | 4457422 | 4457500 | 44 | 32 | 15 | [illegible] |
| RI | 2464 | ENSG00000187091 | PLCD1 | chr3 | - | 38009903 | 38010275 | 38009903 | 38010053 | 38010130 | 38010275 | 18 | 0 | 7 | [illegible] |
| RI | 954 | ENSG00000234912 | SNHG20 | chr17 | + | 77088748 | 77089271 | 77088748 | 77088949 | 77089152 | 77089271 | 45 | 8 | 11 | [illegible] |
| RI | 7671 | ENSG00000129255 | MPDU1 | chr17 | + | 7586898 | 7587271 | 7586898 | 7587017 | 7587160 | 7587271 | 16 | 12 | 36 | [illegible] |
| RI | 5038 | ENSG00000137497 | NUMA1 | chr11 | - | 72007188 | 72008845 | 72007188 | 72007435 | 72008687 | 72008845 | 133 | 62 | 137 | [illegible] |
| RI | 1327 | ENSG00000214078 | CPNE1 | chr20 | - | 35630438 | 35631034 | 35630438 | 35630490 | 35630900 | 35631034 | 59 | 46 | 76 | [illegible] |
| RI | 1640 | ENSG00000140400 | MAN2C1 | chr15 | - | 75356605 | 75356902 | 75356605 | 75356685 | 75356792 | 75356902 | 32 | 6 | 28 | [illegible] |
| RI | 3169 | ENSG00000153914 | SREK1 | chr5 | + | 66162413 | 66163922 | 66162413 | 66162592 | 66163791 | 66163922 | 87 | 88 | 96 | [illegible] |
| RI | 3745 | ENSG00000237943 | PRKCQ-AS1 | chr10 | + | 6583804 | 6584195 | 6583804 | 6583918 | 6584117 | 6584195 | 519 | 606 | 608 | [illegible] |
| RI | 5742 | ENSG00000006062 | MAP3K14 | chr17 | - | 45270412 | 45271221 | 45270412 | 45270563 | 45271057 | 45271221 | 14 | 9 | 14 | [illegible] |
| RI | 3340 | ENSG00000158864 | NDUFS2 | chr1 | + | 161202366 | 161203543 | 161202366 | 161202480 | 161203436 | 161203543 | 3 | 7 | 11 | [illegible] |
| RI | 1243 | ENSG00000260565 | ERVK13-1 | chr16 | - | 2669410 | 2672570 | 2669410 | 2669561 | 2671594 | 2672570 | 79 | 49 | 59 | [illegible] |
| RI | 1886 | ENSG00000213983 | AP1G2 | chr14 | - | 23563583 | 23563856 | 23563583 | 23563638 | 23563715 | 23563856 | 25 | 15 | 21 | [illegible] |
| RI | 3653 | ENSG00000038274 | MAT2B | chr5 | + | 163512001 | 163513669 | 163512001 | 163512196 | 163513554 | 163513669 | 18 | 32 | 27 | [illegible] |
| RI | 1820 | ENSG00000100911 | PSME2 | chr14 | - | 24144399 | 24145155 | 24144399 | 24144468 | 24145057 | 24145155 | 107 | 102 | 93 | [illegible] |
| RI | 5912 | ENSG00000089818 | NECAP1 | chr12 | + | 8089935 | 8090299 | 8089935 | 8090036 | 8090194 | 8090299 | 55 | 51 | 25 | [illegible] |
| RI | 2146 | ENSG00000105698 | USF2 | chr19 | + | 35269802 | 35270597 | 35269802 | 35270003 | 35270446 | 35270597 | 60 | 33 | 46 | [illegible] |

| | | | | | | | | | | | | | | | | | | | | | | | | | | | | | | | | | | | | | | | | | | | | | | |
|---|---|---|---|---|---|---|---|---|---|---|---|---|---|---|---|---|---|---|---|---|---|---|---|---|---|---|---|---|---|---|---|---|---|---|---|---|---|---|---|---|---|---|---|---|---|---|
| RI | 4270 | ENSG00000143224 | PPOX | chr1 | + | 161170906 | 161171181 | 161170906 | 161170949 | 161171033 | 161171181 | 18 | 4 | 10 | 10 | 12 | 27 | 15 | 19 | 32 | 8 | 17 | 12 | 5 | 8 | 16 | 38 | 21 | 7 | 13 | 28 | 233 | 149 | 0.835206807 | 1 | 0.299 | 0.146 | 0.252 | 0.167 | 0.49 | 0.222 | 0.268 | 0.314 | 0.282 | 0.268 | 0 |
| RI | 5568 | ENSG00000115282 | TTC31 | chr2 | + | 74490655 | 74491184 | 74490655 | 74490739 | 74491127 | 74491184 | 43 | 39 | 32 | 46 | 67 | 40 | 43 | 44 | 39 | 44 | 22 | 50 | 57 | 32 | 87 | 27 | 30 | 35 | 60 | 93 | 298 | 149 | 0.836156228 | 1 | 0.35 | 0.312 | 0.267 | 0.371 | 0.432 | 0.289 | 0.455 | 0.449 | 0.211 | 0.319 | 0.002 |
| RI | 1049 | ENSG00000006282 | SPATA20 | chr17 | + | 50549984 | 50550312 | 50549984 | 50550115 | 50550229 | 50550312 | 15 | 1 | 11 | 8 | 4 | 0 | 0 | 0 | 2 | 0 | 8 | 7 | 7 | 15 | 24 | 0 | 3 | 0 | 0 | 0 | 263 | 149 | 0.836341888 | 1 | 1 | 1 | 1 | 0.694 | 1 | 1 | 0.569 | 1 | 1 | 1 | 0.025 |
| RI | 1730 | ENSG00000198513 | ATL1 | chr14 | + | 50590940 | 50591639 | 50590940 | 50591075 | 50591534 | 50591639 | 3 | 0 | 0 | 0 | 0 | 20 | 16 | 8 | 7 | 29 | 2 | 0 | 1 | 4 | 0 | 12 | 26 | 5 | 13 | 31 | 298 | 149 | 0.836494112 | 1 | 0.07 | 0 | 0 | 0 | 0 | 0.077 | 0 | 0.091 | 0.133 | 0 | -0.046 |
| RI | 8138 | ENSG00000107862 | GBF1 | chr10 | + | 102352457 | 102353654 | 102352457 | 102352518 | 102353599 | 102353654 | 11 | 12 | 11 | 18 | 9 | 77 | 54 | 84 | 79 | 165 | 37 | 4 | 31 | 7 | 4 | 193 | 86 | 65 | 119 | 136 | 298 | 149 | 0.836609866 | 1 | 0.067 | 0.1 | 0.061 | 0.102 | 0.027 | 0.087 | 0.023 | 0.193 | 0.029 | 0.014 | 0.002 |
| RI | 2020 | ENSG00000196182 | STK40 | chr1 | - | 36339623 | 36343448 | 36339623 | 36341973 | 36343363 | 36343448 | 6 | 6 | 3 | 5 | 2 | 61 | 30 | 40 | 65 | 28 | 10 | 12 | 4 | 4 | 11 | 100 | 70 | 42 | 51 | 259 | 298 | 149 | 0.837784033 | 1 | 0.047 | 0.091 | 0.036 | 0.037 | 0.034 | 0.048 | 0.079 | 0.045 | 0.038 | 0.021 | 0.003 |
| RI | 7659 | ENSG00000122218 | COPA | chr1 | - | 160332447 | 160333679 | 160332447 | 160332557 | 160333602 | 160333679 | 60 | 40 | 46 | 63 | 35 | 308 | 213 | 213 | 290 | 232 | 43 | 140 | 230 | 159 | 7 | 746 | 421 | 263 | 383 | 453 | 298 | 149 | 0.838019048 | 1 | 0.089 | 0.086 | 0.097 | 0.098 | 0.07 | 0.028 | 0.143 | 0.304 | 0.172 | 0.008 | -0.043 |
| RI | 6510 | ENSG00000122566 | HNRNPA2B1 | chr7 | - | 26193250 | 26193694 | 26193250 | 26193373 | 26193574 | 26193694 | 987 | 1441 | 853 | 1266 | 671 | 513 | 453 | 428 | 580 | 539 | 378 | 1245 | 1725 | 691 | 59 | 140 | 387 | 493 | 369 | 144 | 298 | 149 | 0.838159077 | 1 | 0.49 | 0.614 | 0.499 | 0.522 | 0.384 | 0.574 | 0.617 | 0.636 | 0.484 | 0.17 | 0.006 |
| RI | 6168 | ENSG00000167380 | ZNF226 | chr19 | + | 44165587 | 44165807 | 44165587 | 44165613 | 44165756 | 44165807 | 25 | 16 | 25 | 40 | 31 | 1 | 4 | 1 | 2 | 1 | 41 | 12 | 18 | 6 | 22 | 2 | 0 | 1 | 6 | 0 | 292 | 149 | 0.838661735 | 1 | 0.927 | 0.671 | 0.927 | 0.911 | 0.941 | 0.913 | 1 | 0.902 | 0.338 | 1 | 0.045 |
| RI | 3848 | ENSG00000163608 | NEPRO | chr3 | - | 113012733 | 113013403 | 113012733 | 113012811 | 113013271 | 113013403 | 48 | 43 | 58 | 74 | 43 | 39 | 32 | 37 | 36 | 60 | 0 | 30 | 73 | 16 | 5 | 9 | 17 | 20 | 7 | 7 | 298 | 149 | 0.839101155 | 1 | 0.381 | 0.402 | 0.439 | 0.507 | 0.264 | 0 | 0.469 | 0.646 | 0.533 | 0.263 | 0.016 |
| RI | 4373 | ENSG00000158769 | F11R | chr1 | - | 161001019 | 161001353 | 161001019 | 161001127 | 161001284 | 161001353 | 4 | 0 | 0 | 3 | 3 | 83 | 57 | 50 | 116 | 117 | 4 | 11 | 1 | 4 | 0 | 178 | 119 | 77 | 187 | 843 | 298 | 149 | 0.839148051 | 1 | 0.024 | 0 | 0 | 0.013 | 0.013 | 0.011 | 0.044 | 0.006 | 0.011 | 0 | -0.004 |
| RI | 1139 | ENSG00000008517 | IL32 | chr16 | + | 3067376 | 3067613 | 3067376 | 3067415 | 3067553 | 3067613 | 253 | 41 | 227 | 172 | 145 | 1108 | 240 | 818 | 786 | 661 | 100 | 187 | 76 | 34 | 113 | 422 | 567 | 360 | 200 | 682 | 287 | 149 | 0.841513327 | 1 | 0.106 | 0.081 | 0.126 | 0.102 | 0.102 | 0.11 | 0.146 | 0.099 | 0.081 | 0.079 | 0 |
| RI | 3448 | ENSG00000144591 | GMPPA | chr2 | + | 219501475 | 219502037 | 219501475 | 219501579 | 219501850 | 219502037 | 24 | 23 | 21 | 23 | 16 | 37 | 13 | 52 | 39 | 42 | 25 | 37 | 23 | 31 | 33 | 53 | 38 | 52 | 42 | 75 | 298 | 149 | 0.842748564 | 1 | 0.245 | 0.469 | 0.168 | 0.228 | 0.16 | 0.191 | 0.327 | 0.181 | 0.27 | 0.18 | 0.024 |
| RI | 8020 | ENSG00000198056 | PRIM1 | chr12 | - | 56746780 | 56747032 | 56746780 | 56746854 | 56746925 | 56747032 | 7 | 5 | 9 | 14 | 17 | 22 | 15 | 23 | 27 | 26 | 10 | 3 | 8 | 3 | 0 | 0 | 11 | 11 | 19 | 10 | 220 | 149 | 0.843116127 | 1 | 0.177 | 0.184 | 0.209 | 0.26 | 0.307 | 1 | 0.156 | 0.33 | 0.097 | 0 | -0.089 |
| RI | 2937 | ENSG00000010278 | CD9 | chr12 | + | 6235475 | 6236275 | 6235475 | 6235565 | 6236191 | 6236275 | 2 | 5 | 7 | 6 | 0 | 47 | 89 | 48 | 135 | 83 | 13 | 10 | 6 | 3 | 43 | 355 | 70 | 72 | 85 | 1144 | 298 | 149 | 0.843790562 | 1 | 0.021 | 0.027 | 0.068 | 0.022 | 0 | 0.018 | 0.067 | 0.04 | 0.017 | 0.018 | -0.004 |
| RI | 4266 | ENSG00000198837 | DENND4B | chr1 | - | 153930691 | 153931064 | 153930691 | 153930857 | 153930946 | 153931064 | 8 | 4 | 11 | 11 | 1 | 37 | 5 | 22 | 28 | 5 | 5 | 8 | 8 | 5 | 8 | 22 | 19 | 10 | 7 | 32 | 238 | 149 | 0.843918808 | 1 | 0.119 | 0.334 | 0.238 | 0.197 | 0.111 | 0.125 | 0.209 | 0.334 | 0.309 | 0.135 | -0.023 |
| RI | 8490 | ENSG00000111335 | OAS2 | chr12 | + | 112997519 | 112998410 | 112997519 | 112997755 | 112998265 | 112998410 | 50 | 49 | 46 | 49 | 78 | 216 | 226 | 195 | 301 | 392 | 139 | 87 | 69 | 153 | 21 | 1064 | 509 | 110 | 732 | 195 | 298 | 149 | 0.844419476 | 1 | 0.104 | 0.098 | 0.106 | 0.075 | 0.09 | 0.061 | 0.079 | 0.239 | 0.095 | 0.051 | -0.01 |
| RI | 8045 | ENSG00000156650 | KAT6B | chr10 | + | 74975398 | 74977437 | 74975398 | 74975781 | 74977315 | 74977437 | 90 | 69 | 88 | 94 | 168 | 30 | 8 | 15 | 18 | 14 | 20 | 99 | 87 | 63 | 12 | 10 | 20 | 22 | 5 | 0 | 298 | 149 | 0.844635449 | 1 | 0.6 | 0.812 | 0.746 | 0.723 | 0.857 | 0.5 | 0.712 | 0.664 | 0.863 | 1 | 0 |
| RI | 6093 | ENSG00000112739 | PRPF4B | chr6 | + | 4060412 | 4062299 | 4060412 | 4060686 | 4061654 | 4062299 | 440 | 352 | 383 | 377 | 366 | 0 | 7 | 2 | 0 | 4 | 22 | 149 | 241 | 62 | 7 | 0 | 0 | 1 | 0 | 0 | 298 | 149 | 0.845446263 | 1 | 1 | 0.962 | 0.99 | 1 | 0.979 | 1 | 1 | 0.992 | 1 | 1 | -0.012 |
| RI | 983 | ENSG00000187838 | PLSCR3 | chr17 | - | 7393600 | 7394159 | 7393600 | 7393836 | 7394103 | 7394159 | 14 | 7 | 19 | 17 | 28 | 6 | 2 | 11 | 5 | 9 | 16 | 4 | 10 | 12 | 40 | 19 | 17 | 5 | 0 | 1 | 298 | 149 | 0.845707368 | 1 | 0.538 | 0.636 | 0.463 | 0.63 | 0.609 | 0.296 | 0.105 | 0.5 | 1 | 0.952 | 0.005 |
| RI | 2944 | ENSG00000173540 | GMPPB | chr3 | - | 49722595 | 49723302 | 49722595 | 49723114 | 49723253 | 49723302 | 11 | 4 | 9 | 12 | 1 | 14 | 7 | 17 | 11 | 11 | 0 | 16 | 5 | 5 | 15 | 17 | 31 | 4 | 9 | 2 | 288 | 149 | 0.846060119 | 1 | 0.289 | 0.228 | 0.215 | 0.361 | 0.045 | 0 | 0.211 | 0.393 | 0.223 | 0.795 | -0.097 |
| RI | 128 | ENSG00000198089 | SFI1 | chr22 | + | 31618114 | 31618547 | 31618114 | 31618226 | 31618313 | 31618547 | 8 | 6 | 7 | 7 | 8 | 16 | 9 | 5 | 10 | 1 | 7 | 3 | 2 | 7 | 23 | 0 | 5 | 3 | 18 | 14 | 236 | 149 | 0.847564858 | 1 | 0.24 | 0.296 | 0.469 | 0.306 | 0.835 | 1 | 0.275 | 0.296 | 0.197 | 0.509 | -0.026 |
| RI | 2901 | ENSG00000172053 | QARS1 | chr3 | - | 49099760 | 49100091 | 49099760 | 49099853 | 49099960 | 49100091 | 4 | 2 | 8 | 6 | 2 | 176 | 81 | 167 | 128 | 190 | 1 | 13 | 2 | 4 | 0 | 152 | 150 | 101 | 82 | 215 | 256 | 149 | 0.847941837 | 1 | 0.013 | 0.014 | 0.027 | 0.027 | 0.006 | 0.004 | 0.048 | 0.011 | 0.028 | 0 | -0.001 |
| RI | 8499 | ENSG00000139613 | SMARCC2 | chr12 | - | 56164302 | 56164731 | 56164302 | 56164368 | 56164647 | 56164731 | 71 | 25 | 65 | 58 | 31 | 2 | 0 | 0 | 2 | 4 | 122 | 92 | 63 | 72 | 148 | 0 | 6 | 2 | 3 | 0 | 298 | 149 | 0.848845765 | 1 | 0.947 | 1 | 1 | 0.935 | 0.795 | 1 | 0.885 | 0.94 | 0.923 | 1 | -0.014 |
| RI | 1611 | ENSG00000092439 | TRPM7 | chr15 | - | 50575723 | 50575919 | 50575723 | 50575789 | 50575868 | 50575919 | 17 | 16 | 15 | 17 | 40 | 132 | 124 | 101 | 142 | 185 | 11 | 18 | 20 | 2 | 0 | 47 | 114 | 90 | 30 | 17 | 228 | 149 | 0.848847273 | 1 | 0.078 | 0.078 | 0.088 | 0.073 | 0.124 | 0.133 | 0.094 | 0.127 | 0.042 | 0 | 0.009 |
| RI | 7427 | ENSG00000070061 | ELP1 | chr9 | - | 108896495 | 108897038 | 108896495 | 108896644 | 108896952 | 108897038 | 11 | 8 | 10 | 9 | 14 | 85 | 38 | 51 | 73 | 71 | 3 | 10 | 10 | 10 | 0 | 39 | 23 | 31 | 46 | 33 | 298 | 149 | 0.850703868 | 1 | 0.061 | 0.095 | 0.089 | 0.058 | 0.09 | 0.037 | 0.179 | 0.139 | 0.098 | 0 | -0.012 |
| RI | 2873 | ENSG00000146192 | FGD2 | chr6 | + | 37013608 | 37014100 | 37013608 | 37013765 | 37013961 | 37014100 | 36 | 20 | 34 | 21 | 32 | 146 | 127 | 81 | 152 | 135 | 83 | 25 | 16 | 41 | 104 | 444 | 239 | 67 | 162 | 326 | 298 | 149 | 0.850874883 | 1 | 0.11 | 0.073 | 0.173 | 0.065 | 0.106 | 0.085 | 0.05 | 0.107 | 0.112 | 0.138 | 0.007 |
| RI | 539 | ENSG00000064601 | CTSA | chr20 | + | 45894993 | 45897806 | 45894993 | 45897040 | 45897716 | 45897806 | 17 | 15 | 10 | 20 | 16 | 580 | 396 | 285 | 568 | 465 | 68 | 39 | 37 | 60 | 121 | 1856 | 1281 | 571 | 1060 | 7454 | 298 | 149 | 0.850880344 | 1 | 0.014 | 0.019 | 0.017 | 0.017 | 0.017 | 0.018 | 0.015 | 0.031 | 0.028 | 0.008 | -0.003 |
| RI | 5640 | ENSG00000163516 | ANKZF1 | chr2 | + | 219234132 | 219235875 | 219234132 | 219234288 | 219235707 | 219235875 | 188 | 171 | 222 | 200 | 207 | 0 | 0 | 0 | 1 | 2 | 247 | 331 | 290 | 265 | 277 | 9 | 4 | 0 | 0 | 0 | 298 | 149 | 0.852147591 | 1 | 1 | 1 | 1 | 0.99 | 0.981 | 0.932 | 0.976 | 1 | 1 | 1 | 0.013 |
| RI | 2366 | ENSG00000011243 | AKAP8L | chr19 | - | 15403474 | 15404042 | 15403474 | 15403555 | 15404009 | 15404042 | 225 | 164 | 237 | 265 | 220 | 4 | 0 | 3 | 0 | 1 | 406 | 320 | 262 | 279 | 526 | 0 | 0 | 5 | 3 | 5 | 298 | 149 | 0.852359556 | 1 | 0.966 | 1 | 0.975 | 1 | 0.991 | 1 | 1 | 0.963 | 0.979 | 0.981 | 0.002 |
| RI | 528 | ENSG00000064601 | CTSA | chr20 | + | 45891568 | 45892027 | 45891568 | 45891762 | 45891915 | 45892027 | 10 | 9 | 4 | 8 | 8 | 147 | 78 | 71 | 107 | 84 | 20 | 25 | 7 | 28 | 79 | 290 | 319 | 105 | 216 | 1352 | 298 | 149 | 0.852841246 | 1 | 0.033 | 0.055 | 0.027 | 0.036 | 0.045 | 0.033 | 0.038 | 0.032 | 0.061 | 0.028 | 0.001 |
| RI | 1036 | ENSG00000185813 | PCYT2 | chr17 | - | 81907553 | 81907857 | 81907553 | 81907598 | 81907772 | 81907857 | 3 | 5 | 12 | 10 | 6 | 35 | 15 | 38 | 27 | 24 | 3 | 6 | 1 | 8 | 13 | 50 | 23 | 1 | 4 | 64 | 298 | 149 | 0.853729559 | 1 | 0.041 | 0.143 | 0.136 | 0.156 | 0.111 | 0.029 | 0.115 | 0.333 | 0.5 | 0.092 | -0.096 |
| RI | 4010 | ENSG00000111602 | TIMELESS | chr12 | - | 56420568 | 56420881 | 56420568 | 56420687 | 56420812 | 56420881 | 0 | 0 | 0 | 2 | 3 | 15 | 7 | 7 | 11 | 15 | 0 | 0 | 1 | 2 | 0 | 71 | 15 | 10 | 16 | 47 | 274 | 149 | 0.854177808 | 1 | 0 | 0 | 0 | 0.09 | 0.098 | 0 | 0 | 0.052 | 0.064 | 0 | 0.014 |
| RI | 2015 | ENSG00000135506 | OS9 | chr12 | + | 57718992 | 57720263 | 57718992 | 57719182 | 57720098 | 57720263 | 40 | 26 | 20 | 39 | 52 | 413 | 258 | 285 | 350 | 455 | 111 | 39 | 64 | 48 | 73 | 1073 | 474 | 316 | 560 | 955 | 298 | 149 | 0.854693972 | 1 | 0.046 | 0.048 | 0.034 | 0.053 | 0.054 | 0.049 | 0.04 | 0.092 | 0.041 | 0.037 | -0.005 |
| RI | 2233 | ENSG00000124782 | RREB1 | chr6 | + | 7181123 | 7182082 | 7181123 | 7181289 | 7181869 | 7182082 | 187 | 162 | 207 | 195 | 241 | 2 | 2 | 0 | 0 | 0 | 773 | 362 | 359 | 574 | 488 | 0 | 5 | 0 | 0 | 8 | 298 | 149 | 0.85470244 | 1 | 0.979 | 0.976 | 1 | 1 | 1 | 1 | 0.973 | 1 | 1 | 0.968 | 0.003 |
| RI | 7609 | ENSG00000205220 | PSMB10 | chr16 | - | 67935962 | 67936312 | 67935962 | 67936103 | 67936214 | 67936312 | 68 | 6 | 45 | 57 | 29 | 147 | 51 | 88 | 100 | 45 | 56 | 39 | 29 | 33 | 61 | 68 | 112 | 48 | 111 | 133 | 260 | 149 | 0.854966477 | 1 | 0.21 | 0.063 | 0.227 | 0.246 | 0.27 | 0.321 | 0.166 | 0.257 | 0.146 | 0.208 | -0.016 |
| RI | 3717 | ENSG00000138002 | IFT172 | chr2 | - | 27453629 | 27454418 | 27453629 | 27453739 | 27453981 | 27454418 | 4 | 2 | 4 | 7 | 7 | 10 | 4 | 7 | 10 | 6 | 0 | 6 | 8 | 0 | 28 | 19 | 6 | 9 | 1 | 3 | 298 | 149 | 0.855599371 | 1 | 0.167 | 0.2 | 0.222 | 0.259 | 0.368 | 0 | 0.333 | 0.308 | 0 | 0.824 | -0.05 |
| RI | 2605 | ENSG00000120708 | TGFBI | chr5 | + | 136046334 | 136047015 | 136046334 | 136046495 | 136046850 | 136047015 | 44 | 35 | 37 | 30 | 25 | 483 | 193 | 277 | 253 | 355 | 61 | 14 | 18 | 34 | 81 | 487 | 178 | 186 | 255 | 829 | 298 | 149 | 0.855924777 | 1 | 0.044 | 0.083 | 0.063 | 0.056 | 0.034 | 0.059 | 0.038 | 0.046 | 0.062 | 0.047 | 0.006 |
| RI | 3639 | ENSG00000111674 | ENO2 | chr12 | + | 6916670 | 6917107 | 6916670 | 6916729 | 6917037 | 6917107 | 11 | 17 | 32 | [illegible] | [illegible] | [illegible] | [illegible] | [illegible] | [illegible] | [illegible] | [illegible] | [illegible] | [illegible] | [illegible] | [illegible] | [illegible] | [illegible] | [illegible] | [illegible] | [illegible] | [illegible] | [illegible] | [illegible] | [illegible] | [illegible] | [illegible] | [illegible] | [illegible] | [illegible] | [illegible] | [illegible] | [illegible] | [illegible] | [illegible] | [illegible] | [illegible] | [illegible] |
| RI | 3928 | ENSG00000055609 | KMT2C | chr7 | - | 152202933 | 152205225 | 152202933 | 152203064 | 152205105 | 152205225 | 16 | 15 | 10 | [illegible] | [illegible] | [illegible] | [illegible] | [illegible] | [illegible] | [illegible] | [illegible] | [illegible] | [illegible] | [illegible] | [illegible] | [illegible] | [illegible] | [illegible] | [illegible] | [illegible] | [illegible] | [illegible] | [illegible] | [illegible] | [illegible] | [illegible] | [illegible] | [illegible] | [illegible] | [illegible] | [illegible] | [illegible] | [illegible] | [illegible] | [illegible] | [illegible] | [illegible] |
| RI | 5618 | ENSG00000119912 | IDE | chr10 | - | 92468878 | 92470345 | 92468878 | 92468990 | 92470253 | 92470345 | 11 | 12 | 9 | [illegible] | [illegible] | [illegible] | [illegible] | [illegible] | [illegible] | [illegible] | [illegible] | [illegible] | [illegible] | [illegible] | [illegible] | [illegible] | [illegible] | [illegible] | [illegible] | [illegible] | [illegible] | [illegible] | [illegible] | [illegible] | [illegible] | [illegible] | [illegible] | [illegible] | [illegible] | [illegible] | [illegible] | [illegible] | [illegible] | [illegible] | [illegible] | [illegible] | [illegible] |
| RI | 4907 | ENSG00000115541 | HSPE1 | chr2 | + | 197503038 | 197503381 | 197503038 | 197503128 | 197503208 | 197503381 | 15 | 2 | 8 | [illegible] | [illegible] | [illegible] | [illegible] | [illegible] | [illegible] | [illegible] | [illegible] | [illegible] | [illegible] | [illegible] | [illegible] | [illegible] | [illegible] | [illegible] | [illegible] | [illegible] | [illegible] | [illegible] | [illegible] | [illegible] | [illegible] | [illegible] | [illegible] | [illegible] | [illegible] | [illegible] | [illegible] | [illegible] | [illegible] | [illegible] | [illegible] | [illegible] | [illegible] |
| RI | 3489 | ENSG00000143761 | ARF1 | chr1 | + | 228097077 | 228097452 | 228097077 | 228097262 | 228097341 | 228097452 | 46 | 35 | 39 | [illegible] | [illegible] | [illegible] | [illegible] | [illegible] | [illegible] | [illegible] | [illegible] | [illegible] | [illegible] | [illegible] | [illegible] | [illegible] | [illegible] | [illegible] | [illegible] | [illegible] | [illegible] | [illegible] | [illegible] | [illegible] | [illegible] | [illegible] | [illegible] | [illegible] | [illegible] | [illegible] | [illegible] | [illegible] | [illegible] | [illegible] | [illegible] | [illegible] | [illegible] |
| RI | 4491 | ENSG00000072110 | ACTN1 | chr14 | - | 68877081 | 68878523 | 68877081 | 68877240 | 68878457 | 68878523 | 15 | 15 | 34 | [illegible] | [illegible] | [illegible] | [illegible] | [illegible] | [illegible] | [illegible] | [illegible] | [illegible] | [illegible] | [illegible] | [illegible] | [illegible] | [illegible] | [illegible] | [illegible] | [illegible] | [illegible] | [illegible] | [illegible] | [illegible] | [illegible] | [illegible] | [illegible] | [illegible] | [illegible] | [illegible] | [illegible] | [illegible] | [illegible] | [illegible] | [illegible] | [illegible] | [illegible] |
| RI | 236 | ENSG00000129353 | SLC44A2 | chr19 | + | 10635162 | 10635515 | 10635162 | 10635255 | 10635430 | 10635515 | 82 | 55 | 91 | [illegible] | [illegible] | [illegible] | [illegible] | [illegible] | [illegible] | [illegible] | [illegible] | [illegible] | [illegible] | [illegible] | [illegible] | [illegible] | [illegible] | [illegible] | [illegible] | [illegible] | [illegible] | [illegible] | [illegible] | [illegible] | [illegible] | [illegible] | [illegible] | [illegible] | [illegible] | [illegible] | [illegible] | [illegible] | [illegible] | [illegible] | [illegible] | [illegible] | [illegible] |
| RI | 542 | ENSG00000064601 | CTSA | chr20 | + | 45894993 | 45898109 | 45894993 | 45897806 | 45898004 | 45898109 | 31 | 20 | 24 | [illegible] | [illegible] | [illegible] | [illegible] | [illegible] | [illegible] | [illegible] | [illegible] | [illegible] | [illegible] | [illegible] | [illegible] | [illegible] | [illegible] | [illegible] | [illegible] | [illegible] | [illegible] | [illegible] | [illegible] | [illegible] | [illegible] | [illegible] | [illegible] | [illegible] | [illegible] | [illegible] | [illegible] | [illegible] | [illegible] | [illegible] | [illegible] | [illegible] | [illegible] |
| RI | 5565 | ENSG00000115282 | TTC31 | chr2 | + | 74490024 | 74490473 | 74490024 | 74490127 | 74490243 | 74490473 | 31 | 15 | 20 | [illegible] | [illegible] | [illegible] | [illegible] | [illegible] | [illegible] | [illegible] | [illegible] | [illegible] | [illegible] | [illegible] | [illegible] | [illegible] | [illegible] | [illegible] | [illegible] | [illegible] | [illegible] | [illegible] | [illegible] | [illegible] | [illegible] | [illegible] | [illegible] | [illegible] | [illegible] | [illegible] | [illegible] | [illegible] | [illegible] | [illegible] | [illegible] | [illegible] | [illegible] |
| RI | 1356 | ENSG00000102879 | CORO1A | chr16 | + | 30185208 | 30186720 | 30185208 | 30185407 | 30186597 | 30186720 | 248 | 91 | 197 | [illegible] | [illegible] | [illegible] | [illegible] | [illegible] | [illegible] | [illegible] | [illegible] | [illegible] | [illegible] | [illegible] | [illegible] | [illegible] | [illegible] | [illegible] | [illegible] | [illegible] | [illegible] | [illegible] | [illegible] | [illegible] | [illegible] | [illegible] | [illegible] | [illegible] | [illegible] | [illegible] | [illegible] | [illegible] | [illegible] | [illegible] | [illegible] | [illegible] | [illegible] |
| RI | 1593 | ENSG00000102901 | CENPT | chr16 | - | 67829764 | 67830548 | 67829764 | 67830088 | 67830389 | 67830548 | 68 | 40 | 68 | [illegible] | [illegible] | [illegible] | [illegible] | [illegible] | [illegible] | [illegible] | [illegible] | [illegible] | [illegible] | [illegible] | [illegible] | [illegible] | [illegible] | [illegible] | [illegible] | [illegible] | [illegible] | [illegible] | [illegible] | [illegible] | [illegible] | [illegible] | [illegible] | [illegible] | [illegible] | [illegible] | [illegible] | [illegible] | [illegible] | [illegible] | [illegible] | [illegible] | [illegible] |
| RI | 1954 | ENSG00000112679 | DUSP22 | chr6 | + | 348768 | 351355 | 348768 | 348841 | 350821 | 351355 | 209 | 181 | 193 | [illegible] | [illegible] | [illegible] | [illegible] | [illegible] | [illegible] | [illegible] | [illegible] | [illegible] | [illegible] | [illegible] | [illegible] | [illegible] | [illegible] | [illegible] | [illegible] | [illegible] | [illegible] | [illegible] | [illegible] | [illegible] | [illegible] | [illegible] | [illegible] | [illegible] | [illegible] | [illegible] | [illegible] | [illegible] | [illegible] | [illegible] | [illegible] | [illegible] | [illegible] |
| RI | 6095 | ENSG00000112739 | PRPF4B | chr6 | + | 4061654 | 4062299 | 4061654 | 4061700 | 4062212 | 4062299 | 158 | 128 | 122 | [illegible] | [illegible] | [illegible] | [illegible] | [illegible] | [illegible] | [illegible] | [illegible] | [illegible] | [illegible] | [illegible] | [illegible] | [illegible] | [illegible] | [illegible] | [illegible] | [illegible] | [illegible] | [illegible] | [illegible] | [illegible] | [illegible] | [illegible] | [illegible] | [illegible] | [illegible] | [illegible] | [illegible] | [illegible] | [illegible] | [illegible] | [illegible] | [illegible] | [illegible] |
| RI | 8139 | ENSG00000107862 | GBF1 | chr10 | + | 102358505 | 102359435 | 102358505 | 102358729 | 102359266 | 102359435 | 10 | 5 | 9 | [illegible] | [illegible] | [illegible] | [illegible] | [illegible] | [illegible] | [illegible] | [illegible] | [illegible] | [illegible] | [illegible] | [illegible] | [illegible] | [illegible] | [illegible] | [illegible] | [illegible] | [illegible] | [illegible] | [illegible] | [illegible] | [illegible] | [illegible] | [illegible] | [illegible] | [illegible] | [illegible] | [illegible] | [illegible] | [illegible] | [illegible] | [illegible] | [illegible] | [illegible] |
| RI | 2210 | ENSG00000160201 | U2AF1 | chr21 | - | 43100452 | 43104402 | 43100452 | 43100519 | 43104314 | 43104402 | 2 | 4 | 2 | [illegible] | [illegible] | [illegible] | [illegible] | [illegible] | [illegible] | [illegible] | [illegible] | [illegible] | [illegible] | [illegible] | [illegible] | [illegible] | [illegible] | [illegible] | [illegible] | [illegible] | [illegible] | [illegible] | [illegible] | [illegible] | [illegible] | [illegible] | [illegible] | [illegible] | [illegible] | [illegible] | [illegible] | [illegible] | [illegible] | [illegible] | [illegible] | [illegible] | [illegible] |
| RI | 4180 | ENSG00000143033 | MTF2 | chr1 | + | 93110229 | 93110428 | 93110229 | 93110246 | 93110326 | 93110428 | 136 | 86 | 123 | [illegible] | [illegible] | [illegible] | [illegible] | [illegible] | [illegible] | [illegible] | [illegible] | [illegible] | [illegible] | [illegible] | [illegible] | [illegible] | [illegible] | [illegible] | [illegible] | [illegible] | [illegible] | [illegible] | [illegible] | [illegible] | [illegible] | [illegible] | [illegible] | [illegible] | [illegible] | [illegible] | [illegible] | [illegible] | [illegible] | [illegible] | [illegible] | [illegible] | [illegible] |
| RI | 8279 | ENSG00000241370 | RPP21 | chr6 | + | 30345297 | 30345573 | 30345297 | 30345398 | 30345490 | 30345573 | 8 | 10 | 2 | [illegible] | [illegible] | [illegible] | [illegible] | [illegible] | [illegible] | [illegible] | [illegible] | [illegible] | [illegible] | [illegible] | [illegible] | [illegible] | [illegible] | [illegible] | [illegible] | [illegible] | [illegible] | [illegible] | [illegible] | [illegible] | [illegible] | [illegible] | [illegible] | [illegible] | [illegible] | [illegible] | [illegible] | [illegible] | [illegible] | [illegible] | [illegible] | [illegible] | [illegible] |
| RI | 6919 | ENSG00000248323 | LUCAT1 | chr5 | - | 91314079 | 91314377 | 91314079 | 91314247 | 91314331 | 91314377 | 18 | 12 | 24 | [illegible] | [illegible] | [illegible] | [illegible] | [illegible] | [illegible] | [illegible] | [illegible] | [illegible] | [illegible] | [illegible] | [illegible] | [illegible] | [illegible] | [illegible] | [illegible] | [illegible] | [illegible] | [illegible] | [illegible] | [illegible] | [illegible] | [illegible] | [illegible] | [illegible] | [illegible] | [illegible] | [illegible] | [illegible] | [illegible] | [illegible] | [illegible] | [illegible] | [illegible] |
| RI | 7440 | ENSG00000119139 | TJP2 | chr9 | + | 69252814 | 69255187 | 69252814 | 69252900 | 69254208 | 69255187 | 3 | 2 | 2 | [illegible] | [illegible] | [illegible] | [illegible] | [illegible] | [illegible] | [illegible] | [illegible] | [illegible] | [illegible] | [illegible] | [illegible] | [illegible] | [illegible] | [illegible] | [illegible] | [illegible] | [illegible] | [illegible] | [illegible] | [illegible] | [illegible] | [illegible] | [illegible] | [illegible] | [illegible] | [illegible] | [illegible] | [illegible] | [illegible] | [illegible] | [illegible] | [illegible] | [illegible] |
| RI | 1961 | ENSG00000138081 | FBXO11 | chr2 | - | 47808328 | 47809266 | 47808328 | 47808427 | 47809157 | 47809266 | 41 | 77 | 74 | [illegible] | [illegible] | [illegible] | [illegible] | [illegible] | [illegible] | [illegible] | [illegible] | [illegible] | [illegible] | [illegible] | [illegible] | [illegible] | [illegible] | [illegible] | [illegible] | [illegible] | [illegible] | [illegible] | [illegible] | [illegible] | [illegible] | [illegible] | [illegible] | [illegible] | [illegible] | [illegible] | [illegible] | [illegible] | [illegible] | [illegible] | [illegible] | [illegible] | [illegible] |
| RI | 7846 | ENSG00000174483 | BBS1 | chr11 | + | 66515692 | 66515933 | 66515692 | 66515731 | 66515860 | 66515933 | 17 | 8 | 5 | [illegible] | [illegible] | [illegible] | [illegible] | [illegible] | [illegible] | [illegible] | [illegible] | [illegible] | [illegible] | [illegible] | [illegible] | [illegible] | [illegible] | [illegible] | [illegible] | [illegible] | [illegible] | [illegible] | [illegible] | [illegible] | [illegible] | [illegible] | [illegible] | [illegible] | [illegible] | [illegible] | [illegible] | [illegible] | [illegible] | [illegible] | [illegible] | [illegible] | [illegible] |
| RI | 7886 | ENSG00000072518 | MARK2 | chr11 | + | 63899873 | 63900678 | 63899873 | 63900110 | 63900558 | 63900678 | 20 | 12 | 22 | [illegible] | [illegible] | [illegible] | [illegible] | [illegible] | [illegible] | [illegible] | [illegible] | [illegible] | [illegible] | [illegible] | [illegible] | [illegible] | [illegible] | [illegible] | [illegible] | [illegible] | [illegible] | [illegible] | [illegible] | [illegible] | [illegible] | [illegible] | [illegible] | [illegible] | [illegible] | [illegible] | [illegible] | [illegible] | [illegible] | [illegible] | [illegible] | [illegible] | [illegible] |
| RI | 584 | ENSG00000130589 | HELZ2 | chr20 | - | 63570684 | 63573221 | 63570684 | 63570868 | 63572107 | 63573221 | 15 | 4 | 9 | [illegible] | [illegible] | [illegible] | [illegible] | [illegible] | [illegible] | [illegible] | [illegible] | [illegible] | [illegible] | [illegible] | [illegible] | [illegible] | [illegible] | [illegible] | [illegible] | [illegible] | [illegible] | [illegible] | [illegible] | [illegible] | [illegible] | [illegible] | [illegible] | [illegible] | [illegible] | [illegible] | [illegible] | [illegible] | [illegible] | [illegible] | [illegible] | [illegible] | [illegible] |
| RI | 182 | ENSG00000105607 | GCDH | chr19 | + | 12891159 | 12891395 | 12891159 | 12891202 | 12891270 | 12891395 | 2 | 3 | 0 | [illegible] | [illegible] | [illegible] | [illegible] | [illegible] | [illegible] | [illegible] | [illegible] | [illegible] | [illegible] | [illegible] | [illegible] | [illegible] | [illegible] | [illegible] | [illegible] | [illegible] | [illegible] | [illegible] | [illegible] | [illegible] | [illegible] | [illegible] | [illegible] | [illegible] | [illegible] | [illegible] | [illegible] | [illegible] | [illegible] | [illegible] | [illegible] | [illegible] | [illegible] |
| RI | 2009 | ENSG00000135596 | MICAL1 | chr6 | - | 109451599 | 109452401 | 109451599 | 109451700 | 109452245 | 109452401 | 106 | 83 | 112 | [illegible] | [illegible] | [illegible] | [illegible] | [illegible] | [illegible] | [illegible] | [illegible] | [illegible] | [illegible] | [illegible] | [illegible] | [illegible] | [illegible] | [illegible] | [illegible] | [illegible] | [illegible] | [illegible] | [illegible] | [illegible] | [illegible] | [illegible] | [illegible] | [illegible] | [illegible] | [illegible] | [illegible] | [illegible] | [illegible] | [illegible] | [illegible] | [illegible] | [illegible] |
| RI | 5711 | ENSG00000115053 | NCL | chr2 | - | 231457000 | 231457800 | 231457000 | 231457124 | 231457642 | 231457800 | 308 | 293 | 302 | [illegible] | [illegible] | [illegible] | [illegible] | [illegible] | [illegible] | [illegible] | [illegible] | [illegible] | [illegible] | [illegible] | [illegible] | [illegible] | [illegible] | [illegible] | [illegible] | [illegible] | [illegible] | [illegible] | [illegible] | [illegible] | [illegible] | [illegible] | [illegible] | [illegible] | [illegible] | [illegible] | [illegible] | [illegible] | [illegible] | [illegible] | [illegible] | [illegible] | [illegible] |
| RI | 7589 | ENSG00000108405 | P2RX1 | chr17 | - | 3903550 | 3904024 | 3903550 | 3903631 | 3903927 | 3904024 | 25 | 22 | 23 | [illegible] | [illegible] | [illegible] | [illegible] | [illegible] | [illegible] | [illegible] | [illegible] | [illegible] | [illegible] | [illegible] | [illegible] | [illegible] | [illegible] | [illegible] | [illegible] | [illegible] | [illegible] | [illegible] | [illegible] | [illegible] | [illegible] | [illegible] | [illegible] | [illegible] | [illegible] | [illegible] | [illegible] | [illegible] | [illegible] | [illegible] | [illegible] | [illegible] | [illegible] |
| RI | 4327 | ENSG00000184730 | APOBR | chr16 | + | 28495098 | 28497996 | 28495098 | 28496089 | 28496116 | 28497996 | 53 | 20 | 14 | [illegible] | [illegible] | [illegible] | [illegible] | [illegible] | [illegible] | [illegible] | [illegible] | [illegible] | [illegible] | [illegible] | [illegible] | [illegible] | [illegible] | [illegible] | [illegible] | [illegible] | [illegible] | [illegible] | [illegible] | [illegible] | [illegible] | [illegible] | [illegible] | [illegible] | [illegible] | [illegible] | [illegible] | [illegible] | [illegible] | [illegible] | [illegible] | [illegible] | [illegible] |
| RI | 4332 | ENSG00000115548 | KDM3A | chr2 | + | 86491140 | 86492716 | 86491140 | 86491275 | 86492038 | 86492716 | 40 | 35 | 50 | [illegible] | [illegible] | [illegible] | [illegible] | [illegible] | [illegible] | [illegible] | [illegible] | [illegible] | [illegible] | [illegible] | [illegible] | [illegible] | [illegible] | [illegible] | [illegible] | [illegible] | [illegible] | [illegible] | [illegible] | [illegible] | [illegible] | [illegible] | [illegible] | [illegible] | [illegible] | [illegible] | [illegible] | [illegible] | [illegible] | [illegible] | [illegible] | [illegible] | [illegible] |
| RI | 6115 | ENSG00000125656 | CLPP | chr19 | + | 6361868 | 6362542 | 6361868 | 6361940 | 6362445 | 6362542 | 4 | 4 | 5 | [illegible] | [illegible] | [illegible] | [illegible] | [illegible] | [illegible] | [illegible] | [illegible] | [illegible] | [illegible] | [illegible] | [illegible] | [illegible] | [illegible] | [illegible] | [illegible] | [illegible] | [illegible] | [illegible] | [illegible] | [illegible] | [illegible] | [illegible] | [illegible] | [illegible] | [illegible] | [illegible] | [illegible] | [illegible] | [illegible] | [illegible] | [illegible] | [illegible] | [illegible] |
| RI | 3353 | ENSG00000158864 | NDUFS2 | chr1 | + | 161213863 | 161214299 | 161213863 | 161213921 | 161214155 | 161214299 | 26 | 31 | 38 | [illegible] | [illegible] | [illegible] | [illegible] | [illegible] | [illegible] | [illegible] | [illegible] | [illegible] | [illegible] | [illegible] | [illegible] | [illegible] | [illegible] | [illegible] | [illegible] | [illegible] | [illegible] | [illegible] | [illegible] | [illegible] | [illegible] | [illegible] | [illegible] | [illegible] | [illegible] | [illegible] | [illegible] | [illegible] | [illegible] | [illegible] | [illegible] | [illegible] | [illegible] |
| RI | 8298 | ENSG00000238105 | GOLGA2P5 | chr12 | - | 100157164 | 100157739 | 100157164 | 100157256 | 100157527 | 100157739 | 2 | 7 | 7 | [illegible] | [illegible] | [illegible] | [illegible] | [illegible] | [illegible] | [illegible] | [illegible] | [illegible] | [illegible] | [illegible] | [illegible] | [illegible] | [illegible] | [illegible] | [illegible] | [illegible] | [illegible] | [illegible] | [illegible] | [illegible] | [illegible] | [illegible] | [illegible] | [illegible] | [illegible] | [illegible] | [illegible] | [illegible] | [illegible] | [illegible] | [illegible] | [illegible] | [illegible] |
| RI | 8283 | ENSG00000105221 | AKT2 | chr19 | - | 40236256 | 40238091 | 40236256 | 40236385 | 40237968 | 40238091 | 12 | 1 | 15 | [illegible] | [illegible] | [illegible] | [illegible] | [illegible] | [illegible] | [illegible] | [illegible] | [illegible] | [illegible] | [illegible] | [illegible] | [illegible] | [illegible] | [illegible] | [illegible] | [illegible] | [illegible] | [illegible] | [illegible] | [illegible] | [illegible] | [illegible] | [illegible] | [illegible] | [illegible] | [illegible] | [illegible] | [illegible] | [illegible] | [illegible] | [illegible] | [illegible] | [illegible] |
| RI | 7396 | ENSG00000197070 | ARRDC1 | chr9 | + | 137612895 | 137613510 | 137612895 | 137613006 | 137613459 | 137613510 | 50 | 25 | 27 | [illegible] | [illegible] | [illegible] | [illegible] | [illegible] | [illegible] | [illegible] | [illegible] | [illegible] | [illegible] | [illegible] | [illegible] | [illegible] | [illegible] | [illegible] | [illegible] | [illegible] | [illegible] | [illegible] | [illegible] | [illegible] | [illegible] | [illegible] | [illegible] | [illegible] | [illegible] | [illegible] | [illegible] | [illegible] | [illegible] | [illegible] | [illegible] | [illegible] | [illegible] |
| RI | 3063 | ENSG00000169727 | GPS1 | chr17 | + | 82056839 | 82057470 | 82056839 | 82056974 | 82057052 | 82057470 | 4 | 3 | 1 | [illegible] | [illegible] | [illegible] | [illegible] | [illegible] | [illegible] | [illegible] | [illegible] | [illegible] | [illegible] | [illegible] | [illegible] | [illegible] | [illegible] | [illegible] | [illegible] | [illegible] | [illegible] | [illegible] | [illegible] | [illegible] | [illegible] | [illegible] | [illegible] | [illegible] | [illegible] | [illegible] | [illegible] | [illegible] | [illegible] | [illegible] | [illegible] | [illegible] | [illegible] |
| RI | 6821 | ENSG00000068438 | FTSJ1 | chrX | + | 48481425 | 48481715 | 48481425 | 48481528 | 48481631 | 48481715 | 6 | 4 | 3 | [illegible] | [illegible] | [illegible] | [illegible] | [illegible] | [illegible] | [illegible] | [illegible] | [illegible] | [illegible] | [illegible] | [illegible] | [illegible] | [illegible] | [illegible] | [illegible] | [illegible] | [illegible] | [illegible] | [illegible] | [illegible] | [illegible] | [illegible] | [illegible] | [illegible] | [illegible] | [illegible] | [illegible] | [illegible] | [illegible] | [illegible] | [illegible] | [illegible] | [illegible] |
| RI | 1533 | ENSG00000103876 | FAH | chr15 | + | 80168051 | 80168316 | 80168051 | 80168149 | 80168263 | 80168316 | 2 | 0 | 0 | [illegible] | [illegible] | [illegible] | [illegible] | [illegible] | [illegible] | [illegible] | [illegible] | [illegible] | [illegible] | [illegible] | [illegible] | [illegible] | [illegible] | [illegible] | [illegible] | [illegible] | [illegible] | [illegible] | [illegible] | [illegible] | [illegible] | [illegible] | [illegible] | [illegible] | [illegible] | [illegible] | [illegible] | [illegible] | [illegible] | [illegible] | [illegible] | [illegible] | [illegible] |
| RI | 7322 | ENSG00000139112 | GABARAPL1 | chr12 | + | 10213835 | 10214187 | 10213835 | 10213974 | 10214073 | 10214187 | 95 | 49 | 82 | [illegible] | [illegible] | [illegible] | [illegible] | [illegible] | [illegible] | [illegible] | [illegible] | [illegible] | [illegible] | [illegible] | [illegible] | [illegible] | [illegible] | [illegible] | [illegible] | [illegible] | [illegible] | [illegible] | [illegible] | [illegible] | [illegible] | [illegible] | [illegible] | [illegible] | [illegible] | [illegible] | [illegible] | [illegible] | [illegible] | [illegible] | [illegible] | [illegible] | [illegible] |
| RI | 1797 | ENSG00000119638 | NEK9 | chr14 | - | 75105949 | 75106702 | 75105949 | 75105996 | 75106501 | 75106702 | 34 | 17 | 34 | [illegible] | [illegible] | [illegible] | [illegible] | [illegible] | [illegible] | [illegible] | [illegible] | [illegible] | [illegible] | [illegible] | [illegible] | [illegible] | [illegible] | [illegible] | [illegible] | [illegible] | [illegible] | [illegible] | [illegible] | [illegible] | [illegible] | [illegible] | [illegible] | [illegible] | [illegible] | [illegible] | [illegible] | [illegible] | [illegible] | [illegible] | [illegible] | [illegible] | [illegible] |
| RI | 3235 | ENSG00000095906 | NUBP2 | chr16 | + | 1786536 | 1786955 | 1786536 | 1786655 | 1786756 | 1786955 | 12 | 3 | 2 | [illegible] | [illegible] | [illegible] | [illegible] | [illegible] | [illegible] | [illegible] | [illegible] | [illegible] | [illegible] | [illegible] | [illegible] | [illegible] | [illegible] | [illegible] | [illegible] | [illegible] | [illegible] | [illegible] | [illegible] | [illegible] | [illegible] | [illegible] | [illegible] | [illegible] | [illegible] | [illegible] | [illegible] | [illegible] | [illegible] | [illegible] | [illegible] | [illegible] | [illegible] |
| RI | 8245 | ENSG00000284770 | TBCE | chr1 | + | 235435744 | 235436608 | 235435744 | 235435840 | 235436543 | 235436608 | 23 | 17 | 20 | [illegible] | [illegible] | [illegible] | [illegible] | [illegible] | [illegible] | [illegible] | [illegible] | [illegible] | [illegible] | [illegible] | [illegible] | [illegible] | [illegible] | [illegible] | [illegible] | [illegible] | [illegible] | [illegible] | [illegible] | [illegible] | [illegible] | [illegible] | [illegible] | [illegible] | [illegible] | [illegible] | [illegible] | [illegible] | [illegible] | [illegible] | [illegible] | [illegible] | [illegible] |
| RI | 2367 | ENSG00000011243 | AKAP8L | chr19 | - | 15403474 | 15404042 | 15403474 | 15403715 | 15404009 | 15404042 | 22 | 11 | 19 | [illegible] | [illegible] | [illegible] | [illegible] | [illegible] | [illegible] | [illegible] | [illegible] | [illegible] | [illegible] | [illegible] | [illegible] | [illegible] | [illegible] | [illegible] | [illegible] | [illegible] | [illegible] | [illegible] | [illegible] | [illegible] | [illegible] | [illegible] | [illegible] | [illegible] | [illegible] | [illegible] | [illegible] | [illegible] | [illegible] | [illegible] | [illegible] | [illegible] | [illegible] |
| RI | 144 | ENSG00000274602 | PI4KAP1 | chr22 | - | 18536649 | 18537412 | 18536649 | 18536808 | 18537291 | 18537412 | 7 | 9 | 31 | [illegible] | [illegible] | [illegible] | [illegible] | [illegible] | [illegible] | [illegible] | [illegible] | [illegible] | [illegible] | [illegible] | [illegible] | [illegible] | [illegible] | [illegible] | [illegible] | [illegible] | [illegible] | [illegible] | [illegible] | [illegible] | [illegible] | [illegible] | [illegible] | [illegible] | [illegible] | [illegible] | [illegible] | [illegible] | [illegible] | [illegible] | [illegible] | [illegible] | [illegible] |
| RI | 1800 | ENSG00000119650 | IFT43 | chr14 | + | 76082294 | 76082692 | 76082294 | 76082367 | 76082616 | 76082692 | 15 | 3 | 14 | [illegible] | [illegible] | [illegible] | [illegible] | [illegible] | [illegible] | [illegible] | [illegible] | [illegible] | [illegible] | [illegible] | [illegible] | [illegible] | [illegible] | [illegible] | [illegible] | [illegible] | [illegible] | [illegible] | [illegible] | [illegible] | [illegible] | [illegible] | [illegible] | [illegible] | [illegible] | [illegible] | [illegible] | [illegible] | [illegible] | [illegible] | [illegible] | [illegible] | [illegible] |
| RI | 6116 | ENSG00000044090 | CUL7 | chr6 | - | 43038565 | 43038987 | 43038565 | 43038692 | 43038841 | 43038987 | 12 | 8 | 8 | [illegible] | [illegible] | [illegible] | [illegible] | [illegible] | [illegible] | [illegible] | [illegible] | [illegible] | [illegible] | [illegible] | [illegible] | [illegible] | [illegible] | [illegible] | [illegible] | [illegible] | [illegible] | [illegible] | [illegible] | [illegible] | [illegible] | [illegible] | [illegible] | [illegible] | [illegible] | [illegible] | [illegible] | [illegible] | [illegible] | [illegible] | [illegible] | [illegible] | [illegible] |
| RI | 2385 | ENSG00000013306 | SLC25A39 | chr17 | - | 44321057 | 44321558 | 44321057 | 44321231 | 44321433 | 44321558 | 4 | 1 | 13 | [illegible] | [illegible] | [illegible] | [illegible] | [illegible] | [illegible] | [illegible] | [illegible] | [illegible] | [illegible] | [illegible] | [illegible] | [illegible] | [illegible] | [illegible] | [illegible] | [illegible] | [illegible] | [illegible] | [illegible] | [illegible] | [illegible] | [illegible] | [illegible] | [illegible] | [illegible] | [illegible] | [illegible] | [illegible] | [illegible] | [illegible] | [illegible] | [illegible] | [illegible] |
| RI | 2408 | ENSG00000105875 | WDR91 | chr7 | - | 135204267 | 135206058 | 135204267 | 135204465 | 135205927 | 135206058 | 33 | 25 | 27 | [illegible] | [illegible] | [illegible] | [illegible] | [illegible] | [illegible] | [illegible] | [illegible] | [illegible] | [illegible] | [illegible] | [illegible] | [illegible] | [illegible] | [illegible] | [illegible] | [illegible] | [illegible] | [illegible] | [illegible] | [illegible] | [illegible] | [illegible] | [illegible] | [illegible] | [illegible] | [illegible] | [illegible] | [illegible] | [illegible] | [illegible] | [illegible] | [illegible] | [illegible] |
| RI | 7924 | ENSG00000166340 | TPP1 | chr11 | - | 6616974 | 6617428 | 6616974 | 6617153 | 6617300 | 6617428 | 15 | 16 | 14 | [illegible] | [illegible] | [illegible] | [illegible] | [illegible] | [illegible] | [illegible] | [illegible] | [illegible] | [illegible] | [illegible] | [illegible] | [illegible] | [illegible] | [illegible] | [illegible] | [illegible] | [illegible] | [illegible] | [illegible] | [illegible] | [illegible] | [illegible] | [illegible] | [illegible] | [illegible] | [illegible] | [illegible] | [illegible] | [illegible] | [illegible] | [illegible] | [illegible] | [illegible] |
| RI | 4315 | ENSG00000117308 | GALE | chr1 | - | 23798116 | 23798730 | 23798116 | 23798230 | 23798614 | 23798730 | 4 | 8 | 6 | [illegible] | [illegible] | [illegible] | [illegible] | [illegible] | [illegible] | [illegible] | [illegible] | [illegible] | [illegible] | [illegible] | [illegible] | [illegible] | [illegible] | [illegible] | [illegible] | [illegible] | [illegible] | [illegible] | [illegible] | [illegible] | [illegible] | [illegible] | [illegible] | [illegible] | [illegible] | [illegible] | [illegible] | [illegible] | [illegible] | [illegible] | [illegible] | [illegible] | [illegible] |
| RI | 1638 | ENSG00000140553 | UNC45A | chr15 | + | 90949653 | 90950267 | 90949653 | 90949720 | 90950153 | 90950267 | 11 | 3 | 13 | [illegible] | [illegible] | [illegible] | [illegible] | [illegible] | [illegible] | [illegible] | [illegible] | [illegible] | [illegible] | [illegible] | [illegible] | [illegible] | [illegible] | [illegible] | [illegible] | [illegible] | [illegible] | [illegible] | [illegible] | [illegible] | [illegible] | [illegible] | [illegible] | [illegible] | [illegible] | [illegible] | [illegible] | [illegible] | [illegible] | [illegible] | [illegible] | [illegible] | [illegible] |
| RI | 4117 | ENSG00000116337 | AMPD2 | chr1 | + | 109628082 | 109629235 | 109628082 | 109628495 | 109629108 | 109629235 | 57 | 38 | 58 | [illegible] | [illegible] | [illegible] | [illegible] | [illegible] | [illegible] | [illegible] | [illegible] | [illegible] | [illegible] | [illegible] | [illegible] | [illegible] | [illegible] | [illegible] | [illegible] | [illegible] | [illegible] | [illegible] | [illegible] | [illegible] | [illegible] | [illegible] | [illegible] | [illegible] | [illegible] | [illegible] | [illegible] | [illegible] | [illegible] | [illegible] | [illegible] | [illegible] | [illegible] |
| RI | 3831 | ENSG00000180879 | SSR4 | chrX | + | 153798070 | 153798493 | 153798070 | 153798136 | 153798328 | 153798493 | 43 | 7 | 33 | [illegible] | [illegible] | [illegible] | [illegible] | [illegible] | [illegible] | [illegible] | [illegible] | [illegible] | [illegible] | [illegible] | [illegible] | [illegible] | [illegible] | [illegible] | [illegible] | [illegible] | [illegible] | [illegible] | [illegible] | [illegible] | [illegible] | [illegible] | [illegible] | [illegible] | [illegible] | [illegible] | [illegible] | [illegible] | [illegible] | [illegible] | [illegible] | [illegible] | [illegible] |
| RI | 7300 | ENSG00000165233 | CARD19 | chr9 | + | 93110567 | 93112289 | 93110567 | 93110721 | 93112217 | 93112289 | 23 | 8 | 11 | [illegible] | [illegible] | [illegible] | [illegible] | [illegible] | [illegible] | [illegible] | [illegible] | [illegible] | [illegible] | [illegible] | [illegible] | [illegible] | [illegible] | [illegible] | [illegible] | [illegible] | [illegible] | [illegible] | [illegible] | [illegible] | [illegible] | [illegible] | [illegible] | [illegible] | [illegible] | [illegible] | [illegible] | [illegible] | [illegible] | [illegible] | [illegible] | [illegible] | [illegible] |
| RI | 5560 | ENSG00000120697 | ALG5 | chr13 | - | 36949861 | 36952599 | 36949861 | 36950057 | 36952513 | 36952599 | 14 | 11 | 7 | [illegible] | [illegible] | [illegible] | [illegible] | [illegible] | [illegible] | [illegible] | [illegible] | [illegible] | [illegible] | [illegible] | [illegible] | [illegible] | [illegible] | [illegible] | [illegible] | [illegible] | [illegible] | [illegible] | [illegible] | [illegible] | [illegible] | [illegible] | [illegible] | [illegible] | [illegible] | [illegible] | [illegible] | [illegible] | [illegible] | [illegible] | [illegible] | [illegible] | [illegible] |
| RI | 1569 | ENSG00000168970 | JMJD7-PLA2G4B | chr15 | + | 41836474 | 41836940 | 41836474 | 41836551 | 41836780 | 41836940 | 18 | 17 | 28 | [illegible] | [illegible] | [illegible] | [illegible] | [illegible] | [illegible] | [illegible] | [illegible] | [illegible] | [illegible] | [illegible] | [illegible] | [illegible] | [illegible] | [illegible] | [illegible] | [illegible] | [illegible] | [illegible] | [illegible] | [illegible] | [illegible] | [illegible] | [illegible] | [illegible] | [illegible] | [illegible] | [illegible] | [illegible] | [illegible] | [illegible] | [illegible] | [illegible] | [illegible] |
| RI | 6195 | ENSG00000095015 | MAP3K1 | chr5 | + | 56887377 | 56888357 | 56887377 | 56887520 | 56888225 | 56888357 | 132 | 106 | 85 | [illegible] | [illegible] | [illegible] | [illegible] | [illegible] | [illegible] | [illegible] | [illegible] | [illegible] | [illegible] | [illegible] | [illegible] | [illegible] | [illegible] | [illegible] | [illegible] | [illegible] | [illegible] | [illegible] | [illegible] | [illegible] | [illegible] | [illegible] | [illegible] | [illegible] | [illegible] | [illegible] | [illegible] | [illegible] | [illegible] | [illegible] | [illegible] | [illegible] | [illegible] |
| RI | 1901 | ENSG00000090061 | CCNK | chr14 | + | 99500765 | 99502376 | 99500765 | 99500871 | 99502206 | 99502376 | 90 | 102 | 78 | [illegible] | [illegible] | [illegible] | [illegible] | [illegible] | [illegible] | [illegible] | [illegible] | [illegible] | [illegible] | [illegible] | [illegible] | [illegible] | [illegible] | [illegible] | [illegible] | [illegible] | [illegible] | [illegible] | [illegible] | [illegible] | [illegible] | [illegible] | [illegible] | [illegible] | [illegible] | [illegible] | [illegible] | [illegible] | [illegible] | [illegible] | [illegible] | [illegible] | [illegible] |
| RI | 4082 | ENSG00000116691 | MIIP | chr1 | + | 12029222 | 12030124 | 12029222 | 12029894 | 12030027 | 12030124 | 11 | 6 | 11 | [illegible] | [illegible] | [illegible] | [illegible] | [illegible] | [illegible] | [illegible] | [illegible] | [illegible] | [illegible] | [illegible] | [illegible] | [illegible] | [illegible] | [illegible] | [illegible] | [illegible] | [illegible] | [illegible] | [illegible] | [illegible] | [illegible] | [illegible] | [illegible] | [illegible] | [illegible] | [illegible] | [illegible] | [illegible] | [illegible] | [illegible] | [illegible] | [illegible] | [illegible] |
| RI | 6054 | ENSG00000135486 | HNRNPA1 | chr12 | + | 54283078 | 54283234 | 54283078 | 54283093 | 54283213 | 54283234 | 552 | 485 | 575 | [illegible] | [illegible] | [illegible] | [illegible] | [illegible] | [illegible] | [illegible] | [illegible] | [illegible] | [illegible] | [illegible] | [illegible] | [illegible] | [illegible] | [illegible] | [illegible] | [illegible] | [illegible] | [illegible] | [illegible] | [illegible] | [illegible] | [illegible] | [illegible] | [illegible] | [illegible] | [illegible] | [illegible] | [illegible] | [illegible] | [illegible] | [illegible] | [illegible] | [illegible] |
| RI | 7011 | ENSG00000258388 | PPT2-EGFL8 | chr6 | + | 32166137 | 32166620 | 32166137 | 32166232 | 32166497 | 32166620 | 14 | 3 | 11 | [illegible] | [illegible] | [illegible] | [illegible] | [illegible] | [illegible] | [illegible] | [illegible] | [illegible] | [illegible] | [illegible] | [illegible] | [illegible] | [illegible] | [illegible] | [illegible] | [illegible] | [illegible] | [illegible] | [illegible] | [illegible] | [illegible] | [illegible] | [illegible] | [illegible] | [illegible] | [illegible] | [illegible] | [illegible] | [illegible] | [illegible] | [illegible] | [illegible] | [illegible] |
| RI | 1468 | ENSG00000140983 | RHOT2 | chr16 | + | 669552 | 670175 | 669552 | 669606 | 670122 | 670175 | 9 | 13 | 12 | [illegible] | [illegible] | [illegible] | [illegible] | [illegible] | [illegible] | [illegible] | [illegible] | [illegible] | [illegible] | [illegible] | [illegible] | [illegible] | [illegible] | [illegible] | [illegible] | [illegible] | [illegible] | [illegible] | [illegible] | [illegible] | [illegible] | [illegible] | [illegible] | [illegible] | [illegible] | [illegible] | [illegible] | [illegible] | [illegible] | [illegible] | [illegible] | [illegible] | [illegible] |
| RI | 2631 | ENSG00000100650 | SRSF5 | chr14 | + | 69769181 | 69770540 | 69769181 | 69769251 | 69769510 | 69770540 | 914 | 934 | 835 | [illegible] | [illegible] | [illegible] | [illegible] | [illegible] | [illegible] | [illegible] | [illegible] | [illegible] | [illegible] | [illegible] | [illegible] | [illegible] | [illegible] | [illegible] | [illegible] | [illegible] | [illegible] | [illegible] | [illegible] | [illegible] | [illegible] | [illegible] | [illegible] | [illegible] | [illegible] | [illegible] | [illegible] | [illegible] | [illegible] | [illegible] | [illegible] | [illegible] | [illegible] |
| RI | 4032 | ENSG00000138095 | LRPPRC | chr2 | - | 43894544 | 43896708 | 43894544 | 43894629 | 43896633 | 43896708 | 11 | 19 | 19 | [illegible] | [illegible] | [illegible] | [illegible] | [illegible] | [illegible] | [illegible] | [illegible] | [illegible] | [illegible] | [illegible] | [illegible] | [illegible] | [illegible] | [illegible] | [illegible] | [illegible] | [illegible] | [illegible] | [illegible] | [illegible] | [illegible] | [illegible] | [illegible] | [illegible] | [illegible] | [illegible] | [illegible] | [illegible] | [illegible] | [illegible] | [illegible] | [illegible] | [illegible] |
| RI | 3906 | ENSG00000141556 | TBCD | chr17 | + | 82929361 | 82930643 | 82929361 | 82929500 | 82930521 | 82930643 | 3 | 1 | 6 | [illegible] | [illegible] | [illegible] | [illegible] | [illegible] | [illegible] | [illegible] | [illegible] | [illegible] | [illegible] | [illegible] | [illegible] | [illegible] | [illegible] | [illegible] | [illegible] | [illegible] | [illegible] | [illegible] | [illegible] | [illegible] | [illegible] | [illegible] | [illegible] | [illegible] | [illegible] | [illegible] | [illegible] | [illegible] | [illegible] | [illegible] | [illegible] | [illegible] | [illegible] |
| RI | 6585 | ENSG00000196305 | IARS1 | chr9 | - | 92286535 | 92287910 | 92286535 | 92286618 | 92287790 | 92287910 | 22 | 19 | 36 | [illegible] | [illegible] | [illegible] | [illegible] | [illegible] | [illegible] | [illegible] | [illegible] | [illegible] | [illegible] | [illegible] | [illegible] | [illegible] | [illegible] | [illegible] | [illegible] | [illegible] | [illegible] | [illegible] | [illegible] | [illegible] | [illegible] | [illegible] | [illegible] | [illegible] | [illegible] | [illegible] | [illegible] | [illegible] | [illegible] | [illegible] | [illegible] | [illegible] | [illegible] |
| RI | 5916 | ENSG00000089818 | NECAP1 | chr12 | + | 8090194 | 8092784 | 8090194 | 8090343 | 8092675 | 8092784 | 253 | 186 | 211 | [illegible] | [illegible] | [illegible] | [illegible] | [illegible] | [illegible] | [illegible] | [illegible] | [illegible] | [illegible] | [illegible] | [illegible] | [illegible] | [illegible] | [illegible] | [illegible] | [illegible] | [illegible] | [illegible] | [illegible] | [illegible] | [illegible] | [illegible] | [illegible] | [illegible] | [illegible] | [illegible] | [illegible] | [illegible] | [illegible] | [illegible] | [illegible] | [illegible] | [illegible] |
| RI | 513 | ENSG00000196562 | SULF2 | chr20 | - | 47659398 | 47659730 | 47659398 | 47659452 | 47659696 | 47659730 | 76 | 89 | 50 | [illegible] | [illegible] | [illegible] | [illegible] | [illegible] | [illegible] | [illegible] | [illegible] | [illegible] | [illegible] | [illegible] | [illegible] | [illegible] | [illegible] | [illegible] | [illegible] | [illegible] | [illegible] | [illegible] | [illegible] | [illegible] | [illegible] | [illegible] | [illegible] | [illegible] | [illegible] | [illegible] | [illegible] | [illegible] | [illegible] | [illegible] | [illegible] | [illegible] | [illegible] |
| RI | 7400 | ENSG00000107104 | KANK1 | chr9 | + | 732377 | 734835 | 732377 | 732617 | 734747 | 734835 | 4 | 0 | 2 | [illegible] | [illegible] | [illegible] | [illegible] | [illegible] | [illegible] | [illegible] | [illegible] | [illegible] | [illegible] | [illegible] | [illegible] | [illegible] | [illegible] | [illegible] | [illegible] | [illegible] | [illegible] | [illegible] | [illegible] | [illegible] | [illegible] | [illegible] | [illegible] | [illegible] | [illegible] | [illegible] | [illegible] | [illegible] | [illegible] | [illegible] | [illegible] | [illegible] | [illegible] |
| RI | 4703 | ENSG00000089127 | OAS1 | chr12 | + | 112917546 | 112919903 | 112917546 | 112917700 | 112919388 | 112919903 | 153 | 147 | 142 | [illegible] | [illegible] | [illegible] | [illegible] | [illegible] | [illegible] | [illegible] | [illegible] | [illegible] | [illegible] | [illegible] | [illegible] | [illegible] | [illegible] | [illegible] | [illegible] | [illegible] | [illegible] | [illegible] | [illegible] | [illegible] | [illegible] | [illegible] | [illegible] | [illegible] | [illegible] | [illegible] | [illegible] | [illegible] | [illegible] | [illegible] | [illegible] | [illegible] | [illegible] |
| RI | 1315 | ENSG00000138834 | MAPK8IP3 | chr16 | + | 1767804 | 1768107 | 1767804 | 1767918 | 1768068 | 1768107 | 6 | 2 | 5 | [illegible] | [illegible] | [illegible] | [illegible] | [illegible] | [illegible] | [illegible] | [illegible] | [illegible] | [illegible] | [illegible] | [illegible] | [illegible] | [illegible] | [illegible] | [illegible] | [illegible] | [illegible] | [illegible] | [illegible] | [illegible] | [illegible] | [illegible] | [illegible] | [illegible] | [illegible] | [illegible] | [illegible] | [illegible] | [illegible] | [illegible] | [illegible] | [illegible] | [illegible] |
| RI | 4035 | ENSG00000138095 | LRPPRC | chr2 | - | 43899465 | 43901524 | 43899465 | 43899605 | 43901319 | 43901524 | 33 | 39 | 33 | [illegible] | [illegible] | [illegible] | [illegible] | [illegible] | [illegible] | [illegible] | [illegible] | [illegible] | [illegible] | [illegible] | [illegible] | [illegible] | [illegible] | [illegible] | [illegible] | [illegible] | [illegible] | [illegible] | [illegible] | [illegible] | [illegible] | [illegible] | [illegible] | [illegible] | [illegible] | [illegible] | [illegible] | [illegible] | [illegible] | [illegible] | [illegible] | [illegible] | [illegible] |
| RI | 5484 | ENSG00000144895 | EIF2A | chr3 | + | 150567901 | 150568046 | 150567901 | 150567929 | 150568012 | 150568046 | 352 | 311 | 495 | [illegible] | [illegible] | [illegible] | [illegible] | [illegible] | [illegible] | [illegible] | [illegible] | [illegible] | [illegible] | [illegible] | [illegible] | [illegible] | [illegible] | [illegible] | [illegible] | [illegible] | [illegible] | [illegible] | [illegible] | [illegible] | [illegible] | [illegible] | [illegible] | [illegible] | [illegible] | [illegible] | [illegible] | [illegible] | [illegible] | [illegible] | [illegible] | [illegible] | [illegible] |
| RI | 7957 | ENSG00000103479 | RBL2 | chr16 | + | 53465437 | 53467169 | 53465437 | 53465602 | 53467057 | 53467169 | 507 | 700 | 524 | [illegible] | [illegible] | [illegible] | [illegible] | [illegible] | [illegible] | [illegible] | [illegible] | [illegible] | [illegible] | [illegible] | [illegible] | [illegible] | [illegible] | [illegible] | [illegible] | [illegible] | [illegible] | [illegible] | [illegible] | [illegible] | [illegible] | [illegible] | [illegible] | [illegible] | [illegible] | [illegible] | [illegible] | [illegible] | [illegible] | [illegible] | [illegible] | [illegible] | [illegible] |
| RI | 7931 | ENSG00000166340 | TPP1 | chr11 | - | 6617625 | 6619267 | 6617625 | 6618915 | 6619195 | 6619267 | 5 | 3 | 17 | [illegible] | [illegible] | [illegible] | [illegible] | [illegible] | [illegible] | [illegible] | [illegible] | [illegible] | [illegible] | [illegible] | [illegible] | [illegible] | [illegible] | [illegible] | [illegible] | [illegible] | [illegible] | [illegible] | [illegible] | [illegible] | [illegible] | [illegible] | [illegible] | [illegible] | [illegible] | [illegible] | [illegible] | [illegible] | [illegible] | [illegible] | [illegible] | [illegible] | [illegible] |
| RI | 8204 | ENSG00000148672 | GLUD1 | chr10 | - | 87060160 | 87060825 | 87060160 | 87060241 | 87060687 | 87060825 | 33 | 29 | 20 | [illegible] | [illegible] | [illegible] | [illegible] | [illegible] | [illegible] | [illegible] | [illegible] | [illegible] | [illegible] | [illegible] | [illegible] | [illegible] | [illegible] | [illegible] | [illegible] | [illegible] | [illegible] | [illegible] | [illegible] | [illegible] | [illegible] | [illegible] | [illegible] | [illegible] | [illegible] | [illegible] | [illegible] | [illegible] | [illegible] | [illegible] | [illegible] | [illegible] | [illegible] |
| RI | 5856 | ENSG00000108465 | CDK5RAP3 | chr17 | + | 47975513 | 47976013 | 47975513 | 47975638 | 47975868 | 47976013 | 308 | 201 | 290 | [illegible] | [illegible] | [illegible] | [illegible] | [illegible] | [illegible] | [illegible] | [illegible] | [illegible] | [illegible] | [illegible] | [illegible] | [illegible] | [illegible] | [illegible] | [illegible] | [illegible] | [illegible] | [illegible] | [illegible] | [illegible] | [illegible] | [illegible] | [illegible] | [illegible] | [illegible] | [illegible] | [illegible] | [illegible] | [illegible] | [illegible] | [illegible] | [illegible] | [illegible] |
| RI | 1039 | ENSG00000082641 | NFE2L1 | chr17 | + | 48050606 | 48051628 | 48050606 | 48050819 | 48051116 | 48051628 | 344 | 217 | 220 | [illegible] | [illegible] | [illegible] | [illegible] | [illegible] | [illegible] | [illegible] | [illegible] | [illegible] | [illegible] | [illegible] | [illegible] | [illegible] | [illegible] | [illegible] | [illegible] | [illegible] | [illegible] | [illegible] | [illegible] | [illegible] | [illegible] | [illegible] | [illegible] | [illegible] | [illegible] | [illegible] | [illegible] | [illegible] | [illegible] | [illegible] | [illegible] | [illegible] | [illegible] |
| RI | 6484 | ENSG00000065000 | AP3D1 | chr19 | - | 2109085 | 2109958 | 2109085 | 2109125 | 2109872 | 2109958 | 120 | 68 | 127 | [illegible] | [illegible] | [illegible] | [illegible] | [illegible] | [illegible] | [illegible] | [illegible] | [illegible] | [illegible] | [illegible] | [illegible] | [illegible] | [illegible] | [illegible] | [illegible] | [illegible] | [illegible] | [illegible] | [illegible] | [illegible] | [illegible] | [illegible] | [illegible] | [illegible] | [illegible] | [illegible] | [illegible] | [illegible] | [illegible] | [illegible] | [illegible] | [illegible] | [illegible] |
| RI | 8115 | ENSG00000057608 | GDI2 | chr10 | - | 5766240 | 5766638 | 5766240 | 5766295 | 5766493 | 5766638 | 31 | 44 | 27 | [illegible] | [illegible] | [illegible] | [illegible] | [illegible] | [illegible] | [illegible] | [illegible] | [illegible] | [illegible] | [illegible] | [illegible] | [illegible] | [illegible] | [illegible] | [illegible] | [illegible] | [illegible] | [illegible] | [illegible] | [illegible] | [illegible] | [illegible] | [illegible] | [illegible] | [illegible] | [illegible] | [illegible] | [illegible] | [illegible] | [illegible] | [illegible] | [illegible] | [illegible] |
| RI | 4283 | ENSG00000153187 | HNRNPU | chr1 | - | 244860334 | 244862534 | 244860334 | 244860474 | 244862460 | 244862534 | 279 | 279 | 236 | [illegible] | [illegible] | [illegible] | [illegible] | [illegible] | [illegible] | [illegible] | [illegible] | [illegible] | [illegible] | [illegible] | [illegible] | [illegible] | [illegible] | [illegible] | [illegible] | [illegible] | [illegible] | [illegible] | [illegible] | [illegible] | [illegible] | [illegible] | [illegible] | [illegible] | [illegible] | [illegible] | [illegible] | [illegible] | [illegible] | [illegible] | [illegible] | [illegible] | [illegible] |
| RI | 367 | ENSG00000205744 | DENND1C | chr19 | - | 6471264 | 6471496 | 6471264 | 6471305 | 6471405 | 6471496 | 14 | 2 | 6 | [illegible] | [illegible] | [illegible] | [illegible] | [illegible] | [illegible] | [illegible] | [illegible] | [illegible] | [illegible] | [illegible] | [illegible] | [illegible] | [illegible] | [illegible] | [illegible] | [illegible] | [illegible] | [illegible] | [illegible] | [illegible] | [illegible] | [illegible] | [illegible] | [illegible] | [illegible] | [illegible] | [illegible] | [illegible] | [illegible] | [illegible] | [illegible] | [illegible] | [illegible] |
| RI | 5426 | ENSG00000189046 | ALKBH2 | chr12 | - | 109092506 | 109093472 | 109092506 | 109092787 | 109093246 | 109093472 | 43 | 38 | 56 | [illegible] | [illegible] | [illegible] | [illegible] | [illegible] | [illegible] | [illegible] | [illegible] | [illegible] | [illegible] | [illegible] | [illegible] | [illegible] | [illegible] | [illegible] | [illegible] | [illegible] | [illegible] | [illegible] | [illegible] | [illegible] | [illegible] | [illegible] | [illegible] | [illegible] | [illegible] | [illegible] | [illegible] | [illegible] | [illegible] | [illegible] | [illegible] | [illegible] | [illegible] |
| RI | 3572 | ENSG00000111640 | GAPDH | chr12 | + | 6536683 | 6537010 | 6536683 | 6536790 | 6536919 | 6537010 | 81 | 46 | 88 | [illegible] | [illegible] | [illegible] | [illegible] | [illegible] | [illegible] | [illegible] | [illegible] | [illegible] | [illegible] | [illegible] | [illegible] | [illegible] | [illegible] | [illegible] | [illegible] | [illegible] | [illegible] | [illegible] | [illegible] | [illegible] | [illegible] | [illegible] | [illegible] | [illegible] | [illegible] | [illegible] | [illegible] | [illegible] | [illegible] | [illegible] | [illegible] | [illegible] | [illegible] |
| RI | 5024 | ENSG00000092330 | TINF2 | chr14 | - | 24239642 | 24240155 | 24239642 | 24239931 | 24240063 | 24240155 | 270 | 132 | 210 | [illegible] | [illegible] | [illegible] | [illegible] | [illegible] | [illegible] | [illegible] | [illegible] | [illegible] | [illegible] | [illegible] | [illegible] | [illegible] | [illegible] | [illegible] | [illegible] | [illegible] | [illegible] | [illegible] | [illegible] | [illegible] | [illegible] | [illegible] | [illegible] | [illegible] | [illegible] | [illegible] | [illegible] | [illegible] | [illegible] | [illegible] | [illegible] | [illegible] | [illegible] |
| RI | 3185 | ENSG00000099330 | OCEL1 | chr19 | + | 17228802 | 17229215 | 17228802 | 17228826 | 17228945 | 17229215 | 67 | 58 | 66 | [illegible] | [illegible] | [illegible] | [illegible] | [illegible] | [illegible] | [illegible] | [illegible] | [illegible] | [illegible] | [illegible] | [illegible] | [illegible] | [illegible] | [illegible] | [illegible] | [illegible] | [illegible] | [illegible] | [illegible] | [illegible] | [illegible] | [illegible] | [illegible] | [illegible] | [illegible] | [illegible] | [illegible] | [illegible] | [illegible] | [illegible] | [illegible] | [illegible] | [illegible] |
| RI | 5795 | ENSG00000115307 | AUP1 | chr2 | - | 74528750 | 74529282 | 74528750 | 74528935 | 74529131 | 74529282 | 67 | 23 | 78 | [illegible] | [illegible] | [illegible] | [illegible] | [illegible] | [illegible] | [illegible] | [illegible] | [illegible] | [illegible] | [illegible] | [illegible] | [illegible] | [illegible] | [illegible] | [illegible] | [illegible] | [illegible] | [illegible] | [illegible] | [illegible] | [illegible] | [illegible] | [illegible] | [illegible] | [illegible] | [illegible] | [illegible] | [illegible] | [illegible] | [illegible] | [illegible] | [illegible] | [illegible] |
| RI | 1264 | ENSG00000159713 | TPPP3 | chr16 | - | 67390478 | 67391117 | 67390478 | 67390632 | 67390923 | 67391117 | 4 | 1 | 3 | [illegible] | [illegible] | [illegible] | [illegible] | [illegible] | [illegible] | [illegible] | [illegible] | [illegible] | [illegible] | [illegible] | [illegible] | [illegible] | [illegible] | [illegible] | [illegible] | [illegible] | [illegible] | [illegible] | [illegible] | [illegible] | [illegible] | [illegible] | [illegible] | [illegible] | [illegible] | [illegible] | [illegible] | [illegible] | [illegible] | [illegible] | [illegible] | [illegible] | [illegible] |
| RI | 6299 | ENSG00000145191 | EIF2B5 | chr3 | + | 184143051 | 184144224 | 184143051 | 184143142 | 184144098 | 184144224 | 55 | 66 | 95 | [illegible] | [illegible] | [illegible] | [illegible] | [illegible] | [illegible] | [illegible] | [illegible] | [illegible] | [illegible] | [illegible] | [illegible] | [illegible] | [illegible] | [illegible] | [illegible] | [illegible] | [illegible] | [illegible] | [illegible] | [illegible] | [illegible] | [illegible] | [illegible] | [illegible] | [illegible] | [illegible] | [illegible] | [illegible] | [illegible] | [illegible] | [illegible] | [illegible] | [illegible] |
| RI | 2761 | ENSG00000185000 | DGAT1 | chr8 | - | 144318094 | 144318362 | 144318094 | 144318169 | 144318260 | 144318362 | 5 | 4 | 2 | [illegible] | [illegible] | [illegible] | [illegible] | [illegible] | [illegible] | [illegible] | [illegible] | [illegible] | [illegible] | [illegible] | [illegible] | [illegible] | [illegible] | [illegible] | [illegible] | [illegible] | [illegible] | [illegible] | [illegible] | [illegible] | [illegible] | [illegible] | [illegible] | [illegible] | [illegible] | [illegible] | [illegible] | [illegible] | [illegible] | [illegible] | [illegible] | [illegible] | [illegible] |
| RI | 8590 | ENSG00000148362 | PAXX | chr9 | + | 136992639 | 136993243 | 136992639 | 136992974 | 136993052 | 136993243 | 47 | 12 | 26 | [illegible] | [illegible] | [illegible] | [illegible] | [illegible] | [illegible] | [illegible] | [illegible] | [illegible] | [illegible] | [illegible] | [illegible] | [illegible] | [illegible] | [illegible] | [illegible] | [illegible] | [illegible] | [illegible] | [illegible] | [illegible] | [illegible] | [illegible] | [illegible] | [illegible] | [illegible] | [illegible] | [illegible] | [illegible] | [illegible] | [illegible] | [illegible] | [illegible] | [illegible] |
| RI | 3532 | ENSG00000183258 | DDX41 | chr5 | - | 177512495 | 177512876 | 177512495 | 177512645 | 177512779 | 177512876 | 5 | 10 | 13 | [illegible] | [illegible] | [illegible] | [illegible] | [illegible] | [illegible] | [illegible] | [illegible] | [illegible] | [illegible] | [illegible] | [illegible] | [illegible] | [illegible] | [illegible] | [illegible] | [illegible] | [illegible] | [illegible] | [illegible] | [illegible] | [illegible] | [illegible] | [illegible] | [illegible] | [illegible] | [illegible] | [illegible] | [illegible] | [illegible] | [illegible] | [illegible] | [illegible] | [illegible] |
| RI | 2165 | ENSG00000065357 | DGKA | chr12 | + | 55940081 | 55940433 | 55940081 | 55940168 | 55940313 | 55940433 | 114 | 115 | 236 | [illegible] | [illegible] | [illegible] | [illegible] | [illegible] | [illegible] | [illegible] | [illegible] | [illegible] | [illegible] | [illegible] | [illegible] | [illegible] | [illegible] | [illegible] | [illegible] | [illegible] | [illegible] | [illegible] | [illegible] | [illegible] | [illegible] | [illegible] | [illegible] | [illegible] | [illegible] | [illegible] | [illegible] | [illegible] | [illegible] | [illegible] | [illegible] | [illegible] | [illegible] |
| RI | 6954 | ENSG00000215301 | DDX3X | chrX | + | 41342736 | 41344128 | 41342736 | 41342836 | 41344029 | 41344128 | 316 | 393 | 304 | [illegible] | [illegible] | [illegible] | [illegible] | [illegible] | [illegible] | [illegible] | [illegible] | [illegible] | [illegible] | [illegible] | [illegible] | [illegible] | [illegible] | [illegible] | [illegible] | [illegible] | [illegible] | [illegible] | [illegible] | [illegible] | [illegible] | [illegible] | [illegible] | [illegible] | [illegible] | [illegible] | [illegible] | [illegible] | [illegible] | [illegible] | [illegible] | [illegible] | [illegible] |
| RI | 6904 | ENSG00000090316 | MAEA | chr4 | + | 1338421 | 1340136 | 1338421 | 1338617 | 1339073 | 1340136 | 20 | 5 | 31 | [illegible] | [illegible] | [illegible] | [illegible] | [illegible] | [illegible] | [illegible] | [illegible] | [illegible] | [illegible] | [illegible] | [illegible] | [illegible] | [illegible] | [illegible] | [illegible] | [illegible] | [illegible] | [illegible] | [illegible] | [illegible] | [illegible] | [illegible] | [illegible] | [illegible] | [illegible] | [illegible] | [illegible] | [illegible] | [illegible] | [illegible] | [illegible] | [illegible] | [illegible] |
| RI | 2933 | ENSG00000168591 | TMUB2 | chr17 | + | 44188934 | 44189299 | 44188934 | 44189091 | 44189206 | 44189299 | 111 | 62 | 63 | [illegible] | [illegible] | [illegible] | [illegible] | [illegible] | [illegible] | [illegible] | [illegible] | [illegible] | [illegible] | [illegible] | [illegible] | [illegible] | [illegible] | [illegible] | [illegible] | [illegible] | [illegible] | [illegible] | [illegible] | [illegible] | [illegible] | [illegible] | [illegible] | [illegible] | [illegible] | [illegible] | [illegible] | [illegible] | [illegible] | [illegible] | [illegible] | [illegible] | [illegible] |
| RI | 3576 | ENSG00000163931 | TKT | chr3 | - | 53234982 | 53235174 | 53234982 | 53235116 | 53235148 | 53235174 | 1235 | 546 | 765 | [illegible] | [illegible] | [illegible] | [illegible] | [illegible] | [illegible] | [illegible] | [illegible] | [illegible] | [illegible] | [illegible] | [illegible] | [illegible] | [illegible] | [illegible] | [illegible] | [illegible] | [illegible] | [illegible] | [illegible] | [illegible] | [illegible] | [illegible] | [illegible] | [illegible] | [illegible] | [illegible] | [illegible] | [illegible] | [illegible] | [illegible] | [illegible] | [illegible] | [illegible] |
| RI | 688 | ENSG00000101782 | RIOK3 | chr18 | + | 23477005 | 23477268 | 23477005 | 23477086 | 23477178 | 23477268 | 206 | 170 | 173 | [illegible] | [illegible] | [illegible] | [illegible] | [illegible] | [illegible] | [illegible] | [illegible] | [illegible] | [illegible] | [illegible] | [illegible] | [illegible] | [illegible] | [illegible] | [illegible] | [illegible] | [illegible] | [illegible] | [illegible] | [illegible] | [illegible] | [illegible] | [illegible] | [illegible] | [illegible] | [illegible] | [illegible] | [illegible] | [illegible] | [illegible] | [illegible] | [illegible] | [illegible] |
| RI | 5933 | ENSG00000114388 | NPRL2 | chr3 | - | 50349664 | 50350022 | 50349664 | 50349828 | 50349930 | 50350022 | 18 | 14 | 18 | [illegible] | [illegible] | [illegible] | [illegible] | [illegible] | [illegible] | [illegible] | [illegible] | [illegible] | [illegible] | [illegible] | [illegible] | [illegible] | [illegible] | [illegible] | [illegible] | [illegible] | [illegible] | [illegible] | [illegible] | [illegible] | [illegible] | [illegible] | [illegible] | [illegible] | [illegible] | [illegible] | [illegible] | [illegible] | [illegible] | [illegible] | [illegible] | [illegible] | [illegible] |
| RI | 3142 | ENSG00000041357 | PSMA4 | chr15 | + | 78542176 | 78542645 | 78542176 | 78542219 | 78542575 | 78542645 | 384 | 290 | 364 | [illegible] | [illegible] | [illegible] | [illegible] | [illegible] | [illegible] | [illegible] | [illegible] | [illegible] | [illegible] | [illegible] | [illegible] | [illegible] | [illegible] | [illegible] | [illegible] | [illegible] | [illegible] | [illegible] | [illegible] | [illegible] | [illegible] | [illegible] | [illegible] | [illegible] | [illegible] | [illegible] | [illegible] | [illegible] | [illegible] | [illegible] | [illegible] | [illegible] | [illegible] |
| RI | 5391 | ENSG00000159479 | MED8 | chr1 | - | 43383916 | 43385106 | 43383916 | 43384552 | 43385045 | 43385106 | 125 | 67 | 103 | [illegible] | [illegible] | [illegible] | [illegible] | [illegible] | [illegible] | [illegible] | [illegible] | [illegible] | [illegible] | [illegible] | [illegible] | [illegible] | [illegible] | [illegible] | [illegible] | [illegible] | [illegible] | [illegible] | [illegible] | [illegible] | [illegible] | [illegible] | [illegible] | [illegible] | [illegible] | [illegible] | [illegible] | [illegible] | [illegible] | [illegible] | [illegible] | [illegible] | [illegible] |
| RI | 5740 | ENSG00000186635 | ARAP1 | chr11 | - | 72712437 | 72713243 | 72712437 | 72712568 | 72713175 | 72713243 | 39 | 20 | 37 | [illegible] | [illegible] | [illegible] | [illegible] | [illegible] | [illegible] | [illegible] | [illegible] | [illegible] | [illegible] | [illegible] | [illegible] | [illegible] | [illegible] | [illegible] | [illegible] | [illegible] | [illegible] | [illegible] | [illegible] | [illegible] | [illegible] | [illegible] | [illegible] | [illegible] | [illegible] | [illegible] | [illegible] | [illegible] | [illegible] | [illegible] | [illegible] | [illegible] | [illegible] |
| RI | 7522 | ENSG00000115459 | ELMOD3 | chr2 | + | 85371085 | 85371562 | 85371085 | 85371209 | 85371439 | 85371562 | 20 | 1 | 12 | [illegible] | [illegible] | [illegible] | [illegible] | [illegible] | [illegible] | [illegible] | [illegible] | [illegible] | [illegible] | [illegible] | [illegible] | [illegible] | [illegible] | [illegible] | [illegible] | [illegible] | [illegible] | [illegible] | [illegible] | [illegible] | [illegible] | [illegible] | [illegible] | [illegible] | [illegible] | [illegible] | [illegible] | [illegible] | [illegible] | [illegible] | [illegible] | [illegible] | [illegible] |
| RI | 1253 | ENSG00000188603 | CLN3 | chr16 | - | 28487455 | 28488662 | 28487455 | 28487541 | 28488590 | 28488662 | 8 | 10 | 8 | [illegible] | [illegible] | [illegible] | [illegible] | [illegible] | [illegible] | [illegible] | [illegible] | [illegible] | [illegible] | [illegible] | [illegible] | [illegible] | [illegible] | [illegible] | [illegible] | [illegible] | [illegible] | [illegible] | [illegible] | [illegible] | [illegible] | [illegible] | [illegible] | [illegible] | [illegible] | [illegible] | [illegible] | [illegible] | [illegible] | [illegible] | [illegible] | [illegible] | [illegible] |
| RI | 5954 | ENSG00000141504 | SAT2 | chr17 | - | 7627142 | 7627414 | 7627142 | 7627226 | 7627362 | 7627414 | 43 | 31 | 44 | [illegible] | [illegible] | [illegible] | [illegible] | [illegible] | [illegible] | [illegible] | [illegible] | [illegible] | [illegible] | [illegible] | [illegible] | [illegible] | [illegible] | [illegible] | [illegible] | [illegible] | [illegible] | [illegible] | [illegible] | [illegible] | [illegible] | [illegible] | [illegible] | [illegible] | [illegible] | [illegible] | [illegible] | [illegible] | [illegible] | [illegible] | [illegible] | [illegible] | [illegible] |
| RI | 3177 | ENSG00000182481 | KPNA2 | chr17 | + | 68043099 | 68044071 | 68043099 | 68043363 | 68043837 | 68044071 | 2 | 2 | 1 | [illegible] | [illegible] | [illegible] | [illegible] | [illegible] | [illegible] | [illegible] | [illegible] | [illegible] | [illegible] | [illegible] | [illegible] | [illegible] | [illegible] | [illegible] | [illegible] | [illegible] | [illegible] | [illegible] | [illegible] | [illegible] | [illegible] | [illegible] | [illegible] | [illegible] | [illegible] | [illegible] | [illegible] | [illegible] | [illegible] | [illegible] | [illegible] | [illegible] | [illegible] |
| RI | 7921 | ENSG00000166340 | TPP1 | chr11 | - | 6612781 | 6614991 | 6612781 | 6614686 | 6614865 | 6614991 | 1 | 3 | 3 | [illegible] | [illegible] | [illegible] | [illegible] | [illegible] | [illegible] | [illegible] | [illegible] | [illegible] | [illegible] | [illegible] | [illegible] | [illegible] | [illegible] | [illegible] | [illegible] | [illegible] | [illegible] | [illegible] | [illegible] | [illegible] | [illegible] | [illegible] | [illegible] | [illegible] | [illegible] | [illegible] | [illegible] | [illegible] | [illegible] | [illegible] | [illegible] | [illegible] | [illegible] |
| RI | 2310 | ENSG00000164611 | PTTG1 | chr5 | + | 160422301 | 160422893 | 160422301 | 160422403 | 160422708 | 160422893 | 7 | 4 | 10 | [illegible] | [illegible] | [illegible] | [illegible] | [illegible] | [illegible] | [illegible] | [illegible] | [illegible] | [illegible] | [illegible] | [illegible] | [illegible] | [illegible] | [illegible] | [illegible] | [illegible] | [illegible] | [illegible] | [illegible] | [illegible] | [illegible] | [illegible] | [illegible] | [illegible] | [illegible] | [illegible] | [illegible] | [illegible] | [illegible] | [illegible] | [illegible] | [illegible] | [illegible] |
| RI | 6800 | ENSG00000005302 | MSL3 | chrX | + | 11763779 | 11765729 | 11763779 | 11763938 | 11765469 | 11765729 | 203 | 150 | 210 | [illegible] | [illegible] | [illegible] | [illegible] | [illegible] | [illegible] | [illegible] | [illegible] | [illegible] | [illegible] | [illegible] | [illegible] | [illegible] | [illegible] | [illegible] | [illegible] | [illegible] | [illegible] | [illegible] | [illegible] | [illegible] | [illegible] | [illegible] | [illegible] | [illegible] | [illegible] | [illegible] | [illegible] | [illegible] | [illegible] | [illegible] | [illegible] | [illegible] | [illegible] |
| RI | 6517 | ENSG00000204257 | HLA-DMA | chr6 | - | 32949610 | 32950803 | 32949610 | 32949787 | 32950518 | 32950803 | 280 | 188 | 255 | [illegible] | [illegible] | [illegible] | [illegible] | [illegible] | [illegible] | [illegible] | [illegible] | [illegible] | [illegible] | [illegible] | [illegible] | [illegible] | [illegible] | [illegible] | [illegible] | [illegible] | [illegible] | [illegible] | [illegible] | [illegible] | [illegible] | [illegible] | [illegible] | [illegible] | [illegible] | [illegible] | [illegible] | [illegible] | [illegible] | [illegible] | [illegible] | [illegible] | [illegible] |
| RI | 5738 | ENSG00000186635 | ARAP1 | chr11 | - | 72695541 | 72695865 | 72695541 | 72695628 | 72695717 | 72695865 | 27 | 7 | 3 | [illegible] | [illegible] | [illegible] | [illegible] | [illegible] | [illegible] | [illegible] | [illegible] | [illegible] | [illegible] | [illegible] | [illegible] | [illegible] | [illegible] | [illegible] | [illegible] | [illegible] | [illegible] | [illegible] | [illegible] | [illegible] | [illegible] | [illegible] | [illegible] | [illegible] | [illegible] | [illegible] | [illegible] | [illegible] | [illegible] | [illegible] | [illegible] | [illegible] | [illegible] |
| RI | 6433 | ENSG00000157764 | BRAF | chr7 | - | 140753274 | 140754233 | 140753274 | 140753393 | 140754186 | 140754233 | 15 | 5 | 2 | [illegible] | [illegible] | [illegible] | [illegible] | [illegible] | [illegible] | [illegible] | [illegible] | [illegible] | [illegible] | [illegible] | [illegible] | [illegible] | [illegible] | [illegible] | [illegible] | [illegible] | [illegible] | [illegible] | [illegible] | [illegible] | [illegible] | [illegible] | [illegible] | [illegible] | [illegible] | [illegible] | [illegible] | [illegible] | [illegible] | [illegible] | [illegible] | [illegible] | [illegible] |
| RI | 1701 | ENSG00000166140 | ZFYVE19 | chr15 | + | 40813337 | 40813811 | 40813337 | 40813417 | 40813712 | 40813811 | 1 | 2 | 12 | [illegible] | [illegible] | [illegible] | [illegible] | [illegible] | [illegible] | [illegible] | [illegible] | [illegible] | [illegible] | [illegible] | [illegible] | [illegible] | [illegible] | [illegible] | [illegible] | [illegible] | [illegible] | [illegible] | [illegible] | [illegible] | [illegible] | [illegible] | [illegible] | [illegible] | [illegible] | [illegible] | [illegible] | [illegible] | [illegible] | [illegible] | [illegible] | [illegible] | [illegible] |
| RI | 6064 | ENSG00000204351 | SKIV2L | chr6 | + | 31961040 | 31961675 | 31961040 | 31961094 | 31961546 | 31961675 | 61 | 39 | 36 | [illegible] | [illegible] | [illegible] | [illegible] | [illegible] | [illegible] | [illegible] | [illegible] | [illegible] | [illegible] | [illegible] | [illegible] | [illegible] | [illegible] | [illegible] | [illegible] | [illegible] | [illegible] | [illegible] | [illegible] | [illegible] | [illegible] | [illegible] | [illegible] | [illegible] | [illegible] | [illegible] | [illegible] | [illegible] | [illegible] | [illegible] | [illegible] | [illegible] | [illegible] |
| RI | 6183 | ENSG00000214021 | TTLL3 | chr3 | + | 9817644 | 9820741 | 9817644 | 9818920 | 9820545 | 9820741 | 65 | 50 | 57 | [illegible] | [illegible] | [illegible] | [illegible] | [illegible] | [illegible] | [illegible] | [illegible] | [illegible] | [illegible] | [illegible] | [illegible] | [illegible] | [illegible] | [illegible] | [illegible] | [illegible] | [illegible] | [illegible] | [illegible] | [illegible] | [illegible] | [illegible] | [illegible] | [illegible] | [illegible] | [illegible] | [illegible] | [illegible] | [illegible] | [illegible] | [illegible] | [illegible] | [illegible] |
| RI | 6520 | ENSG00000105835 | NAMPT | chr7 | - | 106269153 | 106271979 | 106269153 | 106269312 | 106269990 | 106271979 | 391 | 709 | 170 | [illegible] | [illegible] | [illegible] | [illegible] | [illegible] | [illegible] | [illegible] | [illegible] | [illegible] | [illegible] | [illegible] | [illegible] | [illegible] | [illegible] | [illegible] | [illegible] | [illegible] | [illegible] | [illegible] | [illegible] | [illegible] | [illegible] | [illegible] | [illegible] | [illegible] | [illegible] | [illegible] | [illegible] | [illegible] | [illegible] | [illegible] | [illegible] | [illegible] | [illegible] |
| RI | 8504 | ENSG00000135473 | PAN2 | chr12 | - | 56322614 | 56323209 | 56322614 | 56322758 | 56323061 | 56323209 | 81 | 109 | 107 | [illegible] | [illegible] | [illegible] | [illegible] | [illegible] | [illegible] | [illegible] | [illegible] | [illegible] | [illegible] | [illegible] | [illegible] | [illegible] | [illegible] | [illegible] | [illegible] | [illegible] | [illegible] | [illegible] | [illegible] | [illegible] | [illegible] | [illegible] | [illegible] | [illegible] | [illegible] | [illegible] | [illegible] | [illegible] | [illegible] | [illegible] | [illegible] | [illegible] | [illegible] |
| RI | 548 | ENSG00000124198 | ARFGEF2 | chr20 | + | 49013563 | 49013960 | 49013563 | 49013694 | 49013830 | 49013960 | 24 | 10 | 14 | [illegible] | [illegible] | [illegible] | [illegible] | [illegible] | [illegible] | [illegible] | [illegible] | [illegible] | [illegible] | [illegible] | [illegible] | [illegible] | [illegible] | [illegible] | [illegible] | [illegible] | [illegible] | [illegible] | [illegible] | [illegible] | [illegible] | [illegible] | [illegible] | [illegible] | [illegible] | [illegible] | [illegible] | [illegible] | [illegible] | [illegible] | [illegible] | [illegible] | [illegible] |
| RI | 1747 | ENSG00000197102 | DYNC1H1 | chr14 | + | 102042227 | 102042507 | 102042227 | 102042288 | 102042383 | 102042507 | 11 | 2 | 13 | [illegible] | [illegible] | [illegible] | [illegible] | [illegible] | [illegible] | [illegible] | [illegible] | [illegible] | [illegible] | [illegible] | [illegible] | [illegible] | [illegible] | [illegible] | [illegible] | [illegible] | [illegible] | [illegible] | [illegible] | [illegible] | [illegible] | [illegible] | [illegible] | [illegible] | [illegible] | [illegible] | [illegible] | [illegible] | [illegible] | [illegible] | [illegible] | [illegible] | [illegible] |
| RI | 3844 | ENSG00000049246 | PER3 | chr1 | + | 7829836 | 7830161 | 7829836 | 7829951 | 7830005 | 7830161 | 27 | 15 | 5 | [illegible] | [illegible] | [illegible] | [illegible] | [illegible] | [illegible] | [illegible] | [illegible] | [illegible] | [illegible] | [illegible] | [illegible] | [illegible] | [illegible] | [illegible] | [illegible] | [illegible] | [illegible] | [illegible] | [illegible] | [illegible] | [illegible] | [illegible] | [illegible] | [illegible] | [illegible] | [illegible] | [illegible] | [illegible] | [illegible] | [illegible] | [illegible] | [illegible] | [illegible] |
| RI | 5584 | ENSG00000117984 | CTSD | chr11 | - | 1753802 | 1754138 | 1753802 | 1753901 | 1753993 | 1754138 | 13 | 5 | 7 | [illegible] | [illegible] | [illegible] | [illegible] | [illegible] | [illegible] | [illegible] | [illegible] | [illegible] | [illegible] | [illegible] | [illegible] | [illegible] | [illegible] | [illegible] | [illegible] | [illegible] | [illegible] | [illegible] | [illegible] | [illegible] | [illegible] | [illegible] | [illegible] | [illegible] | [illegible] | [illegible] | [illegible] | [illegible] | [illegible] | [illegible] | [illegible] | [illegible] | [illegible] |
| RI | 248 | ENSG00000167604 | NFKBID | chr19 | - | 35896725 | 35897058 | 35896725 | 35896831 | 35896912 | 35897058 | 12 | 9 | 12 | [illegible] | [illegible] | [illegible] | [illegible] | [illegible] | [illegible] | [illegible] | [illegible] | [illegible] | [illegible] | [illegible] | [illegible] | [illegible] | [illegible] | [illegible] | [illegible] | [illegible] | [illegible] | [illegible] | [illegible] | [illegible] | [illegible] | [illegible] | [illegible] | [illegible] | [illegible] | [illegible] | [illegible] | [illegible] | [illegible] | [illegible] | [illegible] | [illegible] | [illegible] |
| RI | 6000 | ENSG00000156508 | EEF1A1 | chr6 | - | 73519336 | 73520056 | 73519336 | 73519516 | 73519882 | 73520056 | 445 | 469 | 667 | [illegible] | [illegible] | [illegible] | [illegible] | [illegible] | [illegible] | [illegible] | [illegible] | [illegible] | [illegible] | [illegible] | [illegible] | [illegible] | [illegible] | [illegible] | [illegible] | [illegible] | [illegible] | [illegible] | [illegible] | [illegible] | [illegible] | [illegible] | [illegible] | [illegible] | [illegible] | [illegible] | [illegible] | [illegible] | [illegible] | [illegible] | [illegible] | [illegible] | [illegible] |
| RI | 2789 | ENSG00000206573 | THUMPD3-AS1 | chr3 | - | 9390289 | 9391826 | 9390289 | 9390497 | 9390884 | 9391826 | 104 | 89 | 50 | [illegible] | [illegible] | [illegible] | [illegible] | [illegible] | [illegible] | [illegible] | [illegible] | [illegible] | [illegible] | [illegible] | [illegible] | [illegible] | [illegible] | [illegible] | [illegible] | [illegible] | [illegible] | [illegible] | [illegible] | [illegible] | [illegible] | [illegible] | [illegible] | [illegible] | [illegible] | [illegible] | [illegible] | [illegible] | [illegible] | [illegible] | [illegible] | [illegible] | [illegible] |
| RI | 4751 | ENSG00000159348 | CYB5R1 | chr1 | - | 202965886 | 202966600 | 202965886 | 202966109 | 202966527 | 202966600 | 28 | 22 | 31 | [illegible] | [illegible] | [illegible] | [illegible] | [illegible] | [illegible] | [illegible] | [illegible] | [illegible] | [illegible] | [illegible] | [illegible] | [illegible] | [illegible] | [illegible] | [illegible] | [illegible] | [illegible] | [illegible] | [illegible] | [illegible] | [illegible] | [illegible] | [illegible] | [illegible] | [illegible] | [illegible] | [illegible] | [illegible] | [illegible] | [illegible] | [illegible] | [illegible] | [illegible] |
| RI | 5522 | ENSG00000011132 | APBA3 | chr19 | - | 3751188 | 3751553 | 3751188 | 3751329 | 3751433 | 3751553 | 1 | 1 | 1 | [illegible] | [illegible] | [illegible] | [illegible] | [illegible] | [illegible] | [illegible] | [illegible] | [illegible] | [illegible] | [illegible] | [illegible] | [illegible] | [illegible] | [illegible] | [illegible] | [illegible] | [illegible] | [illegible] | [illegible] | [illegible] | [illegible] | [illegible] | [illegible] | [illegible] | [illegible] | [illegible] | [illegible] | [illegible] | [illegible] | [illegible] | [illegible] | [illegible] | [illegible] |
| RI | 193 | ENSG00000170889 | RPS9 | chr19 | + | 54206275 | 54206724 | 54206275 | 54206462 | 54206552 | 54206724 | 130 | 62 | 91 | [illegible] | [illegible] | [illegible] | [illegible] | [illegible] | [illegible] | [illegible] | [illegible] | [illegible] | [illegible] | [illegible] | [illegible] | [illegible] | [illegible] | [illegible] | [illegible] | [illegible] | [illegible] | [illegible] | [illegible] | [illegible] | [illegible] | [illegible] | [illegible] | [illegible] | [illegible] | [illegible] | [illegible] | [illegible] | [illegible] | [illegible] | [illegible] | [illegible] | [illegible] |
| RI | 4466 | ENSG00000137965 | IFI44 | chr1 | + | 78655013 | 78655511 | 78655013 | 78655209 | 78655361 | 78655511 | 120 | 168 | 47 | [illegible] | [illegible] | [illegible] | [illegible] | [illegible] | [illegible] | [illegible] | [illegible] | [illegible] | [illegible] | [illegible] | [illegible] | [illegible] | [illegible] | [illegible] | [illegible] | [illegible] | [illegible] | [illegible] | [illegible] | [illegible] | [illegible] | [illegible] | [illegible] | [illegible] | [illegible] | [illegible] | [illegible] | [illegible] | [illegible] | [illegible] | [illegible] | [illegible] | [illegible] |
| RI | 4207 | ENSG00000215788 | TNFRSF25 | chr1 | - | 6462862 | 6463127 | 6462862 | 6462970 | 6463071 | 6463127 | 68 | 38 | 122 | [illegible] | [illegible] | [illegible] | [illegible] | [illegible] | [illegible] | [illegible] | [illegible] | [illegible] | [illegible] | [illegible] | [illegible] | [illegible] | [illegible] | [illegible] | [illegible] | [illegible] | [illegible] | [illegible] | [illegible] | [illegible] | [illegible] | [illegible] | [illegible] | [illegible] | [illegible] | [illegible] | [illegible] | [illegible] | [illegible] | [illegible] | [illegible] | [illegible] | [illegible] |
| RI | 2725 | ENSG00000100412 | ACO2 | chr22 | + | 41522829 | 41523278 | 41522829 | 41522987 | 41523204 | 41523278 | 8 | 6 | 16 | [illegible] | [illegible] | [illegible] | [illegible] | [illegible] | [illegible] | [illegible] | [illegible] | [illegible] | [illegible] | [illegible] | [illegible] | [illegible] | [illegible] | [illegible] | [illegible] | [illegible] | [illegible] | [illegible] | [illegible] | [illegible] | [illegible] | [illegible] | [illegible] | [illegible] | [illegible] | [illegible] | [illegible] | [illegible] | [illegible] | [illegible] | [illegible] | [illegible] | [illegible] |
| RI | 4856 | ENSG00000149532 | CPSF7 | chr11 | - | 61416104 | 61416519 | 61416104 | 61416162 | 61416363 | 61416519 | 185 | 140 | 193 | [illegible] | [illegible] | [illegible] | [illegible] | [illegible] | [illegible] | [illegible] | [illegible] | [illegible] | [illegible] | [illegible] | [illegible] | [illegible] | [illegible] | [illegible] | [illegible] | [illegible] | [illegible] | [illegible] | [illegible] | [illegible] | [illegible] | [illegible] | [illegible] | [illegible] | [illegible] | [illegible] | [illegible] | [illegible] | [illegible] | [illegible] | [illegible] | [illegible] | [illegible] |
| RI | 1645 | ENSG00000177082 | WDR73 | chr15 | - | 84645470 | 84646348 | 84645470 | 84645836 | 84646183 | 84646348 | 45 | 37 | 44 | [illegible] | [illegible] | [illegible] | [illegible] | [illegible] | [illegible] | [illegible] | [illegible] | [illegible] | [illegible] | [illegible] | [illegible] | [illegible] | [illegible] | [illegible] | [illegible] | [illegible] | [illegible] | [illegible] | [illegible] | [illegible] | [illegible] | [illegible] | [illegible] | [illegible] | [illegible] | [illegible] | [illegible] | [illegible] | [illegible] | [illegible] | [illegible] | [illegible] | [illegible] |
| RI | 7923 | ENSG00000166340 | TPP1 | chr11 | - | 6616314 | 6616859 | 6616314 | 6616503 | 6616660 | 6616859 | 37 | 9 | 20 | [illegible] | [illegible] | [illegible] | [illegible] | [illegible] | [illegible] | [illegible] | [illegible] | [illegible] | [illegible] | [illegible] | [illegible] | [illegible] | [illegible] | [illegible] | [illegible] | [illegible] | [illegible] | [illegible] | [illegible] | [illegible] | [illegible] | [illegible] | [illegible] | [illegible] | [illegible] | [illegible] | [illegible] | [illegible] | [illegible] | [illegible] | [illegible] | [illegible] | [illegible] |
| RI | 4641 | ENSG00000227057 | WDR46 | chr6 | - | 33279107 | 33279610 | 33279107 | 33279374 | 33279496 | 33279610 | 21 | 16 | 22 | [illegible] | [illegible] | [illegible] | [illegible] | [illegible] | [illegible] | [illegible] | [illegible] | [illegible] | [illegible] | [illegible] | [illegible] | [illegible] | [illegible] | [illegible] | [illegible] | [illegible] | [illegible] | [illegible] | [illegible] | [illegible] | [illegible] | [illegible] | [illegible] | [illegible] | [illegible] | [illegible] | [illegible] | [illegible] | [illegible] | [illegible] | [illegible] | [illegible] | [illegible] |
| RI | 2098 | ENSG00000141279 | NPEPPS | chr17 | + | 47619008 | 47619784 | 47619008 | 47619164 | 47619277 | 47619784 | 23 | 13 | 21 | [illegible] | [illegible] | [illegible] | [illegible] | [illegible] | [illegible] | [illegible] | [illegible] | [illegible] | [illegible] | [illegible] | [illegible] | [illegible] | [illegible] | [illegible] | [illegible] | [illegible] | [illegible] | [illegible] | [illegible] | [illegible] | [illegible] | [illegible] | [illegible] | [illegible] | [illegible] | [illegible] | [illegible] | [illegible] | [illegible] | [illegible] | [illegible] | [illegible] | [illegible] |
| RI | 6975 | ENSG00000147050 | KDM6A | chrX | + | 45082575 | 45082789 | 45082575 | 45082640 | 45082714 | 45082789 | 50 | 86 | 35 | [illegible] | [illegible] | [illegible] | [illegible] | [illegible] | [illegible] | [illegible] | [illegible] | [illegible] | [illegible] | [illegible] | [illegible] | [illegible] | [illegible] | [illegible] | [illegible] | [illegible] | [illegible] | [illegible] | [illegible] | [illegible] | [illegible] | [illegible] | [illegible] | [illegible] | [illegible] | [illegible] | [illegible] | [illegible] | [illegible] | [illegible] | [illegible] | [illegible] | [illegible] |
| RI | 6326 | ENSG00000134905 | CARS2 | chr13 | - | 110645966 | 110647239 | 110645966 | 110646090 | 110647100 | 110647239 | 95 | 61 | 71 | [illegible] | [illegible] | [illegible] | [illegible] | [illegible] | [illegible] | [illegible] | [illegible] | [illegible] | [illegible] | [illegible] | [illegible] | [illegible] | [illegible] | [illegible] | [illegible] | [illegible] | [illegible] | [illegible] | [illegible] | [illegible] | [illegible] | [illegible] | [illegible] | [illegible] | [illegible] | [illegible] | [illegible] | [illegible] | [illegible] | [illegible] | [illegible] | [illegible] | [illegible] |
| RI | 6856 | ENSG00000215021 | PHB2 | chr12 | - | 6970195 | 6970645 | 6970195 | 6970280 | 6970416 | 6970645 | 5 | 1 | 2 | [illegible] | [illegible] | [illegible] | [illegible] | [illegible] | [illegible] | [illegible] | [illegible] | [illegible] | [illegible] | [illegible] | [illegible] | [illegible] | [illegible] | [illegible] | [illegible] | [illegible] | [illegible] | [illegible] | [illegible] | [illegible] | [illegible] | [illegible] | [illegible] | [illegible] | [illegible] | [illegible] | [illegible] | [illegible] | [illegible] | [illegible] | [illegible] | [illegible] | [illegible] |
| RI | 194 | ENSG00000170889 | RPS9 | chr19 | + | 54206552 | 54207647 | 54206552 | 54206724 | 54207397 | 54207647 | 87 | 45 | 51 | [illegible] | [illegible] | [illegible] | [illegible] | [illegible] | [illegible] | [illegible] | [illegible] | [illegible] | [illegible] | [illegible] | [illegible] | [illegible] | [illegible] | [illegible] | [illegible] | [illegible] | [illegible] | [illegible] | [illegible] | [illegible] | [illegible] | [illegible] | [illegible] | [illegible] | [illegible] | [illegible] | [illegible] | [illegible] | [illegible] | [illegible] | [illegible] | [illegible] | [illegible] |
| RI | 2919 | ENSG00000221978 | CCNL2 | chr1 | - | 1390765 | 1392803 | 1390765 | 1391575 | 1392789 | 1392803 | 191 | 184 | 221 | [illegible] | [illegible] | [illegible] | [illegible] | [illegible] | [illegible] | [illegible] | [illegible] | [illegible] | [illegible] | [illegible] | [illegible] | [illegible] | [illegible] | [illegible] | [illegible] | [illegible] | [illegible] | [illegible] | [illegible] | [illegible] | [illegible] | [illegible] | [illegible] | [illegible] | [illegible] | [illegible] | [illegible] | [illegible] | [illegible] | [illegible] | [illegible] | [illegible] | [illegible] |
| RI | 7242 | ENSG00000008988 | RPS20 | chr8 | - | 56074059 | 56074506 | 56074059 | 56074159 | 56074380 | 56074506 | 73 | 51 | 94 | [illegible] | [illegible] | [illegible] | [illegible] | [illegible] | [illegible] | [illegible] | [illegible] | [illegible] | [illegible] | [illegible] | [illegible] | [illegible] | [illegible] | [illegible] | [illegible] | [illegible] | [illegible] | [illegible] | [illegible] | [illegible] | [illegible] | [illegible] | [illegible] | [illegible] | [illegible] | [illegible] | [illegible] | [illegible] | [illegible] | [illegible] | [illegible] | [illegible] | [illegible] |
| RI | 2483 | ENSG00000077232 | DNAJC10 | chr2 | + | 182717015 | 182718290 | 182717015 | 182717072 | 182717940 | 182718290 | 24 | 28 | 15 | [illegible] | [illegible] | [illegible] | [illegible] | [illegible] | [illegible] | [illegible] | [illegible] | [illegible] | [illegible] | [illegible] | [illegible] | [illegible] | [illegible] | [illegible] | [illegible] | [illegible] | [illegible] | [illegible] | [illegible] | [illegible] | [illegible] | [illegible] | [illegible] | [illegible] | [illegible] | [illegible] | [illegible] | [illegible] | [illegible] | [illegible] | [illegible] | [illegible] | [illegible] |
| RI | 2791 | ENSG00000206573 | THUMPD3-AS1 | chr3 | - | 9390289 | 9391826 | 9390289 | 9390497 | 9391678 | 9391826 | 108 | 134 | 89 | [illegible] | [illegible] | [illegible] | [illegible] | [illegible] | [illegible] | [illegible] | [illegible] | [illegible] | [illegible] | [illegible] | [illegible] | [illegible] | [illegible] | [illegible] | [illegible] | [illegible] | [illegible] | [illegible] | [illegible] | [illegible] | [illegible] | [illegible] | [illegible] | [illegible] | [illegible] | [illegible] | [illegible] | [illegible] | [illegible] | [illegible] | [illegible] | [illegible] | [illegible] |
| RI | 5750 | ENSG00000169045 | HNRNPH1 | chr5 | - | 179616868 | 179617110 | 179616868 | 179616958 | 179617050 | 179617110 | 549 | 579 | 545 | [illegible] | [illegible] | [illegible] | [illegible] | [illegible] | [illegible] | [illegible] | [illegible] | [illegible] | [illegible] | [illegible] | [illegible] | [illegible] | [illegible] | [illegible] | [illegible] | [illegible] | [illegible] | [illegible] | [illegible] | [illegible] | [illegible] | [illegible] | [illegible] | [illegible] | [illegible] | [illegible] | [illegible] | [illegible] | [illegible] | [illegible] | [illegible] | [illegible] | [illegible] |
| RI | 687 | ENSG00000100983 | GSS | chr20 | - | 34942487 | 34943006 | 34942487 | 34942627 | 34942930 | 34943006 | 41 | 14 | 41 | [illegible] | [illegible] | [illegible] | [illegible] | [illegible] | [illegible] | [illegible] | [illegible] | [illegible] | [illegible] | [illegible] | [illegible] | [illegible] | [illegible] | [illegible] | [illegible] | [illegible] | [illegible] | [illegible] | [illegible] | [illegible] | [illegible] | [illegible] | [illegible] | [illegible] | [illegible] | [illegible] | [illegible] | [illegible] | [illegible] | [illegible] | [illegible] | [illegible] | [illegible] |
| RI | 3688 | ENSG00000099622 | CIRBP | chr19 | + | 1271980 | 1274440 | 1271980 | 1272051 | 1274306 | 1274440 | 133 | 70 | 132 | [illegible] | [illegible] | [illegible] | [illegible] | [illegible] | [illegible] | [illegible] | [illegible] | [illegible] | [illegible] | [illegible] | [illegible] | [illegible] | [illegible] | [illegible] | [illegible] | [illegible] | [illegible] | [illegible] | [illegible] | [illegible] | [illegible] | [illegible] | [illegible] | [illegible] | [illegible] | [illegible] | [illegible] | [illegible] | [illegible] | [illegible] | [illegible] | [illegible] | [illegible] |
| RI | 6937 | ENSG00000046651 | OFD1 | chrX | + | 13758336 | 13760720 | 13758336 | 13758448 | 13760114 | 13760720 | 63 | 56 | 73 | [illegible] | [illegible] | [illegible] | [illegible] | [illegible] | [illegible] | [illegible] | [illegible] | [illegible] | [illegible] | [illegible] | [illegible] | [illegible] | [illegible] | [illegible] | [illegible] | [illegible] | [illegible] | [illegible] | [illegible] | [illegible] | [illegible] | [illegible] | [illegible] | [illegible] | [illegible] | [illegible] | [illegible] | [illegible] | [illegible] | [illegible] | [illegible] | [illegible] | [illegible] |
| RI | 3060 | ENSG00000169727 | GPS1 | chr17 | + | 82056285 | 82056550 | 82056285 | 82056391 | 82056469 | 82056550 | 17 | 6 | 8 | [illegible] | [illegible] | [illegible] | [illegible] | [illegible] | [illegible] | [illegible] | [illegible] | [illegible] | [illegible] | [illegible] | [illegible] | [illegible] | [illegible] | [illegible] | [illegible] | [illegible] | [illegible] | [illegible] | [illegible] | [illegible] | [illegible] | [illegible] | [illegible] | [illegible] | [illegible] | [illegible] | [illegible] | [illegible] | [illegible] | [illegible] | [illegible] | [illegible] | [illegible] |
| RI | 4737 | ENSG00000164828 | SUN1 | chr7 | + | 852608 | 852952 | 852608 | 852667 | 852809 | 852952 | 57 | 29 | 55 | [illegible] | [illegible] | [illegible] | [illegible] | [illegible] | [illegible] | [illegible] | [illegible] | [illegible] | [illegible] | [illegible] | [illegible] | [illegible] | [illegible] | [illegible] | [illegible] | [illegible] | [illegible] | [illegible] | [illegible] | [illegible] | [illegible] | [illegible] | [illegible] | [illegible] | [illegible] | [illegible] | [illegible] | [illegible] | [illegible] | [illegible] | [illegible] | [illegible] | [illegible] |
| RI | 3510 | ENSG00000269893 | SNHG8 | chr4 | + | 118279388 | 118279820 | 118279388 | 118279431 | 118279623 | 118279820 | 193 | 82 | 170 | [illegible] | [illegible] | [illegible] | [illegible] | [illegible] | [illegible] | [illegible] | [illegible] | [illegible] | [illegible] | [illegible] | [illegible] | [illegible] | [illegible] | [illegible] | [illegible] | [illegible] | [illegible] | [illegible] | [illegible] | [illegible] | [illegible] | [illegible] | [illegible] | [illegible] | [illegible] | [illegible] | [illegible] | [illegible] | [illegible] | [illegible] | [illegible] | [illegible] | [illegible] |
| RI | 4403 | ENSG00000158552 | ZFAND2B | chr2 | + | 219208255 | 219208486 | 219208255 | 219208348 | 219208425 | 219208486 | 26 | 7 | 7 | [illegible] | [illegible] | [illegible] | [illegible] | [illegible] | [illegible] | [illegible] | [illegible] | [illegible] | [illegible] | [illegible] | [illegible] | [illegible] | [illegible] | [illegible] | [illegible] | [illegible] | [illegible] | [illegible] | [illegible] | [illegible] | [illegible] | [illegible] | [illegible] | [illegible] | [illegible] | [illegible] | [illegible] | [illegible] | [illegible] | [illegible] | [illegible] | [illegible] | [illegible] |
| RI | 7605 | ENSG00000152556 | PFKM | chr12 | + | 48141739 | 48142066 | 48141739 | 48141827 | 48141913 | 48142066 | 2 | 2 | 5 | [illegible] | [illegible] | [illegible] | [illegible] | [illegible] | [illegible] | [illegible] | [illegible] | [illegible] | [illegible] | [illegible] | [illegible] | [illegible] | [illegible] | [illegible] | [illegible] | [illegible] | [illegible] | [illegible] | [illegible] | [illegible] | [illegible] | [illegible] | [illegible] | [illegible] | [illegible] | [illegible] | [illegible] | [illegible] | [illegible] | [illegible] | [illegible] | [illegible] | [illegible] |
| RI | 3227 | ENSG00000184014 | DENND5A | chr11 | - | 9142721 | 9144278 | 9142721 | 9142845 | 9144096 | 9144278 | 7 | 22 | 14 | [illegible] | [illegible] | [illegible] | [illegible] | [illegible] | [illegible] | [illegible] | [illegible] | [illegible] | [illegible] | [illegible] | [illegible] | [illegible] | [illegible] | [illegible] | [illegible] | [illegible] | [illegible] | [illegible] | [illegible] | [illegible] | [illegible] | [illegible] | [illegible] | [illegible] | [illegible] | [illegible] | [illegible] | [illegible] | [illegible] | [illegible] | [illegible] | [illegible] | [illegible] |
| RI | 5538 | ENSG00000182004 | SNRPE | chr1 | + | 203861602 | 203862222 | 203861602 | 203861743 | 203862181 | 203862222 | 26 | 16 | 43 | [illegible] | [illegible] | [illegible] | [illegible] | [illegible] | [illegible] | [illegible] | [illegible] | [illegible] | [illegible] | [illegible] | [illegible] | [illegible] | [illegible] | [illegible] | [illegible] | [illegible] | [illegible] | [illegible] | [illegible] | [illegible] | [illegible] | [illegible] | [illegible] | [illegible] | [illegible] | [illegible] | [illegible] | [illegible] | [illegible] | [illegible] | [illegible] | [illegible] | [illegible] |
| RI | 6012 | ENSG00000204420 | MPIG6B | chr6 | + | 31724586 | 31724844 | 31724586 | 31724627 | 31724764 | 31724844 | 10 | 5 | 3 | [illegible] | [illegible] | [illegible] | [illegible] | [illegible] | [illegible] | [illegible] | [illegible] | [illegible] | [illegible] | [illegible] | [illegible] | [illegible] | [illegible] | [illegible] | [illegible] | [illegible] | [illegible] | [illegible] | [illegible] | [illegible] | [illegible] | [illegible] | [illegible] | [illegible] | [illegible] | [illegible] | [illegible] | [illegible] | [illegible] | [illegible] | [illegible] | [illegible] | [illegible] |
| RI | 5964 | ENSG00000168000 | BSCL2 | chr11 | - | 62690786 | 62691421 | 62690786 | 62691141 | 62691279 | 62691421 | 8 | 5 | 3 | [illegible] | [illegible] | [illegible] | [illegible] | [illegible] | [illegible] | [illegible] | [illegible] | [illegible] | [illegible] | [illegible] | [illegible] | [illegible] | [illegible] | [illegible] | [illegible] | [illegible] | [illegible] | [illegible] | [illegible] | [illegible] | [illegible] | [illegible] | [illegible] | [illegible] | [illegible] | [illegible] | [illegible] | [illegible] | [illegible] | [illegible] | [illegible] | [illegible] | [illegible] |
| RI | 594 | ENSG00000160410 | SHKBP1 | chr19 | + | 40577395 | 40577630 | 40577395 | 40577441 | 40577556 | 40577630 | 8 | 2 | 6 | [illegible] | [illegible] | [illegible] | [illegible] | [illegible] | [illegible] | [illegible] | [illegible] | [illegible] | [illegible] | [illegible] | [illegible] | [illegible] | [illegible] | [illegible] | [illegible] | [illegible] | [illegible] | [illegible] | [illegible] | [illegible] | [illegible] | [illegible] | [illegible] | [illegible] | [illegible] | [illegible] | [illegible] | [illegible] | [illegible] | [illegible] | [illegible] | [illegible] | [illegible] |
| RI | 2573 | ENSG00000104972 | LILRB1 | chr19 | + | 54635556 | 54636652 | 54635556 | 54635609 | 54636496 | 54636652 | 23 | 19 | 9 | [illegible] | [illegible] | [illegible] | [illegible] | [illegible] | [illegible] | [illegible] | [illegible] | [illegible] | [illegible] | [illegible] | [illegible] | [illegible] | [illegible] | [illegible] | [illegible] | [illegible] | [illegible] | [illegible] | [illegible] | [illegible] | [illegible] | [illegible] | [illegible] | [illegible] | [illegible] | [illegible] | [illegible] | [illegible] | [illegible] | [illegible] | [illegible] | [illegible] | [illegible] |
| RI | 2099 | ENSG00000141279 | NPEPPS | chr17 | + | 47619008 | 47619784 | 47619008 | 47619164 | 47619736 | 47619784 | 19 | 38 | 36 | [illegible] | [illegible] | [illegible] | [illegible] | [illegible] | [illegible] | [illegible] | [illegible] | [illegible] | [illegible] | [illegible] | [illegible] | [illegible] | [illegible] | [illegible] | [illegible] | [illegible] | [illegible] | [illegible] | [illegible] | [illegible] | [illegible] | [illegible] | [illegible] | [illegible] | [illegible] | [illegible] | [illegible] | [illegible] | [illegible] | [illegible] | [illegible] | [illegible] | [illegible] |
| RI | 933 | ENSG00000261408 | TEN1-CDK3 | chr17 | + | 76002510 | 76003398 | 76002510 | 76002612 | 76003194 | 76003398 | 9 | 3 | 9 | [illegible] | [illegible] | [illegible] | [illegible] | [illegible] | [illegible] | [illegible] | [illegible] | [illegible] | [illegible] | [illegible] | [illegible] | [illegible] | [illegible] | [illegible] | [illegible] | [illegible] | [illegible] | [illegible] | [illegible] | [illegible] | [illegible] | [illegible] | [illegible] | [illegible] | [illegible] | [illegible] | [illegible] | [illegible] | [illegible] | [illegible] | [illegible] | [illegible] | [illegible] |
| RI | 5578 | ENSG00000077150 | NFKB2 | chr10 | + | 102396448 | 102396823 | 102396448 | 102396489 | 102396724 | 102396823 | 33 | 40 | 69 | [illegible] | [illegible] | [illegible] | [illegible] | [illegible] | [illegible] | [illegible] | [illegible] | [illegible] | [illegible] | [illegible] | [illegible] | [illegible] | [illegible] | [illegible] | [illegible] | [illegible] | [illegible] | [illegible] | [illegible] | [illegible] | [illegible] | [illegible] | [illegible] | [illegible] | [illegible] | [illegible] | [illegible] | [illegible] | [illegible] | [illegible] | [illegible] | [illegible] | [illegible] |
| RI | 2992 | ENSG00000160796 | NBEAL2 | chr3 | + | 47007240 | 47007697 | 47007240 | 47007350 | 47007524 | 47007697 | 19 | 11 | 23 | [illegible] | [illegible] | [illegible] | [illegible] | [illegible] | [illegible] | [illegible] | [illegible] | [illegible] | [illegible] | [illegible] | [illegible] | [illegible] | [illegible] | [illegible] | [illegible] | [illegible] | [illegible] | [illegible] | [illegible] | [illegible] | [illegible] | [illegible] | [illegible] | [illegible] | [illegible] | [illegible] | [illegible] | [illegible] | [illegible] | [illegible] | [illegible] | [illegible] | [illegible] |
| RI | 5831 | ENSG00000119707 | RBM25 | chr14 | + | 73099666 | 73103478 | 73099666 | 73099750 | 73103191 | 73103478 | 460 | 649 | 515 | [illegible] | [illegible] | [illegible] | [illegible] | [illegible] | [illegible] | [illegible] | [illegible] | [illegible] | [illegible] | [illegible] | [illegible] | [illegible] | [illegible] | [illegible] | [illegible] | [illegible] | [illegible] | [illegible] | [illegible] | [illegible] | [illegible] | [illegible] | [illegible] | [illegible] | [illegible] | [illegible] | [illegible] | [illegible] | [illegible] | [illegible] | [illegible] | [illegible] | [illegible] |
| RI | 1590 | ENSG00000140521 | POLG | chr15 | - | 89317938 | 89318749 | 89317938 | 89318037 | 89318540 | 89318749 | 42 | 54 | 62 | [illegible] | [illegible] | [illegible] | [illegible] | [illegible] | [illegible] | [illegible] | [illegible] | [illegible] | [illegible] | [illegible] | [illegible] | [illegible] | [illegible] | [illegible] | [illegible] | [illegible] | [illegible] | [illegible] | [illegible] | [illegible] | [illegible] | [illegible] | [illegible] | [illegible] | [illegible] | [illegible] | [illegible] | [illegible] | [illegible] | [illegible] | [illegible] | [illegible] | [illegible] |
| RI | 7360 | ENSG00000089053 | ANAPC5 | chr12 | - | 121318276 | 121318608 | 121318276 | 121318404 | 121318500 | 121318608 | 64 | 60 | 76 | [illegible] | [illegible] | [illegible] | [illegible] | [illegible] | [illegible] | [illegible] | [illegible] | [illegible] | [illegible] | [illegible] | [illegible] | [illegible] | [illegible] | [illegible] | [illegible] | [illegible] | [illegible] | [illegible] | [illegible] | [illegible] | [illegible] | [illegible] | [illegible] | [illegible] | [illegible] | [illegible] | [illegible] | [illegible] | [illegible] | [illegible] | [illegible] | [illegible] | [illegible] |
| RI | [illegible] | [illegible] | RABEPK | chr9 | [illegible] | [illegible] | [illegible] | [illegible] | [illegible] | [illegible] | [illegible] | [illegible] | [illegible] | [illegible] | [illegible] | [illegible] | [illegible] | [illegible] | [illegible] | [illegible] | [illegible] | [illegible] | [illegible] | [illegible] | [illegible] | [illegible] | [illegible] | [illegible] | [illegible] | [illegible] | [illegible] | [illegible] | [illegible] | [illegible] | [illegible] | [illegible] | [illegible] | [illegible] | [illegible] | [illegible] | [illegible] | [illegible] | [illegible] | [illegible] | [illegible] | [illegible] |
| RI | [illegible] | [illegible] | ABCA7 | chr19 | [illegible] | [illegible] | [illegible] | [illegible] | [illegible] | [illegible] | [illegible] | [illegible] | [illegible] | [illegible] | [illegible] | [illegible] | [illegible] | [illegible] | [illegible] | [illegible] | [illegible] | [illegible] | [illegible] | [illegible] | [illegible] | [illegible] | [illegible] | [illegible] | [illegible] | [illegible] | [illegible] | [illegible] | [illegible] | [illegible] | [illegible] | [illegible] | [illegible] | [illegible] | [illegible] | [illegible] | [illegible] | [illegible] | [illegible] | [illegible] | [illegible] | [illegible] |
| RI | [illegible] | [illegible] | CEP120 | chr5 | [illegible] | [illegible] | [illegible] | [illegible] | [illegible] | [illegible] | [illegible] | [illegible] | [illegible] | [illegible] | [illegible] | [illegible] | [illegible] | [illegible] | [illegible] | [illegible] | [illegible] | [illegible] | [illegible] | [illegible] | [illegible] | [illegible] | [illegible] | [illegible] | [illegible] | [illegible] | [illegible] | [illegible] | [illegible] | [illegible] | [illegible] | [illegible] | [illegible] | [illegible] | [illegible] | [illegible] | [illegible] | [illegible] | [illegible] | [illegible] | [illegible] | [illegible] |
| RI | [illegible] | [illegible] | ILKAP | chr2 | [illegible] | [illegible] | [illegible] | [illegible] | [illegible] | [illegible] | [illegible] | [illegible] | [illegible] | [illegible] | [illegible] | [illegible] | [illegible] | [illegible] | [illegible] | [illegible] | [illegible] | [illegible] | [illegible] | [illegible] | [illegible] | [illegible] | [illegible] | [illegible] | [illegible] | [illegible] | [illegible] | [illegible] | [illegible] | [illegible] | [illegible] | [illegible] | [illegible] | [illegible] | [illegible] | [illegible] | [illegible] | [illegible] | [illegible] | [illegible] | [illegible] | [illegible] |
| RI | [illegible] | [illegible] | MED13L | chr12 | [illegible] | [illegible] | [illegible] | [illegible] | [illegible] | [illegible] | [illegible] | [illegible] | [illegible] | [illegible] | [illegible] | [illegible] | [illegible] | [illegible] | [illegible] | [illegible] | [illegible] | [illegible] | [illegible] | [illegible] | [illegible] | [illegible] | [illegible] | [illegible] | [illegible] | [illegible] | [illegible] | [illegible] | [illegible] | [illegible] | [illegible] | [illegible] | [illegible] | [illegible] | [illegible] | [illegible] | [illegible] | [illegible] | [illegible] | [illegible] | [illegible] | [illegible] |

| | | | | | | | | | | | | | | | | | | | | | | | | | | | | | | | | | | | | | | | | | | | | | | |
|---|---|---|---|---|---|---|---|---|---|---|---|---|---|---|---|---|---|---|---|---|---|---|---|---|---|---|---|---|---|---|---|---|---|---|---|---|---|---|---|---|---|---|---|---|---|---|
| RI | 7127 | ENSG00000147576 | ADHFE1 | chr8 | + | 66444366 | 66444748 | 66444366 | 66444420 | 66444593 | 66444748 | 11 | 13 | 25 | 13 | 7 | 26 | 12 | 21 | 20 | 23 | 0 | 11 | 24 | 9 | 0 | 2 | 17 | 8 | 5 | 6 | 298 | 149 | 0.968253694 | 1 | 0.175 | 0.351 | 0.373 | 0.245 | 0.132 | 0 | 0.244 | 0.6 | 0.474 | 0 | -0.008 |
| RI | 6358 | ENSG00000075624 | ACTB | chr7 | - | 5528280 | 5529400 | 5528280 | 5528671 | 5529160 | 5529400 | 4390 | 1825 | 3229 | 3306 | 3261 | 55 | 29 | 47 | 34 | 11 | 4966 | 4017 | 2223 | 3769 | 18303 | 60 | 39 | 28 | 29 | 206 | 298 | 149 | 0.968272291 | 1 | 0.976 | 0.969 | 0.972 | 0.98 | 0.993 | 0.976 | 0.981 | 0.975 | 0.985 | 0.978 | -0.001 |
| RI | 3426 | ENSG00000164088 | PPM1M | chr3 | + | 52249173 | 52250595 | 52249173 | 52249322 | 52249669 | 52250595 | 69 | 22 | 37 | 41 | 52 | 240 | 95 | 136 | 116 | 232 | 95 | 55 | 53 | 59 | 29 | 240 | 237 | 108 | 205 | 310 | 298 | 149 | 0.969581155 | 1 | 0.126 | 0.104 | 0.12 | 0.15 | 0.101 | 0.165 | 0.104 | 0.197 | 0.126 | 0.045 | -0.007 |
| RI | 5331 | ENSG00000064703 | DDX20 | chr1 | + | 111761225 | 111762337 | 111761225 | 111761284 | 111762254 | 111762337 | 15 | 14 | 12 | 34 | 62 | 41 | 43 | 41 | 34 | 67 | 9 | 5 | 22 | 23 | 0 | 17 | 26 | 32 | 5 | 3 | 298 | 149 | 0.970764747 | 1 | 0.155 | 0.14 | 0.128 | 0.333 | 0.316 | 0.209 | 0.088 | 0.256 | 0.697 | 0 | -0.036 |
| RI | 7386 | ENSG00000077463 | SIRT6 | chr19 | - | 4175027 | 4175760 | 4175027 | 4175151 | 4175679 | 4175760 | 8 | 6 | 8 | 11 | 5 | 18 | 2 | 6 | 18 | 22 | 11 | 0 | 5 | 13 | 6 | 6 | 24 | 9 | 1 | 16 | 298 | 149 | 0.971747653 | 1 | 0.182 | 0.6 | 0.4 | 0.234 | 0.102 | 0.478 | 0 | 0.217 | 0.867 | 0.158 | -0.04 |
| RI | 177 | ENSG00000006015 | REX1BD | chr19 | + | 18590853 | 18592336 | 18590853 | 18590933 | 18592107 | 18592336 | 8 | 5 | 10 | 16 | 10 | 43 | 22 | 34 | 30 | 6 | 3 | 8 | 12 | 2 | 38 | 22 | 61 | 8 | 12 | 41 | 298 | 149 | 0.972347869 | 1 | 0.085 | 0.102 | 0.128 | 0.211 | 0.455 | 0.064 | 0.062 | 0.429 | 0.077 | 0.317 | 0.006 |
| RI | 1874 | ENSG00000161914 | ZNF653 | chr19 | - | 11486986 | 11487903 | 11486986 | 11487158 | 11487291 | 11487903 | 9 | 6 | 8 | 15 | 3 | 8 | 0 | 2 | 2 | 5 | 1 | 8 | 11 | 7 | 12 | 14 | 3 | 0 | 5 | 0 | 282 | 149 | 0.97256457 | 1 | 0.373 | 1 | 0.679 | 0.798 | 0.241 | 0.036 | 0.585 | 1 | 0.425 | 1 | 0.009 |
| RI | 5402 | ENSG00000176978 | DPP7 | chr9 | - | 137114242 | 137114576 | 137114242 | 137114382 | 137114462 | 137114576 | 2 | 0 | 10 | 0 | 4 | 22 | 14 | 16 | 16 | 0 | 6 | 3 | 3 | 0 | 0 | 13 | 0 | 13 | 17 | 14 | 229 | 149 | 0.973917658 | 1 | 0.056 | 0 | 0.289 | 0 | 1 | 0.231 | 1 | 0.131 | 0 | 0 | -0.003 |
| RI | 3490 | ENSG00000143578 | CREB3L4 | chr1 | + | 153968929 | 153969455 | 153968929 | 153969176 | 153969333 | 153969455 | 3 | 1 | 7 | 4 | 4 | 17 | 2 | 2 | 13 | 15 | 5 | 2 | 2 | 0 | 14 | 0 | 9 | 14 | 7 | 12 | 298 | 149 | 0.975045607 | 1 | 0.081 | 0.2 | 0.636 | 0.133 | 0.118 | 1 | 0.1 | 0.067 | 0 | 0.368 | -0.073 |
| RI | 3360 | ENSG00000103507 | BCKDK | chr16 | + | 31109672 | 31110124 | 31109672 | 31109783 | 31110076 | 31110124 | 7 | 8 | 7 | 5 | 0 | 22 | 6 | 7 | 21 | 9 | 15 | 3 | 28 | 3 | 12 | 64 | 39 | 5 | 13 | 39 | 298 | 149 | 0.975939658 | 1 | 0.137 | 0.4 | 0.333 | 0.106 | 0 | 0.105 | 0.037 | 0.737 | 0.103 | 0.133 | -0.028 |
| RI | 166 | ENSG00000099624 | ATP5F1D | chr19 | + | 1244314 | 1244825 | 1244314 | 1244440 | 1244736 | 1244825 | 66 | 9 | 34 | 33 | 24 | 51 | 24 | 52 | 35 | 24 | 37 | 27 | 20 | 52 | 79 | 39 | 35 | 27 | 43 | 121 | 298 | 149 | 0.976689282 | 1 | 0.393 | 0.158 | 0.246 | 0.32 | 0.333 | 0.322 | 0.278 | 0.27 | 0.377 | 0.246 | -0.009 |
| RI | 4706 | ENSG00000232956 | SNHG15 | chr7 | - | 44983022 | 44984046 | 44983022 | 44983322 | 44983821 | 44984046 | 26 | 25 | 21 | 38 | 27 | 67 | 25 | 65 | 60 | 79 | 94 | 22 | 35 | 44 | 143 | 173 | 121 | 64 | 157 | 121 | 298 | 149 | 0.977163233 | 1 | 0.163 | 0.333 | 0.139 | 0.241 | 0.146 | 0.214 | 0.083 | 0.215 | 0.123 | 0.371 | 0.003 |
| RI | 7730 | ENSG00000151498 | ACAD8 | chr11 | + | 134259607 | 134261179 | 134259607 | 134259745 | 134261043 | 134261179 | 35 | 19 | 32 | 34 | 32 | 13 | 8 | 12 | 9 | 23 | 48 | 68 | 43 | 65 | 82 | 43 | 7 | 13 | 30 | 56 | 298 | 149 | 0.977620135 | 1 | 0.574 | 0.543 | 0.571 | 0.654 | 0.41 | 0.358 | 0.829 | 0.623 | 0.52 | 0.423 | 0 |
| RI | 5059 | ENSG00000106105 | GARS1 | chr7 | + | 30628559 | 30631541 | 30628559 | 30628669 | 30631447 | 30631541 | 14 | 16 | 20 | 26 | 22 | 160 | 135 | 150 | 146 | 341 | 7 | 21 | 46 | 14 | 0 | 152 | 180 | 133 | 166 | 161 | 298 | 149 | 0.979109985 | 1 | 0.042 | 0.056 | 0.062 | 0.082 | 0.031 | 0.023 | 0.055 | 0.147 | 0.04 | 0 | 0.002 |
| RI | 4348 | ENSG00000169764 | UGP2 | chr2 | + | 63887401 | 63890185 | 63887401 | 63887644 | 63890080 | 63890185 | 23 | 33 | 33 | 34 | 37 | 273 | 179 | 206 | 294 | 322 | 3 | 23 | 20 | 0 | 15 | 70 | 117 | 172 | 84 | 24 | 298 | 149 | 0.979420703 | 1 | 0.04 | 0.084 | 0.074 | 0.055 | 0.054 | 0.021 | 0.089 | 0.055 | 0 | 0.238 | -0.019 |
| RI | 352 | ENSG00000174886 | NDUFA11 | chr19 | - | 5894674 | 5896575 | 5894674 | 5894854 | 5896452 | 5896575 | 13 | 6 | 18 | 13 | 14 | 49 | 26 | 37 | 33 | 30 | 16 | 8 | 24 | 8 | 16 | 85 | 28 | 21 | 25 | 44 | 298 | 149 | 0.979479306 | 1 | 0.117 | 0.103 | 0.196 | 0.165 | 0.189 | 0.086 | 0.125 | 0.364 | 0.138 | 0.154 | -0.019 |
| RI | 1851 | ENSG00000165959 | CLMN | chr14 | - | 95193848 | 95194596 | 95193848 | 95193919 | 95194535 | 95194596 | 25 | 37 | 21 | 20 | 35 | 33 | 21 | 31 | 10 | 60 | 23 | 39 | 37 | 57 | 10 | 22 | 25 | 23 | 29 | 74 | 298 | 149 | 0.980657077 | 1 | 0.275 | 0.468 | 0.253 | 0.5 | 0.226 | 0.343 | 0.438 | 0.446 | 0.496 | 0.063 | -0.013 |
| RI | 4274 | ENSG00000042429 | MED17 | chr11 | + | 93807500 | 93809876 | 93807500 | 93807635 | 93809716 | 93809876 | 24 | 24 | 17 | 39 | 11 | 75 | 46 | 56 | 37 | 66 | 0 | 49 | 6 | 27 | 0 | 2 | 18 | 41 | 20 | 10 | 298 | 149 | 0.98240922 | 1 | 0.138 | 0.207 | 0.132 | 0.345 | 0.077 | 0 | 0.576 | 0.068 | 0.403 | 0 | -0.03 |
| RI | 2227 | ENSG00000169567 | HINT1 | chr5 | - | 131159295 | 131162676 | 131159295 | 131162477 | 131162571 | 131162676 | 159 | 158 | 219 | 197 | 475 | 54 | 25 | 64 | 39 | 104 | 421 | 171 | 184 | 353 | 114 | 52 | 50 | 45 | 65 | 55 | 243 | 149 | 0.983847359 | 1 | 0.644 | 0.795 | 0.677 | 0.756 | 0.737 | 0.832 | 0.677 | 0.715 | 0.769 | 0.56 | 0.011 |
| RI | 456 | ENSG00000130511 | SSBP4 | chr19 | + | 18432558 | 18432735 | 18432558 | 18432604 | 18432699 | 18432735 | 5 | 2 | 2 | 0 | 2 | 67 | 21 | 34 | 79 | 23 | 0 | 3 | 2 | 9 | 0 | 49 | 97 | 27 | 63 | 47 | 244 | 149 | 0.984332029 | 1 | 0.044 | 0.055 | 0.035 | 0 | 0.05 | 0 | 0.019 | 0.043 | 0.08 | 0 | 0.008 |
| RI | 4689 | ENSG00000175634 | RPS6KB2 | chr11 | + | 67433994 | 67434275 | 67433994 | 67434057 | 67434197 | 67434275 | 15 | 7 | 14 | 15 | 2 | 64 | 41 | 68 | 78 | 37 | 10 | 9 | 4 | 6 | 39 | 82 | 60 | 48 | 74 | 96 | 289 | 149 | 0.985343114 | 1 | 0.108 | 0.081 | 0.096 | 0.09 | 0.027 | 0.059 | 0.072 | 0.041 | 0.04 | 0.173 | 0.003 |
| RI | 4171 | ENSG00000165671 | NSD1 | chr5 | + | 177135086 | 177136030 | 177135086 | 177135200 | 177135940 | 177136030 | 143 | 120 | 131 | 151 | 154 | 93 | 70 | 114 | 96 | 107 | 167 | 220 | 180 | 282 | 97 | 117 | 114 | 134 | 187 | 87 | 298 | 149 | 0.987971556 | 1 | 0.435 | 0.462 | 0.365 | 0.44 | 0.418 | 0.416 | 0.491 | 0.402 | 0.43 | 0.358 | 0.005 |
| RI | 2112 | ENSG00000159753 | CARMIL2 | chr16 | + | 67651400 | 67651845 | 67651400 | 67651514 | 67651684 | 67651845 | 3 | 12 | 24 | 26 | 15 | 49 | 32 | 60 | 79 | 22 | 0 | 23 | 22 | 24 | 2 | 35 | 38 | 15 | 33 | 59 | 298 | 149 | 0.988531172 | 1 | 0.03 | 0.158 | 0.167 | 0.141 | 0.254 | 0 | 0.232 | 0.423 | 0.267 | 0.017 | -0.038 |
| RI | 4423 | ENSG00000111653 | ING4 | chr12 | - | 6652661 | 6653050 | 6652661 | 6652770 | 6652944 | 6653050 | 15 | 12 | 12 | 13 | 29 | 10 | 12 | 14 | 9 | 12 | 8 | 11 | 25 | 9 | 6 | 4 | 16 | 11 | 0 | 31 | 298 | 149 | 0.988803107 | 1 | 0.429 | 0.333 | 0.3 | 0.419 | 0.547 | 0.5 | 0.256 | 0.532 | 1 | 0.088 | -0.07 |
| RI | 432 | ENSG00000130726 | TRIM28 | chr19 | + | 58548739 | 58548910 | 58548739 | 58548776 | 58548861 | 58548910 | 52 | 37 | 59 | 56 | 55 | 134 | 53 | 115 | 125 | 136 | 31 | 34 | 33 | 34 | 37 | 74 | 83 | 63 | 61 | 81 | 234 | 149 | 0.989386378 | 1 | 0.198 | 0.308 | 0.246 | 0.222 | 0.205 | 0.211 | 0.207 | 0.25 | 0.262 | 0.225 | 0.005 |
| RI | 7397 | ENSG00000197070 | ARRDC1 | chr9 | + | 137613614 | 137614214 | 137613614 | 137613769 | 137614031 | 137614214 | 38 | 12 | 30 | 17 | 14 | 42 | 20 | 32 | 27 | 19 | 37 | 5 | 17 | 21 | 108 | 10 | 55 | 17 | 54 | 61 | 298 | 149 | 0.991146343 | 1 | 0.311 | 0.231 | 0.319 | 0.239 | 0.269 | 0.649 | 0.043 | 0.333 | 0.163 | 0.47 | -0.058 |
| RI | 2760 | ENSG00000185000 | DGAT1 | chr8 | - | 144317021 | 144317252 | 144317021 | 144317109 | 144317186 | 144317252 | 26 | 10 | 13 | 27 | 14 | 27 | 7 | 16 | 10 | 14 | 8 | 35 | 11 | 17 | 85 | 23 | 8 | 21 | 24 | 20 | 226 | 149 | 0.991194114 | 1 | 0.388 | 0.485 | 0.349 | 0.64 | 0.397 | 0.187 | 0.743 | 0.257 | 0.318 | 0.737 | 0.003 |
| RI | 3356 | ENSG00000213024 | NUP62 | chr19 | - | 49927693 | 49929504 | 49927693 | 49927815 | 49929325 | 49929504 | 25 | 23 | 44 | 60 | 51 | 21 | 17 | 17 | 21 | 16 | 142 | 42 | 49 | 50 | 127 | 50 | 33 | 37 | 12 | 55 | 298 | 149 | 0.991603314 | 1 | 0.373 | 0.404 | 0.564 | 0.588 | 0.614 | 0.587 | 0.389 | 0.398 | 0.676 | 0.536 | -0.009 |
| RI | 7938 | ENSG00000173992 | CCS | chr11 | + | 66599115 | 66599636 | 66599115 | 66599253 | 66599458 | 66599636 | 66 | 44 | 80 | 92 | 32 | 32 | 8 | 22 | 22 | 13 | 128 | 96 | 50 | 45 | 104 | 34 | 19 | 15 | 14 | 49 | 298 | 149 | 0.991823067 | 1 | 0.508 | 0.733 | 0.645 | 0.676 | 0.552 | 0.653 | 0.716 | 0.625 | 0.616 | 0.515 | -0.002 |
| RI | 6822 | ENSG00000068438 | FTSJ1 | chrX | + | 48482596 | 48483027 | 48482596 | 48482794 | 48482985 | 48483027 | 42 | 22 | 21 | 27 | 21 | 38 | 13 | 34 | 33 | 24 | 28 | 33 | 14 | 9 | 61 | 21 | 27 | 17 | 28 | 52 | 298 | 149 | 0.993033537 | 1 | 0.356 | 0.458 | 0.236 | 0.29 | 0.304 | 0.4 | 0.379 | 0.292 | 0.138 | 0.37 | 0.013 |
| RI | 4785 | ENSG00000162909 | CAPN2 | chr1 | + | 223769840 | 223770525 | 223769840 | 223769909 | 223770446 | 223770525 | 98 | 93 | 54 | 62 | 62 | 358 | 251 | 324 | 485 | 780 | 163 | 96 | 93 | 103 | 67 | 815 | 441 | 339 | 545 | 901 | 298 | 149 | 0.994748504 | 1 | 0.12 | 0.156 | 0.077 | 0.06 | 0.038 | 0.091 | 0.098 | 0.121 | 0.086 | 0.036 | 0.004 |
| RI | 7019 | ENSG00000142046 | TMEM91 | chr19 | + | 41382771 | 41384083 | 41382771 | 41382921 | 41383560 | 41384083 | 23 | 19 | 30 | 19 | 9 | 7 | 9 | 9 | 27 | 4 | 80 | 35 | 58 | 47 | 191 | 50 | 18 | 25 | 9 | 177 | 298 | 149 | 0.995995427 | 1 | 0.622 | 0.514 | 0.625 | 0.26 | 0.529 | 0.444 | 0.493 | 0.537 | 0.723 | 0.35 | 0.001 |
| RI | 1099 | ENSG00000125691 | RPL23 | chr17 | - | 38850080 | 38852732 | 38850080 | 38850214 | 38852603 | 38852732 | 14 | 16 | 31 | 29 | 32 | 4 | 2 | 13 | 4 | 6 | 42 | 29 | 22 | 33 | 20 | 11 | 3 | 5 | 9 | 6 | 298 | 149 | 0.996226462 | 1 | 0.636 | 0.8 | 0.544 | 0.784 | 0.727 | 0.656 | 0.829 | 0.688 | 0.647 | 0.625 | 0.009 |
| RI | 16 | ENSG00000265590 | CFAP298-TCP10L | chr21 | - | 32573283 | 32574124 | 32573283 | 32573368 | 32574095 | 32574124 | 0 | 0 | 0 | 0 | 0 | 2 | 0 | 0 | 0 | 0 | 0 | 1 | 0 | 0 | 0 | 0 | 0 | 0 | 0 | 0 | 298 | 149 | 1 | 1 | 0 | NA | NA | NA | NA | NA | 1 | NA | NA | NA | -1 |
| RI | 478 | ENSG00000186204 | CYP4F12 | chr19 | + | 15683492 | 15684882 | 15683492 | 15683763 | 15684815 | 15684882 | 0 | 0 | 0 | 0 | 0 | 0 | 0 | 0 | 1 | 0 | 0 | 2 | 2 | 3 | 0 | 0 | 0 | 0 | 0 | 0 | 298 | 149 | 1 | 1 | NA | NA | NA | 0 | NA | NA | 1 | | 1 | NA | -1 |
| RI | 595 | ENSG00000088340 | FER1L4 | chr20 | - | 35559155 | 35559627 | 35559155 | 35559256 | 35559340 | 35559627 | 0 | 0 | 0 | 0 | 0 | 0 | 0 | 1 | 0 | 0 | 0 | 0 | 1 | 0 | 0 | 0 | 0 | 0 | 0 | 0 | 233 | 149 | 1 | 1 | NA | NA | 0 | NA | NA | NA | NA | 1 | NA | NA | -1 |
| RI | 667 | ENSG00000115257 | PCSK4 | chr19 | - | 1486852 | 1487313 | 1486852 | 1487065 | 1487140 | 1487313 | 0 | 0 | 0 | 0 | 0 | 0 | 0 | 0 | 1 | 0 | 0 | 2 | 0 | 0 | 0 | 0 | 0 | 0 | 0 | 0 | 224 | 149 | 1 | 1 | NA | NA | NA | 0 | NA | NA | 1 | NA | NA | | -1 |
| RI | 1093 | ENSG00000091536 | MYO15A | chr17 | + | 18159274 | 18160017 | 18159274 | 18159347 | 18159934 | 18160017 | 0 | 0 | 0 | 0 | 0 | 1 | 0 | 0 | 0 | 0 | 0 | 0 | 0 | 1 | 0 | 0 | 0 | 0 | 0 | 0 | 298 | 149 | 1 | 1 | 0 | NA | NA | NA | NA | NA | NA | NA | 1 | NA | -1 |
| RI | 1124 | ENSG00000171962 | DRC3 | chr17 | + | 17992764 | 17994418 | 17992764 | 17992911 | 17994298 | 17994418 | 0 | 0 | 0 | 0 | 0 | 1 | 0 | 0 | 0 | 0 | 0 | 2 | 0 | 0 | 0 | 0 | 0 | 0 | 0 | 0 | 298 | 149 | 1 | 1 | 0 | NA | NA | NA | NA | NA | 1 | NA | NA | NA | -1 |
| RI | 1444 | ENSG00000175267 | VWA3A | chr16 | + | 22100193 | 22100493 | 22100193 | 22100318 | 22100415 | 22100493 | 0 | 0 | 0 | 0 | 0 | 0 | 0 | 1 | 0 | 0 | 0 | 0 | 2 | 5 | 4 | 0 | 0 | 0 | 0 | 0 | 246 | 149 | 1 | 1 | NA | NA | 0 | NA | NA | NA | NA | 1 | 1 | 1 | -1 |
| RI | 1541 | ENSG00000103723 | AP3B2 | chr15 | - | 82659844 | 82661922 | 82659844 | 82659983 | 82661824 | 82661922 | 0 | 0 | 0 | 0 | 0 | 0 | 0 | 0 | 0 | 1 | 2 | 0 | 0 | 1 | 0 | 0 | 0 | 0 | 0 | 0 | 298 | 149 | 1 | 1 | NA | NA | NA | NA | 0 | 1 | NA | NA | 1 | NA | -1 |
| RI | 2244 | ENSG00000183010 | PYCR1 | chr17 | - | 81936747 | 81937300 | 81936747 | 81936861 | 81937257 | 81937300 | 0 | 0 | 0 | 0 | 0 | 0 | 0 | 0 | 1 | 0 | 0 | 0 | 0 | 0 | 2 | 0 | 0 | 0 | 0 | 0 | 298 | 149 | 1 | 1 | NA | NA | NA | 0 | NA | NA | NA | NA | NA | 1 | -1 |
| RI | 2816 | ENSG00000163702 | IL17RC | chr3 | + | 9923880 | 9924291 | 9923880 | 9924137 | 9924231 | 9924291 | 0 | 0 | 0 | 0 | 0 | 0 | 3 | 0 | 0 | 0 | 0 | 0 | 0 | 2 | 0 | 0 | 0 | 0 | 0 | 0 | 243 | 149 | 1 | 1 | NA | 0 | NA | NA | NA | NA | NA | NA | 1 | NA | -1 |
| RI | 2830 | ENSG00000108641 | B9D1 | chr17 | - | 19343056 | 19343857 | 19343056 | 19343461 | 19343789 | 19343857 | 0 | 0 | 0 | 0 | 0 | 0 | 0 | 1 | 0 | 0 | 0 | 2 | 2 | 0 | 6 | 0 | 0 | 0 | 0 | 0 | 298 | 149 | 1 | 1 | NA | NA | 0 | NA | NA | NA | 1 | 1 | NA | 1 | -1 |
| RI | 3231 | ENSG00000169908 | TM4SF1 | chr3 | - | 149375442 | 149375769 | 149375442 | 149375588 | 149375679 | 149375769 | 0 | 0 | 0 | 0 | 0 | 4 | 0 | 0 | 0 | 0 | 0 | 0 | 0 | 0 | 1 | 0 | 0 | 0 | 0 | 0 | 240 | 149 | 1 | 1 | 0 | NA | NA | NA | NA | | NA | NA | NA | 1 | -1 |
| RI | 4067 | ENSG00000143416 | SELENBP1 | chr1 | - | 151369003 | 151369769 | 151369003 | 151369554 | 151369712 | 151369769 | 0 | 0 | 0 | 0 | 0 | 0 | 1 | 0 | 1 | 2 | 0 | 0 | 1 | 2 | 0 | 0 | 0 | 0 | 0 | 0 | 298 | 149 | 1 | 1 | NA | 0 | NA | 0 | 0 | NA | NA | 1 | 1 | NA | -1 |
| RI | 4316 | ENSG00000162572 | SCNN1D | chr1 | + | 1287507 | 1287836 | 1287507 | 1287596 | 1287672 | 1287836 | 0 | 0 | 0 | 0 | 0 | 2 | 0 | 0 | 0 | 0 | 0 | 6 | 0 | 0 | 0 | 0 | 0 | 0 | 0 | 0 | 225 | 149 | 1 | 1 | 0 | NA | | NA | NA | NA | 1 | NA | NA | NA | -1 |
| RI | 4517 | ENSG00000143126 | CELSR2 | chr1 | + | 109271216 | 109271722 | 109271216 | 109271296 | 109271385 | 109271722 | 0 | 0 | 0 | 0 | 0 | 2 | 2 | 1 | 0 | 3 | 7 | 0 | 0 | 0 | 0 | 0 | 0 | 0 | 0 | 0 | 238 | 149 | 1 | 1 | 0 | 0 | 0 | NA | 0 | 1 | NA | NA | NA | NA | -1 |
| RI | 5012 | ENSG00000076706 | MCAM | chr11 | - | 119310755 | 119311185 | 119310755 | 119310903 | 119311089 | 119311185 | 0 | 0 | 0 | 0 | 0 | 0 | 0 | 0 | 2 | 0 | 2 | 3 | 1 | 0 | 0 | 0 | 0 | 0 | 0 | 0 | 298 | 149 | 1 | 1 | NA | NA | NA | 0 | NA | 1 | 1 | 1 | NA | NA | -1 |
| RI | 5039 | ENSG00000156113 | KCNMA1 | chr10 | - | 76909965 | 76915049 | 76909965 | 76910096 | 76914935 | 76915049 | 0 | 0 | 0 | 0 | 0 | 0 | 3 | 0 | 2 | 0 | 1 | 0 | 0 | 0 | 2 | 0 | 0 | 0 | 0 | 0 | 298 | 149 | 1 | 1 | NA | 0 | NA | NA | NA | 1 | NA | NA | NA | 1 | -1 |
| RI | 5359 | ENSG00000186567 | CEACAM19 | chr19 | + | 44682566 | 44684359 | 44682566 | 44682620 | 44683433 | 44684359 | 0 | 0 | 0 | 0 | 0 | 0 | 0 | 0 | 0 | 1 | 1 | 0 | 0 | 0 | 0 | 0 | 0 | 0 | 0 | 0 | 298 | 149 | 1 | 1 | NA | NA | NA | NA | 0 | 1 | NA | NA | NA | NA | -1 |
| RI | 6382 | ENSG00000179698 | WDR97 | chr8 | + | 144111100 | 144111781 | 144111100 | 144111486 | 144111631 | 144111781 | 0 | 0 | 0 | 0 | 0 | 0 | 0 | 0 | 1 | 0 | 0 | 2 | 1 | 0 | 9 | 0 | 0 | 0 | 0 | 0 | 294 | 149 | 1 | 1 | NA | NA | NA | 0 | NA | NA | 1 | 1 | NA | 1 | -1 |
| RI | 6605 | ENSG00000055118 | KCNH2 | chr7 | - | 150950920 | 150951835 | 150950920 | 150951120 | 150951447 | 150951835 | 0 | 0 | 0 | 0 | 0 | 0 | 2 | 2 | 1 | 4 | 0 | 0 | 2 | 0 | 0 | 0 | 0 | 0 | 0 | 0 | 298 | 149 | 1 | 1 | NA | 0 | 0 | 0 | 0 | NA | NA | 1 | NA | NA | -1 |
| RI | 7076 | ENSG00000147852 | VLDLR | chr9 | + | 2651873 | 2652949 | 2651873 | 2651954 | 2652779 | 2652949 | 0 | 0 | 0 | 0 | 0 | 0 | 0 | 0 | 3 | 0 | 0 | 0 | 1 | 0 | 0 | 0 | 0 | 0 | 0 | 0 | 298 | 149 | 1 | 1 | NA | NA | NA | 0 | NA | NA | NA | 1 | NA | NA | -1 |
| RI | 8113 | ENSG00000225177 | NA | chr6 | + | 138696596 | 138697288 | 138696596 | 138696755 | 138696902 | 138697288 | 0 | 0 | 0 | 0 | 0 | 0 | 0 | 0 | 0 | 3 | 0 | 0 | 0 | 0 | 12 | 0 | 0 | 0 | 0 | 0 | 296 | 149 | 1 | 1 | NA | NA | NA | NA | 0 | NA | NA | NA | NA | 1 | -1 |
| RI | 8248 | ENSG00000186081 | KRT5 | chr12 | - | 52517896 | 52518163 | 52517896 | 52517992 | 52518102 | 52518163 | 1 | 0 | 1 | 0 | 0 | 9 | 8 | 3 | 9 | 3 | 0 | 0 | 4 | 0 | 0 | 0 | 0 | 0 | 0 | 0 | 259 | 149 | 1 | 1 | 0.06 | 0 | 0.161 | 0 | 0 | NA | NA | 1 | NA | NA | -0.956 |
| RI | 6857 | ENSG00000102038 | SMARCA1 | chrX | - | 129480700 | 129481185 | 129480700 | 129480814 | 129481074 | 129481185 | 0 | 0 | 0 | 3 | 0 | 1 | 2 | 2 | 5 | 3 | 0 | 0 | 0 | 1 | 0 | 0 | 0 | 0 | 0 | 0 | 298 | 149 | 1 | 1 | 0 | 0 | 0 | 0.231 | 0 | NA | NA | NA | 1 | NA | -0.954 |
| RI | 5542 | ENSG00000160766 | GBAP1 | chr1 | - | 155216383 | 155217029 | 155216383 | 155216621 | 155216856 | 155217029 | 2 | 0 | 0 | 0 | 0 | 2 | 1 | 0 | 6 | 10 | 0 | 0 | 0 | 10 | 0 | 0 | 0 | 0 | 0 | 0 | 298 | 149 | 1 | 1 | 0.333 | 0 | NA | 0 | 0 | NA | NA | NA | 1 | NA | -0.917 |
| RI | 4682 | ENSG00000186827 | TNFRSF4 | chr1 | - | 1211339 | 1211832 | 1211339 | 1211625 | 1211703 | 1211832 | 3 | 0 | 0 | 0 | 0 | 3 | 1 | 4 | 1 | 0 | 0 | 0 | 0 | 1 | 0 | 0 | 0 | 0 | 0 | 0 | 227 | 149 | 1 | 1 | 0.396 | 0 | 0 | 0 | NA | NA | NA | NA | 1 | NA | -0.901 |
| RI | 2785 | ENSG00000163913 | IFT122 | chr3 | + | 129514388 | 129515599 | 129514388 | 129514554 | 129515487 | 129515599 | 1 | 0 | 4 | 2 | 0 | 2 | 3 | 7 | 8 | 6 | 0 | 1 | 8 | 1 | 0 | 0 | 0 | 0 | 0 | 0 | 298 | 149 | 1 | 1 | 0.2 | 0 | 0.222 | 0.111 | 0 | NA | 1 | 1 | 1 | NA | -0.893 |
| RI | 4097 | ENSG00000256269 | HMBS | chr11 | + | 119088634 | 119089131 | 119088634 | 119088848 | 119089081 | 119089131 | 0 | 0 | 0 | 0 | 2 | 0 | 0 | 0 | 0 | 8 | 0 | 0 | 0 | 0 | 3 | 0 | 0 | 0 | 0 | 0 | 298 | 149 | 1 | 1 | NA | NA | NA | NA | 0.111 | NA | NA | NA | NA | 1 | -0.889 |
| RI | 4250 | ENSG00000081041 | CXCL2 | chr4 | - | 74098798 | 74099195 | 74098798 | 74098922 | 74099020 | 74099195 | 0 | 0 | 0 | 1 | 0 | 3 | 0 | 0 | 2 | 0 | 2 | 0 | 0 | 0 | 0 | 0 | 0 | 0 | 0 | 0 | 247 | 149 | 1 | 1 | 0 | NA | NA | 0.232 | NA | 1 | NA | NA | NA | NA | -0.884 |
| RI | 5736 | ENSG00000072195 | SPEG | chr2 | + | 219464432 | 219466861 | 219464432 | 219464608 | 219466056 | 219466861 | 0 | 0 | 3 | 2 | 3 | 3 | 1 | 2 | 7 | 5 | 0 | 0 | 0 | 1 | 0 | 0 | 0 | 0 | 0 | 0 | 298 | 149 | 1 | 1 | 0 | 0 | 0.429 | 0.125 | 0.231 | NA | NA | NA | 1 | NA | -0.843 |
| RI | 5324 | ENSG00000237037 | NDUFA6-DT | chr22 | + | 42124046 | 42124562 | 42124046 | 42124130 | 42124519 | 42124562 | 2 | 0 | 0 | 0 | 0 | 1 | 1 | 0 | 0 | 6 | 0 | 0 | 0 | 3 | 0 | 0 | 0 | 0 | 0 | 0 | 298 | 149 | 1 | 1 | 0.5 | 0 | NA | NA | 0 | NA | NA | NA | 1 | NA | -0.833 |
| RI | 6386 | ENSG00000142303 | ADAMTS10 | chr19 | - | 8585131 | 8585660 | 8585131 | 8585308 | 8585455 | 8585660 | 4 | 1 | 0 | 2 | 0 | 10 | 1 | 5 | 3 | 0 | 0 | 0 | 4 | 0 | 0 | 0 | 0 | 0 | 0 | 0 | 296 | 149 | 1 | 1 | 0.168 | 0.335 | 0 | 0.251 | NA | NA | NA | 1 | NA | NA | -0.811 |
| RI | 1552 | ENSG00000103723 | AP3B2 | chr15 | - | 82688735 | 82689232 | 82688735 | 82688831 | 82689157 | 82689232 | 0 | 0 | 0 | 1 | 0 | 0 | 0 | 0 | 2 | 0 | 0 | 0 | 0 | 0 | 9 | 0 | 0 | 0 | 0 | 0 | 298 | 149 | 1 | 1 | NA | NA | NA | 0.2 | NA | NA | NA | NA | NA | 1 | -0.8 |
| RI | 7741 | ENSG00000065717 | TLE2 | chr19 | - | 3019282 | 3019773 | 3019282 | 3019463 | 3019698 | 3019773 | 0 | 4 | 0 | 2 | 14 | 0 | 4 | 6 | 5 | 12 | 0 | 3 | 0 | 0 | 0 | 0 | 0 | 0 | 0 | 0 | 298 | 149 | 1 | 1 | NA | 0.333 | 0 | 0.167 | 0.368 | NA | 1 | NA | NA | NA | -0.783 |
| RI | 7642 | ENSG00000116774 | OLFML3 | chr1 | + | 113980331 | 113982254 | 113980331 | 113980617 | 113980948 | 113982254 | 0 | 0 | 4 | 0 | 0 | 2 | 0 | 1 | 2 | 0 | 0 | 0 | 0 | 1 | 0 | 0 | 0 | 0 | 0 | 0 | 298 | 149 | 1 | 1 | 0 | NA | 0.667 | 0 | NA | NA | NA | NA | 1 | NA | -0.778 |
| RI | 4087 | ENSG00000222009 | BTBD19 | chr1 | + | 44812995 | 44813269 | 44812995 | 44813064 | 44813137 | 44813269 | 0 | 0 | 0 | 2 | 0 | 0 | 0 | 0 | 4 | 0 | 2 | 0 | 3 | 0 | 25 | 0 | 0 | 0 | 0 | 0 | 222 | 149 | 1 | 1 | NA | NA | NA | 0.251 | NA | 1 | NA | 1 | NA | 1 | -0.749 |
| RI | 321 | ENSG00000105655 | ISYNA1 | chr19 | - | 18437598 | 18437988 | 18437598 | 18437760 | 18437859 | 18437988 | 0 | 0 | 4 | 1 | 0 | 2 | 0 | 2 | 2 | 0 | 0 | 0 | 0 | 1 | 0 | 0 | 0 | 0 | 0 | 0 | 248 | 149 | 1 | 1 | 0 | NA | 0.546 | 0.231 | NA | NA | NA | NA | 1 | NA | -0.741 |
| RI | 2128 | ENSG00000154134 | ROBO3 | chr11 | + | 124877160 | 124877309 | 124877160 | 124877184 | 124877266 | 124877309 | 0 | 0 | 5 | 4 | 2 | 2 | 0 | 10 | 3 | 2 | 0 | 3 | 0 | 1 | 2 | 0 | 0 | 0 | 0 | 0 | 231 | 149 | 1 | 1 | 0 | NA | 0.244 | 0.462 | 0.392 | NA | 1 | NA | 1 | 1 | -0.726 |
| RI | 2685 | ENSG00000020577 | SAMD4A | chr14 | + | 54784339 | 54784620 | 54784339 | 54784447 | 54784536 | 54784620 | 0 | 4 | 2 | 0 | 0 | 4 | 0 | 2 | 4 | 3 | 0 | 0 | 2 | 0 | 10 | 0 | 0 | 0 | 0 | 0 | 238 | 149 | 1 | 1 | 0 | 1 | 0.385 | 0 | 0 | NA | NA | 1 | NA | 1 | -0.723 |
| RI | 1871 | ENSG00000105122 | RASAL3 | chr19 | - | 15451623 | 15452815 | 15451623 | 15451938 | 15452657 | 15452815 | 3 | 2 | 3 | 0 | 1 | 2 | 2 | 1 | 3 | 8 | 0 | 4 | 8 | 12 | 12 | 0 | 0 | 0 | 0 | 0 | 298 | 149 | 1 | 1 | 0.429 | 0.333 | 0.6 | 0 | 0.059 | NA | 1 | 1 | 1 | 1 | -0.716 |
| RI | 115 | ENSG00000100124 | ANKRD54 | chr22 | - | 37832957 | 37833206 | 37832957 | 37833082 | 37833158 | 37833206 | 2 | 3 | 1 | 9 | 0 | 5 | 3 | 6 | 2 | 11 | 5 | 17 | 7 | 7 | 0 | 0 | 0 | 0 | 0 | 0 | 225 | 149 | 1 | 1 | 0.209 | 0.398 | 0.099 | 0.749 | 0 | 1 | 1 | 1 | 1 | NA | -0.709 |
| RI | 225 | ENSG00000167733 | HSD11B1L | chr19 | + | 5684818 | 5685119 | 5684818 | 5684905 | 5684988 | 5685119 | 0 | 2 | 0 | 1 | 0 | 3 | 0 | 1 | 3 | 0 | 10 | 1 | 3 | 3 | 0 | 0 | 0 | 0 | 0 | 0 | 232 | 149 | 1 | 1 | 0 | 1 | 0 | 0.176 | NA | 1 | 1 | 1 | 1 | NA | -0.706 |
| RI | 398 | ENSG00000130244 | FAM98C | chr19 | + | 38403337 | 38403694 | 38403337 | 38403486 | 38403559 | 38403694 | 0 | 0 | 0 | 0 | 0 | 1 | 3 | 2 | 2 | 0 | 0 | 0 | 0 | 7 | 0 | 0 | 0 | 0 | 2 | 0 | 222 | 149 | 1 | 1 | 0 | 0 | 0 | 0 | NA | NA | NA | NA | 0.701 | NA | -0.701 |
| RI | 5047 | ENSG00000197841 | ZNF181 | chr19 | + | 34739147 | 34739621 | 34739147 | 34739268 | 34739525 | 34739621 | 8 | 1 | 14 | 16 | 8 | 30 | 5 | 27 | 20 | 16 | 0 | 21 | 13 | 4 | 1 | 0 | 1 | 5 | 0 | 0 | 298 | 149 | 1 | 1 | 0.118 | 0.091 | 0.206 | 0.286 | 0.2 | NA | 0.913 | 0.565 | 1 | 1 | -0.689 |
| RI | 33 | ENSG00000275895 | U2AF1L5 | chr21 | - | 6492130 | 6496080 | 6492130 | 6492197 | 6495992 | 6496080 | 0 | 0 | 1 | 0 | 0 | 0 | 0 | 0 | 1 | 1 | 0 | 0 | 0 | 14 | 0 | 0 | 0 | 0 | 0 | 0 | 298 | 149 | 1 | 1 | NA | NA | 1 | 0 | 0 | NA | NA | NA | 1 | NA | -0.667 |
| RI | 1216 | ENSG00000166743 | ACSM1 | chr16 | - | 20636738 | 20637451 | 20636738 | 20636840 | 20637370 | 20637451 | 0 | 0 | 0 | 0 | 0 | 0 | 2 | 0 | 1 | 0 | 0 | 1 | 2 | 0 | 0 | 0 | 0 | 0 | 0 | 8 | 298 | 149 | 1 | 1 | NA | 0 | NA | 0 | NA | NA | 1 | 1 | NA | 0 | -0.667 |
| RI | 1227 | ENSG00000103034 | NDRG4 | chr16 | + | 58507807 | 58507999 | 58507807 | 58507864 | 58507947 | 58507999 | 0 | 1 | 0 | 0 | 0 | 2 | 0 | 0 | 1 | 0 | 0 | 2 | 2 | 0 | 0 | 0 | 0 | 0 | 0 | 0 | 232 | 149 | 1 | 1 | 0 | 1 | NA | 0 | NA | NA | 1 | 1 | NA | NA | -0.667 |
| RI | 1446 | ENSG00000174628 | IQCK | chr16 | + | 19763847 | 19764112 | 19763847 | 19763900 | 19764034 | 19764112 | 0 | 0 | 5 | 0 | 0 | 3 | 1 | 0 | 0 | 0 | 0 | 5 | 0 | 0 | 0 | 0 | 0 | 0 | 0 | 0 | 283 | 149 | 1 | 1 | 0 | 0 | 1 | NA | NA | NA | 1 | NA | NA | NA | -0.667 |
| RI | 2624 | ENSG00000185684 | EP400P1 | chr12 | + | 132103996 | 132105304 | 132103996 | 132104170 | 132104377 | 132105304 | 0 | 0 | 0 | 0 | 4 | 6 | 0 | 0 | 1 | 0 | 0 | 0 | 0 | 4 | 0 | 0 | 0 | 0 | 0 | 0 | 298 | 149 | 1 | 1 | 0 | NA | NA | 0 | 1 | NA | NA | NA | 1 | NA | -0.667 |
| RI | 3799 | ENSG00000065361 | ERBB3 | chr12 | + | 56094098 | 56094556 | 56094098 | 56094189 | 56094401 | 56094556 | 0 | 0 | 1 | 0 | 0 | 0 | 0 | 1 | 0 | 0 | 1 | 0 | 2 | 0 | 2 | 0 | 0 | 0 | 0 | 0 | 298 | 149 | 1 | 1 | NA | NA | 0.333 | NA | NA | 1 | NA | 1 | NA | 1 | -0.667 |
| RI | 4543 | ENSG00000089685 | BIRC5 | chr17 | + | 78215965 | 78216781 | 78215965 | 78216034 | 78216663 | 78216781 | 1 | 0 | 0 | 0 | 0 | 0 | 0 | 1 | 1 | 0 | 0 | 0 | 0 | 2 | 0 | 0 | 0 | 0 | 0 | 0 | 298 | 149 | 1 | 1 | 1 | NA | 0 | 0 | NA | NA | NA | NA | 1 | NA | -0.667 |
| RI | 4604 | ENSG00000037280 | FLT4 | chr5 | - | 180619252 | 180619769 | 180619252 | 180619366 | 180619664 | 180619769 | 0 | 0 | 0 | 10 | 0 | 1 | 0 | 1 | 0 | 0 | 0 | 1 | 1 | 0 | 0 | 0 | 0 | 0 | 0 | 0 | 298 | 149 | 1 | 1 | 0 | NA | 0 | 1 | NA | NA | 1 | 1 | NA | NA | -0.667 |
| RI | 4934 | ENSG00000119888 | EPCAM | chr2 | + | 47373462 | 47374048 | 47373462 | 47373570 | 47373807 | 47374048 | 0 | 0 | 0 | 0 | 0 | 0 | 0 | 1 | 0 | 0 | 0 | 1 | 0 | 0 | 1 | 0 | 0 | 0 | 6 | 0 | 298 | 149 | 1 | 1 | NA | NA | 0 | NA | NA | NA | 1 | NA | 0 | 1 | -0.667 |
| RI | 747 | ENSG00000161513 | FDXR | chr17 | - | 74863895 | 74864347 | 74863895 | 74864067 | 74864147 | 74864347 | 1 | 0 | 0 | 0 | 0 | 4 | 0 | 7 | 8 | 9 | 9 | 4 | 0 | 3 | 4 | 0 | 0 | 3 | 0 | 3 | 229 | 149 | 1 | 1 | 0.14 | NA | 0 | 0 | 0 | 1 | 1 | 0 | 1 | 0.465 | -0.658 |
| RI | 3985 | ENSG00000188004 | SNHG28 | chr1 | - | 159837916 | 159841235 | 159837916 | 159840866 | 159841125 | 159841235 | 1 | 2 | 0 | 0 | 0 | 5 | 0 | 0 | 1 | 0 | 0 | 0 | 2 | 0 | 2 | 0 | 0 | 0 | 0 | 0 | 298 | 149 | 1 | 1 | 0.091 | 1 | NA | 0 | NA | NA | NA | 1 | NA | 1 | -0.636 |
| RI | 8337 | ENSG00000144061 | NPHP1 | chr2 | - | 110163047 | 110164730 | 110163047 | 110163132 | 110164687 | 110164730 | 1 | 0 | 0 | 4 | 0 | 4 | 0 | 0 | 0 | 2 | 0 | 0 | 0 | 1 | 0 | 0 | 0 | 0 | 0 | 0 | 298 | 149 | 1 | 1 | 0.111 | NA | NA | 1 | 0 | NA | NA | NA | 1 | NA | -0.63 |
| RI | 3800 | ENSG00000065361 | ERBB3 | chr12 | + | 56094098 | 56095310 | 56094098 | 56094556 | 56095256 | 56095310 | 0 | 0 | 0 | 0 | 0 | 0 | 0 | 0 | 2 | 0 | 1 | 0 | 2 | 0 | 3 | 0 | 9 | 1 | 0 | 0 | 298 | 149 | 1 | 1 | NA | NA | NA | 0 | NA | 1 | 0 | 0.5 | NA | 1 | -0.625 |
| RI | 7527 | ENSG00000175287 | PHYHD1 | chr9 | + | 128941444 | 128942038 | 128941444 | 128941571 | 128941667 | 128942038 | 1 | 0 | 1 | 3 | 6 | 1 | 1 | 2 | 1 | 2 | 0 | 0 | 3 | 0 | 10 | 0 | 0 | 0 | 0 | 0 | 245 | 149 | 1 | 1 | 0.378 | 0 | 0.233 | 0.646 | 0.646 | NA | NA | 1 | NA | 1 | -0.619 |
| RI | 8575 | ENSG00000225579 | EDNRB-AS1 | chr13 | + | 77833364 | 77834990 | 77833364 | 77833427 | 77833732 | 77834990 | 2 | 9 | 1 | 2 | 0 | 2 | 0 | 1 | 3 | 2 | 0 | 1 | 7 | 0 | 0 | 0 | 0 | 0 | 0 | 0 | 298 | 149 | 1 | 1 | 0.333 | 1 | 0.333 | 0.25 | 0 | NA | 1 | 1 | NA | NA | -0.617 |
| RI | 11 | ENSG00000154734 | ADAMTS1 | chr21 | - | 26839874 | 26840562 | 26839874 | 26840061 | 26840275 | 26840562 | 3 | 1 | 4 | 3 | 0 | 10 | 3 | 2 | 13 | 0 | 2 | 0 | 4 | 1 | 0 | 0 | 0 | 2 | 0 | 0 | 298 | 149 | 1 | 1 | 0.13 | 0.143 | 0.5 | 0.103 | NA | 1 | NA | 0.5 | 1 | NA | -0.614 |
| RI | 4299 | ENSG00000007402 | CACNA2D2 | chr3 | - | 50364887 | 50365184 | 50364887 | 50364970 | 50365074 | 50365184 | 3 | 1 | 1 | 0 | 0 | 24 | 6 | 4 | 3 | 9 | 0 | 0 | 0 | 2 | 7 | 0 | 3 | 0 | 0 | 0 | 253 | 149 | 1 | 1 | 0.069 | 0.089 | 0.128 | 0 | 0 | NA | 0 | NA | 1 | 1 | -0.609 |
| RI | 8210 | ENSG00000142920 | AZIN2 | chr1 | + | 33117901 | 33120530 | 33117901 | 33118116 | 33120043 | 33120530 | 1 | 2 | 7 | 1 | 4 | 2 | 0 | 5 | 3 | 8 | 0 | 3 | 6 | 1 | 1 | 0 | 0 | 0 | 0 | 0 | 298 | 149 | 1 | 1 | 0.2 | 1 | 0.412 | 0.143 | 0.2 | NA | 1 | 1 | 1 | 1 | -0.609 |
| RI | 3004 | ENSG00000114859 | CLCN2 | chr3 | - | 184355693 | 184357094 | 184355693 | 184355778 | 184356992 | 184357094 | 0 | 0 | 2 | 1 | 0 | 4 | 1 | 5 | 4 | 1 | 0 | 6 | 5 | 4 | 8 | 0 | 0 | 5 | 0 | 9 | 298 | 149 | 1 | 1 | 0 | 0 | 0.167 | 0.111 | 0 | NA | 1 | 0.333 | 1 | 0.308 | -0.605 |
| RI | 1190 | ENSG00000162004 | CCDC78 | chr16 | - | 725077 | 725293 | 725077 | 725145 | 725236 | 725293 | 3 | 0 | 4 | 0 | 0 | 0 | 1 | 0 | 1 | 5 | 0 | 0 | 0 | 8 | 0 | 0 | 0 | 0 | 0 | 0 | 240 | 149 | 1 | 1 | 1 | 0 | 1 | 0 | 0 | NA | NA | NA | 1 | NA | -0.6 |
| RI | 5046 | ENSG00000197841 | ZNF181 | chr19 | + | 34739147 | 34739621 | 34739147 | 34739268 | 34739522 | 34739621 | 9 | 1 | 14 | 15 | 4 | 27 | 10 | 19 | 13 | 31 | 0 | 17 | 13 | 4 | 1 | 0 | 13 | 3 | 0 | 0 | 298 | 149 | 1 | 1 | 0.143 | 0.048 | 0.269 | 0.366 | 0.061 | NA | 0.395 | 0.684 | 1 | 1 | -0.592 |
| RI | 7629 | ENSG00000172927 | MYEOV | chr11 | + | 69295232 | 69297287 | 69295232 | 69295495 | 69295591 | 69297287 | 0 | 0 | 0 | 8 | 0 | 0 | 2 | 0 | 1 | 0 | 15 | 0 | 2 | 0 | 25 | 0 | 0 | 0 | 0 | 0 | 245 | 149 | 1 | 1 | NA | 0 | NA | 0.83 | NA | 1 | NA | 1 | NA | 1 | -0.585 |
| RI | 8084 | ENSG00000235823 | OLMALINC | chr10 | + | 100381215 | 100383372 | 100381215 | 100381448 | 100382427 | 100383372 | 2 | 1 | 1 | 0 | 0 | 0 | 1 | 1 | 0 | 5 | 0 | 1 | 0 | 5 | 0 | 0 | 0 | 0 | 0 | 0 | 298 | 149 | 1 | 1 | 1 | 0.333 | 0.333 | NA | 0 | NA | 1 | NA | 1 | NA | -0.584 |
| RI | 1817 | ENSG00000196497 | IPO4 | chr14 | - | 24188343 | 24188638 | 24188343 | 24188423 | 24188551 | 24188638 | 0 | 0 | 0 | 0 | 0 | 5 | 2 | 3 | 1 | 0 | 0 | 6 | 4 | 6 | 0 | 0 | 6 | 0 | 5 | 0 | 277 | 149 | 1 | 1 | 0 | 0 | 0 | 0 | NA | NA | 0.35 | 1 | 0.392 | NA | -0.581 |
| RI | 5870 | ENSG00000072163 | LIMS2 | chr2 | - | 127640269 | 127640988 | 127640269 | 127640318 | 127640895 | 127640988 | 1 | 0 | 1 | 7 | 3 | 0 | 1 | 2 | 6 | 1 | 0 | 0 | 9 | 0 | 0 | 0 | 3 | 0 | 0 | 0 | 298 | 149 | 1 | 1 | 1 | 0 | 0.2 | 0.368 | 0.6 | NA | NA | 1 | NA | NA | -0.566 |
| RI | 5969 | ENSG00000226979 | LTA | chr6 | + | 31572733 | 31573033 | 31572733 | 31572841 | 31572927 | 31573033 | 0 | 0 | 3 | 0 | 0 | 12 | 4 | 1 | 3 | 0 | 0 | 1 | 2 | 8 | 0 | 0 | 3 | 0 | 0 | 0 | 235 | 149 | 1 | 1 | 0 | 0 | 0.655 | 0 | NA | NA | 0.174 | 1 | 1 | NA | -0.561 |
| RI | 1759 | ENSG00000101384 | JAG1 | chr20 | - | 10650246 | 10651694 | 10650246 | 10650360 | 10651580 | 10651694 | 0 | 0 | 0 | 1 | 0 | 6 | 0 | 2 | 2 | 0 | 2 | 10 | 0 | 8 | 6 | 2 | 0 | 4 | 1 | 0 | 298 | 149 | 1 | 1 | 0 | NA | 0 | 0.2 | NA | 0.333 | 1 | 0 | 0.8 | 1 | -0.56 |
| RI | 5158 | ENSG00000146263 | MMS22L | chr6 | - | 97269901 | 97272881 | 97269901 | 97269992 | 97272703 | 97272881 | 8 | 3 | 13 | 7 | 0 | 24 | 9 | 21 | 11 | 30 | 0 | 27 | 3 | 7 | 0 | 0 | 12 | 1 | 0 | 0 | 298 | 149 | 1 | 1 | 0.143 | 0.143 | 0.236 | 0.241 | 0 | NA | 0.529 | 0.6 | 1 | NA | -0.557 |
| RI | 4324 | ENSG00000103253 | HAGHL | chr16 | + | 729008 | 729715 | 729008 | 729088 | 729287 | 729715 | 2 | 0 | 7 | 5 | 13 | 10 | 0 | 5 | 4 | 8 | 4 | 15 | 0 | 4 | 0 | 0 | 0 | 0 | 1 | 0 | 298 | 149 | 1 | 1 | 0.091 | NA | 0.412 | 0.385 | 0.448 | 1 | 1 | NA | 0.667 | NA | -0.555 |
| RI | 2838 | ENSG00000110844 | PRPF40B | chr12 | + | 49633013 | 49633547 | 49633013 | 49633124 | 49633426 | 49633547 | 5 | 2 | 2 | 6 | 5 | 4 | 9 | 12 | 4 | 2 | 16 | 7 | 4 | 0 | 22 | 0 | 0 | 1 | 0 | 3 | 298 | 149 | 1 | 1 | 0.385 | 0.1 | 0.077 | 0.429 | 0.556 | 1 | 1 | 0.667 | NA | 0.786 | -0.554 |
| RI | 6971 | ENSG00000187325 | TAF9B | chrX | - | 78137748 | 78138098 | 78137748 | 78137883 | 78137961 | 78138098 | 11 | 4 | 16 | 12 | 9 | 12 | 6 | 14 | 16 | 25 | 6 | 32 | 18 | 23 | 0 | 0 | 0 | 11 | 0 | 0 | 227 | 149 | 1 | 1 | 0.376 | 0.304 | 0.429 | 0.33 | 0.191 | 1 | 1 | 0.518 | 1 | NA | -0.553 |
| RI | 3593 | ENSG00000159214 | CCDC24 | chr1 | + | 43993889 | 43995162 | 43993889 | 43993964 | 43995107 | 43995162 | 0 | 1 | 2 | 2 | 2 | 1 | 4 | 5 | 3 | 10 | 12 | 6 | 1 | 2 | 0 | 0 | 0 | 1 | 2 | 0 | 298 | 149 | 1 | 1 | 0 | 0.111 | 0.167 | 0.25 | 0.091 | 1 | 1 | 0.333 | 0.333 | NA | -0.543 |
| RI | 8463 | ENSG00000122966 | CIT | chr12 | - | 119710539 | 119712347 | 119710539 | 119710620 | 119712177 | 119712347 | 0 | 1 | 0 | 0 | 0 | 1 | 1 | 1 | 1 | 2 | 0 | 3 | 1 | 17 | 0 | 0 | 0 | 0 | 11 | 12 | 298 | 149 | 1 | 1 | 0 | 0.333 | 0 | 0 | 0 | NA | 1 | 1 | 0.436 | 0 | -0.542 |
| RI | 6602 | ENSG00000188878 | FBF1 | chr17 | - | 75920006 | 75920429 | 75920006 | 75920107 | 75920273 | 75920429 | 4 | 2 | 0 | 3 | 0 | 2 | 4 | 4 | 8 | 0 | 6 | 0 | 0 | 9 | 6 | 0 | 0 | 1 | 0 | 0 | 298 | 149 | 1 | 1 | 0.5 | 0.2 | 0 | 0.158 | NA | 1 | NA | 0 | 1 | 1 | -0.535 |
| RI | 4770 | ENSG00000075702 | WDR62 | chr19 | + | 36101213 | 36102851 | 36101213 | 36101317 | 36101663 | 36102851 | 1 | 0 | 1 | 0 | 0 | 1 | 1 | 2 | 1 | 0 | 0 | 5 | 0 | 0 | 4 | 12 | 0 | 0 | 0 | 0 | 298 | 149 | 1 | 1 | 0.333 | 0 | 0.2 | 0 | NA | 0 | 1 | NA | NA | 1 | -0.533 |
| RI | 1207 | ENSG00000242612 | DECR2 | chr16 | + | 406684 | 406952 | 406684 | 406833 | 406889 | 406952 | 1 | 1 | 0 | 1 | 0 | 1 | 0 | 2 | 9 | 0 | 0 | 0 | 0 | 10 | 11 | 0 | 0 | 0 | 0 | 2 | 205 | 149 | 1 | 1 | 0.421 | 1 | 0 | 0.075 | NA | NA | NA | NA | 1 | 0.8 | -0.526 |
| RI | 4639 | ENSG00000197580 | BCO2 | chr11 | + | 112193473 | 112193994 | 112193473 | 112193697 | 112193878 | 112193994 | 6 | 7 | 1 | 4 | 0 | 2 | 2 | 3 | 0 | 1 | 0 | 2 | 5 | 0 | 0 | 0 | 0 | 0 | 0 | 0 | 298 | 149 | 1 | 1 | 0.6 | 0.636 | 0.143 | 1 | 0 | NA | 1 | 1 | NA | NA | -0.524 |
| RI | 1653 | ENSG00000255571 | MIR9-3HG | chr15 | + | 89387430 | 89387881 | 89387430 | 89387538 | 89387816 | 89387881 | 2 | 3 | 0 | 2 | 0 | 4 | 1 | 0 | 1 | 0 | 0 | 4 | 8 | 3 | 4 | 0 | 0 | 1 | 0 | 0 | 298 | 149 | 1 | 1 | 0.2 | 0.6 | NA | 0.5 | NA | NA | 1 | 0.8 | 1 | 1 | -0.517 |
| RI | 5157 | ENSG00000146263 | MMS22L | chr6 | - | 97263334 | 97268002 | 97263334 | 97263448 | 97267871 | 97268002 | 11 | 11 | 14 | 8 | 17 | 18 | 8 | 14 | 6 | 34 | 0 | 21 | 13 | 10 | 4 | 0 | 11 | 0 | 1 | 0 | 298 | 149 | 1 | 1 | 0.234 | 0.407 | 0.333 | 0.4 | 0.2 | NA | 0.488 | 1 | 0.833 | 1 | -0.515 |
| RI | 5818 | ENSG00000230006 | ANKRD36BP2 | chr2 | + | 88804379 | 88804880 | 88804379 | 88804450 | 88804786 | 88804880 | 0 | 15 | 22 | 3 | 19 | 2 | 3 | 6 | 2 | 5 | 12 | 0 | 3 | 0 | 0 | 0 | 0 | 0 | 0 | 0 | 298 | 149 | 1 | 1 | 0 | 0.714 | 0.647 | 0.429 | 0.655 | 1 | NA | 1 | NA | NA | -0.511 |
| RI | 890 | ENSG00000141294 | LRRC46 | chr17 | + | 47834424 | 47835399 | 47834424 | 47834533 | 47835352 | 47835399 | 4 | 3 | 2 | 0 | 0 | 3 | 1 | 0 | 4 | 4 | 8 | 0 | 1 | 5 | 10 | 0 | 0 | 0 | 0 | 3 | 298 | 149 | 1 | 1 | 0.4 | 0.6 | 1 | 0 | 0 | 1 | NA | 1 | 1 | 0.625 | -0.506 |
| RI | 69 | ENSG00000128298 | BAIAP2L2 | chr22 | - | 38086241 | 38087264 | 38086241 | 38086449 | 38087123 | 38087264 | 0 | 0 | 0 | 1 | 0 | 0 | 0 | 1 | 0 | 0 | 4 | 0 | 0 | 0 | 0 | 0 | 0 | 0 | 0 | 0 | 298 | 149 | 1 | 1 | NA | NA | 0 | 1 | NA | 1 | NA | NA | NA | NA | -0.5 |
| RI | 202 | ENSG00000174951 | FUT1 | chr19 | - | 48753324 | 48753660 | 48753324 | 48753563 | 48753647 | 48753660 | 0 | 0 | 0 | 0 | 2 | 1 | 0 | 0 | 0 | 0 | 0 | 0 | 0 | 4 | 1 | 0 | 0 | 0 | 0 | 0 | 233 | 149 | 1 | 1 | 0 | NA | NA | NA | 1 | NA | NA | NA | 1 | 1 | -0.5 |
| RI | 328 | ENSG00000187266 | EPOR | chr19 | - | 11382846 | 11383232 | 11382846 | 11382982 | 11383096 | 11383232 | 0 | 0 | 0 | 1 | 0 | 0 | 1 | 0 | 0 | 0 | 0 | 0 | 0 | 8 | 0 | 0 | 0 | 0 | 0 | 0 | 263 | 149 | 1 | 1 | NA | 0 | NA | 1 | NA | NA | NA | NA | 1 | NA | -0.5 |
| RI | 479 | ENSG00000186204 | CYP4F12 | chr19 | + | 15696160 | 15696512 | 15696160 | 15696225 | 15696429 | 15696512 | 0 | 0 | 1 | 0 | 0 | 0 | 1 | 0 | 0 | 0 | 2 | 0 | 0 | 0 | 0 | 0 | 0 | 0 | 0 | 0 | 298 | 149 | 1 | 1 | NA | 0 | 1 | NA | NA | 1 | NA | NA | NA | NA | -0.5 |
| RI | 761 | ENSG00000108823 | SGCA | chr17 | + | 50169091 | 50170351 | 50169091 | 50169254 | 50170142 | 50170351 | 0 | 0 | 1 | 0 | 0 | 0 | 2 | 0 | 0 | 0 | 2 | 0 | 5 | 0 | 0 | 0 | 0 | 0 | 0 | 0 | 298 | 149 | 1 | 1 | NA | 0 | 1 | NA | NA | 1 | NA | 1 | NA | NA | -0.5 |
| RI | 785 | ENSG00000177303 | CASKIN2 | chr17 | - | 75503388 | 75503760 | 75503388 | 75503527 | 75503658 | 75503760 | 0 | 0 | 0 | 0 | 0 | 2 | 0 | 0 | 1 | 0 | 2 | 0 | 0 | 0 | 0 | 0 | 1 | 0 | 0 | 0 | 280 | 149 | 1 | 1 | 0 | NA | NA | 0 | NA | 1 | 0 | NA | NA | NA | -0.5 |
| RI | 829 | ENSG00000108852 | MPP2 | chr17 | - | 43882902 | 43883355 | 43882902 | 43883052 | 43883202 | 43883355 | 0 | 0 | 0 | 0 | 0 | 2 | 0 | 0 | 0 | 0 | 1 | 0 | 0 | 0 | 0 | 0 | 0 | 7 | 0 | 0 | 298 | 149 | 1 | 1 | 0 | NA | NA | NA | NA | 1 | NA | 0 | NA | NA | -0.5 |
| RI | 1405 | ENSG00000103226 | NOMO3 | chr16 | + | 16275011 | 16275443 | 16275011 | 16275171 | 16275316 | 16275443 | 0 | 2 | 0 | 0 | 0 | 0 | 0 | 0 | 0 | 1 | 0 | 5 | 0 | 0 | 0 | 0 | 0 | 0 | 0 | 0 | 294 | 149 | 1 | 1 | NA | 1 | NA | NA | 0 | NA | 1 | NA | NA | NA | -0.5 |
| RI | 1711 | ENSG00000153902 | LGI4 | chr19 | - | 35133692 | 35134104 | 35133692 | 35133764 | 35134032 | 35134104 | 0 | 0 | 0 | 8 | 0 | 0 | 2 | 0 | 0 | 0 | 1 | 0 | 0 | 0 | 0 | 0 | 0 | 0 | 0 | 0 | 298 | 149 | 1 | 1 | NA | 0 | NA | 1 | NA | 1 | NA | NA | NA | NA | -0.5 |
| RI | 2580 | ENSG00000169291 | SHE | chr1 | - | 154479480 | 154486062 | 154479480 | 154484335 | 154485942 | 154486062 | 0 | 0 | 1 | 2 | 0 | 0 | 2 | 0 | 1 | 0 | 1 | 3 | 0 | 0 | 0 | 0 | 0 | 0 | 0 | 0 | 298 | 149 | 1 | 1 | NA | 0 | 1 | 0.5 | NA | 1 | 1 | NA | NA | NA | -0.5 |
| RI | 3791 | ENSG00000164050 | PLXNB1 | chr3 | - | 48411862 | 48412304 | 48411862 | 48412009 | 48412237 | 48412304 | 0 | 0 | 2 | 0 | 0 | 0 | 0 | 0 | 1 | 0 | 0 | 0 | 0 | 2 | 0 | 0 | 0 | 0 | 0 | 0 | 298 | 149 | 1 | 1 | NA | NA | 1 | 0 | NA | NA | NA | NA | 1 | NA | -0.5 |
| RI | 3797 | ENSG00000065361 | ERBB3 | chr12 | + | 56087576 | 56087913 | 56087576 | 56087642 | 56087794 | 56087913 | 0 | 0 | 2 | 0 | 0 | 0 | 0 | 0 | 0 | 5 | 1 | 1 | 8 | 0 | 0 | 0 | 0 | 0 | 0 | 0 | 298 | 149 | 1 | 1 | NA | NA | 1 | NA | 0 | 1 | 1 | 1 | NA | NA | -0.5 |
| RI | 4372 | ENSG00000111344 | RASAL1 | chr12 | - | 113127811 | 113128178 | 113127811 | 113127873 | 113128064 | 113128178 | 0 | 0 | 0 | 1 | 0 | 1 | 0 | 0 | 0 | 0 | 1 | 0 | 0 | 0 | 0 | 0 | 0 | 0 | 0 | 0 | 298 | 149 | 1 | 1 | 0 | NA | NA | 1 | NA | 1 | NA | NA | NA | NA | -0.5 |
| RI | 4417 | ENSG00000146205 | ANO7 | chr2 | + | 241223185 | 241223955 | 241223185 | 241223781 | 241223904 | 241223955 | 0 | 0 | 0 | 1 | 0 | 1 | 0 | 0 | 0 | 0 | 0 | 0 | 0 | 4 | 0 | 0 | 0 | 0 | 0 | 0 | 272 | 149 | 1 | 1 | 0 | NA | NA | 1 | NA | NA | NA | NA | 1 | NA | -0.5 |
| RI | 4768 | ENSG00000075702 | WDR62 | chr19 | + | 36065957 | 36066427 | 36065957 | 36066015 | 36066207 | 36066427 | 0 | 0 | 2 | 0 | 0 | 0 | 0 | 0 | 0 | 1 | 0 | 0 | 4 | 3 | 0 | 0 | 0 | 0 | 0 | 0 | 298 | 149 | 1 | 1 | NA | NA | 1 | NA | 0 | NA | NA | 1 | 1 | NA | -0.5 |
| RI | 5013 | ENSG00000076706 | MCAM | chr11 | - | 119310755 | 119311421 | 119310755 | 119311185 | 119311279 | 119311421 | 0 | 0 | 3 | 0 | 0 | 1 | 0 | 0 | 0 | 0 | 21 | 0 | 0 | 8 | 0 | 0 | 0 | 0 | 0 | 0 | 243 | 149 | 1 | 1 | 0 | NA | 1 | NA | NA | 1 | NA | NA | NA | 1 | -0.5 |
| RI | 5018 | ENSG00000072682 | P4HA2 | chr5 | - | 132217196 | 132217848 | 132217196 | 132217348 | 132217751 | 132217848 | 0 | 0 | 0 | 0 | 0 | 0 | 0 | 0 | 1 | 0 | 0 | 1 | 0 | 0 | 0 | 0 | 3 | 0 | 0 | 0 | 298 | 149 | 1 | 1 | NA | NA | NA | 0 | NA | NA | 1 | 0 | NA | NA | -0.5 |
| RI | 5048 | ENSG00000131037 | EPS8L1 | chr19 | + | 55082091 | 55082349 | 55082091 | 55082180 | 55082274 | 55082349 | 0 | 0 | 0 | 0 | 0 | 1 | 0 | 0 | 7 | 0 | 0 | 2 | 0 | 0 | 0 | 0 | 8 | 0 | 0 | 0 | 243 | 149 | 1 | 1 | 0 | NA | NA | 0 | NA | NA | 1 | 0 | NA | NA | -0.5 |
| RI | 6038 | ENSG00000204301 | NOTCH4 | chr6 | - | 32201116 | 32203882 | 32201116 | 32201500 | 32203769 | 32203882 | 0 | 0 | 4 | 0 | 0 | 0 | 0 | 2 | 0 | 0 | 0 | 7 | 0 | 0 | 0 | 0 | 0 | 0 | 0 | 0 | 298 | 149 | 1 | 1 | NA | NA | 0.5 | NA | NA | NA | 1 | NA | NA | NA | -0.5 |
| RI | 6329 | ENSG00000167103 | PIP5KL1 | chr9 | - | 127927152 | 127927331 | 127927152 | 127927208 | 127927296 | 127927331 | 0 | 4 | 0 | 2 | 0 | 0 | 0 | 1 | 0 | 3 | 0 | 2 | 0 | 0 | 6 | 0 | 0 | 0 | 0 | 0 | 237 | 149 | 1 | 1 | NA | 1 | 0 | 1 | 0 | NA | 1 | NA | NA | 1 | -0.5 |
| RI | 6355 | ENSG00000158525 | CPA5 | chr7 | + | 130367371 | 130367990 | 130367371 | 130367571 | 130367905 | 130367990 | 1 | 0 | 0 | 3 | 0 | 0 | 1 | 0 | 0 | 4 | 0 | 0 | 4 | 2 | 0 | 0 | 0 | 0 | 0 | 0 | 298 | 149 | 1 | 1 | 1 | 0 | NA | 1 | 0 | NA | NA | 1 | 1 | NA | -0.5 |
| RI | 7281 | ENSG00000111664 | GNB3 | chr12 | + | 6841257 | 6841624 | 6841257 | 6841344 | 6841585 | 6841624 | 1 | 0 | 0 | 0 | 0 | 0 | 0 | 0 | 0 | 3 | 0 | 0 | 1 | 18 | 0 | 0 | 0 | 0 | 0 | 0 | 298 | 149 | 1 | 1 | 1 | NA | NA | NA | 0 | NA | NA | 1 | 1 | NA | -0.5 |
| RI | 8034 | ENSG00000151150 | ANK3 | chr10 | - | 60059339 | 60060006 | 60059339 | 60059430 | 60059697 | 60060006 | 6 | 2 | 29 | 6 | 8 | 22 | 9 | 35 | 15 | 24 | 0 | 24 | 31 | 12 | 10 | 0 | 22 | 37 | 0 | 0 | 298 | 149 | 1 | 1 | 0.12 | 0.1 | 0.293 | 0.167 | 0.143 | NA | 0.353 | 0.295 | 1 | 1 | -0.497 |
| RI | 691 | ENSG00000090659 | CD209 | chr19 | - | 7747305 | 7747511 | 7747305 | 7747365 | 7747465 | 7747511 | 0 | 0 | 2 | 2 | 0 | 5 | 0 | 0 | 1 | 0 | 0 | 0 | 2 | 0 | 0 | 0 | 0 | 0 | 0 | 0 | 249 | 149 | 1 | 1 | 0 | NA | 1 | 0.545 | NA | NA | NA | 1 | NA | NA | -0.485 |
| RI | 474 | ENSG00000105135 | ILVBL | chr19 | - | 15122692 | 15123030 | 15122692 | 15122804 | 15122891 | 15123030 | 5 | 4 | 1 | 1 | 1 | 5 | 5 | 4 | 7 | 6 | 0 | 19 | 2 | 13 | 7 | 0 | 0 | 2 | 15 | 0 | 236 | 149 | 1 | 1 | 0.387 | 0.336 | 0.136 | 0.083 | 0.095 | NA | 1 | 0.387 | 0.354 | 1 | -0.478 |
| RI | 1818 | ENSG00000196497 | IPO4 | chr14 | - | 24188551 | 24188797 | 24188551 | 24188638 | 24188718 | 24188797 | 1 | 0 | 0 | 3 | 3 | 2 | 0 | 0 | 2 | 2 | 6 | 9 | 0 | 2 | 12 | 0 | 5 | 0 | 0 | 0 | 229 | 149 | 1 | 1 | 0.245 | NA | NA | 0.494 | 0.494 | 1 | 0.539 | NA | 1 | 1 | -0.474 |
| RI | 2238 | ENSG00000197603 | CPLANE1 | chr5 | - | 37121616 | 37122488 | 37121616 | 37121784 | 37122429 | 37122488 | 9 | 4 | 5 | 5 | 11 | 11 | 7 | 4 | 0 | 13 | 13 | 17 | 7 | 3 | 0 | 0 | 0 | 2 | 0 | 0 | 298 | 149 | 1 | 1 | 0.29 | 0.222 | 0.385 | 1 | 0.297 | 1 | 1 | 0.636 | 1 | NA | -0.47 |
| RI | 4851 | ENSG00000169255 | B3GALNT1 | chr3 | - | 161103426 | 161104406 | 161103426 | 161103517 | 161104318 | 161104406 | 0 | 3 | 0 | 1 | 0 | 0 | 1 | 3 | 0 | 0 | 0 | 0 | 2 | 2 | 0 | 0 | 0 | 0 | 0 | 0 | 298 | 149 | 1 | 1 | NA | 0.6 | 0 | 1 | NA | NA | NA | 1 | 1 | NA | -0.467 |
| RI | 4308 | ENSG00000135541 | AHI1 | chr6 | - | 135463124 | 135466373 | 135463124 | 135463306 | 135465813 | 135466373 | 2 | 1 | 0 | 5 | 3 | 5 | 11 | 7 | 1 | 12 | 0 | 1 | 5 | 0 | 0 | 0 | 0 | 0 | 1 | 0 | 298 | 149 | 1 | 1 | 0.167 | 0.043 | 0 | 0.714 | 0.111 | NA | 1 | 1 | 0 | NA | -0.46 |
| RI | 3467 | ENSG00000162836 | ACP6 | chr1 | - | 147647367 | 147648411 | 147647367 | 147647566 | 147648245 | 147648411 | 6 | 2 | 5 | 8 | 0 | 8 | 1 | 7 | 3 | 0 | 0 | 0 | 10 | 11 | 7 | 0 | 0 | 0 | 4 | 0 | 298 | 149 | 1 | 1 | 0.273 | 0.5 | 0.263 | 0.571 | NA | NA | NA | 1 | 0.579 | 1 | -0.458 |
| RI | 2372 | ENSG00000106624 | AEBP1 | chr7 | + | 44110724 | 44111057 | 44110724 | 44110809 | 44110912 | 44111057 | 1 | 0 | 4 | 2 | 0 | 12 | 10 | 9 | 5 | 12 | 4 | 3 | 0 | 0 | 2 | 0 | 9 | 0 | 1 | 0 | 252 | 149 | 1 | 1 | 0.047 | 0 | 0.208 | 0.191 | 0 | 1 | 0.165 | NA | 0 | 1 | -0.452 |
| RI | 7003 | ENSG00000130158 | DOCK6 | chr19 | - | 11202393 | 11202715 | 11202393 | 11202483 | 11202583 | 11202715 | 0 | 0 | 0 | 1 | 0 | 0 | 1 | 0 | 5 | 0 | 2 | 0 | 0 | 0 | 0 | 0 | 1 | 0 | 0 | 0 | 249 | 149 | 1 | 1 | NA | 0 | NA | 0.107 | NA | 1 | 0 | NA | NA | NA | -0.447 |
| RI | 6078 | ENSG00000269313 | MAGIX | chrX | + | 49164706 | 49165090 | 49164706 | 49164789 | 49164907 | 49165090 | 1 | 0 | 6 | 3 | 0 | 2 | 1 | 0 | 0 | 0 | 4 | 3 | 2 | 5 | 3 | 0 | 0 | 0 | 0 | 0 | 267 | 149 | 1 | 1 | 0.218 | 0 | 1 | 1 | NA | 1 | 1 | 1 | 1 | 1 | -0.446 |
| RI | 7074 | ENSG00000147852 | VLDLR | chr9 | + | 2650369 | 2651954 | 2650369 | 2650516 | 2651873 | 2651954 | 1 | 0 | 1 | 5 | 0 | 4 | 0 | 0 | 2 | 0 | 0 | 2 | 0 | 0 | 0 | 0 | 0 | 0 | 0 | 0 | 298 | 149 | 1 | 1 | 0.111 | NA | 1 | 0.556 | NA | NA | 1 | NA | NA | NA | -0.444 |
| RI | 7308 | ENSG00000177685 | CRACR2B | chr11 | + | 830249 | 830713 | 830249 | 830337 | 830620 | 830713 | 0 | 0 | 0 | 0 | 0 | 1 | 0 | 0 | 5 | 0 | 0 | 1 | 1 | 0 | 0 | 0 | 1 | 0 | 3 | 0 | 298 | 149 | 1 | 1 | 0 | NA | NA | 0 | NA | NA | 0.333 | 1 | 0 | NA | -0.444 |
| RI | 8287 | ENSG00000132141 | CCT6B | chr17 | - | 34939182 | 34939713 | 34939182 | 34939330 | 34939616 | 34939713 | 1 | 0 | 2 | 1 | 0 | 2 | 0 | 4 | 1 | 4 | 2 | 0 | 4 | 2 | 0 | 0 | 7 | 2 | 0 | 0 | 298 | 149 | 1 | 1 | 0.2 | NA | 0.2 | 0.333 | 0 | 1 | 0 | 0.5 | 1 | NA | -0.442 |
| RI | 3892 | ENSG00000139645 | ANKRD52 | chr12 | - | 56257282 | 56257911 | 56257282 | 56257361 | 56257827 | 56257911 | 9 | 5 | 4 | 9 | 8 | 12 | 2 | 10 | 11 | 9 | 5 | 16 | 8 | 0 | 16 | 8 | 2 | 0 | 0 | 0 | 298 | 149 | 1 | 1 | 0.273 | 0.556 | 0.167 | 0.29 | 0.308 | 0.238 | 0.8 | 1 | NA | 1 | -0.441 |
| RI | 1059 | ENSG00000005884 | ITGA3 | chr17 | + | 50075458 | 50075735 | 50075458 | 50075526 | 50075598 | 50075735 | 0 | 0 | 2 | 1 | 0 | 0 | 0 | 3 | 4 | 0 | 0 | 1 | 0 | 0 | 2 | 1 | 0 | 0 | 0 | 0 | 221 | 149 | 1 | 1 | NA | NA | 0.31 | 0.144 | NA | 0 | 1 | NA | NA | 1 | -0.44 |
| RI | 8475 | ENSG00000073060 | SCARB1 | chr12 | - | 124782682 | 124786503 | 124782682 | 124782811 | 124786356 | 124786503 | 9 | 5 | 16 | 10 | 8 | 3 | 5 | 10 | 7 | 18 | 20 | 17 | 0 | 10 | 36 | 0 | 7 | 0 | 0 | 5 | 298 | 149 | 1 | 1 | 0.6 | 0.333 | 0.444 | 0.417 | 0.182 | 1 | 0.548 | NA | 1 | 0.783 | -0.438 |
| RI | 6276 | ENSG00000183735 | TBK1 | chr12 | + | 64495698 | 64496406 | 64495698 | 64495775 | 64496366 | 64496406 | 18 | 15 | 11 | 9 | 11 | 89 | 71 | 72 | 88 | 98 | 2 | 16 | 57 | 10 | 0 | 0 | 24 | 43 | 8 | 0 | 298 | 149 | 1 | 1 | 0.092 | 0.096 | 0.071 | 0.049 | 0.053 | 1 | 0.25 | 0.399 | 0.385 | NA | -0.436 |
| RI | 5680 | ENSG00000169247 | SH3TC2 | chr5 | - | 149006880 | 149009001 | 149006880 | 149007069 | 149008850 | 149009001 | 0 | 1 | 0 | 0 | 0 | 0 | 0 | 0 | 2 | 0 | 18 | 1 | 4 | 4 | 4 | 0 | 0 | 1 | 0 | 0 | 298 | 149 | 1 | 1 | NA | 1 | NA | 0 | NA | 1 | 1 | 0.667 | | 1 | -0.433 |
| RI | 8286 | ENSG00000257337 | NA | chr12 | - | 53045599 | 53046061 | 53045599 | 53045653 | 53045960 | 53046061 | 4 | 0 | 5 | 0 | 0 | 0 | 0 | 1 | 0 | 3 | 0 | 0 | 0 | 1 | 0 | 0 | 0 | 0 | 0 | 0 | 298 | 149 | 1 | 1 | 1 | NA | 0.714 | NA | 0 | NA | NA | NA | 1 | NA | -0.429 |
| RI | 5102 | ENSG00000131748 | STARD3 | chr17 | + | 39662250 | 39663484 | 39662250 | 39662367 | 39662803 | 39663484 | 7 | 0 | 7 | 7 | 0 | 3 | 0 | 1 | 5 | 0 | 27 | 11 | 5 | 6 | 9 | 0 | 0 | 0 | 0 | 0 | 298 | 149 | 1 | 1 | 0.538 | NA | 0.778 | 0.412 | NA | 1 | 1 | 1 | 1 | 1 | -0.424 |
| RI | 6516 | ENSG00000136244 | IL6 | chr7 | + | 22727199 | 22727634 | 22727199 | 22727281 | 22727443 | 22727634 | 0 | 0 | 0 | 0 | 0 | 0 | 0 | 0 | 0 | 1 | 6 | 2 | 0 | 0 | 15 | 0 | 2 | 9 | 26 | 2 | 298 | 149 | 1 | 1 | NA | NA | NA | NA | 0 | 1 | 0.333 | 0 | 0 | 0.789 | -0.424 |
| RI | 1188 | ENSG00000162004 | CCDC78 | chr16 | - | 724321 | 724806 | 724321 | 724509 | 724680 | 724806 | 8 | 5 | 2 | 0 | 0 | 0 | 0 | 2 | 2 | 0 | 0 | 21 | 0 | 5 | 12 | 0 | 0 | 0 | 0 | 0 | 298 | 149 | 1 | 1 | 1 | 1 | 0.333 | 0 | NA | NA | 1 | NA | 1 | 1 | -0.417 |
| RI | 365 | ENSG00000125731 | SH2D3A | chr19 | - | 6754051 | 6754425 | 6754051 | 6754163 | 6754250 | 6754425 | 3 | 3 | 2 | 6 | 0 | 11 | 10 | 3 | 4 | 0 | 2 | 10 | 1 | 0 | 4 | 0 | 4 | 4 | 0 | 0 | 236 | 149 | 1 | 1 | 0.147 | 0.159 | 0.296 | 0.486 | NA | 1 | 0.612 | 0.136 | NA | 1 | -0.415 |
| RI | 7982 | ENSG00000187531 | SIRT7 | chr17 | - | 81917829 | 81918132 | 81917829 | 81917967 | 81918038 | 81918132 | 0 | 0 | 2 | 1 | 0 | 5 | 1 | 4 | 4 | 0 | 0 | 2 | 2 | 1 | 12 | 0 | 0 | 6 | 10 | 2 | 220 | 149 | 1 | 1 | 0 | 0 | 0.253 | 0.145 | NA | NA | 1 | 0.184 | 0.063 | 0.803 | -0.413 |
| RI | 4300 | ENSG00000007402 | CACNA2D2 | chr3 | - | 50364887 | 50365184 | 50364887 | 50364976 | 50365074 | 50365184 | 2 | 1 | 1 | 0 | 0 | 8 | 6 | 2 | 5 | 6 | 0 | 0 | 0 | 2 | 7 | 6 | 0 | 2 | 0 | 0 | 247 | 149 | 1 | 1 | 0.131 | 0.091 | 0.232 | 0 | 0 | 0 | NA | 0 | 1 | 1 | -0.409 |
| RI | 5205 | ENSG00000178038 | ALS2CL | chr3 | - | 46676641 | 46677022 | 46676641 | 46676738 | 46676848 | 46677022 | 4 | 0 | 0 | 2 | 1 | 0 | 0 | 4 | 2 | 0 | 0 | 0 | 4 | 1 | 5 | 0 | 0 | 0 | 0 | 0 | 259 | 149 | 1 | 1 | 1 | NA | 0 | 0.365 | 1 | NA | NA | 1 | 1 | 1 | -0.409 |
| RI | 7417 | ENSG00000107331 | ABCA2 | chr9 | - | 137022940 | 137023840 | 137022940 | 137023052 | 137023837 | 137023840 | 4 | 2 | 1 | 2 | 0 | 1 | 6 | 1 | 2 | 0 | 0 | 8 | 6 | 7 | 1 | 0 | 1 | 0 | 8 | 0 | 298 | 149 | 1 | 1 | 0.667 | 0.143 | 0.333 | 0.333 | NA | NA | 0.8 | 1 | 0.304 | 1 | -0.407 |
| RI | 2396 | ENSG00000198689 | SLC9A6 | chrX | + | 135985434 | 135985827 | 135985434 | 135985477 | 135985602 | 135985827 | 3 | 1 | 1 | 1 | 4 | 1 | 2 | 3 | 0 | 0 | 0 | 0 | 4 | 0 | 0 | 0 | 0 | 0 | 0 | 0 | 274 | 149 | 1 | 1 | 0.62 | 0.214 | 0.153 | 1 | 1 | NA | NA | 1 | NA | NA | -0.403 |
| RI | 4862 | ENSG00000169031 | COL4A3 | chr2 | + | 227304018 | 227305083 | 227304018 | 227304144 | 227304984 | 227305083 | 6 | 6 | 3 | 1 | 4 | 0 | 2 | 4 | 4 | 0 | 0 | 4 | 1 | 0 | 0 | 0 | 0 | 0 | 0 | 0 | 298 | 149 | 1 | 1 | 1 | 0.6 | 0.273 | 0.111 | 1 | NA | 1 | 1 | NA | NA | -0.403 |
| RI | 7416 | ENSG00000107331 | ABCA2 | chr9 | - | 137022701 | 137023052 | 137022701 | 137022865 | 137022940 | 137023052 | 0 | 0 | 1 | 0 | 0 | 25 | 3 | 3 | 14 | 1 | 0 | 7 | 0 | 0 | 2 | 0 | 10 | 6 | 0 | 0 | 224 | 149 | 1 | 1 | 0 | 0 | 0.181 | 0 | 0 | NA | 0.318 | 0 | NA | 1 | -0.403 |
| RI | 2985 | ENSG00000233058 | LINC00884 | chr3 | + | 194487139 | 194488726 | 194487139 | 194487172 | 194488598 | 194488726 | 0 | 1 | 0 | 1 | 4 | 1 | 0 | 0 | 0 | 3 | 0 | 0 | 2 | 0 | 0 | 0 | 0 | 0 | 0 | 0 | 298 | 149 | 1 | 1 | 0 | 1 | NA | 1 | 0.4 | NA | NA | 1 | NA | NA | -0.4 |
| RI | 5131 | ENSG00000186281 | GPAT2 | chr2 | - | 96026182 | 96027106 | 96026182 | 96026305 | 96027006 | 96027106 | 0 | 0 | 1 | 1 | 0 | 0 | 0 | 0 | 2 | 0 | 0 | 0 | 3 | 0 | 2 | 0 | 0 | 0 | 0 | 0 | 298 | 149 | 1 | 1 | NA | NA | 1 | 0.2 | NA | NA | NA | 1 | NA | 1 | -0.4 |
| RI | 1185 | ENSG00000162004 | CCDC78 | chr16 | - | 722921 | 723161 | 722921 | 723022 | 723094 | 723161 | 2 | 1 | 1 | 0 | 0 | 1 | 0 | 3 | 5 | 2 | 0 | 3 | 0 | 10 | 6 | 0 | 0 | 3 | 0 | 0 | 221 | 149 | 1 | 1 | 0.574 | 1 | 0.183 | 0 | 0 | NA | 1 | 0 | 1 | 1 | -0.399 |
| RI | 3522 | ENSG00000270629 | NBPF14 | chr1 | - | 148595542 | 148595890 | 148595542 | 148595752 | 148595849 | 148595890 | 9 | 1 | 10 | 4 | 0 | 1 | 2 | 1 | 5 | 1 | 0 | 0 | 8 | 3 | 1 | 0 | 0 | 4 | 0 | 0 | 246 | 149 | 1 | 1 | 0.845 | 0.232 | 0.858 | 0.326 | 0 | NA | NA | 0.548 | 1 | 1 | -0.397 |
| RI | 7849 | ENSG00000213619 | NDUFS3 | chr11 | + | 47579268 | 47580622 | 47579268 | 47579338 | 47580524 | 47580622 | 8 | 1 | 2 | 8 | 0 | 1 | 1 | 1 | 1 | 0 | 4 | 1 | 5 | 7 | 20 | 0 | 0 | 0 | 0 | 0 | 298 | 149 | 1 | 1 | 0.8 | 0.333 | 0.5 | 0.8 | NA | 1 | 1 | 1 | 1 | 1 | -0.392 |
| RI | 336 | ENSG00000267454 | ZNF582-DT | chr19 | + | 56398276 | 56399168 | 56398276 | 56398383 | 56398893 | 56399168 | 4 | 3 | 3 | 4 | 4 | 2 | 3 | 6 | 6 | 15 | 0 | 5 | 1 | 0 | 0 | 0 | 0 | 0 | 0 | 23 | 298 | 149 | 1 | 1 | 0.5 | 0.333 | 0.2 | 0.25 | 0.118 | NA | 1 | 1 | NA | 0 | -0.386 |
| RI | 4840 | ENSG00000175054 | ATR | chr3 | - | 142467933 | 142469569 | 142467933 | 142468068 | 142469336 | 142469569 | 18 | 6 | 7 | 11 | 2 | 39 | 28 | 39 | 41 | 83 | 4 | 0 | 4 | 0 | 6 | 0 | 3 | 2 | 0 | 4 | 298 | 149 | 1 | 1 | 0.188 | 0.097 | 0.082 | 0.118 | 0.012 | 1 | 0 | 0.5 | NA | 0.429 | -0.383 |
| RI | 3396 | ENSG00000117266 | CDK18 | chr1 | + | 205529520 | 205530349 | 205529520 | 205529563 | 205530258 | 205530349 | 0 | 1 | 3 | 0 | 0 | 0 | 3 | 2 | 0 | 0 | 2 | 1 | 0 | 0 | 0 | 0 | 0 | 0 | 0 | 15 | 298 | 149 | 1 | 1 | NA | 0.143 | 0.429 | NA | NA | 1 | 1 | NA | NA | 0 | -0.381 |
| RI | 3640 | ENSG00000079308 | TNS1 | chr2 | - | 217880897 | 217882411 | 217880897 | 217881014 | 217882345 | 217882411 | 0 | 0 | 0 | 2 | 0 | 1 | 1 | 4 | 2 | 0 | 7 | 3 | 6 | 17 | 8 | 0 | 10 | 3 | 9 | 16 | 298 | 149 | 1 | 1 | 0 | 0 | 0 | 0.333 | NA | 1 | 0.13 | 0.5 | 0.486 | 0.2 | -0.38 |
| RI | 3852 | ENSG00000236432 | MFF-DT | chr2 | - | 227265874 | 227268769 | 227265874 | 227265935 | 227268627 | 227268769 | 0 | 0 | 0 | 2 | 5 | 0 | 0 | 0 | 3 | 0 | 0 | 5 | 0 | 0 | 0 | 0 | 0 | 0 | 0 | 0 | 298 | 149 | 1 | 1 | NA | NA | NA | 0.25 | 1 | NA | 1 | NA | NA | NA | -0.375 |
| RI | 222 | ENSG00000182472 | CAPN12 | chr19 | - | 38735369 | 38735544 | 38735369 | 38735429 | 38735501 | 38735544 | 2 | 0 | 0 | 1 | 0 | 0 | 0 | 0 | 2 | 0 | 0 | 5 | 1 | 0 | 0 | 0 | 0 | 0 | 0 | 0 | 221 | 149 | 1 | 1 | 1 | NA | NA | 0.252 | NA | NA | 1 | 1 | NA | NA | -0.374 |
| RI | 168 | ENSG00000104936 | DMPK | chr19 | - | 45770970 | 45771396 | 45770970 | 45771056 | 45771349 | 45771396 | 4 | 2 | 1 | 3 | 0 | 2 | 3 | 0 | 6 | 2 | 0 | 0 | 3 | 1 | 10 | 0 | 0 | 4 | 0 | 0 | 298 | 149 | 1 | 1 | 0.5 | 0.25 | 1 | 0.2 | 0 | NA | NA | 0.273 | 1 | 1 | -0.368 |
| RI | 1260 | ENSG00000007516 | BAIAP3 | chr16 | + | 1344021 | 1344317 | 1344021 | 1344146 | 1344226 | 1344317 | 1 | 0 | 2 | 2 | 1 | 2 | 2 | 0 | 4 | 2 | 7 | 4 | 1 | 0 | 4 | 7 | 0 | 0 | 0 | 3 | 229 | 149 | 1 | 1 | 0.245 | 0 | 1 | 0.245 | 0.245 | 0.394 | 1 | 1 | NA | 0.465 | -0.368 |
| RI | 4303 | ENSG00000135541 | AHI1 | chr6 | - | 135300499 | 135302817 | 135300499 | 135300558 | 135301407 | 135302817 | 8 | 10 | 6 | 5 | 0 | 4 | 9 | 13 | 3 | 1 | 8 | 6 | 0 | 0 | 0 | 0 | 0 | 3 | 0 | 0 | 298 | 149 | 1 | 1 | 0.5 | 0.357 | 0.188 | 0.455 | 0 | 1 | 1 | 0 | NA | NA | -0.367 |
| RI | 6387 | ENSG00000142303 | ADAMTS10 | chr19 | - | 8596072 | 8596412 | 8596072 | 8596219 | 8596306 | 8596412 | 22 | 8 | 22 | 17 | 5 | 5 | 4 | 5 | 9 | 2 | 0 | 4 | 17 | 5 | 37 | 0 | 0 | 0 | 0 | 0 | 236 | 149 | 1 | 1 | 0.735 | 0.558 | 0.735 | 0.544 | 0.612 | NA | 1 | 1 | 1 | 1 | -0.363 |
| RI | 8259 | ENSG00000182199 | SHMT2 | chr12 | + | 57232210 | 57232575 | 57232210 | 57232292 | 57232424 | 57232575 | 0 | 0 | 5 | 3 | 0 | 1 | 3 | 0 | 0 | 0 | 32 | 4 | 8 | 10 | 9 | 0 | 5 | 0 | 0 | 0 | 281 | 149 | 1 | 1 | 0 | 0 | 1 | 1 | NA | 1 | 0.298 | 1 | 1 | 1 | -0.36 |
| RI | 6175 | ENSG00000109339 | MAPK10 | chr4 | - | 86098420 | 86101217 | 86098420 | 86098595 | 86101051 | 86101217 | 0 | 0 | 0 | 3 | 0 | 0 | 1 | 0 | 2 | 4 | 0 | 4 | 0 | 0 | 0 | 19 | 0 | 0 | 0 | 0 | 298 | 149 | 1 | 1 | NA | 0 | NA | 0.429 | 0 | 0 | 1 | NA | NA | NA | -0.357 |
| RI | 5155 | ENSG00000256223 | ZNF10 | chr12 | + | 133130610 | 133131014 | 133130610 | 133130754 | 133130836 | 133131014 | 8 | 1 | 6 | 10 | 4 | 3 | 1 | 5 | 2 | 0 | 0 | 4 | 8 | 4 | 5 | 0 | 0 | 0 | 0 | 0 | 231 | 149 | 1 | 1 | 0.632 | 0.392 | 0.436 | 0.763 | 1 | NA | 1 | 1 | 1 | 1 | -0.355 |
| RI | 5250 | ENSG00000224078 | SNHG14 | chr15 | + | 25053827 | 25055230 | 25053827 | 25054866 | 25054969 | 25055230 | 7 | 1 | 1 | 2 | 0 | 0 | 0 | 2 | 0 | 1 | 0 | 2 | 0 | 0 | 0 | 0 | 0 | 0 | 0 | 0 | 252 | 149 | 1 | 1 | 1 | 1 | 0.228 | 1 | 0 | NA | 1 | NA | NA | NA | -0.354 |
| RI | 8172 | ENSG00000152582 | SPEF2 | chr5 | + | 35697689 | 35700752 | 35697689 | 35697793 | 35700495 | 35700752 | 4 | 0 | 4 | 4 | 5 | 11 | 6 | 2 | 8 | 1 | 0 | 2 | 0 | 2 | 0 | 0 | 0 | 3 | 0 | 0 | 298 | 149 | 1 | 1 | 0.154 | 0 | 0.5 | 0.2 | 0.714 | NA | 1 | 0 | 1 | NA | -0.353 |
| RI | 1106 | ENSG00000131899 | LLGL1 | chr17 | + | 18235469 | 18236760 | 18235469 | 18235537 | 18236606 | 18236760 | 0 | 2 | 5 | 0 | 0 | 6 | 0 | 3 | 5 | 0 | 0 | 2 | 4 | 2 | 0 | 0 | 6 | 0 | 0 | 0 | 298 | 149 | 1 | 1 | 0 | 1 | 0.455 | 0 | NA | NA | 0.143 | 1 | 1 | NA | -0.351 |
| RI | 1694 | ENSG00000206149 | HERC2P9 | chr15 | + | 28640921 | 28641312 | 28640921 | 28641092 | 28641169 | 28641312 | 9 | 0 | 0 | 5 | 1 | 2 | 3 | 0 | 3 | 12 | 20 | 0 | 11 | 0 | 3 | 5 | 2 | 0 | 0 | 0 | 226 | 149 | 1 | 1 | 0.748 | 0 | NA | 0.524 | 0.052 | 0.725 | 0 | 1 | NA | 1 | -0.35 |
| RI | 5256 | ENSG00000224078 | SNHG14 | chr15 | + | 25093833 | 25095828 | 25093833 | 25093974 | 25095626 | 25095828 | 21 | 15 | 17 | 8 | 11 | 24 | 14 | 5 | 12 | 12 | 31 | 1 | 17 | 9 | 0 | 0 | 0 | 9 | 7 | 0 | 298 | 149 | 1 | 1 | 0.304 | 0.349 | 0.63 | 0.25 | 0.314 | 1 | 1 | 0.486 | 0.391 | NA | -0.35 |
| RI | 3716 | ENSG00000138002 | IFT172 | chr2 | - | 27449293 | 27449562 | 27449293 | 27449380 | 27449498 | 27449562 | 4 | 0 | 2 | 0 | 4 | 5 | 1 | 3 | 5 | 6 | 12 | 2 | 0 | 2 | 0 | 6 | 0 | 5 | 1 | 0 | 267 | 149 | 1 | 1 | 0.309 | 0 | 0.271 | 0 | 0.271 | 0.527 | 1 | 0 | 0.527 | NA | -0.343 |
| RI | 8387 | ENSG00000111271 | ACAD10 | chr12 | + | 111755667 | 111757099 | 111755667 | 111755745 | 111756332 | 111757099 | 4 | 0 | 1 | 7 | 11 | 6 | 0 | 2 | 4 | 5 | 0 | 2 | 2 | 9 | 25 | 0 | 0 | 1 | 10 | 0 | 298 | 149 | 1 | 1 | 0.25 | NA | 0.2 | 0.467 | 0.524 | NA | 1 | 0.5 | 0.31 | 1 | -0.342 |
| RI | 2326 | ENSG00000165810 | BTNL9 | chr5 | + | 181055432 | 181056015 | 181055432 | 181055453 | 181055988 | 181056015 | 2 | 1 | 1 | 6 | 0 | 0 | 3 | 0 | 3 | 0 | 3 | 0 | 0 | 0 | 0 | 0 | 0 | 0 | 0 | 0 | 298 | 149 | 1 | 1 | 1 | 0.143 | 1 | 0.5 | NA | 1 | NA | NA | NA | NA | -0.339 |
| RI | 7612 | ENSG00000062822 | POLD1 | chr19 | + | 50401998 | 50402373 | 50401998 | 50402124 | 50402204 | 50402373 | 7 | 0 | 0 | 5 | 6 | 2 | 0 | 4 | 16 | 2 | 1 | 0 | 2 | 5 | 7 | 0 | 0 | 0 | 2 | 13 | 229 | 149 | 1 | 1 | 0.695 | NA | 0 | 0.169 | 0.661 | 1 | NA | 1 | 0.619 | 0.259 | -0.338 |
| RI | 2718 | ENSG00000172771 | EFCAB12 | chr3 | - | 129411157 | 129415401 | 129411157 | 129411354 | 129415244 | 129415401 | 2 | 0 | 1 | 1 | 1 | 2 | 0 | 0 | 1 | 0 | 0 | 4 | 1 | 2 | 0 | 0 | 0 | 0 | 0 | 0 | 298 | 149 | 1 | 1 | 0.333 | NA | 1 | 0.333 | 1 | NA | 1 | 1 | 1 | NA | -0.334 |
| RI | 8107 | ENSG00000103024 | NME3 | chr16 | - | 1771069 | 1771410 | 1771069 | 1771171 | 1771268 | 1771410 | 10 | 2 | 1 | 12 | 3 | 5 | 1 | 2 | 0 | 0 | 0 | 0 | 1 | 0 | 6 | 0 | 0 | 0 | 0 | 0 | 246 | 149 | 1 | 1 | 0.548 | 0.548 | 0.232 | 1 | 1 | NA | NA | 1 | NA | 1 | -0.334 |
| RI | 44 | ENSG00000206140 | TMEM191C | chr22 | + | 21467893 | 21468431 | 21467893 | 21468017 | 21468353 | 21468431 | 1 | 0 | 3 | 0 | 0 | 0 | 0 | 0 | 2 | 0 | 15 | 0 | 2 | 0 | 0 | 0 | 0 | 0 | 0 | 0 | 298 | 149 | 1 | 1 | 1 | NA | 1 | 0 | NA | 1 | NA | 1 | NA | NA | -0.333 |
| RI | 729 | ENSG00000198720 | ANKRD13B | chr17 | + | 29613353 | 29614761 | 29613353 | 29613719 | 29614262 | 29614761 | 4 | 0 | 0 | 0 | 0 | 1 | 0 | 0 | 0 | 0 | 0 | 0 | 0 | 1 | 2 | 0 | 0 | 0 | 0 | 0 | 298 | 149 | 1 | 1 | 0.667 | NA | NA | NA | NA | NA | NA | NA | 1 | 1 | -0.333 |
| RI | 1108 | ENSG00000131899 | LLGL1 | chr17 | + | 18240577 | 18241999 | 18240577 | 18240873 | 18241884 | 18241999 | 2 | 0 | 1 | 0 | 0 | 0 | 1 | 0 | 0 | 0 | 22 | 4 | 4 | 0 | 23 | 0 | 0 | 0 | 0 | 0 | 298 | 149 | 1 | 1 | 1 | 0 | 1 | NA | NA | 1 | 1 | 1 | NA | 1 | -0.333 |
| RI | 1501 | ENSG00000161996 | WDR90 | chr16 | + | 653745 | 655147 | 653745 | 653803 | 655028 | 655147 | 0 | 0 | 4 | 0 | 3 | 0 | 3 | 0 | 0 | 0 | 0 | 2 | 1 | 5 | 12 | 0 | 0 | 0 | 0 | 0 | 298 | 149 | 1 | 1 | NA | 0 | 1 | NA | 1 | NA | 1 | 1 | 1 | 1 | -0.333 |
| RI | 1780 | ENSG00000129467 | ADCY4 | chr14 | - | 24322065 | 24322708 | 24322065 | 24322224 | 24322623 | 24322708 | 0 | 0 | 1 | 1 | 0 | 0 | 0 | 0 | 0 | 3 | 9 | 0 | 0 | 0 | 0 | 0 | 0 | 0 | 0 | 0 | 298 | 149 | 1 | 1 | NA | NA | 1 | 1 | 0 | 1 | NA | NA | NA | NA | -0.333 |
| RI | 2437 | ENSG00000168824 | NSG1 | chr4 | + | 4385856 | 4387173 | 4385856 | 4386643 | 4387118 | 4387173 | 3 | 0 | 0 | 1 | 0 | 0 | 0 | 1 | 0 | 0 | 0 | 2 | 0 | 0 | 0 | 0 | 0 | 0 | 0 | 0 | 298 | 149 | 1 | 1 | 1 | NA | 0 | 1 | NA | NA | 1 | NA | NA | NA | -0.333 |
| RI | 3005 | ENSG00000114859 | CLCN2 | chr3 | - | 184355693 | 184357094 | 184355693 | 184355844 | 184356992 | 184357094 | 0 | 0 | 2 | 5 | 0 | 0 | 0 | 0 | 0 | 3 | 0 | 12 | 5 | 5 | 0 | 0 | 0 | 0 | 0 | 0 | 298 | 149 | 1 | 1 | NA | NA | 1 | 1 | 0 | NA | 1 | 1 | 1 | NA | -0.333 |
| RI | 3389 | ENSG00000204580 | DDR1 | chr6 | + | 30888687 | 30889010 | 30888687 | 30888814 | 30888907 | 30889010 | 0 | 0 | 1 | 4 | 0 | 3 | 4 | 6 | 9 | 8 | 0 | 1 | 4 | 0 | 1 | 0 | 1 | 1 | 0 | 6 | 242 | 149 | 1 | 1 | 0 | 0 | 0.093 | 0.215 | 0 | NA | 0.381 | 0.711 | NA | 0.093 | -0.333 |
| RI | 3792 | ENSG00000164050 | PLXNB1 | chr3 | - | 48413068 | 48413818 | 48413068 | 48413169 | 48413669 | 48413818 | 0 | 2 | 0 | 1 | 0 | 1 | 0 | 0 | 0 | 0 | 0 | 0 | 0 | 0 | 5 | 0 | 0 | 0 | 0 | 0 | 298 | 149 | 1 | 1 | 0 | 1 | NA | 1 | NA | NA | NA | NA | NA | 1 | -0.333 |

| | | | | | | | | | | | | | | |
|---|---|---|---|---|---|---|---|---|---|---|---|---|---|---|
| RI | 4257 | ENSG00000143801 | PSEN2 | chr1 | + | 226888828 | 226890965 | 226888828 | 226890133 | 226890655 | 226890965 | 2 | 0 | 4 |
| RI | 4518 | ENSG00000143126 | CELSR2 | chr1 | + | 109271385 | 109271722 | 109271385 | 109271512 | 109271599 | 109271722 | 0 | 1 | 0 |
| RI | 4772 | ENSG00000075702 | WDR62 | chr19 | + | 36101663 | 36102851 | 36101663 | 36101774 | 36102013 | 36102851 | 0 | 0 | 0 |
| RI | 5770 | ENSG00000228203 | GRASLND | chr2 | - | 6915822 | 6917224 | 6915822 | 6915967 | 6916815 | 6917224 | 0 | 0 | 3 |
| RI | 6531 | ENSG00000164877 | MICALL2 | chr7 | - | 1436741 | 1437608 | 1436741 | 1436856 | 1437534 | 1437608 | 0 | 0 | 0 |
| RI | 6730 | ENSG00000204632 | HLA-G | chr6 | + | 29829417 | 29829932 | 29829417 | 29829693 | 29829815 | 29829932 | 0 | 0 | 4 |
| RI | 2078 | ENSG00000156345 | CDK20 | chr9 | - | 87969193 | 87970897 | 87969193 | 87969349 | 87970775 | 87970897 | 4 | 3 | 8 |
| RI | 3192 | ENSG00000241163 | LINC00877 | chr3 | - | 72060509 | 72061094 | 72060509 | 72060611 | 72060802 | 72061094 | 4 | 7 | 14 |
| RI | 4358 | ENSG00000099840 | IZUMO4 | chr19 | + | 2098435 | 2099584 | 2098435 | 2098450 | 2099254 | 2099584 | 2 | 1 | 0 |
| RI | 8446 | ENSG00000135476 | ESPL1 | chr12 | + | 53288537 | 53289303 | 53288537 | 53288699 | 53289089 | 53289303 | 0 | 0 | 1 |
| RI | 402 | ENSG00000124466 | LYPD3 | chr19 | - | 43463125 | 43463769 | 43463125 | 43463287 | 43463598 | 43463769 | 2 | 0 | 1 |
| RI | 1153 | ENSG00000102878 | HSF4 | chr16 | + | 67165945 | 67166395 | 67165945 | 67166070 | 67166319 | 67166395 | 5 | 1 | 0 |
| RI | 2717 | ENSG00000172771 | EFCAB12 | chr3 | - | 129402522 | 129404403 | 129402522 | 129402579 | 129404249 | 129404403 | 1 | 0 | 5 |
| RI | 7940 | ENSG00000149582 | TMEM25 | chr11 | + | 118531191 | 118531871 | 118531191 | 118531234 | 118531777 | 118531871 | 1 | 0 | 5 |
| RI | 4628 | ENSG00000268043 | NBPF12 | chr1 | + | 146987953 | 146988126 | 146987953 | 146988008 | 146988012 | 146988126 | 0 | 4 | 4 |
| RI | 211 | ENSG00000132024 | CC2D1A | chr19 | + | 13927892 | 13928188 | 13927892 | 13928030 | 13928126 | 13928188 | 3 | 1 | 1 |
| RI | 3594 | ENSG00000159214 | CCDC24 | chr1 | + | 43995600 | 43995856 | 43995600 | 43995670 | 43995777 | 43995856 | 0 | 0 | 0 |
| RI | 131 | ENSG00000235954 | TTC28-AS1 | chr22 | + | 27997410 | 27998152 | 27997410 | 27997739 | 27997985 | 27998152 | 11 | 7 | 10 |
| RI | 2121 | ENSG00000177595 | PIDD1 | chr11 | - | 799814 | 800244 | 799814 | 800014 | 800130 | 800244 | 30 | 4 | 15 |
| RI | 7166 | ENSG00000271853 | NA | chr1 | - | 153632033 | 153632240 | 153632033 | 153632063 | 153632154 | 153632240 | 5 | 0 | 5 |
| RI | 2089 | ENSG00000124784 | RIOK1 | chr6 | + | 7402602 | 7402897 | 7402602 | 7402715 | 7402816 | 7402897 | 20 | 27 | 24 |
| RI | 7356 | ENSG00000156042 | CFAP70 | chr10 | - | 73293262 | 73297173 | 73293262 | 73293388 | 73297041 | 73297173 | 0 | 3 | 0 |
| RI | 4822 | ENSG00000149231 | CCDC82 | chr11 | - | 96387529 | 96389912 | 96387529 | 96387613 | 96389843 | 96389912 | 22 | 31 | 17 |
| RI | 5869 | ENSG00000072163 | LIMS2 | chr2 | - | 127640069 | 127640318 | 127640069 | 127640145 | 127640269 | 127640318 | 0 | 0 | 1 |
| RI | 2867 | ENSG00000088387 | DOCK9 | chr13 | - | 98855897 | 98860522 | 98855897 | 98856031 | 98860404 | 98860522 | 7 | 9 | 12 |
| RI | 1839 | ENSG00000099797 | TECR | chr19 | + | 14564785 | 14565123 | 14564785 | 14564858 | 14565065 | 14565123 | 10 | 2 | 7 |
| RI | 2177 | ENSG00000079215 | SLC1A3 | chr5 | + | 36671028 | 36674091 | 36671028 | 36671233 | 36674048 | 36674091 | 0 | 0 | 0 |
| RI | 7310 | ENSG00000177685 | CRACR2B | chr11 | + | 830865 | 831295 | 830865 | 831032 | 831223 | 831295 | 0 | 0 | 0 |
| RI | 7993 | ENSG00000167535 | CACNB3 | chr12 | + | 48824668 | 48825243 | 48824668 | 48824733 | 48825162 | 48825243 | 0 | 0 | 0 |
| [illegible] | | | | | | | | | | | | | | |

| | | | | | | | | | | | | | | |
|---|---|---|---|---|---|---|---|---|---|---|---|---|---|---|
| RI | 2510 | ENSG00000109654 | TRIM2 | chr4 | + | 153294304 | 153296036 | 153294304 | 153294485 | 153295312 | 153296036 | 2 | 1 | 0 |
| RI | 5452 | ENSG00000143799 | PARP1 | chr1 | - | 226385503 | 226386442 | 226385503 | 226385680 | 226386218 | 226386442 | 6 | 3 | 6 |
| RI | 3438 | ENSG00000143322 | ABL2 | chr1 | - | 179107717 | 179109441 | 179107717 | 179108898 | 179109207 | 179109441 | 34 | 21 | 28 |
| RI | 2285 | ENSG00000254231 | NA | chr8 | - | 86335702 | 86338387 | 86335702 | 86335761 | 86338119 | 86338387 | 5 | 2 | 5 |
| RI | 5001 | ENSG00000092529 | CAPN3 | chr15 | + | 42409930 | 42410496 | 42409930 | 42409995 | 42410431 | 42410496 | 0 | 1 | 2 |
| RI | 7378 | ENSG00000122435 | TRMT13 | chr1 | + | 100140407 | 100141019 | 100140407 | 100140514 | 100140851 | 100141019 | 58 | 54 | 61 |
| RI | 8077 | ENSG00000180229 | HERC2P3 | chr15 | - | 20438926 | 20439655 | 20438926 | 20439123 | 20439507 | 20439655 | 0 | 2 | 0 |
| RI | 6037 | ENSG00000204301 | NOTCH4 | chr6 | - | 32194842 | 32196421 | 32194842 | 32196150 | 32196323 | 32196421 | 0 | 2 | 6 |
| RI | 8464 | ENSG00000122966 | CIT | chr12 | - | 119712177 | 119712695 | 119712177 | 119712347 | 119712590 | 119712695 | 0 | 0 | 0 |
| RI | 4449 | ENSG00000163781 | TOPBP1 | chr3 | - | 133598174 | 133598630 | 133598174 | 133598462 | 133598555 | 133598630 | 0 | 1 | 2 |
| RI | 4948 | ENSG00000171503 | ETFDH | chr4 | + | 158680123 | 158681614 | 158680123 | 158680182 | 158680466 | 158681614 | 29 | 46 | 52 |
| RI | 2042 | ENSG00000154309 | DISP1 | chr1 | + | 222992012 | 222994982 | 222992012 | 222992110 | 222994884 | 222994982 | 3 | 1 | 0 |
| RI | 5255 | ENSG00000224078 | SNHG14 | chr15 | + | 25089545 | 25091719 | 25089545 | 25089866 | 25091589 | 25091719 | 1 | 5 | 2 |
| RI | 546 | ENSG00000061656 | SPAG4 | chr20 | + | 35618449 | 35618720 | 35618449 | 35618475 | 35618611 | 35618720 | 3 | 0 | 9 |
| RI | 1970 | ENSG00000130545 | CRB3 | chr19 | + | 6466465 | 6467221 | 6466465 | 6466609 | 6466950 | 6467221 | 2 | 2 | 4 |
| RI | 2130 | ENSG00000154134 | ROBO3 | chr11 | + | 124877518 | 124878131 | 124877518 | 124877752 | 124877936 | 124878131 | 9 | 4 | 10 |
| RI | 3387 | ENSG00000085465 | OVGP1 | chr1 | - | 111421275 | 111421673 | 111421275 | 111421461 | 111421564 | 111421673 | 0 | 0 | 6 |
| RI | 6209 | ENSG00000187796 | CARD9 | chr9 | - | 136364482 | 136366845 | 136364482 | 136364559 | 136366799 | 136366845 | 10 | 13 | 4 |
| RI | 8432 | ENSG00000139428 | MMAB | chr12 | - | 109561039 | 109561329 | 109561039 | 109561104 | 109561242 | 109561329 | 1 | 2 | 4 |
| RI | 1557 | ENSG00000062524 | LTK | chr15 | - | 41507094 | 41507657 | 41507094 | 41507290 | 41507513 | 41507657 | 13 | 9 | 20 |
| RI | 95 | ENSG00000198832 | SELENOM | chr22 | - | 31105929 | 31107533 | 31105929 | 31105965 | 31107376 | 31107533 | 2 | 12 | 11 |
| RI | 3694 | ENSG00000018510 | AGPS | chr2 | + | 177440964 | 177442486 | 177440964 | 177442235 | 177442406 | 177442486 | 18 | 25 | 25 |
| RI | 4074 | ENSG00000177943 | MAMDC4 | chr9 | + | 136856463 | 136856826 | 136856463 | 136856575 | 136856709 | 136856826 | 7 | 3 | 18 |
| RI | 6081 | ENSG00000124549 | BTN2A3P | chr6 | + | 26422932 | 26423759 | 26422932 | 26423277 | 26423398 | 26423759 | 13 | 9 | 4 |
| RI | 304 | ENSG00000161265 | U2AF1L4 | chr19 | - | 35743154 | 35743345 | 35743154 | 35743208 | 35743295 | 35743345 | 3 | 7 | 5 |
| RI | 3475 | ENSG00000064687 | ABCA7 | chr19 | + | 1056906 | 1058747 | 1056906 | 1057084 | 1058617 | 1058747 | 2 | 4 | 2 |
| RI | 5401 | ENSG00000176978 | DPP7 | chr9 | - | 137114242 | 137114576 | 137114242 | 137114378 | 137114462 | 137114576 | 6 | 0 | 10 |
| RI | 915 | ENSG00000011143 | MKS1 | chr17 | - | 58207084 | 58208001 | 58207084 | 58207218 | 58207893 | 58208001 | 1 | 1 | 5 |
| RI | 3661 | ENSG00000139372 | TDG | chr12 | + | 103982798 | 103983389 | 103982798 | 103982934 | 103983294 | 103983389 | 17 | 18 | 13 |
| RI | 7901 | ENSG00000247708 | STX18-AS1 | chr4 | - | 4648976 | 4649542 | 4648976 | 4649109 | 4649369 | 4649542 | 6 | 3 | 4 |
| RI | 1629 | ENSG00000174326 | SLC16A11 | chr17 | - | 7041620 | 7042763 | 7041620 | 7041908 | 7041995 | 7042763 | 3 | 0 | 5 |
| RI | 6889 | ENSG00000182484 | WASH6P | chrX | + | 156024119 | 156025666 | 156024119 | 156024272 | 156025240 | 156025666 | 0 | 0 | 0 |
| RI | 774 | ENSG00000073350 | LLGL2 | chr17 | + | 75574212 | 75574668 | 75574212 | 75574260 | 75574609 | 75574668 | 2 | 0 | 1 |
| RI | 1027 | ENSG00000173818 | ENDOV | chr17 | + | 80415452 | 80415821 | 80415452 | 80415518 | 80415649 | 80415821 | 3 | 0 | 5 |
| RI | 1088 | ENSG00000276070 | CCL4L2 | chr17 | + | 36211775 | 36212878 | 36211775 | 36211890 | 36212481 | 36212878 | 58 | 12 | 3 |
| RI | 8494 | ENSG00000204852 | TCTN1 | chr12 | + | 110641541 | 110642389 | 110641541 | 110641627 | 110642248 | 110642389 | 8 | 10 | 16 |
| RI | 460 | ENSG00000064545 | TMEM161A | chr19 | - | 19132351 | 19132754 | 19132351 | 19132508 | 19132656 | 19132754 | 3 | 1 | 8 |
| RI | 348 | ENSG00000133247 | KMT5C | chr19 | + | 55341793 | 55342380 | 55341793 | 55342046 | 55342214 | 55342380 | 3 | 3 | 3 |
| RI | 1179 | ENSG00000125124 | BBS2 | chr16 | - | 56509356 | 56510034 | 56509356 | 56509494 | 56509956 | 56510034 | 13 | 5 | 4 |
| RI | 2079 | ENSG00000156345 | CDK20 | chr9 | - | 87970567 | 87970897 | 87970567 | 87970630 | 87970775 | 87970897 | 1 | 2 | 8 |
| RI | 3668 | ENSG00000167962 | ZNF598 | chr16 | - | 2002520 | 2003728 | 2002520 | 2002732 | 2003615 | 2003728 | 1 | 1 | 6 |
| RI | 5990 | ENSG00000124593 | NA | chr6 | + | 41783461 | 41784238 | 41783461 | 41783605 | 41783733 | 41784238 | 0 | 2 | 0 |
| RI | 7939 | ENSG00000149582 | TMEM25 | chr11 | + | 118531191 | 118531871 | 118531191 | 118531234 | 118531774 | 118531871 | 1 | 0 | 5 |
| RI | 3411 | ENSG00000267575 | NA | chr19 | + | 27805604 | 27806782 | 27805604 | 27806185 | 27806322 | 27806782 | 1 | 1 | 4 |
| RI | 3551 | ENSG00000127054 | INTS11 | chr1 | - | 1312225 | 1312700 | 1312225 | 1312368 | 1312592 | 1312700 | 11 | 13 | 13 |
| RI | 443 | ENSG00000161249 | DMKN | chr19 | - | 35510183 | 35510510 | 35510183 | 35510252 | 35510378 | 35510510 | 2 | 2 | 11 |
| RI | 942 | ENSG00000215041 | NEURL4 | chr17 | - | 7318064 | 7318356 | 7318064 | 7318172 | 7318268 | 7318356 | 14 | 4 | 9 |
| RI | 5203 | ENSG00000178038 | ALS2CL | chr3 | - | 46674565 | 46675976 | 46674565 | 46674739 | 46675617 | 46675976 | 6 | 6 | 13 |
| RI | 7773 | ENSG00000087053 | MTMR2 | chr11 | - | 95862271 | 95865676 | 95862271 | 95862366 | 95865439 | 95865676 | 15 | 8 | 7 |
| RI | 6431 | ENSG00000002933 | TMEM176A | chr7 | + | 150801535 | 150802325 | 150801535 | 150801724 | 150802214 | 150802325 | 7 | 1 | 0 |
| RI | 7544 | ENSG00000130714 | POMT1 | chr9 | + | 131511336 | 131512136 | 131511336 | 131511472 | 131512040 | 131512136 | 6 | 3 | 7 |
| RI | 4582 | ENSG00000077152 | UBE2T | chr1 | - | 202334988 | 202335818 | 202334988 | 202335058 | 202335645 | 202335818 | 2 | 0 | 1 |
| RI | 6446 | ENSG00000115677 | HDLBP | chr2 | - | 241268953 | 241269295 | 241268953 | 241269105 | 241269200 | 241269295 | 16 | 6 | 12 |
| RI | 7385 | ENSG00000077463 | SIRT6 | chr19 | - | 4175027 | 4175760 | 4175027 | 4175111 | 4175679 | 4175760 | 9 | 7 | 6 |
| RI | 2376 | ENSG00000179152 | TCAIM | chr3 | + | 44400354 | 44401334 | 44400354 | 44400587 | 44401202 | 44401334 | 8 | 7 | 8 |
| RI | 7208 | ENSG00000182325 | FBXL6 | chr8 | - | 144356989 | 144357502 | 144356989 | 144357121 | 144357438 | 144357502 | 8 | 0 | 4 |
| RI | 940 | ENSG00000186765 | FSCN2 | chr17 | + | 81536145 | 81536789 | 81536145 | 81536267 | 81536621 | 81536789 | 4 | 4 | 7 |
| RI | 825 | ENSG00000267080 | ASB16-AS1 | chr17 | - | 44181622 | 44183093 | 44181622 | 44181720 | 44182929 | 44183093 | 6 | 2 | 10 |
| RI | 1072 | ENSG00000177369 | FLJ40194 | chr17 | + | 49252245 | 49252918 | 49252245 | 49252674 | 49252853 | 49252918 | 4 | 4 | 2 |
| RI | 4070 | ENSG00000164509 | IL31RA | chr5 | + | 55916643 | 55922850 | 55916643 | 55916834 | 55922395 | 55922850 | 2 | 2 | 2 |
| RI | 4519 | ENSG00000143126 | CELSR2 | chr1 | + | 109272277 | 109272728 | 109272277 | 109272405 | 109272639 | 109272728 | 0 | 0 | 0 |
| RI | 5249 | ENSG00000224078 | SNHG14 | chr15 | + | 25053827 | 25055230 | 25053827 | 25053967 | 25054969 | 25055230 | 4 | 2 | 0 |
| RI | 6633 | ENSG00000196998 | WDR45 | chrX | - | 49100581 | 49101144 | 49100581 | 49100677 | 49101049 | 49101144 | 6 | 2 | 2 |
| RI | 8550 | ENSG00000187535 | IFT140 | chr16 | - | 1525886 | 1526796 | 1525886 | 1526077 | 1526732 | 1526796 | 0 | 3 | 7 |
| RI | 1107 | ENSG00000131899 | LLGL1 | chr17 | + | 18236834 | 18237773 | 18236834 | 18236939 | 18237480 | 18237773 | 5 | 3 | 3 |
| RI | 5189 | ENSG00000048342 | CC2D2A | chr4 | + | 15560530 | 15563522 | 15560530 | 15560622 | 15563354 | 15563522 | 13 | 27 | 2 |
| RI | 6626 | ENSG00000107897 | ACBD5 | chr10 | - | 27240318 | 27240797 | 27240318 | 27240484 | 27240673 | 27240797 | 30 | 12 | 14 |
| RI | 1844 | ENSG00000100722 | ZC3H14 | chr14 | + | 88567857 | 88568153 | 88567857 | 88567916 | 88568038 | 88568153 | 12 | 13 | 24 |
| RI | 2470 | ENSG00000250722 | SELENOP | chr5 | - | 42804655 | 42807108 | 42804655 | 42804773 | 42806895 | 42807108 | 10 | 5 | 4 |
| RI | 5266 | ENSG00000224078 | SNHG14 | chr15 | + | 25111764 | 25117515 | 25111764 | 25111894 | 25117409 | 25117515 | 16 | 10 | 9 |
| RI | 8185 | ENSG00000072694 | FCGR2B | chr1 | + | 161671391 | 161673229 | 161671391 | 161671649 | 161672974 | 161673229 | 13 | 13 | 16 |
| RI | 1834 | ENSG00000099797 | TECR | chr19 | + | 14563657 | 14563903 | 14563657 | 14563702 | 14563770 | 14563903 | 13 | 4 | 6 |
| RI | 4622 | ENSG00000179593 | ALOX15B | chr17 | + | 8042368 | 8042884 | 8042368 | 8042491 | 8042780 | 8042884 | 0 | 0 | 0 |
| RI | 5014 | ENSG00000076706 | MCAM | chr11 | - | 119312769 | 119314576 | 119312769 | 119312949 | 119314488 | 119314576 | 0 | 0 | 0 |
| RI | 4023 | ENSG00000157193 | LRP8 | chr1 | - | 53271226 | 53275753 | 53271226 | 53271346 | 53275630 | 53275753 | 2 | 0 | 6 |
| RI | 5267 | ENSG00000224078 | SNHG14 | chr15 | + | 25111764 | 25117515 | 25111764 | 25115656 | 25117409 | 25117515 | 13 | 7 | 8 |
| RI | 5617 | ENSG00000174915 | PTDSS2 | chr11 | + | 490419 | 491399 | 490419 | 490642 | 491120 | 491399 | 12 | 2 | 7 |
| RI | 6193 | ENSG00000172638 | EFEMP2 | chr11 | - | 65870120 | 65870658 | 65870120 | 65870237 | 65870535 | 65870658 | 5 | 1 | 3 |
| RI | 2155 | ENSG00000106123 | EPHB6 | chr7 | + | 142865959 | 142866605 | 142865959 | 142866316 | 142866480 | 142866605 | 0 | 0 | 3 |
| RI | 5130 | ENSG00000186281 | GPAT2 | chr2 | - | 96024772 | 96025603 | 96024772 | 96024843 | 96025484 | 96025603 | 0 | 1 | 4 |
| RI | 8423 | ENSG00000140682 | TGFB1I1 | chr16 | + | 31474562 | 31476185 | 31474562 | 31474757 | 31476011 | 31476185 | 0 | 2 | 0 |
| RI | 1832 | ENSG00000186648 | CARMIL3 | chr14 | + | 24062708 | 24063183 | 24062708 | 24062846 | 24063119 | 24063183 | 2 | 3 | 6 |
| RI | 310 | ENSG00000115268 | RPS15 | chr19 | + | 1438806 | 1440253 | 1438806 | 1438892 | 1439960 | 1440253 | 17 | 5 | 23 |
| RI | 6712 | ENSG00000142675 | CNKSR1 | chr1 | + | 26188233 | 26188503 | 26188233 | 26188307 | 26188441 | 26188503 | 0 | 0 | 0 |
| RI | 2506 | ENSG00000131591 | C1orf159 | chr1 | - | 1083914 | 1084383 | 1083914 | 1084086 | 1084352 | 1084383 | 5 | 4 | 3 |
| RI | 4603 | ENSG00000037280 | FLT4 | chr5 | - | 180613010 | 180614179 | 180613010 | 180613110 | 180614067 | 180614179 | 1 | 2 | 1 |
| RI | 4623 | ENSG00000269713 | NBPF9 | chr1 | - | 149060522 | 149061383 | 149060522 | 149060695 | 149061331 | 149061383 | 1 | 1 | 0 |
| RI | 3585 | ENSG00000163472 | TMEM79 | chr1 | + | 156285183 | 156286473 | 156285183 | 156285983 | 156286259 | 156286473 | 6 | 3 | 2 |
| RI | 7079 | ENSG00000110697 | PITPNM1 | chr11 | - | 67493687 | 67494070 | 67493687 | 67493837 | 67493921 | 67494070 | 6 | 0 | 11 |
| RI | 1034 | ENSG00000185813 | PCYT2 | chr17 | - | 81905669 | 81906177 | 81905669 | 81905735 | 81906099 | 81906177 | 0 | 3 | 4 |
| RI | 7903 | ENSG00000248712 | CCDC153 | chr11 | - | 119193150 | 119193885 | 119193150 | 119193243 | 119193765 | 119193885 | 2 | 6 | 2 |
| RI | 2664 | ENSG00000154743 | TSEN2 | chr3 | + | 12529761 | 12531659 | 12529761 | 12529873 | 12531538 | 12531659 | 3 | 2 | 18 |
| RI | 3902 | ENSG00000109184 | DCUN1D4 | chr4 | + | 51886567 | 51887244 | 51886567 | 51886638 | 51887122 | 51887244 | 20 | 11 | 18 |
| RI | 7826 | ENSG00000149743 | TRPT1 | chr11 | - | 64224099 | 64224341 | 64224099 | 64224162 | 64224284 | 64224341 | 5 | 4 | 25 |
| RI | 801 | ENSG00000131462 | TUBG1 | chr17 | + | 42613848 | 42614412 | 42613848 | 42613998 | 42614259 | 42614412 | 0 | 0 | 1 |
| RI | 1015 | ENSG00000169738 | DCXR | chr17 | - | 82036190 | 82036448 | 82036190 | 82036322 | 82036383 | 82036448 | 7 | 10 | 6 |
| RI | 2581 | ENSG00000185033 | SEMA4B | chr15 | + | 90224967 | 90225397 | 90224967 | 90225178 | 90225281 | 90225397 | 7 | 0 | 9 |
| RI | 8585 | ENSG00000134901 | POGLUT2 | chr13 | - | 102790900 | 102791430 | 102790900 | 102791138 | 102791257 | 102791430 | 0 | 0 | 0 |
| RI | 2265 | ENSG00000143774 | GUK1 | chr1 | + | 228147405 | 228147699 | 228147405 | 228147544 | 228147620 | 228147699 | 7 | 8 | 15 |
| RI | 4580 | ENSG00000077152 | UBE2T | chr1 | - | 202333009 | 202333335 | 202333009 | 202333093 | 202333236 | 202333335 | 2 | 1 | 6 |
| RI | 6811 | ENSG00000156313 | RPGR | chrX | - | 38287093 | 38288041 | 38287093 | 38287245 | 38287864 | 38288041 | 4 | 17 | 14 |
| RI | 7449 | ENSG00000159899 | NPR2 | chr9 | + | 35808508 | 35808853 | 35808508 | 35808683 | 35808754 | 35808853 | 8 | 5 | 14 |
| RI | 1081 | ENSG00000141698 | NT5C3B | chr17 | - | 41835587 | 41835957 | 41835587 | 41835682 | 41835858 | 41835957 | 0 | 4 | 10 |
| RI | 2991 | ENSG00000114739 | ACVR2B | chr3 | + | 38478140 | 38478518 | 38478140 | 38478292 | 38478374 | 38478518 | 0 | 0 | 0 |
| RI | 4325 | ENSG00000111445 | RFC5 | chr12 | + | 118017689 | 118018053 | 118017689 | 118017815 | 118017994 | 118018053 | 10 | 16 | 7 |
| RI | 8476 | ENSG00000073060 | SCARB1 | chr12 | - | 124782682 | 124786503 | 124782682 | 124782811 | 124786440 | 124786503 | 19 | 8 | 10 |
| RI | 3239 | ENSG00000095906 | NUBP2 | chr16 | + | 1787940 | 1788207 | 1787940 | 1788051 | 1788160 | 1788207 | 2 | 0 | 2 |
| RI | 4352 | ENSG00000169242 | EFNA1 | chr1 | + | 155133502 | 155133780 | 155133502 | 155133568 | 155133729 | 155133780 | 4 | 0 | 5 |
| RI | 6598 | ENSG00000188878 | FBF1 | chr17 | - | 75913912 | 75914298 | 75913912 | 75914050 | 75914121 | 75914298 | 1 | 1 | 3 |
| RI | 8232 | ENSG00000110888 | CAPRIN2 | chr12 | - | 30714958 | 30716676 | 30714958 | 30715144 | 30716507 | 30716676 | 40 | 32 | 41 |
| RI | 1005 | ENSG00000100360 | IFT27 | chr22 | - | 36775673 | 36776125 | 36775673 | 36775794 | 36776030 | 36776125 | 12 | 4 | 12 |
| RI | 1391 | ENSG00000123815 | COQ8B | chr19 | - | 40700309 | 40702693 | 40700309 | 40700447 | 40702599 | 40702693 | 4 | 1 | 9 |
| RI | 3408 | ENSG00000138382 | METTL5 | chr2 | - | 169824488 | 169824905 | 169824488 | 169824640 | 169824867 | 169824905 | 2 | 0 | 4 |
| RI | 3476 | ENSG00000064687 | ABCA7 | chr19 | + | 1056906 | 1058747 | 1056906 | 1057429 | 1058617 | 1058747 | 6 | 3 | 6 |
| RI | 3980 | ENSG00000188004 | SNHG28 | chr1 | - | 159838364 | 159840866 | 159838364 | 159838451 | 159840710 | 159840866 | 8 | 6 | 2 |
| RI | 4231 | ENSG00000132781 | MUTYH | chr1 | - | 45332917 | 45333324 | 45332917 | 45332959 | 45333284 | 45333324 | 5 | 3 | 12 |
| RI | 5040 | ENSG00000285533 | RELA-DT | chr11 | + | 65663426 | 65665543 | 65663426 | 65663683 | 65665420 | 65665543 | 6 | 4 | 6 |
| RI | 5631 | ENSG00000111665 | CDCA3 | chr12 | - | 6850466 | 6851009 | 6850466 | 6850596 | 6850832 | 6851009 | 0 | 1 | 0 |
| RI | 6077 | ENSG00000269313 | MAGIX | chrX | + | 49164706 | 49165090 | 49164706 | 49164789 | 49164862 | 49165090 | 1 | 0 | 1 |
| RI | 6668 | ENSG00000109736 | MFSD10 | chr4 | - | 2931374 | 2931936 | 2931374 | 2931465 | 2931824 | 2931936 | 5 | 2 | 2 |
| RI | 6706 | ENSG00000204271 | SPIN3 | chrX | - | 56978291 | 56978660 | 56978291 | 56978417 | 56978579 | 56978660 | 13 | 2 | 5 |
| RI | 8079 | ENSG00000214279 | SCART1 | chr10 | + | 133457278 | 133458656 | 133457278 | 133457575 | 133458359 | 133458656 | 13 | 1 | 4 |
| RI | 8267 | ENSG00000181588 | MEX3D | chr19 | - | 1554671 | 1556923 | 1554671 | 1555389 | 1555576 | 1556923 | 2 | 1 | 2 |
| RI | 8508 | ENSG00000174233 | ADCY6 | chr12 | - | 48777103 | 48777521 | 48777103 | 48777409 | 48777409 | 48777521 | 0 | 1 | 0 |
| RI | 965 | ENSG00000176170 | SPHK1 | chr17 | + | 76385450 | 76386137 | 76385450 | 76385654 | 76385984 | 76386137 | 3 | 0 | 0 |
| RI | 7516 | ENSG00000165699 | TSC1 | chr9 | - | 132897183 | 132897610 | 132897183 | 132897345 | 132897427 | 132897610 | 6 | 11 | 8 |
| RI | 3337 | ENSG00000159588 | CCDC17 | chr1 | - | 45622750 | 45623652 | 45622750 | 45622834 | 45622954 | 45623652 | 8 | 2 | 0 |
| RI | 5247 | ENSG00000204713 | TRIM27 | chr6 | - | 28903001 | 28908840 | 28903001 | 28904665 | 28908807 | 28908840 | 17 | 29 | 27 |
| RI | 5698 | ENSG00000079785 | DDX1 | chr2 | + | 15602547 | 15603275 | 15602547 | 15602631 | 15603191 | 15603275 | 21 | 32 | 52 |
| RI | 3890 | ENSG00000152527 | PLEKHH2 | chr2 | + | 43710488 | 43712383 | 43710488 | 43710575 | 43712224 | 43712383 | 2 | 0 | 5 |
| RI | 573 | ENSG00000185513 | L3MBTL1 | chr20 | + | 43540752 | 43541189 | 43540752 | 43540815 | 43540933 | 43541189 | 4 | 1 | 3 |
| RI | 1305 | ENSG00000197912 | SPG7 | chr16 | + | 89554485 | 89556085 | 89554485 | 89554563 | 89555920 | 89556085 | 5 | 1 | 19 |
| RI | 5069 | ENSG00000115421 | PAPOLG | chr2 | + | 60793970 | 60795020 | 60793970 | 60794191 | 60794963 | 60795020 | 7 | 8 | 5 |
| RI | 6414 | ENSG00000123191 | ATP7B | chr13 | - | 51957515 | 51960322 | 51957515 | 51957607 | 51960147 | 51960322 | 2 | 1 | 0 |
| RI | 652 | ENSG00000176225 | RTTN | chr18 | - | 70005197 | 70006484 | 70005197 | 70005267 | 70006380 | 70006484 | 4 | 6 | 3 |
| RI | 8233 | ENSG00000110888 | CAPRIN2 | chr12 | - | 30719078 | 30720915 | 30719078 | 30719225 | 30720810 | 30720915 | 15 | 11 | 10 |
| RI | 430 | ENSG00000105204 | DYRK1B | chr19 | - | 39826671 | 39826987 | 39826671 | 39826867 | 39826951 | 39826987 | 17 | 5 | 14 |
| RI | 1187 | ENSG00000162004 | CCDC78 | chr16 | - | 722921 | 723936 | 722921 | 723161 | 723856 | 723936 | 2 | 1 | 3 |
| RI | 2586 | ENSG00000133048 | CHI3L1 | chr1 | - | 203178930 | 203179877 | 203178930 | 203179585 | 203179760 | 203179877 | 0 | 0 | 0 |
| RI | 3629 | ENSG00000160446 | ZDHHC12 | chr9 | - | 128722437 | 128724127 | 128722437 | 128722574 | 128723993 | 128724127 | 2 | 0 | 1 |
| RI | 15 | ENSG00000241945 | PWP2 | chr21 | + | 44124579 | 44126119 | 44124579 | 44124727 | 44126010 | 44126119 | 0 | 0 | 0 |
| RI | 4586 | ENSG00000106462 | EZH2 | chr7 | - | 148816683 | 148818117 | 148816683 | 148816778 | 148817876 | 148818117 | 14 | 17 | 6 |
| RI | 5741 | ENSG00000006062 | MAP3K14 | chr17 | - | 45270412 | 45271221 | 45270412 | 45270563 | 45270851 | 45271221 | 16 | 8 | 9 |
| RI | 5785 | ENSG00000123607 | TTC21B | chr2 | - | 165870796 | 165874832 | 165870796 | 165871020 | 165874678 | 165874832 | 44 | 33 | 22 |
| RI | 2600 | ENSG00000182150 | ERCC6L2 | chr9 | + | 95970575 | 95973088 | 95970575 | 95970656 | 95971932 | 95973088 | 25 | 37 | 15 |
| RI | 5260 | ENSG00000224078 | SNHG14 | chr15 | + | 25107909 | 25109403 | 25107909 | 25108038 | 25108997 | 25109403 | 5 | 7 | 3 |
| RI | 6397 | ENSG00000179818 | PCBP1-AS1 | chr2 | - | 70049195 | 70051305 | 70049195 | 70050252 | 70051202 | 70051305 | 26 | 12 | 10 |
| RI | 1964 | ENSG00000112655 | PTK7 | chr6 | + | 43142171 | 43143620 | 43142171 | 43142299 | 43143416 | 43143620 | 6 | 0 | 1 |
| RI | 4866 | ENSG00000183963 | SMTN | chr22 | + | 31099061 | 31099896 | 31099061 | 31099179 | 31099744 | 31099896 | 2 | 0 | 0 |
| RI | 425 | ENSG00000099326 | MZF1 | chr19 | - | 58570343 | 58571429 | 58570343 | 58570527 | 58570993 | 58571429 | 19 | 10 | 15 |
| RI | 8421 | ENSG00000140682 | TGFB1I1 | chr16 | + | 31473440 | 31473734 | 31473440 | 31473556 | 31473681 | 31473734 | 0 | 0 | 0 |
| RI | 8392 | ENSG00000111077 | TNS2 | chr12 | + | 53060674 | 53061264 | 53060674 | 53060759 | 53060848 | 53061264 | 6 | 3 | 6 |
| RI | 5931 | ENSG00000114388 | NPRL2 | chr3 | - | 50349664 | 50350022 | 50349664 | 50349828 | 50349912 | 50350022 | 18 | 14 | 18 |
| RI | 5071 | ENSG00000067445 | TRO | chrX | + | 54926582 | 54927105 | 54926582 | 54926625 | 54927042 | 54927105 | 2 | 0 | 0 |
| RI | 1757 | ENSG00000101384 | JAG1 | chr20 | - | 10648548 | 10649107 | 10648548 | 10648676 | 10649060 | 10649107 | 4 | 2 | 4 |
| RI | 3062 | ENSG00000169727 | GPS1 | chr17 | + | 82056628 | 82057470 | 82056628 | 82056766 | 82057052 | 82057470 | 6 | 1 | 0 |
| RI | 3452 | ENSG00000144591 | GMPPA | chr2 | + | 219504082 | 219505362 | 219504082 | 219504218 | 219505227 | 219505362 | 15 | 9 | 6 |
| RI | 4503 | ENSG00000115556 | PLCD4 | chr2 | + | 218628028 | 218629663 | 218628028 | 218628230 | 218629518 | 218629663 | 1 | 0 | 4 |
| RI | 7855 | ENSG00000172803 | SNX32 | chr11 | + | 65849919 | 65850271 | 65849919 | 65850030 | 65850149 | 65850271 | 0 | 0 | 0 |
| RI | 8390 | ENSG00000111077 | TNS2 | chr12 | + | 53058315 | 53058637 | 53058315 | 53058445 | 53058571 | 53058637 | 0 | 0 | 2 |
| RI | 8420 | ENSG00000140682 | TGFB1I1 | chr16 | + | 31472170 | 31473556 | 31472170 | 31472201 | 31473440 | 31473556 | 0 | 0 | 0 |
| RI | 5675 | ENSG00000123983 | ACSL3 | chr2 | + | 222927016 | 222928936 | 222927016 | 222927189 | 222928861 | 222928936 | 54 | 68 | 64 |
| RI | 8491 | ENSG00000111335 | OAS2 | chr12 | + | 113004474 | 113005222 | 113004474 | 113004673 | 113004933 | 113005222 | 22 | 17 | 8 |
| RI | 1956 | ENSG00000204536 | CCHCR1 | chr6 | - | 31144686 | 31145073 | 31144686 | 31144788 | 31144884 | 31145073 | 2 | 0 | 4 |
| RI | 7337 | ENSG00000148339 | SLC25A25 | chr9 | + | 128106352 | 128107179 | 128106352 | 128106520 | 128107028 | 128107179 | 1 | 0 | 1 |
| RI | 8007 | ENSG00000196155 | PLEKHG4 | chr16 | + | 67277509 | 67280806 | 67277509 | 67280543 | 67280710 | 67280806 | 1 | 6 | 6 |
| RI | 1089 | ENSG00000276070 | CCL4L2 | chr17 | + | 36211775 | 36212878 | 36211775 | 36211890 | 36212496 | 36212878 | 62 | 12 | 3 |
| RI | 4259 | ENSG00000143801 | PSEN2 | chr1 | + | 226897719 | 226899555 | 226897719 | 226897829 | 226899451 | 226899555 | 3 | 9 | 4 |
| RI | 6022 | ENSG00000112139 | MDGA1 | chr6 | - | 37654799 | 37655896 | 37654799 | 37654932 | 37655699 | 37655896 | 0 | 4 | 4 |
| RI | 7329 | ENSG00000197816 | CCDC180 | chr9 | + | 97307735 | 97308132 | 97307735 | 97307806 | 97307982 | 97308132 | 3 | 1 | 7 |
| RI | 7428 | ENSG00000070061 | ELP1 | chr9 | - | 108897147 | 108898581 | 108897147 | 108897283 | 108898501 | 108898581 | 6 | 10 | 3 |
| RI | 3302 | ENSG00000228794 | LINC01128 | chr1 | + | 851926 | 852766 | 851926 | 852014 | 852670 | 852766 | 18 | 4 | 31 |
| RI | 4841 | ENSG00000155367 | PPM1J | chr1 | - | 112712743 | 112713031 | 112712743 | 112712803 | 112712920 | 112713031 | 3 | 1 | 5 |
| RI | 4009 | ENSG00000214595 | EML6 | chr2 | + | 54866765 | 54869367 | 54866765 | 54866884 | 54869180 | 54869367 | 0 | 0 | 0 |
| RI | 4560 | ENSG00000225470 | JPX | chrX | + | 73998953 | 73999669 | 73998953 | 73999051 | 73999185 | 73999669 | 18 | 3 | 8 |
| RI | 1467 | ENSG00000140983 | RHOT2 | chr16 | + | 668655 | 670175 | 668655 | 668699 | 670122 | 670175 | 7 | 5 | 3 |
| RI | 2811 | ENSG00000160221 | GATD3A | chr21 | + | 44144820 | 44145723 | 44144820 | 44144916 | 44145570 | 44145723 | 0 | 32 | 0 |
| RI | 2445 | ENSG00000109501 | WFS1 | chr4 | + | 6277450 | 6277687 | 6277450 | 6277521 | 6277647 | 6277687 | 2 | 3 | 6 |
| RI | 6672 | ENSG00000172613 | RAD9A | chr11 | + | 67395911 | 67396175 | 67395911 | 67396021 | 67396110 | 67396175 | 0 | 0 | 2 |
| RI | 7799 | ENSG00000132199 | ENOSF1 | chr18 | - | 690548 | 691276 | 690548 | 690745 | 691203 | 691276 | 5 | 12 | 15 |
| RI | 8375 | ENSG00000111752 | PHC1 | chr12 | + | 8932562 | 8933350 | 8932562 | 8933063 | 8933254 | 8933350 | 21 | 11 | 16 |
| RI | 1 | ENSG00000160285 | LSS | chr21 | - | 46188445 | 46191235 | 46188445 | 46188821 | 46191075 | 46191235 | 14 | 4 | 6 |
| RI | 4868 | ENSG00000135924 | DNAJB2 | chr2 | + | 219282489 | 219283489 | 219282836 | 219282929 | 219283418 | 219283489 | 9 | 11 | 12 |
| RI | 6234 | ENSG00000234545 | FAM133B | chr7 | - | 92577654 | 92578182 | 92577654 | 92577676 | 92578149 | 92578182 | 84 | 70 | 69 |
| RI | 6250 | ENSG00000179406 | LINC00174 | chr7 | - | 66491030 | 66491471 | 66491030 | 66491239 | 66491349 | 66491471 | 20 | 7 | 14 |
| RI | 2059 | ENSG00000133706 | LARS1 | chr5 | - | 146140203 | 146143084 | 146140203 | 146140261 | 146142855 | 146143084 | 51 | 59 | 51 |
| RI | 2133 | ENSG00000154153 | RETREG1 | chr5 | - | 16478033 | 16478987 | 16478033 | 16478095 | 16478849 | 16478987 | 5 | 12 | 13 |
| RI | 2435 | ENSG00000160972 | PPP1R16A | chr8 | + | 144500095 | 144500391 | 144500095 | 144500199 | 144500266 | 144500391 | 3 | 2 | 0 |
| RI | 4304 | ENSG00000135541 | AHI1 | chr6 | - | 135300499 | 135302817 | 135300499 | 135300558 | 135302669 | 135302817 | 11 | 8 | 7 |
| RI | 5425 | ENSG00000235703 | LINC00894 | chrX | + | 149945347 | 149948175 | 149945347 | 149945677 | 149945783 | 149948175 | 10 | 2 | 3 |
| RI | 5621 | ENSG00000119912 | IDE | chr10 | - | 92506441 | 92508205 | 92506441 | 92506522 | 92508112 | 92508205 | 18 | 11 | 10 |
| RI | 6417 | ENSG00000164896 | FASTK | chr7 | - | 151076912 | 151077236 | 151076912 | 151077027 | 151077100 | 151077236 | 13 | 5 | 10 |
| RI | 8394 | ENSG00000111077 | TNS2 | chr12 | + | 53062619 | 53063257 | 53062619 | 53062697 | 53063088 | 53063257 | 16 | 8 | 20 |
| RI | 800 | ENSG00000131462 | TUBG1 | chr17 | + | 42613646 | 42613998 | 42613646 | 42613733 | 42613848 | 42613998 | 0 | 0 | 2 |
| RI | 6553 | ENSG00000127914 | AKAP9 | chr7 | + | 92062273 | 92065463 | 92062273 | 92062486 | 92065230 | 92065463 | 9 | 13 | 9 |
| RI | 8534 | ENSG00000185189 | NRBP2 | chr8 | - | 143835819 | 143836030 | 143835819 | 143835875 | 143835966 | 143836030 | 0 | 0 | 0 |
| RI | 4163 | ENSG00000003393 | ALS2 | chr2 | - | 201727211 | 201727775 | 201727211 | 201727278 | 201727704 | 201727775 | 8 | 6 | 4 |
| RI | 5824 | ENSG00000124356 | STAMBP | chr2 | + | 73844812 | 73845262 | 73844812 | 73845036 | 73845166 | 73845262 | 3 | 5 | 3 |
| RI | 7411 | ENSG00000176919 | C8G | chr9 | + | 136946650 | 136946974 | 136946650 | 136946689 | 136946767 | 136946974 | 2 | 0 | 0 |
| RI | 7183 | ENSG00000183250 | LINC01547 | chr21 | - | 44933314 | 44935096 | 44933314 | 44934368 | 44934933 | 44935096 | 23 | 22 | 19 |
| RI | 6179 | ENSG00000105877 | DNAH11 | chr7 | + | 21894622 | 21894999 | 21894622 | 21894805 | 21894883 | 21894999 | 0 | 2 | 7 |
| RI | 4377 | ENSG00000135837 | CEP350 | chr1 | + | 180110996 | 180113380 | 180110996 | 180111195 | 180112436 | 180113380 | 154 | 97 | 106 |
| RI | 6445 | ENSG00000115677 | HDLBP | chr2 | - | 241235120 | 241235594 | 241235120 | 241235275 | 241235489 | 241235594 | 8 | 10 | 25 |
| RI | 634 | ENSG00000196756 | SNHG17 | chr20 | - | 38425934 | 38426503 | 38425934 | 38426052 | 38426418 | 38426503 | 18 | 13 | 6 |
| RI | 1061 | ENSG00000267248 | NA | chr17 | - | 60134755 | 60135693 | 60134755 | 60134851 | 60135096 | 60135693 | 5 | 4 | 4 |
| RI | 1198 | ENSG00000175595 | ERCC4 | chr16 | + | 13924529 | 13926756 | 13924529 | 13924637 | 13926560 | 13926756 | 14 | 8 | 5 |
| RI | 1443 | ENSG00000161999 | JMJD8 | chr16 | - | 683860 | 684155 | 683860 | 683909 | 684065 | 684155 | 2 | 2 | 5 |
| RI | 2769 | ENSG00000229036 | VDAC1P8 | chr6 | + | 143505339 | 143506100 | 143505339 | 143505457 | 143505969 | 143506100 | 8 | 26 | 11 |
| RI | 3318 | ENSG00000223393 | NA | chr1 | + | 230874541 | 230875565 | 230874541 | 230875155 | 230875305 | 230875565 | 1 | 2 | 1 |

| | | | | | | | | | | | | |
|---|---|---|---|---|---|---|---|---|---|---|---|---|
| RI | 3517 | ENSG00000117569 | PTBP2 | chr1 | + | 96804799 | 96806452 | 96804799 | 96804939 | 96806418 | 96806452 | 8 [illegible] |
| RI | 3718 | ENSG00000138002 | IFT172 | chr2 | - | 27453629 | 27454418 | 27453629 | 27453739 | 27454353 | 27454418 | 4 [illegible] |
| RI | 4659 | ENSG00000001497 | LAS1L | chrX | - | 65522430 | 65523707 | 65522430 | 65522493 | 65523559 | 65523707 | 8 [illegible] |
| RI | 4769 | ENSG00000075702 | WDR62 | chr19 | + | 36065957 | 36066427 | 36065957 | 36066015 | 36066256 | 36066427 | 0 [illegible] |
| RI | 5423 | ENSG00000166762 | CATSPER2 | chr15 | - | 43632198 | 43635416 | 43632198 | 43632363 | 43635359 | 43635416 | 8 [illegible] |
| RI | 3089 | ENSG00000164073 | MFSD8 | chr4 | - | 127921523 | 127921963 | 127921523 | 127921771 | 127921859 | 127921963 | 14 [illegible] |
| RI | 6698 | ENSG00000174080 | CTSF | chr11 | - | 66567443 | 66568082 | 66567443 | 66567662 | 66567983 | 66568082 | 0 [illegible] |
| RI | 7220 | ENSG00000185803 | SLC52A2 | chr8 | + | 144359622 | 144360493 | 144359622 | 144359747 | 144360114 | 144360493 | 38 [illegible] |
| RI | 1386 | ENSG00000263072 | ZNF213-AS1 | chr16 | - | 3129247 | 3129653 | 3129247 | 3129310 | 3129460 | 3129653 | 25 [illegible] |
| RI | 6111 | ENSG00000232940 | HCG25 | chr6 | + | 33249533 | 33249889 | 33249533 | 33249655 | 33249789 | 33249889 | 2 [illegible] |
| RI | 7423 | ENSG00000070061 | ELP1 | chr9 | - | 108880051 | 108881765 | 108880051 | 108880165 | 108881662 | 108881765 | 9 [illegible] |
| RI | 350 | ENSG00000271109 | NA | chr19 | - | 34860173 | 34860715 | 34860173 | 34860302 | 34860533 | 34860715 | 33 [illegible] |
| RI | 1466 | ENSG00000140983 | RHOT2 | chr16 | + | 668487 | 668699 | 668487 | 668569 | 668658 | 668699 | 7 [illegible] |
| RI | 3767 | ENSG00000072210 | ALDH3A2 | chr17 | + | 19648780 | 19649124 | 19648780 | 19648843 | 19648934 | 19649124 | 32 [illegible] |
| RI | 5223 | ENSG00000125875 | TBC1D20 | chr20 | - | 440247 | 442043 | 440247 | 440389 | 440964 | 442043 | 22 [illegible] |
| RI | 5475 | ENSG00000223960 | CHROMR | chr2 | + | 178426365 | 178430889 | 178426365 | 178426680 | 178430769 | 178430889 | 42 [illegible] |
| RI | 1529 | ENSG00000198690 | FAN1 | chr15 | + | 30925126 | 30928656 | 30925126 | 30925291 | 30925788 | 30928656 | 12 [illegible] |
| RI | 2398 | ENSG00000213445 | SIPA1 | chr11 | + | 65646455 | 65647065 | 65646455 | 65646663 | 65646969 | 65647065 | 14 [illegible] |
| RI | 2824 | ENSG00000242086 | MUC20-OT1 | chr3 | + | 195708153 | 195708841 | 195708153 | 195708513 | 195708746 | 195708841 | 12 [illegible] |
| RI | 3216 | ENSG00000121410 | A1BG | chr19 | - | 58351390 | 58352555 | 58351390 | 58351687 | 58352282 | 58352555 | 9 [illegible] |
| RI | 5352 | ENSG00000066923 | STAG3 | chr7 | + | 100188452 | 100189016 | 100188452 | 100188529 | 100188811 | 100189016 | 2 [illegible] |
| RI | 7390 | ENSG00000213722 | DDAH2 | chr6 | - | 31728172 | 31728745 | 31728172 | 31728292 | 31728645 | 31728745 | 10 [illegible] |
| RI | 1768 | ENSG00000119682 | AREL1 | chr14 | - | 74669946 | 74670871 | 74669946 | 74670142 | 74670761 | 74670871 | 31 [illegible] |
| RI | 1806 | ENSG00000100888 | CHD8 | chr14 | - | 21400412 | 21401071 | 21400412 | 21400612 | 21400874 | 21401071 | 46 [illegible] |
| RI | 2869 | ENSG00000184470 | TXNRD2 | chr22 | - | 19875521 | 19877234 | 19875521 | 19875807 | 19877108 | 19877234 | 9 [illegible] |
| RI | 3625 | ENSG00000141577 | CEP131 | chr17 | - | 81189596 | 81189975 | 81189596 | 81189843 | 81189914 | 81189975 | 1 [illegible] |
| RI | 4018 | ENSG00000120798 | NR2C1 | chr12 | - | 95057730 | 95058489 | 95057730 | 95057878 | 95058309 | 95058489 | 179 [illegible] |
| RI | 8288 | ENSG00000170486 | KRT72 | chr12 | - | 52585588 | 52586172 | 52585588 | 52585835 | 52585962 | 52586172 | 4 [illegible] |
| RI | 4307 | ENSG00000135541 | AHI1 | chr6 | - | 135301407 | 135302817 | 135301407 | 135301539 | 135302711 | 135302817 | 10 [illegible] |
| RI | 7249 | ENSG00000137563 | GGH | chr8 | - | 63023906 | 63024186 | 63023906 | 63023997 | 63024079 | 63024186 | 0 [illegible] |
| RI | 7391 | ENSG00000130653 | PNPLA7 | chr9 | - | 137460633 | 137461620 | 137460633 | 137460737 | 137461535 | 137461620 | 1 [illegible] |
| RI | 158 | ENSG00000170949 | ZNF160 | chr19 | - | 53091412 | 53091720 | 53091412 | 53091553 | 53091636 | 53091720 | 14 [illegible] |
| RI | 1556 | ENSG00000062524 | LTK | chr15 | - | 41504341 | 41504640 | 41504341 | 41504432 | 41504505 | 41504640 | 1 [illegible] |
| RI | 2114 | ENSG00000131844 | MCCC2 | chr5 | + | 71641002 | 71642727 | 71641002 | 71641075 | 71642663 | 71642727 | 44 [illegible] |
| RI | 4753 | ENSG00000147813 | NAPRT | chr8 | - | 143574784 | 143575345 | 143574784 | 143575093 | 143575229 | 143575345 | 28 [illegible] |
| RI | 7447 | ENSG00000155636 | RBM45 | chr2 | + | 178123521 | 178123912 | 178123521 | 178123651 | 178123827 | 178123912 | 25 [illegible] |
| RI | 3617 | ENSG00000166482 | MFAP4 | chr17 | - | 19385098 | 19385454 | 19385098 | 19385281 | 19385357 | 19385454 | 5 [illegible] |
| RI | 1195 | ENSG00000033011 | ALG1 | chr16 | + | 5073152 | 5075536 | 5073152 | 5073256 | 5075397 | 5075536 | 5 [illegible] |
| RI | 3023 | ENSG00000173145 | NOC3L | chr10 | - | 94357173 | 94358215 | 94357173 | 94357331 | 94358082 | 94358215 | 35 [illegible] |
| RI | 7912 | ENSG00000179532 | DNHD1 | chr11 | + | 6497584 | 6498961 | 6497584 | 6497667 | 6498722 | 6498961 | 11 [illegible] |
| RI | 886 | ENSG00000185359 | HGS | chr17 | + | 81696606 | 81696998 | 81696606 | 81696747 | 81696829 | 81696998 | 6 [illegible] |
| RI | 2684 | ENSG00000020577 | SAMD4A | chr14 | + | 54784339 | 54784620 | 54784339 | 54784381 | 54784536 | 54784620 | 0 [illegible] |
| RI | 2818 | ENSG00000081307 | UBA5 | chr3 | + | 132671776 | 132672177 | 132671776 | 132671881 | 132672049 | 132672177 | 23 [illegible] |
| RI | 3790 | ENSG00000164050 | PLXNB1 | chr3 | - | 48410454 | 48411036 | 48410454 | 48410558 | 48410867 | 48411036 | 0 [illegible] |
| RI | 7540 | ENSG00000130714 | POMT1 | chr9 | + | 131509902 | 131510415 | 131509902 | 131510062 | 131510259 | 131510415 | 11 [illegible] |
| RI | 1421 | ENSG00000167508 | MVD | chr16 | - | 88657435 | 88658029 | 88657435 | 88657582 | 88657734 | 88658029 | 7 [illegible] |
| RI | 2066 | ENSG00000125459 | MSTO1 | chr1 | + | 155611215 | 155611603 | 155611215 | 155611291 | 155611426 | 155611603 | 5 [illegible] |
| RI | 3008 | ENSG00000135083 | CCNJL | chr5 | - | 160254128 | 160255708 | 160254128 | 160254335 | 160255548 | 160255708 | 6 [illegible] |
| RI | 5972 | ENSG00000204410 | MSH5 | chr6 | + | 31743903 | 31744299 | 31743903 | 31744025 | 31744138 | 31744299 | 9 [illegible] |
| RI | 4355 | ENSG00000099840 | IZUMO4 | chr19 | + | 2097928 | 2098127 | 2097928 | 2097955 | 2098051 | 2098127 | 2 [illegible] |
| RI | 7717 | ENSG00000136319 | TTC5 | chr14 | - | 20291982 | 20295526 | 20291982 | 20292127 | 20295311 | 20295526 | 7 [illegible] |
| RI | 524 | ENSG00000026036 | RTEL1-TNFRSF6B | chr20 | + | 63695777 | 63697191 | 63695777 | 63695854 | 63696759 | 63697191 | 6 [illegible] |
| RI | 4393 | ENSG00000145020 | AMT | chr3 | - | 49422103 | 49422451 | 49422103 | 49422271 | 49422360 | 49422451 | 16 [illegible] |
| RI | 5924 | ENSG00000114388 | NPRL2 | chr3 | - | 50347758 | 50348563 | 50347758 | 50347901 | 50348522 | 50348563 | 13 [illegible] |
| RI | 6964 | ENSG00000068366 | ACSL4 | chrX | - | 109663210 | 109665494 | 109663210 | 109663402 | 109665419 | 109665494 | 24 [illegible] |
| RI | 7742 | ENSG00000065717 | TLE2 | chr19 | - | 3019282 | 3019773 | 3019282 | 3019466 | 3019698 | 3019773 | 0 [illegible] |
| RI | 2092 | ENSG00000146670 | CDCA5 | chr11 | - | 65083363 | 65083550 | 65083363 | 65083399 | 65083484 | 65083550 | 1 [illegible] |
| RI | 3991 | ENSG00000233184 | NA | chr1 | + | 101080886 | 101083574 | 101080886 | 101080998 | 101083460 | 101083574 | 22 [illegible] |
| RI | 4119 | ENSG00000116337 | AMPD2 | chr1 | + | 109629326 | 109629916 | 109629326 | 109629486 | 109629795 | 109629916 | 15 [illegible] |
| RI | 5381 | ENSG00000114786 | ABHD14A-ACY1 | chr3 | + | 51986987 | 51987196 | 51986987 | 51987061 | 51987146 | 51987196 | 3 [illegible] |
| RI | 6901 | ENSG00000120509 | PDZD11 | chrX | - | 70289254 | 70289593 | 70289254 | 70289354 | 70289521 | 70289593 | 3 [illegible] |
| RI | 771 | ENSG00000129197 | RPAIN | chr17 | + | 5425970 | 5428211 | 5425970 | 5426082 | 5428070 | 5428211 | 73 [illegible] |
| RI | 2413 | ENSG00000139625 | MAP3K12 | chr12 | - | 53486946 | 53487428 | 53486946 | 53487155 | 53487254 | 53487428 | 13 [illegible] |
| RI | 5932 | ENSG00000114388 | NPRL2 | chr3 | - | 50349664 | 50350022 | 50349664 | 50349833 | 50349912 | 50350022 | 16 [illegible] |
| RI | 6164 | ENSG00000242802 | AP5Z1 | chr7 | + | 4789831 | 4790591 | 4789831 | 4789929 | 4790213 | 4790591 | 18 [illegible] |
| RI | 2837 | ENSG00000110844 | PRPF40B | chr12 | + | 49631859 | 49632880 | 49631859 | 49631925 | 49632854 | 49632880 | 6 [illegible] |
| RI | 3461 | ENSG00000144591 | GMPPA | chr2 | + | 219505457 | 219506072 | 219505457 | 219505555 | 219505979 | 219506072 | 13 [illegible] |
| RI | 3648 | ENSG00000239900 | ADSL | chr22 | + | 40361487 | 40363071 | 40361487 | 40361677 | 40362980 | 40363071 | 22 [illegible] |
| RI | 4265 | ENSG00000198837 | DENND4B | chr1 | - | 153930538 | 153931064 | 153930538 | 153930603 | 153930946 | 153931064 | 8 [illegible] |
| RI | 4989 | ENSG00000080947 | CROCCP3 | chr1 | - | 16491009 | 16491316 | 16491009 | 16491084 | 16491185 | 16491316 | 14 [illegible] |
| RI | 5146 | ENSG00000138035 | PNPT1 | chr2 | - | 55680711 | 55680918 | 55680711 | 55680759 | 55680854 | 55680918 | 6 [illegible] |
| RI | 5629 | ENSG00000111665 | CDCA3 | chr12 | - | 6849322 | 6849858 | 6849322 | 6849413 | 6849564 | 6849858 | 0 [illegible] |
| RI | 5801 | ENSG00000168280 | KIF5C | chr2 | + | 148991009 | 148994538 | 148991009 | 148991198 | 148994420 | 148994538 | 0 [illegible] |
| RI | 6097 | ENSG00000155906 | RMND1 | chr6 | - | 151421244 | 151423631 | 151421244 | 151421321 | 151423524 | 151423631 | 29 [illegible] |
| RI | 7575 | ENSG00000166313 | APBB1 | chr11 | - | 6401573 | 6401982 | 6401573 | 6401688 | 6401976 | 6401982 | 15 [illegible] |
| RI | 7995 | ENSG00000150433 | TMEM218 | chr11 | - | 125102131 | 125102809 | 125102131 | 125102351 | 125102636 | 125102809 | 17 [illegible] |
| RI | 2176 | ENSG00000197375 | SLC22A5 | chr5 | + | 132378377 | 132384301 | 132378377 | 132378481 | 132384146 | 132384301 | 15 [illegible] |
| RI | 5242 | ENSG00000225138 | SLC9A3-AS1 | chr5 | + | 479781 | 480216 | 479781 | 479997 | 480067 | 480216 | 12 [illegible] |
| RI | 3596 | ENSG00000163704 | PRRT3 | chr3 | - | 9945541 | 9948001 | 9945541 | 9945711 | 9946084 | 9948001 | 12 [illegible] |
| RI | 3719 | ENSG00000138002 | IFT172 | chr2 | - | 27453981 | 27454418 | 27453981 | 27454162 | 27454353 | 27454418 | 3 [illegible] |
| RI | 4384 | ENSG00000145020 | AMT | chr3 | - | 49416793 | 49417718 | 49416793 | 49417357 | 49417614 | 49417718 | 31 [illegible] |
| RI | 6267 | ENSG00000183735 | TBK1 | chr12 | + | 64481841 | 64484499 | 64481841 | 64481956 | 64484302 | 64484499 | 82 [illegible] |
| RI | 8357 | ENSG00000135297 | MTO1 | chr6 | + | 73466208 | 73466606 | 73466208 | 73466408 | 73466492 | 73466606 | 19 [illegible] |
| RI | 932 | ENSG00000261408 | TEN1-CDK3 | chr17 | + | 76000140 | 76001541 | 76000140 | 76000389 | 76001411 | 76001541 | 13 [illegible] |
| RI | 1850 | ENSG00000165959 | CLMN | chr14 | - | 95193848 | 95194596 | 95193848 | 95193919 | 95194214 | 95194596 | 15 [illegible] |
| RI | 2262 | ENSG00000143774 | GUK1 | chr1 | + | 228145509 | 228146067 | 228145509 | 228145649 | 228146025 | 228146067 | 23 [illegible] |
| RI | 2560 | ENSG00000145907 | G3BP1 | chr5 | + | 151795478 | 151797428 | 151795478 | 151795575 | 151797196 | 151797428 | 35 [illegible] |
| RI | 4409 | ENSG00000143575 | HAX1 | chr1 | + | 154273335 | 154273961 | 154273335 | 154273598 | 154273749 | 154273961 | 18 [illegible] |
| RI | 7354 | ENSG00000168003 | SLC3A2 | chr11 | + | 62881892 | 62882999 | 62881892 | 62882618 | 62882907 | 62882999 | 12 [illegible] |
| RI | 3128 | ENSG00000165272 | AQP3 | chr9 | - | 33442851 | 33443458 | 33442851 | 33442970 | 33443178 | 33443458 | 0 [illegible] |
| RI | 4025 | ENSG00000162542 | TMCO4 | chr1 | - | 19682239 | 19683444 | 19682239 | 19682714 | 19683287 | 19683444 | 44 [illegible] |
| RI | 4435 | ENSG00000173531 | MST1 | chr3 | - | 49687148 | 49687478 | 49687148 | 49687285 | 49687363 | 49687478 | 2 [illegible] |
| RI | 6693 | ENSG00000174080 | CTSF | chr11 | - | 66567245 | 66567662 | 66567245 | 66567321 | 66567443 | 66567662 | 6 [illegible] |
| RI | 1212 | ENSG00000086504 | MRPL28 | chr16 | - | 368500 | 370225 | 368500 | 369211 | 369930 | 370225 | 7 [illegible] |
| RI | 1659 | ENSG00000140365 | COMMD4 | chr15 | + | 75338061 | 75338420 | 75338061 | 75338138 | 75338354 | 75338420 | 8 [illegible] |
| RI | 1973 | ENSG00000224424 | PRKAR2A-AS1 | chr3 | + | 48848104 | 48851976 | 48848104 | 48848408 | 48848986 | 48851976 | 1 [illegible] |
| RI | 2101 | ENSG00000141279 | NPEPPS | chr17 | + | 47619277 | 47619784 | 47619277 | 47619372 | 47619736 | 47619784 | 29 [illegible] |
| RI | 3361 | ENSG00000103507 | BCKDK | chr16 | + | 31109672 | 31110124 | 31109672 | 31109783 | 31110103 | 31110124 | 19 [illegible] |
| RI | 4663 | ENSG00000001497 | LAS1L | chrX | - | 65523559 | 65524614 | 65523559 | 65523876 | 65524055 | 65524614 | 6 [illegible] |
| RI | 4760 | ENSG00000181409 | AATK | chr17 | - | 81120200 | 81120984 | 81120200 | 81120399 | 81120827 | 81120984 | 4 [illegible] |
| RI | 5458 | ENSG00000164032 | H2AZ1 | chr4 | - | 99948087 | 99948940 | 99948087 | 99948523 | 99948813 | 99948940 | 35 [illegible] |
| RI | 6061 | ENSG00000243649 | CFB | chr6 | + | 31947944 | 31948512 | 31947944 | 31948081 | 31948373 | 31948512 | 4 [illegible] |
| RI | 6277 | ENSG00000183735 | TBK1 | chr12 | + | 64495698 | 64496406 | 64495698 | 64495775 | 64496372 | 64496406 | 21 [illegible] |
| RI | 7371 | ENSG00000167094 | TTC16 | chr9 | + | 127724119 | 127724897 | 127724119 | 127724364 | 127724755 | 127724897 | 15 [illegible] |
| RI | 4158 | ENSG00000003393 | ALS2 | chr2 | - | 201704121 | 201704603 | 201704121 | 201704218 | 201704453 | 201704603 | 12 [illegible] |
| RI | 7677 | ENSG00000168060 | NAALADL1 | chr11 | - | 65046188 | 65046526 | 65046188 | 65046362 | 65046444 | 65046526 | 4 [illegible] |
| RI | 917 | ENSG00000011143 | MKS1 | chr17 | - | 58208104 | 58208583 | 58208104 | 58208174 | 58208512 | 58208583 | 0 [illegible] |
| RI | 3954 | ENSG00000066322 | ELOVL1 | chr1 | - | 43364927 | 43365376 | 43364927 | 43365097 | 43365185 | 43365376 | 15 [illegible] |
| RI | 3994 | ENSG00000113597 | TRAPPC13 | chr5 | + | 65660698 | 65664384 | 65660698 | 65660897 | 65664233 | 65664384 | 31 [illegible] |
| RI | 5261 | ENSG00000224078 | SNHG14 | chr15 | + | 25107909 | 25109403 | 25107909 | 25108038 | 25109197 | 25109403 | 15 [illegible] |
| RI | 7426 | ENSG00000070061 | ELP1 | chr9 | - | 108896495 | 108897038 | 108896495 | 108896644 | 108896890 | 108897038 | 11 [illegible] |
| RI | 7488 | ENSG00000005238 | FAM214B | chr9 | - | 35107380 | 35108349 | 35107380 | 35107568 | 35108127 | 35108349 | 63 [illegible] |
| RI | 7856 | ENSG00000172803 | SNX32 | chr11 | + | 65849919 | 65850554 | 65849919 | 65850271 | 65850430 | 65850554 | 2 [illegible] |
| RI | 2136 | ENSG00000154153 | RETREG1 | chr5 | - | 16478849 | 16483472 | 16478849 | 16478987 | 16483345 | 16483472 | 16 [illegible] |
| RI | 3433 | ENSG00000171680 | PLEKHG5 | chr1 | - | 6474450 | 6475138 | 6474450 | 6474587 | 6475046 | 6475138 | 0 [illegible] |
| RI | 8237 | ENSG00000228784 | LINC00954 | chr2 | + | 19881566 | 19882260 | 19881566 | 19881618 | 19881705 | 19882260 | 15 [illegible] |
| RI | 3542 | ENSG00000247828 | TMEM161B-AS1 | chr5 | + | 88269467 | 88270585 | 88269467 | 88269687 | 88270525 | 88270585 | 26 [illegible] |
| RI | 4354 | ENSG00000099840 | IZUMO4 | chr19 | + | 2097251 | 2097495 | 2097251 | 2097332 | 2097423 | 2097495 | 0 [illegible] |
| RI | 4584 | ENSG00000106462 | EZH2 | chr7 | - | 148813958 | 148815546 | 148813958 | 148814137 | 148815505 | 148815546 | 2 [illegible] |
| RI | 6117 | ENSG00000044090 | CUL7 | chr6 | - | 43045226 | 43045682 | 43045226 | 43045402 | 43045586 | 43045682 | 6 [illegible] |
| RI | 7917 | ENSG00000179532 | DNHD1 | chr11 | + | 6570008 | 6571423 | 6570008 | 6570100 | 6570617 | 6571423 | 2 [illegible] |
| RI | 7963 | ENSG00000133895 | MEN1 | chr11 | - | 64807551 | 64808099 | 64807551 | 64807680 | 64807904 | 64808099 | 13 [illegible] |
| RI | 172 | ENSG00000161551 | ZNF577 | chr19 | - | 51886820 | 51887951 | 51886820 | 51886912 | 51887895 | 51887951 | 19 [illegible] |
| RI | 631 | ENSG00000196756 | SNHG17 | chr20 | - | 38421005 | 38422241 | 38421005 | 38421098 | 38422095 | 38422241 | 24 [illegible] |
| RI | 2522 | ENSG00000128609 | NDUFA5 | chr7 | - | 123557403 | 123557781 | 123557403 | 123557448 | 123557624 | 123557781 | 20 [illegible] |
| RI | 2543 | ENSG00000138796 | HADH | chr4 | + | 108027687 | 108028069 | 108027687 | 108027760 | 108027938 | 108028069 | 34 [illegible] |
| RI | 2878 | ENSG00000240303 | ACAD11 | chr3 | - | 132578795 | 132579558 | 132578795 | 132578881 | 132579491 | 132579558 | 84 [illegible] |
| RI | 5645 | ENSG00000111261 | MANSC1 | chr12 | - | 12343091 | 12343414 | 12343091 | 12343173 | 12343275 | 12343414 | 9 [illegible] |
| RI | 7034 | ENSG00000104626 | ERI1 | chr8 | + | 9029791 | 9033338 | 9029791 | 9030081 | 9032548 | 9033338 | 33 [illegible] |
| RI | 7421 | ENSG00000099958 | DERL3 | chr22 | - | 23834502 | 23837154 | 23834502 | 23836962 | 23837063 | 23837154 | 2 [illegible] |
| RI | 7495 | ENSG00000213930 | GALT | chr9 | + | 34647491 | 34647961 | 34647491 | 34647567 | 34647831 | 34647961 | 38 [illegible] |
| RI | 416 | ENSG00000166770 | ZNF667-AS1 | chr19 | + | 56494162 | 56495437 | 56494162 | 56494342 | 56495047 | 56495437 | 20 [illegible] |
| RI | 7763 | ENSG00000170291 | ELP5 | chr17 | + | 7258587 | 7259940 | 7258587 | 7258683 | 7259570 | 7259940 | 35 [illegible] |
| RI | 181 | ENSG00000105643 | ARRDC2 | chr19 | + | 18008169 | 18008584 | 18008169 | 18008228 | 18008453 | 18008584 | 10 [illegible] |
| RI | 2204 | ENSG00000086189 | DIMT1 | chr5 | - | 62390875 | 62392990 | 62390875 | 62390982 | 62392925 | 62392990 | 63 [illegible] |
| RI | 2345 | ENSG00000184584 | STING1 | chr5 | - | 139481158 | 139482302 | 139481158 | 139481342 | 139482209 | 139482302 | 9 [illegible] |
| RI | 2429 | ENSG00000164889 | SLC4A2 | chr7 | + | 151074038 | 151074841 | 151074038 | 151074293 | 151074674 | 151074841 | 7 [illegible] |
| RI | 2776 | ENSG00000268758 | ADGRE4P | chr19 | - | 6981079 | 6981770 | 6981079 | 6981215 | 6981726 | 6981770 | 10 [illegible] |
| RI | 3336 | ENSG00000113810 | SMC4 | chr3 | + | 160412325 | 160412849 | 160412325 | 160412453 | 160412529 | 160412849 | 32 [illegible] |
| RI | 3897 | ENSG00000178028 | DMAP1 | chr1 | + | 44213470 | 44213858 | 44213470 | 44213662 | 44213748 | 44213858 | 19 [illegible] |
| RI | 4228 | ENSG00000132781 | MUTYH | chr1 | - | 45332762 | 45333170 | 45332762 | 45332834 | 45333096 | 45333170 | 5 [illegible] |
| RI | 4305 | ENSG00000135541 | AHI1 | chr6 | - | 135300499 | 135302817 | 135300499 | 135300558 | 135302711 | 135302817 | 9 [illegible] |
| RI | 4595 | ENSG00000131165 | CHMP1A | chr16 | - | 89646526 | 89647331 | 89646526 | 89646733 | 89647202 | 89647331 | 24 [illegible] |
| RI | 8191 | ENSG00000130640 | TUBGCP2 | chr10 | - | 133283881 | 133285213 | 133283881 | 133284002 | 133285084 | 133285213 | 19 [illegible] |
| RI | 809 | ENSG00000185624 | P4HB | chr17 | - | 81845560 | 81845991 | 81845560 | 81845742 | 81845817 | 81845991 | 15 [illegible] |
| RI | 1491 | ENSG00000183751 | TBL3 | chr16 | + | 1974055 | 1974292 | 1974055 | 1974107 | 1974196 | 1974292 | 0 [illegible] |
| RI | 3264 | ENSG00000113108 | APBB3 | chr5 | - | 140561586 | 140562227 | 140561586 | 140561701 | 140562099 | 140562227 | 41 [illegible] |
| RI | 4431 | ENSG00000197989 | SNHG12 | chr1 | - | 28580559 | 28581871 | 28580559 | 28581229 | 28581589 | 28581871 | 9 [illegible] |
| RI | 5329 | ENSG00000064703 | DDX20 | chr1 | + | 111760705 | 111761284 | 111760705 | 111760848 | 111761225 | 111761284 | 29 [illegible] |
| RI | 5519 | ENSG00000075336 | TIMM21 | chr18 | + | 74155144 | 74155403 | 74155144 | 74155207 | 74155305 | 74155403 | 45 [illegible] |
| RI | 1068 | ENSG00000198496 | NBR2 | chr17 | + | 43138656 | 43140283 | 43138656 | 43138922 | 43139815 | 43140283 | 20 [illegible] |
| RI | 1420 | ENSG00000099999 | RNF215 | chr22 | - | 30379481 | 30379780 | 30379481 | 30379622 | 30379710 | 30379780 | 6 [illegible] |
| RI | 2756 | ENSG00000138031 | ADCY3 | chr2 | - | 24918312 | 24919839 | 24918312 | 24919184 | 24919682 | 24919839 | 0 [illegible] |
| RI | 3009 | ENSG00000151093 | OXSM | chr3 | + | 25790989 | 25791997 | 25790989 | 25791315 | 25791769 | 25791997 | 41 [illegible] |
| RI | 4028 | ENSG00000086015 | MAST2 | chr1 | + | 46032595 | 46032718 | 46032595 | 46032701 | 46032704 | 46032718 | 10 [illegible] |
| RI | 1564 | ENSG00000185880 | TRIM69 | chr15 | + | 44756367 | 44758854 | 44756367 | 44756463 | 44758612 | 44758854 | 9 [illegible] |
| RI | 2797 | ENSG00000163703 | CRELD1 | chr3 | + | 9933856 | 9934612 | 9933856 | 9934080 | 9934419 | 9934612 | 39 [illegible] |
| RI | 3307 | ENSG00000148399 | DPH7 | chr9 | - | 137575708 | 137576167 | 137575708 | 137575892 | 137576079 | 137576167 | 62 [illegible] |
| RI | 4493 | ENSG00000072110 | ACTN1 | chr14 | - | 68877081 | 68879069 | 68877081 | 68878523 | 68878988 | 68879069 | 10 [illegible] |
| RI | 6561 | ENSG00000181513 | ACBD4 | chr17 | + | 45135686 | 45136232 | 45135686 | 45135953 | 45136107 | 45136232 | 5 [illegible] |
| RI | 7872 | ENSG00000078124 | ACER3 | chr11 | + | 77015015 | 77015659 | 77015015 | 77015117 | 77015638 | 77015659 | 20 [illegible] |
| RI | 10 | ENSG00000154734 | ADAMTS1 | chr21 | - | 26838438 | 26839762 | 26838438 | 26838614 | 26839586 | 26839762 | 1 [illegible] |
| RI | 1572 | ENSG00000168970 | JMJD7-PLA2G4B | chr15 | + | 41845202 | 41845775 | 41845202 | 41845320 | 41845637 | 41845775 | 6 [illegible] |
| RI | 2131 | ENSG00000154134 | ROBO3 | chr11 | + | 124878583 | 124879341 | 124878583 | 124878796 | 124879189 | 124879341 | 3 [illegible] |
| RI | 2355 | ENSG00000011243 | AKAP8L | chr19 | - | 15398629 | 15399410 | 15398629 | 15398731 | 15399301 | 15399410 | 37 [illegible] |
| RI | 3977 | ENSG00000188004 | SNHG28 | chr1 | - | 159834522 | 159836155 | 159834522 | 159835777 | 159835978 | 159836155 | 32 [illegible] |
| RI | 4452 | ENSG00000090097 | PCBP4 | chr3 | - | 51960525 | 51960898 | 51960525 | 51960642 | 51960865 | 51960898 | 4 [illegible] |
| RI | 6684 | ENSG00000134668 | SPOCD1 | chr1 | - | 31798206 | 31798631 | 31798206 | 31798323 | 31798441 | 31798631 | 0 [illegible] |
| RI | 545 | ENSG00000061656 | SPAG4 | chr20 | + | 35617519 | 35617840 | 35617519 | 35617586 | 35617778 | 35617840 | 3 [illegible] |
| RI | 2354 | ENSG00000011243 | AKAP8L | chr19 | - | 15398629 | 15399410 | 15398629 | 15398703 | 15399301 | 15399410 | 37 [illegible] |
| RI | 3315 | ENSG00000117054 | ACADM | chr1 | + | 75750450 | 75750891 | 75750450 | 75750546 | 75750769 | 75750891 | 11 [illegible] |
| RI | 3338 | ENSG00000159588 | CCDC17 | chr1 | - | 45622954 | 45623652 | 45622954 | 45623117 | 45623556 | 45623652 | 11 [illegible] |
| RI | 5418 | ENSG00000039650 | PNKP | chr19 | - | 49861448 | 49861881 | 49861448 | 49861510 | 49861771 | 49861881 | 11 [illegible] |
| RI | 5703 | ENSG00000079785 | DDX1 | chr2 | + | 15627053 | 15629697 | 15627053 | 15627145 | 15629601 | 15629697 | 23 [illegible] |
| RI | 759 | ENSG00000004939 | SLC4A1 | chr17 | - | 44262635 | 44262934 | 44262635 | 44262726 | 44262851 | 44262934 | 0 [illegible] |
| RI | 3346 | ENSG00000158864 | NDUFS2 | chr1 | + | 161209482 | 161210710 | 161209482 | 161210188 | 161210590 | 161210710 | 19 [illegible] |
| RI | 3364 | ENSG00000158793 | NIT1 | chr1 | + | 161119133 | 161119612 | 161119133 | 161119388 | 161119492 | 161119612 | 25 [illegible] |
| RI | 8159 | ENSG00000107862 | GBF1 | chr10 | + | 102382055 | 102382899 | 102382055 | 102382210 | 102382307 | 102382899 | 24 [illegible] |
| RI | 210 | ENSG00000132024 | CC2D1A | chr19 | + | 13927892 | 13928188 | 13927892 | 13928030 | 13928123 | 13928188 | 2 [illegible] |
| RI | 843 | ENSG00000175106 | TVP23C | chr17 | - | 15536979 | 15540561 | 15536979 | 15538153 | 15540432 | 15540561 | 19 [illegible] |
| RI | 2804 | ENSG00000163703 | CRELD1 | chr3 | + | 9943076 | 9943515 | 9943076 | 9943229 | 9943380 | 9943515 | 5 [illegible] |
| RI | 2809 | ENSG00000163703 | CRELD1 | chr3 | + | 9943076 | 9944095 | 9943076 | 9943515 | 9943993 | 9944095 | 30 [illegible] |
| RI | 3314 | ENSG00000117054 | ACADM | chr1 | + | 75742167 | 75745914 | 75742167 | 75744391 | 75745834 | 75745914 | 25 [illegible] |
| RI | 5105 | ENSG00000160753 | RUSC1 | chr1 | + | 155325863 | 155327132 | 155325863 | 155325910 | 155326609 | 155327132 | 5 [illegible] |
| RI | 7059 | ENSG00000160781 | PAQR6 | chr1 | - | 156245781 | 156246250 | 156245781 | 156245987 | 156246122 | 156246250 | 4 [illegible] |
| RI | 603 | ENSG00000171703 | TCEA2 | chr20 | + | 64071869 | 64072346 | 64071869 | 64071941 | 64072171 | 64072346 | 29 [illegible] |
| RI | 1583 | ENSG00000114062 | UBE3A | chr15 | - | 25407066 | 25409207 | 25407066 | 25407137 | 25409087 | 25409207 | 53 [illegible] |
| RI | 2768 | ENSG00000229036 | VDAC1P8 | chr6 | + | 143503393 | 143504142 | 143503393 | 143503514 | 143503930 | 143504142 | 27 [illegible] |
| RI | 4666 | ENSG00000001497 | LAS1L | chrX | - | 65528259 | 65529258 | 65528259 | 65528369 | 65528963 | 65529258 | 35 [illegible] |
| RI | 5272 | ENSG00000168890 | TMEM150A | chr2 | - | 85601020 | 85601482 | 85601020 | 85601107 | 85601305 | 85601482 | 8 [illegible] |
| RI | 7040 | ENSG00000147416 | ATP6V1B2 | chr8 | + | 20212781 | 20217324 | 20212781 | 20212905 | 20217219 | 20217324 | 56 [illegible] |
| RI | 7256 | ENSG00000134539 | KLRD1 | chr12 | + | 10307977 | 10309688 | 10307977 | 10308084 | 10309625 | 10309688 | 460 [illegible] |
| RI | 8239 | ENSG00000228784 | LINC00954 | chr2 | + | 19881705 | 19882260 | 19881705 | 19881886 | 19882214 | 19882260 | 28 [illegible] |
| RI | 777 | ENSG00000182173 | TSEN54 | chr17 | + | 75521408 | 75523335 | 75521408 | 75521510 | 75521675 | 75523335 | 20 [illegible] |
| RI | 875 | ENSG00000161381 | PLXDC1 | chr17 | - | 39079103 | 39079528 | 39079103 | 39079164 | 39079501 | 39079528 | 11 [illegible] |
| RI | 3904 | ENSG00000058272 | PPP1R12A | chr12 | - | 79778549 | 79779351 | 79778549 | 79778600 | 79779320 | 79779351 | 56 [illegible] |
| RI | 4161 | ENSG00000003393 | ALS2 | chr2 | - | 201726663 | 201727775 | 201726663 | 201726866 | 201727211 | 201727775 | 2 [illegible] |
| RI | 4657 | ENSG00000001497 | LAS1L | chrX | - | 65520701 | 65523707 | 65520701 | 65520817 | 65523559 | 65523707 | 13 [illegible] |
| RI | 4672 | ENSG00000093010 | COMT | chr22 | + | 19961696 | 19962815 | 19961696 | 19961874 | 19962526 | 19962815 | 13 [illegible] |
| RI | 7137 | ENSG00000158106 | RHPN1 | chr8 | + | 143379828 | 143380175 | 143379828 | 143379985 | 143380061 | 143380175 | 0 [illegible] |
| RI | 7350 | ENSG00000168003 | SLC3A2 | chr11 | + | 62881698 | 62882066 | 62881698 | 62881718 | 62881892 | 62882066 | 13 [illegible] |
| RI | 8022 | ENSG00000183426 | NPIPA1 | chr16 | + | 14935159 | 14935397 | 14935159 | 14935276 | 14935344 | 14935397 | 9 [illegible] |
| RI | 8349 | ENSG00000132341 | RAN | chr12 | + | 130872583 | 130872920 | 130872583 | 130872617 | 130872835 | 130872920 | 41 [illegible] |
| RI | 2207 | ENSG00000086189 | DIMT1 | chr5 | - | 62398645 | 62398881 | 62398645 | 62398739 | 62398823 | 62398881 | 11 [illegible] |
| RI | 3600 | ENSG00000162772 | ATF3 | chr1 | + | 212615021 | 212615261 | 212615021 | 212615060 | 212615151 | 212615261 | 11 [illegible] |
| RI | 4994 | ENSG00000112514 | CUTA | chr6 | - | 33417096 | 33417310 | 33417096 | 33417142 | 33417246 | 33417310 | 30 [illegible] |
| RI | 5611 | ENSG00000064201 | TSPAN32 | chr11 | + | 2314484 | 2316667 | 2314484 | 2314571 | 2316575 | 2316667 | 148 [illegible] |
| RI | 6042 | ENSG00000137312 | FLOT1 | chr6 | - | 30741189 | 30741704 | 30741189 | 30741333 | 30741661 | 30741704 | 64 [illegible] |
| RI | 6206 | ENSG00000223705 | NSUN5P1 | chr7 | + | 75415980 | 75416191 | 75415980 | 75416030 | 75416102 | 75416191 | 22 [illegible] |

| | | | | | | | | | | | | |
|---|---|---|---|---|---|---|---|---|---|---|---|---|
| RI | 6218 | ENSG00000128581 | IFT22 | chr7 | - | 101318123 | 101319032 | 101318123 | 101318213 | 101318955 | 101319032 | 0 | [illegible] |
| RI | 250 | ENSG00000130816 | DNMT1 | chr19 | - | 10139675 | 10140328 | 10139675 | 10139849 | 10140045 | 10140328 | 6 | [illegible] |
| RI | 3480 | ENSG00000239305 | RNF103 | chr2 | - | 86622660 | 86623865 | 86622660 | 86622827 | 86623815 | 86623865 | 41 | [illegible] |
| RI | 4510 | ENSG00000168610 | STAT3 | chr17 | - | 42323259 | 42323625 | 42323259 | 42323354 | 42323452 | 42323625 | 29 | [illegible] |
| RI | 4838 | ENSG00000175054 | ATR | chr3 | - | 142465096 | 142468068 | 142465096 | 142465240 | 142467933 | 142468068 | 14 | [illegible] |
| RI | 5257 | ENSG00000224078 | SNHG14 | chr15 | + | 25093833 | 25095828 | 25093833 | 25093974 | 25095718 | 25095828 | 28 | [illegible] |
| RI | 5646 | ENSG00000135363 | LMO2 | chr11 | - | 33869345 | 33869987 | 33869345 | 33869586 | 33869709 | 33869987 | 16 | [illegible] |
| RI | 5697 | ENSG00000197635 | DPP4 | chr2 | - | 162046701 | 162047006 | 162046701 | 162046805 | 162046914 | 162047006 | 11 | [illegible] |
| RI | 6161 | ENSG00000168944 | CEP120 | chr5 | - | 123389923 | 123391337 | 123389923 | 123390365 | 123391109 | 123391337 | 30 | [illegible] |
| RI | 577 | ENSG00000124104 | SNX21 | chr20 | + | 45840638 | 45842881 | 45840638 | 45840780 | 45842032 | 45842881 | 15 | [illegible] |
| RI | 1181 | ENSG00000140650 | PMM2 | chr16 | + | 8802080 | 8802389 | 8802080 | 8802145 | 8802289 | 8802389 | 29 | [illegible] |
| RI | 1376 | ENSG00000187193 | MT1X | chr16 | + | 56682469 | 56684196 | 56682469 | 56682568 | 56683957 | 56684196 | 5 | [illegible] |
| RI | 2157 | ENSG00000152942 | RAD17 | chr5 | + | 69372033 | 69372217 | 69372033 | 69372054 | 69372156 | 69372217 | 148 | [illegible] |
| RI | 3818 | ENSG00000158887 | MPZ | chr1 | - | 161304734 | 161305977 | 161304734 | 161305289 | 161305533 | 161305977 | 45 | [illegible] |
| RI | 4578 | ENSG00000173947 | PIFO | chr1 | + | 111348515 | 111350223 | 111348515 | 111348646 | 111350105 | 111350223 | 0 | [illegible] |
| RI | 1137 | ENSG00000008517 | IL32 | chr16 | + | 3065368 | 3065826 | 3065368 | 3065494 | 3065780 | 3065826 | 193 | [illegible] |
| RI | 2436 | ENSG00000160972 | PPP1R16A | chr8 | + | 144500266 | 144500612 | 144500266 | 144500391 | 144500486 | 144500612 | 1 | [illegible] |
| RI | 2828 | ENSG00000164053 | ATRIP | chr3 | + | 48465486 | 48467645 | 48465486 | 48466309 | 48466629 | 48467645 | 6 | [illegible] |
| RI | 3308 | ENSG00000162601 | MYSM1 | chr1 | - | 58661405 | 58663522 | 58661405 | 58661511 | 58663449 | 58663522 | 41 | [illegible] |
| RI | 3811 | ENSG00000145833 | DDX46 | chr5 | + | 134816429 | 134817714 | 134816429 | 134816603 | 134817495 | 134817714 | 21 | [illegible] |
| RI | 4267 | ENSG00000143224 | PPOX | chr1 | + | 161167099 | 161167486 | 161167099 | 161167234 | 161167338 | 161167486 | 13 | [illegible] |
| RI | 6533 | ENSG00000164877 | MICALL2 | chr7 | - | 1440590 | 1442484 | 1440590 | 1440684 | 1442191 | 1442484 | 0 | [illegible] |
| RI | 262 | ENSG00000142541 | RPL13A | chr19 | + | 49489849 | 49490576 | 49489849 | 49489922 | 49490474 | 49490576 | 80 | [illegible] |
| RI | 2193 | ENSG00000145604 | SKP2 | chr5 | + | 36163644 | 36166662 | 36163644 | 36164695 | 36166518 | 36166662 | 16 | [illegible] |
| RI | 2386 | ENSG00000127184 | COX7C | chr5 | + | 86619352 | 86619878 | 86619352 | 86619496 | 86619738 | 86619878 | 40 | [illegible] |
| RI | 2691 | ENSG00000108591 | DRG2 | chr17 | + | 18099016 | 18100435 | 18099016 | 18099093 | 18100362 | 18100435 | 12 | [illegible] |
| RI | 3880 | ENSG00000172250 | SERHL | chr22 | + | 42502350 | 42502768 | 42502350 | 42502392 | 42502748 | 42502768 | 10 | [illegible] |
| RI | 4132 | ENSG00000162298 | SYVN1 | chr11 | - | 65132247 | 65132780 | 65132247 | 65132308 | 65132731 | 65132780 | 66 | [illegible] |
| RI | 4949 | ENSG00000171503 | ETFDH | chr4 | + | 158680123 | 158681614 | 158680123 | 158680182 | 158681570 | 158681614 | 24 | [illegible] |
| RI | 4957 | ENSG00000053900 | ANAPC4 | chr4 | + | 25415465 | 25417739 | 25415465 | 25415540 | 25417615 | 25417739 | 34 | [illegible] |
| RI | 6709 | ENSG00000204271 | SPIN3 | chrX | - | 56993786 | 56994949 | 56993786 | 56993888 | 56994722 | 56994949 | 21 | [illegible] |
| RI | 7798 | ENSG00000132199 | ENOSF1 | chr18 | - | 690548 | 691276 | 690548 | 690631 | 691203 | 691276 | 16 | [illegible] |
| RI | 75 | ENSG00000128159 | TUBGCP6 | chr22 | - | 50219956 | 50221874 | 50219956 | 50220015 | 50220191 | 50221874 | 35 | [illegible] |
| RI | 171 | ENSG00000161551 | ZNF577 | chr19 | - | 51886820 | 51887936 | 51886820 | 51886912 | 51887831 | 51887936 | 20 | [illegible] |
| RI | 176 | ENSG00000006015 | REX1BD | chr19 | + | 18590853 | 18592336 | 18590853 | 18590933 | 18592088 | 18592336 | 8 | [illegible] |
| RI | 1147 | ENSG00000168488 | ATXN2L | chr16 | + | 28835932 | 28837237 | 28835932 | 28836122 | 28836747 | 28837237 | 23 | [illegible] |
| RI | 1306 | ENSG00000197912 | SPG7 | chr16 | + | 89554485 | 89556085 | 89554485 | 89554563 | 89555958 | 89556085 | 4 | [illegible] |
| RI | 1580 | ENSG00000089195 | TRMT6 | chr20 | - | 5941052 | 5942036 | 5941052 | 5941139 | 5941950 | 5942036 | 11 | [illegible] |
| RI | 1619 | ENSG00000186654 | PRR5 | chr22 | + | 44702217 | 44702608 | 44702217 | 44702409 | 44702491 | 44702608 | 13 | [illegible] |
| RI | 2542 | ENSG00000138796 | HADH | chr4 | + | 108023473 | 108028069 | 108023473 | 108023563 | 108027938 | 108028069 | 24 | [illegible] |
| RI | 5890 | ENSG00000169884 | WNT10B | chr12 | - | 48967945 | 48968319 | 48967945 | 48968006 | 48968148 | 48968319 | 13 | [illegible] |
| RI | 6591 | ENSG00000234741 | GAS5 | chr1 | - | 173864256 | 173865282 | 173864256 | 173864304 | 173865228 | 173865282 | 115 | [illegible] |
| RI | 8203 | ENSG00000122877 | EGR2 | chr10 | - | 62815860 | 62816366 | 62815860 | 62816069 | 62816252 | 62816366 | 17 | [illegible] |
| RI | 1161 | ENSG00000125124 | BBS2 | chr16 | - | 56496966 | 56497880 | 56496966 | 56497124 | 56497742 | 56497880 | 18 | [illegible] |
| RI | 1502 | ENSG00000161996 | WDR90 | chr16 | + | 655057 | 655468 | 655057 | 655147 | 655306 | 655468 | 1 | [illegible] |
| RI | 1587 | ENSG00000140521 | POLG | chr15 | - | 89317375 | 89318749 | 89317375 | 89317476 | 89318540 | 89318749 | 111 | [illegible] |
| RI | 1721 | ENSG00000165795 | NDRG2 | chr14 | - | 21024029 | 21025061 | 21024029 | 21024096 | 21025028 | 21025061 | 6 | [illegible] |
| RI | 2057 | ENSG00000133706 | LARS1 | chr5 | - | 146130017 | 146133081 | 146130017 | 146130158 | 146132897 | 146133081 | 10 | [illegible] |
| RI | 2393 | ENSG00000181929 | PRKAG1 | chr12 | - | 49005305 | 49005543 | 49005305 | 49005364 | 49005461 | 49005543 | 24 | [illegible] |
| RI | 3058 | ENSG00000169727 | GPS1 | chr17 | + | 82053867 | 82054810 | 82053867 | 82054049 | 82054497 | 82054810 | 3 | [illegible] |
| RI | 4167 | ENSG00000003393 | ALS2 | chr2 | - | 201728511 | 201729183 | 201728511 | 201728640 | 201729051 | 201729183 | 0 | [illegible] |
| RI | 7113 | ENSG00000121879 | PIK3CA | chr3 | + | 179218209 | 179219735 | 179218209 | 179218334 | 179219570 | 179219735 | 23 | [illegible] |
| RI | 7280 | ENSG00000147789 | ZNF7 | chr8 | + | 144829042 | 144829604 | 144829042 | 144829090 | 144829522 | 144829604 | 13 | [illegible] |
| RI | 8419 | ENSG00000105409 | ATP1A3 | chr19 | - | 41966581 | 41967340 | 41966581 | 41966965 | 41967248 | 41967340 | 7 | [illegible] |
| RI | 5227 | ENSG00000125875 | TBC1D20 | chr20 | - | 440247 | 441689 | 440247 | 440389 | 441619 | 441689 | 30 | [illegible] |
| RI | 6301 | ENSG00000145191 | EIF2B5 | chr3 | + | 184143051 | 184144224 | 184143051 | 184143565 | 184144127 | 184144224 | 39 | [illegible] |
| RI | 6579 | ENSG00000196305 | IARS1 | chr9 | - | 92250712 | 92251885 | 92250712 | 92250759 | 92251807 | 92251885 | 69 | [illegible] |
| RI | 6869 | ENSG00000196924 | FLNA | chrX | - | 154350030 | 154351696 | 154350030 | 154350207 | 154351580 | 154351696 | 124 | [illegible] |
| RI | 6940 | ENSG00000147133 | TAF1 | chrX | + | 71423116 | 71424066 | 71423116 | 71423264 | 71423973 | 71424066 | 84 | [illegible] |
| RI | 7022 | ENSG00000142046 | TMEM91 | chr19 | + | 41382771 | 41384083 | 41382771 | 41382921 | 41383873 | 41384083 | 37 | [illegible] |
| RI | 8570 | ENSG00000083844 | ZNF264 | chr19 | + | 57191500 | 57191946 | 57191500 | 57191527 | 57191698 | 57191946 | 21 | [illegible] |
| RI | 615 | ENSG00000124228 | DDX27 | chr20 | + | 49233567 | 49236231 | 49233567 | 49233709 | 49236149 | 49236231 | 25 | [illegible] |
| RI | 4302 | ENSG00000224870 | MRPL20-AS1 | chr1 | + | 1399519 | 1400303 | 1399519 | 1399689 | 1400157 | 1400303 | 40 | [illegible] |
| RI | 4344 | ENSG00000087191 | PSMC5 | chr17 | + | 63827459 | 63827923 | 63827459 | 63827514 | 63827850 | 63827923 | 20 | [illegible] |
| RI | 4911 | ENSG00000160818 | GPATCH4 | chr1 | - | 156596080 | 156596478 | 156596080 | 156596248 | 156596386 | 156596478 | 14 | [illegible] |
| RI | 8167 | ENSG00000168209 | DDIT4 | chr10 | + | 72273923 | 72274421 | 72273923 | 72274060 | 72274224 | 72274421 | 38 | [illegible] |
| RI | 8577 | ENSG00000124496 | TRERF1 | chr6 | - | 42268153 | 42269848 | 42268153 | 42269251 | 42269498 | 42269848 | 118 | [illegible] |
| RI | 859 | ENSG00000132382 | MYBBP1A | chr17 | - | 4545025 | 4545345 | 4545025 | 4545175 | 4545258 | 4545345 | 1 | [illegible] |
| RI | 1810 | ENSG00000100888 | CHD8 | chr14 | - | 21430800 | 21431858 | 21430800 | 21431724 | 21431810 | 21431858 | 55 | [illegible] |
| RI | 3905 | ENSG00000058272 | PPP1R12A | chr12 | - | 79778549 | 79779351 | 79778549 | 79778600 | 79779338 | 79779351 | 57 | [illegible] |
| RI | 4331 | ENSG00000115548 | KDM3A | chr2 | + | 86489519 | 86491057 | 86489519 | 86489659 | 86490880 | 86491057 | 40 | [illegible] |
| RI | 4383 | ENSG00000145020 | AMT | chr3 | - | 49416777 | 49417718 | 49416777 | 49417367 | 49417614 | 49417718 | 31 | [illegible] |
| RI | 4716 | ENSG00000120694 | HSPH1 | chr13 | - | 31140183 | 31141259 | 31140183 | 31140309 | 31141121 | 31141259 | 13 | [illegible] |
| RI | 5709 | ENSG00000115053 | NCL | chr2 | - | 231456009 | 231456764 | 231456009 | 231456159 | 231456630 | 231456764 | 53 | [illegible] |
| RI | 5905 | ENSG00000170364 | SETMAR | chr3 | + | 4312897 | 4313761 | 4312897 | 4313229 | 4313666 | 4313761 | 18 | [illegible] |
| RI | 6314 | ENSG00000101574 | METTL4 | chr18 | - | 2552694 | 2555038 | 2552694 | 2552764 | 2554668 | 2555038 | 2 | [illegible] |
| RI | 7395 | ENSG00000197070 | ARRDC1 | chr9 | + | 137612895 | 137613510 | 137612895 | 137613006 | 137613422 | 137613510 | 64 | [illegible] |
| RI | 7451 | ENSG00000257524 | NA | chr9 | - | 127885185 | 127886788 | 127885185 | 127885343 | 127886635 | 127886788 | 115 | [illegible] |
| RI | 451 | ENSG00000261824 | LINC00662 | chr19 | - | 27792999 | 27793899 | 27792999 | 27793232 | 27793531 | 27793899 | 5 | [illegible] |
| RI | 686 | ENSG00000263753 | LINC00667 | chr18 | + | 5240647 | 5241889 | 5240647 | 5241255 | 5241698 | 5241889 | 43 | [illegible] |
| RI | 1397 | ENSG00000123815 | COQ8B | chr19 | - | 40705095 | 40705447 | 40705095 | 40705181 | 40705349 | 40705447 | 45 | [illegible] |
| RI | 1646 | ENSG00000177082 | WDR73 | chr15 | - | 84645972 | 84646348 | 84645972 | 84646085 | 84646183 | 84646348 | 36 | [illegible] |
| RI | 1845 | ENSG00000100722 | ZC3H14 | chr14 | + | 88609266 | 88609803 | 88609266 | 88609403 | 88609678 | 88609803 | 12 | [illegible] |
| RI | 3520 | ENSG00000068120 | COASY | chr17 | + | 42562147 | 42563322 | 42562147 | 42562219 | 42562932 | 42563322 | 51 | [illegible] |
| RI | 4665 | ENSG00000001497 | LAS1L | chrX | - | 65528259 | 65529258 | 65528259 | 65528369 | 65528875 | 65529258 | 34 | [illegible] |
| RI | 5160 | ENSG00000166801 | FAM111A | chr11 | + | 59144861 | 59145856 | 59144861 | 59144900 | 59145826 | 59145856 | 157 | [illegible] |
| RI | 5201 | ENSG00000178038 | ALS2CL | chr3 | - | 46671487 | 46672033 | 46671487 | 46671584 | 46671883 | 46672033 | 2 | [illegible] |
| RI | 5705 | ENSG00000079785 | DDX1 | chr2 | + | 15628444 | 15629697 | 15628444 | 15628517 | 15629601 | 15629697 | 23 | [illegible] |
| RI | 5727 | ENSG00000035141 | FAM136A | chr2 | - | 70300839 | 70301603 | 70300839 | 70300980 | 70301407 | 70301603 | 25 | [illegible] |
| RI | 5845 | ENSG00000167792 | NDUFV1 | chr11 | + | 67606957 | 67607324 | 67606957 | 67607076 | 67607298 | 67607324 | 23 | [illegible] |
| RI | 7751 | ENSG00000165644 | COMTD1 | chr10 | - | 75234937 | 75235178 | 75234937 | 75234992 | 75235059 | 75235178 | 0 | [illegible] |
| RI | 8576 | ENSG00000166529 | ZSCAN21 | chr7 | + | 100056910 | 100057405 | 100056910 | 100057276 | 100057387 | 100057405 | 13 | [illegible] |
| RI | 681 | ENSG00000263753 | LINC00667 | chr18 | + | 5238080 | 5241889 | 5238080 | 5238893 | 5241698 | 5241889 | 118 | [illegible] |
| RI | 779 | ENSG00000182173 | TSEN54 | chr17 | + | 75521408 | 75522333 | 75521408 | 75521510 | 75522006 | 75522333 | 64 | [illegible] |
| RI | 1038 | ENSG00000108671 | PSMD11 | chr17 | + | 32479250 | 32480197 | 32479250 | 32479376 | 32480145 | 32480197 | 35 | [illegible] |
| RI | 2378 | ENSG00000128694 | OSGEPL1 | chr2 | - | 189761419 | 189761660 | 189761419 | 189761525 | 189761646 | 189761660 | 53 | [illegible] |
| RI | 2872 | ENSG00000075711 | DLG1 | chr3 | - | 197065275 | 197065809 | 197065275 | 197065464 | 197065707 | 197065809 | 91 | [illegible] |
| RI | 5322 | ENSG00000170445 | HARS1 | chr5 | - | 140677907 | 140679127 | 140677907 | 140678015 | 140679061 | 140679127 | 69 | [illegible] |
| RI | 5488 | ENSG00000198830 | HMGN2 | chr1 | + | 26473482 | 26473732 | 26473482 | 26473585 | 26473702 | 26473732 | 108 | [illegible] |
| RI | 5583 | ENSG00000117984 | CTSD | chr11 | - | 1753802 | 1754138 | 1753802 | 1753880 | 1753993 | 1754138 | 112 | [illegible] |
| RI | 6525 | ENSG00000105835 | NAMPT | chr7 | - | 106274945 | 106277179 | 106274945 | 106275043 | 106277083 | 106277179 | 268 | [illegible] |
| RI | 6884 | ENSG00000147140 | NONO | chrX | + | 71297835 | 71298508 | 71297835 | 71297938 | 71298477 | 71298508 | 111 | [illegible] |
| RI | 6949 | ENSG00000215301 | DDX3X | chrX | + | 41337690 | 41339083 | 41337690 | 41338912 | 41339035 | 41339083 | 178 | [illegible] |
| RI | 7181 | ENSG00000100023 | PPIL2 | chr22 | + | 21694936 | 21695496 | 21694936 | 21695070 | 21695397 | 21695496 | 25 | [illegible] |
| RI | 7238 | ENSG00000104408 | EIF3E | chr8 | - | 108201742 | 108203117 | 108201742 | 108201923 | 108202989 | 108203117 | 102 | [illegible] |
| RI | 7352 | ENSG00000168003 | SLC3A2 | chr11 | + | 62881892 | 62882618 | 62881892 | 62882066 | 62882504 | 62882618 | 13 | [illegible] |
| RI | 7996 | ENSG00000150433 | TMEM218 | chr11 | - | 125102131 | 125102809 | 125102131 | 125102317 | 125102729 | 125102809 | 24 | [illegible] |
| RI | 1610 | ENSG00000166411 | IDH3A | chr15 | + | 78156907 | 78157631 | 78156907 | 78157162 | 78157572 | 78157631 | 39 | [illegible] |
| RI | 1654 | ENSG00000255571 | MIR9-3HG | chr15 | + | 89387816 | 89389482 | 89387816 | 89387881 | 89389344 | 89389482 | 7 | [illegible] |
| RI | 2431 | ENSG00000124279 | FASTKD3 | chr5 | - | 7866645 | 7868170 | 7866645 | 7867389 | 7867952 | 7868170 | 35 | [illegible] |
| RI | 2625 | ENSG00000185684 | EP400P1 | chr12 | + | 132103996 | 132105304 | 132103996 | 132104170 | 132104566 | 132105304 | 3 | [illegible] |
| RI | 3359 | ENSG00000213024 | NUP62 | chr19 | - | 49927693 | 49929504 | 49927693 | 49927848 | 49929416 | 49929504 | 26 | [illegible] |
| RI | 3693 | ENSG00000018510 | AGPS | chr2 | + | 177440964 | 177442486 | 177440964 | 177441040 | 177442406 | 177442486 | 16 | [illegible] |
| RI | 3776 | ENSG00000204435 | CSNK2B | chr6 | + | 31666820 | 31668654 | 31666820 | 31666903 | 31668538 | 31668654 | 16 | [illegible] |
| RI | 4036 | ENSG00000138095 | LRPPRC | chr2 | - | 43957384 | 43960634 | 43957384 | 43957451 | 43957725 | 43960634 | 21 | [illegible] |
| RI | 6238 | ENSG00000042317 | SPATA7 | chr14 | + | 88427629 | 88429463 | 88427629 | 88427696 | 88429347 | 88429463 | 2 | [illegible] |
| RI | 6740 | ENSG00000107371 | EXOSC3 | chr9 | - | 37780274 | 37780880 | 37780274 | 37780358 | 37780657 | 37780880 | 174 | [illegible] |
| RI | 146 | ENSG00000274602 | PI4KAP1 | chr22 | - | 18548010 | 18548469 | 18548010 | 18548118 | 18548317 | 18548469 | 0 | [illegible] |
| RI | 272 | ENSG00000011304 | PTBP1 | chr19 | + | 804291 | 804702 | 804291 | 804438 | 804540 | 804702 | 47 | [illegible] |
| RI | 1456 | ENSG00000140694 | PARN | chr16 | - | 14591912 | 14593378 | 14591912 | 14592033 | 14593300 | 14593378 | 16 | [illegible] |
| RI | 2129 | ENSG00000154134 | ROBO3 | chr11 | + | 124877518 | 124878131 | 124877518 | 124877658 | 124877936 | 124878131 | 11 | [illegible] |
| RI | 3044 | ENSG00000053524 | MCF2L2 | chr3 | - | 183224097 | 183228366 | 183224097 | 183224190 | 183228296 | 183228366 | 4 | [illegible] |
| RI | 3414 | ENSG00000170581 | STAT2 | chr12 | - | 56354777 | 56355351 | 56354777 | 56354863 | 56354989 | 56355351 | 37 | [illegible] |
| RI | 4056 | ENSG00000117174 | ZNHIT6 | chr1 | - | 85707628 | 85708433 | 85707628 | 85707936 | 85708053 | 85708433 | 98 | [illegible] |
| RI | 4474 | ENSG00000141456 | PELP1 | chr17 | - | 4671690 | 4673145 | 4671690 | 4671882 | 4672113 | 4673145 | 104 | [illegible] |
| RI | 5937 | ENSG00000085998 | POMGNT1 | chr1 | - | 46188682 | 46189357 | 46188682 | 46188980 | 46189241 | 46189357 | 57 | [illegible] |
| RI | 6089 | ENSG00000168538 | TRAPPC11 | chr4 | + | 183708406 | 183708574 | 183708406 | 183708416 | 183708535 | 183708574 | 86 | [illegible] |
| RI | 6405 | ENSG00000168090 | COPS6 | chr7 | + | 100090591 | 100091152 | 100090591 | 100090654 | 100091037 | 100091152 | 15 | [illegible] |
| RI | 7222 | ENSG00000185803 | SLC52A2 | chr8 | + | 144359622 | 144360493 | 144359622 | 144359747 | 144360171 | 144360493 | 29 | [illegible] |
| RI | 7261 | ENSG00000104365 | IKBKB | chr8 | + | 42319269 | 42320844 | 42319269 | 42319421 | 42320674 | 42320844 | 69 | [illegible] |
| RI | 7497 | ENSG00000213930 | GALT | chr9 | + | 34647088 | 34648171 | 34647088 | 34647258 | 34648047 | 34648171 | 30 | [illegible] |
| RI | 7517 | ENSG00000165699 | TSC1 | chr9 | - | 132910570 | 132911568 | 132910570 | 132910692 | 132911409 | 132911568 | 68 | [illegible] |
| RI | 7868 | ENSG00000171204 | TMEM126B | chr11 | + | 85635666 | 85636536 | 85635666 | 85635778 | 85636045 | 85636536 | 33 | [illegible] |
| RI | 8186 | ENSG00000072694 | FCGR2B | chr1 | + | 161673959 | 161677365 | 161673959 | 161674073 | 161677327 | 161677365 | 4 | [illegible] |
| RI | 684 | ENSG00000263753 | LINC00667 | chr18 | + | 5240647 | 5241889 | 5240647 | 5240819 | 5241698 | 5241889 | 54 | [illegible] |
| RI | 2802 | ENSG00000163703 | CRELD1 | chr3 | + | 9943076 | 9943515 | 9943076 | 9943153 | 9943380 | 9943515 | 12 | [illegible] |
| RI | 3519 | ENSG00000068120 | COASY | chr17 | + | 42562150 | 42562487 | 42562150 | 42562219 | 42562355 | 42562487 | 31 | [illegible] |
| RI | 4478 | ENSG00000156983 | BRPF1 | chr3 | + | 9742024 | 9742541 | 9742024 | 9742171 | 9742351 | 9742541 | 27 | [illegible] |
| RI | 4874 | ENSG00000163534 | FCRL1 | chr1 | - | 157801914 | 157802664 | 157801914 | 157802162 | 157802614 | 157802664 | 105 | [illegible] |
| RI | 4936 | ENSG00000198160 | MIER1 | chr1 | + | 66984571 | 66988619 | 66984571 | 66984592 | 66986402 | 66988619 | 404 | [illegible] |
| RI | 5328 | ENSG00000064703 | DDX20 | chr1 | + | 111760705 | 111761125 | 111760705 | 111760865 | 111760986 | 111761125 | 32 | [illegible] |
| RI | 5543 | ENSG00000160766 | GBAP1 | chr1 | - | 155217239 | 155218049 | 155217239 | 155217373 | 155217963 | 155218049 | 2 | [illegible] |
| RI | 6069 | ENSG00000204348 | DXO | chr6 | - | 31970342 | 31970825 | 31970342 | 31970478 | 31970592 | 31970825 | 32 | [illegible] |
| RI | 7375 | ENSG00000148337 | CIZ1 | chr9 | - | 128178708 | 128179415 | 128178708 | 128178904 | 128178988 | 128179415 | 27 | [illegible] |
| RI | 8596 | ENSG00000127022 | CANX | chr5 | + | 179722803 | 179723779 | 179722803 | 179723019 | 179723663 | 179723779 | 19 | [illegible] |
| RI | 685 | ENSG00000263753 | LINC00667 | chr18 | + | 5240647 | 5241889 | 5240647 | 5240964 | 5241698 | 5241889 | 33 | [illegible] |
| RI | 2186 | ENSG00000165819 | METTL3 | chr14 | - | 21503172 | 21503577 | 21503172 | 21503291 | 21503505 | 21503577 | 176 | [illegible] |
| RI | 2565 | ENSG00000145907 | G3BP1 | chr5 | + | 151799888 | 151800346 | 151799888 | 151800043 | 151800217 | 151800346 | 35 | [illegible] |
| RI | 3076 | ENSG00000188001 | TPRG1 | chr3 | + | 189150655 | 189151136 | 189150655 | 189150805 | 189150903 | 189151136 | 29 | [illegible] |
| RI | 5677 | ENSG00000123983 | ACSL3 | chr2 | + | 222933165 | 222934687 | 222933165 | 222933280 | 222934529 | 222934687 | 6 | [illegible] |
| RI | 5720 | ENSG00000130414 | NDUFA10 | chr2 | - | 240011616 | 240014860 | 240011616 | 240011660 | 240014738 | 240014860 | 130 | [illegible] |
| RI | 6205 | ENSG00000223705 | NSUN5P1 | chr7 | + | 75415591 | 75416191 | 75415591 | 75415770 | 75416102 | 75416191 | 33 | [illegible] |
| RI | 6775 | ENSG00000175463 | TBC1D10C | chr11 | + | 67409406 | 67410089 | 67409406 | 67409454 | 67409871 | 67410089 | 312 | [illegible] |
| RI | 6874 | ENSG00000147140 | NONO | chrX | + | 71290628 | 71291972 | 71290628 | 71290806 | 71291778 | 71291972 | 69 | [illegible] |
| RI | 6923 | ENSG00000186787 | SPIN2B | chrX | - | 57119681 | 57120631 | 57119681 | 57120101 | 57120404 | 57120631 | 28 | [illegible] |
| RI | 7201 | ENSG00000104763 | ASAH1 | chr8 | - | 18064456 | 18067298 | 18064456 | 18064474 | 18067219 | 18067298 | 485 | [illegible] |
| RI | 7936 | ENSG00000166387 | PPFIBP2 | chr11 | + | 7653023 | 7653756 | 7653023 | 7653200 | 7653501 | 7653756 | 74 | [illegible] |
| RI | 27 | ENSG00000141959 | PFKL | chr21 | + | 44324855 | 44326060 | 44324855 | 44324917 | 44325960 | 44326060 | 26 | [illegible] |
| RI | 960 | ENSG00000163597 | SNHG16 | chr17 | + | 76557969 | 76558600 | 76557969 | 76558279 | 76558373 | 76558600 | 46 | [illegible] |
| RI | 1173 | ENSG00000125124 | BBS2 | chr16 | - | 56500853 | 56502456 | 56500853 | 56501025 | 56502316 | 56502456 | 89 | [illegible] |
| RI | 1483 | ENSG00000167965 | MLST8 | chr16 | + | 2208209 | 2208613 | 2208209 | 2208334 | 2208449 | 2208613 | 5 | [illegible] |
| RI | 2192 | ENSG00000145604 | SKP2 | chr5 | + | 36163644 | 36166662 | 36163644 | 36163756 | 36166518 | 36166662 | 15 | [illegible] |
| RI | 3066 | ENSG00000159433 | STARD9 | chr15 | + | 42663280 | 42663917 | 42663280 | 42663490 | 42663819 | 42663917 | 6 | [illegible] |
| RI | 3232 | ENSG00000137135 | ARHGEF39 | chr9 | - | 35660093 | 35662002 | 35660093 | 35660967 | 35661712 | 35662002 | 76 | [illegible] |
| RI | 3664 | ENSG00000111325 | OGFOD2 | chr12 | + | 122978324 | 122978569 | 122978324 | 122978359 | 122978441 | 122978569 | 31 | [illegible] |
| RI | 4884 | ENSG00000158092 | NCK1 | chr3 | + | 136945582 | 136946295 | 136945582 | 136945752 | 136946151 | 136946295 | 453 | [illegible] |
| RI | 4909 | ENSG00000143303 | RRNAD1 | chr1 | + | 156731990 | 156732473 | 156731990 | 156732115 | 156732365 | 156732473 | 40 | [illegible] |
| RI | 5586 | ENSG00000169689 | CENPX | chr17 | - | 82019292 | 82019909 | 82019292 | 82019381 | 82019857 | 82019909 | 1 | [illegible] |
| RI | 5693 | ENSG00000160710 | ADAR | chr1 | - | 154589756 | 154590409 | 154589756 | 154589928 | 154590341 | 154590409 | 330 | [illegible] |
| RI | 5982 | ENSG00000204256 | BRD2 | chr6 | + | 32976561 | 32977570 | 32976561 | 32976936 | 32977259 | 32977570 | 209 | [illegible] |
| RI | 6184 | ENSG00000214021 | TTLL3 | chr3 | + | 9826996 | 9827240 | 9826996 | 9827072 | 9827214 | 9827240 | 91 | [illegible] |
| RI | 6976 | ENSG00000147050 | KDM6A | chrX | + | 45082575 | 45083608 | 45082575 | 45082640 | 45083459 | 45083608 | 60 | [illegible] |
| RI | 7432 | ENSG00000070061 | ELP1 | chr9 | - | 108917546 | 108919349 | 108917546 | 108917662 | 108919252 | 108919349 | 63 | [illegible] |
| RI | 8554 | ENSG00000148444 | COMMD3 | chr10 | + | 22318271 | 22319018 | 22318271 | 22318306 | 22318958 | 22319018 | 157 | [illegible] |
| RI | 198 | ENSG00000239382 | ALKBH6 | chr19 | - | 36013020 | 36013422 | 36013020 | 36013089 | 36013309 | 36013422 | 26 | [illegible] |
| RI | 1169 | ENSG00000125124 | BBS2 | chr16 | - | 56500853 | 56502456 | 56500853 | 56501025 | 56501944 | 56502456 | 53 | [illegible] |
| RI | 1297 | ENSG00000197912 | SPG7 | chr16 | + | 89540943 | 89542279 | 89540943 | 89540977 | 89541152 | 89542279 | 65 | [illegible] |
| RI | 1538 | ENSG00000260916 | CCPG1 | chr15 | - | 55359538 | 55360944 | 55359538 | 55359690 | 55360839 | 55360944 | 675 | [illegible] |
| RI | 2302 | ENSG00000105223 | PLD3 | chr19 | + | 40365552 | 40365930 | 40365552 | 40365585 | 40365717 | 40365930 | 125 | [illegible] |
| RI | 2596 | ENSG00000016864 | GLT8D1 | chr3 | - | 52695189 | 52696040 | 52695189 | 52695302 | 52695927 | 52696040 | 107 | [illegible] |
| RI | 2649 | ENSG00000203875 | SNHG5 | chr6 | - | 85678135 | 85678748 | 85678135 | 85678188 | 85678402 | 85678748 | 207 | [illegible] |
| RI | 2939 | ENSG00000165782 | PIP4P1 | chr14 | - | 20457857 | 20458702 | 20457857 | 20458346 | 20458588 | 20458702 | 181 | [illegible] |
| RI | 3087 | ENSG00000164073 | MFSD8 | chr4 | - | 127917824 | 127920836 | 127917824 | 127919338 | 127920556 | 127920836 | 67 | [illegible] |
| RI | 3156 | ENSG00000100462 | PRMT5 | chr14 | - | 22920528 | 22921056 | 22920528 | 22920731 | 22920839 | 22921056 | 35 | [illegible] |
| RI | 3562 | ENSG00000117862 | TXNDC12 | chr1 | - | 52020130 | 52021012 | 52020130 | 52020405 | 52020925 | 52021012 | 654 | [illegible] |
| RI | 4017 | ENSG00000069424 | KCNAB2 | chr1 | + | 6097268 | 6097555 | 6097268 | 6097357 | 6097516 | 6097555 | 41 | [illegible] |
| RI | 4022 | ENSG00000157193 | LRP8 | chr1 | - | 53255116 | 53257464 | 53255116 | 53255185 | 53257239 | 53257464 | 1 | [illegible] |
| RI | 4152 | ENSG00000001630 | CYP51A1 | chr7 | - | 92117043 | 92117212 | 92117043 | 92117092 | 92117176 | 92117212 | 80 | [illegible] |
| RI | 4972 | ENSG00000231500 | RPS18 | chr6 | + | 33275964 | 33276510 | 33275964 | 33276066 | 33276390 | 33276510 | 193 | [illegible] |
| RI | 5360 | ENSG00000162910 | MRPL55 | chr1 | - | 228108436 | 228109016 | 228108436 | 228108508 | 228108954 | 228109016 | 41 | [illegible] |
| RI | 6188 | ENSG00000106327 | TFR2 | chr7 | - | 100627263 | 100627661 | 100627263 | 100627491 | 100627576 | 100627661 | 0 | [illegible] |
| RI | 6774 | ENSG00000175463 | TBC1D10C | chr11 | + | 67405594 | 67409133 | 67405594 | 67405701 | 67408978 | 67409133 | 57 | [illegible] |
| RI | 7005 | ENSG00000120699 | EXOSC8 | chr13 | + | 37002933 | 37004561 | 37002933 | 37003007 | 37004476 | 37004561 | 68 | [illegible] |
| RI | 7107 | ENSG00000166169 | POLL | chr10 | - | 101584601 | 101584919 | 101584601 | 101584720 | 101584877 | 101584919 | 103 | [illegible] |
| RI | 7359 | ENSG00000089053 | ANAPC5 | chr12 | - | 121318276 | 121318608 | 121318276 | 121318351 | 121318500 | 121318608 | 121 | [illegible] |
| RI | 7415 | ENSG00000107331 | ABCA2 | chr9 | - | 137021391 | 137022001 | 137021391 | 137021610 | 137021890 | 137022001 | 0 | [illegible] |
| RI | 8218 | ENSG00000172243 | CLEC7A | chr12 | - | 10116776 | 10118590 | 10116776 | 10117249 | 10118153 | 10118590 | 186 | [illegible] |
| RI | 213 | ENSG00000104957 | CCDC130 | chr19 | + | 13758867 | 13759272 | 13758867 | 13759010 | 13759110 | 13759272 | 32 | [illegible] |
| RI | 619 | ENSG00000080189 | SLC35C2 | chr20 | - | 46358386 | 46358763 | 46358386 | 46358558 | 46358690 | 46358763 | 105 | [illegible] |
| RI | 624 | ENSG00000174365 | SNHG11 | chr20 | + | 38447465 | 38448730 | 38447465 | 38447623 | 38448665 | 38448730 | 1 | [illegible] |
| RI | 780 | ENSG00000182173 | TSEN54 | chr17 | + | 75521704 | 75523335 | 75521704 | 75522521 | 75523249 | 75523335 | 35 | [illegible] |
| RI | 1051 | ENSG00000068489 | PRR11 | chr17 | + | 59185053 | 59185562 | 59185053 | 59185204 | 59185439 | 59185562 | 1 | [illegible] |
| RI | 1918 | ENSG00000092203 | TOX4 | chr14 | + | 21487450 | 21488850 | 21487450 | 21487602 | 21488589 | 21488850 | 238 | [illegible] |
| RI | 2961 | ENSG00000034533 | ASTE1 | chr3 | - | 131024004 | 131025331 | 131024004 | 131024238 | 131025163 | 131025331 | 168 | [illegible] |
| RI | 3064 | ENSG00000159433 | STARD9 | chr15 | + | 42662793 | 42663917 | 42662793 | 42662891 | 42663280 | 42663917 | 7 | [illegible] |
| RI | 3419 | ENSG00000089682 | RBM41 | chrX | - | 107088435 | 107088839 | 107088435 | 107088573 | 107088734 | 107088839 | 198 | [illegible] |
| RI | 3574 | ENSG00000111640 | GAPDH | chr12 | + | 6536683 | 6537996 | 6536683 | 6537010 | 6537583 | 6537996 | 35 | [illegible] |
| RI | 3927 | ENSG00000055609 | KMT2C | chr7 | - | 152162116 | 152163826 | 152162116 | 152163177 | 152163699 | 152163826 | 200 | [illegible] |
| RI | 4385 | ENSG00000145020 | AMT | chr3 | - | 49416821 | 49417718 | 49416821 | 49417309 | 49417614 | 49417718 | 46 | [illegible] |

| | | | | | | | | | | | | | | |
|---|---|---|---|---|---|---|---|---|---|---|---|---|---|---|
| RI | 4410 | ENSG00000143575 | HAX1 | chr1 | + | 154273335 | 154273961 | 154273335 | 154273534 | 154273773 | 154273961 | 122 | 79 | 171 |
| RI | 5347 | ENSG00000005483 | KMT2E | chr7 | + | 105090008 | 105090273 | 105090008 | 105090126 | 105090127 | 105090273 | 230 | 206 | 242 |
| RI | 5363 | ENSG00000146414 | SHPRH | chr6 | - | 145954689 | 145955354 | 145954689 | 145954853 | 145955186 | 145955354 | 202 | 216 | 148 |
| RI | 5387 | ENSG00000106636 | YKT6 | chr7 | + | 44208132 | 44211124 | 44208132 | 44208198 | 44208794 | 44211124 | 46 | 25 | 18 |
| RI | 7224 | ENSG00000185900 | POMK | chr8 | + | 43122106 | 43125767 | 43122106 | 43123199 | 43123265 | 43125767 | 33 | 16 | 30 |
| RI | 7342 | ENSG00000158856 | DMTN | chr8 | + | 22069015 | 22069518 | 22069015 | 22069060 | 22069418 | 22069518 | 0 | 0 | 0 |
| RI | 7563 | ENSG00000178177 | LCORL | chr4 | - | 17873387 | 17878213 | 17873387 | 17876834 | 17876939 | 17878213 | 50 | 35 | 36 |
| RI | 805 | ENSG00000185624 | P4HB | chr17 | - | 81845143 | 81845742 | 81845143 | 81845230 | 81845459 | 81845742 | 37 | 22 | 32 |
| RI | 1105 | ENSG00000166582 | CENPV | chr17 | - | 16348615 | 16349333 | 16348615 | 16348685 | 16349139 | 16349333 | 35 | 24 | 17 |
| RI | 2088 | ENSG00000174032 | SLC25A30 | chr13 | - | 45393315 | 45399078 | 45393315 | 45396015 | 45398939 | 45399078 | 22 | 19 | 17 |
| RI | 2294 | ENSG00000170144 | HNRNPA3 | chr2 | + | 177215977 | 177216771 | 177215977 | 177216188 | 177216675 | 177216771 | 294 | 418 | 266 |
| RI | 3343 | ENSG00000158864 | NDUFS2 | chr1 | + | 161209482 | 161210188 | 161209482 | 161209595 | 161210110 | 161210188 | 32 | 42 | 59 |
| RI | 3926 | ENSG00000055609 | KMT2C | chr7 | - | 152162116 | 152163826 | 152162116 | 152163177 | 152163273 | 152163826 | 225 | 180 | 198 |
| RI | 3974 | ENSG00000198093 | ZNF649 | chr19 | - | 51899797 | 51900294 | 51899797 | 51899912 | 51900092 | 51900294 | 55 | 45 | 58 |
| RI | 4613 | ENSG00000019582 | CD74 | chr5 | - | 150405084 | 150406321 | 150405084 | 150405671 | 150406258 | 150406321 | 78 | 90 | 79 |
| RI | 4890 | ENSG00000070610 | GBA2 | chr9 | - | 35736866 | 35737939 | 35736866 | 35737604 | 35737747 | 35737939 | 56 | 23 | 51 |
| RI | 5241 | ENSG00000131143 | COX4I1 | chr16 | + | 85804936 | 85805864 | 85804936 | 85805153 | 85805732 | 85805864 | 41 | 43 | 70 |
| RI | 5826 | ENSG00000124356 | STAMBP | chr2 | + | 73847386 | 73847753 | 73847386 | 73847572 | 73847662 | 73847753 | 98 | 74 | 75 |
| RI | 7341 | ENSG00000165525 | NEMF | chr14 | - | 49784924 | 49785320 | 49784924 | 49785004 | 49785219 | 49785320 | 61 | 101 | 63 |
| RI | 8067 | ENSG00000165646 | SLC18A2 | chr10 | + | 117254047 | 117254497 | 117254047 | 117254131 | 117254404 | 117254497 | 0 | 0 | 1 |
| RI | 8190 | ENSG00000099246 | RAB18 | chr10 | + | 27533734 | 27533994 | 27533734 | 27533853 | 27533940 | 27533994 | 95 | 102 | 88 |
| RI | 537 | ENSG00000064601 | CTSA | chr20 | + | 45894993 | 45897040 | 45894993 | 45895133 | 45896964 | 45897040 | 9 | 3 | 14 |
| RI | 725 | ENSG00000166398 | GARRE1 | chr19 | + | 34299678 | 34300968 | 34299678 | 34299923 | 34300754 | 34300968 | 50 | 26 | 65 |
| RI | 838 | ENSG00000108788 | MLX | chr17 | + | 42569506 | 42570183 | 42569506 | 42569606 | 42570096 | 42570183 | 209 | 130 | 163 |
| RI | 975 | ENSG00000179094 | PER1 | chr17 | - | 8144055 | 8144993 | 8144055 | 8144268 | 8144750 | 8144993 | 35 | 22 | 51 |
| RI | 1281 | ENSG00000140854 | KATNB1 | chr16 | + | 57736977 | 57737283 | 57736977 | 57737063 | 57737225 | 57737283 | 91 | 33 | 69 |
| RI | 1799 | ENSG00000100564 | PIGH | chr14 | - | 67593742 | 67593952 | 67593742 | 67593813 | 67593909 | 67593952 | 57 | 29 | 78 |
| RI | 1989 | ENSG00000175061 | SNHG29 | chr17 | + | 16439326 | 16440253 | 16439326 | 16439414 | 16440110 | 16440253 | 74 | 47 | 56 |
| RI | 2339 | ENSG00000126456 | IRF3 | chr19 | - | 49664673 | 49665857 | 49664673 | 49664846 | 49665483 | 49665857 | 98 | 65 | 126 |

| | | | | | | | | | | | | | | | | | | | | | | | | | | | | | | | | | | | | | | | | | | | | | | |
|---|---|---|---|---|---|---|---|---|---|---|---|---|---|---|---|---|---|---|---|---|---|---|---|---|---|---|---|---|---|---|---|---|---|---|---|---|---|---|---|---|---|---|---|---|---|---|
| RI | 3486 | ENSG00000092841 | MYL6 | chr12 | + | 56159586 | 56160148 | 56159586 | 56159622 | 56160024 | 56160148 | 1663 | 1483 | 1094 | 2009 | 2066 | 4 | 3 | 5 | 6 | 3 | 5008 | 2325 | 1410 | 2736 | 9235 | 21 | 7 | 4 | 15 | 14 | 298 | 149 | 1 | 1 | 0.995 | 0.996 | 0.991 | 0.994 | 0.997 | 0.992 | 0.994 | 0.994 | 0.989 | 0.997 | 0.001 |
| RI | 3623 | ENSG00000133226 | SRRM1 | chr1 | + | 24652433 | 24652628 | 24652433 | 24652620 | 24652621 | 24652628 | 242 | 213 | 206 | 210 | 541 | 0 | 0 | 0 | 0 | 0 | 369 | 247 | 239 | 330 | 154 | 0 | 0 | 1 | 0 | 0 | 150 | 149 | 1 | 1 | 1 | 1 | 1 | 1 | 1 | 1 | 1 | 0.996 | 1 | 1 | 0.001 |
| RI | 4044 | ENSG00000142534 | RPS11 | chr19 | + | 49497193 | 49498046 | 49497193 | 49497282 | 49497519 | 49498046 | 4958 | 3179 | 5125 | 4807 | 4747 | 0 | 0 | 0 | 0 | 0 | 7873 | 4955 | 4344 | 4641 | 9195 | 0 | 2 | 4 | 0 | 0 | 298 | 149 | 1 | 1 | 1 | 1 | 1 | 1 | 1 | 1 | 0.999 | 0.998 | 1 | 1 | 0.001 |
| RI | 4789 | ENSG00000141510 | TP53 | chr17 | - | 7676381 | 7676622 | 7676381 | 7676403 | 7676520 | 7676622 | 7 | 0 | 4 | 1 | 1 | 185 | 97 | 169 | 140 | 414 | 0 | 1 | 3 | 0 | 0 | 233 | 170 | 55 | 106 | 187 | 266 | 149 | 1 | 1 | 0.021 | 0 | 0.013 | 0.004 | 0.001 | 0 | 0.003 | 0.03 | 0 | 0 | 0.001 |
| RI | 5076 | ENSG00000265972 | TXNIP | chr1 | - | 145994280 | 145995031 | 145994280 | 145994437 | 145994928 | 145995031 | 1420 | 902 | 1421 | 1162 | 1844 | 0 | 0 | 0 | 0 | 0 | 752 | 1881 | 1231 | 668 | 637 | 0 | 3 | 2 | 0 | 0 | 298 | 149 | 1 | 1 | 1 | 1 | 1 | 1 | 1 | 1 | 0.997 | 0.997 | 1 | 1 | 0.001 |
| RI | 5719 | ENSG00000130414 | NDUFA10 | chr2 | - | 240011616 | 240014860 | 240011616 | 240011696 | 240014694 | 240014860 | 94 | 72 | 132 | 121 | 113 | 0 | 0 | 0 | 1 | 0 | 175 | 124 | 113 | 172 | 61 | 0 | 0 | 0 | 2 | 0 | 298 | 149 | 1 | 1 | 1 | 1 | 1 | 0.984 | 1 | 1 | 1 | 1 | 0.977 | 1 | 0.001 |
| RI | 6091 | ENSG00000112739 | PRPF4B | chr6 | + | 4060412 | 4061147 | 4060412 | 4060644 | 4061014 | 4061147 | 436 | 302 | 358 | 382 | 389 | 0 | 0 | 0 | 0 | 0 | 36 | 141 | 287 | 54 | 3 | 0 | 0 | 1 | 0 | 0 | 298 | 149 | 1 | 1 | 1 | 1 | 1 | 1 | 1 | 1 | 1 | 0.993 | 1 | 1 | 0.001 |
| RI | 6507 | ENSG00000122566 | HNRNPA2B1 | chr7 | - | 26190992 | 26192577 | 26190992 | 26191650 | 26192494 | 26192577 | 687 | 785 | 770 | 896 | 652 | 1 | 1 | 4 | 2 | 2 | 140 | 468 | 679 | 180 | 65 | 0 | 1 | 1 | 2 | 0 | 298 | 149 | 1 | 1 | 0.997 | 0.997 | 0.99 | 0.996 | 0.994 | 1 | 0.996 | 0.997 | 0.978 | 1 | 0.001 |
| RI | 7203 | ENSG00000104763 | ASAH1 | chr8 | - | 18064456 | 18067298 | 18064456 | 18064531 | 18067249 | 18067298 | 520 | 376 | 215 | 345 | 325 | 0 | 0 | 0 | 0 | 0 | 425 | 251 | 385 | 378 | 509 | 0 | 0 | 1 | 0 | 0 | 298 | 149 | 1 | 1 | 1 | 1 | 1 | 1 | 1 | 1 | 1 | 0.995 | 1 | 1 | 0.001 |
| RI | 8000 | ENSG00000144802 | NFKBIZ | chr3 | + | 101853090 | 101853863 | 101853090 | 101853234 | 101853600 | 101853863 | 945 | 976 | 1066 | 1260 | 1639 | 1 | 0 | 1 | 2 | 2 | 1131 | 989 | 941 | 1280 | 657 | 7 | 0 | 1 | 0 | 0 | 298 | 149 | 1 | 1 | 0.998 | 1 | 0.998 | 0.997 | 0.998 | 0.988 | 1 | 0.998 | 1 | 1 | 0.001 |
| RI | 8238 | ENSG00000228784 | LINC00954 | chr2 | + | 19881566 | 19882260 | 19881566 | 19881618 | 19882214 | 19882260 | 22 | 13 | 25 | 13 | 16 | 0 | 0 | 2 | 2 | 3 | 27 | 2 | 19 | 11 | 0 | 0 | 0 | 0 | 6 | 0 | 298 | 149 | 1 | 1 | 1 | 1 | 0.862 | 0.765 | 0.727 | 1 | 1 | 1 | 0.478 | NA | 0.001 |
| RI | 8296 | ENSG00000197971 | MBP | chr18 | - | 77016831 | 77017065 | 77016831 | 77016913 | 77016979 | 77017065 | 489 | 265 | 382 | 509 | 288 | 2 | 0 | 0 | 1 | 0 | 469 | 538 | 312 | 411 | 348 | 0 | 1 | 0 | 3 | 0 | 215 | 149 | 1 | 1 | 0.994 | 1 | 1 | 0.997 | 1 | 1 | 0.997 | 1 | 0.99 | 1 | 0.001 |
| RI | 382 | ENSG00000105640 | RPL18A | chr19 | + | 17861292 | 17861472 | 17861292 | 17861307 | 17861424 | 17861472 | 8069 | 2856 | 8543 | 6308 | 9084 | 1 | 0 | 0 | 0 | 6 | 4489 | 5831 | 3543 | 2879 | 6888 | 0 | 8 | 0 | 13 | 0 | 266 | 149 | 1 | 1 | 1 | 1 | 1 | 1 | 0.999 | 1 | 0.998 | 1 | 0.992 | 1 | 0.002 |
| RI | 1170 | ENSG00000125124 | BBS2 | chr16 | - | 56501352 | 56502456 | 56501352 | 56501497 | 56501944 | 56502456 | 65 | 54 | 92 | 68 | 45 | 0 | 0 | 0 | 1 | 0 | 18 | 50 | 67 | 18 | 12 | 0 | 1 | 0 | 0 | 0 | 298 | 149 | 1 | 1 | 1 | 1 | 1 | 0.971 | 1 | 1 | 0.962 | 1 | 1 | 1 | 0.002 |
| RI | 1771 | ENSG00000100934 | SEC23A | chr14 | - | 39103031 | 39103233 | 39103031 | 39103123 | 39103197 | 39103233 | 8 | 6 | 11 | 10 | 11 | 0 | 0 | 1 | 0 | 0 | 0 | 10 | 13 | 4 | 11 | 0 | 0 | 1 | 0 | 0 | 223 | 149 | 1 | 1 | 1 | 1 | 0.88 | 1 | 1 | NA | 1 | 0.897 | 1 | 1 | 0.002 |
| RI | 2458 | ENSG00000186468 | RPS23 | chr5 | - | 82275208 | 82276255 | 82275208 | 82275295 | 82276177 | 82276255 | 5664 | 2888 | 5862 | 4527 | 18199 | 2 | 3 | 10 | 2 | 19 | 2297 | 2059 | 1988 | 1662 | 2244 | 2 | 0 | 3 | 2 | 13 | 298 | 149 | 1 | 1 | 0.999 | 0.998 | 0.997 | 0.999 | 0.998 | 0.998 | 1 | 0.997 | 0.998 | 0.989 | 0.002 |
| RI | 2795 | ENSG00000113811 | SELENOK | chr3 | - | 53885825 | 53885912 | 53885825 | 53885830 | 53885833 | 53885912 | 160 | 107 | 126 | 151 | 238 | 0 | 0 | 0 | 0 | 0 | 110 | 114 | 151 | 157 | 138 | 0 | 1 | 0 | 0 | 0 | 152 | 149 | 1 | 1 | 1 | 1 | 1 | 1 | 1 | 1 | 0.991 | 1 | 1 | 1 | 0.002 |
| RI | 2861 | ENSG00000168291 | PDHB | chr3 | - | 58430656 | 58430942 | 58430656 | 58430788 | 58430842 | 58430942 | 228 | 146 | 190 | 240 | 222 | 1 | 0 | 0 | 0 | 0 | 251 | 163 | 127 | 186 | 173 | 0 | 0 | 0 | 2 | 0 | 203 | 149 | 1 | 1 | 0.994 | 1 | 1 | 1 | 1 | 1 | 1 | 1 | 0.986 | 1 | 0.002 |
| RI | 3092 | ENSG00000111642 | CHD4 | chr12 | - | 6578028 | 6578546 | 6578028 | 6578135 | 6578408 | 6578546 | 115 | 80 | 80 | 110 | 163 | 0 | 0 | 0 | 0 | 0 | 442 | 203 | 192 | 403 | 219 | 0 | 0 | 1 | 0 | 0 | 298 | 149 | 1 | 1 | 1 | 1 | 1 | 1 | 1 | 1 | 1 | 0.99 | 1 | 1 | 0.002 |
| RI | 3656 | ENSG00000186470 | BTN3A2 | chr6 | + | 26368564 | 26368912 | 26368564 | 26368608 | 26368720 | 26368912 | 484 | 221 | 353 | 228 | 400 | 1 | 0 | 0 | 0 | 0 | 331 | 320 | 132 | 81 | 581 | 0 | 3 | 0 | 0 | 0 | 261 | 149 | 1 | 1 | 0.996 | 1 | 1 | 1 | 1 | 1 | 0.984 | 1 | 1 | 1 | 0.002 |
| RI | 3757 | ENSG00000071894 | CPSF1 | chr8 | - | 144399991 | 144400276 | 144399991 | 144400085 | 144400165 | 144400276 | 0 | 0 | 1 | 0 | 0 | 36 | 23 | 32 | 30 | 24 | 0 | 1 | 0 | 0 | 0 | 16 | 62 | 37 | 15 | 30 | 229 | 149 | 1 | 1 | 0 | 0 | 0.02 | 0 | 0 | 0 | 0.01 | 0 | 0 | 0 | 0.002 |
| RI | 3871 | ENSG00000108654 | DDX5 | chr17 | - | 64502161 | 64503098 | 64502161 | 64502223 | 64502925 | 64503098 | 787 | 621 | 780 | 867 | 747 | 0 | 1 | 0 | 2 | 0 | 347 | 952 | 702 | 631 | 58 | 0 | 0 | 7 | 0 | 0 | 298 | 149 | 1 | 1 | 1 | 0.997 | 1 | 0.995 | 1 | 1 | 1 | 0.98 | 1 | 1 | 0.002 |
| RI | 5920 | ENSG00000128284 | APOL3 | chr22 | - | 36145472 | 36145599 | 36145472 | 36145499 | 36145576 | 36145599 | 351 | 323 | 413 | 308 | 276 | 0 | 1 | 0 | 1 | 2 | 589 | 258 | 230 | 464 | 378 | 0 | 0 | 5 | 0 | 0 | 226 | 149 | 1 | 1 | 1 | 0.995 | 1 | 0.995 | 0.989 | 1 | 1 | 0.968 | 1 | 1 | 0.002 |
| RI | 5984 | ENSG00000204256 | BRD2 | chr6 | + | 32978125 | 32980464 | 32978125 | 32978388 | 32980341 | 32980464 | 78 | 93 | 72 | 134 | 65 | 0 | 1 | 0 | 0 | 0 | 685 | 122 | 148 | 303 | 305 | 0 | 0 | 0 | 0 | 5 | 298 | 149 | 1 | 1 | 1 | 0.979 | 1 | 1 | 1 | 1 | 1 | 1 | 1 | 0.968 | 0.002 |
| RI | 5999 | ENSG00000156508 | EEF1A1 | chr6 | - | 73519336 | 73520056 | 73519336 | 73519507 | 73519882 | 73520056 | 2053 | 1522 | 2391 | 2036 | 4181 | 7 | 1 | 8 | 0 | 3 | 2203 | 1482 | 1136 | 1504 | 1941 | 17 | 5 | 2 | 0 | 0 | 298 | 149 | 1 | 1 | 0.993 | 0.999 | 0.993 | 1 | 0.999 | 0.985 | 0.993 | 0.996 | 1 | 1 | 0.002 |
| RI | 6624 | ENSG00000124222 | STX16 | chr20 | + | 58651271 | 58652087 | 58651271 | 58651414 | 58652072 | 58652087 | 629 | 551 | 675 | 687 | 578 | 0 | 0 | 0 | 0 | 1 | 819 | 721 | 765 | 818 | 604 | 0 | 4 | 0 | 0 | 0 | 298 | 149 | 1 | 1 | 1 | 1 | 1 | 1 | 0.997 | 1 | 0.989 | 1 | 1 | 1 | 0.002 |
| RI | 6657 | ENSG00000102316 | MAGED2 | chrX | + | 54809721 | 54810213 | 54809721 | 54809808 | 54809922 | 54810213 | 422 | 190 | 225 | 327 | 303 | 0 | 0 | 0 | 0 | 1 | 204 | 258 | 117 | 137 | 714 | 0 | 0 | 1 | 0 | 0 | 263 | 149 | 1 | 1 | 1 | 1 | 1 | 1 | 0.994 | 1 | 1 | 0.985 | 1 | 1 | 0.002 |
| RI | 6978 | ENSG00000147050 | KDM6A | chrX | + | 45082575 | 45083608 | 45082575 | 45082640 | 45083518 | 45083608 | 173 | 208 | 140 | 185 | 372 | 0 | 0 | 0 | 0 | 0 | 172 | 117 | 205 | 170 | 34 | 0 | 0 | 1 | 0 | 0 | 298 | 149 | 1 | 1 | 1 | 1 | 1 | 1 | 1 | 1 | 1 | 0.99 | 1 | 1 | 0.002 |
| RI | 7062 | ENSG00000205581 | HMGN1 | chr21 | - | 39345829 | 39347292 | 39345829 | 39345958 | 39347113 | 39347292 | 211 | 219 | 245 | 240 | 340 | 0 | 0 | 0 | 1 | 0 | 16 | 117 | 149 | 47 | 15 | 0 | 1 | 0 | 0 | 0 | 298 | 149 | 1 | 1 | 1 | 1 | 1 | 0.992 | 1 | 1 | 0.983 | 1 | 1 | 1 | 0.002 |
| RI | 7819 | ENSG00000162236 | STX5 | chr11 | - | 62824458 | 62825339 | 62824458 | 62824565 | 62825282 | 62825339 | 71 | 42 | 62 | 97 | 39 | 0 | 0 | 0 | 0 | 0 | 358 | 240 | 181 | 251 | 275 | 0 | 0 | 0 | 1 | 0 | 298 | 149 | 1 | 1 | 1 | 1 | 1 | 1 | 1 | 1 | 1 | 1 | 0.992 | 1 | 0.002 |
| RI | 8132 | ENSG00000107959 | PITRM1 | chr10 | - | 3147571 | 3149753 | 3147571 | 3147737 | 3149620 | 3149753 | 30 | 31 | 39 | 47 | 53 | 0 | 1 | 1 | 0 | 1 | 8 | 56 | 66 | 84 | 34 | 0 | 4 | 1 | 0 | 0 | 298 | 149 | 1 | 1 | 1 | 0.939 | 0.951 | 1 | 0.964 | 1 | 0.875 | 0.971 | 1 | 1 | 0.002 |
| RI | 36 | ENSG00000275464 | NA | chr21 | - | 5135934 | 5137474 | 5135934 | 5136043 | 5137326 | 5137474 | 0 | 0 | 0 | 1 | 0 | 40 | 17 | 57 | 30 | 20 | 0 | 0 | 0 | 0 | 0 | 34 | 27 | 36 | 43 | 55 | 298 | 149 | 1 | 1 | 0 | 0 | 0 | 0.016 | 0 | 0 | 0 | 0 | 0 | 0 | 0.003 |
| RI | 519 | ENSG00000203896 | LIME1 | chr20 | + | 63738007 | 63738499 | 63738007 | 63738060 | 63738182 | 63738499 | 3 | 2 | 3 | 0 | 0 | 57 | 16 | 31 | 26 | 18 | 0 | 2 | 2 | 0 | 0 | 0 | 54 | 12 | 2 | 24 | 271 | 149 | 1 | 1 | 0.028 | 0.064 | 0.051 | 0 | 0 | NA | 0.02 | 0.084 | 0 | 0 | 0.003 |
| RI | 1279 | ENSG00000051108 | HERPUD1 | chr16 | + | 56939894 | 56940245 | 56939894 | 56940088 | 56940176 | 56940245 | 428 | 236 | 332 | 434 | 497 | 0 | 0 | 0 | 3 | 0 | 956 | 670 | 727 | 508 | 635 | 0 | 5 | 2 | 3 | 0 | 237 | 149 | 1 | 1 | 1 | 1 | 1 | 0.989 | 1 | 1 | 0.988 | 0.996 | 0.991 | 1 | 0.003 |
| RI | 1359 | ENSG00000160014 | CALM3 | chr19 | + | 46605826 | 46608340 | 46605826 | 46605857 | 46608198 | 46608340 | 93 | 42 | 66 | 96 | 81 | 0 | 1 | 0 | 0 | 0 | 277 | 122 | 82 | 137 | 379 | 0 | 4 | 0 | 0 | 0 | 298 | 149 | 1 | 1 | 1 | 0.955 | 1 | 1 | 1 | 1 | 0.938 | 1 | 1 | 1 | 0.003 |
| RI | 2226 | ENSG00000169567 | HINT1 | chr5 | - | 131159295 | 131162477 | 131159295 | 131159611 | 131162028 | 131162477 | 111 | 98 | 113 | 157 | 128 | 4 | 0 | 0 | 1 | 1 | 268 | 93 | 125 | 188 | 83 | 0 | 3 | 1 | 3 | 0 | 298 | 149 | 1 | 1 | 0.933 | 1 | 1 | 0.987 | 0.985 | 1 | 0.939 | 0.984 | 0.969 | 1 | 0.003 |
| RI | 2261 | ENSG00000143774 | GUK1 | chr1 | + | 228145510 | 228146067 | 228145510 | 228145624 | 228146012 | 228146067 | 25 | 19 | 12 | 29 | 27 | 1 | 0 | 0 | 0 | 0 | 47 | 34 | 22 | 40 | 64 | 0 | 0 | 0 | 2 | 0 | 298 | 149 | 1 | 1 | 0.926 | 1 | 1 | 1 | 1 | 1 | 1 | 1 | 0.909 | 1 | 0.003 |
| RI | 2476 | ENSG00000070756 | PABPC1 | chr8 | - | 100721390 | 100722087 | 100721390 | 100721452 | 100721728 | 100722087 | 1940 | 927 | 1792 | 1766 | 825 | 0 | 0 | 0 | 0 | 0 | 1715 | 1150 | 1290 | 1367 | 2877 | 6 | 0 | 0 | 4 | 0 | 298 | 149 | 1 | 1 | 1 | 1 | 1 | 1 | 1 | 0.993 | 1 | 1 | 0.994 | 1 | 0.003 |
| RI | 2652 | ENSG00000203875 | SNHG5 | chr6 | - | 85678148 | 85678748 | 85678148 | 85678188 | 85678697 | 85678748 | 120 | 142 | 199 | 158 | 174 | 0 | 1 | 1 | 1 | 0 | 110 | 173 | 186 | 143 | 191 | 0 | 0 | 1 | 3 | 0 | 298 | 149 | 1 | 1 | 1 | 0.986 | 0.99 | 0.988 | 1 | 1 | 1 | 0.989 | 0.96 | 1 | 0.003 |
| RI | 3011 | ENSG00000075785 | RAB7A | chr3 | + | 128806371 | 128807671 | 128806371 | 128806449 | 128807542 | 128807671 | 885 | 631 | 702 | 756 | 1086 | 0 | 1 | 0 | 0 | 0 | 1622 | 903 | 768 | 1182 | 1439 | 0 | 0 | 5 | 0 | 5 | 298 | 149 | 1 | 1 | 1 | 0.997 | 1 | 1 | 1 | 1 | 1 | 0.987 | 1 | 0.993 | 0.003 |
| RI | 4282 | ENSG00000153187 | HNRNPU | chr1 | - | 244859274 | 244862381 | 244859274 | 244859374 | 244860357 | 244862381 | 410 | 414 | 410 | 459 | 478 | 0 | 0 | 0 | 0 | 0 | 196 | 385 | 362 | 476 | 67 | 0 | 0 | 3 | 0 | 0 | 298 | 149 | 1 | 1 | 1 | 1 | 1 | 1 | 1 | 1 | 1 | 0.984 | 1 | 1 | 0.003 |
| RI | 5712 | ENSG00000115053 | NCL | chr2 | - | 231457000 | 231459125 | 231457000 | 231457124 | 231458265 | 231459125 | 293 | 254 | 310 | 326 | 380 | 0 | 0 | 0 | 0 | 0 | 191 | 301 | 221 | 227 | 14 | 0 | 2 | 0 | 0 | 0 | 298 | 149 | 1 | 1 | 1 | 1 | 1 | 1 | 1 | 1 | 0.987 | 1 | 1 | 1 | 0.003 |
| RI | 5713 | ENSG00000115053 | NCL | chr2 | - | 231457000 | 231459125 | 231457000 | 231457124 | 231459000 | 231459125 | 387 | 341 | 372 | 459 | 456 | 0 | 0 | 0 | 0 | 1 | 174 | 380 | 214 | 212 | 19 | 0 | 0 | 2 | 0 | 0 | 298 | 149 | 1 | 1 | 1 | 1 | 1 | 1 | 0.996 | 1 | 1 | 0.982 | 1 | 1 | 0.003 |
| RI | 5724 | ENSG00000180228 | PRKRA | chr2 | - | 178444421 | 178445883 | 178444421 | 178444500 | 178445425 | 178445883 | 27 | 17 | 33 | 24 | 21 | 8 | 5 | 11 | 7 | 7 | 5 | 46 | 44 | 10 | 0 | 4 | 20 | 2 | 3 | 0 | 298 | 149 | 1 | 1 | 0.628 | 0.63 | 0.6 | 0.632 | 0.6 | 0.385 | 0.535 | 0.917 | 0.625 | NA | 0.003 |
| RI | 5881 | ENSG00000111679 | PTPN6 | chr12 | + | 6954804 | 6954994 | 6954804 | 6954853 | 6954953 | 6954994 | 1142 | 565 | 794 | 942 | 664 | 1 | 3 | 0 | 1 | 0 | 1802 | 1327 | 601 | 1155 | 2418 | 0 | 0 | 6 | 4 | 4 | 249 | 149 | 1 | 1 | 0.999 | 0.991 | 1 | 0.998 | 1 | 1 | 1 | 0.984 | 0.994 | 0.997 | 0.003 |
| RI | 6319 | ENSG00000179218 | CALR | chr19 | + | 12943712 | 12944489 | 12943712 | 12943988 | 12944211 | 12944489 | 1320 | 689 | 1326 | 1325 | 2244 | 0 | 0 | 0 | 0 | 0 | 1522 | 1476 | 1013 | 1246 | 1967 | 10 | 0 | 0 | 4 | 0 | 298 | 149 | 1 | 1 | 1 | 1 | 1 | 1 | 1 | 0.987 | 1 | 1 | 1 | 1 | 0.003 |
| RI | 6781 | ENSG00000072778 | ACADVL | chr17 | + | 7223812 | 7224069 | 7223812 | 7223858 | 7223967 | 7224069 | 127 | 111 | 117 | 124 | 293 | 0 | 0 | 0 | 0 | 0 | 351 | 247 | 161 | 225 | 432 | 0 | 0 | 0 | 2 | 0 | 258 | 149 | 1 | 1 | 1 | 1 | 1 | 1 | 1 | 1 | 1 | 1 | 0.985 | 1 | 0.003 |
| RI | 7871 | ENSG00000166333 | ILK | chr11 | + | 6608883 | 6609408 | 6608883 | 6608967 | 6609298 | 6609408 | 65 | 65 | 109 | 109 | 96 | 0 | 0 | 2 | 1 | 2 | 208 | 125 | 128 | 118 | 159 | 0 | 0 | 1 | 6 | 0 | 298 | 149 | 1 | 1 | 1 | 1 | 0.965 | 0.982 | 0.96 | 1 | 1 | 0.985 | 0.908 | 1 | 0.003 |
| RI | 8074 | ENSG00000175197 | DDIT3 | chr12 | - | 57517268 | 57517753 | 57517268 | 57517542 | 57517705 | 57517753 | 52 | 35 | 74 | 51 | 143 | 0 | 0 | 0 | 0 | 0 | 131 | 155 | 136 | 123 | 164 | 0 | 0 | 1 | 0 | 0 | 298 | 149 | 1 | 1 | 1 | 1 | 1 | 1 | 1 | 1 | 1 | 0.986 | 1 | 1 | 0.003 |
| RI | 867 | ENSG00000108298 | RPL19 | chr17 | + | 39202316 | 39203109 | 39202316 | 39202443 | 39202999 | 39203109 | 295 | 214 | 377 | 283 | 415 | 0 | 0 | 0 | 0 | 0 | 301 | 252 | 214 | 196 | 374 | 0 | 0 | 0 | 2 | 0 | 298 | 149 | 1 | 1 | 1 | 1 | 1 | 1 | 1 | 1 | 1 | 1 | 0.98 | 1 | 0.004 |
| RI | 1365 | ENSG00000149923 | PPP4C | chr16 | + | 30082745 | 30083567 | 30082745 | 30082963 | 30083393 | 30083567 | 24 | 10 | 43 | 27 | 18 | 0 | 0 | 0 | 0 | 0 | 172 | 106 | 92 | 135 | 171 | 0 | 0 | 1 | 0 | 0 | 298 | 149 | 1 | 1 | 1 | 1 | 1 | 1 | 1 | 1 | 1 | 0.979 | 1 | 1 | 0.004 |
| RI | 1660 | ENSG00000140365 | COMMD4 | chr15 | + | 75338984 | 75339878 | 75338984 | 75339104 | 75339701 | 75339878 | 8 | 4 | 5 | 17 | 10 | 0 | 1 | 0 | 0 | 0 | 24 | 16 | 11 | 17 | 41 | 0 | 0 | 3 | 0 | 0 | 298 | 149 | 1 | 1 | 1 | 0.667 | 1 | 1 | 1 | 1 | 1 | 0.647 | 1 | 1 | 0.004 |
| RI | 1944 | ENSG00000103811 | CTSH | chr15 | - | 78932371 | 78935082 | 78932371 | 78932458 | 78933730 | 78935082 | 18 | 21 | 17 | 41 | 28 | 1 | 0 | 0 | 0 | 0 | 80 | 33 | 27 | 30 | 59 | 0 | 0 | 0 | 0 | 4 | 298 | 149 | 1 | 1 | 0.9 | 1 | 1 | 1 | 1 | 1 | 1 | 1 | 1 | 0.881 | 0.004 |
| RI | 2168 | ENSG00000065357 | DGKA | chr12 | + | 55941997 | 55942263 | 55941997 | 55942037 | 55942173 | 55942263 | 151 | 192 | 322 | 260 | 228 | 0 | 0 | 0 | 0 | 0 | 187 | 214 | 189 | 230 | 226 | 0 | 0 | 2 | 0 | 0 | 285 | 149 | 1 | 1 | 1 | 1 | 1 | 1 | 1 | 1 | 1 | 0.98 | 1 | 1 | 0.004 |
| RI | 2360 | ENSG00000011243 | AKAP8L | chr19 | - | 15397713 | 15400864 | 15397713 | 15399410 | 15400793 | 15400864 | 35 | 16 | 38 | 49 | 14 | 0 | 0 | 0 | 0 | 0 | 95 | 44 | 42 | 107 | 213 | 0 | 0 | 0 | 1 | 0 | 298 | 149 | 1 | 1 | 1 | 1 | 1 | 1 | 1 | 1 | 1 | 1 | 0.982 | 1 | 0.004 |
| RI | 3391 | ENSG00000132535 | DLG4 | chr17 | - | 7193963 | 7194495 | 7193963 | 7194000 | 7194318 | 7194495 | 1 | 2 | 0 | 0 | 0 | 22 | 11 | 4 | 29 | 8 | 0 | 0 | 4 | 0 | 0 | 13 | 30 | 21 | 16 | 72 | 298 | 149 | 1 | 1 | 0.022 | 0.083 | 0 | 0 | 0 | 0 | 0 | 0.087 | 0 | 0 | 0.004 |
| RI | 3573 | ENSG00000111640 | GAPDH | chr12 | + | 6536683 | 6537010 | 6536683 | 6536817 | 6536919 | 6537010 | 33 | 24 | 26 | 57 | 25 | 0 | 0 | 0 | 0 | 0 | 75 | 162 | 65 | 157 | 97 | 0 | 0 | 0 | 2 | 0 | 251 | 149 | 1 | 1 | 1 | 1 | 1 | 1 | 1 | 1 | 1 | 1 | 0.979 | 1 | 0.004 |
| RI | 3911 | ENSG00000134222 | PSRC1 | chr1 | - | 109282517 | 109282731 | 109282517 | 109282575 | 109282679 | 109282731 | 0 | 0 | 2 | 0 | 0 | 7 | 2 | 6 | 6 | 0 | 3 | 0 | 2 | 0 | 0 | 24 | 2 | 9 | 15 | 20 | 253 | 149 | 1 | 1 | 0 | 0 | 0.164 | 0 | NA | 0.069 | 0 | 0.116 | 0 | 0 | 0.004 |
| RI | 5830 | ENSG00000119707 | RBM25 | chr14 | + | 73099666 | 73101179 | 73099666 | 73099750 | 73100713 | 73101179 | 327 | 418 | 350 | 495 | 848 | 0 | 1 | 0 | 3 | 9 | 91 | 233 | 252 | 129 | 31 | 0 | 0 | 0 | 4 | 0 | 298 | 149 | 1 | 1 | 1 | 0.995 | 1 | 0.988 | 0.979 | 1 | 1 | 1 | 0.942 | 1 | 0.004 |
| RI | 5855 | ENSG00000108465 | CDK5RAP3 | chr17 | + | 47974399 | 47975337 | 47974399 | 47974448 | 47975171 | 47975337 | 380 | 310 | 348 | 400 | 253 | 1 | 0 | 2 | 0 | 0 | 757 | 477 | 458 | 729 | 610 | 0 | 0 | 4 | 8 | 0 | 298 | 149 | 1 | 1 | 0.995 | 1 | 0.989 | 1 | 1 | 1 | 1 | 0.983 | 0.979 | 1 | 0.004 |
| RI | 7443 | ENSG00000244187 | TMEM141 | chr9 | + | 136791946 | 136792358 | 136791946 | 136792030 | 136792250 | 136792358 | 2 | 1 | 4 | 0 | 0 | 62 | 36 | 29 | 47 | 26 | 0 | 4 | 2 | 0 | 0 | 12 | 52 | 24 | 32 | 56 | 298 | 149 | 1 | 1 | 0.016 | 0.014 | 0.065 | 0 | 0 | 0 | 0.037 | 0.04 | 0 | 0 | 0.004 |
| RI | 7445 | ENSG00000213213 | CCDC183 | chr9 | + | 136806077 | 136806867 | 136806077 | 136806672 | 136806756 | 136806867 | 5 | 4 | 9 | 2 | 4 | 3 | 2 | 3 | 2 | 0 | 0 | 2 | 0 | 1 | 0 | 0 | 0 | 0 | 2 | 0 | 233 | 149 | 1 | 1 | 0.516 | 0.561 | 0.657 | 0.39 | 1 | NA | 1 | NA | 0.242 | NA | 0.004 |
| RI | 8295 | ENSG00000197971 | MBP | chr18 | - | 77016831 | 77017065 | 77016831 | 77016846 | 77016913 | 77017065 | 329 | 169 | 260 | 328 | 116 | 0 | 1 | 0 | 1 | 0 | 275 | 356 | 186 | 285 | 179 | 6 | 0 | 0 | 0 | 0 | 216 | 149 | 1 | 1 | 1 | 0.991 | 1 | 0.996 | 1 | 0.969 | 1 | 1 | 1 | 1 | 0.004 |
| RI | 97 | ENSG00000100031 | GGT1 | chr22 | + | 24610249 | 24610531 | 24610249 | 24610336 | 24610454 | 24610531 | 21 | 8 | 20 | 25 | 33 | 0 | 1 | 0 | 1 | 0 | 33 | 27 | 10 | 20 | 50 | 0 | 4 | 0 | 0 | 0 | 267 | 149 | 1 | 1 | 1 | 0.817 | 1 | 1 | 1 | 1 | 0.79 | 1 | 1 | 1 | 0.005 |
| RI | 781 | ENSG00000182173 | TSEN54 | chr17 | + | 75521408 | 75523335 | 75521408 | 75521510 | 75523274 | 75523335 | 33 | 12 | 34 | 29 | 20 | 0 | 0 | 0 | 0 | 0 | 19 | 59 | 76 | 49 | 58 | 0 | 0 | 1 | 0 | 0 | 298 | 149 | 1 | 1 | 1 | 1 | 1 | 1 | 1 | 1 | 1 | 0.974 | 1 | 1 | 0.005 |
| RI | 1316 | ENSG00000130175 | PRKCSH | chr19 | + | 11436040 | 11436505 | 11436040 | 11436196 | 11436388 | 11436505 | 1 | 0 | 3 | 4 | 1 | 220 | 82 | 191 | 208 | 144 | 0 | 0 | 0 | 0 | 0 | 171 | 225 | 144 | 139 | 337 | 298 | 149 | 1 | 1 | 0.002 | 0 | 0.008 | 0.01 | 0.003 | 0 | 0 | 0 | 0 | 0 | 0.005 |
| RI | 5147 | ENSG00000171824 | EXOSC10 | chr1 | - | 11076841 | 11077651 | 11076841 | 11076948 | 11077600 | 11077651 | 29 | 33 | 55 | 38 | 49 | 0 | 0 | 1 | 0 | 0 | 162 | 122 | 86 | 98 | 41 | 0 | 0 | 0 | 3 | 0 | 298 | 149 | 1 | 1 | 1 | 1 | 0.965 | 1 | 1 | 1 | 1 | 1 | 0.942 | 1 | 0.005 |
| RI | 5167 | ENSG00000042493 | CAPG | chr2 | - | 85401528 | 85401683 | 85401528 | 85401588 | 85401650 | 85401683 | 335 | 146 | 189 | 230 | 150 | 0 | 0 | 0 | 0 | 0 | 740 | 593 | 210 | 655 | 1005 | 0 | 2 | 3 | 0 | 0 | 211 | 149 | 1 | 1 | 1 | 1 | 1 | 1 | 1 | 1 | 0.995 | 0.98 | 1 | 1 | 0.005 |
| RI | 5219 | ENSG00000125875 | TBC1D20 | chr20 | - | 435496 | 438841 | 435496 | 435970 | 437425 | 438841 | 210 | 129 | 171 | 157 | 184 | 0 | 1 | 2 | 0 | 1 | 254 | 180 | 95 | 151 | 301 | 0 | 0 | 0 | 6 | 0 | 298 | 149 | 1 | 1 | 1 | 0.985 | 0.977 | 1 | 0.989 | 1 | 1 | 1 | 0.926 | 1 | 0.005 |
| RI | 5228 | ENSG00000125875 | TBC1D20 | chr20 | - | 441587 | 442043 | 441587 | 441689 | 441778 | 442043 | 13 | 12 | 16 | 17 | 16 | 0 | 0 | 0 | 4 | 0 | 67 | 38 | 16 | 15 | 30 | 0 | 0 | 0 | 4 | 0 | 238 | 149 | 1 | 1 | 1 | 1 | 1 | 0.727 | 1 | 1 | 1 | 1 | 0.701 | 1 | 0.005 |
| RI | 5756 | ENSG00000163170 | BOLA3 | chr2 | - | 74145188 | 74147912 | 74145188 | 74145303 | 74147820 | 74147912 | 0 | 0 | 3 | 0 | 1 | 8 | 1 | 5 | 11 | 3 | 7 | 0 | 0 | 0 | 0 | 9 | 1 | 4 | 0 | 9 | 298 | 149 | 1 | 1 | 0 | 0 | 0.231 | 0 | 0.143 | 0.28 | 0 | 0 | NA | 0 | 0.005 |
| RI | 5876 | ENSG00000111011 | RSRC2 | chr12 | - | 122518838 | 122521428 | 122518838 | 122519051 | 122521384 | 122521428 | 204 | 191 | 178 | 256 | 227 | 0 | 0 | 0 | 1 | 2 | 17 | 434 | 250 | 230 | 23 | 0 | 0 | 2 | 4 | 0 | 298 | 149 | 1 | 1 | 1 | 1 | 1 | 0.992 | 0.983 | 1 | 1 | 0.984 | 0.966 | 1 | 0.005 |
| RI | 6399 | ENSG00000179818 | PCBP1-AS1 | chr2 | - | 70085815 | 70086273 | 70085815 | 70085903 | 70086123 | 70086273 | 103 | 80 | 96 | 102 | 124 | 0 | 0 | 1 | 1 | 0 | 121 | 116 | 78 | 149 | 143 | 0 | 0 | 0 | 5 | 0 | 298 | 149 | 1 | 1 | 1 | 1 | 0.98 | 0.981 | 1 | 1 | 1 | 1 | 0.937 | 1 | 0.005 |
| RI | 6664 | ENSG00000204001 | LCN8 | chr9 | - | 136755243 | 136756592 | 136755243 | 136755516 | 136756521 | 136756592 | 0 | 1 | 0 | 2 | 1 | 0 | 0 | 0 | 0 | 7 | 0 | 13 | 0 | 0 | 0 | 0 | 3 | 0 | 0 | 0 | 298 | 149 | 1 | 1 | NA | 1 | NA | 1 | 0.067 | NA | 0.684 | NA | NA | NA | 0.005 |
| RI | 7477 | ENSG00000172270 | BSG | chr19 | + | 580378 | 580782 | 580378 | 580461 | 580645 | 580782 | 6 | 4 | 10 | 8 | 3 | 639 | 331 | 402 | 587 | 472 | 0 | 4 | 5 | 2 | 10 | 829 | 856 | 601 | 991 | 1553 | 298 | 149 | 1 | 1 | 0.005 | 0.006 | 0.012 | 0.007 | 0.003 | 0 | 0.002 | 0.004 | 0.001 | 0.003 | 0.005 |
| RI | 7558 | ENSG00000106397 | PLOD3 | chr7 | - | 101212841 | 101213204 | 101212841 | 101212943 | 101213106 | 101213204 | 2 | 0 | 2 | 2 | 2 | 83 | 48 | 63 | 90 | 128 | 0 | 0 | 1 | 0 | 2 | 66 | 68 | 53 | 74 | 84 | 298 | 149 | 1 | 1 | 0.012 | 0 | 0.016 | 0.011 | 0.008 | 0 | 0 | 0.009 | 0 | 0.012 | 0.005 |
| RI | 8028 | ENSG00000090621 | PABPC4 | chr1 | - | 39562322 | 39564542 | 39562322 | 39562416 | 39564470 | 39564542 | 81 | 46 | 84 | 93 | 87 | 0 | 0 | 0 | 0 | 0 | 84 | 61 | 86 | 90 | 65 | 0 | 0 | 1 | 0 | 0 | 298 | 149 | 1 | 1 | 1 | 1 | 1 | 1 | 1 | 1 | 1 | 0.977 | 1 | 1 | 0.005 |
| RI | 79 | ENSG00000184702 | SEPTIN5 | chr22 | + | 19721821 | 19722339 | 19721821 | 19721957 | 19722236 | 19722339 | 1 | 3 | 0 | 3 | 0 | 57 | 52 | 22 | 135 | 45 | 0 | 0 | 0 | 4 | 17 | 217 | 113 | 82 | 147 | 1419 | 298 | 149 | 1 | 1 | 0.009 | 0.028 | 0 | 0.011 | 0 | 0 | 0 | 0 | 0.013 | 0.006 | 0.006 |
| RI | 203 | ENSG00000167644 | C19orf33 | chr19 | + | 38304250 | 38304490 | 38304250 | 38304283 | 38304374 | 38304490 | 0 | 3 | 0 | 0 | 0 | 19 | 19 | 4 | 62 | 14 | 0 | 0 | 0 | 3 | 0 | 33 | 24 | 16 | 29 | 363 | 240 | 149 | 1 | 1 | 0 | 0.089 | 0 | 0 | 0 | 0 | 0 | 0 | 0.06 | 0 | 0.006 |
| RI | 742 | ENSG00000161939 | RNASEK-C17orf49 | chr17 | + | 7014144 | 7015156 | 7014144 | 7014277 | 7015061 | 7015156 | 254 | 153 | 234 | 245 | 200 | 1 | 0 | 0 | 1 | 0 | 557 | 285 | 207 | 363 | 1096 | 0 | 0 | 2 | 5 | 0 | 298 | 149 | 1 | 1 | 0.992 | 1 | 1 | 0.992 | 1 | 1 | 1 | 0.981 | 0.973 | 1 | 0.006 |
| RI | 1050 | ENSG00000006282 | SPATA20 | chr17 | + | 50550707 | 50551190 | 50550707 | 50550917 | 50550997 | 50551190 | 0 | 0 | 3 | 3 | 1 | 10 | 0 | 13 | 10 | 1 | 2 | 1 | 0 | 1 | 2 | 3 | 14 | 6 | 3 | 3 | 229 | 149 | 1 | 1 | 0 | NA | 0.131 | 0.163 | 0.394 | 0.303 | 0.044 | 0 | 0.178 | 0.303 | 0.006 |
| RI | 1271 | ENSG00000205250 | E2F4 | chr16 | + | 67192760 | 67193170 | 67192760 | 67192824 | 67193008 | 67193170 | 136 | 73 | 68 | 117 | 81 | 1 | 0 | 0 | 0 | 0 | 193 | 136 | 68 | 190 | 169 | 0 | 3 | 0 | 0 | 0 | 298 | 149 | 1 | 1 | 0.986 | 1 | 1 | 1 | 1 | 1 | 0.958 | 1 | 1 | 1 | 0.006 |
| RI | 2254 | ENSG00000161960 | EIF4A1 | chr17 | + | 7576523 | 7577165 | 7576523 | 7576690 | 7577055 | 7577165 | 231 | 155 | 209 | 271 | 136 | 0 | 0 | 0 | 0 | 0 | 547 | 292 | 214 | 363 | 485 | 0 | 0 | 0 | 0 | 8 | 298 | 149 | 1 | 1 | 1 | 1 | 1 | 1 | 1 | 1 | 1 | 1 | 1 | 0.968 | 0.006 |
| RI | 2892 | ENSG00000178035 | IMPDH2 | chr3 | - | 49028430 | 49028806 | 49028430 | 49028532 | 49028721 | 49028806 | 70 | 41 | 78 | 88 | 73 | 0 | 0 | 0 | 0 | 0 | 155 | 120 | 136 | 136 | 164 | 0 | 0 | 2 | 0 | 0 | 298 | 149 | 1 | 1 | 1 | 1 | 1 | 1 | 1 | 1 | 1 | 0.971 | 1 | 1 | 0.006 |
| RI | 5066 | ENSG00000084774 | CAD | chr2 | + | 27242619 | 27242973 | 27242619 | 27242775 | 27242871 | 27242973 | 0 | 0 | 1 | 0 | 0 | 18 | 10 | 19 | 15 | 18 | 0 | 0 | 0 | 0 | 0 | 17 | 0 | 16 | 23 | 23 | 245 | 149 | 1 | 1 | 0 | 0 | 0.031 | 0 | 0 | 0 | NA | 0 | 0 | 0 | 0.006 |
| RI | 6356 | ENSG00000075624 | ACTB | chr7 | - | 5528280 | 5529059 | 5528280 | 5528719 | 5529018 | 5529059 | 158 | 94 | 152 | 163 | 148 | 0 | 0 | 0 | 0 | 0 | 311 | 192 | 175 | 252 | 603 | 0 | 0 | 0 | 0 | 10 | 298 | 149 | 1 | 1 | 1 | 1 | 1 | 1 | 1 | 1 | 1 | 1 | 1 | 0.968 | 0.006 |
| RI | 7025 | ENSG00000241852 | C8orf58 | chr8 | + | 22601971 | 22602312 | 22601971 | 22602080 | 22602199 | 22602312 | 4 | 0 | 7 | 1 | 1 | 1 | 0 | 5 | 2 | 1 | 10 | 1 | 3 | 0 | 19 | 0 | 1 | 4 | 4 | 13 | 268 | 149 | 1 | 1 | 0.69 | NA | 0.438 | 0.218 | 0.357 | 1 | 0.357 | 0.294 | 0 | 0.448 | 0.006 |
| RI | 8327 | ENSG00000089094 | KDM2B | chr12 | - | 121429508 | 121430469 | 121429508 | 121430163 | 121430276 | 121430469 | 128 | 63 | 100 | 123 | 132 | 0 | 0 | 0 | 0 | 0 | 233 | 140 | 124 | 112 | 187 | 0 | 0 | 2 | 0 | 0 | 262 | 149 | 1 | 1 | 1 | 1 | 1 | 1 | 1 | 1 | 1 | 0.972 | 1 | 1 | 0.006 |
| RI | 8457 | ENSG00000123349 | PFDN5 | chr12 | + | 53295838 | 53298150 | 53295838 | 53295941 | 53298044 | 53298150 | 198 | 152 | 201 | 218 | 234 | 0 | 0 | 0 | 0 | 0 | 297 | 255 | 176 | 207 | 275 | 0 | 4 | 0 | 0 | 0 | 298 | 149 | 1 | 1 | 1 | 1 | 1 | 1 | 1 | 1 | 0.97 | 1 | 1 | 1 | 0.006 |
| RI | 329 | ENSG00000076662 | ICAM3 | chr19 | - | 10334159 | 10335976 | 10334159 | 10334408 | 10335670 | 10335976 | 18 | 10 | 17 | 10 | 5 | 0 | 0 | 0 | 1 | 0 | 13 | 16 | 4 | 4 | 4 | 0 | 2 | 0 | 0 | 0 | 298 | 149 | 1 | 1 | 1 | 1 | 1 | 0.833 | 1 | 1 | 0.8 | 1 | 1 | 1 | 0.007 |
| RI | 751 | ENSG00000005961 | ITGA2B | chr17 | - | 44380599 | 44381061 | 44380599 | 44380645 | 44380878 | 44381061 | 2 | 1 | 2 | 4 | 2 | 117 | 71 | 27 | 227 | 178 | 0 | 4 | 4 | 1 | 26 | 315 | 176 | 142 | 104 | 4427 | 298 | 149 | 1 | 1 | 0.008 | 0.007 | 0.036 | 0.009 | 0.006 | 0 | 0.011 | 0.014 | 0.005 | 0.003 | 0.007 |
| RI | 3741 | ENSG00000114626 | ABTB1 | chr3 | + | 127677464 | 127680185 | 127677464 | 127677563 | 127679984 | 127680185 | 14 | 4 | 37 | 22 | 8 | 1 | 0 | 0 | 0 | 0 | 25 | 21 | 21 | 8 | 51 | 0 | 2 | 0 | 0 | 0 | 298 | 149 | 1 | 1 | 0.875 | 1 | 1 | 1 | 1 | 1 | 0.84 | 1 | 1 | 1 | 0.007 |
| RI | 4052 | ENSG00000066279 | ASPM | chr1 | - | 197124869 | 197125191 | 197124869 | 197124955 | 197125045 | 197125191 | 0 | 4 | 9 | 1 | 0 | 3 | 0 | 5 | 3 | 4 | 0 | 0 | 2 | 0 | 0 | 0 | 0 | 0 | 1 | 5 | 239 | 149 | 1 | 1 | 0 | 1 | 0.529 | 0.172 | 0 | NA | NA | 1 | 0 | 0 | 0.007 |
| RI | 4561 | ENSG00000225470 | JPX | chrX | + | 73998767 | 74000496 | 73998767 | 73998844 | 73999354 | 74000496 | 40 | 53 | 51 | 33 | 103 | 0 | 1 | 0 | 0 | 0 | 9 | 26 | 27 | 20 | 8 | 0 | 0 | 1 | 0 | 0 | 298 | 149 | 1 | 1 | 1 | 0.964 | 1 | 1 | 1 | 1 | 1 | 0.931 | 1 | 1 | 0.007 |
| RI | 4849 | ENSG00000137496 | IL18BP | chr11 | + | 71999227 | 71999662 | 71999227 | 71999370 | 71999616 | 71999662 | 60 | 43 | 50 | 45 | 84 | 1 | 1 | 0 | 0 | 0 | 83 | 74 | 32 | 65 | 80 | 0 | 0 | 2 | 0 | 0 | 298 | 149 | 1 | 1 | 0.968 | 0.956 | 1 | 1 | 1 | 1 | 1 | 0.889 | 1 | 1 | 0.007 |
| RI | 5886 | ENSG00000167515 | TRAPPC2L | chr16 | + | 88859344 | 88859750 | 88859344 | 88859443 | 88859572 | 88859750 | 34 | 8 | 32 | 14 | 41 | 0 | 0 | 3 | 0 | 1 | 29 | 41 | 19 | 23 | 73 | 0 | 0 | 3 | 0 | 0 | 278 | 149 | 1 | 1 | 1 | 1 | 0.851 | 1 | 0.956 | 1 | 1 | 0.772 | 1 | 1 | 0.007 |
| RI | 6025 | ENSG00000204264 | PSMB8 | chr6 | - | 32842671 | 32843089 | 32842671 | 32842711 | 32842941 | 32843089 | 187 | 122 | 170 | 220 | 142 | 0 | 0 | 0 | 0 | 0 | 211 | 228 | 192 | 228 | 327 | 0 | 0 | 0 | 4 | 0 | 298 | 149 | 1 | 1 | 1 | 1 | 1 | 1 | 1 | 1 | 1 | 1 | 0.966 | 1 | 0.007 |
| RI | 7974 | ENSG00000187240 | DYNC2H1 | chr11 | + | 103154450 | 103154809 | 103154450 | 103154606 | 103154694 | 103154809 | 8 | 4 | 10 | 5 | 12 | 6 | 6 | 5 | 3 | 3 | 0 | 0 | 0 | 2 | 0 | 0 | 1 | 0 | 0 | 0 | 237 | 149 | 1 | 1 | 0.456 | 0.295 | 0.557 | 0.512 | 0.715 | NA | 0 | NA | 1 | NA | 0.007 |
| RI | 8087 | ENSG00000105287 | PRKD2 | chr19 | - | 46716130 | 46717114 | 46716130 | 46716220 | 46716950 | 46717114 | 53 | 25 | 39 | 44 | 24 | 0 | 0 | 1 | 0 | 0 | 64 | 50 | 22 | 51 | 89 | 0 | 0 | 1 | 0 | 0 | 298 | 149 | 1 | 1 | 1 | 1 | 0.951 | 1 | 1 | 1 | 1 | 0.917 | 1 | 1 | 0.007 |
| RI | 623 | ENSG00000174365 | SNHG11 | chr20 | + | 38447465 | 38448730 | 38447465 | 38447623 | 38448661 | 38448730 | 1 | 5 | 1 | 1 | 0 | 4 | 1 | 8 | 7 | 3 | 0 | 4 | 3 | 0 | 3 | 0 | 9 | 4 | 16 | 4 | 298 | 149 | 1 | 1 | 0.111 | 0.714 | 0.059 | 0.067 | 0 | NA | 0.182 | 0.273 | 0 | 0.273 | 0.008 |
| RI | 1651 | ENSG00000103569 | AQP9 | chr15 | + | 58183960 | 58185911 | 58183960 | 58184440 | 58184486 | 58185911 | 121 | 74 | 56 | 108 | 169 | 0 | 0 | 0 | 0 | 0 | 105 | 216 | 474 | 153 | 55 | 0 | 7 | 0 | 0 | 0 | 195 | 149 | 1 | 1 | 1 | 1 | 1 | 1 | 1 | 1 | 0.959 | 1 | 1 | 1 | 0.008 |
| RI | 2132 | ENSG00000154153 | RETREG1 | chr5 | - | 16477661 | 16478098 | 16477661 | 16477788 | 16478033 | 16478098 | 8 | 8 | 9 | 1 | 23 | 85 | 68 | 155 | 73 | 326 | 0 | 0 | 2 | 2 | 0 | 11 | 26 | 18 | 12 | 8 | 298 | 149 | 1 | 1 | 0.045 | 0.056 | 0.028 | 0.007 | 0.034 | 0 | 0 | 0.053 | 0.077 | 0 | 0.008 |
| RI | 2235 | ENSG00000113716 | HMGXB3 | chr5 | + | 150024261 | 150024680 | 150024261 | 150024408 | 150024504 | 150024680 | 207 | 121 | 111 | 150 | 134 | 1 | 0 | 0 | 1 | 0 | 126 | 263 | 239 | 220 | 213 | 5 | 0 | 0 | 0 | 0 | 245 | 149 | 1 | 1 | 0.992 | 1 | 1 | 0.989 | 1 | 0.939 | 1 | 1 | 1 | 1 | 0.008 |
| RI | 2296 | ENSG00000170144 | HNRNPA3 | chr2 | + | 177215977 | 177216940 | 177215977 | 177216592 | 177216859 | 177216940 | 220 | 411 | 230 | 315 | 143 | 0 | 0 | 0 | 0 | 1 | 294 | 312 | 308 | 398 | 78 | 6 | 0 | 2 | 0 | 0 | 298 | 149 | 1 | 1 | 1 | 1 | 1 | 1 | 0.986 | 0.961 | 1 | 0.987 | 1 | 1 | 0.008 |
| RI | 5678 | ENSG00000144504 | ANKMY1 | chr2 | - | 240523884 | 240524381 | 240523884 | 240524009 | 240524303 | 240524381 | 85 | 58 | 65 | 98 | 69 | 0 | 0 | 0 | 0 | 0 | 55 | 124 | 95 | 57 | 116 | 0 | 0 | 2 | 0 | 0 | 298 | 149 | 1 | 1 | 1 | 1 | 1 | 1 | 1 | 1 | 1 | 0.96 | 1 | 1 | 0.008 |
| RI | 7366 | ENSG00000140988 | RPS2 | chr16 | - | 1962496 | 1963256 | 1962496 | 1962909 | 1962909 | 1963256 | 411 | 186 | 455 | 343 | 245 | 0 | 0 | 0 | 0 | 0 | 254 | 274 | 223 | 196 | 349 | 5 | 0 | 0 | 0 | 0 | 298 | 149 | 1 | 1 | 1 | 1 | 1 | 1 | 1 | 0.962 | 1 | 1 | 1 | 1 | 0.008 |
| RI | 752 | ENSG00000005961 | ITGA2B | chr17 | - | 44383937 | 44384138 | 44383937 | 44383946 | 44384084 | 44384138 | 3 | 0 | 2 | 2 | 0 | 143 | 99 | 22 | 230 | 79 | 10 | 0 | 0 | 0 | 33 | 534 | 192 | 118 | 150 | 5128 | 287 | 149 | 1 | 1 | 0.011 | 0 | 0.045 | 0.004 | 0 | 0.01 | 0 | 0 | 0 | 0.003 | 0.009 |
| RI | 883 | ENSG00000185359 | HGS | chr17 | + | 81696606 | 81696747 | 81696606 | 81696618 | 81696684 | 81696747 | 54 | 19 | 34 | 39 | 18 | 0 | 0 | 0 | 0 | 0 | 62 | 47 | 20 | 29 | 34 | 0 | 0 | 0 | 1 | 0 | 215 | 149 | 1 | 1 | 1 | 1 | 1 | 1 | 1 | 1 | 1 | 1 | 0.953 | 1 | 0.009 |
| RI | 1295 | ENSG00000197912 | SPG7 | chr16 | + | 89530682 | 89532066 | 89530682 | 89530808 | 89531924 | 89532066 | 16 | 9 | 39 | 20 | 14 | 0 | 0 | 0 | 0 | 0 | 30 | 26 | 44 | 30 | 49 | 0 | 0 | 1 | 0 | 0 | 298 | 149 | 1 | 1 | 1 | 1 | 1 | 1 | 1 | 1 | 1 | 0.957 | 1 | 1 | 0.009 |
| RI | 2799 | ENSG00000163703 | CRELD1 | chr3 | + | 9940849 | 9941206 | 9940849 | 9941016 | 9941110 | 9941206 | 2 | 0 | 2 | 1 | 4 | 6 | 4 | 1 | 5 | 0 | 0 | 0 | 0 | 4 | 1 | 4 | 0 | 0 | 0 | 8 | 243 | 149 | 1 | 1 | 0.17 | 0 | 0.551 | 0.109 | 1 | 0 | NA | NA | 1 | 0.071 | 0.009 |
| RI | 3100 | ENSG00000161217 | PCYT1A | chr3 | - | 196247366 | 196248323 | 196247366 | 196247643 | 196248206 | 196248323 | 31 | 34 | 42 | 41 | 14 | 0 | 0 | 0 | 0 | 0 | 37 | 19 | 45 | 44 | 52 | 0 | 0 | 1 | 0 | 0 | 298 | 149 | 1 | 1 | 1 | 1 | 1 | 1 | 1 | 1 | 1 | 0.957 | 1 | 1 | 0.009 |
| RI | 4096 | ENSG00000256269 | HMBS | chr11 | + | 119088634 | 119089131 | 119088634 | 119088707 | 119089081 | 119089131 | 0 | 0 | 1 | 2 | 2 | 29 | 18 | 19 | 23 | 14 | 0 | 1 | 0 | 0 | 3 | 18 | 40 | 19 | 23 | 18 | 298 | 149 | 1 | 1 | 0 | 0 | 0.026 | 0.042 | 0.067 | 0 | 0.012 | 0 | 0 | 0.077 | 0.009 |
| RI | 4366 | ENSG00000144381 | HSPD1 | chr2 | - | 197494156 | 197495376 | 197494156 | 197494250 | 197495293 | 197495376 | 43 | 45 | 50 | 61 | 87 | 0 | 1 | 1 | 0 | 0 | 12 | 21 | 73 | 14 | 19 | 0 | 0 | 0 | 1 | 0 | 298 | 149 | 1 | 1 | 1 | 0.957 | 0.962 | 1 | 1 | 1 | 1 | 1 | 0.875 | 1 | 0.009 |
| RI | 6051 | ENSG00000135486 | HNRNPA1 | chr12 | + | 54282393 | 54282874 | 54282393 | 54282486 | 54282799 | 54282874 | 241 | 242 | 260 | 306 | 245 | 0 | 0 | 0 | 0 | 0 | 332 | 450 | 251 | 585 | 173 | 0 | 0 | 0 | 0 | 4 | 298 | 149 | 1 | 1 | 1 | 1 | 1 | 1 | 1 | 1 | 1 | 1 | 1 | 0.956 | 0.009 |
| RI | 6812 | ENSG00000169359 | SLC33A1 | chr3 | - | 155829687 | 155829903 | 155829687 | 155829755 | 155829855 | 155829903 | 212 | 152 | 148 | 143 | 212 | 1 | 0 | 0 | 0 | 2 | 19 | 87 | 78 | 74 | 160 | 0 | 0 | 1 | 0 | 5 | 249 | 149 | 1 | 1 | 0.992 | 1 | 1 | 1 | 0.984 | 1 | 1 | 0.979 | 1 | 0.95 | 0.009 |
| RI | 8090 | ENSG00000177239 | MAN1B1 | chr9 | + | 137101483 | 137105916 | 137101483 | 137101672 | 137105607 | 137105916 | 24 | 38 | 26 | 40 | 11 | 0 | 0 | 0 | 0 | 0 | 38 | 22 | 43 | 30 | 72 | 0 | 0 | 1 | 0 | 0 | 298 | 149 | 1 | 1 | 1 | 1 | 1 | 1 | 1 | 1 | 1 | 0.956 | 1 | 1 | 0.009 |
| RI | 8131 | ENSG00000107959 | PITRM1 | chr10 | - | 3143388 | 3143731 | 3143388 | 3143501 | 3143605 | 3143731 | 25 | 17 | 36 | 31 | 21 | 0 | 0 | 0 | 0 | 0 | 8 | 38 | 36 | 32 | 33 | 0 | 1 | 0 | 0 | 0 | 253 | 149 | 1 | 1 | 1 | 1 | 1 | 1 | 1 | 1 | 0.957 | 1 | 1 | 1 | 0.009 |
| RI | 8354 | ENSG00000092531 | SNAP23 | chr15 | + | 42513173 | 42513447 | 42513173 | 42513209 | 42513398 | 42513447 | 226 | 261 | 202 | 244 | 206 | 0 | 0 | 1 | 0 | 0 | 194 | 195 | 312 | 143 | 62 | 0 | 5 | 1 | 0 | 0 | 298 | 149 | 1 | 1 | 1 | 1 | 0.99 | 1 | 1 | 1 | 0.951 | 0.994 | 1 | 1 | 0.009 |
| RI | 381 | ENSG00000105483 | CARD8 | chr19 | - | 48232452 | 48234543 | 48232452 | 48232789 | 48234402 | 48234543 | 260 | 266 | 362 | 313 | 238 | 1 | 0 | 0 | 0 | 0 | 127 | 180 | 183 | 125 | 31 | 3 | 0 | 1 | 0 | 0 | 298 | 149 | 1 | 1 | 0.992 | 1 | 1 | 1 | 1 | 0.955 | 1 | 0.989 | 1 | 1 | 0.01 |
| RI | 1054 | ENSG00000073584 | SMARCE1 | chr17 | - | 40630713 | 40632367 | 40630713 | 40630924 | 40632194 | 40632367 | 85 | 92 | 84 | 88 | 75 | 0 | 0 | 0 | 0 | 0 | 0 | 57 | 135 | 14 | 0 | 0 | 0 | 2 | 0 | 0 | 298 | 149 | 1 | 1 | 1 | 1 | 1 | 1 | 1 | NA | 1 | 0.971 | 1 | NA | 0.01 |
| RI | 1378 | ENSG00000196118 | CCDC189 | chr16 | - | 30759011 | 30759247 | 30759011 | 30759101 | 30759177 | 30759247 | 4 | 0 | 0 | 0 | 1 | 0 | 6 | 0 | 0 | 0 | 0 | 7 | 2 | 4 | 15 | 5 | 0 | 1 | 0 | 4 | 225 | 149 | 1 | 1 | 1 | 0 | NA | NA | 1 | 0 | 1 | 0.57 | 1 | 0.713 | 0.01 |
| RI | 2162 | ENSG00000065357 | DGKA | chr12 | + | 55940081 | 55940433 | 55940081 | 55940170 | 55940251 | 55940433 | 93 | 92 | 193 | 130 | 229 | 0 | 0 | 1 | 0 | 5 | 120 | 121 | 137 | 149 | 103 | 0 | 0 | 0 | 1 | 6 | 230 | 149 | 1 | 1 | 1 | 1 | 0.992 | 1 | 0.967 | 1 | 1 | 1 | 0.99 | 0.917 | 0.01 |
| RI | 2250 | ENSG00000111057 | KRT18 | chr12 | + | 52950327 | 52950906 | 52950327 | 52950410 | 52950749 | 52950906 | 0 | 0 | 0 | 1 | 0 | 9 | 7 | 15 | 10 | 14 | 0 | 0 | 0 | 0 | 0 | 4 | 5 | 3 | 11 | 19 | 298 | 149 | 1 | 1 | 0 | 0 | 0 | 0.048 | 0 | 0 | 0 | 0 | 0 | 0 | 0.01 |
| RI | 2274 | ENSG00000104731 | KLHDC4 | chr16 | - | 87755193 | 87756477 | 87755193 | 87755254 | 87756398 | 87756477 | 65 | 24 | 61 | 56 | 35 | 0 | 0 | 0 | 0 | 0 | 58 | 76 | 73 | 97 | 106 | 0 | 0 | 2 | 0 | 0 | 298 | 149 | 1 | 1 | 1 | 1 | 1 | 1 | 1 | 1 | 1 | 0.948 | 1 | 1 | 0.01 |
| RI | 3567 | ENSG00000265241 | RBM8A | chr1 | - | 145926481 | 145927077 | 145926481 | 145926618 | 145927017 | 145927077 | 64 | 78 | 67 | 112 | 78 | 0 | 0 | 0 | 0 | 0 | 243 | 113 | 89 | 223 | 76 | 0 | 0 | 0 | 6 | 0 | 298 | 149 | 1 | 1 | 1 | 1 | 1 | 1 | 1 | 1 | 1 | 1 | 0.949 | 1 | 0.01 |
| RI | 3725 | ENSG00000142208 | AKT1 | chr14 | - | 104772877 | 104773379 | 104772877 | 104772993 | 104773250 | 104773379 | 134 | 47 | 110 | 147 | 117 | 0 | 0 | 0 | 1 | 0 | 251 | 192 | 116 | 90 | 266 | 0 | 0 | 4 | 0 | 0 | 298 | 149 | 1 | 1 | 1 | 1 | 1 | 0.987 | 1 | 1 | 1 | 0.935 | 1 | 1 | 0.01 |
| RI | 4296 | ENSG00000092621 | PHGDH | chr1 | + | 119741766 | 119743044 | 119741766 | 119741897 | 119742806 | 119743044 | 1 | 0 | 0 | 0 | 0 | 3 | 1 | 15 | 4 | 15 | 0 | 0 | 0 | 2 | 0 | 20 | 5 | 11 | 10 | 14 | 298 | 149 | 1 | 1 | 0.143 | 0 | 0 | 0 | 0 | 0 | 0 | 0 | 0.091 | 0 | 0.01 |
| RI | 5262 | ENSG00000224078 | SNHG14 | chr15 | + | 25108997 | 25109403 | 25108997 | 25109099 | 25109197 | 25109403 | 16 | 3 | 2 | 10 | 11 | 3 | 3 | 4 | 1 | 9 | 0 | 5 | 8 | 2 | 0 | 0 | 8 | 12 | 0 | 0 | 247 | 149 | 1 | 1 | 0.763 | 0.376 | 0.232 | 0.858 | 0.424 | NA | 0.274 | 0.287 | 1 | NA | 0.01 |
| RI | 5661 | ENSG00000165280 | VCP | chr9 | - | 35057375 | 35059219 | 35057375 | 35057527 | 35059063 | 35059219 | 81 | 52 | 74 | 88 | 82 | 0 | 0 | 0 | 0 | 0 | 276 | 184 | 95 | 237 | 215 | 0 | 0 | 1 | 0 | 3 | 298 | 149 | 1 | 1 | 1 | 1 | 1 | 1 | 1 | 1 | 1 | 0.979 | 1 | 0.973 | 0.01 |
| RI | 6801 | ENSG00000005302 | MSL3 | chrX | + | 11763779 | 11765729 | 11763779 | 11763938 | 11765521 | 11765729 | 165 | 113 | 184 | 144 | 132 | 0 | 0 | 0 | 0 | 0 | 78 | 112 | 168 | 110 | 44 | 0 | 0 | 0 | 3 | 0 | 298 | 149 | 1 | 1 | 1 | 1 | 1 | 1 | 1 | 1 | 1 | 1 | 0.948 | 1 | 0.01 |
| RI | 7414 | ENSG00000107331 | ABCA2 | chr9 | - | 137011406 | 137011749 | 137011406 | 137011554 | 137011633 | 137011749 | 1 | 0 | 3 | 0 | 0 | 31 | 2 | 25 | 36 | 18 | 0 | 4 | 0 | 0 | 0 | 81 | 54 | 22 | 20 | 62 | 228 | 149 | 1 | 1 | 0.021 | 0 | 0.073 | 0 | 0 | 0 | 0.046 | 0 | 0 | 0 | 0.01 |
| RI | 1296 | ENSG00000197912 | SPG7 | chr16 | + | 89540943 | 89541346 | 89540943 | 89541027 | 89541152 | 89541346 | 63 | 51 | 63 | 77 | 47 | 2 | 0 | 0 | 2 | 0 | 101 | 122 | 74 | 226 | 152 | 0 | 4 | 1 | 10 | 0 | 274 | 149 | 1 | 1 | 0.945 | 1 | 1 | 0.954 | 1 | 1 | 0.943 | 0.976 | 0.925 | 1 | 0.011 |
| RI | 2065 | ENSG00000125459 | MSTO1 | chr1 | + | 155610240 | 155610560 | 155610240 | 155610332 | 155610424 | 155610560 | 2 | 0 | 3 | 1 | 0 | 0 | 0 | 1 | 1 | 0 | 7 | 0 | 0 | 0 | 9 | 0 | 5 | 0 | 0 | 0 | 241 | 149 | 1 | 1 | 1 | NA | 0.65 | 0.382 | NA | 1 | 0 | NA | NA | 1 | 0.011 |
| RI | 2314 | ENSG00000103932 | RPAP1 | chr15 | - | 41517797 | 41518182 | 41517797 | 41517857 | 41518005 | 41518182 | 11 | 5 | 3 | 2 | 5 | 6 | 3 | 2 | 8 | 3 | 11 | 0 | 2 | 0 | 0 | 0 | 7 | 1 | 0 | 3 | 297 | 149 | 1 | 1 | 0.479 | 0.455 | 0.429 | 0.111 | 0.455 | 1 | 0 | 0.501 | NA | 0 | 0.011 |
| RI | 3306 | ENSG00000148399 | DPH7 | chr9 | - | 137564892 | 137565154 | 137564892 | 137564958 | 137565084 | 137565154 | 4 | 4 | 4 | 4 | 2 | 44 | 24 | 25 | 36 | 16 | 0 | 0 | 2 | 6 | 1 | 0 | 11 | 9 | 28 | 50 | 275 | 149 | 1 | 1 | 0.047 | 0.083 | 0.08 | 0.057 | 0.063 | NA | 0 | 0.107 | 0.104 | 0.011 | 0.011 |
| RI | 4709 | ENSG00000232956 | SNHG15 | chr7 | - | 44983821 | 44986097 | 44983821 | 44984046 | 44985881 | 44986097 | 53 | 29 | 62 | 47 | 31 | 0 | 0 | 0 | 0 | 0 | 93 | 70 | 60 | 114 | 191 | 0 | 2 | 0 | 0 | 0 | 298 | 149 | 1 | 1 | 1 | 1 | 1 | 1 | 1 | 1 | 0.946 | 1 | 1 | 1 | 0.011 |
| RI | 5590 | ENSG00000225889 | NA | chr2 | + | 64251118 | 64251834 | 64251118 | 64251337 | 64251688 | 64251834 | 59 | 42 | 58 | 71 | 45 | 0 | 0 | 1 | 0 | 0 | 105 | 61 | 134 | 99 | 86 | 4 | 0 | 1 | 0 | 0 | 298 | 149 | 1 | 1 | 1 | 1 | 0.967 | 1 | 1 | 0.929 | 1 | 0.985 | 1 | 1 | 0.011 |
| RI | 6318 | ENSG00000183723 | CMTM4 | chr16 | - | 66614749 | 66622222 | 66614749 | 66617358 | 66622061 | 66622222 | 6 | 13 | 14 | 8 | 17 | 1 | 1 | 0 | 2 | 0 | 17 | 17 | 15 | 29 | 0 | 0 | 2 | 2 | 4 | 0 | 298 | 149 | 1 | 1 | 0.75 | 0.867 | 1 | 0.667 | 1 | 1 | 0.81 | 0.789 | 0.784 | NA | 0.011 |
| RI | 6470 | ENSG00000169583 | CLIC3 | chr9 | - | 136994929 | 136995292 | 136994929 | 136995105 | 136995185 | 136995292 | 11 | 0 | 3 | 5 | 0 | 93 | 25 | 24 | 22 | 1 | 0 | 0 | 1 | 0 | 0 | 30 | 18 | 3 | 11 | 0 | 229 | 149 | 1 | 1 | 0.071 | 0 | 0.075 | 0.129 | 0 | 0 | 0 | 0.178 | 0 | NA | 0.011 |
| RI | 6654 | ENSG00000182220 | ATP6AP2 | chrX | + | 40599591 | 40600881 | 40599591 | 40599741 | 40600768 | 40600881 | 42 | 12 | 29 | 33 | 29 | 1 | 0 | 2 | 0 | 0 | 12 | 41 | 38 | 47 | 0 | 0 | 0 | 0 | 5 | 0 | 298 | 149 | 1 | 1 | 0.955 | 1 | 0.879 | 1 | 1 | 1 | 1 | 1 | 0.825 | NA | 0.011 |
| RI | 7024 | ENSG00000142046 | TMEM91 | chr19 | + | 41383560 | 41384083 | 41383560 | 41383632 | 41383885 | 41384083 | 21 | 14 | 17 | 15 | 12 | 0 | 0 | 0 | 0 | 0 | 35 | 34 | 29 | 30 | 84 | 0 | 1 | 0 | 0 | 0 | 298 | 149 | 1 | 1 | 1 | 1 | 1 | 1 | 1 | 1 | 0.944 | 1 | 1 | 1 | 0.011 |
| RI | 7250 | ENSG00000137563 | GGH | chr8 | - | 63026157 | 63030217 | 63026157 | 63026296 | 63027180 | 63030217 | 1 | 0 | 3 | 0 | 3 | 15 | 3 | 23 | 9 | 6 | 1 | 1 | 0 | 0 | 0 | 10 | 3 | 9 | 34 | 0 | 298 | 149 | 1 | 1 | 0.032 | 0 | 0.061 | 0 | 0.2 | 0.048 | 0.143 | 0 | 0 | NA | 0.011 |
| RI | 8135 | ENSG00000008300 | CELSR3 | chr3 | - | 48641850 | 48642467 | 48641850 | 48642009 | 48642357 | 48642467 | 1 | 1 | 5 | 1 | 0 | 2 | 0 | 0 | 1 | 0 | 8 | 1 | 0 | 0 | 1 | 2 | 0 | 0 | 0 | 2 | 298 | 149 | 1 | 1 | 0.2 | 1 | 1 | 0.333 | NA | 0.667 | 1 | NA | NA | 0.2 | 0.011 |
| RI | 8386 | ENSG00000175899 | A2M | chr12 | - | 9067711 | 9068224 | 9067711 | 9067839 | 9068182 | 9068224 | 185 | 115 | 137 | 116 | 182 | 0 | 0 | 0 | 1 | 0 | 42 | 140 | 32 | 10 | 0 | 0 | 0 | 1 | 0 | 0 | 298 | 149 | 1 | 1 | 1 | 1 | 1 | 0.983 | 1 | 1 | 1 | 0.941 | 1 | NA | 0.011 |
| RI | 8474 | ENSG00000174106 | LEMD3 | chr12 | + | 65245668 | 65245939 | 65245668 | 65245774 | 65245860 | 65245939 | 37 | 46 | 28 | 37 | 45 | 135 | 82 | 101 | 97 | 360 | 4 | 21 | 27 | 11 | 0 | 36 | 53 | 70 | 38 | 0 | 235 | 149 | 1 | 1 | 0.148 | 0.262 | 0.149 | 0.195 | 0.073 | 0.066 | 0.201 | 0.197 | 0.155 | NA | 0.011 |
| RI | 351 | ENSG00000142453 | CARM1 | chr19 | + | 10920833 | 10921127 | 10920833 | 10920946 | 10921049 | 10921127 | 1 | 0 | 1 | 3 | 0 | 43 | 24 | 26 | 22 | 37 | 0 | 3 | 1 | 0 | 0 | 39 | 73 | 22 | 47 | 101 | 252 | 149 | 1 | 1 | 0.014 | 0 | 0.022 | 0.075 | 0 | 0 | 0.024 | 0.026 | 0 | 0 | 0.012 |
| RI | 4905 | ENSG00000118961 | LDAH | chr2 | - | 20685546 | 20687094 | 20685546 | 20685657 | 20686906 | 20687094 | 55 | 68 | 65 | 88 | 135 | 0 | 0 | 0 | 0 | 0 | 19 | 64 | 46 | 40 | 4 | 0 | 2 | 0 | 0 | 0 | 298 | 149 | 1 | 1 | 1 | 1 | 1 | 1 | 1 | 1 | 0.941 | 1 | 1 | 1 | 0.012 |
| RI | 7908 | ENSG00000102317 | RBM3 | chrX | + | 48575560 | 48576083 | 48575560 | 48575667 | 48575921 | 48576083 | 66 | 35 | 72 | 89 | 53 | 0 | 0 | 1 | 0 | 0 | 148 | 59 | 52 | 83 | 129 | 0 | 0 | 0 | 0 | 6 | 298 | 149 | 1 | 1 | 1 | 1 | 0.973 | 1 | 1 | 1 | 1 | 1 | 1 | 0.915 | 0.012 |
| RI | 257 | ENSG00000142541 | RPL13A | chr19 | + | 49489849 | 49490297 | 49489849 | 49489922 | 49490126 | 49490297 | 81 | 49 | 107 | 100 | 67 | 0 | 0 | 0 | 0 | 0 | 174 | 104 | 59 | 94 | 201 | 0 | 0 | 2 | 0 | 0 | 298 | 149 | 1 | 1 | 1 | 1 | 1 | 1 | 1 | 1 | 1 | 0.937 | 1 | 1 | 0.013 |
| RI | 303 | ENSG00000161265 | U2AF1L4 | chr19 | - | 35743154 | 35743345 | 35743154 | 35743205 | 35743295 | 35743345 | 3 | 7 | 5 | 7 | 4 | 0 | 2 | 0 | 0 | 0 | 10 | 22 | 10 | 10 | 18 | 0 | 0 | 2 | 1 | 0 | 239 | 149 | 1 | 1 | 1 | 0.686 | 1 | 1 | 1 | 1 | 1 | 0.757 | 0.862 | 1 | 0.013 |
| RI | 816 | ENSG00000184009 | ACTG1 | chr17 | - | 81511187 | 81511803 | 81511187 | 81511626 | 81511762 | 81511803 | 28 | 9 | 23 | 33 | 38 | 0 | 0 | 0 | 0 | 0 | 5 | 15 | 36 | 27 | 20 | 0 | 0 | 0 | 1 | 0 | 285 | 149 | 1 | 1 | 1 | 1 | 1 | 1 | 1 | 1 | 1 | 1 | 0.934 | 1 | 0.013 |
| RI | 3263 | ENSG00000113108 | APBB3 | chr5 | - | 140561586 | 140561849 | 140561586 | 140561722 | 140561843 | 140561849 | 20 | 20 | 24 | 28 | 13 | 1 | 0 | 0 | 2 | 0 | 42 | 51 | 35 | 20 | 35 | 0 | 0 | 0 | 3 | 1 | 270 | 149 | 1 | 1 | 0.917 | 1 | 1 | 0.885 | 1 | 1 | 1 | 1 | 0.786 | 0.951 | 0.013 |
| RI | 3850 | ENSG00000073756 | PTGS2 | chr1 | - | 186675897 | 186676713 | 186675897 | 186676184 | 186676466 | 186676713 | 13 | 36 | 41 | 27 | 33 | 118 | 93 | 80 | 225 | 322 | 0 | 4 | 14 | 5 | 0 | 7 | 11 | 37 | 44 | 0 | 298 | 149 | 1 | 1 | 0.052 | 0.162 | 0.204 | 0.057 | 0.049 | 0 | 0.154 | 0.159 | 0.054 | NA | 0.013 |
| RI | 6393 | ENSG00000133216 | EPHB2 | chr1 | + | 22913461 | 22921500 | 22913461 | 22913564 | 22913657 | 22921500 | 0 | 3 | 1 | 16 | 2 | 0 | 0 | 0 | 0 | 0 | 72 | 23 | 8 | 59 | 45 | 3 | 0 | 0 | 0 | 0 | 242 | 149 | 1 | 1 | NA | 1 | 1 | 1 | 1 | 0.937 | 1 | 1 | 1 | 1 | 0.013 |
| RI | 6571 | ENSG00000234456 | MAGI2-AS3 | chr7 | + | 79458846 | 79459640 | 79458846 | 79458999 | 79459535 | 79459640 | 0 | 0 | 1 | 0 | 0 | 4 | 61 | 7 | 23 | 3 | 0 | 0 | 0 | 0 | 0 | 0 | 2 | 44 | 3 | 0 | 298 | 149 | 1 | 1 | 0 | 0 | 0.067 | 0 | 0 | NA | 0 | 0 | 0 | NA | 0.013 |
| RI | 7112 | ENSG00000133740 | E2F5 | chr8 | + | 85213752 | 85214518 | 85213752 | 85213902 | 85214159 | 85214518 | 117 | 103 | 102 | 173 | 309 | 0 | 0 | 0 | 1 | 2 | 61 | 113 | 81 | 110 | 63 | 0 | 0 | 0 | 0 | 3 | 298 | 149 | 1 | 1 | 1 | 1 | 1 | 0.989 | 0.987 | 1 | 1 | 1 | 1 | 0.913 | 0.013 |
| RI | 8023 | ENSG00000090621 | PABPC4 | chr1 | - | 39560815 | 39562203 | 39560815 | 39561122 | 39562072 | 39562203 | 28 | 22 | 25 | 33 | 37 | 0 | 0 | 0 | 0 | 0 | 63 | 16 | 28 | 59 | 1 | 0 | 0 | 0 | 2 | 0 | 298 | 149 | 1 | 1 | 1 | 1 | 1 | 1 | 1 | 1 | 1 | 1 | 0.937 | 1 | 0.013 |
| RI | 8212 | ENSG00000151490 | PTPRO | chr12 | + | 15594936 | 15597399 | 15594936 | 15595057 | 15596089 | 15597399 | 25 | 29 | 28 | 24 | 21 | 7 | 21 | 16 | 10 | 49 | 0 | 9 | 27 | 31 | 0 | 21 | 6 | 2 | 20 | 0 | 298 | 149 | 1 | 1 | 0.641 | 0.408 | 0.467 | 0.545 | 0.176 | 0 | 0.429 | 0.871 | 0.437 | NA | 0.013 |
| RI | 2806 | ENSG00000163703 | CRELD1 | chr3 | + | 9943076 | 9943515 | 9943076 | 9943253 | 9943481 | 9943515 | 48 | 17 | 37 | 56 | 44 | 1 | 0 | 0 | 0 | 0 | 26 | 41 | 13 | 32 | 58 | 0 | 0 | 0 | 2 | 0 | 298 | 149 | 1 | 1 | 0.96 | 1 | 1 | 1 | 1 | 1 | 1 | 1 | 0.889 | 1 | 0.014 |
| RI | 4504 | ENSG00000115556 | PLCD4 | chr2 | + | 218633604 | 218634221 | 218633604 | 218633761 | 218634104 | 218634221 | 27 | 11 | 12 | 27 | 13 | 0 | 0 | 0 | 0 | 0 | 25 | 33 | 26 | 53 | 60 | 0 | 0 | 1 | 0 | 0 | 298 | 149 | 1 | 1 | 1 | 1 | 1 | 1 | 1 | 1 | 1 | 0.929 | 1 | 1 | 0.014 |
| RI | 7243 | ENSG00000283189 | NA | chr3 | - | 49419709 | 49420342 | 49419709 | 49419788 | 49420214 | 49420342 | 23 | 11 | 22 | 15 | 11 | 0 | 0 | 0 | 1 | 0 | 10 | 28 | 26 | 22 | 52 | 0 | 0 | 3 | 0 | 0 | 298 | 149 | 1 | 1 | 1 | 1 | 1 | 0.882 | 1 | 1 | 1 | 0.812 | 1 | 1 | 0.014 |
| RI | 7860 | ENSG00000142102 | PGGHG | chr11 | + | 293162 | 293727 | 293162 | 293235 | 293669 | 293727 | 118 | 45 | 118 | 76 | 51 | 0 | 0 | 0 | 0 | 0 | 13 | 75 | 81 | 58 | 148 | 0 | 0 | 3 | 0 | 0 | 298 | 149 | 1 | 1 | 1 | 1 | 1 | 1 | 1 | 1 | 1 | 0.931 | 1 | 1 | 0.014 |
| RI | 7913 | ENSG00000179532 | DNHD1 | chr11 | + | 6497770 | 6498961 | 6497770 | 6497844 | 6498722 | 6498961 | 13 | 6 | 8 | 3 | 2 | 0 | 0 | 0 | 0 | 0 | 36 | 25 | 26 | 25 | 31 | 0 | 0 | 1 | 0 | 0 | 298 | 149 | 1 | 1 | 1 | 1 | 1 | 1 | 1 | 1 | 1 | 0.929 | 1 | 1 | 0.014 |
| RI | 8114 | ENSG00000196372 | ASB13 | chr10 | - | 5638866 | 5640830 | 5638866 | 5639371 | 5640695 | 5640830 | 29 | 17 | 45 | 14 | 28 | 0 | 0 | 0 | 0 | 0 | 18 | 27 | 8 | 10 | 41 | 0 | 1 | 0 | 0 | 0 | 298 | 149 | 1 | 1 | 1 | 1 | 1 | 1 | 1 | 1 | 0.931 | 1 | 1 | 1 | 0.014 |
| RI | 3199 | ENSG00000010327 | STAB1 | chr3 | + | 52519943 | 52520290 | 52519943 | 52520120 | 52520203 | 52520290 | 0 | 0 | 3 | 1 | 0 | 28 | 3 | 17 | 24 | 1 | 0 | 0 | 1 | 0 | 0 | 64 | 29 | 12 | 22 | 79 | 232 | 149 | 1 | 1 | 0 | 0 | 0.102 | 0.026 | 0 | 0 | 0 | 0.051 | 0 | 0 | 0.015 |
| RI | 4390 | ENSG00000145020 | AMT | chr3 | - | 49418970 | 49420342 | 49418970 | 49419405 | 49420210 | 49420342 | 26 | 15 | 21 | 17 | 14 | 0 | 0 | 0 | 0 | 0 | 12 | 35 | 18 | 12 | 51 | 0 | 0 | 0 | 0 | 2 | 298 | 149 | 1 | 1 | 1 | 1 | 1 | 1 | 1 | 1 | 1 | 1 | 1 | 0.927 | 0.015 |
| RI | 4820 | ENSG00000149231 | CCDC82 | chr11 | - | 96383961 | 96387613 | 96383961 | 96384761 | 96387529 | 96387613 | 91 | 95 | 80 | 95 | 67 | 2 | 0 | 1 | 2 | 2 | 0 | 48 | 44 | 14 | 0 | 0 | 4 | 0 | 0 | 0 | 298 | 149 | 1 | 1 | 0.958 | 1 | 0.976 | 0.96 | 0.944 | NA | 0.857 | 1 | 1 | NA | 0.015 |
| RI | 5339 | ENSG00000084623 | EIF3I | chr1 | + | 32226170 | 32226530 | 32226170 | 32226283 | 32226402 | 32226530 | 39 | 41 | 39 | 56 | 40 | 0 | 0 | 0 | 0 | 0 | 56 | 67 | 28 | 69 | 129 | 0 | 0 | 0 | 3 | 0 | 268 | 149 | 1 | 1 | 1 | 1 | 1 | 1 | 1 | 1 | 1 | 1 | 0.927 | 1 | 0.015 |
| RI | 1923 | ENSG00000125746 | EML2 | chr19 | - | 45632860 | 45633139 | 45632860 | 45632971 | 45633069 | 45633139 | 1 | 1 | 0 | 2 | 0 | 36 | 11 | 21 | 47 | 10 | 0 | 0 | 0 | 1 | 0 | 78 | 37 | 21 | 40 | 94 | 247 | 149 | 1 | 1 | 0.016 | 0.052 | 0 | 0.025 | 0 | 0 | 0 | 0 | 0.015 | 0 | 0.016 |
| RI | 1999 | ENSG00000226067 | LINC00623 | chr1 | + | 120952438 | 120953460 | 120952438 | 120952785 | 120953222 | 120953460 | 131 | 60 | 38 | 89 | 84 | 1 | 0 | 0 | 0 | 0 | 61 | 75 | 40 | 46 | 68 | 0 | 4 | 0 | 0 | 0 | 298 | 149 | 1 | 1 | 0.985 | 1 | 1 | 1 | 1 | 1 | 0.904 | 1 | 1 | 1 | 0.016 |
| RI | 4508 | ENSG00000168610 | STAT3 | chr17 | - | 42316788 | 42317224 | 42316788 | 42316889 | 42317181 | 42317224 | 28 | 21 | 24 | 30 | 22 | 0 | 0 | 0 | 0 | 0 | 90 | 35 | 70 | 92 | 57 | 0 | 0 | 0 | 4 | 0 | 298 | 149 | 1 | 1 | 1 | 1 | 1 | 1 | 1 | 1 | 1 | 1 | 0.92 | 1 | 0.016 |
| RI | 5135 | ENSG00000267796 | LIN37 | chr19 | + | 35753086 | 35754157 | 35753086 | 35753193 | 35754016 | 35754157 | 33 | 28 | 31 | 43 | 14 | 0 | 0 | 3 | 0 | 0 | 72 | 70 | 17 | 19 | 6 | 0 | 0 | 0 | 3 | 0 | 298 | 149 | 1 | 1 | 1 | 1 | 0.838 | 1 | 1 | 1 | 1 | 1 | 0.76 | 1 | 0.016 |
| RI | 7560 | ENSG00000185920 | PTCH1 | chr9 | - | 95443411 | 95447451 | 95443411 | 95446391 | 95446910 | 95447451 | 2 | 0 | 0 | 1 | 0 | 19 | 5 | 5 | 17 | 5 | 0 | 0 | 0 | 0 | 0 | 0 | 0 | 6 | 2 | 4 | 298 | 149 | 1 | 1 | 0.05 | 0 | 0 | 0.029 | 0 | NA | NA | 0 | 0 | 0 | 0.016 |
| RI | 7700 | ENSG00000168067 | MAP4K2 | chr11 | - | 64792171 | 64792422 | 64792171 | 64792275 | 64792363 | 64792422 | 1 | 0 | 7 | 1 | 1 | 9 | 5 | 25 | 21 | 2 | 0 | 3 | 2 | 0 | 5 | 0 | 22 | 17 | 6 | 15 | 237 | 149 | 1 | 1 | 0.065 | 0 | 0.15 | 0.029 | 0.239 | NA | 0.079 | 0.069 | 0 | 0.173 | 0.016 |
| RI | 2337 | ENSG00000183621 | ZNF438 | chr10 | - | 30848530 | 30850367 | 30848530 | 30849443 | 30850221 | 30850367 | 55 | 28 | 48 | 33 | 74 | 1 | 0 | 0 | 0 | 0 | 106 | 46 | 30 | 23 | 34 | 0 | 0 | 2 | 0 | 0 | 298 | 149 | 1 | 1 | 0.965 | 1 | 1 | 1 | 1 | 1 | 1 | 0.882 | 1 | 1 | 0.017 |
| RI | 3144 | ENSG00000168811 | IL12A | chr3 | + | 159993450 | 159993609 | 159993450 | 159993492 | 159993567 | 159993609 | 6 | 2 | 2 | 8 | 9 | 4 | 3 | 7 | 3 | 0 | 0 | 11 | 0 | 7 | 10 | 0 | 8 | 5 | 4 | 0 | 224 | 149 | 1 | 1 | 0.499 | 0.307 | 0.16 | 0.639 | 1 | NA | 0.478 | 0 | 0.538 | 1 | 0.017 |
| RI | 3421 | ENSG00000184432 | COPB2 | chr3 | - | 139373665 | 139374588 | 139373665 | 139373833 | 139374488 | 139374588 | 177 | 129 | 133 | 138 | 135 | 0 | 0 | 0 | 0 | 0 | 49 | 84 | 273 | 54 | 0 | 0 | 3 | 0 | 0 | 0 | 298 | 149 | 1 | 1 | 1 | 1 | 1 | 1 | 1 | 1 | 0.933 | 1 | 1 | NA | 0.017 |
| RI | 4185 | ENSG00000173039 | RELA | chr11 | - | 65653601 | 65655000 | 65653601 | 65654509 | 65654818 | 65655000 | 105 | 39 | 71 | 102 | 66 | 0 | 0 | 0 | 0 | 0 | 174 | 160 | 84 | 95 | 177 | 0 | 0 | 1 | 0 | 6 | 298 | 149 | 1 | 1 | 1 | 1 | 1 | 1 | 1 | 1 | 1 | 0.977 | 1 | 0.937 | 0.017 |
| RI | 4451 | ENSG00000090097 | PCBP4 | chr3 | - | 51960525 | 51960898 | 51960525 | 51960637 | 51960865 | 51960898 | 3 | 0 | 2 | 1 | 3 | 1 | 0 | 1 | 2 | 0 | 0 | 3 | 2 | 1 | 3 | 0 | 12 | 0 | 0 | 11 | 298 | 149 | 1 | 1 | 0.6 | NA | 0.5 | 0.2 | 1 | NA | 0.111 | 1 | 1 | 0.12 | 0.017 |
| RI | 6408 | ENSG00000121716 | PILRB | chr7 | + | 100352563 | 100353123 | 100352563 | 100352667 | 100352996 | 100353123 | 108 | 45 | 108 | 79 | 62 | 0 | 0 | 0 | 0 | 0 | 60 | 86 | 32 | 62 | 108 | 0 | 0 | 0 | 0 | 5 | 298 | 149 | 1 | 1 | 1 | 1 | 1 | 1 | 1 | 1 | 1 | 1 | 1 | 0.915 | 0.017 |
| RI | 2239 | ENSG00000197603 | CPLANE1 | chr5 | - | 37139339 | 37142480 | 37139339 | 37139370 | 37142309 | 37142480 | 2 | 8 | 3 | 4 | 7 | 36 | 23 | 16 | 21 | 25 | 0 | 2 | 5 | 0 | 0 | 0 | 14 | 13 | 2 | 0 | 298 | 149 | 1 | 1 | 0.027 | 0.148 | 0.086 | 0.087 | 0.123 | NA | 0.067 | 0.161 | 0 | NA | 0.018 |
| RI | 4744 | ENSG00000131778 | CHD1L | chr1 | + | 147272170 | 147272630 | 147272170 | 147272281 | 147272446 | 147272630 | 19 | 4 | 8 | 9 | 25 | 0 | 0 | 0 | 0 | 0 | 32 | 25 | 25 | 31 | 0 | 0 | 1 | 0 | 0 | 0 | 298 | 149 | 1 | 1 | 1 | 1 | 1 | 1 | 1 | 1 | 0.926 | 1 | 1 | NA | 0.018 |
| RI | 4819 | ENSG00000149231 | CCDC82 | chr11 | - | 96383961 | 96386293 | 96383961 | 96384761 | 96386253 | 96386293 | 138 | 110 | 99 | 116 | 91 | 98 | 74 | 70 | 99 | 90 | 0 | 42 | 50 | 13 | 0 | 6 | 32 | 7 | 14 | 0 | 298 | 149 | 1 | 1 | 0.413 | 0.426 | 0.414 | 0.369 | 0.336 | 0 | 0.396 | 0.781 | 0.317 | NA | 0.018 |
| RI | 4824 | ENSG00000185163 | DDX51 | chr12 | - | 132140830 | 132141420 | 132140830 | 132141020 | 132141102 | 132141420 | 5 | 5 | 1 | 11 | 1 | 0 | 0 | 1 | 4 | 1 | 0 | 10 | 0 | 0 | 11 | 0 | 0 | 5 | 0 | 0 | 231 | 149 | 1 | 1 | 1 | 1 | 0.392 | 0.639 | 0.392 | NA | 1 | 0 | NA | 1 | 0.018 |
| RI | 5459 | ENSG00000141013 | GAS8 | chr16 | + | 90036380 | 90037399 | 90036380 | 90036586 | 90037231 | 90037399 | 0 | 3 | 3 | 4 | 0 | 4 | 1 | 4 | 5 | 0 | 0 | 6 | 6 | 7 | 14 | 15 | 20 | 4 | 5 | 11 | 298 | 149 | 1 | 1 | 0 | 0.6 | 0.273 | 0.286 | NA | 0 | 0.13 | 0.429 | 0.412 | 0.389 | 0.018 |
| RI | 5974 | ENSG00000204410 | MSH5 | chr6 | + | 31759096 | 31759512 | 31759096 | 31759177 | 31759424 | 31759512 | 10 | 7 | 6 | 2 | 0 | 6 | 4 | 3 | 5 | 8 | 0 | 9 | 1 | 8 | 5 | 0 | 7 | 7 | 3 | 12 | 298 | 149 | 1 | 1 | 0.455 | 0.467 | 0.5 | 0.167 | 0 | NA | 0.391 | 0.067 | 0.571 | 0.172 | 0.018 |
| RI | 208 | ENSG00000132024 | CC2D1A | chr19 | + | 13926977 | 13928030 | 13926977 | 13927077 | 13927892 | 13928030 | 4 | 14 | 8 | 16 | 10 | 0 | 0 | 0 | 0 | 0 | 7 | 11 | 9 | 11 | 38 | 0 | 0 | 0 | 0 | 2 | 298 | 149 | 1 | 1 | 1 | 1 | 1 | 1 | 1 | 1 | 1 | 1 | 1 | 0.905 | 0.019 |
| RI | 3423 | ENSG00000131697 | NPHP4 | chr1 | - | 5862810 | 5866458 | 5862810 | 5864033 | 5864337 | 5866458 | 2 | 0 | 2 | 0 | 0 | 7 | 2 | 4 | 3 | 3 | 0 | 3 | 0 | 0 | 0 | 5 | 5 | 3 | 7 | 29 | 298 | 149 | 1 | 1 | 0.125 | 0 | 0.2 | 0 | 0 | 0 | 0.231 | 0 | 0 | 0 | 0.019 |
| RI | 4101 | ENSG00000256269 | HMBS | chr11 | + | 119092124 | 119092811 | 119092124 | 119092163 | 119092757 | 119092811 | 1 | 3 | 5 | 2 | 4 | 2 | 0 | 2 | 0 | 1 | 15 | 3 | 8 | 3 | 0 | 0 | 22 | 0 | 1 | 0 | 298 | 149 | 1 | 1 | 0.2 | 1 | 0.556 | 1 | 0.667 | 1 | 0.064 | 1 | 0.6 | NA | 0.019 |
| RI | 5351 | ENSG00000213145 | CRIP1 | chr14 | + | 105486316 | 105488388 | 105486316 | 105487299 | 105488330 | 105488388 | 22 | 2 | 11 | 28 | 15 | 0 | 0 | 1 | 0 | 0 | 19 | 14 | 17 | 6 | 52 | 0 | 0 | 0 | 1 | 0 | 298 | 149 | 1 | 1 | 1 | 1 | 0.846 | 1 | 1 | 1 | 1 | 1 | 0.75 | 1 | 0.019 |
| RI | 6895 | ENSG00000102393 | GLA | chrX | - | 101401232 | 101401809 | 101401232 | 101401347 | 101401631 | 101401809 | 34 | 50 | 40 | 65 | 87 | 1 | 0 | 0 | 1 | 2 | 108 | 128 | 92 | 226 | 237 | 0 | 4 | 5 | 3 | 5 | 298 | 149 | 1 | 1 | 0.944 | 1 | 1 | 0.97 | 0.956 | 1 | 0.941 | 0.902 | 0.974 | 0.96 | 0.019 |
| RI | 7288 | ENSG00000120217 | CD274 | chr9 | + | 5462833 | 5463121 | 5462833 | 5462968 | 5463074 | 5463121 | 19 | 31 | 47 | 36 | 53 | 0 | 0 | 0 | 0 | 3 | 34 | 28 | 23 | 17 | 19 | 0 | 0 | 3 | 0 | 0 | 255 | 149 | 1 | 1 | 1 | 1 | 1 | 1 | 0.912 | 1 | 1 | 0.818 | 1 | 1 | 0.019 |
| RI | 298 | ENSG00000063177 | RPL18 | chr19 | - | 48616078 | 48617423 | 48616078 | 48616202 | 48617315 | 48617423 | 17 | 18 | 42 | 31 | 15 | 0 | 0 | 0 | 0 | 0 | 37 | 49 | 32 | 45 | 40 | 2 | 0 | 0 | 0 | 0 | 298 | 149 | 1 | 1 | 1 | 1 | 1 | 1 | 1 | 0.902 | 1 | 1 | 1 | 1 | 0.02 |
| RI | 1353 | ENSG00000007541 | PIGQ | chr16 | + | 580182 | 580972 | 580182 | 580263 | 580857 | 580972 | 0 | 0 | 3 | 0 | 1 | 7 | 0 | 9 | 11 | 9 | 0 | 0 | 0 | 1 | 0 | 11 | 7 | 4 | 3 | 3 | 298 | 149 | 1 | 1 | 0 | NA | 0.143 | 0 | 0.053 | 0 | 0 | 0 | 0.143 | 0 | 0.02 |
| RI | 2267 | ENSG00000124357 | NAGK | chr2 | + | 71072640 | 71073594 | 71072640 | 71072751 | 71072825 | 71073594 | 36 | 29 | 20 | 37 | 12 | 0 | 0 | 0 | 0 | 0 | 108 | 29 | 27 | 34 | 76 | 0 | 0 | 2 | 0 | 0 | 223 | 149 | 1 | 1 | 1 | 1 | 1 | 1 | 1 | 1 | 1 | 0.9 | 1 | 1 | 0.02 |
| RI | 2954 | ENSG00000145029 | NICN1 | chr3 | - | 49424948 | 49425438 | 49424948 | 49425146 | 49425366 | 49425438 | 23 | 16 | 6 | 43 | 11 | 0 | 0 | 0 | 0 | 0 | 22 | 18 | 21 | 17 | 49 | 0 | 1 | 0 | 0 | 0 | 298 | 149 | 1 | 1 | 1 | 1 | 1 | 1 | 1 | 1 | 0.9 | 1 | 1 | 1 | 0.02 |
| RI | 4252 | ENSG00000011009 | LYPLA2 | chr1 | + | 23794223 | 23794600 | 23794223 | 23794325 | 23794551 | 23794600 | 54 | 11 | 47 | 54 | 32 | 0 | 0 | 0 | 0 | 0 | 35 | 66 | 36 | 35 | 82 | 0 | 0 | 2 | 0 | 0 | 298 | 149 | 1 | 1 | 1 | 1 | 1 | 1 | 1 | 1 | 1 | 0.9 | 1 | 1 | 0.02 |
| RI | 8155 | ENSG00000107862 | GBF1 | chr10 | + | 102379852 | 102380362 | 102379852 | 102379954 | 102380332 | 102380362 | 53 | 26 | 43 | 29 | 20 | 0 | 0 | 0 | 0 | 0 | 118 | 89 | 52 | 104 | 183 | 0 | 5 | 0 | 0 | 0 | 298 | 149 | 1 | 1 | 1 | 1 | 1 | 1 | 1 | 1 | 0.899 | 1 | 1 | 1 | 0.02 |
| RI | 3094 | ENSG00000111642 | CHD4 | chr12 | - | 6583026 | 6583378 | 6583026 | 6583113 | 6583195 | 6583378 | 25 | 14 | 20 | 14 | 23 | 1 | 0 | 1 | 0 | 0 | 48 | 48 | 38 | 70 | 62 | 0 | 0 | 0 | 14 | 0 | 231 | 149 | 1 | 1 | 0.942 | 1 | 0.928 | 1 | 1 | 1 | 1 | 1 | 0.763 | 1 | 0.021 |
| RI | 5226 | ENSG00000125875 | TBC1D20 | chr20 | - | 439107 | 442043 | 439107 | 439295 | 441587 | 442043 | 17 | 11 | 20 | 24 | 10 | 0 | 0 | 0 | 0 | 0 | 103 | 51 | 25 | 50 | 51 | 0 | 3 | 0 | 0 | 0 | 298 | 149 | 1 | 1 | 1 | 1 | 1 | 1 | 1 | 1 | 0.895 | 1 | 1 | 1 | 0.021 |
| RI | 6447 | ENSG00000115677 | HDLBP | chr2 | - | 241268953 | 241269295 | 241268953 | 241269105 | 241269205 | 241269295 | 16 | 6 | 12 | 14 | 3 | 0 | 0 | 0 | 0 | 0 | 42 | 14 | 42 | 28 | 27 | 0 | 0 | 0 | 2 | 0 | 249 | 149 | 1 | 1 | 1 | 1 | 1 | 1 | 1 | 1 | 1 | 1 | 0.893 | 1 | 0.021 |
| RI | 773 | ENSG00000073350 | LLGL2 | chr17 | + | 75574212 | 75574668 | 75574212 | 75574260 | 75574452 | 75574668 | 2 | 0 | 0 | 3 | 0 | 30 | 13 | 11 | 18 | 23 | 0 | 0 | 0 | 0 | 0 | 4 | 27 | 7 | 15 | 20 | 298 | 149 | 1 | 1 | 0.032 | 0 | 0 | 0.077 | 0 | 0 | 0 | 0 | 0 | 0 | 0.022 |
| RI | 2373 | ENSG00000106624 | AEBP1 | chr7 | + | 44111506 | 44112050 | 44111506 | 44111630 | 44111853 | 44112050 | 0 | 0 | 1 | 1 | 0 | 8 | 4 | 9 | 8 | 3 | 0 | 0 | 0 | 0 | 0 | 0 | 12 | 2 | 0 | 21 | 298 | 149 | 1 | 1 | 0 | 0 | 0.053 | 0.059 | 0 | NA | 0 | 0 | NA | 0 | 0.022 |
| RI | 3129 | ENSG00000165272 | AQP3 | chr9 | - | 33442851 | 33443458 | 33442851 | 33442970 | 33443320 | 33443458 | 6 | 3 | 9 | 7 | 8 | 50 | 18 | 70 | 25 | 33 | 0 | 1 | 27 | 0 | 0 | 0 | 37 | 43 | 3 | 9 | 298 | 149 | 1 | 1 | 0.057 | 0.077 | 0.06 | 0.123 | 0.108 | NA | 0.013 | 0.239 | 0 | 0 | 0.022 |
| RI | 7104 | ENSG00000094914 | AAAS | chr12 | - | 53315334 | 53315782 | 53315334 | 53315486 | 53315726 | 53315782 | 4 | 12 | 12 | 20 | 7 | 0 | 0 | 0 | 0 | 1 | 23 | 15 | 13 | 4 | 34 | 0 | 0 | 0 | 1 | 0 | 298 | 149 | 1 | 1 | 1 | 1 | 1 | 1 | 0.778 | 1 | 1 | 1 | 0.667 | 1 | 0.022 |
| RI | 8533 | ENSG00000234772 | LINC00412 | chr13 | - | 27236281 | 27236587 | 27236281 | 27236356 | 27236444 | 27236587 | 3 | 8 | 18 | 4 | 3 | 0 | 3 | 0 | 0 | 0 | 3 | 8 | 5 | 5 | 0 | 0 | 0 | 0 | 2 | 0 | 237 | 149 | 1 | 1 | 1 | 0.626 | 1 | 1 | 1 | 1 | 1 | 1 | 0.611 | NA | 0.022 |
| RI | 818 | ENSG00000184009 | ACTG1 | chr17 | - | 81511187 | 81512142 | 81511187 | 81511803 | 81511902 | 81512142 | 10 | 2 | 7 | 14 | 1 | 0 | 0 | 0 | 0 | 0 | 9 | 8 | 15 | 13 | 15 | 0 | 0 | 0 | 1 | 0 | 248 | 149 | 1 | 1 | 1 | 1 | 1 | 1 | 1 | 1 | 1 | 1 | 0.886 | 1 | 0.023 |
| RI | 2898 | ENSG00000172053 | QARS1 | chr3 | - | 49098191 | 49098480 | 49098191 | 49098258 | 49098372 | 49098480 | 17 | 15 | 32 | 29 | 9 | 0 | 0 | 0 | 0 | 0 | 8 | 27 | 21 | 11 | 33 | 0 | 2 | 0 | 0 | 0 | 263 | 149 | 1 | 1 | 1 | 1 | 1 | 1 | 1 | 1 | 0.884 | 1 | 1 | 1 | 0.023 |
| RI | 6692 | ENSG00000174080 | CTSF | chr11 | - | 66565670 | 66565927 | 66565670 | 66565751 | 66565830 | 66565927 | 0 | 1 | 0 | 0 | 0 | 32 | 5 | 37 | 15 | 14 | 0 | 0 | 0 | 0 | 0 | 0 | 16 | 15 | 10 | 7 | 228 | 149 | 1 | 1 | 0 | 0.116 | 0 | 0 | 0 | NA | 0 | 0 | 0 | 0 | 0.023 |

| | | | | | | | | | | | | | | | | | | | | | | | | | | | | | | | | | | | | | | | | | | | | | | |
|---|---|---|---|---|---|---|---|---|---|---|---|---|---|---|---|---|---|---|---|---|---|---|---|---|---|---|---|---|---|---|---|---|---|---|---|---|---|---|---|---|---|---|---|---|---|---|
| RI | 7141 | ENSG00000172269 | DPAGT1 | chr11 | - | 119097141 | 119097551 | 119097141 | 119097297 | 119097444 | 119097551 | 20 | 12 | 4 | 15 | 8 | 0 | 0 | 3 | 1 | 0 | 0 | 0 | 10 | 2 | 20 | 0 | 0 | 0 | 1 | 0 | 296 | 149 | 1 | 1 | 1 | 1 | 0.402 | 0.883 | 1 | NA | NA | 1 | 0.502 | 1 | 0.023 |
| RI | 534 | ENSG00000064601 | CTSA | chr20 | + | 45892724 | 45894741 | 45892724 | 45892880 | 45894649 | 45894741 | 17 | 5 | 13 | 20 | 12 | 0 | 0 | 0 | 0 | 0 | 50 | 22 | 44 | 30 | 57 | 0 | 0 | 0 | 2 | 0 | 298 | 149 | 1 | 1 | 1 | 1 | 1 | 1 | 1 | 1 | 1 | 1 | 0.882 | 1 | 0.024 |
| RI | 1480 | ENSG00000167965 | MLST8 | chr16 | + | 2206357 | 2206659 | 2206357 | 2206409 | 2206483 | 2206659 | 5 | 7 | 2 | 7 | 8 | 0 | 0 | 0 | 0 | 0 | 16 | 3 | 3 | 6 | 33 | 0 | 0 | 0 | 0 | 3 | 223 | 149 | 1 | 1 | 1 | 1 | 1 | 1 | 1 | 1 | 1 | 1 | 1 | 0.88 | 0.024 |
| RI | 6456 | ENSG00000167703 | SLC43A2 | chr17 | - | 1616561 | 1616769 | 1616561 | 1616630 | 1616716 | 1616769 | 73 | 40 | 41 | 33 | 15 | 0 | 0 | 0 | 0 | 0 | 25 | 48 | 51 | 47 | 106 | 0 | 3 | 1 | 0 | 0 | 235 | 149 | 1 | 1 | 1 | 1 | 1 | 1 | 1 | 1 | 0.91 | 0.97 | 1 | 1 | 0.024 |
| RI | 6477 | ENSG00000184216 | IRAK1 | chrX | - | 154019430 | 154019812 | 154019430 | 154019598 | 154019676 | 154019812 | 2 | 0 | 0 | 2 | 0 | 26 | 7 | 18 | 17 | 13 | 0 | 0 | 0 | 0 | 0 | 12 | 31 | 14 | 18 | 32 | 227 | 149 | 1 | 1 | 0.048 | 0 | 0 | 0.072 | 0 | 0 | 0 | 0 | 0 | 0 | 0.024 |
| RI | 6935 | ENSG00000046651 | OFD1 | chrX | + | 13757659 | 13760720 | 13757659 | 13757790 | 13758336 | 13760720 | 16 | 23 | 20 | 26 | 32 | 32 | 39 | 32 | 42 | 42 | 0 | 21 | 32 | 8 | 0 | 10 | 31 | 22 | 19 | 0 | 298 | 149 | 1 | 1 | 0.2 | 0.228 | 0.238 | 0.236 | 0.276 | 0 | 0.253 | 0.421 | 0.174 | NA | 0.024 |
| RI | 7175 | ENSG00000100023 | PPIL2 | chr22 | + | 21670565 | 21671059 | 21670565 | 21670853 | 21670996 | 21671059 | 23 | 13 | 21 | 23 | 10 | 0 | 0 | 0 | 0 | 0 | 89 | 29 | 45 | 29 | 68 | 0 | 0 | 0 | 2 | 0 | 292 | 149 | 1 | 1 | 1 | 1 | 1 | 1 | 1 | 1 | 1 | 1 | 0.881 | 1 | 0.024 |
| RI | 1876 | ENSG00000182809 | CRIP2 | chr14 | + | 105478265 | 105478507 | 105478265 | 105478360 | 105478449 | 105478507 | 0 | 0 | 3 | 0 | 0 | 15 | 5 | 13 | 14 | 3 | 0 | 0 | 0 | 0 | 0 | 2 | 6 | 0 | 0 | 6 | 238 | 149 | 1 | 1 | 0 | 0 | 0.126 | 0 | 0 | 0 | 0 | NA | NA | 0 | 0.025 |
| RI | 5263 | ENSG00000224078 | SNHG14 | chr15 | + | 25108997 | 25109403 | 25108997 | 25109103 | 25109197 | 25109403 | 16 | 3 | 2 | 10 | 11 | 37 | 8 | 25 | 20 | 60 | 0 | 5 | 8 | 2 | 0 | 0 | 31 | 16 | 17 | 0 | 243 | 149 | 1 | 1 | 0.21 | 0.187 | 0.047 | 0.235 | 0.101 | NA | 0.09 | 0.235 | 0.067 | NA | 0.025 |
| RI | 7345 | ENSG00000203780 | FANK1 | chr10 | + | 126009221 | 126009592 | 126009221 | 126009266 | 126009372 | 126009592 | 2 | 3 | 2 | 6 | 2 | 1 | 0 | 0 | 2 | 3 | 0 | 1 | 1 | 0 | 0 | 0 | 0 | 0 | 4 | 0 | 255 | 149 | 1 | 1 | 0.539 | 1 | 1 | 0.637 | 0.28 | NA | 1 | 1 | 0 | NA | 0.025 |
| RI | 7433 | ENSG00000070061 | ELP1 | chr9 | - | 108917546 | 108919349 | 108917546 | 108917670 | 108919252 | 108919349 | 64 | 62 | 65 | 87 | 52 | 0 | 0 | 0 | 0 | 0 | 39 | 32 | 56 | 32 | 12 | 0 | 0 | 4 | 0 | 0 | 298 | 149 | 1 | 1 | 1 | 1 | 1 | 1 | 1 | 1 | 1 | 0.875 | 1 | 1 | 0.025 |
| RI | 291 | ENSG00000142546 | NOSIP | chr19 | - | 49557089 | 49558978 | 49557089 | 49558199 | 49558896 | 49558978 | 27 | 16 | 63 | 25 | 30 | 15 | 6 | 21 | 8 | 13 | 0 | 13 | 10 | 12 | 5 | 0 | 8 | 2 | 1 | 20 | 298 | 149 | 1 | 1 | 0.474 | 0.571 | 0.6 | 0.61 | 0.536 | NA | 0.448 | 0.714 | 0.857 | 0.111 | 0.026 |
| RI | 1463 | ENSG00000140983 | RHOT2 | chr16 | + | 668352 | 668699 | 668352 | 668411 | 668655 | 668699 | 9 | 3 | 12 | 17 | 6 | 1 | 0 | 0 | 0 | 0 | 0 | 24 | 12 | 17 | 45 | 0 | 4 | 0 | 0 | 0 | 298 | 149 | 1 | 1 | 0.818 | 1 | 1 | 1 | 1 | NA | 0.75 | 1 | 1 | 1 | 0.026 |
| RI | 3210 | ENSG00000168615 | ADAM9 | chr8 | + | 39018852 | 39021714 | 39018852 | 39018881 | 39021642 | 39021714 | 15 | 15 | 18 | 28 | 16 | 0 | 0 | 0 | 0 | 0 | 8 | 27 | 74 | 44 | 7 | 0 | 2 | 0 | 0 | 0 | 298 | 149 | 1 | 1 | 1 | 1 | 1 | 1 | 1 | 1 | 0.871 | 1 | 1 | 1 | 0.026 |
| RI | 3598 | ENSG00000162772 | ATF3 | chr1 | + | 212615021 | 212615261 | 212615021 | 212615060 | 212615147 | 212615261 | 11 | 14 | 10 | 22 | 12 | 0 | 0 | 0 | 0 | 0 | 93 | 29 | 21 | 103 | 36 | 0 | 0 | 2 | 0 | 0 | 236 | 149 | 1 | 1 | 1 | 1 | 1 | 1 | 1 | 1 | 1 | 0.869 | 1 | 1 | 0.026 |
| RI | 7590 | ENSG00000122952 | ZWINT | chr10 | - | 56357437 | 56358459 | 56357437 | 56358185 | 56358376 | 56358459 | 1 | 0 | 4 | 5 | 3 | 13 | 6 | 20 | 5 | 9 | 0 | 8 | 1 | 10 | 0 | 9 | 13 | 22 | 36 | 0 | 298 | 149 | 1 | 1 | 0.037 | 0 | 0.091 | 0.333 | 0.143 | 0 | 0.235 | 0.022 | 0.122 | NA | 0.026 |
| RI | 3179 | ENSG00000182481 | KPNA2 | chr17 | + | 68043837 | 68044503 | 68043837 | 68044071 | 68044324 | 68044503 | 31 | 26 | 27 | 21 | 18 | 0 | 0 | 0 | 0 | 0 | 66 | 70 | 13 | 94 | 32 | 0 | 0 | 1 | 0 | 0 | 298 | 149 | 1 | 1 | 1 | 1 | 1 | 1 | 1 | 1 | 1 | 0.867 | 1 | 1 | 0.027 |
| RI | 6471 | ENSG00000169583 | CLIC3 | chr9 | - | 136995441 | 136995757 | 136995441 | 136995567 | 136995647 | 136995757 | 14 | 0 | 3 | 2 | 0 | 75 | 13 | 17 | 34 | 12 | 0 | 0 | 2 | 0 | 0 | 12 | 41 | 13 | 4 | 0 | 229 | 149 | 1 | 1 | 0.108 | 0 | 0.103 | 0.037 | 0 | 0 | 0 | 0.091 | 0 | NA | 0.027 |
| RI | 7909 | ENSG00000149292 | TTC12 | chr11 | + | 113315950 | 113316315 | 113315950 | 113316053 | 113316242 | 113316315 | 59 | 28 | 54 | 54 | 24 | 9 | 1 | 2 | 3 | 0 | 0 | 20 | 13 | 8 | 7 | 0 | 2 | 3 | 0 | 0 | 298 | 149 | 1 | 1 | 0.766 | 0.933 | 0.931 | 0.9 | 1 | NA | 0.833 | 0.684 | 1 | 1 | 0.027 |
| RI | 733 | ENSG00000270379 | HEATR9 | chr17 | - | 35863501 | 35864302 | 35863501 | 35863559 | 35864245 | 35864302 | 8 | 0 | 12 | 4 | 7 | 13 | 1 | 1 | 3 | 8 | 0 | 15 | 9 | 11 | 11 | 0 | 9 | 12 | 9 | 20 | 298 | 149 | 1 | 1 | 0.235 | 0 | 0.857 | 0.4 | 0.304 | NA | 0.455 | 0.273 | 0.379 | 0.216 | 0.028 |
| [illegible] | | | | | | | | | | | | | | | | | | | | | | | | | | | | | | | | | | | | | | | | | | | | |

| | | | | | | | | | | | | | |
|---|---|---|---|---|---|---|---|---|---|---|---|---|---|
| RI | 5466 | ENSG00000237765 | FAM200B | chr4 | + | 15686235 | 15690447 | 15686235 | 15687148 | 15688550 | 15690447 | 47 | 33 |
| RI | 8384 | ENSG00000133641 | C12orf29 | chr12 | + | 88043598 | 88045779 | 88043598 | 88043715 | 88045581 | 88045779 | 16 | 14 |
| RI | 8468 | ENSG00000122966 | CIT | chr12 | - | 119775785 | 119776408 | 119775785 | 119775839 | 119776357 | 119776408 | 2 | 0 |
| RI | 35 | ENSG00000237484 | LINC01684 | chr21 | + | 24488112 | 24488912 | 24488112 | 24488309 | 24488479 | 24488912 | 8 | 11 |
| RI | 7639 | ENSG00000113163 | CERT1 | chr5 | - | 75400204 | 75403058 | 75400204 | 75400291 | 75402971 | 75403058 | 17 | 16 |
| RI | 3173 | ENSG00000135722 | FBXL8 | chr16 | + | 67162847 | 67164174 | 67162847 | 67163852 | 67164018 | 67164174 | 0 | 0 |
| RI | 4216 | ENSG00000147180 | ZNF711 | chrX | + | 85246929 | 85247651 | 85246929 | 85247183 | 85247546 | 85247651 | 2 | 1 |
| RI | 4861 | ENSG00000171848 | RRM2 | chr2 | + | 10122738 | 10123057 | 10122738 | 10122897 | 10122982 | 10123057 | 0 | 0 |
| RI | 7478 | ENSG00000172270 | BSG | chr19 | + | 580378 | 580782 | 580378 | 580461 | 580655 | 580782 | 12 | 9 |
| RI | 8080 | ENSG00000214279 | SCART1 | chr10 | + | 133457278 | 133459326 | 133457278 | 133458656 | 133459020 | 133459326 | 23 | 7 |
| RI | 906 | ENSG00000076382 | SPAG5 | chr17 | - | 28578009 | 28578292 | 28578009 | 28578090 | 28578217 | 28578292 | 2 | 1 |
| RI | 2512 | ENSG00000109654 | TRIM2 | chr4 | + | 153322647 | 153324148 | 153322647 | 153322816 | 153324077 | 153324148 | 8 | 1 |
| RI | 2518 | ENSG00000186431 | FCAR | chr19 | + | 54888006 | 54889067 | 54888006 | 54888642 | 54888975 | 54889067 | 1 | 3 |
| RI | 3201 | ENSG00000010327 | STAB1 | chr3 | + | 52521595 | 52521951 | 52521595 | 52521700 | 52521843 | 52521951 | 9 | 1 |
| RI | 7994 | ENSG00000150433 | TMEM218 | chr11 | - | 125102131 | 125102809 | 125102131 | 125102317 | 125102636 | 125102809 | 12 | 20 |
| RI | 8231 | ENSG00000110888 | CAPRIN2 | chr12 | - | 30714958 | 30716676 | 30714958 | 30715141 | 30716507 | 30716676 | 38 | 32 |
| RI | 2349 | ENSG00000112992 | NNT | chr5 | + | 43647923 | 43649308 | 43647923 | 43648440 | 43649146 | 43649308 | 15 | 15 |
| RI | 7483 | ENSG00000166348 | USP54 | chr10 | - | 73575511 | 73576361 | 73575511 | 73575675 | 73576135 | 73576361 | 8 | 11 |
| RI | 7330 | ENSG00000197816 | CCDC180 | chr9 | + | 97370640 | 97374648 | 97370640 | 97370778 | 97374542 | 97374648 | 4 | 2 |
| RI | 1228 | ENSG00000103034 | NDRG4 | chr16 | + | 58508961 | 58509189 | 58508961 | 58509009 | 58509153 | 58509189 | 0 | 2 |
| RI | 939 | ENSG00000186765 | FSCN2 | chr17 | + | 81536145 | 81536789 | 81536145 | 81536267 | 81536549 | 81536789 | 5 | 8 |
| RI | 446 | ENSG00000126249 | PDCD2L | chr19 | + | 34404648 | 34404990 | 34404648 | 34404815 | 34404929 | 34404990 | 0 | 1 |
| RI | 5541 | ENSG00000160766 | GBAP1 | chr1 | - | 155215562 | 155215787 | 155215562 | 155215651 | 155215761 | 155215787 | 13 | 3 |
| RI | 2127 | ENSG00000177595 | PIDD1 | chr11 | - | 802681 | 803647 | 802681 | 802891 | 803173 | 803647 | 15 | 1 |
| RI | 4339 | ENSG00000198019 | FCGR1B | chr1 | + | 121087820 | 121088908 | 121087820 | 121087841 | 121088653 | 121088908 | 6 | 4 |
| RI | 1134 | ENSG00000162078 | ZG16B | chr16 | + | 2830414 | 2830796 | 2830414 | 2830493 | 2830693 | 2830796 | 1 | 0 |
| RI | 5408 | ENSG00000049768 | FOXP3 | chrX | - | 49256750 | 49257012 | 49256750 | 49256855 | 49256924 | 49257012 | 1 | 1 |
| RI | 7206 | ENSG00000182325 | FBXL6 | chr8 | - | 144356299 | 144356713 | 144356299 | 144356531 | 144356599 | 144356713 | 3 | 0 |
| RI | 1385 | ENSG00000085644 | ZNF213 | chr16 | + | 3137165 | 3142804 | 3137165 | 3137679 | 3140688 | 3142804 | 1 | 2 |

| | | | | | | | | | | | | | | | | | | | | | | | | | | | | | | | | | | | | | | | | | | | | | | |
|---|---|---|---|---|---|---|---|---|---|---|---|---|---|---|---|---|---|---|---|---|---|---|---|---|---|---|---|---|---|---|---|---|---|---|---|---|---|---|---|---|---|---|---|---|---|---|
| RI | 1531 | ENSG00000166147 | FBN1 | chr15 | - | 48427566 | 48428471 | 48427566 | 48427773 | 48428345 | 48428471 | 0 | 0 | 0 | 2 | 0 | 0 | 0 | 1 | 0 | 0 | 0 | 0 | 0 | 0 | 0 | 0 | 0 | 2 | 0 | 0 | 298 | 149 | 1 | 1 | NA | NA | 0 | 1 | NA | NA | NA | 0 | NA | NA | 0.5 |
| RI | 1542 | ENSG00000103723 | AP3B2 | chr15 | - | 82662167 | 82662922 | 82662167 | 82662252 | 82662693 | 82662922 | 1 | 0 | 1 | 0 | 0 | 0 | 0 | 0 | 0 | 0 | 0 | 0 | 4 | 0 | 0 | 0 | 0 | 2 | 0 | 0 | 298 | 149 | 1 | 1 | 1 | NA | 1 | NA | NA | NA | NA | 0.5 | NA | NA | 0.5 |
| RI | 1770 | ENSG00000100749 | VRK1 | chr14 | + | 96945048 | 96945403 | 96945048 | 96945150 | 96945237 | 96945403 | 4 | 1 | 3 | 1 | 1 | 0 | 0 | 0 | 0 | 0 | 5 | 0 | 0 | 0 | 0 | 0 | 1 | 0 | 0 | 0 | 236 | 149 | 1 | 1 | 1 | 1 | 1 | 1 | 1 | 1 | 0 | NA | NA | NA | 0.5 |
| RI | 2427 | ENSG00000136960 | ENPP2 | chr8 | - | 119570704 | 119572250 | 119570704 | 119570841 | 119572175 | 119572250 | 2 | 0 | 0 | 0 | 0 | 1 | 0 | 0 | 0 | 0 | 0 | 0 | 0 | 0 | 0 | 0 | 5 | 0 | 0 | 0 | 298 | 149 | 1 | 1 | 0.5 | NA | NA | NA | NA | NA | 0 | NA | NA | NA | 0.5 |
| RI | 4061 | ENSG00000182983 | ZNF662 | chr3 | + | 42908021 | 42908909 | 42908021 | 42908148 | 42908795 | 42908909 | 0 | 0 | 3 | 5 | 0 | 0 | 1 | 0 | 0 | 1 | 0 | 0 | 0 | 0 | 0 | 0 | 0 | 0 | 1 | 0 | 298 | 149 | 1 | 1 | NA | 0 | 1 | 1 | 0 | NA | NA | NA | 0 | NA | 0.5 |
| RI | 4064 | ENSG00000143416 | SELENBP1 | chr1 | - | 151365767 | 151366453 | 151365767 | 151365846 | 151366274 | 151366453 | 1 | 0 | 0 | 0 | 0 | 0 | 0 | 0 | 0 | 2 | 0 | 0 | 0 | 0 | 0 | 0 | 4 | 0 | 6 | 0 | 298 | 149 | 1 | 1 | 1 | NA | NA | NA | 0 | NA | 0 | NA | 0 | NA | 0.5 |
| RI | 4298 | ENSG00000104327 | CALB1 | chr8 | - | 90060628 | 90063153 | 90060628 | 90060700 | 90063099 | 90063153 | 5 | 11 | 10 | 2 | 2 | 0 | 0 | 0 | 0 | 0 | 0 | 0 | 5 | 0 | 0 | 0 | 2 | 0 | 0 | 0 | 298 | 149 | 1 | 1 | 1 | 1 | 1 | 1 | 1 | NA | 0 | 1 | NA | NA | 0.5 |
| RI | 5707 | ENSG00000229727 | NA | chr2 | + | 7428542 | 7429872 | 7428542 | 7429273 | 7429695 | 7429872 | 0 | 2 | 0 | 3 | 0 | 0 | 0 | 0 | 0 | 0 | 0 | 0 | 1 | 0 | 0 | 0 | 3 | 0 | 0 | 0 | 298 | 149 | 1 | 1 | NA | 1 | NA | 1 | NA | NA | 0 | 1 | NA | NA | 0.5 |
| RI | 6532 | ENSG00000164877 | MICALL2 | chr7 | - | 1438096 | 1438353 | 1438096 | 1438220 | 1438288 | 1438353 | 0 | 0 | 0 | 24 | 0 | 1 | 0 | 0 | 0 | 0 | 0 | 0 | 0 | 0 | 0 | 0 | 0 | 2 | 0 | 0 | 217 | 149 | 1 | 1 | 0 | NA | NA | 1 | NA | NA | NA | 0 | NA | NA | 0.5 |
| RI | 7219 | ENSG00000185803 | SLC52A2 | chr8 | + | 144358594 | 144359423 | 144358594 | 144359065 | 144359187 | 144359423 | 1 | 0 | 0 | 0 | 6 | 0 | 0 | 0 | 0 | 0 | 0 | 2 | 0 | 0 | 0 | 0 | 0 | 6 | 0 | 0 | 271 | 149 | 1 | 1 | 1 | NA | NA | NA | 1 | NA | 1 | 0 | NA | NA | 0.5 |
| RI | 7695 | ENSG00000100867 | DHRS2 | chr14 | + | 23639793 | 23643219 | 23639793 | 23639895 | 23643151 | 23643219 | 0 | 0 | 0 | 0 | 1 | 0 | 0 | 0 | 0 | 0 | 0 | 1 | 1 | 0 | 0 | 1 | 0 | 0 | 1 | 0 | 298 | 149 | 1 | 1 | NA | NA | NA | NA | 1 | 0 | 1 | 1 | 0 | NA | 0.5 |
| RI | 8202 | ENSG00000165731 | RET | chr10 | + | 43113555 | 43114736 | 43113555 | 43113675 | 43114479 | 43114736 | 0 | 0 | 1 | 0 | 0 | 1 | 0 | 0 | 0 | 0 | 0 | 0 | 0 | 0 | 0 | 0 | 0 | 2 | 0 | 0 | 298 | 149 | 1 | 1 | 0 | NA | 1 | NA | NA | NA | NA | 0 | NA | NA | 0.5 |
| RI | 8507 | ENSG00000174233 | ADCY6 | chr12 | - | 48773468 | 48774098 | 48773468 | 48773647 | 48773939 | 48774098 | 0 | 0 | 2 | 0 | 0 | 0 | 0 | 0 | 0 | 0 | 0 | 1 | 0 | 0 | 0 | 0 | 0 | 0 | 0 | 4 | 298 | 149 | 1 | 1 | NA | NA | 1 | NA | NA | NA | 1 | NA | NA | 0 | 0.5 |
| RI | 7861 | ENSG00000182272 | B4GALNT4 | chr11 | + | 375638 | 375956 | 375638 | 375773 | 375846 | 375956 | 3 | 0 | 0 | 3 | 0 | 0 | 0 | 0 | 2 | 1 | 0 | 0 | 0 | 0 | 0 | 0 | 0 | 1 | 0 | 1 | 222 | 149 | 1 | 1 | 1 | NA | NA | 0.502 | 0 | NA | NA | 0 | NA | 0 | 0.501 |
| RI | 1693 | ENSG00000206149 | HERC2P9 | chr15 | + | 28636486 | 28637107 | 28636486 | 28636701 | 28636974 | 28637107 | 3 | 1 | 6 | 2 | 2 | 0 | 2 | 0 | 0 | 0 | 0 | 4 | 4 | 1 | 0 | 0 | 8 | 15 | 0 | 11 | 298 | 149 | 1 | 1 | 1 | 0.2 | 1 | 1 | 1 | NA | 0.2 | 0.118 | 1 | 0 | 0.511 |
| RI | 6547 | ENSG00000159784 | FAM131B | chr7 | - | 143359325 | 143359767 | 143359325 | 143359419 | 143359731 | 143359767 | 0 | 1 | 0 | 1 | 3 | 2 | 0 | 2 | 0 | 1 | 0 | 0 | 0 | 0 | 0 | 0 | 4 | 0 | 0 | 5 | 298 | 149 | 1 | 1 | 0 | 1 | 0 | 1 | 0.6 | NA | 0 | NA | NA | 0 | 0.52 |
| RI | 1565 | ENSG00000185880 | TRIM69 | chr15 | + | 44756367 | 44758854 | 44756367 | 44756463 | 44758620 | 44758854 | 9 | 4 | 2 | 13 | 10 | 0 | 0 | 0 | 0 | 4 | 0 | 9 | 2 | 11 | 0 | 9 | 10 | 1 | 2 | 0 | 298 | 149 | 1 | 1 | 1 | 1 | 1 | 1 | 0.556 | 0 | 0.31 | 0.5 | 0.733 | NA | 0.525 |
| RI | 5551 | ENSG00000249816 | LINC00964 | chr8 | + | 124849274 | 124849953 | 124849274 | 124849336 | 124849744 | 124849953 | 0 | 0 | 0 | 1 | 0 | 0 | 0 | 0 | 0 | 0 | 0 | 0 | 0 | 5 | 0 | 0 | 0 | 0 | 3 | 0 | 298 | 149 | 1 | 1 | NA | NA | NA | 1 | NA | NA | NA | NA | 0.455 | NA | 0.545 |
| RI | 6501 | ENSG00000136247 | ZDHHC4 | chr7 | + | 6577433 | 6577720 | 6577433 | 6577498 | 6577589 | 6577720 | 3 | 0 | 0 | 0 | 0 | 0 | 0 | 0 | 0 | 0 | 0 | 0 | 0 | 0 | 5 | 0 | 0 | 0 | 0 | 4 | 240 | 149 | 1 | 1 | 1 | NA | NA | NA | NA | NA | NA | NA | NA | 0.437 | 0.563 |
| RI | 1092 | ENSG00000091536 | MYO15A | chr17 | + | 18157155 | 18157882 | 18157155 | 18157230 | 18157721 | 18157882 | 0 | 0 | 3 | 0 | 0 | 0 | 0 | 0 | 0 | 0 | 0 | 0 | 0 | 3 | 0 | 0 | 0 | 0 | 2 | 0 | 298 | 149 | 1 | 1 | NA | NA | 1 | NA | NA | NA | NA | NA | 0.429 | NA | 0.571 |
| RI | 6469 | ENSG00000000971 | CFH | chr1 | + | 196740618 | 196742051 | 196740618 | 196740792 | 196741874 | 196742051 | 12 | 10 | 15 | 12 | 18 | 6 | 1 | 4 | 12 | 3 | 0 | 0 | 0 | 0 | 0 | 0 | 0 | 1 | 0 | 0 | 298 | 149 | 1 | 1 | 0.5 | 0.833 | 0.652 | 0.333 | 0.75 | NA | NA | 0 | NA | NA | 0.614 |
| RI | 5687 | ENSG00000135976 | ANKRD36 | chr2 | + | 97249143 | 97249645 | 97249143 | 97249214 | 97249551 | 97249645 | 0 | 2 | 1 | 1 | 0 | 1 | 1 | 0 | 0 | 0 | 0 | 0 | 0 | 0 | 0 | 0 | 0 | 0 | 1 | 0 | 298 | 149 | 1 | 1 | 0 | 0.5 | 1 | 1 | NA | NA | NA | NA | 0 | NA | 0.625 |
| RI | 6145 | ENSG00000170485 | NPAS2 | chr2 | + | 100982230 | 100988276 | 100982230 | 100982377 | 100988078 | 100988276 | 0 | 1 | 1 | 2 | 8 | 2 | 0 | 2 | 0 | 0 | 0 | 0 | 0 | 0 | 0 | 0 | 0 | 0 | 1 | 0 | 298 | 149 | 1 | 1 | 0 | 1 | 0.2 | 1 | 1 | NA | NA | NA | 0 | NA | 0.64 |
| RI | 6430 | ENSG00000198753 | PLXNB3 | chrX | + | 153777527 | 153778095 | 153777527 | 153777688 | 153777947 | 153778095 | 0 | 0 | 3 | 1 | 0 | 2 | 0 | 0 | 0 | 0 | 2 | 0 | 0 | 0 | 1 | 11 | 16 | 6 | 1 | 58 | 298 | 149 | 1 | 1 | 0 | NA | 1 | 1 | NA | 0.083 | 0 | 0 | 0 | 0.009 | 0.648 |
| RI | 3754 | ENSG00000237094 | NA | chr1 | - | 498398 | 498976 | 498398 | 498456 | 498683 | 498976 | 0 | 0 | 1 | 0 | 3 | 0 | 0 | 1 | 0 | 0 | 0 | 0 | 0 | 0 | 0 | 5 | 0 | 0 | 2 | 0 | 298 | 149 | 1 | 1 | NA | NA | 0.333 | NA | 1 | 0 | NA | NA | 0 | NA | 0.666 |
| RI | 8134 | ENSG00000008300 | CELSR3 | chr3 | - | 48639673 | 48641524 | 48639673 | 48640559 | 48641323 | 48641524 | 0 | 0 | 2 | 2 | 0 | 0 | 0 | 0 | 2 | 0 | 0 | 0 | 0 | 0 | 0 | 0 | 3 | 0 | 0 | 0 | 298 | 149 | 1 | 1 | NA | NA | 1 | 0.333 | NA | NA | 0 | NA | NA | NA | 0.666 |
| RI | 2082 | ENSG00000155980 | KIF5A | chr12 | + | 57575639 | 57576862 | 57575639 | 57575757 | 57576086 | 57576862 | 0 | 0 | 1 | 0 | 2 | 0 | 1 | 0 | 0 | 0 | 0 | 0 | 0 | 0 | 0 | 0 | 0 | 0 | 0 | 5 | 298 | 149 | 1 | 1 | NA | 0 | 1 | NA | 1 | NA | NA | NA | NA | 0 | 0.667 |
| RI | 4376 | ENSG00000167799 | NUDT8 | chr11 | - | 67627937 | 67628400 | 67627937 | 67628251 | 67628322 | 67628400 | 3 | 1 | 1 | 0 | 0 | 1 | 0 | 0 | 1 | 0 | 0 | 0 | 0 | 0 | 0 | 0 | 1 | 0 | 0 | 0 | 220 | 149 | 1 | 1 | 0.67 | 1 | 1 | 0 | NA | NA | 0 | NA | NA | NA | 0.667 |
| RI | 5007 | ENSG00000204677 | FAM153CP | chr5 | + | 178038837 | 178039411 | 178038837 | 178038917 | 178039386 | 178039411 | 0 | 8 | 0 | 2 | 0 | 1 | 0 | 0 | 0 | 0 | 0 | 0 | 0 | 0 | 0 | 0 | 0 | 1 | 0 | 0 | 298 | 149 | 1 | 1 | 0 | 1 | NA | 1 | NA | NA | NA | 0 | NA | NA | 0.667 |
| RI | 8347 | ENSG00000267272 | LINC01140 | chr1 | + | 87133613 | 87136669 | 87133613 | 87133718 | 87133964 | 87136669 | 1 | 2 | 0 | 0 | 0 | 0 | 0 | 1 | 0 | 0 | 0 | 0 | 0 | 0 | 0 | 0 | 0 | 2 | 0 | 0 | 298 | 149 | 1 | 1 | 1 | 1 | 0 | NA | NA | NA | NA | 0 | NA | NA | 0.667 |
| RI | 2026 | ENSG00000127990 | SGCE | chr7 | - | 94599696 | 94600857 | 94599696 | 94599723 | 94600645 | 94600857 | 1 | 3 | 4 | 8 | 8 | 3 | 0 | 0 | 0 | 16 | 0 | 0 | 0 | 0 | 0 | 0 | 14 | 8 | 0 | 0 | 298 | 149 | 1 | 1 | 0.143 | 1 | 1 | 1 | 0.2 | NA | 0 | 0 | NA | NA | 0.669 |
| RI | 2178 | ENSG00000079215 | SLC1A3 | chr5 | + | 36676891 | 36679860 | 36676891 | 36677184 | 36679626 | 36679860 | 0 | 1 | 1 | 1 | 0 | 0 | 1 | 0 | 0 | 0 | 0 | 0 | 6 | 0 | 0 | 0 | 2 | 7 | 12 | 0 | 298 | 149 | 1 | 1 | NA | 0.333 | 1 | 1 | NA | NA | 0 | 0.3 | 0 | NA | 0.678 |
| RI | 3131 | ENSG00000197415 | VEPH1 | chr3 | - | 157459621 | 157460355 | 157459621 | 157459926 | 157460180 | 157460355 | 2 | 1 | 2 | 0 | 0 | 0 | 0 | 0 | 4 | 0 | 0 | 0 | 0 | 1 | 0 | 0 | 0 | 0 | 9 | 0 | 298 | 149 | 1 | 1 | 1 | 1 | 1 | 0 | NA | NA | NA | NA | 0.053 | NA | 0.697 |
| RI | 3734 | ENSG00000077782 | FGFR1 | chr8 | - | 38427920 | 38428435 | 38427920 | 38428093 | 38428345 | 38428435 | 1 | 0 | 1 | 0 | 0 | 0 | 0 | 0 | 0 | 0 | 0 | 3 | 0 | 6 | 0 | 0 | 1 | 1 | 8 | 0 | 298 | 149 | 1 | 1 | 1 | NA | 1 | NA | NA | NA | 0.6 | 0 | 0.273 | NA | 0.709 |
| RI | 3712 | ENSG00000182389 | CACNB4 | chr2 | - | 151870530 | 151872493 | 151870530 | 151870611 | 151872416 | 151872493 | 2 | 1 | 5 | 11 | 0 | 0 | 1 | 0 | 0 | 0 | 0 | 0 | 1 | 0 | 0 | 0 | 0 | 2 | 0 | 17 | 298 | 149 | 1 | 1 | 1 | 0.333 | 1 | 1 | NA | NA | NA | 0.2 | NA | 0 | 0.733 |
| RI | 4323 | ENSG00000103253 | HAGHL | chr16 | + | 728315 | 728604 | 728315 | 728424 | 728503 | 728604 | 1 | 0 | 3 | 1 | 0 | 0 | 0 | 5 | 0 | 0 | 0 | 0 | 0 | 0 | 0 | 0 | 12 | 0 | 0 | 0 | 228 | 149 | 1 | 1 | 1 | NA | 0.282 | 1 | NA | NA | 0 | NA | NA | NA | 0.761 |
| RI | 7961 | ENSG00000133895 | MEN1 | chr11 | - | 64806231 | 64807219 | 64806231 | 64806368 | 64807178 | 64807219 | 6 | 4 | 2 | 5 | 2 | 0 | 0 | 0 | 0 | 0 | 5 | 0 | 0 | 0 | 0 | 9 | 0 | 0 | 0 | 0 | 298 | 149 | 1 | 1 | 1 | 1 | 1 | 1 | 1 | 0.217 | NA | NA | NA | NA | 0.783 |
| RI | 7252 | ENSG00000104368 | PLAT | chr8 | - | 42181936 | 42182890 | 42181936 | 42182022 | 42182718 | 42182890 | 0 | 2 | 0 | 0 | 0 | 0 | 0 | 0 | 0 | 0 | 0 | 0 | 2 | 0 | 0 | 0 | 0 | 4 | 0 | 0 | 298 | 149 | 1 | 1 | NA | 1 | NA | NA | NA | NA | NA | 0.2 | NA | NA | 0.8 |
| RI | 7283 | ENSG00000249898 | MCPH1-AS1 | chr8 | - | 6620134 | 6621635 | 6620134 | 6620224 | 6620991 | 6621635 | 6 | 5 | 2 | 1 | 1 | 0 | 0 | 0 | 0 | 0 | 0 | 0 | 1 | 0 | 0 | 0 | 0 | 2 | 0 | 0 | 298 | 149 | 1 | 1 | 1 | 1 | 1 | 1 | 1 | NA | NA | 0.2 | NA | NA | 0.8 |
| RI | 5202 | ENSG00000178038 | ALS2CL | chr3 | - | 46671487 | 46672033 | 46671487 | 46671768 | 46671883 | 46672033 | 2 | 2 | 1 | 2 | 0 | 0 | 1 | 0 | 0 | 0 | 0 | 0 | 0 | 0 | 1 | 0 | 0 | 0 | 0 | 19 | 264 | 149 | 1 | 1 | 1 | 0.53 | 1 | 1 | NA | NA | NA | NA | NA | 0.029 | 0.854 |
| RI | 87 | ENSG00000075275 | CELSR1 | chr22 | - | 46361173 | 46364251 | 46361173 | 46363247 | 46363995 | 46364251 | 0 | 0 | 3 | 1 | 6 | 0 | 0 | 1 | 0 | 0 | 0 | 0 | 0 | 0 | 0 | 0 | 0 | 5 | 0 | 0 | 298 | 149 | 1 | 1 | NA | NA | 0.6 | 1 | 1 | NA | NA | 0 | NA | NA | 0.867 |
| RI | 6255 | ENSG00000105707 | HPN | chr19 | + | 35065867 | 35066558 | 35065867 | 35066032 | 35066248 | 35066558 | 0 | 0 | 1 | 0 | 1 | 0 | 0 | 0 | 0 | 0 | 0 | 0 | 1 | 0 | 0 | 0 | 3 | 2 | 0 | 4 | 298 | 149 | 1 | 1 | NA | NA | 1 | NA | 1 | NA | 0 | 0.2 | NA | 0 | 0.933 |
| RI | 46 | ENSG00000206140 | TMEM191C | chr22 | + | 21468943 | 21469346 | 21468943 | 21469062 | 21469273 | 21469346 | 0 | 0 | 1 | 0 | 0 | 0 | 0 | 0 | 0 | 0 | 0 | 0 | 0 | 0 | 0 | 13 | 0 | 0 | 0 | 0 | 298 | 149 | 1 | 1 | NA | NA | 1 | NA | NA | 0 | NA | NA | NA | NA | 1 |
| RI | 294 | ENSG00000102575 | ACP5 | chr19 | - | 11577592 | 11578194 | 11577592 | 11577679 | 11578023 | 11578194 | 7 | 1 | 7 | 6 | 2 | 0 | 0 | 0 | 0 | 0 | 0 | 0 | 0 | 0 | 0 | 7 | 0 | 0 | 0 | 0 | 298 | 149 | 1 | 1 | 1 | 1 | 1 | 1 | 1 | 0 | NA | NA | NA | NA | 1 |
| RI | 469 | ENSG00000160117 | ANKLE1 | chr19 | + | 17282655 | 17283002 | 17282655 | 17282761 | 17282863 | 17283002 | 0 | 2 | 4 | 1 | 0 | 0 | 0 | 0 | 0 | 0 | 0 | 0 | 0 | 0 | 0 | 0 | 2 | 0 | 0 | 0 | 251 | 149 | 1 | 1 | NA | 1 | 1 | 1 | NA | NA | 0 | NA | NA | NA | 1 |
| RI | 596 | ENSG00000088340 | FER1L4 | chr20 | - | 35559340 | 35560638 | 35559340 | 35559627 | 35560122 | 35560638 | 1 | 0 | 0 | 0 | 0 | 0 | 0 | 0 | 0 | 0 | 0 | 0 | 0 | 0 | 0 | 0 | 0 | 1 | 0 | 0 | 298 | 149 | 1 | 1 | 1 | NA | NA | NA | NA | NA | NA | 0 | NA | NA | 1 |
| RI | 2452 | ENSG00000145362 | ANK2 | chr4 | + | 113369513 | 113373173 | 113369513 | 113369805 | 113373089 | 113373173 | 0 | 0 | 0 | 1 | 0 | 0 | 0 | 0 | 0 | 0 | 0 | 0 | 0 | 0 | 0 | 0 | 0 | 2 | 0 | 0 | 298 | 149 | 1 | 1 | NA | NA | NA | 1 | NA | NA | NA | 0 | NA | NA | 1 |
| RI | 2729 | ENSG00000189337 | KAZN | chr1 | + | 15060171 | 15063622 | 15060171 | 15060302 | 15063571 | 15063622 | 0 | 0 | 0 | 1 | 0 | 0 | 0 | 0 | 0 | 0 | 0 | 0 | 0 | 0 | 0 | 0 | 0 | 0 | 0 | 4 | 298 | 149 | 1 | 1 | NA | NA | NA | 1 | NA | NA | NA | NA | NA | 0 | 1 |
| RI | 3708 | ENSG00000137225 | CAPN11 | chr6 | + | 44179618 | 44180163 | 44179618 | 44179630 | 44179951 | 44180163 | 0 | 0 | 0 | 2 | 0 | 0 | 0 | 0 | 0 | 0 | 0 | 0 | 0 | 0 | 0 | 0 | 2 | 0 | 0 | 20 | 298 | 149 | 1 | 1 | NA | NA | NA | 1 | NA | NA | 0 | NA | NA | 0 | 1 |
| RI | 4515 | ENSG00000004777 | ARHGAP33 | chr19 | + | 35787167 | 35788822 | 35787167 | 35787559 | 35787643 | 35788822 | 1 | 0 | 0 | 4 | 0 | 0 | 0 | 0 | 0 | 0 | 0 | 0 | 0 | 0 | 0 | 0 | 0 | 0 | 3 | 19 | 233 | 149 | 1 | 1 | 1 | NA | NA | 1 | NA | NA | NA | NA | 0 | 0 | 1 |
| RI | 4781 | ENSG00000075702 | WDR62 | chr19 | + | 36102947 | 36104675 | 36102947 | 36103981 | 36104517 | 36104675 | 2 | 0 | 0 | 3 | 0 | 0 | 0 | 0 | 0 | 0 | 0 | 0 | 0 | 0 | 0 | 0 | 2 | 0 | 1 | 0 | 298 | 149 | 1 | 1 | 1 | NA | NA | 1 | NA | NA | 0 | NA | 0 | NA | 1 |
| RI | 5049 | ENSG00000131037 | EPS8L1 | chr19 | + | 55083377 | 55083644 | 55083377 | 55083519 | 55083615 | 55083644 | 0 | 0 | 0 | 1 | 0 | 0 | 0 | 0 | 0 | 0 | 0 | 0 | 0 | 0 | 0 | 0 | 2 | 0 | 0 | 0 | 245 | 149 | 1 | 1 | NA | NA | NA | 1 | NA | NA | 0 | NA | NA | NA | 1 |
| RI | 5284 | ENSG00000063660 | GPC1 | chr2 | + | 240463346 | 240464746 | 240463346 | 240463512 | 240464615 | 240464746 | 0 | 0 | 1 | 7 | 2 | 0 | 0 | 0 | 0 | 0 | 0 | 0 | 0 | 0 | 0 | 10 | 0 | 4 | 0 | 0 | 298 | 149 | 1 | 1 | NA | NA | 1 | 1 | 1 | 0 | NA | 0 | NA | NA | 1 |
| RI | 5491 | ENSG00000096696 | DSP | chr6 | + | 7571382 | 7572068 | 7571382 | 7571584 | 7571841 | 7572068 | 0 | 0 | 1 | 1 | 0 | 0 | 0 | 0 | 0 | 0 | 0 | 0 | 0 | 0 | 0 | 2 | 10 | 1 | 3 | 0 | 298 | 149 | 1 | 1 | NA | NA | 1 | 1 | NA | 0 | 0 | 0 | 0 | NA | 1 |
| RI | 6381 | ENSG00000179698 | WDR97 | chr8 | + | 144111100 | 144111486 | 144111100 | 144111222 | 144111425 | 144111486 | 2 | 0 | 3 | 1 | 0 | 0 | 0 | 0 | 0 | 0 | 0 | 0 | 0 | 0 | 0 | 0 | 0 | 0 | 3 | 0 | 298 | 149 | 1 | 1 | 1 | NA | 1 | 1 | NA | NA | NA | NA | 0 | NA | 1 |
| RI | 6691 | ENSG00000174080 | CTSF | chr11 | - | 66565670 | 66565927 | 66565670 | 66565749 | 66565830 | 66565927 | 0 | 1 | 0 | 0 | 0 | 0 | 0 | 0 | 0 | 0 | 0 | 0 | 0 | 0 | 0 | 0 | 2 | 0 | 0 | 0 | 230 | 149 | 1 | 1 | NA | 1 | NA | NA | NA | NA | 0 | NA | NA | NA | 1 |
| RI | 7089 | ENSG00000164683 | HEY1 | chr8 | - | 79766996 | 79767294 | 79766996 | 79767092 | 79767218 | 79767294 | 0 | 0 | 0 | 1 | 0 | 0 | 0 | 0 | 0 | 0 | 0 | 0 | 0 | 0 | 0 | 0 | 0 | 4 | 0 | 0 | 275 | 149 | 1 | 1 | NA | NA | NA | 1 | NA | NA | NA | 0 | NA | NA | 1 |
| RI | 7247 | ENSG00000197140 | ADAM32 | chr8 | + | 39165029 | 39169997 | 39165029 | 39165196 | 39169915 | 39169997 | 2 | 0 | 0 | 0 | 0 | 0 | 0 | 0 | 0 | 0 | 0 | 0 | 0 | 0 | 0 | 0 | 2 | 0 | 0 | 0 | 298 | 149 | 1 | 1 | 1 | NA | NA | NA | NA | NA | 0 | NA | NA | NA | 1 |

U12_RI_filtered

| | | | | | | | | | | | | | inclusion_counts_HC | | | | | skipping_counts_HC | | | | | Inclusion counts_SLE | | | | | skipping_counts_SLE | | | | | | | | | Inclusion_level(normalized) | | | | | | | | | | |
|---|---|---|---|---|---|---|---|---|---|---|---|---|---|---|---|---|---|---|---|---|---|---|---|---|---|---|---|---|---|---|---|---|---|---|---|---|---|---|---|---|---|---|---|---|---|---|---|
| Splice_type | ID | Gene_ID | Gene_Symbol | Chromosome | Strand | LongExonStart_0base | LongExonEnd | ShortES | ShortEE | FlankingES | FlankingEE | ID | SRR13214146 | SRR13214147 | SRR13214148 | SRR13214149 | SRR13214150 | SRR13214146 | SRR13214147 | SRR13214148 | SRR13214149 | SRR13214150 | SRR13214141 | SRR13214242 | SRR13214143 | SRR13214144 | SRR13214145 | SRR13214141 | SRR13214242 | SRR13214143 | SRR13214144 | SRR13214145 | IncFormLen | SkipFormLen | p value | FDR | SRR13214146 | SRR13214147 | SRR13214148 | SRR13214149 | SRR13214150 | SRR13214141 | SRR13214242 | SRR13214143 | SRR13214144 | SRR13214145 | Inclevel_diff |
| RI | 3093 | ENSG00000111642 | CHD4 | chr12 | - | 6578028 | 6578546 | 6578028 | 6578137 | 6578408 | 6578546 | 3093 | 52 | 59 | 39 | 58 | 56 | 384 | 226 | 344 | 359 | 525 | 383 | 139 | 146 | 347 | 175 | 361 | 239 | 246 | 215 | 405 | 298 | 149 | 0 | 0 | 0.063 | 0.115 | 0.054 | 0.075 | 0.051 | 0.347 | 0.225 | 0.229 | 0.447 | 0.178 | -0.214 |
| RI | 5341 | ENSG00000084623 | EIF3I | chr1 | + | 32228726 | 32229208 | 32228726 | 32228816 | 32229134 | 32229208 | 5341 | 21 | 20 | 21 | 30 | 12 | 136 | 82 | 153 | 117 | 130 | 103 | 69 | 44 | 94 | 110 | 112 | 86 | 84 | 107 | 255 | 298 | 149 | 0 | 0 | 0.072 | 0.109 | 0.064 | 0.114 | 0.044 | 0.315 | 0.286 | 0.208 | 0.305 | 0.177 | -0.178 |
| RI | 5336 | ENSG00000084623 | EIF3I | chr1 | + | 32224033 | 32224475 | 32224033 | 32224121 | 32224409 | 32224475 | 5336 | 26 | 17 | 22 | 48 | 13 | 198 | 124 | 225 | 216 | 151 | 99 | 74 | 61 | 137 | 122 | 245 | 213 | 141 | 163 | 377 | 298 | 149 | 0 | 0 | 0.062 | 0.064 | 0.047 | 0.1 | 0.041 | 0.168 | 0.148 | 0.178 | 0.296 | 0.139 | -0.123 |
| RI | 3097 | ENSG00000111642 | CHD4 | chr12 | - | 6587383 | 6587949 | 6587383 | 6587559 | 6587711 | 6587949 | 3097 | 21 | 27 | 30 | 51 | 17 | 359 | 260 | 345 | 423 | 371 | 148 | 167 | 92 | 192 | 237 | 1058 | 572 | 515 | 874 | 977 | 298 | 149 | 0 | 0 | 0.028 | 0.049 | 0.042 | 0.057 | 0.022 | 0.065 | 0.127 | 0.082 | 0.099 | 0.108 | -0.057 |
| RI | 6850 | ENSG00000215021 | PHB2 | chr12 | - | 6967170 | 6967779 | 6967170 | 6967248 | 6967675 | 6967779 | 6850 | 14 | 18 | 19 | 35 | 64 | 593 | 281 | 561 | 447 | 725 | 51 | 64 | 62 | 62 | 68 | 534 | 279 | 280 | 305 | 534 | 298 | 149 | 0 | 0 | 0.012 | 0.031 | 0.017 | 0.038 | 0.042 | 0.046 | 0.103 | 0.1 | 0.092 | 0.06 | -0.052 |
| RI | 8453 | ENSG00000123349 | PFDN5 | chr12 | + | 53295838 | 53297924 | 53295838 | 53296275 | 53297849 | 53297924 | 8453 | 56 | 52 | 86 | 116 | 142 | 1544 | 881 | 1393 | 1315 | 1587 | 347 | 138 | 115 | 204 | 194 | 2006 | 1172 | 805 | 1133 | 1967 | 298 | 149 | 0 | 0 | 0.018 | 0.029 | 0.03 | 0.042 | 0.043 | 0.08 | 0.056 | 0.067 | 0.083 | 0.047 | -0.034 |
| RI | 5513 | ENSG00000112367 | FIG4 | chr6 | + | 109786301 | 109789677 | 109786301 | 109786449 | 109789593 | 109789677 | 5513 | 19 | 15 | 14 | 24 | 13 | 74 | 62 | 41 | 65 | 151 | 0 | 1 | 0 | 0 | 0 | 76 | 49 | 33 | 69 | 17 | 298 | 149 | 0 | 0 | 0.114 | 0.108 | 0.146 | 0.156 | 0.041 | 0 | 0.01 | 0 | 0 | 0 | 0.111 |
| RI | 3095 | ENSG00000111642 | CHD4 | chr12 | - | 6583026 | 6583378 | 6583026 | 6583113 | 6583197 | 6583378 | 3095 | 25 | 15 | 21 | 17 | 23 | 323 | 203 | 328 | 349 | 402 | 57 | 48 | 42 | 76 | 75 | 407 | 290 | 237 | 413 | 561 | 233 | 149 | 1.41787E-12 | 6.97944E-11 | 0.047 | 0.045 | 0.039 | 0.03 | 0.035 | 0.082 | 0.096 | 0.102 | 0.105 | 0.079 | -0.054 |
| RI | 5450 | ENSG00000143799 | PARP1 | chr1 | - | 226365953 | 226367608 | 226365953 | 226366052 | 226367479 | 226367608 | 5450 | 18 | 8 | 21 | 11 | 9 | 358 | 206 | 282 | 377 | 468 | 13 | 34 | 5 | 42 | 39 | 232 | 223 | 178 | 165 | 280 | 298 | 149 | 7.04425E-12 | 3.1523E-10 | 0.025 | 0.019 | 0.036 | 0.014 | 0.01 | 0.027 | 0.071 | 0.014 | 0.113 | 0.065 | -0.037 |
| RI | 5449 | ENSG00000143799 | PARP1 | chr1 | - | 226363098 | 226365154 | 226363098 | 226364070 | 226365001 | 226365154 | 5449 | 9 | 4 | 9 | 4 | 3 | 132 | 49 | 120 | 100 | 124 | 29 | 15 | 13 | 12 | 27 | 100 | 64 | 65 | 90 | 130 | 298 | 149 | 6.17517E-11 | 2.41568E-09 | 0.033 | 0.039 | 0.036 | 0.02 | 0.012 | 0.127 | 0.105 | 0.091 | 0.062 | 0.094 | -0.068 |
| RI | 338 | ENSG00000013275 | PSMC4 | chr19 | + | 39980285 | 39980717 | 39980285 | 39980454 | 39980661 | 39980717 | 338 | 26 | 7 | 30 | 23 | 24 | 149 | 88 | 115 | 114 | 228 | 91 | 28 | 34 | 56 | 68 | 186 | 149 | 88 | 97 | 161 | 298 | 149 | 1.50653E-10 | 5.52739E-09 | 0.08 | 0.038 | 0.115 | 0.092 | 0.05 | 0.197 | 0.086 | 0.162 | 0.224 | 0.174 | -0.094 |
| RI | 1679 | ENSG00000174444 | RPL4 | chr15 | - | 66502611 | 66503164 | 66502611 | 66502750 | 66503057 | 66503164 | 1679 | 88 | 56 | 98 | 85 | 111 | 1518 | 660 | 1745 | 1247 | 1546 | 106 | 96 | 81 | 141 | 83 | 765 | 811 | 915 | 663 | 1305 | 298 | 149 | 1.64716E-09 | 4.8407E-08 | 0.028 | 0.041 | 0.027 | 0.033 | 0.035 | 0.065 | 0.056 | 0.042 | 0.096 | 0.031 | -0.025 |
| RI | 8017 | ENSG00000198258 | UBL5 | chr19 | + | 9828326 | 9828675 | 9828326 | 9828393 | 9828591 | 9828675 | 8017 | 29 | 8 | 31 | 31 | 25 | 138 | 76 | 100 | 133 | 149 | 22 | 80 | 43 | 67 | 80 | 100 | 106 | 48 | 113 | 163 | 298 | 149 | 3.1146E-09 | 8.47832E-08 | 0.095 | 0.05 | 0.134 | 0.104 | 0.077 | 0.099 | 0.274 | 0.309 | 0.229 | 0.197 | -0.13 |
| RI | 1814 | ENSG00000196497 | IPO4 | chr14 | - | 24183455 | 24183667 | 24183455 | 24183513 | 24183589 | 24183667 | 1814 | 9 | 6 | 14 | 9 | 8 | 27 | 11 | 35 | 43 | 42 | 32 | 20 | 18 | 65 | 37 | 21 | 13 | 27 | 16 | 31 | 225 | 149 | 6.18161E-09 | 1.5876E-07 | 0.181 | 0.265 | 0.209 | 0.122 | 0.112 | 0.502 | 0.505 | 0.306 | 0.729 | 0.441 | -0.319 |
| RI | 7726 | ENSG00000130935 | NOL11 | chr17 | + | 67738121 | 67739010 | 67738121 | 67738355 | 67738931 | 67739010 | 7726 | 84 | 67 | 65 | 64 | 111 | 121 | 63 | 80 | 86 | 179 | 52 | 37 | 78 | 56 | 31 | 6 | 17 | 49 | 5 | 0 | 298 | 149 | 1.01673E-08 | 2.45136E-07 | 0.258 | 0.347 | 0.289 | 0.271 | 0.237 | 0.812 | 0.521 | 0.443 | 0.848 | 1 | -0.444 |
| RI | 1899 | ENSG00000131966 | ACTR10 | chr14 | + | 58223621 | 58223856 | 58223621 | 58223701 | 58223782 | 58223856 | 1899 | 8 | 6 | 4 | 12 | 15 | 107 | 74 | 98 | 95 | 199 | 14 | 29 | 24 | 18 | 0 | 46 | 70 | 49 | 54 | 85 | 230 | 149 | 6.80842E-08 | 1.31001E-06 | 0.046 | 0.05 | 0.026 | 0.076 | 0.047 | 0.165 | 0.212 | 0.241 | 0.178 | 0 | -0.11 |
| RI | 246 | ENSG00000005007 | UPF1 | chr19 | + | 18865288 | 18865778 | 18865288 | 18865450 | 18865560 | 18865778 | 246 | 3 | 0 | 3 | 0 | 0 | 40 | 24 | 40 | 52 | 25 | 8 | 13 | 13 | 2 | 6 | 49 | 50 | 43 | 21 | 81 | 259 | 149 | 9.09003E-08 | 1.71549E-06 | 0.041 | 0 | 0.041 | 0 | 0 | 0.086 | 0.13 | 0.148 | 0.052 | 0.041 | -0.075 |
| RI | 3240 | ENSG00000197386 | HTT | chr4 | + | 3187650 | 3189093 | 3187650 | 3187886 | 3188950 | 3189093 | 3240 | 22 | 15 | 23 | 21 | 30 | 135 | 71 | 75 | 110 | 261 | 60 | 75 | 73 | 89 | 42 | 167 | 95 | 43 | 108 | 285 | 298 | 149 | 1.97511E-07 | 3.46201E-06 | 0.075 | 0.096 | 0.133 | 0.087 | 0.054 | 0.152 | 0.283 | 0.459 | 0.292 | 0.069 | -0.162 |
| RI | 337 | ENSG00000013275 | PSMC4 | chr19 | + | 39980069 | 39980454 | 39980069 | 39980146 | 39980285 | 39980454 | 337 | 23 | 14 | 26 | 25 | 28 | 119 | 58 | 108 | 110 | 134 | 76 | 50 | 37 | 68 | 97 | 237 | 167 | 64 | 97 | 154 | 288 | 149 | 2.35032E-07 | 4.02416E-06 | 0.091 | 0.111 | 0.111 | 0.105 | 0.098 | 0.142 | 0.134 | 0.23 | 0.266 | 0.246 | -0.1 |
| RI | 2678 | ENSG00000184205 | TSPYL2 | chrX | + | 53084531 | 53084866 | 53084531 | 53084622 | 53084754 | 53084866 | 2678 | 7 | 7 | 27 | 29 | 23 | 31 | 25 | 65 | 32 | 52 | 27 | 44 | 116 | 62 | 49 | 24 | 35 | 68 | 16 | 46 | 281 | 149 | 2.53599E-07 | 4.2923E-06 | 0.107 | 0.129 | 0.181 | 0.325 | 0.19 | 0.374 | 0.4 | 0.475 | 0.673 | 0.361 | -0.27 |
| RI | 8458 | ENSG00000123349 | PFDN5 | chr12 | + | 53295838 | 53298150 | 53295838 | 53297924 | 53298044 | 53298150 | 8458 | 58 | 41 | 57 | 75 | 66 | 1917 | 1073 | 1773 | 1559 | 1822 | 161 | 88 | 71 | 103 | 91 | 2468 | 1410 | 1035 | 1314 | 2469 | 269 | 149 | 3.56032E-07 | 5.80961E-06 | 0.016 | 0.021 | 0.017 | 0.026 | 0.02 | 0.035 | 0.033 | 0.037 | 0.042 | 0.02 | -0.013 |
| RI | 5448 | ENSG00000143799 | PARP1 | chr1 | - | 226363098 | 226364070 | 226363098 | 226363160 | 226363942 | 226364070 | 5448 | 14 | 9 | 18 | 19 | 11 | 130 | 79 | 139 | 120 | 145 | 61 | 25 | 10 | 27 | 16 | 72 | 136 | 78 | 62 | 59 | 298 | 149 | 3.60851E-07 | 5.8559E-06 | 0.051 | 0.054 | 0.061 | 0.073 | 0.037 | 0.298 | 0.084 | 0.06 | 0.179 | 0.119 | -0.093 |
| RI | 7817 | ENSG00000156599 | ZDHHC5 | chr11 | + | 57690030 | 57690437 | 57690030 | 57690203 | 57690334 | 57690437 | 7817 | 28 | 14 | 11 | 30 | 27 | 164 | 80 | 140 | 134 | 132 | 61 | 22 | 27 | 71 | 73 | 162 | 161 | 77 | 123 | 158 | 280 | 149 | 4.16157E-07 | 6.60817E-06 | 0.083 | 0.085 | 0.04 | 0.106 | 0.098 | 0.167 | 0.068 | 0.157 | 0.235 | 0.197 | -0.082 |
| RI | 5451 | ENSG00000143799 | PARP1 | chr1 | - | 226379921 | 226381208 | 226379921 | 226380164 | 226381067 | 226381208 | 5451 | 14 | 13 | 33 | 35 | 20 | 353 | 283 | 356 | 451 | 401 | 37 | 42 | 39 | 68 | 42 | 408 | 422 | 312 | 290 | 512 | 298 | 149 | 6.95475E-07 | 1.04004E-05 | 0.019 | 0.022 | 0.044 | 0.037 | 0.024 | 0.043 | 0.047 | 0.059 | 0.105 | 0.039 | -0.029 |
| RI | 6245 | ENSG00000171777 | RASGRP4 | chr19 | - | 38409057 | 38410096 | 38409057 | 38409370 | 38409954 | 38410096 | 6245 | 92 | 43 | 72 | 97 | 128 | 4 | 4 | 0 | 1 | 0 | 360 | 106 | 115 | 247 | 380 | 34 | 3 | 7 | 38 | 16 | 298 | 149 | 7.25616E-07 | 1.07915E-05 | 0.92 | 0.843 | 1 | 0.98 | 1 | 0.841 | 0.946 | 0.891 | 0.765 | 0.922 | 0.076 |
| RI | 1426 | ENSG00000038358 | EDC4 | chr16 | + | 67881264 | 67881533 | 67881264 | 67881417 | 67881496 | 67881533 | 1426 | 5 | 6 | 12 | 16 | 4 | 90 | 32 | 58 | 72 | 63 | 9 | 34 | 32 | 46 | 41 | 89 | 57 | 59 | 76 | 130 | 228 | 149 | 8.66627E-07 | 1.26088E-05 | 0.035 | 0.109 | 0.119 | 0.127 | 0.04 | 0.062 | 0.28 | 0.262 | 0.283 | 0.171 | -0.126 |
| RI | 8175 | ENSG00000119977 | TCTN3 | chr10 | - | 95683100 | 95683629 | 95683100 | 95683180 | 95683521 | 95683629 | 8175 | 17 | 12 | 24 | 12 | 35 | 1 | 2 | 4 | 1 | 4 | 22 | 17 | 3 | 8 | 10 | 0 | 0 | 0 | 0 | 0 | 298 | 149 | 1.38697E-06 | 1.93228E-05 | 0.895 | 0.75 | 0.75 | 0.857 | 0.814 | 1 | 1 | 1 | 1 | 1 | -0.187 |
| RI | 1678 | ENSG00000174444 | RPL4 | chr15 | - | 66500948 | 66501504 | 66500948 | 66501105 | 66501374 | 66501504 | 1678 | 528 | 332 | 659 | 544 | 2649 | 5995 | 3495 | 8096 | 6670 | 18363 | 399 | 366 | 317 | 191 | 451 | 5552 | 4126 | 4448 | 3327 | 7491 | 298 | 149 | 2.76283E-06 | 3.58335E-05 | 0.042 | 0.045 | 0.039 | 0.039 | 0.067 | 0.035 | 0.042 | 0.034 | 0.028 | 0.029 | 0.013 |
| RI | 7196 | ENSG00000174744 | BRMS1 | chr11 | - | 66340773 | 66341333 | 66340773 | 66341046 | 66341205 | 66341333 | 7196 | 12 | 11 | 15 | 23 | 9 | 165 | 117 | 116 | 141 | 148 | 57 | 32 | 17 | 20 | 46 | 187 | 173 | 138 | 147 | 169 | 298 | 149 | 3.39934E-06 | 4.29977E-05 | 0.035 | 0.045 | 0.061 | 0.075 | 0.03 | 0.132 | 0.085 | 0.058 | 0.064 | 0.12 | -0.043 |
| RI | 127 | ENSG00000198089 | SFI1 | chr22 | + | 31604868 | 31606430 | 31604868 | 31604945 | 31606327 | 31606430 | 127 | 20 | 23 | 52 | 50 | 21 | 15 | 13 | 19 | 18 | 21 | 124 | 51 | 65 | 103 | 48 | 25 | 9 | 9 | 2 | 15 | 298 | 149 | 3.84768E-06 | 4.76484E-05 | 0.4 | 0.469 | 0.578 | 0.581 | 0.333 | 0.713 | 0.739 | 0.783 | 0.963 | 0.615 | -0.29 |
| RI | 5345 | ENSG00000084623 | EIF3I | chr1 | + | 32230926 | 32231602 | 32230926 | 32231019 | 32231114 | 32231602 | 5345 | 7 | 12 | 10 | 19 | 15 | 103 | 66 | 135 | 150 | 110 | 46 | 38 | 19 | 22 | 53 | 184 | 111 | 90 | 122 | 270 | 244 | 149 | 4.55102E-06 | 5.45291E-05 | 0.04 | 0.1 | 0.043 | 0.072 | 0.077 | 0.132 | 0.173 | 0.114 | 0.099 | 0.107 | -0.059 |
| RI | 2661 | ENSG00000141452 | RMC1 | chr18 | + | 23530228 | 23530612 | 23530228 | 23530297 | 23530510 | 23530612 | 2661 | 202 | 71 | 105 | 97 | 57 | 9 | 3 | 6 | 6 | 0 | 240 | 168 | 144 | 146 | 97 | 0 | 0 | 1 | 0 | 6 | 298 | 149 | 4.60916E-06 | 5.50027E-05 | 0.918 | 0.922 | 0.897 | 0.89 | 1 | 1 | 1 | 0.986 | 1 | 0.89 | -0.05 |
| RI | 4834 | ENSG00000103168 | TAF1C | chr16 | - | 84181327 | 84181663 | 84181327 | 84181463 | 84181591 | 84181663 | 4834 | 42 | 23 | 27 | 54 | 28 | 8 | 2 | 3 | 7 | 0 | 97 | 75 | 56 | 57 | 152 | 0 | 3 | 3 | 2 | 0 | 277 | 149 | 6.00778E-06 | 6.9448E-05 | 0.738 | 0.861 | 0.829 | 0.806 | 1 | 1 | 0.931 | 0.909 | 0.939 | 1 | -0.109 |
| RI | 789 | ENSG00000214087 | ARL16 | chr17 | - | 81682033 | 81682675 | 81682033 | 81682149 | 81682490 | 81682675 | 789 | 31 | 32 | 49 | 43 | 22 | 10 | 2 | 5 | 6 | 8 | 119 | 107 | 48 | 86 | 100 | 5 | 9 | 0 | 0 | 3 | 298 | 149 | 6.46135E-06 | 7.44E-05 | 0.608 | 0.889 | 0.831 | 0.782 | 0.579 | 0.922 | 0.856 | 1 | 1 | 0.943 | -0.206 |
| RI | 2689 | ENSG00000108591 | DRG2 | chr17 | + | 18099016 | 18100435 | 18099016 | 18099093 | 18099632 | 18100435 | 2689 | 19 | 13 | 26 | 36 | 33 | 3 | 1 | 2 | 2 | 4 | 43 | 37 | 29 | 8 | 49 | 0 | 0 | 1 | 0 | 0 | 298 | 149 | 6.57535E-06 | 7.52725E-05 | 0.76 | 0.867 | 0.867 | 0.9 | 0.805 | 1 | 1 | 0.935 | 1 | 1 | -0.147 |
| RI | 1680 | ENSG00000174444 | RPL4 | chr15 | - | 66503057 | 66503529 | 66503057 | 66503164 | 66503357 | 66503529 | 1680 | 120 | 86 | 126 | 103 | 89 | 1975 | 906 | 2191 | 1665 | 2448 | 71 | 106 | 73 | 152 | 104 | 1019 | 1110 | 1048 | 811 | 1946 | 298 | 149 | 1.3055E-05 | 0.000140519 | 0.029 | 0.045 | 0.028 | 0.03 | 0.018 | 0.034 | 0.046 | 0.034 | 0.086 | 0.026 | -0.015 |
| RI | 2407 | ENSG00000105875 | WDR91 | chr7 | - | 135204267 | 135206058 | 135204267 | 135204433 | 135205927 | 135206058 | 2407 | 36 | 21 | 27 | 27 | 19 | 39 | 10 | 42 | 25 | 29 | 91 | 60 | 34 | 51 | 49 | 20 | 23 | 15 | 32 | 16 | 298 | 149 | 1.44037E-05 | 0.000153399 | 0.316 | 0.512 | 0.243 | 0.351 | 0.247 | 0.695 | 0.566 | 0.531 | 0.443 | 0.605 | -0.234 |
| RI | 2159 | ENSG00000111641 | NOP2 | chr12 | - | 6560446 | 6560787 | 6560446 | 6560569 | 6560697 | 6560787 | 2159 | 0 | 1 | 1 | 7 | 0 | 48 | 22 | 65 | 62 | 43 | 8 | 8 | 6 | 0 | 18 | 66 | 54 | 37 | 48 | 116 | 277 | 149 | 2.28241E-05 | 0.000226973 | 0 | 0.024 | 0.008 | 0.057 | 0 | 0.061 | 0.074 | 0.08 | 0 | 0.077 | -0.041 |
| RI | 5797 | ENSG00000143499 | SMYD2 | chr1 | + | 214330949 | 214334308 | 214330949 | 214331070 | 214332017 | 214334308 | 5797 | 39 | 35 | 47 | 52 | 41 | 30 | 12 | 42 | 46 | 43 | 0 | 24 | 22 | 28 | 14 | 53 | 33 | 19 | 44 | 52 | 298 | 149 | 2.83524E-05 | 0.000274553 | 0.394 | 0.593 | 0.359 | 0.361 | 0.323 | 0 | 0.267 | 0.367 | 0.241 | 0.119 | 0.207 |
| RI | 5212 | ENSG00000149499 | EML3 | chr11 | - | 62602758 | 62603247 | 62602758 | 62602779 | 62603148 | 62603247 | 5212 | 96 | 47 | 79 | 86 | 62 | 8 | 6 | 4 | 5 | 7 | 45 | 99 | 57 | 66 | 135 | 0 | 0 | 2 | 2 | 5 | 298 | 149 | 3.04082E-05 | 0.000292067 | 0.857 | 0.797 | 0.908 | 0.896 | 0.816 | 1 | 1 | 0.934 | 0.943 | 0.931 | -0.107 |
| RI | 3570 | ENSG00000120008 | WDR11 | chr10 | + | 120889077 | 120890009 | 120889077 | 120889184 | 120889894 | 120890009 | 3570 | 4 | 9 | 6 | 5 | 3 | 206 | 168 | 123 | 164 | 272 | 13 | 2 | 16 | 8 | 0 | 96 | 126 | 69 | 113 | 29 | 298 | 149 | 3.07884E-05 | 0.000295239 | 0.01 | 0.026 | 0.024 | 0.015 | 0.005 | 0.063 | 0.008 | 0.104 | 0.034 | 0 | -0.026 |
| RI | 8449 | ENSG00000123349 | PFDN5 | chr12 | + | 53295838 | 53296275 | 53295838 | 53295941 | 53296243 | 53296275 | 8449 | 225 | 175 | 280 | 256 | 294 | 1022 | 540 | 881 | 838 | 1024 | 408 | 309 | 235 | 260 | 366 | 1143 | 630 | 468 | 705 | 1141 | 298 | 149 | 3.11087E-05 | 0.000296967 | 0.099 | 0.139 | 0.137 | 0.133 | 0.126 | 0.151 | 0.197 | 0.201 | 0.156 | 0.138 | -0.042 |
| RI | 4550 | ENSG00000163463 | KRTCAP2 | chr1 | - | 155172564 | 155172892 | 155172564 | 155172628 | 155172737 | 155172892 | 4550 | 6 | 1 | 7 | 11 | 4 | 138 | 47 | 115 | 113 | 54 | 9 | 15 | 6 | 6 | 50 | 71 | 129 | 65 | 116 | 90 | 258 | 149 | 3.14875E-05 | 0.000299995 | 0.024 | 0.012 | 0.034 | 0.053 | 0.041 | 0.068 | 0.063 | 0.051 | 0.029 | 0.243 | -0.058 |
| RI | 6853 | ENSG00000215021 | PHB2 | chr12 | - | 6967170 | 6968021 | 6967170 | 6967779 | 6967891 | 6968021 | 6853 | 17 | 14 | 39 | 41 | 18 | 318 | 194 | 346 | 291 | 328 | 60 | 27 | 40 | 48 | 51 | 389 | 202 | 205 | 287 | 473 | 261 | 149 | 4.98674E-05 | 0.000450408 | 0.03 | 0.04 | 0.06 | 0.074 | 0.03 | 0.081 | 0.071 | 0.1 | 0.087 | 0.058 | -0.033 |
| RI | 1411 | ENSG00000051523 | CYBA | chr16 | - | 88646115 | 88646838 | 88646115 | 88646197 | 88646754 | 88646838 | 1411 | 35 | 23 | 47 | 46 | 13 | 966 | 318 | 481 | 600 | 485 | 31 | 85 | 55 | 58 | 101 | 743 | 721 | 391 | 688 | 1017 | 298 | 149 | 5.415E-05 | 0.000486115 | 0.018 | 0.035 | 0.047 | 0.037 | 0.013 | 0.02 | 0.056 | 0.066 | 0.04 | 0.047 | -0.016 |
| RI | 1924 | ENSG00000125746 | EML2 | chr19 | - | 45638504 | 45638873 | 45638504 | 45638634 | 45638844 | 45638873 | 1924 | 11 | 2 | 19 | 3 | 16 | 50 | 28 | 46 | 44 | 36 | 14 | 1 | 5 | 9 | 0 | 110 | 44 | 52 | 59 | 67 | 298 | 149 | 7.80422E-05 | 0.000664258 | 0.099 | 0.034 | 0.171 | 0.033 | 0.182 | 0.06 | 0.011 | 0.046 | 0.071 | 0 | 0.066 |
| RI | 5453 | ENSG00000143799 | PARP1 | chr1 | - | 226385503 | 226386442 | 226385503 | 226385680 | 226386325 | 226386442 | 5453 | 5 | 3 | 10 | 9 | 0 | 293 | 182 | 269 | 300 | 282 | 0 | 10 | 30 | 32 | 0 | 266 | 349 | 245 | 207 | 449 | 298 | 149 | 8.25958E-05 | 0.000696991 | 0.008 | 0.008 | 0.018 | 0.015 | 0 | 0 | 0.014 | 0.058 | 0.072 | 0 | -0.019 |
| RI | 3297 | ENSG00000198925 | ATG9A | chr2 | - | 219227769 | 219228053 | 219227769 | 219227813 | 219227921 | 219228053 | 3297 | 12 | 7 | 14 | 31 | 6 | 31 | 16 | 26 | 16 | 14 | 8 | 40 | 21 | 38 | 61 | 16 | 9 | 5 | 8 | 22 | 257 | 149 | 0.000176036 | 0.001337788 | 0.183 | 0.202 | 0.238 | 0.529 | 0.199 | 0.225 | 0.72 | 0.709 | 0.734 | 0.616 | -0.331 |
| RI | 931 | ENSG00000180891 | CUEDC1 | chr17 | - | 57866473 | 57867415 | 57866473 | 57866544 | 57867356 | 57867415 | 931 | 9 | 15 | 9 | 5 | 13 | 25 | 17 | 17 | 22 | 13 | 19 | 10 | 5 | 9 | 2 | 63 | 38 | 14 | 48 | 74 | 298 | 149 | 0.000210822 | 0.001560559 | 0.153 | 0.306 | 0.209 | 0.102 | 0.333 | 0.131 | 0.116 | 0.152 | 0.086 | 0.013 | 0.121 |
| RI | 3967 | ENSG00000163479 | SSR2 | chr1 | - | 156011809 | 156012565 | 156011809 | 156011887 | 156012384 | 156012565 | 3967 | 22 | 6 | 8 | 15 | 6 | 2 | 2 | 0 | 0 | 6 | 46 | 16 | 19 | 28 | 70 | 0 | 0 | 2 | 0 | 0 | 298 | 149 | 0.000212166 | 0.001566577 | 0.846 | 0.6 | 1 | 1 | 0.333 | 1 | 1 | 0.826 | 1 | 1 | -0.209 |
| RI | 3478 | ENSG00000160741 | CRTC2 | chr1 | - | 153954872 | 153955166 | 153954872 | 153954989 | 153955064 | 153955166 | 3478 | 39 | 21 | 45 | 29 | 47 | 41 | 33 | 36 | 36 | 28 | 105 | 106 | 81 | 82 | 158 | 56 | 83 | 40 | 16 | 43 | 224 | 149 | 0.000296593 | 0.002062519 | 0.388 | 0.297 | 0.454 | 0.349 | 0.528 | 0.555 | 0.459 | 0.574 | 0.773 | 0.71 | -0.211 |
| RI | 5436 | ENSG00000197858 | GPAA1 | chr8 | + | 144085288 | 144085743 | 144085288 | 144085479 | 144085572 | 144085743 | 5436 | 7 | 0 | 4 | 2 | 5 | 51 | 8 | 42 | 45 | 40 | 1 | 10 | 8 | 8 | 8 | 9 | 56 | 17 | 11 | 27 | 242 | 149 | 0.000306338 | 0.002126367 | 0.078 | 0 | 0.055 | 0.027 | 0.071 | 0.064 | 0.099 | 0.225 | 0.309 | 0.154 | -0.124 |
| RI | 6246 | ENSG00000171777 | RASGRP4 | chr19 | - | 38411114 | 38411381 | 38411114 | 38411249 | 38411344 | 38411381 | 6246 | 2 | 12 | 7 | 7 | 0 | 44 | 25 | 33 | 33 | 23 | 56 | 22 | 8 | 39 | 55 | 62 | 56 | 46 | 117 | 140 | 244 | 149 | 0.000331383 | 0.002281446 | 0.027 | 0.227 | 0.115 | 0.115 | 0 | 0.355 | 0.193 | 0.096 | 0.169 | 0.193 | -0.104 |
| RI | 5079 | ENSG00000112855 | HARS2 | chr5 | + | 140691452 | 140691756 | 140691452 | 140691515 | 140691606 | 140691756 | 5079 | 39 | 18 | 39 | 28 | 22 | 1 | 1 | 4 | 7 | 3 | 8 | 11 | 26 | 15 | 56 | 2 | 0 | 2 | 0 | 0 | 240 | 149 | 0.000332513 | 0.002283899 | 0.96 | 0.918 | 0.858 | 0.713 | 0.82 | 0.713 | 1 | 0.89 | 1 | 1 | -0.067 |
| RI | 5213 | ENSG00000149499 | EML3 | chr11 | - | 62602758 | 62603247 | 62602758 | 62602889 | 62603148 | 62603247 | 5213 | 58 | 25 | 44 | 62 | 50 | 54 | 38 | 65 | 44 | 70 | 33 | 52 | 55 | 55 | 120 | 27 | 38 | 17 | 37 | 52 | 298 | 149 | 0.000423595 | 0.00280828 | 0.349 | 0.248 | 0.253 | 0.413 | 0.263 | 0.379 | 0.406 | 0.618 | 0.426 | 0.536 | -0.168 |
| RI | 5571 | ENSG00000160695 | VPS11 | chr11 | + | 119078172 | 119078997 | 119078172 | 119078334 | 119078794 | 119078997 | 5571 | 6 | 6 | 10 | 20 | 26 | 0 | 0 | 0 | 0 | 0 | 81 | 29 | 28 | 93 | 56 | 10 | 0 | 0 | 3 | 0 | 298 | 149 | 0.000428733 | 0.002835976 | 1 | 1 | 1 | 1 | 1 | 0.802 | 1 | 1 | 0.939 | 1 | 0.052 |
| RI | 788 | ENSG00000214087 | ARL16 | chr17 | - | 81682033 | 81682675 | 81682033 | 81682149 | 81682442 | 81682675 | 788 | 30 | 32 | 59 | 52 | 26 | 11 | 7 | 11 | 6 | 10 | 108 | 102 | 46 | 90 | 95 | 26 | 7 | 3 | 4 | 1 | 298 | 149 | 0.000507544 | 0.003295309 | 0.577 | 0.696 | 0.728 | 0.812 | 0.565 | 0.675 | 0.879 | 0.885 | 0.918 | 0.979 | -0.192 |
| RI | 1427 | ENSG00000038358 | EDC4 | chr16 | + | 67881667 | 67882109 | 67881667 | 67881845 | 67881953 | 67882109 | 1427 | 38 | 19 | 44 | 48 | 12 | 87 | 28 | 53 | 56 | 29 | 87 | 68 | 89 | 98 | 96 | 120 | 64 | 41 | 69 | 78 | 257 | 149 | 0.000514019 | 0.003329291 | 0.202 | 0.282 | 0.325 | 0.332 | 0.193 | 0.296 | 0.381 | 0.557 | 0.452 | 0.416 | -0.154 |
| RI | 3181 | ENSG00000144848 | ATG3 | chr3 | - | 112532509 | 112534337 | 112532509 | 112532780 | 112534268 | 112534337 | 3181 | 24 | 19 | 14 | 22 | 19 | 451 | 366 | 375 | 527 | 868 | 6 | 4 | 4 | 20 | 14 | 764 | 368 | 328 | 555 | 395 | 298 | 149 | 0.000527358 | 0.003404482 | 0.026 | 0.025 | 0.018 | 0.02 | 0.011 | 0.004 | 0.005 | 0.006 | 0.018 | 0.017 | 0.01 |
| RI | 1761 | ENSG00000100949 | RABGGTA | chr14 | - | 24268109 | 24268420 | 24268109 | 24268198 | 24268368 | 24268420 | 1761 | 13 | 14 | 11 | 15 | 8 | 46 | 21 | 16 | 33 | 24 | 30 | 33 | 28 | 22 | 30 | 47 | 24 | 11 | 13 | 52 | 298 | 149 | 0.000579106 | 0.003686184 | 0.124 | 0.25 | 0.256 | 0.185 | 0.143 | 0.242 | 0.407 | 0.56 | 0.458 | 0.224 | -0.187 |
| RI | 7602 | ENSG00000030066 | NUP160 | chr11 | - | 47812063 | 47812429 | 47812063 | 47812224 | 47812301 | 47812429 | 7602 | 13 | 9 | 10 | 6 | 20 | 70 | 29 | 61 | 57 | 89 | 19 | 12 | 8 | 17 | 16 | 13 | 33 | 18 | 49 | 0 | 226 | 149 | 0.000870585 | 0.005290685 | 0.109 | 0.17 | 0.098 | 0.065 | 0.129 | 0.491 | 0.193 | 0.227 | 0.186 | 1 | -0.305 |
| RI | 8599 | ENSG00000134899 | ERCC5 | chr13 | + | 102853756 | 102854374 | 102853756 | 102853872 | 102854287 | 102854374 | 8599 | 118 | 108 | 117 | 122 | 145 | 182 | 142 | 187 | 172 | 302 | 18 | 160 | 170 | 44 | 19 | 16 | 132 | 83 | 60 | 4 | 298 | 149 | 0.001007452 | 0.006027939 | 0.245 | 0.276 | 0.238 | 0.262 | 0.194 | 0.36 | 0.377 | 0.506 | 0.268 | 0.704 | -0.2 |
| RI | 7710 | ENSG00000099783 | HNRNPM | chr19 | + | 8465323 | 8465515 | 8465323 | 8465362 | 8465479 | 8465515 | 7710 | 556 | 426 | 501 | 573 | 727 | 270 | 174 | 205 | 237 | 342 | 793 | 669 | 455 | 725 | 551 | 335 | 192 | 180 | 193 | 145 | 266 | 149 | 0.001131891 | 0.006652814 | 0.536 | 0.578 | 0.578 | 0.575 | 0.544 | 0.57 | 0.661 | 0.586 | 0.678 | 0.68 | -0.073 |
| RI | 7880 | ENSG00000167986 | DDB1 | chr11 | - | 61331542 | 61333105 | 61331542 | 61331691 | 61332907 | 61333105 | 7880 | 16 | 11 | 29 | 7 | 19 | 79 | 31 | 70 | 74 | 42 | 47 | 49 | 32 | 60 | 15 | 52 | 94 | 49 | 52 | 117 | 298 | 149 | 0.001175407 | 0.006888026 | 0.092 | 0.151 | 0.172 | 0.045 | 0.184 | 0.311 | 0.207 | 0.246 | 0.366 | 0.06 | -0.109 |
| RI | 5340 | ENSG00000084623 | EIF3I | chr1 | + | 32226170 | 32226530 | 32226170 | 32226320 | 32226402 | 32226530 | 5340 | 17 | 18 | 12 | 27 | 10 | 277 | 137 | 260 | 256 | 250 | 31 | 39 | 15 | 29 | 74 | 373 | 288 | 216 | 217 | 547 | 231 | 149 | 0.001224396 | 0.007132653 | 0.038 | 0.078 | 0.029 | 0.064 | 0.025 | 0.051 | 0.08 | 0.043 | 0.079 | 0.08 | -0.02 |
| RI | 6476 | ENSG00000104133 | SPG11 | chr15 | - | 44622219 | 44622799 | 44622219 | 44622347 | 44622727 | 44622799 | 6476 | 62 | 121 | 35 | 76 | 50 | 205 | 167 | 138 | 181 | 295 | 43 | 112 | 143 | 66 | 18 | 83 | 181 | 134 | 119 | 5 | 298 | 149 | 0.00123373 | 0.007172878 | 0.131 | 0.266 | 0.113 | 0.174 | 0.078 | 0.206 | 0.236 | 0.348 | 0.217 | 0.643 | -0.178 |
| RI | 7083 | ENSG00000167302 | TEPSIN | chr17 | - | 81236979 | 81237459 | 81236979 | 81237071 | 81237386 | 81237459 | 7083 | 23 | 2 | 24 | 14 | 13 | 45 | 17 | 30 | 39 | 21 | 31 | 48 | 27 | 23 | 47 | 26 | 25 | 19 | 17 | 107 | 298 | 149 | 0.001266708 | 0.007328546 | 0.204 | 0.056 | 0.286 | 0.152 | 0.236 | 0.373 | 0.49 | 0.415 | 0.404 | 0.18 | -0.186 |
| RI | 8455 | ENSG00000123349 | PFDN5 | chr12 | + | 53296429 | 53297924 | 53296429 | 53296551 | 53297849 | 53297924 | 8455 | 52 | 40 | 62 | 82 | 101 | 15 | 13 | 21 | 15 | 12 | 254 | 116 | 102 | 163 | 121 | 14 | 13 | 6 | 24 | 26 | 298 | 149 | 0.00162206 | 0.009084791 | 0.634 | 0.606 | 0.596 | 0.732 | 0.808 | 0.901 | 0.817 | 0.895 | 0.773 | 0.699 | -0.142 |
| RI | 5803 | ENSG00000136699 | SMPD4 | chr2 | - | 130156034 | 130156675 | 130156034 | 130156135 | 130156584 | 130156675 | 5803 | 15 | 5 | 22 | 12 | 10 | 39 | 9 | 27 | 30 | 29 | 49 | 23 | 8 | 40 | 28 | 28 | 36 | 18 | 21 | 25 | 298 | 149 | 0.001685799 | 0.009394352 | 0.161 | 0.217 | 0.289 | 0.167 | 0.147 | 0.467 | 0.242 | 0.182 | 0.488 | 0.359 | -0.151 |
| RI | 4635 | ENSG00000129084 | PSMA1 | chr11 | - | 14513816 | 14514491 | 14513816 | 14513887 | 14514402 | 14514491 | 4635 | 233 | 168 | 143 | 205 | 415 | 356 | 185 | 288 | 292 | 314 | 48 | 133 | 175 | 114 | 34 | 258 | 233 | 230 | 216 | 52 | 298 | 149 | 0.002509661 | 0.013177393 | 0.247 | 0.312 | 0.199 | 0.26 | 0.398 | 0.085 | 0.222 | 0.276 | 0.209 | 0.246 | 0.076 |
| RI | 2780 | ENSG00000178950 | GAK | chr4 | - | 850935 | 851974 | 850935 | 851084 | 851749 | 851974 | 2780 | 26 | 25 | 51 | 29 | 21 | 118 | 67 | 71 | 112 | 40 | 104 | 49 | 53 | 82 | 106 | 153 | 142 | 94 | 102 | 162 | 298 | 149 | 0.002954819 | 0.01511179 | 0.099 | 0.157 | 0.264 | 0.115 | 0.208 | 0.254 | 0.147 | 0.22 | 0.287 | 0.247 | -0.062 |
| RI | 1902 | ENSG00000090061 | CCNK | chr14 | + | 99502718 | 99503644 | 99502718 | 99502984 | 99503610 | 99503644 | 1902 | 29 | 14 | 19 | 29 | 32 | 18 | 17 | 14 | 21 | 26 | 12 | 62 | 39 | 55 | 33 | 61 | 46 | 37 | 58 | 136 | 298 | 149 | 0.003498498 | 0.017395311 | 0.446 | 0.292 | 0.404 | 0.408 | 0.381 | 0.09 | 0.403 | 0.345 | 0.322 | 0.108 | 0.133 |
| RI | 5435 | ENSG00000197858 | GPAA1 | chr8 | + | 144083388 | 144083861 | 144083388 | 144083500 | 144083817 | 144083861 | 5435 | 55 | 23 | 53 | 28 | 19 | 0 | 0 | 0 | 0 | 0 | 54 | 40 | 26 | 26 | 70 | 11 | 0 | 0 | 0 | 0 | 298 | 149 | 0.0053919 | 0.025138084 | 1 | 1 | 1 | 1 | 1 | 0.711 | 1 | 1 | 1 | 1 | 0.058 |
| RI | 8176 | ENSG00000119977 | TCTN3 | chr10 | - | 95683100 | 95683629 | 95683100 | 95683195 | 95683521 | 95683629 | 8176 | 5 | 5 | 14 | 0 | 10 | 34 | 15 | 33 | 33 | 57 | 0 | 13 | 0 | 0 | 0 | 65 | 34 | 18 | 11 | 14 | 298 | 149 | 0.007031508 | 0.031370934 | 0.068 | 0.143 | 0.175 | 0 | 0.081 | 0 | 0.16 | 0 | 0 | 0 | 0.061 |
| RI | 136 | ENSG00000100325 | ASCC2 | chr22 | - | 29793359 | 29793676 | 29793359 | 29793490 | 29793576 | 29793676 | 136 | 9 | 9 | 9 | 17 | 4 | 138 | 79 | 86 | 124 | 106 | 49 | 10 | 8 | 18 | 20 | 150 | 119 | 88 | 158 | 182 | 235 | 149 | 0.007807408 | 0.034442391 | 0.04 | 0.067 | 0.062 | 0.08 | 0.023 | 0.172 | 0.051 | 0.054 | 0.067 | 0.065 | -0.027 |
| RI | 4125 | ENSG00000215305 | VPS16 | chr20 | + | 2863948 | 2864287 | 2863948 | 2864083 | 2864178 | 2864287 | 4125 | 23 | 15 | 27 | 36 | 10 | 63 | 24 | 51 | 85 | 11 | 3 | 40 | 15 | 44 | 57 | 149 | 76 | 83 | 125 | 105 | 244 | 149 | 0.007839363 | 0.03455755 | 0.182 | 0.276 | 0.244 | 0.205 | 0.357 | 0.012 | 0.243 | 0.099 | 0.177 | 0.249 | 0.097 |
| RI | 3126 | ENSG00000130787 | HIP1R | chr12 | + | 122860046 | 122860210 | 122860046 | 122860077 | 122860147 | 122860210 | 3126 | 8 | 1 | 8 | 15 | 10 | 31 | 5 | 19 | 7 | 21 | 24 | 33 | 14 | 15 | 54 | 28 | 19 | 26 | 1 | 22 | 219 | 149 | 0.008295106 | 0.036242004 | 0.149 | 0.12 | 0.223 | 0.593 | 0.245 | 0.368 | 0.542 | 0.268 | 0.911 | 0.625 | -0.277 |
| RI | 4534 | ENSG00000143258 | USP21 | chr1 | + | 161164533 | 161164942 | 161164533 | 161164612 | 161164834 | 161164942 | 4534 | 35 | 19 | 40 | 42 | 47 | 32 | 28 | 27 | 35 | 87 | 44 | 47 | 40 | 24 | 112 | 27 | 34 | 15 | 20 | 77 | 298 | 149 | 0.008837727 | 0.038301138 | 0.354 | 0.253 | 0.426 | 0.375 | 0.213 | 0.449 | 0.409 | 0.571 | 0.375 | 0.421 | -0.121 |
| RI | 2405 | ENSG00000105875 | WDR91 | chr7 | - | 135186971 | 135189452 | 135186971 | 135188545 | 135189343 | 135189452 | 2405 | 14 | 7 | 7 | 2 | 1 | 15 | 13 | 26 | 25 | 11 | 13 | 21 | 4 | 23 | 38 | 10 | 31 | 33 | 21 | 33 | 298 | 149 | 0.008936151 | 0.038589391 | 0.318 | 0.212 | 0.119 | 0.038 | 0.043 | 0.394 | 0.253 | 0.057 | 0.354 | 0.365 | -0.139 |
| RI | 5540 | ENSG00000144635 | DYNC1LI1 | chr3 | - | 32570345 | 32570858 | 32570345 | 32570419 | 32570624 | 32570858 | 5540 | 2 | 3 | 2 | 7 | 0 | 136 | 94 | 108 | 124 | 61 | 10 | 6 | 9 | 10 | 7 | 227 | 172 | 145 | 186 | 108 | 298 | 149 | 0.009030722 | 0.038937574 | 0.007 | 0.016 | 0.009 | 0.027 | 0 | 0.022 | 0.017 | 0.03 | 0.026 | 0.031 | -0.013 |
| RI | 7170 | ENSG00000014216 | CAPN1 | chr11 | + | 65209848 | 65210096 | 65209848 | 65209917 | 65210017 | 65210096 | 7170 | 5 | 7 | 4 | 23 | 1 | 103 | 45 | 88 | 112 | 78 | 0 | 15 | 0 | 5 | 14 | 129 | 135 | 54 | 55 | 350 | 249 | 149 | 0.009347952 | 0.040013298 | 0.028 | 0.085 | 0.026 | 0.109 | 0.008 | 0 | 0.062 | 0 | 0.052 | 0.023 | 0.024 |
| RI | 1603 | ENSG00000140199 | SLC12A6 | chr15 | - | 34336409 | 34338057 | 34336409 | 34336686 | 34337960 | 34338057 | 1603 | 113 | 114 | 138 | 126 | 248 | 4 | 1 | 2 | 4 | 4 | 299 | 142 | 255 | 221 | 162 | 15 | 5 | 12 | 9 | 0 | 298 | 149 | 0.010226493 | 0.04317934 | 0.934 | 0.983 | 0.972 | 0.94 | 0.969 | 0.909 | 0.934 | 0.914 | 0.925 | 1 | 0.023 |
| RI | 8448 | ENSG00000123349 | PFDN5 | chr12 | + | 53295838 | 53296275 | 53295838 | 53295941 | 53296218 | 53296275 | 8448 | 212 | 165 | 266 | 260 | 310 | 2 | 0 | 3 | 2 | 4 | 424 | 288 | 230 | 258 | 299 | 0 | 0 | 4 | 4 | 20 | 298 | 149 | 0.0107447 | 0.044949675 | 0.981 | 1 | 0.978 | 0.985 | 0.975 | 1 | 1 | 0.966 | 0.97 | 0.882 | 0.02 |
| RI | 1763 | ENSG00000100949 | RABGGTA | chr14 | - | 24270836 | 24271169 | 24270836 | 24270947 | 24271112 | 24271169 | 1763 | 7 | 3 | 10 | 14 | 1 | 20 | 4 | 7 | 19 | 1 | 0 | 15 | 1 | 2 | 4 | 25 | 19 | 14 | 3 | 12 | 298 | 149 | 0.011289502 | 0.047095403 | 0.149 | 0.273 | 0.417 | 0.269 | 0.333 | 0 | 0.283 | 0.034 | 0.25 | 0.143 | 0.146 |
| RI | 4181 | ENSG00000087095 | NLK | chr17 | + | 28194581 | 28196381 | 28194581 | 28194708 | 28196179 | 28196381 | 4181 | 61 | 77 | 76 | 76 | 211 | 2 | 2 | 2 | 2 | 0 | 60 | 48 | 62 | 51 | 10 | 0 | 2 | 2 | 2 | 7 | 298 | 149 | 0.015285702 | 0.06134011 | 0.938 | 0.951 | 0.95 | 0.95 | 1 | 1 | 0.923 | 0.939 | 0.927 | 0.417 | 0.117 |
| RI | 8459 | ENSG00000123349 | PFDN5 | chr12 | + | 53297849 | 53298150 | 53297849 | 53297924 | 53298048 | 53298150 | 8459 | 112 | 79 | 101 | 117 | 103 | 9 | 6 | 2 | 6 | 13 | 230 | 124 | 106 | 111 | 130 | 0 | 3 | 5 | 4 | 13 | 273 | 149 | 0.015661841 | 0.062679197 | 0.872 | 0.878 | 0.965 | 0.914 | 0.812 | 1 | 0.958 | 0.92 | 0.938 | 0.845 | -0.044 |
| RI | 5337 | ENSG00000084623 | EIF3I | chr1 | + | 32224409 | 32226530 | 32224409 | 32224475 | 32226170 | 32226530 | 5337 | 43 | 30 | 29 | 56 | 29 | 168 | 96 | 189 | 173 | 102 | 68 | 63 | 31 | 78 | 89 | 189 | 191 | 125 | 124 | 328 | 298 | 149 | 0.016781525 | 0.066262346 | 0.113 | 0.135 | 0.071 | 0.139 | 0.124 | 0.152 | 0.142 | 0.11 | 0.239 | 0.119 | -0.036 |
| RI | 601 | ENSG00000171703 | TCEA2 | chr20 | + | 64069764 | 64070414 | 64069764 | 64069821 | 64070259 | 64070414 | 601 | 3 | 10 | 6 | 12 | 0 | 19 | 2 | 20 | 13 | 4 | 8 | 8 | 7 | 32 | 20 | 0 | 11 | 10 | 16 | 6 | 298 | 149 | 0.017808203 | 0.069434361 | 0.073 | 0.714 | 0.13 | 0.316 | 0 | 1 | 0.267 | 0.259 | 0.5 | 0.625 | -0.284 |
| RI | 1734 | ENSG00000185024 | BRF1 | chr14 | - | 105219150 | 105220130 | 105219150 | 105219232 | 105220068 | 105220130 | 1734 | 13 | 5 | 3 | 11 | 0 | 20 | 3 | 16 | 17 | 4 | 0 | 6 | 2 | 8 | 9 | 18 | 17 | 19 | 39 | 39 | 298 | 149 | 0.018788091 | 0.072442071 | 0.245 | 0.455 | 0.086 | 0.244 | 0 | 0 | 0.15 | 0.05 | 0.093 | 0.103 | 0.127 |
| RI | 5431 | ENSG00000214517 | PPME1 | chr11 | + | 74250953 | 74251715 | 74250953 | 74251018 | 74251647 | 74251715 | 5431 | 2 | 3 | 4 | 5 | 6 | 42 | 12 | 26 | 23 | 70 | 21 | 0 | 3 | 16 | 0 | 22 | 29 | 8 | 5 | 12 | 298 | 149 | 0.019402387 | 0.074567273 | 0.023 | 0.111 | 0.071 | 0.098 | 0.041 | 0.323 | 0 | 0.158 | 0.615 | 0 | -0.15 |
| RI | 5509 | ENSG00000112367 | FIG4 | chr6 | + | 109735149 | 109738453 | 109735149 | 109735298 | 109738324 | 109738453 | 5509 | 54 | 31 | 29 | 25 | 39 | 65 | 29 | 46 | 64 | 91 | 0 | 11 | 26 | 0 | 7 | 75 | 40 | 5 | 93 | 4 | 298 | 149 | 0.020187548 | 0.077033492 | 0.293 | 0.348 | 0.24 | 0.163 | 0.176 | 0 | 0.121 | 0.722 | 0 | 0.467 | -0.018 |
| RI | 7749 | ENSG00000175550 | DRAP1 | chr11 | + | 65919779 | 65920041 | 65919779 | 65919852 | 65919947 | 65920041 | 7749 | 32 | 12 | 17 | 17 | 10 | 195 | 140 | 165 | 239 | 104 | 0 | 11 | 15 | 21 | 16 | 124 | 169 | 95 | 256 | 269 | 244 | 149 | 0.020877667 | 0.07915557 | 0.091 | 0.05 | 0.059 | 0.042 | 0.055 | 0 | 0.038 | 0.088 | 0.048 | 0.035 | 0.018 |
| RI | 2948 | ENSG00000164068 | RNF123 | chr3 | + | 49705533 | 49706065 | 49705533 | 49705679 | 49705981 | 49706065 | 2948 | 8 | 3 | 4 | 10 | 1 | 17 | 10 | 25 | 25 | 6 | 16 | 10 | 6 | 12 | 9 | 4 | 22 | 5 | 16 | 44 | 298 | 149 | 0.021345174 | 0.080566098 | 0.19 | 0.13 | 0.074 | 0.167 | 0.077 | 0.667 | 0.185 | 0.375 | 0.273 | 0.093 | -0.191 |
| RI | 3331 | ENSG00000117228 | GBP1 | chr1 | - | 89055112 | 89056228 | 89055112 | 89055215 | 89056015 | 89056228 | 3331 | 51 | 48 | 34 | 55 | 19 | 187 | 293 | 217 | 265 | 266 | 31 | 8 | 53 | 45 | 5 | 289 | 245 | 144 | 449 | 41 | 298 | 149 | 0.028276183 | 0.102032628 | 0.12 | 0.076 | 0.073 | 0.094 | 0.034 | 0.051 | 0.016 | 0.155 | 0.048 | 0.057 | 0.014 |
| RI | 6834 | ENSG00000177042 | TMEM80 | chr11 | + | 698868 | 700235 | 698868 | 698888 | 700141 | 700235 | 6834 | 3 | 2 | 12 | 4 | 3 | 17 | 10 | 17 | 8 | 11 | 0 | 14 | 5 | 13 | 28 | 6 | 14 | 6 | 20 | 24 | 298 | 149 | 0.029331133 | 0.105005456 | 0.081 | 0.091 | 0.261 | 0.2 | 0.12 | 0 | 0.333 | 0.294 | 0.245 | 0.368 | -0.097 |
| RI | 2949 | ENSG00000164068 | RNF123 | chr3 | + | 49712478 | 49713587 | 49712478 | 49712656 | 49713512 | 49713587 | 2949 | 12 | 2 | 7 | 4 | 3 | 9 | 9 | 11 | 22 | 11 | 28 | 14 | 2 | 6 | 15 | 3 | 2 | 17 | 6 | 25 | 298 | 149 | 0.031160648 | 0.11055012 | 0.4 | 0.1 | 0.241 | 0.083 | 0.12 | 0.824 | 0.778 | 0.056 | 0.333 | 0.231 | -0.256 |
| RI | 7379 | ENSG00000198000 | NOL8 | chr9 | - | 92298256 | 92298954 | 92298256 | 92298336 | 92298883 | 92298954 | 7379 | 73 | 68 | 85 | 57 | 96 | 145 | 153 | 115 | 127 | 341 | 10 | 27 | 70 | 11 | 8 | 0 | 34 | 75 | 6 | 23 | 298 | 149 | 0.031818556 | 0.112478881 | 0.201 | 0.182 | 0.27 | 0.183 | 0.123 | 1 | 0.284 | 0.318 | 0.478 | 0.148 | -0.254 |
| RI | 7195 | ENSG00000174744 | BRMS1 | chr11 | - | 66340773 | 66341333 | 66340773 | 66340870 | 66341205 | 66341333 | 7195 | 12 | 7 | 17 | 16 | 12 | 0 | 0 | 0 | 0 | 0 | 11 | 26 | 4 | 15 | 34 | 10 | 0 | 0 | 0 | 0 | 298 | 149 | 0.034763137 | 0.121424469 | 1 | 1 | 1 | 1 | 1 | 0.355 | 1 | 1 | 1 | 1 | 0.129 |
| RI | 1613 | ENSG00000167077 | MEI1 | chr22 | + | 41793828 | 41794477 | 41793828 | 41793910 | 41794370 | 41794477 | 1613 | 41 | 13 | 33 | 39 | 37 | 13 | 2 | 18 | 13 | 6 | 42 | 39 | 48 | 33 | 62 | 0 | 18 | 4 | 2 | 18 | 298 | 149 | 0.035201404 | 0.122675334 | 0.612 | 0.765 | 0.478 | 0.6 | 0.755 | 1 | 0.52 | 0.857 | 0.892 | 0.633 | -0.138 |
| RI | 7937 | ENSG00000187147 | RNF220 | chr1 | + | 44649882 | 44651723 | 44649882 | 44649957 | 44650703 | 44651723 | 7937 | 10 | 5 | 21 | 13 | 7 | 231 | 106 | 214 | 212 | 209 | 14 | 8 | 12 | 21 | 29 | 255 | 183 | 143 | 103 | 321 | 298 | 149 | 0.035477119 | 0.123345112 | 0.021 | 0.023 | 0.047 | 0.03 | 0.016 | 0.027 | 0.021 | 0.04 | 0.093 | 0.043 | -0.017 |
| RI | 2158 | ENSG00000111641 | NOP2 | chr12 | - | 6560097 | 6560569 | 6560097 | 6560326 | 6560446 | 6560569 | 2158 | 30 | 17 | 41 | 33 | 23 | 24 | 29 | 39 | 53 | 70 | 33 | 39 | 26 | 38 | 45 | 30 | 28 | 27 | 37 | 51 | 269 | 149 | 0.036805899 | 0.126845068 | 0.409 | 0.245 | 0.368 | 0.256 | 0.154 | 0.379 | 0.436 | 0.348 | 0.363 | 0.328 | -0.084 |
| RI | 4099 | ENSG00000256269 | HMBS | chr11 | + | 119089682 | 119090067 | 119089682 | 119089760 | 119089989 | 119090067 | 4099 | 0 | 1 | 2 | 5 | 2 | 28 | 9 | 20 | 13 | 26 | 14 | 0 | 7 | 0 | 0 | 26 | 15 | 8 | 13 | 14 | 298 | 149 | 0.037024372 | 0.127507838 | 0 | 0.053 | 0.048 | 0.161 | 0.037 | 0.212 | 0 | 0.304 | 0 | 0 | -0.043 |
| RI | 4100 | ENSG00000256269 | HMBS | chr11 | + | 119092124 | 119092523 | 119092124 | 119092163 | 119092403 | 119092523 | 4100 | 4 | 7 | 2 | 5 | 7 | 6 | 1 | 9 | 8 | 5 | 1 | 5 | 4 | 1 | 4 | 15 | 10 | 3 | 8 | 10 | 298 | 149 | 0.038072294 | 0.129770942 | 0.25 | 0.778 | 0.1 | 0.238 | 0.412 | 0.032 | 0.2 | 0.4 | 0.059 | 0.167 | 0.184 |
| RI | 5346 | ENSG00000084623 | EIF3I | chr1 | + | 32230926 | 32231602 | 32230926 | 32231019 | 32231259 | 32231602 | 5346 | 103 | 67 | 124 | 113 | 104 | 0 | 0 | 0 | 0 | 0 | 120 | 165 | 95 | 89 | 181 | 0 | 2 | 0 | 0 | 11 | 298 | 149 | 0.038475097 | 0.130883808 | 1 | 1 | 1 | 1 | 1 | 1 | 0.976 | 1 | 1 | 0.892 | 0.026 |
| RI | 5343 | ENSG00000084623 | EIF3I | chr1 | + | 32228726 | 32231019 | 32228726 | 32228816 | 32230926 | 32231019 | 5343 | 21 | 16 | 19 | 25 | 19 | 0 | 0 | 0 | 6 | 3 | 110 | 72 | 39 | 95 | 89 | 13 | 5 | 5 | 11 | 21 | 298 | 149 | 0.045585876 | 0.149680805 | 1 | 1 | 1 | 0.676 | 0.76 | 0.809 | 0.878 | 0.796 | 0.812 | 0.679 | 0.092 |
| RI | 7877 | ENSG00000167986 | DDB1 | chr11 | - | 61302256 | 61302751 | 61302256 | 61302359 | 61302581 | 61302751 | 7877 | 14 | 12 | 10 | 20 | 8 | 227 | 112 | 198 | 172 | 224 | 15 | 21 | 11 | 4 | 38 | 318 | 153 | 104 | 106 | 231 | 298 | 149 | 0.048711201 | 0.157664145 | 0.03 | 0.051 | 0.025 | 0.055 | 0.018 | 0.023 | 0.064 | 0.05 | 0.019 | 0.076 | -0.011 |
| RI | 7875 | ENSG00000166471 | TMEM41B | chr11 | - | 9280653 | 9283593 | 9280653 | 9281041 | 9283370 | 9283593 | 7875 | 286 | 343 | 359 | 366 | 154 | 12 | 6 | 5 | 12 | 16 | 327 | 223 | 374 | 339 | 112 | 0 | 36 | 25 | 24 | 0 | 298 | 149 | 0.049307851 | 0.159507926 | 0.923 | 0.966 | 0.973 | 0.938 | 0.828 | 1 | 0.756 | 0.882 | 0.876 | 1 | 0.023 |
| RI | 6539 | ENSG00000104915 | STX10 | chr19 | - | 13144061 | 13144671 | 13144061 | 13144486 | 13144576 | 13144671 | 6539 | 24 | 20 | 31 | 21 | 32 | 89 | 35 | 65 | 58 | 53 | 71 | 32 | 51 | 73 | 149 | 173 | 111 | 45 | 68 | 207 | 239 | 149 | 0.050613762 | 0.162752037 | 0.144 | 0.263 | 0.229 | 0.184 | 0.273 | 0.204 | 0.152 | 0.414 | 0.401 | 0.31 | -0.078 |
| RI | 1733 | ENSG00000185024 | BRF1 | chr14 | - | 105211121 | 105212164 | 105211121 | 105211293 | 105212112 | 105212164 | 1733 | 20 | 17 | 14 | 21 | 2 | 56 | 14 | 27 | 44 | 29 | 58 | 31 | 22 | 9 | 57 | 32 | 41 | 31 | 31 | 100 | 298 | 149 | 0.053918163 | 0.170957911 | 0.152 | 0.378 | 0.206 | 0.193 | 0.033 | 0.475 | 0.274 | 0.262 | 0.127 | 0.222 | -0.08 |
| RI | 792 | ENSG00000214087 | ARL16 | chr17 | - | 81682033 | 81683594 | 81682033 | 81683126 | 81683535 | 81683594 | 792 | 2 | 0 | 2 | 2 | 1 | 9 | 5 | 8 | 6 | 2 | 8 | 22 | 15 | 0 | 0 | 0 | 20 | 13 | 5 | 31 | 298 | 149 | 0.055142869 | 0.174093493 | 0.1 | 0 | 0.111 | 0.143 | 0.2 | 1 | 0.355 | 0.366 | 0 | 0 | -0.233 |
| RI | 4237 | ENSG00000133392 | MYH11 | chr16 | - | 15720838 | 15721634 | 15720838 | 15721051 | 15721421 | 15721634 | 4237 | 13 | 8 | 14 | 7 | 4 | 0 | 0 | 2 | 0 | 0 | 10 | 16 | 23 | 5 | 74 | 0 | 0 | 0 | 10 | 0 | 298 | 149 | 0.055780714 | 0.175731561 | 1 | 1 | 0.778 | 1 | 1 | 1 | 1 | 1 | 0.2 | 1 | 0.116 |
| RI | 2463 | ENSG00000187091 | PLCD1 | chr3 | - | 38009271 | 38009811 | 38009271 | 38009431 | 38009652 | 38009811 | 2463 | 15 | 5 | 22 | 15 | 9 | 24 | 4 | 13 | 19 | 29 | 12 | 6 | 4 | 0 | 14 | 7 | 23 | 19 | 10 | 26 | 298 | 149 | 0.055842349 | 0.17583196 | 0.238 | 0.385 | 0.458 | 0.283 | 0.134 | 0.462 | 0.115 | 0.095 | 0 | 0.212 | 0.123 |
| RI | 2414 | ENSG00000158290 | CUL4B | chrX | - | 120538123 | 120538770 | 120538123 | 120538209 | 120538659 | 120538770 | 2414 | 96 | 114 | 108 | 133 | 106 | 196 | 156 | 157 | 193 | 341 | 48 | 39 | 73 | 8 | 0 | 33 | 129 | 158 | 65 | 16 | 298 | 149 | 0.05761937 | 0.180274163 | 0.197 | 0.268 | 0.256 | 0.256 | 0.135 | 0.421 | 0.131 | 0.188 | 0.058 | 0 | 0.063 |
| RI | 2677 | ENSG00000177054 | ZDHHC13 | chr11 | + | 19166301 | 19170568 | 19166301 | 19166385 | 19170410 | 19170568 | 2677 | 11 | 18 | 10 | 22 | 9 | 28 | 19 | 24 | 28 | 30 | 3 | 8 | 4 | 0 | 0 | 2 | 11 | 15 | 6 | 9 | 298 | 149 | 0.057989684 | 0.181049189 | 0.164 | 0.321 | 0.172 | 0.282 | 0.13 | 0.429 | 0.267 | 0.118 | 0 | 0 | 0.051 |
| RI | 4876 | ENSG00000154451 | GBP5 | chr1 | - | 89262219 | 89263948 | 89262219 | 89262401 | 89263735 | 89263948 | 4876 | 71 | 89 | 125 | 98 | 108 | 4 | 4 | 5 | 10 | 15 | 40 | 154 | 81 | 59 | 18 | 3 | 5 | 9 | 0 | 0 | 298 | 149 | 0.066450173 | 0.201810372 | 0.899 | 0.918 | 0.926 | 0.831 | 0.783 | 0.87 | 0.939 | 0.818 | 1 | 1 | -0.054 |
| RI | 1132 | ENSG00000072849 | DERL2 | chr17 | - | 5481100 | 5481389 | 5481100 | 5481142 | 5481295 | 5481389 | 1132 | 74 | 63 | 91 | 75 | 87 | 3 | 2 | 2 | 5 | 4 | 46 | 97 | 172 | 123 | 32 | 0 | 7 | 0 | 3 | 0 | 298 | 149 | 0.068213572 | 0.206000802 | 0.925 | 0.94 | 0.958 | 0.882 | 0.916 | 1 | 0.874 | 1 | 0.953 | 1 | -0.041 |
| RI | 6903 | ENSG00000090316 | MAEA | chr4 | + | 1336860 | 1338617 | 1336860 | 1336994 | 1338425 | 1338617 | 6903 | 20 | 12 | 16 | 17 | 10 | 1 | 0 | 2 | 0 | 0 | 8 | 17 | 42 | 9 | 6 | 0 | 9 | 0 | 0 | 0 | 298 | 149 | 0.0691378 | 0.208153407 | 0.909 | 1 | 0.8 | 1 | 1 | 1 | 0.486 | 1 | 1 | 1 | 0.045 |
| RI | 7780 | ENSG00000110274 | CEP164 | chr11 | + | 117394919 | 117395191 | 117394919 | 117395003 | 117395122 | 117395191 | 7780 | 4 | 0 | 5 | 5 | 2 | 32 | 14 | 30 | 34 | 23 | 4 | 7 | 7 | 22 | 3 | 77 | 40 | 22 | 21 | 75 | 268 | 149 | 0.07037787 | 0.209960644 | 0.065 | 0 | 0.085 | 0.076 | 0.046 | 0.028 | 0.089 | 0.15 | 0.368 | 0.022 | -0.077 |
| RI | 8433 | ENSG00000033030 | ZCCHC8 | chr12 | - | 122483278 | 122483563 | 122483278 | 122483344 | 122483459 | 122483563 | 8433 | 26 | 52 | 48 | 75 | 22 | 58 | 43 | 42 | 49 | 97 | 111 | 80 | 48 | 160 | 0 | 30 | 41 | 24 | 13 | 26 | 264 | 149 | 0.077320655 | 0.225102568 | 0.202 | 0.406 | 0.392 | 0.463 | 0.113 | 0.676 | 0.524 | 0.53 | 0.874 | 0 | -0.206 |
| RI | 3330 | ENSG00000117228 | GBP1 | chr1 | - | 89054681 | 89055215 | 89054681 | 89054875 | 89055112 | 89055215 | 3330 | 26 | 45 | 14 | 28 | 41 | 338 | 506 | 381 | 415 | 559 | 11 | 25 | 54 | 57 | 8 | 436 | 346 | 238 | 731 | 34 | 298 | 149 | 0.079686518 | 0.230626293 | 0.037 | 0.043 | 0.018 | 0.033 | 0.035 | 0.012 | 0.035 | 0.102 | 0.038 | 0.105 | -0.025 |
| RI | 7193 | ENSG00000174744 | BRMS1 | chr11 | - | 66338242 | 66338785 | 66338242 | 66338282 | 66338720 | 66338785 | 7193 | 4 | 2 | 12 | 2 | 10 | 81 | 20 | 76 | 56 | 62 | 3 | 13 | 8 | 8 | 6 | 2 | 85 | 37 | 34 | 73 | 298 | 149 | 0.079733586 | 0.230649506 | 0.024 | 0.048 | 0.073 | 0.018 | 0.075 | 0.429 | 0.071 | 0.098 | 0.105 | 0.039 | -0.101 |
| RI | 4921 | ENSG00000130713 | EXOSC2 | chr9 | + | 130697581 | 130698251 | 130697581 | 130697627 | 130698161 | 130698251 | 4921 | 31 | 15 | 34 | 22 | 8 | 89 | 38 | 99 | 83 | 88 | 39 | 42 | 10 | 37 | 49 | 170 | 81 | 59 | 64 | 113 | 298 | 149 | 0.084952789 | 0.242892607 | 0.148 | 0.165 | 0.147 | 0.117 | 0.043 | 0.103 | 0.206 | 0.078 | 0.224 | 0.178 | -0.034 |
| RI | 5802 | ENSG00000136699 | SMPD4 | chr2 | - | 130154276 | 130155259 | 130154276 | 130154482 | 130155095 | 130155259 | 5802 | 10 | 2 | 14 | 8 | 6 | 38 | 11 | 40 | 39 | 30 | 0 | 11 | 14 | 18 | 38 | 59 | 52 | 20 | 6 | 29 | 298 | 149 | 0.086035047 | 0.245156307 | 0.116 | 0.083 | 0.149 | 0.093 | 0.091 | 0 | 0.096 | 0.259 | 0.6 | 0.396 | -0.164 |
| RI | 2751 | ENSG00000099949 | LZTR1 | chr22 | + | 20994103 | 20994727 | 20994103 | 20994269 | 20994557 | 20994727 | 2751 | 13 | 9 | 15 | 23 | 5 | 13 | 7 | 10 | 5 | 0 | 0 | 11 | 0 | 3 | 19 | 4 | 5 | 2 | 0 | 21 | 298 | 149 | 0.089210509 | 0.252130266 | 0.333 | 0.391 | 0.429 | 0.697 | 1 | 0 | 0.524 | 0 | 1 | 0.311 | 0.203 |
| RI | 8126 | ENSG00000148688 | RPP30 | chr10 | + | 90894774 | 90895483 | 90894774 | 90894891 | 90895453 | 90895483 | 8126 | 54 | 54 | 53 | 65 | 53 | 48 | 9 | 61 | 26 | 47 | 28 | 33 | 76 | 16 | 0 | 24 | 29 | 17 | 24 | 30 | 298 | 149 | 0.091383723 | 0.25753991 | 0.36 | 0.75 | 0.303 | 0.556 | 0.361 | 0.368 | 0.363 | 0.691 | 0.25 | 0 | 0.132 |
| RI | 4600 | ENSG00000185946 | RNPC3 | chr1 | + | 103525698 | 103526262 | 103525698 | 103525930 | 103526066 | 103526262 | 4600 | 114 | 65 | 92 | 104 | 86 | 5 | 5 | 5 | 2 | 1 | 101 | 63 | 70 | 36 | 15 | 0 | 5 | 0 | 7 | 16 | 285 | 149 | 0.09173208 | 0.258335395 | 0.923 | 0.872 | 0.906 | 0.965 | 0.978 | 1 | 0.868 | 1 | 0.729 | 0.329 | 0.144 |
| RI | 4835 | ENSG00000103168 | TAF1C | chr16 | - | 84181745 | 84182058 | 84181745 | 84181863 | 84181941 | 84182058 | 4835 | 13 | 10 | 23 | 27 | 5 | 5 | 8 | 13 | 16 | 0 | 61 | 21 | 30 | 52 | 76 | 1 | 15 | 8 | 14 | 16 | 227 | 149 | 0.092622295 | 0.260657407 | 0.631 | 0.451 | 0.537 | 0.526 | 1 | 0.976 | 0.479 | 0.711 | 0.709 | 0.757 | -0.097 |
| RI | 3469 | ENSG00000011485 | PPP5C | chr19 | + | 46387365 | 46388448 | 46387365 | 46387453 | 46388407 | 46388448 | 3469 | 13 | 7 | 2 | 7 | 7 | 161 | 102 | 142 | 167 | 122 | 7 | 15 | 1 | 1 | 0 | 158 | 151 | 59 | 115 | 217 | 298 | 149 | 0.101070847 | 0.280688995 | 0.039 | 0.033 | 0.007 | 0.021 | 0.028 | 0.022 | 0.047 | 0.008 | 0.004 | 0 | 0.009 |
| RI | 7879 | ENSG00000167986 | DDB1 | chr11 | - | 61314046 | 61314486 | 61314046 | 61314210 | 61314307 | 61314486 | 7879 | 16 | 5 | 21 | 9 | 8 | 303 | 135 | 258 | 232 | 380 | 18 | 13 | 17 | 9 | 12 | 287 | 208 | 136 | 130 | 369 | 246 | 149 | 0.105728339 | 0.290888355 | 0.031 | 0.022 | 0.047 | 0.023 | 0.013 | 0.037 | 0.036 | 0.07 | 0.04 | 0.019 | -0.013 |
| RI | 1762 | ENSG00000100949 | RABGGTA | chr14 | - | 24268513 | 24268826 | 24268513 | 24268619 | 24268724 | 24268826 | 1762 | 7 | 15 | 6 | 12 | 6 | 26 | 10 | 18 | 20 | 15 | 30 | 3 | 7 | 5 | 13 | 36 | 28 | 23 | 27 | 64 | 254 | 149 | 0.10872863 | 0.297067538 | 0.136 | 0.468 | 0.164 | 0.26 | 0.19 | 0.328 | 0.059 | 0.151 | 0.098 | 0.106 | 0.095 |
| RI | 6112 | ENSG00000223501 | VPS52 | chr6 | - | 33264373 | 33264900 | 33264373 | 33264497 | 33264781 | 33264900 | 6112 | 62 | 43 | 50 | 66 | 75 | 36 | 9 | 30 | 22 | 47 | 122 | 85 | 76 | 97 | 126 | 56 | 25 | 19 | 27 | 42 | 298 | 149 | 0.111841391 | 0.302771356 | 0.463 | 0.705 | 0.455 | 0.6 | 0.444 | 0.521 | 0.63 | 0.667 | 0.642 | 0.6 | -0.079 |
| RI | 6249 | ENSG00000171777 | RASGRP4 | chr19 | - | 38419859 | 38420262 | 38419859 | 38420013 | 38420130 | 38420262 | 6249 | 7 | 7 | 10 | 7 | 6 | 31 | 4 | 24 | 16 | 18 | 27 | 7 | 22 | 23 | 86 | 35 | 48 | 27 | 26 | 85 | 266 | 149 | 0.112933548 | 0.305104529 | 0.112 | 0.495 | 0.189 | 0.197 | 0.157 | 0.302 | 0.076 | 0.313 | 0.331 | 0.362 | -0.047 |
| RI | 1481 | ENSG00000167965 | MLST8 | chr16 | + | 2206357 | 2206659 | 2206357 | 2206409 | 2206496 | 2206659 | 1481 | 7 | 7 | 2 | 7 | 8 | 6 | 4 | 7 | 8 | 5 | 16 | 3 | 3 | 6 | 36 | 12 | 3 | 8 | 2 | 0 | 236 | 149 | 0.126080959 | 0.33233388 | 0.424 | 0.525 | 0.153 | 0.356 | 0.503 | 0.457 | 0.387 | 0.191 | 0.654 | 1 | -0.146 |
| RI | 2734 | ENSG00000109133 | TMEM33 | chr4 | + | 41935128 | 41935529 | 41935128 | 41935310 | 41935466 | 41935529 | 2734 | 104 | 78 | 94 | 93 | 31 | 2 | 2 | 5 | 0 | 0 | 82 | 60 | 59 | 50 | 78 | 0 | 1 | 11 | 2 | 0 | 298 | 149 | 0.126608586 | 0.333278483 | 0.963 | 0.951 | 0.904 | 1 | 1 | 1 | 0.968 | 0.728 | 0.926 | 1 | 0.039 |
| RI | 3127 | ENSG00000130787 | HIP1R | chr12 | + | 122860422 | 122860784 | 122860422 | 122860523 | 122860678 | 122860784 | 3127 | 1 | 2 | 8 | 9 | 0 | 35 | 14 | 28 | 25 | 34 | 0 | 2 | 13 | 3 | 15 | 8 | 67 | 34 | 10 | 25 | 298 | 149 | 0.127877334 | 0.33601931 | 0.014 | 0.067 | 0.125 | 0.153 | 0 | 0 | 0.015 | 0.16 | 0.13 | 0.231 | -0.035 |
| RI | 1760 | ENSG00000100949 | RABGGTA | chr14 | - | 24266429 | 24266889 | 24266429 | 24266517 | 24266775 | 24266889 | 1760 | 12 | 4 | 7 | 6 | 4 | 20 | 11 | 27 | 31 | 31 | 52 | 5 | 18 | 0 | 20 | 42 | 46 | 17 | 15 | 50 | 298 | 149 | 0.140548471 | 0.362699789 | 0.231 | 0.154 | 0.115 | 0.088 | 0.061 | 0.382 | 0.052 | 0.346 | 0 | 0.167 | -0.06 |
| RI | 5077 | ENSG00000198520 | ARMH1 | chr1 | + | 44724319 | 44724668 | 44724319 | 44724392 | 44724538 | 44724668 | 5077 | 24 | 26 | 22 | 27 | 1 | 4 | 4 | 5 | 1 | 3 | 22 | 7 | 10 | 44 | 14 | 5 | 9 | 2 | 13 | 8 | 295 | 149 | 0.15237815 | 0.388140463 | 0.752 | 0.767 | 0.69 | 0.932 | 0.144 | 0.69 | 0.282 | 0.716 | 0.631 | 0.469 | 0.099 |
| RI | 5355 | ENSG00000162645 | GBP2 | chr1 | - | 89106131 | 89108291 | 89106131 | 89107052 | 89108022 | 89108291 | 5355 | 128 | 139 | 90 | 119 | 186 | 55 | 32 | 33 | 23 | 67 | 115 | 31 | 79 | 53 | 18 | 8 | 21 | 12 | 26 | 0 | 298 | 149 | 0.182271207 | 0.448241474 | 0.538 | 0.685 | 0.577 | 0.721 | 0.581 | 0.878 | 0.425 | 0.767 | 0.505 | 1 | -0.095 |
| RI | 5511 | ENSG00000112367 | FIG4 | chr6 | + | 109743109 | 109743772 | 109743109 | 109743272 | 109743674 | 109743772 | 5511 | 38 | 10 | 7 | 5 | 7 | 51 | 40 | 42 | 44 | 91 | 5 | 23 | 10 | 3 | 0 | 75 | 65 | 18 | 62 | 18 | 298 | 149 | 0.185330674 | 0.453601467 | 0.271 | 0.111 | 0.077 | 0.054 | 0.037 | 0.032 | 0.15 | 0.217 | 0.024 | 0 | 0.025 |
| RI | 7725 | ENSG00000130935 | NOL11 | chr17 | + | 67737507 | 67737972 | 67737507 | 67737692 | 67737846 | 67737972 | 7725 | 62 | 50 | 59 | 52 | 108 | 143 | 90 | 113 | 117 | 160 | 22 | 71 | 65 | 19 | 0 | 7 | 50 | 79 | 27 | 14 | 298 | 149 | 0.187330189 | 0.457444988 | 0.178 | 0.217 | 0.207 | 0.182 | 0.252 | 0.611 | 0.415 | 0.291 | 0.26 | 0 | -0.108 |
| RI | 1272 | ENSG00000205250 | E2F4 | chr16 | + | 67192760 | 67193170 | 67192760 | 67192870 | 67193008 | 67193170 | 1272 | 70 | 37 | 31 | 71 | 30 | 121 | 68 | 124 | 112 | 81 | 123 | 57 | 46 | 138 | 77 | 169 | 141 | 90 | 155 | 208 | 287 | 149 | 0.190431076 | 0.464249429 | 0.231 | 0.22 | 0.115 | 0.248 | 0.161 | 0.274 | 0.173 | 0.21 | 0.316 | 0.161 | -0.032 |
| RI | 1412 | ENSG00000051523 | CYBA | chr16 | - | 88646754 | 88647175 | 88646754 | 88646838 | 88647100 | 88647175 | 1412 | 55 | 29 | 49 | 58 | 13 | 741 | 223 | 390 | 408 | 310 | 42 | 10 | 52 | 50 | 71 | 425 | 612 | 331 | 484 | 844 | 298 | 149 | 0.190569887 | 0.464396174 | 0.036 | 0.061 | 0.059 | 0.066 | 0.021 | 0.047 | 0.008 | 0.073 | 0.049 | 0.04 | 0.005 |
| RI | 2688 | ENSG00000108591 | DRG2 | chr17 | + | 18099016 | 18100435 | 18099016 | 18099077 | 18099632 | 18100435 | 2688 | 19 | 14 | 24 | 40 | 33 | 40 | 17 | 34 | 46 | 38 | 33 | 38 | 33 | 8 | 49 | 75 | 104 | 20 | 40 | 64 | 298 | 149 | 0.207303799 | 0.493767557 | 0.192 | 0.292 | 0.261 | 0.303 | 0.303 | 0.18 | 0.154 | 0.452 | 0.091 | 0.277 | 0.039 |
| RI | 2404 | ENSG00000105875 | WDR91 | chr7 | - | 135186971 | 135189452 | 135186971 | 135187169 | 135189343 | 135189452 | 2404 | 12 | 10 | 7 | 6 | 1 | 5 | 0 | 2 | 2 | 0 | 0 | 12 | 2 | 18 | 3 | 8 | 0 | 14 | 3 | 0 | 298 | 149 | 0.216189574 | 0.509589711 | 0.545 | 1 | 0.636 | 0.6 | 1 | 0 | 1 | 0.067 | 0.75 | 1 | 0.193 |
| RI | 6902 | ENSG00000090316 | MAEA | chr4 | + | 1336860 | 1338617 | 1336860 | 1336994 | 1338421 | 1338617 | 6902 | 15 | 9 | 15 | 13 | 8 | 58 | 15 | 35 | 43 | 30 | 8 | 17 | 34 | 9 | 0 | 4 | 37 | 22 | 27 | 19 | 298 | 149 | 0.223866933 | 0.52433861 | 0.115 | 0.231 | 0.176 | 0.131 | 0.118 | 0.5 | 0.187 | 0.436 | 0.143 | 0 | -0.099 |
| RI | 5942 | ENSG00000112282 | MED23 | chr6 | - | 131592387 | 131593171 | 131592387 | 131592460 | 131593005 | 131593171 | 5942 | 38 | 34 | 44 | 45 | 50 | 71 | 42 | 45 | 48 | 48 | 3 | 19 | 18 | 10 | 6 | 0 | 31 | 33 | 10 | 0 | 298 | 149 | 0.227515136 | 0.530988506 | 0.211 | 0.288 | 0.328 | 0.319 | 0.342 | 1 | 0.235 | 0.214 | 0.333 | 1 | -0.259 |
| RI | 186 | ENSG00000104824 | HNRNPL | chr19 | - | 38840095 | 38840559 | 38840095 | 38840376 | 38840487 | 38840559 | 186 | 66 | 64 | 72 | 104 | 91 | 267 | 162 | 233 | 290 | 407 | 104 | 183 | 102 | 164 | 110 | 443 | 360 | 359 | 323 | 466 | 260 | 149 | 0.227712349 | 0.531037024 | 0.124 | 0.185 | 0.15 | 0.17 | 0.114 | 0.119 | 0.226 | 0.14 | 0.225 | 0.119 | -0.017 |
| RI | 8174 | ENSG00000119977 | TCTN3 | chr10 | - | 95682650 | 95683629 | 95682650 | 95682804 | 95683100 | 95683629 | 8174 | 5 | 17 | 9 | 3 | 19 | 45 | 35 | 49 | 42 | 74 | 6 | 10 | 21 | 15 | 0 | 49 | 45 | 6 | 17 | 21 | 298 | 149 | 0.248990474 | 0.570835702 | 0.053 | 0.195 | 0.084 | 0.034 | 0.114 | 0.058 | 0.1 | 0.636 | 0.306 | 0 | -0.124 |
| RI | 2952 | ENSG00000164068 | RNF123 | chr3 | + | 49720799 | 49721105 | 49720799 | 49720894 | 49721019 | 49721105 | 2952 | 32 | 14 | 49 | 35 | 19 | 20 | 6 | 11 | 19 | 10 | 16 | 44 | 32 | 47 | 40 | 27 | 34 | 9 | 10 | 51 | 274 | 149 | 0.266766107 | 0.600299959 | 0.465 | 0.559 | 0.708 | 0.5 | 0.508 | 0.244 | 0.413 | 0.659 | 0.719 | 0.299 | 0.081 |
| RI | 7045 | ENSG00000221914 | PPP2R2A | chr8 | + | 26362683 | 26363890 | 26362683 | 26362848 | 26363720 | 26363890 | 7045 | 27 | 17 | 18 | 20 | 17 | 99 | 61 | 62 | 73 | 119 | 0 | 42 | 20 | 1 | 7 | 93 | 50 | 65 | 51 | 39 | 298 | 149 | 0.273760112 | 0.610226785 | 0.12 | 0.122 | 0.127 | 0.12 | 0.067 | 0 | 0.296 | 0.133 | 0.01 | 0.082 | 0.007 |
| RI | 8456 | ENSG00000123349 | PFDN5 | chr12 | + | 53297849 | 53298150 | 53297849 | 53297924 | 53298018 | 53298150 | 8456 | 18 | 26 | 28 | 46 | 39 | 3 | 5 | 1 | 1 | 3 | 108 | 57 | 53 | 87 | 60 | 0 | 9 | 0 | 0 | 7 | 243 | 149 | 0.274373474 | 0.611132773 | 0.786 | 0.761 | 0.945 | 0.966 | 0.889 | 1 | 0.795 | 1 | 1 | 0.84 | -0.058 |
| RI | 6852 | ENSG00000215021 | PHB2 | chr12 | - | 6967170 | 6968021 | 6967170 | 6967335 | 6967891 | 6968021 | 6852 | 30 | 19 | 46 | 35 | 47 | 9 | 1 | 3 | 2 | 2 | 59 | 58 | 69 | 44 | 62 | 5 | 4 | 4 | 18 | 7 | 298 | 149 | 0.278742278 | 0.61829915 | 0.625 | 0.905 | 0.885 | 0.897 | 0.922 | 0.855 | 0.879 | 0.896 | 0.55 | 0.816 | 0.048 |
| RI | 8600 | ENSG00000134899 | ERCC5 | chr13 | + | 102872197 | 102873343 | 102872197 | 102872398 | 102873258 | 102873343 | 8600 | 21 | 38 | 16 | 37 | 50 | 267 | 225 | 262 | 240 | 295 | 20 | 26 | 40 | 4 | 0 | 11 | 166 | 300 | 77 | 4 | 298 | 149 | 0.281217428 | 0.622154064 | 0.038 | 0.078 | 0.03 | 0.072 | 0.078 | 0.476 | 0.073 | 0.062 | 0.025 | 0 | -0.068 |
| RI | 3571 | ENSG00000120008 | WDR11 | chr10 | + | 120905875 | 120906855 | 120905875 | 120906021 | 120906775 | 120906855 | 3571 | 60 | 62 | 64 | 39 | 40 | 105 | 69 | 71 | 97 | 100 | 18 | 66 | 57 | 17 | 16 | 26 | 66 | 50 | 63 | 15 | 298 | 149 | 0.28573902 | 0.628860057 | 0.222 | 0.31 | 0.311 | 0.167 | 0.167 | 0.257 | 0.333 | 0.363 | 0.119 | 0.348 | -0.049 |
| RI | 3332 | ENSG00000117228 | GBP1 | chr1 | - | 89057991 | 89059043 | 89057991 | 89058234 | 89058840 | 89059043 | 3332 | 24 | 66 | 37 | 41 | 25 | 153 | 118 | 118 | 172 | 100 | 21 | 31 | 39 | 32 | 15 | 67 | 110 | 50 | 198 | 18 | 298 | 149 | 0.292318332 | 0.639290776 | 0.073 | 0.219 | 0.136 | 0.106 | 0.111 | 0.135 | 0.124 | 0.281 | 0.075 | 0.294 | -0.053 |
| RI | 2426 | ENSG00000032742 | IFT88 | chr13 | + | 20601704 | 20601933 | 20601704 | 20601748 | 20601850 | 20601933 | 2426 | 52 | 56 | 46 | 57 | 96 | 27 | 12 | 4 | 8 | 23 | 7 | 45 | 17 | 29 | 16 | 13 | 1 | 2 | 2 | 2 | 251 | 149 | 0.294882738 | 0.643361418 | 0.533 | 0.735 | 0.872 | 0.809 | 0.712 | 0.242 | 0.964 | 0.835 | 0.896 | 0.826 | -0.02 |
| RI | 8555 | ENSG00000117226 | GBP3 | chr1 | - | 89013184 | 89014279 | 89013184 | 89013427 | 89014082 | 89014279 | 8555 | 59 | 25 | 46 | 72 | 16 | 68 | 65 | 69 | 52 | 94 | 26 | 56 | 47 | 26 | 0 | 6 | 58 | 35 | 33 | 16 | 298 | 149 | 0.294942055 | 0.643361418 | 0.303 | 0.161 | 0.25 | 0.409 | 0.078 | 0.684 | 0.326 | 0.402 | 0.283 | 0 | -0.099 |
| RI | 8382 | ENSG00000135452 | TSPAN31 | chr12 | + | 57746175 | 57746720 | 57746175 | 57746256 | 57746588 | 57746720 | 8382 | 20 | 20 | 23 | 28 | 20 | 62 | 36 | 35 | 31 | 43 | 56 | 14 | 38 | 22 | 20 | 53 | 25 | 14 | 43 | 78 | 298 | 149 | 0.30742123 | 0.662267399 | 0.139 | 0.217 | 0.247 | 0.311 | 0.189 | 0.346 | 0.219 | 0.576 | 0.204 | 0.114 | -0.071 |
| RI | 7698 | ENSG00000068831 | RASGRP2 | chr11 | - | 64735541 | 64735980 | 64735541 | 64735664 | 64735902 | 64735980 | 7698 | 7 | 5 | 6 | 7 | 3 | 174 | 73 | 192 | 171 | 101 | 0 | 6 | 7 | 6 | 0 | 179 | 122 | 71 | 78 | 184 | 298 | 149 | 0.308667035 | 0.664466536 | 0.02 | 0.033 | 0.015 | 0.02 | 0.015 | 0 | 0.024 | 0.047 | 0.037 | 0 | -0.001 |
| RI | 2660 | ENSG00000141452 | RMC1 | chr18 | + | 23530228 | 23530612 | 23530228 | 23530297 | 23530386 | 23530612 | 2660 | 70 | 21 | 38 | 26 | 6 | 112 | 63 | 51 | 95 | 72 | 130 | 64 | 46 | 36 | 10 | 131 | 121 | 61 | 87 | 119 | 238 | 149 | 0.314498442 | 0.6748065 | 0.281 | 0.173 | 0.318 | 0.146 | 0.05 | 0.383 | 0.249 | 0.321 | 0.206 | 0.05 | -0.048 |
| RI | 4465 | ENSG00000070010 | UFD1 | chr22 | - | 19450620 | 19454830 | 19450620 | 19450744 | 19454801 | 19454830 | 4465 | 74 | 72 | 95 | 127 | 89 | 0 | 0 | 0 | 3 | 6 | 229 | 138 | 258 | 261 | 69 | 0 | 0 | 0 | 0 | 0 | 298 | 149 | 0.31765003 | 0.679860696 | 1 | 1 | 1 | 0.955 | 0.881 | 1 | 1 | 1 | 1 | 1 | -0.033 |
| RI | 5273 | ENSG00000168890 | TMEM150A | chr2 | - | 85601020 | 85601482 | 85601020 | 85601107 | 85601434 | 85601482 | 5273 | 12 | 3 | 9 | 10 | 11 | 4 | 3 | 7 | 10 | 1 | 9 | 22 | 14 | 17 | 1 | 7 | 0 | 5 | 0 | 10 | 298 | 149 | 0.317774739 | 0.679860696 | 0.6 | 0.333 | 0.391 | 0.333 | 0.846 | 0.391 | 1 | 0.583 | 1 | 0.048 | -0.104 |
| RI | 6755 | ENSG00000100802 | C14orf93 | chr14 | - | 22998426 | 22999156 | 22998426 | 22998741 | 22999053 | 22999156 | 6755 | 77 | 29 | 57 | 38 | 65 | 4 | 2 | 3 | 4 | 5 | 47 | 59 | 49 | 54 | 92 | 0 | 2 | 9 | 3 | 0 | 298 | 149 | 0.32881789 | 0.699181885 | 0.906 | 0.879 | 0.905 | 0.826 | 0.867 | 1 | 0.937 | 0.731 | 0.9 | 1 | -0.037 |
| RI | 5798 | ENSG00000143499 | SMYD2 | chr1 | + | 214332017 | 214334308 | 214332017 | 214332192 | 214334199 | 214334308 | 5798 | 42 | 21 | 33 | 35 | 30 | 58 | 32 | 45 | 65 | 81 | 22 | 8 | 19 | 16 | 12 | 40 | 26 | 28 | 24 | 38 | 298 | 149 | 0.330566838 | 0.700882381 | 0.266 | 0.247 | 0.268 | 0.212 | 0.156 | 0.216 | 0.133 | 0.253 | 0.25 | 0.136 | 0.032 |
| RI | 5338 | ENSG00000084623 | EIF3I | chr1 | + | 32224409 | 32226530 | 32224409 | 32224475 | 32226402 | 32226530 | 5338 | 42 | 23 | 23 | 43 | 21 | 0 | 0 | 1 | 2 | 0 | 58 | 66 | 19 | 61 | 91 | 0 | 0 | 1 | 0 | 6 | 298 | 149 | 0.333438411 | 0.706210359 | 1 | 1 | 0.92 | 0.915 | 1 | 1 | 1 | 0.905 | 1 | 0.883 | 0.009 |
| RI | 8016 | ENSG00000198258 | UBL5 | chr19 | + | 9828326 | 9828675 | 9828326 | 9828389 | 9828591 | 9828675 | 8016 | 30 | 8 | 31 | 28 | 25 | 1 | 2 | 0 | 0 | 1 | 14 | 77 | 43 | 67 | 80 | 0 | 0 | 0 | 0 | 8 | 298 | 149 | 0.343810899 | 0.721453278 | 0.938 | 0.667 | 1 | 1 | 0.926 | 1 | 1 | 1 | 1 | 0.833 | -0.06 |
| RI | 775 | ENSG00000073350 | LLGL2 | chr17 | + | 75574452 | 75574668 | 75574452 | 75574495 | 75574609 | 75574668 | 775 | 4 | 1 | 1 | 4 | 0 | 25 | 12 | 10 | 17 | 15 | 0 | 0 | 2 | 0 | 4 | 4 | 22 | 2 | 11 | 8 | 263 | 149 | 0.357823877 | 0.741116985 | 0.083 | 0.045 | 0.054 | 0.118 | 0 | 0 | 0 | 0.362 | 0 | 0.221 | -0.057 |
| RI | 7194 | ENSG00000174744 | BRMS1 | chr11 | - | 66340773 | 66341046 | 66340773 | 66340870 | 66340966 | 66341046 | 7194 | 1 | 1 | 13 | 8 | 3 | 124 | 51 | 95 | 119 | 108 | 2 | 21 | 2 | 10 | 8 | 106 | 148 | 110 | 135 | 192 | 245 | 149 | 0.359860602 | 0.744552217 | 0.005 | 0.012 | 0.077 | 0.039 | 0.017 | 0.011 | 0.079 | 0.011 | 0.043 | 0.025 | -0.004 |
| RI | 2692 | ENSG00000108591 | DRG2 | chr17 | + | 18099632 | 18100435 | 18099632 | 18099715 | 18100362 | 18100435 | 2692 | 19 | 11 | 20 | 36 | 25 | 4 | 3 | 0 | 2 | 3 | 50 | 40 | 27 | 16 | 26 | 0 | 0 | 2 | 2 | 7 | 298 | 149 | 0.362668686 | 0.750099414 | 0.704 | 0.647 | 1 | 0.9 | 0.806 | 1 | 1 | 0.871 | 0.8 | 0.65 | -0.053 |
| RI | 2403 | ENSG00000105875 | WDR91 | chr7 | - | 135186971 | 135188545 | 135186971 | 135187169 | 135188432 | 135188545 | 2403 | 10 | 12 | 15 | 9 | 1 | 12 | 5 | 13 | 21 | 5 | 12 | 27 | 10 | 11 | 22 | 22 | 13 | 18 | 10 | 8 | 298 | 149 | 0.366957635 | 0.756321965 | 0.294 | 0.545 | 0.366 | 0.176 | 0.091 | 0.214 | 0.509 | 0.217 | 0.355 | 0.579 | -0.08 |
| RI | 1685 | ENSG00000137841 | PLCB2 | chr15 | - | 40290576 | 40290837 | 40290576 | 40290672 | 40290760 | 40290837 | 1685 | 21 | 10 | 18 | 17 | 3 | 218 | 166 | 145 | 226 | 127 | 46 | 27 | 9 | 13 | 26 | 383 | 314 | 192 | 407 | 584 | 237 | 149 | 0.377944351 | 0.772847911 | 0.057 | 0.036 | 0.072 | 0.045 | 0.015 | 0.07 | 0.051 | 0.029 | 0.02 | 0.027 | 0.006 |
| RI | 6836 | ENSG00000134516 | DOCK2 | chr5 | + | 170078974 | 170081984 | 170078974 | 170079146 | 170081841 | 170081984 | 6836 | 17 | 12 | 12 | 28 | 3 | 2 | 0 | 0 | 4 | 0 | 11 | 22 | 21 | 28 | 15 | 0 | 2 | 2 | 0 | 0 | 298 | 149 | 0.378115873 | 0.772847911 | 0.81 | 1 | 1 | 0.778 | 1 | 1 | 0.846 | 0.84 | 1 | 1 | -0.02 |
| RI | 3091 | ENSG00000111642 | CHD4 | chr12 | - | 6578028 | 6578546 | 6578028 | 6578117 | 6578408 | 6578546 | 3091 | 136 | 93 | 110 | 132 | 184 | 1 | 1 | 4 | 2 | 16 | 511 | 221 | 201 | 409 | 225 | 16 | 9 | 27 | 5 | 5 | 298 | 149 | 0.389814831 | 0.790469003 | 0.986 | 0.979 | 0.932 | 0.971 | 0.852 | 0.941 | 0.925 | 0.788 | 0.976 | 0.957 | 0.027 |
| RI | 2091 | ENSG00000175727 | MLXIP | chr12 | + | 122135488 | 122137590 | 122135488 | 122135666 | 122137468 | 122137590 | 2091 | 11 | 18 | 31 | 27 | 24 | 57 | 16 | 57 | 70 | 33 | 32 | 2 | 11 | 23 | 48 | 22 | 61 | 14 | 50 | 191 | 298 | 149 | 0.397356338 | 0.803005094 | 0.088 | 0.36 | 0.214 | 0.162 | 0.267 | 0.421 | 0.016 | 0.282 | 0.187 | 0.112 | 0.015 |
| RI | 8014 | ENSG00000143702 | CEP170 | chr1 | - | 243185778 | 243186422 | 243185778 | 243186072 | 243186258 | 243186422 | 8014 | 170 | 182 | 126 | 148 | 163 | 71 | 48 | 50 | 57 | 82 | 26 | 118 | 141 | 31 | 11 | 0 | 42 | 19 | 11 | 22 | 298 | 149 | 0.399891706 | 0.806473304 | 0.545 | 0.655 | 0.558 | 0.565 | 0.498 | 1 | 0.584 | 0.788 | 0.585 | 0.2 | -0.067 |
| RI | 4556 | ENSG00000135749 | PCNX2 | chr1 | - | 233217898 | 233218184 | 233217898 | 233217931 | 233218030 | 233218184 | 4556 | 19 | 4 | 19 | 7 | 23 | 79 | 44 | 71 | 85 | 156 | 13 | 3 | 6 | 2 | 10 | 24 | 46 | 73 | 56 | 45 | 248 | 149 | 0.402579019 | 0.810509293 | 0.126 | 0.052 | 0.139 | 0.047 | 0.081 | 0.246 | 0.038 | 0.047 | 0.021 | 0.118 | -0.005 |
| RI | 791 | ENSG00000214087 | ARL16 | chr17 | - | 81682033 | 81683126 | 81682033 | 81682675 | 81683012 | 81683126 | 791 | 15 | 10 | 5 | 12 | 4 | 17 | 9 | 24 | 12 | 10 | 0 | 13 | 34 | 12 | 2 | 5 | 35 | 12 | 11 | 45 | 298 | 149 | 0.413684312 | 0.82764407 | 0.306 | 0.357 | 0.094 | 0.333 | 0.167 | 0 | 0.157 | 0.586 | 0.353 | 0.022 | 0.028 |
| RI | 1815 | ENSG00000196497 | IPO4 | chr14 | - | 24188215 | 24188423 | 24188215 | 24188257 | 24188343 | 24188423 | 1815 | 3 | 2 | 2 | 7 | 7 | 3 | 3 | 4 | 5 | 6 | 0 | 13 | 5 | 19 | 13 | 1 | 7 | 5 | 8 | 12 | 235 | 149 | 0.427112697 | 0.844927898 | 0.388 | 0.297 | 0.241 | 0.47 | 0.425 | 0 | 0.541 | 0.388 | 0.601 | 0.407 | -0.023 |
| RI | 2947 | ENSG00000164068 | RNF123 | chr3 | + | 49704649 | 49705186 | 49704649 | 49704756 | 49705081 | 49705186 | 2947 | 17 | 6 | 25 | 17 | 6 | 4 | 1 | 0 | 0 | 0 | 10 | 23 | 3 | 12 | 11 | 0 | 0 | 0 | 7 | 0 | 298 | 149 | 0.43656097 | 0.859302115 | 0.68 | 0.75 | 1 | 1 | 1 | 1 | 1 | 1 | 0.462 | 1 | -0.006 |
| RI | 2733 | ENSG00000109133 | TMEM33 | chr4 | + | 41935128 | 41935529 | 41935128 | 41935262 | 41935466 | 41935529 | 2733 | 134 | 94 | 122 | 100 | 37 | 4 | 3 | 2 | 5 | 11 | 86 | 76 | 69 | 72 | 101 | 0 | 11 | 5 | 2 | 0 | 298 | 149 | 0.445151105 | 0.871563665 | 0.944 | 0.94 | 0.968 | 0.909 | 0.627 | 1 | 0.776 | 0.873 | 0.947 | 1 | -0.042 |
| RI | 5080 | ENSG00000112855 | HARS2 | chr5 | + | 140693490 | 140693665 | 140693490 | 140693508 | 140693590 | 140693665 | 5080 | 26 | 29 | 26 | 29 | 39 | 0 | 3 | 0 | 0 | 2 | 76 | 46 | 79 | 79 | 71 | 2 | 0 | 1 | 4 | 0 | 231 | 149 | 0.449971301 | 0.877800686 | 1 | 0.862 | 1 | 1 | 0.926 | 0.961 | 1 | 0.981 | 0.927 | 1 | -0.016 |
| RI | 2752 | ENSG00000099949 | LZTR1 | chr22 | + | 20994869 | 20995872 | 20994869 | 20995026 | 20995745 | 20995872 | 2752 | 10 | 6 | 11 | 10 | 10 | 62 | 20 | 32 | 47 | 10 | 9 | 4 | 3 | 7 | 21 | 66 | 20 | 14 | 35 | 48 | 298 | 149 | 0.450664114 | 0.878487297 | 0.075 | 0.13 | 0.147 | 0.096 | 0.333 | 0.064 | 0.091 | 0.097 | 0.091 | 0.179 | 0.052 |
| RI | 4870 | ENSG00000162852 | CNST | chr1 | + | 246633923 | 246634587 | 246633923 | 246634010 | 246634472 | 246634587 | 4870 | 17 | 29 | 23 | 26 | 34 | 168 | 216 | 157 | 237 | 305 | 0 | 14 | 37 | 12 | 25 | 123 | 216 | 172 | 185 | 115 | 298 | 149 | 0.451700861 | 0.879432097 | 0.048 | 0.063 | 0.068 | 0.052 | 0.053 | 0 | 0.031 | 0.097 | 0.031 | 0.098 | 0.005 |
| RI | 5234 | ENSG00000186141 | POLR3C | chr1 | + | 145826453 | 145826709 | 145826453 | 145826492 | 145826605 | 145826709 | 5234 | 117 | 95 | 126 | 146 | 165 | 14 | 12 | 15 | 19 | 28 | 104 | 160 | 91 | 166 | 150 | 13 | 20 | 4 | 30 | 14 | 262 | 149 | 0.456482001 | 0.885239389 | 0.826 | 0.818 | 0.827 | 0.814 | 0.77 | 0.82 | 0.82 | 0.928 | 0.759 | 0.859 | -0.026 |
| RI | 5214 | ENSG00000149499 | EML3 | chr11 | - | 62603943 | 62604201 | 62603943 | 62604041 | 62604112 | 62604201 | 5214 | 14 | 6 | 5 | 26 | 8 | 89 | 41 | 94 | 83 | 85 | 15 | 14 | 11 | 21 | 17 | 104 | 83 | 75 | 78 | 120 | 220 | 149 | 0.469934252 | 0.903319762 | 0.096 | 0.09 | 0.035 | 0.175 | 0.06 | 0.089 | 0.103 | 0.09 | 0.154 | 0.088 | -0.014 |
| RI | 2868 | ENSG00000184470 | TXNRD2 | chr22 | - | 19875521 | 19877234 | 19875521 | 19875807 | 19877039 | 19877234 | 2868 | 0 | 1 | 3 | 9 | 6 | 15 | 1 | 4 | 10 | 1 | 18 | 2 | 2 | 3 | 22 | 5 | 10 | 8 | 2 | 22 | 298 | 149 | 0.4704967 | 0.904106704 | 0 | 0.333 | 0.273 | 0.31 | 0.75 | 0.643 | 0.091 | 0.111 | 0.429 | 0.333 | 0.012 |
| RI | 790 | ENSG00000214087 | ARL16 | chr17 | - | 81682033 | 81683126 | 81682033 | 81682149 | 81683012 | 81683126 | 790 | 32 | 26 | 36 | 45 | 17 | 30 | 8 | 8 | 6 | 34 | 76 | 91 | 54 | 81 | 57 | 34 | 18 | 8 | 43 | 32 | 298 | 149 | 0.473156612 | 0.908331526 | 0.348 | 0.619 | 0.692 | 0.789 | 0.2 | 0.528 | 0.717 | 0.771 | 0.485 | 0.471 | -0.065 |
| RI | 5002 | ENSG00000092529 | CAPN3 | chr15 | + | 42410427 | 42410666 | 42410427 | 42410496 | 42410587 | 42410666 | 5002 | 2 | 5 | 5 | 3 | 2 | 24 | 11 | 20 | 40 | 45 | 21 | 9 | 3 | 8 | 0 | 25 | 64 | 38 | 166 | 19 | 240 | 149 | 0.479537699 | 0.917302198 | 0.049 | 0.22 | 0.134 | 0.044 | 0.027 | 0.343 | 0.08 | 0.047 | 0.029 | 0 | -0.005 |
| RI | 6851 | ENSG00000215021 | PHB2 | chr12 | - | 6967170 | 6968021 | 6967170 | 6967238 | 6967891 | 6968021 | 6851 | 55 | 49 | 81 | 85 | 116 | 14 | 6 | 2 | 5 | 13 | 166 | 93 | 101 | 113 | 114 | 31 | 14 | 22 | 11 | 0 | 298 | 149 | 0.489343042 | 0.930933768 | 0.663 | 0.803 | 0.953 | 0.895 | 0.817 | 0.728 | 0.769 | 0.697 | 0.837 | 1 | 0.02 |
| RI | 1111 | ENSG00000118046 | STK11 | chr19 | + | 1220372 | 1220717 | 1220372 | 1220505 | 1220580 | 1220717 | 1111 | 15 | 4 | 17 | 9 | 16 | 64 | 32 | 70 | 51 | 48 | 28 | 11 | 3 | 1 | 10 | 19 | 68 | 14 | 15 | 42 | 224 | 149 | 0.490133114 | 0.931836597 | 0.135 | 0.077 | 0.139 | 0.105 | 0.181 | 0.495 | 0.097 | 0.125 | 0.042 | 0.137 | -0.052 |
| RI | 4984 | ENSG00000124067 | SLC12A4 | chr16 | - | 67950961 | 67951304 | 67950961 | 67951060 | 67951139 | 67951304 | 4984 | 5 | 8 | 8 | 11 | 1 | 9 | 4 | 5 | 5 | 2 | 0 | 9 | 4 | 6 | 26 | 5 | 7 | 7 | 11 | 4 | 228 | 149 | 0.492531237 | 0.934891394 | 0.266 | 0.567 | 0.511 | 0.59 | 0.246 | 0 | 0.457 | 0.272 | 0.263 | 0.809 | 0.076 |
| RI | 8601 | ENSG00000134899 | ERCC5 | chr13 | + | 102872197 | 102873343 | 102872197 | 102872398 | 102873260 | 102873343 | 8601 | 21 | 39 | 16 | 40 | 56 | 7 | 5 | 6 | 5 | 4 | 20 | 31 | 41 | 4 | 3 | 0 | 3 | 5 | 7 | 0 | 298 | 149 | 0.516131201 | 0.966641409 | 0.6 | 0.796 | 0.571 | 0.8 | 0.875 | 1 | 0.838 | 0.804 | 0.222 | 1 | -0.044 |
| RI | 2260 | ENSG00000143889 | HNRNPLL | chr2 | - | 38602437 | 38603036 | 38602437 | 38602600 | 38602856 | 38603036 | 2260 | 34 | 16 | 14 | 31 | 4 | 1 | 1 | 0 | 0 | 1 | 18 | 19 | 9 | 28 | 12 | 0 | 2 | 3 | 0 | 0 | 298 | 149 | 0.519780301 | 0.971012726 | 0.944 | 0.889 | 1 | 1 | 0.667 | 1 | 0.826 | 0.6 | 1 | 1 | 0.015 |
| RI | 2950 | ENSG00000164068 | RNF123 | chr3 | + | 49712478 | 49713587 | 49712478 | 49712892 | 49713512 | 49713587 | 2950 | 3 | 1 | 4 | 1 | 1 | 3 | 0 | 1 | 0 | 3 | 6 | 11 | 4 | 2 | 12 | 0 | 2 | 2 | 0 | 9 | 298 | 149 | 0.524921023 | 0.974452698 | 0.333 | 1 | 0.667 | 1 | 0.143 | 1 | 0.733 | 0.5 | 1 | 0.4 | -0.098 |
| RI | 8452 | ENSG00000123349 | PFDN5 | chr12 | + | 53295838 | 53297924 | 53295838 | 53295941 | 53297849 | 53297924 | 8452 | 192 | 157 | 217 | 240 | 285 | 3 | 13 | 11 | 3 | 11 | 433 | 302 | 224 | 293 | 324 | 34 | 5 | 5 | 4 | 20 | 298 | 149 | 0.526469632 | 0.976204343 | 0.97 | 0.858 | 0.908 | 0.976 | 0.928 | 0.864 | 0.968 | 0.957 | 0.973 | 0.89 | -0.002 |
| RI | 2753 | ENSG00000099949 | LZTR1 | chr22 | + | 20995962 | 20996801 | 20995962 | 20996112 | 20996695 | 20996801 | 2753 | 10 | 3 | 9 | 12 | 13 | 41 | 5 | 23 | 18 | 6 | 8 | 12 | 2 | 6 | 24 | 44 | 8 | 6 | 19 | 42 | 298 | 149 | 0.52729852 | 0.977330518 | 0.109 | 0.231 | 0.164 | 0.25 | 0.52 | 0.083 | 0.429 | 0.143 | 0.136 | 0.222 | 0.052 |
| RI | 7231 | ENSG00000164941 | INTS8 | chr8 | + | 94867139 | 94867337 | 94867139 | 94867196 | 94867275 | 94867337 | 7231 | 13 | 26 | 16 | 28 | 30 | 213 | 145 | 138 | 116 | 199 | 12 | 12 | 10 | 2 | 0 | 8 | 69 | 59 | 32 | 11 | 228 | 149 | 0.531060799 | 0.983069928 | 0.038 | 0.105 | 0.07 | 0.136 | 0.09 | 0.495 | 0.102 | 0.1 | 0.039 | 0 | -0.059 |
| RI | 8450 | ENSG00000123349 | PFDN5 | chr12 | + | 53295838 | 53296467 | 53295838 | 53295941 | 53296429 | 53296467 | 8450 | 146 | 118 | 157 | 161 | 186 | 0 | 1 | 7 | 1 | 0 | 195 | 197 | 138 | 147 | 203 | 0 | 2 | 0 | 3 | 0 | 298 | 149 | 0.555750848 | 1 | 1 | 0.983 | 0.918 | 0.988 | 1 | 1 | 0.98 | 1 | 0.961 | 1 | -0.01 |
| RI | 1112 | ENSG00000118046 | STK11 | chr19 | + | 1221212 | 1222006 | 1221212 | 1221340 | 1221948 | 1222006 | 1112 | 1 | 4 | 10 | 4 | 0 | 118 | 53 | 79 | 95 | 109 | 11 | 12 | 2 | 3 | 0 | 148 | 108 | 35 | 98 | 212 | 298 | 149 | 0.56003204 | 1 | 0.004 | 0.036 | 0.06 | 0.021 | 0 | 0.036 | 0.053 | 0.028 | 0.015 | 0 | -0.002 |
| RI | 8383 | ENSG00000135452 | TSPAN31 | chr12 | + | 57746175 | 57746720 | 57746175 | 57746256 | 57746592 | 57746720 | 8383 | 31 | 24 | 32 | 33 | 24 | 4 | 1 | 2 | 6 | 0 | 58 | 17 | 47 | 27 | 37 | 10 | 0 | 0 | 4 | 0 | 298 | 149 | 0.561255074 | 1 | 0.795 | 0.923 | 0.889 | 0.733 | 1 | 0.744 | 1 | 1 | 0.771 | 1 | -0.035 |
| RI | 4836 | ENSG00000103168 | TAF1C | chr16 | - | 84183075 | 84183328 | 84183075 | 84183149 | 84183243 | 84183328 | 4836 | 7 | 1 | 12 | 17 | 6 | 28 | 15 | 23 | 30 | 11 | 13 | 27 | 7 | 1 | 9 | 15 | 29 | 32 | 12 | 23 | 243 | 149 | 0.567599424 | 1 | 0.133 | 0.039 | 0.242 | 0.258 | 0.251 | 0.347 | 0.363 | 0.118 | 0.049 | 0.194 | -0.03 |
| RI | 2690 | ENSG00000108591 | DRG2 | chr17 | + | 18099016 | 18100435 | 18099016 | 18099077 | 18100362 | 18100435 | 2690 | 12 | 10 | 19 | 36 | 13 | 3 | 0 | 1 | 1 | 0 | 48 | 25 | 22 | 3 | 39 | 0 | 0 | 0 | 0 | 0 | 298 | 149 | 0.6486741 | 1 | 0.667 | 1 | 0.905 | 0.947 | 1 | 1 | 1 | 1 | 1 | 1 | -0.096 |
| RI | 6248 | ENSG00000171777 | RASGRP4 | chr19 | - | 38419859 | 38420262 | 38419859 | 38419971 | 38420130 | 38420262 | 6248 | 16 | 7 | 18 | 19 | 8 | 0 | 0 | 0 | 0 | 0 | 37 | 14 | 26 | 33 | 116 | 0 | 0 | 0 | 5 | 0 | 298 | 149 | 0.649173485 | 1 | 1 | 1 | 1 | 1 | 1 | 1 | 1 | 1 | 0.767 | 1 | 0.047 |
| RI | 6247 | ENSG00000171777 | RASGRP4 | chr19 | - | 38412930 | 38413297 | 38412930 | 38413049 | 38413192 | 38413297 | 6247 | 11 | 5 | 5 | 8 | 4 | 22 | 13 | 7 | 29 | 1 | 7 | 20 | 14 | 21 | 31 | 34 | 24 | 19 | 92 | 77 | 292 | 149 | 0.651219557 | 1 | 0.203 | 0.164 | 0.267 | 0.123 | 0.671 | 0.095 | 0.298 | 0.273 | 0.104 | 0.17 | 0.098 |
| RI | 2462 | ENSG00000187091 | PLCD1 | chr3 | - | 38008457 | 38009158 | 38008457 | 38008636 | 38009041 | 38009158 | 2462 | 15 | 5 | 6 | 6 | 6 | 23 | 3 | 12 | 14 | 9 | 5 | 8 | 9 | 9 | 9 | 0 | 17 | 25 | 10 | 19 | 298 | 149 | 0.666743373 | 1 | 0.246 | 0.455 | 0.2 | 0.176 | 0.25 | 1 | 0.19 | 0.153 | 0.31 | 0.191 | -0.103 |
| RI | 2754 | ENSG00000099949 | LZTR1 | chr22 | + | 20996695 | 20996966 | 20996695 | 20996801 | 20996885 | 20996966 | 2754 | 14 | 5 | 13 | 12 | 5 | 41 | 13 | 22 | 24 | 6 | 25 | 16 | 4 | 0 | 8 | 18 | 5 | 14 | 20 | 14 | 233 | 149 | 0.669324038 | 1 | 0.179 | 0.197 | 0.274 | 0.242 | 0.348 | 0.47 | 0.672 | 0.154 | 0 | 0.268 | -0.065 |

| | | | | | | | | | | | | | | | | | | | | | | | | | | | | | | | | | | | | | | | | | | | | | | | |
|---|---|---|---|---|---|---|---|---|---|---|---|---|---|---|---|---|---|---|---|---|---|---|---|---|---|---|---|---|---|---|---|---|---|---|---|---|---|---|---|---|---|---|---|---|---|---|---|
| RI | 4549 | ENSG00000163463 | KRTCAP2 | chr1 | - | 155169790 | 155172489 | 155169790 | 155169857 | 155172350 | 155172489 | 4549 | 29 | 23 | 29 | 31 | 23 | 1 | 0 | 0 | 0 | 0 | 54 | 31 | 43 | 34 | 40 | 0 | 3 | 0 | 3 | 0 | 298 | 149 | 0.674179769 | 1 | 0.935 | 1 | 1 | 1 | 1 | 1 | 0.838 | 1 | 0.85 | 1 | 0.049 |
| RI | 5539 | ENSG00000144635 | DYNC1LI1 | chr3 | - | 32530283 | 32533097 | 32530283 | 32530328 | 32532985 | 32533097 | 5539 | 12 | 14 | 7 | 7 | 9 | 0 | 0 | 1 | 0 | 0 | 14 | 13 | 19 | 9 | 0 | 0 | 0 | 0 | 0 | 2 | 298 | 149 | 0.680564883 | 1 | 1 | 1 | 0.778 | 1 | 1 | 1 | 1 | 1 | 1 | 0 | 0.156 |
| RI | 703 | ENSG00000101557 | USP14 | chr18 | + | 209970 | 210493 | 209970 | 210031 | 210385 | 210493 | 703 | 64 | 81 | 59 | 78 | 98 | 173 | 115 | 147 | 153 | 415 | 6 | 101 | 136 | 43 | 0 | 109 | 102 | 71 | 98 | 34 | 298 | 149 | 0.6896282 | 1 | 0.156 | 0.26 | 0.167 | 0.203 | 0.106 | 0.027 | 0.331 | 0.489 | 0.18 | 0 | -0.027 |
| RI | 1035 | ENSG00000185813 | PCYT2 | chr17 | - | 81906099 | 81906546 | 81906099 | 81906177 | 81906463 | 81906546 | 1035 | 8 | 0 | 5 | 8 | 2 | 12 | 7 | 11 | 6 | 10 | 14 | 4 | 0 | 9 | 0 | 13 | 21 | 8 | 11 | 5 | 298 | 149 | 0.720704873 | 1 | 0.25 | 0 | 0.185 | 0.4 | 0.091 | 0.35 | 0.087 | 0 | 0.29 | 0 | 0.04 |
| RI | 5000 | ENSG00000092529 | CAPN3 | chr15 | + | 42409930 | 42410496 | 42409930 | 42409995 | 42410427 | 42410496 | 5000 | 0 | 1 | 2 | 1 | 8 | 24 | 14 | 27 | 37 | 31 | 0 | 4 | 5 | 14 | 0 | 50 | 81 | 31 | 170 | 16 | 298 | 149 | 0.727783528 | 1 | 0 | 0.034 | 0.036 | 0.013 | 0.114 | 0 | 0.024 | 0.075 | 0.04 | 0 | 0.012 |
| RI | 602 | ENSG00000171703 | TCEA2 | chr20 | + | 64069764 | 64070635 | 64069764 | 64070414 | 64070488 | 64070635 | 602 | 5 | 6 | 7 | 5 | 1 | 22 | 10 | 24 | 10 | 30 | 16 | 0 | 2 | 3 | 11 | 33 | 30 | 7 | 27 | 31 | 223 | 149 | 0.738863609 | 1 | 0.132 | 0.286 | 0.163 | 0.25 | 0.022 | 0.245 | 0 | 0.16 | 0.069 | 0.192 | 0.037 |
| RI | 7878 | ENSG00000167986 | DDB1 | chr11 | - | 61314046 | 61314486 | 61314046 | 61314202 | 61314307 | 61314486 | 7878 | 19 | 7 | 22 | 11 | 19 | 9 | 0 | 3 | 7 | 11 | 18 | 24 | 18 | 18 | 19 | 4 | 13 | 6 | 6 | 7 | 254 | 149 | 0.758045711 | 1 | 0.553 | 1 | 0.811 | 0.48 | 0.503 | 0.725 | 0.52 | 0.638 | 0.638 | 0.614 | 0.042 |
| RI | 2693 | ENSG00000108591 | DRG2 | chr17 | + | 18099632 | 18100435 | 18099632 | 18099723 | 18100362 | 18100435 | 2693 | 15 | 11 | 18 | 35 | 14 | 34 | 11 | 26 | 33 | 52 | 39 | 32 | 27 | 12 | 19 | 58 | 50 | 17 | 24 | 56 | 298 | 149 | 0.759417463 | 1 | 0.181 | 0.333 | 0.257 | 0.347 | 0.119 | 0.252 | 0.242 | 0.443 | 0.2 | 0.145 | -0.009 |
| RI | 1686 | ENSG00000137841 | PLCB2 | chr15 | - | 40291264 | 40291924 | 40291264 | 40291487 | 40291848 | 40291924 | 1686 | 81 | 43 | 70 | 63 | 13 | 123 | 76 | 42 | 76 | 25 | 153 | 66 | 48 | 85 | 143 | 98 | 161 | 83 | 123 | 184 | 298 | 149 | 0.7596441 | 1 | 0.248 | 0.221 | 0.455 | 0.293 | 0.206 | 0.438 | 0.17 | 0.224 | 0.257 | 0.28 | 0.011 |
| RI | 2946 | ENSG00000164068 | RNF123 | chr3 | + | 49704649 | 49705186 | 49704649 | 49704756 | 49704983 | 49705186 | 2946 | 9 | 2 | 9 | 10 | 0 | 15 | 6 | 9 | 9 | 17 | 0 | 12 | 4 | 11 | 0 | 14 | 2 | 10 | 4 | 16 | 298 | 149 | 0.771007208 | 1 | 0.231 | 0.143 | 0.333 | 0.357 | 0 | 0 | 0.75 | 0.167 | 0.579 | 0 | -0.086 |
| RI | 5344 | ENSG00000084623 | EIF3I | chr1 | + | 32228726 | 32231019 | 32228726 | 32229208 | 32230926 | 32231019 | 5344 | 8 | 7 | 5 | 15 | 8 | 144 | 102 | 162 | 177 | 124 | 7 | 20 | 0 | 7 | 20 | 208 | 117 | 113 | 146 | 337 | 298 | 149 | 0.773124776 | 1 | 0.027 | 0.033 | 0.015 | 0.041 | 0.031 | 0.017 | 0.079 | 0 | 0.023 | 0.029 | 0 |
| RI | 5836 | ENSG00000144224 | UBXN4 | chr2 | + | 135753538 | 135754277 | 135753538 | 135753567 | 135754158 | 135754277 | 5836 | 193 | 222 | 176 | 196 | 297 | 186 | 103 | 131 | 174 | 430 | 65 | 141 | 231 | 156 | 38 | 55 | 114 | 78 | 80 | 115 | 298 | 149 | 0.817597568 | 1 | 0.342 | 0.519 | 0.402 | 0.36 | 0.257 | 0.371 | 0.382 | 0.597 | 0.494 | 0.142 | -0.021 |
| RI | 1479 | ENSG00000167965 | MLST8 | chr16 | + | 2206357 | 2206659 | 2206357 | 2206409 | 2206478 | 2206659 | 1479 | 5 | 7 | 2 | 7 | 8 | 2 | 0 | 0 | 2 | 0 | 16 | 3 | 3 | 6 | 33 | 0 | 0 | 0 | 0 | 0 | 218 | 149 | 0.818634052 | 1 | 0.631 | 1 | 1 | 0.705 | 1 | 1 | 1 | 1 | 1 | 1 | -0.133 |
| RI | 4982 | ENSG00000124067 | SLC12A4 | chr16 | - | 67945086 | 67945553 | 67945086 | 67945220 | 67945368 | 67945553 | 4982 | 11 | 9 | 8 | 10 | 14 | 10 | 7 | 16 | 7 | 4 | 8 | 5 | 11 | 10 | 31 | 17 | 11 | 5 | 3 | 10 | 297 | 149 | 0.818959886 | 1 | 0.356 | 0.392 | 0.201 | 0.417 | 0.637 | 0.191 | 0.186 | 0.525 | 0.626 | 0.609 | -0.027 |
| RI | 8454 | ENSG00000123349 | PFDN5 | chr12 | + | 53296429 | 53297924 | 53296429 | 53296467 | 53297849 | 53297924 | 8454 | 54 | 44 | 73 | 95 | 103 | 0 | 2 | 5 | 0 | 0 | 272 | 119 | 104 | 164 | 130 | 0 | 2 | 0 | 0 | 0 | 298 | 149 | 0.819733137 | 1 | 1 | 0.917 | 0.88 | 1 | 1 | 1 | 0.967 | 1 | 1 | 1 | -0.034 |
| RI | 5434 | ENSG00000197858 | GPAA1 | chr8 | + | 144083388 | 144083861 | 144083388 | 144083500 | 144083713 | 144083861 | 5434 | 18 | 12 | 22 | 15 | 9 | 47 | 17 | 22 | 14 | 2 | 26 | 18 | 25 | 16 | 8 | 9 | 24 | 3 | 25 | 67 | 298 | 149 | 0.821469205 | 1 | 0.161 | 0.261 | 0.333 | 0.349 | 0.692 | 0.591 | 0.273 | 0.806 | 0.242 | 0.056 | -0.034 |
| RI | 2464 | ENSG00000187091 | PLCD1 | chr3 | - | 38009903 | 38010275 | 38009903 | 38010053 | 38010130 | 38010275 | 2464 | 18 | 0 | 7 | 8 | 17 | 11 | 3 | 4 | 5 | 14 | 14 | 14 | 5 | 5 | 18 | 0 | 26 | 7 | 3 | 23 | 226 | 149 | 0.824669268 | 1 | 0.519 | 0 | 0.536 | 0.513 | 0.445 | 1 | 0.262 | 0.32 | 0.524 | 0.34 | -0.087 |
| RI | 3848 | ENSG00000163608 | NEPRO | chr3 | - | 113012733 | 113013403 | 113012733 | 113012811 | 113013271 | 113013403 | 3848 | 48 | 43 | 58 | 74 | 43 | 39 | 32 | 37 | 36 | 60 | 0 | 30 | 73 | 16 | 5 | 9 | 17 | 20 | 7 | 7 | 298 | 149 | 0.839101155 | 1 | 0.381 | 0.402 | 0.439 | 0.507 | 0.264 | 0 | 0.469 | 0.646 | 0.533 | 0.263 | 0.016 |
| RI | 8020 | ENSG00000198056 | PRIM1 | chr12 | - | 56746780 | 56747032 | 56746780 | 56746854 | 56746925 | 56747032 | 8020 | 7 | 5 | 9 | 14 | 17 | 22 | 15 | 23 | 27 | 26 | 10 | 3 | 8 | 3 | 0 | 0 | 11 | 11 | 19 | 10 | 220 | 149 | 0.843116127 | 1 | 0.177 | 0.184 | 0.209 | 0.26 | 0.307 | 1 | 0.156 | 0.33 | 0.097 | 0 | -0.089 |
| RI | 128 | ENSG00000198089 | SFI1 | chr22 | + | 31618114 | 31618547 | 31618114 | 31618226 | 31618313 | 31618547 | 128 | 8 | 6 | 7 | 7 | 8 | 16 | 9 | 5 | 10 | 1 | 7 | 3 | 2 | 7 | 23 | 0 | 5 | 3 | 18 | 14 | 236 | 149 | 0.847564858 | 1 | 0.24 | 0.296 | 0.469 | 0.306 | 0.835 | 1 | 0.275 | 0.296 | 0.197 | 0.509 | -0.026 |
| RI | 1036 | ENSG00000185813 | PCYT2 | chr17 | - | 81907553 | 81907857 | 81907553 | 81907598 | 81907772 | 81907857 | 1036 | 3 | 5 | 12 | 10 | 6 | 35 | 15 | 38 | 27 | 24 | 3 | 6 | 1 | 8 | 13 | 50 | 23 | 1 | 4 | 64 | 298 | 149 | 0.853729559 | 1 | 0.041 | 0.143 | 0.136 | 0.156 | 0.111 | 0.029 | 0.115 | 0.333 | 0.5 | 0.092 | -0.096 |
| RI | 2408 | ENSG00000105875 | WDR91 | chr7 | - | 135204267 | 135206058 | 135204267 | 135204465 | 135205927 | 135206058 | 2408 | 33 | 25 | 27 | 27 | 20 | 3 | 0 | 3 | 0 | 0 | 75 | 57 | 38 | 54 | 55 | 0 | 4 | 0 | 3 | 0 | 298 | 149 | 0.886449399 | 1 | 0.846 | 1 | 0.818 | 1 | 1 | 1 | 0.877 | 1 | 0.9 | 1 | -0.023 |
| RI | 1901 | ENSG00000090061 | CCNK | chr14 | + | 99500765 | 99502376 | 99500765 | 99500871 | 99502206 | 99502376 | 1901 | 90 | 102 | 78 | 115 | 118 | 0 | 0 | 3 | 6 | 7 | 151 | 129 | 106 | 173 | 84 | 0 | 4 | 11 | 2 | 3 | 298 | 149 | 0.890941827 | 1 | 1 | 1 | 0.929 | 0.906 | 0.894 | 1 | 0.942 | 0.828 | 0.977 | 0.933 | 0.01 |
| RI | 6904 | ENSG00000090316 | MAEA | chr4 | + | 1338421 | 1340136 | 1338421 | 1338617 | 1339073 | 1340136 | 6904 | 20 | 5 | 31 | 21 | 29 | 160 | 78 | 107 | 116 | 70 | 5 | 7 | 37 | 12 | 24 | 82 | 109 | 44 | 60 | 92 | 298 | 149 | 0.90998577 | 1 | 0.059 | 0.031 | 0.127 | 0.083 | 0.172 | 0.03 | 0.031 | 0.296 | 0.091 | 0.115 | -0.018 |
| RI | 6433 | ENSG00000157764 | BRAF | chr7 | - | 140753274 | 140754233 | 140753274 | 140753393 | 140754186 | 140754233 | 6433 | 15 | 5 | 2 | 11 | 9 | 59 | 54 | 57 | 88 | 95 | 0 | 10 | 21 | 3 | 0 | 32 | 62 | 104 | 33 | 32 | 298 | 149 | 0.918908353 | 1 | 0.113 | 0.044 | 0.017 | 0.059 | 0.045 | 0 | 0.075 | 0.092 | 0.043 | 0 | 0.014 |
| RI | 6856 | ENSG00000215021 | PHB2 | chr12 | - | 6970195 | 6970645 | 6970195 | 6970280 | 6970416 | 6970645 | 6856 | 5 | 1 | 2 | 10 | 1 | 81 | 25 | 65 | 63 | 22 | 0 | 3 | 4 | 6 | 4 | 38 | 40 | 20 | 52 | 93 | 285 | 149 | 0.937692054 | 1 | 0.031 | 0.02 | 0.016 | 0.077 | 0.023 | 0 | 0.038 | 0.095 | 0.057 | 0.022 | -0.009 |
| RI | 5538 | ENSG00000182004 | SNRPE | chr1 | + | 203861602 | 203862222 | 203861602 | 203861743 | 203862181 | 203862222 | 5538 | 26 | 16 | 43 | 31 | 23 | 0 | 1 | 2 | 4 | 10 | 37 | 51 | 45 | 38 | 10 | 0 | 7 | 9 | 5 | 0 | 298 | 149 | 0.952615526 | 1 | 1 | 0.889 | 0.915 | 0.795 | 0.535 | 1 | 0.785 | 0.714 | 0.792 | 1 | -0.031 |
| RI | 4785 | ENSG00000162909 | CAPN2 | chr1 | + | 223769840 | 223770525 | 223769840 | 223769909 | 223770446 | 223770525 | 4785 | 98 | 93 | 54 | 62 | 62 | 358 | 251 | 324 | 485 | 780 | 163 | 96 | 93 | 103 | 67 | 815 | 441 | 339 | 545 | 901 | 298 | 149 | 0.994748504 | 1 | 0.12 | 0.156 | 0.077 | 0.06 | 0.038 | 0.091 | 0.098 | 0.121 | 0.086 | 0.036 | 0.004 |
| RI | 1124 | ENSG00000171962 | DRC3 | chr17 | + | 17992764 | 17994418 | 17992764 | 17992911 | 17994298 | 17994418 | 1124 | 0 | 0 | 0 | 0 | 0 | 1 | 0 | 0 | 0 | 0 | 0 | 2 | 0 | 0 | 0 | 0 | 0 | 0 | 0 | 0 | 298 | 149 | 1 | 1 | 0 | NA | NA | NA | NA | NA | 1 | NA | NA | NA | -1 |
| RI | 2785 | ENSG00000163913 | IFT122 | chr3 | + | 129514388 | 129515599 | 129514388 | 129514554 | 129515487 | 129515599 | 2785 | 1 | 0 | 4 | 2 | 0 | 2 | 3 | 7 | 8 | 6 | 0 | 1 | 8 | 1 | 0 | 0 | 0 | 0 | 0 | 0 | 298 | 149 | 1 | 1 | 0.2 | 0 | 0.222 | 0.111 | 0 | NA | 1 | 1 | 1 | NA | -0.893 |
| RI | 4097 | ENSG00000256269 | HMBS | chr11 | + | 119088634 | 119089131 | 119088634 | 119088848 | 119089081 | 119089131 | 4097 | 0 | 0 | 0 | 0 | 2 | 0 | 0 | 0 | 0 | 8 | 0 | 0 | 0 | 0 | 3 | 0 | 0 | 0 | 0 | 0 | 298 | 149 | 1 | 1 | NA | NA | NA | NA | 0.111 | NA | NA | NA | NA | 1 | -0.889 |
| RI | 5736 | ENSG00000072195 | SPEG | chr2 | + | 219464432 | 219466861 | 219464432 | 219464608 | 219466056 | 219466861 | 5736 | 0 | 0 | 3 | 2 | 3 | 3 | 1 | 2 | 7 | 5 | 0 | 0 | 0 | 1 | 0 | 0 | 0 | 0 | 0 | 0 | 298 | 149 | 1 | 1 | 0 | 0 | 0.429 | 0.125 | 0.231 | NA | NA | NA | 1 | NA | -0.843 |
| RI | 4299 | ENSG00000007402 | CACNA2D2 | chr3 | - | 50364887 | 50365184 | 50364887 | 50364970 | 50365074 | 50365184 | 4299 | 3 | 1 | 1 | 0 | 0 | 24 | 6 | 4 | 3 | 9 | 0 | 0 | 0 | 2 | 7 | 0 | 3 | 0 | 0 | 0 | 253 | 149 | 1 | 1 | 0.069 | 0.089 | 0.128 | 0 | 0 | NA | 0 | NA | 1 | 1 | -0.609 |
| RI | 1817 | ENSG00000196497 | IPO4 | chr14 | - | 24188343 | 24188638 | 24188343 | 24188423 | 24188551 | 24188638 | 1817 | 0 | 0 | 0 | 0 | 0 | 5 | 2 | 3 | 1 | 0 | 0 | 6 | 4 | 6 | 0 | 0 | 6 | 0 | 5 | 0 | 277 | 149 | 1 | 1 | 0 | 0 | 0 | 0 | NA | NA | 0.35 | 1 | 0.392 | NA | -0.581 |
| RI | 1818 | ENSG00000196497 | IPO4 | chr14 | - | 24188551 | 24188797 | 24188551 | 24188638 | 24188718 | 24188797 | 1818 | 1 | 0 | 0 | 3 | 3 | 2 | 0 | 0 | 2 | 2 | 6 | 9 | 0 | 2 | 12 | 0 | 5 | 0 | 0 | 0 | 229 | 149 | 1 | 1 | 0.245 | NA | NA | 0.494 | 0.494 | 1 | 0.539 | NA | 1 | 1 | -0.474 |
| RI | 4300 | ENSG00000007402 | CACNA2D2 | chr3 | - | 50364887 | 50365184 | 50364887 | 50364976 | 50365074 | 50365184 | 4300 | 2 | 1 | 1 | 0 | 0 | 8 | 6 | 2 | 5 | 6 | 0 | 0 | 0 | 2 | 7 | 6 | 0 | 2 | 0 | 0 | 247 | 149 | 1 | 1 | 0.131 | 0.091 | 0.232 | 0 | 0 | 0 | NA | 0 | 1 | 1 | -0.409 |
| RI | 2396 | ENSG00000198689 | SLC9A6 | chrX | + | 135985434 | 135985827 | 135985434 | 135985477 | 135985602 | 135985827 | 2396 | 3 | 1 | 1 | 1 | 4 | 1 | 2 | 3 | 0 | 0 | 0 | 0 | 4 | 0 | 0 | 0 | 0 | 0 | 0 | 0 | 274 | 149 | 1 | 1 | 0.62 | 0.214 | 0.153 | 1 | 1 | NA | NA | 1 | NA | NA | -0.403 |
| RI | 6175 | ENSG00000109339 | MAPK10 | chr4 | - | 86098420 | 86101217 | 86098420 | 86098595 | 86101051 | 86101217 | 6175 | 0 | 0 | 0 | 3 | 0 | 0 | 1 | 0 | 2 | 4 | 0 | 4 | 0 | 0 | 0 | 19 | 0 | 0 | 0 | 0 | 298 | 149 | 1 | 1 | NA | 0 | NA | 0.429 | 0 | 0 | 1 | NA | NA | NA | -0.357 |
| RI | 1106 | ENSG00000131899 | LLGL1 | chr17 | + | 18235469 | 18236760 | 18235469 | 18235537 | 18236606 | 18236760 | 1106 | 0 | 2 | 5 | 0 | 0 | 6 | 0 | 3 | 5 | 0 | 0 | 2 | 4 | 2 | 0 | 0 | 6 | 0 | 0 | 0 | 298 | 149 | 1 | 1 | 0 | 1 | 0.455 | 0 | NA | NA | 0.143 | 1 | 1 | NA | -0.351 |
| RI | 1108 | ENSG00000131899 | LLGL1 | chr17 | + | 18240577 | 18241999 | 18240577 | 18240873 | 18241884 | 18241999 | 1108 | 2 | 0 | 1 | 0 | 0 | 0 | 1 | 0 | 0 | 0 | 22 | 4 | 4 | 0 | 23 | 0 | 0 | 0 | 0 | 0 | 298 | 149 | 1 | 1 | 1 | 0 | 1 | NA | NA | 1 | 1 | 1 | NA | 1 | -0.333 |
| RI | 6345 | ENSG00000068885 | IFT80 | chr3 | - | 160383436 | 160384646 | 160383436 | 160383504 | 160384405 | 160384646 | 6345 | 24 | 33 | 28 | 47 | 69 | 3 | 7 | 6 | 13 | 10 | 10 | 8 | 8 | 6 | 0 | 0 | 0 | 0 | 0 | 0 | 298 | 149 | 1 | 1 | 0.8 | 0.702 | 0.7 | 0.644 | 0.775 | 1 | 1 | 1 | 1 | NA | -0.276 |
| RI | 458 | ENSG00000064545 | TMEM161A | chr19 | - | 19121307 | 19121668 | 19121307 | 19121421 | 19121524 | 19121668 | 458 | 2 | 0 | 3 | 6 | 0 | 4 | 1 | 2 | 2 | 4 | 0 | 0 | 0 | 3 | 0 | 0 | 5 | 0 | 0 | 0 | 252 | 149 | 1 | 1 | 0.228 | 0 | 0.47 | 0.639 | 0 | NA | 0 | NA | 1 | NA | -0.233 |
| RI | 4983 | ENSG00000124067 | SLC12A4 | chr16 | - | 67950653 | 67951060 | 67950653 | 67950711 | 67950961 | 67951060 | 4983 | 7 | 4 | 9 | 6 | 0 | 11 | 5 | 8 | 11 | 8 | 0 | 10 | 2 | 2 | 0 | 5 | 3 | 6 | 0 | 0 | 298 | 149 | 1 | 1 | 0.241 | 0.286 | 0.36 | 0.214 | 0 | 0 | 0.625 | 0.143 | 1 | NA | -0.222 |
| RI | 4102 | ENSG00000256269 | HMBS | chr11 | + | 119092124 | 119092811 | 119092124 | 119092523 | 119092757 | 119092811 | 4102 | 9 | 6 | 4 | 3 | 10 | 17 | 3 | 4 | 9 | 5 | 14 | 11 | 7 | 6 | 0 | 0 | 15 | 3 | 5 | 0 | 298 | 149 | 1 | 1 | 0.209 | 0.5 | 0.333 | 0.143 | 0.5 | 1 | 0.268 | 0.538 | 0.375 | NA | -0.208 |
| RI | 2425 | ENSG00000032742 | IFT88 | chr13 | + | 20567669 | 20567997 | 20567669 | 20567785 | 20567884 | 20567997 | 2425 | 11 | 5 | 4 | 9 | 2 | 2 | 2 | 1 | 2 | 2 | 0 | 5 | 12 | 8 | 1 | 0 | 6 | 0 | 0 | 0 | 248 | 149 | 1 | 1 | 0.768 | 0.6 | 0.706 | 0.73 | 0.375 | NA | 0.334 | 1 | 1 | 1 | -0.198 |
| RI | 8388 | ENSG00000133858 | ZFC3H1 | chr12 | - | 71611057 | 71611887 | 71611057 | 71611097 | 71611785 | 71611887 | 8388 | 48 | 60 | 30 | 25 | 62 | 64 | 41 | 63 | 58 | 106 | 0 | 27 | 18 | 3 | 8 | 0 | 56 | 14 | 1 | 3 | 298 | 149 | 1 | 1 | 0.273 | 0.423 | 0.192 | 0.177 | 0.226 | NA | 0.194 | 0.391 | 0.6 | 0.571 | -0.181 |
| RI | 1621 | ENSG00000103852 | TTC23 | chr15 | - | 99218587 | 99219048 | 99218587 | 99218713 | 99218897 | 99219048 | 1621 | 0 | 0 | 0 | 0 | 0 | 3 | 8 | 5 | 0 | 5 | 12 | 4 | 1 | 0 | 0 | 4 | 12 | 3 | 3 | 13 | 298 | 149 | 1 | 1 | 0 | 0 | 0 | NA | 0 | 0.6 | 0.143 | 0.143 | 0 | 0 | -0.177 |
| RI | 4833 | ENSG00000103168 | TAF1C | chr16 | - | 84180169 | 84181186 | 84180169 | 84180344 | 84181042 | 84181186 | 4833 | 26 | 1 | 16 | 17 | 9 | 3 | 1 | 4 | 7 | 2 | 0 | 25 | 26 | 8 | 64 | 0 | 0 | 6 | 2 | 9 | 298 | 149 | 1 | 1 | 0.812 | 0.333 | 0.667 | 0.548 | 0.692 | NA | 1 | 0.684 | 0.667 | 0.78 | -0.172 |
| RI | 5452 | ENSG00000143799 | PARP1 | chr1 | - | 226385503 | 226386442 | 226385503 | 226385680 | 226386218 | 226386442 | 5452 | 6 | 3 | 6 | 10 | 0 | 1 | 0 | 2 | 0 | 0 | 8 | 17 | 29 | 23 | 0 | 0 | 0 | 0 | 0 | 0 | 298 | 149 | 1 | 1 | 0.75 | 1 | 0.6 | 1 | NA | 1 | 1 | 1 | 1 | NA | -0.162 |
| RI | 5001 | ENSG00000092529 | CAPN3 | chr15 | + | 42409930 | 42410496 | 42409930 | 42409995 | 42410431 | 42410496 | 5001 | 0 | 1 | 2 | 1 | 8 | 4 | 0 | 0 | 0 | 0 | 0 | 4 | 5 | 14 | 0 | 0 | 0 | 0 | 1 | 0 | 298 | 149 | 1 | 1 | 0 | 1 | 1 | 1 | 1 | NA | 1 | 1 | 0.875 | NA | -0.158 |
| RI | 95 | ENSG00000198832 | SELENOM | chr22 | - | 31105929 | 31107533 | 31105929 | 31105965 | 31107376 | 31107533 | 95 | 2 | 12 | 11 | 3 | 7 | 6 | 4 | 15 | 10 | 3 | 0 | 4 | 0 | 3 | 14 | 5 | 12 | 0 | 0 | 2 | 298 | 149 | 1 | 1 | 0.143 | 0.6 | 0.268 | 0.13 | 0.538 | 0 | 0.143 | NA | 1 | 0.778 | -0.144 |
| RI | 774 | ENSG00000073350 | LLGL2 | chr17 | + | 75574212 | 75574668 | 75574212 | 75574260 | 75574609 | 75574668 | 774 | 2 | 0 | 1 | 4 | 0 | 3 | 1 | 0 | 8 | 0 | 0 | 0 | 2 | 0 | 4 | 0 | 13 | 0 | 8 | 0 | 298 | 149 | 1 | 1 | 0.25 | 0 | 1 | 0.2 | NA | NA | 0 | 1 | 0 | 1 | -0.138 |
| RI | 8494 | ENSG00000204852 | TCTN1 | chr12 | + | 110641541 | 110642389 | 110641541 | 110641627 | 110642248 | 110642389 | 8494 | 8 | 10 | 16 | 15 | 5 | 8 | 5 | 7 | 4 | 9 | 0 | 4 | 0 | 1 | 6 | 6 | 4 | 0 | 0 | 0 | 298 | 149 | 1 | 1 | 0.333 | 0.5 | 0.533 | 0.652 | 0.217 | 0 | 0.333 | NA | 1 | 1 | -0.136 |
| RI | 460 | ENSG00000064545 | TMEM161A | chr19 | - | 19132351 | 19132754 | 19132351 | 19132508 | 19132656 | 19132754 | 460 | 3 | 1 | 8 | 1 | 2 | 6 | 7 | 5 | 4 | 0 | 8 | 0 | 0 | 0 | 0 | 0 | 0 | 4 | 0 | 0 | 297 | 149 | 1 | 1 | 0.201 | 0.067 | 0.445 | 0.111 | 1 | 1 | NA | 0 | NA | NA | -0.135 |
| RI | 1107 | ENSG00000131899 | LLGL1 | chr17 | + | 18236834 | 18237773 | 18236834 | 18236939 | 18237480 | 18237773 | 1107 | 5 | 3 | 3 | 8 | 0 | 8 | 3 | 7 | 3 | 0 | 0 | 17 | 2 | 2 | 11 | 4 | 3 | 6 | 0 | 9 | 298 | 149 | 1 | 1 | 0.238 | 0.333 | 0.176 | 0.571 | NA | 0 | 0.739 | 0.143 | 1 | 0.379 | -0.123 |
| RI | 1034 | ENSG00000185813 | PCYT2 | chr17 | - | 81905669 | 81906177 | 81905669 | 81905735 | 81906099 | 81906177 | 1034 | 0 | 3 | 4 | 4 | 0 | 15 | 2 | 12 | 14 | 14 | 0 | 0 | 2 | 0 | 0 | 8 | 16 | 0 | 0 | 16 | 298 | 149 | 1 | 1 | 0 | 0.429 | 0.143 | 0.125 | 0 | 0 | 0 | 1 | NA | 0 | -0.111 |
| RI | 4325 | ENSG00000111445 | RFC5 | chr12 | + | 118017689 | 118018053 | 118017689 | 118017815 | 118017994 | 118018053 | 4325 | 10 | 16 | 7 | 7 | 24 | 2 | 0 | 0 | 1 | 0 | 4 | 17 | 21 | 23 | 17 | 0 | 0 | 0 | 0 | 0 | 298 | 149 | 1 | 1 | 0.714 | 1 | 1 | 0.778 | 1 | 1 | 1 | 1 | 1 | 1 | -0.102 |
| RI | 8267 | ENSG00000181588 | MEX3D | chr19 | - | 1554671 | 1556923 | 1554671 | 1555389 | 1555576 | 1556923 | 8267 | 2 | 1 | 2 | 2 | 2 | 0 | 0 | 1 | 0 | 0 | 5 | 1 | 2 | 0 | 0 | 0 | 0 | 0 | 0 | 0 | 298 | 149 | 1 | 1 | 1 | 1 | 0.5 | 1 | 1 | 1 | 1 | 1 | NA | NA | -0.1 |
| RI | 2811 | ENSG00000160221 | GATD3A | chr21 | + | 44144820 | 44145723 | 44144820 | 44144916 | 44145570 | 44145723 | 2811 | 0 | 32 | 0 | 0 | 44 | 0 | 0 | 0 | 0 | 4 | 20 | 0 | 0 | 0 | 0 | 0 | 0 | 0 | 0 | 0 | 298 | 149 | 1 | 1 | NA | 1 | NA | NA | 0.846 | 1 | NA | NA | NA | NA | -0.077 |
| RI | 5423 | ENSG00000166762 | CATSPER2 | chr15 | - | 43632198 | 43635416 | 43632198 | 43632363 | 43635359 | 43635416 | 5423 | 8 | 10 | 8 | 9 | 1 | 2 | 0 | 0 | 0 | 0 | 0 | 0 | 9 | 6 | 0 | 0 | 0 | 0 | 0 | 0 | 298 | 149 | 1 | 1 | 0.667 | 1 | 1 | 1 | 1 | NA | NA | 1 | 1 | NA | -0.067 |
| RI | 2869 | ENSG00000184470 | TXNRD2 | chr22 | - | 19875521 | 19877234 | 19875521 | 19875807 | 19877108 | 19877234 | 2869 | 9 | 9 | 19 | 15 | 12 | 0 | 1 | 0 | 0 | 2 | 29 | 2 | 2 | 15 | 27 | 0 | 0 | 0 | 0 | 2 | 298 | 149 | 1 | 1 | 1 | 0.818 | 1 | 1 | 0.75 | 1 | 1 | 1 | 1 | 0.871 | -0.061 |
| RI | 7421 | ENSG00000099958 | DERL3 | chr22 | - | 23834502 | 23837154 | 23834502 | 23836962 | 23837063 | 23837154 | 7421 | 2 | 2 | 9 | 1 | 6 | 4 | 4 | 11 | 0 | 0 | 0 | 6 | 3 | 3 | 9 | 0 | 3 | 14 | 0 | 2 | 250 | 149 | 1 | 1 | 0.23 | 0.23 | 0.328 | 1 | 1 | NA | 0.544 | 0.113 | 1 | 0.728 | -0.039 |
| RI | 1068 | ENSG00000198496 | NBR2 | chr17 | + | 43138656 | 43140283 | 43138656 | 43138922 | 43139815 | 43140283 | 1068 | 20 | 12 | 31 | 19 | 20 | 2 | 0 | 2 | 0 | 0 | 61 | 33 | 10 | 33 | 16 | 0 | 0 | 0 | 0 | 1 | 298 | 149 | 1 | 1 | 0.833 | 1 | 0.886 | 1 | 1 | 1 | 1 | 1 | 1 | 0.889 | -0.034 |
| RI | 603 | ENSG00000171703 | TCEA2 | chr20 | + | 64071869 | 64072346 | 64071869 | 64071941 | 64072171 | 64072346 | 603 | 29 | 5 | 27 | 10 | 13 | 14 | 4 | 22 | 8 | 7 | 0 | 19 | 22 | 3 | 47 | 0 | 30 | 4 | 3 | 22 | 298 | 149 | 1 | 1 | 0.509 | 0.385 | 0.38 | 0.385 | 0.481 | NA | 0.241 | 0.733 | 0.333 | 0.516 | -0.028 |
| RI | 5272 | ENSG00000168890 | TMEM150A | chr2 | - | 85601020 | 85601482 | 85601020 | 85601107 | 85601305 | 85601482 | 5272 | 8 | 4 | 7 | 5 | 16 | 0 | 0 | 0 | 1 | 0 | 6 | 16 | 15 | 17 | 0 | 0 | 0 | 1 | 0 | 0 | 298 | 149 | 1 | 1 | 1 | 1 | 1 | 0.714 | 1 | 1 | 1 | 0.882 | 1 | NA | -0.028 |
| RI | 3308 | ENSG00000162601 | MYSM1 | chr1 | - | 58661405 | 58663522 | 58661405 | 58661511 | 58663449 | 58663522 | 3308 | 41 | 79 | 45 | 37 | 7 | 1 | 7 | 6 | 15 | 1 | 0 | 28 | 51 | 12 | 9 | 0 | 2 | 6 | 5 | 0 | 298 | 149 | 1 | 1 | 0.953 | 0.849 | 0.789 | 0.552 | 0.778 | NA | 0.875 | 0.81 | 0.545 | 1 | -0.023 |
| RI | 2691 | ENSG00000108591 | DRG2 | chr17 | + | 18099016 | 18100435 | 18099016 | 18099093 | 18100362 | 18100435 | 2691 | 12 | 9 | 21 | 32 | 13 | 0 | 0 | 0 | 2 | 0 | 58 | 24 | 18 | 3 | 39 | 0 | 0 | 0 | 0 | 0 | 298 | 149 | 1 | 1 | 1 | 1 | 1 | 0.889 | 1 | 1 | 1 | 1 | 1 | 1 | -0.022 |
| RI | 1483 | ENSG00000167965 | MLST8 | chr16 | + | 2208209 | 2208613 | 2208209 | 2208334 | 2208449 | 2208613 | 1483 | 5 | 3 | 6 | 2 | 0 | 5 | 5 | 15 | 5 | 7 | 0 | 10 | 1 | 0 | 0 | 0 | 2 | 6 | 1 | 12 | 264 | 149 | 1 | 1 | 0.361 | 0.253 | 0.184 | 0.184 | 0 | NA | 0.738 | 0.086 | 0 | 0 | -0.01 |
| RI | 2088 | ENSG00000174032 | SLC25A30 | chr13 | - | 45393315 | 45399078 | 45393315 | 45396015 | 45398939 | 45399078 | 2088 | 22 | 19 | 17 | 19 | 4 | 0 | 1 | 0 | 0 | 0 | 35 | 24 | 3 | 32 | 11 | 0 | 0 | 0 | 1 | 0 | 298 | 149 | 1 | 1 | 1 | 0.905 | 1 | 1 | 1 | 1 | 1 | 1 | 0.941 | 1 | -0.007 |
| RI | 7792 | ENSG00000074266 | EED | chr11 | + | 86263944 | 86266216 | 86263944 | 86264263 | 86266082 | 86266216 | 7792 | 6 | 1 | 4 | 7 | 2 | 51 | 27 | 61 | 69 | 66 | 0 | 3 | 5 | 1 | 0 | 5 | 17 | 40 | 28 | 0 | 298 | 149 | 1 | 1 | 0.056 | 0.018 | 0.032 | 0.048 | 0.015 | 0 | 0.081 | 0.059 | 0.018 | NA | -0.006 |
| RI | 3877 | ENSG00000145740 | SLC30A5 | chr5 | + | 69121746 | 69121895 | 69121746 | 69121752 | 69121753 | 69121895 | 3877 | 81 | 50 | 79 | 80 | 104 | 0 | 0 | 2 | 0 | 0 | 47 | 23 | 31 | 31 | 15 | 0 | 0 | 0 | 0 | 0 | 150 | 149 | 1 | 1 | 1 | 1 | 0.975 | 1 | 1 | 1 | 1 | 1 | 1 | 1 | -0.005 |
| RI | 2223 | ENSG00000113141 | IK | chr5 | + | 140661619 | 140661708 | 140661619 | 140661631 | 140661632 | 140661708 | 2223 | 256 | 167 | 277 | 315 | 379 | 0 | 2 | 0 | 0 | 0 | 151 | 388 | 298 | 318 | 127 | 0 | 0 | 0 | 0 | 0 | 150 | 149 | 1 | 1 | 1 | 0.988 | 1 | 1 | 1 | 1 | 1 | 1 | 1 | 1 | -0.002 |
| RI | 5311 | ENSG00000132676 | DAP3 | chr1 | + | 155738156 | 155739010 | 155738156 | 155738258 | 155738608 | 155739010 | 5311 | 210 | 135 | 238 | 235 | 180 | 1 | 0 | 0 | 0 | 0 | 188 | 193 | 180 | 146 | 242 | 0 | 0 | 0 | 0 | 0 | 298 | 149 | 1 | 1 | 0.991 | 1 | 1 | 1 | 1 | 1 | 1 | 1 | 1 | 1 | -0.002 |
| RI | 8598 | ENSG00000134899 | ERCC5 | chr13 | + | 102853756 | 102854374 | 102853756 | 102853872 | 102854142 | 102854374 | 8598 | 81 | 91 | 89 | 130 | 95 | 0 | 0 | 0 | 0 | 1 | 17 | 93 | 155 | 46 | 27 | 0 | 0 | 1 | 0 | 0 | 298 | 149 | 1 | 1 | 1 | 1 | 1 | 1 | 0.979 | 1 | 1 | 0.987 | 1 | 1 | -0.002 |
| RI | 7654 | ENSG00000179832 | MROH1 | chr8 | + | 144261113 | 144261349 | 144261113 | 144261216 | 144261283 | 144261349 | 7654 | 2 | 4 | 0 | 2 | 8 | 2 | 6 | 5 | 7 | 2 | 4 | 9 | 4 | 0 | 0 | 8 | 14 | 1 | 2 | 0 | 216 | 149 | 1 | 1 | 0.408 | 0.315 | 0 | 0.165 | 0.734 | 0.256 | 0.307 | 0.734 | 0 | NA | 0 |
| RI | 3092 | ENSG00000111642 | CHD4 | chr12 | - | 6578028 | 6578546 | 6578028 | 6578135 | 6578408 | 6578546 | 3092 | 115 | 80 | 80 | 110 | 163 | 0 | 0 | 0 | 0 | 0 | 442 | 203 | 192 | 403 | 219 | 0 | 0 | 1 | 0 | 0 | 298 | 149 | 1 | 1 | 1 | 1 | 1 | 1 | 1 | 1 | 1 | 0.99 | 1 | 1 | 0.002 |
| RI | 1271 | ENSG00000205250 | E2F4 | chr16 | + | 67192760 | 67193170 | 67192760 | 67192824 | 67193008 | 67193170 | 1271 | 136 | 73 | 68 | 117 | 81 | 1 | 0 | 0 | 0 | 0 | 193 | 136 | 68 | 190 | 169 | 0 | 3 | 0 | 0 | 0 | 298 | 149 | 1 | 1 | 0.986 | 1 | 1 | 1 | 1 | 1 | 0.958 | 1 | 1 | 1 | 0.006 |
| RI | 8457 | ENSG00000123349 | PFDN5 | chr12 | + | 53295838 | 53298150 | 53295838 | 53295941 | 53298044 | 53298150 | 8457 | 198 | 152 | 201 | 218 | 234 | 0 | 0 | 0 | 0 | 0 | 297 | 255 | 176 | 207 | 275 | 0 | 4 | 0 | 0 | 0 | 298 | 149 | 1 | 1 | 1 | 1 | 1 | 1 | 1 | 1 | 0.97 | 1 | 1 | 1 | 0.006 |
| RI | 5886 | ENSG00000167515 | TRAPPC2L | chr16 | + | 88859344 | 88859750 | 88859344 | 88859443 | 88859572 | 88859750 | 5886 | 34 | 8 | 32 | 14 | 41 | 0 | 0 | 3 | 0 | 1 | 29 | 41 | 19 | 23 | 73 | 0 | 0 | 3 | 0 | 0 | 278 | 149 | 1 | 1 | 1 | 1 | 0.851 | 1 | 0.956 | 1 | 1 | 0.772 | 1 | 1 | 0.007 |
| RI | 4096 | ENSG00000256269 | HMBS | chr11 | + | 119088634 | 119089131 | 119088634 | 119088707 | 119089081 | 119089131 | 4096 | 0 | 0 | 1 | 2 | 2 | 29 | 18 | 19 | 23 | 14 | 0 | 1 | 0 | 0 | 3 | 18 | 40 | 19 | 23 | 18 | 298 | 149 | 1 | 1 | 0 | 0 | 0.026 | 0.042 | 0.067 | 0 | 0.012 | 0 | 0 | 0.077 | 0.009 |
| RI | 7112 | ENSG00000133740 | E2F5 | chr8 | + | 85213752 | 85214518 | 85213752 | 85213902 | 85214159 | 85214518 | 7112 | 117 | 103 | 102 | 173 | 309 | 0 | 0 | 0 | 1 | 2 | 61 | 113 | 81 | 110 | 63 | 0 | 0 | 0 | 0 | 3 | 298 | 149 | 1 | 1 | 1 | 1 | 1 | 0.989 | 0.987 | 1 | 1 | 1 | 1 | 0.913 | 0.013 |
| RI | 5339 | ENSG00000084623 | EIF3I | chr1 | + | 32226170 | 32226530 | 32226170 | 32226283 | 32226402 | 32226530 | 5339 | 39 | 41 | 39 | 56 | 40 | 0 | 0 | 0 | 0 | 0 | 56 | 67 | 28 | 69 | 129 | 0 | 0 | 0 | 3 | 0 | 268 | 149 | 1 | 1 | 1 | 1 | 1 | 1 | 1 | 1 | 1 | 1 | 0.927 | 1 | 0.015 |
| RI | 1923 | ENSG00000125746 | EML2 | chr19 | - | 45632860 | 45633139 | 45632860 | 45632971 | 45633069 | 45633139 | 1923 | 1 | 1 | 0 | 2 | 0 | 36 | 11 | 21 | 47 | 10 | 0 | 0 | 0 | 1 | 0 | 78 | 37 | 21 | 40 | 94 | 247 | 149 | 1 | 1 | 0.016 | 0.052 | 0 | 0.025 | 0 | 0 | 0 | 0 | 0.015 | 0 | 0.016 |
| RI | 4744 | ENSG00000131778 | CHD1L | chr1 | + | 147272170 | 147272630 | 147272170 | 147272281 | 147272446 | 147272630 | 4744 | 19 | 4 | 8 | 9 | 25 | 0 | 0 | 0 | 0 | 0 | 32 | 25 | 25 | 31 | 0 | 0 | 1 | 0 | 0 | 0 | 298 | 149 | 1 | 1 | 1 | 1 | 1 | 1 | 1 | 1 | 0.926 | 1 | 1 | NA | 0.018 |
| RI | 4101 | ENSG00000256269 | HMBS | chr11 | + | 119092124 | 119092811 | 119092124 | 119092163 | 119092757 | 119092811 | 4101 | 1 | 3 | 5 | 2 | 4 | 2 | 0 | 2 | 0 | 1 | 15 | 3 | 8 | 3 | 0 | 0 | 22 | 0 | 1 | 0 | 298 | 149 | 1 | 1 | 0.2 | 1 | 0.556 | 1 | 0.667 | 1 | 0.064 | 1 | 0.6 | NA | 0.019 |
| RI | 3094 | ENSG00000111642 | CHD4 | chr12 | - | 6583026 | 6583378 | 6583026 | 6583113 | 6583195 | 6583378 | 3094 | 25 | 14 | 20 | 14 | 23 | 1 | 0 | 1 | 0 | 0 | 48 | 48 | 38 | 70 | 62 | 0 | 0 | 0 | 14 | 0 | 231 | 149 | 1 | 1 | 0.942 | 1 | 0.928 | 1 | 1 | 1 | 1 | 1 | 0.763 | 1 | 0.021 |
| RI | 773 | ENSG00000073350 | LLGL2 | chr17 | + | 75574212 | 75574668 | 75574212 | 75574260 | 75574452 | 75574668 | 773 | 2 | 0 | 0 | 3 | 0 | 30 | 13 | 11 | 18 | 23 | 0 | 0 | 0 | 0 | 0 | 4 | 27 | 7 | 15 | 20 | 298 | 149 | 1 | 1 | 0.032 | 0 | 0 | 0.077 | 0 | 0 | 0 | 0 | 0 | 0 | 0.022 |
| RI | 1480 | ENSG00000167965 | MLST8 | chr16 | + | 2206357 | 2206659 | 2206357 | 2206409 | 2206483 | 2206659 | 1480 | 5 | 7 | 2 | 7 | 8 | 0 | 0 | 0 | 0 | 0 | 16 | 3 | 3 | 6 | 33 | 0 | 0 | 0 | 0 | 3 | 223 | 149 | 1 | 1 | 1 | 1 | 1 | 1 | 1 | 1 | 1 | 1 | 1 | 0.88 | 0.024 |
| RI | 1489 | ENSG00000103249 | CLCN7 | chr16 | - | 1449275 | 1450666 | 1449275 | 1449327 | 1450496 | 1450666 | 1489 | 1 | 1 | 2 | 1 | 0 | 19 | 9 | 6 | 21 | 9 | 0 | 1 | 3 | 0 | 0 | 4 | 30 | 17 | 21 | 45 | 298 | 149 | 1 | 1 | 0.026 | 0.053 | 0.143 | 0.023 | 0 | 0 | 0.016 | 0.081 | 0 | 0 | 0.03 |
| RI | 973 | ENSG00000179029 | TMEM107 | chr17 | - | 8175756 | 8176026 | 8175756 | 8175857 | 8175958 | 8176026 | 973 | 10 | 4 | 8 | 4 | 11 | 6 | 4 | 3 | 7 | 1 | 0 | 4 | 1 | 0 | 20 | 0 | 7 | 0 | 4 | 6 | 250 | 149 | 1 | 1 | 0.498 | 0.373 | 0.614 | 0.254 | 0.868 | NA | 0.254 | 1 | 0 | 0.665 | 0.042 |
| RI | 5510 | ENSG00000112367 | FIG4 | chr6 | + | 109743109 | 109743772 | 109743109 | 109743272 | 109743646 | 109743772 | 5510 | 37 | 11 | 9 | 5 | 7 | 1 | 0 | 0 | 0 | 0 | 5 | 23 | 10 | 3 | 0 | 0 | 3 | 0 | 0 | 0 | 298 | 149 | 1 | 1 | 0.949 | 1 | 1 | 1 | 1 | 1 | 0.793 | 1 | 1 | NA | 0.042 |
| RI | 2415 | ENSG00000158290 | CUL4B | chrX | - | 120538659 | 120540562 | 120538659 | 120538770 | 120540369 | 120540562 | 2415 | 20 | 17 | 17 | 17 | 10 | 0 | 0 | 0 | 3 | 0 | 0 | 5 | 16 | 2 | 0 | 0 | 1 | 0 | 0 | 0 | 298 | 149 | 1 | 1 | 1 | 1 | 1 | 0.739 | 1 | NA | 0.714 | 1 | 1 | NA | 0.043 |
| RI | 7420 | ENSG00000099958 | DERL3 | chr22 | - | 23834502 | 23837154 | 23834502 | 23834932 | 23837063 | 23837154 | 7420 | 5 | 5 | 14 | 3 | 10 | 0 | 0 | 3 | 0 | 2 | 0 | 14 | 8 | 4 | 22 | 0 | 4 | 0 | 0 | 8 | 298 | 149 | 1 | 1 | 1 | 1 | 0.7 | 1 | 0.714 | NA | 0.636 | 1 | 1 | 0.579 | 0.079 |
| RI | 2602 | ENSG00000079337 | RAPGEF3 | chr12 | - | 47748453 | 47748931 | 47748453 | 47748542 | 47748818 | 47748931 | 2602 | 8 | 8 | 3 | 0 | 0 | 0 | 0 | 0 | 1 | 0 | 0 | 6 | 0 | 0 | 7 | 0 | 0 | 0 | 2 | 0 | 298 | 149 | 1 | 1 | 1 | 1 | 1 | 0 | NA | NA | 1 | NA | 0 | 1 | 0.083 |
| RI | 4555 | ENSG00000135749 | PCNX2 | chr1 | - | 232984135 | 232986540 | 232984135 | 232984477 | 232986091 | 232986540 | 4555 | 2 | 9 | 2 | 0 | 0 | 2 | 1 | 4 | 0 | 0 | 0 | 0 | 9 | 0 | 0 | 0 | 5 | 0 | 3 | 0 | 298 | 149 | 1 | 1 | 0.333 | 0.818 | 0.2 | NA | NA | NA | 0 | 1 | 0 | NA | 0.117 |
| RI | 1925 | ENSG00000125746 | EML2 | chr19 | - | 45641917 | 45642352 | 45641917 | 45642041 | 45642133 | 45642352 | 1925 | 3 | 3 | 8 | 2 | 0 | 3 | 0 | 4 | 3 | 0 | 5 | 0 | 3 | 0 | 0 | 7 | 0 | 0 | 0 | 4 | 241 | 149 | 1 | 1 | 0.382 | 1 | 0.553 | 0.292 | NA | 0.306 | NA | 1 | NA | 0 | 0.121 |
| RI | 5389 | ENSG00000156475 | PPP2R2B | chr5 | - | 146878001 | 146878785 | 146878001 | 146878195 | 146878700 | 146878785 | 5389 | 7 | 0 | 8 | 0 | 0 | 1 | 0 | 1 | 0 | 0 | 0 | 0 | 2 | 0 | 6 | 0 | 0 | 0 | 1 | 0 | 298 | 149 | 1 | 1 | 0.778 | NA | 0.8 | NA | NA | NA | NA | 1 | 0 | 1 | 0.122 |
| RI | 2603 | ENSG00000079337 | RAPGEF3 | chr12 | - | 47748453 | 47749536 | 47748453 | 47748931 | 47749389 | 47749536 | 2603 | 2 | 3 | 0 | 1 | 0 | 1 | 0 | 0 | 0 | 1 | 0 | 0 | 0 | 0 | 7 | 0 | 0 | 0 | 2 | 0 | 298 | 149 | 1 | 1 | 0.5 | 1 | NA | 1 | 0 | NA | NA | NA | 0 | 1 | 0.125 |
| RI | 2951 | ENSG00000164068 | RNF123 | chr3 | + | 49713737 | 49713997 | 49713737 | 49713825 | 49713909 | 49713997 | 2951 | 2 | 0 | 3 | 7 | 2 | 14 | 4 | 7 | 7 | 2 | 0 | 0 | 2 | 0 | 0 | 0 | 4 | 4 | 7 | 28 | 233 | 149 | 1 | 1 | 0.084 | 0 | 0.215 | 0.39 | 0.39 | NA | 0 | 0.242 | 0 | 0 | 0.155 |
| RI | 3956 | ENSG00000157445 | CACNA2D3 | chr3 | + | 55073444 | 55073859 | 55073444 | 55073557 | 55073776 | 55073859 | 3956 | 15 | 22 | 10 | 14 | 13 | 34 | 20 | 28 | 10 | 10 | 0 | 0 | 0 | 4 | 0 | 7 | 0 | 0 | 3 | 10 | 298 | 149 | 1 | 1 | 0.181 | 0.355 | 0.152 | 0.412 | 0.394 | 0 | NA | NA | 0.4 | 0 | 0.165 |
| RI | 5271 | ENSG00000135931 | ARMC9 | chr2 | + | 231214830 | 231216793 | 231214830 | 231215001 | 231216637 | 231216793 | 5271 | 0 | 0 | 0 | 1 | 0 | 2 | 2 | 2 | 0 | 2 | 0 | 0 | 0 | 0 | 0 | 0 | 0 | 1 | 0 | 0 | 298 | 149 | 1 | 1 | 0 | 0 | 0 | 1 | 0 | NA | NA | 0 | NA | NA | 0.2 |
| RI | 1754 | ENSG00000174373 | RALGAPA1 | chr14 | - | 35688458 | 35690003 | 35688458 | 35688540 | 35689835 | 35690003 | 1754 | 34 | 26 | 44 | 24 | 80 | 2 | 1 | 5 | 0 | 5 | 0 | 15 | 26 | 0 | 0 | 0 | 2 | 10 | 0 | 0 | 298 | 149 | 1 | 1 | 0.895 | 0.929 | 0.815 | 1 | 0.889 | NA | 0.789 | 0.565 | NA | NA | 0.229 |
| RI | 5512 | ENSG00000112367 | FIG4 | chr6 | + | 109786301 | 109788956 | 109786301 | 109786449 | 109788797 | 109788956 | 5512 | 18 | 9 | 4 | 11 | 7 | 0 | 0 | 2 | 1 | 2 | 0 | 0 | 2 | 2 | 0 | 6 | 3 | 0 | 0 | 0 | 298 | 149 | 1 | 1 | 1 | 1 | 0.5 | 0.846 | 0.636 | 0 | 0 | 1 | 1 | NA | 0.296 |
| RI | 1764 | ENSG00000100949 | RABGGTA | chr14 | - | 24270836 | 24271169 | 24270836 | 24270998 | 24271112 | 24271169 | 1764 | 6 | 1 | 5 | 12 | 1 | 0 | 0 | 0 | 0 | 0 | 0 | 8 | 2 | 2 | 4 | 0 | 4 | 3 | 0 | 0 | 263 | 149 | 1 | 1 | 1 | 1 | 1 | 1 | 1 | NA | 0.531 | 0.274 | 1 | 1 | 0.299 |
| RI | 5514 | ENSG00000112367 | FIG4 | chr6 | + | 109786301 | 109789677 | 109786301 | 109788956 | 109789593 | 109789677 | 5514 | 19 | 21 | 18 | 25 | 12 | 0 | 0 | 0 | 0 | 2 | 0 | 1 | 2 | 2 | 0 | 14 | 0 | 2 | 0 | 0 | 298 | 149 | 1 | 1 | 1 | 1 | 1 | 1 | 0.75 | 0 | 1 | 0.333 | 1 | NA | 0.367 |
| RI | 3975 | ENSG00000154358 | OBSCN | chr1 | + | 228373931 | 228374424 | 228373931 | 228374012 | 228374313 | 228374424 | 3975 | 0 | 0 | 1 | 2 | 3 | 0 | 2 | 3 | 0 | 1 | 0 | 0 | 1 | 0 | 0 | 9 | 2 | 2 | 0 | 0 | 298 | 149 | 1 | 1 | NA | 0 | 0.143 | 1 | 0.6 | 0 | 0 | 0.2 | NA | NA | 0.369 |
| RI | 6827 | ENSG00000184343 | SRPK3 | chrX | + | 153783973 | 153784213 | 153783973 | 153784021 | 153784120 | 153784213 | 6827 | 0 | 0 | 1 | 17 | 0 | 0 | 0 | 0 | 0 | 0 | 2 | 0 | 0 | 2 | 0 | 0 | 0 | 0 | 5 | 0 | 248 | 149 | 1 | 1 | NA | NA | 1 | 1 | NA | 1 | NA | NA | 0.194 | NA | 0.403 |
| RI | 3708 | ENSG00000137225 | CAPN11 | chr6 | + | 44179618 | 44180163 | 44179618 | 44179630 | 44179951 | 44180163 | 3708 | 0 | 0 | 0 | 2 | 0 | 0 | 0 | 0 | 0 | 0 | 0 | 0 | 0 | 0 | 0 | 0 | 2 | 0 | 0 | 20 | 298 | 149 | 1 | 1 | NA | NA | NA | 1 | NA | NA | 0 | NA | NA | 0 | 1 |

MXE

| Splicing_type | ID | Gene_ID | Gene_Symbol | Chromosome | Strand | 1stExonStart_0base | 1stExonEnd | 2ndExonStart_0base | 2ndExonEnd | UpstreamES | UpstreamEE | DownstreamES | DownstreamEE | SRR13214146 |
|---|---|---|---|---|---|---|---|---|---|---|---|---|---|---|
| MXE | 160 | ENSG00000100351 | GRAP2 | chr22 | + | 39947092 | 39947184 | 39960054 | 39960174 | 39901083 | 39901330 | 39965989 | 39966158 | 124 |
| MXE | 380 | ENSG00000170889 | RPS9 | chr19 | + | 54206275 | 54206462 | 54206552 | 54206724 | 54201486 | 54201609 | 54207397 | 54207647 | 4621 |
| MXE | 692 | ENSG00000101347 | SAMHD1 | chr20 | - | 36930759 | 36930875 | 36935028 | 36935189 | 36927181 | 36927252 | 36935340 | 36935403 | 151 |
| MXE | 694 | ENSG00000101347 | SAMHD1 | chr20 | - | 36935028 | 36935189 | 36935340 | 36935403 | 36930759 | 36930875 | 36941038 | 36941111 | 54 |
| MXE | 698 | ENSG00000101347 | SAMHD1 | chr20 | - | 36935340 | 36935403 | 36941038 | 36941111 | 36935028 | 36935189 | 36946737 | 36946804 | 2577 |
| MXE | 1302 | ENSG00000277957 | SENP3-EIF4A1 | chr17 | + | 7574259 | 7574308 | 7574545 | 7574678 | 7572283 | 7572864 | 7575118 | 7575258 | 10 |
| MXE | 2193 | ENSG00000170776 | AKAP13 | chr15 | + | 85658536 | 85658590 | 85662387 | 85662453 | 85655416 | 85655787 | 85664562 | 85664755 | 190 |
| MXE | 2661 | ENSG00000257621 | PSMA3-AS1 | chr14 | - | 58273978 | 58274135 | 58291634 | 58291707 | 58266888 | 58267393 | 58297954 | 58298122 | 68 |
| MXE | 3193 | ENSG00000188994 | ZNF292 | chr6 | + | 87218595 | 87218731 | 87233324 | 87233527 | 87216298 | 87216377 | 87243474 | 87243611 | 215 |
| MXE | 4426 | ENSG00000100650 | SRSF5 | chr14 | + | 69768612 | 69768674 | 69768797 | 69768896 | 69768137 | 69768282 | 69769181 | 69769251 | 10 |
| MXE | 5392 | ENSG00000162736 | NCSTN | chr1 | + | 160351221 | 160351372 | 160351695 | 160351805 | 160350104 | 160350250 | 160352886 | 160352991 | 287 |
| MXE | 10943 | ENSG00000022840 | RNF10 | chr12 | + | 120560725 | 120560886 | 120562944 | 120563070 | 120557545 | 120557682 | 120563331 | 120563623 | 261 |
| MXE | 11459 | ENSG00000043462 | LCP2 | chr5 | - | 170268384 | 170268482 | 170270718 | 170270917 | 170267008 | 170267075 | 170274300 | 170274338 | 515 |
| MXE | 13485 | ENSG00000244754 | N4BP2L2 | chr13 | - | 32522181 | 32522270 | 32527407 | 32527532 | 32521372 | 32521449 | 32535768 | 32537027 | 235 |
| MXE | 14642 | ENSG00000165525 | NEMF | chr14 | - | 49840717 | 49840866 | 49846139 | 49846265 | 49838138 | 49838206 | 49851562 | 49851665 | 207 |
| MXE | 15165 | ENSG00000151465 | CDC123 | chr10 | + | 12238456 | 12238485 | 12246148 | 12246277 | 12237143 | 12237266 | 12250310 | 12250589 | 150 |
| MXE | 15326 | ENSG00000149273 | RPS3 | chr11 | + | 75404019 | 75404207 | 75404671 | 75404868 | 75401639 | 75401733 | 75405613 | 75405692 | 19 |
| MXE | 15327 | ENSG00000149273 | RPS3 | chr11 | + | 75404019 | 75404207 | 75404759 | 75404868 | 75401639 | 75401733 | 75405613 | 75405692 | 19 |
| MXE | 15512 | ENSG00000109920 | FNBP4 | chr11 | - | 47743952 | 47744163 | 47746055 | 47746394 | 47736615 | 47736740 | 47750915 | 47751036 | 280 |
| MXE | 15938 | ENSG00000077420 | APBB1IP | chr10 | + | 26536073 | 26536217 | 26541581 | 26541692 | 26533438 | 26533525 | 26560104 | 26560203 | 466 |
| MXE | 16942 | ENSG00000090376 | IRAK3 | chr12 | + | 66217170 | 66217235 | 66226722 | 66226837 | 66211445 | 66211597 | 66228251 | 66228370 | 332 |
| MXE | 17045 | ENSG00000120690 | ELF1 | chr13 | - | 40958835 | 40959016 | 40981982 | 40982282 | 40951328 | 40951436 | 41019227 | 41019316 | 361 |
| MXE | 406 | ENSG00000129351 | ILF3 | chr19 | + | 10670973 | 10671203 | 10671357 | 10671535 | 10670515 | 10670608 | 10677162 | 10677310 | 1199 |
| MXE | 696 | ENSG00000101347 | SAMHD1 | chr20 | - | 36935080 | 36935189 | 36935340 | 36935403 | 36930759 | 36930875 | 36941038 | 36941111 | 54 |
| MXE | 3877 | ENSG00000140350 | ANP32A | chr15 | - | 68784396 | 68784595 | 68787412 | 68787535 | 68782955 | 68783053 | 68787769 | 68787919 | 1183 |
| MXE | 7383 | ENSG00000111481 | COPZ1 | chr12 | + | 54343224 | 54343316 | 54345459 | 54345515 | 54342205 | 54342287 | 54347766 | 54347844 | 162 |
| MXE | 9333 | ENSG00000172766 | NAA16 | chr13 | + | 41331273 | 41331369 | 41336649 | 41336756 | 41328723 | 41328843 | 41355143 | 41355216 | 100 |
| MXE | 15836 | ENSG00000158545 | ZC3H18 | chr16 | + | 88608933 | 88609051 | 88611267 | 88611536 | 88599790 | 88599948 | 88622196 | 88622388 | 213 |
| MXE | 11549 | ENSG00000064933 | PMS1 | chr2 | + | 189795768 | 189795951 | 189805651 | 189805754 | 189791789 | 189791941 | 189818016 | 189818180 | 123 |
| MXE | 13575 | ENSG00000196924 | FLNA | chrX | - | 154351883 | 154352021 | 154352180 | 154352447 | 154351580 | 154351696 | 154352771 | 154352924 | 19 |
| MXE | 11047 | ENSG00000079263 | SP140 | chr2 | + | 230248884 | 230248968 | 230255451 | 230255532 | 230245862 | 230245940 | 230269531 | 230269618 | 2 |
| MXE | 9240 | ENSG00000163466 | ARPC2 | chr2 | + | 218238663 | 218238850 | 218239390 | 218239484 | 218228737 | 218228850 | 218245419 | 218245457 | 22 |
| MXE | 14676 | ENSG00000116701 | NCF2 | chr1 | - | 183565703 | 183565779 | 183573184 | 183573292 | 183564004 | 183564030 | 183577598 | 183577707 | 4 |
| MXE | 8592 | ENSG00000142875 | PRKACB | chr1 | + | 84197728 | 84197824 | 84214152 | 84214317 | 84196615 | 84196742 | 84235179 | 84235309 | 450 |
| MXE | 4300 | ENSG00000170571 | EMB | chr5 | - | 50399858 | 50399913 | 50403177 | 50403454 | 50396191 | 50399290 | 50405724 | 50405852 | 2434 |
| MXE | 11395 | ENSG00000003402 | CFLAR | chr2 | + | 201133028 | 201133134 | 201135971 | 201136107 | 201116153 | 201116481 | 201140356 | 201140439 | 3 |
| MXE | 3386 | ENSG00000050393 | MCUR1 | chr6 | - | 13791877 | 13791992 | 13798832 | 13798904 | 13786556 | 13790864 | 13800340 | 13800382 | 73 |
| MXE | 6442 | ENSG00000163946 | TASOR | chr3 | - | 56671599 | 56671692 | 56673579 | 56673725 | 56670072 | 56670145 | 56682675 | 56683017 | 178 |
| MXE | 1519 | ENSG00000091592 | NLRP1 | chr17 | - | 5582669 | 5582846 | 5583686 | 5583959 | 5581858 | 5582062 | 5584292 | 5584448 | 24 |
| MXE | 16778 | ENSG00000090686 | USP48 | chr1 | - | 21736445 | 21736625 | 21747066 | 21747149 | 21729703 | 21729832 | 21748137 | 21748271 | 328 |
| MXE | 13344 | ENSG00000136504 | KAT7 | chr17 | + | 49798318 | 49798558 | 49805362 | 49805445 | 49796749 | 49796926 | 49811475 | 49811574 | 266 |
| MXE | 15378 | ENSG00000130592 | LSP1 | chr11 | + | 1880086 | 1880224 | 1881431 | 1881596 | 1853083 | 1853197 | 1883418 | 1883560 | 996 |
| MXE | 5735 | ENSG00000114126 | TFDP2 | chr3 | - | 141995019 | 141995141 | 142000251 | 142000354 | 141993537 | 141993585 | 142005440 | 142005544 | 8 |
| MXE | 159 | ENSG00000100351 | GRAP2 | chr22 | + | 39947092 | 39947184 | 39955818 | 39955910 | 39901083 | 39901330 | 39960054 | 39960174 | 152 |
| MXE | 488 | ENSG00000130164 | LDLR | chr19 | + | 11102663 | 11102786 | 11105219 | 11105600 | 11100222 | 11100345 | 11106564 | 11106687 | 35 |
| MXE | 11059 | ENSG00000064012 | CASP8 | chr2 | + | 201272897 | 201272942 | 201274888 | 201274953 | 201272637 | 201272776 | 201276826 | 201276968 | 227 |
| MXE | 7441 | ENSG00000165119 | HNRNPK | chr9 | - | 83970161 | 83970331 | 83970736 | 83970819 | 83968196 | 83969380 | 83970896 | 83970912 | 1126 |
| MXE | 6352 | ENSG00000089327 | FXYD5 | chr19 | + | 35160708 | 35160801 | 35164155 | 35164245 | 35158343 | 35158400 | 35166141 | 35166171 | 139 |
| MXE | 11058 | ENSG00000064012 | CASP8 | chr2 | + | 201266486 | 201266791 | 201271515 | 201271621 | 201234015 | 201234112 | 201272637 | 201272776 | 52 |
| MXE | 6339 | ENSG00000143222 | UFC1 | chr1 | + | 161157253 | 161157317 | 161157616 | 161157693 | 161156949 | 161157017 | 161158411 | 161158682 | 111 |
| MXE | 3816 | ENSG00000011243 | AKAP8L | chr19 | - | 15398629 | 15399410 | 15400294 | 15400358 | 15397713 | 15397855 | 15400793 | 15400864 | 33 |
| MXE | 11152 | ENSG00000183486 | MX2 | chr21 | + | 41398896 | 41399019 | 41401969 | 41402128 | 41397612 | 41397691 | 41403266 | 41403343 | 19 |
| MXE | 5652 | ENSG00000151576 | QTRT2 | chr3 | + | 114056998 | 114057106 | 114067986 | 114068063 | 114056736 | 114056864 | 114070625 | 114070838 | 9 |
| MXE | 5690 | ENSG00000163681 | SLMAP | chr3 | + | 57890040 | 57890100 | 57896246 | 57896371 | 57864547 | 57864716 | 57896510 | 57896591 | 62 |
| MXE | 5565 | ENSG00000173193 | PARP14 | chr3 | + | 122718358 | 122718958 | 122720254 | 122721653 | 122718070 | 122718277 | 122727811 | 122727986 | 238 |
| MXE | 10471 | ENSG00000162735 | PEX19 | chr1 | - | 160282416 | 160282502 | 160282825 | 160283109 | 160282105 | 160282200 | 160285054 | 160285133 | 26 |
| MXE | 11805 | ENSG00000170275 | CRTAP | chr3 | + | 33132554 | 33132700 | 33134181 | 33134265 | 33124407 | 33124579 | 33142394 | 33143158 | 2 |
| MXE | 12277 | ENSG00000242588 | NA | chr7 | + | 128578889 | 128578933 | 128591849 | 128592017 | 128574750 | 128574791 | 128608117 | 128608213 | 5 |
| MXE | 2791 | ENSG00000089737 | DDX24 | chr14 | - | 94054995 | 94055184 | 94057821 | 94057897 | 94052997 | 94053127 | 94060097 | 94060613 | 94 |
| MXE | 4917 | ENSG00000075711 | DLG1 | chr3 | - | 197119409 | 197119530 | 197130526 | 197130671 | 197115926 | 197116083 | 197136541 | 197136678 | 128 |
| MXE | 10569 | ENSG00000114942 | EEF1B2 | chr2 | + | 206160587 | 206160710 | 206161345 | 206161472 | 206159894 | 206160059 | 206162037 | 206162104 | 1441 |
| MXE | 9566 | ENSG00000143393 | PI4KB | chr1 | - | 151307573 | 151307801 | 151315572 | 151316509 | 151306135 | 151306363 | 151326171 | 151326373 | 71 |
| MXE | 7201 | ENSG00000143363 | PRUNE1 | chr1 | + | 151018556 | 151018669 | 151024610 | 151024795 | 151017811 | 151017904 | 151027232 | 151027327 | 6 |
| MXE | 7306 | ENSG00000117543 | DPH5 | chr1 | - | 100995109 | 100995149 | 101001466 | 101001544 | 100990062 | 100990631 | 101021640 | 101021720 | 3 |
| MXE | 11492 | ENSG00000111679 | PTPN6 | chr12 | + | 6960343 | 6960435 | 6960805 | 6960945 | 6960087 | 6960239 | 6961125 | 6961316 | 264 |
| MXE | 13557 | ENSG00000136938 | ANP32B | chr9 | + | 98004963 | 98005153 | 98011270 | 98011389 | 97998555 | 97998678 | 98012420 | 98012472 | 1257 |
| MXE | 3396 | ENSG00000131504 | DIAPH1 | chr5 | - | 141524142 | 141524229 | 141526037 | 141526173 | 141515020 | 141517008 | 141526296 | 141526461 | 1637 |
| MXE | 3650 | ENSG00000114107 | CEP70 | chr3 | - | 138537177 | 138537347 | 138570317 | 138570498 | 138532513 | 138532570 | 138571033 | 138571157 | 122 |
| MXE | 7629 | ENSG00000074706 | IPCEF1 | chr6 | - | 154247448 | 154247488 | 154265911 | 154265964 | 154246590 | 154246760 | 154289712 | 154289756 | 7 |
| MXE | 5286 | ENSG00000101596 | SMCHD1 | chr18 | + | 2688627 | 2688747 | 2694526 | 2694693 | 2688393 | 2688508 | 2697031 | 2697122 | 649 |
| MXE | 14844 | ENSG00000163527 | STT3B | chr3 | + | 31579808 | 31580096 | 31596797 | 31596863 | 31576395 | 31576504 | 31600359 | 31600459 | 399 |
| MXE | 15525 | ENSG00000110075 | PPP6R3 | chr11 | + | 68544837 | 68545024 | 68564302 | 68564432 | 68537658 | 68537891 | 68569747 | 68569897 | 304 |
| MXE | 12543 | ENSG00000179818 | PCBP1-AS1 | chr2 | - | 70055713 | 70055910 | 70083506 | 70083687 | 70051202 | 70051305 | 70085546 | 70085624 | 36 |
| MXE | 5335 | ENSG00000174891 | RSRC1 | chr3 | + | 158123865 | 158123991 | 158203071 | 158203245 | 158122102 | 158122298 | 158298038 | 158298075 | 202 |
| MXE | 16337 | ENSG00000135655 | USP15 | chr12 | + | 62321463 | 62321609 | 62325871 | 62325933 | 62314789 | 62314916 | 62349220 | 62349307 | 1323 |
| MXE | 12544 | ENSG00000179818 | PCBP1-AS1 | chr2 | - | 70055713 | 70055910 | 70083537 | 70083687 | 70051202 | 70051305 | 70085546 | 70085624 | 37 |
| MXE | 15586 | ENSG00000116898 | MRPS15 | chr1 | - | 36460691 | 36460776 | 36461263 | 36461312 | 36457922 | 36457981 | 36462087 | 36462163 | 13 |
| MXE | 5021 | ENSG00000082996 | RNF13 | chr3 | + | 149895472 | 149895560 | 149921133 | 149921227 | 149872028 | 149872154 | 149960055 | 149960136 | 552 |
| MXE | 8247 | ENSG00000143442 | POGZ | chr1 | - | 151408693 | 151408828 | 151411624 | 151411771 | 151408099 | 151408240 | 151412295 | 151412396 | 152 |
| MXE | 13659 | ENSG00000046651 | OFD1 | chrX | + | 13751248 | 13751368 | 13752703 | 13753441 | 13749426 | 13749533 | 13755150 | 13755242 | 276 |
| MXE | 3980 | ENSG00000138303 | ASCC1 | chr10 | - | 72203426 | 72203524 | 72210731 | 72210831 | 72196937 | 72196989 | 72212187 | 72212285 | 0 |
| MXE | 10228 | ENSG00000000938 | FGR | chr1 | - | 27621558 | 27621657 | 27623690 | 27623929 | 27617192 | 27617296 | 27625088 | 27625151 | 332 |
| MXE | 14166 | ENSG00000213782 | DDX47 | chr12 | + | 12821207 | 12821396 | 12821654 | 12821726 | 12814130 | 12814224 | 12821964 | 12822083 | 123 |
| MXE | 5269 | ENSG00000156030 | MIDEAS | chr14 | - | 73721296 | 73721509 | 73722697 | 73722847 | 73715121 | 73719501 | 73725271 | 73725360 | 201 |
| MXE | 1687 | ENSG00000005844 | ITGAL | chr16 | + | 30494712 | 30494850 | 30496096 | 30496294 | 30494211 | 30494363 | 30496435 | 30496566 | 1259 |
| MXE | 10904 | ENSG00000077150 | NFKB2 | chr10 | + | 102398384 | 102398523 | 102398738 | 102398864 | 102398211 | 102398297 | 102399287 | 102399497 | 181 |
| MXE | 7412 | ENSG00000106610 | STAG3L4 | chr7 | + | 67305986 | 67306083 | 67308576 | 67309139 | 67302620 | 67302885 | 67309563 | 67309715 | 11 |
| MXE | 5213 | ENSG00000206503 | HLA-A | chr6 | + | 29944121 | 29944397 | 29944499 | 29944616 | 29942756 | 29943026 | 29945058 | 29945091 | 43 |
| MXE | 8175 | ENSG00000173575 | CHD2 | chr15 | + | 92991692 | 92991759 | 92992858 | 92992998 | 92991475 | 92991517 | 92996956 | 92997095 | 20 |
| MXE | 2191 | ENSG00000184056 | VPS33B | chr15 | - | 91006371 | 91006445 | 91006651 | 91006729 | 91005972 | 91006059 | 91006949 | 91007046 | 13 |
| MXE | 7411 | ENSG00000106610 | STAG3L4 | chr7 | + | 67305986 | 67306083 | 67308934 | 67309139 | 67302620 | 67302885 | 67309563 | 67309715 | 11 |
| MXE | 10666 | ENSG00000136536 | MARCHF7 | chr2 | + | 159729008 | 159729175 | 159743060 | 159743253 | 159712456 | 159712606 | 159747804 | 159748903 | 90 |
| MXE | 10987 | ENSG00000122026 | RPL21 | chr13 | + | 27254219 | 27254281 | 27255241 | 27255354 | 27253764 | 27253843 | 27256183 | 27256334 | 1230 |
| MXE | 12235 | ENSG00000196792 | STRN3 | chr14 | - | 30913523 | 30913657 | 30929200 | 30929311 | 30912006 | 30912182 | 30935162 | 30935304 | 28 |
| MXE | 4834 | ENSG00000013441 | CLK1 | chr2 | - | 200859679 | 200859746 | 200861237 | 200861466 | 200857972 | 200858089 | 200861701 | 200861862 | 546 |
| MXE | 14683 | ENSG00000116701 | NCF2 | chr1 | - | 183577598 | 183577707 | 183586894 | 183586977 | 183573184 | 183573292 | 183590155 | 183590604 | 1134 |
| MXE | 13943 | ENSG00000075856 | SART3 | chr12 | - | 108543027 | 108543152 | 108544426 | 108544532 | 108538933 | 108539089 | 108545138 | 108545323 | 3 |
| MXE | 2697 | ENSG00000129566 | TEP1 | chr14 | - | 20403722 | 20403884 | 20405450 | 20405585 | 20403376 | 20403448 | 20407872 | 20408463 | 3 |
| MXE | 9795 | ENSG00000138018 | SELENOI | chr2 | + | 26373366 | 26373629 | 26383298 | 26383347 | 26367145 | 26367220 | 26384958 | 26385139 | 23 |
| MXE | 7378 | ENSG00000138095 | LRPPRC | chr2 | - | 43912431 | 43912558 | 43918024 | 43918133 | 43905691 | 43905780 | 43918255 | 43918398 | 272 |
| MXE | 14893 | ENSG00000172716 | SLFN11 | chr17 | - | 35366946 | 35367063 | 35367602 | 35367700 | 35362738 | 35363826 | 35373581 | 35373620 | 2 |
| MXE | 1685 | ENSG00000005844 | ITGAL | chr16 | + | 30494211 | 30494363 | 30496096 | 30496294 | 30489253 | 30489386 | 30496435 | 30496566 | 1344 |
| MXE | 3858 | ENSG00000157107 | FCHO2 | chr5 | + | 72990711 | 72990864 | 73006444 | 73006549 | 72990477 | 72990619 | 73015625 | 73015724 | 150 |
| MXE | 5642 | ENSG00000150457 | LATS2 | chr13 | - | 20979690 | 20979797 | 20981465 | 20981648 | 20973035 | 20975364 | 20983223 | 20983806 | 68 |
| MXE | 623 | ENSG00000268790 | NA | chr19 | - | 16659394 | 16659447 | 16659958 | 16660000 | 16654034 | 16654125 | 16660084 | 16660104 | 4 |
| MXE | 10084 | ENSG00000143870 | PDIA6 | chr2 | - | 10797699 | 10797757 | 10802498 | 10802640 | 10797080 | 10797207 | 10812677 | 10812785 | 66 |
| MXE | 4361 | ENSG00000248672 | LY75-CD302 | chr2 | - | 159783358 | 159783469 | 159806972 | 159807140 | 159780881 | 159780998 | 159808448 | 159808571 | 169 |
| MXE | 10275 | ENSG00000161813 | LARP4 | chr12 | + | 50441589 | 50441643 | 50453459 | 50453672 | 50440438 | 50440549 | 50454313 | 50454417 | 88 |
| MXE | 8636 | ENSG00000019582 | CD74 | chr5 | - | 150402562 | 150402625 | 150403120 | 150404767 | 150401657 | 150402250 | 150405084 | 150405180 | 751 |
| MXE | 13345 | ENSG00000136504 | KAT7 | chr17 | + | 49805362 | 49805445 | 49809118 | 49809208 | 49791885 | 49792033 | 49811475 | 49811574 | 43 |
| MXE | 13836 | ENSG00000145041 | DCAF1 | chr3 | - | 51470928 | 51471005 | 51483718 | 51483836 | 51466802 | 51466876 | 51496733 | 51496780 | 18 |
| MXE | 3055 | ENSG00000153975 | ZUP1 | chr6 | - | 116660735 | 116660846 | 116666633 | 116667207 | 116658802 | 116658924 | 116668565 | 116668766 | 26 |
| MXE | 12581 | ENSG00000175471 | MCTP1 | chr5 | - | 94870416 | 94870491 | 94870871 | 94870973 | 94867221 | 94868452 | 94871314 | 94871417 | 229 |
| MXE | 9605 | ENSG00000168495 | POLR3D | chr8 | + | 22248153 | 22248278 | 22248480 | 22248649 | 22247856 | 22248008 | 22250074 | 22250227 | 39 |
| MXE | 13452 | ENSG00000126756 | UXT | chrX | - | 47657182 | 47657290 | 47657571 | 47657639 | 47652080 | 47652144 | 47657781 | 47657863 | 47 |
| MXE | 3874 | ENSG00000140350 | ANP32A | chr15 | - | 68780409 | 68780473 | 68782955 | 68783053 | 68778534 | 68780142 | 68784396 | 68784595 | 362 |
| MXE | 9074 | ENSG00000140416 | TPM1 | chr15 | + | 63061197 | 63061273 | 63061712 | 63061788 | 63060868 | 63060939 | 63062214 | 63062277 | 59 |
| MXE | 4034 | ENSG00000026297 | RNASET2 | chr6 | - | 166943018 | 166943089 | 166948569 | 166948625 | 166938894 | 166939008 | 166952487 | 166952548 | 103 |
| MXE | 8013 | ENSG00000110852 | CLEC2B | chr12 | - | 9854380 | 9854484 | 9857473 | 9857637 | 9852368 | 9853408 | 9862498 | 9862573 | 833 |
| MXE | 14621 | ENSG00000134077 | THUMPD3 | chr3 | + | 9377818 | 9377888 | 9380502 | 9380618 | 9374515 | 9374646 | 9383198 | 9383309 | 98 |
| MXE | 16856 | ENSG00000177575 | CD163 | chr12 | - | 7495080 | 7495401 | 7496812 | 7497133 | 7487358 | 7488087 | 7498867 | 7499188 | 631 |
| MXE | 1971 | ENSG00000005339 | CREBBP | chr16 | - | 3749626 | 3749683 | 3751725 | 3751806 | 3745276 | 3745354 | 3757287 | 3757291 | 157 |
| MXE | 11204 | ENSG00000223546 | LINC00630 | chrX | + | 102816991 | 102817082 | 102825992 | 102826169 | 102770351 | 102770420 | 102826968 | 102827144 | 2 |
| MXE | 2195 | ENSG00000170776 | AKAP13 | chr15 | + | 85693276 | 85693451 | 85708018 | 85708086 | 85684740 | 85684873 | 85710578 | 85710645 | 120 |
| MXE | 7913 | ENSG00000071054 | MAP4K4 | chr2 | + | 101887777 | 101887937 | 101888795 | 101888935 | 101887087 | 101887237 | 101891165 | 101891249 | 646 |
| MXE | 17041 | ENSG00000198246 | SLC29A3 | chr10 | + | 71344208 | 71344291 | 71351561 | 71351788 | 71322755 | 71323054 | 71356080 | 71356243 | 11 |
| MXE | 5371 | ENSG00000114331 | ACAP2 | chr3 | - | 195392089 | 195392147 | 195410706 | 195411009 | 195381902 | 195382022 | 195412845 | 195412916 | 4 |
| MXE | 9571 | ENSG00000076242 | MLH1 | chr3 | + | 37017505 | 37017599 | 37020309 | 37020463 | 37014431 | 37014544 | 37028783 | 37028932 | 66 |
| MXE | 15269 | ENSG00000095139 | ARCN1 | chr11 | + | 118583178 | 118583358 | 118583808 | 118584014 | 118572389 | 118572550 | 118584479 | 118584644 | 3 |
| MXE | 4661 | ENSG00000025293 | PHF20 | chr20 | + | 35842572 | 35842744 | 35847349 | 35847434 | 35801490 | 35801605 | 35858301 | 35858381 | 113 |
| MXE | 3433 | ENSG00000116459 | ATP5PB | chr1 | + | 111456085 | 111456249 | 111456629 | 111456755 | 111449836 | 111449873 | 111459456 | 111459636 | 11 |
| MXE | 6760 | ENSG00000159592 | GPBP1L1 | chr1 | - | 45633488 | 45633647 | 45634095 | 45634236 | 45630481 | 45630606 | 45640209 | 45640403 | 128 |
| MXE | 2269 | ENSG00000068305 | MEF2A | chr15 | + | 99671322 | 99671454 | 99671519 | 99671657 | 99645560 | 99645764 | 99674392 | 99674612 | 154 |
| MXE | 6415 | ENSG00000070759 | TESK2 | chr1 | - | 45345876 | 45345994 | 45346692 | 45346779 | 45343882 | 45345558 | 45346978 | 45347062 | 6 |
| MXE | 1080 | ENSG00000108953 | YWHAE | chr17 | - | 1361901 | 1362008 | 1364858 | 1365058 | 1361091 | 1361298 | 1400046 | 1400222 | 320 |
| MXE | 4335 | ENSG00000112079 | STK38 | chr6 | - | 36515337 | 36515492 | 36521733 | 36521817 | 36507499 | 36507602 | 36524340 | 36524463 | 675 |
| MXE | 15528 | ENSG00000110075 | PPP6R3 | chr11 | + | 68548066 | 68548204 | 68564302 | 68564432 | 68544837 | 68545024 | 68569747 | 68569897 | 291 |
| MXE | 3358 | ENSG00000127603 | MACF1 | chr1 | + | 39437776 | 39438008 | 39439273 | 39439500 | 39436481 | 39436490 | 39441002 | 39441125 | 48 |
| MXE | 4888 | ENSG00000157601 | MX1 | chr21 | + | 41449136 | 41449295 | 41451190 | 41451243 | 41445999 | 41446141 | 41452620 | 41452869 | 156 |
| MXE | 2341 | ENSG00000140367 | UBE2Q2 | chr15 | + | 75873427 | 75873568 | 75876186 | 75876271 | 75868950 | 75869010 | 75877960 | 75878021 | 224 |
| MXE | 16069 | ENSG00000154124 | OTULIN | chr5 | + | 14687520 | 14687646 | 14690038 | 14690308 | 14681463 | 14681607 | 14692853 | 14699850 | 140 |
| MXE | 7009 | ENSG00000117533 | VAMP4 | chr1 | - | 171719170 | 171719221 | 171728523 | 171728570 | 171710713 | 171710814 | 171738348 | 171738463 | 81 |
| MXE | 3807 | ENSG00000113648 | MACROH2A1 | chr5 | - | 135343259 | 135343434 | 135345967 | 135346057 | 135334416 | 135335141 | 135360496 | 135360607 | 5 |
| MXE | 8588 | ENSG00000134242 | PTPN22 | chr1 | - | 113848539 | 113848626 | 113854906 | 113855049 | 113838543 | 113838620 | 113856381 | 113856441 | 159 |
| MXE | 13749 | ENSG00000143569 | UBAP2L | chr1 | + | 154228614 | 154228725 | 154234590 | 154234759 | 154227281 | 154227359 | 154235195 | 154235291 | 290 |
| MXE | 15703 | ENSG00000102317 | RBM3 | chrX | + | 48575560 | 48575667 | 48576313 | 48576419 | 48575167 | 48575283 | 48577025 | 48577109 | 552 |
| MXE | 11946 | ENSG00000086232 | EIF2AK1 | chr7 | - | 6042932 | 6042993 | 6044561 | 6044661 | 6040891 | 6041219 | 6046070 | 6046151 | 256 |
| MXE | 16402 | ENSG00000100354 | TNRC6B | chr22 | + | 40251178 | 40251200 | 40261831 | 40262173 | 40246014 | 40246102 | 40264687 | 40267036 | 28 |
| MXE | 8880 | ENSG00000117523 | PRRC2C | chr1 | + | 171587647 | 171587751 | 171588378 | 171588505 | 171587002 | 171587221 | 171591586 | 171593511 | 298 |
| MXE | 7672 | ENSG00000165671 | NSD1 | chr5 | + | 177204119 | 177204292 | 177209635 | 177212195 | 177191883 | 177192019 | 177235820 | 177235945 | 322 |
| MXE | 14197 | ENSG00000181396 | OGFOD3 | chr17 | - | 82406417 | 82406482 | 82409367 | 82409410 | 82405323 | 82405380 | 82411454 | 82411530 | 10 |
| MXE | 1498 | ENSG00000029725 | RABEP1 | chr17 | + | 5335183 | 5335344 | 5338018 | 5338138 | 5331948 | 5332152 | 5346789 | 5346925 | 285 |
| MXE | 10822 | ENSG00000064995 | TAF11 | chr6 | - | 34880288 | 34880376 | 34882931 | 34883080 | 34877777 | 34878720 | 34887786 | 34888071 | 21 |
| MXE | 11060 | ENSG00000064012 | CASP8 | chr2 | + | 201272897 | 201272942 | 201274888 | 201274953 | 201272637 | 201272776 | 201284815 | 201285317 | 154 |
| MXE | 15824 | ENSG00000124795 | DEK | chr6 | - | 18256360 | 18256455 | 18257952 | 18258062 | 18255730 | 18255851 | 18258303 | 18258405 | 78 |
| MXE | 8213 | ENSG00000115935 | WIPF1 | chr2 | - | 174581309 | 174581439 | 174585522 | 174585611 | 174575203 | 174575380 | 174597600 | 174597765 | 512 |
| MXE | 13750 | ENSG00000143569 | UBAP2L | chr1 | + | 154228614 | 154228725 | 154234590 | 154234792 | 154227281 | 154227359 | 154235195 | 154235291 | 290 |
| MXE | 16839 | ENSG00000111596 | CNOT2 | chr12 | + | 70310894 | 70311017 | 70319297 | 70319364 | 70278131 | 70278274 | 70329422 | 70329570 | 181 |
| MXE | 14059 | ENSG00000034677 | RNF19A | chr8 | - | 100259853 | 100259997 | 100261541 | 100261755 | 100259062 | 100259246 | 100264670 | 100264785 | 32 |
| MXE | 12847 | ENSG00000146872 | TLK2 | chr17 | + | 62580110 | 62580192 | 62586134 | 62586226 | 62578476 | 62578574 | 62596584 | 62596674 | 87 |
| MXE | 8885 | ENSG00000273136 | NBPF26 | chr1 | + | 120818122 | 120818174 | 120822103 | 120822267 | 120816621 | 120816827 | 120823308 | 120823360 | 23 |
| MXE | 13015 | ENSG00000184863 | RBM33 | chr7 | + | 155700772 | 155700944 | 155706859 | 155707068 | 155680668 | 155680908 | 155711172 | 155711455 | 314 |
| MXE | 3385 | ENSG00000050393 | MCUR1 | chr6 | - | 13791877 | 13791992 | 13793893 | 13793947 | 13786556 | 13790864 | 13798832 | 13798904 | 24 |
| MXE | 2621 | ENSG00000075413 | MARK3 | chr14 | + | 103468085 | 103468186 | 103474992 | 103475210 | 103467078 | 103467191 | 103480386 | 103480490 | 3 |
| MXE | 11657 | ENSG00000100503 | NIN | chr14 | - | 50759856 | 50760359 | 50761789 | 50761911 | 50754741 | 50754867 | 50763825 | 50763964 | 534 |
| MXE | 17161 | ENSG00000270181 | BIVM-ERCC5 | chr13 | + | 102856051 | 102856112 | 102858274 | 102858418 | 102854287 | 102854374 | 102861506 | 102861714 | 91 |
| MXE | 10896 | ENSG00000104218 | CSPP1 | chr8 | + | 67137455 | 67137603 | 67149782 | 67149935 | 67131950 | 67132080 | 67154023 | 67154136 | 41 |
| MXE | 3430 | ENSG00000116459 | ATP5PB | chr1 | + | 111454210 | 111454356 | 111456085 | 111456249 | 111449836 | 111449873 | 111459456 | 111459636 | 237 |
| MXE | 2172 | ENSG00000140199 | SLC12A6 | chr15 | - | 34238964 | 34239160 | 34240660 | 34240829 | 34238231 | 34238331 | 34241232 | 34241337 | 97 |
| MXE | 1888 | ENSG00000077235 | GTF3C1 | chr16 | - | 27462293 | 27462486 | 27463540 | 27463592 | 27460612 | 27461562 | 27464319 | 27464836 | 10 |
| MXE | 13873 | ENSG00000187735 | TCEA1 | chr8 | - | 53999944 | 54000050 | 54010429 | 54010492 | 53993667 | 53993755 | 54022062 | 54022448 | 25 |
| MXE | 14147 | ENSG00000151461 | UPF2 | chr10 | - | 11979056 | 11979165 | 11997671 | 11997757 | 11967340 | 11967454 | 11999905 | 12000009 | 187 |
| MXE | 2194 | ENSG00000170776 | AKAP13 | chr15 | + | 85684740 | 85684873 | 85693276 | 85693451 | 85682157 | 85682212 | 85708018 | 85708086 | 414 |
| MXE | 2007 | ENSG00000213614 | HEXA | chr15 | - | 72351132 | 72351234 | 72353067 | 72353178 | 72350517 | 72350650 | 72355558 | 72355624 | 12 |
| MXE | 7920 | ENSG00000153187 | HNRNPU | chr1 | - | 244858010 | 244858274 | 244858728 | 244858835 | 244857597 | 244857717 | 244859274 | 244859374 | 13 |
| MXE | 8635 | ENSG00000019582 | CD74 | chr5 | - | 150402562 | 150402625 | 150404679 | 150404767 | 150400040 | 150402250 | 150405084 | 150405180 | 731 |
| MXE | 12786 | ENSG00000105819 | PMPCB | chr7 | + | 103303841 | 103304040 | 103304410 | 103304490 | 103300177 | 103300307 | 103307595 | 103307708 | 325 |
| MXE | 12331 | ENSG00000117868 | ESYT2 | chr7 | - | 158741522 | 158741896 | 158743528 | 158743678 | 158739022 | 158739121 | 158748193 | 158748280 | 515 |
| MXE | 5037 | ENSG00000009790 | TRAF3IP3 | chr1 | + | 209779314 | 209779374 | 209780469 | 209780606 | 209778110 | 209778173 | 209781344 | 209781458 | 295 |
| MXE | 4327 | ENSG00000145495 | MARCHF6 | chr5 | + | 10381799 | 10381943 | 10386993 | 10387066 | 10378758 | 10378832 | 10391541 | 10391731 | 238 |
| MXE | 5267 | ENSG00000096384 | HSP90AB1 | chr6 | + | 44248629 | 44248776 | 44249376 | 44249583 | 44247100 | 44247195 | 44249674 | 44249834 | 697 |
| MXE | 7382 | ENSG00000111481 | COPZ1 | chr12 | + | 54342205 | 54342287 | 54345459 | 54345515 | 54340546 | 54340615 | 54347766 | 54347844 | 189 |
| MXE | 17194 | ENSG00000136100 | VPS36 | chr13 | - | 52433661 | 52433748 | 52434792 | 52434882 | 52427186 | 52427219 | 52436289 | 52436404 | 304 |
| MXE | 12965 | ENSG00000127914 | AKAP9 | chr7 | + | 92105677 | 92105763 | 92108493 | 92108633 | 92102593 | 92102826 | 92110121 | 92110571 | 26 |
| MXE | 4858 | ENSG00000113851 | CRBN | chr3 | - | 3167633 | 3167793 | 3172775 | 3172925 | 3153423 | 3156281 | 3174058 | 3174261 | 376 |
| MXE | 4907 | ENSG00000075711 | DLG1 | chr3 | - | 197076585 | 197076685 | 197081050 | 197081117 | 197075836 | 197075870 | 197085579 | 197085756 | 106 |
| MXE | 562 | ENSG00000126261 | UBA2 | chr19 | + | 34434867 | 34434968 | 34438644 | 34438766 | 34428358 | 34428570 | 34443843 | 34443911 | 6 |
| MXE | 6805 | ENSG00000117682 | DHDDS | chr1 | + | 26442707 | 26442873 | 26447558 | 26447660 | 26438167 | 26438284 | 26457790 | 26457905 | 3 |
| MXE | 7182 | ENSG00000166669 | ATF7IP2 | chr16 | + | 10433524 | 10433649 | 10438100 | 10438235 | 10430594 | 10431455 | 10440363 | 10440462 | 147 |
| MXE | 1891 | ENSG00000038532 | CLEC16A | chr16 | + | 10977224 | 10977399 | 10979328 | 10979382 | 10972937 | 10973061 | 10982877 | 10982991 | 71 |
| MXE | 15719 | ENSG00000166387 | PPFIBP2 | chr11 | + | 7648799 | 7648911 | 7649146 | 7649235 | 7648386 | 7648537 | 7649531 | 7649654 | 19 |
| MXE | 11108 | ENSG00000184182 | UBE2F | chr2 | + | 238025341 | 238025412 | 238035877 | 238035940 | 237994743 | 237994809 | 238041287 | 238041692 | 22 |
| MXE | 13710 | ENSG00000125351 | UPF3B | chrX | - | 119843190 | 119843301 | 119845197 | 119845296 | 119841734 | 119841778 | 119851494 | 119851601 | 31 |
| MXE | 2268 | ENSG00000068305 | MEF2A | chr15 | + | 99671322 | 99671454 | 99671519 | 99671657 | 99632977 | 99633173 | 99674392 | 99674612 | 73 |
| MXE | 6568 | ENSG00000198791 | CNOT7 | chr8 | - | 17234715 | 17234860 | 17237211 | 17237373 | 17231569 | 17232537 | 17242991 | 17243185 | 853 |
| MXE | 13833 | ENSG00000145041 | DCAF1 | chr3 | - | 51441384 | 51441897 | 51443765 | 51443903 | 51440969 | 51441071 | 51463113 | 51463227 | 136 |
| MXE | 5374 | ENSG00000054967 | RELT | chr11 | + | 73391143 | 73391223 | 73392210 | 73392468 | 73390754 | 73390921 | 73393836 | 73393917 | 5 |
| MXE | 5710 | ENSG00000082074 | FYB1 | chr5 | - | 39153447 | 39153604 | 39201825 | 39202987 | 39141094 | 39141141 | 39270568 | 39270649 | 109 |
| MXE | 10937 | ENSG00000073849 | ST6GAL1 | chr3 | + | 187038741 | 187038873 | 187051248 | 187051346 | 187021855 | 187022004 | 187072848 | 187072947 | 18 |
| MXE | 638 | ENSG00000123146 | ADGRE5 | chr19 | + | 14388701 | 14388818 | 14390923 | 14391079 | 14388449 | 14388500 | 14397657 | 14397804 | 320 |
| MXE | 11205 | ENSG00000223546 | LINC00630 | chrX | + | 102845856 | 102846020 | 102865041 | 102865152 | 102839800 | 102839863 | 102884142 | 102885406 | 4 |
| MXE | 8171 | ENSG00000173575 | CHD2 | chr15 | + | 92991346 | 92991430 | 92991475 | 92991517 | 92985497 | 92985673 | 92992858 | 92992998 | 13 |
| MXE | 4581 | ENSG00000112893 | MAN2A1 | chr5 | + | 109729341 | 109729513 | 109755328 | 109755456 | 109716119 | 109716264 | 109767534 | 109767708 | 351 |
| MXE | 5984 | ENSG00000141293 | SKAP1 | chr17 | - | 48137228 | 48137337 | 48162468 | 48162569 | 48133441 | 48133816 | 48170608 | 48170659 | 115 |
| MXE | 13694 | ENSG00000215301 | DDX3X | chrX | + | 41347311 | 41347451 | 41347639 | 41347915 | 41346858 | 41347012 | 41357832 | 41358000 | 627 |
| MXE | 13695 | ENSG00000215301 | DDX3X | chrX | + | 41347311 | 41347451 | 41347639 | 41347915 | 41346858 | 41347012 | 41364273 | 41364472 | 627 |
| MXE | 6292 | ENSG00000137776 | SLTM | chr15 | - | 58912562 | 58912610 | 58913498 | 58913696 | 58901259 | 58901287 | 58916934 | 58916999 | 169 |
| MXE | 10797 | ENSG00000184007 | PTP4A2 | chr1 | - | 31910037 | 31910112 | 31915894 | 31915987 | 31906420 | 31908960 | 31918969 | 31919483 | 1248 |
| MXE | 12229 | ENSG00000106245 | BUD31 | chr7 | + | 99410033 | 99410169 | 99411063 | 99411186 | 99408968 | 99409245 | 99416137 | 99416260 | 20 |
| MXE | 4887 | ENSG00000157601 | MX1 | chr21 | + | 41449136 | 41449295 | 41451166 | 41451243 | 41445999 | 41446141 | 41452620 | 41452869 | 156 |
| MXE | 5117 | ENSG00000114867 | EIF4G1 | chr3 | + | 184324877 | 184325114 | 184325268 | 184325373 | 184324200 | 184324347 | 184325479 | 184325639 | 762 |
| MXE | 3199 | ENSG00000112081 | SRSF3 | chr6 | + | 36601151 | 36601190 | 36601707 | 36601794 | 36599820 | 36600276 | 36601961 | 36602040 | 11 |
| MXE | 4572 | ENSG00000152784 | PRDM8 | chr4 | + | 80191471 | 80191569 | 80194112 | 80194239 | 80185364 | 80185518 | 80196351 | 80196601 | 0 |
| MXE | 1234 | ENSG00000132471 | WBP2 | chr17 | - | 75846907 | 75846984 | 75847486 | 75847609 | 75845698 | 75846787 | 75847795 | 75847930 | 181 |
| MXE | 3747 | ENSG00000164308 | ERAP2 | chr5 | + | 96896731 | 96896863 | 96900120 | 96900245 | 96896372 | 96896504 | 96901505 | 96901681 | 193 |
| MXE | 13203 | ENSG00000164062 | APEH | chr3 | + | 49676055 | 49676219 | 49676377 | 49676515 | 49675890 | 49675966 | 49676608 | 49676700 | 315 |
| MXE | 15485 | ENSG00000110321 | EIF4G2 | chr11 | - | 10799551 | 10799756 | 10800089 | 10800348 | 10799212 | 10799424 | 10800431 | 10800647 | 1333 |
| MXE | 1916 | ENSG00000102893 | PHKB | chr16 | + | 47547432 | 47547548 | 47580294 | 47580358 | 47515520 | 47515601 | 47587667 | 47587758 | 66 |
| MXE | 7986 | ENSG00000153283 | CD96 | chr3 | + | 111577501 | 111577549 | 111579026 | 111579234 | 111567522 | 111567647 | 111585322 | 111585378 | 15 |
| MXE | 7687 | ENSG00000151023 | ENKUR | chr10 | - | 24984735 | 24984905 | 24990462 | 24990609 | 24981979 | 24984376 | 24995645 | 24995869 | 96 |
| MXE | 1250 | ENSG00000182446 | NPLOC4 | chr17 | - | 81606690 | 81606814 | 81610209 | 81610258 | 81604547 | 81604727 | 81613317 | 81613494 | 17 |
| MXE | 6291 | ENSG00000119285 | HEATR1 | chr1 | - | 236587401 | 236587490 | 236587947 | 236588043 | 236586240 | 236586452 | 236590846 | 236590954 | 43 |
| MXE | 14822 | ENSG00000106948 | AKNA | chr9 | - | 114355924 | 114356136 | 114357920 | 114358167 | 114350858 | 114351021 | 114359593 | 114359794 | 550 |
| MXE | 5560 | ENSG00000173193 | PARP14 | chr3 | + | 122699389 | 122701635 | 122703741 | 122703978 | 122695425 | 122695662 | 122708189 | 122708268 | 415 |
| MXE | 4785 | ENSG00000163932 | PRKCD | chr3 | + | 53185603 | 53185700 | 53186166 | 53186340 | 53184873 | 53184974 | 53186603 | 53186695 | 209 |
| MXE | 2343 | ENSG00000140367 | UBE2Q2 | chr15 | + | 75873427 | 75873568 | 75879097 | 75879188 | 75868950 | 75869010 | 75883365 | 75883424 | 223 |
| MXE | 6039 | ENSG00000181555 | SETD2 | chr3 | - | 47083719 | 47084382 | 47086194 | 47086314 | 47066991 | 47067118 | 47088112 | 47088249 | 87 |
| MXE | 7746 | ENSG00000156515 | HK1 | chr10 | + | 69392124 | 69392308 | 69394949 | 69395105 | 69389196 | 69389296 | 69398594 | 69398828 | 332 |
| MXE | 15510 | ENSG00000149187 | CELF1 | chr11 | - | 47482694 | 47482856 | 47484388 | 47484523 | 47478876 | 47478952 | 47486749 | 47486798 | 409 |
| MXE | 4044 | ENSG00000062194 | GPBP1 | chr5 | + | 57249408 | 57249576 | 57250953 | 57251141 | 57247074 | 57247215 | 57261179 | 57261282 | 763 |

| | | | | | | | | | | | |
|---|---|---|---|---|---|---|---|---|---|---|---|
| MXE | 9498 | ENSG00000189319 | FAM53B | chr10 | - | 124696157 | 124696212 | 124706635 | 124706887 | 124681606 | 124682379 |
| MXE | 103 | ENSG00000196588 | MRTFA | chr22 | - | 40423535 | 40423685 | 40424205 | 40424381 | 40420846 | 40421100 |
| MXE | 12655 | ENSG00000168092 | PAFAH1B2 | chr11 | + | 117161144 | 117161261 | 117163769 | 117163892 | 117159933 | 117160023 |
| MXE | 4011 | ENSG00000138760 | SCARB2 | chr4 | - | 76174143 | 76174313 | 76175790 | 76175910 | 76169866 | 76169985 |
| MXE | 13690 | ENSG00000215301 | DDX3X | chrX | + | 41346858 | 41347012 | 41347311 | 41347451 | 41346504 | 41346622 |
| MXE | 4050 | ENSG00000070756 | PABPC1 | chr8 | - | 100709458 | 100709731 | 100712361 | 100712457 | 100709132 | 100709223 |
| MXE | 8409 | ENSG00000166938 | DIS3L | chr15 | + | 66323492 | 66323585 | 66325830 | 66326364 | 66322686 | 66322934 |
| MXE | 10396 | ENSG00000153250 | RBMS1 | chr2 | - | 160318168 | 160318227 | 160367215 | 160367391 | 160313155 | 160313247 |
| MXE | 5720 | ENSG00000163507 | CIP2A | chr3 | - | 108582107 | 108582202 | 108582976 | 108583083 | 108581414 | 108581511 |
| MXE | 11853 | ENSG00000168685 | IL7R | chr5 | + | 35867305 | 35867463 | 35871055 | 35871213 | 35860851 | 35860990 |
| MXE | 8016 | ENSG00000136682 | CBWD2 | chr2 | + | 113462407 | 113462494 | 113465128 | 113465173 | 113460672 | 113460719 |
| MXE | 2087 | ENSG00000157823 | AP3S2 | chr15 | - | 89878264 | 89878300 | 89888520 | 89888632 | 89877305 | 89877424 |
| MXE | 16777 | ENSG00000090686 | USP48 | chr1 | - | 21736445 | 21736625 | 21747066 | 21747149 | 21729703 | 21729829 |
| MXE | 15379 | ENSG00000130592 | LSP1 | chr11 | + | 1880086 | 1880224 | 1883418 | 1883560 | 1853083 | 1853197 |
| MXE | 2043 | ENSG00000140479 | PCSK6 | chr15 | - | 101431989 | 101432100 | 101443555 | 101443660 | 101429986 | 101430063 |
| MXE | 2200 | ENSG00000170776 | AKAP13 | chr15 | + | 85735559 | 85735630 | 85736089 | 85736134 | 85734991 | 85735150 |
| MXE | 15820 | ENSG00000124795 | DEK | chr6 | - | 18226173 | 18226242 | 18236451 | 18236600 | 18223859 | 18225730 |
| MXE | 5776 | ENSG00000138750 | NUP54 | chr4 | - | 76132522 | 76132719 | 76134174 | 76134362 | 76131229 | 76131284 |
| MXE | 5038 | ENSG00000009790 | TRAF3IP3 | chr1 | + | 209779314 | 209779374 | 209781344 | 209781458 | 209778110 | 209778173 |
| MXE | 6108 | ENSG00000138382 | METTL5 | chr2 | - | 169819560 | 169819643 | 169821942 | 169822057 | 169812456 | 169812506 |
| MXE | 13253 | ENSG00000288640 | NA | chr7 | + | 112475429 | 112475676 | 112484811 | 112484943 | 112472765 | 112472861 |
| MXE | 7203 | ENSG00000143363 | PRUNE1 | chr1 | + | 151024610 | 151024795 | 151025514 | 151025673 | 151017811 | 151017904 |
| MXE | 13340 | ENSG00000136504 | KAT7 | chr17 | + | 49796749 | 49796926 | 49805362 | 49805445 | 49791885 | 49792033 |
| MXE | 1019 | ENSG00000073605 | GSDMB | chr17 | - | 39908175 | 39908214 | 39908957 | 39909042 | 39906110 | 39906271 |
| [illegible] | [illegible] | [illegible] | [illegible] | [illegible] | [illegible] | [illegible] | [illegible] | [illegible] | [illegible] | [illegible] | [illegible] |

| | | | | | | | | | | | | | | | |
|---|---|---|---|---|---|---|---|---|---|---|---|---|---|---|---|
| MXE | 11507 | ENSG00000137478 | FCHSD2 | chr11 | - | 72849754 | 72849889 | 72867864 | 72868026 | 72843448 | 72843532 | 72887469 | 72887574 | 119 | [illegible] |
| MXE | 7462 | ENSG00000066136 | NFYC | chr1 | + | 40753150 | 40753246 | 40758120 | 40758294 | 40747533 | 40747605 | 40762887 | 40763046 | 0 | [illegible] |
| MXE | 8657 | ENSG00000143398 | PIP5K1A | chr1 | + | 151242437 | 151242567 | 151246919 | 151246965 | 151239954 | 151240039 | 151247862 | 151249531 | 0 | [illegible] |
| MXE | 5888 | ENSG00000117054 | ACADM | chr1 | + | 75749418 | 75749559 | 75750450 | 75750546 | 75745805 | 75745914 | 75761121 | 75761370 | 58 | [illegible] |
| MXE | 13914 | ENSG00000177565 | TBL1XR1 | chr3 | - | 177079826 | 177079945 | 177086964 | 177087041 | 177064919 | 177065022 | 177098465 | 177098541 | 2 | [illegible] |
| MXE | 9992 | ENSG00000262633 | NA | chr17 | + | 46932066 | 46932199 | 46935028 | 46935169 | 46929519 | 46929584 | 46938598 | 46938704 | 3 | [illegible] |
| MXE | 5922 | ENSG00000003400 | CASP10 | chr2 | + | 201192983 | 201193119 | 201195841 | 201195948 | 201187705 | 201187799 | 201208074 | 201208183 | 87 | [illegible] |
| MXE | 1680 | ENSG00000005844 | ITGAL | chr16 | + | 30475305 | 30475400 | 30479090 | 30479208 | 30474195 | 30474298 | 30481438 | 30481584 | 103 | [illegible] |
| MXE | 4415 | ENSG00000182957 | SPATA13 | chr13 | + | 24284134 | 24284271 | 24286213 | 24286393 | 24251693 | 24251862 | 24288998 | 24289178 | 140 | [illegible] |
| MXE | 16607 | ENSG00000196498 | NCOR2 | chr12 | - | 124449814 | 124449867 | 124457105 | 124457162 | 124437929 | 124437996 | 124466172 | 124466286 | 45 | [illegible] |
| MXE | 5759 | ENSG00000215908 | CROCCP2 | chr1 | - | 16633094 | 16633346 | 16642766 | 16642850 | 16632274 | 16632371 | 16643027 | 16643145 | 15 | [illegible] |
| MXE | 3296 | ENSG00000137070 | IL11RA | chr9 | + | 34655604 | 34655665 | 34656738 | 34656908 | 34655217 | 34655317 | 34657034 | 34657149 | 6 | [illegible] |
| MXE | 10433 | ENSG00000159479 | MED8 | chr1 | - | 43385977 | 43386226 | 43386588 | 43386670 | 43383916 | 43385106 | 43386857 | 43386998 | 35 | [illegible] |
| MXE | 13093 | ENSG00000130066 | SAT1 | chrX | + | 23783799 | 23783883 | 23784293 | 23784403 | 23783657 | 23783709 | 23785327 | 23785429 | 911 | [illegible] |
| MXE | 6186 | ENSG00000213390 | ARHGAP19 | chr10 | - | 97243967 | 97244159 | 97246271 | 97246337 | 97235216 | 97235315 | 97259401 | 97259628 | 0 | [illegible] |
| MXE | 12294 | ENSG00000154803 | FLCN | chr17 | - | 17222500 | 17222661 | 17223921 | 17224143 | 17221170 | 17221628 | 17226175 | 17226322 | 8 | [illegible] |
| MXE | 12387 | ENSG00000068885 | IFT80 | chr3 | - | 160307662 | 160307781 | 160319759 | 160319939 | 160303915 | 160303989 | 160343650 | 160343815 | 6 | [illegible] |
| MXE | 238 | ENSG00000100242 | SUN2 | chr22 | - | 38739721 | 38739943 | 38740266 | 38740432 | 38739341 | 38739426 | 38741006 | 38741050 | 681 | [illegible] |
| MXE | 1520 | ENSG00000091592 | NLRP1 | chr17 | - | 5582669 | 5582846 | 5583686 | 5584103 | 5581858 | 5582062 | 5584292 | 5584512 | 37 | [illegible] |
| MXE | 14651 | ENSG00000187239 | FNBP1 | chr9 | - | 129957359 | 129957464 | 129958490 | 129958553 | 129929566 | 129929695 | 129978464 | 129978612 | 300 | [illegible] |
| MXE | 2149 | ENSG00000089195 | TRMT6 | chr20 | - | 5941242 | 5941345 | 5941950 | 5942036 | 5941052 | 5941139 | 5942427 | 5942786 | 13 | [illegible] |
| MXE | 12047 | ENSG00000127947 | PTPN12 | chr7 | + | 77600663 | 77600806 | 77607234 | 77607301 | 77597841 | 77597901 | 77610764 | 77610802 | 352 | [illegible] |
| MXE | 1978 | ENSG00000142192 | APP | chr21 | - | 25997359 | 25997416 | 26000014 | 26000182 | 25982343 | 25982477 | 26021839 | 26022042 | 463 | [illegible] |
| MXE | 3474 | ENSG00000145604 | SKP2 | chr5 | + | 36170343 | 36170442 | 36171602 | 36171733 | 36166518 | 36166662 | 36176964 | 36177016 | 0 | [illegible] |
| MXE | 9315 | ENSG00000154451 | GBP5 | chr1 | - | 89268728 | 89268856 | 89269365 | 89269574 | 89267416 | 89267526 | 89270754 | 89270863 | 257 | [illegible] |
| MXE | 12800 | ENSG00000051825 | MPHOSPH9 | chr12 | - | 123226287 | 123226351 | 123227462 | 123227616 | 123223037 | 123223127 | 123230260 | 123230522 | 177 | [illegible] |
| MXE | 8523 | ENSG00000074201 | CLNS1A | chr11 | - | 77624962 | 77625070 | 77629762 | 77629899 | 77622499 | 77622673 | 77637589 | 77637794 | 54 | [illegible] |
| MXE | 15813 | ENSG00000136167 | LCP1 | chr13 | - | 46146907 | 46147103 | 46148351 | 46148447 | 46144441 | 46144520 | 46150935 | 46151078 | 1851 | [illegible] |
| MXE | 10909 | ENSG00000258984 | UBE2F-SCLY | chr2 | + | 238016565 | 238016633 | 238030555 | 238030613 | 237994743 | 237994809 | 238035877 | 238035940 | 228 | [illegible] |
| MXE | 162 | ENSG00000100351 | GRAP2 | chr22 | + | 39960054 | 39960174 | 39965989 | 39966158 | 39955818 | 39955910 | 39968041 | 39968272 | 492 | [illegible] |
| MXE | 7202 | ENSG00000143363 | PRUNE1 | chr1 | + | 151018556 | 151018669 | 151025514 | 151025673 | 151017811 | 151017904 | 151027232 | 151027327 | 6 | [illegible] |
| MXE | 16336 | ENSG00000135655 | USP15 | chr12 | + | 62314789 | 62314916 | 62321463 | 62321609 | 62302789 | 62302920 | 62325871 | 62325933 | 651 | [illegible] |
| MXE | 16176 | ENSG00000148840 | PPRC1 | chr10 | + | 102144249 | 102144307 | 102145019 | 102145090 | 102143044 | 102143098 | 102146671 | 102147392 | 19 | [illegible] |
| MXE | 15937 | ENSG00000077420 | APBB1IP | chr10 | + | 26500818 | 26501111 | 26511746 | 26511906 | 26496303 | 26496391 | 26513538 | 26513660 | 437 | [illegible] |
| MXE | 3525 | ENSG00000113263 | ITK | chr5 | + | 157238108 | 157238191 | 157240061 | 157240195 | 157232394 | 157232339 | 157241645 | 157241720 | 260 | [illegible] |
| MXE | 2940 | ENSG00000118482 | PHF3 | chr6 | + | 63691736 | 63692043 | 63694580 | 63694764 | 63679999 | 63680161 | 63698222 | 63698367 | 14 | [illegible] |
| MXE | 11308 | ENSG00000116793 | PHTF1 | chr1 | - | 113713278 | 113713438 | 113724758 | 113724893 | 113711939 | 113712113 | 113726417 | 113726574 | 25 | [illegible] |
| MXE | 13682 | ENSG00000215301 | DDX3X | chrX | + | 41344029 | 41344128 | 41344238 | 41344399 | 41343215 | 41343351 | 41345179 | 41345324 | 12 | [illegible] |
| MXE | 2189 | ENSG00000129003 | VPS13C | chr15 | - | 62000563 | 62000626 | 62007307 | 62007479 | 61991672 | 61991802 | 62008654 | 62008761 | 1055 | [illegible] |
| MXE | 16505 | ENSG00000145901 | TNIP1 | chr5 | - | 151064959 | 151065131 | 151080323 | 151080359 | 151063612 | 151063747 | 151080879 | 151081034 | 1 | [illegible] |
| MXE | 8482 | ENSG00000135749 | PCNX2 | chr1 | - | 233227225 | 233227371 | 233236844 | 233236980 | 233218030 | 233218184 | 233250738 | 233250832 | 89 | [illegible] |
| MXE | 3175 | ENSG00000130449 | ZSWIM6 | chr5 | + | 61530051 | 61530198 | 61535483 | 61535619 | 61526249 | 61526396 | 61538813 | 61538971 | 23 | [illegible] |
| MXE | 5359 | ENSG00000051382 | PIK3CB | chr3 | - | 138734633 | 138734804 | 138737706 | 138737886 | 138733360 | 138733438 | 138742557 | 138742781 | 170 | [illegible] |
| MXE | 10377 | ENSG00000121542 | SEC22A | chr3 | + | 123209198 | 123209399 | 123223558 | 123223722 | 123201926 | 123201986 | 123225102 | 123225297 | 17 | [illegible] |
| MXE | 199 | ENSG00000100150 | DEPDC5 | chr22 | + | 31833914 | 31833980 | 31836971 | 31837155 | 31819021 | 31819225 | 31838684 | 31838845 | 6 | [illegible] |
| MXE | 1793 | ENSG00000176953 | NFATC2IP | chr16 | + | 28954564 | 28954682 | 28955977 | 28956058 | 28952131 | 28952204 | 28956150 | 28956337 | 81 | [illegible] |
| MXE | 7515 | ENSG00000115020 | PIKFYVE | chr2 | + | 208276711 | 208276830 | 208288728 | 208288818 | 208273583 | 208273733 | 208298640 | 208298779 | 65 | [illegible] |
| MXE | 3567 | ENSG00000197603 | CPLANE1 | chr5 | - | 37243012 | 37243119 | 37244374 | 37244607 | 37239712 | 37239869 | 37245478 | 37245598 | 30 | [illegible] |
| MXE | 3168 | ENSG00000112234 | FBXL4 | chr6 | - | 98917373 | 98917719 | 98926476 | 98927060 | 98905425 | 98905670 | 98927704 | 98927822 | 56 | [illegible] |
| MXE | 6858 | ENSG00000121774 | KHDRBS1 | chr1 | + | 32036909 | 32037043 | 32039514 | 32039573 | 32033187 | 32033334 | 32042526 | 32042624 | 971 | [illegible] |
| MXE | 168 | ENSG00000100393 | EP300 | chr22 | + | 41169502 | 41169616 | 41170405 | 41170571 | 41168720 | 41168867 | 41172498 | 41172663 | 99 | [illegible] |
| MXE | 7706 | ENSG00000142687 | KIAA0319L | chr1 | - | 35470862 | 35470960 | 35474804 | 35474906 | 35466607 | 35466695 | 35506611 | 35507135 | 8 | [illegible] |
| MXE | 12582 | ENSG00000175471 | MCTP1 | chr5 | - | 94912805 | 94912976 | 94923961 | 94924021 | 94909250 | 94909381 | 94931952 | 94931991 | 25 | [illegible] |
| MXE | 7205 | ENSG00000143363 | PRUNE1 | chr1 | + | 151025514 | 151025673 | 151027232 | 151027327 | 151008477 | 151008671 | 151033805 | 151035207 | 0 | [illegible] |
| MXE | 1964 | ENSG00000197006 | METTL9 | chr16 | + | 21612644 | 21612835 | 21624930 | 21625115 | 21599557 | 21599898 | 21655229 | 21656119 | 193 | [illegible] |
| MXE | 13526 | ENSG00000134516 | DOCK2 | chr5 | + | 169840756 | 169840852 | 169983067 | 169983166 | 169803057 | 169803206 | 169985827 | 169985922 | 278 | [illegible] |
| MXE | 5032 | ENSG00000009790 | TRAF3IP3 | chr1 | + | 209777351 | 209777487 | 209779314 | 209779374 | 209775598 | 209775736 | 209780469 | 209780606 | 782 | [illegible] |
| MXE | 4904 | ENSG00000075711 | DLG1 | chr3 | - | 197075836 | 197075870 | 197076585 | 197076685 | 197066703 | 197066754 | 197081050 | 197081117 | 40 | [illegible] |
| MXE | 2050 | ENSG00000166855 | CLPX | chr15 | - | 65162603 | 65162645 | 65164028 | 65164188 | 65158574 | 65158751 | 65166630 | 65166785 | 133 | [illegible] |
| MXE | 10466 | ENSG00000175215 | CTDSP2 | chr12 | - | 57826345 | 57826402 | 57826995 | 57827097 | 57824226 | 57824319 | 57827533 | 57827590 | 33 | [illegible] |
| MXE | 13933 | ENSG00000075856 | SART3 | chr12 | - | 108531203 | 108531280 | 108535358 | 108535468 | 108530141 | 108530310 | 108536513 | 108536572 | 73 | [illegible] |
| MXE | 3598 | ENSG00000161960 | EIF4A1 | chr17 | + | 7574265 | 7574308 | 7574545 | 7574678 | 7572827 | 7572864 | 7575118 | 7575258 | 6 | [illegible] |
| MXE | 12429 | ENSG00000147065 | MSN | chrX | + | 65716817 | 65716901 | 65727813 | 65727909 | 65588376 | 65588612 | 65729437 | 65729589 | 16 | [illegible] |
| MXE | 1781 | ENSG00000157106 | SMG1 | chr16 | - | 18845428 | 18845651 | 18847452 | 18847607 | 18842207 | 18842454 | 18847815 | 18848033 | 114 | [illegible] |
| MXE | 1472 | ENSG00000276234 | TADA2A | chr17 | + | 37426949 | 37427009 | 37437737 | 37437829 | 37411268 | 37411390 | 37440504 | 37440579 | 0 | [illegible] |
| MXE | 15097 | ENSG00000135362 | PRR5L | chr11 | + | 36437384 | 36437476 | 36446299 | 36446440 | 36431852 | 36431910 | 36451208 | 36451335 | 160 | [illegible] |
| MXE | 9342 | ENSG00000069275 | NUCKS1 | chr1 | - | 205727699 | 205727805 | 205729571 | 205729621 | 205723925 | 205723981 | 205749956 | 205750182 | 17 | [illegible] |
| MXE | 1136 | ENSG00000275700 | AATF | chr17 | + | 36950213 | 36950405 | 36952885 | 36953296 | 36948924 | 36949216 | 36986616 | 36986731 | 87 | [illegible] |
| MXE | 5289 | ENSG00000101596 | SMCHD1 | chr18 | + | 2718314 | 2718434 | 2722518 | 2722663 | 2718157 | 2718235 | 2724898 | 2724995 | 530 | [illegible] |
| MXE | 1258 | ENSG00000129657 | SEC14L1 | chr17 | + | 77213313 | 77213492 | 77213917 | 77214019 | 77211949 | 77212201 | 77216548 | 77216802 | 151 | [illegible] |
| MXE | 2412 | ENSG00000172575 | RASGRP1 | chr15 | - | 38502311 | 38502421 | 38507725 | 38508001 | 38500102 | 38500139 | 38511603 | 38511720 | 261 | [illegible] |
| MXE | 575 | ENSG00000011422 | PLAUR | chr19 | - | 43656478 | 43656640 | 43665315 | 43665459 | 43655438 | 43655573 | 43667580 | 43667691 | 27 | [illegible] |
| MXE | 12267 | ENSG00000269955 | FMC1-LUC7L2 | chr7 | + | 139412550 | 139412580 | 139417537 | 139417729 | 139409562 | 139409654 | 139422162 | 139422599 | 9 | [illegible] |
| MXE | 14273 | ENSG00000089234 | BRAP | chr12 | - | 111681636 | 111681835 | 111683145 | 111683307 | 111679150 | 111679340 | 111685710 | 111685956 | 32 | [illegible] |
| MXE | 5247 | ENSG00000127483 | HP1BP3 | chr1 | - | 20757165 | 20757256 | 20765376 | 20765531 | 20749722 | 20749882 | 20767583 | 20767664 | 554 | [illegible] |
| MXE | 7912 | ENSG00000071054 | MAP4K4 | chr2 | + | 101887087 | 101887237 | 101887777 | 101887937 | 101885186 | 101885287 | 101888795 | 101888935 | 631 | [illegible] |
| MXE | 15412 | ENSG00000087365 | SF3B2 | chr11 | + | 66055075 | 66055315 | 66055534 | 66055585 | 66053026 | 66053104 | 66056837 | 66056955 | 205 | [illegible] |
| MXE | 3141 | ENSG00000164180 | TMEM161B | chr5 | - | 88221673 | 88221760 | 88225768 | 88225866 | 88220562 | 88220719 | 88228444 | 88228528 | 48 | [illegible] |
| MXE | 3449 | ENSG00000065357 | DGKA | chr12 | + | 55937977 | 55938052 | 55938510 | 55938560 | 55937407 | 55937543 | 55938914 | 55938989 | 104 | [illegible] |
| MXE | 10043 | ENSG00000135945 | REV1 | chr2 | - | 99442316 | 99442469 | 99449335 | 99449504 | 99438573 | 99439310 | 99462495 | 99462788 | 177 | [illegible] |
| MXE | 9208 | ENSG00000148660 | CAMK2G | chr10 | - | 73824039 | 73824084 | 73825278 | 73825347 | 73821681 | 73821730 | 73828088 | 73828121 | 1 | [illegible] |
| MXE | 11213 | ENSG00000186635 | ARAP1 | chr11 | - | 72697910 | 72698106 | 72699004 | 72699107 | 72697597 | 72697649 | 72699416 | 72699552 | 103 | [illegible] |
| MXE | 12746 | ENSG00000160856 | FCRL3 | chr1 | - | 157697139 | 157697424 | 157697658 | 157697919 | 157696039 | 157696327 | 157698383 | 157698629 | 253 | [illegible] |
| MXE | 13604 | ENSG00000125354 | SEPTIN6 | chrX | - | 119633359 | 119633492 | 119637026 | 119637195 | 119629317 | 119629508 | 119640691 | 119640788 | 512 | [illegible] |
| MXE | 9573 | ENSG00000076242 | MLH1 | chr3 | + | 37020309 | 37020463 | 37025636 | 37026007 | 37014431 | 37014544 | 37028783 | 37028932 | 5 | [illegible] |
| MXE | 11334 | ENSG00000144233 | AMMECR1L | chr2 | - | 127869453 | 127869544 | 127873827 | 127874272 | 127866899 | 127866996 | 127884202 | 127884312 | 16 | [illegible] |
| MXE | 14804 | ENSG00000106829 | TLE4 | chr9 | + | 79612655 | 79612718 | 79627373 | 79627448 | 79576132 | 79576177 | 79652592 | 79652794 | 76 | [illegible] |
| MXE | 14594 | ENSG00000132334 | PTPRE | chr10 | + | 128073336 | 128073471 | 128076602 | 128076728 | 128072137 | 128072214 | 128077616 | 128077783 | 229 | [illegible] |
| MXE | 5682 | ENSG00000163681 | SLMAP | chr3 | + | 57864806 | 57864857 | 57896246 | 57896371 | 57864547 | 57864716 | 57896510 | 57896591 | 17 | [illegible] |
| MXE | 13691 | ENSG00000215301 | DDX3X | chrX | + | 41346858 | 41347012 | 41347639 | 41347915 | 41346504 | 41346622 | 41364273 | 41364472 | 983 | [illegible] |
| MXE | 5216 | ENSG00000206503 | HLA-A | chr6 | + | 29944121 | 29944397 | 29944499 | 29944616 | 29943267 | 29943543 | 29945058 | 29945091 | 1897 | [illegible] |
| MXE | 3995 | ENSG00000119729 | RHOQ | chr2 | + | 46576086 | 46576251 | 46576560 | 46576656 | 46543753 | 46543812 | 46580927 | 46584688 | 32 | [illegible] |
| MXE | 5752 | ENSG00000160883 | HK3 | chr5 | - | 176890821 | 176890941 | 176891036 | 176891279 | 176890634 | 176890730 | 176891387 | 176891391 | 10 | [illegible] |
| MXE | 13434 | ENSG00000175463 | TBC1D10C | chr11 | + | 67405902 | 67406017 | 67406626 | 67406738 | 67405594 | 67405701 | 67406826 | 67407016 | 170 | [illegible] |
| MXE | 5552 | ENSG00000158062 | UBXN11 | chr1 | - | 26302812 | 26302918 | 26303042 | 26303167 | 26301693 | 26301722 | 26306591 | 26306636 | 3 | [illegible] |
| MXE | 3214 | ENSG00000068796 | KIF2A | chr5 | + | 62366413 | 62366481 | 62373686 | 62373837 | 62365242 | 62365353 | 62377660 | 62377762 | 152 | [illegible] |
| MXE | 12154 | ENSG00000072609 | CHFR | chr12 | - | 132862332 | 132862425 | 132869618 | 132869834 | 132861466 | 132861634 | 132870723 | 132870783 | 110 | [illegible] |
| MXE | 2865 | ENSG00000165516 | KLHDC2 | chr14 | + | 49780212 | 49780322 | 49780702 | 49780775 | 49779594 | 49779675 | 49782369 | 49782457 | 0 | [illegible] |
| MXE | 7816 | ENSG00000143621 | ILF2 | chr1 | - | 153662704 | 153662795 | 153663018 | 153663133 | 153661787 | 153662556 | 153663214 | 153663276 | 196 | [illegible] |
| MXE | 1251 | ENSG00000182446 | NPLOC4 | chr17 | - | 81608727 | 81608822 | 81610209 | 81610258 | 81604547 | 81604727 | 81613317 | 81613494 | 17 | [illegible] |
| MXE | 15889 | ENSG00000138286 | FAM149B1 | chr10 | + | 73210250 | 73210438 | 73228059 | 73228184 | 73208618 | 73208786 | 73230421 | 73230525 | 63 | [illegible] |
| MXE | 11940 | ENSG00000146433 | TMEM181 | chr6 | + | 158605266 | 158605347 | 158607243 | 158607343 | 158589671 | 158589782 | 158608332 | 158608463 | 81 | [illegible] |
| MXE | 2385 | ENSG00000241839 | PLEKHO2 | chr15 | + | 64859893 | 64859998 | 64861476 | 64861575 | 64854920 | 64855037 | 64864898 | 64868000 | 180 | [illegible] |
| MXE | 8258 | ENSG00000143442 | POGZ | chr1 | - | 151430665 | 151430841 | 151440927 | 151441086 | 151428122 | 151428413 | 151442080 | 151442205 | 137 | [illegible] |
| MXE | 10040 | ENSG00000158435 | CNOT11 | chr2 | + | 101257790 | 101257955 | 101262538 | 101262691 | 101252885 | 101253478 | 101264844 | 101265047 | 35 | [illegible] |
| MXE | 8584 | ENSG00000134242 | PTPN22 | chr1 | - | 113834909 | 113834993 | 113837589 | 113838407 | 113834308 | 113834439 | 113838543 | 113838620 | 535 | [illegible] |
| MXE | 3815 | ENSG00000288674 | NA | chr1 | + | 226881887 | 226882048 | 226883704 | 226883919 | 226875364 | 226875550 | 226885537 | 226885679 | 23 | [illegible] |
| MXE | 9622 | ENSG00000119778 | ATAD2B | chr2 | - | 23863380 | 23863555 | 23864808 | 23864924 | 23857414 | 23857503 | 23867834 | 23867887 | 81 | [illegible] |
| MXE | 12259 | ENSG00000160218 | TRAPPC10 | chr21 | + | 44098356 | 44098534 | 44102777 | 44102972 | 44094062 | 44094233 | 44103292 | 44104733 | 164 | [illegible] |
| MXE | 7065 | ENSG00000223745 | CCDC18-AS1 | chr1 | - | 93339648 | 93339679 | 93340457 | 93340510 | 93338130 | 93339536 | 93345723 | 93345846 | 0 | [illegible] |
| MXE | 11658 | ENSG00000100503 | NIN | chr14 | - | 50759856 | 50760359 | 50761789 | 50761972 | 50754741 | 50754867 | 50763825 | 50763964 | 19 | [illegible] |
| MXE | 10224 | ENSG00000000938 | FGR | chr1 | - | 27617192 | 27617296 | 27623041 | 27623144 | 27616856 | 27617006 | 27623690 | 27623929 | 359 | [illegible] |
| MXE | 16844 | ENSG00000111596 | CNOT2 | chr12 | + | 70332766 | 70332846 | 70335437 | 70335563 | 70329422 | 70329570 | 70337388 | 70337513 | 7 | [illegible] |
| MXE | 7514 | ENSG00000115020 | PIKFYVE | chr2 | + | 208276711 | 208276830 | 208285725 | 208285933 | 208273583 | 208273733 | 208288728 | 208288818 | 67 | [illegible] |
| MXE | 682 | ENSG00000132670 | PTPRA | chr20 | + | 3022053 | 3022220 | 3022688 | 3022824 | 3021308 | 3021428 | 3024471 | 3024621 | 126 | [illegible] |
| MXE | 8235 | ENSG00000172667 | ZMAT3 | chr3 | - | 179030879 | 179030999 | 179067482 | 179067809 | 179027645 | 179027812 | 179071594 | 179071809 | 10 | [illegible] |
| MXE | 15462 | ENSG00000288721 | NA | chr6 | - | 41891793 | 41891875 | 41907015 | 41907287 | 41871563 | 41871637 | 41916784 | 41916939 | 9 | [illegible] |
| MXE | 5892 | ENSG00000151466 | SCLT1 | chr4 | - | 128965226 | 128965318 | 128992166 | 128992237 | 128959599 | 128959777 | 128997873 | 128997939 | 106 | [illegible] |
| MXE | 12902 | ENSG00000105939 | ZC3HAV1 | chr7 | - | 139076285 | 139076409 | 139078551 | 139078653 | 139073855 | 139074030 | 139079469 | 139080243 | 196 | [illegible] |
| MXE | 2002 | ENSG00000130703 | OSBPL2 | chr20 | + | 62264628 | 62264718 | 62272124 | 62272259 | 62263615 | 62263691 | 62279156 | 62279339 | 1 | [illegible] |
| MXE | 11211 | ENSG00000186635 | ARAP1 | chr11 | - | 72686094 | 72686191 | 72687438 | 72687502 | 72685643 | 72685681 | 72687687 | 72687738 | 198 | [illegible] |
| MXE | 6852 | ENSG00000107099 | DOCK8 | chr9 | + | 420400 | 420583 | 420948 | 421078 | 418067 | 418207 | 422047 | 422135 | 782 | [illegible] |
| MXE | 12788 | ENSG00000051825 | MPHOSPH9 | chr12 | - | 123161135 | 123161383 | 123163013 | 123163134 | 123159693 | 123160849 | 123163949 | 123164090 | 71 | [illegible] |
| MXE | 16268 | ENSG00000119977 | TCTN3 | chr10 | - | 95687246 | 95687355 | 95687591 | 95687719 | 95687043 | 95687159 | 95692919 | 95693038 | 37 | [illegible] |
| MXE | 5433 | ENSG00000164463 | CREBRF | chr5 | + | 173086500 | 173086626 | 173090314 | 173091401 | 173080584 | 173080784 | 173108623 | 173108818 | 307 | [illegible] |
| MXE | 8095 | ENSG00000138688 | KIAA1109 | chr4 | + | 122189999 | 122190079 | 122192228 | 122192379 | 122187894 | 122188088 | 122196634 | 122196791 | 438 | [illegible] |
| MXE | 13970 | ENSG00000165392 | WRN | chr8 | + | 31068257 | 31068327 | 31076172 | 31076287 | 31067032 | 31067182 | 31080866 | 31081296 | 19 | [illegible] |
| MXE | 14812 | ENSG00000011454 | RABGAP1 | chr9 | + | 123070349 | 123070424 | 123073551 | 123073677 | 123065347 | 123065601 | 123074284 | 123074317 | 0 | [illegible] |
| MXE | 11056 | ENSG00000064012 | CASP8 | chr2 | + | 201266460 | 201266791 | 201271515 | 201271621 | 201234015 | 201234112 | 201272637 | 201272776 | 79 | [illegible] |
| MXE | 11502 | ENSG00000137478 | FCHSD2 | chr11 | - | 72840876 | 72840959 | 72842620 | 72842841 | 72836744 | 72838874 | 72843150 | 72843328 | 182 | [illegible] |
| MXE | 2588 | ENSG00000249624 | NA | chr21 | + | 33262792 | 33263637 | 33266011 | 33266175 | 33260596 | 33260727 | 33268393 | 33268517 | 154 | [illegible] |
| MXE | 2034 | ENSG00000103994 | ZNF106 | chr15 | - | 42449770 | 42451954 | 42456957 | 42457158 | 42448071 | 42448705 | 42466052 | 42466114 | 251 | [illegible] |
| MXE | 17013 | ENSG00000067167 | TRAM1 | chr8 | - | 70594505 | 70594590 | 70596262 | 70596321 | 70587105 | 70587176 | 70597894 | 70598011 | 193 | [illegible] |
| MXE | 8847 | ENSG00000135899 | SP110 | chr2 | - | 230214949 | 230215118 | 230216780 | 230216928 | 230212760 | 230213027 | 230221704 | 230221784 | 23 | [illegible] |
| MXE | 14722 | ENSG00000148337 | CIZ1 | chr9 | - | 128187862 | 128187934 | 128190328 | 128190444 | 128185546 | 128185776 | 128190687 | 128190862 | 0 | [illegible] |
| MXE | 11361 | ENSG00000136699 | SMPD4 | chr2 | - | 130173278 | 130173354 | 130173513 | 130173604 | 130172786 | 130172895 | 130174913 | 130175000 | 0 | [illegible] |
| MXE | 10933 | ENSG00000170266 | GLB1 | chr3 | - | 33065462 | 33065557 | 33072543 | 33072713 | 33058088 | 33058269 | 33097010 | 33097146 | 13 | [illegible] |
| MXE | 4240 | ENSG00000204842 | ATXN2 | chr12 | - | 111516153 | 111516363 | 111518248 | 111518427 | 111513356 | 111513539 | 111520881 | 111520973 | 11 | [illegible] |
| MXE | 4435 | ENSG00000203875 | SNHG5 | chr6 | - | 85677423 | 85677492 | 85677794 | 85677875 | 85666003 | 85666147 | 85677953 | 85678032 | 130 | [illegible] |
| MXE | 1016 | ENSG00000073605 | GSDMB | chr17 | - | 39906960 | 39906987 | 39908175 | 39908214 | 39906110 | 39906271 | 39909755 | 39909924 | 1 | [illegible] |
| MXE | 9451 | ENSG00000067225 | PKM | chr15 | - | 72200473 | 72200655 | 72202453 | 72202620 | 72199028 | 72199756 | 72206727 | 72206880 | 1243 | [illegible] |
| MXE | 13348 | ENSG00000136504 | KAT7 | chr17 | + | 49805362 | 49805445 | 49809118 | 49809208 | 49798318 | 49798558 | 49811475 | 49811574 | 145 | [illegible] |
| MXE | 1681 | ENSG00000005844 | ITGAL | chr16 | + | 30475512 | 30475580 | 30479090 | 30479208 | 30475305 | 30475400 | 30481438 | 30481584 | 436 | [illegible] |
| MXE | 10668 | ENSG00000136536 | MARCHF7 | chr2 | + | 159729008 | 159729175 | 159743060 | 159743253 | 159712522 | 159712661 | 159747804 | 159748126 | 143 | [illegible] |
| MXE | 11576 | ENSG00000205189 | ZBTB10 | chr8 | + | 80499493 | 80500382 | 80513909 | 80514008 | 80486212 | 80487782 | 80518402 | 80518438 | 47 | [illegible] |
| MXE | 6853 | ENSG00000107099 | DOCK8 | chr9 | + | 426884 | 426981 | 429701 | 429854 | 422047 | 422135 | 432165 | 432324 | 222 | [illegible] |
| MXE | 4215 | ENSG00000164190 | NIPBL | chr5 | + | 36984675 | 36986301 | 36995621 | 36995804 | 36975775 | 36976402 | 36996471 | 36997002 | 189 | [illegible] |
| MXE | 9499 | ENSG00000189319 | FAM53B | chr10 | - | 124696157 | 124696212 | 124706635 | 124706887 | 124681606 | 124682379 | 124744012 | 124744269 | 77 | [illegible] |
| MXE | 7200 | ENSG00000143363 | PRUNE1 | chr1 | + | 151018556 | 151018669 | 151024610 | 151024795 | 151017811 | 151017904 | 151025514 | 151025673 | 6 | [illegible] |
| MXE | 12772 | ENSG00000187953 | PMS2CL | chr7 | + | 6741684 | 6741852 | 6746087 | 6746188 | 6737075 | 6737936 | 6747035 | 6747205 | 47 | [illegible] |
| MXE | 5035 | ENSG00000009790 | TRAF3IP3 | chr1 | + | 209778110 | 209778173 | 209781344 | 209781458 | 209777351 | 209777487 | 209782055 | 209782307 | 228 | [illegible] |
| MXE | 7190 | ENSG00000198178 | CLEC4C | chr12 | - | 7737428 | 7737574 | 7741420 | 7741531 | 7730796 | 7730912 | 7746330 | 7746423 | 68 | [illegible] |
| MXE | 7505 | ENSG00000115020 | PIKFYVE | chr2 | + | 208273583 | 208273733 | 208276711 | 208276830 | 208271510 | 208271691 | 208277536 | 208277708 | 58 | [illegible] |
| MXE | 8925 | ENSG00000065413 | ANKRD44 | chr2 | - | 197136591 | 197136662 | 197147026 | 197147105 | 197125836 | 197126037 | 197187022 | 197187106 | 243 | [illegible] |
| MXE | 1976 | ENSG00000005339 | CREBBP | chr16 | - | 3782683 | 3782926 | 3791980 | 3792094 | 3781203 | 3781306 | 3793385 | 3793626 | 147 | [illegible] |
| MXE | 11706 | ENSG00000197081 | IGF2R | chr6 | + | 160062531 | 160062619 | 160063414 | 160063630 | 160061752 | 160061928 | 160064400 | 160064531 | 353 | [illegible] |
| MXE | 7739 | ENSG00000156515 | HK1 | chr10 | + | 69364782 | 69364902 | 69368535 | 69368631 | 69359896 | 69360045 | 69369236 | 69369336 | 273 | [illegible] |
| MXE | 7978 | ENSG00000173209 | AHSA2P | chr2 | + | 61185838 | 61185865 | 61185979 | 61186081 | 61185501 | 61185618 | 61186445 | 61186497 | 5 | [illegible] |
| MXE | 10149 | ENSG00000112078 | KCTD20 | chr6 | + | 36474788 | 36475062 | 36481561 | 36481759 | 36470051 | 36470257 | 36484713 | 36484824 | 138 | [illegible] |
| MXE | 7091 | ENSG00000171103 | TRMT61B | chr2 | - | 28852407 | 28852499 | 28861117 | 28861308 | 28851071 | 28851298 | 28865016 | 28865119 | 86 | [illegible] |
| MXE | 3349 | ENSG00000127603 | MACF1 | chr1 | + | 39302923 | 39303078 | 39309569 | 39309696 | 39300209 | 39300362 | 39310244 | 39310428 | 452 | [illegible] |
| MXE | 6931 | ENSG00000163714 | U2SURP | chr3 | + | 143050938 | 143051049 | 143054942 | 143055119 | 143043116 | 143043276 | 143056311 | 143056500 | 385 | [illegible] |
| MXE | 15652 | ENSG00000081237 | PTPRC | chr1 | + | 198692346 | 198692373 | 198696711 | 198696909 | 198639225 | 198639341 | 198699563 | 198699704 | 1520 | [illegible] |
| MXE | 12594 | ENSG00000136280 | CCM2 | chr7 | + | 45038252 | 45038426 | 45063917 | 45064001 | 45027671 | 45027797 | 45064462 | 45064646 | 80 | [illegible] |
| MXE | 3848 | ENSG00000247774 | PCED1B-AS1 | chr12 | - | 47209280 | 47209458 | 47210639 | 47210761 | 47205899 | 47206585 | 47216164 | 47216264 | 212 | [illegible] |
| MXE | 16492 | ENSG00000114302 | PRKAR2A | chr3 | - | 48756378 | 48756444 | 48765003 | 48765078 | 48752175 | 48752317 | 48772954 | 48773108 | 0 | [illegible] |
| MXE | 809 | ENSG00000170471 | RALGAPB | chr20 | + | 38517783 | 38518000 | 38521496 | 38521698 | 38517505 | 38517654 | 38524777 | 38524945 | 119 | [illegible] |
| MXE | 2678 | ENSG00000100722 | ZC3H14 | chr14 | + | 88596733 | 88596808 | 88601923 | 88602083 | 88577984 | 88578140 | 88607242 | 88607363 | 11 | [illegible] |
| MXE | 6464 | ENSG00000008130 | NADK | chr1 | - | 1761951 | 1762035 | 1765227 | 1765446 | 1757180 | 1757310 | 1779904 | 1779962 | 535 | [illegible] |
| MXE | 6665 | ENSG00000054118 | THRAP3 | chr1 | + | 36291373 | 36291546 | 36292597 | 36292709 | 36289059 | 36289764 | 36293850 | 36293935 | 721 | [illegible] |
| MXE | 1750 | ENSG00000169896 | ITGAM | chr16 | + | 31297744 | 31297954 | 31321240 | 31321371 | 31297513 | 31297654 | 31321463 | 31321627 | 96 | [illegible] |
| MXE | 4815 | ENSG00000163964 | PIGX | chr3 | + | 196717808 | 196717954 | 196721176 | 196721356 | 196716857 | 196716921 | 196722414 | 196722556 | 6 | [illegible] |
| MXE | 740 | ENSG00000125900 | SIRPD | chr20 | - | 1537154 | 1537313 | 1551690 | 1552038 | 1534250 | 1534441 | 1557580 | 1557705 | 4 | [illegible] |
| MXE | 9968 | ENSG00000138363 | ATIC | chr2 | + | 215325986 | 215326138 | 215326821 | 215326978 | 215319664 | 215319731 | 215332381 | 215332507 | 3 | [illegible] |
| MXE | 4216 | ENSG00000164190 | NIPBL | chr5 | + | 36984675 | 36986301 | 36995621 | 36995804 | 36975775 | 36976402 | 37000372 | 37000570 | 191 | [illegible] |
| MXE | 13973 | ENSG00000115419 | GLS | chr2 | + | 190913192 | 190913364 | 190920214 | 190920248 | 190910262 | 190910321 | 190921023 | 190921056 | 98 | [illegible] |
| MXE | 8715 | ENSG00000023902 | PLEKHO1 | chr1 | + | 150156065 | 150156206 | 150156910 | 150157015 | 150150911 | 150151058 | 150158818 | 150159630 | 314 | [illegible] |
| MXE | 10931 | ENSG00000132824 | SERINC3 | chr20 | - | 44509890 | 44510028 | 44512800 | 44512994 | 44506826 | 44506996 | 44513878 | 44514040 | 393 | [illegible] |
| MXE | 15149 | ENSG00000153339 | TRAPPC8 | chr18 | - | 31832083 | 31832173 | 31839311 | 31839457 | 31829196 | 31830989 | 31846715 | 31846817 | 251 | [illegible] |
| MXE | 362 | ENSG00000257103 | LSM14A | chr19 | + | 34196633 | 34196763 | 34208928 | 34209051 | 34194477 | 34194641 | 34215595 | 34215661 | 354 | [illegible] |
| MXE | 5348 | ENSG00000111642 | CHD4 | chr12 | - | 6581648 | 6581814 | 6582136 | 6582281 | 6581290 | 6581388 | 6582614 | 6582748 | 180 | [illegible] |
| MXE | 379 | ENSG00000170889 | RPS9 | chr19 | + | 54201486 | 54201609 | 54206275 | 54206462 | 54201159 | 54201281 | 54207397 | 54207647 | 1633 | [illegible] |
| MXE | 5562 | ENSG00000173193 | PARP14 | chr3 | + | 122703741 | 122703978 | 122704526 | 122704748 | 122695425 | 122695662 | 122708189 | 122708268 | 1 | [illegible] |
| MXE | 5547 | ENSG00000158062 | UBXN11 | chr1 | - | 26294204 | 26294331 | 26296918 | 26296995 | 26285822 | 26286037 | 26297961 | 26298062 | 6 | [illegible] |
| MXE | 9584 | ENSG00000163904 | SENP2 | chr3 | + | 185614563 | 185614740 | 185621825 | 185621905 | 185613344 | 185613408 | 185623997 | 185624082 | 82 | [illegible] |
| MXE | 14472 | ENSG00000135040 | NAA35 | chr9 | + | 85958471 | 85958586 | 85959792 | 85959867 | 85956359 | 85956393 | 85962012 | 85962180 | 20 | [illegible] |
| MXE | 11035 | ENSG00000089123 | TASP1 | chr20 | - | 13587249 | 13587370 | 13623445 | 13623514 | 13580896 | 13580981 | 13625184 | 13625252 | 9 | [illegible] |
| MXE | 1618 | ENSG00000153066 | TXNDC11 | chr16 | - | 11691289 | 11692082 | 11698124 | 11698325 | 11688302 | 11688445 | 11700451 | 11700564 | 234 | [illegible] |
| MXE | 5883 | ENSG00000164307 | ERAP1 | chr5 | - | 96797174 | 96797309 | 96803402 | 96803943 | 96795041 | 96795162 | 96807859 | 96807945 | 105 | [illegible] |
| MXE | 1518 | ENSG00000091592 | NLRP1 | chr17 | - | 5553385 | 5553556 | 5558338 | 5560043 | 5541856 | 5542027 | 5581858 | 5582062 | 176 | [illegible] |
| MXE | 14843 | ENSG00000163527 | STT3B | chr3 | + | 31576395 | 31576504 | 31579808 | 31580096 | 31532637 | 31533312 | 31596797 | 31596863 | 77 | [illegible] |
| MXE | 12801 | ENSG00000051825 | MPHOSPH9 | chr12 | - | 123226287 | 123226351 | 123227545 | 123227616 | 123223037 | 123223127 | 123230260 | 123230522 | 57 | [illegible] |
| MXE | 9627 | ENSG00000172888 | ZNF621 | chr3 | + | 40529318 | 40529445 | 40529494 | 40529575 | 40525778 | 40525864 | 40530208 | 40530316 | 26 | [illegible] |
| MXE | 2520 | ENSG00000100796 | PPP4R3A | chr14 | - | 91481575 | 91482193 | 91485631 | 91485730 | 91476908 | 91476986 | 91490746 | 91490802 | 191 | [illegible] |
| MXE | 3122 | ENSG00000150867 | PIP4K2A | chr10 | - | 22573310 | 22573457 | 22607926 | 22608023 | 22567850 | 22567889 | 22609619 | 22609717 | 428 | [illegible] |
| MXE | 15493 | ENSG00000163131 | CTSS | chr1 | - | 150751780 | 150752008 | 150755000 | 150755150 | 150750005 | 150750171 | 150757857 | 150757980 | 4056 | [illegible] |
| MXE | 9449 | ENSG00000198160 | MIER1 | chr1 | + | 66959678 | 66959743 | 66963087 | 66963160 | 66958860 | 66958983 | 66970807 | 66970959 | 168 | [illegible] |
| MXE | 10593 | ENSG00000163902 | RPN1 | chr3 | - | 128629950 | 128630143 | 128631947 | 128632157 | 128626732 | 128626832 | 128637798 | 128638105 | 471 | [illegible] |
| MXE | 11347 | ENSG00000114956 | DGUOK | chr2 | + | 73938909 | 73939022 | 73957124 | 73957240 | 73926825 | 73927052 | 73958145 | 73958245 | 13 | [illegible] |
| MXE | 16881 | ENSG00000050426 | LETMD1 | chr12 | + | 51053777 | 51053860 | 51056143 | 51056245 | 51052091 | 51052207 | 51056349 | 51056502 | 49 | [illegible] |
| MXE | 8716 | ENSG00000172939 | OXSR1 | chr3 | + | 38236838 | 38236961 | 38242742 | 38242778 | 38230364 | 38230430 | 38246074 | 38246221 | 143 | [illegible] |
| MXE | 5545 | ENSG00000158062 | UBXN11 | chr1 | - | 26294204 | 26294331 | 26296918 | 26296995 | 26285822 | 26286037 | 26297426 | 26297481 | 165 | [illegible] |
| MXE | 17089 | ENSG00000013364 | MVP | chr16 | + | 29835703 | 29835798 | 29836721 | 29836958 | 29834185 | 29834437 | 29840177 | 29840267 | 57 | [illegible] |
| MXE | 1696 | ENSG00000176715 | ACSF3 | chr16 | + | 89112091 | 89112246 | 89120800 | 89120913 | 89102603 | 89102759 | 89133135 | 89133262 | 67 | [illegible] |
| MXE | 12278 | ENSG00000242588 | NA | chr7 | + | 128579998 | 128580094 | 128591849 | 128592017 | 128578889 | 128578933 | 128608117 | 128608213 | 85 | [illegible] |
| MXE | 8061 | ENSG00000197183 | NOL4L | chr20 | - | 32456117 | 32456395 | 32474600 | 32474742 | 32453575 | 32453761 | 32511346 | 32511456 | 79 | [illegible] |
| MXE | 7336 | ENSG00000069424 | KCNAB2 | chr1 | + | 6041830 | 6041872 | 6082194 | 6082274 | 6040516 | 6040645 | 6085203 | 6085248 | 39 | [illegible] |
| MXE | 3756 | ENSG00000153310 | CYRIB | chr8 | - | 129862228 | 129862334 | 129871374 | 129871496 | 129855610 | 129855747 | 129903311 | 129903350 | 21 | [illegible] |
| MXE | 14327 | ENSG00000104687 | GSR | chr8 | - | 30700080 | 30700135 | 30703092 | 30703240 | 30696379 | 30696479 | 30708071 | 30708141 | 316 | [illegible] |
| MXE | 2340 | ENSG00000140367 | UBE2Q2 | chr15 | + | 75868950 | 75869010 | 75876186 | 75876271 | 75859877 | 75859982 | 75877960 | 75878021 | 89 | [illegible] |
| MXE | 8590 | ENSG00000134242 | PTPN22 | chr1 | - | 113852026 | 113852104 | 113854470 | 113854537 | 113848539 | 113848626 | 113858477 | 113858573 | 26 | [illegible] |
| MXE | 3904 | ENSG00000181929 | PRKAG1 | chr12 | - | 49005742 | 49005881 | 49008425 | 49008599 | 49005461 | 49005543 | 49013061 | 49013110 | 2 | [illegible] |
| MXE | 12158 | ENSG00000198900 | TOP1 | chr20 | + | 41061393 | 41061490 | 41076170 | 41076294 | 41029430 | 41029455 | 41077581 | 41077637 | 37 | [illegible] |
| MXE | 6582 | ENSG00000153563 | CD8A | chr2 | - | 86789322 | 86789433 | 86789639 | 86789750 | 86788529 | 86788560 | 86790327 | 86790681 | 135 | [illegible] |
| MXE | 7889 | ENSG00000074603 | DPP8 | chr15 | - | 65451959 | 65452102 | 65456224 | 65456371 | 65450998 | 65451110 | 65463760 | 65463906 | 158 | [illegible] |
| MXE | 12857 | ENSG00000123066 | MED13L | chr12 | - | 116232058 | 116232156 | 116236820 | 116236890 | 116230532 | 116230622 | 116237467 | 116237569 | 0 | [illegible] |
| MXE | 5898 | ENSG00000067066 | SP100 | chr2 | + | 230446818 | 230446902 | 230449087 | 230449150 | 230444177 | 230444346 | 230449560 | 230449710 | 236 | [illegible] |
| MXE | 11538 | ENSG00000116539 | ASH1L | chr1 | - | 155415743 | 155415923 | 155438326 | 155439068 | 155395458 | 155395553 | 155459796 | 155459898 | 86 | [illegible] |
| MXE | 2586 | ENSG00000249624 | NA | chr21 | + | 33260596 | 33260727 | 33266011 | 33266175 | 33252661 | 33252830 | 33268393 | 33268517 | 209 | [illegible] |
| MXE | 11441 | ENSG00000151687 | ANKAR | chr2 | + | 189728659 | 189728821 | 189737682 | 189737841 | 189728266 | 189728420 | 189738564 | 189738682 | 6 | [illegible] |
| MXE | 4553 | ENSG00000086061 | DNAJA1 | chr9 | + | 33026812 | 33026990 | 33029884 | 33029989 | 33026474 | 33026616 | 33030439 | 33030667 | 436 | [illegible] |
| MXE | 6189 | ENSG00000213390 | ARHGAP19 | chr10 | - | 97259401 | 97259628 | 97263419 | 97263629 | 97246271 | 97246337 | 97264825 | 97264906 | 61 | [illegible] |
| MXE | 1496 | ENSG00000177731 | FLII | chr17 | - | 18249325 | 18249408 | 18250837 | 18251017 | 18249126 | 18249201 | 18251264 | 18251477 | 225 | [illegible] |
| MXE | 1320 | ENSG00000141298 | SSH2 | chr17 | - | 29635967 | 29636802 | 29648143 | 29648344 | 29625937 | 29632931 | 29650653 | 29650800 | 511 | [illegible] |
| MXE | 16469 | ENSG00000112297 | CRYBG1 | chr6 | + | 106558485 | 106558625 | 106560802 | 106560926 | 106555767 | 106555897 | 106561341 | 106561500 | 284 | [illegible] |
| MXE | 571 | ENSG00000011422 | PLAUR | chr19 | - | 43652224 | 43652371 | 43655438 | 43655573 | 43648578 | 43649143 | 43665315 | 43665459 | 7 | [illegible] |
| MXE | 15283 | ENSG00000197860 | SGTB | chr5 | - | 65708488 | 65708558 | 65712960 | 65713064 | 65704278 | 65704378 | 65720707 | 65720829 | 85 | [illegible] |
| MXE | 335 | ENSG00000099331 | MYO9B | chr19 | + | 17185920 | 17186001 | 17187934 | 17188045 | 17184864 | 17184987 | 17191096 | 17191219 | 65 | [illegible] |
| MXE | 1805 | ENSG00000140987 | ZSCAN32 | chr16 | - | 3384458 | 3384941 | 3390009 | 3390133 | 3382084 | 3383711 | 3390422 | 3390517 | 2 | [illegible] |
| MXE | 15559 | ENSG00000013725 | CD6 | chr11 | + | 61006573 | 61006642 | 61017478 | 61017550 | 60971679 | 60971914 | 61017758 | 61018013 | 107 | [illegible] |
| MXE | 9838 | ENSG00000166478 | ZNF143 | chr11 | + | 9472676 | 9472769 | 9473776 | 9473849 | 9471301 | 9471420 | 9473940 | 9474024 | 133 | [illegible] |
| MXE | 12848 | ENSG00000146872 | TLK2 | chr17 | + | 62586134 | 62586226 | 62596584 | 62596674 | 62578476 | 62580192 | 62600650 | 62600820 | 125 | [illegible] |
| MXE | 4944 | ENSG00000088726 | TMEM40 | chr3 | - | 12743899 | 12743989 | 12748654 | 12748792 | 12742453 | 12742507 | 12749759 | 12749840 | 14 | [illegible] |

| | | | | | | | | | | | | | | | | | |
|---|---|---|---|---|---|---|---|---|---|---|---|---|---|---|---|---|---|
| MXE | 3011 | ENSG00000115760 | BIRC6 | chr2 | + | 32463193 | 32463381 | 32464508 | 32464823 | 32453807 | 32453942 | 32465064 | 32465164 | 58 | 34 | 59 | 77 |
| MXE | 8596 | ENSG00000185946 | RNPC3 | chr1 | + | 103543295 | 103543447 | 103544940 | 103545102 | 103541349 | 103541475 | 103546247 | 103546342 | 147 | 194 | 106 | 164 |
| MXE | 8455 | ENSG00000137275 | RIPK1 | chr6 | + | 3076763 | 3076987 | 3077778 | 3077935 | 3068493 | 3068661 | 3080978 | 3081116 | 22 | 9 | 14 | 22 |
| MXE | 5819 | ENSG00000143458 | GABPB2 | chr1 | + | 151088157 | 151088297 | 151093191 | 151093386 | 151070803 | 151070934 | 151097851 | 151098002 | 6 | 0 | 5 | 2 |
| MXE | 16026 | ENSG00000077147 | TM9SF3 | chr10 | - | 96533050 | 96533190 | 96544075 | 96544206 | 96530539 | 96530608 | 96547894 | 96547989 | 412 | 263 | 320 | 365 |
| MXE | 14826 | ENSG00000107077 | KDM4C | chr9 | + | 6805598 | 6805774 | 6814630 | 6814745 | 6792971 | 6793132 | 6849506 | 6849700 | 274 | 206 | 285 | 311 |
| MXE | 6450 | ENSG00000172071 | EIF2AK3 | chr2 | - | 88593271 | 88593405 | 88595468 | 88595663 | 88590817 | 88591052 | 88613723 | 88613853 | 30 | 19 | 23 | 15 |
| MXE | 4209 | ENSG00000198478 | SH3BGRL2 | chr6 | + | 79673613 | 79673799 | 79696484 | 79696565 | 79631328 | 79631506 | 79699497 | 79703655 | 100 | 113 | 27 | 251 |
| MXE | 1293 | ENSG00000130402 | ACTN4 | chr19 | + | 38710256 | 38710342 | 38711275 | 38711361 | 38709394 | 38709476 | 38714468 | 38714561 | 329 | 160 | 227 | 295 |
| MXE | 7381 | ENSG00000138095 | LRPPRC | chr2 | - | 43974149 | 43974295 | 43974613 | 43974758 | 43973794 | 43973900 | 43975090 | 43975217 | 130 | 77 | 112 | 128 |
| MXE | 12370 | ENSG00000146963 | LUC7L2 | chr7 | + | 139376061 | 139376156 | 139398598 | 139398697 | 139359893 | 139360322 | 139402136 | 139402247 | 53 | 31 | 29 | 47 |
| MXE | 16145 | ENSG00000172661 | WASHC2C | chr10 | + | 45785508 | 45785631 | 45787034 | 45787247 | 45784820 | 45784901 | 45788870 | 45789491 | 130 | 87 | 93 | 114 |
| MXE | 8725 | ENSG00000178691 | SUZ12 | chr17 | + | 31940421 | 31940486 | 31947616 | 31947685 | 31940285 | 31940332 | 31966146 | 31966196 | 65 | 69 | 94 | 79 |
| MXE | 5022 | ENSG00000082996 | RNF13 | chr3 | + | 149902071 | 149902162 | 149911977 | 149912083 | 149895472 | 149895560 | 149921133 | 149921227 | 311 | 222 | 192 | 271 |
| MXE | 5686 | ENSG00000163681 | SLMAP | chr3 | + | 57871635 | 57871698 | 57896246 | 57896371 | 57864547 | 57864716 | 57896510 | 57896591 | 4 | 4 | 2 | 2 |
| MXE | 6758 | ENSG00000072501 | SMC1A | chrX | - | 53411994 | 53412253 | 53414757 | 53414870 | 53411760 | 53411901 | 53422491 | 53422654 | 0 | 1 | 3 | 3 |
| MXE | 2485 | ENSG00000159110 | IFNAR2 | chr21 | + | 33241839 | 33241977 | 33243672 | 33243714 | 33229937 | 33230216 | 33244950 | 33245074 | 16 | 7 | 15 | 11 |
| MXE | 2294 | ENSG00000069667 | RORA | chr15 | - | 60511225 | 60511621 | 60514615 | 60514757 | 60505507 | 60505629 | 60531765 | 60531851 | 268 | 166 | 204 | 226 |
| MXE | 5290 | ENSG00000101596 | SMCHD1 | chr18 | + | 2722518 | 2722663 | 2724898 | 2724995 | 2718314 | 2718434 | 2726451 | 2726524 | 418 | 357 | 400 | 395 |
| MXE | 3920 | ENSG00000108039 | XPNPEP1 | chr10 | - | 109891721 | 109891826 | 109893011 | 109893075 | 109888502 | 109888595 | 109907690 | 109907815 | 7 | 7 | 10 | 3 |
| MXE | 11078 | ENSG00000145349 | CAMK2D | chr4 | - | 113547643 | 113547716 | 113548656 | 113548729 | 113537340 | 113537443 | 113552030 | 113552096 | 145 | 93 | 131 | 99 |
| MXE | 14859 | ENSG00000266714 | MYO15B | chr17 | + | 75594474 | 75594588 | 75594700 | 75594759 | 75592678 | 75592840 | 75594839 | 75594972 | 3 | 2 | 4 | 14 |
| MXE | 11591 | ENSG00000163939 | PBRM1 | chr3 | - | 52668497 | 52668645 | 52678499 | 52678597 | 52662132 | 52662276 | 52679573 | 52679713 | 80 | 40 | 69 | 65 |
| MXE | 11473 | ENSG00000123600 | METTL8 | chr2 | - | 171326041 | 171326148 | 171337452 | 171337502 | 171325840 | 171325906 | 171339183 | 171339256 | 14 | 7 | 7 | 10 |
| MXE | 8814 | ENSG00000078177 | N4BP2 | chr4 | + | 40117868 | 40118024 | 40119931 | 40122309 | 40113431 | 40113508 | 40123126 | 40123212 | 44 | 27 | 30 | 20 |
| MXE | 11748 | ENSG00000112062 | MAPK14 | chr6 | + | 36095986 | 36096066 | 36100187 | 36100267 | 36076536 | 36076608 | 36102570 | 36102649 | 359 | 181 | 231 | 297 |
| MXE | 12474 | ENSG00000154511 | DIPK1A | chr1 | - | 92850847 | 92850955 | 92876295 | 92876430 | 92847182 | 92847359 | 92961375 | 92961522 | 28 | 7 | 33 | 26 |
| MXE | 12711 | ENSG00000106443 | PHF14 | chr7 | + | 10990702 | 10990847 | 11013746 | 11013906 | 10982371 | 10983159 | 11022867 | 11022979 | 127 | 131 | 125 | 111 |
| MXE | 15665 | ENSG00000081237 | PTPRC | chr1 | + | 198699563 | 198699704 | 198703297 | 198703372 | 198696711 | 198696909 | 198704471 | 198704498 | 924 | 411 | 590 | 657 |
| MXE | 12771 | ENSG00000187953 | PMS2CL | chr7 | + | 6737075 | 6737936 | 6741684 | 6741852 | 6735304 | 6735389 | 6746087 | 6746188 | 10 | 19 | 25 | 31 |
| MXE | 12709 | ENSG00000106443 | PHF14 | chr7 | + | 10982371 | 10983159 | 10990702 | 10990847 | 10974834 | 10974945 | 11013746 | 11013906 | 45 | 40 | 36 | 58 |
| MXE | 4176 | ENSG00000047315 | POLR2B | chr4 | + | 57010360 | 57010504 | 57010747 | 57010887 | 57006815 | 57007002 | 57010988 | 57011100 | 240 | 129 | 167 | 169 |
| MXE | 5034 | ENSG00000009790 | TRAF3IP3 | chr1 | + | 209778110 | 209778173 | 209780469 | 209780606 | 209777351 | 209777487 | 209781344 | 209781458 | 227 | 164 | 229 | 204 |
| MXE | 8291 | ENSG00000257379 | NA | chr12 | + | 53455460 | 53455493 | 53455884 | 53456001 | 53455346 | 53455370 | 53459271 | 53459403 | 1036 | 679 | 843 | 1132 |
| MXE | 6576 | ENSG00000072786 | STK10 | chr5 | - | 172090231 | 172090362 | 172093411 | 172093960 | 172082960 | 172083084 | 172096425 | 172096560 | 482 | 168 | 292 | 293 |
| MXE | 12070 | ENSG00000134250 | NOTCH2 | chr1 | - | 119922235 | 119922446 | 119922635 | 119922778 | 119921712 | 119921809 | 119923636 | 119923984 | 283 | 171 | 194 | 263 |
| MXE | 16698 | ENSG00000139350 | NEDD1 | chr12 | + | 96907603 | 96907856 | 96917620 | 96917737 | 96907237 | 96907300 | 96919984 | 96920125 | 5 | 1 | 1 | 8 |
| MXE | 11185 | ENSG00000130414 | NDUFA10 | chr2 | - | 240011616 | 240014860 | 240018552 | 240018639 | 240007315 | 240007370 | 240021196 | 240021412 | 196 | 134 | 151 | 136 |
| MXE | 12288 | ENSG00000133606 | MKRN1 | chr7 | - | 140455094 | 140455233 | 140455789 | 140455900 | 140453032 | 140454729 | 140456651 | 140456866 | 267 | 241 | 218 | 356 |
| MXE | 16126 | ENSG00000157625 | TAB3 | chrX | - | 30842965 | 30843049 | 30846550 | 30846644 | 30834050 | 30834152 | 30852777 | 30852938 | 51 | 45 | 43 | 74 |
| MXE | 14697 | ENSG00000089053 | ANAPC5 | chr12 | - | 121337290 | 121337392 | 121342002 | 121342069 | 121335516 | 121335723 | 121345838 | 121346031 | 180 | 151 | 143 | 213 |
| MXE | 5537 | ENSG00000169180 | XPO6 | chr16 | - | 28132333 | 28132403 | 28133840 | 28133933 | 28125688 | 28125848 | 28135215 | 28135324 | 117 | 77 | 130 | 115 |
| MXE | 15159 | ENSG00000175216 | CKAP5 | chr11 | - | 46816197 | 46816404 | 46818309 | 46818503 | 46811006 | 46811178 | 46821174 | 46821257 | 119 | 89 | 97 | 158 |
| MXE | 5316 | ENSG00000165288 | BRWD3 | chrX | - | 80707426 | 80707503 | 80709427 | 80709577 | 80704677 | 80704846 | 80716156 | 80716250 | 149 | 147 | 117 | 167 |
| MXE | 15854 | ENSG00000088298 | EDEM2 | chr20 | - | 35123889 | 35124034 | 35126250 | 35126375 | 35118597 | 35118719 | 35131641 | 35131783 | 75 | 30 | 59 | 73 |
| MXE | 5437 | ENSG00000145819 | ARHGAP26 | chr5 | + | 142894237 | 142894348 | 142901934 | 142902039 | 142885297 | 142885399 | 142903539 | 142903669 | 317 | 367 | 219 | 438 |
| MXE | 8844 | ENSG00000135899 | SP110 | chr2 | - | 230170620 | 230170761 | 230171695 | 230171767 | 230165185 | 230169237 | 230172065 | 230172174 | 6 | 11 | 6 | 6 |
| MXE | 5393 | ENSG00000162736 | NCSTN | chr1 | + | 160351287 | 160351372 | 160351695 | 160351805 | 160350104 | 160350250 | 160352886 | 160352991 | 2 | 0 | 0 | 3 |
| MXE | 8576 | ENSG00000124942 | AHNAK | chr11 | - | 62535944 | 62536098 | 62536468 | 62536567 | 62535002 | 62535190 | 62546659 | 62546796 | 76 | 43 | 30 | 41 |
| MXE | 6192 | ENSG00000134248 | LAMTOR5 | chr1 | - | 110403918 | 110404036 | 110406317 | 110406379 | 110401248 | 110401583 | 110407032 | 110407128 | 4 | 2 | 11 | 5 |
| MXE | 7024 | ENSG00000107625 | DDX50 | chr10 | + | 68936005 | 68936079 | 68941059 | 68941194 | 68934798 | 68934918 | 68943212 | 68943257 | 23 | 7 | 16 | 29 |
| MXE | 6191 | ENSG00000143344 | RGL1 | chr1 | + | 183806374 | 183806485 | 183847565 | 183847774 | 183805119 | 183805324 | 183865995 | 183866073 | 0 | 2 | 0 | 0 |
| MXE | 16865 | ENSG00000174106 | LEMD3 | chr12 | + | 65238668 | 65238814 | 65239928 | 65240030 | 65238501 | 65238581 | 65240135 | 65240238 | 220 | 131 | 229 | 197 |
| MXE | 5111 | ENSG00000114867 | EIF4G1 | chr3 | + | 184315762 | 184315856 | 184316131 | 184316218 | 184315488 | 184315545 | 184317716 | 184317816 | 50 | 24 | 28 | 28 |
| MXE | 9345 | ENSG00000159339 | PADI4 | chr1 | + | 17354532 | 17354687 | 17355982 | 17356127 | 17348729 | 17348785 | 17356356 | 17356459 | 38 | 14 | 33 | 27 |
| MXE | 11238 | ENSG00000118495 | PLAGL1 | chr6 | - | 143964786 | 143964818 | 143966157 | 143966198 | 143960468 | 143960542 | 143968906 | 143968978 | 2 | 1 | 0 | 0 |
| MXE | 16622 | ENSG00000135090 | TAOK3 | chr12 | - | 118244893 | 118244965 | 118255447 | 118255655 | 118243414 | 118243516 | 118266654 | 118266759 | 180 | 155 | 114 | 146 |
| MXE | 13195 | ENSG00000115241 | PPM1G | chr2 | - | 27383951 | 27384092 | 27384672 | 27384767 | 27383365 | 27383600 | 27384974 | 27387158 | 1 | 3 | 2 | 7 |
| MXE | 6482 | ENSG00000139842 | CUL4A | chr13 | + | 113233896 | 113233986 | 113235062 | 113235145 | 113233152 | 113233339 | 113236822 | 113236890 | 78 | 65 | 61 | 81 |
| MXE | 1719 | ENSG00000197912 | SPG7 | chr16 | + | 89526328 | 89526468 | 89529134 | 89529245 | 89524005 | 89524247 | 89530682 | 89530808 | 30 | 31 | 43 | 42 |
| MXE | 9837 | ENSG00000122481 | RWDD3 | chr1 | + | 95244210 | 95244698 | 95246541 | 95246657 | 95234209 | 95234315 | 95246755 | 95247225 | 11 | 4 | 9 | 2 |
| MXE | 8792 | ENSG00000162607 | USP1 | chr1 | + | 62443158 | 62443319 | 62444737 | 62445429 | 62442194 | 62442299 | 62447340 | 62447511 | 189 | 128 | 147 | 160 |
| MXE | 16659 | ENSG00000135297 | MTO1 | chr6 | + | 73482448 | 73482620 | 73486597 | 73486761 | 73482028 | 73482244 | 73492233 | 73492352 | 129 | 90 | 138 | 146 |
| MXE | 8403 | ENSG00000135905 | DOCK10 | chr2 | - | 224886062 | 224886185 | 224886458 | 224886526 | 224885670 | 224885805 | 224896294 | 224896377 | 11 | 9 | 3 | 8 |
| MXE | 1751 | ENSG00000169896 | ITGAM | chr16 | + | 31321240 | 31321371 | 31321463 | 31321627 | 31297741 | 31297954 | 31324398 | 31324553 | 332 | 131 | 154 | 188 |
| MXE | 242 | ENSG00000100055 | CYTH4 | chr22 | + | 37292620 | 37292703 | 37294659 | 37294724 | 37282440 | 37282588 | 37295998 | 37296065 | 15 | 5 | 11 | 13 |
| MXE | 16101 | ENSG00000099290 | WASHC2A | chr10 | + | 50125368 | 50125449 | 50127159 | 50127222 | 50125112 | 50125241 | 50127582 | 50127795 | 87 | 36 | 78 | 62 |
| MXE | 10557 | ENSG00000122085 | MTERF4 | chr2 | - | 241089979 | 241090044 | 241094276 | 241094429 | 241089272 | 241089365 | 241097242 | 241097427 | 2 | 1 | 4 | 10 |
| MXE | 5204 | ENSG00000198700 | IPO9 | chr1 | + | 201871160 | 201871327 | 201872827 | 201872961 | 201870582 | 201870858 | 201874249 | 201874372 | 36 | 14 | 39 | 41 |
| MXE | 11628 | ENSG00000198223 | CSF2RA | chrX | + | 1300490 | 1300626 | 1303922 | 1304019 | 1295426 | 1295456 | 1309401 | 1309934 | 42 | 27 | 32 | 25 |
| MXE | 7416 | ENSG00000058804 | NDC1 | chr1 | - | 53807655 | 53807791 | 53809694 | 53809746 | 53806424 | 53806517 | 53818970 | 53819079 | 27 | 14 | 16 | 20 |
| MXE | 9966 | ENSG00000231259 | ANAPC1P2 | chr2 | - | 87047187 | 87047306 | 87049588 | 87049803 | 87043712 | 87043984 | 87052813 | 87052992 | 162 | 96 | 169 | 160 |
| MXE | 1749 | ENSG00000169896 | ITGAM | chr16 | + | 31297741 | 31297954 | 31321240 | 31321371 | 31297513 | 31297654 | 31321463 | 31321627 | 51 | 19 | 14 | 30 |
| MXE | 2937 | ENSG00000118482 | PHF3 | chr6 | + | 63684128 | 63685911 | 63691736 | 63692043 | 63679999 | 63680161 | 63694580 | 63694764 | 426 | 434 | 390 | 547 |
| MXE | 10820 | ENSG00000064995 | TAF11 | chr6 | - | 34879625 | 34879736 | 34879966 | 34880063 | 34877461 | 34878720 | 34880288 | 34880376 | 146 | 96 | 137 | 154 |
| MXE | 8686 | ENSG00000127124 | HIVEP3 | chr1 | - | 41579736 | 41585318 | 41628748 | 41628947 | 41575543 | 41575689 | 41918412 | 41918707 | 1 | 0 | 0 | 2 |
| MXE | 13831 | ENSG00000121316 | PLBD1 | chr12 | - | 14540763 | 14540902 | 14542207 | 14542291 | 14536569 | 14536710 | 14553192 | 14553412 | 337 | 168 | 124 | 187 |
| MXE | 1197 | ENSG00000198909 | MAP3K3 | chr17 | + | 63681765 | 63681899 | 63685516 | 63685590 | 63666939 | 63667060 | 63688526 | 63688594 | 295 | 158 | 190 | 226 |
| MXE | 12153 | ENSG00000072609 | CHFR | chr12 | - | 132862332 | 132862425 | 132869618 | 132869798 | 132861466 | 132861634 | 132870723 | 132870783 | 187 | 165 | 171 | 215 |
| MXE | 13139 | ENSG00000139218 | SCAF11 | chr12 | - | 45961699 | 45961857 | 45964106 | 45964188 | 45951649 | 45951727 | 45990352 | 45990574 | 30 | 16 | 28 | 51 |
| MXE | 17170 | ENSG00000127022 | CANX | chr5 | + | 179722803 | 179723019 | 179723659 | 179723779 | 179720403 | 179720560 | 179724656 | 179724783 | 1112 | 610 | 999 | 893 |
| MXE | 11793 | ENSG00000204305 | AGER | chr6 | - | 32181350 | 32181477 | 32181523 | 32181632 | 32180967 | 32181239 | 32182246 | 32182388 | 24 | 14 | 19 | 20 |
| MXE | 15314 | ENSG00000171466 | ZNF562 | chr19 | - | 9656546 | 9656653 | 9660719 | 9660874 | 9641806 | 9653881 | 9675014 | 9675100 | 18 | 12 | 13 | 21 |
| MXE | 8780 | ENSG00000146574 | CCZ1B | chr7 | - | 6811951 | 6812063 | 6814763 | 6814845 | 6806003 | 6806037 | 6819765 | 6819941 | 125 | 52 | 77 | 84 |
| MXE | 3527 | ENSG00000113263 | ITK | chr5 | + | 157240061 | 157240195 | 157241645 | 157241720 | 157238108 | 157238191 | 157243622 | 157243794 | 419 | 196 | 505 | 342 |
| MXE | 7666 | ENSG00000164306 | PRIMPOL | chr4 | + | 184685411 | 184685498 | 184685575 | 184685684 | 184682247 | 184682336 | 184691498 | 184691581 | 3 | 6 | 1 | 1 |
| MXE | 4912 | ENSG00000075711 | DLG1 | chr3 | - | 197104902 | 197105005 | 197115926 | 197116083 | 197090911 | 197091026 | 197119409 | 197119530 | 152 | 153 | 176 | 182 |
| MXE | 14253 | ENSG00000116604 | MEF2D | chr1 | - | 156479289 | 156479346 | 156479585 | 156479796 | 156477011 | 156477202 | 156480833 | 156480971 | 133 | 47 | 74 | 119 |
| MXE | 5205 | ENSG00000198700 | IPO9 | chr1 | + | 201874151 | 201874936 | 201875151 | 201875228 | 201874249 | 201874372 | 201875943 | 201884291 | 66 | 34 | 76 | 55 |
| MXE | 6106 | ENSG00000138382 | METTL5 | chr2 | - | 169819560 | 169819643 | 169821091 | 169821273 | 169812456 | 169812506 | 169821942 | 169822057 | 88 | 75 | 97 | 110 |
| MXE | 4478 | ENSG00000141562 | NARF | chr17 | + | 82472563 | 82472698 | 82478570 | 82478667 | 82468763 | 82468896 | 82478799 | 82478918 | 233 | 135 | 188 | 188 |

| | | | | | | | | | | | | | | | | | | | | | | | | | | | |
|---|---|---|---|---|---|---|---|---|---|---|---|---|---|---|---|---|---|---|---|---|---|---|---|---|---|---|---|
| MXE | 11121 | ENSG00000135976 | ANKRD36 | chr2 | + | 97146485 | 97146516 | 97149294 | 97149361 | 97144639 | 97144712 | 97151878 | 97151939 | 7 | [illegible] | [illegible] | [illegible] | [illegible] | [illegible] | [illegible] | [illegible] | [illegible] | [illegible] | [illegible] | [illegible] | [illegible] | [illegible] |
| MXE | 6840 | ENSG00000088035 | ALG6 | chr1 | + | 63411145 | 63411331 | 63411681 | 63411806 | 63407061 | 63407126 | 63411925 | 63412061 | 49 | [illegible] | [illegible] | [illegible] | [illegible] | [illegible] | [illegible] | [illegible] | [illegible] | [illegible] | [illegible] | [illegible] | [illegible] | [illegible] |
| MXE | 1744 | ENSG00000103274 | NUBP1 | chr16 | + | 10752609 | 10752678 | 10755720 | 10755753 | 10747142 | 10747276 | 10756689 | 10756780 | 110 | [illegible] | [illegible] | [illegible] | [illegible] | [illegible] | [illegible] | [illegible] | [illegible] | [illegible] | [illegible] | [illegible] | [illegible] | [illegible] |
| MXE | 5238 | ENSG00000136731 | UGGT1 | chr2 | + | 128183674 | 128183789 | 128186682 | 128186799 | 128182129 | 128182290 | 128187448 | 128187614 | 102 | [illegible] | [illegible] | [illegible] | [illegible] | [illegible] | [illegible] | [illegible] | [illegible] | [illegible] | [illegible] | [illegible] | [illegible] | [illegible] |
| MXE | 15914 | ENSG00000119541 | VPS4B | chr18 | - | 63407431 | 63407499 | 63410289 | 63410446 | 63403706 | 63403826 | 63411466 | 63411578 | 206 | 178 | 182 | 187 | 185 | 92 | 75 | 70 | 84 | 173 | 146 | 268 | 230 | 269 |
| MXE | 474 | ENSG00000076928 | ARHGEF1 | chr19 | + | 41896376 | 41896482 | 41897314 | 41897326 | 41895348 | 41895486 | 41897945 | 41898113 | 368 | 220 | 339 | 316 | 180 | 19 | 20 | 18 | 26 | 13 | 479 | 412 | 291 | 347 |
| MXE | 1153 | ENSG00000160551 | TAOK1 | chr17 | + | 29510863 | 29510992 | 29517452 | 29517656 | 29507895 | 29508132 | 29522279 | 29522519 | 294 | 261 | 261 | 313 | 407 | 180 | 136 | 161 | 172 | 186 | 205 | 326 | 279 | 138 |
| MXE | 6044 | ENSG00000181555 | SETD2 | chr3 | - | 47103345 | 47103423 | 47105996 | 47106120 | 47101457 | 47101555 | 47113875 | 47114004 | 160 | 92 | 107 | 154 | 232 | 140 | 134 | 136 | 160 | 210 | 4 | 95 | 131 | 54 |
| MXE | 16667 | ENSG00000198270 | TMEM116 | chr12 | - | 111936691 | 111936830 | 111937159 | 111937243 | 111933885 | 111934030 | 111938160 | 111938210 | 39 | 32 | 40 | 30 | 78 | 25 | 22 | 19 | 30 | 34 | 8 | 26 | 19 | 43 |
| MXE | 7106 | ENSG00000096060 | FKBP5 | chr6 | - | 35620131 | 35620274 | 35637013 | 35637158 | 35619095 | 35619210 | 35642719 | 35642843 | 959 | 715 | 732 | 537 | 1173 | 1983 | 1679 | 1524 | 1214 | 1998 | 673 | 2107 | 2116 | 572 |
| MXE | 1151 | ENSG00000196535 | MYO18A | chr17 | - | 29115663 | 29115840 | 29116443 | 29116455 | 29115350 | 29115441 | 29118044 | 29118189 | 3 | 3 | 1 | 7 | 2 | 59 | 31 | 47 | 69 | 56 | 0 | 3 | 0 | 2 |
| MXE | 6420 | ENSG00000180957 | PITPNB | chr22 | - | 27897129 | 27897137 | 27897800 | 27897892 | 27896551 | 27896626 | 27910963 | 27911109 | 99 | 68 | 87 | 86 | 162 | 15 | 8 | 10 | 15 | 21 | 24 | 44 | 39 | 60 |
| MXE | 11350 | ENSG00000114956 | DGUOK | chr2 | + | 73946718 | 73946906 | 73957124 | 73957240 | 73938909 | 73939022 | 73958145 | 73958245 | 128 | 90 | 122 | 113 | 186 | 81 | 44 | 52 | 68 | 121 | 159 | 157 | 111 | 125 |
| MXE | 639 | ENSG00000123146 | ADGRE5 | chr19 | + | 14390923 | 14391079 | 14397657 | 14397804 | 14388701 | 14388818 | 14397888 | 14397916 | 808 | 335 | 413 | 605 | 204 | 987 | 504 | 634 | 827 | 670 | 1232 | 776 | 801 | 780 |
| MXE | 15261 | ENSG00000122218 | COPA | chr1 | - | 160333602 | 160333679 | 160335241 | 160335322 | 160332447 | 160332557 | 160339908 | 160339982 | 195 | 95 | 117 | 146 | 147 | 196 | 111 | 132 | 183 | 118 | 233 | 206 | 127 | 105 |
| MXE | 12396 | ENSG00000042980 | ADAM28 | chr8 | + | 24313387 | 24313580 | 24320235 | 24320307 | 24311360 | 24311437 | 24321217 | 24321289 | 32 | 28 | 32 | 37 | 75 | 34 | 33 | 32 | 29 | 69 | 29 | 58 | 20 | 13 |
| MXE | 15762 | ENSG00000103479 | RBL2 | chr16 | + | 53447041 | 53447106 | 53451702 | 53451831 | 53442657 | 53442858 | 53453451 | 53453463 | 145 | 78 | 160 | 153 | 281 | 810 | 603 | 848 | 905 | 1461 | 35 | 54 | 53 | 39 |
| MXE | 3775 | ENSG00000153310 | CYRIB | chr8 | - | 129903311 | 129903350 | 129904498 | 129904585 | 129879388 | 129879471 | 129970942 | 129970995 | 18 | 12 | 9 | 9 | 0 | 504 | 358 | 311 | 461 | 350 | 6 | 21 | 15 | 32 |
| MXE | 990 | ENSG00000101019 | UQCC1 | chr20 | - | 35366556 | 35366614 | 35374183 | 35374256 | 35347163 | 35347272 | 35394091 | 35394196 | 0 | 0 | 1 | 0 | 0 | 28 | 13 | 16 | 24 | 22 | 11 | 0 | 2 | 5 |
| MXE | 11645 | ENSG00000156256 | USP16 | chr21 | + | 29036270 | 29036374 | 29037278 | 29037463 | 29034836 | 29034940 | 29038334 | 29038430 | 274 | 240 | 211 | 268 | 312 | 151 | 123 | 145 | 162 | 326 | 35 | 211 | 133 | 35 |
| MXE | 8815 | ENSG00000078177 | N4BP2 | chr4 | + | 40119931 | 40122309 | 40123126 | 40123212 | 40117868 | 40118024 | 40124159 | 40124205 | 78 | 75 | 51 | 71 | 118 | 65 | 47 | 60 | 66 | 65 | 0 | 38 | 33 | 5 |
| MXE | 3552 | ENSG00000280987 | MATR3 | chr5 | + | 139318907 | 139319033 | 139319333 | 139319501 | 139317595 | 139317721 | 139321897 | 139322029 | 666 | 451 | 603 | 689 | 829 | 1444 | 1317 | 1324 | 1701 | 1524 | 17 | 325 | 384 | 138 |
| MXE | 3853 | ENSG00000247774 | PCED1B-AS1 | chr12 | - | 47209280 | 47209482 | 47210639 | 47210761 | 47206510 | 47206585 | 47216164 | 47216200 | 212 | 118 | 315 | 215 | 694 | 62 | 14 | 71 | 56 | 87 | 52 | 148 | 82 | 40 |
| MXE | 3067 | ENSG00000198399 | ITSN2 | chr2 | - | 24299908 | 24300171 | 24301153 | 24301239 | 24298664 | 24298814 | 24301964 | 24302102 | 163 | 118 | 120 | 163 | 203 | 111 | 90 | 77 | 112 | 187 | 71 | 137 | 125 | 83 |
| MXE | 8323 | ENSG00000177409 | SAMD9L | chr7 | - | 93144731 | 93144833 | 93145908 | 93146040 | 93135762 | 93135991 | 93148193 | 93148345 | 45 | 13 | 17 | 27 | 45 | 220 | 206 | 162 | 261 | 115 | 0 | 3 | 1 | 7 |
| MXE | 15709 | ENSG00000166340 | TPP1 | chr11 | - | 6616004 | 6616074 | 6616314 | 6616503 | 6615441 | 6615562 | 6617300 | 6617428 | 0 | 3 | 0 | 0 | 0 | 51 | 31 | 30 | 36 | 33 | 6 | 6 | 0 | 6 |
| MXE | 7029 | ENSG00000093183 | SEC22C | chr3 | - | 42563764 | 42563880 | 42566743 | 42566836 | 42563522 | 42563686 | 42568864 | 42569073 | 1 | 0 | 0 | 0 | 0 | 13 | 7 | 14 | 9 | 7 | 0 | 0 | 5 | 6 |
| MXE | 9711 | ENSG00000197451 | HNRNPAB | chr5 | + | 178209329 | 178209432 | 178210131 | 178210272 | 178207093 | 178207225 | 178210552 | 178211155 | 407 | 315 | 324 | 461 | 320 | 91 | 36 | 73 | 132 | 70 | 992 | 536 | 388 | 712 |
| MXE | 11683 | ENSG00000196262 | PPIA | chr7 | + | 44797519 | 44797621 | 44798747 | 44798814 | 44796680 | 44796793 | 44799246 | 44799277 | 8 | 7 | 9 | 11 | 16 | 117 | 71 | 117 | 109 | 87 | 6 | 0 | 9 | 11 |
| MXE | 8728 | ENSG00000143119 | CD53 | chr1 | + | 110891391 | 110891471 | 110892344 | 110892533 | 110873147 | 110873248 | 110894326 | 110894401 | 1146 | 619 | 729 | 935 | 981 | 1204 | 632 | 981 | 1206 | 1596 | 1160 | 1156 | 738 | 1029 |
| MXE | 3881 | ENSG00000023734 | STRAP | chr12 | + | 15894046 | 15894143 | 15895358 | 15895496 | 15890596 | 15890669 | 15897881 | 15898018 | 72 | 58 | 60 | 86 | 135 | 321 | 284 | 204 | 319 | 442 | 45 | 76 | 71 | 46 |
| MXE | 3892 | ENSG00000005156 | LIG3 | chr17 | + | 34999781 | 34999856 | 35001256 | 35001403 | 34998603 | 34999449 | 35001908 | 35002104 | 32 | 19 | 33 | 35 | 17 | 106 | 66 | 98 | 100 | 58 | 52 | 18 | 9 | 25 |
| MXE | 11254 | ENSG00000115977 | AAK1 | chr2 | - | 69525032 | 69525112 | 69527215 | 69527319 | 69520833 | 69520988 | 69530007 | 69530140 | 444 | 375 | 456 | 457 | 707 | 31 | 19 | 45 | 22 | 50 | 425 | 357 | 468 | 356 |
| MXE | 8701 | ENSG00000003756 | RBM5 | chr3 | + | 50105548 | 50105709 | 50106766 | 50106864 | 50104973 | 50105142 | 50107481 | 50107569 | 369 | 279 | 357 | 430 | 602 | 163 | 112 | 113 | 129 | 261 | 555 | 422 | 409 | 561 |
| MXE | 2239 | ENSG00000285304 | NA | chr22 | + | 37643097 | 37643174 | 37643643 | 37643788 | 37642894 | 37643006 | 37644636 | 37644705 | 81 | 43 | 50 | 66 | 38 | 378 | 226 | 266 | 362 | 214 | 129 | 86 | 56 | 120 |
| MXE | 15341 | ENSG00000099783 | HNRNPM | chr19 | + | 8462528 | 8462589 | 8463496 | 8463504 | 8455404 | 8455574 | 8463592 | 8463686 | 325 | 231 | 176 | 280 | 258 | 9 | 5 | 9 | 6 | 15 | 514 | 252 | 228 | 426 |
| MXE | 6362 | ENSG00000143110 | C1orf162 | chr1 | + | 111477333 | 111477428 | 111477725 | 111477775 | 111476797 | 111476867 | 111477982 | 111478512 | 112 | 77 | 71 | 110 | 147 | 134 | 63 | 60 | 98 | 159 | 283 | 102 | 142 | 211 |
| MXE | 13497 | ENSG00000102054 | RBBP7 | chrX | - | 16852415 | 16852550 | 16852550 | 16852628 | 16852045 | 16852122 | 16852748 | 16852875 | 211 | 119 | 180 | 232 | 271 | 5 | 2 | 3 | 4 | 0 | 126 | 191 | 159 | 199 |
| MXE | 13894 | ENSG00000205581 | HMGN1 | chr21 | - | 39347378 | 39347483 | 39347915 | 39347952 | 39345145 | 39345274 | 39348291 | 39348339 | 0 | 4 | 4 | 3 | 1 | 17 | 8 | 19 | 13 | 14 | 0 | 0 | 3 | 1 |
| MXE | 7339 | ENSG00000229474 | PATL2 | chr15 | - | 44673234 | 44673377 | 44674149 | 44674230 | 44672387 | 44672456 | 44675485 | 44675619 | 33 | 5 | 11 | 15 | 17 | 43 | 15 | 39 | 37 | 31 | 9 | 37 | 11 | 12 |
| MXE | 17124 | ENSG00000275066 | SYNRG | chr17 | - | 37571790 | 37571987 | 37576340 | 37576418 | 37570636 | 37570885 | 37584647 | 37584709 | 40 | 30 | 27 | 37 | 95 | 189 | 116 | 203 | 185 | 314 | 10 | 17 | 9 | 18 |
| MXE | 6893 | ENSG00000163960 | UBXN7 | chr3 | - | 196402951 | 196403019 | 196407245 | 196407393 | 196393553 | 196393619 | 196432326 | 196432427 | 148 | 124 | 119 | 142 | 88 | 309 | 248 | 268 | 285 | 233 | 165 | 150 | 171 | 105 |
| MXE | 2350 | ENSG00000140367 | UBE2Q2 | chr15 | + | 75890918 | 75891014 | 75896994 | 75897061 | 75890434 | 75890483 | 75899426 | 75899650 | 176 | 108 | 117 | 148 | 115 | 51 | 41 | 52 | 43 | 68 | 49 | 80 | 59 | 48 |
| MXE | 16284 | ENSG00000266412 | NCOA4 | chr10 | - | 46009410 | 46009551 | 46010222 | 46011206 | 46005087 | 46006597 | 46012882 | 46013026 | 547 | 301 | 341 | 532 | 326 | 1422 | 928 | 746 | 1512 | 3198 | 424 | 507 | 458 | 519 |
| MXE | 14587 | ENSG00000124702 | KLHDC3 | chr6 | + | 43017133 | 43017346 | 43017852 | 43017968 | 43014248 | 43014348 | 43018144 | 43018216 | 36 | 13 | 23 | 28 | 14 | 111 | 59 | 149 | 160 | 93 | 23 | 15 | 11 | 4 |
| MXE | 4938 | ENSG00000183323 | CCDC125 | chr5 | - | 69292187 | 69292362 | 69294792 | 69294900 | 69285336 | 69285467 | 69300011 | 69300127 | 36 | 15 | 35 | 40 | 56 | 41 | 31 | 51 | 41 | 55 | 83 | 39 | 51 | 144 |
| MXE | 4183 | ENSG00000137601 | NEK1 | chr4 | - | 169477123 | 169477352 | 169477431 | 169477497 | 169463242 | 169463395 | 169479402 | 169479534 | 36 | 18 | 19 | 29 | 49 | 56 | 25 | 34 | 20 | 64 | 25 | 19 | 9 | 22 |
| MXE | 4822 | ENSG00000124570 | SERPINB6 | chr6 | - | 2955523 | 2955670 | 2959167 | 2959342 | 2954591 | 2954709 | 2968778 | 2968913 | 13 | 8 | 13 | 16 | 11 | 347 | 201 | 226 | 246 | 336 | 6 | 4 | 8 | 17 |
| MXE | 4144 | ENSG00000164483 | SAMD3 | chr6 | - | 130175840 | 130176008 | 130184102 | 130184187 | 130154824 | 130155025 | 130184437 | 130184623 | 233 | 69 | 80 | 161 | 174 | 197 | 115 | 117 | 184 | 211 | 23 | 110 | 51 | 44 |
| MXE | 8436 | ENSG00000100612 | DHRS7 | chr14 | - | 60149352 | 60149568 | 60152938 | 60153178 | 60144118 | 60145013 | 60153958 | 60154065 | 467 | 182 | 287 | 308 | 394 | 810 | 493 | 398 | 443 | 546 | 388 | 252 | 221 | 238 |
| MXE | 2037 | ENSG00000137815 | RTF1 | chr15 | + | 41438320 | 41438431 | 41452900 | 41453048 | 41417103 | 41417313 | 41457671 | 41457876 | 33 | 25 | 25 | 51 | 6 | 111 | 52 | 88 | 74 | 96 | 82 | 90 | 110 | 120 |
| MXE | 10150 | ENSG00000112078 | KCTD20 | chr6 | + | 36479120 | 36479223 | 36479590 | 36479711 | 36470051 | 36470257 | 36481561 | 36481759 | 14 | 16 | 6 | 12 | 25 | 201 | 208 | 138 | 241 | 339 | 39 | 20 | 21 | 31 |
| MXE | 7813 | ENSG00000066427 | ATXN3 | chr14 | - | 92093745 | 92093831 | 92096673 | 92096838 | 92093251 | 92093318 | 92106528 | 92106574 | 40 | 25 | 20 | 32 | 15 | 99 | 47 | 55 | 76 | 70 | 40 | 35 | 38 | 23 |
| MXE | 6434 | ENSG00000163946 | TASOR | chr3 | - | 56646521 | 56647223 | 56660730 | 56660834 | 56641348 | 56641752 | 56660913 | 56661017 | 240 | 193 | 140 | 225 | 284 | 227 | 193 | 186 | 241 | 249 | 51 | 177 | 104 | 45 |
| MXE | 12451 | ENSG00000168137 | SETD5 | chr3 | + | 9434589 | 9434630 | 9434823 | 9434882 | 9434333 | 9434485 | 9435727 | 9435906 | 3 | 1 | 0 | 1 | 2 | 97 | 95 | 79 | 85 | 98 | 0 | 14 | 5 | 5 |
| MXE | 2892 | ENSG00000112679 | DUSP22 | chr6 | + | 335113 | 335163 | 345853 | 345928 | 311879 | 311962 | 348102 | 348274 | 13 | 8 | 19 | 14 | 29 | 163 | 167 | 147 | 265 | 246 | 80 | 34 | 23 | 23 |
| MXE | 3395 | ENSG00000111962 | UST | chr6 | + | 148941278 | 148941434 | 148953871 | 148953951 | 148886985 | 148887029 | 148964409 | 148964563 | 11 | 7 | 13 | 18 | 1 | 7 | 4 | 11 | 6 | 14 | 0 | 3 | 2 | 6 |
| MXE | 16853 | ENSG00000177575 | CD163 | chr12 | - | 7479859 | 7479996 | 7481160 | 7481256 | 7471179 | 7471397 | 7482642 | 7482762 | 129 | 72 | 51 | 141 | 119 | 45 | 22 | 16 | 41 | 81 | 490 | 899 | 579 | 349 |
| MXE | 1253 | ENSG00000182446 | NPLOC4 | chr17 | - | 81613317 | 81613494 | 81622165 | 81622278 | 81608727 | 81608822 | 81629724 | 81629805 | 87 | 30 | 79 | 92 | 113 | 1 | 1 | 1 | 7 | 5 | 111 | 87 | 59 | 71 |
| MXE | 333 | ENSG00000099331 | MYO9B | chr19 | + | 17175662 | 17175741 | 17183828 | 17183868 | 17172758 | 17172963 | 17184864 | 17184898 | 81 | 34 | 54 | 74 | 35 | 59 | 21 | 44 | 46 | 51 | 82 | 49 | 76 | 53 |
| MXE | 8598 | ENSG00000031698 | SARS1 | chr1 | + | 109236390 | 109236548 | 109237243 | 109237373 | 109235976 | 109236106 | 109237730 | 109238148 | 230 | 112 | 205 | 228 | 430 | 199 | 112 | 158 | 183 | 280 | 271 | 163 | 96 | 122 |
| MXE | 6185 | ENSG00000213390 | ARHGAP19 | chr10 | - | 97243967 | 97244159 | 97246271 | 97246337 | 97235216 | 97235315 | 97256317 | 97256404 | 25 | 22 | 30 | 20 | 34 | 92 | 42 | 71 | 73 | 70 | 30 | 12 | 10 | 41 |
| MXE | 14088 | ENSG00000171045 | TSNARE1 | chr8 | - | 142345742 | 142345892 | 142354636 | 142354763 | 142343965 | 142344472 | 142403103 | 142403182 | 4 | 3 | 1 | 6 | 2 | 13 | 5 | 15 | 17 | 19 | 0 | 4 | 0 | 0 |
| MXE | 10309 | ENSG00000180488 | MIGA1 | chr1 | + | 77863893 | 77864028 | 77866337 | 77866391 | 77861223 | 77861322 | 77873000 | 77873120 | 62 | 32 | 55 | 46 | 52 | 4 | 3 | 7 | 2 | 5 | 26 | 48 | 60 | 15 |
| MXE | 10878 | ENSG00000120697 | ALG5 | chr13 | - | 36985626 | 36985740 | 36989483 | 36989576 | 36971976 | 36972036 | 36993603 | 36993672 | 92 | 46 | 88 | 50 | 88 | 63 | 21 | 39 | 46 | 115 | 10 | 47 | 89 | 63 |
| MXE | 8177 | ENSG00000173575 | CHD2 | chr15 | + | 93002176 | 93002317 | 93004616 | 93004751 | 93000511 | 93000640 | 93009144 | 93009323 | 148 | 154 | 153 | 177 | 274 | 219 | 145 | 190 | 234 | 274 | 257 | 310 | 352 | 302 |
| MXE | 2309 | ENSG00000128881 | TTBK2 | chr15 | - | 42827927 | 42828032 | 42829937 | 42830078 | 42817001 | 42817097 | 42840359 | 42840433 | 37 | 30 | 37 | 59 | 51 | 46 | 27 | 44 | 33 | 23 | 21 | 12 | 18 | 17 |
| MXE | 10155 | ENSG00000142856 | ITGB3BP | chr1 | - | 63478763 | 63478833 | 63490082 | 63490218 | 63454889 | 63454968 | 63508527 | 63508570 | 84 | 51 | 49 | 82 | 106 | 49 | 38 | 17 | 40 | 72 | 24 | 19 | 24 | 19 |
| MXE | 10916 | ENSG00000125257 | ABCC4 | chr13 | - | 95234609 | 95234834 | 95246974 | 95247095 | 95210691 | 95210781 | 95247642 | 95247753 | 58 | 58 | 6 | 67 | 12 | 32 | 33 | 12 | 47 | 38 | 55 | 58 | 15 | 58 |
| MXE | 10088 | ENSG00000169379 | ARL13B | chr3 | + | 93997079 | 93997136 | 94003658 | 94003908 | 93996547 | 93996684 | 94035330 | 94035436 | 4 | 1 | 5 | 6 | 0 | 61 | 30 | 25 | 50 | 30 | 13 | 6 | 1 | 8 |
| MXE | 4908 | ENSG00000075711 | DLG1 | chr3 | - | 197081050 | 197081117 | 197085579 | 197085756 | 197076585 | 197076683 | 197090911 | 197091026 | 122 | 94 | 136 | 182 | 198 | 19 | 10 | 15 | 9 | 34 | 31 | 79 | 131 | 48 |
| MXE | 14673 | ENSG00000116701 | NCF2 | chr1 | - | 183565703 | 183565779 | 183566919 | 183566988 | 183564004 | 183564030 | 183567203 | 183567345 | 530 | 271 | 289 | 291 | 641 | 200 | 152 | 157 | 137 | 205 | 1056 | 534 | 310 | 582 |
| MXE | 7817 | ENSG00000143621 | ILF2 | chr1 | - | 153662704 | 153662795 | 153663214 | 153663276 | 153661787 | 153662556 | 153664042 | 153664130 | 411 | 319 | 421 | 423 | 1261 | 327 | 193 | 305 | 332 | 416 | 730 | 485 | 501 | 600 |
| MXE | 12899 | ENSG00000105939 | ZC3HAV1 | chr7 | - | 139053964 | 139054095 | 139061035 | 139061138 | 139053450 | 139053581 | 139064878 | 139064999 | 260 | 193 | 272 | 306 | 697 | 276 | 175 | 299 | 271 | 486 | 249 | 283 | 246 | 242 |
| MXE | 3802 | ENSG00000134644 | PUM1 | chr1 | - | 30995053 | 30995220 | 31006993 | 31007102 | 30992389 | 30992660 | 31028795 | 31028864 | 125 | 72 | 112 | 125 | 115 | 266 | 159 | 218 | 271 | 355 | 193 | 123 | 104 | 108 |
| MXE | 15189 | ENSG00000073921 | PICALM | chr11 | - | 86022366 | 86022469 | 86026291 | 86026367 | 86014869 | 86014963 | 86031468 | 86031611 | 1392 | 1115 | 720 | 1045 | 1133 | 887 | 677 | 504 | 750 | 908 | 1395 | 1321 | 1562 | 1499 |
| MXE | 1215 | ENSG00000007202 | KIAA0100 | chr17 | - | 28643163 | 28643311 | 28643609 | 28643659 | 28642868 | 28642987 | 28644057 | 28644187 | 26 | 7 | 17 | 15 | 9 | 71 | 49 | 46 | 70 | 91 | 52 | 28 | 22 | 33 |
| MXE | 13131 | ENSG00000130741 | EIF2S3 | chrX | + | 24071557 | 24071727 | 24073090 | 24073263 | 24067963 | 24068108 | 24076721 | 24078810 | 1126 | 826 | 1137 | 1086 | 1205 | 1677 | 1338 | 1596 | 1405 | 2258 | 795 | 503 | 608 | 982 |
| MXE | 8762 | ENSG00000101040 | ZMYND8 | chr20 | - | 47262287 | 47262428 | 47276313 | 47276795 | 47249286 | 47249439 | 47282101 | 47282217 | 134 | 92 | 112 | 130 | 157 | 50 | 45 | 58 | 60 | 36 | 119 | 113 | 76 | 110 |
| MXE | 14703 | ENSG00000184465 | WDR27 | chr6 | - | 169667135 | 169668776 | 169670568 | 169670693 | 169665485 | 169665556 | 169672254 | 169672396 | 5 | 6 | 9 | 10 | 4 | 2 | 0 | 1 | 1 | 10 | 27 | 6 | 5 | 3 |
| MXE | 4956 | ENSG00000198218 | QRICH1 | chr3 | - | 49044389 | 49044504 | 49046424 | 49046579 | 49033082 | 49033228 | 49047068 | 49047246 | 215 | 83 | 188 | 181 | 166 | 74 | 50 | 119 | 81 | 139 | 134 | 183 | 151 | 140 |
| MXE | 6371 | ENSG00000163931 | TKT | chr3 | - | 53234982 | 53235174 | 53240250 | 53240348 | 53233155 | 53233274 | 53241131 | 53241245 | 257 | 83 | 121 | 116 | 57 | 1234 | 537 | 643 | 699 | 363 | 105 | 223 | 153 | 202 |
| MXE | 10114 | ENSG00000164919 | COX6C | chr8 | - | 99887489 | 99887618 | 99891907 | 99892047 | 99878147 | 99878265 | 99893638 | 99893684 | 16 | 5 | 14 | 12 | 7 | 630 | 125 | 271 | 256 | 345 | 22 | 13 | 11 | 21 |
| [illegible] | | | | | | | | | | | | | | | | | | | | | | | | | | | |

| | | | | | | | | | | | | | | | | | | | | | | | | | | | | | | | | | | | | | | | | | | | | | | | | |
|---|---|---|---|---|---|---|---|---|---|---|---|---|---|---|---|---|---|---|---|---|---|---|---|---|---|---|---|---|---|---|---|---|---|---|---|---|---|---|---|---|---|---|---|---|---|---|---|---|
| MXE | 7610 | ENSG00000163322 | ABRAXAS1 | chr4 | - | 83463493 | 83463608 | 83467453 | 83467538 | 83459516 | 83462902 | 83469031 | 83469151 | 61 | 54 | 50 | 42 | 81 | 24 | 14 | 31 | 18 | 18 | 22 | 14 | 24 | 4 | 31 | 0 | 5 | 3 | 3 | 0 | 234 | 264 | 0.000351785 | 0.00451906 | 0.741 | 0.813 | 0.645 | 0.725 | 0.835 | 1 | 0.76 | 0.9 | 0.601 | 1 | -0.1 |
| MXE | 16593 | ENSG00000007392 | LUC7L | chr16 | - | 190540 | 190570 | 192926 | 193015 | 188968 | 190135 | 199061 | 199238 | 114 | 133 | 113 | 146 | 125 | 24 | 27 | 27 | 20 | 31 | 296 | 135 | 181 | 208 | 175 | 55 | 11 | 23 | 27 | 4 | 238 | 179 | 0.000352387 | 0.004522905 | 0.781 | 0.787 | 0.759 | 0.846 | 0.752 | 0.802 | 0.902 | 0.855 | 0.853 | 0.971 | -0.092 |
| MXE | 10048 | ENSG00000163644 | PPM1K | chr4 | - | 88265000 | 88265135 | 88277142 | 88277243 | 88262576 | 88262726 | 88284405 | 88284631 | 2 | 7 | 8 | 8 | 2 | 193 | 100 | 170 | 178 | 134 | 3 | 0 | 2 | 0 | 0 | 122 | 198 | 68 | 115 | 13 | 250 | 284 | 0.000359327 | 0.004608018 | 0.012 | 0.074 | 0.051 | 0.049 | 0.017 | 0.027 | 0 | 0.032 | 0 | 0 | 0.029 |
| MXE | 11440 | ENSG00000151687 | ANKAR | chr2 | + | 189728659 | 189728821 | 189733106 | 189733229 | 189728266 | 189728420 | 189737682 | 189737841 | 17 | 18 | 17 | 8 | 41 | 13 | 5 | 7 | 6 | 13 | 14 | 9 | 4 | 7 | 0 | 6 | 12 | 5 | 7 | 69 | 298 | 272 | 0.000359979 | 0.004612428 | 0.544 | 0.767 | 0.689 | 0.549 | 0.742 | 0.68 | 0.406 | 0.422 | 0.477 | 0 | 0.261 |
| MXE | 16140 | ENSG00000172661 | WASHC2C | chr10 | + | 45728861 | 45729026 | 45737982 | 45738045 | 45727416 | 45727539 | 45743389 | 45743483 | 41 | 25 | 21 | 30 | 12 | 0 | 5 | 2 | 1 | 0 | 60 | 21 | 23 | 32 | 21 | 12 | 1 | 10 | 6 | 3 | 298 | 212 | 0.000360646 | 0.004617016 | 1 | 0.781 | 0.882 | 0.955 | 1 | 0.781 | 0.937 | 0.621 | 0.791 | 0.833 | 0.131 |
| MXE | 5082 | ENSG00000132153 | DHX30 | chr3 | + | 47827346 | 47827477 | 47829023 | 47829134 | 47819226 | 47819264 | 47840876 | 47841178 | 19 | 10 | 27 | 8 | 22 | 107 | 64 | 136 | 111 | 139 | 15 | 8 | 23 | 8 | 3 | 178 | 122 | 130 | 101 | 202 | 280 | 260 | 0.000367497 | 0.004700688 | 0.142 | 0.127 | 0.156 | 0.063 | 0.128 | 0.073 | 0.057 | 0.141 | 0.069 | 0.014 | 0.052 |
| MXE | 11681 | ENSG00000204256 | BRD2 | chr6 | + | 32977441 | 32977570 | 32979827 | 32980132 | 32976561 | 32976936 | 32980341 | 32980464 | 663 | 418 | 547 | 564 | 713 | 211 | 158 | 164 | 240 | 213 | 962 | 664 | 584 | 611 | 1130 | 516 | 269 | 200 | 446 | 560 | 278 | 298 | 0.000368515 | 0.004709683 | 0.771 | 0.739 | 0.781 | 0.716 | 0.782 | 0.666 | 0.726 | 0.758 | 0.595 | 0.684 | 0.072 |
| MXE | 3455 | ENSG00000176783 | RUFY1 | chr5 | + | 179577074 | 179577136 | 179580946 | 179581012 | 179569301 | 179569425 | 179585795 | 179585865 | 60 | 97 | 46 | 143 | 120 | 145 | 182 | 64 | 275 | 204 | 290 | 112 | 69 | 95 | 567 | 787 | 293 | 156 | 309 | 1494 | 211 | 215 | 0.000369595 | 0.00471945 | 0.297 | 0.352 | 0.423 | 0.346 | 0.375 | 0.273 | 0.28 | 0.311 | 0.239 | 0.279 | 0.082 |
| MXE | 13927 | ENSG00000064115 | TM7SF3 | chr12 | - | 26983492 | 26983596 | 26990449 | 26990627 | 26982772 | 26982859 | 26995236 | 26995408 | 162 | 77 | 104 | 114 | 141 | 5 | 2 | 11 | 6 | 8 | 135 | 111 | 55 | 150 | 213 | 18 | 11 | 6 | 6 | 35 | 298 | 253 | 0.000370293 | 0.004721187 | 0.965 | 0.97 | 0.889 | 0.942 | 0.937 | 0.864 | 0.895 | 0.886 | 0.955 | 0.838 | 0.053 |
| MXE | 14927 | ENSG00000095319 | NUP188 | chr9 | + | 128999171 | 128999317 | 128999623 | 128999805 | 128998537 | 128998623 | 129001528 | 129001729 | 40 | 20 | 46 | 31 | 56 | 91 | 75 | 96 | 135 | 74 | 38 | 6 | 20 | 24 | 37 | 208 | 133 | 62 | 103 | 130 | 295 | 298 | 0.000370364 | 0.004721187 | 0.307 | 0.212 | 0.326 | 0.188 | 0.433 | 0.156 | 0.044 | 0.246 | 0.191 | 0.223 | 0.121 |
| MXE | 2342 | ENSG00000140367 | UBE2Q2 | chr15 | + | 75873427 | 75873568 | 75877960 | 75878021 | 75868950 | 75869010 | 75879097 | 75879188 | 227 | 188 | 181 | 241 | 353 | 116 | 110 | 84 | 103 | 108 | 234 | 182 | 196 | 102 | 132 | 11 | 63 | 58 | 32 | 42 | 290 | 210 | 0.000371647 | 0.004733506 | 0.586 | 0.553 | 0.609 | 0.629 | 0.703 | 0.939 | 0.677 | 0.71 | 0.698 | 0.695 | -0.128 |
| MXE | 3801 | ENSG00000134644 | PUM1 | chr1 | - | 30992389 | 30992660 | 31005852 | 31006031 | 30981311 | 30981405 | 31006993 | 31007102 | 191 | 195 | 222 | 241 | 323 | 111 | 62 | 87 | 108 | 110 | 372 | 299 | 259 | 282 | 165 | 109 | 106 | 62 | 62 | 43 | 298 | 298 | 0.000372493 | 0.004740243 | 0.632 | 0.759 | 0.718 | 0.691 | 0.746 | 0.773 | 0.738 | 0.807 | 0.82 | 0.793 | -0.077 |
| MXE | 14538 | ENSG00000131089 | ARHGEF9 | chrX | - | 63724531 | 63724711 | 63754303 | 63754676 | 63706257 | 63706449 | 63754920 | 63755078 | 8 | 4 | 5 | 6 | 3 | 47 | 21 | 42 | 26 | 43 | 0 | 0 | 8 | 0 | 0 | 45 | 42 | 27 | 20 | 38 | 298 | 298 | 0.000375128 | 0.004769713 | 0.145 | 0.16 | 0.106 | 0.188 | 0.065 | 0 | 0 | 0.229 | 0 | 0 | 0.087 |
| MXE | 7030 | ENSG00000093183 | SEC22C | chr3 | - | 42563764 | 42563980 | 42566743 | 42566836 | 42563522 | 42563686 | 42568864 | 42569073 | 1 | 0 | 0 | 0 | 0 | 21 | 11 | 25 | 15 | 19 | 0 | 0 | 5 | 6 | 1 | 25 | 16 | 34 | 5 | 8 | 242 | 298 | 0.000376196 | 0.004779224 | 0.055 | 0 | 0 | 0 | 0 | 0 | 0 | 0.153 | 0.596 | 0.133 | -0.165 |
| MXE | 13054 | ENSG00000274523 | RCC1L | chr7 | - | 75052710 | 75052796 | 75057528 | 75057616 | 75042182 | 75043109 | 75058587 | 75058769 | 34 | 7 | 29 | 20 | 25 | 9 | 11 | 13 | 31 | 15 | 76 | 41 | 24 | 20 | 39 | 5 | 31 | 4 | 6 | 0 | 237 | 235 | 0.000377716 | 0.004794455 | 0.789 | 0.387 | 0.689 | 0.39 | 0.623 | 0.938 | 0.567 | 0.856 | 0.768 | 1 | -0.25 |
| MXE | 13254 | ENSG00000069509 | FUNDC1 | chrX | - | 44527236 | 44527365 | 44538466 | 44538542 | 44523638 | 44524275 | 44541944 | 44542101 | 34 | 28 | 47 | 34 | 92 | 55 | 18 | 30 | 58 | 64 | 0 | 21 | 11 | 14 | 0 | 63 | 11 | 24 | 42 | 18 | 225 | 278 | 0.000378274 | 0.004797461 | 0.433 | 0.658 | 0.659 | 0.42 | 0.64 | 0 | 0.702 | 0.362 | 0.292 | 0 | 0.291 |
| MXE | 4996 | ENSG00000187742 | SECISBP2 | chr9 | + | 89319651 | 89319797 | 89325426 | 89325676 | 89318499 | 89318612 | 89325896 | 89326038 | 23 | 17 | 30 | 13 | 8 | 135 | 138 | 132 | 132 | 279 | 0 | 34 | 45 | 34 | 28 | 46 | 140 | 141 | 91 | 44 | 295 | 298 | 0.000379234 | 0.004801667 | 0.147 | 0.111 | 0.187 | 0.09 | 0.028 | 0 | 0.197 | 0.244 | 0.274 | 0.391 | -0.109 |
| MXE | 9481 | ENSG00000134058 | CDK7 | chr5 | + | 69258042 | 69258153 | 69259817 | 69259936 | 69255459 | 69255528 | 69262204 | 69262304 | 42 | 20 | 31 | 43 | 91 | 46 | 40 | 50 | 61 | 111 | 4 | 58 | 27 | 5 | 0 | 32 | 84 | 74 | 44 | 5 | 260 | 268 | 0.000379248 | 0.004801667 | 0.485 | 0.34 | 0.39 | 0.421 | 0.458 | 0.114 | 0.416 | 0.273 | 0.105 | 0 | 0.237 |
| MXE | 7685 | ENSG00000134574 | DDB2 | chr11 | + | 47234756 | 47234934 | 47235269 | 47235412 | 47234572 | 47234672 | 47237836 | 47238001 | 47 | 39 | 45 | 35 | 49 | 20 | 7 | 12 | 20 | 17 | 66 | 89 | 52 | 49 | 85 | 10 | 19 | 4 | 12 | 8 | 298 | 292 | 0.000381918 | 0.004831374 | 0.697 | 0.845 | 0.786 | 0.632 | 0.739 | 0.866 | 0.821 | 0.927 | 0.8 | 0.912 | -0.125 |
| MXE | 5647 | ENSG00000174738 | NR1D2 | chr3 | + | 23956036 | 23956125 | 23959670 | 23959924 | 23954625 | 23954803 | 23961976 | 23962334 | 83 | 48 | 51 | 64 | 91 | 9 | 3 | 10 | 2 | 15 | 22 | 15 | 46 | 25 | 4 | 9 | 10 | 7 | 19 | 0 | 238 | 298 | 0.000382376 | 0.004833083 | 0.92 | 0.952 | 0.865 | 0.976 | 0.884 | 0.754 | 0.653 | 0.892 | 0.622 | 1 | 0.135 |
| MXE | 12307 | ENSG00000162600 | OMA1 | chr1 | - | 58536512 | 58536741 | 58538794 | 58539310 | 58534157 | 58534331 | 58546663 | 58546706 | 43 | 14 | 28 | 51 | 26 | 340 | 115 | 182 | 267 | 301 | 0 | 13 | 14 | 11 | 4 | 54 | 206 | 116 | 79 | 123 | 298 | 298 | 0.00038317 | 0.004839025 | 0.112 | 0.109 | 0.133 | 0.16 | 0.08 | 0 | 0.059 | 0.108 | 0.122 | 0.031 | 0.055 |
| MXE | 16802 | ENSG00000110958 | PTGES3 | chr12 | - | 56670274 | 56670364 | 56671748 | 56671847 | 56666203 | 56666266 | 56672739 | 56672809 | 1099 | 886 | 898 | 1009 | 1104 | 635 | 358 | 499 | 512 | 531 | 1111 | 737 | 680 | 788 | 890 | 280 | 400 | 296 | 248 | 264 | 248 | 239 | 0.000384879 | 0.004856493 | 0.625 | 0.705 | 0.634 | 0.655 | 0.667 | 0.793 | 0.64 | 0.689 | 0.754 | 0.765 | -0.071 |
| MXE | 17157 | ENSG00000134900 | TPP2 | chr13 | + | 102651358 | 102651397 | 102657055 | 102657207 | 102649407 | 102649486 | 102663647 | 102663744 | 7 | 9 | 5 | 10 | 9 | 425 | 320 | 378 | 360 | 674 | 8 | 7 | 17 | 3 | 0 | 89 | 257 | 192 | 111 | 45 | 188 | 298 | 0.000385517 | 0.004860437 | 0.025 | 0.043 | 0.021 | 0.042 | 0.021 | 0.125 | 0.041 | 0.123 | 0.041 | 0 | -0.036 |
| MXE | 7549 | ENSG00000132122 | SPATA6 | chr1 | - | 48413109 | 48413151 | 48451551 | 48451600 | 48411463 | 48411588 | 48452993 | 48453131 | 0 | 3 | 2 | 3 | 1 | 6 | 5 | 6 | 1 | 7 | 0 | 0 | 1 | 0 | 0 | 8 | 4 | 2 | 3 | 2 | 198 | 191 | 0.000387088 | 0.004876133 | 0 | 0.367 | 0.243 | 0.743 | 0.121 | 0 | 0 | 0.325 | 0 | 0 | 0.23 |
| MXE | 14650 | ENSG00000187239 | FNBP1 | chr9 | - | 129929566 | 129929695 | 129957359 | 129957464 | 129927194 | 129927341 | 129958490 | 129958553 | 562 | 391 | 530 | 521 | 548 | 406 | 253 | 413 | 426 | 543 | 759 | 484 | 498 | 604 | 549 | 459 | 310 | 285 | 375 | 299 | 254 | 278 | 0.000388759 | 0.00489306 | 0.602 | 0.628 | 0.584 | 0.572 | 0.525 | 0.644 | 0.631 | 0.657 | 0.638 | 0.668 | -0.065 |
| MXE | 8157 | ENSG00000126698 | DNAJC8 | chr1 | - | 28205257 | 28205349 | 28208341 | 28208413 | 28203746 | 28203822 | 28209971 | 28210066 | 272 | 198 | 251 | 330 | 568 | 100 | 100 | 88 | 100 | 260 | 982 | 336 | 276 | 483 | 644 | 222 | 93 | 90 | 150 | 194 | 221 | 241 | 0.000389644 | 0.004900065 | 0.748 | 0.683 | 0.757 | 0.783 | 0.704 | 0.828 | 0.798 | 0.77 | 0.778 | 0.784 | -0.057 |
| MXE | 9860 | ENSG00000112855 | HARS2 | chr5 | + | 140691794 | 140692017 | 140693590 | 140693665 | 140691454 | 140691515 | 140693934 | 140694054 | 0 | 0 | 2 | 5 | 7 | 38 | 18 | 19 | 35 | 38 | 0 | 0 | 0 | 0 | 0 | 27 | 15 | 11 | 23 | 52 | 298 | 224 | 0.000390914 | 0.004911909 | 0 | 0 | 0.073 | 0.097 | 0.122 | 0 | 0 | 0 | 0 | 0 | 0.058 |
| MXE | 6179 | ENSG00000163875 | MEAF6 | chr1 | - | 37496546 | 37496604 | 37496706 | 37496736 | 37495884 | 37495918 | 37501803 | 37501996 | 1 | 2 | 5 | 2 | 8 | 38 | 11 | 21 | 16 | 50 | 0 | 4 | 0 | 0 | 0 | 16 | 9 | 14 | 18 | 33 | 179 | 207 | 0.00039178 | 0.004918086 | 0.03 | 0.174 | 0.216 | 0.126 | 0.156 | 0 | 0.339 | 0 | 0 | 0 | 0.073 |
| MXE | 9360 | ENSG00000115211 | EIF2B4 | chr2 | - | 27367745 | 27367822 | 27368024 | 27368113 | 27367456 | 27367559 | 27368371 | 27368463 | 5 | 1 | 2 | 4 | 11 | 31 | 21 | 32 | 35 | 44 | 0 | 3 | 2 | 0 | 0 | 6 | 30 | 19 | 9 | 26 | 238 | 226 | 0.000392064 | 0.004918086 | 0.133 | 0.043 | 0.056 | 0.098 | 0.192 | 0 | 0.087 | 0.091 | 0 | 0 | 0.069 |
| MXE | 10934 | ENSG00000198162 | MAN1A2 | chr1 | + | 117414712 | 117414831 | 117420568 | 117420649 | 117405548 | 117405645 | 117442230 | 117442325 | 732 | 403 | 403 | 597 | 607 | 168 | 137 | 140 | 184 | 131 | 1045 | 436 | 420 | 488 | 1445 | 51 | 171 | 131 | 96 | 107 | 268 | 230 | 0.000393664 | 0.004934015 | 0.789 | 0.716 | 0.712 | 0.736 | 0.799 | 0.946 | 0.686 | 0.733 | 0.814 | 0.921 | -0.07 |
| MXE | 12325 | ENSG00000146918 | NCAPG2 | chr7 | - | 158644288 | 158644388 | 158645518 | 158645619 | 158631168 | 158631717 | 158646459 | 158646563 | 8 | 21 | 20 | 13 | 29 | 5 | 32 | 27 | 20 | 59 | 14 | 25 | 14 | 21 | 19 | 6 | 11 | 18 | 10 | 0 | 250 | 249 | 0.000396171 | 0.004958869 | 0.614 | 0.395 | 0.425 | 0.393 | 0.329 | 0.699 | 0.694 | 0.437 | 0.677 | 1 | -0.27 |
| MXE | 7662 | ENSG00000164306 | PRIMPOL | chr4 | + | 184678231 | 184678394 | 184685575 | 184685684 | 184672172 | 184672460 | 184691498 | 184691581 | 19 | 17 | 12 | 11 | 23 | 20 | 21 | 18 | 12 | 42 | 6 | 0 | 0 | 2 | 0 | 5 | 4 | 10 | 8 | 6 | 298 | 258 | 0.000396311 | 0.004958869 | 0.451 | 0.412 | 0.366 | 0.442 | 0.322 | 0.51 | 0 | 0 | 0.178 | 0 | 0.261 |
| MXE | 13596 | ENSG00000163682 | RPL9 | chr4 | - | 39454863 | 39454944 | 39456405 | 39456538 | 39454532 | 39454649 | 39457585 | 39457681 | 4120 | 2469 | 2958 | 2355 | 6472 | 1803 | 995 | 1594 | 1174 | 4375 | 1642 | 1364 | 1901 | 1536 | 3188 | 462 | 639 | 768 | 650 | 972 | 282 | 230 | 0.00039763 | 0.004971207 | 0.651 | 0.669 | 0.602 | 0.621 | 0.547 | 0.744 | 0.635 | 0.669 | 0.658 | 0.728 | -0.069 |

[illegible]

| | | | | | | | | | |
|---|---|---|---|---|---|---|---|---|---|
| MXE | 10414 | ENSG00000123338 | NCKAP1L | chr12 | + | 54518650 | 54518732 | 54518913 | 54518972 |
| MXE | 11806 | ENSG00000170275 | CRTAP | chr3 | + | 33132554 | 33132700 | 33134181 | 33134265 |
| MXE | 6860 | ENSG00000025800 | KPNA6 | chr1 | + | 32159399 | 32159531 | 32160614 | 32160703 |
| MXE | 8361 | ENSG00000169991 | IFFO2 | chr1 | - | 18918361 | 18918502 | 18919677 | 18919773 |
| MXE | 16450 | ENSG00000101290 | CDS2 | chr20 | + | 5186686 | 5186839 | 5189066 | 5189186 |
| MXE | 14878 | ENSG00000165282 | PIGO | chr9 | - | 35093420 | 35093580 | 35093900 | 35094024 |
| MXE | 3912 | ENSG00000038427 | VCAN | chr5 | + | 83519348 | 83522309 | 83537006 | 83542268 |
| MXE | 13429 | ENSG00000161791 | FMNL3 | chr12 | - | 49651932 | 49652212 | 49653225 | 49653327 |
| MXE | 3224 | ENSG00000189091 | SF3B3 | chr16 | + | 70565367 | 70565524 | 70567410 | 70567536 |
| MXE | 16177 | ENSG00000148840 | PPRC1 | chr10 | + | 102144249 | 102144307 | 102146671 | 102147392 |
| MXE | 14509 | ENSG00000134333 | LDHA | chr11 | + | 18400836 | 18401010 | 18403737 | 18403811 |
| MXE | 11896 | ENSG00000135317 | SNX14 | chr6 | - | 85547083 | 85547226 | 85547505 | 85547550 |
| MXE | 14661 | ENSG00000198780 | FAM169A | chr5 | - | 74834425 | 74834597 | 74838964 | 74839050 |
| MXE | 3888 | ENSG00000119522 | DENND1A | chr9 | - | 123671290 | 123671371 | 123676719 | 123676789 |
| MXE | 2710 | ENSG00000119689 | DLST | chr14 | + | 74889094 | 74889147 | 74889274 | 74889349 |
| MXE | 12710 | ENSG00000106443 | PHF14 | chr7 | + | 10990702 | 10990847 | 11013746 | 11013906 |
| MXE | 8364 | ENSG00000135842 | NIBAN1 | chr1 | - | 184818637 | 184818825 | 184823166 | 184823329 |
| MXE | 6638 | ENSG00000179335 | CLK3 | chr15 | + | 74625801 | 74625968 | 74627351 | 74627446 |
| MXE | 8678 | ENSG00000118197 | DDX59 | chr1 | - | 200650424 | 200650676 | 200654440 | 200654582 |
| MXE | 12705 | ENSG00000011275 | RNF216 | chr7 | - | 5624055 | 5624125 | 5641153 | 5641376 |
| MXE | 3345 | ENSG00000167881 | SRP68 | chr17 | - | 76040902 | 76040978 | 76043828 | 76043958 |
| MXE | 3574 | ENSG00000196911 | KPNA5 | chr6 | + | 116692292 | 116692392 | 116698703 | 116698798 |
| MXE | 14975 | ENSG00000215126 | CBWD6 | chr9 | - | 41170283 | 41170423 | 41171846 | 41171934 |
| MXE | 1449 | ENSG00000186566 | GPATCH8 | chr17 | - | 44405920 | 44406051 | 44424348 | 44424492 |
| MXE | 4886 | ENSG00000157601 | MX1 | chr21 | + | 41437014 | 41437152 | 41439693 | 41439848 |
| MXE | 4894 | ENSG00000068650 | ATP11A | chr13 | + | 112858144 | 112858290 | 112859392 | 112859452 |
| MXE | 4958 | ENSG00000183726 | TMEM50A | chr1 | + | 25351625 | 25351693 | 25352881 | 25352974 |
| MXE | 16852 | ENSG00000177575 | CD163 | chr12 | - | 7479859 | 7479913 | 7481160 | 7481256 |
| MXE | 11724 | ENSG00000204592 | HLA-E | chr6 | + | 30491536 | 30491653 | 30492403 | 30492436 |
| MXE | 6565 | ENSG00000090863 | GLG1 | chr16 | - | 74492956 | 74493140 | 74494759 | 74494831 |
| MXE | 15908 | ENSG00000197771 | MCMBP | chr10 | - | 119840842 | 119840960 | 119842471 | 119842595 |
| MXE | 2772 | ENSG00000140105 | WARS1 | chr14 | - | 100369086 | 100369258 | 100374135 | 100374265 |

| | | | | | | | | | | |
|---|---|---|---|---|---|---|---|---|---|---|
| MXE | 6145 | ENSG00000163719 | MTMR14 | chr3 | + | 9672684 | 9672758 | 9677316 | 9677387 | 9671047 |
| MXE | 3602 | ENSG00000146282 | RARS2 | chr6 | - | 87524556 | 87524652 | 87529541 | 87529648 | 87521463 |
| MXE | 6911 | ENSG00000091127 | PUS7 | chr7 | - | 105470760 | 105470848 | 105472131 | 105472193 | 105468336 |
| MXE | 1684 | ENSG00000005844 | ITGAL | chr16 | + | 30489253 | 30489386 | 30496435 | 30496566 | 30489081 |
| MXE | 930 | ENSG00000175354 | PTPN2 | chr18 | - | 12817155 | 12817365 | 12819213 | 12819282 | 12814202 |
| MXE | 4336 | ENSG00000112079 | STK38 | chr6 | - | 36517716 | 36517840 | 36521733 | 36521817 | 36515337 |
| MXE | 2185 | ENSG00000129003 | VPS13C | chr15 | - | 61945714 | 61945882 | 61946306 | 61946410 | 61941762 |
| MXE | 5217 | ENSG00000206503 | HLA-A | chr6 | + | 29944121 | 29944397 | 29944499 | 29944634 | 29943267 |
| MXE | 11139 | ENSG00000260238 | PMF1-BGLAP | chr1 | + | 156232319 | 156232425 | 156233627 | 156233728 | 156212981 |
| MXE | 2720 | ENSG00000100605 | ITPK1 | chr14 | - | 92962750 | 92962849 | 92993879 | 92993997 | 92962354 |
| MXE | 577 | ENSG00000105738 | SIPA1L3 | chr19 | + | 38119305 | 38119882 | 38130497 | 38130772 | 38110226 |
| MXE | 13163 | ENSG00000197157 | SND1 | chr7 | + | 127807483 | 127807573 | 127844323 | 127844424 | 127721286 |
| MXE | 2393 | ENSG00000137878 | GCOM1 | chr15 | + | 57674967 | 57675068 | 57711983 | 57712188 | 57661449 |
| MXE | 13323 | ENSG00000170832 | USP32 | chr17 | - | 60271349 | 60271481 | 60288522 | 60288682 | 60269449 |
| MXE | 2387 | ENSG00000137814 | HAUS2 | chr15 | + | 42558197 | 42558290 | 42561269 | 42561402 | 42548827 |
| MXE | 8170 | ENSG00000173575 | CHD2 | chr15 | + | 92956458 | 92956649 | 92967324 | 92967513 | 92955422 |
| MXE | 8712 | ENSG00000162654 | GBP4 | chr1 | - | 89190037 | 89190318 | 89191260 | 89191506 | 89188581 |
| MXE | 2576 | ENSG00000092108 | SCFD1 | chr14 | + | 30633946 | 30634037 | 30638183 | 30638247 | 30630476 |
| MXE | 3502 | ENSG00000153113 | CAST | chr5 | + | 96750586 | 96750682 | 96754059 | 96754161 | 96748517 |
| MXE | 7220 | ENSG00000144357 | UBR3 | chr2 | + | 169896506 | 169896735 | 169905113 | 169905293 | 169895180 |
| MXE | 15385 | ENSG00000140299 | BNIP2 | chr15 | - | 59679591 | 59679768 | 59680240 | 59680308 | 59677812 |
| MXE | 2187 | ENSG00000129003 | VPS13C | chr15 | - | 61962748 | 61962852 | 61963834 | 61963951 | 61962370 |
| MXE | 2945 | ENSG00000111913 | RIPOR2 | chr6 | - | 24832255 | 24832391 | 24835702 | 24835871 | 24830508 |
| MXE | 3490 | ENSG00000147044 | CASK | chrX | - | 41553718 | 41553915 | 41559778 | 41559847 | 41542690 |
| MXE | 5960 | ENSG00000165240 | ATP7A | chrX | + | 77922998 | 77923042 | 77962622 | 77962814 | 77910738 |
| MXE | 9903 | ENSG00000275052 | PPP4R3B | chr2 | - | 55564901 | 55565041 | 55568193 | 55568363 | 55564312 |
| MXE | 10986 | ENSG00000122026 | RPL21 | chr13 | + | 27253764 | 27253843 | 27255241 | 27255349 | 27251560 |
| MXE | 8444 | ENSG00000121964 | GTDC1 | chr2 | - | 144235675 | 144235737 | 144317478 | 144317588 | 144211514 |
| MXE | 15048 | ENSG00000135387 | CAPRIN1 | chr11 | + | 34097195 | 34097296 | 34097697 | 34097761 | 34096478 |
| MXE | 9364 | ENSG00000115211 | EIF2B4 | chr2 | - | 27368371 | 27368463 | 27369005 | 27369209 | 27368024 |
| MXE | 15660 | ENSG00000081237 | PTPRC | chr1 | + | 198696711 | 198696909 | 198702386 | 198702530 | 198692346 |

| | | | | | | | | | | | | | | | | | | | | | | | | | | | | | | | | | | | | | | | | | | | | | | | | |
|---|---|---|---|---|---|---|---|---|---|---|---|---|---|---|---|---|---|---|---|---|---|---|---|---|---|---|---|---|---|---|---|---|---|---|---|---|---|---|---|---|---|---|---|---|---|---|---|---|
| MXE | 8577 | ENSG00000106462 | EZH2 | chr7 | - | 148816683 | 148816778 | 148817221 | 148817391 | 148815505 | 148815546 | 148817876 | 148818117 | 8 | 5 | 19 | 11 | 14 | 10 | 4 | 18 | 7 | 17 | 24 | 30 | 15 | 13 | 20 | 3 | 19 | 15 | 4 | 0 | 298 | 244 | 0.002598566 | 0.020849935 | 0.396 | 0.506 | 0.464 | 0.563 | 0.403 | 0.868 | 0.564 | 0.45 | 0.727 | 1 | -0.255 |
| MXE | 5842 | ENSG00000117298 | ECE1 | chr1 | - | 21227930 | 21228041 | 21233557 | 21233661 | 21227158 | 21227226 | 21235849 | 21235927 | 149 | 117 | 131 | 132 | 322 | 135 | 94 | 154 | 177 | 108 | 256 | 242 | 189 | 228 | 462 | 447 | 273 | 240 | 315 | 710 | 253 | 260 | 0.00260401 | 0.0208824 | 0.531 | 0.561 | 0.466 | 0.434 | 0.754 | 0.37 | 0.477 | 0.447 | 0.427 | 0.401 | 0.125 |
| MXE | 11326 | ENSG00000057252 | SOAT1 | chr1 | + | 179302676 | 179302802 | 179323436 | 179323495 | 179293713 | 179293936 | 179335505 | 179335657 | 83 | 36 | 42 | 45 | 9 | 136 | 81 | 74 | 104 | 153 | 31 | 40 | 40 | 40 | 15 | 31 | 64 | 44 | 59 | 37 | 275 | 208 | 0.002613535 | 0.02094754 | 0.316 | 0.252 | 0.3 | 0.247 | 0.043 | 0.431 | 0.321 | 0.407 | 0.339 | 0.235 | -0.115 |
| MXE | 16579 | ENSG00000111144 | LTA4H | chr12 | - | 96021084 | 96021137 | 96022146 | 96022251 | 96019167 | 96019240 | 96024478 | 96024547 | 941 | 720 | 621 | 884 | 1475 | 500 | 307 | 237 | 413 | 284 | 854 | 672 | 610 | 1699 | 888 | 459 | 362 | 335 | 957 | 446 | 254 | 202 | 0.002619056 | 0.020973211 | 0.599 | 0.651 | 0.676 | 0.63 | 0.805 | 0.597 | 0.596 | 0.592 | 0.585 | 0.613 | 0.076 |
| MXE | 8579 | ENSG00000131013 | PPIL4 | chr6 | - | 149526651 | 149526776 | 149533457 | 149533574 | 149525142 | 149525209 | 149534677 | 149534774 | 170 | 160 | 146 | 171 | 191 | 139 | 87 | 96 | 124 | 153 | 57 | 132 | 183 | 124 | 39 | 6 | 64 | 136 | 53 | 12 | 266 | 274 | 0.002619546 | 0.020973211 | 0.557 | 0.655 | 0.61 | 0.587 | 0.563 | 0.907 | 0.68 | 0.581 | 0.707 | 0.77 | -0.135 |
| MXE | 3896 | ENSG00000181929 | PRKAG1 | chr12 | - | 49003143 | 49003290 | 49003756 | 49003922 | 49002273 | 49003006 | 49004506 | 49004633 | 136 | 90 | 110 | 81 | 227 | 197 | 144 | 129 | 131 | 223 | 149 | 135 | 77 | 64 | 97 | 273 | 200 | 131 | 169 | 200 | 298 | 296 | 0.002624925 | 0.020994057 | 0.407 | 0.383 | 0.459 | 0.38 | 0.503 | 0.352 | 0.401 | 0.369 | 0.273 | 0.325 | 0.082 |
| MXE | 10883 | ENSG00000085365 | SCAMP1 | chr5 | + | 78418774 | 78418903 | 78449932 | 78450034 | 78416540 | 78416649 | 78459244 | 78459362 | 40 | 28 | 25 | 35 | 51 | 30 | 20 | 16 | 31 | 65 | 18 | 25 | 5 | 3 | 8 | 0 | 11 | 12 | 0 | 0 | 278 | 251 | 0.00262496 | 0.020994057 | 0.546 | 0.558 | 0.585 | 0.505 | 0.415 | 1 | 0.672 | 0.273 | 1 | 1 | -0.267 |
| MXE | 16737 | ENSG00000133858 | ZFC3H1 | chr12 | - | 71644876 | 71645075 | 71647748 | 71647813 | 71644094 | 71644318 | 71656884 | 71657301 | 123 | 108 | 107 | 151 | 418 | 165 | 221 | 185 | 191 | 288 | 11 | 132 | 85 | 68 | 8 | 58 | 238 | 158 | 144 | 38 | 214 | 298 | 0.0026291 | 0.02101592 | 0.509 | 0.405 | 0.446 | 0.524 | 0.669 | 0.209 | 0.436 | 0.428 | 0.397 | 0.227 | 0.171 |
| MXE | 15599 | ENSG00000149262 | INTS4 | chr11 | - | 77910969 | 77911152 | 77918030 | 77918264 | 77907716 | 77907810 | 77918820 | 77918978 | 2 | 1 | 0 | 3 | 0 | 18 | 10 | 15 | 10 | 21 | 2 | 4 | 6 | 3 | 0 | 29 | 6 | 11 | 10 | 8 | 298 | 298 | 0.002630539 | 0.021016179 | 0.1 | 0.091 | 0 | 0.231 | 0 | 0.065 | 0.4 | 0.353 | 0.231 | 0 | -0.125 |
| MXE | 15664 | ENSG00000081237 | PTPRC | chr1 | + | 198699563 | 198699704 | 198702386 | 198702530 | 198696711 | 198696909 | 198703297 | 198703372 | 1280 | 759 | 1064 | 1166 | 2732 | 480 | 285 | 404 | 474 | 350 | 150 | 947 | 402 | 116 | 111 | 25 | 212 | 130 | 26 | 0 | 290 | 293 | 0.002634071 | 0.02103315 | 0.729 | 0.729 | 0.727 | 0.713 | 0.887 | 0.858 | 0.819 | 0.758 | 0.818 | 1 | -0.094 |
| MXE | 7278 | ENSG00000233184 | NA | chr1 | + | 101075394 | 101075716 | 101083460 | 101083574 | 101072659 | 101072830 | 101086851 | 101087189 | 12 | 4 | 21 | 13 | 16 | 16 | 8 | 11 | 6 | 20 | 4 | 14 | 5 | 12 | 2 | 0 | 6 | 6 | 0 | 0 | 298 | 263 | 0.00264169 | 0.021082715 | 0.398 | 0.306 | 0.628 | 0.657 | 0.414 | 1 | 0.673 | 0.424 | 1 | 1 | -0.339 |
| MXE | 15812 | ENSG00000136167 | LCP1 | chr13 | - | 46130813 | 46130938 | 46134126 | 46134250 | 46125919 | 46127723 | 46142291 | 46142425 | 3824 | 2649 | 2556 | 3394 | 4625 | 5125 | 3604 | 3309 | 4674 | 6615 | 5461 | 2880 | 2762 | 4477 | 1909 | 9985 | 4664 | 4290 | 7597 | 3010 | 273 | 274 | 0.002668986 | 0.021289185 | 0.428 | 0.425 | 0.437 | 0.422 | 0.412 | 0.354 | 0.383 | 0.393 | 0.372 | 0.389 | 0.047 |
| MXE | 9334 | ENSG00000172766 | NAA16 | chr13 | + | 41336649 | 41336756 | 41355143 | 41355216 | 41331273 | 41331369 | 41358303 | 41358718 | 11 | 16 | 18 | 36 | 45 | 16 | 20 | 16 | 23 | 62 | 10 | 33 | 13 | 5 | 19 | 3 | 7 | 28 | 1 | 0 | 256 | 222 | 0.002684547 | 0.021401885 | 0.374 | 0.41 | 0.494 | 0.576 | 0.386 | 0.743 | 0.803 | 0.287 | 0.813 | 1 | -0.281 |
| MXE | 16757 | ENSG00000175582 | RAB6A | chr11 | - | 73718612 | 73718718 | 73718784 | 73718890 | 73716250 | 73716362 | 73720845 | 73720899 | 648 | 481 | 435 | 698 | 841 | 989 | 681 | 681 | 940 | 1138 | 870 | 531 | 385 | 807 | 950 | 1922 | 958 | 1052 | 1303 | 1336 | 255 | 255 | 0.002693527 | 0.021462022 | 0.396 | 0.414 | 0.39 | 0.426 | 0.425 | 0.312 | 0.357 | 0.268 | 0.382 | 0.416 | 0.063 |
| MXE | 12224 | ENSG00000179406 | LINC00174 | chr7 | - | 66467681 | 66467719 | 66476258 | 66476327 | 66450070 | 66450236 | 66479008 | 66479096 | 9 | 4 | 20 | 11 | 13 | 1 | 0 | 0 | 0 | 1 | 24 | 8 | 8 | 12 | 23 | 0 | 7 | 1 | 2 | 0 | 218 | 187 | 0.002726192 | 0.021710722 | 0.885 | 1 | 1 | 1 | 0.918 | 1 | 0.495 | 0.873 | 0.837 | 1 | 0.12 |
| MXE | 13977 | ENSG00000123124 | WWP1 | chr8 | + | 86435632 | 86435704 | 86438584 | 86438673 | 86435451 | 86435527 | 86442618 | 86442778 | 126 | 101 | 116 | 122 | 141 | 93 | 58 | 74 | 72 | 148 | 5 | 70 | 57 | 72 | 14 | 0 | 32 | 40 | 17 | 0 | 221 | 238 | 0.002730311 | 0.021731941 | 0.593 | 0.652 | 0.628 | 0.646 | 0.506 | 1 | 0.702 | 0.605 | 0.82 | 1 | -0.22 |
| MXE | 6850 | ENSG00000107099 | DOCK8 | chr9 | + | 336581 | 336718 | 339005 | 339099 | 334224 | 334384 | 340158 | 340321 | 481 | 349 | 420 | 473 | 472 | 923 | 703 | 686 | 839 | 1059 | 889 | 707 | 519 | 738 | 464 | 1328 | 992 | 800 | 1028 | 898 | 286 | 243 | 0.002733059 | 0.021742228 | 0.307 | 0.297 | 0.342 | 0.324 | 0.275 | 0.363 | 0.377 | 0.355 | 0.379 | 0.305 | -0.047 |
| MXE | 13436 | ENSG00000072778 | ACADVL | chr17 | + | 7223132 | 7223237 | 7223643 | 7223730 | 7222666 | 7222865 | 7223812 | 7223875 | 188 | 129 | 107 | 158 | 197 | 244 | 139 | 169 | 241 | 243 | 479 | 354 | 210 | 360 | 779 | 429 | 396 | 251 | 357 | 774 | 254 | 236 | 0.002743712 | 0.021815357 | 0.417 | 0.463 | 0.37 | 0.379 | 0.43 | 0.509 | 0.454 | 0.437 | 0.484 | 0.483 | -0.062 |
| MXE | 6994 | ENSG00000126070 | AGO3 | chr1 | + | 35973374 | 35973511 | 36004340 | 36004475 | 35972023 | 35972232 | 36008689 | 36008777 | 81 | 76 | 116 | 99 | 154 | 53 | 45 | 35 | 51 | 83 | 41 | 158 | 149 | 73 | 98 | 31 | 38 | 22 | 29 | 9 | 286 | 284 | 0.002746106 | 0.021821165 | 0.603 | 0.626 | 0.767 | 0.658 | 0.648 | 0.568 | 0.805 | 0.871 | 0.714 | 0.915 | -0.114 |
| MXE | 12869 | ENSG00000122566 | HNRNPA2B1 | chr7 | - | 26193574 | 26193652 | 26195846 | 26195909 | 26193250 | 26193373 | 26196400 | 26196481 | 1032 | 1085 | 897 | 1171 | 1574 | 346 | 296 | 261 | 391 | 319 | 687 | 950 | 1372 | 1325 | 326 | 97 | 278 | 343 | 251 | 83 | 212 | 227 | 0.002747364 | 0.021821165 | 0.762 | 0.797 | 0.786 | 0.762 | 0.841 | 0.883 | 0.785 | 0.811 | 0.85 | 0.808 | -0.038 |
| MXE | 13849 | ENSG00000152601 | MBNL1 | chr3 | + | 152414940 | 152415111 | 152432716 | 152432920 | 152299404 | 152300367 | 152445281 | 152445539 | 710 | 585 | 709 | 710 | 1354 | 942 | 806 | 766 | 1136 | 960 | 357 | 445 | 517 | 287 | 217 | 425 | 1071 | 799 | 452 | 612 | 298 | 298 | 0.002749353 | 0.02182394 | 0.43 | 0.421 | 0.481 | 0.385 | 0.585 | 0.457 | 0.294 | 0.393 | 0.388 | 0.262 | 0.102 |
| MXE | 12334 | ENSG00000117868 | ESYT2 | chr7 | - | 158764676 | 158764853 | 158767653 | 158767774 | 158763082 | 158763165 | 158773340 | 158773396 | 277 | 143 | 298 | 263 | 569 | 1035 | 545 | 920 | 945 | 1572 | 126 | 142 | 125 | 195 | 149 | 859 | 464 | 589 | 614 | 754 | 270 | 298 | 0.002750634 | 0.02182394 | 0.228 | 0.225 | 0.263 | 0.235 | 0.285 | 0.139 | 0.252 | 0.19 | 0.26 | 0.179 | 0.043 |
| MXE | 3007 | ENSG00000115760 | BIRC6 | chr2 | + | 32448794 | 32448928 | 32453807 | 32453942 | 32445520 | 32445668 | 32463193 | 32463381 | 84 | 64 | 93 | 78 | 166 | 62 | 38 | 57 | 77 | 161 | 36 | 87 | 42 | 22 | 10 | 0 | 50 | 33 | 11 | 0 | 283 | 284 | 0.002766104 | 0.021931729 | 0.576 | 0.628 | 0.621 | 0.504 | 0.509 | 1 | 0.636 | 0.561 | 0.667 | 1 | -0.205 |
| MXE | 7647 | ENSG00000100591 | AHSA1 | chr14 | + | 77464597 | 77464686 | 77465538 | 77465667 | 77462641 | 77462759 | 77468082 | 77468184 | 79 | 52 | 67 | 94 | 157 | 180 | 102 | 129 | 170 | 165 | 124 | 33 | 38 | 55 | 90 | 276 | 138 | 109 | 149 | 198 | 238 | 278 | 0.002767156 | 0.021931729 | 0.339 | 0.373 | 0.378 | 0.392 | 0.526 | 0.344 | 0.218 | 0.289 | 0.301 | 0.347 | 0.102 |
| MXE | 15567 | ENSG00000013725 | CD6 | chr11 | + | 61013918 | 61014014 | 61015712 | 61015835 | 61013422 | 61013563 | 61017478 | 61017550 | 155 | 90 | 147 | 214 | 371 | 187 | 140 | 274 | 253 | 333 | 93 | 169 | 82 | 42 | 160 | 162 | 302 | 294 | 70 | 255 | 245 | 272 | 0.002772099 | 0.021959257 | 0.479 | 0.416 | 0.373 | 0.484 | 0.553 | 0.389 | 0.383 | 0.236 | 0.4 | 0.411 | 0.097 |
| MXE | 9715 | ENSG00000140612 | SEC11A | chr15 | - | 84670724 | 84670782 | 84691534 | 84691644 | 84669537 | 84670069 | 84716024 | 84716460 | 167 | 59 | 123 | 166 | 31 | 417 | 249 | 331 | 333 | 437 | 121 | 142 | 117 | 141 | 98 | 284 | 291 | 256 | 375 | 250 | 259 | 207 | 0.002775778 | 0.021968884 | 0.242 | 0.159 | 0.229 | 0.285 | 0.054 | 0.254 | 0.281 | 0.268 | 0.231 | 0.239 | -0.061 |
| MXE | 4718 | ENSG00000155903 | RASA2 | chr3 | + | 141581099 | 141581177 | 141586024 | 141586098 | 141580367 | 141580451 | 141586645 | 141586752 | 122 | 96 | 141 | 105 | 190 | 210 | 129 | 192 | 168 | 139 | 9 | 61 | 63 | 18 | 0 | 29 | 119 | 104 | 40 | 27 | 227 | 223 | 0.002776255 | 0.021968884 | 0.363 | 0.422 | 0.419 | 0.38 | 0.573 | 0.234 | 0.335 | 0.373 | 0.307 | 0 | 0.182 |
| MXE | 1429 | ENSG00000108651 | UTP6 | chr17 | - | 31884423 | 31884505 | 31885979 | 31886061 | 31880180 | 31880754 | 31887235 | 31887313 | 22 | 26 | 26 | 32 | 27 | 101 | 102 | 72 | 95 | 142 | 56 | 20 | 25 | 47 | 15 | 89 | 84 | 45 | 92 | 43 | 231 | 231 | 0.002778319 | 0.021969446 | 0.179 | 0.203 | 0.265 | 0.252 | 0.16 | 0.386 | 0.192 | 0.357 | 0.338 | 0.259 | -0.095 |
| [illegible] | | | | | | | | | | | | | | | | | | | | | | | | | | | | | | | | | | | | | | | | | | | | | | | | |

| | | | | | | | | | | |
|---|---|---|---|---|---|---|---|---|---|---|
| MXE | 11813 | ENSG00000100220 | RTCB | chr22 | - | 32406661 | 32406761 | 32408174 | 32408242 | [illegible] |
| MXE | 11937 | ENSG00000107341 | UBE2R2 | chr9 | + | 33900173 | 33900271 | 33911963 | 33912098 | [illegible] |
| MXE | 7815 | ENSG00000066427 | ATXN3 | chr14 | - | 92096092 | 92096137 | 92096673 | 92096838 | [illegible] |
| MXE | 4447 | ENSG00000205339 | IPO7 | chr11 | + | 9430874 | 9431003 | 9433569 | 9433636 | [illegible] |
| MXE | 4825 | ENSG00000013441 | CLK1 | chr2 | - | 200854615 | 200854695 | 200855003 | 200855086 | [illegible] |
| MXE | 6168 | ENSG00000285708 | NA | chr3 | - | 71299819 | 71299880 | 71359149 | 71359244 | [illegible] |
| MXE | 4159 | ENSG00000177058 | SLC38A9 | chr5 | - | 55633694 | 55635657 | 55645788 | 55645895 | [illegible] |
| MXE | 4127 | ENSG00000132694 | ARHGEF11 | chr1 | - | 156979228 | 156979286 | 156980436 | 156980486 | [illegible] |
| MXE | 2278 | ENSG00000048471 | SNX29 | chr16 | + | 12477718 | 12477859 | 12524701 | 12524841 | [illegible] |
| MXE | 3220 | ENSG00000189091 | SF3B3 | chr16 | + | 70556178 | 70556334 | 70561629 | 70561784 | [illegible] |
| MXE | 13887 | ENSG00000205581 | HMGN1 | chr21 | - | 39345829 | 39345958 | 39347378 | 39347483 | [illegible] |
| MXE | 4926 | ENSG00000114316 | USP4 | chr3 | - | 49280743 | 49280847 | 49283986 | 49284136 | [illegible] |
| MXE | 10369 | ENSG00000172493 | AFF1 | chr4 | + | 87132270 | 87132408 | 87134473 | 87134694 | [illegible] |
| MXE | 11445 | ENSG00000151687 | ANKAR | chr2 | + | 189737682 | 189737841 | 189738564 | 189738682 | [illegible] |
| MXE | 11716 | ENSG00000120802 | TMPO | chr12 | + | 98544950 | 98545061 | 98546358 | 98546447 | [illegible] |
| MXE | 1254 | ENSG00000072818 | ACAP1 | chr17 | + | 7343860 | 7343956 | 7344048 | 7344123 | [illegible] |
| MXE | 1845 | ENSG00000103248 | MTHFSD | chr16 | - | 86546558 | 86546649 | 86548034 | 86548111 | [illegible] |
| MXE | 6309 | ENSG00000071127 | WDR1 | chr4 | - | 10088292 | 10088373 | 10088663 | 10088741 | [illegible] |
| MXE | 7609 | ENSG00000115828 | QPCT | chr2 | + | 37359579 | 37359858 | 37367231 | 37367408 | [illegible] |
| MXE | 14920 | ENSG00000119383 | PTPA | chr9 | + | 129128984 | 129129110 | 129131521 | 129131639 | [illegible] |
| MXE | 14881 | ENSG00000213930 | GALT | chr9 | + | 34647491 | 34647705 | 34647831 | 34647961 | [illegible] |
| MXE | 2030 | ENSG00000103994 | ZNF106 | chr15 | - | 42424833 | 42425025 | 42428017 | 42428134 | [illegible] |
| MXE | 11520 | ENSG00000114166 | KAT2B | chr3 | + | 20146315 | 20146430 | 20147962 | 20147999 | [illegible] |
| MXE | 7621 | ENSG00000132300 | PTCD3 | chr2 | + | 86124994 | 86125082 | 86125454 | 86125515 | [illegible] |
| MXE | 11235 | ENSG00000118495 | PLAGL1 | chr6 | - | 143947984 | 143948460 | 143960468 | 143960542 | [illegible] |
| MXE | 15192 | ENSG00000156110 | ADK | chr10 | + | 74525255 | 74525426 | 74589281 | 74589317 | [illegible] |
| MXE | 711 | ENSG00000173418 | NAA20 | chr20 | + | 20026783 | 20026919 | 20032507 | 20032653 | [illegible] |

[illegible]

| | | | | | | | | | | | | | | | |
|---|---|---|---|---|---|---|---|---|---|---|---|---|---|---|---|
| MXE | 11136 | ENSG00000021574 | SPAST | chr2 | + | 32098795 | 32098891 | 32114637 | 32114825 | 32089521 | 32089605 | 32115701 | 32115835 | 32 | [illegible] |
| MXE | 14445 | ENSG00000100815 | TRIP11 | chr14 | - | 92010985 | 92011072 | 92011754 | 92011795 | 92007639 | 92007852 | 92014214 | 92014577 | 26 | [illegible] |
| MXE | 5737 | ENSG00000135776 | ABCB10 | chr1 | - | 229542236 | 229542371 | 229547498 | 229547701 | 229540605 | 229540752 | 229549233 | 229549434 | 29 | [illegible] |
| MXE | 1172 | ENSG00000125450 | NUP85 | chr17 | + | 75234636 | 75234788 | 75235099 | 75235201 | 75233057 | 75233158 | 75235577 | 75235691 | 15 | [illegible] |
| MXE | 1021 | ENSG00000108840 | HDAC5 | chr17 | - | 44092171 | 44092284 | 44092380 | 44092527 | 44091699 | 44091831 | 44092675 | 44092806 | 130 | [illegible] |
| MXE | 2744 | ENSG00000198604 | BAZ1A | chr14 | - | 34775918 | 34776515 | 34783118 | 34783232 | 34774326 | 34774490 | 34783761 | 34783927 | 98 | [illegible] |
| MXE | 16587 | ENSG00000127314 | RAP1B | chr12 | + | 68651994 | 68652051 | 68654111 | 68654252 | 68650399 | 68650468 | 68656305 | 68656449 | 622 | [illegible] |
| MXE | 722 | ENSG00000124198 | ARFGEF2 | chr20 | + | 49028529 | 49028668 | 49032048 | 49032166 | 49025312 | 49025481 | 49033022 | 49033199 | 33 | [illegible] |
| MXE | 15411 | ENSG00000175294 | CATSPER1 | chr11 | - | 66020316 | 66020389 | 66020563 | 66020627 | 66020139 | 66020200 | 66020810 | 66020954 | 5 | [illegible] |
| MXE | 3470 | ENSG00000145604 | SKP2 | chr5 | + | 36163644 | 36166662 | 36171602 | 36171733 | 36152770 | 36153042 | 36176964 | 36177016 | 23 | [illegible] |
| MXE | 14898 | ENSG00000165699 | TSC1 | chr9 | - | 132902604 | 132902787 | 132903650 | 132903817 | 132901588 | 132901699 | 132904410 | 132904454 | 29 | [illegible] |
| MXE | 5862 | ENSG00000073417 | PDE8A | chr15 | + | 85089337 | 85089416 | 85090717 | 85090915 | 85083555 | 85083644 | 85091043 | 85091181 | 46 | [illegible] |
| MXE | 915 | ENSG00000263917 | NA | chr18 | + | 32037115 | 32037269 | 32042178 | 32042273 | 32018828 | 32019027 | 32045641 | 32045732 | 126 | [illegible] |
| MXE | 10337 | ENSG00000137075 | RNF38 | chr9 | - | 36369718 | 36369932 | 36375933 | 36376127 | 36357774 | 36357942 | 36400553 | 36400872 | 12 | [illegible] |
| MXE | 6631 | ENSG00000162946 | DISC1 | chr1 | + | 231701954 | 231702024 | 231749925 | 231750076 | 231693825 | 231694805 | 231767139 | 231767269 | 15 | [illegible] |
| MXE | 3919 | ENSG00000108039 | XPNPEP1 | chr10 | - | 109870730 | 109870904 | 109871791 | 109871861 | 109869952 | 109870029 | 109873366 | 109873427 | 27 | [illegible] |
| MXE | 7099 | ENSG00000117115 | PADI2 | chr1 | - | 17084598 | 17084702 | 17086520 | 17086699 | 17083725 | 17083837 | 17092407 | 17092533 | 35 | [illegible] |
| MXE | 8879 | ENSG00000117523 | PRRC2C | chr1 | + | 171568246 | 171568339 | 171571319 | 171571421 | 171566591 | 171566843 | 171574926 | 171575128 | 272 | [illegible] |
| MXE | 1061 | ENSG00000030582 | GRN | chr17 | + | 44351549 | 44351795 | 44352014 | 44352248 | 44351362 | 44351460 | 44352340 | 44352571 | 1295 | [illegible] |
| MXE | 11901 | ENSG00000135317 | SNX14 | chr6 | - | 85557975 | 85558060 | 85565331 | 85565419 | 85549722 | 85549879 | 85567533 | 85567577 | 44 | [illegible] |
| MXE | 12856 | ENSG00000123066 | MED13L | chr12 | - | 116111427 | 116111512 | 116237467 | 116237705 | 116096668 | 116096752 | 116277059 | 116277174 | 56 | [illegible] |
| MXE | 10747 | ENSG00000177479 | ARIH2 | chr3 | + | 48927461 | 48927813 | 48944981 | 48945181 | 48922747 | 48922811 | 48961611 | 48961679 | 153 | [illegible] |
| MXE | 564 | ENSG00000126261 | UBA2 | chr19 | + | 34438644 | 34438766 | 34443843 | 34443911 | 34434867 | 34434968 | 34444999 | 34445121 | 421 | [illegible] |
| MXE | 9892 | ENSG00000073614 | KDM5A | chr12 | - | 297040 | 297200 | 306945 | 307089 | 295572 | 295793 | 307453 | 308005 | 163 | [illegible] |
| MXE | 10290 | ENSG00000132676 | DAP3 | chr1 | + | 155717005 | 155717128 | 155721516 | 155721618 | 155709772 | 155709824 | 155725381 | 155725490 | 235 | [illegible] |
| MXE | 3415 | ENSG00000140853 | NLRC5 | chr16 | + | 57051537 | 57051621 | 57054750 | 57054840 | 57047544 | 57047628 | 57055031 | 57055036 | 37 | [illegible] |
| MXE | 6904 | ENSG00000117335 | CD46 | chr1 | + | 207767778 | 207767823 | 207783291 | 207783330 | 207767012 | 207767195 | 207785070 | 207785106 | 115 | [illegible] |
| MXE | 16027 | ENSG00000171307 | ZDHHC16 | chr10 | + | 97455638 | 97455783 | 97455973 | 97456044 | 97454713 | 97454799 | 97456776 | 97457249 | 0 | [illegible] |
| MXE | 42 | ENSG00000156273 | BACH1 | chr21 | + | 29326058 | 29327393 | 29329486 | 29329693 | 29321220 | 29321514 | 29342398 | 29346148 | 290 | [illegible] |
| MXE | 9123 | ENSG00000103365 | GGA2 | chr16 | - | 23493359 | 23493458 | 23494302 | 23494378 | 23491676 | 23491800 | 23495693 | 23495778 | 25 | [illegible] |
| MXE | 12129 | ENSG00000124486 | USP9X | chrX | + | 41140655 | 41140771 | 41140965 | 41141217 | 41136803 | 41137022 | 41141292 | 41141431 | 102 | [illegible] |
| MXE | 1191 | ENSG00000261371 | PECAM1 | chr17 | - | 64360139 | 64360415 | 64363148 | 64363397 | 64356110 | 64356398 | 64369749 | 64370025 | 1215 | [illegible] |
| [illegible] | | | | | | | | | | | | | | | |

| | | | | | | | | | | | | | | |
|---|---|---|---|---|---|---|---|---|---|---|---|---|---|---|
| MXE | 4184 | ENSG00000137601 | NEK1 | chr4 | - | 169537808 | 169537911 | 169555719 | 169555851 | 169508247 | 169508331 | 169555931 | 169556095 | 15 |
| MXE | 12420 | ENSG00000012124 | CD22 | chr19 | + | 35332552 | 35332924 | 35336148 | 35336341 | 35329211 | 35329230 | 35337754 | 35338021 | 10 |
| MXE | 16704 | ENSG00000141376 | BCAS3 | chr17 | + | 61380505 | 61380571 | 61388617 | 61388708 | 61368326 | 61368494 | 61391976 | 61392831 | 4 |
| MXE | 3051 | ENSG00000152969 | JAKMIP1 | chr4 | - | 6078938 | 6078998 | 6080171 | 6080312 | 6064879 | 6065008 | 6081608 | 6081755 | 49 |
| MXE | 6388 | ENSG00000198952 | SMG5 | chr1 | - | 156272319 | 156272398 | 156273360 | 156273450 | 156268289 | 156268415 | 156274596 | 156274686 | 94 |
| MXE | 10954 | ENSG00000188647 | PTAR1 | chr9 | - | 69718649 | 69718684 | 69723325 | 69723630 | 69709524 | 69718568 | 69732138 | 69732352 | 92 |
| MXE | 1314 | ENSG00000083290 | ULK2 | chr17 | - | 19780985 | 19781104 | 19783696 | 19783905 | 19780471 | 19780629 | 19785936 | 19786086 | 10 |
| MXE | 15544 | ENSG00000110075 | PPP6R3 | chr11 | + | 68587926 | 68588024 | 68590659 | 68590714 | 68575957 | 68576043 | 68591575 | 68591706 | 6 |
| MXE | 626 | ENSG00000167460 | TPM4 | chr19 | + | 16086422 | 16086540 | 16088026 | 16088097 | 16081448 | 16082046 | 16089044 | 16089653 | 231 |
| MXE | 3813 | ENSG00000020633 | RUNX3 | chr1 | - | 24907258 | 24907417 | 24919239 | 24919344 | 24899510 | 24902666 | 24927573 | 24927730 | 391 |
| MXE | 11533 | ENSG00000072401 | UBE2D1 | chr10 | + | 58361494 | 58361526 | 58363608 | 58363686 | 58361337 | 58361401 | 58364770 | 58364876 | 39 |
| MXE | 12240 | ENSG00000004059 | ARF5 | chr7 | + | 127589082 | 127589163 | 127589484 | 127589594 | 127588402 | 127588565 | 127590065 | 127590137 | 90 |
| MXE | 15963 | ENSG00000100109 | TFIP11 | chr22 | - | 26506774 | 26506928 | 26510063 | 26510281 | 26506302 | 26506459 | 26512098 | 26512175 | 0 |
| MXE | 15540 | ENSG00000110075 | PPP6R3 | chr11 | + | 68583042 | 68583129 | 68587926 | 68588024 | 68575957 | 68576043 | 68590659 | 68590714 | 83 |
| MXE | 1115 | ENSG00000141252 | VPS53 | chr17 | - | 655837 | 655953 | 661808 | 661895 | 653290 | 653410 | 697417 | 697484 | 22 |
| MXE | 11471 | ENSG00000123600 | METTL8 | chr2 | - | 171326041 | 171326148 | 171330558 | 171330698 | 171325840 | 171325906 | 171331803 | 171331867 | 32 |
| MXE | 10179 | ENSG00000146463 | ZMYM4 | chr1 | + | 35358924 | 35359446 | 35361193 | 35361255 | 35325359 | 35325405 | 35361618 | 35361789 | 68 |
| MXE | 1528 | ENSG00000108559 | NUP88 | chr17 | - | 5387778 | 5387904 | 5388801 | 5388960 | 5387385 | 5387670 | 5391560 | 5391662 | 123 |
| MXE | 15158 | ENSG00000175216 | CKAP5 | chr11 | - | 46783273 | 46783368 | 46784487 | 46784673 | 46780427 | 46780485 | 46788680 | 46788773 | 61 |
| MXE | 10306 | ENSG00000180488 | MIGA1 | chr1 | + | 77806974 | 77807101 | 77813733 | 77813867 | 77803269 | 77803406 | 77815107 | 77815231 | 111 |
| MXE | 8030 | ENSG00000160691 | SHC1 | chr1 | - | 154968770 | 154968834 | 154969377 | 154969448 | 154968494 | 154968614 | 154970031 | 154970199 | 23 |
| MXE | 13223 | ENSG00000188917 | TRMT2B | chrX | - | 101035612 | 101035683 | 101042232 | 101042312 | 101023585 | 101023616 | 101051783 | 101052082 | 12 |
| MXE | 11278 | ENSG00000113407 | TARS1 | chr5 | + | 33453288 | 33453412 | 33454944 | 33455066 | 33448540 | 33448731 | 33455586 | 33455704 | 100 |
| MXE | 12708 | ENSG00000011275 | RNF216 | chr7 | - | 5721032 | 5721172 | 5725323 | 5725438 | 5716715 | 5716766 | 5729431 | 5729596 | 61 |
| MXE | 6353 | ENSG00000089327 | FXYD5 | chr19 | + | 35164155 | 35164245 | 35166141 | 35166171 | 35160708 | 35160801 | 35166250 | 35166325 | 562 |
| MXE | 5928 | ENSG00000003400 | CASP10 | chr2 | + | 201195841 | 201195948 | 201205881 | 201205973 | 201187705 | 201187799 | 201208074 | 201208183 | 21 |
| MXE | 9802 | ENSG00000115825 | PRKD3 | chr2 | - | 37290867 | 37290999 | 37293132 | 37293271 | 37289355 | 37289513 | 37316236 | 37317023 | 65 |
| MXE | 17193 | ENSG00000136100 | VPS36 | chr13 | - | 52426988 | 52427066 | 52427186 | 52427219 | 52425769 | 52426066 | 52429235 | 52429367 | 17 |
| MXE | 2974 | ENSG00000006744 | ELAC2 | chr17 | - | 13010612 | 13010671 | 13011662 | 13011782 | 13005920 | 13005979 | 13013206 | 13013275 | 114 |
| MXE | 2247 | ENSG00000169375 | SIN3A | chr15 | - | 75392241 | 75392815 | 75400039 | 75400156 | 75389651 | 75389821 | 75400729 | 75400940 | 103 |
| MXE | 11668 | ENSG00000024048 | UBR2 | chr6 | + | 42592150 | 42592229 | 42594190 | 42594304 | 42573733 | 42573993 | 42603587 | 42603718 | 68 |
| MXE | 10829 | ENSG00000134987 | WDR36 | chr5 | + | 111104176 | 111104352 | 111104696 | 111104817 | 111103785 | 111103918 | 111105294 | 111105360 | 96 |
| MXE | 7596 | ENSG00000165359 | INTS6L | chrX | + | 135556167 | 135556300 | 135569336 | 135569431 | 135551993 | 135552146 | 135572814 | 135573033 | 71 |
| MXE | 11630 | ENSG00000198223 | CSF2RA | chrX | + | 1303922 | 1304019 | 1305445 | 1305527 | 1295426 | 1295456 | 1309401 | 1309934 | 0 |
| MXE | 14470 | ENSG00000083223 | TUT7 | chr9 | - | 86322324 | 86322475 | 86322872 | 86323960 | 86319583 | 86319670 | 86325333 | 86325514 | 137 |
| MXE | 15291 | ENSG00000137500 | CCDC90B | chr11 | - | 83274638 | 83274740 | 83278725 | 83278829 | 83273950 | 83273992 | 83280140 | 83280260 | 106 |
| MXE | 14035 | ENSG00000215114 | UBXN2B | chr8 | + | 58434394 | 58434504 | 58439632 | 58439770 | 58433159 | 58433243 | 58445906 | 58446065 | 123 |
| MXE | 5110 | ENSG00000114867 | EIF4G1 | chr3 | + | 184315762 | 184315856 | 184316131 | 184316218 | 184315165 | 184315545 | 184317320 | 184317497 | 52 |
| MXE | 14971 | ENSG00000167721 | TSR1 | chr17 | - | 2330514 | 2330625 | 2330946 | 2331109 | 2329342 | 2329475 | 2332168 | 2332359 | 57 |

[illegible]

| | | | | | | | | | | | | | | |
|---|---|---|---|---|---|---|---|---|---|---|---|---|---|---|
| [illegible] | | | | | | | | | | | | | | |
| MXE | 15212 | ENSG00000113163 | CERT1 | chr5 | - | 75400204 | 75400297 | 75402971 | 75403058 | 75389591 | 75389687 | 75411010 | 75411103 | 104 |
| MXE | 3009 | ENSG00000115760 | BIRC6 | chr2 | + | 32453807 | 32453942 | 32463193 | 32463381 | 32448794 | 32448928 | 32464508 | 32464823 | 128 |
| MXE | 16557 | ENSG00000124164 | VAPB | chr20 | + | 58434601 | 58434705 | 58438944 | 58439025 | 58418210 | 58418363 | 58440906 | 58441083 | 34 |
| MXE | 6187 | ENSG00000213390 | ARHGAP19 | chr10 | - | 97246271 | 97246337 | 97256317 | 97256404 | 97243967 | 97244159 | 97259401 | 97259628 | 40 |
| MXE | 5743 | ENSG00000196187 | TMEM63A | chr1 | - | 225859195 | 225859349 | 225860859 | 225860997 | 225856910 | 225857017 | 225862217 | 225862351 | 164 |
| MXE | 11329 | ENSG00000175198 | PCCA | chr13 | + | 100340156 | 100340259 | 100368471 | 100368574 | 100330560 | 100330671 | 100425632 | 100425731 | 27 |
| MXE | 15984 | ENSG00000100225 | FBXO7 | chr22 | + | 32478980 | 32479275 | 32483896 | 32484124 | 32475256 | 32475398 | 32485067 | 32485209 | 24 |
| MXE | 7431 | ENSG00000198408 | OGA | chr10 | - | 101807729 | 101807901 | 101810183 | 101810314 | 101806044 | 101806143 | 101813029 | 101813127 | 56 |
| MXE | 13431 | ENSG00000177646 | ACAD9 | chr3 | + | 128899161 | 128899234 | 128899286 | 128899461 | 128897631 | 128897710 | 128901275 | 128901349 | 0 |
| MXE | 4821 | ENSG00000124570 | SERPINB6 | chr6 | - | 2954591 | 2954709 | 2955523 | 2955670 | 2953043 | 2953186 | 2959167 | 2959342 | 199 |
| MXE | 347 | ENSG00000079805 | DNM2 | chr19 | + | 10802287 | 10802358 | 10805915 | 10805967 | 10798485 | 10798572 | 10812263 | 10812300 | 353 |
| MXE | 10388 | ENSG00000120594 | PLXDC2 | chr10 | + | 20068169 | 20068239 | 20143294 | 20143417 | 20046868 | 20047015 | 20147783 | 20147902 | 232 |
| MXE | 2585 | ENSG00000249624 | NA | chr21 | + | 33260596 | 33260727 | 33262792 | 33263637 | 33252661 | 33252830 | 33268393 | 33268517 | 209 |
| MXE | 16415 | ENSG00000226210 | WASH8P | chr12 | - | 17347 | 17483 | 17722 | 17859 | 16968 | 17170 | 18036 | 18183 | 13 |
| MXE | 9918 | ENSG00000115464 | USP34 | chr2 | - | 61204496 | 61204601 | 61206016 | 61206124 | 61192900 | 61192980 | 61206759 | 61206886 | 167 |
| MXE | 780 | ENSG00000203880 | PCMTD2 | chr20 | + | 64265257 | 64265429 | 64267886 | 64268010 | 64264428 | 64264531 | 64273220 | 64276226 | 214 |
| MXE | 16710 | ENSG00000078674 | PCM1 | chr8 | + | 17969576 | 17969748 | 17972328 | 17972687 | 17966979 | 17967170 | 17985446 | 17985619 | 249 |
| MXE | 2265 | ENSG00000185787 | MORF4L1 | chr15 | + | 78893538 | 78893627 | 78894057 | 78894230 | 78892211 | 78892313 | 78894819 | 78894904 | 1725 |
| MXE | 6062 | ENSG00000144711 | IQSEC1 | chr3 | - | 12915103 | 12915133 | 12915593 | 12915733 | 12913427 | 12913553 | 12920429 | 12920596 | 172 |
| MXE | 1459 | ENSG00000125970 | RALY | chr20 | + | 34073818 | 34073866 | 34075873 | 34076040 | 34073562 | 34073635 | 34076701 | 34076815 | 500 |
| MXE | 1987 | ENSG00000103197 | TSC2 | chr16 | + | 2058746 | 2058873 | 2060669 | 2062008 | 2057104 | 2057178 | 2062496 | 2062600 | 15 |
| MXE | 1502 | ENSG00000074370 | ATP2A3 | chr17 | - | 3951580 | 3951685 | 3953346 | 3953429 | 3951250 | 3951389 | 3953692 | 3953710 | 44 |
| MXE | 11220 | ENSG00000197299 | BLM | chr15 | + | 90769437 | 90769586 | 90782821 | 90782928 | 90769132 | 90769231 | 90784920 | 90785081 | 27 |
| MXE | 6559 | ENSG00000135597 | REPS1 | chr6 | - | 138911275 | 138911371 | 138912764 | 138912950 | 138908667 | 138908816 | 138914696 | 138914761 | 69 |
| MXE | 7822 | ENSG00000265491 | RNF115 | chr1 | - | 145750406 | 145750500 | 145751437 | 145751510 | 145747994 | 145748110 | 145752977 | 145753049 | 54 |
| MXE | 7170 | ENSG00000115239 | ASB3 | chr2 | - | 53714383 | 53714581 | 53716565 | 53716743 | 53700270 | 53700528 | 53728711 | 53728847 | 137 |
| MXE | 6052 | ENSG00000116478 | HDAC1 | chr1 | + | 32316664 | 32316782 | 32324478 | 32324553 | 32302620 | 32302733 | 32326938 | 32327077 | 305 |
| MXE | 4416 | ENSG00000182957 | SPATA13 | chr13 | + | 24286213 | 24286393 | 24288998 | 24289178 | 24284134 | 24284271 | 24290651 | 24290884 | 212 |
| MXE | 14034 | ENSG00000215114 | UBXN2B | chr8 | + | 58433159 | 58433243 | 58434394 | 58434504 | 58430518 | 58430669 | 58439632 | 58439770 | 70 |
| MXE | 12424 | ENSG00000012124 | CD22 | chr19 | + | 35336148 | 35336341 | 35337754 | 35338021 | 35332552 | 35332924 | 35338167 | 35338431 | 1 |
| MXE | 7374 | ENSG00000138095 | LRPPRC | chr2 | - | 43894544 | 43894629 | 43896633 | 43896708 | 43889733 | 43889876 | 43899218 | 43899334 | 32 |
| MXE | 11394 | ENSG00000003402 | CFLAR | chr2 | + | 201129728 | 201130146 | 201133028 | 201133134 | 201118490 | 201118901 | 201135971 | 201136107 | 158 |
| MXE | 2834 | ENSG00000255185 | PDXDC2P | chr16 | - | 70014103 | 70014167 | 70018046 | 70018097 | 70011840 | 70011929 | 70021823 | 70021949 | 14 |
| MXE | 15376 | ENSG00000107036 | RIC1 | chr9 | + | 5753535 | 5753646 | 5754840 | 5754930 | 5753199 | 5753238 | 5756211 | 5756372 | 148 |
| MXE | 10277 | ENSG00000102081 | FMR1 | chrX | + | 147940575 | 147940662 | 147943130 | 147943326 | 147937465 | 147937600 | 147944868 | 147945051 | 84 |
| MXE | 6845 | ENSG00000230124 | ACBD6 | chr1 | - | 180314691 | 180314722 | 180397515 | 180397605 | 180288229 | 180288517 | 180413365 | 180413471 | 54 |
| [illegible] | | | | | | | | | | | | | | |

| | | | | | |
|---|---|---|---|---|---|
| MXE | 15081 | ENSG00000138660 | AP1AR | chr4 | + |
| MXE | 5736 | ENSG00000114126 | TFDP2 | chr3 | - |
| MXE | 6216 | ENSG00000117020 | AKT3 | chr1 | - |
| MXE | 3034 | ENSG00000119899 | SLC17A5 | chr6 | - |
| MXE | 10005 | ENSG00000100644 | HIF1A | chr14 | + |
| MXE | 12469 | ENSG00000085788 | DDHD2 | chr8 | + |
| MXE | 7082 | ENSG00000162614 | NEXN | chr1 | + |
| MXE | 14618 | ENSG00000106771 | TMEM245 | chr9 | - |
| MXE | 3595 | ENSG00000112983 | BRD8 | chr5 | - |
| MXE | 9847 | ENSG00000119844 | AFTPH | chr2 | + |
| MXE | 2339 | ENSG00000140367 | UBE2Q2 | chr15 | + |
| MXE | 11399 | ENSG00000003402 | CFLAR | chr2 | + |
| MXE | 15968 | ENSG00000100109 | TFIP11 | chr22 | - |
| MXE | 8064 | ENSG00000103657 | HERC1 | chr15 | - |
| MXE | 2985 | ENSG00000196367 | TRRAP | chr7 | + |
| MXE | 14665 | ENSG00000197329 | PELI1 | chr2 | - |
| MXE | 13776 | ENSG00000165675 | ENOX2 | chrX | - |
| MXE | 10327 | ENSG00000129292 | PHF20L1 | chr8 | + |
| MXE | 7247 | ENSG00000145687 | SSBP2 | chr5 | - |
| MXE | 12890 | ENSG00000105835 | NAMPT | chr7 | - |
| MXE | 16831 | ENSG00000138777 | PPA2 | chr4 | - |
| MXE | 9857 | ENSG00000152127 | MGAT5 | chr2 | + |
| MXE | 2962 | ENSG00000162711 | NLRP3 | chr1 | + |
| MXE | 13689 | ENSG00000215301 | DDX3X | chrX | + |
| MXE | 11508 | ENSG00000137478 | FCHSD2 | chr11 | - |
| MXE | 48 | ENSG00000241837 | ATP5PO | chr21 | - |
| MXE | 6920 | ENSG00000109323 | MANBA | chr4 | - |
| MXE | 3633 | ENSG00000125846 | ZNF133 | chr20 | + |
| MXE | 10395 | ENSG00000153250 | RBMS1 | chr2 | - |
| MXE | 143 | ENSG00000243156 | MICAL3 | chr22 | - |
| MXE | 7057 | ENSG00000065978 | YBX1 | chr1 | + |
| MXE | 7611 | ENSG00000163322 | ABRAXAS1 | chr4 | - |
| MXE | 16174 | ENSG00000148840 | PPRC1 | chr10 | + |
| MXE | 114 | ENSG00000198911 | SREBF2 | chr22 | + |
| MXE | 8680 | ENSG00000118197 | DDX59 | chr1 | - |
| MXE | 4334 | ENSG00000112079 | STK38 | chr6 | - |
| MXE | 16913 | ENSG00000177119 | ANO6 | chr12 | + |
| MXE | 11305 | ENSG00000182732 | RGS6 | chr14 | + |
| MXE | 3477 | ENSG00000119862 | LGALSL | chr2 | + |
| MXE | 4711 | ENSG00000174227 | PIGG | chr4 | + |
| MXE | 6654 | ENSG00000114098 | ARMC8 | chr3 | + |
| MXE | 16820 | ENSG00000205352 | PRR13 | chr12 | + |
| MXE | 12327 | ENSG00000105778 | AVL9 | chr7 | + |
| MXE | 4927 | ENSG00000114316 | USP4 | chr3 | - |
| MXE | 9797 | ENSG00000115825 | PRKD3 | chr2 | - |
| MXE | 16110 | ENSG00000108055 | SMC3 | chr10 | + |
| MXE | 3750 | ENSG00000172239 | PAIP1 | chr5 | - |
| MXE | 9474 | ENSG00000171503 | ETFDH | chr4 | + |
| MXE | 11950 | ENSG00000178381 | ZFAND2A | chr7 | - |
| MXE | 10497 | ENSG00000123106 | CCDC91 | chr12 | + |
| MXE | 10135 | ENSG00000147526 | TACC1 | chr8 | + |
| MXE | 5740 | ENSG00000196187 | TMEM63A | chr1 | - |
| MXE | 14741 | ENSG00000198000 | NOL8 | chr9 | - |
| MXE | 17035 | ENSG00000136104 | RNASEH2B | chr13 | + |
| MXE | 5383 | ENSG00000175556 | LONRF3 | chrX | + |
| MXE | 6816 | ENSG00000260404 | NA | chr4 | + |
| MXE | 100 | ENSG00000099991 | CABIN1 | chr22 | + |
| MXE | 12162 | ENSG00000169919 | GUSB | chr7 | - |
| MXE | 12320 | ENSG00000147894 | C9orf72 | chr9 | - |
| MXE | 2693 | ENSG00000129566 | TEP1 | chr14 | - |
| MXE | 2129 | ENSG00000137842 | TMEM62 | chr15 | + |
| MXE | 92 | ENSG00000182944 | EWSR1 | chr22 | + |
| MXE | 788 | ENSG00000154930 | ACSS1 | chr20 | - |
| MXE | 10613 | ENSG00000196295 | GARS1-DT | chr7 | - |
| MXE | 8612 | ENSG00000069702 | TGFBR3 | chr1 | - |
| MXE | 10488 | ENSG00000135912 | TTLL4 | chr2 | + |
| MXE | 11779 | ENSG00000164430 | CGAS | chr6 | - |
| MXE | 12631 | ENSG00000100997 | ABHD12 | chr20 | - |
| MXE | 12225 | ENSG00000179406 | LINC00174 | chr7 | - |
| MXE | 1683 | ENSG00000005844 | ITGAL | chr16 | + |
| MXE | 11162 | ENSG00000074054 | CLASP1 | chr2 | - |
| MXE | 4060 | ENSG00000131507 | NDFIP1 | chr5 | + |
| MXE | 8426 | ENSG00000070540 | WIPI1 | chr17 | - |
| MXE | 1609 | ENSG00000125166 | GOT2 | chr16 | - |
| MXE | 5042 | ENSG00000010295 | IFFO1 | chr12 | - |
| MXE | 2138 | ENSG00000159459 | UBR1 | chr15 | - |
| MXE | 1452 | ENSG00000186566 | GPATCH8 | chr17 | - |
| MXE | 15718 | ENSG00000166387 | PPFIBP2 | chr11 | + |
| MXE | 2082 | ENSG00000250021 | ARPIN-AP3S2 | chr15 | - |
| MXE | 10096 | ENSG00000285756 | NA | chrX | - |
| MXE | 11460 | ENSG00000043462 | LCP2 | chr5 | - |
| MXE | 6985 | ENSG00000146007 | ZMAT2 | chr5 | + |
| MXE | 2152 | ENSG00000114062 | UBE3A | chr15 | - |
| MXE | 13456 | ENSG00000118922 | KLF12 | chr13 | - |
| MXE | 17055 | ENSG00000117226 | GBP3 | chr1 | - |
| MXE | 8176 | ENSG00000173575 | CHD2 | chr15 | + |
| MXE | 12579 | ENSG00000175471 | MCTP1 | chr5 | - |
| MXE | 6759 | ENSG00000185344 | ATP6V0A2 | chr12 | + |
| MXE | 8151 | ENSG00000165733 | BMS1 | chr10 | + |
| MXE | 3116 | ENSG00000150867 | PIP4K2A | chr10 | - |
| MXE | 10499 | ENSG00000123106 | CCDC91 | chr12 | + |
| MXE | 9483 | ENSG00000067208 | EVI5 | chr1 | - |
| MXE | 1934 | ENSG00000080603 | SRCAP | chr16 | + |
| MXE | 1942 | ENSG00000047578 | KATNIP | chr16 | + |
| MXE | 5133 | ENSG00000111731 | C2CD5 | chr12 | - |
| MXE | 12472 | ENSG00000154511 | DIPK1A | chr1 | - |
| MXE | 746 | ENSG00000125991 | ERGIC3 | chr20 | + |
| MXE | 9956 | ENSG00000068878 | PSME4 | chr2 | - |
| MXE | 8788 | ENSG00000158985 | CDC42SE2 | chr5 | + |
| MXE | 4820 | ENSG00000241553 | ARPC4 | chr3 | + |
| MXE | 14251 | ENSG00000164941 | INTS8 | chr8 | + |
| MXE | 12101 | ENSG00000146733 | PSPH | chr7 | - |
| MXE | 13727 | ENSG00000147050 | KDM6A | chrX | + |
| MXE | 11844 | ENSG00000136861 | CDK5RAP2 | chr9 | - |
| MXE | 5104 | ENSG00000060971 | ACAA1 | chr3 | - |
| MXE | 4649 | ENSG00000198586 | TLK1 | chr2 | - |
| MXE | 4512 | ENSG00000283782 | NA | chr5 | + |
| MXE | 8667 | ENSG00000117360 | PRPF3 | chr1 | + |
| MXE | 17116 | ENSG00000173473 | SMARCC1 | chr3 | - |
| MXE | 10773 | ENSG00000163412 | EIF4E3 | chr3 | - |
| MXE | 16985 | ENSG00000115310 | RTN4 | chr2 | - |
| MXE | 9781 | ENSG00000118960 | HS1BP3 | chr2 | - |
| MXE | 12976 | ENSG00000122591 | FAM126A | chr7 | - |
| MXE | 8464 | ENSG00000137275 | RIPK1 | chr6 | + |
| MXE | 3413 | ENSG00000140853 | NLRC5 | chr16 | + |
| MXE | 14867 | ENSG00000198887 | SMC5 | chr9 | + |
| MXE | 16911 | ENSG00000177119 | ANO6 | chr12 | + |
| MXE | 7158 | ENSG00000174839 | DENND6A | chr3 | - |
| MXE | 7235 | ENSG00000171148 | TADA3 | chr3 | - |
| MXE | 1188 | ENSG00000261371 | PECAM1 | chr17 | - |
| MXE | 868 | ENSG00000125827 | TMX4 | chr20 | - |
| MXE | 3857 | ENSG00000168288 | MMADHC | chr2 | - |
| MXE | 8840 | ENSG00000173846 | PLK3 | chr1 | + |
| MXE | 4691 | ENSG00000106066 | CPVL | chr7 | - |
| MXE | 2175 | ENSG00000140199 | SLC12A6 | chr15 | - |
| MXE | 10590 | ENSG00000239779 | WBP1 | chr2 | + |
| MXE | 8883 | ENSG00000273136 | NBPF26 | chr1 | + |
| MXE | 14829 | ENSG00000107077 | KDM4C | chr9 | + |
| MXE | 2584 | ENSG00000249624 | NA | chr21 | + |
| MXE | 8328 | ENSG00000177409 | SAMD9L | chr7 | - |
| MXE | 5424 | ENSG00000078070 | MCCC1 | chr3 | - |
| MXE | 5190 | ENSG00000116350 | SRSF4 | chr1 | - |
| MXE | 8796 | ENSG00000142405 | NLRP12 | chr19 | - |
| MXE | 424 | ENSG00000130816 | DNMT1 | chr19 | - |
| MXE | 1243 | ENSG00000161526 | SAP30BP | chr17 | + |
| MXE | 2482 | ENSG00000140563 | MCTP2 | chr15 | + |
| MXE | 6885 | ENSG00000100991 | TRPC4AP | chr20 | - |
| MXE | 9800 | ENSG00000115825 | PRKD3 | chr2 | - |
| MXE | 12985 | ENSG00000146776 | ATXN7L1 | chr7 | - |
| MXE | 16065 | ENSG00000107554 | DNMBP | chr10 | - |
| MXE | 12639 | ENSG00000100997 | ABHD12 | chr20 | - |
| MXE | 3588 | ENSG00000124201 | ZNFX1 | chr20 | - |
| MXE | 9937 | ENSG00000032444 | PNPLA6 | chr19 | + |
| MXE | 7348 | ENSG00000120798 | NR2C1 | chr12 | - |
| MXE | 3653 | ENSG00000145715 | RASA1 | chr5 | + |
| MXE | 4230 | ENSG00000141385 | AFG3L2 | chr18 | - |
| MXE | 9662 | ENSG00000120265 | PCMT1 | chr6 | + |
| MXE | 8288 | ENSG00000112303 | VNN2 | chr6 | - |
| MXE | 12055 | ENSG00000214021 | TTLL3 | chr3 | + |
| MXE | 5953 | ENSG00000134352 | IL6ST | chr5 | - |
| MXE | 9265 | ENSG00000159176 | CSRP1 | chr1 | - |
| MXE | 2935 | ENSG00000118482 | PHF3 | chr6 | + |
| MXE | 4729 | ENSG00000129071 | MBD4 | chr3 | - |
| MXE | 2968 | ENSG00000120925 | RNF170 | chr8 | - |
| MXE | 10144 | ENSG00000112078 | KCTD20 | chr6 | + |
| MXE | 8521 | ENSG00000177200 | CHD9 | chr16 | + |
| MXE | 7309 | ENSG00000117543 | DPH5 | chr1 | - |
| MXE | 6217 | ENSG00000117020 | AKT3 | chr1 | - |
| MXE | 13971 | ENSG00000165392 | WRN | chr8 | + |
| MXE | 4307 | ENSG00000124203 | ZNF831 | chr20 | + |
| MXE | 4653 | ENSG00000198586 | TLK1 | chr2 | - |
| MXE | 4015 | ENSG00000134153 | EMC7 | chr15 | - |
| MXE | 11842 | ENSG00000136861 | CDK5RAP2 | chr9 | - |
| MXE | 8638 | ENSG00000019582 | CD74 | chr5 | - |
| MXE | 543 | ENSG00000105677 | TMEM147 | chr19 | + |
| MXE | 7512 | ENSG00000115020 | PIKFYVE | chr2 | + |
| MXE | 15823 | ENSG00000124795 | DEK | chr6 | - |
| MXE | 7754 | ENSG00000023318 | ERP44 | chr9 | - |
| MXE | 10260 | ENSG00000143207 | COP1 | chr1 | - |
| MXE | 1949 | ENSG00000090905 | TNRC6A | chr16 | + |
| MXE | 13448 | ENSG00000258529 | NA | chr11 | - |
| MXE | 9572 | ENSG00000076242 | MLH1 | chr3 | + |
| MXE | 11811 | ENSG00000136044 | APPL2 | chr12 | - |
| MXE | 13134 | ENSG00000130985 | UBA1 | chrX | + |
| MXE | 14257 | ENSG00000085719 | CPNE3 | chr8 | + |
| MXE | 15608 | ENSG00000159063 | ALG8 | chr11 | - |
| MXE | 2976 | ENSG00000006744 | ELAC2 | chr17 | - |
| MXE | 9979 | ENSG00000264538 | SUZ12P1 | chr17 | + |
| MXE | 9037 | ENSG00000120694 | HSPH1 | chr13 | - |
| MXE | 4952 | ENSG00000172053 | QARS1 | chr3 | - |
| MXE | 10480 | ENSG00000181381 | DDX60L | chr4 | - |
| MXE | 15795 | ENSG00000133816 | MICAL2 | chr11 | + |
| MXE | 13003 | ENSG00000265681 | RPL17 | chr18 | - |
| MXE | 4923 | ENSG00000114127 | XRN1 | chr3 | - |
| MXE | 13988 | ENSG00000156162 | DPY19L4 | chr8 | + |
| MXE | 7680 | ENSG00000001461 | NIPAL3 | chr1 | + |
| MXE | 3217 | ENSG00000189091 | SF3B3 | chr16 | + |
| MXE | 8668 | ENSG00000117360 | PRPF3 | chr1 | + |
| MXE | 10667 | ENSG00000136536 | MARCHF7 | chr2 | + |
| MXE | 2298 | ENSG00000140400 | MAN2C1 | chr15 | - |
| MXE | 6659 | ENSG00000237943 | PRKCQ-AS1 | chr10 | + |
| MXE | 13128 | ENSG00000130741 | EIF2S3 | chrX | + |
| MXE | 5853 | ENSG00000198925 | ATG9A | chr2 | - |
| MXE | 8627 | ENSG00000075151 | EIF4G3 | chr1 | - |
| MXE | 6595 | ENSG00000257315 | ZBED6 | chr1 | + |
| MXE | 374 | ENSG00000104880 | ARHGEF18 | chr19 | + |
| MXE | 16528 | ENSG00000139323 | POC1B | chr12 | - |
| MXE | 4791 | ENSG00000100003 | SEC14L2 | chr22 | + |
| MXE | 5646 | ENSG00000174738 | NR1D2 | chr3 | + |
| MXE | 11977 | ENSG00000141664 | ZCCHC2 | chr18 | + |
| MXE | 3849 | ENSG00000247774 | PCED1B-AS1 | chr12 | - |
| MXE | 5554 | ENSG00000085265 | FCN1 | chr9 | - |
| MXE | 12668 | ENSG00000089289 | IGBP1 | chrX | + |
| MXE | 16011 | ENSG00000108100 | CCNY | chr10 | + |
| MXE | 11055 | ENSG00000121741 | ZMYM2 | chr13 | + |
| MXE | 4150 | ENSG00000152818 | UTRN | chr6 | + |
| MXE | 6112 | ENSG00000115306 | SPTBN1 | chr2 | + |
| MXE | 8698 | ENSG00000268043 | NBPF12 | chr1 | + |
| MXE | 1576 | ENSG00000166747 | AP1G1 | chr16 | - |
| MXE | 16408 | ENSG00000100354 | TNRC6B | chr22 | + |
| MXE | 9945 | ENSG00000064225 | ST3GAL6 | chr3 | + |
| MXE | 2228 | ENSG00000174485 | DENND4A | chr15 | - |
| MXE | 4631 | ENSG00000065882 | TBC1D1 | chr4 | + |
| MXE | 10063 | ENSG00000091409 | ITGA6 | chr2 | + |
| MXE | 9955 | ENSG00000068878 | PSME4 | chr2 | - |
| MXE | 14713 | ENSG00000114030 | KPNA1 | chr3 | - |
| MXE | 16518 | ENSG00000012822 | CALCOCO1 | chr12 | - |
| MXE | 4173 | ENSG00000150977 | RILPL2 | chr12 | - |
| MXE | 13233 | ENSG00000188917 | TRMT2B | chrX | - |
| MXE | 15215 | ENSG00000113163 | CERT1 | chr5 | - |
| MXE | 1110 | ENSG00000141252 | VPS53 | chr17 | - |
| MXE | 11756 | ENSG00000096070 | BRPF3 | chr6 | + |
| MXE | 7874 | ENSG00000087206 | UIMC1 | chr5 | - |
| MXE | 3447 | ENSG00000065357 | DGKA | chr12 | + |
| MXE | 11804 | ENSG00000170275 | CRTAP | chr3 | + |
| MXE | 2739 | ENSG00000015133 | CCDC88C | chr14 | - |
| MXE | 12678 | ENSG00000123505 | AMD1 | chr6 | + |
| MXE | 15834 | ENSG00000134470 | IL15RA | chr10 | - |
| MXE | 14068 | ENSG00000100380 | ST13 | chr22 | - |
| MXE | 16666 | ENSG00000198270 | TMEM116 | chr12 | - |
| MXE | 8795 | ENSG00000142405 | NLRP12 | chr19 | - |
| MXE | 15124 | ENSG00000149516 | MS4A3 | chr11 | + |
| MXE | 3507 | ENSG00000148110 | MFSD14B | chr9 | + |

| | | | | | | | | | | | | | | | |
|---|---|---|---|---|---|---|---|---|---|---|---|---|---|---|---|
| MXE | 3557 | ENSG00000113716 | HMGXB3 | chr5 | + | 150036635 | 150036937 | 150037399 | 150037527 | 150032453 | 150032603 | 150040747 | 150040879 | 36 | [illegible] |
| MXE | 217 | ENSG00000100201 | DDX17 | chr22 | - | 38498084 | 38498150 | 38498439 | 38498573 | 38495795 | 38495937 | 38499399 | 38499499 | 1396 | [illegible] |
| MXE | 6883 | ENSG00000100991 | TRPC4AP | chr20 | - | 35044504 | 35044712 | 35049865 | 35049994 | 35035122 | 35035308 | 35054975 | 35055031 | 303 | [illegible] |
| MXE | 391 | ENSG00000104814 | MAP4K1 | chr19 | - | 38614244 | 38614292 | 38614389 | 38614445 | 38614042 | 38614085 | 38616194 | 38616259 | 110 | [illegible] |
| MXE | 7116 | ENSG00000162894 | FCMR | chr1 | - | 206911729 | 206911952 | 206913758 | 206914094 | 206910209 | 206910340 | 206921817 | 206921941 | 1245 | [illegible] |
| MXE | 6778 | ENSG00000054282 | SDCCAG8 | chr1 | + | 243304712 | 243304777 | 243316754 | 243316893 | 243286382 | 243286397 | 243330539 | 243330692 | 4 | [illegible] |
| MXE | 13092 | ENSG00000169084 | DHRSX | chrX | - | 2243022 | 2243230 | 2266739 | 2266947 | 2219505 | 2221229 | 2291501 | 2291603 | 50 | [illegible] |
| MXE | 10167 | ENSG00000186141 | POLR3C | chr1 | + | 145836493 | 145836574 | 145838055 | 145838206 | 145833489 | 145833582 | 145842338 | 145842493 | 25 | [illegible] |
| MXE | 6636 | ENSG00000179335 | CLK3 | chr15 | + | 74620008 | 74620225 | 74622119 | 74622216 | 74608371 | 74608426 | 74622493 | 74622560 | 17 | [illegible] |
| MXE | 4191 | ENSG00000114738 | MAPKAPK3 | chr3 | + | 50640365 | 50640505 | 50641706 | 50641771 | 50617513 | 50617784 | 50642252 | 50642332 | 167 | [illegible] |
| MXE | 3508 | ENSG00000148110 | MFSD14B | chr9 | + | 94453957 | 94454070 | 94456239 | 94456337 | 94451455 | 94451527 | 94458321 | 94458464 | 149 | [illegible] |
| MXE | 4100 | ENSG00000008513 | ST3GAL1 | chr8 | - | 133507198 | 133507321 | 133539636 | 133539729 | 133499134 | 133499189 | 133545773 | 133545926 | 15 | [illegible] |
| MXE | 14805 | ENSG00000106829 | TLE4 | chr9 | + | 79627373 | 79627448 | 79652592 | 79652794 | 79612655 | 79612718 | 79704782 | 79704902 | 129 | [illegible] |
| MXE | 1341 | ENSG00000062716 | VMP1 | chr17 | + | 59731420 | 59731522 | 59735337 | 59735473 | 59707650 | 59707748 | 59738836 | 59738947 | 86 | [illegible] |
| MXE | 1807 | ENSG00000140987 | ZSCAN32 | chr16 | - | 3390009 | 3390133 | 3390422 | 3390517 | 3383700 | 3384941 | 3393648 | 3393814 | 8 | [illegible] |
| MXE | 16374 | ENSG00000013561 | RNF14 | chr5 | + | 141974803 | 141974955 | 141980122 | 141980351 | 141973582 | 141973742 | 141983379 | 141983552 | 59 | [illegible] |
| MXE | 8704 | ENSG00000003756 | RBM5 | chr3 | + | 50114179 | 50114251 | 50115427 | 50115607 | 50113949 | 50114099 | 50115905 | 50115943 | 139 | [illegible] |
| MXE | 17010 | ENSG00000067167 | TRAM1 | chr8 | - | 70583649 | 70583793 | 70586894 | 70586999 | 70583163 | 70583324 | 70587105 | 70587176 | 674 | [illegible] |
| MXE | 12806 | ENSG00000104133 | SPG11 | chr15 | - | 44570524 | 44570658 | 44572682 | 44572820 | 44569397 | 44569505 | 44573546 | 44573745 | 76 | [illegible] |
| MXE | 3030 | ENSG00000181036 | FCRL6 | chr1 | + | 159813488 | 159813554 | 159815427 | 159815459 | 159810093 | 159810216 | 159815535 | 159816257 | 143 | [illegible] |
| MXE | 5046 | ENSG00000176261 | ZBTB8OS | chr1 | - | 32631826 | 32631879 | 32633644 | 32633727 | 32627507 | 32627544 | 32633950 | 32634072 | 188 | [illegible] |
| MXE | 10519 | ENSG00000091490 | SEL1L3 | chr4 | - | 25832994 | 25833110 | 25833447 | 25833569 | 25830097 | 25830156 | 25835196 | 25835323 | 109 | [illegible] |
| MXE | 4413 | ENSG00000077684 | JADE1 | chr4 | + | 128861703 | 128862225 | 128867855 | 128867973 | 128857337 | 128857454 | 128871354 | 128872334 | 113 | [illegible] |
| MXE | 4386 | ENSG00000254870 | ATP6V1G2-DDX39B | chr6 | - | 31538762 | 31538855 | 31539146 | 31539274 | 31536499 | 31536683 | 31540321 | 31540664 | 283 | [illegible] |
| MXE | 14026 | ENSG00000120992 | LYPLA1 | chr8 | - | 54055059 | 54055133 | 54062253 | 54062324 | 54052654 | 54052756 | 54063327 | 54063375 | 105 | [illegible] |
| MXE | 13528 | ENSG00000134516 | DOCK2 | chr5 | + | 170067509 | 170067686 | 170069136 | 170069220 | 170056437 | 170057666 | 170075946 | 170076084 | 487 | [illegible] |
| MXE | 10261 | ENSG00000143207 | COP1 | chr1 | - | 176136487 | 176136547 | 176149005 | 176149074 | 176135009 | 176135086 | 176162868 | 176162988 | 178 | [illegible] |
| MXE | 6662 | ENSG00000054118 | THRAP3 | chr1 | + | 36259381 | 36259484 | 36282532 | 36282700 | 36224442 | 36224505 | 36286367 | 36287270 | 71 | [illegible] |
| MXE | 15284 | ENSG00000129255 | MPDU1 | chr17 | + | 7585945 | 7586078 | 7586691 | 7586777 | 7585731 | 7585797 | 7586898 | 7587017 | 46 | [illegible] |
| MXE | 2515 | ENSG00000100796 | PPP4R3A | chr14 | - | 91462734 | 91462877 | 91465249 | 91465419 | 91462048 | 91462239 | 91470836 | 91470995 | 249 | [illegible] |
| MXE | 2933 | ENSG00000118482 | PHF3 | chr6 | + | 63679999 | 63680161 | 63691736 | 63692043 | 63635822 | 63636150 | 63694580 | 63694764 | 46 | [illegible] |
| MXE | 13796 | ENSG00000147439 | BIN3 | chr8 | - | 22634412 | 22634525 | 22636524 | 22636586 | 22630441 | 22630578 | 22636921 | 22636962 | 106 | [illegible] |
| MXE | 2165 | ENSG00000138592 | USP8 | chr15 | + | 50465046 | 50465191 | 50471632 | 50471795 | 50462279 | 50462322 | 50476848 | 50476993 | 207 | [illegible] |
| MXE | 8383 | ENSG00000152223 | EPG5 | chr18 | - | 45930675 | 45930830 | 45934808 | 45934966 | 45928868 | 45929009 | 45939599 | 45939755 | 92 | [illegible] |
| MXE | 215 | ENSG00000100201 | DDX17 | chr22 | - | 38494885 | 38495046 | 38495795 | 38495937 | 38494629 | 38494802 | 38498084 | 38498150 | 1175 | [illegible] |
| MXE | 14770 | ENSG00000106868 | SUSD1 | chr9 | - | 112042100 | 112042165 | 112052398 | 112052438 | 112041866 | 112041960 | 112058427 | 112058686 | 76 | [illegible] |
| MXE | 6754 | ENSG00000026508 | CD44 | chr11 | + | 35209964 | 35210054 | 35211245 | 35211449 | 35208104 | 35208206 | 35214851 | 35214914 | 43 | [illegible] |
| MXE | 4619 | ENSG00000017260 | ATP2C1 | chr3 | + | 130969291 | 130969396 | 130975381 | 130975488 | 130967387 | 130967419 | 130980581 | 130980679 | 52 | [illegible] |
| MXE | 1443 | ENSG00000159210 | SNF8 | chr17 | - | 48931642 | 48931717 | 48933204 | 48933346 | 48929315 | 48930612 | 48936169 | 48936242 | 76 | [illegible] |
| MXE | 6462 | ENSG00000168434 | COG7 | chr16 | - | 23416966 | 23417121 | 23418699 | 23418827 | 23413447 | 23413564 | 23424748 | 23424947 | 6 | [illegible] |
| MXE | 11768 | ENSG00000124523 | SIRT5 | chr6 | + | 13588330 | 13588464 | 13591668 | 13591894 | 13584075 | 13584225 | 13595476 | 13595564 | 25 | [illegible] |
| MXE | 15728 | ENSG00000173933 | RBM4 | chr11 | + | 66639699 | 66640123 | 66643449 | 66643528 | 66638704 | 66638752 | 66643858 | 66644089 | 129 | [illegible] |
| MXE | 4440 | ENSG00000168310 | IRF2 | chr4 | - | 184408157 | 184408275 | 184418166 | 184418213 | 184398914 | 184399079 | 184418531 | 184418708 | 112 | [illegible] |
| MXE | 13649 | ENSG00000086758 | HUWE1 | chrX | - | 53603367 | 53603501 | 53604588 | 53604834 | 53602563 | 53602658 | 53607522 | 53607699 | 331 | [illegible] |
| MXE | 7477 | ENSG00000084754 | HADHA | chr2 | - | 26232159 | 26232279 | 26234216 | 26234355 | 26230191 | 26230294 | 26236854 | 26236988 | 681 | [illegible] |
| MXE | 14784 | ENSG00000122490 | SLC66A2 | chr18 | - | 79919183 | 79919400 | 79933968 | 79934022 | 79902419 | 79904183 | 79943328 | 79943462 | 2 | [illegible] |
| MXE | 15694 | ENSG00000054654 | SYNE2 | chr14 | + | 64031015 | 64031357 | 64047999 | 64048155 | 64029894 | 64030059 | 64049610 | 64049876 | 678 | [illegible] |
| MXE | 5317 | ENSG00000165288 | BRWD3 | chrX | - | 80791853 | 80791952 | 80808538 | 80808598 | 80745568 | 80745729 | 80809012 | 80809042 | 42 | [illegible] |
| MXE | 16664 | ENSG00000198270 | TMEM116 | chr12 | - | 111933885 | 111934030 | 111937159 | 111937243 | 111932585 | 111932659 | 111938160 | 111938210 | 29 | [illegible] |
| MXE | 11148 | ENSG00000124491 | F13A1 | chr6 | - | 6250810 | 6250929 | 6266557 | 6266809 | 6248311 | 6248419 | 6305350 | 6305539 | 839 | [illegible] |
| MXE | 8841 | ENSG00000173846 | PLK3 | chr1 | + | 44804338 | 44804501 | 44805265 | 44805379 | 44804162 | 44804246 | 44805486 | 44805950 | 40 | [illegible] |
| MXE | 731 | ENSG00000124207 | CSE1L | chr20 | + | 49085282 | 49085386 | 49088008 | 49088106 | 49084025 | 49084162 | 49089246 | 49089397 | 69 | [illegible] |
| MXE | 1058 | ENSG00000278053 | DDX52 | chr17 | - | 37632112 | 37632298 | 37633287 | 37633418 | 37630029 | 37630173 | 37642109 | 37642308 | 41 | [illegible] |
| MXE | 14595 | ENSG00000056586 | RC3H2 | chr9 | - | 122855731 | 122855878 | 122857922 | 122858093 | 122855183 | 122855397 | 122858668 | 122859102 | 212 | [illegible] |
| MXE | 1335 | ENSG00000055483 | USP36 | chr17 | - | 78803384 | 78803978 | 78806155 | 78806286 | 78802323 | 78802535 | 78806958 | 78807636 | 39 | [illegible] |
| MXE | 2075 | ENSG00000182718 | ANXA2 | chr15 | - | 60393037 | 60393440 | 60396150 | 60396427 | 60386027 | 60386086 | 60397942 | 60397986 | 2 | [illegible] |
| MXE | 1958 | ENSG00000140694 | PARN | chr16 | - | 14586317 | 14586361 | 14591912 | 14593378 | 14584748 | 14584791 | 14599903 | 14599960 | 72 | [illegible] |
| MXE | 14611 | ENSG00000127084 | FGD3 | chr9 | + | 93032768 | 93032873 | 93034540 | 93034681 | 93029873 | 93029996 | 93035337 | 93036222 | 185 | [illegible] |
| MXE | 15121 | ENSG00000149516 | MS4A3 | chr11 | + | 60061489 | 60061554 | 60062467 | 60062605 | 60061145 | 60061316 | 60064261 | 60064318 | 0 | [illegible] |
| MXE | 14620 | ENSG00000160563 | MED27 | chr9 | - | 131939380 | 131939474 | 132014336 | 132014467 | 131893884 | 131893992 | 132077441 | 132077586 | 54 | [illegible] |
| MXE | 14346 | ENSG00000149084 | HSD17B12 | chr11 | + | 43754045 | 43754121 | 43798319 | 43798427 | 43750910 | 43750957 | 43815436 | 43815501 | 3 | [illegible] |
| MXE | 7063 | ENSG00000223745 | CCDC18-AS1 | chr1 | - | 93325768 | 93325895 | 93337376 | 93339274 | 93324634 | 93324729 | 93339356 | 93339536 | 54 | [illegible] |
| MXE | 1712 | ENSG00000103313 | MEFV | chr16 | - | 3244472 | 3244588 | 3246524 | 3246547 | 3244253 | 3244286 | 3247015 | 3247246 | 38 | [illegible] |
| MXE | 3661 | ENSG00000172785 | CBWD1 | chr9 | - | 134978 | 135030 | 152033 | 152078 | 123385 | 123454 | 154708 | 154795 | 14 | [illegible] |
| MXE | 1371 | ENSG00000108671 | PSMD11 | chr17 | + | 32479250 | 32479376 | 32479850 | 32479886 | 32477520 | 32477583 | 32480145 | 32480197 | 153 | [illegible] |
| MXE | 14685 | ENSG00000116580 | GON4L | chr1 | - | 155751959 | 155752587 | 155754374 | 155754488 | 155751766 | 155751869 | 155756957 | 155757077 | 48 | [illegible] |
| MXE | 14060 | ENSG00000034677 | RNF19A | chr8 | - | 100261541 | 100261755 | 100264670 | 100264785 | 100259853 | 100259997 | 100268784 | 100268920 | 78 | [illegible] |
| MXE | 14516 | ENSG00000156471 | PTDSS1 | chr8 | + | 96320245 | 96320345 | 96330212 | 96330281 | 96309556 | 96309622 | 96331025 | 96331095 | 105 | [illegible] |
| MXE | 2199 | ENSG00000170776 | AKAP13 | chr15 | + | 85727380 | 85727463 | 85730512 | 85730707 | 85727065 | 85727247 | 85734991 | 85735150 | 436 | [illegible] |
| MXE | 2416 | ENSG00000248905 | FMN1 | chr15 | - | 33092132 | 33092203 | 33153047 | 33155045 | 33088798 | 33088974 | 33180197 | 33180262 | 0 | [illegible] |
| MXE | 3987 | ENSG00000132952 | USPL1 | chr13 | + | 30630834 | 30631474 | 30637743 | 30637857 | 30621763 | 30621892 | 30642627 | 30642757 | 99 | [illegible] |
| MXE | 2529 | ENSG00000092200 | RPGRIP1 | chr14 | + | 21334604 | 21334705 | 21343035 | 21343228 | 21330248 | 21330387 | 21345112 | 21345197 | 5 | [illegible] |
| MXE | 8268 | ENSG00000170271 | FAXDC2 | chr5 | - | 154822471 | 154822577 | 154823386 | 154823592 | 154821259 | 154821426 | 154830800 | 154830922 | 55 | [illegible] |
| MXE | 6150 | ENSG00000078304 | PPP2R5C | chr14 | + | 101786017 | 101786183 | 101856685 | 101856885 | 101762904 | 101762970 | 101882160 | 101882271 | 285 | [illegible] |
| MXE | 3726 | ENSG00000104320 | NBN | chr8 | - | 89980733 | 89980893 | 89981374 | 89981523 | 89978219 | 89978323 | 89981952 | 89982002 | 32 | [illegible] |
| MXE | 9884 | ENSG00000219626 | FAM228B | chr2 | + | 24135118 | 24135187 | 24146747 | 24146835 | 24095140 | 24095229 | 24146929 | 24147086 | 5 | [illegible] |
| MXE | 5351 | ENSG00000051382 | PIK3CB | chr3 | - | 138656141 | 138656274 | 138657689 | 138657835 | 138652697 | 138655526 | 138663905 | 138664029 | 96 | [illegible] |
| MXE | 11082 | ENSG00000115234 | SNX17 | chr2 | + | 27374688 | 27374758 | 27375060 | 27375153 | 27374345 | 27374433 | 27375505 | 27375709 | 130 | [illegible] |
| MXE | 1587 | ENSG00000187555 | USP7 | chr16 | - | 8895641 | 8895741 | 8896998 | 8897099 | 8895030 | 8895150 | 8898359 | 8898437 | 161 | [illegible] |
| MXE | 2214 | ENSG00000092439 | TRPM7 | chr15 | - | 50580873 | 50580908 | 50583088 | 50583159 | 50578638 | 50578664 | 50586391 | 50586488 | 63 | [illegible] |
| MXE | 1617 | ENSG00000153066 | TXNDC11 | chr16 | - | 11688302 | 11688445 | 11691289 | 11692082 | 11687856 | 11687966 | 11698124 | 11698325 | 101 | [illegible] |
| MXE | 13826 | ENSG00000145868 | FBXO38 | chr5 | + | 148438331 | 148438498 | 148439646 | 148439792 | 148433634 | 148433737 | 148440423 | 148440527 | 121 | [illegible] |
| MXE | 11390 | ENSG00000198721 | ECI2 | chr6 | - | 4125249 | 4125370 | 4126134 | 4126237 | 4119185 | 4119275 | 4127761 | 4127831 | 96 | [illegible] |
| MXE | 489 | ENSG00000130164 | LDLR | chr19 | + | 11120369 | 11120522 | 11123173 | 11123344 | 11120091 | 11120233 | 11128007 | 11128085 | 126 | [illegible] |
| MXE | 12565 | ENSG00000180357 | ZNF609 | chr15 | + | 64622826 | 64623052 | 64670345 | 64670433 | 64499291 | 64500166 | 64673915 | 64676256 | 166 | [illegible] |
| MXE | 15384 | ENSG00000140299 | BNIP2 | chr15 | - | 59671182 | 59671314 | 59672636 | 59672739 | 59669275 | 59669362 | 59677910 | 59678016 | 271 | [illegible] |
| MXE | 4989 | ENSG00000166128 | RAB8B | chr15 | + | 63244755 | 63244816 | 63249644 | 63249705 | 63189605 | 63189748 | 63255506 | 63255584 | 262 | [illegible] |
| MXE | 4273 | ENSG00000102445 | RUBCNL | chr13 | - | 46361440 | 46361573 | 46362537 | 46362598 | 46359485 | 46359631 | 46363114 | 46363213 | 10 | [illegible] |
| MXE | 5249 | ENSG00000127483 | HP1BP3 | chr1 | - | 20765376 | 20765531 | 20767583 | 20767664 | 20757165 | 20757256 | 20770929 | 20771073 | 177 | [illegible] |
| MXE | 8305 | ENSG00000138802 | SEC24B | chr4 | + | 109462900 | 109463644 | 109473003 | 109473186 | 109433812 | 109434002 | 109491326 | 109491407 | 21 | [illegible] |
| MXE | 3844 | ENSG00000247774 | PCED1B-AS1 | chr12 | - | 47206510 | 47206585 | 47209280 | 47209458 | 47205897 | 47206145 | 47210639 | 47210761 | 193 | [illegible] |
| MXE | 14022 | ENSG00000198851 | CD3E | chr11 | + | 118312617 | 118312866 | 118314447 | 118314494 | 118312152 | 118312170 | 118315485 | 118316173 | 2127 | [illegible] |
| MXE | 9042 | ENSG00000100271 | TTLL1 | chr22 | - | 43051800 | 43051887 | 43059383 | 43059527 | 43046409 | 43046573 | 43063812 | 43063921 | 24 | [illegible] |
| MXE | 6594 | ENSG00000257315 | ZBED6 | chr1 | + | 203830122 | 203830203 | 203831660 | 203831771 | 203829779 | 203829896 | 203833790 | 203833853 | 321 | [illegible] |
| MXE | 8128 | ENSG00000136098 | NEK3 | chr13 | - | 52135728 | 52135863 | 52136115 | 52136259 | 52133688 | 52133815 | 52136799 | 52136902 | 35 | [illegible] |
| MXE | 10391 | ENSG00000152117 | SMPD4BP | chr2 | + | 131501357 | 131501508 | 131506311 | 131506442 | 131501180 | 131501233 | 131508536 | 131508608 | 12 | [illegible] |
| MXE | 8071 | ENSG00000103657 | HERC1 | chr15 | - | 63734723 | 63734849 | 63746917 | 63747083 | 63732923 | 63733145 | 63747723 | 63747858 | 85 | [illegible] |
| MXE | 7922 | ENSG00000178685 | PARP10 | chr8 | - | 143984034 | 143984104 | 143984209 | 143984431 | 143983166 | 143983811 | 143984543 | 143985328 | 96 | [illegible] |
| MXE | 12607 | ENSG00000151948 | GLT1D1 | chr12 | + | 128875913 | 128876062 | 128888638 | 128888744 | 128853477 | 128853649 | 128899235 | 128899287 | 12 | [illegible] |
| MXE | 3817 | ENSG00000011243 | AKAP8L | chr19 | - | 15399301 | 15399410 | 15400294 | 15400358 | 15397713 | 15397855 | 15400793 | 15400864 | 33 | [illegible] |
| MXE | 9977 | ENSG00000163219 | ARHGAP25 | chr2 | + | 68782232 | 68782320 | 68787839 | 68787956 | 68775220 | 68775420 | 68807272 | 68807480 | 196 | [illegible] |
| MXE | 2092 | ENSG00000128915 | ICE2 | chr15 | - | 60466593 | 60466713 | 60468060 | 60468162 | 60456656 | 60456794 | 60476062 | 60476167 | 35 | [illegible] |
| MXE | 3347 | ENSG00000167881 | SRP68 | chr17 | - | 76061109 | 76061219 | 76061491 | 76061574 | 76060307 | 76060390 | 76063975 | 76064171 | 105 | [illegible] |
| MXE | 13482 | ENSG00000165609 | NUDT5 | chr10 | - | 12170716 | 12170770 | 12170899 | 12170908 | 12165329 | 12167811 | 12172764 | 12172866 | 7 | [illegible] |
| MXE | 7732 | ENSG00000048052 | HDAC9 | chr7 | + | 18666212 | 18666476 | 18727579 | 18727757 | 18648465 | 18648683 | 18749004 | 18749138 | 97 | [illegible] |
| MXE | 5653 | ENSG00000151576 | QTRT2 | chr3 | + | 114065236 | 114065457 | 114067986 | 114068063 | 114056734 | 114056864 | 114070625 | 114070838 | 43 | [illegible] |
| MXE | 3041 | ENSG00000255819 | KLRC4-KLRK1 | chr12 | - | 10388770 | 10388875 | 10389942 | 10390027 | 10386902 | 10387010 | 10391834 | 10391891 | 49 | [illegible] |
| MXE | 15475 | ENSG00000156599 | ZDHHC5 | chr11 | + | 57695919 | 57696043 | 57696760 | 57696873 | 57693782 | 57693915 | 57698558 | 57698835 | 143 | [illegible] |
| MXE | 7276 | ENSG00000233184 | NA | chr1 | + | 101075394 | 101075716 | 101077362 | 101077508 | 101072659 | 101072830 | 101080388 | 101080613 | 12 | [illegible] |
| MXE | 15342 | ENSG00000099783 | HNRNPM | chr19 | + | 8465323 | 8465362 | 8465479 | 8465515 | 8463592 | 8463686 | 8466234 | 8466388 | 93 | [illegible] |
| MXE | 2815 | ENSG00000100479 | POLE2 | chr14 | - | 49655670 | 49655843 | 49663314 | 49663387 | 49655004 | 49655094 | 49664625 | 49664663 | 11 | [illegible] |
| MXE | 14749 | ENSG00000048828 | FAM120A | chr9 | + | 93471140 | 93471387 | 93476255 | 93476338 | 93451684 | 93452389 | 93497470 | 93497599 | 103 | [illegible] |
| MXE | 13817 | ENSG00000068383 | INPP5A | chr10 | + | 132650417 | 132650505 | 132690391 | 132690455 | 132645867 | 132645968 | 132697815 | 132697919 | 35 | [illegible] |
| MXE | 7797 | ENSG00000081014 | AP4E1 | chr15 | + | 50930804 | 50930971 | 50934623 | 50934697 | 50929008 | 50929168 | 50941441 | 50941564 | 94 | [illegible] |
| MXE | 10240 | ENSG00000087152 | ATXN7L3 | chr17 | - | 44195096 | 44195140 | 44195418 | 44195487 | 44194767 | 44194839 | 44195799 | 44195828 | 77 | [illegible] |
| MXE | 7591 | ENSG00000171793 | CTPS1 | chr1 | + | 41009444 | 41009589 | 41010160 | 41010254 | 41008658 | 41008890 | 41011657 | 41011754 | 62 | [illegible] |
| MXE | 12959 | ENSG00000127914 | AKAP9 | chr7 | + | 91980872 | 91980958 | 91992157 | 91992211 | 91980288 | 91980333 | 91992884 | 91993055 | 8 | [illegible] |
| MXE | 1131 | ENSG00000033627 | ATP6V0A1 | chr17 | + | 42466428 | 42466507 | 42468009 | 42468107 | 42460847 | 42461011 | 42470089 | 42470218 | 62 | [illegible] |
| MXE | 13514 | ENSG00000172534 | HCFC1 | chrX | - | 153956190 | 153956411 | 153956624 | 153956763 | 153954065 | 153955542 | 153956917 | 153957060 | 56 | [illegible] |
| MXE | 8302 | ENSG00000151422 | FER | chr5 | + | 108946129 | 108946222 | 108954728 | 108954932 | 108897647 | 108897848 | 108959224 | 108959347 | 12 | [illegible] |
| MXE | 4530 | ENSG00000168397 | ATG4B | chr2 | + | 241651263 | 241651335 | 241659107 | 241659187 | 241651006 | 241651111 | 241666644 | 241666838 | 109 | [illegible] |
| MXE | 15251 | ENSG00000110367 | DDX6 | chr11 | - | 118779631 | 118779736 | 118781120 | 118781184 | 118768222 | 118768352 | 118786051 | 118786518 | 134 | [illegible] |
| MXE | 16524 | ENSG00000139323 | POC1B | chr12 | - | 89467616 | 89467685 | 89472167 | 89472275 | 89466769 | 89466922 | 89491935 | 89492115 | 48 | [illegible] |
| MXE | 4190 | ENSG00000114738 | MAPKAPK3 | chr3 | + | 50617513 | 50617784 | 50640365 | 50640505 | 50617169 | 50617226 | 50641706 | 50641771 | 15 | [illegible] |
| MXE | 2389 | ENSG00000137814 | HAUS2 | chr15 | + | 42559338 | 42559408 | 42561269 | 42561402 | 42558197 | 42558290 | 42563748 | 42563857 | 8 | [illegible] |
| MXE | 720 | ENSG00000124198 | ARFGEF2 | chr20 | + | 48973144 | 48973284 | 48974765 | 48974874 | 48972325 | 48972425 | 48976015 | 48976199 | 92 | [illegible] |
| MXE | 15667 | ENSG00000081237 | PTPRC | chr1 | + | 198741868 | 198742026 | 198742231 | 198742367 | 198735126 | 198735252 | 198744053 | 198744203 | 3642 | [illegible] |
| MXE | 5277 | ENSG00000284952 | NA | chr3 | - | 155833856 | 155834041 | 155842431 | 155842997 | 155833467 | 155833585 | 155853222 | 155854442 | 1 | [illegible] |
| MXE | 2111 | ENSG00000128739 | SNRPN | chr15 | + | 24974310 | 24974456 | 24976876 | 24977029 | 24967931 | 24968082 | 24977777 | 24977916 | 274 | [illegible] |
| MXE | 5333 | ENSG00000122068 | FYTTD1 | chr3 | + | 197768438 | 197768587 | 197770131 | 197770244 | 197756425 | 197756557 | 197773402 | 197773499 | 449 | [illegible] |
| MXE | 12960 | ENSG00000127914 | AKAP9 | chr7 | + | 92014248 | 92014328 | 92016128 | 92016267 | 92012428 | 92012642 | 92017016 | 92017102 | 87 | [illegible] |
| MXE | 11761 | ENSG00000137200 | CMTR1 | chr6 | + | 37461548 | 37461645 | 37461969 | 37462102 | 37459565 | 37459684 | 37462828 | 37463008 | 124 | [illegible] |
| MXE | 15784 | ENSG00000118276 | B4GALT6 | chr18 | - | 31645354 | 31645479 | 31657975 | 31658089 | 31638643 | 31638760 | 31684311 | 31684571 | 1 | [illegible] |
| MXE | 15145 | ENSG00000183597 | TANGO2 | chr22 | + | 20055942 | 20056013 | 20061529 | 20061683 | 20052464 | 20052584 | 20063337 | 20063442 | 27 | [illegible] |
| MXE | 412 | ENSG00000129353 | SLC44A2 | chr19 | + | 10634973 | 10635073 | 10636322 | 10636585 | 10634755 | 10634887 | 10636661 | 10636756 | 178 | [illegible] |
| MXE | 12941 | ENSG00000141380 | SS18 | chr18 | - | 26035830 | 26035923 | 26038554 | 26038659 | 26035004 | 26035127 | 26039288 | 26039456 | 123 | [illegible] |
| MXE | 15864 | ENSG00000172671 | ZFAND4 | chr10 | - | 45652983 | 45653059 | 45662579 | 45662681 | 45651965 | 45652033 | 45663541 | 45663842 | 5 | [illegible] |
| MXE | 872 | ENSG00000141644 | MBD1 | chr18 | - | 50273563 | 50273863 | 50274185 | 50274353 | 50272823 | 50272955 | 50274976 | 50275045 | 39 | [illegible] |
| MXE | 14588 | ENSG00000124702 | KLHDC3 | chr6 | + | 43017518 | 43017695 | 43017852 | 43017968 | 43017133 | 43017346 | 43018144 | 43018216 | 128 | [illegible] |
| MXE | 2237 | ENSG00000099308 | MAST3 | chr19 | + | 18128399 | 18128458 | 18128865 | 18128951 | 18124641 | 18124774 | 18130493 | 18130702 | 88 | [illegible] |
| MXE | 6908 | ENSG00000091127 | PUS7 | chr7 | - | 105468336 | 105468463 | 105470687 | 105470848 | 105465312 | 105465414 | 105472131 | 105472193 | 8 | [illegible] |
| MXE | 11458 | ENSG00000043462 | LCP2 | chr5 | - | 170258865 | 170258878 | 170261106 | 170261137 | 170258036 | 170258166 | 170262634 | 170262742 | 82 | [illegible] |
| MXE | 12573 | ENSG00000121716 | PILRB | chr7 | + | 100355142 | 100355240 | 100356393 | 100356883 | 100353894 | 100354012 | 100358219 | 100358259 | 65 | [illegible] |
| MXE | 799 | ENSG00000101425 | BPI | chr20 | + | 38307566 | 38307681 | 38308929 | 38309058 | 38304149 | 38304353 | 38310490 | 38310652 | 13 | [illegible] |
| MXE | 6376 | ENSG00000026751 | SLAMF7 | chr1 | + | 160748193 | 160748514 | 160749820 | 160750093 | 160739056 | 160739356 | 160751344 | 160751448 | 183 | [illegible] |
| MXE | 1878 | ENSG00000141101 | NOB1 | chr16 | - | 69749558 | 69749630 | 69752240 | 69752371 | 69749212 | 69749338 | 69754593 | 69754726 | 36 | [illegible] |
| MXE | 8460 | ENSG00000137275 | RIPK1 | chr6 | + | 3082546 | 3083313 | 3085258 | 3085408 | 3080978 | 3081116 | 3089580 | 3089657 | 1 | [illegible] |
| MXE | 8399 | ENSG00000168610 | STAT3 | chr17 | - | 42338730 | 42338812 | 42339313 | 42339409 | 42337762 | 42337857 | 42345558 | 42345657 | 130 | [illegible] |
| MXE | 7926 | ENSG00000043514 | TRIT1 | chr1 | - | 39847953 | 39848097 | 39850118 | 39850261 | 39847547 | 39847660 | 39853969 | 39854068 | 3 | [illegible] |
| MXE | 4651 | ENSG00000198586 | TLK1 | chr2 | - | 171022871 | 171023129 | 171028338 | 171028405 | 171014850 | 171014948 | 171046173 | 171046362 | 556 | [illegible] |
| MXE | 5943 | ENSG00000172175 | MALT1 | chr18 | + | 58741864 | 58742014 | 58744337 | 58744495 | 58735201 | 58735329 | 58745665 | 58745791 | 407 | [illegible] |
| MXE | 2519 | ENSG00000100796 | PPP4R3A | chr14 | - | 91476908 | 91476986 | 91481575 | 91482193 | 91476407 | 91476524 | 91485631 | 91485730 | 411 | [illegible] |
| MXE | 11355 | ENSG00000011566 | MAP4K3 | chr2 | - | 39303073 | 39303117 | 39307942 | 39308005 | 39299742 | 39299801 | 39309460 | 39309519 | 22 | [illegible] |
| MXE | 2864 | ENSG00000165516 | KLHDC2 | chr14 | + | 49779747 | 49779806 | 49780212 | 49780322 | 49779594 | 49779675 | 49780702 | 49780775 | 21 | [illegible] |
| MXE | 15565 | ENSG00000013725 | CD6 | chr11 | + | 61013422 | 61013563 | 61015712 | 61015835 | 61011069 | 61011135 | 61017478 | 61017550 | 184 | [illegible] |
| MXE | 9453 | ENSG00000115806 | GORASP2 | chr2 | + | 170954649 | 170954782 | 170956435 | 170956559 | 170951327 | 170951458 | 170961662 | 170961749 | 156 | [illegible] |
| MXE | 1852 | ENSG00000103091 | WDR59 | chr16 | - | 74887682 | 74887755 | 74889702 | 74889815 | 74886269 | 74886396 | 74892483 | 74892565 | 16 | [illegible] |
| MXE | 1432 | ENSG00000168259 | DNAJC7 | chr17 | - | 41987818 | 41987910 | 41988731 | 41988896 | 41983562 | 41983636 | 41989403 | 41989557 | 257 | [illegible] |
| MXE | 16474 | ENSG00000120253 | NUP43 | chr6 | - | 149742389 | 149742570 | 149743637 | 149743715 | 149738642 | 149738778 | 149745939 | 149746062 | 67 | [illegible] |
| MXE | 2004 | ENSG00000130703 | OSBPL2 | chr20 | + | 62286582 | 62286711 | 62289206 | 62289330 | 62284045 | 62284169 | 62291702 | 62291793 | 114 | [illegible] |
| MXE | 11417 | ENSG00000138380 | CARF | chr2 | + | 202961236 | 202961426 | 202966977 | 202967098 | 202955673 | 202955758 | 202969918 | 202970062 | 42 | [illegible] |
| MXE | 11518 | ENSG00000114166 | KAT2B | chr3 | + | 20072332 | 20072459 | 20095262 | 20095408 | 20040445 | 20040780 | 20099861 | 20099954 | 39 | [illegible] |
| MXE | 16274 | ENSG00000066084 | DIP2B | chr12 | + | 50728547 | 50728678 | 50731368 | 50731537 | 50727702 | 50727812 | 50732365 | 50732536 | 67 | [illegible] |
| MXE | 4867 | ENSG00000078618 | NRDC | chr1 | - | 51825286 | 51825382 | 51827795 | 51827869 | 51823663 | 51823786 | 51834016 | 51834073 | 199 | [illegible] |
| MXE | 3488 | ENSG00000147044 | CASK | chrX | - | 41553718 | 41553915 | 41555599 | 41555635 | 41542690 | 41542806 | 41557031 | 41557100 | 0 | [illegible] |
| MXE | 4961 | ENSG00000004766 | VPS50 | chr7 | + | 93239865 | 93239950 | 93252652 | 93252775 | 93232365 | 93232500 | 93253859 | 93253931 | 44 | [illegible] |
| MXE | 3662 | ENSG00000172785 | CBWD1 | chr9 | - | 146101 | 146158 | 152033 | 152078 | 134978 | 135030 | 154708 | 154795 | 19 | [illegible] |
| MXE | 15293 | ENSG00000137500 | CCDC90B | chr11 | - | 83274638 | 83274763 | 83278725 | 83278829 | 83273950 | 83273992 | 83280140 | 83280260 | 106 | [illegible] |
| MXE | 11336 | ENSG00000144233 | AMMECR1L | chr2 | - | 127870813 | 127870928 | 127871248 | 127871359 | 127869453 | 127869544 | 127873827 | 127874272 | 13 | [illegible] |
| MXE | 4697 | ENSG00000109606 | DHX15 | chr4 | - | 24570653 | 24570847 | 24576242 | 24576678 | 24556250 | 24556410 | 24584322 | 24584554 | 254 | [illegible] |
| MXE | 622 | ENSG00000268790 | NA | chr19 | - | 16654034 | 16654125 | 16659394 | 16659447 | 16647095 | 16647261 | 16659958 | 16660000 | 23 | [illegible] |
| MXE | 11924 | ENSG00000131018 | SYNE1 | chr6 | - | 152244536 | 152244656 | 152249160 | 152249262 | 152242239 | 152242440 | 152254879 | 152255089 | 310 | [illegible] |
| MXE | 11666 | ENSG00000100504 | PYGL | chr14 | - | 50912155 | 50912303 | 50913028 | 50913130 | 50911977 | 50912036 | 50914700 | 50914815 | 68 | [illegible] |
| MXE | 6569 | ENSG00000198791 | CNOT7 | chr8 | - | 17237211 | 17237373 | 17242991 | 17243185 | 17234592 | 17234860 | 17245035 | 17245091 | 729 | [illegible] |
| MXE | 13103 | ENSG00000177324 | BEND2 | chrX | - | 18191000 | 18191108 | 18201814 | 18201940 | 18180509 | 18180650 | 18203500 | 18203915 | 54 | [illegible] |
| MXE | 15841 | ENSG00000148481 | MINDY3 | chr10 | - | 15837203 | 15837318 | 15841425 | 15841599 | 15834542 | 15834616 | 15843211 | 15843272 | 326 | [illegible] |
| MXE | 8397 | ENSG00000054267 | ARID4B | chr1 | - | 235234412 | 235234492 | 235240312 | 235240451 | 235231112 | 235231189 | 235246419 | 235246511 | 97 | [illegible] |
| MXE | 9893 | ENSG00000073614 | KDM5A | chr12 | - | 331818 | 331938 | 333486 | 333649 | 328834 | 329029 | 334240 | 334422 | 728 | [illegible] |
| MXE | 5617 | ENSG00000233452 | STXBP5-AS1 | chr6 | - | 146911551 | 146911665 | 147001440 | 147001543 | 146850824 | 146851331 | 147074804 | 147074847 | 1 | [illegible] |
| MXE | 2488 | ENSG00000100580 | TMED8 | chr14 | - | 77343696 | 77343823 | 77346348 | 77346478 | 77343177 | 77343483 | 77351672 | 77351751 | 12 | [illegible] |
| MXE | 15951 | ENSG00000163249 | CCNYL1 | chr2 | + | 207737410 | 207737446 | 207740654 | 207740706 | 207726841 | 207726876 | 207742222 | 207742342 | 4 | [illegible] |
| MXE | 5767 | ENSG00000143811 | PYCR2 | chr1 | - | 225922203 | 225922383 | 225923700 | 225923771 | 225921857 | 225922079 | 225924043 | 225924250 | 10 | [illegible] |
| MXE | 7008 | ENSG00000185158 | LRRC37B | chr17 | + | 32049101 | 32049394 | 32050002 | 32050107 | 32047166 | 32047901 | 32053265 | 32053413 | 24 | [illegible] |
| MXE | 6359 | ENSG00000143110 | C1orf162 | chr1 | + | 111476797 | 111476867 | 111477333 | 111477428 | 111476017 | 111476065 | 111477650 | 111477775 | 41 | [illegible] |
| MXE | 8568 | ENSG00000213079 | SCAF8 | chr6 | + | 154778000 | 154778045 | 154787860 | 154788022 | 154773988 | 154774072 | 154792822 | 154792858 | 26 | [illegible] |
| MXE | 11773 | ENSG00000124523 | SIRT5 | chr6 | + | 13596962 | 13597016 | 13599031 | 13599155 | 13595476 | 13595564 | 13600833 | 13600949 | 13 | [illegible] |
| MXE | 815 | ENSG00000170471 | RALGAPB | chr20 | + | 38567906 | 38567232 | 38570768 | 38570847 | 38565355 | 38565478 | 38574149 | 38574298 | 143 | [illegible] |
| MXE | 14945 | ENSG00000137073 | UBAP2 | chr9 | - | 33943419 | 33943589 | 33948373 | 33948587 | 33941648 | 33941862 | 33953284 | 33953474 | 78 | [illegible] |
| MXE | 6146 | ENSG00000163719 | MTMR14 | chr3 | + | 9672684 | 9672758 | 9677983 | 9678058 | 9671047 | 9671170 | 9683177 | 9683244 | 145 | [illegible] |
| MXE | 5232 | ENSG00000100034 | PPM1F | chr22 | - | 21939531 | 21939680 | 21945842 | 21946108 | 21934023 | 21934226 | 21952791 | 21952831 | 56 | [illegible] |
| MXE | 8224 | ENSG00000142765 | SYTL1 | chr1 | + | 27349651 | 27349765 | 27349971 | 27350132 | 27349397 | 27349498 | 27350388 | 27350485 | 66 | [illegible] |
| MXE | 12411 | ENSG00000198198 | SZT2 | chr1 | + | 43415081 | 43415213 | 43419733 | 43419944 | 43404379 | 43404550 | 43420152 | 43420323 | 16 | [illegible] |
| MXE | 17138 | ENSG00000102710 | SUPT20H | chr13 | - | 37028147 | 37028305 | 37029764 | 37029836 | 37026789 | 37026816 | 37031566 | 37031623 | 49 | [illegible] |
| MXE | 1625 | ENSG00000153066 | TXNDC11 | chr16 | - | 11721576 | 11721670 | 11730644 | 11730774 | 11700451 | 11700564 | 11733981 | 11734079 | 69 | [illegible] |
| MXE | 7704 | ENSG00000142687 | KIAA0319L | chr1 | - | 35470862 | 35470960 | 35474804 | 35474906 | 35462620 | 35462713 | 35478965 | 35479212 | 58 | [illegible] |
| MXE | 11952 | ENSG00000164587 | RPS14 | chr5 | - | 150446801 | 150446963 | 150447584 | 150447735 | 150445608 | 150445685 | 150449702 | 150449739 | 6339 | [illegible] |
| MXE | 7866 | ENSG00000042429 | MED17 | chr11 | + | 93797534 | 93797719 | 93801834 | 93801972 | 93796409 | 93796540 | 93807517 | 93807635 | 35 | [illegible] |
| MXE | 12751 | ENSG00000113013 | HSPA9 | chr5 | - | 138567648 | 138567722 | 138568924 | 138569049 | 138567454 | 138567561 | 138570959 | 138571141 | 271 | [illegible] |
| MXE | 10714 | ENSG00000168958 | MFF | chr2 | + | 227342744 | 227342819 | 227347225 | 227347384 | 227340291 | 227340380 | 227355676 | 227355761 | 41 | [illegible] |
| MXE | 15232 | ENSG00000109452 | INPP4B | chr4 | - | 142305457 | 142305537 | 142347373 | 142347556 | 142270662 | 142270774 | 142402937 | 142403009 | 1 | [illegible] |
| MXE | 14678 | ENSG00000116701 | NCF2 | chr1 | - | 183570779 | 183570839 | 183573184 | 183573292 | 183564004 | 183564030 | 183577598 | 183577707 | 4 | [illegible] |
| MXE | 9288 | ENSG00000142327 | RNPEPL1 | chr2 | + | 240573774 | 240573891 | 240574112 | 240574348 | 240573109 | 240573261 | 240574514 | 240574628 | 60 | [illegible] |
| MXE | 9825 | ENSG00000151092 | NGLY1 | chr3 | - | 25739576 | 25739799 | 25740402 | 25740450 | 25737333 | 25737455 | 25751097 | 25751263 | 2 | [illegible] |
| MXE | 7833 | ENSG00000176393 | RNPEP | chr1 | + | 201989382 | 201989531 | 201996146 | 201996263 | 201988903 | 201989044 | 201997318 | 201997554 | 120 | [illegible] |
| MXE | 5387 | ENSG00000185507 | IRF7 | chr11 | - | 613950 | 614037 | 614173 | 614399 | 613784 | 613865 | 614475 | 614534 | 21 | [illegible] |
| MXE | 386 | ENSG00000141867 | BRD4 | chr19 | - | 15267415 | 15267551 | 15268904 | 15269042 | 15265353 | 15265643 | 15272814 | 15273133 | 469 | [illegible] |
| MXE | 5683 | ENSG00000163681 | SLMAP | chr3 | + | 57865241 | 57865292 | 57871635 | 57871698 | 57864806 | 57864857 | 57890040 | 57890100 | 29 | [illegible] |
| MXE | 12868 | ENSG00000122566 | HNRNPA2B1 | chr7 | - | 26193574 | 26193598 | 26193598 | 26193694 | 26193250 | 26193373 | 26195846 | 26195909 | 436 | [illegible] |
| MXE | 13606 | ENSG00000125354 | SEPTIN6 | chrX | - | 119649936 | 119650098 | 119663481 | 119663677 | 119640691 | 119640788 | 119675553 | 119675668 | 139 | [illegible] |
| MXE | 6541 | ENSG00000122335 | SERAC1 | chr6 | - | 158128107 | 158128270 | 158130372 | 158130486 | 158120424 | 158120575 | 158143055 | 158143184 | 6 | [illegible] |
| MXE | 279 | ENSG00000228315 | GUSBP11 | chr22 | - | 23694118 | 23695517 | 23700725 | 23700845 | 23683724 | 23683867 | 23705428 | 23705541 | 12 | [illegible] |
| MXE | 3456 | ENSG00000176783 | RUFY1 | chr5 | + | 179589545 | 179589647 | 179591624 | 179591741 | 179585795 | 179585865 | 179593477 | 179593645 | 182 | [illegible] |
| MXE | 14186 | ENSG00000173064 | HECTD4 | chr12 | - | 112273654 | 112273795 | 112274846 | 112274960 | 112270226 | 112270459 | 112279227 | 112279386 | 28 | [illegible] |
| MXE | 15138 | ENSG00000100731 | PCNX1 | chr14 | + | 71009633 | 71009724 | 71011491 | 71011549 | 70976941 | 70978648 | 71012999 | 71013202 | 37 | [illegible] |
| MXE | 4960 | ENSG00000004766 | VPS50 | chr7 | + | 93239865 | 93239934 | 93252652 | 93252775 | 93232339 | 93232500 | 93253859 | 93253931 | 45 | [illegible] |
| MXE | 5654 | ENSG00000168028 | RPSA | chr3 | + | 39408605 | 39408724 | 39410753 | 39410999 | 39407602 | 39407786 | 39411648 | 39411777 | 3326 | [illegible] |
| MXE | 14384 | ENSG00000119397 | CNTRL | chr9 | + | 121096421 | 121096563 | 121098385 | 121098572 | 121094887 | 121095018 | 121107801 | 121107995 | 119 | [illegible] |
| MXE | 10641 | ENSG00000114450 | GNB4 | chr3 | - | 179419398 | 179419505 | 179420888 | 179420927 | 179416492 | 179416556 | 179426143 | 179426200 | 0 | [illegible] |
| MXE | 6427 | ENSG00000113194 | FAF2 | chr5 | + | 176494183 | 176494275 | 176496485 | 176496663 | 176493998 | 176494084 | 176498913 | 176499085 | 42 | [illegible] |
| MXE | 5100 | ENSG00000060971 | ACAA1 | chr3 | - | 38127785 | 38127866 | 38129289 | 38129388 | 38126060 | 38126700 | 38131925 | 38132005 | 23 | [illegible] |
| MXE | 9689 | ENSG00000128815 | WDFY4 | chr10 | + | 48873490 | 48873697 | 48875088 | 48875140 | 48867264 | 48867342 | 48877032 | 48877199 | 100 | [illegible] |
| MXE | 10053 | ENSG00000182973 | CNOT10 | chr3 | + | 32737409 | 32737490 | 32759457 | 32759571 | 32734799 | 32734976 | 32762732 | 32762863 | 17 | [illegible] |
| MXE | 989 | ENSG00000101019 | UQCC1 | chr20 | - | 35366556 | 35366614 | 35374183 | 35374256 | 35347163 | 35347272 | 35381917 | 35382025 | 46 | [illegible] |
| MXE | 6467 | ENSG00000088854 | C20orf194 | chr20 | - | 3324891 | 3324968 | 3326196 | 3326284 | 3322651 | 3322696 | 3330645 | 3330682 | 12 | [illegible] |
| MXE | 14575 | ENSG00000197969 | VPS13A | chr9 | + | 77214328 | 77214386 | 77219953 | 77220081 | 77213233 | 77213314 | 77220276 | 77220383 | 49 | [illegible] |
| MXE | 14419 | ENSG00000132326 | PER2 | chr2 | - | 238261728 | 238261837 | 238262190 | 238262344 | 238260827 | 238260953 | 238262951 | 238263058 | 37 | [illegible] |
| MXE | 2040 | ENSG00000137804 | NUSAP1 | chr15 | + | 41371526 | 41371684 | 41375711 | 41375828 | 41365401 | 41365589 | 41377195 | 41377304 | 9 | [illegible] |
| MXE | 16246 | ENSG00000107863 | ARHGAP21 | chr10 | - | 24629995 | 24630050 | 24633401 | 24633480 | 24619472 | 24621369 | 24635010 | 24635103 | 24 | [illegible] |
| MXE | 9087 | ENSG00000135336 | ORC3 | chr6 | + | 87612088 | 87612248 | 87616313 | 87616427 | 87609095 | 87609229 | 87621353 | 87621487 | 144 | [illegible] |
| MXE | 16998 | ENSG00000135439 | AGAP2 | chr12 | - | 57727680 | 57727771 | 57727936 | 57728085 | 57727359 | 57727582 | 57729638 | 57729767 | 117 | [illegible] |
| MXE | 10323 | ENSG00000155729 | KCTD18 | chr2 | - | 200504769 | 200504959 | 200506856 | 200507091 | 200499001 | 200499084 | 200510010 | 200510068 | 6 | [illegible] |
| MXE | 11120 | ENSG00000135976 | ANKRD36 | chr2 | + | 97144639 | 97144712 | 97146485 | 97146516 | 97144517 | 97144546 | 97149294 | 97149361 | 80 | [illegible] |
| MXE | 14269 | ENSG00000089234 | BRAP | chr12 | - | 111660599 | 111660675 | 111665638 | 111665787 | 111659206 | 111659345 | 111672660 | 111672774 | 56 | [illegible] |

| | | | | | | | | | | | | | | |
|---|---|---|---|---|---|---|---|---|---|---|---|---|---|---|
| MXE | 7405 | ENSG00000131374 | TBC1D5 | chr3 | - | 17586352 | 17586557 | 17623848 | 17623913 | 17576481 | 17576549 | 17742480 | 17742612 | 10 |
| MXE | 8388 | ENSG00000072110 | ACTN1 | chr14 | - | 68893654 | 68893747 | 68902476 | 68902562 | 68892052 | 68892283 | 68904654 | 68904736 | 127 |
| MXE | 2822 | ENSG00000080815 | PSEN1 | chr14 | + | 73206385 | 73206472 | 73211768 | 73211942 | 73192643 | 73192864 | 73217125 | 73217244 | 4 |
| MXE | 15611 | ENSG00000159063 | ALG8 | chr11 | - | 78109441 | 78109581 | 78112649 | 78112770 | 78106806 | 78106912 | 78113885 | 78113989 | 56 |
| MXE | 3216 | ENSG00000189091 | SF3B3 | chr16 | + | 70530744 | 70530917 | 70532478 | 70532620 | 70528872 | 70529199 | 70535307 | 70535420 | 407 |
| MXE | 3333 | ENSG00000259040 | BLOC1S5-TXNDC5 | chr6 | - | 7889494 | 7889581 | 7891620 | 7891736 | 7888704 | 7888848 | 7895105 | 7895202 | 175 |
| MXE | 10561 | ENSG00000168016 | TRANK1 | chr3 | - | 36834761 | 36834907 | 36838612 | 36838716 | 36830872 | 36833919 | 36842621 | 36842710 | 188 |
| MXE | 3533 | ENSG00000133466 | C1QTNF6 | chr22 | - | 37191671 | 37191737 | 37195380 | 37195512 | 37190864 | 37190928 | 37197602 | 37197841 | 2 |
| MXE | 3687 | ENSG00000204120 | GIGYF2 | chr2 | + | 232845731 | 232845886 | 232850261 | 232850409 | 232844368 | 232844574 | 232856792 | 232858608 | 71 |
| MXE | 10827 | ENSG00000169016 | E2F6 | chr2 | - | 11453581 | 11453798 | 11455341 | 11455474 | 11451650 | 11451806 | 11457178 | 11457233 | 5 |
| MXE | 13985 | ENSG00000064313 | TAF2 | chr8 | - | 119791323 | 119791459 | 119793365 | 119793451 | 119789591 | 119789746 | 119795531 | 119795631 | 53 |
| MXE | 12131 | ENSG00000124486 | USP9X | chrX | + | 41187991 | 41188117 | 41189308 | 41189475 | 41186516 | 41186642 | 41196250 | 41196359 | 83 |
| MXE | 651 | ENSG00000141968 | VAV1 | chr19 | + | 6825302 | 6825406 | 6826611 | 6826711 | 6825052 | 6825121 | 6828075 | 6828171 | 166 |
| MXE | 5766 | ENSG00000143811 | PYCR2 | chr1 | - | 225921857 | 225922079 | 225922203 | 225922383 | 225921551 | 225921644 | 225923700 | 225923771 | 38 |
| MXE | 7560 | ENSG00000111331 | OAS3 | chr12 | + | 112941569 | 112941852 | 112944475 | 112944651 | 112938473 | 112938707 | 112946742 | 112946981 | 19 |
| MXE | 5754 | ENSG00000197386 | HTT | chr4 | + | 3086938 | 3087022 | 3099273 | 3099394 | 3074680 | 3075088 | 3103823 | 3103883 | 0 |
| MXE | 1482 | ENSG00000277791 | PSMB3 | chr17 | + | 38760430 | 38760608 | 38762410 | 38762505 | 38755882 | 38755990 | 38764118 | 38764225 | 282 |
| MXE | 13463 | ENSG00000196776 | CD47 | chr3 | - | 108071092 | 108071182 | 108079990 | 108080344 | 108060744 | 108060852 | 108090862 | 108091025 | 112 |
| MXE | 12446 | ENSG00000164535 | DAGLB | chr7 | - | 6425987 | 6426114 | 6430479 | 6430607 | 6424751 | 6424835 | 6432836 | 6432959 | 41 |
| MXE | 7901 | ENSG00000071054 | MAP4K4 | chr2 | + | 101835899 | 101835978 | 101839818 | 101839994 | 101834408 | 101834463 | 101842608 | 101842681 | 185 |
| MXE | 13949 | ENSG00000100403 | ZC3H7B | chr22 | + | 41325563 | 41325597 | 41327205 | 41327364 | 41320654 | 41320713 | 41330022 | 41330103 | 26 |
| MXE | 4162 | ENSG00000177058 | SLC38A9 | chr5 | - | 55649206 | 55649314 | 55652528 | 55652723 | 55645788 | 55645895 | 55656714 | 55656774 | 36 |
| MXE | 13496 | ENSG00000169359 | SLC33A1 | chr3 | - | 155829687 | 155829903 | 155842431 | 155842619 | 155821023 | 155828377 | 155853222 | 155854427 | 147 |
| MXE | 9569 | ENSG00000076242 | MLH1 | chr3 | + | 37014431 | 37014544 | 37017505 | 37017599 | 37012010 | 37012099 | 37020309 | 37020463 | 85 |
| MXE | 15753 | ENSG00000149311 | ATM | chr11 | + | 108331443 | 108331557 | 108331878 | 108332037 | 108330213 | 108330421 | 108332761 | 108332900 | 656 |
| MXE | 6437 | ENSG00000163946 | TASOR | chr3 | - | 56663540 | 56663572 | 56666259 | 56666384 | 56660968 | 56661017 | 56668396 | 56668489 | 138 |
| MXE | 13585 | ENSG00000147140 | NONO | chrX | + | 71294226 | 71294348 | 71296564 | 71296660 | 71291778 | 71291972 | 71296850 | 71297047 | 402 |
| MXE | 1844 | ENSG00000177663 | IL17RA | chr22 | + | 17105590 | 17105602 | 17105852 | 17105954 | 17104725 | 17104810 | 17107726 | 17107768 | 60 |
| MXE | 12449 | ENSG00000164535 | DAGLB | chr7 | - | 6434761 | 6435020 | 6436361 | 6436533 | 6432836 | 6432959 | 6445952 | 6446104 | 10 |
| MXE | 5641 | ENSG00000149480 | MTA2 | chr11 | - | 62598521 | 62598639 | 62600165 | 62600259 | 62598108 | 62598390 | 62600621 | 62600689 | 218 |
| MXE | 965 | ENSG00000134046 | MBD2 | chr18 | - | 54166075 | 54166166 | 54187713 | 54187837 | 54164522 | 54164700 | 54188873 | 54189011 | 12 |
| MXE | 12723 | ENSG00000071462 | BUD23 | chr7 | + | 73693623 | 73693669 | 73693991 | 73694050 | 73693328 | 73693414 | 73697604 | 73697694 | 46 |
| MXE | 9887 | ENSG00000219626 | FAM228B | chr2 | + | 24137908 | 24138100 | 24139369 | 24139450 | 24135118 | 24135187 | 24146747 | 24146835 | 21 |
| MXE | 4551 | ENSG00000055955 | ITIH4 | chr3 | - | 52819753 | 52819807 | 52819939 | 52819990 | 52819392 | 52819518 | 52820290 | 52820317 | 12 |
| MXE | 2614 | ENSG00000075413 | MARK3 | chr14 | + | 103448918 | 103448967 | 103451917 | 103451983 | 103428386 | 103428440 | 103462404 | 103462461 | 57 |
| MXE | 14617 | ENSG00000106771 | TMEM245 | chr9 | - | 109080838 | 109080943 | 109090921 | 109091155 | 109073247 | 109073438 | 109093474 | 109093591 | 38 |
| MXE | 13187 | ENSG00000068354 | TBC1D25 | chrX | + | 48544868 | 48545023 | 48558896 | 48559024 | 48541332 | 48541442 | 48559157 | 48559346 | 40 |
| MXE | 3052 | ENSG00000152969 | JAKMIP1 | chr4 | - | 6085419 | 6085629 | 6105472 | 6105967 | 6084845 | 6084965 | 6112721 | 6112997 | 16 |
| MXE | 549 | ENSG00000157540 | DYRK1A | chr21 | + | 37421773 | 37421866 | 37425689 | 37425781 | 37420311 | 37420384 | 37428866 | 37428908 | 16 |
| MXE | 8065 | ENSG00000103657 | HERC1 | chr15 | - | 63635960 | 63636142 | 63637504 | 63637643 | 63634732 | 63634888 | 63638410 | 63638536 | 96 |
| MXE | 2577 | ENSG00000092108 | SCFD1 | chr14 | + | 30633946 | 30634037 | 30639776 | 30639864 | 30630476 | 30630565 | 30643315 | 30643405 | 53 |
| MXE | 13688 | ENSG00000215301 | DDX3X | chrX | + | 41346858 | 41347012 | 41347314 | 41347454 | 41346504 | 41346622 | 41347639 | 41349884 | 1022 |
| MXE | 15258 | ENSG00000122218 | COPA | chr1 | - | 160332447 | 160332557 | 160333602 | 160333679 | 160325542 | 160325652 | 160335241 | 160335322 | 243 |
| MXE | 10887 | ENSG00000085365 | SCAMP1 | chr5 | + | 78421800 | 78421960 | 78459244 | 78459362 | 78418774 | 78418903 | 78475503 | 78476026 | 73 |
| MXE | 648 | ENSG00000142347 | MYO1F | chr19 | - | 8550159 | 8550356 | 8550561 | 8550694 | 8548236 | 8548317 | 8551739 | 8551874 | 189 |
| MXE | 15052 | ENSG00000184060 | ADAP2 | chr17 | + | 30926826 | 30926918 | 30931888 | 30931968 | 30922939 | 30923070 | 30934184 | 30934297 | 12 |
| MXE | 10181 | ENSG00000146463 | ZMYM4 | chr1 | + | 35387453 | 35387604 | 35388909 | 35389082 | 35387002 | 35387278 | 35389947 | 35390098 | 107 |
| MXE | 3618 | ENSG00000113742 | CPEB4 | chr5 | + | 173932449 | 173932500 | 173943025 | 173943049 | 173910522 | 173910604 | 173944966 | 173945140 | 42 |
| MXE | 12696 | ENSG00000129103 | SUMF2 | chr7 | + | 56072996 | 56073111 | 56074173 | 56074218 | 56068481 | 56068638 | 56076833 | 56076889 | 115 |
| MXE | 7410 | ENSG00000143401 | ANP32E | chr1 | - | 150229071 | 150229237 | 150230570 | 150230693 | 150226607 | 150226795 | 150231776 | 150231926 | 447 |
| MXE | 646 | ENSG00000142347 | MYO1F | chr19 | - | 8544296 | 8544464 | 8545649 | 8545736 | 8541727 | 8541991 | 8548035 | 8548043 | 382 |
| MXE | 10159 | ENSG00000143924 | EML4 | chr2 | + | 42303314 | 42303446 | 42304483 | 42304551 | 42303103 | 42303229 | 42315961 | 42316050 | 209 |
| MXE | 8914 | ENSG00000065413 | ANKRD44 | chr2 | - | 197005693 | 197005910 | 197007805 | 197007923 | 197001752 | 197001840 | 197008943 | 197009031 | 413 |
| MXE | 15950 | ENSG00000163249 | CCNYL1 | chr2 | + | 207733946 | 207734047 | 207737410 | 207737446 | 207726841 | 207726876 | 207740654 | 207740706 | 40 |
| MXE | 5530 | ENSG00000147996 | CBWD5 | chr9 | + | 65690555 | 65690601 | 65690966 | 65691026 | 65682765 | 65682857 | 65692570 | 65692608 | 6 |
| MXE | 2188 | ENSG00000129003 | VPS13C | chr15 | - | 61982458 | 61982573 | 61983819 | 61984012 | 61981341 | 61981478 | 61984856 | 61984999 | 418 |
| MXE | 6294 | ENSG00000106780 | MEGF9 | chr9 | - | 120607740 | 120608010 | 120612395 | 120612539 | 120600810 | 120605641 | 120622615 | 120622755 | 407 |
| MXE | 8222 | ENSG00000124126 | PREX1 | chr20 | - | 48708259 | 48708421 | 48726289 | 48726391 | 48700752 | 48700886 | 48734545 | 48734650 | 196 |
| MXE | 748 | ENSG00000101150 | TPD52L2 | chr20 | + | 63875815 | 63875875 | 63882718 | 63882820 | 63873667 | 63873816 | 63889189 | 63889238 | 30 |
| MXE | 17070 | ENSG00000105352 | CEACAM4 | chr19 | - | 41621650 | 41621768 | 41625600 | 41625960 | 41620574 | 41620627 | 41626899 | 41627074 | 14 |
| MXE | 12005 | ENSG00000160799 | CCDC12 | chr3 | - | 46923328 | 46923363 | 46923606 | 46923668 | 46922235 | 46922312 | 46925455 | 46925535 | 46 |
| MXE | 15739 | ENSG00000149311 | ATM | chr11 | + | 108243952 | 108244118 | 108244787 | 108245026 | 108235669 | 108235834 | 108246963 | 108247127 | 212 |
| MXE | 934 | ENSG00000100983 | GSS | chr20 | - | 34935575 | 34935642 | 34936762 | 34936840 | 34931938 | 34932133 | 34936942 | 34937023 | 24 |
| MXE | 15875 | ENSG00000136935 | GOLGA1 | chr9 | - | 124928187 | 124928285 | 124929215 | 124929290 | 124926708 | 124926741 | 124931315 | 124931406 | 52 |
| MXE | 3985 | ENSG00000006459 | KDM7A | chr7 | - | 140133538 | 140133656 | 140139104 | 140139190 | 140129492 | 140129653 | 140176743 | 140176983 | 4 |
| MXE | 16454 | ENSG00000129315 | CCNT1 | chr12 | - | 48699777 | 48699840 | 48701012 | 48701073 | 48698137 | 48698183 | 48705767 | 48705896 | 7 |
| MXE | 3287 | ENSG00000137177 | KIF13A | chr6 | - | 17808767 | 17808930 | 17817019 | 17817233 | 17805474 | 17805615 | 17825767 | 17825934 | 47 |
| MXE | 5973 | ENSG00000150753 | CCT5 | chr5 | + | 10262480 | 10262618 | 10263133 | 10263314 | 10261559 | 10261745 | 10264655 | 10264902 | 227 |
| MXE | 3811 | ENSG00000113648 | MACROH2A1 | chr5 | - | 135350822 | 135350913 | 135352945 | 135353045 | 135345967 | 135346057 | 135360496 | 135360607 | 379 |
| MXE | 6966 | ENSG00000170248 | PDCD6IP | chr3 | + | 33826479 | 33826580 | 33828837 | 33828969 | 33825186 | 33825340 | 33836043 | 33836266 | 122 |
| MXE | 13922 | ENSG00000171988 | JMJD1C | chr10 | - | 63215552 | 63215696 | 63217206 | 63217331 | 63215262 | 63215455 | 63268662 | 63269187 | 4 |
| MXE | 8849 | ENSG00000110200 | ANAPC15 | chr11 | - | 72111156 | 72111281 | 72111411 | 72111496 | 72110543 | 72110603 | 72112649 | 72112766 | 37 |
| MXE | 6334 | ENSG00000163319 | MRPS18C | chr4 | + | 83458345 | 83458429 | 83459739 | 83459797 | 83456908 | 83456958 | 83460972 | 83461032 | 100 |
| MXE | 15642 | ENSG00000172183 | ISG20 | chr15 | + | 88639342 | 88639594 | 88652109 | 88652310 | 88636152 | 88636292 | 88655414 | 88655621 | 83 |
| MXE | 7154 | ENSG00000095787 | WAC | chr10 | + | 28595732 | 28596041 | 28608185 | 28608431 | 28590719 | 28590832 | 28610698 | 28610821 | 571 |
| MXE | 7225 | ENSG00000008869 | HEATR5B | chr2 | - | 36988645 | 36988859 | 36990647 | 36990799 | 36980894 | 36981794 | 37000585 | 37000813 | 58 |
| MXE | 13231 | ENSG00000188917 | TRMT2B | chrX | - | 101041316 | 101041371 | 101042041 | 101042312 | 101036973 | 101037073 | 101051643 | 101051733 | 3 |
| MXE | 13645 | ENSG00000086758 | HUWE1 | chrX | - | 53558654 | 53558809 | 53558970 | 53559060 | 53557381 | 53557427 | 53559353 | 53559532 | 75 |
| MXE | 16253 | ENSG00000047056 | WDR37 | chr10 | + | 1103601 | 1103836 | 1105125 | 1105267 | 1096169 | 1096246 | 1124217 | 1124352 | 14 |
| MXE | 4761 | ENSG00000204463 | BAG6 | chr6 | - | 31644306 | 31644414 | 31644524 | 31644602 | 31644081 | 31644194 | 31644945 | 31645198 | 116 |
| MXE | 7312 | ENSG00000117543 | DPH5 | chr1 | - | 101001466 | 101001544 | 101021640 | 101021765 | 100995109 | 100995149 | 101025682 | 101025815 | 9 |
| MXE | 10238 | ENSG00000087152 | ATXN7L3 | chr17 | - | 44194767 | 44194839 | 44195096 | 44195140 | 44194516 | 44194674 | 44195418 | 44195442 | 89 |
| MXE | 13234 | ENSG00000188917 | TRMT2B | chrX | - | 101041316 | 101041371 | 101051250 | 101051733 | 101037916 | 101038051 | 101051816 | 101052111 | 21 |
| MXE | 1903 | ENSG00000264772 | NA | chr17 | + | 7580465 | 7580585 | 7580710 | 7580783 | 7577568 | 7580327 | 7580895 | 7581066 | 360 |
| MXE | 9610 | ENSG00000117625 | RCOR3 | chr1 | + | 211295675 | 211295753 | 211304082 | 211304140 | 211289331 | 211289396 | 211312354 | 211312487 | 49 |
| MXE | 382 | ENSG00000239382 | ALKBH6 | chr19 | - | 36011403 | 36011464 | 36013000 | 36013089 | 36010893 | 36011045 | 36013343 | 36013422 | 11 |
| MXE | 1674 | ENSG00000103222 | ABCC1 | chr16 | + | 16079351 | 16079478 | 16083365 | 16083542 | 16076325 | 16076401 | 16090404 | 16090588 | 47 |
| MXE | 7947 | ENSG00000162511 | LAPTM5 | chr1 | - | 30739808 | 30739937 | 30741639 | 30741716 | 30738939 | 30739062 | 30742455 | 30742549 | 714 |
| MXE | 8468 | ENSG00000149474 | KAT14 | chr20 | + | 18150820 | 18150942 | 18159083 | 18159265 | 18145232 | 18145351 | 18161819 | 18162239 | 31 |
| MXE | 11848 | ENSG00000164674 | SYTL3 | chr6 | + | 158762075 | 158762178 | 158763303 | 158763509 | 158760639 | 158760745 | 158764494 | 158764869 | 166 |
| MXE | 16275 | ENSG00000066084 | DIP2B | chr12 | + | 50731368 | 50731537 | 50732365 | 50732536 | 50728547 | 50728678 | 50734134 | 50734196 | 121 |
| MXE | 3239 | ENSG00000012660 | ELOVL5 | chr6 | - | 53287861 | 53287942 | 53290400 | 53290535 | 53276178 | 53276256 | 53291775 | 53291963 | 4 |
| MXE | 13427 | ENSG00000119760 | SUPT7L | chr2 | - | 27657344 | 27657669 | 27660983 | 27661388 | 27655364 | 27655602 | 27662178 | 27662281 | 48 |
| MXE | 7019 | ENSG00000162851 | TFB2M | chr1 | - | 246548544 | 246548607 | 246551212 | 246551302 | 246544520 | 246544681 | 246556572 | 246556721 | 33 |
| MXE | 3743 | ENSG00000164308 | ERAP2 | chr5 | + | 96883791 | 96883930 | 96886654 | 96886789 | 96879563 | 96880260 | 96889184 | 96889305 | 238 |
| MXE | 11437 | ENSG00000151687 | ANKAR | chr2 | + | 189692254 | 189692418 | 189694980 | 189695161 | 189689526 | 189689964 | 189696149 | 189696369 | 7 |
| MXE | 16996 | ENSG00000111300 | NAA25 | chr12 | - | 112060269 | 112060359 | 112061180 | 112061388 | 112054387 | 112054568 | 112068879 | 112068992 | 23 |
| MXE | 5101 | ENSG00000060971 | ACAA1 | chr3 | - | 38127785 | 38127866 | 38129289 | 38129388 | 38126509 | 38126700 | 38131595 | 38131638 | 48 |
| MXE | 7802 | ENSG00000066427 | ATXN3 | chr14 | - | 92088729 | 92088817 | 92093251 | 92093318 | 92083125 | 92083258 | 92096673 | 92096838 | 0 |
| MXE | 9157 | ENSG00000168118 | RAB4A | chr1 | + | 229288728 | 229288843 | 229295847 | 229295910 | 229286485 | 229286566 | 229297481 | 229297636 | 71 |
| MXE | 44 | ENSG00000160294 | MCM3AP | chr21 | - | 46254391 | 46254526 | 46254775 | 46254844 | 46251514 | 46251682 | 46256788 | 46256986 | 14 |
| MXE | 6265 | ENSG00000176890 | TYMS | chr18 | + | 669071 | 669173 | 670691 | 670867 | 662145 | 662320 | 671379 | 671451 | 6 |
| MXE | 11217 | ENSG00000006062 | MAP3K14 | chr17 | - | 45287153 | 45287364 | 45289235 | 45289305 | 45286430 | 45287045 | 45290489 | 45290765 | 40 |
| MXE | 16931 | ENSG00000111540 | RAB5B | chr12 | + | 55989946 | 55990098 | 55990681 | 55990804 | 55986868 | 55987123 | 55991359 | 55991453 | 152 |
| MXE | 17187 | ENSG00000005810 | MYCBP2 | chr13 | - | 77164453 | 77164541 | 77165272 | 77165391 | 77161905 | 77161955 | 77166328 | 77166554 | 165 |
| MXE | 6599 | ENSG00000058673 | ZC3H11A | chr1 | + | 203796292 | 203796496 | 203801574 | 203801628 | 203795698 | 203795775 | 203802956 | 203803016 | 4 |
| MXE | 8826 | ENSG00000118007 | STAG1 | chr3 | - | 136359147 | 136359296 | 136363365 | 136363467 | 136357719 | 136357848 | 136366942 | 136367082 | 73 |
| MXE | 3553 | ENSG00000113716 | HMGXB3 | chr5 | + | 150006472 | 150006647 | 150010110 | 150010608 | 150004850 | 150004989 | 150012254 | 150012353 | 49 |
| MXE | 13339 | ENSG00000136504 | KAT7 | chr17 | + | 49796749 | 49796926 | 49798318 | 49798558 | 49791885 | 49792033 | 49805362 | 49805445 | 195 |
| MXE | 10813 | ENSG00000283761 | NA | chr1 | + | 100015301 | 100015420 | 100043072 | 100043229 | 100011364 | 100011533 | 100049908 | 100050004 | 79 |
| MXE | 4498 | ENSG00000185305 | ARL15 | chr5 | - | 54154579 | 54154639 | 54171783 | 54171928 | 54113201 | 54113410 | 54310294 | 54310415 | 5 |
| MXE | 17142 | ENSG00000102710 | SUPT20H | chr13 | - | 37047877 | 37047936 | 37048563 | 37048599 | 37047534 | 37047601 | 37051487 | 37051583 | 14 |
| MXE | 3498 | ENSG00000153113 | CAST | chr5 | + | 96727488 | 96727530 | 96729152 | 96729209 | 96722638 | 96722698 | 96729611 | 96729725 | 146 |
| MXE | 2694 | ENSG00000129566 | TEP1 | chr14 | - | 20396620 | 20396730 | 20400983 | 20401141 | 20395858 | 20395949 | 20401456 | 20401581 | 71 |
| MXE | 14663 | ENSG00000136828 | RALGPS1 | chr9 | + | 127196473 | 127196631 | 127210694 | 127210778 | 127195090 | 127195217 | 127212130 | 127212236 | 12 |
| MXE | 15924 | ENSG00000181192 | DHTKD1 | chr10 | + | 12094072 | 12094271 | 12097683 | 12097996 | 12091512 | 12091684 | 12100177 | 12100262 | 50 |
| MXE | 8779 | ENSG00000134313 | KIDINS220 | chr2 | - | 8736867 | 8736999 | 8747886 | 8748000 | 8734654 | 8734753 | 8750111 | 8750335 | 98 |
| MXE | 11556 | ENSG00000128284 | APOL3 | chr22 | - | 36149770 | 36149964 | 36153284 | 36153423 | 36149258 | 36149371 | 36160668 | 36160755 | 2 |
| MXE | 593 | ENSG00000105186 | ANKRD27 | chr19 | - | 32644324 | 32644479 | 32646458 | 32646615 | 32643571 | 32643631 | 32649681 | 32649792 | 81 |
| MXE | 5638 | ENSG00000151726 | ACSL1 | chr4 | - | 184770254 | 184770332 | 184770398 | 184770476 | 184768315 | 184768450 | 184773080 | 184773154 | 154 |
| MXE | 8376 | ENSG00000152223 | EPG5 | chr18 | - | 45870566 | 45870742 | 45876235 | 45876342 | 45867562 | 45867748 | 45878375 | 45878448 | 41 |
| MXE | 14961 | ENSG00000165140 | FBP1 | chr9 | - | 94606814 | 94606952 | 94617767 | 94617860 | 94605456 | 94605576 | 94620328 | 94620491 | 78 |
| MXE | 1034 | ENSG00000132388 | UBE2G1 | chr17 | - | 4289229 | 4289408 | 4296716 | 4296814 | 4282797 | 4282921 | 4307020 | 4307123 | 82 |
| MXE | 6967 | ENSG00000170248 | PDCD6IP | chr3 | + | 33826479 | 33826580 | 33828852 | 33828969 | 33825186 | 33825340 | 33836043 | 33836266 | 122 |
| MXE | 5395 | ENSG00000162736 | NCSTN | chr1 | + | 160355862 | 160355958 | 160356259 | 160356347 | 160355654 | 160355757 | 160356599 | 160356754 | 83 |
| MXE | 16962 | ENSG00000121749 | TBC1D15 | chr12 | + | 71872928 | 71873003 | 71880468 | 71880607 | 71872069 | 71872168 | 71884810 | 71885021 | 253 |
| MXE | 6650 | ENSG00000114626 | ABTB1 | chr3 | + | 127677464 | 127677563 | 127677675 | 127677843 | 127677167 | 127677286 | 127679984 | 127680185 | 43 |
| MXE | 11830 | ENSG00000065427 | KARS1 | chr16 | - | 75640183 | 75640349 | 75641563 | 75641723 | 75636453 | 75636547 | 75644282 | 75644462 | 36 |
| MXE | 3286 | ENSG00000137177 | KIF13A | chr6 | - | 17789871 | 17789910 | 17794248 | 17794395 | 17787775 | 17787875 | 17794571 | 17794704 | 84 |
| MXE | 2866 | ENSG00000165516 | KLHDC2 | chr14 | + | 49780212 | 49780322 | 49780702 | 49780775 | 49779747 | 49779806 | 49782369 | 49782457 | 99 |
| MXE | 15413 | ENSG00000087365 | SF3B2 | chr11 | + | 66058313 | 66058405 | 66058829 | 66059045 | 66058053 | 66058150 | 66059200 | 66059338 | 124 |
| MXE | 12569 | ENSG00000086619 | ERO1B | chr1 | - | 236243421 | 236243495 | 236249884 | 236249967 | 236236277 | 236236398 | 236252049 | 236252091 | 69 |
| MXE | 7109 | ENSG00000197780 | TAF13 | chr1 | - | 109066134 | 109066232 | 109074986 | 109075065 | 109064139 | 109064693 | 109075920 | 109076003 | 1 |
| MXE | 16490 | ENSG00000172819 | RARG | chr12 | - | 53215645 | 53215794 | 53227361 | 53227687 | 53215292 | 53215434 | 53231168 | 53231235 | 6 |
| MXE | 11746 | ENSG00000107672 | NSMCE4A | chr10 | - | 121963237 | 121963328 | 121965285 | 121965385 | 121961422 | 121961517 | 121967654 | 121967806 | 78 |
| MXE | 17009 | ENSG00000197111 | PCBP2 | chr12 | + | 53468776 | 53468832 | 53471637 | 53471807 | 53467220 | 53467293 | 53479405 | 53479824 | 77 |
| MXE | 6441 | ENSG00000163946 | TASOR | chr3 | - | 56670072 | 56670145 | 56671599 | 56671692 | 56669699 | 56669791 | 56673579 | 56673725 | 454 |
| MXE | 5543 | ENSG00000169180 | XPO6 | chr16 | - | 28177219 | 28177332 | 28180940 | 28181031 | 28175897 | 28176095 | 28208986 | 28209122 | 16 |
| MXE | 7389 | ENSG00000150681 | RGS18 | chr1 | + | 192159219 | 192159321 | 192160377 | 192160439 | 192158461 | 192158756 | 192181291 | 192181458 | 488 |
| MXE | 11688 | ENSG00000186063 | AIDA | chr1 | - | 222686929 | 222687036 | 222687594 | 222687658 | 222676095 | 222676218 | 222693788 | 222693843 | 104 |
| MXE | 4713 | ENSG00000174227 | PIGG | chr4 | + | 508828 | 508970 | 515972 | 516185 | 507404 | 507593 | 521055 | 521227 | 90 |
| MXE | 15099 | ENSG00000266173 | STRADA | chr17 | - | 63714005 | 63714108 | 63726637 | 63726695 | 63713405 | 63713527 | 63728333 | 63728413 | 4 |
| MXE | 13777 | ENSG00000165675 | ENOX2 | chrX | - | 130679541 | 130679748 | 130688862 | 130689018 | 130669964 | 130670198 | 130703119 | 130703254 | 29 |
| MXE | 2833 | ENSG00000029364 | SLC39A9 | chr14 | + | 69442068 | 69442266 | 69453240 | 69453309 | 69424093 | 69424202 | 69454811 | 69454897 | 122 |
| MXE | 12626 | ENSG00000067596 | DHX8 | chr17 | + | 43508338 | 43508520 | 43513361 | 43513502 | 43507808 | 43508019 | 43517166 | 43517322 | 128 |
| MXE | 296 | ENSG00000100077 | GRK3 | chr22 | + | 25672295 | 25672347 | 25674436 | 25674528 | 25667738 | 25667800 | 25678815 | 25678915 | 69 |
| MXE | 9151 | ENSG00000113615 | SEC24A | chr5 | + | 134708712 | 134708888 | 134715023 | 134715161 | 134705326 | 134705437 | 134718068 | 134718173 | 48 |
| MXE | 16498 | ENSG00000040531 | CTNS | chr17 | + | 3640187 | 3640267 | 3647443 | 3647522 | 3637106 | 3637316 | 3654997 | 3655101 | 27 |
| MXE | 7943 | ENSG00000142599 | RERE | chr1 | - | 8358195 | 8358916 | 8361459 | 8361490 | 8356099 | 8356246 | 8361762 | 8361876 | 20 |
| MXE | 1211 | ENSG00000007202 | KIAA0100 | chr17 | - | 28641888 | 28642078 | 28642274 | 28642335 | 28640559 | 28640683 | 28642868 | 28642987 | 82 |
| MXE | 12161 | ENSG00000169919 | GUSB | chr7 | - | 65976014 | 65976202 | 65979833 | 65979911 | 65974918 | 65975071 | 65980223 | 65980409 | 20 |
| MXE | 9321 | ENSG00000165071 | TMEM71 | chr8 | - | 132757233 | 132757294 | 132758839 | 132758915 | 132751784 | 132751997 | 132760475 | 132760630 | 232 |
| MXE | 16618 | ENSG00000089094 | KDM2B | chr12 | - | 121442991 | 121443030 | 121443679 | 121443793 | 121442156 | 121442836 | 121444011 | 121444272 | 99 |
| MXE | 7620 | ENSG00000132300 | PTCD3 | chr2 | + | 86118920 | 86119044 | 86123700 | 86123762 | 86117054 | 86117159 | 86124994 | 86125082 | 201 |
| MXE | 11892 | ENSG00000135317 | SNX14 | chr6 | - | 85530191 | 85530275 | 85533598 | 85533800 | 85528261 | 85528362 | 85536791 | 85536924 | 225 |
| MXE | 2868 | ENSG00000152240 | HAUS1 | chr18 | + | 46118180 | 46118316 | 46119925 | 46120060 | 46105193 | 46105368 | 46122466 | 46122590 | 31 |
| MXE | 890 | ENSG00000124155 | PIGT | chr20 | + | 45416516 | 45416694 | 45418851 | 45418979 | 45416140 | 45416343 | 45419294 | 45419395 | 98 |
| MXE | 4487 | ENSG00000197142 | ACSL5 | chr10 | + | 112416887 | 112417022 | 112417845 | 112417941 | 112413172 | 112413307 | 112421592 | 112421626 | 158 |
| MXE | 15297 | ENSG00000197798 | FAM118B | chr11 | + | 126234994 | 126235087 | 126240791 | 126241044 | 126229224 | 126229293 | 126250505 | 126250733 | 21 |
| MXE | 15808 | ENSG00000166224 | SGPL1 | chr10 | + | 70871047 | 70871146 | 70873350 | 70873589 | 70869791 | 70869897 | 70875401 | 70875548 | 72 |
| MXE | 11667 | ENSG00000024048 | UBR2 | chr6 | + | 42573733 | 42573993 | 42592150 | 42592229 | 42564028 | 42564397 | 42594190 | 42594304 | 118 |
| MXE | 187 | ENSG00000100150 | DEPDC5 | chr22 | + | 31760655 | 31760702 | 31764974 | 31765060 | 31758526 | 31758633 | 31766584 | 31766668 | 22 |
| MXE | 8839 | ENSG00000173846 | PLK3 | chr1 | + | 44804162 | 44804246 | 44804338 | 44804501 | 44803930 | 44804024 | 44805265 | 44805379 | 29 |
| MXE | 1053 | ENSG00000109118 | PHF12 | chr17 | - | 28923908 | 28924302 | 28926990 | 28927063 | 28921687 | 28921808 | 28950064 | 28950246 | 75 |
| MXE | 13299 | ENSG00000164074 | ABHD18 | chr4 | + | 128008919 | 128008998 | 128017362 | 128017501 | 128001715 | 128001760 | 128020079 | 128020169 | 5 |
| MXE | 10715 | ENSG00000168958 | MFF | chr2 | + | 227342744 | 227342819 | 227352513 | 227352573 | 227340291 | 227340380 | 227355676 | 227355761 | 41 |
| MXE | 2370 | ENSG00000137841 | PLCB2 | chr15 | - | 40298797 | 40298964 | 40299127 | 40299228 | 40298561 | 40298708 | 40301956 | 40302032 | 169 |
| MXE | 10169 | ENSG00000102900 | NUP93 | chr16 | + | 56758537 | 56758655 | 56798475 | 56798538 | 56748233 | 56748426 | 56805503 | 56805552 | 38 |
| MXE | 11222 | ENSG00000138433 | CIR1 | chr2 | - | 174351618 | 174351725 | 174378922 | 174379040 | 174350659 | 174350745 | 174380201 | 174380250 | 190 |
| MXE | 1517 | ENSG00000091592 | NLRP1 | chr17 | - | 5533896 | 5533988 | 5536850 | 5536940 | 5533303 | 5533384 | 5539414 | 5539585 | 31 |
| MXE | 14888 | ENSG00000148300 | REXO4 | chr9 | - | 133407806 | 133407881 | 133408767 | 133408842 | 133406058 | 133407072 | 133410984 | 133411073 | 34 |
| MXE | 3819 | ENSG00000217128 | FNIP1 | chr5 | - | 131677702 | 131677872 | 131679028 | 131679175 | 131671504 | 131672924 | 131698916 | 131699002 | 99 |
| MXE | 9758 | ENSG00000138050 | THUMPD2 | chr2 | - | 39768900 | 39769093 | 39769709 | 39770119 | 39768423 | 39768501 | 39771504 | 39771640 | 30 |
| MXE | 8220 | ENSG00000124126 | PREX1 | chr20 | - | 48649995 | 48650206 | 48650893 | 48651055 | 48649299 | 48649576 | 48651395 | 48651583 | 364 |
| MXE | 2888 | ENSG00000198663 | C6orf89 | chr6 | + | 36914553 | 36914693 | 36916444 | 36916574 | 36914283 | 36914435 | 36919577 | 36919701 | 67 |
| MXE | 2659 | ENSG00000032219 | ARID4A | chr14 | + | 58344694 | 58344767 | 58346410 | 58346505 | 58330002 | 58330169 | 58347019 | 58347117 | 51 |
| MXE | 5670 | ENSG00000166927 | MS4A7 | chr11 | + | 60386716 | 60386773 | 60392684 | 60392786 | 60383128 | 60383288 | 60393786 | 60394522 | 56 |
| MXE | 12075 | ENSG00000134250 | NOTCH2 | chr1 | - | 119967432 | 119967621 | 119969510 | 119969744 | 119966375 | 119966489 | 119986959 | 119987082 | 182 |
| MXE | 17014 | ENSG00000067167 | TRAM1 | chr8 | - | 70594505 | 70594590 | 70597894 | 70598011 | 70587105 | 70587176 | 70600018 | 70600082 | 11 |
| MXE | 16506 | ENSG00000179889 | PDXDC1 | chr16 | + | 15004186 | 15004333 | 15006393 | 15006583 | 14998339 | 14998405 | 15008778 | 15008847 | 8 |
| MXE | 5283 | ENSG00000107758 | PPP3CB | chr10 | - | 73439871 | 73439901 | 73444724 | 73444822 | 73436805 | 73438420 | 73446491 | 73446576 | 160 |
| MXE | 8137 | ENSG00000134198 | TSPAN2 | chr1 | - | 115062134 | 115062232 | 115072904 | 115073007 | 115060463 | 115060538 | 115089363 | 115089414 | 13 |
| MXE | 10900 | ENSG00000133059 | DSTYK | chr1 | - | 205159546 | 205159679 | 205160113 | 205160270 | 205157272 | 205157386 | 205161257 | 205161387 | 33 |
| MXE | 9621 | ENSG00000119778 | ATAD2B | chr2 | - | 23857414 | 23857503 | 23863380 | 23863555 | 23856008 | 23856442 | 23864808 | 23864924 | 96 |
| MXE | 11664 | ENSG00000175575 | PAAF1 | chr11 | + | 73900310 | 73900420 | 73909398 | 73909593 | 73899145 | 73899244 | 73914412 | 73914504 | 1 |
| MXE | 12468 | ENSG00000085788 | DDHD2 | chr8 | + | 38245741 | 38245950 | 38246232 | 38246300 | 38241964 | 38242385 | 38247712 | 38247835 | 109 |
| MXE | 7692 | ENSG00000142784 | WDTC1 | chr1 | + | 27282238 | 27282285 | 27283337 | 27283449 | 27263151 | 27263235 | 27287673 | 27287861 | 213 |
| MXE | 3859 | ENSG00000157107 | FCHO2 | chr5 | + | 73006444 | 73006549 | 73015625 | 73015724 | 72990711 | 72990864 | 73017211 | 73017308 | 43 |
| MXE | 6468 | ENSG00000088854 | C20orf194 | chr20 | - | 3348524 | 3348623 | 3359515 | 3359593 | 3343675 | 3343731 | 3374047 | 3374154 | 13 |
| MXE | 1327 | ENSG00000141298 | SSH2 | chr17 | - | 29684562 | 29684684 | 29695458 | 29695523 | 29677672 | 29677741 | 29702958 | 29703062 | 78 |
| MXE | 244 | ENSG00000100055 | CYTH4 | chr22 | + | 37310987 | 37311064 | 37311455 | 37311527 | 37309211 | 37309323 | 37312019 | 37312174 | 84 |
| MXE | 9495 | ENSG00000102786 | INTS6 | chr13 | - | 51395299 | 51395483 | 51423029 | 51423090 | 51389318 | 51389444 | 51430293 | 51430383 | 16 |
| MXE | 16448 | ENSG00000101290 | CDS2 | chr20 | + | 5184857 | 5184945 | 5186686 | 5186839 | 5183060 | 5183143 | 5189066 | 5189186 | 122 |
| MXE | 6901 | ENSG00000117335 | CD46 | chr1 | + | 207767012 | 207767195 | 207767778 | 207767823 | 207759638 | 207759724 | 207770320 | 207770362 | 60 |
| MXE | 13684 | ENSG00000215301 | DDX3X | chrX | + | 41344029 | 41344128 | 41344238 | 41344399 | 41343736 | 41343822 | 41345179 | 41345324 | 321 |
| MXE | 5149 | ENSG00000108219 | TSPAN14 | chr10 | + | 80512143 | 80512269 | 80514018 | 80514063 | 80509300 | 80509471 | 80516203 | 80516323 | 150 |
| MXE | 7284 | ENSG00000169398 | PTK2 | chr8 | - | 140746759 | 140746860 | 140752231 | 140752316 | 140744651 | 140744767 | 140761164 | 140761262 | 18 |
| MXE | 10253 | ENSG00000065183 | WDR3 | chr1 | + | 117955314 | 117955358 | 117957067 | 117957196 | 117954579 | 117954627 | 117958909 | 117959003 | 27 |
| MXE | 3054 | ENSG00000153975 | ZUP1 | chr6 | - | 116658802 | 116658924 | 116660735 | 116660846 | 116656683 | 116656852 | 116666633 | 116666912 | 30 |
| MXE | 16623 | ENSG00000135090 | TAOK3 | chr12 | - | 118255447 | 118255655 | 118266654 | 118266759 | 118244900 | 118244965 | 118359520 | 118359720 | 13 |
| MXE | 5095 | ENSG00000060971 | ACAA1 | chr3 | - | 38126509 | 38126700 | 38127785 | 38127866 | 38125825 | 38125881 | 38129289 | 38129388 | 115 |
| MXE | 5207 | ENSG00000206503 | HLA-A | chr6 | + | 29942756 | 29943543 | 29944121 | 29944397 | 29942551 | 29942626 | 29944499 | 29944616 | 2941 |
| MXE | 2167 | ENSG00000138592 | USP8 | chr15 | + | 50471632 | 50471795 | 50476848 | 50476997 | 50465046 | 50465191 | 50477275 | 50477499 | 218 |
| MXE | 8540 | ENSG00000121892 | PDS5A | chr4 | - | 39869393 | 39869462 | 39872985 | 39873144 | 39866860 | 39866997 | 39874288 | 39874412 | 272 |
| MXE | 15304 | ENSG00000139697 | SBNO1 | chr12 | - | 123345257 | 123345567 | 123348028 | 123348133 | 123340987 | 123341088 | 123350309 | 123350441 | 96 |
| MXE | 12612 | ENSG00000151948 | GLT1D1 | chr12 | + | 128899235 | 128899287 | 128914932 | 128914989 | 128888638 | 128888744 | 128926358 | 128926454 | 23 |
| MXE | 8751 | ENSG00000116815 | CD58 | chr1 | - | 116519230 | 116519267 | 116521905 | 116521983 | 116514533 | 116514822 | 116535964 | 116536228 | 140 |
| MXE | 9614 | ENSG00000113569 | NUP155 | chr5 | - | 37341089 | 37341242 | 37342548 | 37342646 | 37337817 | 37337918 | 37348504 | 37348596 | 48 |
| MXE | 345 | ENSG00000079805 | DNM2 | chr19 | + | 10786563 | 10786706 | 10793719 | 10793855 | 10782959 | 10783120 | 10795371 | 10795439 | 339 |
| MXE | 16906 | ENSG00000111335 | OAS2 | chr12 | + | 113006412 | 113006600 | 113007704 | 113007943 | 113003304 | 113005222 | 113009086 | 113011138 | 199 |
| MXE | 8159 | ENSG00000126698 | DNAJC8 | chr1 | - | 28205257 | 28205349 | 28214939 | 28214996 | 28203746 | 28203822 | 28228921 | 28229023 | 159 |
| MXE | 1022 | ENSG00000141506 | PIK3R5 | chr17 | - | 8889972 | 8890126 | 8890737 | 8890912 | 8889138 | 8889222 | 8893585 | 8893655 | 138 |
| MXE | 8856 | ENSG00000070882 | OSBPL3 | chr7 | - | 24809806 | 24809951 | 24815058 | 24815203 | 24806775 | 24806902 | 24816609 | 24816688 | 97 |
| MXE | 1120 | ENSG00000120071 | KANSL1 | chr17 | - | 46033402 | 46033460 | 46034160 | 46034282 | 46033079 | 46033192 | 46038537 | 46038686 | 9 |
| MXE | 16244 | ENSG00000125485 | DDX31 | chr9 | - | 132618329 | 132618441 | 132625663 | 132625745 | 132612086 | 132612255 | 132630263 | 132630403 | 14 |
| MXE | 16407 | ENSG00000100354 | TNRC6B | chr22 | + | 40264687 | 40267036 | 40270121 | 40270280 | 40251178 | 40251200 | 40273424 | 40273600 | 124 |
| MXE | 15305 | ENSG00000139697 | SBNO1 | chr12 | - | 123345257 | 123345570 | 123348028 | 123348133 | 123340987 | 123341088 | 123350309 | 123350441 | 96 |
| MXE | 5092 | ENSG00000060971 | ACAA1 | chr3 | - | 38126161 | 38126700 | 38127785 | 38127866 | 38125825 | 38125881 | 38129289 | 38129388 | 115 |
| MXE | 11803 | ENSG00000170275 | CRTAP | chr3 | + | 33129938 | 33130067 | 33132554 | 33132700 | 33124407 | 33124579 | 33134181 | 33134265 | 344 |
| MXE | 13407 | ENSG00000131724 | IL13RA1 | chrX | + | 118766843 | 118766976 | 118773878 | 118773975 | 118766529 | 118766577 | 118776426 | 118776511 | 172 |
| MXE | 3264 | ENSG00000174032 | SLC25A30 | chr13 | - | 45408926 | 45409074 | 45411361 | 45411480 | 45405882 | 45405977 | 45418299 | 45418380 | 29 |
| MXE | 5538 | ENSG00000169180 | XPO6 | chr16 | - | 28133840 | 28133933 | 28135215 | 28135324 | 28132333 | 28132403 | 28146093 | 28146203 | 203 |
| MXE | 2120 | ENSG00000128739 | SNRPN | chr15 | + | 24975357 | 24975509 | 24976304 | 24976416 | 24974287 | 24974456 | 24976876 | 24977029 | 478 |
| MXE | 793 | ENSG00000197586 | ENTPD6 | chr20 | + | 25197104 | 25197226 | 25206521 | 25206590 | 25196153 | 25196254 | 25207078 | 25207397 | 0 |
| MXE | 6578 | ENSG00000072786 | STK10 | chr5 | - | 172127372 | 172127421 | 172156623 | 172156788 | 172117480 | 172117630 | 172187886 | 172188224 | 345 |
| MXE | 7269 | ENSG00000157500 | APPL1 | chr3 | + | 57267741 | 57267792 | 57268397 | 57268487 | 57260627 | 57260774 | 57269540 | 57269751 | 14 |
| MXE | 9684 | ENSG00000015676 | NUDCD3 | chr7 | - | 44404439 | 44404583 | 44427570 | 44427703 | 44392296 | 44392485 | 44484967 | 44485284 | 68 |
| MXE | 8053 | ENSG00000150455 | TIRAP | chr11 | + | 126290461 | 126290585 | 126292476 | 126293055 | 126283092 | 126283153 | 126293667 | 126294933 | 0 |
| MXE | 16744 | ENSG00000284934 | NA | chr12 | - | 122218265 | 122218397 | 122224511 | 122224644 | 122216758 | 122216869 | 122227355 | 122227421 | 5 |
| MXE | 14439 | ENSG00000163378 | EOGT | chr3 | - | 68988924 | 68989017 | 68998010 | 68998114 | 68988505 | 68988577 | 69001607 | 69001714 | 20 |
| MXE | 10512 | ENSG00000039650 | PNKP | chr19 | - | 49861607 | 49861695 | 49861771 | 49861881 | 49861448 | 49861510 | 49862043 | 49862105 | 47 |
| MXE | 695 | ENSG00000101347 | SAMHD1 | chr20 | - | 36935028 | 36935189 | 36941038 | 36941111 | 36930759 | 36930875 | 36946737 | 36946804 | 510 |
| MXE | 4675 | ENSG00000121060 | TRIM25 | chr17 | - | 56899114 | 56899180 | 56904254 | 56904488 | 56895925 | 56895952 | 56908467 | 56908563 | 129 |

| | | | | | | | | | | | | | | | | | | | | | | | | | | | | | | | | | | | | | | | | | | | | | | | | |
|---|---|---|---|---|---|---|---|---|---|---|---|---|---|---|---|---|---|---|---|---|---|---|---|---|---|---|---|---|---|---|---|---|---|---|---|---|---|---|---|---|---|---|---|---|---|---|---|---|
| MXE | 9260 | ENSG00000181817 | LSM10 | chr1 | - | 36392357 | 36392487 | 36393765 | 36394153 | 36391237 | 36391604 | 36397766 | 36397887 | 54 | 24 | 40 | 31 | 22 | 12 | 4 | 13 | 6 | 1 | 58 | 37 | 18 | 40 | 49 | 25 | 10 | 10 | 9 | 15 | 298 | 279 | 0.038361287 | 0.15381579 | 0.808 | 0.849 | 0.742 | 0.829 | 0.954 | 0.685 | 0.776 | 0.628 | 0.806 | 0.754 | 0.107 |
| MXE | 414 | ENSG00000011600 | TYROBP | chr19 | - | 35907217 | 35907264 | 35907445 | 35907580 | 35904402 | 35904634 | 35908167 | 35908275 | 39 | 15 | 21 | 25 | 25 | 1093 | 379 | 405 | 692 | 587 | 81 | 68 | 22 | 39 | 84 | 1475 | 1020 | 554 | 1022 | 1650 | 284 | 196 | 0.038378782 | 0.153844648 | 0.024 | 0.027 | 0.035 | 0.024 | 0.029 | 0.037 | 0.044 | 0.027 | 0.026 | 0.034 | -0.006 |
| MXE | 5361 | ENSG00000175066 | GK5 | chr3 | - | 142187703 | 142187779 | 142198801 | 142198933 | 142186451 | 142186513 | 142204694 | 142204788 | 129 | 65 | 50 | 98 | 113 | 37 | 23 | 16 | 32 | 45 | 31 | 51 | 73 | 61 | 22 | 35 | 16 | 43 | 21 | 12 | 281 | 225 | 0.038428136 | 0.153985128 | 0.736 | 0.694 | 0.714 | 0.71 | 0.668 | 0.415 | 0.718 | 0.576 | 0.699 | 0.595 | 0.104 |
| MXE | 9241 | ENSG00000163466 | ARPC2 | chr2 | + | 218238663 | 218238850 | 218245419 | 218245546 | 218232999 | 218234397 | 218249363 | 218249464 | 2384 | 1668 | 1504 | 2002 | 3165 | 2418 | 1354 | 1685 | 1972 | 2975 | 5150 | 2211 | 1670 | 2764 | 3501 | 2535 | 1966 | 1593 | 2605 | 2784 | 298 | 276 | 0.038434441 | 0.153985128 | 0.477 | 0.533 | 0.453 | 0.485 | 0.496 | 0.653 | 0.51 | 0.493 | 0.496 | 0.538 | -0.049 |
| MXE | 8720 | ENSG00000197323 | TRIM33 | chr1 | - | 114410183 | 114410316 | 114421435 | 114421636 | 114408676 | 114408740 | 114424590 | 114424755 | 104 | 71 | 103 | 131 | 176 | 58 | 27 | 44 | 33 | 24 | 10 | 144 | 74 | 33 | 0 | 13 | 50 | 44 | 17 | 4 | 298 | 282 | 0.038454133 | 0.154022721 | 0.629 | 0.713 | 0.689 | 0.79 | 0.874 | 0.421 | 0.732 | 0.614 | 0.648 | 0 | 0.256 |
| MXE | 1245 | ENSG00000108469 | RECQL5 | chr17 | - | 75630973 | 75631010 | 75631149 | 75631249 | 75630778 | 75630837 | 75631449 | 75631668 | 22 | 8 | 13 | 16 | 4 | 6 | 2 | 3 | 5 | 11 | 7 | 9 | 5 | 9 | 18 | 0 | 5 | 0 | 5 | 0 | 249 | 186 | 0.038478479 | 0.154078925 | 0.733 | 0.749 | 0.764 | 0.705 | 0.214 | 1 | 0.573 | 1 | 0.573 | 1 | -0.196 |
| MXE | 6255 | ENSG00000124406 | ATP8A1 | chr4 | - | 42590810 | 42590884 | 42594288 | 42594362 | 42588259 | 42588329 | 42600477 | 42600518 | 126 | 80 | 129 | 131 | 168 | 4 | 5 | 8 | 2 | 1 | 82 | 91 | 60 | 29 | 38 | 0 | 0 | 1 | 0 | 5 | 223 | 223 | 0.03854609 | 0.154308301 | 0.969 | 0.941 | 0.942 | 0.985 | 0.994 | 1 | 1 | 0.984 | 1 | 0.884 | -0.007 |
| MXE | 2662 | ENSG00000100888 | CHD8 | chr14 | - | 21412912 | 21412996 | 21414300 | 21414418 | 21409850 | 21409988 | 21414937 | 21414993 | 145 | 116 | 130 | 149 | 192 | 70 | 76 | 73 | 102 | 61 | 89 | 157 | 115 | 126 | 68 | 37 | 102 | 153 | 126 | 45 | 267 | 233 | 0.038594076 | 0.154459013 | 0.644 | 0.571 | 0.608 | 0.56 | 0.733 | 0.677 | 0.573 | 0.396 | 0.466 | 0.569 | 0.087 |
| MXE | 10725 | ENSG00000135636 | DYSF | chr2 | + | 71668753 | 71668842 | 71669111 | 71669207 | 71667375 | 71667515 | 71669604 | 71669746 | 32 | 30 | 23 | 23 | 25 | 31 | 41 | 22 | 18 | 44 | 105 | 128 | 58 | 128 | 222 | 119 | 106 | 43 | 89 | 145 | 238 | 245 | 0.038626267 | 0.154546447 | 0.515 | 0.43 | 0.518 | 0.568 | 0.369 | 0.476 | 0.554 | 0.581 | 0.597 | 0.612 | -0.084 |
| MXE | 11550 | ENSG00000064933 | PMS1 | chr2 | + | 189843963 | 189844080 | 189852654 | 189852777 | 189818016 | 189818180 | 189853938 | 189854082 | 151 | 92 | 95 | 114 | 132 | 46 | 28 | 40 | 40 | 86 | 16 | 36 | 8 | 14 | 8 | 0 | 7 | 5 | 3 | 4 | 266 | 272 | 0.038671055 | 0.154668192 | 0.77 | 0.771 | 0.708 | 0.745 | 0.611 | 1 | 0.84 | 0.621 | 0.827 | 0.672 | -0.071 |
| MXE | 13232 | ENSG00000188917 | TRMT2B | chrX | - | 101041316 | 101041371 | 101051250 | 101051366 | 101036973 | 101037073 | 101051643 | 101051733 | 2 | 3 | 0 | 1 | 0 | 5 | 10 | 11 | 7 | 12 | 0 | 4 | 2 | 0 | 5 | 13 | 1 | 6 | 1 | 2 | 265 | 204 | 0.038677401 | 0.154668192 | 0.235 | 0.188 | 0 | 0.099 | 0 | 0 | 0.755 | 0.204 | 0 | 0.658 | -0.219 |
| MXE | 5354 | ENSG00000051382 | PIK3CB | chr3 | - | 138684624 | 138684803 | 138688874 | 138688974 | 138683677 | 138683787 | 138690999 | 138691143 | 74 | 64 | 68 | 95 | 114 | 47 | 50 | 75 | 71 | 88 | 21 | 79 | 111 | 125 | 91 | 45 | 97 | 73 | 117 | 141 | 249 | 298 | 0.038689749 | 0.154676172 | 0.653 | 0.605 | 0.52 | 0.616 | 0.608 | 0.358 | 0.494 | 0.645 | 0.561 | 0.436 | 0.102 |
| MXE | 2039 | ENSG00000137804 | NUSAP1 | chr15 | + | 41365404 | 41365589 | 41371526 | 41371684 | 41358148 | 41358258 | 41375711 | 41375828 | 18 | 13 | 30 | 20 | 18 | 13 | 18 | 24 | 25 | 28 | 111 | 82 | 61 | 103 | 46 | 56 | 14 | 73 | 56 | 57 | 298 | 298 | 0.038743945 | 0.154841754 | 0.581 | 0.419 | 0.556 | 0.444 | 0.391 | 0.665 | 0.854 | 0.455 | 0.648 | 0.447 | -0.136 |
| MXE | 6204 | ENSG00000073792 | IGF2BP2 | chr3 | - | 185649402 | 185649534 | 185652093 | 185652168 | 185647024 | 185647138 | 185657285 | 185657402 | 9 | 15 | 3 | 19 | 8 | 35 | 78 | 37 | 77 | 107 | 45 | 4 | 7 | 22 | 14 | 290 | 81 | 55 | 161 | 350 | 224 | 281 | 0.038751896 | 0.154841754 | 0.244 | 0.194 | 0.092 | 0.236 | 0.086 | 0.163 | 0.058 | 0.138 | 0.146 | 0.048 | 0.06 |
| MXE | 16421 | ENSG00000130119 | GNL3L | chrX | + | 54540134 | 54540242 | 54541272 | 54541389 | 54539039 | 54539101 | 54542954 | 54543038 | 101 | 59 | 75 | 57 | 92 | 103 | 75 | 99 | 102 | 72 | 190 | 89 | 48 | 117 | 174 | 109 | 75 | 63 | 124 | 120 | 257 | 266 | 0.038770417 | 0.154874338 | 0.504 | 0.449 | 0.439 | 0.366 | 0.569 | 0.643 | 0.551 | 0.441 | 0.494 | 0.6 | -0.08 |
| MXE | 14247 | ENSG00000164941 | INTS8 | chr8 | + | 94849915 | 94850091 | 94851552 | 94851686 | 94849461 | 94849532 | 94853804 | 94853915 | 389 | 301 | 260 | 325 | 512 | 122 | 96 | 113 | 91 | 142 | 39 | 206 | 222 | 146 | 35 | 5 | 57 | 48 | 48 | 5 | 298 | 283 | 0.038796274 | 0.154936202 | 0.752 | 0.749 | 0.686 | 0.772 | 0.774 | 0.881 | 0.774 | 0.815 | 0.743 | 0.869 | -0.07 |
| MXE | 7898 | ENSG00000074603 | DPP8 | chr15 | - | 65497863 | 65498032 | 65507242 | 65507355 | 65490188 | 65490299 | 65512294 | 65512564 | 47 | 45 | 53 | 50 | 94 | 105 | 92 | 81 | 105 | 138 | 0 | 48 | 44 | 26 | 18 | 52 | 92 | 109 | 74 | 27 | 262 | 298 | 0.038857705 | 0.155140064 | 0.337 | 0.357 | 0.427 | 0.351 | 0.437 | 0 | 0.372 | 0.315 | 0.286 | 0.431 | 0.101 |
| MXE | 15420 | ENSG00000076043 | REXO2 | chr11 | + | 114445978 | 114446087 | 114447825 | 114447879 | 114440655 | 114440739 | 114449845 | 114449917 | 5 | 5 | 3 | 6 | 2 | 47 | 36 | 35 | 37 | 61 | 8 | 4 | 14 | 1 | 7 | 32 | 25 | 34 | 29 | 70 | 258 | 203 | 0.038880877 | 0.155167308 | 0.077 | 0.099 | 0.063 | 0.113 | 0.025 | 0.164 | 0.112 | 0.245 | 0.026 | 0.073 | -0.049 |
| MXE | 1444 | ENSG00000159210 | SNF8 | chr17 | - | 48931642 | 48931717 | 48937019 | 48937124 | 48930370 | 48930612 | 48940923 | 48941062 | 102 | 85 | 79 | 80 | 89 | 32 | 18 | 26 | 37 | 31 | 291 | 86 | 65 | 127 | 240 | 99 | 29 | 56 | 76 | 100 | 254 | 224 | 0.038885301 | 0.155167308 | 0.738 | 0.806 | 0.728 | 0.656 | 0.717 | 0.722 | 0.723 | 0.506 | 0.596 | 0.679 | 0.084 |
| MXE | 7308 | ENSG00000117543 | DPH5 | chr1 | - | 100995109 | 100995149 | 101001466 | 101001587 | 100992636 | 100992740 | 101013709 | 101013818 | 69 | 41 | 79 | 76 | 151 | 60 | 39 | 62 | 50 | 61 | 40 | 25 | 47 | 22 | 27 | 70 | 40 | 44 | 10 | 29 | 270 | 189 | 0.03891723 | 0.155253248 | 0.446 | 0.424 | 0.471 | 0.516 | 0.634 | 0.286 | 0.304 | 0.428 | 0.606 | 0.395 | 0.094 |
| MXE | 16633 | ENSG00000111450 | STX2 | chr12 | - | 130806981 | 130807090 | 130808630 | 130808704 | 130801414 | 130801488 | 130812956 | 130813031 | 40 | 33 | 40 | 46 | 66 | 13 | 8 | 12 | 13 | 10 | 54 | 41 | 45 | 67 | 87 | 38 | 21 | 17 | 11 | 28 | 223 | 258 | 0.038931711 | 0.155269559 | 0.781 | 0.827 | 0.794 | 0.804 | 0.884 | 0.622 | 0.693 | 0.754 | 0.876 | 0.782 | 0.073 |
| MXE | 8662 | ENSG00000117360 | PRPF3 | chr1 | + | 150324894 | 150325087 | 150325750 | 150325881 | 150321532 | 150321592 | 150328319 | 150328466 | 41 | 25 | 46 | 35 | 39 | 89 | 54 | 76 | 60 | 80 | 91 | 87 | 48 | 71 | 37 | 92 | 74 | 71 | 108 | 93 | 298 | 280 | 0.038972146 | 0.155389342 | 0.302 | 0.303 | 0.363 | 0.354 | 0.314 | 0.482 | 0.525 | 0.388 | 0.382 | 0.272 | -0.083 |

| | | | | | | | | | | |
|---|---|---|---|---|---|---|---|---|---|---|
| MXE | 10058 | ENSG00000167107 | ACSF2 | chr17 | + | 50461241 | 50461370 | 50461632 | 50461686 | [illegible] |
| MXE | 7723 | ENSG00000082512 | TRAF5 | chr1 | + | 211365375 | 211365468 | 211369451 | 211369592 | [illegible] |
| MXE | 9085 | ENSG00000143297 | FCRL5 | chr1 | - | 157546187 | 157546280 | 157546942 | 157547197 | [illegible] |
| MXE | 12441 | ENSG00000169410 | PTPN9 | chr15 | - | 75524208 | 75524298 | 75527117 | 75527261 | [illegible] |
| MXE | 8356 | ENSG00000010803 | SCMH1 | chr1 | - | 41152598 | 41152741 | 41159724 | 41159933 | [illegible] |
| MXE | 778 | ENSG00000080839 | RBL1 | chr20 | - | 37003701 | 37003866 | 37007410 | 37007559 | [illegible] |
| MXE | 11636 | ENSG00000188554 | NBR1 | chr17 | + | 43201685 | 43201780 | 43202654 | 43202712 | [illegible] |
| MXE | 7250 | ENSG00000145687 | SSBP2 | chr5 | - | 81615472 | 81615557 | 81636556 | 81636618 | [illegible] |
| MXE | 15922 | ENSG00000181192 | DHTKD1 | chr10 | + | 12088985 | 12089255 | 12091512 | 12091684 | [illegible] |
| MXE | 6865 | ENSG00000164403 | SHROOM1 | chr5 | - | 132823849 | 132824419 | 132824614 | 132824821 | [illegible] |
| MXE | 11509 | ENSG00000137478 | FCHSD2 | chr11 | - | 72887469 | 72887574 | 72889828 | 72889945 | [illegible] |
| MXE | 14714 | ENSG00000149177 | PTPRJ | chr11 | + | 48144674 | 48144885 | 48146875 | 48146963 | [illegible] |
| MXE | 2094 | ENSG00000091157 | WDR7 | chr18 | + | 56938532 | 56938682 | 56939310 | 56939393 | [illegible] |
| MXE | 13714 | ENSG00000183354 | KIAA2026 | chr9 | - | 5944870 | 5944960 | 5968018 | 5969436 | [illegible] |
| MXE | 15530 | ENSG00000110075 | PPP6R3 | chr11 | + | 68551120 | 68551186 | 68567013 | 68567166 | [illegible] |
| MXE | 12355 | ENSG00000018610 | STEEP1 | chrX | - | 119544352 | 119544491 | 119545462 | 119545504 | [illegible] |
| MXE | 10111 | ENSG00000164172 | MOCS2 | chr5 | - | 53102096 | 53102224 | 53107076 | 53107221 | [illegible] |
| MXE | 13322 | ENSG00000170832 | USP32 | chr17 | - | 60269449 | 60269557 | 60271349 | 60271481 | [illegible] |
| MXE | 9714 | ENSG00000112335 | SNX3 | chr6 | - | 108214497 | 108214622 | 108222949 | 108223045 | [illegible] |
| MXE | 9543 | ENSG00000009307 | CSDE1 | chr1 | - | 114730507 | 114730648 | 114732603 | 114732816 | [illegible] |
| MXE | 7403 | ENSG00000131374 | TBC1D5 | chr3 | - | 17562018 | 17562142 | 17622647 | 17622765 | [illegible] |
| MXE | 1093 | ENSG00000129197 | RPAIN | chr17 | + | 5425970 | 5426299 | 5428070 | 5428211 | [illegible] |
| MXE | 12462 | ENSG00000102531 | FNDC3A | chr13 | + | 49178567 | 49178654 | 49185963 | 49186102 | [illegible] |
| MXE | 4017 | ENSG00000160685 | ZBTB7B | chr1 | + | 155010257 | 155010397 | 155010915 | 155011015 | [illegible] |
| MXE | 10826 | ENSG00000169016 | E2F6 | chr2 | - | 11453581 | 11453722 | 11455341 | 11455474 | [illegible] |
| MXE | 10713 | ENSG00000168958 | MFF | chr2 | + | 227342744 | 227342819 | 227347225 | 227347384 | [illegible] |
| MXE | 11408 | ENSG00000138380 | CARF | chr2 | + | 202924296 | 202924415 | 202942739 | 202942967 | [illegible] |
| MXE | 16373 | ENSG00000013561 | RNF14 | chr5 | + | 141974803 | 141974955 | 141978302 | 141978830 | [illegible] |
| MXE | 15332 | ENSG00000170185 | USP38 | chr4 | + | 143209563 | 143209657 | 143212317 | 143212424 | [illegible] |
| MXE | 6456 | ENSG00000143952 | VPS54 | chr2 | - | 63944599 | 63944655 | 63947382 | 63947490 | [illegible] |
| MXE | 9815 | ENSG00000151092 | NGLY1 | chr3 | - | 25736308 | 25736400 | 25737333 | 25737455 | [illegible] |
| MXE | 1834 | ENSG00000103005 | USB1 | chr16 | + | 58017333 | 58017439 | 58018971 | 58019055 | [illegible] |
| MXE | 9476 | ENSG00000171503 | ETFDH | chr4 | + | 158706188 | 158706371 | 158706628 | 158706850 | [illegible] |
| MXE | 2497 | ENSG00000197102 | DYNC1H1 | chr14 | + | 102036488 | 102036642 | 102038459 | 102038606 | [illegible] |
| MXE | 11819 | ENSG00000164691 | TAGAP | chr6 | - | 159043588 | 159043655 | 159043977 | 159044031 | [illegible] |
| MXE | 12975 | ENSG00000122591 | FAM126A | chr7 | - | 22964414 | 22964531 | 22976209 | 22976305 | [illegible] |
| MXE | 9467 | ENSG00000198873 | GRK5 | chr10 | + | 119396694 | 119396772 | 119423165 | 119423266 | [illegible] |
| MXE | 6494 | ENSG00000152495 | CAMK4 | chr5 | + | 111446685 | 111446776 | 111449128 | 111449203 | [illegible] |
| MXE | 528 | ENSG00000239998 | LILRA2 | chr19 | + | 54573830 | 54573912 | 54574300 | 54574582 | [illegible] |
| MXE | 11396 | ENSG00000003402 | CFLAR | chr2 | + | 201133028 | 201133134 | 201135971 | 201136107 | [illegible] |
| MXE | 14629 | ENSG00000119487 | MAPKAP1 | chr9 | - | 125559632 | 125559809 | 125585554 | 125585727 | [illegible] |
| MXE | 9471 | ENSG00000131408 | NR1H2 | chr19 | + | 50377732 | 50377870 | 50378148 | 50378439 | [illegible] |
| MXE | 13827 | ENSG00000104219 | ZDHHC2 | chr8 | + | 17205654 | 17205775 | 17207959 | 17208092 | [illegible] |
| MXE | 13515 | ENSG00000172534 | HCFC1 | chrX | - | 153963224 | 153963433 | 153964123 | 153964284 | [illegible] |
| MXE | 8121 | ENSG00000145020 | AMT | chr3 | - | 49419709 | 49419788 | 49420210 | 49420342 | [illegible] |
| MXE | 7107 | ENSG00000059691 | GATB | chr4 | - | 151707987 | 151708101 | 151716008 | 151716131 | [illegible] |
| MXE | 16431 | ENSG00000109685 | NSD2 | chr4 | + | 1904215 | 1904378 | 1916870 | 1917037 | [illegible] |
| MXE | 5836 | ENSG00000198793 | MTOR | chr1 | - | 11150125 | 11150226 | 11157151 | 11157291 | [illegible] |
| MXE | 6771 | ENSG00000198265 | HELZ | chr17 | - | 67190156 | 67190355 | 67193966 | 67194042 | [illegible] |
| MXE | 6035 | ENSG00000148290 | SURF1 | chr9 | - | 133352693 | 133352766 | 133353748 | 133353940 | [illegible] |
| MXE | 7916 | ENSG00000135250 | SRPK2 | chr7 | - | 105143083 | 105143330 | 105145282 | 105145308 | [illegible] |
| MXE | 11147 | ENSG00000124491 | F13A1 | chr6 | - | 6174579 | 6174867 | 6181987 | 6182141 | [illegible] |
| MXE | 11956 | ENSG00000122512 | PMS2 | chr7 | - | 5997325 | 5997423 | 5999107 | 5999275 | [illegible] |
| MXE | 12112 | ENSG00000151116 | UEVLD | chr11 | - | 18566346 | 18566482 | 18575346 | 18575412 | [illegible] |
| MXE | 9257 | ENSG00000044459 | CNTLN | chr9 | + | 17298189 | 17298352 | 17309057 | 17309252 | [illegible] |
| MXE | 16703 | ENSG00000141376 | BCAS3 | chr17 | + | 61084466 | 61084564 | 61368326 | 61368494 | [illegible] |
| MXE | 14891 | ENSG00000165219 | GAPVD1 | chr9 | + | 125323797 | 125323923 | 125326415 | 125326589 | [illegible] |
| MXE | 14198 | ENSG00000160948 | VPS28 | chr8 | - | 144424717 | 144424819 | 144424945 | 144425000 | [illegible] |
| MXE | 12143 | ENSG00000062725 | APPBP2 | chr17 | - | 60466290 | 60466459 | 60479147 | 60479271 | [illegible] |
| MXE | 8004 | ENSG00000197943 | PLCG2 | chr16 | + | 81931496 | 81931654 | 81934428 | 81934531 | [illegible] |
| MXE | 4838 | ENSG00000136040 | PLXNC1 | chr12 | + | 94220015 | 94220163 | 94224227 | 94224315 | [illegible] |
| MXE | 12049 | ENSG00000127947 | PTPN12 | chr7 | + | 77637017 | 77637048 | 77638623 | 77638731 | [illegible] |
| MXE | 3356 | ENSG00000127603 | MACF1 | chr1 | + | 39433047 | 39433155 | 39434413 | 39434632 | [illegible] |
| MXE | 15307 | ENSG00000139697 | SBNO1 | chr12 | - | 123348028 | 123348133 | 123350309 | 123350441 | [illegible] |
| MXE | 254 | ENSG00000198089 | SFI1 | chr22 | + | 31550253 | 31550348 | 31556941 | 31557059 | [illegible] |
| MXE | 8240 | ENSG00000143643 | TTC13 | chr1 | - | 230958223 | 230958299 | 230961208 | 230961303 | [illegible] |
| MXE | 2064 | ENSG00000182718 | ANXA2 | chr15 | - | 60390212 | 60390317 | 60393037 | 60393440 | [illegible] |
| MXE | 2961 | ENSG00000162711 | NLRP3 | chr1 | + | 247434102 | 247434273 | 247435969 | 247436140 | [illegible] |
| MXE | 9159 | ENSG00000162909 | CAPN2 | chr1 | + | 223744099 | 223744218 | 223745305 | 223745439 | [illegible] |
| MXE | 6846 | ENSG00000230124 | ACBD6 | chr1 | - | 180397515 | 180397605 | 180413365 | 180413471 | [illegible] |
| MXE | 16665 | ENSG00000198270 | TMEM116 | chr12 | - | 111933885 | 111934030 | 111938160 | 111938210 | [illegible] |
| MXE | 14768 | ENSG00000149308 | NPAT | chr11 | - | 108169927 | 108170043 | 108172198 | 108173851 | [illegible] |
| MXE | 13135 | ENSG00000112561 | TFEB | chr6 | - | 41686089 | 41686237 | 41687093 | 41687169 | [illegible] |
| MXE | 4410 | ENSG00000077684 | JADE1 | chr4 | + | 128842952 | 128843038 | 128846374 | 128846532 | [illegible] |
| MXE | 13295 | ENSG00000107185 | RGP1 | chr9 | + | 35751626 | 35751754 | 35751955 | 35752145 | [illegible] |
| MXE | 4198 | ENSG00000084676 | NCOA1 | chr2 | + | 24673365 | 24673463 | 24682950 | 24683128 | [illegible] |
| MXE | 7794 | ENSG00000174780 | SRP72 | chr4 | + | 56483138 | 56483270 | 56484735 | 56484864 | [illegible] |
| MXE | 540 | ENSG00000105486 | LIG1 | chr19 | - | 48131075 | 48131171 | 48132981 | 48133097 | [illegible] |
| MXE | 7356 | ENSG00000134215 | VAV3 | chr1 | - | 107574046 | 107574198 | 107596211 | 107596341 | [illegible] |
| MXE | 10272 | ENSG00000161813 | LARP4 | chr12 | + | 50430494 | 50430570 | 50435487 | 50435624 | [illegible] |
| MXE | 5924 | ENSG00000003400 | CASP10 | chr2 | + | 201192983 | 201193119 | 201205881 | 201205973 | [illegible] |
| MXE | 3338 | ENSG00000165097 | KDM1B | chr6 | + | 18185771 | 18185810 | 18191196 | 18191381 | [illegible] |
| MXE | 10551 | ENSG00000107643 | MAPK8 | chr10 | + | 48424087 | 48424159 | 48424513 | 48424585 | [illegible] |
| MXE | 6201 | ENSG00000135932 | CAB39 | chr2 | + | 230798728 | 230798897 | 230810262 | 230810322 | [illegible] |
| MXE | 2735 | ENSG00000015133 | CCDC88C | chr14 | - | 91281456 | 91281525 | 91283328 | 91283517 | [illegible] |
| MXE | 16404 | ENSG00000100354 | TNRC6B | chr22 | + | 40261831 | 40262173 | 40264687 | 40267036 | [illegible] |
| MXE | 12812 | ENSG00000104133 | SPG11 | chr15 | - | 44572682 | 44572820 | 44574901 | 44575041 | [illegible] |
| MXE | 12374 | ENSG00000091106 | NLRC4 | chr2 | - | 32249606 | 32251601 | 32252418 | 32252679 | [illegible] |
| MXE | 16452 | ENSG00000111371 | SLC38A1 | chr12 | - | 46201097 | 46201198 | 46203009 | 46203089 | [illegible] |
| MXE | 10345 | ENSG00000031003 | FAM13B | chr5 | - | 137960165 | 137960214 | 137962404 | 137962469 | [illegible] |
| MXE | 5907 | ENSG00000067066 | SP100 | chr2 | + | 230469042 | 230469096 | 230470014 | 230470098 | [illegible] |
| MXE | 4777 | ENSG00000143669 | LYST | chr1 | - | 235773841 | 235773991 | 235774912 | 235775086 | [illegible] |
| MXE | 14724 | ENSG00000136811 | ODF2 | chr9 | + | 128460512 | 128460666 | 128460941 | 128461010 | [illegible] |
| MXE | 2631 | ENSG00000119638 | NEK9 | chr14 | - | 75088479 | 75088641 | 75091269 | 75091478 | [illegible] |
| MXE | 10210 | ENSG00000197429 | IPP | chr1 | - | 45700069 | 45700190 | 45714245 | 45714466 | [illegible] |
| MXE | 10229 | ENSG00000000938 | FGR | chr1 | - | 27621558 | 27621657 | 27625088 | 27625151 | [illegible] |
| MXE | 10186 | ENSG00000138795 | LEF1 | chr4 | - | 108064335 | 108064384 | 108070662 | 108070770 | [illegible] |
| MXE | 9534 | ENSG00000131876 | SNRPA1 | chr15 | - | 101287655 | 101287702 | 101291961 | 101292040 | [illegible] |
| MXE | 10417 | ENSG00000123338 | NCKAP1L | chr12 | + | 54526527 | 54526746 | 54528246 | 54528377 | [illegible] |
| MXE | 17163 | ENSG00000139687 | RB1 | chr13 | + | 48307279 | 48307406 | 48342598 | 48342714 | [illegible] |
| MXE | 10056 | ENSG00000120675 | DNAJC15 | chr13 | + | 43068929 | 43069003 | 43078611 | 43078688 | [illegible] |
| MXE | 4627 | ENSG00000163694 | RBM47 | chr4 | - | 40432650 | 40432862 | 40436440 | 40436647 | [illegible] |
| MXE | 1152 | ENSG00000160551 | TAOK1 | chr17 | + | 29480367 | 29480481 | 29482196 | 29482288 | [illegible] |
| MXE | 13775 | ENSG00000165675 | ENOX2 | chrX | - | 130627957 | 130628043 | 130631467 | 130631576 | [illegible] |
| MXE | 2736 | ENSG00000015133 | CCDC88C | chr14 | - | 91309858 | 91309986 | 91313079 | 91314150 | [illegible] |
| MXE | 4516 | ENSG00000005059 | MCUB | chr4 | + | 109659010 | 109659086 | 109660194 | 109660365 | [illegible] |
| MXE | 12675 | ENSG00000123505 | AMD1 | chr6 | + | 110887504 | 110887591 | 110888856 | 110888983 | [illegible] |
| MXE | 12455 | ENSG00000241973 | PI4KA | chr22 | - | 20798583 | 20798687 | 20799092 | 20799276 | [illegible] |
| MXE | 1029 | ENSG00000161955 | TNFSF13 | chr17 | + | 7559623 | 7559702 | 7560048 | 7560167 | [illegible] |
| MXE | 14116 | ENSG00000249859 | PVT1 | chr8 | + | 128010317 | 128010444 | 128070159 | 128070361 | [illegible] |
| MXE | 10085 | ENSG00000169379 | ARL13B | chr3 | + | 93996547 | 93996684 | 93997246 | 93997301 | [illegible] |
| MXE | 2664 | ENSG00000254505 | CHMP4A | chr14 | - | 24210653 | 24210768 | 24211410 | 24211592 | [illegible] |
| MXE | 8817 | ENSG00000166046 | TCP11L2 | chr12 | + | 106318343 | 106318464 | 106321485 | 106321706 | [illegible] |
| MXE | 13757 | ENSG00000143569 | UBAP2L | chr1 | + | 154251480 | 154251653 | 154253899 | 154254089 | [illegible] |
| MXE | 1090 | ENSG00000126858 | RHOT1 | chr17 | + | 32203889 | 32203973 | 32206909 | 32207029 | [illegible] |
| MXE | 51 | ENSG00000159086 | PAXBP1 | chr21 | - | 32755212 | 32755353 | 32759079 | 32759269 | [illegible] |
| MXE | 5195 | ENSG00000163755 | HPS3 | chr3 | + | 149155106 | 149155215 | 149157349 | 149157531 | [illegible] |
| MXE | 9374 | ENSG00000116984 | MTR | chr1 | + | 236874725 | 236874846 | 236880754 | 236880836 | [illegible] |
| MXE | 9171 | ENSG00000119004 | CYP20A1 | chr2 | + | 203251966 | 203252109 | 203266513 | 203266681 | [illegible] |
| MXE | 2032 | ENSG00000103994 | ZNF106 | chr15 | - | 42442072 | 42442414 | 42444201 | 42444262 | [illegible] |
| MXE | 7078 | ENSG00000141556 | TBCD | chr17 | + | 82900650 | 82900731 | 82905935 | 82906053 | [illegible] |
| MXE | 8600 | ENSG00000151838 | CCDC175 | chr14 | - | 59540674 | 59540746 | 59543343 | 59543454 | [illegible] |
| MXE | 5419 | ENSG00000109861 | CTSC | chr11 | - | 88300529 | 88300645 | 88309162 | 88309318 | [illegible] |
| MXE | 11348 | ENSG00000114956 | DGUOK | chr2 | + | 73938909 | 73939022 | 73950584 | 73950732 | [illegible] |
| MXE | 118 | ENSG00000242259 | C22orf39 | chr22 | - | 19359335 | 19359484 | 19361236 | 19361341 | [illegible] |
| MXE | 6487 | ENSG00000140575 | IQGAP1 | chr15 | + | 90474063 | 90474133 | 90474484 | 90474693 | [illegible] |
| MXE | 8471 | ENSG00000115590 | IL1R2 | chr2 | + | 102022186 | 102022249 | 102026110 | 102026253 | [illegible] |
| MXE | 4707 | ENSG00000155508 | CNOT8 | chr5 | + | 154867691 | 154867792 | 154870660 | 154870822 | [illegible] |
| MXE | 5284 | ENSG00000107758 | PPP3CB | chr10 | - | 73474918 | 73475030 | 73478498 | 73478623 | [illegible] |
| MXE | 12454 | ENSG00000241973 | PI4KA | chr22 | - | 20796145 | 20796314 | 20798583 | 20798687 | [illegible] |
| MXE | 9542 | ENSG00000009307 | CSDE1 | chr1 | - | 114730257 | 114730422 | 114730507 | 114730648 | [illegible] |
| MXE | 12530 | ENSG00000179818 | PCBP1-AS1 | chr2 | - | 70053730 | 70053815 | 70059642 | 70059680 | [illegible] |
| MXE | 7209 | ENSG00000121766 | ZCCHC17 | chr1 | + | 31310043 | 31310164 | 31319108 | 31319166 | [illegible] |
| MXE | 9691 | ENSG00000115525 | ST3GAL5 | chr2 | - | 85851263 | 85851703 | 85861180 | 85861292 | [illegible] |
| MXE | 16288 | ENSG00000266412 | NCOA4 | chr10 | - | 46013549 | 46013639 | 46014443 | 46014552 | [illegible] |
| MXE | 14105 | ENSG00000249859 | PVT1 | chr8 | + | 127969415 | 127969537 | 127983903 | 127984204 | [illegible] |
| MXE | 16849 | ENSG00000128585 | MKLN1 | chr7 | + | 131387119 | 131387262 | 131388883 | 131388972 | [illegible] |
| MXE | 7814 | ENSG00000066427 | ATXN3 | chr14 | - | 92096092 | 92096134 | 92096673 | 92096838 | [illegible] |
| MXE | 1707 | ENSG00000103051 | COG4 | chr16 | - | 70515867 | 70516040 | 70517625 | 70517740 | [illegible] |
| MXE | 16760 | ENSG00000135124 | P2RX4 | chr12 | + | 121228962 | 121229099 | 121232413 | 121232507 | [illegible] |
| MXE | 5209 | ENSG00000206503 | HLA-A | chr6 | + | 29943267 | 29943543 | 29944121 | 29944397 | [illegible] |
| MXE | 15538 | ENSG00000110075 | PPP6R3 | chr11 | + | 68575957 | 68576043 | 68583042 | 68583129 | [illegible] |
| MXE | 3530 | ENSG00000113263 | ITK | chr5 | + | 157244261 | 157244478 | 157245725 | 157245790 | [illegible] |
| MXE | 9055 | ENSG00000141458 | NPC1 | chr18 | - | 23533354 | 23533517 | 23534445 | 23534559 | [illegible] |
| MXE | 15725 | ENSG00000130830 | MPP1 | chrX | - | 154786203 | 154786400 | 154789953 | 154790022 | [illegible] |
| MXE | 13663 | ENSG00000046651 | OFD1 | chrX | + | 13753367 | 13753441 | 13755150 | 13755242 | [illegible] |
| MXE | 971 | ENSG00000101639 | CEP192 | chr18 | + | 12999420 | 12999588 | 13001456 | 13001582 | [illegible] |
| MXE | 3732 | ENSG00000164221 | CCDC112 | chr5 | - | 115269702 | 115269798 | 115271212 | 115271626 | [illegible] |
| MXE | 14711 | ENSG00000114030 | KPNA1 | chr3 | - | 122433660 | 122433788 | 122437169 | 122437295 | [illegible] |
| MXE | 11536 | ENSG00000116209 | TMEM59 | chr1 | - | 54045691 | 54045786 | 54047266 | 54047372 | [illegible] |
| MXE | 10655 | ENSG00000091009 | RBM27 | chr5 | + | 146261509 | 146261806 | 146263490 | 146263631 | [illegible] |
| MXE | 3148 | ENSG00000018189 | RUFY3 | chr4 | + | 70800140 | 70800205 | 70802955 | 70802983 | [illegible] |
| MXE | 2423 | ENSG00000128829 | EIF2AK4 | chr15 | + | 40025976 | 40026089 | 40030358 | 40030456 | [illegible] |
| MXE | 12316 | ENSG00000122482 | ZNF644 | chr1 | - | 90938271 | 90941309 | 90982309 | 90982370 | [illegible] |
| MXE | 3452 | ENSG00000112996 | MRPS30 | chr5 | + | 44811914 | 44812020 | 44813105 | 44813282 | [illegible] |
| MXE | 10083 | ENSG00000010244 | ZNF207 | chr17 | + | 32360891 | 32360967 | 32361467 | 32361515 | [illegible] |
| MXE | 11861 | ENSG00000111670 | GNPTAB | chr12 | - | 101766090 | 101766294 | 101768036 | 101768160 | [illegible] |
| MXE | 2225 | ENSG00000174485 | DENND4A | chr15 | - | 65729071 | 65729247 | 65729533 | 65729678 | [illegible] |
| MXE | 6891 | ENSG00000168894 | RNF181 | chr2 | + | 85596856 | 85596931 | 85597103 | 85597178 | [illegible] |
| MXE | 13220 | ENSG00000188917 | TRMT2B | chrX | - | 101020486 | 101020588 | 101021100 | 101021315 | [illegible] |
| MXE | 8430 | ENSG00000116688 | MFN2 | chr1 | + | 11982522 | 11982590 | 11992554 | 11992690 | [illegible] |
| MXE | 13612 | ENSG00000116128 | BCL9 | chr1 | + | 147611577 | 147611889 | 147612882 | 147613199 | [illegible] |
| MXE | 1986 | ENSG00000103197 | TSC2 | chr16 | + | 2058746 | 2058873 | 2060669 | 2060813 | [illegible] |
| MXE | 8769 | ENSG00000101040 | ZMYND8 | chr20 | - | 47310055 | 47310204 | 47347855 | 47347926 | [illegible] |
| MXE | 9652 | ENSG00000113761 | ZNF346 | chr5 | + | 177041777 | 177041870 | 177044391 | 177044533 | [illegible] |
| MXE | 17005 | ENSG00000110851 | PRDM4 | chr12 | - | 107743196 | 107743282 | 107744542 | 107744661 | [illegible] |
| MXE | 5291 | ENSG00000101596 | SMCHD1 | chr18 | + | 2729274 | 2729409 | 2732264 | 2732492 | [illegible] |
| MXE | 10140 | ENSG00000147526 | TACC1 | chr8 | + | 38831124 | 38831177 | 38836161 | 38836287 | [illegible] |
| MXE | 7642 | ENSG00000063978 | RNF4 | chr4 | + | 2511955 | 2511965 | 2512437 | 2512597 | [illegible] |
| MXE | 4094 | ENSG00000136925 | TSTD2 | chr9 | - | 97617756 | 97617877 | 97625680 | 97625997 | [illegible] |
| MXE | 1372 | ENSG00000108510 | MED13 | chr17 | - | 61960866 | 61961090 | 61961587 | 61961779 | [illegible] |
| MXE | 7482 | ENSG00000120733 | KDM3B | chr5 | + | 138379583 | 138379708 | 138381515 | 138381590 | [illegible] |
| MXE | 1667 | ENSG00000103222 | ABCC1 | chr16 | + | 16014490 | 16014628 | 16033108 | 16033170 | [illegible] |
| MXE | 8168 | ENSG00000173575 | CHD2 | chr15 | + | 92948951 | 92949076 | 92953356 | 92953573 | [illegible] |
| MXE | 3885 | ENSG00000119522 | DENND1A | chr9 | - | 123652012 | 123652123 | 123667025 | 123667079 | [illegible] |
| MXE | 6336 | ENSG00000158882 | TOMM40L | chr1 | + | 161226887 | 161226955 | 161227257 | 161227350 | [illegible] |
| MXE | 7753 | ENSG00000205323 | SARNP | chr12 | - | 55794806 | 55794880 | 55796024 | 55796076 | [illegible] |
| MXE | 5886 | ENSG00000117054 | ACADM | chr1 | + | 75739979 | 75740110 | 75745805 | 75745914 | [illegible] |
| MXE | 1989 | ENSG00000175265 | GOLGA8A | chr15 | - | 34407523 | 34407735 | 34435382 | 34435471 | [illegible] |
| MXE | 12270 | ENSG00000160219 | GAB3 | chrX | - | 154680131 | 154680248 | 154697131 | 154697213 | [illegible] |
| MXE | 2876 | ENSG00000136758 | YME1L1 | chr10 | - | 27123546 | 27123699 | 27126695 | 27126786 | [illegible] |
| MXE | 10674 | ENSG00000148341 | SH3GLB2 | chr9 | - | 129010669 | 129010693 | 129012235 | 129012298 | [illegible] |
| MXE | 10639 | ENSG00000168918 | INPP5D | chr2 | + | 233184407 | 233184521 | 233185842 | 233185925 | [illegible] |
| MXE | 9696 | ENSG00000151779 | NBAS | chr2 | - | 15276850 | 15277101 | 15287072 | 15287183 | [illegible] |
| MXE | 752 | ENSG00000078747 | ITCH | chr20 | + | 34408650 | 34408792 | 34412514 | 34412639 | [illegible] |
| MXE | 4531 | ENSG00000168397 | ATG4B | chr2 | + | 241651263 | 241651335 | 241653511 | 241653610 | [illegible] |
| MXE | 2404 | ENSG00000141564 | RPTOR | chr17 | + | 80643727 | 80643810 | 80707840 | 80707999 | [illegible] |
| MXE | 2378 | ENSG00000125107 | CNOT1 | chr16 | - | 58556846 | 58556993 | 58558472 | 58558674 | [illegible] |
| MXE | 12937 | ENSG00000131373 | HACL1 | chr3 | - | 15585247 | 15585342 | 15586524 | 15586602 | [illegible] |
| MXE | 9402 | ENSG00000235750 | KIAA0040 | chr1 | - | 175166561 | 175166737 | 175192639 | 175192716 | [illegible] |
| MXE | 9207 | ENSG00000160310 | PRMT2 | chr21 | + | 46658744 | 46658920 | 46663382 | 46663554 | [illegible] |
| MXE | 11111 | ENSG00000184182 | UBE2F | chr2 | + | 238025341 | 238025412 | 238035877 | 238035940 | [illegible] |
| MXE | 6130 | ENSG00000271383 | NBPF19 | chr1 | + | 149477944 | 149478047 | 149478879 | 149479094 | [illegible] |
| MXE | 4303 | ENSG00000170571 | EMB | chr5 | - | 50410876 | 50410965 | 50411196 | 50411383 | [illegible] |
| MXE | 2240 | ENSG00000285304 | NA | chr22 | + | 37643643 | 37643788 | 37644636 | 37644705 | [illegible] |
| MXE | 10496 | ENSG00000123106 | CCDC91 | chr12 | + | 28225794 | 28225861 | 28255580 | 28255662 | [illegible] |
| MXE | 9924 | ENSG00000115464 | USP34 | chr2 | - | 61348755 | 61348886 | 61349249 | 61349285 | [illegible] |
| MXE | 16500 | ENSG00000011523 | CEP68 | chr2 | + | 65069398 | 65069801 | 65071453 | 65072980 | [illegible] |
| MXE | 15349 | ENSG00000121895 | TMEM156 | chr4 | - | 38988850 | 38988970 | 38993737 | 38993998 | [illegible] |
| MXE | 2091 | ENSG00000128915 | ICE2 | chr15 | - | 60456656 | 60456794 | 60466593 | 60466713 | [illegible] |
| MXE | 6900 | ENSG00000117335 | CD46 | chr1 | + | 207757013 | 207757202 | 207757539 | 207757642 | [illegible] |
| MXE | 9251 | ENSG00000113368 | LMNB1 | chr5 | + | 126822780 | 126822885 | 126825987 | 126826107 | [illegible] |
| MXE | 13209 | ENSG00000138593 | SECISBP2L | chr15 | - | 49016847 | 49017015 | 49017547 | 49017628 | [illegible] |
| MXE | 11169 | ENSG00000111906 | HDDC2 | chr6 | - | 125292840 | 125292909 | 125298649 | 125298816 | [illegible] |
| MXE | 1075 | ENSG00000153933 | DGKE | chr17 | + | 56856511 | 56856625 | 56858593 | 56858665 | [illegible] |
| MXE | 8933 | ENSG00000154174 | TOMM70 | chr3 | - | 100375017 | 100375152 | 100377704 | 100377912 | [illegible] |
| MXE | 5636 | ENSG00000151726 | ACSL1 | chr4 | - | 184763166 | 184763255 | 184764852 | 184764925 | [illegible] |
| MXE | 4567 | ENSG00000118503 | TNFAIP3 | chr6 | + | 137875687 | 137875835 | 137875995 | 137876166 | [illegible] |
| MXE | 16139 | ENSG00000172661 | WASHC2C | chr10 | + | 45728861 | 45729026 | 45737982 | 45738045 | [illegible] |
| MXE | 14123 | ENSG00000249859 | PVT1 | chr8 | + | 128070159 | 128070272 | 128095691 | 128095817 | [illegible] |
| MXE | 9653 | ENSG00000113761 | ZNF346 | chr5 | + | 177041777 | 177041870 | 177044391 | 177044533 | [illegible] |
| MXE | 3126 | ENSG00000129534 | MIS18BP1 | chr14 | - | 45227662 | 45227814 | 45231140 | 45231298 | [illegible] |
| MXE | 867 | ENSG00000125827 | TMX4 | chr20 | - | 7999731 | 7999822 | 8001495 | 8001541 | [illegible] |
| MXE | 12275 | ENSG00000170632 | ARMC10 | chr7 | + | 103086868 | 103086937 | 103092476 | 103092653 | [illegible] |
| MXE | 4104 | ENSG00000104517 | UBR5 | chr8 | - | 102261145 | 102261259 | 102261914 | 102262070 | [illegible] |
| MXE | 14413 | ENSG00000143198 | MGST3 | chr1 | + | 165649840 | 165649964 | 165651013 | 165651087 | [illegible] |
| MXE | 15092 | ENSG00000277072 | STAG3L2 | chr7 | - | 74885022 | 74885146 | 74887682 | 74887799 | [illegible] |
| MXE | 8031 | ENSG00000160691 | SHC1 | chr1 | - | 154969377 | 154969448 | 154970031 | 154970199 | [illegible] |
| MXE | 5929 | ENSG00000003400 | CASP10 | chr2 | + | 201195841 | 201195948 | 201203729 | 201203766 | [illegible] |
| MXE | 8581 | ENSG00000171960 | PPIH | chr1 | + | 42659521 | 42659566 | 42660861 | 42660904 | [illegible] |
| MXE | 16796 | ENSG00000215039 | CD27-AS1 | chr12 | - | 6448639 | 6448737 | 6450340 | 6450980 | [illegible] |
| MXE | 14417 | ENSG00000177076 | ACER2 | chr9 | + | 19434946 | 19435084 | 19446280 | 19446418 | [illegible] |
| MXE | 8805 | ENSG00000142405 | NLRP12 | chr19 | - | 53801226 | 53801397 | 53803951 | 53804122 | [illegible] |
| MXE | 4452 | ENSG00000287542 | NA | chr4 | + | 88605794 | 88606049 | 88649839 | 88649999 | [illegible] |
| MXE | 13840 | ENSG00000168172 | HOOK3 | chr8 | + | 42997549 | 42997637 | 43002106 | 43002141 | [illegible] |
| MXE | 12155 | ENSG00000072609 | CHFR | chr12 | - | 132877554 | 132877654 | 132887195 | 132887340 | [illegible] |
| MXE | 11959 | ENSG00000164654 | MIOS | chr7 | + | 7589404 | 7589563 | 7594979 | 7595132 | [illegible] |
| MXE | 9417 | ENSG00000160714 | UBE2Q1 | chr1 | - | 154553031 | 154553172 | 154554734 | 154554785 | [illegible] |
| MXE | 16481 | ENSG00000082898 | XPO1 | chr2 | - | 61490641 | 61490776 | 61492034 | 61492198 | [illegible] |
| MXE | 7837 | ENSG00000036549 | ZZZ3 | chr1 | - | 77631849 | 77633405 | 77639448 | 77639602 | [illegible] |
| MXE | 13183 | ENSG00000137642 | SORL1 | chr11 | + | 121543547 | 121543726 | 121545242 | 121545429 | [illegible] |
| MXE | 1853 | ENSG00000103091 | WDR59 | chr16 | - | 74942737 | 74942826 | 74948518 | 74948556 | [illegible] |
| MXE | 128 | ENSG00000100084 | HIRA | chr22 | - | 19353995 | 19354118 | 19355759 | 19355865 | [illegible] |

| | | | | | | | | | | | | | | | | | | | | | | | | | | | | | | | | | | | | | | | | | | | | | | | | |
|---|---|---|---|---|---|---|---|---|---|---|---|---|---|---|---|---|---|---|---|---|---|---|---|---|---|---|---|---|---|---|---|---|---|---|---|---|---|---|---|---|---|---|---|---|---|---|---|---|
| MXE | 5590 | ENSG00000196126 | HLA-DRB1 | chr6 | - | 32580246 | 32580270 | 32580745 | 32580856 | 32578768 | 32579104 | 32581556 | 32581838 | 1141 | 546 | 763 | 689 | 771 | 0 | 192 | 255 | 1 | 0 | 1673 | 1006 | 442 | 872 | 2732 | 0 | 0 | 114 | 232 | 537 | 260 | 173 | 0.058652916 | 0.209034963 | 1 | 0.654 | 0.666 | 0.998 | 1 | 1 | 1 | 0.721 | 0.714 | 0.772 | 0.022 |
| MXE | 15653 | ENSG00000081237 | PTPRC | chr1 | + | 198692346 | 198692373 | 198703297 | 198703372 | 198639225 | 198639341 | 198704471 | 198704485 | 538 | 358 | 408 | 620 | 1270 | 556 | 372 | 381 | 503 | 762 | 44 | 301 | 167 | 42 | 30 | 23 | 199 | 124 | 40 | 7 | 176 | 224 | 0.058677664 | 0.209073289 | 0.552 | 0.551 | 0.577 | 0.611 | 0.68 | 0.709 | 0.658 | 0.632 | 0.572 | 0.845 | -0.089 |
| MXE | 4682 | ENSG00000047365 | ARAP2 | chr4 | - | 36150899 | 36151044 | 36158729 | 36158864 | 36148404 | 36148507 | 36159330 | 36159505 | 245 | 153 | 187 | 261 | 309 | 249 | 146 | 184 | 205 | 190 | 14 | 128 | 43 | 27 | 16 | 49 | 89 | 89 | 18 | 24 | 284 | 294 | 0.058751067 | 0.209284915 | 0.505 | 0.52 | 0.513 | 0.569 | 0.627 | 0.228 | 0.598 | 0.333 | 0.608 | 0.408 | 0.112 |
| MXE | 1400 | ENSG00000108423 | TUBD1 | chr17 | - | 59866608 | 59866749 | 59886082 | 59886230 | 59863663 | 59863847 | 59890830 | 59891041 | 19 | 14 | 33 | 15 | 49 | 27 | 14 | 20 | 14 | 27 | 3 | 12 | 4 | 3 | 3 | 0 | 5 | 1 | 0 | 7 | 297 | 290 | 0.05876672 | 0.209290773 | 0.407 | 0.494 | 0.617 | 0.511 | 0.639 | 1 | 0.701 | 0.796 | 1 | 0.295 | -0.225 |
| MXE | 5420 | ENSG00000109861 | CTSC | chr11 | - | 88309162 | 88309318 | 88312387 | 88312554 | 88296132 | 88296264 | 88334936 | 88335082 | 364 | 218 | 189 | 226 | 213 | 6 | 5 | 3 | 6 | 6 | 574 | 325 | 181 | 317 | 430 | 0 | 23 | 5 | 25 | 4 | 298 | 298 | 0.058804043 | 0.209373785 | 0.984 | 0.978 | 0.984 | 0.974 | 0.973 | 1 | 0.934 | 0.973 | 0.927 | 0.991 | 0.014 |
| MXE | 14902 | ENSG00000165699 | TSC1 | chr9 | - | 132925586 | 132925739 | 132928766 | 132928952 | 132923347 | 132923492 | 132935032 | 132935095 | 73 | 56 | 98 | 83 | 60 | 20 | 8 | 12 | 26 | 35 | 52 | 106 | 62 | 91 | 39 | 24 | 27 | 3 | 1 | 8 | 298 | 298 | 0.058861985 | 0.209515432 | 0.785 | 0.875 | 0.891 | 0.761 | 0.632 | 0.684 | 0.797 | 0.954 | 0.989 | 0.83 | -0.062 |
| MXE | 13853 | ENSG00000138378 | STAT4 | chr2 | - | 191058017 | 191058111 | 191058201 | 191058219 | 191054489 | 191054534 | 191058709 | 191058769 | 7 | 1 | 7 | 3 | 4 | 309 | 185 | 268 | 262 | 447 | 0 | 0 | 1 | 0 | 0 | 53 | 104 | 142 | 55 | 9 | 167 | 243 | 0.058871873 | 0.209515432 | 0.032 | 0.008 | 0.037 | 0.016 | 0.013 | 0 | 0 | 0.01 | 0 | 0 | 0.019 |
| MXE | 10215 | ENSG00000009335 | UBE3C | chr7 | + | 157223253 | 157223351 | 157225406 | 157225539 | 157220688 | 157220776 | 157231079 | 157231327 | 60 | 29 | 45 | 45 | 106 | 170 | 108 | 161 | 117 | 234 | 107 | 59 | 49 | 31 | 42 | 160 | 97 | 107 | 120 | 98 | 247 | 282 | 0.058894862 | 0.20954733 | 0.287 | 0.235 | 0.242 | 0.305 | 0.341 | 0.433 | 0.41 | 0.343 | 0.228 | 0.329 | -0.067 |
| MXE | 8881 | ENSG00000273136 | NBPF26 | chr1 | + | 120811885 | 120812095 | 120813890 | 120813993 | 120810346 | 120810558 | 120814828 | 120815043 | 42 | 45 | 21 | 18 | 50 | 28 | 33 | 20 | 26 | 40 | 56 | 33 | 43 | 58 | 10 | 8 | 11 | 33 | 31 | 14 | 298 | 252 | 0.059006267 | 0.209893723 | 0.559 | 0.536 | 0.47 | 0.369 | 0.514 | 0.855 | 0.717 | 0.524 | 0.613 | 0.377 | -0.128 |
| MXE | 560 | ENSG00000126261 | UBA2 | chr19 | + | 34431860 | 34431931 | 34433347 | 34433412 | 34430575 | 34430659 | 34434867 | 34434968 | 62 | 38 | 71 | 69 | 70 | 345 | 225 | 311 | 356 | 783 | 51 | 44 | 57 | 45 | 28 | 261 | 247 | 206 | 209 | 195 | 220 | 214 | 0.059026224 | 0.209914731 | 0.149 | 0.141 | 0.182 | 0.159 | 0.08 | 0.16 | 0.148 | 0.212 | 0.173 | 0.123 | -0.021 |
| MXE | 2270 | ENSG00000068305 | MEF2A | chr15 | + | 99690240 | 99690428 | 99706728 | 99706855 | 99675398 | 99675458 | 99710633 | 99710760 | 112 | 118 | 116 | 112 | 149 | 141 | 119 | 92 | 117 | 190 | 74 | 149 | 146 | 219 | 43 | 79 | 152 | 130 | 135 | 24 | 298 | 276 | 0.059052443 | 0.209957995 | 0.424 | 0.479 | 0.539 | 0.47 | 0.421 | 0.465 | 0.476 | 0.51 | 0.6 | 0.624 | -0.068 |
| MXE | 5963 | ENSG00000165240 | ATP7A | chrX | + | 78011448 | 78011674 | 78012878 | 78013112 | 78009101 | 78009263 | 78014661 | 78014753 | 0 | 1 | 2 | 10 | 3 | 35 | 18 | 26 | 31 | 27 | 0 | 11 | 6 | 9 | 0 | 16 | 17 | 20 | 7 | 16 | 298 | 298 | 0.059080689 | 0.210008446 | 0 | 0.053 | 0.071 | 0.244 | 0.1 | 0 | 0.393 | 0.231 | 0.562 | 0 | -0.144 |
| MXE | 4781 | ENSG00000143669 | LYST | chr1 | - | 235793502 | 235793612 | 235812970 | 235813061 | 235791572 | 235792125 | 235830225 | 235830424 | 223 | 257 | 135 | 218 | 223 | 216 | 240 | 163 | 171 | 331 | 33 | 179 | 87 | 30 | 21 | 26 | 94 | 55 | 20 | 34 | 240 | 259 | 0.059181476 | 0.210274967 | 0.527 | 0.536 | 0.472 | 0.579 | 0.421 | 0.578 | 0.673 | 0.631 | 0.618 | 0.4 | -0.073 |
| MXE | 2396 | ENSG00000138613 | APH1B | chr15 | + | 63287423 | 63287546 | 63302344 | 63302472 | 63286557 | 63286628 | 63305613 | 63305801 | 43 | 35 | 33 | 36 | 22 | 27 | 16 | 23 | 14 | 24 | 45 | 43 | 18 | 30 | 30 | 25 | 58 | 25 | 28 | 24 | 272 | 277 | 0.059187577 | 0.210274967 | 0.619 | 0.69 | 0.594 | 0.724 | 0.483 | 0.647 | 0.43 | 0.423 | 0.522 | 0.56 | 0.106 |
| MXE | 13301 | ENSG00000157881 | PANK4 | chr1 | - | 2515561 | 2515717 | 2518163 | 2518264 | 2514353 | 2514466 | 2518515 | 2518597 | 16 | 4 | 10 | 14 | 5 | 8 | 7 | 8 | 20 | 12 | 22 | 2 | 11 | 17 | 5 | 0 | 7 | 2 | 11 | 6 | 250 | 298 | 0.059197892 | 0.210274967 | 0.704 | 0.405 | 0.598 | 0.455 | 0.332 | 1 | 0.254 | 0.868 | 0.648 | 0.498 | -0.155 |
| MXE | 7155 | ENSG00000095787 | WAC | chr10 | + | 28608185 | 28608431 | 28610698 | 28610821 | 28590719 | 28590832 | 28611773 | 28611922 | 158 | 68 | 143 | 170 | 278 | 656 | 516 | 548 | 614 | 1191 | 77 | 110 | 74 | 98 | 8 | 238 | 480 | 688 | 534 | 83 | 298 | 272 | 0.059245231 | 0.210393097 | 0.18 | 0.107 | 0.192 | 0.202 | 0.176 | 0.228 | 0.173 | 0.089 | 0.143 | 0.081 | 0.029 |
| MXE | 5500 | ENSG00000168405 | CMAHP | chr6 | - | 25109955 | 25110103 | 25113591 | 25113782 | 25109499 | 25109581 | 25114990 | 25115100 | 24 | 29 | 34 | 43 | 30 | 52 | 43 | 26 | 45 | 63 | 7 | 49 | 24 | 2 | 12 | 0 | 26 | 64 | 8 | 0 | 298 | 297 | 0.059296651 | 0.210520791 | 0.315 | 0.402 | 0.566 | 0.488 | 0.322 | 1 | 0.653 | 0.272 | 0.199 | 1 | -0.206 |
| MXE | 13227 | ENSG00000188917 | TRMT2B | chrX | - | 101037916 | 101038051 | 101042232 | 101042312 | 101036973 | 101037073 | 101051783 | 101052082 | 12 | 3 | 7 | 3 | 2 | 40 | 19 | 30 | 35 | 70 | 28 | 21 | 6 | 5 | 0 | 15 | 42 | 25 | 20 | 49 | 229 | 284 | 0.059309371 | 0.210520791 | 0.271 | 0.164 | 0.224 | 0.096 | 0.034 | 0.698 | 0.383 | 0.229 | 0.237 | 0 | -0.152 |
| MXE | 11958 | ENSG00000164654 | MIOS | chr7 | + | 7583117 | 7583372 | 7585635 | 7585805 | 7574097 | 7574196 | 7588497 | 7588563 | 119 | 95 | 111 | 121 | 221 | 80 | 75 | 76 | 77 | 102 | 50 | 78 | 123 | 97 | 8 | 30 | 50 | 40 | 39 | 5 | 298 | 298 | 0.05937328 | 0.210625363 | 0.598 | 0.559 | 0.594 | 0.611 | 0.684 | 0.625 | 0.609 | 0.755 | 0.713 | 0.615 | -0.054 |
| MXE | 16263 | ENSG00000062650 | WAPL | chr10 | - | 86461175 | 86461287 | 86467278 | 86467506 | 86460398 | 86460496 | 86470991 | 86471103 | 177 | 113 | 161 | 137 | 176 | 228 | 196 | 204 | 230 | 361 | 37 | 103 | 107 | 47 | 2 | 59 | 193 | 214 | 107 | 26 | 298 | 261 | 0.059378724 | 0.210625363 | 0.405 | 0.336 | 0.409 | 0.343 | 0.299 | 0.355 | 0.319 | 0.305 | 0.278 | 0.063 | 0.094 |
| MXE | 3190 | ENSG00000111860 | CEP85L | chr6 | - | 118483705 | 118483858 | 118491685 | 118491865 | 118481778 | 118481933 | 118511297 | 118511415 | 176 | 113 | 209 | 161 | 282 | 147 | 118 | 139 | 113 | 225 | 28 | 69 | 70 | 31 | 21 | 7 | 49 | 46 | 27 | 4 | 298 | 298 | 0.059381127 | 0.210625363 | 0.545 | 0.489 | 0.601 | 0.588 | 0.556 | 0.8 | 0.585 | 0.603 | 0.534 | 0.84 | -0.117 |
| MXE | 1846 | ENSG00000103248 | MTHFSD | chr16 | - | 86551972 | 86552146 | 86552178 | 86552292 | 86548463 | 86548577 | 86554644 | 86554751 | 9 | 21 | 24 | 28 | 35 | 20 | 11 | 23 | 13 | 29 | 47 | 14 | 2 | 16 | 0 | 25 | 24 | 7 | 11 | 29 | 263 | 298 | 0.059409218 | 0.210674987 | 0.338 | 0.684 | 0.542 | 0.709 | 0.578 | 0.681 | 0.398 | 0.245 | 0.622 | 0 | 0.181 |

[illegible]

| | | | | | | | | | | | | | | | | | | | | | | | | | | | | | | | | | | | | | | | | | | | | | | | | |
|---|---|---|---|---|---|---|---|---|---|---|---|---|---|---|---|---|---|---|---|---|---|---|---|---|---|---|---|---|---|---|---|---|---|---|---|---|---|---|---|---|---|---|---|---|---|---|---|---|
| MXE | 5540 | ENSG00000169180 | XPO6 | chr16 | - | 28146093 | 28146203 | 28152658 | 28152785 | 28135215 | 28135324 | 28156073 | 28156527 | 119 | 59 | 135 | 111 | 175 | 148 | 120 | 152 | 146 | 249 | 107 | 62 | 119 | 67 | 124 | 259 | 134 | 182 | 190 | 119 | 276 | 259 | 0.070204829 | 0.237030541 | 0.43 | 0.316 | 0.455 | 0.416 | 0.397 | 0.279 | 0.303 | 0.38 | 0.249 | 0.494 | 0.062 |
| MXE | 8753 | ENSG00000101040 | ZMYND8 | chr20 | - | 47224316 | 47224556 | 47227202 | 47227281 | 47221313 | 47221474 | 47229725 | 47229806 | 3 | 1 | 10 | 20 | 37 | 50 | 33 | 45 | 81 | 114 | 0 | 19 | 5 | 9 | 5 | 41 | 90 | 89 | 78 | 23 | 228 | 298 | 0.07033807 | 0.237421492 | 0.073 | 0.038 | 0.225 | 0.244 | 0.298 | 0 | 0.216 | 0.068 | 0.131 | 0.221 | 0.048 |
| MXE | 5915 | ENSG00000157216 | SSBP3 | chr1 | - | 54257126 | 54257186 | 54258068 | 54258149 | 54251793 | 54251860 | 54281437 | 54281527 | 18 | 3 | 4 | 4 | 9 | 307 | 106 | 177 | 237 | 256 | 8 | 4 | 3 | 12 | 0 | 249 | 231 | 133 | 240 | 382 | 230 | 209 | 0.070352406 | 0.237421492 | 0.051 | 0.025 | 0.02 | 0.015 | 0.031 | 0.028 | 0.015 | 0.02 | 0.043 | 0 | 0.007 |
| MXE | 11845 | ENSG00000164674 | SYTL3 | chr6 | + | 158682924 | 158682989 | 158708321 | 158708391 | 158665394 | 158665613 | 158713799 | 158713878 | 33 | 15 | 7 | 16 | 9 | 56 | 18 | 26 | 22 | 21 | 11 | 28 | 61 | 11 | 21 | 38 | 36 | 10 | 11 | 22 | 214 | 219 | 0.070459444 | 0.237729017 | 0.376 | 0.46 | 0.216 | 0.427 | 0.305 | 0.229 | 0.443 | 0.862 | 0.506 | 0.494 | -0.15 |
| MXE | 5253 | ENSG00000172673 | THEMIS | chr6 | - | 127812882 | 127813931 | 127900841 | 127901081 | 127719687 | 127719823 | 127918427 | 127918540 | 97 | 82 | 86 | 57 | 77 | 88 | 86 | 127 | 107 | 133 | 23 | 64 | 22 | 11 | 3 | 0 | 71 | 24 | 16 | 3 | 298 | 298 | 0.070563081 | 0.238024935 | 0.524 | 0.488 | 0.404 | 0.348 | 0.367 | 1 | 0.474 | 0.478 | 0.407 | 0.5 | -0.146 |
| MXE | 10297 | ENSG00000081320 | STK17B | chr2 | - | 196139619 | 196139799 | 196141248 | 196141297 | 196133582 | 196137729 | 196143559 | 196143686 | 737 | 742 | 736 | 849 | 1971 | 1796 | 1574 | 1619 | 1815 | 5727 | 328 | 601 | 1210 | 589 | 81 | 588 | 1495 | 2320 | 1262 | 159 | 198 | 298 | 0.070588528 | 0.238057023 | 0.382 | 0.415 | 0.406 | 0.413 | 0.341 | 0.456 | 0.377 | 0.44 | 0.413 | 0.434 | -0.033 |
| MXE | 6243 | ENSG00000112144 | CILK1 | chr6 | - | 53019226 | 53019359 | 53031064 | 53031144 | 53018329 | 53018501 | 53032532 | 53032654 | 29 | 16 | 25 | 40 | 79 | 28 | 25 | 41 | 55 | 68 | 0 | 10 | 6 | 12 | 14 | 16 | 17 | 8 | 13 | 31 | 229 | 282 | 0.0707104 | 0.238340339 | 0.561 | 0.441 | 0.429 | 0.472 | 0.589 | 0 | 0.42 | 0.48 | 0.532 | 0.357 | 0.141 |
| MXE | 11784 | ENSG00000188229 | TUBB4B | chr9 | + | 137241720 | 137241829 | 137241910 | 137242021 | 137241286 | 137241417 | 137242495 | 137243707 | 29 | 12 | 24 | 46 | 14 | 113 | 37 | 87 | 116 | 54 | 50 | 35 | 17 | 27 | 226 | 235 | 106 | 52 | 207 | 946 | 258 | 260 | 0.070710609 | 0.238340339 | 0.205 | 0.246 | 0.218 | 0.286 | 0.207 | 0.177 | 0.25 | 0.248 | 0.116 | 0.194 | 0.035 |
| MXE | 16168 | ENSG00000107669 | ATE1 | chr10 | - | 121899865 | 121899994 | 121902390 | 121902620 | 121870005 | 121870038 | 121910905 | 121911151 | 94 | 99 | 89 | 119 | 346 | 18 | 15 | 24 | 23 | 37 | 97 | 86 | 85 | 113 | 109 | 28 | 27 | 19 | 23 | 13 | 298 | 278 | 0.070724235 | 0.238340339 | 0.83 | 0.86 | 0.776 | 0.828 | 0.897 | 0.764 | 0.748 | 0.807 | 0.821 | 0.887 | 0.033 |
| MXE | 1544 | ENSG00000124120 | TTPAL | chr20 | + | 44484336 | 44484530 | 44486595 | 44486706 | 44479984 | 44480444 | 44489262 | 44494603 | 6 | 8 | 18 | 15 | 14 | 40 | 25 | 55 | 39 | 98 | 0 | 25 | 9 | 29 | 34 | 38 | 29 | 28 | 43 | 7 | 298 | 260 | 0.07073635 | 0.238340339 | 0.116 | 0.218 | 0.222 | 0.251 | 0.111 | 0 | 0.429 | 0.219 | 0.37 | 0.809 | -0.182 |
| MXE | 10347 | ENSG00000031003 | FAM13B | chr5 | - | 137985256 | 137985389 | 137987460 | 137987616 | 137962404 | 137962469 | 137988273 | 137988315 | 327 | 273 | 235 | 350 | 378 | 295 | 273 | 172 | 271 | 295 | 105 | 343 | 236 | 196 | 191 | 113 | 179 | 202 | 124 | 57 | 298 | 282 | 0.070787483 | 0.238371939 | 0.512 | 0.486 | 0.564 | 0.55 | 0.548 | 0.468 | 0.645 | 0.525 | 0.599 | 0.76 | -0.067 |
| MXE | 13711 | ENSG00000183354 | KIAA2026 | chr9 | - | 5933750 | 5933881 | 5944870 | 5944960 | 5929043 | 5929159 | 5954015 | 5954095 | 104 | 141 | 112 | 138 | 208 | 18 | 13 | 22 | 18 | 22 | 44 | 166 | 141 | 58 | 19 | 8 | 19 | 15 | 0 | 0 | 239 | 280 | 0.070791425 | 0.238371939 | 0.871 | 0.927 | 0.856 | 0.9 | 0.917 | 0.866 | 0.911 | 0.917 | 1 | 1 | -0.045 |
| MXE | 8608 | ENSG00000162664 | ZNF326 | chr1 | + | 90020791 | 90020922 | 90022249 | 90022345 | 90018684 | 90018784 | 90027353 | 90028226 | 76 | 99 | 102 | 99 | 155 | 57 | 52 | 54 | 83 | 98 | 18 | 34 | 96 | 29 | 12 | 7 | 55 | 70 | 34 | 17 | 280 | 245 | 0.070797926 | 0.238371939 | 0.538 | 0.625 | 0.623 | 0.511 | 0.581 | 0.692 | 0.351 | 0.545 | 0.427 | 0.382 | 0.096 |
| MXE | 1900 | ENSG00000122299 | ZC3H7A | chr16 | - | 11763477 | 11763659 | 11765052 | 11765153 | 11762670 | 11762747 | 11765488 | 11765561 | 70 | 54 | 57 | 71 | 110 | 166 | 116 | 149 | 133 | 232 | 64 | 86 | 86 | 59 | 15 | 114 | 126 | 168 | 117 | 28 | 250 | 298 | 0.070809549 | 0.238371939 | 0.335 | 0.357 | 0.313 | 0.389 | 0.361 | 0.401 | 0.449 | 0.379 | 0.375 | 0.39 | -0.048 |
| MXE | 2752 | ENSG00000196405 | EVL | chr14 | + | 100126706 | 100126771 | 100128518 | 100128748 | 100123538 | 100123602 | 100129562 | 100129684 | 248 | 151 | 210 | 196 | 561 | 273 | 183 | 244 | 273 | 110 | 180 | 127 | 96 | 138 | 192 | 162 | 246 | 113 | 142 | 295 | 214 | 298 | 0.070864006 | 0.23850152 | 0.558 | 0.535 | 0.545 | 0.5 | 0.877 | 0.607 | 0.418 | 0.542 | 0.575 | 0.475 | 0.08 |
| MXE | 7724 | ENSG00000082512 | TRAF5 | chr1 | + | 211369451 | 211369592 | 211371301 | 211371470 | 211365375 | 211365468 | 211372127 | 211372795 | 54 | 46 | 50 | 54 | 99 | 128 | 140 | 192 | 178 | 296 | 9 | 80 | 51 | 11 | 1 | 22 | 167 | 77 | 35 | 1 | 290 | 298 | 0.070895478 | 0.238522807 | 0.302 | 0.252 | 0.211 | 0.238 | 0.256 | 0.296 | 0.33 | 0.405 | 0.244 | 0.507 | -0.105 |
| MXE | 17071 | ENSG00000137822 | TUBGCP4 | chr15 | + | 43401715 | 43401850 | 43403682 | 43403799 | 43400043 | 43400221 | 43404412 | 43404552 | 12 | 22 | 17 | 33 | 24 | 4 | 3 | 9 | 26 | 25 | 12 | 38 | 28 | 27 | 31 | 0 | 2 | 7 | 17 | 20 | 284 | 266 | 0.070902261 | 0.238522807 | 0.738 | 0.873 | 0.639 | 0.543 | 0.473 | 1 | 0.947 | 0.789 | 0.598 | 0.592 | -0.132 |
| MXE | 8234 | ENSG00000164300 | SERINC5 | chr5 | - | 80150881 | 80150948 | 80158835 | 80158962 | 80147244 | 80147284 | 80166382 | 80166478 | 115 | 82 | 149 | 131 | 103 | 39 | 31 | 69 | 61 | 110 | 85 | 81 | 71 | 127 | 27 | 20 | 16 | 61 | 8 | 19 | 276 | 216 | 0.070993882 | 0.238777262 | 0.698 | 0.674 | 0.628 | 0.627 | 0.423 | 0.769 | 0.798 | 0.477 | 0.926 | 0.527 | -0.089 |
| MXE | 15094 | ENSG00000149182 | ARFGAP2 | chr11 | - | 47172280 | 47172333 | 47175850 | 47175923 | 47171663 | 47171800 | 47176515 | 47176634 | 149 | 89 | 142 | 118 | 185 | 21 | 15 | 10 | 9 | 14 | 158 | 128 | 82 | 115 | 225 | 26 | 22 | 16 | 14 | 19 | 222 | 202 | 0.071063291 | 0.238956913 | 0.866 | 0.844 | 0.928 | 0.923 | 0.923 | 0.847 | 0.841 | 0.823 | 0.882 | 0.915 | 0.035 |
| MXE | 14040 | ENSG00000120899 | PTK2B | chr8 | + | 27436250 | 27436348 | 27437121 | 27437206 | 27435742 | 27435793 | 27437395 | 27437496 | 94 | 65 | 81 | 97 | 41 | 104 | 64 | 65 | 106 | 122 | 143 | 183 | 103 | 197 | 273 | 199 | 137 | 107 | 157 | 144 | 247 | 234 | 0.071220024 | 0.239430053 | 0.461 | 0.49 | 0.541 | 0.464 | 0.241 | 0.405 | 0.559 | 0.477 | 0.543 | 0.642 | -0.086 |
| MXE | 4747 | ENSG00000131389 | SLC6A6 | chr3 | + | 14472204 | 14472317 | 14477204 | 14477342 | 14467852 | 14467956 | 14478465 | 14478568 | 3 | 7 | 1 | 6 | 0 | 80 | 59 | 58 | 96 | 77 | 0 | 0 | 6 | 5 | 2 | 141 | 103 | 88 | 77 | 175 | 262 | 287 | 0.071360906 | 0.239782517 | 0.039 | 0.115 | 0.019 | 0.064 | 0 | 0 | 0 | 0.069 | 0.066 | 0.012 | 0.018 |
| MXE | 4613 | ENSG00000109133 | TMEM33 | chr4 | + | 41938601 | 41938696 | 41943746 | 41943814 | 41935128 | 41935529 | 41944792 | 41944926 | 18 | 22 | 20 | 21 | 19 | 97 | 53 | 46 | 74 | 99 | 25 | 18 | 27 | 7 | 15 | 10 | 62 | 58 | 60 | 13 | 244 | 217 | 0.071372388 | 0.239782517 | 0.142 | 0.27 | 0.279 | 0.202 | 0.146 | 0.69 | 0.205 | 0.293 | 0.094 | 0.506 | -0.15 |
| MXE | 14752 | ENSG00000048828 | FAM120A | chr9 | + | 93529352 | 93529580 | 93532154 | 93532329 | 93527154 | 93527242 | 93543221 | 93543471 | 237 | 111 | 203 | 207 | 212 | 254 | 167 | 212 | 268 | 209 | 314 | 220 | 161 | 214 | 242 | 436 | 263 | 187 | 353 | 345 | 298 | 298 | 0.071378537 | 0.239782517 | 0.483 | 0.399 | 0.489 | 0.436 | 0.504 | 0.419 | 0.455 | 0.463 | 0.377 | 0.412 | 0.037 |
| MXE | 12324 | ENSG00000066422 | ZBTB11 | chr3 | - | 101664963 | 101665808 | 101671129 | 101671361 | 101664537 | 101664714 | 101671977 | 101672213 | 94 | 66 | 70 | 66 | 105 | 167 | 113 | 147 | 152 | 152 | 16 | 58 | 23 | 34 | 42 | 26 | 73 | 116 | 22 | 1 | 298 | 298 | 0.071389065 | 0.239782517 | 0.36 | 0.369 | 0.323 | 0.303 | 0.409 | 0.381 | 0.443 | 0.165 | 0.607 | 0.977 | -0.162 |
| MXE | 13286 | ENSG00000187676 | B3GLCT | chr13 | + | 31274508 | 31274628 | 31276701 | 31276771 | 31269213 | 31269277 | 31284647 | 31284761 | 18 | 7 | 10 | 11 | 20 | 24 | 8 | 15 | 10 | 16 | 26 | 2 | 3 | 13 | 4 | 0 | 5 | 9 | 7 | 0 | 269 | 219 | 0.071434732 | 0.239881971 | 0.379 | 0.416 | 0.352 | 0.472 | 0.504 | 1 | 0.246 | 0.213 | 0.602 | 1 | -0.188 |
| MXE | 62 | ENSG00000159131 | GART | chr21 | - | 33530758 | 33530884 | 33531488 | 33531557 | 33528849 | 33528937 | 33532344 | 33532456 | 48 | 27 | 28 | 38 | 48 | 118 | 64 | 80 | 77 | 110 | 58 | 39 | 38 | 52 | 37 | 155 | 87 | 38 | 60 | 71 | 218 | 275 | 0.07154844 | 0.240209817 | 0.339 | 0.347 | 0.306 | 0.384 | 0.355 | 0.321 | 0.361 | 0.558 | 0.522 | 0.397 | -0.086 |
| MXE | 6090 | ENSG00000107566 | ERLIN1 | chr10 | - | 100175944 | 100176070 | 100178132 | 100178194 | 100174207 | 100174281 | 100179200 | 100179247 | 11 | 5 | 9 | 14 | 27 | 78 | 66 | 65 | 69 | 139 | 0 | 15 | 13 | 6 | 13 | 124 | 90 | 85 | 86 | 68 | 211 | 275 | 0.071632803 | 0.24040798 | 0.155 | 0.09 | 0.153 | 0.209 | 0.202 | 0 | 0.178 | 0.166 | 0.083 | 0.199 | 0.037 |
| MXE | 8721 | ENSG00000197323 | TRIM33 | chr1 | - | 114421435 | 114421636 | 114424590 | 114424755 | 114410183 | 114410316 | 114425448 | 114425723 | 211 | 173 | 145 | 222 | 234 | 105 | 79 | 104 | 92 | 44 | 46 | 224 | 162 | 91 | 14 | 10 | 125 | 42 | 23 | 0 | 298 | 298 | 0.071639647 | 0.24040798 | 0.668 | 0.687 | 0.582 | 0.707 | 0.842 | 0.821 | 0.642 | 0.794 | 0.798 | 1 | -0.114 |
| MXE | 9600 | ENSG00000263528 | IKBKE | chr1 | + | 206493022 | 206493119 | 206493265 | 206493378 | 206491647 | 206491749 | 206493919 | 206493991 | 52 | 34 | 54 | 51 | 61 | 36 | 11 | 28 | 36 | 32 | 123 | 91 | 28 | 67 | 139 | 91 | 26 | 7 | 34 | 30 | 246 | 262 | 0.071664442 | 0.240437179 | 0.606 | 0.767 | 0.673 | 0.601 | 0.67 | 0.59 | 0.788 | 0.81 | 0.677 | 0.831 | -0.076 |
| MXE | 5097 | ENSG00000060971 | ACAA1 | chr3 | - | 38126509 | 38126700 | 38129289 | 38129388 | 38125825 | 38125881 | 38131595 | 38131638 | 48 | 49 | 45 | 57 | 45 | 62 | 24 | 43 | 34 | 17 | 109 | 133 | 32 | 86 | 182 | 115 | 72 | 68 | 118 | 166 | 248 | 298 | 0.07177705 | 0.240760918 | 0.482 | 0.71 | 0.557 | 0.668 | 0.761 | 0.532 | 0.689 | 0.361 | 0.467 | 0.568 | 0.112 |
| MXE | 13085 | ENSG00000006757 | PNPLA4 | chrX | - | 7922003 | 7922098 | 7925939 | 7926132 | 7921712 | 7921848 | 7927685 | 7927739 | 2 | 1 | 1 | 2 | 6 | 48 | 22 | 51 | 35 | 97 | 0 | 0 | 0 | 4 | 0 | 38 | 18 | 12 | 35 | 26 | 298 | 244 | 0.071822931 | 0.240860738 | 0.033 | 0.036 | 0.016 | 0.045 | 0.048 | 0 | 0 | 0 | 0.086 | 0 | 0.018 |
| MXE | 5558 | ENSG00000173193 | PARP14 | chr3 | + | 122695425 | 122695662 | 122699389 | 122701635 | 122692300 | 122692543 | 122703741 | 122703978 | 351 | 458 | 250 | 310 | 631 | 54 | 58 | 33 | 50 | 41 | 167 | 330 | 301 | 311 | 57 | 28 | 62 | 19 | 55 | 17 | 298 | 298 | 0.071922277 | 0.241095346 | 0.867 | 0.888 | 0.883 | 0.861 | 0.939 | 0.856 | 0.842 | 0.941 | 0.85 | 0.77 | 0.036 |

[illegible]

| | | | | | | | | | | | | | | |
|---|---|---|---|---|---|---|---|---|---|---|---|---|---|---|
| MXE | 81 | ENSG00000254413 | CHKB-CPT1B | chr22 | - | 50579726 | 50579830 | 50579973 | 50580082 | 50579425 | 50579507 | 50580189 | 50580416 | 184 |
| MXE | 10730 | ENSG00000132912 | DCTN4 | chr5 | - | 150715564 | 150715662 | 150718275 | 150718383 | 150708439 | 150711362 | 150719715 | 150719770 | 72 |
| MXE | 8003 | ENSG00000197943 | PLCG2 | chr16 | + | 81905402 | 81905507 | 81907684 | 81907774 | 81900611 | 81900780 | 81908415 | 81908591 | 79 |
| MXE | 2789 | ENSG00000156414 | TDRD9 | chr14 | + | 104018091 | 104018192 | 104022156 | 104022330 | 104015980 | 104016088 | 104024568 | 104024680 | 4 |
| MXE | 8930 | ENSG00000116667 | C1orf21 | chr1 | + | 184507587 | 184507682 | 184590738 | 184590815 | 184477385 | 184477603 | 184598400 | 184598461 | 174 |
| MXE | 15409 | ENSG00000135365 | PHF21A | chr11 | - | 45971115 | 45971367 | 45979759 | 45979966 | 45969814 | 45969901 | 46076753 | 46076819 | 227 |
| MXE | 5089 | ENSG00000114742 | WDR48 | chr3 | + | 39074725 | 39074950 | 39077138 | 39077213 | 39069642 | 39069744 | 39078136 | 39078239 | 95 |
| MXE | 14899 | ENSG00000165699 | TSC1 | chr9 | - | 132910570 | 132910689 | 132911001 | 132911113 | 132907300 | 132907370 | 132911452 | 132911568 | 15 |
| MXE | 1791 | ENSG00000261740 | BOLA2-SMG1P6 | chr16 | - | 29446801 | 29447026 | 29454015 | 29454113 | 29444397 | 29444499 | 29454190 | 29454651 | 9 |
| MXE | 13259 | ENSG00000132356 | PRKAA1 | chr5 | - | 40771718 | 40771863 | 40775409 | 40775503 | 40767506 | 40767690 | 40777444 | 40777586 | 221 |
| MXE | 1129 | ENSG00000108799 | EZH1 | chr17 | - | 42722794 | 42722915 | 42727634 | 42727763 | 42720272 | 42720449 | 42728824 | 42728952 | 140 |
| MXE | 13250 | ENSG00000128731 | HERC2 | chr15 | - | 28292887 | 28293022 | 28299401 | 28299516 | 28280067 | 28280287 | 28321361 | 28321464 | 15 |
| MXE | 9469 | ENSG00000131408 | NR1H2 | chr19 | + | 50376234 | 50376576 | 50376718 | 50376826 | 50372870 | 50372985 | 50377586 | 50377648 | 6 |
| MXE | 1821 | ENSG00000118900 | UBN1 | chr16 | + | 4852878 | 4853166 | 4857989 | 4858076 | 4847480 | 4848210 | 4858567 | 4858663 | 10 |
| MXE | 7183 | ENSG00000166669 | ATF7IP2 | chr16 | + | 10457371 | 10457529 | 10472109 | 10472183 | 10440363 | 10440462 | 10473478 | 10473534 | 14 |
| MXE | 12002 | ENSG00000089820 | ARHGAP4 | chrX | - | 153933942 | 153934001 | 153934376 | 153934475 | 153932538 | 153932584 | 153934883 | 153934999 | 11 |
| MXE | 13385 | ENSG00000147394 | ZNF185 | chrX | + | 152920709 | 152920748 | 152922172 | 152922256 | 152920327 | 152920411 | 152928574 | 152928661 | 6 |
| MXE | 2024 | ENSG00000182511 | FES | chr15 | + | 90889305 | 90889443 | 90889516 | 90889636 | 90887186 | 90887370 | 90889839 | 90889962 | 75 |
| MXE | 9898 | ENSG00000075884 | ARHGAP15 | chr2 | + | 143202133 | 143202202 | 143216383 | 143216445 | 143155476 | 143155655 | 143228580 | 143228668 | 142 |
| MXE | 13202 | ENSG00000147202 | DIAPH2 | chrX | + | 96763099 | 96763120 | 96881578 | 96881718 | 96758153 | 96758258 | 96912327 | 96912402 | 9 |
| MXE | 6499 | ENSG00000125637 | PSD4 | chr2 | + | 113185364 | 113186255 | 113193047 | 113193128 | 113184956 | 113185073 | 113193257 | 113193370 | 42 |
| MXE | 8292 | ENSG00000257379 | NA | chr12 | + | 53455884 | 53456001 | 53459271 | 53459403 | 53455460 | 53455493 | 53461014 | 53461143 | 2041 |
| MXE | 13752 | ENSG00000143569 | UBAP2L | chr1 | + | 154234590 | 154234792 | 154235195 | 154235291 | 154228614 | 154228725 | 154236565 | 154236611 | 670 |
| MXE | 3293 | ENSG00000141279 | NPEPPS | chr17 | + | 47585499 | 47585745 | 47586147 | 47586244 | 47579389 | 47579511 | 47586364 | 47586398 | 3 |
| MXE | 11102 | ENSG00000184182 | UBE2F | chr2 | + | 237994743 | 237994809 | 238025341 | 238025412 | 237987962 | 237987992 | 238035877 | 238035940 | 69 |
| MXE | 16803 | ENSG00000110958 | PTGES3 | chr12 | - | 56671748 | 56671809 | 56672739 | 56672809 | 56670274 | 56670364 | 56672951 | 56673065 | 591 |
| MXE | 1581 | ENSG00000166747 | AP1G1 | chr16 | - | 71771155 | 71771252 | 71773220 | 71773362 | 71769618 | 71769699 | 71774467 | 71774592 | 131 |
| MXE | 15609 | ENSG00000159063 | ALG8 | chr11 | - | 78106806 | 78106946 | 78109441 | 78109581 | 78104355 | 78104453 | 78112649 | 78112770 | 46 |
| MXE | 14374 | ENSG00000104529 | EEF1D | chr8 | - | 143581228 | 143581328 | 143586218 | 143586290 | 143581053 | 143581154 | 143586728 | 143586849 | 248 |
| MXE | 12900 | ENSG00000105939 | ZC3HAV1 | chr7 | - | 139055204 | 139055295 | 139061035 | 139061138 | 139053964 | 139054095 | 139064878 | 139064999 | 260 |
| MXE | 9845 | ENSG00000115966 | ATF2 | chr2 | - | 175136411 | 175136486 | 175151059 | 175151158 | 175130137 | 175130207 | 175168049 | 175168144 | 61 |
| [illegible] | [illegible] | [illegible] | [illegible] | [illegible] | [illegible] | [illegible] | [illegible] | [illegible] | [illegible] | [illegible] | [illegible] | [illegible] | [illegible] | [illegible] |

[illegible]

| | | | | | | | | | | | | | | |
|---|---|---|---|---|---|---|---|---|---|---|---|---|---|---|
| MXE | 2809 | ENSG00000033170 | FUT8 | chr14 | + | 65561336 | 65561766 | 65615977 | 65616093 | 65455620 | 65455718 | 65616210 | 65616373 | 29 |
| MXE | 16726 | ENSG00000111271 | ACAD10 | chr12 | + | 111721670 | 111721739 | 111727961 | 111728143 | 111715820 | 111715962 | 111729805 | 111729956 | 7 |
| MXE | 6975 | ENSG00000173230 | GOLGB1 | chr3 | - | 121699811 | 121699885 | 121702480 | 121702595 | 121693740 | 121698929 | 121714860 | 121714976 | 55 |
| MXE | 7479 | ENSG00000176148 | TCP11L1 | chr11 | + | 33043749 | 33043936 | 33054592 | 33054725 | 33039571 | 33039792 | 33057114 | 33057235 | 9 |
| MXE | 11231 | ENSG00000047932 | GOPC | chr6 | - | 117569571 | 117569736 | 117570859 | 117570955 | 117566853 | 117567034 | 117573466 | 117573632 | 97 |
| MXE | 8072 | ENSG00000103657 | HERC1 | chr15 | - | 63756436 | 63756748 | 63758174 | 63758369 | 63755228 | 63755325 | 63764095 | 63764191 | 306 |
| MXE | 4391 | ENSG00000121210 | TMEM131L | chr4 | + | 153556965 | 153557082 | 153580825 | 153580903 | 153555786 | 153555910 | 153581406 | 153581560 | 34 |
| MXE | 14163 | ENSG00000166925 | TSC22D4 | chr7 | - | 100467551 | 100467600 | 100474273 | 100474440 | 100466518 | 100467168 | 100477276 | 100478307 | 132 |
| MXE | 10670 | ENSG00000136536 | MARCHF7 | chr2 | + | 159743060 | 159743253 | 159747804 | 159748903 | 159729008 | 159729175 | 159752401 | 159752571 | 437 |
| MXE | 16944 | ENSG00000127311 | HELB | chr12 | + | 66324906 | 66325126 | 66331153 | 66331645 | 66323982 | 66324211 | 66338000 | 66338169 | 1 |
| MXE | 2917 | ENSG00000137210 | TMEM14B | chr6 | + | 10749621 | 10749698 | 10751132 | 10751234 | 10749201 | 10749268 | 10755141 | 10755232 | 28 |
| MXE | 1690 | ENSG00000005844 | ITGAL | chr16 | + | 30496096 | 30496294 | 30496435 | 30496566 | 30494712 | 30494850 | 30499073 | 30499234 | 1111 |
| MXE | 9805 | ENSG00000151092 | NGLY1 | chr3 | - | 25736003 | 25736149 | 25737333 | 25737455 | 25733871 | 25733982 | 25739576 | 25739799 | 82 |
| MXE | 9934 | ENSG00000117643 | MAN1C1 | chr1 | + | 25686439 | 25686536 | 25746667 | 25746783 | 25616790 | 25618337 | 25749254 | 25749335 | 13 |
| MXE | 11065 | ENSG00000122203 | KIAA1191 | chr5 | - | 176350612 | 176350737 | 176352621 | 176352748 | 176348249 | 176348356 | 176355570 | 176355749 | 57 |
| MXE | 1861 | ENSG00000074755 | ZZEF1 | chr17 | - | 4072607 | 4072756 | 4074149 | 4074351 | 4071338 | 4071446 | 4075096 | 4075180 | 134 |
| MXE | 7924 | ENSG00000043514 | TRIT1 | chr1 | - | 39847219 | 39847297 | 39847547 | 39847660 | 39844530 | 39844640 | 39847985 | 39848097 | 49 |
| MXE | 3397 | ENSG00000131504 | DIAPH1 | chr5 | - | 141529171 | 141529273 | 141529602 | 141529697 | 141528701 | 141528941 | 141534334 | 141534433 | 131 |
| MXE | 7136 | ENSG00000141027 | NCOR1 | chr17 | - | 16171802 | 16171995 | 16186553 | 16186687 | 16164870 | 16165161 | 16194461 | 16194639 | 282 |
| MXE | 13920 | ENSG00000171988 | JMJD1C | chr10 | - | 63198512 | 63198727 | 63200475 | 63200677 | 63197410 | 63197563 | 63206594 | 63208801 | 249 |
| MXE | 4749 | ENSG00000131389 | SLC6A6 | chr3 | + | 14472204 | 14472317 | 14477204 | 14477342 | 14468087 | 14468212 | 14478465 | 14478568 | 34 |
| MXE | 7354 | ENSG00000157193 | LRP8 | chr1 | - | 53271027 | 53271153 | 53271226 | 53271346 | 53266472 | 53266647 | 53275630 | 53275753 | 2 |
| MXE | 5058 | ENSG00000176261 | ZBTB8OS | chr1 | - | 32633644 | 32633727 | 32634701 | 32634792 | 32631826 | 32631879 | 32650432 | 32650525 | 10 |
| MXE | 5414 | ENSG00000187862 | TTC24 | chr1 | + | 156583337 | 156583450 | 156583796 | 156583895 | 156583041 | 156583170 | 156584876 | 156584974 | 7 |
| MXE | 344 | ENSG00000079805 | DNM2 | chr19 | + | 10782959 | 10783120 | 10786563 | 10786706 | 10777117 | 10777216 | 10793719 | 10793855 | 394 |
| MXE | 7516 | ENSG00000115020 | PIKFYVE | chr2 | + | 208277536 | 208277708 | 208285725 | 208285933 | 208276711 | 208276830 | 208288728 | 208288818 | 107 |
| MXE | 10631 | ENSG00000144895 | EIF2A | chr3 | + | 150558387 | 150558462 | 150562541 | 150562660 | 150552355 | 150552425 | 150563514 | 150563614 | 181 |
| MXE | 5561 | ENSG00000173193 | PARP14 | chr3 | + | 122699389 | 122701635 | 122704526 | 122704748 | 122695425 | 122695662 | 122708189 | 122708268 | 415 |
| MXE | 16136 | ENSG00000165732 | DDX21 | chr10 | + | 68974669 | 68974743 | 68977528 | 68977688 | 68973544 | 68973664 | 68978841 | 68978976 | 12 |
| MXE | 15943 | ENSG00000156650 | KAT6B | chr10 | + | 74972506 | 74972639 | 74975398 | 74975781 | 74970019 | 74970101 | 74977315 | 74977437 | 137 |
| MXE | 457 | ENSG00000248098 | BCKDHA | chr19 | + | 41410922 | 41411009 | 41414048 | 41414157 | 41410636 | 41410816 | 41419134 | 41419296 | 40 |
| MXE | 2234 | ENSG00000128394 | APOBEC3F | chr22 | + | 39045427 | 39045542 | 39049424 | 39049581 | 39044940 | 39045220 | 39052073 | 39052353 | 24 |
| MXE | 17160 | ENSG00000270181 | BIVM-ERCC5 | chr13 | + | 102834465 | 102834552 | 102838642 | 102838739 | 102831564 | 102831697 | 102839571 | 102839803 | 37 |
| MXE | 16519 | ENSG00000012822 | CALCOCO1 | chr12 | - | 53716259 | 53716415 | 53719738 | 53719829 | 53715792 | 53716047 | 53721466 | 53721615 | 95 |
| MXE | 8233 | ENSG00000164300 | SERINC5 | chr5 | - | 80150881 | 80150948 | 80158835 | 80158956 | 80147244 | 80147284 | 80166382 | 80166478 | 1 |
| MXE | 1599 | ENSG00000187555 | USP7 | chr16 | - | 8921156 | 8921295 | 8923353 | 8923413 | 8920358 | 8920447 | 8930292 | 8930397 | 89 |
| MXE | 9493 | ENSG00000102786 | INTS6 | chr13 | - | 51389318 | 51389444 | 51423029 | 51423090 | 51387385 | 51387540 | 51430293 | 51430383 | 17 |
| MXE | 1879 | ENSG00000167261 | DPEP2 | chr16 | - | 67991114 | 67991184 | 67991837 | 67991979 | 67990820 | 67990939 | 67992063 | 67992193 | 40 |
| MXE | 14018 | ENSG00000198851 | CD3E | chr11 | + | 118308426 | 118308441 | 118312152 | 118312170 | 118307287 | 118307308 | 118312617 | 118312866 | 261 |
| MXE | 15255 | ENSG00000122218 | COPA | chr1 | - | 160325542 | 160325652 | 160332447 | 160332557 | 160323430 | 160323530 | 160333602 | 160333679 | 254 |
| MXE | 15893 | ENSG00000144815 | NXPE3 | chr3 | + | 101785401 | 101785689 | 101801234 | 101801989 | 101782659 | 101782780 | 101807052 | 101807126 | 46 |
| MXE | 11452 | ENSG00000146285 | SCML4 | chr6 | - | 107746688 | 107746889 | 107749683 | 107749813 | 107744948 | 107745143 | 107772171 | 107772386 | 41 |
| MXE | 12453 | ENSG00000168137 | SETD5 | chr3 | + | 9443307 | 9443417 | 9445047 | 9445300 | 9442127 | 9442245 | 9445656 | 9445740 | 89 |
| MXE | 16162 | ENSG00000107669 | ATE1 | chr10 | - | 121870005 | 121870038 | 121898840 | 121898969 | 121841081 | 121841263 | 121902390 | 121902620 | 29 |
| MXE | 1889 | ENSG00000077235 | GTF3C1 | chr16 | - | 27488226 | 27488415 | 27488553 | 27488635 | 27485996 | 27486154 | 27489042 | 27489178 | 44 |
| MXE | 5018 | ENSG00000120686 | UFM1 | chr13 | + | 38354238 | 38354296 | 38358092 | 38358132 | 38349995 | 38350055 | 38359300 | 38359781 | 25 |
| MXE | 13408 | ENSG00000167548 | KMT2D | chr12 | - | 49054306 | 49054416 | 49054527 | 49054751 | 49053977 | 49054140 | 49054899 | 49055026 | 48 |
| MXE | 11177 | ENSG00000115053 | NCL | chr2 | - | 231457642 | 231457800 | 231458265 | 231458389 | 231457000 | 231457124 | 231459000 | 231459125 | 1720 |
| MXE | 3163 | ENSG00000198833 | UBE2J1 | chr6 | - | 89342323 | 89342455 | 89343682 | 89343756 | 89338458 | 89338543 | 89352538 | 89352722 | 50 |
| MXE | 10282 | ENSG00000131051 | RBM39 | chr20 | - | 35729311 | 35729365 | 35729461 | 35729527 | 35725037 | 35725155 | 35731940 | 35732135 | 260 |
| MXE | 7111 | ENSG00000162894 | FCMR | chr1 | - | 206909724 | 206909868 | 206910209 | 206910340 | 206909461 | 206909520 | 206911729 | 206911952 | 303 |
| MXE | 4491 | ENSG00000136051 | WASHC4 | chr12 | + | 105143126 | 105143243 | 105144286 | 105144455 | 105142452 | 105142558 | 105144717 | 105144872 | 125 |
| MXE | 5489 | ENSG00000116191 | RALGPS2 | chr1 | + | 178833423 | 178833550 | 178877497 | 178877635 | 178821611 | 178821704 | 178878901 | 178878992 | 77 |
| MXE | 9862 | ENSG00000110934 | BIN2 | chr12 | - | 51284715 | 51284780 | 51288107 | 51288188 | 51281037 | 51281528 | 51291590 | 51292344 | 136 |
| MXE | 5345 | ENSG00000186001 | LRCH3 | chr3 | + | 197866111 | 197866219 | 197870159 | 197870278 | 197865422 | 197865471 | 197871324 | 197871462 | 86 |
| MXE | 517 | ENSG00000105483 | CARD8 | chr19 | - | 48249522 | 48249633 | 48249750 | 48249847 | 48240961 | 48241063 | 48255791 | 48255898 | 145 |

| | | | | | | | | | | | |
|---|---|---|---|---|---|---|---|---|---|---|---|
| MXE | 16226 | ENSG00000078403 | MLLT10 | chr10 | + | 21670448 | 21670704 | 21673349 | 21673919 | 21651672 | 21651768 |
| MXE | 10717 | ENSG00000123472 | ATPAF1 | chr1 | - | 46643193 | 46643301 | 46645160 | 46645256 | 46632736 | 46635970 |
| MXE | 11632 | ENSG00000188554 | NBR1 | chr17 | + | 43193347 | 43193638 | 43194349 | 43194499 | 43193093 | 43193253 |
| MXE | 13101 | ENSG00000119321 | FKBP15 | chr9 | - | 113207211 | 113207296 | 113211476 | 113211592 | 113206508 | 113206578 |
| MXE | 15708 | ENSG00000166340 | TPP1 | chr11 | - | 6616004 | 6616074 | 6616314 | 6616503 | 6615441 | 6615562 |
| MXE | 6528 | ENSG00000117395 | EBNA1BP2 | chr1 | - | 43168938 | 43169028 | 43170755 | 43170879 | 43167159 | 43167235 |
| MXE | 9356 | ENSG00000070814 | TCOF1 | chr5 | + | 150389886 | 150390023 | 150391543 | 150391657 | 150387901 | 150388088 |
| MXE | 15028 | ENSG00000117751 | PPP1R8 | chr1 | + | 27838698 | 27838852 | 27841013 | 27841234 | 27832755 | 27832816 |
| MXE | 16746 | ENSG00000187109 | NAP1L1 | chr12 | - | 76055018 | 76055090 | 76056032 | 76056161 | 76053769 | 76053909 |
| MXE | 6678 | ENSG00000114982 | KANSL3 | chr2 | - | 96619671 | 96619762 | 96631311 | 96631482 | 96619358 | 96619544 |
| MXE | 10540 | ENSG00000135838 | NPL | chr1 | + | 182818812 | 182818859 | 182822114 | 182822199 | 182816713 | 182816806 |
| MXE | 14284 | ENSG00000100288 | CHKB | chr22 | - | 50579726 | 50579802 | 50579973 | 50580082 | 50579425 | 50579507 |
| MXE | 1494 | ENSG00000205560 | CPT1B | chr22 | - | 50576535 | 50576637 | 50576856 | 50577034 | 50576197 | 50576335 |
| MXE | 13864 | ENSG00000134014 | ELP3 | chr8 | + | 28132277 | 28132404 | 28137697 | 28137891 | 28129501 | 28129663 |
| MXE | 9872 | ENSG00000166986 | MARS1 | chr12 | + | 57500320 | 57500522 | 57504224 | 57504299 | 57498419 | 57498623 |
| MXE | 13349 | ENSG00000136504 | KAT7 | chr17 | + | 49805362 | 49805445 | 49811475 | 49811574 | 49798318 | 49798558 |
| MXE | 12376 | ENSG00000091106 | NLRC4 | chr2 | - | 32252418 | 32252679 | 32256774 | 32256893 | 32249606 | 32251601 |
| MXE | 16331 | ENSG00000132305 | IMMT | chr2 | - | 86166507 | 86166644 | 86170748 | 86170841 | 86161975 | 86162079 |
| MXE | 5905 | ENSG00000067066 | SP100 | chr2 | + | 230450171 | 230450255 | 230461261 | 230461414 | 230449560 | 230449710 |
| MXE | 14375 | ENSG00000104529 | EEF1D | chr8 | - | 143581228 | 143581328 | 143586728 | 143586852 | 143581053 | 143581154 |
| MXE | 11322 | ENSG00000115993 | TRAK2 | chr2 | - | 201392908 | 201393046 | 201394797 | 201394872 | 201389800 | 201389880 |
| MXE | 14810 | ENSG00000011454 | RABGAP1 | chr9 | + | 122997258 | 122997361 | 122998596 | 122998766 | 122996538 | 122996605 |
| MXE | 5731 | ENSG00000139719 | VPS33A | chr12 | - | 122242381 | 122242508 | 122244568 | 122244762 | 122239877 | 122239945 |
| MXE | 9057 | ENSG00000141458 | NPC1 | chr18 | - | 23560230 | 23560480 | 23561359 | 23561527 | 23557116 | 23557190 |
| MXE | 16922 | ENSG00000204852 | TCTN1 | chr12 | + | 110641541 | 110642389 | 110644991 | 110645129 | 110641023 | 110641149 |
| MXE | 13602 | ENSG00000090316 | MAEA | chr4 | + | 1315396 | 1315600 | 1322380 | 1322503 | 1311978 | 1312161 |
| MXE | 15952 | ENSG00000163249 | CCNYL1 | chr2 | + | 207737410 | 207737446 | 207740654 | 207740706 | 207733946 | 207734047 |
| MXE | 11696 | ENSG00000169826 | CSGALNACT2 | chr10 | + | 43163865 | 43164044 | 43167003 | 43167098 | 43160493 | 43160595 |
| MXE | 16968 | ENSG00000186908 | ZDHHC17 | chr12 | + | 76815897 | 76816019 | 76821057 | 76821167 | 76815145 | 76815210 |
| MXE | 5390 | ENSG00000162736 | NCSTN | chr1 | + | 160349548 | 160349670 | 160351695 | 160351805 | 160348998 | 160349122 |
| MXE | 10675 | ENSG00000148341 | SH3GLB2 | chr9 | - | 129012235 | 129012298 | 129014410 | 129014503 | 129010669 | 129010693 |

[illegible]

| | | | | | | | | | | | | | | | | | | | | | | | | | | | | | | | | | | | | | | | | | | | | | | | | |
|---|---|---|---|---|---|---|---|---|---|---|---|---|---|---|---|---|---|---|---|---|---|---|---|---|---|---|---|---|---|---|---|---|---|---|---|---|---|---|---|---|---|---|---|---|---|---|---|---|
| MXE | 2930 | ENSG00000124789 | NUP153 | chr6 | - | 17668974 | 17669028 | 17669429 | 17669546 | 17665238 | 17665385 | 17674904 | 17675033 | 121 | 89 | 130 | 143 | 196 | 30 | 28 | 40 | 26 | 76 | 48 | 111 | 145 | 97 | 40 | 12 | 50 | 54 | 36 | 12 | 266 | 203 | 0.128734923 | 0.359025528 | 0.755 | 0.708 | 0.713 | 0.808 | 0.663 | 0.753 | 0.629 | 0.672 | 0.673 | 0.718 | 0.04 |
| MXE | 17069 | ENSG00000105352 | CEACAM4 | chr19 | - | 41619669 | 41619711 | 41620210 | 41620242 | 41618970 | 41619395 | 41620574 | 41620627 | 3 | 0 | 1 | 8 | 1 | 3 | 1 | 0 | 5 | 1 | 11 | 3 | 3 | 12 | 1 | 5 | 29 | 17 | 23 | 8 | 181 | 191 | 0.128817369 | 0.359109936 | 0.513 | 0 | 1 | 0.628 | 0.513 | 0.699 | 0.098 | 0.157 | 0.355 | 0.117 | 0.246 |
| MXE | 7318 | ENSG00000006831 | ADIPOR2 | chr12 | + | 1777853 | 1778025 | 1780450 | 1780637 | 1772841 | 1772961 | 1780888 | 1781076 | 107 | 85 | 73 | 115 | 194 | 81 | 46 | 44 | 67 | 97 | 125 | 95 | 88 | 148 | 253 | 65 | 136 | 51 | 77 | 282 | 298 | 298 | 0.128824658 | 0.359109936 | 0.569 | 0.649 | 0.624 | 0.632 | 0.667 | 0.658 | 0.411 | 0.633 | 0.658 | 0.473 | 0.062 |
| MXE | 9856 | ENSG00000152127 | MGAT5 | chr2 | + | 134402987 | 134403137 | 134412868 | 134413015 | 134362274 | 134362408 | 134422802 | 134422919 | 135 | 97 | 95 | 153 | 231 | 162 | 73 | 155 | 156 | 169 | 186 | 123 | 102 | 79 | 175 | 92 | 193 | 164 | 189 | 358 | 298 | 296 | 0.1288373 | 0.359109936 | 0.453 | 0.569 | 0.378 | 0.493 | 0.576 | 0.668 | 0.388 | 0.382 | 0.293 | 0.327 | 0.082 |
| MXE | 14835 | ENSG00000107077 | KDM4C | chr9 | + | 7128065 | 7128236 | 7165237 | 7165357 | 7105401 | 7105500 | 7169797 | 7169829 | 21 | 9 | 14 | 20 | 15 | 83 | 67 | 74 | 95 | 111 | 15 | 18 | 3 | 8 | 1 | 77 | 89 | 64 | 74 | 18 | 298 | 269 | 0.128887065 | 0.35915463 | 0.186 | 0.108 | 0.146 | 0.16 | 0.109 | 0.15 | 0.154 | 0.041 | 0.089 | 0.048 | 0.045 |
| MXE | 935 | ENSG00000100983 | GSS | chr20 | - | 34941712 | 34941829 | 34942487 | 34942627 | 34936942 | 34937023 | 34942930 | 34943006 | 29 | 11 | 11 | 28 | 20 | 30 | 21 | 27 | 20 | 22 | 9 | 39 | 23 | 6 | 34 | 30 | 50 | 20 | 44 | 59 | 289 | 266 | 0.128901414 | 0.35915463 | 0.471 | 0.325 | 0.273 | 0.563 | 0.456 | 0.216 | 0.418 | 0.514 | 0.112 | 0.347 | 0.096 |
| MXE | 9487 | ENSG00000067208 | EVI5 | chr1 | - | 92636201 | 92636336 | 92662718 | 92662865 | 92625793 | 92625934 | 92663419 | 92663452 | 0 | 2 | 4 | 0 | 4 | 61 | 74 | 50 | 81 | 105 | 12 | 1 | 1 | 0 | 0 | 71 | 51 | 56 | 44 | 28 | 296 | 284 | 0.129044223 | 0.359485493 | 0 | 0.025 | 0.071 | 0 | 0.035 | 0.14 | 0.018 | 0.017 | 0 | 0 | -0.009 |
| MXE | 2287 | ENSG00000128908 | INO80 | chr15 | - | 41056621 | 41056706 | 41058638 | 41058781 | 41055246 | 41055364 | 41059866 | 41059926 | 81 | 60 | 80 | 101 | 117 | 79 | 57 | 80 | 64 | 106 | 97 | 124 | 71 | 84 | 33 | 119 | 113 | 72 | 76 | 56 | 292 | 234 | 0.129246303 | 0.35994339 | 0.451 | 0.458 | 0.445 | 0.558 | 0.469 | 0.395 | 0.468 | 0.441 | 0.47 | 0.321 | 0.057 |
| MXE | 16877 | ENSG00000050426 | LETMD1 | chr12 | + | 51052091 | 51052207 | 51056143 | 51056245 | 51049033 | 51049185 | 51056349 | 51056502 | 42 | 17 | 44 | 38 | 109 | 50 | 26 | 48 | 41 | 61 | 13 | 35 | 23 | 18 | 64 | 49 | 77 | 45 | 21 | 30 | 265 | 251 | 0.129271667 | 0.35994339 | 0.443 | 0.382 | 0.465 | 0.467 | 0.629 | 0.201 | 0.301 | 0.326 | 0.448 | 0.669 | 0.088 |
| MXE | 13106 | ENSG00000196998 | WDR45 | chrX | - | 49076644 | 49076750 | 49077642 | 49077747 | 49076429 | 49076524 | 49077836 | 49077911 | 27 | 11 | 28 | 34 | 21 | 117 | 27 | 107 | 112 | 55 | 31 | 75 | 33 | 41 | 132 | 151 | 198 | 116 | 93 | 210 | 254 | 255 | 0.129280872 | 0.35994339 | 0.188 | 0.29 | 0.208 | 0.234 | 0.277 | 0.171 | 0.276 | 0.222 | 0.307 | 0.387 | -0.033 |
| MXE | 14689 | ENSG00000116580 | GON4L | chr1 | - | 155763311 | 155763564 | 155764999 | 155766709 | 155762189 | 155762374 | 155767424 | 155767541 | 66 | 46 | 45 | 43 | 93 | 85 | 80 | 86 | 85 | 64 | 101 | 60 | 45 | 66 | 17 | 68 | 65 | 64 | 41 | 31 | 298 | 298 | 0.129465159 | 0.360380572 | 0.437 | 0.365 | 0.344 | 0.336 | 0.592 | 0.598 | 0.48 | 0.413 | 0.617 | 0.354 | -0.078 |
| MXE | 908 | ENSG00000078043 | PIAS2 | chr18 | - | 46846706 | 46846841 | 46855344 | 46855435 | 46844733 | 46844839 | 46855564 | 46855615 | 31 | 27 | 24 | 29 | 26 | 49 | 47 | 47 | 53 | 103 | 13 | 14 | 20 | 7 | 7 | 7 | 55 | 26 | 16 | 0 | 240 | 284 | 0.129486138 | 0.360380572 | 0.428 | 0.405 | 0.377 | 0.393 | 0.23 | 0.687 | 0.231 | 0.477 | 0.341 | 1 | -0.181 |
| MXE | 7695 | ENSG00000142784 | WDTC1 | chr1 | + | 27301225 | 27301461 | 27305000 | 27305193 | 27297937 | 27298111 | 27306185 | 27308244 | 93 | 25 | 70 | 68 | 60 | 97 | 53 | 65 | 107 | 19 | 13 | 66 | 28 | 44 | 109 | 101 | 85 | 39 | 56 | 93 | 298 | 298 | 0.129590769 | 0.360604598 | 0.489 | 0.321 | 0.519 | 0.389 | 0.759 | 0.114 | 0.437 | 0.418 | 0.44 | 0.54 | 0.106 |
| MXE | 16533 | ENSG00000238105 | GOLGA2P5 | chr12 | - | 100168451 | 100168633 | 100170564 | 100170667 | 100167549 | 100167639 | 100170936 | 100170987 | 5 | 6 | 8 | 5 | 4 | 13 | 10 | 10 | 21 | 16 | 0 | 10 | 4 | 3 | 16 | 17 | 32 | 20 | 31 | 27 | 252 | 298 | 0.129628983 | 0.360643762 | 0.313 | 0.415 | 0.486 | 0.22 | 0.228 | 0 | 0.27 | 0.191 | 0.103 | 0.412 | 0.137 |
| MXE | 9077 | ENSG00000164828 | SUN1 | chr7 | + | 848417 | 848615 | 849922 | 850033 | 843340 | 843520 | 851383 | 851482 | 90 | 65 | 68 | 87 | 94 | 30 | 11 | 11 | 17 | 16 | 140 | 55 | 49 | 96 | 113 | 12 | 17 | 13 | 10 | 14 | 298 | 260 | 0.129766548 | 0.360959268 | 0.724 | 0.838 | 0.844 | 0.817 | 0.837 | 0.911 | 0.738 | 0.767 | 0.893 | 0.876 | -0.025 |
| MXE | 3978 | ENSG00000138303 | ASCC1 | chr10 | - | 72133056 | 72133181 | 72152868 | 72152988 | 72096519 | 72097450 | 72161537 | 72161674 | 64 | 39 | 60 | 59 | 51 | 7 | 7 | 9 | 5 | 23 | 39 | 79 | 50 | 50 | 6 | 0 | 21 | 8 | 2 | 0 | 269 | 274 | 0.129816993 | 0.361032368 | 0.903 | 0.85 | 0.872 | 0.923 | 0.693 | 1 | 0.793 | 0.864 | 0.962 | 1 | -0.076 |
| MXE | 3640 | ENSG00000132780 | NASP | chr1 | + | 45602254 | 45602365 | 45606481 | 45606591 | 45591222 | 45591270 | 45613168 | 45613248 | 54 | 33 | 40 | 45 | 23 | 150 | 122 | 88 | 136 | 268 | 79 | 42 | 49 | 99 | 51 | 312 | 110 | 153 | 235 | 145 | 260 | 259 | 0.129859319 | 0.361082863 | 0.264 | 0.212 | 0.312 | 0.248 | 0.079 | 0.201 | 0.276 | 0.242 | 0.296 | 0.259 | -0.032 |
| MXE | 9526 | ENSG00000083520 | DIS3 | chr13 | - | 72775924 | 72776092 | 72777419 | 72777493 | 72775210 | 72775375 | 72778186 | 72778380 | 91 | 55 | 49 | 58 | 68 | 99 | 101 | 55 | 102 | 129 | 18 | 52 | 33 | 9 | 15 | 27 | 79 | 102 | 26 | 24 | 223 | 298 | 0.129967617 | 0.361276193 | 0.551 | 0.421 | 0.543 | 0.432 | 0.413 | 0.471 | 0.468 | 0.302 | 0.316 | 0.455 | 0.07 |
| MXE | 12654 | ENSG00000168092 | PAFAH1B2 | chr11 | + | 117159933 | 117160023 | 117161144 | 117161261 | 117152440 | 117152528 | 117163769 | 117163874 | 87 | 83 | 89 | 108 | 79 | 163 | 109 | 121 | 119 | 189 | 39 | 77 | 86 | 90 | 36 | 191 | 130 | 148 | 132 | 63 | 239 | 266 | 0.129977211 | 0.361276193 | 0.373 | 0.459 | 0.45 | 0.503 | 0.318 | 0.185 | 0.397 | 0.393 | 0.431 | 0.389 | 0.062 |
| MXE | 4047 | ENSG00000002549 | LAP3 | chr4 | + | 17595409 | 17595534 | 17598455 | 17598558 | 17588818 | 17588977 | 17604587 | 17604667 | 107 | 152 | 113 | 184 | 137 | 168 | 197 | 163 | 264 | 91 | 989 | 193 | 126 | 567 | 203 | 1329 | 373 | 224 | 766 | 297 | 274 | 252 | 0.130122445 | 0.361590863 | 0.369 | 0.415 | 0.389 | 0.391 | 0.581 | 0.406 | 0.322 | 0.341 | 0.405 | 0.386 | 0.057 |
| MXE | 1747 | ENSG00000169896 | ITGAM | chr16 | + | 31261691 | 31261797 | 31265394 | 31265498 | 31259966 | 31260092 | 31265810 | 31265881 | 196 | 51 | 94 | 93 | 25 | 261 | 53 | 123 | 141 | 55 | 217 | 236 | 97 | 232 | 284 | 234 | 265 | 126 | 269 | 344 | 255 | 253 | 0.130138827 | 0.361590863 | 0.427 | 0.488 | 0.431 | 0.396 | 0.311 | 0.479 | 0.469 | 0.433 | 0.461 | 0.45 | -0.048 |
| MXE | 13038 | ENSG00000003436 | TFPI | chr2 | - | 187484810 | 187484987 | 187488336 | 187488375 | 187484123 | 187484216 | 187496880 | 187497078 | 6 | 9 | 1 | 17 | 26 | 25 | 65 | 7 | 62 | 46 | 0 | 34 | 8 | 3 | 0 | 26 | 81 | 19 | 34 | 13 | 188 | 298 | 0.130356905 | 0.362129446 | 0.276 | 0.18 | 0.185 | 0.303 | 0.473 | 0 | 0.4 | 0.4 | 0.123 | 0 | 0.099 |
| MXE | 4177 | ENSG00000084092 | NOA1 | chr4 | - | 56966619 | 56966736 | 56968383 | 56968515 | 56964405 | 56964526 | 56973147 | 56973353 | 171 | 94 | 178 | 141 | 376 | 47 | 40 | 62 | 43 | 84 | 169 | 103 | 74 | 66 | 187 | 48 | 46 | 30 | 29 | 62 | 281 | 266 | 0.130422427 | 0.362229184 | 0.775 | 0.69 | 0.731 | 0.756 | 0.809 | 0.769 | 0.679 | 0.7 | 0.683 | 0.741 | 0.038 |
| MXE | 9233 | ENSG00000111801 | BTN3A3 | chr6 | + | 26445703 | 26445985 | 26448616 | 26448637 | 26443956 | 26444304 | 26448727 | 26448754 | 107 | 46 | 81 | 85 | 137 | 108 | 49 | 70 | 113 | 73 | 52 | 96 | 42 | 38 | 115 | 119 | 86 | 27 | 51 | 159 | 298 | 170 | 0.130441299 | 0.362229184 | 0.361 | 0.349 | 0.398 | 0.3 | 0.517 | 0.2 | 0.389 | 0.47 | 0.298 | 0.292 | 0.055 |
| MXE | 15310 | ENSG00000196814 | MVB12B | chr9 | + | 126392065 | 126392195 | 126395574 | 126395697 | 126386561 | 126386658 | 126421853 | 126421948 | 67 | 40 | 56 | 42 | 45 | 20 | 21 | 20 | 13 | 5 | 102 | 121 | 64 | 98 | 143 | 30 | 53 | 29 | 35 | 70 | 279 | 272 | 0.130479952 | 0.362269184 | 0.766 | 0.65 | 0.732 | 0.759 | 0.898 | 0.768 | 0.69 | 0.683 | 0.732 | 0.666 | 0.053 |
| MXE | 5591 | ENSG00000196126 | HLA-DRB1 | chr6 | - | 32580745 | 32580856 | 32581556 | 32581838 | 32580246 | 32580270 | 32584108 | 32584378 | 206 | 401 | 548 | 129 | 174 | 149 | 617 | 726 | 237 | 21 | 577 | 338 | 193 | 744 | 1767 | 18 | 28 | 376 | 860 | 2375 | 298 | 260 | 0.130673516 | 0.362731788 | 0.547 | 0.362 | 0.397 | 0.322 | 0.878 | 0.965 | 0.913 | 0.309 | 0.43 | 0.394 | -0.101 |
| MXE | 15733 | ENSG00000149582 | TMEM25 | chr11 | + | 118532149 | 118532461 | 118532916 | 118533207 | 118531774 | 118531871 | 118533856 | 118533887 | 1 | 2 | 5 | 1 | 0 | 1 | 3 | 13 | 0 | 3 | 0 | 0 | 0 | 5 | 0 | 15 | 8 | 3 | 0 | 3 | 298 | 298 | 0.130739291 | 0.362731788 | 0.5 | 0.4 | 0.278 | 1 | 0 | 0 | 0 | 0 | 1 | 0 | 0.236 |
| MXE | 3493 | ENSG00000147044 | CASK | chrX | - | 41787177 | 41787283 | 41853114 | 41853227 | 41745523 | 41745601 | 41922929 | 41923007 | 28 | 20 | 26 | 22 | 12 | 62 | 49 | 69 | 60 | 96 | 15 | 15 | 14 | 15 | 31 | 28 | 54 | 48 | 24 | 41 | 262 | 255 | 0.130759096 | 0.362731788 | 0.305 | 0.284 | 0.268 | 0.263 | 0.108 | 0.343 | 0.213 | 0.221 | 0.378 | 0.424 | -0.07 |
| MXE | 10754 | ENSG00000177479 | ARIH2 | chr3 | + | 48927461 | 48927813 | 48945135 | 48945181 | 48924717 | 48924839 | 48961611 | 48961679 | 120 | 65 | 102 | 132 | 137 | 15 | 11 | 25 | 25 | 13 | 73 | 143 | 97 | 89 | 184 | 0 | 23 | 14 | 11 | 24 | 298 | 195 | 0.130767308 | 0.362731788 | 0.84 | 0.795 | 0.728 | 0.776 | 0.873 | 1 | 0.803 | 0.819 | 0.841 | 0.834 | -0.057 |
| MXE | 6878 | ENSG00000070190 | DAPP1 | chr4 | + | 99840288 | 99840422 | 99853217 | 99853348 | 99835622 | 99835745 | 99861577 | 99861625 | 115 | 95 | 80 | 143 | 66 | 211 | 326 | 173 | 404 | 365 | 106 | 120 | 99 | 78 | 28 | 418 | 357 | 208 | 455 | 146 | 283 | 280 | 0.130767966 | 0.362731788 | 0.35 | 0.224 | 0.314 | 0.259 | 0.152 | 0.201 | 0.25 | 0.32 | 0.145 | 0.159 | 0.045 |
| MXE | 4802 | ENSG00000084463 | WBP11 | chr12 | - | 14794536 | 14794736 | 14794970 | 14795104 | 14793730 | 14793922 | 14796806 | 14797003 | 300 | 236 | 319 | 305 | 887 | 314 | 206 | 296 | 364 | 327 | 751 | 346 | 377 | 404 | 360 | 484 | 388 | 469 | 487 | 436 | 283 | 298 | 0.130857488 | 0.362912728 | 0.502 | 0.547 | 0.532 | 0.469 | 0.741 | 0.62 | 0.484 | 0.458 | 0.466 | 0.465 | 0.06 |
| MXE | 9316 | ENSG00000163788 | SNRK | chr3 | + | 43332168 | 43332310 | 43340286 | 43340499 | 43303097 | 43303792 | 43343343 | 43343478 | 125 | 64 | 153 | 118 | 244 | 363 | 215 | 352 | 355 | 582 | 62 | 70 | 111 | 65 | 7 | 123 | 244 | 405 | 237 | 29 | 291 | 298 | 0.131030144 | 0.363324119 | 0.261 | 0.234 | 0.308 | 0.254 | 0.3 | 0.34 | 0.227 | 0.219 | 0.219 | 0.198 | 0.031 |
| MXE | 7000 | ENSG00000213593 | TMX2 | chr11 | + | 57737835 | 57738026 | 57738353 | 57738430 | 57737607 | 57737668 | 57738663 | 57738770 | 16 | 16 | 23 | 32 | 24 | 96 | 60 | 73 | 84 | 107 | 13 | 29 | 24 | 34 | 48 | 122 | 57 | 54 | 95 | 98 | 298 | 226 | 0.131073094 | 0.363375769 | 0.112 | 0.168 | 0.193 | 0.224 | 0.145 | 0.075 | 0.278 | 0.252 | 0.213 | 0.271 | -0.049 |
| MXE | 3494 | ENSG00000147044 | CASK | chrX | - | 41787177 | 41787283 | 41853096 | 41853227 | 41745523 | 41745601 | 41922929 | 41923010 | 27 | 22 | 29 | 26 | 14 | 62 | 49 | 69 | 60 | 96 | 15 | 17 | 14 | 21 | 31 | 28 | 54 | 48 | 24 | 41 | 280 | 255 | 0.131205399 | 0.363675076 | 0.284 | 0.29 | 0.277 | 0.283 | 0.117 | 0.328 | 0.223 | 0.21 | 0.443 | 0.408 | -0.072 |
| MXE | 15755 | ENSG00000149311 | ATM | chr11 | + | 108331878 | 108332037 | 108332761 | 108332900 | 108331443 | 108331557 | 108333885 | 108333968 | 802 | 621 | 865 | 799 | 1302 | 745 | 718 | 709 | 797 | 949 | 53 | 238 | 240 | 37 | 35 | 35 | 254 | 245 | 98 | 45 | 298 | 288 | 0.131276348 | 0.363804235 | 0.51 | 0.455 | 0.541 | 0.492 | 0.57 | 0.594 | 0.475 | 0.486 | 0.267 | 0.429 | 0.063 |
| MXE | 9938 | ENSG00000138035 | PNPT1 | chr2 | - | 55656304 | 55656371 | 55660156 | 55660193 | 55656130 | 55656220 | 55661955 | 55662026 | 33 | 28 | 36 | 28 | 65 | 26 | 29 | 38 | 40 | 33 | 23 | 44 | 7 | 34 | 5 | 20 | 23 | 34 | 65 | 10 | 186 | 216 | 0.131484275 | 0.364267314 | 0.596 | 0.529 | 0.524 | 0.448 | 0.696 | 0.572 | 0.69 | 0.193 | 0.378 | 0.367 | 0.119 |
| MXE | 5977 | ENSG00000197948 | FCHSD1 | chr5 | - | 141649886 | 141649954 | 141650358 | 141650404 | 141649394 | 141649530 | 141651019 | 141651117 | 45 | 14 | 20 | 27 | 9 | 38 | 23 | 39 | 23 | 21 | 6 | 36 | 14 | 22 | 24 | 51 | 49 | 12 | 48 | 53 | 195 | 217 | 0.131492211 | 0.364267314 | 0.569 | 0.404 | 0.363 | 0.566 | 0.323 | 0.116 | 0.45 | 0.565 | 0.338 | 0.335 | 0.084 |
| MXE | 4059 | ENSG00000131507 | NDFIP1 | chr5 | + | 142135729 | 142135817 | 142140562 | 142140629 | 142132211 | 142132342 | 142144570 | 142144676 | 235 | 129 | 222 | 237 | 309 | 87 | 76 | 96 | 69 | 131 | 218 | 161 | 114 | 115 | 257 | 45 | 63 | 43 | 59 | 68 | 237 | 216 | 0.131547747 | 0.364353604 | 0.711 | 0.607 | 0.678 | 0.758 | 0.683 | 0.815 | 0.7 | 0.707 | 0.64 | 0.775 | -0.04 |
| MXE | 9580 | ENSG00000163904 | SENP2 | chr3 | + | 185598957 | 185599024 | 185600764 | 185600855 | 185590113 | 185590169 | 185606329 | 185606491 | 0 | 0 | 0 | 0 | 0 | 44 | 36 | 44 | 63 | 72 | 0 | 0 | 0 | 11 | 0 | 76 | 53 | 55 | 94 | 29 | 216 | 240 | 0.131600114 | 0.364428317 | 0 | 0 | 0 | 0 | 0 | 0 | 0 | 0 | 0.115 | 0 | -0.023 |
| MXE | 7086 | ENSG00000198740 | ZNF652 | chr17 | - | 49311311 | 49311456 | 49311926 | 49312042 | 49289205 | 49298924 | 49312697 | 49312845 | 157 | 100 | 146 | 136 | 297 | 95 | 66 | 64 | 109 | 235 | 57 | 112 | 88 | 79 | 44 | 64 | 80 | 96 | 75 | 19 | 265 | 294 | 0.131639821 | 0.364428317 | 0.647 | 0.627 | 0.717 | 0.581 | 0.584 | 0.497 | 0.608 | 0.504 | 0.539 | 0.72 | 0.058 |
| MXE | 8561 | ENSG00000153207 | AHCTF1 | chr1 | - | 246890955 | 246891060 | 246891778 | 246891919 | 246889965 | 246890059 | 246894658 | 246894748 | 90 | 97 | 132 | 172 | 152 | 66 | 55 | 60 | 73 | 110 | 0 | 51 | 106 | 37 | 7 | 4 | 43 | 55 | 17 | 17 | 290 | 254 | 0.1316479 | 0.364428317 | 0.544 | 0.607 | 0.658 | 0.674 | 0.548 | 0 | 0.51 | 0.628 | 0.656 | 0.265 | 0.194 |
| MXE | 3608 | ENSG00000057935 | MTA3 | chr2 | + | 42579106 | 42579200 | 42609457 | 42609584 | 42570436 | 42570504 | 42640172 | 42640236 | 44 | 24 | 53 | 37 | 28 | 24 | 20 | 46 | 30 | 34 | 23 | 22 | 46 | 32 | 1 | 21 | 62 | 36 | 27 | 7 | 243 | 276 | 0.131750701 | 0.364598186 | 0.676 | 0.577 | 0.567 | 0.583 | 0.483 | 0.554 | 0.287 | 0.592 | 0.574 | 0.14 | 0.148 |
| MXE | 16001 | ENSG00000138346 | DNA2 | chr10 | - | 68465666 | 68465812 | 68468122 | 68468306 | 68459103 | 68459235 | 68469980 | 68470163 | 30 | 23 | 24 | 37 | 49 | 45 | 34 | 12 | 42 | 32 | 65 | 47 | 29 | 45 | 79 | 35 | 34 | 33 | 51 | 17 | 298 | 295 | 0.131758073 | 0.364598186 | 0.398 | 0.401 | 0.664 | 0.466 | 0.603 | 0.648 | 0.578 | 0.465 | 0.466 | 0.821 | -0.089 |
| MXE | 10003 | ENSG00000135624 | CCT7 | chr2 | + | 73247761 | 73247926 | 73248990 | 73249179 | 73244543 | 73244715 | 73249818 | 73249844 | 417 | 212 | 396 | 402 | 477 | 352 | 262 | 375 | 412 | 512 | 432 | 363 | 196 | 327 | 360 | 568 | 376 | 226 | 311 | 451 | 298 | 298 | 0.131854377 | 0.364797109 | 0.542 | 0.447 | 0.514 | 0.494 | 0.482 | 0.432 | 0.491 | 0.464 | 0.513 | 0.444 | 0.027 |
| MXE | 11337 | ENSG00000144233 | AMMECR1L | chr2 | - | 127871248 | 127871359 | 127873827 | 127874272 | 127870813 | 127870928 | 127884202 | 127884312 | 21 | 23 | 23 | 10 | 28 | 12 | 3 | 7 | 9 | 2 | 37 | 10 | 22 | 37 | 20 | 0 | 2 | 9 | 0 | 13 | 298 | 260 | 0.131882742 | 0.364808028 | 0.604 | 0.87 | 0.741 | 0.492 | 0.924 | 1 | 0.814 | 0.681 | 1 | 0.573 | -0.087 |
| MXE | 6433 | ENSG00000163946 | TASOR | chr3 | - | 56638705 | 56638765 | 56639985 | 56640130 | 56633043 | 56633966 | 56641348 | 56641752 | 344 | 305 | 234 | 335 | 420 | 83 | 91 | 82 | 90 | 139 | 44 | 199 | 166 | 59 | 16 | 0 | 33 | 45 | 8 | 15 | 294 | 209 | 0.131930718 | 0.364873181 | 0.747 | 0.704 | 0.67 | 0.726 | 0.682 | 1 | 0.811 | 0.724 | 0.84 | 0.431 | -0.055 |
| MXE | 6461 | ENSG00000078140 | UBE2K | chr4 | + | 39755656 | 39755739 | 39774833 | 39774933 | 39745751 | 39745810 | 39778359 | 39779983 | 50 | 26 | 38 | 24 | 84 | 0 | 0 | 5 | 4 | 0 | 23 | 25 | 23 | 24 | 26 | 1 | 0 | 2 | 2 | 6 | 232 | 249 | 0.132077618 | 0.365211849 | 1 | 1 | 0.891 | 0.866 | 1 | 0.961 | 1 | 0.925 | 0.928 | 0.823 | 0.024 |
| MXE | 9547 | ENSG00000163382 | NAXE | chr1 | + | 156592556 | 156592670 | 156593407 | 156593555 | 156592100 | 156592475 | 156593881 | 156594090 | 61 | 26 | 43 | 50 | 52 | 75 | 36 | 52 | 65 | 117 | 71 | 44 | 36 | 38 | 53 | 61 | 46 | 42 | 59 | 67 | 263 | 297 | 0.132155061 | 0.365340512 | 0.479 | 0.449 | 0.483 | 0.465 | 0.334 | 0.568 | 0.519 | 0.492 | 0.421 | 0.472 | -0.052 |
| MXE | 9583 | ENSG00000163904 | SENP2 | chr3 | + | 185609246 | 185609350 | 185611650 | 185611745 | 185606329 | 185606498 | 185612606 | 185612658 | 100 | 56 | 68 | 83 | 195 | 136 | 110 | 115 | 136 | 247 | 152 | 108 | 136 | 149 | 89 | 247 | 183 | 250 | 323 | 137 | 253 | 244 | 0.132173057 | 0.365340512 | 0.415 | 0.329 | 0.363 | 0.371 | 0.432 | 0.372 | 0.363 | 0.344 | 0.308 | 0.385 | 0.028 |
| MXE | 9697 | ENSG00000151779 | NBAS | chr2 | - | 15478225 | 15478289 | 15488893 | 15489022 | 15475686 | 15475880 | 15504144 | 15504213 | 132 | 90 | 84 | 122 | 178 | 73 | 62 | 47 | 69 | 83 | 151 | 103 | 82 | 133 | 178 | 66 | 67 | 54 | 34 | 57 | 278 | 213 | 0.132233523 | 0.365440035 | 0.581 | 0.527 | 0.578 | 0.575 | 0.622 | 0.637 | 0.541 | 0.538 | 0.75 | 0.705 | -0.058 |
| MXE | 1913 | ENSG00000102910 | LONP2 | chr16 | + | 48347506 | 48347714 | 48348099 | 48348290 | 48334215 | 48334358 | 48351580 | 48351851 | 61 | 64 | 57 | 79 | 102 | 162 | 106 | 150 | 156 | 108 | 67 | 69 | 37 | 72 | 51 | 187 | 150 | 149 | 142 | 90 | 298 | 298 | 0.132352116 | 0.365598374 | 0.274 | 0.376 | 0.275 | 0.336 | 0.486 | 0.264 | 0.315 | 0.199 | 0.336 | 0.362 | 0.054 |
| MXE | 13869 | ENSG00000168575 | SLC20A2 | chr8 | - | 42444645 | 42444762 | 42459895 | 42459992 | 42439449 | 42439653 | 42463004 | 42463090 | 14 | 16 | 15 | 25 | 14 | 19 | 15 | 32 | 31 | 18 | 20 | 27 | 22 | 15 | 27 | 13 | 49 | 17 | 11 | 22 | 246 | 266 | 0.132355645 | 0.365598374 | 0.443 | 0.536 | 0.336 | 0.466 | 0.457 | 0.625 | 0.373 | 0.583 | 0.596 | 0.57 | -0.102 |
| MXE | 9432 | ENSG00000138398 | PPIG | chr2 | + | 169607103 | 169607148 | 169608670 | 169608758 | 169606038 | 169606146 | 169614463 | 169614493 | 207 | 125 | 147 | 173 | 207 | 43 | 44 | 32 | 27 | 158 | 39 | 101 | 138 | 41 | 49 | 4 | 25 | 38 | 11 | 7 | 194 | 237 | 0.132364231 | 0.365598374 | 0.855 | 0.776 | 0.849 | 0.887 | 0.615 | 0.923 | 0.832 | 0.816 | 0.82 | 0.895 | -0.061 |
| MXE | 3835 | ENSG00000168394 | TAP1 | chr6 | - | 32850943 | 32851149 | 32852108 | 32852239 | 32850319 | 32850517 | 32852387 | 32852502 | 241 | 95 | 156 | 169 | 95 | 298 | 144 | 185 | 221 | 305 | 232 | 219 | 105 | 291 | 310 | 407 | 281 | 145 | 303 | 276 | 280 | 298 | 0.132507007 | 0.365925082 | 0.463 | 0.413 | 0.473 | 0.449 | 0.249 | 0.378 | 0.453 | 0.435 | 0.505 | 0.545 | -0.054 |
| MXE | 16711 | ENSG00000078674 | PCM1 | chr8 | + | 17972328 | 17972687 | 17980590 | 17980755 | 17969576 | 17969748 | 17985446 | 17985619 | 379 | 324 | 337 | 312 | 531 | 173 | 156 | 133 | 151 | 355 | 35 | 115 | 136 | 19 | 2 | 7 | 41 | 58 | 11 | 0 | 298 | 298 | 0.132566932 | 0.366022909 | 0.687 | 0.675 | 0.717 | 0.674 | 0.599 | 0.833 | 0.737 | 0.701 | 0.633 | 1 | -0.11 |
| MXE | 17165 | ENSG00000153395 | LPCAT1 | chr5 | - | 1473956 | 1474110 | 1474559 | 1474685 | 1470825 | 1470924 | 1477403 | 1477486 | 187 | 135 | 150 | 214 | 113 | 449 | 204 | 286 | 331 | 293 | 168 | 290 | 92 | 211 | 228 | 438 | 277 | 130 | 341 | 377 | 275 | 298 | 0.13259971 | 0.366045763 | 0.311 | 0.418 | 0.362 | 0.412 | 0.295 | 0.294 | 0.532 | 0.434 | 0.401 | 0.396 | -0.052 |
| MXE | 5094 | ENSG00000060971 | ACAA1 | chr3 | - | 38126161 | 38126700 | 38129289 | 38129388 | 38125825 | 38125881 | 38131595 | 38131638 | 48 | 49 | 45 | 57 | 45 | 83 | 45 | 84 | 86 | 34 | 109 | 133 | 32 | 86 | 182 | 120 | 139 | 37 | 99 | 142 | 248 | 298 | 0.132673288 | 0.366181216 | 0.41 | 0.567 | 0.392 | 0.443 | 0.614 | 0.522 | 0.535 | 0.51 | 0.511 | 0.606 | -0.052 |
| MXE | 16764 | ENSG00000127328 | RAB3IP | chr12 | + | 69755570 | 69755659 | 69756404 | 69756663 | 69755381 | 69755478 | 69784719 | 69784815 | 2 | 0 | 1 | 2 | 2 | 90 | 53 | 46 | 64 | 74 | 3 | 6 | 2 | 0 | 0 | 43 | 43 | 58 | 59 | 44 | 238 | 298 | 0.132725224 | 0.366256899 | 0.027 | 0 | 0.026 | 0.038 | 0.033 | 0.08 | 0.149 | 0.041 | 0 | 0 | -0.029 |
| MXE | 6267 | ENSG00000153815 | CMIP | chr16 | + | 81696559 | 81696667 | 81699684 | 81699801 | 81693438 | 81693487 | 81701659 | 81701800 | 53 | 28 | 22 | 65 | 53 | 155 | 116 | 101 | 191 | 122 | 234 | 127 | 64 | 94 | 492 | 590 | 307 | 184 | 282 | 1315 | 257 | 266 | 0.13289804 | 0.366564825 | 0.261 | 0.2 | 0.184 | 0.26 | 0.31 | 0.291 | 0.3 | 0.265 | 0.257 | 0.279 | -0.035 |
| MXE | 6962 | ENSG00000117593 | DARS2 | chr1 | + | 173845228 | 173845291 | 173850326 | 173850479 | 173840865 | 173840973 | 173853348 | 173853567 | 23 | 12 | 7 | 21 | 38 | 16 | 21 | 8 | 30 | 62 | 9 | 10 | 3 | 26 | 6 | 26 | 31 | 12 | 28 | 7 | 212 | 298 | 0.132909232 | 0.366564825 | 0.669 | 0.445 | 0.552 | 0.496 | 0.463 | 0.327 | 0.312 | 0.26 | 0.566 | 0.546 | 0.123 |
| MXE | 3644 | ENSG00000055163 | CYFIP2 | chr5 | + | 157287018 | 157287108 | 157288611 | 157288659 | 157285338 | 157285478 | 157294782 | 157294860 | 518 | 195 | 330 | 314 | 222 | 6 | 13 | 0 | 0 | 0 | 169 | 307 | 149 | 230 | 222 | 0 | 12 | 3 | 7 | 0 | 239 | 197 | 0.132910419 | 0.366564825 | 0.986 | 0.925 | 1 | 1 | 1 | 1 | 0.955 | 0.976 | 0.964 | 1 | 0.003 |
| MXE | 6379 | ENSG00000026751 | SLAMF7 | chr1 | + | 160748193 | 160748514 | 160751344 | 160751448 | 160739056 | 160739356 | 160752185 | 160752248 | 183 | 71 | 137 | 141 | 71 | 150 | 86 | 121 | 123 | 169 | 115 | 229 | 104 | 124 | 117 | 114 | 159 | 84 | 98 | 137 | 298 | 253 | 0.132986832 | 0.366681968 | 0.509 | 0.412 | 0.49 | 0.493 | 0.263 | 0.461 | 0.55 | 0.512 | 0.518 | 0.42 | -0.059 |
| MXE | 15195 | ENSG00000153107 | ANAPC1 | chr2 | - | 111864805 | 111864951 | 111868022 | 111868096 | 111863674 | 111863895 | 111872629 | 111872712 | 39 | 23 | 28 | 37 | 79 | 122 | 120 | 138 | 127 | 284 | 44 | 39 | 26 | 24 | 8 | 104 | 45 | 95 | 68 | 57 | 223 | 295 | 0.13300198 | 0.366681968 | 0.297 | 0.202 | 0.212 | 0.278 | 0.269 | 0.359 | 0.534 | 0.266 | 0.318 | 0.157 | -0.075 |
| MXE | 371 | ENSG00000104880 | ARHGEF18 | chr19 | + | 7378393 | 7378451 | 7379121 | 7379166 | 7376642 | 7376757 | 7380916 | 7380994 | 48 | 30 | 83 | 49 | 59 | 53 | 37 | 69 | 70 | 63 | 29 | 38 | 33 | 29 | 119 | 40 | 63 | 31 | 56 | 151 | 207 | 194 | 0.133096149 | 0.366873887 | 0.459 | 0.432 | 0.53 | 0.396 | 0.467 | 0.405 | 0.361 | 0.499 | 0.327 | 0.425 | 0.053 |
| MXE | 9965 | ENSG00000231259 | ANAPC1P2 | chr2 | - | 87043712 | 87043984 | 87049588 | 87049803 | 87037293 | 87037528 | 87052813 | 87052992 | 160 | 96 | 169 | 158 | 106 | 4 | 4 | 3 | 7 | 2 | 144 | 263 | 225 | 141 | 233 | 0 | 4 | 2 | 11 | 0 | 298 | 298 | 0.133128333 | 0.366894907 | 0.976 | 0.96 | 0.983 | 0.958 | 0.981 | 1 | 0.985 | 0.991 | 0.928 | 1 | -0.009 |
| MXE | 7160 | ENSG00000089012 | SIRPG | chr20 | - | 1635266 | 1635599 | 1636187 | 1636505 | 1630221 | 1630306 | 1649051 | 1649408 | 17 | 7 | 44 | 20 | 47 | 58 | 12 | 71 | 43 | 88 | 6 | 6 | 3 | 11 | 6 | 8 | 38 | 13 | 15 | 27 | 298 | 298 | 0.133154578 | 0.366899556 | 0.227 | 0.368 | 0.383 | 0.317 | 0.348 | 0.429 | 0.136 | 0.188 | 0.423 | 0.182 | 0.057 |
| MXE | 9985 | ENSG00000101577 | LPIN2 | chr18 | - | 2926722 | 2926805 | 2929064 | 2929158 | 2925223 | 2925368 | 2931255 | 2931443 | 147 | 125 | 186 | 189 | 124 | 106 | 69 | 84 | 96 | 151 | 124 | 140 | 99 | 134 | 83 | 77 | 49 | 74 | 41 | 46 | 243 | 232 | 0.133211345 | 0.366988291 | 0.57 | 0.634 | 0.679 | 0.653 | 0.439 | 0.606 | 0.732 | 0.561 | 0.757 | 0.633 | -0.063 |
| MXE | 16845 | ENSG00000111596 | CNOT2 | chr12 | + | 70332766 | 70332846 | 70335437 | 70335563 | 70330286 | 70330469 | 70337388 | 70337513 | 81 | 53 | 53 | 49 | 63 | 341 | 297 | 332 | 367 | 561 | 9 | 57 | 54 | 26 | 5 | 161 | 318 | 308 | 298 | 27 | 229 | 275 | 0.133286337 | 0.367127189 | 0.222 | 0.176 | 0.161 | 0.138 | 0.119 | 0.063 | 0.177 | 0.174 | 0.095 | 0.182 | 0.025 |
| MXE | 9687 | ENSG00000033178 | UBA6 | chr4 | - | 67673696 | 67673777 | 67677610 | 67677722 | 67670469 | 67670592 | 67678438 | 67678533 | 112 | 85 | 92 | 126 | 105 | 133 | 166 | 100 | 150 | 271 | 66 | 82 | 73 | 56 | 23 | 61 | 96 | 110 | 83 | 11 | 261 | 230 | 0.133498213 | 0.367618849 | 0.426 | 0.311 | 0.448 | 0.425 | 0.255 | 0.488 | 0.429 | 0.369 | 0.373 | 0.648 | -0.088 |
| MXE | 5130 | ENSG00000107829 | FBXW4 | chr10 | - | 101624744 | 101624810 | 101667885 | 101667980 | 101612336 | 101612477 | 101672914 | 101673047 | 68 | 30 | 60 | 44 | 79 | 45 | 31 | 50 | 43 | 75 | 48 | 57 | 37 | 18 | 75 | 0 | 61 | 38 | 14 | 55 | 244 | 215 | 0.133514048 | 0.367618849 | 0.571 | 0.46 | 0.514 | 0.474 | 0.481 | 1 | 0.452 | 0.462 | 0.531 | 0.546 | -0.098 |
| MXE | 12877 | ENSG00000257923 | CUX1 | chr7 | + | 102070338 | 102070417 | 102097363 | 102097501 | 102028097 | 102028145 | 102104335 | 102104459 | 115 | 80 | 63 | 114 | 110 | 160 | 134 | 107 | 152 | 204 | 380 | 126 | 127 | 182 | 407 | 570 | 267 | 225 | 297 | 888 | 228 | 287 | 0.133644295 | 0.367909667 | 0.475 | 0.429 | 0.426 | 0.486 | 0.404 | 0.456 | 0.373 | 0.415 | 0.435 | 0.366 | 0.035 |
| MXE | 16133 | ENSG00000165732 | DDX21 | chr10 | + | 68970200 | 68970350 | 68971890 | 68972052 | 68968975 | 68969121 | 68973544 | 68973664 | 305 | 253 | 240 | 292 | 525 | 472 | 361 | 364 | 450 | 304 | 74 | 285 | 329 | 177 | 29 | 78 | 351 | 415 | 141 | 4 | 298 | 298 | 0.133716475 | 0.367998222 | 0.393 | 0.412 | 0.397 | 0.394 | 0.633 | 0.487 | 0.448 | 0.442 | 0.557 | 0.879 | -0.117 |
| MXE | 16344 | ENSG00000155256 | ZFYVE27 | chr10 | + | 97738476 | 97738649 | 97744728 | 97744915 | 97737120 | 97737321 | 97748268 | 97748364 | 5 | 3 | 2 | 15 | 2 | 12 | 8 | 9 | 17 | 30 | 0 | 0 | 4 | 4 | 0 | 5 | 2 | 5 | 6 | 15 | 298 | 298 | 0.133741357 | 0.367998222 | 0.294 | 0.273 | 0.182 | 0.469 | 0.062 | 0 | 0 | 0.444 | 0.4 | 0 | 0.087 |
| MXE | 7612 | ENSG00000163322 | ABRAXAS1 | chr4 | - | 83469031 | 83469151 | 83470202 | 83470396 | 83467453 | 83467538 | 83472221 | 83472288 | 145 | 124 | 109 | 125 | 173 | 52 | 64 | 38 | 49 | 96 | 37 | 51 | 63 | 10 | 20 | 22 | 14 | 32 | 8 | 38 | 298 | 269 | 0.133750358 | 0.367998222 | 0.716 | 0.636 | 0.721 | 0.697 | 0.619 | 0.603 | 0.767 | 0.64 | 0.53 | 0.322 | 0.105 |
| MXE | 10901 | ENSG00000133059 | DSTYK | chr1 | - | 205163722 | 205163955 | 205169162 | 205169832 | 205162922 | 205163006 | 205187417 | 205187806 | 36 | 39 | 42 | 31 | 9 | 55 | 50 | 64 | 57 | 56 | 26 | 19 | 45 | 8 | 29 | 93 | 54 | 96 | 88 | 30 | 298 | 298 | 0.133786893 | 0.368030968 | 0.396 | 0.438 | 0.396 | 0.352 | 0.138 | 0.218 | 0.26 | 0.319 | 0.083 | 0.492 | 0.07 |
| MXE | 9234 | ENSG00000111801 | BTN3A3 | chr6 | + | 26448616 | 26448637 | 26448727 | 26448754 | 26448247 | 26448448 | 26449661 | 26449688 | 3 | 0 | 2 | 3 | 7 | 178 | 112 | 147 | 145 | 188 | 0 | 10 | 3 | 6 | 2 | 220 | 111 | 82 | 108 | 152 | 170 | 176 | 0.133828476 | 0.368077582 | 0.017 | 0 | 0.014 | 0.021 | 0.037 | 0 | 0.085 | 0.036 | 0.054 | 0.013 | -0.02 |
| MXE | 5821 | ENSG00000143458 | GABPB2 | chr1 | + | 151088469 | 151088517 | 151090405 | 151090573 | 151088189 | 151088297 | 151093191 | 151093386 | 0 | 0 | 10 | 2 | 0 | 83 | 67 | 97 | 82 | 100 | 0 | 0 | 0 | 0 | 5 | 99 | 53 | 40 | 53 | 20 | 197 | 298 | 0.13391202 | 0.36823957 | 0 | 0 | 0.135 | 0.036 | 0 | 0 | 0 | 0 | 0 | 0.274 | -0.021 |
| MXE | 4863 | ENSG00000078618 | NRDC | chr1 | - | 51791577 | 51791661 | 51792045 | 51792098 | 51790899 | 51790990 | 51792376 | 51792424 | 60 | 31 | 47 | 54 | 54 | 165 | 140 | 154 | 176 | 260 | 57 | 52 | 57 | 40 | 9 | 61 | 149 | 146 | 170 | 42 | 202 | 233 | 0.133959933 | 0.368277658 | 0.295 | 0.203 | 0.26 | 0.261 | 0.193 | 0.519 | 0.287 | 0.31 | 0.213 | 0.198 | -0.063 |
| MXE | 14892 | ENSG00000165219 | GAPVD1 | chr9 | + | 125323797 | 125323923 | 125326415 | 125326589 | 125321432 | 125321562 | 125330077 | 125330218 | 116 | 76 | 73 | 89 | 93 | 137 | 82 | 80 | 94 | 158 | 73 | 100 | 83 | 117 | 37 | 141 | 107 | 129 | 123 | 116 | 275 | 298 | 0.133987021 | 0.368277658 | 0.478 | 0.501 | 0.497 | 0.506 | 0.389 | 0.359 | 0.503 | 0.411 | 0.508 | 0.257 | 0.067 |
| MXE | 1112 | ENSG00000141252 | VPS53 | chr17 | - | 627173 | 627316 | 631549 | 631628 | 623532 | 623674 | 653290 | 653410 | 30 | 13 | 21 | 35 | 50 | 89 | 60 | 82 | 100 | 110 | 71 | 37 | 25 | 23 | 52 | 171 | 77 | 58 | 88 | 97 | 228 | 292 | 0.133999823 | 0.368277658 | 0.302 | 0.217 | 0.247 | 0.31 | 0.368 | 0.347 | 0.381 | 0.356 | 0.251 | 0.407 | -0.06 |
| MXE | 15231 | ENSG00000109452 | INPP4B | chr4 | - | 142260491 | 142260564 | 142270662 | 142270774 | 142237863 | 142238011 | 142305457 | 142305537 | 105 | 52 | 135 | 64 | 117 | 72 | 85 | 107 | 71 | 54 | 24 | 85 | 77 | 28 | 27 | 28 | 41 | 19 | 16 | 24 | 261 | 222 | 0.134071908 | 0.368379071 | 0.554 | 0.342 | 0.518 | 0.434 | 0.648 | 0.422 | 0.638 | 0.775 | 0.598 | 0.489 | -0.085 |
| MXE | 9787 | ENSG00000068784 | SRBD1 | chr2 | - | 45553622 | 45553730 | 45562652 | 45562756 | 45551124 | 45551282 | 45567935 | 45568021 | 8 | 3 | 3 | 8 | 4 | 76 | 45 | 40 | 43 | 80 | 0 | 18 | 9 | 3 | 0 | 15 | 46 | 25 | 34 | 34 | 253 | 257 | 0.134086037 | 0.368379071 | 0.097 | 0.063 | 0.071 | 0.159 | 0.048 | 0 | 0.284 | 0.268 | 0.082 | 0 | -0.039 |
| MXE | 16983 | ENSG00000139132 | FGD4 | chr12 | + | 32607937 | 32608095 | 32610775 | 32610834 | 32602160 | 32602317 | 32611136 | 32611283 | 2 | 3 | 3 | 2 | 27 | 43 | 39 | 28 | 67 | 78 | 9 | 7 | 5 | 1 | 32 | 49 | 49 | 44 | 33 | 82 | 298 | 208 | 0.134169878 | 0.368482622 | 0.031 | 0.051 | 0.07 | 0.02 | 0.195 | 0.114 | 0.091 | 0.073 | 0.021 | 0.214 | -0.029 |
| MXE | 1613 | ENSG00000089505 | CMTM1 | chr16 | + | 66571136 | 66571270 | 66577103 | 66577202 | 66569935 | 66570094 | 66578830 | 66579135 | 0 | 0 | 0 | 0 | 3 | 3 | 1 | 1 | 6 | 3 | 0 | 0 | 9 | 0 | 0 | 16 | 6 | 10 | 1 | 5 | 283 | 248 | 0.134173056 | 0.368482622 | 0 | 0 | 0 | 0 | 0.467 | 0 | 0 | 0.441 | 0 | 0 | 0.005 |
| MXE | 9409 | ENSG00000187741 | FANCA | chr16 | - | 89739132 | 89739289 | 89739477 | 89739553 | 89738881 | 89738974 | 89739993 | 89740099 | 16 | 5 | 5 | 14 | 2 | 13 | 2 | 7 | 3 | 1 | 33 | 2 | 4 | 1 | 9 | 1 | 8 | 1 | 2 | 0 | 225 | 298 | 0.134329935 | 0.368845658 | 0.62 | 0.768 | 0.486 | 0.861 | 0.726 | 0.978 | 0.249 | 0.841 | 0.398 | 1 | -0.001 |
| MXE | 12397 | ENSG00000042980 | ADAM28 | chr8 | + | 24321217 | 24321289 | 24321314 | 24321448 | 24320235 | 24320307 | 24323833 | 24324003 | 82 | 103 | 111 | 100 | 240 | 9 | 0 | 2 | 1 | 1 | 64 | 161 | 43 | 47 | 9 | 0 | 0 | 0 | 0 | 14 | 221 | 283 | 0.134435571 | 0.369067886 | 0.921 | 1 | 0.986 | 0.992 | 0.997 | 1 | 1 | 1 | 1 | 0.452 | 0.089 |
| MXE | 355 | ENSG00000104824 | HNRNPL | chr19 | - | 38838893 | 38839015 | 38840095 | 38840376 | 38838396 | 38838598 | 38840487 | 38840559 | 270 | 162 | 235 | 291 | 407 | 366 | 187 | 347 | 370 | 464 | 477 | 361 | 363 | 332 | 474 | 379 | 496 | 446 | 331 | 583 | 298 | 271 | 0.13460469 | 0.369412235 | 0.402 | 0.441 | 0.381 | 0.417 | 0.444 | 0.534 | 0.398 | 0.425 | 0.477 | 0.425 | -0.035 |
| MXE | 14542 | ENSG00000108669 | CYTH1 | chr17 | - | 78692416 | 78692493 | 78698819 | 78698968 | 78680970 | 78681042 | 78700330 | 78700443 | 835 | 440 | 792 | 963 | 1065 | 598 | 395 | 519 | 575 | 715 | 1308 | 935 | 861 | 1011 | 1107 | 953 | 697 | 699 | 791 | 913 | 298 | 226 | 0.134610456 | 0.369412235 | 0.514 | 0.458 | 0.536 | 0.559 | 0.53 | 0.51 | 0.504 | 0.483 | 0.492 | 0.479 | 0.026 |
| MXE | 4954 | ENSG00000172053 | QARS1 | chr3 | - | 49102418 | 49102472 | 49103344 | 49103431 | 49102204 | 49102265 | 49103630 | 49103706 | 3 | 1 | 2 | 0 | 3 | 84 | 49 | 75 | 79 | 85 | 12 | 7 | 0 | 0 | 0 | 147 | 56 | 71 | 60 | 194 | 236 | 203 | 0.134849622 | 0.370000617 | 0.03 | 0.017 | 0.022 | 0 | 0.029 | 0.066 | 0.097 | 0 | 0 | 0 | -0.013 |
| MXE | 15588 | ENSG00000101972 | STAG2 | chrX | + | 124037526 | 124037623 | 124042568 | 124042645 | 124030960 | 124031125 | 124045163 | 124045208 | 209 | 132 | 178 | 204 | 329 | 248 | 182 | 214 | 229 | 637 | 9 | 44 | 29 | 6 | 7 | 0 | 102 | 51 | 11 | 0 | 246 | 226 | 0.134956356 | 0.370187897 | 0.436 | 0.4 | 0.433 | 0.45 | 0.322 | 1 | 0.284 | 0.343 | 0.334 | 1 | -0.184 |
| MXE | 8809 | ENSG00000142405 | NLRP12 | chr19 | - | 53807494 | 53807665 | 53809586 | 53811288 | 53805279 | 53805450 | 53814907 | 53814988 | 104 | 50 | 59 | 68 | 88 | 24 | 13 | 22 | 18 | 31 | 84 | 165 | 85 | 124 | 193 | 87 | 43 | 32 | 46 | 48 | 298 | 298 | 0.134967435 | 0.370187897 | 0.812 | 0.794 | 0.728 | 0.791 | 0.739 | 0.491 | 0.793 | 0.726 | 0.729 | 0.801 | 0.065 |
| MXE | 988 | ENSG00000101019 | UQCC1 | chr20 | - | 35347163 | 35347272 | 35366556 | 35366614 | 35314687 | 35314765 | 35374183 | 35374256 | 32 | 30 | 29 | 29 | 24 | 20 | 13 | 22 | 21 | 18 | 22 | 21 | 35 | 37 | 17 | 21 | 35 | 19 | 38 | 25 | 207 | 258 | 0.135009576 | 0.370235511 | 0.666 | 0.63 | 0.563 | 0.633 | 0.624 | 0.566 | 0.428 | 0.697 | 0.548 | 0.459 | 0.084 |
| MXE | 12738 | ENSG00000068976 | PYGM | chr11 | - | 64747223 | 64747358 | 64750375 | 64750583 | 64746920 | 64746987 | 64751324 | 64751466 | 7 | 6 | 8 | 5 | 7 | 16 | 7 | 14 | 9 | 12 | 6 | 5 | 10 | 3 | 3 | 17 | 30 | 14 | 3 | 21 | 298 | 284 | 0.135179465 | 0.370633364 | 0.294 | 0.45 | 0.353 | 0.346 | 0.357 | 0.252 | 0.137 | 0.405 | 0.488 | 0.12 | 0.08 |
| MXE | 13921 | ENSG00000171988 | JMJD1C | chr10 | - | 63215552 | 63215696 | 63264650 | 63264764 | 63215262 | 63215455 | 63268662 | 63269066 | 133 | 143 | 123 | 176 | 288 | 200 | 216 | 185 | 263 | 506 | 29 | 69 | 58 | 55 | 24 | 46 | 144 | 91 | 15 | 18 | 263 | 293 | 0.135273041 | 0.370660585 | 0.426 | 0.424 | 0.426 | 0.427 | 0.388 | 0.413 | 0.348 | 0.415 | 0.803 | 0.598 | -0.097 |
| MXE | 5734 | ENSG00000114126 | TFDP2 | chr3 | - | 141993537 | 141993585 | 141995019 | 141995141 | 141978519 | 141978682 | 142005440 | 142005544 | 83 | 41 | 59 | 63 | 93 | 25 | 12 | 20 | 25 | 45 | 41 | 42 | 35 | 61 | 23 | 7 | 14 | 13 | 14 | 8 | 271 | 197 | 0.135288008 | 0.370660585 | 0.707 | 0.713 | 0.682 | 0.647 | 0.6 | 0.81 | 0.686 | 0.662 | 0.76 | 0.676 | -0.049 |
| MXE | 9412 | ENSG00000187741 | FANCA | chr16 | - | 89749729 | 89749902 | 89752137 | 89752222 | 89748658 | 89748767 | 89758576 | 89758705 | 12 | 12 | 3 | 5 | 7 | 27 | 16 | 12 | 16 | 4 | 33 | 3 | 3 | 4 | 13 | 39 | 21 | 18 | 22 | 54 | 234 | 298 | 0.135288053 | 0.370660585 | 0.361 | 0.489 | 0.241 | 0.285 | 0.69 | 0.519 | 0.154 | 0.175 | 0.188 | 0.235 | 0.159 |
| MXE | 14952 | ENSG00000137073 | UBAP2 | chr9 | - | 33963725 | 33963791 | 33971650 | 33971754 | 33960825 | 33960878 | 33973182 | 33973237 | 43 | 51 | 45 | 54 | 130 | 116 | 60 | 89 | 110 | 299 | 69 | 63 | 40 | 79 | 84 | 122 | 81 | 74 | 103 | 166 | 253 | 215 | 0.135303865 | 0.370660585 | 0.24 | 0.419 | 0.301 | 0.294 | 0.27 | 0.325 | 0.398 | 0.315 | 0.395 | 0.301 | -0.042 |
| MXE | 5639 | ENSG00000033327 | GAB2 | chr11 | - | 78225107 | 78225202 | 78226464 | 78227051 | 78223411 | 78223676 | 78250156 | 78250400 | 47 | 41 | 27 | 49 | 20 | 73 | 37 | 45 | 30 | 41 | 130 | 58 | 109 | 124 | 113 | 98 | 95 | 106 | 70 | 93 | 298 | 244 | 0.135313442 | 0.370660585 | 0.345 | 0.476 | 0.329 | 0.572 | 0.285 | 0.521 | 0.333 | 0.457 | 0.592 | 0.499 | -0.079 |

| | | | | | | | | | | | | | | | | | | | | | | | | | |
|---|---|---|---|---|---|---|---|---|---|---|---|---|---|---|---|---|---|---|---|---|---|---|---|---|---|
| MXE | 13430 | ENSG00000177646 | ACAD9 | chr3 | + | 128895309 | 128895416 | 128896435 | 128896536 | 128893554 | 128893656 | 128897631 | 128897710 | 95 | 80 | 72 | 81 | 178 | 24 | 10 | 13 | 26 | 48 | 213 | 106 |
| MXE | 2238 | ENSG00000099308 | MAST3 | chr19 | + | 18134816 | 18134982 | 18135739 | 18135841 | 18134578 | 18134711 | 18137238 | 18137361 | 88 | 34 | 66 | 95 | 82 | 65 | 27 | 57 | 56 | 29 | 43 | 64 |
| MXE | 8878 | ENSG00000117523 | PRRC2C | chr1 | + | 171545478 | 171545687 | 171557239 | 171558143 | 171539970 | 171542229 | 171561017 | 171561103 | 332 | 297 | 321 | 357 | 681 | 262 | 232 | 314 | 345 | 534 | 133 | 319 |
| MXE | 8706 | ENSG00000176407 | KCMF1 | chr2 | + | 85027888 | 85028056 | 85035015 | 85035155 | 84971160 | 84971467 | 85043563 | 85043665 | 76 | 49 | 59 | 102 | 47 | 249 | 119 | 143 | 206 | 201 | 85 | 85 |
| MXE | 10867 | ENSG00000105953 | OGDH | chr7 | + | 44624316 | 44624565 | 44645326 | 44645518 | 44606587 | 44606653 | 44647656 | 44647759 | 173 | 137 | 139 | 203 | 111 | 130 | 106 | 91 | 132 | 156 | 251 | 204 |
| MXE | 3424 | ENSG00000163468 | CCT3 | chr1 | - | 156333546 | 156333622 | 156335826 | 156335888 | 156324971 | 156325089 | 156338153 | 156338292 | 45 | 15 | 35 | 41 | 1 | 259 | 147 | 241 | 249 | 271 | 42 | 24 |
| MXE | 2620 | ENSG00000075413 | MARK3 | chr14 | + | 103468080 | 103468186 | 103474992 | 103475210 | 103467078 | 103467191 | 103480386 | 103480490 | 17 | 16 | 20 | 12 | 36 | 152 | 125 | 165 | 135 | 203 | 26 | 48 |
| MXE | 15607 | ENSG00000159063 | ALG8 | chr11 | - | 78104355 | 78104453 | 78106806 | 78106946 | 78103979 | 78104052 | 78109441 | 78109581 | 30 | 13 | 52 | 23 | 34 | 14 | 15 | 22 | 23 | 11 | 23 | 24 |
| MXE | 12258 | ENSG00000160218 | TRAPPC10 | chr21 | + | 44084121 | 44084263 | 44087698 | 44087928 | 44082787 | 44083302 | 44089832 | 44089933 | 117 | 46 | 104 | 92 | 101 | 223 | 112 | 156 | 166 | 200 | 80 | 91 |
| MXE | 11167 | ENSG00000197930 | ERO1A | chr14 | - | 52652238 | 52652308 | 52653068 | 52653315 | 52646374 | 52646461 | 52657916 | 52658009 | 597 | 529 | 494 | 677 | 948 | 95 | 93 | 60 | 76 | 118 | 497 | 727 |
| MXE | 10650 | ENSG00000135926 | TMBIM1 | chr2 | - | 218279037 | 218279091 | 218279288 | 218279353 | 218278514 | 218278565 | 218280025 | 218280126 | 116 | 33 | 51 | 93 | 83 | 139 | 66 | 49 | 97 | 144 | 118 | 95 |
| MXE | 2723 | ENSG00000100461 | RBM23 | chr14 | - | 22905335 | 22905453 | 22906194 | 22906368 | 22905093 | 22905246 | 22909482 | 22909595 | 96 | 35 | 47 | 55 | 40 | 81 | 42 | 72 | 74 | 46 | 67 | 64 |
| MXE | 438 | ENSG00000079385 | CEACAM1 | chr19 | - | 42516772 | 42516865 | 42518947 | 42519235 | 42512349 | 42512479 | 42521266 | 42521521 | 2 | 4 | 8 | 10 | 5 | 0 | 0 | 0 | 0 | 0 | 22 | 6 |
| MXE | 12185 | ENSG00000188786 | MTF1 | chr1 | - | 37835078 | 37835215 | 37838624 | 37838756 | 37832244 | 37832322 | 37839919 | 37840158 | 151 | 140 | 109 | 144 | 233 | 136 | 125 | 128 | 148 | 283 | 282 | 163 |
| MXE | 1876 | ENSG00000103047 | TANGO6 | chr16 | + | 68900433 | 68900546 | 68902327 | 68902504 | 68880547 | 68880630 | 68907442 | 68907575 | 93 | 61 | 101 | 66 | 51 | 25 | 16 | 36 | 26 | 41 | 161 | 88 |
| MXE | 15450 | ENSG00000103544 | VPS35L | chr16 | + | 19591789 | 19591874 | 19601663 | 19601723 | 19581524 | 19581653 | 19608177 | 19608274 | 105 | 69 | 72 | 84 | 119 | 10 | 6 | 5 | 9 | 12 | 104 | 82 |
| MXE | 3997 | ENSG00000170142 | UBE2E1 | chr3 | + | 23887566 | 23887699 | 23889111 | 23889259 | 23811457 | 23811510 | 23890508 | 23891316 | 220 | 180 | 164 | 198 | 580 | 214 | 126 | 138 | 136 | 318 | 213 | 132 |
| MXE | 1408 | ENSG00000073584 | SMARCE1 | chr17 | - | 40630713 | 40630924 | 40631591 | 40631693 | 40624961 | 40628993 | 40632194 | 40632367 | 205 | 167 | 204 | 190 | 365 | 175 | 159 | 160 | 166 | 168 | 174 | 243 |
| MXE | 15942 | ENSG00000156650 | KAT6B | chr10 | + | 74972506 | 74972639 | 74975398 | 74975454 | 74970019 | 74970101 | 74977315 | 74977437 | 137 | 114 | 162 | 160 | 198 | 22 | 18 | 12 | 19 | 24 | 35 | 53 |
| MXE | 13540 | ENSG00000035681 | NSMAF | chr8 | - | 58602057 | 58602137 | 58603209 | 58603386 | 58601444 | 58601535 | 58605926 | 58606035 | 130 | 84 | 178 | 135 | 217 | 30 | 20 | 42 | 43 | 49 | 185 | 123 |
| MXE | 9302 | ENSG00000075651 | PLD1 | chr3 | - | 171735491 | 171735637 | 171737531 | 171737659 | 171734864 | 171734970 | 171737891 | 171738082 | 37 | 18 | 24 | 25 | 18 | 28 | 29 | 25 | 19 | 38 | 22 | 32 |
| MXE | 11277 | ENSG00000113407 | TARS1 | chr5 | + | 33448540 | 33448731 | 33453288 | 33453412 | 33445323 | 33445404 | 33454944 | 33455066 | 84 | 50 | 66 | 74 | 94 | 119 | 77 | 103 | 136 | 134 | 33 | 58 |
| MXE | 13039 | ENSG00000003436 | TFPI | chr2 | - | 187513615 | 187513646 | 187529363 | 187529485 | 187503647 | 187503770 | 187554199 | 187554492 | 1 | 8 | 3 | 20 | 2 | 2 | 3 | 3 | 5 | 1 | 27 | 3 |
| MXE | 9821 | ENSG00000151092 | NGLY1 | chr3 | - | 25737333 | 25737455 | 25739576 | 25739799 | 25736003 | 25736149 | 25751097 | 25751263 | 67 | 56 | 50 | 59 | 80 | 121 | 101 | 95 | 133 | 115 | 18 | 45 |
| MXE | 2571 | ENSG00000171469 | ZNF561 | chr19 | - | 9617048 | 9617171 | 9617620 | 9617758 | 9614020 | 9614103 | 9618090 | 9618179 | 4 | 6 | 2 | 2 | 1 | 67 | 45 | 47 | 55 | 55 | 6 | 0 |
| MXE | 11911 | ENSG00000135317 | SNX14 | chr6 | - | 85565331 | 85565419 | 85567533 | 85567577 | 85557975 | 85558060 | 85572136 | 85572215 | 35 | 22 | 17 | 37 | 39 | 91 | 62 | 45 | 74 | 129 | 3 | 2 |
| MXE | 6051 | ENSG00000132716 | DCAF8 | chr1 | - | 160220458 | 160220691 | 160221086 | 160221210 | 160218848 | 160218968 | 160222650 | 160222781 | 2 | 0 | 1 | 0 | 6 | 13 | 8 | 15 | 6 | 28 | 0 | 0 |
| MXE | 2748 | ENSG00000131966 | ACTR10 | chr14 | + | 58215204 | 58215284 | 58219693 | 58219729 | 58208998 | 58209107 | 58223621 | 58223701 | 8 | 14 | 8 | 13 | 28 | 7 | 0 | 6 | 8 | 7 | 6 | 20 |
| MXE | 1562 | ENSG00000102984 | ZNF821 | chr16 | - | 71864902 | 71865048 | 71867911 | 71868037 | 71864137 | 71864242 | 71879906 | 71880023 | 15 | 7 | 7 | 8 | 13 | 12 | 6 | 11 | 15 | 12 | 22 | 22 |
| MXE | 12128 | ENSG00000124486 | USP9X | chrX | + | 41123470 | 41123724 | 41128999 | 41129145 | 41085444 | 41086109 | 41131456 | 41131536 | 128 | 88 | 135 | 130 | 100 | 186 | 147 | 186 | 223 | 96 | 74 | 148 |
| MXE | 15985 | ENSG00000100225 | FBXO7 | chr22 | + | 32479132 | 32479275 | 32483896 | 32484124 | 32475236 | 32475398 | 32485067 | 32485209 | 5 | 1 | 6 | 5 | 0 | 184 | 96 | 166 | 173 | 251 | 0 | 2 |
| MXE | 5465 | ENSG00000168827 | GFM1 | chr3 | + | 158681994 | 158682157 | 158690162 | 158690323 | 158666303 | 158666386 | 158691138 | 158691192 | 60 | 61 | 60 | 73 | 158 | 97 | 45 | 53 | 63 | 86 | 30 | 60 |
| MXE | 2670 | ENSG00000100889 | PCK2 | chr14 | + | 24096891 | 24097137 | 24098202 | 24098387 | 24094158 | 24094434 | 24098474 | 24098678 | 21 | 10 | 28 | 27 | 2 | 18 | 14 | 11 | 13 | 10 | 15 | 20 |
| MXE | 4054 | ENSG00000070756 | PABPC1 | chr8 | - | 100713086 | 100713181 | 100715461 | 100715601 | 100712651 | 100712789 | 100717772 | 100717888 | 7293 | 5502 | 7495 | 7063 | 9301 | 6996 | 5164 | 7402 | 6658 | 8539 | 8011 | 4080 |
| MXE | 8538 | ENSG00000121892 | PDS5A | chr4 | - | 39841947 | 39842056 | 39844655 | 39844801 | 39837011 | 39838208 | 39845817 | 39848561 | 313 | 269 | 248 | 319 | 566 | 332 | 297 | 272 | 348 | 459 | 53 | 143 |
| MXE | 3913 | ENSG00000038427 | VCAN | chr5 | + | 83547970 | 83548084 | 83553363 | 83553522 | 83545536 | 83545650 | 83554955 | 83555038 | 579 | 403 | 320 | 480 | 369 | 1224 | 846 | 661 | 1049 | 1265 | 124 | 623 |
| MXE | 12850 | ENSG00000123066 | MED13L | chr12 | - | 115963406 | 115963519 | 115966081 | 115966243 | 115958575 | 115961398 | 115968939 | 115969097 | 159 | 103 | 138 | 141 | 192 | 134 | 121 | 140 | 140 | 141 | 171 | 171 |
| MXE | 10514 | ENSG00000091490 | SEL1L3 | chr4 | - | 25776276 | 25776360 | 25779075 | 25779203 | 25767739 | 25767830 | 25782241 | 25782418 | 158 | 141 | 272 | 189 | 216 | 88 | 76 | 102 | 120 | 176 | 151 | 350 |
| MXE | 14388 | ENSG00000119397 | CNTRL | chr9 | + | 121162053 | 121162271 | 121164942 | 121165100 | 121161855 | 121161971 | 121166106 | 121166180 | 174 | 182 | 151 | 198 | 201 | 152 | 146 | 144 | 127 | 291 | 22 | 62 |
| [illegible] | | | | | | | | | | | | | | | | | | | | | | | | | |

| | | | | | | | | | | | | | | | | | | | | | | | | | | |
|---|---|---|---|---|---|---|---|---|---|---|---|---|---|---|---|---|---|---|---|---|---|---|---|---|---|---|
| MXE | 10364 | ENSG00000152520 | PAN3 | chr13 | + | 28256291 | 28256539 | 28260446 | 28260551 | 28220230 | 28220378 | 28261400 | 28261458 | 107 | 60 | 84 | 92 | 175 | 182 | 83 | 134 | 150 | 151 | 52 | 80 | [illegible] |
| MXE | 3283 | ENSG00000137177 | KIF13A | chr6 | - | 17778946 | 17779099 | 17779591 | 17779684 | 17777276 | 17777354 | 17780729 | 17780906 | 28 | 25 | 14 | 33 | 29 | 43 | 29 | 14 | 27 | 43 | 38 | 30 | [illegible] |
| MXE | 7510 | ENSG00000115020 | PIKFYVE | chr2 | + | 208274020 | 208274053 | 208277536 | 208277708 | 208273583 | 208273733 | 208285725 | 208285933 | 1 | 8 | 2 | 3 | 1 | 88 | 68 | 80 | 64 | 105 | 1 | 3 | [illegible] |
| MXE | 2983 | ENSG00000226032 | NA | chr6 | - | 159043588 | 159043655 | 159043977 | 159044031 | 159042191 | 159042244 | 159044119 | 159044204 | 145 | 72 | 138 | 146 | 188 | 252 | 185 | 325 | 305 | 414 | 80 | 141 | [illegible] |
| MXE | 16084 | ENSG00000197535 | MYO5A | chr15 | - | 52425829 | 52425974 | 52428397 | 52428569 | 52416144 | 52416301 | 52433174 | 52433285 | 74 | 35 | 38 | 55 | 64 | 102 | 107 | 76 | 108 | 124 | 0 | 53 | [illegible] |
| MXE | 4629 | ENSG00000038219 | BOD1L1 | chr4 | - | 13595859 | 13595944 | 13597103 | 13597168 | 13591922 | 13591966 | 13598945 | 13605084 | 59 | 40 | 45 | 40 | 78 | 72 | 48 | 57 | 61 | 101 | 12 | 22 | [illegible] |
| MXE | 3528 | ENSG00000113263 | ITK | chr5 | + | 157241645 | 157241720 | 157245725 | 157245790 | 157240061 | 157240195 | 157245880 | 157245999 | 154 | 88 | 196 | 150 | 331 | 82 | 63 | 97 | 98 | 101 | 57 | 106 | [illegible] |
| MXE | 13395 | ENSG00000235437 | LINC01278 | chrX | - | 63432363 | 63432491 | 63509943 | 63510079 | 63399780 | 63399903 | 63516909 | 63517082 | 14 | 3 | 19 | 4 | 15 | 8 | 6 | 17 | 10 | 21 | 0 | 5 | [illegible] |
| MXE | 12878 | ENSG00000257923 | CUX1 | chr7 | + | 102097363 | 102097501 | 102104335 | 102104459 | 102070338 | 102070417 | 102111697 | 102111774 | 243 | 173 | 193 | 278 | 332 | 119 | 90 | 88 | 77 | 141 | 843 | 379 | [illegible] |
| MXE | 1354 | ENSG00000172794 | RAB37 | chr17 | + | 74740767 | 74740878 | 74742253 | 74742295 | 74736597 | 74736731 | 74743128 | 74743195 | 3 | 1 | 5 | 2 | 2 | 63 | 37 | 38 | 58 | 43 | 0 | 1 | [illegible] |
| MXE | 2787 | ENSG00000198752 | CDC42BPB | chr14 | - | 103003927 | 103004023 | 103012096 | 103012188 | 102999564 | 102999713 | 103056998 | 103057549 | 2 | 1 | 0 | 2 | 4 | 21 | 12 | 19 | 22 | 8 | 6 | 2 | [illegible] |
| MXE | 3704 | ENSG00000087470 | DNM1L | chr12 | + | 32740063 | 32740240 | 32740408 | 32740518 | 32737104 | 32737161 | 32742588 | 32742748 | 3 | 0 | 3 | 13 | 8 | 123 | 117 | 93 | 153 | 115 | 0 | 4 | [illegible] |
| MXE | 2415 | ENSG00000248905 | FMN1 | chr15 | - | 32910473 | 32910535 | 32926173 | 32926261 | 32908489 | 32908578 | 32964106 | 32964257 | 10 | 53 | 21 | 28 | 22 | 4 | 18 | 13 | 12 | 17 | 39 | 56 | [illegible] |
| MXE | 3863 | ENSG00000113391 | FAM172A | chr5 | - | 93958776 | 93958968 | 93964457 | 93964523 | 93881469 | 93881688 | 94050730 | 94050831 | 39 | 26 | 35 | 20 | 32 | 163 | 141 | 144 | 172 | 289 | 3 | 19 | [illegible] |
| MXE | 1312 | ENSG00000154874 | CCDC144B | chr17 | - | 18587665 | 18587879 | 18594720 | 18594862 | 18583340 | 18583523 | 18595147 | 18595181 | 8 | 17 | 7 | 40 | 26 | 16 | 48 | 9 | 75 | 54 | 0 | 16 | [illegible] |
| MXE | 5135 | ENSG00000111731 | C2CD5 | chr12 | - | 22482556 | 22482743 | 22484696 | 22484888 | 22478027 | 22478477 | 22490122 | 22490218 | 56 | 43 | 28 | 67 | 53 | 141 | 107 | 89 | 110 | 218 | 8 | 10 | [illegible] |
| MXE | 1125 | ENSG00000108799 | EZH1 | chr17 | - | 42706006 | 42706185 | 42707957 | 42708083 | 42705087 | 42705183 | 42708875 | 42708916 | 84 | 52 | 117 | 116 | 136 | 51 | 32 | 57 | 50 | 24 | 180 | 106 | [illegible] |
| MXE | 702 | ENSG00000064601 | CTSA | chr20 | + | 45894649 | 45894741 | 45894822 | 45894901 | 45893987 | 45894072 | 45894993 | 45895133 | 422 | 307 | 243 | 415 | 507 | 484 | 342 | 258 | 467 | 934 | 1100 | 1127 | [illegible] |
| MXE | 10012 | ENSG00000148737 | TCF7L2 | chr10 | + | 113159919 | 113159992 | 113160618 | 113160691 | 113158020 | 113158069 | 113165554 | 113167678 | 10 | 3 | 0 | 0 | 5 | 11 | 9 | 1 | 5 | 16 | 17 | 0 | [illegible] |
| MXE | 10116 | ENSG00000166164 | BRD7 | chr16 | - | 50339975 | 50340086 | 50350022 | 50350167 | 50334710 | 50334895 | 50354424 | 50354482 | 211 | 178 | 159 | 178 | 258 | 123 | 134 | 92 | 150 | 145 | 259 | 174 | [illegible] |
| MXE | 441 | ENSG00000183207 | RUVBL2 | chr19 | + | 49003278 | 49003334 | 49004276 | 49004418 | 48999318 | 48999373 | 49007017 | 49007147 | 8 | 2 | 2 | 8 | 4 | 162 | 64 | 102 | 156 | 92 | 0 | 6 | [illegible] |
| MXE | 516 | ENSG00000105483 | CARD8 | chr19 | - | 48246074 | 48246381 | 48249522 | 48249633 | 48240961 | 48241063 | 48249750 | 48249847 | 311 | 204 | 257 | 343 | 226 | 11 | 0 | 10 | 7 | 11 | 171 | 194 | [illegible] |
| [illegible] | [illegible] | [illegible] | [illegible] | [illegible] | [illegible] | [illegible] | [illegible] | [illegible] | [illegible] | [illegible] | [illegible] | [illegible] | [illegible] | [illegible] | [illegible] | [illegible] | [illegible] | [illegible] | [illegible] | [illegible] | [illegible] | [illegible] | [illegible] | [illegible] | [illegible] | [illegible] |

| | | | | | | | | | | | | | | | | | | | | | |
|---|---|---|---|---|---|---|---|---|---|---|---|---|---|---|---|---|---|---|---|---|---|
| MXE | 4639 | ENSG00000108509 | CAMTA2 | chr17 | - | 4981677 | 4981742 | 4982088 | 4982160 | 4981224 | 4981359 | 4982756 | 4982892 | 21 | 22 | 15 | 29 | 26 | 15 | 20 | 9 |
| MXE | 5119 | ENSG00000114867 | EIF4G1 | chr3 | + | 184326526 | 184326629 | 184326880 | 184326983 | 184325850 | 184325951 | 184327215 | 184327448 | 171 | 109 | 161 | 177 | 120 | 203 | 131 | 155 |
| MXE | 13317 | ENSG00000139436 | GIT2 | chr12 | - | 109967457 | 109967503 | 109980951 | 109981046 | 109965525 | 109965577 | 109983372 | 109983503 | 158 | 131 | 139 | 177 | 105 | 186 | 163 | 159 |
| MXE | 11363 | ENSG00000136699 | SMPD4 | chr2 | - | 130173513 | 130173656 | 130174913 | 130175000 | 130173278 | 130173354 | 130176553 | 130176637 | 23 | 10 | 23 | 31 | 19 | 40 | 20 | 28 |
| MXE | 16271 | ENSG00000066084 | DIP2B | chr12 | + | 50723201 | 50723323 | 50724774 | 50724886 | 50721272 | 50721396 | 50727702 | 50727812 | 32 | 25 | 43 | 39 | 58 | 47 | 34 | 45 |
| MXE | 10880 | ENSG00000124813 | RUNX2 | chr6 | + | 45491940 | 45492114 | 45545216 | 45545282 | 45437946 | 45438051 | 45546826 | 45547209 | 58 | 64 | 68 | 53 | 71 | 4 | 6 | 4 |
| MXE | 10360 | ENSG00000152520 | PAN3 | chr13 | + | 28177864 | 28177935 | 28197184 | 28197346 | 28176492 | 28176559 | 28220230 | 28220378 | 150 | 86 | 108 | 123 | 240 | 58 | 59 | 66 |
| MXE | 6029 | ENSG00000140307 | GTF2A2 | chr15 | - | 59650668 | 59650773 | 59652205 | 59652326 | 59638865 | 59639157 | 59657405 | 59657520 | 132 | 73 | 120 | 120 | 45 | 4 | 4 | 0 |
| MXE | 9877 | ENSG00000152061 | RABGAP1L | chr1 | + | 174370978 | 174371072 | 174393994 | 174394145 | 174304985 | 174305127 | 174637374 | 174637488 | 52 | 27 | 24 | 45 | 20 | 196 | 160 | 131 |
| MXE | 10257 | ENSG00000143207 | COP1 | chr1 | - | 176116623 | 176116681 | 176149005 | 176149074 | 176085775 | 176085890 | 176162868 | 176162988 | 139 | 170 | 98 | 114 | 242 | 95 | 79 | 54 |
| MXE | 3408 | ENSG00000043143 | JADE2 | chr5 | + | 134564493 | 134564610 | 134566115 | 134566580 | 134562199 | 134562367 | 134573644 | 134573762 | 138 | 52 | 148 | 124 | 63 | 166 | 85 | 110 |
| MXE | 10625 | ENSG00000258555 | SPECC1L-ADORA2A | chr22 | + | 24411587 | 24411704 | 24412647 | 24412707 | 24369217 | 24369320 | 24414533 | 24414744 | 19 | 13 | 22 | 33 | 33 | 29 | 10 | 28 |
| MXE | 14710 | ENSG00000114030 | KPNA1 | chr3 | - | 122427537 | 122427716 | 122433660 | 122433788 | 122421901 | 122427172 | 122437169 | 122437295 | 27 | 14 | 21 | 30 | 63 | 160 | 127 | 154 |
| MXE | 11641 | ENSG00000114423 | CBLB | chr3 | - | 105703987 | 105704173 | 105720046 | 105720250 | 105702093 | 105702459 | 105734008 | 105734140 | 54 | 28 | 34 | 28 | 73 | 90 | 86 | 113 |
| MXE | 616 | ENSG00000118620 | ZNF430 | chr19 | + | 21022788 | 21022881 | 21033455 | 21033582 | 21020716 | 21020803 | 21034088 | 21036802 | 22 | 10 | 25 | 20 | 13 | 25 | 26 | 30 |
| MXE | 8900 | ENSG00000172965 | MIR4435-2HG | chr2 | - | 111233627 | 111233739 | 111248147 | 111248241 | 111207775 | 111208661 | 111344060 | 111344237 | 22 | 17 | 6 | 23 | 26 | 3 | 1 | 2 |
| MXE | 16993 | ENSG00000111300 | NAA25 | chr12 | - | 112048291 | 112048443 | 112049482 | 112049611 | 112047664 | 112047790 | 112053557 | 112053657 | 7 | 5 | 1 | 10 | 6 | 67 | 76 | 67 |
| MXE | 4242 | ENSG00000174652 | ZNF266 | chr19 | - | 9415653 | 9415742 | 9417827 | 9417908 | 9412428 | 9414720 | 9418504 | 9418631 | 102 | 74 | 82 | 87 | 118 | 126 | 74 | 70 |
| MXE | 9900 | ENSG00000075884 | ARHGAP15 | chr2 | + | 143436912 | 143437042 | 143487372 | 143487495 | 143435600 | 143435699 | 143519265 | 143519364 | 905 | 746 | 838 | 689 | 1952 | 534 | 392 | 461 |
| MXE | 2718 | ENSG00000063761 | ADCK1 | chr14 | + | 77859075 | 77859279 | 77887090 | 77887249 | 77822434 | 77822518 | 77899099 | 77899258 | 4 | 1 | 6 | 10 | 2 | 11 | 7 | 8 |
| MXE | 6083 | ENSG00000164253 | WDR41 | chr5 | - | 77449759 | 77449860 | 77451290 | 77451353 | 77440812 | 77440997 | 77453618 | 77453928 | 39 | 20 | 22 | 34 | 76 | 86 | 46 | 62 |
| MXE | 10575 | ENSG00000157985 | AGAP1 | chr2 | + | 235908737 | 235908906 | 235930764 | 235930923 | 235883344 | 235883449 | 235968461 | 235968623 | 75 | 23 | 32 | 25 | 66 | 15 | 1 | 8 |
| MXE | 12195 | ENSG00000083168 | KAT6A | chr8 | - | 41987454 | 41987563 | 42048377 | 42049302 | 41981838 | 41981954 | 42051900 | 42051987 | 75 | 31 | 72 | 42 | 19 | 146 | 73 | 179 |
| MXE | 3219 | ENSG00000189091 | SF3B3 | chr16 | + | 70541668 | 70541834 | 70544437 | 70544533 | 70539103 | 70539207 | 70548369 | 70548442 | 145 | 71 | 162 | 136 | 342 | 126 | 80 | 118 |
| MXE | 14361 | ENSG00000133794 | ARNTL | chr11 | + | 13381160 | 13381254 | 13385688 | 13385791 | 13378321 | 13378445 | 13386595 | 13387262 | 52 | 19 | 46 | 50 | 43 | 6 | 3 | 9 |
| MXE | 15210 | ENSG00000113163 | CERT1 | chr5 | - | 75389591 | 75389687 | 75400204 | 75400297 | 75385901 | 75386034 | 75402971 | 75403058 | 139 | 133 | 126 | 166 | 206 | 96 | 68 | 80 |
| MXE | 5079 | ENSG00000171109 | MFN1 | chr3 | + | 179375219 | 179375341 | 179377041 | 179377168 | 179368035 | 179368103 | 179377343 | 179377448 | 132 | 53 | 78 | 96 | 116 | 75 | 49 | 58 |
| MXE | 14574 | ENSG00000164975 | SNAPC3 | chr9 | + | 15451319 | 15451402 | 15453040 | 15453205 | 15447094 | 15447244 | 15457959 | 15458067 | 52 | 64 | 55 | 49 | 134 | 118 | 147 | 115 |
| MXE | 13579 | ENSG00000196924 | FLNA | chrX | - | 154365348 | 154365486 | 154366023 | 154366224 | 154365135 | 154365259 | 154366307 | 154366470 | 1909 | 778 | 1019 | 1420 | 727 | 2310 | 1004 | 1221 |
| MXE | 1752 | ENSG00000169896 | ITGAM | chr16 | + | 31326855 | 31326935 | 31328146 | 31328230 | 31325499 | 31325622 | 31329227 | 31329303 | 58 | 16 | 31 | 25 | 29 | 274 | 84 | 154 |
| MXE | 16417 | ENSG00000011258 | MBTD1 | chr17 | - | 51206887 | 51207005 | 51209313 | 51209430 | 51203790 | 51203925 | 51217335 | 51217416 | 7 | 3 | 2 | 3 | 4 | 68 | 54 | 75 |
| MXE | 13357 | ENSG00000099204 | ABLIM1 | chr10 | - | 114451623 | 114451671 | 114465697 | 114465827 | 114445311 | 114445403 | 114468180 | 114468216 | 336 | 276 | 597 | 504 | 1546 | 13 | 1 | 2 |
| MXE | 7125 | ENSG00000055609 | KMT2C | chr7 | - | 152265037 | 152265209 | 152273704 | 152273867 | 152263015 | 152263130 | 152286299 | 152286376 | 7 | 9 | 7 | 27 | 6 | 183 | 148 | 169 |
| MXE | 13122 | ENSG00000169891 | REPS2 | chrX | + | 17054807 | 17054950 | 17062437 | 17062532 | 17052381 | 17052445 | 17068401 | 17068471 | 16 | 12 | 4 | 9 | 2 | 12 | 16 | 4 |
| MXE | 16643 | ENSG00000089041 | P2RX7 | chr12 | + | 121160901 | 121160974 | 121162270 | 121162520 | 121156078 | 121156147 | 121165356 | 121165437 | 20 | 22 | 20 | 16 | 42 | 36 | 30 | 25 |
| MXE | 7722 | ENSG00000082512 | TRAF5 | chr1 | + | 211361087 | 211361162 | 211365375 | 211365468 | 211360701 | 211360779 | 211369451 | 211369592 | 63 | 44 | 77 | 74 | 76 | 46 | 24 | 35 |
| MXE | 7012 | ENSG00000114439 | BBX | chr3 | + | 107791239 | 107791299 | 107798522 | 107798720 | 107778370 | 107778519 | 107801094 | 107801281 | 26 | 18 | 29 | 24 | 50 | 281 | 211 | 244 |
| MXE | 3414 | ENSG00000140853 | NLRC5 | chr16 | + | 57028071 | 57028155 | 57028301 | 57028385 | 57025367 | 57027018 | 57029772 | 57029856 | 42 | 18 | 63 | 47 | 42 | 144 | 80 | 144 |
| MXE | 9439 | ENSG00000109381 | ELF2 | chr4 | - | 139061864 | 139062057 | 139067683 | 139067770 | 139060323 | 139060674 | 139071865 | 139072039 | 54 | 47 | 49 | 47 | 60 | 175 | 141 | 152 |
| MXE | 90 | ENSG00000182944 | EWSR1 | chr22 | + | 29273737 | 29273864 | 29278130 | 29278216 | 29272379 | 29272431 | 29282389 | 29282557 | 17 | 5 | 19 | 16 | 18 | 154 | 75 | 149 |
| MXE | 14190 | ENSG00000104763 | ASAH1 | chr8 | - | 18069791 | 18069878 | 18071299 | 18071390 | 18067219 | 18067298 | 18075540 | 18075587 | 99 | 43 | 77 | 70 | 23 | 518 | 402 | 285 |
| MXE | 4517 | ENSG00000005059 | MCUB | chr4 | + | 109660194 | 109660365 | 109664289 | 109664394 | 109659010 | 109659086 | 109682581 | 109682742 | 182 | 145 | 209 | 203 | 264 | 420 | 441 | 502 |
| MXE | 13426 | ENSG00000119760 | SUPT7L | chr2 | - | 27655364 | 27655602 | 27657344 | 27657669 | 27650808 | 27653747 | 27660983 | 27661388 | 77 | 52 | 79 | 83 | 78 | 130 | 94 | 77 |
| MXE | 13868 | ENSG00000168575 | SLC20A2 | chr8 | - | 42439449 | 42439653 | 42444645 | 42444762 | 42436988 | 42437577 | 42459895 | 42459992 | 30 | 24 | 40 | 49 | 13 | 16 | 14 | 16 |
| [illegible] | | | | | | | | | | | | | | | | | | | | | |

| | | | | | | | | | | |
|---|---|---|---|---|---|---|---|---|---|---|
| MXE | 10455 | ENSG00000064999 | ANKS1A | chr6 | + | 34982751 | 34982827 | 34983112 | 34983214 | 34981689 | 34981986 |
| MXE | 4578 | ENSG00000189308 | LIN54 | chr4 | - | 82939888 | 82939962 | 82946257 | 82946474 | 82939538 | 82939736 |
| MXE | 7619 | ENSG00000132704 | FCRL2 | chr1 | - | 157766854 | 157766971 | 157770408 | 157770666 | 157749649 | 157749677 |
| MXE | 12777 | ENSG00000177189 | RPS6KA3 | chrX | - | 20164898 | 20165060 | 20169401 | 20169491 | 20162963 | 20163040 |
| MXE | 13814 | ENSG00000136279 | DBNL | chr7 | + | 44052866 | 44052941 | 44056756 | 44056903 | 44051829 | 44051942 |
| MXE | 278 | ENSG00000228315 | GUSBP11 | chr22 | - | 23640307 | 23640423 | 23652478 | 23652576 | 23638486 | 23638942 |
| MXE | 8074 | ENSG00000006756 | ARSD | chrX | - | 2909816 | 2909979 | 2910658 | 2910793 | 2908720 | 2908842 |
| MXE | 9785 | ENSG00000068784 | SRBD1 | chr2 | - | 45547521 | 45547612 | 45551124 | 45551282 | 45546731 | 45546839 |
| MXE | 10620 | ENSG00000163872 | YEATS2 | chr3 | + | 183724418 | 183724531 | 183728689 | 183728851 | 183721890 | 183722136 |
| MXE | 11702 | ENSG00000074842 | MYDGF | chr19 | - | 4660668 | 4660750 | 4664875 | 4664937 | 4659687 | 4660003 |
| MXE | 11101 | ENSG00000184182 | UBE2F | chr2 | + | 237994743 | 237994809 | 238025341 | 238025412 | 237987962 | 237987992 |
| MXE | 5829 | ENSG00000147251 | DOCK11 | chrX | + | 118561382 | 118561517 | 118566004 | 118566182 | 118546020 | 118546116 |
| MXE | 636 | ENSG00000127507 | ADGRE2 | chr19 | - | 14773937 | 14774054 | 14774255 | 14774306 | 14771879 | 14772497 |
| MXE | 3833 | ENSG00000151292 | CSNK1G3 | chr5 | + | 123553106 | 123553147 | 123557494 | 123557564 | 123545414 | 123545841 |
| MXE | 5761 | ENSG00000105856 | HBP1 | chr7 | + | 107179878 | 107180062 | 107182372 | 107182601 | 107168960 | 107169027 |
| MXE | 7593 | ENSG00000162976 | SLC66A3 | chr2 | + | 11164203 | 11164261 | 11171924 | 11172045 | 11160624 | 11160694 |
| MXE | 15434 | ENSG00000087053 | MTMR2 | chr11 | - | 95882373 | 95882496 | 95887611 | 95887759 | 95865600 | 95865676 |
| MXE | 4706 | ENSG00000155508 | CNOT8 | chr5 | + | 154865191 | 154865385 | 154870660 | 154870822 | 154863206 | 154863395 |
| MXE | 14693 | ENSG00000116580 | GON4L | chr1 | - | 155820605 | 155820656 | 155821473 | 155821548 | 155815804 | 155816262 |
| MXE | 5459 | ENSG00000168811 | IL12A | chr3 | + | 159993011 | 159993125 | 159993450 | 159993492 | 159990166 | 159990312 |
| MXE | 15076 | ENSG00000143549 | TPM3 | chr1 | - | 154170648 | 154170711 | 154172028 | 154172104 | 154170399 | 154170469 |
| MXE | 13460 | ENSG00000196776 | CD47 | chr3 | - | 108049618 | 108049651 | 108051938 | 108051970 | 108043090 | 108047292 |
| MXE | 12974 | ENSG00000122591 | FAM126A | chr7 | - | 22961234 | 22961322 | 22964414 | 22964531 | 22960255 | 22960415 |
| MXE | 16554 | ENSG00000061987 | MON2 | chr12 | + | 62571391 | 62571582 | 62578444 | 62578505 | 62566321 | 62566450 |
| MXE | 1951 | ENSG00000090905 | TNRC6A | chr16 | + | 24805673 | 24805733 | 24806205 | 24806283 | 24805013 | 24805151 |
| MXE | 11820 | ENSG00000132424 | PNISR | chr6 | - | 99409172 | 99409344 | 99410740 | 99410964 | 99408080 | 99408271 |
| MXE | 8478 | ENSG00000135749 | PCNX2 | chr1 | - | 233200153 | 233200264 | 233208517 | 233208689 | 233198938 | 233199030 |
| MXE | 16864 | ENSG00000135677 | GNS | chr12 | - | 64739380 | 64739499 | 64740605 | 64740688 | 64737003 | 64737107 |
| MXE | 9524 | ENSG00000018699 | TTC27 | chr2 | + | 32811023 | 32811221 | 32812503 | 32812615 | 32786983 | 32787149 |
| MXE | 3691 | ENSG00000081791 | DELE1 | chr5 | + | 141929581 | 141929740 | 141933258 | 141933401 | 141928150 | 141928298 |
| MXE | 16504 | ENSG00000145901 | TNIP1 | chr5 | - | 151056765 | 151056957 | 151060317 | 151060395 | 151052164 | 151052259 |
| MXE | 1279 | ENSG00000141030 | COPS3 | chr17 | - | 17254858 | 17254945 | 17261965 | 17262106 | 17248925 | 17249039 |
| MXE | 10506 | ENSG00000122025 | FLT3 | chr13 | - | 28035934 | 28036043 | 28037184 | 28037288 | 28035494 | 28035673 |
| MXE | 6440 | ENSG00000163946 | TASOR | chr3 | - | 56666259 | 56666384 | 56668396 | 56668558 | 56663540 | 56663572 |
| MXE | 1242 | ENSG00000161526 | SAP30BP | chr17 | + | 75693439 | 75693482 | 75699782 | 75699871 | 75671815 | 75671863 |
| MXE | 15860 | ENSG00000088298 | EDEM2 | chr20 | - | 35137879 | 35138005 | 35142372 | 35142478 | 35134737 | 35134949 |
| MXE | 9324 | ENSG00000182400 | TRAPPC6B | chr14 | - | 39154210 | 39154294 | 39158284 | 39158402 | 39151745 | 39151839 |
| MXE | 14922 | ENSG00000119383 | PTPA | chr9 | + | 129134794 | 129134894 | 129136470 | 129136595 | 129131521 | 129131639 |
| MXE | 13150 | ENSG00000111581 | NUP107 | chr12 | + | 68691967 | 68692112 | 68696850 | 68696922 | 68690630 | 68690746 |
| MXE | 5688 | ENSG00000163681 | SLMAP | chr3 | + | 57871635 | 57871698 | 57890040 | 57890100 | 57864806 | 57864857 |
| MXE | 1239 | ENSG00000108395 | TRIM37 | chr17 | - | 59062566 | 59062648 | 59064354 | 59064405 | 59061031 | 59061108 |
| MXE | 859 | ENSG00000125844 | RRBP1 | chr20 | - | 17627347 | 17627382 | 17627503 | 17627682 | 17625511 | 17625602 |
| MXE | 13630 | ENSG00000152332 | UHMK1 | chr1 | + | 162499954 | 162500247 | 162503753 | 162503848 | 162497804 | 162498268 |
| MXE | 1237 | ENSG00000108395 | TRIM37 | chr17 | - | 59047682 | 59047819 | 59049177 | 59049393 | 59041812 | 59041898 |
| MXE | 4774 | ENSG00000143669 | LYST | chr1 | - | 235751209 | 235751362 | 235753043 | 235753274 | 235746335 | 235746527 |
| MXE | 9072 | ENSG00000077254 | USP33 | chr1 | - | 77736055 | 77736158 | 77739264 | 77739417 | 77734346 | 77734416 |
| MXE | 1764 | ENSG00000214078 | CPNE1 | chr20 | - | 35630438 | 35630490 | 35630900 | 35631034 | 35627279 | 35627413 |
| MXE | 9520 | ENSG00000096080 | MRPS18A | chr6 | - | 43675201 | 43675617 | 43678517 | 43678625 | 43671783 | 43671906 |
| MXE | 16779 | ENSG00000090686 | USP48 | chr1 | - | 21747066 | 21747149 | 21748137 | 21748271 | 21736445 | 21736625 |
| MXE | 13352 | ENSG00000140332 | TLE3 | chr15 | - | 70064453 | 70064470 | 70065996 | 70066218 | 70060529 | 70060649 |
| MXE | 7970 | ENSG00000048707 | VPS13D | chr1 | + | 12455997 | 12456130 | 12460200 | 12460396 | 12416659 | 12416827 |
| MXE | 15897 | ENSG00000173915 | ATP5MK | chr10 | - | 103392190 | 103392283 | 103392370 | 103392466 | 103389049 | 103389166 |
| MXE | 4716 | ENSG00000155903 | RASA2 | chr3 | + | 141540532 | 141540609 | 141553856 | 141553940 | 141529707 | 141529802 |
| MXE | 3103 | ENSG00000272325 | NUDT3 | chr6 | - | 34293450 | 34293535 | 34295640 | 34295685 | 34279678 | 34288931 |
| MXE | 16316 | ENSG00000148672 | GLUD1 | chr10 | - | 87060687 | 87060825 | 87060914 | 87061052 | 87060160 | 87060241 |
| MXE | 5114 | ENSG00000114867 | EIF4G1 | chr3 | + | 184316131 | 184316218 | 184317320 | 184317431 | 184315762 | 184315856 |
| MXE | 10366 | ENSG00000152520 | PAN3 | chr13 | + | 28281314 | 28281379 | 28287983 | 28288122 | 28280411 | 28280541 |
| MXE | 13718 | ENSG00000271147 | ARMCX5-GPRASP2 | chrX | + | 102601463 | 102601509 | 102601879 | 102601977 | 102599525 | 102599847 |
| MXE | 5753 | ENSG00000160883 | HK3 | chr5 | - | 176890821 | 176890941 | 176891036 | 176891191 | 176890634 | 176890730 |
| MXE | 10101 | ENSG00000285756 | NA | chrX | - | 3825087 | 3825231 | 3826600 | 3826729 | 3822373 | 3822580 |
| MXE | 8778 | ENSG00000134313 | KIDINS220 | chr2 | - | 8734654 | 8734753 | 8736867 | 8736999 | 8733443 | 8733680 |
| MXE | 16369 | ENSG00000149100 | EIF3M | chr11 | + | 32594913 | 32595013 | 32595965 | 32596047 | 32593768 | 32593949 |
| MXE | 4731 | ENSG00000183624 | HMCES | chr3 | + | 129290678 | 129290804 | 129298353 | 129298535 | 129288853 | 129288997 |
| MXE | 10917 | ENSG00000115652 | UXS1 | chr2 | - | 106096717 | 106096821 | 106098715 | 106098773 | 106093307 | 106094157 |
| MXE | 14846 | ENSG00000163527 | STT3B | chr3 | + | 31624913 | 31625085 | 31625953 | 31626127 | 31623673 | 31623861 |
| MXE | 2196 | ENSG00000170776 | AKAP13 | chr15 | + | 85708018 | 85708086 | 85710578 | 85710645 | 85693098 | 85693451 |
| MXE | 2830 | ENSG00000119723 | COQ6 | chr14 | + | 73959414 | 73959522 | 73961736 | 73961903 | 73959161 | 73959224 |
| MXE | 12127 | ENSG00000136273 | HUS1 | chr7 | - | 47965299 | 47965438 | 47967805 | 47967925 | 47964319 | 47964632 |
| MXE | 10221 | ENSG00000071073 | MGAT4A | chr2 | - | 98658217 | 98658264 | 98663045 | 98663179 | 98656351 | 98656465 |
| MXE | 10203 | ENSG00000123240 | OPTN | chr10 | + | 13123994 | 13124110 | 13125417 | 13125567 | 13122384 | 13122487 |
| MXE | 11358 | ENSG00000136699 | SMPD4 | chr2 | - | 130158183 | 130158265 | 130161185 | 130161272 | 130157250 | 130157396 |
| MXE | 825 | ENSG00000196756 | SNHG17 | chr20 | - | 38422091 | 38422241 | 38422290 | 38422368 | 38421005 | 38421098 |
| MXE | 2220 | ENSG00000092439 | TRPM7 | chr15 | - | 50614122 | 50614263 | 50619744 | 50619798 | 50613706 | 50613841 |
| MXE | 15913 | ENSG00000119541 | VPS4B | chr18 | - | 63397033 | 63397253 | 63400047 | 63400196 | 63393408 | 63393549 |
| MXE | 14483 | ENSG00000150776 | NKAPD1 | chr11 | + | 112078214 | 112078315 | 112080408 | 112080558 | 112075566 | 112075643 |
| MXE | 10492 | ENSG00000004455 | AK2 | chr1 | - | 33021592 | 33021703 | 33024441 | 33024567 | 33021366 | 33021461 |
| MXE | 5572 | ENSG00000058262 | SEC61A1 | chr3 | + | 128064876 | 128065037 | 128066953 | 128067151 | 128060507 | 128060661 |
| MXE | 5787 | ENSG00000137074 | APTX | chr9 | - | 32987543 | 32987731 | 32988082 | 32988124 | 32985970 | 32986030 |
| MXE | 15207 | ENSG00000110218 | PANX1 | chr11 | + | 94178368 | 94178592 | 94179601 | 94180257 | 94153490 | 94153630 |
| MXE | 6742 | ENSG00000285215 | NA | chr1 | + | 120972785 | 120972910 | 120973107 | 120973246 | 120969118 | 120969354 |
| MXE | 7699 | ENSG00000142687 | KIAA0319L | chr1 | - | 35444160 | 35444303 | 35451627 | 35451776 | 35442905 | 35443028 |
| MXE | 14882 | ENSG00000213930 | GALT | chr9 | + | 34647491 | 34647705 | 34647831 | 34647961 | 34647088 | 34647258 |
| MXE | 1511 | ENSG00000091592 | NLRP1 | chr17 | - | 5517745 | 5517887 | 5521523 | 5521786 | 5515472 | 5515517 |
| MXE | 16843 | ENSG00000111596 | CNOT2 | chr12 | + | 70330286 | 70330469 | 70332766 | 70332846 | 70329422 | 70329570 |
| MXE | 13380 | ENSG00000054983 | GALC | chr14 | - | 87982204 | 87982243 | 87984393 | 87984533 | 87976357 | 87976488 |
| MXE | 4418 | ENSG00000109756 | RAPGEF2 | chr4 | + | 159193199 | 159193256 | 159210499 | 159210583 | 159186641 | 159186712 |
| MXE | 11389 | ENSG00000198721 | ECI2 | chr6 | - | 4119185 | 4119275 | 4126134 | 4126237 | 4117307 | 4117451 |
| MXE | 15046 | ENSG00000110422 | HIPK3 | chr11 | + | 33348521 | 33348818 | 33349146 | 33349287 | 33347851 | 33348013 |
| MXE | 4149 | ENSG00000152818 | UTRN | chr6 | + | 144482208 | 144482388 | 144485384 | 144485519 | 144479811 | 144479982 |
| MXE | 13555 | ENSG00000101966 | XIAP | chrX | + | 123891237 | 123891316 | 123892730 | 123892773 | 123888618 | 123888718 |
| MXE | 8852 | ENSG00000160746 | ANO10 | chr3 | - | 43565652 | 43565727 | 43574808 | 43574864 | 43561219 | 43561402 |
| MXE | 6957 | ENSG00000160058 | BSDC1 | chr1 | - | 32383829 | 32383997 | 32386778 | 32386895 | 32381213 | 32381268 |
| MXE | 15751 | ENSG00000149311 | ATM | chr11 | + | 108307896 | 108307984 | 108310159 | 108310315 | 108304674 | 108304852 |
| MXE | 10973 | ENSG00000102125 | TAZ | chrX | + | 154414100 | 154414190 | 154419545 | 154419623 | 154413481 | 154413567 |
| MXE | 7128 | ENSG00000141027 | NCOR1 | chr17 | - | 16064067 | 16064187 | 16064869 | 16065019 | 16062104 | 16062270 |
| MXE | 6944 | ENSG00000160058 | BSDC1 | chr1 | - | 32378214 | 32378283 | 32378723 | 32378839 | 32377969 | 32378048 |
| MXE | 8190 | ENSG00000188641 | DPYD | chr1 | - | 97373560 | 97373644 | 97382392 | 97382461 | 97306176 | 97306297 |
| MXE | 3409 | ENSG00000043143 | JADE2 | chr5 | + | 134566115 | 134566580 | 134573644 | 134573762 | 134564493 | 134564610 |
| MXE | 2524 | ENSG00000100749 | VRK1 | chr14 | + | 96837761 | 96837817 | 96847256 | 96847344 | 96833466 | 96833631 |
| MXE | 9587 | ENSG00000163904 | SENP2 | chr3 | + | 185617479 | 185617611 | 185623997 | 185624082 | 185614563 | 185614740 |
| MXE | 16857 | ENSG00000177575 | CD163 | chr12 | - | 7501138 | 7501462 | 7502477 | 7502564 | 7498867 | 7499188 |
| MXE | 15534 | ENSG00000110075 | PPP6R3 | chr11 | + | 68574108 | 68574224 | 68583042 | 68583129 | 68571039 | 68571104 |
| MXE | 2207 | ENSG00000092470 | WDR76 | chr15 | + | 43851086 | 43851245 | 43856945 | 43857163 | 43843900 | 43844054 |
| MXE | 2355 | ENSG00000137764 | MAP2K5 | chr15 | + | 67727915 | 67727945 | 67748230 | 67748257 | 67703336 | 67703408 |
| MXE | 8293 | ENSG00000257379 | NA | chr12 | + | 53464759 | 53464852 | 53465931 | 53465973 | 53462492 | 53462567 |
| MXE | 14727 | ENSG00000136811 | ODF2 | chr9 | + | 128469182 | 128469353 | 128471307 | 128471468 | 128460941 | 128461010 |
| MXE | 9842 | ENSG00000166478 | ZNF143 | chr11 | + | 9516200 | 9516362 | 9525239 | 9525386 | 9512447 | 9512596 |
| MXE | 15453 | ENSG00000103544 | VPS35L | chr16 | + | 19642395 | 19642476 | 19644885 | 19644949 | 19640014 | 19640100 |
| MXE | 1426 | ENSG00000108960 | MMD | chr17 | - | 55403766 | 55403868 | 55411256 | 55411417 | 55401468 | 55401538 |
| MXE | 5426 | ENSG00000078070 | MCCC1 | chr3 | - | 183092408 | 183092545 | 183094558 | 183094605 | 183090935 | 183091031 |
| MXE | 17031 | ENSG00000136169 | SETDB2 | chr13 | + | 49485629 | 49485723 | 49488289 | 49488630 | 49483463 | 49483563 |
| MXE | 13782 | ENSG00000101974 | ATP11C | chrX | - | 139774689 | 139774953 | 139782546 | 139782728 | 139768259 | 139768434 |
| MXE | 10279 | ENSG00000135966 | TGFBRAP1 | chr2 | - | 105272854 | 105273014 | 105273543 | 105273690 | 105269271 | 105269705 |
| MXE | 4520 | ENSG00000177054 | ZDHHC13 | chr11 | + | 19152157 | 19152320 | 19152558 | 19152684 | 19150726 | 19150791 |
| MXE | 7483 | ENSG00000120733 | KDM3B | chr5 | + | 138381515 | 138381590 | 138386021 | 138386190 | 138379583 | 138379708 |
| MXE | 5632 | ENSG00000121578 | B4GALT4 | chr3 | - | 119218649 | 119218772 | 119224057 | 119224245 | 119211739 | 119216344 |
| MXE | 136 | ENSG00000100418 | DESI1 | chr22 | - | 41604043 | 41604153 | 41607839 | 41607861 | 41603258 | 41603381 |
| MXE | 5732 | ENSG00000114126 | TFDP2 | chr3 | - | 141963811 | 141963963 | 141970072 | 141970141 | 141959673 | 141959840 |
| MXE | 4978 | ENSG00000043093 | DCUN1D1 | chr3 | - | 182963880 | 182964049 | 182965536 | 182965753 | 182961225 | 182961356 |
| MXE | 3129 | ENSG00000129534 | MIS18BP1 | chr14 | - | 45231140 | 45231298 | 45232732 | 45232820 | 45227662 | 45227814 |
| MXE | 7767 | ENSG00000112029 | FBXO5 | chr6 | - | 152972271 | 152972454 | 152973045 | 152973136 | 152970528 | 152971414 |
| MXE | 14102 | ENSG00000249859 | PVT1 | chr8 | + | 127939507 | 127939676 | 127983903 | 127984204 | 127890588 | 127890998 |
| MXE | 4768 | ENSG00000066777 | ARFGEF1 | chr8 | - | 67232854 | 67232945 | 67238342 | 67238493 | 67228223 | 67228264 |
| MXE | 9878 | ENSG00000141429 | GALNT1 | chr18 | + | 35677590 | 35677757 | 35687015 | 35687186 | 35663627 | 35663802 |
| MXE | 14199 | ENSG00000160948 | VPS28 | chr8 | - | 144424717 | 144424819 | 144424945 | 144425051 | 144424214 | 144424268 |
| MXE | 16783 | ENSG00000110906 | KCTD10 | chr12 | - | 109457991 | 109458078 | 109460635 | 109460805 | 109457629 | 109457682 |
| MXE | 6457 | ENSG00000143952 | VPS54 | chr2 | - | 63944599 | 63944655 | 63947382 | 63947490 | 63942464 | 63942561 |
| MXE | 4614 | ENSG00000109133 | TMEM33 | chr4 | + | 41939195 | 41939383 | 41939720 | 41939828 | 41938601 | 41938696 |
| MXE | 8599 | ENSG00000151838 | CCDC175 | chr14 | - | 59525281 | 59525434 | 59531771 | 59531910 | 59521573 | 59521676 |
| MXE | 13419 | ENSG00000147010 | SH3KBP1 | chrX | - | 19631863 | 19631958 | 19645399 | 19645475 | 19607937 | 19608045 |
| MXE | 8098 | ENSG00000178537 | SLC25A20 | chr3 | - | 48879357 | 48879448 | 48883996 | 48884124 | 48862541 | 48862659 |
| MXE | 14113 | ENSG00000249859 | PVT1 | chr8 | + | 128009589 | 128009718 | 128082752 | 128082848 | 127989161 | 127989291 |
| MXE | 6961 | ENSG00000117593 | DARS2 | chr1 | + | 173831534 | 173831630 | 173833375 | 173833499 | 173830659 | 173830761 |
| MXE | 12367 | ENSG00000164808 | SPIDR | chr8 | + | 47727199 | 47727293 | 47728932 | 47729047 | 47713488 | 47713641 |
| MXE | 3889 | ENSG00000119522 | DENND1A | chr9 | - | 123676719 | 123676789 | 123757702 | 123757822 | 123671290 | 123671371 |
| MXE | 7664 | ENSG00000164306 | PRIMPOL | chr4 | + | 184685408 | 184685498 | 184685575 | 184685684 | 184682247 | 184682336 |
| MXE | 13224 | ENSG00000188917 | TRMT2B | chrX | - | 101036973 | 101037073 | 101041316 | 101041371 | 101035612 | 101035683 |
| MXE | 12206 | ENSG00000274272 | NA | chr7 | - | 100577272 | 100577389 | 100577496 | 100577558 | 100577085 | 100577154 |
| MXE | 2886 | ENSG00000112584 | FAM120B | chr6 | + | 170388286 | 170388493 | 170391012 | 170391121 | 170358225 | 170358318 |
| MXE | 1081 | ENSG00000198862 | LTN1 | chr21 | - | 28932469 | 28932664 | 28935108 | 28935329 | 28931154 | 28931322 |
| MXE | 10910 | ENSG00000258984 | UBE2F-SCLY | chr2 | + | 238083247 | 238083354 | 238091217 | 238091254 | 238082044 | 238082209 |
| MXE | 2561 | ENSG00000197249 | SERPINA1 | chr14 | - | 94382591 | 94383241 | 94388559 | 94388663 | 94380870 | 94381141 |
| MXE | 13387 | ENSG00000147394 | ZNF185 | chrX | + | 152922172 | 152922256 | 152928574 | 152928661 | 152920706 | 152920748 |
| MXE | 8857 | ENSG00000070882 | OSBPL3 | chr7 | - | 24815058 | 24815203 | 24816609 | 24816688 | 24809806 | 24809951 |
| MXE | 7253 | ENSG00000154358 | OBSCN | chr1 | + | 228245412 | 228245688 | 228246527 | 228246803 | 228244295 | 228244571 |
| MXE | 13819 | ENSG00000147471 | PLPBP | chr8 | + | 37775338 | 37775481 | 37775917 | 37776016 | 37772754 | 37772889 |
| MXE | 15906 | ENSG00000197771 | MCMBP | chr10 | - | 119835539 | 119835704 | 119836895 | 119837029 | 119832011 | 119832100 |
| MXE | 13077 | ENSG00000161203 | AP2M1 | chr3 | + | 184181084 | 184181226 | 184181695 | 184181815 | 184180848 | 184180984 |
| MXE | 5894 | ENSG00000151466 | SCLT1 | chr4 | - | 129039040 | 129039096 | 129043394 | 129043467 | 129003740 | 129003876 |
| MXE | 12054 | ENSG00000214021 | TTLL3 | chr3 | + | 9816073 | 9816202 | 9818821 | 9818920 | 9813247 | 9813345 |
| MXE | 15061 | ENSG00000157837 | SPPL3 | chr12 | - | 120784473 | 120784593 | 120791468 | 120791557 | 120783673 | 120783752 |
| MXE | 7291 | ENSG00000010810 | FYN | chr6 | - | 111700103 | 111700268 | 111702884 | 111703034 | 111696276 | 111696456 |
| MXE | 11558 | ENSG00000114388 | NPRL2 | chr3 | - | 50349385 | 50349494 | 50349664 | 50349833 | 50348873 | 50349010 |
| MXE | 12572 | ENSG00000121716 | PILRB | chr7 | + | 100355142 | 100355240 | 100356749 | 100356883 | 100353209 | 100353270 |
| MXE | 5281 | ENSG00000288520 | NA | chr6 | - | 154221256 | 154221328 | 154223169 | 154223243 | 154214217 | 154214276 |
| MXE | 14706 | ENSG00000144524 | COPS7B | chr2 | + | 231788554 | 231788732 | 231794262 | 231794351 | 231786433 | 231786538 |
| MXE | 14473 | ENSG00000090060 | PAPOLA | chr14 | + | 96527429 | 96527539 | 96527952 | 96528006 | 96525309 | 96525391 |
| MXE | 3886 | ENSG00000119522 | DENND1A | chr9 | - | 123667025 | 123667079 | 123671290 | 123671371 | 123652012 | 123652123 |
| MXE | 3536 | ENSG00000132842 | AP3B1 | chr5 | - | 78175625 | 78175697 | 78175783 | 78175838 | 78165609 | 78165672 |
| MXE | 15828 | ENSG00000124795 | DEK | chr6 | - | 18258303 | 18258405 | 18263842 | 18263996 | 18257952 | 18258062 |
| MXE | 8671 | ENSG00000117360 | PRPF3 | chr1 | + | 150346017 | 150346136 | 150346407 | 150346491 | 150344375 | 150344547 |
| MXE | 13559 | ENSG00000109111 | SUPT6H | chr17 | + | 28678544 | 28678634 | 28678820 | 28678963 | 28678075 | 28678192 |
| MXE | 878 | ENSG00000060069 | CTDP1 | chr18 | + | 79728906 | 79729069 | 79736354 | 79736521 | 79714490 | 79715528 |
| MXE | 5902 | ENSG00000067066 | SP100 | chr2 | + | 230449087 | 230449150 | 230461261 | 230461414 | 230446818 | 230446902 |
| MXE | 6365 | ENSG00000163374 | YY1AP1 | chr1 | - | 155670319 | 155670464 | 155672559 | 155672731 | 155668626 | 155668777 |
| MXE | 2232 | ENSG00000167004 | PDIA3 | chr15 | + | 43766727 | 43766910 | 43768488 | 43768597 | 43765886 | 43766012 |
| MXE | 11225 | ENSG00000169045 | HNRNPH1 | chr5 | - | 179618144 | 179618323 | 179620891 | 179621035 | 179617988 | 179618060 |
| MXE | 3061 | ENSG00000135596 | MICAL1 | chr6 | - | 109448731 | 109448879 | 109449399 | 109449481 | 109448202 | 109448393 |
| MXE | 9809 | ENSG00000151092 | NGLY1 | chr3 | - | 25736003 | 25736149 | 25739576 | 25739799 | 25733871 | 25733982 |
| MXE | 1693 | ENSG00000005844 | ITGAL | chr16 | + | 30513770 | 30513846 | 30516972 | 30517086 | 30511049 | 30511136 |
| MXE | 11223 | ENSG00000125503 | PPP1R12C | chr19 | - | 55092441 | 55092544 | 55092621 | 55092662 | 55092221 | 55092326 |
| MXE | 3375 | ENSG00000131844 | MCCC2 | chr5 | + | 71636103 | 71636167 | 71641002 | 71641075 | 71635150 | 71635246 |
| MXE | 12366 | ENSG00000164808 | SPIDR | chr8 | + | 47595810 | 47596006 | 47598945 | 47599196 | 47440322 | 47440542 |
| MXE | 3460 | ENSG00000215421 | ZNF407 | chr18 | + | 74781427 | 74781502 | 74877196 | 74877363 | 74641007 | 74641122 |
| MXE | 10125 | ENSG00000140386 | SCAPER | chr15 | - | 76857808 | 76857879 | 76862415 | 76862533 | 76841733 | 76841931 |
| MXE | 5951 | ENSG00000134352 | IL6ST | chr5 | - | 55951463 | 55951604 | 55951928 | 55952075 | 55947492 | 55947589 |
| MXE | 7447 | ENSG00000188739 | RBM34 | chr1 | - | 235152705 | 235152765 | 235154880 | 235155112 | 235148403 | 235148447 |
| MXE | 16592 | ENSG00000122965 | RBM19 | chr12 | - | 113948832 | 113949036 | 113950082 | 113950154 | 113947333 | 113947464 |
| MXE | 8734 | ENSG00000143119 | CD53 | chr1 | + | 110894326 | 110894401 | 110896652 | 110896733 | 110892344 | 110892533 |
| MXE | 4392 | ENSG00000121210 | TMEM131L | chr4 | + | 153558257 | 153558368 | 153580825 | 153580903 | 153556965 | 153557082 |
| MXE | 7592 | ENSG00000162976 | SLC66A3 | chr2 | + | 11164203 | 11164261 | 11171924 | 11172045 | 11160624 | 11160694 |
| MXE | 98 | ENSG00000099991 | CABIN1 | chr22 | + | 24041138 | 24041273 | 24042903 | 24043084 | 24038347 | 24038461 |
| MXE | 2359 | ENSG00000136381 | IREB2 | chr15 | + | 78466270 | 78466489 | 78470531 | 78470601 | 78465250 | 78465388 |
| MXE | 7060 | ENSG00000223745 | CCDC18-AS1 | chr1 | - | 93311489 | 93311608 | 93318167 | 93318228 | 93305387 | 93305470 |
| MXE | 3407 | ENSG00000043143 | JADE2 | chr5 | + | 134562199 | 134562367 | 134564493 | 134564610 | 134560745 | 134560957 |
| MXE | 10578 | ENSG00000101104 | PABPC1L | chr20 | + | 44916755 | 44916871 | 44918905 | 44919045 | 44912659 | 44912853 |
| MXE | 574 | ENSG00000011422 | PLAUR | chr19 | - | 43655438 | 43655573 | 43656478 | 43656640 | 43652224 | 43652371 |
| MXE | 12884 | ENSG00000257923 | CUX1 | chr7 | + | 102196633 | 102197305 | 102200070 | 102200172 | 102195506 | 102195603 |
| MXE | 777 | ENSG00000124209 | RAB22A | chr20 | + | 58343717 | 58343799 | 58353272 | 58353344 | 58311042 | 58311122 |
| MXE | 7860 | ENSG00000143224 | PPOX | chr1 | + | 161168992 | 161169183 | 161170408 | 161170519 | 161168412 | 161168576 |
| MXE | 1346 | ENSG00000062716 | VMP1 | chr17 | + | 59738836 | 59738947 | 59764970 | 59765138 | 59737452 | 59737543 |
| MXE | 1843 | ENSG00000158805 | ZNF276 | chr16 | + | 89723136 | 89723709 | 89727278 | 89727357 | 89722530 | 89722834 |
| MXE | 11797 | ENSG00000204305 | AGER | chr6 | - | 32182246 | 32182388 | 32182538 | 32182698 | 32181605 | 32181632 |
| MXE | 2013 | ENSG00000159322 | ADPGK | chr15 | - | 72755555 | 72755654 | 72760406 | 72760527 | 72751368 | 72752892 |
| MXE | 1495 | ENSG00000091542 | ALKBH5 | chr17 | + | 18194954 | 18195035 | 18206814 | 18206970 | 18183827 | 18185013 |
| MXE | 13667 | ENSG00000147133 | TAF1 | chrX | + | 71392874 | 71392994 | 71393300 | 71393476 | 71392568 | 71392718 |
| MXE | 4666 | ENSG00000025293 | PHF20 | chr20 | + | 35847349 | 35847434 | 35863012 | 35863400 | 35842572 | 35842744 |
| MXE | 13990 | ENSG00000156162 | DPY19L4 | chr8 | + | 94761699 | 94761834 | 94765182 | 94765314 | 94756035 | 94756159 |
| MXE | 1509 | ENSG00000091592 | NLRP1 | chr17 | - | 5517745 | 5517887 | 5521523 | 5521786 | 5501399 | 5501872 |
| MXE | 440 | ENSG00000099860 | GADD45B | chr19 | + | 2476528 | 2476630 | 2477028 | 2477251 | 2476126 | 2476402 |
| MXE | 15044 | ENSG00000110422 | HIPK3 | chr11 | + | 33347851 | 33348013 | 33348521 | 33348818 | 33347628 | 33347753 |
| MXE | 6915 | ENSG00000288208 | NA | chr1 | - | 45355302 | 45355449 | 45385911 | 45385960 | 45334390 | 45334511 |
| MXE | 9341 | ENSG00000069275 | NUCKS1 | chr1 | - | 205723925 | 205723981 | 205727699 | 205727805 | 205720500 | 205720653 |
| MXE | 359 | ENSG00000142002 | DPP9 | chr19 | - | 4697550 | 4697651 | 4700215 | 4700277 | 4695377 | 4695555 |
| MXE | 9522 | ENSG00000018699 | TTC27 | chr2 | + | 32633875 | 32634005 | 32640269 | 32640410 | 32630522 | 32630700 |
| MXE | 16860 | ENSG00000135677 | GNS | chr12 | - | 64723005 | 64723113 | 64728955 | 64729057 | 64721594 | 64721705 |
| MXE | 3183 | ENSG00000133706 | LARS1 | chr5 | - | 146142855 | 146143084 | 146143411 | 146143550 | 146140203 | 146140261 |
| MXE | 7491 | ENSG00000120029 | ARMH3 | chr10 | - | 102009976 | 102010023 | 102011722 | 102011783 | 102009373 | 102009449 |
| MXE | 5481 | ENSG00000173889 | PHC3 | chr3 | - | 170136418 | 170136665 | 170145422 | 170145521 | 170128683 | 170129552 |
| MXE | 6144 | ENSG00000163719 | MTMR14 | chr3 | + | 9662266 | 9662375 | 9669431 | 9669492 | 9653620 | 9653769 |
| MXE | 15098 | ENSG00000135362 | PRR5L | chr11 | + | 36437384 | 36437476 | 36446299 | 36446440 | 36431852 | 36431910 |
| MXE | 12653 | ENSG00000168092 | PAFAH1B2 | chr11 | + | 117152440 | 117152528 | 117159933 | 117160023 | 117144283 | 117144418 |
| MXE | 15978 | ENSG00000078369 | GNB1 | chr1 | - | 1790394 | 1790596 | 1793244 | 1793311 | 1789052 | 1789269 |
| MXE | 12418 | ENSG00000198198 | SZT2 | chr1 | + | 43426714 | 43426809 | 43427055 | 43427179 | 43426037 | 43426151 |
| MXE | 16081 | ENSG00000197535 | MYO5A | chr15 | - | 52353870 | 52354014 | 52359967 | 52360081 | 52353604 | 52353658 |
| MXE | 960 | ENSG00000152229 | PSTPIP2 | chr18 | - | 45992105 | 45992202 | 45993604 | 45993703 | 45991901 | 45991983 |
| MXE | 6181 | ENSG00000085433 | WDR47 | chr1 | - | 108981732 | 108981864 | 108982608 | 108982779 | 108974535 | 108974754 |
| MXE | 13413 | ENSG00000139083 | ETV6 | chr12 | + | 11839139 | 11839304 | 11853426 | 11853561 | 11752449 | 11752579 |
| MXE | 1222 | ENSG00000136436 | CALCOCO2 | chr17 | + | 48852925 | 48853012 | 48856091 | 48856187 | 48852339 | 48852628 |
| MXE | 17150 | ENSG00000134900 | TPP2 | chr13 | + | 102643221 | 102643376 | 102644556 | 102644673 | 102640269 | 102640376 |
| MXE | 13105 | ENSG00000196998 | WDR45 | chrX | - | 49076429 | 49076524 | 49077642 | 49077747 | 49075865 | 49075945 |
| MXE | 8709 | ENSG00000124571 | XPO5 | chr6 | - | 43572505 | 43572578 | 43573479 | 43573601 | 43570856 | 43570994 |
| MXE | 1 | ENSG00000155313 | USP25 | chr21 | + | 15805120 | 15805258 | 15811136 | 15811210 | 15799756 | 15799843 |
| MXE | 12444 | ENSG00000164535 | DAGLB | chr7 | - | 6424751 | 6424835 | 6425987 | 6426114 | 6421726 | 6421804 |
| MXE | 6320 | ENSG00000117640 | MTFR1L | chr1 | + | 25823237 | 25823323 | 25823345 | 25823439 | 25823018 | 25823128 |
| MXE | 10622 | ENSG00000163872 | YEATS2 | chr3 | + | 183756527 | 183756689 | 183758861 | 183758965 | 183754125 | 183754365 |
| MXE | 13792 | ENSG00000198373 | WWP2 | chr16 | + | 69840125 | 69840263 | 69871803 | 69871931 | 69799173 | 69799295 |
| MXE | 16087 | ENSG00000213047 | DENND1B | chr1 | - | 197672036 | 197672156 | 197674119 | 197674169 | 197658299 | 197658369 |
| MXE | 14222 | ENSG00000168522 | FNTA | chr8 | + | 43072180 | 43072307 | 43077215 | 43077364 | 43069554 | 43069659 |
| MXE | 481 | ENSG00000012061 | ERCC1 | chr19 | - | 45413962 | 45414034 | 45414860 | 45414960 | 45413676 | 45413745 |
| MXE | 9564 | ENSG00000143393 | PI4KB | chr1 | - | 151306135 | 151306363 | 151307573 | 151307801 | 151303540 | 151303650 |
| MXE | 8862 | ENSG00000082701 | GSK3B | chr3 | - | 119876412 | 119876508 | 120002045 | 120002239 | 119863418 | 119863605 |
| MXE | 11754 | ENSG00000096070 | BRPF3 | chr6 | + | 36217916 | 36218010 | 36222167 | 36222265 | 36213879 | 36214386 |

| | | | | | |
|---|---|---|---|---|---|
| MXE | 9431 | ENSG00000138398 | PPIG | chr2 | + |
| MXE | 567 | ENSG00000105379 | ETFB | chr19 | - |
| MXE | 12600 | ENSG00000136280 | CCM2 | chr7 | + |
| MXE | 2634 | ENSG00000100554 | ATP6V1D | chr14 | - |
| MXE | 6164 | ENSG00000285708 | NA | chr3 | - |
| MXE | 5689 | ENSG00000163681 | SLMAP | chr3 | + |
| MXE | 13418 | ENSG00000104870 | FCGRT | chr19 | + |
| MXE | 6887 | ENSG00000100991 | TRPC4AP | chr20 | - |
| MXE | 2104 | ENSG00000128739 | SNRPN | chr15 | + |
| MXE | 10734 | ENSG00000187446 | CHP1 | chr15 | + |
| MXE | 15058 | ENSG00000157837 | SPPL3 | chr12 | - |
| MXE | 769 | ENSG00000101294 | HM13 | chr20 | + |
| MXE | 15519 | ENSG00000110075 | PPP6R3 | chr11 | + |
| MXE | 10576 | ENSG00000115365 | LANCL1 | chr2 | - |
| MXE | 14518 | ENSG00000130958 | SLC35D2 | chr9 | - |
| MXE | 6881 | ENSG00000100991 | TRPC4AP | chr20 | - |
| MXE | 15732 | ENSG00000162408 | NOL9 | chr1 | - |
| MXE | 15654 | ENSG00000081237 | PTPRC | chr1 | + |
| MXE | 3707 | ENSG00000158987 | RAPGEF6 | chr5 | - |
| MXE | 9198 | ENSG00000116127 | ALMS1 | chr2 | + |
| MXE | 9242 | ENSG00000163466 | ARPC2 | chr2 | + |
| MXE | 12831 | ENSG00000086730 | LAT2 | chr7 | + |
| MXE | 13366 | ENSG00000147905 | ZCCHC7 | chr9 | + |
| MXE | 14476 | ENSG00000180776 | ZDHHC20 | chr13 | - |
| MXE | 3464 | ENSG00000112851 | ERBIN | chr5 | + |
| MXE | 5697 | ENSG00000130313 | PGLS | chr19 | + |
| MXE | 2448 | ENSG00000226777 | FAM30A | chr14 | + |
| MXE | 14940 | ENSG00000130714 | POMT1 | chr9 | + |
| MXE | 10796 | ENSG00000065675 | PRKCQ | chr10 | - |
| MXE | 1829 | ENSG00000140718 | FTO | chr16 | + |
| MXE | 10494 | ENSG00000036257 | CUL3 | chr2 | - |
| MXE | 1265 | ENSG00000178971 | CTC1 | chr17 | - |
| MXE | 17056 | ENSG00000125304 | TM9SF2 | chr13 | + |
| MXE | 14228 | ENSG00000104164 | BLOC1S6 | chr15 | + |
| MXE | 12745 | ENSG00000160856 | FCRL3 | chr1 | - |
| MXE | 688 | ENSG00000101347 | SAMHD1 | chr20 | - |
| MXE | 11041 | ENSG00000079263 | SP140 | chr2 | + |
| MXE | 12280 | ENSG00000242588 | NA | chr7 | + |
| MXE | 10882 | ENSG00000085365 | SCAMP1 | chr5 | + |
| MXE | 9186 | ENSG00000213064 | SFT2D2 | chr1 | + |
| MXE | 738 | ENSG00000101421 | CHMP4B | chr20 | + |
| MXE | 2730 | ENSG00000100461 | RBM23 | chr14 | - |
| MXE | 1832 | ENSG00000140718 | FTO | chr16 | + |
| MXE | 6322 | ENSG00000119185 | ITGB1BP1 | chr2 | - |
| MXE | 9328 | ENSG00000168924 | LETM1 | chr4 | - |
| MXE | 6941 | ENSG00000162434 | JAK1 | chr1 | - |
| MXE | 6274 | ENSG00000145287 | PLAC8 | chr4 | - |
| MXE | 5744 | ENSG00000196187 | TMEM63A | chr1 | - |
| MXE | 5494 | ENSG00000100462 | PRMT5 | chr14 | - |
| MXE | 7120 | ENSG00000055609 | KMT2C | chr7 | - |
| MXE | 4883 | ENSG00000168297 | PXK | chr3 | + |
| MXE | 3044 | ENSG00000255819 | KLRC4-KLRK1 | chr12 | - |
| MXE | 3928 | ENSG00000059378 | PARP12 | chr7 | - |
| MXE | 6769 | ENSG00000138069 | RAB1A | chr2 | - |
| MXE | 6238 | ENSG00000064687 | ABCA7 | chr19 | + |
| MXE | 13081 | ENSG00000124222 | STX16 | chr20 | + |
| MXE | 16025 | ENSG00000203791 | EEF1AKMT2 | chr10 | - |
| MXE | 13850 | ENSG00000152601 | MBNL1 | chr3 | + |
| MXE | 9843 | ENSG00000115966 | ATF2 | chr2 | - |
| MXE | 5799 | ENSG00000134684 | YARS1 | chr1 | - |
| MXE | 5982 | ENSG00000130695 | CEP85 | chr1 | + |
| MXE | 10098 | ENSG00000285756 | NA | chrX | - |
| MXE | 4782 | ENSG00000143669 | LYST | chr1 | - |
| MXE | 14991 | ENSG00000172340 | SUCLG2 | chr3 | - |
| MXE | 2256 | ENSG00000140577 | CRTC3 | chr15 | + |
| MXE | 6175 | ENSG00000285708 | NA | chr3 | - |
| MXE | 2811 | ENSG00000033170 | FUT8 | chr14 | + |
| MXE | 6780 | ENSG00000054282 | SDCCAG8 | chr1 | + |
| MXE | 10501 | ENSG00000186625 | KATNA1 | chr6 | - |
| MXE | 10404 | ENSG00000124151 | NCOA3 | chr20 | + |
| MXE | 4174 | ENSG00000118564 | FBXL5 | chr4 | - |
| MXE | 15194 | ENSG00000153107 | ANAPC1 | chr2 | - |
| MXE | 14997 | ENSG00000256591 | NA | chr11 | + |
| MXE | 2409 | ENSG00000172575 | RASGRP1 | chr15 | - |
| MXE | 6041 | ENSG00000181555 | SETD2 | chr3 | - |
| MXE | 14905 | ENSG00000115459 | ELMOD3 | chr2 | + |
| MXE | 15358 | ENSG00000025796 | SEC63 | chr6 | - |
| MXE | 6343 | ENSG00000078142 | PIK3C3 | chr18 | + |
| MXE | 15091 | ENSG00000096401 | CDC5L | chr6 | + |
| MXE | 10219 | ENSG00000059804 | SLC2A3 | chr12 | - |
| MXE | 13928 | ENSG00000064115 | TM7SF3 | chr12 | - |
| MXE | 1899 | ENSG00000122299 | ZC3H7A | chr16 | - |
| MXE | 7914 | ENSG00000123636 | BAZ2B | chr2 | - |
| MXE | 1389 | ENSG00000154229 | PRKCA | chr17 | + |
| MXE | 7622 | ENSG00000132300 | PTCD3 | chr2 | + |
| MXE | 15140 | ENSG00000100731 | PCNX1 | chr14 | + |
| MXE | 5961 | ENSG00000165240 | ATP7A | chrX | + |
| MXE | 13841 | ENSG00000168172 | HOOK3 | chr8 | + |
| MXE | 14636 | ENSG00000165525 | NEMF | chr14 | - |
| MXE | 14023 | ENSG00000120992 | LYPLA1 | chr8 | - |
| MXE | 6485 | ENSG00000140575 | IQGAP1 | chr15 | + |
| MXE | 14184 | ENSG00000173064 | HECTD4 | chr12 | - |
| MXE | 6392 | ENSG00000134318 | ROCK2 | chr2 | - |
| MXE | 12434 | ENSG00000010318 | PHF7 | chr3 | + |
| MXE | 5262 | ENSG00000105993 | DNAJB6 | chr7 | + |
| MXE | 1270 | ENSG00000133030 | MPRIP | chr17 | + |
| MXE | 5773 | ENSG00000198087 | CD2AP | chr6 | + |
| MXE | 11442 | ENSG00000151687 | ANKAR | chr2 | + |
| MXE | 1092 | ENSG00000129197 | RPAIN | chr17 | + |
| MXE | 7277 | ENSG00000233184 | NA | chr1 | + |
| MXE | 15451 | ENSG00000103544 | VPS35L | chr16 | + |
| MXE | 5630 | ENSG00000127481 | UBR4 | chr1 | - |
| MXE | 9631 | ENSG00000112031 | MTRF1L | chr6 | - |
| MXE | 11044 | ENSG00000079263 | SP140 | chr2 | + |
| MXE | 2704 | ENSG00000119684 | MLH3 | chr14 | - |
| MXE | 15678 | ENSG00000072518 | MARK2 | chr11 | + |
| MXE | 467 | ENSG00000188033 | ZNF490 | chr19 | - |
| MXE | 8677 | ENSG00000269713 | NBPF9 | chr1 | - |
| MXE | 10481 | ENSG00000181381 | DDX60L | chr4 | - |
| MXE | 6648 | ENSG00000108443 | RPS6KB1 | chr17 | + |
| MXE | 7932 | ENSG00000158006 | PAFAH2 | chr1 | - |
| MXE | 16158 | ENSG00000122958 | VPS26A | chr10 | + |
| MXE | 11426 | ENSG00000147443 | DOK2 | chr8 | - |
| MXE | 9702 | ENSG00000134243 | SORT1 | chr1 | - |
| MXE | 16899 | ENSG00000188906 | LRRK2 | chr12 | + |
| MXE | 6946 | ENSG00000160058 | BSDC1 | chr1 | - |
| MXE | 2446 | ENSG00000197555 | SIPA1L1 | chr14 | + |
| MXE | 10583 | ENSG00000198589 | LRBA | chr4 | - |
| MXE | 14351 | ENSG00000143748 | NVL | chr1 | - |
| MXE | 1374 | ENSG00000108510 | MED13 | chr17 | - |
| MXE | 11993 | ENSG00000139154 | AEBP2 | chr12 | + |
| MXE | 10976 | ENSG00000040933 | INPP4A | chr2 | + |
| MXE | 5815 | ENSG00000162777 | DENND2D | chr1 | - |
| MXE | 15055 | ENSG00000197563 | PIGN | chr18 | - |
| MXE | 12903 | ENSG00000105939 | ZC3HAV1 | chr7 | - |
| MXE | 179 | ENSG00000100239 | PPP6R2 | chr22 | + |
| MXE | 12335 | ENSG00000117868 | ESYT2 | chr7 | - |
| MXE | 5275 | ENSG00000284952 | NA | chr3 | - |
| MXE | 4737 | ENSG00000111321 | LTBR | chr12 | + |
| MXE | 7460 | ENSG00000066136 | NFYC | chr1 | + |
| MXE | 1466 | ENSG00000177302 | TOP3A | chr17 | - |
| MXE | 7290 | ENSG00000010810 | FYN | chr6 | - |
| MXE | 8093 | ENSG00000138688 | KIAA1109 | chr4 | + |
| MXE | 6552 | ENSG00000139372 | TDG | chr12 | + |
| MXE | 119 | ENSG00000242259 | C22orf39 | chr22 | - |
| MXE | 3271 | ENSG00000124784 | RIOK1 | chr6 | + |
| MXE | 10361 | ENSG00000152520 | PAN3 | chr13 | + |
| MXE | 7637 | ENSG00000143167 | GPA33 | chr1 | - |
| MXE | 9942 | ENSG00000285542 | NA | chr2 | + |
| MXE | 15745 | ENSG00000149311 | ATM | chr11 | + |
| MXE | 8641 | ENSG00000270231 | NBPF8 | chr1 | + |
| MXE | 1169 | ENSG00000125450 | NUP85 | chr17 | + |
| MXE | 2388 | ENSG00000137814 | HAUS2 | chr15 | + |
| MXE | 2675 | ENSG00000100722 | ZC3H14 | chr14 | + |
| MXE | 7938 | ENSG00000197312 | DDI2 | chr1 | + |
| MXE | 410 | ENSG00000129351 | ILF3 | chr19 | + |
| MXE | 9210 | ENSG00000148660 | CAMK2G | chr10 | - |
| MXE | 16070 | ENSG00000107679 | PLEKHA1 | chr10 | + |
| MXE | 14764 | ENSG00000021776 | AQR | chr15 | - |
| MXE | 8691 | ENSG00000268043 | NBPF12 | chr1 | + |
| MXE | 9071 | ENSG00000077254 | USP33 | chr1 | - |
| MXE | 14912 | ENSG00000148356 | LRSAM1 | chr9 | + |
| MXE | 2442 | ENSG00000197555 | SIPA1L1 | chr14 | + |
| MXE | 13945 | ENSG00000147649 | MTDH | chr8 | + |
| MXE | 14959 | ENSG00000132589 | FLOT2 | chr17 | - |
| MXE | 6797 | ENSG00000162889 | MAPKAPK2 | chr1 | + |
| MXE | 14265 | ENSG00000089234 | BRAP | chr12 | - |
| MXE | 15988 | ENSG00000052749 | RRP12 | chr10 | - |
| MXE | 3609 | ENSG00000057935 | MTA3 | chr2 | + |
| MXE | 10935 | ENSG00000073849 | ST6GAL1 | chr3 | + |
| MXE | 63 | ENSG00000159131 | GART | chr21 | - |
| MXE | 15980 | ENSG00000078369 | GNB1 | chr1 | - |
| MXE | 4921 | ENSG00000114127 | XRN1 | chr3 | - |
| MXE | 17177 | ENSG00000005810 | MYCBP2 | chr13 | - |
| MXE | 8207 | ENSG00000180353 | HCLS1 | chr3 | - |
| MXE | 834 | ENSG00000101310 | SEC23B | chr20 | + |
| MXE | 9361 | ENSG00000115211 | EIF2B4 | chr2 | - |
| MXE | 14937 | ENSG00000187764 | SEMA4D | chr9 | - |
| MXE | 11589 | ENSG00000163939 | PBRM1 | chr3 | - |
| MXE | 1695 | ENSG00000170100 | ZNF778 | chr16 | + |
| MXE | 7721 | ENSG00000082512 | TRAF5 | chr1 | + |
| MXE | 17182 | ENSG00000005810 | MYCBP2 | chr13 | - |
| MXE | 15627 | ENSG00000078124 | ACER3 | chr11 | + |
| MXE | 5407 | ENSG00000143612 | C1orf43 | chr1 | - |
| MXE | 9546 | ENSG00000163382 | NAXE | chr1 | + |
| MXE | 12667 | ENSG00000089289 | IGBP1 | chrX | + |
| MXE | 9705 | ENSG00000138646 | HERC5 | chr4 | + |
| MXE | 7387 | ENSG00000126091 | ST3GAL3 | chr1 | + |
| MXE | 1231 | ENSG00000161533 | ACOX1 | chr17 | - |
| MXE | 1319 | ENSG00000108604 | SMARCD2 | chr17 | - |
| MXE | 6416 | ENSG00000070759 | TESK2 | chr1 | - |
| MXE | 23 | ENSG00000185917 | SETD4 | chr21 | - |
| MXE | 1210 | ENSG00000007202 | KIAA0100 | chr17 | - |
| MXE | 6602 | ENSG00000163659 | TIPARP | chr3 | + |
| MXE | 7034 | ENSG00000095564 | BTAF1 | chr10 | + |
| MXE | 9636 | ENSG00000185811 | IKZF1 | chr7 | + |
| MXE | 8732 | ENSG00000143119 | CD53 | chr1 | + |
| MXE | 2765 | ENSG00000259316 | NA | chr15 | - |
| MXE | 9916 | ENSG00000115464 | USP34 | chr2 | - |
| MXE | 1164 | ENSG00000141699 | RETREG3 | chr17 | - |
| MXE | 5563 | ENSG00000173193 | PARP14 | chr3 | + |
| MXE | 11016 | ENSG00000131781 | FMO5 | chr1 | - |
| MXE | 15254 | ENSG00000122218 | COPA | chr1 | - |
| MXE | 10349 | ENSG00000031003 | FAM13B | chr5 | - |
| MXE | 12332 | ENSG00000117868 | ESYT2 | chr7 | - |
| MXE | 3603 | ENSG00000146282 | RARS2 | chr6 | - |
| MXE | 15442 | ENSG00000110274 | CEP164 | chr11 | + |
| MXE | 7614 | ENSG00000163322 | ABRAXAS1 | chr4 | - |
| MXE | 9964 | ENSG00000101336 | HCK | chr20 | + |
| MXE | 12964 | ENSG00000127914 | AKAP9 | chr7 | + |
| MXE | 6445 | ENSG00000173083 | HPSE | chr4 | - |
| MXE | 5992 | ENSG00000116514 | RNF19B | chr1 | - |
| MXE | 14793 | ENSG00000070061 | ELP1 | chr9 | - |
| MXE | 7907 | ENSG00000071054 | MAP4K4 | chr2 | + |
| MXE | 7442 | ENSG00000165119 | HNRNPK | chr9 | - |
| MXE | 3165 | ENSG00000075568 | TMEM131 | chr2 | - |
| MXE | 6143 | ENSG00000163719 | MTMR14 | chr3 | + |
| MXE | 10683 | ENSG00000169499 | PLEKHA2 | chr8 | + |
| MXE | 14021 | ENSG00000198851 | CD3E | chr11 | + |
| MXE | 15878 | ENSG00000165322 | ARHGAP12 | chr10 | - |
| MXE | 16163 | ENSG00000107669 | ATE1 | chr10 | - |
| MXE | 17074 | ENSG00000125457 | MIF4GD | chr17 | - |
| MXE | 15822 | ENSG00000124795 | DEK | chr6 | - |
| MXE | 2248 | ENSG00000169375 | SIN3A | chr15 | - |
| MXE | 189 | ENSG00000100150 | DEPDC5 | chr22 | + |
| MXE | 16699 | ENSG00000139350 | NEDD1 | chr12 | + |
| MXE | 2951 | ENSG00000111913 | RIPOR2 | chr6 | - |
| MXE | 6693 | ENSG00000102606 | ARHGEF7 | chr13 | + |
| MXE | 2655 | ENSG00000100938 | GMPR2 | chr14 | + |
| MXE | 16341 | ENSG00000135655 | USP15 | chr12 | + |
| MXE | 9634 | ENSG00000185811 | IKZF1 | chr7 | + |
| MXE | 6071 | ENSG00000143457 | GOLPH3L | chr1 | - |
| MXE | 13127 | ENSG00000130741 | EIF2S3 | chrX | + |
| MXE | 16212 | ENSG00000150093 | ITGB1 | chr10 | - |
| MXE | 14834 | ENSG00000107077 | KDM4C | chr9 | + |
| MXE | 12165 | ENSG00000161405 | IKZF3 | chr17 | - |
| MXE | 6764 | ENSG00000135999 | EPC2 | chr2 | + |
| MXE | 14226 | ENSG00000168522 | FNTA | chr8 | + |
| MXE | 13181 | ENSG00000137642 | SORL1 | chr11 | + |
| MXE | 2360 | ENSG00000136381 | IREB2 | chr15 | + |
| MXE | 9376 | ENSG00000155275 | TRMT44 | chr4 | + |
| MXE | 10623 | ENSG00000163872 | YEATS2 | chr3 | + |
| MXE | 16010 | ENSG00000108100 | CCNY | chr10 | + |

| | | | | | | | | | | | | | | | | | | | | | | | | | |
|---|---|---|---|---|---|---|---|---|---|---|---|---|---|---|---|---|---|---|---|---|---|---|---|---|---|
| MXE | 8624 | ENSG00000075151 | EIF4G3 | chr1 | - | 20981047 | 20981227 | 21001198 | 21001324 | 20980333 | 20980448 | 21002712 | 21002808 | 0 | [illegible] | [illegible] | [illegible] | [illegible] | [illegible] | [illegible] | [illegible] | [illegible] | [illegible] | [illegible] | [illegible] |
| MXE | 13940 | ENSG00000075856 | SART3 | chr12 | - | 108538933 | 108539089 | 108543027 | 108543152 | 108538064 | 108538203 | 108544426 | 108544478 | 128 | [illegible] | [illegible] | [illegible] | [illegible] | [illegible] | [illegible] | [illegible] | [illegible] | [illegible] | [illegible] | [illegible] |
| MXE | 16685 | ENSG00000112200 | ZNF451 | chr6 | + | 57133041 | 57133192 | 57134743 | 57134870 | 57128728 | 57128840 | 57141301 | 57141368 | 132 | [illegible] | [illegible] | [illegible] | [illegible] | [illegible] | [illegible] | [illegible] | [illegible] | [illegible] | [illegible] | [illegible] |
| MXE | 14753 | ENSG00000048828 | FAM120A | chr9 | + | 93558580 | 93558718 | 93561108 | 93561250 | 93557826 | 93558010 | 93562207 | 93562304 | 68 | 27 | 39 | 64 | 22 | 172 | 83 | 109 | 170 | 56 | 63 | 52 |
| MXE | 7820 | ENSG00000143621 | ILF2 | chr1 | - | 153664042 | 153664130 | 153664395 | 153664474 | 153663214 | 153663276 | 153665219 | 153665336 | 193 | 175 | 234 | 218 | 572 | 156 | 75 | 155 | 143 | 256 | 389 | 238 |
| MXE | 4108 | ENSG00000104517 | UBR5 | chr8 | - | 102287432 | 102287564 | 102289452 | 102289576 | 102286359 | 102286617 | 102293604 | 102293844 | 212 | 226 | 218 | 257 | 286 | 471 | 405 | 408 | 455 | 652 | 93 | 264 |
| MXE | 974 | ENSG00000101639 | CEP192 | chr18 | + | 13038369 | 13038579 | 13040829 | 13040956 | 13037236 | 13037301 | 13042203 | 13042334 | 74 | 24 | 49 | 62 | 87 | 161 | 130 | 128 | 80 | 277 | 73 | 38 |
| MXE | 1329 | ENSG00000173821 | RNF213 | chr17 | + | 80361733 | 80361888 | 80363101 | 80363314 | 80360060 | 80360206 | 80363608 | 80363790 | 301 | 170 | 206 | 274 | 285 | 382 | 166 | 267 | 359 | 144 | 499 | 288 |
| MXE | 7643 | ENSG00000100591 | AHSA1 | chr14 | + | 77462159 | 77462242 | 77464597 | 77464686 | 77459615 | 77459806 | 77465538 | 77465667 | 57 | 34 | 36 | 65 | 92 | 151 | 112 | 139 | 211 | 208 | 92 | 59 |
| MXE | 12256 | ENSG00000160218 | TRAPPC10 | chr21 | + | 44063537 | 44063785 | 44074323 | 44074470 | 44059102 | 44059214 | 44075038 | 44075153 | 83 | 36 | 70 | 59 | 66 | 142 | 75 | 90 | 107 | 101 | 28 | 82 |
| MXE | 11579 | ENSG00000163939 | PBRM1 | chr3 | - | 52550746 | 52550817 | 52554723 | 52554879 | 52550420 | 52550637 | 52561766 | 52561968 | 48 | 32 | 49 | 47 | 72 | 30 | 13 | 22 | 35 | 52 | 0 | 17 |
| MXE | 11006 | ENSG00000119912 | IDE | chr10 | - | 92483254 | 92483337 | 92487195 | 92487318 | 92479276 | 92479421 | 92490492 | 92490595 | 102 | 92 | 116 | 117 | 61 | 9 | 7 | 7 | 8 | 15 | 8 | 75 |
| MXE | 7744 | ENSG00000156515 | HK1 | chr10 | + | 69386322 | 69386418 | 69389196 | 69389296 | 69384795 | 69384915 | 69392124 | 69392308 | 369 | 293 | 285 | 411 | 286 | 107 | 59 | 70 | 100 | 84 | 1144 | 556 |
| MXE | 8463 | ENSG00000137275 | RIPK1 | chr6 | + | 3083084 | 3083313 | 3085258 | 3085408 | 3080978 | 3081116 | 3104224 | 3104315 | 144 | 78 | 83 | 99 | 103 | 11 | 8 | 5 | 7 | 3 | 47 | 90 |
| MXE | 3974 | ENSG00000112685 | EXOC2 | chr6 | - | 619429 | 619543 | 629834 | 629961 | 617710 | 617835 | 632940 | 633117 | 64 | 43 | 63 | 68 | 90 | 93 | 57 | 108 | 109 | 89 | 21 | 30 |
| MXE | 16617 | ENSG00000120805 | ARL1 | chr12 | - | 101402864 | 101402946 | 101405843 | 101405981 | 101401061 | 101401173 | 101407641 | 101407746 | 101 | 46 | 79 | 97 | 49 | 105 | 96 | 94 | 79 | 162 | 4 | 40 |
| MXE | 1470 | ENSG00000108306 | FBXL20 | chr17 | - | 39282728 | 39282855 | 39285477 | 39285573 | 39281388 | 39281463 | 39297126 | 39297195 | 33 | 21 | 23 | 26 | 44 | 47 | 33 | 49 | 47 | 56 | 40 | 51 |
| MXE | 85 | ENSG00000100226 | GTPBP1 | chr22 | + | 38716651 | 38717000 | 38721741 | 38721865 | 38715906 | 38716087 | 38724296 | 38724411 | 92 | 85 | 50 | 101 | 82 | 72 | 31 | 55 | 64 | 93 | 140 | 104 |
| MXE | 2773 | ENSG00000140105 | WARS1 | chr14 | - | 100369086 | 100369258 | 100374135 | 100374265 | 100361729 | 100361921 | 100375510 | 100375607 | 0 | 0 | 16 | 0 | 7 | 268 | 276 | 329 | 277 | 153 | 0 | 0 |
| MXE | 1094 | ENSG00000129197 | RPAIN | chr17 | + | 5426235 | 5426299 | 5428070 | 5428211 | 5422768 | 5422829 | 5432541 | 5432876 | 11 | 3 | 5 | 8 | 3 | 57 | 32 | 44 | 69 | 99 | 8 | 2 |
| MXE | 8329 | ENSG00000177409 | SAMD9L | chr7 | - | 93145384 | 93145532 | 93146882 | 93147146 | 93144731 | 93144833 | 93148193 | 93148345 | 133 | 85 | 97 | 177 | 109 | 154 | 215 | 133 | 220 | 166 | 38 | 50 |
| MXE | 3350 | ENSG00000127603 | MACF1 | chr1 | + | 39327217 | 39327353 | 39331202 | 39336653 | 39324645 | 39324734 | 39337181 | 39337331 | 295 | 247 | 248 | 296 | 394 | 238 | 143 | 204 | 225 | 193 | 89 | 425 |
| MXE | 8315 | ENSG00000109667 | SLC2A9 | chr4 | - | 9941912 | 9942045 | 9980591 | 9980737 | 9920384 | 9920572 | 9985668 | 9985793 | 9 | 6 | 8 | 9 | 2 | 8 | 1 | 5 | 2 | 7 | 11 | 10 |
| MXE | 5356 | ENSG00000051382 | PIK3CB | chr3 | - | 138688874 | 138688974 | 138690999 | 138691143 | 138684624 | 138684803 | 138694785 | 138694907 | 108 | 99 | 94 | 147 | 80 | 95 | 86 | 86 | 124 | 117 | 67 | 144 |
| MXE | 2169 | ENSG00000138592 | USP8 | chr15 | + | 50495847 | 50496084 | 50497088 | 50497231 | 50493184 | 50494280 | 50498595 | 50498728 | 112 | 95 | 94 | 96 | 162 | 64 | 54 | 66 | 80 | 123 | 15 | 44 |
| MXE | 16097 | ENSG00000099290 | WASHC2A | chr10 | + | 50092161 | 50092233 | 50093267 | 50093386 | 50091430 | 50091518 | 50093859 | 50093917 | 57 | 50 | 44 | 46 | 81 | 25 | 29 | 14 | 31 | 26 | 82 | 41 |
| MXE | 7053 | ENSG00000287725 | NA | chr11 | + | 69078891 | 69079020 | 69079833 | 69079883 | 69078733 | 69078793 | 69081399 | 69081499 | 10 | 6 | 7 | 5 | 6 | 1 | 1 | 4 | 8 | 2 | 8 | 9 |
| MXE | 6118 | ENSG00000197746 | PSAP | chr10 | - | 71818616 | 71818724 | 71819030 | 71819111 | 71816297 | 71817476 | 71819464 | 71819622 | 1974 | 1520 | 1241 | 1722 | 1207 | 4657 | 3071 | 2563 | 3410 | 2822 | 5929 | 3041 |
| MXE | 8374 | ENSG00000152223 | EPG5 | chr18 | - | 45867562 | 45867748 | 45870566 | 45870742 | 45866797 | 45867007 | 45876235 | 45876342 | 49 | 22 | 48 | 55 | 54 | 117 | 54 | 69 | 83 | 106 | 14 | 45 |
| MXE | 8555 | ENSG00000060688 | SNRNP40 | chr1 | - | 31261528 | 31261632 | 31267870 | 31267932 | 31259567 | 31260121 | 31269157 | 31269240 | 57 | 32 | 49 | 40 | 78 | 89 | 45 | 61 | 62 | 118 | 26 | 62 |
| MXE | 2549 | ENSG00000197249 | SERPINA1 | chr14 | - | 94379463 | 94379611 | 94380870 | 94381141 | 94376746 | 94378640 | 94382591 | 94383241 | 2249 | 1512 | 1058 | 1654 | 2598 | 1273 | 743 | 568 | 743 | 1125 | 6418 | 4249 |
| MXE | 13288 | ENSG00000158201 | ABHD3 | chr18 | - | 21656860 | 21657026 | 21657103 | 21657152 | 21650900 | 21651763 | 21659169 | 21659343 | 54 | 36 | 48 | 78 | 58 | 60 | 30 | 45 | 39 | 69 | 18 | 41 |
| MXE | 8130 | ENSG00000138668 | HNRNPD | chr4 | - | 82358658 | 82358820 | 82371527 | 82371584 | 82357312 | 82357444 | 82373445 | 82373991 | 19 | 10 | 14 | 20 | 3 | 691 | 677 | 642 | 802 | 1043 | 5 | 8 |
| MXE | 14448 | ENSG00000216937 | CCDC7 | chr10 | + | 32462682 | 32462703 | 32463016 | 32463049 | 32456250 | 32456334 | 32472480 | 32472542 | 23 | 17 | 6 | 24 | 24 | 1 | 3 | 3 | 6 | 7 | 21 | 12 |
| MXE | 16170 | ENSG00000107669 | ATE1 | chr10 | - | 121902390 | 121902620 | 121910905 | 121911151 | 121898840 | 121898969 | 121913789 | 121913893 | 165 | 149 | 125 | 177 | 447 | 30 | 42 | 65 | 49 | 44 | 125 | 133 |
| MXE | 12825 | ENSG00000104133 | SPG11 | chr15 | - | 44621758 | 44621934 | 44622219 | 44622347 | 44620124 | 44620403 | 44622727 | 44622799 | 188 | 153 | 127 | 168 | 277 | 302 | 278 | 235 | 269 | 443 | 75 | 176 |
| MXE | 14449 | ENSG00000216937 | CCDC7 | chr10 | + | 32462682 | 32462703 | 32471063 | 32471230 | 32456250 | 32456334 | 32472480 | 32472542 | 23 | 17 | 6 | 24 | 24 | 53 | 43 | 24 | 54 | 138 | 21 | 12 |
| MXE | 9199 | ENSG00000116127 | ALMS1 | chr2 | + | 73534823 | 73534949 | 73550266 | 73550437 | 73519774 | 73520016 | 73557219 | 73557354 | 69 | 49 | 63 | 43 | 58 | 74 | 39 | 47 | 43 | 84 | 8 | 27 |
| MXE | 6503 | ENSG00000125637 | PSD4 | chr2 | + | 113192379 | 113192589 | 113193047 | 113193128 | 113185364 | 113186255 | 113193257 | 113193370 | 173 | 128 | 170 | 239 | 314 | 193 | 160 | 180 | 256 | 329 | 260 | 205 |
| MXE | 7639 | ENSG00000131115 | ZNF227 | chr19 | + | 44217790 | 44217852 | 44228445 | 44228572 | 44213086 | 44213244 | 44229732 | 44229816 | 19 | 16 | 14 | 14 | 35 | 16 | 14 | 23 | 20 | 30 | 20 | 6 |
| MXE | 7587 | ENSG00000144455 | SUMF1 | chr3 | - | 4420063 | 4420146 | 4449265 | 4449340 | 4418009 | 4418132 | 4452875 | 4453049 | 58 | 30 | 56 | 46 | 20 | 89 | 71 | 82 | 71 | 30 | 89 | 94 |
| MXE | 10161 | ENSG00000143924 | EML4 | chr2 | + | 42315961 | 42316050 | 42317426 | 42317524 | 42304483 | 42304551 | 42325466 | 42325554 | 80 | 71 | 104 | 103 | 131 | 95 | 62 | 122 | 104 | 92 | 27 | 40 |
| MXE | 12130 | ENSG00000124486 | USP9X | chrX | + | 41186516 | 41186642 | 41187991 | 41188117 | 41184396 | 41184675 | 41189308 | 41189475 | 77 | 61 | 93 | 58 | 105 | 231 | 129 | 142 | 171 | 243 | 4 | 40 |
| MXE | 10213 | ENSG00000009335 | UBE3C | chr7 | + | 157220688 | 157220776 | 157223253 | 157223351 | 157216866 | 157216971 | 157225406 | 157225539 | 81 | 60 | 79 | 115 | 162 | 67 | 51 | 67 | 52 | 99 | 156 | 61 |
| MXE | 15838 | ENSG00000148481 | MINDY3 | chr10 | - | 15782154 | 15782226 | 15786560 | 15786648 | 15778169 | 15779141 | 15789246 | 15789319 | 29 | 13 | 13 | 20 | 18 | 15 | 14 | 20 | 17 | 25 | 12 | 22 |
| MXE | 5435 | ENSG00000164463 | CREBRF | chr5 | + | 173112305 | 173112379 | 173123079 | 173123202 | 173110521 | 173110711 | 173133629 | 173139284 | 226 | 189 | 190 | 241 | 285 | 96 | 67 | 58 | 99 | 155 | 27 | 116 |
| MXE | 16045 | ENSG00000175029 | CTBP2 | chr10 | - | 125038996 | 125039155 | 125110989 | 125111093 | 125003337 | 125003492 | 125160826 | 125161026 | 4 | 0 | 0 | 5 | 1 | 108 | 41 | 54 | 100 | 42 | 0 | 3 |
| MXE | 7731 | ENSG00000048052 | HDAC9 | chr7 | + | 18629349 | 18629481 | 18634626 | 18634742 | 18593907 | 18594029 | 18644670 | 18644793 | 47 | 21 | 18 | 34 | 55 | 97 | 76 | 49 | 90 | 154 | 32 | 22 |
| MXE | 10632 | ENSG00000144895 | EIF2A | chr3 | + | 150562541 | 150562660 | 150564298 | 150564381 | 150558387 | 150558462 | 150567692 | 150567766 | 253 | 196 | 252 | 256 | 524 | 172 | 142 | 217 | 183 | 409 | 47 | 171 |
| MXE | 12105 | ENSG00000136250 | AOAH | chr7 | - | 36632035 | 36632106 | 36637850 | 36637910 | 36623189 | 36623250 | 36659165 | 36659265 | 404 | 307 | 272 | 267 | 259 | 342 | 260 | 252 | 277 | 289 | 871 | 572 |
| MXE | 14522 | ENSG00000130958 | SLC35D2 | chr9 | - | 96336716 | 96336784 | 96360153 | 96360221 | 96324090 | 96324169 | 96364463 | 96364550 | 30 | 21 | 20 | 19 | 24 | 26 | 30 | 34 | 35 | 69 | 14 | 23 |
| MXE | 2514 | ENSG00000092036 | HAUS4 | chr14 | - | 22950313 | 22950410 | 22951554 | 22951689 | 22947600 | 22947731 | 22952327 | 22952459 | 82 | 39 | 80 | 72 | 112 | 14 | 12 | 14 | 3 | 20 | 96 | 101 |
| MXE | 11489 | ENSG00000255587 | RAB44 | chr6 | + | 36725861 | 36725943 | 36727576 | 36727691 | 36721150 | 36722733 | 36728699 | 36728801 | 13 | 1 | 7 | 8 | 0 | 19 | 12 | 5 | 8 | 3 | 8 | 11 |
| MXE | 12880 | ENSG00000257923 | CUX1 | chr7 | + | 102158559 | 102158608 | 102170445 | 102170550 | 102115206 | 102115273 | 102178468 | 102178657 | 51 | 44 | 29 | 37 | 52 | 72 | 62 | 50 | 56 | 32 | 107 | 46 |
| MXE | 3132 | ENSG00000137055 | PLAA | chr9 | - | 26910337 | 26910439 | 26913878 | 26913947 | 26907802 | 26907998 | 26917096 | 26917165 | 114 | 96 | 82 | 82 | 130 | 46 | 45 | 51 | 42 | 73 | 81 | 92 |
| MXE | 1478 | ENSG00000108599 | AKAP10 | chr17 | - | 19924407 | 19924517 | 19936285 | 19936430 | 19920035 | 19920118 | 19939712 | 19939849 | 145 | 104 | 119 | 141 | 140 | 54 | 35 | 44 | 57 | 93 | 115 | 119 |
| MXE | 10835 | ENSG00000108883 | EFTUD2 | chr17 | - | 44868286 | 44868350 | 44872445 | 44872570 | 44867806 | 44867897 | 44875933 | 44876100 | 185 | 85 | 106 | 133 | 140 | 63 | 44 | 66 | 59 | 65 | 166 | 150 |
| MXE | 5750 | ENSG00000138964 | PARVG | chr22 | + | 44181741 | 44181917 | 44183317 | 44183408 | 44180924 | 44181185 | 44185807 | 44185872 | 10 | 1 | 0 | 7 | 3 | 122 | 73 | 106 | 137 | 67 | 6 | 7 |
| MXE | 1780 | ENSG00000157106 | SMG1 | chr16 | - | 18829925 | 18830115 | 18830218 | 18830369 | 18829285 | 18829755 | 18832939 | 18833166 | 86 | 79 | 83 | 95 | 100 | 204 | 174 | 195 | 154 | 286 | 6 | 65 |
| MXE | 83 | ENSG00000184979 | USP18 | chr22 | + | 18161789 | 18161935 | 18167254 | 18167334 | 18160171 | 18160268 | 18167889 | 18168036 | 13 | 17 | 10 | 28 | 8 | 9 | 8 | 8 | 23 | 15 | 96 | 128 |
| MXE | 12747 | ENSG00000171681 | ATF7IP | chr12 | + | 14423908 | 14425473 | 14434336 | 14434423 | 14385063 | 14385400 | 14436105 | 14436251 | 59 | 43 | 46 | 48 | 25 | 1056 | 1018 | 840 | 1104 | 1472 | 1 | 81 |
| MXE | 13311 | ENSG00000139436 | GIT2 | chr12 | - | 109945259 | 109945349 | 109951166 | 109951316 | 109939164 | 109939247 | 109953091 | 109953234 | 117 | 92 | 96 | 113 | 109 | 23 | 19 | 21 | 23 | 24 | 118 | 110 |
| MXE | 11615 | ENSG00000079819 | EPB41L2 | chr6 | - | 130867458 | 130867581 | 130869811 | 130870126 | 130865535 | 130865634 | 130885095 | 130885268 | 17 | 9 | 16 | 20 | 37 | 37 | 23 | 32 | 21 | 52 | 17 | 1 |
| MXE | 11360 | ENSG00000136699 | SMPD4 | chr2 | - | 130167457 | 130167663 | 130174913 | 130175000 | 130164373 | 130164445 | 130176553 | 130176637 | 23 | 10 | 23 | 31 | 19 | 56 | 27 | 58 | 52 | 80 | 26 | 27 |
| MXE | 4914 | ENSG00000075711 | DLG1 | chr3 | - | 197104902 | 197105005 | 197119409 | 197119530 | 197090911 | 197091026 | 197130526 | 197130671 | 118 | 115 | 159 | 170 | 255 | 74 | 67 | 68 | 109 | 125 | 46 | 96 |
| MXE | 16735 | ENSG00000111271 | ACAD10 | chr12 | + | 111749010 | 111749103 | 111749172 | 111749345 | 111748316 | 111748475 | 111755667 | 111755745 | 0 | 0 | 0 | 0 | 20 | 10 | 6 | 24 | 32 | 14 | 0 | 0 |
| MXE | 7923 | ENSG00000043514 | TRIT1 | chr1 | - | 39844530 | 39844640 | 39847219 | 39847297 | 39844100 | 39844218 | 39847547 | 39847660 | 17 | 11 | 21 | 20 | 29 | 34 | 19 | 29 | 36 | 60 | 19 | 17 |
| MXE | 15070 | ENSG00000110514 | MADD | chr11 | + | 47311731 | 47311842 | 47315219 | 47315327 | 47309506 | 47309612 | 47323670 | 47323835 | 172 | 123 | 122 | 239 | 238 | 120 | 35 | 95 | 99 | 100 | 284 | 195 |
| MXE | 5008 | ENSG00000156976 | EIF4A2 | chr3 | + | 186784863 | 186784696 | 186785882 | 186786051 | 186784428 | 186784477 | 186786163 | 186786273 | 1814 | 1081 | 1758 | 1836 | 1602 | 1067 | 770 | 1099 | 1134 | 2154 | 389 | 953 |

| | | | | | | | | | | | | | |
|---|---|---|---|---|---|---|---|---|---|---|---|---|---|
| MXE | 14539 | ENSG00000131089 | ARHGEF9 | chrX | - | 63724531 | 63724711 | 63754307 | 63754676 | 63706257 | 63706449 | 63755018 | 63755113 | 0 |
| MXE | 9674 | ENSG00000144028 | SNRNP200 | chr2 | - | 96279450 | 96279559 | 96281813 | 96281922 | 96278808 | 96278998 | 96283200 | 96283352 | 100 |
| MXE | 1073 | ENSG00000153933 | DGKE | chr17 | + | 56845689 | 56845809 | 56847921 | 56848065 | 56844018 | 56844178 | 56848695 | 56848853 | 41 |
| MXE | 2912 | ENSG00000090615 | GOLGA3 | chr12 | - | 132816539 | 132816812 | 132821995 | 132822311 | 132813306 | 132813419 | 132828802 | 132828840 | 18 |
| MXE | 2590 | ENSG00000249624 | NA | chr21 | + | 33268393 | 33268517 | 33276595 | 33276753 | 33252661 | 33252830 | 33279751 | 33279918 | 27 |
| MXE | 13548 | ENSG00000131381 | RBSN | chr3 | - | 15077061 | 15077164 | 15078074 | 15078161 | 15075605 | 15075710 | 15080731 | 15080802 | 17 |
| MXE | 7589 | ENSG00000037474 | NSUN2 | chr5 | - | 6607199 | 6607384 | 6609825 | 6609922 | 6606819 | 6606912 | 6610954 | 6611085 | 46 |
| MXE | 1524 | ENSG00000083454 | P2RX5 | chr17 | - | 3688625 | 3688759 | 3689491 | 3689627 | 3688011 | 3688105 | 3690069 | 3690150 | 32 |
| MXE | 8246 | ENSG00000143442 | POGZ | chr1 | - | 151408099 | 151408240 | 151408408 | 151408581 | 151407234 | 151407291 | 151408693 | 151408828 | 79 |
| MXE | 14919 | ENSG00000119383 | PTPA | chr9 | + | 129128984 | 129129110 | 129131521 | 129131639 | 129123051 | 129123138 | 129134794 | 129134894 | 47 |
| MXE | 2972 | ENSG00000204623 | ZNRD1ASP | chr6 | - | 30034907 | 30035110 | 30035915 | 30035983 | 30021666 | 30021761 | 30058114 | 30058190 | 13 |
| MXE | 9043 | ENSG00000134759 | ELP2 | chr18 | + | 36138269 | 36138426 | 36138794 | 36138872 | 36136306 | 36136377 | 36139418 | 36139613 | 64 |
| MXE | 13229 | ENSG00000188917 | TRMT2B | chrX | - | 101037916 | 101038051 | 101051250 | 101051733 | 101036973 | 101037073 | 101051816 | 101052111 | 21 |
| MXE | 3600 | ENSG00000172081 | MOB3A | chr19 | - | 2078139 | 2078679 | 2085174 | 2085328 | 2076810 | 2077013 | 2096225 | 2096326 | 65 |
| MXE | 16289 | ENSG00000266412 | NCOA4 | chr10 | - | 46013549 | 46013639 | 46014443 | 46014552 | 46012882 | 46013026 | 46015125 | 46015266 | 16 |
| MXE | 5681 | ENSG00000163681 | SLMAP | chr3 | + | 57864806 | 57864857 | 57890040 | 57890100 | 57864547 | 57864716 | 57896510 | 57896591 | 17 |
| MXE | 11332 | ENSG00000144233 | AMMECR1L | chr2 | - | 127869453 | 127869544 | 127870813 | 127870928 | 127866899 | 127866996 | 127873827 | 127874272 | 4 |
| MXE | 12828 | ENSG00000104133 | SPG11 | chr15 | - | 44633504 | 44633637 | 44648865 | 44649011 | 44629232 | 44629388 | 44651490 | 44651939 | 174 |
| MXE | 4023 | ENSG00000172869 | DMXL1 | chr5 | + | 119101934 | 119102006 | 119110150 | 119110283 | 119097978 | 119098104 | 119114474 | 119114541 | 35 |
| MXE | 7058 | ENSG00000223745 | CCDC18-AS1 | chr1 | - | 93305387 | 93305470 | 93318167 | 93318228 | 93264644 | 93264772 | 93324634 | 93324729 | 0 |
| MXE | 13935 | ENSG00000075856 | SART3 | chr12 | - | 108537487 | 108537595 | 108538064 | 108538203 | 108536707 | 108536785 | 108538933 | 108539089 | 129 |
| MXE | 10689 | ENSG00000132530 | XAF1 | chr17 | + | 6759661 | 6759718 | 6760405 | 6760601 | 6758088 | 6758224 | 6762154 | 6762240 | 33 |
| MXE | 14272 | ENSG00000089234 | BRAP | chr12 | - | 111679150 | 111679340 | 111683145 | 111683307 | 111672660 | 111672774 | 111685710 | 111685956 | 32 |
| MXE | 2440 | ENSG00000119537 | KDSR | chr18 | - | 63355203 | 63355299 | 63355497 | 63355563 | 63350887 | 63351079 | 63359735 | 63359792 | 20 |
| MXE | 12883 | ENSG00000257923 | CUX1 | chr7 | + | 102178474 | 102178657 | 102189812 | 102189871 | 102170445 | 102170550 | 102193841 | 102193890 | 15 |
| MXE | 15022 | ENSG00000115084 | SLC35F5 | chr2 | - | 113750424 | 113750568 | 113755164 | 113755306 | 113746276 | 113746339 | 113755453 | 113755544 | 44 |
| MXE | 14416 | ENSG00000114209 | PDCD10 | chr3 | - | 167704841 | 167704895 | 167720061 | 167720273 | 167697008 | 167697126 | 167734805 | 167734930 | 180 |
| MXE | 6598 | ENSG00000257315 | ZBED6 | chr1 | + | 203850514 | 203850681 | 203851056 | 203851124 | 203849710 | 203850026 | 203852140 | 203854999 | 269 |
| MXE | 16943 | ENSG00000127311 | HELB | chr12 | + | 66323982 | 66324211 | 66324982 | 66325126 | 66322723 | 66322783 | 66331153 | 66331645 | 56 |
| MXE | 12006 | ENSG00000106415 | GLCCI1 | chr7 | + | 8022482 | 8022569 | 8055435 | 8055549 | 8003907 | 8004059 | 8060095 | 8060248 | 28 |
| MXE | 8558 | ENSG00000160679 | CHTOP | chr1 | + | 153638294 | 153638448 | 153642242 | 153642429 | 153636571 | 153636653 | 153643226 | 153643364 | 160 |
| MXE | 5388 | ENSG00000087269 | NOP14 | chr4 | - | 2949933 | 2950213 | 2951174 | 2951245 | 2948277 | 2948408 | 2952274 | 2952397 | 96 |

| | | | | | | | | | | | | | | | |
|---|---|---|---|---|---|---|---|---|---|---|---|---|---|---|---|
| MXE | 16432 | ENSG00000109685 | NSD2 | chr4 | + | 1912090 | 1912194 | 1916870 | 1917037 | 1904215 | 1904378 | 1918140 | 1918623 | 10 | [illegible] |
| MXE | 2060 | ENSG00000182718 | ANXA2 | chr15 | - | 60390212 | 60390317 | 60393012 | 60393440 | 60386027 | 60386086 | 60397257 | 60397338 | 0 | [illegible] |
| MXE | 7628 | ENSG00000185808 | PIGP | chr21 | - | 37067261 | 37067380 | 37069551 | 37069624 | 37065373 | 37065712 | 37072433 | 37072537 | 40 | [illegible] |
| MXE | 14911 | ENSG00000198431 | TXNRD1 | chr12 | + | 104319469 | 104319585 | 104321090 | 104321316 | 104318912 | 104319055 | 104325336 | 104325429 | 126 | [illegible] |
| MXE | 7728 | ENSG00000117481 | NSUN4 | chr1 | + | 46346920 | 46347075 | 46352867 | 46353028 | 46344800 | 46345144 | 46360703 | 46360828 | 14 | [illegible] |
| MXE | 4990 | ENSG00000166128 | RAB8B | chr15 | + | 63244755 | 63244816 | 63249644 | 63249705 | 63223847 | 63223979 | 63255506 | 63255584 | 5 | [illegible] |
| MXE | 15139 | ENSG00000100731 | PCNX1 | chr14 | + | 71036064 | 71036157 | 71045132 | 71045283 | 71033930 | 71034036 | 71046963 | 71047105 | 54 | [illegible] |
| MXE | 7268 | ENSG00000146247 | PHIP | chr6 | - | 78970780 | 78970888 | 78978591 | 78978711 | 78970048 | 78970173 | 78982885 | 78982943 | 53 | [illegible] |
| MXE | 15248 | ENSG00000110367 | DDX6 | chr11 | - | 118758773 | 118758902 | 118759921 | 118760044 | 118757170 | 118757287 | 118763211 | 118763306 | 739 | [illegible] |
| MXE | 14988 | ENSG00000166199 | ALKBH3 | chr11 | + | 43892040 | 43892129 | 43897194 | 43897354 | 43889724 | 43889828 | 43901515 | 43901725 | 148 | [illegible] |
| MXE | 11604 | ENSG00000277147 | LINC00869 | chr1 | + | 149661301 | 149661419 | 149664520 | 149664646 | 149660853 | 149661089 | 149665081 | 149665133 | 13 | [illegible] |
| MXE | 16473 | ENSG00000120253 | NUP43 | chr6 | - | 149738642 | 149738778 | 149742389 | 149742570 | 149736470 | 149736622 | 149743637 | 149743715 | 171 | [illegible] |
| MXE | 3425 | ENSG00000198612 | COPS8 | chr2 | + | 237089861 | 237089994 | 237094089 | 237094197 | 237088604 | 237088653 | 237095821 | 237095884 | 67 | [illegible] |
| MXE | 12342 | ENSG00000139641 | ESYT1 | chr12 | + | 56133780 | 56133873 | 56134341 | 56134428 | 56133587 | 56133674 | 56136743 | 56136893 | 236 | [illegible] |
| MXE | 4106 | ENSG00000104517 | UBR5 | chr8 | - | 102270038 | 102270183 | 102271133 | 102271245 | 102268977 | 102269108 | 102272528 | 102272772 | 135 | [illegible] |
| MXE | 7486 | ENSG00000120733 | KDM3B | chr5 | + | 138420705 | 138420962 | 138424074 | 138424341 | 138418952 | 138419232 | 138425410 | 138425582 | 256 | [illegible] |
| MXE | 7681 | ENSG00000139636 | LMBR1L | chr12 | - | 49101249 | 49101263 | 49101450 | 49101549 | 49100555 | 49100646 | 49102119 | 49102196 | 2 | [illegible] |
| MXE | 14053 | ENSG00000137770 | CTDSPL2 | chr15 | + | 44496379 | 44496458 | 44497026 | 44497138 | 44490783 | 44490999 | 44499726 | 44499813 | 109 | [illegible] |
| MXE | 4357 | ENSG00000163625 | WDFY3 | chr4 | - | 84831412 | 84831605 | 84836928 | 84837090 | 84829003 | 84829190 | 84841153 | 84841263 | 60 | [illegible] |
| MXE | 4019 | ENSG00000138772 | ANXA3 | chr4 | + | 78573179 | 78573267 | 78579026 | 78579121 | 78554435 | 78554488 | 78582176 | 78582290 | 2 | [illegible] |
| MXE | 10547 | ENSG00000047230 | CTPS2 | chrX | - | 16670579 | 16670674 | 16678361 | 16678450 | 16667661 | 16667724 | 16683093 | 16683226 | 34 | [illegible] |
| MXE | 579 | ENSG00000269858 | EGLN2 | chr19 | + | 40807137 | 40807274 | 40807483 | 40807551 | 40801289 | 40801415 | 40807808 | 40807936 | 7 | [illegible] |
| MXE | 9539 | ENSG00000009307 | CSDE1 | chr1 | - | 114725220 | 114725333 | 114726210 | 114726386 | 114723882 | 114724002 | 114726982 | 114727090 | 1846 | [illegible] |
| MXE | 16964 | ENSG00000121749 | TBC1D15 | chr12 | + | 71880468 | 71880607 | 71884810 | 71885021 | 71872928 | 71873003 | 71893221 | 71893324 | 389 | [illegible] |
| MXE | 2861 | ENSG00000197776 | KLHDC1 | chr14 | + | 49725685 | 49725769 | 49728925 | 49729009 | 49723873 | 49723952 | 49729489 | 49729548 | 20 | [illegible] |
| MXE | 66 | ENSG00000142207 | URB1 | chr21 | - | 32373646 | 32373772 | 32378444 | 32378541 | 32372506 | 32372631 | 32383421 | 32383554 | 29 | [illegible] |
| MXE | 16965 | ENSG00000121749 | TBC1D15 | chr12 | + | 71907021 | 71907138 | 71913825 | 71913926 | 71897846 | 71897941 | 71917697 | 71917775 | 128 | [illegible] |
| MXE | 12881 | ENSG00000257923 | CUX1 | chr7 | + | 102170445 | 102170550 | 102178468 | 102178657 | 102158559 | 102158608 | 102189812 | 102189871 | 92 | [illegible] |
| MXE | 149 | ENSG00000100330 | MTMR3 | chr22 | + | 30012367 | 30012563 | 30013355 | 30013541 | 30009017 | 30009129 | 30016527 | 30016698 | 129 | [illegible] |
| MXE | 14529 | ENSG00000135018 | UBQLN1 | chr9 | - | 83665029 | 83665145 | 83686003 | 83686155 | 83663874 | 83664043 | 83707499 | 83707934 | 123 | [illegible] |
| MXE | 13603 | ENSG00000090316 | MAEA | chr4 | + | 1322380 | 1322503 | 1327626 | 1327703 | 1315396 | 1315600 | 1332756 | 1332865 | 59 | [illegible] |
| MXE | 9981 | ENSG00000264538 | SUZ12P1 | chr17 | + | 30768833 | 30768867 | 30769000 | 30769064 | 30766502 | 30766588 | 30769871 | 30770034 | 14 | [illegible] |
| MXE | 4192 | ENSG00000114738 | MAPKAPK3 | chr3 | + | 50646139 | 50646264 | 50646739 | 50646825 | 50645709 | 50645785 | 50647122 | 50647203 | 225 | [illegible] |
| MXE | 7519 | ENSG00000115020 | PIKFYVE | chr2 | + | 208329841 | 208329913 | 208330522 | 208330694 | 208328179 | 208328280 | 208333314 | 208333493 | 13 | [illegible] |
| MXE | 16772 | ENSG00000090686 | USP48 | chr1 | - | 21706466 | 21706589 | 21706743 | 21706868 | 21706125 | 21706187 | 21715388 | 21715457 | 79 | [illegible] |
| MXE | 14173 | ENSG00000168404 | MLKL | chr16 | - | 74678898 | 74678980 | 74682650 | 74682786 | 74675612 | 74675764 | 74685485 | 74685583 | 143 | [illegible] |
| MXE | 1387 | ENSG00000141540 | TTYH2 | chr17 | + | 74243976 | 74244049 | 74252233 | 74252376 | 74243373 | 74243469 | 74253080 | 74253266 | 3 | [illegible] |
| MXE | 195 | ENSG00000100150 | DEPDC5 | chr22 | + | 31804841 | 31804915 | 31806121 | 31806191 | 31804161 | 31804223 | 31809610 | 31809647 | 24 | [illegible] |
| MXE | 16376 | ENSG00000013561 | RNF14 | chr5 | + | 141978302 | 141978830 | 141980122 | 141980351 | 141974803 | 141974955 | 141983379 | 141983552 | 37 | [illegible] |
| MXE | 15074 | ENSG00000080546 | SESN1 | chr6 | - | 108994461 | 108994609 | 108998512 | 108998755 | 108992786 | 108992899 | 109000490 | 109000673 | 78 | [illegible] |
| MXE | 5826 | ENSG00000132549 | VPS13B | chr8 | + | 99819411 | 99819582 | 99819920 | 99820122 | 99818712 | 99818888 | 99821293 | 99821482 | 93 | [illegible] |
| MXE | 3204 | ENSG00000198625 | MDM4 | chr1 | + | 204532190 | 204532246 | 204537429 | 204537497 | 204530683 | 204530817 | 204538208 | 204538308 | 237 | [illegible] |
| MXE | 12197 | ENSG00000168014 | C2CD3 | chr11 | - | 74054606 | 74054671 | 74057405 | 74057544 | 74049336 | 74049542 | 74074252 | 74074505 | 8 | [illegible] |
| MXE | 16416 | ENSG00000122694 | GLIPR2 | chr9 | + | 36147785 | 36147894 | 36148546 | 36148650 | 36136732 | 36136791 | 36150871 | 36150949 | 31 | [illegible] |
| MXE | 8694 | ENSG00000268043 | NBPF12 | chr1 | + | 146970646 | 146970719 | 146976928 | 146977001 | 146969381 | 146969596 | 146977465 | 146977671 | 29 | [illegible] |
| MXE | 12041 | ENSG00000127947 | PTPN12 | chr7 | + | 77592184 | 77592256 | 77597841 | 77597901 | 77585542 | 77585581 | 77600663 | 77600806 | 91 | [illegible] |
| MXE | 11327 | ENSG00000057252 | SOAT1 | chr1 | + | 179344946 | 179345076 | 179347599 | 179347697 | 179343589 | 179343635 | 179348843 | 179348942 | 105 | [illegible] |
| MXE | 17023 | ENSG00000108349 | CASC3 | chr17 | + | 40167370 | 40167612 | 40167849 | 40167948 | 40166796 | 40166861 | 40168202 | 40168255 | 9 | [illegible] |
| MXE | 10996 | ENSG00000119912 | IDE | chr10 | - | 92465675 | 92465843 | 92468878 | 92468990 | 92463730 | 92464003 | 92470253 | 92470345 | 28 | [illegible] |
| MXE | 8896 | ENSG00000172965 | MIR4435-2HG | chr2 | - | 111208755 | 111208977 | 111233627 | 111233739 | 111207775 | 111208661 | 111248147 | 111248241 | 1 | [illegible] |
| MXE | 373 | ENSG00000104880 | ARHGEF18 | chr19 | + | 7444203 | 7444454 | 7447042 | 7447168 | 7441911 | 7442052 | 7451148 | 7451266 | 328 | [illegible] |
| MXE | 8080 | ENSG00000071205 | ARHGAP10 | chr4 | + | 147847150 | 147847222 | 147857552 | 147857654 | 147822895 | 147822957 | 147864845 | 147864956 | 41 | [illegible] |
| MXE | 3696 | ENSG00000120705 | ETF1 | chr5 | - | 138513567 | 138513706 | 138517560 | 138517700 | 138512763 | 138512954 | 138518691 | 138518867 | 248 | [illegible] |
| MXE | 1691 | ENSG00000005844 | ITGAL | chr16 | + | 30504174 | 30504264 | 30505243 | 30505300 | 30499337 | 30499489 | 30505388 | 30505462 | 530 | [illegible] |
| MXE | 15782 | ENSG00000118276 | B4GALT6 | chr18 | - | 31638643 | 31638760 | 31645354 | 31645479 | 31630958 | 31631146 | 31657975 | 31658089 | 39 | [illegible] |
| MXE | 13866 | ENSG00000134014 | ELP3 | chr8 | + | 28160228 | 28160456 | 28161996 | 28162078 | 28158567 | 28158633 | 28189648 | 28189992 | 144 | [illegible] |
| MXE | 2527 | ENSG00000100934 | SEC23A | chr14 | - | 39055142 | 39055296 | 39063323 | 39063413 | 39048651 | 39048729 | 39064912 | 39064993 | 33 | [illegible] |
| MXE | 2825 | ENSG00000053770 | AP5M1 | chr14 | + | 57280194 | 57280422 | 57282088 | 57282228 | 57274243 | 57274889 | 57282933 | 57283019 | 144 | [illegible] |
| MXE | 14070 | ENSG00000153317 | ASAP1 | chr8 | - | 130079901 | 130079971 | 130091972 | 130092143 | 130076347 | 130076406 | 130112093 | 130112322 | 19 | [illegible] |
| MXE | 4668 | ENSG00000025293 | PHF20 | chr20 | + | 35858301 | 35858381 | 35899369 | 35899648 | 35847349 | 35847434 | 35913248 | 35913316 | 64 | [illegible] |
| MXE | 5260 | ENSG00000117133 | RPF1 | chr1 | + | 84495881 | 84496063 | 84496243 | 84496370 | 84495372 | 84495455 | 84497428 | 84498352 | 256 | [illegible] |
| MXE | 2035 | ENSG00000103994 | ZNF106 | chr15 | - | 42456957 | 42457158 | 42466052 | 42466114 | 42449770 | 42451954 | 42472235 | 42472321 | 135 | [illegible] |
| MXE | 8842 | ENSG00000164924 | YWHAZ | chr8 | - | 100923954 | 100924050 | 100924134 | 100924298 | 100916522 | 100920752 | 100924915 | 100925039 | 3596 | [illegible] |
| MXE | 7229 | ENSG00000110619 | CARS1 | chr11 | - | 3026675 | 3026797 | 3028995 | 3029084 | 3020219 | 3020332 | 3029302 | 3029443 | 50 | [illegible] |
| MXE | 15794 | ENSG00000133816 | MICAL2 | chr11 | + | 12236176 | 12236245 | 12239435 | 12239585 | 12227024 | 12227131 | 12256784 | 12256971 | 45 | [illegible] |
| MXE | 3500 | ENSG00000153113 | CAST | chr5 | + | 96742654 | 96742756 | 96746341 | 96746425 | 96741493 | 96741580 | 96747344 | 96747392 | 309 | [illegible] |
| MXE | 4218 | ENSG00000164190 | NIPBL | chr5 | + | 37057185 | 37057332 | 37058890 | 37059165 | 37052365 | 37052566 | 37060843 | 37061018 | 231 | [illegible] |
| MXE | 9890 | ENSG00000219626 | FAM228B | chr2 | + | 24139369 | 24139450 | 24146747 | 24146835 | 24137908 | 24138100 | 24146929 | 24147086 | 55 | [illegible] |
| MXE | 8087 | ENSG00000131828 | PDHA1 | chrX | + | 19353081 | 19353173 | 19354490 | 19354583 | 19351280 | 19351407 | 19355348 | 19355504 | 60 | [illegible] |
| MXE | 11263 | ENSG00000153094 | BCL2L11 | chr2 | + | 111153779 | 111153904 | 111161418 | 111161550 | 111150043 | 111150147 | 111164132 | 111168445 | 12 | [illegible] |
| MXE | 16655 | ENSG00000135297 | MTO1 | chr6 | + | 73482039 | 73482244 | 73482448 | 73482620 | 73480674 | 73480805 | 73497735 | 73497896 | 70 | [illegible] |
| MXE | 16350 | ENSG00000155256 | ZFYVE27 | chr10 | + | 97744728 | 97744915 | 97748268 | 97748364 | 97743093 | 97743164 | 97749473 | 97749586 | 18 | [illegible] |
| MXE | 6438 | ENSG00000163946 | TASOR | chr3 | - | 56663540 | 56663572 | 56666259 | 56666384 | 56662384 | 56662490 | 56668396 | 56668558 | 138 | [illegible] |
| MXE | 7561 | ENSG00000111331 | OAS3 | chr12 | + | 112944475 | 112944651 | 112946742 | 112946981 | 112941569 | 112941852 | 112947945 | 112948099 | 51 | [illegible] |
| MXE | 7603 | ENSG00000053254 | FOXN3 | chr14 | - | 89180700 | 89180806 | 89280949 | 89281014 | 89156171 | 89162969 | 89350671 | 89350808 | 164 | [illegible] |
| MXE | 4331 | ENSG00000145495 | MARCHF6 | chr5 | + | 10417269 | 10417404 | 10429892 | 10430028 | 10415467 | 10415669 | 10433593 | 10433684 | 198 | [illegible] |
| MXE | 14599 | ENSG00000056586 | RC3H2 | chr9 | - | 122883203 | 122883379 | 122890311 | 122890545 | 122880593 | 122880794 | 122892908 | 122893026 | 147 | [illegible] |
| MXE | 3495 | ENSG00000153113 | CAST | chr5 | + | 96695835 | 96695907 | 96722638 | 96722698 | 96675538 | 96675601 | 96727488 | 96727530 | 23 | [illegible] |
| MXE | 3013 | ENSG00000115760 | BIRC6 | chr2 | + | 32464508 | 32464823 | 32465064 | 32465164 | 32463193 | 32463381 | 32467524 | 32467739 | 79 | [illegible] |
| MXE | 4444 | ENSG00000137656 | BUD13 | chr11 | - | 116757765 | 116757950 | 116759073 | 116759179 | 116757145 | 116757227 | 116760734 | 116760952 | 78 | [illegible] |
| MXE | 15573 | ENSG00000110713 | NUP98 | chr11 | - | 3771747 | 3771928 | 3773631 | 3773739 | 3768580 | 3768744 | 3775881 | 3776021 | 210 | [illegible] |
| MXE | 8583 | ENSG00000134242 | PTPN22 | chr1 | - | 113819576 | 113819654 | 113825141 | 113825172 | 113813810 | 113814969 | 113829591 | 113829707 | 67 | [illegible] |
| MXE | 14363 | ENSG00000104365 | IKBKB | chr8 | + | 42272082 | 42272205 | 42288633 | 42288728 | 42271317 | 42271469 | 42290155 | 42290273 | 39 | [illegible] |
| MXE | 10256 | ENSG00000143207 | COP1 | chr1 | - | 176116623 | 176116681 | 176135009 | 176135086 | 176085775 | 176085890 | 176149005 | 176149074 | 23 | [illegible] |
| MXE | 14320 | ENSG00000135679 | MDM2 | chr12 | + | 68828770 | 68828931 | 68835828 | 68835984 | 68824554 | 68824651 | 68836671 | 68836749 | 273 | [illegible] |
| MXE | 13036 | ENSG00000205903 | ZNF316 | chr7 | + | 6637845 | 6638009 | 6639041 | 6639141 | 6637317 | 6637447 | 6641824 | 6641962 | 3 | [illegible] |
| MXE | 11112 | ENSG00000184182 | UBE2F | chr2 | + | 238030555 | 238030613 | 238032221 | 238032254 | 238025341 | 238025412 | 238035877 | 238035940 | 119 | [illegible] |
| MXE | 12097 | ENSG00000106635 | BCL7B | chr7 | - | 73539881 | 73540052 | 73543547 | 73543644 | 73537933 | 73538013 | 73552166 | 73552242 | 86 | [illegible] |
| MXE | 15060 | ENSG00000157837 | SPPL3 | chr12 | - | 120783673 | 120783752 | 120784473 | 120784593 | 120782654 | 120782767 | 120791468 | 120791557 | 58 | [illegible] |
| MXE | 10663 | ENSG00000184613 | NELL2 | chr12 | - | 44703725 | 44703854 | 44711291 | 44711394 | 44665435 | 44665609 | 44714649 | 44714741 | 163 | [illegible] |
| MXE | 4641 | ENSG00000108509 | CAMTA2 | chr17 | - | 4981677 | 4981742 | 4982088 | 4982227 | 4981224 | 4981359 | 4982756 | 4982892 | 12 | [illegible] |
| MXE | 10255 | ENSG00000166439 | RNF169 | chr11 | + | 74817595 | 74817714 | 74834675 | 74834775 | 74810183 | 74810330 | 74835545 | 74842413 | 136 | [illegible] |
| MXE | 1590 | ENSG00000187555 | USP7 | chr16 | - | 8899120 | 8899188 | 8900529 | 8900630 | 8898530 | 8898639 | 8900989 | 8901057 | 213 | [illegible] |
| MXE | 7586 | ENSG00000138756 | BMP2K | chr4 | + | 78844927 | 78845049 | 78847187 | 78847269 | 78842384 | 78842527 | 78850923 | 78851056 | 104 | [illegible] |
| MXE | 15535 | ENSG00000110075 | PPP6R3 | chr11 | + | 68575957 | 68576043 | 68583042 | 68583129 | 68571039 | 68571104 | 68587926 | 68588024 | 15 | [illegible] |
| MXE | 5795 | ENSG00000113108 | APBB3 | chr5 | - | 140560312 | 140560504 | 140561017 | 140561101 | 140558267 | 140558821 | 140561364 | 140561449 | 8 | [illegible] |
| MXE | 11709 | ENSG00000197081 | IGF2R | chr6 | + | 160078200 | 160078362 | 160079579 | 160079787 | 160075846 | 160075996 | 160080128 | 160080275 | 421 | [illegible] |
| MXE | 4110 | ENSG00000104517 | UBR5 | chr8 | - | 102305084 | 102305295 | 102311298 | 102311496 | 102304042 | 102304169 | 102311734 | 102311889 | 150 | [illegible] |
| MXE | 1411 | ENSG00000073584 | SMARCE1 | chr17 | - | 40631591 | 40631693 | 40632194 | 40632367 | 40630713 | 40630924 | 40635930 | 40636102 | 212 | [illegible] |
| MXE | 16485 | ENSG00000161638 | ITGA5 | chr12 | - | 54402179 | 54402330 | 54402982 | 54403050 | 54402000 | 54402093 | 54403186 | 54403324 | 29 | [illegible] |
| MXE | 10555 | ENSG00000122085 | MTERF4 | chr2 | - | 241089979 | 241090044 | 241094276 | 241094432 | 241087224 | 241089365 | 241097242 | 241097427 | 8 | [illegible] |
| MXE | 15240 | ENSG00000198879 | SFMBT2 | chr10 | - | 7276891 | 7276989 | 7283903 | 7283959 | 7248547 | 7248649 | 7285865 | 7285954 | 5 | [illegible] |
| MXE | 10449 | ENSG00000162923 | WDR26 | chr1 | - | 224411426 | 224411565 | 224418259 | 224418416 | 224404429 | 224404570 | 224419517 | 224419615 | 379 | [illegible] |
| MXE | 15235 | ENSG00000110721 | CHKA | chr11 | - | 68074716 | 68074830 | 68081403 | 68081457 | 68070723 | 68070857 | 68097018 | 68097130 | 9 | [illegible] |
| MXE | 14347 | ENSG00000134539 | KLRD1 | chr12 | + | 10309625 | 10309688 | 10311463 | 10311615 | 10309387 | 10309480 | 10313409 | 10313513 | 1598 | [illegible] |
| MXE | 10807 | ENSG00000144635 | DYNC1LI1 | chr3 | - | 32530460 | 32530520 | 32532985 | 32533097 | 32530283 | 32530328 | 32534510 | 32534646 | 130 | [illegible] |
| MXE | 4975 | ENSG00000248487 | ABHD14A | chr3 | + | 51978258 | 51978374 | 51980392 | 51980628 | 51977870 | 51978082 | 51980835 | 51981196 | 9 | [illegible] |
| MXE | 16809 | ENSG00000123349 | PFDN5 | chr12 | + | 53295838 | 53296275 | 53296429 | 53296551 | 53295524 | 53295639 | 53297849 | 53297924 | 1395 | [illegible] |
| MXE | 12779 | ENSG00000177189 | RPS6KA3 | chrX | - | 20169401 | 20169491 | 20172745 | 20172868 | 20167588 | 20167747 | 20175163 | 20175288 | 20 | [illegible] |
| MXE | 11535 | ENSG00000138376 | BARD1 | chr2 | - | 214780559 | 214781509 | 214792296 | 214792445 | 214769231 | 214769312 | 214797060 | 214797117 | 15 | [illegible] |
| MXE | 4710 | ENSG00000174227 | PIGG | chr4 | + | 501100 | 501170 | 505717 | 505927 | 500395 | 500601 | 507404 | 507593 | 10 | [illegible] |
| MXE | 9873 | ENSG00000152061 | RABGAP1L | chr1 | + | 174231144 | 174231355 | 174241482 | 174241657 | 174220971 | 174221164 | 174250474 | 174250632 | 302 | [illegible] |
| MXE | 12140 | ENSG00000164715 | LMTK2 | chr7 | + | 98171540 | 98171674 | 98185050 | 98185135 | 98159337 | 98159425 | 98186876 | 98186998 | 63 | [illegible] |
| MXE | 3411 | ENSG00000043143 | JADE2 | chr5 | + | 134566115 | 134566580 | 134573644 | 134573762 | 134564493 | 134564610 | 134578496 | 134583230 | 128 | [illegible] |
| MXE | 1026 | ENSG00000167851 | CD300A | chr17 | + | 74474531 | 74474685 | 74477435 | 74477530 | 74466657 | 74466743 | 74481288 | 74481326 | 20 | [illegible] |
| MXE | 14868 | ENSG00000198887 | SMC5 | chr9 | + | 70347918 | 70348038 | 70350113 | 70350293 | 70347612 | 70347717 | 70350375 | 70350471 | 62 | [illegible] |
| MXE | 7851 | ENSG00000081154 | PCNP | chr3 | + | 101585436 | 101585511 | 101590214 | 101590270 | 101579588 | 101580004 | 101592626 | 101592771 | 181 | [illegible] |
| MXE | 820 | ENSG00000130684 | ZNF337 | chr20 | - | 25685566 | 25685662 | 25685995 | 25686249 | 25676948 | 25677037 | 25686390 | 25686466 | 5 | [illegible] |
| MXE | 10919 | ENSG00000115652 | UXS1 | chr2 | - | 106101057 | 106101118 | 106104793 | 106104837 | 106098715 | 106098773 | 106112645 | 106112765 | 15 | [illegible] |
| MXE | 14910 | ENSG00000198431 | TXNRD1 | chr12 | + | 104311289 | 104311412 | 104313244 | 104313317 | 104288930 | 104289040 | 104315776 | 104315896 | 84 | [illegible] |
| MXE | 3870 | ENSG00000100385 | IL2RB | chr22 | - | 37139116 | 37139222 | 37142433 | 37142512 | 37137586 | 37137735 | 37143520 | 37143635 | 169 | [illegible] |
| MXE | 2038 | ENSG00000137804 | NUSAP1 | chr15 | + | 41365401 | 41365589 | 41371526 | 41371684 | 41358148 | 41358258 | 41375711 | 41375828 | 10 | [illegible] |
| MXE | 14030 | ENSG00000120992 | LYPLA1 | chr8 | - | 54063327 | 54063375 | 54065747 | 54065813 | 54062253 | 54062324 | 54100907 | 54100939 | 13 | [illegible] |
| MXE | 8153 | ENSG00000136238 | RAC1 | chr7 | + | 6400125 | 6400188 | 6401867 | 6402027 | 6391923 | 6392041 | 6402315 | 6402481 | 97 | [illegible] |
| MXE | 608 | ENSG00000089639 | GMIP | chr19 | - | 19635414 | 19635569 | 19635643 | 19635721 | 19635024 | 19635213 | 19636706 | 19636796 | 20 | [illegible] |
| MXE | 2731 | ENSG00000197324 | LRP10 | chr14 | + | 22873310 | 22873446 | 22875054 | 22875245 | 22872737 | 22872782 | 22875354 | 22876372 | 64 | [illegible] |
| MXE | 2469 | ENSG00000133961 | NUMB | chr14 | - | 73297210 | 73297285 | 73323129 | 73323204 | 73292733 | 73292874 | 73355625 | 73355766 | 145 | [illegible] |
| MXE | 7601 | ENSG00000102763 | VWA8 | chr13 | - | 41727193 | 41727313 | 41732079 | 41732155 | 41721369 | 41721575 | 41761127 | 41761204 | 25 | [illegible] |
| MXE | 7122 | ENSG00000055609 | KMT2C | chr7 | - | 152224014 | 152224179 | 152224434 | 152224616 | 152222572 | 152222682 | 152229922 | 152230027 | 95 | [illegible] |
| MXE | 6581 | ENSG00000018510 | AGPS | chr2 | + | 177513818 | 177513908 | 177521268 | 177521368 | 177507969 | 177508031 | 177523747 | 177523805 | 97 | [illegible] |
| MXE | 865 | ENSG00000088812 | ATRN | chr20 | + | 3572730 | 3572951 | 3575826 | 3575948 | 3565347 | 3565432 | 3576858 | 3576997 | 59 | [illegible] |
| MXE | 10370 | ENSG00000286239 | NA | chr2 | - | 223880123 | 223880232 | 223881941 | 223882072 | 223878528 | 223878730 | 223884647 | 223884749 | 182 | [illegible] |
| MXE | 8580 | ENSG00000171960 | PPIH | chr1 | + | 42659521 | 42659566 | 42660861 | 42660904 | 42659227 | 42659251 | 42664862 | 42664955 | 45 | [illegible] |
| MXE | 9148 | ENSG00000113615 | SEC24A | chr5 | + | 134676022 | 134676125 | 134679601 | 134679728 | 134675044 | 134675217 | 134682372 | 134682482 | 41 | [illegible] |
| MXE | 15065 | ENSG00000110514 | MADD | chr11 | + | 47290148 | 47290299 | 47290609 | 47290816 | 47289866 | 47290053 | 47293882 | 47293983 | 97 | [illegible] |
| MXE | 7579 | ENSG00000204149 | AGAP6 | chr10 | + | 50004684 | 50004720 | 50008024 | 50008076 | 50001995 | 50002096 | 50008710 | 50010407 | 15 | [illegible] |
| MXE | 138 | ENSG00000188677 | PARVB | chr22 | + | 44100052 | 44100123 | 44119037 | 44119140 | 44093927 | 44094017 | 44131486 | 44131627 | 16 | [illegible] |
| MXE | 16463 | ENSG00000119801 | YPEL5 | chr2 | + | 30148244 | 30148541 | 30155849 | 30155910 | 30146940 | 30147062 | 30156627 | 30156792 | 67 | [illegible] |
| MXE | 1147 | ENSG00000159259 | CHAF1B | chr21 | + | 36387597 | 36387730 | 36391550 | 36391668 | 36386059 | 36386262 | 36394546 | 36394650 | 14 | [illegible] |
| MXE | 7326 | ENSG00000007237 | GAS7 | chr17 | - | 9934165 | 9934244 | 9940125 | 9940200 | 9926640 | 9926769 | 9943120 | 9943236 | 103 | [illegible] |
| MXE | 8567 | ENSG00000120948 | TARDBP | chr1 | + | 11015523 | 11015646 | 11016122 | 11016254 | 11013715 | 11013965 | 11016843 | 11017007 | 3 | [illegible] |
| MXE | 8438 | ENSG00000121964 | GTDC1 | chr2 | - | 143952772 | 143952843 | 143957194 | 143957339 | 143951981 | 143952073 | 143970652 | 143970762 | 26 | [illegible] |
| MXE | 6892 | ENSG00000163960 | UBXN7 | chr3 | - | 196368027 | 196368155 | 196369420 | 196369511 | 196362293 | 196362687 | 196371895 | 196372042 | 126 | [illegible] |
| MXE | 17054 | ENSG00000117226 | GBP3 | chr1 | - | 89009391 | 89009494 | 89011746 | 89012027 | 89008946 | 89009140 | 89013184 | 89013427 | 71 | [illegible] |
| MXE | 3613 | ENSG00000086589 | RBM22 | chr5 | - | 150698498 | 150698631 | 150699255 | 150699271 | 150696790 | 150696891 | 150700443 | 150700497 | 3 | [illegible] |
| MXE | 16102 | ENSG00000099290 | WASHC2A | chr10 | + | 50125368 | 50125449 | 50127582 | 50127795 | 50125112 | 50125241 | 50129418 | 50129784 | 85 | [illegible] |
| MXE | 4274 | ENSG00000102445 | RUBCNL | chr13 | - | 46371940 | 46372597 | 46377889 | 46378005 | 46368732 | 46368815 | 46387676 | 46387879 | 0 | [illegible] |
| MXE | 11271 | ENSG00000172765 | TMCC1 | chr3 | - | 129838897 | 129839060 | 129860939 | 129861012 | 129832773 | 129832826 | 129880308 | 129880562 | 7 | [illegible] |
| MXE | 7149 | ENSG00000196419 | XRCC6 | chr22 | + | 41653528 | 41653690 | 41656902 | 41657032 | 41650722 | 41650891 | 41658251 | 41658352 | 552 | [illegible] |
| MXE | 13043 | ENSG00000161010 | MRNIP | chr5 | - | 179844151 | 179844227 | 179853196 | 179853276 | 179841906 | 179842064 | 179853377 | 179853437 | 0 | [illegible] |
| MXE | 3924 | ENSG00000059378 | PARP12 | chr7 | - | 140034234 | 140034331 | 140041643 | 140041839 | 140028612 | 140028688 | 140046883 | 140047007 | 136 | [illegible] |
| MXE | 1675 | ENSG00000103222 | ABCC1 | chr16 | + | 16079351 | 16079478 | 16086823 | 16086991 | 16076325 | 16076401 | 16090404 | 16090588 | 47 | [illegible] |
| MXE | 11893 | ENSG00000135317 | SNX14 | chr6 | - | 85533598 | 85533800 | 85536791 | 85536924 | 85530191 | 85530275 | 85543181 | 85543306 | 6 | [illegible] |
| MXE | 14746 | ENSG00000213722 | DDAH2 | chr6 | - | 31728172 | 31728292 | 31728450 | 31728524 | 31727542 | 31727692 | 31728645 | 31728745 | 13 | [illegible] |
| MXE | 12808 | ENSG00000104133 | SPG11 | chr15 | - | 44570524 | 44570658 | 44574901 | 44575041 | 44569397 | 44569505 | 44583813 | 44584558 | 44 | [illegible] |
| MXE | 14586 | ENSG00000110455 | ACCS | chr11 | + | 44081178 | 44081320 | 44083168 | 44083311 | 44081017 | 44081065 | 44083694 | 44083893 | 28 | [illegible] |
| MXE | 11988 | ENSG00000099968 | BCL2L13 | chr22 | + | 17688985 | 17689142 | 17696140 | 17696210 | 17683213 | 17683321 | 17702242 | 17702386 | 200 | [illegible] |
| MXE | 3213 | ENSG00000068796 | KIF2A | chr5 | + | 62365242 | 62365353 | 62366413 | 62366481 | 62363694 | 62363899 | 62373686 | 62373837 | 263 | [illegible] |
| MXE | 3518 | ENSG00000113558 | SKP1 | chr5 | - | 134160986 | 134161130 | 134167169 | 134167243 | 134157337 | 134158595 | 134173925 | 134174022 | 236 | [illegible] |
| MXE | 3926 | ENSG00000059378 | PARP12 | chr7 | - | 140037714 | 140037856 | 140041643 | 140041839 | 140034234 | 140034331 | 140046883 | 140047007 | 138 | [illegible] |
| MXE | 12136 | ENSG00000187796 | CARD9 | chr9 | - | 136365140 | 136365217 | 136366799 | 136366845 | 136364482 | 136364559 | 136367215 | 136367257 | 0 | [illegible] |
| MXE | 12823 | ENSG00000104133 | SPG11 | chr15 | - | 44620124 | 44620403 | 44621758 | 44621934 | 44615362 | 44615566 | 44622219 | 44622347 | 277 | [illegible] |
| MXE | 10983 | ENSG00000064201 | TSPAN32 | chr11 | + | 2316228 | 2316312 | 2316575 | 2316667 | 2314484 | 2314571 | 2317343 | 2317525 | 23 | [illegible] |
| MXE | 5785 | ENSG00000137074 | APTX | chr9 | - | 32987543 | 32987729 | 32987759 | 32987846 | 32985970 | 32986030 | 32988082 | 32988129 | 1 | [illegible] |
| MXE | 8836 | ENSG00000067057 | PFKP | chr10 | + | 3118781 | 3118869 | 3129818 | 3129983 | 3116775 | 3116846 | 3132379 | 3132441 | 20 | [illegible] |
| MXE | 11388 | ENSG00000198721 | ECI2 | chr6 | - | 4119185 | 4119275 | 4125249 | 4126237 | 4117307 | 4117451 | 4127761 | 4127831 | 116 | [illegible] |
| MXE | 9976 | ENSG00000163219 | ARHGAP25 | chr2 | + | 68775220 | 68775420 | 68782232 | 68782320 | 68734810 | 68735260 | 68787839 | 68787956 | 155 | [illegible] |
| MXE | 16732 | ENSG00000111271 | ACAD10 | chr12 | + | 111729805 | 111729956 | 111736830 | 111737004 | 111727961 | 111728143 | 111744642 | 111745043 | 27 | [illegible] |
| MXE | 15999 | ENSG00000156671 | SAMD8 | chr10 | + | 75164644 | 75164740 | 75168540 | 75168658 | 75150513 | 75151106 | 75176065 | 75176216 | 35 | [illegible] |
| MXE | 9993 | ENSG00000172116 | CD8B | chr2 | - | 86846683 | 86846773 | 86852996 | 86853086 | 86844921 | 86844958 | 86858056 | 86858416 | 124 | [illegible] |
| MXE | 13098 | ENSG00000166579 | NDEL1 | chr17 | + | 8454795 | 8454887 | 8460008 | 8460160 | 8450780 | 8450953 | 8490246 | 8490411 | 286 | [illegible] |
| MXE | 14520 | ENSG00000130958 | SLC35D2 | chr9 | - | 96336716 | 96336784 | 96343903 | 96343996 | 96324090 | 96324169 | 96345298 | 96345401 | 49 | [illegible] |
| MXE | 14797 | ENSG00000107164 | FUBP3 | chr9 | + | 130614287 | 130614345 | 130616354 | 130616517 | 130612955 | 130613027 | 130617796 | 130617895 | 56 | [illegible] |
| MXE | 13687 | ENSG00000215301 | DDX3X | chrX | + | 41346858 | 41347012 | 41347311 | 41347451 | 41346493 | 41346622 | 41347639 | 41349867 | 1022 | [illegible] |
| MXE | 13793 | ENSG00000198373 | WWP2 | chr16 | + | 69930129 | 69930258 | 69931151 | 69931227 | 69929447 | 69929529 | 69931508 | 69931580 | 95 | [illegible] |
| MXE | 1766 | ENSG00000087263 | OGFOD1 | chr16 | + | 56466875 | 56466967 | 56467164 | 56467293 | 56466151 | 56466268 | 56467904 | 56468018 | 44 | [illegible] |
| MXE | 9867 | ENSG00000110934 | BIN2 | chr12 | - | 51295795 | 51295878 | 51297088 | 51297164 | 51291590 | 51292344 | 51299202 | 51299288 | 437 | [illegible] |
| MXE | 14313 | ENSG00000135679 | MDM2 | chr12 | + | 68813553 | 68813628 | 68816811 | 68816945 | 68809211 | 68809292 | 68820324 | 68820374 | 77 | [illegible] |
| MXE | 17105 | ENSG00000124496 | TRERF1 | chr6 | - | 42265750 | 42265797 | 42268153 | 42269848 | 42264703 | 42264854 | 42300637 | 42300749 | 37 | [illegible] |
| MXE | 8 | ENSG00000159267 | HLCS | chr21 | - | 36946008 | 36946154 | 36947332 | 36948287 | 36938831 | 36938994 | 36962035 | 36962170 | 1 | [illegible] |
| MXE | 17078 | ENSG00000139668 | WDFY2 | chr13 | + | 51675169 | 51675243 | 51703595 | 51703650 | 51660595 | 51660663 | 51719197 | 51719348 | 32 | [illegible] |
| MXE | 2068 | ENSG00000182718 | ANXA2 | chr15 | - | 60390212 | 60390317 | 60397257 | 60397338 | 60386027 | 60386086 | 60397942 | 60397986 | 117 | [illegible] |
| MXE | 2759 | ENSG00000259316 | NA | chr15 | - | 64204864 | 64204949 | 64207508 | 64207594 | 64204440 | 64204589 | 64213889 | 64214124 | 23 | [illegible] |
| MXE | 7915 | ENSG00000123636 | BAZ2B | chr2 | - | 159448241 | 159448409 | 159453612 | 159453801 | 159446781 | 159446975 | 159478574 | 159478721 | 58 | [illegible] |
| MXE | 7625 | ENSG00000132300 | PTCD3 | chr2 | + | 86134838 | 86134987 | 86136520 | 86136562 | 86134291 | 86134377 | 86136981 | 86137140 | 120 | [illegible] |
| MXE | 8422 | ENSG00000178607 | ERN1 | chr17 | - | 64052779 | 64052979 | 64053271 | 64053371 | 64049054 | 64049202 | 64054249 | 64054439 | 64 | [illegible] |
| MXE | 9492 | ENSG00000102786 | INTS6 | chr13 | - | 51389318 | 51389444 | 51395299 | 51395483 | 51387385 | 51387540 | 51430293 | 51430383 | 118 | [illegible] |
| MXE | 12925 | ENSG00000013374 | NUB1 | chr7 | + | 151349072 | 151349240 | 151351423 | 151351482 | 151345344 | 151345466 | 151352811 | 151352882 | 334 | [illegible] |
| MXE | 17079 | ENSG00000139668 | WDFY2 | chr13 | + | 51703595 | 51703650 | 51719197 | 51719348 | 51675169 | 51675243 | 51727677 | 51727790 | 36 | [illegible] |
| MXE | 14046 | ENSG00000122882 | ECD | chr10 | - | 73156274 | 73156453 | 73156567 | 73156655 | 73154255 | 73154448 | 73160433 | 73160551 | 89 | [illegible] |
| MXE | 16150 | ENSG00000134452 | FBH1 | chr10 | + | 5918341 | 5918478 | 5921257 | 5921357 | 5917589 | 5917676 | 5921447 | 5921569 | 112 | [illegible] |
| MXE | 9637 | ENSG00000185811 | IKZF1 | chr7 | + | 50376532 | 50376793 | 50382539 | 50382707 | 50327637 | 50327757 | 50391728 | 50391863 | 355 | [illegible] |
| MXE | 10784 | ENSG00000184708 | EIF4ENIF1 | chr22 | - | 31463680 | 31463967 | 31468174 | 31468302 | 31462931 | 31463133 | 31471843 | 31471917 | 39 | [illegible] |
| MXE | 11486 | ENSG00000111011 | RSRC2 | chr12 | - | 122511108 | 122511188 | 122515104 | 122515227 | 122508217 | 122508450 | 122517226 | 122517430 | 17 | [illegible] |
| MXE | 16384 | ENSG00000150045 | KLRF1 | chr12 | + | 9833302 | 9833452 | 9841811 | 9841951 | 9832315 | 9832414 | 9842320 | 9842433 | 1199 | [illegible] |
| MXE | 12322 | ENSG00000121680 | PEX16 | chr11 | - | 45915467 | 45915568 | 45916226 | 45916303 | 45914603 | 45914684 | 45917457 | 45917493 | 22 | [illegible] |
| MXE | 13513 | ENSG00000171311 | EXOSC1 | chr10 | - | 97441170 | 97441259 | 97443236 | 97443311 | 97437699 | 97437750 | 97445731 | 97445847 | 16 | [illegible] |
| MXE | 16238 | ENSG00000148634 | HERC4 | chr10 | - | 68038092 | 68038169 | 68044403 | 68044563 | 68033964 | 68034186 | 68072882 | 68073186 | 128 | [illegible] |
| MXE | 12291 | ENSG00000133606 | MKRN1 | chr7 | - | 140459706 | 140459936 | 140471882 | 140472011 | 140459006 | 140459233 | 140479159 | 140479336 | 23 | [illegible] |
| MXE | 5450 | ENSG00000169221 | TBC1D10B | chr16 | - | 30359171 | 30359361 | 30359537 | 30359603 | 30358662 | 30358817 | 30359726 | 30359841 | 19 | [illegible] |
| MXE | 8334 | ENSG00000083720 | OXCT1 | chr5 | - | 41803068 | 41803163 | 41807330 | 41807438 | 41801021 | 41801070 | 41840450 | 41840511 | 61 | [illegible] |
| MXE | 1938 | ENSG00000047578 | KATNIP | chr16 | + | 27631064 | 27631162 | 27648603 | 27648735 | 27628660 | 27628830 | 27677728 | 27677996 | 52 | [illegible] |
| MXE | 16841 | ENSG00000111596 | CNOT2 | chr12 | + | 70319297 | 70319364 | 70335437 | 70335563 | 70310894 | 70311017 | 70337388 | 70337513 | 262 | [illegible] |
| MXE | 757 | ENSG00000118705 | RPN2 | chr20 | + | 37232295 | 37232391 | 37234019 | 37234095 | 37229972 | 37230059 | 37236579 | 37236709 | 216 | [illegible] |
| MXE | 6813 | ENSG00000117682 | DHDDS | chr1 | + | 26457790 | 26457905 | 26460036 | 26460144 | 26447558 | 26447660 | 26468894 | 26469226 | 19 | [illegible] |
| MXE | 14387 | ENSG00000119397 | CNTRL | chr9 | + | 121150169 | 121150483 | 121152484 | 121152693 | 121148671 | 121148861 | 121154720 | 121154913 | 146 | [illegible] |
| MXE | 7903 | ENSG00000071054 | MAP4K4 | chr2 | + | 101855976 | 101856138 | 101858995 | 101859082 | 101844100 | 101844311 | 101859735 | 101859864 | 190 | [illegible] |
| MXE | 6342 | ENSG00000078142 | PIK3C3 | chr18 | + | 41993269 | 41993341 | 41995889 | 41995994 | 41990458 | 41990554 | 41996637 | 41996730 | 129 | [illegible] |
| MXE | 8656 | ENSG00000143398 | PIP5K1A | chr1 | + | 151239954 | 151240039 | 151242122 | 151242269 | 151239129 | 151239178 | 151242437 | 151242567 | 36 | [illegible] |
| MXE | 3005 | ENSG00000115760 | BIRC6 | chr2 | + | 32406498 | 32406557 | 32414768 | 32416163 | 32401462 | 32401623 | 32429145 | 32429295 | 53 | [illegible] |
| MXE | 15749 | ENSG00000149311 | ATM | chr11 | + | 108293312 | 108293477 | 108294926 | 108296393 | 108292618 | 108292793 | 108297286 | 108297382 | 246 | [illegible] |
| MXE | 656 | ENSG00000196365 | LONP1 | chr19 | - | 5708341 | 5708403 | 5711770 | 5712002 | 5707696 | 5707826 | 5713133 | 5713253 | 129 | [illegible] |
| MXE | 15603 | ENSG00000149262 | INTS4 | chr11 | - | 77955941 | 77956062 | 77958745 | 77958834 | 77941179 | 77941251 | 77960340 | 77960391 | 71 | [illegible] |
| MXE | 2838 | ENSG00000203546 | NA | chr14 | - | 31398672 | 31398781 | 31400300 | 31400512 | 31395198 | 31395348 | 31402849 | 31403049 | 65 | [illegible] |
| MXE | 16525 | ENSG00000139323 | POC1B | chr12 | - | 89467616 | 89467685 | 89491935 | 89492115 | 89466769 | 89466922 | 89525119 | 89525204 | 42 | [illegible] |
| MXE | 5219 | ENSG00000166501 | PRKCB | chr16 | + | 23837374 | 23837406 | 23988507 | 23988590 | 23835982 | 23836348 | 24032135 | 24032247 | 13 | [illegible] |
| MXE | 16480 | ENSG00000082898 | XPO1 | chr2 | - | 61488587 | 61488771 | 61490641 | 61490776 | 61488164 | 61488271 | 61492034 | 61492198 | 599 | [illegible] |
| MXE | 11393 | ENSG00000143374 | TARS2 | chr1 | + | 150496809 | 150496908 | 150497529 | 150497747 | 150496481 | 150496628 | 150498501 | 150498664 | 16 | [illegible] |
| MXE | 3957 | ENSG00000138744 | NAAA | chr4 | - | 75931213 | 75931304 | 75936108 | 75936235 | 75925734 | 75925811 | 75940000 | 75940165 | 54 | [illegible] |
| MXE | 16153 | ENSG00000108094 | CUL2 | chr10 | - | 35011847 | 35011964 | 35013698 | 35013800 | 35008550 | 35010442 | 35016191 | 35016394 | 83 | [illegible] |
| MXE | 4738 | ENSG00000111321 | LTBR | chr12 | + | 6388799 | 6388825 | 6390111 | 6390340 | 6388253 | 6388505 | 6390659 | 6390735 | 9 | [illegible] |
| MXE | 15083 | ENSG00000108854 | SMURF2 | chr17 | - | 64547599 | 64547801 | 64551583 | 64551704 | 64546262 | 64546338 | 64554855 | 64554993 | 84 | [illegible] |
| MXE | 5334 | ENSG00000122068 | FYTTD1 | chr3 | + | 197770131 | 197770244 | 197774148 | 197774210 | 197768438 | 197768587 | 197776926 | 197777001 | 557 | [illegible] |
| MXE | 4039 | ENSG00000137802 | MAPKBP1 | chr15 | + | 41814549 | 41814739 | 41815258 | 41815405 | 41812918 | 41813101 | 41815623 | 41815799 | 4 | [illegible] |
| MXE | 9980 | ENSG00000264538 | SUZ12P1 | chr17 | + | 30735032 | 30735097 | 30743300 | 30743369 | 30734896 | 30734943 | 30758171 | 30758335 | 23 | [illegible] |
| MXE | 13297 | ENSG00000094631 | HDAC6 | chrX | + | 48823921 | 48824068 | 48824165 | 48824294 | 48823671 | 48823785 | 48824543 | 48824860 | 67 | [illegible] |
| MXE | 11634 | ENSG00000188554 | NBR1 | chr17 | + | 43196941 | 43197106 | 43200166 | 43200608 | 43196480 | 43196591 | 43201685 | 43201780 | 193 | [illegible] |
| MXE | 13629 | ENSG00000071553 | ATP6AP1 | chrX | + | 154432265 | 154432376 | 154433659 | 154433720 | 154431829 | 154431904 | 154434207 | 154434446 | 95 | [illegible] |
| MXE | 9959 | ENSG00000068878 | PSME4 | chr2 | - | 53925538 | 53925689 | 53925958 | 53926023 | 53923320 | 53923419 | 53927393 | 53927483 | 40 | [illegible] |
| MXE | 16975 | ENSG00000111269 | CREBL2 | chr12 | + | 12635776 | 12635974 | 12637569 | 12637714 | 12611875 | 12612187 | 12641993 | 12645108 | 71 | [illegible] |
| MXE | 4704 | ENSG00000155508 | CNOT8 | chr5 | + | 154865191 | 154865385 | 154867691 | 154867792 | 154858626 | 154858768 | 154870660 | 154870822 | 283 | [illegible] |
| MXE | 2777 | ENSG00000100784 | RPS6KA5 | chr14 | - | 90943077 | 90943185 | 90947434 | 90947550 | 90923112 | 90923196 | 90978305 | 90978524 | 45 | [illegible] |
| MXE | 14036 | ENSG00000215114 | UBXN2B | chr8 | + | 58434857 | 58434948 | 58439632 | 58439770 | 58434394 | 58434504 | 58445906 | 58446068 | 12 | [illegible] |

[illegible]

| | | | | | | | | | | |
|---|---|---|---|---|---|---|---|---|---|---|
| MXE | 16741 | ENSG00000136521 | NDUFB5 | chr3 | + | 179615982 | 179616049 | 179616982 | 179617044 | [illegible] |
| MXE | 11227 | ENSG00000138835 | RGS3 | chr9 | + | 113536795 | 113536918 | 113583449 | 113584427 | [illegible] |
| MXE | 1523 | ENSG00000083454 | P2RX5 | chr17 | - | 3688625 | 3688676 | 3690069 | 3690150 | [illegible] |
| MXE | 7083 | ENSG00000126749 | EMG1 | chr12 | + | 6975228 | 6975378 | 6983001 | 6983206 | [illegible] |
| MXE | 13503 | ENSG00000197713 | RPE | chr2 | + | 210003437 | 210003468 | 210009656 | 210009736 | [illegible] |
| MXE | 16137 | ENSG00000165732 | DDX21 | chr10 | + | 68977528 | 68977688 | 68978841 | 68978976 | [illegible] |
| MXE | 10963 | ENSG00000168214 | RBPJ | chr4 | + | 26420550 | 26420725 | 26424341 | 26424479 | [illegible] |
| MXE | 4988 | ENSG00000166128 | RAB8B | chr15 | + | 63223847 | 63223979 | 63244755 | 63244816 | [illegible] |
| MXE | 13834 | ENSG00000145041 | DCAF1 | chr3 | - | 51466802 | 51466876 | 51470928 | 51471005 | [illegible] |
| MXE | 11366 | ENSG00000115970 | THADA | chr2 | - | 43344121 | 43344237 | 43397970 | 43398139 | [illegible] |
| MXE | 10745 | ENSG00000177479 | ARIH2 | chr3 | + | 48924717 | 48924839 | 48964918 | 48964982 | [illegible] |
| MXE | 7825 | ENSG00000132604 | TERF2 | chr16 | - | 69370482 | 69370629 | 69372268 | 69372355 | [illegible] |
| MXE | 5831 | ENSG00000147251 | DOCK11 | chrX | + | 118651963 | 118652077 | 118654601 | 118654817 | [illegible] |
| MXE | 11362 | ENSG00000136699 | SMPD4 | chr2 | - | 130173278 | 130173354 | 130173513 | 130173656 | [illegible] |
| MXE | 10662 | ENSG00000132405 | TBC1D14 | chr4 | + | 7006631 | 7006726 | 7009876 | 7009948 | [illegible] |
| MXE | 1666 | ENSG00000103222 | ABCC1 | chr16 | + | 16014490 | 16014628 | 16016495 | 16016621 | [illegible] |
| MXE | 16879 | ENSG00000050426 | LETMD1 | chr12 | + | 51053777 | 51053860 | 51055986 | 51056021 | [illegible] |
| MXE | 14634 | ENSG00000095383 | TBC1D2 | chr9 | - | 98233415 | 98233549 | 98243993 | 98244129 | [illegible] |
| MXE | 1760 | ENSG00000214078 | CPNE1 | chr20 | - | 35626566 | 35626803 | 35627279 | 35627413 | [illegible] |
| MXE | 15024 | ENSG00000110756 | HPS5 | chr11 | - | 18281949 | 18282220 | 18283794 | 18283901 | [illegible] |
| MXE | 2695 | ENSG00000129566 | TEP1 | chr14 | - | 20403376 | 20403448 | 20403722 | 20403884 | [illegible] |
| MXE | 11040 | ENSG00000079263 | SP140 | chr2 | + | 230237082 | 230237260 | 230243730 | 230243811 | [illegible] |
| MXE | 352 | ENSG00000189068 | VSTM1 | chr19 | - | 54042276 | 54042369 | 54051409 | 54051448 | [illegible] |
| MXE | 5975 | ENSG00000197948 | FCHSD1 | chr5 | - | 141645770 | 141645821 | 141646086 | 141646191 | [illegible] |
| MXE | 17121 | ENSG00000275066 | SYNRG | chr17 | - | 37535978 | 37536127 | 37539191 | 37539245 | [illegible] |
| MXE | 6399 | ENSG00000116001 | TIA1 | chr2 | - | 70215370 | 70215494 | 70216207 | 70216292 | [illegible] |
| MXE | 2747 | ENSG00000198604 | BAZ1A | chr14 | - | 34807450 | 34807538 | 34810934 | 34811036 | [illegible] |
| [illegible] | | | | | | | | | | |

| | | | | | | | | | | | | | | | |
|---|---|---|---|---|---|---|---|---|---|---|---|---|---|---|---|
| MXE | 12138 | ENSG00000173715 | C11orf80 | chr11 | + | 66788159 | 66788269 | 66796280 | 66796364 | 66759042 | 66759096 | 66800640 | 66800696 | 2 | [illegible] |
| MXE | 16901 | ENSG00000111335 | OAS2 | chr12 | + | 112997519 | 112998410 | 113002931 | 113003102 | 112995295 | 112995474 | 113004933 | 113005222 | 215 | [illegible] |
| MXE | 8519 | ENSG00000184178 | SCFD2 | chr4 | - | 53273825 | 53274001 | 53313635 | 53313763 | 53145332 | 53145582 | 53352597 | 53352766 | 48 | [illegible] |
| MXE | 4202 | ENSG00000084676 | NCOA1 | chr2 | + | 24705085 | 24705233 | 24706567 | 24707888 | 24697657 | 24697798 | 24710930 | 24711111 | 202 | [illegible] |
| MXE | 10552 | ENSG00000114054 | PCCB | chr3 | + | 136256554 | 136256623 | 136260478 | 136260535 | 136255855 | 136255975 | 136261951 | 136262065 | 52 | [illegible] |
| MXE | 9131 | ENSG00000047410 | TPR | chr1 | - | 186361621 | 186361709 | 186361788 | 186361869 | 186360764 | 186360905 | 186362287 | 186362380 | 154 | [illegible] |
| MXE | 7370 | ENSG00000116171 | SCP2 | chr1 | + | 53027968 | 53028071 | 53038916 | 53039046 | 53014965 | 53015043 | 53047857 | 53047937 | 302 | [illegible] |
| MXE | 743 | ENSG00000088970 | KIZ | chr20 | + | 21145564 | 21145654 | 21161870 | 21162507 | 21136389 | 21136552 | 21162849 | 21163159 | 56 | [illegible] |
| MXE | 561 | ENSG00000126261 | UBA2 | chr19 | + | 34433347 | 34433412 | 34434867 | 34434968 | 34431860 | 34431931 | 34438644 | 34438764 | 94 | [illegible] |
| MXE | 15068 | ENSG00000110514 | MADD | chr11 | + | 47293882 | 47293983 | 47295495 | 47295576 | 47290609 | 47290816 | 47295896 | 47296055 | 97 | [illegible] |
| MXE | 13522 | ENSG00000134516 | DOCK2 | chr5 | + | 169689251 | 169689333 | 169695802 | 169695938 | 169684195 | 169684350 | 169698373 | 169698449 | 223 | [illegible] |
| MXE | 12104 | ENSG00000136250 | AOAH | chr7 | - | 36540318 | 36540491 | 36549438 | 36549475 | 36532285 | 36532344 | 36576573 | 36576656 | 92 | [illegible] |
| MXE | 6952 | ENSG00000160058 | BSDC1 | chr1 | - | 32381213 | 32381268 | 32383829 | 32383997 | 32378723 | 32378839 | 32394079 | 32394140 | 1 | [illegible] |
| MXE | 6909 | ENSG00000091127 | PUS7 | chr7 | - | 105468336 | 105468463 | 105472131 | 105472193 | 105465312 | 105465414 | 105481051 | 105481177 | 11 | [illegible] |
| MXE | 11670 | ENSG00000024048 | UBR2 | chr6 | + | 42603587 | 42603718 | 42605720 | 42605859 | 42594190 | 42594304 | 42606588 | 42606651 | 247 | [illegible] |
| MXE | 3867 | ENSG00000113391 | FAM172A | chr5 | - | 93964457 | 93964523 | 94053125 | 94053227 | 93958776 | 93958968 | 94111510 | 94111676 | 28 | [illegible] |
| MXE | 8674 | ENSG00000125686 | MED1 | chr17 | - | 39431941 | 39432016 | 39434248 | 39434320 | 39431114 | 39431188 | 39439164 | 39439193 | 84 | [illegible] |
| MXE | 11961 | ENSG00000136295 | TTYH3 | chr7 | + | 2658285 | 2658459 | 2658939 | 2659015 | 2656397 | 2656534 | 2661667 | 2664800 | 44 | [illegible] |
| MXE | 13547 | ENSG00000131381 | RBSN | chr3 | - | 15075605 | 15075710 | 15077061 | 15077164 | 15070072 | 15074930 | 15078074 | 15078161 | 35 | [illegible] |
| MXE | 15539 | ENSG00000110075 | PPP6R3 | chr11 | + | 68575957 | 68576043 | 68587926 | 68588024 | 68574108 | 68574224 | 68590659 | 68590714 | 153 | [illegible] |
| MXE | 9701 | ENSG00000162961 | DPY30 | chr2 | - | 31891991 | 31892134 | 31932066 | 31932135 | 31883385 | 31883462 | 31943301 | 31943383 | 30 | [illegible] |
| MXE | 14334 | ENSG00000104695 | PPP2CB | chr8 | - | 30797580 | 30797754 | 30799545 | 30799755 | 30794191 | 30794281 | 30812319 | 30812818 | 72 | [illegible] |
| MXE | 11034 | ENSG00000089123 | TASP1 | chr20 | - | 13580896 | 13580981 | 13587249 | 13587370 | 13569506 | 13569586 | 13623445 | 13623514 | 10 | [illegible] |
| MXE | 10656 | ENSG00000091009 | RBM27 | chr5 | + | 146263490 | 146263631 | 146267648 | 146267768 | 146261509 | 146261806 | 146269206 | 146269281 | 79 | [illegible] |
| MXE | 2417 | ENSG00000206149 | HERC2P9 | chr15 | + | 28618218 | 28618378 | 28629265 | 28629441 | 28612141 | 28612173 | 28632636 | 28632807 | 17 | [illegible] |
| MXE | 1917 | ENSG00000102893 | PHKB | chr16 | + | 47663676 | 47663734 | 47665941 | 47666032 | 47661718 | 47661800 | 47669214 | 47669417 | 26 | [illegible] |
| MXE | 9094 | ENSG00000131778 | CHD1L | chr1 | + | 147268781 | 147268878 | 147272170 | 147272630 | 147267425 | 147267518 | 147275353 | 147275468 | 25 | [illegible] |
| MXE | 13351 | ENSG00000140332 | TLE3 | chr15 | - | 70064453 | 70064470 | 70066013 | 70066218 | 70060529 | 70060649 | 70074502 | 70074607 | 31 | [illegible] |
| MXE | 12674 | ENSG00000123505 | AMD1 | chr6 | + | 110887504 | 110887591 | 110890253 | 110890356 | 110874783 | 110875215 | 110892160 | 110892203 | 110 | [illegible] |
| MXE | 4356 | ENSG00000163625 | WDFY3 | chr4 | - | 84756925 | 84757161 | 84765809 | 84766027 | 84755265 | 84755400 | 84766251 | 84766372 | 53 | [illegible] |
| MXE | 16377 | ENSG00000103966 | EHD4 | chr15 | - | 41919209 | 41919622 | 41943066 | 41943164 | 41909698 | 41909863 | 41953763 | 41953940 | 59 | [illegible] |
| MXE | 5353 | ENSG00000051382 | PIK3CB | chr3 | - | 138683677 | 138683787 | 138694785 | 138694907 | 138681966 | 138682045 | 138698906 | 138699095 | 107 | [illegible] |
| MXE | 2049 | ENSG00000128989 | ARPP19 | chr15 | - | 52564189 | 52564246 | 52568847 | 52568900 | 52557099 | 52557222 | 52569115 | 52569167 | 87 | [illegible] |
| MXE | 13958 | ENSG00000166169 | POLL | chr10 | - | 101582762 | 101582891 | 101583507 | 101583681 | 101580247 | 101580416 | 101584601 | 101584720 | 31 | [illegible] |
| MXE | 6621 | ENSG00000148019 | CEP78 | chr9 | + | 78253231 | 78253277 | 78254835 | 78254964 | 78251904 | 78252043 | 78262906 | 78262984 | 31 | [illegible] |
| MXE | 13474 | ENSG00000111203 | ITFG2 | chr12 | + | 2821261 | 2821359 | 2821691 | 2821792 | 2820723 | 2820872 | 2822793 | 2822911 | 16 | [illegible] |
| MXE | 2650 | ENSG00000171723 | GPHN | chr14 | + | 66776463 | 66776521 | 66879938 | 66880033 | 66681106 | 66681185 | 66916002 | 66916034 | 12 | [illegible] |
| MXE | 12141 | ENSG00000164715 | LMTK2 | chr7 | + | 98190727 | 98190877 | 98191613 | 98194572 | 98186876 | 98186998 | 98203573 | 98203706 | 26 | [illegible] |
| MXE | 5968 | ENSG00000150753 | CCT5 | chr5 | + | 10254673 | 10254838 | 10255954 | 10256153 | 10254144 | 10254205 | 10258110 | 10258303 | 387 | [illegible] |
| MXE | 11482 | ENSG00000166436 | TRIM66 | chr11 | - | 8620049 | 8620124 | 8621031 | 8621237 | 8619382 | 8619535 | 8621644 | 8621819 | 4 | [illegible] |
| MXE | 6721 | ENSG00000204435 | CSNK2B | chr6 | + | 31669096 | 31669172 | 31669327 | 31669508 | 31668538 | 31668654 | 31669835 | 31670074 | 148 | [illegible] |
| MXE | 2810 | ENSG00000033170 | FUT8 | chr14 | + | 65721774 | 65722021 | 65724146 | 65724323 | 65669242 | 65669480 | 65733230 | 65733381 | 48 | [illegible] |
| MXE | 15218 | ENSG00000066294 | CD84 | chr1 | - | 160550937 | 160551035 | 160553359 | 160553497 | 160549916 | 160549979 | 160553894 | 160554146 | 120 | [illegible] |
| MXE | 5812 | ENSG00000162777 | DENND2D | chr1 | - | 111189211 | 111189253 | 111192139 | 111192317 | 111188786 | 111188916 | 111194577 | 111194726 | 483 | [illegible] |
| MXE | 2431 | ENSG00000159128 | IFNGR2 | chr21 | + | 33432176 | 33432336 | 33432713 | 33432871 | 33426883 | 33427032 | 33436827 | 33436944 | 1011 | [illegible] |
| MXE | 10508 | ENSG00000164164 | OTUD4 | chr4 | - | 145144310 | 145144434 | 145146266 | 145146429 | 145143945 | 145144001 | 145150512 | 145150699 | 142 | [illegible] |
| MXE | 7827 | ENSG00000108021 | TASOR2 | chr10 | + | 5712822 | 5712918 | 5717658 | 5717750 | 5684837 | 5685175 | 5720543 | 5720668 | 15 | [illegible] |
| MXE | 801 | ENSG00000124228 | DDX27 | chr20 | + | 49226429 | 49226535 | 49228714 | 49228888 | 49225112 | 49225199 | 49230198 | 49230349 | 82 | [illegible] |
| MXE | 15059 | ENSG00000157837 | SPPL3 | chr12 | - | 120782654 | 120782767 | 120783673 | 120783752 | 120768952 | 120769059 | 120784473 | 120784593 | 95 | [illegible] |
| MXE | 8907 | ENSG00000089127 | OAS1 | chr12 | + | 112908535 | 112908824 | 112911050 | 112911235 | 112906972 | 112907219 | 112916508 | 112916738 | 288 | [illegible] |
| MXE | 6737 | ENSG00000145703 | IQGAP2 | chr5 | + | 76673951 | 76674036 | 76677217 | 76677350 | 76673448 | 76673589 | 76683114 | 76683217 | 154 | [illegible] |
| MXE | 7873 | ENSG00000113812 | ACTR8 | chr3 | - | 53878356 | 53878467 | 53879938 | 53880109 | 53877646 | 53877751 | 53881978 | 53882152 | 31 | [illegible] |
| MXE | 9411 | ENSG00000187741 | FANCA | chr16 | - | 89742799 | 89742938 | 89744958 | 89745071 | 89740803 | 89740866 | 89746583 | 89746688 | 9 | [illegible] |
| MXE | 9284 | ENSG00000170606 | HSPA4 | chr5 | + | 133067416 | 133067557 | 133070073 | 133070496 | 133064979 | 133065037 | 133073229 | 133073329 | 45 | [illegible] |
| MXE | 16442 | ENSG00000170456 | DENND5B | chr12 | - | 31451939 | 31452476 | 31460193 | 31460381 | 31447537 | 31447769 | 31479588 | 31480255 | 14 | [illegible] |
| MXE | 13609 | ENSG00000122122 | SASH3 | chrX | + | 129792327 | 129792476 | 129792626 | 129792836 | 129790939 | 129791081 | 129792988 | 129793139 | 659 | [illegible] |
| MXE | 11546 | ENSG00000116539 | ASH1L | chr1 | - | 155477885 | 155482449 | 155521099 | 155521618 | 155459898 | 155459974 | 155562778 | 155562841 | 23 | [illegible] |
| MXE | 4195 | ENSG00000084676 | NCOA1 | chr2 | + | 24584475 | 24584560 | 24658660 | 24658766 | 24564294 | 24564430 | 24665748 | 24665915 | 103 | [illegible] |
| MXE | 16875 | ENSG00000050426 | LETMD1 | chr12 | + | 51052091 | 51052207 | 51053777 | 51053860 | 51049033 | 51049185 | 51056349 | 51056502 | 42 | [illegible] |
| MXE | 16330 | ENSG00000132305 | IMMT | chr2 | - | 86161975 | 86162079 | 86166507 | 86166644 | 86159246 | 86159671 | 86170748 | 86170844 | 83 | [illegible] |
| MXE | 15445 | ENSG00000149136 | SSRP1 | chr11 | - | 57330854 | 57330927 | 57331667 | 57331889 | 57330290 | 57330429 | 57332151 | 57332280 | 169 | [illegible] |
| MXE | 12554 | ENSG00000179818 | PCBP1-AS1 | chr2 | - | 70059642 | 70059680 | 70083537 | 70083687 | 70053730 | 70053792 | 70085546 | 70085624 | 33 | [illegible] |
| MXE | 9269 | ENSG00000138709 | LARP1B | chr4 | + | 128091000 | 128091144 | 128098185 | 128098330 | 128082164 | 128082305 | 128107138 | 128107313 | 35 | [illegible] |
| MXE | 10211 | ENSG00000091436 | MAP3K20 | chr2 | + | 173209728 | 173209835 | 173217114 | 173217250 | 173203795 | 173203870 | 173221149 | 173222204 | 38 | [illegible] |
| MXE | 6377 | ENSG00000026751 | SLAMF7 | chr1 | + | 160748193 | 160748514 | 160749820 | 160750093 | 160739056 | 160739356 | 160752185 | 160752248 | 183 | [illegible] |
| MXE | 7179 | ENSG00000133106 | EPSTI1 | chr13 | - | 42889210 | 42889242 | 42895008 | 42895108 | 42886387 | 42888502 | 42900309 | 42900383 | 237 | [illegible] |
| MXE | 12350 | ENSG00000167081 | PBX3 | chr9 | + | 125960683 | 125960849 | 125963011 | 125963101 | 125935471 | 125935607 | 125965830 | 125965923 | 46 | [illegible] |
| MXE | 6084 | ENSG00000164253 | WDR41 | chr5 | - | 77464760 | 77464809 | 77489456 | 77489572 | 77463094 | 77463226 | 77492169 | 77492215 | 33 | [illegible] |
| MXE | 5849 | ENSG00000130024 | PHF10 | chr6 | - | 169717822 | 169717906 | 169718787 | 169718918 | 169715954 | 169716088 | 169721004 | 169721111 | 121 | [illegible] |
| MXE | 9550 | ENSG00000198836 | OPA1 | chr3 | + | 193626091 | 193626202 | 193631611 | 193631665 | 193618868 | 193618936 | 193635417 | 193635522 | 64 | [illegible] |
| MXE | 4618 | ENSG00000017260 | ATP2C1 | chr3 | + | 130969291 | 130969396 | 130980581 | 130980679 | 130967329 | 130967419 | 130992950 | 130993001 | 51 | [illegible] |
| MXE | 14657 | ENSG00000198780 | FAM169A | chr5 | - | 74813839 | 74814019 | 74834425 | 74834597 | 74805155 | 74805284 | 74838964 | 74839050 | 104 | [illegible] |
| MXE | 11989 | ENSG00000099968 | BCL2L13 | chr22 | + | 17696140 | 17696210 | 17702242 | 17702386 | 17688985 | 17689142 | 17726676 | 17730855 | 57 | [illegible] |
| MXE | 8738 | ENSG00000174123 | TLR10 | chr4 | - | 38775774 | 38775963 | 38776109 | 38776426 | 38772238 | 38775652 | 38782920 | 38782990 | 30 | [illegible] |
| MXE | 16111 | ENSG00000108055 | SMC3 | chr10 | + | 110590411 | 110590572 | 110590990 | 110591132 | 110589891 | 110589991 | 110593072 | 110593223 | 118 | [illegible] |
| MXE | 8919 | ENSG00000065413 | ANKRD44 | chr2 | - | 197025195 | 197025267 | 197187022 | 197187106 | 197013510 | 197013712 | 197310577 | 197310646 | 157 | [illegible] |
| MXE | 16733 | ENSG00000111271 | ACAD10 | chr12 | + | 111733922 | 111734068 | 111736830 | 111737004 | 111729805 | 111729956 | 111744642 | 111745043 | 10 | [illegible] |
| MXE | 14475 | ENSG00000180776 | ZDHHC20 | chr13 | - | 21387507 | 21387634 | 21391721 | 21391854 | 21382919 | 21383009 | 21400372 | 21400493 | 198 | [illegible] |
| MXE | 14275 | ENSG00000120868 | APAF1 | chr12 | + | 98671534 | 98671719 | 98677424 | 98677551 | 98670972 | 98671086 | 98680276 | 98680402 | 96 | [illegible] |
| MXE | 14656 | ENSG00000158467 | AHCYL2 | chr7 | + | 129405835 | 129405899 | 129406377 | 129406466 | 129405096 | 129405213 | 129409475 | 129409546 | 17 | [illegible] |
| MXE | 294 | ENSG00000100077 | GRK3 | chr22 | + | 25667738 | 25667800 | 25672295 | 25672347 | 25663629 | 25663704 | 25674436 | 25674528 | 154 | [illegible] |
| MXE | 2617 | ENSG00000075413 | MARK3 | chr14 | + | 103451917 | 103451983 | 103457141 | 103457212 | 103448918 | 103448967 | 103462404 | 103462461 | 68 | [illegible] |
| MXE | 7017 | ENSG00000135148 | TRAFD1 | chr12 | + | 112145585 | 112145662 | 112148073 | 112148304 | 112142088 | 112142295 | 112149750 | 112149808 | 43 | [illegible] |
| MXE | 3516 | ENSG00000163635 | ATXN7 | chr3 | + | 63988058 | 63988324 | 63990175 | 63990374 | 63982938 | 63983021 | 63990737 | 63990859 | 283 | [illegible] |
| MXE | 2115 | ENSG00000128739 | SNRPN | chr15 | + | 24974397 | 24974456 | 24974641 | 24974957 | 24962113 | 24962209 | 24975357 | 24975509 | 302 | [illegible] |
| MXE | 3789 | ENSG00000164219 | PGGT1B | chr5 | - | 115230975 | 115231021 | 115236389 | 115236522 | 115221823 | 115222008 | 115237857 | 115238009 | 91 | [illegible] |
| MXE | 10907 | ENSG00000258984 | UBE2F-SCLY | chr2 | + | 237987962 | 237987992 | 238016565 | 238016633 | 237973091 | 237973225 | 238035877 | 238035940 | 27 | [illegible] |
| MXE | 3796 | ENSG00000005100 | DHX33 | chr17 | - | 5453820 | 5453980 | 5455159 | 5455271 | 5453579 | 5453668 | 5455996 | 5456182 | 15 | [illegible] |
| MXE | 12681 | ENSG00000155096 | AZIN1 | chr8 | - | 102833055 | 102833218 | 102834188 | 102834263 | 102829820 | 102829936 | 102834665 | 102834747 | 345 | [illegible] |
| MXE | 1425 | ENSG00000108960 | MMD | chr17 | - | 55403766 | 55403868 | 55407745 | 55407820 | 55401468 | 55401538 | 55411256 | 55411417 | 50 | [illegible] |
| MXE | 4043 | ENSG00000171497 | PPID | chr4 | - | 158713118 | 158713260 | 158715296 | 158715403 | 158710761 | 158710848 | 158715561 | 158715668 | 63 | [illegible] |
| MXE | 17101 | ENSG00000197442 | MAP3K5 | chr6 | - | 136642519 | 136642569 | 136650983 | 136651091 | 136639542 | 136639638 | 136656306 | 136656460 | 82 | [illegible] |
| MXE | 6034 | ENSG00000090989 | EXOC1 | chr4 | + | 55892634 | 55892711 | 55893551 | 55893780 | 55891314 | 55891422 | 55896716 | 55896900 | 43 | [illegible] |
| MXE | 12570 | ENSG00000121716 | PILRB | chr7 | + | 100353372 | 100353483 | 100353894 | 100354012 | 100353209 | 100353270 | 100355142 | 100355240 | 22 | [illegible] |
| MXE | 3549 | ENSG00000124782 | RREB1 | chr6 | + | 7228996 | 7231907 | 7246423 | 7247221 | 7226466 | 7226656 | 7248510 | 7249274 | 82 | [illegible] |
| MXE | 8311 | ENSG00000131791 | PRKAB2 | chr1 | - | 147166845 | 147166939 | 147167766 | 147167933 | 147166497 | 147166618 | 147171988 | 147172167 | 12 | [illegible] |
| MXE | 7421 | ENSG00000153922 | CHD1 | chr5 | - | 98881974 | 98882123 | 98883087 | 98883237 | 98881246 | 98881375 | 98885577 | 98885649 | 288 | [illegible] |
| MXE | 12400 | ENSG00000118058 | KMT2A | chr11 | + | 118507528 | 118507609 | 118509135 | 118509200 | 118502397 | 118506646 | 118509947 | 118510118 | 92 | [illegible] |
| MXE | 14092 | ENSG00000066855 | MTFR1 | chr8 | + | 65682352 | 65682451 | 65693643 | 65693759 | 65669872 | 65670018 | 65704693 | 65704929 | 13 | [illegible] |
| MXE | 5226 | ENSG00000187514 | PTMA | chr2 | + | 231711886 | 231711983 | 231712442 | 231712516 | 231711347 | 231711419 | 231712803 | 231713536 | 103 | [illegible] |
| MXE | 188 | ENSG00000100150 | DEPDC5 | chr22 | + | 31764974 | 31765060 | 31766584 | 31766668 | 31760656 | 31760702 | 31768813 | 31768863 | 21 | [illegible] |
| MXE | 9840 | ENSG00000166478 | ZNF143 | chr11 | + | 9474552 | 9474633 | 9478389 | 9478586 | 9473940 | 9474024 | 9479471 | 9479546 | 16 | [illegible] |
| MXE | 11695 | ENSG00000169826 | CSGALNACT2 | chr10 | + | 43160493 | 43160595 | 43167003 | 43167098 | 43158714 | 43158931 | 43175950 | 43176032 | 140 | [illegible] |
| MXE | 8032 | ENSG00000160691 | SHC1 | chr1 | - | 154969377 | 154969448 | 154970031 | 154970199 | 154968770 | 154968834 | 154974247 | 154974336 | 45 | [illegible] |
| MXE | 13454 | ENSG00000118922 | KLF12 | chr13 | - | 73715367 | 73715525 | 73764937 | 73765000 | 73686088 | 73695671 | 73813151 | 73813287 | 182 | [illegible] |
| MXE | 13832 | ENSG00000145041 | DCAF1 | chr3 | - | 51403142 | 51403395 | 51412378 | 51412480 | 51397841 | 51398827 | 51412992 | 51413066 | 124 | [illegible] |
| MXE | 3837 | ENSG00000128872 | TMOD2 | chr15 | + | 51768261 | 51768418 | 51773711 | 51773834 | 51766372 | 51766567 | 51776931 | 51777018 | 31 | [illegible] |
| MXE | 10584 | ENSG00000143799 | PARP1 | chr1 | - | 226381067 | 226381208 | 226383102 | 226383183 | 226379921 | 226380164 | 226385503 | 226385680 | 297 | [illegible] |
| MXE | 12984 | ENSG00000181513 | ACBD4 | chr17 | + | 45137018 | 45137139 | 45137367 | 45137454 | 45136691 | 45136776 | 45137759 | 45137830 | 21 | [illegible] |
| MXE | 248 | ENSG00000198089 | SFI1 | chr22 | + | 31528689 | 31528863 | 31550253 | 31550348 | 31508254 | 31508376 | 31556941 | 31557059 | 105 | [illegible] |
| MXE | 14721 | ENSG00000148337 | CIZ1 | chr9 | - | 128178708 | 128178904 | 128179072 | 128179415 | 128178368 | 128178490 | 128180414 | 128180523 | 91 | [illegible] |
| MXE | 6664 | ENSG00000054118 | THRAP3 | chr1 | + | 36259381 | 36259484 | 36282532 | 36282700 | 36225411 | 36225463 | 36286367 | 36287270 | 8 | [illegible] |
| MXE | 12853 | ENSG00000123066 | MED13L | chr12 | - | 115980749 | 115980938 | 115982383 | 115982603 | 115975514 | 115975738 | 115983116 | 115983540 | 83 | [illegible] |
| MXE | 13765 | ENSG00000242574 | HLA-DMB | chr6 | - | 32935535 | 32935652 | 32937171 | 32937456 | 32934628 | 32934987 | 32938683 | 32938688 | 223 | [illegible] |
| MXE | 13648 | ENSG00000086758 | HUWE1 | chrX | - | 53563745 | 53563821 | 53564573 | 53564722 | 53562830 | 53562929 | 53565066 | 53565239 | 491 | [illegible] |
| MXE | 6082 | ENSG00000164253 | WDR41 | chr5 | - | 77436260 | 77436394 | 77438239 | 77438361 | 77430932 | 77433287 | 77440812 | 77440997 | 83 | [illegible] |
| MXE | 11418 | ENSG00000138380 | CARF | chr2 | + | 202974333 | 202974496 | 202977268 | 202977332 | 202971504 | 202971738 | 202981554 | 202981685 | 68 | [illegible] |
| MXE | 7868 | ENSG00000042429 | MED17 | chr11 | + | 93807517 | 93807635 | 93809716 | 93809876 | 93801834 | 93801972 | 93811852 | 93812378 | 73 | [illegible] |
| MXE | 837 | ENSG00000101310 | SEC23B | chr20 | + | 18542295 | 18542402 | 18543018 | 18543172 | 18535652 | 18535742 | 18545955 | 18546033 | 160 | [illegible] |
| MXE | 1705 | ENSG00000103051 | COG4 | chr16 | - | 70481358 | 70481487 | 70481763 | 70481865 | 70480567 | 70481144 | 70482091 | 70482175 | 31 | [illegible] |
| MXE | 9275 | ENSG00000175054 | ATR | chr3 | - | 142497012 | 142497192 | 142498596 | 142498774 | 142496360 | 142496520 | 142499626 | 142499718 | 99 | [illegible] |
| MXE | 11841 | ENSG00000136861 | CDK5RAP2 | chr9 | - | 120453455 | 120453873 | 120467859 | 120467997 | 120447894 | 120448126 | 120477349 | 120477450 | 0 | [illegible] |
| MXE | 2855 | ENSG00000100714 | MTHFD1 | chr14 | + | 64417887 | 64418024 | 64424803 | 64424931 | 64415638 | 64415739 | 64425729 | 64425827 | 32 | [illegible] |
| MXE | 11371 | ENSG00000115970 | THADA | chr2 | - | 43556344 | 43556555 | 43560233 | 43560385 | 43552203 | 43552339 | 43566697 | 43566821 | 48 | [illegible] |
| MXE | 1785 | ENSG00000007541 | PIGQ | chr16 | + | 580182 | 580263 | 580857 | 580972 | 579068 | 579180 | 582247 | 582309 | 7 | [illegible] |
| MXE | 7741 | ENSG00000156515 | HK1 | chr10 | + | 69376933 | 69377089 | 69379861 | 69380095 | 69369440 | 69369624 | 69382486 | 69382791 | 253 | [illegible] |
| MXE | 283 | ENSG00000100325 | ASCC2 | chr22 | - | 29800990 | 29801110 | 29801993 | 29802208 | 29793359 | 29793676 | 29804637 | 29804824 | 70 | [illegible] |
| MXE | 7897 | ENSG00000074603 | DPP8 | chr15 | - | 65490188 | 65490299 | 65497863 | 65498032 | 65487689 | 65487818 | 65500605 | 65500779 | 189 | [illegible] |
| MXE | 14185 | ENSG00000173064 | HECTD4 | chr12 | - | 112270226 | 112270459 | 112273654 | 112273795 | 112269703 | 112269849 | 112274846 | 112274960 | 36 | [illegible] |
| MXE | 5847 | ENSG00000130024 | PHF10 | chr6 | - | 169717822 | 169717900 | 169718787 | 169718918 | 169715954 | 169716088 | 169721004 | 169721111 | 121 | [illegible] |
| MXE | 6188 | ENSG00000213390 | ARHGAP19 | chr10 | - | 97246271 | 97246337 | 97259401 | 97259628 | 97243967 | 97244159 | 97263419 | 97263629 | 77 | [illegible] |
| MXE | 13446 | ENSG00000075539 | FRYL | chr4 | - | 48540352 | 48540960 | 48542026 | 48542121 | 48539970 | 48540068 | 48543806 | 48543997 | 83 | [illegible] |
| MXE | 8831 | ENSG00000118007 | STAG1 | chr3 | - | 136623145 | 136623248 | 136630869 | 136630981 | 136604308 | 136604473 | 136752194 | 136752366 | 113 | [illegible] |
| MXE | 5244 | ENSG00000198034 | RPS4X | chrX | - | 72273231 | 72273389 | 72273800 | 72273972 | 72272041 | 72272772 | 72275052 | 72275150 | 7810 | [illegible] |
| MXE | 11726 | ENSG00000204568 | MRPS18B | chr6 | + | 30622831 | 30622898 | 30624882 | 30624942 | 30619492 | 30619989 | 30625501 | 30626395 | 39 | [illegible] |
| MXE | 11938 | ENSG00000146433 | TMEM181 | chr6 | + | 158580939 | 158580995 | 158583953 | 158584044 | 158573419 | 158573523 | 158585303 | 158585425 | 9 | [illegible] |
| MXE | 8736 | ENSG00000143119 | CD53 | chr1 | + | 110894959 | 110895055 | 110896652 | 110896733 | 110894326 | 110894401 | 110897808 | 110897892 | 596 | [illegible] |
| MXE | 13734 | ENSG00000147050 | KDM6A | chrX | + | 45078399 | 45078505 | 45079145 | 45079351 | 45076696 | 45076826 | 45082575 | 45082640 | 104 | [illegible] |
| MXE | 16583 | ENSG00000127314 | RAP1B | chr12 | + | 68624670 | 68624873 | 68641033 | 68641153 | 68610907 | 68611043 | 68648698 | 68648781 | 35 | [illegible] |
| MXE | 15452 | ENSG00000103544 | VPS35L | chr16 | + | 19640014 | 19640100 | 19642395 | 19642476 | 19637593 | 19637656 | 19644885 | 19644949 | 32 | [illegible] |
| MXE | 1824 | ENSG00000118900 | UBN1 | chr16 | + | 4859864 | 4859968 | 4860663 | 4861102 | 4859024 | 4859159 | 4868832 | 4868903 | 191 | [illegible] |
| MXE | 163 | ENSG00000100387 | RBX1 | chr22 | + | 40953554 | 40953633 | 40964046 | 40964117 | 40951377 | 40951476 | 40967798 | 40967884 | 185 | [illegible] |
| MXE | 11198 | ENSG00000112799 | LY86 | chr6 | + | 6626292 | 6626421 | 6649624 | 6649677 | 6624925 | 6625012 | 6654543 | 6654982 | 158 | [illegible] |
| MXE | 9358 | ENSG00000115211 | EIF2B4 | chr2 | - | 27367073 | 27367201 | 27367456 | 27367559 | 27366758 | 27366936 | 27367745 | 27367802 | 27 | [illegible] |
| MXE | 14852 | ENSG00000128973 | CLN6 | chr15 | - | 68211555 | 68211643 | 68211648 | 68211863 | 68211262 | 68211318 | 68214289 | 68214388 | 16 | [illegible] |
| MXE | 14615 | ENSG00000165121 | NA | chr9 | + | 85822879 | 85823080 | 85829462 | 85829606 | 85815965 | 85816263 | 85837758 | 85837965 | 37 | [illegible] |
| MXE | 16335 | ENSG00000096717 | SIRT1 | chr10 | + | 67888881 | 67889123 | 67891401 | 67891554 | 67887416 | 67887533 | 67906789 | 67906937 | 108 | [illegible] |
| MXE | 3112 | ENSG00000055208 | TAB2 | chr6 | + | 149397968 | 149398062 | 149399103 | 149399184 | 149397603 | 149397764 | 149409576 | 149411395 | 206 | [illegible] |
| MXE | 13704 | ENSG00000134982 | APC | chr5 | + | 112821895 | 112821991 | 112827107 | 112827247 | 112818965 | 112819344 | 112827928 | 112828006 | 57 | [illegible] |
| MXE | 16552 | ENSG00000061987 | MON2 | chr12 | + | 62493914 | 62494042 | 62495015 | 62495147 | 62484169 | 62484233 | 62498918 | 62499048 | 78 | [illegible] |
| MXE | 2354 | ENSG00000137764 | MAP2K5 | chr15 | + | 67703336 | 67703408 | 67727915 | 67727945 | 67693517 | 67693568 | 67748230 | 67748257 | 20 | [illegible] |
| MXE | 7648 | ENSG00000060339 | CCAR1 | chr10 | + | 68737844 | 68737889 | 68740628 | 68740661 | 68736875 | 68737048 | 68742375 | 68742569 | 21 | [illegible] |
| MXE | 8571 | ENSG00000094975 | SUCO | chr1 | + | 172575517 | 172575623 | 172577538 | 172577559 | 172573890 | 172573998 | 172577763 | 172577819 | 77 | [illegible] |
| MXE | 10982 | ENSG00000064201 | TSPAN32 | chr11 | + | 2304106 | 2304204 | 2308735 | 2308810 | 2302843 | 2302958 | 2313653 | 2313755 | 53 | [illegible] |
| MXE | 3798 | ENSG00000126456 | IRF3 | chr19 | - | 49664442 | 49664673 | 49664673 | 49664846 | 49663342 | 49663514 | 49665630 | 49665705 | 113 | [illegible] |
| MXE | 15066 | ENSG00000110514 | MADD | chr11 | + | 47290609 | 47290816 | 47293882 | 47293983 | 47290148 | 47290299 | 47295495 | 47295567 | 53 | [illegible] |
| MXE | 1783 | ENSG00000157106 | SMG1 | chr16 | - | 18885540 | 18885666 | 18889371 | 18889585 | 18885089 | 18885162 | 18890862 | 18890921 | 152 | [illegible] |
| MXE | 7624 | ENSG00000132300 | PTCD3 | chr2 | + | 86127940 | 86127991 | 86130647 | 86130737 | 86127160 | 86127305 | 86131077 | 86131106 | 29 | [illegible] |
| MXE | 14814 | ENSG00000011454 | RABGAP1 | chr9 | + | 123071411 | 123071501 | 123074284 | 123074428 | 123070349 | 123070424 | 123076244 | 123076286 | 0 | [illegible] |
| MXE | 15662 | ENSG00000081237 | PTPRC | chr1 | + | 198699563 | 198699704 | 198703297 | 198703372 | 198692346 | 198692373 | 198704471 | 198704498 | 994 | [illegible] |
| MXE | 11796 | ENSG00000204305 | AGER | chr6 | - | 32182246 | 32182388 | 32182567 | 32182698 | 32181523 | 32181632 | 32182840 | 32183023 | 6 | [illegible] |
| MXE | 7880 | ENSG00000074603 | DPP8 | chr15 | - | 65450998 | 65451110 | 65451959 | 65452102 | 65442462 | 65447006 | 65454262 | 65454415 | 90 | [illegible] |
| MXE | 10672 | ENSG00000144029 | MRPS5 | chr2 | - | 95100836 | 95100894 | 95101676 | 95101723 | 95100473 | 95100536 | 95104639 | 95104730 | 43 | [illegible] |
| MXE | 11765 | ENSG00000137200 | CMTR1 | chr6 | + | 37471020 | 37471077 | 37471846 | 37471904 | 37462828 | 37463008 | 37472418 | 37472487 | 47 | [illegible] |
| MXE | 8238 | ENSG00000143643 | TTC13 | chr1 | - | 230940439 | 230940556 | 230945388 | 230945454 | 230933778 | 230933861 | 230954332 | 230954403 | 104 | [illegible] |
| MXE | 16083 | ENSG00000197535 | MYO5A | chr15 | - | 52416144 | 52416301 | 52425829 | 52425974 | 52410332 | 52410476 | 52428397 | 52428569 | 89 | [illegible] |
| MXE | 4717 | ENSG00000155903 | RASA2 | chr3 | + | 141553856 | 141553940 | 141555839 | 141555912 | 141540532 | 141540609 | 141558885 | 141558962 | 214 | [illegible] |
| MXE | 2535 | ENSG00000165929 | TC2N | chr14 | - | 91792366 | 91792558 | 91798298 | 91798399 | 91787512 | 91787627 | 91798988 | 91798998 | 202 | [illegible] |
| MXE | 15312 | ENSG00000135249 | RINT1 | chr7 | + | 105536564 | 105536612 | 105536657 | 105536749 | 105532823 | 105532869 | 105542407 | 105542649 | 18 | [illegible] |
| MXE | 14478 | ENSG00000180776 | ZDHHC20 | chr13 | - | 21400372 | 21400493 | 21401652 | 21401685 | 21391721 | 21391854 | 21402796 | 21402866 | 45 | [illegible] |
| MXE | 7607 | ENSG00000053254 | FOXN3 | chr14 | - | 89280949 | 89281014 | 89350671 | 89350808 | 89190449 | 89190490 | 89411933 | 89412490 | 289 | [illegible] |
| MXE | 6745 | ENSG00000076321 | KLHL20 | chr1 | + | 173753212 | 173753307 | 173755922 | 173756038 | 173751763 | 173751922 | 173756975 | 173757159 | 46 | [illegible] |
| MXE | 14003 | ENSG00000095485 | CWF19L1 | chr10 | - | 100256261 | 100256476 | 100260965 | 100261044 | 100253420 | 100253539 | 100261978 | 100262063 | 50 | [illegible] |
| MXE | 336 | ENSG00000099331 | MYO9B | chr19 | + | 17197791 | 17197858 | 17200292 | 17200426 | 17194555 | 17195473 | 17200638 | 17200829 | 23 | [illegible] |
| MXE | 13536 | ENSG00000135048 | CEMIP2 | chr9 | - | 71740062 | 71740232 | 71745017 | 71745579 | 71734805 | 71734994 | 71746200 | 71746341 | 262 | [illegible] |
| MXE | 16872 | ENSG00000050426 | LETMD1 | chr12 | + | 51052091 | 51052207 | 51055834 | 51056021 | 51049033 | 51049185 | 51056143 | 51056237 | 36 | [illegible] |
| MXE | 8338 | ENSG00000138642 | HERC6 | chr4 | + | 88417424 | 88417579 | 88423859 | 88423973 | 88413076 | 88413266 | 88428579 | 88428750 | 35 | [illegible] |
| MXE | 12415 | ENSG00000198198 | SZT2 | chr1 | + | 43426367 | 43426538 | 43427055 | 43427179 | 43426037 | 43426151 | 43427280 | 43427445 | 21 | [illegible] |
| MXE | 16550 | ENSG00000111142 | METAP2 | chr12 | + | 95494956 | 95495138 | 95496003 | 95496098 | 95494055 | 95494217 | 95504064 | 95504161 | 133 | [illegible] |
| MXE | 13027 | ENSG00000055130 | CUL1 | chr7 | + | 148759547 | 148759638 | 148760332 | 148760496 | 148759303 | 148759354 | 148766560 | 148766723 | 104 | [illegible] |
| MXE | 6880 | ENSG00000100991 | TRPC4AP | chr20 | - | 35006434 | 35006575 | 35007549 | 35007640 | 35005694 | 35005803 | 35008663 | 35008747 | 54 | [illegible] |
| MXE | 14728 | ENSG00000136811 | ODF2 | chr9 | + | 128469182 | 128469353 | 128471307 | 128471468 | 128460941 | 128461067 | 128472912 | 128473042 | 61 | [illegible] |
| MXE | 14141 | ENSG00000123908 | AGO2 | chr8 | - | 140558484 | 140558572 | 140559394 | 140559529 | 140557088 | 140557236 | 140560373 | 140560510 | 75 | [illegible] |
| MXE | 7555 | ENSG00000263956 | NBPF11 | chr1 | - | 148127140 | 148127288 | 148135123 | 148135247 | 148126828 | 148127038 | 148135671 | 148135813 | 0 | [illegible] |
| MXE | 8358 | ENSG00000030419 | IKZF2 | chr2 | - | 213013790 | 213013934 | 213021992 | 213022130 | 212999690 | 213008084 | 213049712 | 213049880 | 77 | [illegible] |
| MXE | 11043 | ENSG00000079263 | SP140 | chr2 | + | 230245862 | 230245940 | 230248884 | 230248968 | 230244987 | 230245071 | 230255451 | 230255532 | 15 | [illegible] |
| MXE | 3843 | ENSG00000179152 | TCAIM | chr3 | + | 44392854 | 44392977 | 44396399 | 44396497 | 44367455 | 44367708 | 44396742 | 44396834 | 20 | [illegible] |
| MXE | 12765 | ENSG00000089006 | SNX5 | chr20 | - | 17950290 | 17950396 | 17951499 | 17951595 | 17950131 | 17950207 | 17952586 | 17952710 | 247 | [illegible] |
| MXE | 3459 | ENSG00000261556 | SMG1P7 | chr16 | - | 70235312 | 70235443 | 70241035 | 70241260 | 70234996 | 70235158 | 70243626 | 70243799 | 5 | [illegible] |
| MXE | 16986 | ENSG00000115310 | RTN4 | chr2 | - | 54973820 | 54973867 | 54974694 | 54974764 | 54973512 | 54973621 | 54982514 | 54982653 | 237 | [illegible] |
| MXE | 11884 | ENSG00000204227 | RING1 | chr6 | + | 33209914 | 33210130 | 33211157 | 33211547 | 33209559 | 33209786 | 33211728 | 33212002 | 66 | [illegible] |
| MXE | 9661 | ENSG00000120265 | PCMT1 | chr6 | + | 149771161 | 149771266 | 149773137 | 149773169 | 149749694 | 149749956 | 149789953 | 149790058 | 69 | [illegible] |
| MXE | 7341 | ENSG00000229474 | PATL2 | chr15 | - | 44710095 | 44710214 | 44710870 | 44710929 | 44676474 | 44676565 | 44711019 | 44711323 | 16 | [illegible] |
| MXE | 13201 | ENSG00000147202 | DIAPH2 | chrX | + | 96758153 | 96758258 | 96763099 | 96763120 | 96738585 | 96738762 | 96881578 | 96881718 | 110 | [illegible] |
| MXE | 8367 | ENSG00000117859 | OSBPL9 | chr1 | + | 51750144 | 51750195 | 51756319 | 51756358 | 51748368 | 51748398 | 51760689 | 51760780 | 64 | [illegible] |
| MXE | 12343 | ENSG00000139641 | ESYT1 | chr12 | + | 56138926 | 56139013 | 56142284 | 56142425 | 56138767 | 56138839 | 56142577 | 56142732 | 160 | [illegible] |
| MXE | 9281 | ENSG00000175054 | ATR | chr3 | - | 142562231 | 142563109 | 142566120 | 142566261 | 142561242 | 142561421 | 142568062 | 142568154 | 54 | [illegible] |
| MXE | 669 | ENSG00000104885 | DOT1L | chr19 | + | 2191083 | 2191240 | 2193688 | 2193783 | 2189731 | 2189795 | 2194514 | 2194577 | 0 | [illegible] |
| MXE | 1803 | ENSG00000122390 | NAA60 | chr16 | + | 3448470 | 3448540 | 3479470 | 3479600 | 3443705 | 3443827 | 3482501 | 3482598 | 2 | [illegible] |
| MXE | 6233 | ENSG00000156136 | DCK | chr4 | + | 71022366 | 71022560 | 71023558 | 71023706 | 70998066 | 70998182 | 71025815 | 71025931 | 225 | [illegible] |
| MXE | 1931 | ENSG00000103426 | CORO7-PAM16 | chr16 | - | 4412355 | 4412430 | 4413307 | 4413404 | 4408180 | 4408251 | 4416458 | 4416587 | 36 | [illegible] |
| MXE | 2164 | ENSG00000138592 | USP8 | chr15 | + | 50462279 | 50462322 | 50465046 | 50465191 | 50458999 | 50459162 | 50471632 | 50471735 | 58 | [illegible] |
| MXE | 3965 | ENSG00000083312 | TNPO1 | chr5 | + | 72893615 | 72893703 | 72896457 | 72896556 | 72893376 | 72893535 | 72897055 | 72897151 | 83 | [illegible] |
| MXE | 5863 | ENSG00000073417 | PDE8A | chr15 | + | 85089337 | 85089416 | 85091043 | 85091181 | 85083555 | 85083644 | 85097947 | 85098036 | 25 | [illegible] |
| MXE | 1163 | ENSG00000078687 | TNRC6C | chr17 | + | 78091439 | 78091607 | 78092932 | 78093124 | 78086852 | 78087093 | 78093619 | 78093763 | 16 | [illegible] |
| MXE | 6560 | ENSG00000135597 | REPS1 | chr6 | - | 138915857 | 138915976 | 138917554 | 138917624 | 138914696 | 138914761 | 138920214 | 138920316 | 8 | [illegible] |
| MXE | 2562 | ENSG00000197249 | SERPINA1 | chr14 | - | 94382591 | 94383241 | 94388559 | 94388663 | 94380870 | 94381141 | 94388818 | 94389010 | 24 | [illegible] |
| MXE | 9620 | ENSG00000119778 | ATAD2B | chr2 | - | 23857414 | 23857503 | 23864808 | 23864924 | 23833918 | 23834078 | 23867834 | 23867946 | 79 | [illegible] |
| MXE | 6436 | ENSG00000163946 | TASOR | chr3 | - | 56662384 | 56662490 | 56668396 | 56668558 | 56660913 | 56661017 | 56669699 | 56669791 | 101 | [illegible] |
| MXE | 9557 | ENSG00000196975 | ANXA4 | chr2 | + | 69788053 | 69788141 | 69803392 | 69803515 | 69781519 | 69781574 | 69804532 | 69804627 | 62 | [illegible] |
| MXE | 6623 | ENSG00000148019 | CEP78 | chr9 | + | 78254835 | 78254964 | 78262906 | 78262984 | 78253231 | 78253277 | 78264149 | 78264316 | 71 | [illegible] |
| MXE | 6998 | ENSG00000126070 | AGO3 | chr1 | + | 36034173 | 36034333 | 36036176 | 36036267 | 36027113 | 36027298 | 36039789 | 36039984 | 44 | [illegible] |
| MXE | 10833 | ENSG00000108883 | EFTUD2 | chr17 | - | 44857074 | 44857157 | 44859079 | 44859181 | 44854917 | 44855004 | 44859904 | 44860045 | 112 | [illegible] |
| MXE | 4758 | ENSG00000047849 | MAP4 | chr3 | - | 47915950 | 47917174 | 47918718 | 47918841 | 47914816 | 47914939 | 47921764 | 47921878 | 158 | [illegible] |
| MXE | 6272 | ENSG00000130826 | DKC1 | chrX | + | 154765443 | 154765530 | 154765906 | 154765998 | 154764898 | 154764966 | 154766215 | 154766400 | 25 | [illegible] |
| MXE | 9309 | ENSG00000135924 | DNAJB2 | chr2 | + | 219282836 | 219282929 | 219283132 | 219283235 | 219281938 | 219282061 | 219283418 | 219283489 | 46 | [illegible] |
| MXE | 12735 | ENSG00000253729 | PRKDC | chr8 | - | 47885943 | 47886147 | 47887546 | 47887705 | 47881911 | 47882097 | 47888517 | 47888650 | 78 | [illegible] |
| MXE | 8606 | ENSG00000106665 | CLIP2 | chr7 | + | 74353879 | 74354004 | 74357279 | 74357477 | 74338447 | 74339004 | 74360174 | 74360278 | 8 | [illegible] |
| MXE | 16047 | ENSG00000107938 | EDRF1 | chr10 | + | 125749411 | 125749565 | 125753693 | 125753845 | 125747862 | 125748012 | 125763300 | 125763524 | 25 | [illegible] |
| MXE | 7969 | ENSG00000048707 | VPS13D | chr1 | + | 12416659 | 12416827 | 12455997 | 12456130 | 12415086 | 12415221 | 12460200 | 12460396 | 131 | [illegible] |
| MXE | 15947 | ENSG00000156650 | KAT6B | chr10 | + | 74981786 | 74981928 | 74985079 | 74985241 | 74979223 | 74979339 | 74989018 | 74989112 | 75 | [illegible] |
| MXE | 9278 | ENSG00000175054 | ATR | chr3 | - | 142538481 | 142538625 | 142540903 | 142541034 | 142536107 | 142536201 | 142542664 | 142542757 | 76 | [illegible] |
| MXE | 10801 | ENSG00000163803 | PLB1 | chr2 | + | 28604654 | 28604759 | 28605852 | 28605948 | 28603965 | 28604047 | 28606495 | 28606567 | 5 | [illegible] |
| MXE | 7296 | ENSG00000266028 | SRGAP2 | chr1 | + | 206454877 | 206455024 | 206458622 | 206458947 | 206453198 | 206453380 | 206461036 | 206461227 | 21 | [illegible] |
| MXE | 2802 | ENSG00000151445 | VIPAS39 | chr14 | - | 77437807 | 77437881 | 77441065 | 77441093 | 77435843 | 77435919 | 77442559 | 77442662 | 9 | [illegible] |
| MXE | 11097 | ENSG00000184182 | UBE2F | chr2 | + | 237987962 | 237987992 | 238016565 | 238016633 | 237973091 | 237973225 | 238025341 | 238025412 | 27 | [illegible] |
| MXE | 15193 | ENSG00000156110 | ADK | chr10 | + | 74589281 | 74589317 | 74600378 | 74600493 | 74525255 | 74525426 | 74670182 | 74670269 | 25 | [illegible] |
| MXE | 1085 | ENSG00000126561 | STAT5A | chr17 | + | 42289401 | 42289539 | 42291971 | 42292061 | 42288055 | 42288598 | 42295618 | 42295793 | 22 | [illegible] |
| MXE | 13744 | ENSG00000117676 | RPS6KA1 | chr1 | + | 26555536 | 26555625 | 26556653 | 26556718 | 26555150 | 26555221 | 26556997 | 26557100 | 112 | [illegible] |
| MXE | 9852 | ENSG00000206562 | METTL6 | chr3 | - | 15414020 | 15414162 | 15415771 | 15415942 | 15409759 | 15411437 | 15424954 | 15425089 | 60 | [illegible] |
| MXE | 12915 | ENSG00000164815 | ORC5 | chr7 | - | 104136780 | 104136893 | 104161071 | 104161182 | 104126340 | 104126893 | 104165234 | 104165282 | 50 | [illegible] |
| MXE | 65 | ENSG00000142207 | URB1 | chr21 | - | 32352709 | 32352906 | 32353932 | 32354103 | 32350703 | 32350922 | 32354858 | 32354997 | 16 | [illegible] |
| MXE | 647 | ENSG00000142347 | MYO1F | chr19 | - | 8545649 | 8545736 | 8548035 | 8548122 | 8544296 | 8544464 | 8548236 | 8548317 | 234 | [illegible] |

| | | | | | | | | | | | | | | | | | | | | |
|---|---|---|---|---|---|---|---|---|---|---|---|---|---|---|---|---|---|---|---|---|
| MXE | 6531 | ENSG00000101474 | APMAP | chr20 | - | 24978766 | 24978882 | 24983902 | 24984019 | [illegible] | [illegible] | [illegible] | [illegible] | [illegible] | [illegible] | [illegible] | [illegible] | [illegible] | [illegible] | [illegible] |
| MXE | 6765 | ENSG00000135999 | EPC2 | chr2 | + | 148753926 | 148754133 | 148761781 | 148761930 | [illegible] | [illegible] | [illegible] | [illegible] | [illegible] | [illegible] | [illegible] | [illegible] | [illegible] | [illegible] | [illegible] |
| MXE | 12193 | ENSG00000088930 | XRN2 | chr20 | + | 21356585 | 21356665 | 21357735 | 21357792 | [illegible] | [illegible] | [illegible] | [illegible] | [illegible] | [illegible] | [illegible] | [illegible] | [illegible] | [illegible] | [illegible] |
| MXE | 7847 | ENSG00000204217 | BMPR2 | chr2 | + | 202518821 | 202519052 | 202520086 | 202520201 | 202514887 | 202514979 | 202530793 | 202530954 | 62 | 31 | 53 | 61 | 71 | 100 | 105 |
| MXE | 12326 | ENSG00000105778 | AVL9 | chr7 | + | 32558928 | 32559464 | 32570019 | 32570154 | 32558558 | 32558628 | 32573198 | 32573418 | 105 | 69 | 117 | 118 | 119 | 85 | 83 |
| MXE | 8673 | ENSG00000125686 | MED1 | chr17 | - | 39431114 | 39431188 | 39431941 | 39432016 | 39427700 | 39427790 | 39434248 | 39434320 | 25 | 19 | 29 | 28 | 48 | 31 | 27 |
| MXE | 12450 | ENSG00000168137 | SETD5 | chr3 | + | 9429826 | 9430051 | 9433844 | 9433950 | 9428822 | 9429009 | 9434276 | 9434485 | 25 | 21 | 22 | 33 | 26 | 57 | 20 |
| MXE | 15075 | ENSG00000080546 | SESN1 | chr6 | - | 108998512 | 108998755 | 109000490 | 109000673 | 108994461 | 108994609 | 109001287 | 109001488 | 102 | 67 | 103 | 80 | 206 | 97 | 71 |
| MXE | 551 | ENSG00000157540 | DYRK1A | chr21 | + | 37425689 | 37425781 | 37428866 | 37428908 | 37421773 | 37421866 | 37430268 | 37430400 | 30 | 24 | 17 | 34 | 55 | 12 | 0 |
| MXE | 13442 | ENSG00000075539 | FRYL | chr4 | - | 48500029 | 48500220 | 48501622 | 48501733 | 48497540 | 48499680 | 48505546 | 48505615 | 73 | 51 | 70 | 67 | 62 | 223 | 173 |
| MXE | 2299 | ENSG00000140400 | MAN2C1 | chr15 | - | 75363209 | 75363290 | 75363998 | 75364188 | 75362641 | 75362748 | 75364487 | 75364665 | 128 | 53 | 154 | 132 | 58 | 4 | 1 |
| MXE | 13537 | ENSG00000135048 | CEMIP2 | chr9 | - | 71745017 | 71745579 | 71746200 | 71746341 | 71740062 | 71740232 | 71750042 | 71750385 | 287 | 157 | 124 | 160 | 215 | 197 | 89 |
| MXE | 10431 | ENSG00000106636 | YKT6 | chr7 | + | 44204567 | 44204650 | 44206384 | 44206485 | 44200958 | 44201239 | 44207387 | 44207492 | 31 | 8 | 19 | 18 | 5 | 104 | 77 |
| MXE | 14280 | ENSG00000167632 | TRAPPC9 | chr8 | - | 140426614 | 140426641 | 140435111 | 140435240 | 140405576 | 140405698 | 140439051 | 140439197 | 86 | 40 | 72 | 105 | 50 | 12 | 15 |
| MXE | 6919 | ENSG00000109323 | MANBA | chr4 | - | 102690879 | 102690957 | 102714437 | 102714561 | 102690595 | 102690771 | 102722870 | 102723041 | 176 | 93 | 153 | 180 | 241 | 44 | 36 |
| MXE | 6125 | ENSG00000170581 | STAT2 | chr12 | - | 56348751 | 56348851 | 56348923 | 56349059 | 56348528 | 56348623 | 56349162 | 56349261 | 42 | 25 | 34 | 57 | 34 | 39 | 29 |
| MXE | 13686 | ENSG00000215301 | DDX3X | chrX | + | 41346858 | 41347012 | 41347314 | 41347451 | 41346228 | 41346622 | 41347639 | 41347694 | 1022 | 660 | 846 | 1038 | 1493 | 424 | 284 |
| MXE | 14916 | ENSG00000119383 | PTPA | chr9 | + | 129123051 | 129123138 | 129128984 | 129129110 | 129120512 | 129120610 | 129131521 | 129131639 | 37 | 36 | 46 | 47 | 58 | 85 | 32 |
| MXE | 555 | ENSG00000105383 | CD33 | chr19 | + | 51235156 | 51235253 | 51235594 | 51235676 | 51226308 | 51226356 | 51239517 | 51239942 | 48 | 20 | 4 | 27 | 12 | 29 | 14 |
| MXE | 5827 | ENSG00000132549 | VPS13B | chr8 | + | 99868288 | 99868465 | 99870784 | 99870887 | 99861775 | 99861946 | 99871447 | 99871697 | 26 | 11 | 26 | 19 | 17 | 11 | 6 |
| MXE | 10868 | ENSG00000105953 | OGDH | chr7 | + | 44645326 | 44645518 | 44647656 | 44647759 | 44624316 | 44624565 | 44666735 | 44666851 | 344 | 246 | 252 | 324 | 418 | 200 | 87 |
| MXE | 13635 | ENSG00000104613 | INTS10 | chr8 | + | 19819572 | 19819676 | 19820378 | 19820518 | 19818274 | 19818342 | 19822438 | 19822520 | 33 | 9 | 21 | 22 | 12 | 181 | 98 |
| MXE | 6012 | ENSG00000137393 | RNF144B | chr6 | + | 18439683 | 18439744 | 18459606 | 18459751 | 18427580 | 18427685 | 18463290 | 18463380 | 12 | 19 | 10 | 7 | 27 | 44 | 99 |
| MXE | 6535 | ENSG00000239900 | ADSL | chr22 | + | 40353072 | 40353117 | 40354247 | 40354327 | 40349831 | 40350035 | 40358863 | 40359035 | 24 | 11 | 20 | 7 | 19 | 44 | 43 |
| MXE | 16109 | ENSG00000114013 | CD86 | chr3 | + | 122106197 | 122106500 | 122109264 | 122109408 | 122103511 | 122103847 | 122118047 | 122118093 | 212 | 167 | 133 | 172 | 410 | 341 | 196 |
| MXE | 7072 | ENSG00000273189 | NA | chr19 | + | 49478908 | 49479047 | 49480297 | 49480476 | 49476422 | 49476566 | 49480551 | 49480620 | 41 | 27 | 70 | 49 | 29 | 31 | 7 |
| MXE | 3284 | ENSG00000137177 | KIF13A | chr6 | - | 17785514 | 17785641 | 17787775 | 17787875 | 17783645 | 17783701 | 17789871 | 17789910 | 43 | 31 | 22 | 32 | 41 | 10 | 11 |
| MXE | 17053 | ENSG00000081189 | MEF2C | chr5 | - | 88760987 | 88761125 | 88761184 | 88761328 | 88751856 | 88752043 | 88804597 | 88804801 | 404 | 332 | 360 | 428 | 714 | 20 | 27 |
| MXE | 9346 | ENSG00000077157 | PPP1R12B | chr1 | + | 202416786 | 202416917 | 202422619 | 202422738 | 202348727 | 202349142 | 202425565 | 202425725 | 4 | 14 | 10 | 6 | 1 | 33 | 34 |
| MXE | 6295 | ENSG00000204152 | TIMM23B | chr10 | + | 49952125 | 49952219 | 49955001 | 49955060 | 49945031 | 49945090 | 49958369 | 49958480 | 31 | 31 | 42 | 40 | 31 | 32 | 13 |
| MXE | 8690 | ENSG00000268043 | NBPF12 | chr1 | + | 146951813 | 146952037 | 146959858 | 146960006 | 146951347 | 146951489 | 146960108 | 146960318 | 2 | 0 | 6 | 0 | 8 | 3 | 7 |
| MXE | 2651 | ENSG00000171723 | GPHN | chr14 | + | 66824473 | 66824566 | 66842677 | 66842716 | 66776463 | 66776521 | 66879938 | 66880033 | 68 | 26 | 48 | 29 | 50 | 3 | 0 |
| MXE | 1236 | ENSG00000092929 | UNC13D | chr17 | - | 75842856 | 75842923 | 75843013 | 75843073 | 75842432 | 75842613 | 75843158 | 75843248 | 35 | 14 | 25 | 18 | 6 | 150 | 75 |
| MXE | 11166 | ENSG00000197930 | ERO1A | chr14 | - | 52646374 | 52646461 | 52652238 | 52652308 | 52646153 | 52646287 | 52653068 | 52653315 | 195 | 190 | 184 | 222 | 291 | 267 | 242 |
| MXE | 5486 | ENSG00000276291 | FRG1HP | chr9 | + | 41008880 | 41009006 | 41011616 | 41011730 | 40994632 | 40994699 | 41032833 | 41033012 | 1 | 13 | 2 | 5 | 10 | 28 | 2 |
| MXE | 16706 | ENSG00000105810 | CDK6 | chr7 | - | 92623035 | 92623086 | 92671425 | 92671535 | 92618071 | 92618207 | 92725625 | 92725793 | 54 | 45 | 57 | 35 | 89 | 14 | 13 |
| MXE | 827 | ENSG00000196756 | SNHG17 | chr20 | - | 38422091 | 38422241 | 38425934 | 38426052 | 38421005 | 38421098 | 38426418 | 38426503 | 0 | 0 | 0 | 3 | 0 | 2 | 5 |
| MXE | 5655 | ENSG00000168028 | RPSA | chr3 | + | 39410753 | 39410999 | 39411648 | 39411777 | 39408203 | 39408724 | 39411895 | 39412061 | 5437 | 2689 | 6417 | 5176 | 11295 | 1409 | 550 |
| MXE | 8822 | ENSG00000001497 | LAS1L | chrX | - | 65531356 | 65531438 | 65532560 | 65532630 | 65529593 | 65529878 | 65533609 | 65533735 | 42 | 27 | 51 | 50 | 55 | 47 | 35 |
| MXE | 15940 | ENSG00000156650 | KAT6B | chr10 | + | 74959969 | 74960078 | 74969659 | 74969775 | 74842599 | 74843478 | 74970019 | 74970101 | 36 | 19 | 32 | 32 | 28 | 66 | 43 |
| MXE | 16164 | ENSG00000107669 | ATE1 | chr10 | - | 121898840 | 121898969 | 121899865 | 121899994 | 121841081 | 121841263 | 121902390 | 121902620 | 52 | 34 | 46 | 50 | 109 | 29 | 42 |
| MXE | 1395 | ENSG00000108406 | DHX40 | chr17 | + | 59573115 | 59573235 | 59573739 | 59573967 | 59570517 | 59570663 | 59574187 | 59574254 | 119 | 58 | 93 | 96 | 163 | 69 | 39 |
| MXE | 1322 | ENSG00000141298 | SSH2 | chr17 | - | 29650653 | 29650800 | 29655560 | 29655607 | 29647693 | 29648344 | 29666866 | 29666995 | 192 | 131 | 101 | 136 | 189 | 512 | 359 |
| MXE | 7682 | ENSG00000139636 | LMBR1L | chr12 | - | 49101249 | 49101323 | 49101450 | 49101549 | 49100555 | 49100646 | 49102119 | 49102196 | 2 | 3 | 10 | 6 | 8 | 3 | 0 |
| MXE | 5239 | ENSG00000115827 | DCAF17 | chr2 | + | 171457970 | 171458075 | 171468887 | 171469030 | 171453123 | 171453213 | 171473865 | 171473975 | 24 | 28 | 22 | 18 | 47 | 43 | 20 |
| MXE | 6305 | ENSG00000138674 | SEC31A | chr4 | - | 82875726 | 82875822 | 82878729 | 82878928 | 82874610 | 82874751 | 82880798 | 82880922 | 326 | 260 | 293 | 348 | 430 | 203 | 136 |
| MXE | 830 | ENSG00000196756 | SNHG17 | chr20 | - | 38422095 | 38422241 | 38425934 | 38426052 | 38421005 | 38421098 | 38426418 | 38426503 | 0 | 0 | 0 | 3 | 0 | 1 | 3 |
| MXE | 11203 | ENSG00000164902 | PHAX | chr5 | + | 126608363 | 126608484 | 126617249 | 126617333 | 126603569 | 126604183 | 126624574 | 126627252 | 143 | 104 | 103 | 125 | 168 | 56 | 59 |
| MXE | 5678 | ENSG00000163681 | SLMAP | chr3 | + | 57864806 | 57864857 | 57865241 | 57865292 | 57864547 | 57864637 | 57871635 | 57871698 | 51 | 54 | 49 | 76 | 81 | 6 | 8 |
| MXE | 5203 | ENSG00000198700 | IPO9 | chr1 | + | 201855782 | 201855934 | 201857095 | 201857194 | 201855123 | 201855182 | 201858446 | 201858553 | 65 | 43 | 48 | 66 | 135 | 103 | 34 |
| MXE | 15856 | ENSG00000088298 | EDEM2 | chr20 | - | 35126250 | 35126375 | 35137879 | 35138005 | 35123889 | 35124034 | 35142372 | 35142478 | 32 | 20 | 20 | 27 | 32 | 91 | 17 |
| MXE | 11838 | ENSG00000085491 | SLC25A24 | chr1 | - | 108139057 | 108139208 | 108143542 | 108143710 | 108134042 | 108136837 | 108148278 | 108148386 | 103 | 93 | 72 | 73 | 128 | 82 | 52 |
| MXE | 14078 | ENSG00000156787 | TBC1D31 | chr8 | + | 123082701 | 123082817 | 123084161 | 123084340 | 123077110 | 123077257 | 123093590 | 123093742 | 59 | 28 | 43 | 60 | 78 | 181 | 70 |
| MXE | 4227 | ENSG00000138443 | ABI2 | chr2 | + | 203391045 | 203391143 | 203394699 | 203394846 | 203380207 | 203380384 | 203395655 | 203395780 | 63 | 47 | 43 | 67 | 83 | 16 | 15 |
| MXE | 12866 | ENSG00000122566 | HNRNPA2B1 | chr7 | - | 26190992 | 26192338 | 26192494 | 26192577 | 26189665 | 26190460 | 26193250 | 26193373 | 290 | 250 | 229 | 315 | 332 | 267 | 294 |
| MXE | 2977 | ENSG00000143379 | SETDB1 | chr1 | + | 150960562 | 150961191 | 150962586 | 150962719 | 150959177 | 150959347 | 150962973 | 150963139 | 120 | 80 | 77 | 90 | 86 | 141 | 84 |
| MXE | 15976 | ENSG00000178440 | TIMM23B-AGAP6 | chr10 | + | 49987164 | 49987234 | 49988750 | 49988938 | 49973011 | 49973142 | 49989307 | 49989376 | 5 | 0 | 3 | 0 | 0 | 29 | 17 |
| MXE | 11587 | ENSG00000163939 | PBRM1 | chr3 | - | 52644703 | 52644789 | 52648343 | 52648442 | 52641953 | 52642045 | 52651741 | 52651810 | 144 | 125 | 125 | 144 | 262 | 4 | 3 |
| MXE | 15465 | ENSG00000110079 | MS4A4A | chr11 | + | 60301000 | 60301057 | 60306099 | 60306201 | 60297196 | 60297325 | 60308106 | 60308960 | 3 | 1 | 5 | 5 | 8 | 35 | 27 |
| MXE | 111 | ENSG00000198911 | SREBF2 | chr22 | + | 41877228 | 41877421 | 41877941 | 41878123 | 41875542 | 41875724 | 41880715 | 41880992 | 156 | 49 | 118 | 147 | 202 | 33 | 2 |
| MXE | 2749 | ENSG00000131966 | ACTR10 | chr14 | + | 58219693 | 58219729 | 58223621 | 58223701 | 58215204 | 58215284 | 58223782 | 58223827 | 28 | 13 | 14 | 12 | 42 | 45 | 31 |
| MXE | 13361 | ENSG00000099204 | ABLIM1 | chr10 | - | 114468180 | 114468216 | 114472976 | 114473132 | 114465697 | 114465827 | 114473878 | 114473956 | 220 | 202 | 411 | 377 | 731 | 60 | 37 |
| MXE | 16799 | ENSG00000100599 | RIN3 | chr14 | + | 92615406 | 92615479 | 92641237 | 92641329 | 92577359 | 92577477 | 92651581 | 92653075 | 26 | 15 | 31 | 21 | 21 | 92 | 57 |
| MXE | 12286 | ENSG00000196511 | TPK1 | chr7 | - | 144648806 | 144648879 | 144682908 | 144682978 | 144623165 | 144623261 | 144765879 | 144765951 | 22 | 15 | 12 | 23 | 10 | 39 | 30 |

| | | | | | | | | | | | | | | | | | | | | | | | | | | | | | | | | | | | | | | | | | | | | | | | | |
|---|---|---|---|---|---|---|---|---|---|---|---|---|---|---|---|---|---|---|---|---|---|---|---|---|---|---|---|---|---|---|---|---|---|---|---|---|---|---|---|---|---|---|---|---|---|---|---|---|
| MXE | 12598 | ENSG00000136280 | CCM2 | chr7 | + | 45063917 | 45064001 | 45073459 | 45073571 | 45038252 | 45038426 | 45074269 | 45074408 | 48 | 24 | 42 | 44 | 58 | 52 | 14 | 33 | 54 | 33 | 7 | 28 | 8 | 27 | 35 | 3 | 26 | 19 | 23 | 37 | 233 | 261 | 0.4709915 | 0.83909051 | 0.508 | 0.658 | 0.588 | 0.477 | 0.663 | 0.723 | 0.547 | 0.32 | 0.568 | 0.514 | 0.044 |
| MXE | 6940 | ENSG00000145740 | SLC30A5 | chr5 | + | 69121693 | 69121895 | 69123198 | 69123425 | 69118498 | 69118628 | 69128003 | 69128132 | 54 | 34 | 68 | 63 | 96 | 107 | 110 | 105 | 99 | 110 | 39 | 19 | 14 | 19 | 19 | 27 | 57 | 56 | 44 | 40 | 298 | 298 | 0.471199119 | 0.8393603 | 0.335 | 0.236 | 0.393 | 0.389 | 0.466 | 0.591 | 0.25 | 0.2 | 0.302 | 0.322 | 0.031 |
| MXE | 15366 | ENSG00000136527 | TRA2B | chr3 | - | 185922010 | 185922126 | 185923795 | 185923984 | 185921103 | 185921187 | 185925463 | 185925626 | 697 | 444 | 699 | 770 | 766 | 610 | 518 | 622 | 795 | 807 | 731 | 856 | 680 | 911 | 476 | 800 | 969 | 720 | 1031 | 392 | 298 | 265 | 0.471530549 | 0.83985055 | 0.504 | 0.433 | 0.5 | 0.463 | 0.458 | 0.448 | 0.44 | 0.456 | 0.44 | 0.519 | 0.011 |
| MXE | 3506 | ENSG00000148110 | MFSD14B | chr9 | + | 94440984 | 94441099 | 94444963 | 94445182 | 94438400 | 94438528 | 94446822 | 94446936 | 36 | 18 | 30 | 48 | 44 | 71 | 36 | 60 | 63 | 46 | 68 | 66 | 28 | 33 | 74 | 100 | 88 | 45 | 74 | 55 | 264 | 298 | 0.471628648 | 0.83992514 | 0.364 | 0.361 | 0.361 | 0.462 | 0.519 | 0.434 | 0.458 | 0.413 | 0.335 | 0.603 | -0.035 |
| MXE | 2005 | ENSG00000182054 | IDH2 | chr15 | - | 90087111 | 90087263 | 90087438 | 90087575 | 90085274 | 90085387 | 90088358 | 90088502 | 154 | 54 | 139 | 152 | 192 | 145 | 69 | 153 | 121 | 153 | 228 | 331 | 183 | 197 | 205 | 235 | 151 | 119 | 115 | 407 | 286 | 298 | 0.471772996 | 0.840082069 | 0.525 | 0.449 | 0.486 | 0.567 | 0.567 | 0.503 | 0.695 | 0.616 | 0.641 | 0.344 | -0.041 |
| MXE | 9264 | ENSG00000159176 | CSRP1 | chr1 | - | 201490175 | 201490344 | 201496191 | 201496261 | 201488854 | 201488984 | 201507069 | 201507123 | 0 | 0 | 0 | 0 | 1 | 45 | 7 | 29 | 24 | 37 | 0 | 0 | 0 | 0 | 8 | 18 | 40 | 21 | 33 | 145 | 219 | 298 | 0.471877908 | 0.840154068 | 0 | 0 | 0 | 0 | 0.035 | 0 | 0 | 0 | 0 | 0.07 | -0.007 |
| MXE | 10258 | ENSG00000143207 | COP1 | chr1 | - | 176116623 | 176116681 | 176149005 | 176149074 | 176085775 | 176085890 | 176175909 | 176176007 | 6 | 7 | 3 | 1 | 9 | 92 | 79 | 54 | 68 | 133 | 3 | 8 | 3 | 3 | 0 | 71 | 53 | 72 | 56 | 73 | 218 | 207 | 0.471925899 | 0.840154068 | 0.058 | 0.078 | 0.05 | 0.014 | 0.06 | 0.039 | 0.125 | 0.038 | 0.048 | 0 | 0.002 |
| MXE | 17141 | ENSG00000102710 | SUPT20H | chr13 | - | 37045246 | 37045376 | 37047345 | 37047408 | 37044077 | 37044181 | 37047534 | 37047601 | 7 | 1 | 3 | 3 | 3 | 235 | 162 | 183 | 245 | 444 | 0 | 0 | 3 | 3 | 0 | 81 | 116 | 99 | 134 | 49 | 212 | 279 | 0.471998787 | 0.840183709 | 0.038 | 0.008 | 0.021 | 0.016 | 0.009 | 0 | 0 | 0.038 | 0.029 | 0 | 0.005 |
| MXE | 15264 | ENSG00000170145 | SIK2 | chr11 | + | 111700885 | 111701010 | 111701451 | 111701575 | 111688000 | 111688162 | 111703202 | 111703423 | 99 | 40 | 69 | 72 | 125 | 55 | 24 | 41 | 46 | 61 | 76 | 80 | 32 | 57 | 136 | 52 | 58 | 29 | 42 | 49 | 274 | 273 | 0.472109434 | 0.84026942 | 0.642 | 0.624 | 0.626 | 0.609 | 0.671 | 0.593 | 0.579 | 0.524 | 0.575 | 0.734 | 0.033 |
| [illegible] | | | | | | | | | | | | | | | | | | | | | | | | | | | | | | | | | | | | | | | | | | | | | | | | |

| | | | | | | | |
|---|---|---|---|---|---|---|---|
| MXE | 2349 | ENSG00000140367 | UBE2Q2 | chr15 | + | 75890434 | [illegible] |
| MXE | 1997 | ENSG00000167202 | TBC1D2B | chr15 | - | 78016545 | [illegible] |
| MXE | 1360 | ENSG00000109066 | TMEM104 | chr17 | + | 74777263 | [illegible] |
| MXE | 961 | ENSG00000152229 | PSTPIP2 | chr18 | - | 46024608 | [illegible] |
| MXE | 13292 | ENSG00000137801 | THBS1 | chr15 | + | 39589804 | [illegible] |
| MXE | 14808 | ENSG00000204394 | VARS1 | chr6 | - | 31783095 | [illegible] |
| MXE | 6323 | ENSG00000079277 | MKNK1 | chr1 | - | 46562648 | [illegible] |
| MXE | 2132 | ENSG00000159459 | UBR1 | chr15 | - | 42977879 | [illegible] |
| MXE | 9923 | ENSG00000115464 | USP34 | chr2 | - | 61314874 | [illegible] |
| MXE | 1490 | ENSG00000006125 | AP2B1 | chr17 | + | 35641876 | [illegible] |
| MXE | 10148 | ENSG00000112078 | KCTD20 | chr6 | + | 36474788 | [illegible] |
| MXE | 5938 | ENSG00000003400 | CASP10 | chr2 | + | 201205881 | [illegible] |
| MXE | 16997 | ENSG00000111300 | NAA25 | chr12 | - | 112081059 | [illegible] |
| MXE | 16269 | ENSG00000066084 | DIP2B | chr12 | + | 50691048 | [illegible] |
| MXE | 15089 | ENSG00000152556 | PFKM | chr12 | + | 48141913 | [illegible] |
| MXE | 5679 | ENSG00000163681 | SLMAP | chr3 | + | 57864806 | [illegible] |
| MXE | 8047 | ENSG00000159023 | EPB41 | chr1 | + | 29112367 | [illegible] |
| MXE | 15892 | ENSG00000144815 | NXPE3 | chr3 | + | 101782659 | [illegible] |
| MXE | 8768 | ENSG00000101040 | ZMYND8 | chr20 | - | 47291795 | [illegible] |
| MXE | 6302 | ENSG00000138674 | SEC31A | chr4 | - | 82828999 | [illegible] |
| MXE | 13646 | ENSG00000086758 | HUWE1 | chrX | - | 53562830 | [illegible] |
| MXE | 12548 | ENSG00000179818 | PCBP1-AS1 | chr2 | - | 70055713 | [illegible] |
| MXE | 9246 | ENSG00000152683 | SLC30A6 | chr2 | + | 32184272 | [illegible] |
| MXE | 13391 | ENSG00000014257 | ACP3 | chr3 | + | 132331646 | [illegible] |
| MXE | 7139 | ENSG00000122417 | ODF2L | chr1 | - | 86371017 | [illegible] |
| MXE | 7752 | ENSG00000205323 | SARNP | chr12 | - | 55757382 | [illegible] |
| MXE | 9870 | ENSG00000162191 | UBXN1 | chr11 | - | 62676812 | [illegible] |
| MXE | 15236 | ENSG00000139722 | VPS37B | chr12 | - | 122868479 | [illegible] |
| MXE | 8035 | ENSG00000159023 | EPB41 | chr1 | + | 28993329 | [illegible] |
| MXE | 12490 | ENSG00000179818 | PCBP1-AS1 | chr2 | - | 69996685 | [illegible] |
| MXE | 6008 | ENSG00000137393 | RNF144B | chr6 | + | 18427580 | [illegible] |
| MXE | 16122 | ENSG00000011376 | LARS2 | chr3 | + | 45500441 | [illegible] |
| MXE | 10158 | ENSG00000143924 | EML4 | chr2 | + | 42256500 | [illegible] |
| MXE | 9654 | ENSG00000113761 | ZNF346 | chr5 | + | 177041777 | [illegible] |
| MXE | 10166 | ENSG00000186141 | POLR3C | chr1 | + | 145836493 | [illegible] |
| MXE | 10608 | ENSG00000196295 | GARS1-DT | chr7 | - | 30552099 | [illegible] |
| MXE | 8669 | ENSG00000117360 | PRPF3 | chr1 | + | 150344433 | [illegible] |
| MXE | 8263 | ENSG00000176095 | IP6K1 | chr3 | - | 49738211 | [illegible] |
| MXE | 6018 | ENSG00000010322 | NISCH | chr3 | + | 52489335 | [illegible] |
| MXE | 13415 | ENSG00000008294 | SPAG9 | chr17 | - | 51047374 | [illegible] |
| MXE | 15198 | ENSG00000139626 | ITGB7 | chr12 | - | 53200242 | [illegible] |
| MXE | 16322 | ENSG00000139620 | KANSL2 | chr12 | - | 48679654 | [illegible] |
| MXE | 4671 | ENSG00000121060 | TRIM25 | chr17 | - | 56895342 | [illegible] |
| MXE | 1428 | ENSG00000123159 | GIPC1 | chr19 | - | 14482688 | [illegible] |
| MXE | 1189 | ENSG00000261371 | PECAM1 | chr17 | - | 64350379 | [illegible] |
| MXE | 6757 | ENSG00000164331 | ANKRA2 | chr5 | - | 73554860 | [illegible] |
| MXE | 10336 | ENSG00000137075 | RNF38 | chr9 | - | 36369718 | [illegible] |
| MXE | 13924 | ENSG00000160588 | MPZL3 | chr11 | - | 118235423 | [illegible] |
| MXE | 8191 | ENSG00000188641 | DPYD | chr1 | - | 97382392 | [illegible] |
| MXE | 9084 | ENSG00000105402 | NAPA | chr19 | - | 47493415 | [illegible] |
| MXE | 300 | ENSG00000100077 | GRK3 | chr22 | + | 25678815 | [illegible] |
| MXE | 8045 | ENSG00000159023 | EPB41 | chr1 | + | 29097806 | [illegible] |
| MXE | 13848 | ENSG00000152601 | MBNL1 | chr3 | + | 152299404 | [illegible] |
| MXE | 5597 | ENSG00000092098 | RNF31 | chr14 | + | 24157831 | [illegible] |
| MXE | 6332 | ENSG00000247828 | TMEM161B-AS1 | chr5 | + | 88283009 | [illegible] |
| MXE | 7908 | ENSG00000071054 | MAP4K4 | chr2 | + | 101882550 | [illegible] |
| MXE | 1233 | ENSG00000161533 | ACOX1 | chr17 | - | 75960214 | [illegible] |
| MXE | 16585 | ENSG00000127314 | RAP1B | chr12 | + | 68624674 | [illegible] |
| MXE | 16265 | ENSG00000059573 | ALDH18A1 | chr10 | - | 95642991 | [illegible] |
| MXE | 2646 | ENSG00000134001 | EIF2S1 | chr14 | + | 67374467 | [illegible] |
| MXE | 10791 | ENSG00000144228 | SPOPL | chr2 | + | 138550902 | [illegible] |
| MXE | 4098 | ENSG00000008513 | ST3GAL1 | chr8 | - | 133475718 | [illegible] |
| MXE | 2672 | ENSG00000203485 | INF2 | chr14 | + | 104706909 | [illegible] |
| MXE | 2673 | ENSG00000203485 | INF2 | chr14 | + | 104706909 | [illegible] |
| MXE | 182 | ENSG00000100376 | FAM118A | chr22 | + | 45332424 | [illegible] |
| MXE | 10676 | ENSG00000148341 | SH3GLB2 | chr9 | - | 129021090 | [illegible] |
| MXE | 1037 | ENSG00000070413 | DGCR2 | chr22 | - | 19041057 | [illegible] |
| MXE | 11750 | ENSG00000096070 | BRPF3 | chr6 | + | 36210215 | [illegible] |
| MXE | 14584 | ENSG00000110455 | ACCS | chr11 | + | 44071255 | [illegible] |
| MXE | 11073 | ENSG00000122203 | KIAA1191 | chr5 | - | 176352621 | [illegible] |
| MXE | 14552 | ENSG00000107242 | PIP5K1B | chr9 | + | 68876676 | [illegible] |
| MXE | 5838 | ENSG00000198793 | MTOR | chr1 | - | 11167441 | [illegible] |
| MXE | 3229 | ENSG00000112624 | BICRAL | chr6 | + | 42855855 | [illegible] |
| MXE | 9331 | ENSG00000070785 | EIF2B3 | chr1 | - | 44897354 | [illegible] |
| MXE | 16372 | ENSG00000013561 | RNF14 | chr5 | + | 141974803 | [illegible] |
| MXE | 10630 | ENSG00000144895 | EIF2A | chr3 | + | 150558387 | [illegible] |
| MXE | 10834 | ENSG00000108883 | EFTUD2 | chr17 | - | 44867806 | [illegible] |
| MXE | 11927 | ENSG00000131018 | SYNE1 | chr6 | - | 152409558 | [illegible] |
| MXE | 16660 | ENSG00000135297 | MTO1 | chr6 | + | 73482448 | [illegible] |
| MXE | 4705 | ENSG00000155508 | CNOT8 | chr5 | + | 154865191 | [illegible] |
| MXE | 5059 | ENSG00000176261 | ZBTB8OS | chr1 | - | 32633644 | [illegible] |
| MXE | 267 | ENSG00000244625 | MIATNB | chr22 | + | 26717252 | [illegible] |
| MXE | 6832 | ENSG00000007341 | ST7L | chr1 | - | 112583971 | [illegible] |
| MXE | 2223 | ENSG00000174485 | DENND4A | chr15 | - | 65701050 | [illegible] |
| MXE | 5733 | ENSG00000114126 | TFDP2 | chr3 | - | 141993537 | [illegible] |
| MXE | 2594 | ENSG00000100450 | GZMH | chr14 | - | 24607148 | [illegible] |
| MXE | 11671 | ENSG00000024048 | UBR2 | chr6 | + | 42612170 | [illegible] |
| MXE | 5394 | ENSG00000162736 | NCSTN | chr1 | + | 160355654 | [illegible] |
| MXE | 3348 | ENSG00000127603 | MACF1 | chr1 | + | 39293457 | [illegible] |
| MXE | 8284 | ENSG00000169554 | ZEB2 | chr2 | - | 144429768 | [illegible] |
| MXE | 5115 | ENSG00000114867 | EIF4G1 | chr3 | + | 184316131 | [illegible] |
| MXE | 16912 | ENSG00000177119 | ANO6 | chr12 | + | 45388160 | [illegible] |
| MXE | 9434 | ENSG00000138078 | PREPL | chr2 | - | 44328936 | [illegible] |
| MXE | 3378 | ENSG00000131844 | MCCC2 | chr5 | + | 71646210 | [illegible] |
| MXE | 11822 | ENSG00000112237 | CCNC | chr6 | - | 99549507 | [illegible] |
| MXE | 14684 | ENSG00000129675 | ARHGEF6 | chrX | - | 136677935 | [illegible] |
| MXE | 1761 | ENSG00000214078 | CPNE1 | chr20 | - | 35626734 | [illegible] |
| MXE | 3323 | ENSG00000140455 | USP3 | chr15 | + | 63558105 | [illegible] |
| MXE | 13917 | ENSG00000177565 | TBL1XR1 | chr3 | - | 177098465 | [illegible] |
| MXE | 3852 | ENSG00000247774 | PCED1B-AS1 | chr12 | - | 47209280 | [illegible] |
| MXE | 13104 | ENSG00000177324 | BEND2 | chrX | - | 18195295 | [illegible] |
| MXE | 13996 | ENSG00000110074 | FOXRED1 | chr11 | + | 126274926 | [illegible] |
| MXE | 16649 | ENSG00000135297 | MTO1 | chr6 | + | 73481000 | [illegible] |
| MXE | 9435 | ENSG00000138078 | PREPL | chr2 | - | 44332458 | [illegible] |
| MXE | 10694 | ENSG00000113712 | CSNK1A1 | chr5 | - | 149515668 | [illegible] |
| MXE | 13504 | ENSG00000068438 | FTSJ1 | chrX | + | 48478448 | [illegible] |
| MXE | 6997 | ENSG00000126070 | AGO3 | chr1 | + | 36027113 | [illegible] |
| MXE | 6653 | ENSG00000114098 | ARMC8 | chr3 | + | 138223388 | [illegible] |
| MXE | 13026 | ENSG00000055130 | CUL1 | chr7 | + | 148759303 | [illegible] |
| MXE | 13673 | ENSG00000215301 | DDX3X | chrX | + | 41342736 | [illegible] |
| MXE | 9504 | ENSG00000151893 | CACUL1 | chr10 | - | 118686597 | [illegible] |
| MXE | 6501 | ENSG00000125637 | PSD4 | chr2 | + | 113185876 | [illegible] |
| MXE | 6799 | ENSG00000162739 | SLAMF6 | chr1 | - | 160490574 | [illegible] |
| MXE | 12200 | ENSG00000168014 | C2CD3 | chr11 | - | 74100524 | [illegible] |
| MXE | 14889 | ENSG00000165219 | GAPVD1 | chr9 | + | 125298889 | [illegible] |
| MXE | 9865 | ENSG00000110934 | BIN2 | chr12 | - | 51288107 | [illegible] |
| MXE | 5959 | ENSG00000134352 | IL6ST | chr5 | - | 55969549 | [illegible] |
| MXE | 5448 | ENSG00000113269 | RNF130 | chr5 | - | 179980128 | [illegible] |
| MXE | 11944 | ENSG00000125779 | PANK2 | chr20 | + | 3912457 | [illegible] |
| MXE | 13479 | ENSG00000111203 | ITFG2 | chr12 | + | 2830833 | [illegible] |
| MXE | 10795 | ENSG00000065675 | PRKCQ | chr10 | - | 6430809 | [illegible] |
| MXE | 14164 | ENSG00000166925 | TSC22D4 | chr7 | - | 100474273 | [illegible] |
| MXE | 10239 | ENSG00000087152 | ATXN7L3 | chr17 | - | 44194767 | [illegible] |
| MXE | 9313 | ENSG00000163534 | FCRL1 | chr1 | - | 157802376 | [illegible] |
| MXE | 5036 | ENSG00000009790 | TRAF3IP3 | chr1 | + | 209778110 | [illegible] |
| MXE | 10776 | ENSG00000163412 | EIF4E3 | chr3 | - | 71710411 | [illegible] |
| MXE | 6079 | ENSG00000271723 | MROH7-TTC4 | chr1 | + | 54733628 | [illegible] |
| MXE | 8381 | ENSG00000152223 | EPG5 | chr18 | - | 45916437 | [illegible] |
| MXE | 14051 | ENSG00000137770 | CTDSPL2 | chr15 | + | 44458990 | [illegible] |
| MXE | 5946 | ENSG00000109320 | NFKB1 | chr4 | + | 102529835 | [illegible] |
| MXE | 15270 | ENSG00000095139 | ARCN1 | chr11 | + | 118592708 | [illegible] |
| MXE | 16294 | ENSG00000151923 | TIAL1 | chr10 | - | 119582458 | [illegible] |
| MXE | 15461 | ENSG00000133318 | RTN3 | chr11 | + | 63752506 | [illegible] |
| MXE | 1317 | ENSG00000266967 | AARSD1 | chr17 | - | 42955841 | [illegible] |
| MXE | 7834 | ENSG00000176393 | RNPEP | chr1 | + | 201989382 | [illegible] |
| MXE | 10586 | ENSG00000131236 | CAP1 | chr1 | + | 40041428 | [illegible] |
| MXE | 8082 | ENSG00000124574 | ABCC10 | chr6 | + | 43441861 | [illegible] |
| MXE | 2967 | ENSG00000120925 | RNF170 | chr8 | - | 42870003 | [illegible] |
| MXE | 9262 | ENSG00000166377 | ATP9B | chr18 | + | 79253380 | [illegible] |
| MXE | 3027 | ENSG00000181036 | FCRL6 | chr1 | + | 159808960 | [illegible] |
| MXE | 4025 | ENSG00000172869 | DMXL1 | chr5 | + | 119110150 | [illegible] |
| MXE | 13495 | ENSG00000164031 | DNAJB14 | chr4 | - | 99908710 | [illegible] |
| MXE | 17120 | ENSG00000275066 | SYNRG | chr17 | - | 37535978 | [illegible] |
| MXE | 6033 | ENSG00000090989 | EXOC1 | chr4 | + | 55891314 | [illegible] |
| MXE | 6750 | ENSG00000183020 | AP2A2 | chr11 | + | 970168 | [illegible] |
| MXE | 8239 | ENSG00000143643 | TTC13 | chr1 | - | 230954332 | [illegible] |
| MXE | 11323 | ENSG00000115993 | TRAK2 | chr2 | - | 201399376 | [illegible] |
| MXE | 7049 | ENSG00000136628 | EPRS1 | chr1 | - | 220030385 | [illegible] |
| MXE | 4568 | ENSG00000118503 | TNFAIP3 | chr6 | + | 137875995 | [illegible] |
| MXE | 13237 | ENSG00000188917 | TRMT2B | chrX | - | 101042041 | [illegible] |
| MXE | 11669 | ENSG00000024048 | UBR2 | chr6 | + | 42594190 | [illegible] |
| MXE | 10115 | ENSG00000166164 | BRD7 | chr16 | - | 50334710 | [illegible] |
| MXE | 11400 | ENSG00000003402 | CFLAR | chr2 | + | 201140356 | [illegible] |
| MXE | 14271 | ENSG00000089234 | BRAP | chr12 | - | 111665638 | [illegible] |
| MXE | 8352 | ENSG00000189362 | NEMP2 | chr2 | - | 190517519 | [illegible] |
| MXE | 16770 | ENSG00000186815 | TPCN1 | chr12 | + | 113284580 | [illegible] |
| MXE | 7678 | ENSG00000115685 | PPP1R7 | chr2 | + | 241158483 | [illegible] |
| MXE | 2492 | ENSG00000100813 | ACIN1 | chr14 | - | 23090521 | [illegible] |
| MXE | 3531 | ENSG00000116906 | GNPAT | chr1 | + | 231267679 | [illegible] |
| MXE | 6055 | ENSG00000132185 | FCRLA | chr1 | + | 161710759 | [illegible] |
| MXE | 12930 | ENSG00000197448 | GSTK1 | chr7 | + | 143264085 | [illegible] |
| MXE | 8726 | ENSG00000143815 | LBR | chr1 | - | 225410290 | [illegible] |
| MXE | 13746 | ENSG00000143569 | UBAP2L | chr1 | + | 154225083 | [illegible] |
| MXE | 10703 | ENSG00000249141 | NA | chr6 | - | 166948569 | [illegible] |
| MXE | 2827 | ENSG00000198718 | TOGARAM1 | chr14 | + | 45025782 | [illegible] |
| MXE | 2660 | ENSG00000032219 | ARID4A | chr14 | + | 58346410 | [illegible] |
| MXE | 13172 | ENSG00000140543 | DET1 | chr15 | - | 88527598 | [illegible] |
| MXE | 13472 | ENSG00000005302 | MSL3 | chrX | + | 11768572 | [illegible] |
| MXE | 6010 | ENSG00000137393 | RNF144B | chr6 | + | 18427580 | [illegible] |
| MXE | 16064 | ENSG00000148700 | ADD3 | chr10 | + | 110105999 | [illegible] |
| MXE | 6980 | ENSG00000158517 | NCF1 | chr7 | + | 74783524 | [illegible] |
| MXE | 3223 | ENSG00000189091 | SF3B3 | chr16 | + | 70565064 | [illegible] |
| MXE | 13915 | ENSG00000177565 | TBL1XR1 | chr3 | - | 177079826 | [illegible] |
| MXE | 8066 | ENSG00000103657 | HERC1 | chr15 | - | 63637504 | [illegible] |
| MXE | 15740 | ENSG00000149311 | ATM | chr11 | + | 108244787 | [illegible] |
| MXE | 2500 | ENSG00000197102 | DYNC1H1 | chr14 | + | 102040597 | [illegible] |
| MXE | 11299 | ENSG00000080345 | RIF1 | chr2 | + | 151433076 | [illegible] |
| MXE | 3120 | ENSG00000150867 | PIP4K2A | chr10 | - | 22567850 | [illegible] |
| MXE | 4811 | ENSG00000113971 | NPHP3 | chr3 | - | 132682702 | [illegible] |
| MXE | 3688 | ENSG00000204120 | GIGYF2 | chr2 | + | 232847347 | [illegible] |
| MXE | 6299 | ENSG00000270629 | NBPF14 | chr1 | - | 148585140 | [illegible] |
| MXE | 2994 | ENSG00000069974 | RAB27A | chr15 | - | 55234781 | [illegible] |
| MXE | 3467 | ENSG00000112851 | ERBIN | chr5 | + | 66075023 | [illegible] |
| MXE | 14044 | ENSG00000122882 | ECD | chr10 | - | 73154255 | [illegible] |
| MXE | 6178 | ENSG00000163875 | MEAF6 | chr1 | - | 37495884 | [illegible] |
| MXE | 4062 | ENSG00000131507 | NDFIP1 | chr5 | + | 142140562 | [illegible] |
| MXE | 6600 | ENSG00000058673 | ZC3H11A | chr1 | + | 203801574 | [illegible] |
| MXE | 9367 | ENSG00000116984 | MTR | chr1 | + | 236803427 | [illegible] |
| MXE | 14278 | ENSG00000167632 | TRAPPC9 | chr8 | - | 140311247 | [illegible] |
| MXE | 8511 | ENSG00000225470 | JPX | chrX | + | 73994839 | [illegible] |
| MXE | 13535 | ENSG00000135048 | CEMIP2 | chr9 | - | 71740062 | [illegible] |
| MXE | 4167 | ENSG00000109814 | UGDH | chr4 | - | 39510660 | [illegible] |
| MXE | 10470 | ENSG00000162735 | PEX19 | chr1 | - | 160282416 | [illegible] |
| MXE | 2181 | ENSG00000129003 | VPS13C | chr15 | - | 61884127 | [illegible] |
| MXE | 1077 | ENSG00000108953 | YWHAE | chr17 | - | 1354210 | [illegible] |
| MXE | 12052 | ENSG00000214021 | TTLL3 | chr3 | + | 9813247 | [illegible] |
| MXE | 12083 | ENSG00000198815 | FOXJ3 | chr1 | - | 42205761 | [illegible] |
| MXE | 12042 | ENSG00000127947 | PTPN12 | chr7 | + | 77592184 | [illegible] |
| MXE | 17020 | ENSG00000065615 | CYB5R4 | chr6 | + | 83909008 | [illegible] |
| MXE | 10495 | ENSG00000036257 | CUL3 | chr2 | - | 224511353 | [illegible] |
| MXE | 15744 | ENSG00000149311 | ATM | chr11 | + | 108258985 | [illegible] |
| MXE | 15477 | ENSG00000156599 | ZDHHC5 | chr11 | + | 57696760 | [illegible] |
| MXE | 13478 | ENSG00000111203 | ITFG2 | chr12 | + | 2830216 | [illegible] |
| MXE | 16149 | ENSG00000196233 | LCOR | chr10 | + | 96944112 | [illegible] |
| MXE | 8070 | ENSG00000103657 | HERC1 | chr15 | - | 63723181 | [illegible] |
| MXE | 4687 | ENSG00000047365 | ARAP2 | chr4 | - | 36212395 | [illegible] |
| MXE | 7691 | ENSG00000142784 | WDTC1 | chr1 | + | 27263151 | [illegible] |
| MXE | 1104 | ENSG00000141252 | VPS53 | chr17 | - | 586269 | [illegible] |
| MXE | 14146 | ENSG00000151461 | UPF2 | chr10 | - | 11952065 | [illegible] |
| MXE | 4638 | ENSG00000065882 | TBC1D1 | chr4 | + | 38118032 | [illegible] |
| MXE | 17082 | ENSG00000198176 | TFDP1 | chr13 | + | 113623179 | [illegible] |
| MXE | 7976 | ENSG00000173209 | AHSA2P | chr2 | + | 61181319 | [illegible] |
| MXE | 1362 | ENSG00000109066 | TMEM104 | chr17 | + | 74777266 | [illegible] |
| MXE | 10684 | ENSG00000169499 | PLEKHA2 | chr8 | + | 38957322 | [illegible] |
| MXE | 4547 | ENSG00000064886 | CHI3L2 | chr1 | + | 111236023 | [illegible] |
| MXE | 16482 | ENSG00000082898 | XPO1 | chr2 | - | 61492914 | [illegible] |
| MXE | 4584 | ENSG00000132466 | ANKRD17 | chr4 | - | 73090666 | [illegible] |
| MXE | 10284 | ENSG00000131051 | RBM39 | chr20 | - | 35738967 | [illegible] |
| MXE | 9780 | ENSG00000175662 | TOM1L2 | chr17 | - | 17884633 | [illegible] |
| MXE | 9236 | ENSG00000163466 | ARPC2 | chr2 | + | 218225919 | [illegible] |
| MXE | 4348 | ENSG00000182150 | ERCC6L2 | chr9 | + | 95928078 | [illegible] |
| MXE | 7877 | ENSG00000087206 | UIMC1 | chr5 | - | 176975395 | [illegible] |
| MXE | 12712 | ENSG00000106443 | PHF14 | chr7 | + | 11013746 | [illegible] |
| MXE | 10094 | ENSG00000149499 | EML3 | chr11 | - | 62605854 | [illegible] |
| MXE | 4733 | ENSG00000082805 | ERC1 | chr12 | + | 1180539 | [illegible] |
| MXE | 9384 | ENSG00000070961 | ATP2B1 | chr12 | - | 89621548 | [illegible] |
| MXE | 16030 | ENSG00000120438 | TCP1 | chr6 | - | 159783940 | [illegible] |
| MXE | 3514 | ENSG00000104497 | SNX16 | chr8 | - | 81823791 | [illegible] |
| MXE | 8936 | ENSG00000110090 | CPT1A | chr11 | - | 68794803 | [illegible] |
| MXE | 8938 | ENSG00000135404 | CD63 | chr12 | - | 55726120 | [illegible] |
| MXE | 12547 | ENSG00000179818 | PCBP1-AS1 | chr2 | - | 70055713 | [illegible] |
| MXE | 15583 | ENSG00000154310 | TNIK | chr3 | - | 171228164 | [illegible] |
| MXE | 16651 | ENSG00000135297 | MTO1 | chr6 | + | 73482028 | [illegible] |

| | | | | | | | | | | | | | | |
|---|---|---|---|---|---|---|---|---|---|---|---|---|---|---|
| MXE | 15405 | ENSG00000149091 | DGKZ | chr11 | + | 46378456 | 46378500 | 46378990 | 46379111 | 46378197 | 46378229 | 46379202 | 46379236 | 9 |
| MXE | 6476 | ENSG00000182473 | EXOC7 | chr17 | - | 76090328 | 76090397 | 76091142 | 76091235 | 76089174 | 76089320 | 76094413 | 76094581 | 17 |
| MXE | 10171 | ENSG00000102900 | NUP93 | chr16 | + | 56838951 | 56839069 | 56839520 | 56839604 | 56837607 | 56837726 | 56841704 | 56841833 | 62 |
| MXE | 9792 | ENSG00000013810 | TACC3 | chr4 | + | 1723727 | 1723870 | 1727707 | 1728787 | 1723420 | 1723583 | 1730886 | 1730962 | 88 |
| MXE | 11994 | ENSG00000089820 | ARHGAP4 | chrX | - | 153910921 | 153910999 | 153911128 | 153911189 | 153910699 | 153910834 | 153912699 | 153912802 | 72 |
| MXE | 13216 | ENSG00000230590 | FTX | chrX | - | 74280393 | 74280494 | 74280930 | 74281085 | 74274259 | 74274410 | 74281701 | 74281848 | 100 |
| MXE | 14364 | ENSG00000104365 | IKBKB | chr8 | + | 42306342 | 42306432 | 42308900 | 42309025 | 42305186 | 42305275 | 42314321 | 42314429 | 56 |
| MXE | 432 | ENSG00000076944 | STXBP2 | chr19 | + | 7639018 | 7639100 | 7639730 | 7639807 | 7638725 | 7638775 | 7640730 | 7640809 | 71 |
| MXE | 14288 | ENSG00000158473 | CD1D | chr1 | + | 158182877 | 158183156 | 158183935 | 158184035 | 158182031 | 158182310 | 158184128 | 158186427 | 131 |
| MXE | 15556 | ENSG00000172543 | CTSW | chr11 | + | 65882429 | 65882526 | 65882608 | 65882689 | 65882174 | 65882329 | 65882778 | 65882904 | 544 |
| MXE | 16334 | ENSG00000132305 | IMMT | chr2 | - | 86173649 | 86173761 | 86179432 | 86179622 | 86171207 | 86171312 | 86181298 | 86181372 | 83 |
| MXE | 3692 | ENSG00000120705 | ETF1 | chr5 | - | 138508668 | 138508816 | 138510564 | 138510629 | 138506094 | 138508387 | 138511044 | 138511200 | 108 |
| MXE | 203 | ENSG00000100150 | DEPDC5 | chr22 | + | 31836998 | 31837155 | 31838684 | 31838845 | 31833865 | 31833980 | 31843094 | 31843212 | 17 |
| MXE | 15143 | ENSG00000183597 | TANGO2 | chr22 | + | 20053436 | 20053551 | 20055942 | 20056013 | 20052464 | 20052584 | 20063337 | 20063442 | 33 |
| MXE | 7310 | ENSG00000117543 | DPH5 | chr1 | - | 100995109 | 100995149 | 101001466 | 101001587 | 100992636 | 100992740 | 101021640 | 101021765 | 10 |
| MXE | 2046 | ENSG00000185418 | TARS3 | chr15 | - | 101685895 | 101686062 | 101699330 | 101699435 | 101684074 | 101684237 | 101701085 | 101701184 | 22 |
| MXE | 1982 | ENSG00000172366 | MCRIP2 | chr16 | + | 647416 | 647544 | 647782 | 647884 | 642119 | 642249 | 648119 | 648474 | 3 |
| MXE | 10579 | ENSG00000101104 | PABPC1L | chr20 | + | 44921593 | 44921731 | 44924160 | 44924256 | 44919182 | 44919277 | 44930517 | 44930726 | 47 |
| MXE | 16245 | ENSG00000139517 | LNX2 | chr13 | - | 27559841 | 27559985 | 27562412 | 27562781 | 27556235 | 27556413 | 27567639 | 27567839 | 17 |
| MXE | 14000 | ENSG00000095485 | CWF19L1 | chr10 | - | 100236849 | 100236969 | 100238021 | 100238231 | 100235666 | 100235764 | 100243697 | 100243772 | 59 |
| MXE | 3263 | ENSG00000174032 | SLC25A30 | chr13 | - | 45405882 | 45405977 | 45408926 | 45409074 | 45404326 | 45404412 | 45418299 | 45418455 | 4 |
| MXE | 4369 | ENSG00000129116 | PALLD | chr4 | + | 168894578 | 168894677 | 168898492 | 168898714 | 168890921 | 168891057 | 168903756 | 168903906 | 28 |
| MXE | 2907 | ENSG00000078319 | PMS2P1 | chr7 | - | 100328835 | 100329019 | 100330181 | 100330284 | 100325690 | 100325779 | 100330462 | 100330549 | 68 |
| MXE | 5830 | ENSG00000147251 | DOCK11 | chrX | + | 118618549 | 118618728 | 118624538 | 118624655 | 118615599 | 118615711 | 118627503 | 118627579 | 221 |
| MXE | 9149 | ENSG00000113615 | SEC24A | chr5 | + | 134679601 | 134679728 | 134682372 | 134682482 | 134676022 | 134676125 | 134686789 | 134686902 | 47 |
| MXE | 4495 | ENSG00000143624 | INTS3 | chr1 | + | 153763232 | 153763362 | 153764689 | 153764734 | 153762727 | 153762847 | 153764943 | 153765063 | 79 |
| MXE | 15279 | ENSG00000196323 | ZBTB44 | chr11 | - | 130239811 | 130239896 | 130260855 | 130261929 | 130238443 | 130238607 | 130314374 | 130314612 | 26 |
| MXE | 7287 | ENSG00000154710 | RABGEF1 | chr7 | + | 66783674 | 66783841 | 66795510 | 66795592 | 66775226 | 66775393 | 66797373 | 66797506 | 70 |
| MXE | 4550 | ENSG00000055955 | ITIH4 | chr3 | - | 52819753 | 52819792 | 52819939 | 52819990 | 52819392 | 52819518 | 52820290 | 52820317 | 12 |
| MXE | 7726 | ENSG00000117481 | NSUN4 | chr1 | + | 46344800 | 46345144 | 46352867 | 46353028 | 46340788 | 46340919 | 46360703 | 46360828 | 27 |
| MXE | 12470 | ENSG00000085788 | DDHD2 | chr8 | + | 38246232 | 38246300 | 38247712 | 38247835 | 38245741 | 38245950 | 38249707 | 38249747 | 24 |
| MXE | 9270 | ENSG00000138709 | LARP1B | chr4 | + | 128091346 | 128091512 | 128098185 | 128098330 | 128091000 | 128091144 | 128107138 | 128107313 | 25 |

[illegible]

| | | | | | | | | | | | | | | |
|---|---|---|---|---|---|---|---|---|---|---|---|---|---|---|
| MXE | 16558 | ENSG00000124164 | VAPB | chr20 | + | 58438944 | 58439025 | 58440906 | 58441083 | 58434601 | 58434705 | 58444076 | 58445552 | 78 [illegible] |
| MXE | 1269 | ENSG00000179094 | PER1 | chr17 | - | 8148643 | 8148786 | 8149461 | 8149663 | 8148180 | 8148259 | 8149754 | 8149876 | 66 [illegible] |
| MXE | 16509 | ENSG00000135720 | DYNC1LI2 | chr16 | - | 66727687 | 66727805 | 66728200 | 66728242 | 66725827 | 66725944 | 66729039 | 66729099 | 29 [illegible] |
| MXE | 13538 | ENSG00000074966 | TXK | chr4 | - | 48110537 | 48110603 | 48112306 | 48112512 | 48104900 | 48104955 | 48113206 | 48113309 | 211 [illegible] |
| MXE | 11258 | ENSG00000153094 | BCL2L11 | chr2 | + | 111123732 | 111124139 | 111150043 | 111150147 | 111120913 | 111121188 | 111153779 | 111153904 | 79 [illegible] |
| MXE | 4746 | ENSG00000131389 | SLC6A6 | chr3 | + | 14468087 | 14468212 | 14479084 | 14479185 | 14467852 | 14467956 | 14481670 | 14481841 | 23 [illegible] |
| MXE | 14696 | ENSG00000089053 | ANAPC5 | chr12 | - | 121320384 | 121320459 | 121327095 | 121327231 | 121319696 | 121319818 | 121328315 | 121328458 | 146 [illegible] |
| MXE | 852 | ENSG00000282826 | FRG1CP | chr20 | - | 28588642 | 28588757 | 28590556 | 28590614 | 28586298 | 28586403 | 28591393 | 28591519 | 45 [illegible] |
| MXE | 2036 | ENSG00000140264 | SERF2 | chr15 | + | 43801131 | 43801187 | 43801517 | 43801569 | 43799357 | 43800784 | 43801710 | 43804427 | 18 [illegible] |
| MXE | 15444 | ENSG00000110274 | CEP164 | chr11 | + | 117395122 | 117395191 | 117395546 | 117395722 | 117394919 | 117395003 | 117396053 | 117396180 | 14 [illegible] |
| MXE | 12603 | ENSG00000136280 | CCM2 | chr7 | + | 45070388 | 45071872 | 45072725 | 45072783 | 45070093 | 45070168 | 45073459 | 45073571 | 0 [illegible] |
| MXE | 16714 | ENSG00000100982 | PCIF1 | chr20 | + | 45939214 | 45939339 | 45940474 | 45940612 | 45938980 | 45939123 | 45940808 | 45940939 | 69 [illegible] |
| MXE | 11190 | ENSG00000102034 | ELF4 | chrX | - | 130071039 | 130071232 | 130071335 | 130071419 | 130069299 | 130069677 | 130072225 | 130072417 | 271 [illegible] |
| MXE | 7035 | ENSG00000095564 | BTAF1 | chr10 | + | 91964076 | 91964201 | 91966636 | 91966757 | 91962537 | 91962678 | 91980453 | 91980558 | 290 [illegible] |
| MXE | 14014 | ENSG00000172172 | MRPL13 | chr8 | - | 120413990 | 120414112 | 120419851 | 120419938 | 120395436 | 120396125 | 120425305 | 120425366 | 40 [illegible] |
| MXE | 1583 | ENSG00000040199 | PHLPP2 | chr16 | - | 71702597 | 71702731 | 71714511 | 71714801 | 71690518 | 71690709 | 71723768 | 71723812 | 7 [illegible] |
| MXE | 10175 | ENSG00000138073 | PREB | chr2 | - | 27132842 | 27132923 | 27133116 | 27133721 | 27132510 | 27132727 | 27134286 | 27134666 | 42 [illegible] |
| MXE | 8442 | ENSG00000121964 | GTDC1 | chr2 | - | 143970652 | 143970762 | 144007181 | 144007535 | 143951981 | 143952073 | 144141881 | 144142055 | 59 [illegible] |
| MXE | 14726 | ENSG00000136811 | ODF2 | chr9 | + | 128460941 | 128461010 | 128469182 | 128469353 | 128460512 | 128460666 | 128471307 | 128471468 | 21 [illegible] |
| MXE | 12181 | ENSG00000180233 | ZNRF2 | chr7 | + | 30323641 | 30323737 | 30355727 | 30355833 | 30284596 | 30285826 | 30362376 | 30362456 | 7 [illegible] |
| MXE | 6718 | ENSG00000204435 | CSNK2B | chr6 | + | 31667867 | 31667970 | 31668538 | 31668654 | 31666820 | 31666903 | 31669096 | 31669172 | 136 [illegible] |
| MXE | 10131 | ENSG00000155097 | ATP6V1C1 | chr8 | + | 103064713 | 103064811 | 103066320 | 103066447 | 103063134 | 103063228 | 103068651 | 103069076 | 55 [illegible] |
| MXE | 11249 | ENSG00000068724 | TTC7A | chr2 | + | 47051745 | 47051880 | 47060768 | 47060971 | 47049948 | 47050046 | 47073701 | 47073923 | 26 [illegible] |
| MXE | 13672 | ENSG00000215301 | DDX3X | chrX | + | 41342736 | 41342836 | 41343736 | 41343822 | 41340011 | 41342653 | 41344029 | 41344128 | 642 [illegible] |
| MXE | 13011 | ENSG00000196305 | IARS1 | chr9 | - | 92253361 | 92253453 | 92255534 | 92255604 | 92251807 | 92251885 | 92256679 | 92256800 | 15 [illegible] |
| MXE | 5780 | ENSG00000138750 | NUP54 | chr4 | - | 76136185 | 76136412 | 76144148 | 76144292 | 76134174 | 76134362 | 76144389 | 76144473 | 84 [illegible] |
| MXE | 1038 | ENSG00000070413 | DGCR2 | chr22 | - | 19048439 | 19048643 | 19056985 | 19057162 | 19041806 | 19041959 | 19063201 | 19063278 | 113 [illegible] |
| MXE | 5772 | ENSG00000064961 | HMG20B | chr19 | + | 3575539 | 3575660 | 3576260 | 3576307 | 3573691 | 3574586 | 3576552 | 3576625 | 29 [illegible] |
| MXE | 5813 | ENSG00000162777 | DENND2D | chr1 | - | 111192139 | 111192317 | 111194577 | 111194726 | 111189211 | 111189253 | 111195915 | 111196056 | 372 [illegible] |
| MXE | 11321 | ENSG00000123607 | TTC21B | chr2 | - | 165945523 | 165945690 | 165949393 | 165949504 | 165943218 | 165943341 | 165949594 | 165949724 | 113 [illegible] |
| MXE | 1902 | ENSG00000122299 | ZC3H7A | chr16 | - | 11776750 | 11776909 | 11779165 | 11779363 | 11776451 | 11776532 | 11781424 | 11781464 | 177 [illegible] |
| MXE | 5062 | ENSG00000172046 | USP19 | chr3 | - | 49116450 | 49116607 | 49116726 | 49116943 | 49116279 | 49116351 | 49117656 | 49117711 | 2 [illegible] |
| MXE | 16832 | ENSG00000138777 | PPA2 | chr4 | - | 105424195 | 105424322 | 105446382 | 105446502 | 105399036 | 105399164 | 105449349 | 105449403 | 118 [illegible] |
| MXE | 12911 | ENSG00000119471 | HSDL2 | chr9 | + | 112438430 | 112438625 | 112441698 | 112441770 | 112418859 | 112418958 | 112454012 | 112454162 | 59 [illegible] |
| MXE | 10429 | ENSG00000163728 | TTC14 | chr3 | + | 180604224 | 180604309 | 180604851 | 180605007 | 180603123 | 180603323 | 180605765 | 180605837 | 188 [illegible] |
| MXE | 12750 | ENSG00000113013 | HSPA9 | chr5 | - | 138558552 | 138558657 | 138559863 | 138560091 | 138557868 | 138557986 | 138561579 | 138561789 | 313 [illegible] |
| MXE | 9615 | ENSG00000113569 | NUP155 | chr5 | - | 37342548 | 37342646 | 37348504 | 37348596 | 37341089 | 37341242 | 37349171 | 37349245 | 26 [illegible] |
| MXE | 4353 | ENSG00000182150 | ERCC6L2 | chr9 | + | 95966561 | 95966714 | 95970575 | 95973088 | 95955913 | 95956013 | 95978060 | 95978215 | 42 [illegible] |
| MXE | 2205 | ENSG00000169032 | MAP2K1 | chr15 | + | 66481754 | 66481879 | 66484989 | 66485191 | 66444655 | 66444707 | 66487227 | 66487292 | 90 [illegible] |
| MXE | 6380 | ENSG00000026751 | SLAMF7 | chr1 | + | 160749820 | 160750093 | 160751344 | 160751448 | 160739056 | 160739356 | 160752185 | 160752248 | 36 [illegible] |
| MXE | 12084 | ENSG00000198815 | FOXJ3 | chr1 | - | 42227882 | 42227966 | 42265114 | 42265189 | 42199101 | 42199230 | 42278347 | 42278672 | 101 [illegible] |
| MXE | 13738 | ENSG00000112159 | MDN1 | chr6 | - | 89684875 | 89684985 | 89685826 | 89685973 | 89683830 | 89683904 | 89686901 | 89687023 | 53 [illegible] |
| MXE | 14984 | ENSG00000148396 | SEC16A | chr9 | - | 136443822 | 136443900 | 136445644 | 136445719 | 136440100 | 136441823 | 136446854 | 136446949 | 53 [illegible] |
| MXE | 1595 | ENSG00000187555 | USP7 | chr16 | - | 8915444 | 8915525 | 8916501 | 8916556 | 8915253 | 8915344 | 8917025 | 8917156 | 146 [illegible] |
| MXE | 10478 | ENSG00000181381 | DDX60L | chr4 | - | 168416681 | 168416797 | 168419301 | 168419397 | 168415656 | 168415799 | 168420260 | 168420380 | 20 [illegible] |
| MXE | 12087 | ENSG00000106609 | TMEM248 | chr7 | + | 66944975 | 66945261 | 66948543 | 66948694 | 66941847 | 66942024 | 66950951 | 66951135 | 212 [illegible] |
| MXE | 8556 | ENSG00000160679 | CHTOP | chr1 | + | 153636571 | 153636653 | 153638294 | 153638448 | 153633981 | 153634343 | 153642242 | 153642429 | 108 [illegible] |
| MXE | 7755 | ENSG00000023318 | ERP44 | chr9 | - | 100022041 | 100022226 | 100052416 | 100052532 | 100020615 | 100020731 | 100057819 | 100057859 | 409 [illegible] |
| MXE | 6154 | ENSG00000078304 | PPP2R5C | chr14 | + | 101882160 | 101882271 | 101883256 | 101883349 | 101856685 | 101856885 | 101883431 | 101883562 | 573 [illegible] |
| MXE | 14776 | ENSG00000106868 | SUSD1 | chr9 | - | 112124256 | 112124436 | 112142319 | 112142499 | 112112770 | 112112868 | 112143470 | 112143623 | 78 [illegible] |
| MXE | 11387 | ENSG00000198721 | ECI2 | chr6 | - | 4119185 | 4119275 | 4125249 | 4125370 | 4117307 | 4117451 | 4126134 | 4126237 | 25 [illegible] |
| MXE | 3029 | ENSG00000181036 | FCRL6 | chr1 | + | 159813488 | 159813554 | 159814220 | 159814292 | 159810093 | 159810216 | 159815535 | 159815802 | 143 [illegible] |
| MXE | 16688 | ENSG00000106133 | NSUN5P2 | chr7 | - | 72952151 | 72952295 | 72953325 | 72953430 | 72950072 | 72950196 | 72953510 | 72953677 | 14 [illegible] |
| MXE | 15108 | ENSG00000183431 | SF3A3 | chr1 | - | 37987783 | 37987836 | 37989547 | 37989595 | 37987572 | 37987678 | 37989869 | 37989973 | 62 [illegible] |
| MXE | 7044 | ENSG00000160049 | DFFA | chr1 | - | 10463057 | 10463209 | 10467189 | 10467332 | 10456521 | 10461702 | 10469176 | 10469338 | 72 [illegible] |
| MXE | 6444 | ENSG00000173083 | HPSE | chr4 | - | 83306202 | 83306317 | 83308844 | 83308951 | 83302149 | 83302268 | 83309401 | 83309495 | 80 [illegible] |
| MXE | 2154 | ENSG00000114062 | UBE3A | chr15 | - | 25407066 | 25407137 | 25408619 | 25408684 | 25405460 | 25405502 | 25409087 | 25409207 | 89 [illegible] |
| MXE | 8858 | ENSG00000196865 | NHLRC2 | chr10 | + | 113898109 | 113898209 | 113901665 | 113901897 | 113884250 | 113884380 | 113902470 | 113902593 | 41 [illegible] |
| MXE | 96 | ENSG00000182944 | EWSR1 | chr22 | + | 29299233 | 29299331 | 29299598 | 29299851 | 29298732 | 29298895 | 29300121 | 29300336 | 316 [illegible] |
| MXE | 14506 | ENSG00000134333 | LDHA | chr11 | + | 18399430 | 18399548 | 18402839 | 18403013 | 18396818 | 18396968 | 18403693 | 18403811 | 624 [illegible] |
| MXE | 2537 | ENSG00000165929 | TC2N | chr14 | - | 91797784 | 91797901 | 91798298 | 91798399 | 91787512 | 91787627 | 91798988 | 91799064 | 202 [illegible] |
| MXE | 1066 | ENSG00000108819 | PPP1R9B | chr17 | - | 50139428 | 50139581 | 50140092 | 50140228 | 50139262 | 50139316 | 50141268 | 50141373 | 514 [illegible] |
| MXE | 4264 | ENSG00000241258 | CRCP | chr7 | + | 66134279 | 66134374 | 66145442 | 66145500 | 66127703 | 66127740 | 66152207 | 66152870 | 0 [illegible] |
| MXE | 4003 | ENSG00000136709 | WDR33 | chr2 | - | 127726650 | 127726777 | 127763061 | 127763159 | 127725060 | 127725215 | 127764827 | 127764979 | 81 [illegible] |
| MXE | 8194 | ENSG00000188641 | DPYD | chr1 | - | 97593217 | 97593387 | 97595058 | 97595166 | 97573759 | 97573970 | 97679094 | 97679182 | 554 [illegible] |
| MXE | 9070 | ENSG00000077254 | USP33 | chr1 | - | 77734346 | 77734416 | 77736055 | 77736158 | 77730617 | 77730731 | 77739264 | 77739417 | 142 [illegible] |
| MXE | 9773 | ENSG00000175662 | TOM1L2 | chr17 | - | 17850892 | 17850952 | 17861475 | 17861551 | 17848822 | 17848859 | 17866295 | 17866419 | 1 [illegible] |
| MXE | 134 | ENSG00000100209 | HSCB | chr22 | + | 28745863 | 28746008 | 28751240 | 28751288 | 28744614 | 28744704 | 28757077 | 28757210 | 8 [illegible] |
| MXE | 4805 | ENSG00000198964 | SGMS1 | chr10 | - | 50466889 | 50466932 | 50519830 | 50519921 | 50460672 | 50460814 | 50590152 | 50590247 | 36 [illegible] |
| MXE | 14969 | ENSG00000178177 | LCORL | chr4 | - | 17909093 | 17909345 | 17960264 | 17960311 | 17886067 | 17886161 | 17961902 | 17962032 | 1 [illegible] |
| MXE | 9866 | ENSG00000110934 | BIN2 | chr12 | - | 51291590 | 51292344 | 51295795 | 51295878 | 51288107 | 51288188 | 51297088 | 51297164 | 435 [illegible] |
| MXE | 12514 | ENSG00000179818 | PCBP1-AS1 | chr2 | - | 70051202 | 70051305 | 70083537 | 70083687 | 70032132 | 70032255 | 70085546 | 70085624 | 33 [illegible] |
| MXE | 11257 | ENSG00000153094 | BCL2L11 | chr2 | + | 111123732 | 111123869 | 111150043 | 111150147 | 111120913 | 111121188 | 111164132 | 111165048 | 71 [illegible] |
| MXE | 5628 | ENSG00000127481 | UBR4 | chr1 | - | 19128210 | 19128318 | 19128977 | 19129074 | 19127622 | 19127739 | 19138006 | 19138181 | 38 [illegible] |
| MXE | 15499 | ENSG00000163131 | CTSS | chr1 | - | 150757857 | 150757980 | 150764637 | 150764764 | 150755000 | 150755150 | 150765697 | 150765778 | 3551 [illegible] |
| MXE | 6370 | ENSG00000137996 | RTCA | chr1 | + | 100267165 | 100267245 | 100267499 | 100267538 | 100266523 | 100266624 | 100268151 | 100268295 | 2 [illegible] |
| MXE | 14268 | ENSG00000089234 | BRAP | chr12 | - | 111659206 | 111659345 | 111665638 | 111665787 | 111658735 | 111658845 | 111672660 | 111672774 | 56 [illegible] |
| MXE | 1963 | ENSG00000130731 | METTL26 | chr16 | - | 634888 | 634956 | 635280 | 635340 | 634718 | 634797 | 635611 | 635774 | 6 [illegible] |
| MXE | 11929 | ENSG00000271793 | NA | chr6 | - | 85677423 | 85677492 | 85677953 | 85678032 | 85666003 | 85666147 | 85678135 | 85678733 | 67 [illegible] |
| MXE | 3651 | ENSG00000114107 | CEP70 | chr3 | - | 138570317 | 138570498 | 138571033 | 138571157 | 138536918 | 138537347 | 138571265 | 138571356 | 133 [illegible] |
| MXE | 14894 | ENSG00000172716 | SLFN11 | chr17 | - | 35366946 | 35367063 | 35367540 | 35367700 | 35363567 | 35363826 | 35373581 | 35373620 | 2 [illegible] |
| MXE | 17179 | ENSG00000005810 | MYCBP2 | chr13 | - | 77061661 | 77061790 | 77062595 | 77062697 | 77061168 | 77061301 | 77064614 | 77064734 | 187 [illegible] |
| MXE | 2513 | ENSG00000174373 | RALGAPA1 | chr14 | - | 35762709 | 35762753 | 35770941 | 35770999 | 35760828 | 35761006 | 35775005 | 35775055 | 28 [illegible] |
| MXE | 3032 | ENSG00000119899 | SLC17A5 | chr6 | - | 73600350 | 73600441 | 73610399 | 73610547 | 73593378 | 73595214 | 73615314 | 73615447 | 30 [illegible] |
| MXE | 8205 | ENSG00000180353 | HCLS1 | chr3 | - | 121647318 | 121647448 | 121657278 | 121657352 | 121644229 | 121644928 | 121658263 | 121658347 | 598 [illegible] |
| MXE | 6736 | ENSG00000145703 | IQGAP2 | chr5 | + | 76606178 | 76606303 | 76611019 | 76611183 | 76600817 | 76600972 | 76627409 | 76627500 | 541 [illegible] |
| MXE | 15553 | ENSG00000214787 | MS4A4E | chr11 | - | 60224964 | 60225093 | 60228593 | 60228627 | 60214570 | 60214614 | 60229911 | 60230055 | 0 [illegible] |
| MXE | 1775 | ENSG00000196712 | NF1 | chr17 | + | 31327498 | 31327839 | 31330295 | 31330498 | 31325819 | 31326252 | 31334837 | 31335031 | 112 [illegible] |
| MXE | 11193 | ENSG00000115904 | SOS1 | chr2 | - | 39007030 | 39007193 | 39010583 | 39010703 | 39006411 | 39006529 | 39012125 | 39012348 | 89 [illegible] |
| MXE | 15895 | ENSG00000144815 | NXPE3 | chr3 | + | 101807052 | 101807126 | 101816794 | 101817001 | 101801234 | 101801989 | 101821403 | 101822823 | 28 [illegible] |
| MXE | 8202 | ENSG00000180353 | HCLS1 | chr3 | - | 121637145 | 121637256 | 121642926 | 121642981 | 121636433 | 121636489 | 121644817 | 121644929 | 353 [illegible] |
| MXE | 10006 | ENSG00000100644 | HIF1A | chr14 | + | 61738086 | 61738373 | 61740754 | 61741188 | 61736888 | 61737109 | 61744704 | 61744813 | 394 [illegible] |
| MXE | 14383 | ENSG00000119397 | CNTRL | chr9 | + | 121094887 | 121095018 | 121096421 | 121096563 | 121090274 | 121090405 | 121098385 | 121098572 | 46 [illegible] |
| MXE | 9330 | ENSG00000070785 | EIF2B3 | chr1 | - | 44881611 | 44881739 | 44897354 | 44897444 | 44879817 | 44880008 | 44926627 | 44926655 | 35 [illegible] |
| MXE | 5235 | ENSG00000136152 | COG3 | chr13 | + | 45492158 | 45492250 | 45493346 | 45493486 | 45491411 | 45491538 | 45496151 | 45496312 | 45 [illegible] |
| MXE | 10118 | ENSG00000125875 | TBC1D20 | chr20 | - | 445049 | 445130 | 447888 | 448074 | 441856 | 442043 | 462335 | 462516 | 20 [illegible] |
| MXE | 16716 | ENSG00000123329 | ARHGAP9 | chr12 | - | 57474041 | 57474176 | 57474422 | 57474476 | 57473602 | 57473708 | 57474625 | 57474703 | 119 [illegible] |
| MXE | 13326 | ENSG00000101901 | ALG13 | chrX | + | 111689086 | 111689197 | 111711674 | 111711725 | 111684964 | 111685103 | 111712483 | 111712530 | 0 [illegible] |
| MXE | 5379 | ENSG00000138246 | DNAJC13 | chr3 | + | 132463695 | 132463817 | 132465994 | 132466070 | 132462466 | 132462523 | 132466298 | 132466394 | 76 [illegible] |
| MXE | 3823 | ENSG00000082397 | EPB41L3 | chr18 | - | 5394676 | 5394793 | 5395066 | 5395147 | 5392380 | 5393478 | 5395608 | 5395707 | 57 [illegible] |
| MXE | 8664 | ENSG00000117360 | PRPF3 | chr1 | + | 150328319 | 150328466 | 150332683 | 150332767 | 150325750 | 150325881 | 150332978 | 150333199 | 91 [illegible] |
| MXE | 2894 | ENSG00000120688 | WBP4 | chr13 | + | 41076043 | 41076237 | 41080645 | 41080809 | 41072781 | 41072857 | 41082703 | 41084006 | 65 [illegible] |
| MXE | 16094 | ENSG00000099290 | WASHC2A | chr10 | + | 50078674 | 50078737 | 50080757 | 50080931 | 50069546 | 50069711 | 50084071 | 50084165 | 36 [illegible] |
| MXE | 14877 | ENSG00000165282 | PIGO | chr9 | - | 35093029 | 35093209 | 35093420 | 35093580 | 35091239 | 35092767 | 35093900 | 35094024 | 28 [illegible] |
| MXE | 12305 | ENSG00000162600 | OMA1 | chr1 | - | 58530600 | 58530729 | 58533952 | 58534060 | 58527260 | 58527335 | 58534157 | 58534331 | 384 [illegible] |
| MXE | 7645 | ENSG00000100591 | AHSA1 | chr14 | + | 77462641 | 77462759 | 77465538 | 77465667 | 77462159 | 77462242 | 77468082 | 77468184 | 105 [illegible] |
| MXE | 12228 | ENSG00000106245 | BUD31 | chr7 | + | 99410033 | 99410169 | 99411063 | 99411186 | 99408640 | 99409004 | 99416137 | 99416260 | 36 [illegible] |
| MXE | 13610 | ENSG00000122122 | SASH3 | chrX | + | 129792626 | 129792836 | 129792988 | 129793139 | 129792327 | 129792476 | 129793641 | 129795201 | 290 [illegible] |
| MXE | 16686 | ENSG00000112200 | ZNF451 | chr6 | + | 57134743 | 57134870 | 57141301 | 57141455 | 57133041 | 57133192 | 57141947 | 57142095 | 67 [illegible] |
| MXE | 5857 | ENSG00000156875 | MFSD14A | chr1 | + | 100068472 | 100068586 | 100069613 | 100069685 | 100067976 | 100068195 | 100077164 | 100077277 | 102 [illegible] |
| MXE | 11571 | ENSG00000197548 | ATG7 | chr3 | + | 11340644 | 11340735 | 11342134 | 11342279 | 11332971 | 11333093 | 11347876 | 11348035 | 56 [illegible] |
| MXE | 8363 | ENSG00000135842 | NIBAN1 | chr1 | - | 184803584 | 184803692 | 184818637 | 184818825 | 184798078 | 184798190 | 184823166 | 184823329 | 125 [illegible] |
| MXE | 14953 | ENSG00000137073 | UBAP2 | chr9 | - | 33996222 | 33996333 | 33998786 | 33998864 | 33988972 | 33989126 | 34011626 | 34011664 | 0 [illegible] |
| MXE | 950 | ENSG00000101782 | RIOK3 | chr18 | + | 23477178 | 23477268 | 23479316 | 23479424 | 23477005 | 23477086 | 23481171 | 23481580 | 173 [illegible] |
| MXE | 13780 | ENSG00000156504 | PABIR2 | chrX | - | 134785885 | 134785950 | 134787471 | 134787533 | 134781820 | 134781917 | 134788729 | 134788831 | 49 [illegible] |
| MXE | 14515 | ENSG00000156471 | PTDSS1 | chr8 | + | 96284108 | 96284153 | 96287021 | 96287146 | 96273298 | 96273390 | 96295097 | 96295256 | 36 [illegible] |
| MXE | 16124 | ENSG00000011376 | LARS2 | chr3 | + | 45520218 | 45520296 | 45523996 | 45524108 | 45517902 | 45518072 | 45541828 | 45541885 | 10 [illegible] |
| MXE | 4830 | ENSG00000013441 | CLK1 | chr2 | - | 200856681 | 200856811 | 200856890 | 200857884 | 200854615 | 200854695 | 200857972 | 200858089 | 561 [illegible] |
| MXE | 5741 | ENSG00000196187 | TMEM63A | chr1 | - | 225856651 | 225856738 | 225856910 | 225857017 | 225855877 | 225855940 | 225859195 | 225859349 | 31 [illegible] |
| MXE | 1556 | ENSG00000140829 | DHX38 | chr16 | + | 72097676 | 72097781 | 72098644 | 72098792 | 72096821 | 72097009 | 72098926 | 72099045 | 60 [illegible] |
| MXE | 8250 | ENSG00000143442 | POGZ | chr1 | - | 151423948 | 151424286 | 151427822 | 151428041 | 151423396 | 151423551 | 151428122 | 151428413 | 37 [illegible] |
| MXE | 15633 | ENSG00000078124 | ACER3 | chr11 | + | 77011329 | 77011375 | 77015015 | 77015659 | 76998762 | 76998821 | 77016674 | 77016779 | 1 [illegible] |
| MXE | 6340 | ENSG00000162521 | RBBP4 | chr1 | + | 32657426 | 32657572 | 32668738 | 32668854 | 32651913 | 32652061 | 32668971 | 32669132 | 594 [illegible] |
| MXE | 15695 | ENSG00000054654 | SYNE2 | chr14 | + | 64101931 | 64102042 | 64107490 | 64107607 | 64098724 | 64098821 | 64113340 | 64113572 | 231 [illegible] |
| MXE | 10549 | ENSG00000020922 | MRE11 | chr11 | - | 94464112 | 94464239 | 94476288 | 94476403 | 94460935 | 94461036 | 94478734 | 94478876 | 108 [illegible] |
| MXE | 15961 | ENSG00000138190 | EXOC6 | chr10 | + | 92934309 | 92934430 | 92935813 | 92935885 | 92934143 | 92934190 | 92940726 | 92940824 | 84 [illegible] |
| MXE | 4056 | ENSG00000197020 | ZNF100 | chr19 | - | 21744016 | 21744112 | 21744940 | 21745067 | 21726724 | 21727989 | 21765693 | 21765783 | 11 [illegible] |
| MXE | 7002 | ENSG00000213593 | TMX2 | chr11 | + | 57737912 | 57738026 | 57738663 | 57738770 | 57737607 | 57737668 | 57738973 | 57739039 | 46 [illegible] |
| MXE | 3708 | ENSG00000158987 | RAPGEF6 | chr5 | - | 131505363 | 131505522 | 131508070 | 131508207 | 131504625 | 131504778 | 131510313 | 131510491 | 160 [illegible] |
| MXE | 3654 | ENSG00000145715 | RASA1 | chr5 | + | 87380508 | 87380595 | 87383712 | 87383780 | 87379734 | 87379850 | 87385300 | 87385389 | 50 [illegible] |
| MXE | 9990 | ENSG00000145241 | CENPC | chr4 | - | 67495158 | 67495212 | 67505204 | 67505284 | 67493404 | 67493988 | 67506787 | 67506934 | 88 [illegible] |
| MXE | 7965 | ENSG00000156642 | NPTN | chr15 | - | 73573661 | 73573795 | 73587523 | 73587618 | 73570149 | 73570423 | 73591965 | 73592137 | 222 [illegible] |
| MXE | 2925 | ENSG00000124789 | NUP153 | chr6 | - | 17624560 | 17624833 | 17625807 | 17626164 | 17616526 | 17616695 | 17628654 | 17629539 | 196 [illegible] |
| MXE | 14341 | ENSG00000105339 | DENND3 | chr8 | + | 141168003 | 141168525 | 141175199 | 141175459 | 141166189 | 141166389 | 141176590 | 141176761 | 75 [illegible] |
| MXE | 16674 | ENSG00000185482 | STAC3 | chr12 | - | 57245144 | 57245211 | 57246803 | 57246901 | 57244915 | 57244965 | 57248125 | 57248198 | 29 [illegible] |
| MXE | 2845 | ENSG00000258989 | NA | chr14 | + | 61530406 | 61530595 | 61547742 | 61547886 | 61529127 | 61529213 | 61657604 | 61657964 | 357 [illegible] |
| MXE | 15371 | ENSG00000107036 | RIC1 | chr9 | + | 5732387 | 5732479 | 5742868 | 5743013 | 5720613 | 5720750 | 5743688 | 5743737 | 73 [illegible] |
| MXE | 15726 | ENSG00000130830 | MPP1 | chrX | - | 154789953 | 154790022 | 154790982 | 154791068 | 154786203 | 154786400 | 154791768 | 154791847 | 82 [illegible] |
| MXE | 15321 | ENSG00000171634 | BPTF | chr17 | + | 67886127 | 67886316 | 67891843 | 67892034 | 67875555 | 67875744 | 67893369 | 67893725 | 28 [illegible] |
| MXE | 7961 | ENSG00000142197 | DOP1B | chr21 | + | 36247516 | 36247628 | 36248379 | 36248568 | 36245047 | 36246677 | 36251161 | 36251284 | 34 [illegible] |
| MXE | 15722 | ENSG00000109917 | ZPR1 | chr11 | - | 116782157 | 116782244 | 116782918 | 116783029 | 116779771 | 116779837 | 116783529 | 116783619 | 129 [illegible] |
| MXE | 1388 | ENSG00000154229 | PRKCA | chr17 | + | 66496200 | 66496283 | 66641354 | 66641466 | 66306095 | 66306127 | 66645382 | 66645511 | 17 [illegible] |
| MXE | 11703 | ENSG00000197081 | IGF2R | chr6 | + | 160027184 | 160027314 | 160029549 | 160029655 | 160024571 | 160024704 | 160032550 | 160032713 | 332 [illegible] |
| MXE | 16903 | ENSG00000111335 | OAS2 | chr12 | + | 113002931 | 113003102 | 113004933 | 113005222 | 112997519 | 112998410 | 113006412 | 113006600 | 477 [illegible] |
| MXE | 2027 | ENSG00000069966 | GNB5 | chr15 | - | 52147458 | 52147535 | 52149883 | 52149925 | 52141139 | 52141272 | 52153939 | 52154076 | 4 [illegible] |
| MXE | 446 | ENSG00000063176 | SPHK2 | chr19 | + | 48625890 | 48626362 | 48627691 | 48627841 | 48620401 | 48620553 | 48628161 | 48628277 | 7 [illegible] |
| MXE | 13671 | ENSG00000215301 | DDX3X | chrX | + | 41339038 | 41339083 | 41340635 | 41340814 | 41337407 | 41337465 | 41341483 | 41341616 | 262 [illegible] |
| MXE | 14073 | ENSG00000153317 | ASAP1 | chr8 | - | 130214555 | 130214701 | 130236921 | 130236994 | 130188108 | 130188183 | 130358016 | 130358143 | 40 [illegible] |
| MXE | 11052 | ENSG00000155749 | FLACC1 | chr2 | - | 201330735 | 201330833 | 201342369 | 201342431 | 201330469 | 201330522 | 201344169 | 201344263 | 6 [illegible] |
| MXE | 11091 | ENSG00000129691 | ASH2L | chr8 | + | 38116649 | 38116725 | 38119269 | 38119363 | 38114904 | 38115000 | 38120931 | 38121149 | 147 [illegible] |
| MXE | 15926 | ENSG00000181192 | DHTKD1 | chr10 | + | 12100177 | 12100262 | 12101041 | 12101181 | 12097683 | 12097996 | 12106245 | 12106396 | 20 [illegible] |
| MXE | 8744 | ENSG00000181722 | ZBTB20 | chr3 | - | 114710267 | 114710313 | 114801100 | 114801174 | 114693527 | 114693575 | 114900303 | 114900342 | 141 [illegible] |
| MXE | 5739 | ENSG00000196187 | TMEM63A | chr1 | - | 225853628 | 225853791 | 225855877 | 225855940 | 225852663 | 225852769 | 225856651 | 225856738 | 69 [illegible] |
| MXE | 12782 | ENSG00000177189 | RPS6KA3 | chrX | - | 20176995 | 20177084 | 20186295 | 20186366 | 20176433 | 20176498 | 20188496 | 20188534 | 0 [illegible] |
| MXE | 5063 | ENSG00000172046 | USP19 | chr3 | - | 49116726 | 49116943 | 49117656 | 49117785 | 49116450 | 49116607 | 49117946 | 49117978 | 7 [illegible] |
| MXE | 3198 | ENSG00000112081 | SRSF3 | chr6 | + | 36601151 | 36601190 | 36601707 | 36601794 | 36598848 | 36598983 | 36601961 | 36602197 | 98 [illegible] |
| MXE | 10756 | ENSG00000177479 | ARIH2 | chr3 | + | 48927461 | 48927813 | 48961611 | 48961679 | 48924717 | 48924839 | 48967124 | 48967275 | 36 [illegible] |
| MXE | 6833 | ENSG00000007341 | ST7L | chr1 | - | 112583971 | 112584126 | 112597970 | 112598086 | 112582374 | 112582472 | 112610840 | 112611003 | 4 [illegible] |
| MXE | 7385 | ENSG00000126091 | ST3GAL3 | chr1 | + | 43814890 | 43814933 | 43838218 | 43838311 | 43736262 | 43736380 | 43894382 | 43894477 | 0 [illegible] |
| MXE | 7796 | ENSG00000144136 | SLC20A1 | chr2 | + | 112647323 | 112647464 | 112647652 | 112647738 | 112646562 | 112647162 | 112652701 | 112652798 | 75 [illegible] |
| MXE | 6808 | ENSG00000117682 | DHDDS | chr1 | + | 26442730 | 26442873 | 26447558 | 26447660 | 26438167 | 26438284 | 26457790 | 26457905 | 43 [illegible] |
| MXE | 11699 | ENSG00000244482 | LILRA6 | chr19 | - | 54240830 | 54241127 | 54241661 | 54241878 | 54240273 | 54240576 | 54242025 | 54242310 | 9 [illegible] |
| MXE | 1649 | ENSG00000188603 | CLN3 | chr16 | - | 28487661 | 28487741 | 28488590 | 28488662 | 28487455 | 28487541 | 28489289 | 28489386 | 36 [illegible] |
| MXE | 15470 | ENSG00000162231 | NXF1 | chr11 | - | 62803418 | 62803572 | 62803791 | 62803978 | 62802176 | 62802260 | 62805328 | 62805440 | 134 [illegible] |
| MXE | 1559 | ENSG00000102984 | ZNF821 | chr16 | - | 71864137 | 71864242 | 71867911 | 71868037 | 71859679 | 71860672 | 71879906 | 71880023 | 15 [illegible] |
| MXE | 14177 | ENSG00000173064 | HECTD4 | chr12 | - | 112179173 | 112179397 | 112183058 | 112183266 | 112178930 | 112179082 | 112184186 | 112185493 | 59 [illegible] |
| MXE | 8242 | ENSG00000011405 | PIK3C2A | chr11 | - | 17135110 | 17135159 | 17136481 | 17136625 | 17134818 | 17135029 | 17145667 | 17145731 | 54 [illegible] |
| MXE | 15560 | ENSG00000013725 | CD6 | chr11 | + | 61009571 | 61009874 | 61011069 | 61011135 | 61008533 | 61008845 | 61013422 | 61013563 | 76 [illegible] |
| MXE | 15395 | ENSG00000157600 | TMEM164 | chrX | + | 110067346 | 110067396 | 110109079 | 110109146 | 110003500 | 110004164 | 110144797 | 110144876 | 2 [illegible] |
| MXE | 6317 | ENSG00000118308 | IRAG2 | chr12 | + | 25063722 | 25063816 | 25066364 | 25066512 | 25062819 | 25062900 | 25069349 | 25069431 | 47 [illegible] |
| MXE | 17158 | ENSG00000134900 | TPP2 | chr13 | + | 102657055 | 102657207 | 102663647 | 102663744 | 102649407 | 102649486 | 102664794 | 102664925 | 176 [illegible] |
| MXE | 12509 | ENSG00000179818 | PCBP1-AS1 | chr2 | - | 70051202 | 70051305 | 70053730 | 70053792 | 69996685 | 69996888 | 70059642 | 70059680 | 0 [illegible] |
| MXE | 4329 | ENSG00000145495 | MARCHF6 | chr5 | + | 10415472 | 10415669 | 10417269 | 10417404 | 10414432 | 10414502 | 10423734 | 10423835 | 5 [illegible] |
| MXE | 5438 | ENSG00000145819 | ARHGAP26 | chr5 | + | 142901934 | 142902039 | 142903539 | 142903669 | 142894237 | 142894348 | 142907703 | 142907804 | 167 [illegible] |
| MXE | 12364 | ENSG00000164808 | SPIDR | chr8 | + | 47407860 | 47407961 | 47440322 | 47440542 | 47291032 | 47291137 | 47595810 | 47596006 | 0 [illegible] |
| MXE | 5052 | ENSG00000176261 | ZBTB8OS | chr1 | - | 32633644 | 32633727 | 32633950 | 32634072 | 32627507 | 32627544 | 32634767 | 32634792 | 63 [illegible] |
| MXE | 16545 | ENSG00000028203 | VEZT | chr12 | + | 95257149 | 95257239 | 95262905 | 95263081 | 95256549 | 95256622 | 95266356 | 95266632 | 17 [illegible] |
| MXE | 8772 | ENSG00000042445 | RETSAT | chr2 | - | 85343638 | 85343798 | 85344238 | 85344348 | 85341954 | 85343381 | 85344593 | 85344732 | 15 [illegible] |
| MXE | 117 | ENSG00000242259 | C22orf39 | chr22 | - | 19356889 | 19357051 | 19359335 | 19359484 | 19356229 | 19356288 | 19361236 | 19361341 | 56 [illegible] |
| MXE | 9030 | ENSG00000072858 | SIDT1 | chr3 | + | 113611007 | 113611144 | 113616099 | 113616176 | 113608418 | 113608536 | 113619679 | 113619726 | 25 [illegible] |
| MXE | 7542 | ENSG00000076685 | NT5C2 | chr10 | - | 103111744 | 103111805 | 103136248 | 103136372 | 103106588 | 103106706 | 103139405 | 103139479 | 7 [illegible] |
| MXE | 13852 | ENSG00000138378 | STAT4 | chr2 | - | 191054489 | 191054534 | 191058017 | 191058111 | 191041064 | 191041148 | 191058201 | 191058219 | 160 [illegible] |
| MXE | 11708 | ENSG00000197081 | IGF2R | chr6 | + | 160075846 | 160075996 | 160078200 | 160078362 | 160073756 | 160073975 | 160079579 | 160079787 | 273 [illegible] |
| MXE | 16128 | ENSG00000165732 | DDX21 | chr10 | + | 68962081 | 68962157 | 68963290 | 68963469 | 68959805 | 68960249 | 68965376 | 68965494 | 268 [illegible] |
| MXE | 14553 | ENSG00000107242 | PIP5K1B | chr9 | + | 68934889 | 68935045 | 68940645 | 68940790 | 68923301 | 68923386 | 68991139 | 68991257 | 46 [illegible] |
| MXE | 1910 | ENSG00000102910 | LONP2 | chr16 | + | 48296014 | 48296165 | 48299661 | 48299788 | 48277337 | 48277479 | 48303171 | 48303305 | 124 [illegible] |
| MXE | 15566 | ENSG00000013725 | CD6 | chr11 | + | 61013918 | 61014014 | 61015712 | 61015835 | 61011069 | 61011135 | 61017478 | 61017550 | 0 [illegible] |
| MXE | 6767 | ENSG00000138069 | RAB1A | chr2 | - | 65088938 | 65089070 | 65090982 | 65091078 | 65086853 | 65088690 | 65097970 | 65098066 | 200 [illegible] |
| MXE | 4950 | ENSG00000172053 | QARS1 | chr3 | - | 49101354 | 49101441 | 49101619 | 49101705 | 49100574 | 49100674 | 49101827 | 49101899 | 139 [illegible] |
| MXE | 1082 | ENSG00000198862 | LTN1 | chr21 | - | 28943666 | 28943904 | 28944382 | 28944596 | 28943261 | 28943336 | 28945806 | 28945951 | 52 [illegible] |
| MXE | 5416 | ENSG00000131323 | TRAF3 | chr14 | + | 102886188 | 102886269 | 102889559 | 102889634 | 102876357 | 102876525 | 102891324 | 102891417 | 35 [illegible] |
| MXE | 7801 | ENSG00000066427 | ATXN3 | chr14 | - | 92088729 | 92088817 | 92093251 | 92093318 | 92083125 | 92083258 | 92093745 | 92093831 | 31 [illegible] |
| MXE | 542 | ENSG00000090372 | STRN4 | chr19 | - | 46733038 | 46733236 | 46736822 | 46736901 | 46730731 | 46730873 | 46738163 | 46738172 | 57 [illegible] |
| MXE | 15793 | ENSG00000175224 | ATG13 | chr11 | + | 46665382 | 46665519 | 46667772 | 46667887 | 46659334 | 46659485 | 46668498 | 46668576 | 79 [illegible] |
| MXE | 15579 | ENSG00000073910 | FRY | chr13 | + | 32147281 | 32147385 | 32147838 | 32147947 | 32136870 | 32136972 | 32149747 | 32149834 | 5 [illegible] |
| MXE | 11818 | ENSG00000164691 | TAGAP | chr6 | - | 159042077 | 159042244 | 159043588 | 159043655 | 159041353 | 159041515 | 159043977 | 159044031 | 398 [illegible] |
| MXE | 4397 | ENSG00000121210 | TMEM131L | chr4 | + | 153632717 | 153632838 | 153634191 | 153634280 | 153627604 | 153627687 | 153635431 | 153635571 | 88 [illegible] |
| MXE | 6836 | ENSG00000007341 | ST7L | chr1 | - | 112597970 | 112598086 | 112600793 | 112600848 | 112591524 | 112591603 | 112610840 | 112611003 | 18 [illegible] |
| MXE | 4600 | ENSG00000100412 | ACO2 | chr22 | + | 41522829 | 41522982 | 41523204 | 41523278 | 41520170 | 41520276 | 41523829 | 41523941 | 197 [illegible] |
| MXE | 4156 | ENSG00000152818 | UTRN | chr6 | + | 144839172 | 144839284 | 144840739 | 144840832 | 144836300 | 144836541 | 144846804 | 144846827 | 282 [illegible] |
| MXE | 2186 | ENSG00000129003 | VPS13C | chr15 | - | 61951823 | 61951980 | 61954420 | 61954554 | 61950944 | 61951024 | 61958607 | 61958653 | 268 [illegible] |
| MXE | 1339 | ENSG00000154265 | ABCA5 | chr17 | - | 69306724 | 69306954 | 69308279 | 69308368 | 69304093 | 69304810 | 69309261 | 69309320 | 12 [illegible] |
| MXE | 15971 | ENSG00000065613 | SLK | chr10 | + | 104001443 | 104001572 | 104002171 | 104003527 | 103999866 | 103999948 | 104005560 | 104005691 | 236 [illegible] |
| MXE | 15505 | ENSG00000174483 | BBS1 | chr11 | + | 66514405 | 66514678 | 66515539 | 66515586 | 66511204 | 66511239 | 66515692 | 66515731 | 7 [illegible] |
| MXE | 11626 | ENSG00000198223 | CSF2RA | chrX | + | 1285777 | 1285920 | 1288518 | 1288642 | 1282677 | 1282779 | 1288758 | 1288888 | 72 [illegible] |
| MXE | 3222 | ENSG00000189091 | SF3B3 | chr16 | + | 70560468 | 70560591 | 70561629 | 70561784 | 70556885 | 70557029 | 70563875 | 70564050 | 109 [illegible] |
| MXE | 627 | ENSG00000131355 | ADGRE3 | chr19 | - | 14654981 | 14655165 | 14658479 | 14658550 | 14651084 | 14651204 | 14661962 | 14662118 | 38 [illegible] |
| MXE | 14929 | ENSG00000137145 | DENND4C | chr9 | + | 19300186 | 19300331 | 19305351 | 19305527 | 19299228 | 19299287 | 19316416 | 19316517 | 183 [illegible] |
| MXE | 11092 | ENSG00000166197 | NOLC1 | chr10 | + | 102157448 | 102157555 | 102158048 | 102158214 | 102157018 | 102157074 | 102159192 | 102159308 | 0 [illegible] |
| MXE | 5083 | ENSG00000157036 | EXOG | chr3 | + | 38499680 | 38499747 | 38500089 | 38500164 | 38497628 | 38497778 | 38501354 | 38501494 | 0 [illegible] |
| MXE | 1407 | ENSG00000011260 | UTP18 | chr17 | + | 51288547 | 51288591 | 51293902 | 51294045 | 51288028 | 51288203 | 51296964 | 51297003 | 1 [illegible] |
| MXE | 6098 | ENSG00000103342 | GSPT1 | chr16 | - | 11877406 | 11877580 | 11883014 | 11883095 | 11876085 | 11876175 | 11885180 | 11885274 | 204 [illegible] |
| MXE | 5011 | ENSG00000164327 | RICTOR | chr5 | - | 38975536 | 38975604 | 38978582 | 38978650 | 38969686 | 38971959 | 38981866 | 38982036 | 106 [illegible] |
| MXE | 14182 | ENSG00000173064 | HECTD4 | chr12 | - | 112246900 | 112247076 | 112247461 | 112247550 | 112243873 | 112244009 | 112248066 | 112248171 | 42 [illegible] |
| MXE | 5136 | ENSG00000111731 | C2CD5 | chr12 | - | 22484696 | 22484888 | 22490122 | 22490218 | 22482556 | 22482743 | 22493222 | 22493337 | 89 [illegible] |
| MXE | 4200 | ENSG00000084676 | NCOA1 | chr2 | + | 24693251 | 24693347 | 24697657 | 24697798 | 24691480 | 24691660 | 24705085 | 24705233 | 190 [illegible] |
| MXE | 12887 | ENSG00000105835 | NAMPT | chr7 | - | 106254363 | 106254504 | 106261587 | 106261707 | 106253016 | 106253151 | 106263391 | 106263617 | 234 [illegible] |
| MXE | 1880 | ENSG00000167261 | DPEP2 | chr16 | - | 67991114 | 67991184 | 67991837 | 67991979 | 67990820 | 67990997 | 67992063 | 67992193 | 41 [illegible] |
| MXE | 12431 | ENSG00000147065 | MSN | chrX | + | 65727813 | 65727909 | 65729437 | 65729712 | 65716817 | 65716901 | 65731106 | 65731190 | 646 [illegible] |
| MXE | 6884 | ENSG00000100991 | TRPC4AP | chr20 | - | 35049865 | 35049994 | 35054975 | 35055031 | 35044504 | 35044712 | 35057513 | 35057571 | 85 [illegible] |
| MXE | 14737 | ENSG00000090054 | SPTLC1 | chr9 | - | 92076797 | 92077035 | 92079433 | 92079566 | 92067965 | 92068098 | 92080015 | 92080088 | 33 [illegible] |
| MXE | 2011 | ENSG00000185596 | WASH3P | chr15 | + | 101973219 | 101973352 | 101973591 | 101973727 | 101972900 | 101973047 | 101973904 | 101973939 | 48 [illegible] |
| MXE | 11723 | ENSG00000135316 | SYNCRIP | chr6 | - | 85636966 | 85637143 | 85637242 | 85637356 | 85623976 | 85624112 | 85640220 | 85640328 | 77 [illegible] |
| MXE | 3559 | ENSG00000153071 | DAB2 | chr5 | - | 39381453 | 39381616 | 39382617 | 39383271 | 39376649 | 39377282 | 39388798 | 39388852 | 6 [illegible] |
| MXE | 4168 | ENSG00000109814 | UGDH | chr4 | - | 39514082 | 39514184 | 39521350 | 39521519 | 39510660 | 39510861 | 39527282 | 39527439 | 42 [illegible] |
| MXE | 2579 | ENSG00000092108 | SCFD1 | chr14 | + | 30638124 | 30638247 | 30639776 | 30639864 | 30633946 | 30634037 | 30643315 | 30643405 | 141 [illegible] |
| MXE | 16235 | ENSG00000148634 | HERC4 | chr10 | - | 68025545 | 68025676 | 68032777 | 68032869 | 68014025 | 68014186 | 68033964 | 68034186 | 55 [illegible] |
| MXE | 17065 | ENSG00000180370 | PAK2 | chr3 | + | 196814450 | 196814568 | 196818056 | 196818156 | 196812738 | 196812851 | 196820370 | 196820567 | 501 [illegible] |
| MXE | 2960 | ENSG00000162711 | NLRP3 | chr1 | + | 247429584 | 247429755 | 247435969 | 247436140 | 247423846 | 247425599 | 247443971 | 247444142 | 20 [illegible] |
| MXE | 13021 | ENSG00000234741 | GAS5 | chr1 | - | 173864674 | 173864704 | 173865228 | 173865282 | 173864483 | 173864506 | 173865470 | 173865547 | 147 [illegible] |
| MXE | 297 | ENSG00000100077 | GRK3 | chr22 | + | 25672295 | 25672347 | 25678815 | 25678915 | 25667738 | 25667800 | 25685169 | 25685248 | 59 [illegible] |
| MXE | 16516 | ENSG00000188342 | GTF2F2 | chr13 | + | 45149769 | 45149788 | 45151686 | 45151831 | 45136732 | 45136806 | 45207423 | 45207505 | 8 [illegible] |
| MXE | 14376 | ENSG00000104529 | EEF1D | chr8 | - | 143581228 | 143581328 | 143586728 | 143586849 | 143581053 | 143581154 | 143597347 | 143597399 | 37 [illegible] |

| | | | | | | |
|---|---|---|---|---|---|---|
| MXE | 9387 | ENSG00000135829 | DHX9 | chr1 | + | [illegible] |
| MXE | 2054 | ENSG00000090487 | SPG21 | chr15 | - | [illegible] |
| MXE | 13215 | ENSG00000230590 | FTX | chrX | - | [illegible] |
| MXE | 3185 | ENSG00000133706 | LARS1 | chr5 | - | [illegible] |
| MXE | 15460 | ENSG00000086848 | ALG9 | chr11 | - | [illegible] |
| MXE | 931 | ENSG00000175354 | PTPN2 | chr18 | - | [illegible] |
| MXE | 7244 | ENSG00000145687 | SSBP2 | chr5 | - | [illegible] |
| MXE | 10065 | ENSG00000111275 | ALDH2 | chr12 | + | [illegible] |
| MXE | 15441 | ENSG00000110274 | CEP164 | chr11 | + | [illegible] |
| MXE | 229 | ENSG00000054611 | TBC1D22A | chr22 | + | [illegible] |
| MXE | 2326 | ENSG00000104043 | ATP8B4 | chr15 | - | [illegible] |
| MXE | 15289 | ENSG00000149547 | EI24 | chr11 | + | [illegible] |
| MXE | 8167 | ENSG00000173575 | CHD2 | chr15 | + | [illegible] |
| MXE | 812 | ENSG00000170471 | RALGAPB | chr20 | + | [illegible] |
| MXE | 14260 | ENSG00000154114 | TBCEL | chr11 | + | [illegible] |
| MXE | 16155 | ENSG00000108094 | CUL2 | chr10 | - | [illegible] |
| MXE | 15700 | ENSG00000054654 | SYNE2 | chr14 | + | [illegible] |
| MXE | 10236 | ENSG00000088179 | PTPN4 | chr2 | + | [illegible] |
| MXE | 4663 | ENSG00000025293 | PHF20 | chr20 | + | [illegible] |
| MXE | 2495 | ENSG00000197102 | DYNC1H1 | chr14 | + | [illegible] |
| MXE | 15214 | ENSG00000113163 | CERT1 | chr5 | - | [illegible] |
| MXE | 21 | ENSG00000154727 | GABPA | chr21 | + | [illegible] |
| MXE | 4126 | ENSG00000132694 | ARHGEF11 | chr1 | - | [illegible] |
| MXE | 9619 | ENSG00000119778 | ATAD2B | chr2 | - | [illegible] |
| MXE | 9820 | ENSG00000151092 | NGLY1 | chr3 | - | [illegible] |
| MXE | 3179 | ENSG00000135341 | MAP3K7 | chr6 | - | [illegible] |
| MXE | 2943 | ENSG00000111913 | RIPOR2 | chr6 | - | [illegible] |
| MXE | 2315 | ENSG00000169371 | SNUPN | chr15 | - | [illegible] |
| MXE | 9178 | ENSG00000119004 | CYP20A1 | chr2 | + | [illegible] |
| MXE | 11652 | ENSG00000072121 | ZFYVE26 | chr14 | - | [illegible] |
| MXE | 16816 | ENSG00000288663 | NA | chr12 | + | [illegible] |
| MXE | 4924 | ENSG00000114127 | XRN1 | chr3 | - | [illegible] |
| MXE | 13480 | ENSG00000111203 | ITFG2 | chr12 | + | [illegible] |
| MXE | 15199 | ENSG00000070718 | AP3M2 | chr8 | + | [illegible] |
| MXE | 14745 | ENSG00000077463 | SIRT6 | chr19 | - | [illegible] |
| MXE | 1406 | ENSG00000011260 | UTP18 | chr17 | + | [illegible] |
| MXE | 2319 | ENSG00000140365 | COMMD4 | chr15 | + | [illegible] |
| MXE | 3426 | ENSG00000198612 | COPS8 | chr2 | + | [illegible] |
| MXE | 3181 | ENSG00000135341 | MAP3K7 | chr6 | - | [illegible] |
| MXE | 4994 | ENSG00000188313 | PLSCR1 | chr3 | - | [illegible] |
| MXE | 375 | ENSG00000104880 | ARHGEF18 | chr19 | + | [illegible] |
| MXE | 11794 | ENSG00000204305 | AGER | chr6 | - | [illegible] |
| MXE | 5882 | ENSG00000122674 | CCZ1 | chr7 | + | [illegible] |
| MXE | 399 | ENSG00000197961 | ZNF121 | chr19 | - | [illegible] |
| MXE | 1315 | ENSG00000083290 | ULK2 | chr17 | - | [illegible] |
| MXE | 6835 | ENSG00000007341 | ST7L | chr1 | - | [illegible] |
| MXE | 13747 | ENSG00000143569 | UBAP2L | chr1 | + | [illegible] |
| MXE | 8661 | ENSG00000173744 | AGFG1 | chr2 | + | [illegible] |
| MXE | 8401 | ENSG00000135905 | DOCK10 | chr2 | - | [illegible] |
| MXE | 3486 | ENSG00000147044 | CASK | chrX | - | [illegible] |
| MXE | 293 | ENSG00000100077 | GRK3 | chr22 | + | [illegible] |
| MXE | 9165 | ENSG00000119004 | CYP20A1 | chr2 | + | [illegible] |
| MXE | 2301 | ENSG00000140598 | EFL1 | chr15 | - | [illegible] |
| MXE | 14917 | ENSG00000119383 | PTPA | chr9 | + | [illegible] |
| MXE | 1728 | ENSG00000197912 | SPG7 | chr16 | + | [illegible] |
| MXE | 1659 | ENSG00000121281 | ADCY7 | chr16 | + | [illegible] |
| MXE | 7716 | ENSG00000137198 | GMPR | chr6 | + | [illegible] |
| MXE | 2222 | ENSG00000092439 | TRPM7 | chr15 | - | [illegible] |
| MXE | 14289 | ENSG00000164733 | CTSB | chr8 | - | [illegible] |
| MXE | 594 | ENSG00000105186 | ANKRD27 | chr19 | - | [illegible] |
| MXE | 4776 | ENSG00000143669 | LYST | chr1 | - | [illegible] |
| MXE | 13665 | ENSG00000147133 | TAF1 | chrX | + | [illegible] |
| MXE | 16785 | ENSG00000151148 | UBE3B | chr12 | + | [illegible] |
| MXE | 16237 | ENSG00000148634 | HERC4 | chr10 | - | [illegible] |
| MXE | 1529 | ENSG00000108559 | NUP88 | chr17 | - | [illegible] |
| MXE | 9338 | ENSG00000242247 | ARFGAP3 | chr22 | - | [illegible] |
| MXE | 8195 | ENSG00000188641 | DPYD | chr1 | - | [illegible] |
| MXE | 4515 | ENSG00000150961 | SEC24D | chr4 | - | [illegible] |
| MXE | 4479 | ENSG00000141562 | NARF | chr17 | + | [illegible] |
| MXE | 16172 | ENSG00000107669 | ATE1 | chr10 | - | [illegible] |
| MXE | 3410 | ENSG00000043143 | JADE2 | chr5 | + | [illegible] |
| MXE | 11661 | ENSG00000065883 | CDK13 | chr7 | + | [illegible] |
| MXE | 4076 | ENSG00000136813 | ECPAS | chr9 | - | [illegible] |
| MXE | 12580 | ENSG00000175471 | MCTP1 | chr5 | - | [illegible] |
| MXE | 13381 | ENSG00000147394 | ZNF185 | chrX | + | [illegible] |
| MXE | 14954 | ENSG00000137073 | UBAP2 | chr9 | - | [illegible] |
| MXE | 4939 | ENSG00000183323 | CCDC125 | chr5 | - | [illegible] |
| MXE | 15621 | ENSG00000078124 | ACER3 | chr11 | + | [illegible] |
| MXE | 16080 | ENSG00000197535 | MYO5A | chr15 | - | [illegible] |
| MXE | 5714 | ENSG00000168615 | ADAM9 | chr8 | + | [illegible] |
| MXE | 5081 | ENSG00000132153 | DHX30 | chr3 | + | [illegible] |
| MXE | 131 | ENSG00000183765 | CHEK2 | chr22 | - | [illegible] |
| MXE | 5423 | ENSG00000078070 | MCCC1 | chr3 | - | [illegible] |
| MXE | 8458 | ENSG00000137275 | RIPK1 | chr6 | + | [illegible] |
| MXE | 2371 | ENSG00000214013 | GANC | chr15 | + | [illegible] |
| MXE | 5310 | ENSG00000103064 | SLC7A6 | chr16 | + | [illegible] |
| MXE | 8316 | ENSG00000109667 | SLC2A9 | chr4 | - | [illegible] |
| MXE | 16938 | ENSG00000101160 | CTSZ | chr20 | - | [illegible] |
| MXE | 7359 | ENSG00000134215 | VAV3 | chr1 | - | [illegible] |
| MXE | 13643 | ENSG00000086758 | HUWE1 | chrX | - | [illegible] |
| MXE | 13657 | ENSG00000046651 | OFD1 | chrX | + | [illegible] |
| MXE | 3352 | ENSG00000127603 | MACF1 | chr1 | + | [illegible] |
| MXE | 11386 | ENSG00000177463 | NR2C2 | chr3 | + | [illegible] |
| MXE | 4598 | ENSG00000100412 | ACO2 | chr22 | + | [illegible] |
| MXE | 1246 | ENSG00000108469 | RECQL5 | chr17 | - | [illegible] |
| MXE | 15418 | ENSG00000255663 | NA | chr11 | + | [illegible] |
| MXE | 4632 | ENSG00000065882 | TBC1D1 | chr4 | + | [illegible] |
| MXE | 8105 | ENSG00000237188 | NA | chr1 | + | [illegible] |
| MXE | 4324 | ENSG00000164597 | COG5 | chr7 | - | [illegible] |
| MXE | 10286 | ENSG00000131051 | RBM39 | chr20 | - | [illegible] |
| MXE | 12114 | ENSG00000151116 | UEVLD | chr11 | - | [illegible] |
| MXE | 12931 | ENSG00000145375 | SPATA5 | chr4 | + | [illegible] |
| MXE | 15632 | ENSG00000078124 | ACER3 | chr11 | + | [illegible] |
| MXE | 11242 | ENSG00000072415 | MPP5 | chr14 | + | [illegible] |
| MXE | 16068 | ENSG00000163947 | ARHGEF3 | chr3 | - | [illegible] |
| MXE | 13975 | ENSG00000123124 | WWP1 | chr8 | + | [illegible] |
| MXE | 8850 | ENSG00000110200 | ANAPC15 | chr11 | - | [illegible] |
| MXE | 9122 | ENSG00000103365 | GGA2 | chr16 | - | [illegible] |
| MXE | 6117 | ENSG00000132768 | DPH2 | chr1 | + | [illegible] |
| MXE | 4346 | ENSG00000123352 | SPATS2 | chr12 | + | [illegible] |
| MXE | 3841 | ENSG00000179152 | TCAIM | chr3 | + | [illegible] |
| MXE | 1240 | ENSG00000161526 | SAP30BP | chr17 | + | [illegible] |
| MXE | 4797 | ENSG00000242086 | MUC20-OT1 | chr3 | + | [illegible] |
| MXE | 15925 | ENSG00000181192 | DHTKD1 | chr10 | + | [illegible] |
| MXE | 7101 | ENSG00000167257 | RNF214 | chr11 | + | [illegible] |
| MXE | 2114 | ENSG00000128739 | SNRPN | chr15 | + | [illegible] |
| MXE | 15663 | ENSG00000081237 | PTPRC | chr1 | + | [illegible] |
| MXE | 13382 | ENSG00000147394 | ZNF185 | chrX | + | [illegible] |
| MXE | 14735 | ENSG00000090054 | SPTLC1 | chr9 | - | [illegible] |
| MXE | 3780 | ENSG00000153310 | CYRIB | chr8 | - | [illegible] |
| MXE | 1024 | ENSG00000268434 | NA | chr19 | - | [illegible] |
| MXE | 9497 | ENSG00000189319 | FAM53B | chr10 | - | [illegible] |
| MXE | 15905 | ENSG00000122033 | MTIF3 | chr13 | - | [illegible] |
| MXE | 7958 | ENSG00000142197 | DOP1B | chr21 | + | [illegible] |
| MXE | 33 | ENSG00000157617 | C2CD2 | chr21 | - | [illegible] |
| MXE | 5107 | ENSG00000159147 | DONSON | chr21 | - | [illegible] |
| MXE | 2303 | ENSG00000077238 | IL4R | chr16 | + | [illegible] |
| MXE | 15888 | ENSG00000138286 | FAM149B1 | chr10 | + | [illegible] |
| MXE | 3930 | ENSG00000059378 | PARP12 | chr7 | - | [illegible] |
| MXE | 7172 | ENSG00000113360 | DROSHA | chr5 | - | [illegible] |
| MXE | 8562 | ENSG00000153207 | AHCTF1 | chr1 | - | [illegible] |
| MXE | 8264 | ENSG00000049618 | ARID1B | chr6 | + | [illegible] |
| MXE | 11852 | ENSG00000112739 | PRPF4B | chr6 | + | [illegible] |
| MXE | 361 | ENSG00000257103 | LSM14A | chr19 | + | [illegible] |
| MXE | 9050 | ENSG00000134759 | ELP2 | chr18 | + | [illegible] |
| MXE | 14432 | ENSG00000080503 | SMARCA2 | chr9 | + | [illegible] |
| MXE | 6368 | ENSG00000120008 | WDR11 | chr10 | + | [illegible] |
| MXE | 4910 | ENSG00000075711 | DLG1 | chr3 | - | [illegible] |
| MXE | 6969 | ENSG00000170248 | PDCD6IP | chr3 | + | [illegible] |
| MXE | 2880 | ENSG00000112584 | FAM120B | chr6 | + | [illegible] |
| MXE | 4457 | ENSG00000287542 | NA | chr4 | + | [illegible] |
| MXE | 16086 | ENSG00000213047 | DENND1B | chr1 | - | [illegible] |
| MXE | 14926 | ENSG00000095319 | NUP188 | chr9 | + | [illegible] |
| MXE | 15073 | ENSG00000080546 | SESN1 | chr6 | - | [illegible] |
| MXE | 16119 | ENSG00000186020 | ZNF529 | chr19 | - | [illegible] |
| MXE | 7295 | ENSG00000266028 | SRGAP2 | chr1 | + | [illegible] |
| MXE | 3971 | ENSG00000214655 | ZSWIM8 | chr10 | + | [illegible] |
| MXE | 16871 | ENSG00000050426 | LETMD1 | chr12 | + | [illegible] |
| MXE | 13598 | ENSG00000005889 | ZFX | chrX | + | [illegible] |
| MXE | 1333 | ENSG00000087157 | PGS1 | chr17 | + | [illegible] |
| MXE | 15393 | ENSG00000107290 | SETX | chr9 | - | [illegible] |
| MXE | 814 | ENSG00000170471 | RALGAPB | chr20 | + | [illegible] |
| MXE | 12940 | ENSG00000131373 | HACL1 | chr3 | - | [illegible] |
| MXE | 3972 | ENSG00000038382 | TRIO | chr5 | + | [illegible] |
| MXE | 8566 | ENSG00000163001 | CFAP36 | chr2 | + | [illegible] |
| MXE | 1944 | ENSG00000047578 | KATNIP | chr16 | + | [illegible] |
| MXE | 5367 | ENSG00000120685 | PROSER1 | chr13 | - | [illegible] |
| MXE | 9033 | ENSG00000186854 | TRABD2A | chr2 | - | [illegible] |
| MXE | 16807 | ENSG00000123349 | PFDN5 | chr12 | + | [illegible] |
| MXE | 15275 | ENSG00000155158 | TTC39B | chr9 | - | [illegible] |
| MXE | 350 | ENSG00000189068 | VSTM1 | chr19 | - | [illegible] |
| MXE | 9393 | ENSG00000147099 | HDAC8 | chrX | - | [illegible] |
| MXE | 861 | ENSG00000089063 | TMEM230 | chr20 | - | [illegible] |
| MXE | 7233 | ENSG00000162869 | PPP1R21 | chr2 | + | [illegible] |
| MXE | 14619 | ENSG00000160563 | MED27 | chr9 | - | [illegible] |
| MXE | 4911 | ENSG00000075711 | DLG1 | chr3 | - | [illegible] |
| MXE | 1074 | ENSG00000153933 | DGKE | chr17 | + | [illegible] |
| MXE | 11583 | ENSG00000163939 | PBRM1 | chr3 | - | [illegible] |
| MXE | 13156 | ENSG00000110848 | CD69 | chr12 | - | [illegible] |
| MXE | 11451 | ENSG00000162419 | GMEB1 | chr1 | + | [illegible] |
| MXE | 15025 | ENSG00000110756 | HPS5 | chr11 | - | [illegible] |
| MXE | 2408 | ENSG00000154237 | LRRK1 | chr15 | + | [illegible] |
| MXE | 16950 | ENSG00000075089 | ACTR6 | chr12 | + | [illegible] |
| MXE | 2874 | ENSG00000288564 | NA | chr6 | - | [illegible] |
| MXE | 4877 | ENSG00000058056 | USP13 | chr3 | + | [illegible] |
| MXE | 6512 | ENSG00000114861 | FOXP1 | chr3 | - | [illegible] |
| MXE | 3188 | ENSG00000178385 | PLEKHM3 | chr2 | - | [illegible] |
| MXE | 13786 | ENSG00000104361 | NIPAL2 | chr8 | - | [illegible] |
| MXE | 16058 | ENSG00000198825 | INPP5F | chr10 | + | [illegible] |
| MXE | 10295 | ENSG00000132676 | DAP3 | chr1 | + | [illegible] |
| MXE | 1380 | ENSG00000262879 | NA | chr17 | - | [illegible] |
| MXE | 6877 | ENSG00000070190 | DAPP1 | chr4 | + | [illegible] |
| MXE | 14873 | ENSG00000148358 | GPR107 | chr9 | + | [illegible] |
| MXE | 15689 | ENSG00000054654 | SYNE2 | chr14 | + | [illegible] |
| MXE | 12885 | ENSG00000257923 | CUX1 | chr7 | + | [illegible] |
| MXE | 13083 | ENSG00000124222 | STX16 | chr20 | + | [illegible] |
| MXE | 1957 | ENSG00000140694 | PARN | chr16 | - | [illegible] |
| MXE | 7234 | ENSG00000162869 | PPP1R21 | chr2 | + | [illegible] |
| MXE | 2870 | ENSG00000152240 | HAUS1 | chr18 | + | [illegible] |
| MXE | 10653 | ENSG00000091009 | RBM27 | chr5 | + | [illegible] |
| MXE | 9162 | ENSG00000162909 | CAPN2 | chr1 | + | [illegible] |
| MXE | 14245 | ENSG00000164941 | INTS8 | chr8 | + | [illegible] |
| MXE | 236 | ENSG00000100307 | CBX7 | chr22 | - | [illegible] |
| MXE | 2733 | ENSG00000197324 | LRP10 | chr14 | + | [illegible] |
| MXE | 10507 | ENSG00000018280 | SLC11A1 | chr2 | + | [illegible] |
| MXE | 14218 | ENSG00000254673 | NA | chr8 | + | [illegible] |
| MXE | 5336 | ENSG00000174891 | RSRC1 | chr3 | + | [illegible] |
| MXE | 6831 | ENSG00000007341 | ST7L | chr1 | - | [illegible] |
| MXE | 16848 | ENSG00000198015 | MRPL42 | chr12 | + | [illegible] |
| MXE | 14720 | ENSG00000148337 | CIZ1 | chr9 | - | [illegible] |
| MXE | 1819 | ENSG00000167264 | DUS2 | chr16 | + | [illegible] |
| MXE | 1437 | ENSG00000131467 | PSME3 | chr17 | + | [illegible] |
| MXE | 15537 | ENSG00000110075 | PPP6R3 | chr11 | + | [illegible] |
| MXE | 17166 | ENSG00000132965 | ALOX5AP | chr13 | + | [illegible] |
| MXE | 11501 | ENSG00000137478 | FCHSD2 | chr11 | - | [illegible] |
| MXE | 14831 | ENSG00000107077 | KDM4C | chr9 | + | [illegible] |
| MXE | 3324 | ENSG00000140455 | USP3 | chr15 | + | [illegible] |
| MXE | 15290 | ENSG00000149547 | EI24 | chr11 | + | [illegible] |
| MXE | 5030 | ENSG00000009790 | TRAF3IP3 | chr1 | + | [illegible] |
| MXE | 10311 | ENSG00000170445 | HARS1 | chr5 | - | [illegible] |
| MXE | 2407 | ENSG00000141564 | RPTOR | chr17 | + | [illegible] |
| MXE | 11209 | ENSG00000162885 | B3GALNT2 | chr1 | - | [illegible] |
| MXE | 7324 | ENSG00000141446 | ESCO1 | chr18 | - | [illegible] |
| MXE | 14795 | ENSG00000070061 | ELP1 | chr9 | - | [illegible] |
| MXE | 14751 | ENSG00000048828 | FAM120A | chr9 | + | [illegible] |
| MXE | 14719 | ENSG00000099381 | SETD1A | chr16 | + | [illegible] |
| MXE | 16488 | ENSG00000123268 | ATF1 | chr12 | + | [illegible] |
| MXE | 13533 | ENSG00000135048 | CEMIP2 | chr9 | - | [illegible] |
| MXE | 1984 | ENSG00000103126 | AXIN1 | chr16 | - | [illegible] |
| MXE | 5346 | ENSG00000186001 | LRCH3 | chr3 | + | [illegible] |
| MXE | 14798 | ENSG00000198771 | RCSD1 | chr1 | + | [illegible] |
| MXE | 7084 | ENSG00000134222 | PSRC1 | chr1 | - | [illegible] |
| MXE | 16287 | ENSG00000266412 | NCOA4 | chr10 | - | [illegible] |
| MXE | 8454 | ENSG00000185379 | RAD51D | chr17 | - | [illegible] |
| MXE | 3537 | ENSG00000132842 | AP3B1 | chr5 | - | [illegible] |
| MXE | 5918 | ENSG00000003400 | CASP10 | chr2 | + | [illegible] |
| MXE | 4769 | ENSG00000066777 | ARFGEF1 | chr8 | - | [illegible] |
| MXE | 2860 | ENSG00000154001 | PPP2R5E | chr14 | - | [illegible] |
| MXE | 3427 | ENSG00000116459 | ATP5PB | chr1 | + | [illegible] |
| MXE | 14219 | ENSG00000168522 | FNTA | chr8 | + | [illegible] |
| MXE | 7360 | ENSG00000134215 | VAV3 | chr1 | - | [illegible] |
| MXE | 6959 | ENSG00000117593 | DARS2 | chr1 | + | [illegible] |
| MXE | 3432 | ENSG00000116459 | ATP5PB | chr1 | + | [illegible] |

| | | | | | | | | | | | | | | |
|---|---|---|---|---|---|---|---|---|---|---|---|---|---|---|
| MXE | 8685 | ENSG00000127124 | HIVEP3 | chr1 | - | 41524734 | 41524910 | 41579736 | 41585318 | 41518401 | 41518488 | 41628748 | 41628947 | 20 |
| MXE | 13520 | ENSG00000134516 | DOCK2 | chr5 | + | 169669287 | 169669328 | 169671077 | 169671174 | 169654402 | 169654486 | 169674296 | 169674445 | 62 |
| MXE | 9775 | ENSG00000175662 | TOM1L2 | chr17 | - | 17861475 | 17861551 | 17862730 | 17862848 | 17848822 | 17848859 | 17866295 | 17866419 | 16 |
| MXE | 17130 | ENSG00000139597 | N4BP2L1 | chr13 | - | 32404202 | 32404271 | 32404320 | 32404397 | 32400722 | 32403200 | 32407644 | 32407772 | 41 |
| MXE | 16493 | ENSG00000114302 | PRKAR2A | chr3 | - | 48765247 | 48765349 | 48772954 | 48773108 | 48765003 | 48765078 | 48782985 | 48783092 | 153 |
| MXE | 372 | ENSG00000104880 | ARHGEF18 | chr19 | + | 7379121 | 7379166 | 7380916 | 7380994 | 7378393 | 7378451 | 7382791 | 7382894 | 73 |
| MXE | 1405 | ENSG00000011260 | UTP18 | chr17 | + | 51266181 | 51266280 | 51268836 | 51268904 | 51263273 | 51263386 | 51273361 | 51273450 | 49 |
| MXE | 174 | ENSG00000100083 | GGA1 | chr22 | + | 37614189 | 37614274 | 37616921 | 37616997 | 37608833 | 37608903 | 37618447 | 37618546 | 0 |
| MXE | 3735 | ENSG00000072364 | AFF4 | chr5 | - | 132904367 | 132904404 | 132927120 | 132927207 | 132902441 | 132902487 | 132932177 | 132932222 | 67 |
| MXE | 12912 | ENSG00000067182 | TNFRSF1A | chr12 | - | 6329777 | 6330066 | 6330266 | 6330295 | 6328756 | 6329622 | 6330597 | 6330711 | 112 |
| MXE | 8759 | ENSG00000101040 | ZMYND8 | chr20 | - | 47246007 | 47246517 | 47249286 | 47249439 | 47236325 | 47236516 | 47262287 | 47262428 | 50 |
| MXE | 4370 | ENSG00000129116 | PALLD | chr4 | + | 168921533 | 168921741 | 168924254 | 168924420 | 168915894 | 168916027 | 168924944 | 168925078 | 68 |
| MXE | 3332 | ENSG00000259040 | BLOC1S5-TXNDC5 | chr6 | - | 7888704 | 7888848 | 7889494 | 7889581 | 7885960 | 7886043 | 7891620 | 7891736 | 64 |
| MXE | 12368 | ENSG00000164808 | SPIDR | chr8 | + | 47728932 | 47729047 | 47729411 | 47729465 | 47727195 | 47727293 | 47735306 | 47735374 | 37 |
| MXE | 13549 | ENSG00000196092 | PAX5 | chr9 | - | 36846842 | 36846929 | 36882003 | 36882105 | 36840550 | 36840636 | 36923354 | 36923484 | 7 |
| MXE | 6050 | ENSG00000198734 | F5 | chr1 | - | 169552556 | 169552734 | 169555181 | 169555347 | 169550639 | 169550739 | 169556645 | 169556867 | 36 |
| MXE | 541 | ENSG00000105486 | LIG1 | chr19 | - | 48151231 | 48151339 | 48153871 | 48153967 | 48150087 | 48150210 | 48157013 | 48157140 | 81 |
| MXE | 5382 | ENSG00000175556 | LONRF3 | chrX | + | 118978344 | 118978463 | 118989407 | 118989672 | 118974613 | 118975597 | 118990469 | 118990560 | 6 |
| MXE | 12583 | ENSG00000147535 | PLPP5 | chr8 | - | 38267266 | 38267391 | 38267896 | 38267960 | 38266140 | 38266311 | 38268370 | 38268407 | 25 |
| MXE | 7384 | ENSG00000186716 | BCR | chr22 | + | 23292540 | 23292638 | 23295023 | 23295155 | 23290338 | 23290413 | 23309423 | 23309483 | 115 |
| MXE | 161 | ENSG00000100351 | GRAP2 | chr22 | + | 39955818 | 39955910 | 39960054 | 39960174 | 39946816 | 39947184 | 39965989 | 39966158 | 303 |
| MXE | 8005 | ENSG00000203705 | TATDN3 | chr1 | + | 212797111 | 212797196 | 212802700 | 212802763 | 212796516 | 212796590 | 212804319 | 212804429 | 19 |
| MXE | 7992 | ENSG00000133805 | AMPD3 | chr11 | + | 10487234 | 10487364 | 10493348 | 10493543 | 10484819 | 10485039 | 10494898 | 10495030 | 26 |
| MXE | 9919 | ENSG00000115464 | USP34 | chr2 | - | 61206016 | 61206124 | 61206759 | 61206886 | 61204496 | 61204601 | 61208898 | 61208977 | 157 |
| MXE | 6088 | ENSG00000107566 | ERLIN1 | chr10 | - | 100156144 | 100156234 | 100164003 | 100164095 | 100154859 | 100154939 | 100167347 | 100167406 | 21 |
| MXE | 14365 | ENSG00000104365 | IKBKB | chr8 | + | 42308900 | 42309025 | 42314321 | 42314429 | 42306342 | 42306432 | 42316209 | 42316339 | 55 |
| MXE | 16835 | ENSG00000111596 | CNOT2 | chr12 | + | 70285399 | 70285586 | 70294236 | 70294294 | 70278131 | 70278274 | 70310894 | 70311017 | 10 |
| MXE | 13960 | ENSG00000166169 | POLL | chr10 | - | 101582762 | 101582891 | 101584601 | 101584720 | 101580247 | 101580416 | 101584877 | 101584919 | 4 |
| MXE | 7504 | ENSG00000136068 | FLNB | chr3 | + | 58155959 | 58156075 | 58159553 | 58159686 | 58154790 | 58154928 | 58163153 | 58163330 | 32 |
| MXE | 16850 | ENSG00000128585 | MKLN1 | chr7 | + | 131443480 | 131443702 | 131445773 | 131445903 | 131437784 | 131437997 | 131463216 | 131463364 | 134 |
| MXE | 1471 | ENSG00000108306 | FBXL20 | chr17 | - | 39301000 | 39301075 | 39303584 | 39303639 | 39298989 | 39299084 | 39343179 | 39343241 | 19 |
| MXE | 7994 | ENSG00000133805 | AMPD3 | chr11 | + | 10496760 | 10496938 | 10500085 | 10500249 | 10495569 | 10495733 | 10501469 | 10501590 | 1 |
| [illegible] | [illegible] | [illegible] | [illegible] | [illegible] | [illegible] | [illegible] | [illegible] | [illegible] | [illegible] | [illegible] | [illegible] | [illegible] | [illegible] | [illegible] |

| | | | | | | | | | |
|---|---|---|---|---|---|---|---|---|---|
| MXE | 13989 | ENSG00000156162 | DPY19L4 | chr8 | + | 94756035 | 94756159 | 94761699 | 94761834 |
| MXE | 1030 | ENSG00000161955 | TNFSF13 | chr17 | + | 7559623 | 7559702 | 7559845 | 7559893 |
| MXE | 2801 | ENSG00000151445 | VIPAS39 | chr14 | - | 77435843 | 77435919 | 77437807 | 77437881 |
| MXE | 9947 | ENSG00000175104 | TRAF6 | chr11 | - | 36498489 | 36498640 | 36501219 | 36501537 |
| MXE | 5834 | ENSG00000198793 | MTOR | chr1 | - | 11144967 | 11145045 | 11146675 | 11146791 |
| MXE | 4383 | ENSG00000157800 | SLC37A3 | chr7 | - | 140369589 | 140369682 | 140380281 | 140380390 |
| MXE | 14340 | ENSG00000105339 | DENND3 | chr8 | + | 141163332 | 141163429 | 141165185 | 141165289 |
| MXE | 8828 | ENSG00000118007 | STAG1 | chr3 | - | 136369107 | 136369282 | 136377659 | 136377752 |
| MXE | 12401 | ENSG00000084070 | SMAP2 | chr1 | + | 40362299 | 40362436 | 40406735 | 40406869 |
| MXE | 2821 | ENSG00000080815 | PSEN1 | chr14 | + | 73192643 | 73192864 | 73206385 | 73206472 |
| MXE | 7153 | ENSG00000095787 | WAC | chr10 | + | 28535909 | 28536059 | 28537583 | 28537815 |
| MXE | 7702 | ENSG00000142687 | KIAA0319L | chr1 | - | 35456012 | 35456241 | 35462620 | 35462713 |
| MXE | 5428 | ENSG00000258653 | NA | chr14 | + | 73893796 | 73893932 | 73896315 | 73896534 |
| MXE | 14596 | ENSG00000056586 | RC3H2 | chr9 | - | 122859916 | 122860131 | 122865348 | 122865541 |
| MXE | 282 | ENSG00000228315 | GUSBP11 | chr22 | - | 23705428 | 23705541 | 23709129 | 23709241 |
| MXE | 4521 | ENSG00000163811 | WDR43 | chr2 | + | 28901986 | 28902124 | 28906459 | 28906581 |
| MXE | 4544 | ENSG00000108591 | DRG2 | chr17 | + | 18099016 | 18099077 | 18099632 | 18099723 |
| MXE | 10358 | ENSG00000213145 | CRIP1 | chr14 | + | 105487212 | 105487299 | 105488165 | 105488260 |
| MXE | 11851 | ENSG00000168538 | TRAPPC11 | chr4 | + | 183686617 | 183686748 | 183691315 | 183691471 |
| MXE | 2109 | ENSG00000128739 | SNRPN | chr15 | + | 24974310 | 24974456 | 24974641 | 24974957 |
| MXE | 1025 | ENSG00000167851 | CD300A | chr17 | + | 74469945 | 74470242 | 74473535 | 74473874 |
| MXE | 16806 | ENSG00000123349 | PFDN5 | chr12 | + | 53295838 | 53295941 | 53297849 | 53297924 |
| [illegible] | | | | | | | | | |

| | | | | | | | | | | |
|---|---|---|---|---|---|---|---|---|---|---|
| MXE | 11898 | ENSG00000135317 | SNX14 | chr6 | - | 85547115 | 85547226 | 85547505 | 85547550 | [illegible] |
| MXE | 5945 | ENSG00000109320 | NFKB1 | chr4 | + | 102525511 | 102525557 | 102529835 | 102529914 | [illegible] |
| MXE | 356 | ENSG00000180448 | ARHGAP45 | chr19 | + | 1073673 | 1073746 | 1073947 | 1074014 | [illegible] |
| MXE | 10832 | ENSG00000108883 | EFTUD2 | chr17 | - | 44852408 | 44852562 | 44853295 | 44853390 | [illegible] |
| MXE | 12053 | ENSG00000214021 | TTLL3 | chr3 | + | 9816073 | 9816202 | 9817644 | 9818920 | [illegible] |
| MXE | 11201 | ENSG00000059588 | TARBP1 | chr1 | - | 234430086 | 234430301 | 234433409 | 234433571 | [illegible] |
| MXE | 14974 | ENSG00000215126 | CBWD6 | chr9 | - | 41170278 | 41170423 | 41171846 | 41171934 | [illegible] |
| MXE | 490 | ENSG00000130164 | LDLR | chr19 | + | 11123173 | 11123344 | 11128007 | 11128085 | [illegible] |
| MXE | 1833 | ENSG00000103005 | USB1 | chr16 | + | 58014272 | 58014326 | 58017333 | 58017439 | [illegible] |
| MXE | 2208 | ENSG00000092470 | WDR76 | chr15 | + | 43856945 | 43857163 | 43858670 | 43858823 | [illegible] |
| MXE | 6873 | ENSG00000143164 | DCAF6 | chr1 | + | 168065589 | 168065746 | 168066376 | 168066465 | [illegible] |
| MXE | 3268 | ENSG00000130363 | RSPH3 | chr6 | - | 158983661 | 158983807 | 158986279 | 158986421 | [illegible] |
| MXE | 12391 | ENSG00000150768 | DLAT | chr11 | + | 112033403 | 112033530 | 112037272 | 112037460 | [illegible] |
| MXE | 13461 | ENSG00000196776 | CD47 | chr3 | - | 108059451 | 108059544 | 108060744 | 108060852 | [illegible] |
| MXE | 11567 | ENSG00000197548 | ATG7 | chr3 | + | 11299361 | 11299416 | 11306942 | 11307060 | [illegible] |
| MXE | 12613 | ENSG00000151948 | GLT1D1 | chr12 | + | 128899235 | 128899287 | 128926358 | 128926454 | [illegible] |
| MXE | 9671 | ENSG00000183943 | PRKX | chrX | - | 3626418 | 3626514 | 3641851 | 3641971 | [illegible] |
| MXE | 16753 | ENSG00000118971 | CCND2 | chr12 | + | 4278759 | 4278919 | 4288841 | 4288990 | [illegible] |
| MXE | 11590 | ENSG00000163939 | PBRM1 | chr3 | - | 52651741 | 52651810 | 52658198 | 52658315 | [illegible] |
| MXE | 15990 | ENSG00000165669 | FAM204A | chr10 | - | 118335113 | 118335213 | 118335395 | 118335426 | [illegible] |
| MXE | 5539 | ENSG00000169180 | XPO6 | chr16 | - | 28135215 | 28135324 | 28146093 | 28146203 | [illegible] |
| MXE | 16684 | ENSG00000112200 | ZNF451 | chr6 | + | 57128728 | 57128840 | 57133041 | 57133192 | [illegible] |
| MXE | 2395 | ENSG00000138613 | APH1B | chr15 | + | 63286557 | 63286628 | 63302344 | 63302472 | [illegible] |
| MXE | 6011 | ENSG00000137393 | RNF144B | chr6 | + | 18439683 | 18439744 | 18457154 | 18457359 | [illegible] |
| MXE | 3781 | ENSG00000153310 | CYRIB | chr8 | - | 129904498 | 129904585 | 129970942 | 129970995 | [illegible] |
| MXE | 8761 | ENSG00000101040 | ZMYND8 | chr20 | - | 47249286 | 47249439 | 47262287 | 47262428 | [illegible] |
| MXE | 9079 | ENSG00000164828 | SUN1 | chr7 | + | 848417 | 848615 | 851383 | 851482 | [illegible] |
| MXE | 10537 | ENSG00000135838 | NPL | chr1 | + | 182818540 | 182818689 | 182818812 | 182818859 | [illegible] |
| MXE | 4001 | ENSG00000138767 | CNOT6L | chr4 | - | 77773080 | 77773166 | 77774529 | 77774716 | [illegible] |
| MXE | 8918 | ENSG00000065413 | ANKRD44 | chr2 | - | 197025195 | 197025267 | 197078702 | 197078814 | [illegible] |
| MXE | 2545 | ENSG00000092199 | HNRNPC | chr14 | - | 21236372 | 21236460 | 21262600 | 21262768 | [illegible] |
| MXE | 5468 | ENSG00000075420 | FNDC3B | chr3 | + | 172247532 | 172247776 | 172251259 | 172251541 | [illegible] |
| MXE | 16882 | ENSG00000050426 | LETMD1 | chr12 | + | 51053811 | 51053860 | 51056143 | 51056245 | [illegible] |
| [illegible] | | | | | | | | | | |

| | | | | | | | | | | | | | | | | | | | | | | | | | | | | | | | | | | | | | | | | | | | | | | | | |
|---|---|---|---|---|---|---|---|---|---|---|---|---|---|---|---|---|---|---|---|---|---|---|---|---|---|---|---|---|---|---|---|---|---|---|---|---|---|---|---|---|---|---|---|---|---|---|---|---|
| MXE | 219 | ENSG00000100347 | SAMM50 | chr22 | + | 43973235 | 43973323 | 43976054 | 43976183 | 43972870 | 43973001 | 43976749 | 43976821 | 61 | 42 | 63 | 71 | 91 | 101 | 49 | 99 | 107 | 97 | 128 | 57 | 22 | 48 | 94 | 145 | 71 | 67 | 82 | 104 | 237 | 278 | 0.75015291 | 1 | 0.415 | 0.501 | 0.427 | 0.438 | 0.524 | 0.509 | 0.485 | 0.278 | 0.407 | 0.515 | 0.022 |
| MXE | 17102 | ENSG00000197442 | MAP3K5 | chr6 | - | 136705109 | 136705133 | 136720449 | 136720589 | 136698488 | 136698682 | 136791709 | 136792477 | 112 | 43 | 99 | 146 | 61 | 30 | 26 | 32 | 50 | 41 | 73 | 86 | 99 | 54 | 58 | 28 | 35 | 36 | 30 | 13 | 289 | 173 | 0.750356329 | 1 | 0.691 | 0.497 | 0.649 | 0.636 | 0.471 | 0.609 | 0.595 | 0.622 | 0.519 | 0.728 | -0.026 |
| MXE | 14813 | ENSG00000011454 | RABGAP1 | chr9 | + | 123071411 | 123071501 | 123073551 | 123073677 | 123070349 | 123070424 | 123076244 | 123076286 | 0 | 7 | 3 | 1 | 0 | 95 | 64 | 59 | 84 | 125 | 0 | 7 | 1 | 0 | 0 | 23 | 64 | 58 | 69 | 54 | 239 | 275 | 0.750474956 | 1 | 0 | 0.112 | 0.055 | 0.014 | 0 | 0 | 0.112 | 0.019 | 0 | 0 | 0.01 |
| MXE | 13179 | ENSG00000137642 | SORL1 | chr11 | + | 121470006 | 121470123 | 121478117 | 121478243 | 121452313 | 121452616 | 121488031 | 121488193 | 251 | 103 | 194 | 168 | 156 | 446 | 311 | 440 | 406 | 1158 | 103 | 60 | 78 | 100 | 82 | 229 | 192 | 300 | 299 | 276 | 266 | 275 | 0.75060291 | 1 | 0.368 | 0.255 | 0.313 | 0.3 | 0.122 | 0.317 | 0.244 | 0.212 | 0.257 | 0.235 | 0.019 |
| MXE | 10664 | ENSG00000184613 | NELL2 | chr12 | - | 44711291 | 44711394 | 44714649 | 44714741 | 44703725 | 44703854 | 44774746 | 44774849 | 79 | 61 | 100 | 77 | 226 | 189 | 191 | 288 | 158 | 394 | 14 | 10 | 68 | 9 | 16 | 40 | 32 | 142 | 11 | 26 | 241 | 252 | 0.750617434 | 1 | 0.304 | 0.25 | 0.266 | 0.338 | 0.375 | 0.268 | 0.246 | 0.334 | 0.461 | 0.392 | -0.034 |
| MXE | 16429 | ENSG00000205356 | TECPR1 | chr7 | - | 98236775 | 98236921 | 98238508 | 98238610 | 98233420 | 98233911 | 98240850 | 98240951 | 53 | 38 | 67 | 50 | 63 | 29 | 24 | 86 | 57 | 35 | 66 | 73 | 58 | 32 | 98 | 44 | 79 | 27 | 19 | 91 | 251 | 295 | 0.750829043 | 1 | 0.682 | 0.65 | 0.478 | 0.508 | 0.679 | 0.638 | 0.521 | 0.716 | 0.664 | 0.559 | -0.02 |
| MXE | 9080 | ENSG00000164828 | SUN1 | chr7 | + | 849519 | 849603 | 849922 | 850033 | 848417 | 848615 | 851383 | 851482 | 81 | 98 | 84 | 98 | 122 | 42 | 13 | 13 | 17 | 20 | 178 | 92 | 58 | 107 | 127 | 40 | 19 | 28 | 14 | 23 | 233 | 260 | 0.750880482 | 1 | 0.683 | 0.894 | 0.878 | 0.865 | 0.872 | 0.832 | 0.844 | 0.698 | 0.895 | 0.86 | 0.013 |
| MXE | 1174 | ENSG00000067191 | CACNB1 | chr17 | - | 39184783 | 39184864 | 39185130 | 39185150 | 39184324 | 39184383 | 39186495 | 39186572 | 0 | 2 | 0 | 0 | 0 | 36 | 11 | 10 | 23 | 18 | 0 | 5 | 0 | 0 | 0 | 25 | 41 | 34 | 13 | 26 | 169 | 230 | 0.751017603 | 1 | 0 | 0.198 | 0 | 0 | 0 | 0 | 0.142 | 0 | 0 | 0 | 0.011 |
| MXE | 8424 | ENSG00000163041 | H3-3A | chr1 | + | 226064328 | 226064479 | 226065655 | 226065809 | 226062715 | 226062811 | 226071350 | 226072019 | 743 | 372 | 552 | 744 | 873 | 2548 | 1517 | 1883 | 3291 | 3405 | 890 | 968 | 502 | 806 | 2191 | 4515 | 2938 | 2569 | 3127 | 6847 | 298 | 298 | 0.751246906 | 1 | 0.226 | 0.197 | 0.227 | 0.184 | 0.204 | 0.165 | 0.248 | 0.163 | 0.205 | 0.242 | 0.003 |
| MXE | 13681 | ENSG00000215301 | DDX3X | chrX | + | 41343736 | 41344128 | 41344238 | 41344564 | 41343215 | 41343351 | 41345179 | 41345324 | 195 | 131 | 169 | 252 | 253 | 6 | 0 | 4 | 6 | 1 | 29 | 72 | 175 | 109 | 5 | 0 | 1 | 1 | 3 | 1 | 298 | 298 | 0.751290574 | 1 | 0.97 | 1 | 0.977 | 0.977 | 0.996 | 1 | 0.986 | 0.994 | 0.973 | 0.833 | 0.027 |
| MXE | 13249 | ENSG00000128731 | HERC2 | chr15 | - | 28182400 | 28182512 | 28186576 | 28186752 | 28179141 | 28179223 | 28190964 | 28191056 | 37 | 33 | 40 | 62 | 49 | 32 | 28 | 42 | 28 | 21 | 79 | 70 | 21 | 33 | 29 | 0 | 33 | 34 | 50 | 57 | 298 | 261 | 0.751571723 | 1 | 0.503 | 0.508 | 0.455 | 0.66 | 0.671 | 1 | 0.65 | 0.351 | 0.366 | 0.308 | 0.024 |
| MXE | 3947 | ENSG00000113300 | CNOT6 | chr5 | + | 180569109 | 180569340 | 180571229 | 180571432 | 180567848 | 180568003 | 180573987 | 180578347 | 99 | 60 | 70 | 77 | 177 | 138 | 87 | 95 | 98 | 364 | 24 | 77 | 35 | 40 | 8 | 41 | 84 | 58 | 71 | 24 | 298 | 298 | 0.751669024 | 1 | 0.418 | 0.408 | 0.424 | 0.44 | 0.327 | 0.369 | 0.478 | 0.376 | 0.36 | 0.25 | 0.037 |
| MXE | 6735 | ENSG00000145703 | IQGAP2 | chr5 | + | 76562395 | 76562552 | 76570579 | 76570657 | 76461569 | 76461669 | 76575692 | 76575769 | 379 | 226 | 240 | 331 | 415 | 393 | 270 | 269 | 413 | 392 | 128 | 290 | 272 | 76 | 66 | 97 | 305 | 316 | 92 | 81 | 298 | 227 | 0.751671206 | 1 | 0.424 | 0.389 | 0.405 | 0.379 | 0.446 | 0.501 | 0.42 | 0.396 | 0.386 | 0.383 | -0.009 |
| MXE | 14517 | ENSG00000130958 | SLC35D2 | chr9 | - | 96321997 | 96322080 | 96324090 | 96324169 | 96320997 | 96321341 | 96336716 | 96336784 | 52 | 50 | 62 | 58 | 91 | 20 | 26 | 31 | 30 | 33 | 199 | 85 | 53 | 78 | 348 | 29 | 47 | 41 | 60 | 176 | 228 | 232 | 0.751715126 | 1 | 0.726 | 0.662 | 0.671 | 0.663 | 0.737 | 0.875 | 0.648 | 0.568 | 0.569 | 0.668 | 0.026 |
| MXE | 9214 | ENSG00000148660 | CAMK2G | chr10 | - | 73847987 | 73848082 | 73848525 | 73848609 | 73847224 | 73847347 | 73849012 | 73849115 | 13 | 9 | 22 | 10 | 18 | 52 | 24 | 39 | 40 | 59 | 6 | 26 | 8 | 18 | 26 | 79 | 41 | 36 | 47 | 61 | 233 | 244 | 0.751863807 | 1 | 0.207 | 0.282 | 0.371 | 0.207 | 0.242 | 0.074 | 0.399 | 0.189 | 0.286 | 0.309 | 0.01 |
| MXE | 4489 | ENSG00000136051 | WASHC4 | chr12 | + | 105137885 | 105138011 | 105140293 | 105140401 | 105133769 | 105133896 | 105140898 | 105141045 | 247 | 265 | 196 | 249 | 632 | 147 | 126 | 90 | 128 | 143 | 6 | 66 | 36 | 11 | 22 | 0 | 29 | 21 | 4 | 10 | 275 | 257 | 0.751892794 | 1 | 0.611 | 0.663 | 0.671 | 0.645 | 0.805 | 1 | 0.68 | 0.616 | 0.72 | 0.673 | -0.059 |
| MXE | 2156 | ENSG00000114062 | UBE3A | chr15 | - | 25407066 | 25407212 | 25408619 | 25408684 | 25405460 | 25405502 | 25409087 | 25409207 | 89 | 78 | 86 | 144 | 159 | 21 | 13 | 11 | 14 | 26 | 137 | 93 | 78 | 75 | 258 | 43 | 26 | 3 | 19 | 12 | 214 | 295 | 0.75190987 | 1 | 0.854 | 0.892 | 0.915 | 0.934 | 0.894 | 0.815 | 0.831 | 0.973 | 0.845 | 0.967 | 0.012 |
| MXE | 1475 | ENSG00000276234 | TADA2A | chr17 | + | 37444695 | 37444768 | 37458523 | 37458587 | 37442563 | 37442652 | 37462077 | 37462121 | 55 | 29 | 41 | 51 | 13 | 12 | 2 | 13 | 4 | 6 | 30 | 15 | 6 | 23 | 53 | 16 | 8 | 1 | 5 | 0 | 222 | 213 | 0.752438705 | 1 | 0.815 | 0.933 | 0.752 | 0.924 | 0.675 | 0.643 | 0.643 | 0.852 | 0.815 | 1 | 0.029 |
| MXE | 8783 | ENSG00000135913 | USP37 | chr2 | - | 218560819 | 218560883 | 218562672 | 218562813 | 218558497 | 218558677 | 218568177 | 218568326 | 17 | 7 | 12 | 14 | 5 | 14 | 17 | 10 | 8 | 25 | 14 | 10 | 11 | 2 | 0 | 8 | 18 | 7 | 11 | 2 | 290 | 213 | 0.75265838 | 1 | 0.471 | 0.232 | 0.468 | 0.562 | 0.128 | 0.562 | 0.29 | 0.536 | 0.118 | 0 | 0.071 |
| MXE | 7475 | ENSG00000084754 | HADHA | chr2 | - | 26192309 | 26192424 | 26193576 | 26193772 | 26190654 | 26191628 | 26194569 | 26194638 | 425 | 224 | 324 | 406 | 381 | 755 | 524 | 605 | 676 | 913 | 638 | 391 | 226 | 381 | 729 | 1263 | 754 | 466 | 771 | 1565 | 298 | 264 | 0.753007203 | 1 | 0.333 | 0.275 | 0.322 | 0.347 | 0.27 | 0.309 | 0.315 | 0.301 | 0.304 | 0.292 | 0.005 |
| MXE | 13449 | ENSG00000258529 | NA | chr11 | - | 111838248 | 111838378 | 111840654 | 111840809 | 111837467 | 111837615 | 111844600 | 111844723 | 46 | 25 | 38 | 24 | 55 | 56 | 31 | 52 | 61 | 73 | 87 | 43 | 39 | 16 | 18 | 47 | 37 | 82 | 55 | 64 | 298 | 279 | 0.753114073 | 1 | 0.435 | 0.43 | 0.406 | 0.269 | 0.414 | 0.634 | 0.521 | 0.308 | 0.214 | 0.208 | 0.014 |
| MXE | 17122 | ENSG00000275066 | SYNRG | chr17 | - | 37570636 | 37570885 | 37571790 | 37571987 | 37568790 | 37568924 | 37576340 | 37576418 | 343 | 218 | 372 | 324 | 632 | 133 | 82 | 167 | 119 | 170 | 155 | 307 | 146 | 195 | 105 | 45 | 93 | 69 | 74 | 36 | 298 | 298 | 0.753743923 | 1 | 0.721 | 0.727 | 0.69 | 0.731 | 0.788 | 0.775 | 0.767 | 0.679 | 0.725 | 0.745 | -0.007 |
| MXE | 15201 | ENSG00000070718 | AP3M2 | chr8 | + | 42165426 | 42165560 | 42167163 | 42167371 | 42165070 | 42165156 | 42167665 | 42167810 | 19 | 21 | 20 | 17 | 29 | 39 | 18 | 44 | 37 | 56 | 8 | 9 | 66 | 37 | 5 | 26 | 34 | 61 | 17 | 44 | 283 | 298 | 0.753806119 | 1 | 0.339 | 0.551 | 0.324 | 0.326 | 0.353 | 0.245 | 0.218 | 0.533 | 0.696 | 0.107 | 0.019 |
| MXE | 3384 | ENSG00000050393 | MCUR1 | chr6 | - | 13788645 | 13788704 | 13791877 | 13791992 | 13787133 | 13787355 | 13793893 | 13793947 | 72 | 72 | 57 | 200 | 122 | 0 | 0 | 0 | 0 | 0 | 222 | 116 | 60 | 94 | 602 | 0 | 0 | 5 | 5 | 3 | 264 | 208 | 0.753961932 | 1 | 1 | 1 | 1 | 1 | 1 | 1 | 1 | 0.904 | 0.937 | 0.994 | 0.033 |
| MXE | 17136 | ENSG00000123104 | ITPR2 | chr12 | - | 26711172 | 26711268 | 26715298 | 26715445 | 26695605 | 26695650 | 26715751 | 26715835 | 81 | 113 | 123 | 97 | 139 | 97 | 130 | 104 | 112 | 146 | 41 | 70 | 74 | 46 | 46 | 105 | 56 | 68 | 49 | 20 | 296 | 245 | 0.754075106 | 1 | 0.409 | 0.418 | 0.495 | 0.418 | 0.441 | 0.244 | 0.509 | 0.474 | 0.437 | 0.656 | -0.028 |
| MXE | 7600 | ENSG00000137628 | DDX60 | chr4 | - | 168287047 | 168287203 | 168288173 | 168288315 | 168285392 | 168285498 | 168291747 | 168291906 | 52 | 53 | 61 | 85 | 136 | 42 | 58 | 53 | 55 | 106 | 10 | 51 | 17 | 13 | 6 | 11 | 29 | 13 | 25 | 0 | 291 | 298 | 0.754250332 | 1 | 0.559 | 0.483 | 0.541 | 0.613 | 0.568 | 0.482 | 0.643 | 0.572 | 0.347 | 1 | -0.056 |
| MXE | 15288 | ENSG00000149547 | EI24 | chr11 | + | 125578948 | 125579068 | 125581210 | 125581322 | 125578132 | 125578257 | 125582345 | 125582420 | 28 | 17 | 41 | 36 | 62 | 78 | 49 | 60 | 52 | 100 | 28 | 35 | 16 | 50 | 39 | 84 | 82 | 38 | 34 | 57 | 269 | 261 | 0.754830092 | 1 | 0.258 | 0.252 | 0.399 | 0.402 | 0.376 | 0.244 | 0.293 | 0.29 | 0.588 | 0.399 | -0.025 |
| MXE | 8225 | ENSG00000142765 | SYTL1 | chr1 | + | 27350793 | 27350952 | 27351257 | 27351336 | 27350388 | 27350485 | 27351455 | 27351555 | 115 | 42 | 68 | 71 | 89 | 87 | 25 | 38 | 54 | 33 | 64 | 51 | 36 | 81 | 105 | 44 | 17 | 18 | 63 | 92 | 298 | 228 | 0.754854914 | 1 | 0.503 | 0.562 | 0.578 | 0.501 | 0.674 | 0.527 | 0.697 | 0.605 | 0.496 | 0.466 | 0.005 |
| MXE | 15989 | ENSG00000075945 | KIFAP3 | chr1 | - | 170039232 | 170039288 | 170046711 | 170046866 | 170038289 | 170038431 | 170055304 | 170055436 | 90 | 74 | 71 | 67 | 208 | 54 | 45 | 40 | 45 | 132 | 33 | 88 | 36 | 21 | 79 | 7 | 59 | 30 | 7 | 74 | 298 | 205 | 0.755468903 | 1 | 0.534 | 0.531 | 0.55 | 0.506 | 0.52 | 0.764 | 0.506 | 0.452 | 0.674 | 0.423 | -0.036 |
| MXE | 1554 | ENSG00000172831 | CES2 | chr16 | + | 66942104 | 66942249 | 66942647 | 66942785 | 66941767 | 66941848 | 66943298 | 66943371 | 57 | 40 | 63 | 47 | 54 | 60 | 45 | 69 | 70 | 42 | 62 | 64 | 45 | 55 | 77 | 88 | 37 | 45 | 99 | 99 | 294 | 287 | 0.755644515 | 1 | 0.481 | 0.465 | 0.471 | 0.396 | 0.557 | 0.408 | 0.628 | 0.494 | 0.352 | 0.432 | 0.011 |
| MXE | 15414 | ENSG00000145649 | GZMA | chr5 | + | 55105473 | 55105618 | 55107793 | 55107935 | 55102645 | 55102752 | 55108124 | 55108394 | 1316 | 411 | 569 | 738 | 752 | 914 | 274 | 375 | 582 | 709 | 501 | 809 | 389 | 449 | 300 | 460 | 477 | 279 | 315 | 231 | 294 | 291 | 0.755938479 | 1 | 0.588 | 0.598 | 0.6 | 0.557 | 0.512 | 0.519 | 0.627 | 0.58 | 0.585 | 0.562 | -0.004 |
| MXE | 15316 | ENSG00000171466 | ZNF562 | chr19 | - | 9658008 | 9658135 | 9660719 | 9660874 | 9653707 | 9653881 | 9675014 | 9675052 | 18 | 12 | 13 | 21 | 6 | 12 | 20 | 22 | 4 | 7 | 6 | 16 | 12 | 9 | 6 | 30 | 4 | 6 | 21 | 0 | 298 | 276 | 0.755980179 | 1 | 0.581 | 0.357 | 0.354 | 0.829 | 0.443 | 0.156 | 0.787 | 0.649 | 0.284 | 1 | -0.062 |
| MXE | 8067 | ENSG00000103657 | HERC1 | chr15 | - | 63648068 | 63648199 | 63649724 | 63649925 | 63645482 | 63645682 | 63651252 | 63651380 | 78 | 67 | 69 | 102 | 105 | 242 | 189 | 229 | 281 | 172 | 18 | 84 | 124 | 52 | 20 | 110 | 166 | 221 | 174 | 64 | 298 | 280 | 0.756117725 | 1 | 0.232 | 0.25 | 0.221 | 0.254 | 0.365 | 0.133 | 0.322 | 0.345 | 0.219 | 0.227 | 0.015 |
| MXE | 418 | ENSG00000130816 | DNMT1 | chr19 | - | 10163325 | 10163360 | 10166597 | 10166685 | 10162666 | 10162748 | 10168329 | 10168364 | 50 | 45 | 53 | 88 | 71 | 6 | 4 | 4 | 3 | 5 | 295 | 50 | 70 | 135 | 223 | 24 | 1 | 9 | 7 | 11 | 237 | 184 | 0.756484573 | 1 | 0.866 | 0.897 | 0.911 | 0.958 | 0.917 | 0.905 | 0.975 | 0.858 | 0.937 | 0.94 | -0.013 |
| MXE | 1476 | ENSG00000276234 | TADA2A | chr17 | + | 37458523 | 37458587 | 37462077 | 37462121 | 37444695 | 37444768 | 37465430 | 37465471 | 14 | 13 | 19 | 10 | 3 | 7 | 11 | 22 | 12 | 13 | 36 | 2 | 1 | 14 | 12 | 4 | 7 | 7 | 11 | 11 | 213 | 193 | 0.75657306 | 1 | 0.644 | 0.517 | 0.439 | 0.43 | 0.173 | 0.891 | 0.206 | 0.115 | 0.536 | 0.497 | -0.008 |
| MXE | 1508 | ENSG00000091592 | NLRP1 | chr17 | - | 5515472 | 5515517 | 5517745 | 5517867 | 5514121 | 5515073 | 5521523 | 5521786 | 51 | 66 | 76 | 51 | 52 | 32 | 32 | 43 | 61 | 22 | 173 | 87 | 94 | 61 | 201 | 102 | 55 | 45 | 78 | 119 | 291 | 194 | 0.756565585 | 1 | 0.515 | 0.579 | 0.541 | 0.358 | 0.612 | 0.531 | 0.513 | 0.582 | 0.343 | 0.53 | 0.021 |
| MXE | 10274 | ENSG00000161813 | LARP4 | chr12 | + | 50435487 | 50435624 | 50437734 | 50437838 | 50430494 | 50430570 | 50440438 | 50440549 | 153 | 199 | 139 | 169 | 400 | 101 | 121 | 146 | 153 | 377 | 297 | 129 | 128 | 123 | 124 | 170 | 164 | 83 | 107 | 92 | 286 | 253 | 0.756778196 | 1 | 0.573 | 0.593 | 0.457 | 0.494 | 0.484 | 0.607 | 0.41 | 0.577 | 0.504 | 0.544 | -0.008 |
| MXE | 95 | ENSG00000182944 | EWSR1 | chr22 | + | 29288605 | 29288786 | 29291564 | 29291599 | 29286922 | 29287134 | 29292136 | 29292169 | 194 | 75 | 135 | 115 | 103 | 72 | 53 | 73 | 56 | 18 | 143 | 169 | 80 | 155 | 259 | 101 | 56 | 32 | 106 | 75 | 298 | 184 | 0.756839988 | 1 | 0.625 | 0.466 | 0.533 | 0.559 | 0.779 | 0.466 | 0.651 | 0.607 | 0.474 | 0.681 | 0.017 |
| MXE | 15874 | ENSG00000136935 | GOLGA1 | chr9 | - | 124921128 | 124921240 | 124921711 | 124921892 | 124911900 | 124912026 | 124923094 | 124923223 | 39 | 38 | 18 | 41 | 44 | 23 | 34 | 32 | 25 | 26 | 51 | 45 | 59 | 147 | 0 | 27 | 22 | 58 | 20 | 44 | 298 | 261 | 0.756843736 | 1 | 0.598 | 0.495 | 0.33 | 0.59 | 0.597 | 0.623 | 0.642 | 0.471 | 0.866 | 0 | 0.002 |
| MXE | 5057 | ENSG00000176261 | ZBTB8OS | chr1 | - | 32633644 | 32633727 | 32633950 | 32634110 | 32631826 | 32631879 | 32650401 | 32650525 | 1 | 2 | 3 | 1 | 0 | 44 | 35 | 38 | 38 | 77 | 0 | 0 | 0 | 3 | 0 | 9 | 21 | 13 | 18 | 8 | 298 | 232 | 0.756935246 | 1 | 0.017 | 0.043 | 0.058 | 0.02 | 0 | 0 | 0 | 0 | 0.115 | 0 | 0.005 |
| MXE | 3057 | ENSG00000135596 | MICAL1 | chr6 | - | 109447356 | 109447440 | 109447680 | 109447722 | 109447072 | 109447229 | 109447874 | 109447963 | 2 | 7 | 1 | 7 | 2 | 49 | 25 | 27 | 43 | 13 | 2 | 5 | 7 | 13 | 6 | 75 | 57 | 20 | 49 | 80 | 191 | 233 | 0.756956175 | 1 | 0.047 | 0.255 | 0.043 | 0.166 | 0.158 | 0.032 | 0.097 | 0.299 | 0.245 | 0.084 | -0.018 |
| MXE | 16788 | ENSG00000033030 | ZCCHC8 | chr12 | - | 122482634 | 122482695 | 122483278 | 122483563 | 122481944 | 122482087 | 122489385 | 122489463 | 60 | 52 | 51 | 64 | 113 | 75 | 60 | 64 | 95 | 109 | 42 | 20 | 29 | 24 | 6 | 50 | 57 | 54 | 72 | 0 | 298 | 210 | 0.757040984 | 1 | 0.361 | 0.379 | 0.36 | 0.322 | 0.422 | 0.372 | 0.198 | 0.275 | 0.19 | 1 | -0.038 |
| MXE | 9730 | ENSG00000137497 | NUMA1 | chr11 | - | 72010785 | 72010854 | 72012894 | 72016260 | 72009267 | 72009387 | 72016407 | 72016530 | 155 | 71 | 123 | 117 | 87 | 74 | 59 | 74 | 87 | 88 | 85 | 139 | 83 | 91 | 144 | 93 | 106 | 57 | 45 | 105 | 298 | 218 | 0.757181632 | 1 | 0.605 | 0.468 | 0.549 | 0.496 | 0.42 | 0.401 | 0.49 | 0.516 | 0.597 | 0.501 | 0.007 |
| MXE | 9756 | ENSG00000259080 | NA | chr2 | + | 27601355 | 27601442 | 27603139 | 27603307 | 27599969 | 27600053 | 27607844 | 27608039 | 2 | 0 | 2 | 7 | 3 | 97 | 95 | 126 | 139 | 187 | 0 | 4 | 1 | 0 | 6 | 125 | 84 | 90 | 100 | 33 | 236 | 298 | 0.757400406 | 1 | 0.025 | 0 | 0.02 | 0.06 | 0.02 | 0 | 0.057 | 0.014 | 0 | 0.187 | -0.027 |
| MXE | 145 | ENSG00000284554 | NA | chr22 | + | 39025556 | 39025671 | 39029362 | 39029519 | 39025069 | 39025349 | 39031693 | 39031973 | 22 | 4 | 7 | 12 | 20 | 60 | 24 | 45 | 45 | 42 | 5 | 23 | 11 | 14 | 3 | 48 | 65 | 14 | 39 | 36 | 264 | 298 | 0.757455289 | 1 | 0.293 | 0.158 | 0.149 | 0.231 | 0.35 | 0.105 | 0.285 | 0.47 | 0.288 | 0.086 | -0.011 |
| MXE | 9982 | ENSG00000101577 | LPIN2 | chr18 | - | 2922046 | 2922199 | 2923774 | 2923861 | 2921532 | 2921647 | 2924397 | 2924546 | 132 | 74 | 121 | 92 | 110 | 141 | 83 | 137 | 163 | 190 | 57 | 53 | 61 | 42 | 77 | 63 | 87 | 104 | 72 | 79 | 236 | 298 | 0.7575488 | 1 | 0.542 | 0.53 | 0.527 | 0.416 | 0.422 | 0.533 | 0.435 | 0.425 | 0.424 | 0.552 | 0.014 |
| MXE | 5941 | ENSG00000172175 | MALT1 | chr18 | + | 58700440 | 58700591 | 58714082 | 58714109 | 58696365 | 58696487 | 58715934 | 58715967 | 158 | 98 | 140 | 134 | 217 | 67 | 55 | 50 | 61 | 83 | 19 | 32 | 60 | 40 | 41 | 2 | 22 | 72 | 31 | 7 | 298 | 176 | 0.75771085 | 1 | 0.582 | 0.513 | 0.623 | 0.565 | 0.607 | 0.849 | 0.462 | 0.33 | 0.432 | 0.776 | 0.008 |
| MXE | 10611 | ENSG00000196295 | GARS1-DT | chr7 | - | 30579230 | 30579314 | 30584710 | 30584808 | 30568832 | 30568989 | 30585329 | 30585479 | 15 | 12 | 7 | 10 | 4 | 24 | 6 | 18 | 28 | 17 | 28 | 24 | 7 | 12 | 21 | 20 | 30 | 18 | 44 | 46 | 247 | 233 | 0.757911441 | 1 | 0.371 | 0.654 | 0.268 | 0.252 | 0.182 | 0.569 | 0.43 | 0.268 | 0.205 | 0.301 | -0.009 |
| MXE | 16555 | ENSG00000125651 | GTF2F1 | chr19 | - | 6383310 | 6383495 | 6387388 | 6387559 | 6381696 | 6381850 | 6389443 | 6389637 | 245 | 93 | 207 | 277 | 223 | 93 | 60 | 77 | 86 | 93 | 509 | 333 | 261 | 351 | 610 | 166 | 151 | 113 | 97 | 284 | 298 | 298 | 0.758189292 | 1 | 0.725 | 0.608 | 0.729 | 0.763 | 0.706 | 0.754 | 0.688 | 0.698 | 0.783 | 0.682 | -0.015 |
| MXE | 16637 | ENSG00000076513 | ANKRD13A | chr12 | + | 110024045 | 110024112 | 110025741 | 110025823 | 110019138 | 110019328 | 110027704 | 110027766 | 260 | 177 | 250 | 278 | 599 | 193 | 83 | 148 | 148 | 426 | 701 | 330 | 280 | 341 | 534 | 403 | 164 | 186 | 199 | 357 | 216 | 231 | 0.758305355 | 1 | 0.59 | 0.695 | 0.644 | 0.668 | 0.601 | 0.65 | 0.683 | 0.617 | 0.647 | 0.615 | -0.003 |
| MXE | 7633 | ENSG00000148187 | MRRF | chr9 | + | 122291748 | 122291840 | 122313226 | 122313386 | 122285168 | 122285287 | 122322539 | 122323461 | 60 | 53 | 37 | 53 | 79 | 37 | 37 | 52 | 25 | 75 | 73 | 70 | 68 | 70 | 37 | 78 | 44 | 42 | 44 | 29 | 241 | 298 | 0.758547415 | 1 | 0.667 | 0.639 | 0.468 | 0.724 | 0.566 | 0.536 | 0.663 | 0.667 | 0.663 | 0.612 | -0.015 |
| MXE | 11197 | ENSG00000115904 | SOS1 | chr2 | - | 39058672 | 39058804 | 39067627 | 39067753 | 39056701 | 39056866 | 39120335 | 39120450 | 47 | 15 | 28 | 28 | 17 | 193 | 185 | 146 | 201 | 222 | 18 | 48 | 22 | 13 | 0 | 85 | 178 | 157 | 175 | 85 | 275 | 281 | 0.759843676 | 1 | 0.199 | 0.077 | 0.164 | 0.125 | 0.073 | 0.178 | 0.216 | 0.125 | 0.071 | 0 | 0.01 |
| MXE | 9378 | ENSG00000155275 | TRMT44 | chr4 | + | 8451959 | 8452028 | 8452881 | 8452989 | 8449668 | 8449888 | 8454741 | 8454813 | 15 | 15 | 15 | 15 | 27 | 23 | 21 | 14 | 13 | 50 | 21 | 3 | 15 | 23 | 23 | 19 | 15 | 12 | 27 | 24 | 218 | 257 | 0.760278065 | 1 | 0.435 | 0.457 | 0.558 | 0.576 | 0.389 | 0.566 | 0.191 | 0.596 | 0.501 | 0.53 | 0.006 |
| MXE | 15580 | ENSG00000073910 | FRY | chr13 | + | 32187545 | 32187656 | 32194142 | 32194297 | 32186259 | 32186420 | 32201940 | 32202040 | 27 | 26 | 36 | 39 | 12 | 48 | 29 | 29 | 38 | 59 | 27 | 9 | 22 | 33 | 5 | 33 | 23 | 29 | 36 | 5 | 260 | 298 | 0.760305118 | 1 | 0.392 | 0.507 | 0.587 | 0.541 | 0.189 | 0.484 | 0.31 | 0.465 | 0.512 | 0.534 | -0.018 |
| MXE | 7711 | ENSG00000113318 | MSH3 | chr5 | + | 80775693 | 80775758 | 80778719 | 80778836 | 80768834 | 80769003 | 80787564 | 80787672 | 35 | 17 | 14 | 16 | 33 | 41 | 29 | 31 | 33 | 80 | 0 | 27 | 44 | 19 | 10 | 14 | 22 | 43 | 27 | 16 | 214 | 266 | 0.760586614 | 1 | 0.515 | 0.422 | 0.36 | 0.376 | 0.339 | 0 | 0.604 | 0.56 | 0.467 | 0.437 | -0.011 |
| MXE | 1276 | ENSG00000141030 | COPS3 | chr17 | - | 17248925 | 17249039 | 17254858 | 17254945 | 17247479 | 17247560 | 17260300 | 17260474 | 167 | 122 | 98 | 100 | 135 | 259 | 224 | 172 | 206 | 253 | 292 | 93 | 173 | 131 | 97 | 370 | 232 | 291 | 332 | 173 | 236 | 263 | 0.760708025 | 1 | 0.418 | 0.378 | 0.388 | 0.351 | 0.373 | 0.468 | 0.309 | 0.399 | 0.305 | 0.385 | 0.008 |
| MXE | 16648 | ENSG00000135297 | MTO1 | chr6 | + | 73480674 | 73480805 | 73482448 | 73482620 | 73479935 | 73480367 | 73492233 | 73492352 | 16 | 10 | 17 | 16 | 15 | 88 | 65 | 103 | 103 | 170 | 5 | 1 | 2 | 8 | 10 | 26 | 84 | 47 | 25 | 20 | 280 | 298 | 0.760896185 | 1 | 0.162 | 0.141 | 0.149 | 0.142 | 0.086 | 0.17 | 0.013 | 0.043 | 0.254 | 0.347 | -0.029 |
| MXE | 1285 | ENSG00000132383 | RPA1 | chr17 | + | 1888674 | 1888851 | 1891832 | 1891940 | 1883811 | 1883944 | 1895008 | 1895095 | 160 | 100 | 129 | 181 | 279 | 136 | 96 | 145 | 124 | 235 | 130 | 120 | 52 | 107 | 175 | 150 | 108 | 81 | 78 | 102 | 298 | 257 | 0.760950459 | 1 | 0.504 | 0.473 | 0.434 | 0.557 | 0.506 | 0.428 | 0.489 | 0.356 | 0.542 | 0.597 | 0.012 |
| MXE | 6139 | ENSG00000163719 | MTMR14 | chr3 | + | 9653620 | 9653769 | 9662266 | 9662375 | 9649504 | 9649742 | 9669431 | 9669492 | 95 | 46 | 117 | 101 | 37 | 3 | 2 | 1 | 2 | 7 | 116 | 177 | 124 | 142 | 258 | 8 | 0 | 9 | 7 | 5 | 298 | 258 | 0.760983252 | 1 | 0.965 | 0.952 | 0.99 | 0.978 | 0.821 | 0.926 | 1 | 0.923 | 0.946 | 0.978 | -0.013 |
| MXE | 11351 | ENSG00000114956 | DGUOK | chr2 | + | 73950600 | 73950732 | 73957124 | 73957240 | 73946718 | 73946906 | 73958145 | 73958245 | 15 | 9 | 20 | 20 | 48 | 82 | 44 | 52 | 67 | 118 | 13 | 32 | 18 | 12 | 9 | 73 | 25 | 33 | 58 | 65 | 281 | 265 | 0.761443796 | 1 | 0.147 | 0.162 | 0.266 | 0.22 | 0.277 | 0.144 | 0.547 | 0.34 | 0.163 | 0.115 | -0.047 |
| MXE | 11919 | ENSG00000001167 | NFYA | chr6 | + | 41091527 | 41091694 | 41092911 | 41093085 | 41090203 | 41090309 | 41094395 | 41094497 | 74 | 50 | 72 | 75 | 68 | 88 | 58 | 79 | 87 | 105 | 72 | 49 | 66 | 77 | 11 | 41 | 54 | 52 | 88 | 47 | 298 | 298 | 0.761493723 | 1 | 0.457 | 0.463 | 0.477 | 0.463 | 0.393 | 0.637 | 0.476 | 0.559 | 0.467 | 0.19 | -0.015 |
| MXE | 6489 | ENSG00000152495 | CAMK4 | chr5 | + | 111344023 | 111344102 | 111374849 | 111374912 | 111224383 | 111224644 | 111376859 | 111376942 | 17 | 25 | 20 | 38 | 10 | 51 | 70 | 105 | 51 | 258 | 9 | 4 | 8 | 6 | 0 | 16 | 21 | 38 | 29 | 5 | 228 | 212 | 0.761553592 | 1 | 0.237 | 0.249 | 0.15 | 0.409 | 0.035 | 0.343 | 0.15 | 0.164 | 0.161 | 0 | 0.052 |
| MXE | 10539 | ENSG00000135838 | NPL | chr1 | + | 182818540 | 182818689 | 182822114 | 182822199 | 182816713 | 182816806 | 182825780 | 182825820 | 84 | 62 | 88 | 85 | 61 | 35 | 18 | 29 | 45 | 50 | 135 | 113 | 80 | 128 | 85 | 62 | 58 | 44 | 89 | 8 | 298 | 234 | 0.761681007 | 1 | 0.653 | 0.73 | 0.704 | 0.597 | 0.489 | 0.631 | 0.605 | 0.588 | 0.53 | 0.893 | -0.015 |
| MXE | 9442 | ENSG00000066027 | PPP2R5A | chr1 | + | 212342187 | 212342280 | 212345933 | 212345993 | 212333496 | 212333598 | 212347346 | 212347406 | 253 | 115 | 144 | 181 | 302 | 216 | 125 | 159 | 232 | 492 | 91 | 168 | 127 | 167 | 72 | 142 | 126 | 157 | 234 | 61 | 298 | 280 | 0.761780663 | 1 | 0.575 | 0.516 | 0.512 | 0.474 | 0.415 | 0.426 | 0.607 | 0.483 | 0.452 | 0.577 | -0.011 |
| MXE | 264 | ENSG00000198089 | SFI1 | chr22 | + | 31602606 | 31602785 | 31606327 | 31606430 | 31602211 | 31602293 | 31607936 | 31608033 | 84 | 87 | 91 | 116 | 110 | 18 | 15 | 29 | 17 | 37 | 87 | 106 | 71 | 65 | 187 | 21 | 12 | 11 | 23 | 77 | 298 | 252 | 0.761870015 | 1 | 0.798 | 0.831 | 0.726 | 0.852 | 0.715 | 0.778 | 0.882 | 0.845 | 0.705 | 0.673 | 0.008 |
| MXE | 13302 | ENSG00000157881 | PANK4 | chr1 | - | 2518163 | 2518264 | 2518515 | 2518597 | 2515561 | 2515717 | 2519142 | 2519295 | 16 | 4 | 10 | 17 | 8 | 14 | 3 | 9 | 6 | 13 | 13 | 18 | 7 | 0 | 39 | 25 | 4 | 10 | 14 | 0 | 231 | 250 | 0.761986557 | 1 | 0.553 | 0.591 | 0.546 | 0.492 | 0.591 | 0.36 | 0.83 | 0.431 | 0 | 1 | 0.03 |
| MXE | 7323 | ENSG00000141446 | ESCO1 | chr18 | - | 21560858 | 21560990 | 21564202 | 21564317 | 21540576 | 21540700 | 21566145 | 21566206 | 92 | 36 | 66 | 66 | 113 | 2 | 3 | 5 | 7 | 22 | 8 | 18 | 26 | 12 | 17 | 5 | 1 | 0 | 0 | 8 | 264 | 281 | 0.762087135 | 1 | 0.98 | 0.927 | 0.934 | 0.909 | 0.845 | 0.63 | 0.95 | 1 | 1 | 0.693 | 0.064 |
| MXE | 3115 | ENSG00000155111 | CDK19 | chr6 | - | 110670430 | 110670541 | 110746125 | 110746201 | 110667433 | 110667574 | 110773875 | 110773937 | 9 | 4 | 3 | 9 | 7 | 49 | 30 | 19 | 48 | 69 | 4 | 6 | 7 | 3 | 0 | 1 | 35 | 34 | 72 | 11 | 225 | 260 | 0.762109215 | 1 | 0.175 | 0.134 | 0.154 | 0.178 | 0.105 | 0.822 | 0.165 | 0.192 | 0.046 | 0 | -0.096 |
| MXE | 4463 | ENSG00000141452 | RMC1 | chr18 | + | 23506969 | 23507054 | 23509192 | 23509279 | 23504370 | 23504447 | 23515855 | 23515996 | 32 | 31 | 46 | 49 | 14 | 131 | 102 | 85 | 102 | 143 | 58 | 51 | 24 | 60 | 27 | 182 | 140 | 95 | 151 | 211 | 234 | 236 | 0.762181928 | 1 | 0.198 | 0.235 | 0.353 | 0.326 | 0.09 | 0.243 | 0.269 | 0.203 | 0.286 | 0.114 | 0.017 |
| MXE | 6128 | ENSG00000170581 | STAT2 | chr12 | - | 56349588 | 56349636 | 56350096 | 56350190 | 56349344 | 56349509 | 56350411 | 56350432 | 149 | 66 | 87 | 183 | 76 | 52 | 21 | 27 | 49 | 69 | 433 | 222 | 78 | 292 | 257 | 150 | 27 | 38 | 132 | 147 | 243 | 197 | 0.762280682 | 1 | 0.699 | 0.718 | 0.723 | 0.752 | 0.472 | 0.701 | 0.87 | 0.625 | 0.642 | 0.586 | -0.012 |
| MXE | 14571 | ENSG00000079387 | SENP1 | chr12 | - | 48096342 | 48096427 | 48097993 | 48098124 | 48088552 | 48088960 | 48101468 | 48101516 | 54 | 28 | 44 | 39 | 63 | 42 | 42 | 37 | 42 | 81 | 13 | 30 | 39 | 38 | 16 | 27 | 49 | 34 | 34 | 3 | 280 | 234 | 0.762326359 | 1 | 0.518 | 0.358 | 0.498 | 0.437 | 0.394 | 0.287 | 0.338 | 0.489 | 0.483 | 0.817 | -0.042 |
| MXE | 15934 | ENSG00000155629 | PIK3AP1 | chr10 | - | 96609711 | 96609867 | 96616638 | 96616711 | 96603739 | 96604049 | 96620351 | 96620411 | 339 | 266 | 203 | 276 | 428 | 203 | 133 | 112 | 143 | 196 | 588 | 378 | 221 | 570 | 171 | 352 | 204 | 108 | 317 | 76 | 222 | 298 | 0.762402858 | 1 | 0.692 | 0.729 | 0.709 | 0.722 | 0.746 | 0.692 | 0.713 | 0.733 | 0.707 | 0.751 | 0 |
| MXE | 2401 | ENSG00000273173 | SNURF | chr15 | + | 24975357 | 24975509 | 24976304 | 24976416 | 24967931 | 24968082 | 24976876 | 24977029 | 1 | 1 | 4 | 8 | 10 | 75 | 68 | 98 | 96 | 128 | 5 | 10 | 1 | 3 | 7 | 51 | 99 | 40 | 22 | 291 | 298 | 261 | 0.762599241 | 1 | 0.012 | 0.013 | 0.035 | 0.068 | 0.064 | 0.079 | 0.081 | 0.021 | 0.107 | 0.021 | -0.023 |
| MXE | 16209 | ENSG00000107862 | GBF1 | chr10 | + | 102359269 | 102359435 | 102360183 | 102360395 | 102358505 | 102358729 | 102361021 | 102361120 | 117 | 55 | 108 | 63 | 60 | 74 | 37 | 67 | 85 | 76 | 200 | 79 | 73 | 142 | 164 | 119 | 90 | 80 | 69 | 157 | 298 | 298 | 0.762715437 | 1 | 0.613 | 0.598 | 0.617 | 0.426 | 0.441 | 0.627 | 0.467 | 0.477 | 0.673 | 0.511 | -0.012 |
| MXE | 14453 | ENSG00000216937 | CCDC7 | chr10 | + | 32574428 | 32574526 | 32583033 | 32583307 | 32571858 | 32571893 | 32584231 | 32584304 | 26 | 10 | 9 | 19 | 45 | 291 | 222 | 167 | 181 | 774 | 0 | 23 | 6 | 1 | 22 | 128 | 227 | 151 | 90 | 130 | 247 | 298 | 0.762924462 | 1 | 0.097 | 0.052 | 0.061 | 0.112 | 0.066 | 0 | 0.109 | 0.046 | 0.013 | 0.17 | 0.01 |
| MXE | 16937 | ENSG00000101160 | CTSZ | chr20 | - | 58997854 | 58997954 | 58998597 | 58998690 | 58997602 | 58997753 | 59001464 | 59001644 | 4 | 1 | 0 | 0 | 0 | 11 | 15 | 4 | 4 | 36 | 0 | 0 | 1 | 0 | 4 | 69 | 26 | 11 | 24 | 32 | 242 | 249 | 0.762938411 | 1 | 0.272 | 0.064 | 0 | 0 | 0 | 0 | 0 | 0.086 | 0 | 0.114 | 0.027 |

| | | | | | | | | | | | | | | | |
|---|---|---|---|---|---|---|---|---|---|---|---|---|---|---|---|
| MXE | 8597 | ENSG00000185946 | RNPC3 | chr1 | + | 103543298 | 103543447 | 103544940 | 103545102 | 103541349 | 103541475 | 103546247 | 103546342 | 35 | [illegible] |
| MXE | 6451 | ENSG00000169231 | THBS3 | chr1 | - | 155203512 | 155203539 | 155204854 | 155204957 | 155203222 | 155203305 | 155205059 | 155205316 | 6 | [illegible] |
| MXE | 15267 | ENSG00000119318 | RAD23B | chr9 | + | 107300140 | 107300222 | 107302034 | 107302114 | 107283278 | 107283695 | 107306378 | 107306647 | 13 | [illegible] |
| MXE | 7748 | ENSG00000123064 | DDX54 | chr12 | - | 113163131 | 113163274 | 113164066 | 113164285 | 113162931 | 113163045 | 113165643 | 113165717 | 65 | [illegible] |
| MXE | 7667 | ENSG00000114933 | INO80D | chr2 | - | 206062798 | 206063045 | 206063150 | 206063244 | 206056197 | 206056943 | 206085900 | 206086174 | 11 | [illegible] |
| MXE | 10905 | ENSG00000258984 | UBE2F-SCLY | chr2 | + | 237973091 | 237973225 | 237987962 | 237987992 | 237967013 | 237967132 | 237994743 | 237994809 | 34 | [illegible] |
| MXE | 9202 | ENSG00000112339 | HBS1L | chr6 | - | 135039572 | 135039767 | 135042000 | 135042126 | 135002733 | 135002842 | 135050581 | 135050647 | 130 | [illegible] |
| MXE | 15503 | ENSG00000188690 | UROS | chr10 | - | 125815033 | 125815130 | 125816176 | 125816260 | 125812213 | 125812288 | 125816436 | 125816525 | 18 | [illegible] |
| MXE | 5143 | ENSG00000108219 | TSPAN14 | chr10 | + | 80463087 | 80463127 | 80468546 | 80468687 | 80459301 | 80459507 | 80489216 | 80489314 | 9 | [illegible] |
| MXE | 4980 | ENSG00000123737 | EXOSC9 | chr4 | + | 121804621 | 121804759 | 121811582 | 121811671 | 121802914 | 121803017 | 121813233 | 121813293 | 89 | [illegible] |
| MXE | 15859 | ENSG00000088298 | EDEM2 | chr20 | - | 35134737 | 35134949 | 35137879 | 35138005 | 35131641 | 35131783 | 35142372 | 35142478 | 34 | [illegible] |
| MXE | 38 | ENSG00000182670 | TTC3 | chr21 | + | 37094004 | 37094090 | 37095349 | 37095444 | 37091292 | 37091413 | 37096580 | 37096643 | 126 | [illegible] |
| MXE | 8462 | ENSG00000137275 | RIPK1 | chr6 | + | 3083084 | 3083313 | 3085258 | 3085408 | 3080978 | 3081116 | 3089580 | 3089657 | 145 | [illegible] |
| MXE | 3722 | ENSG00000288643 | NA | chr5 | + | 62526141 | 62526257 | 62530708 | 62530785 | 62515387 | 62515501 | 62536701 | 62536781 | 13 | [illegible] |
| MXE | 3787 | ENSG00000164219 | PGGT1B | chr5 | - | 115221823 | 115222008 | 115230975 | 115231021 | 115216864 | 115216973 | 115236389 | 115236522 | 78 | [illegible] |
| MXE | 1042 | ENSG00000170004 | CHD3 | chr17 | + | 7907890 | 7908019 | 7908696 | 7908829 | 7907600 | 7907702 | 7909142 | 7909338 | 14 | [illegible] |
| MXE | 6630 | ENSG00000162946 | DISC1 | chr1 | + | 231693825 | 231694805 | 231749925 | 231750076 | 231626814 | 231626934 | 231767139 | 231767269 | 6 | [illegible] |
| MXE | 10966 | ENSG00000115762 | PLEKHB2 | chr2 | + | 131130720 | 131130760 | 131132901 | 131133015 | 131126683 | 131126786 | 131140166 | 131140275 | 41 | [illegible] |
| MXE | 7679 | ENSG00000001461 | NIPAL3 | chr1 | + | 24449480 | 24449626 | 24453407 | 24453504 | 24445184 | 24445244 | 24456137 | 24456273 | 14 | [illegible] |
| MXE | 17100 | ENSG00000197442 | MAP3K5 | chr6 | - | 136639542 | 136639638 | 136650983 | 136651091 | 136637324 | 136637406 | 136656306 | 136656460 | 79 | [illegible] |
| MXE | 16173 | ENSG00000107669 | ATE1 | chr10 | - | 121913789 | 121913893 | 121922348 | 121922411 | 121910905 | 121911151 | 121924265 | 121924329 | 9 | [illegible] |
| MXE | 3182 | ENSG00000133706 | LARS1 | chr5 | - | 146140203 | 146143084 | 146143411 | 146143550 | 146135600 | 146135664 | 146144266 | 146144349 | 108 | [illegible] |
| MXE | 11431 | ENSG00000119707 | RBM25 | chr14 | + | 73077368 | 73077536 | 73083493 | 73083551 | 73076318 | 73076368 | 73088000 | 73088161 | 126 | [illegible] |
| MXE | 4347 | ENSG00000182150 | ERCC6L2 | chr9 | + | 95897848 | 95897971 | 95907077 | 95907271 | 95880868 | 95881293 | 95915667 | 95915829 | 66 | [illegible] |
| MXE | 4759 | ENSG00000211897 | IGHG3 | chr14 | - | 105770104 | 105770149 | 105770292 | 105770337 | 105769656 | 105769986 | 105770480 | 105770525 | 11 | [illegible] |
| MXE | 8089 | ENSG00000083544 | TDRD3 | chr13 | + | 60444686 | 60444748 | 60460379 | 60460540 | 60439687 | 60439772 | 60467237 | 60467379 | 3 | [illegible] |
| MXE | 16165 | ENSG00000107669 | ATE1 | chr10 | - | 121898840 | 121898969 | 121899865 | 121899994 | 121870005 | 121870038 | 121902390 | 121902620 | 72 | [illegible] |
| MXE | 9353 | ENSG00000070814 | TCOF1 | chr5 | + | 150372005 | 150372236 | 150374173 | 150374386 | 150369528 | 150369602 | 150374616 | 150374811 | 87 | [illegible] |
| MXE | 6679 | ENSG00000155115 | GTF3C6 | chr6 | + | 110960571 | 110960616 | 110962391 | 110962505 | 110960413 | 110960477 | 110967509 | 110967872 | 120 | [illegible] |
| MXE | 5871 | ENSG00000228794 | LINC01128 | chr1 | + | 841199 | 841373 | 847653 | 847806 | 829002 | 829104 | 851926 | 852110 | 5 | [illegible] |
| MXE | 8660 | ENSG00000173744 | AGFG1 | chr2 | + | 227491546 | 227491640 | 227519947 | 227520063 | 227472151 | 227472588 | 227523762 | 227523925 | 27 | [illegible] |
| MXE | 16589 | ENSG00000088992 | TESC | chr12 | - | 117046771 | 117046838 | 117049018 | 117049158 | 117046558 | 117046666 | 117056805 | 117056886 | 44 | [illegible] |
| MXE | 8028 | ENSG00000160691 | SHC1 | chr1 | - | 154965542 | 154965781 | 154965945 | 154966080 | 154962297 | 154963931 | 154966161 | 154966231 | 25 | [illegible] |
| MXE | 15699 | ENSG00000054654 | SYNE2 | chr14 | + | 64190070 | 64190237 | 64202800 | 64202963 | 64188549 | 64188708 | 64208757 | 64208945 | 170 | [illegible] |
| MXE | 877 | ENSG00000060069 | CTDP1 | chr18 | + | 79717809 | 79718016 | 79728906 | 79729069 | 79714490 | 79715528 | 79736354 | 79736521 | 14 | [illegible] |
| MXE | 16017 | ENSG00000119979 | DENND10 | chr10 | + | 119129514 | 119129622 | 119132514 | 119132609 | 119123468 | 119123569 | 119136470 | 119137826 | 104 | [illegible] |
| MXE | 7850 | ENSG00000132128 | LRRC41 | chr1 | - | 46280191 | 46280290 | 46281124 | 46281385 | 46279181 | 46279614 | 46285361 | 46286499 | 33 | [illegible] |
| MXE | 9831 | ENSG00000122507 | BBS9 | chr7 | + | 33351218 | 33351323 | 33352858 | 33352873 | 33349067 | 33349170 | 33357854 | 33357995 | 57 | [illegible] |
| MXE | 10201 | ENSG00000123240 | OPTN | chr10 | + | 13116266 | 13116340 | 13118887 | 13119040 | 13112452 | 13112635 | 13122384 | 13122487 | 72 | [illegible] |
| MXE | 15526 | ENSG00000110075 | PPP6R3 | chr11 | + | 68548066 | 68548204 | 68551120 | 68551186 | 68544837 | 68545024 | 68554144 | 68554220 | 291 | [illegible] |
| MXE | 6387 | ENSG00000198952 | SMG5 | chr1 | - | 156250857 | 156250996 | 156251402 | 156251477 | 156249223 | 156250670 | 156252413 | 156252504 | 44 | [illegible] |
| MXE | 7809 | ENSG00000066427 | ATXN3 | chr14 | - | 92093745 | 92093831 | 92096092 | 92096134 | 92093251 | 92093318 | 92096673 | 92096838 | 9 | [illegible] |
| MXE | 13477 | ENSG00000111203 | ITFG2 | chr12 | + | 2830216 | 2830885 | 2836744 | 2836956 | 2824089 | 2824252 | 2858011 | 2858331 | 5 | [illegible] |
| MXE | 9763 | ENSG00000115761 | NOL10 | chr2 | - | 10675793 | 10675871 | 10681970 | 10682069 | 10673519 | 10673557 | 10684566 | 10684612 | 48 | [illegible] |
| MXE | 2358 | ENSG00000136381 | IREB2 | chr15 | + | 78466270 | 78466489 | 78470531 | 78470601 | 78462921 | 78463087 | 78471740 | 78471924 | 5 | [illegible] |
| MXE | 1771 | ENSG00000090857 | PDPR | chr16 | + | 70136193 | 70136386 | 70138898 | 70139023 | 70132150 | 70132300 | 70142233 | 70142389 | 102 | [illegible] |
| MXE | 1727 | ENSG00000197912 | SPG7 | chr16 | + | 89536764 | 89536948 | 89540943 | 89542279 | 89532462 | 89532636 | 89544647 | 89544772 | 17 | [illegible] |
| MXE | 9280 | ENSG00000175054 | ATR | chr3 | - | 142556382 | 142556575 | 142558623 | 142558776 | 142555876 | 142556139 | 142559250 | 142559441 | 56 | [illegible] |
| MXE | 11921 | ENSG00000167483 | NIBAN3 | chr19 | + | 17533586 | 17533701 | 17537375 | 17537543 | 17532262 | 17532388 | 17539149 | 17539265 | 22 | [illegible] |
| MXE | 10441 | ENSG00000152359 | POC5 | chr5 | - | 75677773 | 75677950 | 75685206 | 75685484 | 75674430 | 75674578 | 75689011 | 75689165 | 42 | [illegible] |
| MXE | 2875 | ENSG00000136758 | YME1L1 | chr10 | - | 27117575 | 27117727 | 27119293 | 27119449 | 27116218 | 27116345 | 27120434 | 27120547 | 400 | [illegible] |
| MXE | 12408 | ENSG00000198198 | SZT2 | chr1 | + | 43403600 | 43403774 | 43404379 | 43404550 | 43403176 | 43403302 | 43415081 | 43415213 | 55 | [illegible] |
| MXE | 13166 | ENSG00000197892 | KIF13B | chr8 | - | 29075280 | 29075343 | 29092744 | 29092878 | 29071619 | 29072316 | 29099132 | 29099241 | 50 | [illegible] |
| MXE | 5278 | ENSG00000234857 | HNRNPUL2-BSCL2 | chr11 | - | 62723913 | 62723990 | 62724290 | 62724426 | 62723586 | 62723726 | 62726618 | 62727384 | 83 | [illegible] |
| MXE | 15343 | ENSG00000099783 | HNRNPM | chr19 | + | 8471325 | 8471427 | 8473663 | 8473708 | 8468773 | 8468834 | 8474166 | 8474244 | 304 | [illegible] |
| MXE | 11302 | ENSG00000168763 | CNNM3 | chr2 | + | 96828098 | 96828195 | 96828566 | 96828700 | 96827730 | 96827900 | 96828995 | 96829134 | 34 | [illegible] |
| MXE | 6486 | ENSG00000140575 | IQGAP1 | chr15 | + | 90467449 | 90467592 | 90472839 | 90473010 | 90466268 | 90466436 | 90473714 | 90473798 | 918 | [illegible] |
| MXE | 15179 | ENSG00000147316 | MCPH1 | chr8 | + | 6442066 | 6442156 | 6444392 | 6445547 | 6438952 | 6439096 | 6455142 | 6455252 | 50 | [illegible] |
| MXE | 5991 | ENSG00000141293 | SKAP1 | chr17 | - | 48363788 | 48363814 | 48396679 | 48396785 | 48345904 | 48346006 | 48430074 | 48430214 | 110 | [illegible] |
| MXE | 5910 | ENSG00000067066 | SP100 | chr2 | + | 230503065 | 230503110 | 230504185 | 230504290 | 230498460 | 230498535 | 230506302 | 230506445 | 40 | [illegible] |
| MXE | 1031 | ENSG00000161955 | TNFSF13 | chr17 | + | 7559623 | 7559702 | 7559850 | 7559893 | 7559067 | 7559297 | 7560048 | 7560167 | 78 | [illegible] |
| MXE | 7134 | ENSG00000141027 | NCOR1 | chr17 | - | 16164978 | 16165161 | 16171802 | 16172020 | 16158759 | 16158873 | 16186553 | 16186687 | 4 | [illegible] |
| MXE | 10923 | ENSG00000132323 | ILKAP | chr2 | - | 238185180 | 238185287 | 238188130 | 238188257 | 238184019 | 238184113 | 238189852 | 238189972 | 34 | [illegible] |
| MXE | 1890 | ENSG00000077235 | GTF3C1 | chr16 | - | 27528597 | 27528721 | 27533290 | 27533387 | 27511748 | 27511901 | 27537783 | 27537927 | 71 | [illegible] |
| MXE | 3351 | ENSG00000127603 | MACF1 | chr1 | + | 39340803 | 39340953 | 39346976 | 39347210 | 39340501 | 39340717 | 39349477 | 39349627 | 362 | [illegible] |
| MXE | 9686 | ENSG00000033178 | UBA6 | chr4 | - | 67663138 | 67663215 | 67663884 | 67663947 | 67661662 | 67662255 | 67665188 | 67665292 | 44 | [illegible] |
| MXE | 251 | ENSG00000198089 | SFI1 | chr22 | + | 31546860 | 31546971 | 31550253 | 31550348 | 31508258 | 31508376 | 31556941 | 31557059 | 16 | [illegible] |
| MXE | 10723 | ENSG00000135636 | DYSF | chr2 | + | 71516988 | 71517039 | 71520177 | 71520208 | 71516179 | 71516242 | 71520788 | 71520904 | 9 | [illegible] |
| MXE | 17192 | ENSG00000136100 | VPS36 | chr13 | - | 52417991 | 52418056 | 52423573 | 52423639 | 52417056 | 52417141 | 52425931 | 52426066 | 248 | [illegible] |
| MXE | 3522 | ENSG00000135074 | ADAM19 | chr5 | - | 157513433 | 157513505 | 157518822 | 157518888 | 157509300 | 157509467 | 157519838 | 157520031 | 29 | [illegible] |
| MXE | 10782 | ENSG00000184708 | EIF4ENIF1 | chr22 | - | 31462931 | 31463133 | 31463680 | 31463967 | 31458474 | 31458650 | 31468174 | 31468302 | 55 | [illegible] |
| MXE | 13895 | ENSG00000205581 | HMGN1 | chr21 | - | 39347378 | 39347483 | 39347915 | 39347952 | 39345829 | 39345958 | 39348291 | 39348339 | 3 | [illegible] |
| MXE | 7566 | ENSG00000077549 | CAPZB | chr1 | - | 19356634 | 19356751 | 19357421 | 19357563 | 19344357 | 19345252 | 19378539 | 19378653 | 756 | [illegible] |
| MXE | 9083 | ENSG00000164828 | SUN1 | chr7 | + | 849922 | 850033 | 851383 | 851482 | 849519 | 849603 | 851949 | 852043 | 55 | [illegible] |
| MXE | 7097 | ENSG00000163125 | RPRD2 | chr1 | + | 150460059 | 150460317 | 150464526 | 150464727 | 150457287 | 150457570 | 150470560 | 150473556 | 203 | [illegible] |
| MXE | 13638 | ENSG00000104613 | INTS10 | chr8 | + | 19832027 | 19832110 | 19837051 | 19837160 | 19830405 | 19830559 | 19842847 | 19842927 | 88 | [illegible] |
| MXE | 363 | ENSG00000257103 | LSM14A | chr19 | + | 34208928 | 34209051 | 34215123 | 34215300 | 34196633 | 34196763 | 34215595 | 34215661 | 316 | [illegible] |
| MXE | 8209 | ENSG00000092847 | AGO1 | chr1 | + | 35893591 | 35893810 | 35894036 | 35894171 | 35893096 | 35893278 | 35894314 | 35894402 | 5 | [illegible] |
| MXE | 5904 | ENSG00000067066 | SP100 | chr2 | + | 230449560 | 230449710 | 230461261 | 230461414 | 230449087 | 230449150 | 230462434 | 230462518 | 305 | [illegible] |
| MXE | 2703 | ENSG00000119684 | MLH3 | chr14 | - | 75030542 | 75030702 | 75032067 | 75032179 | 75022994 | 75023018 | 75038339 | 75038412 | 1 | [illegible] |
| MXE | 11544 | ENSG00000116539 | ASH1L | chr1 | - | 155477885 | 155482449 | 155521099 | 155521618 | 155459796 | 155459898 | 155562152 | 155562291 | 27 | [illegible] |
| MXE | 5321 | ENSG00000164073 | MFSD8 | chr4 | - | 127921523 | 127921771 | 127930682 | 127930817 | 127917798 | 127920836 | 127932984 | 127933093 | 76 | [illegible] |
| MXE | 14773 | ENSG00000106868 | SUSD1 | chr9 | - | 112078537 | 112078724 | 112080073 | 112080165 | 112062936 | 112063033 | 112098469 | 112098662 | 136 | [illegible] |
| MXE | 3803 | ENSG00000112992 | NNT | chr5 | + | 43609515 | 43609551 | 43612907 | 43613137 | 43609142 | 43609346 | 43624031 | 43624120 | 1 | [illegible] |
| MXE | 11338 | ENSG00000144233 | AMMECR1L | chr2 | - | 127873827 | 127874272 | 127884202 | 127884312 | 127871248 | 127871359 | 127885809 | 127885883 | 9 | [illegible] |
| MXE | 1181 | ENSG00000271605 | MILR1 | chr17 | + | 64452596 | 64452866 | 64457399 | 64457684 | 64449313 | 64449355 | 64460821 | 64460932 | 70 | [illegible] |
| MXE | 4271 | ENSG00000102445 | RUBCNL | chr13 | - | 46345446 | 46345600 | 46349285 | 46349347 | 46344740 | 46344831 | 46350112 | 46350351 | 15 | [illegible] |
| MXE | 6471 | ENSG00000153561 | RMND5A | chr2 | + | 86751895 | 86752030 | 86765026 | 86765193 | 86740926 | 86741069 | 86765858 | 86766024 | 123 | [illegible] |
| MXE | 3434 | ENSG00000116459 | ATP5PB | chr1 | + | 111456085 | 111456249 | 111456629 | 111456755 | 111454210 | 111454356 | 111459456 | 111459636 | 814 | [illegible] |
| MXE | 12281 | ENSG00000242588 | NA | chr7 | + | 128623574 | 128623781 | 128625796 | 128625855 | 128622478 | 128622694 | 128626294 | 128626473 | 66 | [illegible] |
| MXE | 12832 | ENSG00000086730 | LAT2 | chr7 | + | 74221636 | 74221692 | 74223723 | 74223783 | 74220703 | 74220734 | 74224017 | 74224197 | 79 | [illegible] |
| MXE | 8474 | ENSG00000163463 | KRTCAP2 | chr1 | - | 155169790 | 155169857 | 155172564 | 155172892 | 155169408 | 155169560 | 155173220 | 155173251 | 94 | [illegible] |
| MXE | 1106 | ENSG00000141252 | VPS53 | chr17 | - | 623532 | 623674 | 631549 | 631628 | 586269 | 586364 | 653290 | 653410 | 29 | [illegible] |
| MXE | 5881 | ENSG00000229180 | NA | chr7 | - | 66576754 | 66576929 | 66588511 | 66588580 | 66573384 | 66573550 | 66592256 | 66592397 | 0 | [illegible] |
| MXE | 15372 | ENSG00000107036 | RIC1 | chr9 | + | 5738449 | 5738538 | 5742868 | 5743013 | 5732387 | 5732479 | 5743688 | 5743737 | 31 | [illegible] |
| MXE | 897 | ENSG00000074657 | ZNF532 | chr18 | + | 58934432 | 58934614 | 58939444 | 58939621 | 58918270 | 58920633 | 58948066 | 58948229 | 26 | [illegible] |
| MXE | 15702 | ENSG00000054654 | SYNE2 | chr14 | + | 64216247 | 64216424 | 64218397 | 64218512 | 64214193 | 64214470 | 64219207 | 64219410 | 95 | [illegible] |
| MXE | 2831 | ENSG00000029364 | SLC39A9 | chr14 | + | 69424093 | 69424202 | 69442068 | 69442266 | 69398691 | 69399465 | 69453240 | 69453309 | 58 | [illegible] |
| MXE | 12578 | ENSG00000175471 | MCTP1 | chr5 | - | 94714776 | 94714886 | 94779109 | 94779163 | 94710817 | 94710927 | 94799012 | 94799132 | 26 | [illegible] |
| MXE | 1729 | ENSG00000197912 | SPG7 | chr16 | + | 89549080 | 89549336 | 89549499 | 89549713 | 89548002 | 89548113 | 89550493 | 89550609 | 8 | [illegible] |
| MXE | 8026 | ENSG00000111348 | ARHGDIB | chr12 | - | 14949801 | 14949885 | 14950531 | 14950724 | 14947688 | 14947949 | 14961536 | 14961591 | 3587 | [illegible] |
| MXE | 10485 | ENSG00000135912 | TTLL4 | chr2 | + | 218747002 | 218747194 | 218747289 | 218747372 | 218746154 | 218746231 | 218747596 | 218747725 | 30 | [illegible] |
| MXE | 8532 | ENSG00000136161 | RCBTB2 | chr13 | - | 48515585 | 48515741 | 48521897 | 48521962 | 48515204 | 48515355 | 48522307 | 48522403 | 8 | [illegible] |
| MXE | 13551 | ENSG00000196092 | PAX5 | chr9 | - | 36923354 | 36923484 | 36930887 | 36930944 | 36882003 | 36882105 | 36966548 | 36966724 | 0 | [illegible] |
| MXE | 2325 | ENSG00000214331 | PDPR2P | chr16 | - | 74350616 | 74350738 | 74352039 | 74352203 | 74349741 | 74349859 | 74352296 | 74352378 | 42 | [illegible] |
| MXE | 12671 | ENSG00000176105 | YES1 | chr18 | - | 747919 | 748018 | 751704 | 751804 | 745947 | 746051 | 756556 | 756835 | 66 | [illegible] |
| MXE | 11557 | ENSG00000114388 | NPRL2 | chr3 | - | 50349385 | 50349494 | 50349664 | 50349828 | 50348873 | 50349010 | 50349930 | 50350022 | 28 | [illegible] |
| MXE | 4721 | ENSG00000163913 | IFT122 | chr3 | + | 129451913 | 129451972 | 129458598 | 129458677 | 129449870 | 129449937 | 129461227 | 129461304 | 13 | [illegible] |
| MXE | 5765 | ENSG00000143811 | PYCR2 | chr1 | - | 225921551 | 225921644 | 225921857 | 225922079 | 225921207 | 225921371 | 225922203 | 225922383 | 66 | [illegible] |
| MXE | 11915 | ENSG00000146842 | TMEM209 | chr7 | - | 130178401 | 130178527 | 130181622 | 130181719 | 130175511 | 130175609 | 130184183 | 130184255 | 78 | [illegible] |
| MXE | 12849 | ENSG00000128595 | CALU | chr7 | + | 128754261 | 128754455 | 128754528 | 128754722 | 128748572 | 128748804 | 128758870 | 128759037 | 226 | [illegible] |
| MXE | 16897 | ENSG00000188906 | LRRK2 | chr12 | + | 40348408 | 40348509 | 40351538 | 40351733 | 40346752 | 40346923 | 40354298 | 40354492 | 186 | [illegible] |
| MXE | 6525 | ENSG00000121940 | CLCC1 | chr1 | - | 108943474 | 108943615 | 108943835 | 108944057 | 108941404 | 108941498 | 108947610 | 108947718 | 61 | [illegible] |
| MXE | 14660 | ENSG00000198780 | FAM169A | chr5 | - | 74834425 | 74834597 | 74838964 | 74839050 | 74805155 | 74805284 | 74840073 | 74840173 | 61 | [illegible] |
| MXE | 16115 | ENSG00000095585 | BLNK | chr10 | - | 96227409 | 96227566 | 96230793 | 96230834 | 96223825 | 96223989 | 96242734 | 96242784 | 12 | [illegible] |
| MXE | 507 | ENSG00000130755 | GMFG | chr19 | - | 39333076 | 39333126 | 39335434 | 39335531 | 39329543 | 39329626 | 39335973 | 39336005 | 187 | [illegible] |
| MXE | 2211 | ENSG00000166411 | IDH3A | chr15 | + | 78157547 | 78157631 | 78160091 | 78160206 | 78155212 | 78155275 | 78161580 | 78161768 | 30 | [illegible] |
| MXE | 1051 | ENSG00000109118 | PHF12 | chr17 | - | 28921687 | 28921808 | 28923908 | 28924302 | 28919142 | 28919275 | 28926990 | 28927063 | 249 | [illegible] |
| MXE | 11208 | ENSG00000152348 | ATG10 | chr5 | + | 82164398 | 82164537 | 82178489 | 82178587 | 82058494 | 82058602 | 82252561 | 82252659 | 13 | [illegible] |
| MXE | 3289 | ENSG00000163933 | RFT1 | chr3 | - | 53092368 | 53092618 | 53099380 | 53099486 | 53088482 | 53092070 | 53103952 | 53104097 | 25 | [illegible] |
| MXE | 10699 | ENSG00000115484 | CCT4 | chr2 | - | 61876919 | 61877052 | 61877392 | 61877514 | 61876094 | 61876234 | 61878868 | 61879011 | 363 | [illegible] |
| MXE | 6149 | ENSG00000078304 | PPP2R5C | chr14 | + | 101786017 | 101786183 | 101818991 | 101819085 | 101762904 | 101762970 | 101856685 | 101856885 | 507 | [illegible] |
| MXE | 12672 | ENSG00000110497 | AMBRA1 | chr11 | - | 46508190 | 46508370 | 46512726 | 46512813 | 46494123 | 46494204 | 46541944 | 46543398 | 18 | [illegible] |
| MXE | 15911 | ENSG00000119541 | VPS4B | chr18 | - | 63393408 | 63393549 | 63397033 | 63397253 | 63389189 | 63391076 | 63399241 | 63399323 | 304 | [illegible] |
| MXE | 1105 | ENSG00000141252 | VPS53 | chr17 | - | 601794 | 601896 | 623532 | 623674 | 586269 | 586364 | 627173 | 627316 | 91 | [illegible] |
| MXE | 5993 | ENSG00000116514 | RNF19B | chr1 | - | 32938396 | 32938528 | 32942251 | 32942459 | 32936453 | 32937309 | 32944018 | 32944159 | 113 | [illegible] |
| MXE | 113 | ENSG00000198911 | SREBF2 | chr22 | + | 41877941 | 41878123 | 41878658 | 41878758 | 41875542 | 41875724 | 41880715 | 41880992 | 33 | [illegible] |
| MXE | 15222 | ENSG00000149289 | ZC3H12C | chr11 | + | 110159255 | 110159490 | 110163272 | 110163379 | 110152918 | 110153058 | 110164340 | 110165934 | 15 | [illegible] |
| MXE | 675 | ENSG00000132670 | PTPRA | chr20 | + | 2947981 | 2948024 | 2955608 | 2955698 | 2923206 | 2923285 | 2964271 | 2964350 | 677 | [illegible] |
| MXE | 17184 | ENSG00000005810 | MYCBP2 | chr13 | - | 77156057 | 77156202 | 77157936 | 77158109 | 77150733 | 77150949 | 77161905 | 77161955 | 159 | [illegible] |
| MXE | 670 | ENSG00000064932 | SBNO2 | chr19 | - | 1122893 | 1123045 | 1123533 | 1123639 | 1122657 | 1122791 | 1123941 | 1124022 | 47 | [illegible] |
| MXE | 5224 | ENSG00000166501 | PRKCB | chr16 | + | 24174517 | 24174580 | 24180789 | 24180928 | 24172269 | 24172361 | 24185110 | 24185191 | 117 | [illegible] |
| MXE | 11881 | ENSG00000135503 | ACVR1B | chr12 | + | 51983998 | 51984166 | 51985191 | 51985348 | 51980968 | 51981199 | 51986817 | 51986942 | 35 | [illegible] |
| MXE | 10460 | ENSG00000076003 | MCM6 | chr2 | - | 135865012 | 135865163 | 135866131 | 135866277 | 135862606 | 135862748 | 135866562 | 135866728 | 89 | [illegible] |
| MXE | 1328 | ENSG00000173821 | RNF213 | chr17 | + | 80349769 | 80349906 | 80350300 | 80350396 | 80344677 | 80348286 | 80351684 | 80351803 | 356 | [illegible] |
| MXE | 8199 | ENSG00000027001 | MIPEP | chr13 | - | 23841334 | 23841488 | 23858859 | 23858912 | 23839648 | 23839726 | 23862301 | 23862362 | 6 | [illegible] |
| MXE | 1772 | ENSG00000090857 | PDPR | chr16 | + | 70146133 | 70146228 | 70148463 | 70148553 | 70144420 | 70144533 | 70156474 | 70156859 | 25 | [illegible] |
| MXE | 9187 | ENSG00000213064 | SFT2D2 | chr1 | + | 168235100 | 168235182 | 168236588 | 168236624 | 168231833 | 168231919 | 168236711 | 168236770 | 133 | [illegible] |
| MXE | 16396 | ENSG00000144579 | CTDSP1 | chr2 | + | 218402110 | 218402215 | 218403034 | 218403127 | 218401563 | 218401712 | 218403231 | 218403415 | 47 | [illegible] |
| MXE | 1244 | ENSG00000161526 | SAP30BP | chr17 | + | 75703310 | 75703371 | 75704755 | 75704814 | 75702479 | 75702571 | 75706007 | 75706092 | 38 | [illegible] |
| MXE | 1438 | ENSG00000197381 | ADARB1 | chr21 | + | 45128401 | 45128573 | 45171609 | 45171684 | 45074579 | 45074793 | 45175729 | 45176664 | 8 | [illegible] |
| MXE | 16601 | ENSG00000204482 | LST1 | chr6 | + | 31587568 | 31587733 | 31587943 | 31587966 | 31587199 | 31587318 | 31588517 | 31588909 | 176 | [illegible] |
| MXE | 13363 | ENSG00000099204 | ABLIM1 | chr10 | - | 114472976 | 114473132 | 114473878 | 114473956 | 114468180 | 114468216 | 114487957 | 114488016 | 91 | [illegible] |
| MXE | 14821 | ENSG00000106948 | AKNA | chr9 | - | 114343707 | 114343803 | 114346668 | 114346784 | 114342008 | 114342125 | 114347723 | 114347900 | 177 | [illegible] |
| MXE | 3952 | ENSG00000048028 | USP28 | chr11 | - | 113830866 | 113830943 | 113833419 | 113833557 | 113829196 | 113829345 | 113834248 | 113834335 | 243 | [illegible] |
| MXE | 7243 | ENSG00000145687 | SSBP2 | chr5 | - | 81440557 | 81440636 | 81446867 | 81446922 | 81437429 | 81437458 | 81448789 | 81448825 | 11 | [illegible] |
| MXE | 9875 | ENSG00000152061 | RABGAP1L | chr1 | + | 174252479 | 174252590 | 174272413 | 174272480 | 174250474 | 174250632 | 174275832 | 174275935 | 218 | [illegible] |
| MXE | 12576 | ENSG00000175471 | MCTP1 | chr5 | - | 94708511 | 94708609 | 94710817 | 94710927 | 94703740 | 94705203 | 94714776 | 94714886 | 25 | [illegible] |
| MXE | 7836 | ENSG00000176393 | RNPEP | chr1 | + | 201999901 | 202000015 | 202001375 | 202001488 | 201996146 | 201996263 | 202001658 | 202001767 | 2 | [illegible] |
| MXE | 1238 | ENSG00000108395 | TRIM37 | chr17 | - | 59061031 | 59061108 | 59064354 | 59064405 | 59056874 | 59057054 | 59070822 | 59070947 | 21 | [illegible] |
| MXE | 2026 | ENSG00000069966 | GNB5 | chr15 | - | 52133377 | 52133469 | 52135612 | 52135756 | 52128195 | 52128244 | 52141139 | 52141272 | 30 | [illegible] |
| MXE | 16812 | ENSG00000123349 | PFDN5 | chr12 | + | 53296218 | 53296275 | 53296429 | 53296551 | 53295838 | 53295941 | 53297849 | 53297924 | 626 | [illegible] |
| MXE | 12430 | ENSG00000147065 | MSN | chrX | + | 65716817 | 65716901 | 65727813 | 65727909 | 65667653 | 65667853 | 65729437 | 65729712 | 946 | [illegible] |
| MXE | 4078 | ENSG00000136813 | ECPAS | chr9 | - | 111436939 | 111437108 | 111440371 | 111440521 | 111433232 | 111433372 | 111442305 | 111442424 | 75 | [illegible] |
| MXE | 5584 | ENSG00000175792 | RUVBL1 | chr3 | - | 128101558 | 128101648 | 128104772 | 128104924 | 128100594 | 128100744 | 128112887 | 128113020 | 92 | [illegible] |
| MXE | 1206 | ENSG00000073969 | NSF | chr17 | + | 46693846 | 46693921 | 46694474 | 46694662 | 46692902 | 46693068 | 46704758 | 46704854 | 44 | [illegible] |
| MXE | 4101 | ENSG00000008513 | ST3GAL1 | chr8 | - | 133507198 | 133507321 | 133545773 | 133545926 | 133499134 | 133499189 | 133571692 | 133571887 | 164 | [illegible] |
| MXE | 1726 | ENSG00000197912 | SPG7 | chr16 | + | 89536764 | 89536948 | 89540943 | 89541384 | 89532462 | 89532636 | 89544647 | 89544772 | 17 | [illegible] |
| MXE | 10718 | ENSG00000171720 | HDAC3 | chr5 | - | 141626012 | 141626071 | 141626193 | 141626283 | 141625684 | 141625764 | 141627892 | 141627957 | 34 | [illegible] |
| MXE | 9850 | ENSG00000163898 | LIPH | chr3 | - | 185524070 | 185524160 | 185527483 | 185527585 | 185519141 | 185519309 | 185533570 | 185533679 | 1 | [illegible] |
| MXE | 11922 | ENSG00000167483 | NIBAN3 | chr19 | + | 17543523 | 17543631 | 17546685 | 17546797 | 17543316 | 17543433 | 17549443 | 17549527 | 15 | [illegible] |
| MXE | 15855 | ENSG00000088298 | EDEM2 | chr20 | - | 35126250 | 35126375 | 35131641 | 35131783 | 35123889 | 35124034 | 35134737 | 35134949 | 51 | [illegible] |
| MXE | 4143 | ENSG00000186431 | FCAR | chr19 | + | 54885234 | 54885525 | 54888006 | 54888228 | 54875329 | 54875365 | 54889648 | 54889890 | 63 | [illegible] |
| MXE | 14840 | ENSG00000165275 | TRMT10B | chr9 | + | 37763628 | 37763753 | 37768075 | 37768228 | 37762576 | 37762685 | 37769940 | 37770019 | 14 | [illegible] |
| MXE | 16605 | ENSG00000196498 | NCOR2 | chr12 | - | 124372021 | 124372610 | 124378236 | 124378384 | 124363678 | 124363799 | 124385744 | 124385887 | 269 | [illegible] |
| MXE | 13732 | ENSG00000147050 | KDM6A | chrX | + | 45053828 | 45053955 | 45059005 | 45059104 | 45051708 | 45051802 | 45059246 | 45059466 | 157 | [illegible] |
| MXE | 4199 | ENSG00000084676 | NCOA1 | chr2 | + | 24673365 | 24673463 | 24682950 | 24683128 | 24665748 | 24665915 | 24691480 | 24691660 | 271 | [illegible] |
| MXE | 2020 | ENSG00000140157 | NIPA2 | chr15 | + | 22853211 | 22853268 | 22858539 | 22858630 | 22851638 | 22851870 | 22860628 | 22860789 | 27 | [illegible] |
| MXE | 12439 | ENSG00000077454 | LRCH4 | chr7 | - | 100577272 | 100577496 | 100577389 | 100577558 | 100576901 | 100577005 | 100577663 | 100577740 | 88 | [illegible] |
| MXE | 7890 | ENSG00000074603 | DPP8 | chr15 | - | 65454262 | 65454415 | 65456224 | 65456371 | 65450998 | 65451110 | 65463760 | 65463906 | 158 | [illegible] |
| MXE | 4203 | ENSG00000084676 | NCOA1 | chr2 | + | 24762686 | 24762776 | 24768078 | 24768135 | 24757972 | 24758156 | 24768220 | 24770702 | 287 | [illegible] |
| MXE | 2130 | ENSG00000137842 | TMEM62 | chr15 | + | 43169592 | 43169677 | 43181180 | 43181299 | 43160393 | 43160794 | 43184259 | 43185089 | 31 | [illegible] |
| MXE | 7228 | ENSG00000008869 | HEATR5B | chr2 | - | 37079119 | 37079331 | 37083288 | 37083436 | 37076910 | 37077019 | 37084268 | 37084372 | 140 | [illegible] |
| MXE | 7849 | ENSG00000181523 | SGSH | chr17 | - | 80214614 | 80214765 | 80215032 | 80215138 | 80214171 | 80214328 | 80217031 | 80217192 | 8 | [illegible] |
| MXE | 16936 | ENSG00000150990 | DHX37 | chr12 | - | 124980489 | 124980838 | 124982510 | 124982623 | 124977341 | 124977490 | 124986095 | 124986265 | 51 | [illegible] |
| MXE | 3962 | ENSG00000083312 | TNPO1 | chr5 | + | 72855773 | 72855923 | 72861807 | 72861914 | 72851243 | 72851319 | 72865595 | 72865729 | 200 | [illegible] |
| MXE | 896 | ENSG00000074657 | ZNF532 | chr18 | + | 58918270 | 58920633 | 58939444 | 58939621 | 58865469 | 58865579 | 58948066 | 58948229 | 22 | [illegible] |
| MXE | 3071 | ENSG00000126214 | KLC1 | chr14 | + | 103687080 | 103687211 | 103692358 | 103692425 | 103679383 | 103679518 | 103700654 | 103700727 | 16 | [illegible] |
| MXE | 15152 | ENSG00000153339 | TRAPPC8 | chr18 | - | 31855659 | 31855807 | 31857539 | 31857982 | 31853848 | 31853945 | 31864626 | 31864781 | 63 | [illegible] |
| MXE | 1067 | ENSG00000108819 | PPP1R9B | chr17 | - | 50139428 | 50139581 | 50141268 | 50141373 | 50139262 | 50139316 | 50143597 | 50143718 | 286 | [illegible] |
| MXE | 4922 | ENSG00000114127 | XRN1 | chr3 | - | 142412543 | 142412663 | 142414134 | 142414291 | 142404906 | 142405076 | 142417139 | 142417229 | 261 | [illegible] |
| MXE | 2372 | ENSG00000214013 | GANC | chr15 | + | 42329305 | 42329449 | 42330575 | 42330672 | 42327362 | 42327442 | 42338388 | 42338490 | 31 | [illegible] |
| MXE | 1804 | ENSG00000122390 | NAA60 | chr16 | + | 3479470 | 3479600 | 3482501 | 3482598 | 3476221 | 3476337 | 3483362 | 3483597 | 145 | [illegible] |
| MXE | 4502 | ENSG00000155744 | FAM126B | chr2 | - | 200992886 | 200993003 | 200997444 | 200997540 | 200992280 | 200992368 | 201008972 | 201009088 | 27 | [illegible] |
| MXE | 16116 | ENSG00000139651 | ZNF740 | chr12 | + | 53184890 | 53185040 | 53185386 | 53185476 | 53181676 | 53181992 | 53185953 | 53186077 | 107 | [illegible] |
| MXE | 13567 | ENSG00000012048 | BRCA1 | chr17 | - | 43094743 | 43094860 | 43095845 | 43095922 | 43090943 | 43091032 | 43097243 | 43097289 | 4 | [illegible] |
| MXE | 8326 | ENSG00000177409 | SAMD9L | chr7 | - | 93145384 | 93145532 | 93145908 | 93146040 | 93144731 | 93144833 | 93146882 | 93147146 | 72 | [illegible] |
| MXE | 13294 | ENSG00000107185 | RGP1 | chr9 | + | 35751265 | 35751412 | 35751955 | 35752145 | 35750657 | 35750989 | 35752650 | 35752802 | 48 | [illegible] |
| MXE | 12952 | ENSG00000141380 | SS18 | chr18 | - | 26082362 | 26082528 | 26084051 | 26084175 | 26078075 | 26078160 | 26087500 | 26087577 | 1 | [illegible] |
| MXE | 7204 | ENSG00000143363 | PRUNE1 | chr1 | + | 151024610 | 151024795 | 151025514 | 151025673 | 151018466 | 151018669 | 151027232 | 151027327 | 54 | [illegible] |
| MXE | 7478 | ENSG00000084754 | HADHA | chr2 | - | 26234216 | 26234355 | 26236854 | 26236988 | 26232159 | 26232279 | 26238933 | 26239004 | 567 | [illegible] |
| MXE | 2459 | ENSG00000258461 | NA | chr15 | + | 42403740 | 42403777 | 42408210 | 42408324 | 42402123 | 42403002 | 42409302 | 42409380 | 13 | [illegible] |
| MXE | 7878 | ENSG00000087206 | UIMC1 | chr5 | - | 176975395 | 176975480 | 176982468 | 176982623 | 176970741 | 176970866 | 177006649 | 177006715 | 53 | [illegible] |
| MXE | 8528 | ENSG00000187605 | TET3 | chr2 | + | 74003109 | 74003166 | 74046277 | 74048411 | 73985979 | 73986706 | 74073548 | 74073639 | 45 | [illegible] |
| MXE | 2553 | ENSG00000197249 | SERPINA1 | chr14 | - | 94379463 | 94379611 | 94388559 | 94388663 | 94378448 | 94378640 | 94390456 | 94390650 | 72 | [illegible] |
| MXE | 4145 | ENSG00000083896 | YTHDC1 | chr4 | - | 68332793 | 68332847 | 68333307 | 68333397 | 68332102 | 68332197 | 68337026 | 68337450 | 129 | [illegible] |
| MXE | 12375 | ENSG00000091106 | NLRC4 | chr2 | - | 32252418 | 32252679 | 32256774 | 32256893 | 32241032 | 32241125 | 32265586 | 32265732 | 35 | [illegible] |
| MXE | 12471 | ENSG00000085788 | DDHD2 | chr8 | + | 38247712 | 38247835 | 38249707 | 38249803 | 38246183 | 38246300 | 38251911 | 38252028 | 64 | [illegible] |
| MXE | 11889 | ENSG00000112511 | PHF1 | chr6 | + | 33414965 | 33415144 | 33415260 | 33415329 | 33414724 | 33414829 | 33415589 | 33415670 | 49 | [illegible] |
| MXE | 16248 | ENSG00000136738 | STAM | chr10 | + | 17684831 | 17684927 | 17688026 | 17688173 | 17684674 | 17684750 | 17693221 | 17693281 | 47 | [illegible] |
| MXE | 12707 | ENSG00000011275 | RNF216 | chr7 | - | 5652412 | 5652510 | 5711760 | 5711839 | 5641153 | 5641376 | 5712714 | 5712863 | 35 | [illegible] |
| MXE | 2230 | ENSG00000167004 | PDIA3 | chr15 | + | 43756648 | 43756766 | 43761423 | 43761531 | 43753823 | 43753902 | 43763076 | 43763206 | 308 | [illegible] |
| MXE | 3178 | ENSG00000135341 | MAP3K7 | chr6 | - | 90518446 | 90518562 | 90519257 | 90519319 | 90513572 | 90516681 | 90523677 | 90523783 | 98 | [illegible] |
| MXE | 8390 | ENSG00000119725 | ZNF410 | chr14 | + | 73896347 | 73896534 | 73898070 | 73898262 | 73893796 | 73893932 | 73903959 | 73904110 | 16 | [illegible] |
| MXE | 8557 | ENSG00000160679 | CHTOP | chr1 | + | 153636571 | 153636653 | 153638294 | 153638448 | 153633981 | 153634343 | 153645063 | 153646305 | 108 | [illegible] |
| MXE | 3420 | ENSG00000140853 | NLRC5 | chr16 | + | 57077718 | 57077802 | 57077942 | 57078020 | 57077295 | 57077379 | 57079049 | 57079133 | 74 | [illegible] |
| MXE | 13321 | ENSG00000170832 | USP32 | chr17 | - | 60211375 | 60211514 | 60212023 | 60212098 | 60211012 | 60211118 | 60213580 | 60213662 | 106 | [illegible] |
| MXE | 13851 | ENSG00000138378 | STAT4 | chr2 | - | 191036163 | 191036299 | 191039198 | 191039297 | 191034547 | 191034597 | 191041064 | 191041148 | 161 | [illegible] |
| MXE | 2202 | ENSG00000170776 | AKAP13 | chr15 | + | 85740221 | 85740272 | 85741045 | 85741495 | 85736089 | 85736134 | 85743491 | 85743825 | 75 | [illegible] |
| MXE | 13524 | ENSG00000134516 | DOCK2 | chr5 | + | 169714027 | 169714211 | 169714359 | 169714457 | 169712119 | 169712223 | 169716212 | 169716302 | 1366 | [illegible] |
| MXE | 16730 | ENSG00000111271 | ACAD10 | chr12 | + | 111729805 | 111729956 | 111733922 | 111734068 | 111721670 | 111721739 | 111736830 | 111737004 | 3 | [illegible] |
| MXE | 6394 | ENSG00000116266 | STXBP3 | chr1 | + | 108800219 | 108800305 | 108807400 | 108807549 | 108798144 | 108798237 | 108808782 | 108809523 | 124 | [illegible] |
| MXE | 7973 | ENSG00000072803 | FBXW11 | chr5 | - | 171878010 | 171878129 | 171891466 | 171891604 | 171876284 | 171876534 | 171899003 | 171899094 | 133 | [illegible] |
| MXE | 3060 | ENSG00000135596 | MICAL1 | chr6 | - | 109447680 | 109447722 | 109448202 | 109448393 | 109447356 | 109447440 | 109448731 | 109448870 | 56 | [illegible] |
| MXE | 10007 | ENSG00000170802 | FOXN2 | chr2 | + | 48359046 | 48359147 | 48362642 | 48362707 | 48346200 | 48346751 | 48373291 | 48373328 | 94 | [illegible] |
| MXE | 9164 | ENSG00000119004 | CYP20A1 | chr2 | + | 203245845 | 203245895 | 203251966 | 203252109 | 203238981 | 203239134 | 203266513 | 203266681 | 20 | [illegible] |
| MXE | 1606 | ENSG00000179583 | CIITA | chr16 | + | 10922406 | 10922490 | 10923227 | 10923325 | 10922166 | 10922250 | 10935115 | 10935200 | 40 | [illegible] |
| MXE | 14839 | ENSG00000165275 | TRMT10B | chr9 | + | 37762576 | 37762685 | 37763628 | 37763753 | 37761902 | 37762117 | 37768075 | 37768228 | 26 | [illegible] |
| MXE | 937 | ENSG00000100983 | GSS | chr20 | - | 34942487 | 34942627 | 34942930 | 34943006 | 34941712 | 34941829 | 34945952 | 34946098 | 18 | [illegible] |
| MXE | 4358 | ENSG00000109762 | SNX25 | chr4 | + | 185266968 | 185267155 | 185288011 | 185288082 | 185264437 | 185264610 | 185310634 | 185310816 | 73 | [illegible] |
| MXE | 5858 | ENSG00000073417 | PDE8A | chr15 | + | 85076732 | 85076787 | 85083555 | 85083644 | 85075861 | 85075918 | 85089337 | 85089416 | 14 | [illegible] |
| MXE | 45 | ENSG00000160294 | MCM3AP | chr21 | - | 46261279 | 46261411 | 46264116 | 46264217 | 46260792 | 46260906 | 46265320 | 46265523 | 30 | [illegible] |
| MXE | 12039 | ENSG00000127947 | PTPN12 | chr7 | + | 77583554 | 77583650 | 77585542 | 77585581 | 77581426 | 77581503 | 77592184 | 77592256 | 509 | [illegible] |
| MXE | 13476 | ENSG00000111203 | ITFG2 | chr12 | + | 2824089 | 2824252 | 2836744 | 2836956 | 2823769 | 2823943 | 2840841 | 2840995 | 87 | [illegible] |
| MXE | 7883 | ENSG00000074603 | DPP8 | chr15 | - | 65450998 | 65451110 | 65451959 | 65452102 | 65446620 | 65447006 | 65456224 | 65456371 | 28 | [illegible] |
| MXE | 15163 | ENSG00000151465 | CDC123 | chr10 | + | 12210289 | 12210322 | 12215739 | 12215835 | 12209966 | 12210024 | 12217360 | 12217467 | 37 | [illegible] |
| MXE | 15649 | ENSG00000167986 | DDB1 | chr11 | - | 61316491 | 61316567 | 61321594 | 61321697 | 61314883 | 61316393 | 61322295 | 61322412 | 170 | [illegible] |
| MXE | 9408 | ENSG00000187741 | FANCA | chr16 | - | 89739128 | 89739289 | 89739477 | 89739553 | 89738743 | 89738974 | 89739993 | 89740099 | 16 | [illegible] |
| MXE | 9445 | ENSG00000066027 | PPP2R5A | chr1 | + | 212347346 | 212347406 | 212349188 | 212349242 | 212345802 | 212345933 | 212356625 | 212356676 | 96 | [illegible] |
| MXE | 7415 | ENSG00000076650 | GPATCH1 | chr19 | + | 33101494 | 33101574 | 33106694 | 33106899 | 33100172 | 33100274 | 33109716 | 33109877 | 0 | [illegible] |
| MXE | 2057 | ENSG00000159216 | RUNX1 | chr21 | - | 34880556 | 34880713 | 34892924 | 34892963 | 34859473 | 34859578 | 35048841 | 35048958 | 21 | [illegible] |
| MXE | 11019 | ENSG00000225830 | ERCC6 | chr10 | - | 49505883 | 49506012 | 49524032 | 49524777 | 49500537 | 49500696 | 49528416 | 49528525 | 121 | [illegible] |
| MXE | 14385 | ENSG00000119397 | CNTRL | chr9 | + | 121125715 | 121125936 | 121135805 | 121135982 | 121123930 | 121124084 | 121138544 | 121138679 | 184 | [illegible] |

| | | | | | | | | | | | | | | | | | | | | | | | | | |
|---|---|---|---|---|---|---|---|---|---|---|---|---|---|---|---|---|---|---|---|---|---|---|---|---|---|
| MXE | 1382 | ENSG00000109062 | SLC9A3R1 | chr17 | + | 74762011 | 74762173 | 74766935 | 74766976 | 74748627 | 74749287 | 74768146 | 74768236 | 137 | 79 | 111 | 112 | 41 | 122 | 43 | 101 | 110 | 172 | 89 | 109 |
| MXE | 8275 | ENSG00000141127 | PRPSAP2 | chr17 | + | 18911102 | 18911251 | 18923913 | 18923984 | 18889821 | 18889877 | 18928810 | 18928957 | 68 | 37 | 85 | 64 | 67 | 13 | 6 | 17 | 15 | 37 | 90 | 42 |
| MXE | 3152 | ENSG00000137413 | TAF8 | chr6 | + | 42057388 | 42057513 | 42066311 | 42066459 | 42055951 | 42056014 | 42068464 | 42068607 | 143 | 98 | 127 | 130 | 259 | 92 | 44 | 82 | 79 | 68 | 224 | 147 |
| MXE | 12695 | ENSG00000129103 | SUMF2 | chr7 | + | 56072996 | 56073111 | 56074173 | 56074218 | 56068481 | 56068638 | 56074585 | 56074736 | 115 | 64 | 109 | 81 | 159 | 21 | 7 | 30 | 24 | 41 | 59 | 135 |
| MXE | 10244 | ENSG00000137817 | PARP6 | chr15 | - | 72250019 | 72250092 | 72250844 | 72250954 | 72242619 | 72242699 | 72251206 | 72251255 | 56 | 19 | 58 | 50 | 96 | 29 | 33 | 46 | 38 | 87 | 38 | 52 |
| MXE | 14431 | ENSG00000080503 | SMARCA2 | chr9 | + | 2104002 | 2104169 | 2110253 | 2110417 | 2101569 | 2101616 | 2115821 | 2116049 | 203 | 119 | 202 | 177 | 243 | 203 | 108 | 220 | 214 | 201 | 84 | 154 |
| MXE | 12402 | ENSG00000091732 | ZC3HC1 | chr7 | - | 130023510 | 130023723 | 130024262 | 130024506 | 130022318 | 130022525 | 130026157 | 130026312 | 24 | 12 | 17 | 14 | 40 | 10 | 2 | 22 | 10 | 14 | 15 | 17 |
| MXE | 15693 | ENSG00000054654 | SYNE2 | chr14 | + | 64029894 | 64030059 | 64031015 | 64031357 | 64027483 | 64027793 | 64047999 | 64048155 | 439 | 284 | 266 | 320 | 206 | 191 | 147 | 101 | 161 | 113 | 28 | 103 |
| MXE | 9038 | ENSG00000132906 | CASP9 | chr1 | - | 15495272 | 15495452 | 15504610 | 15504758 | 15493891 | 15494001 | 15505989 | 15506079 | 13 | 14 | 13 | 19 | 20 | 18 | 6 | 20 | 31 | 46 | 18 | 31 |
| MXE | 14610 | ENSG00000127084 | FGD3 | chr9 | + | 93020316 | 93020424 | 93022326 | 93022389 | 93019830 | 93019861 | 93029873 | 93029996 | 193 | 128 | 171 | 158 | 377 | 48 | 18 | 40 | 63 | 39 | 173 | 229 |
| MXE | 3290 | ENSG00000163933 | RFT1 | chr3 | - | 53119883 | 53120021 | 53121698 | 53121800 | 53111829 | 53111908 | 53122373 | 53122563 | 27 | 29 | 41 | 24 | 9 | 117 | 82 | 157 | 121 | 98 | 31 | 44 |
| MXE | 633 | ENSG00000127507 | ADGRE2 | chr19 | - | 14752328 | 14752526 | 14756237 | 14756345 | 14751435 | 14751671 | 14764432 | 14764610 | 72 | 57 | 43 | 76 | 60 | 27 | 21 | 26 | 29 | 21 | 121 | 71 |
| MXE | 3058 | ENSG00000135596 | MICAL1 | chr6 | - | 109447356 | 109447440 | 109448202 | 109448393 | 109447072 | 109447229 | 109448731 | 109448879 | 56 | 40 | 51 | 73 | 7 | 49 | 25 | 27 | 44 | 13 | 66 | 111 |
| MXE | 8133 | ENSG00000145979 | TBC1D7 | chr6 | - | 13307599 | 13307745 | 13316570 | 13316708 | 13306397 | 13306527 | 13320907 | 13321095 | 24 | 15 | 26 | 21 | 43 | 37 | 38 | 23 | 40 | 29 | 11 | 32 |
| MXE | 13847 | ENSG00000221914 | PPP2R2A | chr8 | + | 26354467 | 26354633 | 26360168 | 26360281 | 26338889 | 26338987 | 26360973 | 26361001 | 163 | 136 | 155 | 134 | 186 | 151 | 109 | 113 | 169 | 296 | 83 | 118 |
| MXE | 4280 | ENSG00000002586 | CD99 | chrX | + | 2726259 | 2726373 | 2738202 | 2738256 | 2723313 | 2723364 | 2740778 | 2740804 | 569 | 245 | 326 | 431 | 525 | 604 | 294 | 373 | 467 | 465 | 961 | 833 |
| MXE | 2083 | ENSG00000250021 | ARPIN-AP3S2 | chr15 | - | 89871474 | 89871546 | 89888520 | 89888632 | 89837614 | 89837722 | 89889048 | 89889140 | 55 | 37 | 49 | 28 | 25 | 78 | 50 | 81 | 51 | 69 | 78 | 69 |
| MXE | 14371 | ENSG00000254995 | STX16-NPEPL1 | chr20 | + | 58691033 | 58691227 | 58691404 | 58691562 | 58673630 | 58673711 | 58691723 | 58691780 | 9 | 6 | 12 | 13 | 32 | 17 | 28 | 24 | 38 | 26 | 14 | 11 |
| MXE | 751 | ENSG00000078747 | ITCH | chr20 | + | 34393790 | 34393881 | 34408650 | 34408792 | 34369393 | 34369470 | 34412514 | 34412639 | 143 | 62 | 126 | 173 | 155 | 108 | 93 | 115 | 117 | 134 | 13 | 235 |
| MXE | 8710 | ENSG00000162654 | GBP4 | chr1 | - | 89186999 | 89187102 | 89188581 | 89188794 | 89186332 | 89186526 | 89190037 | 89190318 | 258 | 183 | 203 | 236 | 210 | 176 | 106 | 87 | 85 | 190 | 705 | 255 |
| MXE | 14403 | ENSG00000099219 | ERMP1 | chr9 | - | 5812888 | 5813035 | 5823895 | 5824001 | 5812124 | 5812217 | 5830726 | 5831028 | 4 | 11 | 8 | 4 | 4 | 87 | 42 | 40 | 64 | 35 | 0 | 2 |
| MXE | 5809 | ENSG00000164330 | EBF1 | chr5 | - | 158796344 | 158796475 | 158823175 | 158823317 | 158777412 | 158777539 | 158840028 | 158840110 | 17 | 19 | 18 | 26 | 49 | 19 | 15 | 27 | 27 | 44 | 13 | 27 |
| MXE | 16267 | ENSG00000119977 | TCTN3 | chr10 | - | 95680471 | 95680609 | 95682650 | 95682804 | 95663400 | 95664300 | 95683100 | 95683180 | 46 | 35 | 49 | 45 | 74 | 24 | 12 | 34 | 17 | 31 | 49 | 45 |
| MXE | 17093 | ENSG00000185989 | RASA3 | chr13 | - | 114024278 | 114024355 | 114052051 | 114052155 | 114021403 | 114021508 | 114073719 | 114073837 | 228 | 124 | 175 | 173 | 229 | 247 | 157 | 240 | 212 | 272 | 215 | 176 |
| MXE | 4293 | ENSG00000147548 | NSD3 | chr8 | - | 38299443 | 38299590 | 38304586 | 38304757 | 38295795 | 38295952 | 38305247 | 38305445 | 53 | 42 | 55 | 70 | 87 | 60 | 26 | 58 | 42 | 84 | 24 | 23 |
| MXE | 10202 | ENSG00000123240 | OPTN | chr10 | + | 13116266 | 13116340 | 13118905 | 13119040 | 13112452 | 13112635 | 13122384 | 13122487 | 72 | 43 | 89 | 80 | 58 | 140 | 104 | 122 | 144 | 80 | 39 | 48 |
| MXE | 11823 | ENSG00000146070 | PLA2G7 | chr6 | - | 46714459 | 46714553 | 46716383 | 46716528 | 46712268 | 46712337 | 46716974 | 46717096 | 53 | 38 | 54 | 36 | 52 | 56 | 35 | 42 | 25 | 42 | 102 | 2 |
| MXE | 8307 | ENSG00000138802 | SEC24B | chr4 | + | 109473003 | 109473186 | 109481676 | 109481781 | 109462900 | 109463644 | 109491326 | 109491407 | 227 | 150 | 152 | 163 | 285 | 22 | 16 | 14 | 16 | 31 | 136 | 226 |
| MXE | 10276 | ENSG00000102081 | FMR1 | chrX | + | 147937465 | 147937600 | 147940575 | 147940662 | 147936503 | 147936613 | 147943130 | 147943326 | 158 | 140 | 130 | 170 | 235 | 156 | 129 | 105 | 147 | 199 | 97 | 83 |
| MXE | 12990 | ENSG00000122643 | NT5C3A | chr7 | - | 33024038 | 33024108 | 33026816 | 33026915 | 33022052 | 33022099 | 33035933 | 33035983 | 92 | 139 | 67 | 112 | 247 | 134 | 159 | 105 | 233 | 271 | 57 | 153 |
| MXE | 15775 | ENSG00000100600 | LGMN | chr14 | - | 92714375 | 92714451 | 92716135 | 92716221 | 92713822 | 92713885 | 92717379 | 92717461 | 19 | 18 | 22 | 46 | 19 | 12 | 6 | 17 | 43 | 58 | 16 | 21 |
| MXE | 16141 | ENSG00000172661 | WASHC2C | chr10 | + | 45737982 | 45738045 | 45740072 | 45740246 | 45728861 | 45729026 | 45743389 | 45743483 | 32 | 51 | 38 | 55 | 34 | 97 | 103 | 109 | 138 | 175 | 146 | 48 |
| MXE | 11054 | ENSG00000121741 | ZMYM2 | chr13 | + | 19993825 | 19993919 | 20002849 | 20003135 | 19993062 | 19993564 | 20005073 | 20005239 | 32 | 49 | 38 | 43 | 120 | 100 | 81 | 96 | 110 | 144 | 10 | 29 |
| MXE | 2286 | ENSG00000128908 | INO80 | chr15 | - | 41053928 | 41054014 | 41056621 | 41056706 | 41049934 | 41050102 | 41058638 | 41058781 | 92 | 80 | 79 | 91 | 119 | 145 | 120 | 86 | 111 | 374 | 158 | 108 |
| MXE | 10491 | ENSG00000004455 | AK2 | chr1 | - | 33021366 | 33021461 | 33021592 | 33021703 | 33014521 | 33014594 | 33024441 | 33024567 | 199 | 101 | 150 | 136 | 122 | 189 | 123 | 162 | 180 | 480 | 189 | 164 |
| MXE | 12673 | ENSG00000101307 | SIRPB1 | chr20 | - | 1566152 | 1566267 | 1570804 | 1571137 | 1561384 | 1565497 | 1571719 | 1572037 | 30 | 16 | 18 | 30 | 26 | 115 | 71 | 59 | 56 | 75 | 107 | 52 |
| MXE | 6308 | ENSG00000071127 | WDR1 | chr4 | - | 10087706 | 10087940 | 10088292 | 10088373 | 10084423 | 10084530 | 10088663 | 10088741 | 395 | 367 | 360 | 374 | 294 | 830 | 653 | 641 | 907 | 525 | 731 | 503 |
| MXE | 2271 | ENSG00000068305 | MEF2A | chr15 | + | 99706728 | 99706855 | 99710633 | 99710760 | 99690240 | 99690428 | 99712389 | 99713782 | 88 | 90 | 69 | 124 | 143 | 158 | 131 | 114 | 173 | 245 | 64 | 140 |
| MXE | 5585 | ENSG00000175792 | RUVBL1 | chr3 | - | 128104772 | 128104924 | 128112887 | 128113020 | 128101558 | 128101648 | 128119327 | 128119414 | 86 | 51 | 81 | 86 | 82 | 132 | 73 | 120 | 96 | 124 | 86 | 74 |
| MXE | 15211 | ENSG00000113163 | CERT1 | chr5 | - | 75389591 | 75389687 | 75400204 | 75400297 | 75385901 | 75386034 | 75411010 | 75411103 | 1 | 0 | 1 | 0 | 0 | 96 | 69 | 80 | 69 | 95 | 0 | 7 |
| MXE | 12038 | ENSG00000127947 | PTPN12 | chr7 | + | 77571077 | 77571186 | 77585542 | 77585581 | 77537294 | 77537645 | 77592184 | 77592256 | 243 | 212 | 127 | 220 | 192 | 24 | 26 | 37 | 35 | 55 | 149 | 228 |
| MXE | 13386 | ENSG00000147394 | ZNF185 | chrX | + | 152920709 | 152920748 | 152922172 | 152922256 | 152920327 | 152920411 | 152931671 | 152931776 | 6 | 10 | 6 | 11 | 11 | 7 | 8 | 3 | 20 | 5 | 26 | 13 |
| MXE | 15419 | ENSG00000076043 | REXO2 | chr11 | + | 114443855 | 114443933 | 114445978 | 114446087 | 114440655 | 114440739 | 114447825 | 114447879 | 39 | 20 | 39 | 66 | 57 | 136 | 74 | 143 | 136 | 232 | 7 | 42 |
| MXE | 13702 | ENSG00000134982 | APC | chr5 | + | 112815494 | 112815593 | 112818965 | 112819344 | 112801278 | 112801383 | 112821895 | 112821991 | 50 | 39 | 61 | 39 | 103 | 92 | 104 | 57 | 111 | 105 | 0 | 19 |
| MXE | 8913 | ENSG00000065413 | ANKRD44 | chr2 | - | 197001752 | 197001840 | 197005693 | 197005910 | 197000418 | 197000502 | 197007805 | 197007923 | 304 | 199 | 247 | 257 | 253 | 115 | 57 | 102 | 84 | 103 | 131 | 215 |
| MXE | 4443 | ENSG00000137656 | BUD13 | chr11 | - | 116757765 | 116757950 | 116758268 | 116758407 | 116757145 | 116757227 | 116759073 | 116759179 | 68 | 61 | 57 | 61 | 62 | 60 | 38 | 43 | 42 | 29 | 82 | 47 |
| MXE | 733 | ENSG00000175063 | UBE2C | chr20 | + | 45814383 | 45814470 | 45815540 | 45815745 | 45813436 | 45813464 | 45815853 | 45815913 | 1 | 8 | 3 | 2 | 0 | 3 | 6 | 11 | 7 | 3 | 1 | 0 |
| MXE | 3914 | ENSG00000285000 | NA | chr5 | + | 77036555 | 77036720 | 77039530 | 77039719 | 77035540 | 77035743 | 77046346 | 77046677 | 172 | 131 | 109 | 123 | 202 | 140 | 132 | 143 | 139 | 387 | 15 | 65 |
| MXE | 15119 | ENSG00000152270 | PDE3B | chr11 | + | 14834981 | 14835095 | 14843826 | 14844026 | 14832721 | 14832833 | 14859042 | 14859246 | 167 | 187 | 392 | 235 | 380 | 60 | 61 | 157 | 94 | 128 | 10 | 74 |
| MXE | 13871 | ENSG00000187735 | TCEA1 | chr8 | - | 53993667 | 53993755 | 53999944 | 54000050 | 53988113 | 53988259 | 54010429 | 54010492 | 387 | 287 | 315 | 375 | 405 | 461 | 404 | 378 | 474 | 604 | 69 | 307 |
| MXE | 8022 | ENSG00000087191 | PSMC5 | chr17 | + | 63829851 | 63829949 | 63830132 | 63830189 | 63829493 | 63829563 | 63830270 | 63830501 | 228 | 130 | 207 | 236 | 185 | 109 | 90 | 114 | 125 | 184 | 249 | 202 |
| MXE | 3629 | ENSG00000104671 | DCTN6 | chr8 | + | 30175084 | 30175190 | 30177125 | 30177214 | 30164110 | 30164175 | 30179407 | 30179455 | 43 | 27 | 37 | 40 | 32 | 99 | 76 | 71 | 75 | 160 | 5 | 43 |
| MXE | 15956 | ENSG00000138190 | EXOC6 | chr10 | + | 92909426 | 92909631 | 92915757 | 92915913 | 92899598 | 92899644 | 92919981 | 92920050 | 137 | 84 | 88 | 127 | 223 | 106 | 114 | 77 | 70 | 116 | 36 | 81 |
| MXE | 11934 | ENSG00000149657 | LSM14B | chr20 | + | 62126303 | 62126439 | 62129784 | 62129952 | 62124616 | 62124780 | 62130529 | 62130605 | 31 | 12 | 21 | 50 | 21 | 9 | 7 | 1 | 12 | 2 | 16 | 13 |
| MXE | 12810 | ENSG00000104133 | SPG11 | chr15 | - | 44572682 | 44572820 | 44573546 | 44573745 | 44570524 | 44570658 | 44574901 | 44575041 | 73 | 50 | 59 | 59 | 99 | 101 | 44 | 66 | 99 | 85 | 16 | 53 |
| MXE | 617 | ENSG00000130475 | FCHO1 | chr19 | + | 17776246 | 17776271 | 17776634 | 17776686 | 17775982 | 17776161 | 17778136 | 17778228 | 14 | 14 | 11 | 15 | 12 | 21 | 7 | 11 | 25 | 29 | 3 | 16 |
| MXE | 630 | ENSG00000131355 | ADGRE3 | chr19 | - | 14658512 | 14658550 | 14661962 | 14662118 | 14654981 | 14655165 | 14663417 | 14663540 | 68 | 62 | 46 | 53 | 141 | 15 | 7 | 7 | 13 | 19 | 33 | 11 |
| MXE | 7032 | ENSG00000095564 | BTAF1 | chr10 | + | 91939951 | 91940066 | 91942421 | 91942568 | 91935651 | 91935780 | 91951402 | 91951566 | 126 | 79 | 89 | 84 | 55 | 269 | 280 | 297 | 286 | 294 | 72 | 38 |
| MXE | 3066 | ENSG00000198399 | ITSN2 | chr2 | - | 24286211 | 24286351 | 24293687 | 24293775 | 24275712 | 24275849 | 24295663 | 24295804 | 154 | 160 | 139 | 173 | 238 | 218 | 234 | 194 | 248 | 404 | 11 | 87 |
| MXE | 9523 | ENSG00000018699 | TTC27 | chr2 | + | 32678855 | 32678922 | 32702806 | 32702920 | 32672271 | 32672384 | 32733827 | 32733923 | 35 | 34 | 25 | 37 | 36 | 35 | 22 | 51 | 43 | 72 | 4 | 17 |
| MXE | 11774 | ENSG00000124523 | SIRT5 | chr6 | + | 13596962 | 13597016 | 13599031 | 13599155 | 13595476 | 13595564 | 13611789 | 13614558 | 12 | 6 | 20 | 20 | 5 | 0 | 2 | 0 | 3 | 0 | 21 | 12 |
| MXE | 7911 | ENSG00000071054 | MAP4K4 | chr2 | + | 101885106 | 101885287 | 101887087 | 101887261 | 101882550 | 101882685 | 101887777 | 101887937 | 153 | 156 | 158 | 200 | 329 | 127 | 101 | 73 | 121 | 249 | 372 | 202 |
| MXE | 6772 | ENSG00000198265 | HELZ | chr17 | - | 67203318 | 67203443 | 67215898 | 67215935 | 67200612 | 67201185 | 67218594 | 67218822 | 41 | 16 | 33 | 44 | 31 | 295 | 176 | 212 | 267 | 162 | 17 | 57 |
| MXE | 1011 | ENSG00000073605 | GSDMB | chr17 | - | 39906110 | 39906271 | 39906960 | 39906987 | 39905846 | 39905985 | 39908175 | 39908214 | 6 | 0 | 1 | 7 | 0 | 52 | 45 | 30 | 75 | 26 | 0 | 5 |
| MXE | 10285 | ENSG00000131051 | RBM39 | chr20 | - | 35738967 | 35739017 | 35740524 | 35740597 | 35734185 | 35734257 | 35740823 | 35740887 | 77 | 45 | 99 | 53 | 35 | 42 | 27 | 24 | 53 | 29 | 19 | 43 |
| MXE | 3977 | ENSG00000204859 | ZBTB48 | chr1 | + | 6585918 | 6586030 | 6586694 | 6586787 | 6582057 | 6582299 | 6587204 | 6587291 | 7 | 12 | 4 | 8 | 24 | 22 | 6 | 18 | 13 | 30 | 9 | 19 |
| MXE | 5939 | ENSG00000172175 | MALT1 | chr18 | + | 58700440 | 58700591 | 58709377 | 58709556 | 58696365 | 58696487 | 58709975 | 58710072 | 165 | 107 | 145 | 140 | 229 | 155 | 113 | 170 | 181 | 313 | 19 | 32 |
| MXE | 9213 | ENSG00000148660 | CAMK2G | chr10 | - | 73847224 | 73847347 | 73848525 | 73848609 | 73842457 | 73842541 | 73849012 | 73849115 | 12 | 6 | 18 | 7 | 15 | 107 | 74 | 127 | 138 | 147 | 6 | 16 |
| MXE | 9238 | ENSG00000163466 | ARPC2 | chr2 | + | 218234351 | 218234397 | 218239390 | 218239484 | 218228737 | 218228850 | 218245419 | 218245546 | 852 | 640 | 627 | 766 | 725 | 2152 | 1726 | 1491 | 1889 | 3162 | 1480 | 818 |
| MXE | 2564 | ENSG00000197249 | SERPINA1 | chr14 | - | 94388559 | 94388663 | 94388818 | 94389010 | 94382591 | 94383241 | 94390419 | 94390692 | 7 | 1 | 2 | 4 | 2 | 330 | 189 | 135 | 186 | 136 | 0 | 4 |
| MXE | 6953 | ENSG00000160058 | BSDC1 | chr1 | - | 32381213 | 32381268 | 32386778 | 32386895 | 32378723 | 32378839 | 32394079 | 32394140 | 83 | 43 | 76 | 63 | 84 | 14 | 11 | 16 | 15 | 6 | 77 | 37 |
| MXE | 14955 | ENSG00000137073 | UBAP2 | chr9 | - | 33998786 | 33998864 | 34011626 | 34011664 | 33996222 | 33996333 | 34017049 | 34017189 | 7 | 0 | 0 | 4 | 2 | 100 | 85 | 97 | 121 | 188 | 9 | 8 |
| MXE | 3824 | ENSG00000082397 | EPB41L3 | chr18 | - | 5395066 | 5395147 | 5395608 | 5395707 | 5394676 | 5394793 | 5396200 | 5396332 | 76 | 89 | 36 | 65 | 92 | 51 | 58 | 25 | 52 | 88 | 115 | 93 |
| MXE | 16157 | ENSG00000176986 | SEC24C | chr10 | + | 73767053 | 73767170 | 73767836 | 73768007 | 73766759 | 73766853 | 73768809 | 73768906 | 198 | 114 | 173 | 124 | 213 | 80 | 46 | 69 | 79 | 85 | 343 | 200 |
| MXE | 14074 | ENSG00000147654 | EBAG9 | chr8 | + | 109554728 | 109554887 | 109556934 | 109557042 | 109553864 | 109553943 | 109560837 | 109560929 | 166 | 102 | 135 | 125 | 362 | 116 | 113 | 151 | 113 | 287 | 60 | 80 |
| MXE | 3102 | ENSG00000150403 | TMCO3 | chr13 | + | 113503495 | 113503588 | 113520615 | 113520733 | 113501751 | 113501880 | 113534044 | 113534240 | 90 | 55 | 71 | 73 | 94 | 31 | 14 | 22 | 38 | 16 | 225 | 98 |
| MXE | 14179 | ENSG00000173064 | HECTD4 | chr12 | - | 112203635 | 112203772 | 112204485 | 112204623 | 112200637 | 112200798 | 112207873 | 112208000 | 58 | 38 | 39 | 71 | 58 | 71 | 50 | 31 | 63 | 26 | 61 | 69 |
| MXE | 15584 | ENSG00000185187 | SIGIRR | chr11 | - | 408072 | 408206 | 408694 | 408893 | 407816 | 407957 | 409867 | 410027 | 51 | 14 | 38 | 43 | 9 | 61 | 20 | 57 | 35 | 39 | 5 | 17 |
| MXE | 3401 | ENSG00000173692 | PSMD1 | chr2 | + | 231087116 | 231087181 | 231138735 | 231138850 | 231085018 | 231085114 | 231146239 | 231146356 | 68 | 39 | 59 | 53 | 88 | 59 | 34 | 44 | 60 | 75 | 55 | 32 |
| MXE | 15581 | ENSG00000073910 | FRY | chr13 | + | 32231178 | 32231300 | 32234573 | 32234761 | 32228455 | 32228654 | 32236077 | 32236172 | 20 | 4 | 20 | 16 | 8 | 21 | 23 | 27 | 13 | 15 | 10 | 14 |
| MXE | 11248 | ENSG00000068724 | TTC7A | chr2 | + | 47023407 | 47023465 | 47024286 | 47024359 | 47021861 | 47021979 | 47029223 | 47029384 | 30 | 7 | 23 | 32 | 18 | 41 | 11 | 22 | 18 | 15 | 12 | 20 |
| MXE | 11829 | ENSG00000065427 | KARS1 | chr16 | - | 75636453 | 75636547 | 75640183 | 75640349 | 75635911 | 75636098 | 75641563 | 75641723 | 297 | 175 | 264 | 273 | 313 | 199 | 133 | 178 | 202 | 306 | 323 | 249 |
| MXE | 1290 | ENSG00000105053 | VRK3 | chr19 | - | 50016023 | 50016163 | 50020584 | 50020647 | 50007568 | 50007826 | 50025266 | 50025372 | 3 | 0 | 1 | 6 | 4 | 20 | 14 | 8 | 21 | 23 | 0 | 0 |
| MXE | 11131 | ENSG00000179051 | RCC2 | chr1 | - | 17422704 | 17422836 | 17425540 | 17425684 | 17422202 | 17422291 | 17429105 | 17429199 | 170 | 88 | 117 | 139 | 73 | 168 | 92 | 164 | 170 | 105 | 404 | 211 |
| MXE | 6345 | ENSG00000136059 | VILL | chr3 | + | 38002395 | 38002575 | 38003167 | 38003313 | 38001701 | 38001860 | 38004254 | 38004399 | 29 | 2 | 40 | 22 | 27 | 15 | 7 | 14 | 15 | 15 | 24 | 38 |
| MXE | 8391 | ENSG00000119725 | ZNF410 | chr14 | + | 73909340 | 73909430 | 73920979 | 73921105 | 73904901 | 73905083 | 73922065 | 73922206 | 26 | 24 | 27 | 19 | 68 | 25 | 16 | 18 | 21 | 20 | 24 | 39 |
| MXE | 11134 | ENSG00000021574 | SPAST | chr2 | + | 32087494 | 32087578 | 32089521 | 32089605 | 32063781 | 32064246 | 32098795 | 32098891 | 8 | 4 | 7 | 9 | 1 | 41 | 45 | 43 | 43 | 56 | 0 | 3 |
| MXE | 11364 | ENSG00000115970 | THADA | chr2 | - | 43232712 | 43232882 | 43279764 | 43279896 | 43230850 | 43231343 | 43286907 | 43287061 | 13 | 8 | 21 | 9 | 10 | 24 | 14 | 15 | 10 | 26 | 0 | 6 |
| MXE | 11465 | ENSG00000110080 | ST3GAL4 | chr11 | + | 126408306 | 126408496 | 126409267 | 126409411 | 126408098 | 126408194 | 126413504 | 126413648 | 33 | 13 | 22 | 32 | 15 | 39 | 27 | 22 | 46 | 52 | 19 | 38 |
| MXE | 4018 | ENSG00000160613 | PCSK7 | chr11 | - | 117207066 | 117207160 | 117207653 | 117207735 | 117206680 | 117206798 | 117207969 | 117208064 | 4 | 2 | 3 | 8 | 8 | 133 | 60 | 100 | 110 | 87 | 3 | 6 |
| MXE | 11439 | ENSG00000151687 | ANKAR | chr2 | + | 189728266 | 189728420 | 189728659 | 189728821 | 189720618 | 189720787 | 189737682 | 189737841 | 8 | 7 | 3 | 4 | 8 | 15 | 15 | 11 | 6 | 37 | 0 | 2 |
| MXE | 7432 | ENSG00000088832 | FKBP1A | chr20 | - | 1372075 | 1372240 | 1375490 | 1375603 | 1368977 | 1370072 | 1392833 | 1392881 | 142 | 67 | 78 | 116 | 32 | 1006 | 522 | 472 | 809 | 1045 | 213 | 103 |
| MXE | 3354 | ENSG00000127603 | MACF1 | chr1 | + | 39427960 | 39428287 | 39429241 | 39429326 | 39427454 | 39427614 | 39429826 | 39430068 | 165 | 119 | 156 | 198 | 149 | 207 | 105 | 107 | 154 | 127 | 12 | 144 |

| | | | | | | | | | | | | | | | |
|---|---|---|---|---|---|---|---|---|---|---|---|---|---|---|---|
| MXE | 8806 | ENSG00000142405 | NLRP12 | chr19 | - | 53801226 | 53801397 | 53805279 | 53805450 | 53795858 | 53796029 | 53807494 | 53807665 | 24 | [illegible] |
| MXE | 435 | ENSG00000076944 | STXBP2 | chr19 | + | 7640899 | 7641003 | 7641704 | 7641853 | 7639730 | 7639807 | 7642033 | 7642118 | 0 | [illegible] |
| MXE | 13641 | ENSG00000133687 | TMTC1 | chr12 | - | 29572104 | 29572218 | 29583406 | 29583574 | 29556856 | 29557000 | 29604177 | 29604299 | 2 | [illegible] |
| MXE | 6563 | ENSG00000113119 | TMCO6 | chr5 | + | 140644099 | 140644194 | 140644572 | 140644740 | 140643779 | 140643966 | 140644984 | 140645404 | 3 | [illegible] |
| MXE | 16979 | ENSG00000139132 | FGD4 | chr12 | + | 32576265 | 32576449 | 32581959 | 32582467 | 32564136 | 32564289 | 32598496 | 32598586 | 21 | [illegible] |
| MXE | 13058 | ENSG00000174428 | GTF2IRD2B | chr7 | + | 75126341 | 75126385 | 75133134 | 75133212 | 75125915 | 75125970 | 75135000 | 75135057 | 8 | [illegible] |
| MXE | 12413 | ENSG00000198198 | SZT2 | chr1 | + | 43422082 | 43422225 | 43422479 | 43422632 | 43421173 | 43421303 | 43422768 | 43422883 | 28 | [illegible] |
| MXE | 3278 | ENSG00000156639 | ZFAND3 | chr6 | + | 38082391 | 38082457 | 38116571 | 38116739 | 38061592 | 38061775 | 38152234 | 38154624 | 127 | [illegible] |
| MXE | 8480 | ENSG00000135749 | PCNX2 | chr1 | - | 233208517 | 233208689 | 233217898 | 233217931 | 233200153 | 233200264 | 233218030 | 233218184 | 46 | [illegible] |
| MXE | 6269 | ENSG00000164506 | STXBP5 | chr6 | + | 147364000 | 147364166 | 147373730 | 147373842 | 147363334 | 147363704 | 147382777 | 147382998 | 103 | [illegible] |
| MXE | 4786 | ENSG00000278259 | MYO19 | chr17 | - | 36510745 | 36510917 | 36511364 | 36511455 | 36509061 | 36509135 | 36513428 | 36513505 | 9 | [illegible] |
| MXE | 13588 | ENSG00000128654 | MTX2 | chr2 | + | 176329300 | 176329426 | 176330583 | 176330660 | 176328873 | 176328912 | 176337492 | 176337679 | 30 | [illegible] |
| MXE | 1737 | ENSG00000188559 | RALGAPA2 | chr20 | - | 20536655 | 20536784 | 20546703 | 20546832 | 20535744 | 20535803 | 20571457 | 20571613 | 61 | [illegible] |
| MXE | 10317 | ENSG00000005483 | KMT2E | chr7 | + | 105106521 | 105106772 | 105107165 | 105107222 | 105105858 | 105106003 | 105107361 | 105107925 | 351 | [illegible] |
| MXE | 1166 | ENSG00000125450 | NUP85 | chr17 | + | 75211991 | 75212062 | 75213075 | 75213119 | 75209822 | 75209985 | 75215753 | 75215775 | 66 | [illegible] |
| MXE | 3260 | ENSG00000174032 | SLC25A30 | chr13 | - | 45405882 | 45405977 | 45408926 | 45409074 | 45402274 | 45402370 | 45411361 | 45411480 | 51 | [illegible] |
| MXE | 16397 | ENSG00000144579 | CTDSP1 | chr2 | + | 218402113 | 218402215 | 218403034 | 218403127 | 218401166 | 218401712 | 218403231 | 218403417 | 80 | [illegible] |
| MXE | 3614 | ENSG00000103496 | STX4 | chr16 | + | 31034203 | 31034325 | 31034461 | 31034536 | 31034012 | 31034114 | 31034969 | 31035040 | 0 | [illegible] |
| MXE | 8387 | ENSG00000072110 | ACTN1 | chr14 | - | 68878457 | 68878523 | 68878988 | 68879069 | 68877081 | 68877240 | 68879961 | 68880108 | 218 | [illegible] |
| MXE | 589 | ENSG00000105186 | ANKRD27 | chr19 | - | 32643430 | 32643484 | 32643571 | 32643631 | 32643286 | 32643352 | 32644324 | 32644479 | 33 | [illegible] |
| MXE | 1930 | ENSG00000131797 | CLUHP3 | chr16 | + | 31705671 | 31705792 | 31705889 | 31706034 | 31705348 | 31705472 | 31706890 | 31707422 | 22 | [illegible] |
| MXE | 5845 | ENSG00000117298 | ECE1 | chr1 | - | 21247220 | 21247363 | 21255946 | 21256138 | 21244988 | 21245103 | 21257524 | 21257590 | 176 | [illegible] |
| MXE | 9323 | ENSG00000082212 | ME2 | chr18 | + | 50921073 | 50921187 | 50924097 | 50924212 | 50920660 | 50920758 | 50925755 | 50925898 | 232 | [illegible] |
| MXE | 5483 | ENSG00000173889 | PHC3 | chr3 | - | 170145422 | 170145521 | 170149085 | 170149244 | 170136365 | 170136665 | 170171389 | 170171450 | 17 | [illegible] |
| MXE | 3934 | ENSG00000141084 | RANBP10 | chr16 | - | 67729679 | 67729828 | 67729937 | 67730046 | 67729279 | 67729484 | 67731471 | 67731584 | 6 | [illegible] |
| MXE | 14652 | ENSG00000187239 | FNBP1 | chr9 | - | 129958490 | 129958553 | 129978464 | 129978612 | 129957359 | 129957464 | 129979317 | 129979374 | 821 | [illegible] |
| MXE | 5880 | ENSG00000072422 | RHOBTB1 | chr10 | - | 60892809 | 60892995 | 60910886 | 60910990 | 60888211 | 60889185 | 60911350 | 60911552 | 6 | [illegible] |
| MXE | 2844 | ENSG00000119688 | ABCD4 | chr14 | - | 74297929 | 74298069 | 74300149 | 74300268 | 74296332 | 74296449 | 74302874 | 74302934 | 28 | [illegible] |
| MXE | 12352 | ENSG00000186591 | UBE2H | chr7 | - | 129839206 | 129839335 | 129857510 | 129857563 | 129830731 | 129835061 | 129858901 | 129858941 | 43 | [illegible] |
| MXE | 563 | ENSG00000126261 | UBA2 | chr19 | + | 34434867 | 34434968 | 34438644 | 34438766 | 34433347 | 34433412 | 34443843 | 34443911 | 316 | [illegible] |
| MXE | 15404 | ENSG00000149091 | DGKZ | chr11 | + | 46369940 | 46370009 | 46371312 | 46371384 | 46369493 | 46369550 | 46371486 | 46371603 | 203 | [illegible] |
| MXE | 6167 | ENSG00000285708 | NA | chr3 | - | 71299819 | 71299880 | 71359149 | 71359244 | 71198201 | 71198392 | 71467250 | 71467315 | 8 | [illegible] |
| MXE | 2648 | ENSG00000100596 | SPTLC2 | chr14 | - | 77562395 | 77562489 | 77570383 | 77570508 | 77557040 | 77557146 | 77576766 | 77576915 | 110 | [illegible] |
| MXE | 16616 | ENSG00000120805 | ARL1 | chr12 | - | 101401061 | 101401173 | 101402864 | 101402946 | 101396398 | 101396577 | 101407641 | 101407746 | 1 | [illegible] |
| MXE | 14242 | ENSG00000164941 | INTS8 | chr8 | + | 94831991 | 94832174 | 94836523 | 94836631 | 94828974 | 94829026 | 94838493 | 94838618 | 325 | [illegible] |
| MXE | 7177 | ENSG00000121957 | GPSM2 | chr1 | + | 108922416 | 108922576 | 108923999 | 108924214 | 108918612 | 108918789 | 108929700 | 108930377 | 19 | [illegible] |
| MXE | 4388 | ENSG00000069869 | NEDD4 | chr15 | - | 55834037 | 55834145 | 55837788 | 55837849 | 55833007 | 55833104 | 55838106 | 55838180 | 8 | [illegible] |
| MXE | 207 | ENSG00000100150 | DEPDC5 | chr22 | + | 31897481 | 31897653 | 31901741 | 31901802 | 31893581 | 31893751 | 31905983 | 31906066 | 10 | [illegible] |
| MXE | 1057 | ENSG00000278053 | DDX52 | chr17 | - | 37630029 | 37630173 | 37632112 | 37632298 | 37628560 | 37628672 | 37633287 | 37633418 | 76 | [illegible] |
| MXE | 3862 | ENSG00000113391 | FAM172A | chr5 | - | 93881469 | 93881688 | 93958776 | 93958968 | 93824147 | 93824267 | 93964457 | 93964523 | 359 | [illegible] |
| MXE | 11978 | ENSG00000168944 | CEP120 | chr5 | - | 123391109 | 123391337 | 123393299 | 123393497 | 123389923 | 123390140 | 123399135 | 123399284 | 96 | [illegible] |
| MXE | 6635 | ENSG00000133067 | LGR6 | chr1 | + | 202301163 | 202301235 | 202303278 | 202303347 | 202300848 | 202300920 | 202304558 | 202304630 | 11 | [illegible] |
| MXE | 7245 | ENSG00000145687 | SSBP2 | chr5 | - | 81442652 | 81442723 | 81446867 | 81446922 | 81440537 | 81440636 | 81448789 | 81448825 | 11 | [illegible] |
| MXE | 5039 | ENSG00000009790 | TRAF3IP3 | chr1 | + | 209780469 | 209780606 | 209781344 | 209781458 | 209779314 | 209779374 | 209782055 | 209782320 | 124 | [illegible] |
| MXE | 2055 | ENSG00000090487 | SPG21 | chr15 | - | 64976474 | 64976555 | 64980863 | 64981025 | 64974601 | 64974747 | 64983506 | 64983593 | 110 | [illegible] |
| MXE | 7640 | ENSG00000179632 | MAF1 | chr8 | + | 144106554 | 144106674 | 144106834 | 144106963 | 144106342 | 144106464 | 144107087 | 144107217 | 178 | [illegible] |
| MXE | 14229 | ENSG00000165102 | HGSNAT | chr8 | + | 43173712 | 43173743 | 43178073 | 43178234 | 43172309 | 43172386 | 43182144 | 43182260 | 9 | [illegible] |
| MXE | 5430 | ENSG00000258653 | NA | chr14 | + | 73920979 | 73921105 | 73923394 | 73923522 | 73909340 | 73909430 | 73931502 | 73932278 | 51 | [illegible] |
| MXE | 7811 | ENSG00000066427 | ATXN3 | chr14 | - | 92093745 | 92093831 | 92096092 | 92096134 | 92093251 | 92093318 | 92106528 | 92106574 | 8 | [illegible] |
| MXE | 596 | ENSG00000187994 | RINL | chr19 | - | 38871077 | 38871227 | 38871646 | 38871711 | 38870569 | 38870992 | 38871797 | 38871870 | 31 | [illegible] |
| MXE | 16790 | ENSG00000168010 | ATG16L2 | chr11 | + | 72826517 | 72826589 | 72826702 | 72826823 | 72826172 | 72826243 | 72827187 | 72827272 | 35 | [illegible] |
| MXE | 2992 | ENSG00000069974 | RAB27A | chr15 | - | 55228608 | 55228712 | 55230400 | 55230486 | 55223888 | 55224012 | 55234781 | 55234956 | 278 | [illegible] |
| MXE | 17110 | ENSG00000173473 | SMARCC1 | chr3 | - | 47670657 | 47670717 | 47675474 | 47675588 | 47662333 | 47662592 | 47676628 | 47676782 | 55 | [illegible] |
| MXE | 5582 | ENSG00000168038 | ULK4 | chr3 | - | 41800131 | 41800293 | 41819422 | 41819506 | 41789660 | 41789843 | 41835863 | 41835971 | 12 | [illegible] |
| MXE | 9625 | ENSG00000197043 | ANXA6 | chr5 | - | 151109752 | 151109846 | 151110626 | 151110644 | 151108454 | 151108550 | 151117126 | 151117180 | 25 | [illegible] |
| MXE | 15928 | ENSG00000181192 | DHTKD1 | chr10 | + | 12107908 | 12108015 | 12112899 | 12113064 | 12106245 | 12106396 | 12117672 | 12117755 | 34 | [illegible] |
| MXE | 16285 | ENSG00000266412 | NCOA4 | chr10 | - | 46009410 | 46009551 | 46010222 | 46011206 | 46009123 | 46009212 | 46012882 | 46013026 | 544 | [illegible] |
| MXE | 9854 | ENSG00000285641 | NA | chr1 | - | 153982718 | 153982808 | 153983218 | 153983296 | 153982534 | 153982600 | 153983520 | 153983581 | 11 | [illegible] |
| MXE | 9185 | ENSG00000213064 | SFT2D2 | chr1 | + | 168231833 | 168231919 | 168235100 | 168235182 | 168231513 | 168231600 | 168236588 | 168236624 | 70 | [illegible] |
| MXE | 1096 | ENSG00000182173 | TSEN54 | chr17 | + | 75521704 | 75522521 | 75523274 | 75523335 | 75521408 | 75521510 | 75523662 | 75523779 | 76 | [illegible] |
| MXE | 2368 | ENSG00000140464 | PML | chr15 | + | 74032571 | 74032715 | 74033155 | 74033414 | 74024856 | 74024927 | 74034477 | 74034530 | 17 | [illegible] |
| MXE | 10944 | ENSG00000022840 | RNF10 | chr12 | + | 120562944 | 120563070 | 120563331 | 120563623 | 120560725 | 120560886 | 120563809 | 120563943 | 133 | [illegible] |
| MXE | 4281 | ENSG00000002586 | CD99 | chrX | + | 2726259 | 2726373 | 2738199 | 2738256 | 2723313 | 2723364 | 2740778 | 2741101 | 569 | [illegible] |
| MXE | 12977 | ENSG00000122591 | FAM126A | chr7 | - | 22978268 | 22978448 | 22983943 | 22984045 | 22977340 | 22977421 | 22991060 | 22991139 | 107 | [illegible] |
| MXE | 6023 | ENSG00000143515 | ATP8B2 | chr1 | + | 154330395 | 154330454 | 154331047 | 154331146 | 154328104 | 154328172 | 154331443 | 154331505 | 10 | [illegible] |
| MXE | 5947 | ENSG00000109320 | NFKB1 | chr4 | + | 102533844 | 102533885 | 102537857 | 102537956 | 102529835 | 102529914 | 102566986 | 102567135 | 28 | [illegible] |
| MXE | 6367 | ENSG00000120008 | WDR11 | chr10 | + | 120866568 | 120866764 | 120867065 | 120867169 | 120865629 | 120865744 | 120871169 | 120871346 | 145 | [illegible] |
| MXE | 9308 | ENSG00000135924 | DNAJB2 | chr2 | + | 219280577 | 219280687 | 219281717 | 219281771 | 219279515 | 219279898 | 219281938 | 219282061 | 16 | [illegible] |
| MXE | 13508 | ENSG00000171311 | EXOSC1 | chr10 | - | 97437699 | 97437750 | 97438669 | 97438703 | 97437190 | 97437275 | 97441170 | 97441259 | 25 | [illegible] |
| MXE | 13186 | ENSG00000137642 | SORL1 | chr11 | + | 121625084 | 121625277 | 121627554 | 121627767 | 121622161 | 121622268 | 121629495 | 121629678 | 658 | [illegible] |
| MXE | 331 | ENSG00000099331 | MYO9B | chr19 | + | 17159344 | 17159484 | 17162349 | 17162466 | 17156908 | 17157038 | 17162987 | 17163122 | 193 | [illegible] |
| MXE | 102 | ENSG00000196588 | MRTFA | chr22 | - | 40420846 | 40421100 | 40423535 | 40423685 | 40420404 | 40420576 | 40424205 | 40424381 | 154 | [illegible] |
| MXE | 2958 | ENSG00000162711 | NLRP3 | chr1 | + | 247429584 | 247429755 | 247434162 | 247434273 | 247423846 | 247425599 | 247435969 | 247436140 | 21 | [illegible] |
| MXE | 8611 | ENSG00000143313 | TMCO1 | chr1 | - | 165743166 | 165743311 | 165752101 | 165752169 | 165724292 | 165728121 | 165754227 | 165754274 | 95 | [illegible] |
| MXE | 7226 | ENSG00000008869 | HEATR5B | chr2 | - | 37005631 | 37005759 | 37007049 | 37007304 | 37003541 | 37003686 | 37008610 | 37008848 | 139 | [illegible] |
| MXE | 3611 | ENSG00000164574 | GALNT10 | chr5 | + | 154376276 | 154376462 | 154380447 | 154380631 | 154329571 | 154329738 | 154386312 | 154386430 | 78 | [illegible] |
| MXE | 13165 | ENSG00000102316 | MAGED2 | chrX | + | 54809302 | 54809376 | 54809721 | 54810213 | 54807598 | 54807802 | 54810820 | 54811129 | 0 | [illegible] |
| MXE | 2144 | ENSG00000101266 | CSNK2A1 | chr20 | - | 500030 | 500171 | 503064 | 503281 | 499832 | 499934 | 505117 | 505229 | 2 | [illegible] |
| MXE | 12191 | ENSG00000088930 | XRN2 | chr20 | + | 21328558 | 21328670 | 21330480 | 21330539 | 21326489 | 21326601 | 21330615 | 21330705 | 584 | [illegible] |
| MXE | 7968 | ENSG00000048707 | VPS13D | chr1 | + | 12415086 | 12415221 | 12416659 | 12416827 | 12403824 | 12403973 | 12455997 | 12456071 | 81 | [illegible] |
| MXE | 10588 | ENSG00000138442 | WDR12 | chr2 | - | 202897299 | 202897415 | 202899530 | 202899637 | 202895997 | 202896219 | 202901024 | 202901120 | 91 | [illegible] |
| MXE | 996 | ENSG00000154889 | MPPE1 | chr18 | - | 11897525 | 11897620 | 11905737 | 11905953 | 11896983 | 11897356 | 11906202 | 11906309 | 5 | [illegible] |
| MXE | 12265 | ENSG00000269955 | FMC1-LUC7L2 | chr7 | + | 139402136 | 139402247 | 139405643 | 139405787 | 139398598 | 139398697 | 139407173 | 139407350 | 289 | [illegible] |
| MXE | 8479 | ENSG00000135749 | PCNX2 | chr1 | - | 233200153 | 233200264 | 233208517 | 233208610 | 233199019 | 233199030 | 233217898 | 233217931 | 2 | [illegible] |
| MXE | 12139 | ENSG00000173715 | C11orf80 | chr11 | + | 66821642 | 66821764 | 66822563 | 66822674 | 66816052 | 66816199 | 66828265 | 66828364 | 18 | [illegible] |
| MXE | 1537 | ENSG00000008517 | IL32 | chr16 | + | 3067376 | 3067415 | 3067442 | 3067613 | 3065783 | 3065826 | 3067983 | 3068010 | 74 | [illegible] |
| MXE | 3200 | ENSG00000198625 | MDM4 | chr1 | + | 204526359 | 204526434 | 204530683 | 204530817 | 204525483 | 204525596 | 204532190 | 204532246 | 49 | [illegible] |
| MXE | 5783 | ENSG00000137074 | APTX | chr9 | - | 32984630 | 32984857 | 32985970 | 32986030 | 32974457 | 32974561 | 32987543 | 32987846 | 20 | [illegible] |
| MXE | 6419 | ENSG00000132773 | TOE1 | chr1 | + | 45341948 | 45342107 | 45342383 | 45342643 | 45341472 | 45341569 | 45342842 | 45343002 | 25 | [illegible] |
| MXE | 147 | ENSG00000100330 | MTMR3 | chr22 | + | 30002879 | 30002993 | 30007113 | 30007319 | 29998760 | 29998857 | 30007900 | 30008032 | 68 | [illegible] |
| MXE | 14944 | ENSG00000137073 | UBAP2 | chr9 | - | 33943419 | 33943589 | 33944364 | 33944639 | 33941648 | 33941862 | 33953284 | 33953474 | 1 | [illegible] |
| MXE | 12824 | ENSG00000104133 | SPG11 | chr15 | - | 44620189 | 44620403 | 44621758 | 44621934 | 44615362 | 44615566 | 44622219 | 44622347 | 277 | [illegible] |
| MXE | 10241 | ENSG00000157483 | MYO1E | chr15 | - | 59163156 | 59163303 | 59171896 | 59172042 | 59161072 | 59161230 | 59173745 | 59173915 | 27 | [illegible] |
| MXE | 8011 | ENSG00000196235 | SUPT5H | chr19 | + | 39445881 | 39445965 | 39453355 | 39453521 | 39445557 | 39445637 | 39457674 | 39457740 | 1 | [illegible] |
| MXE | 9967 | ENSG00000166949 | SMAD3 | chr15 | + | 67170553 | 67170604 | 67181240 | 67181453 | 67166778 | 67166853 | 67187364 | 67187447 | 10 | [illegible] |
| MXE | 13020 | ENSG00000234741 | GAS5 | chr1 | - | 173864483 | 173864704 | 173865228 | 173865282 | 173864256 | 173864304 | 173865470 | 173865547 | 147 | [illegible] |
| MXE | 16842 | ENSG00000111596 | CNOT2 | chr12 | + | 70329422 | 70329570 | 70332766 | 70332846 | 70319297 | 70319364 | 70335437 | 70335563 | 217 | [illegible] |
| MXE | 6246 | ENSG00000160741 | CRTC2 | chr1 | - | 153953265 | 153953369 | 153953537 | 153953606 | 153952804 | 153952834 | 153954254 | 153954316 | 19 | [illegible] |
| MXE | 5288 | ENSG00000101596 | SMCHD1 | chr18 | + | 2718157 | 2718235 | 2718314 | 2718434 | 2707806 | 2707920 | 2722518 | 2722550 | 225 | [illegible] |
| MXE | 1860 | ENSG00000074755 | ZZEF1 | chr17 | - | 4072607 | 4072756 | 4074149 | 4074351 | 4070683 | 4070924 | 4075096 | 4075178 | 141 | [illegible] |
| MXE | 6091 | ENSG00000077097 | TOP2B | chr3 | - | 25637214 | 25637312 | 25638164 | 25638310 | 25635935 | 25636148 | 25642321 | 25642385 | 359 | [illegible] |
| MXE | 15153 | ENSG00000153339 | TRAPPC8 | chr18 | - | 31874479 | 31874704 | 31890734 | 31890866 | 31873429 | 31873538 | 31897785 | 31897891 | 51 | [illegible] |
| MXE | 2536 | ENSG00000165929 | TC2N | chr14 | - | 91792366 | 91792558 | 91797784 | 91797901 | 91787512 | 91787627 | 91798988 | 91799064 | 14 | [illegible] |
| MXE | 5381 | ENSG00000132591 | ERAL1 | chr17 | + | 28856504 | 28856582 | 28857938 | 28857985 | 28856263 | 28856391 | 28858145 | 28858207 | 15 | [illegible] |
| MXE | 16440 | ENSG00000170456 | DENND5B | chr12 | - | 31442774 | 31442925 | 31447537 | 31447769 | 31432374 | 31433248 | 31451939 | 31452476 | 9 | [illegible] |
| MXE | 3754 | ENSG00000153310 | CYRIB | chr8 | - | 129846803 | 129846874 | 129849242 | 129849369 | 129839592 | 129842205 | 129850834 | 129850914 | 598 | [illegible] |
| MXE | 4027 | ENSG00000026297 | RNASET2 | chr6 | - | 166931043 | 166931118 | 166934090 | 166934136 | 166922112 | 166929791 | 166938894 | 166939008 | 68 | [illegible] |
| MXE | 11422 | ENSG00000146576 | C7orf26 | chr7 | + | 6594434 | 6594600 | 6599828 | 6600356 | 6591645 | 6591867 | 6602100 | 6602140 | 86 | [illegible] |
| MXE | 6826 | ENSG00000007341 | ST7L | chr1 | - | 112555867 | 112556018 | 112576985 | 112577088 | 112550600 | 112550693 | 112578344 | 112578417 | 10 | [illegible] |
| MXE | 16775 | ENSG00000090686 | USP48 | chr1 | - | 21721649 | 21721764 | 21723897 | 21724095 | 21721035 | 21721166 | 21728569 | 21728719 | 84 | [illegible] |
| MXE | 6017 | ENSG00000010322 | NISCH | chr3 | + | 52476446 | 52476599 | 52477573 | 52477642 | 52473733 | 52473829 | 52478096 | 52478282 | 97 | [illegible] |
| MXE | 5484 | ENSG00000173889 | PHC3 | chr3 | - | 170145422 | 170145521 | 170149085 | 170149244 | 170136418 | 170136665 | 170171372 | 170171450 | 48 | [illegible] |
| MXE | 16441 | ENSG00000170456 | DENND5B | chr12 | - | 31447537 | 31447769 | 31451939 | 31452476 | 31442774 | 31442925 | 31460193 | 31460381 | 16 | [illegible] |
| MXE | 1027 | ENSG00000167851 | CD300A | chr17 | + | 74477435 | 74477530 | 74481288 | 74481833 | 74474531 | 74474685 | 74484000 | 74484168 | 154 | [illegible] |
| MXE | 2182 | ENSG00000129003 | VPS13C | chr15 | - | 61890164 | 61890400 | 61907263 | 61907390 | 61884127 | 61884269 | 61908991 | 61909125 | 501 | [illegible] |
| MXE | 9052 | ENSG00000125633 | CCDC93 | chr2 | - | 117939028 | 117939111 | 117945528 | 117945582 | 117936701 | 117936739 | 117946810 | 117946882 | 18 | [illegible] |
| MXE | 14281 | ENSG00000167632 | TRAPPC9 | chr8 | - | 140435111 | 140435240 | 140439051 | 140439197 | 140405576 | 140405698 | 140450789 | 140450930 | 59 | [illegible] |
| MXE | 678 | ENSG00000132670 | PTPRA | chr20 | + | 2964860 | 2965202 | 2975214 | 2975241 | 2964271 | 2964350 | 2986764 | 2986849 | 1127 | [illegible] |
| MXE | 9179 | ENSG00000119004 | CYP20A1 | chr2 | + | 203285611 | 203285732 | 203289764 | 203289876 | 203280058 | 203280113 | 203292261 | 203292326 | 99 | [illegible] |
| MXE | 10316 | ENSG00000005483 | KMT2E | chr7 | + | 105066726 | 105066807 | 105068474 | 105068545 | 105063350 | 105063580 | 105073618 | 105073677 | 431 | [illegible] |
| MXE | 89 | ENSG00000186575 | NF2 | chr22 | + | 29639089 | 29639212 | 29642201 | 29642285 | 29636750 | 29636876 | 29654656 | 29654725 | 106 | [illegible] |
| MXE | 9554 | ENSG00000143727 | ACP1 | chr2 | + | 272036 | 272150 | 272191 | 272305 | 271865 | 271939 | 276979 | 277085 | 55 | [illegible] |
| MXE | 9576 | ENSG00000076242 | MLH1 | chr3 | + | 37047518 | 37047683 | 37048903 | 37049017 | 37042267 | 37042331 | 37050485 | 37050844 | 59 | [illegible] |
| MXE | 263 | ENSG00000198089 | SFI1 | chr22 | + | 31602606 | 31602785 | 31604868 | 31606430 | 31602211 | 31602293 | 31607936 | 31608033 | 84 | [illegible] |
| MXE | 7905 | ENSG00000071054 | MAP4K4 | chr2 | + | 101860824 | 101860986 | 101864929 | 101865036 | 101859642 | 101859864 | 101866427 | 101866579 | 9 | [illegible] |
| MXE | 6138 | ENSG00000163719 | MTMR14 | chr3 | + | 9653620 | 9653769 | 9662266 | 9662375 | 9649432 | 9649742 | 9668718 | 9668794 | 95 | [illegible] |
| MXE | 7840 | ENSG00000163564 | PYHIN1 | chr1 | + | 158938396 | 158938542 | 158939079 | 158939247 | 158936890 | 158937175 | 158941976 | 158942399 | 414 | [illegible] |
| MXE | 13073 | ENSG00000161203 | AP2M1 | chr3 | + | 184176950 | 184177067 | 184178856 | 184179191 | 184174904 | 184174959 | 184180168 | 184180212 | 89 | [illegible] |
| MXE | 1759 | ENSG00000214078 | CPNE1 | chr20 | - | 35626562 | 35626803 | 35627279 | 35627413 | 35626043 | 35626381 | 35630438 | 35630456 | 226 | [illegible] |
| MXE | 10302 | ENSG00000133872 | SARAF | chr8 | - | 30066027 | 30066139 | 30066776 | 30066918 | 30063125 | 30063913 | 30069641 | 30070059 | 1446 | [illegible] |
| MXE | 2454 | ENSG00000100429 | HDAC10 | chr22 | - | 50248830 | 50248859 | 50249327 | 50249454 | 50248661 | 50248751 | 50249634 | 50249703 | 20 | [illegible] |
| MXE | 8923 | ENSG00000065413 | ANKRD44 | chr2 | - | 197122649 | 197122792 | 197125836 | 197126037 | 197121331 | 197121544 | 197136591 | 197136662 | 496 | [illegible] |
| MXE | 11834 | ENSG00000124508 | BTN2A2 | chr6 | + | 26385014 | 26385362 | 26388012 | 26388294 | 26383791 | 26383915 | 26390004 | 26390211 | 27 | [illegible] |
| MXE | 554 | ENSG00000157540 | DYRK1A | chr21 | + | 37486466 | 37486614 | 37496117 | 37496258 | 37480637 | 37480826 | 37505282 | 37505357 | 413 | [illegible] |
| MXE | 14174 | ENSG00000168404 | MLKL | chr16 | - | 74678898 | 74678980 | 74682650 | 74682786 | 74676300 | 74676425 | 74685485 | 74685583 | 138 | [illegible] |
| MXE | 8029 | ENSG00000160691 | SHC1 | chr1 | - | 154966318 | 154966517 | 154967670 | 154967797 | 154966161 | 154966231 | 154967979 | 154968031 | 156 | [illegible] |
| MXE | 5015 | ENSG00000125037 | EMC3 | chr3 | - | 9973627 | 9973709 | 9974383 | 9974488 | 9970581 | 9970661 | 9976956 | 9977050 | 198 | [illegible] |
| MXE | 15927 | ENSG00000181192 | DHTKD1 | chr10 | + | 12101041 | 12101181 | 12106245 | 12106396 | 12100177 | 12100262 | 12107908 | 12108015 | 27 | [illegible] |
| MXE | 16048 | ENSG00000180229 | HERC2P3 | chr15 | - | 20466194 | 20466370 | 20483402 | 20483517 | 20462830 | 20463001 | 20505390 | 20505493 | 16 | [illegible] |
| MXE | 6242 | ENSG00000112144 | CILK1 | chr6 | - | 53018329 | 53018501 | 53019226 | 53019359 | 53016082 | 53016250 | 53031064 | 53031144 | 80 | [illegible] |
| MXE | 527 | ENSG00000239998 | LILRA2 | chr19 | + | 54573697 | 54573912 | 54574300 | 54574582 | 54573027 | 54573435 | 54574730 | 54575033 | 14 | [illegible] |
| MXE | 1881 | ENSG00000038358 | EDC4 | chr16 | + | 67881496 | 67881533 | 67881667 | 67882109 | 67881264 | 67881417 | 67882211 | 67882327 | 13 | [illegible] |
| MXE | 14140 | ENSG00000123908 | AGO2 | chr8 | - | 140547467 | 140547627 | 140549113 | 140549298 | 140544212 | 140544303 | 140551302 | 140551436 | 137 | [illegible] |
| MXE | 3006 | ENSG00000115760 | BIRC6 | chr2 | + | 32436052 | 32436184 | 32439507 | 32439686 | 32433643 | 32433804 | 32441328 | 32441462 | 134 | [illegible] |
| MXE | 3693 | ENSG00000120705 | ETF1 | chr5 | - | 138510564 | 138510629 | 138511044 | 138511200 | 138508668 | 138508816 | 138511474 | 138511604 | 304 | [illegible] |
| MXE | 2817 | ENSG00000100479 | POLE2 | chr14 | - | 49669523 | 49669598 | 49674122 | 49674216 | 49666329 | 49666413 | 49674349 | 49674427 | 0 | [illegible] |
| MXE | 17038 | ENSG00000136147 | PHF11 | chr13 | + | 49520893 | 49520940 | 49522042 | 49522107 | 49518017 | 49518151 | 49523174 | 49523207 | 24 | [illegible] |
| MXE | 3719 | ENSG00000273247 | NA | chr5 | - | 131561977 | 131562047 | 131592382 | 131592466 | 131548046 | 131548190 | 131603270 | 131603327 | 63 | [illegible] |
| MXE | 13510 | ENSG00000171311 | EXOSC1 | chr10 | - | 97438662 | 97438703 | 97443236 | 97443311 | 97437699 | 97437750 | 97445731 | 97445847 | 16 | [illegible] |
| MXE | 6657 | ENSG00000114098 | ARMC8 | chr3 | + | 138272966 | 138273116 | 138274448 | 138274544 | 138270039 | 138270132 | 138284430 | 138284526 | 123 | [illegible] |
| MXE | 5440 | ENSG00000145819 | ARHGAP26 | chr5 | + | 142932046 | 142932125 | 143014079 | 143014116 | 142913198 | 142913293 | 143037195 | 143037223 | 105 | [illegible] |
| MXE | 15982 | ENSG00000078369 | GNB1 | chr1 | - | 1806474 | 1806538 | 1815755 | 1815862 | 1804418 | 1804581 | 1825396 | 1825499 | 9 | [illegible] |
| MXE | 2107 | ENSG00000128739 | SNRPN | chr15 | + | 24974287 | 24974456 | 24976876 | 24977029 | 24967931 | 24968082 | 24977777 | 24977916 | 2 | [illegible] |
| MXE | 14772 | ENSG00000106868 | SUSD1 | chr9 | - | 112062936 | 112063033 | 112078537 | 112078724 | 112058427 | 112058686 | 112080073 | 112080165 | 194 | [illegible] |
| MXE | 8435 | ENSG00000258644 | SYNJ2BP-COX16 | chr14 | - | 70342657 | 70342729 | 70373032 | 70373131 | 70329173 | 70329236 | 70375675 | 70375771 | 39 | [illegible] |
| MXE | 1622 | ENSG00000153066 | TXNDC11 | chr16 | - | 11698124 | 11698325 | 11700451 | 11700564 | 11692049 | 11692082 | 11721313 | 11721435 | 0 | [illegible] |
| MXE | 4308 | ENSG00000124203 | ZNF831 | chr20 | + | 59206904 | 59207056 | 59252977 | 59253138 | 59195868 | 59196005 | 59253897 | 59259113 | 170 | [illegible] |
| MXE | 12914 | ENSG00000128563 | PRKRIP1 | chr7 | + | 102404597 | 102404683 | 102407433 | 102407498 | 102399547 | 102399648 | 102425013 | 102425446 | 60 | [illegible] |
| MXE | 15180 | ENSG00000119392 | GLE1 | chr9 | + | 128508875 | 128509097 | 128515528 | 128515639 | 128504654 | 128504904 | 128522667 | 128522816 | 27 | [illegible] |
| MXE | 4885 | ENSG00000168297 | PXK | chr3 | + | 58397604 | 58397722 | 58399298 | 58399377 | 58397038 | 58397200 | 58403861 | 58403910 | 112 | [illegible] |
| MXE | 10772 | ENSG00000163412 | EIF4E3 | chr3 | - | 71693874 | 71693941 | 71710411 | 71710484 | 71690009 | 71690165 | 71728475 | 71728623 | 4 | [illegible] |
| MXE | 10648 | ENSG00000135926 | TMBIM1 | chr2 | - | 218277934 | 218277974 | 218278514 | 218278565 | 218277632 | 218277670 | 218279037 | 218279091 | 24 | [illegible] |
| MXE | 8197 | ENSG00000188641 | DPYD | chr1 | - | 97828113 | 97828196 | 97883263 | 97883374 | 97740391 | 97740479 | 97920883 | 97921034 | 192 | [illegible] |
| MXE | 12345 | ENSG00000048392 | RRM2B | chr8 | - | 102218813 | 102218947 | 102224045 | 102224140 | 102214053 | 102214158 | 102224884 | 102225018 | 53 | [illegible] |
| MXE | 1078 | ENSG00000108953 | YWHAE | chr17 | - | 1361091 | 1361298 | 1361901 | 1362008 | 1354210 | 1354347 | 1364858 | 1365058 | 329 | [illegible] |
| MXE | 7118 | ENSG00000055609 | KMT2C | chr7 | - | 152139185 | 152139259 | 152139674 | 152139791 | 152134921 | 152138904 | 152144712 | 152144881 | 206 | [illegible] |
| MXE | 8185 | ENSG00000138614 | INTS14 | chr15 | - | 65605128 | 65605236 | 65607158 | 65607442 | 65599773 | 65599929 | 65611097 | 65611289 | 22 | [illegible] |
| MXE | 8327 | ENSG00000177409 | SAMD9L | chr7 | - | 93145384 | 93145532 | 93145908 | 93146040 | 93144731 | 93144833 | 93147087 | 93147146 | 40 | [illegible] |
| MXE | 11769 | ENSG00000124523 | SIRT5 | chr6 | + | 13588330 | 13588464 | 13591668 | 13591894 | 13584986 | 13585127 | 13595476 | 13595564 | 0 | [illegible] |
| MXE | 4266 | ENSG00000119414 | PPP6C | chr9 | - | 125158240 | 125158382 | 125160840 | 125160906 | 125153905 | 125153985 | 125171084 | 125171180 | 230 | [illegible] |
| MXE | 16409 | ENSG00000100354 | TNRC6B | chr22 | + | 40264687 | 40267036 | 40273424 | 40273600 | 40261831 | 40262173 | 40277076 | 40277151 | 95 | [illegible] |
| MXE | 3014 | ENSG00000115760 | BIRC6 | chr2 | + | 32465064 | 32465164 | 32467524 | 32467739 | 32464508 | 32464823 | 32467902 | 32468111 | 139 | [illegible] |
| MXE | 28 | ENSG00000160208 | RRP1B | chr21 | + | 43686803 | 43686935 | 43690287 | 43690440 | 43685769 | 43685789 | 43691438 | 43691502 | 88 | [illegible] |
| MXE | 13736 | ENSG00000147050 | KDM6A | chrX | + | 45082575 | 45082640 | 45082714 | 45082789 | 45079145 | 45079351 | 45083459 | 45083608 | 80 | [illegible] |
| MXE | 8009 | ENSG00000196235 | SUPT5H | chr19 | + | 39445803 | 39445965 | 39453355 | 39453521 | 39445440 | 39445637 | 39457674 | 39457740 | 19 | [illegible] |
| MXE | 3748 | ENSG00000164308 | ERAP2 | chr5 | + | 96900120 | 96900189 | 96901505 | 96901681 | 96896731 | 96896863 | 96902273 | 96902353 | 93 | [illegible] |
| MXE | 14406 | ENSG00000122692 | SMU1 | chr9 | - | 33060464 | 33060584 | 33062048 | 33062177 | 33057597 | 33057714 | 33068823 | 33068934 | 157 | [illegible] |
| MXE | 16313 | ENSG00000023041 | ZDHHC6 | chr10 | - | 112440533 | 112440695 | 112441728 | 112441812 | 112438335 | 112438389 | 112442191 | 112442351 | 0 | [illegible] |
| MXE | 4616 | ENSG00000134852 | CLOCK | chr4 | - | 55453676 | 55453824 | 55455896 | 55456003 | 55453053 | 55453129 | 55456217 | 55456300 | 46 | [illegible] |
| MXE | 1547 | ENSG00000067955 | CBFB | chr16 | + | 67036638 | 67036755 | 67066681 | 67066798 | 67029726 | 67030177 | 67082212 | 67082339 | 2 | [illegible] |
| MXE | 16304 | ENSG00000099246 | RAB18 | chr10 | + | 27526827 | 27526889 | 27531528 | 27531615 | 27509874 | 27509930 | 27532506 | 27532579 | 408 | [illegible] |
| MXE | 16138 | ENSG00000165732 | DDX21 | chr10 | + | 68978841 | 68978976 | 68981536 | 68981581 | 68977528 | 68977688 | 68982542 | 68985067 | 131 | [illegible] |
| MXE | 6019 | ENSG00000010322 | NISCH | chr3 | + | 52490074 | 52490231 | 52490704 | 52490833 | 52489335 | 52489678 | 52491351 | 52491513 | 43 | [illegible] |
| MXE | 1139 | ENSG00000275700 | AATF | chr17 | + | 36953769 | 36953907 | 36986616 | 36986731 | 36952885 | 36953296 | 36988518 | 36988720 | 269 | [illegible] |
| MXE | 3890 | ENSG00000119522 | DENND1A | chr9 | - | 123757702 | 123757822 | 123769513 | 123769563 | 123676719 | 123676789 | 123792586 | 123792630 | 27 | [illegible] |
| MXE | 729 | ENSG00000124214 | STAU1 | chr20 | - | 49153932 | 49154071 | 49174194 | 49174269 | 49151581 | 49151747 | 49188115 | 49188367 | 4 | [illegible] |
| MXE | 3439 | ENSG00000152942 | RAD17 | chr5 | + | 69371453 | 69371557 | 69372033 | 69372217 | 69371034 | 69371171 | 69373829 | 69374087 | 7 | [illegible] |
| MXE | 6562 | ENSG00000135597 | REPS1 | chr6 | - | 138915857 | 138915976 | 138917554 | 138917646 | 138914696 | 138914761 | 138920214 | 138920316 | 5 | [illegible] |
| MXE | 544 | ENSG00000105677 | TMEM147 | chr19 | + | 35546525 | 35546585 | 35546671 | 35547029 | 35545887 | 35545957 | 35547118 | 35547240 | 12 | [illegible] |
| MXE | 10446 | ENSG00000162923 | WDR26 | chr1 | - | 224398514 | 224398593 | 224398888 | 224399034 | 224397726 | 224398226 | 224400949 | 224401019 | 658 | [illegible] |
| MXE | 12558 | ENSG00000179818 | PCBP1-AS1 | chr2 | - | 70077300 | 70077381 | 70083537 | 70083687 | 70055655 | 70055910 | 70085546 | 70085624 | 33 | [illegible] |
| MXE | 17162 | ENSG00000121644 | DESI2 | chr1 | + | 244686596 | 244686669 | 244689248 | 244689342 | 244653125 | 244653355 | 244691878 | 244692020 | 7 | [illegible] |
| MXE | 10087 | ENSG00000169379 | ARL13B | chr3 | + | 93996562 | 93996684 | 94003658 | 94003908 | 93995873 | 93995944 | 94035330 | 94035436 | 5 | [illegible] |
| MXE | 8386 | ENSG00000072110 | ACTN1 | chr14 | - | 68878457 | 68878520 | 68878988 | 68879069 | 68877081 | 68877240 | 68879961 | 68880108 | 218 | [illegible] |
| MXE | 9355 | ENSG00000070814 | TCOF1 | chr5 | + | 150387942 | 150388088 | 150389886 | 150390023 | 150379531 | 150379732 | 150391543 | 150391795 | 2 | [illegible] |
| MXE | 14227 | ENSG00000168522 | FNTA | chr8 | + | 43083117 | 43083180 | 43084709 | 43084881 | 43077215 | 43080441 | 43085159 | 43085788 | 15 | [illegible] |
| MXE | 1970 | ENSG00000005339 | CREBBP | chr16 | - | 3745276 | 3745354 | 3749626 | 3749683 | 3744893 | 3744961 | 3751725 | 3751806 | 60 | [illegible] |
| MXE | 12616 | ENSG00000151948 | GLT1D1 | chr12 | + | 128912423 | 128912460 | 128914932 | 128914989 | 128899235 | 128899287 | 128926358 | 128926454 | 7 | [illegible] |
| MXE | 8088 | ENSG00000083544 | TDRD3 | chr13 | + | 60444682 | 60444748 | 60460379 | 60460540 | 60439667 | 60439772 | 60467237 | 60467379 | 14 | [illegible] |
| MXE | 7646 | ENSG00000100591 | AHSA1 | chr14 | + | 77462641 | 77462759 | 77468082 | 77468184 | 77462159 | 77462242 | 77469076 | 77469470 | 105 | [illegible] |
| MXE | 7171 | ENSG00000115239 | ASB3 | chr2 | - | 53716565 | 53716743 | 53728711 | 53728847 | 53714383 | 53714581 | 53729457 | 53729570 | 115 | [illegible] |
| MXE | 10789 | ENSG00000144228 | SPOPL | chr2 | + | 138550482 | 138550604 | 138550973 | 138551054 | 138550156 | 138550294 | 138552553 | 138552582 | 129 | [illegible] |
| MXE | 1039 | ENSG00000070413 | DGCR2 | chr22 | - | 19056985 | 19057162 | 19063201 | 19063278 | 19048439 | 19048643 | 19064847 | 19065058 | 56 | [illegible] |
| MXE | 4164 | ENSG00000177058 | SLC38A9 | chr5 | - | 55664692 | 55664863 | 55669227 | 55669321 | 55656714 | 55656774 | 55669556 | 55669620 | 42 | [illegible] |
| MXE | 3177 | ENSG00000130449 | ZSWIM6 | chr5 | + | 61538813 | 61538971 | 61539595 | 61539759 | 61535483 | 61535619 | 61541883 | 61541965 | 91 | [illegible] |
| MXE | 2481 | ENSG00000140563 | MCTP2 | chr15 | + | 94440175 | 94440298 | 94442918 | 94442960 | 94401899 | 94402019 | 94458136 | 94458246 | 27 | [illegible] |
| MXE | 10340 | ENSG00000137075 | RNF38 | chr9 | - | 36375933 | 36376127 | 36390466 | 36390616 | 36369718 | 36369932 | 36401097 | 36401196 | 1 | [illegible] |
| MXE | 3523 | ENSG00000113263 | ITK | chr5 | + | 157211286 | 157211368 | 157214190 | 157214319 | 157208888 | 157208993 | 157217866 | 157217907 | 184 | [illegible] |
| MXE | 11268 | ENSG00000172765 | TMCC1 | chr3 | - | 129827802 | 129828508 | 129832773 | 129832826 | 129670329 | 129671264 | 129880308 | 129880559 | 6 | [illegible] |
| MXE | 7088 | ENSG00000117090 | SLAMF1 | chr1 | - | 160619775 | 160619849 | 160634612 | 160634897 | 160612487 | 160612580 | 160637190 | 160637529 | 75 | [illegible] |
| MXE | 2541 | ENSG00000151327 | FAM177A1 | chr14 | + | 35053277 | 35053451 | 35077149 | 35077216 | 35046271 | 35046628 | 35078926 | 35079024 | 12 | [illegible] |
| MXE | 9936 | ENSG00000032444 | PNPLA6 | chr19 | + | 7553874 | 7554015 | 7554208 | 7554272 | 7551361 | 7551437 | 7554554 | 7554723 | 26 | [illegible] |
| MXE | 15555 | ENSG00000172543 | CTSW | chr11 | + | 65881406 | 65881520 | 65882174 | 65882329 | 65880201 | 65880286 | 65882608 | 65882689 | 324 | [illegible] |
| MXE | 16846 | ENSG00000111596 | CNOT2 | chr12 | + | 70344127 | 70344228 | 70346179 | 70346324 | 70342257 | 70342307 | 70353828 | 70353913 | 178 | [illegible] |
| MXE | 7999 | ENSG00000197872 | CYRIA | chr2 | - | 16563988 | 16564094 | 16565645 | 16565767 | 16562004 | 16562141 | 16588049 | 16588129 | 102 | [illegible] |
| MXE | 14446 | ENSG00000100815 | TRIP11 | chr14 | - | 92011754 | 92011795 | 92014214 | 92014577 | 92007639 | 92007852 | 92015695 | 92015751 | 126 | [illegible] |
| MXE | 6691 | ENSG00000102606 | ARHGEF7 | chr13 | + | 111205288 | 111205373 | 111217678 | 111217880 | 111153713 | 111153991 | 111233204 | 111233293 | 81 | [illegible] |
| MXE | 10950 | ENSG00000164548 | TRA2A | chr7 | - | 23522131 | 23522432 | 23531180 | 23531270 | 23521706 | 23521840 | 23531788 | 23532041 | 16 | [illegible] |
| MXE | 12876 | ENSG00000257923 | CUX1 | chr7 | + | 102028097 | 102028145 | 102070338 | 102070417 | 101916114 | 101916225 | 102097363 | 102097501 | 52 | [illegible] |
| MXE | 7562 | ENSG00000111331 | OAS3 | chr12 | + | 112944475 | 112944651 | 112946742 | 112946981 | 112941569 | 112941852 | 112961070 | 112961246 | 51 | [illegible] |
| MXE | 8335 | ENSG00000083720 | OXCT1 | chr5 | - | 41805566 | 41805681 | 41807330 | 41807438 | 41803068 | 41803163 | 41840450 | 41840511 | 65 | [illegible] |
| MXE | 1925 | ENSG00000069345 | DNAJA2 | chr16 | - | 46967512 | 46967646 | 46968083 | 46968164 | 46964610 | 46964807 | 46971348 | 46971572 | 139 | [illegible] |
| MXE | 7853 | ENSG00000136643 | RPS6KC1 | chr1 | + | 213077695 | 213077816 | 213104453 | 213104569 | 213071005 | 213071041 | 213117316 | 213117410 | 22 | [illegible] |
| MXE | 1516 | ENSG00000091592 | NLRP1 | chr17 | - | 5533896 | 5533988 | 5536850 | 5536940 | 5532821 | 5532984 | 5539414 | 5539585 | 40 | [illegible] |
| MXE | 10711 | ENSG00000112367 | FIG4 | chr6 | + | 109715077 | 109715176 | 109727108 | 109727265 | 109691295 | 109691501 | 109732636 | 109732687 | 28 | [illegible] |
| MXE | 43 | ENSG00000160294 | MCM3AP | chr21 | - | 46251528 | 46251682 | 46254391 | 46254526 | 46246306 | 46246886 | 46254775 | 46254844 | 62 | [illegible] |
| MXE | 4844 | ENSG00000131375 | CAPN7 | chr3 | + | 15240753 | 15240853 | 15245525 | 15245671 | 15240472 | 15240617 | 15246731 | 15246794 | 68 | [illegible] |
| MXE | 2383 | ENSG00000125107 | CNOT1 | chr16 | - | 58588798 | 58588906 | 58599235 | 58599511 | 58587779 | 58587878 | 58629727 | 58629826 | 74 | [illegible] |
| MXE | 2786 | ENSG00000099899 | TRMT2A | chr22 | - | 20115265 | 20115447 | 20115671 | 20115780 | 20114964 | 20115079 | 20116037 | 20116612 | 67 | [illegible] |
| MXE | 4854 | ENSG00000261796 | ISY1-RAB43 | chr3 | - | 129130549 | 129130636 | 129134073 | 129134195 | 129130087 | 129130188 | 129134831 | 129134954 | 120 | [illegible] |
| MXE | 5567 | ENSG00000163539 | CLASP2 | chr3 | - | 33644756 | 33644903 | 33663444 | 33663515 | 33632291 | 33632371 | 33684358 | 33684456 | 52 | [illegible] |
| MXE | 12557 | ENSG00000179818 | PCBP1-AS1 | chr2 | - | 70059642 | 70059680 | 70083506 | 70083687 | 70055713 | 70055910 | 70085546 | 70085624 | 33 | [illegible] |
| MXE | 3745 | ENSG00000164308 | ERAP2 | chr5 | + | 96889184 | 96889305 | 96892298 | 96892453 | 96886654 | 96886789 | 96895245 | 96895359 | 442 | [illegible] |

| | | | | | | | | | | | | | | | | | | | | | | | | | | | | | | | | | | | | | | | | | | | | | | | | |
|---|---|---|---|---|---|---|---|---|---|---|---|---|---|---|---|---|---|---|---|---|---|---|---|---|---|---|---|---|---|---|---|---|---|---|---|---|---|---|---|---|---|---|---|---|---|---|---|---|
| MXE | 5368 | ENSG00000095951 | HIVEP1 | chr6 | + | 12089183 | 12089237 | 12119889 | 12125870 | 12015525 | 12015668 | 12129758 | 12129892 | 23 | 21 | 27 | 29 | 29 | 60 | 88 | 70 | 76 | 86 | 21 | 8 | 25 | 14 | 0 | 0 | 62 | 75 | 50 | 15 | 203 | 298 | 0.886404492 | 1 | 0.36 | 0.259 | 0.362 | 0.359 | 0.331 | 1 | 0.159 | 0.329 | 0.291 | 0 | -0.022 |
| MXE | 5147 | ENSG00000108219 | TSPAN14 | chr10 | + | 80507227 | 80507374 | 80509300 | 80509471 | 80504727 | 80504778 | 80512143 | 80512269 | 98 | 29 | 91 | 86 | 62 | 169 | 78 | 141 | 160 | 84 | 99 | 142 | 67 | 51 | 124 | 173 | 162 | 178 | 84 | 217 | 296 | 298 | 0.886507437 | 1 | 0.369 | 0.272 | 0.394 | 0.351 | 0.426 | 0.366 | 0.469 | 0.275 | 0.379 | 0.365 | -0.008 |
| MXE | 14561 | ENSG00000113924 | HGD | chr3 | - | 120638454 | 120638581 | 120641588 | 120641693 | 120633146 | 120633328 | 120646972 | 120647052 | 13 | 25 | 6 | 22 | 6 | 3 | 7 | 5 | 16 | 5 | 17 | 4 | 4 | 15 | 51 | 40 | 2 | 0 | 1 | 66 | 254 | 276 | 0.886746133 | 1 | 0.825 | 0.795 | 0.566 | 0.599 | 0.566 | 0.316 | 0.685 | 1 | 0.942 | 0.456 | -0.01 |
| MXE | 12433 | ENSG00000153814 | JAZF1 | chr7 | - | 27840697 | 27840867 | 27895219 | 27895416 | 27830572 | 27832976 | 27991908 | 27992056 | 96 | 56 | 31 | 74 | 86 | 123 | 43 | 37 | 81 | 40 | 51 | 81 | 75 | 56 | 62 | 50 | 47 | 69 | 53 | 68 | 298 | 298 | 0.886762618 | 1 | 0.438 | 0.566 | 0.456 | 0.477 | 0.683 | 0.505 | 0.633 | 0.521 | 0.514 | 0.477 | -0.006 |
| MXE | 15986 | ENSG00000052749 | RRP12 | chr10 | - | 97390422 | 97390539 | 97390738 | 97390844 | 97388488 | 97388624 | 97393683 | 97393760 | 48 | 21 | 25 | 29 | 13 | 33 | 14 | 18 | 27 | 24 | 107 | 45 | 42 | 71 | 59 | 68 | 60 | 46 | 68 | 42 | 255 | 266 | 0.886778934 | 1 | 0.603 | 0.61 | 0.592 | 0.528 | 0.361 | 0.621 | 0.439 | 0.488 | 0.521 | 0.594 | 0.006 |
| MXE | 14890 | ENSG00000165219 | GAPVD1 | chr9 | + | 125301982 | 125302826 | 125305062 | 125305149 | 125298885 | 125299106 | 125307412 | 125307547 | 108 | 102 | 97 | 109 | 177 | 20 | 9 | 31 | 41 | 30 | 210 | 139 | 113 | 214 | 40 | 41 | 22 | 34 | 25 | 18 | 298 | 236 | 0.886783091 | 1 | 0.81 | 0.9 | 0.712 | 0.678 | 0.824 | 0.802 | 0.833 | 0.725 | 0.871 | 0.638 | 0.011 |
| MXE | 12680 | ENSG00000155096 | AZIN1 | chr8 | - | 102829820 | 102829936 | 102833055 | 102833218 | 102829271 | 102829486 | 102834188 | 102834263 | 255 | 142 | 215 | 286 | 398 | 494 | 277 | 338 | 442 | 634 | 113 | 180 | 207 | 211 | 128 | 272 | 275 | 414 | 386 | 116 | 298 | 265 | 0.886871717 | 1 | 0.315 | 0.313 | 0.361 | 0.365 | 0.358 | 0.27 | 0.368 | 0.308 | 0.327 | 0.495 | -0.011 |
| MXE | 14862 | ENSG00000060237 | WNK1 | chr12 | + | 859244 | 859464 | 861012 | 861343 | 857160 | 857249 | 862082 | 862270 | 514 | 454 | 405 | 473 | 750 | 393 | 284 | 274 | 313 | 303 | 512 | 685 | 573 | 530 | 345 | 291 | 415 | 285 | 440 | 222 | 298 | 298 | 0.88692545 | 1 | 0.567 | 0.615 | 0.596 | 0.602 | 0.712 | 0.638 | 0.623 | 0.668 | 0.546 | 0.608 | 0.002 |
| MXE | 16702 | ENSG00000141376 | BCAS3 | chr17 | + | 60889694 | 60889771 | 60902619 | 60902703 | 60874661 | 60874738 | 60910531 | 60910702 | 25 | 20 | 21 | 31 | 25 | 14 | 25 | 32 | 35 | 28 | 20 | 33 | 25 | 24 | 48 | 51 | 32 | 30 | 26 | 28 | 226 | 233 | 0.886954296 | 1 | 0.648 | 0.452 | 0.404 | 0.477 | 0.479 | 0.288 | 0.515 | 0.462 | 0.488 | 0.639 | 0.014 |
| MXE | 17112 | ENSG00000173473 | SMARCC1 | chr3 | - | 47676628 | 47676782 | 47678197 | 47678311 | 47675474 | 47675588 | 47680436 | 47680508 | 243 | 199 | 242 | 213 | 524 | 77 | 36 | 60 | 43 | 82 | 191 | 211 | 159 | 182 | 113 | 20 | 75 | 41 | 32 | 20 | 263 | 298 | 0.887016096 | 1 | 0.781 | 0.862 | 0.82 | 0.849 | 0.879 | 0.915 | 0.761 | 0.815 | 0.866 | 0.865 | -0.006 |
| MXE | 4981 | ENSG00000123737 | EXOSC9 | chr4 | + | 121807539 | 121807622 | 121809966 | 121810099 | 121804621 | 121804759 | 121811582 | 121811641 | 61 | 51 | 92 | 83 | 131 | 78 | 74 | 70 | 102 | 84 | 42 | 61 | 79 | 13 | 15 | 26 | 67 | 114 | 86 | 0 | 232 | 282 | 0.887187374 | 1 | 0.487 | 0.456 | 0.615 | 0.497 | 0.655 | 0.663 | 0.525 | 0.457 | 0.155 | 1 | -0.018 |
| MXE | 3705 | ENSG00000087470 | DNM1L | chr12 | + | 32740408 | 32740518 | 32742588 | 32742748 | 32740063 | 32740240 | 32743353 | 32743581 | 70 | 77 | 53 | 85 | 71 | 154 | 187 | 138 | 183 | 180 | 10 | 23 | 80 | 32 | 12 | 33 | 92 | 136 | 52 | 36 | 259 | 298 | 0.887192666 | 1 | 0.343 | 0.321 | 0.306 | 0.348 | 0.312 | 0.259 | 0.223 | 0.404 | 0.415 | 0.277 | 0.01 |
| MXE | 17076 | ENSG00000125457 | MIF4GD | chr17 | - | 75268082 | 75268192 | 75269346 | 75269469 | 75267745 | 75267901 | 75270113 | 75270245 | 9 | 5 | 11 | 3 | 7 | 68 | 46 | 50 | 51 | 56 | 20 | 4 | 13 | 0 | 5 | 56 | 109 | 23 | 63 | 99 | 272 | 259 | 0.887363743 | 1 | 0.112 | 0.094 | 0.173 | 0.053 | 0.106 | 0.254 | 0.034 | 0.35 | 0 | 0.046 | -0.029 |
| MXE | 2986 | ENSG00000196367 | TRRAP | chr7 | + | 98906176 | 98906255 | 98908727 | 98908962 | 98903378 | 98903517 | 98910055 | 98910419 | 43 | 39 | 42 | 49 | 73 | 65 | 59 | 68 | 67 | 82 | 59 | 49 | 52 | 55 | 40 | 93 | 61 | 45 | 72 | 81 | 228 | 298 | 0.887425519 | 1 | 0.464 | 0.464 | 0.447 | 0.489 | 0.538 | 0.453 | 0.512 | 0.602 | 0.5 | 0.392 | -0.011 |
| MXE | 2302 | ENSG00000077238 | IL4R | chr16 | + | 27344868 | 27345020 | 27346466 | 27346618 | 27342120 | 27342259 | 27352539 | 27352696 | 150 | 64 | 147 | 130 | 150 | 86 | 54 | 92 | 75 | 135 | 152 | 229 | 148 | 174 | 178 | 95 | 154 | 151 | 100 | 119 | 298 | 298 | 0.887562327 | 1 | 0.636 | 0.542 | 0.615 | 0.634 | 0.526 | 0.615 | 0.598 | 0.495 | 0.635 | 0.599 | 0.002 |
| MXE | 16125 | ENSG00000011376 | LARS2 | chr3 | + | 45523996 | 45524108 | 45541828 | 45541956 | 45520218 | 45520296 | 45547350 | 45547404 | 12 | 8 | 10 | 15 | 9 | 10 | 5 | 10 | 6 | 18 | 9 | 50 | 9 | 23 | 4 | 15 | 8 | 6 | 4 | 48 | 261 | 277 | 0.887720337 | 1 | 0.56 | 0.629 | 0.515 | 0.726 | 0.347 | 0.389 | 0.869 | 0.614 | 0.859 | 0.081 | -0.007 |
| MXE | 3976 | ENSG00000140471 | LINS1 | chr15 | - | 100580443 | 100580736 | 100580800 | 100580945 | 100580262 | 100580352 | 100602120 | 100602174 | 82 | 53 | 114 | 69 | 150 | 87 | 65 | 72 | 60 | 130 | 26 | 21 | 65 | 23 | 1 | 12 | 31 | 33 | 22 | 4 | 294 | 298 | 0.8877468 | 1 | 0.489 | 0.452 | 0.616 | 0.538 | 0.539 | 0.687 | 0.407 | 0.666 | 0.514 | 0.202 | 0.032 |
| MXE | 15467 | ENSG00000101017 | CD40 | chr20 | + | 46123125 | 46123219 | 46126639 | 46126701 | 46122609 | 46122756 | 46128137 | 46128224 | 35 | 6 | 18 | 28 | 30 | 19 | 17 | 18 | 44 | 63 | 26 | 32 | 11 | 16 | 27 | 45 | 55 | 5 | 20 | 33 | 243 | 211 | 0.887887916 | 1 | 0.615 | 0.235 | 0.465 | 0.356 | 0.293 | 0.334 | 0.336 | 0.656 | 0.41 | 0.415 | -0.037 |
| MXE | 13797 | ENSG00000156171 | DRAM2 | chr1 | - | 111120515 | 111120693 | 111124741 | 111124881 | 111119876 | 111119959 | 111126226 | 111126294 | 18 | 8 | 6 | 20 | 38 | 292 | 195 | 264 | 343 | 520 | 23 | 16 | 3 | 5 | 4 | 217 | 233 | 182 | 163 | 82 | 289 | 298 | 0.88812027 | 1 | 0.06 | 0.041 | 0.023 | 0.057 | 0.07 | 0.099 | 0.066 | 0.017 | 0.031 | 0.048 | -0.002 |
| MXE | 170 | ENSG00000100124 | ANKRD54 | chr22 | - | 37832957 | 37833109 | 37833158 | 37833206 | 37832636 | 37832744 | 37833683 | 37833755 | 3 | 2 | 1 | 5 | 7 | 13 | 6 | 12 | 13 | 8 | 5 | 0 | 2 | 0 | 9 | 6 | 7 | 2 | 8 | 10 | 197 | 298 | 0.888149441 | 1 | 0.259 | 0.335 | 0.112 | 0.368 | 0.57 | 0.558 | 0 | 0.602 | 0 | 0.577 | -0.019 |
| MXE | 16650 | ENSG00000135297 | MTO1 | chr6 | + | 73481000 | 73481116 | 73482039 | 73482244 | 73480674 | 73480805 | 73482448 | 73482620 | 1 | 0 | 0 | 5 | 5 | 111 | 55 | 71 | 93 | 122 | 0 | 0 | 8 | 0 | 0 | 110 | 51 | 59 | 24 | 19 | 265 | 298 | 0.888164678 | 1 | 0.01 | 0 | 0 | 0.057 | 0.044 | 0 | 0 | 0.132 | 0 | 0 | -0.004 |
| MXE | 14589 | ENSG00000136937 | NCBP1 | chr9 | + | 97643203 | 97643360 | 97645116 | 97645224 | 97641561 | 97641662 | 97647491 | 97647561 | 49 | 40 | 45 | 48 | 40 | 6 | 4 | 2 | 6 | 8 | 44 | 21 | 15 | 7 | 5 | 0 | 5 | 6 | 0 | 0 | 298 | 257 | 0.888384777 | 1 | 0.876 | 0.896 | 0.951 | 0.873 | 0.812 | 1 | 0.784 | 0.683 | 1 | 1 | -0.012 |
| MXE | 7676 | ENSG00000115685 | PPP1R7 | chr2 | + | 241158465 | 241158549 | 241159212 | 241159343 | 241157806 | 241157862 | 241160331 | 241160433 | 1 | 1 | 0 | 0 | 4 | 364 | 239 | 276 | 360 | 735 | 7 | 4 | 7 | 1 | 0 | 703 | 327 | 252 | 343 | 353 | 233 | 280 | 0.888422844 | 1 | 0.003 | 0.005 | 0 | 0 | 0.006 | 0.012 | 0.014 | 0.032 | 0.003 | 0 | -0.009 |
| MXE | 3015 | ENSG00000115760 | BIRC6 | chr2 | + | 32468436 | 32468783 | 32469394 | 32469614 | 32467902 | 32468111 | 32470167 | 32470301 | 132 | 116 | 141 | 148 | 251 | 149 | 137 | 121 | 122 | 106 | 42 | 99 | 141 | 39 | 17 | 31 | 119 | 94 | 31 | 9 | 298 | 298 | 0.888707719 | 1 | 0.47 | 0.458 | 0.538 | 0.548 | 0.703 | 0.575 | 0.454 | 0.6 | 0.557 | 0.654 | -0.025 |
| MXE | 3372 | ENSG00000131844 | MCCC2 | chr5 | + | 71599658 | 71599760 | 71602505 | 71602633 | 71596279 | 71596364 | 71604355 | 71604468 | 59 | 32 | 42 | 44 | 30 | 38 | 29 | 51 | 33 | 49 | 67 | 46 | 25 | 39 | 67 | 53 | 49 | 44 | 20 | 63 | 251 | 277 | 0.888861241 | 1 | 0.631 | 0.549 | 0.476 | 0.595 | 0.403 | 0.582 | 0.509 | 0.385 | 0.683 | 0.54 | -0.009 |
| MXE | 4445 | ENSG00000137656 | BUD13 | chr11 | - | 116758268 | 116758407 | 116759073 | 116759179 | 116757765 | 116757950 | 116760734 | 116760952 | 82 | 34 | 41 | 40 | 34 | 51 | 65 | 61 | 53 | 57 | 41 | 35 | 38 | 54 | 4 | 66 | 43 | 18 | 26 | 36 | 255 | 288 | 0.889053567 | 1 | 0.645 | 0.371 | 0.432 | 0.46 | 0.403 | 0.412 | 0.479 | 0.705 | 0.701 | 0.111 | -0.019 |
| MXE | 8421 | ENSG00000178607 | ERN1 | chr17 | - | 64049054 | 64049202 | 64052779 | 64052979 | 64047857 | 64047985 | 64053271 | 64053371 | 97 | 46 | 83 | 73 | 36 | 154 | 85 | 123 | 154 | 33 | 39 | 35 | 114 | 78 | 74 | 84 | 52 | 186 | 119 | 86 | 298 | 297 | 0.889117835 | 1 | 0.386 | 0.35 | 0.402 | 0.321 | 0.521 | 0.316 | 0.401 | 0.379 | 0.395 | 0.462 | 0.005 |
| MXE | 16606 | ENSG00000196498 | NCOR2 | chr12 | - | 124378236 | 124378384 | 124385744 | 124385887 | 124372021 | 124372610 | 124398118 | 124398181 | 103 | 72 | 72 | 90 | 113 | 35 | 36 | 36 | 42 | 63 | 430 | 143 | 104 | 126 | 297 | 97 | 70 | 73 | 88 | 119 | 292 | 297 | 0.889188525 | 1 | 0.75 | 0.67 | 0.67 | 0.685 | 0.646 | 0.818 | 0.675 | 0.592 | 0.593 | 0.717 | 0.005 |
| MXE | 10106 | ENSG00000153214 | TMEM87B | chr2 | + | 112091711 | 112091783 | 112095227 | 112095299 | 112089624 | 112089718 | 112097043 | 112097152 | 199 | 175 | 148 | 201 | 98 | 1 | 3 | 0 | 2 | 0 | 105 | 246 | 139 | 108 | 52 | 0 | 6 | 4 | 0 | 0 | 221 | 221 | 0.889354448 | 1 | 0.995 | 0.983 | 1 | 0.99 | 1 | 1 | 0.976 | 0.972 | 1 | 1 | 0.004 |
| MXE | 9991 | ENSG00000262633 | NA | chr17 | + | 46929519 | 46929584 | 46931098 | 46931207 | 46923132 | 46923221 | 46932066 | 46932199 | 12 | 8 | 19 | 10 | 16 | 127 | 94 | 109 | 141 | 134 | 5 | 9 | 1 | 16 | 23 | 104 | 95 | 69 | 70 | 126 | 214 | 258 | 0.889367594 | 1 | 0.102 | 0.093 | 0.174 | 0.079 | 0.126 | 0.055 | 0.103 | 0.017 | 0.216 | 0.18 | 0.001 |
| MXE | 14314 | ENSG00000135679 | MDM2 | chr12 | + | 68814602 | 68814689 | 68815627 | 68815720 | 68813553 | 68813628 | 68816811 | 68816945 | 19 | 6 | 6 | 32 | 23 | 10 | 12 | 17 | 5 | 2 | 5 | 30 | 26 | 21 | 6 | 0 | 15 | 37 | 33 | 0 | 236 | 242 | 0.889382229 | 1 | 0.661 | 0.339 | 0.266 | 0.868 | 0.922 | 1 | 0.672 | 0.419 | 0.395 | 1 | -0.086 |
| MXE | 2963 | ENSG00000120925 | RNF170 | chr8 | - | 42865415 | 42865489 | 42873930 | 42874006 | 42861329 | 42861855 | 42887727 | 42887871 | 29 | 28 | 56 | 32 | 41 | 40 | 40 | 48 | 27 | 44 | 36 | 18 | 29 | 19 | 33 | 46 | 27 | 17 | 20 | 33 | 225 | 223 | 0.889432887 | 1 | 0.418 | 0.41 | 0.536 | 0.54 | 0.48 | 0.437 | 0.398 | 0.628 | 0.485 | 0.498 | -0.012 |
| MXE | 13862 | ENSG00000134014 | ELP3 | chr8 | + | 28107912 | 28107976 | 28110369 | 28110438 | 28106712 | 28106783 | 28113018 | 28113173 | 110 | 50 | 92 | 105 | 119 | 75 | 62 | 68 | 89 | 116 | 117 | 61 | 23 | 38 | 27 | 43 | 79 | 31 | 37 | 20 | 213 | 218 | 0.890126304 | 1 | 0.6 | 0.452 | 0.581 | 0.547 | 0.512 | 0.736 | 0.441 | 0.432 | 0.512 | 0.58 | -0.002 |
| MXE | 364 | ENSG00000257103 | LSM14A | chr19 | + | 34208928 | 34209051 | 34215595 | 34215661 | 34196633 | 34196763 | 34219390 | 34219573 | 267 | 185 | 274 | 309 | 509 | 9 | 11 | 7 | 15 | 9 | 121 | 254 | 312 | 309 | 144 | 9 | 2 | 13 | 14 | 0 | 272 | 215 | 0.890398866 | 1 | 0.959 | 0.93 | 0.969 | 0.942 | 0.978 | 0.914 | 0.99 | 0.95 | 0.946 | 1 | -0.004 |
| MXE | 8303 | ENSG00000151422 | FER | chr5 | + | 108946129 | 108946222 | 108954728 | 108954932 | 108924605 | 108924789 | 108959224 | 108959347 | 2 | 3 | 1 | 3 | 0 | 93 | 76 | 47 | 90 | 59 | 0 | 0 | 0 | 0 | 0 | 6 | 46 | 26 | 12 | 50 | 242 | 298 | 0.890435673 | 1 | 0.026 | 0.046 | 0.026 | 0.039 | 0 | 0 | 0 | 0 | 0 | 0 | 0.027 |
| MXE | 4694 | ENSG00000005436 | GCFC2 | chr2 | - | 75690637 | 75690719 | 75691976 | 75692100 | 75689968 | 75690081 | 75694240 | 75694427 | 78 | 66 | 59 | 70 | 170 | 22 | 23 | 23 | 30 | 17 | 9 | 38 | 23 | 3 | 18 | 7 | 3 | 7 | 14 | 0 | 273 | 231 | 0.890550721 | 1 | 0.75 | 0.708 | 0.685 | 0.664 | 0.894 | 0.521 | 0.915 | 0.735 | 0.153 | 1 | 0.075 |
| MXE | 15629 | ENSG00000078124 | ACER3 | chr11 | + | 76994155 | 76994226 | 77011329 | 77011375 | 76990538 | 76990574 | 77016674 | 77016779 | 4 | 3 | 0 | 7 | 4 | 0 | 0 | 1 | 1 | 2 | 17 | 11 | 2 | 9 | 5 | 0 | 0 | 3 | 0 | 0 | 220 | 195 | 0.890657798 | 1 | 1 | 1 | 0 | 0.861 | 0.639 | 1 | 1 | 0.371 | 1 | 1 | -0.174 |
| MXE | 8908 | ENSG00000089127 | OAS1 | chr12 | + | 112908535 | 112908824 | 112911097 | 112911235 | 112906972 | 112907219 | 112916508 | 112916738 | 288 | 230 | 174 | 294 | 300 | 246 | 175 | 148 | 210 | 176 | 1004 | 595 | 154 | 953 | 378 | 831 | 480 | 132 | 524 | 271 | 298 | 287 | 0.890813608 | 1 | 0.53 | 0.559 | 0.531 | 0.574 | 0.621 | 0.538 | 0.544 | 0.529 | 0.637 | 0.573 | -0.001 |
| MXE | 5350 | ENSG00000175110 | MRPS22 | chr3 | + | 139350178 | 139351060 | 139352646 | 139352792 | 139348159 | 139348324 | 139355681 | 139355790 | 79 | 66 | 77 | 82 | 62 | 233 | 167 | 171 | 166 | 286 | 141 | 80 | 82 | 99 | 20 | 274 | 177 | 232 | 257 | 122 | 298 | 295 | 0.890844251 | 1 | 0.251 | 0.281 | 0.308 | 0.328 | 0.177 | 0.337 | 0.309 | 0.259 | 0.276 | 0.14 | 0.005 |
| MXE | 6825 | ENSG00000007341 | ST7L | chr1 | - | 112555867 | 112556018 | 112576985 | 112577088 | 112541950 | 112542090 | 112578344 | 112578417 | 10 | 12 | 25 | 13 | 37 | 0 | 5 | 0 | 0 | 0 | 20 | 13 | 10 | 24 | 23 | 0 | 0 | 0 | 4 | 0 | 252 | 298 | 0.890850659 | 1 | 1 | 0.739 | 1 | 1 | 1 | 1 | 1 | 1 | 0.876 | 1 | -0.027 |
| MXE | 8457 | ENSG00000137275 | RIPK1 | chr6 | + | 3080978 | 3081116 | 3083084 | 3083313 | 3077778 | 3077935 | 3085258 | 3085408 | 133 | 60 | 86 | 104 | 88 | 60 | 33 | 43 | 78 | 48 | 88 | 79 | 73 | 48 | 37 | 22 | 44 | 36 | 44 | 36 | 287 | 298 | 0.891011843 | 1 | 0.697 | 0.654 | 0.675 | 0.581 | 0.656 | 0.806 | 0.651 | 0.678 | 0.531 | 0.516 | 0.016 |
| MXE | 3429 | ENSG00000116459 | ATP5PB | chr1 | + | 111454210 | 111454356 | 111456226 | 111456249 | 111449836 | 111449873 | 111456629 | 111456755 | 238 | 111 | 185 | 183 | 218 | 64 | 41 | 59 | 46 | 119 | 71 | 134 | 160 | 74 | 56 | 13 | 45 | 32 | 44 | 29 | 295 | 172 | 0.891510704 | 1 | 0.684 | 0.612 | 0.646 | 0.699 | 0.516 | 0.761 | 0.635 | 0.745 | 0.495 | 0.53 | -0.002 |
| MXE | 2178 | ENSG00000137812 | KNL1 | chr15 | + | 40629272 | 40629371 | 40640911 | 40641027 | 40628610 | 40628678 | 40644996 | 40645087 | 8 | 15 | 13 | 23 | 11 | 22 | 22 | 29 | 34 | 74 | 48 | 11 | 10 | 13 | 6 | 76 | 22 | 28 | 41 | 37 | 248 | 265 | 0.892198497 | 1 | 0.28 | 0.421 | 0.324 | 0.42 | 0.137 | 0.403 | 0.348 | 0.276 | 0.253 | 0.148 | 0.031 |
| MXE | 841 | ENSG00000171456 | ASXL1 | chr20 | + | 32431582 | 32431679 | 32432879 | 32432985 | 32428763 | 32431484 | 32433283 | 32438415 | 69 | 42 | 100 | 97 | 57 | 133 | 100 | 139 | 113 | 182 | 118 | 106 | 73 | 110 | 127 | 243 | 190 | 158 | 170 | 258 | 246 | 255 | 0.892384373 | 1 | 0.35 | 0.303 | 0.427 | 0.471 | 0.245 | 0.335 | 0.366 | 0.324 | 0.401 | 0.338 | 0.006 |
| MXE | 16321 | ENSG00000139620 | KANSL2 | chr12 | - | 48679035 | 48679150 | 48679654 | 48679833 | 48671798 | 48671962 | 48681381 | 48681641 | 87 | 53 | 84 | 101 | 86 | 64 | 37 | 76 | 73 | 82 | 111 | 121 | 60 | 98 | 136 | 94 | 83 | 75 | 91 | 63 | 298 | 264 | 0.89245057 | 1 | 0.546 | 0.559 | 0.495 | 0.551 | 0.482 | 0.511 | 0.564 | 0.415 | 0.488 | 0.657 | 0 |
| MXE | 2420 | ENSG00000102921 | N4BP1 | chr16 | - | 48546146 | 48546254 | 48551385 | 48551482 | 48538725 | 48543261 | 48553538 | 48553669 | 76 | 49 | 58 | 68 | 122 | 135 | 103 | 129 | 123 | 283 | 39 | 54 | 37 | 43 | 19 | 30 | 135 | 132 | 71 | 47 | 246 | 257 | 0.892464162 | 1 | 0.37 | 0.332 | 0.32 | 0.366 | 0.311 | 0.576 | 0.295 | 0.227 | 0.388 | 0.297 | -0.017 |
| MXE | 5919 | ENSG00000003400 | CASP10 | chr2 | + | 201187705 | 201187799 | 201192983 | 201193119 | 201185770 | 201186124 | 201203729 | 201203766 | 57 | 21 | 33 | 41 | 48 | 4 | 1 | 4 | 0 | 8 | 32 | 20 | 29 | 79 | 5 | 0 | 2 | 4 | 8 | 0 | 243 | 285 | 0.892892862 | 1 | 0.944 | 0.961 | 0.906 | 1 | 0.876 | 1 | 0.921 | 0.895 | 0.921 | 1 | -0.01 |
| MXE | 2929 | ENSG00000124789 | NUP153 | chr6 | - | 17665238 | 17665385 | 17668974 | 17669028 | 17662017 | 17662070 | 17669292 | 17669337 | 104 | 78 | 110 | 121 | 127 | 210 | 183 | 236 | 292 | 393 | 69 | 118 | 186 | 132 | 32 | 217 | 325 | 374 | 248 | 99 | 203 | 296 | 0.893115056 | 1 | 0.419 | 0.383 | 0.405 | 0.377 | 0.32 | 0.317 | 0.346 | 0.42 | 0.437 | 0.32 | 0.013 |
| MXE | 11849 | ENSG00000164116 | GUCY1A1 | chr4 | + | 155703931 | 155703993 | 155708235 | 155708294 | 155696755 | 155697122 | 155710541 | 155711251 | 4 | 30 | 4 | 43 | 28 | 2 | 11 | 2 | 13 | 24 | 31 | 18 | 12 | 52 | 180 | 12 | 16 | 4 | 13 | 91 | 211 | 208 | 0.893203562 | 1 | 0.663 | 0.729 | 0.663 | 0.765 | 0.535 | 0.718 | 0.526 | 0.747 | 0.798 | 0.661 | -0.019 |
| MXE | 4488 | ENSG00000136051 | WASHC4 | chr12 | + | 105133769 | 105133896 | 105137885 | 105138011 | 105127128 | 105127289 | 105140293 | 105140401 | 221 | 168 | 148 | 188 | 327 | 276 | 302 | 251 | 285 | 553 | 3 | 55 | 31 | 12 | 3 | 6 | 90 | 24 | 10 | 25 | 276 | 275 | 0.893382876 | 1 | 0.444 | 0.357 | 0.37 | 0.397 | 0.371 | 0.333 | 0.378 | 0.563 | 0.545 | 0.107 | 0.003 |
| MXE | 2275 | ENSG00000048471 | SNX29 | chr16 | + | 12002990 | 12003043 | 12027319 | 12027444 | 11999296 | 11999358 | 12042896 | 12043077 | 8 | 5 | 13 | 3 | 0 | 34 | 20 | 26 | 31 | 47 | 10 | 12 | 20 | 9 | 10 | 97 | 41 | 62 | 120 | 91 | 202 | 274 | 0.893486325 | 1 | 0.242 | 0.253 | 0.404 | 0.116 | 0 | 0.123 | 0.284 | 0.304 | 0.092 | 0.13 | 0.016 |
| MXE | 8119 | ENSG00000145020 | AMT | chr3 | - | 49418970 | 49419151 | 49419259 | 49421572 | 49417817 | 49417973 | 49422103 | 49422271 | 17 | 0 | 8 | 5 | 0 | 32 | 19 | 25 | 16 | 11 | 0 | 14 | 13 | 7 | 8 | 41 | 31 | 25 | 22 | 74 | 298 | 298 | 0.893647764 | 1 | 0.347 | 0 | 0.242 | 0.238 | 0 | 0 | 0.311 | 0.342 | 0.241 | 0.098 | -0.033 |
| MXE | 4355 | ENSG00000163625 | WDFY3 | chr4 | - | 84688085 | 84688265 | 84690505 | 84690664 | 84683942 | 84684125 | 84691630 | 84691785 | 46 | 21 | 25 | 25 | 57 | 40 | 23 | 19 | 16 | 21 | 45 | 27 | 28 | 78 | 11 | 11 | 35 | 43 | 37 | 5 | 298 | 298 | 0.894086811 | 1 | 0.535 | 0.477 | 0.568 | 0.61 | 0.731 | 0.804 | 0.435 | 0.394 | 0.678 | 0.688 | -0.016 |
| MXE | 1763 | ENSG00000214078 | CPNE1 | chr20 | - | 35630438 | 35630490 | 35630740 | 35630795 | 35627279 | 35627413 | 35630900 | 35631034 | 50 | 34 | 45 | 39 | 40 | 99 | 38 | 110 | 31 | 55 | 42 | 33 | 19 | 44 | 86 | 15 | 103 | 47 | 70 | 148 | 204 | 201 | 0.894101404 | 1 | 0.332 | 0.469 | 0.287 | 0.553 | 0.417 | 0.734 | 0.24 | 0.285 | 0.382 | 0.364 | 0.011 |
| MXE | 13527 | ENSG00000134516 | DOCK2 | chr5 | + | 170034398 | 170034555 | 170036514 | 170036555 | 170027862 | 170027948 | 170041054 | 170041145 | 551 | 411 | 483 | 570 | 659 | 95 | 80 | 76 | 98 | 175 | 1185 | 561 | 637 | 751 | 369 | 148 | 148 | 108 | 133 | 96 | 298 | 190 | 0.894166965 | 1 | 0.787 | 0.766 | 0.802 | 0.788 | 0.706 | 0.836 | 0.707 | 0.79 | 0.783 | 0.71 | 0.005 |
| MXE | 15682 | ENSG00000165916 | PSMC3 | chr11 | - | 47420263 | 47420409 | 47422573 | 47422722 | 47419115 | 47419197 | 47422829 | 47422973 | 56 | 16 | 51 | 48 | 31 | 175 | 98 | 166 | 153 | 96 | 64 | 44 | 45 | 47 | 35 | 236 | 157 | 95 | 139 | 223 | 298 | 295 | 0.89425153 | 1 | 0.241 | 0.139 | 0.233 | 0.237 | 0.242 | 0.212 | 0.217 | 0.319 | 0.251 | 0.134 | -0.008 |
| MXE | 8777 | ENSG00000134313 | KIDINS220 | chr2 | - | 8733443 | 8733680 | 8734654 | 8734753 | 8728857 | 8731982 | 8736867 | 8736999 | 80 | 82 | 65 | 77 | 115 | 96 | 98 | 104 | 133 | 186 | 45 | 67 | 41 | 49 | 12 | 17 | 113 | 111 | 108 | 14 | 248 | 298 | 0.894280364 | 1 | 0.5 | 0.501 | 0.429 | 0.41 | 0.426 | 0.761 | 0.416 | 0.307 | 0.353 | 0.507 | -0.016 |
| MXE | 8441 | ENSG00000121964 | GTDC1 | chr2 | - | 143957194 | 143957339 | 144007181 | 144007535 | 143952772 | 143952843 | 144141881 | 144142055 | 61 | 53 | 65 | 57 | 102 | 53 | 30 | 33 | 33 | 55 | 26 | 29 | 48 | 29 | 38 | 28 | 4 | 18 | 56 | 16 | 298 | 294 | 0.89438391 | 1 | 0.532 | 0.635 | 0.66 | 0.63 | 0.647 | 0.478 | 0.877 | 0.725 | 0.338 | 0.701 | -0.003 |
| MXE | 8318 | ENSG00000177409 | SAMD9L | chr7 | - | 93144731 | 93144833 | 93146882 | 93147146 | 93130055 | 93135991 | 93148193 | 93148385 | 131 | 89 | 97 | 179 | 111 | 220 | 206 | 162 | 261 | 115 | 38 | 50 | 15 | 43 | 0 | 23 | 90 | 51 | 23 | 4 | 298 | 251 | 0.894430878 | 1 | 0.334 | 0.267 | 0.335 | 0.366 | 0.448 | 0.582 | 0.319 | 0.199 | 0.612 | 0 | 0.008 |
| MXE | 7846 | ENSG00000204217 | BMPR2 | chr2 | + | 202467518 | 202467689 | 202513718 | 202513829 | 202464808 | 202464979 | 202514887 | 202514979 | 91 | 59 | 54 | 76 | 170 | 49 | 42 | 28 | 47 | 69 | 49 | 82 | 24 | 65 | 134 | 68 | 19 | 28 | 13 | 61 | 298 | 260 | 0.894783086 | 1 | 0.618 | 0.551 | 0.627 | 0.585 | 0.682 | 0.386 | 0.79 | 0.428 | 0.814 | 0.657 | -0.002 |
| MXE | 8754 | ENSG00000101040 | ZMYND8 | chr20 | - | 47227202 | 47227281 | 47229725 | 47229806 | 47224316 | 47224556 | 47236325 | 47236516 | 35 | 17 | 26 | 25 | 21 | 29 | 24 | 31 | 39 | 86 | 2 | 49 | 17 | 32 | 32 | 45 | 27 | 48 | 51 | 6 | 230 | 228 | 0.895033878 | 1 | 0.545 | 0.413 | 0.454 | 0.389 | 0.195 | 0.042 | 0.643 | 0.26 | 0.383 | 0.841 | -0.035 |
| MXE | 8891 | ENSG00000160688 | FLAD1 | chr1 | + | 154992564 | 154992673 | 154992712 | 154992786 | 154990158 | 154990257 | 154992901 | 154993111 | 2 | 0 | 3 | 0 | 0 | 26 | 6 | 21 | 7 | 7 | 0 | 0 | 2 | 1 | 1 | 2 | 41 | 12 | 16 | 54 | 258 | 223 | 0.89553943 | 1 | 0.062 | 0 | 0.11 | 0 | 0 | 0 | 0 | 0.126 | 0.051 | 0.016 | -0.004 |
| MXE | 12460 | ENSG00000102531 | FNDC3A | chr13 | + | 49075288 | 49075364 | 49114654 | 49114731 | 49006151 | 49006289 | 49131136 | 49131374 | 20 | 24 | 23 | 38 | 41 | 79 | 73 | 75 | 93 | 128 | 31 | 33 | 21 | 5 | 2 | 29 | 87 | 51 | 41 | 20 | 225 | 226 | 0.895650956 | 1 | 0.203 | 0.248 | 0.235 | 0.291 | 0.243 | 0.518 | 0.276 | 0.293 | 0.109 | 0.091 | -0.013 |
| MXE | 4806 | ENSG00000089220 | PEBP1 | chr12 | + | 118138038 | 118138148 | 118139450 | 118139551 | 118136123 | 118136344 | 118144585 | 118145584 | 48 | 7 | 50 | 42 | 23 | 162 | 119 | 209 | 169 | 263 | 18 | 41 | 48 | 16 | 38 | 143 | 210 | 140 | 116 | 278 | 259 | 250 | 0.895687848 | 1 | 0.222 | 0.054 | 0.188 | 0.193 | 0.078 | 0.108 | 0.159 | 0.249 | 0.117 | 0.117 | -0.003 |
| MXE | 13670 | ENSG00000215301 | DDX3X | chrX | + | 41339035 | 41339083 | 41340635 | 41340814 | 41337407 | 41337465 | 41341483 | 41341616 | 1368 | 729 | 1093 | 1282 | 615 | 8 | 19 | 14 | 26 | 16 | 215 | 628 | 801 | 511 | 208 | 7 | 9 | 17 | 13 | 0 | 197 | 298 | 0.895838429 | 1 | 0.996 | 0.983 | 0.992 | 0.987 | 0.983 | 0.979 | 0.991 | 0.986 | 0.983 | 1 | 0 |
| MXE | 4421 | ENSG00000109756 | RAPGEF2 | chr4 | + | 159304341 | 159304473 | 159314590 | 159314768 | 159243773 | 159243791 | 159322346 | 159322363 | 77 | 113 | 80 | 91 | 111 | 43 | 35 | 33 | 40 | 51 | 46 | 115 | 175 | 83 | 13 | 22 | 46 | 76 | 43 | 3 | 281 | 298 | 0.896168651 | 1 | 0.655 | 0.774 | 0.72 | 0.707 | 0.698 | 0.689 | 0.726 | 0.709 | 0.672 | 0.821 | -0.013 |
| MXE | 839 | ENSG00000105576 | TNPO2 | chr19 | - | 12705491 | 12705599 | 12705681 | 12705768 | 12705239 | 12705398 | 12706195 | 12706367 | 8 | 5 | 11 | 25 | 5 | 29 | 15 | 28 | 20 | 22 | 23 | 5 | 6 | 3 | 57 | 41 | 14 | 20 | 19 | 84 | 236 | 257 | 0.896212217 | 1 | 0.231 | 0.266 | 0.3 | 0.576 | 0.198 | 0.379 | 0.28 | 0.246 | 0.147 | 0.425 | 0.019 |
| MXE | 16216 | ENSG00000136754 | ABI1 | chr10 | - | 26755654 | 26755741 | 26759061 | 26759235 | 26751597 | 26751783 | 26765217 | 26765318 | 217 | 216 | 177 | 212 | 438 | 1 | 7 | 1 | 0 | 5 | 242 | 405 | 431 | 245 | 129 | 0 | 1 | 1 | 0 | 0 | 298 | 236 | 0.896252934 | 1 | 0.994 | 0.961 | 0.993 | 1 | 0.986 | 1 | 0.997 | 0.997 | 1 | 1 | -0.012 |
| MXE | 6844 | ENSG00000230124 | ACBD6 | chr1 | - | 180288269 | 180288517 | 180314691 | 180314722 | 180281305 | 180281386 | 180397515 | 180397605 | 20 | 14 | 24 | 19 | 42 | 0 | 9 | 1 | 7 | 0 | 14 | 6 | 17 | 8 | 10 | 0 | 15 | 2 | 0 | 0 | 180 | 298 | 0.896434094 | 1 | 1 | 0.72 | 0.975 | 0.818 | 1 | 1 | 0.398 | 0.934 | 1 | 1 | 0.036 |
| MXE | 7991 | ENSG00000153283 | CD96 | chr3 | + | 111600725 | 111600914 | 111606699 | 111606792 | 111598119 | 111598210 | 111623753 | 111623822 | 498 | 414 | 477 | 553 | 1008 | 167 | 139 | 204 | 177 | 465 | 53 | 215 | 256 | 90 | 50 | 9 | 97 | 107 | 28 | 25 | 298 | 242 | 0.896955181 | 1 | 0.708 | 0.707 | 0.655 | 0.717 | 0.638 | 0.827 | 0.643 | 0.66 | 0.723 | 0.619 | -0.009 |
| MXE | 14709 | ENSG00000196116 | TDRD7 | chr9 | + | 97464841 | 97465028 | 97472292 | 97472495 | 97460177 | 97460764 | 97473491 | 97473626 | 58 | 63 | 37 | 45 | 104 | 50 | 45 | 35 | 48 | 69 | 48 | 75 | 37 | 80 | 1 | 31 | 56 | 16 | 37 | 10 | 298 | 298 | 0.896979833 | 1 | 0.537 | 0.583 | 0.514 | 0.484 | 0.601 | 0.608 | 0.573 | 0.698 | 0.684 | 0.091 | 0.013 |
| MXE | 1929 | ENSG00000131797 | CLUHP3 | chr16 | + | 31705348 | 31705472 | 31705671 | 31705792 | 31704219 | 31704390 | 31705889 | 31706034 | 10 | 5 | 9 | 16 | 14 | 15 | 6 | 27 | 38 | 34 | 10 | 4 | 11 | 10 | 11 | 14 | 6 | 30 | 10 | 44 | 273 | 270 | 0.897223918 | 1 | 0.397 | 0.452 | 0.248 | 0.294 | 0.289 | 0.414 | 0.397 | 0.266 | 0.497 | 0.198 | -0.018 |
| MXE | 483 | ENSG00000012061 | ERCC1 | chr19 | - | 45419097 | 45419197 | 45420323 | 45420427 | 45416490 | 45416897 | 45421177 | 45421393 | 39 | 27 | 46 | 21 | 17 | 64 | 44 | 54 | 52 | 49 | 59 | 32 | 15 | 33 | 39 | 110 | 70 | 31 | 51 | 67 | 253 | 249 | 0.897399672 | 1 | 0.375 | 0.377 | 0.456 | 0.284 | 0.255 | 0.345 | 0.31 | 0.323 | 0.389 | 0.364 | 0.003 |
| MXE | 1272 | ENSG00000108439 | PNPO | chr17 | + | 47945558 | 47945612 | 47945860 | 47945989 | 47944615 | 47944715 | 47946322 | 47946393 | 17 | 5 | 17 | 5 | 4 | 24 | 10 | 31 | 29 | 18 | 4 | 29 | 12 | 17 | 12 | 23 | 42 | 25 | 24 | 56 | 203 | 278 | 0.897734327 | 1 | 0.492 | 0.406 | 0.429 | 0.191 | 0.233 | 0.192 | 0.486 | 0.397 | 0.492 | 0.227 | -0.009 |
| MXE | 5422 | ENSG00000078070 | MCCC1 | chr3 | - | 183017265 | 183017337 | 183020129 | 183020237 | 183015217 | 183015566 | 183022416 | 183022554 | 60 | 34 | 39 | 63 | 62 | 26 | 12 | 22 | 21 | 56 | 14 | 51 | 52 | 45 | 19 | 24 | 38 | 17 | 16 | 2 | 257 | 221 | 0.897740749 | 1 | 0.665 | 0.709 | 0.604 | 0.721 | 0.488 | 0.334 | 0.536 | 0.725 | 0.707 | 0.891 | -0.001 |
| MXE | 2155 | ENSG00000114062 | UBE3A | chr15 | - | 25407066 | 25407212 | 25408571 | 25408684 | 25405460 | 25405502 | 25409087 | 25409207 | 79 | 73 | 81 | 127 | 151 | 21 | 13 | 11 | 14 | 26 | 148 | 90 | 78 | 62 | 249 | 43 | 26 | 3 | 19 | 12 | 262 | 295 | 0.898602732 | 1 | 0.809 | 0.863 | 0.892 | 0.911 | 0.867 | 0.795 | 0.796 | 0.967 | 0.786 | 0.959 | 0.008 |
| MXE | 14967 | ENSG00000070476 | ZXDC | chr3 | - | 126441764 | 126441946 | 126459652 | 126459737 | 126439631 | 126439727 | 126461534 | 126462220 | 31 | 16 | 12 | 26 | 14 | 8 | 5 | 11 | 6 | 24 | 50 | 21 | 25 | 22 | 20 | 31 | 22 | 11 | 6 | 7 | 234 | 298 | 0.898722301 | 1 | 0.832 | 0.803 | 0.581 | 0.847 | 0.426 | 0.673 | 0.549 | 0.743 | 0.824 | 0.784 | -0.017 |
| MXE | 8280 | ENSG00000169554 | ZEB2 | chr2 | - | 144404835 | 144405024 | 144424795 | 144424864 | 144403915 | 144404130 | 144429768 | 144430026 | 0 | 0 | 2 | 2 | 0 | 1100 | 855 | 469 | 718 | 803 | 0 | 5 | 5 | 11 | 3 | 437 | 989 | 1138 | 479 | 304 | 218 | 298 | 0.89872589 | 1 | 0 | 0 | 0.006 | 0.004 | 0 | 0 | 0.007 | 0.006 | 0.03 | 0.013 | -0.009 |
| MXE | 11180 | ENSG00000115053 | NCL | chr2 | - | 231457642 | 231457800 | 231459000 | 231459125 | 231457000 | 231457124 | 231460151 | 231460293 | 792 | 599 | 756 | 748 | 1214 | 667 | 459 | 600 | 604 | 816 | 550 | 574 | 611 | 647 | 103 | 430 | 343 | 450 | 413 | 134 | 274 | 298 | 0.898790556 | 1 | 0.564 | 0.587 | 0.578 | 0.574 | 0.618 | 0.582 | 0.645 | 0.596 | 0.63 | 0.455 | 0.003 |
| MXE | 13225 | ENSG00000188917 | TRMT2B | chrX | - | 101036973 | 101037073 | 101041316 | 101041371 | 101035612 | 101035683 | 101042232 | 101042312 | 8 | 3 | 5 | 12 | 4 | 59 | 29 | 62 | 50 | 74 | 0 | 3 | 12 | 2 | 14 | 45 | 50 | 22 | 41 | 68 | 204 | 249 | 0.898813153 | 1 | 0.142 | 0.112 | 0.09 | 0.227 | 0.062 | 0 | 0.068 | 0.4 | 0.056 | 0.201 | -0.018 |
| MXE | 5672 | ENSG00000143862 | ARL8A | chr1 | - | 202135149 | 202135220 | 202135458 | 202135526 | 202133403 | 202134516 | 202135706 | 202135800 | 69 | 25 | 39 | 67 | 50 | 83 | 52 | 72 | 86 | 182 | 80 | 64 | 83 | 105 | 104 | 257 | 91 | 143 | 201 | 160 | 217 | 220 | 0.898824717 | 1 | 0.457 | 0.328 | 0.354 | 0.441 | 0.218 | 0.24 | 0.416 | 0.37 | 0.346 | 0.397 | 0.006 |
| MXE | 7417 | ENSG00000153922 | CHD1 | chr5 | - | 98858179 | 98858390 | 98858963 | 98859015 | 98853984 | 98856725 | 98859971 | 98860068 | 143 | 129 | 113 | 137 | 264 | 123 | 128 | 100 | 126 | 194 | 23 | 44 | 73 | 10 | 14 | 16 | 54 | 80 | 2 | 12 | 201 | 298 | 0.899162662 | 1 | 0.633 | 0.599 | 0.626 | 0.617 | 0.669 | 0.681 | 0.547 | 0.575 | 0.881 | 0.634 | -0.035 |
| MXE | 11672 | ENSG00000024048 | UBR2 | chr6 | + | 42615070 | 42615178 | 42616001 | 42616090 | 42612170 | 42612291 | 42617408 | 42617507 | 111 | 104 | 114 | 91 | 202 | 57 | 66 | 57 | 62 | 90 | 14 | 132 | 129 | 77 | 30 | 28 | 46 | 72 | 33 | 12 | 257 | 238 | 0.89918927 | 1 | 0.643 | 0.593 | 0.649 | 0.576 | 0.675 | 0.316 | 0.727 | 0.624 | 0.684 | 0.698 | 0.017 |
| MXE | 4080 | ENSG00000157869 | RAB28 | chr4 | - | 13381490 | 13381594 | 13460698 | 13460828 | 13376544 | 13376622 | 13474317 | 13474406 | 67 | 53 | 63 | 80 | 70 | 71 | 35 | 49 | 60 | 106 | 58 | 31 | 56 | 35 | 17 | 30 | 45 | 33 | 25 | 35 | 279 | 253 | 0.89967342 | 1 | 0.461 | 0.579 | 0.538 | 0.547 | 0.375 | 0.637 | 0.384 | 0.606 | 0.559 | 0.306 | 0.002 |
| MXE | 11721 | ENSG00000135316 | SYNCRIP | chr6 | - | 85623976 | 85624112 | 85636966 | 85637143 | 85622481 | 85622687 | 85637242 | 85637356 | 459 | 407 | 459 | 546 | 1047 | 616 | 524 | 558 | 669 | 1174 | 170 | 368 | 355 | 256 | 35 | 251 | 403 | 427 | 316 | 50 | 298 | 285 | 0.90002063 | 1 | 0.416 | 0.426 | 0.44 | 0.438 | 0.46 | 0.393 | 0.466 | 0.443 | 0.437 | 0.401 | 0.008 |
| MXE | 112 | ENSG00000198911 | SREBF2 | chr22 | + | 41877228 | 41877421 | 41878658 | 41878758 | 41875542 | 41875724 | 41880715 | 41880992 | 156 | 49 | 118 | 147 | 202 | 2 | 0 | 4 | 3 | 4 | 164 | 119 | 66 | 143 | 381 | 0 | 4 | 3 | 1 | 7 | 298 | 249 | 0.900022196 | 1 | 0.985 | 1 | 0.961 | 0.976 | 0.977 | 1 | 0.961 | 0.948 | 0.992 | 0.978 | 0.004 |
| MXE | 12119 | ENSG00000196923 | PDLIM7 | chr5 | - | 177491403 | 177491420 | 177491976 | 177491976 | 177491009 | 177491146 | 177492404 | 177492435 | 18 | 2 | 12 | 13 | 20 | 5 | 0 | 1 | 1 | 8 | 59 | 34 | 19 | 42 | 89 | 21 | 2 | 2 | 2 | 28 | 298 | 166 | 0.900073484 | 1 | 0.667 | 1 | 0.87 | 0.879 | 0.582 | 0.61 | 0.904 | 0.841 | 0.921 | 0.639 | 0.017 |
| MXE | 376 | ENSG00000104880 | ARHGEF18 | chr19 | + | 7459902 | 7459994 | 7462151 | 7462334 | 7458511 | 7458690 | 7463817 | 7463955 | 172 | 130 | 226 | 186 | 196 | 199 | 125 | 181 | 173 | 87 | 163 | 219 | 172 | 178 | 246 | 117 | 192 | 89 | 128 | 333 | 241 | 298 | 0.900089333 | 1 | 0.517 | 0.563 | 0.607 | 0.571 | 0.736 | 0.633 | 0.585 | 0.705 | 0.632 | 0.477 | -0.008 |
| MXE | 16154 | ENSG00000108094 | CUL2 | chr10 | - | 35044771 | 35044868 | 35049682 | 35049765 | 35044565 | 35044676 | 35054433 | 35054539 | 41 | 39 | 28 | 31 | 63 | 94 | 63 | 72 | 91 | 145 | 10 | 35 | 29 | 11 | 0 | 11 | 62 | 66 | 38 | 5 | 232 | 246 | 0.900212049 | 1 | 0.316 | 0.396 | 0.292 | 0.265 | 0.315 | 0.491 | 0.374 | 0.318 | 0.235 | 0 | 0.033 |
| MXE | 2153 | ENSG00000114062 | UBE3A | chr15 | - | 25407066 | 25407137 | 25408571 | 25408684 | 25405460 | 25405502 | 25409087 | 25409207 | 79 | 73 | 81 | 127 | 151 | 8 | 4 | 7 | 9 | 8 | 148 | 90 | 78 | 62 | 249 | 27 | 13 | 3 | 4 | 0 | 262 | 220 | 0.900428539 | 1 | 0.892 | 0.939 | 0.907 | 0.922 | 0.941 | 0.822 | 0.853 | 0.956 | 0.929 | 1 | 0.008 |
| MXE | 5974 | ENSG00000197948 | FCHSD1 | chr5 | - | 141642396 | 141642503 | 141643088 | 141643200 | 141641701 | 141641757 | 141644217 | 141644438 | 98 | 28 | 99 | 93 | 121 | 7 | 1 | 2 | 2 | 16 | 68 | 108 | 38 | 55 | 119 | 0 | 1 | 6 | 0 | 20 | 237 | 256 | 0.9005343 | 1 | 0.938 | 0.968 | 0.982 | 0.98 | 0.891 | 1 | 0.992 | 0.872 | 1 | 0.865 | 0.006 |
| MXE | 3660 | ENSG00000113719 | ERGIC1 | chr5 | + | 172924004 | 172924109 | 172926508 | 172926569 | 172914564 | 172914838 | 172932435 | 172932478 | 105 | 61 | 86 | 104 | 75 | 42 | 16 | 15 | 21 | 6 | 72 | 193 | 134 | 79 | 122 | 17 | 49 | 28 | 12 | 23 | 254 | 210 | 0.900553041 | 1 | 0.674 | 0.759 | 0.826 | 0.804 | 0.912 | 0.778 | 0.765 | 0.798 | 0.845 | 0.814 | -0.005 |
| MXE | 8108 | ENSG00000130939 | UBE4B | chr1 | + | 10144939 | 10145039 | 10146962 | 10147090 | 10137066 | 10137205 | 10149183 | 10149282 | 27 | 18 | 29 | 28 | 69 | 40 | 20 | 26 | 35 | 32 | 9 | 23 | 32 | 15 | 24 | 12 | 25 | 8 | 37 | 23 | 249 | 277 | 0.900696434 | 1 | 0.429 | 0.5 | 0.554 | 0.471 | 0.706 | 0.455 | 0.506 | 0.817 | 0.311 | 0.537 | 0.007 |
| MXE | 16636 | ENSG00000076513 | ANKRD13A | chr12 | + | 110021107 | 110021301 | 110024045 | 110024112 | 110019138 | 110019328 | 110025741 | 110025823 | 6 | 4 | 3 | 10 | 4 | 1025 | 795 | 967 | 1154 | 1958 | 4 | 4 | 16 | 31 | 0 | 2994 | 1309 | 1270 | 1490 | 2344 | 298 | 216 | 0.901042474 | 1 | 0.004 | 0.004 | 0.002 | 0.006 | 0.001 | 0.001 | 0.002 | 0.009 | 0.015 | 0 | -0.002 |
| MXE | 7509 | ENSG00000115020 | PIKFYVE | chr2 | + | 208274020 | 208274053 | 208276711 | 208276830 | 208273583 | 208273733 | 208285725 | 208285933 | 1 | 8 | 2 | 3 | 1 | 73 | 53 | 53 | 80 | 71 | 1 | 3 | 7 | 4 | 0 | 77 | 89 | 72 | 50 | 64 | 182 | 268 | 0.901297338 | 1 | 0.02 | 0.182 | 0.053 | 0.052 | 0.02 | 0.019 | 0.047 | 0.125 | 0.105 | 0 | 0.006 |
| MXE | 790 | ENSG00000154930 | ACSS1 | chr20 | - | 25022939 | 25023092 | 25030758 | 25030958 | 25021388 | 25021536 | 25048084 | 25048181 | 42 | 27 | 35 | 35 | 19 | 104 | 42 | 91 | 74 | 44 | 47 | 39 | 15 | 25 | 69 | 104 | 52 | 68 | 53 | 139 | 298 | 298 | 0.901539251 | 1 | 0.288 | 0.391 | 0.278 | 0.321 | 0.302 | 0.311 | 0.429 | 0.181 | 0.321 | 0.332 | 0.001 |
| MXE | 16159 | ENSG00000087338 | GMCL1 | chr2 | + | 69847542 | 69847627 | 69849651 | 69849742 | 69844130 | 69844196 | 69854822 | 69854860 | 22 | 13 | 42 | 32 | 51 | 74 | 33 | 47 | 64 | 81 | 8 | 20 | 27 | 3 | 0 | 0 | 19 | 25 | 36 | 8 | 234 | 240 | 0.901646563 | 1 | 0.234 | 0.288 | 0.478 | 0.339 | 0.392 | 1 | 0.519 | 0.526 | 0.079 | 0 | -0.079 |
| MXE | 8286 | ENSG00000112303 | VNN2 | chr6 | - | 132752460 | 132752577 | 132753417 | 132753491 | 132751144 | 132751518 | 132755842 | 132756035 | 0 | 0 | 1 | 5 | 0 | 233 | 90 | 220 | 244 | 539 | 0 | 8 | 4 | 0 | 0 | 111 | 377 | 251 | 291 | 19 | 223 | 266 | 0.901719434 | 1 | 0 | 0 | 0.005 | 0.024 | 0 | 0 | 0.025 | 0.019 | 0 | 0 | -0.003 |
| MXE | 2291 | ENSG00000128908 | INO80 | chr15 | - | 41074369 | 41074565 | 41079700 | 41079904 | 41073427 | 41073495 | 41081019 | 41081073 | 127 | 68 | 110 | 105 | 178 | 92 | 70 | 76 | 102 | 141 | 123 | 145 | 120 | 171 | 113 | 171 | 120 | 124 | 122 | 41 | 298 | 298 | 0.901774245 | 1 | 0.58 | 0.493 | 0.591 | 0.507 | 0.558 | 0.418 | 0.547 | 0.492 | 0.584 | 0.734 | -0.009 |
| MXE | 14505 | ENSG00000134333 | LDHA | chr11 | + | 18399430 | 18399548 | 18400836 | 18401010 | 18396818 | 18396968 | 18402839 | 18403013 | 626 | 471 | 537 | 639 | 888 | 646 | 332 | 699 | 692 | 806 | 1547 | 866 | 928 | 1322 | 797 | 1324 | 1052 | 752 | 1566 | 802 | 267 | 298 | 0.902028948 | 1 | 0.52 | 0.613 | 0.462 | 0.508 | 0.551 | 0.566 | 0.479 | 0.579 | 0.485 | 0.526 | 0.004 |
| MXE | 2700 | ENSG00000100603 | SNW1 | chr14 | - | 77734946 | 77735012 | 77735936 | 77736006 | 77732484 | 77732601 | 77736970 | 77737007 | 238 | 227 | 235 | 289 | 255 | 61 | 58 | 64 | 56 | 82 | 125 | 291 | 361 | 269 | 60 | 71 | 60 | 107 | 63 | 3 | 219 | 215 | 0.902398594 | 1 | 0.793 | 0.793 | 0.783 | 0.835 | 0.753 | 0.633 | 0.826 | 0.768 | 0.807 | 0.952 | -0.006 |
| MXE | 16898 | ENSG00000188906 | LRRK2 | chr12 | + | 40351538 | 40351733 | 40354298 | 40354492 | 40348408 | 40348509 | 40356114 | 40356187 | 213 | 208 | 103 | 162 | 293 | 121 | 112 | 67 | 82 | 204 | 38 | 125 | 107 | 13 | 5 | 4 | 66 | 67 | 7 | 27 | 298 | 298 | 0.90264046 | 1 | 0.638 | 0.65 | 0.606 | 0.664 | 0.59 | 0.905 | 0.654 | 0.615 | 0.65 | 0.156 | 0.034 |
| MXE | 15244 | ENSG00000147457 | CHMP7 | chr8 | + | 23255246 | 23255432 | 23256459 | 23256593 | 23249209 | 23249381 | 23258032 | 23258081 | 84 | 56 | 111 | 82 | 86 | 73 | 56 | 109 | 102 | 61 | 32 | 88 | 40 | 55 | 95 | 63 | 54 | 34 | 40 | 92 | 298 | 283 | 0.902677979 | 1 | 0.522 | 0.487 | 0.492 | 0.433 | 0.572 | 0.325 | 0.607 | 0.528 | 0.566 | 0.495 | -0.003 |
| MXE | 10759 | ENSG00000177479 | ARIH2 | chr3 | + | 48945135 | 48945181 | 48961611 | 48961679 | 48927461 | 48927813 | 48964918 | 48964982 | 20 | 5 | 15 | 18 | 18 | 212 | 98 | 187 | 252 | 347 | 23 | 17 | 11 | 5 | 16 | 244 | 187 | 142 | 136 | 307 | 195 | 217 | 0.902750688 | 1 | 0.095 | 0.054 | 0.082 | 0.074 | 0.055 | 0.095 | 0.092 | 0.079 | 0.039 | 0.055 | 0 |
| MXE | 13040 | ENSG00000182487 | NCF1B | chr7 | + | 73229566 | 73230260 | 73231810 | 73231890 | 73227408 | 73227464 | 73234613 | 73234718 | 93 | 46 | 74 | 30 | 101 | 31 | 11 | 12 | 21 | 7 | 211 | 73 | 66 | 174 | 348 | 53 | 42 | 11 | 47 | 42 | 298 | 229 | 0.902767578 | 1 | 0.697 | 0.763 | 0.826 | 0.523 | 0.917 | 0.754 | 0.572 | 0.822 | 0.74 | 0.864 | -0.005 |
| MXE | 16889 | ENSG00000188906 | LRRK2 | chr12 | + | 40240482 | 40240617 | 40243549 | 40243681 | 40237968 | 40238103 | 40249825 | 40249945 | 306 | 272 | 144 | 169 | 334 | 199 | 204 | 106 | 139 | 397 | 81 | 170 | 179 | 44 | 14 | 52 | 153 | 120 | 42 | 22 | 284 | 281 | 0.902953518 | 1 | 0.603 | 0.569 | 0.573 | 0.546 | 0.454 | 0.606 | 0.524 | 0.596 | 0.509 | 0.386 | 0.025 |
| MXE | 5436 | ENSG00000145819 | ARHGAP26 | chr5 | + | 142885297 | 142885399 | 142894237 | 142894348 | 142879373 | 142879445 | 142901934 | 142902039 | 172 | 164 | 95 | 150 | 130 | 176 | 167 | 143 | 236 | 192 | 245 | 224 | 301 | 232 | 78 | 273 | 303 | 277 | 298 | 186 | 251 | 260 | 0.903248929 | 1 | 0.503 | 0.504 | 0.408 | 0.397 | 0.412 | 0.482 | 0.434 | 0.53 | 0.446 | 0.303 | 0.006 |
| MXE | 8014 | ENSG00000143970 | ASXL2 | chr2 | - | 25753533 | 25753639 | 25756017 | 25756114 | 25749695 | 25750413 | 25759481 | 25759645 | 93 | 61 | 63 | 70 | 83 | 149 | 110 | 116 | 86 | 194 | 12 | 132 | 42 | 85 | 20 | 80 | 136 | 77 | 103 | 22 | 246 | 255 | 0.903348215 | 1 | 0.393 | 0.365 | 0.36 | 0.458 | 0.307 | 0.135 | 0.502 | 0.361 | 0.461 | 0.485 | -0.012 |
| MXE | 16736 | ENSG00000133858 | ZFC3H1 | chr12 | - | 71619314 | 71619409 | 71619925 | 71620124 | 71615205 | 71615316 | 71620209 | 71620315 | 209 | 158 | 179 | 221 | 151 | 60 | 72 | 58 | 60 | 104 | 6 | 90 | 80 | 11 | 0 | 1 | 24 | 23 | 2 | 8 | 298 | 244 | 0.903754525 | 1 | 0.74 | 0.642 | 0.716 | 0.751 | 0.543 | 0.831 | 0.754 | 0.74 | 0.818 | 0 | 0.05 |
| MXE | 13519 | ENSG00000134516 | DOCK2 | chr5 | + | 169669287 | 169669328 | 169670541 | 169670597 | 169654402 | 169654486 | 169671077 | 169671174 | 61 | 20 | 42 | 28 | 6 | 252 | 182 | 159 | 173 | 207 | 14 | 63 | 21 | 61 | 45 | 315 | 247 | 294 | 299 | 181 | 190 | 205 | 0.903943933 | 1 | 0.207 | 0.106 | 0.222 | 0.149 | 0.03 | 0.046 | 0.216 | 0.072 | 0.18 | 0.212 | -0.002 |
| MXE | 8362 | ENSG00000135842 | NIBAN1 | chr1 | - | 184803584 | 184803692 | 184805945 | 184806056 | 184798078 | 184798190 | 184808073 | 184808235 | 66 | 51 | 73 | 92 | 121 | 198 | 179 | 169 | 203 | 392 | 118 | 112 | 115 | 99 | 52 | 335 | 238 | 386 | 291 | 138 | 260 | 257 | 0.903960765 | 1 | 0.248 | 0.22 | 0.299 | 0.309 | 0.234 | 0.258 | 0.317 | 0.227 | 0.252 | 0.271 | -0.003 |
| MXE | 12 | ENSG00000160255 | ITGB2 | chr21 | - | 44910724 | 44910785 | 44920820 | 44921050 | 44910283 | 44910372 | 44931813 | 44931989 | 0 | 0 | 0 | 0 | 0 | 75 | 18 | 17 | 35 | 9 | 0 | 0 | 0 | 8 | 0 | 51 | 86 | 24 | 66 | 87 | 298 | 210 | 0.903988599 | 1 | 0 | 0 | 0 | 0 | 0 | 0 | 0 | 0 | 0.079 | 0 | -0.016 |
| MXE | 7338 | ENSG00000113532 | ST8SIA4 | chr5 | - | 100886342 | 100886600 | 100895653 | 100895785 | 100856102 | 100856396 | 100902842 | 100903282 | 49 | 40 | 70 | 80 | 82 | 192 | 127 | 123 | 180 | 284 | 10 | 48 | 48 | 19 | 23 | 22 | 133 | 132 | 64 | 49 | 281 | 298 | 0.904285454 | 1 | 0.213 | 0.25 | 0.376 | 0.32 | 0.234 | 0.325 | 0.277 | 0.278 | 0.239 | 0.332 | -0.012 |
| MXE | 4866 | ENSG00000078618 | NRDC | chr1 | - | 51823663 | 51823786 | 51825286 | 51825382 | 51821497 | 51821555 | 51827795 | 51827869 | 78 | 49 | 61 | 68 | 107 | 342 | 285 | 267 | 311 | 646 | 32 | 61 | 55 | 57 | 18 | 128 | 334 | 293 | 263 | 88 | 245 | 272 | 0.90430957 | 1 | 0.202 | 0.16 | 0.202 | 0.195 | 0.155 | 0.217 | 0.169 | 0.172 | 0.194 | 0.185 | -0.005 |
| MXE | 4884 | ENSG00000168297 | PXK | chr3 | + | 58391146 | 58391220 | 58391772 | 58391847 | 58390581 | 58390659 | 58394997 | 58395102 | 59 | 30 | 33 | 63 | 125 | 89 | 71 | 45 | 62 | 131 | 118 | 57 | 44 | 96 | 51 | 140 | 90 | 59 | 84 | 99 | 223 | 224 | 0.90483046 | 1 | 0.4 | 0.298 | 0.424 | 0.505 | 0.489 | 0.458 | 0.389 | 0.428 | 0.534 | 0.341 | -0.007 |
| MXE | 5401 | ENSG00000173226 | IQCB1 | chr3 | - | 121795456 | 121795566 | 121799195 | 121799374 | 121790072 | 121790215 | 121807343 | 121807443 | 18 | 26 | 25 | 38 | 97 | 84 | 74 | 82 | 109 | 154 | 0 | 38 | 20 | 8 | 10 | 30 | 44 | 61 | 23 | 4 | 298 | 259 | 0.904838555 | 1 | 0.157 | 0.234 | 0.209 | 0.233 | 0.354 | 0 | 0.429 | 0.222 | 0.232 | 0.685 | -0.076 |
| MXE | 7759 | ENSG00000175581 | MRPL48 | chr11 | + | 73825707 | 73825796 | 73844806 | 73844976 | 73808312 | 73808350 | 73863171 | 73863261 | 26 | 20 | 26 | 54 | 51 | 0 | 1 | 2 | 0 | 7 | 16 | 34 | 17 | 14 | 10 | 6 | 0 | 1 | 0 | 0 | 238 | 298 | 0.904881187 | 1 | 1 | 0.962 | 0.942 | 1 | 0.901 | 0.77 | 1 | 0.955 | 1 | 1 | 0.016 |
| MXE | 2521 | ENSG00000119682 | AREL1 | chr14 | - | 74669946 | 74670126 | 74670761 | 74670871 | 74669648 | 74669774 | 74671407 | 74671483 | 90 | 54 | 68 | 69 | 91 | 108 | 62 | 85 | 104 | 92 | 45 | 68 | 110 | 97 | 86 | 112 | 94 | 66 | 116 | 84 | 259 | 298 | 0.904957774 | 1 | 0.489 | 0.501 | 0.479 | 0.433 | 0.532 | 0.316 | 0.454 | 0.657 | 0.49 | 0.541 | -0.005 |
| MXE | 5102 | ENSG00000060971 | ACAA1 | chr3 | - | 38129289 | 38129388 | 38131595 | 38131638 | 38126161 | 38126700 | 38131925 | 38132005 | 18 | 14 | 18 | 28 | 10 | 23 | 10 | 20 | 14 | 12 | 12 | 55 | 23 | 68 | 44 | 60 | 13 | 26 | 39 | 42 | 192 | 248 | 0.905372547 | 1 | 0.503 | 0.644 | 0.538 | 0.721 | 0.518 | 0.205 | 0.845 | 0.533 | 0.693 | 0.575 | 0.015 |
| MXE | 1173 | ENSG00000189159 | JPT1 | chr17 | - | 75146665 | 75146684 | 75147555 | 75147653 | 75135306 | 75136250 | 75148528 | 75148671 | 232 | 116 | 136 | 173 | 194 | 14 | 3 | 9 | 20 | 12 | 190 | 244 | 124 | 245 | 423 | 5 | 25 | 22 | 10 | 18 | 247 | 168 | 0.905498725 | 1 | 0.919 | 0.963 | 0.911 | 0.855 | 0.917 | 0.963 | 0.869 | 0.793 | 0.943 | 0.941 | 0.011 |
| MXE | 10716 | ENSG00000168958 | MFF | chr2 | + | 227347225 | 227347384 | 227352513 | 227352573 | 227340291 | 227340380 | 227355676 | 227355761 | 22 | 18 | 26 | 20 | 68 | 30 | 11 | 49 | 45 | 52 | 30 | 23 | 30 | 10 | 62 | 36 | 23 | 33 | 48 | 30 | 298 | 209 | 0.905661204 | 1 | 0.34 | 0.534 | 0.271 | 0.238 | 0.478 | 0.369 | 0.412 | 0.389 | 0.127 | 0.592 | -0.006 |
| MXE | 13870 | ENSG00000178234 | GALNT11 | chr7 | + | 152103111 | 152103278 | 152105244 | 152105370 | 152094189 | 152094522 | 152108037 | 152108287 | 11 | 4 | 7 | 5 | 8 | 36 | 14 | 34 | 55 | 38 | 11 | 0 | 0 | 10 | 8 | 20 | 42 | 11 | 17 | 26 | 298 | 275 | 0.905699004 | 1 | 0.22 | 0.209 | 0.16 | 0.077 | 0.163 | 0.337 | 0 | 0 | 0.352 | 0.221 | -0.016 |
| MXE | 14077 | ENSG00000140525 | FANCI | chr15 | + | 89268398 | 89268525 | 89273376 | 89273469 | 89264521 | 89264607 | 89274167 | 89274304 | 7 | 0 | 14 | 9 | 11 | 14 | 7 | 13 | 12 | 22 | 5 | 7 | 4 | 8 | 0 | 0 | 5 | 20 | 26 | 5 | 276 | 242 | 0.905754249 | 1 | 0.305 | 0 | 0.486 | 0.397 | 0.305 | 1 | 0.551 | 0.149 | 0.212 | 0 | -0.084 |
| MXE | 13116 | ENSG00000173674 | EIF1AX | chrX | - | 20132181 | 20132263 | 20133956 | 20134007 | 20130515 | 20130607 | 20135737 | 20135841 | 68 | 50 | 61 | 43 | 116 | 53 | 63 | 89 | 55 | 169 | 6 | 30 | 25 | 26 | 5 | 16 | 5 | 34 | 61 | 7 | 200 | 231 | 0.905949783 | 1 | 0.597 | 0.478 | 0.442 | 0.475 | 0.442 | 0.302 | 0.874 | 0.459 | 0.33 | 0.452 | 0.003 |
| MXE | 4477 | ENSG00000154743 | TSEN2 | chr3 | + | 12519058 | 12519197 | 12528887 | 12528924 | 12516610 | 12516661 | 12529761 | 12529873 | 12 | 11 | 12 | 14 | 10 | 3 | 2 | 11 | 16 | 3 | 16 | 21 | 2 | 4 | 0 | 0 | 5 | 3 | 3 | 4 | 288 | 186 | 0.906052471 | 1 | 0.721 | 0.78 | 0.413 | 0.361 | 0.683 | 1 | 0.731 | 0.301 | 0.463 | 0 | 0.093 |
| MXE | 8298 | ENSG00000179912 | R3HDM2 | chr12 | - | 57303175 | 57303217 | 57310263 | 57310463 | 57300094 | 57300181 | 57395748 | 57395818 | 160 | 110 | 164 | 160 | 127 | 94 | 69 | 64 | 79 | 113 | 282 | 170 | 157 | 223 | 220 | 180 | 105 | 95 | 115 | 124 | 298 | 191 | 0.906598572 | 1 | 0.522 | 0.505 | 0.622 | 0.565 | 0.419 | 0.501 | 0.509 | 0.514 | 0.554 | 0.532 | 0.005 |
| MXE | 7476 | ENSG00000084754 | HADHA | chr2 | - | 26201148 | 26201320 | 26204061 | 26204196 | 26197690 | 26197777 | 26209779 | 26209889 | 269 | 163 | 238 | 184 | 283 | 1039 | 634 | 798 | 759 | 2007 | 427 | 215 | 114 | 214 | 343 | 1502 | 911 | 549 | 975 | 1643 | 284 | 298 | 0.907084646 | 1 | 0.214 | 0.212 | 0.238 | 0.203 | 0.129 | 0.23 | 0.198 | 0.179 | 0.187 | 0.18 | 0.004 |
| MXE | 423 | ENSG00000130816 | DNMT1 | chr19 | - | 10168329 | 10168364 | 10173089 | 10173174 | 10166591 | 10166685 | 10173870 | 10173905 | 58 | 54 | 59 | 92 | 107 | 8 | 0 | 6 | 3 | 8 | 172 | 76 | 54 | 139 | 126 | 17 | 4 | 7 | 6 | 5 | 234 | 184 | 0.907213336 | 1 | 0.851 | 1 | 0.885 | 0.96 | 0.913 | 0.888 | 0.937 | 0.858 | 0.948 | 0.952 | 0.005 |
| MXE | 5674 | ENSG00000143862 | ARL8A | chr1 | - | 202135706 | 202135800 | 202138367 | 202138448 | 202135458 | 202135526 | 202144449 | 202144737 | 60 | 16 | 42 | 37 | 35 | 117 | 70 | 67 | 112 | 206 | 63 | 76 | 41 | 85 | 99 | 314 | 132 | 187 | 175 | 262 | 230 | 243 | 0.907312178 | 1 | 0.351 | 0.195 | 0.398 | 0.259 | 0.152 | 0.175 | 0.378 | 0.188 | 0.339 | 0.285 | -0.002 |
| MXE | 12142 | ENSG00000062725 | APPBP2 | chr17 | - | 60460662 | 60460787 | 60461993 | 60462061 | 60456295 | 60456381 | 60464020 | 60464110 | 105 | 70 | 78 | 104 | 119 | 89 | 62 | 56 | 81 | 170 | 0 | 78 | 65 | 41 | 4 | 9 | 30 | 42 | 17 | 7 | 217 | 274 | 0.907461843 | 1 | 0.598 | 0.588 | 0.638 | 0.618 | 0.469 | 0 | 0.767 | 0.661 | 0.753 | 0.419 | 0.062 |
| MXE | 10002 | ENSG00000135624 | CCT7 | chr2 | + | 73243003 | 73243129 | 73244543 | 73244715 | 73234235 | 73234384 | 73247761 | 73247926 | 0 | 0 | 0 | 0 | 0 | 418 | 211 | 396 | 399 | 476 | 0 | 0 | 4 | 8 | 0 | 432 | 363 | 193 | 326 | 356 | 275 | 298 | 0.907677305 | 1 | 0 | 0 | 0 | 0 | 0 | 0 | 0 | 0.022 | 0.026 | 0 | -0.01 |
| MXE | 6935 | ENSG00000062485 | CS | chr12 | - | 56283791 | 56283857 | 56285915 | 56286023 | 56282859 | 56282991 | 56286594 | 56286645 | 92 | 41 | 39 | 76 | 46 | 123 | 87 | 120 | 156 | 191 | 86 | 51 | 28 | 72 | 100 | 152 | 132 | 86 | 226 | 199 | 257 | 215 | 0.907738186 | 1 | 0.385 | 0.283 | 0.214 | 0.29 | 0.168 | 0.321 | 0.244 | 0.214 | 0.21 | 0.296 | 0.011 |
| MXE | 15242 | ENSG00000198879 | SFMBT2 | chr10 | - | 7276891 | 7276989 | 7285865 | 7285954 | 7248649 | 7248768 | 7367648 | 7367889 | 18 | 33 | 28 | 25 | 30 | 34 | 28 | 38 | 23 | 31 | 6 | 34 | 38 | 26 | 0 | 3 | 39 | 35 | 16 | 11 | 238 | 247 | 0.908038741 | 1 | 0.355 | 0.55 | 0.433 | 0.53 | 0.501 | 0.675 | 0.475 | 0.53 | 0.628 | 0 | 0.012 |
| MXE | 15780 | ENSG00000118276 | B4GALT6 | chr18 | - | 31626998 | 31627121 | 31630958 | 31631146 | 31626282 | 31626384 | 31638643 | 31638760 | 56 | 24 | 21 | 25 | 10 | 27 | 17 | 13 | 10 | 10 | 10 | 15 | 2 | 2 | 10 | 0 | 5 | 12 | 4 | 8 | 298 | 272 | 0.908100554 | 1 | 0.654 | 0.563 | 0.596 | 0.416 | 0.477 | 1 | 0.732 | 0.132 | 0.313 | 0.533 | -0.001 |
| MXE | 11572 | ENSG00000197548 | ATG7 | chr3 | + | 11342134 | 11342279 | 11347876 | 11348035 | 11340644 | 11340735 | 11358417 | 11358610 | 75 | 43 | 49 | 49 | 82 | 33 | 19 | 27 | 26 | 34 | 88 | 92 | 24 | 114 | 128 | 51 | 30 | 16 | 48 | 56 | 294 | 298 | 0.908520949 | 1 | 0.697 | 0.696 | 0.648 | 0.656 | 0.71 | 0.636 | 0.757 | 0.603 | 0.707 | 0.699 | 0.001 |
| MXE | 2794 | ENSG00000119720 | NRDE2 | chr14 | - | 90290220 | 90290607 | 90292696 | 90292872 | 90288216 | 90289145 | 90298259 | 90298380 | 16 | 13 | 15 | 34 | 18 | 24 | 11 | 24 | 19 | 24 | 19 | 48 | 37 | 21 | 6 | 19 | 20 | 29 | 38 | 24 | 298 | 298 | 0.908648177 | 1 | 0.4 | 0.542 | 0.385 | 0.642 | 0.429 | 0.5 | 0.706 | 0.561 | 0.356 | 0.2 | 0.015 |
| MXE | 1423 | ENSG00000108960 | MMD | chr17 | - | 55401468 | 55401538 | 55403766 | 55403868 | 55392621 | 55394534 | 55407745 | 55407820 | 224 | 309 | 100 | 406 | 234 | 27 | 34 | 17 | 81 | 59 | 384 | 244 | 182 | 385 | 373 | 65 | 74 | 20 | 54 | 57 | 251 | 219 | 0.908708391 | 1 | 0.879 | 0.888 | 0.837 | 0.814 | 0.776 | 0.838 | 0.742 | 0.888 | 0.862 | 0.851 | 0.003 |
| MXE | 16624 | ENSG00000089154 | GCN1 | chr12 | - | 120138322 | 120138415 | 120138694 | 120138856 | 120137900 | 120138044 | 120140858 | 120141023 | 71 | 34 | 103 | 90 | 115 | 36 | 17 | 46 | 24 | 18 | 15 | 101 | 52 | 34 | 84 | 12 | 13 | 24 | 5 | 45 | 298 | 242 | 0.908742501 | 1 | 0.616 | 0.619 | 0.645 | 0.753 | 0.838 | 0.504 | 0.863 | 0.638 | 0.847 | 0.603 | 0.003 |
| MXE | 13902 | ENSG00000174197 | MGA | chr15 | + | 41760322 | 41760529 | 41761738 | 41761850 | 41757787 | 41757839 | 41762128 | 41762194 | 111 | 93 | 91 | 89 | 171 | 153 | 112 | 128 | 111 | 246 | 11 | 62 | 38 | 21 | 0 | 0 | 93 | 76 | 39 | 6 | 298 | 261 | 0.908821779 | 1 | 0.389 | 0.421 | 0.384 | 0.413 | 0.378 | 1 | 0.369 | 0.305 | 0.32 | 0 | -0.002 |
| MXE | 6156 | ENSG00000078304 | PPP2R5C | chr14 | + | 101883431 | 101883562 | 101890236 | 101890296 | 101883256 | 101883349 | 101892999 | 101893108 | 520 | 366 | 432 | 506 | 893 | 304 | 144 | 210 | 243 | 586 | 91 | 371 | 311 | 206 | 60 | 86 | 180 | 165 | 102 | 18 | 280 | 209 | 0.909349071 | 1 | 0.561 | 0.655 | 0.606 | 0.609 | 0.532 | 0.441 | 0.606 | 0.585 | 0.601 | 0.713 | 0.003 |
| MXE | 2799 | ENSG00000151445 | VIPAS39 | chr14 | - | 77434261 | 77434303 | 77435258 | 77435393 | 77433841 | 77433931 | 77435843 | 77435919 | 26 | 13 | 15 | 22 | 82 | 14 | 7 | 12 | 12 | 11 | 103 | 36 | 16 | 17 | 22 | 19 | 15 | 11 | 8 | 14 | 284 | 191 | 0.909447828 | 1 | 0.555 | 0.555 | 0.457 | 0.552 | 0.834 | 0.785 | 0.617 | 0.494 | 0.588 | 0.514 | -0.009 |
| MXE | 5449 | ENSG00000123609 | NMI | chr2 | - | 151275757 | 151275864 | 151278827 | 151278990 | 151275483 | 151275670 | 151281947 | 151282043 | 418 | 430 | 275 | 463 | 765 | 210 | 292 | 204 | 321 | 321 | 363 | 568 | 436 | 534 | 117 | 313 | 277 | 265 | 312 | 46 | 298 | 256 | 0.909742968 | 1 | 0.631 | 0.559 | 0.537 | 0.553 | 0.672 | 0.499 | 0.638 | 0.586 | 0.595 | 0.686 | -0.01 |
| MXE | 8515 | ENSG00000144218 | AFF3 | chr2 | - | 100112385 | 100112524 | 100129223 | 100129306 | 100109844 | 100110333 | 100142483 | 100142595 | 5 | 6 | 6 | 4 | 15 | 3 | 1 | 1 | 1 | 6 | 8 | 1 | 3 | 4 | 6 | 0 | 7 | 2 | 3 | 0 | 232 | 288 | 0.909777578 | 1 | 0.674 | 0.882 | 0.882 | 0.832 | 0.756 | 1 | 0.151 | 0.651 | 0.623 | 1 | 0.12 |
| MXE | 12390 | ENSG00000150768 | DLAT | chr11 | + | 112028791 | 112028945 | 112037272 | 112037460 | 112028514 | 112028639 | 112039243 | 112039397 | 57 | 23 | 56 | 26 | 66 | 43 | 24 | 48 | 38 | 72 | 30 | 54 | 26 | 30 | 7 | 18 | 28 | 23 | 31 | 27 | 298 | 298 | 0.909884863 | 1 | 0.57 | 0.489 | 0.538 | 0.406 | 0.478 | 0.625 | 0.659 | 0.531 | 0.492 | 0.206 | -0.006 |
| MXE | 17099 | ENSG00000197442 | MAP3K5 | chr6 | - | 136639542 | 136639638 | 136642519 | 136642569 | 136637324 | 136637406 | 136650983 | 136651091 | 30 | 14 | 14 | 22 | 40 | 135 | 69 | 83 | 120 | 96 | 13 | 34 | 27 | 27 | 21 | 75 | 113 | 129 | 105 | 106 | 199 | 245 | 0.909979409 | 1 | 0.215 | 0.2 | 0.172 | 0.184 | 0.339 | 0.176 | 0.27 | 0.205 | 0.24 | 0.196 | 0.005 |
| MXE | 3461 | ENSG00000112851 | ERBIN | chr5 | + | 66046352 | 66046538 | 66048666 | 66048781 | 66044136 | 66044310 | 66050782 | 66050966 | 351 | 304 | 309 | 354 | 588 | 199 | 148 | 350 | 395 | 723 | 27 | 170 | 200 | 31 | 14 | 8 | 164 | 141 | 39 | 18 | 298 | 264 | 0.910000346 | 1 | 0.61 | 0.645 | 0.439 | 0.443 | 0.419 | 0.749 | 0.479 | 0.557 | 0.413 | 0.408 | -0.01 |
| MXE | 141 | ENSG00000015475 | BID | chr22 | - | 17743802 | 17744013 | 17750104 | 17750174 | 17739348 | 17739488 | 17774380 | 17774495 | 21 | 7 | 8 | 21 | 11 | 76 | 45 | 41 | 65 | 49 | 22 | 35 | 5 | 23 | 28 | 92 | 87 | 48 | 84 | 165 | 219 | 298 | 0.910250885 | 1 | 0.273 | 0.175 | 0.21 | 0.305 | 0.234 | 0.246 | 0.354 | 0.124 | 0.271 | 0.188 | 0.003 |
| MXE | 12186 | ENSG00000188786 | MTF1 | chr1 | - | 37835670 | 37835744 | 37838624 | 37838756 | 37835078 | 37835215 | 37839919 | 37840158 | 150 | 140 | 109 | 144 | 233 | 78 | 60 | 71 | 67 | 88 | 282 | 163 | 189 | 463 | 140 | 124 | 88 | 105 | 214 | 60 | 281 | 223 | 0.910739332 | 1 | 0.604 | 0.649 | 0.549 | 0.63 | 0.678 | 0.643 | 0.595 | 0.588 | 0.632 | 0.649 | 0.001 |
| MXE | 16104 | ENSG00000099290 | WASHC2A | chr10 | + | 50126056 | 50126179 | 50127582 | 50127795 | 50125368 | 50125449 | 50129418 | 50130039 | 113 | 42 | 84 | 95 | 172 | 41 | 34 | 42 | 51 | 37 | 127 | 48 | 108 | 86 | 35 | 53 | 35 | 31 | 33 | 21 | 272 | 298 | 0.910939943 | 1 | 0.751 | 0.575 | 0.687 | 0.671 | 0.836 | 0.724 | 0.6 | 0.792 | 0.741 | 0.646 | 0.003 |
| MXE | 1840 | ENSG00000102974 | CTCF | chr16 | + | 67616744 | 67616878 | 67620696 | 67620817 | 67611950 | 67612121 | 67621441 | 67621591 | 324 | 237 | 298 | 317 | 607 | 188 | 117 | 200 | 211 | 275 | 202 | 239 | 164 | 181 | 91 | 95 | 134 | 98 | 150 | 32 | 283 | 270 | 0.91144643 | 1 | 0.622 | 0.659 | 0.587 | 0.589 | 0.678 | 0.67 | 0.63 | 0.615 | 0.535 | 0.731 | -0.009 |
| MXE | 10898 | ENSG00000116750 | UCHL5 | chr1 | - | 193029531 | 193029657 | 193049745 | 193049851 | 193029387 | 193029449 | 193051753 | 193051817 | 60 | 43 | 43 | 47 | 56 | 136 | 108 | 77 | 141 | 203 | 15 | 11 | 19 | 12 | 0 | 9 | 33 | 66 | 14 | 14 | 255 | 275 | 0.911627696 | 1 | 0.322 | 0.3 | 0.376 | 0.264 | 0.229 | 0.643 | 0.264 | 0.237 | 0.48 | 0 | -0.027 |
| MXE | 16029 | ENSG00000171307 | ZDHHC16 | chr10 | + | 97455673 | 97455783 | 97455973 | 97456044 | 97454713 | 97454799 | 97456776 | 97456924 | 2 | 2 | 1 | 1 | 0 | 38 | 9 | 25 | 33 | 34 | 4 | 1 | 1 | 0 | 0 | 43 | 42 | 5 | 31 | 35 | 219 | 220 | 0.911775039 | 1 | 0.05 | 0.182 | 0.039 | 0.03 | 0 | 0.085 | 0.023 | 0.167 | 0 | 0 | 0.005 |
| MXE | 11264 | ENSG00000113387 | SUB1 | chr5 | + | 32588511 | 32588584 | 32591562 | 32591685 | 32571090 | 32571174 | 32598960 | 32599069 | 4 | 9 | 7 | 3 | 11 | 849 | 527 | 860 | 808 | 1007 | 0 | 26 | 0 | 2 | 0 | 651 | 1081 | 1029 | 676 | 435 | 222 | 272 | 0.911947724 | 1 | 0.006 | 0.02 | 0.01 | 0.005 | 0.013 | 0 | 0.029 | 0 | 0.004 | 0 | 0.004 |
| MXE | 4217 | ENSG00000164190 | NIPBL | chr5 | + | 36995621 | 36995804 | 37000372 | 37000570 | 36984675 | 36986301 | 37000816 | 37000888 | 419 | 388 | 286 | 382 | 686 | 176 | 144 | 105 | 151 | 280 | 11 | 93 | 156 | 27 | 0 | 0 | 53 | 23 | 8 | 16 | 298 | 298 | 0.91210624 | 1 | 0.704 | 0.729 | 0.731 | 0.717 | 0.71 | 1 | 0.637 | 0.872 | 0.771 | 0 | 0.062 |
| MXE | 5413 | ENSG00000197415 | VEPH1 | chr3 | - | 157381155 | 157381376 | 157413880 | 157414090 | 157364302 | 157364512 | 157428321 | 157428488 | 7 | 10 | 0 | 10 | 10 | 9 | 14 | 4 | 17 | 4 | 20 | 7 | 6 | 1 | 18 | 29 | 27 | 1 | 21 | 7 | 298 | 298 | 0.912353922 | 1 | 0.437 | 0.417 | 0 | 0.37 | 0.714 | 0.408 | 0.206 | 0.857 | 0.045 | 0.72 | -0.06 |
| MXE | 7108 | ENSG00000059691 | GATB | chr4 | - | 151716008 | 151716131 | 151716875 | 151717074 | 151707987 | 151708101 | 151719424 | 151719538 | 45 | 43 | 50 | 34 | 49 | 31 | 14 | 30 | 19 | 27 | 91 | 67 | 40 | 67 | 64 | 85 | 46 | 13 | 30 | 21 | 298 | 272 | 0.91239907 | 1 | 0.57 | 0.737 | 0.603 | 0.62 | 0.624 | 0.494 | 0.571 | 0.737 | 0.671 | 0.736 | -0.011 |
| MXE | 2010 | ENSG00000185596 | WASH3P | chr15 | + | 101966069 | 101966223 | 101972595 | 101972694 | 101961617 | 101961641 | 101972900 | 101973047 | 19 | 2 | 2 | 16 | 0 | 101 | 21 | 55 | 48 | 43 | 9 | 10 | 13 | 3 | 13 | 61 | 73 | 35 | 91 | 118 | 298 | 248 | 0.912517866 | 1 | 0.135 | 0.073 | 0.029 | 0.217 | 0 | 0.109 | 0.102 | 0.236 | 0.027 | 0.084 | -0.021 |
| MXE | 17155 | ENSG00000134900 | TPP2 | chr13 | + | 102649407 | 102649486 | 102657055 | 102657207 | 102648906 | 102649151 | 102663647 | 102663744 | 152 | 102 | 148 | 156 | 167 | 249 | 204 | 231 | 215 | 337 | 23 | 107 | 94 | 60 | 7 | 39 | 150 | 112 | 67 | 43 | 228 | 298 | 0.912745994 | 1 | 0.444 | 0.395 | 0.456 | 0.487 | 0.393 | 0.435 | 0.482 | 0.523 | 0.539 | 0.175 | 0.004 |
| MXE | 16486 | ENSG00000157895 | C12orf43 | chr12 | - | 121006320 | 121006394 | 121010827 | 121010926 | 121005002 | 121005093 | 121011103 | 121011146 | 27 | 3 | 16 | 21 | 8 | 16 | 13 | 20 | 19 | 20 | 9 | 33 | 11 | 8 | 21 | 28 | 20 | 11 | 18 | 21 | 248 | 223 | 0.912827342 | 1 | 0.603 | 0.172 | 0.418 | 0.498 | 0.265 | 0.224 | 0.597 | 0.473 | 0.286 | 0.473 | -0.019 |
| MXE | 11627 | ENSG00000198223 | CSF2RA | chrX | + | 1300490 | 1300626 | 1303922 | 1304019 | 1295426 | 1295456 | 1305445 | 1305527 | 48 | 36 | 38 | 28 | 70 | 46 | 39 | 31 | 34 | 51 | 146 | 58 | 45 | 130 | 121 | 178 | 79 | 27 | 58 | 147 | 285 | 246 | 0.913767403 | 1 | 0.474 | 0.443 | 0.514 | 0.415 | 0.542 | 0.415 | 0.388 | 0.59 | 0.659 | 0.415 | -0.016 |
| MXE | 4154 | ENSG00000152818 | UTRN | chr6 | + | 144678405 | 144678495 | 144700086 | 144700243 | 144577098 | 144577288 | 144730356 | 144730486 | 381 | 249 | 266 | 281 | 239 | 520 | 610 | 455 | 626 | 917 | 76 | 211 | 174 | 37 | 28 | 201 | 382 | 376 | 115 | 45 | 298 | 298 | 0.913783605 | 1 | 0.423 | 0.29 | 0.369 | 0.31 | 0.207 | 0.274 | 0.356 | 0.316 | 0.243 | 0.384 | 0.005 |
| MXE | 10440 | ENSG00000214960 | CRPPA | chr7 | - | 16308522 | 16308627 | 16376091 | 16376241 | 16301420 | 16301466 | 16406060 | 16406337 | 13 | 8 | 10 | 5 | 3 | 4 | 7 | 0 | 1 | 0 | 9 | 8 | 4 | 15 | 22 | 0 | 4 | 3 | 0 | 10 | 298 | 254 | 0.91381219 | 1 | 0.735 | 0.493 | 1 | 0.81 | 1 | 1 | 0.63 | 0.532 | 1 | 0.652 | 0.045 |
| MXE | 5837 | ENSG00000198793 | MTOR | chr1 | - | 11157151 | 11157291 | 11167441 | 11167517 | 11150125 | 11150226 | 11199257 | 11199403 | 17 | 29 | 19 | 23 | 36 | 60 | 33 | 7 | 48 | 17 | 56 | 12 | 41 | 44 | 14 | 70 | 30 | 27 | 55 | 57 | 225 | 289 | 0.913997923 | 1 | 0.267 | 0.53 | 0.332 | 0.381 | 0.731 | 0.507 | 0.339 | 0.661 | 0.507 | 0.24 | -0.003 |
| MXE | 7036 | ENSG00000095564 | BTAF1 | chr10 | + | 91966636 | 91966757 | 91980453 | 91980558 | 91964076 | 91964201 | 91981642 | 91981792 | 238 | 203 | 283 | 297 | 317 | 162 | 108 | 164 | 202 | 150 | 54 | 104 | 188 | 67 | 16 | 12 | 78 | 125 | 31 | 19 | 270 | 254 | 0.914009751 | 1 | 0.58 | 0.639 | 0.619 | 0.58 | 0.665 | 0.809 | 0.556 | 0.586 | 0.67 | 0.442 | 0.004 |
| MXE | 4411 | ENSG00000077684 | JADE1 | chr4 | + | 128855629 | 128855797 | 128857337 | 128857454 | 128852056 | 128852268 | 128861703 | 128862225 | 72 | 55 | 83 | 76 | 99 | 86 | 36 | 90 | 56 | 38 | 40 | 100 | 64 | 30 | 13 | 12 | 41 | 40 | 39 | 31 | 298 | 266 | 0.914150915 | 1 | 0.428 | 0.577 | 0.452 | 0.548 | 0.699 | 0.748 | 0.685 | 0.588 | 0.407 | 0.272 | 0.001 |
| MXE | 1734 | ENSG00000072864 | NDE1 | chr16 | + | 15691143 | 15691323 | 15694164 | 15694256 | 15687374 | 15687511 | 15696708 | 15696860 | 202 | 165 | 229 | 236 | 246 | 45 | 16 | 38 | 20 | 18 | 469 | 235 | 234 | 225 | 459 | 33 | 38 | 39 | 39 | 51 | 298 | 241 | 0.91421893 | 1 | 0.784 | 0.893 | 0.83 | 0.905 | 0.917 | 0.92 | 0.833 | 0.829 | 0.823 | 0.879 | 0.009 |
| MXE | 4734 | ENSG00000181789 | COPG1 | chr3 | + | 129268494 | 129268620 | 129268931 | 129269000 | 129267936 | 129268040 | 129271766 | 129271909 | 57 | 35 | 35 | 45 | 66 | 88 | 79 | 82 | 83 | 39 | 80 | 77 | 48 | 45 | 93 | 114 | 97 | 89 | 95 | 119 | 275 | 218 | 0.914288807 | 1 | 0.339 | 0.26 | 0.253 | 0.301 | 0.573 | 0.357 | 0.386 | 0.299 | 0.273 | 0.383 | 0.006 |
| MXE | 10977 | ENSG00000040933 | INPP4A | chr2 | + | 98545968 | 98546073 | 98546585 | 98546694 | 98543876 | 98544007 | 98552785 | 98552969 | 179 | 82 | 160 | 166 | 141 | 178 | 90 | 146 | 192 | 163 | 43 | 206 | 73 | 118 | 152 | 98 | 128 | 126 | 114 | 103 | 254 | 258 | 0.914354301 | 1 | 0.505 | 0.481 | 0.527 | 0.468 | 0.468 | 0.308 | 0.62 | 0.37 | 0.513 | 0.6 | 0.008 |
| MXE | 2879 | ENSG00000112584 | FAM120B | chr6 | + | 170317369 | 170319124 | 170323078 | 170323259 | 170306757 | 170306842 | 170330448 | 170330550 | 33 | 8 | 21 | 18 | 5 | 96 | 67 | 99 | 82 | 88 | 0 | 32 | 17 | 10 | 7 | 58 | 64 | 69 | 52 | 47 | 298 | 298 | 0.914510599 | 1 | 0.256 | 0.107 | 0.175 | 0.18 | 0.054 | 0 | 0.333 | 0.198 | 0.161 | 0.13 | -0.01 |
| MXE | 8295 | ENSG00000161929 | SCIMP | chr17 | - | 5221286 | 5221350 | 5223332 | 5223456 | 5214924 | 5214998 | 5234734 | 5234860 | 136 | 82 | 65 | 77 | 87 | 100 | 69 | 63 | 55 | 77 | 136 | 112 | 21 | 50 | 155 | 65 | 104 | 13 | 48 | 168 | 273 | 213 | 0.914725751 | 1 | 0.515 | 0.481 | 0.446 | 0.522 | 0.469 | 0.62 | 0.457 | 0.558 | 0.448 | 0.419 | -0.014 |
| MXE | 6905 | ENSG00000117335 | CD46 | chr1 | + | 207767778 | 207767823 | 207785070 | 207785106 | 207767012 | 207767195 | 207785618 | 207785682 | 115 | 96 | 115 | 130 | 255 | 138 | 98 | 108 | 131 | 171 | 54 | 105 | 71 | 90 | 56 | 52 | 122 | 82 | 52 | 47 | 194 | 185 | 0.915157777 | 1 | 0.443 | 0.483 | 0.504 | 0.486 | 0.587 | 0.498 | 0.451 | 0.452 | 0.623 | 0.532 | -0.011 |
| MXE | 7465 | ENSG00000066136 | NFYC | chr1 | + | 40766595 | 40766703 | 40769355 | 40769415 | 40762887 | 40763046 | 40770708 | 40770940 | 24 | 10 | 37 | 31 | 10 | 21 | 8 | 8 | 26 | 11 | 65 | 14 | 17 | 16 | 68 | 33 | 14 | 10 | 16 | 47 | 257 | 209 | 0.915271497 | 1 | 0.482 | 0.504 | 0.79 | 0.492 | 0.425 | 0.616 | 0.448 | 0.58 | 0.448 | 0.541 | 0.012 |
| MXE | 8813 | ENSG00000144535 | DIS3L2 | chr2 | + | 232130618 | 232130943 | 232136471 | 232136719 | 232087486 | 232087721 | 232163458 | 232163632 | 56 | 51 | 77 | 62 | 78 | 61 | 54 | 52 | 72 | 67 | 97 | 76 | 49 | 74 | 78 | 71 | 80 | 49 | 77 | 71 | 298 | 298 | 0.915288534 | 1 | 0.479 | 0.486 | 0.597 | 0.463 | 0.538 | 0.577 | 0.487 | 0.5 | 0.49 | 0.523 | -0.003 |
| MXE | 15026 | ENSG00000110756 | HPS5 | chr11 | - | 18291441 | 18292019 | 18292898 | 18292976 | 18287892 | 18288013 | 18295019 | 18295169 | 21 | 21 | 31 | 20 | 41 | 40 | 32 | 41 | 58 | 68 | 1 | 41 | 13 | 14 | 3 | 39 | 29 | 15 | 19 | 4 | 227 | 298 | 0.915450108 | 1 | 0.408 | 0.463 | 0.498 | 0.312 | 0.442 | 0.033 | 0.65 | 0.532 | 0.492 | 0.496 | -0.016 |
| MXE | 13909 | ENSG00000198919 | DZIP3 | chr3 | - | 108644163 | 108644781 | 108646616 | 108646649 | 108642437 | 108642514 | 108647942 | 108648112 | 20 | 19 | 14 | 24 | 35 | 8 | 6 | 12 | 1 | 9 | 6 | 8 | 4 | 3 | 0 | 0 | 1 | 3 | 10 | 4 | 298 | 182 | 0.915518247 | 1 | 0.604 | 0.659 | 0.416 | 0.936 | 0.704 | 1 | 0.83 | 0.449 | 1 | 0 | 0.008 |
| MXE | 3846 | ENSG00000247774 | PCED1B-AS1 | chr12 | - | 47206510 | 47206588 | 47216164 | 47216264 | 47205897 | 47206069 | 47216383 | 47216434 | 46 | 8 | 41 | 31 | 42 | 52 | 29 | 122 | 70 | 147 | 28 | 9 | 22 | 1 | 9 | 37 | 30 | 41 | 26 | 6 | 249 | 227 | 0.915543128 | 1 | 0.446 | 0.201 | 0.235 | 0.288 | 0.207 | 0.408 | 0.215 | 0.328 | 0.034 | 0.578 | -0.037 |
| MXE | 13481 | ENSG00000111203 | ITFG2 | chr12 | + | 2836744 | 2836956 | 2840841 | 2840995 | 2824089 | 2824252 | 2858011 | 2858331 | 0 | 4 | 2 | 7 | 9 | 4 | 1 | 4 | 1 | 2 | 9 | 4 | 11 | 9 | 5 | 8 | 2 | 2 | 3 | 12 | 298 | 298 | 0.916026132 | 1 | 0 | 0.8 | 0.333 | 0.875 | 0.818 | 0.529 | 0.667 | 0.846 | 0.75 | 0.294 | -0.052 |
| MXE | 8289 | ENSG00000151414 | NEK7 | chr1 | + | 198232552 | 198232637 | 198253039 | 198253180 | 198156997 | 198157276 | 198262574 | 198262637 | 57 | 18 | 23 | 30 | 2 | 198 | 171 | 164 | 154 | 294 | 0 | 17 | 8 | 4 | 7 | 84 | 63 | 54 | 27 | 37 | 234 | 290 | 0.916205994 | 1 | 0.263 | 0.115 | 0.148 | 0.194 | 0.008 | 0 | 0.251 | 0.155 | 0.155 | 0.19 | -0.005 |
| MXE | 11297 | ENSG00000129158 | SERGEF | chr11 | - | 17992930 | 17992993 | 17995795 | 17995909 | 17988596 | 17988755 | 18000496 | 18000557 | 120 | 30 | 75 | 79 | 87 | 11 | 10 | 8 | 8 | 19 | 29 | 93 | 47 | 53 | 72 | 6 | 15 | 5 | 0 | 21 | 263 | 212 | 0.916743851 | 1 | 0.898 | 0.707 | 0.883 | 0.888 | 0.787 | 0.796 | 0.833 | 0.883 | 1 | 0.734 | -0.017 |
| MXE | 719 | ENSG00000124198 | ARFGEF2 | chr20 | + | 48963829 | 48963898 | 48965871 | 48966023 | 48953555 | 48953790 | 48969146 | 48969277 | 22 | 15 | 37 | 27 | 51 | 86 | 55 | 66 | 66 | 99 | 14 | 14 | 7 | 17 | 14 | 27 | 49 | 44 | 25 | 19 | 218 | 298 | 0.916807414 | 1 | 0.259 | 0.272 | 0.434 | 0.359 | 0.413 | 0.415 | 0.281 | 0.179 | 0.482 | 0.502 | -0.024 |
| MXE | 5663 | ENSG00000100439 | ABHD4 | chr14 | + | 22603924 | 22604079 | 22605786 | 22605866 | 22603389 | 22603762 | 22606421 | 22606533 | 80 | 40 | 32 | 76 | 41 | 0 | 0 | 6 | 1 | 5 | 80 | 60 | 42 | 56 | 266 | 0 | 9 | 6 | 2 | 0 | 298 | 229 | 0.91680894 | 1 | 1 | 1 | 0.804 | 0.983 | 0.863 | 1 | 0.837 | 0.843 | 0.956 | 1 | 0.003 |
| MXE | 1926 | ENSG00000069345 | DNAJA2 | chr16 | - | 46968003 | 46968164 | 46971348 | 46971572 | 46967512 | 46967646 | 46971895 | 46971955 | 214 | 158 | 160 | 283 | 245 | 313 | 182 | 286 | 358 | 487 | 357 | 201 | 281 | 248 | 175 | 515 | 352 | 313 | 448 | 289 | 298 | 230 | 0.917119819 | 1 | 0.345 | 0.401 | 0.302 | 0.379 | 0.28 | 0.349 | 0.306 | 0.409 | 0.299 | 0.319 | 0.005 |
| MXE | 15814 | ENSG00000136167 | LCP1 | chr13 | - | 46148351 | 46148447 | 46150935 | 46151078 | 46146905 | 46147103 | 46152779 | 46152945 | 4032 | 2267 | 2808 | 3378 | 3658 | 1771 | 1109 | 1102 | 1414 | 1705 | 4905 | 3376 | 2802 | 4493 | 1825 | 1963 | 1333 | 1240 | 2071 | 905 | 292 | 245 | 0.917132576 | 1 | 0.656 | 0.632 | 0.681 | 0.667 | 0.643 | 0.677 | 0.68 | 0.655 | 0.645 | 0.629 | -0.001 |
| MXE | 8916 | ENSG00000065413 | ANKRD44 | chr2 | - | 197008943 | 197009031 | 197013510 | 197013712 | 197007805 | 197007923 | 197025195 | 197025267 | 469 | 369 | 400 | 401 | 944 | 392 | 303 | 383 | 350 | 310 | 265 | 419 | 325 | 212 | 230 | 196 | 337 | 286 | 233 | 78 | 298 | 298 | 0.917145014 | 1 | 0.488 | 0.492 | 0.454 | 0.477 | 0.708 | 0.518 | 0.497 | 0.475 | 0.42 | 0.701 | 0.002 |
| MXE | 592 | ENSG00000105186 | ANKRD27 | chr19 | - | 32643571 | 32643631 | 32644324 | 32644479 | 32643430 | 32643484 | 32649681 | 32649792 | 10 | 15 | 6 | 7 | 3 | 47 | 15 | 20 | 45 | 27 | 13 | 7 | 10 | 4 | 21 | 36 | 26 | 29 | 35 | 96 | 298 | 209 | 0.917190957 | 1 | 0.13 | 0.412 | 0.174 | 0.098 | 0.072 | 0.202 | 0.159 | 0.195 | 0.074 | 0.133 | 0.025 |
| MXE | 12096 | ENSG00000173200 | PARP15 | chr3 | + | 122613040 | 122613268 | 122615778 | 122615857 | 122610493 | 122610730 | 122617014 | 122617164 | 72 | 27 | 18 | 68 | 42 | 124 | 85 | 81 | 120 | 81 | 19 | 54 | 32 | 7 | 0 | 14 | 74 | 68 | 16 | 41 | 298 | 228 | 0.91814798 | 1 | 0.308 | 0.196 | 0.145 | 0.302 | 0.284 | 0.509 | 0.358 | 0.265 | 0.251 | 0 | -0.03 |
| MXE | 16291 | ENSG00000266412 | NCOA4 | chr10 | + | 46014443 | 46014552 | 46015125 | 46015266 | 46013549 | 46013639 | 46016539 | 46016694 | 625 | 369 | 464 | 606 | 710 | 591 | 354 | 378 | 505 | 652 | 527 | 610 | 450 | 497 | 456 | 530 | 482 | 466 | 541 | 298 | 290 | 258 | 0.918193982 | 1 | 0.485 | 0.481 | 0.522 | 0.516 | 0.492 | 0.469 | 0.53 | 0.462 | 0.45 | 0.577 | 0.002 |
| MXE | 12552 | ENSG00000179818 | PCBP1-AS1 | chr2 | - | 70059642 | 70059680 | 70083537 | 70083687 | 70051202 | 70051305 | 70085566 | 70085624 | 37 | 19 | 27 | 23 | 43 | 8 | 8 | 12 | 10 | 7 | 12 | 72 | 7 | 64 | 20 | 9 | 14 | 21 | 11 | 0 | 298 | 187 | 0.918599667 | 1 | 0.744 | 0.598 | 0.585 | 0.591 | 0.794 | 0.456 | 0.763 | 0.173 | 0.785 | 1 | 0.027 |
| MXE | 11186 | ENSG00000180228 | PRKRA | chr2 | - | 178436144 | 178436319 | 178441609 | 178441704 | 178431439 | 178432254 | 178443266 | 178443384 | 46 | 25 | 41 | 67 | 46 | 115 | 102 | 136 | 112 | 233 | 65 | 24 | 15 | 23 | 12 | 98 | 77 | 121 | 79 | 30 | 244 | 298 | 0.918699543 | 1 | 0.328 | 0.23 | 0.269 | 0.422 | 0.194 | 0.448 | 0.276 | 0.131 | 0.262 | 0.328 | 0 |
| MXE | 6802 | ENSG00000138138 | ATAD1 | chr10 | - | 87776320 | 87776427 | 87784469 | 87784670 | 87770951 | 87771041 | 87790309 | 87790430 | 238 | 130 | 218 | 196 | 257 | 144 | 98 | 130 | 125 | 182 | 38 | 159 | 109 | 45 | 25 | 50 | 65 | 54 | 42 | 11 | 298 | 256 | 0.918877753 | 1 | 0.587 | 0.533 | 0.59 | 0.574 | 0.548 | 0.395 | 0.678 | 0.634 | 0.479 | 0.661 | -0.003 |
| MXE | 3556 | ENSG00000113716 | HMGXB3 | chr5 | + | 150032453 | 150032603 | 150037399 | 150037527 | 150030740 | 150030839 | 150040747 | 150040879 | 49 | 41 | 79 | 54 | 57 | 39 | 23 | 41 | 42 | 33 | 96 | 88 | 107 | 92 | 110 | 82 | 57 | 66 | 61 | 44 | 298 | 277 | 0.919397738 | 1 | 0.539 | 0.624 | 0.642 | 0.544 | 0.616 | 0.521 | 0.589 | 0.601 | 0.584 | 0.699 | -0.006 |
| MXE | 11931 | ENSG00000149657 | LSM14B | chr20 | + | 62126225 | 62126439 | 62129784 | 62129952 | 62124616 | 62124780 | 62130218 | 62130296 | 13 | 5 | 7 | 8 | 1 | 57 | 27 | 34 | 35 | 18 | 14 | 19 | 2 | 17 | 12 | 74 | 18 | 60 | 82 | 118 | 298 | 298 | 0.919632556 | 1 | 0.186 | 0.156 | 0.171 | 0.186 | 0.053 | 0.159 | 0.514 | 0.032 | 0.172 | 0.092 | -0.043 |
| MXE | 5550 | ENSG00000158062 | UBXN11 | chr1 | - | 26301693 | 26301722 | 26302812 | 26302918 | 26297961 | 26298062 | 26303042 | 26303167 | 4 | 12 | 1 | 9 | 4 | 10 | 4 | 4 | 0 | 13 | 13 | 7 | 6 | 2 | 26 | 0 | 7 | 17 | 12 | 3 | 255 | 178 | 0.919948909 | 1 | 0.218 | 0.677 | 0.149 | 0.611 | 1 | 1 | 0.411 | 0.198 | 0.104 | 0.858 | 0.017 |
| MXE | 13919 | ENSG00000171988 | JMJD1C | chr10 | - | 63186214 | 63186383 | 63189167 | 63189446 | 63185562 | 63185653 | 63190893 | 63191108 | 327 | 272 | 320 | 343 | 523 | 234 | 231 | 202 | 285 | 375 | 24 | 95 | 46 | 29 | 23 | 42 | 33 | 48 | 7 | 35 | 298 | 298 | 0.920086931 | 1 | 0.583 | 0.541 | 0.613 | 0.546 | 0.582 | 0.364 | 0.742 | 0.489 | 0.806 | 0.397 | 0.013 |
| MXE | 2682 | ENSG00000165959 | CLMN | chr14 | - | 95213218 | 95213409 | 95215640 | 95215733 | 95210685 | 95210879 | 95221690 | 95221774 | 30 | 17 | 17 | 11 | 17 | 20 | 11 | 32 | 6 | 13 | 18 | 22 | 6 | 14 | 19 | 10 | 11 | 13 | 10 | 20 | 242 | 298 | 0.920332229 | 1 | 0.649 | 0.656 | 0.395 | 0.693 | 0.617 | 0.689 | 0.711 | 0.362 | 0.633 | 0.539 | 0.015 |
| MXE | 2807 | ENSG00000125746 | EML2 | chr19 | - | 45626704 | 45626839 | 45629950 | 45630046 | 45624718 | 45624818 | 45632860 | 45632971 | 20 | 7 | 21 | 22 | 8 | 44 | 27 | 46 | 50 | 19 | 22 | 35 | 14 | 23 | 24 | 70 | 66 | 22 | 48 | 78 | 245 | 284 | 0.92044016 | 1 | 0.345 | 0.231 | 0.346 | 0.338 | 0.328 | 0.267 | 0.381 | 0.425 | 0.357 | 0.263 | -0.021 |
| MXE | 3751 | ENSG00000172239 | PAIP1 | chr5 | - | 43543003 | 43543116 | 43547727 | 43547913 | 43538923 | 43539035 | 43555829 | 43555999 | 171 | 72 | 128 | 124 | 152 | 141 | 127 | 116 | 171 | 198 | 28 | 82 | 101 | 57 | 44 | 24 | 111 | 114 | 116 | 27 | 298 | 262 | 0.920936942 | 1 | 0.516 | 0.333 | 0.492 | 0.389 | 0.403 | 0.506 | 0.394 | 0.438 | 0.302 | 0.589 | -0.019 |
| MXE | 2769 | ENSG00000211893 | IGHG2 | chr14 | - | 105643622 | 105643949 | 105644067 | 105644103 | 105643067 | 105643525 | 105644495 | 105644790 | 28 | 11 | 123 | 43 | 35 | 649 | 252 | 3544 | 1021 | 801 | 3 | 123 | 98 | 18 | 19 | 243 | 2354 | 3457 | 312 | 375 | 185 | 298 | 0.921166548 | 1 | 0.065 | 0.066 | 0.053 | 0.064 | 0.066 | 0.019 | 0.078 | 0.044 | 0.085 | 0.075 | 0.003 |
| MXE | 3953 | ENSG00000048028 | USP28 | chr11 | - | 113852500 | 113852633 | 113854257 | 113854335 | 113841662 | 113841768 | 113875444 | 113875528 | 3 | 0 | 5 | 4 | 2 | 111 | 19 | 35 | 59 | 44 | 0 | 9 | 0 | 3 | 0 | 68 | 31 | 34 | 33 | 3 | 227 | 282 | 0.921234455 | 1 | 0.032 | 0 | 0.151 | 0.078 | 0.053 | 0 | 0.265 | 0 | 0.099 | 0 | -0.01 |
| MXE | 6587 | ENSG00000010292 | NCAPD2 | chr12 | + | 6509716 | 6509792 | 6510074 | 6510133 | 6495075 | 6495225 | 6510628 | 6510810 | 60 | 24 | 46 | 50 | 109 | 42 | 28 | 39 | 32 | 54 | 20 | 45 | 33 | 47 | 91 | 64 | 28 | 8 | 19 | 83 | 225 | 208 | 0.921409734 | 1 | 0.569 | 0.442 | 0.522 | 0.591 | 0.651 | 0.224 | 0.598 | 0.792 | 0.696 | 0.503 | -0.008 |
| MXE | 16766 | ENSG00000127328 | RAB3IP | chr12 | + | 69784719 | 69784815 | 69794436 | 69794514 | 69756404 | 69756663 | 69795140 | 69795216 | 36 | 34 | 25 | 33 | 48 | 54 | 37 | 42 | 52 | 36 | 27 | 22 | 28 | 33 | 34 | 41 | 29 | 42 | 68 | 17 | 245 | 227 | 0.921713049 | 1 | 0.382 | 0.46 | 0.355 | 0.37 | 0.553 | 0.379 | 0.413 | 0.382 | 0.31 | 0.649 | -0.003 |
| MXE | 5364 | ENSG00000182606 | TRAK1 | chr3 | + | 42189015 | 42189124 | 42191557 | 42191636 | 42188044 | 42188145 | 42193074 | 42193205 | 85 | 41 | 51 | 87 | 60 | 90 | 72 | 54 | 99 | 98 | 137 | 64 | 56 | 83 | 125 | 161 | 96 | 77 | 145 | 101 | 258 | 228 | 0.921848061 | 1 | 0.455 | 0.335 | 0.455 | 0.437 | 0.351 | 0.429 | 0.371 | 0.391 | 0.336 | 0.522 | -0.003 |
| MXE | 10463 | ENSG00000008083 | JARID2 | chr6 | + | 15374116 | 15374252 | 15452005 | 15452175 | 15246068 | 15246584 | 15468541 | 15468718 | 144 | 122 | 120 | 154 | 192 | 138 | 123 | 121 | 124 | 165 | 420 | 266 | 289 | 377 | 263 | 445 | 160 | 267 | 291 | 347 | 285 | 298 | 0.921861327 | 1 | 0.522 | 0.509 | 0.509 | 0.565 | 0.549 | 0.497 | 0.635 | 0.531 | 0.575 | 0.442 | -0.005 |
| MXE | 16425 | ENSG00000136404 | TM6SF1 | chr15 | + | 83121920 | 83122003 | 83126754 | 83126847 | 83119577 | 83119681 | 83127357 | 83127477 | 10 | 13 | 12 | 14 | 13 | 20 | 20 | 19 | 28 | 24 | 0 | 7 | 14 | 20 | 14 | 26 | 17 | 12 | 27 | 3 | 232 | 242 | 0.921942883 | 1 | 0.343 | 0.404 | 0.397 | 0.343 | 0.361 | 0 | 0.3 | 0.549 | 0.436 | 0.83 | -0.053 |
| MXE | 16709 | ENSG00000078674 | PCM1 | chr8 | + | 17969576 | 17969748 | 17972328 | 17972687 | 17966979 | 17967170 | 17980590 | 17980755 | 249 | 199 | 251 | 223 | 410 | 209 | 235 | 159 | 200 | 323 | 8 | 57 | [illegible] | [illegible] | [illegible] | [illegible] | [illegible] | [illegible] | [illegible] | [illegible] | [illegible] | [illegible] | [illegible] | [illegible] | [illegible] | [illegible] | [illegible] | [illegible] | [illegible] | [illegible] | [illegible] | [illegible] | [illegible] | [illegible] | [illegible] |
| MXE | 6738 | ENSG00000145703 | IQGAP2 | chr5 | + | 76697986 | 76698147 | 76701075 | 76701213 | 76695453 | 76695666 | 76702481 | 76702590 | 353 | 239 | 203 | 261 | 312 | 106 | 72 | 61 | 111 | 89 | 18 | 204 | [illegible] | [illegible] | [illegible] | [illegible] | [illegible] | [illegible] | [illegible] | [illegible] | [illegible] | [illegible] | [illegible] | [illegible] | [illegible] | [illegible] | [illegible] | [illegible] | [illegible] | [illegible] | [illegible] | [illegible] | [illegible] | [illegible] | [illegible] |
| MXE | 13804 | ENSG00000111364 | DDX55 | chr12 | + | 123616510 | 123616603 | 123617757 | 123617872 | 123615184 | 123615316 | 123618668 | 123618837 | 15 | 4 | 11 | 16 | 18 | 14 | 13 | 21 | 19 | 16 | 7 | 11 | [illegible] | [illegible] | [illegible] | [illegible] | [illegible] | [illegible] | [illegible] | [illegible] | [illegible] | [illegible] | [illegible] | [illegible] | [illegible] | [illegible] | [illegible] | [illegible] | [illegible] | [illegible] | [illegible] | [illegible] | [illegible] | [illegible] | [illegible] |
| MXE | 12254 | ENSG00000160218 | TRAPPC10 | chr21 | + | 44052279 | 44052476 | 44055697 | 44055893 | 44037791 | 44037927 | 44059102 | 44059214 | 284 | 143 | 177 | 232 | 378 | 214 | 111 | 124 | 163 | 200 | 347 | 246 | [illegible] | [illegible] | [illegible] | [illegible] | [illegible] | [illegible] | [illegible] | [illegible] | [illegible] | [illegible] | [illegible] | [illegible] | [illegible] | [illegible] | [illegible] | [illegible] | [illegible] | [illegible] | [illegible] | [illegible] | [illegible] | [illegible] | [illegible] |
| MXE | 47 | ENSG00000241837 | ATP5PO | chr21 | - | 33906448 | 33907453 | 33912288 | 33912399 | 33903934 | 33904021 | 33914449 | 33915768 | 254 | 131 | 289 | 197 | 387 | 5 | 3 | 1 | 0 | 0 | 247 | 188 | [illegible] | [illegible] | [illegible] | [illegible] | [illegible] | [illegible] | [illegible] | [illegible] | [illegible] | [illegible] | [illegible] | [illegible] | [illegible] | [illegible] | [illegible] | [illegible] | [illegible] | [illegible] | [illegible] | [illegible] | [illegible] | [illegible] | [illegible] |
| MXE | 6330 | ENSG00000247828 | TMEM161B-AS1 | chr5 | + | 88282842 | 88282866 | 88283009 | 88283086 | 88282041 | 88282106 | 88285740 | 88285851 | 29 | 37 | 26 | 17 | 35 | 5 | 5 | 4 | 4 | 7 | 4 | 33 | [illegible] | [illegible] | [illegible] | [illegible] | [illegible] | [illegible] | [illegible] | [illegible] | [illegible] | [illegible] | [illegible] | [illegible] | [illegible] | [illegible] | [illegible] | [illegible] | [illegible] | [illegible] | [illegible] | [illegible] | [illegible] | [illegible] | [illegible] |
| MXE | 4243 | ENSG00000174652 | ZNF266 | chr19 | - | 9434074 | 9434235 | 9434797 | 9434859 | 9420064 | 9420218 | 9435107 | 9435193 | 7 | 2 | 26 | 9 | 22 | 6 | 4 | 3 | 3 | 6 | 0 | 24 | [illegible] | [illegible] | [illegible] | [illegible] | [illegible] | [illegible] | [illegible] | [illegible] | [illegible] | [illegible] | [illegible] | [illegible] | [illegible] | [illegible] | [illegible] | [illegible] | [illegible] | [illegible] | [illegible] | [illegible] | [illegible] | [illegible] | [illegible] |
| MXE | 15317 | ENSG00000171466 | ZNF562 | chr19 | - | 9658008 | 9658135 | 9659378 | 9659467 | 9656546 | 9656653 | 9660719 | 9660874 | 36 | 12 | 22 | 16 | 16 | 77 | 77 | 49 | 72 | 77 | 37 | 29 | [illegible] | [illegible] | [illegible] | [illegible] | [illegible] | [illegible] | [illegible] | [illegible] | [illegible] | [illegible] | [illegible] | [illegible] | [illegible] | [illegible] | [illegible] | [illegible] | [illegible] | [illegible] | [illegible] | [illegible] | [illegible] | [illegible] | [illegible] |
| [illegible] | | | | | | | | | | | | | | | | | | | | | | | | | | | | | | | | | | | | | | | | | | | | | | | | |

| | | | | | | | | | | | | | | | |
|---|---|---|---|---|---|---|---|---|---|---|---|---|---|---|---|
| MXE | 2376 | ENSG00000125107 | CNOT1 | chr16 | - | 58547185 | 58547296 | 58547565 | 58547682 | 58546671 | 58546749 | 58549718 | 58549898 | 239 | [illegible] |
| MXE | 12822 | ENSG00000104133 | SPG11 | chr15 | - | 44615362 | 44615566 | 44620189 | 44620403 | 44613429 | 44613536 | 44621758 | 44621934 | 302 | [illegible] |
| MXE | 9032 | ENSG00000186854 | TRABD2A | chr2 | - | 84832054 | 84832145 | 84839148 | 84839323 | 84823952 | 84824204 | 84841860 | 84842007 | 64 | [illegible] |
| MXE | 2425 | ENSG00000206190 | ATP10A | chr15 | - | 25727159 | 25727266 | 25736055 | 25736141 | 25725950 | 25726082 | 25781018 | 25781223 | 23 | [illegible] |
| MXE | 1907 | ENSG00000130733 | YIPF2 | chr19 | - | 10925685 | 10925773 | 10927629 | 10927716 | 10924075 | 10924192 | 10927798 | 10927959 | 11 | [illegible] |
| MXE | 16459 | ENSG00000284770 | TBCE | chr1 | + | 235414432 | 235414618 | 235427139 | 235427239 | 235380018 | 235380149 | 235430704 | 235430804 | 7 | [illegible] |
| MXE | 9565 | ENSG00000143393 | PI4KB | chr1 | - | 151307573 | 151307801 | 151315572 | 151316509 | 151306135 | 151306363 | 151324744 | 151324917 | 46 | [illegible] |
| MXE | 2506 | ENSG00000012983 | MAP4K5 | chr14 | - | 50434393 | 50434571 | 50434961 | 50435065 | 50429191 | 50429260 | 50437475 | 50437534 | 39 | [illegible] |
| MXE | 2306 | ENSG00000103569 | AQP9 | chr15 | + | 58174917 | 58175036 | 58179127 | 58179345 | 58173067 | 58173205 | 58183960 | 58184440 | 77 | [illegible] |
| MXE | 2595 | ENSG00000075413 | MARK3 | chr14 | + | 103428386 | 103428440 | 103448918 | 103448967 | 103405075 | 103405267 | 103451917 | 103451983 | 84 | [illegible] |
| MXE | 3604 | ENSG00000146282 | RARS2 | chr6 | - | 87548590 | 87548646 | 87555407 | 87555505 | 87545615 | 87545699 | 87562701 | 87562785 | 105 | [illegible] |
| MXE | 13078 | ENSG00000171115 | GIMAP8 | chr7 | + | 150470828 | 150470874 | 150474011 | 150474638 | 150466670 | 150467334 | 150477091 | 150479393 | 226 | [illegible] |
| MXE | 4188 | ENSG00000120519 | SLC10A7 | chr4 | - | 146509912 | 146510049 | 146517037 | 146517120 | 146503848 | 146503924 | 146521617 | 146521830 | 69 | [illegible] |
| MXE | 11524 | ENSG00000131378 | RFTN1 | chr3 | - | 16326772 | 16326876 | 16357931 | 16358047 | 16323375 | 16323457 | 16370075 | 16370279 | 146 | [illegible] |
| MXE | 10669 | ENSG00000136536 | MARCHF7 | chr2 | + | 159743060 | 159743253 | 159745769 | 159745937 | 159729008 | 159729175 | 159747804 | 159748903 | 458 | [illegible] |
| MXE | 8348 | ENSG00000204138 | PHACTR4 | chr1 | + | 28476106 | 28476291 | 28480450 | 28480604 | 28473553 | 28474151 | 28489169 | 28489225 | 161 | [illegible] |
| MXE | 10072 | ENSG00000115355 | CCDC88A | chr2 | - | 55334093 | 55335164 | 55336680 | 55336818 | 55332130 | 55332693 | 55339463 | 55339648 | 134 | [illegible] |
| MXE | 3035 | ENSG00000119899 | SLC17A5 | chr6 | - | 73636620 | 73636707 | 73638411 | 73638499 | 73635232 | 73635500 | 73641690 | 73641924 | 10 | [illegible] |
| MXE | 7787 | ENSG00000146376 | ARHGAP18 | chr6 | - | 129605876 | 129605959 | 129607892 | 129608052 | 129600641 | 129600848 | 129611532 | 129611610 | 160 | [illegible] |
| MXE | 5220 | ENSG00000166501 | PRKCB | chr16 | + | 24035418 | 24035547 | 24092790 | 24092947 | 24032135 | 24032247 | 24094162 | 24094297 | 447 | [illegible] |
| MXE | 10465 | ENSG00000008083 | JARID2 | chr6 | + | 15410223 | 15410365 | 15452005 | 15452175 | 15374116 | 15374252 | 15468541 | 15468718 | 189 | [illegible] |
| MXE | 10280 | ENSG00000186318 | BACE1 | chr11 | - | 117290899 | 117291049 | 117291711 | 117291813 | 117290487 | 117290659 | 117293053 | 117293188 | 1 | [illegible] |
| MXE | 12204 | ENSG00000274272 | NA | chr7 | - | 100577085 | 100577154 | 100577272 | 100577389 | 100576693 | 100577005 | 100577496 | 100577558 | 166 | [illegible] |
| MXE | 9779 | ENSG00000175662 | TOM1L2 | chr17 | - | 17882704 | 17882863 | 17893660 | 17893810 | 17879626 | 17879743 | 17898595 | 17898674 | 44 | [illegible] |
| MXE | 13616 | ENSG00000258890 | CEP95 | chr17 | + | 64521401 | 64521527 | 64522701 | 64522895 | 64519320 | 64519436 | 64525769 | 64525882 | 38 | [illegible] |
| MXE | 14591 | ENSG00000148908 | RGS10 | chr10 | - | 119515508 | 119515652 | 119526031 | 119526118 | 119499816 | 119500259 | 119527305 | 119527424 | 33 | [illegible] |
| MXE | 15741 | ENSG00000149311 | ATM | chr11 | + | 108244787 | 108245026 | 108246963 | 108247136 | 108243952 | 108244118 | 108248932 | 108249102 | 355 | [illegible] |
| [illegible] | | | | | | | | | | | | | | | |

| | | | | | | | | | | | | | | |
|---|---|---|---|---|---|---|---|---|---|---|---|---|---|---|
| MXE | 39 | ENSG00000182670 | TTC3 | chr21 | + | 37152950 | 37153277 | 37156654 | 37156906 | 37151892 | 37152029 | 37159698 | 37159745 | [illegible] |
| MXE | 6417 | ENSG00000070759 | TESK2 | chr1 | - | 45346978 | 45347062 | 45347608 | 45347693 | 45346692 | 45346779 | 45347917 | 45348000 | [illegible] |
| MXE | 5279 | ENSG00000266338 | NBPF15 | chr1 | - | 144424689 | 144424862 | 144426277 | 144426450 | 144423869 | 144423975 | 144427046 | 144427098 | [illegible] |
| MXE | 17178 | ENSG00000005810 | MYCBP2 | chr13 | - | 77061168 | 77061301 | 77061661 | 77061790 | 77059522 | 77059626 | 77062595 | 77062697 | [illegible] |
| MXE | 16042 | ENSG00000175029 | CTBP2 | chr10 | - | 124992694 | 124992812 | 124993201 | 124993329 | 124984316 | 124989698 | 124993854 | 124993985 | [illegible] |
| MXE | 10804 | ENSG00000144635 | DYNC1LI1 | chr3 | - | 32528445 | 32528601 | 32529539 | 32529660 | 32525973 | 32526908 | 32530283 | 32530328 | [illegible] |
| MXE | 9987 | ENSG00000101577 | LPIN2 | chr18 | - | 2931255 | 2931443 | 2934350 | 2934450 | 2929064 | 2929158 | 2937691 | 2938037 | [illegible] |
| MXE | 13296 | ENSG00000121390 | PSPC1 | chr13 | - | 19741564 | 19741649 | 19751270 | 19751467 | 19730238 | 19730344 | 19759322 | 19759418 | [illegible] |
| MXE | 9448 | ENSG00000143093 | STRIP1 | chr1 | + | 110039394 | 110039515 | 110040634 | 110040703 | 110039171 | 110039306 | 110041535 | 110041642 | [illegible] |
| MXE | 7830 | ENSG00000108021 | TASOR2 | chr10 | + | 5714134 | 5714207 | 5717658 | 5717750 | 5712822 | 5712918 | 5720543 | 5720668 | [illegible] |
| MXE | 16920 | ENSG00000204852 | TCTN1 | chr12 | + | 110641023 | 110641149 | 110644966 | 110645129 | 110640382 | 110640517 | 110647210 | 110647336 | [illegible] |
| MXE | 4328 | ENSG00000145495 | MARCHF6 | chr5 | + | 10386993 | 10387066 | 10390331 | 10390500 | 10381799 | 10381943 | 10391541 | 10391731 | [illegible] |
| MXE | 9044 | ENSG00000134759 | ELP2 | chr18 | + | 36138269 | 36138426 | 36138794 | 36138872 | 36136306 | 36136377 | 36141136 | 36141201 | [illegible] |
| MXE | 7641 | ENSG00000063978 | RNF4 | chr4 | + | 2497006 | 2497121 | 2500658 | 2500738 | 2490336 | 2490502 | 2511955 | 2511965 | [illegible] |
| MXE | 2511 | ENSG00000174373 | RALGAPA1 | chr14 | - | 35756792 | 35756908 | 35760828 | 35761006 | 35752023 | 35752162 | 35762709 | 35762753 | [illegible] |
| MXE | 6697 | ENSG00000115808 | STRN | chr2 | - | 36855211 | 36855352 | 36857855 | 36858023 | 36850999 | 36851107 | 36861131 | 36861253 | [illegible] |
| MXE | 1192 | ENSG00000261371 | PECAM1 | chr17 | - | 64375050 | 64375356 | 64377823 | 64378117 | 64369749 | 64370025 | 64390488 | 64390515 | [illegible] |
| MXE | 16619 | ENSG00000089094 | KDM2B | chr12 | - | 121453119 | 121453344 | 121494578 | 121494665 | 121444449 | 121445418 | 121509566 | 121510039 | [illegible] |
| MXE | 6554 | ENSG00000134698 | AGO4 | chr1 | + | 35832056 | 35832185 | 35832436 | 35832570 | 35831811 | 35831931 | 35833989 | 35834174 | [illegible] |
| MXE | 10408 | ENSG00000124151 | NCOA3 | chr20 | + | 47639007 | 47639202 | 47639576 | 47639822 | 47637647 | 47637783 | 47639924 | 47640051 | [illegible] |
| MXE | 6304 | ENSG00000138674 | SEC31A | chr4 | - | 82870324 | 82870424 | 82871943 | 82872086 | 82867154 | 82867316 | 82874610 | 82874751 | [illegible] |
| MXE | 16762 | ENSG00000135108 | FBXO21 | chr12 | - | 117166897 | 117167077 | 117172470 | 117172607 | 117165484 | 117165617 | 117174204 | 117174341 | [illegible] |
| MXE | 6003 | ENSG00000174437 | ATP2A2 | chr12 | + | 110345248 | 110345382 | 110346000 | 110346118 | 110344885 | 110344971 | 110346200 | 110346321 | [illegible] |
| MXE | 4862 | ENSG00000078618 | NRDC | chr1 | - | 51789567 | 51789657 | 51790532 | 51790649 | 51789190 | 51789433 | 51790899 | 51790990 | [illegible] |
| MXE | 2203 | ENSG00000170776 | AKAP13 | chr15 | + | 85740221 | 85740272 | 85743491 | 85743825 | 85736089 | 85736134 | 85744627 | 85744677 | [illegible] |
| MXE | 13060 | ENSG00000175274 | TP53I11 | chr11 | - | 44937303 | 44937352 | 44937554 | 44937613 | 44936802 | 44936899 | 44938206 | 44938366 | [illegible] |
| MXE | 7745 | ENSG00000156515 | HK1 | chr10 | + | 69389196 | 69389296 | 69392124 | 69392308 | 69386322 | 69386418 | 69394949 | 69395105 | [illegible] |
| MXE | 11384 | ENSG00000115942 | ORC2 | chr2 | - | 200942684 | 200942777 | 200949553 | 200949643 | 200941247 | 200941279 | 200957400 | 200957544 | [illegible] |
| MXE | 588 | ENSG00000105186 | ANKRD27 | chr19 | - | 32607634 | 32607832 | 32615657 | 32615780 | 32605834 | 32605954 | 32617588 | 32617633 | [illegible] |
| MXE | 4052 | ENSG00000070756 | PABPC1 | chr8 | - | 100712651 | 100712789 | 100713086 | 100713181 | 100712361 | 100712457 | 100715461 | 100715601 | [illegible] |
| MXE | 11897 | ENSG00000135317 | SNX14 | chr6 | - | 85547111 | 85547226 | 85547505 | 85547550 | 85543604 | 85543760 | 85548300 | 85548376 | [illegible] |
| MXE | 13000 | ENSG00000186088 | GSAP | chr7 | - | 77311840 | 77311940 | 77312100 | 77312202 | 77310750 | 77311449 | 77313487 | 77313549 | [illegible] |
| MXE | 7803 | ENSG00000066427 | ATXN3 | chr14 | - | 92088729 | 92088817 | 92096092 | 92096137 | 92083125 | 92083258 | 92096673 | 92096838 | [illegible] |
| MXE | 9329 | ENSG00000070785 | EIF2B3 | chr1 | - | 44879817 | 44880008 | 44881611 | 44881739 | 44875617 | 44875695 | 44897354 | 44897444 | [illegible] |
| MXE | 9054 | ENSG00000125633 | CCDC93 | chr2 | - | 117977993 | 117978030 | 117985968 | 117986069 | 117975187 | 117975280 | 117995445 | 117995502 | [illegible] |
| MXE | 727 | ENSG00000088888 | MAVS | chr20 | + | 3857634 | 3857809 | 3861331 | 3861504 | 3854557 | 3854741 | 3862253 | 3862413 | [illegible] |
| MXE | 476 | ENSG00000076928 | ARHGEF1 | chr19 | + | 41904948 | 41905036 | 41905174 | 41905261 | 41904215 | 41904383 | 41905759 | 41905827 | [illegible] |
| MXE | 13144 | ENSG00000068400 | GRIPAP1 | chrX | - | 48997249 | 48997357 | 48998153 | 48998180 | 48993427 | 48993578 | 48999237 | 48999299 | [illegible] |
| MXE | 11030 | ENSG00000198363 | ASPH | chr8 | - | 61583943 | 61584029 | 61618977 | 61619019 | 61576771 | 61576858 | 61633682 | 61633727 | [illegible] |
| MXE | 4724 | ENSG00000163913 | IFT122 | chr3 | + | 129492140 | 129492194 | 129495445 | 129495607 | 129488256 | 129488397 | 129499901 | 129500068 | [illegible] |
| MXE | 1941 | ENSG00000047578 | KATNIP | chr16 | + | 27677728 | 27677996 | 27681398 | 27681530 | 27648603 | 27648735 | 27698327 | 27698500 | [illegible] |
| MXE | 8924 | ENSG00000065413 | ANKRD44 | chr2 | - | 197125836 | 197126037 | 197136591 | 197136662 | 197125380 | 197125468 | 197147026 | 197147105 | [illegible] |
| MXE | 11975 | ENSG00000064419 | TNPO3 | chr7 | - | 128991998 | 128992090 | 128993806 | 128993914 | 128989960 | 128990100 | 128997388 | 128997535 | [illegible] |
| MXE | 14397 | ENSG00000226752 | CUTALP | chr9 | + | 120844992 | 120845196 | 120847293 | 120847353 | 120844866 | 120844899 | 120851237 | 120851497 | [illegible] |
| MXE | 1095 | ENSG00000129197 | RPAIN | chr17 | + | 5426235 | 5426299 | 5428070 | 5428211 | 5425970 | 5426082 | 5432541 | 5432876 | [illegible] |
| MXE | 2854 | ENSG00000100714 | MTHFD1 | chr14 | + | 64400792 | 64400877 | 64411089 | 64411149 | 64388310 | 64388468 | 64412471 | 64412525 | [illegible] |
| MXE | 4333 | ENSG00000112079 | STK38 | chr6 | - | 36498362 | 36498486 | 36499872 | 36499990 | 36497779 | 36497875 | 36506582 | 36506644 | [illegible] |
| MXE | 9352 | ENSG00000146409 | SLC18B1 | chr6 | - | 132787433 | 132787581 | 132789763 | 132789837 | 132783932 | 132784089 | 132790176 | 132790272 | [illegible] |
| MXE | 10589 | ENSG00000140368 | PSTPIP1 | chr15 | + | 77032297 | 77032428 | 77032861 | 77032952 | 77031179 | 77031278 | 77035507 | 77035563 | [illegible] |
| MXE | 15148 | ENSG00000254772 | EEF1G | chr11 | - | 62570964 | 62571108 | 62571539 | 62571682 | 62567398 | 62567528 | 62571837 | 62571901 | [illegible] |
| MXE | 5403 | ENSG00000173226 | IQCB1 | chr3 | - | 121797117 | 121797227 | 121799195 | 121799374 | 121795456 | 121795566 | 121807343 | 121807443 | [illegible] |
| MXE | 8309 | ENSG00000140992 | PDPK1 | chr16 | + | 2595792 | 2595850 | 2597122 | 2597275 | 2586675 | 2586893 | 2597650 | 2597928 | [illegible] |
| MXE | 12855 | ENSG00000123066 | MED13L | chr12 | - | 116096668 | 116096752 | 116111427 | 116111512 | 116022455 | 116022601 | 116237467 | 116237687 | [illegible] |
| MXE | 16487 | ENSG00000123268 | ATF1 | chr12 | + | 50780139 | 50780238 | 50795908 | 50796009 | 50764077 | 50764307 | 50809455 | 50809475 | [illegible] |
| MXE | 10381 | ENSG00000169087 | HSPBAP1 | chr3 | - | 122768700 | 122768882 | 122777720 | 122777906 | 122759223 | 122759360 | 122793616 | 122793714 | [illegible] |
| MXE | 8294 | ENSG00000161929 | SCIMP | chr17 | - | 5221286 | 5221350 | 5223353 | 5223456 | 5214924 | 5214998 | 5234734 | 5234823 | [illegible] |
| MXE | 1004 | ENSG00000101557 | USP14 | chr18 | + | 166786 | 166819 | 178932 | 179037 | 163307 | 163453 | 180235 | 180339 | [illegible] |
| MXE | 1506 | ENSG00000091592 | NLRP1 | chr17 | - | 5501736 | 5501872 | 5521523 | 5521786 | 5499426 | 5499950 | 5530480 | 5530704 | [illegible] |
| MXE | 7419 | ENSG00000153922 | CHD1 | chr5 | - | 98872050 | 98872201 | 98872416 | 98872555 | 98870686 | 98870803 | 98873592 | 98873723 | [illegible] |
| MXE | 2713 | ENSG00000119689 | DLST | chr14 | + | 74889276 | 74889349 | 74889896 | 74889952 | 74889094 | 74889147 | 74891055 | 74891167 | [illegible] |
| MXE | 15306 | ENSG00000139697 | SBNO1 | chr12 | - | 123348028 | 123348133 | 123350309 | 123350441 | 123345257 | 123345567 | 123364700 | 123364843 | [illegible] |
| MXE | 7810 | ENSG00000066427 | ATXN3 | chr14 | - | 92093745 | 92093831 | 92096092 | 92096137 | 92093251 | 92093318 | 92096673 | 92096838 | [illegible] |
| MXE | 11914 | ENSG00000146842 | TMEM209 | chr7 | - | 130175511 | 130175609 | 130181622 | 130181719 | 130173824 | 130173939 | 130184183 | 130184255 | [illegible] |
| MXE | 11953 | ENSG00000122512 | PMS2 | chr7 | - | 5986758 | 5987620 | 5989799 | 5989955 | 5982823 | 5982991 | 5991972 | 5992057 | [illegible] |
| MXE | 3583 | ENSG00000124201 | ZNFX1 | chr20 | - | 49251526 | 49251622 | 49252719 | 49252830 | 49245896 | 49249711 | 49253665 | 49253811 | [illegible] |
| MXE | 5722 | ENSG00000285162 | NA | chr7 | + | 29367931 | 29367987 | 29393678 | 29393710 | 29354624 | 29354663 | 29398372 | 29398486 | [illegible] |
| MXE | 13031 | ENSG00000183773 | AIFM3 | chr22 | + | 20977031 | 20977095 | 20977699 | 20977776 | 20976866 | 20976938 | 20977887 | 20978005 | [illegible] |
| MXE | 10218 | ENSG00000183495 | EP400 | chr12 | + | 131991406 | 131991456 | 131994866 | 131994956 | 131990635 | 131990714 | 132005076 | 132005184 | [illegible] |
| MXE | 10077 | ENSG00000138801 | PAPSS1 | chr4 | - | 107687038 | 107687177 | 107693770 | 107694006 | 107682014 | 107682133 | 107701170 | 107701285 | [illegible] |
| MXE | 16722 | ENSG00000145050 | MANF | chr3 | + | 51386207 | 51386335 | 51387736 | 51387878 | 51385290 | 51385436 | 51388904 | 51389397 | [illegible] |
| MXE | 13052 | ENSG00000166716 | ZNF592 | chr15 | + | 84764706 | 84764815 | 84778182 | 84778312 | 84748591 | 84748664 | 84782656 | 84784895 | [illegible] |
| MXE | 2957 | ENSG00000162711 | NLRP3 | chr1 | + | 247429584 | 247429755 | 247434102 | 247434273 | 247423846 | 247425599 | 247435969 | 247436140 | [illegible] |
| MXE | 12829 | ENSG00000086730 | LAT2 | chr7 | + | 74216824 | 74216864 | 74219743 | 74219787 | 74215946 | 74216069 | 74219959 | 74219972 | [illegible] |
| MXE | 15309 | ENSG00000066926 | FECH | chr18 | - | 57551314 | 57551374 | 57554259 | 57554424 | 57544376 | 57550846 | 57554844 | 57554952 | [illegible] |
| MXE | 15206 | ENSG00000110218 | PANX1 | chr11 | + | 94153490 | 94153630 | 94178368 | 94178592 | 94128840 | 94129493 | 94179601 | 94180257 | [illegible] |
| MXE | 7939 | ENSG00000143157 | POGK | chr1 | + | 166846611 | 166846738 | 166847493 | 166847592 | 166840954 | 166841088 | 166848937 | 166849208 | [illegible] |
| MXE | 12988 | ENSG00000122643 | NT5C3A | chr7 | - | 33019634 | 33019724 | 33021271 | 33021357 | 33017203 | 33017601 | 33022052 | 33022099 | [illegible] |
| MXE | 645 | ENSG00000142347 | MYO1F | chr19 | - | 8541905 | 8541991 | 8544296 | 8544464 | 8539946 | 8540028 | 8545649 | 8545736 | [illegible] |
| MXE | 7863 | ENSG00000114841 | DNAH1 | chr3 | + | 52351961 | 52352103 | 52352551 | 52352707 | 52350507 | 52350590 | 52353102 | 52353301 | [illegible] |
| MXE | 11195 | ENSG00000115904 | SOS1 | chr2 | - | 39012125 | 39012348 | 39013459 | 39013563 | 39010583 | 39010703 | 39013866 | 39013989 | [illegible] |
| MXE | 1171 | ENSG00000125450 | NUP85 | chr17 | + | 75218184 | 75218306 | 75225102 | 75225237 | 75215753 | 75215823 | 75225341 | 75225464 | [illegible] |
| MXE | 4163 | ENSG00000177058 | SLC38A9 | chr5 | - | 55649206 | 55649314 | 55656714 | 55656774 | 55645788 | 55645895 | 55664692 | 55664863 | [illegible] |
| MXE | 7461 | ENSG00000066136 | NFYC | chr1 | + | 40749572 | 40749686 | 40753150 | 40753246 | 40747533 | 40747605 | 40758120 | 40758294 | [illegible] |
| MXE | 7998 | ENSG00000197872 | CYRIA | chr2 | - | 16562004 | 16562141 | 16565645 | 16565767 | 16561455 | 16561533 | 16588049 | 16588129 | [illegible] |
| MXE | 9274 | ENSG00000175054 | ATR | chr3 | - | 142496360 | 142496520 | 142497012 | 142497192 | 142491142 | 142493311 | 142498596 | 142498774 | [illegible] |
| MXE | 12926 | ENSG00000013374 | NUB1 | chr7 | + | 151355767 | 151355950 | 151356127 | 151356222 | 151352811 | 151352882 | 151360140 | 151360209 | [illegible] |
| MXE | 7684 | ENSG00000134574 | DDB2 | chr11 | + | 47216857 | 47217049 | 47232813 | 47232959 | 47216335 | 47216472 | 47234572 | 47234672 | [illegible] |
| MXE | 12989 | ENSG00000122643 | NT5C3A | chr7 | - | 33019634 | 33019724 | 33024038 | 33024108 | 33017203 | 33017601 | 33026816 | 33026915 | [illegible] |
| MXE | 10428 | ENSG00000163728 | TTC14 | chr3 | + | 180603123 | 180603323 | 180604851 | 180605007 | 180602890 | 180603015 | 180605765 | 180605837 | [illegible] |
| MXE | 15760 | ENSG00000149311 | ATM | chr11 | + | 108347278 | 108347365 | 108353765 | 108353880 | 108345742 | 108345908 | 108354810 | 108354874 | [illegible] |
| MXE | 16171 | ENSG00000107669 | ATE1 | chr10 | - | 121902390 | 121902620 | 121910905 | 121911151 | 121899865 | 121899994 | 121913789 | 121913893 | [illegible] |
| MXE | 2800 | ENSG00000151445 | VIPAS39 | chr14 | - | 77435258 | 77435393 | 77435843 | 77435919 | 77434261 | 77434303 | 77437807 | 77437881 | [illegible] |
| MXE | 9153 | ENSG00000117153 | KLHL12 | chr1 | - | 202895521 | 202895717 | 202896853 | 202896960 | 202894590 | 202894749 | 202909009 | 202909124 | [illegible] |
| MXE | 1600 | ENSG00000187555 | USP7 | chr16 | - | 8923214 | 8923413 | 8930292 | 8930397 | 8921156 | 8921295 | 8963206 | 8963906 | [illegible] |
| MXE | 15136 | ENSG00000100731 | PCNX1 | chr14 | + | 70988566 | 70988699 | 70995740 | 70995925 | 70976941 | 70978648 | 71009633 | 71009724 | [illegible] |
| MXE | 4103 | ENSG00000104517 | UBR5 | chr8 | - | 102258984 | 102259129 | 102261145 | 102261259 | 102257631 | 102257717 | 102261914 | 102262070 | [illegible] |
| MXE | 10790 | ENSG00000144228 | SPOPL | chr2 | + | 138550482 | 138550604 | 138550902 | 138551054 | 138550156 | 138550294 | 138552553 | 138552681 | [illegible] |
| MXE | 7314 | ENSG00000117543 | DPH5 | chr1 | - | 101001466 | 101001587 | 101013709 | 101013818 | 100995109 | 100995149 | 101021640 | 101021765 | [illegible] |
| MXE | 4441 | ENSG00000155189 | AGPAT5 | chr8 | + | 6741660 | 6741751 | 6747669 | 6747828 | 6732560 | 6732650 | 6755050 | 6755174 | [illegible] |
| MXE | 755 | ENSG00000118705 | RPN2 | chr20 | + | 37198396 | 37198492 | 37203884 | 37203960 | 37184179 | 37184373 | 37204766 | 37204901 | [illegible] |
| MXE | 2984 | ENSG00000226032 | NA | chr6 | - | 159043977 | 159044031 | 159044119 | 159044204 | 159043588 | 159043655 | 159044878 | 159044920 | [illegible] |
| MXE | 11494 | ENSG00000116698 | SMG7 | chr1 | + | 183537144 | 183537215 | 183538379 | 183538440 | 183533675 | 183533832 | 183540983 | 183541103 | [illegible] |
| MXE | 1418 | ENSG00000178852 | EFCAB13 | chr17 | + | 47345015 | 47345098 | 47370436 | 47370508 | 47344161 | 47344292 | 47374471 | 47374639 | [illegible] |
| MXE | 17119 | ENSG00000173473 | SMARCC1 | chr3 | - | 47745907 | 47745993 | 47772816 | 47772936 | 47738028 | 47738110 | 47781602 | 47781893 | [illegible] |
| MXE | 16561 | ENSG00000111785 | RIC8B | chr12 | + | 106825725 | 106825820 | 106842588 | 106842817 | 106814695 | 106815304 | 106843851 | 106843947 | [illegible] |
| MXE | 16405 | ENSG00000100354 | TNRC6B | chr22 | + | 40261831 | 40262173 | 40264687 | 40267036 | 40251178 | 40251200 | 40273424 | 40273600 | [illegible] |
| MXE | 7773 | ENSG00000165458 | INPPL1 | chr11 | + | 72230992 | 72231189 | 72231497 | 72231615 | 72230795 | 72230898 | 72232239 | 72232336 | [illegible] |
| MXE | 5461 | ENSG00000138735 | PDE5A | chr4 | - | 119501169 | 119501253 | 119502580 | 119502655 | 119494396 | 119498738 | 119504535 | 119504599 | [illegible] |
| MXE | 7438 | ENSG00000134453 | RBM17 | chr10 | + | 6101270 | 6101387 | 6104930 | 6105097 | 6096909 | 6097188 | 6106140 | 6106238 | [illegible] |
| MXE | 1912 | ENSG00000102910 | LONP2 | chr16 | + | 48299661 | 48299788 | 48303171 | 48303305 | 48296014 | 48296165 | 48334215 | 48334358 | [illegible] |
| MXE | 17164 | ENSG00000139687 | RB1 | chr13 | + | 48362814 | 48362957 | 48364893 | 48364971 | 48360016 | 48360127 | 48367493 | 48367603 | [illegible] |
| MXE | 5497 | ENSG00000124596 | OARD1 | chr6 | - | 41070075 | 41070134 | 41071131 | 41071276 | 41068840 | 41068953 | 41071595 | 41071675 | [illegible] |
| MXE | 13632 | ENSG00000152332 | UHMK1 | chr1 | + | 162500912 | 162501104 | 162503753 | 162503848 | 162499954 | 162500247 | 162512499 | 162512576 | [illegible] |
| MXE | 11157 | ENSG00000064666 | CNN2 | chr19 | + | 1032558 | 1032669 | 1036129 | 1036246 | 1032391 | 1032458 | 1036415 | 1036472 | [illegible] |
| MXE | 8564 | ENSG00000163001 | CFAP36 | chr2 | + | 55522101 | 55522166 | 55523720 | 55523822 | 55519645 | 55519916 | 55528877 | 55528992 | [illegible] |
| MXE | 5005 | ENSG00000135870 | RC3H1 | chr1 | - | 173972508 | 173972627 | 173978487 | 173978620 | 173970504 | 173970617 | 173980808 | 173981009 | [illegible] |
| MXE | 13238 | ENSG00000188917 | TRMT2B | chrX | - | 101042041 | 101042312 | 101051250 | 101051733 | 101041316 | 101041371 | 101051816 | 101052111 | [illegible] |
| MXE | 13828 | ENSG00000104219 | ZDHHC2 | chr8 | + | 17210387 | 17210480 | 17215236 | 17215349 | 17209931 | 17210058 | 17217171 | 17217246 | [illegible] |
| MXE | 7658 | ENSG00000109536 | FRG1 | chr4 | + | 189952161 | 189952287 | 189955036 | 189955151 | 189943201 | 189943272 | 189957397 | 189957502 | [illegible] |
| MXE | 2534 | ENSG00000165929 | TC2N | chr14 | - | 91792366 | 91792558 | 91797784 | 91797901 | 91787512 | 91787627 | 91798298 | 91798399 | [illegible] |
| MXE | 13207 | ENSG00000138593 | SECISBP2L | chr15 | - | 49009215 | 49009378 | 49011730 | 49011863 | 49000876 | 49001097 | 49012647 | 49012817 | [illegible] |
| MXE | 12100 | ENSG00000146733 | PSPH | chr7 | - | 56015022 | 56015171 | 56017233 | 56017379 | 56011063 | 56011869 | 56019599 | 56019734 | [illegible] |
| MXE | 7208 | ENSG00000121766 | ZCCHC17 | chr1 | + | 31310043 | 31310164 | 31318166 | 31318236 | 31296981 | 31297075 | 31319108 | 31319166 | [illegible] |
| MXE | 3736 | ENSG00000150712 | MTMR12 | chr5 | - | 32234961 | 32235129 | 32239000 | 32239173 | 32233772 | 32233934 | 32242056 | 32242127 | [illegible] |
| MXE | 4693 | ENSG00000157426 | AASDH | chr4 | - | 56378147 | 56378464 | 56382476 | 56382597 | 56371450 | 56371643 | 56384069 | 56384341 | [illegible] |
| MXE | 5347 | ENSG00000186001 | LRCH3 | chr3 | + | 197871324 | 197871462 | 197875697 | 197875775 | 197870159 | 197870278 | 197883540 | 197883772 | [illegible] |
| MXE | 14013 | ENSG00000101190 | TCFL5 | chr20 | - | 62857394 | 62857638 | 62859363 | 62859526 | 62854015 | 62854157 | 62860124 | 62860308 | [illegible] |
| MXE | 14641 | ENSG00000165525 | NEMF | chr14 | - | 49838138 | 49838206 | 49840717 | 49840866 | 49834362 | 49834449 | 49846139 | 49846265 | [illegible] |
| MXE | 10393 | ENSG00000153250 | RBMS1 | chr2 | - | 160300650 | 160300730 | 160303329 | 160303487 | 160286968 | 160287084 | 160313155 | 160313247 | [illegible] |
| MXE | 9635 | ENSG00000185811 | IKZF1 | chr7 | + | 50376532 | 50376793 | 50387344 | 50387470 | 50327378 | 50327757 | 50391728 | 50391863 | [illegible] |
| MXE | 6981 | ENSG00000068097 | HEATR6 | chr17 | - | 60049579 | 60049702 | 60050842 | 60050977 | 60048263 | 60048388 | 60055514 | 60055601 | [illegible] |
| MXE | 16210 | ENSG00000107862 | GBF1 | chr10 | + | 102360183 | 102360395 | 102361717 | 102361912 | 102359269 | 102359435 | 102362474 | 102362664 | [illegible] |
| MXE | 14807 | ENSG00000106829 | TLE4 | chr9 | + | 79722450 | 79722601 | 79722958 | 79723035 | 79721740 | 79721888 | 79725036 | 79725233 | [illegible] |
| MXE | 2767 | ENSG00000183576 | SETD3 | chr14 | - | 99406515 | 99406590 | 99412950 | 99413065 | 99405204 | 99405371 | 99413875 | 99413934 | [illegible] |
| MXE | 130 | ENSG00000183765 | CHEK2 | chr22 | - | 28719394 | 28719485 | 28725242 | 28725367 | 28711908 | 28712017 | 28734402 | 28734721 | [illegible] |
| MXE | 2361 | ENSG00000138629 | UBL7 | chr15 | - | 74451435 | 74451520 | 74452295 | 74452378 | 74450801 | 74450859 | 74456551 | 74456671 | [illegible] |
| MXE | 14322 | ENSG00000070501 | POLB | chr8 | + | 42350006 | 42350065 | 42352518 | 42352568 | 42349015 | 42349090 | 42355515 | 42355567 | [illegible] |
| MXE | 3992 | ENSG00000124279 | FASTKD3 | chr5 | - | 7861582 | 7861652 | 7862822 | 7862997 | 7861148 | 7861263 | 7865897 | 7865983 | [illegible] |
| MXE | 8219 | ENSG00000124126 | PREX1 | chr20 | - | 48639765 | 48639894 | 48642167 | 48642258 | 48637710 | 48637752 | 48642406 | 48642489 | [illegible] |
| MXE | 16427 | ENSG00000205356 | TECPR1 | chr7 | - | 98229038 | 98229166 | 98230960 | 98231118 | 98228013 | 98228116 | 98231223 | 98231373 | [illegible] |
| MXE | 16435 | ENSG00000142166 | IFNAR1 | chr21 | + | 33340998 | 33341174 | 33343534 | 33343676 | 33335523 | 33335647 | 33345245 | 33345360 | [illegible] |
| MXE | 7394 | ENSG00000131374 | TBC1D5 | chr3 | - | 17372074 | 17372247 | 17374470 | 17374540 | 17307991 | 17308134 | 17374628 | 17374679 | [illegible] |
| MXE | 5742 | ENSG00000196187 | TMEM63A | chr1 | - | 225856910 | 225857017 | 225859195 | 225859349 | 225856651 | 225856738 | 225860859 | 225860997 | [illegible] |
| MXE | 4366 | ENSG00000106052 | TAX1BP1 | chr7 | + | 27816348 | 27816394 | 27816889 | 27817038 | 27799964 | 27800090 | 27827737 | 27827820 | [illegible] |
| MXE | 1596 | ENSG00000187555 | USP7 | chr16 | - | 8916501 | 8916556 | 8917025 | 8917156 | 8915444 | 8915525 | 8919030 | 8919139 | [illegible] |
| MXE | 7123 | ENSG00000055609 | KMT2C | chr7 | - | 152224434 | 152224616 | 152229922 | 152230027 | 152224014 | 152224179 | 152230219 | 152230321 | [illegible] |
| MXE | 8312 | ENSG00000151694 | ADAM17 | chr2 | - | 9491100 | 9491151 | 9492897 | 9492986 | 9488485 | 9490518 | 9493746 | 9493825 | [illegible] |
| MXE | 773 | ENSG00000272897 | NA | chr20 | - | 35632794 | 35632923 | 35655201 | 35655344 | 35632516 | 35632696 | 35658929 | 35659014 | [illegible] |
| MXE | 2307 | ENSG00000128881 | TTBK2 | chr15 | - | 42794643 | 42794801 | 42810613 | 42810739 | 42783418 | 42783635 | 42811687 | 42811780 | [illegible] |
| MXE | 6928 | ENSG00000285053 | TBCE | chr1 | + | 235438768 | 235438922 | 235441813 | 235441882 | 235437321 | 235437474 | 235442851 | 235442911 | [illegible] |
| MXE | 14484 | ENSG00000150776 | NKAPD1 | chr11 | + | 112080408 | 112080558 | 112081981 | 112082035 | 112078214 | 112078315 | 112082464 | 112082822 | [illegible] |
| MXE | 4832 | ENSG00000013441 | CLK1 | chr2 | - | 200856890 | 200856985 | 200857717 | 200857884 | 200856681 | 200856811 | 200857972 | 200858089 | [illegible] |
| MXE | 10 | ENSG00000186866 | POFUT2 | chr21 | - | 45283382 | 45283527 | 45285268 | 45285328 | 45282348 | 45282459 | 45285677 | 45285928 | [illegible] |
| MXE | 13 | ENSG00000160191 | PDE9A | chr21 | + | 42686191 | 42686262 | 42687916 | 42687994 | 42653778 | 42653883 | 42698967 | 42699011 | [illegible] |
| MXE | 18 | ENSG00000160191 | PDE9A | chr21 | + | 42731769 | 42731949 | 42732069 | 42732124 | 42698967 | 42699011 | 42733355 | 42733426 | [illegible] |
| MXE | 22 | ENSG00000154734 | ADAMTS1 | chr21 | - | 26840275 | 26840562 | 26840997 | 26841165 | 26839874 | 26840061 | 26841857 | 26841990 | [illegible] |
| MXE | 26 | ENSG00000160209 | PDXK | chr21 | + | 43743668 | 43743807 | 43746078 | 43746125 | 43741666 | 43741771 | 43750499 | 43750545 | [illegible] |
| MXE | 27 | ENSG00000160209 | PDXK | chr21 | + | 43743723 | 43743807 | 43746078 | 43746125 | 43741666 | 43741771 | 43750499 | 43750545 | [illegible] |
| MXE | 30 | ENSG00000265590 | CFAP298-TCP10L | chr21 | - | 32584160 | 32584305 | 32602271 | 32602367 | 32582199 | 32582415 | 32603160 | 32603292 | [illegible] |
| MXE | 40 | ENSG00000182670 | TTC3 | chr21 | + | 37157160 | 37157214 | 37158130 | 37158203 | 37156654 | 37156906 | 37159698 | 37159745 | [illegible] |
| MXE | 41 | ENSG00000156273 | BACH1 | chr21 | + | 29218860 | 29218926 | 29270356 | 29270456 | 29194146 | 29194260 | 29321220 | 29321355 | [illegible] |
| MXE | 50 | ENSG00000241837 | ATP5PO | chr21 | - | 33912288 | 33912399 | 33914449 | 33914500 | 33907340 | 33907453 | 33915727 | 33915761 | [illegible] |
| MXE | 54 | ENSG00000275895 | U2AF1L5 | chr21 | - | 6492130 | 6492197 | 6493043 | 6493110 | 6487362 | 6487412 | 6495992 | 6496080 | [illegible] |
| MXE | 57 | ENSG00000237484 | LINC01684 | chr21 | + | 24485622 | 24485685 | 24488112 | 24488309 | 24441794 | 24441913 | 24526342 | 24526892 | [illegible] |
| MXE | 58 | ENSG00000237484 | LINC01684 | chr21 | + | 24485622 | 24485685 | 24522888 | 24522999 | 24441794 | 24441913 | 24526342 | 24526892 | [illegible] |
| MXE | 59 | ENSG00000237484 | LINC01684 | chr21 | + | 24488112 | 24488309 | 24522888 | 24522999 | 24441794 | 24441913 | 24526342 | 24526892 | [illegible] |
| MXE | 64 | ENSG00000243927 | MRPS6 | chr21 | + | 34103496 | 34103519 | 34125340 | 34125480 | 34102834 | 34102908 | 34142407 | 34143034 | [illegible] |
| MXE | 84 | ENSG00000100427 | MLC1 | chr22 | - | 50080343 | 50080397 | 50083083 | 50083173 | 50079917 | 50080019 | 50084725 | 50084961 | [illegible] |
| MXE | 86 | ENSG00000100211 | CBY1 | chr22 | + | 38656955 | 38657143 | 38665545 | 38665685 | 38656675 | 38656750 | 38668016 | 38668132 | [illegible] |
| MXE | 93 | ENSG00000182944 | EWSR1 | chr22 | + | 29288605 | 29288786 | 29288986 | 29289062 | 29286922 | 29287134 | 29291564 | 29291599 | [illegible] |
| MXE | 101 | ENSG00000099991 | CABIN1 | chr22 | + | 24160329 | 24160406 | 24162065 | 24162215 | 24156532 | 24156576 | 24164399 | 24164489 | [illegible] |
| MXE | 104 | ENSG00000133424 | LARGE1 | chr22 | - | 33283201 | 33283348 | 33304228 | 33304507 | 33277059 | 33277255 | 33316084 | 33316248 | [illegible] |
| MXE | 105 | ENSG00000133424 | LARGE1 | chr22 | - | 33304228 | 33304507 | 33316084 | 33316248 | 33283201 | 33283348 | 33337645 | 33337801 | [illegible] |
| MXE | 106 | ENSG00000133424 | LARGE1 | chr22 | - | 33316084 | 33316248 | 33337645 | 33337801 | 33304228 | 33304507 | 33381918 | 33382044 | [illegible] |
| MXE | 107 | ENSG00000133424 | LARGE1 | chr22 | - | 33337645 | 33337801 | 33381918 | 33382044 | 33316084 | 33316248 | 33384191 | 33384304 | [illegible] |
| MXE | 115 | ENSG00000198911 | SREBF2 | chr22 | + | 41899236 | 41899553 | 41900019 | 41900122 | 41898648 | 41898781 | 41900329 | 41900498 | [illegible] |
| MXE | 116 | ENSG00000242259 | C22orf39 | chr22 | - | 19356229 | 19356288 | 19359335 | 19359484 | 19353995 | 19354118 | 19361236 | 19361341 | [illegible] |
| MXE | 121 | ENSG00000188130 | MAPK12 | chr22 | - | 50248365 | 50248472 | 50248661 | 50248751 | 50248224 | 50248292 | 50248830 | 50248890 | [illegible] |
| MXE | 123 | ENSG00000188130 | MAPK12 | chr22 | - | 50248661 | 50248751 | 50248830 | 50248890 | 50248365 | 50248472 | 50249102 | 50249454 | [illegible] |
| MXE | 124 | ENSG00000232855 | NA | chr21 | - | 28482956 | 28483069 | 28577274 | 28577380 | 28446289 | 28446472 | 28626584 | 28626604 | [illegible] |
| MXE | 146 | ENSG00000100099 | HPS4 | chr22 | - | 26465454 | 26465551 | 26466050 | 26466104 | 26463916 | 26464826 | 26466225 | 26466262 | [illegible] |
| MXE | 151 | ENSG00000100030 | MAPK1 | chr22 | - | 21788693 | 21788808 | 21799011 | 21799128 | 21772872 | 21772982 | 21805849 | 21806039 | [illegible] |
| MXE | 171 | ENSG00000100124 | ANKRD54 | chr22 | - | 37833683 | 37833755 | 37838499 | 37838593 | 37833158 | 37833206 | 37840186 | 37840234 | [illegible] |
| MXE | 172 | ENSG00000100124 | ANKRD54 | chr22 | - | 37833683 | 37833755 | 37838499 | 37838598 | 37833158 | 37833206 | 37840186 | 37840234 | [illegible] |
| MXE | 177 | ENSG00000100095 | SEZ6L | chr22 | + | 26351051 | 26351243 | 26365371 | 26365566 | 26347718 | 26347913 | 26375574 | 26375689 | [illegible] |
| MXE | 180 | ENSG00000138942 | RNF185 | chr22 | + | 31195468 | 31195581 | 31196935 | 31197157 | 31192683 | 31192702 | 31201497 | 31201615 | [illegible] |
| MXE | 181 | ENSG00000100376 | FAM118A | chr22 | + | 45321601 | 45321793 | 45322041 | 45322215 | 45318479 | 45318640 | 45322370 | 45322426 | [illegible] |
| MXE | 185 | ENSG00000100150 | DEPDC5 | chr22 | + | 31758526 | 31758633 | 31760655 | 31760702 | 31754861 | 31754979 | 31764974 | 31765060 | [illegible] |
| MXE | 197 | ENSG00000100150 | DEPDC5 | chr22 | + | 31821501 | 31821637 | 31831063 | 31831225 | 31819021 | 31819225 | 31833914 | 31833980 | [illegible] |
| MXE | 208 | ENSG00000100106 | TRIOBP | chr22 | + | 37713209 | 37713411 | 37715762 | 37715934 | 37710426 | 37710566 | 37723184 | 37726503 | [illegible] |
| MXE | 210 | ENSG00000100372 | SLC25A17 | chr22 | - | 40779757 | 40779894 | 40792524 | 40792676 | 40779008 | 40779125 | 40794513 | 40794580 | [illegible] |
| MXE | 211 | ENSG00000100372 | SLC25A17 | chr22 | - | 40792524 | 40792676 | 40794513 | 40794580 | 40779008 | 40779125 | 40799022 | 40799083 | [illegible] |
| MXE | 212 | ENSG00000100372 | SLC25A17 | chr22 | - | 40792524 | 40792676 | 40797283 | 40797336 | 40779008 | 40779125 | 40799022 | 40799083 | [illegible] |
| MXE | 213 | ENSG00000100372 | SLC25A17 | chr22 | - | 40794513 | 40794580 | 40797283 | 40797336 | 40779008 | 40779125 | 40799022 | 40799083 | [illegible] |
| MXE | 214 | ENSG00000100372 | SLC25A17 | chr22 | - | 40794513 | 40794580 | 40797253 | 40797336 | 40792524 | 40792676 | 40799022 | 40799083 | [illegible] |
| MXE | 221 | ENSG00000248405 | PRR5-ARHGAP8 | chr22 | + | 44802076 | 44802164 | 44808306 | 44808438 | 44786456 | 44786606 | 44814671 | 44814758 | [illegible] |
| MXE | 231 | ENSG00000054611 | TBC1D22A | chr22 | + | 46913333 | 46913446 | 46974289 | 46974399 | 46912073 | 46912188 | 46997633 | 46997709 | [illegible] |
| MXE | 237 | ENSG00000100242 | SUN2 | chr22 | - | 38739721 | 38739943 | 38740266 | 38740432 | 38739341 | 38739426 | 38740888 | 38741050 | [illegible] |
| MXE | 270 | ENSG00000244625 | MIATNB | chr22 | + | 26717252 | 26717459 | 26726643 | 26726720 | 26711829 | 26712008 | 26738061 | 26738240 | [illegible] |
| MXE | 271 | ENSG00000244625 | MIATNB | chr22 | + | 26718139 | 26718231 | 26726643 | 26726720 | 26717252 | 26717459 | 26774155 | 26774217 | [illegible] |
| MXE | 272 | ENSG00000244625 | MIATNB | chr22 | + | 26718139 | 26718231 | 26738061 | 26738240 | 26717252 | 26717459 | 26774155 | 26774217 | [illegible] |
| MXE | 273 | ENSG00000244625 | MIATNB | chr22 | + | 26726643 | 26726720 | 26738061 | 26738240 | 26717252 | 26717459 | 26774155 | 26774217 | [illegible] |
| MXE | 274 | ENSG00000235954 | TTC28-AS1 | chr22 | + | 27940019 | 27940171 | 27994472 | 27994653 | 27935150 | 27935230 | 27997410 | 27998838 | [illegible] |
| MXE | 275 | ENSG00000100281 | HMGXB4 | chr22 | + | 35283961 | 35284043 | 35285996 | 35286061 | 35264647 | 35265603 | 35287346 | 35287452 | [illegible] |
| MXE | 284 | ENSG00000100325 | ASCC2 | chr22 | - | 29822334 | 29822464 | 29825086 | 29825257 | 29816005 | 29816073 | 29832244 | 29832342 | [illegible] |
| MXE | 285 | ENSG00000100325 | ASCC2 | chr22 | - | 29822334 | 29822464 | 29825086 | 29825257 | 29816005 | 29816073 | 29838177 | 29838274 | [illegible] |
| MXE | 286 | ENSG00000100325 | ASCC2 | chr22 | - | 29825086 | 29825257 | 29832244 | 29832342 | 29822334 | 29822464 | 29838177 | 29838274 | [illegible] |
| MXE | 288 | ENSG00000100325 | ASCC2 | chr22 | - | 29827476 | 29827600 | 29833540 | 29833614 | 29825621 | 29825780 | 29834488 | 29834550 | [illegible] |
| MXE | 289 | ENSG00000100325 | ASCC2 | chr22 | - | 29832244 | 29832342 | 29833540 | 29833614 | 29825621 | 29825780 | 29834488 | 29834550 | [illegible] |
| MXE | 303 | ENSG00000099917 | MED15 | chr22 | + | 20551435 | 20551487 | 20554935 | 20555148 | 20537116 | 20537215 | 20564449 | 20564688 | [illegible] |
| MXE | 305 | ENSG00000274602 | PI4KAP1 | chr22 | - | 18537291 | 18537412 | 18538432 | 18538558 | 18536649 | 18536808 | 18538639 | 18538744 | [illegible] |
| MXE | 306 | ENSG00000274602 | PI4KAP1 | chr22 | - | 18537294 | 18537412 | 18538432 | 18538558 | 18536649 | 18536808 | 18538639 | 18538744 | [illegible] |
| MXE | 324 | ENSG00000160336 | ZNF761 | chr19 | + | 53445000 | 53445219 | 53446226 | 53446337 | 53431983 | 53432028 | 53447195 | 53447283 | [illegible] |
| MXE | 326 | ENSG00000270011 | ZNF559-ZNF177 | chr19 | + | 9333014 | 9333113 | 9337795 | 9338017 | 9324694 | 9324780 | 9338493 | 9338582 | [illegible] |
| MXE | 334 | ENSG00000099331 | MYO9B | chr19 | + | 17177796 | 17177886 | 17180926 | 17181040 | 17175662 | 17175741 | 17183828 | 17183868 | [illegible] |
| MXE | 342 | ENSG00000213889 | PPM1N | chr19 | + | 45499948 | 45500066 | 45500455 | 45500561 | 45499609 | 45499725 | 45500649 | 45500710 | [illegible] |
| MXE | 348 | ENSG00000105701 | FKBP8 | chr19 | - | 18537600 | 18537773 | 18539370 | 18539459 | 18533269 | 18533347 | 18541678 | 18541995 | [illegible] |
| MXE | 349 | ENSG00000105701 | FKBP8 | chr19 | - | 18538215 | 18538436 | 18539370 | 18539459 | 18537600 | 18537773 | 18541678 | 18541995 | [illegible] |
| MXE | 353 | ENSG00000189068 | VSTM1 | chr19 | - | 54042276 | 54042369 | 54051409 | 54051448 | 54042168 | 54042196 | 54063743 | 54063953 | [illegible] |
| MXE | 354 | ENSG00000105607 | GCDH | chr19 | + | 12892115 | 12892178 | 12892316 | 12892434 | 12891830 | 12891974 | 12893482 | 12893653 | [illegible] |
| MXE | 357 | ENSG00000188321 | ZNF559 | chr19 | + | 9333014 | 9333113 | 9337795 | 9337865 | 9324694 | 9324780 | 9338493 | 9338582 | [illegible] |
| MXE | 358 | ENSG00000188321 | ZNF559 | chr19 | + | 9338493 | 9338582 | 9339192 | 9339319 | 9337795 | 9337858 | 9341101 | 9341145 | [illegible] |
| MXE | 366 | ENSG00000189144 | ZNF573 | chr19 | - | 37758475 | 37758596 | 37759046 | 37759139 | 37738286 | 37740194 | 37770004 | 37770097 | [illegible] |
| MXE | 367 | ENSG00000189144 | ZNF573 | chr19 | - | 37759046 | 37759139 | 37770004 | 37770097 | 37758475 | 37758596 | 37771563 | 37771696 | [illegible] |
| MXE | 387 | ENSG00000141867 | BRD4 | chr19 | - | 15268904 | 15269042 | 15272814 | 15273133 | 15267415 | 15267551 | 15332289 | 15332539 | [illegible] |
| MXE | 389 | ENSG00000182472 | CAPN12 | chr19 | - | 38733702 | 38733781 | 38735369 | 38735429 | 38731106 | 38731223 | 38735501 | 38735544 | [illegible] |
| MXE | 390 | ENSG00000104814 | MAP4K1 | chr19 | - | 38589177 | 38589275 | 38592449 | 38592497 | 38587640 | 38587817 | 38593281 | 38593337 | [illegible] |
| MXE | 411 | ENSG00000129353 | SLC44A2 | chr19 | + | 10627919 | 10628004 | 10631056 | 10631141 | 10626252 | 10626301 | 10631274 | 10631385 | [illegible] |
| MXE | 417 | ENSG00000011600 | TYROBP | chr19 | - | 35907445 | 35907580 | 35907729 | 35907762 | 35907217 | 35907264 | 35908174 | 35908295 | [illegible] |
| MXE | 431 | ENSG00000196268 | ZNF493 | chr19 | + | 21405128 | 21405255 | 21413465 | 21413508 | 21397118 | 21397267 | 21422912 | 21423040 | [illegible] |
| MXE | 437 | ENSG00000079385 | CEACAM1 | chr19 | - | 42515010 | 42515103 | 42518947 | 42519235 | 42512349 | 42512479 | 42521266 | 42521521 | [illegible] |
| MXE | 439 | ENSG00000079435 | LIPE | chr19 | - | 42408231 | 42408322 | 42410306 | 42410842 | 42408086 | 42408121 | 42423418 | 42424123 | [illegible] |
| MXE | 447 | ENSG00000063176 | SPHK2 | chr19 | + | 48626067 | 48626362 | 48627691 | 48627841 | 48620401 | 48620553 | 48628161 | 48628277 | [illegible] |

| | | | | | | |
|---|---|---|---|---|---|---|
| MXE | 449 | ENSG00000063177 | RPL18 | chr19 | - | [illegible] |
| MXE | 451 | ENSG00000090661 | CERS4 | chr19 | + | [illegible] |
| MXE | 452 | ENSG00000090661 | CERS4 | chr19 | + | [illegible] |
| MXE | 453 | ENSG00000090661 | CERS4 | chr19 | + | [illegible] |
| MXE | 455 | ENSG00000105655 | ISYNA1 | chr19 | - | [illegible] |
| MXE | 456 | ENSG00000105492 | SIGLEC6 | chr19 | - | [illegible] |
| MXE | 460 | ENSG00000188227 | ZNF793 | chr19 | + | [illegible] |
| MXE | 461 | ENSG00000188227 | ZNF793 | chr19 | + | [illegible] |
| MXE | 462 | ENSG00000188227 | ZNF793 | chr19 | + | [illegible] |
| MXE | 463 | ENSG00000121417 | ZNF211 | chr19 | + | [illegible] |
| MXE | 465 | ENSG00000213020 | ZNF611 | chr19 | - | [illegible] |
| MXE | 466 | ENSG00000213020 | ZNF611 | chr19 | - | [illegible] |
| MXE | 469 | ENSG00000071655 | MBD3 | chr19 | - | [illegible] |
| MXE | 472 | ENSG00000245680 | ZNF585B | chr19 | - | [illegible] |
| MXE | 477 | ENSG00000198521 | ZNF43 | chr19 | - | [illegible] |
| MXE | 478 | ENSG00000198521 | ZNF43 | chr19 | - | [illegible] |
| MXE | 479 | ENSG00000198521 | ZNF43 | chr19 | - | [illegible] |
| MXE | 480 | ENSG00000198521 | ZNF43 | chr19 | - | [illegible] |
| MXE | 484 | ENSG00000012061 | ERCC1 | chr19 | - | [illegible] |
| MXE | 486 | ENSG00000187912 | CLEC17A | chr19 | + | [illegible] |
| MXE | 491 | ENSG00000177025 | C19orf18 | chr19 | - | [illegible] |
| MXE | 494 | ENSG00000125731 | SH2D3A | chr19 | - | [illegible] |
| MXE | 498 | ENSG00000105197 | TIMM50 | chr19 | + | [illegible] |
| MXE | 502 | ENSG00000130755 | GMFG | chr19 | - | [illegible] |
| MXE | 505 | ENSG00000130755 | GMFG | chr19 | - | [illegible] |
| MXE | 512 | ENSG00000105483 | CARD8 | chr19 | - | [illegible] |
| MXE | 520 | ENSG00000105640 | RPL18A | chr19 | + | [illegible] |
| MXE | 530 | ENSG00000142230 | SAE1 | chr19 | + | [illegible] |
| MXE | 531 | ENSG00000142227 | EMP3 | chr19 | + | [illegible] |
| MXE | 535 | ENSG00000268027 | NA | chr19 | + | [illegible] |
| MXE | 536 | ENSG00000268027 | NA | chr19 | + | [illegible] |
| MXE | 546 | ENSG00000022556 | NLRP2 | chr19 | + | [illegible] |
| MXE | 548 | ENSG00000131400 | NAPSA | chr19 | - | [illegible] |
| MXE | 559 | ENSG00000240403 | KIR3DL2 | chr19 | + | [illegible] |
| MXE | 570 | ENSG00000249115 | HAUS5 | chr19 | + | [illegible] |
| MXE | 586 | ENSG00000248101 | NA | chr19 | - | [illegible] |
| MXE | 597 | ENSG00000161249 | DMKN | chr19 | - | [illegible] |
| MXE | 598 | ENSG00000161249 | DMKN | chr19 | - | [illegible] |
| MXE | 599 | ENSG00000161249 | DMKN | chr19 | - | [illegible] |
| MXE | 600 | ENSG00000161249 | DMKN | chr19 | - | [illegible] |
| MXE | 601 | ENSG00000161249 | DMKN | chr19 | - | [illegible] |
| MXE | 605 | ENSG00000161249 | DMKN | chr19 | - | [illegible] |
| MXE | 607 | ENSG00000089335 | ZNF302 | chr19 | + | [illegible] |
| MXE | 610 | ENSG00000261824 | LINC00662 | chr19 | - | [illegible] |
| MXE | 615 | ENSG00000197013 | ZNF429 | chr19 | + | [illegible] |
| MXE | 618 | ENSG00000105662 | CRTC1 | chr19 | + | [illegible] |
| MXE | 620 | ENSG00000268790 | NA | chr19 | - | [illegible] |
| MXE | 621 | ENSG00000268790 | NA | chr19 | - | [illegible] |
| MXE | 637 | ENSG00000123146 | ADGRE5 | chr19 | + | [illegible] |
| MXE | 657 | ENSG00000196365 | LONP1 | chr19 | - | [illegible] |
| MXE | 661 | ENSG00000031823 | RANBP3 | chr19 | - | [illegible] |
| MXE | 665 | ENSG00000031823 | RANBP3 | chr19 | - | [illegible] |
| MXE | 683 | ENSG00000080845 | DLGAP4 | chr20 | + | [illegible] |
| MXE | 697 | ENSG00000101347 | SAMHD1 | chr20 | - | [illegible] |
| MXE | 705 | ENSG00000000419 | DPM1 | chr20 | - | [illegible] |
| MXE | 710 | ENSG00000173418 | NAA20 | chr20 | + | [illegible] |
| MXE | 717 | ENSG00000087586 | AURKA | chr20 | - | [illegible] |
| MXE | 739 | ENSG00000100979 | PLTP | chr20 | - | [illegible] |
| MXE | 742 | ENSG00000088970 | KIZ | chr20 | + | [illegible] |
| MXE | 763 | ENSG00000125885 | MCM8 | chr20 | + | [illegible] |
| MXE | 764 | ENSG00000125885 | MCM8 | chr20 | + | [illegible] |
| MXE | 771 | ENSG00000244687 | UBE2V1 | chr20 | - | [illegible] |
| MXE | 781 | ENSG00000064787 | BCAS1 | chr20 | - | [illegible] |
| MXE | 787 | ENSG00000198276 | UCKL1 | chr20 | - | [illegible] |
| MXE | 792 | ENSG00000197586 | ENTPD6 | chr20 | + | [illegible] |
| MXE | 795 | ENSG00000197586 | ENTPD6 | chr20 | + | [illegible] |
| MXE | 797 | ENSG00000101391 | CDK5RAP1 | chr20 | - | [illegible] |
| MXE | 822 | ENSG00000196756 | SNHG17 | chr20 | - | [illegible] |
| MXE | 823 | ENSG00000196756 | SNHG17 | chr20 | - | [illegible] |
| MXE | 824 | ENSG00000196756 | SNHG17 | chr20 | - | [illegible] |
| MXE | 831 | ENSG00000196756 | SNHG17 | chr20 | - | [illegible] |
| MXE | 832 | ENSG00000196756 | SNHG17 | chr20 | - | [illegible] |
| MXE | 840 | ENSG00000105576 | TNPO2 | chr19 | - | [illegible] |
| MXE | 855 | ENSG00000088766 | CRLS1 | chr20 | + | [illegible] |
| MXE | 856 | ENSG00000088766 | CRLS1 | chr20 | + | [illegible] |
| MXE | 857 | ENSG00000125772 | GPCPD1 | chr20 | - | [illegible] |
| MXE | 863 | ENSG00000132623 | ANKEF1 | chr20 | + | [illegible] |
| MXE | 875 | ENSG00000134755 | DSC2 | chr18 | - | [illegible] |
| MXE | 886 | ENSG00000176225 | RTTN | chr18 | - | [illegible] |
| MXE | 888 | ENSG00000134758 | RNF138 | chr18 | + | [illegible] |
| MXE | 900 | ENSG00000134444 | RELCH | chr18 | + | [illegible] |
| MXE | 901 | ENSG00000154655 | L3MBTL4 | chr18 | - | [illegible] |
| MXE | 910 | ENSG00000186111 | PIP5K1C | chr19 | - | [illegible] |
| MXE | 925 | ENSG00000141469 | SLC14A1 | chr18 | + | [illegible] |
| MXE | 926 | ENSG00000141469 | SLC14A1 | chr18 | + | [illegible] |
| MXE | 927 | ENSG00000141469 | SLC14A1 | chr18 | + | [illegible] |
| MXE | 936 | ENSG00000100983 | GSS | chr20 | - | [illegible] |
| MXE | 941 | ENSG00000166479 | TMX3 | chr18 | - | [illegible] |
| MXE | 969 | ENSG00000101773 | RBBP8 | chr18 | + | [illegible] |
| MXE | 970 | ENSG00000101773 | RBBP8 | chr18 | + | [illegible] |
| MXE | 975 | ENSG00000101639 | CEP192 | chr18 | + | [illegible] |
| MXE | 995 | ENSG00000101019 | UQCC1 | chr20 | - | [illegible] |
| MXE | 1001 | ENSG00000166974 | MAPRE2 | chr18 | + | [illegible] |
| MXE | 1005 | ENSG00000141447 | OSBPL1A | chr18 | - | [illegible] |
| MXE | 1007 | ENSG00000141447 | OSBPL1A | chr18 | - | [illegible] |
| MXE | 1008 | ENSG00000141447 | OSBPL1A | chr18 | - | [illegible] |
| MXE | 1009 | ENSG00000167525 | PROCA1 | chr17 | - | [illegible] |
| MXE | 1012 | ENSG00000073605 | GSDMB | chr17 | - | [illegible] |
| MXE | 1013 | ENSG00000073605 | GSDMB | chr17 | - | [illegible] |
| MXE | 1044 | ENSG00000136492 | BRIP1 | chr17 | - | [illegible] |
| MXE | 1045 | ENSG00000136492 | BRIP1 | chr17 | - | [illegible] |
| MXE | 1059 | ENSG00000278053 | DDX52 | chr17 | - | [illegible] |
| MXE | 1060 | ENSG00000030582 | GRN | chr17 | + | [illegible] |
| MXE | 1069 | ENSG00000267060 | PTGES3L | chr17 | - | [illegible] |
| MXE | 1083 | ENSG00000198862 | LTN1 | chr21 | - | [illegible] |
| MXE | 1084 | ENSG00000126561 | STAT5A | chr17 | + | [illegible] |
| MXE | 1086 | ENSG00000126561 | STAT5A | chr17 | + | [illegible] |
| MXE | 1097 | ENSG00000182173 | TSEN54 | chr17 | + | [illegible] |
| MXE | 1098 | ENSG00000182173 | TSEN54 | chr17 | + | [illegible] |
| MXE | 1099 | ENSG00000196704 | AMZ2 | chr17 | + | [illegible] |
| MXE | 1100 | ENSG00000196704 | AMZ2 | chr17 | + | [illegible] |
| MXE | 1109 | ENSG00000141252 | VPS53 | chr17 | - | [illegible] |
| MXE | 1111 | ENSG00000141252 | VPS53 | chr17 | - | [illegible] |
| MXE | 1119 | ENSG00000120071 | KANSL1 | chr17 | - | [illegible] |
| MXE | 1124 | ENSG00000167699 | GLOD4 | chr17 | - | [illegible] |
| MXE | 1132 | ENSG00000033627 | ATP6V0A1 | chr17 | + | [illegible] |
| MXE | 1157 | ENSG00000078687 | TNRC6C | chr17 | + | [illegible] |
| MXE | 1159 | ENSG00000078687 | TNRC6C | chr17 | + | [illegible] |
| MXE | 1175 | ENSG00000067191 | CACNB1 | chr17 | - | [illegible] |
| MXE | 1176 | ENSG00000161381 | PLXDC1 | chr17 | - | [illegible] |
| MXE | 1177 | ENSG00000161381 | PLXDC1 | chr17 | - | [illegible] |
| MXE | 1195 | ENSG00000198909 | MAP3K3 | chr17 | + | [illegible] |
| MXE | 1218 | ENSG00000136436 | CALCOCO2 | chr17 | + | [illegible] |
| MXE | 1223 | ENSG00000153944 | MSI2 | chr17 | + | [illegible] |
| MXE | 1224 | ENSG00000153944 | MSI2 | chr17 | + | [illegible] |
| MXE | 1225 | ENSG00000153944 | MSI2 | chr17 | + | [illegible] |
| MXE | 1229 | ENSG00000121058 | COIL | chr17 | - | [illegible] |
| MXE | 1230 | ENSG00000007168 | PAFAH1B1 | chr17 | + | [illegible] |
| MXE | 1268 | ENSG00000179094 | PER1 | chr17 | - | [illegible] |
| MXE | 1277 | ENSG00000141030 | COPS3 | chr17 | - | [illegible] |
| MXE | 1280 | ENSG00000121067 | SPOP | chr17 | - | [illegible] |
| MXE | 1282 | ENSG00000121067 | SPOP | chr17 | - | [illegible] |
| MXE | 1283 | ENSG00000121067 | SPOP | chr17 | - | [illegible] |
| MXE | 1294 | ENSG00000130402 | ACTN4 | chr19 | + | [illegible] |
| MXE | 1297 | ENSG00000174238 | PITPNA | chr17 | - | [illegible] |
| MXE | 1299 | ENSG00000277957 | SENP3-EIF4A1 | chr17 | + | [illegible] |
| MXE | 1301 | ENSG00000277957 | SENP3-EIF4A1 | chr17 | + | [illegible] |
| MXE | 1309 | ENSG00000273018 | FAM106A | chr17 | - | [illegible] |
| MXE | 1310 | ENSG00000273018 | FAM106A | chr17 | - | [illegible] |
| MXE | 1311 | ENSG00000154874 | CCDC144B | chr17 | - | [illegible] |
| MXE | 1344 | ENSG00000062716 | VMP1 | chr17 | + | [illegible] |
| MXE | 1345 | ENSG00000062716 | VMP1 | chr17 | + | [illegible] |
| MXE | 1349 | ENSG00000062716 | VMP1 | chr17 | + | [illegible] |
| MXE | 1353 | ENSG00000108384 | RAD51C | chr17 | + | [illegible] |
| MXE | 1356 | ENSG00000267123 | SCAT1 | chr17 | - | [illegible] |
| MXE | 1363 | ENSG00000109066 | TMEM104 | chr17 | + | [illegible] |
| MXE | 1364 | ENSG00000173818 | ENDOV | chr17 | + | [illegible] |
| MXE | 1365 | ENSG00000173818 | ENDOV | chr17 | + | [illegible] |
| MXE | 1366 | ENSG00000173818 | ENDOV | chr17 | + | [illegible] |
| MXE | 1367 | ENSG00000173818 | ENDOV | chr17 | + | [illegible] |
| MXE | 1369 | ENSG00000185813 | PCYT2 | chr17 | - | [illegible] |
| MXE | 1370 | ENSG00000185813 | PCYT2 | chr17 | - | [illegible] |
| MXE | 1392 | ENSG00000189050 | RNFT1 | chr17 | - | [illegible] |
| MXE | 1398 | ENSG00000108423 | TUBD1 | chr17 | - | [illegible] |
| MXE | 1399 | ENSG00000108423 | TUBD1 | chr17 | - | [illegible] |
| MXE | 1409 | ENSG00000073584 | SMARCE1 | chr17 | - | [illegible] |
| MXE | 1410 | ENSG00000073584 | SMARCE1 | chr17 | - | [illegible] |
| MXE | 1413 | ENSG00000073584 | SMARCE1 | chr17 | - | [illegible] |
| MXE | 1415 | ENSG00000005243 | COPZ2 | chr17 | - | [illegible] |
| MXE | 1416 | ENSG00000178852 | EFCAB13 | chr17 | + | [illegible] |
| MXE | 1419 | ENSG00000178852 | EFCAB13 | chr17 | + | [illegible] |
| MXE | 1422 | ENSG00000178852 | EFCAB13 | chr17 | + | [illegible] |
| MXE | 1431 | ENSG00000176208 | ATAD5 | chr17 | + | [illegible] |
| MXE | 1436 | ENSG00000188825 | LINC00910 | chr17 | - | [illegible] |
| MXE | 1439 | ENSG00000197381 | ADARB1 | chr21 | + | [illegible] |
| MXE | 1441 | ENSG00000264058 | NA | chr17 | - | [illegible] |
| MXE | 1447 | ENSG00000187607 | ZNF286A | chr17 | + | [illegible] |
| MXE | 1456 | ENSG00000141696 | P3H4 | chr17 | - | [illegible] |
| MXE | 1474 | ENSG00000276234 | TADA2A | chr17 | + | [illegible] |
| MXE | 1481 | ENSG00000108599 | AKAP10 | chr17 | - | [illegible] |
| MXE | 1485 | ENSG00000161395 | PGAP3 | chr17 | - | [illegible] |
| MXE | 1492 | ENSG00000215067 | ALOX12-AS1 | chr17 | - | [illegible] |
| MXE | 1497 | ENSG00000142494 | SLC47A1 | chr17 | + | [illegible] |
| MXE | 1501 | ENSG00000198920 | KIAA0753 | chr17 | - | [illegible] |
| MXE | 1503 | ENSG00000074370 | ATP2A3 | chr17 | - | [illegible] |
| MXE | 1527 | ENSG00000004660 | CAMKK1 | chr17 | - | [illegible] |
| MXE | 1546 | ENSG00000102878 | HSF4 | chr16 | + | [illegible] |
| MXE | 1558 | ENSG00000257017 | HP | chr16 | + | [illegible] |
| MXE | 1566 | ENSG00000140650 | PMM2 | chr16 | + | [illegible] |
| MXE | 1568 | ENSG00000140650 | PMM2 | chr16 | + | [illegible] |
| MXE | 1569 | ENSG00000140650 | PMM2 | chr16 | + | [illegible] |
| MXE | 1574 | ENSG00000175595 | ERCC4 | chr16 | + | [illegible] |
| MXE | 1575 | ENSG00000186153 | WWOX | chr16 | + | [illegible] |
| MXE | 1577 | ENSG00000166747 | AP1G1 | chr16 | - | [illegible] |
| MXE | 1578 | ENSG00000166747 | AP1G1 | chr16 | - | [illegible] |
| MXE | 1579 | ENSG00000166747 | AP1G1 | chr16 | - | [illegible] |
| MXE | 1580 | ENSG00000166747 | AP1G1 | chr16 | - | [illegible] |
| MXE | 1592 | ENSG00000187555 | USP7 | chr16 | - | [illegible] |
| MXE | 1602 | ENSG00000166743 | ACSM1 | chr16 | - | [illegible] |
| MXE | 1608 | ENSG00000125814 | NAPB | chr20 | - | [illegible] |
| MXE | 1610 | ENSG00000103042 | SLC38A7 | chr16 | - | [illegible] |
| MXE | 1611 | ENSG00000039068 | CDH1 | chr16 | + | [illegible] |
| MXE | 1612 | ENSG00000089505 | CMTM1 | chr16 | + | [illegible] |
| MXE | 1614 | ENSG00000089505 | CMTM1 | chr16 | + | [illegible] |
| MXE | 1623 | ENSG00000153066 | TXNDC11 | chr16 | - | [illegible] |
| MXE | 1624 | ENSG00000153066 | TXNDC11 | chr16 | - | [illegible] |
| MXE | 1629 | ENSG00000162066 | AMDHD2 | chr16 | + | [illegible] |
| MXE | 1631 | ENSG00000140995 | DEF8 | chr16 | + | [illegible] |
| MXE | 1632 | ENSG00000140995 | DEF8 | chr16 | + | [illegible] |
| MXE | 1633 | ENSG00000140995 | DEF8 | chr16 | + | [illegible] |
| MXE | 1634 | ENSG00000140995 | DEF8 | chr16 | + | [illegible] |
| MXE | 1636 | ENSG00000260565 | ERVK13-1 | chr16 | - | [illegible] |
| MXE | 1640 | ENSG00000166548 | TK2 | chr16 | - | [illegible] |
| MXE | 1642 | ENSG00000166548 | TK2 | chr16 | - | [illegible] |
| MXE | 1648 | ENSG00000188603 | CLN3 | chr16 | - | [illegible] |
| MXE | 1653 | ENSG00000121274 | TENT4B | chr16 | + | [illegible] |
| MXE | 1669 | ENSG00000103222 | ABCC1 | chr16 | + | [illegible] |
| MXE | 1672 | ENSG00000103222 | ABCC1 | chr16 | + | [illegible] |
| MXE | 1679 | ENSG00000070770 | CSNK2A2 | chr16 | - | [illegible] |
| MXE | 1698 | ENSG00000183889 | NA | chr16 | + | [illegible] |
| MXE | 1699 | ENSG00000159720 | ATP6V0D1 | chr16 | - | [illegible] |
| MXE | 1713 | ENSG00000103313 | MEFV | chr16 | - | [illegible] |
| MXE | 1714 | ENSG00000103313 | MEFV | chr16 | - | [illegible] |
| MXE | 1715 | ENSG00000103313 | MEFV | chr16 | - | [illegible] |
| MXE | 1717 | ENSG00000103194 | USP10 | chr16 | + | [illegible] |
| MXE | 1718 | ENSG00000185324 | CDK10 | chr16 | + | [illegible] |
| MXE | 1720 | ENSG00000197912 | SPG7 | chr16 | + | [illegible] |
| MXE | 1723 | ENSG00000197912 | SPG7 | chr16 | + | [illegible] |
| MXE | 1724 | ENSG00000197912 | SPG7 | chr16 | + | [illegible] |
| MXE | 1725 | ENSG00000197912 | SPG7 | chr16 | + | [illegible] |
| MXE | 1730 | ENSG00000072864 | NDE1 | chr16 | + | [illegible] |
| MXE | 1740 | ENSG00000130175 | PRKCSH | chr19 | + | [illegible] |
| MXE | 1758 | ENSG00000103148 | NPRL3 | chr16 | - | [illegible] |
| MXE | 1768 | ENSG00000157353 | FCSK | chr16 | + | [illegible] |
| MXE | 1774 | ENSG00000196712 | NF1 | chr17 | + | [illegible] |
| MXE | 1777 | ENSG00000196712 | NF1 | chr17 | + | [illegible] |
| MXE | 1778 | ENSG00000196712 | NF1 | chr17 | + | [illegible] |
| MXE | 1784 | ENSG00000089280 | FUS | chr16 | + | [illegible] |
| MXE | 1792 | ENSG00000181031 | RPH3AL | chr17 | - | [illegible] |
| MXE | 1802 | ENSG00000103415 | HMOX2 | chr16 | + | [illegible] |
| MXE | 1806 | ENSG00000140987 | ZSCAN32 | chr16 | - | [illegible] |
| MXE | 1810 | ENSG00000140987 | ZSCAN32 | chr16 | - | [illegible] |
| MXE | 1815 | ENSG00000123815 | COQ8B | chr19 | - | [illegible] |

| | | | | | | | | | | |
|---|---|---|---|---|---|---|---|---|---|---|
| MXE | 1816 | ENSG00000123815 | COQ8B | chr19 | - | 40713739 | 40713869 | 40714066 | 40714153 | [illegible] |
| MXE | 1823 | ENSG00000118900 | UBN1 | chr16 | + | 4859864 | 4859968 | 4860663 | 4860953 | [illegible] |
| MXE | 1826 | ENSG00000103121 | CMC2 | chr16 | - | 80988539 | 80988596 | 80991753 | 80991957 | [illegible] |
| MXE | 1827 | ENSG00000103121 | CMC2 | chr16 | - | 80988539 | 80988596 | 80996922 | 80997020 | [illegible] |
| MXE | 1828 | ENSG00000103121 | CMC2 | chr16 | - | 80991753 | 80991957 | 80996922 | 80997020 | [illegible] |
| MXE | 1847 | ENSG00000103248 | MTHFSD | chr16 | - | 86551972 | 86552146 | 86554644 | 86554748 | [illegible] |
| MXE | 1848 | ENSG00000103248 | MTHFSD | chr16 | - | 86551972 | 86552146 | 86554644 | 86554751 | [illegible] |
| MXE | 1863 | ENSG00000140941 | MAP1LC3B | chr16 | + | 87398814 | 87398870 | 87402174 | 87402281 | [illegible] |
| MXE | 1866 | ENSG00000135698 | MPHOSPH6 | chr16 | - | 82149308 | 82149403 | 82151423 | 82151514 | [illegible] |
| MXE | 1871 | ENSG00000159593 | NAE1 | chr16 | - | 66821449 | 66821559 | 66823226 | 66823262 | [illegible] |
| MXE | 1872 | ENSG00000159593 | NAE1 | chr16 | - | 66823226 | 66823306 | 66823528 | 66823600 | [illegible] |
| MXE | 1873 | ENSG00000159593 | NAE1 | chr16 | - | 66823528 | 66823600 | 66824854 | 66824885 | [illegible] |
| MXE | 1883 | ENSG00000102898 | NUTF2 | chr16 | + | 67847277 | 67847456 | 67865101 | 67865229 | [illegible] |
| MXE | 1885 | ENSG00000197170 | PSMD12 | chr17 | - | 67350228 | 67350336 | 67357302 | 67357431 | [illegible] |
| MXE | 1886 | ENSG00000090565 | RAB11FIP3 | chr16 | + | 491133 | 491268 | 496823 | 496859 | [illegible] |
| MXE | 1919 | ENSG00000174628 | IQCK | chr16 | + | 19735352 | 19735450 | 19763847 | 19764112 | [illegible] |
| MXE | 1927 | ENSG00000069345 | DNAJA2 | chr16 | - | 46968083 | 46968164 | 46971348 | 46971659 | [illegible] |
| MXE | 1933 | ENSG00000141646 | SMAD4 | chr18 | + | 51059865 | 51059916 | 51065422 | 51065606 | [illegible] |
| MXE | 1960 | ENSG00000140694 | PARN | chr16 | - | 14588097 | 14588172 | 14591912 | 14592033 | [illegible] |
| MXE | 1961 | ENSG00000140694 | PARN | chr16 | - | 14617589 | 14617650 | 14623923 | 14623953 | [illegible] |
| MXE | 1965 | ENSG00000102897 | LYRM1 | chr16 | + | 20911182 | 20911264 | 20915555 | 20915714 | [illegible] |
| MXE | 1972 | ENSG00000005339 | CREBBP | chr16 | - | 3749626 | 3749683 | 3751725 | 3751836 | [illegible] |
| MXE | 1980 | ENSG00000168096 | ANKS3 | chr16 | - | 4714050 | 4714186 | 4724749 | 4724831 | [illegible] |
| MXE | 1983 | ENSG00000167972 | ABCA3 | chr16 | - | 2308449 | 2308623 | 2317282 | 2317403 | [illegible] |
| MXE | 1993 | ENSG00000173517 | PEAK1 | chr15 | - | 77252366 | 77252526 | 77286422 | 77286504 | [illegible] |
| MXE | 2003 | ENSG00000130703 | OSBPL2 | chr20 | + | 62285535 | 62286217 | 62286336 | 62286488 | [illegible] |
| MXE | 2008 | ENSG00000213614 | HEXA | chr15 | - | 72362375 | 72362466 | 72370430 | 72370694 | [illegible] |
| MXE | 2012 | ENSG00000284626 | NA | chr15 | + | 90240899 | 90241724 | 90244117 | 90244217 | [illegible] |
| MXE | 2014 | ENSG00000140463 | BBS4 | chr15 | + | 72695176 | 72695228 | 72704423 | 72704481 | [illegible] |
| MXE | 2017 | ENSG00000034053 | APBA2 | chr15 | + | 28995752 | 28995806 | 29053844 | 29054835 | [illegible] |
| MXE | 2018 | ENSG00000137871 | ZNF280D | chr15 | - | 56654197 | 56654234 | 56654384 | 56654503 | [illegible] |
| MXE | 2022 | ENSG00000273025 | NA | chr15 | - | 72288537 | 72288618 | 72288867 | 72288930 | [illegible] |
| MXE | 2028 | ENSG00000185658 | BRWD1 | chr21 | - | 39218528 | 39218660 | 39225085 | 39225197 | [illegible] |
| MXE | 2041 | ENSG00000285920 | NA | chr15 | + | 41332155 | 41333050 | 41342385 | 41342454 | [illegible] |
| MXE | 2044 | ENSG00000185418 | TARS3 | chr15 | - | 101684074 | 101684237 | 101685895 | 101686062 | [illegible] |
| MXE | 2052 | ENSG00000090487 | SPG21 | chr15 | - | 64970113 | 64970222 | 64974601 | 64974747 | [illegible] |
| MXE | 2058 | ENSG00000182718 | ANXA2 | chr15 | - | 60364428 | 60364523 | 60386027 | 60386086 | [illegible] |
| MXE | 2067 | ENSG00000182718 | ANXA2 | chr15 | - | 60390212 | 60390317 | 60396372 | 60396427 | [illegible] |
| MXE | 2069 | ENSG00000182718 | ANXA2 | chr15 | - | 60393012 | 60393440 | 60396150 | 60396427 | [illegible] |
| MXE | 2070 | ENSG00000182718 | ANXA2 | chr15 | - | 60393012 | 60393440 | 60396279 | 60396427 | [illegible] |
| MXE | 2072 | ENSG00000182718 | ANXA2 | chr15 | - | 60393012 | 60393440 | 60396279 | 60396427 | [illegible] |
| MXE | 2073 | ENSG00000182718 | ANXA2 | chr15 | - | 60393012 | 60393440 | 60396372 | 60396427 | [illegible] |
| MXE | 2074 | ENSG00000182718 | ANXA2 | chr15 | - | 60393012 | 60393440 | 60397257 | 60397338 | [illegible] |
| MXE | 2077 | ENSG00000182718 | ANXA2 | chr15 | - | 60393037 | 60393440 | 60396372 | 60396427 | [illegible] |
| MXE | 2078 | ENSG00000182718 | ANXA2 | chr15 | - | 60393037 | 60393440 | 60397257 | 60397338 | [illegible] |
| MXE | 2081 | ENSG00000182718 | ANXA2 | chr15 | - | 60396372 | 60396427 | 60397257 | 60397338 | [illegible] |
| MXE | 2086 | ENSG00000157823 | AP3S2 | chr15 | - | 89878264 | 89878300 | 89888520 | 89888632 | [illegible] |
| MXE | 2089 | ENSG00000128915 | ICE2 | chr15 | - | 60431933 | 60431984 | 60436142 | 60436227 | [illegible] |
| MXE | 2090 | ENSG00000128915 | ICE2 | chr15 | - | 60436142 | 60436227 | 60442415 | 60442545 | [illegible] |
| MXE | 2106 | ENSG00000128739 | SNRPN | chr15 | + | 24974287 | 24974456 | 24975357 | 24975509 | [illegible] |
| MXE | 2108 | ENSG00000128739 | SNRPN | chr15 | + | 24974310 | 24974456 | 24974641 | 24974957 | [illegible] |
| MXE | 2113 | ENSG00000128739 | SNRPN | chr15 | + | 24974386 | 24974456 | 24975357 | 24975509 | [illegible] |
| MXE | 2117 | ENSG00000128739 | SNRPN | chr15 | + | 24974397 | 24974456 | 24975357 | 24975509 | [illegible] |
| MXE | 2119 | ENSG00000128739 | SNRPN | chr15 | + | 24974641 | 24974957 | 24975357 | 24975509 | [illegible] |
| MXE | 2124 | ENSG00000104093 | DMXL2 | chr15 | - | 51463378 | 51463496 | 51465565 | 51465651 | [illegible] |
| MXE | 2125 | ENSG00000104093 | DMXL2 | chr15 | - | 51464674 | 51464876 | 51465565 | 51465651 | [illegible] |
| MXE | 2133 | ENSG00000159459 | UBR1 | chr15 | - | 42983896 | 42983993 | 42984886 | 42984942 | [illegible] |
| MXE | 2135 | ENSG00000159459 | UBR1 | chr15 | - | 43043214 | 43043395 | 43047160 | 43047289 | [illegible] |
| MXE | 2136 | ENSG00000159459 | UBR1 | chr15 | - | 43047160 | 43047289 | 43048391 | 43048491 | [illegible] |
| MXE | 2137 | ENSG00000159459 | UBR1 | chr15 | - | 43047160 | 43047289 | 43048391 | 43048491 | [illegible] |
| MXE | 2140 | ENSG00000101266 | CSNK2A1 | chr20 | - | 489779 | 489881 | 492253 | 492364 | [illegible] |
| MXE | 2143 | ENSG00000101266 | CSNK2A1 | chr20 | - | 500030 | 500115 | 503064 | 503281 | [illegible] |
| MXE | 2146 | ENSG00000196705 | ZNF431 | chr19 | + | 21143550 | 21143643 | 21166334 | 21166461 | [illegible] |
| MXE | 2190 | ENSG00000129003 | VPS13C | chr15 | - | 62023779 | 62023845 | 62028357 | 62028420 | [illegible] |
| MXE | 2212 | ENSG00000092439 | TRPM7 | chr15 | - | 50578638 | 50578664 | 50580873 | 50580908 | [illegible] |
| MXE | 2215 | ENSG00000092439 | TRPM7 | chr15 | - | 50583088 | 50583156 | 50586391 | 50586488 | [illegible] |
| MXE | 2216 | ENSG00000092439 | TRPM7 | chr15 | - | 50583088 | 50583159 | 50586391 | 50586488 | [illegible] |
| MXE | 2219 | ENSG00000092439 | TRPM7 | chr15 | - | 50596254 | 50596381 | 50599121 | 50599296 | [illegible] |
| MXE | 2241 | ENSG00000103978 | TMEM87A | chr15 | - | 42231191 | 42231260 | 42231843 | 42231915 | [illegible] |
| MXE | 2243 | ENSG00000178226 | PRSS36 | chr16 | - | 31140293 | 31140415 | 31141468 | 31141610 | [illegible] |
| MXE | 2244 | ENSG00000178226 | PRSS36 | chr16 | - | 31140678 | 31140757 | 31141468 | 31141610 | [illegible] |
| MXE | 2251 | ENSG00000104177 | MYEF2 | chr15 | - | 48151873 | 48151942 | 48152233 | 48152284 | [illegible] |
| MXE | 2252 | ENSG00000186654 | PRR5 | chr22 | + | 44702491 | 44702608 | 44714590 | 44714671 | [illegible] |
| MXE | 2253 | ENSG00000186654 | PRR5 | chr22 | + | 44731734 | 44731821 | 44732250 | 44732391 | [illegible] |
| MXE | 2260 | ENSG00000260661 | NA | chr15 | - | 92157154 | 92157241 | 92313040 | 92313185 | [illegible] |
| MXE | 2263 | ENSG00000137845 | ADAM10 | chr15 | - | 58682195 | 58682314 | 58717576 | 58717727 | [illegible] |
| MXE | 2274 | ENSG00000048471 | SNX29 | chr16 | + | 11999296 | 11999358 | 12002990 | 12003043 | [illegible] |
| MXE | 2284 | ENSG00000128908 | INO80 | chr15 | - | 41027595 | 41027736 | 41044903 | 41045075 | [illegible] |
| MXE | 2293 | ENSG00000267002 | NA | chr17 | - | 43159422 | 43159727 | 43166979 | 43167849 | [illegible] |
| MXE | 2311 | ENSG00000255571 | MIR9-3HG | chr15 | + | 89379047 | 89379269 | 89387430 | 89387538 | [illegible] |
| MXE | 2312 | ENSG00000140374 | ETFA | chr15 | - | 76283756 | 76283825 | 76285636 | 76285738 | [illegible] |
| MXE | 2313 | ENSG00000140374 | ETFA | chr15 | - | 76284364 | 76284460 | 76284804 | 76284927 | [illegible] |
| MXE | 2322 | ENSG00000156958 | GALK2 | chr15 | + | 49225204 | 49225349 | 49235624 | 49235739 | [illegible] |
| MXE | 2328 | ENSG00000104043 | ATP8B4 | chr15 | - | 50018888 | 50018976 | 50038767 | 50038829 | [illegible] |
| MXE | 2329 | ENSG00000166035 | LIPC | chr15 | + | 58538332 | 58538517 | 58541784 | 58541967 | [illegible] |
| MXE | 2331 | ENSG00000140474 | ULK3 | chr15 | - | 74837750 | 74837798 | 74838151 | 74838192 | [illegible] |
| MXE | 2332 | ENSG00000140474 | ULK3 | chr15 | - | 74837750 | 74837798 | 74838151 | 74838192 | [illegible] |
| MXE | 2334 | ENSG00000140474 | ULK3 | chr15 | - | 74837756 | 74837798 | 74838151 | 74838192 | [illegible] |
| MXE | 2335 | ENSG00000140474 | ULK3 | chr15 | - | 74837756 | 74837798 | 74838151 | 74838192 | [illegible] |
| MXE | 2344 | ENSG00000140367 | UBE2Q2 | chr15 | + | 75873427 | 75873568 | 75883365 | 75883424 | [illegible] |
| MXE | 2362 | ENSG00000138629 | UBL7 | chr15 | - | 74452295 | 74452378 | 74456551 | 74456671 | [illegible] |
| MXE | 2365 | ENSG00000248508 | SRP14-AS1 | chr15 | + | 40056452 | 40056641 | 40059132 | 40059289 | [illegible] |
| MXE | 2374 | ENSG00000075131 | TIPIN | chr15 | - | 66349314 | 66349437 | 66351524 | 66351600 | [illegible] |
| MXE | 2402 | ENSG00000170113 | NIPA1 | chr15 | + | 22812162 | 22812253 | 22814082 | 22814153 | [illegible] |
| MXE | 2403 | ENSG00000170113 | NIPA1 | chr15 | + | 22813632 | 22813714 | 22814082 | 22814153 | [illegible] |
| MXE | 2411 | ENSG00000172575 | RASGRP1 | chr15 | - | 38498793 | 38498946 | 38500102 | 38500139 | [illegible] |
| MXE | 2426 | ENSG00000159128 | IFNGR2 | chr21 | + | 33411444 | 33411575 | 33414887 | 33415020 | [illegible] |
| MXE | 2427 | ENSG00000159128 | IFNGR2 | chr21 | + | 33414887 | 33415020 | 33421479 | 33421685 | [illegible] |
| MXE | 2432 | ENSG00000275835 | TUBGCP5 | chr15 | - | 23006051 | 23006172 | 23006267 | 23006352 | [illegible] |
| MXE | 2441 | ENSG00000119537 | KDSR | chr18 | - | 63355218 | 63355299 | 63355497 | 63355563 | [illegible] |
| MXE | 2443 | ENSG00000197555 | SIPA1L1 | chr14 | + | 71413947 | 71414014 | 71503827 | 71503946 | [illegible] |
| MXE | 2444 | ENSG00000197555 | SIPA1L1 | chr14 | + | 71418129 | 71418196 | 71497853 | 71497889 | [illegible] |
| MXE | 2450 | ENSG00000100429 | HDAC10 | chr22 | - | 50248365 | 50248472 | 50249102 | 50249168 | [illegible] |
| MXE | 2451 | ENSG00000100429 | HDAC10 | chr22 | - | 50248661 | 50248751 | 50249102 | 50249168 | [illegible] |
| MXE | 2453 | ENSG00000100429 | HDAC10 | chr22 | - | 50248661 | 50248751 | 50249102 | 50249168 | [illegible] |
| MXE | 2455 | ENSG00000100429 | HDAC10 | chr22 | - | 50248830 | 50248859 | 50249327 | 50249516 | [illegible] |
| MXE | 2456 | ENSG00000100429 | HDAC10 | chr22 | - | 50248830 | 50248890 | 50249102 | 50249168 | [illegible] |
| MXE | 2461 | ENSG00000140043 | PTGR2 | chr14 | + | 73876997 | 73877168 | 73880054 | 73880176 | [illegible] |
| MXE | 2467 | ENSG00000133961 | NUMB | chr14 | - | 73297210 | 73297285 | 73323129 | 73323204 | [illegible] |
| MXE | 2468 | ENSG00000133961 | NUMB | chr14 | - | 73297210 | 73297285 | 73316389 | 73316422 | [illegible] |
| MXE | 2471 | ENSG00000079134 | THOC1 | chr18 | - | 254272 | 254355 | 259179 | 259275 | [illegible] |
| MXE | 2472 | ENSG00000079134 | THOC1 | chr18 | - | 259179 | 259275 | 259681 | 259730 | [illegible] |
| MXE | 2483 | ENSG00000140563 | MCTP2 | chr15 | + | 94458136 | 94458246 | 94470332 | 94470442 | [illegible] |
| MXE | 2494 | ENSG00000197102 | DYNC1H1 | chr14 | + | 102024284 | 102024349 | 102024475 | 102024580 | [illegible] |
| MXE | 2496 | ENSG00000197102 | DYNC1H1 | chr14 | + | 102034334 | 102034452 | 102036488 | 102036642 | [illegible] |
| MXE | 2499 | ENSG00000197102 | DYNC1H1 | chr14 | + | 102040597 | 102040673 | 102041573 | 102041734 | [illegible] |
| MXE | 2501 | ENSG00000012983 | MAP4K5 | chr14 | - | 50423120 | 50423176 | 50425906 | 50425977 | [illegible] |
| MXE | 2502 | ENSG00000012983 | MAP4K5 | chr14 | - | 50425906 | 50425977 | 50428661 | 50428754 | [illegible] |
| MXE | 2503 | ENSG00000012983 | MAP4K5 | chr14 | - | 50428661 | 50428754 | 50429191 | 50429256 | [illegible] |
| MXE | 2504 | ENSG00000012983 | MAP4K5 | chr14 | - | 50428661 | 50428754 | 50429191 | 50429260 | [illegible] |
| MXE | 2526 | ENSG00000100749 | VRK1 | chr14 | + | 96876029 | 96876120 | 96931330 | 96931556 | [illegible] |
| MXE | 2540 | ENSG00000155465 | SLC7A7 | chr14 | - | 22819637 | 22819756 | 22821121 | 22821232 | [illegible] |
| MXE | 2543 | ENSG00000092199 | HNRNPC | chr14 | - | 21234768 | 21234831 | 21236372 | 21236460 | [illegible] |
| MXE | 2544 | ENSG00000092199 | HNRNPC | chr14 | - | 21234768 | 21234831 | 21262600 | 21262768 | [illegible] |
| MXE | 2546 | ENSG00000092199 | HNRNPC | chr14 | - | 21263310 | 21263336 | 21268622 | 21268668 | [illegible] |
| MXE | 2560 | ENSG00000197249 | SERPINA1 | chr14 | - | 94382591 | 94383241 | 94388559 | 94388660 | [illegible] |
| MXE | 2563 | ENSG00000197249 | SERPINA1 | chr14 | - | 94382591 | 94383241 | 94388559 | 94388663 | [illegible] |
| MXE | 2565 | ENSG00000171469 | ZNF561 | chr19 | - | 9614020 | 9614103 | 9618090 | 9618179 | [illegible] |
| MXE | 2566 | ENSG00000171469 | ZNF561 | chr19 | - | 9614020 | 9614103 | 9618090 | 9618166 | [illegible] |
| MXE | 2572 | ENSG00000092108 | SCFD1 | chr14 | + | 30628208 | 30628279 | 30630476 | 30630565 | [illegible] |
| MXE | 2574 | ENSG00000092108 | SCFD1 | chr14 | + | 30633929 | 30634037 | 30643315 | 30643405 | [illegible] |
| MXE | 2582 | ENSG00000249624 | NA | chr21 | + | 33252661 | 33252830 | 33262792 | 33263637 | [illegible] |
| MXE | 2583 | ENSG00000249624 | NA | chr21 | + | 33252661 | 33252830 | 33262792 | 33263637 | [illegible] |
| MXE | 2598 | ENSG00000075413 | MARK3 | chr14 | + | 103429192 | 103429373 | 103438024 | 103438333 | [illegible] |
| MXE | 2599 | ENSG00000075413 | MARK3 | chr14 | + | 103429192 | 103429373 | 103441469 | 103441563 | [illegible] |
| MXE | 2600 | ENSG00000075413 | MARK3 | chr14 | + | 103429192 | 103429373 | 103438020 | 103438333 | [illegible] |
| MXE | 2601 | ENSG00000075413 | MARK3 | chr14 | + | 103429192 | 103429373 | 103438024 | 103438333 | [illegible] |
| MXE | 2602 | ENSG00000075413 | MARK3 | chr14 | + | 103429192 | 103429373 | 103438184 | 103438333 | [illegible] |
| MXE | 2604 | ENSG00000075413 | MARK3 | chr14 | + | 103429192 | 103429373 | 103448918 | 103448967 | [illegible] |
| MXE | 2606 | ENSG00000075413 | MARK3 | chr14 | + | 103438020 | 103438333 | 103448918 | 103448967 | [illegible] |
| MXE | 2607 | ENSG00000075413 | MARK3 | chr14 | + | 103438024 | 103438333 | 103441469 | 103441563 | [illegible] |
| MXE | 2609 | ENSG00000075413 | MARK3 | chr14 | + | 103438024 | 103438333 | 103448918 | 103448967 | [illegible] |
| MXE | 2611 | ENSG00000075413 | MARK3 | chr14 | + | 103438184 | 103438333 | 103448918 | 103448967 | [illegible] |
| MXE | 2622 | ENSG00000075413 | MARK3 | chr14 | + | 103487378 | 103487707 | 103489052 | 103491107 | [illegible] |
| MXE | 2623 | ENSG00000075413 | MARK3 | chr14 | + | 103487378 | 103487707 | 103490951 | 103491107 | [illegible] |
| MXE | 2624 | ENSG00000075413 | MARK3 | chr14 | + | 103487378 | 103487707 | 103490977 | 103491107 | [illegible] |
| MXE | 2628 | ENSG00000075413 | MARK3 | chr14 | + | 103498501 | 103498528 | 103500155 | 103500200 | [illegible] |
| MXE | 2630 | ENSG00000075413 | MARK3 | chr14 | + | 103498501 | 103498528 | 103500155 | 103500200 | [illegible] |
| MXE | 2632 | ENSG00000100554 | ATP6V1D | chr14 | - | 67337386 | 67337509 | 67338074 | 67338152 | [illegible] |
| MXE | 2633 | ENSG00000100554 | ATP6V1D | chr14 | - | 67338560 | 67338762 | 67340439 | 67340518 | [illegible] |
| MXE | 2635 | ENSG00000100554 | ATP6V1D | chr14 | - | 67347408 | 67347453 | 67349036 | 67349104 | [illegible] |
| MXE | 2647 | ENSG00000185347 | TEDC1 | chr14 | + | 105492578 | 105492734 | 105493834 | 105493933 | [illegible] |
| MXE | 2654 | ENSG00000100938 | GMPR2 | chr14 | + | 24233218 | 24233340 | 24233478 | 24233586 | [illegible] |
| MXE | 2686 | ENSG00000256053 | COA8 | chr14 | + | 103574106 | 103574170 | 103581568 | 103581656 | [illegible] |
| MXE | 2687 | ENSG00000256053 | COA8 | chr14 | + | 103579370 | 103579491 | 103581568 | 103581656 | [illegible] |
| MXE | 2705 | ENSG00000119684 | MLH3 | chr14 | - | 75033418 | 75033490 | 75038339 | 75038412 | [illegible] |
| MXE | 2706 | ENSG00000119684 | MLH3 | chr14 | - | 75033418 | 75033490 | 75039910 | 75040015 | [illegible] |
| MXE | 2707 | ENSG00000119689 | DLST | chr14 | + | 74885585 | 74885634 | 74889094 | 74889147 | [illegible] |
| MXE | 2708 | ENSG00000119689 | DLST | chr14 | + | 74885585 | 74885634 | 74889094 | 74889147 | [illegible] |
| MXE | 2709 | ENSG00000119689 | DLST | chr14 | + | 74885585 | 74885634 | 74889094 | 74889147 | [illegible] |
| MXE | 2711 | ENSG00000119689 | DLST | chr14 | + | 74889274 | 74889349 | 74889896 | 74889952 | [illegible] |
| MXE | 2722 | ENSG00000215277 | RNF212B | chr14 | + | 23260658 | 23260687 | 23270599 | 23270661 | [illegible] |
| MXE | 2726 | ENSG00000100461 | RBM23 | chr14 | - | 22911785 | 22911903 | 22913723 | 22913821 | [illegible] |
| MXE | 2727 | ENSG00000100461 | RBM23 | chr14 | - | 22911785 | 22911903 | 22913723 | 22913845 | [illegible] |
| MXE | 2728 | ENSG00000100461 | RBM23 | chr14 | - | 22911785 | 22911903 | 22917027 | 22917166 | [illegible] |
| MXE | 2729 | ENSG00000100461 | RBM23 | chr14 | - | 22913723 | 22913821 | 22917027 | 22917166 | [illegible] |
| MXE | 2754 | ENSG00000165914 | TTC7B | chr14 | - | 90593485 | 90593626 | 90610741 | 90610839 | [illegible] |
| MXE | 2770 | ENSG00000140105 | WARS1 | chr14 | - | 100365474 | 100365576 | 100365953 | 100366067 | [illegible] |
| MXE | 2771 | ENSG00000140105 | WARS1 | chr14 | - | 100369086 | 100369258 | 100374135 | 100374179 | [illegible] |
| MXE | 2784 | ENSG00000259084 | NA | chr14 | + | 95669550 | 95669617 | 95670441 | 95670577 | [illegible] |
| MXE | 2785 | ENSG00000259084 | NA | chr14 | + | 95669550 | 95669617 | 95670441 | 95670577 | [illegible] |
| MXE | 2795 | ENSG00000100811 | YY1 | chr14 | + | 100262303 | 100262466 | 100274697 | 100274758 | [illegible] |
| MXE | 2796 | ENSG00000070367 | EXOC5 | chr14 | - | 57233987 | 57234032 | 57235710 | 57235820 | [illegible] |
| MXE | 2803 | ENSG00000151445 | VIPAS39 | chr14 | - | 77449292 | 77449357 | 77449713 | 77449752 | [illegible] |
| MXE | 2804 | ENSG00000119636 | BBOF1 | chr14 | + | 74049701 | 74050195 | 74055583 | 74055685 | [illegible] |
| MXE | 2808 | ENSG00000139946 | PELI2 | chr14 | + | 56178334 | 56178464 | 56279675 | 56279777 | [illegible] |
| MXE | 2812 | ENSG00000161558 | TMEM143 | chr19 | - | 48345159 | 48345354 | 48360071 | 48360176 | [illegible] |
| MXE | 2823 | ENSG00000187790 | FANCM | chr14 | + | 45155372 | 45155459 | 45164358 | 45164565 | [illegible] |
| MXE | 2824 | ENSG00000187790 | FANCM | chr14 | + | 45159095 | 45159280 | 45164358 | 45164565 | [illegible] |
| MXE | 2841 | ENSG00000081177 | EXD2 | chr14 | + | 69203916 | 69204000 | 69209423 | 69209803 | [illegible] |
| MXE | 2842 | ENSG00000119688 | ABCD4 | chr14 | - | 74293153 | 74293248 | 74295147 | 74295198 | [illegible] |
| MXE | 2843 | ENSG00000119688 | ABCD4 | chr14 | - | 74297638 | 74298069 | 74299547 | 74299675 | [illegible] |
| MXE | 2846 | ENSG00000103811 | CTSH | chr15 | - | 78926174 | 78926254 | 78926781 | 78926830 | [illegible] |
| MXE | 2847 | ENSG00000103811 | CTSH | chr15 | - | 78926174 | 78926254 | 78926781 | 78926868 | [illegible] |
| MXE | 2848 | ENSG00000103811 | CTSH | chr15 | - | 78926174 | 78926254 | 78926781 | 78926939 | [illegible] |
| MXE | 2849 | ENSG00000103811 | CTSH | chr15 | - | 78926174 | 78926254 | 78927367 | 78927436 | [illegible] |
| MXE | 2851 | ENSG00000103811 | CTSH | chr15 | - | 78926781 | 78926868 | 78927367 | 78927436 | [illegible] |
| MXE | 2852 | ENSG00000103811 | CTSH | chr15 | - | 78926781 | 78926939 | 78927367 | 78927436 | [illegible] |
| MXE | 2859 | ENSG00000198252 | STYX | chr14 | + | 52730458 | 52730531 | 52744851 | 52744884 | [illegible] |
| MXE | 2869 | ENSG00000152240 | HAUS1 | chr18 | + | 46118180 | 46118316 | 46119925 | 46120060 | [illegible] |
| MXE | 2871 | ENSG00000152240 | HAUS1 | chr18 | + | 46122466 | 46122590 | 46123298 | 46123364 | [illegible] |
| MXE | 2873 | ENSG00000092140 | G2E3 | chr14 | + | 30581075 | 30581116 | 30586717 | 30586815 | [illegible] |
| MXE | 2895 | ENSG00000139974 | SLC38A6 | chr14 | + | 61034018 | 61034113 | 61037058 | 61037141 | [illegible] |
| MXE | 2908 | ENSG00000129493 | HEATR5A | chr14 | - | 31293890 | 31294104 | 31295908 | 31296063 | [illegible] |
| MXE | 2909 | ENSG00000129493 | HEATR5A | chr14 | - | 31302294 | 31302519 | 31304904 | 31305177 | [illegible] |
| MXE | 2913 | ENSG00000090615 | GOLGA3 | chr12 | - | 132827606 | 132827713 | 132827807 | 132827902 | [illegible] |
| MXE | 2915 | ENSG00000137210 | TMEM14B | chr6 | + | 10749201 | 10749268 | 10770076 | 10770204 | [illegible] |
| MXE | 2918 | ENSG00000111845 | PAK1IP1 | chr6 | + | 10702564 | 10702639 | 10703404 | 10703457 | [illegible] |
| MXE | 2931 | ENSG00000118482 | PHF3 | chr6 | + | 63646526 | 63646795 | 63691736 | 63692043 | [illegible] |
| MXE | 2932 | ENSG00000118482 | PHF3 | chr6 | + | 63646618 | 63646795 | 63691736 | 63692043 | [illegible] |
| MXE | 2956 | ENSG00000162711 | NLRP3 | chr1 | + | 247418228 | 247418290 | 247418756 | 247419077 | [illegible] |
| MXE | 2969 | ENSG00000213066 | CEP43 | chr6 | + | 167026546 | 167026615 | 167032602 | 167032642 | [illegible] |
| MXE | 2970 | ENSG00000204623 | ZNRD1ASP | chr6 | - | 30021666 | 30021761 | 30034907 | 30035110 | [illegible] |
| MXE | 2973 | ENSG00000125871 | MGME1 | chr20 | + | 17975683 | 17975903 | 17988165 | 17988298 | [illegible] |
| MXE | 2981 | ENSG00000135446 | CDK4 | chr12 | - | 57750441 | 57750563 | 57750655 | 57750765 | [illegible] |
| MXE | 2989 | ENSG00000130779 | CLIP1 | chr12 | - | 122347374 | 122347479 | 122351110 | 122351143 | [illegible] |
| MXE | 2990 | ENSG00000234911 | TEX21P | chr14 | - | 64347522 | 64347648 | 64353952 | 64354074 | [illegible] |
| MXE | 2996 | ENSG00000166024 | R3HCC1L | chr10 | + | 98156092 | 98156147 | 98162882 | 98162975 | [illegible] |
| MXE | 2997 | ENSG00000166024 | R3HCC1L | chr10 | + | 98162882 | 98162975 | 98163292 | 98163397 | [illegible] |
| MXE | 3001 | ENSG00000166024 | R3HCC1L | chr10 | + | 98211306 | 98211348 | 98231511 | 98231687 | [illegible] |
| MXE | 3002 | ENSG00000165025 | SYK | chr9 | + | 90865047 | 90865097 | 90867130 | 90867199 | [illegible] |
| MXE | 3018 | ENSG00000175048 | ZDHHC14 | chr6 | + | 157592987 | 157593146 | 157628348 | 157628374 | [illegible] |
| MXE | 3020 | ENSG00000026652 | AGPAT4 | chr6 | - | 161154148 | 161154310 | 161166247 | 161166417 | [illegible] |
| MXE | 3025 | ENSG00000112290 | WASF1 | chr6 | - | 110160634 | 110160732 | 110178597 | 110178742 | [illegible] |
| MXE | 3031 | ENSG00000181036 | FCRL6 | chr1 | + | 159814220 | 159814292 | 159815427 | 159815459 | [illegible] |
| MXE | 3042 | ENSG00000255819 | KLRC4-KLRK1 | chr12 | - | 10388770 | 10388875 | 10389942 | 10390027 | [illegible] |
| MXE | 3043 | ENSG00000255819 | KLRC4-KLRK1 | chr12 | - | 10389942 | 10390027 | 10393809 | 10393867 | [illegible] |
| MXE | 3069 | ENSG00000198399 | ITSN2 | chr2 | - | 24321813 | 24321880 | 24326659 | 24326681 | [illegible] |
| MXE | 3078 | ENSG00000127990 | SGCE | chr7 | - | 94599696 | 94599723 | 94600645 | 94600857 | [illegible] |
| MXE | 3086 | ENSG00000229956 | ZRANB2-AS2 | chr1 | + | 71093132 | 71093251 | 71223056 | 71223114 | [illegible] |
| MXE | 3100 | ENSG00000143252 | SDHC | chr1 | + | 161323613 | 161323670 | 161328395 | 161328497 | [illegible] |
| MXE | 3117 | ENSG00000150867 | PIP4K2A | chr10 | - | 22550658 | 22550772 | 22567850 | 22567889 | [illegible] |
| MXE | 3139 | ENSG00000154309 | DISP1 | chr1 | + | 222872270 | 222872428 | 222899780 | 222899865 | [illegible] |
| MXE | 3145 | ENSG00000126777 | KTN1 | chr14 | + | 55663954 | 55664041 | 55667240 | 55667330 | [illegible] |
| MXE | 3150 | ENSG00000164414 | SLC35A1 | chr6 | + | 87506381 | 87506448 | 87508419 | 87508596 | [illegible] |
| MXE | 3155 | ENSG00000108468 | CBX1 | chr17 | - | 48075988 | 48076178 | 48076864 | 48077041 | [illegible] |
| MXE | 3158 | ENSG00000093134 | VNN3 | chr6 | - | 132726721 | 132727012 | 132728740 | 132728979 | [illegible] |
| MXE | 3160 | ENSG00000085382 | HACE1 | chr6 | - | 104811310 | 104811393 | 104843222 | 104843298 | [illegible] |
| MXE | 3166 | ENSG00000075568 | TMEM131 | chr2 | - | 97818612 | 97818721 | 97833364 | 97833426 | [illegible] |
| MXE | 3171 | ENSG00000129515 | SNX6 | chr14 | - | 34603347 | 34603471 | 34604134 | 34604174 | [illegible] |
| MXE | 3180 | ENSG00000135341 | MAP3K7 | chr6 | - | 90561621 | 90561667 | 90568557 | 90568623 | [illegible] |
| MXE | 3186 | ENSG00000133706 | LARS1 | chr5 | - | 146164309 | 146164471 | 146168127 | 146168265 | [illegible] |
| MXE | 3196 | ENSG00000125459 | MSTO1 | chr1 | + | 155611215 | 155611241 | 155611548 | 155611603 | [illegible] |

[illegible]

| | | | | | | | | | | | | | | |
|---|---|---|---|---|---|---|---|---|---|---|---|---|---|---|
| MXE | 4450 | ENSG00000182621 | PLCB1 | chr20 | + | 8760406 | 8760460 | 8765138 | 8765358 | 8757045 | 8757178 | 8774538 | 8774719 | 0 |
| MXE | 4462 | ENSG00000141452 | RMC1 | chr18 | + | 23506969 | 23507054 | 23507984 | 23508041 | 23504370 | 23504447 | 23515855 | 23515996 | 32 |
| MXE | 4464 | ENSG00000141452 | RMC1 | chr18 | + | 23507984 | 23508041 | 23509192 | 23509279 | 23504370 | 23504447 | 23515855 | 23515996 | 0 |
| MXE | 4469 | ENSG00000154743 | TSEN2 | chr3 | + | 12505153 | 12505231 | 12516610 | 12516661 | 12503261 | 12503784 | 12519058 | 12519197 | 9 |
| MXE | 4470 | ENSG00000154743 | TSEN2 | chr3 | + | 12505153 | 12505231 | 12516610 | 12516661 | 12503261 | 12503784 | 12528887 | 12528924 | 5 |
| MXE | 4471 | ENSG00000154743 | TSEN2 | chr3 | + | 12505153 | 12505231 | 12519058 | 12519197 | 12503261 | 12503784 | 12528887 | 12528924 | 5 |
| MXE | 4472 | ENSG00000154743 | TSEN2 | chr3 | + | 12505153 | 12505231 | 12528887 | 12528924 | 12503261 | 12503784 | 12529761 | 12529873 | 5 |
| MXE | 4473 | ENSG00000154743 | TSEN2 | chr3 | + | 12506707 | 12506835 | 12516610 | 12516661 | 12505153 | 12505231 | 12528887 | 12528924 | 0 |
| MXE | 4474 | ENSG00000154743 | TSEN2 | chr3 | + | 12506707 | 12506835 | 12519058 | 12519197 | 12505153 | 12505231 | 12528887 | 12528924 | 0 |
| MXE | 4475 | ENSG00000154743 | TSEN2 | chr3 | + | 12516610 | 12516661 | 12519058 | 12519197 | 12503261 | 12503784 | 12528887 | 12528924 | 0 |
| MXE | 4476 | ENSG00000154743 | TSEN2 | chr3 | + | 12516610 | 12516661 | 12519058 | 12519197 | 12505153 | 12505231 | 12528887 | 12528924 | 2 |
| MXE | 4481 | ENSG00000106070 | GRB10 | chr7 | - | 50703820 | 50703908 | 50710874 | 50710990 | 50674435 | 50674658 | 50732271 | 50732368 | 0 |
| MXE | 4482 | ENSG00000106070 | GRB10 | chr7 | - | 50703820 | 50703908 | 50710874 | 50710990 | 50674435 | 50674658 | 50755886 | 50756056 | 2 |
| MXE | 4483 | ENSG00000106070 | GRB10 | chr7 | - | 50705139 | 50705323 | 50710874 | 50710990 | 50703820 | 50703908 | 50755886 | 50756056 | 4 |
| MXE | 4484 | ENSG00000106070 | GRB10 | chr7 | - | 50705139 | 50705323 | 50732271 | 50732368 | 50703820 | 50703908 | 50755886 | 50756056 | 0 |
| MXE | 4485 | ENSG00000106070 | GRB10 | chr7 | - | 50710874 | 50710990 | 50732271 | 50732368 | 50703820 | 50703908 | 50755886 | 50756056 | 0 |
| MXE | 4499 | ENSG00000248049 | UBA6-AS1 | chr4 | + | 67730073 | 67730170 | 67731505 | 67732345 | 67728539 | 67728686 | 67745826 | 67746366 | 20 |
| MXE | 4501 | ENSG00000144290 | SLC4A10 | chr2 | + | 161950686 | 161950848 | 161958486 | 161958555 | 161949147 | 161949261 | 161964134 | 161964308 | 49 |
| MXE | 4508 | ENSG00000113522 | RAD50 | chr5 | + | 132589630 | 132589837 | 132591223 | 132591406 | 132587923 | 132588089 | 132591876 | 132592034 | 0 |
| MXE | 4511 | ENSG00000113522 | RAD50 | chr5 | + | 132592763 | 132592965 | 132594561 | 132594677 | 132591876 | 132592034 | 132595103 | 132595201 | 3 |
| MXE | 4513 | ENSG00000113522 | RAD50 | chr5 | + | 132594868 | 132595044 | 132595103 | 132595201 | 132594561 | 132594677 | 132595572 | 132595810 | 113 |
| MXE | 4523 | ENSG00000163811 | WDR43 | chr2 | + | 28937930 | 28937994 | 28941460 | 28941574 | 28936921 | 28936953 | 28942311 | 28942381 | 22 |
| MXE | 4532 | ENSG00000168397 | ATG4B | chr2 | + | 241651883 | 241651974 | 241653306 | 241653369 | 241651263 | 241651335 | 241653511 | 241653610 | 2 |
| MXE | 4536 | ENSG00000005700 | IBTK | chr6 | - | 82212706 | 82212793 | 82214226 | 82214829 | 82204856 | 82204958 | 82216075 | 82216250 | 51 |
| MXE | 4537 | ENSG00000005700 | IBTK | chr6 | - | 82214226 | 82214829 | 82216075 | 82216250 | 82212706 | 82212793 | 82217959 | 82218137 | 59 |
| MXE | 4539 | ENSG00000005700 | IBTK | chr6 | - | 82234158 | 82234255 | 82240165 | 82240843 | 82231717 | 82231842 | 82247561 | 82247671 | 31 |
| MXE | 4540 | ENSG00000178104 | PDE4DIP | chr1 | + | 148889783 | 148889894 | 148929196 | 148929273 | 148868470 | 148868622 | 148931799 | 148931923 | 1 |
| MXE | 4542 | ENSG00000135299 | ANKRD6 | chr6 | + | 89606006 | 89606105 | 89612271 | 89612370 | 89603028 | 89603127 | 89623409 | 89623544 | 4 |
| MXE | 4555 | ENSG00000196209 | SIRPB2 | chr20 | - | 1477337 | 1477403 | 1478265 | 1478607 | 1473791 | 1473870 | 1479699 | 1480065 | 55 |
| MXE | 4557 | ENSG00000014824 | SLC30A9 | chr4 | + | 42070525 | 42070691 | 42078211 | 42078325 | 42067084 | 42067192 | 42086081 | 42086369 | 56 |
| MXE | 4558 | ENSG00000163138 | PACRGL | chr4 | + | 20707802 | 20707870 | 20709682 | 20709773 | 20704659 | 20704814 | 20712787 | 20712922 | 9 |
| MXE | 4559 | ENSG00000163138 | PACRGL | chr4 | + | 20709682 | 20709773 | 20712787 | 20712922 | 20704659 | 20704814 | 20713431 | 20713539 | 0 |
| MXE | 4560 | ENSG00000163138 | PACRGL | chr4 | + | 20709682 | 20709773 | 20712787 | 20712922 | 20707802 | 20707870 | 20713431 | 20713539 | 3 |
| MXE | 4569 | ENSG00000288695 | NA | chr4 | + | 55361116 | 55361250 | 55367587 | 55367722 | 55359345 | 55359488 | 55369831 | 55370534 | 0 |
| MXE | 4570 | ENSG00000152784 | PRDM8 | chr4 | + | 80185364 | 80185518 | 80191471 | 80191569 | 80184649 | 80184753 | 80194112 | 80194239 | 2 |
| MXE | 4573 | ENSG00000165533 | TTC8 | chr14 | + | 88841424 | 88841557 | 88843805 | 88843850 | 88841036 | 88841196 | 88852970 | 88853056 | 0 |
| MXE | 4579 | ENSG00000189308 | LIN54 | chr4 | - | 82946257 | 82946474 | 82970326 | 82970469 | 82939888 | 82939962 | 82984160 | 82984876 | 0 |
| MXE | 4587 | ENSG00000132466 | ANKRD17 | chr4 | - | 73113791 | 73113908 | 73115820 | 73115916 | 73113083 | 73113311 | 73118687 | 73118850 | 125 |
| MXE | 4596 | ENSG00000100412 | ACO2 | chr22 | + | 41502629 | 41502771 | 41507445 | 41507606 | 41499725 | 41499862 | 41507790 | 41508049 | 0 |
| MXE | 4597 | ENSG00000100412 | ACO2 | chr22 | + | 41517942 | 41518408 | 41518480 | 41518572 | 41517526 | 41517631 | 41520170 | 41520276 | 1 |
| MXE | 4599 | ENSG00000100412 | ACO2 | chr22 | + | 41520170 | 41520276 | 41522829 | 41522987 | 41517526 | 41517631 | 41523204 | 41523278 | 0 |
| MXE | 4601 | ENSG00000187621 | TCL6 | chr14 | + | 95654508 | 95654626 | 95662479 | 95662664 | 95652376 | 95652489 | 95663255 | 95663635 | 11 |
| MXE | 4602 | ENSG00000187621 | TCL6 | chr14 | + | 95654508 | 95654626 | 95662479 | 95662664 | 95652376 | 95652489 | 95669550 | 95669617 | 13 |
| MXE | 4603 | ENSG00000187621 | TCL6 | chr14 | + | 95662479 | 95662664 | 95663255 | 95663635 | 95654508 | 95654626 | 95669550 | 95669617 | 0 |
| MXE | 4604 | ENSG00000187621 | TCL6 | chr14 | + | 95668125 | 95669617 | 95670441 | 95670577 | 95663255 | 95665076 | 95671330 | 95671487 | 4 |
| MXE | 4615 | ENSG00000134852 | CLOCK | chr4 | - | 55453053 | 55453129 | 55455896 | 55456003 | 55450090 | 55450232 | 55456217 | 55456300 | 46 |
| MXE | 4622 | ENSG00000137807 | KIF23 | chr15 | + | 69440767 | 69441079 | 69444789 | 69445041 | 69440307 | 69440487 | 69446008 | 69446091 | 4 |
| MXE | 4623 | ENSG00000137807 | KIF23 | chr15 | + | 69446282 | 69446364 | 69446430 | 69446587 | 69446008 | 69446091 | 69446870 | 69446941 | 14 |
| MXE | 4624 | ENSG00000137807 | KIF23 | chr15 | + | 69446285 | 69446364 | 69446430 | 69446587 | 69446008 | 69446091 | 69446870 | 69446941 | 6 |
| MXE | 4634 | ENSG00000065882 | TBC1D1 | chr4 | + | 38051898 | 38052060 | 38053105 | 38053225 | 38049617 | 38049898 | 38054198 | 38054338 | 0 |
| MXE | 4656 | ENSG00000008311 | AASS | chr7 | - | 122098744 | 122098866 | 122101370 | 122101438 | 122098449 | 122098576 | 122101620 | 122101680 | 2 |
| MXE | 4657 | ENSG00000163104 | SMARCAD1 | chr4 | + | 94226118 | 94226296 | 94233953 | 94234122 | 94208345 | 94208584 | 94236951 | 94237018 | 64 |
| MXE | 4662 | ENSG00000025293 | PHF20 | chr20 | + | 35842572 | 35842744 | 35847349 | 35847434 | 35801490 | 35801605 | 35863012 | 35863400 | 108 |
| MXE | 4664 | ENSG00000025293 | PHF20 | chr20 | + | 35847349 | 35847434 | 35858301 | 35858381 | 35801490 | 35801605 | 35863012 | 35863400 | 2 |
| MXE | 4678 | ENSG00000047365 | ARAP2 | chr4 | - | 36015385 | 36015632 | 36015899 | 36015958 | 36012562 | 36012831 | 36019143 | 36019286 | 0 |
| MXE | 4679 | ENSG00000047365 | ARAP2 | chr4 | - | 36015385 | 36015632 | 36019143 | 36019286 | 36012562 | 36012831 | 36045978 | 36046076 | 1 |
| MXE | 4680 | ENSG00000047365 | ARAP2 | chr4 | - | 36046709 | 36046849 | 36052005 | 36052053 | 36045978 | 36046076 | 36057969 | 36058143 | 0 |
| MXE | 4692 | ENSG00000157426 | AASDH | chr4 | - | 56342834 | 56342966 | 56343561 | 56343684 | 56338604 | 56338791 | 56349262 | 56350058 | 21 |
| MXE | 4695 | ENSG00000268758 | ADGRE4P | chr19 | - | 6954603 | 6954699 | 6957043 | 6957224 | 6952499 | 6953318 | 6960296 | 6960388 | 8 |
| MXE | 4698 | ENSG00000155508 | CNOT8 | chr5 | + | 154858638 | 154858768 | 154865191 | 154865385 | 154858516 | 154858543 | 154870660 | 154870822 | 1 |
| MXE | 4699 | ENSG00000155508 | CNOT8 | chr5 | + | 154858638 | 154858801 | 154865191 | 154865385 | 154858516 | 154858543 | 154870660 | 154870822 | 2 |
| MXE | 4700 | ENSG00000155508 | CNOT8 | chr5 | + | 154858638 | 154858816 | 154865191 | 154865385 | 154858261 | 154858525 | 154870660 | 154870747 | 1 |
| MXE | 4703 | ENSG00000155508 | CNOT8 | chr5 | + | 154863206 | 154863395 | 154865191 | 154865385 | 154858688 | 154858816 | 154867691 | 154867792 | 9 |
| MXE | 4708 | ENSG00000178950 | GAK | chr4 | - | 851749 | 851974 | 859361 | 859493 | 850935 | 851084 | 859605 | 859722 | 0 |
| MXE | 4712 | ENSG00000174227 | PIGG | chr4 | + | 508828 | 508970 | 515972 | 516185 | 500395 | 500601 | 521055 | 521273 | 0 |
| MXE | 4720 | ENSG00000171608 | PIK3CD | chr1 | + | 9717063 | 9717108 | 9717536 | 9717626 | 9716439 | 9716619 | 9718693 | 9718915 | 0 |
| MXE | 4723 | ENSG00000163913 | IFT122 | chr3 | + | 129456113 | 129456264 | 129457921 | 129458015 | 129451913 | 129451998 | 129458598 | 129458677 | 2 |
| MXE | 4726 | ENSG00000116668 | SWT1 | chr1 | + | 185181945 | 185182057 | 185184242 | 185184344 | 185180390 | 185180450 | 185184742 | 185184931 | 23 |
| MXE | 4735 | ENSG00000168026 | TTC21A | chr3 | + | 39130247 | 39130358 | 39130700 | 39130839 | 39130078 | 39130151 | 39130991 | 39131095 | 4 |
| MXE | 4736 | ENSG00000168026 | TTC21A | chr3 | + | 39130247 | 39130358 | 39130991 | 39131095 | 39130078 | 39130151 | 39133051 | 39133240 | 4 |
| MXE | 4740 | ENSG00000131389 | SLC6A6 | chr3 | + | 14443627 | 14443863 | 14445716 | 14445851 | 14416411 | 14416453 | 14447581 | 14447816 | 1 |
| MXE | 4754 | ENSG00000137171 | KLC4 | chr6 | + | 43073222 | 43073338 | 43073901 | 43073965 | 43072823 | 43072964 | 43074621 | 43074908 | 12 |
| MXE | 4755 | ENSG00000174776 | WDR49 | chr3 | - | 167575917 | 167576151 | 167626851 | 167627292 | 167566829 | 167566894 | 167653260 | 167653499 | 5 |
| MXE | 4757 | ENSG00000047849 | MAP4 | chr3 | - | 47902949 | 47903000 | 47909037 | 47912421 | 47877416 | 47877523 | 47914816 | 47914939 | 7 |
| MXE | 4763 | ENSG00000152133 | GPATCH11 | chr2 | + | 37089639 | 37089866 | 37090680 | 37090722 | 37088368 | 37088440 | 37091915 | 37092036 | 38 |
| MXE | 4780 | ENSG00000143669 | LYST | chr1 | - | 235788700 | 235788845 | 235791698 | 235792125 | 235781926 | 235782087 | 235793502 | 235793612 | 249 |
| MXE | 4790 | ENSG00000100003 | SEC14L2 | chr22 | + | 30407414 | 30407603 | 30409186 | 30409282 | 30406341 | 30406385 | 30409425 | 30409486 | 0 |
| MXE | 4792 | ENSG00000163702 | IL17RC | chr3 | + | 9930027 | 9930149 | 9930399 | 9930459 | 9929851 | 9929897 | 9930894 | 9930943 | 1 |
| MXE | 4795 | ENSG00000182568 | SATB1 | chr3 | - | 18415110 | 18415234 | 18416006 | 18416133 | 18410797 | 18411054 | 18416901 | 18417393 | 285 |
| MXE | 4796 | ENSG00000242086 | MUC20-OT1 | chr3 | + | 195685817 | 195685939 | 195686603 | 195686718 | 195683749 | 195683865 | 195688532 | 195688797 | 22 |
| MXE | 4799 | ENSG00000242086 | MUC20-OT1 | chr3 | + | 195701379 | 195701495 | 195708133 | 195708435 | 195699029 | 195699207 | 195708746 | 195708807 | 8 |
| MXE | 4807 | ENSG00000253438 | PCAT1 | chr8 | + | 126589191 | 126589290 | 126713285 | 126713674 | 126557892 | 126557931 | 126877671 | 126877790 | 2 |
| MXE | 4808 | ENSG00000253438 | PCAT1 | chr8 | + | 126878374 | 126878480 | 126926317 | 126926399 | 126877671 | 126877790 | 127006554 | 127006618 | 0 |
| MXE | 4809 | ENSG00000253438 | PCAT1 | chr8 | + | 126878374 | 126878480 | 126938255 | 126938310 | 126877671 | 126877790 | 127006554 | 127006618 | 0 |
| MXE | 4810 | ENSG00000253438 | PCAT1 | chr8 | + | 126926317 | 126926399 | 126938255 | 126938310 | 126877671 | 126877790 | 127006554 | 127006618 | 0 |
| MXE | 4824 | ENSG00000068745 | IP6K2 | chr3 | - | 48708119 | 48708237 | 48711366 | 48711459 | 48695089 | 48695418 | 48717156 | 48717214 | 0 |
| MXE | 4827 | ENSG00000013441 | CLK1 | chr2 | - | 200855003 | 200855086 | 200856681 | 200856811 | 200854615 | 200854695 | 200857717 | 200857884 | 2 |
| MXE | 4829 | ENSG00000013441 | CLK1 | chr2 | - | 200856681 | 200856811 | 200856890 | 200856985 | 200854615 | 200854695 | 200857717 | 200857884 | 268 |
| MXE | 4842 | ENSG00000131370 | SH3BP5 | chr3 | - | 15273115 | 15273405 | 15304102 | 15304231 | 15271600 | 15271671 | 15330503 | 15330566 | 65 |
| MXE | 4845 | ENSG00000131375 | CAPN7 | chr3 | + | 15242177 | 15242253 | 15245525 | 15245671 | 15240753 | 15240853 | 15246731 | 15246794 | 23 |
| MXE | 4853 | ENSG00000261796 | ISY1-RAB43 | chr3 | - | 129130087 | 129130188 | 129130549 | 129130636 | 129094985 | 129095169 | 129134073 | 129134195 | 50 |
| MXE | 4861 | ENSG00000072756 | TRNT1 | chr3 | + | 3139588 | 3139710 | 3140509 | 3140648 | 3137259 | 3137453 | 3144583 | 3144710 | 0 |
| MXE | 4874 | ENSG00000150995 | ITPR1 | chr3 | + | 4725545 | 4725581 | 4727122 | 4727173 | 4711756 | 4711868 | 4733087 | 4733220 | 0 |
| MXE | 4878 | ENSG00000058056 | USP13 | chr3 | + | 179690243 | 179690301 | 179701007 | 179701129 | 179681877 | 179682003 | 179706933 | 179707076 | 0 |
| MXE | 4881 | ENSG00000288698 | NA | chr3 | + | 179757051 | 179757078 | 179761111 | 179761255 | 179754731 | 179754854 | 179764001 | 179764168 | 1 |
| MXE | 4882 | ENSG00000168297 | PXK | chr3 | + | 58369430 | 58369478 | 58382513 | 58382700 | 58365873 | 58365924 | 58394997 | 58395102 | 2 |
| MXE | 4889 | ENSG00000088387 | DOCK9 | chr13 | - | 98845328 | 98845372 | 98845923 | 98845991 | 98837493 | 98837609 | 98848591 | 98848639 | 18 |
| MXE | 4890 | ENSG00000088387 | DOCK9 | chr13 | - | 98845328 | 98845372 | 98845923 | 98846060 | 98837493 | 98837609 | 98848591 | 98848639 | 92 |
| MXE | 4900 | ENSG00000075711 | DLG1 | chr3 | - | 197069218 | 197069260 | 197075836 | 197075870 | 197066703 | 197066754 | 197076585 | 197076685 | 0 |
| MXE | 4901 | ENSG00000075711 | DLG1 | chr3 | - | 197069218 | 197069260 | 197075836 | 197075870 | 197066703 | 197066754 | 197081050 | 197081117 | 8 |
| MXE | 4903 | ENSG00000075711 | DLG1 | chr3 | - | 197069218 | 197069260 | 197075836 | 197075870 | 197068501 | 197068537 | 197081050 | 197081117 | 8 |
| MXE | 4909 | ENSG00000075711 | DLG1 | chr3 | - | 197085295 | 197085756 | 197090911 | 197091026 | 197081050 | 197081117 | 197104902 | 197105005 | 174 |
| MXE | 4913 | ENSG00000075711 | DLG1 | chr3 | - | 197104902 | 197105005 | 197115926 | 197116083 | 197090911 | 197091026 | 197130526 | 197130671 | 0 |
| MXE | 4920 | ENSG00000075711 | DLG1 | chr3 | - | 197155668 | 197155763 | 197161639 | 197161738 | 197149742 | 197149796 | 197194424 | 197194589 | 58 |
| MXE | 4928 | ENSG00000114316 | USP4 | chr3 | - | 49302383 | 49302542 | 49305714 | 49305888 | 49298551 | 49298635 | 49327685 | 49327816 | 0 |
| MXE | 4945 | ENSG00000144741 | SLC25A26 | chr3 | + | 66346363 | 66346408 | 66362859 | 66362929 | 66263331 | 66263379 | 66369477 | 66369542 | 10 |
| MXE | 4946 | ENSG00000144741 | SLC25A26 | chr3 | + | 66356030 | 66356084 | 66362859 | 66362929 | 66346363 | 66346408 | 66369477 | 66369542 | 2 |
| MXE | 4955 | ENSG00000172053 | QARS1 | chr3 | - | 49102418 | 49102472 | 49103344 | 49103466 | 49102204 | 49102265 | 49103630 | 49103706 | 0 |
| MXE | 4963 | ENSG00000004766 | VPS50 | chr7 | + | 93334116 | 93334197 | 93340597 | 93340870 | 93311165 | 93311272 | 93341426 | 93341575 | 27 |
| MXE | 4967 | ENSG00000221978 | CCNL2 | chr1 | - | 1388491 | 1388510 | 1388625 | 1388743 | 1387953 | 1388065 | 1390229 | 1390371 | 2 |
| MXE | 4969 | ENSG00000221978 | CCNL2 | chr1 | - | 1388625 | 1388743 | 1388949 | 1389047 | 1387953 | 1388065 | 1390229 | 1390371 | 0 |
| MXE | 4971 | ENSG00000243989 | ACY1 | chr3 | + | 51985206 | 51985271 | 51985360 | 51985465 | 51984046 | 51984158 | 51985851 | 51985946 | 1 |
| MXE | 4972 | ENSG00000243989 | ACY1 | chr3 | + | 51985206 | 51985271 | 51985400 | 51985465 | 51984046 | 51984158 | 51985851 | 51985946 | 1 |
| MXE | 4973 | ENSG00000243989 | ACY1 | chr3 | + | 51985360 | 51985465 | 51985851 | 51985946 | 51985206 | 51985271 | 51986254 | 51986331 | 4 |
| MXE | 4982 | ENSG00000123737 | EXOSC9 | chr4 | + | 121807539 | 121807622 | 121809966 | 121810073 | 121804621 | 121804759 | 121811582 | 121811671 | 61 |
| MXE | 5007 | ENSG00000156976 | EIF4A2 | chr3 | + | 186784431 | 186784477 | 186784563 | 186784696 | 186783571 | 186783639 | 186785882 | 186786051 | 1 |
| MXE | 5012 | ENSG00000164327 | RICTOR | chr5 | - | 38975536 | 38975604 | 38978582 | 38978648 | 38971876 | 38971959 | 38981866 | 38982036 | 0 |
| MXE | 5014 | ENSG00000164327 | RICTOR | chr5 | - | 39021038 | 39021136 | 39074110 | 39074158 | 39003557 | 39003622 | 39074328 | 39074389 | 0 |
| MXE | 5020 | ENSG00000175455 | CCDC14 | chr3 | - | 123955842 | 123955965 | 123956354 | 123956427 | 123948895 | 123949132 | 123956739 | 123956795 | 2 |
| MXE | 5024 | ENSG00000164081 | TEX264 | chr3 | + | 51672209 | 51672301 | 51672340 | 51672392 | 51671174 | 51671288 | 51674270 | 51674562 | 2 |
| MXE | 5045 | ENSG00000176261 | ZBTB8OS | chr1 | - | 32631826 | 32631879 | 32633261 | 32633354 | 32627507 | 32627544 | 32633644 | 32633727 | 1 |
| MXE | 5051 | ENSG00000176261 | ZBTB8OS | chr1 | - | 32631826 | 32631879 | 32634767 | 32634792 | 32627571 | 32627865 | 32650432 | 32650535 | 2 |
| MXE | 5060 | ENSG00000176261 | ZBTB8OS | chr1 | - | 32633950 | 32634072 | 32634767 | 32634792 | 32633644 | 32633727 | 32650432 | 32650535 | 2 |
| MXE | 5065 | ENSG00000128513 | POT1 | chr7 | - | 124863349 | 124863640 | 124870910 | 124871041 | 124858956 | 124859112 | 124892265 | 124892380 | 73 |
| MXE | 5066 | ENSG00000128513 | POT1 | chr7 | - | 124865128 | 124865299 | 124870910 | 124871041 | 124863349 | 124863640 | 124892265 | 124892380 | 136 |
| MXE | 5067 | ENSG00000180376 | CCDC66 | chr3 | + | 56559568 | 56559594 | 56560861 | 56561012 | 56558845 | 56558910 | 56563683 | 56564125 | 128 |
| MXE | 5068 | ENSG00000180376 | CCDC66 | chr3 | + | 56560861 | 56561012 | 56561256 | 56561378 | 56559568 | 56559594 | 56563683 | 56564125 | 22 |
| MXE | 5069 | ENSG00000180376 | CCDC66 | chr3 | + | 56563683 | 56564125 | 56566949 | 56567053 | 56559568 | 56559594 | 56592969 | 56593101 | 129 |
| MXE | 5073 | ENSG00000009830 | POMT2 | chr14 | - | 77291513 | 77291664 | 77292769 | 77293290 | 77291313 | 77291380 | 77296163 | 77296273 | 0 |
| MXE | 5075 | ENSG00000009830 | POMT2 | chr14 | - | 77301089 | 77301249 | 77302834 | 77302943 | 77299454 | 77299561 | 77304691 | 77304800 | 5 |
| MXE | 5076 | ENSG00000138182 | KIF20B | chr10 | + | 89709166 | 89709253 | 89709926 | 89710065 | 89705293 | 89705441 | 89710960 | 89711090 | 11 |
| MXE | 5077 | ENSG00000138182 | KIF20B | chr10 | + | 89709344 | 89709461 | 89709926 | 89710065 | 89709166 | 89709253 | 89710960 | 89711090 | 33 |
| MXE | 5093 | ENSG00000060971 | ACAA1 | chr3 | - | 38126161 | 38126700 | 38127785 | 38127866 | 38125825 | 38125881 | 38131595 | 38131638 | 1 |
| MXE | 5096 | ENSG00000060971 | ACAA1 | chr3 | - | 38126509 | 38126700 | 38127785 | 38127866 | 38125825 | 38125881 | 38131595 | 38131638 | 1 |
| MXE | 5098 | ENSG00000060971 | ACAA1 | chr3 | - | 38127785 | 38127866 | 38129289 | 38129388 | 38125825 | 38125881 | 38131595 | 38131638 | 48 |
| MXE | 5106 | ENSG00000114859 | CLCN2 | chr3 | - | 184352012 | 184352117 | 184352292 | 184352331 | 184346934 | 184347021 | 184352442 | 184352496 | 1 |
| MXE | 5108 | ENSG00000114867 | EIF4G1 | chr3 | + | 184315488 | 184315545 | 184316131 | 184316218 | 184314568 | 184314674 | 184317716 | 184317816 | 8 |
| MXE | 5112 | ENSG00000114867 | EIF4G1 | chr3 | + | 184316071 | 184316218 | 184317320 | 184317497 | 184315762 | 184315856 | 184317746 | 184317816 | 2 |
| MXE | 5131 | ENSG00000173145 | NOC3L | chr10 | - | 94344414 | 94344515 | 94344852 | 94344933 | 94341672 | 94341745 | 94346424 | 94346556 | 29 |
| MXE | 5132 | ENSG00000111731 | C2CD5 | chr12 | - | 22458483 | 22458585 | 22459491 | 22459542 | 22456970 | 22457161 | 22469708 | 22469795 | 11 |
| MXE | 5138 | ENSG00000111731 | C2CD5 | chr12 | - | 22506710 | 22506819 | 22513293 | 22513379 | 22493222 | 22493337 | 22523425 | 22523624 | 0 |
| MXE | 5140 | ENSG00000111731 | C2CD5 | chr12 | - | 22513293 | 22513379 | 22517985 | 22518137 | 22493222 | 22493337 | 22523425 | 22523624 | 80 |
| MXE | 5192 | ENSG00000178802 | MPI | chr15 | + | 74890437 | 74890654 | 74890978 | 74891035 | 74890041 | 74890089 | 74891378 | 74891579 | 10 |
| MXE | 5193 | ENSG00000178802 | MPI | chr15 | + | 74890526 | 74890654 | 74890978 | 74891035 | 74890041 | 74890089 | 74891378 | 74891579 | 10 |
| MXE | 5194 | ENSG00000178802 | MPI | chr15 | + | 74890555 | 74890654 | 74890978 | 74891035 | 74890041 | 74890089 | 74891378 | 74891579 | 7 |
| MXE | 5196 | ENSG00000163755 | HPS3 | chr3 | + | 149162689 | 149162878 | 149163841 | 149163949 | 149162147 | 149162333 | 149167064 | 149167240 | 60 |
| MXE | 5230 | ENSG00000236548 | RNF217-AS1 | chr6 | - | 124941389 | 124941524 | 124945298 | 124945439 | 124935007 | 124935117 | 124947930 | 124948098 | 0 |
| MXE | 5231 | ENSG00000236548 | RNF217-AS1 | chr6 | - | 124941389 | 124941604 | 124945298 | 124945439 | 124935007 | 124935117 | 124947930 | 124948098 | 0 |
| MXE | 5241 | ENSG00000115827 | DCAF17 | chr2 | + | 171473865 | 171473975 | 171476859 | 171476950 | 171468887 | 171469030 | 171477986 | 171478070 | 26 |
| MXE | 5242 | ENSG00000115827 | DCAF17 | chr2 | + | 171473865 | 171473975 | 171477986 | 171478070 | 171468887 | 171469030 | 171480037 | 171480193 | 26 |
| MXE | 5243 | ENSG00000115827 | DCAF17 | chr2 | + | 171477986 | 171478070 | 171480037 | 171480193 | 171468887 | 171469030 | 171480973 | 171485048 | 0 |
| MXE | 5245 | ENSG00000159433 | STARD9 | chr15 | + | 42665252 | 42665330 | 42665785 | 42665848 | 42662793 | 42663917 | 42669157 | 42673114 | 2 |
| MXE | 5254 | ENSG00000172673 | THEMIS | chr6 | - | 127812882 | 127813931 | 127855029 | 127855188 | 127719687 | 127719823 | 127918427 | 127918597 | 42 |
| MXE | 5265 | ENSG00000238121 | LINC00426 | chr13 | - | 30358253 | 30358376 | 30363160 | 30363226 | 30357741 | 30357889 | 30372284 | 30372374 | 1 |
| MXE | 5266 | ENSG00000096384 | HSP90AB1 | chr6 | + | 44248616 | 44248776 | 44249376 | 44249583 | 44247086 | 44247195 | 44249674 | 44249834 | 7 |
| MXE | 5274 | ENSG00000284952 | NA | chr3 | - | 155829687 | 155829903 | 155842431 | 155842997 | 155828181 | 155828377 | 155853222 | 155854442 | 0 |
| MXE | 5276 | ENSG00000284952 | NA | chr3 | - | 155833467 | 155833585 | 155842431 | 155842997 | 155829687 | 155829903 | 155853222 | 155854442 | 7 |
| MXE | 5280 | ENSG00000266338 | NBPF15 | chr1 | - | 144440401 | 144440581 | 144442669 | 144442920 | 144440140 | 144440295 | 144448774 | 144448916 | 6 |
| MXE | 5285 | ENSG00000188001 | TPRG1 | chr3 | + | 189147583 | 189147647 | 189207375 | 189207594 | 189132398 | 189132697 | 189215291 | 189215383 | 22 |
| MXE | 5314 | ENSG00000145016 | RUBCN | chr3 | - | 197696953 | 197697049 | 197699186 | 197699231 | 197695865 | 197695981 | 197700612 | 197701146 | 0 |
| MXE | 5318 | ENSG00000165288 | BRWD3 | chrX | - | 80809012 | 80809042 | 80809245 | 80809304 | 80808538 | 80808598 | 80809440 | 80809683 | 1 |
| MXE | 5330 | ENSG00000164073 | MFSD8 | chr4 | - | 127957500 | 127957590 | 127965071 | 127965482 | 127949803 | 127949847 | 127965896 | 127965995 | 0 |
| MXE | 5331 | ENSG00000164073 | MFSD8 | chr4 | - | 127957500 | 127957592 | 127965071 | 127965208 | 127949803 | 127949847 | 127965896 | 127965949 | 0 |
| MXE | 5338 | ENSG00000174891 | RSRC1 | chr3 | + | 158298038 | 158298075 | 158354856 | 158354908 | 158203071 | 158203245 | 158460934 | 158461003 | 18 |
| MXE | 5339 | ENSG00000174891 | RSRC1 | chr3 | + | 158354856 | 158354908 | 158460934 | 158461003 | 158298038 | 158298075 | 158537091 | 158537198 | 17 |
| MXE | 5340 | ENSG00000174891 | RSRC1 | chr3 | + | 158354856 | 158354908 | 158460934 | 158461009 | 158298038 | 158298075 | 158537091 | 158537198 | 17 |
| MXE | 5352 | ENSG00000051382 | PIK3CB | chr3 | - | 138657689 | 138657835 | 138663905 | 138664029 | 138656141 | 138656274 | 138665035 | 138665203 | 37 |
| MXE | 5355 | ENSG00000051382 | PIK3CB | chr3 | - | 138684624 | 138684803 | 138688874 | 138688974 | 138683677 | 138683787 | 138694785 | 138694907 | 2 |
| MXE | 5360 | ENSG00000166323 | C11orf65 | chr11 | - | 108405428 | 108405559 | 108406762 | 108406963 | 108393207 | 108393378 | 108461478 | 108461568 | 4 |
| MXE | 5378 | ENSG00000138246 | DNAJC13 | chr3 | + | 132462466 | 132462523 | 132463695 | 132463817 | 132461049 | 132461205 | 132465994 | 132466070 | 35 |
| MXE | 5385 | ENSG00000175556 | LONRF3 | chrX | + | 118989407 | 118989672 | 118990469 | 118990560 | 118978344 | 118978463 | 119006120 | 119006235 | 4 |
| MXE | 5386 | ENSG00000175556 | LONRF3 | chrX | + | 118990469 | 118990560 | 119006120 | 119006235 | 118989407 | 118989672 | 119009125 | 119009247 | 13 |
| MXE | 5396 | ENSG00000188687 | SLC4A5 | chr2 | - | 74241993 | 74242052 | 74247035 | 74247307 | 74239334 | 74239535 | 74248352 | 74248486 | 4 |
| MXE | 5397 | ENSG00000188687 | SLC4A5 | chr2 | - | 74304488 | 74304680 | 74328119 | 74328186 | 74285772 | 74285902 | 74334026 | 74334177 | 2 |
| MXE | 5398 | ENSG00000163687 | DNASE1L3 | chr3 | - | 58204768 | 58204881 | 58205470 | 58205560 | 58200996 | 58201109 | 58208217 | 58208306 | 0 |
| MXE | 5402 | ENSG00000173226 | IQCB1 | chr3 | - | 121797117 | 121797227 | 121799195 | 121799374 | 121790072 | 121790215 | 121807343 | 121807443 | 18 |
| MXE | 5406 | ENSG00000143612 | C1orf43 | chr1 | - | 154213181 | 154213303 | 154213892 | 154213946 | 154212457 | 154212624 | 154214456 | 154214574 | 173 |
| MXE | 5409 | ENSG00000143612 | C1orf43 | chr1 | - | 154213892 | 154213946 | 154214456 | 154214574 | 154213181 | 154213303 | 154219835 | 154219937 | 132 |
| MXE | 5410 | ENSG00000143612 | C1orf43 | chr1 | - | 154214456 | 154214574 | 154218715 | 154218785 | 154213892 | 154213946 | 154219835 | 154219937 | 0 |
| MXE | 5427 | ENSG00000145428 | RNF175 | chr4 | - | 153728206 | 153728361 | 153748644 | 153748786 | 153723350 | 153723458 | 153751437 | 153751475 | 0 |
| MXE | 5432 | ENSG00000164463 | CREBRF | chr5 | + | 173080588 | 173080784 | 173086500 | 173086626 | 173056351 | 173056479 | 173090314 | 173092330 | 1 |
| MXE | 5442 | ENSG00000145819 | ARHGAP26 | chr5 | + | 143207197 | 143207308 | 143207382 | 143207473 | 143147230 | 143147381 | 143213996 | 143214088 | 41 |
| MXE | 5444 | ENSG00000100578 | KIAA0586 | chr14 | + | 58440069 | 58440160 | 58441220 | 58441343 | 58432387 | 58432457 | 58442705 | 58442880 | 6 |
| MXE | 5446 | ENSG00000100578 | KIAA0586 | chr14 | + | 58477122 | 58477241 | 58482512 | 58482712 | 58474606 | 58474797 | 58487006 | 58487166 | 23 |
| MXE | 5474 | ENSG00000259033 | NA | chr14 | + | 70275295 | 70275360 | 70291476 | 70291566 | 70255632 | 70256189 | 70342671 | 70343388 | 0 |
| MXE | 5491 | ENSG00000116191 | RALGPS2 | chr1 | + | 178902105 | 178902211 | 178906775 | 178906867 | 178897663 | 178897756 | 178916329 | 178916375 | 74 |
| MXE | 5498 | ENSG00000168405 | CMAHP | chr6 | - | 25084479 | 25084576 | 25086783 | 25086899 | 25081108 | 25081879 | 25091895 | 25092012 | 27 |
| MXE | 5499 | ENSG00000168405 | CMAHP | chr6 | - | 25091895 | 25092012 | 25093513 | 25093688 | 25086783 | 25086899 | 25097060 | 25097201 | 36 |
| MXE | 5502 | ENSG00000169230 | PRELID1 | chr5 | + | 177305914 | 177305984 | 177306097 | 177306176 | 177304624 | 177304850 | 177306421 | 177306669 | 1 |
| MXE | 5532 | ENSG00000147996 | CBWD5 | chr9 | + | 65700250 | 65700337 | 65702969 | 65703014 | 65698514 | 65698561 | 65720040 | 65720092 | 16 |
| MXE | 5533 | ENSG00000147996 | CBWD5 | chr9 | + | 65704330 | 65704414 | 65707012 | 65707152 | 65702969 | 65703014 | 65708901 | 65708958 | 0 |
| MXE | 5534 | ENSG00000147996 | CBWD5 | chr9 | + | 65704330 | 65704547 | 65707012 | 65707152 | 65702969 | 65703014 | 65708901 | 65708958 | 2 |
| MXE | 5541 | ENSG00000169180 | XPO6 | chr16 | - | 28152658 | 28152785 | 28155701 | 28155771 | 28146093 | 28146203 | 28155945 | 28155975 | 2 |
| MXE | 5542 | ENSG00000169180 | XPO6 | chr16 | - | 28152929 | 28152964 | 28155701 | 28155771 | 28152658 | 28152785 | 28155945 | 28155975 | 2 |
| MXE | 5544 | ENSG00000214922 | HLA-F-AS1 | chr6 | - | 29738550 | 29738886 | 29745472 | 29745727 | 29738334 | 29738455 | 29748899 | 29748993 | 12 |
| MXE | 5580 | ENSG00000168038 | ULK4 | chr3 | - | 41463086 | 41463253 | 41566024 | 41566130 | 41455496 | 41455595 | 41615668 | 41615717 | 3 |
| MXE | 5592 | ENSG00000184106 | TREML3P | chr6 | - | 41215761 | 41215819 | 41217126 | 41217177 | 41209598 | 41209730 | 41217652 | 41217947 | 4 |
| MXE | 5594 | ENSG00000135622 | SEMA4F | chr2 | + | 74662731 | 74662825 | 74673456 | 74673576 | 74657852 | 74657951 | 74674497 | 74674676 | 0 |
| MXE | 5602 | ENSG00000233452 | STXBP5-AS1 | chr6 | - | 146850824 | 146851331 | 146911551 | 146911665 | 146844318 | 146844712 | 147001440 | 147001543 | 1 |
| MXE | 5603 | ENSG00000233452 | STXBP5-AS1 | chr6 | - | 146860617 | 146860743 | 146911551 | 146911665 | 146850824 | 146851331 | 146972044 | 146972070 | 1 |
| MXE | 5604 | ENSG00000233452 | STXBP5-AS1 | chr6 | - | 146860617 | 146860743 | 146972044 | 146972070 | 146850824 | 146851331 | 147001440 | 147001543 | 0 |
| MXE | 5605 | ENSG00000233452 | STXBP5-AS1 | chr6 | - | 146860617 | 146860743 | 146911551 | 146911665 | 146850824 | 146851331 | 147074804 | 147074847 | 0 |
| MXE | 5606 | ENSG00000233452 | STXBP5-AS1 | chr6 | - | 146860617 | 146860743 | 147001440 | 147001543 | 146850824 | 146851331 | 147074804 | 147074847 | 1 |
| MXE | 5607 | ENSG00000233452 | STXBP5-AS1 | chr6 | - | 146860617 | 146860743 | 146911551 | 146911663 | 146851117 | 146851331 | 146972044 | 146972070 | 0 |
| MXE | 5608 | ENSG00000233452 | STXBP5-AS1 | chr6 | - | 146860617 | 146860743 | 146911551 | 146911663 | 146851117 | 146851331 | 147001440 | 147001543 | 0 |
| MXE | 5609 | ENSG00000233452 | STXBP5-AS1 | chr6 | - | 146860617 | 146860743 | 146911551 | 146911665 | 146851117 | 146851331 | 147001440 | 147001543 | 0 |
| MXE | 5610 | ENSG00000233452 | STXBP5-AS1 | chr6 | - | 146860617 | 146860743 | 146911551 | 146911663 | 146851117 | 146851331 | 147074804 | 147074847 | 0 |
| MXE | 5611 | ENSG00000233452 | STXBP5-AS1 | chr6 | - | 146860617 | 146860743 | 146972044 | 146972070 | 146851117 | 146851331 | 147074804 | 147074847 | 0 |
| MXE | 5612 | ENSG00000233452 | STXBP5-AS1 | chr6 | - | 146860617 | 146860743 | 147074804 | 147074847 | 146851117 | 146851331 | 147119416 | 147119477 | 0 |
| MXE | 5613 | ENSG00000233452 | STXBP5-AS1 | chr6 | - | 146864383 | 146864602 | 146911551 | 146911663 | 146860617 | 146860743 | 146972044 | 146972070 | 0 |
| MXE | 5614 | ENSG00000233452 | STXBP5-AS1 | chr6 | - | 146864383 | 146864602 | 146911551 | 146911665 | 146860617 | 146860743 | 146972044 | 146972070 | 1 |
| MXE | 5615 | ENSG00000233452 | STXBP5-AS1 | chr6 | - | 146911551 | 146911663 | 146972044 | 146972070 | 146851117 | 146851331 | 147001440 | 147001543 | 0 |
| MXE | 5616 | ENSG00000233452 | STXBP5-AS1 | chr6 | - | 146911551 | 146911663 | 146972044 | 146972070 | 146851117 | 146851331 | 147074804 | 147074847 | 0 |
| MXE | 5618 | ENSG00000233452 | STXBP5-AS1 | chr6 | - | 146911551 | 146911665 | 146972044 | 146972070 | 146851117 | 146851331 | 147001440 | 147001543 | 0 |
| MXE | 5619 | ENSG00000233452 | STXBP5-AS1 | chr6 | - | 146911551 | 146911665 | 146972044 | 146972070 | 146851117 | 146851331 | 147074804 | 147074847 | 0 |
| MXE | 5620 | ENSG00000233452 | STXBP5-AS1 | chr6 | - | 146911551 | 146911665 | 146972044 | 146972070 | 146860617 | 146860743 | 147001440 | 147001543 | 0 |
| MXE | 5621 | ENSG00000233452 | STXBP5-AS1 | chr6 | - | 146911551 | 146911665 | 146972044 | 146972070 | 146860617 | 146860743 | 147074804 | 147074847 | 0 |
| MXE | 5622 | ENSG00000233452 | STXBP5-AS1 | chr6 | - | 146911551 | 146911665 | 147001440 | 147001543 | 146860617 | 146860743 | 147074804 | 147074847 | 1 |
| MXE | 5624 | ENSG00000233452 | STXBP5-AS1 | chr6 | - | 146972044 | 146972070 | 147001440 | 147001543 | 146860617 | 146860743 | 147074804 | 147074847 | 1 |
| MXE | 5626 | ENSG00000233452 | STXBP5-AS1 | chr6 | - | 147108471 | 147108610 | 147119416 | 147119477 | 147074804 | 147074847 | 147173592 | 147173647 | 3 |
| MXE | 5643 | ENSG00000189283 | FHIT | chr3 | - | 60011370 | 60011400 | 60077369 | 60077526 | 59922345 | 59922414 | 60080790 | 60080840 | 4 |
| MXE | 5656 | ENSG00000163606 | CD200R1 | chr3 | - | 112924489 | 112924535 | 112925084 | 112925193 | 112921204 | 112923799 | 112928815 | 112929064 | 12 |
| MXE | 5657 | ENSG00000163606 | CD200R1 | chr3 | - | 112931105 | 112931171 | 112947855 | 112947924 | 112928917 | 112929507 | 112974790 | 112974912 | 51 |
| MXE | 5664 | ENSG00000075826 | SEC31B | chr10 | - | 100489251 | 100489398 | 100489702 | 100489761 | 100488857 | 100488974 | 100490705 | 100490883 | 4 |
| MXE | 5684 | ENSG00000163681 | SLMAP | chr3 | + | 57865241 | 57865292 | 57883960 | 57884026 | 57864806 | 57864857 | 57890040 | 57890100 | 29 |
| MXE | 5687 | ENSG00000163681 | SLMAP | chr3 | + | 57871635 | 57871698 | 57883960 | 57884026 | 57864806 | 57864857 | 57890040 | 57890100 | 43 |
| MXE | 5693 | ENSG00000165115 | KIF27 | chr9 | - | 83859155 | 83859371 | 83867683 | 83867860 | 83853628 | 83853835 | 83870518 | 83870632 | 6 |
| MXE | 5694 | ENSG00000165115 | KIF27 | chr9 | - | 83859155 | 83859371 | 83870518 | 83870632 | 83853628 | 83853835 | 83880296 | 83880494 | 1 |
| MXE | 5695 | ENSG00000165115 | KIF27 | chr9 | - | 83870518 | 83870632 | 83880296 | 83880494 | 83859155 | 83859371 | 83883812 | 83884018 | 10 |
| MXE | 5712 | ENSG00000168615 | ADAM9 | chr8 | + | 39041945 | 39042117 | 39046001 | 39046058 | 39026676 | 39026810 | 39054480 | 39054573 | 193 |
| MXE | 5713 | ENSG00000168615 | ADAM9 | chr8 | + | 39046001 | 39046058 | 39046953 | 39047046 | 39041945 | 39042117 | 39054480 | 39054573 | 0 |
| MXE | 5719 | ENSG00000163507 | CIP2A | chr3 | - | 108582107 | 108582138 | 108582976 | 108583083 | 108581414 | 108581511 | 108585064 | 108585212 | 0 |
| MXE | 5721 | ENSG00000285162 | NA | chr7 | + | 29354624 | 29354663 | 29367931 | 29367987 | 29146848 | 29146960 | 29393678 | 29393710 | 2 |

| | | | | | | | |
|---|---|---|---|---|---|---|---|
| MXE | 5727 | ENSG00000186469 | GNG2 | chr14 | + | 51913008 | [illegible] |
| MXE | 5728 | ENSG00000170954 | ZNF415 | chr19 | - | 53115209 | [illegible] |
| MXE | 5768 | ENSG00000110871 | COQ5 | chr12 | - | 120510016 | [illegible] |
| MXE | 5771 | ENSG00000064961 | HMG20B | chr19 | + | 3575501 | [illegible] |
| MXE | 5774 | ENSG00000138750 | NUP54 | chr4 | - | 76131229 | [illegible] |
| MXE | 5775 | ENSG00000138750 | NUP54 | chr4 | - | 76132522 | [illegible] |
| MXE | 5779 | ENSG00000138750 | NUP54 | chr4 | - | 76136185 | [illegible] |
| MXE | 5793 | ENSG00000121897 | LIAS | chr4 | + | 39463524 | [illegible] |
| MXE | 5817 | ENSG00000162777 | DENND2D | chr1 | - | 111197919 | [illegible] |
| MXE | 5832 | ENSG00000173930 | SLCO4C1 | chr5 | - | 102270623 | [illegible] |
| MXE | 5839 | ENSG00000117298 | ECE1 | chr1 | - | 21225249 | [illegible] |
| MXE | 5848 | ENSG00000130024 | PHF10 | chr6 | - | 169717822 | [illegible] |
| MXE | 5859 | ENSG00000073417 | PDE8A | chr15 | + | 85083555 | [illegible] |
| MXE | 5860 | ENSG00000073417 | PDE8A | chr15 | + | 85083555 | [illegible] |
| MXE | 5861 | ENSG00000073417 | PDE8A | chr15 | + | 85083555 | [illegible] |
| MXE | 5866 | ENSG00000143365 | RORC | chr1 | - | 151813238 | [illegible] |
| MXE | 5867 | ENSG00000143365 | RORC | chr1 | - | 151813238 | [illegible] |
| MXE | 5872 | ENSG00000228794 | LINC01128 | chr1 | + | 847653 | [illegible] |
| MXE | 5879 | ENSG00000162601 | MYSM1 | chr1 | - | 58668429 | [illegible] |
| MXE | 5884 | ENSG00000117054 | ACADM | chr1 | + | 75728400 | [illegible] |
| MXE | 5885 | ENSG00000117054 | ACADM | chr1 | + | 75733050 | [illegible] |
| MXE | 5887 | ENSG00000117054 | ACADM | chr1 | + | 75745805 | [illegible] |
| MXE | 5890 | ENSG00000111843 | TMEM14C | chr6 | + | 10725906 | [illegible] |
| MXE | 5897 | ENSG00000067066 | SP100 | chr2 | + | 230446818 | [illegible] |
| MXE | 5912 | ENSG00000067066 | SP100 | chr2 | + | 230507992 | [illegible] |
| MXE | 5923 | ENSG00000003400 | CASP10 | chr2 | + | 201192983 | [illegible] |
| MXE | 5927 | ENSG00000003400 | CASP10 | chr2 | + | 201195841 | [illegible] |
| MXE | 5933 | ENSG00000003400 | CASP10 | chr2 | + | 201203729 | [illegible] |
| MXE | 5965 | ENSG00000165240 | ATP7A | chrX | + | 78031399 | [illegible] |
| MXE | 5969 | ENSG00000150753 | CCT5 | chr5 | + | 10254765 | [illegible] |
| MXE | 5997 | ENSG00000145388 | METTL14 | chr4 | + | 118697181 | [illegible] |
| MXE | 6001 | ENSG00000158864 | NDUFS2 | chr1 | + | 161212591 | [illegible] |
| MXE | 6002 | ENSG00000158864 | NDUFS2 | chr1 | + | 161212591 | [illegible] |
| MXE | 6028 | ENSG00000144554 | FANCD2 | chr3 | + | 10049373 | [illegible] |
| MXE | 6030 | ENSG00000135951 | TSGA10 | chr2 | - | 99020279 | [illegible] |
| MXE | 6032 | ENSG00000090989 | EXOC1 | chr4 | + | 55890222 | [illegible] |
| MXE | 6040 | ENSG00000181555 | SETD2 | chr3 | - | 47086194 | [illegible] |
| MXE | 6054 | ENSG00000132185 | FCRLA | chr1 | + | 161710474 | [illegible] |
| MXE | 6058 | ENSG00000072274 | TFRC | chr3 | - | 196075158 | [illegible] |
| MXE | 6061 | ENSG00000116299 | ELAPOR1 | chr1 | + | 109173687 | [illegible] |
| MXE | 6064 | ENSG00000143452 | HORMAD1 | chr1 | - | 150711544 | [illegible] |
| MXE | 6066 | ENSG00000143452 | HORMAD1 | chr1 | - | 150711832 | [illegible] |
| MXE | 6067 | ENSG00000143452 | HORMAD1 | chr1 | - | 150714084 | [illegible] |
| MXE | 6068 | ENSG00000143452 | HORMAD1 | chr1 | - | 150714084 | [illegible] |
| MXE | 6069 | ENSG00000143452 | HORMAD1 | chr1 | - | 150714614 | [illegible] |
| MXE | 6078 | ENSG00000271723 | MROH7-TTC4 | chr1 | + | 54731654 | [illegible] |
| MXE | 6085 | ENSG00000164253 | WDR41 | chr5 | - | 77480080 | [illegible] |
| MXE | 6087 | ENSG00000135537 | AFG1L | chr6 | + | 108401995 | [illegible] |
| MXE | 6092 | ENSG00000143190 | POU2F1 | chr1 | + | 167303473 | [illegible] |
| MXE | 6093 | ENSG00000143190 | POU2F1 | chr1 | + | 167303473 | [illegible] |
| MXE | 6107 | ENSG00000138382 | METTL5 | chr2 | - | 169819560 | [illegible] |
| MXE | 6109 | ENSG00000138382 | METTL5 | chr2 | - | 169821091 | [illegible] |
| MXE | 6122 | ENSG00000170581 | STAT2 | chr12 | - | 56343303 | [illegible] |
| MXE | 6124 | ENSG00000170581 | STAT2 | chr12 | - | 56348751 | [illegible] |
| MXE | 6126 | ENSG00000170581 | STAT2 | chr12 | - | 56348751 | [illegible] |
| MXE | 6132 | ENSG00000234663 | LINC01934 | chr2 | + | 181226470 | [illegible] |
| MXE | 6133 | ENSG00000234663 | LINC01934 | chr2 | + | 181226470 | [illegible] |
| MXE | 6135 | ENSG00000234663 | LINC01934 | chr2 | + | 181238250 | [illegible] |
| MXE | 6136 | ENSG00000234663 | LINC01934 | chr2 | + | 181238250 | [illegible] |
| MXE | 6140 | ENSG00000163719 | MTMR14 | chr3 | + | 9653620 | [illegible] |
| MXE | 6153 | ENSG00000078304 | PPP2R5C | chr14 | + | 101856685 | [illegible] |
| MXE | 6157 | ENSG00000078304 | PPP2R5C | chr14 | + | 101889981 | [illegible] |
| MXE | 6170 | ENSG00000285708 | NA | chr3 | - | 71335159 | [illegible] |
| MXE | 6171 | ENSG00000285708 | NA | chr3 | - | 71359149 | [illegible] |
| MXE | 6172 | ENSG00000285708 | NA | chr3 | - | 71690009 | [illegible] |
| MXE | 6173 | ENSG00000285708 | NA | chr3 | - | 71690009 | [illegible] |
| MXE | 6176 | ENSG00000162627 | SNX7 | chr1 | + | 98701816 | [illegible] |
| MXE | 6194 | ENSG00000162994 | CLHC1 | chr2 | - | 55180509 | [illegible] |
| MXE | 6195 | ENSG00000162994 | CLHC1 | chr2 | - | 55212672 | [illegible] |
| MXE | 6205 | ENSG00000163870 | TPRA1 | chr3 | - | 127575402 | [illegible] |
| MXE | 6206 | ENSG00000163870 | TPRA1 | chr3 | - | 127575748 | [illegible] |
| MXE | 6212 | ENSG00000126016 | AMOT | chrX | - | 112822254 | [illegible] |
| MXE | 6223 | ENSG00000067829 | IDH3G | chrX | - | 153790563 | [illegible] |
| MXE | 6224 | ENSG00000151883 | PARP8 | chr5 | + | 50738979 | [illegible] |
| MXE | 6234 | ENSG00000156136 | DCK | chr4 | + | 71023558 | [illegible] |
| MXE | 6252 | ENSG00000115523 | GNLY | chr2 | + | 85696987 | [illegible] |
| MXE | 6270 | ENSG00000143498 | TAF1A | chr1 | - | 222579758 | [illegible] |
| MXE | 6271 | ENSG00000143498 | TAF1A | chr1 | - | 222579758 | [illegible] |
| MXE | 6282 | ENSG00000117569 | PTBP2 | chr1 | + | 96804784 | [illegible] |
| MXE | 6283 | ENSG00000117569 | PTBP2 | chr1 | + | 96804799 | [illegible] |
| MXE | 6284 | ENSG00000054392 | HHAT | chr1 | + | 210348932 | [illegible] |
| MXE | 6285 | ENSG00000054392 | HHAT | chr1 | + | 210404463 | [illegible] |
| MXE | 6293 | ENSG00000137776 | SLTM | chr15 | - | 58916934 | [illegible] |
| MXE | 6301 | ENSG00000138674 | SEC31A | chr4 | - | 82828999 | [illegible] |
| MXE | 6307 | ENSG00000138674 | SEC31A | chr4 | - | 82878729 | [illegible] |
| MXE | 6327 | ENSG00000247828 | TMEM161B-AS1 | chr5 | + | 88282041 | [illegible] |
| MXE | 6328 | ENSG00000247828 | TMEM161B-AS1 | chr5 | + | 88282041 | [illegible] |
| MXE | 6329 | ENSG00000247828 | TMEM161B-AS1 | chr5 | + | 88282714 | [illegible] |
| MXE | 6348 | ENSG00000172456 | FGGY | chr1 | + | 59378748 | [illegible] |
| MXE | 6350 | ENSG00000172456 | FGGY | chr1 | + | 59512310 | [illegible] |
| MXE | 6355 | ENSG00000089327 | FXYD5 | chr19 | + | 35166141 | [illegible] |
| MXE | 6375 | ENSG00000092871 | RFFL | chr17 | - | 35021370 | [illegible] |
| MXE | 6397 | ENSG00000258465 | NA | chr1 | - | 160263264 | [illegible] |
| MXE | 6404 | ENSG00000156398 | SFXN2 | chr10 | + | 102715222 | [illegible] |
| MXE | 6411 | ENSG00000185842 | DNAH14 | chr1 | + | 225043885 | [illegible] |
| MXE | 6413 | ENSG00000185842 | DNAH14 | chr1 | + | 225252300 | [illegible] |
| MXE | 6428 | ENSG00000104381 | GDAP1 | chr8 | + | 74351273 | [illegible] |
| MXE | 6448 | ENSG00000162687 | KCNT2 | chr1 | - | 196343055 | [illegible] |
| MXE | 6458 | ENSG00000198380 | GFPT1 | chr2 | - | 69354258 | [illegible] |
| MXE | 6469 | ENSG00000116711 | PLA2G4A | chr1 | + | 186911247 | [illegible] |
| MXE | 6473 | ENSG00000066557 | LRRC40 | chr1 | - | 70159329 | [illegible] |
| MXE | 6474 | ENSG00000066557 | LRRC40 | chr1 | - | 70173464 | [illegible] |
| MXE | 6475 | ENSG00000066557 | LRRC40 | chr1 | - | 70187264 | [illegible] |
| MXE | 6477 | ENSG00000119965 | C10orf88 | chr10 | - | 122937704 | [illegible] |
| MXE | 6478 | ENSG00000119965 | C10orf88 | chr10 | - | 122937704 | [illegible] |
| MXE | 6488 | ENSG00000123728 | RAP2C | chrX | - | 132216995 | [illegible] |
| MXE | 6492 | ENSG00000152495 | CAMK4 | chr5 | + | 111394709 | [illegible] |
| MXE | 6497 | ENSG00000187010 | RHD | chr1 | + | 25306595 | [illegible] |
| MXE | 6498 | ENSG00000187010 | RHD | chr1 | + | 25307646 | [illegible] |
| MXE | 6500 | ENSG00000125637 | PSD4 | chr2 | + | 113185661 | [illegible] |
| MXE | 6502 | ENSG00000125637 | PSD4 | chr2 | + | 113185960 | [illegible] |
| MXE | 6507 | ENSG00000164037 | SLC9B1 | chr4 | - | 102945192 | [illegible] |
| MXE | 6508 | ENSG00000164037 | SLC9B1 | chr4 | - | 102946646 | [illegible] |
| MXE | 6510 | ENSG00000114861 | FOXP1 | chr3 | - | 70972010 | [illegible] |
| MXE | 6511 | ENSG00000114861 | FOXP1 | chr3 | - | 71112535 | [illegible] |
| MXE | 6516 | ENSG00000121940 | CLCC1 | chr1 | - | 108941404 | [illegible] |
| MXE | 6517 | ENSG00000121940 | CLCC1 | chr1 | - | 108941404 | [illegible] |
| MXE | 6518 | ENSG00000121940 | CLCC1 | chr1 | - | 108941404 | [illegible] |
| MXE | 6520 | ENSG00000121940 | CLCC1 | chr1 | - | 108941404 | [illegible] |
| MXE | 6523 | ENSG00000121940 | CLCC1 | chr1 | - | 108943474 | [illegible] |
| MXE | 6524 | ENSG00000121940 | CLCC1 | chr1 | - | 108943474 | [illegible] |
| MXE | 6527 | ENSG00000076258 | FMO4 | chr1 | + | 171323003 | [illegible] |
| MXE | 6533 | ENSG00000213160 | KLHL23 | chr2 | + | 169697892 | [illegible] |
| MXE | 6547 | ENSG00000218739 | CEBPZOS | chr2 | + | 37199703 | [illegible] |
| MXE | 6550 | ENSG00000181666 | ZNF875 | chr19 | + | 37344609 | [illegible] |
| MXE | 6558 | ENSG00000009413 | REV3L | chr6 | - | 111311068 | [illegible] |
| MXE | 6573 | ENSG00000095261 | PSMD5 | chr9 | - | 120829098 | [illegible] |
| MXE | 6579 | ENSG00000018510 | AGPS | chr2 | + | 177442119 | [illegible] |
| MXE | 6586 | ENSG00000163565 | IFI16 | chr1 | + | 159045296 | [illegible] |
| MXE | 6590 | ENSG00000110172 | CHORDC1 | chr11 | - | 90206201 | [illegible] |
| MXE | 6592 | ENSG00000110172 | CHORDC1 | chr11 | - | 90211214 | [illegible] |
| MXE | 6601 | ENSG00000058673 | ZC3H11A | chr1 | + | 203850494 | [illegible] |
| MXE | 6610 | ENSG00000163291 | PAQR3 | chr4 | - | 78920533 | [illegible] |
| MXE | 6614 | ENSG00000182389 | CACNB4 | chr2 | - | 151860710 | [illegible] |
| MXE | 6615 | ENSG00000182389 | CACNB4 | chr2 | - | 151860710 | [illegible] |
| MXE | 6616 | ENSG00000182389 | CACNB4 | chr2 | - | 151860710 | [illegible] |
| MXE | 6617 | ENSG00000182389 | CACNB4 | chr2 | - | 151860710 | [illegible] |
| MXE | 6618 | ENSG00000182389 | CACNB4 | chr2 | - | 151870530 | [illegible] |
| MXE | 6619 | ENSG00000182389 | CACNB4 | chr2 | - | 151870530 | [illegible] |
| MXE | 6625 | ENSG00000138002 | IFT172 | chr2 | - | 27484027 | [illegible] |
| MXE | 6626 | ENSG00000138002 | IFT172 | chr2 | - | 27484226 | [illegible] |
| MXE | 6634 | ENSG00000133067 | LGR6 | chr1 | + | 202276305 | [illegible] |
| MXE | 6641 | ENSG00000165029 | ABCA1 | chr9 | - | 104817225 | [illegible] |
| MXE | 6643 | ENSG00000077782 | FGFR1 | chr8 | - | 38421796 | [illegible] |
| MXE | 6647 | ENSG00000115165 | CYTIP | chr2 | - | 157443846 | [illegible] |
| MXE | 6658 | ENSG00000173166 | RAPH1 | chr2 | - | 203470238 | [illegible] |
| MXE | 6660 | ENSG00000237943 | PRKCQ-AS1 | chr10 | + | 6585797 | [illegible] |
| MXE | 6661 | ENSG00000237943 | PRKCQ-AS1 | chr10 | + | 6613933 | [illegible] |
| MXE | 6663 | ENSG00000054118 | THRAP3 | chr1 | + | 36259381 | [illegible] |
| MXE | 6669 | ENSG00000182923 | CEP63 | chr3 | + | 134546384 | [illegible] |
| MXE | 6670 | ENSG00000182923 | CEP63 | chr3 | + | 134546384 | [illegible] |
| MXE | 6671 | ENSG00000182923 | CEP63 | chr3 | + | 134546384 | [illegible] |
| MXE | 6672 | ENSG00000182923 | CEP63 | chr3 | + | 134558141 | [illegible] |
| MXE | 6680 | ENSG00000183826 | BTBD9 | chr6 | - | 38402816 | [illegible] |
| MXE | 6682 | ENSG00000136717 | BIN1 | chr2 | - | 127051153 | [illegible] |
| MXE | 6684 | ENSG00000136717 | BIN1 | chr2 | - | 127052254 | [illegible] |
| MXE | 6689 | ENSG00000102606 | ARHGEF7 | chr13 | + | 111153904 | [illegible] |
| MXE | 6703 | ENSG00000230426 | LINC01036 | chr1 | + | 187254972 | [illegible] |
| MXE | 6704 | ENSG00000230426 | LINC01036 | chr1 | + | 187269395 | [illegible] |
| MXE | 6705 | ENSG00000197721 | CR1L | chr1 | + | 207708177 | [illegible] |
| MXE | 6710 | ENSG00000163517 | HDAC11 | chr3 | + | 13483463 | [illegible] |
| MXE | 6711 | ENSG00000163517 | HDAC11 | chr3 | + | 13483463 | [illegible] |
| MXE | 6712 | ENSG00000163517 | HDAC11 | chr3 | + | 13501870 | [illegible] |
| MXE | 6719 | ENSG00000204435 | CSNK2B | chr6 | + | 31667867 | [illegible] |
| MXE | 6720 | ENSG00000204435 | CSNK2B | chr6 | + | 31668538 | [illegible] |
| MXE | 6744 | ENSG00000076321 | KLHL20 | chr1 | + | 173716002 | [illegible] |
| MXE | 6746 | ENSG00000076321 | KLHL20 | chr1 | + | 173766145 | [illegible] |
| MXE | 6748 | ENSG00000076321 | KLHL20 | chr1 | + | 173775633 | [illegible] |
| MXE | 6768 | ENSG00000138069 | RAB1A | chr2 | - | 65090982 | [illegible] |
| MXE | 6773 | ENSG00000175137 | SH3BP5L | chr1 | - | 248820402 | [illegible] |
| MXE | 6774 | ENSG00000145833 | DDX46 | chr5 | + | 134784365 | [illegible] |
| MXE | 6784 | ENSG00000143493 | INTS7 | chr1 | - | 211946606 | [illegible] |
| MXE | 6789 | ENSG00000134744 | TUT4 | chr1 | - | 52465069 | [illegible] |
| MXE | 6793 | ENSG00000134744 | TUT4 | chr1 | - | 52481803 | [illegible] |
| MXE | 6794 | ENSG00000134744 | TUT4 | chr1 | - | 52493610 | [illegible] |
| MXE | 6800 | ENSG00000138138 | ATAD1 | chr10 | - | 87759700 | [illegible] |
| MXE | 6804 | ENSG00000117682 | DHDDS | chr1 | + | 26442707 | [illegible] |
| MXE | 6807 | ENSG00000117682 | DHDDS | chr1 | + | 26442730 | [illegible] |
| MXE | 6811 | ENSG00000117682 | DHDDS | chr1 | + | 26447558 | [illegible] |
| MXE | 6817 | ENSG00000275131 | PDE4DIPP2 | chr1 | - | 120494233 | [illegible] |
| MXE | 6818 | ENSG00000275131 | PDE4DIPP2 | chr1 | - | 120495029 | [illegible] |
| MXE | 6819 | ENSG00000275131 | PDE4DIPP2 | chr1 | - | 120505710 | [illegible] |
| MXE | 6822 | ENSG00000007341 | ST7L | chr1 | - | 112541950 | [illegible] |
| MXE | 6823 | ENSG00000007341 | ST7L | chr1 | - | 112541950 | [illegible] |
| MXE | 6824 | ENSG00000007341 | ST7L | chr1 | - | 112550600 | [illegible] |
| MXE | 6827 | ENSG00000007341 | ST7L | chr1 | - | 112576985 | [illegible] |
| MXE | 6828 | ENSG00000007341 | ST7L | chr1 | - | 112578344 | [illegible] |
| MXE | 6834 | ENSG00000007341 | ST7L | chr1 | - | 112591524 | [illegible] |
| MXE | 6839 | ENSG00000088035 | ALG6 | chr1 | + | 63411145 | [illegible] |
| MXE | 6842 | ENSG00000133731 | IMPA1 | chr8 | - | 81676233 | [illegible] |
| MXE | 6857 | ENSG00000121774 | KHDRBS1 | chr1 | + | 32036909 | [illegible] |
| MXE | 6867 | ENSG00000205236 | NA | chr7 | - | 102602323 | [illegible] |
| MXE | 6895 | ENSG00000186231 | KLHL32 | chr6 | + | 97064627 | [illegible] |
| MXE | 6896 | ENSG00000186231 | KLHL32 | chr6 | + | 97064627 | [illegible] |
| MXE | 6898 | ENSG00000186231 | KLHL32 | chr6 | + | 97085125 | [illegible] |
| MXE | 6899 | ENSG00000197134 | ZNF257 | chr19 | + | 22072808 | [illegible] |
| MXE | 6902 | ENSG00000117335 | CD46 | chr1 | + | 207767012 | [illegible] |
| MXE | 6914 | ENSG00000288208 | NA | chr1 | - | 45332917 | [illegible] |
| MXE | 6923 | ENSG00000188672 | RHCE | chr1 | - | 25375348 | [illegible] |
| MXE | 6937 | ENSG00000168646 | AXIN2 | chr17 | - | 65535625 | [illegible] |
| MXE | 6951 | ENSG00000160058 | BSDC1 | chr1 | - | 32381213 | [illegible] |
| MXE | 6954 | ENSG00000160058 | BSDC1 | chr1 | - | 32383829 | [illegible] |
| MXE | 6956 | ENSG00000160058 | BSDC1 | chr1 | - | 32383829 | [illegible] |
| MXE | 6958 | ENSG00000101353 | MROH8 | chr20 | - | 37137809 | [illegible] |
| MXE | 6976 | ENSG00000173230 | GOLGB1 | chr3 | - | 121699811 | [illegible] |
| MXE | 6978 | ENSG00000158517 | NCF1 | chr7 | + | 74782938 | [illegible] |
| MXE | 6988 | ENSG00000106714 | CNTNAP3 | chr9 | - | 39099910 | [illegible] |
| MXE | 6989 | ENSG00000181315 | ZNF322 | chr6 | - | 26643658 | [illegible] |
| MXE | 7004 | ENSG00000213593 | TMX2 | chr11 | + | 57738663 | [illegible] |
| MXE | 7005 | ENSG00000100280 | AP1B1 | chr22 | - | 29363000 | [illegible] |
| MXE | 7010 | ENSG00000114439 | BBX | chr3 | + | 107638501 | [illegible] |
| MXE | 7022 | ENSG00000107625 | DDX50 | chr10 | + | 68936005 | [illegible] |
| MXE | 7031 | ENSG00000093183 | SEC22C | chr3 | - | 42565775 | [illegible] |
| MXE | 7039 | ENSG00000028310 | BRD9 | chr5 | - | 868606 | [illegible] |
| MXE | 7040 | ENSG00000028310 | BRD9 | chr5 | - | 881106 | [illegible] |
| MXE | 7043 | ENSG00000160049 | DFFA | chr1 | - | 10462987 | [illegible] |
| MXE | 7050 | ENSG00000248932 | NA | chr3 | + | 139459930 | [illegible] |
| MXE | 7055 | ENSG00000185864 | NPIPB4 | chr16 | - | 21851769 | [illegible] |
| MXE | 7059 | ENSG00000223745 | CCDC18-AS1 | chr1 | - | 93309714 | [illegible] |
| MXE | 7061 | ENSG00000223745 | CCDC18-AS1 | chr1 | - | 93315148 | [illegible] |
| MXE | 7069 | ENSG00000058272 | PPP1R12A | chr12 | - | 79788647 | [illegible] |
| MXE | 7075 | ENSG00000141556 | TBCD | chr17 | + | 82893546 | [illegible] |
| MXE | 7079 | ENSG00000141556 | TBCD | chr17 | + | 82900650 | [illegible] |
| MXE | 7080 | ENSG00000141556 | TBCD | chr17 | + | 82903404 | [illegible] |
| MXE | 7087 | ENSG00000117090 | SLAMF1 | chr1 | - | 160619775 | [illegible] |
| MXE | 7089 | ENSG00000171103 | TRMT61B | chr2 | - | 28851071 | [illegible] |
| MXE | 7090 | ENSG00000171103 | TRMT61B | chr2 | - | 28852407 | [illegible] |
| MXE | 7126 | ENSG00000055609 | KMT2C | chr7 | - | 152273704 | [illegible] |
| MXE | 7144 | ENSG00000175309 | PHYKPL | chr5 | - | 178229939 | [illegible] |
| MXE | 7147 | ENSG00000087301 | TXNDC16 | chr14 | - | 52511239 | [illegible] |
| MXE | 7161 | ENSG00000164124 | TMEM144 | chr4 | + | 158232900 | [illegible] |
| MXE | 7163 | ENSG00000242808 | SOX2-OT | chr3 | + | 181003774 | [illegible] |
| MXE | 7164 | ENSG00000242808 | SOX2-OT | chr3 | + | 181030569 | [illegible] |

| | | | | | | | | | | | |
|---|---|---|---|---|---|---|---|---|---|---|---|
| MXE | 7176 | ENSG00000121957 | GPSM2 | chr1 | + | 108918612 | 108918789 | 108922416 | 108922576 | 108914337 | 108914408 |
| MXE | 7181 | ENSG00000166669 | ATF7IP2 | chr16 | + | 10419580 | 10419623 | 10428867 | 10429016 | 10414573 | 10414612 |
| MXE | 7185 | ENSG00000164136 | IL15 | chr4 | + | 141656185 | 141656307 | 141688821 | 141689035 | 141636725 | 141636748 |
| MXE | 7186 | ENSG00000144426 | NBEAL1 | chr2 | + | 203151464 | 203151589 | 203157698 | 203157825 | 203148990 | 203149148 |
| MXE | 7187 | ENSG00000144426 | NBEAL1 | chr2 | + | 203193794 | 203193911 | 203197301 | 203197391 | 203190291 | 203190389 |
| MXE | 7191 | ENSG00000198178 | CLEC4C | chr12 | - | 7737428 | 7737574 | 7741420 | 7741531 | 7730796 | 7730912 |
| MXE | 7195 | ENSG00000183023 | SLC8A1 | chr2 | - | 40175260 | 40175281 | 40177751 | 40177855 | 40174824 | 40174842 |
| MXE | 7207 | ENSG00000135845 | PIGC | chr1 | - | 172376553 | 172376636 | 172399818 | 172399916 | 172370188 | 172370311 |
| MXE | 7213 | ENSG00000121766 | ZCCHC17 | chr1 | + | 31338956 | 31339048 | 31346639 | 31346818 | 31319108 | 31319166 |
| MXE | 7215 | ENSG00000121766 | ZCCHC17 | chr1 | + | 31338956 | 31339048 | 31346639 | 31346818 | 31337174 | 31337275 |
| MXE | 7221 | ENSG00000144357 | UBR3 | chr2 | + | 170073403 | 170073607 | 170079813 | 170080023 | 170061317 | 170061443 |
| MXE | 7222 | ENSG00000144357 | UBR3 | chr2 | + | 170073427 | 170073607 | 170079813 | 170080023 | 170061317 | 170061443 |
| MXE | 7238 | ENSG00000153560 | UBP1 | chr3 | - | 33425589 | 33425741 | 33439735 | 33439925 | 33416757 | 33416834 |
| MXE | 7240 | ENSG00000145687 | SSBP2 | chr5 | - | 81437429 | 81437458 | 81440537 | 81440636 | 81428584 | 81428683 |
| MXE | 7254 | ENSG00000154358 | OBSCN | chr1 | + | 228246527 | 228246803 | 228249964 | 228250240 | 228245412 | 228245688 |
| MXE | 7256 | ENSG00000154358 | OBSCN | chr1 | + | 228268529 | 228268805 | 228273271 | 228273547 | 228264115 | 228264391 |
| MXE | 7260 | ENSG00000187554 | TLR5 | chr1 | - | 223134681 | 223134864 | 223137177 | 223137263 | 223132474 | 223132639 |
| MXE | 7261 | ENSG00000187554 | TLR5 | chr1 | - | 223137177 | 223137263 | 223141647 | 223141763 | 223132474 | 223132639 |
| MXE | 7264 | ENSG00000188004 | SNHG28 | chr1 | - | 159838364 | 159838451 | 159840710 | 159840866 | 159837916 | 159838027 |
| MXE | 7266 | ENSG00000188004 | SNHG28 | chr1 | - | 159838364 | 159838451 | 159840768 | 159840866 | 159837916 | 159838027 |
| MXE | 7270 | ENSG00000157500 | APPL1 | chr3 | + | 57268397 | 57268487 | 57269540 | 57269751 | 57267741 | 57267792 |
| MXE | 7272 | ENSG00000233184 | NA | chr1 | + | 101072659 | 101072830 | 101075275 | 101075394 | 101025877 | 101026061 |
| MXE | 7273 | ENSG00000233184 | NA | chr1 | + | 101072659 | 101072830 | 101075394 | 101075716 | 101025886 | 101026061 |
| MXE | 7279 | ENSG00000233184 | NA | chr1 | + | 101077362 | 101077508 | 101080388 | 101080613 | 101075394 | 101075716 |
| MXE | 7280 | ENSG00000233184 | NA | chr1 | + | 101077362 | 101077508 | 101083460 | 101083574 | 101075394 | 101075716 |
| MXE | 7281 | ENSG00000233184 | NA | chr1 | + | 101077362 | 101077508 | 101080388 | 101083574 | 101075394 | 101075716 |
| MXE | 7282 | ENSG00000233184 | NA | chr1 | + | 101080388 | 101080613 | 101080886 | 101080998 | 101077362 | 101077508 |
| MXE | 7286 | ENSG00000154710 | RABGEF1 | chr7 | + | 66739995 | 66740792 | 66771882 | 66772078 | 66712166 | 66712224 |
| MXE | 7293 | ENSG00000170476 | MZB1 | chr5 | - | 139388020 | 139388131 | 139388460 | 139388585 | 139387784 | 139387921 |
| MXE | 7300 | ENSG00000113597 | TRAPPC13 | chr5 | + | 65655635 | 65655653 | 65658367 | 65658501 | 65652500 | 65652545 |
| MXE | 7302 | ENSG00000137821 | LRRC49 | chr15 | + | 70895848 | 70895936 | 70900921 | 70901024 | 70893583 | 70893640 |
| MXE | 7303 | ENSG00000197530 | MIB2 | chr1 | + | 1625545 | 1625653 | 1626649 | 1626754 | 1625285 | 1625428 |

[illegible]

| | | | | | | | | | | | |
|---|---|---|---|---|---|---|---|---|---|---|---|
| MXE | 8508 | ENSG00000225470 | JPX | chrX | + | 73962734 | 73962801 | 73994839 | 73994961 | 73946998 | 73947374 |
| MXE | 8509 | ENSG00000225470 | JPX | chrX | + | 73984443 | 73984541 | 73994839 | 73994961 | 73944583 | 73944662 |
| MXE | 8510 | ENSG00000225470 | JPX | chrX | + | 73984443 | 73984541 | 73994863 | 73994961 | 73944583 | 73944662 |
| MXE | 8512 | ENSG00000225470 | JPX | chrX | + | 73994839 | 73994961 | 73998767 | 73998844 | 73944583 | 73944662 |
| MXE | 8513 | ENSG00000225470 | JPX | chrX | + | 73994839 | 73994961 | 74062035 | 74062148 | 73944583 | 73944662 |
| MXE | 8522 | ENSG00000177200 | CHD9 | chr16 | + | 53296955 | 53297158 | 53303719 | 53304625 | 53292832 | 53293052 |
| MXE | 8526 | ENSG00000122678 | POLM | chr7 | - | 44074397 | 44074530 | 44076508 | 44076629 | 44073782 | 44074025 |
| MXE | 8531 | ENSG00000136161 | RCBTB2 | chr13 | - | 48512015 | 48512122 | 48512728 | 48512895 | 48502723 | 48502914 |
| MXE | 8534 | ENSG00000136161 | RCBTB2 | chr13 | - | 48521897 | 48521962 | 48524658 | 48524757 | 48515585 | 48515741 |
| MXE | 8535 | ENSG00000136161 | RCBTB2 | chr13 | - | 48522307 | 48522403 | 48524658 | 48524757 | 48515585 | 48515741 |
| MXE | 8536 | ENSG00000136161 | RCBTB2 | chr13 | - | 48522307 | 48522403 | 48524658 | 48524757 | 48521897 | 48521962 |
| MXE | 8539 | ENSG00000121892 | PDS5A | chr4 | - | 39866860 | 39866997 | 39869393 | 39869462 | 39863335 | 39863459 |
| MXE | 8543 | ENSG00000186409 | CCDC30 | chr1 | + | 42473121 | 42473303 | 42480460 | 42480566 | 42463285 | 42463657 |
| MXE | 8545 | ENSG00000186409 | CCDC30 | chr1 | + | 42506946 | 42507069 | 42534888 | 42534989 | 42498817 | 42498916 |
| MXE | 8546 | ENSG00000186409 | CCDC30 | chr1 | + | 42506946 | 42507069 | 42536475 | 42536601 | 42498817 | 42498916 |
| MXE | 8547 | ENSG00000186409 | CCDC30 | chr1 | + | 42521162 | 42521256 | 42534888 | 42534989 | 42498817 | 42498916 |
| MXE | 8548 | ENSG00000186409 | CCDC30 | chr1 | + | 42521162 | 42521256 | 42536475 | 42536601 | 42498817 | 42498916 |
| MXE | 8549 | ENSG00000186409 | CCDC30 | chr1 | + | 42534888 | 42534989 | 42536475 | 42536601 | 42498817 | 42498916 |
| MXE | 8550 | ENSG00000186409 | CCDC30 | chr1 | + | 42534888 | 42534989 | 42536475 | 42536601 | 42498817 | 42498916 |
| MXE | 8552 | ENSG00000186409 | CCDC30 | chr1 | + | 42545409 | 42545579 | 42566295 | 42566475 | 42536475 | 42536601 |
| MXE | 8553 | ENSG00000186409 | CCDC30 | chr1 | + | 42566295 | 42566475 | 42577019 | 42577229 | 42545409 | 42545579 |
| MXE | 8559 | ENSG00000153207 | AHCTF1 | chr1 | - | 246863923 | 246864116 | 246865254 | 246865333 | 246861958 | 246862153 |
| MXE | 8560 | ENSG00000153207 | AHCTF1 | chr1 | - | 246863923 | 246864116 | 246867243 | 246867351 | 246861958 | 246862153 |
| MXE | 8578 | ENSG00000106462 | EZH2 | chr7 | - | 148816683 | 148816778 | 148817221 | 148817412 | 148815505 | 148815546 |
| MXE | 8594 | ENSG00000085999 | RAD54L | chr1 | + | 46258685 | 46258746 | 46259963 | 46260099 | 46249999 | 46250119 |
| MXE | 8595 | ENSG00000085999 | RAD54L | chr1 | + | 46273354 | 46273465 | 46273623 | 46273747 | 46272671 | 46272802 |
| MXE | 8614 | ENSG00000069702 | TGFBR3 | chr1 | - | 91770841 | 91770970 | 91786185 | 91786227 | 91758612 | 91758750 |
| MXE | 8622 | ENSG00000075151 | EIF4G3 | chr1 | - | 20981047 | 20981227 | 21001198 | 21001307 | 20980333 | 20980448 |
| MXE | 8626 | ENSG00000075151 | EIF4G3 | chr1 | - | 20981047 | 20981227 | 21002712 | 21002808 | 20980333 | 20980448 |
| MXE | 8630 | ENSG00000075151 | EIF4G3 | chr1 | - | 20997600 | 20997633 | 21001198 | 21001312 | 20981047 | 20981227 |
| MXE | 8632 | ENSG00000075151 | EIF4G3 | chr1 | - | 21002712 | 21002808 | 21050865 | 21050994 | 20980333 | 20980448 |
| MXE | 8639 | ENSG00000019582 | CD74 | chr5 | - | 150403120 | 150403312 | 150405084 | 150405671 | 150402562 | 150402625 |
| MXE | 8640 | ENSG00000174827 | PDZK1 | chr1 | - | 145682499 | 145682636 | 145686476 | 145686726 | 145680911 | 145681107 |
| MXE | 8644 | ENSG00000228956 | SATB1-AS1 | chr3 | + | 18472361 | 18472459 | 18516044 | 18516178 | 18465982 | 18466039 |
| MXE | 8647 | ENSG00000228956 | SATB1-AS1 | chr3 | + | 18527950 | 18528139 | 18548063 | 18548193 | 18527173 | 18527326 |
| MXE | 8649 | ENSG00000228956 | SATB1-AS1 | chr3 | + | 18529980 | 18530185 | 18548063 | 18548193 | 18527173 | 18527326 |
| MXE | 8651 | ENSG00000228956 | SATB1-AS1 | chr3 | + | 18529980 | 18530185 | 18548063 | 18548193 | 18527173 | 18527326 |
| MXE | 8652 | ENSG00000228956 | SATB1-AS1 | chr3 | + | 18529980 | 18530185 | 18585386 | 18585442 | 18527173 | 18527326 |
| MXE | 8655 | ENSG00000119950 | MXI1 | chr10 | + | 110245742 | 110245818 | 110260998 | 110261145 | 110244827 | 110244857 |
| MXE | 8659 | ENSG00000133619 | KRBA1 | chr7 | + | 149729187 | 149729370 | 149730146 | 149730341 | 149728522 | 149728675 |
| MXE | 8676 | ENSG00000269713 | NBPF9 | chr1 | - | 149058924 | 149059097 | 149062092 | 149062265 | 149058163 | 149058215 |
| MXE | 8687 | ENSG00000259673 | IQCH-AS1 | chr15 | - | 67430320 | 67430400 | 67459691 | 67459975 | 67421334 | 67421498 |
| MXE | 8693 | ENSG00000268043 | NBPF12 | chr1 | + | 146970646 | 146970719 | 146976928 | 146977001 | 146968447 | 146968550 |
| MXE | 8695 | ENSG00000268043 | NBPF12 | chr1 | + | 146984812 | 146984985 | 146986351 | 146986524 | 146984133 | 146984185 |
| MXE | 8697 | ENSG00000268043 | NBPF12 | chr1 | + | 146988840 | 146988949 | 146990465 | 146990517 | 146987953 | 146988126 |
| MXE | 8708 | ENSG00000124571 | XPO5 | chr6 | - | 43551297 | 43551453 | 43555835 | 43555964 | 43549892 | 43549934 |
| MXE | 8717 | ENSG00000172939 | OXSR1 | chr3 | + | 38249965 | 38250018 | 38251402 | 38251471 | 38247667 | 38247732 |
| MXE | 8724 | ENSG00000223519 | KIF28P | chr1 | - | 246780294 | 246780569 | 246780948 | 246781057 | 246777273 | 246777440 |
| MXE | 8727 | ENSG00000143815 | LBR | chr1 | - | 225411336 | 225411440 | 225412453 | 225412645 | 225410290 | 225410416 |
| MXE | 8741 | ENSG00000181722 | ZBTB20 | chr3 | - | 114582113 | 114582266 | 114599250 | 114599346 | 114500351 | 114500391 |
| MXE | 8742 | ENSG00000181722 | ZBTB20 | chr3 | - | 114693527 | 114693575 | 114801100 | 114801174 | 114500351 | 114500391 |
| MXE | 8743 | ENSG00000181722 | ZBTB20 | chr3 | - | 114710267 | 114710313 | 114713975 | 114714015 | 114693527 | 114693575 |
| MXE | 8745 | ENSG00000181722 | ZBTB20 | chr3 | - | 114713975 | 114714015 | 114743711 | 114743779 | 114693527 | 114693575 |
| MXE | 8746 | ENSG00000181722 | ZBTB20 | chr3 | - | 114713975 | 114714015 | 114801100 | 114801174 | 114693527 | 114693575 |
| MXE | 8749 | ENSG00000181722 | ZBTB20 | chr3 | - | 114801100 | 114801174 | 114900303 | 114900342 | 114713975 | 114714015 |
| MXE | 8750 | ENSG00000163430 | FSTL1 | chr3 | - | 120415922 | 120416027 | 120450683 | 120450746 | 120411853 | 120411983 |
| MXE | 8765 | ENSG00000101040 | ZMYND8 | chr20 | - | 47290186 | 47290274 | 47291795 | 47291888 | 47287228 | 47287284 |
| MXE | 8767 | ENSG00000101040 | ZMYND8 | chr20 | - | 47291795 | 47291888 | 47294665 | 47294779 | 47287228 | 47287284 |
| MXE | 8770 | ENSG00000101040 | ZMYND8 | chr20 | - | 47310055 | 47310204 | 47347855 | 47347926 | 47298728 | 47298947 |
| MXE | 8771 | ENSG00000101040 | ZMYND8 | chr20 | - | 47349860 | 47349988 | 47355449 | 47355520 | 47347855 | 47347926 |
| MXE | 8773 | ENSG00000180881 | CAPS2 | chr12 | - | 75323176 | 75323222 | 75325238 | 75325288 | 75321399 | 75321576 |
| MXE | 8785 | ENSG00000161040 | FBXL13 | chr7 | - | 102832839 | 102832974 | 102854776 | 102854860 | 102822039 | 102822203 |
| MXE | 8786 | ENSG00000161040 | FBXL13 | chr7 | - | 102931880 | 102931933 | 102963532 | 102963665 | 102926273 | 102926374 |
| MXE | 8787 | ENSG00000161040 | FBXL13 | chr7 | - | 103046808 | 103047020 | 103055087 | 103055178 | 103029350 | 103029418 |
| MXE | 8793 | ENSG00000249825 | THBS4-AS1 | chr5 | - | 80072085 | 80072223 | 80072507 | 80072665 | 80071731 | 80071891 |
| MXE | 8797 | ENSG00000142405 | NLRP12 | chr19 | - | 53795858 | 53796029 | 53798239 | 53798413 | 53793742 | 53794136 |
| MXE | 8798 | ENSG00000142405 | NLRP12 | chr19 | - | 53795858 | 53796029 | 53798239 | 53798413 | 53793742 | 53794136 |
| MXE | 8800 | ENSG00000142405 | NLRP12 | chr19 | - | 53798239 | 53798413 | 53801226 | 53801397 | 53793742 | 53794136 |
| MXE | 8801 | ENSG00000142405 | NLRP12 | chr19 | - | 53798239 | 53798413 | 53803948 | 53804122 | 53793742 | 53794136 |
| MXE | 8821 | ENSG00000001497 | LAS1L | chrX | - | 65520701 | 65522285 | 65522430 | 65522493 | 65517986 | 65518073 |
| MXE | 8825 | ENSG00000118007 | STAG1 | chr3 | - | 136359147 | 136359296 | 136363365 | 136363467 | 136349157 | 136349363 |
| MXE | 8827 | ENSG00000118007 | STAG1 | chr3 | - | 136363365 | 136363467 | 136366942 | 136367082 | 136359147 | 136359296 |
| MXE | 8830 | ENSG00000118007 | STAG1 | chr3 | - | 136586801 | 136586926 | 136591444 | 136591549 | 136568764 | 136568861 |
| MXE | 8843 | ENSG00000164924 | YWHAZ | chr8 | - | 100923954 | 100924050 | 100924134 | 100924322 | 100920715 | 100920752 |
| MXE | 8853 | ENSG00000160746 | ANO10 | chr3 | - | 43565652 | 43565727 | 43574808 | 43574864 | 43561219 | 43561402 |
| MXE | 8859 | ENSG00000082701 | GSK3B | chr3 | - | 119866589 | 119866628 | 119875079 | 119875185 | 119863418 | 119863605 |
| MXE | 8863 | ENSG00000082701 | GSK3B | chr3 | - | 119880244 | 119880498 | 119881160 | 119881270 | 119876412 | 119876508 |
| MXE | 8864 | ENSG00000082701 | GSK3B | chr3 | - | 119880244 | 119880498 | 119882636 | 119882834 | 119876412 | 119876508 |
| MXE | 8865 | ENSG00000082701 | GSK3B | chr3 | - | 119880244 | 119880498 | 119900557 | 119900613 | 119876412 | 119876508 |
| MXE | 8866 | ENSG00000082701 | GSK3B | chr3 | - | 119880244 | 119880510 | 119881160 | 119881270 | 119876412 | 119876508 |
| MXE | 8867 | ENSG00000082701 | GSK3B | chr3 | - | 119880244 | 119880510 | 119882636 | 119882834 | 119876412 | 119876508 |
| MXE | 8868 | ENSG00000082701 | GSK3B | chr3 | - | 119880244 | 119880510 | 119900557 | 119900613 | 119876412 | 119876508 |
| MXE | 8869 | ENSG00000082701 | GSK3B | chr3 | - | 119881160 | 119881270 | 119882636 | 119882834 | 119876412 | 119876508 |
| MXE | 8870 | ENSG00000082701 | GSK3B | chr3 | - | 119881160 | 119881270 | 119900557 | 119900613 | 119876412 | 119876508 |
| MXE | 8871 | ENSG00000082701 | GSK3B | chr3 | - | 119882636 | 119882834 | 119900557 | 119900613 | 119876412 | 119876508 |
| MXE | 8872 | ENSG00000082701 | GSK3B | chr3 | - | 119976484 | 119976545 | 119981956 | 119982125 | 119947267 | 119947351 |
| MXE | 8873 | ENSG00000082701 | GSK3B | chr3 | - | 119976484 | 119976545 | 120000551 | 120000723 | 119947267 | 119947351 |
| MXE | 8874 | ENSG00000082701 | GSK3B | chr3 | - | 119981956 | 119982125 | 120000551 | 120000723 | 119947267 | 119947351 |
| MXE | 8884 | ENSG00000273136 | NBPF26 | chr1 | + | 120818122 | 120818174 | 120822103 | 120822156 | 120816621 | 120816827 |
| MXE | 8889 | ENSG00000273136 | NBPF26 | chr1 | + | 120833602 | 120833775 | 120834629 | 120834802 | 120832874 | 120832983 |
| MXE | 8904 | ENSG00000172965 | MIR4435-2HG | chr2 | - | 111247429 | 111247558 | 111248147 | 111248241 | 111239801 | 111239996 |
| MXE | 8906 | ENSG00000172965 | MIR4435-2HG | chr2 | - | 111447211 | 111447349 | 111479906 | 111480061 | 111367486 | 111367575 |
| MXE | 8926 | ENSG00000117262 | GPR89A | chr1 | + | 145608347 | 145608675 | 145612201 | 145612247 | 145608047 | 145608175 |
| MXE | 8927 | ENSG00000117262 | GPR89A | chr1 | + | 145645646 | 145645743 | 145646183 | 145646272 | 145643868 | 145643978 |
| MXE | 8929 | ENSG00000170089 | NA | chr5 | + | 177876531 | 177876688 | 177878643 | 177878848 | 177875521 | 177875860 |
| MXE | 8931 | ENSG00000116667 | C1orf21 | chr1 | + | 184507597 | 184507682 | 184590738 | 184590815 | 184477385 | 184477603 |
| MXE | 8945 | ENSG00000226674 | TEX41 | chr2 | + | 144671601 | 144671664 | 144766369 | 144766530 | 144668011 | 144668108 |
| MXE | 8949 | ENSG00000226674 | TEX41 | chr2 | + | 144691038 | 144691110 | 144766369 | 144766530 | 144667977 | 144668108 |
| MXE | 9031 | ENSG00000186854 | TRABD2A | chr2 | - | 84823952 | 84824204 | 84832054 | 84832145 | 84821649 | 84822100 |
| MXE | 9039 | ENSG00000115594 | IL1R1 | chr2 | + | 102165114 | 102165304 | 102166112 | 102166281 | 102164773 | 102165008 |
| MXE | 9041 | ENSG00000115594 | IL1R1 | chr2 | + | 102166112 | 102166281 | 102168597 | 102168663 | 102165114 | 102165304 |
| MXE | 9046 | ENSG00000134759 | ELP2 | chr18 | + | 36141136 | 36141201 | 36142825 | 36142966 | 36138269 | 36138426 |
| MXE | 9047 | ENSG00000134759 | ELP2 | chr18 | + | 36141136 | 36141201 | 36142825 | 36142966 | 36138269 | 36138426 |
| MXE | 9049 | ENSG00000134759 | ELP2 | chr18 | + | 36145947 | 36146048 | 36146249 | 36146381 | 36142825 | 36142966 |
| MXE | 9058 | ENSG00000173409 | ARV1 | chr1 | + | 230988319 | 230988439 | 230990109 | 230990263 | 230979093 | 230979279 |
| MXE | 9064 | ENSG00000162415 | ZSWIM5 | chr1 | - | 45040391 | 45040538 | 45043218 | 45043395 | 45038935 | 45039073 |
| MXE | 9065 | ENSG00000163157 | TMOD4 | chr1 | - | 151170919 | 151171063 | 151171632 | 151171763 | 151170518 | 151170663 |
| MXE | 9073 | ENSG00000140416 | TPM1 | chr15 | + | 63043705 | 63043831 | 63044026 | 63044152 | 63042757 | 63042943 |
| MXE | 9075 | ENSG00000164828 | SUN1 | chr7 | + | 832504 | 832601 | 838797 | 838986 | 816614 | 816673 |
| MXE | 9078 | ENSG00000164828 | SUN1 | chr7 | + | 848417 | 848615 | 849922 | 850033 | 843340 | 843520 |
| MXE | 9082 | ENSG00000164828 | SUN1 | chr7 | + | 849922 | 850033 | 851383 | 851482 | 843340 | 843520 |
| MXE | 9088 | ENSG00000135336 | ORC3 | chr6 | + | 87634844 | 87634961 | 87636406 | 87636486 | 87621949 | 87622013 |
| MXE | 9089 | ENSG00000135336 | ORC3 | chr6 | + | 87636406 | 87636486 | 87653115 | 87653249 | 87634844 | 87634961 |
| MXE | 9090 | ENSG00000135336 | ORC3 | chr6 | + | 87653115 | 87653249 | 87656902 | 87656982 | 87636406 | 87636486 |
| MXE | 9125 | ENSG00000282246 | NA | chr10 | + | 13611614 | 13611683 | 13614014 | 13614086 | 13610046 | 13610185 |
| MXE | 9132 | ENSG00000168421 | RHOH | chr4 | + | 40197120 | 40197300 | 40242713 | 40242834 | 40192966 | 40193123 |
| MXE | 9133 | ENSG00000168421 | RHOH | chr4 | + | 40200511 | 40200565 | 40212663 | 40212791 | 40196906 | 40197300 |
| MXE | 9135 | ENSG00000168421 | RHOH | chr4 | + | 40200511 | 40200565 | 40235192 | 40235358 | 40196906 | 40197300 |
| MXE | 9137 | ENSG00000168421 | RHOH | chr4 | + | 40212663 | 40212791 | 40235192 | 40235358 | 40196906 | 40197300 |
| MXE | 9144 | ENSG00000115204 | MPV17 | chr2 | - | 27312993 | 27313109 | 27322447 | 27322522 | 27312679 | 27312772 |
| MXE | 9147 | ENSG00000075702 | WDR62 | chr19 | + | 36091199 | 36091311 | 36091401 | 36091465 | 36090444 | 36090520 |
| MXE | 9174 | ENSG00000119004 | CYP20A1 | chr2 | + | 203272645 | 203272748 | 203278572 | 203278688 | 203266513 | 203266681 |
| MXE | 9180 | ENSG00000187922 | LCN10 | chr9 | - | 136740835 | 136741014 | 136741251 | 136741361 | 136739949 | 136740048 |
| MXE | 9181 | ENSG00000058453 | CROCC | chr1 | + | 16931290 | 16931397 | 16936636 | 16936873 | 16930428 | 16930594 |
| MXE | 9188 | ENSG00000168216 | LMBRD1 | chr6 | - | 69705500 | 69705619 | 69708849 | 69709231 | 69701888 | 69701953 |
| MXE | 9189 | ENSG00000168216 | LMBRD1 | chr6 | - | 69705500 | 69705619 | 69709156 | 69709231 | 69701888 | 69701953 |
| MXE | 9191 | ENSG00000168216 | LMBRD1 | chr6 | - | 69708849 | 69709231 | 69710424 | 69710444 | 69701888 | 69701953 |
| MXE | 9192 | ENSG00000168216 | LMBRD1 | chr6 | - | 69709156 | 69709231 | 69710424 | 69710444 | 69701888 | 69701953 |
| MXE | 9197 | ENSG00000116127 | ALMS1 | chr2 | + | 73432128 | 73432291 | 73447959 | 73454067 | 73426452 | 73426553 |
| MXE | 9211 | ENSG00000148660 | CAMK2G | chr10 | - | 73825278 | 73825347 | 73828088 | 73828121 | 73824039 | 73824084 |
| MXE | 9217 | ENSG00000087903 | RFX2 | chr19 | - | 6042043 | 6042123 | 6044192 | 6044282 | 6039979 | 6040241 |
| MXE | 9218 | ENSG00000135605 | TEC | chr4 | - | 48141354 | 48141419 | 48145078 | 48145295 | 48138903 | 48139022 |
| MXE | 9219 | ENSG00000135605 | TEC | chr4 | - | 48170247 | 48170376 | 48171367 | 48171408 | 48168585 | 48168626 |
| MXE | 9232 | ENSG00000165996 | HACD1 | chr10 | - | 17599289 | 17599411 | 17603929 | 17604047 | 17594204 | 17594383 |
| MXE | 9245 | ENSG00000152683 | SLC30A6 | chr2 | + | 32175318 | 32175361 | 32184272 | 32184338 | 32174062 | 32174147 |
| MXE | 9247 | ENSG00000152683 | SLC30A6 | chr2 | + | 32193888 | 32193983 | 32197343 | 32197392 | 32192917 | 32192953 |
| MXE | 9248 | ENSG00000152683 | SLC30A6 | chr2 | + | 32204589 | 32204692 | 32206885 | 32206933 | 32197706 | 32197826 |
| MXE | 9249 | ENSG00000152683 | SLC30A6 | chr2 | + | 32204589 | 32204692 | 32206885 | 32206933 | 32197706 | 32197826 |
| MXE | 9254 | ENSG00000149231 | CCDC82 | chr11 | - | 96383268 | 96383473 | 96386253 | 96386293 | 96373374 | 96373467 |
| MXE | 9255 | ENSG00000149231 | CCDC82 | chr11 | - | 96385936 | 96386020 | 96386253 | 96386293 | 96383961 | 96384761 |
| MXE | 9256 | ENSG00000149231 | CCDC82 | chr11 | - | 96386253 | 96386293 | 96387529 | 96387613 | 96383961 | 96384761 |
| MXE | 9271 | ENSG00000138709 | LARP1B | chr4 | + | 128178430 | 128178642 | 128179405 | 128179512 | 128122116 | 128122188 |
| MXE | 9272 | ENSG00000103168 | TAF1C | chr16 | - | 84180867 | 84180995 | 84181042 | 84181186 | 84180169 | 84180344 |
| MXE | 9273 | ENSG00000175054 | ATR | chr3 | - | 142465096 | 142465240 | 142466323 | 142466533 | 142461939 | 142462090 |
| MXE | 9285 | ENSG00000164144 | ARFIP1 | chr4 | + | 152807203 | 152807248 | 152809672 | 152809733 | 152780108 | 152780226 |
| MXE | 9289 | ENSG00000117280 | RAB29 | chr1 | - | 205771471 | 205771653 | 205772495 | 205772567 | 205770732 | 205770854 |
| MXE | 9297 | ENSG00000096093 | EFHC1 | chr6 | + | 52481545 | 52481662 | 52488556 | 52488675 | 52479639 | 52479787 |
| MXE | 9301 | ENSG00000075651 | PLD1 | chr3 | - | 171686684 | 171686798 | 171687370 | 171687584 | 171677565 | 171677694 |
| MXE | 9307 | ENSG00000099250 | NRP1 | chr10 | - | 33202890 | 33202995 | 33207571 | 33207716 | 33197620 | 33197709 |
| MXE | 9319 | ENSG00000165071 | TMEM71 | chr8 | - | 132687461 | 132687655 | 132722039 | 132722115 | 132685006 | 132685105 |
| MXE | 9336 | ENSG00000172766 | NAA16 | chr13 | + | 41373767 | 41373880 | 41374741 | 41374839 | 41372814 | 41372830 |
| MXE | 9375 | ENSG00000100410 | PHF5A | chr22 | - | 41467447 | 41467614 | 41468123 | 41468147 | 41459716 | 41460487 |
| MXE | 9379 | ENSG00000288637 | NA | chr4 | + | 153394261 | 153394336 | 153397218 | 153397333 | 153358473 | 153358622 |
| MXE | 9380 | ENSG00000288637 | NA | chr4 | + | 153397218 | 153397333 | 153408970 | 153409015 | 153358473 | 153358622 |
| MXE | 9392 | ENSG00000147099 | HDAC8 | chrX | - | 72488932 | 72489041 | 72495155 | 72495268 | 72473415 | 72474729 |
| MXE | 9394 | ENSG00000147099 | HDAC8 | chrX | - | 72495155 | 72495268 | 72567888 | 72568030 | 72488932 | 72489041 |
| MXE | 9401 | ENSG00000235750 | KIAA0040 | chr1 | - | 175166561 | 175166737 | 175177610 | 175177684 | 175156986 | 175161146 |
| MXE | 9403 | ENSG00000106346 | USP42 | chr7 | + | 6116794 | 6116939 | 6135840 | 6135951 | 6115322 | 6115523 |
| MXE | 9405 | ENSG00000152380 | FAM151B | chr5 | + | 80495052 | 80495283 | 80499884 | 80499985 | 80487985 | 80488148 |
| MXE | 9406 | ENSG00000152380 | FAM151B | chr5 | + | 80505942 | 80506076 | 80510713 | 80511086 | 80501791 | 80501917 |
| MXE | 9414 | ENSG00000187741 | FANCA | chr16 | - | 89810706 | 89810802 | 89810928 | 89811071 | 89808293 | 89808367 |
| MXE | 9419 | ENSG00000237298 | TTN-AS1 | chr2 | + | 178531313 | 178531578 | 178535731 | 178535828 | 178528723 | 178528813 |
| MXE | 9420 | ENSG00000237298 | TTN-AS1 | chr2 | + | 178535731 | 178535828 | 178537603 | 178537691 | 178523291 | 178523636 |
| MXE | 9421 | ENSG00000237298 | TTN-AS1 | chr2 | + | 178535731 | 178535828 | 178584572 | 178584993 | 178523291 | 178523636 |
| MXE | 9422 | ENSG00000237298 | TTN-AS1 | chr2 | + | 178536333 | 178536462 | 178537360 | 178537447 | 178535731 | 178535828 |
| MXE | 9423 | ENSG00000237298 | TTN-AS1 | chr2 | + | 178537603 | 178537691 | 178584572 | 178584993 | 178523291 | 178523636 |
| MXE | 9427 | ENSG00000144674 | GOLGA4 | chr3 | + | 37286013 | 37286061 | 37289234 | 37289289 | 37281957 | 37282272 |
| MXE | 9436 | ENSG00000138078 | PREPL | chr2 | - | 44346267 | 44346390 | 44359496 | 44360038 | 44344519 | 44344586 |
| MXE | 9441 | ENSG00000066027 | PPP2R5A | chr1 | + | 212306782 | 212306900 | 212313766 | 212313816 | 212301862 | 212302092 |
| MXE | 9455 | ENSG00000198039 | ZNF273 | chr7 | + | 64917580 | 64917707 | 64918196 | 64918292 | 64917029 | 64917118 |
| MXE | 9456 | ENSG00000111711 | GOLT1B | chr12 | + | 21507116 | 21507151 | 21507943 | 21508109 | 21506884 | 21506976 |
| MXE | 9457 | ENSG00000111711 | GOLT1B | chr12 | + | 21507116 | 21507151 | 21507943 | 21508109 | 21506884 | 21506976 |
| MXE | 9458 | ENSG00000111711 | GOLT1B | chr12 | + | 21507116 | 21507151 | 21508382 | 21508561 | 21506884 | 21506976 |
| MXE | 9461 | ENSG00000028528 | SNX1 | chr15 | + | 64118787 | 64118854 | 64123502 | 64123546 | 64118116 | 64118244 |
| MXE | 9470 | ENSG00000131408 | NR1H2 | chr19 | + | 50377732 | 50377870 | 50378148 | 50378312 | 50377586 | 50377648 |
| MXE | 9475 | ENSG00000171503 | ETFDH | chr4 | + | 158703689 | 158706371 | 158706628 | 158706850 | 158703422 | 158703591 |
| MXE | 9478 | ENSG00000134058 | CDK7 | chr5 | + | 69252417 | 69252451 | 69254601 | 69254669 | 69235393 | 69235453 |
| MXE | 9479 | ENSG00000134058 | CDK7 | chr5 | + | 69254601 | 69254669 | 69255459 | 69255528 | 69252417 | 69252451 |
| MXE | 9486 | ENSG00000067208 | EVI5 | chr1 | - | 92636201 | 92636336 | 92663419 | 92663452 | 92607580 | 92607727 |
| MXE | 9488 | ENSG00000067208 | EVI5 | chr1 | - | 92636201 | 92636336 | 92663419 | 92663452 | 92625793 | 92625934 |
| MXE | 9489 | ENSG00000067208 | EVI5 | chr1 | - | 92662718 | 92662865 | 92663419 | 92663452 | 92636201 | 92636336 |
| MXE | 9491 | ENSG00000102786 | INTS6 | chr13 | - | 51389318 | 51389444 | 51421111 | 51421334 | 51387385 | 51387540 |
| MXE | 9500 | ENSG00000035115 | SH3YL1 | chr2 | - | 242797 | 242871 | 243502 | 243562 | 234159 | 234272 |
| MXE | 9501 | ENSG00000035115 | SH3YL1 | chr2 | - | 242797 | 242871 | 244626 | 244738 | 234159 | 234272 |
| MXE | 9503 | ENSG00000035115 | SH3YL1 | chr2 | - | 243502 | 243562 | 244626 | 244738 | 242797 | 242871 |
| MXE | 9510 | ENSG00000053900 | ANAPC4 | chr4 | + | 25413644 | 25413742 | 25414323 | 25414385 | 25409697 | 25409791 |
| MXE | 9527 | ENSG00000138757 | G3BP2 | chr4 | - | 75657556 | 75657730 | 75657800 | 75657893 | 75656923 | 75657014 |
| MXE | 9544 | ENSG00000150938 | CRIM1 | chr2 | + | 36522091 | 36522313 | 36537351 | 36537546 | 36517326 | 36517542 |
| MXE | 9551 | ENSG00000198836 | OPA1 | chr3 | + | 193642974 | 193643049 | 193643372 | 193643444 | 193642764 | 193642845 |
| MXE | 9552 | ENSG00000198836 | OPA1 | chr3 | + | 193668314 | 193668484 | 193689068 | 193689170 | 193667169 | 193667280 |
| MXE | 9556 | ENSG00000104343 | UBE2W | chr8 | - | 73830380 | 73830472 | 73870268 | 73870301 | 73825146 | 73825249 |
| MXE | 9562 | ENSG00000162929 | KIAA1841 | chr2 | + | 61071605 | 61071792 | 61073457 | 61073551 | 61070325 | 61070500 |
| MXE | 9577 | ENSG00000080947 | CROCCP3 | chr1 | - | 16486425 | 16486661 | 16489684 | 16489791 | 16485568 | 16485665 |
| MXE | 9582 | ENSG00000163904 | SENP2 | chr3 | + | 185598991 | 185599024 | 185600764 | 185600855 | 185598411 | 185598545 |
| MXE | 9590 | ENSG00000044115 | CTNNA1 | chr5 | + | 138783176 | 138783372 | 138810037 | 138810204 | 138781922 | 138782029 |
| MXE | 9594 | ENSG00000182885 | ADGRG3 | chr16 | + | 57680504 | 57680617 | 57684389 | 57684483 | 57680264 | 57680365 |
| MXE | 9597 | ENSG00000144040 | SFXN5 | chr2 | - | 72956979 | 72957103 | 72961130 | 72961248 | 72944739 | 72945099 |
| MXE | 9598 | ENSG00000144040 | SFXN5 | chr2 | - | 73041606 | 73041661 | 73042487 | 73042585 | 73040853 | 73040931 |
| MXE | 9599 | ENSG00000270800 | RPS10-NUDT3 | chr6 | - | 34295640 | 34295685 | 34341861 | 34341972 | 34293450 | 34293535 |
| MXE | 9617 | ENSG00000119778 | ATAD2B | chr2 | - | 23799824 | 23799993 | 23808988 | 23809176 | 23798137 | 23798323 |
| MXE | 9633 | ENSG00000185811 | IKZF1 | chr7 | + | 50319047 | 50319101 | 50376532 | 50376793 | 50304781 | 50304922 |
| MXE | 9641 | ENSG00000185811 | IKZF1 | chr7 | + | 50387344 | 50387470 | 50391728 | 50391863 | 50319047 | 50319101 |
| MXE | 9642 | ENSG00000072682 | P4HA2 | chr5 | - | 132198205 | 132198271 | 132198320 | 132198380 | 132195411 | 132195480 |
| MXE | 9643 | ENSG00000214193 | SH3D21 | chr1 | + | 36319259 | 36319312 | 36319674 | 36320798 | 36319070 | 36319164 |
| MXE | 9665 | ENSG00000146856 | AGBL3 | chr7 | + | 134989249 | 134989310 | 134993492 | 134993678 | 134987888 | 134987996 |
| MXE | 9672 | ENSG00000049167 | ERCC8 | chr5 | - | 60892036 | 60893643 | 60896458 | 60896515 | 60890888 | 60891086 |
| MXE | 9673 | ENSG00000049167 | ERCC8 | chr5 | - | 60909647 | 60909957 | 60915185 | 60915286 | 60904791 | 60904873 |
| MXE | 9675 | ENSG00000107614 | TRDMT1 | chr10 | - | 17160304 | 17160374 | 17161482 | 17161548 | 17159145 | 17159229 |
| MXE | 9676 | ENSG00000107614 | TRDMT1 | chr10 | - | 17160304 | 17160374 | 17162165 | 17162237 | 17159145 | 17159229 |
| MXE | 9678 | ENSG00000107614 | TRDMT1 | chr10 | - | 17161482 | 17161548 | 17162165 | 17162237 | 17160304 | 17160374 |
| MXE | 9679 | ENSG00000107614 | TRDMT1 | chr10 | - | 17161482 | 17161548 | 17168840 | 17168917 | 17160304 | 17160374 |
| MXE | 9680 | ENSG00000107614 | TRDMT1 | chr10 | - | 17162165 | 17162237 | 17168840 | 17168917 | 17160304 | 17160374 |
| MXE | 9681 | ENSG00000173597 | SULT1B1 | chr4 | - | 69749720 | 69749818 | 69755069 | 69755261 | 69734137 | 69734264 |
| MXE | 9695 | ENSG00000049247 | UTS2 | chr1 | - | 7847803 | 7847882 | 7849639 | 7849683 | 7846691 | 7846766 |
| MXE | 9717 | ENSG00000140612 | SEC11A | chr15 | - | 84680712 | 84680832 | 84691534 | 84691644 | 84669543 | 84670069 |
| MXE | 9719 | ENSG00000114120 | SLC25A36 | chr3 | + | 140966347 | 140966434 | 140966933 | 140967036 | 140963126 | 140963227 |
| MXE | 9720 | ENSG00000114120 | SLC25A36 | chr3 | + | 140966347 | 140966434 | 140966999 | 140967036 | 140963126 | 140963227 |
| MXE | 9725 | ENSG00000130595 | TNNT3 | chr11 | + | 1923047 | 1923061 | 1923554 | 1923572 | 1922856 | 1922891 |
| MXE | 9729 | ENSG00000228590 | MIR4432HG | chr2 | - | 60351808 | 60352308 | 60359269 | 60359848 | 60343087 | 60343168 |
| MXE | 9732 | ENSG00000137497 | NUMA1 | chr11 | - | 72049418 | 72049539 | 72062588 | 72062686 | 72035901 | 72035975 |
| MXE | 9734 | ENSG00000137497 | NUMA1 | chr11 | - | 72049418 | 72049539 | 72068495 | 72068617 | 72035901 | 72035975 |
| MXE | 9735 | ENSG00000137497 | NUMA1 | chr11 | - | 72050661 | 72050833 | 72062588 | 72062686 | 72035901 | 72035975 |
| MXE | 9736 | ENSG00000137497 | NUMA1 | chr11 | - | 72050661 | 72050833 | 72068078 | 72068231 | 72035901 | 72035975 |
| MXE | 9737 | ENSG00000137497 | NUMA1 | chr11 | - | 72050661 | 72050833 | 72068495 | 72068617 | 72035901 | 72035975 |
| MXE | 9738 | ENSG00000137497 | NUMA1 | chr11 | - | 72062588 | 72062686 | 72068078 | 72068231 | 72035901 | 72035975 |
| MXE | 9740 | ENSG00000137497 | NUMA1 | chr11 | - | 72068078 | 72068231 | 72068495 | 72068617 | 72035901 | 72035975 |
| MXE | 9748 | ENSG00000285533 | RELA-DT | chr11 | + | 65663426 | 65663683 | 65665420 | 65665543 | 65663005 | 65663148 |
| MXE | 9749 | ENSG00000285533 | RELA-DT | chr11 | + | 65665420 | 65665543 | 65666130 | 65666217 | 65663426 | 65663683 |
| MXE | 9754 | ENSG00000132953 | XPO4 | chr13 | - | 20800830 | 20800990 | 20807456 | 20807634 | 20800155 | 20800325 |
| MXE | 9757 | ENSG00000093217 | XYLB | chr3 | + | 38373114 | 38373279 | 38374461 | 38374502 | 38372654 | 38372736 |
| MXE | 9765 | ENSG00000164134 | NAA15 | chr4 | + | 139354025 | 139354098 | 139357385 | 139357555 | 139351504 | 139351611 |
| MXE | 9766 | ENSG00000164134 | NAA15 | chr4 | + | 139370210 | 139370404 | 139376364 | 139376473 | 139361723 | 139361937 |
| MXE | 9767 | ENSG00000164134 | NAA15 | chr4 | + | 139376364 | 139376473 | 139378755 | 139378854 | 139370210 | 139370404 |

| | | | | | | | | | | | | | | | |
|---|---|---|---|---|---|---|---|---|---|---|---|---|---|---|---|
| MXE | 9768 | ENSG00000164134 | NAA15 | chr4 | + | 139378755 | 139378854 | 139384831 | 139384978 | 139376364 | 139376473 | 139386132 | 139386230 | 50 | [illegible] |
| MXE | 9769 | ENSG00000164134 | NAA15 | chr4 | + | 139384831 | 139384978 | 139386132 | 139386230 | 139378755 | 139378854 | 139387883 | 139391384 | 146 | [illegible] |
| MXE | 9770 | ENSG00000164134 | NAA15 | chr4 | + | 139384834 | 139384978 | 139386132 | 139386230 | 139378755 | 139378854 | 139387883 | 139391368 | 31 | [illegible] |
| MXE | 9783 | ENSG00000068784 | SRBD1 | chr2 | - | 45419787 | 45419894 | 45488239 | 45488331 | 45418364 | 45418541 | 45546731 | 45546839 | 24 | [illegible] |
| MXE | 9784 | ENSG00000068784 | SRBD1 | chr2 | - | 45476992 | 45477075 | 45488239 | 45488331 | 45419787 | 45419894 | 45546731 | 45546839 | 27 | [illegible] |
| MXE | 9789 | ENSG00000106105 | GARS1 | chr7 | + | 30606325 | 30606397 | 30608407 | 30608483 | 30603495 | 30603572 | 30609584 | 30609730 | 0 | [illegible] |
| MXE | 9793 | ENSG00000095002 | MSH2 | chr2 | + | 47466657 | 47466808 | 47470964 | 47471062 | 47463030 | 47463154 | 47475024 | 47475270 | 76 | [illegible] |
| MXE | 9798 | ENSG00000115825 | PRKD3 | chr2 | - | 37256661 | 37256929 | 37259582 | 37259681 | 37254203 | 37254289 | 37260222 | 37260384 | 53 | [illegible] |
| MXE | 9808 | ENSG00000151092 | NGLY1 | chr3 | - | 25736003 | 25736149 | 25737333 | 25737455 | 25733871 | 25733982 | 25751097 | 25751263 | 2 | [illegible] |
| MXE | 9811 | ENSG00000151092 | NGLY1 | chr3 | - | 25736308 | 25736400 | 25737333 | 25737455 | 25733871 | 25733982 | 25751097 | 25751263 | 2 | [illegible] |
| MXE | 9812 | ENSG00000151092 | NGLY1 | chr3 | - | 25736308 | 25736400 | 25739576 | 25739799 | 25733871 | 25733982 | 25751097 | 25751263 | 65 | [illegible] |
| MXE | 9813 | ENSG00000151092 | NGLY1 | chr3 | - | 25736308 | 25736400 | 25740402 | 25740450 | 25733871 | 25733982 | 25751097 | 25751263 | 2 | [illegible] |
| MXE | 9816 | ENSG00000151092 | NGLY1 | chr3 | - | 25736308 | 25736400 | 25739576 | 25739799 | 25736003 | 25736149 | 25751097 | 25751263 | 67 | [illegible] |
| MXE | 9817 | ENSG00000151092 | NGLY1 | chr3 | - | 25736308 | 25736400 | 25740402 | 25740450 | 25736003 | 25736149 | 25751097 | 25751263 | 2 | [illegible] |
| MXE | 9818 | ENSG00000151092 | NGLY1 | chr3 | - | 25737333 | 25737455 | 25739576 | 25739799 | 25733871 | 25733982 | 25751097 | 25751263 | 65 | [illegible] |
| MXE | 9819 | ENSG00000151092 | NGLY1 | chr3 | - | 25737333 | 25737455 | 25740402 | 25740450 | 25733871 | 25733982 | 25751097 | 25751263 | 2 | [illegible] |
| MXE | 9823 | ENSG00000151092 | NGLY1 | chr3 | - | 25739576 | 25739799 | 25740402 | 25740450 | 25733871 | 25733982 | 25751097 | 25751263 | 2 | [illegible] |
| MXE | 9824 | ENSG00000151092 | NGLY1 | chr3 | - | 25739576 | 25739799 | 25740402 | 25740450 | 25736003 | 25736149 | 25751097 | 25751263 | 2 | [illegible] |
| MXE | 9828 | ENSG00000118263 | KLF7 | chr2 | - | 207123975 | 207124404 | 207165466 | 207165706 | 207088457 | 207088581 | 207166756 | 207166847 | 1 | [illegible] |
| MXE | 9829 | ENSG00000084774 | CAD | chr2 | + | 27239330 | 27239471 | 27239696 | 27239798 | 27239041 | 27239232 | 27240264 | 27240361 | 33 | [illegible] |
| MXE | 9830 | ENSG00000250299 | MRPS31P4 | chr13 | + | 52629218 | 52629359 | 52632421 | 52632494 | 52627437 | 52627578 | 52637304 | 52637446 | 5 | [illegible] |
| MXE | 9833 | ENSG00000115421 | PAPOLG | chr2 | + | 60794963 | 60795020 | 60795289 | 60795336 | 60794709 | 60794775 | 60797061 | 60798466 | 104 | [illegible] |
| MXE | 9834 | ENSG00000138658 | ZGRF1 | chr4 | - | 112587279 | 112587929 | 112589723 | 112589874 | 112586444 | 112586583 | 112603523 | 112603697 | 0 | [illegible] |
| MXE | 9835 | ENSG00000138658 | ZGRF1 | chr4 | - | 112589723 | 112589874 | 112603523 | 112603697 | 112587279 | 112587929 | 112606007 | 112606091 | 0 | [illegible] |
| MXE | 9836 | ENSG00000138658 | ZGRF1 | chr4 | - | 112628620 | 112628851 | 112629758 | 112630015 | 112623816 | 112623876 | 112631929 | 112632010 | 2 | [illegible] |
| MXE | 9844 | ENSG00000115966 | ATF2 | chr2 | - | 175136411 | 175136486 | 175147945 | 175147980 | 175130137 | 175130207 | 175151059 | 175151158 | 1 | [illegible] |
| MXE | 9902 | ENSG00000108557 | RAI1 | chr17 | + | 17724027 | 17724159 | 17792932 | 17798513 | 17681457 | 17681793 | 17803755 | 17803849 | 8 | [illegible] |
| MXE | 9907 | ENSG00000109332 | UBE2D3 | chr4 | - | 102809791 | 102809855 | 102826484 | 102826844 | 102809671 | 102809703 | 102827943 | 102828064 | 0 | [illegible] |
| MXE | 9912 | ENSG00000115464 | USP34 | chr2 | - | 61190517 | 61190658 | 61192900 | 61192980 | 61190270 | 61190414 | 61204496 | 61204601 | 3 | [illegible] |
| MXE | 9921 | ENSG00000115464 | USP34 | chr2 | - | 61222618 | 61222663 | 61223247 | 61223296 | 61221501 | 61221606 | 61227066 | 61227218 | 165 | [illegible] |
| MXE | 9929 | ENSG00000283050 | GTF2IP12 | chr4 | - | 119440559 | 119441054 | 119442157 | 119442217 | 119434739 | 119434975 | 119444367 | 119444433 | 2 | [illegible] |
| MXE | 9930 | ENSG00000115919 | KYNU | chr2 | + | 142881344 | 142881430 | 142885348 | 142885536 | 142877663 | 142877736 | 142918608 | 142918729 | 1 | [illegible] |
| MXE | 9931 | ENSG00000115919 | KYNU | chr2 | + | 143029626 | 143029679 | 143033235 | 143033321 | 142985947 | 142986021 | 143040427 | 143040658 | 9 | [illegible] |
| MXE | 9933 | ENSG00000283602 | NA | chr5 | - | 140591073 | 140591150 | 140591669 | 140591752 | 140590874 | 140590991 | 140592023 | 140592101 | 6 | [illegible] |
| MXE | 9948 | ENSG00000185499 | MUC1 | chr1 | - | 155187454 | 155187544 | 155187721 | 155187858 | 155187224 | 155187374 | 155188007 | 155188063 | 3 | [illegible] |
| MXE | 9950 | ENSG00000185499 | MUC1 | chr1 | - | 155187454 | 155187576 | 155187721 | 155187858 | 155187224 | 155187374 | 155188007 | 155188063 | 3 | [illegible] |
| MXE | 9951 | ENSG00000185499 | MUC1 | chr1 | - | 155187721 | 155187858 | 155188007 | 155188063 | 155187454 | 155187576 | 155188162 | 155188212 | 1 | [illegible] |
| MXE | 9962 | ENSG00000101336 | HCK | chr20 | + | 32054215 | 32054281 | 32069687 | 32069751 | 32052241 | 32052486 | 32071661 | 32071782 | 18 | [illegible] |
| MXE | 9994 | ENSG00000166801 | FAM111A | chr11 | + | 59143496 | 59143695 | 59144808 | 59144896 | 59143082 | 59143304 | 59145826 | 59145856 | 76 | [illegible] |
| MXE | 9997 | ENSG00000066455 | GOLGA5 | chr14 | + | 92833121 | 92833347 | 92835558 | 92835664 | 92824545 | 92824644 | 92837385 | 92837449 | 72 | [illegible] |
| MXE | 9999 | ENSG00000144036 | EXOC6B | chr2 | - | 72513131 | 72513252 | 72514633 | 72514680 | 72499900 | 72499972 | 72515042 | 72515126 | 8 | [illegible] |
| [illegible] | | | | | | | | | | | | | | | |

| | | | | | | | | | | |
|---|---|---|---|---|---|---|---|---|---|---|
| MXE | 11046 | ENSG00000079263 | SP140 | chr2 | + | 230248884 | 230248968 | 230253315 | 230253417 | [illegible] |
| MXE | 11049 | ENSG00000079263 | SP140 | chr2 | + | 230253315 | 230253417 | 230255451 | 230255532 | [illegible] |
| MXE | 11062 | ENSG00000165280 | VCP | chr9 | - | 35065250 | 35065381 | 35066674 | 35066900 | [illegible] |
| MXE | 11067 | ENSG00000122203 | KIAA1191 | chr5 | - | 176350612 | 176350737 | 176352621 | 176352748 | [illegible] |
| MXE | 11069 | ENSG00000122203 | KIAA1191 | chr5 | - | 176352621 | 176352748 | 176355570 | 176355749 | [illegible] |
| MXE | 11080 | ENSG00000163596 | ICA1L | chr2 | - | 202796889 | 202796964 | 202811745 | 202811789 | [illegible] |
| MXE | 11081 | ENSG00000115234 | SNX17 | chr2 | + | 27372622 | 27372740 | 27373039 | 27373113 | [illegible] |
| MXE | 11083 | ENSG00000123983 | ACSL3 | chr2 | + | 222892263 | 222892301 | 222900673 | 222900780 | [illegible] |
| MXE | 11088 | ENSG00000144504 | ANKMY1 | chr2 | - | 240507559 | 240507691 | 240509347 | 240509455 | [illegible] |
| MXE | 11089 | ENSG00000144504 | ANKMY1 | chr2 | - | 240507559 | 240507691 | 240511860 | 240512001 | [illegible] |
| MXE | 11090 | ENSG00000144504 | ANKMY1 | chr2 | - | 240509347 | 240509455 | 240511860 | 240512001 | [illegible] |
| MXE | 11093 | ENSG00000164114 | MAP9 | chr4 | - | 155374936 | 155375021 | 155375775 | 155375914 | [illegible] |
| MXE | 11095 | ENSG00000184182 | UBE2F | chr2 | + | 237973648 | 237973747 | 237986161 | 237986292 | [illegible] |
| MXE | 11096 | ENSG00000184182 | UBE2F | chr2 | + | 237973648 | 237973747 | 237987962 | 237987992 | [illegible] |
| MXE | 11099 | ENSG00000184182 | UBE2F | chr2 | + | 237994743 | 237994809 | 238025341 | 238025412 | [illegible] |
| MXE | 11103 | ENSG00000184182 | UBE2F | chr2 | + | 238003367 | 238003527 | 238016532 | 238016633 | [illegible] |
| MXE | 11104 | ENSG00000184182 | UBE2F | chr2 | + | 238003367 | 238003527 | 238016565 | 238016633 | [illegible] |
| MXE | 11105 | ENSG00000184182 | UBE2F | chr2 | + | 238016565 | 238016633 | 238025341 | 238025412 | [illegible] |
| MXE | 11106 | ENSG00000184182 | UBE2F | chr2 | + | 238016565 | 238016633 | 238025341 | 238025412 | [illegible] |
| MXE | 11110 | ENSG00000184182 | UBE2F | chr2 | + | 238025341 | 238025412 | 238030555 | 238030613 | [illegible] |
| MXE | 11123 | ENSG00000135976 | ANKRD36 | chr2 | + | 97187197 | 97187226 | 97189086 | 97189115 | [illegible] |
| MXE | 11126 | ENSG00000135976 | ANKRD36 | chr2 | + | 97202201 | 97202230 | 97207810 | 97207839 | [illegible] |
| MXE | 11127 | ENSG00000135976 | ANKRD36 | chr2 | + | 97205939 | 97205968 | 97209680 | 97209709 | [illegible] |
| MXE | 11128 | ENSG00000135976 | ANKRD36 | chr2 | + | 97207810 | 97207839 | 97209680 | 97209709 | [illegible] |
| MXE | 11132 | ENSG00000071909 | MYO3B | chr2 | + | 170501784 | 170501865 | 170514920 | 170515022 | [illegible] |
| MXE | 11135 | ENSG00000021574 | SPAST | chr2 | + | 32089521 | 32089605 | 32098795 | 32098891 | [illegible] |
| MXE | 11140 | ENSG00000260238 | PMF1-BGLAP | chr1 | + | 156233627 | 156233728 | 156236287 | 156236483 | [illegible] |
| MXE | 11155 | ENSG00000064666 | CNN2 | chr19 | + | 1032391 | 1032458 | 1032558 | 1032663 | [illegible] |
| MXE | 11156 | ENSG00000064666 | CNN2 | chr19 | + | 1032391 | 1032458 | 1032558 | 1032669 | [illegible] |
| MXE | 11159 | ENSG00000074054 | CLASP1 | chr2 | - | 121444938 | 121444962 | 121445448 | 121445496 | [illegible] |
| MXE | 11170 | ENSG00000079785 | DDX1 | chr2 | + | 15613223 | 15613284 | 15617243 | 15617342 | [illegible] |
| MXE | 11173 | ENSG00000115053 | NCL | chr2 | - | 231455400 | 231455624 | 231456009 | 231456159 | [illegible] |
| MXE | 11176 | ENSG00000115053 | NCL | chr2 | - | 231457642 | 231457800 | 231458265 | 231458344 | [illegible] |
| MXE | 11178 | ENSG00000115053 | NCL | chr2 | - | [illegible] | [illegible] | [illegible] | [illegible] | [illegible] |
| MXE | 11181 | ENSG00000272980 | NA | chr6 | + | [illegible] | [illegible] | [illegible] | [illegible] | [illegible] |
| MXE | 11183 | ENSG00000130414 | NDUFA10 | chr2 | - | [illegible] | [illegible] | [illegible] | [illegible] | [illegible] |
| MXE | 11184 | ENSG00000130414 | NDUFA10 | chr2 | - | [illegible] | [illegible] | [illegible] | [illegible] | [illegible] |
| MXE | 11191 | ENSG00000251569 | NA | chr2 | + | [illegible] | [illegible] | [illegible] | [illegible] | [illegible] |
| MXE | 11200 | ENSG00000059588 | TARBP1 | chr1 | - | [illegible] | [illegible] | [illegible] | [illegible] | [illegible] |
| MXE | 11207 | ENSG00000152348 | ATG10 | chr5 | + | [illegible] | [illegible] | [illegible] | [illegible] | [illegible] |
| MXE | 11215 | ENSG00000131368 | MRPS25 | chr3 | - | [illegible] | [illegible] | [illegible] | [illegible] | [illegible] |
| MXE | 11218 | ENSG00000059377 | TBXAS1 | chr7 | + | [illegible] | [illegible] | [illegible] | [illegible] | [illegible] |
| MXE | 11237 | ENSG00000118495 | PLAGL1 | chr6 | - | [illegible] | [illegible] | [illegible] | [illegible] | [illegible] |
| MXE | 11240 | ENSG00000177842 | ZNF620 | chr3 | + | [illegible] | [illegible] | [illegible] | [illegible] | [illegible] |
| MXE | 11241 | ENSG00000177842 | ZNF620 | chr3 | + | [illegible] | [illegible] | [illegible] | [illegible] | [illegible] |
| MXE | 11250 | ENSG00000189350 | TOGARAM2 | chr2 | + | [illegible] | [illegible] | [illegible] | [illegible] | [illegible] |
| MXE | 11251 | ENSG00000189350 | TOGARAM2 | chr2 | + | [illegible] | [illegible] | [illegible] | [illegible] | [illegible] |
| MXE | 11252 | ENSG00000189350 | TOGARAM2 | chr2 | + | [illegible] | [illegible] | [illegible] | [illegible] | [illegible] |
| MXE | 11265 | ENSG00000113387 | SUB1 | chr5 | + | [illegible] | [illegible] | [illegible] | [illegible] | [illegible] |
| MXE | 11267 | ENSG00000172765 | TMCC1 | chr3 | - | [illegible] | [illegible] | [illegible] | [illegible] | [illegible] |
| MXE | 11269 | ENSG00000172765 | TMCC1 | chr3 | - | [illegible] | [illegible] | [illegible] | [illegible] | [illegible] |
| MXE | 11274 | ENSG00000240498 | CDKN2B-AS1 | chr9 | + | [illegible] | [illegible] | [illegible] | [illegible] | [illegible] |
| MXE | 11275 | ENSG00000113407 | TARS1 | chr5 | + | [illegible] | [illegible] | [illegible] | [illegible] | [illegible] |
| MXE | 11281 | ENSG00000105372 | RPS19 | chr19 | + | [illegible] | [illegible] | [illegible] | [illegible] | [illegible] |
| MXE | 11282 | ENSG00000188428 | BLOC1S5 | chr6 | - | [illegible] | [illegible] | [illegible] | [illegible] | [illegible] |
| MXE | 11284 | ENSG00000162383 | SLC1A7 | chr1 | - | [illegible] | [illegible] | [illegible] | [illegible] | [illegible] |
| MXE | 11292 | ENSG00000129158 | SERGEF | chr11 | - | [illegible] | [illegible] | [illegible] | [illegible] | [illegible] |
| MXE | 11294 | ENSG00000129158 | SERGEF | chr11 | - | [illegible] | [illegible] | [illegible] | [illegible] | [illegible] |
| MXE | 11300 | ENSG00000080345 | RIF1 | chr2 | + | [illegible] | [illegible] | [illegible] | [illegible] | [illegible] |
| MXE | 11301 | ENSG00000272752 | STAG3L5P-PVRIG2P-PILRB | chr7 | + | [illegible] | [illegible] | [illegible] | [illegible] | [illegible] |
| MXE | 11303 | ENSG00000124831 | LRRFIP1 | chr2 | + | [illegible] | [illegible] | [illegible] | [illegible] | [illegible] |
| MXE | 11304 | ENSG00000124831 | LRRFIP1 | chr2 | + | [illegible] | [illegible] | [illegible] | [illegible] | [illegible] |
| MXE | 11309 | ENSG00000239665 | NA | chr10 | + | [illegible] | [illegible] | [illegible] | [illegible] | [illegible] |
| MXE | 11312 | ENSG00000239665 | NA | chr10 | + | [illegible] | [illegible] | [illegible] | [illegible] | [illegible] |
| MXE | 11319 | ENSG00000123607 | TTC21B | chr2 | - | [illegible] | [illegible] | [illegible] | [illegible] | [illegible] |
| MXE | 11328 | ENSG00000175198 | PCCA | chr13 | + | [illegible] | [illegible] | [illegible] | [illegible] | [illegible] |
| MXE | 11330 | ENSG00000175198 | PCCA | chr13 | + | [illegible] | [illegible] | [illegible] | [illegible] | [illegible] |
| MXE | 11335 | ENSG00000144233 | AMMECR1L | chr2 | - | [illegible] | [illegible] | [illegible] | [illegible] | [illegible] |
| MXE | 11342 | ENSG00000183091 | NEB | chr2 | - | [illegible] | [illegible] | [illegible] | [illegible] | [illegible] |
| MXE | 11343 | ENSG00000183091 | NEB | chr2 | - | [illegible] | [illegible] | [illegible] | [illegible] | [illegible] |
| MXE | 11353 | ENSG00000168280 | KIF5C | chr2 | + | [illegible] | [illegible] | [illegible] | [illegible] | [illegible] |
| MXE | 11354 | ENSG00000168280 | KIF5C | chr2 | + | [illegible] | [illegible] | [illegible] | [illegible] | [illegible] |
| MXE | 11372 | ENSG00000115604 | IL18R1 | chr2 | + | [illegible] | [illegible] | [illegible] | [illegible] | [illegible] |
| MXE | 11374 | ENSG00000115604 | IL18R1 | chr2 | + | [illegible] | [illegible] | [illegible] | [illegible] | [illegible] |
| MXE | 11375 | ENSG00000115604 | IL18R1 | chr2 | + | [illegible] | [illegible] | [illegible] | [illegible] | [illegible] |
| MXE | 11376 | ENSG00000115604 | IL18R1 | chr2 | + | [illegible] | [illegible] | [illegible] | [illegible] | [illegible] |
| MXE | 11377 | ENSG00000092208 | GEMIN2 | chr14 | + | [illegible] | [illegible] | [illegible] | [illegible] | [illegible] |
| MXE | 11378 | ENSG00000240344 | PPIL3 | chr2 | - | [illegible] | [illegible] | [illegible] | [illegible] | [illegible] |
| MXE | 11379 | ENSG00000240344 | PPIL3 | chr2 | - | [illegible] | [illegible] | [illegible] | [illegible] | [illegible] |
| MXE | 11382 | ENSG00000115942 | ORC2 | chr2 | - | [illegible] | [illegible] | [illegible] | [illegible] | [illegible] |
| MXE | 11402 | ENSG00000215559 | ANKRD20A11P | chr21 | - | [illegible] | [illegible] | [illegible] | [illegible] | [illegible] |
| MXE | 11403 | ENSG00000215559 | ANKRD20A11P | chr21 | - | [illegible] | [illegible] | [illegible] | [illegible] | [illegible] |
| MXE | 11405 | ENSG00000138380 | CARF | chr2 | + | [illegible] | [illegible] | [illegible] | [illegible] | [illegible] |
| MXE | 11406 | ENSG00000138380 | CARF | chr2 | + | [illegible] | [illegible] | [illegible] | [illegible] | [illegible] |
| MXE | 11409 | ENSG00000138380 | CARF | chr2 | + | [illegible] | [illegible] | [illegible] | [illegible] | [illegible] |
| MXE | 11411 | ENSG00000138380 | CARF | chr2 | + | [illegible] | [illegible] | [illegible] | [illegible] | [illegible] |
| MXE | 11413 | ENSG00000138380 | CARF | chr2 | + | [illegible] | [illegible] | [illegible] | [illegible] | [illegible] |
| MXE | 11415 | ENSG00000138380 | CARF | chr2 | + | [illegible] | [illegible] | [illegible] | [illegible] | [illegible] |
| MXE | 11421 | ENSG00000146576 | C7orf26 | chr7 | + | [illegible] | [illegible] | [illegible] | [illegible] | [illegible] |
| MXE | 11427 | ENSG00000124356 | STAMBP | chr2 | + | [illegible] | [illegible] | [illegible] | [illegible] | [illegible] |
| MXE | 11428 | ENSG00000124356 | STAMBP | chr2 | + | [illegible] | [illegible] | [illegible] | [illegible] | [illegible] |
| MXE | 11432 | ENSG00000158417 | EIF5B | chr2 | + | [illegible] | [illegible] | [illegible] | [illegible] | [illegible] |
| MXE | 11434 | ENSG00000115526 | CHST10 | chr2 | - | [illegible] | [illegible] | [illegible] | [illegible] | [illegible] |
| MXE | 11435 | ENSG00000178075 | GRAMD1C | chr3 | + | [illegible] | [illegible] | [illegible] | [illegible] | [illegible] |
| MXE | 11436 | ENSG00000178075 | GRAMD1C | chr3 | + | [illegible] | [illegible] | [illegible] | [illegible] | [illegible] |
| MXE | 11466 | ENSG00000108465 | CDK5RAP3 | chr17 | + | [illegible] | [illegible] | [illegible] | [illegible] | [illegible] |
| MXE | 11467 | ENSG00000164609 | SLU7 | chr5 | - | [illegible] | [illegible] | [illegible] | [illegible] | [illegible] |
| MXE | 11468 | ENSG00000237499 | WAKMAR2 | chr6 | - | [illegible] | [illegible] | [illegible] | [illegible] | [illegible] |
| MXE | 11469 | ENSG00000237499 | WAKMAR2 | chr6 | - | [illegible] | [illegible] | [illegible] | [illegible] | [illegible] |
| MXE | 11470 | ENSG00000237499 | WAKMAR2 | chr6 | - | [illegible] | [illegible] | [illegible] | [illegible] | [illegible] |
| MXE | 11474 | ENSG00000123600 | METTL8 | chr2 | - | [illegible] | [illegible] | [illegible] | [illegible] | [illegible] |
| MXE | 11477 | ENSG00000123600 | METTL8 | chr2 | - | [illegible] | [illegible] | [illegible] | [illegible] | [illegible] |
| MXE | 11479 | ENSG00000123600 | METTL8 | chr2 | - | [illegible] | [illegible] | [illegible] | [illegible] | [illegible] |
| MXE | 11490 | ENSG00000093167 | LRRFIP2 | chr3 | - | [illegible] | [illegible] | [illegible] | [illegible] | [illegible] |
| MXE | 11491 | ENSG00000093167 | LRRFIP2 | chr3 | - | [illegible] | [illegible] | [illegible] | [illegible] | [illegible] |
| MXE | 11498 | ENSG00000235257 | ITGA9-AS1 | chr3 | - | [illegible] | [illegible] | [illegible] | [illegible] | [illegible] |
| MXE | 11500 | ENSG00000235257 | ITGA9-AS1 | chr3 | - | [illegible] | [illegible] | [illegible] | [illegible] | [illegible] |
| MXE | 11522 | ENSG00000187118 | CMC1 | chr3 | + | [illegible] | [illegible] | [illegible] | [illegible] | [illegible] |
| MXE | 11545 | ENSG00000116539 | ASH1L | chr1 | - | [illegible] | [illegible] | [illegible] | [illegible] | [illegible] |
| MXE | 11547 | ENSG00000123219 | CENPK | chr5 | - | [illegible] | [illegible] | [illegible] | [illegible] | [illegible] |
| MXE | 11551 | ENSG00000064933 | PMS1 | chr2 | + | [illegible] | [illegible] | [illegible] | [illegible] | [illegible] |
| MXE | 11552 | ENSG00000064933 | PMS1 | chr2 | + | [illegible] | [illegible] | [illegible] | [illegible] | [illegible] |
| MXE | 11554 | ENSG00000090975 | PITPNM2 | chr12 | - | [illegible] | [illegible] | [illegible] | [illegible] | [illegible] |
| MXE | 11563 | ENSG00000055957 | ITIH1 | chr3 | + | [illegible] | [illegible] | [illegible] | [illegible] | [illegible] |
| MXE | 11568 | ENSG00000197548 | ATG7 | chr3 | + | [illegible] | [illegible] | [illegible] | [illegible] | [illegible] |
| MXE | 11574 | ENSG00000248592 | STIMATE-MUSTN1 | chr3 | - | [illegible] | [illegible] | [illegible] | [illegible] | [illegible] |
| MXE | 11575 | ENSG00000205189 | ZBTB10 | chr8 | + | [illegible] | [illegible] | [illegible] | [illegible] | [illegible] |
| MXE | 11580 | ENSG00000163939 | PBRM1 | chr3 | - | [illegible] | [illegible] | [illegible] | [illegible] | [illegible] |
| MXE | 11593 | ENSG00000011198 | ABHD5 | chr3 | + | [illegible] | [illegible] | [illegible] | [illegible] | [illegible] |
| MXE | 11594 | ENSG00000011198 | ABHD5 | chr3 | + | [illegible] | [illegible] | [illegible] | [illegible] | [illegible] |
| MXE | 11595 | ENSG00000146063 | TRIM41 | chr5 | + | [illegible] | [illegible] | [illegible] | [illegible] | [illegible] |
| MXE | 11606 | ENSG00000277147 | LINC00869 | chr1 | + | [illegible] | [illegible] | [illegible] | [illegible] | [illegible] |
| MXE | 11608 | ENSG00000272305 | NA | chr3 | - | [illegible] | [illegible] | [illegible] | [illegible] | [illegible] |
| MXE | 11609 | ENSG00000272305 | NA | chr3 | - | [illegible] | [illegible] | [illegible] | [illegible] | [illegible] |
| MXE | 11610 | ENSG00000272305 | NA | chr3 | - | [illegible] | [illegible] | [illegible] | [illegible] | [illegible] |
| MXE | 11612 | ENSG00000186017 | ZNF566 | chr19 | - | [illegible] | [illegible] | [illegible] | [illegible] | [illegible] |
| MXE | 11614 | ENSG00000079819 | EPB41L2 | chr6 | - | [illegible] | [illegible] | [illegible] | [illegible] | [illegible] |
| MXE | 11618 | ENSG00000052802 | MSMO1 | chr4 | + | [illegible] | [illegible] | [illegible] | [illegible] | [illegible] |
| MXE | 11621 | ENSG00000100814 | CCNB1IP1 | chr14 | - | [illegible] | [illegible] | [illegible] | [illegible] | [illegible] |
| MXE | 11622 | ENSG00000100814 | CCNB1IP1 | chr14 | - | [illegible] | [illegible] | [illegible] | [illegible] | [illegible] |
| MXE | 11638 | ENSG00000114423 | CBLB | chr3 | - | [illegible] | [illegible] | [illegible] | [illegible] | [illegible] |
| MXE | 11639 | ENSG00000114423 | CBLB | chr3 | - | [illegible] | [illegible] | [illegible] | [illegible] | [illegible] |
| MXE | 11653 | ENSG00000204410 | MSH5 | chr6 | + | [illegible] | [illegible] | [illegible] | [illegible] | [illegible] |
| MXE | 11654 | ENSG00000204410 | MSH5 | chr6 | + | [illegible] | [illegible] | [illegible] | [illegible] | [illegible] |
| MXE | 11655 | ENSG00000204410 | MSH5 | chr6 | + | [illegible] | [illegible] | [illegible] | [illegible] | [illegible] |
| MXE | 11701 | ENSG00000135049 | AGTPBP1 | chr9 | - | [illegible] | [illegible] | [illegible] | [illegible] | [illegible] |
| MXE | 11714 | ENSG00000120802 | TMPO | chr12 | + | [illegible] | [illegible] | [illegible] | [illegible] | [illegible] |
| MXE | 11718 | ENSG00000135316 | SYNCRIP | chr6 | - | [illegible] | [illegible] | [illegible] | [illegible] | [illegible] |
| MXE | 11719 | ENSG00000135316 | SYNCRIP | chr6 | - | [illegible] | [illegible] | [illegible] | [illegible] | [illegible] |
| MXE | 11720 | ENSG00000135316 | SYNCRIP | chr6 | - | [illegible] | [illegible] | [illegible] | [illegible] | [illegible] |
| MXE | 11727 | ENSG00000124786 | SLC35B3 | chr6 | - | [illegible] | [illegible] | [illegible] | [illegible] | [illegible] |
| MXE | 11738 | ENSG00000130340 | SNX9 | chr6 | + | [illegible] | [illegible] | [illegible] | [illegible] | [illegible] |
| MXE | 11740 | ENSG00000107890 | ANKRD26 | chr10 | - | [illegible] | [illegible] | [illegible] | [illegible] | [illegible] |
| MXE | 11742 | ENSG00000107890 | ANKRD26 | chr10 | - | [illegible] | [illegible] | [illegible] | [illegible] | [illegible] |
| MXE | 11743 | ENSG00000107890 | ANKRD26 | chr10 | - | [illegible] | [illegible] | [illegible] | [illegible] | [illegible] |
| MXE | 11749 | ENSG00000144580 | CNOT9 | chr2 | + | [illegible] | [illegible] | [illegible] | [illegible] | [illegible] |
| MXE | 11755 | ENSG00000096070 | BRPF3 | chr6 | + | [illegible] | [illegible] | [illegible] | [illegible] | [illegible] |
| MXE | 11758 | ENSG00000096070 | BRPF3 | chr6 | + | [illegible] | [illegible] | [illegible] | [illegible] | [illegible] |
| MXE | 11759 | ENSG00000010030 | ETV7 | chr6 | - | [illegible] | [illegible] | [illegible] | [illegible] | [illegible] |
| MXE | 11775 | ENSG00000124523 | SIRT5 | chr6 | + | [illegible] | [illegible] | [illegible] | [illegible] | [illegible] |
| MXE | 11783 | ENSG00000010704 | HFE | chr6 | + | [illegible] | [illegible] | [illegible] | [illegible] | [illegible] |
| MXE | 11788 | ENSG00000118873 | RAB3GAP2 | chr1 | - | [illegible] | [illegible] | [illegible] | [illegible] | [illegible] |
| MXE | 11790 | ENSG00000251364 | NA | chr11 | - | [illegible] | [illegible] | [illegible] | [illegible] | [illegible] |
| MXE | 11795 | ENSG00000204305 | AGER | chr6 | - | [illegible] | [illegible] | [illegible] | [illegible] | [illegible] |
| MXE | 11801 | ENSG00000137312 | FLOT1 | chr6 | - | [illegible] | [illegible] | [illegible] | [illegible] | [illegible] |
| MXE | 11824 | ENSG00000083097 | DOP1A | chr6 | + | [illegible] | [illegible] | [illegible] | [illegible] | [illegible] |
| MXE | 11825 | ENSG00000083097 | DOP1A | chr6 | + | [illegible] | [illegible] | [illegible] | [illegible] | [illegible] |
| MXE | 11826 | ENSG00000083097 | DOP1A | chr6 | + | [illegible] | [illegible] | [illegible] | [illegible] | [illegible] |
| MXE | 11831 | ENSG00000065427 | KARS1 | chr16 | - | [illegible] | [illegible] | [illegible] | [illegible] | [illegible] |
| MXE | 11833 | ENSG00000124508 | BTN2A2 | chr6 | + | [illegible] | [illegible] | [illegible] | [illegible] | [illegible] |
| MXE | 11843 | ENSG00000136861 | CDK5RAP2 | chr9 | - | [illegible] | [illegible] | [illegible] | [illegible] | [illegible] |
| MXE | 11855 | ENSG00000155906 | RMND1 | chr6 | - | [illegible] | [illegible] | [illegible] | [illegible] | [illegible] |
| MXE | 11857 | ENSG00000120262 | CCDC170 | chr6 | + | [illegible] | [illegible] | [illegible] | [illegible] | [illegible] |
| MXE | 11863 | ENSG00000170919 | TPT1-AS1 | chr13 | + | [illegible] | [illegible] | [illegible] | [illegible] | [illegible] |
| MXE | 11864 | ENSG00000170919 | TPT1-AS1 | chr13 | + | [illegible] | [illegible] | [illegible] | [illegible] | [illegible] |
| MXE | 11865 | ENSG00000170919 | TPT1-AS1 | chr13 | + | [illegible] | [illegible] | [illegible] | [illegible] | [illegible] |
| MXE | 11866 | ENSG00000170919 | TPT1-AS1 | chr13 | + | [illegible] | [illegible] | [illegible] | [illegible] | [illegible] |
| MXE | 11870 | ENSG00000170919 | TPT1-AS1 | chr13 | + | [illegible] | [illegible] | [illegible] | [illegible] | [illegible] |
| MXE | 11871 | ENSG00000170919 | TPT1-AS1 | chr13 | + | [illegible] | [illegible] | [illegible] | [illegible] | [illegible] |
| MXE | 11872 | ENSG00000170919 | TPT1-AS1 | chr13 | + | [illegible] | [illegible] | [illegible] | [illegible] | [illegible] |
| MXE | 11875 | ENSG00000170919 | TPT1-AS1 | chr13 | + | [illegible] | [illegible] | [illegible] | [illegible] | [illegible] |
| MXE | 11876 | ENSG00000170919 | TPT1-AS1 | chr13 | + | [illegible] | [illegible] | [illegible] | [illegible] | [illegible] |
| MXE | 11899 | ENSG00000135317 | SNX14 | chr6 | - | [illegible] | [illegible] | [illegible] | [illegible] | [illegible] |
| MXE | 11903 | ENSG00000135317 | SNX14 | chr6 | - | [illegible] | [illegible] | [illegible] | [illegible] | [illegible] |
| MXE | 11906 | ENSG00000135317 | SNX14 | chr6 | - | [illegible] | [illegible] | [illegible] | [illegible] | [illegible] |
| MXE | 11907 | ENSG00000135317 | SNX14 | chr6 | - | [illegible] | [illegible] | [illegible] | [illegible] | [illegible] |
| MXE | 11910 | ENSG00000135317 | SNX14 | chr6 | - | [illegible] | [illegible] | [illegible] | [illegible] | [illegible] |
| MXE | 11928 | ENSG00000173511 | VEGFB | chr11 | + | [illegible] | [illegible] | [illegible] | [illegible] | [illegible] |
| MXE | 11957 | ENSG00000164654 | MIOS | chr7 | + | [illegible] | [illegible] | [illegible] | [illegible] | [illegible] |
| MXE | 11960 | ENSG00000164654 | MIOS | chr7 | + | [illegible] | [illegible] | [illegible] | [illegible] | [illegible] |
| MXE | 11979 | ENSG00000168944 | CEP120 | chr5 | - | [illegible] | [illegible] | [illegible] | [illegible] | [illegible] |
| MXE | 11982 | ENSG00000117000 | RLF | chr1 | + | [illegible] | [illegible] | [illegible] | [illegible] | [illegible] |
| MXE | 11992 | ENSG00000086827 | ZW10 | chr11 | - | [illegible] | [illegible] | [illegible] | [illegible] | [illegible] |
| MXE | 11996 | ENSG00000089820 | ARHGAP4 | chrX | - | [illegible] | [illegible] | [illegible] | [illegible] | [illegible] |
| MXE | 11998 | ENSG00000089820 | ARHGAP4 | chrX | - | [illegible] | [illegible] | [illegible] | [illegible] | [illegible] |
| MXE | 11999 | ENSG00000089820 | ARHGAP4 | chrX | - | [illegible] | [illegible] | [illegible] | [illegible] | [illegible] |
| MXE | 12001 | ENSG00000089820 | ARHGAP4 | chrX | - | [illegible] | [illegible] | [illegible] | [illegible] | [illegible] |
| MXE | 12004 | ENSG00000229358 | DPY19L1P1 | chr7 | - | [illegible] | [illegible] | [illegible] | [illegible] | [illegible] |
| MXE | 12007 | ENSG00000109339 | MAPK10 | chr4 | - | [illegible] | [illegible] | [illegible] | [illegible] | [illegible] |
| MXE | 12008 | ENSG00000109339 | MAPK10 | chr4 | - | [illegible] | [illegible] | [illegible] | [illegible] | [illegible] |
| MXE | 12012 | ENSG00000109339 | MAPK10 | chr4 | - | [illegible] | [illegible] | [illegible] | [illegible] | [illegible] |
| MXE | 12015 | ENSG00000109339 | MAPK10 | chr4 | - | [illegible] | [illegible] | [illegible] | [illegible] | [illegible] |
| MXE | 12046 | ENSG00000127947 | PTPN12 | chr7 | + | [illegible] | [illegible] | [illegible] | [illegible] | [illegible] |
| MXE | 12090 | ENSG00000106524 | ANKMY2 | chr7 | - | [illegible] | [illegible] | [illegible] | [illegible] | [illegible] |
| MXE | 12099 | ENSG00000146729 | NIPSNAP2 | chr7 | + | [illegible] | [illegible] | [illegible] | [illegible] | [illegible] |
| MXE | 12109 | ENSG00000151116 | UEVLD | chr11 | - | [illegible] | [illegible] | [illegible] | [illegible] | [illegible] |
| MXE | 12113 | ENSG00000151116 | UEVLD | chr11 | - | [illegible] | [illegible] | [illegible] | [illegible] | [illegible] |
| MXE | 12134 | ENSG00000124486 | USP9X | chrX | + | [illegible] | [illegible] | [illegible] | [illegible] | [illegible] |
| MXE | 12144 | ENSG00000062725 | APPBP2 | chr17 | - | [illegible] | [illegible] | [illegible] | [illegible] | [illegible] |
| MXE | 12148 | ENSG00000228409 | CCT6P1 | chr7 | + | [illegible] | [illegible] | [illegible] | [illegible] | [illegible] |
| MXE | 12152 | ENSG00000072609 | CHFR | chr12 | - | [illegible] | [illegible] | [illegible] | [illegible] | [illegible] |
| MXE | 12163 | ENSG00000127980 | PEX1 | chr7 | - | [illegible] | [illegible] | [illegible] | [illegible] | [illegible] |
| MXE | 12172 | ENSG00000255036 | SUGT1P4-STRA6LP-CCDC180 | chr9 | + | [illegible] | [illegible] | [illegible] | [illegible] | [illegible] |
| MXE | 12173 | ENSG00000255036 | SUGT1P4-STRA6LP-CCDC180 | chr9 | + | [illegible] | [illegible] | [illegible] | [illegible] | [illegible] |
| MXE | 12175 | ENSG00000255036 | SUGT1P4-STRA6LP-CCDC180 | chr9 | + | [illegible] | [illegible] | [illegible] | [illegible] | [illegible] |
| MXE | 12176 | ENSG00000255036 | SUGT1P4-STRA6LP-CCDC180 | chr9 | + | [illegible] | [illegible] | [illegible] | [illegible] | [illegible] |
| MXE | 12188 | ENSG00000169902 | TPST1 | chr7 | + | [illegible] | [illegible] | [illegible] | [illegible] | [illegible] |
| MXE | 12212 | ENSG00000083642 | PDS5B | chr13 | + | [illegible] | [illegible] | [illegible] | [illegible] | [illegible] |
| MXE | 12218 | ENSG00000127995 | CASD1 | chr7 | + | [illegible] | [illegible] | [illegible] | [illegible] | [illegible] |
| MXE | 12221 | ENSG00000179406 | LINC00174 | chr7 | - | [illegible] | [illegible] | [illegible] | [illegible] | [illegible] |
| MXE | 12222 | ENSG00000179406 | LINC00174 | chr7 | - | [illegible] | [illegible] | [illegible] | [illegible] | [illegible] |
| MXE | 12232 | ENSG00000224897 | POT1-AS1 | chr7 | + | [illegible] | [illegible] | [illegible] | [illegible] | [illegible] |
| MXE | 12233 | ENSG00000224897 | POT1-AS1 | chr7 | + | [illegible] | [illegible] | [illegible] | [illegible] | [illegible] |
| MXE | 12249 | ENSG00000183735 | TBK1 | chr12 | + | [illegible] | [illegible] | [illegible] | [illegible] | [illegible] |
| MXE | 12251 | ENSG00000183735 | TBK1 | chr12 | + | [illegible] | [illegible] | [illegible] | [illegible] | [illegible] |
| MXE | 12273 | ENSG00000169660 | HEXD | chr17 | + | [illegible] | [illegible] | [illegible] | [illegible] | [illegible] |
| MXE | 12282 | ENSG00000132823 | OSER1 | chr20 | - | [illegible] | [illegible] | [illegible] | [illegible] | [illegible] |
| MXE | 12285 | ENSG00000196511 | TPK1 | chr7 | - | [illegible] | [illegible] | [illegible] | [illegible] | [illegible] |
| MXE | 12289 | ENSG00000133606 | MKRN1 | chr7 | - | [illegible] | [illegible] | [illegible] | [illegible] | [illegible] |
| MXE | 12292 | ENSG00000133606 | MKRN1 | chr7 | - | [illegible] | [illegible] | [illegible] | [illegible] | [illegible] |
| MXE | 12301 | ENSG00000001631 | KRIT1 | chr7 | - | [illegible] | [illegible] | [illegible] | [illegible] | [illegible] |
| MXE | 12309 | ENSG00000231721 | LINC-PINT | chr7 | - | [illegible] | [illegible] | [illegible] | [illegible] | [illegible] |
| MXE | 12310 | ENSG00000231721 | LINC-PINT | chr7 | - | [illegible] | [illegible] | [illegible] | [illegible] | [illegible] |
| MXE | 12311 | ENSG00000231721 | LINC-PINT | chr7 | - | [illegible] | [illegible] | [illegible] | [illegible] | [illegible] |
| MXE | 12312 | ENSG00000231721 | LINC-PINT | chr7 | - | [illegible] | [illegible] | [illegible] | [illegible] | [illegible] |
| MXE | 12315 | ENSG00000122482 | ZNF644 | chr1 | - | [illegible] | [illegible] | [illegible] | [illegible] | [illegible] |
| MXE | 12336 | ENSG00000119616 | FCF1 | chr14 | + | [illegible] | [illegible] | [illegible] | [illegible] | [illegible] |
| MXE | 12337 | ENSG00000119616 | FCF1 | chr14 | + | [illegible] | [illegible] | [illegible] | [illegible] | [illegible] |
| MXE | 12338 | ENSG00000119616 | FCF1 | chr14 | + | [illegible] | [illegible] | [illegible] | [illegible] | [illegible] |
| MXE | 12339 | ENSG00000119616 | FCF1 | chr14 | + | [illegible] | [illegible] | [illegible] | [illegible] | [illegible] |
| MXE | 12344 | ENSG00000048392 | RRM2B | chr8 | - | [illegible] | [illegible] | [illegible] | [illegible] | [illegible] |
| MXE | 12346 | ENSG00000048392 | RRM2B | chr8 | - | [illegible] | [illegible] | [illegible] | [illegible] | [illegible] |
| MXE | 12351 | ENSG00000128512 | DOCK4 | chr7 | - | [illegible] | [illegible] | [illegible] | [illegible] | [illegible] |

| | | | | | | | | | | |
|---|---|---|---|---|---|---|---|---|---|---|
| MXE | 12354 | ENSG00000134905 | CARS2 | chr13 | - | 110688784 | 110688904 | 110697917 | 110697967 | [illegible] |
| MXE | 12356 | ENSG00000230189 | NA | chr7 | - | 66429883 | 66430017 | 66439134 | 66439302 | [illegible] |
| MXE | 12369 | ENSG00000146963 | LUC7L2 | chr7 | + | 139360204 | 139360322 | 139376061 | 139376156 | [illegible] |
| MXE | 12386 | ENSG00000188312 | CENPP | chr9 | + | 92611313 | 92611393 | 92612522 | 92612614 | [illegible] |
| MXE | 12393 | ENSG00000006530 | AGK | chr7 | + | 141593145 | 141593185 | 141594674 | 141594928 | [illegible] |
| MXE | 12399 | ENSG00000118058 | KMT2A | chr11 | + | 118507507 | 118507609 | 118509135 | 118509200 | [illegible] |
| MXE | 12403 | ENSG00000091732 | ZC3HC1 | chr7 | - | 130043386 | 130043431 | 130049032 | 130049144 | [illegible] |
| MXE | 12409 | ENSG00000198198 | SZT2 | chr1 | + | 43403606 | 43403774 | 43404379 | 43404550 | [illegible] |
| MXE | 12421 | ENSG00000012124 | CD22 | chr19 | + | 35332652 | 35332924 | 35336148 | 35336341 | [illegible] |
| MXE | 12423 | ENSG00000012124 | CD22 | chr19 | + | 35336035 | 35336341 | 35337754 | 35338021 | [illegible] |
| MXE | 12425 | ENSG00000012124 | CD22 | chr19 | + | 35336250 | 35336341 | 35337754 | 35338021 | [illegible] |
| MXE | 12427 | ENSG00000179454 | KLHL28 | chr14 | - | 44931332 | 44931541 | 44934114 | 44934558 | [illegible] |
| MXE | 12435 | ENSG00000113303 | BTNL8 | chr5 | + | 180947511 | 180947625 | 180948354 | 180948375 | [illegible] |
| MXE | 12436 | ENSG00000113303 | BTNL8 | chr5 | + | 180947511 | 180947755 | 180948354 | 180948375 | [illegible] |
| MXE | 12467 | ENSG00000117906 | RCN2 | chr15 | + | 76938725 | 76938815 | 76941636 | 76941690 | [illegible] |
| MXE | 12473 | ENSG00000154511 | DIPK1A | chr1 | - | 92847182 | 92847359 | 92850847 | 92850955 | [illegible] |
| MXE | 12487 | ENSG00000179818 | PCBP1-AS1 | chr2 | - | 69988445 | 69988505 | 69988741 | 69988838 | [illegible] |
| MXE | 12492 | ENSG00000179818 | PCBP1-AS1 | chr2 | - | 70003045 | 70003315 | 70015447 | 70015580 | [illegible] |
| MXE | 12493 | ENSG00000179818 | PCBP1-AS1 | chr2 | - | 70003045 | 70003315 | 70018014 | 70018131 | [illegible] |
| MXE | 12494 | ENSG00000179818 | PCBP1-AS1 | chr2 | - | 70003045 | 70003315 | 70032132 | 70032255 | [illegible] |
| MXE | 12499 | ENSG00000179818 | PCBP1-AS1 | chr2 | - | 70015447 | 70015580 | 70018014 | 70018131 | [illegible] |
| MXE | 12500 | ENSG00000179818 | PCBP1-AS1 | chr2 | - | 70015447 | 70015580 | 70032132 | 70032255 | [illegible] |
| MXE | 12501 | ENSG00000179818 | PCBP1-AS1 | chr2 | - | 70015447 | 70015580 | 70018014 | 70018131 | [illegible] |
| MXE | 12502 | ENSG00000179818 | PCBP1-AS1 | chr2 | - | 70018014 | 70018131 | 70032132 | 70032255 | [illegible] |
| MXE | 12504 | ENSG00000179818 | PCBP1-AS1 | chr2 | - | 70018014 | 70018131 | 70053730 | 70053792 | [illegible] |
| MXE | 12505 | ENSG00000179818 | PCBP1-AS1 | chr2 | - | 70032132 | 70032255 | 70053730 | 70053792 | [illegible] |
| MXE | 12506 | ENSG00000179818 | PCBP1-AS1 | chr2 | - | 70032132 | 70032255 | 70059642 | 70059680 | [illegible] |
| MXE | 12511 | ENSG00000179818 | PCBP1-AS1 | chr2 | - | 70051202 | 70051305 | 70059642 | 70059680 | [illegible] |
| MXE | 12512 | ENSG00000179818 | PCBP1-AS1 | chr2 | - | 70051202 | 70051305 | 70055713 | 70055910 | [illegible] |
| MXE | 12515 | ENSG00000179818 | PCBP1-AS1 | chr2 | - | 70053730 | 70053792 | 70059642 | 70059680 | [illegible] |
| MXE | 12516 | ENSG00000179818 | PCBP1-AS1 | chr2 | - | 70053730 | 70053792 | 70055655 | 70055910 | [illegible] |
| MXE | 12517 | ENSG00000179818 | PCBP1-AS1 | chr2 | - | 70053730 | 70053792 | 70055713 | 70055910 | [illegible] |
| MXE | 12518 | ENSG00000179818 | PCBP1-AS1 | chr2 | - | 70053730 | 70053792 | 70055713 | 70055910 | [illegible] |
| MXE | 12519 | ENSG00000179818 | PCBP1-AS1 | chr2 | - | 70053730 | 70053792 | 70059642 | 70059680 | [illegible] |
| MXE | 12520 | ENSG00000179818 | PCBP1-AS1 | chr2 | - | 70053730 | 70053792 | 70077300 | 70077381 | [illegible] |
| MXE | 12521 | ENSG00000179818 | PCBP1-AS1 | chr2 | - | 70053730 | 70053792 | 70055713 | 70055910 | [illegible] |
| MXE | 12523 | ENSG00000179818 | PCBP1-AS1 | chr2 | - | 70053730 | 70053792 | 70077300 | 70077381 | [illegible] |
| MXE | 12524 | ENSG00000179818 | PCBP1-AS1 | chr2 | - | 70053730 | 70053792 | 70055655 | 70055910 | [illegible] |
| MXE | 12525 | ENSG00000179818 | PCBP1-AS1 | chr2 | - | 70053730 | 70053792 | 70055713 | 70055910 | [illegible] |
| MXE | 12529 | ENSG00000179818 | PCBP1-AS1 | chr2 | - | 70053730 | 70053815 | 70055713 | 70055910 | [illegible] |
| MXE | 12531 | ENSG00000179818 | PCBP1-AS1 | chr2 | - | 70053730 | 70053815 | 70077300 | 70077381 | [illegible] |
| MXE | 12532 | ENSG00000179818 | PCBP1-AS1 | chr2 | - | 70055655 | 70055910 | 70059642 | 70059680 | [illegible] |
| MXE | 12533 | ENSG00000179818 | PCBP1-AS1 | chr2 | - | 70055655 | 70055910 | 70059642 | 70059680 | [illegible] |
| MXE | 12536 | ENSG00000179818 | PCBP1-AS1 | chr2 | - | 70055655 | 70055910 | 70059642 | 70059680 | [illegible] |
| MXE | 12537 | ENSG00000179818 | PCBP1-AS1 | chr2 | - | 70055713 | 70055910 | 70059642 | 70059680 | [illegible] |
| MXE | 12538 | ENSG00000179818 | PCBP1-AS1 | chr2 | - | 70055713 | 70055910 | 70059642 | 70059680 | [illegible] |
| MXE | 12539 | ENSG00000179818 | PCBP1-AS1 | chr2 | - | 70055713 | 70055910 | 70077300 | 70077381 | [illegible] |
| MXE | 12540 | ENSG00000179818 | PCBP1-AS1 | chr2 | - | 70055713 | 70055910 | 70059642 | 70059680 | [illegible] |
| MXE | 12541 | ENSG00000179818 | PCBP1-AS1 | chr2 | - | 70055713 | 70055910 | 70077300 | 70077381 | [illegible] |
| MXE | 12542 | ENSG00000179818 | PCBP1-AS1 | chr2 | - | 70055713 | 70055910 | 70059642 | 70059680 | [illegible] |
| MXE | 12545 | ENSG00000179818 | PCBP1-AS1 | chr2 | - | 70055713 | 70055910 | 70059642 | 70059680 | [illegible] |
| MXE | 12546 | ENSG00000179818 | PCBP1-AS1 | chr2 | - | 70055713 | 70055910 | 70059642 | 70059680 | [illegible] |
| MXE | 12549 | ENSG00000179818 | PCBP1-AS1 | chr2 | - | 70059642 | 70059680 | 70077300 | 70077381 | [illegible] |
| MXE | 12550 | ENSG00000179818 | PCBP1-AS1 | chr2 | - | 70059642 | 70059680 | 70077300 | 70077381 | [illegible] |
| MXE | 12556 | ENSG00000179818 | PCBP1-AS1 | chr2 | - | 70059642 | 70059680 | 70077300 | 70077381 | [illegible] |
| MXE | 12559 | ENSG00000179818 | PCBP1-AS1 | chr2 | - | 70077300 | 70077381 | 70081369 | 70081490 | [illegible] |
| MXE | 12560 | ENSG00000179818 | PCBP1-AS1 | chr2 | - | 70081369 | 70081490 | 70083506 | 70083687 | [illegible] |
| MXE | 12561 | ENSG00000179818 | PCBP1-AS1 | chr2 | - | 70081369 | 70081490 | 70083537 | 70083687 | [illegible] |
| MXE | 12571 | ENSG00000121716 | PILRB | chr7 | + | 100353894 | 100354012 | 100355142 | 100355240 | [illegible] |
| MXE | 12574 | ENSG00000121716 | PILRB | chr7 | + | 100356393 | 100356883 | 100358219 | 100358366 | [illegible] |
| MXE | 12584 | ENSG00000147535 | PLPP5 | chr8 | - | 38267896 | 38267960 | 38268370 | 38268407 | [illegible] |
| MXE | 12585 | ENSG00000147535 | PLPP5 | chr8 | - | 38267896 | 38267960 | 38268370 | 38268461 | [illegible] |
| MXE | 12587 | ENSG00000285953 | NA | chr7 | - | 92244001 | 92244149 | 92244901 | 92245134 | [illegible] |
| MXE | 12588 | ENSG00000086300 | SNX10 | chr7 | + | 26346419 | 26346466 | 26360974 | 26361061 | [illegible] |
| MXE | 12604 | ENSG00000123191 | ATP7B | chr13 | - | 51944108 | 51944291 | 51946283 | 51946478 | [illegible] |
| MXE | 12605 | ENSG00000123191 | ATP7B | chr13 | - | 51946283 | 51946478 | 51949661 | 51949796 | [illegible] |
| MXE | 12606 | ENSG00000123191 | ATP7B | chr13 | - | 51958310 | 51958544 | 51960147 | 51960322 | [illegible] |
| MXE | 12611 | ENSG00000151948 | GLT1D1 | chr12 | + | 128899235 | 128899287 | 128912423 | 128912460 | [illegible] |
| MXE | 12615 | ENSG00000151948 | GLT1D1 | chr12 | + | 128912423 | 128912460 | 128914932 | 128914989 | [illegible] |
| MXE | 12623 | ENSG00000126883 | NUP214 | chr9 | + | 131144576 | 131144754 | 131146128 | 131146304 | [illegible] |
| MXE | 12634 | ENSG00000100997 | ABHD12 | chr20 | - | 25314924 | 25314970 | 25317047 | 25317078 | [illegible] |
| MXE | 12637 | ENSG00000100997 | ABHD12 | chr20 | - | 25317047 | 25317078 | 25320198 | 25320318 | [illegible] |
| MXE | 12640 | ENSG00000130429 | ARPC1B | chr7 | + | 99376432 | 99376574 | 99377337 | 99377439 | [illegible] |
| MXE | 12641 | ENSG00000130429 | ARPC1B | chr7 | + | 99376432 | 99376574 | 99377357 | 99377439 | [illegible] |
| MXE | 12642 | ENSG00000130429 | ARPC1B | chr7 | + | 99376432 | 99376574 | 99377357 | 99377455 | [illegible] |
| MXE | 12643 | ENSG00000130429 | ARPC1B | chr7 | + | 99376443 | 99376574 | 99377337 | 99377439 | [illegible] |
| MXE | 12644 | ENSG00000130429 | ARPC1B | chr7 | + | 99376443 | 99376574 | 99377357 | 99377439 | [illegible] |
| MXE | 12658 | ENSG00000157764 | BRAF | chr7 | - | 140747365 | 140747447 | 140748931 | 140749052 | [illegible] |
| MXE | 12661 | ENSG00000152207 | CYSLTR2 | chr13 | + | 48693437 | 48693510 | 48696525 | 48696626 | [illegible] |
| MXE | 12662 | ENSG00000152207 | CYSLTR2 | chr13 | + | 48693437 | 48693510 | 48696525 | 48696626 | [illegible] |
| MXE | 12688 | ENSG00000135164 | DMTF1 | chr7 | + | 87188038 | 87188301 | 87190944 | 87191027 | [illegible] |
| MXE | 12689 | ENSG00000135164 | DMTF1 | chr7 | + | 87188091 | 87188301 | 87190944 | 87191027 | [illegible] |
| MXE | 12690 | ENSG00000135164 | DMTF1 | chr7 | + | 87188197 | 87188301 | 87190944 | 87191027 | [illegible] |
| MXE | 12693 | ENSG00000258733 | LINC02328 | chr14 | + | 86035713 | 86035815 | 86041725 | 86041833 | [illegible] |
| MXE | 12697 | ENSG00000129103 | SUMF2 | chr7 | + | 56072996 | 56073111 | 56074173 | 56074218 | [illegible] |
| MXE | 12699 | ENSG00000129103 | SUMF2 | chr7 | + | 56074173 | 56074218 | 56076833 | 56076889 | [illegible] |
| MXE | 12726 | ENSG00000106348 | IMPDH1 | chr7 | - | 128409288 | 128409352 | 128409440 | 128409484 | [illegible] |
| MXE | 12727 | ENSG00000106348 | IMPDH1 | chr7 | - | 128409288 | 128409366 | 128409440 | 128409484 | [illegible] |
| MXE | 12740 | ENSG00000130948 | HSD17B3 | chr9 | - | 96252802 | 96252910 | 96254867 | 96254943 | [illegible] |
| MXE | 12743 | ENSG00000181090 | EHMT1 | chr9 | + | 137716625 | 137717182 | 137728348 | 137728593 | [illegible] |
| MXE | 12752 | ENSG00000105793 | GTPBP10 | chr7 | + | 90352815 | 90353009 | 90372154 | 90372228 | [illegible] |
| MXE | 12753 | ENSG00000105793 | GTPBP10 | chr7 | + | 90354457 | 90354549 | 90355085 | 90355230 | [illegible] |
| MXE | 12754 | ENSG00000105793 | GTPBP10 | chr7 | + | 90354457 | 90354549 | 90355085 | 90355230 | [illegible] |
| MXE | 12755 | ENSG00000105793 | GTPBP10 | chr7 | + | 90354457 | 90354549 | 90372154 | 90372228 | [illegible] |
| MXE | 12756 | ENSG00000105793 | GTPBP10 | chr7 | + | 90355085 | 90355230 | 90372154 | 90372228 | [illegible] |
| MXE | 12757 | ENSG00000105793 | GTPBP10 | chr7 | + | 90372154 | 90372228 | 90374301 | 90374354 | [illegible] |
| MXE | 12758 | ENSG00000105793 | GTPBP10 | chr7 | + | 90372154 | 90372228 | 90374301 | 90374354 | [illegible] |
| MXE | 12759 | ENSG00000105793 | GTPBP10 | chr7 | + | 90372154 | 90372228 | 90374301 | 90374354 | [illegible] |
| MXE | 12761 | ENSG00000058091 | CDK14 | chr7 | + | 90608529 | 90608575 | 90666339 | 90666411 | [illegible] |
| MXE | 12762 | ENSG00000058091 | CDK14 | chr7 | + | 90608529 | 90608575 | 90716339 | 90716394 | [illegible] |
| MXE | 12763 | ENSG00000058091 | CDK14 | chr7 | + | 90666339 | 90666411 | 90716339 | 90716394 | [illegible] |
| MXE | 12774 | ENSG00000177189 | RPS6KA3 | chrX | - | 20162963 | 20163040 | 20164898 | 20164961 | [illegible] |
| MXE | 12790 | ENSG00000051825 | MPHOSPH9 | chr12 | - | 123202623 | 123203084 | 123203113 | 123203375 | [illegible] |
| MXE | 12791 | ENSG00000051825 | MPHOSPH9 | chr12 | - | 123203249 | 123203375 | 123210055 | 123210162 | [illegible] |
| MXE | 12792 | ENSG00000051825 | MPHOSPH9 | chr12 | - | 123210055 | 123210162 | 123214743 | 123214834 | [illegible] |
| MXE | 12794 | ENSG00000051825 | MPHOSPH9 | chr12 | - | 123218375 | 123218499 | 123221371 | 123221723 | [illegible] |
| MXE | 12796 | ENSG00000051825 | MPHOSPH9 | chr12 | - | 123218398 | 123218499 | 123221371 | 123221723 | [illegible] |
| MXE | 12797 | ENSG00000051825 | MPHOSPH9 | chr12 | - | 123218398 | 123218499 | 123221371 | 123221895 | [illegible] |
| MXE | 12804 | ENSG00000104133 | SPG11 | chr15 | - | 44567423 | 44567592 | 44572682 | 44572820 | [illegible] |
| MXE | 12807 | ENSG00000104133 | SPG11 | chr15 | - | 44570524 | 44570658 | 44572682 | 44572820 | [illegible] |
| MXE | 12809 | ENSG00000104133 | SPG11 | chr15 | - | 44572682 | 44572820 | 44574901 | 44575041 | [illegible] |
| MXE | 12811 | ENSG00000104133 | SPG11 | chr15 | - | 44572682 | 44572820 | 44573546 | 44573745 | [illegible] |
| MXE | 12813 | ENSG00000104133 | SPG11 | chr15 | - | 44573546 | 44573745 | 44574901 | 44575041 | [illegible] |
| MXE | 12814 | ENSG00000104133 | SPG11 | chr15 | - | 44573546 | 44573745 | 44574901 | 44575041 | [illegible] |
| MXE | 12816 | ENSG00000104133 | SPG11 | chr15 | - | 44601738 | 44601824 | 44604104 | 44604197 | [illegible] |
| MXE | 12817 | ENSG00000104133 | SPG11 | chr15 | - | 44601738 | 44601824 | 44604802 | 44604927 | [illegible] |
| MXE | 12818 | ENSG00000104133 | SPG11 | chr15 | - | 44604104 | 44604197 | 44604802 | 44604927 | [illegible] |
| MXE | 12819 | ENSG00000104133 | SPG11 | chr15 | - | 44604104 | 44604197 | 44606024 | 44606091 | [illegible] |
| MXE | 12830 | ENSG00000086730 | LAT2 | chr7 | + | 74220583 | 74220619 | 74220703 | 74220734 | [illegible] |
| MXE | 12836 | ENSG00000128891 | CCDC32 | chr15 | - | 40539238 | 40539355 | 40557215 | 40557372 | [illegible] |
| MXE | 12840 | ENSG00000188002 | NA | chr5 | - | 1628754 | 1629630 | 1632643 | 1632751 | [illegible] |
| MXE | 12852 | ENSG00000123066 | MED13L | chr12 | - | 115980749 | 115980938 | 115982383 | 115982595 | [illegible] |
| MXE | 12863 | ENSG00000223482 | NUTM2A-AS1 | chr10 | - | 87241181 | 87245855 | 87246114 | 87246212 | [illegible] |
| MXE | 12865 | ENSG00000223482 | NUTM2A-AS1 | chr10 | - | 87291935 | 87292533 | 87307956 | 87308060 | [illegible] |
| MXE | 12871 | ENSG00000197150 | ABCB8 | chr7 | + | 151033442 | 151033604 | 151033604 | 151033917 | [illegible] |
| MXE | 12874 | ENSG00000076555 | ACACB | chr12 | + | 109260064 | 109260149 | 109260479 | 109260657 | [illegible] |
| MXE | 12875 | ENSG00000228649 | SNHG26 | chr7 | + | 22856972 | 22857016 | 22858570 | 22858976 | [illegible] |
| MXE | 12888 | ENSG00000105835 | NAMPT | chr7 | - | 106261587 | 106261707 | 106263391 | 106263606 | [illegible] |
| MXE | 12891 | ENSG00000105835 | NAMPT | chr7 | - | 106265009 | 106265171 | 106266690 | 106266769 | [illegible] |
| MXE | 12892 | ENSG00000105835 | NAMPT | chr7 | - | 106272529 | 106272658 | 106274945 | 106275040 | [illegible] |
| MXE | 12894 | ENSG00000105835 | NAMPT | chr7 | - | 106272529 | 106272658 | 106274945 | 106275068 | [illegible] |
| MXE | 12895 | ENSG00000105835 | NAMPT | chr7 | - | 106272529 | 106272658 | 106274945 | 106275043 | [illegible] |
| MXE | 12896 | ENSG00000105835 | NAMPT | chr7 | - | 106272529 | 106272658 | 106274945 | 106275049 | [illegible] |
| MXE | 12897 | ENSG00000157191 | NECAP2 | chr1 | + | 16450124 | 16450241 | 16451837 | 16452015 | [illegible] |
| MXE | 12909 | ENSG00000119471 | HSDL2 | chr9 | + | 112418859 | 112418958 | 112438430 | 112438625 | [illegible] |
| MXE | 12910 | ENSG00000119471 | HSDL2 | chr9 | + | 112418859 | 112418958 | 112438430 | 112438625 | [illegible] |
| MXE | 12916 | ENSG00000164815 | ORC5 | chr7 | - | 104161071 | 104161182 | 104165234 | 104165282 | [illegible] |
| MXE | 12922 | ENSG00000127824 | TUBA4A | chr2 | - | 219251564 | 219251713 | 219252007 | 219252230 | [illegible] |
| MXE | 12923 | ENSG00000127824 | TUBA4A | chr2 | - | 219251564 | 219251713 | 219252007 | 219252230 | [illegible] |
| MXE | 12933 | ENSG00000159784 | FAM131B | chr7 | - | 143359325 | 143359419 | 143359731 | 143359767 | [illegible] |
| MXE | 12934 | ENSG00000229153 | EPHA1-AS1 | chr7 | + | 143504773 | 143504830 | 143518281 | 143518517 | [illegible] |
| MXE | 12936 | ENSG00000131373 | HACL1 | chr3 | - | 15579909 | 15580045 | 15582876 | 15582989 | [illegible] |
| MXE | 12942 | ENSG00000141380 | SS18 | chr18 | - | 26079010 | 26079125 | 26080304 | 26080423 | [illegible] |
| MXE | 12944 | ENSG00000141380 | SS18 | chr18 | - | 26079010 | 26079125 | 26082362 | 26082528 | [illegible] |
| MXE | 12945 | ENSG00000141380 | SS18 | chr18 | - | 26079010 | 26079125 | 26084051 | 26084175 | [illegible] |
| MXE | 12946 | ENSG00000141380 | SS18 | chr18 | - | 26079010 | 26079146 | 26082362 | 26082528 | [illegible] |
| MXE | 12948 | ENSG00000141380 | SS18 | chr18 | - | 26080304 | 26080423 | 26082362 | 26082528 | [illegible] |
| MXE | 12949 | ENSG00000141380 | SS18 | chr18 | - | 26080304 | 26080423 | 26084051 | 26084175 | [illegible] |
| MXE | 12953 | ENSG00000141380 | SS18 | chr18 | - | 26082362 | 26082528 | 26084051 | 26084175 | [illegible] |
| MXE | 12962 | ENSG00000127914 | AKAP9 | chr7 | + | 92070904 | 92071009 | 92076854 | 92077007 | [illegible] |
| MXE | 12968 | ENSG00000196873 | CBWD3 | chr9 | + | 68259346 | 68259434 | 68260856 | 68261001 | [illegible] |
| MXE | 12969 | ENSG00000196873 | CBWD3 | chr9 | + | 68266219 | 68266306 | 68268933 | 68268978 | [illegible] |
| MXE | 12970 | ENSG00000196873 | CBWD3 | chr9 | + | 68269599 | 68269722 | 68270301 | 68270518 | [illegible] |
| MXE | 12971 | ENSG00000196873 | CBWD3 | chr9 | + | 68269599 | 68269722 | 68274881 | 68274938 | [illegible] |
| MXE | 12980 | ENSG00000106682 | EIF4H | chr7 | + | 74178655 | 74178826 | 74179337 | 74179418 | [illegible] |
| MXE | 12986 | ENSG00000161048 | NAPEPLD | chr7 | - | 103115059 | 103115174 | 103119576 | 103120223 | [illegible] |
| MXE | 12991 | ENSG00000285106 | NA | chr7 | - | 130912109 | 130912152 | 130945719 | 130945835 | [illegible] |
| MXE | 12992 | ENSG00000234456 | MAGI2-AS3 | chr7 | + | 79455889 | 79456180 | 79458658 | 79458730 | [illegible] |
| MXE | 12997 | ENSG00000008277 | ADAM22 | chr7 | + | 88131268 | 88131435 | 88132866 | 88132951 | [illegible] |
| MXE | 12998 | ENSG00000008277 | ADAM22 | chr7 | + | 88131268 | 88131435 | 88132866 | 88132982 | [illegible] |
| MXE | 12999 | ENSG00000008277 | ADAM22 | chr7 | + | 88132866 | 88132951 | 88134328 | 88134419 | [illegible] |
| MXE | 13002 | ENSG00000186088 | GSAP | chr7 | - | 77397345 | 77397415 | 77406028 | 77406105 | [illegible] |
| MXE | 13004 | ENSG00000265681 | RPL17 | chr18 | - | 49490792 | 49490927 | 49491404 | 49491445 | [illegible] |
| MXE | 13008 | ENSG00000196305 | IARS1 | chr9 | - | 92229000 | 92229126 | 92240642 | 92240691 | [illegible] |
| MXE | 13009 | ENSG00000196305 | IARS1 | chr9 | - | 92229000 | 92229126 | 92240855 | 92240961 | [illegible] |
| MXE | 13010 | ENSG00000196305 | IARS1 | chr9 | - | 92253361 | 92253446 | 92255534 | 92255604 | [illegible] |
| MXE | 13016 | ENSG00000126653 | NSRP1 | chr17 | + | 30156539 | 30156662 | 30163066 | 30163162 | [illegible] |
| MXE | 13018 | ENSG00000126653 | NSRP1 | chr17 | + | 30163066 | 30163162 | 30168813 | 30168925 | [illegible] |
| MXE | 13024 | ENSG00000234741 | GAS5 | chr1 | - | 173866527 | 173866567 | 173866990 | 173867043 | [illegible] |
| MXE | 13037 | ENSG00000106077 | ABHD11 | chr7 | - | 73736928 | 73737110 | 73737220 | 73737391 | [illegible] |
| MXE | 13045 | ENSG00000161010 | MRNIP | chr5 | - | 179847977 | 179848066 | 179851242 | 179851337 | [illegible] |
| MXE | 13046 | ENSG00000161010 | MRNIP | chr5 | - | 179847977 | 179848066 | 179851242 | 179851424 | [illegible] |
| MXE | 13048 | ENSG00000161010 | MRNIP | chr5 | - | 179851242 | 179851337 | 179853196 | 179853276 | [illegible] |
| MXE | 13050 | ENSG00000161010 | MRNIP | chr5 | - | 179851242 | 179851424 | 179853196 | 179853276 | [illegible] |
| MXE | 13057 | ENSG00000174428 | GTF2IRD2B | chr7 | + | 75123135 | 75123319 | 75123487 | 75123516 | [illegible] |
| MXE | 13059 | ENSG00000091073 | DTX2 | chr7 | + | 76470299 | 76470419 | 76477113 | 76477178 | [illegible] |
| MXE | 13066 | ENSG00000106603 | COA1 | chr7 | - | 43656032 | 43656153 | 43657215 | 43657269 | [illegible] |
| MXE | 13067 | ENSG00000106603 | COA1 | chr7 | - | 43656032 | 43656153 | 43665657 | 43665722 | [illegible] |
| MXE | 13068 | ENSG00000106603 | COA1 | chr7 | - | 43657215 | 43657269 | 43665657 | 43665722 | [illegible] |
| MXE | 13090 | ENSG00000107897 | ACBD5 | chr10 | - | 27220372 | 27220464 | 27223206 | 27223239 | [illegible] |
| MXE | 13091 | ENSG00000234129 | NA | chrX | - | 11038602 | 11038655 | 11055935 | 11056036 | [illegible] |
| MXE | 13123 | ENSG00000248309 | MEF2C-AS1 | chr5 | + | 89094958 | 89095029 | 89150532 | 89150595 | [illegible] |
| MXE | 13126 | ENSG00000204837 | FGF7P3 | chr9 | - | 39830012 | 39830110 | 39863832 | 39863942 | [illegible] |
| MXE | 13132 | ENSG00000116106 | EPHA4 | chr2 | - | 221455546 | 221455658 | 221456612 | 221456772 | [illegible] |
| MXE | 13137 | ENSG00000139218 | SCAF11 | chr12 | - | 45922935 | 45923154 | 45924727 | 45925074 | [illegible] |
| MXE | 13140 | ENSG00000139218 | SCAF11 | chr12 | - | 45961699 | 45961857 | 45964106 | 45964188 | [illegible] |
| MXE | 13145 | ENSG00000149809 | TM7SF2 | chr11 | + | 65112810 | 65112865 | 65113219 | 65113414 | [illegible] |
| MXE | 13146 | ENSG00000149809 | TM7SF2 | chr11 | + | 65112810 | 65112865 | 65113368 | 65113414 | [illegible] |
| MXE | 13147 | ENSG00000149809 | TM7SF2 | chr11 | + | 65113219 | 65113414 | 65113490 | 65113594 | [illegible] |
| MXE | 13149 | ENSG00000112715 | VEGFA | chr6 | + | 43780731 | 43780803 | 43781955 | 43782052 | [illegible] |
| MXE | 13173 | ENSG00000165259 | HDX | chrX | - | 84440531 | 84440585 | 84468471 | 84469575 | [illegible] |
| MXE | 13174 | ENSG00000165259 | HDX | chrX | - | 84440531 | 84440585 | 84468471 | 84469575 | [illegible] |
| MXE | 13175 | ENSG00000165259 | HDX | chrX | - | 84468471 | 84469575 | 84475250 | 84475397 | [illegible] |
| MXE | 13176 | ENSG00000165259 | HDX | chrX | - | 84475250 | 84475397 | 84488023 | 84488132 | [illegible] |
| MXE | 13177 | ENSG00000165259 | HDX | chrX | - | 84475250 | 84475397 | 84488023 | 84488132 | [illegible] |
| MXE | 13188 | ENSG00000214783 | POLR2J4 | chr7 | - | 43986600 | 43986677 | 44013592 | 44013679 | [illegible] |
| MXE | 13213 | ENSG00000230590 | FTX | chrX | - | 74274341 | 74274410 | 74280393 | 74280494 | [illegible] |
| MXE | 13218 | ENSG00000136143 | SUCLA2 | chr13 | - | 47956839 | 47956878 | 47965577 | 47965640 | [illegible] |
| MXE | 13235 | ENSG00000188917 | TRMT2B | chrX | - | 101042041 | 101042312 | 101051250 | 101051366 | [illegible] |
| MXE | 13241 | ENSG00000137522 | RNF121 | chr11 | + | 71962151 | 71962289 | 71978085 | 71978235 | [illegible] |
| MXE | 13246 | ENSG00000148985 | PGAP2 | chr11 | + | 3811249 | 3811424 | 3823612 | 3823740 | [illegible] |
| MXE | 13247 | ENSG00000148985 | PGAP2 | chr11 | + | 3811249 | 3811424 | 3823882 | 3824135 | [illegible] |
| MXE | 13248 | ENSG00000148985 | PGAP2 | chr11 | + | 3817352 | 3817535 | 3822908 | 3822993 | [illegible] |
| MXE | 13251 | ENSG00000177853 | ZNF518A | chr10 | + | 96151415 | 96151566 | 96155325 | 96155398 | [illegible] |
| MXE | 13255 | ENSG00000156170 | NDUFAF6 | chr8 | + | 95036301 | 95036494 | 95040705 | 95040927 | [illegible] |
| MXE | 13256 | ENSG00000156170 | NDUFAF6 | chr8 | + | 95036370 | 95036494 | 95040705 | 95040927 | [illegible] |
| MXE | 13257 | ENSG00000166432 | ZMAT1 | chrX | - | 101887197 | 101887304 | 101892704 | 101892754 | [illegible] |
| MXE | 13262 | ENSG00000245958 | SEPTIN7P14 | chr4 | + | 119462895 | 119462979 | 119493332 | 119493437 | [illegible] |
| MXE | 13271 | ENSG00000258867 | LINC01146 | chr14 | + | 88025406 | 88025481 | 88044685 | 88044848 | [illegible] |
| MXE | 13275 | ENSG00000258867 | LINC01146 | chr14 | + | 88025406 | 88025481 | 88044614 | 88044848 | [illegible] |
| MXE | 13278 | ENSG00000258867 | LINC01146 | chr14 | + | 88044614 | 88044848 | 88079103 | 88079368 | [illegible] |
| MXE | 13279 | ENSG00000258867 | LINC01146 | chr14 | + | 88044614 | 88044848 | 88079103 | 88079368 | [illegible] |
| MXE | 13280 | ENSG00000258867 | LINC01146 | chr14 | + | 88044685 | 88044848 | 88079103 | 88079368 | [illegible] |
| MXE | 13283 | ENSG00000258867 | LINC01146 | chr14 | + | 88044685 | 88044848 | 88084036 | 88084155 | [illegible] |
| MXE | 13284 | ENSG00000258867 | LINC01146 | chr14 | + | 88079103 | 88079368 | 88086827 | 88087023 | [illegible] |
| MXE | 13285 | ENSG00000187676 | B3GLCT | chr13 | + | 31269213 | 31269277 | 31274508 | 31274628 | [illegible] |
| MXE | 13287 | ENSG00000130734 | ATG4D | chr19 | + | 10546838 | 10547115 | 10547188 | 10547253 | [illegible] |
| MXE | 13289 | ENSG00000173681 | BCLAF3 | chrX | - | 19929784 | 19929940 | 19935808 | 19935898 | [illegible] |
| MXE | 13290 | ENSG00000173681 | BCLAF3 | chrX | - | 19929784 | 19929940 | 19935808 | 19935898 | [illegible] |
| MXE | 13318 | ENSG00000139436 | GIT2 | chr12 | - | 109989689 | 109989802 | 109991626 | 109991760 | [illegible] |
| MXE | 13324 | ENSG00000101901 | ALG13 | chrX | + | 111682131 | 111682235 | 111682235 | 111682294 | [illegible] |
| MXE | 13325 | ENSG00000101901 | ALG13 | chrX | + | 111689086 | 111689197 | 111708964 | 111709048 | [illegible] |
| MXE | 13331 | ENSG00000101901 | ALG13 | chrX | + | 111736713 | 111736879 | 111744667 | 111744904 | [illegible] |
| MXE | 13342 | ENSG00000136504 | KAT7 | chr17 | + | 49796749 | 49796926 | 49805362 | 49805445 | [illegible] |
| MXE | 13346 | ENSG00000136504 | KAT7 | chr17 | + | 49805362 | 49805445 | 49811475 | 49811574 | [illegible] |
| MXE | 13350 | ENSG00000162669 | HFM1 | chr1 | - | 91316390 | 91316476 | 91319077 | 91319209 | [illegible] |
| MXE | 13355 | ENSG00000110583 | NAA40 | chr11 | + | 63946724 | 63947003 | 63952237 | 63952333 | [illegible] |
| MXE | 13356 | ENSG00000110583 | NAA40 | chr11 | + | 63946950 | 63947003 | 63952237 | 63952333 | [illegible] |
| MXE | 13372 | ENSG00000176124 | DLEU1 | chr13 | + | 50390197 | 50390291 | 50415088 | 50415202 | [illegible] |
| MXE | 13373 | ENSG00000176124 | DLEU1 | chr13 | + | 50390197 | 50390291 | 50415088 | 50415202 | [illegible] |
| MXE | 13374 | ENSG00000176124 | DLEU1 | chr13 | + | 50415088 | 50415202 | 50433412 | 50433550 | [illegible] |
| MXE | 13375 | ENSG00000176124 | DLEU1 | chr13 | + | 50433412 | 50433550 | 50492650 | 50492761 | [illegible] |
| MXE | 13376 | ENSG00000176124 | DLEU1 | chr13 | + | 50433412 | 50433550 | 50527865 | 50528064 | [illegible] |

| | | | | | | | | | | |
|---|---|---|---|---|---|---|---|---|---|---|
| MXE | 13388 | ENSG00000147394 | ZNF185 | chrX | + | 152928574 | 152928661 | 152931671 | 152931776 | [illegible] |
| MXE | 13393 | ENSG00000102181 | CD99L2 | chrX | - | 150816278 | 150816455 | 150828305 | 150829604 | [illegible] |
| MXE | 13398 | ENSG00000155085 | AK9 | chr6 | - | 109516429 | 109516642 | 109529010 | 109529073 | [illegible] |
| MXE | 13399 | ENSG00000155085 | AK9 | chr6 | - | 109529010 | 109529073 | 109533250 | 109533470 | [illegible] |
| MXE | 13403 | ENSG00000137216 | TMEM63B | chr6 | + | 44150223 | 44150310 | 44150563 | 44150629 | [illegible] |
| MXE | 13404 | ENSG00000285547 | NA | chrX | - | 72495155 | 72495268 | 72567888 | 72568030 | [illegible] |
| MXE | 13405 | ENSG00000116791 | CRYZ | chr1 | - | 74707102 | 74707204 | 74710097 | 74710247 | [illegible] |
| MXE | 13423 | ENSG00000147010 | SH3KBP1 | chrX | - | 19687616 | 19687691 | 19694994 | 19695051 | [illegible] |
| MXE | 13440 | ENSG00000151413 | NUBPL | chr14 | + | 31654069 | 31654134 | 31667820 | 31667979 | [illegible] |
| MXE | 13441 | ENSG00000151413 | NUBPL | chr14 | + | 31673483 | 31673574 | 31787779 | 31787873 | [illegible] |
| MXE | 13443 | ENSG00000075539 | FRYL | chr4 | - | 48501622 | 48501733 | 48505546 | 48505615 | [illegible] |
| MXE | 13444 | ENSG00000075539 | FRYL | chr4 | - | 48505546 | 48505615 | 48510058 | 48510157 | [illegible] |
| MXE | 13447 | ENSG00000256861 | NA | chr12 | - | 122242381 | 122242508 | 122249870 | 122250045 | [illegible] |
| MXE | 13468 | ENSG00000106976 | DNM1 | chr9 | + | 128224250 | 128224389 | 128226034 | 128226173 | [illegible] |
| MXE | 13484 | ENSG00000244754 | N4BP2L2 | chr13 | - | 32446358 | 32446480 | 32478227 | 32478275 | [illegible] |
| MXE | 13486 | ENSG00000244754 | N4BP2L2 | chr13 | - | 32522181 | 32522270 | 32527407 | 32527532 | [illegible] |
| MXE | 13487 | ENSG00000244754 | N4BP2L2 | chr13 | - | 32522181 | 32522270 | 32527407 | 32527532 | [illegible] |
| MXE | 13488 | ENSG00000244754 | N4BP2L2 | chr13 | - | 32527407 | 32527532 | 32537005 | 32537027 | [illegible] |
| MXE | 13489 | ENSG00000244754 | N4BP2L2 | chr13 | - | 32527457 | 32527532 | 32537005 | 32537027 | [illegible] |
| MXE | 13490 | ENSG00000244754 | N4BP2L2 | chr13 | - | 32537005 | 32537027 | 32538617 | 32538684 | [illegible] |
| MXE | 13491 | ENSG00000156313 | RPGR | chrX | - | 38287093 | 38287245 | 38287860 | 38288041 | [illegible] |
| MXE | 13492 | ENSG00000156313 | RPGR | chrX | - | 38287093 | 38287245 | 38287864 | 38288041 | [illegible] |
| MXE | 13493 | ENSG00000156313 | RPGR | chrX | - | 38304634 | 38304790 | 38310614 | 38310773 | [illegible] |
| MXE | 13494 | ENSG00000156313 | RPGR | chrX | - | 38321026 | 38321089 | 38322852 | 38322945 | [illegible] |
| MXE | 13509 | ENSG00000171311 | EXOSC1 | chr10 | - | 97438662 | 97438703 | 97441170 | 97441259 | [illegible] |
| MXE | 13511 | ENSG00000171311 | EXOSC1 | chr10 | - | 97438669 | 97438703 | 97441170 | 97441259 | [illegible] |
| MXE | 13543 | ENSG00000035681 | NSMAF | chr8 | - | 58637233 | 58637389 | 58639967 | 58640102 | [illegible] |
| MXE | 13546 | ENSG00000131381 | RBSN | chr3 | - | 15075046 | 15075116 | 15075289 | 15075417 | [illegible] |
| MXE | 13563 | ENSG00000134575 | ACP2 | chr11 | - | 47244734 | 47244867 | 47245304 | 47245394 | [illegible] |
| MXE | 13564 | ENSG00000134575 | ACP2 | chr11 | - | 47244734 | 47244867 | 47245304 | 47245422 | [illegible] |
| MXE | 13565 | ENSG00000134575 | ACP2 | chr11 | - | 47244734 | 47244867 | 47245304 | 47245427 | [illegible] |
| MXE | 13568 | ENSG00000012048 | BRCA1 | chr17 | - | 43094743 | 43094860 | 43095845 | 43095922 | [illegible] |
| MXE | 13569 | ENSG00000012048 | BRCA1 | chr17 | - | 43095845 | 43095922 | 43097243 | 43097289 | [illegible] |
| MXE | 13573 | ENSG00000158578 | ALAS2 | chrX | - | 55021051 | 55021274 | 55024717 | 55024840 | [illegible] |
| MXE | 13574 | ENSG00000158578 | ALAS2 | chrX | - | 55024717 | 55024840 | 55025819 | 55026015 | [illegible] |
| MXE | 13580 | ENSG00000196924 | FLNA | chrX | - | 154365744 | 154365801 | 154366307 | 154366470 | [illegible] |
| MXE | 13584 | ENSG00000147140 | NONO | chrX | + | 71284983 | 71285097 | 71288426 | 71288534 | [illegible] |
| MXE | 13587 | ENSG00000147140 | NONO | chrX | + | 71297839 | 71297938 | 71298468 | 71298508 | [illegible] |
| MXE | 13589 | ENSG00000128654 | MTX2 | chr2 | + | 176329369 | 176329426 | 176330583 | 176330660 | [illegible] |
| MXE | 13590 | ENSG00000198814 | GK | chrX | + | 30694399 | 30694537 | 30695082 | 30695132 | [illegible] |
| MXE | 13591 | ENSG00000101868 | POLA1 | chrX | + | 24742001 | 24742121 | 24743229 | 24743329 | [illegible] |
| MXE | 13594 | ENSG00000151689 | INPP1 | chr2 | + | 190348887 | 190349031 | 190359645 | 190359698 | [illegible] |
| MXE | 13595 | ENSG00000169057 | MECP2 | chrX | - | 154057742 | 154057842 | 154058398 | 154058522 | [illegible] |
| MXE | 13613 | ENSG00000116128 | BCL9 | chr1 | + | 147611577 | 147611889 | 147612882 | 147612977 | [illegible] |
| MXE | 13614 | ENSG00000155008 | APOOL | chrX | + | 85046445 | 85046550 | 85051388 | 85051508 | [illegible] |
| MXE | 13618 | ENSG00000248323 | LUCAT1 | chr5 | - | 91244819 | 91245010 | 91264662 | 91264713 | [illegible] |
| MXE | 13619 | ENSG00000248323 | LUCAT1 | chr5 | - | 91305679 | 91305771 | 91306031 | 91306096 | [illegible] |
| MXE | 13620 | ENSG00000248323 | LUCAT1 | chr5 | - | 91305679 | 91305771 | 91312294 | 91312413 | [illegible] |
| MXE | 13621 | ENSG00000248323 | LUCAT1 | chr5 | - | 91305679 | 91305771 | 91312294 | 91312495 | [illegible] |
| MXE | 13623 | ENSG00000248323 | LUCAT1 | chr5 | - | 91306031 | 91306096 | 91312294 | 91312413 | [illegible] |
| MXE | 13624 | ENSG00000248323 | LUCAT1 | chr5 | - | 91306031 | 91306096 | 91312294 | 91312495 | [illegible] |
| MXE | 13626 | ENSG00000248323 | LUCAT1 | chr5 | - | 91311645 | 91311737 | 91312294 | 91312413 | [illegible] |
| MXE | 13627 | ENSG00000248323 | LUCAT1 | chr5 | - | 91313077 | 91313229 | 91313637 | 91313777 | [illegible] |
| MXE | 13628 | ENSG00000248323 | LUCAT1 | chr5 | - | 91313637 | 91313781 | 91313961 | 91314052 | [illegible] |
| MXE | 13633 | ENSG00000101935 | AMMECR1 | chrX | - | 110202445 | 110202536 | 110216517 | 110216632 | [illegible] |
| MXE | 13664 | ENSG00000046651 | OFD1 | chrX | + | 13763744 | 13763855 | 13767126 | 13767284 | [illegible] |
| MXE | 13678 | ENSG00000215301 | DDX3X | chrX | + | 41343220 | 41343351 | 41344029 | 41344128 | [illegible] |
| MXE | 13683 | ENSG00000215301 | DDX3X | chrX | + | 41344029 | 41344128 | 41344238 | 41344564 | [illegible] |
| MXE | 13692 | ENSG00000215301 | DDX3X | chrX | + | 41347311 | 41347451 | 41347639 | 41347915 | [illegible] |
| MXE | 13693 | ENSG00000215301 | DDX3X | chrX | + | 41347311 | 41347451 | 41347639 | 41347915 | [illegible] |
| MXE | 13701 | ENSG00000134982 | APC | chr5 | + | 112754872 | 112755025 | 112766325 | 112766410 | [illegible] |
| MXE | 13703 | ENSG00000134982 | APC | chr5 | + | 112818965 | 112819344 | 112821895 | 112821991 | [illegible] |
| MXE | 13708 | ENSG00000186376 | ZNF75D | chrX | - | 135291471 | 135291563 | 135295767 | 135296039 | [illegible] |
| MXE | 13719 | ENSG00000068366 | ACSL4 | chrX | - | 109683135 | 109683372 | 109696143 | 109696196 | [illegible] |
| MXE | 13723 | ENSG00000198954 | KIFBP | chr10 | + | 69006841 | 69006929 | 69007626 | 69007751 | [illegible] |
| MXE | 13724 | ENSG00000198954 | KIFBP | chr10 | + | 69007626 | 69007751 | 69008840 | 69008925 | [illegible] |
| MXE | 13725 | ENSG00000198954 | KIFBP | chr10 | + | 69008840 | 69008925 | 69010899 | 69011015 | [illegible] |
| MXE | 13728 | ENSG00000147050 | KDM6A | chrX | + | 44974665 | 44974715 | 45010960 | 45011019 | [illegible] |
| MXE | 13729 | ENSG00000147050 | KDM6A | chrX | + | 44974665 | 44974715 | 45010960 | 45011019 | [illegible] |
| MXE | 13737 | ENSG00000147050 | KDM6A | chrX | + | 45082575 | 45082640 | 45082714 | 45082789 | [illegible] |
| MXE | 13758 | ENSG00000229807 | XIST | chrX | - | 73831065 | 73831206 | 73833237 | 73833374 | [illegible] |
| MXE | 13759 | ENSG00000229807 | XIST | chrX | - | 73831065 | 73831206 | 73837439 | 73837503 | [illegible] |
| MXE | 13760 | ENSG00000229807 | XIST | chrX | - | 73831065 | 73831274 | 73833237 | 73833374 | [illegible] |
| MXE | 13761 | ENSG00000229807 | XIST | chrX | - | 73831065 | 73831274 | 73837439 | 73837503 | [illegible] |
| MXE | 13762 | ENSG00000229807 | XIST | chrX | - | 73833237 | 73833374 | 73837439 | 73837503 | [illegible] |
| MXE | 13763 | ENSG00000229807 | XIST | chrX | - | 73833237 | 73833374 | 73837439 | 73837503 | [illegible] |
| MXE | 13764 | ENSG00000229807 | XIST | chrX | - | 73837439 | 73837503 | 73841381 | 73841473 | [illegible] |
| MXE | 13766 | ENSG00000115896 | PLCL1 | chr2 | + | 198387852 | 198388005 | 198571353 | 198571477 | [illegible] |
| MXE | 13767 | ENSG00000107815 | TWNK | chr10 | + | 100989643 | 100989884 | 100990868 | 100991010 | [illegible] |
| MXE | 13770 | ENSG00000063601 | MTMR1 | chrX | + | 150712341 | 150712365 | 150714478 | 150714529 | [illegible] |
| MXE | 13772 | ENSG00000063601 | MTMR1 | chrX | + | 150712943 | 150712970 | 150714478 | 150714529 | [illegible] |
| MXE | 13779 | ENSG00000198932 | GPRASP1 | chrX | + | 102652174 | 102652265 | 102652830 | 102652898 | [illegible] |
| MXE | 13781 | ENSG00000101974 | ATP11C | chrX | - | 139743558 | 139743624 | 139745721 | 139745857 | [illegible] |
| MXE | 13785 | ENSG00000147408 | CSGALNACT1 | chr8 | - | 19585165 | 19585317 | 19591159 | 19591278 | [illegible] |
| MXE | 13788 | ENSG00000120699 | EXOSC8 | chr13 | + | 37001748 | 37001801 | 37002272 | 37002309 | [illegible] |
| MXE | 13801 | ENSG00000197265 | GTF2E2 | chr8 | - | 30636929 | 30636960 | 30637244 | 30637274 | [illegible] |
| MXE | 13802 | ENSG00000158483 | FAM86C1P | chr11 | + | 71791735 | 71791819 | 71793379 | 71793481 | [illegible] |
| MXE | 13806 | ENSG00000198042 | MAK16 | chr8 | + | 33488987 | 33489139 | 33490284 | 33490339 | [illegible] |
| MXE | 13807 | ENSG00000036565 | SLC18A1 | chr8 | - | 20148006 | 20148070 | 20148414 | 20148516 | [illegible] |
| MXE | 13810 | ENSG00000227678 | NA | chr6 | - | 130142683 | 130142813 | 130143749 | 130144051 | [illegible] |
| MXE | 13822 | ENSG00000197467 | COL13A1 | chr10 | + | 69930399 | 69930552 | 69935349 | 69935391 | [illegible] |
| MXE | 13830 | ENSG00000215548 | FRG1JP | chr9 | - | 63837730 | 63837832 | 63842823 | 63842949 | [illegible] |
| MXE | 13835 | ENSG00000145041 | DCAF1 | chr3 | - | 51466805 | 51466876 | 51470928 | 51471005 | [illegible] |
| MXE | 13872 | ENSG00000187735 | TCEA1 | chr8 | - | 53993667 | 53993755 | 53999944 | 54000050 | [illegible] |
| MXE | 13874 | ENSG00000151090 | THRB | chr3 | - | 24228937 | 24229001 | 24297225 | 24297371 | [illegible] |
| MXE | 13875 | ENSG00000151090 | THRB | chr3 | - | 24237990 | 24238081 | 24297225 | 24297371 | [illegible] |
| MXE | 13877 | ENSG00000151090 | THRB | chr3 | - | 24297225 | 24297371 | 24318357 | 24318494 | [illegible] |
| MXE | 13879 | ENSG00000183077 | AFMID | chr17 | + | 78204827 | 78204900 | 78205439 | 78205518 | [illegible] |
| MXE | 13880 | ENSG00000183077 | AFMID | chr17 | + | 78204827 | 78204900 | 78205602 | 78205738 | [illegible] |
| MXE | 13882 | ENSG00000183077 | AFMID | chr17 | + | 78205439 | 78205518 | 78205602 | 78205738 | [illegible] |
| MXE | 13883 | ENSG00000183077 | AFMID | chr17 | + | 78205439 | 78205518 | 78205602 | 78205753 | [illegible] |
| MXE | 13885 | ENSG00000205581 | HMGN1 | chr21 | - | 39345829 | 39345958 | 39347246 | 39347292 | [illegible] |
| MXE | 13888 | ENSG00000205581 | HMGN1 | chr21 | - | 39345829 | 39345958 | 39347915 | 39347952 | [illegible] |
| MXE | 13889 | ENSG00000205581 | HMGN1 | chr21 | - | 39347246 | 39347292 | 39347378 | 39347478 | [illegible] |
| MXE | 13891 | ENSG00000205581 | HMGN1 | chr21 | - | 39347246 | 39347292 | 39347915 | 39347952 | [illegible] |
| MXE | 13892 | ENSG00000205581 | HMGN1 | chr21 | - | 39347378 | 39347478 | 39347915 | 39347952 | [illegible] |
| MXE | 13893 | ENSG00000205581 | HMGN1 | chr21 | - | 39347378 | 39347478 | 39347915 | 39347952 | [illegible] |
| MXE | 13901 | ENSG00000165061 | ZMAT4 | chr8 | - | 40786688 | 40786782 | 40808508 | 40808571 | [illegible] |
| MXE | 13903 | ENSG00000174197 | MGA | chr15 | + | 41761738 | 41761850 | 41762128 | 41762362 | [illegible] |
| MXE | 13910 | ENSG00000198919 | DZIP3 | chr3 | + | 108646616 | 108646649 | 108647942 | 108648112 | [illegible] |
| MXE | 13911 | ENSG00000214954 | LRRC69 | chr8 | + | 91135667 | 91135739 | 91189521 | 91189623 | [illegible] |
| MXE | 13912 | ENSG00000214954 | LRRC69 | chr8 | + | 91189521 | 91189623 | 91200612 | 91200792 | [illegible] |
| MXE | 13916 | ENSG00000177565 | TBL1XR1 | chr3 | - | 177086964 | 177087041 | 177091150 | 177091180 | [illegible] |
| MXE | 13926 | ENSG00000064115 | TM7SF3 | chr12 | - | 26980565 | 26980646 | 26990449 | 26990627 | [illegible] |
| MXE | 13930 | ENSG00000154589 | LY96 | chr8 | + | 74010000 | 74010129 | 74026788 | 74026841 | [illegible] |
| MXE | 13941 | ENSG00000075856 | SART3 | chr12 | - | 108538944 | 108539089 | 108543027 | 108543152 | [illegible] |
| MXE | 13950 | ENSG00000164024 | METAP1 | chr4 | + | 99035953 | 99036011 | 99036042 | 99036089 | [illegible] |
| MXE | 13951 | ENSG00000164024 | METAP1 | chr4 | + | 99035953 | 99036011 | 99037766 | 99037941 | [illegible] |
| MXE | 13952 | ENSG00000164024 | METAP1 | chr4 | + | 99036042 | 99036089 | 99037766 | 99037941 | [illegible] |
| MXE | 13956 | ENSG00000104325 | DECR1 | chr8 | + | 90005447 | 90005845 | 90006035 | 90006277 | [illegible] |
| MXE | 13957 | ENSG00000104325 | DECR1 | chr8 | + | 90005447 | 90005845 | 90017123 | 90017326 | [illegible] |
| MXE | 13962 | ENSG00000166169 | POLL | chr10 | - | 101583507 | 101583681 | 101584601 | 101584720 | [illegible] |
| MXE | 13963 | ENSG00000166169 | POLL | chr10 | - | 101583507 | 101583681 | 101584601 | 101584720 | [illegible] |
| MXE | 13967 | ENSG00000166169 | POLL | chr10 | - | 101583507 | 101583681 | 101587121 | 101587200 | [illegible] |
| MXE | 13968 | ENSG00000166169 | POLL | chr10 | - | 101584601 | 101584919 | 101587121 | 101587200 | [illegible] |
| MXE | 13969 | ENSG00000165392 | WRN | chr8 | + | 31067032 | 31067182 | 31068257 | 31068327 | [illegible] |
| MXE | 13972 | ENSG00000165392 | WRN | chr8 | + | 31120242 | 31120424 | 31124521 | 31124623 | [illegible] |
| MXE | 13980 | ENSG00000253394 | LINC00534 | chr8 | + | 90454408 | 90454523 | 90669039 | 90669133 | [illegible] |
| MXE | 13982 | ENSG00000133739 | LRRCC1 | chr8 | + | 85115099 | 85115275 | 85123412 | 85123606 | [illegible] |
| MXE | 13983 | ENSG00000133739 | LRRCC1 | chr8 | + | 85115374 | 85115584 | 85123412 | 85123606 | [illegible] |
| MXE | 13984 | ENSG00000064313 | TAF2 | chr8 | - | 119744880 | 119745036 | 119746704 | 119746934 | [illegible] |
| MXE | 13987 | ENSG00000156162 | DPY19L4 | chr8 | + | 94734629 | 94734754 | 94738368 | 94738459 | [illegible] |
| MXE | 13993 | ENSG00000108239 | TBC1D12 | chr10 | + | 94500220 | 94500327 | 94507266 | 94507347 | [illegible] |
| MXE | 13994 | ENSG00000108239 | TBC1D12 | chr10 | + | 94500220 | 94500327 | 94510090 | 94510179 | [illegible] |
| MXE | 13995 | ENSG00000108239 | TBC1D12 | chr10 | + | 94522343 | 94522453 | 94531201 | 94531460 | [illegible] |
| MXE | 13997 | ENSG00000110074 | FOXRED1 | chr11 | + | 126274930 | 126275021 | 126275326 | 126275428 | [illegible] |
| MXE | 14006 | ENSG00000174799 | CEP135 | chr4 | + | 55959681 | 55959766 | 55964273 | 55964402 | [illegible] |
| MXE | 14008 | ENSG00000174799 | CEP135 | chr4 | + | 55991933 | 55992085 | 55999301 | 55999417 | [illegible] |
| MXE | 14020 | ENSG00000198851 | CD3E | chr11 | + | 118312152 | 118312170 | 118312617 | 118312866 | [illegible] |
| MXE | 14024 | ENSG00000120992 | LYPLA1 | chr8 | - | 54055059 | 54055133 | 54062253 | 54062324 | [illegible] |
| MXE | 14028 | ENSG00000120992 | LYPLA1 | chr8 | - | 54062253 | 54062324 | 54063327 | 54063375 | [illegible] |
| MXE | 14033 | ENSG00000215114 | UBXN2B | chr8 | + | 58433159 | 58433243 | 58439632 | 58439770 | [illegible] |
| MXE | 14043 | ENSG00000122882 | ECD | chr10 | - | 73154255 | 73154448 | 73156274 | 73156453 | [illegible] |
| MXE | 14055 | ENSG00000137492 | THAP12 | chr11 | - | 76362881 | 76362975 | 76364640 | 76364720 | [illegible] |
| MXE | 14069 | ENSG00000100380 | ST13 | chr22 | - | 40829625 | 40829674 | 40830839 | 40830956 | [illegible] |
| MXE | 14076 | ENSG00000078487 | ZCWPW1 | chr7 | - | 100405012 | 100405093 | 100406693 | 100406798 | [illegible] |
| MXE | 14079 | ENSG00000156787 | TBC1D31 | chr8 | + | 123134113 | 123134206 | 123140760 | 123140901 | [illegible] |
| MXE | 14084 | ENSG00000035928 | RFC1 | chr4 | - | 39321286 | 39321374 | 39323339 | 39323417 | [illegible] |
| MXE | 14099 | ENSG00000249859 | PVT1 | chr8 | + | 127902455 | 127902539 | 127932027 | 127932198 | [illegible] |
| MXE | 14100 | ENSG00000249859 | PVT1 | chr8 | + | 127902455 | 127902539 | 127932052 | 127932198 | [illegible] |
| MXE | 14101 | ENSG00000249859 | PVT1 | chr8 | + | 127939507 | 127939676 | 127969415 | 127969537 | [illegible] |
| MXE | 14103 | ENSG00000249859 | PVT1 | chr8 | + | 127969415 | 127969537 | 127983903 | 127984204 | [illegible] |
| MXE | 14104 | ENSG00000249859 | PVT1 | chr8 | + | 127969415 | 127969537 | 127982359 | 127982479 | [illegible] |
| MXE | 14106 | ENSG00000249859 | PVT1 | chr8 | + | 127982359 | 127982479 | 127983903 | 127984204 | [illegible] |
| MXE | 14109 | ENSG00000249859 | PVT1 | chr8 | + | 128009589 | 128009718 | 128048447 | 128048609 | [illegible] |
| MXE | 14117 | ENSG00000249859 | PVT1 | chr8 | + | 128010317 | 128010444 | 128070159 | 128070361 | [illegible] |
| MXE | 14119 | ENSG00000249859 | PVT1 | chr8 | + | 128031958 | 128032272 | 128070159 | 128070361 | [illegible] |
| MXE | 14124 | ENSG00000249859 | PVT1 | chr8 | + | 128070159 | 128070361 | 128082752 | 128082848 | [illegible] |
| MXE | 14126 | ENSG00000249859 | PVT1 | chr8 | + | 128070159 | 128070361 | 128095691 | 128095817 | [illegible] |
| MXE | 14127 | ENSG00000249859 | PVT1 | chr8 | + | 128070161 | 128070272 | 128082752 | 128082848 | [illegible] |
| MXE | 14130 | ENSG00000249859 | PVT1 | chr8 | + | 128074069 | 128074291 | 128082752 | 128082848 | [illegible] |
| MXE | 14131 | ENSG00000249859 | PVT1 | chr8 | + | 128074072 | 128074291 | 128082752 | 128082848 | [illegible] |
| MXE | 14133 | ENSG00000249859 | PVT1 | chr8 | + | 128080302 | 128080430 | 128082752 | 128082848 | [illegible] |
| MXE | 14135 | ENSG00000249859 | PVT1 | chr8 | + | 128082752 | 128082848 | 128090247 | 128090288 | [illegible] |
| MXE | 14136 | ENSG00000170633 | RNF34 | chr12 | + | 121402741 | 121402807 | 121417503 | 121417911 | [illegible] |
| MXE | 14138 | ENSG00000047249 | ATP6V1H | chr8 | - | 53801798 | 53801896 | 53811163 | 53811217 | [illegible] |
| MXE | 14142 | ENSG00000165923 | AGBL2 | chr11 | - | 47677270 | 47677401 | 47681968 | 47682095 | [illegible] |
| MXE | 14143 | ENSG00000165923 | AGBL2 | chr11 | - | 47679972 | 47680073 | 47685892 | 47686049 | [illegible] |
| MXE | 14144 | ENSG00000165923 | AGBL2 | chr11 | - | 47705863 | 47705917 | 47710376 | 47710511 | [illegible] |
| MXE | 14145 | ENSG00000070081 | NUCB2 | chr11 | + | 17330901 | 17330983 | 17337347 | 17337534 | [illegible] |
| MXE | 14161 | ENSG00000164953 | TMEM67 | chr8 | + | 93759859 | 93759993 | 93761984 | 93762023 | [illegible] |
| MXE | 14175 | ENSG00000079459 | FDFT1 | chr8 | + | 11803321 | 11803518 | 11808793 | 11808891 | [illegible] |
| MXE | 14176 | ENSG00000079459 | FDFT1 | chr8 | + | 11803321 | 11803518 | 11809791 | 11809850 | [illegible] |
| MXE | 14188 | ENSG00000104763 | ASAH1 | chr8 | - | 18062278 | 18062367 | 18063184 | 18063230 | [illegible] |
| MXE | 14191 | ENSG00000185728 | YTHDF3 | chr8 | + | 63169386 | 63169411 | 63186146 | 63187745 | [illegible] |
| MXE | 14192 | ENSG00000185728 | YTHDF3 | chr8 | + | 63172652 | 63172798 | 63173650 | 63173681 | [illegible] |
| MXE | 14193 | ENSG00000107020 | PLGRKT | chr9 | - | 5361757 | 5361888 | 5431896 | 5431983 | [illegible] |
| MXE | 14196 | ENSG00000107020 | PLGRKT | chr9 | - | 5431896 | 5431983 | 5436568 | 5436694 | [illegible] |
| MXE | 14200 | ENSG00000185803 | SLC52A2 | chr8 | + | 144359622 | 144359747 | 144360114 | 144360493 | [illegible] |
| MXE | 14202 | ENSG00000185803 | SLC52A2 | chr8 | + | 144360171 | 144360493 | 144360589 | 144360713 | [illegible] |
| MXE | 14203 | ENSG00000171428 | NAT1 | chr8 | + | 18212390 | 18212774 | 18216876 | 18216994 | [illegible] |
| MXE | 14204 | ENSG00000166225 | FRS2 | chr12 | + | 69537915 | 69537983 | 69557799 | 69557856 | [illegible] |
| MXE | 14205 | ENSG00000132669 | RIN2 | chr20 | + | 19970837 | 19970929 | 19974653 | 19975787 | [illegible] |
| MXE | 14214 | ENSG00000154582 | ELOC | chr8 | - | 73955910 | 73956054 | 73959764 | 73959818 | [illegible] |
| MXE | 14215 | ENSG00000154582 | ELOC | chr8 | - | 73955910 | 73956054 | 73959764 | 73959818 | [illegible] |
| MXE | 14236 | ENSG00000173848 | NET1 | chr10 | + | 5452857 | 5452920 | 5453249 | 5453346 | [illegible] |
| MXE | 14240 | ENSG00000164941 | INTS8 | chr8 | + | 94827721 | 94827793 | 94828974 | 94829026 | [illegible] |
| MXE | 14252 | ENSG00000288596 | C8orf44 | chr8 | + | 66676744 | 66676902 | 66677641 | 66677954 | [illegible] |
| MXE | 14254 | ENSG00000116604 | MEF2D | chr1 | - | 156480642 | 156480777 | 156480833 | 156480971 | [illegible] |
| MXE | 14256 | ENSG00000085719 | CPNE3 | chr8 | + | 86521603 | 86521714 | 86524876 | 86524997 | [illegible] |
| MXE | 14261 | ENSG00000154114 | TBCEL | chr11 | + | 121055051 | 121055308 | 121057507 | 121057614 | [illegible] |
| MXE | 14263 | ENSG00000120526 | NUDCD1 | chr8 | - | 109293343 | 109293524 | 109296383 | 109296569 | [illegible] |
| MXE | 14264 | ENSG00000120526 | NUDCD1 | chr8 | - | 109296383 | 109296569 | 109322308 | 109322463 | [illegible] |
| MXE | 14283 | ENSG00000189298 | ZKSCAN3 | chr6 | + | 28350271 | 28350388 | 28361323 | 28361471 | [illegible] |
| MXE | 14285 | ENSG00000104408 | EIF3E | chr8 | - | 108219749 | 108219830 | 108225711 | 108225833 | [illegible] |
| MXE | 14286 | ENSG00000104408 | EIF3E | chr8 | - | 108242111 | 108242256 | 108243292 | 108243463 | [illegible] |
| MXE | 14287 | ENSG00000104408 | EIF3E | chr8 | - | 108242111 | 108242346 | 108243292 | 108243463 | [illegible] |
| MXE | 14290 | ENSG00000164733 | CTSB | chr8 | - | 11854679 | 11854914 | 11857857 | 11857940 | [illegible] |
| MXE | 14292 | ENSG00000164733 | CTSB | chr8 | - | 11854679 | 11854914 | 11861405 | 11861479 | [illegible] |
| MXE | 14293 | ENSG00000164733 | CTSB | chr8 | - | 11854679 | 11854914 | 11862344 | 11862478 | [illegible] |
| MXE | 14298 | ENSG00000164733 | CTSB | chr8 | - | 11857857 | 11857940 | 11861405 | 11861479 | [illegible] |
| MXE | 14299 | ENSG00000164733 | CTSB | chr8 | - | 11857857 | 11857940 | 11862344 | 11862478 | [illegible] |
| MXE | 14300 | ENSG00000164733 | CTSB | chr8 | - | 11857857 | 11857940 | 11861405 | 11861479 | [illegible] |
| MXE | 14303 | ENSG00000164733 | CTSB | chr8 | - | 11861405 | 11861479 | 11862344 | 11862478 | [illegible] |
| MXE | 14305 | ENSG00000164733 | CTSB | chr8 | - | 11864375 | 11864463 | 11865463 | 11865584 | [illegible] |
| MXE | 14307 | ENSG00000197140 | ADAM32 | chr8 | + | 39169915 | 39169997 | 39186908 | 39187045 | [illegible] |
| MXE | 14309 | ENSG00000197140 | ADAM32 | chr8 | + | 39186908 | 39187045 | 39211143 | 39211324 | [illegible] |
| MXE | 14311 | ENSG00000147601 | TERF1 | chr8 | + | 73030335 | 73030395 | 73030794 | 73030955 | [illegible] |
| MXE | 14317 | ENSG00000135679 | MDM2 | chr12 | + | 68820884 | 68820910 | 68823414 | 68823514 | [illegible] |
| MXE | 14318 | ENSG00000135679 | MDM2 | chr12 | + | 68824554 | 68824651 | 68828770 | 68828931 | [illegible] |
| MXE | 14319 | ENSG00000135679 | MDM2 | chr12 | + | 68828770 | 68828931 | 68835828 | 68835984 | [illegible] |
| MXE | 14329 | ENSG00000104687 | GSR | chr8 | - | 30704646 | 30704864 | 30706989 | 30707104 | [illegible] |
| MXE | 14331 | ENSG00000184445 | KNTC1 | chr12 | + | 122549765 | 122549864 | 122551318 | 122551362 | [illegible] |
| MXE | 14332 | ENSG00000184445 | KNTC1 | chr12 | + | 122597730 | 122597938 | 122601535 | 122601625 | [illegible] |
| MXE | 14333 | ENSG00000184445 | KNTC1 | chr12 | + | 122615469 | 122615526 | 122618342 | 122618397 | [illegible] |
| MXE | 14342 | ENSG00000205913 | SRRM2-AS1 | chr16 | - | 2737479 | 2737610 | 2749544 | 2749659 | [illegible] |
| MXE | 14349 | ENSG00000176623 | RMDN1 | chr8 | - | 86494828 | 86494966 | 86505299 | 86505379 | [illegible] |
| MXE | 14350 | ENSG00000143748 | NVL | chr1 | - | 224232088 | 224232182 | 224232894 | 224232972 | [illegible] |
| MXE | 14352 | ENSG00000143748 | NVL | chr1 | - | 224313049 | 224313151 | 224317693 | 224317793 | [illegible] |
| MXE | 14356 | ENSG00000133794 | ARNTL | chr11 | + | 13354989 | 13355010 | 13355172 | 13355296 | [illegible] |
| MXE | 14357 | ENSG00000133794 | ARNTL | chr11 | + | 13354989 | 13355010 | 13355225 | 13355296 | [illegible] |
| MXE | 14358 | ENSG00000133794 | ARNTL | chr11 | + | 13354989 | 13355010 | 13355232 | 13355296 | [illegible] |
| MXE | 14380 | ENSG00000104529 | EEF1D | chr8 | - | 143590607 | 143590738 | 143592714 | 143592893 | [illegible] |
| MXE | 14382 | ENSG00000253320 | MAILR | chr8 | + | 102937623 | 102937757 | 102967596 | 102967673 | [illegible] |
| MXE | 14390 | ENSG00000135315 | CEP162 | chr6 | - | 84217798 | 84217860 | 84219102 | 84219214 | [illegible] |
| MXE | 14391 | ENSG00000196150 | ZNF250 | chr8 | - | 144886839 | 144886902 | 144889580 | 144889694 | [illegible] |
| MXE | 14399 | ENSG00000036448 | MYOM2 | chr8 | + | 2141140 | 2141177 | 2142374 | 2142397 | [illegible] |
| MXE | 14400 | ENSG00000076554 | TPD52 | chr8 | - | 80044166 | 80044208 | 80050444 | 80050471 | [illegible] |
| MXE | 14401 | ENSG00000232104 | RFX3-AS1 | chr9 | + | 3568793 | 3568889 | 3602522 | 3602584 | [illegible] |
| MXE | 14405 | ENSG00000122692 | SMU1 | chr9 | - | 33056112 | 33056239 | 33056836 | 33056964 | [illegible] |
| MXE | 14409 | ENSG00000164904 | ALDH7A1 | chr5 | - | 126554286 | 126554393 | 126559239 | 126559334 | [illegible] |
| MXE | 14410 | ENSG00000164904 | ALDH7A1 | chr5 | - | 126582850 | 126582974 | 126583931 | 126584012 | [illegible] |
| MXE | 14412 | ENSG00000143198 | MGST3 | chr1 | + | 165649754 | 165649964 | 165651013 | 165651087 | [illegible] |

| | | | | | | | | | |
|---|---|---|---|---|---|---|---|---|---|
| MXE | 14415 | ENSG00000143198 | MGST3 | chr1 | + | 165651013 | 165651087 | 165654278 | 165654351 |
| MXE | 14427 | ENSG00000137509 | PRCP | chr11 | - | 82839260 | 82839425 | 82843183 | 82843304 |
| MXE | 14429 | ENSG00000165271 | NOL6 | chr9 | - | 33467393 | 33467516 | 33467690 | 33467868 |
| MXE | 14444 | ENSG00000100815 | TRIP11 | chr14 | - | 92007639 | 92007852 | 92010985 | 92011072 |
| MXE | 14451 | ENSG00000216937 | CCDC7 | chr10 | + | 32491921 | 32491997 | 32517944 | 32517975 |
| MXE | 14454 | ENSG00000216937 | CCDC7 | chr10 | + | 32623718 | 32623817 | 32675664 | 32675813 |
| MXE | 14456 | ENSG00000216937 | CCDC7 | chr10 | + | 32685969 | 32686080 | 32711619 | 32711730 |
| MXE | 14457 | ENSG00000216937 | CCDC7 | chr10 | + | 32689052 | 32689163 | 32694878 | 32694992 |
| MXE | 14458 | ENSG00000216937 | CCDC7 | chr10 | + | 32689052 | 32689163 | 32711619 | 32711730 |
| MXE | 14459 | ENSG00000216937 | CCDC7 | chr10 | + | 32694878 | 32694992 | 32711619 | 32711730 |
| MXE | 14498 | ENSG00000228340 | MIR646HG | chr20 | + | 60101690 | 60101798 | 60117743 | 60117807 |
| MXE | 14499 | ENSG00000228340 | MIR646HG | chr20 | + | 60101690 | 60101798 | 60121027 | 60121121 |
| MXE | 14500 | ENSG00000228340 | MIR646HG | chr20 | + | 60101690 | 60101798 | 60138493 | 60138644 |
| MXE | 14501 | ENSG00000228340 | MIR646HG | chr20 | + | 60121027 | 60121121 | 60138493 | 60138644 |
| MXE | 14508 | ENSG00000134333 | LDHA | chr11 | + | 18400836 | 18401010 | 18402839 | 18403013 |
| MXE | 14510 | ENSG00000134333 | LDHA | chr11 | + | 18402839 | 18403013 | 18403693 | 18403811 |
| MXE | 14511 | ENSG00000134333 | LDHA | chr11 | + | 18402839 | 18403013 | 18403737 | 18403811 |
| MXE | 14521 | ENSG00000130958 | SLC35D2 | chr9 | - | 96336716 | 96336784 | 96351102 | 96351171 |
| MXE | 14537 | ENSG00000131089 | ARHGEF9 | chrX | - | 63715987 | 63716119 | 63722761 | 63722897 |
| MXE | 14543 | ENSG00000108669 | CYTH1 | chr17 | - | 78696006 | 78696009 | 78698268 | 78698380 |
| MXE | 14545 | ENSG00000108669 | CYTH1 | chr17 | - | 78696006 | 78696009 | 78698819 | 78698968 |
| MXE | 14546 | ENSG00000108669 | CYTH1 | chr17 | - | 78698268 | 78698380 | 78698819 | 78698968 |
| MXE | 14563 | ENSG00000113924 | HGD | chr3 | - | 120644318 | 120644443 | 120646266 | 120646366 |
| MXE | 14573 | ENSG00000079387 | SENP1 | chr12 | - | 48097993 | 48098124 | 48101468 | 48101516 |
| MXE | 14577 | ENSG00000137757 | CASP5 | chr11 | - | 105003273 | 105003383 | 105007082 | 105007334 |
| MXE | 14579 | ENSG00000163558 | PRKCI | chr3 | + | 170263378 | 170263429 | 170267914 | 170268000 |
| MXE | 14580 | ENSG00000163558 | PRKCI | chr3 | + | 170291853 | 170291941 | 170293382 | 170293508 |

[illegible]

| | | | | | | | | | | |
|---|---|---|---|---|---|---|---|---|---|---|
| MXE | 15821 | ENSG00000124795 | DEK | chr6 | - | 18226173 | 18226242 | 18236451 | 18236596 | [illegible] |
| MXE | 15827 | ENSG00000124795 | DEK | chr6 | - | 18258303 | 18258405 | 18263842 | 18263996 | [illegible] |
| MXE | 15830 | ENSG00000170525 | PFKFB3 | chr10 | + | 6224148 | 6224213 | 6226191 | 6226365 | [illegible] |
| MXE | 15833 | ENSG00000151657 | KIN | chr10 | - | 7775750 | 7775799 | 7778837 | 7779019 | [illegible] |
| MXE | 15846 | ENSG00000165309 | ARMC3 | chr10 | + | 22968305 | 22968489 | 22981339 | 22981492 | [illegible] |
| MXE | 15847 | ENSG00000095794 | CREM | chr10 | + | 35179133 | 35179276 | 35188199 | 35188388 | [illegible] |
| MXE | 15848 | ENSG00000095794 | CREM | chr10 | + | 35179133 | 35179276 | 35188199 | 35188388 | [illegible] |
| MXE | 15849 | ENSG00000095794 | CREM | chr10 | + | 35188199 | 35188388 | 35201450 | 35201486 | [illegible] |
| MXE | 15862 | ENSG00000088298 | EDEM2 | chr20 | - | 35142372 | 35142478 | 35144978 | 35145018 | [illegible] |
| MXE | 15863 | ENSG00000172671 | ZFAND4 | chr10 | - | 45651965 | 45652033 | 45652983 | 45653059 | [illegible] |
| MXE | 15865 | ENSG00000143702 | CEP170 | chr1 | - | 243142285 | 243142463 | 243156220 | 243156347 | [illegible] |
| MXE | 15866 | ENSG00000143702 | CEP170 | chr1 | - | 243142285 | 243142463 | 243156220 | 243156455 | [illegible] |
| MXE | 15867 | ENSG00000143702 | CEP170 | chr1 | - | 243142315 | 243142463 | 243156220 | 243156347 | [illegible] |
| MXE | 15868 | ENSG00000143702 | CEP170 | chr1 | - | 243142315 | 243142463 | 243156220 | 243156455 | [illegible] |
| MXE | 15869 | ENSG00000143702 | CEP170 | chr1 | - | 243142366 | 243142463 | 243156220 | 243156347 | [illegible] |
| MXE | 15876 | ENSG00000176928 | GCNT4 | chr5 | - | 75047896 | 75048037 | 75052168 | 75052227 | [illegible] |
| MXE | 15877 | ENSG00000165322 | ARHGAP12 | chr10 | - | 31817787 | 31817886 | 31820386 | 31820488 | [illegible] |
| MXE | 15890 | ENSG00000151150 | ANK3 | chr10 | - | 60059339 | 60059430 | 60059697 | 60060006 | [illegible] |
| MXE | 15898 | ENSG00000164347 | GFM2 | chr5 | - | 74730259 | 74730398 | 74730601 | 74730775 | [illegible] |
| MXE | 15903 | ENSG00000122033 | MTIF3 | chr13 | - | 27439988 | 27440449 | 27445087 | 27445137 | [illegible] |
| MXE | 15904 | ENSG00000122033 | MTIF3 | chr13 | - | 27439988 | 27440449 | 27445087 | 27445156 | [illegible] |
| MXE | 15915 | ENSG00000176155 | CCDC57 | chr17 | - | 82134072 | 82134194 | 82151559 | 82151773 | [illegible] |
| MXE | 15948 | ENSG00000156650 | KAT6B | chr10 | + | 74985079 | 74985241 | 74989018 | 74989112 | [illegible] |
| MXE | 15953 | ENSG00000196531 | NACA | chr12 | - | 56715870 | 56716110 | 56719536 | 56719596 | [illegible] |
| MXE | 15972 | ENSG00000085274 | MYNN | chr3 | + | 169783476 | 169783560 | 169784621 | 169784708 | [illegible] |
| MXE | 15973 | ENSG00000178440 | TIMM23B-AGAP6 | chr10 | + | 49973011 | 49973142 | 49987164 | 49987234 | [illegible] |
| MXE | 15974 | ENSG00000178440 | TIMM23B-AGAP6 | chr10 | + | 49976083 | 49976185 | 49981188 | 49981281 | [illegible] |
| MXE | 15975 | ENSG00000178440 | TIMM23B-AGAP6 | chr10 | + | 49987164 | 49987234 | 49988750 | 49988938 | [illegible] |
| MXE | 15996 | ENSG00000069956 | MAPK6 | chr15 | + | 52045829 | 52047015 | 52049992 | 52050137 | [illegible] |
| MXE | 16000 | ENSG00000255624 | C10orf88B | chr10 | + | 122883724 | 122883931 | 122884716 | 122884852 | [illegible] |
| MXE | 16007 | ENSG00000108100 | CCNY | chr10 | + | 35501500 | 35501535 | 35530123 | 35530243 | [illegible] |
| MXE | 16031 | ENSG00000138161 | CUZD1 | chr10 | - | 122834705 | 122835097 | 122839016 | 122839231 | [illegible] |
| MXE | 16039 | ENSG00000185515 | BRCC3 | chrX | + | 155099314 | 155099389 | 155116056 | 155116188 | [illegible] |
| MXE | 16040 | ENSG00000185515 | BRCC3 | chrX | + | 155116056 | 155116188 | 155116710 | 155116754 | [illegible] |
| MXE | 16041 | ENSG00000185515 | BRCC3 | chrX | + | 155116056 | 155116188 | 155116710 | 155116754 | [illegible] |
| MXE | 16049 | ENSG00000214279 | SCART1 | chr10 | + | 133464605 | 133465174 | 133466234 | 133466381 | [illegible] |
| MXE | 16052 | ENSG00000040341 | STAU2 | chr8 | - | 73688653 | 73688813 | 73709031 | 73709177 | [illegible] |
| MXE | 16055 | ENSG00000105287 | PRKD2 | chr19 | - | 46678365 | 46678663 | 46681649 | 46681748 | [illegible] |
| MXE | 16057 | ENSG00000198825 | INPP5F | chr10 | + | 119757891 | 119757977 | 119781060 | 119781242 | [illegible] |
| MXE | 16059 | ENSG00000198825 | INPP5F | chr10 | + | 119757894 | 119757977 | 119781060 | 119781242 | [illegible] |
| MXE | 16066 | ENSG00000182010 | RTKN2 | chr10 | - | 62218194 | 62218301 | 62223237 | 62223332 | [illegible] |
| MXE | 16072 | ENSG00000157554 | ERG | chr21 | - | 38392375 | 38392444 | 38400573 | 38400645 | [illegible] |
| MXE | 16077 | ENSG00000269609 | C10orf95-AS1 | chr10 | + | 102451398 | 102451555 | 102452089 | 102452187 | [illegible] |
| MXE | 16078 | ENSG00000269609 | C10orf95-AS1 | chr10 | + | 102451398 | 102451555 | 102452089 | 102452187 | [illegible] |
| MXE | 16079 | ENSG00000138111 | MFSD13A | chr10 | + | 102468286 | 102468332 | 102468752 | 102468842 | [illegible] |
| MXE | 16090 | ENSG00000156374 | PCGF6 | chr10 | - | 103333924 | 103333952 | 103345023 | 103345132 | [illegible] |
| MXE | 16092 | ENSG00000156374 | PCGF6 | chr10 | - | 103345023 | 103345132 | 103347237 | 103347297 | [illegible] |
| MXE | 16106 | ENSG00000138311 | ZNF365 | chr10 | + | 62388395 | 62388576 | 62459740 | 62459797 | [illegible] |
| MXE | 16114 | ENSG00000095585 | BLNK | chr10 | - | 96223825 | 96223989 | 96227409 | 96227566 | [illegible] |
| MXE | 16118 | ENSG00000120539 | MASTL | chr10 | + | 27173120 | 27173259 | 27180952 | 27181066 | [illegible] |
| MXE | 16123 | ENSG00000011376 | LARS2 | chr3 | + | 45520218 | 45520296 | 45523996 | 45524108 | [illegible] |
| MXE | 16147 | ENSG00000128805 | ARHGAP22 | chr10 | - | 48459683 | 48459891 | 48479635 | 48479764 | [illegible] |
| MXE | 16169 | ENSG00000107669 | ATE1 | chr10 | - | 121902390 | 121902620 | 121910905 | 121911151 | [illegible] |
| MXE | 16179 | ENSG00000099284 | MACROH2A2 | chr10 | + | 70092336 | 70092642 | 70093019 | 70093507 | [illegible] |
| MXE | 16180 | ENSG00000099284 | MACROH2A2 | chr10 | + | 70092336 | 70092642 | 70093222 | 70093507 | [illegible] |
| MXE | 16183 | ENSG00000107862 | GBF1 | chr10 | + | 102284952 | 102285070 | 102314807 | 102314860 | [illegible] |
| MXE | 16185 | ENSG00000107862 | GBF1 | chr10 | + | 102284952 | 102285070 | 102326359 | 102326420 | [illegible] |
| MXE | 16187 | ENSG00000107862 | GBF1 | chr10 | + | 102284952 | 102285070 | 102332407 | 102332526 | [illegible] |
| MXE | 16188 | ENSG00000107862 | GBF1 | chr10 | + | 102284952 | 102285070 | 102335562 | 102335743 | [illegible] |
| MXE | 16195 | ENSG00000107862 | GBF1 | chr10 | + | 102314807 | 102314860 | 102326359 | 102326420 | [illegible] |
| MXE | 16197 | ENSG00000107862 | GBF1 | chr10 | + | 102314807 | 102314860 | 102332407 | 102332526 | [illegible] |
| MXE | 16199 | ENSG00000107862 | GBF1 | chr10 | + | 102314810 | 102314860 | 102326359 | 102326420 | [illegible] |
| MXE | 16201 | ENSG00000107862 | GBF1 | chr10 | + | 102314810 | 102314860 | 102332407 | 102332526 | [illegible] |
| MXE | 16202 | ENSG00000107862 | GBF1 | chr10 | + | 102314810 | 102314860 | 102335562 | 102335743 | [illegible] |
| MXE | 16204 | ENSG00000107862 | GBF1 | chr10 | + | 102326359 | 102326420 | 102332407 | 102332526 | [illegible] |
| MXE | 16205 | ENSG00000107862 | GBF1 | chr10 | + | 102326359 | 102326420 | 102335562 | 102335743 | [illegible] |
| MXE | 16208 | ENSG00000107862 | GBF1 | chr10 | + | 102332407 | 102332526 | 102335562 | 102335743 | [illegible] |
| MXE | 16211 | ENSG00000107862 | GBF1 | chr10 | + | 102363255 | 102363396 | 102363709 | 102363798 | [illegible] |
| MXE | 16213 | ENSG00000150093 | ITGB1 | chr10 | - | 32936241 | 32936423 | 32951674 | 32951758 | [illegible] |
| MXE | 16214 | ENSG00000150093 | ITGB1 | chr10 | - | 32936349 | 32936423 | 32951674 | 32951758 | [illegible] |
| MXE | 16215 | ENSG00000150093 | ITGB1 | chr10 | - | 32936349 | 32936423 | 32938553 | 32938730 | [illegible] |
| MXE | 16217 | ENSG00000136754 | ABI1 | chr10 | - | 26755654 | 26755741 | 26759061 | 26759238 | [illegible] |
| MXE | 16218 | ENSG00000136754 | ABI1 | chr10 | - | 26755654 | 26755741 | 26763879 | 26763960 | [illegible] |
| MXE | 16219 | ENSG00000136754 | ABI1 | chr10 | - | 26759061 | 26759235 | 26763879 | 26763960 | [illegible] |
| MXE | 16220 | ENSG00000136754 | ABI1 | chr10 | - | 26759061 | 26759238 | 26763879 | 26763960 | [illegible] |
| MXE | 16221 | ENSG00000078403 | MLLT10 | chr10 | + | 21541262 | 21541361 | 21557620 | 21557824 | [illegible] |
| MXE | 16222 | ENSG00000078403 | MLLT10 | chr10 | + | 21541262 | 21541361 | 21571974 | 21572026 | [illegible] |
| MXE | 16225 | ENSG00000078403 | MLLT10 | chr10 | + | 21663749 | 21663843 | 21668916 | 21669052 | [illegible] |
| MXE | 16228 | ENSG00000012232 | EXTL3 | chr8 | + | 28713456 | 28713550 | 28715584 | 28718207 | [illegible] |
| MXE | 16229 | ENSG00000120616 | EPC1 | chr10 | - | 32273013 | 32273281 | 32273921 | 32274038 | [illegible] |
| MXE | 16230 | ENSG00000120616 | EPC1 | chr10 | - | 32273162 | 32273281 | 32273921 | 32274038 | [illegible] |
| MXE | 16247 | ENSG00000136738 | STAM | chr10 | + | 17672974 | 17673078 | 17684674 | 17684750 | [illegible] |
| MXE | 16254 | ENSG00000118518 | RNF146 | chr6 | + | 127280515 | 127280604 | 127282286 | 127282395 | [illegible] |
| MXE | 16255 | ENSG00000118518 | RNF146 | chr6 | + | 127280515 | 127280604 | 127285207 | 127285348 | [illegible] |
| MXE | 16256 | ENSG00000118518 | RNF146 | chr6 | + | 127282286 | 127282395 | 127285207 | 127285348 | [illegible] |
| MXE | 16258 | ENSG00000107736 | CDH23 | chr10 | + | 71730468 | 71730604 | 71731986 | 71732375 | [illegible] |
| MXE | 16277 | ENSG00000188234 | AGAP4 | chr10 | - | 45847859 | 45847954 | 45848842 | 45848912 | [illegible] |
| MXE | 16295 | ENSG00000120049 | KCNIP2 | chr10 | - | 101828150 | 101828258 | 101828388 | 101828459 | [illegible] |
| MXE | 16296 | ENSG00000257242 | LINC01619 | chr12 | - | 91987532 | 91987643 | 91989096 | 91989193 | [illegible] |
| MXE | 16297 | ENSG00000257242 | LINC01619 | chr12 | - | 91987555 | 91987643 | 91989096 | 91989193 | [illegible] |
| MXE | 16298 | ENSG00000257242 | LINC01619 | chr12 | - | 91989096 | 91989193 | 91993369 | 91993529 | [illegible] |
| MXE | 16299 | ENSG00000139734 | DIAPH3 | chr13 | - | 60093627 | 60093732 | 60112009 | 60112186 | [illegible] |
| MXE | 16309 | ENSG00000130640 | TUBGCP2 | chr10 | - | 133295321 | 133295439 | 133297320 | 133297404 | [illegible] |
| MXE | 16312 | ENSG00000023041 | ZDHHC6 | chr10 | - | 112439505 | 112439626 | 112441728 | 112441812 | [illegible] |
| MXE | 16314 | ENSG00000172731 | LRRC20 | chr10 | - | 70323862 | 70324030 | 70340552 | 70340702 | [illegible] |
| MXE | 16317 | ENSG00000148672 | GLUD1 | chr10 | - | 87074550 | 87074614 | 87078483 | 87078543 | [illegible] |
| MXE | 16318 | ENSG00000148672 | GLUD1 | chr10 | - | 87089561 | 87089788 | 87091570 | 87091623 | [illegible] |
| MXE | 16323 | ENSG00000128487 | SPECC1 | chr17 | + | 20124899 | 20124997 | 20204332 | 20205912 | [illegible] |
| MXE | 16325 | ENSG00000128487 | SPECC1 | chr17 | + | 20232199 | 20232405 | 20245925 | 20246574 | [illegible] |
| MXE | 16327 | ENSG00000183230 | CTNNA3 | chr10 | - | 67539502 | 67539669 | 67606856 | 67607049 | [illegible] |
| MXE | 16342 | ENSG00000122884 | P4HA1 | chr10 | - | 73030270 | 73030370 | 73044980 | 73045051 | [illegible] |
| MXE | 16345 | ENSG00000155256 | ZFYVE27 | chr10 | + | 97738476 | 97738649 | 97744728 | 97744915 | [illegible] |
| MXE | 16347 | ENSG00000155256 | ZFYVE27 | chr10 | + | 97738476 | 97738674 | 97744728 | 97744915 | [illegible] |
| MXE | 16351 | ENSG00000155256 | ZFYVE27 | chr10 | + | 97748268 | 97748364 | 97749473 | 97749586 | [illegible] |
| MXE | 16361 | ENSG00000204179 | PTPN20 | chr10 | + | 46932376 | 46932533 | 46940581 | 46940717 | [illegible] |
| MXE | 16362 | ENSG00000204179 | PTPN20 | chr10 | + | 46932376 | 46932533 | 46940622 | 46940717 | [illegible] |
| MXE | 16364 | ENSG00000204179 | PTPN20 | chr10 | + | 46967956 | 46968047 | 46984229 | 46984564 | [illegible] |
| MXE | 16365 | ENSG00000204179 | PTPN20 | chr10 | + | 46967956 | 46968047 | 46987339 | 46987555 | [illegible] |
| MXE | 16367 | ENSG00000184047 | DIABLO | chr12 | - | 122218265 | 122218397 | 122225964 | 122226653 | [illegible] |
| MXE | 16370 | ENSG00000205846 | CLEC6A | chr12 | + | 8465483 | 8465629 | 8476124 | 8476240 | [illegible] |
| MXE | 16378 | ENSG00000151490 | PTPRO | chr12 | + | 15551550 | 15551671 | 15557454 | 15557523 | [illegible] |
| MXE | 16379 | ENSG00000151490 | PTPRO | chr12 | + | 15560192 | 15560276 | 15565592 | 15565628 | [illegible] |
| MXE | 16380 | ENSG00000120647 | CCDC77 | chr12 | + | 428768 | 428865 | 430663 | 430736 | [illegible] |
| MXE | 16383 | ENSG00000150045 | KLRF1 | chr12 | + | 9832315 | 9832414 | 9833302 | 9833452 | [illegible] |
| MXE | 16385 | ENSG00000150045 | KLRF1 | chr12 | + | 9841811 | 9841951 | 9842320 | 9842433 | [illegible] |
| MXE | 16395 | ENSG00000060982 | BCAT1 | chr12 | - | 24878529 | 24878649 | 24881300 | 24881411 | [illegible] |
| MXE | 16403 | ENSG00000100354 | TNRC6B | chr22 | + | 40251178 | 40251200 | 40261831 | 40262173 | [illegible] |
| MXE | 16412 | ENSG00000211455 | STK38L | chr12 | + | 27319327 | 27319423 | 27322142 | 27322234 | [illegible] |
| MXE | 16413 | ENSG00000110888 | CAPRIN2 | chr12 | - | 30719078 | 30719225 | 30720810 | 30720915 | [illegible] |
| MXE | 16443 | ENSG00000064102 | INTS13 | chr12 | - | 26913973 | 26914128 | 26914407 | 26914578 | [illegible] |
| MXE | 16447 | ENSG00000101290 | CDS2 | chr20 | + | 5175182 | 5175279 | 5178816 | 5178956 | [illegible] |
| MXE | 16471 | ENSG00000161970 | RPL26 | chr17 | - | 8379795 | 8379936 | 8382142 | 8382338 | [illegible] |
| MXE | 16478 | ENSG00000076067 | RBMS2 | chr12 | + | 56571697 | 56571855 | 56580277 | 56580375 | [illegible] |
| MXE | 16501 | ENSG00000011523 | CEP68 | chr2 | + | 65069398 | 65069801 | 65071453 | 65072569 | [illegible] |
| MXE | 16503 | ENSG00000145901 | TNIP1 | chr5 | - | 151033479 | 151033579 | 151033607 | 151033799 | [illegible] |
| MXE | 16511 | ENSG00000111145 | ELK3 | chr12 | + | 96247602 | 96247734 | 96259730 | 96259853 | [illegible] |
| MXE | 16513 | ENSG00000152256 | PDK1 | chr2 | + | 172562740 | 172562800 | 172564051 | 172564130 | [illegible] |
| MXE | 16538 | ENSG00000028203 | VEZT | chr12 | + | 95224179 | 95224236 | 95242163 | 95242264 | [illegible] |
| MXE | 16543 | ENSG00000028203 | VEZT | chr12 | + | 95257149 | 95257239 | 95258244 | 95258282 | [illegible] |
| MXE | 16544 | ENSG00000028203 | VEZT | chr12 | + | 95257149 | 95257239 | 95258244 | 95258287 | [illegible] |
| MXE | 16546 | ENSG00000028203 | VEZT | chr12 | + | 95257153 | 95257239 | 95258244 | 95258282 | [illegible] |
| MXE | 16547 | ENSG00000028203 | VEZT | chr12 | + | 95257153 | 95257239 | 95258244 | 95258287 | [illegible] |
| MXE | 16556 | ENSG00000013503 | POLR3B | chr12 | + | 106430272 | 106430473 | 106432317 | 106432480 | [illegible] |
| MXE | 16560 | ENSG00000111785 | RIC8B | chr12 | + | 106814695 | 106815304 | 106825725 | 106825820 | [illegible] |
| MXE | 16563 | ENSG00000111785 | RIC8B | chr12 | + | 106868300 | 106868423 | 106870822 | 106870942 | [illegible] |
| MXE | 16564 | ENSG00000111785 | RIC8B | chr12 | + | 106870822 | 106870942 | 106878986 | 106879125 | [illegible] |
| MXE | 16565 | ENSG00000111785 | RIC8B | chr12 | + | 106874480 | 106874567 | 106878986 | 106879125 | [illegible] |
| MXE | 16566 | ENSG00000111785 | RIC8B | chr12 | + | 106878986 | 106879125 | 106879774 | 106879947 | [illegible] |
| MXE | 16568 | ENSG00000111727 | HCFC2 | chr12 | + | 104086846 | 104087014 | 104087985 | 104088038 | [illegible] |
| MXE | 16569 | ENSG00000111727 | HCFC2 | chr12 | + | 104098342 | 104098480 | 104101967 | 104102153 | [illegible] |
| MXE | 16581 | ENSG00000111144 | LTA4H | chr12 | - | 96024478 | 96024547 | 96027443 | 96027503 | [illegible] |
| MXE | 16584 | ENSG00000127314 | RAP1B | chr12 | + | 68624670 | 68624873 | 68648705 | 68648781 | [illegible] |
| MXE | 16586 | ENSG00000127314 | RAP1B | chr12 | + | 68624674 | 68624873 | 68648698 | 68648781 | [illegible] |
| MXE | 16598 | ENSG00000163312 | HELQ | chr4 | - | 83443516 | 83443614 | 83446013 | 83446086 | [illegible] |
| MXE | 16599 | ENSG00000163312 | HELQ | chr4 | - | 83446013 | 83446086 | 83446834 | 83447035 | [illegible] |
| MXE | 16603 | ENSG00000130177 | CDC16 | chr13 | + | 114265149 | 114265240 | 114269562 | 114269620 | [illegible] |
| MXE | 16631 | ENSG00000111837 | MAK | chr6 | - | 10808809 | 10808942 | 10813643 | 10813723 | [illegible] |
| MXE | 16639 | ENSG00000112053 | SLC26A8 | chr6 | - | 35977203 | 35977351 | 35982120 | 35982203 | [illegible] |
| MXE | 16640 | ENSG00000112053 | SLC26A8 | chr6 | - | 35982120 | 35982203 | 35991658 | 35991808 | [illegible] |
| MXE | 16653 | ENSG00000135297 | MTO1 | chr6 | + | 73482028 | 73482244 | 73482448 | 73482620 | [illegible] |
| MXE | 16654 | ENSG00000135297 | MTO1 | chr6 | + | 73482028 | 73482244 | 73482448 | 73482620 | [illegible] |
| MXE | 16657 | ENSG00000135297 | MTO1 | chr6 | + | 73482039 | 73482244 | 73482448 | 73482620 | [illegible] |
| MXE | 16661 | ENSG00000198270 | TMEM116 | chr12 | - | 111932585 | 111932659 | 111933885 | 111934030 | [illegible] |
| MXE | 16662 | ENSG00000198270 | TMEM116 | chr12 | - | 111932585 | 111932659 | 111933885 | 111934030 | [illegible] |
| MXE | 16668 | ENSG00000198270 | TMEM116 | chr12 | - | 111936741 | 111936830 | 111937159 | 111937243 | [illegible] |
| MXE | 16669 | ENSG00000198270 | TMEM116 | chr12 | - | 111938160 | 111938210 | 111943264 | 111943369 | [illegible] |
| MXE | 16678 | ENSG00000111696 | NT5DC3 | chr12 | - | 103787440 | 103787527 | 103788837 | 103788919 | [illegible] |
| MXE | 16680 | ENSG00000134532 | SOX5 | chr12 | - | 24213342 | 24213417 | 24277215 | 24277312 | [illegible] |
| MXE | 16681 | ENSG00000166012 | TAF1D | chr11 | - | 93733349 | 93733397 | 93734624 | 93734660 | [illegible] |
| MXE | 16682 | ENSG00000166012 | TAF1D | chr11 | - | 93733349 | 93733397 | 93734624 | 93734660 | [illegible] |
| MXE | 16687 | ENSG00000112200 | ZNF451 | chr6 | + | 57147089 | 57148693 | 57150718 | 57150862 | [illegible] |
| MXE | 16692 | ENSG00000135968 | GCC2 | chr2 | + | 108481709 | 108481816 | 108482286 | 108482451 | [illegible] |
| MXE | 16694 | ENSG00000135968 | GCC2 | chr2 | + | 108486434 | 108486648 | 108487698 | 108487820 | [illegible] |
| MXE | 16695 | ENSG00000135968 | GCC2 | chr2 | + | 108486510 | 108486648 | 108487698 | 108487820 | [illegible] |
| MXE | 16700 | ENSG00000120662 | MTRF1 | chr13 | - | 41261690 | 41261825 | 41262214 | 41262331 | [illegible] |
| MXE | 16701 | ENSG00000141376 | BCAS3 | chr17 | + | 60688019 | 60688195 | 60688931 | 60689046 | [illegible] |
| MXE | 16707 | ENSG00000215483 | LINC00598 | chr13 | - | 40466506 | 40466677 | 40469995 | 40470162 | [illegible] |
| MXE | 16727 | ENSG00000111271 | ACAD10 | chr12 | + | 111721670 | 111721739 | 111727961 | 111728143 | [illegible] |
| MXE | 16728 | ENSG00000111271 | ACAD10 | chr12 | + | 111727961 | 111728143 | 111729805 | 111729956 | [illegible] |
| MXE | 16731 | ENSG00000111271 | ACAD10 | chr12 | + | 111729805 | 111729956 | 111736830 | 111737004 | [illegible] |
| MXE | 16734 | ENSG00000111271 | ACAD10 | chr12 | + | 111747587 | 111747725 | 111748316 | 111748475 | [illegible] |
| MXE | 16745 | ENSG00000187109 | NAP1L1 | chr12 | - | 76055018 | 76055090 | 76056032 | 76056158 | [illegible] |
| MXE | 16749 | ENSG00000187109 | NAP1L1 | chr12 | - | 76056032 | 76056161 | 76074202 | 76074239 | [illegible] |
| MXE | 16750 | ENSG00000187109 | NAP1L1 | chr12 | - | 76056573 | 76056672 | 76059776 | 76059878 | [illegible] |
| MXE | 16751 | ENSG00000187109 | NAP1L1 | chr12 | - | 76056573 | 76056672 | 76059797 | 76059878 | [illegible] |
| MXE | 16756 | ENSG00000175582 | RAB6A | chr11 | - | 73716250 | 73716362 | 73718784 | 73718890 | [illegible] |
| MXE | 16763 | ENSG00000135108 | FBXO21 | chr12 | - | 117188758 | 117188793 | 117189226 | 117189362 | [illegible] |
| MXE | 16773 | ENSG00000090686 | USP48 | chr1 | - | 21706466 | 21706589 | 21706743 | 21706907 | [illegible] |
| MXE | 16782 | ENSG00000134545 | KLRC1 | chr12 | - | 10449913 | 10449967 | 10450483 | 10450579 | [illegible] |
| MXE | 16787 | ENSG00000033030 | ZCCHC8 | chr12 | - | 122482634 | 122482695 | 122483278 | 122483344 | [illegible] |
| MXE | 16793 | ENSG00000157538 | VPS26C | chr21 | - | 37229144 | 37229222 | 37231492 | 37231975 | [illegible] |
| MXE | 16795 | ENSG00000166037 | CEP57 | chr11 | + | 95821870 | 95821978 | 95822498 | 95822576 | [illegible] |
| MXE | 16798 | ENSG00000111615 | KRR1 | chr12 | - | 75505197 | 75505254 | 75506315 | 75506399 | [illegible] |
| MXE | 16804 | ENSG00000285625 | NA | chr12 | - | 56724451 | 56724523 | 56725262 | 56725340 | [illegible] |
| MXE | 16805 | ENSG00000123349 | PFDN5 | chr12 | + | 53295838 | 53295941 | 53296429 | 53296551 | [illegible] |
| MXE | 16808 | ENSG00000123349 | PFDN5 | chr12 | + | 53295838 | 53295941 | 53296453 | 53296551 | [illegible] |
| MXE | 16811 | ENSG00000123349 | PFDN5 | chr12 | + | 53296218 | 53296275 | 53296429 | 53296467 | [illegible] |
| MXE | 16813 | ENSG00000123349 | PFDN5 | chr12 | + | 53296243 | 53296275 | 53296429 | 53296467 | [illegible] |
| MXE | 16814 | ENSG00000123349 | PFDN5 | chr12 | + | 53296243 | 53296275 | 53296429 | 53296551 | [illegible] |
| MXE | 16815 | ENSG00000123349 | PFDN5 | chr12 | + | 53296429 | 53296551 | 53297849 | 53297924 | [illegible] |
| MXE | 16817 | ENSG00000288663 | NA | chr12 | + | 53296243 | 53296275 | 53296453 | 53296551 | [illegible] |
| MXE | 16818 | ENSG00000288663 | NA | chr12 | + | 53296453 | 53296551 | 53297849 | 53297924 | [illegible] |
| MXE | 16824 | ENSG00000134283 | PPHLN1 | chr12 | + | 42355160 | 42355222 | 42374862 | 42375074 | [illegible] |
| MXE | 16827 | ENSG00000122966 | CIT | chr12 | - | 119776357 | 119776408 | 119776671 | 119776842 | [illegible] |
| MXE | 16828 | ENSG00000089693 | MLF2 | chr12 | - | 6750144 | 6750305 | 6750712 | 6750766 | [illegible] |
| MXE | 16830 | ENSG00000120837 | NFYB | chr12 | - | 104126113 | 104126244 | 104128423 | 104128517 | [illegible] |
| MXE | 16833 | ENSG00000138777 | PPA2 | chr4 | - | 105424195 | 105424322 | 105437949 | 105438036 | [illegible] |
| MXE | 16834 | ENSG00000111596 | CNOT2 | chr12 | + | 70285399 | 70285586 | 70294070 | 70294294 | [illegible] |
| MXE | 16836 | ENSG00000111596 | CNOT2 | chr12 | + | 70285399 | 70285586 | 70301704 | 70301865 | [illegible] |
| MXE | 16837 | ENSG00000111596 | CNOT2 | chr12 | + | 70294070 | 70294294 | 70301704 | 70301865 | [illegible] |
| MXE | 16838 | ENSG00000111596 | CNOT2 | chr12 | + | 70294236 | 70294294 | 70301704 | 70301865 | [illegible] |
| MXE | 16847 | ENSG00000111247 | RAD51AP1 | chr12 | + | 4545218 | 4545312 | 4545781 | 4545832 | [illegible] |
| MXE | 16851 | ENSG00000129317 | PUS7L | chr12 | - | 43738309 | 43738391 | 43742456 | 43742555 | [illegible] |
| MXE | 16862 | ENSG00000135677 | GNS | chr12 | - | 64728955 | 64729057 | 64737003 | 64737107 | [illegible] |
| MXE | 16866 | ENSG00000174106 | LEMD3 | chr12 | + | 65245668 | 65245774 | 65245860 | 65245939 | [illegible] |
| MXE | 16869 | ENSG00000073060 | SCARB1 | chr12 | - | 124800123 | 124800242 | 124807760 | 124807927 | [illegible] |
| MXE | 16885 | ENSG00000166598 | HSP90B1 | chr12 | + | 103932278 | 103932418 | 103932825 | 103932942 | [illegible] |
| MXE | 16886 | ENSG00000166598 | HSP90B1 | chr12 | + | 103932280 | 103932418 | 103932825 | 103932942 | [illegible] |
| MXE | 16892 | ENSG00000188906 | LRRK2 | chr12 | + | 40259479 | 40259604 | 40263788 | 40263901 | [illegible] |
| MXE | 16907 | ENSG00000111335 | OAS2 | chr12 | + | 113006417 | 113006600 | 113007704 | 113007943 | [illegible] |
| MXE | 16910 | ENSG00000177119 | ANO6 | chr12 | + | 45302013 | 45302093 | 45331294 | 45331423 | [illegible] |
| MXE | 16917 | ENSG00000168778 | TCTN2 | chr12 | + | 123694841 | 123694976 | 123695219 | 123695297 | [illegible] |
| MXE | 16918 | ENSG00000204852 | TCTN1 | chr12 | + | 110634395 | 110634463 | 110634669 | 110634779 | [illegible] |
| MXE | 16919 | ENSG00000204852 | TCTN1 | chr12 | + | 110641023 | 110641149 | 110641541 | 110642389 | [illegible] |
| MXE | 16924 | ENSG00000204852 | TCTN1 | chr12 | + | 110642248 | 110642389 | 110644879 | 110645143 | [illegible] |
| MXE | 16932 | ENSG00000123411 | IKZF4 | chr12 | + | 56019317 | 56019415 | 56021430 | 56021580 | [illegible] |
| MXE | 16933 | ENSG00000123411 | IKZF4 | chr12 | + | 56023670 | 56023764 | 56024661 | 56024905 | [illegible] |
| MXE | 16939 | ENSG00000101160 | CTSZ | chr20 | - | 58998597 | 58998690 | 58999234 | 58999580 | [illegible] |
| MXE | 16947 | ENSG00000075089 | ACTR6 | chr12 | + | 100205675 | 100205744 | 100207662 | 100207786 | [illegible] |
| MXE | 16948 | ENSG00000075089 | ACTR6 | chr12 | + | 100205675 | 100205744 | 100207662 | 100207786 | [illegible] |
| MXE | 16949 | ENSG00000075089 | ACTR6 | chr12 | + | 100205675 | 100205757 | 100207662 | 100207786 | [illegible] |
| MXE | 16951 | ENSG00000075089 | ACTR6 | chr12 | + | 100207662 | 100207786 | 100210072 | 100210208 | [illegible] |
| MXE | 16953 | ENSG00000075089 | ACTR6 | chr12 | + | 100207662 | 100207786 | 100210294 | 100210351 | [illegible] |
| MXE | 16956 | ENSG00000121749 | TBC1D15 | chr12 | + | 71841100 | 71841231 | 71854562 | 71854602 | [illegible] |
| MXE | 16957 | ENSG00000121749 | TBC1D15 | chr12 | + | 71841100 | 71841231 | 71854784 | 71854853 | [illegible] |
| MXE | 16958 | ENSG00000121749 | TBC1D15 | chr12 | + | 71841100 | 71841231 | 71861404 | 71861496 | [illegible] |
| MXE | 16959 | ENSG00000121749 | TBC1D15 | chr12 | + | 71854562 | 71854602 | 71854784 | 71854853 | [illegible] |
| MXE | 16960 | ENSG00000121749 | TBC1D15 | chr12 | + | 71854562 | 71854602 | 71861404 | 71861496 | [illegible] |
| MXE | 16961 | ENSG00000121749 | TBC1D15 | chr12 | + | 71854784 | 71854853 | 71861404 | 71861496 | [illegible] |
| MXE | 16971 | ENSG00000205809 | KLRC2 | chr12 | - | 10434485 | 10434530 | 10434994 | 10435039 | [illegible] |
| MXE | 16972 | ENSG00000184752 | NDUFA12 | chr12 | - | 94996903 | 94996976 | 94997307 | 94998733 | [illegible] |
| MXE | 16973 | ENSG00000075415 | SLC25A3 | chr12 | + | 98595432 | 98595557 | 98595726 | 98595848 | [illegible] |
| MXE | 16976 | ENSG00000118454 | ANKRD13C | chr1 | - | 70270855 | 70270956 | 70274719 | 70274818 | [illegible] |

| | | | | | | | | | | | | | | | | | | | | | | | | | | | | | | | | | | | | | | | | | | | | | | | | |
|---|---|---|---|---|---|---|---|---|---|---|---|---|---|---|---|---|---|---|---|---|---|---|---|---|---|---|---|---|---|---|---|---|---|---|---|---|---|---|---|---|---|---|---|---|---|---|---|---|
| MXE | 16977 | ENSG00000118454 | ANKRD13C | chr1 | - | 70270855 | 70270956 | 70274719 | 70274818 | 70262273 | 70262803 | 70276764 | 70276843 | 41 | 46 | 45 | 53 | 65 | 3 | 1 | 8 | 8 | 12 | 3 | 23 | 34 | 9 | 0 | 0 | 7 | 2 | 3 | 0 | 248 | 250 | 1 | 1 | 0.932 | 0.979 | 0.85 | 0.87 | 0.845 | 1 | 0.768 | 0.945 | 0.752 | NA | 0.029 |
| MXE | 16981 | ENSG00000139132 | FGD4 | chr12 | + | 32602160 | 32602317 | 32607956 | 32608095 | 32598496 | 32598586 | 32610775 | 32610834 | 0 | 0 | 0 | 0 | 0 | 165 | 147 | 75 | 209 | 241 | 0 | 2 | 0 | 4 | 0 | 300 | 159 | 176 | 208 | 301 | 298 | 288 | 1 | 1 | 0 | 0 | 0 | 0 | 0 | 0 | 0.012 | 0 | 0.018 | 0 | -0.006 |
| MXE | 16992 | ENSG00000111300 | NAA25 | chr12 | - | 112047664 | 112047790 | 112048291 | 112048443 | 112043624 | 112043868 | 112049482 | 112049611 | 11 | 7 | 14 | 14 | 8 | 21 | 29 | 35 | 34 | 47 | 15 | 19 | 13 | 15 | 0 | 14 | 28 | 31 | 29 | 0 | 298 | 275 | 1 | 1 | 0.326 | 0.182 | 0.27 | 0.275 | 0.136 | 0.497 | 0.385 | 0.279 | 0.323 | NA | -0.133 |
| MXE | 16994 | ENSG00000111300 | NAA25 | chr12 | - | 112049482 | 112049611 | 112053557 | 112053657 | 112048291 | 112048443 | 112060269 | 112060359 | 15 | 10 | 16 | 11 | 16 | 9 | 8 | 12 | 16 | 4 | 0 | 19 | 13 | 4 | 0 | 15 | 11 | 14 | 17 | 0 | 249 | 278 | 1 | 1 | 0.65 | 0.583 | 0.598 | 0.434 | 0.817 | 0 | 0.659 | 0.509 | 0.208 | NA | 0.272 |
| MXE | 16999 | ENSG00000123415 | SMUG1 | chr12 | - | 54187818 | 54187905 | 54188510 | 54188596 | 54183655 | 54183959 | 54188725 | 54188957 | 1 | 0 | 0 | 0 | 0 | 3 | 2 | 1 | 2 | 3 | 0 | 0 | 0 | 0 | 0 | 0 | 7 | 0 | 0 | 0 | 235 | 236 | 1 | 1 | 0.251 | 0 | 0 | 0 | 0 | NA | 0 | NA | NA | NA | 0.05 |
| MXE | 17000 | ENSG00000111554 | MDM1 | chr12 | - | 68316077 | 68316253 | 68316580 | 68316610 | 68314947 | 68315265 | 68321346 | 68321446 | 18 | 5 | 4 | 13 | 13 | 64 | 74 | 77 | 137 | 138 | 0 | 20 | 3 | 3 | 5 | 0 | 93 | 68 | 83 | 13 | 179 | 298 | 1 | 1 | 0.319 | 0.101 | 0.08 | 0.136 | 0.136 | NA | 0.264 | 0.068 | 0.057 | 0.39 | -0.04 |
| MXE | 17007 | ENSG00000197111 | PCBP2 | chr12 | + | 53455884 | 53456001 | 53459271 | 53459403 | 53455460 | 53455493 | 53460330 | 53460425 | 2022 | 1172 | 1748 | 1905 | 1984 | 3 | 0 | 0 | 8 | 3 | 2663 | 1601 | 1367 | 1747 | 3801 | 0 | 8 | 4 | 1 | 0 | 266 | 281 | 1 | 1 | 0.999 | 1 | 1 | 0.996 | 0.999 | 1 | 0.995 | 0.997 | 0.999 | 1 | 0.001 |
| MXE | 17008 | ENSG00000197111 | PCBP2 | chr12 | + | 53459271 | 53459403 | 53460330 | 53460425 | 53455884 | 53456001 | 53461014 | 53461143 | 3504 | 1903 | 3101 | 3313 | 4481 | 0 | 1 | 0 | 1 | 0 | 4467 | 3131 | 2724 | 2995 | 6963 | 0 | 0 | 0 | 0 | 0 | 281 | 244 | 1 | 1 | 1 | 0.999 | 1 | 1 | 1 | 1 | 1 | 1 | 1 | 1 | 0 |
| MXE | 17016 | ENSG00000092978 | GPATCH2 | chr1 | - | 217491679 | 217491750 | 217498355 | 217498395 | 217449248 | 217449337 | 217514821 | 217514889 | 4 | 10 | 1 | 5 | 8 | 4 | 3 | 4 | 5 | 10 | 0 | 0 | 5 | 0 | 0 | 0 | 4 | 0 | 0 | 0 | 189 | 220 | 1 | 1 | 0.538 | 0.795 | 0.225 | 0.538 | 0.482 | NA | 0 | 1 | NA | NA | 0.016 |
| MXE | 17034 | ENSG00000136104 | RNASEH2B | chr13 | + | 50929460 | 50929582 | 50934884 | 50934999 | 50927406 | 50927478 | 50943320 | 50943394 | 2 | 0 | 4 | 2 | 0 | 463 | 304 | 434 | 434 | 698 | 0 | 1 | 1 | 0 | 0 | 400 | 359 | 247 | 326 | 292 | 271 | 264 | 1 | 1 | 0.004 | 0 | 0.009 | 0.004 | 0 | 0 | 0.003 | 0.004 | 0 | 0 | 0.002 |
| MXE | 17037 | ENSG00000136104 | RNASEH2B | chr13 | + | 50929515 | 50929582 | 50934884 | 50934999 | 50927406 | 50927478 | 50943320 | 50943352 | 0 | 0 | 1 | 0 | 0 | 463 | 304 | 434 | 434 | 698 | 0 | 0 | 0 | 0 | 0 | 400 | 359 | 247 | 326 | 292 | 216 | 264 | 1 | 1 | 0 | 0 | 0.003 | 0 | 0 | 0 | 0 | 0 | 0 | 0 | 0.001 |
| MXE | 17047 | ENSG00000111684 | LPCAT3 | chr12 | - | 6981003 | 6981182 | 6981594 | 6981632 | 6978602 | 6978689 | 6981810 | 6981904 | 10 | 11 | 9 | 5 | 9 | 1 | 0 | 1 | 2 | 0 | 20 | 23 | 27 | 47 | 62 | 0 | 0 | 3 | 1 | 0 | 187 | 298 | 1 | 1 | 0.941 | 1 | 0.935 | 0.799 | 1 | 1 | 1 | 0.935 | 0.987 | 1 | -0.049 |
| MXE | 17049 | ENSG00000227354 | RBM26-AS1 | chr13 | + | 79415110 | 79415364 | 79417834 | 79417916 | 79412681 | 79412907 | 79420759 | 79420815 | 1 | 0 | 1 | 0 | 0 | 3 | 1 | 1 | 0 | 6 | 0 | 0 | 2 | 0 | 0 | 0 | 0 | 0 | 1 | 0 | 298 | 231 | 1 | 1 | 0.205 | 0 | 0.437 | NA | 0 | NA | NA | 1 | 0 | NA | -0.34 |
| MXE | 17050 | ENSG00000227354 | RBM26-AS1 | chr13 | + | 79415261 | 79415364 | 79417834 | 79417916 | 79412681 | 79412907 | 79420759 | 79420815 | 1 | 1 | 2 | 0 | 0 | 3 | 1 | 1 | 0 | 6 | 0 | 0 | 0 | 0 | 0 | 0 | 0 | 0 | 1 | 0 | 252 | 231 | 1 | 1 | 0.234 | 0.478 | 0.647 | NA | 0 | NA | NA | NA | 0 | NA | 0.34 |
| MXE | 17059 | ENSG00000286330 | NA | chr13 | + | 74356146 | 74356322 | 74365024 | 74365087 | 74331951 | 74332013 | 74399995 | 74400168 | 1 | 3 | 2 | 0 | 3 | 4 | 0 | 7 | 1 | 18 | 0 | 0 | 0 | 0 | 0 | 0 | 0 | 2 | 1 | 0 | 298 | 212 | 1 | 1 | 0.151 | 1 | 0.169 | 0 | 0.106 | NA | NA | 0 | 0 | NA | 0.285 |
| MXE | 17060 | ENSG00000286330 | NA | chr13 | + | 74356146 | 74356322 | 74365024 | 74365087 | 74331951 | 74332013 | 74406314 | 74408791 | 0 | 4 | 2 | 0 | 4 | 4 | 5 | 10 | 3 | 19 | 0 | 0 | 0 | 0 | 0 | 0 | 0 | 8 | 1 | 0 | 298 | 212 | 1 | 1 | 0 | 0.363 | 0.125 | 0 | 0.13 | NA | NA | 0 | 0 | NA | 0.124 |
| MXE | 17061 | ENSG00000286330 | NA | chr13 | + | 74365024 | 74365061 | 74399995 | 74400168 | 74356146 | 74356322 | 74406314 | 74407540 | 0 | 1 | 3 | 1 | 2 | 5 | 9 | 11 | 3 | 17 | 0 | 0 | 5 | 0 | 0 | 0 | 4 | 5 | 0 | 0 | 186 | 298 | 1 | 1 | 0 | 0.151 | 0.304 | 0.348 | 0.159 | NA | 0 | 0.616 | NA | NA | -0.116 |
| MXE | 17064 | ENSG00000139496 | NUP58 | chr13 | + | 25319325 | 25319350 | 25320529 | 25320595 | 25315356 | 25315467 | 25320918 | 25321093 | 45 | 31 | 34 | 14 | 26 | 48 | 28 | 38 | 20 | 88 | 35 | 22 | 16 | 32 | 0 | 8 | 49 | 44 | 23 | 0 | 174 | 215 | 1 | 1 | 0.537 | 0.578 | 0.525 | 0.464 | 0.267 | 0.844 | 0.357 | 0.31 | 0.632 | NA | -0.062 |
| MXE | 17067 | ENSG00000133103 | COG6 | chr13 | + | 39694633 | 39694725 | 39699500 | 39699618 | 39689759 | 39689824 | 39719235 | 39719367 | 42 | 26 | 53 | 42 | 120 | 40 | 3 | 22 | 12 | 31 | 0 | 21 | 12 | 8 | 0 | 0 | 3 | 0 | 0 | 0 | 241 | 267 | 1 | 1 | 0.538 | 0.906 | 0.727 | 0.795 | 0.811 | NA | 0.886 | 1 | 1 | NA | -0.207 |
| MXE | 17123 | ENSG00000275066 | SYNRG | chr17 | - | 37571790 | 37571987 | 37576340 | 37576418 | 37568790 | 37568924 | 37584647 | 37584759 | 40 | 31 | 27 | 37 | 95 | 0 | 0 | 4 | 1 | 6 | 10 | 17 | 9 | 18 | 0 | 0 | 5 | 0 | 4 | 0 | 227 | 298 | 1 | 1 | 1 | 1 | 0.899 | 0.98 | 0.954 | 1 | 0.817 | 1 | 0.855 | NA | 0.049 |
| MXE | 17125 | ENSG00000055147 | FAM114A2 | chr5 | - | 154029488 | 154029580 | 154038387 | 154038434 | 154028148 | 154028283 | 154038856 | 154038921 | 0 | 0 | 1 | 0 | 0 | 28 | 35 | 32 | 29 | 60 | 0 | 0 | 0 | 0 | 0 | 2 | 13 | 17 | 28 | 0 | 196 | 241 | 1 | 1 | 0 | 0 | 0.037 | 0 | 0 | 0 | 0 | 0 | 0 | NA | 0.007 |
| MXE | 17139 | ENSG00000102710 | SUPT20H | chr13 | - | 37029764 | 37029836 | 37031566 | 37031623 | 37028147 | 37028305 | 37031738 | 37031895 | 35 | 20 | 27 | 30 | 6 | 88 | 49 | 80 | 94 | 124 | 14 | 28 | 17 | 26 | 0 | 12 | 78 | 50 | 46 | 0 | 206 | 221 | 1 | 1 | 0.299 | 0.305 | 0.266 | 0.255 | 0.049 | 0.556 | 0.278 | 0.267 | 0.377 | NA | -0.135 |
| MXE | 17140 | ENSG00000102710 | SUPT20H | chr13 | - | 37045246 | 37045373 | 37047345 | 37047408 | 37044077 | 37044181 | 37047534 | 37047601 | 7 | 1 | 3 | 3 | 3 | 367 | 305 | 310 | 393 | 682 | 0 | 0 | 3 | 3 | 0 | 209 | 235 | 186 | 223 | 75 | 212 | 276 | 1 | 1 | 0.024 | 0.004 | 0.012 | 0.01 | 0.006 | 0 | 0 | 0.021 | 0.017 | 0 | 0.004 |
| MXE | 17147 | ENSG00000167186 | COQ7 | chr16 | + | 19075720 | 19075860 | 19077305 | 19077374 | 19071927 | 19072106 | 19078080 | 19078283 | 0 | 1 | 0 | 0 | 0 | 1 | 2 | 0 | 5 | 0 | 0 | 0 | 0 | 0 | 0 | 0 | 2 | 0 | 0 | 0 | 289 | 218 | 1 | 1 | 0 | 0.274 | NA | 0 | NA | NA | 0 | NA | NA | NA | 0.091 |
| MXE | 17153 | ENSG00000134900 | TPP2 | chr13 | + | 102644908 | 102646390 | 102647206 | 102647344 | 102644556 | 102644673 | 102649407 | 102649486 | 205 | 136 | 179 | 234 | 183 | 0 | 0 | 0 | 0 | 0 | 25 | 95 | 140 | 51 | 29 | 0 | 0 | 3 | 3 | 0 | 298 | 287 | 1 | 1 | 1 | 1 | 1 | 1 | 1 | 1 | 1 | 0.978 | 0.942 | 1 | 0.016 |
| MXE | 17156 | ENSG00000134900 | TPP2 | chr13 | + | 102651338 | 102651397 | 102657055 | 102657207 | 102649407 | 102649486 | 102663647 | 102663744 | 4 | 1 | 1 | 0 | 1 | 425 | 320 | 378 | 360 | 674 | 0 | 0 | 0 | 0 | 0 | 89 | 257 | 192 | 111 | 45 | 208 | 298 | 1 | 1 | 0.013 | 0.004 | 0.004 | 0 | 0.002 | 0 | 0 | 0 | 0 | 0 | 0.005 |
| MXE | 17167 | ENSG00000163519 | TRAT1 | chr3 | + | 108838933 | 108838967 | 108847067 | 108847129 | 108822771 | 108822934 | 108849165 | 108849254 | 0 | 0 | 1 | 0 | 0 | 119 | 97 | 139 | 133 | 384 | 0 | 0 | 0 | 0 | 0 | 0 | 44 | 47 | 12 | 0 | 183 | 211 | 1 | 1 | 0 | 0 | 0.008 | 0 | 0 | NA | 0 | 0 | 0 | NA | 0.002 |
| MXE | 17169 | ENSG00000127022 | CANX | chr5 | + | 179708238 | 179708380 | 179709872 | 179710065 | 179698933 | 179699102 | 179716104 | 179716294 | 0 | 1 | 0 | 0 | 0 | 2134 | 1461 | 1885 | 1828 | 3349 | 0 | 0 | 0 | 0 | 0 | 1344 | 1590 | 1685 | 1939 | 469 | 291 | 298 | 1 | 1 | 0 | 0.001 | 0 | 0 | 0 | 0 | 0 | 0 | 0 | 0 | 0 |
| MXE | 17172 | ENSG00000127022 | CANX | chr5 | + | 179723663 | 179723779 | 179724656 | 179724783 | 179722803 | 179723019 | 179726679 | 179726759 | 0 | 0 | 1 | 0 | 0 | 1033 | 763 | 850 | 822 | 1280 | 0 | 0 | 2 | 0 | 0 | 523 | 813 | 1087 | 1344 | 149 | 265 | 276 | 1 | 1 | 0 | 0 | 0.001 | 0 | 0 | 0 | 0 | 0.002 | 0 | 0 | 0 |
| MXE | 17173 | ENSG00000133027 | PEMT | chr17 | - | 17506226 | 17506301 | 17509433 | 17509545 | 17505588 | 17505848 | 17576919 | 17577027 | 1 | 0 | 0 | 1 | 0 | 4 | 0 | 1 | 2 | 0 | 0 | 0 | 0 | 0 | 0 | 16 | 5 | 7 | 2 | 7 | 261 | 224 | 1 | 1 | 0.177 | NA | 0 | 0.3 | NA | 0 | 0 | 0 | 0 | 0 | 0.159 |
| MXE | 17189 | ENSG00000136100 | VPS36 | chr13 | - | 52417056 | 52417141 | 52417436 | 52417679 | 52416016 | 52416093 | 52417991 | 52418056 | 0 | 1 | 0 | 0 | 0 | 169 | 142 | 142 | 148 | 146 | 0 | 1 | 0 | 0 | 0 | 43 | 73 | 86 | 83 | 18 | 298 | 234 | 1 | 1 | 0 | 0.005 | 0 | 0 | 0 | 0 | 0.011 | 0 | 0 | 0 | -0.001 |

U12_MXE_filtered

| Splicing type | ID | Gene_ID | Gene_name | Chromosome | Strand | 1stExonStart_0base | 1stExonEnd | 2ndExonStart_0base | 2ndExonEnd | UpstreamES | UpstreamEE | DownstreamES | DownstreamEE |
|---|---|---|---|---|---|---|---|---|---|---|---|---|---|
| MXE | 9333 | ENSG00000172766 | NAA16 | chr13 | + | 41331273 | 41331369 | 41336649 | 41336756 | 41328723 | 41328843 | 41355143 | 41355216 |
| MXE | 3396 | ENSG00000131504 | DIAPH1 | chr5 | - | 141524142 | 141524229 | 141526037 | 141526173 | 141515020 | 141517008 | 141526296 | 141526461 |
| MXE | 3858 | ENSG00000157107 | FCHO2 | chr5 | + | 72990711 | 72990864 | 73006444 | 73006549 | 72990477 | 72990619 | 73015625 | 73015724 |
| MXE | 13836 | ENSG00000145041 | DCAF1 | chr3 | - | 51470928 | 51471005 | 51483718 | 51483836 | 51466802 | 51466876 | 51496733 | 51496780 |
| MXE | 2172 | ENSG00000140199 | SLC12A6 | chr15 | - | 34238964 | 34239160 | 34240660 | 34240829 | 34238231 | 34238331 | 34241232 | 34241337 |
| MXE | 13873 | ENSG00000187735 | TCEA1 | chr8 | - | 53999944 | 54000050 | 54010429 | 54010492 | 53993667 | 53993755 | 54022062 | 54022448 |
| MXE | 13833 | ENSG00000145041 | DCAF1 | chr3 | - | 51441384 | 51441897 | 51443765 | 51443903 | 51440969 | 51441071 | 51463113 | 51463227 |
| MXE | 5720 | ENSG00000163507 | CIP2A | chr3 | - | 108582107 | 108582202 | 108582976 | 108583083 | 108581414 | 108581511 | 108585064 | 108585212 |
| MXE | 1922 | ENSG00000069329 | VPS35 | chr16 | - | 46677314 | 46677398 | 46678942 | 46679156 | 46676504 | 46676692 | 46680670 | 46680853 |
| MXE | 3346 | ENSG00000167881 | SRP68 | chr17 | - | 76060307 | 76060390 | 76061109 | 76061219 | 76057402 | 76057543 | 76061491 | 76061574 |
| MXE | 8313 | ENSG00000070010 | UFD1 | chr22 | - | 19471686 | 19471808 | 19475067 | 19475100 | 19467872 | 19468003 | 19475469 | 19475602 |
| MXE | 10373 | ENSG00000162645 | GBP2 | chr1 | - | 89114015 | 89114296 | 89116991 | 89117234 | 89112471 | 89112684 | 89117576 | 89117773 |
| MXE | 13523 | ENSG00000134516 | DOCK2 | chr5 | + | 169712119 | 169712223 | 169714027 | 169714211 | 169711934 | 169712007 | 169714359 | 169714457 |
| MXE | 2410 | ENSG00000172575 | RASGRP1 | chr15 | - | 38494381 | 38494767 | 38498793 | 38498946 | 38490553 | 38490688 | 38502311 | 38502421 |
| [illegible] | | | | | | | | | | | | | |

| ID | Inclusion_counts_HC | | | | | Skipping_counts_HC | | | | | Inclusion counts_SLE | | | | | Skipping_counts_SLE | | | | |
|---|---|---|---|---|---|---|---|---|---|---|---|---|---|---|---|---|---|---|---|---|
| | SRR13214146 | SRR13214147 | SRR13214148 | SRR13214149 | SRR13214150 | SRR13214146 | SRR13214147 | SRR13214148 | SRR13214149 | SRR13214150 | SRR13214141 | SRR13214242 | SRR13214143 | SRR13214144 | SRR13214145 | SRR13214141 | SRR13214242 | SRR13214143 | SRR13214144 | SRR13214145 |
| 9333 | 100 | 101 | 97 | 100 | 134 | 38 | 35 | 45 | 38 | 91 | 17 | 84 | 93 | 42 | 37 | 0 | 12 | 7 | 3 | 0 |
| 3396 | 1637 | 1193 | 973 | 1557 | 2060 | 386 | 178 | 269 | 282 | 510 | 3482 | 1489 | 1131 | 2075 | 4041 | 294 | 264 | 187 | 205 | 485 |
| 3858 | 150 | 143 | 77 | 175 | 123 | 60 | 44 | 35 | 51 | 48 | 183 | 62 | 175 | 81 | 29 | 0 | 17 | 29 | 6 | 0 |
| 13836 | 18 | 16 | 20 | 24 | 21 | 146 | 92 | 136 | 158 | 132 | 13 | 11 | 9 | 14 | 2 | 353 | 156 | 159 | 245 | 158 |
| 2172 | 97 | 52 | 78 | 78 | 187 | 9 | 0 | 2 | 1 | 3 | 122 | 50 | 55 | 69 | 11 | 18 | 8 | 13 | 10 | 0 |
| 13873 | 25 | 9 | 16 | 21 | 1 | 323 | 270 | 303 | 386 | 621 | 16 | 22 | 26 | 21 | 5 | 116 | 221 | 261 | 203 | 34 |
| 13833 | 136 | 93 | 147 | 109 | 123 | 49 | 42 | 65 | 32 | 52 | 146 | 119 | 160 | 169 | 130 | 11 | 24 | 41 | 41 | 7 |
| 5720 | 0 | 0 | 13 | 5 | 2 | 13 | 11 | 20 | 8 | 6 | 0 | 0 | 0 | 0 | 0 | 1 | 11 | 16 | 12 | 11 |
| 1922 | 568 | 588 | 468 | 589 | 1235 | 184 | 133 | 162 | 201 | 319 | 378 | 488 | 448 | 568 | 111 | 51 | 136 | 54 | 91 | 8 |
| 3346 | 179 | 143 | 170 | 194 | 279 | 77 | 50 | 76 | 90 | 127 | 359 | 270 | 246 | 217 | 223 | 64 | 50 | 51 | 55 | 86 |
| 8313 | 6 | 0 | 3 | 1 | 3 | 184 | 98 | 190 | 180 | 281 | 4 | 14 | 8 | 7 | 35 | 365 | 243 | 211 | 202 | 381 |
| 10373 | 342 | 177 | 331 | 306 | 288 | 528 | 478 | 520 | 528 | 544 | 258 | 116 | 188 | 119 | 84 | 1038 | 327 | 369 | 656 | 285 |
| 13523 | 867 | 654 | 682 | 845 | 1176 | 703 | 550 | 500 | 608 | 584 | 1171 | 1069 | 962 | 1029 | 740 | 1569 | 991 | 1022 | 1081 | 711 |
| 2410 | 6 | 7 | 4 | 17 | 12 | 173 | 127 | 157 | 144 | 288 | 0 | 0 | 2 | 0 | 0 | 37 | 113 | 93 | 19 | 26 |
| [illegible] | | | | | | | | | | | | | | | | | | | | |

| ID | IncFormLen | SkipFormLen | p value | FDR | Inclusion_level(normalized) | | | | | | | | | | Inclevel_diff |
|---|---|---|---|---|---|---|---|---|---|---|---|---|---|---|---|
| | | | | | SRR13214146 | SRR13214147 | SRR13214148 | SRR13214149 | SRR13214150 | SRR13214141 | SRR13214242 | SRR13214143 | SRR13214144 | SRR13214145 | |
| 9333 | 245 | 256 | 4.44089E-16 | 2.36953E-13 | 0.733 | 0.751 | 0.693 | 0.733 | 0.606 | 1 | 0.88 | 0.933 | 0.936 | 1 | -0.247 |
| 3396 | 285 | 236 | 1.94611E-12 | 4.33953E-10 | 0.778 | 0.847 | 0.75 | 0.821 | 0.77 | 0.907 | 0.824 | 0.834 | 0.893 | 0.873 | -0.073 |
| 3858 | 298 | 254 | 7.87284E-11 | 1.14194E-08 | 0.681 | 0.735 | 0.652 | 0.745 | 0.686 | 1 | 0.757 | 0.837 | 0.92 | 1 | -0.203 |
| 13836 | 267 | 226 | 1.49509E-10 | 2.01231E-08 | 0.094 | 0.128 | 0.111 | 0.114 | 0.119 | 0.03 | 0.056 | 0.046 | 0.046 | 0.011 | 0.075 |
| 2172 | 298 | 298 | 5.47522E-09 | 4.84023E-07 | 0.915 | 1 | 0.975 | 0.987 | 0.984 | 0.871 | 0.862 | 0.809 | 0.873 | 1 | 0.089 |
| 13873 | 212 | 255 | 6.55344E-09 | 5.72563E-07 | 0.085 | 0.039 | 0.06 | 0.061 | 0.002 | 0.142 | 0.107 | 0.107 | 0.111 | 0.15 | -0.074 |
| 13833 | 287 | 298 | 1.4196E-08 | 1.08208E-06 | 0.742 | 0.697 | 0.701 | 0.78 | 0.711 | 0.932 | 0.837 | 0.802 | 0.811 | 0.951 | -0.14 |
| 5720 | 256 | 244 | 4.81776E-08 | 3.0116E-06 | 0 | 0 | 0.383 | 0.373 | 0.241 | 0 | 0 | 0 | 0 | 0 | 0.199 |
| 1922 | 298 | 233 | 1.56526E-07 | 8.53467E-06 | 0.707 | 0.776 | 0.693 | 0.696 | 0.752 | 0.853 | 0.737 | 0.866 | 0.83 | 0.916 | -0.116 |
| 3346 | 259 | 232 | 2.51968E-07 | 1.259E-05 | 0.676 | 0.719 | 0.667 | 0.659 | 0.663 | 0.834 | 0.829 | 0.812 | 0.779 | 0.699 | -0.114 |
| 8313 | 182 | 271 | 4.11117E-07 | 1.93147E-05 | 0.046 | 0 | 0.023 | 0.008 | 0.016 | 0.016 | 0.079 | 0.053 | 0.049 | 0.12 | -0.045 |
| 10373 | 298 | 298 | 5.67824E-07 | 2.53991E-05 | 0.393 | 0.27 | 0.389 | 0.367 | 0.346 | 0.199 | 0.262 | 0.338 | 0.154 | 0.228 | 0.117 |
| 13523 | 253 | 298 | 6.13693E-07 | 2.70459E-05 | 0.592 | 0.583 | 0.616 | 0.621 | 0.703 | 0.468 | 0.56 | 0.526 | 0.529 | 0.551 | 0.096 |
| 2410 | 298 | 298 | 1.30739E-06 | 4.9489E-05 | 0.034 | 0.052 | 0.025 | 0.106 | 0.04 | 0 | 0 | 0.021 | 0 | 0 | 0.047 |
| [illegible] | | | | | | | | | | | | | | | |

| | | | | | | | | | | | | | | |
|---|---|---|---|---|---|---|---|---|---|---|---|---|---|---|
| MXE | 13798 | ENSG00000156171 | DRAM2 | chr1 | - | 111122615 | 111122643 | 111124741 | 111124881 | 111120515 | 111120693 | 111126226 | 111126294 | 236 |
| MXE | 5676 | ENSG00000114850 | SSR3 | chr3 | - | 156548865 | 156549003 | 156553654 | 156553781 | 156544307 | 156544439 | 156554956 | 156555149 | 116 |
| MXE | 10160 | ENSG00000143924 | EML4 | chr2 | + | 42304483 | 42304551 | 42315961 | 42316050 | 42303314 | 42303446 | 42317426 | 42317524 | 131 |
| MXE | 12388 | ENSG00000068885 | IFT80 | chr3 | - | 160319759 | 160319939 | 160343650 | 160343815 | 160307662 | 160307781 | 160356012 | 160356150 | 3 |
| MXE | 16921 | ENSG00000204852 | TCTN1 | chr12 | + | 110641541 | 110642389 | 110644966 | 110645129 | 110641023 | 110641149 | 110647210 | 110647336 | 7 |
| MXE | 13955 | ENSG00000137574 | TGS1 | chr8 | + | 55795977 | 55796152 | 55798913 | 55799220 | 55792697 | 55792784 | 55802456 | 55802606 | 56 |
| MXE | 12815 | ENSG00000104133 | SPG11 | chr15 | - | 44573546 | 44573745 | 44574901 | 44575041 | 44572682 | 44572820 | 44583813 | 44584558 | 44 |
| MXE | 13242 | ENSG00000137522 | RNF121 | chr11 | + | 71982760 | 71982915 | 71987003 | 71987111 | 71960749 | 71960891 | 71990596 | 71990717 | 12 |
| MXE | 8561 | ENSG00000153207 | AHCTF1 | chr1 | - | 246890955 | 246891060 | 246891778 | 246891919 | 246889965 | 246890059 | 246894658 | 246894748 | 90 |
| MXE | 16001 | ENSG00000138346 | DNA2 | chr10 | - | 68465666 | 68465812 | 68468122 | 68468306 | 68459103 | 68459235 | 68469980 | 68470163 | 30 |
| MXE | 355 | ENSG00000104824 | HNRNPL | chr19 | - | 38838893 | 38839015 | 38840095 | 38840376 | 38838396 | 38838598 | 38840487 | 38840559 | 270 |
| MXE | 16967 | ENSG00000186908 | ZDHHC17 | chr12 | + | 76815856 | 76816019 | 76821057 | 76821167 | 76815145 | 76815210 | 76822405 | 76822531 | 180 |
| MXE | 7771 | ENSG00000233355 | CHRM3-AS2 | chr1 | - | 239706321 | 239706496 | 239707533 | 239707760 | 239703891 | 239705715 | 239714176 | 239714264 | 38 |
| MXE | 13529 | ENSG00000134516 | DOCK2 | chr5 | + | 170069136 | 170069220 | 170075946 | 170076084 | 170067509 | 170067686 | 170077709 | 170077837 | 70 |
| MXE | 16491 | ENSG00000175782 | SLC35E3 | chr12 | + | 68747929 | 68748040 | 68748075 | 68748245 | 68746124 | 68746779 | 68752031 | 68752190 | 47 |
| MXE | 122 | ENSG00000188130 | MAPK12 | chr22 | - | 50248365 | 50248472 | 50249102 | 50249454 | 50248224 | 50248292 | 50249634 | 50249703 | 21 |
| MXE | 7925 | ENSG00000043514 | TRIT1 | chr1 | - | 39847547 | 39847660 | 39847985 | 39848097 | 39847219 | 39847297 | 39850118 | 39850261 | 29 |
| MXE | 15135 | ENSG00000100731 | PCNX1 | chr14 | + | 70988566 | 70988699 | 70995740 | 70995843 | 70976941 | 70978648 | 71009633 | 71009724 | 39 |
| MXE | 15449 | ENSG00000103544 | VPS35L | chr16 | + | 19573118 | 19573241 | 19575097 | 19575122 | 19569423 | 19569591 | 19579051 | 19579128 | 72 |
| MXE | 8706 | ENSG00000176407 | KCMF1 | chr2 | + | 85027888 | 85028056 | 85035015 | 85035155 | 84971160 | 84971467 | 85043563 | 85043665 | 76 |
| MXE | 15607 | ENSG00000159063 | ALG8 | chr11 | - | 78104355 | 78104453 | 78106806 | 78106946 | 78103979 | 78104052 | 78109441 | 78109581 | 30 |
| MXE | 12258 | ENSG00000160218 | TRAPPC10 | chr21 | + | 44084121 | 44084263 | 44087698 | 44087928 | 44082787 | 44083302 | 44089832 | 44089933 | 117 |
| MXE | 15450 | ENSG00000103544 | VPS35L | chr16 | + | 19591789 | 19591874 | 19601663 | 19601723 | 19581524 | 19581653 | 19608177 | 19608274 | 105 |
| MXE | 2748 | ENSG00000131966 | ACTR10 | chr14 | + | 58215204 | 58215284 | 58219693 | 58219729 | 58208998 | 58209107 | 58223621 | 58223701 | 8 |
| MXE | 3259 | ENSG00000174032 | SLC25A30 | chr13 | - | 45404326 | 45404412 | 45405882 | 45405977 | 45402274 | 45402370 | 45408926 | 45409074 | 72 |
| MXE | 10093 | ENSG00000149499 | EML3 | chr11 | - | 62605641 | 62605773 | 62605854 | 62605980 | 62605112 | 62605180 | 62606062 | 62606212 | 45 |
| MXE | 13986 | ENSG00000064313 | TAF2 | chr8 | - | 119801793 | 119802025 | 119803877 | 119804019 | 119797661 | 119797846 | 119806282 | 119806401 | 44 |
| MXE | 13636 | ENSG00000104613 | INTS10 | chr8 | + | 19830405 | 19830559 | 19832027 | 19832110 | 19826425 | 19826559 | 19833168 | 19833321 | 85 |
| MXE | 2242 | ENSG00000103978 | TMEM87A | chr15 | - | 42233212 | 42233306 | 42236319 | 42236419 | 42231191 | 42231260 | 42237431 | 42237615 | 44 |
| MXE | 16697 | ENSG00000139350 | NEDD1 | chr12 | + | 96907603 | 96907624 | 96917620 | 96917737 | 96907229 | 96907300 | 96919984 | 96920118 | 0 |
| MXE | 108 | ENSG00000100147 | CCDC134 | chr22 | + | 41810206 | 41810291 | 41813263 | 41813445 | 41809878 | 41810000 | 41813750 | 41813822 | 19 |
| MXE | 5755 | ENSG00000197386 | HTT | chr4 | + | 3099273 | 3099394 | 3103823 | 3103883 | 3086938 | 3087022 | 3105356 | 3105436 | 137 |
| MXE | 15057 | ENSG00000197563 | PIGN | chr18 | - | 62162252 | 62162329 | 62163530 | 62163656 | 62161132 | 62161385 | 62186843 | 62187038 | 28 |
| MXE | 10888 | ENSG00000160695 | VPS11 | chr11 | + | 119071595 | 119071843 | 119073197 | 119073399 | 119070233 | 119070397 | 119073799 | 119073951 | 55 |
| MXE | 13030 | ENSG00000055130 | CUL1 | chr7 | + | 148790309 | 148790441 | 148792725 | 148792818 | 148789749 | 148789826 | 148797811 | 148797859 | 89 |
| MXE | 15476 | ENSG00000156599 | ZDHHC5 | chr11 | + | 57695919 | 57696043 | 57698558 | 57699418 | 57693782 | 57693915 | 57699865 | 57700363 | 123 |
| MXE | 13799 | ENSG00000156171 | DRAM2 | chr1 | - | 111124741 | 111124881 | 111126226 | 111126294 | 111120515 | 111120693 | 111131423 | 111131568 | 46 |
| MXE | 12803 | ENSG00000104133 | SPG11 | chr15 | - | 44567423 | 44567592 | 44572682 | 44572820 | 44566216 | 44566305 | 44573546 | 44573745 | 77 |
| MXE | 7917 | ENSG00000135250 | SRPK2 | chr7 | - | 105268805 | 105268869 | 105297420 | 105297533 | 105203693 | 105203785 | 105306435 | 105306571 | 0 |
| MXE | 11363 | ENSG00000136699 | SMPD4 | chr2 | - | 130173513 | 130173656 | 130174913 | 130175000 | 130173278 | 130173354 | 130176553 | 130176637 | 23 |

| | | | | | | | | | | | | | | | | |
|---|---|---|---|---|---|---|---|---|---|---|---|---|---|---|---|---|
| MXE | 16030 | ENSG00000120438 | TCP1 | chr6 | - | 159783940 | 159784067 | 159785899 | 159785997 | 159780934 | 159781110 | 159787742 | 159787871 | 125 | [illegible] | [illegible] | [illegible] |
| MXE | 3263 | ENSG00000174032 | SLC25A30 | chr13 | - | 45405882 | 45405977 | 45408926 | 45409074 | 45404326 | 45404412 | 45418299 | 45418455 | 4 | [illegible] | [illegible] | [illegible] |
| MXE | 14248 | ENSG00000164941 | INTS8 | chr8 | + | 94856776 | 94856978 | 94859510 | 94859632 | 94853804 | 94853915 | 94865505 | 94865690 | 135 | [illegible] | [illegible] | [illegible] |
| MXE | 3106 | ENSG00000136986 | DERL1 | chr8 | - | 123019194 | 123019245 | 123021446 | 123021499 | 123014148 | 123015585 | 123022683 | 123022779 | 122 | 65 | 89 | 86 |
| MXE | 4545 | ENSG00000145348 | TBCK | chr4 | - | 106247138 | 106247287 | 106248244 | 106248306 | 106244625 | 106244764 | 106248920 | 106248982 | 22 | 15 | 9 | 18 |
| MXE | 14461 | ENSG00000137054 | POLR1E | chr9 | + | 37498090 | 37498224 | 37500839 | 37500921 | 37495889 | 37495986 | 37501712 | 37501844 | 53 | 31 | 65 | 62 |
| MXE | 11481 | ENSG00000144134 | RABL2A | chr2 | + | 113635050 | 113635130 | 113640893 | 113641005 | 113634152 | 113634232 | 113641355 | 113641450 | 15 | 1 | 3 | 1 |
| MXE | 13637 | ENSG00000104613 | INTS10 | chr8 | + | 19832027 | 19832110 | 19833168 | 19833321 | 19830405 | 19830559 | 19837051 | 19837160 | 89 | 74 | 108 | 101 |
| MXE | 14249 | ENSG00000164941 | INTS8 | chr8 | + | 94859510 | 94859632 | 94865505 | 94865690 | 94856776 | 94856978 | 94866157 | 94866191 | 108 | 77 | 68 | 109 |
| MXE | 5756 | ENSG00000197386 | HTT | chr4 | + | 3154292 | 3154419 | 3157071 | 3157199 | 3148004 | 3148207 | 3160281 | 3160392 | 49 | 21 | 37 | 37 |
| MXE | 10287 | ENSG00000132676 | DAP3 | chr1 | + | 155689094 | 155689174 | 155709772 | 155709824 | 155687984 | 155688192 | 155717005 | 155717128 | 1 | 0 | 0 | 0 |
| MXE | 8051 | ENSG00000144744 | UBA3 | chr3 | - | 69062076 | 69062179 | 69062981 | 69063137 | 69061582 | 69061927 | 69063438 | 69063503 | 278 | 167 | 213 | 191 |
| MXE | 13336 | ENSG00000147459 | DOCK5 | chr8 | + | 25368191 | 25368250 | 25368570 | 25368725 | 25366869 | 25366970 | 25369555 | 25369641 | 46 | 48 | 45 | 49 |
| MXE | 12255 | ENSG00000160218 | TRAPPC10 | chr21 | + | 44059102 | 44059214 | 44074323 | 44074470 | 44055697 | 44055893 | 44075038 | 44075153 | 190 | 99 | 116 | 157 |
| MXE | 10242 | ENSG00000157483 | MYO1E | chr15 | - | 59178392 | 59178537 | 59188117 | 59188216 | 59174125 | 59174240 | 59195460 | 59195567 | 21 | 18 | 14 | 6 |
| MXE | 9160 | ENSG00000162909 | CAPN2 | chr1 | + | 223745305 | 223745439 | 223746996 | 223747165 | 223744099 | 223744218 | 223749038 | 223749122 | 161 | 83 | 101 | 165 |
| MXE | 16509 | ENSG00000135720 | DYNC1LI2 | chr16 | - | 66727687 | 66727805 | 66728200 | 66728242 | 66725827 | 66725944 | 66729039 | 66729099 | 29 | 34 | 26 | 19 |
| MXE | 15444 | ENSG00000110274 | CEP164 | chr11 | + | 117395122 | 117395191 | 117395546 | 117395722 | 117394919 | 117395003 | 117396053 | 117396180 | 14 | 11 | 14 | 26 |
| MXE | 7035 | ENSG00000095564 | BTAF1 | chr10 | + | 91964076 | 91964201 | 91966636 | 91966757 | 91962537 | 91962678 | 91980453 | 91980558 | 290 | 237 | 273 | 309 |
| MXE | 2513 | ENSG00000174373 | RALGAPA1 | chr14 | - | 35762709 | 35762753 | 35770941 | 35770999 | 35760828 | 35761006 | 35775005 | 35775055 | 28 | 17 | 37 | 40 |
| MXE | 15722 | ENSG00000109917 | ZPR1 | chr11 | - | 116782157 | 116782244 | 116782918 | 116783029 | 116779771 | 116779837 | 116783529 | 116783619 | 129 | 85 | 102 | 143 |
| MXE | 16545 | ENSG00000028203 | VEZT | chr12 | + | 95257149 | 95257239 | 95262905 | 95263081 | 95256549 | 95256622 | 95266356 | 95266632 | 17 | 17 | 17 | 12 |
| MXE | 15441 | ENSG00000110274 | CEP164 | chr11 | + | 117373750 | 117373831 | 117375707 | 117375791 | 117371079 | 117371466 | 117380613 | 117380705 | 30 | 26 | 20 | 24 |
| MXE | 16155 | ENSG00000108094 | CUL2 | chr10 | - | 35049682 | 35049765 | 35054433 | 35054539 | 35044771 | 35044868 | 35060873 | 35060924 | 156 | 132 | 124 | 134 |
| MXE | 2301 | ENSG00000140598 | EFL1 | chr15 | - | 82151464 | 82152423 | 82157712 | 82157860 | 82138657 | 82138842 | 82163852 | 82163984 | 35 | 32 | 29 | 19 |
| MXE | 5107 | ENSG00000159147 | DONSON | chr21 | - | 33583487 | 33583666 | 33584589 | 33584768 | 33582164 | 33582246 | 33585977 | 33586181 | 10 | 17 | 23 | 20 |
| MXE | 8562 | ENSG00000153207 | AHCTF1 | chr1 | - | 246891778 | 246891919 | 246894658 | 246894748 | 246890955 | 246891060 | 246895834 | 246895925 | 47 | 37 | 63 | 57 |
| MXE | 6368 | ENSG00000120008 | WDR11 | chr10 | + | 120890715 | 120890887 | 120900028 | 120900137 | 120889894 | 120890009 | 120901035 | 120901098 | 129 | 92 | 94 | 88 |
| MXE | 16807 | ENSG00000123349 | PFDN5 | chr12 | + | 53295838 | 53295941 | 53296453 | 53296551 | 53295541 | 53295639 | 53297849 | 53297924 | 671 | 420 | 540 | 556 |
| MXE | 10295 | ENSG00000132676 | DAP3 | chr1 | + | 155727607 | 155727738 | 155729041 | 155729122 | 155725926 | 155726019 | 155729207 | 155729366 | 223 | 195 | 195 | 210 |
| MXE | 9162 | ENSG00000162909 | CAPN2 | chr1 | + | 223752795 | 223752956 | 223755479 | 223755649 | 223751996 | 223752071 | 223757368 | 223757380 | 527 | 383 | 398 | 479 |
| MXE | 14245 | ENSG00000164941 | INTS8 | chr8 | + | 94838493 | 94838618 | 94841490 | 94841591 | 94836523 | 94836631 | 94842346 | 94842488 | 6 | 3 | 6 | 7 |
| MXE | 14719 | ENSG00000099381 | SETD1A | chr16 | + | 30967500 | 30967588 | 30969304 | 30969462 | 30966883 | 30967060 | 30969601 | 30969689 | 56 | 34 | 74 | 68 |
| MXE | 13520 | ENSG00000134516 | DOCK2 | chr5 | + | 169669287 | 169669328 | 169671077 | 169671174 | 169654402 | 169654486 | 169674296 | 169674445 | 62 | 21 | 41 | 27 |
| MXE | 10296 | ENSG00000132676 | DAP3 | chr1 | + | 155731943 | 155732033 | 155736945 | 155737063 | 155731355 | 155731415 | 155738156 | 155738529 | 134 | 60 | 185 | 129 |
| MXE | 15610 | ENSG00000159063 | ALG8 | chr11 | - | 78106806 | 78106946 | 78112649 | 78112770 | 78104355 | 78104453 | 78113885 | 78113989 | 57 | 33 | 31 | 31 |
| MXE | 14243 | ENSG00000164941 | INTS8 | chr8 | + | 94836523 | 94836631 | 94838493 | 94838618 | 94831991 | 94832174 | 94841490 | 94841591 | 432 | 372 | 332 | 432 |
| MXE | 8049 | ENSG00000144744 | UBA3 | chr3 | - | 69056611 | 69056693 | 69056778 | 69056815 | 69056182 | 69056283 | 69057255 | 69057309 | 23 | 19 | 15 | 22 |
| MXE | 12198 | ENSG00000168014 | C2CD3 | chr11 | - | 74085521 | 74085886 | 74090812 | 74090936 | 74084880 | 74084970 | 74092415 | 74092588 | 32 | 20 | 15 | 23 |
| MXE | 15615 | ENSG00000159063 | ALG8 | chr11 | - | 78109441 | 78109581 | 78113885 | 78113989 | 78106806 | 78106946 | 78114265 | 78114392 | 66 | 55 | 75 | 59 |
| MXE | 6466 | ENSG00000088854 | C20orf194 | chr20 | - | 3298028 | 3298175 | 3304439 | 3304543 | 3295493 | 3296949 | 3315032 | 3315120 | 9 | 12 | 14 | 8 |
| MXE | 1801 | ENSG00000197818 | SLC9A8 | chr20 | + | 49874704 | 49874821 | 49877980 | 49878063 | 49864738 | 49864844 | 49880923 | 49881035 | 54 | 33 | 51 | 38 |
| MXE | 2816 | ENSG00000100479 | POLE2 | chr14 | - | 49666329 | 49666413 | 49669523 | 49669598 | 49665093 | 49665163 | 49674122 | 49674216 | 21 | 19 | 11 | 22 |
| MXE | 12401 | ENSG00000084070 | SMAP2 | chr1 | + | 40362299 | 40362436 | 40406735 | 40406869 | 40344849 | 40344910 | 40408652 | 40408738 | 61 | 45 | 19 | 43 |
| MXE | 4544 | ENSG00000108591 | DRG2 | chr17 | + | 18099016 | 18099077 | 18099632 | 18099723 | 18098269 | 18098359 | 18100362 | 18100435 | 28 | 9 | 20 | 23 |
| MXE | 16806 | ENSG00000123349 | PFDN5 | chr12 | + | 53295838 | 53295941 | 53297849 | 53297924 | 53295524 | 53295639 | 53298044 | 53298150 | 669 | 407 | 530 | 553 |
| MXE | 9883 | ENSG00000148690 | FRA10AC1 | chr10 | - | 93692008 | 93692093 | 93692645 | 93692729 | 93687403 | 93687449 | 93694860 | 93694937 | 37 | 60 | 43 | 68 |
| MXE | 12347 | ENSG00000198522 | GPN1 | chr2 | + | 27640042 | 27640125 | 27641239 | 27641279 | 27638884 | 27639031 | 27642428 | 27642519 | 29 | 18 | 27 | 35 |
| MXE | 4722 | ENSG00000163913 | IFT122 | chr3 | + | 129451913 | 129451998 | 129458598 | 129458677 | 129449870 | 129449937 | 129461227 | 129461304 | 20 | 11 | 14 | 11 |
| MXE | 13512 | ENSG00000171311 | EXOSC1 | chr10 | - | 97438669 | 97438703 | 97443236 | 97443311 | 97437699 | 97437750 | 97445731 | 97445847 | 16 | 9 | 14 | 10 |
| MXE | 2508 | ENSG00000174373 | RALGAPA1 | chr14 | - | 35627089 | 35627951 | 35634573 | 35634757 | 35625360 | 35625432 | 35635463 | 35635598 | 65 | 43 | 64 | 68 |
| MXE | 256 | ENSG00000198089 | SFI1 | chr22 | + | 31556941 | 31557059 | 31561289 | 31561392 | 31550253 | 31550348 | 31573057 | 31573214 | 67 | 47 | 74 | 69 |
| MXE | 7255 | ENSG00000154358 | OBSCN | chr1 | + | 228249964 | 228250240 | 228256650 | 228256926 | 228246527 | 228246803 | 228264115 | 228264391 | 10 | 14 | 46 | 3 |
| MXE | 12199 | ENSG00000168014 | C2CD3 | chr11 | - | 74085617 | 74085886 | 74090812 | 74090936 | 74084880 | 74084970 | 74092415 | 74092588 | 32 | 20 | 15 | 23 |
| MXE | 9161 | ENSG00000162909 | CAPN2 | chr1 | + | 223746996 | 223747165 | 223749038 | 223749122 | 223745305 | 223745439 | 223750889 | 223750975 | 324 | 165 | 214 | 326 |
| MXE | 3107 | ENSG00000136986 | DERL1 | chr8 | - | 123022683 | 123022779 | 123024985 | 123025050 | 123021446 | 123021499 | 123030604 | 123030716 | 60 | 29 | 56 | 48 |
| MXE | 15455 | ENSG00000103544 | VPS35L | chr16 | + | 19691352 | 19691471 | 19699501 | 19699648 | 19682224 | 19682390 | 19700377 | 19700687 | 45 | 47 | 53 | 58 |
| MXE | 12826 | ENSG00000104133 | SPG11 | chr15 | - | 44622219 | 44622347 | 44622727 | 44622799 | 44621758 | 44621934 | 44626330 | 44626507 | 135 | 116 | 129 | 134 |
| MXE | 9723 | ENSG00000185420 | SMYD3 | chr1 | - | 246335366 | 246335474 | 246355030 | 246355094 | 246330479 | 246330537 | 246507053 | 246507144 | 5 | 1 | 4 | 4 |
| MXE | 15613 | ENSG00000159063 | ALG8 | chr11 | - | 78109441 | 78109581 | 78112649 | 78112770 | 78106806 | 78106946 | 78113885 | 78113989 | 56 | 33 | 31 | 31 |
| MXE | 15989 | ENSG00000075945 | KIFAP3 | chr1 | - | 170039232 | 170039288 | 170046711 | 170046866 | 170038289 | 170038431 | 170055304 | 170055436 | 90 | 74 | 71 | 67 |
| MXE | 16788 | ENSG00000033030 | ZCCHC8 | chr12 | - | 122482634 | 122482695 | 122483278 | 122483563 | 122481944 | 122482087 | 122489385 | 122489463 | 60 | 52 | 51 | 64 |
| MXE | 7711 | ENSG00000113318 | MSH3 | chr5 | + | 80775693 | 80775758 | 80778719 | 80778836 | 80768834 | 80769003 | 80787564 | 80787672 | 35 | 17 | 14 | 16 |
| MXE | 264 | ENSG00000198089 | SFI1 | chr22 | + | 31602606 | 31602785 | 31606327 | 31606430 | 31602211 | 31602293 | 31607936 | 31608033 | 84 | 87 | 91 | 116 |
| MXE | 4463 | ENSG00000141452 | RMC1 | chr18 | + | 23506969 | 23507054 | 23509192 | 23509279 | 23504370 | 23504447 | 23515855 | 23515996 | 32 | 31 | 46 | 49 |
| MXE | 126 | ENSG00000093009 | CDC45 | chr22 | + | 19514748 | 19514887 | 19514964 | 19515048 | 19508529 | 19508691 | 19516526 | 19516645 | 1 | 2 | 4 | 6 |
| MXE | 13846 | ENSG00000221914 | PPP2R2A | chr8 | + | 26338889 | 26338987 | 26354467 | 26354633 | 26293571 | 26293740 | 26360168 | 26360281 | 49 | 54 | 56 | 42 |
| MXE | 15454 | ENSG00000103544 | VPS35L | chr16 | + | 19644885 | 19644949 | 19647783 | 19647882 | 19642395 | 19642476 | 19651975 | 19652090 | 30 | 27 | 38 | 26 |
| MXE | 266 | ENSG00000198089 | SFI1 | chr22 | + | 31604308 | 31604404 | 31604868 | 31604945 | 31603743 | 31603819 | 31606327 | 31606430 | 29 | 22 | 39 | 44 |
| MXE | 1716 | ENSG00000103194 | USP10 | chr16 | + | 84744632 | 84745673 | 84759890 | 84759946 | 84733434 | 84733503 | 84760171 | 84760275 | 4 | 2 | 2 | 0 |
| MXE | 7252 | ENSG00000154358 | OBSCN | chr1 | + | 228219323 | 228219599 | 228224458 | 228224734 | 228217012 | 228217288 | 228243182 | 228243458 | 15 | 3 | 32 | 11 |
| MXE | 8597 | ENSG00000185946 | RNPC3 | chr1 | + | 103543298 | 103543447 | 103544940 | 103545102 | 103541349 | 103541475 | 103546247 | 103546342 | 35 | 63 | 31 | 48 |
| MXE | 7748 | ENSG00000123064 | DDX54 | chr12 | - | 113163131 | 113163274 | 113164066 | 113164285 | 113162931 | 113163045 | 113165643 | 113165717 | 65 | 30 | 61 | 71 |
| MXE | 7667 | ENSG00000114933 | INO80D | chr2 | - | 206062798 | 206063045 | 206063150 | 206063244 | 206056197 | 206056943 | 206085900 | 206086174 | 11 | 6 | 8 | 9 |
| MXE | 1042 | ENSG00000170004 | CHD3 | chr17 | + | 7907890 | 7908019 | 7908696 | 7908829 | 7907600 | 7907702 | 7909142 | 7909338 | 14 | 9 | 33 | 32 |
| MXE | 6630 | ENSG00000162946 | DISC1 | chr1 | + | 231693825 | 231694805 | 231749925 | 231750076 | 231626814 | 231626934 | 231767139 | 231767269 | 6 | 4 | 11 | 0 |
| MXE | 15343 | ENSG00000099783 | HNRNPM | chr19 | + | 8471325 | 8471427 | 8473663 | 8473708 | 8468773 | 8468834 | 8474166 | 8474244 | 304 | 217 | 234 | 317 |
| MXE | 251 | ENSG00000198089 | SFI1 | chr22 | + | 31546860 | 31546971 | 31550253 | 31550348 | 31508258 | 31508376 | 31556941 | 31557059 | 16 | 19 | 24 | 18 |
| MXE | 13638 | ENSG00000104613 | INTS10 | chr8 | + | 19832027 | 19832110 | 19837051 | 19837160 | 19830405 | 19830559 | 19842847 | 19842927 | 88 | 74 | 109 | 100 |
| MXE | 8474 | ENSG00000163463 | KRTCAP2 | chr1 | - | 155169790 | 155169857 | 155172564 | 155172892 | 155169408 | 155169560 | 155173220 | 155173251 | 94 | 17 | 62 | 76 |
| MXE | 4721 | ENSG00000163913 | IFT122 | chr3 | + | 129451913 | 129451998 | 129458598 | 129458677 | 129449870 | 129449937 | 129461227 | 129461304 | 13 | 5 | 8 | 7 |
| MXE | 670 | ENSG00000064932 | SBNO2 | chr19 | - | 1122893 | 1123045 | 1123533 | 1123639 | 1122657 | 1122791 | 1123941 | 1124022 | 47 | 21 | 35 | 42 |
| MXE | 16812 | ENSG00000123349 | PFDN5 | chr12 | + | 53296218 | 53296275 | 53296429 | 53296551 | 53295838 | 53295941 | 53297849 | 53297924 | 626 | 367 | 549 | 516 |
| MXE | 3962 | ENSG00000083312 | TNPO1 | chr5 | + | 72855773 | 72855923 | 72861807 | 72861914 | 72851243 | 72851319 | 72865595 | 72865729 | 200 | 136 | 132 | 150 |
| MXE | 15152 | ENSG00000153339 | TRAPPC8 | chr18 | - | 31855659 | 31855807 | 31857539 | 31857982 | 31853848 | 31853945 | 31864626 | 31864781 | 63 | 40 | 43 | 37 |
| MXE | 1804 | ENSG00000122390 | NAA60 | chr16 | + | 3479470 | 3479600 | 3482501 | 3482598 | 3476221 | 3476337 | 3483362 | 3483597 | 145 | 76 | 116 | 145 |
| MXE | 13524 | ENSG00000134516 | DOCK2 | chr5 | + | 169714027 | 169714211 | 169714359 | 169714457 | 169712119 | 169712223 | 169716212 | 169716302 | 1366 | 943 | 1025 | 1220 |
| MXE | 15649 | ENSG00000167986 | DDB1 | chr11 | - | 61316491 | 61316567 | 61321594 | 61321697 | 61314883 | 61316393 | 61322295 | 61322412 | 170 | 100 | 132 | 160 |
| MXE | 12402 | ENSG00000091732 | ZC3HC1 | chr7 | - | 130023510 | 130023723 | 130024262 | 130024506 | 130022318 | 130022525 | 130026157 | 130026312 | 24 | 12 | 17 | 14 |

| | | | | | | | | | | | | | | | | | | | | | | | | | | | | | | | | | | | | | | | | | | | | | | | | |
|---|---|---|---|---|---|---|---|---|---|---|---|---|---|---|---|---|---|---|---|---|---|---|---|---|---|---|---|---|---|---|---|---|---|---|---|---|---|---|---|---|---|---|---|---|---|---|---|---|
| MXE | 15474 | ENSG00000156599 | ZDHHC5 | chr11 | + | 57682421 | 57682543 | 57688507 | 57688665 | 57681478 | 57681698 | 57690030 | 57690437 | 0 | 1 | 2 | 2 | 0 | 163 | 58 | 157 | 127 | 151 | 0 | 0 | 0 | 0 | 0 | 178 | 152 | 94 | 107 | 141 | 271 | 298 | 1 | 1 | 0 | 0.019 | 0.014 | 0.017 | 0 | 0 | 0 | 0 | 0 | 0 | 0.01 |
| MXE | 13509 | ENSG00000171311 | EXOSC1 | chr10 | - | 97438662 | 97438703 | 97441170 | 97441259 | 97437699 | 97437750 | 97445731 | 97445847 | 1 | 1 | 1 | 2 | 0 | 7 | 1 | 0 | 0 | 4 | 0 | 1 | 0 | 0 | 2 | 11 | 0 | 0 | 2 | 0 | 238 | 190 | 1 | 1 | 0.102 | 0.444 | 1 | 1 | 0 | 0 | 1 | NA | 0 | 1 | 0.009 |
| MXE | 16787 | ENSG00000033030 | ZCCHC8 | chr12 | - | 122482634 | 122482695 | 122483278 | 122483344 | 122481944 | 122482087 | 122489385 | 122489463 | 2 | 0 | 0 | 0 | 2 | 75 | 60 | 64 | 95 | 109 | 0 | 0 | 0 | 0 | 0 | 50 | 57 | 54 | 72 | 0 | 215 | 210 | 1 | 1 | 0.025 | 0 | 0 | 0 | 0.018 | 0 | 0 | 0 | 0 | NA | 0.009 |
| MXE | 9754 | ENSG00000132953 | XPO4 | chr13 | - | 20800830 | 20800990 | 20807456 | 20807634 | 20800155 | 20800325 | 20808435 | 20808581 | 57 | 49 | 65 | 36 | 74 | 69 | 60 | 77 | 60 | 124 | 10 | 38 | 27 | 8 | 0 | 9 | 43 | 68 | 13 | 0 | 298 | 298 | 1 | 1 | 0.452 | 0.45 | 0.458 | 0.375 | 0.374 | 0.526 | 0.469 | 0.284 | 0.381 | NA | 0.007 |
| MXE | 9272 | ENSG00000103168 | TAF1C | chr16 | - | 84180867 | 84180995 | 84181042 | 84181186 | 84180169 | 84180344 | 84181327 | 84181663 | 34 | 12 | 29 | 36 | 14 | 0 | 0 | 0 | 0 | 0 | 36 | 35 | 41 | 18 | 50 | 0 | 1 | 0 | 0 | 0 | 293 | 277 | 1 | 1 | 1 | 1 | 1 | 1 | 1 | 1 | 0.971 | 1 | 1 | 1 | 0.006 |
| MXE | 840 | ENSG00000105576 | TNPO2 | chr19 | - | 12705491 | 12705599 | 12705681 | 12705830 | 12705239 | 12705398 | 12706195 | 12706367 | 2 | 0 | 0 | 0 | 0 | 29 | 15 | 28 | 20 | 22 | 0 | 0 | 0 | 0 | 3 | 41 | 14 | 20 | 19 | 84 | 298 | 257 | 1 | 1 | 0.056 | 0 | 0 | 0 | 0 | 0 | 0 | 0 | 0 | 0.03 | 0.005 |
| MXE | 1717 | ENSG00000103194 | USP10 | chr16 | + | 84744632 | 84745673 | 84762988 | 84763088 | 84733434 | 84733503 | 84764085 | 84764263 | 5 | 0 | 0 | 0 | 0 | 103 | 68 | 70 | 108 | 126 | 0 | 0 | 1 | 0 | 0 | 89 | 81 | 71 | 86 | 149 | 298 | 249 | 1 | 1 | 0.039 | 0 | 0 | 0 | 0 | 0 | 0 | 0.012 | 0 | 0 | 0.005 |
| MXE | 13872 | ENSG00000187735 | TCEA1 | chr8 | - | 53993667 | 53993755 | 53999944 | 54000050 | 53988113 | 53988259 | 54022062 | 54022448 | 6 | 3 | 3 | 3 | 1 | 461 | 403 | 378 | 474 | 604 | 0 | 4 | 0 | 0 | 0 | 176 | 371 | 484 | 256 | 53 | 255 | 237 | 1 | 1 | 0.012 | 0.007 | 0.007 | 0.006 | 0.002 | 0 | 0.01 | 0 | 0 | 0 | 0.005 |
| MXE | 12811 | ENSG00000104133 | SPG11 | chr15 | - | 44572682 | 44572820 | 44573546 | 44573745 | 44570524 | 44570658 | 44583813 | 44584558 | 3 | 0 | 0 | 0 | 1 | 101 | 44 | 66 | 98 | 85 | 0 | 0 | 1 | 0 | 0 | 21 | 92 | 55 | 39 | 14 | 298 | 287 | 1 | 1 | 0.028 | 0 | 0 | 0 | 0.011 | 0 | 0 | 0.017 | 0 | 0 | 0.004 |
| MXE | 7256 | ENSG00000154358 | OBSCN | chr1 | + | 228268529 | 228268805 | 228273271 | 228273547 | 228264115 | 228264391 | 228273769 | 228274048 | 10 | 7 | 10 | 8 | 6 | 1 | 2 | 9 | 2 | 2 | 7 | 5 | 1 | 0 | 0 | 0 | 0 | 0 | 0 | 26 | 298 | 298 | 1 | 1 | 0.909 | 0.778 | 0.526 | 0.8 | 0.75 | 1 | 1 | 1 | NA | 0 | 0.003 |
| MXE | 8559 | ENSG00000153207 | AHCTF1 | chr1 | - | 246863923 | 246864116 | 246865254 | 246865333 | 246861958 | 246862153 | 246867243 | 246867351 | 1 | 1 | 0 | 0 | 0 | 218 | 154 | 198 | 194 | 340 | 0 | 0 | 0 | 0 | 0 | 24 | 47 | 119 | 7 | 18 | 228 | 298 | 1 | 1 | 0.006 | 0.008 | 0 | 0 | 0 | 0 | 0 | 0 | 0 | 0 | 0.003 |
| MXE | 16805 | ENSG00000123349 | PFDN5 | chr12 | + | 53295838 | 53295941 | 53296429 | 53296551 | 53295524 | 53295639 | 53298044 | 53298150 | 669 | 407 | 530 | 553 | 332 | 3 | 1 | 4 | 2 | 0 | 701 | 454 | 363 | 357 | 826 | 12 | 3 | 0 | 0 | 9 | 252 | 271 | 1 | 1 | 0.996 | 0.998 | 0.993 | 0.997 | 1 | 0.984 | 0.994 | 1 | 1 | 0.99 | 0.003 |
| MXE | 286 | ENSG00000100325 | ASCC2 | chr22 | - | 29825086 | 29825257 | 29832244 | 29832342 | 29822334 | 29822464 | 29838177 | 29838274 | 0 | 0 | 1 | 0 | 0 | 108 | 49 | 102 | 98 | 104 | 0 | 0 | 0 | 0 | 0 | 97 | 117 | 88 | 116 | 86 | 247 | 298 | 1 | 1 | 0 | 0 | 0.012 | 0 | 0 | 0 | 0 | 0 | 0 | 0 | 0.002 |
| MXE | 3628 | ENSG00000104671 | DCTN6 | chr8 | + | 30165715 | 30165901 | 30175084 | 30175190 | 30164110 | 30164175 | 30177125 | 30177214 | 0 | 0 | 1 | 0 | 0 | 100 | 89 | 80 | 110 | 120 | 0 | 0 | 0 | 0 | 0 | 28 | 145 | 193 | 112 | 12 | 298 | 255 | 1 | 1 | 0 | 0 | 0.011 | 0 | 0 | 0 | 0 | 0 | 0 | 0 | 0.002 |
| MXE | 12804 | ENSG00000104133 | SPG11 | chr15 | - | 44567423 | 44567592 | 44572682 | 44572820 | 44566216 | 44566305 | 44583813 | 44584558 | 1 | 0 | 0 | 0 | 0 | 107 | 58 | 60 | 74 | 58 | 0 | 0 | 0 | 0 | 0 | 37 | 55 | 52 | 60 | 38 | 287 | 298 | 1 | 1 | 0.01 | 0 | 0 | 0 | 0 | 0 | 0 | 0 | 0 | 0 | 0.002 |
| MXE | 284 | ENSG00000100325 | ASCC2 | chr22 | - | 29822334 | 29822464 | 29825086 | 29825257 | 29816005 | 29816073 | 29832244 | 29832342 | 1 | 0 | 3 | 1 | 3 | 103 | 44 | 77 | 93 | 138 | 6 | 0 | 1 | 0 | 0 | 109 | 81 | 48 | 109 | 87 | 298 | 279 | 1 | 1 | 0.009 | 0 | 0.035 | 0.01 | 0.02 | 0.049 | 0 | 0.019 | 0 | 0 | 0.001 |
| MXE | 7792 | ENSG00000174780 | SRP72 | chr4 | + | 56478591 | 56478649 | 56483138 | 56483270 | 56476670 | 56476702 | 56484735 | 56484864 | 2 | 3 | 0 | 1 | 1 | 445 | 235 | 332 | 351 | 551 | 0 | 0 | 5 | 0 | 0 | 33 | 307 | 281 | 141 | 41 | 207 | 281 | 1 | 1 | 0.006 | 0.017 | 0 | 0.004 | 0.002 | 0 | 0 | 0.024 | 0 | 0 | 0.001 |
| MXE | 10634 | ENSG00000082258 | CCNT2 | chr2 | + | 134942611 | 134942674 | 134946100 | 134946146 | 134939001 | 134939062 | 134947735 | 134947899 | 30 | 13 | 40 | 48 | 74 | 3 | 9 | 12 | 5 | 0 | 0 | 23 | 29 | 16 | 8 | 0 | 2 | 2 | 0 | 9 | 212 | 195 | 1 | 1 | 0.902 | 0.571 | 0.754 | 0.898 | 1 | NA | 0.914 | 0.93 | 1 | 0.45 | 0.001 |
| MXE | 14240 | ENSG00000164941 | INTS8 | chr8 | + | 94827721 | 94827793 | 94828974 | 94829026 | 94824892 | 94825067 | 94831991 | 94832001 | 0 | 0 | 1 | 0 | 0 | 180 | 174 | 151 | 157 | 126 | 0 | 0 | 0 | 0 | 0 | 51 | 126 | 109 | 78 | 18 | 221 | 201 | 1 | 1 | 0 | 0 | 0.006 | 0 | 0 | 0 | 0 | 0 | 0 | 0 | 0.001 |
| MXE | 16814 | ENSG00000123349 | PFDN5 | chr12 | + | 53296243 | 53296275 | 53296429 | 53296551 | 53295838 | 53295941 | 53297849 | 53297924 | 1622 | 887 | 1400 | 1338 | 1648 | 14 | 8 | 19 | 8 | 11 | 1972 | 1168 | 788 | 1165 | 1995 | 6 | 12 | 6 | 26 | 22 | 181 | 271 | 1 | 1 | 0.994 | 0.994 | 0.991 | 0.996 | 0.996 | 0.998 | 0.993 | 0.995 | 0.985 | 0.993 | 0.001 |
| MXE | 8927 | ENSG00000117262 | GPR89A | chr1 | + | 145645646 | 145645743 | 145646183 | 145646272 | 145643868 | 145643978 | 145647174 | 145647267 | 1 | 0 | 0 | 2 | 0 | 0 | 3 | 3 | 0 | 0 | 0 | 7 | 0 | 2 | 0 | 13 | 0 | 1 | 0 | 0 | 246 | 238 | 1 | 1 | 1 | 0 | 0 | 1 | NA | 0 | 1 | 0 | 1 | NA | 0 |
| MXE | 16756 | ENSG00000175582 | RAB6A | chr11 | - | 73716250 | 73716362 | 73718784 | 73718890 | 73679653 | 73679720 | 73720845 | 73720899 | 178 | 127 | 107 | 160 | 217 | 1 | 2 | 1 | 2 | 0 | 210 | 139 | 114 | 187 | 213 | 0 | 0 | 3 | 3 | 0 | 255 | 261 | 1 | 1 | 0.995 | 0.985 | 0.991 | 0.988 | 1 | 1 | 1 | 0.975 | 0.985 | 1 | 0 |
| MXE | 151 | ENSG00000100030 | MAPK1 | chr22 | - | 21788693 | 21788808 | 21799011 | 21799128 | 21772872 | 21772982 | 21805849 | 21806039 | 300 | 159 | 221 | 232 | 216 | 1 | 0 | 0 | 0 | 0 | 289 | 207 | 206 | 223 | 307 | 0 | 0 | 0 | 0 | 0 | 266 | 264 | 1 | 1 | 0.997 | 1 | 1 | 1 | 1 | 1 | 1 | 1 | 1 | 1 | -0.001 |
| MXE | 16813 | ENSG00000123349 | PFDN5 | chr12 | + | 53296243 | 53296275 | 53296429 | 53296467 | 53295838 | 53295941 | 53297849 | 53297924 | 1622 | 887 | 1400 | 1338 | 1648 | 0 | 1 | 8 | 0 | 0 | 1972 | 1168 | 788 | 1165 | 1995 | 0 | 2 | 0 | 0 | 0 | 181 | 187 | 1 | 1 | 1 | 0.999 | 0.994 | 1 | 1 | 1 | 0.998 | 1 | 1 | 1 | -0.001 |
| MXE | 15340 | ENSG00000099783 | HNRNPM | chr19 | + | 8462528 | 8462589 | 8463496 | 8463500 | 8455404 | 8455574 | 8463592 | 8463686 | 325 | 231 | 176 | 280 | 258 | 0 | 0 | 2 | 0 | 2 | 514 | 252 | 228 | 426 | 378 | 0 | 1 | 0 | 4 | 0 | 210 | 153 | 1 | 1 | 1 | 1 | 0.985 | 1 | 0.989 | 1 | 0.995 | 1 | 0.987 | 1 | -0.002 |
| MXE | 16815 | ENSG00000123349 | PFDN5 | chr12 | + | 53296429 | 53296551 | 53297849 | 53297924 | 53295524 | 53295639 | 53298044 | 53298150 | 3 | 1 | 4 | 2 | 0 | 916 | 481 | 863 | 714 | 745 | 12 | 3 | 0 | 0 | 9 | 1102 | 587 | 534 | 583 | 1073 | 271 | 224 | 1 | 1 | 0.003 | 0.002 | 0.004 | 0.002 | 0 | 0.009 | 0.004 | 0 | 0 | 0.007 | -0.002 |
| MXE | 13835 | ENSG00000145041 | DCAF1 | chr3 | - | 51466805 | 51466876 | 51470928 | 51471005 | 51463113 | 51463227 | 51483718 | 51483836 | 74 | 38 | 63 | 63 | 45 | 1 | 0 | 0 | 0 | 0 | 132 | 43 | 47 | 82 | 43 | 0 | 0 | 0 | 0 | 0 | 226 | 220 | 1 | 1 | 0.986 | 1 | 1 | 1 | 1 | 1 | 1 | 1 | 1 | 1 | -0.003 |
| MXE | 15650 | ENSG00000167986 | DDB1 | chr11 | - | 61326778 | 61326893 | 61329362 | 61329584 | 61323978 | 61324137 | 61329957 | 61330074 | 127 | 75 | 122 | 143 | 141 | 0 | 0 | 0 | 2 | 0 | 149 | 165 | 91 | 100 | 207 | 0 | 0 | 0 | 0 | 0 | 298 | 264 | 1 | 1 | 1 | 1 | 1 | 0.984 | 1 | 1 | 1 | 1 | 1 | 1 | -0.003 |
| MXE | 16811 | ENSG00000123349 | PFDN5 | chr12 | + | 53296218 | 53296275 | 53296429 | 53296467 | 53295838 | 53295941 | 53297849 | 53297924 | 626 | 367 | 549 | 516 | 649 | 0 | 1 | 8 | 0 | 0 | 848 | 569 | 331 | 482 | 930 | 0 | 2 | 0 | 0 | 0 | 206 | 187 | 1 | 1 | 1 | 0.997 | 0.984 | 1 | 1 | 1 | 0.996 | 1 | 1 | 1 | -0.003 |
| MXE | 3444 | ENSG00000111641 | NOP2 | chr12 | - | 6564149 | 6564248 | 6566100 | 6566336 | 6563890 | 6563946 | 6566528 | 6566617 | 197 | 146 | 227 | 162 | 162 | 0 | 0 | 2 | 0 | 6 | 258 | 139 | 219 | 166 | 317 | 0 | 1 | 2 | 1 | 0 | 298 | 248 | 1 | 1 | 1 | 1 | 0.99 | 1 | 0.957 | 1 | 0.991 | 0.989 | 0.993 | 1 | -0.005 |
| MXE | 12807 | ENSG00000104133 | SPG11 | chr15 | - | 44570524 | 44570658 | 44572682 | 44572820 | 44569397 | 44569505 | 44583813 | 44584558 | 0 | 0 | 1 | 0 | 0 | 77 | 49 | 59 | 73 | 68 | 0 | 0 | 4 | 0 | 0 | 30 | 91 | 87 | 51 | 37 | 287 | 283 | 1 | 1 | 0 | 0 | 0.016 | 0 | 0 | 0 | 0 | 0.043 | 0 | 0 | -0.005 |
| MXE | 12813 | ENSG00000104133 | SPG11 | chr15 | - | 44573546 | 44573745 | 44574901 | 44575041 | 44567423 | 44567592 | 44583813 | 44584558 | 44 | 17 | 28 | 39 | 25 | 1 | 0 | 0 | 0 | 1 | 14 | 26 | 29 | 33 | 12 | 0 | 0 | 1 | 0 | 0 | 289 | 298 | 1 | 1 | 0.978 | 1 | 1 | 1 | 0.963 | 1 | 1 | 0.968 | 1 | 1 | -0.005 |
| MXE | 16808 | ENSG00000123349 | PFDN5 | chr12 | + | 53295838 | 53295941 | 53296453 | 53296551 | 53295541 | 53295639 | 53298044 | 53298150 | 669 | 407 | 530 | 553 | 332 | 3 | 0 | 7 | 10 | 0 | 701 | 454 | 363 | 357 | 826 | 0 | 3 | 2 | 0 | 0 | 252 | 247 | 1 | 1 | 0.995 | 1 | 0.987 | 0.982 | 1 | 1 | 0.993 | 0.994 | 1 | 1 | -0.005 |
| MXE | 710 | ENSG00000173418 | NAA20 | chr20 | + | 20025668 | 20025767 | 20026783 | 20026919 | 20017358 | 20017449 | 20032507 | 20032653 | 0 | 0 | 0 | 0 | 0 | 88 | 60 | 70 | 97 | 91 | 0 | 0 | 0 | 4 | 0 | 144 | 70 | 41 | 125 | 114 | 248 | 285 | 1 | 1 | 0 | 0 | 0 | 0 | 0 | 0 | 0 | 0 | 0.035 | 0 | -0.007 |
| MXE | 3443 | ENSG00000111641 | NOP2 | chr12 | - | 6564149 | 6564248 | 6566100 | 6566324 | 6563890 | 6563946 | 6566528 | 6566617 | 149 | 115 | 188 | 126 | 105 | 0 | 0 | 2 | 0 | 6 | 201 | 112 | 163 | 122 | 215 | 0 | 1 | 2 | 1 | 0 | 298 | 248 | 1 | 1 | 1 | 1 | 0.987 | 1 | 0.936 | 1 | 0.989 | 0.985 | 0.99 | 1 | -0.008 |
| MXE | 4708 | ENSG00000178950 | GAK | chr4 | - | 851749 | 851974 | 859361 | 859493 | 850935 | 851084 | 859605 | 859722 | 0 | 0 | 1 | 1 | 0 | 162 | 86 | 107 | 146 | 53 | 0 | 0 | 0 | 8 | 0 | 221 | 167 | 127 | 138 | 197 | 281 | 298 | 1 | 1 | 0 | 0 | 0.01 | 0.007 | 0 | 0 | 0 | 0 | 0.058 | 0 | -0.008 |
| MXE | 7927 | ENSG00000043514 | TRIT1 | chr1 | - | 39847985 | 39848097 | 39850118 | 39850261 | 39847547 | 39847660 | 39883317 | 39883511 | 0 | 0 | 0 | 1 | 0 | 52 | 35 | 29 | 35 | 45 | 0 | 0 | 0 | 3 | 0 | 26 | 27 | 19 | 36 | 21 | 292 | 261 | 1 | 1 | 0 | 0 | 0 | 0.025 | 0 | 0 | 0 | 0 | 0.069 | 0 | -0.009 |
| MXE | 285 | ENSG00000100325 | ASCC2 | chr22 | - | 29822334 | 29822464 | 29825086 | 29825257 | 29816005 | 29816073 | 29838177 | 29838274 | 0 | 0 | 0 | 0 | 0 | 103 | 45 | 76 | 93 | 138 | 6 | 0 | 0 | 0 | 0 | 109 | 81 | 48 | 109 | 87 | 298 | 279 | 1 | 1 | 0 | 0 | 0 | 0 | 0 | 0.049 | 0 | 0 | 0 | 0 | -0.01 |
| MXE | 3967 | ENSG00000083312 | TNPO1 | chr5 | + | 72896457 | 72896556 | 72897055 | 72897151 | 72893615 | 72893703 | 72900005 | 72900081 | 172 | 126 | 151 | 174 | 220 | 81 | 60 | 60 | 73 | 66 | 0 | 87 | 88 | 36 | 0 | 0 | 34 | 44 | 10 | 0 | 248 | 245 | 1 | 1 | 0.677 | 0.675 | 0.713 | 0.702 | 0.767 | NA | 0.717 | 0.664 | 0.781 | NA | -0.014 |
| MXE | 12814 | ENSG00000104133 | SPG11 | chr15 | - | 44573546 | 44573745 | 44574901 | 44575041 | 44570524 | 44570658 | 44583813 | 44584558 | 44 | 17 | 28 | 37 | 25 | 3 | 0 | 0 | 0 | 1 | 14 | 28 | 30 | 33 | 12 | 0 | 0 | 1 | 0 | 0 | 289 | 298 | 1 | 1 | 0.938 | 1 | 1 | 1 | 0.963 | 1 | 1 | 0.969 | 1 | 1 | -0.014 |
| MXE | 12403 | ENSG00000091732 | ZC3HC1 | chr7 | - | 130043386 | 130043431 | 130049032 | 130049144 | 130040950 | 130041101 | 130051220 | 130051374 | 42 | 16 | 46 | 31 | 53 | 0 | 1 | 0 | 0 | 0 | 0 | 37 | 12 | 18 | 24 | 0 | 0 | 0 | 0 | 0 | 261 | 194 | 1 | 1 | 1 | 0.922 | 1 | 1 | 1 | NA | 1 | 1 | 1 | 1 | -0.016 |
| MXE | 7552 | ENSG00000256269 | HMBS | chr11 | + | 119091412 | 119091526 | 119091860 | 119091874 | 119090189 | 119090265 | 119092124 | 119092163 | 16 | 4 | 22 | 17 | 7 | 0 | 0 | 0 | 1 | 0 | 34 | 38 | 10 | 29 | 32 | 0 | 0 | 0 | 0 | 0 | 263 | 163 | 1 | 1 | 1 | 1 | 1 | 0.913 | 1 | 1 | 1 | 1 | 1 | 1 | -0.017 |
| MXE | 13984 | ENSG00000064313 | TAF2 | chr8 | - | 119744880 | 119745036 | 119746704 | 119746934 | 119744287 | 119744393 | 119756005 | 119756115 | 94 | 79 | 50 | 78 | 135 | 2 | 2 | 0 | 8 | 2 | 0 | 63 | 47 | 19 | 12 | 0 | 0 | 2 | 0 | 0 | 298 | 298 | 1 | 1 | 0.979 | 0.975 | 1 | 0.907 | 0.985 | NA | 1 | 0.959 | 1 | 1 | -0.021 |
| MXE | 15302 | ENSG00000139697 | SBNO1 | chr12 | - | 123304581 | 123304719 | 123309309 | 123309402 | 123302823 | 123302900 | 123309488 | 123309583 | 50 | 42 | 51 | 65 | 108 | 119 | 86 | 89 | 111 | 121 | 10 | 59 | 44 | 29 | 0 | 30 | 69 | 78 | 30 | 0 | 242 | 287 | 1 | 1 | 0.333 | 0.367 | 0.405 | 0.41 | 0.514 | 0.283 | 0.503 | 0.401 | 0.534 | NA | -0.024 |
| MXE | 16919 | ENSG00000204852 | TCTN1 | chr12 | + | 110641023 | 110641149 | 110641541 | 110642389 | 110640382 | 110640517 | 110644991 | 110645129 | 6 | 11 | 5 | 6 | 16 | 0 | 0 | 0 | 1 | 0 | 6 | 3 | 6 | 7 | 5 | 0 | 0 | 0 | 0 | 0 | 275 | 298 | 1 | 1 | 1 | 1 | 1 | 0.867 | 1 | 1 | 1 | 1 | 1 | 1 | -0.027 |
| MXE | 3968 | ENSG00000083312 | TNPO1 | chr5 | + | 72896457 | 72896556 | 72900005 | 72900081 | 72893615 | 72893703 | 72900973 | 72901073 | 167 | 126 | 150 | 171 | 216 | 17 | 7 | 9 | 14 | 3 | 0 | 87 | 88 | 36 | 0 | 0 | 4 | 5 | 0 | 0 | 248 | 225 | 1 | 1 | 0.899 | 0.942 | 0.938 | 0.917 | 0.985 | NA | 0.952 | 0.941 | 1 | NA | -0.028 |
| MXE | 7553 | ENSG00000256269 | HMBS | chr11 | + | 119091412 | 119091526 | 119091860 | 119091874 | 119090189 | 119090265 | 119092403 | 119092523 | 11 | 2 | 11 | 10 | 3 | 0 | 0 | 0 | 1 | 0 | 19 | 15 | 4 | 14 | 22 | 0 | 0 | 0 | 0 | 0 | 263 | 163 | 1 | 1 | 1 | 1 | 1 | 0.861 | 1 | 1 | 1 | 1 | 1 | 1 | -0.028 |
| MXE | 7683 | ENSG00000139636 | LMBR1L | chr12 | - | 49104718 | 49104885 | 49105923 | 49105957 | 49104447 | 49104551 | 49106960 | 49107045 | 10 | 7 | 9 | 12 | 19 | 0 | 2 | 0 | 0 | 0 | 5 | 14 | 3 | 11 | 25 | 0 | 0 | 0 | 0 | 0 | 183 | 298 | 1 | 1 | 1 | 0.851 | 1 | 1 | 1 | 1 | 1 | 1 | 1 | 1 | -0.03 |
| MXE | 121 | ENSG00000188130 | MAPK12 | chr22 | - | 50248365 | 50248472 | 50248661 | 50248751 | 50248224 | 50248292 | 50248830 | 50248890 | 11 | 1 | 3 | 16 | 3 | 4 | 4 | 6 | 3 | 1 | 29 | 11 | 4 | 10 | 0 | 10 | 3 | 5 | 13 | 0 | 239 | 256 | 1 | 1 | 0.747 | 0.211 | 0.349 | 0.851 | 0.763 | 0.756 | 0.797 | 0.461 | 0.452 | NA | -0.032 |
| MXE | 1369 | ENSG00000185813 | PCYT2 | chr17 | - | 81905381 | 81905447 | 81905669 | 81905735 | 81905065 | 81905154 | 81906099 | 81906177 | 8 | 0 | 7 | 5 | 5 | 3 | 1 | 3 | 3 | 13 | 0 | 15 | 0 | 0 | 5 | 0 | 0 | 2 | 2 | 0 | 215 | 215 | 1 | 1 | 0.727 | 0 | 0.7 | 0.625 | 0.278 | NA | 1 | 0 | 0 | 1 | -0.034 |
| MXE | 16544 | ENSG00000028203 | VEZT | chr12 | + | 95257149 | 95257239 | 95258244 | 95258287 | 95251939 | 95252071 | 95262905 | 95263081 | 73 | 68 | 50 | 67 | 106 | 4 | 3 | 0 | 0 | 6 | 0 | 36 | 36 | 30 | 6 | 0 | 0 | 0 | 0 | 0 | 239 | 192 | 1 | 1 | 0.936 | 0.948 | 1 | 1 | 0.934 | NA | 1 | 1 | 1 | 1 | -0.036 |
| MXE | 4462 | ENSG00000141452 | RMC1 | chr18 | + | 23506969 | 23507054 | 23507984 | 23508041 | 23504370 | 23504447 | 23515855 | 23515996 | 32 | 31 | 46 | 49 | 14 | 0 | 0 | 0 | 3 | 2 | 58 | 51 | 24 | 60 | 27 | 0 | 0 | 0 | 0 | 0 | 234 | 206 | 1 | 1 | 1 | 1 | 1 | 0.935 | 0.86 | 1 | 1 | 1 | 1 | 1 | -0.041 |
| MXE | 2241 | ENSG00000103978 | TMEM87A | chr15 | - | 42231191 | 42231260 | 42231843 | 42231915 | 42228711 | 42228820 | 42233212 | 42233306 | 14 | 2 | 6 | 15 | 15 | 10 | 8 | 7 | 12 | 22 | 0 | 19 | 24 | 9 | 0 | 0 | 20 | 8 | 4 | 11 | 221 | 218 | 1 | 1 | 0.58 | 0.198 | 0.458 | 0.552 | 0.402 | NA | 0.484 | 0.747 | 0.689 | 0 | -0.042 |
| MXE | 12819 | ENSG00000104133 | SPG11 | chr15 | - | 44604104 | 44604197 | 44606024 | 44606091 | 44600466 | 44600632 | 44608443 | 44608605 | 81 | 51 | 53 | 62 | 42 | 5 | 4 | 6 | 3 | 0 | 8 | 58 | 19 | 58 | 0 | 0 | 0 | 0 | 2 | 0 | 216 | 242 | 1 | 1 | 0.948 | 0.935 | 0.908 | 0.959 | 1 | 1 | 1 | 1 | 0.97 | NA | -0.042 |
| MXE | 3018 | ENSG00000175048 | ZDHHC14 | chr6 | + | 157592987 | 157593146 | 157628348 | 157628374 | 157542584 | 157542745 | 157632833 | 157632882 | 12 | 6 | 9 | 13 | 16 | 1 | 0 | 0 | 0 | 1 | 15 | 16 | 13 | 12 | 33 | 0 | 0 | 0 | 0 | 0 | 298 | 175 | 1 | 1 | 0.876 | 1 | 1 | 1 | 0.904 | 1 | 1 | 1 | 1 | 1 | -0.044 |
| MXE | 16543 | ENSG00000028203 | VEZT | chr12 | + | 95257149 | 95257239 | 95258244 | 95258282 | 95251939 | 95252071 | 95262905 | 95263081 | 73 | 68 | 50 | 67 | 106 | 4 | 0 | 3 | 1 | 6 | 0 | 36 | 36 | 30 | 6 | 0 | 0 | 0 | 0 | 0 | 239 | 187 | 1 | 1 | 0.935 | 1 | 0.929 | 0.981 | 0.933 | NA | 1 | 1 | 1 | 1 | -0.044 |
| MXE | 16547 | ENSG00000028203 | VEZT | chr12 | + | 95257153 | 95257239 | 95258244 | 95258287 | 95251939 | 95252071 | 95262905 | 95263081 | 60 | 53 | 33 | 44 | 77 | 4 | 3 | 0 | 0 | 6 | 0 | 33 | 27 | 17 | 6 | 0 | 0 | 0 | 0 | 0 | 235 | 192 | 1 | 1 | 0.925 | 0.935 | 1 | 1 | 0.913 | NA | 1 | 1 | 1 | 1 | -0.045 |
| MXE | 15866 | ENSG00000143702 | CEP170 | chr1 | - | 243142285 | 243142463 | 243156220 | 243156455 | 243139936 | 243140107 | 243164283 | 243166116 | 58 | 71 | 45 | 74 | 153 | 4 | 3 | 5 | 5 | 9 | 8 | 35 | 36 | 0 | 7 | 0 | 0 | 2 | 0 | 0 | 298 | 298 | 1 | 1 | 0.935 | 0.959 | 0.9 | 0.937 | 0.944 | 1 | 1 | 0.947 | NA | 1 | -0.052 |
| MXE | 15865 | ENSG00000143702 | CEP170 | chr1 | - | 243142285 | 243142463 | 243156220 | 243156347 | 243139936 | 243140107 | 243164283 | 243166116 | 10 | 9 | 10 | 2 | 14 | 4 | 3 | 5 | 5 | 9 | 0 | 5 | 1 | 0 | 0 | 0 | 0 | 2 | 0 | 0 | 276 | 298 | 1 | 1 | 0.73 | 0.764 | 0.683 | 0.302 | 0.627 | NA | 1 | 0.351 | NA | NA | -0.054 |
| MXE | 16546 | ENSG00000028203 | VEZT | chr12 | + | 95257153 | 95257239 | 95258244 | 95258282 | 95251939 | 95252071 | 95262905 | 95263081 | 60 | 53 | 33 | 44 | 77 | 4 | 0 | 3 | 1 | 6 | 0 | 33 | 27 | 17 | 6 | 0 | 0 | 0 | 0 | 0 | 235 | 187 | 1 | 1 | 0.923 | 1 | 0.897 | 0.972 | 0.911 | NA | 1 | 1 | 1 | 1 | -0.059 |
| MXE | 16924 | ENSG00000204852 | TCTN1 | chr12 | + | 110642248 | 110642389 | 110644879 | 110645143 | 110641541 | 110641627 | 110647210 | 110647336 | 8 | 6 | 7 | 4 | 8 | 0 | 0 | 0 | 2 | 0 | 6 | 4 | 0 | 0 | 0 | 0 | 0 | 0 | 0 | 0 | 290 | 298 | 1 | 1 | 1 | 1 | 1 | 0.673 | 1 | 1 | 1 | NA | NA | NA | -0.065 |
| MXE | 5879 | ENSG00000162601 | MYSM1 | chr1 | - | 58668429 | 58668566 | 58668631 | 58668682 | 58667846 | 58667921 | 58668983 | 58669038 | 91 | 79 | 94 | 87 | 159 | 39 | 31 | 24 | 49 | 47 | 0 | 52 | 40 | 7 | 0 | 0 | 6 | 8 | 3 | 0 | 200 | 286 | 1 | 1 | 0.769 | 0.785 | 0.849 | 0.717 | 0.829 | NA | 0.925 | 0.877 | 0.769 | NA | -0.067 |
| MXE | 3969 | ENSG00000083312 | TNPO1 | chr5 | + | 72897055 | 72897151 | 72900005 | 72900081 | 72896457 | 72896556 | 72900973 | 72901073 | 169 | 136 | 156 | 173 | 252 | 23 | 7 | 13 | 21 | 9 | 15 | 119 | 121 | 41 | 0 | 0 | 4 | 2 | 0 | 0 | 245 | 225 | 1 | 1 | 0.871 | 0.947 | 0.917 | 0.883 | 0.963 | 1 | 0.965 | 0.982 | 1 | NA | -0.071 |
| MXE | 7918 | ENSG00000135250 | SRPK2 | chr7 | - | 105297420 | 105297533 | 105306435 | 105306571 | 105268805 | 105268869 | 105388647 | 105388702 | 1 | 4 | 3 | 1 | 19 | 1 | 2 | 1 | 1 | 12 | 0 | 1 | 4 | 0 | 0 | 0 | 0 | 8 | 0 | 0 | 285 | 262 | 1 | 1 | 0.479 | 0.648 | 0.734 | 0.479 | 0.593 | NA | 1 | 0.315 | NA | NA | -0.071 |
| MXE | 11960 | ENSG00000164654 | MIOS | chr7 | + | 7594979 | 7595132 | 7596256 | 7596461 | 7589404 | 7589563 | 7605941 | 7606071 | 40 | 18 | 38 | 31 | 85 | 64 | 41 | 73 | 82 | 89 | 0 | 30 | 16 | 19 | 3 | 0 | 49 | 32 | 27 | 2 | 298 | 298 | 1 | 1 | 0.385 | 0.305 | 0.342 | 0.274 | 0.489 | NA | 0.38 | 0.333 | 0.413 | 0.6 | -0.073 |
| MXE | 15151 | ENSG00000153339 | TRAPPC8 | chr18 | - | 31855659 | 31855807 | 31857539 | 31857720 | 31853848 | 31853945 | 31864626 | 31864781 | 0 | 2 | 4 | 1 | 2 | 178 | 158 | 120 | 172 | 270 | 4 | 0 | 0 | 0 | 0 | 7 | 50 | 89 | 13 | 0 | 298 | 297 | 1 | 1 | 0 | 0.012 | 0.032 | 0.006 | 0.007 | 0.363 | 0 | 0 | 0 | NA | -0.079 |
| MXE | 15087 | ENSG00000030066 | NUP160 | chr11 | - | 47818055 | 47818124 | 47819373 | 47819458 | 47815945 | 47816029 | 47821723 | 47821821 | 25 | 21 | 12 | 39 | 38 | 33 | 26 | 31 | 27 | 82 | 28 | 9 | 9 | 19 | 0 | 10 | 19 | 16 | 15 | 0 | 234 | 218 | 1 | 1 | 0.414 | 0.429 | 0.265 | 0.574 | 0.302 | 0.723 | 0.306 | 0.344 | 0.541 | NA | -0.082 |
| MXE | 14702 | ENSG00000184465 | WDR27 | chr6 | - | 169667135 | 169667187 | 169670568 | 169670693 | 169665485 | 169665556 | 169672254 | 169672396 | 5 | 6 | 9 | 10 | 4 | 2 | 0 | 0 | 1 | 1 | 27 | 6 | 5 | 3 | 1 | 0 | 2 | 0 | 0 | 0 | 274 | 201 | 1 | 1 | 0.647 | 1 | 1 | 0.88 | 0.746 | 1 | 0.688 | 1 | 1 | 1 | -0.083 |
| MXE | 7221 | ENSG00000144357 | UBR3 | chr2 | + | 170073403 | 170073607 | 170079813 | 170080023 | 170061317 | 170061443 | 170080544 | 170080684 | 3 | 1 | 0 | 1 | 0 | 52 | 39 | 61 | 58 | 79 | 0 | 0 | 2 | 1 | 0 | 0 | 25 | 19 | 2 | 6 | 298 | 298 | 1 | 1 | 0.055 | 0.025 | 0 | 0.017 | 0 | NA | 0 | 0.095 | 0.333 | 0 | -0.088 |
| MXE | 15868 | ENSG00000143702 | CEP170 | chr1 | - | 243142315 | 243142463 | 243156220 | 243156455 | 243139936 | 243140107 | 243164283 | 243166116 | 58 | 71 | 45 | 74 | 153 | 14 | 13 | 6 | 14 | 22 | 8 | 35 | 36 | 0 | 7 | 0 | 3 | 7 | 0 | 0 | 298 | 297 | 1 | 1 | 0.805 | 0.845 | 0.882 | 0.84 | 0.874 | 1 | 0.921 | 0.837 | NA | 1 | -0.09 |
| MXE | 12008 | ENSG00000109339 | MAPK10 | chr4 | - | 86089205 | 86089277 | 86098523 | 86098595 | 86067772 | 86067955 | 86101051 | 86101217 | 0 | 1 | 0 | 2 | 4 | 0 | 0 | 0 | 1 | 0 | 19 | 0 | 0 | 1 | 2 | 0 | 0 | 0 | 0 | 0 | 221 | 221 | 1 | 1 | NA | 1 | NA | 0.667 | 1 | 1 | NA | NA | 1 | 1 | -0.111 |
| MXE | 9766 | ENSG00000164134 | NAA15 | chr4 | + | 139370210 | 139370404 | 139376364 | 139376473 | 139361723 | 139361937 | 139378755 | 139378854 | 129 | 123 | 93 | 143 | 131 | 39 | 22 | 31 | 37 | 54 | 0 | 101 | 142 | 13 | 0 | 0 | 17 | 13 | 1 | 0 | 298 | 258 | 1 | 1 | 0.741 | 0.829 | 0.722 | 0.77 | 0.677 | NA | 0.837 | 0.904 | 0.918 | NA | -0.139 |
| MXE | 11957 | ENSG00000164654 | MIOS | chr7 | + | 7567606 | 7567688 | 7568025 | 7568123 | 7567052 | 7567484 | 7572435 | 7572667 | 4 | 0 | 0 | 1 | 1 | 23 | 12 | 13 | 27 | 38 | 0 | 1 | 3 | 6 | 0 | 0 | 18 | 4 | 14 | 5 | 231 | 247 | 1 | 1 | 0.157 | 0 | 0 | 0.038 | 0.027 | NA | 0.056 | 0.445 | 0.314 | 0 | -0.159 |
| MXE | 13241 | ENSG00000137522 | RNF121 | chr11 | + | 71962151 | 71962289 | 71978085 | 71978235 | 71960749 | 71960891 | 71982760 | 71982915 | 2 | 0 | 5 | 0 | 1 | 9 | 1 | 2 | 2 | 1 | 0 | 16 | 0 | 0 | 0 | 10 | 2 | 0 | 0 | 0 | 287 | 298 | 1 | 1 | 0.187 | 0 | 0.722 | 0 | 0.509 | 0 | 0.893 | NA | NA | NA | -0.163 |
| MXE | 2251 | ENSG00000104177 | MYEF2 | chr15 | - | 48151873 | 48151942 | 48152233 | 48152284 | 48151472 | 48151571 | 48153791 | 48153893 | 10 | 2 | 9 | 8 | 4 | 13 | 6 | 16 | 8 | 20 | 0 | 5 | 5 | 3 | 0 | 0 | 13 | 12 | 0 | 0 | 200 | 218 | 1 | 1 | 0.456 | 0.267 | 0.38 | 0.522 | 0.179 | NA | 0.295 | 0.312 | 1 | NA | -0.175 |
| MXE | 15150 | ENSG00000153339 | TRAPPC8 | chr18 | - | 31853848 | 31853945 | 31855659 | 31855807 | 31852594 | 31852663 | 31857539 | 31857720 | 141 | 107 | 84 | 105 | 136 | 130 | 68 | 65 | 111 | 121 | 4 | 85 | 67 | 14 | 0 | 0 | 35 | 43 | 14 | 0 | 297 | 246 | 1 | 1 | 0.473 | 0.566 | 0.517 | 0.439 | 0.482 | 1 | 0.668 | 0.563 | 0.453 | NA | -0.176 |
| MXE | 8926 | ENSG00000117262 | GPR89A | chr1 | + | 145608347 | 145608675 | 145612201 | 145612247 | 145608047 | 145608175 | 145616233 | 145616293 | 3 | 0 | 1 | 0 | 0 | 21 | 2 | 26 | 6 | 18 | 0 | 5 | 11 | 8 | 0 | 12 | 23 | 15 | 8 | 0 | 298 | 195 | 1 | 1 | 0.085 | 0 | 0.025 | 0 | 0 | 0 | 0.125 | 0.324 | 0.396 | NA | -0.189 |
| MXE | 8333 | ENSG00000083720 | OXCT1 | chr5 | - | 41749526 | 41749607 | 41762110 | 41762200 | 41739389 | 41739491 | 41794002 | 41794078 | 25 | 12 | 35 | 39 | 24 | 26 | 14 | 27 | 24 | 16 | 7 | 41 | 50 | 13 | 0 | 0 | 10 | 17 | 21 | 0 | 239 | 230 | 1 | 1 | 0.481 | 0.452 | 0.555 | 0.61 | 0.591 | 1 | 0.798 | 0.739 | 0.373 | NA | -0.19 |
| MXE | 7254 | ENSG00000154358 | OBSCN | chr1 | + | 228246527 | 228246803 | 228249964 | 228250240 | 228245412 | 228245688 | 228256650 | 228256926 | 14 | 18 | 25 | 11 | 7 | 14 | 1 | 14 | 13 | 1 | 0 | 7 | 17 | 7 | 46 | 0 | 0 | 0 | 3 | 11 | 298 | 298 | 1 | 1 | 0.5 | 0.947 | 0.641 | 0.458 | 0.875 | NA | 1 | 1 | 0.7 | 0.807 | -0.193 |
| MXE | 8560 | ENSG00000153207 | AHCTF1 | chr1 | - | 246863923 | 246864116 | 246867243 | 246867351 | 246861958 | 246862153 | 246867660 | 246867811 | 64 | 28 | 40 | 30 | 83 | 136 | 92 | 135 | 122 | 230 | 18 | 27 | 22 | 0 | 2 | 5 | 26 | 64 | 0 | 7 | 257 | 298 | 1 | 1 | 0.353 | 0.261 | 0.256 | 0.222 | 0.295 | 0.807 | 0.546 | 0.285 | NA | 0.249 | -0.194 |
| MXE | 9767 | ENSG00000164134 | NAA15 | chr4 | + | 139376364 | 139376473 | 139378755 | 139378854 | 139370210 | 139370404 | 139384831 | 139384978 | 19 | 12 | 20 | 29 | 42 | 127 | 100 | 102 | 125 | 201 | 0 | 6 | 21 | 1 | 5 | 0 | 50 | 72 | 8 | 0 | 258 | 248 | 1 | 1 | 0.126 | 0.103 | 0.159 | 0.182 | 0.167 | NA | 0.103 | 0.219 | 0.107 | 1 | -0.21 |
| MXE | 9088 | ENSG00000135336 | ORC3 | chr6 | + | 87634844 | 87634961 | 87636406 | 87636486 | 87621949 | 87622013 | 87653115 | 87653249 | 78 | 42 | 84 | 57 | 83 | 36 | 38 | 45 | 39 | 114 | 7 | 34 | 15 | 0 | 9 | 0 | 17 | 18 | 0 | 0 | 266 | 229 | 1 | 1 | 0.651 | 0.488 | 0.616 | 0.557 | 0.385 | 1 | 0.633 | 0.418 | NA | 1 | -0.223 |
| MXE | 14701 | ENSG00000184465 | WDR27 | chr6 | - | 169665485 | 169665556 | 169667135 | 169668776 | 169664165 | 169664286 | 169670568 | 169670693 | 3 | 1 | 6 | 10 | 1 | 0 | 2 | 1 | 2 | 0 | 10 | 17 | 0 | 5 | 0 | 0 | 0 | 0 | 0 | 0 | 298 | 220 | 1 | 1 | 1 | 0.27 | 0.816 | 0.787 | 1 | 1 | 1 | NA | 1 | NA | -0.225 |
| MXE | 15352 | ENSG00000130935 | NOL11 | chr17 | + | 67734362 | 67734439 | 67736665 | 67736754 | 67726459 | 67726648 | 67737070 | 67737145 | 64 | 26 | 77 | 55 | 164 | 92 | 30 | 88 | 78 | 190 | 0 | 32 | 47 | 8 | 7 | 0 | 3 | 34 | 25 | 0 | 226 | 238 | 1 | 1 | 0.423 | 0.477 | 0.48 | 0.426 | 0.476 | NA | 0.918 | 0.593 | 0.252 | 1 | -0.234 |
| MXE | 7554 | ENSG00000256269 | HMBS | chr11 | + | 119092124 | 119092163 | 119092403 | 119092523 | 119091412 | 119091526 | 119092757 | 119092811 | 3 | 1 | 2 | 0 | 2 | 17 | 3 | 4 | 9 | 5 | 0 | 13 | 1 | 7 | 0 | 0 | 15 | 3 | 5 | 0 | 188 | 269 | 1 | 1 | 0.202 | 0.323 | 0.417 | 0 | 0.364 | NA | 0.554 | 0.323 | 0.667 | NA | -0.253 |
| MXE | 11037 | ENSG00000153253 | SCN3A | chr2 | - | 165163617 | 165163709 | 165163801 | 165163893 | 165162555 | 165162828 | 165164391 | 165164520 | 5 | 1 | 2 | 4 | 6 | 2 | 2 | 2 | 0 | 0 | 0 | 8 | 4 | 3 | 0 | 0 | 0 | 0 | 0 | 0 | 241 | 241 | 1 | 1 | 0.714 | 0.333 | 0.5 | 1 | 1 | NA | 1 | 1 | 1 | NA | -0.291 |
| MXE | 12007 | ENSG00000109339 | MAPK10 | chr4 | - | 86064265 | 86064390 | 86107222 | 86107352 | 86031367 | 86031431 | 86159297 | 86159467 | 0 | 0 | 0 | 0 | 0 | 2 | 0 | 1 | 1 | 1 | 7 | 2 | 0 | 0 | 1 | 7 | 3 | 0 | 4 | 2 | 279 | 274 | 1 | 1 | 0 | NA | 0 | 0 | 0 | 0.495 | 0.396 | NA | 0 | 0.329 | -0.305 |
| MXE | 1370 | ENSG00000185813 | PCYT2 | chr17 | - | 81905669 | 81905735 | 81906099 | 81906177 | 81905381 | 81905447 | 81906463 | 81906546 | 7 | 7 | 5 | 4 | 5 | 11 | 3 | 8 | 6 | 3 | 5 | 6 | 9 | 11 | 0 | 7 | 0 | 0 | 2 | 0 | 227 | 215 | 1 | 1 | 0.376 | 0.688 | 0.372 | 0.387 | 0.612 | 0.404 | 1 | 1 | 0.839 | NA | -0.324 |
| MXE | 13350 | ENSG00000162669 | HFM1 | chr1 | - | 91316390 | 91316476 | 91319077 | 91319209 | 91316100 | 91316184 | 91319292 | 91319390 | 0 | 3 | 0 | 0 | 4 | 1 | 0 | 0 | 0 | 0 | 0 | 0 | 0 | 0 | 6 | 0 | 0 | 0 | 0 | 0 | 281 | 235 | 1 | 1 | 0 | 1 | NA | NA | 1 | NA | NA | NA | NA | 1 | -0.333 |
| MXE | 12817 | ENSG00000104133 | SPG11 | chr15 | - | 44601738 | 44601824 | 44604802 | 44604927 | 44600466 | 44600632 | 44606024 | 44606091 | 4 | 0 | 2 | 0 | 0 | 0 | 2 | 3 | 1 | 0 | 0 | 2 | 3 | 0 | 0 | 0 | 3 | 0 | 0 | 0 | 274 | 235 | 1 | 1 | 1 | 0 | 0.364 | 0 | NA | NA | 0.364 | 1 | NA | NA | -0.341 |
| MXE | 5427 | ENSG00000145428 | RNF175 | chr4 | - | 153728206 | 153728361 | 153748644 | 153748786 | 153723350 | 153723458 | 153751437 | 153751475 | 0 | 0 | 0 | 1 | 0 | 0 | 4 | 1 | 5 | 0 | 0 | 3 | 0 | 0 | 0 | 3 | 0 | 0 | 0 | 0 | 291 | 298 | 1 | 1 | NA | 0 | 0 | 0.17 | NA | 0 | 1 | NA | NA | NA | -0.443 |
| MXE | 12818 | ENSG00000104133 | SPG11 | chr15 | - | 44604104 | 44604197 | 44604802 | 44604927 | 44600466 | 44600632 | 44606024 | 44606091 | 4 | 0 | 2 | 0 | 0 | 6 | 4 | 9 | 4 | 3 | 0 | 2 | 3 | 0 | 0 | 0 | 0 | 0 | 2 | 0 | 274 | 242 | 1 | 1 | 0.371 | 0 | 0.164 | 0 | 0 | NA | 1 | 1 | 0 | NA | -0.56 |
| MXE | 12015 | ENSG00000109339 | MAPK10 | chr4 | - | 86194308 | 86194335 | 86194335 | 86194407 | 86159297 | 86159467 | 86354529 | 86354644 | 1 | 0 | 0 | 0 | 0 | 1 | 1 | 0 | 1 | 0 | 23 | 3 | 0 | 0 | 8 | 0 | 0 | 0 | 0 | 0 | 221 | 176 | 1 | 1 | 0.443 | 0 | NA | 0 | NA | 1 | 1 | NA | NA | 1 | -0.852 |

**A5SS**

| | | | | | | | | | | | | Inclusion_counts_HC | | | | | Skipping_counts_HC | | | | | Inclusion counts_SLE | | | | | Skipping_counts_SLE | | | | | | | | | Inclusion_level(normalized) | | | | | | | | | | |
|---|---|---|---|---|---|---|---|---|---|---|---|---|---|---|---|---|---|---|---|---|---|---|---|---|---|---|---|---|---|---|---|---|---|---|---|---|---|---|---|---|---|---|---|---|---|---|
| Splicing_event | ID | Gene_ID | Gene_symbol | Chromosome | Strand | LongExonStart_0base | LongExonEnd | ShortES | ShortEE | FlankingES | FlankingEE | SRR13214146 | SRR13214147 | SRR13214148 | SRR13214149 | SRR13214150 | SRR13214146 | SRR13214147 | SRR13214148 | SRR13214149 | SRR13214150 | SRR13214141 | SRR13214242 | SRR13214143 | SRR13214144 | SRR13214145 | SRR13214141 | SRR13214242 | SRR13214143 | SRR13214144 | SRR13214145 | IncFormLen | SkipFormLen | p value | FDR | SRR13214146 | SRR13214147 | SRR13214148 | SRR13214149 | SRR13214150 | SRR13214141 | SRR13214242 | SRR13214143 | SRR13214144 | SRR13214145 | Inclevel_diff |
| A5SS | 7793 | ENSG00000130714 | POMT1 | chr9 | + | 131509902 | 131510141 | 131509902 | 131509996 | 131510259 | 131510415 | 5 | 2 | 12 | 8 | 0 | 8 | 1 | 12 | 6 | 4 | 9 | 11 | 3 | 2 | 23 | 0 | 0 | 0 | 0 | 0 | 294 | 149 | 0 | 0 | 0.241 | 0.503 | 0.336 | 0.403 | 0 | 1 | 1 | 1 | 1 | 1 | -0.703 |
| A5SS | 5360 | ENSG00000084623 | EIF3I | chr1 | + | 32228726 | 32229024 | 32228726 | 32228816 | 32229134 | 32229208 | 17 | 11 | 18 | 19 | 12 | 136 | 82 | 153 | 117 | 130 | 103 | 60 | 39 | 89 | 77 | 112 | 86 | 84 | 107 | 255 | 298 | 149 | 0 | 0 | 0.059 | 0.063 | 0.056 | 0.075 | 0.044 | 0.315 | 0.259 | 0.188 | 0.294 | 0.131 | -0.178 |
| A5SS | 2907 | ENSG00000120686 | UFM1 | chr13 | + | 38349998 | 38350227 | 38349998 | 38350055 | 38354238 | 38354296 | 13 | 17 | 22 | 10 | 9 | 218 | 141 | 177 | 241 | 118 | 24 | 34 | 12 | 15 | 14 | 24 | 74 | 63 | 29 | 23 | 298 | 149 | 0 | 0 | 0.029 | 0.057 | 0.059 | 0.02 | 0.037 | 0.333 | 0.187 | 0.087 | 0.205 | 0.233 | -0.169 |
| A5SS | 7585 | ENSG00000132334 | PTPRE | chr10 | + | 128070807 | 128071741 | 128070807 | 128070901 | 128072137 | 128072214 | 59 | 40 | 45 | 61 | 38 | 281 | 191 | 195 | 275 | 283 | 325 | 114 | 251 | 373 | 70 | 369 | 240 | 304 | 442 | 237 | 298 | 149 | 0 | 0 | 0.095 | 0.095 | 0.103 | 0.1 | 0.063 | 0.306 | 0.192 | 0.292 | 0.297 | 0.129 | -0.152 |
| A5SS | 5148 | ENSG00000181163 | NPM1 | chr5 | + | 171391304 | 171391609 | 171391304 | 171391424 | 171391705 | 171391799 | 102 | 115 | 141 | 162 | 106 | 4105 | 3548 | 5449 | 4941 | 8235 | 134 | 198 | 172 | 210 | 76 | 1944 | 3624 | 4980 | 3284 | 503 | 298 | 149 | 0 | 0 | 0.012 | 0.016 | 0.013 | 0.016 | 0.006 | 0.033 | 0.027 | 0.017 | 0.031 | 0.07 | -0.023 |
| A5SS | 3217 | ENSG00000082074 | FYB1 | chr5 | - | 39125971 | 39126135 | 39125997 | 39126135 | 39124252 | 39124278 | 30 | 69 | 70 | 53 | 121 | 357 | 504 | 230 | 407 | 539 | 0 | 9 | 32 | 4 | 0 | 258 | 414 | 426 | 443 | 91 | 175 | 149 | 0 | 0 | 0.067 | 0.104 | 0.206 | 0.1 | 0.16 | 0 | 0.018 | 0.06 | 0.008 | 0 | 0.11 |
| A5SS | 7039 | ENSG00000147168 | IL2RG | chrX | - | 71110155 | 71110688 | 71110503 | 71110688 | 71109227 | 71109390 | 1416 | 674 | 1100 | 1133 | 1404 | 32 | 12 | 29 | 29 | 24 | 721 | 1166 | 592 | 750 | 1002 | 31 | 66 | 109 | 79 | 79 | 298 | 149 | 0 | 0 | 0.957 | 0.966 | 0.95 | 0.951 | 0.967 | 0.921 | 0.898 | 0.731 | 0.826 | 0.864 | 0.11 |
| A5SS | 7697 | ENSG00000106948 | AKNA | chr9 | - | 114343707 | 114346009 | 114345862 | 114346009 | 114342008 | 114342125 | 735 | 302 | 535 | 515 | 371 | 43 | 24 | 41 | 32 | 28 | 406 | 646 | 291 | 348 | 579 | 103 | 177 | 164 | 103 | 205 | 298 | 149 | 0 | 0 | 0.895 | 0.863 | 0.867 | 0.889 | 0.869 | 0.663 | 0.646 | 0.47 | 0.628 | 0.585 | 0.278 |
| A5SS | 7686 | ENSG00000096746 | HNRNPH3 | chr10 | + | 68337198 | 68338687 | 68337198 | 68337333 | 68339139 | 68339226 | 219 | 237 | 189 | 270 | 273 | 9 | 11 | 11 | 21 | 7 | 139 | 181 | 270 | 113 | 40 | 24 | 51 | 139 | 88 | 12 | 298 | 149 | 0 | 0 | 0.924 | 0.915 | 0.896 | 0.865 | 0.951 | 0.743 | 0.64 | 0.493 | 0.391 | 0.625 | 0.332 |
| A5SS | 3773 | ENSG00000104419 | NDRG1 | chr8 | - | 133256725 | 133256863 | 133256776 | 133256863 | 133254538 | 133254595 | 15 | 5 | 4 | 9 | 8 | 226 | 118 | 191 | 201 | 147 | 54 | 9 | 18 | 29 | 41 | 196 | 129 | 121 | 134 | 166 | 200 | 149 | 1.22E-15 | 6.65E-13 | 0.047 | 0.031 | 0.015 | 0.032 | 0.039 | 0.17 | 0.049 | 0.1 | 0.139 | 0.155 | -0.09 |
| A5SS | 1589 | ENSG00000125107 | CNOT1 | chr16 | - | 58536967 | 58537220 | 58536988 | 58537220 | 58534146 | 58534395 | 36 | 38 | 45 | 68 | 65 | 216 | 113 | 170 | 178 | 293 | 27 | 89 | 85 | 101 | 21 | 30 | 133 | 111 | 114 | 17 | 170 | 149 | 2.00E-15 | 9.89E-13 | 0.127 | 0.228 | 0.188 | 0.251 | 0.163 | 0.441 | 0.37 | 0.402 | 0.437 | 0.52 | -0.243 |
| A5SS | 8846 | ENSG00000061936 | SFSWAP | chr12 | + | 131714070 | 131714240 | 131714070 | 131714193 | 131714821 | 131714953 | 87 | 71 | 69 | 70 | 53 | 24 | 17 | 21 | 28 | 24 | 282 | 139 | 164 | 213 | 198 | 6 | 19 | 22 | 22 | 0 | 196 | 149 | 3.22E-15 | 1.46E-12 | 0.734 | 0.76 | 0.714 | 0.655 | 0.627 | 0.973 | 0.848 | 0.85 | 0.88 | 1 | -0.212 |
| A5SS | 4862 | ENSG00000149532 | CPSF7 | chr11 | - | 61420469 | 61421608 | 61421389 | 61421608 | 61419921 | 61420094 | 225 | 123 | 202 | 202 | 336 | 10 | 0 | 3 | 10 | 9 | 295 | 218 | 95 | 161 | 203 | 18 | 33 | 22 | 43 | 32 | 298 | 149 | 3.89E-15 | 1.63E-12 | 0.918 | 1 | 0.971 | 0.91 | 0.949 | 0.891 | 0.768 | 0.683 | 0.652 | 0.76 | 0.199 |
| A5SS | 2521 | ENSG00000147548 | NSD3 | chr8 | - | 38281466 | 38281583 | 38281499 | 38281583 | 38279539 | 38279681 | 81 | 79 | 82 | 108 | 138 | 71 | 36 | 43 | 59 | 80 | 14 | 51 | 28 | 32 | 4 | 34 | 84 | 102 | 62 | 26 | 182 | 149 | 8.99E-15 | 3.50E-12 | 0.483 | 0.642 | 0.61 | 0.6 | 0.585 | 0.252 | 0.332 | 0.183 | 0.297 | 0.112 | 0.349 |
| A5SS | 1941 | ENSG00000112306 | RPS12 | chr6 | + | 132814971 | 132815224 | 132814971 | 132815088 | 132816460 | 132816563 | 280 | 184 | 338 | 329 | 262 | 5461 | 3259 | 6954 | 5226 | 7029 | 290 | 319 | 350 | 253 | 111 | 3030 | 2748 | 2792 | 2477 | 3528 | 285 | 149 | 5.01E-14 | 1.82E-11 | 0.026 | 0.029 | 0.025 | 0.032 | 0.019 | 0.048 | 0.057 | 0.062 | 0.051 | 0.016 | -0.021 |
| A5SS | 5825 | ENSG00000139531 | SUOX | chr12 | + | 55999332 | 55999590 | 55999332 | 55999458 | 56002211 | 56002271 | 6 | 0 | 1 | 1 | 1 | 8 | 4 | 1 | 0 | 3 | 11 | 1 | 1 | 2 | 6 | 0 | 0 | 0 | 0 | 0 | 281 | 149 | 5.34E-14 | 1.82E-11 | 0.285 | 0 | 0.347 | 1 | 0.15 | 1 | 1 | 1 | 1 | 1 | -0.644 |
| A5SS | 3719 | ENSG00000134419 | RPS15A | chr16 | - | 18790243 | 18790334 | 18790309 | 18790334 | 18788980 | 18789118 | 369 | 98 | 356 | 311 | 296 | 2327 | 936 | 2530 | 1992 | 2105 | 189 | 370 | 234 | 164 | 311 | 854 | 1075 | 833 | 768 | 1443 | 215 | 149 | 6.24E-14 | 2.00E-11 | 0.099 | 0.068 | 0.089 | 0.098 | 0.089 | 0.133 | 0.193 | 0.163 | 0.129 | 0.13 | -0.061 |
| A5SS | 6633 | ENSG00000104133 | SPG11 | chr15 | - | 44596059 | 44596355 | 44596082 | 44596355 | 44595258 | 44595459 | 42 | 23 | 22 | 27 | 37 | 353 | 293 | 229 | 306 | 276 | 28 | 71 | 77 | 48 | 30 | 93 | 320 | 143 | 175 | 124 | 172 | 149 | 9.26E-14 | 2.80E-11 | 0.093 | 0.064 | 0.077 | 0.071 | 0.104 | 0.207 | 0.161 | 0.318 | 0.192 | 0.173 | -0.128 |
| A5SS | 2266 | ENSG00000126456 | IRF3 | chr19 | - | 49665483 | 49665857 | 49665630 | 49665857 | 49664673 | 49664846 | 108 | 137 | 111 | 126 | 131 | 16 | 7 | 21 | 17 | 11 | 355 | 133 | 154 | 271 | 385 | 0 | 10 | 7 | 2 | 7 | 296 | 149 | 3.66E-13 | 1.05E-10 | 0.773 | 0.908 | 0.727 | 0.789 | 0.857 | 1 | 0.87 | 0.917 | 0.986 | 0.965 | -0.137 |
| A5SS | 2145 | ENSG00000100284 | TOM1 | chr22 | + | 35321958 | 35322037 | 35321958 | 35322019 | 35323027 | 35323177 | 208 | 89 | 144 | 177 | 171 | 3 | 0 | 3 | 2 | 3 | 702 | 346 | 269 | 468 | 566 | 65 | 21 | 18 | 28 | 39 | 167 | 149 | 5.23E-13 | 1.42E-10 | 0.984 | 1 | 0.977 | 0.987 | 0.981 | 0.906 | 0.936 | 0.93 | 0.937 | 0.928 | 0.058 |
| A5SS | 8456 | ENSG00000107862 | GBF1 | chr10 | + | 102362474 | 102363396 | 102362474 | 102362664 | 102363709 | 102363798 | 156 | 110 | 87 | 154 | 140 | 6 | 4 | 5 | 8 | 1 | 239 | 158 | 106 | 149 | 319 | 120 | 31 | 29 | 85 | 22 | 298 | 149 | 2.00E-12 | 5.19E-10 | 0.929 | 0.932 | 0.897 | 0.906 | 0.986 | 0.499 | 0.718 | 0.646 | 0.467 | 0.879 | 0.288 |
| A5SS | 4656 | ENSG00000117616 | RSRP1 | chr1 | - | 25244176 | 25245301 | 25245149 | 25245301 | 25243549 | 25243633 | 201 | 139 | 144 | 160 | 163 | 18 | 22 | 26 | 38 | 29 | 121 | 114 | 77 | 117 | 93 | 8 | 4 | 3 | 3 | 0 | 298 | 149 | 2.72E-12 | 6.73E-10 | 0.848 | 0.76 | 0.735 | 0.678 | 0.738 | 0.883 | 0.934 | 0.928 | 0.951 | 1 | -0.187 |
| A5SS | 4271 | ENSG00000176393 | RNPEP | chr1 | + | 201996146 | 201996409 | 201996146 | 201996263 | 201997318 | 201997554 | 11 | 8 | 10 | 5 | 6 | 85 | 43 | 77 | 57 | 102 | 85 | 22 | 26 | 22 | 91 | 120 | 85 | 66 | 94 | 228 | 295 | 149 | 5.06E-12 | 1.20E-09 | 0.061 | 0.086 | 0.062 | 0.042 | 0.029 | 0.263 | 0.116 | 0.166 | 0.106 | 0.168 | -0.108 |
| A5SS | 8844 | ENSG00000050426 | LETMD1 | chr12 | + | 51052091 | 51052246 | 51052091 | 51052207 | 51053777 | 51053860 | 11 | 11 | 11 | 13 | 24 | 67 | 39 | 62 | 39 | 135 | 17 | 20 | 19 | 19 | 29 | 22 | 22 | 32 | 20 | 32 | 188 | 149 | 5.47E-12 | 1.24E-09 | 0.115 | 0.183 | 0.123 | 0.209 | 0.123 | 0.38 | 0.419 | 0.32 | 0.43 | 0.418 | -0.243 |
| A5SS | 3008 | ENSG00000206503 | HLA-A | chr6 | + | 29942756 | 29943543 | 29942756 | 29943026 | 29944499 | 29944616 | 91 | 85 | 176 | 170 | 47 | 1 | 0 | 2 | 0 | 2 | 150 | 82 | 264 | 245 | 213 | 24 | 4 | 2 | 14 | 46 | 298 | 149 | 5.72E-12 | 1.25E-09 | 0.978 | 1 | 0.978 | 1 | 0.922 | 0.758 | 0.911 | 0.985 | 0.897 | 0.698 | 0.126 |
| A5SS | 1266 | ENSG00000149923 | PPP4C | chr16 | + | 30082745 | 30082963 | 30082745 | 30082847 | 30083393 | 30083567 | 4 | 2 | 12 | 7 | 7 | 102 | 54 | 79 | 116 | 106 | 50 | 23 | 25 | 37 | 33 | 155 | 143 | 84 | 110 | 191 | 265 | 149 | 7.28E-12 | 1.52E-09 | 0.022 | 0.02 | 0.079 | 0.033 | 0.036 | 0.154 | 0.083 | 0.143 | 0.159 | 0.089 | -0.088 |
| [illegible] | | | | | | | | | | | | | | | | | | | | | | | | | | | | | | | | | | | | | | | | | | | | | | |

| | | | | | | | | | | | | | | | | | | | | | | | | | | | | | | | | | | | | | | | | | | | | | | |
|---|---|---|---|---|---|---|---|---|---|---|---|---|---|---|---|---|---|---|---|---|---|---|---|---|---|---|---|---|---|---|---|---|---|---|---|---|---|---|---|---|---|---|---|---|---|---|
| A5SS | 1377 | ENSG00000140983 | RHOT2 | chr16 | + | 668104 | 668236 | 668104 | 668154 | 668352 | 668411 | 55 | 15 | 42 | 56 | 8 | 0 | 0 | 0 | 0 | 0 | 16 | 49 | 52 | 31 | 97 | 0 | 0 | 1 | 0 | 16 | 231 | 149 | 9.20E-06 | 0.000241922 | 1 | 1 | 1 | 1 | 1 | 1 | 1 | 0.971 | 1 | 0.796 | 0.047 |
| A5SS | 8849 | ENSG00000166598 | HSP90B1 | chr12 | + | 103943073 | 103943573 | 103943073 | 103943319 | 103943737 | 103943874 | 36 | 23 | 29 | 55 | 23 | 921 | 643 | 1177 | 908 | 1181 | 0 | 4 | 17 | 0 | 0 | 118 | 518 | 1329 | 230 | 31 | 298 | 149 | 9.25E-06 | 0.000241942 | 0.019 | 0.018 | 0.012 | 0.029 | 0.01 | 0 | 0.004 | 0.006 | 0 | 0 | 0.016 |
| A5SS | 4159 | ENSG00000148335 | NTMT1 | chr9 | + | 129632649 | 129632865 | 129632649 | 129632801 | 129634053 | 129634306 | 19 | 8 | 18 | 16 | 14 | 1 | 3 | 0 | 1 | 7 | 57 | 27 | 28 | 39 | 71 | 0 | 0 | 0 | 0 | 0 | 213 | 149 | 9.64E-06 | 0.00025091 | 0.93 | 0.651 | 1 | 0.918 | 0.583 | 1 | 1 | 1 | 1 | 1 | -0.184 |
| A5SS | 6097 | ENSG00000135316 | SYNCRIP | chr6 | - | 85637242 | 85637356 | 85637268 | 85637356 | 85636966 | 85637143 | 501 | 434 | 507 | 578 | 1164 | 5 | 15 | 13 | 13 | 24 | 170 | 390 | 381 | 267 | 35 | 14 | 23 | 15 | 7 | 9 | 175 | 149 | 9.73E-06 | 0.000252212 | 0.988 | 0.961 | 0.971 | 0.974 | 0.976 | 0.912 | 0.935 | 0.956 | 0.97 | 0.768 | 0.066 |
| A5SS | 892 | ENSG00000276231 | PIK3R6 | chr17 | - | 8837802 | 8837871 | 8837818 | 8837871 | 8836790 | 8836923 | 9 | 3 | 3 | 7 | 4 | 0 | 0 | 0 | 0 | 0 | 8 | 13 | 11 | 18 | 45 | 0 | 0 | 0 | 1 | 14 | 165 | 149 | 9.85E-06 | 0.000254026 | 1 | 1 | 1 | 1 | 1 | 1 | 1 | 1 | 0.942 | 0.744 | 0.063 |
| A5SS | 1018 | ENSG00000177302 | TOP3A | chr17 | - | 18308312 | 18308424 | 18308412 | 18308424 | 18306890 | 18306966 | 35 | 23 | 15 | 29 | 34 | 0 | 0 | 0 | 0 | 0 | 86 | 53 | 43 | 23 | 54 | 9 | 0 | 0 | 0 | 7 | 249 | 149 | 9.90E-06 | 0.00025407 | 1 | 1 | 1 | 1 | 1 | 0.851 | 1 | 1 | 1 | 0.822 | 0.065 |
| A5SS | 5741 | ENSG00000064012 | CASP8 | chr2 | + | 201276826 | 201277104 | 201276826 | 201276968 | 201284815 | 201285317 | 917 | 264 | 611 | 907 | 389 | 270 | 163 | 205 | 181 | 231 | 710 | 1195 | 1838 | 907 | 267 | 162 | 115 | 109 | 104 | 98 | 285 | 149 | 1.07E-05 | 0.000273075 | 0.64 | 0.459 | 0.609 | 0.724 | 0.468 | 0.696 | 0.845 | 0.898 | 0.82 | 0.588 | -0.189 |
| A5SS | 1025 | ENSG00000276234 | TADA2A | chr17 | + | 37437737 | 37437922 | 37437737 | 37437829 | 37440504 | 37440662 | 12 | 1 | 2 | 7 | 5 | 76 | 50 | 74 | 69 | 93 | 13 | 17 | 1 | 5 | 8 | 49 | 45 | 25 | 31 | 17 | 242 | 149 | 1.10E-05 | 0.000280172 | 0.089 | 0.012 | 0.016 | 0.059 | 0.032 | 0.14 | 0.189 | 0.024 | 0.09 | 0.225 | -0.092 |
| A5SS | 5561 | ENSG00000168958 | MFF | chr2 | + | 227328677 | 227328899 | 227328677 | 227328789 | 227329605 | 227329783 | 13 | 8 | 24 | 20 | 18 | 5 | 6 | 9 | 5 | 2 | 3 | 8 | 12 | 8 | 0 | 18 | 11 | 8 | 7 | 12 | 259 | 149 | 1.12E-05 | 0.000282315 | 0.599 | 0.434 | 0.605 | 0.697 | 0.838 | 0.087 | 0.295 | 0.463 | 0.397 | 0 | 0.386 |
| A5SS | 5466 | ENSG00000143933 | CALM2 | chr2 | - | 47176235 | 47176511 | 47176440 | 47176511 | 47170733 | 47170764 | 50 | 50 | 68 | 73 | 13 | 1458 | 1149 | 1026 | 1183 | 887 | 30 | 36 | 53 | 63 | 36 | 513 | 958 | 526 | 537 | 466 | 298 | 149 | 1.12E-05 | 0.000282315 | 0.017 | 0.021 | 0.032 | 0.03 | 0.007 | 0.028 | 0.018 | 0.048 | 0.055 | 0.037 | -0.016 |
| A5SS | 7456 | ENSG00000120868 | APAF1 | chr12 | + | 98648625 | 98648815 | 98648625 | 98648782 | 98649486 | 98649684 | 173 | 136 | 113 | 140 | 168 | 13 | 7 | 12 | 9 | 2 | 163 | 153 | 82 | 264 | 32 | 34 | 14 | 24 | 43 | 0 | 182 | 149 | 1.13E-05 | 0.000282971 | 0.916 | 0.941 | 0.885 | 0.927 | 0.986 | 0.797 | 0.899 | 0.737 | 0.834 | 1 | 0.078 |
| A5SS | 1596 | ENSG00000154237 | LRRK1 | chr15 | + | 101022382 | 101022597 | 101022382 | 101022478 | 101024802 | 101024967 | 15 | 34 | 48 | 26 | 11 | 0 | 0 | 2 | 5 | 0 | 80 | 72 | 27 | 98 | 130 | 20 | 1 | 9 | 24 | 38 | 268 | 149 | 1.24E-05 | 0.000310628 | 1 | 1 | 0.93 | 0.743 | 1 | 0.69 | 0.976 | 0.625 | 0.694 | 0.655 | 0.207 |
| A5SS | 6700 | ENSG00000140396 | NCOA2 | chr8 | - | 70141178 | 70141399 | 70141183 | 70141399 | 70138202 | 70138332 | 1 | 0 | 0 | 2 | 0 | 97 | 91 | 87 | 111 | 101 | 0 | 9 | 2 | 11 | 0 | 42 | 115 | 114 | 81 | 21 | 154 | 149 | 1.27E-05 | 0.000314409 | 0.01 | 0 | 0 | 0.017 | 0 | 0 | 0.07 | 0.017 | 0.116 | 0 | -0.035 |
| A5SS | 6041 | ENSG00000042088 | TDP1 | chr14 | + | 89980539 | 89980679 | 89980539 | 89980632 | 89985131 | 89985210 | 2 | 2 | 1 | 4 | 2 | 1 | 1 | 0 | 2 | 0 | 0 | 1 | 8 | 5 | 0 | 23 | 1 | 3 | 10 | 1 | 196 | 149 | 1.30E-05 | 0.000320685 | 0.603 | 0.603 | 1 | 0.603 | 1 | 0 | 0.432 | 0.67 | 0.275 | 0 | 0.486 |
| A5SS | 6008 | ENSG00000148291 | SURF2 | chr9 | + | 133356913 | 133357234 | 133356913 | 133357068 | 133357710 | 133357814 | 10 | 0 | 0 | 8 | 0 | 9 | 5 | 13 | 19 | 1 | 0 | 1 | 0 | 1 | 0 | 21 | 5 | 4 | 10 | 11 | 298 | 149 | 1.38E-05 | 0.000338595 | 0.357 | 0 | 0 | 0.174 | 0 | 0 | 0.091 | 0 | 0.048 | 0 | 0.078 |
| A5SS | 7073 | ENSG00000102103 | PQBP1 | chrX | + | 48902232 | 48902517 | 48902232 | 48902384 | 48902731 | 48902793 | 244 | 138 | 192 | 158 | 177 | 2 | 5 | 5 | 6 | 9 | 216 | 166 | 119 | 126 | 304 | 0 | 2 | 4 | 0 | 0 | 282 | 149 | 1.46E-05 | 0.000358829 | 0.985 | 0.936 | 0.953 | 0.933 | 0.912 | 1 | 0.978 | 0.94 | 1 | 1 | -0.04 |
| A5SS | 3167 | ENSG00000173193 | PARP14 | chr3 | + | 122720254 | 122721653 | 122720254 | 122720388 | 122727811 | 122727986 | 26 | 141 | 58 | 14 | 54 | 185 | 139 | 119 | 182 | 109 | 25 | 86 | 56 | 124 | 16 | 21 | 107 | 103 | 133 | 13 | 298 | 149 | 1.49E-05 | 0.000363747 | 0.066 | 0.337 | 0.196 | 0.037 | 0.199 | 0.373 | 0.287 | 0.214 | 0.318 | 0.381 | -0.148 |
| A5SS | 655 | ENSG00000101782 | RIOK3 | chr18 | + | 23473428 | 23473626 | 23473428 | 23473483 | 23474947 | 23475107 | 411 | 448 | 247 | 482 | 738 | 6 | 3 | 5 | 4 | 9 | 511 | 544 | 564 | 797 | 208 | 7 | 1 | 18 | 38 | 13 | 292 | 149 | 1.50E-05 | 0.000364203 | 0.972 | 0.987 | 0.962 | 0.984 | 0.977 | 0.974 | 0.996 | 0.941 | 0.915 | 0.891 | 0.033 |
| A5SS | 5359 | ENSG00000084623 | EIF3I | chr1 | + | 32226170 | 32226530 | 32226170 | 32226320 | 32228498 | 32228609 | 236 | 145 | 231 | 223 | 356 | 1 | 2 | 2 | 5 | 7 | 296 | 325 | 177 | 195 | 473 | 6 | 11 | 15 | 14 | 12 | 298 | 149 | 1.52E-05 | 0.000367428 | 0.992 | 0.973 | 0.983 | 0.957 | 0.962 | 0.961 | 0.937 | 0.855 | 0.874 | 0.952 | 0.058 |
| A5SS | 505 | ENSG00000185513 | L3MBTL1 | chr20 | + | 43530283 | 43530889 | 43530283 | 43530419 | 43532772 | 43532924 | 3 | 3 | 3 | 1 | 2 | 1 | 2 | 2 | 2 | 4 | 5 | 0 | 4 | 18 | 7 | 0 | 4 | 0 | 0 | 0 | 298 | 149 | 1.54E-05 | 0.00037075 | 0.6 | 0.429 | 0.429 | 0.2 | 0.2 | 1 | 0 | 1 | 1 | 1 | -0.428 |
| A5SS | 7671 | ENSG00000021776 | AQR | chr15 | - | 34906518 | 34906712 | 34906544 | 34906712 | 34904335 | 34904505 | 6 | 2 | 6 | 0 | 0 | 246 | 264 | 183 | 293 | 294 | 11 | 17 | 3 | 1 | 0 | 81 | 229 | 193 | 181 | 10 | 175 | 149 | 1.59E-05 | 0.000381863 | 0.02 | 0.006 | 0.027 | 0 | 0 | 0.104 | 0.059 | 0.013 | 0.005 | 0 | -0.026 |
| A5SS | 4707 | ENSG00000144535 | DIS3L2 | chr2 | + | 232130618 | 232130943 | 232130618 | 232130719 | 232163458 | 232163632 | 27 | 33 | 20 | 10 | 13 | 14 | 10 | 5 | 7 | 14 | 10 | 7 | 21 | 14 | 4 | 20 | 27 | 20 | 17 | 22 | 298 | 149 | 1.61E-05 | 0.000385158 | 0.491 | 0.623 | 0.667 | 0.417 | 0.317 | 0.2 | 0.115 | 0.344 | 0.292 | 0.083 | 0.296 |
| A5SS | 2866 | ENSG00000178035 | IMPDH2 | chr3 | - | 49028198 | 49028322 | 49028247 | 49028322 | 49027709 | 49027916 | 12 | 6 | 12 | 16 | 5 | 100 | 66 | 112 | 82 | 106 | 22 | 22 | 30 | 7 | 26 | 95 | 60 | 34 | 66 | 103 | 198 | 149 | 1.62E-05 | 0.000385158 | 0.083 | 0.064 | 0.075 | 0.128 | 0.034 | 0.148 | 0.216 | 0.399 | 0.074 | 0.16 | -0.123 |
| A5SS | 2845 | ENSG00000157601 | MX1 | chr21 | + | 41441715 | 41443866 | 41441715 | 41441914 | 41445447 | 41445570 | 296 | 515 | 296 | 575 | 412 | 0 | 0 | 0 | 4 | 0 | 2882 | 1095 | 452 | 2113 | 468 | 25 | 13 | 1 | 38 | 6 | 298 | 149 | 1.73E-05 | 0.000408443 | 1 | 1 | 1 | 0.986 | 1 | 0.983 | 0.977 | 0.996 | 0.965 | 0.975 | 0.018 |
| A5SS | 1875 | ENSG00000090615 | GOLGA3 | chr12 | - | 132775065 | 132775305 | 132775140 | 132775305 | 132774156 | 132774320 | 4 | 4 | 3 | 5 | 0 | 29 | 20 | 23 | 23 | 18 | 0 | 0 | 2 | 0 | 0 | 56 | 15 | 6 | 29 | 87 | 224 | 149 | 1.75E-05 | 0.000410539 | 0.084 | 0.117 | 0.08 | 0.126 | 0 | 0 | 0 | 0.181 | 0 | 0 | 0.045 |
| A5SS | 3465 | ENSG00000119782 | FKBP1B | chr2 | + | 24053901 | 24054616 | 24053901 | 24053949 | 24060813 | 24060926 | 0 | 0 | 0 | 1 | 0 | 2 | 1 | 5 | 6 | 6 | 0 | 0 | 2 | 0 | 18 | 12 | 8 | 7 | 4 | 33 | 298 | 149 | 1.75E-05 | 0.000410539 | 0 | 0 | 0 | 0.077 | 0 | 0 | 0 | 0.125 | 0 | 0.214 | -0.052 |
| A5SS | 7275 | ENSG00000183077 | AFMID | chr17 | + | 78205602 | 78205753 | 78205602 | 78205738 | 78205945 | 78206050 | 0 | 0 | 0 | 0 | 3 | 4 | 1 | 5 | 4 | 8 | 0 | 5 | 2 | 0 | 11 | 8 | 3 | 0 | 1 | 0 | 164 | 149 | 1.90E-05 | 0.000443051 | 0 | 0 | 0 | 0 | 0.254 | 0 | 0.602 | 1 | 0 | 1 | -0.47 |
| A5SS | 7431 | ENSG00000185122 | HSF1 | chr8 | + | 144309771 | 144311249 | 144309771 | 144309896 | 144311320 | 144311382 | 122 | 53 | 93 | 93 | 150 | 0 | 0 | 2 | 0 | 3 | 187 | 123 | 79 | 92 | 154 | 8 | 2 | 5 | 7 | 3 | 298 | 149 | 2.02E-05 | 0.000468328 | 1 | 1 | 0.959 | 1 | 0.962 | 0.921 | 0.969 | 0.888 | 0.868 | 0.963 | 0.062 |
| A5SS | 2504 | ENSG00000155229 | MMS19 | chr10 | - | 97461495 | 97461896 | 97461827 | 97461896 | 97460906 | 97461007 | 24 | 21 | 38 | 34 | 23 | 4 | 2 | 4 | 5 | 1 | 27 | 18 | 19 | 17 | 19 | 0 | 0 | 1 | 2 | 0 | 298 | 149 | 2.02E-05 | 0.000468328 | 0.75 | 0.84 | 0.826 | 0.773 | 0.92 | 1 | 1 | 0.905 | 0.81 | 1 | -0.121 |
| A5SS | 8952 | ENSG00000148362 | PAXX | chr9 | + | 136992639 | 136992974 | 136992639 | 136992700 | 136993052 | 136993243 | 24 | 4 | 11 | 12 | 8 | 0 | 0 | 0 | 0 | 0 | 10 | 23 | 11 | 1 | 12 | 9 | 5 | 0 | 0 | 0 | 298 | 149 | 2.03E-05 | 0.000469087 | 1 | 1 | 1 | 1 | 1 | 0.357 | 0.697 | 1 | 1 | 1 | 0.189 |
| A5SS | 1645 | ENSG00000126822 | PLEKHG3 | chr14 | + | 64731360 | 64731806 | 64731360 | 64731543 | 64732094 | 64732181 | 143 | 19 | 63 | 68 | 43 | 1 | 0 | 0 | 1 | 0 | 54 | 21 | 40 | 69 | 75 | 12 | 0 | 0 | 0 | 4 | 298 | 149 | 2.10E-05 | 0.000481624 | 0.986 | 1 | 1 | 0.971 | 1 | 0.692 | 1 | 1 | 1 | 0.904 | 0.072 |
| A5SS | 1777 | ENSG00000105122 | RASAL3 | chr19 | - | 15452044 | 15452815 | 15452657 | 15452815 | 15451623 | 15451938 | 118 | 32 | 104 | 80 | 80 | 2 | 2 | 1 | 3 | 8 | 35 | 138 | 49 | 89 | 125 | 0 | 0 | 0 | 0 | 0 | 298 | 149 | 2.31E-05 | 0.000527935 | 0.967 | 0.889 | 0.981 | 0.93 | 0.833 | 1 | 1 | 1 | 1 | 1 | -0.08 |
| A5SS | 356 | ENSG00000090013 | BLVRB | chr19 | - | 40457498 | 40458244 | 40458173 | 40458244 | 40451363 | 40451492 | 16 | 2 | 5 | 16 | 11 | 3 | 0 | 0 | 0 | 0 | 49 | 10 | 7 | 0 | 53 | 34 | 3 | 0 | 2 | 3 | 298 | 149 | 2.37E-05 | 0.000540511 | 0.727 | 1 | 1 | 1 | 1 | 0.419 | 0.625 | 1 | 0 | 0.898 | 0.357 |
| A5SS | 4808 | ENSG00000168421 | RHOH | chr4 | + | 40242713 | 40242846 | 40242713 | 40242834 | 40243177 | 40244764 | 1 | 0 | 3 | 1 | 1 | 310 | 144 | 391 | 281 | 508 | 0 | 6 | 6 | 19 | 0 | 64 | 306 | 181 | 157 | 148 | 161 | 149 | 2.43E-05 | 0.000551871 | 0.003 | 0 | 0.007 | 0.003 | 0.002 | 0 | 0.018 | 0.03 | 0.101 | 0 | -0.027 |
| A5SS | 7771 | ENSG00000148356 | LRSAM1 | chr9 | + | 127451514 | 127452084 | 127451514 | 127451669 | 127454495 | 127454599 | 30 | 12 | 17 | 18 | 1 | 0 | 0 | 0 | 0 | 0 | 17 | 27 | 5 | 40 | 93 | 9 | 0 | 1 | 0 | 5 | 298 | 149 | 2.50E-05 | 0.000564707 | 1 | 1 | 1 | 1 | 1 | 0.486 | 1 | 0.714 | 1 | 0.903 | 0.179 |
| A5SS | 5329 | ENSG00000197037 | ZSCAN25 | chr7 | + | 99619560 | 99621574 | 99619560 | 99619997 | 99622548 | 99622640 | 49 | 28 | 56 | 52 | 31 | 1 | 0 | 0 | 0 | 0 | 36 | 27 | 44 | 25 | 12 | 0 | 9 | 1 | 3 | 0 | 298 | 149 | 2.69E-05 | 0.000604466 | 0.961 | 1 | 1 | 1 | 1 | 1 | 0.6 | 0.957 | 0.806 | 1 | 0.12 |
| A5SS | 6050 | ENSG00000141504 | SAT2 | chr17 | - | 7627362 | 7627802 | 7627552 | 7627802 | 7627142 | 7627226 | 113 | 43 | 76 | 83 | 100 | 1 | 0 | 1 | 2 | 0 | 195 | 141 | 122 | 136 | 225 | 12 | 3 | 1 | 2 | 11 | 298 | 149 | 2.70E-05 | 0.000604805 | 0.983 | 1 | 0.974 | 0.954 | 1 | 0.89 | 0.959 | 0.984 | 0.971 | 0.911 | 0.039 |
| A5SS | 2359 | ENSG00000114744 | COMMD2 | chr3 | - | 149751322 | 149751485 | 149751402 | 149751485 | 149750677 | 149750851 | 6 | 1 | 9 | 9 | 9 | 91 | 59 | 72 | 80 | 138 | 23 | 23 | 4 | 9 | 0 | 39 | 59 | 75 | 29 | 16 | 229 | 149 | 2.71E-05 | 0.000605133 | 0.041 | 0.011 | 0.075 | 0.068 | 0.041 | 0.277 | 0.202 | 0.034 | 0.168 | 0 | -0.089 |
| A5SS | 6144 | ENSG00000204305 | AGER | chr6 | - | 32183275 | 32183388 | 32183323 | 32183388 | 32183113 | 32183201 | 8 | 7 | 9 | 12 | 6 | 25 | 20 | 28 | 19 | 32 | 59 | 14 | 10 | 10 | 14 | 25 | 4 | 15 | 18 | 6 | 197 | 149 | 2.74E-05 | 0.000607596 | 0.195 | 0.209 | 0.196 | 0.323 | 0.124 | 0.641 | 0.726 | 0.335 | 0.296 | 0.638 | -0.318 |
| A5SS | 2404 | ENSG00000070756 | PABPC1 | chr8 | - | 100718086 | 100718280 | 100718182 | 100718280 | 100717772 | 100717888 | 7361 | 4907 | 7451 | 7158 | 6468 | 25 | 11 | 45 | 18 | 16 | 7727 | 4321 | 5055 | 4738 | 8288 | 77 | 48 | 35 | 46 | 72 | 245 | 149 | 2.81E-05 | 0.000621646 | 0.994 | 0.996 | 0.99 | 0.996 | 0.996 | 0.984 | 0.982 | 0.989 | 0.984 | 0.986 | 0.009 |
| A5SS | 5498 | ENSG00000087087 | SRRT | chr7 | + | 100885700 | 100885774 | 100885700 | 100885762 | 100885862 | 100885941 | 2 | 2 | 6 | 3 | 1 | 110 | 87 | 120 | 145 | 77 | 35 | 1 | 7 | 13 | 8 | 165 | 182 | 117 | 159 | 238 | 161 | 149 | 3.02E-05 | 0.000664751 | 0.017 | 0.021 | 0.044 | 0.019 | 0.012 | 0.164 | 0.005 | 0.052 | 0.07 | 0.03 | -0.042 |
| A5SS | 3373 | ENSG00000125347 | IRF1 | chr5 | - | 132486521 | 132486686 | 132486556 | 132486686 | 132483827 | 132484075 | 19 | 18 | 24 | 14 | 20 | 10 | 0 | 6 | 5 | 2 | 75 | 14 | 17 | 34 | 52 | 0 | 2 | 2 | 4 | 0 | 184 | 149 | 3.12E-05 | 0.000684883 | 0.606 | 1 | 0.764 | 0.694 | 0.89 | 1 | 0.85 | 0.873 | 0.873 | 1 | -0.128 |
| A5SS | 6416 | ENSG00000170632 | ARMC10 | chr7 | + | 103083681 | 103084062 | 103083681 | 103083830 | 103086629 | 103086764 | 16 | 20 | 18 | 5 | 15 | 57 | 56 | 70 | 83 | 147 | 12 | 16 | 16 | 39 | 16 | 35 | 74 | 12 | 68 | 21 | 298 | 149 | 3.22E-05 | 0.00070401 | 0.123 | 0.152 | 0.114 | 0.029 | 0.049 | 0.146 | 0.098 | 0.4 | 0.223 | 0.276 | -0.135 |
| A5SS | 8847 | ENSG00000166598 | HSP90B1 | chr12 | + | 103931520 | 103931748 | 103931520 | 103931623 | 103932276 | 103932418 | 67 | 42 | 60 | 74 | 68 | 498 | 400 | 617 | 485 | 709 | 47 | 84 | 72 | 41 | 6 | 71 | 347 | 879 | 174 | 22 | 274 | 149 | 3.32E-05 | 0.000723752 | 0.068 | 0.054 | 0.05 | 0.077 | 0.05 | 0.265 | 0.116 | 0.043 | 0.114 | 0.129 | -0.074 |
| A5SS | 8629 | ENSG00000040531 | CTNS | chr17 | + | 3655220 | 3655352 | 3655220 | 3655326 | 3656486 | 3656586 | 35 | 35 | 44 | 35 | 92 | 2 | 1 | 0 | 0 | 0 | 55 | 61 | 28 | 43 | 160 | 0 | 4 | 0 | 10 | 8 | 175 | 149 | 3.39E-05 | 0.000734042 | 0.937 | 0.968 | 1 | 1 | 1 | 1 | 0.928 | 1 | 0.785 | 0.945 | 0.049 |
| A5SS | 6003 | ENSG00000112787 | FBRSL1 | chr12 | + | 132572287 | 132572344 | 132572287 | 132572338 | 132572526 | 132572622 | 19 | 4 | 19 | 16 | 13 | 0 | 0 | 3 | 0 | 0 | 10 | 17 | 7 | 10 | 58 | 0 | 0 | 1 | 1 | 16 | 155 | 149 | 3.49E-05 | 0.00075262 | 1 | 1 | 0.859 | 1 | 1 | 1 | 1 | 0.871 | 0.906 | 0.777 | 0.061 |
| A5SS | 1913 | ENSG00000196655 | TRAPPC4 | chr11 | + | 119019142 | 119019424 | 119019142 | 119019317 | 119020149 | 119020253 | 14 | 5 | 14 | 22 | 11 | 21 | 9 | 15 | 31 | 17 | 33 | 17 | 17 | 20 | 35 | 0 | 14 | 11 | 6 | 11 | 256 | 149 | 3.54E-05 | 0.000761141 | 0.28 | 0.244 | 0.352 | 0.292 | 0.274 | 1 | 0.414 | 0.474 | 0.66 | 0.649 | -0.351 |
| A5SS | 5759 | ENSG00000130038 | CRACR2A | chr12 | - | 3648359 | 3648613 | 3648541 | 3648613 | 3644594 | 3644640 | 1 | 0 | 3 | 1 | 1 | 57 | 33 | 50 | 64 | 27 | 7 | 15 | 1 | 1 | 0 | 33 | 49 | 49 | 61 | 2 | 298 | 149 | 3.58E-05 | 0.000766733 | 0.009 | 0 | 0.029 | 0.008 | 0.018 | 0.096 | 0.133 | 0.01 | 0.008 | 0 | -0.037 |
| A5SS | 6006 | ENSG00000177889 | UBE2N | chr12 | - | 93411052 | 93411299 | 93411125 | 93411299 | 93410733 | 93410874 | 625 | 387 | 541 | 623 | 1022 | 11 | 3 | 7 | 5 | 7 | 465 | 499 | 404 | 382 | 248 | 20 | 7 | 10 | 8 | 8 | 222 | 149 | 3.67E-05 | 0.000783614 | 0.974 | 0.989 | 0.981 | 0.988 | 0.99 | 0.94 | 0.98 | 0.964 | 0.97 | 0.954 | 0.023 |
| A5SS | 6979 | ENSG00000175463 | TBC1D10C | chr11 | + | 67406626 | 67406738 | 67406626 | 67406692 | 67406826 | 67407016 | 64 | 17 | 56 | 69 | 22 | 96 | 29 | 94 | 73 | 26 | 5 | 64 | 49 | 29 | 173 | 5 | 60 | 22 | 19 | 44 | 195 | 149 | 3.70E-05 | 0.000783614 | 0.337 | 0.309 | 0.313 | 0.419 | 0.393 | 0.433 | 0.449 | 0.63 | 0.538 | 0.75 | -0.206 |
| A5SS | 357 | ENSG00000090013 | BLVRB | chr19 | - | 40458154 | 40458244 | 40458173 | 40458244 | 40451363 | 40451492 | 179 | 49 | 96 | 119 | 45 | 3 | 0 | 0 | 0 | 0 | 324 | 216 | 112 | 192 | 416 | 34 | 3 | 0 | 2 | 3 | 168 | 149 | 3.70E-05 | 0.000783614 | 0.981 | 1 | 1 | 1 | 1 | 0.894 | 0.985 | 1 | 0.988 | 0.992 | 0.024 |
| A5SS | 863 | ENSG00000007202 | KIAA0100 | chr17 | - | 28643163 | 28643659 | 28643609 | 28643659 | 28642868 | 28642987 | 91 | 71 | 65 | 90 | 111 | 7 | 3 | 1 | 1 | 0 | 156 | 92 | 84 | 126 | 183 | 6 | 11 | 10 | 5 | 22 | 298 | 149 | 3.82E-05 | 0.000806165 | 0.867 | 0.922 | 0.97 | 0.978 | 1 | 0.929 | 0.807 | 0.808 | 0.926 | 0.806 | 0.092 |
| A5SS | 3824 | ENSG00000108654 | DDX5 | chr17 | - | 64500986 | 64501143 | 64501056 | 64501143 | 64500548 | 64500773 | 365 | 196 | 363 | 369 | 452 | 126 | 80 | 88 | 99 | 101 | 48 | 246 | 220 | 140 | 26 | 9 | 40 | 12 | 9 | 10 | 219 | 149 | 3.85E-05 | 0.000809871 | 0.663 | 0.625 | 0.737 | 0.717 | 0.753 | 0.784 | 0.807 | 0.926 | 0.914 | 0.639 | -0.115 |
| A5SS | 3912 | ENSG00000055609 | KMT2C | chr7 | - | 152251938 | 152252090 | 152251971 | 152252090 | 152250852 | 152250966 | 194 | 166 | 140 | 192 | 271 | 18 | 6 | 14 | 8 | 28 | 62 | 113 | 169 | 88 | 72 | 0 | 5 | 3 | 2 | 5 | 182 | 149 | 3.91E-05 | 0.000818296 | 0.898 | 0.958 | 0.891 | 0.952 | 0.888 | 1 | 0.949 | 0.979 | 0.973 | 0.922 | -0.047 |
| A5SS | 7659 | ENSG00000157954 | WIPI2 | chr7 | + | 5193117 | 5193345 | 5193117 | 5193171 | 5199575 | 5199658 | 26 | 40 | 18 | 28 | 37 | 21 | 8 | 18 | 11 | 18 | 141 | 62 | 57 | 112 | 148 | 22 | 21 | 11 | 4 | 17 | 298 | 149 | 4.01E-05 | 0.000833113 | 0.382 | 0.714 | 0.333 | 0.56 | 0.507 | 0.762 | 0.596 | 0.722 | 0.933 | 0.813 | -0.266 |
| A5SS | 951 | ENSG00000108671 | PSMD11 | chr17 | + | 32473800 | 32474081 | 32473800 | 32473945 | 32474763 | 32474824 | 4 | 7 | 8 | 14 | 11 | 149 | 92 | 153 | 144 | 163 | 26 | 28 | 14 | 38 | 0 | 129 | 168 | 99 | 147 | 260 | 285 | 149 | 4.01E-05 | 0.000833113 | 0.014 | 0.038 | 0.027 | 0.048 | 0.034 | 0.095 | 0.08 | 0.069 | 0.119 | 0 | -0.04 |
| A5SS | 6735 | ENSG00000122783 | CYREN | chr7 | - | 135168744 | 135169060 | 135168968 | 135169060 | 135167731 | 135167807 | 122 | 92 | 90 | 124 | 113 | 7 | 1 | 6 | 2 | 1 | 132 | 122 | 113 | 118 | 265 | 0 | 0 | 0 | 0 | 0 | 298 | 149 | 4.18E-05 | 0.000864577 | 0.897 | 0.979 | 0.882 | 0.969 | 0.983 | 1 | 1 | 1 | 1 | 1 | -0.058 |
| A5SS | 1555 | ENSG00000177082 | WDR73 | chr15 | - | 84654226 | 84654280 | 84654233 | 84654280 | 84653631 | 84653699 | 6 | 4 | 9 | 1 | 8 | 17 | 11 | 20 | 16 | 17 | 0 | 3 | 2 | 3 | 0 | 28 | 19 | 20 | 18 | 14 | 156 | 149 | 4.22E-05 | 0.000869394 | 0.252 | 0.258 | 0.301 | 0.056 | 0.31 | 0 | 0.131 | 0.087 | 0.137 | 0 | 0.164 |
| A5SS | 5025 | ENSG00000143486 | EIF2D | chr1 | - | 206611147 | 206611374 | 206611183 | 206611374 | 206609375 | 206609459 | 1 | 1 | 3 | 6 | 0 | 79 | 68 | 75 | 93 | 171 | 17 | 0 | 2 | 11 | 0 | 71 | 45 | 42 | 47 | 141 | 185 | 149 | 4.23E-05 | 0.000869394 | 0.01 | 0.012 | 0.031 | 0.049 | 0 | 0.162 | 0 | 0.037 | 0.159 | 0 | -0.051 |
| A5SS | 2283 | ENSG00000011243 | AKAP8L | chr19 | - | 15398629 | 15400358 | 15400294 | 15400358 | 15397713 | 15397855 | 42 | 17 | 32 | 39 | 14 | 0 | 0 | 0 | 1 | 0 | 44 | 12 | 21 | 48 | 54 | 25 | 1 | 0 | 1 | 0 | 298 | 149 | 4.46E-05 | 0.000911977 | 1 | 1 | 1 | 0.951 | 1 | 0.468 | 0.857 | 1 | 0.96 | 1 | 0.133 |
| A5SS | 5742 | ENSG00000163754 | GYG1 | chr3 | + | 148996741 | 148996904 | 148996741 | 148996892 | 149009275 | 149009402 | 319 | 184 | 222 | 244 | 292 | 4 | 2 | 2 | 5 | 3 | 291 | 451 | 244 | 504 | 361 | 0 | 4 | 3 | 14 | 42 | 161 | 149 | 4.55E-05 | 0.000927589 | 0.987 | 0.988 | 0.99 | 0.978 | 0.989 | 1 | 0.991 | 0.987 | 0.971 | 0.888 | 0.019 |
| A5SS | 4414 | ENSG00000125538 | IL1B | chr2 | - | 112833116 | 112833575 | 112833373 | 112833575 | 112832661 | 112832826 | 37 | 33 | 18 | 23 | 13 | 52 | 51 | 27 | 37 | 31 | 0 | 0 | 16 | 12 | 10 | 68 | 6 | 22 | 37 | 32 | 298 | 149 | 4.60E-05 | 0.000933517 | 0.262 | 0.244 | 0.25 | 0.237 | 0.173 | 0 | 0 | 0.267 | 0.14 | 0.135 | 0.125 |
| A5SS | 1902 | ENSG00000250462 | LRRC37BP1 | chr17 | + | 30631754 | 30632099 | 30631754 | 30631895 | 30633097 | 30633390 | 18 | 11 | 12 | 12 | 7 | 2 | 0 | 0 | 0 | 2 | 19 | 16 | 9 | 10 | 14 | 5 | 2 | 3 | 0 | 13 | 298 | 149 | 4.65E-05 | 0.000939712 | 0.818 | 1 | 1 | 1 | 0.636 | 0.655 | 0.8 | 0.6 | 1 | 0.35 | 0.21 |
| A5SS | 2284 | ENSG00000011243 | AKAP8L | chr19 | - | 15399301 | 15400358 | 15400294 | 15400358 | 15397713 | 15397855 | 175 | 87 | 112 | 177 | 176 | 0 | 0 | 0 | 1 | 0 | 358 | 144 | 169 | 227 | 355 | 25 | 1 | 0 | 1 | 0 | 298 | 149 | 4.68E-05 | 0.000942852 | 1 | 1 | 1 | 0.989 | 1 | 0.877 | 0.986 | 1 | 0.991 | 1 | 0.027 |
| A5SS | 1384 | ENSG00000125877 | ITPA | chr20 | + | 3209512 | 3209617 | 3209512 | 3209566 | 3213168 | 3213226 | 91 | 58 | 82 | 86 | 61 | 0 | 0 | 0 | 0 | 0 | 203 | 115 | 39 | 90 | 113 | 0 | 14 | 2 | 1 | 4 | 200 | 149 | 4.71E-05 | 0.000946485 | 1 | 1 | 1 | 1 | 1 | 1 | 0.86 | 0.936 | 0.985 | 0.955 | 0.053 |
| A5SS | 5273 | ENSG00000138434 | ITPRID2 | chr2 | + | 181915215 | 181917005 | 181915215 | 181916427 | 181918597 | 181918675 | 57 | 60 | 37 | 49 | 63 | 37 | 31 | 26 | 31 | 27 | 12 | 24 | 53 | 22 | 4 | 31 | 24 | 46 | 34 | 6 | 298 | 149 | 4.81E-05 | 0.000961429 | 0.435 | 0.492 | 0.416 | 0.441 | 0.538 | 0.162 | 0.333 | 0.366 | 0.244 | 0.25 | 0.193 |
| A5SS | 749 | ENSG00000126561 | STAT5A | chr17 | + | 42304529 | 42304652 | 42304529 | 42304646 | 42305609 | 42305702 | 80 | 41 | 68 | 84 | 93 | 6 | 1 | 3 | 0 | 10 | 90 | 64 | 43 | 54 | 168 | 0 | 3 | 0 | 0 | 0 | 155 | 149 | 4.87E-05 | 0.000971513 | 0.928 | 0.975 | 0.956 | 1 | 0.899 | 1 | 0.954 | 1 | 1 | 1 | -0.039 |
| A5SS | 6134 | ENSG00000160185 | UBASH3A | chr21 | + | 42434831 | 42435509 | 42434831 | 42434954 | 42437487 | 42437580 | 16 | 10 | 24 | 35 | 42 | 52 | 47 | 47 | 60 | 64 | 2 | 3 | 8 | 9 | 16 | 36 | 65 | 11 | 19 | 114 | 298 | 149 | 5.04E-05 | 0.000998826 | 0.133 | 0.096 | 0.203 | 0.226 | 0.247 | 0.027 | 0.023 | 0.267 | 0.191 | 0.066 | 0.066 |
| A5SS | 6048 | ENSG00000116857 | TMEM9 | chr1 | - | 201153782 | 201153959 | 201153857 | 201153959 | 201151760 | 201151852 | 0 | 3 | 1 | 1 | 0 | 18 | 8 | 31 | 23 | 12 | 0 | 4 | 2 | 0 | 17 | 20 | 12 | 10 | 10 | 20 | 224 | 149 | 5.05E-05 | 0.000998826 | 0 | 0.2 | 0.021 | 0.028 | 0 | 0 | 0.181 | 0.117 | 0 | 0.361 | -0.082 |
| A5SS | 4122 | ENSG00000116337 | AMPD2 | chr1 | + | 109629326 | 109629916 | 109629326 | 109629490 | 109630232 | 109630406 | 88 | 32 | 46 | 46 | 46 | 5 | 2 | 2 | 4 | 2 | 64 | 45 | 38 | 64 | 121 | 5 | 5 | 8 | 33 | 21 | 298 | 149 | 5.48E-05 | 0.001080717 | 0.898 | 0.889 | 0.92 | 0.852 | 0.92 | 0.865 | 0.818 | 0.704 | 0.492 | 0.742 | 0.172 |
| A5SS | 62 | ENSG00000182944 | EWSR1 | chr22 | + | 29298732 | 29299280 | 29298732 | 29298895 | 29299598 | 29299851 | 27 | 24 | 38 | 29 | 24 | 0 | 0 | 0 | 2 | 0 | 124 | 105 | 70 | 125 | 94 | 17 | 0 | 5 | 0 | 2 | 298 | 149 | 5.53E-05 | 0.001085762 | 1 | 1 | 1 | 0.879 | 1 | 0.785 | 1 | 0.875 | 1 | 0.959 | 0.052 |
| A5SS | 5511 | ENSG00000223960 | CHROMR | chr2 | + | 178426365 | 178426661 | 178426365 | 178426505 | 178430694 | 178430889 | 30 | 9 | 20 | 20 | 10 | 11 | 10 | 5 | 9 | 11 | 48 | 32 | 12 | 12 | 11 | 0 | 8 | 5 | 0 | 0 | 298 | 149 | 6.01E-05 | 0.001175738 | 0.577 | 0.31 | 0.667 | 0.526 | 0.312 | 1 | 0.667 | 0.545 | 1 | 1 | -0.364 |
| A5SS | 534 | ENSG00000215440 | NPEPL1 | chr20 | + | 58691723 | 58691830 | 58691723 | 58691780 | 58693736 | 58693922 | 0 | 0 | 5 | 0 | 1 | 10 | 4 | 14 | 12 | 8 | 3 | 13 | 5 | 3 | 5 | 0 | 10 | 13 | 12 | 6 | 199 | 149 | 6.03E-05 | 0.001177067 | 0 | 0 | 0.211 | 0 | 0.086 | 1 | 0.493 | 0.224 | 0.158 | 0.384 | -0.392 |
| A5SS | 7854 | ENSG00000110514 | MADD | chr11 | + | 47284940 | 47285194 | 47284940 | 47285065 | 47289390 | 47289493 | 27 | 7 | 26 | 49 | 64 | 0 | 0 | 0 | 1 | 0 | 60 | 94 | 50 | 82 | 129 | 0 | 7 | 2 | 8 | 0 | 278 | 149 | 6.08E-05 | 0.001181679 | 1 | 1 | 1 | 0.963 | 1 | 1 | 0.878 | 0.931 | 0.846 | 1 | 0.062 |
| A5SS | 4096 | ENSG00000076685 | NT5C2 | chr10 | - | 103097290 | 103098984 | 103098930 | 103098984 | 103095938 | 103095980 | 271 | 224 | 201 | 292 | 380 | 0 | 0 | 0 | 3 | 0 | 478 | 286 | 388 | 439 | 172 | 0 | 14 | 4 | 5 | 7 | 298 | 149 | 6.19E-05 | 0.001199244 | 1 | 1 | 1 | 0.98 | 1 | 1 | 0.911 | 0.98 | 0.978 | 0.925 | 0.037 |
| A5SS | 6632 | ENSG00000051825 | MPHOSPH9 | chr12 | - | 123227462 | 123227616 | 123227545 | 123227616 | 123226287 | 123226351 | 77 | 57 | 31 | 60 | 107 | 3 | 0 | 0 | 0 | 0 | 79 | 70 | 54 | 56 | 15 | 0 | 0 | 6 | 10 | 0 | 232 | 149 | 6.27E-05 | 0.001210165 | 0.943 | 1 | 1 | 1 | 1 | 1 | 1 | 0.853 | 0.782 | 1 | 0.062 |
| A5SS | 2131 | ENSG00000153113 | CAST | chr5 | + | 96722638 | 96722718 | 96722638 | 96722698 | 96726793 | 96726859 | 4 | 2 | 4 | 10 | 39 | 25 | 12 | 13 | 8 | 59 | 0 | 0 | 13 | 3 | 0 | 36 | 55 | 18 | 48 | 15 | 169 | 149 | 6.48E-05 | 0.00124624 | 0.124 | 0.128 | 0.213 | 0.524 | 0.368 | 0 | 0 | 0.389 | 0.052 | 0 | 0.183 |
| A5SS | 369 | ENSG00000105369 | CD79A | chr19 | + | 41878989 | 41879289 | 41878989 | 41879175 | 41879534 | 41879653 | 327 | 97 | 257 | 303 | 311 | 14 | 10 | 12 | 20 | 5 | 209 | 636 | 143 | 159 | 543 | 0 | 12 | 0 | 2 | 20 | 263 | 149 | 6.82E-05 | 0.001299404 | 0.93 | 0.846 | 0.924 | 0.896 | 0.972 | 1 | 0.968 | 1 | 0.978 | 0.939 | -0.063 |
| A5SS | 8468 | ENSG00000107862 | GBF1 | chr10 | + | 102376934 | 102377140 | 102376934 | 102377128 | 102379283 | 102379434 | 12 | 6 | 14 | 16 | 13 | 30 | 14 | 18 | 17 | 21 | 14 | 31 | 27 | 39 | 35 | 5 | 27 | 6 | 27 | 22 | 161 | 149 | 6.82E-05 | 0.001299404 | 0.27 | 0.284 | 0.419 | 0.466 | 0.364 | 0.722 | 0.515 | 0.806 | 0.572 | 0.596 | -0.282 |
| A5SS | 8057 | ENSG00000159352 | PSMD4 | chr1 | + | 151266003 | 151266121 | 151266003 | 151266112 | 151266307 | 151266439 | 9 | 3 | 6 | 6 | 4 | 200 | 75 | 171 | 152 | 168 | 25 | 12 | 6 | 15 | 21 | 147 | 234 | 93 | 163 | 320 | 158 | 149 | 6.84E-05 | 0.001299404 | 0.041 | 0.036 | 0.032 | 0.036 | 0.022 | 0.138 | 0.046 | 0.057 | 0.08 | 0.058 | -0.042 |
| A5SS | 6740 | ENSG00000122783 | CYREN | chr7 | - | 135168785 | 135169060 | 135168968 | 135169060 | 135167731 | 135167807 | 132 | 95 | 96 | 132 | 117 | 7 | 1 | 6 | 2 | 1 | 132 | 126 | 116 | 120 | 287 | 0 | 0 | 0 | 0 | 0 | 298 | 149 | 6.86E-05 | 0.001299404 | 0.904 | 0.979 | 0.889 | 0.971 | 0.983 | 1 | 1 | 1 | 1 | 1 | -0.055 |
| A5SS | 1305 | ENSG00000074755 | ZZEF1 | chr17 | - | 4070683 | 4070924 | 4070810 | 4070924 | 4067162 | 4067242 | 254 | 184 | 186 | 233 | 210 | 19 | 10 | 17 | 7 | 33 | 291 | 278 | 144 | 267 | 455 | 0 | 0 | 12 | 19 | 19 | 276 | 149 | 6.88E-05 | 0.001299404 | 0.878 | 0.909 | 0.855 | 0.947 | 0.775 | 1 | 1 | 0.866 | 0.884 | 0.928 | -0.063 |
| A5SS | 6165 | ENSG00000135486 | HNRNPA1 | chr12 | + | 54282572 | 54282662 | 54282572 | 54282641 | 54283826 | 54283967 | 100 | 97 | 83 | 71 | 129 | 1 | 0 | 0 | 1 | 0 | 61 | 115 | 130 | 140 | 52 | 33 | 0 | 4 | 0 | 0 | 170 | 149 | 6.91E-05 | 0.001300409 | 0.989 | 1 | 1 | 0.984 | 1 | 0.618 | 1 | 0.966 | 1 | 1 | 0.078 |
| A5SS | 3091 | ENSG00000205981 | DNAJC19 | chr3 | - | 180986942 | 180987022 | 180986997 | 180987022 | 180985925 | 180985996 | 68 | 37 | 59 | 76 | 55 | 5 | 5 | 10 | 5 | 2 | 33 | 33 | 31 | 16 | 22 | 0 | 0 | 0 | 4 | 0 | 204 | 149 | 7.17E-05 | 0.001344946 | 0.909 | 0.844 | 0.812 | 0.917 | 0.953 | 1 | 1 | 1 | 0.745 | 1 | -0.062 |
| A5SS | 113 | ENSG00000100242 | SUN2 | chr22 | - | 38740888 | 38741050 | 38741006 | 38741050 | 38740266 | 38740432 | 38 | 16 | 33 | 33 | 11 | 680 | 277 | 476 | 569 | 344 | 42 | 39 | 29 | 59 | 80 | 665 | 613 | 375 | 369 | 715 | 267 | 149 | 7.39E-05 | 0.001381164 | 0.03 | 0.031 | 0.037 | 0.031 | 0.018 | 0.034 | 0.034 | 0.041 | 0.082 | 0.059 | -0.021 |
| A5SS | 5706 | ENSG00000127081 | ZNF484 | chr9 | - | 92877819 | 92878038 | 92877889 | 92878038 | 92875014 | 92875059 | 14 | 17 | 21 | 9 | 13 | 32 | 11 | 29 | 23 | 19 | 50 | 49 | 28 | 23 | 6 | 16 | 21 | 8 | 7 | 9 | 219 | 149 | 7.45E-05 | 0.00138778 | 0.229 | 0.513 | 0.33 | 0.21 | 0.318 | 0.68 | 0.614 | 0.704 | 0.691 | 0.312 | -0.28 |
| A5SS | 4842 | ENSG00000167671 | UBXN6 | chr19 | - | 4446048 | 4446413 | 4446282 | 4446413 | 4445009 | 4445623 | 53 | 20 | 24 | 29 | 16 | 0 | 0 | 0 | 0 | 0 | 43 | 44 | 12 | 19 | 140 | 0 | 0 | 0 | 0 | 14 | 298 | 149 | 7.51E-05 | 0.0013946 | 1 | 1 | 1 | 1 | 1 | 1 | 1 | 1 | 1 | 0.833 | 0.033 |
| A5SS | 7753 | ENSG00000165699 | TSC1 | chr9 | - | 132901588 | 132901699 | 132901630 | 132901699 | 132900714 | 132900837 | 85 | 37 | 61 | 63 | 77 | 0 | 0 | 2 | 0 | 4 | 17 | 53 | 34 | 39 | 2 | 0 | 3 | 4 | 4 | 9 | 191 | 149 | 7.57E-05 | 0.001399778 | 1 | 1 | 0.96 | 1 | 0.938 | 1 | 0.932 | 0.869 | 0.884 | 0.148 | 0.213 |
| A5SS | 3810 | ENSG00000049541 | RFC2 | chr7 | - | 74237288 | 74237442 | 74237361 | 74237442 | 74235531 | 74235645 | 1 | 1 | 1 | 2 | 0 | 84 | 42 | 75 | 74 | 55 | 0 | 5 | 5 | 4 | 15 | 98 | 83 | 55 | 76 | 132 | 222 | 149 | 7.59E-05 | 0.001399778 | 0.008 | 0.016 | 0.009 | 0.018 | 0 | 0 | 0.039 | 0.058 | 0.034 | 0.071 | -0.03 |
| A5SS | 5826 | ENSG00000172936 | MYD88 | chr3 | + | 38140387 | 38140568 | 38140387 | 38140549 | 38140756 | 38140848 | 169 | 93 | 112 | 145 | 153 | 12 | 8 | 17 | 9 | 4 | 375 | 232 | 100 | 229 | 370 | 69 | 18 | 20 | 20 | 69 | 168 | 149 | 7.88E-05 | 0.001449237 | 0.926 | 0.912 | 0.854 | 0.935 | 0.971 | 0.828 | 0.92 | 0.816 | 0.91 | 0.826 | 0.06 |
| A5SS | 1512 | ENSG00000102901 | CENPT | chr16 | - | 67832227 | 67832545 | 67832454 | 67832545 | 67832011 | 67832108 | 97 | 81 | 91 | 115 | 116 | 0 | 0 | 1 | 0 | 0 | 79 | 127 | 77 | 70 | 322 | 6 | 0 | 0 | 6 | 7 | 298 | 149 | 8.30E-05 | 0.001519966 | 1 | 1 | 0.978 | 1 | 1 | 0.868 | 1 | 1 | 0.854 | 0.958 | 0.06 |
| A5SS | 7023 | ENSG00000169359 | SLC33A1 | chr3 | - | 155853107 | 155854456 | 155853222 | 155854456 | 155845012 | 155845076 | 37 | 28 | 55 | 44 | 32 | 4 | 2 | 7 | 1 | 0 | 118 | 70 | 73 | 112 | 64 | 0 | 4 | 1 | 0 | 0 | 264 | 149 | 8.38E-05 | 0.001530227 | 0.839 | 0.888 | 0.816 | 0.961 | 1 | 1 | 0.908 | 0.976 | 1 | 1 | -0.076 |
| A5SS | 4663 | ENSG00000174446 | SNAPC5 | chr15 | - | 66494902 | 66495419 | 66495329 | 66495419 | 66493467 | 66494552 | 8 | 6 | 8 | 9 | 33 | 98 | 76 | 118 | 120 | 116 | 0 | 2 | 10 | 8 | 4 | 42 | 168 | 82 | 133 | 66 | 298 | 149 | 8.57E-05 | 0.001560463 | 0.039 | 0.038 | 0.033 | 0.036 | 0.125 | 0 | 0.006 | 0.057 | 0.029 | 0.029 | 0.03 |
| A5SS | 3230 | ENSG00000184014 | DENND5A | chr11 | - | 9142721 | 9144278 | 9144096 | 9144278 | 9141939 | 9142108 | 196 | 146 | 180 | 204 | 289 | 1 | 4 | 2 | 1 | 0 | 423 | 187 | 211 | 344 | 327 | 7 | 3 | 6 | 16 | 4 | 298 | 149 | 8.75E-05 | 0.001587261 | 0.99 | 0.948 | 0.978 | 0.99 | 1 | 0.968 | 0.969 | 0.946 | 0.915 | 0.976 | 0.026 |
| A5SS | 5506 | ENSG00000237765 | FAM200B | chr4 | + | 15681755 | 15681968 | 15681755 | 15681784 | 15686235 | 15686462 | 40 | 17 | 21 | 36 | 24 | 71 | 46 | 69 | 49 | 106 | 55 | 33 | 29 | 40 | 33 | 45 | 41 | 46 | 38 | 9 | 298 | 149 | 8.86E-05 | 0.00160224 | 0.22 | 0.156 | 0.132 | 0.269 | 0.102 | 0.379 | 0.287 | 0.24 | 0.345 | 0.647 | -0.204 |
| A5SS | 236 | ENSG00000142541 | RPL13A | chr19 | + | 49489849 | 49490090 | 49489849 | 49489912 | 49490231 | 49490297 | 63 | 26 | 71 | 55 | 45 | 176 | 53 | 212 | 108 | 197 | 100 | 40 | 41 | 54 | 137 | 48 | 123 | 87 | 53 | 103 | 298 | 149 | 8.99E-05 | 0.001619149 | 0.152 | 0.197 | 0.143 | 0.203 | 0.103 | 0.51 | 0.14 | 0.191 | 0.337 | 0.399 | -0.156 |
| A5SS | 826 | ENSG00000136448 | NMT1 | chr17 | + | 45081643 | 45081995 | 45081643 | 45081752 | 45086507 | 45086652 | 14 | 6 | 4 | 6 | 12 | 184 | 128 | 146 | 174 | 229 | 40 | 10 | 10 | 7 | 34 | 293 | 159 | 106 | 141 | 293 | 298 | 149 | 9.24E-05 | 0.001655088 | 0.037 | 0.023 | 0.014 | 0.017 | 0.026 | 0.064 | 0.03 | 0.045 | 0.024 | 0.055 | -0.02 |
| A5SS | 5562 | ENSG00000168958 | MFF | chr2 | + | 227328677 | 227330846 | 227328677 | 227328789 | 227332418 | 227332588 | 238 | 118 | 240 | 184 | 321 | 1 | 0 | 2 | 1 | 2 | 282 | 161 | 173 | 166 | 161 | 0 | 3 | 9 | 6 | 5 | 298 | 149 | 9.25E-05 | 0.001655088 | 0.992 | 1 | 0.984 | 0.989 | 0.988 | 1 | 0.964 | 0.906 | 0.933 | 0.942 | 0.042 |
| A5SS | 3913 | ENSG00000141027 | NCOR1 | chr17 | - | 16046950 | 16047093 | 16046989 | 16047093 | 16040440 | 16040494 | 174 | 109 | 151 | 162 | 206 | 6 | 5 | 7 | 1 | 10 | 80 | 163 | 126 | 123 | 71 | 13 | 8 | 23 | 4 | 5 | 188 | 149 | 9.49E-05 | 0.001694101 | 0.958 | 0.945 | 0.945 | 0.992 | 0.942 | 0.83 | 0.942 | 0.813 | 0.961 | 0.918 | 0.064 |
| A5SS | 2081 | ENSG00000146094 | DOK3 | chr5 | - | 177510149 | 177510382 | 177510259 | 177510382 | 177509474 | 177509657 | 4 | 0 | 0 | 6 | 0 | 2 | 3 | 1 | 4 | 2 | 19 | 3 | 1 | 7 | 21 | 4 | 3 | 0 | 2 | 0 | 259 | 149 | 9.66E-05 | 0.001717868 | 0.535 | 0 | 0 | 0.463 | 0 | 0.732 | 0.365 | 1 | 0.668 | 1 | -0.553 |
| A5SS | 7785 | ENSG00000180104 | EXOC3 | chr5 | + | 462156 | 462529 | 462156 | 462307 | 464289 | 465272 | 9 | 21 | 24 | 29 | 19 | 177 | 89 | 135 | 141 | 253 | 46 | 24 | 24 | 27 | 76 | 159 | 183 | 71 | 189 | 257 | 298 | 149 | 9.90E-05 | 0.001754902 | 0.025 | 0.106 | 0.082 | 0.093 | 0.036 | 0.126 | 0.062 | 0.145 | 0.067 | 0.129 | -0.037 |
| A5SS | 5666 | ENSG00000115762 | PLEKHB2 | chr2 | + | 131132901 | 131133015 | 131132901 | 131132991 | 131140169 | 131140275 | 26 | 13 | 13 | 13 | 23 | 28 | 14 | 31 | 33 | 46 | 0 | 9 | 1 | 13 | 8 | 58 | 15 | 26 | 39 | 29 | 173 | 149 | 0.000102136 | 0.00180463 | 0.444 | 0.444 | 0.265 | 0.253 | 0.301 | 0 | 0.341 | 0.032 | 0.223 | 0.192 | 0.184 |
| A5SS | 1888 | ENSG00000006744 | ELAC2 | chr17 | - | 13002440 | 13002579 | 13002445 | 13002579 | 13002273 | 13002359 | 106 | 61 | 83 | 121 | 128 | 1 | 2 | 2 | 0 | 0 | 146 | 104 | 78 | 111 | 161 | 11 | 5 | 6 | 4 | 0 | 154 | 149 | 0.000104695 | 0.001843852 | 0.99 | 0.967 | 0.976 | 1 | 1 | 0.928 | 0.953 | 0.926 | 0.964 | 1 | 0.032 |
| A5SS | 6166 | ENSG00000135486 | HNRNPA1 | chr12 | + | 54282572 | 54282665 | 54282572 | 54282641 | 54283826 | 54283967 | 98 | 93 | 83 | 74 | 131 | 1 | 0 | 0 | 1 | 0 | 54 | 111 | 128 | 135 | 35 | 33 | 0 | 4 | 0 | 0 | 173 | 149 | 0.000106123 | 0.001862966 | 0.988 | 1 | 1 | 0.985 | 1 | 0.585 | 1 | 0.965 | 1 | 1 | 0.085 |
| A5SS | 3477 | ENSG00000137776 | SLTM | chr15 | - | 58893820 | 58893987 | 58893901 | 58893987 | 58893278 | 58893364 | 129 | 175 | 111 | 162 | 164 | 8 | 10 | 12 | 16 | 1 | 12 | 47 | 82 | 71 | 34 | 0 | 1 | 4 | 1 | 0 | 230 | 149 | 0.000111069 | 0.001943528 | 0.913 | 0.919 | 0.857 | 0.868 | 0.991 | 1 | 0.968 | 0.93 | 0.979 | 1 | -0.066 |
| A5SS | 7400 | ENSG00000100023 | PPIL2 | chr22 | + | 21670565 | 21670853 | 21670565 | 21670611 | 21670996 | 21671059 | 26 | 15 | 8 | 6 | 23 | 72 | 51 | 81 | 58 | 60 | 56 | 33 | 43 | 40 | 20 | 105 | 70 | 44 | 59 | 105 | 298 | 149 | 0.0001115 | 0.001944824 | 0.153 | 0.128 | 0.047 | 0.049 | 0.161 | 0.211 | 0.191 | 0.328 | 0.253 | 0.087 | -0.106 |
| A5SS | 5592 | ENSG00000143390 | RFX5 | chr1 | - | 151346484 | 151346606 | 151346488 | 151346606 | 151346204 | 151346333 | 3 | 0 | 0 | 4 | 1 | 282 | 159 | 221 | 208 | 341 | 6 | 5 | 3 | 14 | 8 | 255 | 266 | 128 | 271 | 296 | 153 | 149 | 0.000112354 | 0.001953458 | 0.01 | 0 | 0 | 0.018 | 0.003 | 0.022 | 0.018 | 0.022 | 0.048 | 0.026 | -0.021 |
| A5SS | 814 | ENSG00000141503 | MINK1 | chr17 | + | 4886450 | 4886626 | 4886450 | 4886617 | 4887109 | 4887179 | 165 | 95 | 168 | 178 | 138 | 10 | 1 | 5 | 7 | 2 | 216 | 216 | 163 | 210 | 794 | 20 | 9 | 12 | 20 | 51 | 158 | 149 | 0.000116533 | 0.00201965 | 0.94 | 0.989 | 0.969 | 0.96 | 0.985 | 0.911 | 0.958 | 0.928 | 0.908 | 0.936 | 0.04 |
| A5SS | 5763 | ENSG00000211460 | TSN | chr2 | + | 121757239 | 121757434 | 121757239 | 121757333 | 121758709 | 121758806 | 41 | 21 | 25 | 42 | 17 | 200 | 136 | 173 | 184 | 312 | 35 | 44 | 33 | 59 | 13 | 146 | 223 | 119 | 180 | 69 | 250 | 149 | 0.000117008 | 0.002021452 | 0.109 | 0.084 | 0.079 | 0.12 | 0.031 | 0.125 | 0.105 | 0.142 | 0.163 | 0.101 | -0.043 |
| A5SS | 6778 | ENSG00000087274 | ADD1 | chr4 | + | 2908514 | 2909431 | 2908514 | 2908604 | 2914883 | 2915040 | 45 | 24 | 30 | 29 | 52 | 539 | 329 | 447 | 617 | 489 | 36 | 29 | 15 | 41 | 60 | 846 | 749 | 411 | 651 | 1335 | 298 | 149 | 0.000118862 | 0.002046675 | 0.04 | 0.035 | 0.032 | 0.023 | 0.05 | 0.021 | 0.019 | 0.018 | 0.031 | 0.022 | 0.014 |
| A5SS | 1585 | ENSG00000137841 | PLCB2 | chr15 | - | 40296745 | 40296908 | 40296757 | 40296908 | 40296295 | 40296392 | 12 | 5 | 3 | 1 | 1 | 1 | 0 | 0 | 0 | 0 | 10 | 1 | 7 | 6 | 19 | 0 | 0 | 1 | 0 | 15 | 161 | 149 | 0.00011922 | 0.002046675 | 0.917 | 1 | 1 | 1 | 1 | 1 | 1 | 0.866 | 1 | 0.54 | 0.102 |
| A5SS | 2224 | ENSG00000105223 | PLD3 | chr19 | + | 40365552 | 40365930 | 40365552 | 40365585 | 40366418 | 40366510 | 114 | 72 | 65 | 84 | 91 | 5 | 2 | 3 | 0 | 0 | 132 | 121 | 109 | 95 | 102 | 10 | 10 | 0 | 3 | 20 | 298 | 149 | 0.000120813 | 0.002067496 | 0.919 | 0.947 | 0.915 | 1 | 1 | 0.868 | 0.858 | 1 | 0.941 | 0.718 | 0.079 |
| A5SS | 5692 | ENSG00000243970 | PPIEL | chr1 | - | 39541656 | 39541841 | 39541719 | 39541841 | 39529054 | 39529164 | 0 | 1 | 0 | 3 | 2 | 15 | 18 | 13 | 35 | 2 | 9 | 7 | 4 | 9 | 2 | 19 | 13 | 25 | 16 | 52 | 212 | 149 | 0.000123009 | 0.002096528 | 0 | 0.038 | 0 | 0.057 | 0.413 | 0.25 | 0.275 | 0.101 | 0.283 | 0.026 | -0.085 |
| A5SS | 8552 | ENSG00000213809 | KLRK1 | chr12 | - | 10379446 | 10379792 | 10379699 | 10379792 | 10378553 | 10378705 | 1815 | 981 | 1082 | 1464 | 2039 | 20 | 7 | 12 | 25 | 20 | 173 | 729 | 561 | 299 | 60 | 7 | 0 | 7 | 28 | 6 | 298 | 149 | 0.00012328 | 0.002096528 | 0.978 | 0.986 | 0.978 | 0.967 | 0.981 | 0.925 | 1 | 0.976 | 0.842 | 0.833 | 0.063 |
| A5SS | 971 | ENSG00000108433 | GOSR2 | chr17 | + | 46923159 | 46923632 | 46923159 | 46923221 | 46929519 | 46929584 | 4 | 0 | 3 | 0 | 1 | 119 | 58 | 98 | 110 | 54 | 7 | 5 | 2 | 15 | 3 | 64 | 98 | 56 | 77 | 166 | 298 | 149 | 0.00012451 | 0.002107533 | 0.017 | 0 | 0.015 | 0 | 0.009 | 0.052 | 0.025 | 0.018 | 0.089 | 0.009 | -0.03 |
| A5SS | 5436 | ENSG00000175215 | CTDSP2 | chr12 | - | 57827533 | 57827590 | 57827551 | 57827590 | 57826995 | 57827097 | 63 | 50 | 75 | 105 | 60 | 304 | 136 | 231 | 246 | 113 | 217 | 123 | 79 | 72 | 273 | 241 | 220 | 144 | 147 | 562 | 167 | 149 | 0.000124701 | 0.002107533 | 0.156 | 0.247 | 0.225 | 0.276 | 0.321 | 0.445 | 0.333 | 0.329 | 0.304 | 0.302 | -0.098 |
| A5SS | 332 | ENSG00000142227 | EMP3 | chr19 | + | 48327520 | 48327664 | 48327520 | 48327623 | 48329351 | 48329492 | 3 | 5 | 5 | 11 | 2 | 583 | 234 | 315 | 404 | 251 | 40 | 17 | 7 | 3 | 41 | 772 | 635 | 303 | 783 | 987 | 190 | 149 | 0.000126226 | 0.002122759 | 0.004 | 0.016 | 0.012 | 0.021 | 0.006 | 0.039 | 0.021 | 0.018 | 0.003 | 0.032 | -0.011 |
| A5SS | 573 | ENSG00000105576 | TNPO2 | chr19 | - | 12723768 | 12723932 | 12723772 | 12723932 | 12723248 | 12723365 | 3 | 1 | 3 | 8 | 20 | 1 | 0 | 1 | 0 | 2 | 9 | 7 | 4 | 3 | 0 | 5 | 2 | 2 | 0 | 4 | 153 | 149 | 0.000126383 | 0.002122759 | 0.745 | 1 | 0.745 | 1 | 0.907 | 0.637 | 0.773 | 0.661 | 1 | 0 | 0.265 |
| A5SS | 3076 | ENSG00000111642 | CHD4 | chr12 | - | 6583195 | 6583378 | 6583197 | 6583378 | 6583026 | 6583113 | 1 | 1 | 2 | 3 | 0 | 323 | 203 | 328 | 349 | 402 | 9 | 0 | 4 | 20 | 13 | 407 | 290 | 237 | 413 | 561 | 151 | 149 | 0.000133032 | 0.002227569 | 0.003 | 0.005 | 0.006 | 0.008 | 0 | 0.021 | 0 | 0.016 | 0.046 | 0.022 | -0.017 |
| A5SS | 7109 | ENSG00000163682 | RPL9 | chr4 | - | 39457585 | 39457681 | 39457596 | 39457681 | 39454863 | 39454944 | 157 | 76 | 67 | 61 | 69 | 3 | 6 | 4 | 4 | 3 | 16 | 40 | 96 | 62 | 132 | 0 | 0 | 2 | 0 | 0 | 160 | 149 | 0.000137431 | 0.002294167 | 0.98 | 0.922 | 0.94 | 0.934 | 0.955 | 1 | 1 | 0.978 | 1 | 1 | -0.049 |
| A5SS | 1205 | ENSG00000197912 | SPG7 | chr16 | + | 89540943 | 89542279 | 89540943 | 89541384 | 89544647 | 89544772 | 100 | 121 | 92 | 105 | 67 | 2 | 2 | 2 | 11 | 2 | 273 | 167 | 148 | 318 | 243 | 0 | 0 | 3 | 5 | 2 | 298 | 149 | 0.000145977 | 0.002429373 | 0.962 | 0.968 | 0.958 | 0.827 | 0.944 | 1 | 1 | 0.961 | 0.97 | 0.984 | -0.051 |
| A5SS | 1926 | ENSG00000085511 | MAP3K4 | chr6 | + | 161093772 | 161093851 | 161093772 | 161093839 | 161097079 | 161097176 | 20 | 21 | 29 | 32 | 75 | 30 | 34 | 42 | 22 | 113 | 21 | 21 | 20 | 7 | 3 | 0 | 4 | 12 | 9 | 0 | 161 | 149 | 0.000151586 | 0.002515035 | 0.382 | 0.364 | 0.39 | 0.574 | 0.381 | 1 | 0.829 | 0.607 | 0.419 | 1 | -0.353 |
| A5SS | 63 | ENSG00000099991 | CABIN1 | chr22 | + | 24098013 | 24098192 | 24098013 | 24098105 | 24113565 | 24113748 | 43 | 22 | 40 | 35 | 41 | 5 | 8 | 6 | 3 | 0 | 107 | 45 | 56 | 82 | 138 | 0 | 9 | 0 | 0 | 2 | 236 | 149 | 0.000154389 | 0.002553759 | 0.844 | 0.635 | 0.808 | 0.88 | 1 | 1 | 0.759 | 1 | 1 | 0.978 | -0.114 |
| A5SS | 4176 | ENSG00000115685 | PPP1R7 | chr2 | + | 241158483 | 241158549 | 241158483 | 241158531 | 241159212 | 241159343 | 376 | 239 | 320 | 333 | 673 | 17 | 14 | 10 | 20 | 44 | 588 | 303 | 212 | 333 | 413 | 9 | 6 | 9 | 6 | 21 | 167 | 149 | 0.000157165 | 0.00259179 | 0.952 | 0.938 | 0.966 | 0.937 | 0.932 | 0.983 | 0.978 | 0.955 | 0.98 | 0.946 | -0.023 |
| A5SS | 4152 | ENSG00000148187 | MRRF | chr9 | + | 122265461 | 122265780 | 122265461 | 122265545 | 122270863 | 122271075 | 26 | 17 | 23 | 15 | 31 | 0 | 1 | 3 | 2 | 0 | 0 | 42 | 19 | 36 | 0 | 14 | 4 | 2 | 1 | 3 | 298 | 149 | 0.00016182 | 0.002660079 | 1 | 0.895 | 0.793 | 0.789 | 1 | 0 | 0.84 | 0.826 | 0.947 | 0 | 0.373 |
| A5SS | 7871 | ENSG00000198561 | CTNND1 | chr11 | + | 57761796 | 57762119 | 57761796 | 57762046 | 57789036 | 57789155 | 12 | 35 | 6 | 23 | 10 | 18 | 23 | 13 | 27 | 43 | 85 | 24 | 28 | 86 | 39 | 79 | 20 | 5 | 28 | 11 | 222 | 149 | 0.000162459 | 0.002660079 | 0.309 | 0.505 | 0.237 | 0.364 | 0.135 | 0.419 | 0.446 | 0.79 | 0.673 | 0.704 | -0.296 |
| A5SS | 4632 | ENSG00000131165 | CHMP1A | chr16 | - | 89653798 | 89653923 | 89653903 | 89653923 | 89651568 | 89651646 | 10 | 9 | 7 | 8 | 3 | 96 | 50 | 90 | 69 | 56 | 66 | 22 | 18 | 18 | 21 | 49 | 94 | 92 | 79 | 221 | 254 | 149 | 0.000162772 | 0.002660079 | 0.058 | 0.096 | 0.044 | 0.064 | 0.03 | 0.441 | 0.121 | 0.103 | 0.118 | 0.053 | -0.109 |
| A5SS | 5931 | ENSG00000124356 | STAMBP | chr2 | + | 73828982 | 73829510 | 73828982 | 73829070 | 73830844 | 73831059 | 30 | 10 | 12 | 21 | 29 | 1 | 2 | 1 | 0 | 0 | 35 | 15 | 11 | 30 | 15 | 8 | 0 | 3 | 4 | 0 | 298 | 149 | 0.000171436 | 0.00279328 | 0.938 | 0.714 | 0.857 | 1 | 1 | 0.686 | 1 | 0.647 | 0.789 | 1 | 0.077 |
| A5SS | 5032 | ENSG00000162650 | ATXN7L2 | chr1 | + | 109486056 | 109486304 | 109486056 | 109486122 | 109486505 | 109486610 | 8 | 2 | 4 | 9 | 3 | 3 | 3 | 6 | 3 | 3 | 24 | 12 | 14 | 4 | 18 | 10 | 0 | 2 | 0 | 0 | 298 | 149 | 0.000180455 | 0.002931452 | 0.571 | 0.25 | 0.25 | 0.6 | 0.333 | 0.545 | 1 | 0.778 | 1 | 1 | -0.464 |
| A5SS | 2962 | ENSG00000060971 | ACAA1 | chr3 | - | 38126161 | 38126700 | 38126509 | 38126700 | 38125825 | 38125881 | 96 | 58 | 108 | 105 | 49 | 62 | 24 | 43 | 34 | 17 | 133 | 160 | 60 | 130 | 185 | 114 | 72 | 68 | 118 | 166 | 298 | 149 | 0.000184071 | 0.002981293 | 0.436 | 0.547 | 0.557 | 0.607 | 0.59 | 0.368 | 0.526 | 0.306 | 0.355 | 0.358 | 0.165 |
| A5SS | 4010 | ENSG00000069424 | KCNAB2 | chr1 | + | 6094399 | 6094962 | 6094399 | 6094485 | 6095322 | 6095443 | 20 | 13 | 16 | 24 | 2 | 120 | 53 | 75 | 102 | 65 | 22 | 34 | 42 | 21 | 101 | 74 | 146 | 55 | 102 | 248 | 298 | 149 | 0.000186387 | 0.003000816 | 0.077 | 0.109 | 0.096 | 0.105 | 0.015 | 0.129 | 0.104 | 0.276 | 0.093 | 0.169 | -0.074 |
| A5SS | 2899 | ENSG00000254087 | LYN | chr8 | + | 55941854 | 55941991 | 55941854 | 55941928 | 55946447 | 55946493 | 984 | 839 | 463 | 956 | 1210 | 513 | 610 | 323 | 532 | 737 | 3486 | 1143 | 948 | 1833 | 1965 | 2426 | 818 | 1064 | 1461 | 1801 | 212 | 149 | 0.000186417 | 0.003000816 | 0.574 | 0.492 | 0.502 | 0.558 | 0.536 | 0.502 | 0.495 | 0.385 | 0.469 | 0.434 | 0.075 |
| A5SS | 4032 | ENSG00000126091 | ST3GAL3 | chr1 | + | 43736232 | 43736425 | 43736232 | 43736380 | 43792101 | 43792149 | 3 | 2 | 5 | 4 | 0 | 28 | 13 | 20 | 33 | 24 | 0 | 3 | 0 | 0 | 1 | 14 | 25 | 16 | 10 | 158 | 194 | 149 | 0.000186931 | 0.003000816 | 0.076 | 0.106 | 0.161 | 0.085 | 0 | 0 | 0.084 | 0 | 0 | 0.005 | 0.068 |
| A5SS | 4033 | ENSG00000126091 | ST3GAL3 | chr1 | + | 43736232 | 43736425 | 43736232 | 43736380 | 43814890 | 43814933 | 2 | 2 | 6 | 4 | 0 | 1 | 8 | 1 | 8 | 10 | 0 | 1 | 0 | 0 | 0 | 10 | 0 | 4 | 7 | 5 | 194 | 149 | 0.000190497 | 0.003049071 | 0.606 | 0.161 | 0.822 | 0.277 | 0 | 0 | 1 | 0 | 0 | 0 | 0.173 |
| A5SS | 8401 | ENSG00000103024 | NME3 | chr16 | - | 1771268 | 1771410 | 1771279 | 1771410 | 1771069 | 1771171 | 5 | 1 | 2 | 0 | 3 | 22 | 2 | 7 | 7 | 1 | 1 | 0 | 0 | 0 | 0 | 7 | 6 | 1 | 9 | 1 | 160 | 149 | 0.000194946 | 0.003111136 | 0.175 | 0.318 | 0.21 | 0 | 0.736 | 0.117 | 0 | 0 | 0 | 0 | 0.264 |
| A5SS | 6007 | ENSG00000215375 | MYL5 | chr4 | + | 678957 | 679173 | 678957 | 679033 | 679913 | 680018 | 1 | 0 | 3 | 6 | 7 | 16 | 9 | 21 | 8 | 8 | 0 | 0 | 0 | 3 | 2 | 10 | 18 | 22 | 11 | 12 | 289 | 149 | 0.000199932 | 0.003181373 | 0.031 | 0 | 0.069 | 0.279 | 0.311 | 0 | 0 | 0 | 0.123 | 0.079 | 0.098 |
| A5SS | 3391 | ENSG00000132768 | DPH2 | chr1 | + | 43970595 | 43970817 | 43970595 | 43970708 | 43970965 | 43971189 | 5 | 4 | 0 | 9 | 4 | 2 | 0 | 3 | 5 | 3 | 0 | 1 | 0 | 0 | 3 | 4 | 6 | 3 | 7 | 0 | 258 | 149 | 0.000201099 | 0.003190106 | 0.591 | 1 | 0 | 0.51 | 0.435 | 0 | 0.088 | 0 | 0 | 1 | 0.29 |
| A5SS | 1477 | ENSG00000182718 | ANXA2 | chr15 | - | 60396279 | 60396427 | 60396372 | 60396427 | 60386027 | 60386086 | 11 | 7 | 5 | 13 | 13 | 0 | 1 | 0 | 0 | 0 | 29 | 7 | 7 | 27 | 16 | 2 | 2 | 2 | 1 | 3 | 242 | 149 | 0.000201653 | 0.003190106 | 1 | 0.812 | 1 | 1 | 1 | 0.899 | 0.683 | 0.683 | 0.943 | 0.767 | 0.167 |
| A5SS | 3659 | ENSG00000163565 | IFI16 | chr1 | + | 159051678 | 159053724 | 159051678 | 159052098 | 159054820 | 159055151 | 638 | 457 | 436 | 547 | 813 | 15 | 12 | 13 | 15 | 25 | 51 | 271 | 245 | 86 | 12 | 8 | 8 | 11 | 11 | 7 | 298 | 149 | 0.000204087 | 0.003219248 | 0.955 | 0.95 | 0.944 | 0.948 | 0.942 | 0.761 | 0.944 | 0.918 | 0.796 | 0.462 | 0.172 |
| A5SS | 3778 | ENSG00000139921 | TMX1 | chr14 | + | 51245312 | 51245454 | 51245312 | 51245358 | 51247091 | 51247220 | 15 | 14 | 6 | 15 | 11 | 213 | 116 | 148 | 203 | 259 | 11 | 30 | 14 | 1 | 0 | 36 | 133 | 107 | 34 | 25 | 245 | 149 | 0.000208153 | 0.003273895 | 0.041 | 0.068 | 0.024 | 0.043 | 0.025 | 0.157 | 0.121 | 0.074 | 0.018 | 0 | -0.034 |
| A5SS | 349 | ENSG00000105486 | LIG1 | chr19 | - | 48120166 | 48121322 | 48121169 | 48121322 | 48119136 | 48119190 | 26 | 6 | 19 | 10 | 8 | 7 | 9 | 9 | 15 | 6 | 6 | 6 | 9 | 2 | 0 | 33 | 7 | 7 | 14 | 12 | 298 | 149 | 0.000209816 | 0.003290549 | 0.65 | 0.25 | 0.514 | 0.25 | 0.4 | 0.083 | 0.3 | 0.391 | 0.067 | 0 | 0.245 |
| A5SS | 471 | ENSG00000064601 | CTSA | chr20 | + | 45892724 | 45893311 | 45892724 | 45892880 | 45893987 | 45894072 | 545 | 424 | 314 | 578 | 775 | 1 | 1 | 0 | 2 | 0 | 1562 | 1477 | 601 | 1032 | 7037 | 21 | 7 | 14 | 34 | 21 | 298 | 149 | 0.000212471 | 0.003322606 | 0.996 | 0.995 | 1 | 0.993 | 1 | 0.974 | 0.991 | 0.955 | 0.938 | 0.994 | 0.026 |
| A5SS | 5716 | ENSG00000110931 | CAMKK2 | chr12 | - | 121274055 | 121274370 | 121274063 | 121274370 | 121270897 | 121270945 | 29 | 21 | 22 | 14 | 4 | 0 | 0 | 0 | 0 | 0 | 40 | 27 | 12 | 42 | 90 | 0 | 0 | 0 | 5 | 10 | 157 | 149 | 0.000217614 | 0.003393286 | 1 | 1 | 1 | 1 | 1 | 1 | 1 | 1 | 0.889 | 0.895 | 0.043 |
| A5SS | 4712 | ENSG00000001497 | LAS1L | chrX | - | 65520701 | 65523707 | 65523559 | 65523707 | 65517986 | 65518465 | 4 | 9 | 9 | 6 | 3 | 41 | 35 | 60 | 48 | 68 | 14 | 8 | 14 | 4 | 48 | 44 | 40 | 55 | 46 | 42 | 298 | 149 | 0.000218607 | 0.003399019 | 0.047 | 0.114 | 0.07 | 0.059 | 0.022 | 0.137 | 0.091 | 0.113 | 0.042 | 0.364 | -0.087 |
| A5SS | 8359 | ENSG00000146826 | TRAPPC14 | chr7 | - | 100157292 | 100157459 | 100157372 | 100157459 | 100157093 | 100157214 | 0 | 1 | 2 | 6 | 2 | 52 | 34 | 52 | 46 | 33 | 0 | 14 | 17 | 8 | 0 | 58 | 39 | 23 | 39 | 68 | 229 | 149 | 0.000222227 | 0.003440476 | 0 | 0.019 | 0.024 | 0.078 | 0.038 | 0 | 0.189 | 0.325 | 0.118 | 0 | -0.095 |
| A5SS | 7745 | ENSG00000172716 | SLFN11 | chr17 | - | 35367540 | 35367700 | 35367602 | 35367700 | 35366946 | 35367063 | 11 | 13 | 4 | 2 | 8 | 42 | 16 | 29 | 44 | 60 | 11 | 1 | 0 | 3 | 0 | 94 | 69 | 17 | 40 | 23 | 211 | 149 | 0.000222537 | 0.003440476 | 0.156 | 0.365 | 0.089 | 0.031 | 0.086 | 0.076 | 0.01 | 0 | 0.05 | 0 | 0.118 |
| A5SS | 1918 | ENSG00000115760 | BIRC6 | chr2 | + | 32453807 | 32453954 | 32453807 | 32453942 | 32463193 | 32463381 | 35 | 20 | 33 | 26 | 67 | 62 | 37 | 57 | 79 | 165 | 20 | 26 | 24 | 9 | 33 | 0 | 50 | 30 | 11 | 0 | 161 | 149 | 0.000227073 | 0.003500656 | 0.343 | 0.333 | 0.349 | 0.233 | 0.273 | 1 | 0.325 | 0.425 | 0.431 | 1 | -0.33 |
| A5SS | 2793 | ENSG00000114745 | GORASP1 | chr3 | - | 39100787 | 39101102 | 39101015 | 39101102 | 39100304 | 39100454 | 10 | 5 | 11 | 14 | 1 | 0 | 0 | 0 | 1 | 0 | 9 | 9 | 10 | 12 | 38 | 0 | 9 | 2 | 1 | 0 | 298 | 149 | 0.000228191 | 0.003507951 | 1 | 1 | 1 | 0.875 | 1 | 1 | 0.333 | 0.714 | 0.857 | 1 | 0.194 |
| A5SS | 1664 | ENSG00000100796 | PPP4R3A | chr14 | - | 91475810 | 91475966 | 91475849 | 91475966 | 91473238 | 91473370 | 224 | 193 | 190 | 208 | 416 | 153 | 100 | 91 | 120 | 307 | 88 | 139 | 313 | 190 | 30 | 70 | 42 | 46 | 43 | 6 | 188 | 149 | 0.00023243 | 0.003563058 | 0.537 | 0.605 | 0.623 | 0.579 | 0.518 | 0.499 | 0.724 | 0.844 | 0.778 | 0.798 | -0.156 |
| A5SS | 1757 | ENSG00000165410 | CFL2 | chr14 | - | 34713396 | 34713561 | 34713507 | 34713561 | 34713059 | 34713136 | 40 | 35 | 52 | 72 | 94 | 2 | 2 | 3 | 0 | 7 | 9 | 28 | 26 | 59 | 24 | 0 | 0 | 0 | 1 | 0 | 260 | 149 | 0.000237008 | 0.003623025 | 0.92 | 0.909 | 0.909 | 1 | 0.885 | 1 | 1 | 1 | 0.971 | 1 | -0.07 |
| A5SS | 2338 | ENSG00000137449 | CPEB2 | chr4 | + | 15007304 | 15007595 | 15007304 | 15007586 | 15008337 | 15008427 | 1 | 0 | 2 | 0 | 0 | 15 | 12 | 8 | 13 | 18 | 12 | 0 | 10 | 0 | 0 | 14 | 14 | 8 | 14 | 54 | 158 | 149 | 0.000238994 | 0.003643147 | 0.059 | 0 | 0.191 | 0 | 0 | 0.447 | 0 | 0.541 | 0 | 0 | -0.148 |
| A5SS | 4011 | ENSG00000069424 | KCNAB2 | chr1 | + | 6097268 | 6097555 | 6097268 | 6097357 | 6098484 | 6101180 | 21 | 17 | 21 | 31 | 16 | 210 | 104 | 142 | 224 | 176 | 16 | 53 | 55 | 23 | 85 | 233 | 205 | 93 | 259 | 198 | 298 | 149 | 0.000247908 | 0.003768473 | 0.048 | 0.076 | 0.069 | 0.065 | 0.043 | 0.033 | 0.114 | 0.228 | 0.043 | 0.177 | -0.059 |
| A5SS | 1755 | ENSG00000165410 | CFL2 | chr14 | - | 34713253 | 34713561 | 34713396 | 34713561 | 34713059 | 34713136 | 122 | 95 | 114 | 174 | 317 | 6 | 4 | 5 | 1 | 3 | 131 | 107 | 101 | 150 | 46 | 0 | 0 | 0 | 0 | 0 | 292 | 149 | 0.000253427 | 0.003841642 | 0.912 | 0.924 | 0.921 | 0.989 | 0.982 | 1 | 1 | 1 | 1 | 1 | -0.054 |
| A5SS | 8580 | ENSG00000284770 | TBCE | chr1 | + | 235438768 | 235438988 | 235438768 | 235438922 | 235441813 | 235441882 | 0 | 4 | 0 | 3 | 0 | 55 | 68 | 46 | 59 | 87 | 16 | 0 | 3 | 3 | 0 | 95 | 62 | 33 | 44 | 26 | 215 | 149 | 0.00025957 | 0.003923837 | 0 | 0.039 | 0 | 0.034 | 0 | 0.105 | 0 | 0.059 | 0.045 | 0 | -0.027 |
| A5SS | 1070 | ENSG00000067955 | CBFB | chr16 | + | 67029726 | 67030177 | 67029726 | 67029813 | 67036638 | 67036755 | 5 | 3 | 6 | 7 | 3 | 160 | 121 | 162 | 208 | 132 | 10 | 0 | 13 | 13 | 18 | 112 | 169 | 181 | 214 | 81 | 298 | 149 | 0.000260737 | 0.003930555 | 0.015 | 0.012 | 0.018 | 0.017 | 0.011 | 0.043 | 0 | 0.035 | 0.029 | 0.1 | -0.027 |
| A5SS | 3974 | ENSG00000198093 | ZNF649 | chr19 | - | 51899797 | 51900294 | 51900092 | 51900294 | 51896851 | 51896978 | 46 | 38 | 39 | 44 | 50 | 29 | 35 | 40 | 28 | 50 | 74 | 61 | 42 | 116 | 71 | 43 | 31 | 35 | 2 | 0 | 298 | 149 | 0.000263294 | 0.003958143 | 0.442 | 0.352 | 0.328 | 0.44 | 0.333 | 0.463 | 0.496 | 0.375 | 0.967 | 1 | -0.281 |
| A5SS | 1661 | ENSG00000174373 | RALGAPA1 | chr14 | - | 35688458 | 35690003 | 35689835 | 35690003 | 35686541 | 35686666 | 20 | 11 | 30 | 11 | 57 | 62 | 37 | 74 | 77 | 122 | 0 | 6 | 13 | 0 | 0 | 19 | 37 | 59 | 4 | 4 | 298 | 149 | 0.000264304 | 0.003962377 | 0.139 | 0.129 | 0.169 | 0.067 | 0.189 | 0 | 0.075 | 0.099 | 0 | 0 | 0.104 |
| A5SS | 7355 | ENSG00000078487 | ZCWPW1 | chr7 | - | 100419001 | 100419189 | 100419110 | 100419189 | 100417065 | 100417183 | 10 | 8 | 5 | 7 | 10 | 4 | 4 | 3 | 9 | 13 | 20 | 16 | 9 | 10 | 5 | 5 | 0 | 2 | 4 | 0 | 258 | 149 | 0.000281589 | 0.004209917 | 0.591 | 0.536 | 0.49 | 0.31 | 0.308 | 0.698 | 1 | 0.722 | 0.591 | 1 | -0.355 |
| A5SS | 3808 | ENSG00000163608 | NEPRO | chr3 | - | 113013948 | 113014032 | 113013952 | 113014032 | 113013271 | 113013403 | 2 | 2 | 3 | 6 | 11 | 21 | 22 | 29 | 15 | 23 | 0 | 0 | 4 | 0 | 0 | 4 | 15 | 9 | 1 | 10 | 153 | 149 | 0.000282901 | 0.004217941 | 0.085 | 0.081 | 0.092 | 0.28 | 0.318 | 0 | 0 | 0.302 | 0 | 0 | 0.111 |
| A5SS | 414 | ENSG00000105705 | SUGP1 | chr19 | - | 19316246 | 19316593 | 19316421 | 19316593 | 19310096 | 19310200 | 4 | 3 | 8 | 13 | 19 | 104 | 64 | 85 | 98 | 367 | 43 | 20 | 19 | 17 | 10 | 196 | 153 | 93 | 217 | 201 | 298 | 149 | 0.000284225 | 0.004219777 | 0.019 | 0.023 | 0.045 | 0.062 | 0.025 | 0.099 | 0.061 | 0.093 | 0.038 | 0.024 | -0.028 |
| A5SS | 126 | ENSG00000100325 | ASCC2 | chr22 | - | 29834441 | 29834550 | 29834488 | 29834550 | 29825621 | 29825780 | 5 | 1 | 2 | 4 | 9 | 4 | 0 | 4 | 2 | 0 | 1 | 2 | 11 | 11 | 12 | 0 | 1 | 2 | 0 | 0 | 196 | 149 | 0.00028489 | 0.004219777 | 0.487 | 1 | 0.275 | 0.603 | 1 | 1 | 0.603 | 0.807 | 1 | 1 | -0.209 |
| A5SS | 5132 | ENSG00000110934 | BIN2 | chr12 | - | 51302019 | 51302780 | 51302685 | 51302780 | 51299606 | 51299714 | 1523 | 1226 | 620 | 1668 | 2319 | 7 | 3 | 6 | 16 | 10 | 2952 | 1914 | 1536 | 2095 | 3371 | 36 | 32 | 24 | 8 | 25 | 298 | 149 | 0.000285351 | 0.004219777 | 0.991 | 0.995 | 0.981 | 0.981 | 0.991 | 0.976 | 0.968 | 0.97 | 0.992 | 0.985 | 0.01 |
| A5SS | 5752 | ENSG00000122203 | KIAA1191 | chr5 | - | 176359461 | 176359567 | 176359480 | 176359567 | 176355570 | 176355749 | 4 | 0 | 1 | 2 | 0 | 46 | 43 | 46 | 59 | 62 | 0 | 2 | 1 | 6 | 31 | 87 | 82 | 74 | 75 | 218 | 168 | 149 | 0.000293526 | 0.004328918 | 0.072 | 0 | 0.019 | 0.029 | 0 | 0 | 0.021 | 0.012 | 0.066 | 0.112 | -0.018 |
| A5SS | 7345 | ENSG00000137492 | THAP12 | chr11 | - | 76365779 | 76365972 | 76365851 | 76365972 | 76364337 | 76364432 | 4 | 3 | 3 | 1 | 1 | 43 | 63 | 76 | 65 | 100 | 15 | 6 | 4 | 5 | 0 | 13 | 79 | 70 | 47 | 9 | 221 | 149 | 0.000295142 | 0.004340975 | 0.059 | 0.031 | 0.026 | 0.01 | 0.007 | 0.438 | 0.049 | 0.037 | 0.067 | 0 | -0.092 |
| A5SS | 1419 | ENSG00000213614 | HEXA | chr15 | - | 72347685 | 72348134 | 72348047 | 72348134 | 72346526 | 72346710 | 213 | 126 | 147 | 159 | 189 | 8 | 6 | 11 | 7 | 14 | 348 | 205 | 185 | 255 | 304 | 45 | 25 | 25 | 19 | 20 | 298 | 149 | 0.000299892 | 0.00439895 | 0.93 | 0.913 | 0.87 | 0.919 | 0.871 | 0.795 | 0.804 | 0.787 | 0.87 | 0.884 | 0.073 |
| A5SS | 6348 | ENSG00000127980 | PEX1 | chr7 | - | 92489713 | 92489911 | 92489718 | 92489911 | 92489292 | 92489423 | 91 | 51 | 74 | 86 | 177 | 8 | 3 | 4 | 2 | 13 | 20 | 49 | 60 | 15 | 4 | 0 | 0 | 4 | 0 | 0 | 154 | 149 | 0.000306792 | 0.004481297 | 0.917 | 0.943 | 0.947 | 0.977 | 0.929 | 1 | 1 | 0.936 | 1 | 1 | -0.045 |
| A5SS | 1997 | ENSG00000068796 | KIF2A | chr5 | + | 62352587 | 62352710 | 62352587 | 62352653 | 62353274 | 62353375 | 425 | 522 | 323 | 720 | 820 | 117 | 184 | 92 | 186 | 199 | 643 | 431 | 464 | 380 | 968 | 250 | 222 | 149 | 127 | 370 | 206 | 149 | 0.000307153 | 0.004481297 | 0.724 | 0.672 | 0.717 | 0.737 | 0.749 | 0.65 | 0.584 | 0.693 | 0.684 | 0.654 | 0.067 |
| A5SS | 5048 | ENSG00000033867 | SLC4A7 | chr3 | - | 27433915 | 27434104 | 27433954 | 27434104 | 27424036 | 27424152 | 71 | 29 | 64 | 67 | 103 | 12 | 3 | 8 | 9 | 19 | 25 | 36 | 26 | 3 | 7 | 10 | 7 | 11 | 8 | 11 | 188 | 149 | 0.000314825 | 0.004580953 | 0.824 | 0.885 | 0.864 | 0.855 | 0.811 | 0.665 | 0.803 | 0.652 | 0.229 | 0.335 | 0.311 |
| A5SS | 8713 | ENSG00000135297 | MTO1 | chr6 | + | 73461736 | 73462173 | 73461736 | 73462071 | 73466208 | 73466408 | 7 | 2 | 5 | 11 | 0 | 53 | 26 | 52 | 77 | 33 | 32 | 14 | 0 | 8 | 7 | 60 | 48 | 33 | 47 | 21 | 251 | 149 | 0.000319109 | 0.004630915 | 0.073 | 0.044 | 0.054 | 0.078 | 0 | 0.24 | 0.148 | 0 | 0.092 | 0.165 | -0.079 |
| A5SS | 7344 | ENSG00000137492 | THAP12 | chr11 | - | 76365779 | 76365972 | 76365851 | 76365972 | 76360955 | 76361063 | 5 | 3 | 3 | 1 | 4 | 241 | 180 | 175 | 196 | 689 | 7 | 4 | 8 | 5 | 0 | 11 | 168 | 157 | 70 | 25 | 221 | 149 | 0.000321729 | 0.004652948 | 0.014 | 0.011 | 0.011 | 0.003 | 0.004 | 0.3 | 0.016 | 0.033 | 0.046 | 0 | -0.07 |
| A5SS | 3333 | ENSG00000158864 | NDUFS2 | chr1 | + | 161213863 | 161214032 | 161213863 | 161213921 | 161214155 | 161214299 | 19 | 19 | 19 | 19 | 19 | 127 | 76 | 116 | 142 | 142 | 9 | 16 | 12 | 21 | 5 | 173 | 161 | 113 | 118 | 197 | 260 | 149 | 0.000323035 | 0.004652948 | 0.079 | 0.125 | 0.086 | 0.071 | 0.071 | 0.029 | 0.054 | 0.057 | 0.093 | 0.014 | 0.037 |
| A5SS | 7847 | ENSG00000197601 | FAR1 | chr11 | + | 13711927 | 13712055 | 13711927 | 13712046 | 13712965 | 13713033 | 17 | 10 | 4 | 10 | 13 | 248 | 201 | 157 | 208 | 231 | 0 | 4 | 0 | 0 | 0 | 13 | 75 | 32 | 30 | 16 | 158 | 149 | 0.000323193 | 0.004652948 | 0.061 | 0.045 | 0.023 | 0.043 | 0.05 | 0 | 0.048 | 0 | 0 | 0 | 0.035 |
| A5SS | 7135 | ENSG00000248323 | LUCAT1 | chr5 | - | 91311645 | 91312413 | 91312294 | 91312413 | 91310640 | 91311032 | 14 | 6 | 12 | 36 | 15 | 4 | 0 | 0 | 2 | 1 | 14 | 10 | 78 | 50 | 11 | 6 | 3 | 15 | 4 | 35 | 298 | 149 | 0.000325728 | 0.00467707 | 0.636 | 1 | 1 | 0.9 | 0.882 | 0.538 | 0.625 | 0.722 | 0.862 | 0.136 | 0.307 |
| A5SS | 5957 | ENSG00000167792 | NDUFV1 | chr11 | + | 67606957 | 67607324 | 67606957 | 67607076 | 67608395 | 67608478 | 8 | 2 | 4 | 3 | 8 | 65 | 35 | 66 | 54 | 30 | 0 | 1 | 0 | 5 | 6 | 115 | 96 | 42 | 49 | 99 | 298 | 149 | 0.000335218 | 0.004800674 | 0.058 | 0.028 | 0.029 | 0.027 | 0.118 | 0 | 0.005 | 0 | 0.049 | 0.029 | 0.035 |
| A5SS | 8702 | ENSG00000125458 | NT5C | chr17 | - | 75130964 | 75131281 | 75131180 | 75131281 | 75130752 | 75130867 | 25 | 9 | 25 | 40 | 22 | 2 | 0 | 0 | 0 | 0 | 13 | 24 | 19 | 6 | 38 | 0 | 2 | 5 | 1 | 2 | 298 | 149 | 0.000343956 | 0.00491288 | 0.862 | 1 | 1 | 1 | 1 | 1 | 0.857 | 0.655 | 0.75 | 0.905 | 0.139 |
| A5SS | 4745 | ENSG00000117523 | PRRC2C | chr1 | + | 171514535 | 171514645 | 171514535 | 171514630 | 171515733 | 171515859 | 61 | 60 | 64 | 85 | 61 | 1 | 5 | 8 | 5 | 4 | 6 | 72 | 60 | 42 | 14 | 0 | 3 | 2 | 0 | 0 | 164 | 149 | 0.000352763 | 0.005025481 | 0.982 | 0.916 | 0.879 | 0.939 | 0.933 | 1 | 0.956 | 0.965 | 1 | 1 | -0.054 |
| A5SS | 1857 | ENSG00000100519 | PSMC6 | chr14 | + | 52707201 | 52707529 | 52707201 | 52707304 | 52708307 | 52708388 | 21 | 8 | 14 | 20 | 11 | 145 | 90 | 120 | 129 | 99 | 0 | 41 | 23 | 25 | 0 | 23 | 89 | 83 | 32 | 9 | 298 | 149 | 0.0003689 | 0.005232008 | 0.068 | 0.043 | 0.055 | 0.072 | 0.053 | 0 | 0.187 | 0.122 | 0.281 | 0 | -0.06 |
| A5SS | 8781 | ENSG00000127328 | RAB3IP | chr12 | + | 69800208 | 69800337 | 69800208 | 69800332 | 69801608 | 69801721 | 40 | 30 | 26 | 28 | 60 | 13 | 0 | 1 | 2 | 4 | 13 | 20 | 36 | 48 | 21 | 0 | 3 | 2 | 0 | 0 | 154 | 149 | 0.000369182 | 0.005232008 | 0.749 | 1 | 0.962 | 0.931 | 0.936 | 1 | 0.866 | 0.946 | 1 | 1 | -0.047 |
| A5SS | 5707 | ENSG00000100027 | YPEL1 | chr22 | - | 21710486 | 21710908 | 21710627 | 21710908 | 21703838 | 21703882 | 13 | 4 | 5 | 13 | 0 | 37 | 8 | 16 | 32 | 8 | 13 | 25 | 14 | 16 | 19 | 16 | 13 | 14 | 20 | 35 | 290 | 149 | 0.000373465 | 0.005278958 | 0.153 | 0.204 | 0.138 | 0.173 | 0 | 0.295 | 0.497 | 0.339 | 0.291 | 0.218 | -0.194 |
| A5SS | 1775 | ENSG00000100823 | APEX1 | chr14 | + | 20455577 | 20455773 | 20455577 | 20455703 | 20455913 | 20456101 | 6 | 0 | 6 | 7 | 7 | 121 | 69 | 139 | 102 | 115 | 12 | 4 | 8 | 7 | 16 | 68 | 58 | 52 | 59 | 177 | 219 | 149 | 0.00038427 | 0.005417616 | 0.033 | 0 | 0.029 | 0.045 | 0.04 | 0.107 | 0.045 | 0.095 | 0.075 | 0.058 | -0.047 |
| A5SS | 7233 | ENSG00000111364 | DDX55 | chr12 | + | 123605930 | 123605994 | 123605930 | 123605981 | 123606072 | 123606159 | 1 | 1 | 0 | 0 | 0 | 29 | 19 | 16 | 19 | 14 | 0 | 3 | 4 | 0 | 6 | 16 | 14 | 12 | 34 | 42 | 162 | 149 | 0.000393229 | 0.005529586 | 0.031 | 0.046 | 0 | 0 | 0 | 0 | 0.165 | 0.235 | 0 | 0.116 | -0.088 |
| A5SS | 3878 | ENSG00000109184 | DCUN1D4 | chr4 | + | 51886567 | 51887244 | 51886567 | 51886638 | 51891759 | 51891851 | 21 | 17 | 23 | 13 | 27 | 74 | 65 | 44 | 60 | 108 | 0 | 1 | 5 | 1 | 11 | 21 | 29 | 18 | 9 | 57 | 298 | 149 | 0.000419569 | 0.005884786 | 0.124 | 0.116 | 0.207 | 0.098 | 0.111 | 0 | 0.017 | 0.122 | 0.053 | 0.088 | 0.075 |
| A5SS | 1100 | ENSG00000166747 | AP1G1 | chr16 | - | 71808504 | 71808832 | 71808762 | 71808832 | 71789278 | 71789482 | 7 | 3 | 10 | 10 | 6 | 20 | 14 | 24 | 23 | 26 | 31 | 15 | 13 | 9 | 28 | 5 | 13 | 27 | 20 | 1 | 298 | 149 | 0.000421711 | 0.005899613 | 0.149 | 0.097 | 0.172 | 0.179 | 0.103 | 0.756 | 0.366 | 0.194 | 0.184 | 0.933 | -0.347 |
| A5SS | 8149 | ENSG00000185187 | SIGIRR | chr11 | - | 409859 | 410027 | 409867 | 410027 | 408072 | 408206 | 6 | 1 | 1 | 5 | 0 | 18 | 5 | 20 | 6 | 9 | 4 | 0 | 0 | 0 | 0 | 14 | 8 | 6 | 9 | 8 | 157 | 149 | 0.000433337 | 0.006046719 | 0.24 | 0.16 | 0.045 | 0.442 | 0 | 0.213 | 0 | 0 | 0 | 0 | 0.135 |
| A5SS | 8666 | ENSG00000127314 | RAP1B | chr12 | + | 68610928 | 68611052 | 68610928 | 68611043 | 68648698 | 68648781 | 242 | 201 | 151 | 229 | 124 | 635 | 465 | 365 | 613 | 429 | 159 | 226 | 173 | 130 | 58 | 157 | 485 | 366 | 251 | 95 | 158 | 149 | 0.000435861 | 0.006066384 | 0.264 | 0.29 | 0.281 | 0.261 | 0.214 | 0.489 | 0.305 | 0.308 | 0.328 | 0.365 | -0.097 |
| A5SS | 1487 | ENSG00000166783 | MARF1 | chr16 | - | 15634756 | 15636342 | 15635655 | 15636342 | 15633616 | 15633843 | 205 | 141 | 174 | 206 | 213 | 2 | 4 | 2 | 1 | 2 | 247 | 195 | 206 | 229 | 66 | 24 | 4 | 4 | 1 | 0 | 298 | 149 | 0.00044719 | 0.006208184 | 0.981 | 0.946 | 0.978 | 0.99 | 0.982 | 0.837 | 0.961 | 0.963 | 0.991 | 1 | 0.025 |
| A5SS | 1726 | ENSG00000179364 | PACS2 | chr14 | + | 105368458 | 105368539 | 105368458 | 105368515 | 105369840 | 105369900 | 44 | 55 | 58 | 81 | 45 | 2 | 3 | 8 | 9 | 0 | 102 | 96 | 70 | 56 | 186 | 0 | 5 | 0 | 6 | 0 | 173 | 149 | 0.000456815 | 0.006325666 | 0.95 | 0.94 | 0.862 | 0.886 | 1 | 1 | 0.943 | 1 | 0.889 | 1 | -0.039 |
| A5SS | 4485 | ENSG00000156983 | BRPF1 | chr3 | + | 9742024 | 9742541 | 9742024 | 9742153 | 9742943 | 9743253 | 13 | 13 | 15 | 21 | 9 | 34 | 33 | 40 | 20 | 42 | 32 | 33 | 19 | 45 | 34 | 42 | 26 | 13 | 34 | 63 | 298 | 149 | 0.000482734 | 0.006654408 | 0.16 | 0.165 | 0.158 | 0.344 | 0.097 | 0.276 | 0.388 | 0.422 | 0.398 | 0.212 | -0.154 |
| A5SS | 6237 | ENSG00000135317 | SNX14 | chr6 | - | 85547083 | 85547226 | 85547111 | 85547226 | 85543604 | 85543760 | 5 | 8 | 5 | 2 | 7 | 144 | 86 | 79 | 124 | 194 | 0 | 1 | 0 | 0 | 1 | 41 | 63 | 61 | 53 | 26 | 177 | 149 | 0.000483001 | 0.006654408 | 0.028 | 0.073 | 0.051 | 0.013 | 0.029 | 0 | 0.013 | 0 | 0 | 0.031 | 0.03 |
| A5SS | 5934 | ENSG00000124356 | STAMBP | chr2 | + | 73844812 | 73845036 | 73844812 | 73844888 | 73845166 | 73845262 | 13 | 8 | 18 | 25 | 37 | 122 | 92 | 88 | 135 | 210 | 14 | 17 | 6 | 35 | 0 | 227 | 137 | 125 | 171 | 185 | 297 | 149 | 0.000484809 | 0.006662456 | 0.051 | 0.042 | 0.093 | 0.085 | 0.081 | 0.03 | 0.059 | 0.024 | 0.093 | 0 | 0.029 |
| A5SS | 437 | ENSG00000130254 | SAFB2 | chr19 | - | 5616127 | 5616335 | 5616131 | 5616335 | 5613464 | 5613527 | 23 | 21 | 27 | 19 | 6 | 207 | 140 | 186 | 256 | 188 | 47 | 39 | 36 | 75 | 35 | 408 | 196 | 172 | 222 | 425 | 153 | 149 | 0.000490232 | 0.006720001 | 0.098 | 0.127 | 0.124 | 0.067 | 0.03 | 0.101 | 0.162 | 0.169 | 0.248 | 0.074 | -0.062 |
| A5SS | 8236 | ENSG00000149311 | ATM | chr11 | + | 108294926 | 108296393 | 108294926 | 108295059 | 108297286 | 108297382 | 149 | 129 | 124 | 109 | 177 | 293 | 267 | 353 | 386 | 654 | 18 | 98 | 43 | 11 | 7 | 29 | 55 | 77 | 1 | 2 | 298 | 149 | 0.00050165 | 0.00684496 | 0.203 | 0.195 | 0.149 | 0.124 | 0.119 | 0.237 | 0.471 | 0.218 | 0.846 | 0.636 | -0.324 |
| A5SS | 596 | ENSG00000141644 | MBD1 | chr18 | - | 50272823 | 50272955 | 50272847 | 50272955 | 50272676 | 50272738 | 110 | 66 | 116 | 120 | 80 | 1 | 0 | 0 | 1 | 4 | 156 | 92 | 111 | 102 | 138 | 12 | 6 | 6 | 0 | 0 | 173 | 149 | 0.000501863 | 0.00684496 | 0.99 | 1 | 1 | 0.99 | 0.945 | 0.918 | 0.93 | 0.941 | 1 | 1 | 0.027 |
| A5SS | 1192 | ENSG00000141002 | TCF25 | chr16 | + | 89908965 | 89909107 | 89908965 | 89909083 | 89910590 | 89910663 | 23 | 10 | 6 | 22 | 10 | 8 | 5 | 1 | 2 | 1 | 11 | 16 | 18 | 8 | 31 | 0 | 0 | 0 | 0 | 6 | 173 | 149 | 0.0005139 | 0.0069885 | 0.712 | 0.633 | 0.838 | 0.905 | 0.896 | 1 | 1 | 1 | 1 | 0.817 | -0.167 |
| A5SS | 4400 | ENSG00000145979 | TBC1D7 | chr6 | - | 13328315 | 13328531 | 13328455 | 13328531 | 13326786 | 13326906 | 3 | 2 | 5 | 2 | 4 | 4 | 2 | 3 | 7 | 6 | 0 | 0 | 3 | 0 | 6 | 9 | 17 | 5 | 11 | 23 | 289 | 149 | 0.000514956 | 0.0069885 | 0.279 | 0.34 | 0.462 | 0.128 | 0.256 | 0 | 0 | 0.236 | 0 | 0.119 | 0.222 |
| A5SS | 1558 | ENSG00000196547 | MAN2A2 | chr15 | + | 90913272 | 90913406 | 90913272 | 90913374 | 90913613 | 90913755 | 37 | 18 | 23 | 30 | 3 | 1 | 0 | 1 | 3 | 0 | 53 | 78 | 36 | 87 | 244 | 9 | 8 | 5 | 18 | 0 | 181 | 149 | 0.000530198 | 0.007176612 | 0.968 | 1 | 0.95 | 0.892 | 1 | 0.829 | 0.889 | 0.856 | 0.799 | 1 | 0.087 |
| A5SS | 7604 | ENSG00000106771 | TMEM245 | chr9 | - | 109073247 | 109073438 | 109073355 | 109073438 | 109064476 | 109064567 | 18 | 42 | 18 | 20 | 11 | 117 | 100 | 123 | 134 | 124 | 50 | 38 | 18 | 21 | 6 | 30 | 71 | 47 | 45 | 75 | 257 | 149 | 0.000531454 | 0.007176612 | 0.082 | 0.196 | 0.078 | 0.08 | 0.049 | 0.491 | 0.237 | 0.182 | 0.213 | 0.044 | -0.136 |
| A5SS | 1785 | ENSG00000119689 | DLST | chr14 | + | 74891055 | 74891193 | 74891055 | 74891167 | 74892833 | 74892986 | 3 | 8 | 3 | 3 | 6 | 70 | 37 | 42 | 40 | 125 | 32 | 5 | 13 | 9 | 14 | 73 | 54 | 55 | 52 | 207 | 175 | 149 | 0.000536374 | 0.007225123 | 0.035 | 0.155 | 0.057 | 0.06 | 0.039 | 0.272 | 0.073 | 0.168 | 0.128 | 0.054 | -0.07 |
| A5SS | 3520 | ENSG00000175203 | DCTN2 | chr12 | - | 57538482 | 57538524 | 57538518 | 57538524 | 57535748 | 57535845 | 0 | 3 | 1 | 0 | 0 | 12 | 19 | 26 | 15 | 12 | 0 | 0 | 0 | 1 | 17 | 16 | 19 | 10 | 20 | 37 | 185 | 149 | 0.00055377 | 0.0074325 | 0 | 0.113 | 0.03 | 0 | 0 | 0 | 0 | 0 | 0.039 | 0.27 | -0.033 |
| A5SS | 3377 | ENSG00000095321 | CRAT | chr9 | - | 129101743 | 129102057 | 129101882 | 129102057 | 129100510 | 129100689 | 3 | 0 | 2 | 1 | 0 | 24 | 24 | 22 | 32 | 19 | 0 | 6 | 3 | 11 | 13 | 48 | 33 | 13 | 38 | 150 | 288 | 149 | 0.000554501 | 0.0074325 | 0.061 | 0 | 0.045 | 0.016 | 0 | 0 | 0.086 | 0.107 | 0.13 | 0.043 | -0.049 |
| A5SS | 754 | ENSG00000182173 | TSEN54 | chr17 | + | 75521704 | 75522521 | 75521704 | 75522333 | 75523274 | 75523335 | 23 | 6 | 17 | 31 | 3 | 11 | 4 | 18 | 9 | 5 | 18 | 42 | 14 | 16 | 8 | 0 | 7 | 7 | 0 | 2 | 298 | 149 | 0.000561844 | 0.007512415 | 0.511 | 0.429 | 0.321 | 0.633 | 0.231 | 1 | 0.75 | 0.5 | 1 | 0.667 | -0.358 |
| A5SS | 5907 | ENSG00000285437 | POLR2J3 | chr7 | - | 102572442 | 102572583 | 102572471 | 102572583 | 102569834 | 102569924 | 1 | 6 | 5 | 6 | 2 | 18 | 21 | 38 | 33 | 20 | 3 | 0 | 3 | 0 | 6 | 63 | 74 | 48 | 67 | 45 | 178 | 149 | 0.000576753 | 0.007692869 | 0.044 | 0.193 | 0.099 | 0.132 | 0.077 | 0.038 | 0 | 0.05 | 0 | 0.1 | 0.071 |
| A5SS | 4360 | ENSG00000103657 | HERC1 | chr15 | - | 63625941 | 63626154 | 63625984 | 63626154 | 63624157 | 63624327 | 4 | 3 | 8 | 6 | 3 | 158 | 89 | 127 | 147 | 134 | 0 | 4 | 6 | 3 | 17 | 30 | 34 | 101 | 48 | 30 | 192 | 149 | 0.000578952 | 0.007703311 | 0.019 | 0.025 | 0.047 | 0.031 | 0.017 | 0 | 0.084 | 0.044 | 0.046 | 0.305 | -0.068 |
| A5SS | 5512 | ENSG00000223960 | CHROMR | chr2 | + | 178426365 | 178426680 | 178426365 | 178426505 | 178430694 | 178430889 | 37 | 12 | 24 | 22 | 18 | 11 | 10 | 5 | 9 | 11 | 48 | 24 | 13 | 13 | 11 | 0 | 8 | 5 | 0 | 0 | 298 | 149 | 0.000584307 | 0.007755608 | 0.627 | 0.375 | 0.706 | 0.55 | 0.45 | 1 | 0.6 | 0.565 | 1 | 1 | -0.291 |
| A5SS | 6202 | ENSG00000168538 | TRAPPC11 | chr4 | + | 183674712 | 183674870 | 183674712 | 183674812 | 183675163 | 183675237 | 6 | 3 | 4 | 3 | 3 | 191 | 144 | 131 | 129 | 304 | 9 | 7 | 2 | 8 | 0 | 57 | 143 | 87 | 103 | 5 | 207 | 149 | 0.000601505 | 0.007964452 | 0.022 | 0.015 | 0.022 | 0.016 | 0.007 | 0.102 | 0.034 | 0.016 | 0.053 | 0 | -0.025 |
| A5SS | 7082 | ENSG00000012048 | BRCA1 | chr17 | - | 43106455 | 43106533 | 43106477 | 43106533 | 43104867 | 43104956 | 15 | 11 | 14 | 14 | 27 | 0 | 4 | 0 | 0 | 0 | 32 | 23 | 9 | 22 | 30 | 0 | 8 | 1 | 6 | 0 | 171 | 149 | 0.000622765 | 0.008225942 | 1 | 0.706 | 1 | 1 | 1 | 1 | 0.715 | 0.887 | 0.762 | 1 | 0.068 |
| A5SS | 7156 | ENSG00000046651 | OFD1 | chrX | + | 13746779 | 13746956 | 13746779 | 13746953 | 13749426 | 13749533 | 3 | 4 | 2 | 7 | 1 | 232 | 213 | 193 | 181 | 403 | 0 | 17 | 7 | 1 | 0 | 71 | 119 | 167 | 144 | 47 | 152 | 149 | 0.000630373 | 0.008306273 | 0.013 | 0.018 | 0.01 | 0.037 | 0.002 | 0 | 0.123 | 0.039 | 0.007 | 0 | -0.018 |
| A5SS | 3914 | ENSG00000141027 | NCOR1 | chr17 | - | 16061400 | 16061894 | 16061757 | 16061894 | 16058470 | 16058599 | 272 | 183 | 230 | 282 | 272 | 11 | 6 | 14 | 9 | 10 | 143 | 317 | 167 | 260 | 108 | 0 | 10 | 3 | 5 | 0 | 298 | 149 | 0.00064565 | 0.008487018 | 0.925 | 0.938 | 0.891 | 0.94 | 0.932 | 1 | 0.941 | 0.965 | 0.963 | 1 | -0.049 |
| A5SS | 98 | ENSG00000138942 | RNF185 | chr22 | + | 31196935 | 31197158 | 31196935 | 31196990 | 31201497 | 31201615 | 12 | 7 | 12 | 12 | 2 | 14 | 4 | 11 | 11 | 15 | 30 | 15 | 29 | 31 | 37 | 17 | 13 | 9 | 8 | 6 | 298 | 149 | 0.000655326 | 0.008575001 | 0.3 | 0.467 | 0.353 | 0.353 | 0.062 | 0.469 | 0.366 | 0.617 | 0.66 | 0.755 | -0.266 |

| | | | | | | | | | | | | | | |
|---|---|---|---|---|---|---|---|---|---|---|---|---|---|---|
| A5SS | 8115 | ENSG00000162236 | STX5 | chr11 | - | 62825024 | 62825117 | 62825035 | 62825117 | 62824458 | 62824565 | 9 | 6 | 7 [illegible] |
| A5SS | 6487 | ENSG00000006634 | DBF4 | chr7 | + | 87878052 | 87878225 | 87878052 | 87878133 | 87884978 | 87885158 | 66 | 55 | 97 [illegible] |
| A5SS | 6071 | ENSG00000100503 | NIN | chr14 | - | 50759856 | 50760359 | 50759993 | 50760359 | 50754741 | 50754867 | 332 | 274 | 402 [illegible] |
| A5SS | 3229 | ENSG00000184014 | DENND5A | chr11 | - | 9142721 | 9143485 | 9143402 | 9143485 | 9141939 | 9142108 | 198 | 149 | 184 [illegible] |
| A5SS | 1673 | ENSG00000100749 | VRK1 | chr14 | + | 96855223 | 96855352 | 96855223 | 96855350 | 96856129 | 96856250 | 7 | 6 | 6 [illegible] |
| A5SS | 6021 | ENSG00000089818 | NECAP1 | chr12 | + | 8090194 | 8091850 | 8090194 | 8090299 | 8092675 | 8092784 | 140 | 76 | 127 [illegible] |
| A5SS | 8426 | ENSG00000148688 | RPP30 | chr10 | + | 90894774 | 90895483 | 90894774 | 90894891 | 90895879 | 90895917 | 95 | 50 | 105 [illegible] |
| A5SS | 8611 | ENSG00000082898 | XPO1 | chr2 | - | 61534422 | 61534584 | 61534557 | 61534584 | 61533771 | 61533903 | 11 | 4 | 19 [illegible] |
| A5SS | 8209 | ENSG00000102317 | RBM3 | chrX | + | 48575560 | 48576083 | 48575560 | 48575667 | 48576313 | 48576419 | 30 | 24 | 36 [illegible] |
| A5SS | 5859 | ENSG00000074582 | BCS1L | chr2 | + | 218662196 | 218662679 | 218662196 | 218662260 | 218662882 | 218663000 | 24 | 13 | 30 [illegible] |
| A5SS | 8086 | ENSG00000110274 | CEP164 | chr11 | + | 117408889 | 117409028 | 117408889 | 117409004 | 117409617 | 117409965 | 25 | 15 | 16 [illegible] |
| A5SS | 1693 | ENSG00000092199 | HNRNPC | chr14 | - | 21230957 | 21231072 | 21230996 | 21231072 | 21230318 | 21230366 | 543 | 306 | 499 [illegible] |
| A5SS | 5192 | ENSG00000148737 | TCF7L2 | chr10 | + | 113143922 | 113144025 | 113143922 | 113144013 | 113146010 | 113146097 | 20 | 8 | 13 [illegible] |
| A5SS | 1574 | ENSG00000247240 | UBL7-AS1 | chr15 | + | 74461882 | 74462031 | 74461882 | 74462027 | 74466035 | 74466130 | 2 | 1 | 3 [illegible] |
| A5SS | 3401 | ENSG00000170581 | STAT2 | chr12 | - | 56354418 | 56354614 | 56354465 | 56354614 | 56351291 | 56351450 | 14 | 13 | 6 [illegible] |
| A5SS | 1491 | ENSG00000159459 | UBR1 | chr15 | - | 43025373 | 43025429 | 43025380 | 43025429 | 43024828 | 43024983 | 4 | 5 | 8 [illegible] |
| A5SS | 4316 | ENSG00000166825 | ANPEP | chr15 | - | 89804945 | 89805217 | 89805077 | 89805217 | 89804490 | 89804617 | 14 | 16 | 15 [illegible] |
| A5SS | 3772 | ENSG00000104419 | NDRG1 | chr8 | - | 133246595 | 133246663 | 133246615 | 133246663 | 133244354 | 133244390 | 0 | 3 | 3 [illegible] |
| A5SS | 3396 | ENSG00000170581 | STAT2 | chr12 | - | 56349344 | 56349509 | 56349425 | 56349509 | 56348923 | 56349059 | 35 | 34 | 35 [illegible] |
| A5SS | 2960 | ENSG00000165195 | PIGA | chrX | - | 15331589 | 15331992 | 15331873 | 15331992 | 15325913 | 15326046 | 33 | 25 | 32 [illegible] |
| A5SS | 6815 | ENSG00000167778 | SPRYD3 | chr12 | - | 53074648 | 53075219 | 53075094 | 53075219 | 53073285 | 53073471 | 124 | 54 | 94 [illegible] |
| A5SS | 3583 | ENSG00000169241 | SLC50A1 | chr1 | + | 155136298 | 155136396 | 155136298 | 155136376 | 155137560 | 155137722 | 0 | 0 | 0 [illegible] |
| A5SS | 6738 | ENSG00000122783 | CYREN | chr7 | - | 135168748 | 135169060 | 135168968 | 135169060 | 135167731 | 135167807 | 174 | 135 | 138 [illegible] |
| A5SS | 3048 | ENSG00000182899 | RPL35A | chr3 | + | 197950189 | 197950299 | 197950189 | 197950221 | 197950935 | 197950978 | 8 | 4 | 9 [illegible] |
| A5SS | 7278 | ENSG00000205581 | HMGN1 | chr21 | - | 39345829 | 39347292 | 39347246 | 39347292 | 39345145 | 39345274 | 409 | 500 | 525 [illegible] |
| A5SS | 976 | ENSG00000073584 | SMARCE1 | chr17 | - | 40642117 | 40642559 | 40642454 | 40642559 | 40637491 | 40637572 | 21 | 17 | 28 [illegible] |
| A5SS | 2961 | ENSG00000165195 | PIGA | chrX | - | 15331819 | 15331992 | 15331873 | 15331992 | 15325913 | 15326046 | 26 | 23 | 25 [illegible] |
| A5SS | 8609 | ENSG00000082898 | XPO1 | chr2 | - | 61525491 | 61526521 | 61526419 | 61526521 | 61522610 | 61522683 | 27 | 46 | 24 [illegible] |
| A5SS | 4916 | ENSG00000128191 | DGCR8 | chr22 | + | 20106591 | 20106698 | 20106591 | 20106685 | 20107270 | 20107398 | 93 | 70 | 70 [illegible] |
| A5SS | 7806 | ENSG00000070476 | ZXDC | chr3 | - | 126461269 | 126462220 | 126461534 | 126462220 | 126441764 | 126441946 | 63 | 21 | 55 [illegible] |
| A5SS | 1214 | ENSG00000197912 | SPG7 | chr16 | + | 89554485 | 89556085 | 89554485 | 89554563 | 89556886 | 89557173 | 2 | 1 | 26 [illegible] |
| A5SS | 97 | ENSG00000138942 | RNF185 | chr22 | + | 31196935 | 31197157 | 31196935 | 31196990 | 31201497 | 31201615 | 12 | 7 | 12 [illegible] |
| A5SS | 1796 | ENSG00000150527 | MIA2 | chr14 | + | 39276933 | 39277065 | 39276933 | 39276955 | 39279336 | 39279358 | 168 | 102 | 137 [illegible] |
| A5SS | 6581 | ENSG00000115677 | HDLBP | chr2 | - | 241268953 | 241269295 | 241269200 | 241269295 | 241268476 | 241268541 | 8 | 5 | 12 [illegible] |
| A5SS | 3071 | ENSG00000111642 | CHD4 | chr12 | - | 6573070 | 6573269 | 6573073 | 6573269 | 6570868 | 6571032 | 13 | 2 | 8 [illegible] |
| A5SS | 1277 | ENSG00000108798 | ABI3 | chr17 | + | 49216530 | 49216698 | 49216530 | 49216680 | 49217738 | 49217915 | 231 | 100 | 91 [illegible] |
| A5SS | 624 | ENSG00000099956 | SMARCB1 | chr22 | + | 23800943 | 23801964 | 23800943 | 23801081 | 23803294 | 23803422 | 0 | 2 | 9 [illegible] |
| A5SS | 4153 | ENSG00000148187 | MRRF | chr9 | + | 122285168 | 122285946 | 122285168 | 122285287 | 122286038 | 122286166 | 41 | 23 | 49 [illegible] |
| A5SS | 5717 | ENSG00000161202 | DVL3 | chr3 | + | 184164795 | 184165206 | 184164795 | 184164931 | 184165421 | 184165491 | 48 | 13 | 40 [illegible] |
| A5SS | 6067 | ENSG00000072121 | ZFYVE26 | chr14 | - | 67777513 | 67777735 | 67777558 | 67777735 | 67775859 | 67776106 | 3 | 1 | 2 [illegible] |
| A5SS | 744 | ENSG00000159842 | ABR | chr17 | - | 1057843 | 1058045 | 1057969 | 1058045 | 1056997 | 1057102 | 4 | 7 | 3 [illegible] |
| A5SS | 2132 | ENSG00000153113 | CAST | chr5 | + | 96722638 | 96722718 | 96722638 | 96722698 | 96727488 | 96727530 | 4 | 4 | 5 [illegible] |
| A5SS | 3201 | ENSG00000010327 | STAB1 | chr3 | + | 52523426 | 52523756 | 52523426 | 52523576 | 52523870 | 52524017 | 24 | 11 | 6 [illegible] |
| A5SS | 3322 | ENSG00000113810 | SMC4 | chr3 | + | 160412325 | 160412849 | 160412325 | 160412453 | 160413472 | 160413613 | 26 | 22 | 20 [illegible] |
| A5SS | 3623 | ENSG00000113161 | HMGCR | chr5 | + | 75359397 | 75359556 | 75359397 | 75359511 | 75359984 | 75360139 | 180 | 82 | 159 [illegible] |
| A5SS | 2732 | ENSG00000174227 | PIGG | chr4 | + | 521055 | 521273 | 521055 | 521249 | 521659 | 521941 | 26 | 21 | 30 [illegible] |
| A5SS | 2842 | ENSG00000186577 | SMIM29 | chr6 | - | 34247680 | 34247864 | 34247740 | 34247864 | 34247466 | 34247492 | 24 | 10 | 26 [illegible] |
| A5SS | 2333 | ENSG00000196605 | ZNF846 | chr19 | - | 9764809 | 9765035 | 9764935 | 9765035 | 9763281 | 9763408 | 14 | 10 | 13 [illegible] |
| A5SS | 5007 | ENSG00000196975 | ANXA4 | chr2 | + | 69810593 | 69811456 | 69810593 | 69810673 | 69812652 | 69812709 | 10 | 6 | 8 [illegible] |
| A5SS | 750 | ENSG00000129197 | RPAIN | chr17 | + | 5425970 | 5426299 | 5425970 | 5426082 | 5428070 | 5428211 | 72 | 47 | 88 [illegible] |
| A5SS | 7674 | ENSG00000159069 | FBXW5 | chr9 | - | 136941754 | 136942466 | 136942045 | 136942466 | 136941536 | 136941684 | 6 | 12 | 7 [illegible] |
| A5SS | 8387 | ENSG00000177239 | MAN1B1 | chr9 | + | 137101483 | 137105916 | 137101483 | 137101672 | 137106124 | 137106315 | 6 | 7 | 8 [illegible] |
| A5SS | 3767 | ENSG00000121774 | KHDRBS1 | chr1 | + | 32037834 | 32038119 | 32037834 | 32038036 | 32038551 | 32038619 | 5 | 11 | 5 [illegible] |
| A5SS | 8201 | ENSG00000054654 | SYNE2 | chr14 | + | 63997300 | 63997576 | 63997300 | 63997391 | 63998218 | 63998328 | 66 | 40 | 28 [illegible] |
| A5SS | 3871 | ENSG00000178028 | DMAP1 | chr1 | + | 44220016 | 44220443 | 44220016 | 44220309 | 44220558 | 44220681 | 1 | 1 | 2 [illegible] |
| A5SS | 3481 | ENSG00000270629 | NBPF14 | chr1 | - | 148595849 | 148595890 | 148595854 | 148595890 | 148595542 | 148595752 | 2 | 2 | 1 [illegible] |
| A5SS | 1200 | ENSG00000197912 | SPG7 | chr16 | + | 89528631 | 89529579 | 89528631 | 89529524 | 89530682 | 89530808 | 45 | 44 | 67 [illegible] |
| A5SS | 1988 | ENSG00000079616 | KIF22 | chr16 | + | 29803997 | 29804107 | 29803997 | 29804065 | 29804813 | 29805026 | 13 | 10 | 14 [illegible] |
| A5SS | 1476 | ENSG00000182718 | ANXA2 | chr15 | - | 60396150 | 60396427 | 60396372 | 60396427 | 60386027 | 60386086 | 23 | 21 | 10 [illegible] |
| A5SS | 2025 | ENSG00000141279 | NPEPPS | chr17 | + | 47585499 | 47585745 | 47585499 | 47585700 | 47586364 | 47586398 | 8 | 6 | 12 [illegible] |
| A5SS | 443 | ENSG00000031823 | RANBP3 | chr19 | - | 5957906 | 5957973 | 5957917 | 5957973 | 5951392 | 5951596 | 1 | 0 | 2 [illegible] |
| A5SS | 2903 | ENSG00000164068 | RNF123 | chr3 | + | 49689556 | 49689822 | 49689556 | 49689606 | 49691129 | 49691247 | 2 | 0 | 0 [illegible] |
| A5SS | 7777 | ENSG00000119383 | PTPA | chr9 | + | 129120512 | 129120628 | 129120512 | 129120610 | 129123051 | 129123138 | 1 | 0 | 1 [illegible] |
| A5SS | 2958 | ENSG00000165195 | PIGA | chrX | - | 15331215 | 15331992 | 15331873 | 15331992 | 15325913 | 15326046 | 46 | 42 | 49 [illegible] |
| A5SS | 3715 | ENSG00000115524 | SF3B1 | chr2 | - | 197418895 | 197420542 | 197420427 | 197420542 | 197418508 | 197418588 | 778 | 630 | 836 [illegible] |
| A5SS | 181 | ENSG00000239382 | ALKBH6 | chr19 | - | 36013309 | 36013422 | 36013343 | 36013422 | 36013020 | 36013089 | 0 | 0 | 2 [illegible] |
| A5SS | 6764 | ENSG00000160408 | ST6GALNAC6 | chr9 | - | 127890636 | 127891043 | 127890655 | 127891043 | 127887483 | 127887591 | 54 | 16 | 39 [illegible] |
| A5SS | 8684 | ENSG00000135090 | TAOK3 | chr12 | - | 118359505 | 118359720 | 118359520 | 118359720 | 118266654 | 118266759 | 106 | 75 | 75 [illegible] |
| A5SS | 5809 | ENSG00000130414 | NDUFA10 | chr2 | - | 240014694 | 240014860 | 240014738 | 240014860 | 240011616 | 240011696 | 5 | 0 | 5 [illegible] |
| A5SS | 8767 | ENSG00000182796 | TMEM198B | chr12 | + | 55832958 | 55833038 | 55832958 | 55833014 | 55833446 | 55833541 | 0 | 0 | 1 [illegible] |
| A5SS | 5235 | ENSG00000115355 | CCDC88A | chr2 | - | 55299838 | 55301277 | 55301205 | 55301277 | 55296257 | 55296523 | 91 | 87 | 65 [illegible] |
| A5SS | 3445 | ENSG00000144591 | GMPPA | chr2 | + | 219504082 | 219505362 | 219504082 | 219504213 | 219505457 | 219505555 | 71 | 67 | 67 [illegible] |
| A5SS | 7513 | ENSG00000167325 | RRM1 | chr11 | + | 4119844 | 4119970 | 4119844 | 4119928 | 4121603 | 4121765 | 0 | 1 | 4 [illegible] |
| A5SS | 6267 | ENSG00000168944 | CEP120 | chr5 | - | 123388303 | 123388606 | 123388431 | 123388606 | 123386517 | 123386667 | 13 | 14 | 12 [illegible] |
| A5SS | 422 | ENSG00000127527 | EPS15L1 | chr19 | - | 16402314 | 16402485 | 16402320 | 16402485 | 16395343 | 16395467 | 0 | 0 | 0 [illegible] |
| A5SS | 2545 | ENSG00000106052 | TAX1BP1 | chr7 | + | 27816348 | 27816520 | 27816348 | 27816394 | 27816889 | 27817038 | 84 | 64 | 74 [illegible] |
| A5SS | 2032 | ENSG00000130811 | EIF3G | chr19 | - | 10119066 | 10119171 | 10119087 | 10119171 | 10118867 | 10118956 | 0 | 1 | 0 [illegible] |
| A5SS | 5509 | ENSG00000085978 | ATG16L1 | chr2 | + | 233272965 | 233273777 | 233272965 | 233273052 | 233274675 | 233274778 | 43 | 50 | 62 [illegible] |
| A5SS | 1291 | ENSG00000272578 | NA | chr22 | - | 23686789 | 23686995 | 23686837 | 23686995 | 23683743 | 23683867 | 1 | 1 | 2 [illegible] |
| A5SS | 4290 | ENSG00000087206 | UIMC1 | chr5 | - | 176968554 | 176969291 | 176969052 | 176969291 | 176958092 | 176958154 | 123 | 77 | 107 [illegible] |
| A5SS | 6309 | ENSG00000169490 | TM2D2 | chr8 | - | 38996212 | 38996509 | 38996418 | 38996509 | 38995317 | 38995405 | 18 | 7 | 6 [illegible] |
| A5SS | 2339 | ENSG00000137449 | CPEB2 | chr4 | + | 15007304 | 15007595 | 15007304 | 15007586 | 15017187 | 15017278 | 8 | 9 | 9 [illegible] |
| A5SS | 6380 | ENSG00000158321 | AUTS2 | chr7 | + | 70777102 | 70777217 | 70777102 | 70777174 | 70781614 | 70781756 | 5 | 0 | 7 [illegible] |
| A5SS | 2058 | ENSG00000131844 | MCCC2 | chr5 | + | 71641002 | 71641083 | 71641002 | 71641075 | 71643818 | 71643895 | 0 | 0 | 7 [illegible] |
| A5SS | 1298 | ENSG00000103005 | USB1 | chr16 | + | 58002478 | 58002663 | 58002478 | 58002645 | 58009928 | 58010112 | 1 | 0 | 0 [illegible] |
| A5SS | 6651 | ENSG00000123066 | MED13L | chr12 | - | 115975170 | 115975738 | 115975514 | 115975738 | 115972077 | 115972236 | 108 | 105 | 138 [illegible] |
| A5SS | 7154 | ENSG00000086758 | HUWE1 | chrX | - | 53543840 | 53543968 | 53543885 | 53543968 | 53542442 | 53542539 | 159 | 70 | 117 [illegible] |
| A5SS | 8699 | ENSG00000229117 | RPL41 | chr12 | + | 56116774 | 56116930 | 56116774 | 56116799 | 56117188 | 56117211 | 42 | 41 | 34 [illegible] |
| A5SS | 1545 | ENSG00000128928 | IVD | chr15 | + | 40407635 | 40407812 | 40407635 | 40407725 | 40407938 | 40407990 | 28 | 41 | 24 [illegible] |
| A5SS | 1907 | ENSG00000196367 | TRRAP | chr7 | + | 99030594 | 99030764 | 99030594 | 99030757 | 99030884 | 99031000 | 6 | 8 | 9 [illegible] |
| A5SS | 5447 | ENSG00000123106 | CCDC91 | chr12 | + | 28305648 | 28305806 | 28305648 | 28305779 | 28306741 | 28306945 | 302 | 157 | 220 [illegible] |
| A5SS | 1853 | ENSG00000126773 | PCNX4 | chr14 | + | 60124217 | 60125251 | 60124217 | 60124310 | 60125636 | 60125823 | 218 | 164 | 237 [illegible] |
| A5SS | 442 | ENSG00000031823 | RANBP3 | chr19 | - | 5957906 | 5957973 | 5957917 | 5957973 | 5941798 | 5941835 | 4 | 2 | 1 [illegible] |
| A5SS | 7386 | ENSG00000110344 | UBE4A | chr11 | + | 118372506 | 118372687 | 118372506 | 118372666 | 118373085 | 118373288 | 5 | 1 | 1 [illegible] |
| A5SS | 6839 | ENSG00000124333 | VAMP7 | chrX | + | 155889457 | 155889633 | 155889457 | 155889547 | 155898111 | 155898249 | 39 | 17 | 34 [illegible] |
| A5SS | 8874 | ENSG00000110911 | SLC11A2 | chr12 | - | 50991486 | 50991672 | 50991598 | 50991672 | 50990794 | 50990948 | 1 | 8 | 1 [illegible] |
| A5SS | 6221 | ENSG00000112695 | COX7A2 | chr6 | - | 75240276 | 75240385 | 75240300 | 75240385 | 75237786 | 75237988 | 25 | 9 | 16 [illegible] |
| A5SS | 7673 | ENSG00000196502 | SULT1A1 | chr16 | - | 28609058 | 28610191 | 28609930 | 28610191 | 28608707 | 28608859 | 8 | 1 | 14 [illegible] |
| A5SS | 2667 | ENSG00000189308 | LIN54 | chr4 | - | 82984160 | 82984876 | 82984823 | 82984876 | 82978882 | 82979006 | 44 | 43 | 95 [illegible] |
| A5SS | 4715 | ENSG00000001497 | LAS1L | chrX | - | 65528875 | 65529258 | 65529211 | 65529258 | 65528259 | 65528369 | 26 | 15 | 22 [illegible] |
| A5SS | 4386 | ENSG00000145020 | AMT | chr3 | - | 49419259 | 49420342 | 49419709 | 49420342 | 49418970 | 49419151 | 11 | 6 | 12 [illegible] |
| A5SS | 3471 | ENSG00000114796 | KLHL24 | chr3 | + | 183635622 | 183635793 | 183635622 | 183635747 | 183636220 | 183636311 | 7 | 9 | 13 [illegible] |
| A5SS | 6189 | ENSG00000225791 | TRAM2-AS1 | chr6 | + | 52578049 | 52578594 | 52578049 | 52578393 | 52579374 | 52579609 | 10 | 6 | 14 [illegible] |
| A5SS | 1054 | ENSG00000135709 | KIAA0513 | chr16 | + | 85027750 | 85027858 | 85027750 | 85027851 | 85066899 | 85067400 | 6 | 12 | 16 [illegible] |
| A5SS | 5266 | ENSG00000125875 | TBC1D20 | chr20 | - | 441778 | 442043 | 441856 | 442043 | 441587 | 441689 | 9 | 5 | 6 [illegible] |
| A5SS | 435 | ENSG00000142347 | MYO1F | chr19 | - | 8555636 | 8555796 | 8555658 | 8555796 | 8554653 | 8554743 | 5 | 2 | 3 [illegible] |
| A5SS | 2956 | ENSG00000165195 | PIGA | chrX | - | 15331215 | 15331992 | 15331589 | 15331992 | 15325913 | 15326046 | 64 | 40 | 65 [illegible] |
| A5SS | 562 | ENSG00000224397 | PELATON | chr20 | + | 50277490 | 50277980 | 50277490 | 50277613 | 50278176 | 50278421 | 21 | 17 | 4 [illegible] |
| A5SS | 3763 | ENSG00000088035 | ALG6 | chr1 | + | 63411145 | 63411331 | 63411145 | 63411155 | 63411681 | 63411806 | 30 | 25 | 36 [illegible] |
| A5SS | 821 | ENSG00000125454 | SLC25A19 | chr17 | - | 75288387 | 75288591 | 75288501 | 75288591 | 75286632 | 75286802 | 17 | 7 | 8 [illegible] |
| A5SS | 8607 | ENSG00000076067 | RBMS2 | chr12 | + | 56587553 | 56587664 | 56587553 | 56587659 | 56588293 | 56588374 | 20 | 15 | 3 [illegible] |
| A5SS | 1794 | ENSG00000213983 | AP1G2 | chr14 | - | 23567596 | 23567783 | 23567738 | 23567783 | 23567110 | 23567319 | 18 | 7 | 17 [illegible] |
| A5SS | 3683 | ENSG00000179335 | CLK3 | chr15 | + | 74622119 | 74622560 | 74622119 | 74622216 | 74624901 | 74625018 | 209 | 93 | 166 [illegible] |
| A5SS | 972 | ENSG00000108433 | GOSR2 | chr17 | + | 46923159 | 46923387 | 46923159 | 46923269 | 46929519 | 46929584 | 4 | 0 | 3 [illegible] |
| A5SS | 8406 | ENSG00000269609 | C10orf95-AS1 | chr10 | + | 102450186 | 102450360 | 102450186 | 102450330 | 102455336 | 102456297 | 0 | 0 | 8 [illegible] |
| A5SS | 4476 | ENSG00000177409 | SAMD9L | chr7 | - | 93146882 | 93147146 | 93147087 | 93147146 | 93145384 | 93146040 | 264 | 252 | 174 [illegible] |
| A5SS | 5319 | ENSG00000137817 | PARP6 | chr15 | - | 72265269 | 72265473 | 72265412 | 72265473 | 72265080 | 72265171 | 8 | 6 | 7 [illegible] |
| A5SS | 3883 | ENSG00000223745 | CCDC18-AS1 | chr1 | - | 93345539 | 93345846 | 93345723 | 93345846 | 93340457 | 93340510 | 7 | 10 | 7 [illegible] |
| A5SS | 7502 | ENSG00000104529 | EEF1D | chr8 | - | 143581228 | 143581328 | 143581285 | 143581328 | 143581053 | 143581154 | 2319 | 1173 | 2184 [illegible] |
| A5SS | 2332 | ENSG00000048140 | TSPAN17 | chr5 | + | 176656894 | 176656956 | 176656894 | 176656924 | 176657517 | 176659053 | 34 | 18 | 45 [illegible] |
| A5SS | 6964 | ENSG00000100532 | CGRRF1 | chr14 | + | 54530048 | 54530763 | 54530048 | 54530226 | 54530902 | 54531050 | 13 | 2 | 5 [illegible] |
| A5SS | 5552 | ENSG00000132530 | XAF1 | chr17 | + | 6758088 | 6758263 | 6758088 | 6758224 | 6759661 | 6759718 | 5 | 16 | 3 [illegible] |
| A5SS | 3582 | ENSG00000169241 | SLC50A1 | chr1 | + | 155136298 | 155136396 | 155136298 | 155136376 | 155136827 | 155136951 | 3 | 1 | 0 [illegible] |
| A5SS | 3358 | ENSG00000143420 | ENSA | chr1 | - | 150627311 | 150627592 | 150627466 | 150627592 | 150626478 | 150626526 | 55 | 55 | 80 [illegible] |
| A5SS | 312 | ENSG00000186818 | LILRB4 | chr19 | + | 54666398 | 54666456 | 54666398 | 54666436 | 54666693 | 54667390 | 0 | 1 | 3 [illegible] |
| A5SS | 1422 | ENSG00000213614 | HEXA | chr15 | - | 72353067 | 72355624 | 72355558 | 72355624 | 72351132 | 72351234 | 150 | 76 | 137 [illegible] |
| A5SS | 7371 | ENSG00000172269 | DPAGT1 | chr11 | - | 119097463 | 119098043 | 119097854 | 119098043 | 119097141 | 119097297 | 38 | 25 | 60 [illegible] |
| A5SS | 5439 | ENSG00000176978 | DPP7 | chr9 | - | 137111872 | 137112029 | 137111940 | 137112029 | 137111689 | 137111754 | 92 | 43 | 73 [illegible] |
| A5SS | 8235 | ENSG00000149311 | ATM | chr11 | + | 108246963 | 108247136 | 108246963 | 108247127 | 108248932 | 108249102 | 28 | 12 | 14 [illegible] |
| A5SS | 4716 | ENSG00000001497 | LAS1L | chrX | - | 65528963 | 65529258 | 65529211 | 65529258 | 65528259 | 65528369 | 26 | 15 | 24 [illegible] |
| A5SS | 645 | ENSG00000263753 | LINC00667 | chr18 | + | 5241698 | 5241889 | 5241698 | 5241812 | 5243593 | 5243784 | 47 | 51 | 26 [illegible] |
| A5SS | 3516 | ENSG00000089327 | FXYD5 | chr19 | + | 35157420 | 35157617 | 35157420 | 35157501 | 35158343 | 35158400 | 70 | 24 | 39 [illegible] |
| A5SS | 3762 | ENSG00000088035 | ALG6 | chr1 | + | 63370770 | 63371287 | 63370770 | 63371059 | 63396512 | 63396597 | 11 | 7 | 13 [illegible] |
| A5SS | 1745 | ENSG00000213928 | IRF9 | chr14 | + | 24164062 | 24164955 | 24164062 | 24164134 | 24165846 | 24165962 | 254 | 142 | 236 [illegible] |
| A5SS | 1674 | ENSG00000100749 | VRK1 | chr14 | + | 96855223 | 96855356 | 96855223 | 96855350 | 96856129 | 96856250 | 126 | 77 | 99 [illegible] |
| A5SS | 4499 | ENSG00000104894 | CD37 | chr19 | + | 49335713 | 49335786 | 49335713 | 49335782 | 49335885 | 49336029 | 16 | 5 | 9 [illegible] |
| A5SS | 4081 | ENSG00000087589 | CASS4 | chr20 | + | 56437163 | 56437586 | 56437163 | 56437424 | 56445899 | 56446001 | 45 | 30 | 65 [illegible] |
| A5SS | 7631 | ENSG00000100296 | THOC5 | chr22 | - | 29528273 | 29528466 | 29528425 | 29528466 | 29528077 | 29528177 | 11 | 9 | 11 [illegible] |
| A5SS | 6516 | ENSG00000179818 | PCBP1-AS1 | chr2 | - | 70085815 | 70086273 | 70086054 | 70086273 | 70085546 | 70085624 | 57 | 48 | 50 [illegible] |
| A5SS | 2532 | ENSG00000145495 | MARCHF6 | chr5 | + | 10417269 | 10417410 | 10417269 | 10417404 | 10423734 | 10423835 | 4 | 8 | 6 [illegible] |
| A5SS | 2475 | ENSG00000205937 | RNPS1 | chr16 | - | 2267174 | 2267237 | 2267188 | 2267237 | 2264572 | 2264760 | 9 | 5 | 6 [illegible] |
| A5SS | 7111 | ENSG00000090316 | MAEA | chr4 | + | 1311978 | 1312238 | 1311978 | 1312161 | 1315396 | 1315600 | 2 | 5 | 8 [illegible] |
| A5SS | 1740 | ENSG00000254505 | CHMP4A | chr14 | - | 24211410 | 24211592 | 24211414 | 24211592 | 24210653 | 24210768 | 86 | 12 | 39 [illegible] |
| A5SS | 2075 | ENSG00000043143 | JADE2 | chr5 | + | 134576767 | 134577038 | 134576767 | 134576896 | 134578496 | 134579475 | 11 | 9 | 10 [illegible] |
| A5SS | 7618 | ENSG00000116580 | GON4L | chr1 | - | 155774962 | 155775173 | 155775001 | 155775173 | 155773065 | 155773210 | 22 | 12 | 30 [illegible] |
| A5SS | 6838 | ENSG00000124333 | VAMP7 | chrX | + | 155889457 | 155889612 | 155889457 | 155889547 | 155898111 | 155898249 | 42 | 24 | 38 [illegible] |
| A5SS | 5240 | ENSG00000010244 | ZNF207 | chr17 | + | 32369598 | 32369782 | 32369598 | 32369745 | 32373398 | 32373457 | 190 | 138 | 201 [illegible] |
| A5SS | 7718 | ENSG00000135047 | CTSL | chr9 | + | 87726118 | 87726398 | 87726118 | 87726253 | 87727593 | 87727729 | 8 | 6 | 3 [illegible] |
| A5SS | 6360 | ENSG00000083168 | KAT6A | chr8 | - | 41942786 | 41943000 | 41942792 | 41943000 | 41940841 | 41941444 | 3 | 1 | 1 [illegible] |
| A5SS | 6256 | ENSG00000125779 | PANK2 | chr20 | + | 3907925 | 3908278 | 3907925 | 3908135 | 3910576 | 3910830 | 192 | 105 | 135 [illegible] |
| A5SS | 5193 | ENSG00000198755 | RPL10A | chr6 | + | 35468946 | 35469285 | 35468946 | 35469027 | 35469380 | 35469529 | 93 | 37 | 102 [illegible] |
| A5SS | 1646 | ENSG00000126822 | PLEKHG3 | chr14 | + | 64736852 | 64737375 | 64736852 | 64736891 | 64738741 | 64738855 | 204 | 75 | 81 [illegible] |
| A5SS | 8151 | ENSG00000185101 | ANO9 | chr11 | - | 429473 | 429652 | 429569 | 429652 | 428721 | 428826 | 14 | 0 | 6 [illegible] |
| A5SS | 336 | ENSG00000167613 | LAIR1 | chr19 | - | 54360915 | 54361209 | 54361009 | 54361209 | 54356927 | 54356966 | 309 | 109 | 116 [illegible] |
| A5SS | 3769 | ENSG00000025800 | KPNA6 | chr1 | + | 32156852 | 32156945 | 32156852 | 32156927 | 32157345 | 32157445 | 130 | 80 | 103 [illegible] |
| A5SS | 6582 | ENSG00000115677 | HDLBP | chr2 | - | 241268957 | 241269295 | 241269200 | 241269295 | 241268476 | 241268541 | 7 | 5 | 11 [illegible] |
| A5SS | 8480 | ENSG00000120616 | EPC1 | chr10 | - | 32273013 | 32273281 | 32273162 | 32273281 | 32272025 | 32272167 | 74 | 54 | 90 [illegible] |
| A5SS | 4847 | ENSG00000174748 | RPL15 | chr3 | + | 23917131 | 23917239 | 23917131 | 23917173 | 23917849 | 23918031 | 84 | 20 | 50 [illegible] |
| A5SS | 4675 | ENSG00000162734 | PEA15 | chr1 | + | 160211542 | 160211716 | 160211542 | 160211650 | 160213109 | 160213265 | 297 | 287 | 225 [illegible] |
| A5SS | 294 | ENSG00000178935 | ZNF552 | chr19 | - | 57814532 | 57814913 | 57814710 | 57814913 | 57813293 | 57813420 | 7 | 2 | 5 [illegible] |
| A5SS | 2908 | ENSG00000175455 | CCDC14 | chr3 | - | 123946802 | 123947319 | 123947279 | 123947319 | 123944848 | 123944990 | 236 | 228 | 229 [illegible] |
| A5SS | 2738 | ENSG00000181789 | COPG1 | chr3 | + | 129250681 | 129250894 | 129250681 | 129250734 | 129252280 | 129252361 | 22 | 5 | 13 [illegible] |
| A5SS | 286 | ENSG00000142453 | CARM1 | chr19 | + | 10908549 | 10908747 | 10908549 | 10908668 | 10909102 | 10909207 | 8 | 2 | 3 [illegible] |
| A5SS | 347 | ENSG00000160439 | RDH13 | chr19 | - | 55047104 | 55047488 | 55047386 | 55047488 | 55044342 | 55045309 | 12 | 3 | 17 [illegible] |
| A5SS | 8724 | ENSG00000166012 | TAF1D | chr11 | - | 93736693 | 93737239 | 93737063 | 93737239 | 93736140 | 93736304 | 455 | 397 | 440 [illegible] |
| A5SS | 2328 | ENSG00000108039 | XPNPEP1 | chr10 | - | 109875527 | 109877867 | 109877789 | 109877867 | 109873366 | 109873427 | 147 | 89 | 92 [illegible] |
| A5SS | 8321 | ENSG00000138286 | FAM149B1 | chr10 | + | 73228059 | 73228184 | 73228059 | 73228160 | 73230421 | 73230525 | 24 | 23 | 25 [illegible] |
| A5SS | 8939 | ENSG00000124496 | TRERF1 | chr6 | - | 42362946 | 42363079 | 42362996 | 42363079 | 42300637 | 42300749 | 44 | 20 | 52 [illegible] |
| A5SS | 8072 | ENSG00000160593 | JAML | chr11 | - | 118197967 | 118198091 | 118197997 | 118198091 | 118196734 | 118196821 | 84 | 62 | 56 [illegible] |
| A5SS | 822 | ENSG00000125454 | SLC25A19 | chr17 | - | 75288497 | 75288591 | 75288501 | 75288591 | 75286632 | 75286802 | 3 | 2 | 1 [illegible] |
| A5SS | 4671 | ENSG00000003756 | RBM5 | chr3 | + | 50088953 | 50089057 | 50088953 | 50089029 | 50090381 | 50090451 | 3 | 12 | 5 [illegible] |
| A5SS | 5344 | ENSG00000132676 | DAP3 | chr1 | + | 155689090 | 155689243 | 155689090 | 155689174 | 155709772 | 155709824 | 77 | 39 | 87 [illegible] |
| A5SS | 8424 | ENSG00000196233 | LCOR | chr10 | + | 96832263 | 96832399 | 96832263 | 96832395 | 96833405 | 96833479 | 6 | 6 | 13 [illegible] |
| A5SS | 5789 | ENSG00000160710 | ADAR | chr1 | - | 154590183 | 154590409 | 154590261 | 154590409 | 154589756 | 154589928 | 421 | 273 | 374 [illegible] |
| A5SS | 7746 | ENSG00000172716 | SLFN11 | chr17 | - | 35367540 | 35367700 | 35367649 | 35367700 | 35366946 | 35367063 | 25 | 17 | 10 [illegible] |
| A5SS | 5624 | ENSG00000115282 | TTC31 | chr2 | + | 74483321 | 74483563 | 74483321 | 74483410 | 74490024 | 74490473 | 8 | 10 | 9 [illegible] |
| A5SS | 5391 | ENSG00000162910 | MRPL55 | chr1 | - | 228108436 | 228109021 | 228108954 | 228109021 | 228108234 | 228108318 | 27 | 16 | 27 [illegible] |
| A5SS | 4097 | ENSG00000076685 | NT5C2 | chr10 | - | 103111692 | 103111805 | 103111744 | 103111805 | 103106588 | 103106706 | 5 | 3 | 3 [illegible] |
| A5SS | 8084 | ENSG00000110274 | CEP164 | chr11 | + | 117380613 | 117380714 | 117380613 | 117380705 | 117381700 | 117381868 | 3 | 3 | 6 [illegible] |
| A5SS | 5850 | ENSG00000169045 | HNRNPH1 | chr5 | - | 179616868 | 179616958 | 179616898 | 179616958 | 179616125 | 179616218 | 1826 | 1924 | 1666 [illegible] |
| A5SS | 3971 | ENSG00000163479 | SSR2 | chr1 | - | 156018131 | 156018368 | 156018269 | 156018368 | 156014960 | 156015069 | 11 | 4 | 9 [illegible] |
| A5SS | 5358 | ENSG00000084623 | EIF3I | chr1 | + | 32226170 | 32226530 | 32226170 | 32226283 | 32228498 | 32228609 | 258 | 167 | 258 [illegible] |
| A5SS | 4769 | ENSG00000129968 | ABHD17A | chr19 | - | 1881234 | 1881804 | 1881261 | 1881804 | 1879920 | 1880115 | 97 | 11 | 40 [illegible] |
| A5SS | 4464 | ENSG00000106028 | SSBP1 | chr7 | + | 141738357 | 141738503 | 141738357 | 141738410 | 141739123 | 141739190 | 29 | 24 | 39 [illegible] |
| A5SS | 7071 | ENSG00000102103 | PQBP1 | chrX | + | 48897911 | 48898093 | 48897911 | 48898082 | 48898491 | 48898576 | 22 | 7 | 28 [illegible] |
| A5SS | 1774 | ENSG00000100823 | APEX1 | chr14 | + | 20455225 | 20455394 | 20455225 | 20455389 | 20455577 | 20455703 | 210 | 87 | 172 [illegible] |
| A5SS | 2635 | ENSG00000055955 | ITIH4 | chr3 | - | 52819753 | 52819990 | 52819939 | 52819990 | 52819392 | 52819518 | 44 | 15 | 38 [illegible] |
| A5SS | 8471 | ENSG00000107862 | GBF1 | chr10 | + | 102379520 | 102379729 | 102379520 | 102379651 | 102379852 | 102379954 | 4 | 1 | 4 [illegible] |
| A5SS | 7433 | ENSG00000181396 | OGFOD3 | chr17 | - | 82415397 | 82415627 | 82415405 | 82415627 | 82411454 | 82411530 | 17 | 7 | 12 [illegible] |
| A5SS | 874 | ENSG00000108587 | GOSR1 | chr17 | + | 30481142 | 30481263 | 30481142 | 30481257 | 30484213 | 30484301 | 46 | 33 | 43 [illegible] |
| A5SS | 5696 | ENSG00000255857 | PXN-AS1 | chr12 | + | 120206544 | 120206712 | 120206544 | 120206621 | 120209838 | 120209934 | 4 | 2 | 2 [illegible] |
| A5SS | 887 | ENSG00000163597 | SNHG16 | chr17 | + | 76558373 | 76559043 | 76558373 | 76558600 | 76561287 | 76561398 | 142 | 58 | 153 [illegible] |
| A5SS | 2979 | ENSG00000163660 | CCNL1 | chr3 | - | 157157006 | 157157083 | 157157010 | 157157083 | 157153035 | 157153156 | 3 | 5 | 7 [illegible] |
| A5SS | 2985 | ENSG00000169714 | CNBP | chr3 | - | 129171633 | 129171771 | 129171654 | 129171771 | 129171445 | 129171538 | 393 | 225 | 411 [illegible] |
| A5SS | 6375 | ENSG00000105854 | PON2 | chr7 | - | 95416171 | 95416297 | 95416241 | 95416297 | 95412311 | 95412477 | 8 | 18 | 5 [illegible] |
| A5SS | 3246 | ENSG00000137074 | APTX | chr9 | - | 32988082 | 32988124 | 32988091 | 32988124 | 32987543 | 32987731 | 16 | 4 | 16 [illegible] |
| A5SS | 1080 | ENSG00000102984 | ZNF821 | chr16 | - | 71867892 | 71868037 | 71867911 | 71868037 | 71864902 | 71865048 | 5 | 9 | 27 [illegible] |
| A5SS | 5114 | ENSG00000118263 | KLF7 | chr2 | - | 207123975 | 207124404 | 207124343 | 207124404 | 207088457 | 207088581 | 43 | 27 | 41 [illegible] |
| A5SS | 6837 | ENSG00000124333 | VAMP7 | chrX | + | 155889457 | 155889608 | 155889457 | 155889547 | 155898111 | 155898249 | 31 | 17 | 32 [illegible] |
| A5SS | 7836 | ENSG00000133943 | DGLUCY | chr14 | + | 91204705 | 91204867 | 91204705 | 91204825 | 91215404 | 91215793 | 2 | 3 | 2 [illegible] |
| A5SS | 7101 | ENSG00000102393 | GLA | chrX | - | 101398718 | 101398946 | 101398784 | 101398946 | 101398369 | 101398567 | 5 | 1 | 5 [illegible] |
| A5SS | 4816 | ENSG00000162909 | CAPN2 | chr1 | + | 223761580 | 223761739 | 223761580 | 223761617 | 223762185 | 223762251 | 37 | 27 | 34 [illegible] |
| A5SS | 3460 | ENSG00000115523 | GNLY | chr2 | + | 85694752 | 85694996 | 85694752 | 85694923 | 85695319 | 85695423 | 481 | 99 | 94 [illegible] |
| A5SS | 1846 | ENSG00000103811 | CTSH | chr15 | - | 78944777 | 78945022 | 78944890 | 78945022 | 78939139 | 78939171 | 10 | 7 | 7 [illegible] |
| A5SS | 4259 | ENSG00000106479 | ZNF862 | chr7 | + | 149838374 | 149838635 | 149838374 | 149838461 | 149844624 | 149844736 | 16 | 10 | 8 [illegible] |
| A5SS | 4438 | ENSG00000180353 | HCLS1 | chr3 | - | 121657201 | 121657352 | 121657278 | 121657352 | 121647318 | 121647448 | 29 | 32 | 32 [illegible] |
| A5SS | 4548 | ENSG00000137275 | RIPK1 | chr6 | + | 3083084 | 3085408 | 3083084 | 3083313 | 3089580 | 3089657 | 106 | 102 | 96 [illegible] |
| A5SS | 7901 | ENSG00000183597 | TANGO2 | chr22 | + | 20036759 | 20037094 | 20036759 | 20036854 | 20043354 | 20043443 | 21 | 16 | 13 [illegible] |
| A5SS | 695 | ENSG00000072310 | SREBF1 | chr17 | - | 17819537 | 17819725 | 17819543 | 17819725 | 17819319 | 17819454 | 26 | 3 | 20 [illegible] |

| | | | | | | | | | | | | | |
|---|---|---|---|---|---|---|---|---|---|---|---|---|---|
| A5SS | 8428 | ENSG00000108094 | CUL2 | chr10 | - | 35090178 | 35090642 | 35090429 | 35090642 | 35071198 | 35071339 | 14 | 11 |
| A5SS | 5581 | ENSG00000197622 | CDC42SE1 | chr1 | - | 151055015 | 151055993 | 151055676 | 151055993 | 151054230 | 151054321 | 1271 | 546 |
| A5SS | 856 | ENSG00000108963 | DPH1 | chr17 | + | 2036528 | 2036686 | 2036528 | 2036653 | 2036834 | 2036956 | 38 | 23 |
| A5SS | 7316 | ENSG00000166169 | POLL | chr10 | - | 101584601 | 101584919 | 101584877 | 101584919 | 101582762 | 101582891 | 50 | 18 |
| A5SS | 7002 | ENSG00000111203 | ITFG2 | chr12 | + | 2823769 | 2824013 | 2823769 | 2823943 | 2824089 | 2824252 | 14 | 5 |
| A5SS | 6541 | ENSG00000164896 | FASTK | chr7 | - | 151078841 | 151079021 | 151078922 | 151079021 | 151078561 | 151078701 | 50 | 17 |
| A5SS | 3158 | ENSG00000107404 | DVL1 | chr1 | - | 1339286 | 1339439 | 1339361 | 1339439 | 1338521 | 1338653 | 4 | 2 |
| A5SS | 7473 | ENSG00000135679 | MDM2 | chr12 | + | 68835828 | 68835984 | 68835828 | 68835843 | 68836671 | 68836749 | 577 | 481 |
| A5SS | 1692 | ENSG00000155465 | SLC7A7 | chr14 | - | 22815313 | 22815451 | 22815319 | 22815451 | 22812899 | 22813440 | 31 | 21 |
| A5SS | 7413 | ENSG00000168404 | MLKL | chr16 | - | 74675323 | 74675404 | 74675354 | 74675404 | 74674959 | 74675100 | 19 | 13 |
| A5SS | 4220 | ENSG00000137106 | GRHPR | chr9 | + | 37422695 | 37422833 | 37422695 | 37422805 | 37424844 | 37424975 | 47 | 21 |
| A5SS | 2799 | ENSG00000068745 | IP6K2 | chr3 | - | 48694693 | 48695421 | 48695089 | 48695421 | 48694458 | 48694525 | 36 | 9 |
| A5SS | 1460 | ENSG00000137876 | RSL24D1 | chr15 | - | 55192402 | 55192833 | 55192719 | 55192833 | 55190974 | 55191047 | 27 | 13 |
| A5SS | 2794 | ENSG00000114745 | GORASP1 | chr3 | - | 39100746 | 39100877 | 39100787 | 39100877 | 39100304 | 39100503 | 21 | 5 |
| A5SS | 8060 | ENSG00000101158 | NELFCD | chr20 | + | 58989857 | 58990232 | 58989857 | 58989988 | 58990909 | 58991075 | 15 | 11 |
| A5SS | 8701 | ENSG00000125458 | NT5C | chr17 | - | 75130964 | 75131105 | 75131044 | 75131105 | 75130752 | 75130867 | 26 | 17 |
| A5SS | 659 | ENSG00000134030 | CTIF | chr18 | + | 48757918 | 48758411 | 48757918 | 48758405 | 48761389 | 48761689 | 1 | 1 |
| A5SS | 1507 | ENSG00000138592 | USP8 | chr15 | + | 50424381 | 50424654 | 50424381 | 50424514 | 50439008 | 50439177 | 35 | 42 |
| A5SS | 8762 | ENSG00000002016 | RAD52 | chr12 | - | 929972 | 930144 | 930050 | 930144 | 929818 | 929886 | 3 | 3 |
| A5SS | 6536 | ENSG00000163636 | PSMD6 | chr3 | - | 64022642 | 64022801 | 64022738 | 64022801 | 64022317 | 64022523 | 43 | 55 |
| A5SS | 5275 | ENSG00000148303 | RPL7A | chr9 | + | 133348251 | 133348817 | 133348251 | 133348388 | 133348921 | 133349042 | 7 | 1 |
| A5SS | 1584 | ENSG00000140464 | PML | chr15 | + | 74033155 | 74033422 | 74033155 | 74033414 | 74034477 | 74034530 | 3 | 0 |
| A5SS | 572 | ENSG00000101310 | SEC23B | chr20 | + | 18507939 | 18508297 | 18507939 | 18507972 | 18510821 | 18511056 | 8 | 0 |
| A5SS | 2398 | ENSG00000137802 | MAPKBP1 | chr15 | + | 41812515 | 41813781 | 41812515 | 41813101 | 41814549 | 41814739 | 39 | 11 |
| A5SS | 3009 | ENSG00000206503 | HLA-A | chr6 | + | 29944499 | 29944634 | 29944499 | 29944616 | 29945058 | 29945091 | 91 | 6 |
| A5SS | 8489 | ENSG00000119943 | PYROXD2 | chr10 | - | 98390870 | 98391082 | 98391009 | 98391082 | 98390597 | 98390754 | 7 | 1 |
| A5SS | 4724 | ENSG00000204498 | NFKBIL1 | chr6 | + | 31557627 | 31557849 | 31557627 | 31557804 | 31558021 | 31558829 | 17 | 6 |
| A5SS | 3034 | ENSG00000127483 | HP1BP3 | chr1 | - | 20780344 | 20780540 | 20780349 | 20780540 | 20779811 | 20779911 | 422 | 272 |
| A5SS | 1166 | ENSG00000103275 | UBE2I | chr16 | + | 1315653 | 1315730 | 1315653 | 1315726 | 1320173 | 1320283 | 20 | 20 |
| A5SS | 6939 | ENSG00000140332 | TLE3 | chr15 | - | 70074502 | 70074607 | 70074532 | 70074607 | 70066013 | 70066218 | 31 | 25 |
| A5SS | 5277 | ENSG00000131143 | COX4I1 | chr16 | + | 85804936 | 85805153 | 85804936 | 85805104 | 85805732 | 85805864 | 12 | 4 |
| A5SS | 4423 | ENSG00000166523 | CLEC4E | chr12 | - | 8539141 | 8539306 | 8539216 | 8539306 | 8536089 | 8536205 | 68 | 46 |
| A5SS | 7921 | ENSG00000106290 | TAF6 | chr7 | - | 100114053 | 100114268 | 100114083 | 100114268 | 100113867 | 100113954 | 125 | 66 |
| A5SS | 8548 | ENSG00000079974 | RABL2B | chr22 | - | 50783038 | 50783286 | 50783044 | 50783286 | 50782187 | 50782351 | 3 | 4 |
| A5SS | 3765 | ENSG00000119203 | CPSF3 | chr2 | + | 9467706 | 9467975 | 9467706 | 9467776 | 9471342 | 9471439 | 2 | 3 |
| A5SS | 417 | ENSG00000268790 | NA | chr19 | - | 16647095 | 16647261 | 16647119 | 16647261 | 16578334 | 16578409 | 11 | 4 |
| A5SS | 2443 | ENSG00000137494 | ANKRD42 | chr11 | + | 83198478 | 83198642 | 83198478 | 83198558 | 83206057 | 83206165 | 21 | 20 |
| A5SS | 5869 | ENSG00000153094 | BCL2L11 | chr2 | + | 111123732 | 111124139 | 111123732 | 111123869 | 111150043 | 111150147 | 173 | 87 |
| A5SS | 6970 | ENSG00000173660 | UQCRH | chr1 | + | 46309100 | 46309143 | 46309100 | 46309127 | 46310154 | 46310316 | 5 | 4 |
| A5SS | 7385 | ENSG00000160299 | PCNT | chr21 | + | 46431528 | 46432215 | 46431528 | 46431978 | 46435903 | 46436148 | 50 | 35 |
| A5SS | 7102 | ENSG00000102393 | GLA | chrX | - | 101401232 | 101401809 | 101401631 | 101401809 | 101400665 | 101400757 | 10 | 26 |
| A5SS | 1059 | ENSG00000008517 | IL32 | chr16 | + | 3067376 | 3067613 | 3067376 | 3067415 | 3067983 | 3068010 | 854 | 170 |
| A5SS | 6937 | ENSG00000140332 | TLE3 | chr15 | - | 70074502 | 70074607 | 70074532 | 70074607 | 70060529 | 70060649 | 4 | 9 |
| A5SS | 3096 | ENSG00000132591 | ERAL1 | chr17 | + | 28856504 | 28856685 | 28856504 | 28856582 | 28857938 | 28857985 | 3 | 0 |
| A5SS | 5405 | ENSG00000003509 | NDUFAF7 | chr2 | + | 37237756 | 37237867 | 37237756 | 37237835 | 37241577 | 37241637 | 21 | 10 |
| A5SS | 4389 | ENSG00000249915 | PDCD6 | chr5 | + | 304176 | 306754 | 304176 | 304221 | 311292 | 311402 | 46 | 21 |
| A5SS | 1311 | ENSG00000167508 | MVD | chr16 | - | 88657734 | 88658029 | 88657914 | 88658029 | 88657435 | 88657582 | 4 | 0 |
| A5SS | 6028 | ENSG00000114388 | NPRL2 | chr3 | - | 50348684 | 50348782 | 50348710 | 50348782 | 50348526 | 50348563 | 38 | 18 |
| A5SS | 4335 | ENSG00000173209 | AHSA2P | chr2 | + | 61184711 | 61184924 | 61184711 | 61184800 | 61185501 | 61185618 | 15 | 13 |
| A5SS | 1927 | ENSG00000135596 | MICAL1 | chr6 | - | 109451599 | 109452401 | 109452245 | 109452401 | 109446695 | 109446772 | 54 | 47 |
| A5SS | 103 | ENSG00000100150 | DEPDC5 | chr22 | + | 31810520 | 31810665 | 31810520 | 31810641 | 31814991 | 31815212 | 0 | 0 |
| A5SS | 6341 | ENSG00000173715 | C11orf80 | chr11 | + | 66801030 | 66801161 | 66801030 | 66801102 | 66803994 | 66804194 | 1 | 0 |
| A5SS | 4768 | ENSG00000129968 | ABHD17A | chr19 | - | 1878663 | 1880115 | 1879920 | 1880115 | 1877507 | 1877687 | 21 | 7 |
| [illegible] | | | | | | | | | | | | | |

| | | | | | | | | | | | | | | | | | | | | | | | | |
|---|---|---|---|---|---|---|---|---|---|---|---|---|---|---|---|---|---|---|---|---|---|---|---|---|
| A5SS | 4637 | ENSG00000160654 | CD3G | chr11 | + | 118349026 | 118349171 | 118349026 | 118349050 | 118349742 | 118349970 | 4 | 2 | 7 | 10 | 7 | 389 | 193 | 389 | 415 | 582 | 0 | 28 | [illegible] |
| A5SS | 1367 | ENSG00000140694 | PARN | chr16 | - | 14591912 | 14593378 | 14593300 | 14593378 | 14586317 | 14586361 | 19 | 8 | 11 | 7 | 13 | 166 | 140 | 180 | 179 | 409 | 0 | 6 | [illegible] |
| A5SS | 7173 | ENSG00000215301 | DDX3X | chrX | + | 41344238 | 41344564 | 41344238 | 41344399 | 41345179 | 41345324 | 19 | 6 | 27 | 14 | 11 | 648 | 431 | 503 | 698 | 631 | 0 | 5 | [illegible] |
| A5SS | 149 | ENSG00000270011 | ZNF559-ZNF177 | chr19 | + | 9337795 | 9338017 | 9337795 | 9337858 | 9338493 | 9338582 | 33 | 19 | 30 | 27 | 27 | 29 | 18 | 36 | 34 | 18 | 0 | 22 | [illegible] |
| A5SS | 1932 | ENSG00000126214 | KLC1 | chr14 | + | 103679383 | 103679545 | 103679383 | 103679518 | 103687080 | 103687211 | 70 | 54 | 54 | 64 | 75 | 10 | 8 | 7 | 13 | 19 | 135 | 36 | [illegible] |
| A5SS | 5665 | ENSG00000115762 | PLEKHB2 | chr2 | + | 131132901 | 131133015 | 131132901 | 131132991 | 131140166 | 131140275 | 52 | 43 | 30 | 74 | 70 | 225 | 149 | 193 | 239 | 385 | 37 | 60 | [illegible] |
| A5SS | 2282 | ENSG00000011243 | AKAP8L | chr19 | - | 15398629 | 15399410 | 15399301 | 15399410 | 15397713 | 15397855 | 49 | 38 | 42 | 30 | 33 | 152 | 82 | 94 | 146 | 170 | 32 | 37 | [illegible] |
| A5SS | 5502 | ENSG00000127831 | VIL1 | chr2 | + | 218425614 | 218425811 | 218425614 | 218425799 | 218427964 | 218428073 | 6 | 13 | 2 | 16 | 2 | 0 | 3 | 0 | 5 | 0 | 13 | 20 | [illegible] |
| A5SS | 2136 | ENSG00000166888 | STAT6 | chr12 | - | 57109940 | 57110110 | 57110042 | 57110110 | 57108162 | 57108299 | 64 | 36 | 76 | 68 | 80 | 15 | 17 | 19 | 33 | 46 | 216 | 122 | [illegible] |
| A5SS | 7469 | ENSG00000164733 | CTSB | chr8 | - | 11861360 | 11861479 | 11861405 | 11861479 | 11853328 | 11853479 | 13 | 11 | 12 | 15 | 7 | 0 | 0 | 1 | 1 | 3 | 48 | 18 | [illegible] |
| A5SS | 1935 | ENSG00000135506 | OS9 | chr12 | + | 57695961 | 57696038 | 57695961 | 57695976 | 57696274 | 57696373 | 509 | 297 | 373 | 411 | 582 | 2 | 5 | 4 | 7 | 0 | 862 | 572 | [illegible] |
| A5SS | 233 | ENSG00000130816 | DNMT1 | chr19 | - | 10142027 | 10142220 | 10142079 | 10142220 | 10141104 | 10141189 | 70 | 24 | 52 | 68 | 45 | 2 | 0 | 1 | 7 | 2 | 53 | 42 | [illegible] |
| A5SS | 1138 | ENSG00000010361 | FUZ | chr19 | - | 49812614 | 49812736 | 49812637 | 49812736 | 49812250 | 49812335 | 8 | 3 | 8 | 7 | 7 | 1 | 0 | 1 | 0 | 2 | 8 | 5 | [illegible] |
| A5SS | 5880 | ENSG00000129158 | SERGEF | chr11 | - | 18010070 | 18010139 | 18010074 | 18010139 | 18007940 | 18008076 | 17 | 18 | 14 | 19 | 19 | 18 | 6 | 8 | 18 | 24 | 19 | 6 | [illegible] |
| A5SS | 7620 | ENSG00000116580 | GON4L | chr1 | - | 155857146 | 155857295 | 155857158 | 155857295 | 155853275 | 155853806 | 22 | 15 | 17 | 15 | 22 | 5 | 9 | 6 | 6 | 4 | 12 | 3 | [illegible] |
| A5SS | 1063 | ENSG00000168488 | ATXN2L | chr16 | + | 28826861 | 28826986 | 28826861 | 28826923 | 28829400 | 28829492 | 666 | 455 | 640 | 762 | 706 | 3 | 4 | 2 | 1 | 8 | 1206 | 869 | [illegible] |
| A5SS | 3955 | ENSG00000066322 | ELOVL1 | chr1 | - | 43364541 | 43364647 | 43364614 | 43364647 | 43364323 | 43364460 | 245 | 99 | 147 | 170 | 233 | 12 | 3 | 7 | 4 | 19 | 290 | 227 | [illegible] |
| A5SS | 3692 | ENSG00000136450 | SRSF1 | chr17 | - | 58006906 | 58007261 | 58006943 | 58007261 | 58006342 | 58006527 | 14 | 9 | 21 | 13 | 4 | 216 | 111 | 199 | 203 | 51 | 13 | 15 | [illegible] |
| A5SS | 6334 | ENSG00000183696 | UPP1 | chr7 | + | 48101823 | 48101982 | 48101823 | 48101870 | 48103296 | 48103411 | 175 | 127 | 114 | 131 | 130 | 5 | 8 | 3 | 5 | 1 | 466 | 211 | [illegible] |
| A5SS | 2518 | ENSG00000182093 | GET1 | chr21 | + | 39390697 | 39391476 | 39390697 | 39390863 | 39391768 | 39391836 | 4 | 3 | 5 | 7 | 0 | 34 | 23 | 11 | 36 | 64 | 15 | 0 | [illegible] |
| A5SS | 7336 | ENSG00000120899 | PTK2B | chr8 | + | 27433434 | 27433552 | 27433434 | 27433513 | 27434092 | 27434132 | 305 | 117 | 236 | 256 | 197 | 21 | 9 | 16 | 8 | 12 | 458 | 426 | [illegible] |
| A5SS | 5711 | ENSG00000163516 | ANKZF1 | chr2 | + | 219230227 | 219230405 | 219230227 | 219230350 | 219232259 | 219232362 | 9 | 8 | 19 | 14 | 3 | 4 | 2 | 0 | 1 | 0 | 2 | 6 | [illegible] |
| A5SS | 6752 | ENSG00000234741 | GAS5 | chr1 | - | 173864483 | 173864704 | 173864674 | 173864704 | 173864256 | 173864304 | 721 | 489 | 882 | 678 | 1210 | 9 | 5 | 26 | 22 | 40 | 318 | 322 | [illegible] |
| A5SS | 3124 | ENSG00000100578 | KIAA0586 | chr14 | + | 58459842 | 58460070 | 58459842 | 58459909 | 58460985 | 58461160 | 187 | 213 | 186 | 220 | 399 | 6 | 12 | 0 | 1 | 0 | 17 | 151 | [illegible] |
| A5SS | 973 | ENSG00000108433 | GOSR2 | chr17 | + | 46923159 | 46923632 | 46923159 | 46923269 | 46929519 | 46929584 | 7 | 0 | 4 | 0 | 1 | 0 | 1 | 1 | 2 | 1 | 7 | 6 | [illegible] |
| A5SS | 5505 | ENSG00000237765 | FAM200B | chr4 | + | 15681755 | 15681901 | 15681755 | 15681784 | 15686235 | 15686462 | 33 | 23 | 31 | 44 | 28 | 71 | 46 | 69 | 49 | 106 | 47 | 14 | [illegible] |
| A5SS | 8833 | ENSG00000129317 | PUS7L | chr12 | - | 43754335 | 43755261 | 43755216 | 43755261 | 43748449 | 43748609 | 129 | 89 | 104 | 111 | 137 | 3 | 2 | 2 | 5 | 5 | 26 | 56 | [illegible] |
| A5SS | 3525 | ENSG00000163374 | YY1AP1 | chr1 | - | 155672520 | 155672731 | 155672559 | 155672731 | 155670319 | 155670464 | 21 | 22 | 16 | 21 | 30 | 184 | 101 | 152 | 169 | 162 | 39 | 46 | [illegible] |
| A5SS | 4515 | ENSG00000168610 | STAT3 | chr17 | - | 42322281 | 42322494 | 42322284 | 42322494 | 42317181 | 42317224 | 250 | 149 | 150 | 247 | 343 | 67 | 31 | 57 | 42 | 81 | 352 | 346 | [illegible] |
| A5SS | 6989 | ENSG00000223959 | AFG3L1P | chr16 | + | 89984530 | 89984617 | 89984530 | 89984599 | 89990865 | 89991003 | 33 | 23 | 20 | 34 | 10 | 0 | 0 | 3 | 1 | 0 | 83 | 60 | [illegible] |
| A5SS | 6135 | ENSG00000160185 | UBASH3A | chr21 | + | 42434831 | 42435509 | 42434831 | 42434954 | 42437541 | 42437596 | 16 | 10 | 18 | 32 | 41 | 0 | 0 | 0 | 0 | 0 | 2 | 2 | [illegible] |
| A5SS | 7468 | ENSG00000164733 | CTSB | chr8 | - | 11861299 | 11861479 | 11861405 | 11861479 | 11853328 | 11853479 | 40 | 40 | 28 | 32 | 20 | 0 | 0 | 1 | 1 | 3 | 94 | 35 | [illegible] |
| A5SS | 2393 | ENSG00000026297 | RNASET2 | chr6 | - | 166946540 | 166946739 | 166946681 | 166946739 | 166943018 | 166943089 | 161 | 237 | 171 | 223 | 491 | 375 | 207 | 275 | 455 | 625 | 728 | 226 | [illegible] |
| A5SS | 4241 | ENSG00000227372 | TP73-AS1 | chr1 | - | 3739695 | 3740387 | 3739835 | 3740387 | 3737165 | 3739407 | 18 | 13 | 23 | 27 | 25 | 1 | 1 | 3 | 1 | 10 | 3 | 5 | [illegible] |
| A5SS | 3845 | ENSG00000117620 | SLC35A3 | chr1 | + | 99993536 | 99993745 | 99993536 | 99993741 | 99999260 | 99999415 | 5 | 4 | 2 | 2 | 1 | 92 | 33 | 74 | 71 | 93 | 0 | 2 | [illegible] |
| A5SS | 1140 | ENSG00000260565 | ERVK13-1 | chr16 | - | 2671023 | 2672570 | 2671344 | 2672570 | 2669410 | 2669561 | 35 | 22 | 30 | 36 | 15 | 1 | 4 | 4 | 1 | 0 | 20 | 37 | [illegible] |
| A5SS | 123 | ENSG00000228315 | GUSBP11 | chr22 | - | 23694118 | 23695517 | 23695172 | 23695517 | 23683724 | 23683867 | 24 | 18 | 26 | 27 | 11 | 5 | 0 | 0 | 1 | 0 | 41 | 29 | [illegible] |
| A5SS | 5463 | ENSG00000172890 | NADSYN1 | chr11 | + | 71474394 | 71475622 | 71474394 | 71474526 | 71476200 | 71477225 | 17 | 3 | 14 | 6 | 1 | 1 | 0 | 1 | 3 | 2 | 14 | 11 | [illegible] |
| A5SS | 6689 | ENSG00000013374 | NUB1 | chr7 | + | 151341811 | 151341965 | 151341811 | 151341846 | 151345347 | 151345466 | 3 | 2 | 4 | 1 | 1 | 37 | 22 | 19 | 37 | 14 | 14 | 6 | [illegible] |
| A5SS | 4308 | ENSG00000161835 | TAMALIN | chr12 | + | 52011038 | 52011207 | 52011038 | 52011141 | 52013686 | 52013780 | 10 | 6 | 14 | 7 | 1 | 10 | 3 | 15 | 16 | 19 | 37 | 16 | [illegible] |
| A5SS | 4434 | ENSG00000197989 | SNHG12 | chr1 | - | 28580424 | 28580646 | 28580559 | 28580646 | 28579911 | 28579981 | 58 | 34 | 68 | 59 | 57 | 54 | 23 | 52 | 37 | 58 | 32 | 37 | [illegible] |
| A5SS | 2766 | ENSG00000100003 | SEC14L2 | chr22 | + | 30415758 | 30416087 | 30415758 | 30415865 | 30416233 | 30416403 | 8 | 13 | 32 | 30 | 20 | 0 | 0 | 0 | 0 | 2 | 0 | 20 | [illegible] |
| A5SS | 759 | ENSG00000214087 | ARL16 | chr17 | - | 81682442 | 81682675 | 81682490 | 81682675 | 81682033 | 81682149 | 16 | 11 | 19 | 6 | 15 | 10 | 2 | 5 | 6 | 8 | 39 | 18 | [illegible] |
| A5SS | 1883 | ENSG00000175768 | TOMM5 | chr9 | - | 37592309 | 37592597 | 37592411 | 37592597 | 37588412 | 37588932 | 6 | 8 | 3 | 7 | 0 | 68 | 20 | 55 | 60 | 43 | 0 | 8 | [illegible] |
| A5SS | 3523 | ENSG00000143110 | C1orf162 | chr1 | + | 111476797 | 111477078 | 111476797 | 111476867 | 111477725 | 111477775 | 73 | 43 | 33 | 43 | 48 | 2 | 2 | 1 | 13 | 5 | 122 | 32 | [illegible] |
| A5SS | 2230 | ENSG00000204628 | RACK1 | chr5 | - | 181242169 | 181242345 | 181242173 | 181242345 | 181239486 | 181239582 | 30 | 15 | 27 | 16 | 6 | 1 | 0 | 3 | 4 | 0 | 4 | 12 | [illegible] |
| A5SS | 4402 | ENSG00000158552 | ZFAND2B | chr2 | + | 219207647 | 219207779 | 219207647 | 219207774 | 219207886 | 219208038 | 50 | 17 | 59 | 25 | 32 | 0 | 0 | 0 | 0 | 3 | 56 | 33 | [illegible] |
| A5SS | 3127 | ENSG00000134897 | BIVM | chr13 | + | 102816427 | 102817128 | 102816427 | 102816554 | 102821036 | 102821132 | 7 | 6 | 6 | 8 | 7 | 26 | 21 | 39 | 55 | 46 | 7 | 4 | [illegible] |
| A5SS | 6450 | ENSG00000136286 | MYO1G | chr7 | - | 44975475 | 44975649 | 44975483 | 44975649 | 44975173 | 44975227 | 14 | 26 | 9 | 16 | 7 | 760 | 519 | 534 | 672 | 462 | 81 | 24 | [illegible] |
| A5SS | 3385 | ENSG00000231607 | DLEU2 | chr13 | - | 50044639 | 50044768 | 50044650 | 50044768 | 50029277 | 50029492 | 3 | 0 | 0 | 3 | 7 | 40 | 53 | 31 | 46 | 117 | 0 | 0 | [illegible] |
| A5SS | 8732 | ENSG00000145781 | COMMD10 | chr5 | + | 116092544 | 116092700 | 116092544 | 116092695 | 116134067 | 116134178 | 120 | 77 | 80 | 100 | 192 | 1 | 2 | 0 | 2 | 2 | 15 | 60 | [illegible] |
| A5SS | 8769 | ENSG00000111424 | VDR | chr12 | - | 47878815 | 47879115 | 47878967 | 47879115 | 47865046 | 47865177 | 2 | 2 | 3 | 5 | 1 | 24 | 18 | 16 | 25 | 14 | 15 | 0 | [illegible] |
| A5SS | 8703 | ENSG00000125458 | NT5C | chr17 | - | 75131044 | 75131281 | 75131180 | 75131281 | 75130752 | 75130867 | 58 | 26 | 51 | 60 | 44 | 2 | 0 | 0 | 0 | 0 | 47 | 92 | [illegible] |
| A5SS | 8313 | ENSG00000090621 | PABPC4 | chr1 | - | 39563526 | 39563741 | 39563613 | 39563741 | 39562322 | 39562416 | 7 | 3 | 4 | 6 | 3 | 124 | 62 | 135 | 142 | 71 | 0 | 2 | [illegible] |
| A5SS | 7419 | ENSG00000111678 | C12orf57 | chr12 | + | 6944475 | 6944652 | 6944475 | 6944565 | 6945770 | 6946002 | 321 | 142 | 338 | 279 | 207 | 68 | 28 | 77 | 52 | 56 | 107 | 238 | [illegible] |
| A5SS | 7925 | ENSG00000139626 | ITGB7 | chr12 | - | 53201071 | 53201194 | 53201146 | 53201194 | 53200242 | 53200446 | 652 | 280 | 580 | 601 | 588 | 9 | 0 | 2 | 0 | 11 | 432 | 404 | [illegible] |
| A5SS | 2594 | ENSG00000129187 | DCTD | chr4 | - | 182916285 | 182916539 | 182916418 | 182916539 | 182915880 | 182915888 | 12 | 6 | 28 | 19 | 2 | 5 | 0 | 2 | 1 | 0 | 6 | 13 | [illegible] |
| A5SS | 4574 | ENSG00000126709 | IFI6 | chr1 | - | 27669232 | 27669346 | 27669244 | 27669346 | 27668457 | 27668535 | 4 | 7 | 4 | 12 | 0 | 34 | 24 | 41 | 67 | 8 | 35 | 47 | [illegible] |
| A5SS | 2606 | ENSG00000113522 | RAD50 | chr5 | + | 132579316 | 132580066 | 132579316 | 132579502 | 132587561 | 132587690 | 107 | 126 | 75 | 125 | 202 | 4 | 0 | 1 | 2 | 7 | 12 | 50 | [illegible] |
| A5SS | 6499 | ENSG00000197694 | SPTAN1 | chr9 | + | 128624327 | 128624824 | 128624327 | 128624487 | 128625102 | 128625179 | 21 | 11 | 18 | 9 | 19 | 259 | 144 | 213 | 221 | 350 | 21 | 16 | [illegible] |
| A5SS | 3822 | ENSG00000118217 | ATF6 | chr1 | + | 161778243 | 161778320 | 161778243 | 161778316 | 161781911 | 161781999 | 69 | 54 | 67 | 60 | 128 | 1 | 0 | 4 | 1 | 1 | 79 | 67 | [illegible] |
| A5SS | 1772 | ENSG00000100823 | APEX1 | chr14 | + | 20455225 | 20455389 | 20455225 | 20455328 | 20455577 | 20455703 | 29 | 18 | 40 | 19 | 14 | 13 | 6 | 22 | 10 | 2 | 24 | 35 | [illegible] |
| A5SS | 6611 | ENSG00000133195 | SLC39A11 | chr17 | - | 72947527 | 72947875 | 72947751 | 72947875 | 72849633 | 72849804 | 3 | 2 | 2 | 4 | 2 | 25 | 11 | 18 | 15 | 11 | 7 | 18 | [illegible] |
| A5SS | 6136 | ENSG00000196730 | DAPK1 | chr9 | + | 87497680 | 87498107 | 87497680 | 87497772 | 87498969 | 87499139 | 14 | 15 | 4 | 10 | 9 | 5 | 0 | 0 | 4 | 0 | 29 | 4 | [illegible] |
| A5SS | 6055 | ENSG00000168000 | BSCL2 | chr11 | - | 62705300 | 62705617 | 62705467 | 62705617 | 62702467 | 62702549 | 69 | 15 | 30 | 50 | 36 | 2 | 0 | 0 | 2 | 0 | 16 | 26 | [illegible] |
| A5SS | 6080 | ENSG00000171453 | POLR1C | chr6 | + | 43519705 | 43519905 | 43519705 | 43519838 | 43520065 | 43520185 | 7 | 11 | 17 | 18 | 22 | 68 | 46 | 40 | 61 | 114 | 12 | 10 | [illegible] |
| A5SS | 5138 | ENSG00000166986 | MARS1 | chr12 | + | 57512007 | 57512353 | 57512007 | 57512103 | 57512750 | 57512964 | 121 | 61 | 116 | 129 | 117 | 2 | 0 | 0 | 2 | 0 | 116 | 137 | [illegible] |
| A5SS | 451 | ENSG00000080845 | DLGAP4 | chr20 | + | 36525850 | 36526077 | 36525850 | 36526006 | 36526812 | 36527231 | 1 | 1 | 3 | 0 | 0 | 12 | 9 | 19 | 24 | 12 | 0 | 0 | [illegible] |
| A5SS | 600 | ENSG00000101751 | POLI | chr18 | + | 54283921 | 54284065 | 54283921 | 54284013 | 54287280 | 54287411 | 10 | 8 | 1 | 13 | 1 | 92 | 48 | 44 | 51 | 62 | 0 | 2 | [illegible] |
| A5SS | 4902 | ENSG00000166822 | TMEM170A | chr16 | - | 75451668 | 75451839 | 75451737 | 75451839 | 75443053 | 75447688 | 38 | 9 | 19 | 28 | 12 | 4 | 3 | 7 | 5 | 1 | 18 | 23 | [illegible] |
| A5SS | 157 | ENSG00000104936 | DMPK | chr19 | - | 45778492 | 45779359 | 45779263 | 45779359 | 45778126 | 45778220 | 18 | 11 | 20 | 29 | 5 | 0 | 0 | 1 | 1 | 0 | 3 | 28 | [illegible] |
| A5SS | 2231 | ENSG00000204628 | RACK1 | chr5 | - | 181242169 | 181242345 | 181242173 | 181242345 | 181241491 | 181241563 | 32 | 17 | 28 | 16 | 15 | 3 | 0 | 0 | 5 | 1 | 4 | 14 | [illegible] |
| A5SS | 2317 | ENSG00000181929 | PRKAG1 | chr12 | - | 49018523 | 49018748 | 49018731 | 49018748 | 49013061 | 49013110 | 3 | 5 | 4 | 4 | 0 | 28 | 11 | 10 | 8 | 22 | 3 | 3 | [illegible] |
| A5SS | 2625 | ENSG00000108591 | DRG2 | chr17 | + | 18099016 | 18099093 | 18099016 | 18099077 | 18099632 | 18100435 | 4 | 4 | 2 | 6 | 5 | 40 | 17 | 34 | 46 | 38 | 5 | 1 | [illegible] |
| A5SS | 3415 | ENSG00000184432 | COPB2 | chr3 | - | 139373665 | 139374588 | 139374488 | 139374588 | 139373212 | 139373412 | 298 | 204 | 215 | 258 | 203 | 8 | 10 | 7 | 11 | 6 | 187 | 187 | [illegible] |
| A5SS | 3253 | ENSG00000178922 | HYI | chr1 | - | 43453385 | 43453497 | 43453412 | 43453497 | 43452204 | 43452319 | 4 | 2 | 14 | 9 | 2 | 0 | 0 | 1 | 1 | 6 | 3 | 1 | [illegible] |
| A5SS | 8582 | ENSG00000284770 | TBCE | chr1 | + | 235442851 | 235442961 | 235442851 | 235442911 | 235448348 | 235448440 | 1 | 9 | 5 | 1 | 0 | 73 | 65 | 51 | 73 | 47 | 0 | 3 | [illegible] |
| A5SS | 4103 | ENSG00000256269 | HMBS | chr11 | + | 119088634 | 119088848 | 119088634 | 119088707 | 119089081 | 119089131 | 0 | 0 | 1 | 2 | 8 | 29 | 18 | 19 | 23 | 14 | 0 | 1 | [illegible] |
| A5SS | 6790 | ENSG00000124222 | STX16 | chr20 | + | 58667489 | 58667608 | 58667489 | 58667597 | 58667986 | 58668127 | 8 | 13 | 9 | 14 | 19 | 400 | 355 | 489 | 457 | 425 | 25 | 16 | [illegible] |
| A5SS | 2097 | ENSG00000261556 | SMG1P7 | chr16 | - | 70231322 | 70231523 | 70231436 | 70231523 | 70230993 | 70231158 | 13 | 5 | 10 | 12 | 13 | 18 | 8 | 12 | 27 | 36 | 0 | 0 | [illegible] |
| A5SS | 2272 | ENSG00000134644 | PUM1 | chr1 | - | 30950126 | 30950261 | 30950132 | 30950261 | 30945345 | 30945483 | 139 | 69 | 93 | 129 | 127 | 103 | 50 | 71 | 93 | 79 | 147 | 185 | [illegible] |
| A5SS | 8698 | ENSG00000229117 | RPL41 | chr12 | + | 56116774 | 56116829 | 56116774 | 56116799 | 56117188 | 56117211 | 7 | 10 | 16 | 16 | 7 | 346 | 172 | 306 | 342 | 316 | 0 | 13 | [illegible] |
| A5SS | 6020 | ENSG00000089818 | NECAP1 | chr12 | + | 8090194 | 8090343 | 8090194 | 8090299 | 8092675 | 8092784 | 13 | 9 | 17 | 22 | 23 | 13 | 5 | 9 | 13 | 38 | 0 | 18 | [illegible] |
| A5SS | 249 | ENSG00000104852 | SNRNP70 | chr19 | + | 49102113 | 49102185 | 49102113 | 49102173 | 49104633 | 49104735 | 34 | 21 | 35 | 32 | 64 | 23 | 37 | 35 | 41 | 47 | 19 | 11 | [illegible] |
| A5SS | 1219 | ENSG00000130175 | PRKCSH | chr19 | + | 11447438 | 11447618 | 11447438 | 11447597 | 11447692 | 11447789 | 24 | 6 | 33 | 34 | 25 | 48 | 19 | 58 | 40 | 82 | 28 | 28 | [illegible] |
| A5SS | 5846 | ENSG00000059377 | TBXAS1 | chr7 | + | 139875584 | 139875666 | 139875584 | 139875637 | 139911224 | 139911321 | 18 | 5 | 3 | 9 | 10 | 247 | 119 | 153 | 178 | 196 | 61 | 30 | [illegible] |
| A5SS | 6029 | ENSG00000114388 | NPRL2 | chr3 | - | 50349385 | 50349828 | 50349664 | 50349828 | 50348873 | 50349010 | 68 | 34 | 59 | 76 | 48 | 4 | 0 | 0 | 2 | 2 | 101 | 75 | [illegible] |
| A5SS | 4114 | ENSG00000111331 | OAS3 | chr12 | + | 112967935 | 112968180 | 112967935 | 112968174 | 112969607 | 112969755 | 5 | 4 | 3 | 10 | 0 | 76 | 81 | 32 | 91 | 50 | 56 | 8 | [illegible] |
| A5SS | 5930 | ENSG00000124356 | STAMBP | chr2 | + | 73828966 | 73829510 | 73828966 | 73828996 | 73830844 | 73831059 | 12 | 4 | 8 | 12 | 23 | 13 | 7 | 10 | 13 | 18 | 1 | 14 | [illegible] |
| A5SS | 105 | ENSG00000100150 | DEPDC5 | chr22 | + | 31905983 | 31906094 | 31905983 | 31906066 | 31906204 | 31906975 | 0 | 0 | 0 | 1 | 1 | 17 | 6 | 27 | 29 | 16 | 0 | 0 | [illegible] |
| A5SS | 6235 | ENSG00000168036 | CTNNB1 | chr3 | + | 41199469 | 41199670 | 41199469 | 41199622 | 41224020 | 41224081 | 108 | 44 | 85 | 78 | 38 | 2 | 2 | 0 | 8 | 0 | 55 | 86 | [illegible] |
| A5SS | 1003 | ENSG00000159210 | SNF8 | chr17 | - | 48936130 | 48936242 | 48936169 | 48936242 | 48933204 | 48933346 | 6 | 5 | 3 | 1 | 8 | 83 | 59 | 72 | 91 | 84 | 0 | 6 | [illegible] |
| A5SS | 2169 | ENSG00000153071 | DAB2 | chr5 | - | 39392203 | 39392463 | 39392364 | 39392463 | 39390443 | 39390575 | 8 | 28 | 9 | 10 | 25 | 202 | 317 | 63 | 201 | 219 | 138 | 53 | [illegible] |
| A5SS | 5547 | ENSG00000081059 | TCF7 | chr5 | + | 134138058 | 134138237 | 134138058 | 134138164 | 134138950 | 134139038 | 10 | 15 | 34 | 13 | 19 | 101 | 91 | 167 | 105 | 203 | 21 | 15 | [illegible] |
| A5SS | 4399 | ENSG00000145979 | TBC1D7 | chr6 | - | 13328295 | 13328531 | 13328455 | 13328531 | 13326786 | 13326906 | 4 | 6 | 3 | 6 | 5 | 4 | 2 | 3 | 7 | 6 | 3 | 7 | [illegible] |
| A5SS | 1087 | ENSG00000125124 | BBS2 | chr16 | - | 56501944 | 56502456 | 56502316 | 56502456 | 56501352 | 56501497 | 84 | 93 | 83 | 111 | 48 | 52 | 34 | 50 | 37 | 95 | 32 | 114 | [illegible] |
| A5SS | 8869 | ENSG00000139613 | SMARCC2 | chr12 | - | 56178403 | 56178847 | 56178809 | 56178847 | 56178021 | 56178093 | 282 | 168 | 299 | 305 | 491 | 0 | 0 | 0 | 0 | 0 | 383 | 293 | [illegible] |
| A5SS | 1756 | ENSG00000165410 | CFL2 | chr14 | - | 34713253 | 34713561 | 34713507 | 34713561 | 34713059 | 34713136 | 111 | 97 | 110 | 173 | 290 | 2 | 2 | 3 | 0 | 7 | 71 | 107 | [illegible] |
| A5SS | 4398 | ENSG00000145979 | TBC1D7 | chr6 | - | 13328295 | 13328531 | 13328315 | 13328531 | 13326786 | 13326906 | 4 | 6 | 1 | 6 | 4 | 3 | 2 | 3 | 2 | 3 | 3 | 7 | [illegible] |
| A5SS | 7211 | ENSG00000111237 | VPS29 | chr12 | - | 110499456 | 110499546 | 110499534 | 110499546 | 110498819 | 110498859 | 3 | 3 | 6 | 2 | 6 | 19 | 10 | 13 | 14 | 22 | 0 | 1 | [illegible] |
| A5SS | 2922 | ENSG00000176261 | ZBTB8OS | chr1 | - | 32650428 | 32650550 | 32650432 | 32650550 | 32634767 | 32634792 | 6 | 8 | 7 | 2 | 6 | 86 | 45 | 76 | 52 | 99 | 0 | 10 | [illegible] |
| A5SS | 1654 | ENSG00000139998 | RAB15 | chr14 | - | 64950578 | 64951151 | 64951073 | 64951151 | 64950324 | 64950414 | 6 | 10 | 15 | 4 | 2 | 22 | 9 | 15 | 6 | 5 | 10 | 12 | [illegible] |
| A5SS | 4688 | ENSG00000227057 | WDR46 | chr6 | - | 33288563 | 33288694 | 33288613 | 33288694 | 33288357 | 33288470 | 0 | 0 | 0 | 0 | 1 | 57 | 33 | 60 | 50 | 79 | 1 | 12 | [illegible] |
| A5SS | 8794 | ENSG00000090686 | USP48 | chr1 | - | 21736429 | 21736625 | 21736445 | 21736625 | 21729703 | 21729832 | 18 | 8 | 11 | 12 | 21 | 70 | 49 | 65 | 68 | 98 | 0 | 18 | [illegible] |
| A5SS | 176 | ENSG00000105258 | POLR2I | chr19 | - | 36114150 | 36114251 | 36114176 | 36114251 | 36114014 | 36114066 | 0 | 2 | 2 | 0 | 0 | 26 | 10 | 20 | 53 | 18 | 0 | 0 | [illegible] |
| A5SS | 215 | ENSG00000105723 | GSK3A | chr19 | - | 42241900 | 42242602 | 42242182 | 42242602 | 42239954 | 42240142 | 7 | 9 | 4 | 4 | 9 | 48 | 15 | 35 | 51 | 1 | 0 | 5 | [illegible] |
| A5SS | 6748 | ENSG00000196305 | IARS1 | chr9 | - | 92293443 | 92293695 | 92293610 | 92293695 | 92289300 | 92289426 | 26 | 48 | 41 | 44 | 44 | 42 | 30 | 41 | 38 | 15 | 21 | 51 | [illegible] |
| A5SS | 4582 | ENSG00000122678 | POLM | chr7 | - | 44074130 | 44074530 | 44074397 | 44074530 | 44073782 | 44074025 | 24 | 8 | 25 | 21 | 25 | 4 | 3 | 6 | 1 | 9 | 14 | 19 | [illegible] |
| A5SS | 2906 | ENSG00000164068 | RNF123 | chr3 | + | 49712478 | 49712892 | 49712478 | 49712656 | 49713512 | 49713587 | 14 | 2 | 5 | 3 | 6 | 9 | 9 | 11 | 22 | 11 | 22 | 16 | [illegible] |
| A5SS | 199 | ENSG00000196684 | HSH2D | chr19 | + | 16148723 | 16148875 | 16148723 | 16148800 | 16151446 | 16151578 | 17 | 5 | 17 | 19 | 1 | 0 | 0 | 0 | 0 | 0 | 8 | 21 | [illegible] |
| A5SS | 8312 | ENSG00000090621 | PABPC4 | chr1 | - | 39561684 | 39561787 | 39561787 | 39561787 | 39560815 | 39561122 | 223 | 118 | 217 | 223 | 381 | 2 | 2 | 1 | 1 | 14 | 226 | 143 | [illegible] |
| A5SS | 1228 | ENSG00000214078 | CPNE1 | chr20 | - | 35626562 | 35626803 | 35626566 | 35626803 | 35626043 | 35626381 | 1 | 0 | 0 | 1 | 0 | 131 | 44 | 120 | 102 | 80 | 0 | 0 | [illegible] |
| A5SS | 4342 | ENSG00000111445 | RFC5 | chr12 | + | 118017689 | 118018053 | 118017689 | 118017815 | 118019071 | 118019136 | 9 | 14 | 14 | 8 | 29 | 2 | 1 | 1 | 2 | 0 | 12 | 17 | [illegible] |
| A5SS | 6424 | ENSG00000147894 | C9orf72 | chr9 | - | 27558490 | 27560299 | 27560226 | 27560299 | 27556560 | 27556796 | 407 | 337 | 227 | 282 | 652 | 0 | 0 | 0 | 0 | 4 | 13 | 67 | [illegible] |
| A5SS | 8749 | ENSG00000110987 | BCL7A | chr12 | + | 122054804 | 122054989 | 122054804 | 122054926 | 122059091 | 122062044 | 10 | 12 | 12 | 7 | 23 | 14 | 6 | 27 | 28 | 56 | 6 | 15 | [illegible] |
| A5SS | 1721 | ENSG00000174837 | ADGRE1 | chr19 | + | 6928144 | 6928310 | 6928144 | 6928211 | 6934986 | 6935078 | 8 | 0 | 6 | 7 | 0 | 32 | 24 | 18 | 21 | 52 | 0 | 6 | [illegible] |
| A5SS | 587 | ENSG00000282826 | FRG1CP | chr20 | - | 28601602 | 28601756 | 28601656 | 28601756 | 28600391 | 28600461 | 3 | 8 | 6 | 1 | 6 | 8 | 16 | 6 | 18 | 25 | 0 | 0 | [illegible] |
| A5SS | 5992 | ENSG00000197345 | MRPL21 | chr11 | - | 68896468 | 68896678 | 68896514 | 68896678 | 68893402 | 68893455 | 3 | 1 | 0 | 13 | 0 | 71 | 29 | 80 | 62 | 139 | 5 | 0 | [illegible] |
| A5SS | 912 | ENSG00000141030 | COPS3 | chr17 | - | 17270757 | 17271008 | 17270895 | 17271008 | 17267884 | 17267977 | 220 | 74 | 147 | 116 | 147 | 5 | 0 | 0 | 0 | 1 | 233 | 184 | [illegible] |
| A5SS | 2910 | ENSG00000082996 | RNF13 | chr3 | + | 149895472 | 149895570 | 149895472 | 149895560 | 149902071 | 149902162 | 255 | 173 | 160 | 255 | 238 | 650 | 511 | 462 | 641 | 725 | 219 | 232 | [illegible] |
| A5SS | 8774 | ENSG00000063046 | EIF4B | chr12 | + | 53034611 | 53034724 | 53034611 | 53034709 | 53039237 | 53039343 | 16 | 8 | 27 | 17 | 10 | 3 | 0 | 2 | 0 | 3 | 17 | 11 | [illegible] |
| A5SS | 6898 | ENSG00000107185 | RGP1 | chr9 | + | 35750657 | 35750989 | 35750657 | 35750741 | 35751265 | 35751412 | 50 | 19 | 46 | 57 | 62 | 0 | 0 | 0 | 0 | 0 | 77 | 62 | [illegible] |
| A5SS | 8812 | ENSG00000123349 | PFDN5 | chr12 | + | 53295838 | 53296275 | 53295838 | 53295941 | 53297849 | 53297924 | 1689 | 998 | 1549 | 1475 | 1772 | 3 | 13 | 11 | 3 | 11 | 2190 | 1364 | [illegible] |
| A5SS | 6012 | ENSG00000116539 | ASH1L | chr1 | - | 155440489 | 155440593 | 155440502 | 155440593 | 155438326 | 155439068 | 20 | 19 | 24 | 25 | 56 | 18 | 21 | 27 | 43 | 49 | 14 | 18 | [illegible] |
| A5SS | 5427 | ENSG00000162923 | WDR26 | chr1 | - | 224431476 | 224431581 | 224431524 | 224431581 | 224424517 | 224424654 | 166 | 133 | 142 | 233 | 173 | 178 | 102 | 114 | 138 | 160 | 128 | 266 | [illegible] |
| A5SS | 250 | ENSG00000104852 | SNRNP70 | chr19 | + | 49102113 | 49103587 | 49102113 | 49102173 | 49104633 | 49104735 | 433 | 280 | 515 | 479 | 518 | 23 | 37 | 35 | 41 | 47 | 300 | 290 | [illegible] |
| A5SS | 2986 | ENSG00000169714 | CNBP | chr3 | - | 129171651 | 129171771 | 129171654 | 129171771 | 129171445 | 129171538 | 37 | 5 | 18 | 20 | 24 | 410 | 222 | 355 | 396 | 261 | 9 | 6 | [illegible] |
| A5SS | 6373 | ENSG00000106245 | BUD31 | chr7 | + | 99408968 | 99409245 | 99408968 | 99409004 | 99411063 | 99411186 | 7 | 2 | 6 | 25 | 1 | 0 | 2 | 0 | 1 | 0 | 10 | 8 | [illegible] |
| A5SS | 8880 | ENSG00000075089 | ACTR6 | chr12 | + | 100205675 | 100205757 | 100205675 | 100205744 | 100207662 | 100207786 | 38 | 21 | 17 | 17 | 36 | 165 | 71 | 132 | 115 | 164 | 18 | 28 | [illegible] |
| A5SS | 1773 | ENSG00000100823 | APEX1 | chr14 | + | 20455225 | 20455394 | 20455225 | 20455328 | 20455577 | 20455703 | 234 | 101 | 206 | 165 | 102 | 13 | 6 | 22 | 10 | 2 | 203 | 118 | [illegible] |
| A5SS | 1198 | ENSG00000185324 | CDK10 | chr16 | + | 89689251 | 89689324 | 89689251 | 89689306 | 89690552 | 89690624 | 37 | 26 | 52 | 45 | 32 | 5 | 0 | 0 | 3 | 10 | 26 | 32 | [illegible] |
| A5SS | 5900 | ENSG00000123607 | TTC21B | chr2 | - | 165931740 | 165931856 | 165931757 | 165931856 | 165930171 | 165930364 | 2 | 7 | 6 | 6 | 0 | 51 | 31 | 39 | 60 | 44 | 0 | 5 | [illegible] |
| A5SS | 3399 | ENSG00000170581 | STAT2 | chr12 | - | 56350096 | 56350432 | 56350411 | 56350432 | 56349588 | 56349636 | 291 | 190 | 243 | 383 | 255 | 6 | 1 | 0 | 1 | 3 | 1416 | 539 | [illegible] |
| A5SS | 3297 | ENSG00000067066 | SP100 | chr2 | + | 230467119 | 230467215 | 230467119 | 230467206 | 230469042 | 230469096 | 363 | 260 | 335 | 408 | 428 | 8 | 6 | 8 | 7 | 3 | 153 | 351 | [illegible] |
| A5SS | 7724 | ENSG00000148358 | GPR107 | chr9 | + | 130077298 | 130077378 | 130077298 | 130077374 | 130079629 | 130079769 | 142 | 85 | 91 | 130 | 180 | 6 | 8 | 2 | 7 | 6 | 323 | 180 | [illegible] |
| A5SS | 4495 | ENSG00000117859 | OSBPL9 | chr1 | + | 51669433 | 51669814 | 51669433 | 51669512 | 51714002 | 51714079 | 3 | 7 | 2 | 4 | 4 | 69 | 34 | 55 | 62 | 82 | 0 | 2 | [illegible] |
| A5SS | 1980 | ENSG00000133706 | LARS1 | chr5 | - | 146144623 | 146144709 | 146144662 | 146144709 | 146144471 | 146144537 | 157 | 160 | 147 | 178 | 245 | 6 | 4 | 6 | 1 | 7 | 33 | 129 | [illegible] |
| A5SS | 8485 | ENSG00000225484 | NUTM2B-AS1 | chr10 | - | 79808758 | 79808832 | 79808762 | 79808832 | 79806038 | 79806122 | 110 | 66 | 110 | 123 | 145 | 25 | 9 | 18 | 29 | 18 | 272 | 159 | [illegible] |
| A5SS | 8415 | ENSG00000180198 | RCC1 | chr1 | + | 28530522 | 28530615 | 28530522 | 28530573 | 28531802 | 28531990 | 0 | 1 | 10 | 1 | 0 | 13 | 11 | 14 | 22 | 8 | 6 | 0 | [illegible] |
| A5SS | 873 | ENSG00000181038 | METTL23 | chr17 | + | 76733292 | 76733381 | 76733292 | 76733377 | 76733520 | 76733880 | 4 | 1 | 4 | 11 | 0 | 163 | 63 | 115 | 93 | 277 | 0 | 4 | [illegible] |
| A5SS | 201 | ENSG00000196684 | HSH2D | chr19 | + | 16152551 | 16153208 | 16152551 | 16152641 | 16154398 | 16154491 | 101 | 57 | 69 | 89 | 27 | 0 | 0 | 0 | 0 | 0 | 209 | 133 | [illegible] |
| A5SS | 3904 | ENSG00000055609 | KMT2C | chr7 | - | 152145152 | 152145295 | 152145164 | 152145295 | 152144712 | 152144881 | 151 | 113 | 134 | 158 | 178 | 69 | 46 | 63 | 78 | 121 | 46 | 141 | [illegible] |
| A5SS | 5514 | ENSG00000223960 | CHROMR | chr2 | + | 178426365 | 178426661 | 178426365 | 178426505 | 178430769 | 178430889 | 33 | 12 | 23 | 24 | 16 | 5 | 4 | 8 | 10 | 7 | 55 | 32 | [illegible] |
| A5SS | 2973 | ENSG00000115129 | TP53I3 | chr2 | - | 24082884 | 24083152 | 24083000 | 24083152 | 24080818 | 24081031 | 13 | 10 | 23 | 10 | 8 | 0 | 1 | 0 | 0 | 0 | 18 | 33 | [illegible] |
| A5SS | 5774 | ENSG00000163832 | ELP6 | chr3 | - | 47510148 | 47510254 | 47510183 | 47510254 | 47504329 | 47504448 | 2 | 0 | 5 | 3 | 0 | 52 | 31 | 38 | 31 | 24 | 0 | 0 | [illegible] |
| A5SS | 6774 | ENSG00000161010 | MRNIP | chr5 | - | 179853196 | 179853437 | 179853377 | 179853437 | 179847977 | 179848066 | 18 | 11 | 12 | 19 | 13 | 25 | 15 | 24 | 22 | 8 | 50 | 34 | [illegible] |
| A5SS | 4143 | ENSG00000053254 | FOXN3 | chr14 | - | 89190383 | 89190449 | 89190386 | 89190449 | 89180700 | 89180806 | 59 | 29 | 76 | 60 | 61 | 12 | 17 | 20 | 21 | 35 | 112 | 63 | [illegible] |
| A5SS | 8737 | ENSG00000173442 | EHBP1L1 | chr11 | + | 65581210 | 65581373 | 65581210 | 65581251 | 65584240 | 65584398 | 141 | 68 | 73 | 90 | 95 | 4 | 4 | 5 | 7 | 14 | 232 | 123 | [illegible] |
| A5SS | 6476 | ENSG00000198198 | SZT2 | chr1 | + | 43419733 | 43419961 | 43419733 | 43419944 | 43420152 | 43420323 | 4 | 1 | 2 | 6 | 3 | 26 | 22 | 38 | 26 | 18 | 32 | 5 | [illegible] |
| A5SS | 5969 | ENSG00000164609 | SLU7 | chr5 | - | 160418595 | 160419081 | 160419022 | 160419081 | 160415124 | 160415310 | 22 | 10 | 38 | 25 | 43 | 72 | 69 | 49 | 94 | 98 | 21 | 21 | [illegible] |
| A5SS | 4668 | ENSG00000171603 | CLSTN1 | chr1 | - | 9731121 | 9731390 | 9731205 | 9731390 | 9729027 | 9730705 | 13 | 15 | 30 | 21 | 24 | 233 | 122 | 179 | 188 | 149 | 56 | 44 | [illegible] |
| A5SS | 5065 | ENSG00000162961 | DPY30 | chr2 | - | 32039592 | 32039805 | 32039732 | 32039805 | 32039420 | 32039492 | 3 | 2 | 0 | 4 | 3 | 19 | 22 | 11 | 10 | 14 | 0 | 5 | [illegible] |
| A5SS | 3386 | ENSG00000267575 | NA | chr19 | + | 27793451 | 27793650 | 27793451 | 27793609 | 27797753 | 27798176 | 9 | 3 | 6 | 7 | 17 | 5 | 0 | 0 | 4 | 0 | 6 | 10 | [illegible] |
| A5SS | 3483 | ENSG00000136877 | FPGS | chr9 | + | 127807585 | 127807775 | 127807585 | 127807688 | 127808233 | 127808311 | 12 | 2 | 17 | 10 | 0 | 18 | 4 | 14 | 20 | 12 | 11 | 3 | [illegible] |
| A5SS | 5516 | ENSG00000223960 | CHROMR | chr2 | + | 178426365 | 178426703 | 178426365 | 178426505 | 178430769 | 178430889 | 26 | 9 | 18 | 18 | 10 | 5 | 4 | 8 | 10 | 7 | 48 | 24 | [illegible] |
| A5SS | 8308 | ENSG00000075407 | ZNF37A | chr10 | + | 38095537 | 38095633 | 38095537 | 38095628 | 38095743 | 38095804 | 1 | 2 | 4 | 3 | 2 | 6 | 7 | 15 | 7 | 9 | 0 | 1 | [illegible] |
| A5SS | 2867 | ENSG00000178035 | IMPDH2 | chr3 | - | 49028247 | 49028532 | 49028430 | 49028532 | 49027709 | 49027916 | 125 | 75 | 136 | 117 | 136 | 0 | 0 | 3 | 8 | 1 | 159 | 111 | [illegible] |
| A5SS | 3654 | ENSG00000018510 | AGPS | chr2 | + | 177440964 | 177442235 | 177440964 | 177441036 | 177442406 | 177442486 | 13 | 22 | 24 | 12 | 22 | 218 | 249 | 175 | 240 | 316 | 0 | 3 | [illegible] |
| A5SS | 8452 | ENSG00000107862 | GBF1 | chr10 | + | 102358505 | 102359435 | 102358505 | 102358729 | 102360183 | 102360395 | 69 | 32 | 79 | 58 | 59 | 0 | 7 | 0 | 6 | 3 | 103 | 79 | [illegible] |
| A5SS | 2159 | ENSG00000124782 | RREB1 | chr6 | + | 7181123 | 7181289 | 7181123 | 7181246 | 7181869 | 7182082 | 13 | 14 | 12 | 15 | 9 | 32 | 42 | 54 | 44 | 87 | 3 | 41 | [illegible] |
| A5SS | 8028 | ENSG00000177666 | PNPLA2 | chr11 | + | 823693 | 824130 | 823693 | 823855 | 824313 | 824436 | 56 | 20 | 44 | 36 | 14 | 0 | 0 | 0 | 3 | 0 | 42 | 63 | [illegible] |
| A5SS | 7326 | ENSG00000166946 | CCNDBP1 | chr15 | + | 43185819 | 43186233 | 43185819 | 43185879 | 43189198 | 43189280 | 193 | 77 | 160 | 141 | 157 | 10 | 4 | 8 | 3 | 2 | 92 | 206 | [illegible] |
| A5SS | 8411 | ENSG00000139651 | ZNF740 | chr12 | + | 53181676 | 53182190 | 53181676 | 53181992 | 53185386 | 53185476 | 45 | 19 | 46 | 48 | 40 | 9 | 6 | 8 | 16 | 7 | 83 | 71 | [illegible] |
| A5SS | 2700 | ENSG00000099949 | LZTR1 | chr22 | + | 20983026 | 20983122 | 20983026 | 20983089 | 20985840 | 20985897 | 5 | 0 | 2 | 2 | 0 | 36 | 11 | 26 | 38 | 19 | 4 | 0 | [illegible] |
| A5SS | 1914 | ENSG00000196655 | TRAPPC4 | chr11 | + | 119019142 | 119019424 | 119019142 | 119019317 | 119021759 | 119021886 | 14 | 5 | 15 | 22 | 10 | 6 | 6 | 0 | 12 | 10 | 33 | 14 | [illegible] |
| A5SS | 1216 | ENSG00000138834 | MAPK8IP3 | chr16 | + | 1761223 | 1761305 | 1761223 | 1761278 | 1762350 | 1762481 | 167 | 99 | 139 | 167 | 210 | 13 | 24 | 17 | 41 | 30 | 320 | 241 | [illegible] |
| A5SS | 261 | ENSG00000130159 | ECSIT | chr19 | - | 11507701 | 11508048 | 11507990 | 11508048 | 11507456 | 11507562 | 55 | 22 | 34 | 32 | 21 | 1 | 2 | 2 | 1 | 0 | 38 | 61 | [illegible] |
| A5SS | 6501 | ENSG00000126746 | ZNF384 | chr12 | - | 6689097 | 6689510 | 6689367 | 6689510 | 6688166 | 6688226 | 21 | 4 | 7 | 10 | 17 | 7 | 0 | 6 | 6 | 11 | 0 | 9 | [illegible] |
| A5SS | 7856 | ENSG00000110514 | MADD | chr11 | + | 47295495 | 47295579 | 47295495 | 47295567 | 47295896 | 47296055 | 140 | 94 | 106 | 120 | 163 | 41 | 20 | 20 | 63 | 33 | 163 | 186 | [illegible] |
| A5SS | 5177 | ENSG00000048991 | R3HDM1 | chr2 | + | 135641535 | 135641790 | 135641535 | 135641604 | 135645378 | 135645527 | 12 | 3 | 8 | 6 | 8 | 3 | 6 | 3 | 7 | 3 | 32 | 0 | [illegible] |
| A5SS | 4492 | ENSG00000030419 | IKZF2 | chr2 | - | 213056832 | 213057099 | 213056910 | 213057099 | 213049712 | 213049880 | 69 | 30 | 44 | 54 | 121 | 27 | 13 | 34 | 25 | 50 | 0 | 21 | [illegible] |
| A5SS | 5441 | ENSG00000163848 | ZNF148 | chr3 | - | 125323308 | 125323444 | 125323387 | 125323444 | 125313307 | 125313656 | 283 | 168 | 210 | 268 | 318 | 32 | 18 | 23 | 26 | 43 | 29 | 148 | [illegible] |
| A5SS | 886 | ENSG00000129657 | SEC14L1 | chr17 | + | 77088748 | 77088985 | 77088748 | 77088949 | 77089152 | 77089271 | 6 | 1 | 3 | 1 | 3 | 6 | 5 | 24 | 22 | 27 | 0 | 5 | [illegible] |
| A5SS | 3889 | ENSG00000058272 | PPP1R12A | chr12 | - | 79786776 | 79788783 | 79788647 | 79788783 | 79786373 | 79786478 | 33 | 53 | 42 | 47 | 37 | 694 | 541 | 434 | 615 | 1025 | 0 | 24 | [illegible] |
| A5SS | 345 | ENSG00000268027 | NA | chr19 | + | 41551179 | 41551494 | 41551179 | 41551302 | 41553705 | 41553746 | 52 | 25 | 62 | 49 | 30 | 36 | 17 | 51 | 12 | 63 | 3 | 32 | [illegible] |
| A5SS | 3549 | ENSG00000116001 | TIA1 | chr2 | - | 70236072 | 70236175 | 70236078 | 70236175 | 70230755 | 70230854 | 10 | 1 | 0 | 6 | 15 | 241 | 181 | 189 | 217 | 323 | 0 | 5 | [illegible] |
| A5SS | 2601 | ENSG00000171298 | GAA | chr17 | + | 80101810 | 80101925 | 80101810 | 80101890 | 80104554 | 80105132 | 3 | 0 | 2 | 0 | 0 | 18 | 12 | 13 | 16 | 16 | 0 | 9 | [illegible] |
| A5SS | 2445 | ENSG00000177058 | SLC38A9 | chr5 | - | 55711351 | 55711499 | 55711351 | 55711499 | 55697845 | 55697992 | 5 | 0 | 1 | 1 | 6 | 45 | 28 | 43 | 50 | 43 | 0 | 0 | [illegible] |
| A5SS | 5258 | ENSG00000204344 | STK19 | chr6 | + | 31972620 | 31972919 | 31972620 | 31972757 | 31978902 | 31978998 | 2 | 2 | 1 | 4 | 4 | 24 | 9 | 15 | 14 | 13 | 19 | 1 | [illegible] |
| A5SS | 4422 | ENSG00000166523 | CLEC4E | chr12 | - | 8539141 | 8539306 | 8539199 | 8539306 | 8536089 | 8536205 | 56 | 40 | 57 | 30 | 65 | 15 | 16 | 5 | 8 | 16 | 18 | 41 | [illegible] |
| A5SS | 7252 | ENSG00000221914 | PPP2R2A | chr8 | + | 26293665 | 26293860 | 26293665 | 26293740 | 26338889 | 26338987 | 24 | 20 | 18 | 25 | 9 | 95 | 95 | 104 | 93 | 90 | 11 | 27 | [illegible] |
| A5SS | 8668 | ENSG00000127314 | RAP1B | chr12 | + | 68610928 | 68611056 | 68610928 | 68611052 | 68648698 | 68648781 | 2 | 2 | 3 | 3 | 1 | 242 | 201 | 151 | 229 | 124 | 5 | 0 | [illegible] |
| A5SS | 3500 | ENSG00000247828 | TMEM161B-AS1 | chr5 | + | 88269467 | 88282106 | 88269467 | 88270585 | 88282714 | 88282866 | 57 | 61 | 62 | 52 | 30 | 0 | 0 | 0 | 0 | 0 | 27 | 52 | [illegible] |
| A5SS | 1716 | ENSG00000100564 | PIGH | chr14 | - | 67593727 | 67593952 | 67593742 | 67593952 | 67592634 | 67592718 | 1 | 1 | 0 | 5 | 6 | 37 | 11 | 30 | 37 | 42 | 3 | 0 | [illegible] |
| A5SS | 3555 | ENSG00000048740 | CELF2 | chr10 | + | 11314138 | 11314270 | 11314138 | 11314258 | 11321206 | 11321386 | 72 | 43 | 69 | 76 | 31 | 174 | 91 | 168 | 170 | 107 | 33 | 96 | [illegible] |
| A5SS | 1404 | ENSG00000103197 | TSC2 | chr16 | + | 2081594 | 2082504 | 2081594 | 2081798 | 2083694 | 2083816 | 6 | 3 | 0 | 4 | 0 | 31 | 11 | 27 | 12 | 8 | 0 | 4 | [illegible] |
| A5SS | 4753 | ENSG00000089127 | OAS1 | chr12 | + | 112911050 | 112911258 | 112911050 | 112911235 | 112916508 | 112916738 | 8 | 1 | 1 | 2 | 1 | 293 | 196 | 166 | 239 | 184 | 7 | 15 | [illegible] |
| A5SS | 4390 | ENSG00000249915 | PDCD6 | chr5 | + | 306601 | 306760 | 306601 | 306754 | 311292 | 311402 | 154 | 76 | 103 | 141 | 126 | 40 | 14 | 21 | 18 | 11 | 200 | 101 | [illegible] |
| A5SS | 7079 | ENSG00000083750 | RRAGB | chrX | + | 55717748 | 55718419 | 55717748 | 55717788 | 55719313 | 55719347 | 63 | 66 | 79 | 108 | 55 | 6 | 3 | 1 | 1 | 0 | 47 | 63 | [illegible] |
| A5SS | 6911 | ENSG00000101901 | ALG13 | chrX | + | 111681169 | 111681424 | 111681169 | 111681299 | 111682131 | 111682294 | 14 | 7 | 28 | 17 | 20 | 25 | 13 | 11 | 35 | 3 | 13 | 10 | [illegible] |
| A5SS | 3129 | ENSG00000136603 | SKIL | chr3 | + | 170384532 | 170384765 | 170384532 | 170384627 | 170390222 | 170390464 | 117 | 121 | 84 | 104 | 173 | 5 | 0 | 6 | 1 | 0 | 9 | 45 | [illegible] |
| A5SS | 5668 | ENSG00000095059 | DHPS | chr19 | - | 12680160 | 12680325 | 12680195 | 12680325 | 12679622 | 12679719 | 28 | 21 | 28 | 38 | 23 | 0 | 0 | 0 | 0 | 0 | 21 | 48 | [illegible] |
| A5SS | 8101 | ENSG00000168002 | POLR2G | chr11 | + | 62761794 | 62762184 | 62761794 | 62761904 | 62762866 | 62763026 | 18 | 4 | 16 | 21 | 7 | 176 | 70 | 136 | 167 | 105 | 8 | 29 | [illegible] |
| A5SS | 7642 | ENSG00000136811 | ODF2 | chr9 | + | 128460941 | 128461067 | 128460941 | 128461010 | 128471307 | 128471468 | 29 | 14 | 15 | 19 | 13 | 2 | 3 | 2 | 4 | 1 | 14 | 10 | [illegible] |
| A5SS | 3448 | ENSG00000168710 | AHCYL1 | chr1 | + | 109984764 | 109985543 | 109984764 | 109985172 | 110009033 | 110009145 | 0 | 2 | 1 | 0 | 2 | 65 | 31 | 34 | 52 | 32 | 8 | 0 | [illegible] |
| A5SS | 2381 | ENSG00000138760 | SCARB2 | chr4 | - | 76179516 | 76181101 | 76180953 | 76181101 | 76176436 | 76176528 | 224 | 188 | 166 | 253 | 376 | 5 | 1 | 0 | 6 | 5 | 370 | 248 | [illegible] |
| A5SS | 2180 | ENSG00000205309 | NT5M | chr17 | + | 17344793 | 17344926 | 17344793 | 17344908 | 17346804 | 17347663 | 5 | 3 | 1 | 13 | 10 | 7 | 7 | 2 | 23 | 14 | 19 | 20 | [illegible] |
| A5SS | 5599 | ENSG00000163803 | PLB1 | chr2 | + | 28626427 | 28626548 | 28626427 | 28626508 | 28628562 | 28628628 | 0 | 0 | 0 | 1 | 0 | 33 | 12 | 19 | 21 | 17 | 10 | 0 | [illegible] |
| A5SS | 654 | ENSG00000166562 | SEC11C | chr18 | + | 59152535 | 59152685 | 59152535 | 59152612 | 59155687 | 59155807 | 334 | 181 | 489 | 315 | 325 | 7 | 6 | 6 | 3 | 3 | 300 | 235 | [illegible] |
| A5SS | 7951 | ENSG00000173120 | KDM2A | chr11 | + | 67250085 | 67250798 | 67250085 | 67250346 | 67252693 | 67252857 | 343 | 224 | 378 | 316 | 287 | 6 | 10 | 2 | 1 | 0 | 411 | 455 | [illegible] |
| A5SS | 7183 | ENSG00000100483 | VCPKMT | chr14 | - | 50116068 | 50116179 | 50116079 | 50116179 | 50115838 | 50115911 | 14 | 17 | 19 | 15 | 23 | 11 | 19 | 13 | 21 | 9 | 1 | 17 | [illegible] |
| A5SS | 8670 | ENSG00000102225 | CDK16 | chrX | + | 47222260 | 47222380 | 47222260 | 47222342 | 47223551 | 47223759 | 6 | 9 | 4 | 8 | 12 | 10 | 3 | 5 | 2 | 1 | 42 | 17 | [illegible] |
| A5SS | 2608 | ENSG00000070423 | RNF126 | chr19 | - | 651610 | 651855 | 651691 | 651855 | 650233 | 650296 | 101 | 29 | 72 | 105 | 72 | 6 | 4 | 4 | 4 | 4 | 26 | 80 | [illegible] |
| A5SS | 5854 | ENSG00000169045 | HNRNPH1 | chr5 | - | 179623594 | 179623669 | 179623635 | 179623669 | 179623036 | 179623164 | 171 | 124 | 181 | 223 | 75 | 90 | 61 | 97 | 104 | 14 | 75 | 213 | [illegible] |
| A5SS | 2378 | ENSG00000174720 | LARP7 | chr4 | + | 112644667 | 112644871 | 112644667 | 112644768 | 112646350 | 112646451 | 261 | 160 | 172 | 217 | 383 | 7 | 7 | 22 | 19 | 14 | 37 | 77 | [illegible] |
| A5SS | 3269 | ENSG00000165801 | ARHGEF40 | chr14 | + | 21075331 | 21075765 | 21075331 | 21075499 | 21076359 | 21076456 | 19 | 6 | 13 | 11 | 11 | 4 | 2 | 1 | 4 | 0 | 73 | 18 | [illegible] |
| A5SS | 7589 | ENSG00000196781 | TLE1 | chr9 | - | 81613376 | 81613521 | 81613421 | 81613521 | 81611842 | 81611956 | 37 | 18 | 14 | 22 | 19 | 2 | 0 | 1 | 1 | 1 | 17 | 33 | [illegible] |
| A5SS | 720 | ENSG00000170004 | CHD3 | chr17 | + | 7909142 | 7909384 | 7909142 | 7909338 | 7910427 | 7910591 | 7 | 3 | 23 | 24 | 30 | 78 | 57 | 77 | 68 | 205 | 1 | 13 | [illegible] |
| A5SS | 8432 | ENSG00000198728 | LDB1 | chr10 | - | 102109028 | 102109177 | 102109034 | 102109177 | 102107559 | 102108323 | 27 | 21 | 35 | 36 | 33 | 25 | 8 | 9 | 15 | 13 | 22 | 26 | [illegible] |

| | | | | | | |
|---|---|---|---|---|---|---|
| A5SS | 2146 | ENSG00000100284 | TOM1 | chr22 | + | [illegible] |
| A5SS | 8695 | ENSG00000116198 | CEP104 | chr1 | - | [illegible] |
| A5SS | 4224 | ENSG00000104231 | ZFAND1 | chr8 | - | [illegible] |
| A5SS | 4179 | ENSG00000139636 | LMBR1L | chr12 | - | [illegible] |
| A5SS | 6842 | ENSG00000124333 | VAMP7 | chrX | + | [illegible] |
| A5SS | 1071 | ENSG00000067955 | CBFB | chr16 | + | [illegible] |
| A5SS | 6421 | ENSG00000101574 | METTL4 | chr18 | - | [illegible] |
| A5SS | 3306 | ENSG00000159692 | CTBP1 | chr4 | - | [illegible] |
| A5SS | 338 | ENSG00000167613 | LAIR1 | chr19 | - | [illegible] |
| A5SS | 6387 | ENSG00000183735 | TBK1 | chr12 | + | [illegible] |
| A5SS | 7641 | ENSG00000136811 | ODF2 | chr9 | + | [illegible] |
| A5SS | 1335 | ENSG00000038532 | CLEC16A | chr16 | + | [illegible] |
| A5SS | 1556 | ENSG00000196547 | MAN2A2 | chr15 | + | [illegible] |
| A5SS | 7429 | ENSG00000104763 | ASAH1 | chr8 | - | [illegible] |
| A5SS | 1414 | ENSG00000166145 | SPINT1 | chr15 | + | [illegible] |
| A5SS | 3993 | ENSG00000198912 | C1orf174 | chr1 | - | [illegible] |
| A5SS | 8302 | ENSG00000136935 | GOLGA1 | chr9 | - | [illegible] |
| A5SS | 3783 | ENSG00000244274 | DBNDD2 | chr20 | + | [illegible] |
| A5SS | 6777 | ENSG00000087274 | ADD1 | chr4 | + | [illegible] |
| A5SS | 7159 | ENSG00000147133 | TAF1 | chrX | + | [illegible] |
| A5SS | 1165 | ENSG00000103275 | UBE2I | chr16 | + | [illegible] |
| A5SS | 1275 | ENSG00000187193 | MT1X | chr16 | + | [illegible] |
| A5SS | 3938 | ENSG00000133106 | EPSTI1 | chr13 | - | [illegible] |
| A5SS | 8759 | ENSG00000002016 | RAD52 | chr12 | - | [illegible] |
| A5SS | 8011 | ENSG00000160226 | CFAP410 | chr21 | - | [illegible] |
| A5SS | 8317 | ENSG00000090621 | PABPC4 | chr1 | - | [illegible] |
| A5SS | 5686 | ENSG00000105325 | FZR1 | chr19 | + | [illegible] |
| A5SS | 6284 | ENSG00000164620 | RELL2 | chr5 | + | [illegible] |
| A5SS | 6534 | ENSG00000147535 | PLPP5 | chr8 | - | [illegible] |
| A5SS | 350 | ENSG00000160193 | WDR4 | chr21 | - | [illegible] |
| A5SS | 6630 | ENSG00000051825 | MPHOSPH9 | chr12 | - | [illegible] |
| A5SS | 6983 | ENSG00000146701 | MDH2 | chr7 | + | [illegible] |
| A5SS | 6096 | ENSG00000135316 | SYNCRIP | chr6 | - | [illegible] |
| A5SS | 3059 | ENSG00000165288 | BRWD3 | chrX | - | [illegible] |
| A5SS | 1403 | ENSG00000103197 | TSC2 | chr16 | + | [illegible] |
| A5SS | 1499 | ENSG00000114062 | UBE3A | chr15 | - | [illegible] |
| A5SS | 1231 | ENSG00000214078 | CPNE1 | chr20 | - | [illegible] |
| A5SS | 1052 | ENSG00000172530 | BANP | chr16 | + | [illegible] |
| A5SS | 5191 | ENSG00000166801 | FAM111A | chr11 | + | [illegible] |
| A5SS | 8339 | ENSG00000156650 | KAT6B | chr10 | + | [illegible] |
| A5SS | 2100 | ENSG00000112851 | ERBIN | chr5 | + | [illegible] |
| A5SS | 4185 | ENSG00000143033 | MTF2 | chr1 | + | [illegible] |
| A5SS | 6556 | ENSG00000115274 | INO80B | chr2 | + | [illegible] |
| A5SS | 408 | ENSG00000130511 | SSBP4 | chr19 | + | [illegible] |
| A5SS | 7879 | ENSG00000152556 | PFKM | chr12 | + | [illegible] |
| A5SS | 6467 | ENSG00000042980 | ADAM28 | chr8 | + | [illegible] |
| A5SS | 2495 | ENSG00000174652 | ZNF266 | chr19 | - | [illegible] |
| A5SS | 3494 | ENSG00000119185 | ITGB1BP1 | chr2 | - | [illegible] |
| A5SS | 7222 | ENSG00000168066 | SF1 | chr11 | - | [illegible] |
| A5SS | 6854 | ENSG00000188917 | TRMT2B | chrX | - | [illegible] |
| A5SS | 5963 | ENSG00000108465 | CDK5RAP3 | chr17 | + | [illegible] |
| A5SS | 3049 | ENSG00000182899 | RPL35A | chr3 | + | [illegible] |
| A5SS | 5199 | ENSG00000167895 | TMC8 | chr17 | + | [illegible] |
| A5SS | 6704 | ENSG00000136271 | DDX56 | chr7 | - | [illegible] |
| A5SS | 7497 | ENSG00000104365 | IKBKB | chr8 | + | [illegible] |
| A5SS | 4345 | ENSG00000198860 | TSEN15 | chr1 | + | [illegible] |
| A5SS | 3064 | ENSG00000114473 | IQCG | chr3 | - | [illegible] |
| A5SS | 904 | ENSG00000133030 | MPRIP | chr17 | + | [illegible] |
| A5SS | 3694 | ENSG00000237943 | PRKCQ-AS1 | chr10 | + | [illegible] |
| A5SS | 7646 | ENSG00000139644 | TMBIM6 | chr12 | + | [illegible] |
| A5SS | 1303 | ENSG00000103248 | MTHFSD | chr16 | - | [illegible] |
| A5SS | 8741 | ENSG00000123329 | ARHGAP9 | chr12 | - | [illegible] |
| A5SS | 7808 | ENSG00000178177 | LCORL | chr4 | - | [illegible] |
| A5SS | 8714 | ENSG00000135297 | MTO1 | chr6 | + | [illegible] |
| A5SS | 3638 | ENSG00000113119 | TMCO6 | chr5 | + | [illegible] |
| A5SS | 876 | ENSG00000002919 | SNX11 | chr17 | + | [illegible] |
| A5SS | 3295 | ENSG00000117054 | ACADM | chr1 | + | [illegible] |
| A5SS | 834 | ENSG00000185359 | HGS | chr17 | + | [illegible] |
| A5SS | 2346 | ENSG00000113300 | CNOT6 | chr5 | + | [illegible] |
| A5SS | 2919 | ENSG00000176261 | ZBTB8OS | chr1 | - | [illegible] |
| A5SS | 2440 | ENSG00000186431 | FCAR | chr19 | + | [illegible] |
| A5SS | 7095 | ENSG00000124275 | MTRR | chr5 | + | [illegible] |
| A5SS | 6107 | ENSG00000112146 | FBXO9 | chr6 | + | [illegible] |
| A5SS | 6869 | ENSG00000245958 | SEPTIN7P14 | chr4 | + | [illegible] |
| A5SS | 3947 | ENSG00000156873 | PHKG2 | chr16 | + | [illegible] |
| A5SS | 3226 | ENSG00000015479 | MATR3 | chr5 | + | [illegible] |
| A5SS | 2653 | ENSG00000163161 | ERCC3 | chr2 | - | [illegible] |
| A5SS | 2654 | ENSG00000163161 | ERCC3 | chr2 | - | [illegible] |
| A5SS | 2795 | ENSG00000114745 | GORASP1 | chr3 | - | [illegible] |
| A5SS | 7684 | ENSG00000204394 | VARS1 | chr6 | - | [illegible] |
| A5SS | 8362 | ENSG00000186529 | CYP4F3 | chr19 | + | [illegible] |
| A5SS | 1019 | ENSG00000177302 | TOP3A | chr17 | - | [illegible] |
| A5SS | 7634 | ENSG00000148337 | CIZ1 | chr9 | - | [illegible] |
| A5SS | 1163 | ENSG00000103275 | UBE2I | chr16 | + | [illegible] |
| A5SS | 4145 | ENSG00000167528 | ZNF641 | chr12 | - | [illegible] |
| A5SS | 3620 | ENSG00000177728 | TMEM94 | chr17 | + | [illegible] |
| A5SS | 8956 | ENSG00000127022 | CANX | chr5 | + | [illegible] |
| A5SS | 7428 | ENSG00000104763 | ASAH1 | chr8 | - | [illegible] |
| A5SS | 7603 | ENSG00000165526 | RPUSD4 | chr11 | - | [illegible] |
| A5SS | 7 | ENSG00000186866 | POFUT2 | chr21 | - | [illegible] |
| A5SS | 6849 | ENSG00000164062 | APEH | chr3 | + | [illegible] |
| A5SS | 1623 | ENSG00000226777 | FAM30A | chr14 | + | [illegible] |
| A5SS | 3045 | ENSG00000092931 | MFSD11 | chr17 | + | [illegible] |
| A5SS | 7969 | ENSG00000166452 | AKIP1 | chr11 | + | [illegible] |
| A5SS | 122 | ENSG00000228315 | GUSBP11 | chr22 | - | [illegible] |
| A5SS | 2582 | ENSG00000168310 | IRF2 | chr4 | - | [illegible] |
| A5SS | 6108 | ENSG00000150672 | DLG2 | chr11 | - | [illegible] |
| A5SS | 8221 | ENSG00000166340 | TPP1 | chr11 | - | [illegible] |
| A5SS | 5966 | ENSG00000108465 | CDK5RAP3 | chr17 | + | [illegible] |
| A5SS | 7774 | ENSG00000148356 | LRSAM1 | chr9 | + | [illegible] |
| A5SS | 893 | ENSG00000276231 | PIK3R6 | chr17 | - | [illegible] |
| A5SS | 520 | ENSG00000101294 | HM13 | chr20 | + | [illegible] |
| A5SS | 544 | ENSG00000101365 | IDH3B | chr20 | - | [illegible] |
| A5SS | 4583 | ENSG00000122678 | POLM | chr7 | - | [illegible] |
| A5SS | 5515 | ENSG00000223960 | CHROMR | chr2 | + | [illegible] |
| A5SS | 5325 | ENSG00000143207 | COP1 | chr1 | - | [illegible] |
| A5SS | 2726 | ENSG00000136261 | BZW2 | chr7 | + | [illegible] |
| A5SS | 4486 | ENSG00000156983 | BRPF1 | chr3 | + | [illegible] |
| A5SS | 1095 | ENSG00000033011 | ALG1 | chr16 | + | [illegible] |
| A5SS | 257 | ENSG00000104872 | PIH1D1 | chr19 | - | [illegible] |
| A5SS | 7873 | ENSG00000198561 | CTNND1 | chr11 | + | [illegible] |
| A5SS | 7169 | ENSG00000215301 | DDX3X | chrX | + | [illegible] |
| A5SS | 880 | ENSG00000161533 | ACOX1 | chr17 | - | [illegible] |
| A5SS | 4972 | ENSG00000134058 | CDK7 | chr5 | + | [illegible] |
| A5SS | 158 | ENSG00000079432 | CIC | chr19 | + | [illegible] |
| A5SS | 8946 | ENSG00000167186 | COQ7 | chr16 | + | [illegible] |
| A5SS | 8519 | ENSG00000139620 | KANSL2 | chr12 | - | [illegible] |
| A5SS | 3531 | ENSG00000111640 | GAPDH | chr12 | + | [illegible] |
| A5SS | 8363 | ENSG00000203791 | EEF1AKMT2 | chr10 | - | [illegible] |
| A5SS | 6641 | ENSG00000086730 | LAT2 | chr7 | + | [illegible] |
| A5SS | 4418 | ENSG00000111653 | ING4 | chr12 | - | [illegible] |
| A5SS | 3954 | ENSG00000121766 | ZCCHC17 | chr1 | + | [illegible] |
| A5SS | 8340 | ENSG00000156650 | KAT6B | chr10 | + | [illegible] |
| A5SS | 8114 | ENSG00000172663 | TMEM134 | chr11 | - | [illegible] |
| A5SS | 8530 | ENSG00000150054 | MPP7 | chr10 | - | [illegible] |
| A5SS | 4147 | ENSG00000156482 | RPL30 | chr8 | - | [illegible] |
| A5SS | 521 | ENSG00000101294 | HM13 | chr20 | + | [illegible] |
| A5SS | 851 | ENSG00000198909 | MAP3K3 | chr17 | + | [illegible] |
| A5SS | 3633 | ENSG00000135597 | REPS1 | chr6 | - | [illegible] |
| A5SS | 8112 | ENSG00000172663 | TMEM134 | chr11 | - | [illegible] |
| A5SS | 4120 | ENSG00000116337 | AMPD2 | chr1 | + | [illegible] |
| A5SS | 6645 | ENSG00000188002 | NA | chr5 | - | [illegible] |
| A5SS | 916 | ENSG00000105053 | VRK3 | chr19 | - | [illegible] |
| A5SS | 6988 | ENSG00000223959 | AFG3L1P | chr16 | + | [illegible] |
| A5SS | 3589 | ENSG00000126088 | UROD | chr1 | + | [illegible] |
| A5SS | 4457 | ENSG00000112303 | VNN2 | chr6 | - | [illegible] |
| A5SS | 845 | ENSG00000256525 | POLG2 | chr17 | - | [illegible] |
| A5SS | 8017 | ENSG00000105254 | TBCB | chr19 | + | [illegible] |
| A5SS | 6765 | ENSG00000160408 | ST6GALNAC6 | chr9 | - | [illegible] |
| A5SS | 3682 | ENSG00000179335 | CLK3 | chr15 | + | [illegible] |
| A5SS | 7391 | ENSG00000136518 | ACTL6A | chr3 | + | [illegible] |
| A5SS | 7636 | ENSG00000148337 | CIZ1 | chr9 | - | [illegible] |
| A5SS | 5918 | ENSG00000092208 | GEMIN2 | chr14 | + | [illegible] |
| A5SS | 297 | ENSG00000127452 | FBXL12 | chr19 | - | [illegible] |
| A5SS | 7184 | ENSG00000271147 | ARMCX5-GPRASP2 | chrX | + | [illegible] |
| A5SS | 8341 | ENSG00000156650 | KAT6B | chr10 | + | [illegible] |
| A5SS | 7811 | ENSG00000167721 | TSR1 | chr17 | - | [illegible] |
| A5SS | 3677 | ENSG00000138002 | IFT172 | chr2 | - | [illegible] |
| A5SS | 3900 | ENSG00000162894 | FCMR | chr1 | - | [illegible] |
| A5SS | 6463 | ENSG00000150768 | DLAT | chr11 | + | [illegible] |
| A5SS | 6473 | ENSG00000118058 | KMT2A | chr11 | + | [illegible] |
| A5SS | 6963 | ENSG00000100532 | CGRRF1 | chr14 | + | [illegible] |
| A5SS | 2902 | ENSG00000173540 | GMPPB | chr3 | - | [illegible] |
| A5SS | 5524 | ENSG00000137309 | HMGA1 | chr6 | + | [illegible] |
| A5SS | 6422 | ENSG00000133624 | ZNF767P | chr7 | - | [illegible] |
| A5SS | 4980 | ENSG00000053900 | ANAPC4 | chr4 | + | [illegible] |
| A5SS | 3457 | ENSG00000092841 | MYL6 | chr12 | + | [illegible] |
| A5SS | 5396 | ENSG00000089060 | SLC8B1 | chr12 | - | [illegible] |
| A5SS | 6532 | ENSG00000121716 | PILRB | chr7 | + | [illegible] |
| A5SS | 1517 | ENSG00000129003 | VPS13C | chr15 | - | [illegible] |
| A5SS | 5577 | ENSG00000177479 | ARIH2 | chr3 | + | [illegible] |
| A5SS | 5921 | ENSG00000198721 | ECI2 | chr6 | - | [illegible] |
| A5SS | 6887 | ENSG00000158201 | ABHD3 | chr18 | - | [illegible] |
| A5SS | 4573 | ENSG00000126709 | IFI6 | chr1 | - | [illegible] |
| A5SS | 7872 | ENSG00000198561 | CTNND1 | chr11 | + | [illegible] |
| A5SS | 5597 | ENSG00000173559 | NABP1 | chr2 | + | [illegible] |
| A5SS | 1164 | ENSG00000103275 | UBE2I | chr16 | + | [illegible] |
| A5SS | 5004 | ENSG00000065989 | PDE4A | chr19 | + | [illegible] |
| A5SS | 2743 | ENSG00000047849 | MAP4 | chr3 | - | [illegible] |
| A5SS | 2878 | ENSG00000172053 | QARS1 | chr3 | - | [illegible] |
| A5SS | 2323 | ENSG00000164181 | ELOVL7 | chr5 | - | [illegible] |
| A5SS | 5471 | ENSG00000020922 | MRE11 | chr11 | - | [illegible] |
| A5SS | 3893 | ENSG00000108179 | PPIF | chr10 | + | [illegible] |
| A5SS | 1776 | ENSG00000100823 | APEX1 | chr14 | + | [illegible] |
| A5SS | 2662 | ENSG00000131042 | LILRB2 | chr19 | - | [illegible] |
| A5SS | 1501 | ENSG00000114062 | UBE3A | chr15 | - | [illegible] |
| A5SS | 3074 | ENSG00000111642 | CHD4 | chr12 | - | [illegible] |
| A5SS | 8953 | ENSG00000127022 | CANX | chr5 | + | [illegible] |
| A5SS | 143 | ENSG00000170949 | ZNF160 | chr19 | - | [illegible] |
| A5SS | 3542 | ENSG00000163472 | TMEM79 | chr1 | + | [illegible] |
| A5SS | 8131 | ENSG00000254986 | DPP3 | chr11 | + | [illegible] |
| A5SS | 771 | ENSG00000108799 | EZH1 | chr17 | - | [illegible] |
| A5SS | 4639 | ENSG00000114812 | VIPR1 | chr3 | + | [illegible] |
| A5SS | 568 | ENSG00000196756 | SNHG17 | chr20 | - | [illegible] |
| A5SS | 5684 | ENSG00000064201 | TSPAN32 | chr11 | + | [illegible] |
| A5SS | 1450 | ENSG00000104064 | GABPB1 | chr15 | - | [illegible] |
| A5SS | 2264 | ENSG00000126456 | IRF3 | chr19 | - | [illegible] |
| A5SS | 4518 | ENSG00000168610 | STAT3 | chr17 | - | [illegible] |
| A5SS | 6593 | ENSG00000106477 | CEP41 | chr7 | - | [illegible] |
| A5SS | 2269 | ENSG00000134644 | PUM1 | chr1 | - | [illegible] |
| A5SS | 5290 | ENSG00000224078 | SNHG14 | chr15 | + | [illegible] |
| A5SS | 5019 | ENSG00000112514 | CUTA | chr6 | - | [illegible] |
| A5SS | 4309 | ENSG00000151692 | RNF144A | chr2 | + | [illegible] |
| A5SS | 635 | ENSG00000101544 | ADNP2 | chr18 | + | [illegible] |
| A5SS | 6873 | ENSG00000131196 | NFATC1 | chr18 | + | [illegible] |
| A5SS | 7799 | ENSG00000106397 | PLOD3 | chr7 | - | [illegible] |
| A5SS | 7584 | ENSG00000197816 | CCDC180 | chr9 | + | [illegible] |
| A5SS | 4049 | ENSG00000116670 | MAD2L2 | chr1 | - | [illegible] |
| A5SS | 5154 | ENSG00000179115 | FARSA | chr19 | - | [illegible] |
| A5SS | 1416 | ENSG00000130703 | OSBPL2 | chr20 | + | [illegible] |
| A5SS | 7251 | ENSG00000221914 | PPP2R2A | chr8 | + | [illegible] |
| A5SS | 4621 | ENSG00000106462 | EZH2 | chr7 | - | [illegible] |
| A5SS | 5499 | ENSG00000140368 | PSTPIP1 | chr15 | + | [illegible] |
| A5SS | 2975 | ENSG00000163660 | CCNL1 | chr3 | - | [illegible] |
| A5SS | 6354 | ENSG00000108825 | PTGES3L-AARSD1 | chr17 | - | [illegible] |
| A5SS | 7978 | ENSG00000134910 | STT3A | chr11 | + | [illegible] |
| A5SS | 8285 | ENSG00000158545 | ZC3H18 | chr16 | + | [illegible] |
| A5SS | 8175 | ENSG00000162144 | CYB561A3 | chr11 | - | [illegible] |
| A5SS | 7870 | ENSG00000198561 | CTNND1 | chr11 | + | [illegible] |

| | | | | | | | | | | | | | | | | | | | | | | | | | | | | | | | | | | | | | | | | | | | | | | |
|---|---|---|---|---|---|---|---|---|---|---|---|---|---|---|---|---|---|---|---|---|---|---|---|---|---|---|---|---|---|---|---|---|---|---|---|---|---|---|---|---|---|---|---|---|---|---|
| A5SS | 884 | ENSG00000215041 | NEURL4 | chr17 | - | 7322947 | 7323123 | 7322953 | 7323123 | 7322734 | 7322866 | 13 | 15 | 33 | 26 | 21 | 7 | 0 | 4 | 2 | 8 | 35 | 32 | 5 | 10 | 19 | 20 | 3 | 0 | 4 | 17 | 155 | 149 | 0.117634634 | 0.511316036 | 0.641 | 1 | 0.888 | 0.926 | 0.716 | 0.627 | 0.911 | 1 | 0.706 | 0.518 | 0.082 |
| A5SS | 1232 | ENSG00000214078 | CPNE1 | chr20 | - | 35664358 | 35664900 | 35664759 | 35664900 | 35658929 | 35659014 | 9 | 5 | 6 | 15 | 0 | 195 | 88 | 197 | 165 | 73 | 0 | 9 | 16 | 0 | 28 | 118 | 182 | 151 | 130 | 191 | 298 | 149 | 0.117765092 | 0.511474567 | 0.023 | 0.028 | 0.015 | 0.043 | 0 | 0 | 0.024 | 0.05 | 0 | 0.068 | -0.007 |
| A5SS | 8780 | ENSG00000135108 | FBXO21 | chr12 | - | 117174650 | 117174797 | 117174654 | 117174797 | 117174204 | 117174341 | 81 | 33 | 125 | 76 | 94 | 0 | 0 | 0 | 0 | 0 | 52 | 77 | 79 | 92 | 61 | 5 | 5 | 2 | 0 | 0 | 153 | 149 | 0.118110891 | 0.512567362 | 1 | 1 | 1 | 1 | 1 | 0.91 | 0.937 | 0.975 | 1 | 1 | 0.036 |
| A5SS | 6682 | ENSG00000105939 | ZC3HAV1 | chr7 | - | 139079103 | 139080243 | 139079469 | 139080243 | 139078551 | 139078653 | 145 | 128 | 163 | 180 | 254 | 249 | 179 | 290 | 293 | 396 | 148 | 102 | 98 | 135 | 65 | 195 | 209 | 209 | 282 | 126 | 298 | 149 | 0.118227732 | 0.512665591 | 0.226 | 0.263 | 0.219 | 0.235 | 0.243 | 0.275 | 0.196 | 0.19 | 0.193 | 0.205 | 0.025 |
| A5SS | 6411 | ENSG00000145191 | EIF2B5 | chr3 | + | 184142236 | 184142394 | 184142236 | 184142378 | 184142501 | 184142603 | 11 | 3 | 8 | 7 | 5 | 58 | 66 | 85 | 80 | 108 | 26 | 5 | 13 | 11 | 42 | 311 | 124 | 103 | 165 | 129 | 165 | 149 | 0.118538815 | 0.51360528 | 0.146 | 0.039 | 0.078 | 0.073 | 0.04 | 0.07 | 0.035 | 0.102 | 0.057 | 0.227 | -0.023 |
| A5SS | 3452 | ENSG00000186205 | MTARC1 | chr1 | + | 220797873 | 220798065 | 220797873 | 220798014 | 220805051 | 220805113 | 0 | 3 | 1 | 4 | 0 | 38 | 21 | 32 | 26 | 16 | 0 | 27 | 8 | 7 | 1 | 87 | 152 | 105 | 148 | 180 | 200 | 149 | 0.119777927 | 0.518561241 | 0 | 0.096 | 0.023 | 0.103 | 0 | 0 | 0.117 | 0.054 | 0.034 | 0.004 | 0.003 |
| A5SS | 7637 | ENSG00000148337 | CIZ1 | chr9 | - | 128178988 | 128179415 | 128179072 | 128179415 | 128178708 | 128178904 | 27 | 5 | 16 | 10 | 4 | 10 | 6 | 17 | 7 | 1 | 13 | 16 | 4 | 17 | 34 | 0 | 10 | 2 | 6 | 15 | 233 | 149 | 0.119959909 | 0.518936266 | 0.633 | 0.348 | 0.376 | 0.477 | 0.719 | 1 | 0.506 | 0.561 | 0.644 | 0.592 | -0.15 |
| A5SS | 4964 | ENSG00000131408 | NR1H2 | chr19 | + | 50376472 | 50376826 | 50376472 | 50376576 | 50377586 | 50377648 | 120 | 85 | 76 | 81 | 103 | 6 | 0 | 2 | 0 | 0 | 199 | 117 | 100 | 175 | 255 | 12 | 0 | 2 | 6 | 0 | 298 | 149 | 0.120281188 | 0.519912808 | 0.909 | 1 | 0.95 | 1 | 1 | 0.892 | 1 | 0.962 | 0.936 | 1 | 0.014 |
| A5SS | 7710 | ENSG00000266714 | MYO15B | chr17 | + | 75624542 | 75624688 | 75624542 | 75624639 | 75624770 | 75624916 | 1 | 1 | 4 | 4 | 0 | 42 | 23 | 36 | 42 | 24 | 0 | 0 | 0 | 2 | 0 | 125 | 26 | 21 | 55 | 63 | 198 | 149 | 0.120478016 | 0.520343039 | 0.018 | 0.032 | 0.077 | 0.067 | 0 | 0 | 0 | 0 | 0.027 | 0 | 0.033 |
| A5SS | 6367 | ENSG00000168014 | C2CD3 | chr11 | - | 74085521 | 74085886 | 74085617 | 74085886 | 74084880 | 74084970 | 10 | 3 | 21 | 13 | 10 | 66 | 73 | 94 | 89 | 95 | 27 | 11 | 15 | 21 | 4 | 30 | 113 | 87 | 134 | 41 | 245 | 149 | 0.120571954 | 0.520343039 | 0.084 | 0.024 | 0.12 | 0.082 | 0.06 | 0.354 | 0.056 | 0.095 | 0.087 | 0.056 | -0.056 |
| A5SS | 7729 | ENSG00000213930 | GALT | chr9 | + | 34647088 | 34647961 | 34647088 | 34647258 | 34648333 | 34648456 | 17 | 4 | 15 | 8 | 8 | 0 | 0 | 0 | 0 | 0 | 16 | 26 | 12 | 21 | 3 | 0 | 0 | 0 | 0 | 5 | 298 | 149 | 0.12077139 | 0.520790733 | 1 | 1 | 1 | 1 | 1 | 1 | 1 | 1 | 1 | 0.231 | 0.154 |
| A5SS | 8437 | ENSG00000107959 | PITRM1 | chr10 | - | 3156929 | 3157531 | 3157434 | 3157531 | 3155590 | 3155729 | 233 | 181 | 282 | 261 | 305 | 3 | 0 | 0 | 3 | 1 | 366 | 338 | 299 | 293 | 308 | 0 | 6 | 1 | 1 | 10 | 298 | 149 | 0.12105277 | 0.521590793 | 0.975 | 1 | 1 | 0.978 | 0.993 | 1 | 0.966 | 0.993 | 0.993 | 0.939 | 0.011 |
| A5SS | 5587 | ENSG00000143390 | RFX5 | chr1 | - | 151346421 | 151346606 | 151346488 | 151346606 | 151346204 | 151346333 | 12 | 15 | 15 | 22 | 25 | 282 | 159 | 221 | 208 | 341 | 15 | 32 | 17 | 30 | 20 | 255 | 266 | 128 | 271 | 296 | 216 | 149 | 0.121783695 | 0.524325056 | 0.029 | 0.061 | 0.045 | 0.068 | 0.048 | 0.039 | 0.077 | 0.084 | 0.071 | 0.045 | -0.013 |
| A5SS | 8315 | ENSG00000090621 | PABPC4 | chr1 | - | 39564422 | 39564542 | 39564470 | 39564542 | 39562322 | 39563922 | 101 | 53 | 92 | 112 | 78 | 38 | 14 | 42 | 27 | 20 | 70 | 51 | 62 | 62 | 92 | 9 | 14 | 6 | 17 | 39 | 197 | 149 | 0.122681533 | 0.527773044 | 0.668 | 0.741 | 0.624 | 0.758 | 0.747 | 0.855 | 0.734 | 0.887 | 0.734 | 0.641 | -0.063 |
| A5SS | 1766 | ENSG00000185215 | TNFAIP2 | chr14 | + | 103123460 | 103123951 | 103123460 | 103123483 | 103124374 | 103124506 | 57 | 35 | 28 | 28 | 9 | 1 | 0 | 0 | 0 | 0 | 59 | 24 | 11 | 20 | 79 | 0 | 0 | 0 | 0 | 8 | 298 | 149 | 0.122990982 | 0.528686355 | 0.966 | 1 | 1 | 1 | 1 | 1 | 1 | 1 | 1 | 0.832 | 0.027 |
| A5SS | 6899 | ENSG00000094631 | HDAC6 | chrX | + | 48805630 | 48805779 | 48805630 | 48805671 | 48806367 | 48806464 | 5 | 4 | 5 | 4 | 4 | 127 | 90 | 116 | 151 | 136 | 0 | 7 | 2 | 0 | 6 | 320 | 123 | 117 | 175 | 266 | 257 | 149 | 0.123442403 | 0.530208015 | 0.022 | 0.025 | 0.024 | 0.015 | 0.017 | 0 | 0.032 | 0.01 | 0 | 0.013 | 0.01 |
| A5SS | 1239 | ENSG00000090861 | AARS1 | chr16 | - | 70258032 | 70259186 | 70258979 | 70259186 | 70255727 | 70255836 | 26 | 16 | 29 | 14 | 30 | 9 | 1 | 1 | 12 | 5 | 37 | 29 | 6 | 30 | 56 | 0 | 0 | 0 | 7 | 17 | 298 | 149 | 0.124807173 | 0.535647188 | 0.591 | 0.889 | 0.935 | 0.368 | 0.75 | 1 | 1 | 1 | 0.682 | 0.622 | -0.154 |
| A5SS | 3634 | ENSG00000135597 | REPS1 | chr6 | - | 138926400 | 138930098 | 138929976 | 138930098 | 138921036 | 138921124 | 102 | 99 | 105 | 108 | 157 | 31 | 20 | 21 | 16 | 39 | 74 | 71 | 75 | 47 | 33 | 37 | 16 | 11 | 13 | 22 | 298 | 149 | 0.125115509 | 0.536112104 | 0.622 | 0.712 | 0.714 | 0.771 | 0.668 | 0.5 | 0.689 | 0.773 | 0.644 | 0.429 | 0.09 |
| A5SS | 1042 | ENSG00000177731 | FLII | chr17 | - | 18245743 | 18246062 | 18245837 | 18246062 | 18245554 | 18245660 | 367 | 183 | 243 | 335 | 328 | 0 | 0 | 0 | 0 | 2 | 456 | 382 | 213 | 293 | 737 | 24 | 0 | 0 | 0 | 0 | 243 | 149 | 0.125210165 | 0.536112104 | 1 | 1 | 1 | 1 | 0.99 | 0.921 | 1 | 1 | 1 | 1 | 0.014 |
| A5SS | 3073 | ENSG00000111642 | CHD4 | chr12 | - | 6578337 | 6578546 | 6578408 | 6578546 | 6578028 | 6578117 | 15 | 17 | 5 | 7 | 4 | 1 | 1 | 4 | 2 | 16 | 214 | 46 | 38 | 103 | 77 | 16 | 9 | 27 | 5 | 5 | 220 | 149 | 0.125211041 | 0.536112104 | 0.91 | 0.92 | 0.458 | 0.703 | 0.145 | 0.901 | 0.776 | 0.488 | 0.933 | 0.913 | -0.175 |
| A5SS | 1057 | ENSG00000008517 | IL32 | chr16 | + | 3065609 | 3065710 | 3065609 | 3065655 | 3065783 | 3065826 | 111 | 21 | 81 | 70 | 79 | 5 | 3 | 8 | 5 | 10 | 83 | 59 | 29 | 9 | 84 | 6 | 14 | 9 | 3 | 3 | 204 | 149 | 0.12714604 | 0.543594982 | 0.942 | 0.836 | 0.881 | 0.911 | 0.852 | 0.91 | 0.755 | 0.702 | 0.687 | 0.953 | 0.083 |
| A5SS | 1062 | ENSG00000124120 | TTPAL | chr20 | + | 44484336 | 44484530 | 44484336 | 44484428 | 44486595 | 44486706 | 48 | 42 | 70 | 62 | 123 | 0 | 2 | 3 | 2 | 3 | 102 | 42 | 40 | 48 | 23 | 0 | 4 | 1 | 0 | 0 | 251 | 149 | 0.127160769 | 0.543594982 | 1 | 0.926 | 0.933 | 0.948 | 0.961 | 1 | 0.862 | 0.96 | 1 | 1 | -0.011 |
| A5SS | 1282 | ENSG00000122390 | NAA60 | chr16 | + | 3484698 | 3485083 | 3484698 | 3485061 | 3485466 | 3486952 | 8 | 0 | 4 | 6 | 0 | 11 | 4 | 6 | 5 | 5 | 5 | 20 | 13 | 20 | 47 | 11 | 26 | 23 | 26 | 38 | 171 | 149 | 0.127278584 | 0.543594982 | 0.388 | 0 | 0.367 | 0.511 | 0 | 0.284 | 0.401 | 0.33 | 0.401 | 0.519 | -0.134 |
| A5SS | 3375 | ENSG00000125347 | IRF1 | chr5 | - | 132486521 | 132486686 | 132486556 | 132486686 | 132485666 | 132485716 | 20 | 17 | 25 | 15 | 20 | 29 | 21 | 44 | 29 | 45 | 75 | 14 | 23 | 37 | 52 | 134 | 91 | 39 | 142 | 71 | 184 | 149 | 0.127358251 | 0.543594982 | 0.358 | 0.396 | 0.315 | 0.295 | 0.265 | 0.312 | 0.111 | 0.323 | 0.174 | 0.372 | 0.067 |
| A5SS | 3536 | ENSG00000106400 | ZNHIT1 | chr7 | + | 101218318 | 101221796 | 101218318 | 101218485 | 101222603 | 101222774 | 3 | 1 | 7 | 14 | 4 | 1 | 0 | 0 | 1 | 3 | 15 | 4 | 13 | 6 | 21 | 0 | 12 | 0 | 4 | 0 | 298 | 149 | 0.127556086 | 0.544012712 | 0.6 | 1 | 1 | 0.875 | 0.4 | 1 | 0.143 | 1 | 0.429 | 1 | 0.061 |
| A5SS | 1578 | ENSG00000136381 | IREB2 | chr15 | + | 78439794 | 78439881 | 78439794 | 78439876 | 78462921 | 78463087 | 240 | 183 | 215 | 184 | 281 | 6 | 7 | 4 | 8 | 1 | 159 | 236 | 236 | 174 | 81 | 0 | 6 | 4 | 3 | 19 | 154 | 149 | 0.128047496 | 0.54568087 | 0.975 | 0.962 | 0.981 | 0.957 | 0.996 | 1 | 0.974 | 0.983 | 0.982 | 0.805 | 0.025 |
| A5SS | 2367 | ENSG00000138767 | CNOT6L | chr4 | - | 77819232 | 77819390 | 77819303 | 77819390 | 77776270 | 77776392 | 35 | 23 | 46 | 61 | 33 | 18 | 19 | 42 | 42 | 32 | 11 | 52 | 32 | 17 | 16 | 0 | 15 | 54 | 4 | 13 | 220 | 149 | 0.128182567 | 0.545829052 | 0.568 | 0.451 | 0.426 | 0.496 | 0.411 | 1 | 0.701 | 0.286 | 0.742 | 0.455 | -0.166 |
| A5SS | 3121 | ENSG00000145819 | ARHGAP26 | chr5 | + | 143014079 | 143014293 | 143014079 | 143014116 | 143037195 | 143037261 | 10 | 21 | 13 | 19 | 10 | 134 | 112 | 93 | 119 | 94 | 6 | 38 | 30 | 34 | 0 | 128 | 90 | 114 | 187 | 65 | 298 | 149 | 0.12869753 | 0.5475934 | 0.036 | 0.086 | 0.065 | 0.074 | 0.051 | 0.023 | 0.174 | 0.116 | 0.083 | 0 | -0.017 |
| A5SS | 2458 | ENSG00000138698 | RAP1GDS1 | chr4 | + | 98343138 | 98343319 | 98343138 | 98343261 | 98352475 | 98352601 | 30 | 17 | 27 | 28 | 8 | 103 | 84 | 79 | 83 | 67 | 8 | 7 | 14 | 9 | 24 | 118 | 55 | 52 | 82 | 36 | 207 | 149 | 0.128820693 | 0.547689229 | 0.173 | 0.127 | 0.197 | 0.195 | 0.079 | 0.047 | 0.084 | 0.162 | 0.073 | 0.324 | 0.016 |
| A5SS | 6639 | ENSG00000086730 | LAT2 | chr7 | + | 74214821 | 74215010 | 74214821 | 74214940 | 74215946 | 74216069 | 113 | 34 | 83 | 86 | 46 | 24 | 11 | 26 | 7 | 20 | 218 | 121 | 53 | 151 | 139 | 23 | 23 | 14 | 45 | 11 | 219 | 149 | 0.129581788 | 0.550494996 | 0.762 | 0.678 | 0.685 | 0.893 | 0.61 | 0.866 | 0.782 | 0.72 | 0.695 | 0.896 | -0.066 |
| A5SS | 6959 | ENSG00000133703 | KRAS | chr12 | - | 25250750 | 25250929 | 25250763 | 25250929 | 25245273 | 25245395 | 65 | 47 | 48 | 80 | 48 | 4 | 1 | 5 | 5 | 0 | 5 | 70 | 55 | 33 | 17 | 0 | 1 | 2 | 4 | 0 | 162 | 149 | 0.130117157 | 0.551447615 | 0.937 | 0.977 | 0.898 | 0.936 | 1 | 1 | 0.985 | 0.962 | 0.884 | 1 | -0.017 |
| A5SS | 701 | ENSG00000183011 | NAA38 | chr17 | - | 7856984 | 7857198 | 7857014 | 7857198 | 7856684 | 7856843 | 12 | 1 | 8 | 8 | 7 | 57 | 21 | 35 | 39 | 26 | 6 | 11 | 6 | 4 | 26 | 108 | 93 | 35 | 40 | 132 | 179 | 149 | 0.130125462 | 0.551447615 | 0.149 | 0.038 | 0.16 | 0.146 | 0.183 | 0.044 | 0.09 | 0.125 | 0.077 | 0.141 | 0.04 |
| A5SS | 1554 | ENSG00000177082 | WDR73 | chr15 | - | 84653631 | 84654000 | 84653699 | 84654000 | 84652713 | 84652802 | 51 | 31 | 50 | 55 | 69 | 1 | 0 | 3 | 0 | 4 | 78 | 48 | 42 | 47 | 64 | 0 | 2 | 2 | 4 | 6 | 217 | 149 | 0.130184043 | 0.551447615 | 0.972 | 1 | 0.92 | 1 | 0.922 | 1 | 0.943 | 0.935 | 0.89 | 0.88 | 0.033 |
| A5SS | 6836 | ENSG00000124333 | VAMP7 | chrX | + | 155889457 | 155889633 | 155889457 | 155889612 | 155895622 | 155895680 | 23 | 17 | 8 | 16 | 11 | 280 | 187 | 200 | 265 | 349 | 10 | 18 | 6 | 10 | 18 | 118 | 206 | 173 | 133 | 143 | 170 | 149 | 0.130211353 | 0.551447615 | 0.067 | 0.074 | 0.034 | 0.05 | 0.027 | 0.069 | 0.071 | 0.03 | 0.062 | 0.099 | -0.016 |
| A5SS | 7857 | ENSG00000110514 | MADD | chr11 | + | 47295495 | 47295579 | 47295495 | 47295576 | 47295896 | 47296055 | 138 | 93 | 105 | 118 | 159 | 44 | 20 | 32 | 43 | 35 | 151 | 183 | 100 | 138 | 352 | 34 | 34 | 22 | 31 | 95 | 152 | 149 | 0.13076592 | 0.553029095 | 0.755 | 0.82 | 0.763 | 0.729 | 0.817 | 0.813 | 0.841 | 0.817 | 0.814 | 0.784 | -0.037 |
| A5SS | 7798 | ENSG00000132589 | FLOT2 | chr17 | - | 28883107 | 28883231 | 28883176 | 28883231 | 28882572 | 28882691 | 534 | 289 | 422 | 444 | 435 | 9 | 1 | 1 | 1 | 8 | 664 | 751 | 420 | 843 | 1425 | 0 | 6 | 12 | 16 | 17 | 218 | 149 | 0.130820666 | 0.553029095 | 0.976 | 0.995 | 0.997 | 0.997 | 0.974 | 1 | 0.988 | 0.96 | 0.973 | 0.983 | 0.007 |
| A5SS | 227 | ENSG00000005007 | UPF1 | chr19 | + | 18852986 | 18853104 | 18852986 | 18853071 | 18853251 | 18853350 | 44 | 17 | 46 | 51 | 37 | 99 | 50 | 88 | 73 | 124 | 28 | 45 | 71 | 53 | 90 | 86 | 68 | 74 | 74 | 196 | 182 | 149 | 0.13088965 | 0.553029095 | 0.267 | 0.218 | 0.3 | 0.364 | 0.196 | 0.21 | 0.351 | 0.44 | 0.37 | 0.273 | -0.06 |
| A5SS | 5271 | ENSG00000146556 | WASH2P | chr2 | + | 113596380 | 113596582 | 113596380 | 113596578 | 113596670 | 113596829 | 1 | 1 | 1 | 0 | 4 | 9 | 8 | 6 | 3 | 26 | 0 | 0 | 0 | 0 | 0 | 13 | 10 | 6 | 7 | 1 | 153 | 149 | 0.131389268 | 0.554709383 | 0.098 | 0.109 | 0.14 | 0 | 0.13 | 0 | 0 | 0 | 0 | 0 | 0.095 |
| A5SS | 1334 | ENSG00000038532 | CLEC16A | chr16 | + | 11003073 | 11003305 | 11003073 | 11003257 | 11020192 | 11020325 | 50 | 42 | 36 | 33 | 57 | 22 | 24 | 43 | 42 | 53 | 161 | 97 | 70 | 108 | 125 | 53 | 58 | 49 | 91 | 81 | 197 | 149 | 0.131688282 | 0.555426652 | 0.632 | 0.57 | 0.388 | 0.373 | 0.449 | 0.697 | 0.558 | 0.519 | 0.473 | 0.539 | -0.075 |
| A5SS | 1055 | ENSG00000178188 | SH2B1 | chr16 | + | 28872533 | 28872758 | 28872533 | 28872705 | 28873169 | 28873269 | 19 | 16 | 36 | 19 | 12 | 54 | 31 | 64 | 46 | 22 | 35 | 16 | 14 | 17 | 23 | 69 | 70 | 21 | 51 | 101 | 202 | 149 | 0.131763287 | 0.555426652 | 0.206 | 0.276 | 0.293 | 0.234 | 0.287 | 0.272 | 0.144 | 0.33 | 0.197 | 0.144 | 0.042 |
| A5SS | 4853 | ENSG00000159363 | ATP13A2 | chr1 | - | 17004691 | 17004821 | 17004706 | 17004821 | 17004331 | 17004408 | 8 | 2 | 5 | 6 | 3 | 2 | 3 | 0 | 0 | 0 | 5 | 4 | 1 | 3 | 14 | 0 | 2 | 1 | 6 | 0 | 164 | 149 | 0.131872332 | 0.555456063 | 0.784 | 0.377 | 1 | 1 | 1 | 1 | 0.645 | 0.476 | 0.312 | 1 | 0.146 |
| A5SS | 4341 | ENSG00000111445 | RFC5 | chr12 | + | 118017689 | 118017852 | 118017689 | 118017815 | 118019071 | 118019136 | 1 | 6 | 1 | 5 | 15 | 2 | 1 | 1 | 2 | 0 | 4 | 11 | 3 | 0 | 8 | 9 | 0 | 2 | 7 | 0 | 186 | 149 | 0.132400353 | 0.557248816 | 0.286 | 0.828 | 0.445 | 0.667 | 1 | 0.263 | 1 | 0.546 | 0 | 1 | 0.083 |
| A5SS | 2745 | ENSG00000144034 | TPRKB | chr2 | - | 73734428 | 73734591 | 73734527 | 73734591 | 73732162 | 73732285 | 95 | 63 | 81 | 96 | 49 | 6 | 3 | 1 | 2 | 0 | 115 | 77 | 67 | 113 | 69 | 9 | 3 | 0 | 0 | 9 | 248 | 149 | 0.132836142 | 0.558650914 | 0.905 | 0.927 | 0.98 | 0.966 | 1 | 0.885 | 0.939 | 1 | 1 | 0.822 | 0.026 |
| A5SS | 3289 | ENSG00000117054 | ACADM | chr1 | + | 75732643 | 75732745 | 75732643 | 75732741 | 75732852 | 75732922 | 8 | 4 | 8 | 5 | 9 | 209 | 205 | 190 | 261 | 221 | 24 | 0 | 2 | 1 | 0 | 51 | 134 | 85 | 38 | 56 | 153 | 149 | 0.13306097 | 0.559164325 | 0.036 | 0.019 | 0.039 | 0.018 | 0.038 | 0.314 | 0 | 0.022 | 0.025 | 0 | -0.042 |
| A5SS | 1421 | ENSG00000213614 | HEXA | chr15 | - | 72348030 | 72348134 | 72348047 | 72348134 | 72347685 | 72347758 | 6 | 1 | 1 | 3 | 8 | 194 | 134 | 165 | 150 | 215 | 2 | 0 | 8 | 13 | 22 | 473 | 189 | 200 | 281 | 370 | 166 | 149 | 0.135374307 | 0.567974862 | 0.027 | 0.007 | 0.005 | 0.018 | 0.032 | 0.004 | 0 | 0.035 | 0.04 | 0.051 | -0.008 |
| A5SS | 251 | ENSG00000104852 | SNRNP70 | chr19 | + | 49102113 | 49103587 | 49102113 | 49102185 | 49104633 | 49104735 | 403 | 265 | 483 | 459 | 464 | 3 | 6 | 3 | 10 | 10 | 281 | 285 | 206 | 245 | 559 | 0 | 6 | 0 | 7 | 4 | 298 | 149 | 0.135479486 | 0.567974862 | 0.985 | 0.957 | 0.988 | 0.958 | 0.959 | 1 | 0.96 | 1 | 0.946 | 0.986 | -0.009 |
| A5SS | 8097 | ENSG00000070047 | PHRF1 | chr11 | + | 581491 | 581606 | 581491 | 581597 | 581961 | 582081 | 65 | 22 | 42 | 34 | 16 | 13 | 7 | 5 | 2 | 10 | 8 | 37 | 19 | 28 | 37 | 10 | 21 | 8 | 4 | 5 | 158 | 149 | 0.135503966 | 0.567974862 | 0.825 | 0.748 | 0.888 | 0.941 | 0.601 | 0.43 | 0.624 | 0.691 | 0.868 | 0.875 | 0.103 |
| A5SS | 6961 | ENSG00000167548 | KMT2D | chr12 | - | 49044190 | 49044304 | 49044199 | 49044304 | 49043867 | 49043998 | 117 | 88 | 86 | 119 | 59 | 141 | 81 | 115 | 117 | 83 | 197 | 111 | 82 | 146 | 137 | 290 | 136 | 159 | 135 | 230 | 158 | 149 | 0.135605913 | 0.567974862 | 0.439 | 0.506 | 0.414 | 0.49 | 0.401 | 0.39 | 0.435 | 0.327 | 0.505 | 0.36 | 0.047 |
| A5SS | 6398 | ENSG00000169660 | HEXD | chr17 | + | 82439630 | 82440334 | 82439630 | 82439713 | 82440996 | 82441075 | 15 | 7 | 25 | 26 | 9 | 14 | 2 | 13 | 2 | 1 | 12 | 12 | 10 | 4 | 3 | 4 | 4 | 1 | 6 | 22 | 298 | 149 | 0.135679405 | 0.567974862 | 0.349 | 0.636 | 0.49 | 0.867 | 0.818 | 0.6 | 0.6 | 0.833 | 0.25 | 0.064 | 0.163 |
| A5SS | 5453 | ENSG00000018280 | SLC11A1 | chr2 | + | 218385146 | 218385485 | 218385146 | 218385266 | 218386634 | 218387230 | 29 | 13 | 7 | 33 | 23 | 74 | 40 | 24 | 60 | 22 | 39 | 46 | 64 | 34 | 70 | 112 | 172 | 131 | 129 | 177 | 298 | 149 | 0.135938297 | 0.568621224 | 0.164 | 0.14 | 0.127 | 0.216 | 0.343 | 0.148 | 0.118 | 0.196 | 0.116 | 0.165 | 0.049 |
| A5SS | 3533 | ENSG00000100142 | POLR2F | chr22 | + | 37956772 | 37956901 | 37956772 | 37956842 | 37959345 | 37959476 | 10 | 0 | 8 | 3 | 24 | 56 | 29 | 33 | 44 | 55 | 2 | 14 | 4 | 1 | 17 | 111 | 52 | 32 | 53 | 60 | 208 | 149 | 0.137378361 | 0.574203564 | 0.113 | 0 | 0.148 | 0.047 | 0.238 | 0.013 | 0.162 | 0.082 | 0.013 | 0.169 | 0.021 |
| A5SS | 230 | ENSG00000011600 | TYROBP | chr19 | - | 35908167 | 35908295 | 35908174 | 35908295 | 35907729 | 35907762 | 877 | 278 | 381 | 556 | 489 | 83 | 19 | 39 | 62 | 26 | 1274 | 952 | 572 | 915 | 1445 | 123 | 88 | 54 | 97 | 166 | 156 | 149 | 0.137541036 | 0.574442302 | 0.91 | 0.933 | 0.903 | 0.895 | 0.947 | 0.908 | 0.912 | 0.91 | 0.9 | 0.893 | 0.013 |
| A5SS | 3632 | ENSG00000167962 | ZNF598 | chr16 | - | 2001582 | 2001700 | 2001591 | 2001700 | 2001003 | 2001191 | 58 | 28 | 57 | 55 | 24 | 8 | 9 | 2 | 2 | 2 | 97 | 57 | 59 | 53 | 80 | 4 | 10 | 0 | 7 | 0 | 158 | 149 | 0.137714868 | 0.574727234 | 0.872 | 0.746 | 0.964 | 0.963 | 0.919 | 0.958 | 0.843 | 1 | 0.877 | 1 | -0.043 |
| A5SS | 6617 | ENSG00000160856 | FCRL3 | chr1 | - | 157697139 | 157697919 | 157697658 | 157697919 | 157696039 | 157696327 | 209 | 94 | 96 | 122 | 118 | 9 | 2 | 2 | 2 | 11 | 78 | 132 | 63 | 55 | 57 | 4 | 9 | 14 | 1 | 0 | 298 | 149 | 0.138682197 | 0.578320703 | 0.921 | 0.959 | 0.96 | 0.968 | 0.843 | 0.907 | 0.88 | 0.692 | 0.965 | 1 | 0.041 |
| A5SS | 956 | ENSG00000108370 | RGS9 | chr17 | + | 65177731 | 65177803 | 65177731 | 65177794 | 65189285 | 65189315 | 36 | 12 | 20 | 33 | 20 | 2 | 5 | 2 | 5 | 7 | 14 | 21 | 23 | 13 | 33 | 0 | 0 | 1 | 3 | 8 | 158 | 149 | 0.1391847 | 0.579971772 | 0.944 | 0.694 | 0.904 | 0.862 | 0.729 | 1 | 1 | 0.956 | 0.803 | 0.796 | -0.084 |
| A5SS | 5507 | ENSG00000237765 | FAM200B | chr4 | + | 15681755 | 15681968 | 15681755 | 15681901 | 15686235 | 15686462 | 31 | 18 | 19 | 23 | 24 | 22 | 16 | 26 | 28 | 25 | 48 | 41 | 22 | 59 | 33 | 40 | 9 | 19 | 33 | 46 | 216 | 149 | 0.139746336 | 0.581070527 | 0.493 | 0.437 | 0.335 | 0.362 | 0.398 | 0.453 | 0.759 | 0.444 | 0.552 | 0.331 | -0.103 |
| A5SS | 1921 | ENSG00000115760 | BIRC6 | chr2 | + | 32471013 | 32471151 | 32471013 | 32471124 | 32473111 | 32473239 | 244 | 221 | 211 | 217 | 316 | 60 | 43 | 44 | 47 | 81 | 32 | 249 | 260 | 69 | 14 | 21 | 32 | 36 | 6 | 0 | 176 | 149 | 0.139853757 | 0.581070527 | 0.775 | 0.813 | 0.802 | 0.796 | 0.768 | 0.563 | 0.868 | 0.859 | 0.907 | 1 | -0.049 |
| A5SS | 492 | ENSG00000125843 | AP5S1 | chr20 | + | 3820546 | 3820854 | 3820546 | 3820758 | 3822101 | 3822293 | 2 | 0 | 1 | 3 | 0 | 15 | 5 | 13 | 10 | 6 | 3 | 1 | 4 | 0 | 3 | 0 | 25 | 4 | 15 | 24 | 245 | 149 | 0.139867918 | 0.581070527 | 0.075 | 0 | 0.045 | 0.154 | 0 | 1 | 0.024 | 0.378 | 0 | 0.071 | -0.24 |
| A5SS | 858 | ENSG00000076604 | TRAF4 | chr17 | + | 28747212 | 28747391 | 28747212 | 28747264 | 28747842 | 28747947 | 6 | 3 | 6 | 6 | 8 | 7 | 1 | 10 | 9 | 9 | 31 | 22 | 11 | 12 | 36 | 17 | 17 | 20 | 14 | 7 | 276 | 149 | 0.139875485 | 0.581070527 | 0.316 | 0.618 | 0.245 | 0.265 | 0.324 | 0.496 | 0.411 | 0.229 | 0.316 | 0.735 | -0.084 |
| A5SS | 5867 | ENSG00000135426 | TESPA1 | chr12 | - | 54963741 | 54963950 | 54963784 | 54963950 | 54963142 | 54963242 | 173 | 180 | 246 | 245 | 350 | 2 | 0 | 0 | 1 | 4 | 46 | 94 | 117 | 68 | 214 | 0 | 0 | 1 | 0 | 12 | 192 | 149 | 0.140403999 | 0.582821175 | 0.985 | 1 | 1 | 0.995 | 0.985 | 1 | 1 | 0.989 | 1 | 0.933 | 0.009 |
| A5SS | 194 | ENSG00000178982 | EIF3K | chr19 | + | 38619195 | 38619327 | 38619195 | 38619317 | 38620336 | 38620435 | 1266 | 498 | 1045 | 932 | 508 | 24 | 19 | 46 | 31 | 8 | 719 | 889 | 625 | 768 | 1395 | 18 | 24 | 23 | 23 | 78 | 159 | 149 | 0.140528535 | 0.582893513 | 0.98 | 0.961 | 0.955 | 0.966 | 0.983 | 0.974 | 0.972 | 0.962 | 0.969 | 0.944 | 0.005 |
| A5SS | 6549 | ENSG00000002933 | TMEM176A | chr7 | + | 150804361 | 150804485 | 150804361 | 150804472 | 150804826 | 150805118 | 22 | 11 | 0 | 1 | 3 | 257 | 76 | 4 | 11 | 17 | 14 | 21 | 0 | 0 | 3 | 202 | 222 | 8 | 123 | 4 | 162 | 149 | 0.141232992 | 0.585095274 | 0.073 | 0.117 | 0 | 0.077 | 0.14 | 0.06 | 0.08 | 0 | 0 | 0.408 | -0.028 |
| A5SS | 6788 | ENSG00000124222 | STX16 | chr20 | + | 58652072 | 58652138 | 58652072 | 58652087 | 58659622 | 58659634 | 66 | 48 | 65 | 61 | 56 | 46 | 46 | 64 | 78 | 95 | 40 | 50 | 41 | 68 | 71 | 34 | 35 | 50 | 80 | 20 | 200 | 149 | 0.141274383 | 0.585095274 | 0.517 | 0.437 | 0.431 | 0.368 | 0.305 | 0.467 | 0.516 | 0.379 | 0.388 | 0.726 | -0.084 |
| A5SS | 7398 | ENSG00000213782 | DDX47 | chr12 | + | 12823202 | 12824016 | 12823202 | 12823319 | 12824539 | 12824677 | 148 | 80 | 136 | 169 | 240 | 2 | 1 | 2 | 8 | 19 | 93 | 140 | 117 | 67 | 30 | 0 | 0 | 7 | 6 | 0 | 298 | 149 | 0.141516848 | 0.585653756 | 0.974 | 0.976 | 0.971 | 0.914 | 0.863 | 1 | 1 | 0.893 | 0.848 | 1 | -0.009 |
| A5SS | 3884 | ENSG00000138386 | NAB1 | chr2 | + | 190655976 | 190656159 | 190655976 | 190656153 | 190659157 | 190659995 | 3 | 2 | 1 | 3 | 0 | 24 | 2 | 17 | 34 | 25 | 3 | 5 | 4 | 0 | 0 | 0 | 2 | 14 | 22 | 53 | 155 | 149 | 0.141868784 | 0.586664076 | 0.107 | 0.49 | 0.054 | 0.078 | 0 | 1 | 0.706 | 0.215 | 0 | 0 | -0.238 |
| A5SS | 5267 | ENSG00000140386 | SCAPER | chr15 | - | 76771723 | 76771954 | 76771741 | 76771954 | 76766917 | 76767088 | 16 | 15 | 26 | 17 | 19 | 42 | 33 | 61 | 45 | 68 | 0 | 9 | 15 | 1 | 0 | 11 | 30 | 24 | 5 | 6 | 167 | 149 | 0.142783345 | 0.588955928 | 0.254 | 0.289 | 0.276 | 0.252 | 0.2 | 0 | 0.211 | 0.358 | 0.151 | 0 | 0.11 |
| A5SS | 4472 | ENSG00000137965 | IFI44 | chr1 | + | 78660553 | 78660691 | 78660553 | 78660654 | 78662703 | 78662878 | 14 | 25 | 15 | 35 | 27 | 128 | 212 | 90 | 229 | 168 | 28 | 38 | 21 | 8 | 8 | 45 | 227 | 124 | 199 | 8 | 186 | 149 | 0.142789288 | 0.588955928 | 0.081 | 0.086 | 0.118 | 0.109 | 0.114 | 0.333 | 0.118 | 0.119 | 0.031 | 0.445 | -0.108 |
| A5SS | 3693 | ENSG00000160271 | RALGDS | chr9 | - | 133100495 | 133101762 | 133101519 | 133101762 | 133100267 | 133100382 | 27 | 10 | 4 | 21 | 4 | 75 | 20 | 33 | 45 | 18 | 6 | 23 | 1 | 7 | 7 | 27 | 51 | 41 | 10 | 51 | 298 | 149 | 0.142828931 | 0.588955928 | 0.153 | 0.2 | 0.057 | 0.189 | 0.1 | 0.1 | 0.184 | 0.012 | 0.259 | 0.064 | 0.016 |
| A5SS | 6366 | ENSG00000262246 | CORO7 | chr16 | - | 4407500 | 4407684 | 4407561 | 4407684 | 4405490 | 4405567 | 191 | 73 | 106 | 72 | 48 | 9 | 0 | 7 | 9 | 0 | 218 | 183 | 105 | 242 | 255 | 10 | 14 | 1 | 6 | 42 | 210 | 149 | 0.142855903 | 0.588955928 | 0.938 | 1 | 0.915 | 0.85 | 1 | 0.939 | 0.903 | 0.987 | 0.966 | 0.812 | 0.019 |
| A5SS | 4890 | ENSG00000182400 | TRAPPC6B | chr14 | - | 39159449 | 39159550 | 39159482 | 39159550 | 39158284 | 39158402 | 40 | 30 | 30 | 24 | 39 | 111 | 86 | 79 | 79 | 95 | 10 | 38 | 15 | 25 | 43 | 66 | 55 | 39 | 45 | 28 | 182 | 149 | 0.143616924 | 0.591645192 | 0.228 | 0.222 | 0.237 | 0.199 | 0.252 | 0.11 | 0.361 | 0.239 | 0.313 | 0.557 | -0.088 |
| A5SS | 3658 | ENSG00000145916 | RMND5B | chr5 | + | 178138107 | 178138396 | 178138107 | 178138258 | 178142582 | 178142728 | 8 | 3 | 7 | 4 | 2 | 8 | 0 | 8 | 2 | 0 | 16 | 0 | 13 | 1 | 2 | 7 | 13 | 3 | 0 | 9 | 287 | 149 | 0.143816028 | 0.592017265 | 0.342 | 1 | 0.312 | 0.509 | 1 | 0.543 | 0 | 0.692 | 1 | 0.103 | 0.165 |
| A5SS | 5127 | ENSG00000112855 | HARS2 | chr5 | + | 140691794 | 140692017 | 140691794 | 140691937 | 140693590 | 140693665 | 7 | 11 | 4 | 10 | 4 | 2 | 4 | 0 | 0 | 0 | 10 | 17 | 17 | 19 | 5 | 0 | 0 | 0 | 0 | 0 | 229 | 149 | 0.144456007 | 0.59420226 | 0.695 | 0.641 | 1 | 1 | 1 | 1 | 1 | 1 | 1 | 1 | -0.133 |
| A5SS | 5190 | ENSG00000166801 | FAM111A | chr11 | + | 59142855 | 59142969 | 59142855 | 59142946 | 59143082 | 59143304 | 89 | 96 | 70 | 91 | 61 | 46 | 44 | 52 | 49 | 47 | 27 | 49 | 56 | 59 | 9 | 30 | 44 | 39 | 30 | 8 | 172 | 149 | 0.145472254 | 0.59793052 | 0.626 | 0.654 | 0.538 | 0.617 | 0.529 | 0.438 | 0.491 | 0.554 | 0.63 | 0.494 | 0.071 |
| A5SS | 2191 | ENSG00000178338 | ZNF354B | chr5 | + | 178885613 | 178885737 | 178885613 | 178885733 | 178887620 | 178888122 | 0 | 5 | 0 | 0 | 0 | 2 | 1 | 1 | 4 | 8 | 0 | 0 | 0 | 0 | 0 | 5 | 1 | 1 | 7 | 4 | 153 | 149 | 0.145993831 | 0.599621455 | 0 | 0.83 | 0 | 0 | 0 | 0 | 0 | 0 | 0 | 0 | 0.166 |
| A5SS | 3673 | ENSG00000148019 | CEP78 | chr9 | + | 78236079 | 78236659 | 78236079 | 78236603 | 78240022 | 78240195 | 18 | 2 | 7 | 4 | 3 | 75 | 39 | 45 | 72 | 21 | 0 | 0 | 7 | 1 | 4 | 3 | 45 | 50 | 16 | 6 | 205 | 149 | 0.146383255 | 0.600767478 | 0.149 | 0.036 | 0.102 | 0.039 | 0.094 | 0 | 0 | 0.092 | 0.043 | 0.326 | -0.008 |
| A5SS | 3234 | ENSG00000197386 | HTT | chr4 | + | 3172907 | 3173218 | 3172907 | 3173131 | 3174720 | 3174799 | 4 | 4 | 4 | 9 | 6 | 64 | 38 | 72 | 62 | 29 | 20 | 13 | 9 | 12 | 0 | 98 | 82 | 50 | 54 | 106 | 236 | 149 | 0.147045445 | 0.603030379 | 0.038 | 0.062 | 0.034 | 0.084 | 0.116 | 0.114 | 0.091 | 0.102 | 0.123 | 0 | -0.019 |
| A5SS | 8820 | ENSG00000089693 | MLF2 | chr12 | - | 6750144 | 6750305 | 6750176 | 6750305 | 6749847 | 6750007 | 1 | 0 | 3 | 4 | 5 | 149 | 54 | 73 | 119 | 110 | 0 | 0 | 3 | 3 | 23 | 243 | 116 | 121 | 117 | 201 | 181 | 149 | 0.147372006 | 0.603858053 | 0.005 | 0 | 0.033 | 0.027 | 0.036 | 0 | 0 | 0.02 | 0.021 | 0.086 | -0.005 |
| A5SS | 8264 | ENSG00000133816 | MICAL2 | chr11 | + | 12138389 | 12138464 | 12138389 | 12138460 | 12162078 | 12162419 | 14 | 15 | 3 | 15 | 17 | 23 | 24 | 20 | 25 | 22 | 17 | 3 | 9 | 2 | 21 | 31 | 24 | 16 | 17 | 50 | 153 | 149 | 0.147469194 | 0.603858053 | 0.372 | 0.378 | 0.127 | 0.369 | 0.429 | 0.348 | 0.109 | 0.354 | 0.103 | 0.29 | 0.094 |
| A5SS | 1290 | ENSG00000123154 | WDR83 | chr19 | + | 12669976 | 12670097 | 12669976 | 12670080 | 12670179 | 12670285 | 22 | 2 | 17 | 27 | 1 | 1 | 0 | 2 | 4 | 0 | 56 | 23 | 10 | 14 | 8 | 0 | 0 | 1 | 0 | 0 | 166 | 149 | 0.149589446 | 0.612079522 | 0.952 | 1 | 0.884 | 0.858 | 1 | 1 | 1 | 0.9 | 1 | 1 | -0.041 |
| A5SS | 4570 | ENSG00000187231 | SESTD1 | chr2 | - | 179116573 | 179116790 | 179116667 | 179116790 | 179115064 | 179115256 | 6 | 0 | 2 | 7 | 0 | 56 | 63 | 61 | 82 | 96 | 0 | 1 | 1 | 0 | 0 | 20 | 60 | 116 | 86 | 31 | 243 | 149 | 0.149869285 | 0.612763825 | 0.062 | 0 | 0.02 | 0.05 | 0 | 0 | 0.01 | 0.005 | 0 | 0 | 0.023 |
| A5SS | 7759 | ENSG00000165699 | TSC1 | chr9 | - | 132944542 | 132944619 | 132944571 | 132944619 | 132935032 | 132935095 | 27 | 18 | 47 | 33 | 22 | 16 | 10 | 17 | 21 | 14 | 24 | 55 | 18 | 37 | 5 | 12 | 25 | 16 | 30 | 17 | 178 | 149 | 0.150370163 | 0.614350171 | 0.586 | 0.601 | 0.698 | 0.568 | 0.568 | 0.626 | 0.648 | 0.485 | 0.508 | 0.198 | 0.111 |
| A5SS | 8136 | ENSG00000131626 | PPFIA1 | chr11 | + | 70338373 | 70338453 | 70338373 | 70338434 | 70339170 | 70339306 | 61 | 48 | 37 | 59 | 85 | 6 | 0 | 4 | 6 | 5 | 125 | 57 | 44 | 95 | 29 | 22 | 3 | 7 | 6 | 0 | 168 | 149 | 0.150622675 | 0.614550727 | 0.9 | 1 | 0.891 | 0.897 | 0.938 | 0.834 | 0.944 | 0.848 | 0.934 | 1 | 0.013 |
| A5SS | 2464 | ENSG00000125386 | FAM193A | chr4 | + | 2694945 | 2695142 | 2694945 | 2695082 | 2696362 | 2696593 | 28 | 14 | 23 | 23 | 2 | 7 | 0 | 0 | 0 | 0 | 45 | 31 | 25 | 19 | 20 | 0 | 0 | 1 | 1 | 0 | 209 | 149 | 0.150645107 | 0.614550727 | 0.74 | 1 | 1 | 1 | 1 | 1 | 1 | 0.947 | 0.931 | 1 | -0.028 |
| A5SS | 3495 | ENSG00000079277 | MKNK1 | chr1 | - | 46604164 | 46604255 | 46604226 | 46604255 | 46594112 | 46594280 | 1 | 0 | 2 | 0 | 1 | 3 | 5 | 10 | 6 | 0 | 1 | 4 | 2 | 2 | 4 | 39 | 18 | 11 | 8 | 34 | 211 | 149 | 0.150769501 | 0.614597473 | 0.191 | 0 | 0.124 | 0 | 1 | 0.018 | 0.136 | 0.114 | 0.15 | 0.077 | 0.164 |
| A5SS | 7738 | ENSG00000113356 | POLR3G | chr5 | + | 90497655 | 90497735 | 90497655 | 90497706 | 90501905 | 90501988 | 1 | 0 | 0 | 3 | 7 | 7 | 10 | 8 | 8 | 1 | 0 | 1 | 4 | 0 | 1 | 24 | 4 | 1 | 17 | 13 | 178 | 149 | 0.151646733 | 0.617493143 | 0.107 | 0 | 0 | 0.239 | 0.854 | 0 | 0.173 | 0.77 | 0 | 0.06 | 0.039 |
| A5SS | 7747 | ENSG00000172716 | SLFN11 | chr17 | - | 35367602 | 35367700 | 35367649 | 35367700 | 35366946 | 35367063 | 56 | 20 | 35 | 55 | 67 | 0 | 0 | 0 | 3 | 6 | 102 | 73 | 42 | 42 | 25 | 0 | 0 | 0 | 1 | 0 | 196 | 149 | 0.151706787 | 0.617493143 | 1 | 1 | 1 | 0.933 | 0.895 | 1 | 1 | 1 | 0.97 | 1 | -0.028 |
| A5SS | 564 | ENSG00000196756 | SNHG17 | chr20 | - | 38422086 | 38422241 | 38422091 | 38422241 | 38421005 | 38421098 | 8 | 2 | 5 | 3 | 1 | 1 | 2 | 1 | 0 | 4 | 0 | 12 | 5 | 0 | 21 | 39 | 0 | 5 | 6 | 0 | 154 | 149 | 0.152992358 | 0.622260399 | 0.886 | 0.492 | 0.829 | 1 | 0.195 | 0 | 1 | 0.492 | 0 | 1 | 0.182 |
| A5SS | 2227 | ENSG00000120705 | ETF1 | chr5 | - | 138542704 | 138542936 | 138542832 | 138542936 | 138518691 | 138518867 | 27 | 23 | 34 | 18 | 10 | 75 | 53 | 68 | 108 | 89 | 13 | 33 | 17 | 63 | 25 | 39 | 107 | 117 | 113 | 9 | 277 | 149 | 0.154008837 | 0.62592688 | 0.162 | 0.189 | 0.212 | 0.082 | 0.057 | 0.152 | 0.142 | 0.072 | 0.231 | 0.599 | -0.099 |
| A5SS | 1417 | ENSG00000130703 | OSBPL2 | chr20 | + | 62279156 | 62279519 | 62279156 | 62279349 | 62280008 | 62280221 | 6 | 5 | 3 | 10 | 2 | 0 | 0 | 0 | 6 | 0 | 7 | 16 | 9 | 11 | 7 | 0 | 0 | 0 | 0 | 0 | 298 | 149 | 0.154408331 | 0.627082191 | 1 | 1 | 1 | 0.455 | 1 | 1 | 1 | 1 | 1 | 1 | -0.109 |
| A5SS | 1727 | ENSG00000179364 | PACS2 | chr14 | + | 105381913 | 105382058 | 105381913 | 105382046 | 105382476 | 105382581 | 2 | 2 | 2 | 1 | 0 | 25 | 6 | 13 | 10 | 6 | 0 | 2 | 0 | 0 | 0 | 21 | 24 | 10 | 13 | 47 | 161 | 149 | 0.154639311 | 0.627240837 | 0.069 | 0.236 | 0.125 | 0.085 | 0 | 0 | 0.072 | 0 | 0 | 0 | 0.089 |
| A5SS | 2697 | ENSG00000113504 | SLC12A7 | chr5 | - | 1075890 | 1076236 | 1076137 | 1076236 | 1075370 | 1075490 | 10 | 4 | 0 | 2 | 0 | 29 | 7 | 7 | 18 | 6 | 3 | 6 | 5 | 5 | 6 | 8 | 25 | 11 | 19 | 8 | 298 | 149 | 0.154677913 | 0.627240837 | 0.147 | 0.222 | 0 | 0.053 | 0 | 0.158 | 0.107 | 0.185 | 0.116 | 0.273 | -0.083 |
| A5SS | 2840 | ENSG00000163349 | HIPK1 | chr1 | + | 113957123 | 113957286 | 113957123 | 113957184 | 113958065 | 113958291 | 597 | 400 | 454 | 542 | 1167 | 4 | 9 | 15 | 8 | 17 | 482 | 600 | 501 | 528 | 232 | 7 | 19 | 20 | 6 | 2 | 251 | 149 | 0.157287915 | 0.637120541 | 0.989 | 0.963 | 0.947 | 0.976 | 0.976 | 0.976 | 0.949 | 0.937 | 0.981 | 0.986 | 0.004 |
| A5SS | 7787 | ENSG00000109220 | CHIC2 | chr4 | - | 54048954 | 54049110 | 54049038 | 54049110 | 54014062 | 54014119 | 388 | 241 | 315 | 453 | 699 | 20 | 14 | 19 | 9 | 14 | 205 | 415 | 325 | 179 | 162 | 10 | 19 | 28 | 2 | 5 | 233 | 149 | 0.157348403 | 0.637120541 | 0.925 | 0.917 | 0.914 | 0.97 | 0.97 | 0.929 | 0.933 | 0.881 | 0.983 | 0.954 | 0.003 |
| A5SS | 7896 | ENSG00000062822 | POLD1 | chr19 | + | 50409518 | 50409666 | 50409518 | 50409594 | 50413425 | 50413521 | 24 | 7 | 21 | 26 | 15 | 2 | 0 | 5 | 0 | 0 | 5 | 27 | 19 | 17 | 44 | 0 | 0 | 3 | 0 | 0 | 221 | 149 | 0.157547036 | 0.637450536 | 0.89 | 1 | 0.739 | 1 | 1 | 1 | 1 | 0.81 | 1 | 1 | -0.036 |
| A5SS | 4210 | ENSG00000048052 | HDAC9 | chr7 | + | 18591515 | 18591642 | 18591515 | 18591636 | 18593907 | 18594029 | 70 | 46 | 32 | 55 | 141 | 17 | 4 | 4 | 8 | 7 | 54 | 42 | 22 | 45 | 37 | 0 | 5 | 8 | 8 | 18 | 155 | 149 | 0.15853028 | 0.64095229 | 0.798 | 0.917 | 0.885 | 0.869 | 0.951 | 1 | 0.89 | 0.726 | 0.844 | 0.664 | 0.059 |
| A5SS | 7709 | ENSG00000266714 | MYO15B | chr17 | + | 75624174 | 75624269 | 75624174 | 75624249 | 75624369 | 75624447 | 23 | 15 | 10 | 39 | 8 | 4 | 2 | 3 | 2 | 0 | 24 | 25 | 13 | 66 | 44 | 0 | 2 | 5 | 0 | 2 | 169 | 149 | 0.15869049 | 0.641123718 | 0.835 | 0.869 | 0.746 | 0.945 | 1 | 1 | 0.917 | 0.696 | 1 | 0.951 | -0.034 |
| A5SS | 7356 | ENSG00000140525 | FANCI | chr15 | + | 89247628 | 89247731 | 89247628 | 89247726 | 89258703 | 89258776 | 10 | 5 | 17 | 3 | 13 | 6 | 0 | 1 | 1 | 0 | 1 | 9 | 33 | 46 | 7 | 0 | 0 | 5 | 1 | 0 | 154 | 149 | 0.158987051 | 0.641530148 | 0.617 | 1 | 0.943 | 0.744 | 1 | 1 | 1 | 0.865 | 0.978 | 1 | -0.108 |
| A5SS | 5769 | ENSG00000166197 | NOLC1 | chr10 | + | 102159192 | 102159338 | 102159192 | 102159308 | 102159432 | 102159568 | 9 | 8 | 12 | 8 | 18 | 22 | 17 | 31 | 32 | 26 | 0 | 20 | 10 | 43 | 9 | 15 | 11 | 22 | 34 | 0 | 179 | 149 | 0.15902686 | 0.641530148 | 0.254 | 0.281 | 0.244 | 0.172 | 0.366 | 0 | 0.602 | 0.275 | 0.513 | 1 | -0.215 |
| A5SS | 2093 | ENSG00000113240 | CLK4 | chr5 | - | 178617343 | 178618778 | 178618555 | 178618778 | 178616881 | 178616948 | 23 | 23 | 36 | 26 | 46 | 32 | 20 | 36 | 29 | 21 | 4 | 40 | 21 | 11 | 0 | 0 | 5 | 7 | 3 | 10 | 298 | 149 | 0.159850405 | 0.644374744 | 0.264 | 0.365 | 0.333 | 0.31 | 0.523 | 1 | 0.8 | 0.6 | 0.647 | 0 | -0.25 |
| A5SS | 8163 | ENSG00000159063 | ALG8 | chr11 | - | 78138709 | 78139624 | 78139493 | 78139624 | 78127357 | 78127436 | 4 | 4 | 2 | 2 | 10 | 13 | 9 | 3 | 12 | 9 | 5 | 14 | 4 | 2 | 5 | 0 | 5 | 15 | 4 | 9 | 298 | 149 | 0.160067 | 0.644770253 | 0.133 | 0.182 | 0.25 | 0.077 | 0.357 | 1 | 0.583 | 0.118 | 0.2 | 0.217 | -0.224 |
| A5SS | 4116 | ENSG00000084764 | MAPRE3 | chr2 | + | 27024095 | 27024297 | 27024095 | 27024252 | 27025582 | 27025737 | 20 | 6 | 12 | 13 | 19 | 4 | 2 | 1 | 3 | 1 | 2 | 24 | 2 | 12 | 8 | 5 | 0 | 1 | 7 | 2 | 194 | 149 | 0.160414434 | 0.645691828 | 0.793 | 0.697 | 0.902 | 0.769 | 0.936 | 0.235 | 1 | 0.606 | 0.568 | 0.754 | 0.187 |
| A5SS | 6934 | ENSG00000140332 | TLE3 | chr15 | - | 70058149 | 70058291 | 70058185 | 70058291 | 70057458 | 70057658 | 44 | 13 | 36 | 23 | 43 | 19 | 2 | 15 | 12 | 20 | 66 | 38 | 94 | 76 | 55 | 42 | 25 | 39 | 35 | 41 | 185 | 149 | 0.160932834 | 0.647299693 | 0.651 | 0.84 | 0.659 | 0.607 | 0.634 | 0.559 | 0.55 | 0.66 | 0.636 | 0.519 | 0.093 |
| A5SS | 5906 | ENSG00000144233 | AMMECR1L | chr2 | - | 127870761 | 127870928 | 127870813 | 127870928 | 127869453 | 127869544 | 7 | 2 | 3 | 5 | 1 | 54 | 25 | 34 | 33 | 34 | 0 | 7 | 0 | 2 | 0 | 32 | 36 | 15 | 13 | 25 | 201 | 149 | 0.161204288 | 0.647912655 | 0.088 | 0.056 | 0.061 | 0.101 | 0.021 | 0 | 0.126 | 0 | 0.102 | 0 | 0.02 |
| A5SS | 2021 | ENSG00000141279 | NPEPPS | chr17 | + | 47579389 | 47579511 | 47579389 | 47579455 | 47582741 | 47582849 | 270 | 178 | 199 | 265 | 602 | 1 | 5 | 12 | 4 | 7 | 334 | 339 | 208 | 276 | 507 | 0 | 2 | 4 | 2 | 14 | 205 | 149 | 0.16161094 | 0.649067701 | 0.995 | 0.963 | 0.923 | 0.98 | 0.984 | 1 | 0.992 | 0.974 | 0.99 | 0.963 | -0.015 |
| A5SS | 4178 | ENSG00000139636 | LMBR1L | chr12 | - | 49104718 | 49104885 | 49104745 | 49104885 | 49104447 | 49104551 | 0 | 3 | 2 | 1 | 2 | 24 | 28 | 64 | 38 | 51 | 0 | 4 | 1 | 8 | 0 | 29 | 35 | 16 | 20 | 83 | 176 | 149 | 0.161938678 | 0.649904341 | 0 | 0.083 | 0.026 | 0.022 | 0.032 | 0 | 0.088 | 0.05 | 0.253 | 0 | -0.046 |
| A5SS | 614 | ENSG00000124155 | PIGT | chr20 | + | 45419503 | 45419638 | 45419503 | 45419590 | 45420135 | 45420223 | 4 | 1 | 3 | 6 | 3 | 187 | 74 | 117 | 111 | 102 | 4 | 18 | 14 | 0 | 0 | 190 | 231 | 132 | 127 | 236 | 197 | 149 | 0.162064506 | 0.649930023 | 0.016 | 0.01 | 0.019 | 0.039 | 0.022 | 0.016 | 0.056 | 0.074 | 0 | 0 | -0.008 |
| A5SS | 4786 | ENSG00000131778 | CHD1L | chr1 | + | 147270931 | 147271113 | 147270931 | 147271005 | 147272170 | 147272281 | 22 | 14 | 18 | 10 | 10 | 42 | 17 | 41 | 38 | 38 | 3 | 19 | 12 | 5 | 23 | 27 | 31 | 21 | 33 | 100 | 257 | 149 | 0.162630765 | 0.651720636 | 0.233 | 0.323 | 0.203 | 0.132 | 0.132 | 0.061 | 0.262 | 0.249 | 0.081 | 0.118 | 0.05 |
| A5SS | 4740 | ENSG00000070882 | OSBPL3 | chr7 | - | 24834086 | 24834736 | 24834485 | 24834736 | 24830767 | 24830905 | 19 | 14 | 19 | 15 | 1 | 17 | 11 | 29 | 37 | 16 | 0 | 14 | 26 | 0 | 1 | 10 | 9 | 22 | 14 | 11 | 298 | 149 | 0.163611852 | 0.654741526 | 0.358 | 0.389 | 0.247 | 0.169 | 0.03 | 0 | 0.437 | 0.371 | 0 | 0.043 | 0.068 |
| A5SS | 1121 | ENSG00000099364 | FBXL19 | chr16 | + | 30928466 | 30928628 | 30928466 | 30928499 | 30930072 | 30930584 | 11 | 2 | 20 | 28 | 12 | 31 | 12 | 14 | 27 | 8 | 7 | 25 | 5 | 13 | 26 | 7 | 18 | 5 | 20 | 11 | 278 | 149 | 0.163709772 | 0.654741526 | 0.16 | 0.082 | 0.434 | 0.357 | 0.446 | 0.349 | 0.427 | 0.349 | 0.258 | 0.559 | -0.093 |
| A5SS | 7407 | ENSG00000182872 | RBM10 | chrX | + | 47179879 | 47180040 | 47179879 | 47180037 | 47180211 | 47180309 | 28 | 8 | 21 | 11 | 3 | 18 | 7 | 13 | 16 | 5 | 12 | 29 | 11 | 18 | 11 | 0 | 8 | 6 | 15 | 19 | 152 | 149 | 0.163745538 | 0.654741526 | 0.604 | 0.528 | 0.613 | 0.403 | 0.37 | 1 | 0.78 | 0.642 | 0.541 | 0.362 | -0.161 |
| A5SS | 7741 | ENSG00000165219 | GAPVD1 | chr9 | + | 125331925 | 125332060 | 125331925 | 125332017 | 125332509 | 125332629 | 83 | 49 | 67 | 78 | 84 | 8 | 4 | 24 | 15 | 12 | 109 | 98 | 116 | 101 | 54 | 0 | 8 | 32 | 29 | 1 | 192 | 149 | 0.16484965 | 0.658672392 | 0.89 | 0.905 | 0.684 | 0.801 | 0.845 | 1 | 0.905 | 0.738 | 0.73 | 0.977 | -0.045 |
| A5SS | 2287 | ENSG00000011243 | AKAP8L | chr19 | - | 15403652 | 15403715 | 15403657 | 15403715 | 15403474 | 15403555 | 12 | 11 | 9 | 13 | 8 | 10 | 15 | 24 | 22 | 12 | 18 | 15 | 11 | 29 | 1 | 0 | 34 | 10 | 6 | 14 | 154 | 149 | 0.165360152 | 0.660227401 | 0.537 | 0.415 | 0.266 | 0.364 | 0.392 | 1 | 0.299 | 0.516 | 0.824 | 0.065 | -0.146 |
| A5SS | 5632 | ENSG00000104218 | CSPP1 | chr8 | + | 67113804 | 67115264 | 67113804 | 67113862 | 67115913 | 67116122 | 4 | 7 | 2 | 5 | 9 | 20 | 16 | 24 | 13 | 38 | 0 | 0 | 5 | 6 | 0 | 6 | 3 | 21 | 10 | 18 | 298 | 149 | 0.166451057 | 0.664095784 | 0.091 | 0.179 | 0.04 | 0.161 | 0.106 | 0 | 0 | 0.106 | 0.231 | 0 | 0.048 |
| A5SS | 2329 | ENSG00000108039 | XPNPEP1 | chr10 | - | 109915010 | 109915099 | 109915022 | 109915099 | 109907690 | 109907815 | 62 | 47 | 47 | 66 | 54 | 12 | 0 | 0 | 0 | 0 | 88 | 69 | 40 | 119 | 49 | 0 | 0 | 0 | 8 | 0 | 161 | 149 | 0.167556755 | 0.667427533 | 0.827 | 1 | 1 | 1 | 1 | 1 | 1 | 1 | 0.932 | 1 | -0.021 |
| A5SS | 6687 | ENSG00000139737 | SLAIN1 | chr13 | + | 77744282 | 77744516 | 77744282 | 77744432 | 77746513 | 77746855 | 3 | 5 | 7 | 5 | 3 | 105 | 62 | 140 | 162 | 327 | 0 | 0 | 0 | 0 | 0 | 83 | 100 | 82 | 56 | 73 | 233 | 149 | 0.167620623 | 0.667427533 | 0.018 | 0.049 | 0.031 | 0.019 | 0.006 | 0 | 0 | 0 | 0 | 0 | 0.025 |
| A5SS | 3869 | ENSG00000095564 | BTAF1 | chr10 | + | 91959064 | 91959282 | 91959064 | 91959154 | 91959784 | 91959880 | 380 | 388 | 386 | 376 | 512 | 55 | 37 | 43 | 49 | 86 | 100 | 275 | 449 | 131 | 31 | 10 | 17 | 29 | 8 | 14 | 277 | 149 | 0.167654068 | 0.667427533 | 0.788 | 0.849 | 0.828 | 0.805 | 0.762 | 0.843 | 0.897 | 0.893 | 0.898 | 0.544 | -0.009 |
| A5SS | 4739 | ENSG00000175634 | RPS6KB2 | chr11 | + | 67432951 | 67433194 | 67432951 | 67433042 | 67433339 | 67433447 | 26 | 6 | 21 | 16 | 7 | 5 | 2 | 3 | 6 | 2 | 12 | 30 | 27 | 21 | 28 | 5 | 9 | 14 | 4 | 11 | 298 | 149 | 0.168004304 | 0.66833291 | 0.722 | 0.6 | 0.778 | 0.571 | 0.636 | 0.545 | 0.625 | 0.491 | 0.724 | 0.56 | 0.072 |
| A5SS | 1590 | ENSG00000125107 | CNOT1 | chr16 | - | 58556846 | 58556993 | 58556861 | 58556993 | 58555783 | 58555908 | 208 | 198 | 165 | 207 | 200 | 79 | 69 | 61 | 82 | 54 | 115 | 202 | 132 | 198 | 24 | 26 | 97 | 50 | 142 | 11 | 164 | 149 | 0.169322717 | 0.673085629 | 0.705 | 0.723 | 0.711 | 0.696 | 0.771 | 0.801 | 0.654 | 0.706 | 0.559 | 0.665 | 0.044 |
| A5SS | 8813 | ENSG00000123349 | PFDN5 | chr12 | + | 53296429 | 53296551 | 53296429 | 53296467 | 53297849 | 53297924 | 21 | 17 | 32 | 28 | 14 | 0 | 2 | 5 | 0 | 0 | 32 | 16 | 8 | 27 | 35 | 0 | 2 | 0 | 0 | 0 | 233 | 149 | 0.170451687 | 0.677078889 | 1 | 0.845 | 0.804 | 1 | 1 | 1 | 0.836 | 1 | 1 | 1 | -0.037 |
| A5SS | 7199 | ENSG00000143569 | UBAP2L | chr1 | + | 154234590 | 154234792 | 154234590 | 154234738 | 154235195 | 154235291 | 69 | 40 | 72 | 64 | 108 | 1 | 6 | 4 | 5 | 5 | 158 | 63 | 28 | 71 | 130 | 5 | 6 | 0 | 0 | 6 | 203 | 149 | 0.171024309 | 0.678318522 | 0.981 | 0.83 | 0.93 | 0.904 | 0.941 | 0.959 | 0.885 | 1 | 1 | 0.941 | -0.04 |
| A5SS | 4462 | ENSG00000161929 | SCIMP | chr17 | - | 5223332 | 5223456 | 5223353 | 5223456 | 5214924 | 5214998 | 5 | 6 | 3 | 15 | 2 | 2 | 2 | 0 | 0 | 0 | 15 | 11 | 3 | 4 | 3 | 0 | 0 | 0 | 0 | 8 | 170 | 149 | 0.171073118 | 0.678318522 | 0.687 | 0.724 | 1 | 1 | 1 | 1 | 1 | 1 | 1 | 0.247 | 0.033 |
| A5SS | 423 | ENSG00000127527 | EPS15L1 | chr19 | - | 16440861 | 16440909 | 16440884 | 16440909 | 16437769 | 16437865 | 73 | 29 | 69 | 116 | 101 | 3 | 0 | 4 | 5 | 5 | 108 | 90 | 79 | 125 | 260 | 4 | 0 | 8 | 2 | 25 | 172 | 149 | 0.171137694 | 0.678318522 | 0.955 | 1 | 0.937 | 0.953 | 0.946 | 0.959 | 1 | 0.895 | 0.982 | 0.9 | 0.011 |
| A5SS | 8394 | ENSG00000148498 | PARD3 | chr10 | - | 34337274 | 34337426 | 34337277 | 34337426 | 34336198 | 34336243 | 6 | 11 | 3 | 22 | 10 | 0 | 4 | 3 | 2 | 1 | 45 | 18 | 12 | 26 | 24 | 25 | 0 | 5 | 2 | 14 | 152 | 149 | 0.171388248 | 0.678817211 | 1 | 0.729 | 0.495 | 0.915 | 0.907 | 0.638 | 1 | 0.702 | 0.927 | 0.627 | 0.03 |
| A5SS | 4806 | ENSG00000159658 | EFCAB14 | chr1 | - | 46716290 | 46716443 | 46716294 | 46716443 | 46707905 | 46708051 | 5 | 0 | 1 | 3 | 2 | 219 | 103 | 187 | 197 | 331 | 11 | 6 | 0 | 0 | 0 | 166 | 175 | 150 | 147 | 99 | 153 | 149 | 0.172073745 | 0.681036595 | 0.022 | 0 | 0.005 | 0.015 | 0.006 | 0.061 | 0.032 | 0 | 0 | 0 | -0.009 |
| A5SS | 5968 | ENSG00000132196 | HSD17B7 | chr1 | + | 162797801 | 162798935 | 162797801 | 162797916 | 162799742 | 162799937 | 4 | 12 | 13 | 4 | 4 | 38 | 7 | 10 | 12 | 41 | 35 | 0 | 11 | 7 | 10 | 21 | 26 | 15 | 0 | 8 | 298 | 149 | 0.17294028 | 0.683968754 | 0.05 | 0.462 | 0.394 | 0.143 | 0.047 | 0.455 | 0 | 0.268 | 1 | 0.385 | -0.202 |
| A5SS | 1196 | ENSG00000103313 | MEFV | chr16 | - | 3244135 | 3244286 | 3244253 | 3244286 | 3243859 | 3243892 | 25 | 33 | 13 | 24 | 33 | 369 | 191 | 167 | 172 | 471 | 41 | 27 | 15 | 18 | 47 | 771 | 266 | 196 | 318 | 387 | 267 | 149 | 0.173214827 | 0.684557073 | 0.036 | 0.088 | 0.042 | 0.072 | 0.038 | 0.029 | 0.054 | 0.041 | 0.031 | 0.063 | 0.012 |
| A5SS | 6239 | ENSG00000135317 | SNX14 | chr6 | - | 85547111 | 85547226 | 85547115 | 85547226 | 85543604 | 85543760 | 148 | 90 | 81 | 136 | 197 | 13 | 2 | 8 | 12 | 15 | 41 | 65 | 61 | 53 | 26 | 5 | 17 | 5 | 4 | 0 | 153 | 149 | 0.173719038 | 0.686051526 | 0.917 | 0.978 | 0.908 | 0.917 | 0.927 | 0.889 | 0.788 | 0.922 | 0.928 | 1 | 0.024 |
| A5SS | 1410 | ENSG00000047346 | FAM214A | chr15 | - | 52587055 | 52587219 | 52587150 | 52587219 | 52584744 | 52584944 | 26 | 19 | 21 | 28 | 24 | 64 | 60 | 59 | 68 | 73 | 0 | 34 | 19 | 7 | 0 | 28 | 53 | 60 | 3 | 41 | 244 | 149 | 0.175363401 | 0.692043239 | 0.199 | 0.162 | 0.179 | 0.201 | 0.167 | 0 | 0.281 | 0.162 | 0.588 | 0 | -0.025 |
| A5SS | 8603 | ENSG00000139624 | CERS5 | chr12 | - | 50134399 | 50134702 | 50134545 | 50134702 | 50129798 | 50130694 | 22 | 8 | 8 | 22 | 22 | 16 | 16 | 16 | 27 | 20 | 37 | 31 | 54 | 62 | 11 | 24 | 31 | 19 | 15 | 51 | 295 | 149 | 0.17615315 | 0.694656119 | 0.41 | 0.202 | 0.202 | 0.292 | 0.357 | 0.438 | 0.336 | 0.589 | 0.676 | 0.098 | -0.135 |
| A5SS | 8711 | ENSG00000135297 | MTO1 | chr6 | + | 73461730 | 73462071 | 73461730 | 73461794 | 73466208 | 73466408 | 75 | 33 | 93 | 122 | 55 | 7 | 8 | 6 | 9 | 14 | 87 | 90 | 57 | 84 | 25 | 0 | 5 | 14 | 4 | 5 | 298 | 149 | 0.176349488 | 0.694926801 | 0.843 | 0.673 | 0.886 | 0.871 | 0.663 | 1 | 0.9 | 0.671 | 0.913 | 0.714 | -0.052 |
| A5SS | 2026 | ENSG00000141279 | NPEPPS | chr17 | + | 47586147 | 47586249 | 47586147 | 47586244 | 47586364 | 47586398 | 4 | 7 | 2 | 7 | 2 | 255 | 175 | 200 | 279 | 295 | 6 | 5 | 4 | 4 | 23 | 484 | 261 | 168 | 242 | 545 | 154 | 149 | 0.177206811 | 0.697799904 | 0.015 | 0.037 | 0.01 | 0.024 | 0.007 | 0.012 | 0.018 | 0.023 | 0.016 | 0.039 | -0.003 |
| A5SS | 3730 | ENSG00000119402 | FBXW2 | chr9 | - | 120792886 | 120793257 | 120793148 | 120793257 | 120787768 | 120788278 | 26 | 14 | 20 | 27 | 13 | 24 | 15 | 21 | 29 | 10 | 31 | 28 | 57 | 53 | 6 | 4 | 35 | 13 | 24 | 22 | 298 | 149 | 0.178054626 | 0.700631433 | 0.351 | 0.318 | 0.323 | 0.318 | 0.394 | 0.795 | 0.286 | 0.687 | 0.525 | 0.12 | -0.142 |
| A5SS | 1663 | ENSG00000092036 | HAUS4 | chr14 | - | 22956745 | 22957094 | 22956915 | 22957094 | 22955099 | 22955176 | 9 | 12 | 14 | 12 | 18 | 24 | 6 | 20 | 9 | 8 | 52 | 17 | 14 | 1 | 15 | 0 | 7 | 10 | 4 | 20 | 298 | 149 | 0.180011148 | 0.707818403 | 0.158 | 0.5 | 0.259 | 0.4 | 0.529 | 1 | 0.548 | 0.412 | 0.111 | 0.273 | -0.1 |
| A5SS | 7832 | ENSG00000110768 | GTF2H1 | chr11 | + | 18325946 | 18326153 | 18325946 | 18326019 | 18333059 | 18333228 | 19 | 5 | 16 | 13 | 37 | 0 | 0 | 0 | 0 | 0 | 11 | 36 | 16 | 37 | 4 | 6 | 0 | 0 | 0 | 0 | 283 | 149 | 0.18065317 | 0.709830003 | 1 | 1 | 1 | 1 | 1 | 0.491 | 1 | 1 | 1 | 1 | 0.102 |
| A5SS | 902 | ENSG00000187838 | PLSCR3 | chr17 | - | 7393143 | 7393509 | 7393466 | 7393509 | 7392790 | 7392952 | 41 | 17 | 47 | 27 | 51 | 10 | 5 | 22 | 9 | 14 | 41 | 28 | 16 | 6 | 21 | 22 | 8 | 6 | 4 | 12 | 298 | 149 | 0.18201903 | 0.714680779 | 0.672 | 0.63 | 0.516 | 0.6 | 0.646 | 0.482 | 0.636 | 0.571 | 0.429 | 0.467 | 0.096 |
| A5SS | 365 | ENSG00000104859 | CLASRP | chr19 | + | 45057749 | 45057898 | 45057749 | 45057882 | 45059267 | 45059364 | 179 | 162 | 207 | 219 | 235 | 25 | 20 | 25 | 19 | 24 | 696 | 353 | 246 | 429 | 672 | 67 | 41 | 22 | 17 | 76 | 165 | 149 | 0.182292434 | 0.715133736 | 0.866 | 0.88 | 0.882 | 0.912 | 0.898 | 0.904 | 0.886 | 0.91 | 0.958 | 0.889 | -0.022 |
| A5SS | 8712 | ENSG00000135297 | MTO1 | chr6 | + | 73461736 | 73462173 | 73461736 | 73461794 | 73466208 | 73466408 | 24 | 7 | 41 | 45 | 22 | 7 | 8 | 6 | 9 | 14 | 39 | 46 | 24 | 37 | 4 | 0 | 5 | 14 | 4 | 5 | 298 | 149 | 0.182397212 | 0.715133736 | 0.632 | 0.304 | 0.774 | 0.714 | 0.44 | 1 | 0.821 | 0.462 | 0.822 | 0.286 | -0.105 |
| A5SS | 1795 | ENSG00000160256 | FAM207A | chr21 | + | 44943683 | 44943837 | 44943683 | 44943792 | 44960099 | 44960168 | 4 | 5 | 8 | 6 | 9 | 15 | 0 | 19 | 6 | 4 | 0 | 13 | 5 | 19 | 9 | 6 | 24 | 7 | 11 | 45 | 194 | 149 | 0.183345097 | 0.718332625 | 0.17 | 1 | 0.244 | 0.434 | 0.633 | 0 | 0.294 | 0.354 | 0.57 | 0.133 | 0.226 |
| A5SS | 3351 | ENSG00000072274 | TFRC | chr3 | - | 196081830 | 196082090 | 196082042 | 196082090 | 196077063 | 196077122 | 0 | 1 | 0 | 3 | 2 | 63 | 67 | 98 | 81 | 29 | 0 | 0 | 0 | 2 | 12 | 38 | 136 | 184 | 83 | 19 | 298 | 149 | 0.18495386 | 0.723263237 | 0 | 0.007 | 0 | 0.018 | 0.033 | 0 | 0 | 0 | 0.012 | 0.24 | -0.039 |
| A5SS | 1729 | ENSG00000100938 | GMPR2 | chr14 | + | 24232421 | 24232847 | 24232421 | 24232788 | 24233218 | 24233340 | 21 | 6 | 11 | 24 | 12 | 22 | 7 | 29 | 25 | 31 | 5 | 25 | 26 | 12 | 27 | 9 | 36 | 14 | 29 | 6 | 208 | 149 | 0.184999715 | 0.723263237 | 0.406 | 0.38 | 0.214 | 0.407 | 0.217 | 0.285 | 0.332 | 0.571 | 0.229 | 0.763 | -0.111 |
| A5SS | 5117 | ENSG00000167595 | PROSER3 | chr19 | + | 35762252 | 35762477 | 35762252 | 35762356 | 35764853 | 35764936 | 7 | 3 | 3 | 2 | 6 | 13 | 4 | 5 | 5 | 8 | 9 | 1 | 4 | 3 | 5 | 13 | 22 | 11 | 14 | 7 | 270 | 149 | 0.185108508 | 0.723263237 | 0.229 | 0.293 | 0.249 | 0.181 | 0.293 | 0.276 | 0.024 | 0.167 | 0.106 | 0.283 | 0.078 |
| A5SS | 5081 | ENSG00000137497 | NUMA1 | chr11 | - | 72068078 | 72068231 | 72068104 | 72068231 | 72035901 | 72035975 | 1 | 0 | 0 | 0 | 0 | 6 | 1 | 2 | 2 | 3 | 0 | 0 | 4 | 0 | 0 | 8 | 5 | 0 | 6 | 5 | 175 | 149 | 0.185218043 | 0.723263237 | 0.124 | 0 | 0 | 0 | 0 | 0 | 0 | 1 | 0 | 0 | -0.175 |
| A5SS | 5659 | ENSG00000164548 | TRA2A | chr7 | - | 23522131 | 23522432 | 23522353 | 23522432 | 23521706 | 23521840 | 267 | 274 | 233 | 420 | 331 | 3 | 4 | 4 | 2 | 11 | 261 | 479 | 674 | 546 | 62 | 5 | 1 | 6 | 2 | 6 | 298 | 149 | 0.185268091 | 0.723263237 | 0.978 | 0.972 | 0.967 | 0.991 | 0.938 | 0.963 | 0.996 | 0.983 | 0.993 | 0.838 | 0.015 |
| A5SS | 2226 | ENSG00000120705 | ETF1 | chr5 | - | 138542633 | 138542936 | 138542832 | 138542936 | 138518691 | 138518867 | 30 | 23 | 35 | 18 | 12 | 75 | 53 | 68 | 108 | 89 | 13 | 33 | 18 | 64 | 25 | 39 | 107 | 117 | 113 | 9 | 298 | 149 | 0.185590055 | 0.724000774 | 0.167 | 0.178 | 0.205 | 0.077 | 0.063 | 0.143 | 0.134 | 0.071 | 0.221 | 0.581 | -0.092 |
| A5SS | 3303 | ENSG00000172175 | MALT1 | chr18 | + | 58734306 | 58734566 | 58734306 | 58734381 | 58735201 | 58735329 | 24 | 25 | 31 | 26 | 32 | 204 | 192 | 242 | 223 | 398 | 0 | 13 | 25 | 12 | 20 | 51 | 88 | 175 | 68 | 0 | 298 | 149 | 0.186500034 | 0.727029504 | 0.056 | 0.061 | 0.06 | 0.055 | 0.039 | 0 | 0.069 | 0.067 | 0.081 | 1 | -0.189 |
| A5SS | 2138 | ENSG00000166888 | STAT6 | chr12 | - | 57110042 | 57110110 | 57110076 | 57110110 | 57108162 | 57108299 | 37 | 31 | 36 | 64 | 66 | 1 | 2 | 5 | 1 | 0 | 58 | 93 | 54 | 64 | 140 | 0 | 1 | 0 | 0 | 0 | 183 | 149 | 0.18667093 | 0.727174803 | 0.968 | 0.927 | 0.854 | 0.981 | 1 | 1 | 0.987 | 1 | 1 | 1 | -0.051 |
| A5SS | 3251 | ENSG00000178922 | HYI | chr1 | - | 43451409 | 43451544 | 43451427 | 43451544 | 43451153 | 43451311 | 3 | 3 | 4 | 1 | 3 | 1 | 2 | 9 | 6 | 1 | 0 | 1 | 5 | 0 | 2 | 7 | 16 | 0 | 1 | 0 | 167 | 149 | 0.187244802 | 0.728888563 | 0.728 | 0.572 | 0.284 | 0.129 | 0.728 | 0 | 0.053 | 1 | 0 | 1 | 0.078 |
| A5SS | 5239 | ENSG00000152689 | RASGRP3 | chr2 | + | 33539093 | 33539223 | 33539093 | 33539210 | 33543511 | 33543627 | 1 | 3 | 6 | 10 | 2 | 13 | 3 | 14 | 7 | 10 | 20 | 31 | 2 | 8 | 2 | 22 | 29 | 14 | 14 | 3 | 262 | 149 | 0.187546059 | 0.729173322 | 0.042 | 0.363 | 0.196 | 0.448 | 0.102 | 0.341 | 0.378 | 0.075 | 0.245 | 0.275 | -0.033 |
| A5SS | 1151 | ENSG00000188603 | CLN3 | chr16 | - | 28491481 | 28491835 | 28491713 | 28491835 | 28489289 | 28489386 | 57 | 24 | 35 | 37 | 39 | 2 | 0 | 6 | 4 | 1 | 20 | 79 | 33 | 36 | 112 | 9 | 3 | 4 | 1 | 10 | 298 | 149 | 0.187585934 | 0.729173322 | 0.934 | 1 | 0.745 | 0.822 | 0.951 | 0.526 | 0.929 | 0.805 | 0.947 | 0.848 | 0.079 |
| A5SS | 329 | ENSG00000142230 | SAE1 | chr19 | + | 47130888 | 47131112 | 47130888 | 47131028 | 47143493 | 47143605 | 6 | 0 | 5 | 4 | 0 | 86 | 57 | 103 | 94 | 46 | 0 | 4 | 5 | 3 | 17 | 154 | 82 | 63 | 118 | 136 | 233 | 149 | 0.188106454 | 0.73067475 | 0.043 | 0 | 0.03 | 0.026 | 0 | 0 | 0.03 | 0.048 | 0.016 | 0.074 | -0.014 |
| A5SS | 3774 | ENSG00000104419 | NDRG1 | chr8 | - | 133279965 | 133280267 | 133280231 | 133280267 | 133264546 | 133264652 | 11 | 11 | 7 | 6 | 13 | 49 | 29 | 47 | 37 | 26 | 18 | 12 | 9 | 11 | 23 | 34 | 41 | 33 | 59 | 48 | 298 | 149 | 0.188575302 | 0.73197346 | 0.101 | 0.159 | 0.069 | 0.075 | 0.2 | 0.209 | 0.128 | 0.12 | 0.085 | 0.193 | -0.026 |
| A5SS | 1765 | ENSG00000100664 | EIF5 | chr14 | + | 103339171 | 103339803 | 103339171 | 103339333 | 103340426 | 103340561 | 237 | 228 | 300 | 373 | 543 | 7 | 2 | 2 | 3 | 2 | 178 | 266 | 337 | 190 | 25 | 1 | 0 | 2 | 6 | 7 | 298 | 149 | 0.188848635 | 0.732511955 | 0.944 | 0.983 | 0.987 | 0.984 | 0.993 | 0.989 | 1 | 0.988 | 0.941 | 0.641 | 0.066 |
| A5SS | 7703 | ENSG00000188283 | ZNF383 | chr19 | + | 37235548 | 37235758 | 37235548 | 37235675 | 37235978 | 37236074 | 7 | 3 | 5 | 10 | 20 | 28 | 26 | 18 | 38 | 22 | 21 | 12 | 8 | 5 | 0 | 5 | 25 | 20 | 7 | 7 | 232 | 149 | 0.190443542 | 0.738172191 | 0.138 | 0.069 | 0.151 | 0.145 | 0.369 | 0.73 | 0.236 | 0.204 | 0.314 | 0 | -0.122 |
| A5SS | 416 | ENSG00000181896 | ZNF101 | chr19 | + | 19677863 | 19677990 | 19677863 | 19677986 | 19678725 | 19678786 | 77 | 48 | 147 | 118 | 141 | 8 | 13 | 19 | 9 | 18 | 40 | 99 | 45 | 56 | 33 | 12 | 20 | 5 | 8 | 4 | 153 | 149 | 0.19149117 | 0.741704588 | 0.904 | 0.782 | 0.883 | 0.927 | 0.884 | 0.764 | 0.828 | 0.898 | 0.872 | 0.889 | 0.026 |
| A5SS | 1272 | ENSG00000261740 | BOLA2-SMG1P6 | chr16 | - | 29452108 | 29454113 | 29454015 | 29454113 | 29446801 | 29447026 | 0 | 0 | 2 | 0 | 0 | 13 | 8 | 9 | 16 | 10 | 0 | 0 | 1 | 7 | 0 | 9 | 1 | 14 | 10 | 28 | 298 | 149 | 0.192420086 | 0.744772479 | 0 | 0 | 0.1 | 0 | 0 | 0 | 0 | 0.034 | 0.259 | 0 | -0.039 |
| A5SS | 7795 | ENSG00000130714 | POMT1 | chr9 | + | 131511336 | 131511472 | 131511336 | 131511467 | 131512040 | 131512136 | 5 | 0 | 0 | 5 | 6 | 19 | 9 | 20 | 12 | 10 | 0 | 0 | 2 | 2 | 10 | 10 | 36 | 27 | 28 | 9 | 154 | 149 | 0.193336737 | 0.747788572 | 0.203 | 0 | 0 | 0.287 | 0.367 | 0 | 0 | 0.067 | 0.065 | 0.518 | 0.041 |
| A5SS | 6093 | ENSG00000118418 | HMGN3 | chr6 | - | 79202275 | 79202389 | 79202316 | 79202389 | 79201244 | 79201726 | 99 | 79 | 69 | 59 | 51 | 35 | 29 | 31 | 35 | 39 | 2 | 38 | 70 | 12 | 22 | 7 | 12 | 20 | 38 | 13 | 190 | 149 | 0.194004999 | 0.749840344 | 0.689 | 0.681 | 0.636 | 0.569 | 0.506 | 0.183 | 0.713 | 0.733 | 0.198 | 0.57 | 0.137 |
| A5SS | 7596 | ENSG00000095303 | PTGS1 | chr9 | + | 122386445 | 122386732 | 122386445 | 122386621 | 122390197 | 122390345 | 110 | 64 | 20 | 150 | 42 | 2 | 3 | 1 | 11 | 2 | 271 | 196 | 67 | 128 | 2151 | 41 | 10 | 0 | 0 | 122 | 260 | 149 | 0.195013102 | 0.752681049 | 0.969 | 0.924 | 0.92 | 0.887 | 0.923 | 0.791 | 0.918 | 1 | 1 | 0.91 | 0.001 |
| A5SS | 7855 | ENSG00000110514 | MADD | chr11 | + | 47295495 | 47295576 | 47295495 | 47295567 | 47295896 | 47296055 | 46 | 21 | 33 | 45 | 39 | 41 | 20 | 20 | 63 | 33 | 46 | 37 | 30 | 37 | 120 | 39 | 54 | 14 | 9 | 84 | 158 | 149 | 0.195016589 | 0.752681049 | 0.514 | 0.498 | 0.609 | 0.402 | 0.527 | 0.527 | 0.393 | 0.669 | 0.795 | 0.574 | -0.082 |
| A5SS | 1154 | ENSG00000121281 | ADCY7 | chr16 | + | 50266550 | 50266680 | 50266550 | 50266587 | 50287911 | 50288350 | 19 | 10 | 8 | 24 | 5 | 16 | 9 | 9 | 9 | 13 | 0 | 6 | 9 | 8 | 0 | 6 | 4 | 1 | 7 | 14 | 242 | 149 | 0.19752018 | 0.761803559 | 0.422 | 0.406 | 0.354 | 0.621 | 0.191 | 0 | 0.48 | 0.847 | 0.413 | 0 | 0.051 |
| A5SS | 2024 | ENSG00000141279 | NPEPPS | chr17 | + | 47585499 | 47585745 | 47585499 | 47585700 | 47586147 | 47586244 | 19 | 15 | 28 | 22 | 21 | 237 | 178 | 183 | 253 | 303 | 18 | 41 | 52 | 32 | 31 | 509 | 243 | 149 | 263 | 551 | 194 | 149 | 0.197768149 | 0.762219737 | 0.058 | 0.061 | 0.105 | 0.063 | 0.051 | 0.026 | 0.115 | 0.211 | 0.085 | 0.041 | -0.028 |
| A5SS | 5318 | ENSG00000137817 | PARP6 | chr15 | - | 72260477 | 72260688 | 72260537 | 72260688 | 72259607 | 72259661 | 21 | 10 | 26 | 18 | 42 | 30 | 12 | 40 | 44 | 22 | 67 | 54 | 36 | 50 | 17 | 42 | 56 | 29 | 46 | 29 | 209 | 149 | 0.198399754 | 0.763168643 | 0.333 | 0.373 | 0.317 | 0.226 | 0.576 | 0.532 | 0.407 | 0.469 | 0.437 | 0.295 | -0.063 |
| A5SS | 7443 | ENSG00000168522 | FNTA | chr8 | + | 43077215 | 43080441 | 43077215 | 43077364 | 43084709 | 43084881 | 111 | 90 | 79 | 90 | 129 | 3 | 1 | 0 | 1 | 5 | 49 | 74 | 97 | 91 | 11 | 0 | 0 | 2 | 0 | 0 | 298 | 149 | 0.198413406 | 0.763168643 | 0.949 | 0.978 | 1 | 0.978 | 0.928 | 1 | 1 | 0.96 | 1 | 1 | -0.025 |
| A5SS | 2761 | ENSG00000163703 | CRELD1 | chr3 | + | 9943076 | 9943253 | 9943076 | 9943172 | 9943380 | 9943515 | 2 | 1 | 2 | 1 | 2 | 32 | 12 | 30 | 41 | 14 | 0 | 0 | 0 | 5 | 0 | 39 | 22 | 18 | 51 | 82 | 230 | 149 | 0.198435066 | 0.763168643 | 0.039 | 0.051 | 0.041 | 0.016 | 0.085 | 0 | 0 | 0 | 0.06 | 0 | 0.034 |
| A5SS | 2375 | ENSG00000125089 | SH3TC1 | chr4 | + | 8219334 | 8219530 | 8219334 | 8219449 | 8222839 | 8222970 | 56 | 36 | 19 | 35 | 12 | 5 | 1 | 0 | 8 | 1 | 97 | 56 | 58 | 49 | 184 | 7 | 7 | 8 | 6 | 23 | 230 | 149 | 0.199080684 | 0.765110933 | 0.879 | 0.959 | 1 | 0.739 | 0.886 | 0.9 | 0.838 | 0.824 | 0.841 | 0.838 | 0.044 |
| A5SS | 6141 | ENSG00000100316 | RPL3 | chr22 | - | 39319389 | 39319623 | 39319594 | 39319623 | 39318399 | 39318592 | 13 | 4 | 9 | 10 | 14 | 1132 | 450 | 1434 | 1307 | 1245 | 42 | 2 | 16 | 6 | 5 | 522 | 792 | 620 | 407 | 1327 | 298 | 149 | 0.201714588 | 0.77468651 | 0.006 | 0.004 | 0.003 | 0.004 | 0.006 | 0.039 | 0.001 | 0.013 | 0.007 | 0.002 | -0.008 |
| A5SS | 5710 | ENSG00000163516 | ANKZF1 | chr2 | + | 219230227 | 219230405 | 219230227 | 219230350 | 219231927 | 219232040 | 58 | 39 | 74 | 66 | 25 | 14 | 4 | 21 | 8 | 11 | 28 | 43 | 22 | 23 | 26 | 0 | 17 | 1 | 13 | 0 | 204 | 149 | 0.202480298 | 0.777078832 | 0.752 | 0.877 | 0.72 | 0.858 | 0.624 | 1 | 0.649 | 0.941 | 0.564 | 1 | -0.065 |
| A5SS | 3548 | ENSG00000116001 | TIA1 | chr2 | - | 70216752 | 70216994 | 70216885 | 70216994 | 70216403 | 70216499 | 98 | 110 | 115 | 90 | 112 | 109 | 92 | 128 | 138 | 175 | 14 | 55 | 71 | 51 | 2 | 31 | 121 | 38 | 49 | 29 | 282 | 149 | 0.203158458 | 0.779132015 | 0.322 | 0.387 | 0.322 | 0.256 | 0.253 | 0.193 | 0.194 | 0.497 | 0.355 | 0.035 | 0.053 |
| A5SS | 5483 | ENSG00000114942 | EEF1B2 | chr2 | + | 206159933 | 206160059 | 206159933 | 206160018 | 206160556 | 206160710 | 34 | 10 | 14 | 29 | 15 | 0 | 0 | 0 | 0 | 0 | 47 | 48 | 15 | 11 | 56 | 0 | 0 | 1 | 6 | 0 | 190 | 149 | 0.204806682 | 0.784538128 | 1 | 1 | 1 | 1 | 1 | 1 | 1 | 0.922 | 0.59 | 1 | 0.098 |
| A5SS | 206 | ENSG00000105723 | GSK3A | chr19 | - | 42232929 | 42233205 | 42233109 | 42233205 | 42232495 | 42232682 | 9 | 12 | 23 | 10 | 4 | 157 | 95 | 115 | 177 | 344 | 30 | 26 | 7 | 25 | 10 | 212 | 241 | 145 | 171 | 440 | 298 | 149 | 0.204955096 | 0.784538128 | 0.028 | 0.059 | 0.091 | 0.027 | 0.006 | 0.066 | 0.051 | 0.024 | 0.068 | 0.011 | -0.002 |
| A5SS | 5452 | ENSG00000018280 | SLC11A1 | chr2 | + | 218382959 | 218383102 | 218382959 | 218383070 | 218384318 | 218384365 | 29 | 20 | 6 | 19 | 27 | 0 | 0 | 0 | 0 | 0 | 77 | 56 | 58 | 46 | 82 | 10 | 0 | 0 | 0 | 0 | 181 | 149 | 0.205000591 | 0.784538128 | 1 | 1 | 1 | 1 | 1 | 0.864 | 1 | 1 | 1 | 1 | 0.027 |
| A5SS | 5715 | ENSG00000187736 | NHEJ1 | chr2 | - | 219078073 | 219078206 | 219078088 | 219078206 | 219077245 | 219077364 | 7 | 3 | 1 | 1 | 5 | 86 | 50 | 31 | 49 | 63 | 0 | 4 | 0 | 0 | 5 | 27 | 53 | 24 | 46 | 42 | 164 | 149 | 0.205264018 | 0.784994226 | 0.069 | 0.052 | 0.028 | 0.018 | 0.067 | 0 | 0.064 | 0 | 0 | 0.098 | 0.014 |
| A5SS | 3496 | ENSG00000079277 | MKNK1 | chr1 | - | 46604184 | 46604294 | 46604226 | 46604294 | 46594112 | 46594280 | 14 | 8 | 15 | 10 | 1 | 3 | 5 | 10 | 6 | 0 | 43 | 31 | 18 | 18 | 33 | 39 | 18 | 11 | 8 | 34 | 191 | 149 | 0.20634595 | 0.788577712 | 0.785 | 0.555 | 0.539 | 0.565 | 1 | 0.462 | 0.573 | 0.561 | 0.637 | 0.431 | 0.156 |
| A5SS | 5451 | ENSG00000018280 | SLC11A1 | chr2 | + | 218382959 | 218383102 | 218382959 | 218383070 | 218384242 | 218384365 | 105 | 66 | 31 | 81 | 84 | 1 | 2 | 2 | 1 | 5 | 131 | 173 | 136 | 106 | 221 | 0 | 7 | 0 | 0 | 4 | 181 | 149 | 0.206775642 | 0.789665293 | 0.989 | 0.964 | 0.927 | 0.985 | 0.933 | 1 | 0.953 | 1 | 1 | 0.978 | -0.027 |
| A5SS | 6754 | ENSG00000234741 | GAS5 | chr1 | - | 173865228 | 173865547 | 173865509 | 173865547 | 173864256 | 173864304 | 122 | 87 | 149 | 110 | 215 | 1 | 0 | 9 | 6 | 3 | 48 | 85 | 94 | 82 | 34 | 0 | 0 | 0 | 0 | 8 | 298 | 149 | 0.207082511 | 0.790282625 | 0.984 | 1 | 0.892 | 0.902 | 0.973 | 1 | 1 | 1 | 1 | 0.68 | 0.014 |
| A5SS | 2554 | ENSG00000127419 | TMEM175 | chr4 | + | 948115 | 948615 | 948115 | 948154 | 950420 | 950518 | 22 | 10 | 24 | 36 | 16 | 12 | 8 | 13 | 8 | 1 | 37 | 41 | 10 | 45 | 30 | 38 | 24 | 13 | 9 | 16 | 298 | 149 | 0.208027475 | 0.79333253 | 0.478 | 0.385 | 0.48 | 0.692 | 0.889 | 0.327 | 0.461 | 0.278 | 0.714 | 0.484 | 0.132 |
| A5SS | 1 | ENSG00000160285 | LSS | chr21 | - | 46222629 | 46222738 | 46222662 | 46222738 | 46221853 | 46221975 | 39 | 1 | 18 | 17 | 23 | 4 | 5 | 3 | 3 | 0 | 20 | 25 | 27 | 5 | 28 | 0 | 0 | 1 | 4 | 5 | 182 | 149 | 0.208570355 | 0.794845848 | 0.889 | 0.141 | 0.831 | 0.823 | 1 | 1 | 1 | 0.957 | 0.506 | 0.821 | -0.12 |
| A5SS | 7644 | ENSG00000136811 | ODF2 | chr9 | + | 128496040 | 128496297 | 128496040 | 128496141 | 128498412 | 128498575 | 14 | 10 | 11 | 6 | 10 | 100 | 62 | 67 | 67 | 67 | 9 | 20 | 11 | 9 | 11 | 200 | 124 | 36 | 85 | 145 | 298 | 149 | 0.208860177 | 0.79539334 | 0.065 | 0.075 | 0.076 | 0.043 | 0.069 | 0.022 | 0.075 | 0.133 | 0.05 | 0.037 | 0.002 |
| A5SS | 899 | ENSG00000063244 | U2AF2 | chr19 | + | 55669082 | 55669181 | 55669082 | 55669169 | 55669443 | 55669692 | 17 | 18 | 21 | 31 | 7 | 157 | 60 | 132 | 129 | 69 | 63 | 31 | 26 | 19 | 8 | 151 | 139 | 62 | 111 | 166 | 161 | 149 | 0.209256509 | 0.796345401 | 0.091 | 0.217 | 0.128 | 0.182 | 0.086 | 0.279 | 0.171 | 0.28 | 0.137 | 0.043 | -0.041 |
| A5SS | 1791 | ENSG00000213983 | AP1G2 | chr14 | - | 23565815 | 23566160 | 23566059 | 23566160 | 23565605 | 23565701 | 152 | 63 | 121 | 166 | 94 | 5 | 0 | 5 | 2 | 3 | 230 | 154 | 139 | 157 | 276 | 0 | 2 | 2 | 0 | 10 | 298 | 149 | 0.209521689 | 0.796797366 | 0.938 | 1 | 0.924 | 0.976 | 0.94 | 1 | 0.975 | 0.972 | 1 | 0.932 | -0.02 |
| A5SS | 3363 | ENSG00000132535 | DLG4 | chr17 | - | 7208074 | 7208239 | 7208173 | 7208239 | 7204198 | 7204252 | 2 | 1 | 0 | 4 | 0 | 7 | 3 | 2 | 2 | 11 | 20 | 0 | 0 | 1 | 10 | 14 | 14 | 13 | 23 | 6 | 248 | 149 | 0.209725295 | 0.797014703 | 0.147 | 0.167 | 0 | 0.546 | 0 | 0.462 | 0 | 0 | 0.025 | 0.5 | -0.025 |
| A5SS | 6170 | ENSG00000135486 | HNRNPA1 | chr12 | + | 54282799 | 54282841 | 54282799 | 54282823 | 54283826 | 54283967 | 150 | 120 | 154 | 154 | 160 | 1 | 1 | 1 | 1 | 1 | 111 | 149 | 233 | 184 | 58 | 14 | 0 | 1 | 0 | 0 | 167 | 149 | 0.211242471 | 0.802177373 | 0.993 | 0.991 | 0.993 | 0.993 | 0.993 | 0.876 | 1 | 0.995 | 1 | 1 | 0.018 |
| A5SS | 4051 | ENSG00000066279 | ASPM | chr1 | - | 197121866 | 197122043 | 197121914 | 197122043 | 197117788 | 197117983 | 0 | 0 | 2 | 0 | 8 | 1 | 6 | 8 | 14 | 13 | 0 | 7 | 0 | 0 | 0 | 2 | 0 | 3 | 1 | 12 | 197 | 149 | 0.211378602 | 0.802177373 | 0 | 0 | 0.159 | 0 | 0.318 | 0 | 1 | 0 | 0 | 0 | -0.105 |
| A5SS | 7474 | ENSG00000139546 | TARBP2 | chr12 | + | 53503026 | 53503212 | 53503026 | 53503129 | 53503712 | 53503808 | 4 | 0 | 8 | 8 | 0 | 11 | 1 | 12 | 9 | 6 | 7 | 7 | 5 | 6 | 0 | 9 | 13 | 1 | 0 | 11 | 232 | 149 | 0.211859784 | 0.803443168 | 0.189 | 0 | 0.3 | 0.363 | 0 | 0.333 | 0.257 | 0.763 | 1 | 0 | -0.3 |
| A5SS | 7012 | ENSG00000244754 | N4BP2L2 | chr13 | - | 32536932 | 32537027 | 32537005 | 32537027 | 32527407 | 32527532 | 280 | 276 | 236 | 279 | 297 | 11 | 6 | 8 | 0 | 0 | 75 | 178 | 185 | 82 | 37 | 0 | 0 | 2 | 1 | 6 | 222 | 149 | 0.212364572 | 0.803729754 | 0.945 | 0.969 | 0.952 | 1 | 1 | 1 | 1 | 0.984 | 0.982 | 0.805 | 0.019 |
| A5SS | 6439 | ENSG00000111667 | USP5 | chr12 | + | 6863185 | 6863377 | 6863185 | 6863308 | 6863829 | 6863973 | 12 | 10 | 9 | 6 | 7 | 29 | 14 | 53 | 41 | 16 | 0 | 12 | 17 | 2 | 4 | 22 | 45 | 22 | 11 | 60 | 218 | 149 | 0.212365367 | 0.803729754 | 0.22 | 0.328 | 0.104 | 0.091 | 0.23 | 0 | 0.154 | 0.346 | 0.111 | 0.044 | 0.064 |
| A5SS | 6199 | ENSG00000164674 | SYTL3 | chr6 | + | 158662749 | 158663378 | 158662749 | 158662812 | 158665394 | 158665613 | 153 | 44 | 65 | 82 | 94 | 17 | 24 | 9 | 5 | 1 | 120 | 175 | 264 | 91 | 110 | 9 | 13 | 19 | 3 | 14 | 298 | 149 | 0.212378425 | 0.803729754 | 0.818 | 0.478 | 0.783 | 0.891 | 0.979 | 0.87 | 0.871 | 0.874 | 0.938 | 0.797 | -0.08 |
| A5SS | 1160 | ENSG00000005844 | ITGAL | chr16 | + | 30489253 | 30492069 | 30489253 | 30489386 | 30494211 | 30494363 | 35 | 23 | 26 | 18 | 6 | 1344 | 772 | 783 | 1052 | 799 | 42 | 39 | 14 | 36 | 74 | 1956 | 1590 | 937 | 1197 | 1906 | 298 | 149 | 0.21267019 | 0.804274616 | 0.013 | 0.015 | 0.016 | 0.008 | 0.004 | 0.011 | 0.012 | 0.007 | 0.015 | 0.019 | -0.002 |
| A5SS | 4726 | ENSG00000173846 | PLK3 | chr1 | + | 44804338 | 44804779 | 44804338 | 44804501 | 44805265 | 44805379 | 33 | 8 | 24 | 22 | 22 | 8 | 2 | 19 | 18 | 8 | 60 | 33 | 52 | 42 | 44 | 47 | 16 | 27 | 45 | 26 | 298 | 149 | 0.213106516 | 0.805365043 | 0.673 | 0.667 | 0.387 | 0.379 | 0.579 | 0.39 | 0.508 | 0.491 | 0.318 | 0.458 | 0.104 |
| A5SS | 714 | ENSG00000004975 | DVL2 | chr17 | - | 7230045 | 7230155 | 7230057 | 7230155 | 7229807 | 7229943 | 17 | 14 | 13 | 29 | 23 | 24 | 11 | 24 | 11 | 5 | 33 | 26 | 4 | 21 | 13 | 22 | 12 | 18 | 19 | 39 | 161 | 149 | 0.214047159 | 0.808358528 | 0.396 | 0.541 | 0.334 | 0.709 | 0.81 | 0.581 | 0.667 | 0.171 | 0.506 | 0.236 | 0.126 |
| A5SS | 789 | ENSG00000185624 | P4HB | chr17 | - | 81859810 | 81860535 | 81860326 | 81860535 | 81859180 | 81859387 | 3 | 4 | 5 | 1 | 1 | 109 | 39 | 66 | 69 | 47 | 7 | 2 | 6 | 3 | 10 | 94 | 120 | 64 | 47 | 182 | 298 | 149 | 0.214841254 | 0.810794802 | 0.014 | 0.049 | 0.036 | 0.007 | 0.011 | 0.036 | 0.008 | 0.045 | 0.031 | 0.027 | -0.006 |
| A5SS | 1938 | ENSG00000135506 | OS9 | chr12 | + | 57718145 | 57718421 | 57718145 | 57718376 | 57718992 | 57719182 | 746 | 539 | 624 | 766 | 1251 | 78 | 60 | 52 | 50 | 161 | 2330 | 1129 | 1024 | 1523 | 2108 | 146 | 137 | 58 | 182 | 192 | 194 | 149 | 0.217453903 | 0.819678815 | 0.88 | 0.873 | 0.902 | 0.922 | 0.856 | 0.925 | 0.864 | 0.931 | 0.865 | 0.894 | -0.009 |
| A5SS | 6362 | ENSG00000083168 | KAT6A | chr8 | - | 42049575 | 42049707 | 42049619 | 42049707 | 42048377 | 42049302 | 119 | 97 | 92 | 74 | 66 | 13 | 18 | 18 | 7 | 13 | 57 | 152 | 112 | 121 | 42 | 15 | 18 | 22 | 26 | 11 | 193 | 149 | 0.21760764 | 0.819678815 | 0.876 | 0.806 | 0.798 | 0.891 | 0.797 | 0.746 | 0.867 | 0.797 | 0.782 | 0.747 | 0.046 |
| A5SS | 2744 | ENSG00000047849 | MAP4 | chr3 | - | 47870812 | 47871105 | 47870857 | 47871105 | 47867245 | 47867338 | 14 | 4 | 7 | 11 | 8 | 0 | 0 | 0 | 0 | 0 | 7 | 12 | 5 | 8 | 18 | 0 | 4 | 2 | 0 | 0 | 194 | 149 | 0.217647168 | 0.819678815 | 1 | 1 | 1 | 1 | 1 | 1 | 0.697 | 0.658 | 1 | 1 | 0.129 |
| A5SS | 6413 | ENSG00000134291 | TMEM106C | chr12 | + | 47963595 | 47963704 | 47963595 | 47963633 | 47964208 | 47964423 | 8 | 5 | 3 | 1 | 2 | 8 | 13 | 8 | 11 | 7 | 7 | 2 | 0 | 0 | 23 | 20 | 11 | 11 | 48 | 37 | 220 | 149 | 0.218014281 | 0.820493581 | 0.404 | 0.207 | 0.203 | 0.058 | 0.162 | 0.192 | 0.11 | 0 | 0 | 0.296 | 0.087 |
| A5SS | 2685 | ENSG00000258315 | C17orf49 | chr17 | + | 7016494 | 7016685 | 7016494 | 7016525 | 7016909 | 7017014 | 368 | 189 | 276 | 299 | 346 | 5 | 3 | 7 | 3 | 8 | 410 | 356 | 192 | 234 | 454 | 0 | 3 | 3 | 5 | 28 | 298 | 149 | 0.218322282 | 0.821084907 | 0.974 | 0.969 | 0.952 | 0.98 | 0.956 | 1 | 0.983 | 0.97 | 0.959 | 0.89 | 0.006 |
| A5SS | 1102 | ENSG00000074356 | NCBP3 | chr17 | - | 3823792 | 3825050 | 3824941 | 3825050 | 3821952 | 3822052 | 25 | 14 | 23 | 14 | 20 | 116 | 124 | 90 | 115 | 173 | 46 | 6 | 15 | 63 | 3 | 185 | 76 | 115 | 226 | 59 | 298 | 149 | 0.219622067 | 0.825040646 | 0.097 | 0.053 | 0.113 | 0.057 | 0.055 | 0.111 | 0.038 | 0.061 | 0.122 | 0.025 | 0.004 |
| A5SS | 1546 | ENSG00000140326 | CDAN1 | chr15 | - | 42727620 | 42727769 | 42727667 | 42727769 | 42726309 | 42726417 | 8 | 2 | 7 | 12 | 4 | 1 | 1 | 1 | 4 | 0 | 21 | 3 | 8 | 4 | 25 | 0 | 0 | 2 | 0 | 11 | 196 | 149 | 0.219677305 | 0.825040646 | 0.859 | 0.603 | 0.842 | 0.695 | 1 | 1 | 1 | 0.753 | 1 | 0.633 | -0.077 |
| A5SS | 529 | ENSG00000101052 | IFT52 | chr20 | + | 43605001 | 43605216 | 43605001 | 43605073 | 43613849 | 43613976 | 4 | 4 | 5 | 4 | 15 | 26 | 32 | 39 | 36 | 70 | 4 | 0 | 5 | 3 | 3 | 12 | 43 | 21 | 38 | 11 | 292 | 149 | 0.219864092 | 0.825172683 | 0.073 | 0.06 | 0.061 | 0.054 | 0.099 | 0.145 | 0 | 0.108 | 0.039 | 0.122 | -0.013 |
| A5SS | 712 | ENSG00000270647 | TAF15 | chr17 | + | 35820163 | 35820248 | 35820163 | 35820239 | 35820331 | 35820437 | 531 | 332 | 458 | 459 | 505 | 83 | 78 | 77 | 85 | 129 | 327 | 430 | 348 | 502 | 302 | 125 | 89 | 42 | 136 | 62 | 158 | 149 | 0.220772239 | 0.828010011 | 0.858 | 0.801 | 0.849 | 0.836 | 0.787 | 0.712 | 0.82 | 0.887 | 0.777 | 0.821 | 0.023 |
| A5SS | 3932 | ENSG00000182134 | TDRKH | chr1 | - | 151779950 | 151780140 | 151780050 | 151780140 | 151779102 | 151779242 | 49 | 47 | 74 | 77 | 88 | 3 | 0 | 3 | 11 | 0 | 96 | 44 | 34 | 56 | 87 | 7 | 1 | 4 | 0 | 0 | 249 | 149 | 0.221009037 | 0.82832726 | 0.907 | 1 | 0.937 | 0.807 | 1 | 0.891 | 0.963 | 0.836 | 1 | 1 | -0.008 |
| A5SS | 2572 | ENSG00000203875 | SNHG5 | chr6 | - | 85678402 | 85678748 | 85678693 | 85678748 | 85678135 | 85678188 | 9 | 1 | 20 | 7 | 3 | 43 | 62 | 47 | 32 | 27 | 12 | 2 | 8 | 4 | 10 | 23 | 35 | 55 | 75 | 90 | 298 | 149 | 0.221417929 | 0.829288624 | 0.095 | 0.008 | 0.175 | 0.099 | 0.053 | 0.207 | 0.028 | 0.068 | 0.026 | 0.053 | 0.01 |
| A5SS | 5128 | ENSG00000112855 | HARS2 | chr5 | + | 140691794 | 140692329 | 140691794 | 140691937 | 140693590 | 140693665 | 19 | 14 | 10 | 22 | 11 | 2 | 4 | 0 | 0 | 0 | 20 | 30 | 26 | 25 | 7 | 0 | 0 | 0 | 0 | 0 | 298 | 149 | 0.22283513 | 0.83381688 | 0.826 | 0.636 | 1 | 1 | 1 | 1 | 1 | 1 | 1 | 1 | -0.108 |
| A5SS | 4652 | ENSG00000019582 | CD74 | chr5 | - | 150405084 | 150406321 | 150406258 | 150406321 | 150403120 | 150403312 | 19 | 15 | 12 | 12 | 2 | 0 | 1 | 0 | 0 | 0 | 77 | 69 | 20 | 73 | 155 | 0 | 0 | 5 | 3 | 0 | 298 | 149 | 0.222943022 | 0.83381688 | 1 | 0.882 | 1 | 1 | 1 | 1 | 1 | 0.667 | 0.924 | 1 | 0.058 |
| A5SS | 4352 | ENSG00000116138 | DNAJC16 | chr1 | + | 15566081 | 15566410 | 15566081 | 15566180 | 15567098 | 15567269 | 11 | 11 | 7 | 17 | 15 | 15 | 16 | 18 | 24 | 14 | 6 | 19 | 13 | 23 | 8 | 32 | 13 | 7 | 17 | 3 | 298 | 149 | 0.223086618 | 0.83381688 | 0.268 | 0.256 | 0.163 | 0.262 | 0.349 | 0.086 | 0.422 | 0.481 | 0.404 | 0.571 | -0.133 |
| A5SS | 4986 | ENSG00000138757 | G3BP2 | chr4 | - | 75656854 | 75657014 | 75656923 | 75657014 | 75655767 | 75655870 | 14 | 16 | 19 | 32 | 62 | 327 | 292 | 326 | 412 | 766 | 4 | 29 | 22 | 27 | 0 | 168 | 317 | 546 | 349 | 83 | 218 | 149 | 0.223268044 | 0.833922234 | 0.028 | 0.036 | 0.038 | 0.05 | 0.052 | 0.016 | 0.059 | 0.027 | 0.05 | 0 | 0.01 |
| A5SS | 924 | ENSG00000171928 | TVP23B | chr17 | + | 18790895 | 18791040 | 18790895 | 18790960 | 18797578 | 18797668 | 107 | 46 | 70 | 83 | 223 | 0 | 1 | 4 | 5 | 1 | 16 | 16 | 34 | 13 | 24 | 0 | 0 | 0 | 0 | 0 | 229 | 149 | 0.223524022 | 0.83430571 | 1 | 0.968 | 0.919 | 0.915 | 0.993 | 1 | 1 | 1 | 1 | 1 | -0.041 |
| A5SS | 8948 | ENSG00000134900 | TPP2 | chr13 | + | 102664794 | 102664936 | 102664794 | 102664925 | 102674282 | 102674490 | 39 | 32 | 40 | 42 | 51 | 228 | 170 | 260 | 253 | 283 | 7 | 13 | 24 | 0 | 0 | 0 | 196 | 174 | 82 | 22 | 160 | 149 | 0.224469573 | 0.837260739 | 0.137 | 0.149 | 0.125 | 0.134 | 0.144 | 1 | 0.058 | 0.114 | 0 | 0 | -0.097 |
| A5SS | 8526 | ENSG00000155256 | ZFYVE27 | chr10 | + | 97738476 | 97738674 | 97738476 | 97738649 | 97743093 | 97743164 | 14 | 3 | 13 | 18 | 12 | 0 | 0 | 0 | 0 | 0 | 20 | 2 | 12 | 8 | 30 | 0 | 0 | 0 | 6 | 0 | 174 | 149 | 0.226666731 | 0.844876951 | 1 | 1 | 1 | 1 | 1 | 1 | 1 | 1 | 0.533 | 1 | 0.093 |

| | | | | | | |
|---|---|---|---|---|---|---|
| A5SS | 1325 | ENSG00000159593 | NAE1 | chr16 | - | [illegible] |
| A5SS | 1072 | ENSG00000067955 | CBFB | chr16 | + | [illegible] |
| A5SS | 4058 | ENSG00000204165 | CXorf65 | chrX | - | [illegible] |
| A5SS | 1081 | ENSG00000125124 | BBS2 | chr16 | - | [illegible] |
| A5SS | 8047 | ENSG00000051009 | FAM160A2 | chr11 | - | [illegible] |
| A5SS | 479 | ENSG00000000419 | DPM1 | chr20 | - | [illegible] |
| A5SS | 1508 | ENSG00000138592 | USP8 | chr15 | + | [illegible] |
| A5SS | 5281 | ENSG00000116521 | SCAMP3 | chr1 | - | [illegible] |
| A5SS | 5595 | ENSG00000184708 | EIF4ENIF1 | chr22 | - | [illegible] |
| A5SS | 7788 | ENSG00000130714 | POMT1 | chr9 | + | [illegible] |
| A5SS | 6320 | ENSG00000149564 | ESAM | chr11 | - | [illegible] |
| A5SS | 6857 | ENSG00000188917 | TRMT2B | chrX | - | [illegible] |
| A5SS | 7108 | ENSG00000068308 | OTUD5 | chrX | - | [illegible] |
| A5SS | 6507 | ENSG00000179818 | PCBP1-AS1 | chr2 | - | [illegible] |
| A5SS | 1624 | ENSG00000100429 | HDAC10 | chr22 | - | [illegible] |
| A5SS | 6184 | ENSG00000204348 | DXO | chr6 | - | [illegible] |
| A5SS | 6298 | ENSG00000214021 | TTLL3 | chr3 | + | [illegible] |
| A5SS | 846 | ENSG00000256525 | POLG2 | chr17 | - | [illegible] |
| A5SS | 6657 | ENSG00000136247 | ZDHHC4 | chr7 | + | [illegible] |
| A5SS | 4674 | ENSG00000124571 | XPO5 | chr6 | - | [illegible] |
| A5SS | 5517 | ENSG00000223960 | CHROMR | chr2 | + | [illegible] |
| A5SS | 3172 | ENSG00000122779 | TRIM24 | chr7 | + | [illegible] |
| A5SS | 1565 | ENSG00000104129 | DNAJC17 | chr15 | - | [illegible] |
| A5SS | 5590 | ENSG00000143390 | RFX5 | chr1 | - | [illegible] |
| A5SS | 7749 | ENSG00000172716 | SLFN11 | chr17 | - | [illegible] |
| A5SS | 3676 | ENSG00000138002 | IFT172 | chr2 | - | [illegible] |
| A5SS | 5768 | ENSG00000145216 | FIP1L1 | chr4 | + | [illegible] |
| A5SS | 8246 | ENSG00000074319 | TSG101 | chr11 | - | [illegible] |
| A5SS | 5140 | ENSG00000152061 | RABGAP1L | chr1 | + | [illegible] |
| A5SS | 5108 | ENSG00000151092 | NGLY1 | chr3 | - | [illegible] |
| A5SS | 1906 | ENSG00000196367 | TRRAP | chr7 | + | [illegible] |
| A5SS | 6044 | ENSG00000183569 | SERHL2 | chr22 | + | [illegible] |
| A5SS | 7074 | ENSG00000215021 | PHB2 | chr12 | - | [illegible] |
| A5SS | 746 | ENSG00000108953 | YWHAE | chr17 | - | [illegible] |
| A5SS | 7418 | ENSG00000173064 | HECTD4 | chr12 | - | [illegible] |
| A5SS | 4134 | ENSG00000109180 | OCIAD1 | chr4 | + | [illegible] |
| A5SS | 8893 | ENSG00000170653 | ATF7 | chr12 | - | [illegible] |
| A5SS | 4659 | ENSG00000119950 | MXI1 | chr10 | + | [illegible] |
| A5SS | 3416 | ENSG00000164088 | PPM1M | chr3 | + | [illegible] |
| A5SS | 3440 | ENSG00000144591 | GMPPA | chr2 | + | [illegible] |
| A5SS | 4831 | ENSG00000148660 | CAMK2G | chr10 | - | [illegible] |
| A5SS | 8615 | ENSG00000241370 | RPP21 | chr6 | + | [illegible] |
| A5SS | 862 | ENSG00000007202 | KIAA0100 | chr17 | - | [illegible] |
| A5SS | 66 | ENSG00000196588 | MRTFA | chr22 | - | [illegible] |
| A5SS | 6133 | ENSG00000173588 | CEP83 | chr12 | - | [illegible] |
| A5SS | 15 | ENSG00000160209 | PDXK | chr21 | + | [illegible] |
| A5SS | 5827 | ENSG00000172936 | MYD88 | chr3 | + | [illegible] |
| A5SS | 2480 | ENSG00000145907 | G3BP1 | chr5 | + | [illegible] |
| A5SS | 2496 | ENSG00000174652 | ZNF266 | chr19 | - | [illegible] |
| A5SS | 5544 | ENSG00000107521 | HPS1 | chr10 | - | [illegible] |
| A5SS | 6210 | ENSG00000153406 | NMRAL1 | chr16 | - | [illegible] |
| A5SS | 2668 | ENSG00000101844 | ATG4A | chrX | + | [illegible] |
| A5SS | 2978 | ENSG00000163660 | CCNL1 | chr3 | - | [illegible] |
| A5SS | 3233 | ENSG00000197386 | HTT | chr4 | + | [illegible] |
| A5SS | 1881 | ENSG00000120925 | RNF170 | chr8 | - | [illegible] |
| A5SS | 2277 | ENSG00000112992 | NNT | chr5 | + | [illegible] |
| A5SS | 1475 | ENSG00000182718 | ANXA2 | chr15 | - | [illegible] |
| A5SS | 5681 | ENSG00000102125 | TAZ | chrX | + | [illegible] |
| A5SS | 803 | ENSG00000108788 | MLX | chr17 | + | [illegible] |
| A5SS | 2059 | ENSG00000131844 | MCCC2 | chr5 | + | [illegible] |
| A5SS | 5280 | ENSG00000116521 | SCAMP3 | chr1 | - | [illegible] |
| A5SS | 6968 | ENSG00000163389 | POGLUT1 | chr3 | + | [illegible] |
| A5SS | 4426 | ENSG00000166523 | CLEC4E | chr12 | - | [illegible] |
| A5SS | 512 | ENSG00000078747 | ITCH | chr20 | + | [illegible] |
| A5SS | 235 | ENSG00000142541 | RPL13A | chr19 | + | [illegible] |
| A5SS | 178 | ENSG00000239382 | ALKBH6 | chr19 | - | [illegible] |
| A5SS | 7312 | ENSG00000094914 | AAAS | chr12 | - | [illegible] |
| A5SS | 1879 | ENSG00000162711 | NLRP3 | chr1 | + | [illegible] |
| A5SS | 6933 | ENSG00000140332 | TLE3 | chr15 | - | [illegible] |
| A5SS | 7238 | ENSG00000136279 | DBNL | chr7 | + | [illegible] |
| A5SS | 4078 | ENSG00000249096 | LINC02362 | chr4 | - | [illegible] |
| A5SS | 8334 | ENSG00000197771 | MCMBP | chr10 | - | [illegible] |
| A5SS | 7918 | ENSG00000153827 | TRIP12 | chr2 | - | [illegible] |
| A5SS | 7935 | ENSG00000113163 | CERT1 | chr5 | - | [illegible] |
| A5SS | 6799 | ENSG00000118513 | MYB | chr6 | + | [illegible] |
| A5SS | 8945 | ENSG00000139597 | N4BP2L1 | chr13 | - | [illegible] |
| A5SS | 5973 | ENSG00000123600 | METTL8 | chr2 | - | [illegible] |
| A5SS | 5625 | ENSG00000115282 | TTC31 | chr2 | + | [illegible] |
| A5SS | 3400 | ENSG00000170581 | STAT2 | chr12 | - | [illegible] |
| A5SS | 2868 | ENSG00000178035 | IMPDH2 | chr3 | - | [illegible] |
| A5SS | 5270 | ENSG00000147526 | TACC1 | chr8 | + | [illegible] |
| A5SS | 1301 | ENSG00000158805 | ZNF276 | chr16 | + | [illegible] |
| A5SS | 301 | ENSG00000125734 | GPR108 | chr19 | - | [illegible] |
| A5SS | 5689 | ENSG00000243970 | PPIEL | chr1 | - | [illegible] |
| A5SS | 4690 | ENSG00000101040 | ZMYND8 | chr20 | - | [illegible] |
| A5SS | 5314 | ENSG00000136718 | IMP4 | chr2 | + | [illegible] |
| A5SS | 8184 | ENSG00000132275 | RRP8 | chr11 | - | [illegible] |
| A5SS | 2987 | ENSG00000204310 | AGPAT1 | chr6 | - | [illegible] |
| A5SS | 1215 | ENSG00000103245 | CIAO3 | chr16 | - | [illegible] |
| A5SS | 5041 | ENSG00000134686 | PHC2 | chr1 | - | [illegible] |
| A5SS | 130 | ENSG00000099917 | MED15 | chr22 | + | [illegible] |
| A5SS | 480 | ENSG00000101181 | MTG2 | chr20 | + | [illegible] |
| A5SS | 223 | ENSG00000104969 | SGTA | chr19 | - | [illegible] |
| A5SS | 4430 | ENSG00000138614 | INTS14 | chr15 | - | [illegible] |
| A5SS | 2181 | ENSG00000258466 | NA | chr14 | - | [illegible] |
| A5SS | 6200 | ENSG00000204520 | MICA | chr6 | + | [illegible] |
| A5SS | 618 | ENSG00000124155 | PIGT | chr20 | + | [illegible] |
| A5SS | 793 | ENSG00000275700 | AATF | chr17 | + | [illegible] |
| A5SS | 8144 | ENSG00000110713 | NUP98 | chr11 | - | [illegible] |
| A5SS | 687 | ENSG00000141447 | OSBPL1A | chr18 | - | [illegible] |
| A5SS | 7430 | ENSG00000104763 | ASAH1 | chr8 | - | [illegible] |
| A5SS | 7784 | ENSG00000165948 | IFI27L1 | chr14 | + | [illegible] |
| A5SS | 3764 | ENSG00000088035 | ALG6 | chr1 | + | [illegible] |
| A5SS | 5986 | ENSG00000111679 | PTPN6 | chr12 | + | [illegible] |
| A5SS | 2049 | ENSG00000228903 | RASA4CP | chr7 | - | [illegible] |
| A5SS | 6448 | ENSG00000136286 | MYO1G | chr7 | - | [illegible] |
| A5SS | 2719 | ENSG00000155508 | CNOT8 | chr5 | + | [illegible] |
| A5SS | 7209 | ENSG00000111237 | VPS29 | chr12 | - | [illegible] |
| A5SS | 928 | ENSG00000108604 | SMARCD2 | chr17 | - | [illegible] |
| A5SS | 7941 | ENSG00000066294 | CD84 | chr1 | - | [illegible] |
| A5SS | 6635 | ENSG00000104133 | SPG11 | chr15 | - | [illegible] |
| A5SS | 1651 | ENSG00000100577 | GSTZ1 | chr14 | + | [illegible] |
| A5SS | 7681 | ENSG00000070061 | ELP1 | chr9 | - | [illegible] |
| A5SS | 7683 | ENSG00000132950 | ZMYM5 | chr13 | - | [illegible] |
| A5SS | 592 | ENSG00000089063 | TMEM230 | chr20 | - | [illegible] |
| A5SS | 7452 | ENSG00000164941 | INTS8 | chr8 | + | [illegible] |
| A5SS | 7232 | ENSG00000139197 | PEX5 | chr12 | + | [illegible] |
| A5SS | 95 | ENSG00000100124 | ANKRD54 | chr22 | - | [illegible] |
| A5SS | 8523 | ENSG00000132305 | IMMT | chr2 | - | [illegible] |
| A5SS | 4315 | ENSG00000166825 | ANPEP | chr15 | - | [illegible] |
| A5SS | 5051 | ENSG00000120265 | PCMT1 | chr6 | + | [illegible] |
| A5SS | 4500 | ENSG00000104894 | CD37 | chr19 | + | [illegible] |
| A5SS | 3678 | ENSG00000162946 | DISC1 | chr1 | + | [illegible] |
| A5SS | 6681 | ENSG00000105939 | ZC3HAV1 | chr7 | - | [illegible] |
| A5SS | 522 | ENSG00000101294 | HM13 | chr20 | + | [illegible] |
| A5SS | 6676 | ENSG00000105835 | NAMPT | chr7 | - | [illegible] |
| A5SS | 4994 | ENSG00000061273 | HDAC7 | chr12 | - | [illegible] |
| A5SS | 6351 | ENSG00000255036 | SUGT1P4-STRA6LP-CCDC180 | chr9 | + | [illegible] |
| A5SS | 6091 | ENSG00000244482 | LILRA6 | chr19 | - | [illegible] |
| A5SS | 1822 | ENSG00000080815 | PSEN1 | chr14 | + | [illegible] |
| A5SS | 7608 | ENSG00000148153 | INIP | chr9 | - | [illegible] |
| A5SS | 91 | ENSG00000100099 | HPS4 | chr22 | - | [illegible] |
| A5SS | 5481 | ENSG00000122085 | MTERF4 | chr2 | - | [illegible] |
| A5SS | 413 | ENSG00000130475 | FCHO1 | chr19 | + | [illegible] |
| A5SS | 2232 | ENSG00000204628 | RACK1 | chr5 | - | [illegible] |
| A5SS | 1448 | ENSG00000182511 | FES | chr15 | + | [illegible] |
| A5SS | 716 | ENSG00000161944 | ASGR2 | chr17 | - | [illegible] |
| A5SS | 7508 | ENSG00000147789 | ZNF7 | chr8 | + | [illegible] |
| A5SS | 6352 | ENSG00000177156 | TALDO1 | chr11 | + | [illegible] |
| A5SS | 7850 | ENSG00000159314 | ARHGAP27 | chr17 | - | [illegible] |
| A5SS | 6207 | ENSG00000155906 | RMND1 | chr6 | - | [illegible] |
| A5SS | 234 | ENSG00000130816 | DNMT1 | chr19 | - | [illegible] |
| A5SS | 1979 | ENSG00000133706 | LARS1 | chr5 | - | [illegible] |
| A5SS | 3461 | ENSG00000013573 | DDX11 | chr12 | + | [illegible] |
| A5SS | 2382 | ENSG00000101182 | PSMA7 | chr20 | - | [illegible] |
| A5SS | 5626 | ENSG00000115282 | TTC31 | chr2 | + | [illegible] |
| A5SS | 2513 | ENSG00000058063 | ATP11B | chr3 | + | [illegible] |
| A5SS | 2380 | ENSG00000138760 | SCARB2 | chr4 | - | [illegible] |
| A5SS | 5642 | ENSG00000132323 | ILKAP | chr2 | - | [illegible] |
| A5SS | 8775 | ENSG00000198715 | GLMP | chr1 | - | [illegible] |
| A5SS | 7499 | ENSG00000154188 | ANGPT1 | chr8 | - | [illegible] |
| A5SS | 4350 | ENSG00000169764 | UGP2 | chr2 | + | [illegible] |
| A5SS | 3852 | ENSG00000158517 | NCF1 | chr7 | + | [illegible] |
| A5SS | 8594 | ENSG00000119801 | YPEL5 | chr2 | + | [illegible] |
| A5SS | 836 | ENSG00000185359 | HGS | chr17 | + | [illegible] |
| A5SS | 3389 | ENSG00000084693 | AGBL5 | chr2 | + | [illegible] |
| A5SS | 2928 | ENSG00000180376 | CCDC66 | chr3 | + | [illegible] |
| A5SS | 7874 | ENSG00000198561 | CTNND1 | chr11 | + | [illegible] |
| A5SS | 2349 | ENSG00000139625 | MAP3K12 | chr12 | - | [illegible] |
| A5SS | 7764 | ENSG00000115459 | ELMOD3 | chr2 | + | [illegible] |
| A5SS | 7622 | ENSG00000089053 | ANAPC5 | chr12 | - | [illegible] |
| A5SS | 6728 | ENSG00000186088 | GSAP | chr7 | - | [illegible] |
| A5SS | 5879 | ENSG00000163938 | GNL3 | chr3 | + | [illegible] |
| A5SS | 2451 | ENSG00000213468 | FIRRE | chrX | - | [illegible] |
| A5SS | 6910 | ENSG00000170832 | USP32 | chr17 | - | [illegible] |
| A5SS | 7987 | ENSG00000182240 | BACE2 | chr21 | + | [illegible] |
| A5SS | 7731 | ENSG00000213930 | GALT | chr9 | + | [illegible] |
| A5SS | 2564 | ENSG00000203875 | SNHG5 | chr6 | - | [illegible] |
| A5SS | 1670 | ENSG00000119682 | AREL1 | chr14 | - | [illegible] |
| A5SS | 1989 | ENSG00000125459 | MSTO1 | chr1 | + | [illegible] |
| A5SS | 2981 | ENSG00000175193 | PARL | chr3 | - | [illegible] |
| A5SS | 445 | ENSG00000198858 | R3HDM4 | chr19 | - | [illegible] |
| A5SS | 2473 | ENSG00000204084 | INPP5B | chr1 | - | [illegible] |
| A5SS | 1934 | ENSG00000126214 | KLC1 | chr14 | + | [illegible] |
| A5SS | 514 | ENSG00000198053 | SIRPA | chr20 | + | [illegible] |
| A5SS | 7924 | ENSG00000139626 | ITGB7 | chr12 | - | [illegible] |
| A5SS | 2589 | ENSG00000251022 | THAP9-AS1 | chr4 | - | [illegible] |
| A5SS | 3911 | ENSG00000055609 | KMT2C | chr7 | - | [illegible] |
| A5SS | 7455 | ENSG00000111530 | CAND1 | chr12 | + | [illegible] |
| A5SS | 1743 | ENSG00000196497 | IPO4 | chr14 | - | [illegible] |
| A5SS | 7770 | ENSG00000198431 | TXNRD1 | chr12 | + | [illegible] |
| A5SS | 5658 | ENSG00000164548 | TRA2A | chr7 | - | [illegible] |
| A5SS | 3426 | ENSG00000213390 | ARHGAP19 | chr10 | - | [illegible] |
| A5SS | 6759 | ENSG00000234741 | GAS5 | chr1 | - | [illegible] |
| A5SS | 2314 | ENSG00000005156 | LIG3 | chr17 | + | [illegible] |
| A5SS | 1253 | ENSG00000067836 | ROGDI | chr16 | - | [illegible] |
| A5SS | 8917 | ENSG00000105983 | LMBR1 | chr7 | - | [illegible] |
| A5SS | 5079 | ENSG00000117222 | RBBP5 | chr1 | - | [illegible] |
| A5SS | 5656 | ENSG00000169689 | CENPX | chr17 | - | [illegible] |
| A5SS | 3729 | ENSG00000119402 | FBXW2 | chr9 | - | [illegible] |
| A5SS | 4557 | ENSG00000114491 | UMPS | chr3 | + | [illegible] |
| A5SS | 6601 | ENSG00000071462 | BUD23 | chr7 | + | [illegible] |
| A5SS | 2137 | ENSG00000166888 | STAT6 | chr12 | - | [illegible] |
| A5SS | 5990 | ENSG00000116698 | SMG7 | chr1 | + | [illegible] |
| A5SS | 3580 | ENSG00000169231 | THBS3 | chr1 | - | [illegible] |
| A5SS | 8678 | ENSG00000196498 | NCOR2 | chr12 | - | [illegible] |
| A5SS | 2835 | ENSG00000150995 | ITPR1 | chr3 | + | [illegible] |
| A5SS | 4204 | ENSG00000117481 | NSUN4 | chr1 | + | [illegible] |
| A5SS | 2756 | ENSG00000163703 | CRELD1 | chr3 | + | [illegible] |
| A5SS | 6539 | ENSG00000151948 | GLT1D1 | chr12 | + | [illegible] |
| A5SS | 1026 | ENSG00000171475 | WIPF2 | chr17 | + | [illegible] |
| A5SS | 5820 | ENSG00000180228 | PRKRA | chr2 | - | [illegible] |
| A5SS | 3517 | ENSG00000089327 | FXYD5 | chr19 | + | [illegible] |
| A5SS | 4245 | ENSG00000164022 | AIMP1 | chr4 | + | [illegible] |
| A5SS | 2859 | ENSG00000183323 | CCDC125 | chr5 | - | [illegible] |
| A5SS | 6433 | ENSG00000119616 | FCF1 | chr14 | + | [illegible] |
| A5SS | 4118 | ENSG00000116337 | AMPD2 | chr1 | + | [illegible] |
| A5SS | 3276 | ENSG00000112096 | SOD2 | chr6 | - | [illegible] |
| A5SS | 5677 | ENSG00000102125 | TAZ | chrX | + | [illegible] |

| | | | | | | | | | | | | | | | | | | | | | | | | | | | | | | | | | | | | | | | | | | | | | | |
|---|---|---|---|---|---|---|---|---|---|---|---|---|---|---|---|---|---|---|---|---|---|---|---|---|---|---|---|---|---|---|---|---|---|---|---|---|---|---|---|---|---|---|---|---|---|---|
| A5SS | 7040 | ENSG00000184634 | MED12 | chrX | + | 71137186 | 71137392 | 71137186 | 71137383 | 71137557 | 71137635 | 27 | 8 | 24 | 34 | 3 | 23 | 2 | 15 | 7 | 3 | 1 | 6 | 6 | 36 | 22 | 0 | 13 | 4 | 25 | 12 | 158 | 149 | 0.35331209 | 1 | 0.525 | 0.79 | 0.601 | 0.821 | 0.485 | 1 | 0.303 | 0.586 | 0.576 | 0.634 | 0.025 |
| A5SS | 7834 | ENSG00000133943 | DGLUCY | chr14 | + | 91173288 | 91173439 | 91173288 | 91173403 | 91175933 | 91176056 | 31 | 12 | 40 | 57 | 6 | 1 | 0 | 5 | 5 | 0 | 59 | 24 | 17 | 19 | 51 | 2 | 21 | 0 | 0 | 4 | 185 | 149 | 0.353336217 | 1 | 0.961 | 1 | 0.866 | 0.902 | 1 | 0.96 | 0.479 | 1 | 1 | 0.911 | 0.076 |
| A5SS | 29 | ENSG00000159086 | PAXBP1 | chr21 | - | 32755212 | 32755353 | 32755229 | 32755353 | 32751118 | 32751218 | 69 | 118 | 136 | 101 | 106 | 62 | 32 | 39 | 72 | 65 | 8 | 55 | 37 | 7 | 14 | 18 | 43 | 15 | 7 | 3 | 166 | 149 | 0.353572121 | 1 | 0.5 | 0.768 | 0.758 | 0.557 | 0.594 | 0.285 | 0.534 | 0.689 | 0.473 | 0.807 | 0.078 |
| A5SS | 7760 | ENSG00000115459 | ELMOD3 | chr2 | + | 85355554 | 85355716 | 85355554 | 85355598 | 85356601 | 85356667 | 20 | 11 | 11 | 20 | 14 | 0 | 0 | 0 | 0 | 0 | 6 | 24 | 14 | 24 | 9 | 0 | 3 | 0 | 3 | 0 | 267 | 149 | 0.353787556 | 1 | 1 | 1 | 1 | 1 | 1 | 1 | 0.817 | 1 | 0.817 | 1 | 0.073 |
| A5SS | 1366 | ENSG00000140694 | PARN | chr16 | - | 14591912 | 14593378 | 14593300 | 14593378 | 14584748 | 14584791 | 15 | 6 | 3 | 4 | 12 | 2 | 12 | 1 | 0 | 6 | 0 | 4 | 6 | 12 | 13 | 24 | 7 | 1 | 5 | 0 | 298 | 149 | 0.354211747 | 1 | 0.789 | 0.2 | 0.6 | 1 | 0.5 | 0 | 0.222 | 0.75 | 0.545 | 1 | 0.114 |
| A5SS | 5974 | ENSG00000155506 | LARP1 | chr5 | + | 154793799 | 154794000 | 154793799 | 154793922 | 154794099 | 154794262 | 186 | 116 | 227 | 192 | 174 | 6 | 3 | 2 | 1 | 3 | 175 | 214 | 124 | 176 | 191 | 0 | 0 | 0 | 3 | 8 | 227 | 149 | 0.354773286 | 1 | 0.953 | 0.962 | 0.987 | 0.992 | 0.974 | 1 | 1 | 1 | 0.975 | 0.94 | -0.009 |
| A5SS | 5559 | ENSG00000112367 | FIG4 | chr6 | + | 109786301 | 109788956 | 109786301 | 109786449 | 109789593 | 109789677 | 7 | 0 | 0 | 6 | 3 | 74 | 62 | 41 | 65 | 151 | 14 | 0 | 2 | 0 | 0 | 76 | 49 | 33 | 69 | 17 | 298 | 149 | 0.355552056 | 1 | 0.045 | 0 | 0 | 0.044 | 0.01 | 0.084 | 0 | 0.029 | 0 | 0 | -0.003 |
| A5SS | 935 | ENSG00000108592 | FTSJ3 | chr17 | - | 63826546 | 63826672 | 63826566 | 63826672 | 63826257 | 63826304 | 0 | 3 | 0 | 1 | 4 | 36 | 18 | 35 | 31 | 27 | 0 | 0 | 1 | 2 | 0 | 54 | 26 | 6 | 12 | 34 | 169 | 149 | 0.356340083 | 1 | 0 | 0.128 | 0 | 0.028 | 0.116 | 0 | 0 | 0.128 | 0.128 | 0 | 0.003 |
| A5SS | 3384 | ENSG00000114544 | SLC41A3 | chr3 | - | 126050897 | 126051050 | 126050942 | 126051050 | 126033606 | 126033678 | 5 | 0 | 1 | 0 | 2 | 11 | 10 | 25 | 17 | 3 | 0 | 2 | 4 | 0 | 0 | 11 | 15 | 5 | 16 | 21 | 194 | 149 | 0.35788432 | 1 | 0.259 | 0 | 0.03 | 0 | 0.339 | 0 | 0.093 | 0.381 | 0 | 0 | 0.031 |
| A5SS | 4015 | ENSG00000162542 | TMCO4 | chr1 | - | 19798070 | 19798215 | 19798136 | 19798215 | 19787025 | 19787117 | 5 | 2 | 1 | 5 | 4 | 13 | 2 | 3 | 2 | 4 | 15 | 6 | 1 | 0 | 0 | 0 | 10 | 2 | 4 | 16 | 215 | 149 | 0.358830678 | 1 | 0.21 | 0.409 | 0.188 | 0.634 | 0.409 | 1 | 0.294 | 0.257 | 0 | 0 | 0.06 |
| A5SS | 6733 | ENSG00000122783 | CYREN | chr7 | - | 135168744 | 135169060 | 135168785 | 135169060 | 135167731 | 135167807 | 18 | 12 | 19 | 21 | 21 | 10 | 3 | 6 | 8 | 4 | 39 | 10 | 10 | 17 | 25 | 0 | 4 | 3 | 2 | 22 | 190 | 149 | 0.359402903 | 1 | 0.585 | 0.758 | 0.713 | 0.673 | 0.805 | 1 | 0.662 | 0.723 | 0.87 | 0.471 | -0.038 |
| A5SS | 2927 | ENSG00000180376 | CCDC66 | chr3 | + | 56563683 | 56564125 | 56563683 | 56563825 | 56566593 | 56566759 | 487 | 489 | 475 | 491 | 967 | 0 | 0 | 0 | 0 | 2 | 135 | 217 | 255 | 210 | 82 | 0 | 0 | 0 | 0 | 10 | 298 | 149 | 0.36102474 | 1 | 1 | 1 | 1 | 1 | 0.996 | 1 | 1 | 1 | 1 | 0.804 | 0.038 |
| A5SS | 1027 | ENSG00000273559 | CWC25 | chr17 | - | 38810275 | 38810595 | 38810467 | 38810595 | 38809701 | 38809765 | 43 | 48 | 56 | 79 | 77 | 15 | 15 | 14 | 45 | 48 | 164 | 58 | 88 | 145 | 93 | 39 | 27 | 30 | 62 | 32 | 298 | 149 | 0.361056155 | 1 | 0.589 | 0.615 | 0.667 | 0.467 | 0.445 | 0.678 | 0.518 | 0.595 | 0.539 | 0.592 | -0.028 |
| A5SS | 1440 | ENSG00000137871 | ZNF280D | chr15 | - | 56666678 | 56666986 | 56666799 | 56666986 | 56666394 | 56666535 | 139 | 102 | 135 | 140 | 193 | 2 | 7 | 5 | 4 | 17 | 0 | 38 | 13 | 12 | 20 | 3 | 0 | 1 | 0 | 1 | 270 | 149 | 0.361164128 | 1 | 0.975 | 0.889 | 0.937 | 0.951 | 0.862 | 0 | 1 | 0.878 | 1 | 0.917 | 0.164 |
| A5SS | 2560 | ENSG00000100650 | SRSF5 | chr14 | + | 69767092 | 69767380 | 69767092 | 69767255 | 69768137 | 69768282 | 305 | 195 | 291 | 385 | 164 | 584 | 377 | 529 | 643 | 341 | 117 | 249 | 255 | 260 | 192 | 281 | 553 | 439 | 492 | 377 | 274 | 149 | 0.362599162 | 1 | 0.221 | 0.22 | 0.23 | 0.246 | 0.207 | 0.185 | 0.197 | 0.24 | 0.223 | 0.217 | 0.012 |
| A5SS | 8036 | ENSG00000177830 | CHID1 | chr11 | - | 883147 | 883303 | 883199 | 883303 | 870418 | 870499 | 15 | 5 | 9 | 19 | 4 | 0 | 0 | 0 | 1 | 3 | 5 | 3 | 5 | 18 | 24 | 0 | 6 | 0 | 0 | 0 | 201 | 149 | 0.36387392 | 1 | 1 | 1 | 1 | 0.934 | 0.497 | 1 | 0.27 | 1 | 1 | 1 | 0.032 |
| A5SS | 4144 | ENSG00000167528 | ZNF641 | chr12 | - | 48347148 | 48347343 | 48347251 | 48347343 | 48344598 | 48344712 | 13 | 2 | 18 | 36 | 21 | 31 | 20 | 34 | 22 | 44 | 22 | 38 | 6 | 11 | 30 | 74 | 47 | 54 | 47 | 34 | 252 | 149 | 0.364400135 | 1 | 0.199 | 0.056 | 0.238 | 0.492 | 0.22 | 0.15 | 0.323 | 0.062 | 0.122 | 0.343 | 0.041 |
| A5SS | 127 | ENSG00000100325 | ASCC2 | chr22 | - | 29834441 | 29834550 | 29834488 | 29834550 | 29832244 | 29832342 | 10 | 1 | 3 | 4 | 9 | 11 | 3 | 6 | 2 | 5 | 8 | 2 | 11 | 11 | 15 | 0 | 8 | 6 | 9 | 10 | 196 | 149 | 0.36441363 | 1 | 0.409 | 0.202 | 0.275 | 0.603 | 0.578 | 1 | 0.16 | 0.582 | 0.482 | 0.533 | -0.138 |
| A5SS | 1600 | ENSG00000248334 | WHAMMP2 | chr15 | + | 28751487 | 28751678 | 28751487 | 28751674 | 28754468 | 28754555 | 0 | 0 | 1 | 0 | 0 | 57 | 25 | 45 | 145 | 113 | 2 | 0 | 0 | 6 | 0 | 113 | 15 | 60 | 12 | 169 | 153 | 149 | 0.36555969 | 1 | 0 | 0 | 0.021 | 0 | 0 | 0.017 | 0 | 0 | 0.327 | 0 | -0.065 |
| A5SS | 8957 | ENSG00000005810 | MYCBP2 | chr13 | - | 77076750 | 77076849 | 77076759 | 77076849 | 77070630 | 77070711 | 64 | 30 | 47 | 37 | 121 | 61 | 48 | 55 | 57 | 111 | 16 | 29 | 26 | 13 | 0 | 3 | 14 | 32 | 5 | 6 | 158 | 149 | 0.365943706 | 1 | 0.497 | 0.371 | 0.446 | 0.38 | 0.507 | 0.834 | 0.661 | 0.434 | 0.71 | 0 | -0.088 |
| A5SS | 8348 | ENSG00000179988 | PSTK | chr10 | + | 122983271 | 122983514 | 122983271 | 122983470 | 122986299 | 122986375 | 2 | 0 | 0 | 7 | 0 | 8 | 6 | 8 | 9 | 12 | 9 | 0 | 0 | 1 | 0 | 8 | 2 | 15 | 0 | 3 | 193 | 149 | 0.365956622 | 1 | 0.162 | 0 | 0 | 0.375 | 0 | 0.465 | 0 | 0 | 1 | 0 | -0.186 |
| A5SS | 6863 | ENSG00000148985 | PGAP2 | chr11 | + | 3824269 | 3824455 | 3824269 | 3824376 | 3825019 | 3825128 | 4 | 6 | 7 | 5 | 7 | 13 | 5 | 11 | 9 | 8 | 15 | 1 | 6 | 12 | 12 | 11 | 3 | 9 | 9 | 21 | 228 | 149 | 0.366017785 | 1 | 0.167 | 0.44 | 0.294 | 0.266 | 0.364 | 0.471 | 0.179 | 0.303 | 0.466 | 0.272 | -0.032 |
| A5SS | 6671 | ENSG00000204257 | HLA-DMA | chr6 | - | 32949270 | 32949399 | 32949273 | 32949399 | 32948612 | 32948868 | 324 | 165 | 204 | 279 | 216 | 16 | 6 | 10 | 15 | 20 | 548 | 249 | 166 | 205 | 971 | 15 | 18 | 6 | 18 | 83 | 152 | 149 | 0.366308714 | 1 | 0.952 | 0.964 | 0.952 | 0.948 | 0.914 | 0.973 | 0.931 | 0.964 | 0.918 | 0.92 | 0.005 |
| A5SS | 6783 | ENSG00000106603 | COA1 | chr7 | - | 43650492 | 43650712 | 43650611 | 43650712 | 43648599 | 43648652 | 22 | 30 | 21 | 21 | 23 | 24 | 23 | 24 | 22 | 20 | 36 | 19 | 10 | 18 | 12 | 8 | 9 | 14 | 13 | 22 | 268 | 149 | 0.367985173 | 1 | 0.338 | 0.42 | 0.327 | 0.347 | 0.39 | 0.714 | 0.54 | 0.284 | 0.435 | 0.233 | -0.077 |
| A5SS | 4425 | ENSG00000166523 | CLEC4E | chr12 | - | 8539141 | 8539306 | 8539199 | 8539306 | 8537114 | 8537266 | 64 | 49 | 51 | 33 | 21 | 2 | 0 | 2 | 1 | 3 | 64 | 40 | 184 | 40 | 7 | 0 | 0 | 3 | 4 | 0 | 207 | 149 | 0.367995021 | 1 | 0.958 | 1 | 0.948 | 0.96 | 0.834 | 1 | 1 | 0.978 | 0.878 | 1 | -0.031 |
| A5SS | 121 | ENSG00000228315 | GUSBP11 | chr22 | - | 23686830 | 23686995 | 23686837 | 23686995 | 23683724 | 23683867 | 4 | 2 | 6 | 12 | 2 | 1 | 2 | 3 | 10 | 3 | 5 | 0 | 8 | 11 | 7 | 0 | 5 | 0 | 4 | 7 | 156 | 149 | 0.369436824 | 1 | 0.793 | 0.489 | 0.656 | 0.534 | 0.389 | 1 | 0 | 1 | 0.724 | 0.489 | -0.07 |
| A5SS | 993 | ENSG00000168259 | DNAJC7 | chr17 | - | 42016586 | 42017151 | 42017051 | 42017151 | 42000481 | 42000570 | 17 | 4 | 23 | 14 | 19 | 1 | 0 | 2 | 4 | 0 | 67 | 61 | 17 | 38 | 47 | 0 | 3 | 0 | 0 | 0 | 298 | 149 | 0.369615344 | 1 | 0.895 | 1 | 0.852 | 0.636 | 1 | 1 | 0.91 | 1 | 1 | 1 | -0.105 |
| A5SS | 4952 | ENSG00000067225 | PKM | chr15 | - | 72230899 | 72231386 | 72231115 | 72231386 | 72218943 | 72219110 | 5 | 5 | 1 | 1 | 9 | 156 | 70 | 114 | 211 | 71 | 0 | 4 | 0 | 8 | 25 | 634 | 591 | 204 | 429 | 1420 | 298 | 149 | 0.371666135 | 1 | 0.016 | 0.034 | 0.004 | 0.002 | 0.06 | 0 | 0.003 | 0 | 0.009 | 0.009 | 0.019 |
| A5SS | 4268 | ENSG00000160087 | UBE2J2 | chr1 | - | 1267732 | 1267992 | 1267861 | 1267992 | 1263345 | 1263386 | 11 | 8 | 7 | 13 | 0 | 65 | 46 | 52 | 56 | 69 | 11 | 3 | 3 | 7 | 24 | 106 | 58 | 20 | 78 | 196 | 278 | 149 | 0.372854461 | 1 | 0.083 | 0.085 | 0.067 | 0.111 | 0 | 0.053 | 0.027 | 0.074 | 0.046 | 0.062 | 0.017 |
| A5SS | 6694 | ENSG00000204316 | MRPL38 | chr17 | - | 75904225 | 75904719 | 75904539 | 75904719 | 75902019 | 75902154 | 6 | 3 | 11 | 3 | 4 | 26 | 12 | 33 | 27 | 13 | 0 | 16 | 6 | 12 | 18 | 44 | 35 | 12 | 23 | 60 | 298 | 149 | 0.375875235 | 1 | 0.103 | 0.111 | 0.143 | 0.053 | 0.133 | 0 | 0.186 | 0.2 | 0.207 | 0.13 | -0.036 |
| A5SS | 5755 | ENSG00000115234 | SNX17 | chr2 | + | 27371268 | 27371468 | 27371268 | 27371343 | 27373246 | 27373311 | 22 | 16 | 14 | 10 | 3 | 2 | 1 | 3 | 7 | 0 | 39 | 18 | 26 | 28 | 38 | 21 | 5 | 3 | 0 | 18 | 274 | 149 | 0.375937083 | 1 | 0.857 | 0.897 | 0.717 | 0.437 | 1 | 0.502 | 0.662 | 0.825 | 1 | 0.534 | 0.077 |
| A5SS | 7451 | ENSG00000116852 | KIF21B | chr1 | - | 201004540 | 201004933 | 201004765 | 201004933 | 201004339 | 201004455 | 5 | 4 | 1 | 1 | 6 | 91 | 47 | 138 | 87 | 101 | 7 | 2 | 3 | 3 | 7 | 82 | 127 | 47 | 73 | 131 | 298 | 149 | 0.376041398 | 1 | 0.027 | 0.041 | 0.004 | 0.006 | 0.029 | 0.041 | 0.008 | 0.031 | 0.02 | 0.026 | -0.004 |
| A5SS | 3535 | ENSG00000026751 | SLAMF7 | chr1 | + | 160749820 | 160750132 | 160749820 | 160750093 | 160752185 | 160752248 | 12 | 4 | 1 | 3 | 0 | 15 | 8 | 10 | 11 | 9 | 0 | 0 | 7 | 7 | 4 | 11 | 16 | 10 | 14 | 30 | 188 | 149 | 0.37672462 | 1 | 0.388 | 0.284 | 0.073 | 0.178 | 0 | 0 | 0 | 0.357 | 0.284 | 0.096 | 0.037 |
| A5SS | 5260 | ENSG00000163541 | SUCLG1 | chr2 | - | 84443220 | 84443400 | 84443283 | 84443400 | 84441246 | 84441459 | 15 | 14 | 8 | 10 | 4 | 46 | 34 | 68 | 29 | 81 | 7 | 20 | 12 | 13 | 0 | 25 | 24 | 41 | 53 | 40 | 212 | 149 | 0.376866319 | 1 | 0.186 | 0.224 | 0.076 | 0.195 | 0.034 | 0.164 | 0.369 | 0.171 | 0.147 | 0 | -0.027 |
| A5SS | 6932 | ENSG00000140332 | TLE3 | chr15 | - | 70057458 | 70057658 | 70057473 | 70057658 | 70056297 | 70056374 | 45 | 10 | 30 | 33 | 5 | 37 | 8 | 20 | 27 | 5 | 80 | 28 | 41 | 46 | 57 | 40 | 55 | 36 | 50 | 53 | 164 | 149 | 0.377235721 | 1 | 0.525 | 0.532 | 0.577 | 0.526 | 0.476 | 0.645 | 0.316 | 0.509 | 0.455 | 0.494 | 0.043 |
| A5SS | 482 | ENSG00000177410 | ZFAS1 | chr20 | + | 49278641 | 49278738 | 49278641 | 49278716 | 49279103 | 49279208 | 31 | 17 | 30 | 24 | 13 | 193 | 115 | 232 | 207 | 103 | 63 | 36 | 38 | 43 | 42 | 417 | 171 | 219 | 250 | 446 | 171 | 149 | 0.377478464 | 1 | 0.123 | 0.114 | 0.101 | 0.092 | 0.099 | 0.116 | 0.155 | 0.131 | 0.13 | 0.076 | -0.016 |
| A5SS | 3255 | ENSG00000164330 | EBF1 | chr5 | - | 158823175 | 158823317 | 158823199 | 158823317 | 158796344 | 158796475 | 9 | 16 | 16 | 17 | 33 | 2 | 3 | 0 | 4 | 4 | 16 | 32 | 6 | 13 | 17 | 8 | 0 | 0 | 0 | 0 | 173 | 149 | 0.378735206 | 1 | 0.795 | 0.821 | 1 | 0.785 | 0.877 | 0.633 | 1 | 1 | 1 | 1 | -0.071 |
| A5SS | 7372 | ENSG00000172269 | DPAGT1 | chr11 | - | 119097854 | 119098487 | 119098402 | 119098487 | 119097463 | 119097551 | 27 | 10 | 20 | 36 | 27 | 3 | 1 | 2 | 2 | 0 | 27 | 12 | 11 | 24 | 30 | 9 | 1 | 2 | 0 | 0 | 298 | 149 | 0.379303062 | 1 | 0.818 | 0.833 | 0.833 | 0.9 | 1 | 0.6 | 0.857 | 0.733 | 1 | 1 | 0.039 |
| A5SS | 1007 | ENSG00000126581 | BECN1 | chr17 | - | 42824132 | 42824282 | 42824154 | 42824282 | 42823747 | 42823879 | 80 | 33 | 69 | 75 | 36 | 118 | 52 | 115 | 108 | 116 | 138 | 115 | 59 | 74 | 77 | 74 | 159 | 106 | 108 | 248 | 171 | 149 | 0.379855475 | 1 | 0.371 | 0.356 | 0.343 | 0.377 | 0.213 | 0.619 | 0.387 | 0.327 | 0.374 | 0.213 | -0.052 |
| A5SS | 5888 | ENSG00000143436 | MRPL9 | chr1 | - | 151762954 | 151763146 | 151762989 | 151763146 | 151762375 | 151762500 | 11 | 3 | 8 | 8 | 8 | 67 | 71 | 83 | 98 | 46 | 16 | 0 | 7 | 4 | 3 | 86 | 81 | 59 | 55 | 75 | 184 | 149 | 0.380414803 | 1 | 0.117 | 0.033 | 0.072 | 0.062 | 0.123 | 0.131 | 0 | 0.088 | 0.056 | 0.031 | 0.02 |
| A5SS | 8547 | ENSG00000079974 | RABL2B | chr22 | - | 50769424 | 50769549 | 50769454 | 50769549 | 50769040 | 50769124 | 2 | 1 | 3 | 11 | 2 | 25 | 21 | 27 | 33 | 45 | 0 | 0 | 4 | 3 | 6 | 46 | 26 | 14 | 22 | 29 | 179 | 149 | 0.381036965 | 1 | 0.062 | 0.038 | 0.085 | 0.217 | 0.036 | 0 | 0 | 0.192 | 0.102 | 0.147 | -0.001 |
| A5SS | 7286 | ENSG00000174197 | MGA | chr15 | + | 41736180 | 41736698 | 41736180 | 41736368 | 41740052 | 41740203 | 166 | 105 | 116 | 150 | 127 | 6 | 0 | 0 | 3 | 8 | 27 | 112 | 92 | 15 | 12 | 0 | 0 | 0 | 7 | 0 | 298 | 149 | 0.382424899 | 1 | 0.933 | 1 | 1 | 0.962 | 0.888 | 1 | 1 | 1 | 0.517 | 1 | 0.053 |
| A5SS | 8181 | ENSG00000072518 | MARK2 | chr11 | + | 63903985 | 63904354 | 63903985 | 63904147 | 63904785 | 63905043 | 9 | 2 | 4 | 1 | 0 | 58 | 34 | 33 | 39 | 25 | 0 | 10 | 4 | 0 | 13 | 44 | 48 | 41 | 78 | 67 | 298 | 149 | 0.38278794 | 1 | 0.072 | 0.029 | 0.057 | 0.013 | 0 | 0 | 0.094 | 0.047 | 0 | 0.088 | -0.012 |
| A5SS | 3484 | ENSG00000138674 | SEC31A | chr4 | - | 82842139 | 82842481 | 82842184 | 82842481 | 82828999 | 82829058 | 57 | 44 | 58 | 69 | 54 | 18 | 4 | 6 | 7 | 2 | 53 | 33 | 33 | 59 | 11 | 9 | 3 | 2 | 6 | 0 | 194 | 149 | 0.383335092 | 1 | 0.709 | 0.894 | 0.881 | 0.883 | 0.954 | 0.819 | 0.894 | 0.927 | 0.883 | 1 | -0.04 |
| A5SS | 622 | ENSG00000099956 | SMARCB1 | chr22 | + | 23791755 | 23791894 | 23791755 | 23791867 | 23793558 | 23793688 | 70 | 30 | 57 | 55 | 100 | 83 | 67 | 82 | 75 | 147 | 101 | 110 | 45 | 72 | 104 | 150 | 162 | 39 | 103 | 122 | 176 | 149 | 0.384107813 | 1 | 0.417 | 0.275 | 0.37 | 0.383 | 0.365 | 0.363 | 0.365 | 0.494 | 0.372 | 0.419 | -0.041 |
| A5SS | 8544 | ENSG00000172243 | CLEC7A | chr12 | - | 10126570 | 10127845 | 10127746 | 10127845 | 10125296 | 10125448 | 245 | 369 | 194 | 213 | 262 | 127 | 136 | 94 | 77 | 123 | 139 | 180 | 604 | 247 | 95 | 260 | 80 | 92 | 90 | 119 | 298 | 149 | 0.384414259 | 1 | 0.491 | 0.576 | 0.508 | 0.58 | 0.516 | 0.211 | 0.529 | 0.766 | 0.578 | 0.285 | 0.06 |
| A5SS | 1553 | ENSG00000177082 | WDR73 | chr15 | - | 84645972 | 84646348 | 84646183 | 84646348 | 84645470 | 84645836 | 20 | 20 | 33 | 30 | 11 | 17 | 11 | 18 | 8 | 12 | 33 | 33 | 42 | 22 | 32 | 23 | 17 | 14 | 9 | 13 | 298 | 149 | 0.384449647 | 1 | 0.37 | 0.476 | 0.478 | 0.652 | 0.314 | 0.418 | 0.493 | 0.6 | 0.55 | 0.552 | -0.065 |
| A5SS | 5794 | ENSG00000197635 | DPP4 | chr2 | - | 162046701 | 162047006 | 162046914 | 162047006 | 162045531 | 162045612 | 6 | 6 | 6 | 10 | 16 | 121 | 143 | 183 | 107 | 286 | 0 | 2 | 5 | 6 | 4 | 22 | 34 | 82 | 55 | 0 | 298 | 149 | 0.384862077 | 1 | 0.024 | 0.021 | 0.016 | 0.045 | 0.027 | 0 | 0.029 | 0.03 | 0.052 | 1 | -0.196 |
| A5SS | 777 | ENSG00000185624 | P4HB | chr17 | - | 81845459 | 81845742 | 81845626 | 81845742 | 81845143 | 81845230 | 286 | 184 | 253 | 251 | 323 | 6 | 0 | 0 | 1 | 0 | 763 | 372 | 310 | 570 | 745 | 0 | 2 | 18 | 2 | 0 | 298 | 149 | 0.385845443 | 1 | 0.96 | 1 | 1 | 0.992 | 1 | 1 | 0.989 | 0.896 | 0.993 | 1 | 0.015 |
| A5SS | 1653 | ENSG00000139998 | RAB15 | chr14 | - | 64950578 | 64951151 | 64950942 | 64951151 | 64950324 | 64950414 | 16 | 5 | 5 | 8 | 8 | 3 | 0 | 1 | 1 | 0 | 12 | 6 | 21 | 12 | 21 | 24 | 0 | 1 | 0 | 0 | 298 | 149 | 0.386113004 | 1 | 0.727 | 1 | 0.714 | 0.8 | 1 | 0.2 | 1 | 0.913 | 1 | 1 | 0.026 |
| A5SS | 7427 | ENSG00000104763 | ASAH1 | chr8 | - | 18083980 | 18084980 | 18084675 | 18084980 | 18075540 | 18075587 | 119 | 50 | 88 | 64 | 24 | 3 | 6 | 11 | 15 | 4 | 20 | 74 | 102 | 105 | 102 | 0 | 16 | 4 | 5 | 13 | 298 | 149 | 0.386928579 | 1 | 0.952 | 0.806 | 0.8 | 0.681 | 0.75 | 1 | 0.698 | 0.927 | 0.913 | 0.797 | -0.069 |
| A5SS | 2912 | ENSG00000009790 | TRAF3IP3 | chr1 | + | 209759981 | 209760384 | 209759981 | 209760324 | 209762514 | 209762662 | 324 | 204 | 248 | 235 | 217 | 17 | 16 | 20 | 26 | 15 | 248 | 395 | 207 | 148 | 277 | 8 | 22 | 6 | 20 | 28 | 209 | 149 | 0.38710597 | 1 | 0.931 | 0.901 | 0.898 | 0.866 | 0.912 | 0.957 | 0.928 | 0.961 | 0.841 | 0.876 | -0.011 |
| A5SS | 7687 | ENSG00000096746 | HNRNPH3 | chr10 | + | 68338502 | 68338687 | 68338502 | 68338642 | 68339139 | 68339226 | 214 | 152 | 166 | 228 | 340 | 41 | 44 | 71 | 53 | 98 | 134 | 104 | 161 | 116 | 43 | 19 | 53 | 53 | 53 | 12 | 194 | 149 | 0.387924667 | 1 | 0.8 | 0.726 | 0.642 | 0.768 | 0.727 | 0.844 | 0.601 | 0.7 | 0.627 | 0.733 | 0.032 |
| A5SS | 7033 | ENSG00000102030 | NAA10 | chrX | - | 153933942 | 153934001 | 153933960 | 153934001 | 153932538 | 153932584 | 161 | 94 | 141 | 121 | 246 | 4 | 5 | 2 | 2 | 1 | 183 | 176 | 106 | 176 | 334 | 0 | 0 | 6 | 8 | 8 | 167 | 149 | 0.388941185 | 1 | 0.973 | 0.944 | 0.984 | 0.982 | 0.995 | 1 | 1 | 0.94 | 0.952 | 0.974 | 0.002 |
| A5SS | 7141 | ENSG00000071553 | ATP6AP1 | chrX | + | 154429047 | 154431657 | 154429047 | 154429174 | 154431829 | 154431904 | 16 | 8 | 11 | 17 | 6 | 100 | 38 | 64 | 73 | 23 | 19 | 11 | 15 | 50 | 41 | 157 | 119 | 61 | 62 | 202 | 298 | 149 | 0.389133223 | 1 | 0.074 | 0.095 | 0.079 | 0.104 | 0.115 | 0.057 | 0.044 | 0.109 | 0.287 | 0.092 | -0.024 |
| A5SS | 264 | ENSG00000161265 | U2AF1L4 | chr19 | - | 35744011 | 35744145 | 35744069 | 35744145 | 35743808 | 35743904 | 6 | 3 | 3 | 6 | 24 | 88 | 32 | 89 | 71 | 64 | 20 | 1 | 15 | 2 | 2 | 113 | 101 | 55 | 90 | 56 | 207 | 149 | 0.389142032 | 1 | 0.047 | 0.063 | 0.024 | 0.057 | 0.213 | 0.113 | 0.007 | 0.164 | 0.016 | 0.025 | 0.016 |
| A5SS | 5426 | ENSG00000162923 | WDR26 | chr1 | - | 224431476 | 224431581 | 224431524 | 224431581 | 224419517 | 224419615 | 51 | 47 | 38 | 91 | 30 | 3 | 0 | 0 | 2 | 3 | 16 | 82 | 87 | 56 | 48 | 0 | 0 | 6 | 0 | 0 | 197 | 149 | 0.389565882 | 1 | 0.928 | 1 | 1 | 0.972 | 0.883 | 1 | 1 | 0.916 | 1 | 1 | -0.027 |
| A5SS | 1220 | ENSG00000130175 | PRKCSH | chr19 | + | 11447438 | 11447618 | 11447438 | 11447597 | 11447722 | 11448018 | 30 | 13 | 15 | 30 | 33 | 1 | 0 | 0 | 0 | 6 | 6 | 17 | 24 | 33 | 37 | 0 | 0 | 0 | 0 | 0 | 170 | 149 | 0.389587252 | 1 | 0.963 | 1 | 1 | 1 | 0.828 | 1 | 1 | 1 | 1 | 1 | -0.042 |
| A5SS | 5156 | ENSG00000197062 | ZSCAN26 | chr6 | + | 28271853 | 28272336 | 28271853 | 28271934 | 28272669 | 28272787 | 47 | 16 | 47 | 27 | 32 | 34 | 22 | 29 | 27 | 21 | 1 | 23 | 52 | 11 | 8 | 22 | 11 | 18 | 2 | 0 | 298 | 149 | 0.390608126 | 1 | 0.409 | 0.267 | 0.448 | 0.333 | 0.432 | 0.022 | 0.511 | 0.591 | 0.733 | 1 | -0.194 |
| A5SS | 7442 | ENSG00000168522 | FNTA | chr8 | + | 43077215 | 43080441 | 43077215 | 43077364 | 43083117 | 43083180 | 162 | 132 | 111 | 149 | 188 | 198 | 153 | 166 | 158 | 464 | 60 | 102 | 133 | 138 | 11 | 89 | 168 | 90 | 119 | 30 | 298 | 149 | 0.391147276 | 1 | 0.29 | 0.301 | 0.251 | 0.32 | 0.168 | 0.252 | 0.233 | 0.425 | 0.367 | 0.155 | -0.02 |
| A5SS | 2403 | ENSG00000070756 | PABPC1 | chr8 | - | 100715461 | 100715601 | 100715470 | 100715601 | 100713086 | 100713181 | 9003 | 7553 | 9069 | 8757 | 16337 | 138 | 130 | 188 | 166 | 340 | 11306 | 5135 | 6451 | 6443 | 11758 | 200 | 77 | 156 | 113 | 160 | 158 | 149 | 0.39172759 | 1 | 0.984 | 0.982 | 0.978 | 0.98 | 0.978 | 0.982 | 0.984 | 0.975 | 0.982 | 0.986 | -0.001 |
| A5SS | 2723 | ENSG00000155508 | CNOT8 | chr5 | + | 154858638 | 154858816 | 154858638 | 154858801 | 154865191 | 154865385 | 29 | 27 | 10 | 10 | 9 | 1 | 0 | 0 | 0 | 0 | 19 | 3 | 15 | 14 | 2 | 7 | 0 | 0 | 0 | 0 | 164 | 149 | 0.392124866 | 1 | 0.963 | 1 | 1 | 1 | 1 | 0.711 | 1 | 1 | 1 | 1 | 0.05 |
| A5SS | 7446 | ENSG00000107864 | CPEB3 | chr10 | - | 92192476 | 92192636 | 92192545 | 92192636 | 92180962 | 92181019 | 12 | 27 | 12 | 27 | 17 | 24 | 16 | 12 | 13 | 14 | 13 | 50 | 12 | 18 | 16 | 31 | 15 | 17 | 28 | 21 | 218 | 149 | 0.392138216 | 1 | 0.255 | 0.536 | 0.406 | 0.587 | 0.454 | 0.223 | 0.695 | 0.325 | 0.305 | 0.342 | 0.07 |
| A5SS | 3815 | ENSG00000175390 | EIF3F | chr11 | + | 7991780 | 7991971 | 7991780 | 7991851 | 7992083 | 7992163 | 50 | 29 | 35 | 33 | 29 | 712 | 338 | 742 | 607 | 802 | 5 | 46 | 38 | 59 | 30 | 573 | 583 | 314 | 434 | 1062 | 269 | 149 | 0.392324605 | 1 | 0.037 | 0.045 | 0.025 | 0.029 | 0.02 | 0.005 | 0.042 | 0.063 | 0.07 | 0.015 | -0.008 |
| A5SS | 5935 | ENSG00000124356 | STAMBP | chr2 | + | 73847386 | 73847753 | 73847386 | 73847572 | 73849362 | 73849487 | 147 | 96 | 100 | 129 | 177 | 0 | 3 | 5 | 4 | 4 | 78 | 169 | 90 | 110 | 97 | 15 | 6 | 0 | 0 | 0 | 298 | 149 | 0.392538613 | 1 | 1 | 0.941 | 0.909 | 0.942 | 0.957 | 0.722 | 0.934 | 1 | 1 | 1 | 0.019 |
| A5SS | 7926 | ENSG00000139626 | ITGB7 | chr12 | - | 53201117 | 53201194 | 53201146 | 53201194 | 53200242 | 53200446 | 68 | 28 | 76 | 64 | 86 | 9 | 0 | 2 | 0 | 11 | 52 | 45 | 21 | 25 | 35 | 0 | 6 | 0 | 0 | 21 | 178 | 149 | 0.393292227 | 1 | 0.863 | 1 | 0.97 | 1 | 0.867 | 1 | 0.863 | 1 | 1 | 0.582 | 0.051 |
| A5SS | 6893 | ENSG00000183401 | CCDC159 | chr19 | + | 11346530 | 11346634 | 11346530 | 11346627 | 11349653 | 11349687 | 10 | 8 | 8 | 8 | 4 | 9 | 12 | 18 | 15 | 10 | 19 | 16 | 8 | 14 | 12 | 52 | 11 | 29 | 51 | 28 | 156 | 149 | 0.393530074 | 1 | 0.515 | 0.389 | 0.298 | 0.337 | 0.276 | 0.259 | 0.581 | 0.209 | 0.208 | 0.29 | 0.054 |
| A5SS | 6629 | ENSG00000182165 | TP53TG1 | chr7 | - | 87345042 | 87345528 | 87345304 | 87345528 | 87341522 | 87341684 | 36 | 18 | 32 | 38 | 24 | 4 | 6 | 4 | 2 | 1 | 11 | 27 | 11 | 40 | 27 | 0 | 1 | 1 | 7 | 2 | 298 | 149 | 0.394417775 | 1 | 0.818 | 0.6 | 0.8 | 0.905 | 0.923 | 1 | 0.931 | 0.846 | 0.741 | 0.871 | -0.069 |
| A5SS | 7244 | ENSG00000145868 | FBXO38 | chr5 | + | 148427212 | 148427947 | 148427212 | 148427722 | 148433423 | 148433524 | 90 | 73 | 76 | 104 | 61 | 12 | 8 | 23 | 25 | 23 | 30 | 74 | 38 | 21 | 12 | 9 | 15 | 8 | 2 | 0 | 298 | 149 | 0.39487144 | 1 | 0.789 | 0.82 | 0.623 | 0.675 | 0.57 | 0.625 | 0.712 | 0.704 | 0.84 | 1 | -0.081 |
| A5SS | 7645 | ENSG00000139644 | TMBIM6 | chr12 | + | 49741956 | 49742280 | 49741956 | 49742115 | 49752463 | 49752549 | 13 | 2 | 7 | 9 | 0 | 26 | 7 | 4 | 14 | 7 | 0 | 5 | 7 | 31 | 18 | 8 | 11 | 7 | 11 | 32 | 298 | 149 | 0.394943346 | 1 | 0.2 | 0.125 | 0.467 | 0.243 | 0 | 0 | 0.185 | 0.333 | 0.585 | 0.22 | -0.058 |
| A5SS | 1643 | ENSG00000092148 | HECTD1 | chr14 | - | 31122913 | 31123047 | 31122919 | 31123047 | 31121382 | 31121505 | 166 | 151 | 126 | 152 | 238 | 33 | 33 | 23 | 42 | 57 | 9 | 112 | 151 | 43 | 4 | 0 | 24 | 31 | 11 | 0 | 155 | 149 | 0.395993699 | 1 | 0.829 | 0.815 | 0.84 | 0.777 | 0.801 | 1 | 0.818 | 0.824 | 0.79 | 1 | -0.074 |
| A5SS | 5159 | ENSG00000141367 | CLTC | chr17 | + | 59644275 | 59644495 | 59644275 | 59644483 | 59647397 | 59647666 | 1 | 2 | 2 | 1 | 4 | 247 | 168 | 155 | 175 | 310 | 0 | 12 | 2 | 1 | 0 | 74 | 151 | 194 | 169 | 89 | 161 | 149 | 0.396690864 | 1 | 0.004 | 0.011 | 0.012 | 0.005 | 0.012 | 0 | 0.069 | 0.009 | 0.005 | 0 | -0.008 |
| A5SS | 6515 | ENSG00000179818 | PCBP1-AS1 | chr2 | - | 70085815 | 70086272 | 70086123 | 70086272 | 70085546 | 70085624 | 50 | 39 | 50 | 52 | 67 | 4 | 9 | 10 | 6 | 10 | 36 | 57 | 32 | 95 | 61 | 10 | 2 | 9 | 13 | 0 | 298 | 149 | 0.396861795 | 1 | 0.862 | 0.684 | 0.714 | 0.812 | 0.77 | 0.643 | 0.934 | 0.64 | 0.785 | 1 | -0.032 |
| A5SS | 1829 | ENSG00000119688 | ABCD4 | chr14 | - | 74297638 | 74298069 | 74297929 | 74298069 | 74296332 | 74296449 | 11 | 6 | 9 | 16 | 9 | 17 | 15 | 20 | 23 | 11 | 29 | 4 | 9 | 13 | 15 | 19 | 30 | 7 | 10 | 24 | 298 | 149 | 0.397087649 | 1 | 0.244 | 0.167 | 0.184 | 0.258 | 0.29 | 0.433 | 0.062 | 0.391 | 0.394 | 0.238 | -0.075 |
| A5SS | 2248 | ENSG00000104951 | IL4I1 | chr19 | - | 49929325 | 49929496 | 49929416 | 49929496 | 49927693 | 49927815 | 34 | 22 | 28 | 32 | 19 | 3 | 2 | 0 | 1 | 7 | 50 | 40 | 45 | 18 | 77 | 0 | 14 | 0 | 2 | 0 | 240 | 149 | 0.398356686 | 1 | 0.876 | 0.872 | 1 | 0.952 | 0.628 | 1 | 0.639 | 1 | 0.848 | 1 | -0.032 |
| A5SS | 7335 | ENSG00000198851 | CD3E | chr11 | + | 118312152 | 118313874 | 118312152 | 118312170 | 118314447 | 118314494 | 3293 | 2329 | 3506 | 3746 | 9711 | 11 | 8 | 8 | 4 | 23 | 1889 | 3453 | 2135 | 1499 | 3094 | 15 | 28 | 16 | 5 | 0 | 298 | 149 | 0.398423553 | 1 | 0.993 | 0.993 | 0.995 | 0.998 | 0.995 | 0.984 | 0.984 | 0.985 | 0.993 | 1 | 0.006 |
| A5SS | 2438 | ENSG00000156253 | RWDD2B | chr21 | - | 29008239 | 29008621 | 29008394 | 29008621 | 29007760 | 29008123 | 28 | 5 | 17 | 19 | 31 | 5 | 9 | 7 | 8 | 8 | 20 | 7 | 8 | 4 | 16 | 3 | 3 | 2 | 8 | 0 | 298 | 149 | 0.399508512 | 1 | 0.737 | 0.217 | 0.548 | 0.543 | 0.66 | 0.769 | 0.538 | 0.667 | 0.2 | 1 | -0.094 |
| A5SS | 2921 | ENSG00000176261 | ZBTB8OS | chr1 | - | 32634701 | 32634792 | 32634767 | 32634792 | 32633950 | 32634110 | 13 | 15 | 7 | 14 | 10 | 12 | 4 | 7 | 6 | 17 | 25 | 10 | 12 | 16 | 20 | 0 | 21 | 11 | 18 | 4 | 215 | 149 | 0.401745688 | 1 | 0.429 | 0.722 | 0.409 | 0.618 | 0.29 | 1 | 0.248 | 0.431 | 0.381 | 0.776 | -0.074 |
| A5SS | 4648 | ENSG00000060138 | YBX3 | chr12 | - | 10704050 | 10704148 | 10704059 | 10704148 | 10701959 | 10702134 | 234 | 106 | 122 | 187 | 232 | 14 | 9 | 2 | 13 | 25 | 324 | 405 | 214 | 270 | 958 | 44 | 36 | 9 | 18 | 61 | 158 | 149 | 0.402577667 | 1 | 0.94 | 0.917 | 0.983 | 0.931 | 0.897 | 0.874 | 0.914 | 0.957 | 0.934 | 0.937 | 0.01 |
| A5SS | 4333 | ENSG00000117308 | GALE | chr1 | - | 23796508 | 23796782 | 23796696 | 23796782 | 23796150 | 23796265 | 22 | 13 | 15 | 18 | 10 | 2 | 0 | 0 | 0 | 0 | 45 | 17 | 7 | 7 | 23 | 0 | 0 | 4 | 0 | 0 | 298 | 149 | 0.403789227 | 1 | 0.846 | 1 | 1 | 1 | 1 | 1 | 1 | 0.467 | 1 | 1 | 0.076 |
| A5SS | 5033 | ENSG00000162650 | ATXN7L2 | chr1 | + | 109488846 | 109490128 | 109488846 | 109489100 | 109490270 | 109490392 | 17 | 5 | 13 | 20 | 10 | 0 | 0 | 0 | 0 | 0 | 6 | 38 | 5 | 7 | 13 | 0 | 0 | 0 | 4 | 0 | 298 | 149 | 0.403798107 | 1 | 1 | 1 | 1 | 1 | 1 | 1 | 1 | 1 | 0.467 | 1 | 0.107 |
| A5SS | 6255 | ENSG00000165568 | AKR1E2 | chr10 | + | 4847147 | 4847299 | 4847147 | 4847230 | 4847487 | 4848062 | 7 | 2 | 4 | 2 | 18 | 1 | 0 | 1 | 1 | 11 | 6 | 2 | 8 | 7 | 15 | 0 | 5 | 9 | 0 | 0 | 218 | 149 | 0.405427034 | 1 | 0.827 | 1 | 0.732 | 0.578 | 0.528 | 1 | 0.215 | 0.378 | 1 | 1 | 0.014 |
| A5SS | 5675 | ENSG00000102125 | TAZ | chrX | + | 154413206 | 154413339 | 154413206 | 154413252 | 154413481 | 154413567 | 3 | 1 | 7 | 6 | 2 | 20 | 9 | 16 | 12 | 11 | 11 | 4 | 4 | 8 | 9 | 21 | 36 | 11 | 19 | 24 | 236 | 149 | 0.406109338 | 1 | 0.087 | 0.066 | 0.216 | 0.24 | 0.103 | 0.249 | 0.066 | 0.187 | 0.21 | 0.191 | -0.038 |
| A5SS | 1520 | ENSG00000092439 | TRPM7 | chr15 | - | 50578558 | 50578664 | 50578638 | 50578664 | 50575868 | 50575919 | 7 | 13 | 13 | 12 | 32 | 105 | 97 | 85 | 116 | 146 | 0 | 28 | 9 | 0 | 0 | 26 | 80 | 76 | 30 | 43 | 229 | 149 | 0.406120396 | 1 | 0.042 | 0.08 | 0.091 | 0.063 | 0.125 | 0 | 0.185 | 0.072 | 0 | 0 | 0.029 |
| A5SS | 2048 | ENSG00000228903 | RASA4CP | chr7 | - | 44033932 | 44034311 | 44034175 | 44034311 | 44030780 | 44030901 | 2 | 6 | 6 | 4 | 8 | 1 | 1 | 14 | 15 | 9 | 9 | 0 | 3 | 0 | 7 | 0 | 14 | 2 | 3 | 15 | 298 | 149 | 0.406456249 | 1 | 0.5 | 0.75 | 0.176 | 0.118 | 0.308 | 1 | 0 | 0.429 | 0 | 0.189 | 0.047 |
| A5SS | 8187 | ENSG00000204356 | NELFE | chr6 | - | 31958307 | 31958454 | 31958371 | 31958454 | 31956940 | 31957010 | 3 | 6 | 7 | 6 | 5 | 54 | 43 | 47 | 58 | 59 | 29 | 4 | 8 | 7 | 6 | 114 | 74 | 54 | 88 | 108 | 213 | 149 | 0.406817878 | 1 | 0.037 | 0.089 | 0.094 | 0.067 | 0.056 | 0.151 | 0.036 | 0.094 | 0.053 | 0.037 | -0.006 |
| A5SS | 7370 | ENSG00000172269 | DPAGT1 | chr11 | - | 119097444 | 119097551 | 119097463 | 119097551 | 119097141 | 119097297 | 4 | 0 | 3 | 1 | 0 | 36 | 25 | 57 | 57 | 61 | 0 | 0 | 2 | 3 | 0 | 45 | 31 | 17 | 16 | 57 | 168 | 149 | 0.409268804 | 1 | 0.09 | 0 | 0.045 | 0.015 | 0 | 0 | 0 | 0.094 | 0.143 | 0 | -0.017 |
| A5SS | 6349 | ENSG00000127980 | PEX1 | chr7 | - | 92518959 | 92519078 | 92518994 | 92519078 | 92518140 | 92518255 | 20 | 13 | 13 | 22 | 42 | 63 | 44 | 92 | 58 | 127 | 0 | 13 | 2 | 13 | 8 | 10 | 28 | 43 | 35 | 21 | 184 | 149 | 0.410160888 | 1 | 0.205 | 0.193 | 0.103 | 0.235 | 0.211 | 0 | 0.273 | 0.036 | 0.231 | 0.236 | 0.034 |
| A5SS | 1627 | ENSG00000165795 | NDRG2 | chr14 | - | 21019613 | 21019742 | 21019638 | 21019742 | 21019115 | 21019160 | 0 | 0 | 0 | 0 | 0 | 31 | 24 | 65 | 51 | 51 | 7 | 0 | 0 | 0 | 0 | 42 | 19 | 9 | 8 | 10 | 174 | 149 | 0.410291656 | 1 | 0 | 0 | 0 | 0 | 0 | 0.125 | 0 | 0 | 0 | 0 | -0.025 |
| A5SS | 939 | ENSG00000108384 | RAD51C | chr17 | + | 58692572 | 58692788 | 58692572 | 58692685 | 58694930 | 58695189 | 29 | 38 | 54 | 41 | 19 | 0 | 2 | 8 | 1 | 0 | 41 | 32 | 48 | 30 | 40 | 0 | 11 | 6 | 0 | 0 | 252 | 149 | 0.411177792 | 1 | 1 | 0.918 | 0.8 | 0.96 | 1 | 1 | 0.632 | 0.825 | 1 | 1 | 0.044 |
| A5SS | 4669 | ENSG00000153208 | MERTK | chr2 | + | 111929119 | 111929540 | 111929119 | 111929304 | 111944959 | 111945060 | 28 | 29 | 27 | 32 | 22 | 3 | 0 | 4 | 1 | 0 | 78 | 39 | 33 | 93 | 34 | 32 | 0 | 0 | 0 | 0 | 298 | 149 | 0.411261283 | 1 | 0.824 | 1 | 0.771 | 0.941 | 1 | 0.549 | 1 | 1 | 1 | 1 | -0.003 |
| A5SS | 4758 | ENSG00000232956 | SNHG15 | chr7 | - | 44985881 | 44986097 | 44986020 | 44986097 | 44984356 | 44984426 | 12 | 8 | 21 | 13 | 17 | 14 | 7 | 20 | 13 | 9 | 25 | 35 | 16 | 15 | 40 | 15 | 5 | 21 | 33 | 17 | 288 | 149 | 0.41163604 | 1 | 0.307 | 0.372 | 0.352 | 0.341 | 0.494 | 0.463 | 0.784 | 0.283 | 0.19 | 0.549 | -0.081 |
| A5SS | 1923 | ENSG00000175048 | ZDHHC14 | chr6 | + | 157628348 | 157628486 | 157628348 | 157628374 | 157632833 | 157632882 | 27 | 36 | 24 | 50 | 55 | 1 | 0 | 0 | 4 | 2 | 41 | 32 | 14 | 28 | 23 | 0 | 1 | 0 | 0 | 0 | 261 | 149 | 0.411750453 | 1 | 0.939 | 1 | 1 | 0.877 | 0.94 | 1 | 0.948 | 1 | 1 | 1 | -0.038 |
| A5SS | 7657 | ENSG00000197070 | ARRDC1 | chr9 | + | 137614558 | 137615000 | 137614558 | 137614898 | 137615073 | 137615360 | 31 | 13 | 13 | 24 | 12 | 0 | 0 | 0 | 0 | 0 | 25 | 18 | 14 | 16 | 31 | 0 | 0 | 0 | 0 | 7 | 251 | 149 | 0.41317805 | 1 | 1 | 1 | 1 | 1 | 1 | 1 | 1 | 1 | 1 | 0.724 | 0.055 |
| A5SS | 4638 | ENSG00000106665 | CLIP2 | chr7 | + | 74386520 | 74386604 | 74386520 | 74386597 | 74389102 | 74389259 | 19 | 21 | 8 | 18 | 23 | 0 | 2 | 0 | 0 | 0 | 69 | 9 | 4 | 6 | 94 | 0 | 0 | 0 | 0 | 8 | 156 | 149 | 0.413708249 | 1 | 1 | 0.909 | 1 | 1 | 1 | 1 | 1 | 1 | 1 | 0.918 | -0.002 |
| A5SS | 5909 | ENSG00000115307 | AUP1 | chr2 | - | 74528750 | 74529282 | 74529131 | 74529282 | 74528416 | 74528489 | 210 | 96 | 137 | 215 | 204 | 4 | 2 | 13 | 10 | 16 | 345 | 279 | 143 | 250 | 570 | 28 | 13 | 13 | 27 | 9 | 298 | 149 | 0.414006217 | 1 | 0.963 | 0.96 | 0.84 | 0.915 | 0.864 | 0.86 | 0.915 | 0.846 | 0.822 | 0.969 | 0.026 |
| A5SS | 4539 | ENSG00000143258 | USP21 | chr1 | + | 161163877 | 161163981 | 161163877 | 161163922 | 161164163 | 161164250 | 32 | 19 | 45 | 42 | 50 | 8 | 4 | 4 | 9 | 3 | 39 | 24 | 26 | 17 | 78 | 9 | 1 | 6 | 1 | 22 | 208 | 149 | 0.41414724 | 1 | 0.741 | 0.773 | 0.89 | 0.77 | 0.923 | 0.756 | 0.945 | 0.756 | 0.924 | 0.717 | 0 |
| A5SS | 7800 | ENSG00000106397 | PLOD3 | chr7 | - | 101216162 | 101216326 | 101216196 | 101216326 | 101215907 | 101216020 | 91 | 48 | 40 | 53 | 94 | 4 | 2 | 3 | 3 | 4 | 85 | 94 | 34 | 42 | 140 | 0 | 3 | 0 | 4 | 7 | 183 | 149 | 0.414166348 | 1 | 0.949 | 0.951 | 0.916 | 0.935 | 0.95 | 1 | 0.962 | 1 | 0.895 | 0.942 | -0.02 |
| A5SS | 5596 | ENSG00000198901 | PRC1 | chr15 | - | 90969446 | 90969623 | 90969523 | 90969623 | 90969078 | 90969120 | 5 | 8 | 2 | 3 | 4 | 0 | 0 | 0 | 0 | 0 | 16 | 15 | 12 | 33 | 12 | 0 | 0 | 5 | 0 | 0 | 226 | 149 | 0.414355653 | 1 | 1 | 1 | 1 | 1 | 1 | 1 | 1 | 0.613 | 1 | 1 | 0.077 |
| A5SS | 8667 | ENSG00000127314 | RAP1B | chr12 | + | 68610928 | 68611056 | 68610928 | 68611043 | 68648698 | 68648781 | 2 | 2 | 3 | 3 | 1 | 635 | 465 | 365 | 613 | 429 | 5 | 0 | 0 | 4 | 12 | 157 | 485 | 366 | 251 | 95 | 162 | 149 | 0.41644629 | 1 | 0.003 | 0.004 | 0.008 | 0.004 | 0.002 | 0.028 | 0 | 0 | 0.014 | 0.104 | -0.025 |
| A5SS | 2897 | ENSG00000105373 | NOP53 | chr19 | + | 47752511 | 47752559 | 47752511 | 47752544 | 47755473 | 47755523 | 22 | 11 | 55 | 86 | 32 | 4 | 1 | 2 | 1 | 0 | 33 | 29 | 7 | 10 | 58 | 0 | 0 | 0 | 0 | 0 | 164 | 149 | 0.416670386 | 1 | 0.833 | 0.909 | 0.962 | 0.987 | 1 | 1 | 1 | 1 | 1 | 1 | -0.062 |
| A5SS | 8306 | ENSG00000198056 | PRIM1 | chr12 | - | 56752186 | 56752323 | 56752195 | 56752323 | 56751037 | 56751195 | 0 | 1 | 2 | 0 | 0 | 64 | 35 | 70 | 61 | 59 | 0 | 2 | 0 | 5 | 0 | 4 | 27 | 18 | 34 | 16 | 158 | 149 | 0.416772532 | 1 | 0 | 0.026 | 0.026 | 0 | 0 | 0 | 0.065 | 0 | 0.122 | 0 | -0.027 |
| A5SS | 1295 | ENSG00000103005 | USB1 | chr16 | + | 58001406 | 58001588 | 58001406 | 58001581 | 58002478 | 58002645 | 1 | 0 | 1 | 0 | 2 | 37 | 23 | 31 | 21 | 3 | 0 | 0 | 5 | 0 | 0 | 19 | 34 | 5 | 26 | 13 | 156 | 149 | 0.417199755 | 1 | 0.025 | 0 | 0.03 | 0 | 0.389 | 0 | 0 | 0.489 | 0 | 0 | -0.009 |
| A5SS | 148 | ENSG00000160392 | C19orf47 | chr19 | - | 40341785 | 40341890 | 40341838 | 40341890 | 40336319 | 40336407 | 3 | 1 | 8 | 2 | 0 | 12 | 9 | 10 | 11 | 11 | 0 | 8 | 2 | 0 | 3 | 24 | 11 | 7 | 14 | 22 | 202 | 149 | 0.417364521 | 1 | 0.156 | 0.076 | 0.371 | 0.118 | 0 | 0 | 0.349 | 0.174 | 0 | 0.091 | 0.021 |
| A5SS | 7317 | ENSG00000166169 | POLL | chr10 | - | 101584601 | 101584919 | 101584877 | 101584919 | 101583507 | 101583681 | 79 | 29 | 36 | 52 | 67 | 9 | 3 | 6 | 5 | 10 | 21 | 37 | 40 | 56 | 89 | 24 | 8 | 0 | 3 | 24 | 298 | 149 | 0.418051555 | 1 | 0.814 | 0.829 | 0.75 | 0.839 | 0.77 | 0.304 | 0.698 | 1 | 0.903 | 0.65 | 0.089 |
| A5SS | 4803 | ENSG00000103365 | GGA2 | chr16 | - | 23495688 | 23495778 | 23495693 | 23495778 | 23494302 | 23494378 | 3 | 2 | 2 | 1 | 0 | 43 | 22 | 33 | 23 | 21 | 0 | 0 | 0 | 0 | 10 | 27 | 34 | 15 | 33 | 62 | 154 | 149 | 0.418294612 | 1 | 0.063 | 0.081 | 0.055 | 0.04 | 0 | 0 | 0 | 0 | 0 | 0.135 | 0.021 |
| A5SS | 5683 | ENSG00000040933 | INPP4A | chr2 | + | 98555552 | 98555808 | 98555552 | 98555691 | 98559462 | 98559495 | 163 | 73 | 156 | 189 | 130 | 3 | 0 | 0 | 8 | 0 | 67 | 167 | 112 | 52 | 135 | 0 | 2 | 0 | 0 | 0 | 266 | 149 | 0.418353966 | 1 | 0.968 | 1 | 1 | 0.93 | 1 | 1 | 0.979 | 1 | 1 | 1 | -0.016 |
| A5SS | 7757 | ENSG00000165699 | TSC1 | chr9 | - | 132911409 | 132911568 | 132911452 | 132911568 | 132911001 | 132911113 | 3 | 2 | 3 | 4 | 0 | 24 | 12 | 14 | 22 | 50 | 0 | 0 | 7 | 0 | 0 | 7 | 11 | 18 | 9 | 18 | 192 | 149 | 0.41859732 | 1 | 0.088 | 0.115 | 0.143 | 0.124 | 0 | 0 | 0 | 0.232 | 0 | 0 | 0.048 |
| A5SS | 1703 | ENSG00000171469 | ZNF561 | chr19 | - | 9620915 | 9621231 | 9621161 | 9621231 | 9619431 | 9619582 | 6 | 3 | 20 | 6 | 6 | 9 | 14 | 7 | 13 | 4 | 1 | 4 | 6 | 3 | 0 | 4 | 0 | 6 | 8 | 2 | 298 | 149 | 0.418778205 | 1 | 0.25 | 0.097 | 0.588 | 0.188 | 0.429 | 0.111 | 1 | 0.333 | 0.158 | 0 | -0.01 |
| A5SS | 2064 | ENSG00000177595 | PIDD1 | chr11 | - | 800332 | 800451 | 800383 | 800451 | 800130 | 800244 | 32 | 5 | 10 | 16 | 11 | 8 | 2 | 2 | 4 | 1 | 19 | 30 | 11 | 18 | 18 | 0 | 0 | 8 | 0 | 19 | 200 | 149 | 0.418950156 | 1 | 0.749 | 0.651 | 0.788 | 0.749 | 0.891 | 1 | 1 | 0.506 | 1 | 0.414 | -0.018 |
| A5SS | 6187 | ENSG00000124508 | BTN2A2 | chr6 | + | 26383125 | 26383299 | 26383125 | 26383181 | 26383791 | 26383915 | 37 | 12 | 14 | 21 | 4 | 14 | 7 | 8 | 18 | 0 | 8 | 24 | 19 | 15 | 31 | 4 | 13 | 11 | 0 | 8 | 267 | 149 | 0.41949238 | 1 | 0.596 | 0.489 | 0.494 | 0.394 | 1 | 0.527 | 0.507 | 0.491 | 1 | 0.684 | -0.047 |
| A5SS | 5644 | ENSG00000132323 | ILKAP | chr2 | - | 238194781 | 238194870 | 238194804 | 238194870 | 238194274 | 238194331 | 11 | 0 | 18 | 11 | 7 | 20 | 5 | 27 | 13 | 16 | 0 | 24 | 17 | 26 | 0 | 7 | 33 | 16 | 18 | 62 | 172 | 149 | 0.421191345 | 1 | 0.323 | 0 | 0.366 | 0.423 | 0.275 | 0 | 0.387 | 0.479 | 0.556 | 0 | -0.007 |
| A5SS | 2084 | ENSG00000065357 | DGKA | chr12 | + | 55939414 | 55940170 | 55939414 | 55939529 | 55940313 | 55940433 | 194 | 176 | 351 | 309 | 267 | 14 | 8 | 20 | 9 | 12 | 196 | 182 | 174 | 215 | 153 | 0 | 8 | 23 | 18 | 7 | 298 | 149 | 0.421351516 | 1 | 0.874 | 0.917 | 0.898 | 0.945 | 0.918 | 1 | 0.919 | 0.791 | 0.857 | 0.916 | 0.014 |
| A5SS | 1346 | ENSG00000103319 | EEF2K | chr16 | + | 22258493 | 22258827 | 22258493 | 22258695 | 22260461 | 22260529 | 6 | 4 | 3 | 1 | 4 | 25 | 4 | 16 | 9 | 10 | 19 | 3 | 5 | 0 | 8 | 0 | 14 | 11 | 12 | 37 | 281 | 149 | 0.42433003 | 1 | 0.113 | 0.347 | 0.09 | 0.056 | 0.175 | 1 | 0.102 | 0.194 | 0 | 0.103 | -0.124 |
| A5SS | 3595 | ENSG00000133226 | SRRM1 | chr1 | + | 24654854 | 24655129 | 24654854 | 24655123 | 24660718 | 24660799 | 151 | 131 | 120 | 161 | 215 | 32 | 44 | 45 | 54 | 28 | 370 | 245 | 218 | 253 | 166 | 138 | 36 | 97 | 83 | 39 | 155 | 149 | 0.424400803 | 1 | 0.819 | 0.741 | 0.719 | 0.741 | 0.881 | 0.72 | 0.867 | 0.684 | 0.746 | 0.804 | 0.016 |
| A5SS | 848 | ENSG00000141295 | SCRN2 | chr17 | - | 47838269 | 47838450 | 47838335 | 47838450 | 47837719 | 47838002 | 16 | 12 | 29 | 15 | 11 | 1 | 4 | 9 | 0 | 2 | 9 | 17 | 13 | 5 | 20 | 0 | 2 | 4 | 3 | 0 | 215 | 149 | 0.424444996 | 1 | 0.917 | 0.675 | 0.691 | 1 | 0.792 | 1 | 0.855 | 0.693 | 0.536 | 1 | -0.002 |
| A5SS | 2976 | ENSG00000163660 | CCNL1 | chr3 | - | 157151728 | 157151930 | 157151916 | 157151930 | 157150281 | 157150381 | 586 | 485 | 611 | 699 | 795 | 7 | 6 | 4 | 2 | 1 | 131 | 415 | 410 | 100 | 67 | 3 | 4 | 6 | 0 | 0 | 298 | 149 | 0.424531211 | 1 | 0.977 | 0.976 | 0.987 | 0.994 | 0.997 | 0.956 | 0.981 | 0.972 | 1 | 1 | 0.004 |
| A5SS | 510 | ENSG00000101084 | RAB5IF | chr20 | + | 36609600 | 36609758 | 36609600 | 36609730 | 36612009 | 36612557 | 21 | 6 | 14 | 17 | 15 | 519 | 218 | 313 | 493 | 698 | 18 | 14 | 16 | 14 | 26 | 784 | 552 | 410 | 666 | 747 | 177 | 149 | 0.424627213 | 1 | 0.033 | 0.023 | 0.036 | 0.028 | 0.018 | 0.019 | 0.021 | 0.032 | 0.017 | 0.028 | 0.004 |
| A5SS | 1122 | ENSG00000099364 | FBXL19 | chr16 | + | 30928466 | 30928630 | 30928466 | 30928499 | 30930072 | 30930584 | 7 | 0 | 10 | 11 | 5 | 31 | 12 | 14 | 27 | 8 | 0 | 7 | 3 | 6 | 22 | 7 | 18 | 5 | 20 | 11 | 280 | 149 | 0.424801413 | 1 | 0.107 | 0 | 0.275 | 0.178 | 0.25 | 0 | 0.171 | 0.242 | 0.138 | 0.516 | -0.051 |
| A5SS | 2288 | ENSG00000082397 | EPB41L3 | chr18 | - | 5398020 | 5401043 | 5400968 | 5401043 | 5397057 | 5397426 | 7 | 6 | 5 | 3 | 3 | 0 | 4 | 0 | 5 | 5 | 0 | 2 | 5 | 1 | 3 | 4 | 0 | 0 | 4 | 0 | 298 | 149 | 0.426058643 | 1 | 1 | 0.429 | 1 | 0.231 | 0.231 | 0 | 1 | 1 | 0.111 | 1 | -0.044 |
| A5SS | 773 | ENSG00000131462 | TUBG1 | chr17 | + | 42610422 | 42610629 | 42610422 | 42610590 | 42612074 | 42612143 | 0 | 1 | 2 | 2 | 0 | 14 | 17 | 19 | 15 | 22 | 0 | 3 | 3 | 0 | 2 | 24 | 32 | 12 | 6 | 33 | 188 | 149 | 0.426135926 | 1 | 0 | 0.045 | 0.077 | 0.096 | 0 | 0 | 0.069 | 0.165 | 0 | 0.046 | -0.012 |
| A5SS | 6935 | ENSG00000140332 | TLE3 | chr15 | - | 70058158 | 70058291 | 70058185 | 70058291 | 70057458 | 70057658 | 125 | 43 | 117 | 97 | 126 | 19 | 2 | 15 | 12 | 20 | 289 | 178 | 261 | 226 | 291 | 42 | 25 | 39 | 35 | 41 | 176 | 149 | 0.426505224 | 1 | 0.848 | 0.948 | 0.868 | 0.873 | 0.842 | 0.853 | 0.858 | 0.85 | 0.845 | 0.857 | 0.023 |
| A5SS | 3334 | ENSG00000220804 | LINC01881 | chr2 | + | 242130654 | 242130866 | 242130654 | 242130812 | 242138783 | 242138888 | 14 | 11 | 17 | 9 | 13 | 9 | 5 | 9 | 7 | 16 | 18 | 11 | 11 | 10 | 8 | 43 | 9 | 0 | 15 | 0 | 203 | 149 | 0.42739237 | 1 | 0.533 | 0.618 | 0.581 | 0.486 | 0.374 | 0.235 | 0.473 | 1 | 0.329 | 1 | -0.089 |
| A5SS | 7478 | ENSG00000147687 | TATDN1 | chr8 | - | 124518741 | 124518881 | 124518817 | 124518881 | 124515886 | 124516030 | 26 | 25 | 30 | 37 | 54 | 118 | 102 | 110 | 114 | 125 | 0 | 23 | 18 | 21 | 0 | 55 | 66 | 53 | 12 | 14 | 225 | 149 | 0.428651477 | 1 | 0.127 | 0.14 | 0.153 | 0.177 | 0.222 | 0 | 0.188 | 0.184 | 0.537 | 0 | -0.018 |
| A5SS | 7761 | ENSG00000115459 | ELMOD3 | chr2 | + | 85355554 | 85355896 | 85355554 | 85355598 | 85356601 | 85356667 | 23 | 13 | 12 | 22 | 16 | 0 | 0 | 0 | 0 | 0 | 21 | 24 | 20 | 30 | 15 | 0 | 3 | 0 | 3 | 0 | 298 | 149 | 0.432073552 | 1 | 1 | 1 | 1 | 1 | 1 | 1 | 0.8 | 1 | 0.833 | 1 | 0.073 |
| A5SS | 5676 | ENSG00000102125 | TAZ | chrX | + | 154413206 | 154413352 | 154413206 | 154413252 | 154413481 | 154413567 | 3 | 1 | 5 | 6 | 1 | 20 | 9 | 16 | 12 | 11 | 11 | 4 | 4 | 2 | 9 | 21 | 36 | 11 | 19 | 24 | 249 | 149 | 0.432219262 | 1 | 0.082 | 0.062 | 0.158 | 0.23 | 0.052 | 0.239 | 0.062 | 0.179 | 0.059 | 0.183 | -0.028 |
| A5SS | 5459 | ENSG00000214517 | PPME1 | chr11 | + | 74250953 | 74251715 | 74250953 | 74251018 | 74253491 | 74254703 | 23 | 14 | 20 | 23 | 55 | 0 | 0 | 0 | 0 | 0 | 51 | 20 | 6 | 17 | 28 | 0 | 0 | 0 | 5 | 0 | 298 | 149 | 0.433224028 | 1 | 1 | 1 | 1 | 1 | 1 | 1 | 1 | 1 | 0.63 | 1 | 0.074 |
| A5SS | 6519 | ENSG00000179818 | PCBP1-AS1 | chr2 | - | 70086054 | 70086271 | 70086123 | 70086271 | 70085546 | 70085624 | 22 | 9 | 21 | 11 | 0 | 4 | 9 | 10 | 6 | 10 | 11 | 14 | 7 | 16 | 4 | 10 | 2 | 9 | 13 | 0 | 218 | 149 | 0.435510697 | 1 | 0.79 | 0.406 | 0.589 | 0.556 | 0 | 0.429 | 0.827 | 0.347 | 0.457 | 1 | -0.144 |
| A5SS | 4008 | ENSG00000111602 | TIMELESS | chr12 | - | 56423583 | 56423707 | 56423624 | 56423707 | 56423273 | 56423475 | 24 | 14 | 27 | 21 | 7 | 0 | 2 | 0 | 0 | 0 | 21 | 46 | 15 | 28 | 4 | 6 | 0 | 0 | 0 | 0 | 190 | 149 | 0.435733954 | 1 | 1 | 0.846 | 1 | 1 | 1 | 0.733 | 1 | 1 | 1 | 1 | 0.023 |
| A5SS | 7221 | ENSG00000168066 | SF1 | chr11 | - | 64767566 | 64767844 | 64767570 | 64767844 | 64767191 | 64767246 | 0 | 0 | 0 | 0 | 0 | 17 | 1 | 15 | 10 | 7 | 4 | 3 | 0 | 0 | 0 | 59 | 44 | 16 | 41 | 29 | 153 | 149 | 0.436596901 | 1 | 0 | 0 | 0 | 0 | 0 | 0.062 | 0.062 | 0 | 0 | 0 | -0.025 |
| A5SS | 4542 | ENSG00000082014 | SMARCD3 | chr7 | - | 151242464 | 151242603 | 151242480 | 151242603 | 151242136 | 151242232 | 4 | 1 | 0 | 1 | 0 | 38 | 22 | 26 | 20 | 43 | 0 | 0 | 0 | 0 | 0 | 85 | 54 | 41 | 41 | 127 | 165 | 149 | 0.437286296 | 1 | 0.087 | 0.039 | 0 | 0.043 | 0 | 0 | 0 | 0 | 0 | 0 | 0.034 |
| A5SS | 3829 | ENSG00000163714 | U2SURP | chr3 | + | 143021349 | 143021385 | 143021349 | 143021382 | 143021472 | 143021555 | 114 | 119 | 125 | 131 | 234 | 171 | 97 | 119 | 142 | 269 | 11 | 67 | 82 | 29 | 3 | 24 | 78 | 89 | 33 | 4 | 152 | 149 | 0.438407641 | 1 | 0.395 | 0.546 | 0.507 | 0.475 | 0.46 | 0.31 | 0.457 | 0.475 | 0.463 | 0.424 | 0.051 |
| A5SS | 8453 | ENSG00000107862 | GBF1 | chr10 | + | 102362474 | 102362664 | 102362474 | 102362628 | 102363255 | 102363396 | 140 | 106 | 97 | 167 | 133 | 7 | 3 | 0 | 3 | 10 | 289 | 143 | 105 | 105 | 292 | 16 | 1 | 3 | 8 | 10 | 185 | 149 | 0.440381916 | 1 | 0.942 | 0.966 | 1 | 0.978 | 0.915 | 0.936 | 0.991 | 0.966 | 0.914 | 0.959 | 0.007 |
| A5SS | 429 | ENSG00000167664 | TMIGD2 | chr19 | - | 4294566 | 4294680 | 4294578 | 4294680 | 4292231 | 4292885 | 9 | 4 | 13 | 18 | 31 | 36 | 4 | 20 | 24 | 28 | 12 | 8 | 3 | 6 | 0 | 0 | 3 | 9 | 4 | 12 | 161 | 149 | 0.440423533 | 1 | 0.188 | 0.481 | 0.376 | 0.41 | 0.506 | 1 | 0.712 | 0.236 | 0.581 | 0 | -0.114 |
| A5SS | 1462 | ENSG00000069943 | PIGB | chr15 | + | 55350698 | 55350912 | 55350698 | 55350803 | 55354797 | 55354978 | 93 | 68 | 60 | 67 | 138 | 0 | 0 | 1 | 0 | 0 | 29 | 54 | 63 | 55 | 15 | 0 | 7 | 0 | 0 | 0 | 258 | 149 | 0.441002028 | 1 | 1 | 1 | 0.972 | 1 | 1 | 1 | 0.817 | 1 | 1 | 1 | 0.031 |
| A5SS | 917 | ENSG00000105053 | VRK3 | chr19 | - | 50025266 | 50025372 | 50025345 | 50025372 | 50020584 | 50020647 | 9 | 1 | 2 | 10 | 12 | 75 | 54 | 51 | 56 | 89 | 5 | 2 | 8 | 2 | 21 | 105 | 94 | 55 | 59 | 141 | 228 | 149 | 0.441062373 | 1 | 0.073 | 0.012 | 0.025 | 0.105 | 0.081 | 0.03 | 0.014 | 0.087 | 0.022 | 0.089 | 0.011 |
| A5SS | 6866 | ENSG00000156170 | NDUFAF6 | chr8 | + | 95031994 | 95032098 | 95031994 | 95032094 | 95035453 | 95035576 | 10 | 3 | 12 | 10 | 19 | 7 | 4 | 5 | 14 | 14 | 4 | 15 | 4 | 4 | 1 | 0 | 11 | 9 | 16 | 2 | 153 | 149 | 0.441968679 | 1 | 0.582 | 0.422 | 0.7 | 0.41 | 0.569 | 1 | 0.57 | 0.302 | 0.196 | 0.327 | 0.058 |
| A5SS | 96 | ENSG00000100083 | GGA1 | chr22 | + | 37616921 | 37617048 | 37616921 | 37616997 | 37618447 | 37618546 | 9 | 6 | 9 | 4 | 1 | 37 | 17 | 34 | 22 | 10 | 11 | 10 | 8 | 23 | 9 | 38 | 50 | 22 | 37 | 82 | 200 | 149 | 0.443305302 | 1 | 0.153 | 0.208 | 0.165 | 0.119 | 0.069 | 0.177 | 0.13 | 0.213 | 0.317 | 0.076 | -0.04 |
| A5SS | 3967 | ENSG00000110619 | CARS1 | chr11 | - | 3001961 | 3002053 | 3001969 | 3002053 | 3000928 | 3001248 | 22 | 12 | 8 | 23 | 40 | 52 | 48 | 60 | 61 | 59 | 21 | 13 | 28 | 45 | 27 | 109 | 66 | 28 | 54 | 59 | 157 | 149 | 0.443588756 | 1 | 0.286 | 0.192 | 0.112 | 0.264 | 0.392 | 0.155 | 0.157 | 0.487 | 0.442 | 0.303 | -0.06 |
| A5SS | 8683 | ENSG00000089022 | MAPKAPK5 | chr12 | + | 111888884 | 111889006 | 111888884 | 111889000 | 111890039 | 111890144 | 59 | 40 | 39 | 57 | 72 | 129 | 78 | 109 | 99 | 164 | 111 | 42 | 26 | 44 | 46 | 130 | 58 | 108 | 86 | 96 | 155 | 149 | 0.444361977 | 1 | 0.305 | 0.33 | 0.256 | 0.356 | 0.297 | 0.451 | 0.41 | 0.188 | 0.33 | 0.315 | -0.03 |
| A5SS | 2869 | ENSG00000185909 | KLHDC8B | chr3 | + | 49172635 | 49173145 | 49172635 | 49172801 | 49174238 | 49174403 | 14 | 5 | 3 | 3 | 4 | 0 | 0 | 0 | 2 | 2 | 4 | 7 | 3 | 8 | 206 | 0 | 0 | 0 | 0 | 0 | 298 | 149 | 0.444581142 | 1 | 1 | 1 | 1 | 0.429 | 0.5 | 1 | 1 | 1 | 1 | 1 | -0.214 |
| A5SS | 7765 | ENSG00000115459 | ELMOD3 | chr2 | + | 85355554 | 85355896 | 85355554 | 85355598 | 85356966 | 85357252 | 25 | 15 | 12 | 27 | 20 | 18 | 4 | 7 | 8 | 2 | 21 | 24 | 20 | 30 | 13 | 9 | 10 | 11 | 3 | 0 | 298 | 149 | 0.445616371 | 1 | 0.41 | 0.652 | 0.462 | 0.628 | 0.833 | 0.538 | 0.545 | 0.476 | 0.833 | 1 | -0.081 |
| A5SS | 1904 | ENSG00000226067 | LINC00623 | chr1 | + | 120952438 | 120952785 | 120952438 | 120952625 | 120953222 | 120953460 | 116 | 43 | 38 | 59 | 106 | 3 | 0 | 1 | 0 | 3 | 40 | 38 | 35 | 45 | 44 | 0 | 0 | 0 | 0 | 0 | 298 | 149 | 0.445894879 | 1 | 0.951 | 1 | 0.95 | 1 | 0.946 | 1 | 1 | 1 | 1 | 1 | -0.031 |
| A5SS | 5631 | ENSG00000104218 | CSPP1 | chr8 | + | 67111971 | 67112075 | 67111971 | 67112065 | 67113804 | 67113862 | 3 | 7 | 7 | 5 | 15 | 9 | 3 | 8 | 8 | 14 | 10 | 2 | 11 | 0 | 4 | 0 | 2 | 3 | 4 | 8 | 159 | 149 | 0.446338204 | 1 | 0.238 | 0.686 | 0.451 | 0.369 | 0.501 | 1 | 0.484 | 0.775 | 0 | 0.319 | -0.067 |
| A5SS | 23 | ENSG00000157557 | ETS2 | chr21 | + | 38819502 | 38819871 | 38819502 | 38819766 | 38821585 | 38821704 | 6 | 3 | 0 | 6 | 3 | 80 | 50 | 63 | 54 | 41 | 10 | 14 | 10 | 5 | 0 | 58 | 86 | 197 | 143 | 84 | 254 | 149 | 0.446427626 | 1 | 0.042 | 0.034 | 0 | 0.061 | 0.041 | 0.092 | 0.087 | 0.029 | 0.02 | 0 | -0.01 |
| A5SS | 7542 | ENSG00000163378 | EOGT | chr3 | - | 69012514 | 69012799 | 69012602 | 69012799 | 69011935 | 69011991 | 3 | 3 | 7 | 6 | 4 | 0 | 0 | 0 | 0 | 3 | 23 | 4 | 2 | 0 | 2 | 0 | 0 | 0 | 3 | 0 | 237 | 149 | 0.447242615 | 1 | 1 | 1 | 1 | 1 | 0.456 | 1 | 1 | 1 | 0 | 1 | 0.091 |
| A5SS | 441 | ENSG00000031823 | RANBP3 | chr19 | - | 5931403 | 5931531 | 5931418 | 5931531 | 5927967 | 5928087 | 54 | 18 | 42 | 27 | 25 | 93 | 22 | 47 | 52 | 24 | 74 | 46 | 33 | 49 | 68 | 96 | 51 | 98 | 84 | 105 | 164 | 149 | 0.44795904 | 1 | 0.345 | 0.426 | 0.448 | 0.321 | 0.486 | 0.412 | 0.45 | 0.234 | 0.346 | 0.37 | 0.043 |
| A5SS | 8625 | ENSG00000110719 | TCIRG1 | chr11 | + | 68041267 | 68041388 | 68041267 | 68041346 | 68041752 | 68041831 | 67 | 16 | 23 | 41 | 13 | 0 | 4 | 0 | 3 | 0 | 109 | 52 | 40 | 84 | 108 | 6 | 0 | 0 | 0 | 0 | 191 | 149 | 0.448354712 | 1 | 1 | 0.757 | 1 | 0.914 | 1 | 0.934 | 1 | 1 | 1 | 1 | -0.053 |
| A5SS | 1333 | ENSG00000077235 | GTF3C1 | chr16 | - | 27464319 | 27464836 | 27464394 | 27464836 | 27463540 | 27463592 | 30 | 12 | 13 | 21 | 3 | 5 | 1 | 3 | 3 | 1 | 1 | 15 | 2 | 9 | 4 | 0 | 1 | 1 | 5 | 0 | 224 | 149 | 0.448564351 | 1 | 0.8 | 0.889 | 0.742 | 0.823 | 0.666 | 1 | 0.909 | 0.571 | 0.545 | 1 | -0.021 |
| A5SS | 8012 | ENSG00000160226 | CFAP410 | chr21 | - | 44334060 | 44335804 | 44335757 | 44335804 | 44333032 | 44333262 | 11 | 7 | 6 | 15 | 5 | 6 | 8 | 6 | 14 | 3 | 11 | 2 | 7 | 2 | 12 | 12 | 10 | 13 | 2 | 2 | 298 | 149 | 0.448579831 | 1 | 0.478 | 0.304 | 0.333 | 0.349 | 0.455 | 0.314 | 0.091 | 0.212 | 0.333 | 0.75 | 0.044 |
| A5SS | 4424 | ENSG00000166523 | CLEC4E | chr12 | - | 8539199 | 8539306 | 8539216 | 8539306 | 8536089 | 8536205 | 27 | 22 | 12 | 9 | 18 | 6 | 11 | 7 | 6 | 14 | 49 | 22 | 40 | 33 | 12 | 22 | 17 | 41 | 10 | 9 | 166 | 149 | 0.449786399 | 1 | 0.802 | 0.642 | 0.606 | 0.574 | 0.536 | 0.667 | 0.537 | 0.467 | 0.748 | 0.545 | 0.039 |
| A5SS | 8364 | ENSG00000175470 | PPP2R2D | chr10 | + | 131940581 | 131940841 | 131940581 | 131940694 | 131943967 | 131944145 | 37 | 66 | 28 | 35 | 31 | 90 | 55 | 67 | 58 | 147 | 76 | 18 | 96 | 45 | 18 | 66 | 81 | 39 | 86 | 82 | 296 | 149 | 0.450091152 | 1 | 0.171 | 0.377 | 0.174 | 0.233 | 0.096 | 0.367 | 0.101 | 0.553 | 0.208 | 0.1 | -0.056 |
| A5SS | 3032 | ENSG00000159433 | STARD9 | chr15 | + | 42585520 | 42585637 | 42585520 | 42585634 | 42634855 | 42634972 | 7 | 16 | 11 | 15 | 11 | 0 | 0 | 0 | 0 | 0 | 10 | 31 | 11 | 15 | 7 | 0 | 3 | 0 | 0 | 3 | 152 | 149 | 0.450156165 | 1 | 1 | 1 | 1 | 1 | 1 | 1 | 0.91 | 1 | 1 | 0.696 | 0.079 |
| A5SS | 5444 | ENSG00000114735 | HEMK1 | chr3 | + | 50570930 | 50571332 | 50570930 | 50571294 | 50571709 | 50571801 | 31 | 17 | 86 | 37 | 26 | 8 | 3 | 6 | 9 | 8 | 60 | 38 | 26 | 40 | 32 | 5 | 5 | 1 | 3 | 16 | 187 | 149 | 0.451216413 | 1 | 0.755 | 0.819 | 0.919 | 0.766 | 0.721 | 0.905 | 0.858 | 0.954 | 0.914 | 0.614 | -0.053 |
| A5SS | 5993 | ENSG00000197345 | MRPL21 | chr11 | - | 68900534 | 68900605 | 68900547 | 68900605 | 68897926 | 68898012 | 7 | 1 | 13 | 0 | 12 | 24 | 1 | 21 | 14 | 22 | 0 | 1 | 7 | 3 | 14 | 20 | 17 | 10 | 15 | 19 | 162 | 149 | 0.452930455 | 1 | 0.212 | 0.479 | 0.363 | 0 | 0.334 | 0 | 0.051 | 0.392 | 0.155 | 0.404 | 0.077 |
| A5SS | 6755 | ENSG00000234741 | GAS5 | chr1 | - | 173865470 | 173865547 | 173865509 | 173865547 | 173864256 | 173864304 | 33 | 33 | 62 | 39 | 99 | 1 | 0 | 9 | 6 | 3 | 4 | 31 | 26 | 23 | 14 | 0 | 0 | 0 | 0 | 8 | 188 | 149 | 0.453053085 | 1 | 0.963 | 1 | 0.845 | 0.837 | 0.963 | 1 | 1 | 1 | 1 | 0.581 | 0.005 |
| A5SS | 7403 | ENSG00000100023 | PPIL2 | chr22 | + | 21694754 | 21695070 | 21694754 | 21694817 | 21695393 | 21695496 | 43 | 27 | 21 | 27 | 7 | 0 | 0 | 0 | 0 | 0 | 46 | 18 | 12 | 12 | 23 | 0 | 5 | 0 | 0 | 0 | 298 | 149 | 0.453700957 | 1 | 1 | 1 | 1 | 1 | 1 | 1 | 0.643 | 1 | 1 | 1 | 0.071 |
| A5SS | 118 | ENSG00000100365 | NCF4 | chr22 | + | 36875652 | 36875787 | 36875652 | 36875783 | 36876028 | 36876094 | 3 | 0 | 5 | 1 | 8 | 40 | 35 | 33 | 41 | 46 | 20 | 3 | 5 | 10 | 15 | 94 | 104 | 77 | 262 | 79 | 153 | 149 | 0.453827328 | 1 | 0.068 | 0 | 0.129 | 0.023 | 0.145 | 0.172 | 0.027 | 0.059 | 0.036 | 0.156 | -0.017 |
| A5SS | 2881 | ENSG00000004766 | VPS50 | chr7 | + | 93239865 | 93239950 | 93239865 | 93239934 | 93252652 | 93252775 | 1 | 2 | 1 | 0 | 3 | 68 | 66 | 49 | 66 | 37 | 0 | 0 | 0 | 0 | 6 | 1 | 24 | 20 | 10 | 19 | 165 | 149 | 0.453956166 | 1 | 0.013 | 0.027 | 0.018 | 0 | 0.068 | 0 | 0 | 0 | 0 | 0.222 | -0.019 |
| A5SS | 2076 | ENSG00000154153 | RETREG1 | chr5 | - | 16477661 | 16478098 | 16478033 | 16478098 | 16473070 | 16475234 | 85 | 79 | 184 | 86 | 342 | 2 | 0 | 0 | 0 | 8 | 11 | 15 | 28 | 8 | 10 | 0 | 4 | 0 | 0 | 0 | 298 | 149 | 0.454886875 | 1 | 0.955 | 1 | 1 | 1 | 0.955 | 1 | 0.652 | 1 | 1 | 1 | 0.052 |
| A5SS | 4789 | ENSG00000131778 | CHD1L | chr1 | + | 147272170 | 147272630 | 147272170 | 147272281 | 147275353 | 147275468 | 13 | 2 | 6 | 9 | 23 | 49 | 28 | 53 | 41 | 46 | 12 | 29 | 20 | 16 | 0 | 61 | 52 | 23 | 34 | 121 | 298 | 149 | 0.455670131 | 1 | 0.117 | 0.034 | 0.054 | 0.099 | 0.2 | 0.09 | 0.218 | 0.303 | 0.19 | 0 | -0.059 |
| A5SS | 8569 | ENSG00000101290 | CDS2 | chr20 | + | 5183060 | 5183178 | 5183060 | 5183143 | 5184857 | 5184945 | 0 | 2 | 7 | 3 | 11 | 148 | 70 | 185 | 156 | 444 | 8 | 0 | 4 | 5 | 0 | 133 | 166 | 147 | 152 | 324 | 184 | 149 | 0.456923715 | 1 | 0 | 0.023 | 0.03 | 0.015 | 0.02 | 0.046 | 0 | 0.022 | 0.026 | 0 | -0.001 |
| A5SS | 5861 | ENSG00000163482 | STK36 | chr2 | + | 218672740 | 218673017 | 218672740 | 218672913 | 218673624 | 218673765 | 3 | 1 | 1 | 1 | 0 | 10 | 19 | 13 | 15 | 26 | 0 | 0 | 2 | 1 | 5 | 6 | 20 | 7 | 25 | 24 | 253 | 149 | 0.456980137 | 1 | 0.15 | 0.03 | 0.043 | 0.038 | 0 | 0 | 0 | 0.144 | 0.023 | 0.109 | -0.003 |
| A5SS | 5038 | ENSG00000112031 | MTRF1L | chr6 | - | 152995135 | 152995319 | 152995243 | 152995319 | 152994512 | 152994676 | 214 | 200 | 159 | 215 | 288 | 3 | 3 | 8 | 7 | 2 | 238 | 368 | 202 | 296 | 75 | 0 | 5 | 2 | 7 | 6 | 257 | 149 | 0.457498128 | 1 | 0.976 | 0.975 | 0.92 | 0.947 | 0.988 | 1 | 0.977 | 0.983 | 0.961 | 0.879 | 0.001 |
| A5SS | 3746 | ENSG00000145833 | DDX46 | chr5 | + | 134816429 | 134816606 | 134816429 | 134816603 | 134817495 | 134817714 | 249 | 235 | 254 | 243 | 246 | 103 | 79 | 62 | 81 | 50 | 28 | 164 | 189 | 44 | 22 | 23 | 28 | 62 | 10 | 0 | 152 | 149 | 0.458068486 | 1 | 0.703 | 0.745 | 0.801 | 0.746 | 0.828 | 0.544 | 0.852 | 0.749 | 0.812 | 1 | -0.027 |
| A5SS | 5669 | ENSG00000095059 | DHPS | chr19 | - | 12680149 | 12680325 | 12680160 | 12680325 | 12679800 | 12679922 | 10 | 4 | 17 | 5 | 4 | 97 | 44 | 85 | 80 | 58 | 19 | 15 | 6 | 4 | 23 | 108 | 77 | 79 | 84 | 193 | 160 | 149 | 0.458838953 | 1 | 0.088 | 0.078 | 0.157 | 0.055 | 0.06 | 0.141 | 0.154 | 0.066 | 0.042 | 0.1 | -0.013 |
| A5SS | 5904 | ENSG00000198001 | IRAK4 | chr12 | + | 43772911 | 43773072 | 43772911 | 43772986 | 43773964 | 43774029 | 169 | 167 | 114 | 134 | 176 | 11 | 7 | 13 | 7 | 19 | 82 | 89 | 132 | 50 | 12 | 7 | 0 | 24 | 5 | 3 | 235 | 149 | 0.459546172 | 1 | 0.907 | 0.938 | 0.848 | 0.924 | 0.855 | 0.881 | 1 | 0.777 | 0.864 | 0.717 | 0.047 |
| A5SS | 8766 | ENSG00000187109 | NAP1L1 | chr12 | - | 76059776 | 76059878 | 76059797 | 76059878 | 76056032 | 76056161 | 78 | 70 | 82 | 122 | 6 | 1609 | 1843 | 1604 | 2562 | 3146 | 9 | 77 | 58 | 28 | 59 | 1190 | 1355 | 1462 | 949 | 2393 | 170 | 149 | 0.460086246 | 1 | 0.041 | 0.032 | 0.043 | 0.04 | 0.002 | 0.007 | 0.047 | 0.034 | 0.025 | 0.021 | 0.005 |
| A5SS | 1957 | ENSG00000143252 | SDHC | chr1 | + | 161314380 | 161314874 | 161314380 | 161314425 | 161323613 | 161323670 | 7 | 2 | 5 | 6 | 0 | 79 | 34 | 70 | 78 | 51 | 0 | 16 | 10 | 4 | 0 | 87 | 84 | 54 | 76 | 84 | 298 | 149 | 0.460429531 | 1 | 0.042 | 0.029 | 0.034 | 0.037 | 0 | 0 | 0.087 | 0.085 | 0.026 | 0 | -0.011 |
| A5SS | 8158 | ENSG00000171202 | TMEM126A | chr11 | + | 85654062 | 85654256 | 85654062 | 85654234 | 85655593 | 85655708 | 107 | 46 | 89 | 76 | 192 | 4 | 1 | 3 | 3 | 0 | 36 | 80 | 77 | 82 | 59 | 0 | 0 | 0 | 3 | 0 | 171 | 149 | 0.460765948 | 1 | 0.959 | 0.976 | 0.963 | 0.957 | 1 | 1 | 1 | 1 | 0.96 | 1 | -0.021 |
| A5SS | 2628 | ENSG00000108591 | DRG2 | chr17 | + | 18099632 | 18099723 | 18099632 | 18099715 | 18100362 | 18100435 | 38 | 11 | 28 | 34 | 63 | 4 | 3 | 0 | 2 | 3 | 69 | 58 | 17 | 28 | 63 | 0 | 0 | 2 | 2 | 7 | 157 | 149 | 0.460771093 | 1 | 0.9 | 0.777 | 1 | 0.942 | 0.952 | 1 | 1 | 0.89 | 0.93 | 0.895 | -0.029 |
| A5SS | 6882 | ENSG00000130734 | ATG4D | chr19 | + | 10546838 | 10547115 | 10546838 | 10547071 | 10547188 | 10547253 | 15 | 9 | 15 | 20 | 10 | 1 | 0 | 4 | 1 | 0 | 11 | 27 | 22 | 18 | 32 | 0 | 0 | 0 | 0 | 0 | 193 | 149 | 0.461264917 | 1 | 0.921 | 1 | 0.743 | 0.939 | 1 | 1 | 1 | 1 | 1 | 1 | -0.079 |
| A5SS | 5432 | ENSG00000226419 | SLC16A1-AS1 | chr1 | + | 112956414 | 112957722 | 112956414 | 112956956 | 112960476 | 112960575 | 7 | 10 | 9 | 16 | 1 | 3 | 5 | 4 | 5 | 3 | 24 | 12 | 20 | 6 | 10 | 12 | 6 | 0 | 1 | 13 | 298 | 149 | 0.46221608 | 1 | 0.538 | 0.5 | 0.529 | 0.615 | 0.143 | 0.5 | 0.5 | 1 | 0.75 | 0.278 | -0.141 |
| A5SS | 8490 | ENSG00000119943 | PYROXD2 | chr10 | - | 98410612 | 98410958 | 98410938 | 98410958 | 98407903 | 98407997 | 1 | 2 | 3 | 4 | 3 | 6 | 0 | 7 | 13 | 9 | 2 | 13 | 3 | 0 | 0 | 0 | 2 | 0 | 4 | 11 | 298 | 149 | 0.462516274 | 1 | 0.077 | 1 | 0.176 | 0.133 | 0.143 | 1 | 0.765 | 1 | 0 | 0 | -0.247 |
| A5SS | 3345 | ENSG00000174547 | MRPL11 | chr11 | - | 66438631 | 66438848 | 66438709 | 66438848 | 66438163 | 66438259 | 29 | 14 | 28 | 32 | 13 | 0 | 0 | 4 | 3 | 3 | 62 | 41 | 28 | 33 | 57 | 6 | 3 | 2 | 6 | 3 | 227 | 149 | 0.462771859 | 1 | 1 | 1 | 0.821 | 0.875 | 0.74 | 0.872 | 0.9 | 0.902 | 0.783 | 0.926 | 0.011 |
| A5SS | 7727 | ENSG00000213930 | GALT | chr9 | + | 34647088 | 34647961 | 34647088 | 34647258 | 34648114 | 34648171 | 115 | 61 | 141 | 75 | 61 | 3 | 3 | 2 | 1 | 2 | 126 | 158 | 68 | 91 | 144 | 0 | 1 | 1 | 5 | 11 | 298 | 149 | 0.463582604 | 1 | 0.95 | 0.91 | 0.972 | 0.974 | 0.938 | 1 | 0.988 | 0.971 | 0.901 | 0.867 | 0.003 |
| A5SS | 3616 | ENSG00000239900 | ADSL | chr22 | + | 40361487 | 40361677 | 40361487 | 40361635 | 40362980 | 40363071 | 0 | 3 | 2 | 6 | 2 | 81 | 49 | 92 | 79 | 97 | 0 | 2 | 4 | 3 | 0 | 76 | 71 | 64 | 54 | 64 | 191 | 149 | 0.463847943 | 1 | 0 | 0.046 | 0.017 | 0.056 | 0.016 | 0 | 0.022 | 0.046 | 0.042 | 0 | 0.005 |
| A5SS | 7016 | ENSG00000244754 | N4BP2L2 | chr13 | - | 32538617 | 32538825 | 32538682 | 32538825 | 32533544 | 32537027 | 294 | 300 | 259 | 277 | 304 | 164 | 147 | 146 | 150 | 142 | 48 | 139 | 121 | 70 | 27 | 18 | 124 | 100 | 20 | 23 | 214 | 149 | 0.463953854 | 1 | 0.555 | 0.587 | 0.553 | 0.563 | 0.598 | 0.65 | 0.438 | 0.457 | 0.709 | 0.45 | 0.03 |
| A5SS | 5779 | ENSG00000183486 | MX2 | chr21 | + | 41363268 | 41363416 | 41363268 | 41363389 | 41376835 | 41377155 | 3 | 1 | 5 | 9 | 0 | 1 | 10 | 0 | 10 | 6 | 10 | 1 | 1 | 3 | 3 | 13 | 9 | 6 | 21 | 0 | 176 | 149 | 0.464077552 | 1 | 0.717 | 0.078 | 1 | 0.432 | 0 | 0.394 | 0.086 | 0.124 | 0.108 | 1 | 0.103 |
| A5SS | 4140 | ENSG00000165359 | INTS6L | chrX | + | 135577192 | 135577427 | 135577192 | 135577295 | 135579787 | 135580162 | 72 | 60 | 50 | 81 | 125 | 39 | 17 | 47 | 59 | 94 | 14 | 47 | 68 | 17 | 10 | 1 | 42 | 36 | 9 | 5 | 281 | 149 | 0.464163406 | 1 | 0.495 | 0.652 | 0.361 | 0.421 | 0.414 | 0.881 | 0.372 | 0.5 | 0.5 | 0.515 | -0.085 |
| A5SS | 7763 | ENSG00000115459 | ELMOD3 | chr2 | + | 85355554 | 85355716 | 85355554 | 85355598 | 85356966 | 85357252 | 21 | 13 | 14 | 24 | 16 | 18 | 4 | 7 | 8 | 2 | 17 | 24 | 20 | 25 | 9 | 9 | 10 | 11 | 3 | 0 | 267 | 149 | 0.464459248 | 1 | 0.394 | 0.645 | 0.527 | 0.626 | 0.817 | 0.513 | 0.573 | 0.504 | 0.823 | 1 | -0.081 |
| A5SS | 3718 | ENSG00000072210 | ALDH3A2 | chr17 | + | 19663332 | 19665047 | 19663332 | 19663499 | 19671720 | 19671956 | 64 | 48 | 61 | 70 | 106 | 1 | 0 | 0 | 1 | 3 | 70 | 46 | 29 | 78 | 82 | 0 | 7 | 0 | 0 | 0 | 298 | 149 | 0.464525015 | 1 | 0.97 | 1 | 1 | 0.972 | 0.946 | 1 | 0.767 | 1 | 1 | 1 | 0.024 |
| A5SS | 6638 | ENSG00000086730 | LAT2 | chr7 | + | 74214821 | 74215099 | 74214821 | 74214917 | 74215946 | 74216069 | 51 | 16 | 40 | 39 | 21 | 0 | 0 | 2 | 0 | 0 | 48 | 60 | 18 | 77 | 56 | 0 | 0 | 0 | 0 | 7 | 298 | 149 | 0.46485496 | 1 | 1 | 1 | 0.909 | 1 | 1 | 1 | 1 | 1 | 1 | 0.8 | 0.022 |
| A5SS | 3788 | ENSG00000170791 | CHCHD7 | chr8 | + | 56211788 | 56212174 | 56211788 | 56211837 | 56214597 | 56214667 | 7 | 1 | 6 | 6 | 0 | 23 | 19 | 22 | 28 | 19 | 1 | 2 | 2 | 7 | 0 | 2 | 11 | 11 | 10 | 8 | 298 | 149 | 0.464949955 | 1 | 0.132 | 0.026 | 0.12 | 0.097 | 0 | 0.2 | 0.083 | 0.083 | 0.259 | 0 | -0.05 |
| A5SS | 3961 | ENSG00000163743 | RCHY1 | chr4 | - | 75508819 | 75508935 | 75508823 | 75508935 | 75494100 | 75494179 | 187 | 114 | 109 | 131 | 254 | 16 | 12 | 15 | 16 | 22 | 44 | 153 | 89 | 76 | 31 | 0 | 15 | 9 | 10 | 3 | 153 | 149 | 0.465195445 | 1 | 0.919 | 0.902 | 0.876 | 0.889 | 0.918 | 1 | 0.909 | 0.906 | 0.881 | 0.91 | -0.02 |
| A5SS | 5828 | ENSG00000172936 | MYD88 | chr3 | + | 38140387 | 38140592 | 38140387 | 38140568 | 38140756 | 38140848 | 3 | 3 | 3 | 9 | 18 | 145 | 83 | 98 | 121 | 144 | 0 | 15 | 3 | 26 | 6 | 326 | 199 | 75 | 173 | 289 | 173 | 149 | 0.467853741 | 1 | 0.018 | 0.03 | 0.026 | 0.06 | 0.097 | 0 | 0.061 | 0.033 | 0.115 | 0.018 | 0.001 |
| A5SS | 7325 | ENSG00000121879 | PIK3CA | chr3 | + | 179229271 | 179230121 | 179229271 | 179229442 | 179230224 | 179230376 | 127 | 82 | 98 | 115 | 148 | 0 | 0 | 0 | 0 | 0 | 6 | 47 | 18 | 16 | 0 | 0 | 4 | 0 | 0 | 1 | 298 | 149 | 0.468664901 | 1 | 1 | 1 | 1 | 1 | 1 | 1 | 0.855 | 1 | 1 | 0 | 0.229 |
| A5SS | 5151 | ENSG00000181163 | NPM1 | chr5 | + | 171400152 | 171400925 | 171400152 | 171400210 | 171405301 | 171405403 | 916 | 716 | 1224 | 1018 | 2400 | 30 | 9 | 23 | 19 | 50 | 208 | 513 | 658 | 455 | 59 | 1 | 14 | 7 | 11 | 4 | 298 | 149 | 0.469800562 | 1 | 0.939 | 0.975 | 0.964 | 0.964 | 0.96 | 0.99 | 0.948 | 0.979 | 0.954 | 0.881 | 0.01 |
| A5SS | 7666 | ENSG00000132740 | IGHMBP2 | chr11 | + | 68906068 | 68906242 | 68906068 | 68906238 | 68908144 | 68908337 | 1 | 0 | 0 | 1 | 4 | 31 | 11 | 17 | 15 | 7 | 0 | 0 | 1 | 0 | 0 | 41 | 52 | 21 | 39 | 12 | 153 | 149 | 0.470273094 | 1 | 0.03 | 0 | 0 | 0.061 | 0.358 | 0 | 0 | 0.044 | 0 | 0 | 0.081 |
| A5SS | 6590 | ENSG00000129103 | SUMF2 | chr7 | + | 56068481 | 56068647 | 56068481 | 56068638 | 56072996 | 56073111 | 7 | 2 | 0 | 5 | 2 | 175 | 88 | 162 | 122 | 260 | 0 | 0 | 6 | 4 | 0 | 109 | 161 | 96 | 95 | 221 | 158 | 149 | 0.470599209 | 1 | 0.036 | 0.021 | 0 | 0.037 | 0.007 | 0 | 0 | 0.056 | 0.038 | 0 | 0.001 |
| A5SS | 1199 | ENSG00000185324 | CDK10 | chr16 | + | 89694930 | 89695070 | 89694930 | 89695052 | 89695292 | 89695345 | 52 | 32 | 56 | 54 | 56 | 2 | 1 | 0 | 4 | 2 | 81 | 33 | 40 | 34 | 68 | 0 | 0 | 0 | 0 | 14 | 167 | 149 | 0.470985253 | 1 | 0.959 | 0.966 | 1 | 0.923 | 0.962 | 1 | 1 | 1 | 1 | 0.813 | -0.001 |
| A5SS | 3662 | ENSG00000058673 | ZC3H11A | chr1 | + | 203796292 | 203796601 | 203796292 | 203796496 | 203801574 | 203801628 | 33 | 12 | 45 | 39 | 24 | 13 | 18 | 13 | 16 | 17 | 23 | 35 | 13 | 24 | 10 | 6 | 16 | 19 | 10 | 0 | 254 | 149 | 0.471958464 | 1 | 0.598 | 0.281 | 0.67 | 0.588 | 0.453 | 0.692 | 0.562 | 0.286 | 0.585 | 1 | -0.107 |
| A5SS | 5962 | ENSG00000108465 | CDK5RAP3 | chr17 | + | 47973518 | 47973693 | 47973518 | 47973650 | 47973930 | 47974031 | 14 | 13 | 13 | 9 | 0 | 125 | 77 | 120 | 111 | 116 | 12 | 16 | 16 | 21 | 23 | 255 | 150 | 97 | 149 | 245 | 192 | 149 | 0.472187229 | 1 | 0.08 | 0.116 | 0.078 | 0.059 | 0 | 0.035 | 0.076 | 0.113 | 0.099 | 0.068 | -0.012 |
| A5SS | 1226 | ENSG00000132600 | PRMT7 | chr16 | + | 68311055 | 68311111 | 68311055 | 68311099 | 68315896 | 68316074 | 0 | 3 | 1 | 5 | 2 | 18 | 9 | 14 | 20 | 0 | 0 | 0 | 2 | 11 | 24 | 19 | 25 | 12 | 15 | 6 | 161 | 149 | 0.472264637 | 1 | 0 | 0.236 | 0.062 | 0.188 | 1 | 0 | 0 | 0.134 | 0.404 | 0.787 | 0.032 |
| A5SS | 4906 | ENSG00000100410 | PHF5A | chr22 | - | 41468019 | 41468147 | 41468123 | 41468147 | 41467447 | 41467614 | 21 | 4 | 17 | 17 | 14 | 71 | 23 | 62 | 68 | 56 | 15 | 17 | 14 | 17 | 54 | 82 | 76 | 61 | 86 | 59 | 253 | 149 | 0.474028197 | 1 | 0.148 | 0.093 | 0.139 | 0.128 | 0.128 | 0.097 | 0.116 | 0.119 | 0.104 | 0.35 | -0.03 |
| A5SS | 6753 | ENSG00000234741 | GAS5 | chr1 | - | 173865228 | 173865547 | 173865470 | 173865547 | 173864256 | 173864304 | 95 | 58 | 105 | 87 | 142 | 1 | 0 | 1 | 0 | 5 | 51 | 60 | 70 | 60 | 20 | 0 | 0 | 0 | 0 | 0 | 298 | 149 | 0.475308371 | 1 | 0.979 | 1 | 0.981 | 1 | 0.934 | 1 | 1 | 1 | 1 | 1 | -0.021 |
| A5SS | 2151 | ENSG00000135074 | ADAM19 | chr5 | - | 157513423 | 157513505 | 157513433 | 157513505 | 157509300 | 157509467 | 2 | 7 | 4 | 8 | 7 | 82 | 43 | 81 | 56 | 103 | 8 | 17 | 5 | 1 | 6 | 92 | 232 | 126 | 95 | 74 | 159 | 149 | 0.47534741 | 1 | 0.022 | 0.132 | 0.044 | 0.118 | 0.06 | 0.075 | 0.064 | 0.036 | 0.01 | 0.071 | 0.024 |
| A5SS | 7554 | ENSG00000150776 | NKAPD1 | chr11 | + | 112075566 | 112075684 | 112075566 | 112075643 | 112078214 | 112078315 | 1 | 6 | 3 | 6 | 10 | 67 | 49 | 69 | 86 | 72 | 0 | 6 | 6 | 4 | 0 | 29 | 69 | 63 | 44 | 25 | 190 | 149 | 0.475569978 | 1 | 0.012 | 0.088 | 0.033 | 0.052 | 0.098 | 0 | 0.064 | 0.069 | 0.067 | 0 | 0.017 |
| A5SS | 8600 | ENSG00000243335 | KCTD7 | chr7 | + | 66638252 | 66638431 | 66638252 | 66638391 | 66638855 | 66638929 | 20 | 10 | 15 | 19 | 12 | 5 | 0 | 1 | 0 | 0 | 11 | 29 | 10 | 9 | 4 | 0 | 0 | 0 | 0 | 0 | 189 | 149 | 0.476187961 | 1 | 0.759 | 1 | 0.922 | 1 | 1 | 1 | 1 | 1 | 1 | 1 | -0.064 |
| A5SS | 5233 | ENSG00000091409 | ITGA6 | chr2 | + | 172489484 | 172489658 | 172489484 | 172489613 | 172491023 | 172491122 | 274 | 248 | 382 | 337 | 583 | 15 | 4 | 13 | 20 | 11 | 31 | 117 | 89 | 134 | 34 | 0 | 0 | 1 | 14 | 0 | 194 | 149 | 0.476574187 | 1 | 0.933 | 0.979 | 0.958 | 0.928 | 0.976 | 1 | 1 | 0.986 | 0.88 | 1 | -0.018 |
| A5SS | 372 | ENSG00000249115 | HAUS5 | chr19 | + | 35614034 | 35614231 | 35614034 | 35614059 | 35615041 | 35615147 | 2 | 0 | 2 | 1 | 0 | 18 | 1 | 6 | 4 | 7 | 0 | 4 | 0 | 4 | 0 | 4 | 18 | 6 | 2 | 9 | 298 | 149 | 0.477184874 | 1 | 0.053 | 0 | 0.143 | 0.111 | 0 | 0 | 0.1 | 0 | 0.5 | 0 | -0.059 |
| A5SS | 5213 | ENSG00000104907 | TRMT1 | chr19 | - | 13112895 | 13113011 | 13112908 | 13113011 | 13112704 | 13112817 | 51 | 28 | 48 | 52 | 23 | 6 | 2 | 8 | 3 | 4 | 70 | 38 | 37 | 31 | 40 | 14 | 4 | 2 | 4 | 6 | 162 | 149 | 0.477559436 | 1 | 0.887 | 0.928 | 0.847 | 0.941 | 0.841 | 0.821 | 0.897 | 0.944 | 0.877 | 0.86 | 0.009 |
| A5SS | 56 | ENSG00000185339 | TCN2 | chr22 | + | 30612872 | 30613042 | 30612872 | 30612961 | 30614348 | 30614501 | 9 | 12 | 11 | 27 | 9 | 0 | 0 | 0 | 0 | 0 | 142 | 423 | 132 | 153 | 301 | 0 | 13 | 2 | 0 | 0 | 230 | 149 | 0.4794042 | 1 | 1 | 1 | 1 | 1 | 1 | 1 | 0.955 | 0.977 | 1 | 1 | 0.014 |
| A5SS | 4297 | ENSG00000071054 | MAP4K4 | chr2 | + | 101887087 | 101887261 | 101887087 | 101887237 | 101887777 | 101887937 | 164 | 132 | 91 | 148 | 335 | 646 | 586 | 646 | 795 | 1628 | 362 | 151 | 196 | 242 | 174 | 1598 | 914 | 1309 | 1326 | 750 | 173 | 149 | 0.479677247 | 1 | 0.179 | 0.162 | 0.108 | 0.138 | 0.151 | 0.163 | 0.125 | 0.114 | 0.136 | 0.167 | 0.007 |
| A5SS | 8879 | ENSG00000110911 | SLC11A2 | chr12 | - | 51026309 | 51026384 | 51026314 | 51026384 | 51010694 | 51010766 | 1 | 3 | 8 | 4 | 4 | 12 | 4 | 10 | 15 | 1 | 0 | 8 | 26 | 3 | 4 | 37 | 14 | 8 | 20 | 16 | 154 | 149 | 0.480960344 | 1 | 0.075 | 0.421 | 0.436 | 0.205 | 0.795 | 0 | 0.356 | 0.759 | 0.127 | 0.195 | 0.099 |
| A5SS | 8269 | ENSG00000150433 | TMEM218 | chr11 | - | 125102636 | 125102809 | 125102733 | 125102809 | 125102131 | 125102317 | 16 | 15 | 11 | 11 | 8 | 6 | 4 | 2 | 7 | 8 | 7 | 11 | 3 | 4 | 36 | 11 | 2 | 0 | 0 | 17 | 246 | 149 | 0.482137919 | 1 | 0.618 | 0.694 | 0.769 | 0.488 | 0.377 | 0.278 | 0.769 | 1 | 1 | 0.562 | -0.133 |
| A5SS | 2471 | ENSG00000138796 | HADH | chr4 | + | 108023473 | 108028069 | 108023473 | 108027760 | 108033175 | 108033292 | 22 | 22 | 13 | 20 | 14 | 46 | 40 | 64 | 54 | 39 | 12 | 14 | 11 | 20 | 0 | 59 | 25 | 15 | 27 | 56 | 298 | 149 | 0.483596775 | 1 | 0.193 | 0.216 | 0.092 | 0.156 | 0.152 | 0.092 | 0.219 | 0.268 | 0.27 | 0 | -0.008 |
| A5SS | 7705 | ENSG00000128973 | CLN6 | chr15 | - | 68211555 | 68211863 | 68211674 | 68211863 | 68211262 | 68211318 | 9 | 2 | 5 | 7 | 0 | 54 | 12 | 25 | 37 | 26 | 0 | 4 | 5 | 0 | 6 | 30 | 26 | 7 | 17 | 37 | 268 | 149 | 0.483869368 | 1 | 0.085 | 0.085 | 0.1 | 0.095 | 0 | 0 | 0.079 | 0.284 | 0 | 0.083 | -0.016 |
| A5SS | 3499 | ENSG00000247828 | TMEM161B-AS1 | chr5 | + | 88270525 | 88270741 | 88270525 | 88270585 | 88282041 | 88282106 | 31 | 46 | 48 | 38 | 12 | 29 | 3 | 26 | 11 | 4 | 23 | 29 | 24 | 2 | 2 | 4 | 19 | 10 | 0 | 10 | 298 | 149 | 0.484061732 | 1 | 0.348 | 0.885 | 0.48 | 0.633 | 0.6 | 0.742 | 0.433 | 0.545 | 1 | 0.091 | 0.027 |

| | | | | | | | | | | | | | | | | | | | | | | | | | | | | | | | | | | | | | | | | | | | | | | |
|---|---|---|---|---|---|---|---|---|---|---|---|---|---|---|---|---|---|---|---|---|---|---|---|---|---|---|---|---|---|---|---|---|---|---|---|---|---|---|---|---|---|---|---|---|---|---|
| A5SS | 5518 | ENSG00000223960 | CHROMR | chr2 | + | 178426365 | 178426703 | 178426365 | 178426661 | 178430769 | 178430889 | 5 | 8 | 2 | 2 | 0 | 7 | 3 | 5 | 6 | 6 | 9 | 1 | 3 | 1 | 0 | 7 | 8 | 1 | 8 | 4 | 191 | 149 | 0.484181965 | 1 | 0.358 | 0.675 | 0.238 | 0.206 | 0 | 0.501 | 0.089 | 0.701 | 0.089 | 0 | 0.019 |
| A5SS | 788 | ENSG00000185624 | P4HB | chr17 | - | 81859766 | 81860524 | 81860326 | 81860524 | 81859180 | 81859387 | 3 | 4 | 5 | 1 | 1 | 109 | 39 | 66 | 69 | 47 | 3 | 2 | 6 | 3 | 10 | 94 | 120 | 64 | 47 | 182 | 298 | 149 | 0.48461183 | 1 | 0.014 | 0.049 | 0.036 | 0.007 | 0.011 | 0.016 | 0.008 | 0.045 | 0.031 | 0.027 | -0.002 |
| A5SS | 790 | ENSG00000185624 | P4HB | chr17 | - | 81859884 | 81860523 | 81860326 | 81860523 | 81859180 | 81859387 | 3 | 4 | 5 | 1 | 1 | 109 | 39 | 66 | 69 | 47 | 3 | 2 | 6 | 3 | 10 | 94 | 120 | 64 | 47 | 182 | 298 | 149 | 0.48461183 | 1 | 0.014 | 0.049 | 0.036 | 0.007 | 0.011 | 0.016 | 0.008 | 0.045 | 0.031 | 0.027 | -0.002 |
| A5SS | 7700 | ENSG00000119314 | PTBP3 | chr9 | - | 112297831 | 112297916 | 112297840 | 112297916 | 112275843 | 112276013 | 112 | 89 | 98 | 94 | 174 | 91 | 29 | 58 | 74 | 112 | 49 | 61 | 86 | 85 | 18 | 12 | 71 | 80 | 14 | 12 | 158 | 149 | 0.485624709 | 1 | 0.537 | 0.743 | 0.614 | 0.545 | 0.594 | 0.794 | 0.448 | 0.503 | 0.851 | 0.586 | -0.03 |
| A5SS | 1750 | ENSG00000100889 | PCK2 | chr14 | + | 24094310 | 24094845 | 24094310 | 24094434 | 24096891 | 24097137 | 16 | 8 | 12 | 18 | 3 | 21 | 10 | 27 | 27 | 2 | 32 | 14 | 3 | 6 | 6 | 15 | 20 | 3 | 15 | 77 | 298 | 149 | 0.485682574 | 1 | 0.276 | 0.286 | 0.182 | 0.25 | 0.429 | 0.516 | 0.259 | 0.333 | 0.167 | 0.037 | 0.022 |
| A5SS | 2201 | ENSG00000197321 | SVIL | chr10 | - | 29480536 | 29480813 | 29480549 | 29480813 | 29473837 | 29473989 | 43 | 47 | 77 | 61 | 46 | 0 | 1 | 2 | 2 | 2 | 93 | 56 | 66 | 104 | 74 | 0 | 0 | 0 | 3 | 0 | 162 | 149 | 0.485764281 | 1 | 1 | 0.977 | 0.973 | 0.966 | 0.955 | 1 | 1 | 1 | 0.97 | 1 | -0.02 |
| A5SS | 1737 | ENSG00000129465 | RIPK3 | chr14 | - | 24338210 | 24338295 | 24338248 | 24338295 | 24337872 | 24338040 | 12 | 1 | 10 | 6 | 10 | 14 | 7 | 15 | 12 | 13 | 26 | 12 | 6 | 13 | 10 | 29 | 30 | 7 | 16 | 9 | 187 | 149 | 0.485980474 | 1 | 0.406 | 0.102 | 0.347 | 0.285 | 0.38 | 0.417 | 0.242 | 0.406 | 0.393 | 0.47 | -0.082 |
| A5SS | 5585 | ENSG00000143390 | RFX5 | chr1 | - | 151346421 | 151346606 | 151346481 | 151346606 | 151346204 | 151346333 | 9 | 11 | 15 | 18 | 19 | 61 | 57 | 54 | 44 | 112 | 9 | 27 | 14 | 13 | 5 | 69 | 61 | 33 | 50 | 41 | 209 | 149 | 0.486263229 | 1 | 0.095 | 0.121 | 0.165 | 0.226 | 0.108 | 0.085 | 0.24 | 0.232 | 0.156 | 0.08 | -0.016 |
| A5SS | 3862 | ENSG00000117533 | VAMP4 | chr1 | - | 171710713 | 171710814 | 171710759 | 171710814 | 171709664 | 171709744 | 236 | 169 | 186 | 184 | 273 | 6 | 5 | 4 | 1 | 1 | 146 | 152 | 128 | 99 | 148 | 5 | 0 | 2 | 0 | 3 | 195 | 149 | 0.486363759 | 1 | 0.968 | 0.963 | 0.973 | 0.993 | 0.995 | 0.957 | 1 | 0.98 | 1 | 0.974 | -0.004 |
| A5SS | 699 | ENSG00000169733 | RFNG | chr17 | - | 82051011 | 82051342 | 82051293 | 82051342 | 82050661 | 82050764 | 9 | 0 | 2 | 0 | 0 | 14 | 1 | 6 | 10 | 4 | 4 | 0 | 0 | 4 | 6 | 6 | 13 | 5 | 3 | 10 | 298 | 149 | 0.486536235 | 1 | 0.243 | 0 | 0.143 | 0 | 0 | 0.25 | 0 | 0 | 0.4 | 0.231 | -0.099 |
| A5SS | 5315 | ENSG00000136718 | IMP4 | chr2 | + | 130344628 | 130344725 | 130344628 | 130344712 | 130345375 | 130345485 | 0 | 1 | 1 | 2 | 3 | 47 | 28 | 54 | 52 | 50 | 0 | 0 | 0 | 0 | 0 | 39 | 71 | 45 | 46 | 91 | 162 | 149 | 0.486632758 | 1 | 0 | 0.032 | 0.017 | 0.034 | 0.052 | 0 | 0 | 0 | 0 | 0 | 0.027 |
| A5SS | 637 | ENSG00000263753 | LINC00667 | chr18 | + | 5238119 | 5239196 | 5238119 | 5239049 | 5240647 | 5241255 | 64 | 36 | 46 | 61 | 86 | 15 | 2 | 12 | 15 | 20 | 29 | 31 | 12 | 7 | 45 | 7 | 5 | 2 | 15 | 6 | 296 | 149 | 0.486962392 | 1 | 0.682 | 0.901 | 0.659 | 0.672 | 0.684 | 0.676 | 0.757 | 0.751 | 0.19 | 0.791 | 0.087 |
| A5SS | 229 | ENSG00000011600 | TYROBP | chr19 | - | 35908167 | 35908295 | 35908174 | 35908295 | 35907445 | 35907580 | 38 | 15 | 18 | 28 | 25 | 66 | 15 | 25 | 42 | 20 | 54 | 72 | 24 | 39 | 85 | 59 | 85 | 12 | 66 | 101 | 156 | 149 | 0.487168068 | 1 | 0.355 | 0.489 | 0.407 | 0.389 | 0.544 | 0.466 | 0.447 | 0.656 | 0.361 | 0.446 | -0.038 |
| A5SS | 7676 | ENSG00000159069 | FBXW5 | chr9 | - | 136942045 | 136942466 | 136942099 | 136942466 | 136941536 | 136941684 | 130 | 65 | 85 | 102 | 69 | 6 | 0 | 1 | 4 | 0 | 98 | 209 | 78 | 131 | 269 | 0 | 9 | 2 | 1 | 0 | 203 | 149 | 0.488183005 | 1 | 0.941 | 1 | 0.984 | 0.949 | 1 | 1 | 0.945 | 0.966 | 0.99 | 1 | -0.005 |
| A5SS | 5115 | ENSG00000122507 | BBS9 | chr7 | + | 33152700 | 33152855 | 33152700 | 33152851 | 33155637 | 33155702 | 0 | 2 | 0 | 0 | 1 | 49 | 29 | 56 | 46 | 46 | 0 | 0 | 4 | 0 | 3 | 23 | 34 | 15 | 18 | 39 | 153 | 149 | 0.488537117 | 1 | 0 | 0.063 | 0 | 0 | 0.021 | 0 | 0 | 0.206 | 0 | 0.07 | -0.038 |
| A5SS | 5304 | ENSG00000168890 | TMEM150A | chr2 | - | 85601305 | 85601482 | 85601434 | 85601482 | 85601020 | 85601107 | 9 | 2 | 5 | 7 | 9 | 4 | 3 | 7 | 10 | 1 | 6 | 7 | 8 | 7 | 1 | 7 | 0 | 5 | 0 | 10 | 278 | 149 | 0.488556771 | 1 | 0.547 | 0.263 | 0.277 | 0.273 | 0.828 | 0.315 | 1 | 0.462 | 1 | 0.051 | -0.128 |
| A5SS | 4698 | ENSG00000115170 | ACVR1 | chr2 | - | 157818268 | 157818559 | 157818384 | 157818559 | 157799426 | 157799500 | 4 | 8 | 3 | 3 | 2 | 19 | 15 | 9 | 9 | 30 | 16 | 4 | 3 | 0 | 18 | 24 | 20 | 6 | 26 | 61 | 265 | 149 | 0.489188741 | 1 | 0.106 | 0.231 | 0.158 | 0.158 | 0.036 | 0.273 | 0.101 | 0.219 | 0 | 0.142 | -0.009 |
| A5SS | 4035 | ENSG00000126091 | ST3GAL3 | chr1 | + | 43814890 | 43815004 | 43814890 | 43814933 | 43838218 | 43838311 | 2 | 0 | 0 | 2 | 2 | 30 | 21 | 18 | 28 | 28 | 0 | 0 | 0 | 0 | 0 | 44 | 26 | 18 | 31 | 202 | 220 | 149 | 0.489384227 | 1 | 0.043 | 0 | 0 | 0.046 | 0.046 | 0 | 0 | 0 | 0 | 0 | 0.027 |
| A5SS | 8522 | ENSG00000128487 | SPECC1 | chr17 | + | 20245925 | 20246574 | 20245925 | 20246071 | 20247218 | 20247319 | 1 | 7 | 3 | 0 | 0 | 13 | 6 | 7 | 12 | 27 | 0 | 10 | 2 | 2 | 0 | 2 | 15 | 10 | 6 | 25 | 298 | 149 | 0.492175069 | 1 | 0.037 | 0.368 | 0.176 | 0 | 0 | 0 | 0.25 | 0.091 | 0.143 | 0 | 0.019 |
| A5SS | 4307 | ENSG00000043514 | TRIT1 | chr1 | - | 39850118 | 39850261 | 39850122 | 39850261 | 39847985 | 39848097 | 31 | 13 | 18 | 14 | 22 | 6 | 9 | 11 | 3 | 9 | 31 | 24 | 7 | 17 | 16 | 6 | 0 | 14 | 11 | 44 | 153 | 149 | 0.492424969 | 1 | 0.834 | 0.584 | 0.614 | 0.82 | 0.704 | 0.834 | 1 | 0.327 | 0.601 | 0.262 | 0.106 |
| A5SS | 6415 | ENSG00000134291 | TMEM106C | chr12 | + | 47966088 | 47966229 | 47966088 | 47966172 | 47966682 | 47966732 | 35 | 18 | 27 | 39 | 44 | 22 | 10 | 31 | 7 | 25 | 36 | 34 | 24 | 41 | 161 | 46 | 11 | 37 | 32 | 51 | 206 | 149 | 0.492899737 | 1 | 0.535 | 0.566 | 0.386 | 0.801 | 0.56 | 0.361 | 0.691 | 0.319 | 0.481 | 0.695 | 0.06 |
| A5SS | 6053 | ENSG00000079819 | EPB41L2 | chr6 | - | 130869562 | 130870126 | 130869811 | 130870126 | 130867458 | 130867581 | 26 | 19 | 31 | 15 | 36 | 0 | 2 | 0 | 0 | 4 | 12 | 13 | 8 | 9 | 3 | 0 | 0 | 1 | 0 | 0 | 298 | 149 | 0.495172618 | 1 | 1 | 0.826 | 1 | 1 | 0.818 | 1 | 1 | 0.8 | 1 | 1 | -0.031 |
| A5SS | 940 | ENSG00000108384 | RAD51C | chr17 | + | 58692601 | 58693650 | 58692601 | 58692788 | 58694930 | 58695189 | 4 | 5 | 3 | 4 | 7 | 20 | 25 | 40 | 34 | 19 | 0 | 2 | 5 | 10 | 13 | 41 | 19 | 43 | 18 | 22 | 298 | 149 | 0.495776643 | 1 | 0.091 | 0.091 | 0.036 | 0.056 | 0.156 | 0 | 0.05 | 0.055 | 0.217 | 0.228 | -0.024 |
| A5SS | 7093 | ENSG00000147140 | NONO | chrX | + | 71297376 | 71297938 | 71297376 | 71297461 | 71298468 | 71298508 | 265 | 157 | 254 | 275 | 229 | 0 | 0 | 0 | 0 | 0 | 447 | 256 | 220 | 261 | 369 | 0 | 2 | 1 | 12 | 0 | 298 | 149 | 0.496467326 | 1 | 1 | 1 | 1 | 1 | 1 | 1 | 0.985 | 0.991 | 0.916 | 1 | 0.022 |
| A5SS | 6871 | ENSG00000262919 | CCNQ | chrX | - | 153598961 | 153599117 | 153598965 | 153599117 | 153596003 | 153596187 | 11 | 2 | 11 | 9 | 2 | 1 | 1 | 4 | 1 | 4 | 6 | 7 | 6 | 0 | 5 | 0 | 0 | 4 | 2 | 21 | 153 | 149 | 0.496576944 | 1 | 0.915 | 0.661 | 0.728 | 0.898 | 0.327 | 1 | 1 | 0.594 | 0 | 0.188 | 0.149 |
| A5SS | 7491 | ENSG00000134539 | KLRD1 | chr12 | + | 10311463 | 10311619 | 10311463 | 10311615 | 10313406 | 10313513 | 1 | 3 | 0 | 1 | 1 | 254 | 58 | 63 | 98 | 108 | 0 | 2 | 5 | 0 | 0 | 19 | 61 | 58 | 23 | 16 | 153 | 149 | 0.496964659 | 1 | 0.004 | 0.048 | 0 | 0.01 | 0.009 | 0 | 0.031 | 0.077 | 0 | 0 | -0.007 |
| A5SS | 7319 | ENSG00000103266 | STUB1 | chr16 | + | 681792 | 681903 | 681792 | 681880 | 682019 | 682076 | 9 | 10 | 4 | 11 | 8 | 96 | 33 | 59 | 81 | 87 | 17 | 23 | 9 | 8 | 0 | 59 | 109 | 73 | 56 | 114 | 172 | 149 | 0.49905188 | 1 | 0.075 | 0.208 | 0.055 | 0.105 | 0.074 | 0.2 | 0.155 | 0.096 | 0.11 | 0 | -0.009 |
| A5SS | 6437 | ENSG00000119616 | FCF1 | chr14 | + | 74713484 | 74713620 | 74713484 | 74713552 | 74714871 | 74714943 | 18 | 9 | 14 | 16 | 25 | 150 | 123 | 152 | 186 | 143 | 10 | 21 | 19 | 8 | 0 | 242 | 87 | 98 | 137 | 48 | 217 | 149 | 0.499233107 | 1 | 0.076 | 0.048 | 0.059 | 0.056 | 0.107 | 0.028 | 0.142 | 0.117 | 0.039 | 0 | 0.004 |
| A5SS | 6656 | ENSG00000136247 | ZDHHC4 | chr7 | + | 6577467 | 6577720 | 6577467 | 6577498 | 6578565 | 6578746 | 15 | 1 | 7 | 4 | 3 | 1 | 0 | 2 | 4 | 0 | 31 | 1 | 1 | 10 | 23 | 0 | 2 | 3 | 0 | 14 | 298 | 149 | 0.501260224 | 1 | 0.882 | 1 | 0.636 | 0.333 | | 1 | 0.2 | 0.143 | 1 | 0.451 | 0.211 |
| A5SS | 4435 | ENSG00000197989 | SNHG12 | chr1 | - | 28581093 | 28581229 | 28581110 | 28581229 | 28580559 | 28580646 | 15 | 11 | 13 | 12 | 6 | 48 | 25 | 71 | 66 | 39 | 37 | 2 | 18 | 6 | 9 | 14 | 41 | 61 | 35 | 39 | 166 | 149 | 0.503537375 | 1 | 0.219 | 0.283 | 0.141 | 0.14 | 0.121 | 0.703 | 0.042 | 0.209 | 0.133 | 0.172 | -0.071 |
| A5SS | 6409 | ENSG00000145191 | EIF2B5 | chr3 | + | 184140417 | 184140730 | 184140417 | 184140615 | 184141924 | 184142070 | 78 | 34 | 91 | 102 | 60 | 0 | 0 | 0 | 2 | 8 | 94 | 113 | 66 | 62 | 127 | 9 | 2 | 4 | 0 | 0 | 264 | 149 | 0.503944379 | 1 | 1 | 1 | 1 | 0.966 | 0.809 | 0.855 | 0.97 | 0.903 | 1 | 1 | 0.009 |
| A5SS | 6636 | ENSG00000086730 | LAT2 | chr7 | + | 74214821 | 74214940 | 74214821 | 74214917 | 74215946 | 74216069 | 41 | 17 | 36 | 10 | 32 | 0 | 0 | 2 | 0 | 0 | 55 | 42 | 15 | 73 | 28 | 0 | 0 | 0 | 0 | 7 | 172 | 149 | 0.505364023 | 1 | 1 | 1 | 0.94 | 1 | 1 | 1 | 1 | 1 | 1 | 0.776 | 0.033 |
| A5SS | 6909 | ENSG00000139436 | GIT2 | chr12 | - | 109996098 | 109996368 | 109996172 | 109996368 | 109991626 | 109991760 | 3 | 0 | 4 | 1 | 0 | 61 | 22 | 47 | 56 | 8 | 2 | 0 | 0 | 0 | 0 | 50 | 84 | 47 | 52 | 24 | 223 | 149 | 0.506254344 | 1 | 0.032 | 0 | 0.054 | 0.012 | 0 | 0.026 | 0 | | 0 | 0 | 0.014 |
| A5SS | 3390 | ENSG00000132768 | DPH2 | chr1 | + | 43970009 | 43970322 | 43970009 | 43970270 | 43970595 | 43970708 | 16 | 3 | 13 | 26 | 16 | 0 | 2 | 1 | 1 | 1 | 19 | 5 | 7 | 7 | 7 | 0 | 0 | 0 | 0 | 5 | 201 | 149 | 0.506559588 | 1 | 1 | 0.527 | 0.906 | 0.951 | 0.922 | 1 | 1 | | 1 | 0.509 | -0.041 |
| A5SS | 7032 | ENSG00000102030 | NAA10 | chrX | - | 153932070 | 153932431 | 153932315 | 153932431 | 153930762 | 153930847 | 146 | 86 | 124 | 144 | 77 | 1 | 4 | 1 | 2 | 0 | 114 | 153 | 86 | 123 | 210 | 0 | 0 | 1 | 3 | 0 | 298 | 149 | 0.506862639 | 1 | 0.986 | 0.915 | 0.984 | 0.973 | 1 | 1 | 1 | 0.977 | 0.953 | 1 | -0.014 |
| A5SS | 2509 | ENSG00000105202 | FBL | chr19 | - | 39840616 | 39840787 | 39840655 | 39840787 | 39840413 | 39840515 | 30 | 13 | 28 | 27 | 11 | 6 | 0 | 8 | 3 | 0 | 25 | 3 | 14 | 23 | 42 | 5 | 0 | 1 | 0 | 7 | 188 | 149 | 0.506882218 | 1 | 0.798 | 1 | 0.735 | 0.877 | 1 | 0.798 | 1 | 0.917 | 1 | 0.826 | -0.026 |
| A5SS | 4642 | ENSG00000069702 | TGFBR3 | chr1 | - | 91885877 | 91886151 | 91886029 | 91886151 | 91861470 | 91861644 | 201 | 19 | 63 | 150 | 14 | 17 | 1 | 3 | 5 | 1 | 32 | 95 | 87 | 34 | 10 | 0 | 14 | 10 | 0 | 0 | 298 | 149 | 0.507196022 | 1 | 0.855 | 0.905 | 0.913 | 0.938 | 0.875 | 1 | 0.772 | 0.813 | 1 | 1 | -0.02 |
| A5SS | 4132 | ENSG00000109180 | OCIAD1 | chr4 | + | 48831226 | 48831496 | 48831226 | 48831398 | 48832619 | 48832682 | 18 | 7 | 17 | 12 | 21 | 0 | 1 | 1 | 4 | 3 | 6 | 9 | 9 | 11 | 31 | 0 | 0 | 5 | 2 | 0 | 247 | 149 | 0.507799637 | 1 | 1 | 0.809 | 0.911 | 0.644 | 0.809 | 1 | 1 | 0.521 | 0.768 | 1 | -0.023 |
| A5SS | 2666 | ENSG00000189308 | LIN54 | chr4 | - | 82984160 | 82984876 | 82984823 | 82984876 | 82970326 | 82970469 | 21 | 18 | 53 | 40 | 27 | 0 | 0 | 0 | 0 | 0 | 9 | 55 | 19 | 42 | 21 | 0 | 0 | 0 | 6 | 0 | 298 | 149 | 0.508257986 | 1 | 1 | 1 | 1 | 1 | 1 | 1 | 1 | 1 | 0.778 | 1 | 0.044 |
| A5SS | 5555 | ENSG00000115694 | STK25 | chr2 | - | 241501477 | 241501698 | 241501527 | 241501698 | 241500739 | 241500796 | 101 | 52 | 84 | 85 | 63 | 2 | 0 | 3 | 0 | 3 | 68 | 90 | 74 | 67 | 211 | 0 | 2 | 0 | 4 | 6 | 199 | 149 | 0.509728789 | 1 | 0.974 | 1 | 0.954 | 1 | 0.94 | 1 | 0.971 | 1 | 0.926 | 0.963 | 0.002 |
| A5SS | 1647 | ENSG00000126822 | PLEKHG3 | chr14 | + | 64736852 | 64737375 | 64736852 | 64736891 | 64741035 | 64742455 | 14 | 7 | 3 | 12 | 0 | 6 | 0 | 0 | 4 | 1 | 38 | 7 | 17 | 15 | 35 | 8 | 3 | 0 | 4 | 7 | 298 | 149 | 0.511673786 | 1 | 0.538 | 1 | 1 | 0.6 | 0 | 0.704 | 0.538 | 1 | 0.652 | 0.714 | -0.094 |
| A5SS | 3078 | ENSG00000161217 | PCYT1A | chr3 | - | 196287531 | 196287726 | 196287614 | 196287726 | 196270414 | 196270541 | 5 | 4 | 6 | 3 | 2 | 28 | 6 | 26 | 47 | 17 | 0 | 6 | 9 | 3 | 8 | 41 | 23 | 17 | 14 | 34 | 232 | 149 | 0.5119763 | 1 | 0.103 | 0.3 | 0.129 | 0.039 | 0.07 | 0 | 0.143 | 0.254 | 0.121 | 0.131 | -0.002 |
| A5SS | 1586 | ENSG00000137841 | PLCB2 | chr15 | - | 40296745 | 40296908 | 40296757 | 40296908 | 40296521 | 40296634 | 190 | 99 | 129 | 156 | 79 | 25 | 37 | 26 | 45 | 17 | 294 | 151 | 141 | 197 | 336 | 34 | 38 | 23 | 74 | 61 | 161 | 149 | 0.512468038 | 1 | 0.876 | 0.712 | 0.821 | 0.762 | 0.811 | 0.889 | 0.786 | 0.85 | 0.711 | 0.836 | -0.018 |
| A5SS | 7420 | ENSG00000104763 | ASAH1 | chr8 | - | 18062247 | 18062423 | 18062278 | 18062423 | 18061685 | 18061740 | 10 | 8 | 5 | 18 | 5 | 677 | 617 | 429 | 639 | 890 | 10 | 9 | 17 | 2 | 0 | 373 | 553 | 647 | 459 | 140 | 180 | 149 | 0.512620612 | 1 | 0.012 | 0.011 | 0.01 | 0.023 | 0.005 | 0.022 | 0.013 | 0.021 | 0.004 | 0 | 0 |
| A5SS | 4706 | ENSG00000144535 | DIS3L2 | chr2 | + | 232130618 | 232130943 | 232130618 | 232130719 | 232136471 | 232136719 | 27 | 33 | 21 | 10 | 13 | 47 | 42 | 69 | 76 | 72 | 10 | 7 | 22 | 14 | 4 | 63 | 29 | 19 | 50 | 33 | 298 | 149 | 0.512825029 | 1 | 0.223 | 0.282 | 0.132 | 0.062 | 0.083 | 0.074 | 0.108 | 0.367 | 0.123 | 0.057 | 0.011 |
| A5SS | 1739 | ENSG00000254505 | CHMP4A | chr14 | - | 24211410 | 24211592 | 24211414 | 24211592 | 24210347 | 24210483 | 4 | 2 | 4 | 3 | 0 | 0 | 5 | 2 | 4 | 3 | 9 | 5 | 4 | 8 | 8 | 22 | 0 | 18 | 8 | 0 | 153 | 149 | 0.513653304 | 1 | 1 | 0.28 | 0.661 | 0.422 | 0 | 0.285 | 1 | 0.178 | 0.493 | 1 | -0.119 |
| A5SS | 1111 | ENSG00000187555 | USP7 | chr16 | - | 8923214 | 8923413 | 8923353 | 8923413 | 8921156 | 8921295 | 820 | 575 | 798 | 790 | 883 | 6 | 6 | 5 | 15 | 2 | 999 | 859 | 926 | 1045 | 488 | 11 | 5 | 7 | 9 | 0 | 288 | 149 | 0.513810381 | 1 | 0.986 | 0.98 | 0.988 | 0.965 | 0.996 | 0.979 | 0.989 | 0.986 | 0.984 | 1 | -0.005 |
| A5SS | 6242 | ENSG00000135317 | SNX14 | chr6 | - | 85593578 | 85593857 | 85593833 | 85593857 | 85574257 | 85574378 | 50 | 20 | 33 | 48 | 22 | 1 | 0 | 2 | 3 | 0 | 11 | 29 | 9 | 21 | 10 | 0 | 0 | 2 | 4 | 0 | 298 | 149 | 0.51539174 | 1 | 0.962 | 1 | 0.892 | 0.889 | 1 | 1 | 1 | 0.692 | 0.724 | 1 | 0.065 |
| A5SS | 4828 | ENSG00000115042 | FAHD2A | chr2 | + | 95410526 | 95411026 | 95410526 | 95410586 | 95412433 | 95412542 | 55 | 36 | 31 | 52 | 35 | 8 | 1 | 2 | 3 | 0 | 45 | 26 | 28 | 58 | 68 | 13 | 0 | 1 | 2 | 4 | 298 | 149 | 0.516219096 | 1 | 0.775 | 0.947 | 0.886 | 0.897 | 1 | 0.634 | 1 | 0.933 | 0.935 | 0.895 | 0.022 |
| A5SS | 2983 | ENSG00000169714 | CNBP | chr3 | - | 129171633 | 129171771 | 129171654 | 129171771 | 129171445 | 129171508 | 48 | 33 | 41 | 56 | 30 | 1 | 2 | 3 | 0 | 0 | 38 | 52 | 30 | 65 | 65 | 0 | 6 | 0 | 0 | 0 | 170 | 149 | 0.517729984 | 1 | 0.977 | 0.935 | 0.923 | 1 | 1 | 1 | 0.884 | 1 | 1 | 1 | -0.01 |
| A5SS | 1447 | ENSG00000182511 | FES | chr15 | + | 90884503 | 90884639 | 90884503 | 90884544 | 90885036 | 90885258 | 19 | 2 | 8 | 6 | 0 | 23 | 5 | 10 | 15 | 1 | 32 | 22 | 7 | 22 | 17 | 77 | 51 | 15 | 42 | 50 | 244 | 149 | 0.517772448 | 1 | 0.335 | 0.196 | 0.328 | 0.196 | 0 | 0.202 | 0.208 | 0.222 | 0.242 | 0.172 | 0.002 |
| A5SS | 3398 | ENSG00000170581 | STAT2 | chr12 | - | 56350096 | 56350432 | 56350411 | 56350432 | 56349425 | 56349509 | 28 | 30 | 28 | 50 | 29 | 0 | 0 | 0 | 0 | 0 | 239 | 69 | 48 | 98 | 111 | 9 | 0 | 0 | 0 | 0 | 298 | 149 | 0.520533186 | 1 | 1 | 1 | 1 | 1 | 1 | 0.93 | 1 | 1 | 1 | 1 | 0.014 |
| A5SS | 6895 | ENSG00000173681 | BCLAF3 | chrX | - | 19937417 | 19937532 | 19937504 | 19937532 | 19935808 | 19935898 | 20 | 8 | 9 | 14 | 27 | 20 | 29 | 37 | 38 | 34 | 3 | 0 | 7 | 2 | 7 | 4 | 25 | 29 | 7 | 0 | 236 | 149 | 0.520709759 | 1 | 0.387 | 0.148 | 0.133 | 0.189 | 0.334 | 0.321 | 0 | 0.132 | 0.153 | 1 | -0.083 |
| A5SS | 3442 | ENSG00000144591 | GMPPA | chr2 | + | 219504082 | 219504218 | 219504082 | 219504213 | 219505227 | 219505362 | 0 | 1 | 3 | 2 | 2 | 48 | 41 | 35 | 34 | 51 | 0 | 0 | 3 | 6 | 0 | 73 | 49 | 25 | 47 | 67 | 154 | 149 | 0.521681026 | 1 | 0 | 0.023 | 0.077 | 0.054 | 0.037 | 0 | 0 | 0.104 | 0.11 | 0 | -0.005 |
| A5SS | 5226 | ENSG00000167107 | ACSF2 | chr17 | + | 50461241 | 50461370 | 50461241 | 50461287 | 50461632 | 50461686 | 35 | 10 | 18 | 23 | 11 | 0 | 1 | 0 | 0 | 0 | 17 | 8 | 2 | 16 | 54 | 0 | 4 | 0 | 0 | 0 | 232 | 149 | 0.52216418 | 1 | 1 | 0.865 | 1 | 1 | 1 | 1 | 0.562 | 1 | 1 | 1 | 0.061 |
| A5SS | 2491 | ENSG00000204842 | ATXN2 | chr12 | - | 111485264 | 111485331 | 111485270 | 111485331 | 111482786 | 111482831 | 2 | 5 | 6 | 2 | 0 | 2 | 5 | 4 | 2 | 1 | 9 | 10 | 4 | 16 | 2 | 3 | 3 | 0 | 10 | 28 | 155 | 149 | 0.522228259 | 1 | 0.49 | 0.49 | 0.59 | 0.49 | 0 | 0.743 | 0.762 | 1 | 0.606 | 0.064 | -0.223 |
| A5SS | 5530 | ENSG00000082258 | CCNT2 | chr2 | + | 134946100 | 134946465 | 134946100 | 134946146 | 134947735 | 134947899 | 99 | 65 | 85 | 115 | 122 | 22 | 28 | 28 | 27 | 12 | 18 | 87 | 81 | 18 | 0 | 0 | 20 | 21 | 5 | 11 | 298 | 149 | 0.522641877 | 1 | 0.692 | 0.537 | 0.603 | 0.68 | 0.836 | 1 | 0.685 | 0.659 | 0.643 | 0 | 0.072 |
| A5SS | 7379 | ENSG00000249859 | PVT1 | chr8 | + | 128070159 | 128070361 | 128070159 | 128070272 | 128096517 | 128096654 | 13 | 3 | 13 | 2 | 7 | 12 | 8 | 14 | 4 | 6 | 0 | 7 | 7 | 0 | 7 | 35 | 4 | 6 | 1 | 0 | 238 | 149 | 0.523260942 | 1 | 0.404 | 0.19 | 0.368 | 0.238 | 0.422 | 0 | 0.523 | 0.422 | 0 | 1 | -0.065 |
| A5SS | 2408 | ENSG00000159202 | UBE2Z | chr17 | + | 48912833 | 48913021 | 48912833 | 48912965 | 48916075 | 48916187 | 130 | 86 | 127 | 129 | 174 | 4 | 0 | 5 | 5 | 1 | 179 | 146 | 86 | 145 | 161 | 0 | 5 | 2 | 2 | 4 | 205 | 149 | 0.523349414 | 1 | 0.959 | 1 | 0.949 | 0.949 | 0.992 | 1 | 0.955 | 0.969 | 0.981 | 0.967 | -0.005 |
| A5SS | 6552 | ENSG00000157764 | BRAF | chr7 | - | 140808882 | 140808995 | 140808891 | 140808995 | 140807959 | 140808062 | 1 | 0 | 2 | 1 | 4 | 79 | 96 | 80 | 66 | 271 | 0 | 4 | 4 | 0 | 0 | 47 | 66 | 49 | 27 | 61 | 158 | 149 | 0.524997416 | 1 | 0.012 | 0 | 0.023 | 0.014 | 0.014 | 0 | 0.054 | 0.071 | 0 | 0 | -0.012 |
| A5SS | 5671 | ENSG00000095059 | DHPS | chr19 | - | 12680160 | 12680325 | 12680195 | 12680325 | 12679800 | 12679922 | 125 | 61 | 113 | 117 | 81 | 6 | 2 | 5 | 4 | 0 | 129 | 125 | 100 | 114 | 266 | 0 | 4 | 4 | 9 | 11 | 184 | 149 | 0.52534788 | 1 | 0.944 | 0.961 | 0.948 | 0.959 | 1 | 1 | 0.962 | 0.953 | 0.911 | 0.951 | 0.007 |
| A5SS | 2369 | ENSG00000160179 | ABCG1 | chr21 | + | 42287888 | 42288073 | 42287888 | 42288037 | 42288210 | 42288312 | 12 | 10 | 19 | 18 | 24 | 5 | 3 | 15 | 2 | 0 | 15 | 22 | 58 | 28 | 0 | 3 | 8 | 1 | 0 | 5 | 185 | 149 | 0.527971581 | 1 | 0.659 | 0.729 | 0.505 | 0.879 | 1 | 0.801 | 0.689 | 0.979 | 1 | 0 | 0.061 |
| A5SS | 5012 | ENSG00000076242 | MLH1 | chr3 | + | 37000954 | 37001053 | 37000954 | 37001048 | 37004400 | 37004474 | 134 | 122 | 146 | 145 | 153 | 11 | 1 | 0 | 3 | 0 | 193 | 149 | 156 | 188 | 39 | 10 | 0 | 3 | 6 | 0 | 154 | 149 | 0.528163377 | 1 | 0.922 | 0.992 | 1 | 0.979 | 1 | 0.949 | 1 | 0.981 | 0.968 | 1 | -0.001 |
| A5SS | 6716 | ENSG00000170852 | KBTBD2 | chr7 | - | 32879434 | 32879942 | 32879700 | 32879942 | 32874991 | 32875157 | 462 | 321 | 399 | 460 | 678 | 21 | 4 | 16 | 6 | 5 | 378 | 558 | 750 | 551 | 250 | 10 | 14 | 12 | 12 | 1 | 298 | 149 | 0.52837522 | 1 | 0.917 | 0.976 | 0.926 | 0.975 | 0.985 | 0.95 | 0.952 | 0.969 | 0.958 | 0.992 | -0.008 |
| A5SS | 1056 | ENSG00000142512 | SIGLEC10 | chr19 | - | 51415897 | 51416357 | 51416309 | 51416357 | 51415567 | 51415615 | 42 | 30 | 17 | 46 | 19 | 32 | 32 | 4 | 9 | 0 | 3 | 24 | 10 | 31 | 50 | 14 | 8 | 3 | 3 | 22 | 298 | 149 | 0.52902483 | 1 | 0.396 | 0.319 | 0.68 | 0.719 | 1 | 0.097 | 0.6 | 0.625 | 0.838 | 0.532 | 0.084 |
| A5SS | 4984 | ENSG00000138757 | G3BP2 | chr4 | - | 75653982 | 75655246 | 75655065 | 75655246 | 75648638 | 75648741 | 67 | 63 | 61 | 56 | 146 | 450 | 348 | 421 | 563 | 623 | 52 | 88 | 69 | 59 | 38 | 341 | 397 | 777 | 502 | 120 | 298 | 149 | 0.529625225 | 1 | 0.069 | 0.083 | 0.068 | 0.047 | 0.105 | 0.071 | 0.1 | 0.043 | 0.056 | 0.137 | -0.007 |
| A5SS | 8631 | ENSG00000145901 | TNIP1 | chr5 | - | 151080072 | 151080359 | 151080323 | 151080359 | 151064959 | 151065131 | 42 | 33 | 48 | 48 | 18 | 0 | 2 | 5 | 0 | 1 | 211 | 83 | 94 | 178 | 130 | 0 | 0 | 0 | 2 | 10 | 298 | 149 | 0.530349555 | 1 | 1 | 0.892 | 0.828 | 1 | 0.9 | 1 | 1 | 1 | 0.978 | 0.867 | -0.045 |
| A5SS | 5718 | ENSG00000148297 | MED22 | chr9 | - | 133347921 | 133348117 | 133347941 | 133348117 | 133346539 | 133346700 | 8 | 5 | 17 | 17 | 4 | 0 | 0 | 3 | 0 | 4 | 16 | 19 | 2 | 14 | 29 | 0 | 0 | 0 | 5 | 0 | 169 | 149 | 0.530804288 | 1 | 1 | 1 | 0.833 | 1 | 0.469 | 1 | 1 | 1 | 0.712 | 1 | -0.082 |
| A5SS | 8161 | ENSG00000159063 | ALG8 | chr11 | - | 78124020 | 78124214 | 78124024 | 78124214 | 78121064 | 78121174 | 70 | 45 | 39 | 57 | 63 | 0 | 0 | 0 | 2 | 7 | 28 | 28 | 29 | 33 | 22 | 0 | 0 | 0 | 0 | 0 | 153 | 149 | 0.531217314 | 1 | 1 | 1 | 1 | 0.965 | 0.898 | 1 | 1 | 1 | 1 | 1 | -0.027 |
| A5SS | 5574 | ENSG00000196576 | PLXNB2 | chr22 | - | 50294718 | 50294778 | 50294725 | 50294778 | 50289516 | 50290597 | 24 | 8 | 0 | 21 | 15 | 15 | 9 | 1 | 9 | 2 | 6 | 6 | 2 | 1 | 19 | 8 | 1 | 2 | 1 | 17 | 156 | 149 | 0.531493402 | 1 | 0.604 | 0.459 | 0 | 0.69 | 0.878 | 0.417 | 0.851 | 0.489 | 0.489 | 0.516 | -0.026 |
| A5SS | 7577 | ENSG00000137207 | YIPF3 | chr6 | - | 43515594 | 43515701 | 43515605 | 43515701 | 43513587 | 43513633 | 78 | 44 | 54 | 78 | 76 | 1 | 6 | 3 | 3 | 1 | 172 | 89 | 49 | 87 | 146 | 0 | 3 | 1 | 5 | 7 | 160 | 149 | 0.531575656 | 1 | 0.986 | 0.872 | 0.944 | 0.96 | 0.986 | 1 | 0.965 | 0.979 | 0.942 | 0.951 | -0.018 |
| A5SS | 1732 | ENSG00000100938 | GMPR2 | chr14 | + | 24233478 | 24233598 | 24233478 | 24233593 | 24235736 | 24235820 | 76 | 37 | 84 | 90 | 113 | 0 | 0 | 6 | 2 | 1 | 132 | 100 | 47 | 92 | 136 | 0 | 0 | 1 | 1 | 5 | 154 | 149 | 0.531732964 | 1 | 1 | 1 | 0.931 | 0.978 | 0.991 | 1 | 1 | 0.978 | 0.989 | 0.963 | -0.006 |
| A5SS | 4463 | ENSG00000161929 | SCIMP | chr17 | - | 5223332 | 5223456 | 5223353 | 5223456 | 5221286 | 5221350 | 95 | 89 | 46 | 72 | 66 | 33 | 9 | 19 | 21 | 37 | 94 | 85 | 18 | 31 | 115 | 21 | 28 | 10 | 23 | 35 | 170 | 149 | 0.532633205 | 1 | 0.716 | 0.897 | 0.68 | 0.75 | 0.61 | 0.797 | 0.727 | 0.612 | 0.542 | 0.742 | 0.047 |
| A5SS | 808 | ENSG00000078687 | TNRC6C | chr17 | + | 78048844 | 78051457 | 78048844 | 78051448 | 78064721 | 78064937 | 3 | 5 | 3 | 8 | 4 | 66 | 22 | 55 | 59 | 18 | 2 | 5 | 3 | 8 | 0 | 6 | 35 | 13 | 35 | 31 | 158 | 149 | 0.532709997 | 1 | 0.041 | 0.176 | 0.049 | 0.113 | 0.173 | 0.239 | 0.119 | 0.179 | 0.177 | 0 | -0.032 |
| A5SS | 7254 | ENSG00000197217 | ENTPD4 | chr8 | - | 23441568 | 23441723 | 23441592 | 23441723 | 23439748 | 23439915 | 72 | 30 | 66 | 76 | 88 | 44 | 30 | 72 | 40 | 88 | 54 | 66 | 34 | 52 | 53 | 72 | 21 | 72 | 40 | 57 | 173 | 149 | 0.533360008 | 1 | 0.585 | 0.463 | 0.441 | 0.621 | 0.463 | 0.392 | 0.73 | 0.289 | 0.528 | 0.445 | 0.038 |
| A5SS | 4782 | ENSG00000105402 | NAPA | chr19 | - | 47503392 | 47503502 | 47503422 | 47503502 | 47500632 | 47500749 | 2 | 1 | 1 | 1 | 0 | 248 | 139 | 199 | 267 | 173 | 23 | 3 | 4 | 2 | 3 | 521 | 358 | 214 | 488 | 801 | 179 | 149 | 0.533555752 | 1 | 0.007 | 0.006 | 0.004 | 0.003 | 0 | 0.035 | 0.007 | 0.015 | 0.003 | 0.003 | -0.009 |
| A5SS | 7354 | ENSG00000104783 | KCNN4 | chr19 | - | 43769332 | 43769560 | 43769441 | 43769560 | 43768962 | 43769032 | 3 | 1 | 0 | 4 | 0 | 47 | 18 | 31 | 51 | 46 | 0 | 0 | 0 | 0 | 0 | 43 | 15 | 17 | 37 | 18 | 258 | 149 | 0.534228386 | 1 | 0.036 | 0.031 | 0 | 0.043 | 0 | 0 | 0 | 0 | 0 | 0 | 0.022 |
| A5SS | 2028 | ENSG00000141279 | NPEPPS | chr17 | + | 47619277 | 47619495 | 47619277 | 47619322 | 47619736 | 47619784 | 16 | 5 | 7 | 4 | 3 | 1 | 0 | 1 | 0 | 3 | 13 | 2 | 3 | 6 | 8 | 9 | 0 | 0 | 0 | 0 | 298 | 149 | 0.534711853 | 1 | 0.889 | 1 | 0.778 | 1 | 0.333 | 0.419 | 1 | | 1 | 1 | -0.084 |
| A5SS | 1821 | ENSG00000125746 | EML2 | chr19 | - | 45639294 | 45639397 | 45639356 | 45639397 | 45638844 | 45638873 | 1 | 1 | 0 | 5 | 0 | 31 | 26 | 40 | 34 | 26 | 0 | 0 | 0 | 0 | 5 | 93 | 24 | 42 | 37 | 42 | 211 | 149 | 0.535885045 | 1 | 0.022 | 0.026 | 0 | 0.094 | 0 | 0 | 0 | | 0 | 0.078 | 0.013 |
| A5SS | 5580 | ENSG00000184922 | FMNL1 | chr17 | + | 45232366 | 45232480 | 45232366 | 45232450 | 45233223 | 45233297 | 259 | 133 | 246 | 209 | 160 | 167 | 54 | 155 | 155 | 87 | 336 | 242 | 176 | 339 | 376 | 201 | 181 | 120 | 159 | 280 | 179 | 149 | 0.539343375 | 1 | 0.564 | 0.672 | 0.569 | 0.529 | 0.605 | 0.582 | 0.527 | 0.55 | 0.64 | 0.528 | 0.022 |
| A5SS | 2002 | ENSG00000109775 | UFSP2 | chr4 | - | 185418586 | 185418770 | 185418611 | 185418770 | 185418440 | 185418507 | 80 | 72 | 76 | 72 | 104 | 2 | 2 | 1 | 2 | 1 | 5 | 63 | 83 | 56 | 13 | 0 | 0 | 0 | 0 | 0 | 174 | 149 | 0.539429154 | 1 | 0.972 | 0.969 | 0.985 | 0.969 | 0.989 | 1 | 1 | 1 | 1 | 1 | -0.023 |
| A5SS | 2830 | ENSG00000240682 | ISY1 | chr3 | - | 129134827 | 129134954 | 129134831 | 129134954 | 129134073 | 129134195 | 1 | 1 | 4 | 5 | 0 | 147 | 134 | 124 | 144 | 156 | 0 | 2 | 2 | 4 | 12 | 370 | 126 | 234 | 187 | 298 | 153 | 149 | 0.540118631 | 1 | 0.007 | 0.007 | 0.03 | 0.033 | 0 | 0 | 0.015 | 0.008 | 0.02 | 0.038 | -0.001 |
| A5SS | 2030 | ENSG00000130811 | EIF3G | chr19 | - | 10116799 | 10116989 | 10116823 | 10116989 | 10115966 | 10116074 | 1025 | 629 | 889 | 908 | 865 | 1 | 5 | 2 | 3 | 7 | 1763 | 1346 | 762 | 1087 | 1606 | 37 | 3 | 7 | 4 | 0 | 173 | 149 | 0.540571783 | 1 | 0.999 | 0.991 | 0.997 | 0.996 | 0.991 | 0.976 | 0.997 | 0.989 | 0.996 | 1 | 0.003 |
| A5SS | 3421 | ENSG00000008128 | CDK11A | chr1 | - | 1722587 | 1722831 | 1722707 | 1722831 | 1721595 | 1721711 | 24 | 42 | 39 | 79 | 42 | 84 | 65 | 20 | 91 | 101 | 160 | 87 | 52 | 145 | 89 | 282 | 50 | 119 | 149 | 141 | 269 | 149 | 0.541097753 | 1 | 0.137 | 0.264 | 0.519 | 0.325 | 0.187 | 0.239 | 0.491 | 0.195 | 0.35 | 0.259 | -0.02 |
| A5SS | 486 | ENSG00000101138 | CSTF1 | chr20 | + | 56395520 | 56395759 | 56395520 | 56395721 | 56397206 | 56397484 | 2 | 3 | 4 | 0 | 1 | 62 | 45 | 57 | 56 | 95 | 0 | 0 | 6 | 0 | 0 | 15 | 63 | 48 | 13 | 11 | 187 | 149 | 0.541472869 | 1 | 0.025 | 0.05 | 0.053 | 0 | 0.008 | 0 | 0 | 0.091 | 0 | 0 | 0.009 |
| A5SS | 1142 | ENSG00000260565 | ERVK13-1 | chr16 | - | 2671344 | 2672570 | 2671594 | 2672570 | 2669410 | 2669561 | 55 | 36 | 48 | 40 | 18 | 1 | 1 | 2 | 0 | 2 | 2 | 36 | 41 | 14 | 21 | 0 | 0 | 3 | 4 | 0 | 298 | 149 | 0.542601755 | 1 | 0.965 | 0.947 | 0.923 | 1 | 0.818 | 1 | 1 | 0.872 | 0.636 | 1 | 0.029 |
| A5SS | 1548 | ENSG00000140553 | UNC45A | chr15 | + | 90935284 | 90935375 | 90935284 | 90935330 | 90935543 | 90935705 | 3 | 0 | 7 | 0 | 0 | 8 | 2 | 12 | 6 | 1 | 6 | 0 | 0 | 0 | 10 | 7 | 1 | 1 | 11 | 20 | 194 | 149 | 0.542988393 | 1 | 0.224 | 0 | 0.309 | 0 | 0 | 1 | 0 | 0 | 0 | 0.277 | -0.149 |
| A5SS | 1738 | ENSG00000129559 | NEDD8 | chr14 | - | 24218058 | 24218215 | 24218132 | 24218215 | 24216856 | 24217223 | 9 | 2 | 12 | 15 | 9 | 251 | 88 | 184 | 128 | 210 | 9 | 11 | 17 | 14 | 17 | 270 | 219 | 155 | 152 | 337 | 223 | 149 | 0.543849351 | 1 | 0.023 | 0.015 | 0.042 | 0.073 | 0.028 | 0.022 | 0.032 | 0.068 | 0.058 | 0.033 | -0.006 |
| A5SS | 6906 | ENSG00000139436 | GIT2 | chr12 | - | 109951166 | 109951316 | 109951211 | 109951316 | 109939164 | 109939247 | 37 | 17 | 23 | 28 | 16 | 0 | 4 | 0 | 3 | 0 | 30 | 18 | 27 | 34 | 17 | 0 | 0 | 6 | 0 | 0 | 194 | 149 | 0.546142361 | 1 | 1 | 0.765 | 1 | 0.878 | 1 | 1 | 1 | 0.776 | 1 | 1 | -0.027 |
| A5SS | 6729 | ENSG00000265681 | RPL17 | chr18 | - | 49491351 | 49491445 | 49491404 | 49491445 | 49490792 | 49490927 | 44 | 38 | 57 | 38 | 17 | 2834 | 1463 | 3224 | 2743 | 3592 | 5 | 26 | 24 | 32 | 11 | 1446 | 1639 | 1703 | 1334 | 1154 | 202 | 149 | 0.546580509 | 1 | 0.011 | 0.019 | 0.013 | 0.01 | 0.003 | 0.003 | 0.012 | 0.01 | 0.017 | 0.007 | 0.001 |
| A5SS | 1939 | ENSG00000135506 | OS9 | chr12 | + | 57718992 | 57720263 | 57718992 | 57719182 | 57720405 | 57720518 | 354 | 191 | 237 | 296 | 406 | 297 | 252 | 253 | 278 | 438 | 798 | 462 | 266 | 515 | 687 | 798 | 486 | 375 | 558 | 671 | 298 | 149 | 0.546697581 | 1 | 0.373 | 0.275 | 0.319 | 0.347 | 0.317 | 0.333 | 0.322 | 0.262 | 0.316 | 0.339 | 0.012 |
| A5SS | 4704 | ENSG00000101361 | NOP56 | chr20 | + | 2655933 | 2656214 | 2655933 | 2656034 | 2656400 | 2656549 | 23 | 25 | 41 | 31 | 9 | 91 | 48 | 119 | 86 | 84 | 21 | 20 | 21 | 22 | 37 | 112 | 84 | 44 | 28 | 109 | 298 | 149 | 0.546960666 | 1 | 0.112 | 0.207 | 0.147 | 0.153 | 0.051 | 0.086 | 0.106 | 0.193 | 0.282 | 0.145 | -0.028 |
| A5SS | 5097 | ENSG00000204564 | C6orf136 | chr6 | + | 30650993 | 30651100 | 30650993 | 30651082 | 30651265 | 30651466 | 23 | 15 | 15 | 15 | 44 | 55 | 26 | 56 | 59 | 120 | 20 | 10 | 18 | 20 | 10 | 21 | 34 | 21 | 45 | 79 | 167 | 149 | 0.547494873 | 1 | 0.272 | 0.34 | 0.193 | 0.185 | 0.247 | 0.459 | 0.208 | 0.433 | 0.284 | 0.101 | -0.05 |
| A5SS | 6157 | ENSG00000135486 | HNRNPA1 | chr12 | + | 54281794 | 54281908 | 54281794 | 54281896 | 54283826 | 54283967 | 220 | 132 | 179 | 152 | 164 | 46 | 28 | 39 | 57 | 11 | 151 | 169 | 212 | 143 | 58 | 12 | 40 | 67 | 43 | 0 | 161 | 149 | 0.547680753 | 1 | 0.816 | 0.814 | 0.809 | 0.712 | 0.932 | 0.921 | 0.796 | 0.745 | 0.755 | 1 | -0.027 |
| A5SS | 4650 | ENSG00000019582 | CD74 | chr5 | - | 150403120 | 150404767 | 150404679 | 150404767 | 150402562 | 150402625 | 892 | 507 | 544 | 619 | 945 | 5535 | 3125 | 3624 | 4094 | 5839 | 1495 | 826 | 365 | 693 | 2407 | 8085 | 5785 | 2264 | 5104 | 12057 | 298 | 149 | 0.548657287 | 1 | 0.075 | 0.075 | 0.07 | 0.07 | 0.075 | 0.085 | 0.067 | 0.075 | 0.064 | 0.091 | -0.003 |
| A5SS | 7699 | ENSG00000119314 | PTBP3 | chr9 | - | 112297831 | 112297916 | 112297840 | 112297916 | 112268048 | 112268195 | 37 | 26 | 22 | 30 | 49 | 25 | 4 | 34 | 9 | 16 | 8 | 19 | 17 | 25 | 2 | 6 | 13 | 27 | 8 | 0 | 158 | 149 | 0.549101028 | 1 | 0.583 | 0.86 | 0.379 | 0.759 | 0.743 | 0.557 | 0.58 | 0.373 | 0.747 | 1 | 0.013 |
| A5SS | 5503 | ENSG00000239779 | WBP1 | chr2 | + | 74458831 | 74459117 | 74458831 | 74458919 | 74459642 | 74459745 | 20 | 12 | 26 | 35 | 14 | 2 | 0 | 0 | 0 | 3 | 10 | 8 | 15 | 22 | 27 | 0 | 0 | 0 | 0 | 0 | 298 | 149 | 0.549926177 | 1 | 0.833 | 1 | 1 | 1 | 0.7 | 1 | 1 | 1 | 1 | 1 | -0.093 |
| A5SS | 1564 | ENSG00000104129 | DNAJC17 | chr15 | - | 40774967 | 40775108 | 40775030 | 40775108 | 40774355 | 40774436 | 2 | 6 | 7 | 3 | 1 | 81 | 53 | 74 | 63 | 87 | 10 | 0 | 3 | 12 | 0 | 112 | 59 | 45 | 85 | 83 | 212 | 149 | 0.550107345 | 1 | 0.017 | 0.074 | 0.062 | 0.032 | 0.008 | 0.059 | 0 | 0.045 | 0.09 | 0 | 0 |
| A5SS | 7200 | ENSG00000143569 | UBAP2L | chr1 | + | 154234590 | 154234792 | 154234590 | 154234759 | 154235195 | 154235291 | 36 | 22 | 39 | 44 | 64 | 238 | 157 | 271 | 290 | 400 | 81 | 25 | 18 | 35 | 70 | 310 | 262 | 155 | 257 | 387 | 182 | 149 | 0.550507145 | 1 | 0.11 | 0.103 | 0.105 | 0.11 | 0.116 | 0.176 | 0.072 | 0.087 | 0.1 | 0.129 | -0.004 |
| A5SS | 17 | ENSG00000184900 | SUMO3 | chr21 | - | 44813861 | 44814104 | 44813975 | 44814104 | 44809046 | 44809118 | 2 | 4 | 8 | 8 | 13 | 318 | 243 | 299 | 326 | 307 | 0 | 2 | 2 | 13 | 19 | 1003 | 479 | 336 | 723 | 1509 | 263 | 149 | 0.550515674 | 1 | 0.004 | 0.009 | 0.015 | 0.014 | 0.023 | 0 | 0.002 | 0.003 | 0.01 | 0.007 | 0.009 |
| A5SS | 8073 | ENSG00000160593 | JAML | chr11 | - | 118205881 | 118205991 | 118205914 | 118205991 | 118203427 | 118203665 | 521 | 450 | 366 | 394 | 513 | 4 | 6 | 10 | 2 | 21 | 800 | 270 | 224 | 484 | 732 | 15 | 1 | 5 | 9 | 11 | 182 | 149 | 0.551754023 | 1 | 0.991 | 0.984 | 0.968 | 0.994 | 0.952 | 0.978 | 0.995 | 0.973 | 0.978 | 0.982 | -0.003 |
| A5SS | 2573 | ENSG00000203875 | SNHG5 | chr6 | - | 85678402 | 85678748 | 85678697 | 85678748 | 85678135 | 85678188 | 8 | 0 | 16 | 6 | 3 | 0 | 1 | 1 | 1 | 0 | 12 | 2 | 8 | 4 | 10 | 0 | 0 | 1 | 3 | 0 | 298 | 149 | 0.551754054 | 1 | 1 | 0 | 0.889 | 0.75 | 1 | 1 | 1 | 0.8 | 0.4 | 1 | -0.112 |
| A5SS | 7393 | ENSG00000184575 | XPOT | chr12 | + | 64425037 | 64425182 | 64425037 | 64425138 | 64425337 | 64425457 | 93 | 56 | 89 | 95 | 142 | 3 | 1 | 2 | 2 | 4 | 22 | 27 | 50 | 48 | 4 | 0 | 0 | 3 | 3 | 0 | 193 | 149 | 0.55265846 | 1 | 0.96 | 0.977 | 0.972 | 0.973 | 0.965 | 1 | 1 | 0.928 | 0.925 | 1 | -0.001 |
| A5SS | 460 | ENSG00000273154 | NA | chr20 | + | 63733586 | 63733775 | 63733586 | 63733739 | 63734704 | 63734824 | 2 | 2 | 8 | 1 | 2 | 29 | 4 | 9 | 17 | 0 | 19 | 0 | 3 | 4 | 33 | 33 | 39 | 15 | 7 | 16 | 185 | 149 | 0.55316475 | 1 | 0.053 | 0.287 | 0.417 | 0.045 | 1 | 0.317 | 0 | 0.139 | 0.315 | 0.624 | 0.081 |
| A5SS | 6040 | ENSG00000042088 | TDP1 | chr14 | + | 89980539 | 89980679 | 89980539 | 89980632 | 89984515 | 89984683 | 2 | 2 | 1 | 4 | 2 | 58 | 21 | 49 | 53 | 62 | 0 | 1 | 8 | 1 | 0 | 9 | 78 | 30 | 25 | 42 | 196 | 149 | 0.553556892 | 1 | 0.026 | 0.068 | 0.015 | 0.054 | 0.024 | 0 | 0.01 | 0.169 | 0.03 | 0 | -0.004 |
| A5SS | 8188 | ENSG00000186166 | CENATAC | chr11 | + | 119011220 | 119011311 | 119011220 | 119011283 | 119011938 | 119012003 | 34 | 31 | 59 | 46 | 57 | 12 | 7 | 6 | 7 | 13 | 76 | 45 | 75 | 79 | 30 | 2 | 12 | 8 | 16 | 12 | 177 | 149 | 0.554121398 | 1 | 0.705 | 0.788 | 0.892 | 0.847 | 0.787 | 0.97 | 0.759 | 0.888 | 0.806 | 0.678 | -0.016 |
| A5SS | 7716 | ENSG00000135047 | CTSL | chr9 | + | 87726118 | 87726253 | 87726118 | 87726249 | 87727593 | 87727729 | 12 | 10 | 11 | 31 | 9 | 4 | 1 | 0 | 0 | 0 | 23 | 5 | 18 | 19 | 41 | 0 | 1 | 0 | 0 | 5 | 153 | 149 | 0.555314023 | 1 | 0.745 | 0.907 | 1 | 1 | 1 | 1 | 0.83 | 1 | 1 | 0.889 | -0.013 |
| A5SS | 6027 | ENSG00000114388 | NPRL2 | chr3 | - | 50348482 | 50348563 | 50348526 | 50348563 | 50348316 | 50348410 | 10 | 6 | 6 | 6 | 2 | 37 | 8 | 43 | 31 | 43 | 10 | 0 | 3 | 20 | 0 | 35 | 32 | 34 | 23 | 34 | 153 | 149 | 0.555554261 | 1 | 0.208 | 0.422 | 0.12 | 0.159 | 0.043 | 0.218 | 0 | 0.079 | 0.459 | 0 | 0.039 |
| A5SS | 2129 | ENSG00000147044 | CASK | chrX | - | 41853096 | 41853227 | 41853114 | 41853227 | 41787177 | 41787283 | 3 | 12 | 16 | 12 | 21 | 81 | 50 | 67 | 88 | 115 | 5 | 4 | 13 | 6 | 12 | 46 | 42 | 43 | 33 | 41 | 167 | 149 | 0.555825325 | 1 | 0.032 | 0.176 | 0.176 | 0.108 | 0.14 | 0.088 | 0.078 | 0.212 | 0.14 | 0.207 | -0.019 |
| A5SS | 6736 | ENSG00000122783 | CYREN | chr7 | - | 135168748 | 135169060 | 135168785 | 135169060 | 135167731 | 135167807 | 69 | 54 | 67 | 52 | 56 | 10 | 3 | 6 | 8 | 4 | 84 | 48 | 31 | 52 | 120 | 0 | 4 | 3 | 2 | 22 | 186 | 149 | 0.557087705 | 1 | 0.847 | 0.935 | 0.899 | 0.839 | 0.918 | 1 | 0.906 | 0.892 | 0.954 | 0.814 | -0.026 |
| A5SS | 2574 | ENSG00000203875 | SNHG5 | chr6 | - | 85678497 | 85678748 | 85678693 | 85678748 | 85678135 | 85678188 | 9 | 1 | 17 | 7 | 0 | 43 | 62 | 47 | 32 | 27 | 12 | 2 | 8 | 4 | 10 | 23 | 35 | 55 | 75 | 90 | 298 | 149 | 0.55713982 | 1 | 0.095 | 0.008 | 0.153 | 0.099 | 0 | 0.207 | 0.028 | 0.068 | 0.026 | 0.053 | -0.005 |
| A5SS | 1141 | ENSG00000260565 | ERVK13-1 | chr16 | - | 2671023 | 2672570 | 2671594 | 2672570 | 2669410 | 2669561 | 54 | 32 | 45 | 39 | 18 | 1 | 1 | 2 | 0 | 2 | 2 | 30 | 41 | 14 | 21 | 0 | 0 | 3 | 4 | 0 | 298 | 149 | 0.557360574 | 1 | 0.964 | 0.941 | 0.918 | 1 | 0.818 | 1 | 1 | 0.872 | 0.636 | 1 | 0.027 |
| A5SS | 2492 | ENSG00000204842 | ATXN2 | chr12 | - | 111519831 | 111520076 | 111519878 | 111520076 | 111518248 | 111518427 | 20 | 8 | 11 | 26 | 15 | 53 | 28 | 66 | 46 | 97 | 22 | 5 | 23 | 6 | 25 | 66 | 45 | 38 | 55 | 26 | 196 | 149 | 0.557651962 | 1 | 0.223 | 0.178 | 0.112 | 0.301 | 0.105 | 0.202 | 0.078 | 0.315 | 0.077 | 0.422 | -0.035 |
| A5SS | 8265 | ENSG00000150433 | TMEM218 | chr11 | - | 125102636 | 125102809 | 125102733 | 125102809 | 125101200 | 125101256 | 16 | 9 | 10 | 12 | 6 | 0 | 0 | 0 | 0 | 0 | 7 | 12 | 3 | 4 | 27 | 0 | 0 | 1 | 3 | 0 | 246 | 149 | 0.558036841 | 1 | 1 | 1 | 1 | 1 | 1 | 1 | 1 | 0.645 | 0.447 | 1 | 0.182 |
| A5SS | 3681 | ENSG00000170222 | ADPRM | chr17 | + | 10704909 | 10705615 | 10704909 | 10705527 | 10706437 | 10706554 | 91 | 117 | 149 | 86 | 141 | 54 | 23 | 67 | 44 | 92 | 26 | 33 | 107 | 45 | 0 | 14 | 22 | 34 | 11 | 5 | 237 | 149 | 0.558136961 | 1 | 0.514 | 0.762 | 0.583 | 0.551 | 0.491 | 0.539 | 0.485 | 0.664 | 0.72 | 0 | 0.099 |
| A5SS | 3402 | ENSG00000170581 | STAT2 | chr12 | - | 56354458 | 56354614 | 56354465 | 56354614 | 56351291 | 56351450 | 1 | 2 | 3 | 3 | 4 | 134 | 120 | 109 | 188 | 142 | 2 | 7 | 3 | 0 | 10 | 349 | 209 | 51 | 250 | 264 | 156 | 149 | 0.558532831 | 1 | 0.007 | 0.016 | 0.026 | 0.015 | 0.026 | 0.005 | 0.031 | 0.053 | 0 | 0.035 | -0.007 |
| A5SS | 4491 | ENSG00000156983 | BRPF1 | chr3 | + | 9745572 | 9745930 | 9745572 | 9745709 | 9746299 | 9746454 | 80 | 42 | 81 | 92 | 104 | 0 | 0 | 0 | 0 | 0 | 111 | 81 | 75 | 145 | 227 | 9 | 0 | 0 | 0 | 0 | 298 | 149 | 0.559366305 | 1 | 1 | 1 | 1 | 1 | 1 | 0.86 | 1 | 1 | 1 | 1 | 0.028 |
| A5SS | 5740 | ENSG00000121741 | ZMYM2 | chr13 | + | 20031318 | 20031439 | 20031318 | 20031435 | 20034253 | 20034404 | 5 | 2 | 2 | 1 | 1 | 82 | 96 | 66 | 90 | 115 | 0 | 0 | 4 | 0 | 0 | 15 | 30 | 54 | 1 | 38 | 153 | 149 | 0.561079568 | 1 | 0.056 | 0.02 | 0.029 | 0.011 | 0.008 | 0 | 0 | 0.067 | 0 | 0 | 0.011 |
| A5SS | 5670 | ENSG00000095059 | DHPS | chr19 | - | 12680149 | 12680325 | 12680195 | 12680325 | 12679800 | 12679922 | 38 | 21 | 45 | 42 | 27 | 6 | 2 | 5 | 4 | 0 | 40 | 63 | 27 | 34 | 96 | 0 | 4 | 4 | 9 | 11 | 195 | 149 | 0.56117411 | 1 | 0.829 | 0.889 | 0.873 | 0.889 | 1 | 1 | 0.923 | 0.838 | 0.743 | 0.87 | 0.021 |
| A5SS | 911 | ENSG00000167280 | ENGASE | chr17 | + | 79081897 | 79082877 | 79081897 | 79082063 | 79083019 | 79083123 | 62 | 35 | 50 | 64 | 52 | 8 | 12 | 22 | 18 | 8 | 9 | 21 | 50 | 21 | 60 | 54 | 9 | 8 | 0 | 14 | 298 | 149 | 0.561419531 | 1 | 0.795 | 0.593 | 0.532 | 0.64 | 0.765 | 0.077 | 0.538 | 0.758 | 1 | 0.682 | 0.054 |
| A5SS | 8266 | ENSG00000150433 | TMEM218 | chr11 | - | 125102636 | 125102809 | 125102733 | 125102809 | 125101200 | 125101303 | 17 | 10 | 11 | 13 | 7 | 6 | 2 | 4 | 7 | 9 | 7 | 12 | 3 | 4 | 28 | 0 | 13 | 2 | 14 | 0 | 246 | 149 | 0.562769693 | 1 | 0.632 | 0.752 | 0.625 | 0.529 | 0.32 | 1 | 0.359 | 0.476 | 0.148 | 1 | -0.025 |
| A5SS | 6914 | ENSG00000101901 | ALG13 | chrX | + | 111682131 | 111682300 | 111682131 | 111682294 | 111684964 | 111685103 | 1 | 1 | 2 | 3 | 2 | 87 | 65 | 60 | 84 | 101 | 0 | 0 | 1 | 0 | 0 | 3 | 27 | 34 | 42 | 12 | 155 | 149 | 0.565031441 | 1 | 0.011 | 0.015 | 0.031 | 0.033 | 0.019 | 0 | 0 | 0.027 | 0 | 0 | 0.016 |
| A5SS | 5197 | ENSG00000093144 | ECHDC1 | chr6 | - | 127326997 | 127327144 | 127327001 | 127327144 | 127314815 | 127314896 | 1 | 2 | 0 | 4 | 7 | 71 | 55 | 40 | 64 | 62 | 0 | 1 | 0 | 0 | 6 | 20 | 29 | 22 | 9 | 17 | 153 | 149 | 0.566066255 | 1 | 0.014 | 0.034 | 0 | 0.057 | 0.099 | 0 | 0.032 | 0 | 0 | 0.256 | -0.017 |
| A5SS | 7472 | ENSG00000224597 | SVIL-AS1 | chr10 | + | 29409556 | 29409683 | 29409556 | 29409658 | 29415356 | 29415412 | 12 | 3 | 6 | 12 | 23 | 7 | 13 | 9 | 12 | 6 | 6 | 8 | 2 | 7 | 11 | 18 | 0 | 5 | 2 | 7 | 174 | 149 | 0.566612319 | 1 | 0.595 | 0.165 | 0.363 | 0.461 | 0.766 | 0.222 | 1 | 0.255 | 0.75 | 0.574 | -0.09 |
| A5SS | 3841 | ENSG00000141076 | UTP4 | chr16 | + | 69137800 | 69137977 | 69137800 | 69137885 | 69139824 | 69139914 | 3 | 2 | 1 | 2 | 0 | 46 | 22 | 43 | 57 | 58 | 0 | 1 | 7 | 0 | 2 | 85 | 31 | 31 | 29 | 25 | 241 | 149 | 0.56716749 | 1 | 0.039 | 0.053 | 0.014 | 0.021 | 0 | 0 | 0.02 | 0.123 | 0 | 0.047 | -0.013 |
| A5SS | 5633 | ENSG00000104218 | CSPP1 | chr8 | + | 67118247 | 67118385 | 67118247 | 67118369 | 67118742 | 67118821 | 2 | 0 | 1 | 0 | 3 | 31 | 24 | 31 | 28 | 68 | 0 | 0 | 0 | 0 | 0 | 8 | 14 | 25 | 28 | 29 | 165 | 149 | 0.567861557 | 1 | 0.055 | 0 | 0.028 | 0 | 0.038 | 0 | 0 | 0 | 0 | 0 | 0.024 |
| A5SS | 5802 | ENSG00000115053 | NCL | chr2 | - | 231456009 | 231456764 | 231456630 | 231456764 | 231455400 | 231455624 | 859 | 633 | 848 | 869 | 792 | 32 | 21 | 28 | 38 | 47 | 631 | 607 | 717 | 806 | 156 | 5 | 71 | 28 | 50 | 0 | 298 | 149 | 0.569181709 | 1 | 0.931 | 0.938 | 0.938 | 0.92 | 0.894 | 0.984 | 0.81 | 0.928 | 0.89 | 1 | 0.002 |
| A5SS | 1538 | ENSG00000168904 | LRRC28 | chr15 | + | 99255897 | 99256125 | 99255897 | 99255973 | 99276575 | 99276616 | 49 | 46 | 56 | 54 | 74 | 0 | 0 | 0 | 1 | 0 | 33 | 77 | 18 | 58 | 126 | 5 | 0 | 0 | 1 | 0 | 298 | 149 | 0.57049219 | 1 | 1 | 1 | 1 | 0.964 | 1 | 0.767 | 1 | 1 | 0.967 | 1 | 0.046 |
| A5SS | 2984 | ENSG00000169714 | CNBP | chr3 | - | 129171633 | 129171771 | 129171651 | 129171771 | 129171445 | 129171538 | 392 | 225 | 410 | 373 | 244 | 36 | 5 | 17 | 19 | 21 | 477 | 276 | 212 | 308 | 322 | 9 | 6 | 15 | 19 | 37 | 167 | 149 | 0.570958759 | 1 | 0.907 | 0.976 | 0.956 | 0.946 | 0.912 | 0.979 | 0.976 | 0.927 | 0.935 | 0.886 | -0.001 |
| A5SS | 964 | ENSG00000108984 | MAP2K6 | chr17 | + | 69517499 | 69517613 | 69517499 | 69517551 | 69519312 | 69519432 | 78 | 30 | 89 | 85 | 89 | 2 | 2 | 2 | 1 | 2 | 64 | 88 | 126 | 125 | 33 | 0 | 1 | 0 | 1 | 0 | 211 | 149 | 0.571455853 | 1 | 0.965 | 0.914 | 0.969 | 0.984 | 0.969 | 1 | 0.984 | 1 | 0.989 | 1 | -0.034 |
| A5SS | 7953 | ENSG00000109452 | INPP4B | chr4 | - | 142208424 | 142208529 | 142208446 | 142208529 | 142193086 | 142193195 | 80 | 102 | 98 | 80 | 126 | 1 | 1 | 0 | 4 | 3 | 3 | 9 | 31 | 21 | 5 | 0 | 0 | 1 | 0 | 0 | 171 | 149 | 0.571601529 | 1 | 0.986 | 0.989 | 1 | 0.946 | 0.973 | 1 | 1 | 0.964 | 1 | 1 | -0.014 |
| A5SS | 7075 | ENSG00000215021 | PHB2 | chr12 | - | 6969497 | 6969577 | 6969506 | 6969577 | 6968410 | 6968587 | 5 | 0 | 3 | 5 | 0 | 5 | 4 | 7 | 8 | 10 | 6 | 10 | 5 | 1 | 0 | 16 | 5 | 1 | 16 | 8 | 158 | 149 | 0.572538252 | 1 | 0.485 | 0 | 0.288 | 0.371 | 0 | 0.261 | 0.654 | 0.825 | 0.056 | 0 | -0.13 |
| A5SS | 3524 | ENSG00000140403 | DNAJA4 | chr15 | + | 78274196 | 78274424 | 78274196 | 78274408 | 78275497 | 78275728 | 33 | 27 | 28 | 73 | 37 | 0 | 1 | 0 | 0 | 2 | 47 | 43 | 14 | 40 | 19 | 0 | 0 | 0 | 0 | 6 | 165 | 149 | 0.572924881 | 1 | 1 | 0.961 | 1 | 1 | 0.944 | 1 | 1 | 1 | 1 | 0.741 | 0.033 |
| A5SS | 4959 | ENSG00000175866 | BAIAP2 | chr17 | + | 81085653 | 81085725 | 81085653 | 81085715 | 81086442 | 81086580 | 11 | 9 | 9 | 10 | 4 | 0 | 3 | 2 | 0 | 0 | 20 | 8 | 7 | 7 | 19 | 0 | 0 | 0 | 0 | 0 | 159 | 149 | 0.573505454 | 1 | 1 | 0.738 | 0.808 | 1 | 1 | 1 | 1 | 1 | 1 | 1 | -0.091 |
| A5SS | 5682 | ENSG00000102125 | TAZ | chrX | + | 154420031 | 154420264 | 154420031 | 154420094 | 154420657 | 154420735 | 81 | 40 | 112 | 99 | 81 | 2 | 0 | 0 | 2 | 3 | 47 | 82 | 39 | 73 | 164 | 0 | 0 | 0 | 4 | 0 | 298 | 149 | 0.574146674 | 1 | 0.953 | 1 | 1 | 0.961 | 0.931 | 1 | 1 | 1 | 0.901 | 1 | -0.011 |
| A5SS | 2182 | ENSG00000112983 | BRD8 | chr5 | - | 138171344 | 138171410 | 138171360 | 138171410 | 138171037 | 138171160 | 4 | 2 | 0 | 2 | 1 | 96 | 58 | 76 | 102 | 55 | 0 | 0 | 3 | 0 | 0 | 59 | 60 | 66 | 53 | 25 | 165 | 149 | 0.574564341 | 1 | 0.036 | 0.03 | 0 | 0.017 | 0.016 | 0 | 0 | 0.039 | 0 | 0 | 0.012 |
| A5SS | 7880 | ENSG00000152556 | PFKM | chr12 | + | 48139284 | 48139504 | 48139284 | 48139349 | 48139848 | 48139912 | 3 | 3 | 0 | 0 | 4 | 12 | 8 | 7 | 17 | 13 | 4 | 6 | 5 | 3 | 8 | 59 | 22 | 11 | 18 | 73 | 298 | 149 | 0.574659076 | 1 | 0.111 | 0.158 | 0 | 0 | 0.133 | 0.033 | 0.12 | 0.185 | 0.077 | 0.052 | -0.013 |
| A5SS | 5806 | ENSG00000130414 | NDUFA10 | chr2 | - | 240012331 | 240014860 | 240014738 | 240014860 | 240011616 | 240011696 | 35 | 21 | 53 | 52 | 66 | 60 | 23 | 97 | 69 | 99 | 89 | 44 | 39 | 59 | 7 | 114 | 70 | 45 | 66 | 71 | 298 | 149 | 0.574747636 | 1 | 0.226 | 0.313 | 0.215 | 0.274 | 0.25 | 0.281 | 0.239 | 0.302 | 0.309 | 0.047 | 0.02 |
| A5SS | 6206 | ENSG00000198492 | YTHDF2 | chr1 | + | 28738258 | 28738338 | 28738258 | 28738329 | 28768928 | 28769775 | 25 | 8 | 13 | 15 | 18 | 1 | 0 | 0 | 0 | 5 | 30 | 17 | 16 | 17 | 37 | 0 | 0 | 0 | 0 | 0 | 158 | 149 | 0.57496858 | 1 | 0.959 | 1 | 1 | 1 | 0.772 | 1 | 1 | 1 | 1 | 1 | -0.054 |
| A5SS | 6198 | ENSG00000136861 | CDK5RAP2 | chr9 | - | 120571973 | 120572041 | 120571976 | 120572041 | 120568320 | 120568388 | 55 | 32 | 47 | 61 | 49 | 0 | 0 | 6 | 4 | 0 | 72 | 54 | 43 | 81 | 69 | 4 | 0 | 3 | 0 | 7 | 152 | 149 | 0.575290078 | 1 | 1 | 1 | 0.885 | 0.937 | 1 | 0.946 | 1 | 0.934 | 1 | 0.906 | 0.007 |
| A5SS | 7198 | ENSG00000143569 | UBAP2L | chr1 | + | 154234590 | 154234759 | 154234590 | 154234738 | 154235195 | 154235291 | 271 | 175 | 304 | 310 | 444 | 1 | 6 | 4 | 5 | 5 | 387 | 300 | 165 | 293 | 447 | 5 | 6 | 0 | 0 | 6 | 170 | 149 | 0.576137255 | 1 | 0.996 | 0.962 | 0.985 | 0.982 | 0.987 | 0.985 | 0.978 | 1 | 1 | 0.985 | -0.007 |
| A5SS | 1269 | ENSG00000169592 | INO80E | chr16 | + | 30003649 | 30003714 | 30003649 | 30003670 | 30004567 | 30004657 | 7 | 3 | 8 | 2 | 1 | 1 | 2 | 4 | 0 | 0 | 12 | 1 | 1 | 5 | 5 | 2 | 1 | 4 | 0 | 1 | 193 | 149 | 0.57618846 | 1 | 0.844 | 0.537 | 0.607 | 1 | 1 | 0.822 | 0.436 | 0.162 | 1 | 0.794 | 0.155 |
| A5SS | 4374 | ENSG00000138688 | KIAA1109 | chr4 | + | 122334343 | 122334545 | 122334343 | 122334440 | 122336191 | 122336353 | 46 | 49 | 41 | 69 | 119 | 73 | 65 | 58 | 76 | 103 | 9 | 11 | 7 | 2 | 4 | 14 | 7 | 20 | 5 | 0 | 254 | 149 | 0.578678676 | 1 | 0.27 | 0.307 | 0.293 | 0.348 | 0.404 | 0.569 | 0.48 | 0.17 | 0.19 | 1 | -0.157 |
| A5SS | 7155 | ENSG00000086758 | HUWE1 | chrX | - | 53602563 | 53602658 | 53602590 | 53602658 | 53600117 | 53600309 | 357 | 204 | 312 | 349 | 245 | 8 | 5 | 12 | 8 | 3 | 559 | 362 | 272 | 308 | 288 | 10 | 12 | 2 | 2 | 11 | 176 | 149 | 0.57907423 | 1 | 0.974 | 0.972 | 0.957 | 0.974 | 0.986 | 0.979 | 0.962 | 0.991 | 0.992 | 0.957 | -0.004 |
| A5SS | 4713 | ENSG00000001497 | LAS1L | chrX | - | 65528259 | 65529258 | 65529211 | 65529258 | 65524964 | 65525050 | 75 | 35 | 66 | 61 | 82 | 4 | 3 | 2 | 1 | 0 | 136 | 76 | 48 | 71 | 104 | 0 | 8 | 3 | 4 | 0 | 298 | 149 | 0.580106146 | 1 | 0.904 | 0.854 | 0.943 | 0.968 | 1 | 1 | 0.826 | 0.889 | 0.899 | 1 | 0.011 |
| A5SS | 2442 | ENSG00000186431 | FCAR | chr19 | + | 54888006 | 54888294 | 54888006 | 54888228 | 54889648 | 54889863 | 91 | 67 | 63 | 125 | 77 | 7 | 5 | 9 | 6 | 6 | 249 | 205 | 244 | 574 | 294 | 18 | 15 | 35 | 50 | 10 | 215 | 149 | 0.582440263 | 1 | 0.9 | 0.903 | 0.829 | 0.935 | 0.899 | 0.906 | 0.905 | 0.829 | 0.888 | 0.953 | -0.003 |
| A5SS | 3330 | ENSG00000158864 | NDUFS2 | chr1 | + | 161209482 | 161212480 | 161209482 | 161210710 | 161213379 | 161213475 | 131 | 68 | 126 | 136 | 118 | 1 | 2 | 1 | 3 | 0 | 117 | 164 | 88 | 100 | 131 | 0 | 0 | 0 | 5 | 0 | 298 | 149 | 0.582806896 | 1 | 0.985 | 0.944 | 0.984 | 0.958 | 1 | 1 | 1 | 1 | 0.909 | 1 | -0.008 |
| A5SS | 1511 | ENSG00000102901 | CENPT | chr16 | - | 67831753 | 67831890 | 67831840 | 67831890 | 67831215 | 67831358 | 23 | 19 | 11 | 22 | 34 | 0 | 0 | 1 | 1 | 0 | 12 | 15 | 10 | 25 | 35 | 0 | 0 | 4 | 0 | 0 | 236 | 149 | 0.585615783 | 1 | 1 | 1 | 0.874 | 0.933 | 1 | 1 | 1 | 0.612 | 1 | 1 | 0.039 |
| A5SS | 6835 | ENSG00000124333 | VAMP7 | chrX | + | 155889457 | 155889633 | 155889457 | 155889608 | 155895622 | 155895680 | 28 | 17 | 9 | 16 | 13 | 0 | 2 | 0 | 4 | 0 | 10 | 18 | 6 | 10 | 18 | 0 | 0 | 3 | 0 | 4 | 174 | 149 | 0.585672756 | 1 | 1 | 0.879 | 1 | 0.774 | 1 | 1 | 1 | 0.631 | 1 | 0.794 | 0.046 |
| A5SS | 3438 | ENSG00000144591 | GMPPA | chr2 | + | 219498900 | 219499254 | 219498900 | 219498938 | 219499960 | 219500015 | 17 | 5 | 20 | 8 | 24 | 4 | 0 | 5 | 2 | 1 | 34 | 33 | 2 | 8 | 19 | 0 | 1 | 3 | 4 | 10 | 298 | 149 | 0.586233518 | 1 | 0.68 | 1 | 0.667 | 0.667 | 0.923 | 1 | 0.943 | 0.25 | 0.5 | 0.487 | 0.151 |
| A5SS | 2688 | ENSG00000109133 | TMEM33 | chr4 | + | 41935128 | 41935310 | 41935128 | 41935262 | 41935466 | 41935529 | 51 | 35 | 48 | 25 | 13 | 4 | 3 | 2 | 5 | 11 | 27 | 17 | 25 | 26 | 23 | 0 | 11 | 5 | 2 | 0 | 197 | 149 | 0.587083815 | 1 | 0.906 | 0.898 | 0.948 | 0.791 | 0.472 | 1 | 0.539 | 0.791 | 0.908 | 1 | -0.045 |
| A5SS | 1748 | ENSG00000100897 | DCAF11 | chr14 | + | 24115381 | 24115749 | 24115381 | 24115726 | 24116916 | 24117044 | 99 | 55 | 82 | 95 | 96 | 7 | 3 | 4 | 14 | 7 | 181 | 190 | 63 | 98 | 203 | 23 | 9 | 3 | 3 | 27 | 172 | 149 | 0.587234613 | 1 | 0.925 | 0.941 | 0.947 | 0.855 | 0.922 | 0.872 | 0.948 | 0.948 | 0.966 | 0.867 | -0.002 |
| A5SS | 6584 | ENSG00000155096 | AZIN1 | chr8 | - | 102849986 | 102850144 | 102849998 | 102850144 | 102843550 | 102843747 | 24 | 31 | 26 | 26 | 39 | 99 | 54 | 103 | 104 | 79 | 35 | 33 | 62 | 50 | 37 | 86 | 116 | 142 | 185 | 70 | 161 | 149 | 0.589149777 | 1 | 0.183 | 0.347 | 0.189 | 0.188 | 0.314 | 0.274 | 0.208 | 0.288 | 0.2 | 0.328 | -0.015 |
| A5SS | 8906 | ENSG00000185189 | NRBP2 | chr8 | - | 143839508 | 143840030 | 143839928 | 143840030 | 143839313 | 143839408 | 25 | 14 | 9 | 9 | 6 | 0 | 0 | 0 | 0 | 0 | 9 | 8 | 13 | 8 | 24 | 0 | 0 | 4 | 0 | 0 | 298 | 149 | 0.589813699 | 1 | 1 | 1 | 1 | 1 | 1 | 1 | 1 | 0.619 | 1 | 1 | 0.076 |
| A5SS | 6852 | ENSG00000230590 | FTX | chrX | - | 74280376 | 74280494 | 74280393 | 74280494 | 74274341 | 74274410 | 12 | 13 | 6 | 17 | 6 | 11 | 9 | 4 | 18 | 10 | 0 | 14 | 2 | 3 | 21 | 8 | 6 | 17 | 1 | 5 | 166 | 149 | 0.589845474 | 1 | 0.495 | 0.565 | 0.574 | 0.459 | 0.35 | 0 | 0.677 | 0.096 | 0.729 | 0.79 | 0.03 |
| A5SS | 2514 | ENSG00000185033 | SEMA4B | chr15 | + | 90225281 | 90225427 | 90225281 | 90225397 | 90225660 | 90225827 | 1 | 0 | 0 | 0 | 0 | 13 | 8 | 14 | 10 | 15 | 0 | 0 | 0 | 0 | 6 | 6 | 14 | 8 | 3 | 93 | 179 | 149 | 0.590563954 | 1 | 0.06 | 0 | 0 | 0 | 0 | 0 | 0 | 0 | 0 | 0.051 | 0.002 |
| A5SS | 305 | ENSG00000105193 | RPS16 | chr19 | - | 39435606 | 39435708 | 39435657 | 39435708 | 39433664 | 39433761 | 3942 | 1388 | 4758 | 3666 | 2855 | 88 | 39 | 135 | 88 | 52 | 1598 | 2594 | 1769 | 1277 | 3817 | 64 | 49 | 51 | 33 | 80 | 200 | 149 | 0.590761466 | 1 | 0.971 | 0.964 | 0.963 | 0.969 | 0.976 | 0.949 | 0.975 | 0.963 | 0.966 | 0.973 | 0.003 |
| A5SS | 1498 | ENSG00000114062 | UBE3A | chr15 | - | 25407066 | 25409207 | 25409087 | 25409207 | 25405460 | 25405502 | 50 | 51 | 36 | 37 | 48 | 80 | 69 | 73 | 78 | 103 | 43 | 38 | 14 | 18 | 21 | 16 | 71 | 124 | 40 | 59 | 298 | 149 | 0.59209359 | 1 | 0.238 | 0.27 | 0.198 | 0.192 | 0.189 | 0.573 | 0.211 | 0.053 | 0.184 | 0.151 | -0.017 |
| A5SS | 10 | ENSG00000160255 | ITGB2 | chr21 | - | 44910599 | 44910785 | 44910724 | 44910785 | 44910283 | 44910415 | 18 | 6 | 12 | 12 | 1 | 3 | 1 | 0 | 1 | 0 | 12 | 29 | 6 | 12 | 37 | 0 | 0 | 1 | 4 | 0 | 274 | 149 | 0.593577228 | 1 | 0.765 | 0.765 | 1 | 0.867 | 1 | 1 | 1 | 0.765 | 0.62 | 1 | 0.002 |
| A5SS | 4139 | ENSG00000162298 | SYVN1 | chr11 | - | 65132247 | 65132780 | 65132731 | 65132780 | 65131469 | 65131596 | 78 | 9 | 63 | 59 | 33 | 0 | 4 | 0 | 1 | 0 | 52 | 93 | 44 | 50 | 173 | 0 | 1 | 2 | 3 | 0 | 298 | 149 | 0.594430687 | 1 | 1 | 0.529 | 1 | 0.967 | 1 | 1 | 0.979 | 0.917 | 0.893 | 1 | -0.059 |
| A5SS | 6410 | ENSG00000145191 | EIF2B5 | chr3 | + | 184140417 | 184140740 | 184140417 | 184140615 | 184141924 | 184142070 | 36 | 10 | 36 | 32 | 30 | 0 | 0 | 0 | 2 | 8 | 40 | 39 | 28 | 41 | 59 | 9 | 2 | 4 | 0 | 0 | 274 | 149 | 0.594510729 | 1 | 1 | 1 | 1 | 0.897 | 0.671 | 0.707 | 0.914 | 0.792 | 1 | 1 | 0.031 |
| A5SS | 997 | ENSG00000188825 | LINC00910 | chr17 | - | 43387935 | 43388312 | 43388288 | 43388312 | 43381272 | 43381660 | 3 | 0 | 3 | 0 | 0 | 8 | 3 | 4 | 5 | 15 | 0 | 0 | 0 | 2 | 0 | 3 | 8 | 5 | 0 | 35 | 298 | 149 | 0.594749872 | 1 | 0.158 | 0 | 0.273 | 0 | 0 | 0 | 0 | 0 | 1 | 0 | -0.114 |
| A5SS | 3534 | ENSG00000026751 | SLAMF7 | chr1 | + | 160749820 | 160750132 | 160749820 | 160750093 | 160750303 | 160750423 | 16 | 4 | 4 | 4 | 1 | 152 | 89 | 138 | 119 | 100 | 0 | 3 | 8 | 11 | 15 | 173 | 163 | 131 | 90 | 129 | 188 | 149 | 0.594756146 | 1 | 0.077 | 0.034 | 0.022 | 0.026 | 0.008 | 0 | 0.014 | 0.046 | 0.088 | 0.084 | -0.013 |
| A5SS | 1672 | ENSG00000100749 | VRK1 | chr14 | + | 96852830 | 96852943 | 96852830 | 96852939 | 96853073 | 96853166 | 5 | 5 | 2 | 2 | 2 | 257 | 142 | 193 | 244 | 286 | 0 | 0 | 1 | 7 | 7 | 121 | 223 | 133 | 143 | 177 | 153 | 149 | 0.594793054 | 1 | 0.019 | 0.033 | 0.01 | 0.008 | 0.007 | 0 | 0 | 0.007 | 0.046 | 0.037 | -0.003 |
| A5SS | 3731 | ENSG00000166526 | ZNF3 | chr7 | - | 100077228 | 100077433 | 100077302 | 100077433 | 100075541 | 100075630 | 2 | 4 | 12 | 17 | 3 | 13 | 7 | 20 | 27 | 8 | 3 | 7 | 2 | 9 | 16 | 8 | 14 | 22 | 19 | 25 | 223 | 149 | 0.595568795 | 1 | 0.093 | 0.276 | 0.286 | 0.296 | 0.2 | 0.2 | 0.25 | 0.057 | 0.24 | 0.3 | 0.021 |
| A5SS | 5060 | ENSG00000143622 | RIT1 | chr1 | - | 155911242 | 155911381 | 155911271 | 155911381 | 155910655 | 155910804 | 137 | 98 | 79 | 127 | 61 | 1 | 3 | 0 | 2 | 0 | 35 | 125 | 134 | 96 | 131 | 0 | 0 | 4 | 5 | 0 | 178 | 149 | 0.595703692 | 1 | 0.991 | 0.965 | 1 | 0.982 | 1 | 1 | 1 | 0.966 | 0.941 | 1 | 0.006 |
| A5SS | 1628 | ENSG00000165795 | NDRG2 | chr14 | - | 21021746 | 21021879 | 21021816 | 21021879 | 21020783 | 21020844 | 1 | 0 | 3 | 4 | 0 | 22 | 34 | 50 | 39 | 78 | 0 | 2 | 3 | 0 | 2 | 27 | 6 | 6 | 16 | 22 | 219 | 149 | 0.596124132 | 1 | 0.03 | 0 | 0.039 | 0.065 | 0 | 0 | 0.185 | 0.254 | 0 | 0.058 | -0.073 |
| A5SS | 2849 | ENSG00000075711 | DLG1 | chr3 | - | 197138214 | 197138391 | 197138221 | 197138391 | 197136541 | 197136678 | 5 | 7 | 2 | 10 | 9 | 106 | 115 | 116 | 93 | 113 | 0 | 5 | 6 | 8 | 3 | 55 | 72 | 54 | 53 | 43 | 156 | 149 | 0.596428715 | 1 | 0.043 | 0.055 | 0.016 | 0.093 | 0.071 | 0 | 0.062 | 0.096 | 0.126 | 0.062 | -0.014 |
| A5SS | 2748 | ENSG00000206573 | THUMPD3-AS1 | chr3 | - | 9391426 | 9391826 | 9391678 | 9391826 | 9388873 | 9390497 | 68 | 103 | 89 | 72 | 113 | 13 | 15 | 4 | 22 | 23 | 56 | 76 | 65 | 62 | 27 | 5 | 6 | 0 | 10 | 43 | 298 | 149 | 0.597054346 | 1 | 0.723 | 0.774 | 0.918 | 0.621 | 0.711 | 0.848 | 0.864 | 1 | 0.756 | 0.239 | 0.008 |
| A5SS | 2463 | ENSG00000125386 | FAM193A | chr4 | + | 2694945 | 2695129 | 2694945 | 2695082 | 2696362 | 2696593 | 107 | 52 | 90 | 78 | 51 | 7 | 0 | 0 | 0 | 0 | 243 | 109 | 103 | 104 | 150 | 0 | 0 | 1 | 1 | 0 | 196 | 149 | 0.597604525 | 1 | 0.921 | 1 | 1 | 1 | 1 | 1 | 1 | 0.987 | 0.988 | 1 | -0.011 |
| A5SS | 2150 | ENSG00000113558 | SKP1 | chr5 | - | 134176685 | 134176950 | 134176854 | 134176950 | 134173925 | 134174022 | 7 | 3 | 6 | 3 | 7 | 223 | 108 | 144 | 211 | 112 | 8 | 1 | 5 | 11 | 0 | 137 | 157 | 92 | 123 | 126 | 298 | 149 | 0.597770654 | 1 | 0.015 | 0.014 | 0.02 | 0.007 | 0.03 | 0.028 | 0.003 | 0.026 | 0.043 | 0 | -0.003 |
| A5SS | 7036 | ENSG00000171311 | EXOSC1 | chr10 | - | 97438662 | 97438703 | 97438669 | 97438703 | 97437699 | 97437750 | 7 | 2 | 2 | 6 | 6 | 91 | 58 | 98 | 108 | 83 | 11 | 0 | 2 | 4 | 0 | 83 | 53 | 37 | 95 | 101 | 156 | 149 | 0.598147655 | 1 | 0.068 | 0.032 | 0.019 | 0.05 | 0.065 | 0.112 | 0 | 0.049 | 0.039 | 0 | 0.007 |
| A5SS | 7754 | ENSG00000165699 | TSC1 | chr9 | - | 132909806 | 132910692 | 132910570 | 132910692 | 132907300 | 132907370 | 2 | 1 | 7 | 2 | 9 | 18 | 19 | 19 | 22 | 16 | 0 | 11 | 1 | 8 | 5 | 3 | 32 | 22 | 16 | 20 | 298 | 149 | 0.598794227 | 1 | 0.053 | 0.026 | 0.156 | 0.043 | 0.22 | 0 | 0.147 | 0.022 | 0.2 | 0.111 | 0.004 |
| A5SS | 809 | ENSG00000131470 | PSMC3IP | chr17 | - | 42573818 | 42574210 | 42574098 | 42574210 | 42573477 | 42573623 | 15 | 9 | 15 | 10 | 10 | 3 | 0 | 2 | 6 | 0 | 22 | 20 | 21 | 7 | 29 | 22 | 0 | 4 | 5 | 0 | 298 | 149 | 0.600122948 | 1 | 0.714 | 1 | 0.789 | 0.455 | 1 | 0.333 | 1 | 0.724 | 0.412 | 1 | 0.098 |
| A5SS | 7886 | ENSG00000266173 | STRADA | chr17 | - | 63704340 | 63704582 | 63704431 | 63704582 | 63704004 | 63704047 | 57 | 31 | 54 | 49 | 80 | 7 | 0 | 9 | 5 | 4 | 71 | 42 | 37 | 46 | 63 | 11 | 6 | 4 | 0 | 0 | 240 | 149 | 0.60154898 | 1 | 0.835 | 1 | 0.788 | 0.859 | 0.925 | 0.8 | 0.813 | 0.852 | 1 | 1 | -0.012 |
| A5SS | 4788 | ENSG00000131778 | CHD1L | chr1 | + | 147270931 | 147271195 | 147270931 | 147271113 | 147272170 | 147272281 | 32 | 29 | 9 | 23 | 12 | 4 | 0 | 7 | 0 | 1 | 3 | 9 | 13 | 16 | 10 | 0 | 11 | 0 | 0 | 4 | 231 | 149 | 0.601706976 | 1 | 0.838 | 1 | 0.453 | 1 | 0.886 | 1 | 0.345 | 1 | 1 | 0.617 | 0.043 |
| A5SS | 5621 | ENSG00000120697 | ALG5 | chr13 | - | 36994988 | 36995596 | 36995426 | 36995596 | 36993603 | 36993672 | 120 | 76 | 93 | 83 | 128 | 2 | 0 | 0 | 3 | 0 | 17 | 66 | 100 | 58 | 24 | 0 | 1 | 6 | 0 | 0 | 298 | 149 | 0.6040321 | 1 | 0.968 | 1 | 1 | 0.933 | 1 | 1 | 0.971 | 0.893 | 1 | 1 | 0.007 |
| A5SS | 7766 | ENSG00000115459 | ELMOD3 | chr2 | + | 85355554 | 85355784 | 85355554 | 85355716 | 85356966 | 85357252 | 12 | 14 | 13 | 14 | 7 | 1 | 2 | 3 | 4 | 2 | 46 | 15 | 19 | 40 | 12 | 11 | 1 | 6 | 1 | 0 | 217 | 149 | 0.604194903 | 1 | 0.892 | 0.828 | 0.748 | 0.706 | 0.706 | 0.742 | 0.912 | 0.685 | 0.965 | 1 | -0.085 |
| A5SS | 5276 | ENSG00000138073 | PREB | chr2 | - | 27133116 | 27133721 | 27133531 | 27133721 | 27132842 | 27132923 | 203 | 98 | 178 | 188 | 197 | 0 | 0 | 0 | 0 | 1 | 196 | 170 | 125 | 169 | 232 | 8 | 0 | 1 | 0 | 0 | 298 | 149 | 0.604550449 | 1 | 1 | 1 | 1 | 1 | 0.99 | 0.925 | 1 | 0.984 | 1 | 1 | 0.016 |
| A5SS | 4651 | ENSG00000019582 | CD74 | chr5 | - | 150405084 | 150406321 | 150406258 | 150406321 | 150402562 | 150402625 | 21 | 15 | 17 | 13 | 2 | 4 | 0 | 1 | 0 | 0 | 82 | 68 | 20 | 66 | 136 | 0 | 0 | 0 | 3 | 0 | 298 | 149 | 0.605326591 | 1 | 0.724 | 1 | 0.895 | 1 | 1 | 1 | 1 | 1 | 0.917 | 1 | -0.06 |
| A5SS | 674 | ENSG00000152234 | ATP5F1A | chr18 | - | 46094979 | 46095131 | 46095052 | 46095131 | 46093179 | 46093365 | 13 | 9 | 15 | 18 | 6 | 0 | 0 | 0 | 0 | 0 | 36 | 16 | 24 | 38 | 53 | 0 | 0 | 0 | 6 | 0 | 222 | 149 | 0.607847371 | 1 | 1 | 1 | 1 | 1 | 1 | 1 | 1 | 1 | 0.81 | 1 | 0.038 |
| A5SS | 6222 | ENSG00000112695 | COX7A2 | chr6 | - | 75241122 | 75241265 | 75241175 | 75241265 | 75240300 | 75240385 | 25 | 11 | 13 | 16 | 10 | 489 | 254 | 444 | 433 | 464 | 18 | 13 | 20 | 5 | 12 | 423 | 395 | 329 | 391 | 553 | 202 | 149 | 0.608155342 | 1 | 0.036 | 0.031 | 0.021 | 0.027 | 0.016 | 0.03 | 0.024 | 0.043 | 0.009 | 0.016 | 0.002 |
| A5SS | 5053 | ENSG00000049167 | ERCC8 | chr5 | - | 60903566 | 60903716 | 60903647 | 60903716 | 60902441 | 60902508 | 2 | 2 | 3 | 2 | 4 | 30 | 11 | 12 | 18 | 50 | 5 | 6 | 6 | 0 | 0 | 0 | 40 | 22 | 26 | 9 | 230 | 149 | 0.608221072 | 1 | 0.041 | 0.105 | 0.139 | 0.067 | 0.049 | 1 | 0.089 | 0.15 | 0 | 0 | -0.168 |
| A5SS | 3374 | ENSG00000125347 | IRF1 | chr5 | - | 132486521 | 132486686 | 132486556 | 132486686 | 132484361 | 132484497 | 19 | 17 | 24 | 14 | 20 | 14 | 16 | 17 | 22 | 5 | 75 | 14 | 16 | 32 | 52 | 46 | 17 | 11 | 76 | 24 | 184 | 149 | 0.608231902 | 1 | 0.524 | 0.462 | 0.533 | 0.34 | 0.764 | 0.569 | 0.4 | 0.541 | 0.254 | 0.637 | 0.044 |
| A5SS | 6969 | ENSG00000108679 | LGALS3BP | chr17 | - | 78976994 | 78977214 | 78977139 | 78977214 | 78975964 | 78976156 | 4 | 2 | 3 | 1 | 7 | 28 | 14 | 27 | 32 | 9 | 0 | 14 | 6 | 19 | 16 | 126 | 108 | 38 | 201 | 70 | 294 | 149 | 0.608939631 | 1 | 0.068 | 0.068 | 0.053 | 0.016 | 0.283 | 0 | 0.062 | 0.074 | 0.046 | 0.104 | 0.04 |
| A5SS | 1002 | ENSG00000108773 | KAT2A | chr17 | - | 42119053 | 42119436 | 42119244 | 42119436 | 42118296 | 42118403 | 41 | 14 | 61 | 36 | 41 | 14 | 13 | 22 | 13 | 25 | 10 | 60 | 20 | 34 | 47 | 7 | 35 | 5 | 8 | 18 | 298 | 149 | 0.609797665 | 1 | 0.594 | 0.35 | 0.581 | 0.581 | 0.451 | 0.417 | 0.462 | 0.667 | 0.68 | 0.566 | -0.047 |
| A5SS | 6840 | ENSG00000124333 | VAMP7 | chrX | + | 155889457 | 155889612 | 155889457 | 155889608 | 155898111 | 155898249 | 11 | 9 | 6 | 10 | 6 | 0 | 2 | 0 | 0 | 0 | 21 | 7 | 2 | 16 | 15 | 0 | 0 | 3 | 0 | 0 | 153 | 149 | 0.610073202 | 1 | 1 | 0.814 | 1 | 1 | 1 | 1 | 1 | 0.394 | 1 | 1 | 0.084 |
| A5SS | 3025 | ENSG00000136731 | UGGT1 | chr2 | + | 128091199 | 128091415 | 128091199 | 128091382 | 128097428 | 128097564 | 30 | 5 | 11 | 14 | 6 | 3 | 3 | 6 | 6 | 0 | 2 | 37 | 16 | 15 | 3 | 0 | 4 | 5 | 3 | 3 | 182 | 149 | 0.61112544 | 1 | 0.891 | 0.577 | 0.6 | 0.656 | 1 | 1 | 0.883 | 0.724 | 0.804 | 0.45 | -0.027 |
| A5SS | 5381 | ENSG00000196504 | PRPF40A | chr2 | - | 152717229 | 152717457 | 152717355 | 152717457 | 152715994 | 152716126 | 28 | 19 | 22 | 33 | 20 | 68 | 25 | 73 | 62 | 15 | 0 | 19 | 21 | 20 | 0 | 15 | 46 | 20 | 8 | 5 | 275 | 149 | 0.611824382 | 1 | 0.182 | 0.292 | 0.14 | 0.224 | 0.419 | 0 | 0.183 | 0.363 | 0.575 | 0 | 0.027 |
| A5SS | 102 | ENSG00000100150 | DEPDC5 | chr22 | + | 31797599 | 31797703 | 31797599 | 31797699 | 31798581 | 31798656 | 84 | 59 | 66 | 70 | 80 | 7 | 8 | 12 | 7 | 22 | 124 | 94 | 59 | 69 | 132 | 27 | 26 | 17 | 8 | 4 | 153 | 149 | 0.612521708 | 1 | 0.921 | 0.878 | 0.843 | 0.907 | 0.78 | 0.817 | 0.779 | 0.772 | 0.894 | 0.97 | 0.019 |
| A5SS | 2612 | ENSG00000132376 | INPP5K | chr17 | - | 1509167 | 1509353 | 1509177 | 1509353 | 1508114 | 1508226 | 5 | 1 | 5 | 8 | 0 | 114 | 58 | 109 | 111 | 129 | 5 | 16 | 2 | 4 | 0 | 129 | 125 | 103 | 94 | 218 | 159 | 149 | 0.612961742 | 1 | 0.039 | 0.016 | 0.041 | 0.063 | 0 | 0.035 | 0.107 | 0.018 | 0.038 | 0 | -0.008 |
| A5SS | 820 | ENSG00000125454 | SLC25A19 | chr17 | - | 75288387 | 75288591 | 75288497 | 75288591 | 75286632 | 75286802 | 16 | 7 | 8 | 17 | 16 | 2 | 2 | 1 | 0 | 0 | 39 | 19 | 23 | 21 | 18 | 0 | 0 | 0 | 0 | 0 | 259 | 149 | 0.613288454 | 1 | 0.822 | 0.668 | 0.822 | 1 | 1 | 1 | 1 | 1 | 1 | 1 | -0.138 |
| A5SS | 8484 | ENSG00000049323 | LTBP1 | chr2 | + | 33262721 | 33262824 | 33262721 | 33262821 | 33263293 | 33263392 | 12 | 5 | 0 | 14 | 4 | 33 | 54 | 14 | 102 | 44 | 37 | 15 | 5 | 6 | 129 | 283 | 32 | 71 | 77 | 811 | 152 | 149 | 0.614714249 | 1 | 0.263 | 0.083 | 0 | 0.119 | 0.082 | 0.114 | 0.315 | 0.065 | 0.071 | 0.135 | -0.031 |
| A5SS | 6861 | ENSG00000148985 | PGAP2 | chr11 | + | 3824269 | 3824455 | 3824269 | 3824364 | 3825019 | 3825128 | 4 | 6 | 8 | 5 | 8 | 10 | 1 | 8 | 10 | 5 | 15 | 1 | 7 | 16 | 17 | 12 | 6 | 7 | 16 | 1 | 240 | 149 | 0.61552552 | 1 | 0.199 | 0.788 | 0.383 | 0.237 | 0.498 | 0.437 | 0.094 | 0.383 | 0.383 | 0.913 | -0.021 |
| A5SS | 8319 | ENSG00000090621 | PABPC4 | chr1 | - | 39575758 | 39576790 | 39576551 | 39576790 | 39572392 | 39572586 | 368 | 100 | 216 | 273 | 83 | 0 | 0 | 3 | 1 | 1 | 300 | 196 | 207 | 132 | 224 | 0 | 0 | 9 | 2 | 1 | 298 | 149 | 0.615743996 | 1 | 1 | 1 | 0.973 | 0.993 | 0.976 | 1 | 1 | 0.92 | 0.971 | 0.991 | 0.012 |
| A5SS | 1701 | ENSG00000197249 | SERPINA1 | chr14 | - | 94388800 | 94389010 | 94388818 | 94389010 | 94388559 | 94388663 | 84 | 39 | 21 | 38 | 20 | 41 | 22 | 14 | 29 | 16 | 63 | 76 | 38 | 63 | 127 | 21 | 64 | 33 | 41 | 99 | 167 | 149 | 0.615812517 | 1 | 0.646 | 0.613 | 0.572 | 0.539 | 0.527 | 0.728 | 0.514 | 0.507 | 0.578 | 0.534 | 0.007 |
| A5SS | 1489 | ENSG00000128739 | SNRPN | chr15 | + | 24967028 | 24967240 | 24967028 | 24967152 | 24967931 | 24968082 | 3 | 3 | 10 | 4 | 4 | 5 | 6 | 8 | 5 | 9 | 14 | 12 | 2 | 12 | 10 | 22 | 5 | 0 | 16 | 41 | 237 | 149 | 0.616134826 | 1 | 0.274 | 0.239 | 0.44 | 0.335 | 0.218 | 0.286 | 0.601 | 1 | 0.32 | 0.133 | -0.167 |
| A5SS | 6032 | ENSG00000114388 | NPRL2 | chr3 | - | 50349912 | 50350022 | 50349930 | 50350022 | 50349664 | 50349833 | 1 | 0 | 0 | 3 | 0 | 70 | 24 | 47 | 76 | 42 | 0 | 0 | 0 | 7 | 0 | 55 | 56 | 28 | 24 | 52 | 167 | 149 | 0.616679487 | 1 | 0.013 | 0 | 0 | 0.034 | 0 | 0 | 0 | 0 | 0.206 | 0 | -0.032 |
| A5SS | 7590 | ENSG00000196781 | TLE1 | chr9 | - | 81613376 | 81613521 | 81613421 | 81613521 | 81612291 | 81612393 | 36 | 13 | 15 | 17 | 17 | 0 | 0 | 0 | 0 | 0 | 17 | 23 | 12 | 9 | 5 | 0 | 0 | 0 | 4 | 0 | 194 | 149 | 0.616688328 | 1 | 1 | 1 | 1 | 1 | 1 | 1 | 1 | 1 | 0.633 | 1 | 0.073 |
| A5SS | 3992 | ENSG00000198912 | C1orf174 | chr1 | - | 3892740 | 3892996 | 3892882 | 3892996 | 3890568 | 3891057 | 13 | 12 | 21 | 22 | 10 | 21 | 12 | 11 | 23 | 0 | 21 | 11 | 29 | 27 | 70 | 39 | 5 | 8 | 51 | 47 | 291 | 149 | 0.617325481 | 1 | 0.241 | 0.339 | 0.494 | 0.329 | 1 | 0.216 | 0.53 | 0.65 | 0.213 | 0.433 | 0.072 |
| A5SS | 5256 | ENSG00000129083 | COPB1 | chr11 | - | 14499706 | 14499811 | 14499741 | 14499811 | 14498837 | 14498985 | 45 | 24 | 41 | 34 | 10 | 11 | 2 | 7 | 4 | 7 | 12 | 12 | 24 | 5 | 5 | 0 | 2 | 6 | 5 | 0 | 184 | 149 | 0.617986121 | 1 | 0.768 | 0.907 | 0.826 | 0.873 | 0.536 | 1 | 0.829 | 0.764 | 0.447 | 1 | -0.026 |
| A5SS | 7436 | ENSG00000160948 | VPS28 | chr8 | - | 144425874 | 144426063 | 144425921 | 144426063 | 144424945 | 144425772 | 10 | 9 | 12 | 6 | 21 | 4 | 0 | 1 | 1 | 7 | 15 | 10 | 16 | 4 | 21 | 0 | 8 | 3 | 16 | 0 | 196 | 149 | 0.618018452 | 1 | 0.655 | 1 | 0.901 | 0.82 | 0.695 | 1 | 0.487 | 0.802 | 0.16 | 1 | 0.124 |
| A5SS | 5733 | ENSG00000079263 | SP140 | chr2 | + | 230244987 | 230245080 | 230244987 | 230245071 | 230245862 | 230245940 | 140 | 137 | 179 | 114 | 181 | 30 | 35 | 38 | 42 | 24 | 130 | 294 | 262 | 170 | 72 | 54 | 61 | 32 | 37 | 31 | 158 | 149 | 0.618288183 | 1 | 0.815 | 0.787 | 0.816 | 0.719 | 0.877 | 0.694 | 0.82 | 0.885 | 0.812 | 0.687 | 0.023 |
| A5SS | 8354 | ENSG00000137752 | CASP1 | chr11 | - | 105034207 | 105034474 | 105034423 | 105034474 | 105031164 | 105031280 | 449 | 393 | 229 | 327 | 389 | 0 | 0 | 1 | 0 | 0 | 149 | 210 | 196 | 174 | 19 | 0 | 0 | 8 | 0 | 0 | 298 | 149 | 0.619756433 | 1 | 1 | 1 | 0.991 | 1 | 1 | 1 | 1 | 0.925 | 1 | 1 | 0.013 |
| A5SS | 5478 | ENSG00000204387 | SNHG32 | chr6 | + | 31834907 | 31835200 | 31834907 | 31835057 | 31835410 | 31835451 | 713 | 478 | 1031 | 504 | 454 | 12 | 5 | 25 | 13 | 18 | 447 | 548 | 615 | 526 | 857 | 6 | 13 | 4 | 21 | 19 | 292 | 149 | 0.619866728 | 1 | 0.968 | 0.98 | 0.955 | 0.952 | 0.928 | 0.974 | 0.956 | 0.987 | 0.927 | 0.958 | -0.004 |
| A5SS | 4838 | ENSG00000137101 | CD72 | chr9 | - | 35618221 | 35618413 | 35618271 | 35618413 | 35618013 | 35618121 | 24 | 14 | 21 | 41 | 28 | 8 | 6 | 11 | 10 | 15 | 8 | 42 | 5 | 39 | 27 | 0 | 19 | 8 | 4 | 12 | 199 | 149 | 0.619889284 | 1 | 0.692 | 0.636 | 0.588 | 0.754 | 0.583 | 1 | 0.623 | 0.319 | 0.88 | 0.628 | -0.039 |
| A5SS | 4154 | ENSG00000126012 | KDM5C | chrX | - | 53193772 | 53193851 | 53193781 | 53193851 | 53193436 | 53193636 | 30 | 20 | 35 | 26 | 10 | 41 | 19 | 42 | 32 | 26 | 17 | 31 | 14 | 18 | 47 | 30 | 30 | 19 | 29 | 39 | 158 | 149 | 0.622177164 | 1 | 0.408 | 0.498 | 0.44 | 0.434 | 0.266 | 0.348 | 0.494 | 0.41 | 0.369 | 0.532 | -0.021 |
| A5SS | 1714 | ENSG00000072042 | RDH11 | chr14 | - | 67692933 | 67693052 | 67692972 | 67693052 | 67692437 | 67692593 | 174 | 169 | 106 | 191 | 80 | 0 | 9 | 0 | 0 | 2 | 400 | 254 | 157 | 325 | 567 | 6 | 0 | 0 | 3 | 6 | 188 | 149 | 0.622589284 | 1 | 1 | 0.937 | 1 | 1 | 0.969 | 0.981 | 1 | 1 | 0.988 | 0.987 | -0.01 |
| A5SS | 5851 | ENSG00000169045 | HNRNPH1 | chr5 | - | 179616868 | 179617110 | 179617020 | 179617110 | 179616125 | 179616218 | 1459 | 1414 | 1321 | 1327 | 1195 | 128 | 111 | 62 | 109 | 80 | 326 | 848 | 1373 | 686 | 173 | 66 | 40 | 38 | 57 | 10 | 298 | 149 | 0.62358009 | 1 | 0.851 | 0.864 | 0.914 | 0.859 | 0.882 | 0.712 | 0.914 | 0.948 | 0.858 | 0.896 | 0.008 |

| | | | | | | | | | | | | | | | |
|---|---|---|---|---|---|---|---|---|---|---|---|---|---|---|---|
| A5SS | 6599 | ENSG00000071462 | BUD23 | chr7 | + | 73683766 | 73683901 | 73683766 | 73683804 | 73686635 | 73686731 | 0 | 3 | 4 | [illegible] |
| A5SS | 335 | ENSG00000167613 | LAIR1 | chr19 | - | 54359842 | 54360072 | 54360021 | 54360072 | 54356927 | 54356966 | 11 | 6 | 2 | [illegible] |
| A5SS | 5196 | ENSG00000167220 | HDHD2 | chr18 | - | 47136301 | 47136449 | 47136338 | 47136449 | 47134495 | 47134704 | 18 | 17 | 8 | [illegible] |
| A5SS | 6447 | ENSG00000164808 | SPIDR | chr8 | + | 47279861 | 47280017 | 47279861 | 47279980 | 47284027 | 47284094 | 98 | 47 | 68 | [illegible] |
| A5SS | 6867 | ENSG00000134668 | SPOCD1 | chr1 | - | 31796589 | 31796715 | 31796592 | 31796715 | 31794123 | 31794235 | 5 | 10 | 2 | [illegible] |
| A5SS | 4921 | ENSG00000106346 | USP42 | chr7 | + | 6139091 | 6140195 | 6139091 | 6139194 | 6140913 | 6140984 | 122 | 59 | 73 | [illegible] |
| A5SS | 524 | ENSG00000244687 | UBE2V1 | chr20 | - | 50096482 | 50096820 | 50096671 | 50096820 | 50084128 | 50084254 | 41 | 26 | 36 | [illegible] |
| A5SS | 2356 | ENSG00000214655 | ZSWIM8 | chr10 | + | 73800010 | 73800194 | 73800010 | 73800170 | 73800295 | 73800472 | 32 | 4 | 10 | [illegible] |
| A5SS | 267 | ENSG00000007255 | TRAPPC6A | chr19 | - | 45178092 | 45178237 | 45178134 | 45178237 | 45165126 | 45165194 | 44 | 1 | 43 | [illegible] |
| A5SS | 2874 | ENSG00000172053 | QARS1 | chr3 | - | 49099329 | 49099431 | 49099343 | 49099431 | 49099109 | 49099253 | 0 | 0 | 2 | [illegible] |
| A5SS | 1667 | ENSG00000022976 | ZNF839 | chr14 | + | 102339093 | 102339229 | 102339093 | 102339223 | 102341322 | 102342367 | 0 | 0 | 0 | [illegible] |
| A5SS | 2489 | ENSG00000204842 | ATXN2 | chr12 | - | 111485264 | 111485331 | 111485270 | 111485331 | 111470557 | 111470742 | 3 | 7 | 4 | [illegible] |
| A5SS | 8467 | ENSG00000107862 | GBF1 | chr10 | + | 102376559 | 102376804 | 102376559 | 102376800 | 102376934 | 102377140 | 1 | 2 | 10 | [illegible] |
| A5SS | 6908 | ENSG00000139436 | GIT2 | chr12 | - | 109951166 | 109951316 | 109951211 | 109951316 | 109947255 | 109947504 | 95 | 45 | 52 | [illegible] |
| A5SS | 6016 | ENSG00000064933 | PMS1 | chr2 | + | 189863742 | 189864228 | 189863742 | 189863974 | 189867798 | 189867929 | 142 | 82 | 120 | [illegible] |
| A5SS | 4670 | ENSG00000268043 | NBPF12 | chr1 | + | 146959970 | 146960011 | 146959970 | 146960006 | 146960108 | 146960318 | 0 | 2 | 3 | [illegible] |
| A5SS | 784 | ENSG00000185624 | P4HB | chr17 | - | 81858494 | 81859387 | 81859180 | 81859387 | 81855452 | 81855586 | 10 | 6 | 15 | [illegible] |
| A5SS | 7722 | ENSG00000148334 | PTGES2 | chr9 | - | 128123701 | 128123851 | 128123725 | 128123851 | 128122933 | 128123134 | 55 | 54 | 67 | [illegible] |
| A5SS | 1497 | ENSG00000035664 | DAPK2 | chr15 | - | 63983532 | 63983754 | 63983624 | 63983754 | 63971422 | 63971561 | 15 | 9 | 11 | [illegible] |
| A5SS | 7721 | ENSG00000060237 | WNK1 | chr12 | + | 890452 | 890513 | 890452 | 890510 | 894561 | 894635 | 282 | 238 | 235 | [illegible] |
| A5SS | 537 | ENSG00000184402 | SS18L1 | chr20 | + | 62158671 | 62158942 | 62158671 | 62158748 | 62159876 | 62159961 | 5 | 1 | 9 | [illegible] |
| A5SS | 1083 | ENSG00000125124 | BBS2 | chr16 | - | 56501352 | 56502456 | 56502316 | 56502456 | 56498436 | 56499907 | 90 | 94 | 87 | [illegible] |
| A5SS | 4933 | ENSG00000182362 | YBEY | chr21 | + | 46286869 | 46287123 | 46286869 | 46286988 | 46296161 | 46296230 | 154 | 70 | 46 | [illegible] |
| A5SS | 1744 | ENSG00000213928 | IRF9 | chr14 | + | 24162143 | 24162324 | 24162143 | 24162280 | 24162965 | 24163149 | 271 | 159 | 251 | [illegible] |
| A5SS | 8673 | ENSG00000088992 | TESC | chr12 | - | 117046771 | 117046838 | 117046776 | 117046838 | 117046558 | 117046666 | 12 | 4 | 4 | [illegible] |
| A5SS | 3935 | ENSG00000121957 | GPSM2 | chr1 | + | 108918612 | 108919150 | 108918612 | 108918789 | 108922416 | 108922576 | 22 | 21 | 11 | [illegible] |
| A5SS | 1383 | ENSG00000140990 | NDUFB10 | chr16 | + | 1961152 | 1961291 | 1961152 | 1961258 | 1961496 | 1961636 | 195 | 123 | 147 | [illegible] |
| A5SS | 2195 | ENSG00000103496 | STX4 | chr16 | + | 31034203 | 31034536 | 31034203 | 31034325 | 31034969 | 31035040 | 81 | 57 | 94 | [illegible] |
| A5SS | 6512 | ENSG00000179818 | PCBP1-AS1 | chr2 | - | 70083506 | 70083687 | 70083537 | 70083687 | 70059642 | 70059680 | 5 | 1 | 2 | [illegible] |
| A5SS | 7888 | ENSG00000266173 | STRADA | chr17 | - | 63728333 | 63728413 | 63728357 | 63728413 | 63714005 | 63714108 | 17 | 7 | 20 | [illegible] |
| A5SS | 4576 | ENSG00000188186 | LAMTOR4 | chr7 | + | 100149498 | 100149579 | 100149498 | 100149565 | 100153399 | 100153517 | 70 | 40 | 49 | [illegible] |
| A5SS | 6243 | ENSG00000255339 | NA | chr10 | - | 100529365 | 100529506 | 100529379 | 100529506 | 100526974 | 100527074 | 17 | 7 | 17 | [illegible] |
| A5SS | 3691 | ENSG00000114626 | ABTB1 | chr3 | + | 127674390 | 127674453 | 127674390 | 127674418 | 127674544 | 127674600 | 51 | 28 | 51 | [illegible] |
| A5SS | 8694 | ENSG00000116198 | CEP104 | chr1 | - | 3836494 | 3837519 | 3837291 | 3837519 | 3834924 | 3835092 | 59 | 38 | 62 | [illegible] |
| A5SS | 1393 | ENSG00000183751 | TBL3 | chr16 | + | 1976210 | 1976314 | 1976210 | 1976257 | 1976813 | 1976961 | 54 | 29 | 44 | [illegible] |
| A5SS | 8809 | ENSG00000139192 | TAPBPL | chr12 | + | 6452041 | 6452312 | 6452041 | 6452150 | 6453066 | 6453297 | 207 | 74 | 140 | [illegible] |
| A5SS | 7938 | ENSG00000113163 | CERT1 | chr5 | - | 75425291 | 75425499 | 75425360 | 75425499 | 75419340 | 75419424 | 3 | 4 | 2 | [illegible] |
| A5SS | 3618 | ENSG00000177728 | TMEM94 | chr17 | + | 75471799 | 75471955 | 75471799 | 75471929 | 75485427 | 75485547 | 3 | 2 | 2 | [illegible] |
| A5SS | 2095 | ENSG00000082213 | C5orf22 | chr5 | + | 31534271 | 31534417 | 31534271 | 31534325 | 31535743 | 31535893 | 135 | 61 | 90 | [illegible] |
| A5SS | 1299 | ENSG00000103005 | USB1 | chr16 | + | 58014272 | 58014326 | 58014272 | 58014294 | 58017333 | 58017439 | 82 | 38 | 36 | [illegible] |
| A5SS | 3075 | ENSG00000111642 | CHD4 | chr12 | - | 6581037 | 6581173 | 6581043 | 6581173 | 6580796 | 6580929 | 0 | 0 | 1 | [illegible] |
| A5SS | 5348 | ENSG00000115687 | PASK | chr2 | - | 241115021 | 241115177 | 241115042 | 241115177 | 241112239 | 241112439 | 5 | 4 | 3 | [illegible] |
| A5SS | 6479 | ENSG00000197024 | ZNF398 | chr7 | + | 149153944 | 149154340 | 149153944 | 149154317 | 149166157 | 149166284 | 87 | 62 | 93 | [illegible] |
| A5SS | 1788 | ENSG00000119705 | SLIRP | chr14 | + | 77710837 | 77710896 | 77710837 | 77710880 | 77715771 | 77715879 | 96 | 61 | 114 | [illegible] |
| A5SS | 2249 | ENSG00000196189 | SEMA4A | chr1 | + | 156154549 | 156154985 | 156154549 | 156154717 | 156156413 | 156156574 | 0 | 0 | 0 | [illegible] |
| A5SS | 4912 | ENSG00000147121 | KRBOX4 | chrX | + | 46463197 | 46463308 | 46463197 | 46463293 | 46471141 | 46471185 | 4 | 3 | 3 | [illegible] |
| A5SS | 3365 | ENSG00000164253 | WDR41 | chr5 | - | 77453801 | 77453928 | 77453816 | 77453928 | 77451323 | 77451353 | 2 | 4 | 1 | [illegible] |
| A5SS | 7161 | ENSG00000147133 | TAF1 | chrX | + | 71458240 | 71458366 | 71458240 | 71458357 | 71459551 | 71459708 | 92 | 72 | 104 | [illegible] |
| A5SS | 6412 | ENSG00000145191 | EIF2B5 | chr3 | + | 184143051 | 184143565 | 184143051 | 184143142 | 184144098 | 184144224 | 127 | 100 | 146 | [illegible] |
| A5SS | 5408 | ENSG00000152056 | AP1S3 | chr2 | - | 223777688 | 223777869 | 223777690 | 223777869 | 223775900 | 223776009 | 5 | 1 | 0 | [illegible] |
| A5SS | 4439 | ENSG00000180353 | HCLS1 | chr3 | - | 121657201 | 121657352 | 121657278 | 121657352 | 121656058 | 121656266 | 29 | 30 | 30 | [illegible] |
| A5SS | 3270 | ENSG00000165801 | ARHGEF40 | chr14 | + | 21087000 | 21087463 | 21087000 | 21087105 | 21087967 | 21088098 | 84 | 39 | 43 | [illegible] |
| A5SS | 2675 | ENSG00000102796 | DHRS12 | chr13 | - | 51773920 | 51774034 | 51773929 | 51774034 | 51771820 | 51771911 | 9 | 1 | 3 | [illegible] |
| A5SS | 5428 | ENSG00000162923 | WDR26 | chr1 | - | 224431476 | 224431581 | 224431524 | 224431581 | 224424517 | 224424719 | 51 | 48 | 40 | [illegible] |
| A5SS | 7971 | ENSG00000196323 | ZBTB44 | chr11 | - | 130236792 | 130237009 | 130237093 | 130237093 | 130233307 | 130233382 | 110 | 50 | 85 | [illegible] |
| A5SS | 7717 | ENSG00000135047 | CTSL | chr9 | + | 87726118 | 87726398 | 87726118 | 87726249 | 87727593 | 87727729 | 10 | 6 | 3 | [illegible] |
| A5SS | 1143 | ENSG00000260565 | ERVK13-1 | chr16 | - | 2671344 | 2672273 | 2671594 | 2672273 | 2670863 | 2670974 | 54 | 32 | 44 | [illegible] |
| A5SS | 5435 | ENSG00000175215 | CTDSP2 | chr12 | - | 57827533 | 57827590 | 57827551 | 57827590 | 57826345 | 57826402 | 21 | 2 | 13 | [illegible] |
| A5SS | 4765 | ENSG00000100271 | TTLL1 | chr22 | - | 43069603 | 43069844 | 43069635 | 43069844 | 43068409 | 43068590 | 0 | 0 | 0 | [illegible] |
| A5SS | 6330 | ENSG00000136273 | HUS1 | chr7 | - | 47979420 | 47979553 | 47979467 | 47979553 | 47978688 | 47978816 | 3 | 1 | 2 | [illegible] |
| A5SS | 1942 | ENSG00000196182 | STK40 | chr1 | - | 36367847 | 36368084 | 36367851 | 36368084 | 36361220 | 36361340 | 12 | 7 | 7 | [illegible] |
| A5SS | 2392 | ENSG00000187091 | PLCD1 | chr3 | - | 38008457 | 38009158 | 38009041 | 38009158 | 38008234 | 38008367 | 27 | 3 | 18 | [illegible] |
| A5SS | 6518 | ENSG00000179818 | PCBP1-AS1 | chr2 | - | 70085815 | 70086273 | 70086251 | 70086273 | 70085546 | 70085624 | 27 | 24 | 25 | [illegible] |
| A5SS | 623 | ENSG00000099956 | SMARCB1 | chr22 | + | 23800943 | 23801135 | 23800943 | 23801081 | 23803294 | 23803422 | 0 | 0 | 1 | [illegible] |
| A5SS | 8355 | ENSG00000137752 | CASP1 | chr11 | - | 105034207 | 105034474 | 105034423 | 105034474 | 105033063 | 105033126 | 784 | 589 | 399 | [illegible] |
| A5SS | 6690 | ENSG00000013374 | NUB1 | chr7 | + | 151374096 | 151374243 | 151374096 | 151374201 | 151375847 | 151375943 | 34 | 29 | 51 | [illegible] |
| A5SS | 7769 | ENSG00000115459 | ELMOD3 | chr2 | + | 85371085 | 85371562 | 85371085 | 85371209 | 85377343 | 85377474 | 25 | 9 | 23 | [illegible] |
| A5SS | 7882 | ENSG00000277072 | STAG3L2 | chr7 | - | 74890270 | 74890566 | 74890387 | 74890566 | 74888311 | 74888397 | 15 | 18 | 20 | [illegible] |
| A5SS | 2838 | ENSG00000058056 | USP13 | chr3 | + | 179721401 | 179721735 | 179721401 | 179721589 | 179730188 | 179730260 | 1 | 0 | 1 | [illegible] |
| A5SS | 246 | ENSG00000076944 | STXBP2 | chr19 | + | 7647161 | 7647511 | 7647161 | 7647247 | 7647724 | 7647873 | 215 | 93 | 97 | [illegible] |
| A5SS | 7742 | ENSG00000165219 | GAPVD1 | chr9 | + | 125346818 | 125346995 | 125346818 | 125346941 | 125349389 | 125349519 | 6 | 6 | 1 | [illegible] |
| A5SS | 7530 | ENSG00000157045 | NTAN1 | chr16 | - | 15047417 | 15047550 | 15047441 | 15047550 | 15044333 | 15044407 | 14 | 7 | 12 | [illegible] |
| A5SS | 2847 | ENSG00000068650 | ATP11A | chr13 | + | 112857817 | 112859452 | 112857817 | 112857920 | 112860286 | 112860798 | 130 | 105 | 87 | [illegible] |
| A5SS | 3786 | ENSG00000142330 | CAPN10 | chr2 | + | 240590814 | 240591011 | 240590814 | 240591000 | 240591932 | 240592150 | 26 | 7 | 27 | [illegible] |
| A5SS | 5584 | ENSG00000143390 | RFX5 | chr1 | - | 151346204 | 151346333 | 151346214 | 151346333 | 151345927 | 151345961 | 310 | 197 | 264 | [illegible] |
| A5SS | 810 | ENSG00000131470 | PSMC3IP | chr17 | - | 42573818 | 42574210 | 42574195 | 42574210 | 42573477 | 42573623 | 12 | 4 | 12 | [illegible] |
| A5SS | 6612 | ENSG00000133195 | SLC39A11 | chr17 | - | 72947730 | 72947875 | 72947751 | 72947875 | 72849633 | 72849804 | 13 | 2 | 7 | [illegible] |
| A5SS | 4099 | ENSG00000081760 | AACS | chr12 | + | 125124704 | 125125024 | 125124704 | 125124769 | 125128160 | 125128274 | 8 | 6 | 20 | [illegible] |
| A5SS | 6975 | ENSG00000177646 | ACAD9 | chr3 | + | 128902552 | 128902632 | 128902552 | 128902628 | 128904061 | 128904132 | 6 | 0 | 3 | [illegible] |
| A5SS | 5660 | ENSG00000225889 | NA | chr2 | + | 64251118 | 64251983 | 64251118 | 64251834 | 64252419 | 64252783 | 15 | 41 | 21 | [illegible] |
| A5SS | 4489 | ENSG00000156983 | BRPF1 | chr3 | + | 9742943 | 9743471 | 9742943 | 9743253 | 9743577 | 9743901 | 5 | 2 | 9 | [illegible] |
| A5SS | 303 | ENSG00000105197 | TIMM50 | chr19 | + | 39482884 | 39482963 | 39482884 | 39482916 | 39483134 | 39483156 | 0 | 0 | 2 | [illegible] |
| A5SS | 6974 | ENSG00000177646 | ACAD9 | chr3 | + | 128879622 | 128880017 | 128879622 | 128879841 | 128884652 | 128884746 | 18 | 9 | 28 | [illegible] |
| A5SS | 2022 | ENSG00000141279 | NPEPPS | chr17 | + | 47579389 | 47582849 | 47579389 | 47579511 | 47585499 | 47585700 | 256 | 174 | 201 | [illegible] |
| A5SS | 4747 | ENSG00000273136 | NBPF26 | chr1 | + | 120805417 | 120805458 | 120805417 | 120805453 | 120805555 | 120805765 | 49 | 49 | 23 | [illegible] |
| A5SS | 791 | ENSG00000185624 | P4HB | chr17 | - | 81860326 | 81860569 | 81860346 | 81860569 | 81859180 | 81859387 | 111 | 41 | 67 | [illegible] |
| A5SS | 2352 | ENSG00000158290 | CUL4B | chrX | - | 120538117 | 120538209 | 120538123 | 120538209 | 120536926 | 120537034 | 20 | 14 | 16 | [illegible] |
| A5SS | 6789 | ENSG00000124222 | STX16 | chr20 | + | 58652072 | 58652138 | 58652072 | 58652087 | 58667986 | 58668127 | 24 | 20 | 25 | [illegible] |
| A5SS | 5591 | ENSG00000143390 | RFX5 | chr1 | - | 151346481 | 151346606 | 151346578 | 151346606 | 151346204 | 151346333 | 118 | 88 | 132 | [illegible] |
| A5SS | 4896 | ENSG00000115875 | SRSF7 | chr2 | - | 38748046 | 38748157 | 38748055 | 38748157 | 38746693 | 38746747 | 263 | 145 | 223 | [illegible] |
| A5SS | 6670 | ENSG00000228649 | SNHG26 | chr7 | + | 22858570 | 22858976 | 22858570 | 22858830 | 22860546 | 22860638 | 10 | 5 | 7 | [illegible] |
| A5SS | 1092 | ENSG00000125124 | BBS2 | chr16 | - | 56519745 | 56520049 | 56519956 | 56520049 | 56514452 | 56514680 | 74 | 18 | 41 | [illegible] |
| A5SS | 7853 | ENSG00000110514 | MADD | chr11 | + | 47284940 | 47285194 | 47284940 | 47285065 | 47285450 | 47285590 | 116 | 57 | 88 | [illegible] |
| A5SS | 3863 | ENSG00000114439 | BBX | chr3 | + | 107645835 | 107646087 | 107645835 | 107645909 | 107710451 | 107710622 | 6 | 12 | 17 | [illegible] |
| A5SS | 295 | ENSG00000178980 | SELENOW | chr19 | + | 47780863 | 47781000 | 47780863 | 47780917 | 47781107 | 47781182 | 0 | 4 | 9 | [illegible] |
| A5SS | 256 | ENSG00000104872 | PIH1D1 | chr19 | - | 49449339 | 49449654 | 49449474 | 49449654 | 49448000 | 49448062 | 7 | 0 | 12 | [illegible] |
| A5SS | 823 | ENSG00000189159 | JPT1 | chr17 | - | 75147519 | 75147653 | 75147555 | 75147653 | 75146665 | 75146684 | 0 | 3 | 1 | [illegible] |
| A5SS | 2213 | ENSG00000170144 | HNRNPA3 | chr2 | + | 177212740 | 177212871 | 177212740 | 177212805 | 177215538 | 177215661 | 446 | 334 | 403 | [illegible] |
| A5SS | 4056 | ENSG00000158773 | USF1 | chr1 | - | 161042554 | 161042670 | 161042585 | 161042670 | 161042115 | 161042217 | 196 | 120 | 149 | [illegible] |
| A5SS | 6773 | ENSG00000161010 | MRNIP | chr5 | - | 179853196 | 179853437 | 179853377 | 179853437 | 179844151 | 179844227 | 8 | 6 | 5 | [illegible] |
| A5SS | 1090 | ENSG00000125124 | BBS2 | chr16 | - | 56509356 | 56510034 | 56509956 | 56510034 | 56506119 | 56506224 | 9 | 3 | 3 | [illegible] |
| A5SS | 6523 | ENSG00000179818 | PCBP1-AS1 | chr2 | - | 70086123 | 70086273 | 70086251 | 70086273 | 70085546 | 70085624 | 12 | 19 | 20 | [illegible] |
| A5SS | 8238 | ENSG00000133895 | MEN1 | chr11 | - | 64806082 | 64806368 | 64806231 | 64806368 | 64805634 | 64805770 | 12 | 1 | 1 | [illegible] |
| A5SS | 3171 | ENSG00000133422 | MORC2 | chr22 | - | 30928018 | 30928207 | 30928027 | 30928207 | 30926610 | 30926871 | 45 | 46 | 76 | [illegible] |
| A5SS | 4950 | ENSG00000198160 | MIER1 | chr1 | + | 66984571 | 66984652 | 66984571 | 66984592 | 66986402 | 66988619 | 181 | 126 | 163 | [illegible] |
| A5SS | 5403 | ENSG00000144118 | RALB | chr2 | + | 120278617 | 120278778 | 120278617 | 120278712 | 120280834 | 120280907 | 58 | 36 | 32 | [illegible] |
| A5SS | 1381 | ENSG00000162104 | ADCY9 | chr16 | - | 3993348 | 3993510 | 3993405 | 3993510 | 3992145 | 3992363 | 15 | 6 | 3 | [illegible] |
| A5SS | 8590 | ENSG00000133316 | WDR74 | chr11 | - | 62835918 | 62836036 | 62835960 | 62836036 | 62835694 | 62835841 | 2 | 3 | 2 | [illegible] |
| A5SS | 6606 | ENSG00000071462 | BUD23 | chr7 | + | 73692595 | 73693414 | 73692595 | 73692646 | 73697604 | 73697694 | 17 | 10 | 10 | [illegible] |
| A5SS | 7758 | ENSG00000165699 | TSC1 | chr9 | - | 132925586 | 132927304 | 132927200 | 132927304 | 132923347 | 132923492 | 24 | 15 | 17 | [illegible] |
| A5SS | 1675 | ENSG00000100749 | VRK1 | chr14 | + | 96855223 | 96855356 | 96855223 | 96855352 | 96856129 | 96856250 | 126 | 77 | 98 | [illegible] |
| A5SS | 2487 | ENSG00000145907 | G3BP1 | chr5 | + | 151803884 | 151804162 | 151803884 | 151803996 | 151804405 | 151812785 | 165 | 72 | 156 | [illegible] |
| A5SS | 1136 | ENSG00000010361 | FUZ | chr19 | - | 49812446 | 49812736 | 49812614 | 49812736 | 49812250 | 49812335 | 4 | 4 | 12 | [illegible] |
| A5SS | 4243 | ENSG00000122986 | HVCN1 | chr12 | - | 110661163 | 110661448 | 110661279 | 110661448 | 110655233 | 110655338 | 245 | 131 | 190 | [illegible] |
| A5SS | 5641 | ENSG00000132323 | ILKAP | chr2 | - | 238189852 | 238189972 | 238189858 | 238189972 | 238188130 | 238188257 | 33 | 21 | 39 | [illegible] |
| A5SS | 7807 | ENSG00000070476 | ZXDC | chr3 | - | 126461269 | 126462220 | 126461534 | 126462220 | 126459652 | 126459737 | 63 | 22 | 58 | [illegible] |
| A5SS | 2486 | ENSG00000145907 | G3BP1 | chr5 | + | 151803884 | 151804141 | 151803884 | 151803996 | 151804405 | 151812785 | 162 | 72 | 152 | [illegible] |
| A5SS | 2125 | ENSG00000086189 | DIMT1 | chr5 | - | 62398428 | 62398560 | 62398510 | 62398560 | 62394483 | 62394607 | 2 | 4 | 2 | [illegible] |
| A5SS | 8311 | ENSG00000204161 | TMEM273 | chr10 | - | 49166444 | 49167009 | 49166868 | 49167009 | 49165765 | 49165796 | 2 | 1 | 4 | [illegible] |
| A5SS | 1218 | ENSG00000130175 | PRKCSH | chr19 | + | 11435634 | 11435710 | 11435634 | 11435706 | 11436040 | 11436196 | 26 | 7 | 23 | [illegible] |
| A5SS | 2078 | ENSG00000140853 | NLRC5 | chr16 | + | 57028071 | 57028203 | 57028071 | 57028155 | 57028301 | 57028385 | 13 | 14 | 28 | [illegible] |
| A5SS | 6228 | ENSG00000223501 | VPS52 | chr6 | - | 33271136 | 33271876 | 33271585 | 33271876 | 33270198 | 33270283 | 8 | 5 | 7 | [illegible] |
| A5SS | 6858 | ENSG00000189376 | C8orf76 | chr8 | - | 123231299 | 123231757 | 123231721 | 123231757 | 123226499 | 123226632 | 134 | 96 | 112 | [illegible] |
| A5SS | 1845 | ENSG00000103811 | CTSH | chr15 | - | 78944537 | 78945035 | 78944890 | 78945035 | 78939139 | 78939171 | 10 | 5 | 4 | [illegible] |
| A5SS | 204 | ENSG00000160570 | DEDD2 | chr19 | - | 42215132 | 42215252 | 42215147 | 42215252 | 42209699 | 42209840 | 65 | 31 | 30 | [illegible] |
| A5SS | 8723 | ENSG00000166012 | TAF1D | chr11 | - | 93736140 | 93736304 | 93736289 | 93736304 | 93734624 | 93734660 | 668 | 612 | 671 | [illegible] |
| A5SS | 4762 | ENSG00000172500 | FIBP | chr11 | - | 65885508 | 65885663 | 65885529 | 65885663 | 65885077 | 65885186 | 13 | 2 | 12 | [illegible] |
| A5SS | 161 | ENSG00000006015 | REX1BD | chr19 | + | 18588994 | 18589683 | 18588994 | 18589077 | 18590853 | 18590933 | 82 | 33 | 52 | [illegible] |
| A5SS | 872 | ENSG00000181038 | METTL23 | chr17 | + | 76726829 | 76727213 | 76726829 | 76726879 | 76729689 | 76729794 | 9 | 3 | 15 | [illegible] |
| A5SS | 2395 | ENSG00000026297 | RNASET2 | chr6 | - | 166955466 | 166955627 | 166955483 | 166955627 | 166952487 | 166952548 | 5 | 5 | 3 | [illegible] |
| A5SS | 6494 | ENSG00000204371 | EHMT2 | chr6 | - | 31884890 | 31885016 | 31884911 | 31885016 | 31884644 | 31884799 | 2 | 2 | 4 | [illegible] |
| A5SS | 6747 | ENSG00000196305 | IARS1 | chr9 | - | 92286535 | 92287910 | 92287790 | 92287910 | 92285721 | 92285839 | 171 | 146 | 185 | [illegible] |
| A5SS | 1798 | ENSG00000150527 | MIA2 | chr14 | + | 39291018 | 39291111 | 39291018 | 39291096 | 39293270 | 39293381 | 9 | 9 | 1 | [illegible] |
| A5SS | 2694 | ENSG00000105443 | CYTH2 | chr19 | + | 48474837 | 48474953 | 48474837 | 48474949 | 48478068 | 48478145 | 4 | 1 | 3 | [illegible] |
| A5SS | 1157 | ENSG00000005844 | ITGAL | chr16 | + | 30474195 | 30474298 | 30474195 | 30474287 | 30475288 | 30475400 | 31 | 18 | 10 | [illegible] |
| A5SS | 5720 | ENSG00000089123 | TASP1 | chr20 | - | 13528432 | 13528511 | 13528445 | 13528511 | 13483226 | 13483337 | 24 | 9 | 15 | [illegible] |
| A5SS | 6592 | ENSG00000106477 | CEP41 | chr7 | - | 130400697 | 130400821 | 130400706 | 130400821 | 130400038 | 130400254 | 2 | 0 | 0 | [illegible] |
| A5SS | 3488 | ENSG00000084072 | PPIE | chr1 | + | 39741894 | 39742031 | 39741894 | 39741921 | 39743215 | 39743297 | 14 | 4 | 15 | [illegible] |
| A5SS | 3036 | ENSG00000127483 | HP1BP3 | chr1 | - | 20786252 | 20786652 | 20786648 | 20786652 | 20780344 | 20780540 | 3 | 0 | 0 | [illegible] |
| A5SS | 2200 | ENSG00000113580 | NR3C1 | chr5 | - | 143313998 | 143314168 | 143314001 | 143314168 | 143310096 | 143310213 | 38 | 22 | 20 | [illegible] |
| A5SS | 1420 | ENSG00000213614 | HEXA | chr15 | - | 72348030 | 72348134 | 72348047 | 72348134 | 72346526 | 72346710 | 6 | 1 | 1 | [illegible] |
| A5SS | 308 | ENSG00000130669 | PAK4 | chr19 | + | 39168229 | 39168312 | 39168229 | 39168302 | 39169531 | 39169757 | 0 | 1 | 3 | [illegible] |
| A5SS | 1077 | ENSG00000172831 | CES2 | chr16 | + | 66942647 | 66942785 | 66942647 | 66942737 | 66943298 | 66943371 | 83 | 62 | 110 | [illegible] |
| A5SS | 2273 | ENSG00000134644 | PUM1 | chr1 | - | 30967098 | 30967310 | 30967166 | 30967310 | 30965981 | 30966197 | 13 | 7 | 14 | [illegible] |
| A5SS | 5785 | ENSG00000115514 | TXNDC9 | chr2 | - | 99332983 | 99333242 | 99333021 | 99333242 | 99327534 | 99327653 | 13 | 9 | 6 | [illegible] |
| A5SS | 3420 | ENSG00000008128 | CDK11A | chr1 | - | 1719327 | 1719455 | 1719357 | 1719455 | 1716345 | 1716478 | 19 | 22 | 10 | [illegible] |
| A5SS | 8653 | ENSG00000028203 | VEZT | chr12 | + | 95282312 | 95282644 | 95282312 | 95282359 | 95287663 | 95287857 | 238 | 176 | 212 | [illegible] |
| A5SS | 8391 | ENSG00000154124 | OTULIN | chr5 | + | 14681463 | 14681624 | 14681463 | 14681607 | 14687520 | 14687646 | 26 | 17 | 31 | [illegible] |
| A5SS | 2259 | ENSG00000153310 | CYRIB | chr8 | - | 129896671 | 129896854 | 129896748 | 129896854 | 129879388 | 129879471 | 15 | 15 | 17 | [illegible] |
| A5SS | 8734 | ENSG00000078674 | PCM1 | chr8 | + | 17991541 | 17991700 | 17991541 | 17991676 | 17993482 | 17993619 | 152 | 105 | 123 | [illegible] |
| A5SS | 2085 | ENSG00000065357 | DGKA | chr12 | + | 55940081 | 55940170 | 55940081 | 55940168 | 55940313 | 55940433 | 112 | 94 | 195 | [illegible] |
| A5SS | 104 | ENSG00000100150 | DEPDC5 | chr22 | + | 31838684 | 31838845 | 31838684 | 31838721 | 31843094 | 31843212 | 27 | 15 | 24 | [illegible] |
| A5SS | 2720 | ENSG00000155508 | CNOT8 | chr5 | + | 154858638 | 154858816 | 154858638 | 154858768 | 154863206 | 154863395 | 46 | 49 | 32 | [illegible] |
| A5SS | 3205 | ENSG00000114850 | SSR3 | chr3 | - | 156548865 | 156549003 | 156548904 | 156549003 | 156544307 | 156544439 | 11 | 11 | 12 | [illegible] |
| A5SS | 7298 | ENSG00000176731 | RBIS | chr8 | - | 85219235 | 85219363 | 85219322 | 85219363 | 85217385 | 85217502 | 162 | 19 | 11 | [illegible] |
| A5SS | 6972 | ENSG00000166881 | NEMP1 | chr12 | - | 57070673 | 57070893 | 57070819 | 57070893 | 57069233 | 57069306 | 54 | 55 | 36 | [illegible] |
| A5SS | 3826 | ENSG00000108654 | DDX5 | chr17 | - | 64500986 | 64501143 | 64501056 | 64501143 | 64500548 | 64500882 | 363 | 198 | 363 | [illegible] |
| A5SS | 1082 | ENSG00000125124 | BBS2 | chr16 | - | 56500853 | 56502456 | 56502316 | 56502456 | 56498436 | 56499907 | 119 | 126 | 150 | [illegible] |
| A5SS | 868 | ENSG00000011143 | MKS1 | chr17 | - | 58214258 | 58216243 | 58216087 | 58216243 | 58213764 | 58213869 | 14 | 13 | 11 | [illegible] |
| A5SS | 7989 | ENSG00000066926 | FECH | chr18 | - | 57580054 | 57580199 | 57580072 | 57580199 | 57573245 | 57573365 | 1 | 3 | 0 | [illegible] |
| A5SS | 4903 | ENSG00000143382 | ADAMTSL4 | chr1 | + | 150549821 | 150550276 | 150549821 | 150549895 | 150552204 | 150552308 | 3 | 2 | 3 | [illegible] |
| A5SS | 5815 | ENSG00000130414 | NDUFA10 | chr2 | - | 240018552 | 240018639 | 240018554 | 240018639 | 240011616 | 240014860 | 391 | 277 | 332 | [illegible] |
| A5SS | 1506 | ENSG00000140521 | POLG | chr15 | - | 89325449 | 89325686 | 89325489 | 89325686 | 89324106 | 89324227 | 106 | 67 | 72 | [illegible] |
| A5SS | 6495 | ENSG00000182518 | FAM104B | chrX | - | 55159125 | 55159228 | 55159128 | 55159228 | 55146180 | 55146311 | 3 | 1 | 1 | [illegible] |
| A5SS | 3304 | ENSG00000173171 | MTX1 | chr1 | + | 155210345 | 155210415 | 155210345 | 155210415 | 155210547 | 155210627 | 1 | 0 | 1 | [illegible] |
| A5SS | 101 | ENSG00000100150 | DEPDC5 | chr22 | + | 31784813 | 31785107 | 31784813 | 31784875 | 31792032 | 31792102 | 21 | 40 | 12 | [illegible] |
| A5SS | 6382 | ENSG00000173875 | ZNF791 | chr19 | + | 12612225 | 12612365 | 12612225 | 12612316 | 12623699 | 12623826 | 4 | 3 | 1 | [illegible] |
| A5SS | 4954 | ENSG00000129933 | MAU2 | chr19 | + | 19342775 | 19342875 | 19342775 | 19342866 | 19343836 | 19343940 | 0 | 0 | 0 | [illegible] |
| A5SS | 3249 | ENSG00000121897 | LIAS | chr4 | + | 39467517 | 39468427 | 39467517 | 39467646 | 39470018 | 39470164 | 18 | 13 | 16 | [illegible] |
| A5SS | 1814 | ENSG00000182979 | MTA1 | chr14 | + | 105450057 | 105450324 | 105450057 | 105450184 | 105454192 | 105454310 | 76 | 52 | 105 | [illegible] |
| A5SS | 7042 | ENSG00000134516 | DOCK2 | chr5 | + | 170078974 | 170081984 | 170078974 | 170079146 | 170082795 | 170083377 | 209 | 95 | 167 | [illegible] |
| A5SS | 3325 | ENSG00000144559 | TAMM41 | chr3 | - | 11817079 | 11817337 | 11817191 | 11817337 | 11810775 | 11810897 | 34 | 33 | 32 | [illegible] |
| A5SS | 3443 | ENSG00000144591 | GMPPA | chr2 | + | 219504082 | 219504868 | 219504082 | 219504213 | 219505227 | 219505362 | 15 | 5 | 7 | [illegible] |
| A5SS | 6441 | ENSG00000186591 | UBE2H | chr7 | - | 129880878 | 129880971 | 129880894 | 129880971 | 129879567 | 129879642 | 4 | 2 | 0 | [illegible] |
| A5SS | 4601 | ENSG00000166311 | SMPD1 | chr11 | + | 6391383 | 6392196 | 6391383 | 6392156 | 6393215 | 6393387 | 8 | 5 | 6 | [illegible] |
| A5SS | 5139 | ENSG00000166986 | MARS1 | chr12 | + | 57515919 | 57516158 | 57515919 | 57515991 | 57516244 | 57516337 | 38 | 27 | 36 | [illegible] |
| A5SS | 1358 | ENSG00000171208 | NETO2 | chr16 | - | 47122867 | 47122912 | 47122888 | 47122912 | 47122656 | 47122784 | 15 | 8 | 11 | [illegible] |
| A5SS | 2217 | ENSG00000170144 | HNRNPA3 | chr2 | + | 177219036 | 177219159 | 177219036 | 177219156 | 177219246 | 177219314 | 475 | 503 | 441 | [illegible] |
| A5SS | 7858 | ENSG00000108405 | P2RX1 | chr17 | - | 3903550 | 3904024 | 3903927 | 3904024 | 3903201 | 3903343 | 33 | 36 | 25 | [illegible] |
| A5SS | 6688 | ENSG00000104915 | STX10 | chr19 | - | 13144763 | 13144870 | 13144767 | 13144870 | 13144576 | 13144671 | 111 | 72 | 100 | [illegible] |
| A5SS | 5489 | ENSG00000020129 | NCDN | chr1 | + | 35558155 | 35558484 | 35558155 | 35558223 | 35559106 | 35559247 | 4 | 2 | 5 | [illegible] |
| A5SS | 447 | ENSG00000282508 | LINC01002 | chr19 | - | 203870 | 204065 | 203874 | 204065 | 201755 | 201813 | 1 | 1 | 1 | [illegible] |
| A5SS | 4993 | ENSG00000061273 | HDAC7 | chr12 | - | 47784078 | 47785471 | 47785386 | 47785471 | 47782725 | 47783886 | 116 | 56 | 158 | [illegible] |
| A5SS | 2365 | ENSG00000164889 | SLC4A2 | chr7 | + | 151074038 | 151074293 | 151074038 | 151074099 | 151074674 | 151074841 | 10 | 13 | 13 | [illegible] |
| A5SS | 5901 | ENSG00000119535 | CSF3R | chr1 | - | 36473718 | 36473887 | 36473763 | 36473887 | 36472516 | 36472686 | 0 | 3 | 1 | [illegible] |
| A5SS | 2402 | ENSG00000070756 | PABPC1 | chr8 | - | 100715461 | 100715601 | 100715470 | 100715601 | 100712651 | 100712789 | 384 | 329 | 399 | [illegible] |
| A5SS | 444 | ENSG00000099817 | POLR2E | chr19 | - | 1093903 | 1095375 | 1095258 | 1095375 | 1091791 | 1091907 | 211 | 101 | 136 | [illegible] |
| A5SS | 5817 | ENSG00000130414 | NDUFA10 | chr2 | - | 240021086 | 240021412 | 240021196 | 240021412 | 240018552 | 240018639 | 23 | 15 | 12 | [illegible] |
| A5SS | 4703 | ENSG00000158985 | CDC42SE2 | chr5 | + | 131359208 | 131359558 | 131359208 | 131359547 | 131385542 | 131385644 | 14 | 2 | 5 | [illegible] |
| A5SS | 2275 | ENSG00000134644 | PUM1 | chr1 | - | 31059203 | 31059577 | 31059298 | 31059577 | 31028795 | 31028864 | 370 | 189 | 310 | [illegible] |
| A5SS | 5887 | ENSG00000143436 | MRPL9 | chr1 | - | 151762949 | 151763146 | 151762989 | 151763146 | 151762375 | 151762500 | 10 | 3 | 9 | [illegible] |
| A5SS | 8157 | ENSG00000171204 | TMEM126B | chr11 | + | 85634085 | 85634279 | 85634085 | 85634176 | 85635666 | 85635778 | 348 | 137 | 309 | [illegible] |
| A5SS | 5964 | ENSG00000108465 | CDK5RAP3 | chr17 | + | 47973518 | 47973696 | 47973518 | 47973693 | 47973930 | 47974031 | 0 | 0 | 1 | [illegible] |
| A5SS | 4471 | ENSG00000070010 | UFD1 | chr22 | - | 19475469 | 19475602 | 19475495 | 19475602 | 19475067 | 19475100 | 79 | 50 | 64 | [illegible] |
| A5SS | 6897 | ENSG00000233093 | LINC00892 | chrX | + | 136640773 | 136640873 | 136640773 | 136640840 | 136642102 | 136642426 | 22 | 24 | 36 | [illegible] |
| A5SS | 8037 | ENSG00000177830 | CHID1 | chr11 | - | 902147 | 902330 | 902197 | 902330 | 900935 | 900980 | 2 | 2 | 2 | [illegible] |
| A5SS | 120 | ENSG00000235954 | TTC28-AS1 | chr22 | + | 27997410 | 27998838 | 27997410 | 27997739 | 27999429 | 28001658 | 15 | 4 | 14 | [illegible] |
| A5SS | 1571 | ENSG00000214331 | PDPR2P | chr16 | - | 74352039 | 74352378 | 74352296 | 74352378 | 74350616 | 74350738 | 40 | 29 | 52 | [illegible] |
| A5SS | 1306 | ENSG00000135686 | KLHL36 | chr16 | + | 84656870 | 84657944 | 84656870 | 84657913 | 84659759 | 84659917 | 53 | 17 | 42 | [illegible] |
| A5SS | 5157 | ENSG00000197062 | ZSCAN26 | chr6 | + | 28271853 | 28272339 | 28271853 | 28271934 | 28272669 | 28272787 | 53 | 23 | 53 | [illegible] |
| A5SS | 4526 | ENSG00000135241 | PNPLA8 | chr7 | - | 108487758 | 108487953 | 108487944 | 108487953 | 108479183 | 108479379 | 200 | 158 | 141 | [illegible] |
| A5SS | 5473 | ENSG00000114054 | PCCB | chr3 | + | 136326802 | 136327255 | 136326802 | 136326910 | 136327633 | 136327732 | 27 | 13 | 27 | [illegible] |
| A5SS | 5845 | ENSG00000006062 | MAP3K14 | chr17 | - | 45270851 | 45271221 | 45271057 | 45271221 | 45270412 | 45270563 | 6 | 4 | 9 | [illegible] |
| A5SS | 4854 | ENSG00000159363 | ATP13A2 | chr1 | - | 17004691 | 17004821 | 17004706 | 17004821 | 17004331 | 17004411 | 17 | 4 | 19 | [illegible] |
| A5SS | 6905 | ENSG00000179195 | ZNF664 | chr12 | + | 123973218 | 123973352 | 123973218 | 123973303 | 123973885 | 123974553 | 13 | 2 | 5 | [illegible] |
| A5SS | 7675 | ENSG00000159069 | FBXW5 | chr9 | - | 136941754 | 136942466 | 136942099 | 136942466 | 136941536 | 136941684 | 62 | 30 | 39 | [illegible] |
| A5SS | 8148 | ENSG00000185187 | SIGIRR | chr11 | - | 407816 | 407957 | 407851 | 407957 | 407424 | 407568 | 84 | 37 | 68 | [illegible] |
| A5SS | 5620 | ENSG00000120697 | ALG5 | chr13 | - | 36994988 | 36995596 | 36995424 | 36995596 | 36993603 | 36993672 | 119 | 76 | 89 | [illegible] |
| A5SS | 1876 | ENSG00000146457 | WTAP | chr6 | + | 159748190 | 159750010 | 159748190 | 159748369 | 159753459 | 159753614 | 653 | 451 | 525 | [illegible] |

| | | | | | | | | | | | | | | | |
|---|---|---|---|---|---|---|---|---|---|---|---|---|---|---|---|
| A5SS | 8397 | ENSG00000148498 | PARD3 | chr10 | - | 34347955 | 34348115 | 34347964 | 34348115 | 34341626 | 34341816 | 4 | 7 | 0 | [illegible] |
| A5SS | 1115 | ENSG00000166451 | CENPN | chr16 | + | 81014135 | 81014391 | 81014135 | 81014181 | 81017325 | 81017385 | 2 | 1 | 4 | [illegible] |
| A5SS | 4876 | ENSG00000162852 | CNST | chr1 | + | 246633923 | 246634587 | 246633923 | 246634010 | 246641748 | 246641770 | 214 | 297 | 174 | [illegible] |
| A5SS | 7138 | ENSG00000248323 | LUCAT1 | chr5 | - | 91313961 | 91314404 | 91314079 | 91314404 | 91313637 | 91313777 | 17 | 10 | 16 | [illegible] |
| A5SS | 4427 | ENSG00000166523 | CLEC4E | chr12 | - | 8539199 | 8539306 | 8539216 | 8539306 | 8537114 | 8537266 | 14 | 6 | 9 | [illegible] |
| A5SS | 1823 | ENSG00000165898 | ISCA2 | chr14 | + | 74494274 | 74494404 | 74494274 | 74494390 | 74494825 | 74497106 | 8 | 4 | 13 | [illegible] |
| A5SS | 2051 | ENSG00000181991 | MRPS11 | chr15 | + | 88467907 | 88469145 | 88467907 | 88468024 | 88472626 | 88472725 | 9 | 6 | 8 | [illegible] |
| A5SS | 6628 | ENSG00000105819 | PMPCB | chr7 | + | 103299442 | 103299533 | 103299442 | 103299529 | 103300177 | 103300307 | 3 | 2 | 2 | [illegible] |
| A5SS | 8803 | ENSG00000168010 | ATG16L2 | chr11 | + | 72822477 | 72823797 | 72822477 | 72822543 | 72824059 | 72824122 | 26 | 19 | 14 | [illegible] |
| A5SS | 5027 | ENSG00000092529 | CAPN3 | chr15 | + | 42408210 | 42408437 | 42408210 | 42408324 | 42409302 | 42409380 | 2 | 3 | 0 | [illegible] |
| A5SS | 7986 | ENSG00000085117 | CD82 | chr11 | + | 44605057 | 44605429 | 44605057 | 44605182 | 44615271 | 44615373 | 28 | 10 | 29 | [illegible] |
| A5SS | 8182 | ENSG00000072518 | MARK2 | chr11 | + | 63906087 | 63907567 | 63906087 | 63906114 | 63908259 | 63908304 | 1 | 7 | 3 | [illegible] |
| A5SS | 7829 | ENSG00000083799 | CYLD | chr16 | + | 50742761 | 50742841 | 50742761 | 50742822 | 50749575 | 50750202 | 161 | 141 | 202 | [illegible] |
| A5SS | 1032 | ENSG00000214719 | NA | chr17 | + | 30631754 | 30631979 | 30631754 | 30631927 | 30633097 | 30633390 | 17 | 9 | 9 | [illegible] |
| A5SS | 2718 | ENSG00000115073 | ACTR1B | chr2 | - | 97658426 | 97659003 | 97658878 | 97659003 | 97658223 | 97658316 | 60 | 13 | 70 | [illegible] |
| A5SS | 7567 | ENSG00000135018 | UBQLN1 | chr9 | - | 83666349 | 83669327 | 83669184 | 83669327 | 83665029 | 83665145 | 408 | 260 | 341 | [illegible] |
| A5SS | 2750 | ENSG00000206573 | THUMPD3-AS1 | chr3 | - | 9390884 | 9391826 | 9391678 | 9391826 | 9390289 | 9390497 | 67 | 99 | 88 | [illegible] |
| A5SS | 6122 | ENSG00000124523 | SIRT5 | chr6 | + | 13599031 | 13600949 | 13599031 | 13599155 | 13611789 | 13615158 | 31 | 15 | 31 | [illegible] |
| A5SS | 3614 | ENSG00000239900 | ADSL | chr22 | + | 40349831 | 40353117 | 40349831 | 40350035 | 40354247 | 40354327 | 95 | 58 | 95 | [illegible] |
| A5SS | 3554 | ENSG00000048740 | CELF2 | chr10 | + | 11314138 | 11314270 | 11314138 | 11314258 | 11321188 | 11321386 | 5 | 6 | 4 | [illegible] |
| A5SS | 6831 | ENSG00000124333 | VAMP7 | chrX | + | 155889457 | 155889608 | 155889457 | 155889547 | 155895622 | 155895680 | 31 | 17 | 32 | [illegible] |
| A5SS | 5494 | ENSG00000143799 | PARP1 | chr1 | - | 226386218 | 226386442 | 226386325 | 226386442 | 226385503 | 226385680 | 3 | 1 | 6 | [illegible] |
| A5SS | 3418 | ENSG00000008128 | CDK11A | chr1 | - | 1707408 | 1707584 | 1707524 | 1707584 | 1705641 | 1705732 | 39 | 17 | 55 | [illegible] |
| A5SS | 8408 | ENSG00000095585 | BLNK | chr10 | - | 96227409 | 96227566 | 96227421 | 96227566 | 96223825 | 96223989 | 72 | 70 | 99 | [illegible] |
| A5SS | 8665 | ENSG00000110801 | PSMD9 | chr12 | + | 121899633 | 121899889 | 121899633 | 121899845 | 121903005 | 121903107 | 0 | 0 | 0 | [illegible] |
| A5SS | 7701 | ENSG00000165275 | TRMT10B | chr9 | + | 37763628 | 37763865 | 37763628 | 37763753 | 37768075 | 37768228 | 30 | 27 | 38 | [illegible] |
| A5SS | 4305 | ENSG00000178685 | PARP10 | chr8 | - | 143985720 | 143985975 | 143985724 | 143985975 | 143985568 | 143985648 | 60 | 19 | 35 | [illegible] |
| A5SS | 1734 | ENSG00000100938 | GMPR2 | chr14 | + | 24238588 | 24238803 | 24238588 | 24238748 | 24238957 | 24239240 | 95 | 52 | 82 | [illegible] |
| A5SS | 4184 | ENSG00000133275 | CSNK1G2 | chr19 | + | 1979494 | 1979643 | 1979494 | 1979618 | 1979751 | 1979835 | 183 | 85 | 148 | [illegible] |
| A5SS | 61 | ENSG00000182944 | EWSR1 | chr22 | + | 29288605 | 29291599 | 29288605 | 29288786 | 29292136 | 29292169 | 255 | 169 | 230 | [illegible] |
| A5SS | 7884 | ENSG00000149182 | ARFGAP2 | chr11 | - | 47171425 | 47171557 | 47171452 | 47171557 | 47168122 | 47168251 | 206 | 142 | 213 | [illegible] |
| A5SS | 2400 | ENSG00000070756 | PABPC1 | chr8 | - | 100704279 | 100704390 | 100704296 | 100704390 | 100702915 | 100703359 | 68 | 67 | 83 | [illegible] |
| A5SS | 1993 | ENSG00000215190 | LINC00680 | chr6 | - | 57948872 | 57949068 | 57948917 | 57949068 | 57946077 | 57947177 | 27 | 10 | 37 | [illegible] |
| A5SS | 613 | ENSG00000124155 | PIGT | chr20 | + | 45416516 | 45418979 | 45416516 | 45416694 | 45419294 | 45419395 | 55 | 15 | 27 | [illegible] |
| A5SS | 6249 | ENSG00000131018 | SYNE1 | chr6 | - | 152135025 | 152135232 | 152135103 | 152135232 | 152133275 | 152133488 | 2 | 1 | 1 | [illegible] |
| A5SS | 262 | ENSG00000105520 | PLPPR2 | chr19 | + | 11363712 | 11363835 | 11363712 | 11363819 | 11364160 | 11364212 | 11 | 2 | 3 | [illegible] |
| A5SS | 3645 | ENSG00000160789 | LMNA | chr1 | + | 156136213 | 156136549 | 156136213 | 156136436 | 156136920 | 156137028 | 3 | 0 | 1 | [illegible] |
| A5SS | 3467 | ENSG00000127948 | POR | chr7 | + | 75972412 | 75972764 | 75972412 | 75972461 | 75979450 | 75979579 | 0 | 0 | 1 | [illegible] |
| A5SS | 3430 | ENSG00000117448 | AKR1A1 | chr1 | + | 45568926 | 45569006 | 45568926 | 45568999 | 45569142 | 45569229 | 2 | 0 | 0 | [illegible] |
| A5SS | 6683 | ENSG00000198315 | ZKSCAN8 | chr6 | + | 28148315 | 28148838 | 28148315 | 28148824 | 28149482 | 28149624 | 7 | 5 | 9 | [illegible] |
| A5SS | 3891 | ENSG00000198740 | ZNF652 | chr17 | - | 49316825 | 49317983 | 49317365 | 49317983 | 49312697 | 49312845 | 283 | 250 | 289 | [illegible] |
| A5SS | 3079 | ENSG00000172830 | SSH3 | chr11 | + | 67306837 | 67307113 | 67306837 | 67306962 | 67307370 | 67307436 | 9 | 9 | 3 | [illegible] |
| A5SS | 7001 | ENSG00000111203 | ITFG2 | chr12 | + | 2818105 | 2818353 | 2818105 | 2818277 | 2820085 | 2820225 | 8 | 3 | 10 | [illegible] |
| A5SS | 4086 | ENSG00000136068 | FLNB | chr3 | + | 58141857 | 58141929 | 58141857 | 58141896 | 58142649 | 58142752 | 27 | 5 | 8 | [illegible] |
| A5SS | 7488 | ENSG00000178209 | PLEC | chr8 | - | 143935010 | 143935117 | 143935022 | 143935117 | 143934809 | 143934929 | 130 | 58 | 126 | [illegible] |
| A5SS | 3148 | ENSG00000100462 | PRMT5 | chr14 | - | 22920839 | 22921056 | 22920987 | 22921056 | 22920528 | 22920731 | 31 | 8 | 26 | [illegible] |
| A5SS | 4622 | ENSG00000106462 | EZH2 | chr7 | - | 148846469 | 148846598 | 148846496 | 148846598 | 148832633 | 148832750 | 24 | 12 | 22 | [illegible] |
| A5SS | 646 | ENSG00000263753 | LINC00667 | chr18 | + | 5238080 | 5239196 | 5238080 | 5239049 | 5243593 | 5246494 | 63 | 35 | 42 | [illegible] |
| A5SS | 861 | ENSG00000109113 | RAB34 | chr17 | - | 28715194 | 28715276 | 28715218 | 28715276 | 28715031 | 28715122 | 37 | 13 | 28 | [illegible] |
| A5SS | 4928 | ENSG00000130713 | EXOSC2 | chr9 | + | 130697581 | 130698251 | 130697581 | 130697627 | 130699328 | 130699394 | 124 | 69 | 150 | [illegible] |
| A5SS | 3896 | ENSG00000145860 | RNF145 | chr5 | - | 159168872 | 159169055 | 159168991 | 159169055 | 159162931 | 159163079 | 67 | 50 | 52 | [illegible] |
| A5SS | 7342 | ENSG00000137492 | THAP12 | chr11 | - | 76365319 | 76365972 | 76365779 | 76365972 | 76360955 | 76361063 | 9 | 8 | 4 | [illegible] |
| A5SS | 5965 | ENSG00000108465 | CDK5RAP3 | chr17 | + | 47973930 | 47974035 | 47973930 | 47974031 | 47974399 | 47974448 | 1 | 4 | 1 | [illegible] |
| A5SS | 8385 | ENSG00000177239 | MAN1B1 | chr9 | + | 137088074 | 137088398 | 137088074 | 137088183 | 137088868 | 137089005 | 11 | 9 | 13 | [illegible] |
| A5SS | 5259 | ENSG00000204344 | STK19 | chr6 | + | 31978902 | 31979010 | 31978902 | 31978998 | 31979413 | 31979553 | 5 | 5 | 0 | [illegible] |
| A5SS | 903 | ENSG00000187838 | PLSCR3 | chr17 | - | 7393600 | 7394267 | 7394103 | 7394267 | 7393466 | 7393509 | 42 | 25 | 50 | [illegible] |
| A5SS | 4664 | ENSG00000269713 | NBPF9 | chr1 | - | 149065525 | 149065689 | 149065636 | 149065689 | 149064430 | 149064482 | 6 | 3 | 6 | [illegible] |
| A5SS | 6654 | ENSG00000128604 | IRF5 | chr7 | + | 128945844 | 128946034 | 128945844 | 128946006 | 128946500 | 128946562 | 148 | 71 | 82 | [illegible] |
| A5SS | 8488 | ENSG00000059573 | ALDH18A1 | chr10 | - | 95633490 | 95633649 | 95633496 | 95633649 | 95632958 | 95633049 | 66 | 29 | 67 | [illegible] |
| A5SS | 2905 | ENSG00000164068 | RNF123 | chr3 | + | 49704983 | 49705186 | 49704983 | 49705182 | 49705533 | 49705679 | 3 | 0 | 0 | [illegible] |
| A5SS | 7285 | ENSG00000174197 | MGA | chr15 | + | 41736180 | 41736698 | 41736180 | 41736368 | 41739905 | 41740203 | 139 | 93 | 95 | [illegible] |
| A5SS | 3274 | ENSG00000132879 | FBXO44 | chr1 | + | 11658266 | 11658393 | 11658266 | 11658364 | 11658735 | 11658871 | 0 | 5 | 5 | [illegible] |
| A5SS | 6145 | ENSG00000137312 | FLOT1 | chr6 | - | 30741189 | 30741704 | 30741613 | 30741704 | 30740678 | 30740798 | 360 | 183 | 274 | [illegible] |
| A5SS | 952 | ENSG00000262879 | NA | chr17 | - | 47033277 | 47033381 | 47033319 | 47033381 | 47021581 | 47021721 | 8 | 3 | 18 | [illegible] |
| A5SS | 1819 | ENSG00000092203 | TOX4 | chr14 | + | 21477495 | 21477592 | 21477495 | 21477564 | 21487450 | 21487693 | 3 | 0 | 2 | [illegible] |
| A5SS | 7253 | ENSG00000221914 | PPP2R2A | chr8 | + | 26362683 | 26363890 | 26362683 | 26362848 | 26366314 | 26366406 | 111 | 121 | 109 | [illegible] |
| A5SS | 2558 | ENSG00000173020 | GRK2 | chr11 | + | 67280731 | 67281184 | 67280731 | 67280783 | 67281458 | 67281558 | 466 | 220 | 414 | [illegible] |
| A5SS | 220 | ENSG00000129353 | SLC44A2 | chr19 | + | 10634755 | 10635073 | 10634755 | 10634887 | 10636322 | 10636585 | 48 | 20 | 44 | [illegible] |
| A5SS | 6640 | ENSG00000086730 | LAT2 | chr7 | + | 74214821 | 74215099 | 74214821 | 74214940 | 74215946 | 74216069 | 35 | 10 | 30 | [illegible] |
| A5SS | 2806 | ENSG00000225697 | SLC26A6 | chr3 | - | 48631223 | 48631306 | 48631252 | 48631306 | 48630992 | 48631140 | 6 | 4 | 2 | [illegible] |
| A5SS | 3823 | ENSG00000108654 | DDX5 | chr17 | - | 64500548 | 64502069 | 64502009 | 64502069 | 64498256 | 64500326 | 5950 | 4529 | 5160 | [illegible] |
| A5SS | 2262 | ENSG00000183621 | ZNF438 | chr10 | - | 30848530 | 30850367 | 30850221 | 30850367 | 30835566 | 30835640 | 41 | 20 | 38 | [illegible] |
| A5SS | 8061 | ENSG00000101158 | NELFCD | chr20 | + | 58993623 | 58993764 | 58993623 | 58993675 | 58994109 | 58994239 | 117 | 71 | 85 | [illegible] |
| A5SS | 8561 | ENSG00000136404 | TM6SF1 | chr15 | + | 83119577 | 83119734 | 83119577 | 83119681 | 83124671 | 83124776 | 16 | 11 | 4 | [illegible] |
| A5SS | 1371 | ENSG00000130731 | METTL26 | chr16 | - | 635517 | 635774 | 635611 | 635774 | 635280 | 635340 | 4 | 5 | 0 | [illegible] |
| A5SS | 569 | ENSG00000196756 | SNHG17 | chr20 | - | 38425934 | 38426503 | 38426418 | 38426503 | 38422290 | 38422368 | 19 | 12 | 6 | [illegible] |
| A5SS | 6069 | ENSG00000204410 | MSH5 | chr6 | + | 31743903 | 31744299 | 31743903 | 31744025 | 31744545 | 31744581 | 22 | 25 | 20 | [illegible] |
| A5SS | 7739 | ENSG00000165219 | GAPVD1 | chr9 | + | 125321432 | 125321562 | 125321432 | 125321499 | 125323797 | 125323923 | 147 | 99 | 101 | [illegible] |
| A5SS | 1587 | ENSG00000166233 | ARIH1 | chr15 | + | 72580730 | 72580991 | 72580730 | 72580877 | 72582074 | 72582187 | 236 | 142 | 202 | [illegible] |
| A5SS | 1575 | ENSG00000140474 | ULK3 | chr15 | - | 74837750 | 74837798 | 74837756 | 74837798 | 74837368 | 74837435 | 30 | 8 | 0 | [illegible] |
| A5SS | 1285 | ENSG00000131652 | THOC6 | chr16 | + | 3025923 | 3026166 | 3025923 | 3025988 | 3026250 | 3026288 | 47 | 32 | 39 | [illegible] |
| A5SS | 6823 | ENSG00000112715 | VEGFA | chr6 | + | 43781955 | 43782087 | 43781955 | 43782052 | 43784540 | 43784562 | 18 | 4 | 10 | [illegible] |
| A5SS | 4059 | ENSG00000011295 | TTC19 | chr17 | + | 16000117 | 16000382 | 16000117 | 16000245 | 16001914 | 16002025 | 22 | 24 | 34 | [illegible] |
| A5SS | 5732 | ENSG00000226950 | DANCR | chr4 | + | 52713493 | 52713702 | 52713493 | 52713542 | 52714198 | 52714238 | 155 | 72 | 160 | [illegible] |
| A5SS | 334 | ENSG00000167613 | LAIR1 | chr19 | - | 54356711 | 54356966 | 54356927 | 54356966 | 54356490 | 54356619 | 15 | 10 | 19 | [illegible] |
| A5SS | 2550 | ENSG00000123213 | NLN | chr5 | + | 65792453 | 65792655 | 65792453 | 65792601 | 65809514 | 65809701 | 39 | 27 | 20 | [illegible] |
| A5SS | 3615 | ENSG00000239900 | ADSL | chr22 | + | 40361487 | 40361666 | 40361487 | 40361635 | 40362980 | 40363071 | 2 | 3 | 2 | [illegible] |
| A5SS | 5141 | ENSG00000131748 | STARD3 | chr17 | + | 39657774 | 39657852 | 39657774 | 39657839 | 39657972 | 39658026 | 91 | 20 | 76 | [illegible] |
| A5SS | 811 | ENSG00000131470 | PSMC3IP | chr17 | - | 42574098 | 42574210 | 42574195 | 42574210 | 42573477 | 42573623 | 7 | 3 | 3 | [illegible] |
| A5SS | 6116 | ENSG00000137200 | CMTR1 | chr6 | + | 37475320 | 37475916 | 37475320 | 37475412 | 37476125 | 37476194 | 18 | 17 | 16 | [illegible] |
| A5SS | 5342 | ENSG00000131051 | RBM39 | chr20 | - | 35738967 | 35740887 | 35740823 | 35740887 | 35731940 | 35732135 | 872 | 788 | 802 | [illegible] |
| A5SS | 1016 | ENSG00000177302 | TOP3A | chr17 | - | 18290539 | 18290686 | 18290556 | 18290686 | 18285406 | 18285520 | 1 | 0 | 0 | [illegible] |
| A5SS | 3046 | ENSG00000007923 | DNAJC11 | chr1 | - | 6638082 | 6638364 | 6638294 | 6638364 | 6637446 | 6637504 | 2 | 0 | 10 | [illegible] |
| A5SS | 1482 | ENSG00000037757 | MRI1 | chr19 | + | 13765953 | 13766129 | 13765953 | 13766065 | 13768560 | 13768870 | 34 | 6 | 22 | [illegible] |
| A5SS | 5903 | ENSG00000198001 | IRAK4 | chr12 | + | 43758950 | 43759592 | 43758950 | 43759016 | 43768102 | 43768272 | 6 | 1 | 0 | [illegible] |
| A5SS | 3664 | ENSG00000058673 | ZC3H11A | chr1 | + | 203801574 | 203803016 | 203801574 | 203801628 | 203816925 | 203817124 | 398 | 275 | 371 | [illegible] |
| A5SS | 6234 | ENSG00000112511 | PHF1 | chr6 | + | 33414724 | 33415144 | 33414724 | 33414829 | 33415234 | 33415329 | 66 | 39 | 77 | [illegible] |
| A5SS | 5308 | ENSG00000155366 | RHOC | chr1 | - | 112705052 | 112705168 | 112705099 | 112705168 | 112703643 | 112703806 | 8 | 0 | 2 | [illegible] |
| A5SS | 1053 | ENSG00000172530 | BANP | chr16 | + | 87975047 | 87975207 | 87975047 | 87975203 | 87981035 | 87981127 | 11 | 5 | 12 | [illegible] |
| A5SS | 2737 | ENSG00000082805 | ERC1 | chr12 | + | 1112214 | 1112307 | 1112214 | 1112298 | 1115865 | 1116033 | 8 | 0 | 0 | [illegible] |
| A5SS | 6329 | ENSG00000136273 | HUS1 | chr7 | - | 47979409 | 47979553 | 47979467 | 47979553 | 47978688 | 47978816 | 3 | 0 | 7 | [illegible] |
| A5SS | 8020 | ENSG00000149503 | INCENP | chr11 | + | 62123997 | 62124163 | 62123997 | 62124055 | 62128150 | 62128301 | 29 | 25 | 16 | [illegible] |
| A5SS | 786 | ENSG00000185624 | P4HB | chr17 | - | 81858934 | 81859387 | 81859180 | 81859387 | 81855452 | 81855586 | 9 | 7 | 17 | [illegible] |
| A5SS | 2777 | ENSG00000242086 | MUC20-OT1 | chr3 | + | 195708133 | 195708387 | 195708133 | 195708268 | 195708746 | 195708807 | 10 | 3 | 15 | [illegible] |
| A5SS | 4746 | ENSG00000117523 | PRRC2C | chr1 | + | 171584418 | 171584541 | 171584418 | 171584526 | 171587002 | 171587221 | 181 | 106 | 163 | [illegible] |
| A5SS | 7122 | ENSG00000232119 | MCTS1 | chrX | + | 120608224 | 120608463 | 120608224 | 120608358 | 120611010 | 120611078 | 9 | 8 | 7 | [illegible] |
| A5SS | 6547 | ENSG00000130429 | ARPC1B | chr7 | + | 99377337 | 99377455 | 99377337 | 99377439 | 99385701 | 99385778 | 19 | 4 | 10 | [illegible] |
| A5SS | 5122 | ENSG00000119844 | AFTPH | chr2 | + | 64579485 | 64581273 | 64579485 | 64579546 | 64585421 | 64585545 | 16 | 19 | 25 | [illegible] |
| A5SS | 4612 | ENSG00000204406 | MBD5 | chr2 | + | 148489385 | 148490787 | 148489385 | 148490594 | 148502435 | 148502509 | 11 | 8 | 15 | [illegible] |
| A5SS | 5143 | ENSG00000131748 | STARD3 | chr17 | + | 39662250 | 39662367 | 39662250 | 39662344 | 39662803 | 39663484 | 3 | 0 | 2 | [illegible] |
| A5SS | 7467 | ENSG00000164733 | CTSB | chr8 | - | 11861299 | 11861479 | 11861360 | 11861479 | 11853328 | 11853479 | 27 | 29 | 16 | [illegible] |
| A5SS | 1495 | ENSG00000089195 | TRMT6 | chr20 | - | 5943826 | 5944031 | 5943947 | 5944031 | 5943558 | 5943683 | 11 | 5 | 3 | [illegible] |
| A5SS | 6987 | ENSG00000223959 | AFG3L1P | chr16 | + | 89981772 | 89981923 | 89981772 | 89981918 | 89984530 | 89984599 | 47 | 35 | 37 | [illegible] |
| A5SS | 4310 | ENSG00000234684 | SDCBP2-AS1 | chr20 | + | 1325420 | 1325782 | 1325420 | 1325529 | 1373077 | 1373178 | 15 | 2 | 6 | [illegible] |
| A5SS | 1634 | ENSG00000100445 | SDR39U1 | chr14 | - | 24441673 | 24441795 | 24441677 | 24441795 | 24440782 | 24440926 | 61 | 24 | 75 | [illegible] |
| A5SS | 910 | ENSG00000068079 | IFI35 | chr17 | + | 43013266 | 43013379 | 43013266 | 43013373 | 43013475 | 43013662 | 68 | 32 | 43 | [illegible] |
| A5SS | 2565 | ENSG00000203875 | SNHG5 | chr6 | - | 85677790 | 85677875 | 85677794 | 85677875 | 85677423 | 85677492 | 0 | 2 | 0 | [illegible] |
| A5SS | 1171 | ENSG00000168701 | TMEM208 | chr16 | + | 67228494 | 67228657 | 67228494 | 67228631 | 67228796 | 67228881 | 9 | 1 | 7 | [illegible] |
| A5SS | 5655 | ENSG00000169689 | CENPX | chr17 | - | 82019292 | 82019381 | 82019307 | 82019381 | 82018702 | 82019219 | 27 | 8 | 20 | [illegible] |
| A5SS | 2507 | ENSG00000105202 | FBL | chr19 | - | 39840682 | 39840787 | 39840721 | 39840787 | 39840413 | 39840509 | 26 | 14 | 38 | [illegible] |
| A5SS | 6693 | ENSG00000204316 | MRPL38 | chr17 | - | 75904221 | 75904719 | 75904539 | 75904719 | 75902019 | 75902154 | 3 | 3 | 11 | [illegible] |
| A5SS | 6022 | ENSG00000089818 | NECAP1 | chr12 | + | 8090194 | 8091850 | 8090194 | 8090343 | 8092675 | 8092784 | 154 | 87 | 150 | [illegible] |
| A5SS | 4979 | ENSG00000128463 | EMC4 | chr15 | + | 34225535 | 34225658 | 34225535 | 34225650 | 34227692 | 34227846 | 5 | 1 | 3 | [illegible] |
| A5SS | 8743 | ENSG00000133641 | C12orf29 | chr12 | + | 88043598 | 88045779 | 88043598 | 88043715 | 88046806 | 88046976 | 97 | 52 | 107 | [illegible] |
| A5SS | 1038 | ENSG00000274487 | NPEPPSP1 | chr17 | - | 38202735 | 38202981 | 38202780 | 38202981 | 38202237 | 38202334 | 0 | 0 | 0 | [illegible] |
| A5SS | 8870 | ENSG00000135473 | PAN2 | chr12 | - | 56324036 | 56324182 | 56324048 | 56324182 | 56323806 | 56323913 | 13 | 6 | 9 | [illegible] |
| A5SS | 8224 | ENSG00000187147 | RNF220 | chr1 | + | 44635544 | 44635666 | 44635544 | 44635588 | 44636029 | 44636162 | 159 | 107 | 121 | [illegible] |
| A5SS | 6300 | ENSG00000214021 | TTLL3 | chr3 | + | 9826996 | 9827240 | 9826996 | 9827072 | 9828959 | 9829395 | 71 | 28 | 51 | [illegible] |
| A5SS | 412 | ENSG00000130475 | FCHO1 | chr19 | + | 17754316 | 17754466 | 17754316 | 17754361 | 17754591 | 17754685 | 28 | 6 | 23 | [illegible] |
| A5SS | 7972 | ENSG00000196323 | ZBTB44 | chr11 | - | 130236792 | 130237093 | 130237009 | 130237093 | 130236109 | 130236216 | 112 | 52 | 99 | [illegible] |
| A5SS | 1004 | ENSG00000126581 | BECN1 | chr17 | - | 42818543 | 42819609 | 42819547 | 42819609 | 42818220 | 42818415 | 277 | 180 | 195 | [illegible] |
| A5SS | 5370 | ENSG00000155729 | KCTD18 | chr2 | - | 200504747 | 200504959 | 200504769 | 200504959 | 200498890 | 200499084 | 123 | 85 | 99 | [illegible] |
| A5SS | 3376 | ENSG00000125347 | IRF1 | chr5 | - | 132486521 | 132486686 | 132486556 | 132486686 | 132486250 | 132486373 | 20 | 17 | 26 | [illegible] |
| A5SS | 3273 | ENSG00000132879 | FBXO44 | chr1 | + | 11658266 | 11658393 | 11658266 | 11658364 | 11658532 | 11658628 | 14 | 9 | 28 | [illegible] |
| A5SS | 5414 | ENSG00000163728 | TTC14 | chr3 | + | 180604224 | 180604338 | 180604224 | 180604309 | 180604477 | 180604607 | 10 | 10 | 5 | [illegible] |
| A5SS | 7688 | ENSG00000011454 | RABGAP1 | chr9 | + | 123065347 | 123065601 | 123065347 | 123065461 | 123070349 | 123070424 | 4 | 11 | 16 | [illegible] |
| A5SS | 6832 | ENSG00000124333 | VAMP7 | chrX | + | 155889457 | 155889612 | 155889457 | 155889547 | 155895622 | 155895680 | 316 | 202 | 233 | [illegible] |
| A5SS | 7011 | ENSG00000244754 | N4BP2L2 | chr13 | - | 32535768 | 32537027 | 32537005 | 32537027 | 32527407 | 32527532 | 825 | 699 | 702 | [illegible] |
| A5SS | 1091 | ENSG00000125124 | BBS2 | chr16 | - | 56511132 | 56511284 | 56511158 | 56511284 | 56510858 | 56510921 | 1 | 2 | 5 | [illegible] |
| A5SS | 2577 | ENSG00000171566 | PLRG1 | chr4 | - | 154547010 | 154547064 | 154547037 | 154547064 | 154546122 | 154546213 | 198 | 117 | 160 | [illegible] |
| A5SS | 8700 | ENSG00000229117 | RPL41 | chr12 | + | 56116774 | 56116930 | 56116774 | 56116829 | 56117188 | 56117211 | 39 | 35 | 23 | [illegible] |
| A5SS | 3395 | ENSG00000170581 | STAT2 | chr12 | - | 56349344 | 56349509 | 56349425 | 56349509 | 56348751 | 56348851 | 37 | 39 | 37 | [illegible] |
| A5SS | 5467 | ENSG00000135838 | NPL | chr1 | + | 182806144 | 182806531 | 182806144 | 182806232 | 182812155 | 182812213 | 17 | 8 | 4 | [illegible] |
| A5SS | 7783 | ENSG00000137145 | DENND4C | chr9 | + | 19331977 | 19332188 | 19331977 | 19332184 | 19334976 | 19335105 | 6 | 5 | 3 | [illegible] |
| A5SS | 8322 | ENSG00000138286 | FAM149B1 | chr10 | + | 73232938 | 73233197 | 73232938 | 73233163 | 73234816 | 73234940 | 5 | 6 | 5 | [illegible] |
| A5SS | 3874 | ENSG00000164010 | ERMAP | chr1 | + | 42835731 | 42835764 | 42835731 | 42835760 | 42837157 | 42837190 | 15 | 13 | 18 | [illegible] |
| A5SS | 88 | ENSG00000100413 | POLR3H | chr22 | - | 41532603 | 41532745 | 41532658 | 41532745 | 41532093 | 41532157 | 25 | 1 | 20 | [illegible] |
| A5SS | 6833 | ENSG00000124333 | VAMP7 | chrX | + | 155889457 | 155889633 | 155889457 | 155889547 | 155895622 | 155895680 | 59 | 32 | 41 | [illegible] |
| A5SS | 825 | ENSG00000136448 | NMT1 | chr17 | + | 45081643 | 45081995 | 45081643 | 45081752 | 45083657 | 45083755 | 14 | 6 | 5 | [illegible] |
| A5SS | 2779 | ENSG00000242086 | MUC20-OT1 | chr3 | + | 195708153 | 195708513 | 195708153 | 195708268 | 195708746 | 195708841 | 11 | 4 | 18 | [illegible] |
| A5SS | 7831 | ENSG00000110756 | HPS5 | chr11 | - | 18321945 | 18322174 | 18322132 | 18322174 | 18311913 | 18312024 | 15 | 8 | 9 | [illegible] |
| A5SS | 8662 | ENSG00000137955 | RABGGTB | chr1 | + | 75787496 | 75790517 | 75787496 | 75787604 | 75791284 | 75791337 | 206 | 202 | 200 | [illegible] |
| A5SS | 8000 | ENSG00000171634 | BPTF | chr17 | + | 67945408 | 67946325 | 67945408 | 67945896 | 67947725 | 67947808 | 122 | 96 | 90 | [illegible] |
| A5SS | 4383 | ENSG00000183918 | SH2D1A | chrX | + | 124370175 | 124370320 | 124370175 | 124370311 | 124371350 | 124371488 | 94 | 30 | 147 | [illegible] |
| A5SS | 8706 | ENSG00000132341 | RAN | chr12 | + | 130872835 | 130872920 | 130872835 | 130872912 | 130873002 | 130873128 | 621 | 392 | 578 | [illegible] |
| A5SS | 567 | ENSG00000196756 | SNHG17 | chr20 | - | 38422095 | 38422984 | 38422290 | 38422984 | 38421005 | 38421098 | 22 | 15 | 28 | [illegible] |
| A5SS | 337 | ENSG00000167613 | LAIR1 | chr19 | - | 54359842 | 54360072 | 54360021 | 54360072 | 54358005 | 54358625 | 11 | 5 | 2 | [illegible] |
| A5SS | 768 | ENSG00000167699 | GLOD4 | chr17 | - | 778621 | 778744 | 778694 | 778744 | 776867 | 776988 | 24 | 12 | 34 | [illegible] |
| A5SS | 3514 | ENSG00000171163 | ZNF692 | chr1 | - | 248856287 | 248856422 | 248856372 | 248856422 | 248855557 | 248855946 | 52 | 35 | 49 | [illegible] |
| A5SS | 1636 | ENSG00000100445 | SDR39U1 | chr14 | - | 24442097 | 24442260 | 24442177 | 24442260 | 24440782 | 24440926 | 35 | 13 | 22 | [illegible] |
| A5SS | 6110 | ENSG00000111846 | GCNT2 | chr6 | + | 10528630 | 10529836 | 10528630 | 10528841 | 10621350 | 10621443 | 37 | 45 | 30 | [illegible] |
| A5SS | 2257 | ENSG00000172239 | PAIP1 | chr5 | - | 43556581 | 43557077 | 43556818 | 43557077 | 43555829 | 43555999 | 9 | 3 | 12 | [illegible] |
| A5SS | 7552 | ENSG00000257365 | FNTB | chr14 | + | 65032358 | 65032521 | 65032358 | 65032503 | 65032609 | 65032696 | 1 | 1 | 0 | [illegible] |
| A5SS | 2192 | ENSG00000143774 | GUK1 | chr1 | + | 228141123 | 228141507 | 228141123 | 228141288 | 228145510 | 228145624 | 19 | 3 | 31 | [illegible] |
| A5SS | 1437 | ENSG00000034053 | APBA2 | chr15 | + | 28921639 | 28921749 | 28921639 | 28921718 | 28995752 | 28995806 | 94 | 44 | 62 | [illegible] |
| A5SS | 4338 | ENSG00000029534 | ANK1 | chr8 | - | 41661800 | 41661941 | 41661875 | 41661941 | 41655300 | 41655753 | 15 | 17 | 11 | [illegible] |
| A5SS | 5542 | ENSG00000147454 | SLC25A37 | chr8 | + | 23566107 | 23566813 | 23566107 | 23566336 | 23568321 | 23568378 | 60 | 55 | 80 | [illegible] |
| A5SS | 5643 | ENSG00000132323 | ILKAP | chr2 | - | 238189858 | 238189972 | 238189862 | 238189972 | 238188130 | 238188257 | 5 | 2 | 4 | [illegible] |
| A5SS | 719 | ENSG00000219200 | RNASEK | chr17 | + | 7013318 | 7013535 | 7013318 | 7013516 | 7013665 | 7013742 | 11 | 4 | 5 | [illegible] |
| A5SS | 2139 | ENSG00000113282 | CLINT1 | chr5 | - | 157858929 | 157859145 | 157858952 | 157859145 | 157817442 | 157817547 | 94 | 52 | 65 | [illegible] |
| A5SS | 6872 | ENSG00000131196 | NFATC1 | chr18 | + | 79450953 | 79451126 | 79450953 | 79451041 | 79451675 | 79451816 | 53 | 36 | 77 | [illegible] |
| A5SS | 7995 | ENSG00000135249 | RINT1 | chr7 | + | 105536564 | 105536749 | 105536564 | 105536612 | 105542407 | 105542649 | 96 | 62 | 81 | [illegible] |
| A5SS | 5397 | ENSG00000153250 | RBMS1 | chr2 | - | 160286968 | 160287084 | 160286977 | 160287084 | 160284994 | 160285044 | 226 | 176 | 136 | [illegible] |
| A5SS | 112 | ENSG00000100242 | SUN2 | chr22 | - | 38740888 | 38741050 | 38741006 | 38741050 | 38739721 | 38739943 | 33 | 16 | 30 | [illegible] |
| A5SS | 7843 | ENSG00000149428 | HYOU1 | chr11 | - | 119057019 | 119057137 | 119057026 | 119057137 | 119056069 | 119056167 | 26 | 14 | 39 | [illegible] |
| A5SS | 1637 | ENSG00000100445 | SDR39U1 | chr14 | - | 24442108 | 24442260 | 24442177 | 24442260 | 24440782 | 24440926 | 33 | 11 | 13 | [illegible] |
| A5SS | 2848 | ENSG00000075711 | DLG1 | chr3 | - | 197085295 | 197085756 | 197085579 | 197085756 | 197081050 | 197081117 | 17 | 13 | 7 | [illegible] |
| A5SS | 3125 | ENSG00000041357 | PSMA4 | chr15 | + | 78540426 | 78540539 | 78540426 | 78540463 | 78541904 | 78541930 | 547 | 349 | 494 | [illegible] |
| A5SS | 8055 | ENSG00000175294 | CATSPER1 | chr11 | - | 66018787 | 66018902 | 66018826 | 66018902 | 66017059 | 66017174 | 19 | 11 | 9 | [illegible] |
| A5SS | 6862 | ENSG00000148985 | PGAP2 | chr11 | + | 3824269 | 3824449 | 3824269 | 3824376 | 3825019 | 3825128 | 4 | 6 | 8 | [illegible] |
| A5SS | 1505 | ENSG00000140521 | POLG | chr15 | - | 89317938 | 89318749 | 89318540 | 89318749 | 89317375 | 89317536 | 43 | 26 | 47 | [illegible] |
| A5SS | 5533 | ENSG00000168918 | INPP5D | chr2 | + | 233163706 | 233163926 | 233163706 | 233163903 | 233164306 | 233164424 | 3 | 0 | 3 | [illegible] |
| A5SS | 5657 | ENSG00000164548 | TRA2A | chr7 | - | 23522120 | 23522432 | 23522131 | 23522432 | 23521706 | 23521840 | 74 | 80 | 60 | [illegible] |
| A5SS | 593 | ENSG00000069399 | BCL3 | chr19 | + | 44758245 | 44758473 | 44758245 | 44758413 | 44758723 | 44758841 | 0 | 1 | 8 | [illegible] |
| A5SS | 6408 | ENSG00000145191 | EIF2B5 | chr3 | + | 184140079 | 184140740 | 184140079 | 184140730 | 184141924 | 184142070 | 2 | 3 | 4 | [illegible] |
| A5SS | 2508 | ENSG00000105202 | FBL | chr19 | - | 39840616 | 39840787 | 39840619 | 39840787 | 39840413 | 39840515 | 29 | 12 | 24 | [illegible] |
| A5SS | 3493 | ENSG00000183258 | DDX41 | chr5 | - | 177514553 | 177514837 | 177514700 | 177514837 | 177513684 | 177513847 | 3 | 3 | 2 | [illegible] |
| A5SS | 7767 | ENSG00000115459 | ELMOD3 | chr2 | + | 85355554 | 85355896 | 85355554 | 85355716 | 85356966 | 85357252 | 19 | 18 | 15 | [illegible] |
| A5SS | 4138 | ENSG00000241878 | PISD | chr22 | - | 31621333 | 31621885 | 31621648 | 31621885 | 31620995 | 31621142 | 109 | 75 | 109 | [illegible] |
| A5SS | 5582 | ENSG00000197622 | CDC42SE1 | chr1 | - | 151056650 | 151056786 | 151056655 | 151056786 | 151055676 | 151055993 | 142 | 86 | 113 | [illegible] |
| A5SS | 2308 | ENSG00000127184 | COX7C | chr5 | + | 86619352 | 86619878 | 86619352 | 86619496 | 86620667 | 86620766 | 20 | 16 | 23 | [illegible] |
| A5SS | 8342 | ENSG00000156650 | KAT6B | chr10 | + | 74975398 | 74976330 | 74975398 | 74975781 | 74977315 | 74977437 | 94 | 77 | 91 | [illegible] |
| A5SS | 3631 | ENSG00000167962 | ZNF598 | chr16 | - | 2000390 | 2000481 | 2000408 | 2000481 | 1999495 | 2000256 | 19 | 12 | 7 | [illegible] |
| A5SS | 6578 | ENSG00000115677 | HDLBP | chr2 | - | 241230068 | 241230269 | 241230152 | 241230269 | 241229832 | 241229961 | 0 | 3 | 6 | [illegible] |
| A5SS | 7778 | ENSG00000180773 | SLC36A4 | chr11 | - | 93182805 | 93182894 | 93182834 | 93182894 | 93181690 | 93181786 | 83 | 98 | 50 | [illegible] |
| A5SS | 931 | ENSG00000173821 | RNF213 | chr17 | + | 80388611 | 80389367 | 80388611 | 80388689 | 80389827 | 80389917 | 545 | 335 | 461 | [illegible] |
| A5SS | 4111 | ENSG00000111331 | OAS3 | chr12 | + | 112948860 | 112949205 | 112948860 | 112949130 | 112950692 | 112950975 | 95 | 85 | 64 | [illegible] |
| A5SS | 4692 | ENSG00000042445 | RETSAT | chr2 | - | 85345512 | 85346094 | 85345974 | 85346094 | 85344593 | 85344732 | 0 | 5 | 5 | [illegible] |
| A5SS | 6946 | ENSG00000102181 | CD99L2 | chrX | - | 150777335 | 150777482 | 150777443 | 150777482 | 150776173 | 150776293 | 0 | 0 | 2 | [illegible] |
| A5SS | 4608 | ENSG00000165689 | ENTR1 | chr9 | - | 136405090 | 136405202 | 136405117 | 136405202 | 136404630 | 136404693 | 60 | 38 | 77 | [illegible] |
| A5SS | 5402 | ENSG00000124151 | NCOA3 | chr20 | + | 47649004 | 47649109 | 47649004 | 47649097 | 47650981 | 47651276 | 9 | 17 | 7 | [illegible] |
| A5SS | 3144 | ENSG00000276291 | FRG1HP | chr9 | + | 40992260 | 40992775 | 40992260 | 40992479 | 40993252 | 40993434 | 51 | 36 | 77 | [illegible] |
| A5SS | 571 | ENSG00000196756 | SNHG17 | chr20 | - | 38435026 | 38435352 | 38435196 | 38435352 | 38434503 | 38434626 | 3 | 1 | 8 | [illegible] |
| A5SS | 8773 | ENSG00000063046 | EIF4B | chr12 | + | 53034611 | 53034724 | 53034611 | 53034709 | 53037408 | 53037622 | 224 | 130 | 259 | [illegible] |
| A5SS | 7376 | ENSG00000249859 | PVT1 | chr8 | + | 127795345 | 127796071 | 127795345 | 127796008 | 127890588 | 127890998 | 10 | 4 | 4 | [illegible] |
| A5SS | 6180 | ENSG00000132424 | PNISR | chr6 | - | 99403828 | 99404702 | 99404602 | 99404702 | 99402539 | 99402710 | 408 | 449 | 436 | [illegible] |
| A5SS | 1764 | ENSG00000165959 | CLMN | chr14 | - | 95194214 | 95194596 | 95194535 | 95194596 | 95193848 | 95193919 | 16 | 26 | 17 | [illegible] |
| A5SS | 243 | ENSG00000011304 | PTBP1 | chr19 | + | 805012 | 807902 | 805012 | 805187 | 808359 | 808452 | 199 | 81 | 158 | [illegible] |
| A5SS | 2776 | ENSG00000242086 | MUC20-OT1 | chr3 | + | 195708133 | 195708435 | 195708133 | 195708268 | 195708746 | 195708801 | 12 | 4 | 21 | [illegible] |

| | | | | | | |
|---|---|---|---|---|---|---|
| A5SS | 1051 | ENSG00000172530 | BANP | chr16 | + | [illegible] |
| A5SS | 4272 | ENSG00000176393 | RNPEP | chr1 | + | [illegible] |
| A5SS | 395 | ENSG00000089335 | ZNF302 | chr19 | + | [illegible] |
| A5SS | 2307 | ENSG00000127184 | COX7C | chr5 | + | [illegible] |
| A5SS | 6814 | ENSG00000196123 | KIAA0895L | chr16 | - | [illegible] |
| A5SS | 7306 | ENSG00000165806 | CASP7 | chr10 | + | [illegible] |
| A5SS | 3856 | ENSG00000068097 | HEATR6 | chr17 | - | [illegible] |
| A5SS | 3459 | ENSG00000143578 | CREB3L4 | chr1 | + | [illegible] |
| A5SS | 8007 | ENSG00000168071 | CCDC88B | chr11 | + | [illegible] |
| A5SS | 6658 | ENSG00000122566 | HNRNPA2B1 | chr7 | - | [illegible] |
| A5SS | 7550 | ENSG00000165233 | CARD19 | chr9 | + | [illegible] |
| A5SS | 3526 | ENSG00000163374 | YY1AP1 | chr1 | - | [illegible] |
| A5SS | 1903 | ENSG00000226067 | LINC00623 | chr1 | + | [illegible] |
| A5SS | 366 | ENSG00000104859 | CLASRP | chr19 | + | [illegible] |
| A5SS | 8310 | ENSG00000183426 | NPIPA1 | chr16 | + | [illegible] |
| A5SS | 1467 | ENSG00000166855 | CLPX | chr15 | - | [illegible] |
| A5SS | 2634 | ENSG00000055955 | ITIH4 | chr3 | - | [illegible] |
| A5SS | 4420 | ENSG00000173575 | CHD2 | chr15 | + | [illegible] |
| A5SS | 6072 | ENSG00000100503 | NIN | chr14 | - | [illegible] |
| A5SS | 3204 | ENSG00000114850 | SSR3 | chr3 | - | [illegible] |
| A5SS | 7408 | ENSG00000110077 | MS4A6A | chr11 | - | [illegible] |
| A5SS | 8478 | ENSG00000150093 | ITGB1 | chr10 | - | [illegible] |
| A5SS | 8564 | ENSG00000205356 | TECPR1 | chr7 | - | [illegible] |
| A5SS | 6 | ENSG00000186866 | POFUT2 | chr21 | - | [illegible] |
| A5SS | 3290 | ENSG00000117054 | ACADM | chr1 | + | [illegible] |
| A5SS | 4824 | ENSG00000139631 | CSAD | chr12 | - | [illegible] |
| A5SS | 1101 | ENSG00000166747 | AP1G1 | chr16 | - | [illegible] |
| A5SS | 6943 | ENSG00000197620 | EOLA1 | chrX | + | [illegible] |
| A5SS | 6231 | ENSG00000044090 | CUL7 | chr6 | - | [illegible] |
| A5SS | 2160 | ENSG00000124782 | RREB1 | chr6 | + | [illegible] |
| A5SS | 3828 | ENSG00000163714 | U2SURP | chr3 | + | [illegible] |
| A5SS | 311 | ENSG00000077312 | SNRPA | chr19 | + | [illegible] |
| A5SS | 2894 | ENSG00000123737 | EXOSC9 | chr4 | + | [illegible] |
| A5SS | 2891 | ENSG00000043093 | DCUN1D1 | chr3 | - | [illegible] |
| A5SS | 5500 | ENSG00000140368 | PSTPIP1 | chr15 | + | [illegible] |
| A5SS | 3207 | ENSG00000189180 | ZNF33A | chr10 | + | [illegible] |
| A5SS | 5593 | ENSG00000143390 | RFX5 | chr1 | - | [illegible] |
| A5SS | 7123 | ENSG00000258890 | CEP95 | chr17 | + | [illegible] |
| A5SS | 543 | ENSG00000101365 | IDH3B | chr20 | - | [illegible] |
| A5SS | 343 | ENSG00000167555 | ZNF528 | chr19 | + | [illegible] |
| A5SS | 7288 | ENSG00000174197 | MGA | chr15 | + | [illegible] |
| A5SS | 4270 | ENSG00000160087 | UBE2J2 | chr1 | - | [illegible] |
| A5SS | 6531 | ENSG00000242294 | STAG3L5P | chr7 | + | [illegible] |
| A5SS | 5289 | ENSG00000224078 | SNHG14 | chr15 | + | [illegible] |
| A5SS | 5780 | ENSG00000064666 | CNN2 | chr19 | + | [illegible] |
| A5SS | 5987 | ENSG00000111679 | PTPN6 | chr12 | + | [illegible] |
| A5SS | 5588 | ENSG00000143390 | RFX5 | chr1 | - | [illegible] |
| A5SS | 2896 | ENSG00000188313 | PLSCR1 | chr3 | - | [illegible] |
| A5SS | 2320 | ENSG00000198689 | SLC9A6 | chrX | + | [illegible] |
| A5SS | 1047 | ENSG00000091592 | NLRP1 | chr17 | - | [illegible] |
| A5SS | 3850 | ENSG00000173230 | GOLGB1 | chr3 | - | [illegible] |
| A5SS | 3122 | ENSG00000145819 | ARHGAP26 | chr5 | + | [illegible] |
| A5SS | 5336 | ENSG00000151623 | NR3C2 | chr4 | - | [illegible] |
| A5SS | 4451 | ENSG00000170271 | FAXDC2 | chr5 | - | [illegible] |
| A5SS | 2477 | ENSG00000214063 | TSPAN4 | chr11 | + | [illegible] |
| A5SS | 6204 | ENSG00000112739 | PRPF4B | chr6 | + | [illegible] |
| A5SS | 7174 | ENSG00000215301 | DDX3X | chrX | + | [illegible] |
| A5SS | 1274 | ENSG00000103472 | RRN3P2 | chr16 | + | [illegible] |
| A5SS | 4955 | ENSG00000129933 | MAU2 | chr19 | + | [illegible] |
| A5SS | 683 | ENSG00000178127 | NDUFV2 | chr18 | + | [illegible] |
| A5SS | 1373 | ENSG00000007376 | RPUSD1 | chr16 | - | [illegible] |
| A5SS | 5034 | ENSG00000119778 | ATAD2B | chr2 | - | [illegible] |
| A5SS | 4419 | ENSG00000111653 | ING4 | chr12 | - | [illegible] |
| A5SS | 6860 | ENSG00000148985 | PGAP2 | chr11 | + | [illegible] |
| A5SS | 8166 | ENSG00000078124 | ACER3 | chr11 | + | [illegible] |
| A5SS | 4861 | ENSG00000149532 | CPSF7 | chr11 | - | [illegible] |
| A5SS | 346 | ENSG00000268027 | NA | chr19 | + | [illegible] |
| A5SS | 1736 | ENSG00000100888 | CHD8 | chr14 | - | [illegible] |
| A5SS | 156 | ENSG00000104936 | DMPK | chr19 | - | [illegible] |
| A5SS | 7838 | ENSG00000133943 | DGLUCY | chr14 | + | [illegible] |
| A5SS | 8096 | ENSG00000140749 | IGSF6 | chr16 | - | [illegible] |
| A5SS | 7017 | ENSG00000244754 | N4BP2L2 | chr13 | - | [illegible] |
| A5SS | 3117 | ENSG00000109861 | CTSC | chr11 | - | [illegible] |
| A5SS | 5772 | ENSG00000169247 | SH3TC2 | chr5 | - | [illegible] |
| A5SS | 4501 | ENSG00000072110 | ACTN1 | chr14 | - | [illegible] |
| A5SS | 8316 | ENSG00000090621 | PABPC4 | chr1 | - | [illegible] |
| A5SS | 2638 | ENSG00000196209 | SIRPB2 | chr20 | - | [illegible] |
| A5SS | 530 | ENSG00000160410 | SHKBP1 | chr19 | + | [illegible] |
| A5SS | 7593 | ENSG00000196781 | TLE1 | chr9 | - | [illegible] |
| A5SS | 6414 | ENSG00000134291 | TMEM106C | chr12 | + | [illegible] |
| A5SS | 3665 | ENSG00000196628 | TCF4 | chr18 | - | [illegible] |
| A5SS | 2483 | ENSG00000145907 | G3BP1 | chr5 | + | [illegible] |
| A5SS | 6031 | ENSG00000114388 | NPRL2 | chr3 | - | [illegible] |
| A5SS | 8758 | ENSG00000002016 | RAD52 | chr12 | - | [illegible] |
| A5SS | 1502 | ENSG00000114062 | UBE3A | chr15 | - | [illegible] |
| A5SS | 2505 | ENSG00000155229 | MMS19 | chr10 | - | [illegible] |
| A5SS | 8542 | ENSG00000069493 | CLEC2D | chr12 | + | [illegible] |
| A5SS | 7934 | ENSG00000162300 | ZFPL1 | chr11 | + | [illegible] |
| A5SS | 1385 | ENSG00000125877 | ITPA | chr20 | + | [illegible] |
| A5SS | 1638 | ENSG00000100445 | SDR39U1 | chr14 | - | [illegible] |
| A5SS | 4356 | ENSG00000150455 | TIRAP | chr11 | + | [illegible] |
| A5SS | 3958 | ENSG00000144357 | UBR3 | chr2 | + | [illegible] |
| A5SS | 2604 | ENSG00000113522 | RAD50 | chr5 | + | [illegible] |
| A5SS | 839 | ENSG00000185359 | HGS | chr17 | + | [illegible] |
| A5SS | 6859 | ENSG00000148985 | PGAP2 | chr11 | + | [illegible] |
| A5SS | 2752 | ENSG00000278259 | MYO19 | chr17 | - | [illegible] |
| A5SS | 3734 | ENSG00000026508 | CD44 | chr11 | + | [illegible] |
| A5SS | 7890 | ENSG00000266173 | STRADA | chr17 | - | [illegible] |
| A5SS | 8932 | ENSG00000198176 | TFDP1 | chr13 | + | [illegible] |
| A5SS | 6064 | ENSG00000114423 | CBLB | chr3 | - | [illegible] |
| A5SS | 8117 | ENSG00000110321 | EIF4G2 | chr11 | - | [illegible] |
| A5SS | 3599 | ENSG00000137409 | MTCH1 | chr6 | - | [illegible] |
| A5SS | 8135 | ENSG00000110075 | PPP6R3 | chr11 | + | [illegible] |
| A5SS | 1230 | ENSG00000214078 | CPNE1 | chr20 | - | [illegible] |
| A5SS | 7231 | ENSG00000182196 | ARL6IP4 | chr12 | + | [illegible] |
| A5SS | 7624 | ENSG00000080802 | CNOT4 | chr7 | - | [illegible] |
| A5SS | 2735 | ENSG00000129071 | MBD4 | chr3 | - | [illegible] |
| A5SS | 4223 | ENSG00000104231 | ZFAND1 | chr8 | - | [illegible] |
| A5SS | 2061 | ENSG00000172262 | ZNF131 | chr5 | + | [illegible] |
| A5SS | 307 | ENSG00000130669 | PAK4 | chr19 | + | [illegible] |
| A5SS | 487 | ENSG00000087586 | AURKA | chr20 | - | [illegible] |
| A5SS | 1127 | ENSG00000167207 | NOD2 | chr16 | + | [illegible] |
| A5SS | 1402 | ENSG00000161996 | WDR90 | chr16 | + | [illegible] |
| A5SS | 1552 | ENSG00000214432 | VPS33B-DT | chr15 | + | [illegible] |
| A5SS | 2252 | ENSG00000249738 | NA | chr5 | + | [illegible] |
| A5SS | 3302 | ENSG00000012171 | SEMA3B | chr3 | + | [illegible] |
| A5SS | 3860 | ENSG00000106714 | CNTNAP3 | chr9 | - | [illegible] |
| A5SS | 4693 | ENSG00000158528 | PPP1R9A | chr7 | + | [illegible] |
| A5SS | 5080 | ENSG00000143036 | SLC44A3 | chr1 | + | [illegible] |
| A5SS | 5947 | ENSG00000162723 | SLAMF9 | chr1 | - | [illegible] |
| A5SS | 6844 | ENSG00000158813 | EDA | chrX | + | [illegible] |
| A5SS | 7311 | ENSG00000164024 | METAP1 | chr4 | + | [illegible] |
| A5SS | 7611 | ENSG00000168003 | SLC3A2 | chr11 | + | [illegible] |
| A5SS | 7864 | ENSG00000245573 | BDNF-AS | chr11 | + | [illegible] |
| A5SS | 7865 | ENSG00000245573 | BDNF-AS | chr11 | + | [illegible] |
| A5SS | 8690 | ENSG00000144061 | NPHP1 | chr2 | - | [illegible] |
| A5SS | 8691 | ENSG00000144061 | NPHP1 | chr2 | - | [illegible] |
| A5SS | 6317 | ENSG00000106524 | ANKMY2 | chr7 | - | [illegible] |
| A5SS | 3342 | ENSG00000158485 | CD1B | chr1 | - | [illegible] |
| A5SS | 4006 | ENSG00000214595 | EML6 | chr2 | + | [illegible] |
| A5SS | 2418 | ENSG00000125744 | RTN2 | chr19 | - | [illegible] |
| A5SS | 4219 | ENSG00000175262 | C1orf127 | chr1 | - | [illegible] |
| A5SS | 942 | ENSG00000267123 | SCAT1 | chr17 | - | [illegible] |
| A5SS | 4948 | ENSG00000173214 | MFSD4B | chr6 | + | [illegible] |
| A5SS | 1273 | ENSG00000261740 | BOLA2-SMG1P6 | chr16 | - | [illegible] |
| A5SS | 4100 | ENSG00000136895 | GARNL3 | chr9 | + | [illegible] |
| A5SS | 8791 | ENSG00000167371 | PRRT2 | chr16 | + | [illegible] |
| A5SS | 4924 | ENSG00000187741 | FANCA | chr16 | - | [illegible] |
| A5SS | 2765 | ENSG00000122970 | IFT81 | chr12 | + | [illegible] |
| A5SS | 8352 | ENSG00000255624 | C10orf88B | chr10 | + | [illegible] |
| A5SS | 2954 | ENSG00000101049 | SGK2 | chr20 | + | [illegible] |
| A5SS | 3934 | ENSG00000133116 | KL | chr13 | + | [illegible] |
| A5SS | 3630 | ENSG00000167550 | RHEBL1 | chr12 | - | [illegible] |
| A5SS | 4007 | ENSG00000214595 | EML6 | chr2 | + | [illegible] |
| A5SS | 7792 | ENSG00000130714 | POMT1 | chr9 | + | [illegible] |
| A5SS | 5195 | ENSG00000167220 | HDHD2 | chr18 | - | [illegible] |
| A5SS | 5722 | ENSG00000153253 | SCN3A | chr2 | - | [illegible] |
| A5SS | 3543 | ENSG00000197343 | ZNF655 | chr7 | + | [illegible] |
| A5SS | 459 | ENSG00000258366 | RTEL1 | chr20 | + | [illegible] |
| A5SS | 2221 | ENSG00000204120 | GIGYF2 | chr2 | + | [illegible] |
| A5SS | 2871 | ENSG00000172053 | QARS1 | chr3 | - | [illegible] |
| A5SS | 3784 | ENSG00000260271 | NA | chr6 | + | [illegible] |
| A5SS | 4016 | ENSG00000102048 | ASB9 | chrX | - | [illegible] |
| A5SS | 6896 | ENSG00000142675 | CNKSR1 | chr1 | + | [illegible] |
| A5SS | 7520 | ENSG00000147862 | NFIB | chr9 | - | [illegible] |
| A5SS | 8912 | ENSG00000198246 | SLC29A3 | chr10 | + | [illegible] |
| A5SS | 6864 | ENSG00000148985 | PGAP2 | chr11 | + | [illegible] |
| A5SS | 4764 | ENSG00000115594 | IL1R1 | chr2 | + | [illegible] |
| A5SS | 4267 | ENSG00000160087 | UBE2J2 | chr1 | - | [illegible] |
| A5SS | 1307 | ENSG00000260630 | SNAI3-AS1 | chr16 | + | [illegible] |
| A5SS | 2412 | ENSG00000163812 | ZDHHC3 | chr3 | - | [illegible] |
| A5SS | 7476 | ENSG00000147687 | TATDN1 | chr8 | - | [illegible] |
| A5SS | 6602 | ENSG00000071462 | BUD23 | chr7 | + | [illegible] |
| A5SS | 4065 | ENSG00000143416 | SELENBP1 | chr1 | - | [illegible] |
| A5SS | 7923 | ENSG00000254536 | NA | chr10 | + | [illegible] |
| A5SS | 994 | ENSG00000168259 | DNAJC7 | chr17 | - | [illegible] |
| A5SS | 743 | ENSG00000141560 | FN3KRP | chr17 | + | [illegible] |
| A5SS | 354 | ENSG00000022556 | NLRP2 | chr19 | + | [illegible] |
| A5SS | 6100 | ENSG00000221838 | AP4M1 | chr7 | + | [illegible] |
| A5SS | 4034 | ENSG00000126091 | ST3GAL3 | chr1 | + | [illegible] |
| A5SS | 4306 | ENSG00000043514 | TRIT1 | chr1 | - | [illegible] |
| A5SS | 370 | ENSG00000161298 | ZNF382 | chr19 | + | [illegible] |
| A5SS | 6555 | ENSG00000152207 | CYSLTR2 | chr13 | + | [illegible] |
| A5SS | 8916 | ENSG00000227354 | RBM26-AS1 | chr13 | + | [illegible] |
| A5SS | 4002 | ENSG00000197530 | MIB2 | chr1 | + | [illegible] |
| A5SS | 4448 | ENSG00000090097 | PCBP4 | chr3 | - | [illegible] |
| A5SS | 2502 | ENSG00000106692 | FKTN | chr9 | + | [illegible] |
| A5SS | 3018 | ENSG00000187609 | EXD3 | chr9 | - | [illegible] |
| A5SS | 193 | ENSG00000182472 | CAPN12 | chr19 | - | [illegible] |
| A5SS | 390 | ENSG00000161249 | DMKN | chr19 | - | [illegible] |
| A5SS | 428 | ENSG00000131355 | ADGRE3 | chr19 | - | [illegible] |
| A5SS | 1515 | ENSG00000138606 | SHF | chr15 | - | [illegible] |
| A5SS | 1712 | ENSG00000172717 | FAM71D | chr14 | + | [illegible] |
| A5SS | 2011 | ENSG00000155980 | KIF5A | chr12 | + | [illegible] |
| A5SS | 2108 | ENSG00000242110 | AMACR | chr5 | - | [illegible] |
| A5SS | 2727 | ENSG00000161013 | MGAT4B | chr5 | - | [illegible] |
| A5SS | 3861 | ENSG00000152527 | PLEKHH2 | chr2 | + | [illegible] |
| A5SS | 5163 | ENSG00000283050 | GTF2IP12 | chr4 | - | [illegible] |
| A5SS | 5598 | ENSG00000109805 | NCAPG | chr4 | + | [illegible] |
| A5SS | 5886 | ENSG00000143436 | MRPL9 | chr1 | - | [illegible] |
| A5SS | 6066 | ENSG00000059758 | CDK17 | chr12 | - | [illegible] |
| A5SS | 6772 | ENSG00000168078 | PBK | chr8 | - | [illegible] |
| A5SS | 6791 | ENSG00000006757 | PNPLA4 | chrX | - | [illegible] |
| A5SS | 7269 | ENSG00000160781 | PAQR6 | chr1 | - | [illegible] |
| A5SS | 7544 | ENSG00000148057 | IDNK | chr9 | + | [illegible] |
| A5SS | 7991 | ENSG00000239523 | MYLK-AS1 | chr3 | + | [illegible] |
| A5SS | 8751 | ENSG00000110881 | ASIC1 | chr12 | + | [illegible] |
| A5SS | 5187 | ENSG00000145241 | CENPC | chr4 | - | [illegible] |
| A5SS | 968 | ENSG00000108423 | TUBD1 | chr17 | - | [illegible] |
| A5SS | 7629 | ENSG00000167094 | TTC16 | chr9 | + | [illegible] |
| A5SS | 1524 | ENSG00000167077 | MEI1 | chr22 | + | [illegible] |
| A5SS | 3751 | ENSG00000167384 | ZNF180 | chr19 | - | [illegible] |
| A5SS | 6506 | ENSG00000179818 | PCBP1-AS1 | chr2 | - | [illegible] |
| A5SS | 8793 | ENSG00000090686 | USP48 | chr1 | - | [illegible] |
| A5SS | 8832 | ENSG00000198015 | MRPL42 | chr12 | + | [illegible] |
| A5SS | 6379 | ENSG00000158321 | AUTS2 | chr7 | + | [illegible] |
| A5SS | 6188 | ENSG00000269313 | MAGIX | chrX | + | [illegible] |
| A5SS | 5840 | ENSG00000072195 | SPEG | chr2 | + | [illegible] |
| A5SS | 6417 | ENSG00000236753 | MKLN1-AS | chr7 | - | [illegible] |

| | | | | | | | | | | | | | | |
|---|---|---|---|---|---|---|---|---|---|---|---|---|---|---|
| A5SS | 6215 | ENSG00000197279 | ZNF165 | chr6 | + | 28080567 | 28081530 | 28080567 | 28080970 | 28085480 | 28085891 | 0 | 0 | 2 |
| A5SS | 3849 | ENSG00000173230 | GOLGB1 | chr3 | - | 121669196 | 121669355 | 121669211 | 121669355 | 121668313 | 121668508 | 2 | 4 | 2 |
| A5SS | 3690 | ENSG00000077782 | FGFR1 | chr8 | - | 38428345 | 38428435 | 38428351 | 38428435 | 38427920 | 38428093 | 0 | 0 | 0 |
| A5SS | 7083 | ENSG00000012048 | BRCA1 | chr17 | - | 43125270 | 43125370 | 43125276 | 43125370 | 43124016 | 43124115 | 2 | 1 | 3 |
| A5SS | 8333 | ENSG00000197771 | MCMBP | chr10 | - | 119843247 | 119843426 | 119843253 | 119843426 | 119840842 | 119840960 | 3 | 1 | 3 |
| A5SS | 8581 | ENSG00000284770 | TBCE | chr1 | + | 235438768 | 235438988 | 235438768 | 235438926 | 235441813 | 235441882 | 0 | 4 | 0 |
| A5SS | 1778 | ENSG00000141428 | C18orf21 | chr18 | + | 35972624 | 35972840 | 35972624 | 35972805 | 35974895 | 35975095 | 1 | 0 | 0 |
| A5SS | 3798 | ENSG00000049246 | PER3 | chr1 | + | 7810437 | 7810588 | 7810437 | 7810567 | 7819284 | 7819420 | 10 | 6 | 7 |
| A5SS | 5332 | ENSG00000185480 | PARPBP | chr12 | + | 102153868 | 102153989 | 102153868 | 102153976 | 102165728 | 102165883 | 2 | 0 | 0 |
| A5SS | 1456 | ENSG00000062524 | LTK | chr15 | - | 41507513 | 41507657 | 41507561 | 41507657 | 41507094 | 41507290 | 12 | 2 | 6 |
| A5SS | 4131 | ENSG00000109180 | OCIAD1 | chr4 | + | 48831226 | 48831433 | 48831226 | 48831398 | 48832619 | 48832682 | 0 | 0 | 0 |
| A5SS | 7609 | ENSG00000168003 | SLC3A2 | chr11 | + | 62856108 | 62856453 | 62856108 | 62856381 | 62870744 | 62870837 | 0 | 0 | 1 |
| A5SS | 6211 | ENSG00000153406 | NMRAL1 | chr16 | - | 4463641 | 4463850 | 4463659 | 4463850 | 4463419 | 4463512 | 1 | 0 | 4 |
| A5SS | 7215 | ENSG00000198932 | GPRASP1 | chrX | + | 102651686 | 102651908 | 102651686 | 102651733 | 102652174 | 102652265 | 17 | 5 | 15 |
| A5SS | 4330 | ENSG00000135541 | AHI1 | chr6 | - | 135495312 | 135495898 | 135495813 | 135495898 | 135492227 | 135492291 | 13 | 8 | 5 |
| A5SS | 4371 | ENSG00000149527 | PLCH2 | chr1 | + | 2502111 | 2502910 | 2502111 | 2502409 | 2503921 | 2505532 | 2 | 1 | 1 |
| A5SS | 7647 | ENSG00000198000 | NOL8 | chr9 | - | 92324905 | 92325350 | 92325305 | 92325350 | 92324022 | 92324209 | 7 | 2 | 6 |
| A5SS | 7096 | ENSG00000124275 | MTRR | chr5 | + | 7869147 | 7869215 | 7869147 | 7869189 | 7875257 | 7875375 | 3 | 0 | 3 |
| A5SS | 3123 | ENSG00000100578 | KIAA0586 | chr14 | + | 58427727 | 58427857 | 58427727 | 58427813 | 58429362 | 58429433 | 10 | 2 | 13 |
| A5SS | 914 | ENSG00000185504 | FAAP100 | chr17 | - | 81551731 | 81552052 | 81551927 | 81552052 | 81550247 | 81551203 | 5 | 0 | 0 |
| A5SS | 7349 | ENSG00000120903 | CHRNA2 | chr8 | - | 27469760 | 27469981 | 27469805 | 27469981 | 27469334 | 27469379 | 0 | 0 | 0 |
| A5SS | 2265 | ENSG00000126456 | IRF3 | chr19 | - | 49664442 | 49664673 | 49664552 | 49664673 | 49663342 | 49663514 | 8 | 2 | 16 |
| A5SS | 6880 | ENSG00000130734 | ATG4D | chr19 | + | 10546838 | 10547075 | 10546838 | 10547071 | 10547188 | 10547253 | 1 | 1 | 1 |
| A5SS | 3410 | ENSG00000234663 | LINC01934 | chr2 | + | 181226470 | 181226539 | 181226470 | 181226535 | 181262828 | 181262977 | 1 | 3 | 0 |
| A5SS | 4203 | ENSG00000146540 | C7orf50 | chr7 | - | 1138188 | 1138257 | 1138217 | 1138257 | 1127256 | 1127386 | 1 | 2 | 1 |
| A5SS | 2499 | ENSG00000174652 | ZNF266 | chr19 | - | 9434064 | 9434235 | 9434074 | 9434235 | 9433667 | 9433783 | 0 | 4 | 0 |
| A5SS | 2627 | ENSG00000108591 | DRG2 | chr17 | + | 18099016 | 18099093 | 18099016 | 18099077 | 18100362 | 18100435 | 1 | 3 | 0 |
| A5SS | 282 | ENSG00000221946 | FXYD7 | chr19 | + | 35151439 | 35151482 | 35151439 | 35151473 | 35151632 | 35151673 | 2 | 4 | 13 |
| A5SS | 5915 | ENSG00000136699 | SMPD4 | chr2 | - | 130161090 | 130161272 | 130161185 | 130161272 | 130157250 | 130157396 | 1 | 0 | 1 |
| A5SS | 6956 | ENSG00000101928 | MOSPD1 | chrX | - | 134896813 | 134897034 | 134896816 | 134897034 | 134891478 | 134891640 | 1 | 0 | 0 |
| A5SS | 782 | ENSG00000185624 | P4HB | chr17 | - | 81858494 | 81859387 | 81858934 | 81859387 | 81855452 | 81855586 | 6 | 2 | 4 |
| A5SS | 8481 | ENSG00000118518 | RNF146 | chr6 | + | 127266863 | 127267044 | 127266863 | 127266925 | 127280230 | 127280340 | 2 | 7 | 3 |
| A5SS | 8622 | ENSG00000151746 | BICD1 | chr12 | + | 32337498 | 32337816 | 32337498 | 32337706 | 32367669 | 32367745 | 2 | 3 | 7 |
| A5SS | 3977 | ENSG00000154358 | OBSCN | chr1 | + | 228356150 | 228356344 | 228356150 | 228356193 | 228359596 | 228361250 | 4 | 2 | 5 |
| A5SS | 6301 | ENSG00000214021 | TTLL3 | chr3 | + | 9833103 | 9833273 | 9833103 | 9833245 | 9834426 | 9834503 | 2 | 1 | 6 |
| A5SS | 1126 | ENSG00000167207 | NOD2 | chr16 | + | 50712193 | 50712401 | 50712193 | 50712373 | 50719924 | 50720008 | 0 | 0 | 1 |
| A5SS | 2925 | ENSG00000128513 | POT1 | chr7 | - | 124827200 | 124827305 | 124827213 | 124827305 | 124825251 | 124825357 | 4 | 2 | 0 |
| A5SS | 2676 | ENSG00000102796 | DHRS12 | chr13 | - | 51773920 | 51774034 | 51773929 | 51774034 | 51772765 | 51773033 | 4 | 1 | 2 |
| A5SS | 2455 | ENSG00000128609 | NDUFA5 | chr7 | - | 123546582 | 123546817 | 123546657 | 123546817 | 123545610 | 123545676 | 9 | 3 | 9 |
| A5SS | 7227 | ENSG00000255052 | FAM66D | chr8 | + | 12121986 | 12122160 | 12121986 | 12122050 | 12145422 | 12145546 | 1 | 0 | 10 |
| A5SS | 7380 | ENSG00000249859 | PVT1 | chr8 | + | 128070159 | 128070361 | 128070159 | 128070272 | 128096520 | 128096654 | 2 | 2 | 3 |
| A5SS | 7794 | ENSG00000130714 | POMT1 | chr9 | + | 131509902 | 131510141 | 131509902 | 131510062 | 131510259 | 131510415 | 0 | 1 | 2 |
| A5SS | 2699 | ENSG00000198931 | APRT | chr16 | - | 88811496 | 88811656 | 88811549 | 88811656 | 88810069 | 88810148 | 4 | 0 | 0 |
| A5SS | 1435 | ENSG00000172345 | STARD5 | chr15 | - | 81323801 | 81324141 | 81324000 | 81324141 | 81322898 | 81322948 | 6 | 2 | 12 |
| A5SS | 4458 | ENSG00000112303 | VNN2 | chr6 | - | 132753827 | 132753949 | 132753875 | 132753949 | 132752723 | 132752749 | 4 | 3 | 0 |
| A5SS | 2407 | ENSG00000137806 | NDUFAF1 | chr15 | - | 41394104 | 41394358 | 41394264 | 41394358 | 41388447 | 41388522 | 3 | 1 | 2 |
| A5SS | 2626 | ENSG00000108591 | DRG2 | chr17 | + | 18093812 | 18093973 | 18093812 | 18093968 | 18100362 | 18100435 | 5 | 0 | 0 |
| A5SS | 2547 | ENSG00000023287 | RB1CC1 | chr8 | - | 52628031 | 52628168 | 52628040 | 52628168 | 52624716 | 52624787 | 46 | 15 | 30 |
| A5SS | 2290 | ENSG00000129595 | EPB41L4A | chr5 | - | 112170255 | 112170369 | 112170300 | 112170369 | 112168994 | 112169105 | 0 | 0 | 0 |
| A5SS | 3380 | ENSG00000095321 | CRAT | chr9 | - | 129108898 | 129109277 | 129109178 | 129109277 | 129107813 | 129108077 | 2 | 1 | 1 |
| A5SS | 8284 | ENSG00000095066 | HOOK2 | chr19 | - | 12774634 | 12774741 | 12774668 | 12774741 | 12772993 | 12773044 | 0 | 2 | 1 |
| A5SS | 2544 | ENSG00000061918 | GUCY1B1 | chr4 | + | 155772601 | 155772753 | 155772601 | 155772697 | 155774967 | 155775068 | 0 | 1 | 0 |
| A5SS | 4465 | ENSG00000106028 | SSBP1 | chr7 | + | 141738357 | 141738503 | 141738357 | 141738410 | 141742168 | 141742229 | 0 | 0 | 0 |
| A5SS | 7598 | ENSG00000182919 | C11orf54 | chr11 | + | 93741962 | 93742094 | 93741962 | 93742066 | 93747296 | 93747448 | 2 | 0 | 0 |
| A5SS | 1916 | ENSG00000157551 | KCNJ15 | chr21 | + | 38296925 | 38297023 | 38296925 | 38297011 | 38298143 | 38298282 | 0 | 0 | 1 |
| A5SS | 1096 | ENSG00000167083 | GNGT2 | chr17 | - | 49209242 | 49209400 | 49209352 | 49209400 | 49208844 | 49208954 | 4 | 1 | 2 |
| A5SS | 7072 | ENSG00000102103 | PQBP1 | chrX | + | 48897911 | 48898093 | 48897911 | 48898082 | 48901189 | 48901301 | 4 | 0 | 3 |
| A5SS | 84 | ENSG00000100209 | HSCB | chr22 | + | 28744614 | 28744709 | 28744614 | 28744704 | 28751240 | 28751288 | 2 | 1 | 2 |
| A5SS | 5881 | ENSG00000158158 | CNNM4 | chr2 | + | 96788589 | 96788894 | 96788589 | 96788665 | 96797011 | 96797155 | 2 | 0 | 1 |
| A5SS | 7187 | ENSG00000198954 | KIFBP | chr10 | + | 69005731 | 69007751 | 69005731 | 69005915 | 69008840 | 69008925 | 22 | 7 | 10 |
| A5SS | 1465 | ENSG00000128833 | MYO5C | chr15 | - | 52260861 | 52261127 | 52260901 | 52261127 | 52256638 | 52256720 | 1 | 0 | 0 |
| A5SS | 1869 | ENSG00000204536 | CCHCR1 | chr6 | - | 31157357 | 31157500 | 31157384 | 31157500 | 31157022 | 31157089 | 0 | 0 | 0 |
| A5SS | 1943 | ENSG00000127990 | SGCE | chr7 | - | 94587777 | 94588732 | 94588688 | 94588732 | 94585263 | 94585515 | 3 | 1 | 6 |
| A5SS | 2149 | ENSG00000116991 | SIPA1L2 | chr1 | - | 232608826 | 232608988 | 232608911 | 232608988 | 232574173 | 232574222 | 0 | 0 | 0 |
| A5SS | 3451 | ENSG00000186205 | MTARC1 | chr1 | + | 220797873 | 220798065 | 220797873 | 220798014 | 220798151 | 220798275 | 0 | 0 | 1 |
| A5SS | 3736 | ENSG00000164050 | PLXNB1 | chr3 | - | 48415175 | 48415347 | 48415211 | 48415347 | 48414798 | 48415041 | 0 | 0 | 3 |
| A5SS | 3744 | ENSG00000171595 | DNAI2 | chr17 | + | 74312002 | 74312230 | 74312002 | 74312075 | 74314120 | 74314271 | 0 | 1 | 5 |
| A5SS | 3928 | ENSG00000243701 | DUBR | chr3 | + | 107246007 | 107246137 | 107246007 | 107246052 | 107248577 | 107248831 | 0 | 3 | 0 |
| A5SS | 4001 | ENSG00000197530 | MIB2 | chr1 | + | 1623388 | 1623699 | 1623388 | 1623482 | 1623773 | 1623945 | 12 | 2 | 5 |
| A5SS | 6088 | ENSG00000169314 | C22orf15 | chr22 | + | 23764259 | 23764402 | 23764259 | 23764397 | 23764638 | 23764713 | 0 | 0 | 0 |
| A5SS | 6684 | ENSG00000164877 | MICALL2 | chr7 | - | 1437433 | 1437608 | 1437534 | 1437608 | 1436741 | 1436856 | 0 | 0 | 0 |
| A5SS | 6798 | ENSG00000118513 | MYB | chr6 | + | 135194355 | 135194460 | 135194355 | 135194451 | 135194959 | 135195081 | 0 | 0 | 1 |
| A5SS | 7005 | ENSG00000244754 | N4BP2L2 | chr13 | - | 32538682 | 32538825 | 32538777 | 32538825 | 32480589 | 32480637 | 0 | 1 | 0 |
| A5SS | 7067 | ENSG00000102001 | CACNA1F | chrX | - | 49226936 | 49227127 | 49226969 | 49227127 | 49226609 | 49226702 | 2 | 0 | 0 |
| A5SS | 7931 | ENSG00000261645 | DISC1FP1 | chr11 | + | 90844879 | 90844957 | 90844879 | 90844949 | 90852615 | 90852707 | 0 | 0 | 0 |
| A5SS | 7958 | ENSG00000189164 | ZNF527 | chr19 | + | 37379119 | 37379246 | 37379119 | 37379236 | 37379486 | 37379692 | 0 | 0 | 0 |
| A5SS | 8305 | ENSG00000121903 | ZSCAN20 | chr1 | + | 33472644 | 33472691 | 33472644 | 33472687 | 33479178 | 33479705 | 1 | 0 | 0 |
| A5SS | 8763 | ENSG00000002016 | RAD52 | chr12 | - | 949601 | 949665 | 949605 | 949665 | 932974 | 933076 | 3 | 0 | 6 |
| A5SS | 8858 | ENSG00000168778 | TCTN2 | chr12 | + | 123671112 | 123671348 | 123671112 | 123671322 | 123671506 | 123671614 | 0 | 0 | 0 |
| A5SS | 8866 | ENSG00000135451 | TROAP | chr12 | + | 49323844 | 49324037 | 49323844 | 49323946 | 49325500 | 49325658 | 0 | 0 | 12 |
| A5SS | 4895 | ENSG00000172766 | NAA16 | chr13 | + | 41362030 | 41362873 | 41362030 | 41362159 | 41367438 | 41367652 | 37 | 37 | 38 |
| A5SS | 2742 | ENSG00000174776 | WDR49 | chr3 | - | 167554649 | 167554798 | 167554682 | 167554798 | 167536869 | 167537000 | 12 | 5 | 1 |
| A5SS | 5313 | ENSG00000088179 | PTPN4 | chr2 | + | 119915178 | 119916690 | 119915178 | 119915242 | 119932423 | 119932549 | 10 | 22 | 14 |
| A5SS | 8799 | ENSG00000130396 | AFDN | chr6 | + | 167965756 | 167966270 | 167965756 | 167966045 | 167969113 | 167969198 | 0 | 0 | 1 |
| A5SS | 2616 | ENSG00000178104 | PDE4DIP | chr1 | + | 149001590 | 149002026 | 149001590 | 149001696 | 149002849 | 149003042 | 0 | 1 | 6 |
| A5SS | 2648 | ENSG00000163138 | PACRGL | chr4 | + | 20700458 | 20700747 | 20700458 | 20700617 | 20704465 | 20704533 | 4 | 0 | 2 |
| A5SS | 6881 | ENSG00000130734 | ATG4D | chr19 | + | 10546838 | 10547082 | 10546838 | 10547071 | 10547188 | 10547253 | 2 | 1 | 2 |
| A5SS | 943 | ENSG00000267123 | SCAT1 | chr17 | - | 78631918 | 78632064 | 78631989 | 78632064 | 78630031 | 78630430 | 0 | 0 | 1 |
| A5SS | 6278 | ENSG00000236039 | LINC02889 | chr7 | - | 17457339 | 17457589 | 17457458 | 17457589 | 17456677 | 17456762 | 1 | 0 | 0 |
| A5SS | 3887 | ENSG00000058272 | PPP1R12A | chr12 | - | 79779320 | 79779351 | 79779338 | 79779351 | 79778549 | 79778600 | 1 | 3 | 1 |
| A5SS | 8800 | ENSG00000130396 | AFDN | chr6 | + | 167965756 | 167966276 | 167965756 | 167966045 | 167969113 | 167969198 | 1 | 0 | 1 |
| A5SS | 775 | ENSG00000131462 | TUBG1 | chr17 | + | 42613848 | 42614136 | 42613848 | 42613998 | 42614259 | 42614412 | 0 | 0 | 0 |
| A5SS | 2170 | ENSG00000115661 | STK16 | chr2 | + | 219245478 | 219245796 | 219245478 | 219245522 | 219245895 | 219246085 | 8 | 1 | 3 |
| A5SS | 5278 | ENSG00000225138 | SLC9A3-AS1 | chr5 | + | 477243 | 477465 | 477243 | 477336 | 477805 | 477849 | 4 | 4 | 3 |
| A5SS | 4913 | ENSG00000147121 | KRBOX4 | chrX | + | 46463197 | 46463308 | 46463197 | 46463293 | 46472752 | 46473594 | 1 | 0 | 1 |
| A5SS | 707 | ENSG00000270379 | HEATR9 | chr17 | - | 35862950 | 35863125 | 35862994 | 35863125 | 35858887 | 35859070 | 3 | 4 | 2 |
| A5SS | 6268 | ENSG00000168944 | CEP120 | chr5 | - | 123403216 | 123403344 | 123403255 | 123403344 | 123399135 | 123399284 | 8 | 0 | 4 |
| A5SS | 7536 | ENSG00000080503 | SMARCA2 | chr9 | + | 2158455 | 2158978 | 2158455 | 2158587 | 2159813 | 2159909 | 1 | 0 | 1 |
| A5SS | 6250 | ENSG00000131018 | SYNE1 | chr6 | - | 152309830 | 152310017 | 152309834 | 152310017 | 152308488 | 152308628 | 2 | 1 | 1 |
| A5SS | 5023 | ENSG00000218510 | LINC00339 | chr1 | + | 22025510 | 22026119 | 22025510 | 22025639 | 22030279 | 22030617 | 1 | 3 | 4 |
| A5SS | 5268 | ENSG00000140386 | SCAPER | chr15 | - | 76795240 | 76795440 | 76795279 | 76795440 | 76774854 | 76775117 | 0 | 1 | 1 |
| A5SS | 1535 | ENSG00000128923 | MINDY2 | chr15 | + | 58821716 | 58821823 | 58821716 | 58821819 | 58831773 | 58831916 | 1 | 6 | 0 |
| A5SS | 3991 | ENSG00000198912 | C1orf174 | chr1 | - | 3891534 | 3891779 | 3891625 | 3891779 | 3890568 | 3891057 | 4 | 0 | 8 |
| A5SS | 4384 | ENSG00000135953 | MFSD9 | chr2 | - | 102736529 | 102736888 | 102736645 | 102736888 | 102732321 | 102732409 | 1 | 2 | 4 |
| A5SS | 4314 | ENSG00000085185 | BCORL1 | chrX | + | 129981672 | 129981783 | 129981672 | 129981752 | 130005187 | 130005317 | 1 | 0 | 0 |
| A5SS | 2649 | ENSG00000163138 | PACRGL | chr4 | + | 20700458 | 20700787 | 20700458 | 20700617 | 20704465 | 20704533 | 7 | 0 | 4 |
| A5SS | 3004 | ENSG00000178802 | MPI | chr15 | + | 74890041 | 74890201 | 74890041 | 74890089 | 74890526 | 74890654 | 5 | 0 | 4 |
| A5SS | 2039 | ENSG00000151612 | ZNF827 | chr4 | - | 145764974 | 145765165 | 145764987 | 145765165 | 145763089 | 145763122 | 0 | 0 | 1 |
| A5SS | 8779 | ENSG00000139116 | KIF21A | chr12 | - | 39321347 | 39322882 | 39322667 | 39322882 | 39319905 | 39320013 | 0 | 2 | 2 |
| A5SS | 4483 | ENSG00000156983 | BRPF1 | chr3 | + | 9742024 | 9742171 | 9742024 | 9742153 | 9742351 | 9742541 | 2 | 2 | 2 |
| A5SS | 5300 | ENSG00000143537 | ADAM15 | chr1 | + | 155061903 | 155062117 | 155061903 | 155061975 | 155062244 | 155062369 | 10 | 2 | 1 |
| A5SS | 816 | ENSG00000129219 | PLD2 | chr17 | + | 4819428 | 4819582 | 4819428 | 4819549 | 4821792 | 4821907 | 1 | 5 | 12 |
| A5SS | 1562 | ENSG00000255571 | MIR9-3HG | chr15 | + | 89388503 | 89388746 | 89388503 | 89388581 | 89389344 | 89389482 | 3 | 0 | 1 |
| A5SS | 3099 | ENSG00000136449 | MYCBPAP | chr17 | + | 50517292 | 50517497 | 50517292 | 50517452 | 50517594 | 50517698 | 0 | 0 | 1 |
| A5SS | 3458 | ENSG00000143761 | ARF1 | chr1 | + | 228083149 | 228083374 | 228083149 | 228083309 | 228097077 | 228097262 | 2 | 0 | 1 |
| A5SS | 1958 | ENSG00000228624 | HDAC2-AS2 | chr6 | + | 113991892 | 113992018 | 113991892 | 113992014 | 113993342 | 113993487 | 3 | 0 | 0 |
| A5SS | 1254 | ENSG00000103227 | LMF1 | chr16 | - | 934189 | 934254 | 934243 | 934254 | 910930 | 911079 | 0 | 0 | 0 |
| A5SS | 8450 | ENSG00000107862 | GBF1 | chr10 | + | 102354369 | 102354468 | 102354369 | 102354444 | 102358038 | 102358186 | 0 | 1 | 0 |
| A5SS | 1128 | ENSG00000167207 | NOD2 | chr16 | + | 50712193 | 50712637 | 50712193 | 50712373 | 50722621 | 50722705 | 2 | 6 | 1 |
| A5SS | 8482 | ENSG00000118518 | RNF146 | chr6 | + | 127266882 | 127267095 | 127266882 | 127266925 | 127280230 | 127280340 | 8 | 6 | 7 |
| A5SS | 79 | ENSG00000183765 | CHEK2 | chr22 | - | 28730419 | 28730548 | 28730455 | 28730548 | 28725242 | 28725367 | 1 | 1 | 0 |
| A5SS | 5527 | ENSG00000204634 | TBC1D8 | chr2 | - | 101090208 | 101090364 | 101090253 | 101090364 | 101059420 | 101059539 | 0 | 1 | 0 |
| A5SS | 3474 | ENSG00000050327 | ARHGEF5 | chr7 | + | 144362657 | 144365823 | 144362657 | 144362853 | 144366354 | 144366412 | 2 | 0 | 7 |
| A5SS | 6333 | ENSG00000124486 | USP9X | chrX | + | 41229566 | 41229827 | 41229566 | 41229779 | 41230500 | 41230596 | 34 | 25 | 18 |
| A5SS | 8245 | ENSG00000133895 | MEN1 | chr11 | - | 64810513 | 64810551 | 64810525 | 64810551 | 64809664 | 64810132 | 0 | 0 | 1 |
| A5SS | 4821 | ENSG00000116127 | ALMS1 | chr2 | + | 73489633 | 73491498 | 73489633 | 73490158 | 73519774 | 73520016 | 63 | 47 | 62 |
| A5SS | 7348 | ENSG00000120903 | CHRNA2 | chr8 | - | 27469528 | 27469981 | 27469805 | 27469981 | 27469334 | 27469379 | 0 | 0 | 0 |
| A5SS | 6149 | ENSG00000262943 | ALOX12P2 | chr17 | + | 6894317 | 6894519 | 6894317 | 6894513 | 6894883 | 6895006 | 2 | 0 | 1 |
| A5SS | 1908 | ENSG00000196367 | TRRAP | chr7 | + | 99030594 | 99030764 | 99030594 | 99030757 | 99031344 | 99031465 | 1 | 1 | 0 |
| A5SS | 2040 | ENSG00000151612 | ZNF827 | chr4 | - | 145764978 | 145765165 | 145764987 | 145765165 | 145763089 | 145763122 | 0 | 2 | 0 |
| A5SS | 2172 | ENSG00000111879 | FAM184A | chr6 | - | 118961760 | 118961963 | 118961928 | 118961963 | 118959828 | 118960184 | 0 | 0 | 7 |
| A5SS | 2629 | ENSG00000108591 | DRG2 | chr17 | + | 18093812 | 18093973 | 18093812 | 18093968 | 18103800 | 18103889 | 5 | 0 | 0 |
| A5SS | 2640 | ENSG00000163138 | PACRGL | chr4 | + | 20700458 | 20700747 | 20700458 | 20700617 | 20701757 | 20701884 | 2 | 0 | 2 |
| A5SS | 2661 | ENSG00000165533 | TTC8 | chr14 | + | 88841036 | 88841514 | 88841036 | 88841196 | 88843805 | 88843850 | 0 | 1 | 3 |
| A5SS | 3231 | ENSG00000114670 | NEK11 | chr3 | + | 131026878 | 131028002 | 131026878 | 131027006 | 131029612 | 131029878 | 0 | 0 | 2 |
| A5SS | 3515 | ENSG00000171163 | ZNF692 | chr1 | - | 248858917 | 248859085 | 248859014 | 248859085 | 248858130 | 248858321 | 3 | 4 | 7 |
| A5SS | 4260 | ENSG00000106479 | ZNF862 | chr7 | + | 149838374 | 149838635 | 149838374 | 149838461 | 149846150 | 149846255 | 3 | 6 | 0 |
| A5SS | 4401 | ENSG00000145979 | TBC1D7 | chr6 | - | 13328295 | 13328531 | 13328455 | 13328531 | 13326786 | 13326963 | 1 | 0 | 2 |
| A5SS | 5661 | ENSG00000231672 | DIRC3 | chr2 | - | 217433756 | 217434116 | 217433947 | 217434116 | 217432922 | 217433021 | 1 | 3 | 2 |
| A5SS | 5693 | ENSG00000243970 | PPIEL | chr1 | - | 39541656 | 39541841 | 39541719 | 39541841 | 39537190 | 39537343 | 0 | 1 | 1 |
| A5SS | 5893 | ENSG00000239665 | NA | chr10 | + | 13652845 | 13652929 | 13652845 | 13652882 | 13654352 | 13654427 | 3 | 0 | 1 |
| A5SS | 6566 | ENSG00000074527 | NTN4 | chr12 | - | 95790254 | 95790764 | 95790727 | 95790764 | 95786938 | 95787468 | 2 | 0 | 1 |
| A5SS | 7940 | ENSG00000066294 | CD84 | chr1 | - | 160553359 | 160553497 | 160553377 | 160553497 | 160549916 | 160549979 | 1 | 0 | 5 |
| A5SS | 8287 | ENSG00000233673 | ANAPC1P1 | chr2 | + | 86889066 | 86889276 | 86889066 | 86889179 | 86893087 | 86893181 | 2 | 0 | 1 |
| A5SS | 8422 | ENSG00000128805 | ARHGAP22 | chr10 | - | 48455001 | 48455134 | 48455053 | 48455134 | 48454087 | 48454161 | 5 | 1 | 0 |
| A5SS | 8797 | ENSG00000130396 | AFDN | chr6 | + | 167965756 | 167966270 | 167965756 | 167965853 | 167969113 | 167969198 | 6 | 5 | 6 |
| A5SS | 8798 | ENSG00000130396 | AFDN | chr6 | + | 167965756 | 167966276 | 167965756 | 167965853 | 167969113 | 167969198 | 7 | 5 | 6 |
| A5SS | 3060 | ENSG00000105717 | PBX4 | chr19 | - | 19570098 | 19570299 | 19570108 | 19570299 | 19569448 | 19569584 | 2 | 3 | 0 |
| A5SS | 3194 | ENSG00000284707 | NA | chr7 | - | 97936946 | 97937160 | 97937079 | 97937160 | 97928709 | 97928777 | 0 | 0 | 1 |
| A5SS | 2497 | ENSG00000174652 | ZNF266 | chr19 | - | 9419050 | 9419299 | 9419216 | 9419299 | 9418765 | 9418895 | 69 | 33 | 20 |
| A5SS | 2826 | ENSG00000206053 | JPT2 | chr16 | + | 1683508 | 1683620 | 1683508 | 1683592 | 1685438 | 1685587 | 2 | 0 | 1 |
| A5SS | 5695 | ENSG00000243970 | PPIEL | chr1 | - | 39548669 | 39548746 | 39548682 | 39548746 | 39547578 | 39547634 | 4 | 2 | 6 |
| A5SS | 1125 | ENSG00000167207 | NOD2 | chr16 | + | 50712193 | 50712637 | 50712193 | 50712373 | 50716586 | 50716670 | 2 | 6 | 4 |
| A5SS | 953 | ENSG00000262879 | NA | chr17 | - | 47056455 | 47056753 | 47056459 | 47056753 | 47033277 | 47033381 | 0 | 0 | 0 |
| A5SS | 4544 | ENSG00000287514 | NA | chr9 | + | 36781469 | 36781897 | 36781469 | 36781893 | 36793993 | 36796516 | 0 | 0 | 0 |
| A5SS | 977 | ENSG00000005884 | ITGA3 | chr17 | + | 50074143 | 50074292 | 50074143 | 50074280 | 50074447 | 50074534 | 0 | 0 | 0 |
| A5SS | 5651 | ENSG00000021300 | PLEKHB1 | chr11 | + | 73647654 | 73647895 | 73647654 | 73647738 | 73649011 | 73649087 | 1 | 0 | 0 |
| A5SS | 1389 | ENSG00000172366 | MCRIP2 | chr16 | + | 647416 | 647884 | 647416 | 647544 | 648119 | 648474 | 14 | 0 | 13 |
| A5SS | 1602 | ENSG00000127586 | CHTF18 | chr16 | + | 788623 | 788775 | 788623 | 788756 | 788930 | 789125 | 8 | 5 | 13 |
| A5SS | 238 | ENSG00000142541 | RPL13A | chr19 | + | 49489849 | 49489922 | 49489849 | 49489912 | 49490474 | 49490576 | 4 | 0 | 4 |
| A5SS | 2672 | ENSG00000125375 | DMAC2L | chr14 | + | 50322510 | 50322806 | 50322510 | 50322719 | 50323944 | 50324116 | 5 | 9 | 5 |
| A5SS | 2424 | ENSG00000136925 | TSTD2 | chr9 | - | 97627305 | 97627612 | 97627397 | 97627612 | 97625680 | 97625997 | 8 | 18 | 7 |
| A5SS | 76 | ENSG00000232855 | NA | chr21 | - | 28577218 | 28577380 | 28577274 | 28577380 | 28446289 | 28446472 | 2 | 9 | 6 |
| A5SS | 241 | ENSG00000142541 | RPL13A | chr19 | + | 49490231 | 49490301 | 49490231 | 49490297 | 49490479 | 49490576 | 6 | 3 | 1 |
| A5SS | 420 | ENSG00000160117 | ANKLE1 | chr19 | + | 17283224 | 17283962 | 17283224 | 17283884 | 17284088 | 17284266 | 4 | 2 | 2 |
| A5SS | 3799 | ENSG00000164023 | SGMS2 | chr4 | + | 107858471 | 107858584 | 107858471 | 107858553 | 107899574 | 107899692 | 1 | 0 | 1 |
| A5SS | 6533 | ENSG00000121716 | PILRB | chr7 | + | 100356749 | 100356938 | 100356749 | 100356883 | 100358263 | 100358366 | 5 | 2 | 4 |
| A5SS | 7411 | ENSG00000127415 | IDUA | chr4 | + | 987808 | 987949 | 987808 | 987911 | 1000611 | 1000697 | 2 | 6 | 2 |
| A5SS | 561 | ENSG00000174365 | SNHG11 | chr20 | + | 38446696 | 38448093 | 38446696 | 38447623 | 38448661 | 38448730 | 1 | 1 | 1 |
| A5SS | 1559 | ENSG00000169609 | C15orf40 | chr15 | - | 82991931 | 82992046 | 82991955 | 82992046 | 82989105 | 82989157 | 1 | 0 | 3 |
| A5SS | 6524 | ENSG00000179818 | PCBP1-AS1 | chr2 | - | 70086221 | 70086273 | 70086251 | 70086273 | 70085546 | 70085624 | 1 | 2 | 3 |
| A5SS | 155 | ENSG00000160111 | CPAMD8 | chr19 | - | 16975086 | 16975258 | 16975096 | 16975258 | 16970890 | 16971033 | 0 | 0 | 0 |
| A5SS | 5392 | ENSG00000162910 | MRPL55 | chr1 | - | 228109148 | 228109299 | 228109260 | 228109299 | 228108954 | 228109021 | 6 | 0 | 4 |
| A5SS | 2852 | ENSG00000075711 | DLG1 | chr3 | - | 197298417 | 197298612 | 197298535 | 197298612 | 197297185 | 197297235 | 2 | 0 | 3 |
| A5SS | 3960 | ENSG00000163743 | RCHY1 | chr4 | - | 75508819 | 75508935 | 75508823 | 75508935 | 75491888 | 75491933 | 0 | 1 | 6 |
| A5SS | 6947 | ENSG00000155966 | AFF2 | chrX | + | 148651998 | 148652131 | 148651998 | 148652119 | 148661907 | 148662768 | 0 | 0 | 0 |
| A5SS | 2500 | ENSG00000174652 | ZNF266 | chr19 | - | 9434064 | 9434235 | 9434074 | 9434235 | 9433667 | 9433786 | 0 | 2 | 0 |
| A5SS | 4354 | ENSG00000159023 | EPB41 | chr1 | + | 29068732 | 29068783 | 29068732 | 29068768 | 29097806 | 29097935 | 0 | 3 | 0 |
| A5SS | 1616 | ENSG00000276550 | HERC2P2 | chr15 | + | 22561558 | 22561934 | 22561558 | 22561789 | 22562948 | 22563087 | 36 | 16 | 38 |
| A5SS | 2222 | ENSG00000204120 | GIGYF2 | chr2 | + | 232731121 | 232731258 | 232731121 | 232731234 | 232735154 | 232735238 | 2 | 2 | 0 |
| A5SS | 3454 | ENSG00000064687 | ABCA7 | chr19 | + | 1056906 | 1057429 | 1056906 | 1057084 | 1058617 | 1058747 | 0 | 2 | 3 |
| A5SS | 8357 | ENSG00000119979 | DENND10 | chr10 | + | 119104763 | 119105022 | 119104763 | 119105006 | 119107967 | 119108164 | 0 | 1 | 6 |
| A5SS | 7210 | ENSG00000111237 | VPS29 | chr12 | - | 110499456 | 110499546 | 110499534 | 110499546 | 110496822 | 110497002 | 3 | 4 | 6 |
| A5SS | 6890 | ENSG00000204271 | SPIN3 | chrX | - | 56995215 | 56995530 | 56995241 | 56995530 | 56994722 | 56994949 | 2 | 1 | 3 |
| A5SS | 6931 | ENSG00000140332 | TLE3 | chr15 | - | 70057458 | 70057658 | 70057473 | 70057658 | 70055048 | 70055298 | 0 | 0 | 2 |
| A5SS | 6962 | ENSG00000100802 | C14orf93 | chr14 | - | 23010016 | 23010143 | 23010100 | 23010143 | 23000958 | 23001037 | 0 | 0 | 0 |
| A5SS | 1615 | ENSG00000276550 | HERC2P2 | chr15 | + | 22561558 | 22561934 | 22561558 | 22561789 | 22562642 | 22562731 | 23 | 11 | 22 |
| A5SS | 6111 | ENSG00000078269 | SYNJ2 | chr6 | + | 158059256 | 158059537 | 158059256 | 158059353 | 158061991 | 158062164 | 3 | 0 | 2 |
| A5SS | 7194 | ENSG00000166788 | SAAL1 | chr11 | - | 18086856 | 18087054 | 18086865 | 18087054 | 18083534 | 18083728 | 0 | 0 | 1 |
| A5SS | 7195 | ENSG00000166788 | SAAL1 | chr11 | - | 18086861 | 18087054 | 18086865 | 18087054 | 18083534 | 18083728 | 0 | 0 | 1 |
| A5SS | 4977 | ENSG00000035115 | SH3YL1 | chr2 | - | 262200 | 262786 | 262630 | 262786 | 260084 | 261130 | 22 | 3 | 8 |
| A5SS | 1861 | ENSG00000152240 | HAUS1 | chr18 | + | 46105193 | 46105408 | 46105193 | 46105368 | 46119925 | 46120060 | 1 | 0 | 1 |
| A5SS | 6702 | ENSG00000122591 | FAM126A | chr7 | - | 22983943 | 22984045 | 22983948 | 22984045 | 22977340 | 22977421 | 7 | 4 | 1 |
| A5SS | 3607 | ENSG00000144120 | TMEM177 | chr2 | + | 119679166 | 119679617 | 119679166 | 119679225 | 119680831 | 119682118 | 5 | 2 | 4 |
| A5SS | 7614 | ENSG00000156042 | CFAP70 | chr10 | - | 73291225 | 73291444 | 73291349 | 73291444 | 73278178 | 73278337 | 2 | 3 | 3 |
| A5SS | 755 | ENSG00000182173 | TSEN54 | chr17 | + | 75521704 | 75522535 | 75521704 | 75522521 | 75523274 | 75523335 | 6 | 6 | 6 |
| A5SS | 5373 | ENSG00000169432 | SCN9A | chr2 | - | 166284452 | 166284824 | 166284485 | 166284824 | 166281678 | 166281808 | 0 | 1 | 2 |
| A5SS | 448 | ENSG00000197766 | CFD | chr19 | + | 859663 | 859765 | 859663 | 859744 | 860616 | 860773 | 0 | 0 | 0 |
| A5SS | 2361 | ENSG00000136960 | ENPP2 | chr8 | - | 119582417 | 119582602 | 119582429 | 119582602 | 119580115 | 119580167 | 4 | 12 | 5 |
| A5SS | 5371 | ENSG00000155729 | KCTD18 | chr2 | - | 200509627 | 200510068 | 200510010 | 200510068 | 200506856 | 200507091 | 12 | 17 | 10 |
| A5SS | 7494 | ENSG00000110841 | PPFIBP1 | chr12 | + | 27664361 | 27664446 | 27664361 | 27664437 | 27667165 | 27667320 | 5 | 5 | 6 |
| A5SS | 8134 | ENSG00000110075 | PPP6R3 | chr11 | + | 68505144 | 68505271 | 68505144 | 68505249 | 68515027 | 68515166 | 1 | 3 | 5 |
| A5SS | 4391 | ENSG00000136098 | NEK3 | chr13 | - | 52143914 | 52143987 | 52143915 | 52143987 | 52136799 | 52136902 | 1 | 0 | 0 |
| A5SS | 3118 | ENSG00000109861 | CTSC | chr11 | - | 88337546 | 88337723 | 88337707 | 88337723 | 88334936 | 88335082 | 9 | 1 | 2 |
| A5SS | 4382 | ENSG00000111358 | GTF2H3 | chr12 | + | 123650993 | 123651156 | 123650993 | 123651056 | 123652523 | 123652561 | 2 | 1 | 3 |
| A5SS | 2029 | ENSG00000141279 | NPEPPS | chr17 | + | 47619277 | 47619495 | 47619277 | 47619372 | 47619736 | 47619784 | 14 | 5 | 6 |
| A5SS | 5841 | ENSG00000072195 | SPEG | chr2 | + | 219443107 | 219443440 | 219443107 | 219443150 | 219444029 | 219444059 | 2 | 0 | 3 |
| A5SS | 75 | ENSG00000232855 | NA | chr21 | - | 28577152 | 28577380 | 28577274 | 28577380 | 28446289 | 28446472 | 8 | 19 | 12 |
| A5SS | 7213 | ENSG00000067840 | PDZD4 | chrX | - | 153807850 | 153808139 | 153807896 | 153808139 | 153807278 | 153807369 | 1 | 0 | 0 |

| | | | | | | | | | | | | | | | | | | | | | | | | | | | | | | | | | | | | | | | | | | | | | | |
|---|---|---|---|---|---|---|---|---|---|---|---|---|---|---|---|---|---|---|---|---|---|---|---|---|---|---|---|---|---|---|---|---|---|---|---|---|---|---|---|---|---|---|---|---|---|---|
| A5SS | 566 | ENSG00000196756 | SNHG17 | chr20 | - | 38422091 | 38422241 | 38422095 | 38422241 | 38421005 | 38421098 | 1 | 2 | 1 | 0 | 4 | 0 | 0 | 3 | 0 | 1 | 39 | 0 | 5 | 6 | 5 | 0 | 0 | 0 | 1 | 0 | 153 | 149 | 1 | 1 | 1 | 1 | 0.245 | NA | 0.796 | 1 | NA | 1 | 0.854 | 1 | -0.203 |
| A5SS | 3061 | ENSG00000105717 | PBX4 | chr19 | - | 19570585 | 19570833 | 19570686 | 19570833 | 19570108 | 19570299 | 9 | 6 | 13 | 6 | 2 | 3 | 2 | 3 | 9 | 0 | 0 | 12 | 9 | 2 | 13 | 0 | 9 | 0 | 0 | 0 | 250 | 149 | 1 | 1 | 0.641 | 0.641 | 0.721 | 0.284 | 1 | NA | 0.443 | 1 | 1 | 1 | -0.203 |
| A5SS | 4340 | ENSG00000111445 | RFC5 | chr12 | + | 118017689 | 118017852 | 118017689 | 118017815 | 118017994 | 118018053 | 1 | 6 | 1 | 3 | 15 | 2 | 0 | 0 | 1 | 0 | 4 | 11 | 3 | 0 | 8 | 0 | 0 | 0 | 0 | 0 | 186 | 149 | 1 | 1 | 0.286 | 1 | 1 | 0.706 | 1 | 1 | 1 | 1 | NA | 1 | -0.202 |
| A5SS | 8655 | ENSG00000090006 | LTBP4 | chr19 | + | 40611858 | 40611984 | 40611858 | 40611976 | 40612072 | 40612192 | 24 | 4 | 7 | 9 | 13 | 5 | 1 | 1 | 11 | 0 | 0 | 3 | 23 | 0 | 11 | 0 | 0 | 1 | 0 | 0 | 157 | 149 | 1 | 1 | 0.82 | 0.792 | 0.869 | 0.437 | 1 | NA | 1 | 0.956 | NA | 1 | -0.202 |
| A5SS | 6703 | ENSG00000106268 | NUDT1 | chr7 | + | 2242904 | 2243048 | 2242904 | 2242975 | 2244562 | 2244726 | 0 | 0 | 1 | 3 | 0 | 10 | 0 | 0 | 2 | 1 | 8 | 2 | 0 | 22 | 0 | 0 | 0 | 4 | 2 | 14 | 222 | 149 | 1 | 1 | 0 | NA | 1 | 0.502 | 0 | 1 | 1 | 0 | 0.881 | 0 | -0.201 |
| A5SS | 275 | ENSG00000266916 | ZNF793-AS1 | chr19 | - | 37505845 | 37506040 | 37505971 | 37506040 | 37497158 | 37497592 | 2 | 4 | 0 | 2 | 3 | 0 | 0 | 1 | 0 | 0 | 0 | 0 | 2 | 0 | 7 | 0 | 0 | 0 | 0 | 0 | 275 | 149 | 1 | 1 | 1 | 1 | 0 | 1 | 1 | NA | NA | 1 | NA | 1 | -0.2 |
| A5SS | 461 | ENSG00000273154 | NA | chr20 | + | 63733586 | 63733775 | 63733586 | 63733739 | 63737532 | 63737647 | 2 | 2 | 8 | 0 | 2 | 0 | 0 | 0 | 1 | 0 | 19 | 0 | 3 | 4 | 33 | 0 | 0 | 0 | 0 | 0 | 185 | 149 | 1 | 1 | 1 | 1 | 1 | 0 | 1 | 1 | NA | 1 | 1 | 1 | -0.2 |
| A5SS | 5194 | ENSG00000167220 | HDHD2 | chr18 | - | 47136301 | 47136449 | 47136338 | 47136449 | 47130243 | 47130328 | 3 | 1 | 0 | 0 | 1 | 0 | 0 | 1 | 0 | 0 | 0 | 10 | 1 | 7 | 0 | 0 | 0 | 0 | 1 | 0 | 186 | 149 | 1 | 1 | 1 | 1 | 0 | NA | 1 | NA | 1 | 1 | 0.849 | NA | -0.2 |
| A5SS | 5230 | ENSG00000157184 | CPT2 | chr1 | + | 53210014 | 53211369 | 53210014 | 53211319 | 53212780 | 53212816 | 2 | 0 | 1 | 2 | 2 | 1 | 0 | 0 | 1 | 0 | 0 | 0 | 1 | 0 | 0 | 0 | 0 | 0 | 0 | 0 | 199 | 149 | 1 | 1 | 0.6 | NA | 1 | 0.6 | 1 | NA | NA | 1 | NA | NA | -0.2 |
| A5SS | 5485 | ENSG00000185621 | LMLN | chr3 | + | 197960199 | 197960440 | 197960199 | 197960319 | 197974376 | 197974474 | 7 | 2 | 8 | 3 | 0 | 0 | 0 | 0 | 0 | 1 | 0 | 1 | 1 | 0 | 7 | 0 | 0 | 0 | 0 | 0 | 270 | 149 | 1 | 1 | 1 | 1 | 1 | 1 | 0 | NA | 1 | 1 | NA | 1 | -0.2 |
| A5SS | 5570 | ENSG00000127249 | ATP13A4 | chr3 | - | 193440489 | 193440637 | 193440557 | 193440637 | 193439022 | 193439065 | 0 | 0 | 4 | 3 | 8 | 1 | 2 | 0 | 1 | 5 | 1 | 0 | 1 | 4 | 0 | 0 | 0 | 3 | 1 | 0 | 217 | 149 | 1 | 1 | 0 | 0 | 1 | 0.673 | 0.523 | 1 | NA | 0.186 | 0.733 | NA | -0.2 |
| A5SS | 6060 | ENSG00000198223 | CSF2RA | chrX | + | 1274754 | 1274842 | 1274754 | 1274818 | 1282408 | 1282560 | 4 | 2 | 1 | 0 | 1 | 0 | 0 | 0 | 2 | 0 | 0 | 0 | 0 | 7 | 0 | 0 | 0 | 0 | 0 | 0 | 173 | 149 | 1 | 1 | 1 | 1 | 1 | 0 | 1 | NA | NA | NA | 1 | NA | -0.2 |
| A5SS | 3310 | ENSG00000165240 | ATP7A | chrX | + | 77922998 | 77923052 | 77922998 | 77923042 | 77971620 | 77971761 | 0 | 2 | 1 | 0 | 0 | 3 | 2 | 4 | 10 | 5 | 0 | 0 | 2 | 0 | 0 | 0 | 2 | 0 | 2 | 0 | 159 | 149 | 1 | 1 | 0 | 0.484 | 0.19 | 0 | 0 | NA | 0 | 1 | 0 | NA | -0.199 |
| A5SS | 7226 | ENSG00000258388 | PPT2-EGFL8 | chr6 | + | 32164578 | 32164681 | 32164578 | 32164657 | 32166137 | 32166620 | 1 | 1 | 7 | 3 | 0 | 1 | 1 | 0 | 2 | 0 | 0 | 2 | 4 | 0 | 1 | 0 | 0 | 4 | 0 | 0 | 173 | 149 | 1 | 1 | 0.463 | 0.463 | 1 | 0.564 | NA | NA | 1 | 0.463 | NA | 1 | -0.199 |
| A5SS | 6525 | ENSG00000179818 | PCBP1-AS1 | chr2 | - | 70086251 | 70086485 | 70086430 | 70086485 | 70085546 | 70085624 | 3 | 0 | 8 | 7 | 2 | 2 | 0 | 0 | 1 | 0 | 0 | 5 | 4 | 15 | 4 | 0 | 0 | 0 | 0 | 0 | 298 | 149 | 1 | 1 | 0.429 | NA | 1 | 0.778 | 1 | NA | 1 | 1 | 1 | 1 | -0.198 |
| A5SS | 6047 | ENSG00000116857 | TMEM9 | chr1 | - | 201153782 | 201153959 | 201153848 | 201153959 | 201151760 | 201151852 | 0 | 2 | 1 | 1 | 0 | 0 | 0 | 1 | 0 | 0 | 0 | 4 | 2 | 0 | 17 | 0 | 0 | 0 | 0 | 0 | 215 | 149 | 1 | 1 | NA | 1 | 0.409 | 1 | NA | NA | 1 | 1 | NA | 1 | -0.197 |
| A5SS | 2452 | ENSG00000213468 | FIRRE | chrX | - | 131794296 | 131794466 | 131794323 | 131794466 | 131783886 | 131784039 | 0 | 1 | 2 | 2 | 2 | 0 | 3 | 0 | 0 | 0 | 0 | 6 | 3 | 0 | 0 | 0 | 0 | 0 | 0 | 0 | 176 | 149 | 1 | 1 | NA | 0.22 | 1 | 1 | 1 | NA | 1 | 1 | NA | NA | -0.195 |
| A5SS | 1882 | ENSG00000250510 | GPR162 | chr12 | + | 6823467 | 6824765 | 6823467 | 6823763 | 6825483 | 6825673 | 16 | 3 | 8 | 3 | 2 | 0 | 2 | 0 | 1 | 0 | 34 | 10 | 1 | 21 | 4 | 0 | 0 | 0 | 0 | 0 | 298 | 149 | 1 | 1 | 1 | 0.429 | 1 | 0.6 | 1 | 1 | 1 | 1 | 1 | 1 | -0.194 |
| A5SS | 4392 | ENSG00000136098 | NEK3 | chr13 | - | 52143914 | 52143987 | 52143915 | 52143987 | 52141019 | 52141069 | 1 | 0 | 7 | 0 | 9 | 6 | 1 | 0 | 0 | 3 | 0 | 3 | 12 | 0 | 0 | 0 | 4 | 0 | 4 | 0 | 150 | 149 | 1 | 1 | 0.142 | 0 | 1 | NA | 0.749 | NA | 1 | 1 | 0 | NA | -0.194 |
| A5SS | 7041 | ENSG00000184634 | MED12 | chrX | + | 71137186 | 71137392 | 71137186 | 71137383 | 71137560 | 71137635 | 2 | 3 | 3 | 1 | 0 | 2 | 2 | 1 | 7 | 4 | 0 | 0 | 0 | 6 | 2 | 0 | 4 | 0 | 2 | 0 | 158 | 149 | 1 | 1 | 0.485 | 0.586 | 0.739 | 0.119 | 0 | NA | 0 | NA | 0.739 | 1 | -0.194 |
| A5SS | 4718 | ENSG00000114354 | TFG | chr3 | + | 100736575 | 100736716 | 100736575 | 100736704 | 100744837 | 100744931 | 1 | 1 | 1 | 7 | 0 | 0 | 3 | 0 | 0 | 0 | 0 | 12 | 0 | 0 | 0 | 0 | 0 | 0 | 0 | 0 | 161 | 149 | 1 | 1 | 1 | 0.236 | 1 | 1 | NA | NA | 1 | NA | NA | NA | -0.191 |
| A5SS | 5876 | ENSG00000228203 | GRASLND | chr2 | - | 6918284 | 6918680 | 6918401 | 6918680 | 6916815 | 6917224 | 0 | 4 | 0 | 4 | 2 | 0 | 0 | 0 | 3 | 0 | 0 | 0 | 0 | 1 | 6 | 0 | 0 | 0 | 0 | 0 | 266 | 149 | 1 | 1 | NA | 1 | NA | 0.428 | 1 | NA | NA | NA | 1 | 1 | -0.191 |
| A5SS | 6888 | ENSG00000125676 | THOC2 | chrX | - | 123620374 | 123620965 | 123620906 | 123620965 | 123619400 | 123619436 | 25 | 24 | 16 | 36 | 12 | 49 | 37 | 36 | 31 | 49 | 1 | 0 | 11 | 0 | 0 | 0 | 14 | 3 | 0 | 1 | 298 | 149 | 1 | 1 | 0.203 | 0.245 | 0.182 | 0.367 | 0.109 | 1 | 0 | 0.647 | NA | 0 | -0.191 |
| A5SS | 1093 | ENSG00000140650 | PMM2 | chr16 | + | 8849848 | 8851050 | 8849848 | 8849993 | 8855033 | 8855217 | 6 | 5 | 10 | 9 | 0 | 4 | 3 | 0 | 0 | 0 | 18 | 6 | 17 | 32 | 16 | 0 | 0 | 0 | 1 | 5 | 298 | 149 | 1 | 1 | 0.429 | 0.455 | 1 | 1 | NA | 1 | 1 | 1 | 0.941 | 0.615 | -0.19 |
| A5SS | 8890 | ENSG00000123415 | SMUG1 | chr12 | - | 54188676 | 54188957 | 54188725 | 54188957 | 54187818 | 54187905 | 2 | 2 | 0 | 2 | 0 | 2 | 0 | 0 | 0 | 0 | 0 | 0 | 1 | 0 | 0 | 0 | 0 | 0 | 0 | 0 | 198 | 149 | 1 | 1 | 0.429 | 1 | NA | 1 | NA | NA | NA | 1 | NA | NA | -0.19 |
| A5SS | 3933 | ENSG00000182134 | TDRKH | chr1 | - | 151790379 | 151790488 | 151790398 | 151790488 | 151782898 | 151783049 | 7 | 2 | 5 | 2 | 7 | 0 | 0 | 6 | 1 | 0 | 0 | 4 | 1 | 0 | 0 | 0 | 0 | 0 | 0 | 0 | 168 | 149 | 1 | 1 | 1 | 1 | 0.425 | 0.639 | 1 | NA | 1 | 1 | NA | NA | -0.187 |
| A5SS | 4919 | ENSG00000118961 | LDAH | chr2 | - | 20801291 | 20801465 | 20801309 | 20801465 | 20774809 | 20774979 | 2 | 3 | 6 | 3 | 6 | 3 | 3 | 2 | 4 | 1 | 0 | 0 | 2 | 3 | 7 | 0 | 3 | 0 | 0 | 0 | 167 | 149 | 1 | 1 | 0.373 | 0.472 | 0.728 | 0.401 | 0.843 | NA | 0 | 1 | 1 | 1 | -0.187 |
| A5SS | 8637 | ENSG00000224429 | LINC00539 | chr13 | - | 21312145 | 21312365 | 21312278 | 21312365 | 21311041 | 21311118 | 4 | 1 | 9 | 4 | 0 | 1 | 0 | 1 | 4 | 1 | 0 | 9 | 2 | 0 | 0 | 0 | 0 | 1 | 0 | 0 | 282 | 149 | 1 | 1 | 0.679 | 1 | 0.826 | 0.346 | 0 | NA | 1 | 0.514 | NA | NA | -0.187 |
| A5SS | 2575 | ENSG00000203875 | SNHG5 | chr6 | - | 85678497 | 85678748 | 85678697 | 85678748 | 85678135 | 85678188 | 8 | 0 | 13 | 6 | 0 | 0 | 1 | 1 | 1 | 0 | 12 | 2 | 8 | 4 | 10 | 0 | 0 | 1 | 3 | 0 | 298 | 149 | 1 | 1 | 1 | 0 | 0.867 | 0.75 | NA | 1 | 1 | 0.8 | 0.4 | 1 | -0.186 |
| A5SS | 4407 | ENSG00000143851 | PTPN7 | chr1 | - | 202148553 | 202148699 | 202148580 | 202148699 | 202148212 | 202148460 | 1 | 1 | 2 | 10 | 2 | 4 | 2 | 1 | 8 | 0 | 0 | 21 | 5 | 2 | 9 | 0 | 3 | 0 | 1 | 14 | 176 | 149 | 1 | 1 | 0.175 | 0.297 | 0.629 | 0.514 | 1 | NA | 0.856 | 1 | 0.629 | 0.352 | -0.186 |
| A5SS | 4872 | ENSG00000236810 | ELOA-AS1 | chr1 | - | 23772267 | 23772426 | 23772271 | 23772426 | 23772110 | 23772166 | 4 | 1 | 3 | 0 | 0 | 4 | 0 | 3 | 4 | 9 | 0 | 3 | 0 | 2 | 3 | 0 | 4 | 0 | 4 | 0 | 153 | 149 | 1 | 1 | 0.493 | 1 | 0.493 | 0 | 0 | NA | 0.422 | NA | 0.327 | 1 | -0.186 |
| A5SS | 7648 | ENSG00000198000 | NOL8 | chr9 | - | 92324905 | 92325350 | 92325317 | 92325350 | 92324022 | 92324209 | 3 | 2 | 1 | 6 | 8 | 3 | 0 | 2 | 0 | 2 | 0 | 11 | 2 | 5 | 0 | 0 | 6 | 0 | 0 | 0 | 298 | 149 | 1 | 1 | 0.333 | 1 | 0.2 | 1 | 0.667 | NA | 0.478 | 1 | 1 | NA | -0.186 |
| A5SS | 7698 | ENSG00000119408 | NEK6 | chr9 | + | 124261472 | 124261583 | 124261472 | 124261579 | 124301935 | 124302054 | 5 | 0 | 3 | 2 | 5 | 0 | 0 | 1 | 6 | 0 | 41 | 2 | 19 | 18 | 0 | 3 | 0 | 1 | 3 | 0 | 153 | 149 | 1 | 1 | 1 | NA | 0.745 | 0.245 | 1 | 0.93 | 1 | 0.949 | 0.854 | NA | -0.186 |
| A5SS | 8718 | ENSG00000134532 | SOX5 | chr12 | - | 23895792 | 23896024 | 23895897 | 23896024 | 23845982 | 23846193 | 2 | 9 | 0 | 14 | 6 | 3 | 1 | 0 | 0 | 3 | 0 | 16 | 3 | 0 | 0 | 0 | 4 | 0 | 0 | 0 | 254 | 149 | 1 | 1 | 0.281 | 0.841 | NA | 1 | 0.54 | NA | 0.701 | 1 | NA | NA | -0.185 |
| A5SS | 1064 | ENSG00000168488 | ATXN2L | chr16 | + | 28835932 | 28836176 | 28835932 | 28836122 | 28836325 | 28836490 | 0 | 3 | 0 | 1 | 0 | 0 | 0 | 1 | 2 | 3 | 0 | 1 | 2 | 0 | 0 | 1 | 0 | 0 | 0 | 8 | 203 | 149 | 1 | 1 | NA | 1 | 0 | 0.268 | 0 | 0 | 1 | 1 | NA | 0 | -0.183 |
| A5SS | 4164 | ENSG00000003393 | ALS2 | chr2 | - | 201741570 | 201741854 | 201741673 | 201741854 | 201738669 | 201738735 | 4 | 1 | 5 | 3 | 1 | 13 | 9 | 5 | 9 | 11 | 0 | 2 | 8 | 0 | 0 | 4 | 2 | 0 | 11 | 0 | 252 | 149 | 1 | 1 | 0.154 | 0.062 | 0.372 | 0.165 | 0.051 | 0 | 0.372 | 1 | 0 | NA | -0.182 |
| A5SS | 5890 | ENSG00000239665 | NA | chr10 | + | 13646614 | 13646780 | 13646614 | 13646686 | 13647132 | 13647276 | 8 | 8 | 4 | 1 | 1 | 0 | 0 | 1 | 1 | 0 | 0 | 11 | 4 | 8 | 0 | 0 | 0 | 0 | 0 | 0 | 243 | 149 | 1 | 1 | 1 | 1 | 0.71 | 0.38 | 1 | NA | 1 | 1 | 1 | NA | -0.182 |
| A5SS | 2052 | ENSG00000181991 | MRPS11 | chr15 | + | 88467907 | 88469145 | 88467907 | 88468024 | 88475109 | 88475239 | 9 | 3 | 7 | 2 | 5 | 0 | 1 | 0 | 1 | 0 | 0 | 8 | 9 | 7 | 0 | 0 | 0 | 0 | 0 | 0 | 298 | 149 | 1 | 1 | 1 | 0.6 | 1 | 0.5 | 1 | NA | 1 | 1 | 1 | NA | -0.18 |
| A5SS | 8376 | ENSG00000235823 | OLMALINC | chr10 | + | 100373567 | 100374134 | 100373567 | 100374012 | 100381215 | 100381448 | 1 | 0 | 0 | 2 | 0 | 6 | 2 | 6 | 3 | 4 | 0 | 0 | 1 | 0 | 0 | 0 | 7 | 0 | 2 | 6 | 271 | 149 | 1 | 1 | 0.084 | 0 | 0 | 0.268 | 0 | NA | 0 | 1 | 0 | 0 | -0.18 |
| A5SS | 1886 | ENSG00000066933 | MYO9A | chr15 | - | 71951701 | 71951896 | 71951776 | 71951896 | 71938851 | 71938927 | 4 | 2 | 6 | 7 | 0 | 45 | 24 | 35 | 50 | 80 | 0 | 0 | 2 | 0 | 7 | 0 | 21 | 9 | 5 | 1 | 224 | 149 | 1 | 1 | 0.056 | 0.053 | 0.102 | 0.085 | 0 | NA | 0 | 0.129 | 0 | 0.823 | -0.179 |
| A5SS | 2895 | ENSG00000125247 | TMTC4 | chr13 | - | 100670282 | 100670569 | 100670359 | 100670569 | 100668578 | 100668794 | 6 | 5 | 2 | 3 | 1 | 25 | 6 | 22 | 20 | 6 | 0 | 6 | 3 | 12 | 0 | 8 | 7 | 1 | 20 | 0 | 226 | 149 | 1 | 1 | 0.137 | 0.355 | 0.057 | 0.09 | 0.099 | 0 | 0.361 | 0.664 | 0.283 | NA | -0.179 |
| A5SS | 4586 | ENSG00000122678 | POLM | chr7 | - | 44078739 | 44078811 | 44078755 | 44078811 | 44078321 | 44078438 | 2 | 1 | 4 | 2 | 0 | 1 | 0 | 0 | 1 | 0 | 0 | 0 | 0 | 0 | 8 | 0 | 0 | 0 | 0 | 0 | 165 | 149 | 1 | 1 | 0.644 | 1 | 1 | 0.644 | NA | NA | NA | NA | NA | 1 | -0.178 |
| A5SS | 396 | ENSG00000089335 | ZNF302 | chr19 | + | 34677646 | 34677856 | 34677646 | 34677796 | 34678736 | 34678813 | 14 | 19 | 21 | 17 | 18 | 1 | 4 | 8 | 2 | 1 | 14 | 17 | 9 | 2 | 0 | 0 | 0 | 0 | 0 | 0 | 209 | 149 | 1 | 1 | 0.909 | 0.772 | 0.652 | 0.858 | 0.928 | 1 | 1 | 1 | 1 | NA | -0.176 |
| A5SS | 3988 | ENSG00000169398 | PTK2 | chr8 | - | 140925561 | 140925749 | 140925660 | 140925749 | 140890542 | 140890769 | 0 | 1 | 0 | 0 | 0 | 2 | 1 | 2 | 5 | 19 | 8 | 0 | 0 | 0 | 0 | 0 | 3 | 1 | 12 | 0 | 248 | 149 | 1 | 1 | 0 | 0.375 | 0 | 0 | 0 | 1 | 0 | 0 | 0 | NA | -0.175 |
| A5SS | 5059 | ENSG00000255028 | NA | chr11 | - | 109512578 | 109512725 | 109512590 | 109512725 | 109415923 | 109416171 | 0 | 0 | 0 | 1 | 2 | 0 | 0 | 0 | 0 | 1 | 0 | 0 | 0 | 2 | 0 | 0 | 0 | 0 | 0 | 0 | 161 | 149 | 1 | 1 | NA | NA | NA | 1 | 0.649 | NA | NA | NA | 1 | NA | -0.175 |
| A5SS | 276 | ENSG00000251369 | ZNF550 | chr19 | - | 57543650 | 57544583 | 57544236 | 57544583 | 57541839 | 57543243 | 9 | 11 | 21 | 13 | 21 | 0 | 0 | 4 | 3 | 4 | 0 | 4 | 3 | 0 | 0 | 0 | 0 | 0 | 0 | 0 | 298 | 149 | 1 | 1 | 1 | 1 | 0.724 | 0.684 | 0.724 | NA | 1 | 1 | NA | NA | -0.174 |
| A5SS | 3848 | ENSG00000173230 | GOLGB1 | chr3 | - | 121669196 | 121669355 | 121669211 | 121669355 | 121668060 | 121668158 | 18 | 19 | 22 | 13 | 2 | 41 | 14 | 34 | 26 | 30 | 0 | 13 | 10 | 1 | 0 | 3 | 3 | 0 | 5 | 0 | 164 | 149 | 1 | 1 | 0.285 | 0.552 | 0.37 | 0.312 | 0.057 | 0 | 0.797 | 1 | 0.154 | NA | -0.173 |
| A5SS | 4404 | ENSG00000115756 | HPCAL1 | chr2 | + | 10422982 | 10425981 | 10422982 | 10423088 | 10426219 | 10426495 | 0 | 6 | 9 | 12 | 0 | 0 | 0 | 1 | 3 | 0 | 6 | 7 | 5 | 10 | 13 | 0 | 0 | 0 | 0 | 0 | 298 | 149 | 1 | 1 | NA | 1 | 0.818 | 0.667 | NA | 1 | 1 | 1 | 1 | 1 | -0.172 |
| A5SS | 245 | ENSG00000196268 | ZNF493 | chr19 | + | 21397132 | 21397422 | 21397132 | 21397267 | 21405128 | 21405255 | 3 | 8 | 3 | 10 | 3 | 12 | 6 | 11 | 5 | 7 | 0 | 20 | 3 | 1 | 0 | 0 | 10 | 5 | 0 | 10 | 298 | 149 | 1 | 1 | 0.111 | 0.4 | 0.12 | 0.5 | 0.176 | NA | 0.5 | 0.231 | 1 | 0 | -0.171 |
| A5SS | 2411 | ENSG00000163812 | ZDHHC3 | chr3 | - | 44959126 | 44959460 | 44959130 | 44959460 | 44933887 | 44933984 | 0 | 0 | 0 | 0 | 10 | 0 | 0 | 2 | 0 | 0 | 0 | 4 | 8 | 12 | 0 | 0 | 1 | 1 | 0 | 8 | 153 | 149 | 1 | 1 | NA | NA | 0 | NA | 1 | NA | 0.796 | 0.886 | 1 | 0 | -0.17 |
| A5SS | 8199 | ENSG00000247708 | STX18-AS1 | chr4 | + | 4542142 | 4542625 | 4542142 | 4542475 | 4544737 | 4544907 | 33 | 3 | 30 | 11 | 30 | 4 | 0 | 1 | 8 | 0 | 0 | 21 | 18 | 7 | 3 | 0 | 0 | 0 | 0 | 0 | 298 | 149 | 1 | 1 | 0.805 | 1 | 0.938 | 0.407 | 1 | NA | 1 | 1 | 1 | 1 | -0.17 |
| A5SS | 2597 | ENSG00000141562 | NARF | chr17 | + | 82481081 | 82481218 | 82481081 | 82481211 | 82483173 | 82483304 | 3 | 0 | 1 | 1 | 1 | 0 | 0 | 0 | 2 | 0 | 0 | 0 | 3 | 5 | 3 | 0 | 0 | 0 | 0 | 0 | 156 | 149 | 1 | 1 | 1 | NA | 1 | 0.323 | 1 | NA | NA | 1 | 1 | 1 | -0.169 |
| A5SS | 4298 | ENSG00000123636 | BAZ2B | chr2 | - | 159337461 | 159337772 | 159337566 | 159337772 | 159332539 | 159332686 | 49 | 38 | 32 | 36 | 29 | 6 | 2 | 3 | 5 | 10 | 0 | 19 | 25 | 2 | 0 | 0 | 0 | 1 | 0 | 0 | 254 | 149 | 1 | 1 | 0.827 | 0.918 | 0.862 | 0.809 | 0.63 | NA | 1 | 0.936 | 1 | NA | -0.169 |
| A5SS | 8039 | ENSG00000121895 | TMEM156 | chr4 | - | 38998622 | 38998909 | 38998639 | 38998909 | 38993737 | 38993998 | 6 | 12 | 4 | 9 | 29 | 16 | 35 | 50 | 28 | 65 | 0 | 8 | 9 | 7 | 0 | 0 | 18 | 25 | 0 | 4 | 166 | 149 | 1 | 1 | 0.252 | 0.235 | 0.067 | 0.224 | 0.286 | NA | 0.285 | 0.244 | 1 | 0 | -0.169 |
| A5SS | 3605 | ENSG00000121940 | CLCC1 | chr1 | - | 108963342 | 108963484 | 108963360 | 108963484 | 108962308 | 108962469 | 0 | 0 | 0 | 0 | 0 | 6 | 5 | 2 | 2 | 1 | 0 | 8 | 0 | 0 | 0 | 0 | 7 | 4 | 9 | 0 | 167 | 149 | 1 | 1 | 0 | 0 | 0 | 0 | 0 | NA | 0.505 | 0 | 0 | NA | -0.168 |
| A5SS | 1537 | ENSG00000103852 | TTC23 | chr15 | - | 99249170 | 99249799 | 99249621 | 99249799 | 99228532 | 99228732 | 0 | 2 | 2 | 0 | 1 | 0 | 0 | 1 | 0 | 0 | 0 | 3 | 0 | 0 | 8 | 0 | 0 | 0 | 0 | 0 | 298 | 149 | 1 | 1 | NA | 1 | 0.5 | NA | 1 | NA | 1 | NA | NA | 1 | -0.167 |
| A5SS | 3035 | ENSG00000127483 | HP1BP3 | chr1 | - | 20786252 | 20786652 | 20786648 | 20786652 | 20779811 | 20779911 | 3 | 0 | 0 | 0 | 0 | 0 | 0 | 2 | 0 | 0 | 0 | 5 | 0 | 0 | 1 | 8 | 0 | 0 | 0 | 0 | 298 | 149 | 1 | 1 | 1 | NA | 0 | NA | NA | 0 | 1 | NA | NA | 1 | -0.167 |
| A5SS | 4554 | ENSG00000132681 | ATP1A4 | chr1 | + | 160167277 | 160167429 | 160167277 | 160167412 | 160171250 | 160171440 | 0 | 0 | 0 | 0 | 3 | 2 | 0 | 0 | 0 | 0 | 0 | 0 | 3 | 0 | 2 | 0 | 0 | 0 | 2 | 0 | 166 | 149 | 1 | 1 | 0 | NA | NA | NA | 1 | NA | NA | 1 | 0 | 1 | -0.167 |
| A5SS | 4635 | ENSG00000104714 | ERICH1 | chr8 | - | 625674 | 627238 | 627002 | 627238 | 614745 | 615284 | 2 | 1 | 0 | 1 | 2 | 0 | 1 | 0 | 0 | 0 | 7 | 5 | 1 | 13 | 4 | 0 | 0 | 0 | 0 | 0 | 298 | 149 | 1 | 1 | 1 | 0.333 | NA | 1 | 1 | 1 | 1 | 1 | 1 | 1 | -0.167 |
| A5SS | 7188 | ENSG00000198954 | KIFBP | chr10 | + | 69005731 | 69007751 | 69005731 | 69005915 | 69010899 | 69011015 | 4 | 3 | 2 | 2 | 3 | 1 | 0 | 0 | 1 | 0 | 0 | 13 | 0 | 0 | 0 | 0 | 0 | 0 | 0 | 0 | 298 | 149 | 1 | 1 | 0.667 | 1 | 1 | 0.5 | 1 | NA | 1 | NA | NA | NA | -0.167 |
| A5SS | 8375 | ENSG00000235823 | OLMALINC | chr10 | + | 100373567 | 100374080 | 100373567 | 100374012 | 100381215 | 100381448 | 1 | 0 | 0 | 2 | 0 | 6 | 2 | 6 | 3 | 4 | 0 | 0 | 1 | 0 | 0 | 0 | 7 | 0 | 2 | 6 | 217 | 149 | 1 | 1 | 0.103 | 0 | 0 | 0.314 | 0 | NA | 0 | 1 | 0 | 0 | -0.167 |
| A5SS | 4376 | ENSG00000256128 | LINC00944 | chr12 | - | 126732310 | 126732407 | 126732342 | 126732407 | 126730697 | 126730803 | 0 | 0 | 0 | 1 | 1 | 7 | 1 | 3 | 12 | 6 | 0 | 5 | 0 | 0 | 0 | 0 | 6 | 3 | 0 | 0 | 181 | 149 | 1 | 1 | 0 | 0 | 0 | 0.064 | 0.121 | NA | 0.407 | 0 | NA | NA | -0.166 |
| A5SS | 5407 | ENSG00000003509 | NDUFAF7 | chr2 | + | 37241577 | 37241791 | 37241577 | 37241637 | 37243862 | 37243973 | 19 | 23 | 19 | 28 | 48 | 3 | 2 | 8 | 5 | 6 | 15 | 14 | 19 | 13 | 0 | 0 | 0 | 6 | 0 | 0 | 298 | 149 | 1 | 1 | 0.76 | 0.852 | 0.543 | 0.737 | 0.8 | 1 | 1 | 0.613 | 1 | NA | -0.165 |
| A5SS | 629 | ENSG00000166401 | SERPINB8 | chr18 | + | 63970028 | 63970187 | 63970028 | 63970170 | 63979800 | 63979938 | 2 | 1 | 1 | 1 | 0 | 1 | 4 | 2 | 2 | 0 | 0 | 3 | 0 | 3 | 0 | 3 | 2 | 0 | 0 | 0 | 166 | 149 | 1 | 1 | 0.642 | 0.183 | 0.31 | 0.31 | NA | 0 | 0.574 | NA | 1 | NA | -0.163 |
| A5SS | 1050 | ENSG00000172530 | BANP | chr16 | + | 87975047 | 87975203 | 87975047 | 87975185 | 87978439 | 87978692 | 8 | 4 | 2 | 2 | 0 | 1 | 0 | 2 | 0 | 0 | 0 | 6 | 6 | 0 | 5 | 0 | 0 | 0 | 0 | 0 | 167 | 149 | 1 | 1 | 0.877 | 1 | 0.472 | 1 | NA | NA | 1 | 1 | NA | 1 | -0.163 |
| A5SS | 7129 | ENSG00000197114 | ZGPAT | chr20 | + | 63733586 | 63733775 | 63733586 | 63733739 | 63734644 | 63734824 | 2 | 2 | 8 | 0 | 2 | 0 | 3 | 0 | 0 | 2 | 19 | 0 | 3 | 4 | 33 | 0 | 0 | 0 | 4 | 0 | 185 | 149 | 1 | 1 | 1 | 0.349 | 1 | NA | 0.446 | 1 | NA | 1 | 0.446 | 1 | -0.163 |
| A5SS | 1286 | ENSG00000123815 | COQ8B | chr19 | - | 40703368 | 40703622 | 40703540 | 40703622 | 40702599 | 40702693 | 2 | 0 | 1 | 2 | 1 | 14 | 7 | 15 | 6 | 2 | 3 | 0 | 0 | 0 | 0 | 0 | 4 | 5 | 6 | 0 | 298 | 149 | 1 | 1 | 0.067 | 0 | 0.032 | 0.143 | 0.2 | 1 | 0 | 0 | 0 | NA | -0.162 |
| A5SS | 7748 | ENSG00000172716 | SLFN11 | chr17 | - | 35373473 | 35373620 | 35373581 | 35373620 | 35366946 | 35367063 | 9 | 4 | 12 | 9 | 2 | 0 | 1 | 3 | 0 | 2 | 0 | 15 | 1 | 9 | 0 | 0 | 0 | 3 | 0 | 0 | 257 | 149 | 1 | 1 | 1 | 0.699 | 0.699 | 1 | 0.367 | NA | 0.744 | 1 | 1 | NA | -0.162 |
| A5SS | 74 | ENSG00000232855 | NA | chr21 | - | 28577152 | 28577380 | 28577218 | 28577380 | 28446289 | 28446472 | 8 | 10 | 7 | 11 | 0 | 2 | 0 | 1 | 2 | 0 | 0 | 5 | 3 | 1 | 0 | 0 | 0 | 0 | 0 | 0 | 215 | 149 | 1 | 1 | 0.735 | 1 | 0.829 | 0.792 | NA | NA | 1 | 1 | 1 | NA | -0.161 |
| A5SS | 2715 | ENSG00000087266 | SH3BP2 | chr4 | + | 2823394 | 2823573 | 2823394 | 2823569 | 2824612 | 2824730 | 6 | 0 | 1 | 2 | 0 | 10 | 6 | 5 | 4 | 0 | 1 | 0 | 6 | 9 | 14 | 21 | 7 | 4 | 10 | 4 | 153 | 149 | 1 | 1 | 0.369 | 0 | 0.163 | 0.327 | NA | 0.044 | 0 | 0.594 | 0.467 | 0.773 | -0.161 |
| A5SS | 5456 | ENSG00000188282 | RUFY4 | chr2 | + | 218073815 | 218073945 | 218073815 | 218073885 | 218075092 | 218075740 | 1 | 0 | 0 | 1 | 0 | 5 | 1 | 0 | 1 | 0 | 18 | 1 | 2 | 7 | 7 | 19 | 11 | 12 | 31 | 0 | 209 | 149 | 1 | 1 | 0.125 | 0 | NA | 0.416 | NA | 0.403 | 0.061 | 0.106 | 0.139 | 1 | -0.161 |
| A5SS | 3937 | ENSG00000073282 | TP63 | chr3 | + | 189868579 | 189868716 | 189868579 | 189868704 | 189869323 | 189869406 | 0 | 1 | 6 | 3 | 0 | 1 | 1 | 1 | 0 | 1 | 4 | 0 | 1 | 0 | 0 | 11 | 0 | 0 | 0 | 0 | 161 | 149 | 1 | 1 | 0 | 0.481 | 0.847 | 1 | 0 | 0.252 | NA | 1 | NA | NA | -0.16 |
| A5SS | 3100 | ENSG00000162736 | NCSTN | chr1 | + | 160343703 | 160343888 | 160343703 | 160343802 | 160344721 | 160344826 | 7 | 6 | 10 | 12 | 5 | 1 | 0 | 0 | 0 | 5 | 90 | 32 | 25 | 28 | 0 | 0 | 0 | 0 | 0 | 0 | 235 | 149 | 1 | 1 | 0.816 | 1 | 1 | 1 | 0.388 | 1 | 1 | 1 | 1 | NA | -0.159 |
| A5SS | 6220 | ENSG00000170919 | TPT1-AS1 | chr13 | + | 45383759 | 45383843 | 45383759 | 45383838 | 45389734 | 45389820 | 1 | 0 | 0 | 0 | 2 | 5 | 8 | 9 | 3 | 5 | 0 | 2 | 5 | 0 | 0 | 0 | 21 | 7 | 0 | 0 | 154 | 149 | 1 | 1 | 0.162 | 0 | 0 | 0 | 0.279 | NA | 0.084 | 0.409 | NA | NA | -0.158 |
| A5SS | 8938 | ENSG00000124496 | TRERF1 | chr6 | - | 42263259 | 42263568 | 42263319 | 42263568 | 42259338 | 42259723 | 3 | 7 | 5 | 7 | 2 | 1 | 3 | 4 | 1 | 0 | 0 | 0 | 5 | 6 | 17 | 0 | 0 | 2 | 0 | 0 | 209 | 149 | 1 | 1 | 0.681 | 0.625 | 0.471 | 0.833 | 1 | NA | NA | 0.641 | 1 | 1 | -0.158 |
| A5SS | 6420 | ENSG00000001631 | KRIT1 | chr7 | - | 92245789 | 92245914 | 92245843 | 92245914 | 92244901 | 92245171 | 2 | 0 | 7 | 10 | 11 | 1 | 0 | 0 | 2 | 0 | 0 | 3 | 2 | 0 | 3 | 0 | 0 | 0 | 0 | 0 | 203 | 149 | 1 | 1 | 0.595 | NA | 1 | 0.786 | 1 | NA | 1 | 1 | NA | 1 | -0.155 |
| A5SS | 1990 | ENSG00000125459 | MSTO1 | chr1 | + | 155611688 | 155611852 | 155611688 | 155611827 | 155611982 | 155612100 | 5 | 0 | 0 | 10 | 3 | 17 | 0 | 8 | 14 | 5 | 26 | 9 | 0 | 0 | 0 | 6 | 6 | 4 | 7 | 0 | 174 | 149 | 1 | 1 | 0.201 | 0 | 0 | 0.38 | 0.339 | 0.788 | 0.562 | 0 | 0 | NA | -0.154 |
| A5SS | 3444 | ENSG00000144591 | GMPPA | chr2 | + | 219504082 | 219504868 | 219504082 | 219504218 | 219505227 | 219505362 | 15 | 5 | 4 | 13 | 2 | 0 | 1 | 0 | 2 | 1 | 0 | 5 | 8 | 19 | 11 | 0 | 0 | 1 | 0 | 0 | 298 | 149 | 1 | 1 | 1 | 0.714 | 1 | 0.765 | 0.5 | NA | 1 | 0.8 | 1 | 1 | -0.154 |
| A5SS | 5458 | ENSG00000235703 | LINC00894 | chrX | - | 149945783 | 149948175 | 149945783 | 149947347 | 149948305 | 149948394 | 19 | 9 | 7 | 13 | 3 | 1 | 4 | 5 | 4 | 10 | 22 | 9 | 5 | 0 | 0 | 0 | 2 | 0 | 6 | 0 | 298 | 149 | 1 | 1 | 0.905 | 0.529 | 0.412 | 0.619 | 0.13 | 1 | 0.692 | 1 | 0 | NA | -0.154 |
| A5SS | 8024 | ENSG00000005801 | ZNF195 | chr11 | - | 3370942 | 3371070 | 3370974 | 3371070 | 3361742 | 3361889 | 2 | 9 | 8 | 5 | 6 | 3 | 1 | 3 | 0 | 3 | 0 | 2 | 10 | 5 | 7 | 0 | 0 | 5 | 0 | 0 | 181 | 149 | 1 | 1 | 0.354 | 0.881 | 0.687 | 1 | 0.622 | NA | 1 | 1 | 0.452 | 1 | -0.154 |
| A5SS | 4351 | ENSG00000186049 | KRT73 | chr12 | - | 52610614 | 52611329 | 52611203 | 52611329 | 52609246 | 52609281 | 10 | 7 | 36 | 31 | 17 | 7 | 0 | 4 | 0 | 0 | 0 | 19 | 11 | 6 | 10 | 0 | 0 | 0 | 0 | 0 | 298 | 149 | 1 | 1 | 0.417 | 1 | 0.818 | 1 | 1 | NA | 1 | 1 | 1 | 1 | -0.153 |
| A5SS | 6388 | ENSG00000183735 | TBK1 | chr12 | + | 64480011 | 64480131 | 64480011 | 64480122 | 64481851 | 64482021 | 2 | 1 | 1 | 0 | 2 | 3 | 0 | 0 | 0 | 0 | 0 | 3 | 2 | 0 | 0 | 0 | 0 | 0 | 0 | 0 | 158 | 149 | 1 | 1 | 0.386 | 1 | 1 | NA | 1 | NA | 1 | 1 | NA | NA | -0.153 |
| A5SS | 7592 | ENSG00000196781 | TLE1 | chr9 | - | 81652183 | 81652288 | 81652213 | 81652288 | 81633347 | 81633364 | 5 | 3 | 4 | 2 | 0 | 2 | 1 | 0 | 0 | 0 | 0 | 3 | 5 | 5 | 4 | 0 | 0 | 0 | 0 | 0 | 179 | 149 | 1 | 1 | 0.675 | 0.714 | 1 | 1 | NA | NA | 1 | 1 | 1 | 1 | -0.153 |
| A5SS | 4163 | ENSG00000003393 | ALS2 | chr2 | - | 201727700 | 201727775 | 201727704 | 201727775 | 201727211 | 201727278 | 0 | 2 | 1 | 0 | 0 | 6 | 1 | 3 | 6 | 8 | 0 | 6 | 0 | 0 | 0 | 3 | 0 | 2 | 0 | 0 | 153 | 149 | 1 | 1 | 0 | 0.661 | 0.245 | 0 | 0 | 0 | 1 | 0 | NA | NA | -0.152 |
| A5SS | 6327 | ENSG00000223705 | NSUN5P1 | chr7 | + | 75415591 | 75415778 | 75415591 | 75415770 | 75415980 | 75416191 | 1 | 3 | 0 | 6 | 0 | 3 | 1 | 5 | 2 | 5 | 3 | 0 | 5 | 0 | 0 | 0 | 0 | 5 | 0 | 7 | 157 | 149 | 1 | 1 | 0.24 | 0.74 | 0 | 0.74 | 0 | 1 | NA | 0.487 | NA | 0 | -0.152 |
| A5SS | 4156 | ENSG00000131115 | ZNF227 | chr19 | + | 44228445 | 44228633 | 44228445 | 44228572 | 44229732 | 44229816 | 5 | 1 | 4 | 4 | 9 | 17 | 17 | 24 | 15 | 34 | 0 | 2 | 12 | 4 | 0 | 6 | 16 | 11 | 2 | 0 | 210 | 149 | 1 | 1 | 0.173 | 0.04 | 0.106 | 0.159 | 0.158 | 0 | 0.081 | 0.436 | 0.587 | NA | -0.149 |
| A5SS | 8 | ENSG00000186866 | POFUT2 | chr21 | - | 45283351 | 45283527 | 45283382 | 45283527 | 45282803 | 45282938 | 4 | 0 | 0 | 0 | 0 | 3 | 0 | 3 | 4 | 0 | 0 | 1 | 0 | 3 | 0 | 0 | 2 | 2 | 0 | 1 | 180 | 149 | 1 | 1 | 0.525 | NA | 0 | 0 | NA | NA | 0.293 | 0 | 1 | 0 | -0.148 |
| A5SS | 2841 | ENSG00000169696 | ASPSCR1 | chr17 | + | 82016940 | 82017113 | 82016940 | 82017088 | 82017308 | 82017403 | 2 | 4 | 10 | 2 | 2 | 1 | 0 | 0 | 0 | 1 | 1 | 15 | 1 | 0 | 33 | 0 | 0 | 0 | 0 | 0 | 174 | 149 | 1 | 1 | 0.631 | 1 | 1 | 1 | 0.631 | 1 | 1 | 1 | NA | 1 | -0.148 |
| A5SS | 6340 | ENSG00000173715 | C11orf80 | chr11 | + | 66788159 | 66788273 | 66788159 | 66788269 | 66796280 | 66796364 | 1 | 0 | 1 | 1 | 0 | 7 | 0 | 5 | 7 | 10 | 7 | 0 | 0 | 0 | 0 | 0 | 0 | 6 | 9 | 18 | 153 | 149 | 1 | 1 | 0.122 | NA | 0.163 | 0.122 | 0 | 1 | NA | 0 | 0 | 0 | -0.148 |
| A5SS | 3877 | ENSG00000185864 | NPIPB4 | chr16 | - | 21879337 | 21879457 | 21879377 | 21879457 | 21857579 | 21857658 | 4 | 4 | 4 | 0 | 5 | 2 | 0 | 0 | 0 | 1 | 2 | 2 | 3 | 8 | 0 | 0 | 0 | 0 | 0 | 0 | 189 | 149 | 1 | 1 | 0.612 | 1 | 1 | NA | 0.798 | 1 | 1 | 1 | 1 | NA | -0.147 |
| A5SS | 5648 | ENSG00000151718 | WWC2 | chr4 | + | 183280775 | 183280969 | 183280775 | 183280897 | 183282707 | 183282906 | 0 | 2 | 0 | 2 | 2 | 10 | 8 | 14 | 4 | 10 | 0 | 0 | 1 | 0 | 0 | 14 | 6 | 0 | 11 | 0 | 221 | 149 | 1 | 1 | 0 | 0.144 | 0 | 0.252 | 0.119 | 0 | 0 | 1 | 0 | NA | -0.147 |
| A5SS | 6086 | ENSG00000247134 | NA | chr8 | + | 32996200 | 32996355 | 32996200 | 32996316 | 33041634 | 33041771 | 0 | 0 | 0 | 0 | 0 | 1 | 0 | 2 | 3 | 0 | 0 | 0 | 0 | 0 | 1 | 0 | 0 | 3 | 2 | 1 | 188 | 149 | 1 | 1 | 0 | NA | 0 | 0 | NA | NA | NA | 0 | 0 | 0.442 | -0.147 |
| A5SS | 3381 | ENSG00000095321 | CRAT | chr9 | - | 129109102 | 129109277 | 129109178 | 129109277 | 129107813 | 129108077 | 0 | 1 | 1 | 3 | 0 | 7 | 3 | 5 | 8 | 0 | 0 | 6 | 0 | 3 | 4 | 0 | 3 | 0 | 10 | 32 | 225 | 149 | 1 | 1 | 0 | 0.181 | 0.117 | 0.199 | NA | NA | 0.57 | NA | 0.166 | 0.076 | -0.146 |
| A5SS | 7128 | ENSG00000046647 | GEMIN8 | chrX | - | 14026051 | 14026221 | 14026139 | 14026221 | 14021463 | 14021511 | 12 | 15 | 14 | 9 | 14 | 8 | 5 | 9 | 15 | 18 | 0 | 4 | 18 | 9 | 0 | 0 | 4 | 4 | 3 | 0 | 237 | 149 | 1 | 1 | 0.485 | 0.654 | 0.494 | 0.274 | 0.328 | NA | 0.386 | 0.739 | 0.654 | NA | -0.146 |
| A5SS | 7338 | ENSG00000156795 | NTAQ1 | chr8 | + | 123416773 | 123416932 | 123416773 | 123416856 | 123427923 | 123428023 | 12 | 0 | 5 | 12 | 5 | 1 | 3 | 2 | 2 | 8 | 0 | 1 | 3 | 0 | 4 | 11 | 0 | 1 | 0 | 0 | 225 | 149 | 1 | 1 | 0.888 | 0 | 0.623 | 0.799 | 0.293 | 0 | 1 | 0.665 | NA | 1 | -0.146 |
| A5SS | 6018 | ENSG00000183960 | KCNH8 | chr3 | + | 19450105 | 19451404 | 19450105 | 19450305 | 19456767 | 19456982 | 14 | 6 | 5 | 9 | 6 | 2 | 0 | 0 | 0 | 3 | 0 | 13 | 6 | 3 | 2 | 0 | 0 | 0 | 0 | 0 | 298 | 149 | 1 | 1 | 0.778 | 1 | 1 | 1 | 0.5 | NA | 1 | 1 | 1 | 1 | -0.144 |
| A5SS | 3285 | ENSG00000111961 | SASH1 | chr6 | + | 148519546 | 148520382 | 148519546 | 148519893 | 148525290 | 148525365 | 0 | 3 | 2 | 6 | 0 | 6 | 7 | 12 | 4 | 7 | 5 | 0 | 0 | 1 | 6 | 27 | 6 | 0 | 15 | 0 | 298 | 149 | 1 | 1 | 0 | 0.176 | 0.077 | 0.429 | 0 | 0.085 | 0 | NA | 0.032 | 1 | -0.143 |
| A5SS | 8534 | ENSG00000066651 | TRMT11 | chr6 | + | 125986552 | 125986773 | 125986552 | 125986690 | 125993756 | 125993822 | 10 | 0 | 4 | 5 | 3 | 1 | 0 | 2 | 0 | 0 | 0 | 2 | 7 | 0 | 4 | 0 | 0 | 0 | 0 | 0 | 232 | 149 | 1 | 1 | 0.865 | NA | 0.562 | 1 | 1 | NA | 1 | 1 | NA | 1 | -0.143 |
| A5SS | 3357 | ENSG00000085465 | OVGP1 | chr1 | - | 111421275 | 111421673 | 111421564 | 111421673 | 111419609 | 111419726 | 4 | 6 | 14 | 8 | 15 | 0 | 1 | 1 | 2 | 0 | 0 | 6 | 11 | 1 | 15 | 0 | 0 | 0 | 0 | 0 | 298 | 149 | 1 | 1 | 1 | 0.75 | 0.875 | 0.667 | 1 | NA | 1 | 1 | 1 | 1 | -0.142 |
| A5SS | 8105 | ENSG00000132199 | ENOSF1 | chr18 | - | 712377 | 712630 | 712503 | 712630 | 706469 | 706578 | 1 | 1 | 0 | 0 | 0 | 1 | 6 | 8 | 10 | 0 | 0 | 0 | 0 | 0 | 5 | 0 | 3 | 3 | 1 | 0 | 275 | 149 | 1 | 1 | 0.351 | 0.083 | 0 | 0 | NA | NA | 0 | 0 | 0 | 1 | -0.142 |
| A5SS | 226 | ENSG00000005007 | UPF1 | chr19 | + | 18849007 | 18849076 | 18849007 | 18849031 | 18850074 | 18850242 | 3 | 4 | 2 | 5 | 0 | 3 | 0 | 0 | 0 | 0 | 39 | 21 | 6 | 9 | 14 | 0 | 0 | 0 | 0 | 0 | 194 | 149 | 1 | 1 | 0.434 | 1 | 1 | 1 | NA | 1 | 1 | 1 | 1 | 1 | -0.141 |
| A5SS | 8118 | ENSG00000105808 | RASA4 | chr7 | - | 102594446 | 102594559 | 102594491 | 102594559 | 102594049 | 102594242 | 4 | 2 | 0 | 1 | 3 | 0 | 2 | 0 | 0 | 0 | 20 | 0 | 0 | 0 | 1 | 0 | 0 | 0 | 0 | 0 | 194 | 149 | 1 | 1 | 1 | 0.434 | NA | 1 | 1 | 1 | NA | NA | 1 | NA | -0.141 |
| A5SS | 8256 | ENSG00000261008 | LINC01572 | chr16 | - | 72570434 | 72570515 | 72570450 | 72570515 | 72555264 | 72555326 | 2 | 7 | 1 | 0 | 0 | 1 | 1 | 5 | 2 | 4 | 0 | 8 | 4 | 0 | 0 | 13 | 0 | 5 | 0 | 0 | 165 | 149 | 1 | 1 | 0.644 | 0.863 | 0.153 | 0 | 0 | 0 | 1 | 0.419 | NA | NA | -0.141 |
| A5SS | 64 | ENSG00000161179 | YDJC | chr22 | - | 21629542 | 21629761 | 21629601 | 21629761 | 21629307 | 21629407 | 19 | 8 | 10 | 13 | 2 | 2 | 0 | 2 | 1 | 1 | 0 | 13 | 10 | 3 | 5 | 0 | 0 | 1 | 0 | 0 | 208 | 149 | 1 | 1 | 0.872 | 1 | 0.782 | 0.903 | 0.589 | NA | 1 | 0.878 | 1 | 1 | -0.14 |
| A5SS | 362 | ENSG00000159917 | ZNF235 | chr19 | - | 44298807 | 44298903 | 44298819 | 44298903 | 44236109 | 44236729 | 0 | 0 | 3 | 0 | 0 | 0 | 2 | 2 | 3 | 0 | 0 | 0 | 0 | 8 | 0 | 0 | 0 | 2 | 0 | 1 | 161 | 149 | 1 | 1 | NA | 0 | 0.581 | 0 | NA | NA | NA | 0 | 1 | 0 | -0.14 |
| A5SS | 5015 | ENSG00000124067 | SLC12A4 | chr16 | - | 67962243 | 67963559 | 67963464 | 67963559 | 67961574 | 67961706 | 4 | 0 | 4 | 1 | 1 | 3 | 2 | 2 | 1 | 5 | 0 | 1 | 4 | 3 | 0 | 0 | 0 | 5 | 3 | 18 | 298 | 149 | 1 | 1 | 0.4 | 0 | 0.5 | 0.333 | 0.091 | NA | 1 | 0.286 | 0.333 | 0 | -0.14 |
| A5SS | 2341 | ENSG00000150756 | ATPSCKMT | chr5 | - | 10236477 | 10236615 | 10236521 | 10236615 | 10235210 | 10235261 | 10 | 9 | 11 | 6 | 24 | 5 | 0 | 0 | 2 | 0 | 10 | 0 | 6 | 9 | 0 | 0 | 0 | 0 | 0 | 0 | 193 | 149 | 1 | 1 | 0.607 | 1 | 1 | 0.698 | 1 | 1 | NA | 1 | 1 | NA | -0.139 |
| A5SS | 5691 | ENSG00000243970 | PPIEL | chr1 | - | 39529015 | 39529164 | 39529054 | 39529164 | 39525698 | 39525848 | 5 | 2 | 1 | 10 | 0 | 0 | 2 | 0 | 0 | 0 | 2 | 4 | 3 | 5 | 0 | 0 | 0 | 0 | 0 | 0 | 188 | 149 | 1 | 1 | 1 | 0.442 | 1 | 1 | NA | 1 | 1 | 1 | 1 | NA | -0.139 |
| A5SS | 5252 | ENSG00000105948 | TTC26 | chr7 | + | 139134654 | 139134792 | 139134654 | 139134721 | 139137846 | 139137939 | 8 | 5 | 2 | 5 | 3 | 0 | 0 | 3 | 0 | 0 | 0 | 0 | 6 | 7 | 0 | 0 | 0 | 0 | 0 | 0 | 220 | 149 | 1 | 1 | 1 | 1 | 0.311 | 1 | 1 | NA | NA | 1 | 1 | NA | -0.138 |
| A5SS | 708 | ENSG00000270379 | HEATR9 | chr17 | - | 35864757 | 35864890 | 35864877 | 35864890 | 35864496 | 35864553 | 3 | 5 | 4 | 3 | 0 | 2 | 0 | 0 | 0 | 0 | 0 | 3 | 2 | 6 | 0 | 0 | 0 | 0 | 0 | 0 | 269 | 149 | 1 | 1 | 0.454 | 1 | 1 | 1 | NA | NA | 1 | 1 | 1 | NA | -0.137 |
| A5SS | 2885 | ENSG00000147231 | RADX | chrX | + | 106637759 | 106637924 | 106637759 | 106637794 | 106640551 | 106640721 | 18 | 21 | 22 | 20 | 11 | 2 | 0 | 2 | 3 | 1 | 16 | 7 | 21 | 3 | 0 | 0 | 0 | 0 | 0 | 0 | 279 | 149 | 1 | 1 | 0.828 | 1 | 0.855 | 0.781 | 0.855 | 1 | 1 | 1 | 1 | NA | -0.136 |
| A5SS | 5089 | ENSG00000138050 | THUMPD2 | chr2 | - | 39771504 | 39771640 | 39771514 | 39771640 | 39768423 | 39768501 | 1 | 1 | 5 | 3 | 1 | 2 | 0 | 0 | 0 | 0 | 0 | 0 | 4 | 0 | 0 | 0 | 0 | 0 | 0 | 0 | 159 | 149 | 1 | 1 | 0.319 | 1 | 1 | 1 | 1 | NA | NA | 1 | NA | NA | -0.136 |
| A5SS | 5296 | ENSG00000224078 | SNHG14 | chr15 | + | 25111764 | 25115656 | 25111764 | 25111894 | 25117409 | 25117515 | 15 | 17 | 10 | 12 | 14 | 1 | 3 | 0 | 0 | 3 | 0 | 18 | 10 | 3 | 0 | 0 | 0 | 0 | 0 | 0 | 298 | 149 | 1 | 1 | 0.882 | 0.739 | 1 | 1 | 0.7 | NA | 1 | 1 | 1 | NA | -0.136 |
| A5SS | 7852 | ENSG00000167554 | ZNF610 | chr19 | + | 52347706 | 52347944 | 52347706 | 52347831 | 52349153 | 52349235 | 2 | 5 | 1 | 3 | 5 | 0 | 0 | 0 | 0 | 6 | 0 | 5 | 0 | 2 | 0 | 0 | 0 | 0 | 0 | 0 | 262 | 149 | 1 | 1 | 1 | 1 | 1 | 1 | 0.322 | NA | 1 | NA | 1 | NA | -0.136 |
| A5SS | 1304 | ENSG00000103248 | MTHFSD | chr16 | - | 86555168 | 86555235 | 86555172 | 86555235 | 86554644 | 86554748 | 8 | 2 | 3 | 4 | 3 | 0 | 4 | 0 | 0 | 0 | 0 | 8 | 1 | 0 | 5 | 0 | 0 | 0 | 0 | 0 | 153 | 149 | 1 | 1 | 1 | 0.327 | 1 | 1 | 1 | NA | 1 | 1 | NA | 1 | -0.135 |
| A5SS | 1597 | ENSG00000206149 | HERC2P9 | chr15 | + | 28641169 | 28641312 | 28641169 | 28641226 | 28641429 | 28641581 | 8 | 1 | 1 | 4 | 5 | 1 | 3 | 2 | 4 | 17 | 30 | 0 | 7 | 0 | 0 | 5 | 2 | 2 | 0 | 0 | 235 | 149 | 1 | 1 | 0.835 | 0.174 | 0.241 | 0.388 | 0.157 | 0.792 | 0 | 0.689 | NA | NA | -0.135 |
| A5SS | 7801 | ENSG00000185920 | PTCH1 | chr9 | - | 95449840 | 95453620 | 95453477 | 95453620 | 95449068 | 95449323 | 41 | 7 | 23 | 21 | 11 | 6 | 1 | 4 | 6 | 0 | 0 | 35 | 3 | 10 | 20 | 0 | 8 | 0 | 0 | 0 | 298 | 149 | 1 | 1 | 0.774 | 0.778 | 0.742 | 0.636 | 1 | NA | 0.686 | 1 | 1 | 1 | -0.135 |
| A5SS | 7942 | ENSG00000066294 | CD84 | chr1 | - | 160553359 | 160553497 | 160553377 | 160553497 | 160552704 | 160552737 | 0 | 0 | 5 | 0 | 4 | 1 | 5 | 3 | 3 | 12 | 0 | 5 | 4 | 0 | 0 | 18 | 0 | 14 | 16 | 0 | 167 | 149 | 1 | 1 | 0 | 0 | 0.598 | 0 | 0.229 | 0 | 1 | 0.203 | 0 | NA | -0.135 |
| A5SS | 532 | ENSG00000088340 | FER1L4 | chr20 | - | 35560122 | 35560638 | 35560539 | 35560638 | 35559340 | 35559627 | 0 | 0 | 1 | 0 | 0 | 0 | 0 | 1 | 0 | 0 | 0 | 2 | 4 | 0 | 0 | 0 | 0 | 3 | 1 | 0 | 298 | 149 | 1 | 1 | NA | NA | 0.333 | NA | NA | NA | 1 | 0.4 | 0 | NA | -0.134 |
| A5SS | 1135 | ENSG00000010361 | FUZ | chr19 | - | 49812614 | 49812736 | 49812637 | 49812736 | 49811630 | 49811699 | 1 | 0 | 3 | 1 | 2 | 0 | 0 | 0 | 1 | 0 | 6 | 0 | 1 | 13 | 7 | 0 | 0 | 0 | 0 | 0 | 172 | 149 | 1 | 1 | 1 | NA | 1 | 0.464 | 1 | 1 | NA | 1 | 1 | 1 | -0.134 |
| A5SS | 2420 | ENSG00000167995 | BEST1 | chr11 | + | 61958145 | 61958501 | 61958145 | 61958298 | 61959497 | 61959578 | 6 | 3 | 4 | 5 | 1 | 1 | 0 | 2 | 1 | 3 | 0 | 3 | 5 | 3 | 7 | 0 | 0 | 2 | 0 | 4 | 298 | 149 | 1 | 1 | 0.75 | 1 | 0.5 | 0.714 | 0.143 | NA | 1 | 0.556 | 1 | 0.467 | -0.134 |
| A5SS | 4968 | ENSG00000171503 | ETFDH | chr4 | + | 158680466 | 158681614 | 158680466 | 158680607 | 158682194 | 158682424 | 15 | 21 | 23 | 18 | 17 | 9 | 24 | 11 | 17 | 16 | 12 | 30 | 32 | 13 | 0 | 19 | 0 | 8 | 26 | 0 | 298 | 149 | 1 | 1 | 0.455 | 0.304 | 0.511 | 0.346 | 0.347 | 0.24 | 1 | 0.667 | 0.2 | NA | -0.134 |
| A5SS | 7236 | ENSG00000227678 | NA | chr6 | - | 130142683 | 130142813 | 130142751 | 130142813 | 130133565 | 130133719 | 14 | 10 | 24 | 18 | 6 | 7 | 3 | 0 | 4 | 6 | 0 | 7 | 15 | 5 | 10 | 0 | 12 | 0 | 0 | 0 | 217 | 149 | 1 | 1 | 0.579 | 0.696 | 1 | 0.755 | 0.407 | NA | 0.286 | 1 | 1 | 1 | -0.134 |
| A5SS | 1411 | ENSG00000047346 | FAM214A | chr15 | - | 52678827 | 52679372 | 52679275 | 52679372 | 52678005 | 52678122 | 13 | 4 | 7 | 2 | 3 | 0 | 0 | 0 | 0 | 3 | 0 | 0 | 8 | 0 | 2 | 0 | 0 | 0 | 0 | 0 | 298 | 149 | 1 | 1 | 1 | 1 | 1 | 1 | 0.333 | NA | NA | 1 | NA | 1 | -0.133 |
| A5SS | 2158 | ENSG00000164398 | ACSL6 | chr5 | - | 131994030 | 131994251 | 131994135 | 131994251 | 131990852 | 131990967 | 1 | 1 | 1 | 0 | 9 | 0 | 0 | 0 | 0 | 6 | 0 | 0 | 0 | 6 | 0 | 0 | 0 | 0 | 0 | 0 | 254 | 149 | 1 | 1 | 1 | 1 | 1 | NA | 0.468 | NA | NA | NA | 1 | NA | -0.133 |
| A5SS | 3628 | ENSG00000111325 | OGFOD2 | chr12 | + | 122976870 | 122977166 | 122976870 | 122976963 | 122978324 | 122978569 | 11 | 2 | 7 | 9 | 1 | 0 | 2 | 0 | 0 | 0 | 0 | 3 | 10 | 7 | 7 | 0 | 0 | 0 | 0 | 0 | 298 | 149 | 1 | 1 | 1 | 0.333 | 1 | 1 | 1 | NA | 1 | 1 | 1 | 1 | -0.133 |
| A5SS | 5563 | ENSG00000171720 | HDAC3 | chr5 | - | 141629859 | 141630125 | 141630043 | 141630125 | 141629172 | 141629306 | 6 | 2 | 3 | 17 | 2 | 0 | 0 | 0 | 0 | 2 | 9 | 15 | 7 | 22 | 18 | 0 | 0 | 0 | 0 | 0 | 298 | 149 | 1 | 1 | 1 | 1 | 1 | 1 | 0.333 | 1 | 1 | 1 | 1 | 1 | -0.133 |
| A5SS | 6929 | ENSG00000241489 | NA | chrX | - | 149526529 | 149527068 | 149526935 | 149527068 | 149502086 | 149502223 | 14 | 6 | 2 | 5 | 15 | 0 | 0 | 2 | 0 | 0 | 32 | 6 | 11 | 21 | 6 | 0 | 0 | 0 | 0 | 0 | 298 | 149 | 1 | 1 | 1 | 1 | 0.333 | 1 | 1 | 1 | 1 | 1 | 1 | 1 | -0.133 |
| A5SS | 7106 | ENSG00000169057 | MECP2 | chrX | - | 154033438 | 154034174 | 154033923 | 154034174 | 154032458 | 154032557 | 2 | 4 | 5 | 6 | 5 | 2 | 0 | 0 | 0 | 0 | 40 | 20 | 29 | 34 | 42 | 0 | 0 | 0 | 0 | 0 | 298 | 149 | 1 | 1 | 0.333 | 1 | 1 | 1 | 1 | 1 | 1 | 1 | 1 | 1 | -0.133 |
| A5SS | 7599 | ENSG00000182919 | C11orf54 | chr11 | + | 93741962 | 93742141 | 93741962 | 93742066 | 93747296 | 93747448 | 5 | 7 | 4 | 8 | 26 | 0 | 3 | 0 | 2 | 0 | 0 | 2 | 13 | 7 | 0 | 0 | 0 | 0 | 0 | 0 | 224 | 149 | 1 | 1 | 1 | 0.608 | 1 | 0.727 | 1 | NA | 1 | 1 | 1 | NA | -0.133 |
| A5SS | 4372 | ENSG00000052723 | SIKE1 | chr1 | - | 114780436 | 114780685 | 114780448 | 114780685 | 114780109 | 114780215 | 1 | 1 | 1 | 1 | 0 | 16 | 9 | 15 | 18 | 7 | 0 | 5 | 6 | 3 | 0 | 4 | 23 | 5 | 10 | 6 | 161 | 149 | 1 | 1 | 0.055 | 0.093 | 0.058 | 0.049 | 0 | 0 | 0.167 | 0.526 | 0.217 | 0 | -0.131 |
| A5SS | 8275 | ENSG00000134460 | IL2RA | chr10 | - | 6021477 | 6021693 | 6021504 | 6021693 | 6019869 | 6019941 | 10 | 20 | 17 | 17 | 27 | 0 | 12 | 2 | 0 | 3 | 0 | 18 | 12 | 4 | 0 | 0 | 0 | 0 | 0 | 0 | 176 | 149 | 1 | 1 | 1 | 0.585 | 0.878 | 1 | 0.884 | NA | 1 | 1 | 1 | NA | -0.131 |
| A5SS | 667 | ENSG00000166845 | C18orf54 | chr18 | + | 54361642 | 54361948 | 54361642 | 54361891 | 54362770 | 54362921 | 8 | 4 | 0 | 2 | 9 | 1 | 2 | 3 | 1 | 1 | 0 | 1 | 0 | 2 | 0 | 0 | 1 | 0 | 0 | 0 | 206 | 149 | 1 | 1 | 0.853 | 0.591 | 0 | 0.591 | 0.867 | NA | 0.42 | NA | 1 | NA | -0.13 |
| A5SS | 7842 | ENSG00000110717 | NDUFS8 | chr11 | + | 68032285 | 68032523 | 68032285 | 68032336 | 68032922 | 68033012 | 10 | 1 | 0 | 2 | 4 | 11 | 4 | 11 | 20 | 5 | 0 | 7 | 6 | 13 | 10 | 0 | 10 | 6 | 19 | 13 | 298 | 149 | 1 | 1 | 0.312 | 0.111 | 0 | 0.048 | 0.286 | NA | 0.259 | 0.333 | 0.255 | 0.278 | -0.13 |
| A5SS | 2827 | ENSG00000197808 | ZNF461 | chr19 | - | 36658298 | 36658425 | 36658335 | 36658425 | 36656447 | 36656543 | 14 | 9 | 15 | 20 | 28 | 1 | 5 | 0 | 2 | 6 | 0 | 15 | 19 | 16 | 20 | 0 | 0 | 0 | 2 | 0 | 186 | 149 | 1 | 1 | 0.918 | 0.59 | 1 | 0.889 | 0.789 | NA | 1 | 1 | 0.865 | 1 | -0.129 |
| A5SS | 2950 | ENSG00000157036 | EXOG | chr3 | + | 38497628 | 38497778 | 38497628 | 38497689 | 38498925 | 38498989 | 4 | 4 | 7 | 3 | 0 | 0 | 0 | 0 | 2 | 0 | 0 | 3 | 1 | 0 | 6 | 0 | 0 | 0 | 0 | 0 | 238 | 149 | 1 | 1 | 1 | 1 | 1 | 0.484 | NA | NA | 1 | 1 | NA | 1 | -0.129 |
| A5SS | 2991 | ENSG00000111731 | C2CD5 | chr12 | - | 22519039 | 22519239 | 22519114 | 22519239 | 22518095 | 22518137 | 16 | 1 | 10 | 16 | 5 | 0 | 0 | 0 | 0 | 6 | 2 | 14 | 6 | 7 | 0 | 0 | 0 | 0 | 0 | 0 | 224 | 149 | 1 | 1 | 1 | 1 | 1 | 1 | 0.357 | 1 | 1 | 1 | 1 | NA | -0.129 |
| A5SS | 6176 | ENSG00000135486 | HNRNPA1 | chr12 | + | 54283078 | 54283234 | 54283078 | 54283228 | 54283826 | 54283967 | 3 | 0 | 1 | 0 | 0 | 1 | 0 | 0 | 0 | 0 | 0 | 0 | 2 | 0 | 0 | 0 | 0 | 0 | 0 | 0 | 155 | 149 | 1 | 1 | 0.743 | NA | 1 | NA | NA | NA | NA | 1 | NA | NA | -0.129 |
| A5SS | 5077 | ENSG00000177683 | THAP5 | chr7 | - | 108565829 | 108566022 | 108565951 | 108566022 | 108564992 | 108565105 | 88 | 60 | 72 | 58 | 131 | 8 | 10 | 7 | 14 | 59 | 20 | 36 | 55 | 33 | 0 | 0 | 7 | 10 | 0 | 0 | 271 | 149 | 1 | 1 | 0.858 | 0.767 | 0.85 | 0.695 | 0.55 | 1 | 0.739 | 0.751 | 1 | NA | -0.128 |
| A5SS | 5788 | ENSG00000144362 | PHOSPHO2 | chr2 | + | 169694453 | 169694741 | 169694453 | 169694622 | 169695214 | 169695247 | 1 | 1 | 0 | 2 | 0 | 4 | 2 | 5 | 3 | 4 | 0 | 0 | 0 | 0 | 4 | 0 | 4 | 1 | 1 | 0 | 268 | 149 | 1 | 1 | 0.122 | 0.218 | 0 | 0.27 | 0 | NA | 0 | 0 | 0 | 1 | -0.128 |
| A5SS | 8371 | ENSG00000180229 | HERC2P3 | chr15 | - | 20443338 | 20443521 | 20443370 | 20443521 | 20440473 | 20440642 | 0 | 0 | 0 | 2 | 0 | 3 | 5 | 3 | 20 | 3 | 0 | 0 | 0 | 21 | 0 | 0 | 7 | 0 | 23 | 6 | 181 | 149 | 1 | 1 | 0 | 0 | 0 | 0.076 | 0 | NA | 0 | NA | 0.429 | 0 | -0.128 |
| A5SS | 1920 | ENSG00000115760 | BIRC6 | chr2 | + | 32453807 | 32453954 | 32453807 | 32453942 | 32467524 | 32467739 | 3 | 4 | 3 | 0 | 2 | 1 | 0 | 0 | 2 | 3 | 4 | 4 | 4 | 0 | 0 | 0 | 0 | 0 | 6 | 0 | 161 | 149 | 1 | 1 | 0.735 | 1 | 1 | 0 | 0.382 | 1 | 1 | 1 | 0 | NA | -0.127 |
| A5SS | 5690 | ENSG00000243970 | PPIEL | chr1 | - | 39529015 | 39529164 | 39529054 | 39529164 | 39523806 | 39523896 | 5 | 2 | 1 | 9 | 0 | 2 | 4 | 0 | 7 | 0 | 2 | 4 | 3 | 5 | 0 | 0 | 0 | 1 | 0 | 15 | 188 | 149 | 1 | 1 | 0.665 | 0.284 | 1 | 0.505 | NA | 1 | 1 | 0.704 | 1 | 0 | -0.127 |
| A5SS | 6806 | ENSG00000118513 | MYB | chr6 | + | 135200289 | 135200557 | 135200289 | 135200415 | 135201638 | 135201749 | 8 | 3 | 4 | 12 | 5 | 14 | 15 | 33 | 20 | 32 | 28 | 13 | 17 | 24 | 0 | 30 | 29 | 12 | 85 | 0 | 291 | 149 | 1 | 1 | 0.226 | 0.093 | 0.058 | 0.235 | 0.074 | 0.323 | 0.187 | 0.42 | 0.126 | NA | -0.127 |
| A5SS | 1592 | ENSG00000151575 | TEX9 | chr15 | + | 56365446 | 56365893 | 56365446 | 56365670 | 56373440 | 56373504 | 2 | 4 | 0 | 0 | 16 | 5 | 8 | 7 | 2 | 4 | 0 | 0 | 1 | 0 | 0 | 0 | 0 | 1 | 0 | 0 | 298 | 149 | 1 | 1 | 0.167 | 0.2 | 0 | 0 | 0.667 | NA | NA | 0.333 | NA | NA | -0.126 |
| A5SS | 2010 | ENSG00000272168 | CASC15 | chr6 | + | 22110746 | 22110920 | 22110746 | 22110818 | 22191566 | 22191656 | 1 | 3 | 8 | 15 | 1 | 1 | 0 | 0 | 0 | 0 | 9 | 5 | 0 | 7 | 28 | 0 | 0 | 0 | 0 | 0 | 251 | 149 | 1 | 1 | 0.372 | 1 | 1 | 1 | 1 | 1 | 1 | NA | 1 | 1 | -0.126 |
| A5SS | 8960 | ENSG00000071564 | TCF3 | chr19 | - | 1631950 | 1632116 | 1632037 | 1632116 | 1627358 | 1627426 | 2 | 0 | 1 | 1 | 1 | 6 | 1 | 10 | 7 | 1 | 0 | 0 | 1 | 6 | 0 | 0 | 15 | 9 | 0 | 1 | 236 | 149 | 1 | 1 | 0.174 | 0 | 0.059 | 0.083 | 0.387 | NA | 0 | 0.066 | 1 | 0 | -0.126 |
| A5SS | 891 | ENSG00000141576 | RNF157 | chr17 | - | 76154279 | 76154328 | 76154282 | 76154328 | 76152354 | 76152465 | 0 | 0 | 2 | 1 | 0 | 7 | 8 | 13 | 3 | 12 | 0 | 0 | 0 | 1 | 0 | 20 | 12 | 8 | 0 | 3 | 152 | 149 | 1 | 1 | 0 | 0 | 0.131 | 0.246 | 0 | 0 | 0 | 0 | 1 | 0 | -0.125 |
| A5SS | 4917 | ENSG00000248333 | CDK11B | chr1 | - | 1642346 | 1642508 | 1642385 | 1642508 | 1641905 | 1642001 | 0 | 0 | 2 | 0 | 2 | 0 | 3 | 5 | 4 | 0 | 0 | 3 | 11 | 10 | 0 | 18 | 6 | 7 | 10 | 15 | 188 | 149 | 1 | 1 | NA | 0 | 0.241 | 0 | 0.284 | 0 | 0.284 | 0.555 | 0.442 | 0 | -0.125 |
| A5SS | 7314 | ENSG00000188343 | CIBAR1 | chr8 | + | 93701223 | 93702543 | 93701223 | 93701458 | 93703619 | 93703688 | 0 | 0 | 0 | 0 | 0 | 2 | 2 | 3 | 5 | 2 | 8 | 0 | 0 | 0 | 0 | 4 | 6 | 1 | 9 | 0 | 298 | 149 | 1 | 1 | 0 | 0 | 0 | 0 | 0 | 0.5 | 0 | 0 | 0 | NA | -0.125 |
| A5SS | 7976 | ENSG00000245498 | NA | chr11 | + | 124800447 | 124800894 | 124800447 | 124800718 | 124801983 | 124802157 | 6 | 4 | 2 | 0 | 4 | 0 | 2 | 0 | 0 | 0 | 0 | 0 | 0 | 5 | 0 | 0 | 0 | 0 | 0 | 0 | 298 | 149 | 1 | 1 | 1 | 0.5 | 1 | NA | 1 | NA | NA | NA | 1 | NA | -0.125 |
| A5SS | 8652 | ENSG00000028203 | VEZT | chr12 | + | 95258244 | 95258287 | 95258244 | 95258282 | 95262905 | 95263081 | 5 | 3 | 1 | 0 | 16 | 8 | 1 | 3 | 2 | 15 | 0 | 0 | 0 | 4 | 0 | 0 | 0 | 1 | 0 | 0 | 154 | 149 | 1 | 1 | 0.377 | 0.744 | 0.244 | 0 | 0.508 | NA | NA | 0 | 1 | NA | -0.125 |
| A5SS | 6668 | ENSG00000228649 | SNHG26 | chr7 | + | 22858570 | 22858830 | 22858570 | 22858795 | 22860546 | 22860638 | 4 | 2 | 17 | 7 | 14 | 0 | 2 | 1 | 0 | 0 | 0 | 21 | 13 | 0 | 21 | 0 | 0 | 1 | 0 | 0 | 184 | 149 | 1 | 1 | 1 | 0.447 | 0.932 | 1 | 1 | NA | 1 | 1 | NA | 1 | -0.124 |
| A5SS | 1324 | ENSG00000159593 | NAE1 | chr16 | - | 66826667 | 66826780 | 66826676 | 66826780 | 66826522 | 66826583 | 1 | 0 | 1 | 0 | 3 | 14 | 34 | 28 | 25 | 29 | 0 | 4 | 2 | 0 | 0 | 21 | 11 | 3 | 9 | 0 | 158 | 149 | 1 | 1 | 0.063 | 0 | 0.033 | 0 | 0.089 | 0 | 0.255 | 0.386 | 0 | NA | -0.123 |
| A5SS | 6926 | ENSG00000285171 | NA | chrX | - | 71106353 | 71106426 | 71106358 | 71106426 | 71105999 | 71106137 | 0 | 0 | 0 | 0 | 0 | 0 | 2 | 7 | 6 | 12 | 1 | 0 | 0 | 3 | 0 | 0 | 7 | 6 | 5 | 0 | 154 | 149 | 1 | 1 | 0 | 0 | 0 | 0 | 0 | NA | 0 | 0 | 0.367 | NA | -0.122 |
| A5SS | 812 | ENSG00000131470 | PSMC3IP | chr17 | - | 42577079 | 42577302 | 42577212 | 42577302 | 42574098 | 42574210 | 8 | 0 | 1 | 4 | 10 | 1 | 0 | 2 | 5 | 0 | 28 | 5 | 10 | 4 | 20 | 20 | 0 | 1 | 7 | 0 | 282 | 149 | 1 | 1 | 0.809 | NA | 0.209 | 0.297 | 1 | 0.425 | 1 | 0.841 | 0.232 | 1 | -0.121 |
| A5SS | 2783 | ENSG00000242086 | MUC20-OT1 | chr3 | + | 195708428 | 195708555 | 195708428 | 195708513 | 195708746 | 195708841 | 4 | 2 | 4 | 2 | 0 | 1 | 0 | 1 | 0 | 0 | 9 | 0 | 6 | 0 | 0 | 0 | 0 | 0 | 0 | 0 | 191 | 149 | 1 | 1 | 0.757 | 1 | 0.757 | 1 | NA | 1 | NA | 1 | NA | NA | -0.121 |
| A5SS | 3835 | ENSG00000158488 | CD1E | chr1 | + | 158355826 | 158356001 | 158355826 | 158355940 | 158356497 | 158356591 | 3 | 2 | 0 | 4 | 5 | 2 | 0 | 0 | 0 | 0 | 21 | 1 | 0 | 0 | 0 | 0 | 0 | 0 | 0 | 0 | 210 | 149 | 1 | 1 | 0.516 | 1 | NA | 1 | 1 | 1 | 1 | NA | NA | NA | -0.121 |
| A5SS | 8729 | ENSG00000120662 | MTRF1 | chr13 | - | 41262047 | 41262331 | 41262214 | 41262331 | 41260492 | 41260915 | 4 | 2 | 4 | 1 | 2 | 3 | 0 | 0 | 0 | 0 | 8 | 7 | 5 | 10 | 0 | 0 | 0 | 0 | 0 | 0 | 298 | 149 | 1 | 1 | 0.4 | 1 | 1 | 1 | 1 | 1 | 1 | 1 | 1 | NA | -0.12 |
| A5SS | 8796 | ENSG00000130396 | AFDN | chr6 | + | 167965756 | 167966045 | 167965756 | 167965853 | 167969113 | 167969198 | 10 | 7 | 9 | 5 | 4 | 0 | 0 | 0 | 0 | 3 | 15 | 1 | 7 | 11 | 0 | 0 | 0 | 0 | 0 | 0 | 298 | 149 | 1 | 1 | 1 | 1 | 1 | 1 | 0.4 | 1 | 1 | 1 | 1 | NA | -0.12 |
| A5SS | 2824 | ENSG00000178700 | DHFR2 | chr3 | - | 94062745 | 94062916 | 94062868 | 94062916 | 94057921 | 94061606 | 23 | 4 | 14 | 23 | 3 | 1 | 1 | 6 | 4 | 0 | 7 | 5 | 6 | 7 | 0 | 0 | 0 | 2 | 0 | 0 | 272 | 149 | 1 | 1 | 0.926 | 0.687 | 0.561 | 0.759 | 1 | 1 | 1 | 0.622 | 1 | NA | -0.119 |
| A5SS | 4694 | ENSG00000180881 | CAPS2 | chr12 | - | 75316283 | 75316434 | 75316311 | 75316434 | 75312847 | 75312915 | 3 | 0 | 0 | 6 | 0 | 5 | 4 | 1 | 3 | 0 | 0 | 2 | 0 | 0 | 0 | 0 | 3 | 0 | 0 | 0 | 177 | 149 | 1 | 1 | 0.336 | 0 | 0 | 0.627 | NA | NA | 0.359 | NA | NA | NA | -0.118 |
| A5SS | 4361 | ENSG00000166352 | IFTAP | chr11 | + | 36594501 | 36594641 | 36594501 | 36594592 | 36610080 | 36610239 | 4 | 2 | 3 | 2 | 13 | 4 | 3 | 7 | 2 | 9 | 4 | 6 | 2 | 4 | 0 | 3 | 2 | 3 | 3 | 0 | 198 | 149 | 1 | 1 | 0.429 | 0.334 | 0.244 | 0.429 | 0.521 | 0.501 | 0.693 | 0.334 | 0.501 | NA | -0.116 |
| A5SS | 6120 | ENSG00000124523 | SIRT5 | chr6 | + | 13574600 | 13574744 | 13574600 | 13574624 | 13579450 | 13579609 | 0 | 0 | 1 | 0 | 0 | 6 | 1 | 2 | 7 | 2 | 0 | 2 | 0 | 6 | 0 | 2 | 9 | 2 | 3 | 0 | 269 | 149 | 1 | 1 | 0 | 0 | 0.217 | 0 | 0 | 0 | 0.11 | 0 | 0.526 | NA | -0.116 |
| A5SS | 3864 | ENSG00000251131 | NA | chr5 | + | 43023918 | 43024109 | 43023918 | 43024097 | 43024198 | 43024607 | 2 | 0 | 3 | 0 | 3 | 2 | 2 | 4 | 2 | 0 | 0 | 0 | 2 | 1 | 0 | 3 | 0 | 0 | 1 | 0 | 161 | 149 | 1 | 1 | 0.481 | 0 | 0.41 | 0 | 1 | 0 | NA | 1 | 0.481 | NA | -0.115 |
| A5SS | 800 | ENSG00000100038 | TOP3B | chr22 | - | 21962127 | 21962602 | 21962428 | 21962602 | 21960320 | 21960449 | 9 | 2 | 1 | 4 | 7 | 7 | 3 | 2 | 9 | 9 | 0 | 3 | 7 | 1 | 20 | 0 | 16 | 1 | 6 | 8 | 298 | 149 | 1 | 1 | 0.391 | 0.25 | 0.2 | 0.182 | 0.28 | NA | 0.086 | 0.778 | 0.077 | 0.556 | -0.114 |
| A5SS | 2429 | ENSG00000109654 | TRIM2 | chr4 | + | 153222242 | 153223039 | 153222242 | 153222759 | 153270334 | 153270519 | 1 | 3 | 5 | 1 | 3 | 0 | 0 | 0 | 0 | 2 | 0 | 0 | 1 | 10 | 0 | 0 | 0 | 0 | 0 | 0 | 298 | 149 | 1 | 1 | 1 | 1 | 1 | 1 | 0.429 | NA | NA | 1 | 1 | NA | -0.114 |
| A5SS | 5288 | ENSG00000224078 | SNHG14 | chr15 | + | 25087466 | 25089866 | 25087466 | 25087599 | 25091589 | 25091719 | 14 | 6 | 20 | 6 | 10 | 0 | 4 | 0 | 0 | 0 | 26 | 21 | 6 | 16 | 0 | 0 | 0 | 0 | 0 | 0 | 298 | 149 | 1 | 1 | 1 | 0.429 | 1 | 1 | 1 | 1 | 1 | 1 | 1 | NA | -0.114 |
| A5SS | 4242 | ENSG00000144747 | TMF1 | chr3 | - | 69047348 | 69048562 | 69047357 | 69048562 | 69044491 | 69044595 | 16 | 12 | 10 | 17 | 12 | 155 | 133 | 113 | 123 | 180 | 0 | 13 | 16 | 7 | 0 | 0 | 80 | 76 | 16 | 0 | 158 | 149 | 1 | 1 | 0.089 | 0.078 | 0.077 | 0.115 | 0.059 | NA | 0.133 | 0.166 | 0.292 | NA | -0.113 |
| A5SS | 726 | ENSG00000161513 | FDXR | chr17 | - | 74872768 | 74872968 | 74872768 | 74872968 | 74872035 | 74872133 | 0 | 0 | 0 | 2 | 0 | 1 | 0 | 2 | 2 | 3 | 0 | 0 | 3 | 7 | 0 | 0 | 6 | 4 | 4 | 22 | 246 | 149 | 1 | 1 | 0 | NA | 1 | 0.377 | 0 | NA | 0 | 0.312 | 0.515 | 0 | -0.112 |
| A5SS | 922 | ENSG00000129245 | FXR2 | chr17 | - | 7606015 | 7606149 | 7606096 | 7606149 | 7605644 | 7605738 | 2 | 0 | 0 | 1 | 2 | 29 | 26 | 27 | 22 | 48 | 7 | 6 | 4 | 0 | 17 | 16 | 37 | 21 | 21 | 36 | 230 | 149 | 1 | 1 | 0.043 | 0 | 0 | 0.029 | 0.026 | 0.221 | 0.095 | 0.11 | 0 | 0.234 | -0.112 |
| A5SS | 4874 | ENSG00000099250 | NRP1 | chr10 | - | 33202890 | 33202995 | 33202911 | 33202995 | 33197649 | 33197709 | 11 | 10 | 9 | 5 | 13 | 0 | 0 | 0 | 4 | 1 | 0 | 0 | 0 | 5 | 0 | 0 | 0 | 0 | 0 | 0 | 170 | 149 | 1 | 1 | 1 | 1 | 1 | 0.523 | 0.919 | NA | NA | NA | 1 | NA | -0.112 |
| A5SS | 4781 | ENSG00000140416 | TPM1 | chr15 | + | 63064063 | 63064190 | 63064063 | 63064142 | 63065895 | 63066093 | 3 | 4 | 4 | 5 | 5 | 1 | 1 | 0 | 0 | 0 | 4 | 0 | 4 | 4 | 0 | 0 | 0 | 0 | 0 | 0 | 197 | 149 | 1 | 1 | 0.694 | 0.752 | 1 | 1 | 1 | 1 | NA | 1 | 1 | NA | -0.111 |
| A5SS | 6669 | ENSG00000228649 | SNHG26 | chr7 | + | 22858570 | 22858976 | 22858570 | 22858795 | 22860546 | 22860638 | 11 | 5 | 16 | 6 | 12 | 0 | 2 | 1 | 0 | 0 | 8 | 8 | 19 | 2 | 0 | 0 | 0 | 0 | 0 | 0 | 298 | 149 | 1 | 1 | 1 | 0.556 | 0.889 | 1 | 1 | 1 | 1 | 1 | 1 | NA | -0.111 |
| A5SS | 7089 | ENSG00000254685 | FPGT | chr1 | + | 74199663 | 74199845 | 74199663 | 74199831 | 74201317 | 74201410 | 10 | 11 | 1 | 12 | 7 | 67 | 31 | 37 | 32 | 64 | 0 | 7 | 2 | 4 | 0 | 0 | 12 | 7 | 4 | 4 | 163 | 149 | 1 | 1 | 0.12 | 0.245 | 0.024 | 0.255 | 0.091 | NA | 0.348 | 0.207 | 0.478 | 0 | -0.111 |
| A5SS | 177 | ENSG00000239382 | ALKBH6 | chr19 | - | 36013000 | 36013089 | 36013020 | 36013089 | 36010893 | 36011045 | 0 | 1 | 1 | 1 | 0 | 1 | 0 | 0 | 3 | 0 | 0 | 2 | 1 | 0 | 0 | 0 | 0 | 0 | 1 | 0 | 169 | 149 | 1 | 1 | 0 | 1 | 1 | 0.227 | NA | NA | 1 | 1 | 0 | NA | -0.11 |
| A5SS | 1992 | ENSG00000125459 | MSTO1 | chr1 | + | 155613656 | 155613766 | 155613656 | 155613733 | 155614061 | 155614933 | 3 | 5 | 3 | 1 | 5 | 0 | 0 | 0 | 1 | 0 | 0 | 5 | 6 | 1 | 4 | 0 | 0 | 0 | 0 | 0 | 182 | 149 | 1 | 1 | 1 | 1 | 1 | 0.45 | 1 | NA | 1 | 1 | 1 | 1 | -0.11 |
| A5SS | 6208 | ENSG00000143314 | MRPL24 | chr1 | - | 156740930 | 156741088 | 156741011 | 156741088 | 156738521 | 156738764 | 0 | 0 | 1 | 0 | 0 | 11 | 4 | 9 | 7 | 17 | 0 | 7 | 2 | 0 | 0 | 26 | 19 | 3 | 2 | 0 | 230 | 149 | 1 | 1 | 0 | 0 | 0.067 | 0 | 0 | 0 | 0.193 | 0.302 | 0 | NA | -0.11 |
| A5SS | 7730 | ENSG00000213930 | GALT | chr9 | + | 34648761 | 34648907 | 34648761 | 34648894 | 34648997 | 34649081 | 0 | 0 | 2 | 3 | 1 | 24 | 11 | 24 | 26 | 20 | 0 | 4 | 0 | 0 | 14 | 0 | 6 | 6 | 9 | 43 | 162 | 149 | 1 | 1 | 0 | 0 | 0.071 | 0.096 | 0.044 | NA | 0.38 | 0 | 0 | 0.23 | -0.11 |
| A5SS | 4192 | ENSG00000142687 | KIAA0319L | chr1 | - | 35557206 | 35557403 | 35557361 | 35557403 | 35554349 | 35554519 | 19 | 12 | 5 | 16 | 0 | 10 | 6 | 8 | 10 | 0 | 16 | 19 | 4 | 2 | 26 | 4 | 6 | 5 | 14 | 0 | 298 | 149 | 1 | 1 | 0.487 | 0.5 | 0.238 | 0.444 | NA | 0.667 | 0.613 | 0.286 | 0.067 | 1 | -0.109 |
| A5SS | 8486 | ENSG00000152582 | SPEF2 | chr5 | + | 35694287 | 35694363 | 35694287 | 35694348 | 35695734 | 35695796 | 0 | 3 | 1 | 1 | 1 | 12 | 4 | 8 | 11 | 2 | 0 | 3 | 0 | 0 | 0 | 0 | 2 | 3 | 0 | 0 | 164 | 149 | 1 | 1 | 0 | 0.405 | 0.102 | 0.076 | 0.312 | NA | 0.577 | 0 | NA | NA | -0.109 |
| A5SS | 8926 | ENSG00000228889 | UBAC2-AS1 | chr13 | - | 99200365 | 99200710 | 99200637 | 99200710 | 99196376 | 99197802 | 5 | 1 | 6 | 10 | 8 | 3 | 0 | 0 | 0 | 0 | 0 | 15 | 3 | 3 | 3 | 0 | 0 | 0 | 0 | 0 | 298 | 149 | 1 | 1 | 0.455 | 1 | 1 | 1 | 1 | NA | 1 | 1 | 1 | 1 | -0.109 |
| A5SS | 7162 | ENSG00000147144 | CCDC120 | chrX | + | 49061958 | 49062104 | 49061958 | 49062058 | 49062234 | 49062325 | 0 | 1 | 0 | 1 | 4 | 1 | 0 | 0 | 0 | 0 | 0 | 1 | 5 | 10 | 4 | 0 | 1 | 0 | 0 | 0 | 195 | 149 | 1 | 1 | 0 | 1 | NA | 1 | 1 | NA | 0.433 | 1 | 1 | 1 | -0.108 |
| A5SS | 8381 | ENSG00000198825 | INPP5F | chr10 | + | 119792156 | 119792213 | 119792156 | 119792211 | 119796714 | 119796913 | 46 | 40 | 36 | 44 | 27 | 3 | 2 | 3 | 6 | 8 | 8 | 32 | 34 | 33 | 0 | 0 | 0 | 0 | 0 | 0 | 151 | 149 | 1 | 1 | 0.938 | 0.952 | 0.922 | 0.879 | 0.769 | 1 | 1 | 1 | 1 | NA | -0.108 |
| A5SS | 4005 | ENSG00000226688 | ENTPD1-AS1 | chr10 | - | 95907662 | 95907903 | 95907777 | 95907903 | 95904829 | 95904938 | 12 | 2 | 4 | 9 | 4 | 0 | 0 | 4 | 6 | 0 | 0 | 5 | 7 | 8 | 11 | 0 | 3 | 0 | 0 | 0 | 264 | 149 | 1 | 1 | 1 | 1 | 0.361 | 0.458 | 1 | NA | 0.485 | 1 | 1 | 1 | -0.107 |
| A5SS | 4146 | ENSG00000163322 | ABRAXAS1 | chr4 | - | 83484823 | 83485100 | 83484985 | 83485100 | 83482153 | 83482244 | 1 | 0 | 0 | 0 | 0 | 28 | 15 | 18 | 26 | 3 | 0 | 0 | 0 | 0 | 1 | 0 | 8 | 22 | 0 | 1 | 298 | 149 | 1 | 1 | 0.018 | 0 | 0 | 0 | 0 | NA | 0 | 0 | NA | 0.333 | -0.107 |
| A5SS | 4901 | ENSG00000196440 | ARMCX4 | chrX | + | 101486022 | 101486098 | 101486022 | 101486092 | 101487192 | 101487272 | 0 | 0 | 0 | 0 | 0 | 6 | 4 | 14 | 11 | 0 | 0 | 0 | 2 | 0 | 0 | 0 | 1 | 7 | 0 | 0 | 155 | 149 | 1 | 1 | 0 | 0 | 0 | 0 | NA | NA | 0 | 0.215 | NA | NA | -0.107 |
| A5SS | 6062 | ENSG00000198223 | CSF2RA | chrX | + | 1274754 | 1274842 | 1274754 | 1274818 | 1285777 | 1285920 | 0 | 1 | 1 | 0 | 1 | 0 | 0 | 0 | 0 | 2 | 0 | 22 | 2 | 10 | 11 | 0 | 3 | 1 | 0 | 0 | 173 | 149 | 1 | 1 | NA | 1 | 1 | NA | 0.301 | NA | 0.863 | 0.633 | 1 | 1 | -0.107 |
| A5SS | 8592 | ENSG00000105370 | LIM2 | chr19 | - | 51387142 | 51387449 | 51387268 | 51387449 | 51382417 | 51382567 | 8 | 6 | 6 | 3 | 4 | 5 | 0 | 0 | 0 | 0 | 0 | 3 | 4 | 1 | 0 | 0 | 0 | 0 | 0 | 0 | 275 | 149 | 1 | 1 | 0.464 | 1 | 1 | 1 | 1 | NA | 1 | 1 | 1 | NA | -0.107 |
| A5SS | 6059 | ENSG00000198223 | CSF2RA | chrX | + | 1274754 | 1274842 | 1274754 | 1274818 | 1281870 | 1282220 | 1 | 1 | 1 | 0 | 4 | 0 | 1 | 0 | 0 | 0 | 12 | 4 | 3 | 9 | 0 | 0 | 0 | 0 | 1 | 0 | 173 | 149 | 1 | 1 | 1 | 0.463 | 1 | NA | 1 | 1 | 1 | 1 | 0.886 | NA | -0.106 |
| A5SS | 7117 | ENSG00000133142 | TCEAL4 | chrX | + | 103586218 | 103586301 | 103586218 | 103586291 | 103586648 | 103587518 | 0 | 0 | 0 | 0 | 0 | 10 | 8 | 1 | 7 | 5 | 0 | 0 | 1 | 0 | 0 | 14 | 0 | 2 | 7 | 0 | 159 | 149 | 1 | 1 | 0 | 0 | 0 | 0 | 0 | 0 | NA | 0.319 | 0 | NA | -0.106 |
| A5SS | 8113 | ENSG00000172663 | TMEM134 | chr11 | - | 67468016 | 67468092 | 67468027 | 67468092 | 67467500 | 67467563 | 1 | 0 | 0 | 3 | 8 | 4 | 0 | 3 | 3 | 3 | 0 | 0 | 7 | 3 | 8 | 0 | 4 | 4 | 1 | 9 | 160 | 149 | 1 | 1 | 0.189 | NA | 0 | 0.482 | 0.713 | NA | 0 | 0.62 | 0.736 | 0.453 | -0.106 |
| A5SS | 89 | ENSG00000183506 | PI4KAP2 | chr22 | - | 21475991 | 21476160 | 21476001 | 21476160 | 21475210 | 21475300 | 0 | 0 | 3 | 0 | 0 | 1 | 0 | 1 | 0 | 0 | 0 | 1 | 0 | 7 | 0 | 0 | 0 | 0 | 9 | 5 | 159 | 149 | 1 | 1 | 0 | NA | 0.738 | NA | NA | NA | 1 | NA | 0.422 | 0 | -0.105 |
| A5SS | 1454 | ENSG00000140264 | SERF2 | chr15 | + | 43792367 | 43792611 | 43792367 | 43792578 | 43792974 | 43793083 | 4 | 1 | 2 | 1 | 3 | 1 | 0 | 0 | 0 | 1 | 10 | 2 | 4 | 5 | 33 | 0 | 0 | 0 | 0 | 0 | 182 | 149 | 1 | 1 | 0.766 | 1 | 1 | 1 | 0.711 | 1 | 1 | 1 | 1 | 1 | -0.105 |
| A5SS | 8344 | ENSG00000156650 | KAT6B | chr10 | + | 74981786 | 74985241 | 74981786 | 74981928 | 74989018 | 74989112 | 84 | 60 | 75 | 72 | 133 | 11 | 5 | 2 | 3 | 11 | 8 | 55 | 59 | 11 | 5 | 0 | 1 | 2 | 0 | 0 | 298 | 149 | 1 | 1 | 0.792 | 0.857 | 0.949 | 0.923 | 0.858 | 1 | 0.965 | 0.937 | 1 | 1 | -0.105 |
| A5SS | 897 | ENSG00000178999 | AURKB | chr17 | - | 8207175 | 8207367 | 8207317 | 8207367 | 8206749 | 8206888 | 9 | 7 | 22 | 23 | 7 | 0 | 1 | 1 | 0 | 1 | 52 | 27 | 31 | 46 | 0 | 0 | 0 | 0 | 0 | 0 | 291 | 149 | 1 | 1 | 1 | 0.782 | 0.918 | 1 | 0.782 | 1 | 1 | 1 | 1 | NA | -0.104 |
| A5SS | 4263 | ENSG00000066427 | ATXN3 | chr14 | - | 92080952 | 92081061 | 92080964 | 92081061 | 92070937 | 92071053 | 0 | 1 | 0 | 3 | 0 | 26 | 22 | 3 | 37 | 35 | 0 | 2 | 2 | 3 | 0 | 0 | 10 | 9 | 51 | 0 | 161 | 149 | 1 | 1 | 0 | 0.04 | 0 | 0.07 | 0 | NA | 0.156 | 0.171 | 0.052 | NA | -0.104 |
| A5SS | 5724 | ENSG00000153253 | SCN3A | chr2 | - | 165140650 | 165140998 | 165140788 | 165140998 | 165139475 | 165139608 | 13 | 6 | 6 | 6 | 1 | 1 | 0 | 0 | 2 | 0 | 0 | 19 | 6 | 1 | 0 | 0 | 0 | 0 | 0 | 0 | 287 | 149 | 1 | 1 | 0.871 | 1 | 1 | 0.609 | 1 | NA | 1 | 1 | 1 | NA | -0.104 |
| A5SS | 7204 | ENSG00000107815 | TWNK | chr10 | + | 100990435 | 100990663 | 100990435 | 100990543 | 100990868 | 100991010 | 2 | 2 | 4 | 2 | 1 | 24 | 9 | 18 | 14 | 30 | 0 | 0 | 0 | 4 | 0 | 0 | 11 | 0 | 2 | 3 | 269 | 149 | 1 | 1 | 0.044 | 0.11 | 0.11 | 0.073 | 0.018 | NA | 0 | NA | 0.526 | 0 | -0.104 |
| A5SS | 3797 | ENSG00000091127 | PUS7 | chr7 | - | 105470687 | 105470848 | 105470760 | 105470848 | 105468336 | 105468463 | 2 | 10 | 17 | 24 | 13 | 6 | 0 | 0 | 2 | 9 | 8 | 18 | 4 | 8 | 0 | 15 | 0 | 0 | 0 | 0 | 222 | 149 | 1 | 1 | 0.183 | 1 | 1 | 0.89 | 0.492 | 0.264 | 1 | 1 | 1 | NA | -0.103 |
| A5SS | 6696 | ENSG00000097033 | SH3GLB1 | chr1 | + | 86728402 | 86728489 | 86728402 | 86728441 | 86730087 | 86730111 | 10 | 5 | 7 | 5 | 6 | 0 | 0 | 0 | 4 | 0 | 0 | 0 | 7 | 0 | 0 | 0 | 0 | 0 | 0 | 0 | 197 | 149 | 1 | 1 | 1 | 1 | 1 | 0.486 | 1 | NA | NA | 1 | NA | NA | -0.103 |
| A5SS | 6876 | ENSG00000174080 | CTSF | chr11 | - | 66567890 | 66568082 | 66567983 | 66568082 | 66567443 | 66567662 | 0 | 0 | 0 | 1 | 0 | 4 | 5 | 2 | 8 | 0 | 0 | 0 | 3 | 0 | 0 | 0 | 4 | 2 | 2 | 17 | 242 | 149 | 1 | 1 | 0 | 0 | 0 | 0.071 | NA | NA | 0 | 0.48 | 0 | 0 | -0.102 |
| [illegible] | | | | | | | | | | | | | | | | | | | | | | | | | | | | | | | | | | | | | | | | | | | | | | |

| | | | | | | | | | | | |
|---|---|---|---|---|---|---|---|---|---|---|---|
| A5SS | 8129 | ENSG00000260923 | LINC02193 | chr16 | + | 90215292 | 90216198 | 90215292 | 90215556 | 90222243 | 90222292 |
| A5SS | 857 | ENSG00000108963 | DPH1 | chr17 | + | 2041767 | 2042231 | 2041767 | 2041875 | 2042604 | 2043898 |
| A5SS | 4613 | ENSG00000120948 | TARDBP | chr1 | + | 11022123 | 11022250 | 11022123 | 11022241 | 11023192 | 11023329 |
| A5SS | 7098 | ENSG00000182484 | WASH6P | chrX | + | 156022322 | 156022539 | 156022322 | 156022459 | 156022698 | 156022834 |
| A5SS | 536 | ENSG00000124205 | EDN3 | chr20 | + | 59322371 | 59322452 | 59322371 | 59322417 | 59324330 | 59325992 |
| A5SS | 4193 | ENSG00000165046 | LETM2 | chr8 | + | 38386500 | 38387182 | 38386500 | 38386568 | 38387949 | 38388030 |
| A5SS | 4275 | ENSG00000143801 | PSEN2 | chr1 | + | 226891742 | 226891849 | 226891742 | 226891844 | 226894006 | 226894125 |
| A5SS | 2006 | ENSG00000120318 | ARAP3 | chr5 | - | 141671264 | 141671400 | 141671324 | 141671400 | 141670511 | 141670628 |
| A5SS | 3696 | ENSG00000182923 | CEP63 | chr3 | + | 134495295 | 134495364 | 134495295 | 134495360 | 134507108 | 134507286 |
| A5SS | 6469 | ENSG00000288558 | DUS4L-BCAP29 | chr7 | + | 107563970 | 107564247 | 107563970 | 107564209 | 107567049 | 107567186 |
| A5SS | 2254 | ENSG00000164308 | ERAP2 | chr5 | + | 96909579 | 96909809 | 96909579 | 96909764 | 96912636 | 96912798 |
| A5SS | 2470 | ENSG00000138796 | HADH | chr4 | + | 108023473 | 108028069 | 108023473 | 108023563 | 108033175 | 108033292 |
| A5SS | 3388 | ENSG00000167315 | ACAA2 | chr18 | - | 49813681 | 49813953 | 49813816 | 49813953 | 49802686 | 49802853 |
| A5SS | 5734 | ENSG00000079263 | SP140 | chr2 | + | 230244987 | 230245080 | 230244987 | 230245071 | 230247915 | 230248065 |
| A5SS | 8911 | ENSG00000187535 | IFT140 | chr16 | - | 1526618 | 1526796 | 1526732 | 1526796 | 1525886 | 1526077 |
| A5SS | 493 | ENSG00000101220 | C20orf27 | chr20 | - | 3760006 | 3760186 | 3760081 | 3760186 | 3758535 | 3758678 |
| A5SS | 3485 | ENSG00000138674 | SEC31A | chr4 | - | 82842139 | 82842481 | 82842184 | 82842481 | 82830936 | 82830975 |
| A5SS | 6521 | ENSG00000179818 | PCBP1-AS1 | chr2 | - | 70086054 | 70086273 | 70086251 | 70086273 | 70085546 | 70085624 |
| A5SS | 1008 | ENSG00000141698 | NT5C3B | chr17 | - | 41835587 | 41835957 | 41835858 | 41835957 | 41835202 | 41835272 |
| A5SS | 3068 | ENSG00000174891 | RSRC1 | chr3 | + | 158203071 | 158203269 | 158203071 | 158203245 | 158298038 | 158298075 |
| A5SS | 6856 | ENSG00000188917 | TRMT2B | chrX | - | 101051250 | 101051733 | 101051643 | 101051733 | 101041316 | 101041371 |
| A5SS | 8132 | ENSG00000165915 | SLC39A13 | chr11 | + | 47414776 | 47414909 | 47414776 | 47414905 | 47415059 | 47415159 |
| A5SS | 203 | ENSG00000197256 | KANK2 | chr19 | - | 11195555 | 11195820 | 11195621 | 11195820 | 11194474 | 11194590 |
| A5SS | 1260 | ENSG00000186260 | MRTFB | chr16 | + | 14246472 | 14247507 | 14246472 | 14247357 | 14248925 | 14249081 |
| A5SS | 3641 | ENSG00000213071 | LPAL2 | chr6 | - | 160482661 | 160482819 | 160482687 | 160482819 | 160478507 | 160478688 |
| A5SS | 7018 | ENSG00000156313 | RPGR | chrX | - | 38287093 | 38288041 | 38287860 | 38288041 | 38276586 | 38276772 |
| A5SS | 2309 | ENSG00000171462 | DLK2 | chr6 | - | 43453004 | 43453135 | 43453022 | 43453135 | 43451939 | 43452084 |
| A5SS | 2387 | ENSG00000180316 | PNPLA1 | chr6 | + | 36291319 | 36291555 | 36291319 | 36291552 | 36293060 | 36293126 |
| A5SS | 6118 | ENSG00000112139 | MDGA1 | chr6 | - | 37646197 | 37646375 | 37646267 | 37646375 | 37644496 | 37644649 |
| A5SS | 1691 | ENSG00000092200 | RPGRIP1 | chr14 | + | 21324617 | 21325070 | 21324617 | 21324725 | 21325231 | 21325383 |
| A5SS | 3836 | ENSG00000158488 | CD1E | chr1 | + | 158355826 | 158356105 | 158355826 | 158355940 | 158356497 | 158356591 |
| A5SS | 4343 | ENSG00000111445 | RFC5 | chr12 | + | 118017689 | 118018053 | 118017689 | 118017852 | 118019071 | 118019136 |
| A5SS | 4357 | ENSG00000150455 | TIRAP | chr11 | + | 126289664 | 126289833 | 126289664 | 126289817 | 126290802 | 126290961 |
| A5SS | 3280 | ENSG00000228794 | LINC01128 | chr1 | + | 851926 | 852094 | 851926 | 852014 | 852670 | 852766 |
| A5SS | 5227 | ENSG00000167107 | ACSF2 | chr17 | + | 50461241 | 50461370 | 50461241 | 50461287 | 50462183 | 50462302 |
| A5SS | 5855 | ENSG00000074582 | BCS1L | chr2 | + | 218660036 | 218660218 | 218660036 | 218660166 | 218660938 | 218661307 |
| A5SS | 6744 | ENSG00000144840 | RABL3 | chr3 | - | 120738526 | 120738667 | 120738549 | 120738667 | 120730695 | 120730787 |
| A5SS | 7601 | ENSG00000182919 | C11orf54 | chr11 | + | 93741962 | 93742094 | 93741962 | 93742066 | 93747301 | 93747448 |
| A5SS | 2800 | ENSG00000068745 | IP6K2 | chr3 | - | 48715314 | 48715527 | 48715388 | 48715527 | 48695089 | 48695418 |
| A5SS | 3236 | ENSG00000197386 | HTT | chr4 | + | 3172907 | 3173331 | 3172907 | 3173218 | 3174720 | 3174799 |
| A5SS | 1283 | ENSG00000263072 | ZNF213-AS1 | chr16 | - | 3129247 | 3129653 | 3129460 | 3129653 | 3114967 | 3116519 |
| A5SS | 287 | ENSG00000065268 | WDR18 | chr19 | + | 989761 | 990103 | 989761 | 989895 | 990222 | 990364 |
| A5SS | 807 | ENSG00000175106 | TVP23C | chr17 | - | 15553684 | 15553829 | 15553764 | 15553829 | 15547058 | 15547148 |
| A5SS | 6848 | ENSG00000172613 | RAD9A | chr11 | + | 67396110 | 67396175 | 67396110 | 67396171 | 67396262 | 67396400 |
| A5SS | 8030 | ENSG00000177697 | CD151 | chr11 | + | 832951 | 833026 | 832951 | 833022 | 836062 | 836153 |
| A5SS | 1860 | ENSG00000152240 | HAUS1 | chr18 | + | 46105193 | 46105408 | 46105193 | 46105368 | 46118180 | 46118316 |
| A5SS | 3092 | ENSG00000205981 | DNAJC19 | chr3 | - | 180986947 | 180987022 | 180986997 | 180987022 | 180985925 | 180985996 |
| A5SS | 8413 | ENSG00000180198 | RCC1 | chr1 | + | 28508829 | 28508905 | 28508829 | 28508901 | 28529857 | 28529939 |
| A5SS | 8707 | ENSG00000112053 | SLC26A8 | chr6 | - | 35954819 | 35955520 | 35955151 | 35955520 | 35951444 | 35951499 |
| A5SS | 5189 | ENSG00000256223 | ZNF10 | chr12 | + | 133130626 | 133131014 | 133130626 | 133130754 | 133144433 | 133144525 |
| A5SS | 6807 | ENSG00000118513 | MYB | chr6 | + | 135200289 | 135200674 | 135200289 | 135200415 | 135201638 | 135201749 |
| A5SS | 7163 | ENSG00000147144 | CCDC120 | chrX | + | 49061958 | 49062117 | 49061958 | 49062058 | 49062234 | 49062325 |
| A5SS | 7694 | ENSG00000184939 | ZFP90 | chr16 | + | 68539290 | 68539479 | 68539290 | 68539469 | 68539757 | 68539825 |
| A5SS | 8819 | ENSG00000122966 | CIT | chr12 | - | 119784188 | 119785065 | 119784959 | 119785065 | 119783907 | 119784051 |
| A5SS | 2684 | ENSG00000187621 | TCL6 | chr14 | + | 95668125 | 95669617 | 95668125 | 95668580 | 95670441 | 95670577 |
| A5SS | 3563 | ENSG00000104381 | GDAP1 | chr8 | + | 74351273 | 74351466 | 74351273 | 74351321 | 74360136 | 74360310 |
| A5SS | 4102 | ENSG00000132122 | SPATA6 | chr1 | - | 48355669 | 48355769 | 48355717 | 48355769 | 48305786 | 48305878 |
| A5SS | 4846 | ENSG00000185163 | DDX51 | chr12 | - | 132141102 | 132141420 | 132141274 | 132141420 | 132140830 | 132141020 |
| A5SS | 5727 | ENSG00000153253 | SCN3A | chr2 | - | 165140701 | 165140998 | 165140788 | 165140998 | 165139475 | 165139608 |
| A5SS | 5856 | ENSG00000074582 | BCS1L | chr2 | + | 218660036 | 218660245 | 218660036 | 218660166 | 218660938 | 218661307 |
| A5SS | 7839 | ENSG00000139324 | TMTC3 | chr12 | + | 88142295 | 88142487 | 88142295 | 88142396 | 88148287 | 88148504 |
| A5SS | 8764 | ENSG00000111199 | TRPV4 | chr12 | - | 109793484 | 109793600 | 109793526 | 109793600 | 109792651 | 109792817 |
| A5SS | 1428 | ENSG00000198690 | FAN1 | chr15 | + | 30903911 | 30904024 | 30903911 | 30904011 | 30904511 | 30905897 |
| A5SS | 2548 | ENSG00000166004 | CEP295 | chr11 | + | 93695496 | 93695673 | 93695496 | 93695634 | 93696319 | 93696417 |
| A5SS | 4585 | ENSG00000122678 | POLM | chr7 | - | 44078553 | 44078811 | 44078755 | 44078811 | 44078321 | 44078438 |
| A5SS | 6879 | ENSG00000258867 | LINC01146 | chr14 | + | 88084036 | 88084483 | 88084036 | 88084155 | 88096498 | 88097053 |
| A5SS | 281 | ENSG00000221946 | FXYD7 | chr19 | + | 35151253 | 35151482 | 35151253 | 35151328 | 35151632 | 35151673 |
| A5SS | 757 | ENSG00000182173 | TSEN54 | chr17 | + | 75521408 | 75523335 | 75521408 | 75521510 | 75524261 | 75524723 |
| A5SS | 6218 | ENSG00000170919 | TPT1-AS1 | chr13 | + | 45379194 | 45380001 | 45379194 | 45379335 | 45389734 | 45389820 |
| A5SS | 2162 | ENSG00000281398 | SNHG4 | chr5 | + | 139278326 | 139278496 | 139278326 | 139278408 | 139279050 | 139279129 |
| A5SS | 379 | ENSG00000159882 | ZNF230 | chr19 | + | 44008789 | 44008991 | 44008789 | 44008916 | 44009083 | 44009170 |
| A5SS | 7549 | ENSG00000165233 | CARD19 | chr9 | + | 93107673 | 93107827 | 93107673 | 93107816 | 93108990 | 93109092 |
| A5SS | 1500 | ENSG00000114062 | UBE3A | chr15 | - | 25408571 | 25408684 | 25408619 | 25408684 | 25407066 | 25407212 |
| A5SS | 7225 | ENSG00000288550 | NA | chr3 | + | 9923880 | 9924020 | 9923880 | 9923981 | 9924231 | 9924291 |
| A5SS | 8443 | ENSG00000107959 | PITRM1 | chr10 | - | 3172181 | 3172782 | 3172716 | 3172782 | 3170103 | 3170206 |
| A5SS | 2557 | ENSG00000163629 | PTPN13 | chr4 | + | 86764592 | 86764739 | 86764592 | 86764724 | 86765394 | 86765488 |
| A5SS | 8138 | ENSG00000172977 | KAT5 | chr11 | + | 65712921 | 65713058 | 65712921 | 65713055 | 65713347 | 65713503 |
| A5SS | 403 | ENSG00000261824 | LINC00662 | chr19 | - | 27771543 | 27771989 | 27771663 | 27771989 | 27746097 | 27746340 |
| A5SS | 5287 | ENSG00000224078 | SNHG14 | chr15 | + | 25056482 | 25057088 | 25056482 | 25056622 | 25059154 | 25059294 |
| A5SS | 6505 | ENSG00000179818 | PCBP1-AS1 | chr2 | - | 69996685 | 69996888 | 69996857 | 69996888 | 69988741 | 69988838 |
| A5SS | 8491 | ENSG00000119977 | TCTN3 | chr10 | - | 95682650 | 95683629 | 95683100 | 95683629 | 95680471 | 95680609 |
| A5SS | 4572 | ENSG00000126709 | IFI6 | chr1 | - | 27669220 | 27669346 | 27669232 | 27669346 | 27668457 | 27668535 |
| A5SS | 4823 | ENSG00000139631 | CSAD | chr12 | - | 53171176 | 53171441 | 53171272 | 53171441 | 53170422 | 53170502 |
| A5SS | 4825 | ENSG00000139631 | CSAD | chr12 | - | 53171272 | 53171441 | 53171325 | 53171441 | 53170422 | 53170502 |
| A5SS | 2027 | ENSG00000141279 | NPEPPS | chr17 | + | 47619277 | 47619372 | 47619277 | 47619322 | 47619736 | 47619784 |
| A5SS | 2949 | ENSG00000157036 | EXOG | chr3 | + | 38497628 | 38497778 | 38497628 | 38497684 | 38498925 | 38498989 |
| A5SS | 3473 | ENSG00000117569 | PTBP2 | chr1 | + | 96804799 | 96806452 | 96804799 | 96804939 | 96806865 | 96806958 |
| A5SS | 3640 | ENSG00000092010 | PSME1 | chr14 | + | 24138344 | 24138403 | 24138344 | 24138399 | 24138735 | 24138959 |
| A5SS | 4004 | ENSG00000197530 | MIB2 | chr1 | + | 1625545 | 1626754 | 1625545 | 1625653 | 1626836 | 1626999 |
| A5SS | 4069 | ENSG00000153363 | LINC00467 | chr1 | + | 211396766 | 211396841 | 211396766 | 211396831 | 211426210 | 211426283 |
| A5SS | 8760 | ENSG00000002016 | RAD52 | chr12 | - | 929763 | 930144 | 930050 | 930144 | 927144 | 927263 |
| A5SS | 1885 | ENSG00000106686 | SPATA6L | chr9 | - | 4625216 | 4625566 | 4625326 | 4625566 | 4622407 | 4622510 |
| A5SS | 2128 | ENSG00000147044 | CASK | chrX | - | 41853096 | 41853227 | 41853114 | 41853227 | 41745523 | 41745601 |
| A5SS | 2292 | ENSG00000105341 | DMAC2 | chr19 | - | 41438217 | 41438414 | 41438234 | 41438414 | 41433536 | 41433673 |
| A5SS | 4479 | ENSG00000204138 | PHACTR4 | chr1 | + | 28457831 | 28457859 | 28457831 | 28457855 | 28459084 | 28459258 |
| A5SS | 4490 | ENSG00000156983 | BRPF1 | chr3 | + | 9744223 | 9744541 | 9744223 | 9744508 | 9745007 | 9745155 |
| A5SS | 6181 | ENSG00000132424 | PNISR | chr6 | - | 99415568 | 99415747 | 99415635 | 99415747 | 99414571 | 99414690 |
| A5SS | 6443 | ENSG00000134905 | CARS2 | chr13 | - | 110650636 | 110651100 | 110651033 | 110651100 | 110648317 | 110649297 |
| A5SS | 8909 | ENSG00000136104 | RNASEH2B | chr13 | + | 50956357 | 50956441 | 50956357 | 50956421 | 50969931 | 50970433 |
| A5SS | 5480 | ENSG00000122085 | MTERF4 | chr2 | - | 241090277 | 241090449 | 241090302 | 241090449 | 241089979 | 241090044 |
| A5SS | 2622 | ENSG00000164039 | BDH2 | chr4 | - | 103085348 | 103086540 | 103086479 | 103086540 | 103082870 | 103082929 |
| A5SS | 4743 | ENSG00000109445 | ZNF330 | chr4 | + | 141220929 | 141221077 | 141220929 | 141221016 | 141222365 | 141222491 |
| A5SS | 6297 | ENSG00000214021 | TTLL3 | chr3 | + | 9809739 | 9809871 | 9809739 | 9809848 | 9810620 | 9810709 |
| A5SS | 8427 | ENSG00000148688 | RPP30 | chr10 | + | 90900569 | 90900668 | 90900569 | 90900666 | 90902242 | 90902344 |
| A5SS | 668 | ENSG00000166845 | C18orf54 | chr18 | + | 54361642 | 54362431 | 54361642 | 54361891 | 54362770 | 54362921 |
| A5SS | 1245 | ENSG00000157353 | FCSK | chr16 | + | 70475349 | 70475493 | 70475349 | 70475415 | 70475647 | 70475767 |
| A5SS | 2673 | ENSG00000125375 | DMAC2L | chr14 | + | 50322510 | 50322806 | 50322510 | 50322719 | 50325608 | 50327957 |
| A5SS | 381 | ENSG00000073008 | PVR | chr19 | + | 44658741 | 44658900 | 44658741 | 44658765 | 44661291 | 44661323 |
| A5SS | 3966 | ENSG00000213672 | NCKIPSD | chr3 | - | 48682868 | 48683012 | 48682902 | 48683012 | 48682347 | 48682552 |
| A5SS | 6508 | ENSG00000179818 | PCBP1-AS1 | chr2 | - | 70055655 | 70055910 | 70055713 | 70055910 | 70053730 | 70053792 |
| A5SS | 405 | ENSG00000187187 | ZNF546 | chr19 | + | 39998247 | 39998410 | 39998247 | 39998332 | 40006095 | 40006182 |
| A5SS | 709 | ENSG00000270379 | HEATR9 | chr17 | - | 35865214 | 35865396 | 35865218 | 35865396 | 35864496 | 35864553 |
| A5SS | 1947 | ENSG00000127990 | SGCE | chr7 | - | 94599696 | 94600857 | 94600645 | 94600857 | 94598774 | 94598963 |
| A5SS | 4014 | ENSG00000157193 | LRP8 | chr1 | - | 53271226 | 53275753 | 53275630 | 53275753 | 53271027 | 53271153 |
| A5SS | 4337 | ENSG00000231551 | PDE4DIPP6 | chr1 | + | 148424626 | 148424678 | 148424626 | 148424638 | 148425505 | 148425664 |
| A5SS | 4447 | ENSG00000090097 | PCBP4 | chr3 | - | 51959894 | 51960320 | 51960188 | 51960320 | 51959576 | 51959651 |
| A5SS | 8222 | ENSG00000166340 | TPP1 | chr11 | - | 6617625 | 6618915 | 6618775 | 6618915 | 6616974 | 6617153 |
| A5SS | 7375 | ENSG00000249859 | PVT1 | chr8 | + | 127795893 | 127796071 | 127795893 | 127796008 | 127855154 | 127855319 |
| A5SS | 2846 | ENSG00000184470 | TXNRD2 | chr22 | - | 19877039 | 19877234 | 19877108 | 19877234 | 19875521 | 19875807 |
| A5SS | 895 | ENSG00000178971 | CTC1 | chr17 | - | 8236057 | 8236342 | 8236082 | 8236342 | 8235052 | 8235285 |
| A5SS | 5860 | ENSG00000163481 | RNF25 | chr2 | - | 218668238 | 218668341 | 218668278 | 218668341 | 218668078 | 218668146 |
| A5SS | 7509 | ENSG00000135315 | CEP162 | chr6 | - | 84175209 | 84175347 | 84175213 | 84175347 | 84174726 | 84174954 |
| A5SS | 1169 | ENSG00000176715 | ACSF3 | chr16 | + | 89093851 | 89093996 | 89093851 | 89093992 | 89098590 | 89098763 |
| A5SS | 2785 | ENSG00000164053 | ATRIP | chr3 | + | 48446736 | 48447092 | 48446736 | 48446799 | 48450036 | 48450170 |
| A5SS | 2917 | ENSG00000176261 | ZBTB8OS | chr1 | - | 32634701 | 32634792 | 32634767 | 32634792 | 32631826 | 32631879 |
| A5SS | 2932 | ENSG00000009830 | POMT2 | chr14 | - | 77279536 | 77279928 | 77279822 | 77279928 | 77278728 | 77278869 |
| A5SS | 3041 | ENSG00000251287 | ALG1L2 | chr3 | + | 130094402 | 130094513 | 130094402 | 130094467 | 130097174 | 130097250 |
| A5SS | 4731 | ENSG00000171791 | BCL2 | chr18 | - | 63317529 | 63318666 | 63318081 | 63318666 | 63280886 | 63281214 |
| A5SS | 5842 | ENSG00000072195 | SPEG | chr2 | + | 219443107 | 219443637 | 219443107 | 219443150 | 219444652 | 219444742 |
| A5SS | 7677 | ENSG00000107317 | PTGDS | chr9 | + | 136979222 | 136979401 | 136979222 | 136979299 | 136979945 | 136980062 |
| A5SS | 5331 | ENSG00000185480 | PARPBP | chr12 | + | 102153868 | 102153989 | 102153868 | 102153976 | 102164437 | 102164608 |
| A5SS | 3905 | ENSG00000055609 | KMT2C | chr7 | - | 152158856 | 152159072 | 152158862 | 152159072 | 152157760 | 152157854 |
| A5SS | 4767 | ENSG00000213963 | NA | chr2 | - | 177323238 | 177323396 | 177323303 | 177323396 | 177311232 | 177311328 |
| A5SS | 8102 | ENSG00000168002 | POLR2G | chr11 | + | 62761794 | 62762866 | 62761794 | 62761904 | 62765181 | 62765232 |
| A5SS | 8421 | ENSG00000175482 | POLD4 | chr11 | - | 67353210 | 67353534 | 67353363 | 67353534 | 67352987 | 67353077 |
| A5SS | 1761 | ENSG00000100722 | ZC3H14 | chr14 | + | 88602827 | 88603060 | 88602827 | 88603045 | 88607242 | 88607363 |
| A5SS | 8338 | ENSG00000166987 | MBD6 | chr12 | + | 57529159 | 57529329 | 57529159 | 57529325 | 57529610 | 57529674 |
| A5SS | 3907 | ENSG00000055609 | KMT2C | chr7 | - | 152158856 | 152159072 | 152158862 | 152159072 | 152157760 | 152157925 |
| A5SS | 6403 | ENSG00000145191 | EIF2B5 | chr3 | + | 184140079 | 184140221 | 184140079 | 184140157 | 184141924 | 184142070 |
| A5SS | 8837 | ENSG00000073060 | SCARB1 | chr12 | - | 124786356 | 124786503 | 124786440 | 124786503 | 124782682 | 124782811 |
| A5SS | 1438 | ENSG00000034053 | APBA2 | chr15 | + | 28921639 | 28921749 | 28921639 | 28921718 | 29053844 | 29054835 |
| A5SS | 1641 | ENSG00000140043 | PTGR2 | chr14 | + | 73876997 | 73880176 | 73876997 | 73877168 | 73881204 | 73881292 |
| A5SS | 2683 | ENSG00000187621 | TCL6 | chr14 | + | 95663255 | 95665076 | 95663255 | 95663635 | 95669550 | 95669617 |
| A5SS | 4108 | ENSG00000188732 | FAM221A | chr7 | + | 23692182 | 23694913 | 23692182 | 23692253 | 23698191 | 23698299 |
| A5SS | 4584 | ENSG00000122678 | POLM | chr7 | - | 44078553 | 44078811 | 44078739 | 44078811 | 44078321 | 44078438 |
| A5SS | 6383 | ENSG00000173598 | NUDT4 | chr12 | + | 93378577 | 93378790 | 93378577 | 93378693 | 93394608 | 93394719 |
| A5SS | 8634 | ENSG00000174206 | C12orf66 | chr12 | - | 64222002 | 64222296 | 64222161 | 64222296 | 64215677 | 64215963 |
| A5SS | 1978 | ENSG00000139433 | GLTP | chr12 | - | 109857854 | 109858193 | 109858139 | 109858193 | 109857525 | 109857659 |
| A5SS | 2310 | ENSG00000013375 | PGM3 | chr6 | - | 83169062 | 83169323 | 83169241 | 83169323 | 83161149 | 83161242 |
| A5SS | 3069 | ENSG00000174891 | RSRC1 | chr3 | + | 158460934 | 158461009 | 158460934 | 158461003 | 158537091 | 158537198 |
| A5SS | 3155 | ENSG00000162302 | RPS6KA4 | chr11 | + | 64359377 | 64359449 | 64359377 | 64359403 | 64360162 | 64360381 |
| A5SS | 5942 | ENSG00000154328 | NEIL2 | chr8 | + | 11769709 | 11770335 | 11769709 | 11769791 | 11771445 | 11771585 |
| A5SS | 7140 | ENSG00000248323 | LUCAT1 | chr5 | - | 91314079 | 91314402 | 91314164 | 91314402 | 91313637 | 91313781 |
| A5SS | 8283 | ENSG00000095066 | HOOK2 | chr19 | - | 12774560 | 12774741 | 12774668 | 12774741 | 12772993 | 12773044 |
| A5SS | 425 | ENSG00000105135 | ILVBL | chr19 | - | 15116000 | 15116321 | 15116151 | 15116321 | 15115849 | 15115921 |
| A5SS | 2240 | ENSG00000102100 | SLC35A2 | chrX | - | 48911468 | 48911646 | 48911545 | 48911646 | 48909813 | 48909996 |
| A5SS | 4186 | ENSG00000162971 | TYW5 | chr2 | - | 199948317 | 199948472 | 199948336 | 199948472 | 199943764 | 199943834 |
| A5SS | 4604 | ENSG00000160229 | ZNF66 | chr19 | + | 20776303 | 20776481 | 20776303 | 20776450 | 20792511 | 20792638 |
| A5SS | 5054 | ENSG00000049167 | ERCC8 | chr5 | - | 60904787 | 60904873 | 60904791 | 60904873 | 60903647 | 60903716 |
| A5SS | 5726 | ENSG00000153253 | SCN3A | chr2 | - | 165140701 | 165140998 | 165140784 | 165140998 | 165139475 | 165139608 |
| A5SS | 7505 | ENSG00000104529 | EEF1D | chr8 | - | 143593874 | 143594071 | 143593892 | 143594071 | 143592714 | 143592893 |
| A5SS | 8210 | ENSG00000110171 | TRIM3 | chr11 | - | 6458064 | 6458296 | 6458069 | 6458296 | 6457695 | 6457847 |
| A5SS | 8727 | ENSG00000139350 | NEDD1 | chr12 | + | 96907603 | 96907856 | 96907603 | 96907624 | 96917620 | 96917737 |
| A5SS | 1280 | ENSG00000103415 | HMOX2 | chr16 | + | 4483636 | 4483754 | 4483636 | 4483738 | 4505483 | 4505610 |
| A5SS | 4937 | ENSG00000177542 | SLC25A22 | chr11 | - | 794452 | 794513 | 794469 | 794513 | 793528 | 793619 |
| A5SS | 5134 | ENSG00000131061 | ZNF341 | chr20 | + | 33757147 | 33757343 | 33757147 | 33757322 | 33758715 | 33758806 |
| A5SS | 5713 | ENSG00000115486 | GGCX | chr2 | - | 85558439 | 85558662 | 85558538 | 85558662 | 85556181 | 85556260 |
| A5SS | 6653 | ENSG00000128604 | IRF5 | chr7 | + | 128941278 | 128941424 | 128941278 | 128941391 | 128942070 | 128942276 |
| A5SS | 8839 | ENSG00000073060 | SCARB1 | chr12 | - | 124807760 | 124807927 | 124807765 | 124807927 | 124800123 | 124800242 |
| A5SS | 133 | ENSG00000226287 | TMEM191A | chr22 | + | 20703632 | 20703751 | 20703632 | 20703692 | 20703967 | 20704040 |
| A5SS | 2605 | ENSG00000113522 | RAD50 | chr5 | + | 132575776 | 132575928 | 132575776 | 132575863 | 132587561 | 132587690 |
| A5SS | 5326 | ENSG00000197037 | ZSCAN25 | chr7 | + | 99619560 | 99619997 | 99619560 | 99619993 | 99621372 | 99621574 |
| A5SS | 5536 | ENSG00000166435 | XRRA1 | chr11 | - | 74933800 | 74933872 | 74933824 | 74933872 | 74930299 | 74930372 |
| A5SS | 5743 | ENSG00000163754 | GYG1 | chr3 | + | 148996741 | 148996904 | 148996741 | 148996892 | 149024052 | 149024272 |
| A5SS | 7917 | ENSG00000153827 | TRIP12 | chr2 | - | 229807704 | 229807864 | 229807707 | 229807864 | 229805729 | 229805802 |
| A5SS | 8098 | ENSG00000070047 | PHRF1 | chr11 | + | 581491 | 581606 | 581491 | 581597 | 581964 | 582081 |
| A5SS | 477 | ENSG00000000419 | DPM1 | chr20 | - | 50941104 | 50941209 | 50941128 | 50941209 | 50940864 | 50940933 |
| A5SS | 1730 | ENSG00000100938 | GMPR2 | chr14 | + | 24233478 | 24233593 | 24233478 | 24233586 | 24235736 | 24235820 |
| A5SS | 2706 | ENSG00000008311 | AASS | chr7 | - | 122078861 | 122079712 | 122079596 | 122079712 | 122077837 | 122078014 |
| A5SS | 3106 | ENSG00000173226 | IQCB1 | chr3 | - | 121834248 | 121834479 | 121834390 | 121834479 | 121828860 | 121828972 |
| A5SS | 4421 | ENSG00000126226 | PCID2 | chr13 | - | 113208023 | 113208650 | 113208598 | 113208650 | 113200426 | 113200516 |
| A5SS | 8921 | ENSG00000117226 | GBP3 | chr1 | - | 89013184 | 89014279 | 89014082 | 89014279 | 89011746 | 89012027 |
| A5SS | 2953 | ENSG00000157036 | EXOG | chr3 | + | 38497628 | 38497778 | 38497628 | 38497689 | 38501354 | 38501494 |
| A5SS | 3970 | ENSG00000163479 | SSR2 | chr1 | - | 156018131 | 156018368 | 156018251 | 156018368 | 156011809 | 156011887 |
| A5SS | 5013 | ENSG00000076242 | MLH1 | chr3 | + | 37000954 | 37001053 | 37000954 | 37001048 | 37006990 | 37007063 |
| A5SS | 5892 | ENSG00000239665 | NA | chr10 | + | 13644916 | 13645136 | 13644916 | 13645074 | 13654352 | 13654427 |
| A5SS | 6708 | ENSG00000181513 | ACBD4 | chr17 | + | 45135686 | 45136232 | 45135686 | 45135953 | 45136499 | 45136620 |
| A5SS | 1233 | ENSG00000087263 | OGFOD1 | chr16 | + | 56451520 | 56451760 | 56451520 | 56451567 | 56453262 | 56453408 |
| A5SS | 1694 | ENSG00000187097 | ENTPD5 | chr14 | - | 74015823 | 74015930 | 74015847 | 74015930 | 74011090 | 74011150 |
| A5SS | 3777 | ENSG00000164403 | SHROOM1 | chr5 | - | 132823849 | 132824419 | 132824280 | 132824419 | 132823622 | 132823764 |
| A5SS | 3840 | ENSG00000229961 | NA | chr1 | + | 157182859 | 157183252 | 157182859 | 157183014 | 157185119 | 157185413 |
| A5SS | 4236 | ENSG00000233355 | CHRM3-AS2 | chr1 | - | 239707533 | 239707760 | 239707614 | 239707760 | 239706424 | 239706496 |
| A5SS | 5101 | ENSG00000106105 | GARS1 | chr7 | + | 30603495 | 30609730 | 30603495 | 30603572 | 30615895 | 30616058 |
| A5SS | 6471 | ENSG00000288558 | DUS4L-BCAP29 | chr7 | + | 107563970 | 107564676 | 107563970 | 107564247 | 107567049 | 107567186 |
| A5SS | 6710 | ENSG00000267673 | FDX2 | chr19 | - | 10315385 | 10315759 | 10315704 | 10315759 | 10310852 | 10310940 |
| A5SS | 7876 | ENSG00000198561 | CTNND1 | chr11 | + | 57761796 | 57762119 | 57761796 | 57762068 | 57794009 | 57794081 |
| A5SS | 8215 | ENSG00000179532 | DNHD1 | chr11 | + | 6570008 | 6570396 | 6570008 | 6570100 | 6570617 | 6571423 |
| A5SS | 3066 | ENSG00000174891 | RSRC1 | chr3 | + | 158110088 | 158110223 | 158110088 | 158110211 | 158122102 | 158122298 |
| A5SS | 5619 | ENSG00000010219 | DYRK4 | chr12 | + | 4590329 | 4590554 | 4590329 | 4590440 | 4591159 | 4591298 |
| A5SS | 6087 | ENSG00000247134 | NA | chr8 | + | 33041634 | 33041771 | 33041634 | 33041743 | 33044366 | 33044855 |
| A5SS | 1131 | ENSG00000140995 | DEF8 | chr16 | + | 89949416 | 89949638 | 89949416 | 89949513 | 89954242 | 89954376 |
| A5SS | 2241 | ENSG00000102100 | SLC35A2 | chrX | - | 48911506 | 48911646 | 48911545 | 48911646 | 48909813 | 48909996 |
| A5SS | 3120 | ENSG00000164342 | TLR3 | chr4 | + | 186082319 | 186084172 | 186082319 | 186082550 | 186084644 | 186088073 |
| A5SS | 5016 | ENSG00000124067 | SLC12A4 | chr16 | - | 67963436 | 67963559 | 67963464 | 67963559 | 67961574 | 67961706 |
| A5SS | 5723 | ENSG00000153253 | SCN3A | chr2 | - | 165140650 | 165140998 | 165140784 | 165140998 | 165139475 | 165139608 |
| A5SS | 6801 | ENSG00000118513 | MYB | chr6 | + | 135195747 | 135195954 | 135195747 | 135195917 | 135196960 | 135197323 |
| A5SS | 7617 | ENSG00000116580 | GON4L | chr1 | - | 155774962 | 155775173 | 155775001 | 155775173 | 155771066 | 155771217 |
| A5SS | 54 | ENSG00000100139 | MICALL1 | chr22 | + | 37927410 | 37927826 | 37927410 | 37927820 | 37931798 | 37931933 |
| A5SS | 1105 | ENSG00000242612 | DECR2 | chr16 | + | 410242 | 410404 | 410242 | 410367 | 410690 | 410784 |

| | | | | | | | | | | | | | | |
|---|---|---|---|---|---|---|---|---|---|---|---|---|---|---|
| A5SS | 7180 | ENSG00000234665 | LERFS | chr9 | - | 62897418 | 62898095 | 62897528 | 62898095 | 62869189 | 62869304 | 38 | 0 | 0 |
| A5SS | 7364 | ENSG00000241560 | ZBTB20-AS1 | chr3 | + | 114366966 | 114367221 | 114366966 | 114367028 | 114387422 | 114387579 | 13 | 17 | 17 |
| A5SS | 7623 | ENSG00000080802 | CNOT4 | chr7 | - | 135410514 | 135410648 | 135410523 | 135410648 | 135395633 | 135395883 | 7 | 3 | 7 |
| A5SS | 7894 | ENSG00000239445 | ST3GAL6-AS1 | chr3 | - | 98718418 | 98718521 | 98718423 | 98718521 | 98715649 | 98715777 | 3 | 0 | 0 |
| A5SS | 1678 | ENSG00000100749 | VRK1 | chr14 | + | 96855223 | 96855356 | 96855223 | 96855352 | 96856527 | 96856586 | 5 | 1 | 0 |
| A5SS | 4346 | ENSG00000163166 | IWS1 | chr2 | - | 127526174 | 127526489 | 127526382 | 127526489 | 127523675 | 127523791 | 40 | 32 | 30 |
| A5SS | 232 | ENSG00000130816 | DNMT1 | chr19 | - | 10137075 | 10137280 | 10137084 | 10137280 | 10136120 | 10136287 | 0 | 1 | 1 |
| A5SS | 341 | ENSG00000181027 | FKRP | chr19 | + | 46747233 | 46747305 | 46747233 | 46747294 | 46748026 | 46748088 | 1 | 2 | 0 |
| A5SS | 5352 | ENSG00000064703 | DDX20 | chr1 | + | 111760705 | 111760865 | 111760705 | 111760848 | 111760986 | 111761125 | 0 | 0 | 1 |
| A5SS | 7661 | ENSG00000132740 | IGHMBP2 | chr11 | + | 68906068 | 68906200 | 68906068 | 68906135 | 68908144 | 68908337 | 4 | 1 | 5 |
| A5SS | 7868 | ENSG00000110429 | FBXO3 | chr11 | - | 33753110 | 33754500 | 33754454 | 33754500 | 33751522 | 33751607 | 7 | 9 | 15 |
| A5SS | 319 | ENSG00000104866 | PPP1R37 | chr19 | + | 45140505 | 45140661 | 45140505 | 45140606 | 45141321 | 45141386 | 3 | 1 | 1 |
| A5SS | 4003 | ENSG00000197530 | MIB2 | chr1 | + | 1623430 | 1623699 | 1623430 | 1623679 | 1623773 | 1623945 | 7 | 2 | 4 |
| A5SS | 5757 | ENSG00000123983 | ACSL3 | chr2 | + | 222927016 | 222927242 | 222927016 | 222927189 | 222928861 | 222928936 | 3 | 7 | 3 |
| A5SS | 5919 | ENSG00000196290 | NIF3L1 | chr2 | + | 200889368 | 200889652 | 200889368 | 200889391 | 200890522 | 200890585 | 8 | 7 | 9 |
| A5SS | 6470 | ENSG00000288558 | DUS4L-BCAP29 | chr7 | + | 107563970 | 107564676 | 107563970 | 107564209 | 107567049 | 107567186 | 14 | 11 | 15 |
| A5SS | 8871 | ENSG00000101160 | CTSZ | chr20 | - | 58996638 | 58997753 | 58997602 | 58997753 | 58992761 | 58992821 | 14 | 11 | 3 |
| A5SS | 2060 | ENSG00000131844 | MCCC2 | chr5 | + | 71641002 | 71642727 | 71641002 | 71641083 | 71643818 | 71643895 | 19 | 25 | 22 |
| A5SS | 7230 | ENSG00000182196 | ARL6IP4 | chr12 | + | 122981128 | 122981299 | 122981128 | 122981266 | 122981570 | 122981879 | 11 | 5 | 3 |
| A5SS | 681 | ENSG00000154889 | MPPE1 | chr18 | - | 11905737 | 11905953 | 11905911 | 11905953 | 11897525 | 11897620 | 25 | 15 | 9 |
| A5SS | 3139 | ENSG00000173889 | PHC3 | chr3 | - | 170181686 | 170181733 | 170181701 | 170181733 | 170178772 | 170178938 | 1 | 0 | 5 |
| A5SS | 3431 | ENSG00000135932 | CAB39 | chr2 | + | 230713547 | 230714058 | 230713547 | 230713802 | 230759958 | 230760115 | 9 | 13 | 6 |
| A5SS | 4375 | ENSG00000157796 | WDR19 | chr4 | + | 39232161 | 39232308 | 39232161 | 39232272 | 39234765 | 39234875 | 0 | 0 | 4 |
| A5SS | 7881 | ENSG00000152556 | PFKM | chr12 | + | 48141739 | 48142066 | 48141739 | 48141827 | 48142781 | 48142946 | 14 | 9 | 15 |
| A5SS | 8442 | ENSG00000107959 | PITRM1 | chr10 | - | 3171760 | 3172782 | 3172716 | 3172782 | 3170103 | 3170206 | 0 | 0 | 0 |
| A5SS | 8761 | ENSG00000002016 | RAD52 | chr12 | - | 929818 | 930144 | 930050 | 930144 | 927144 | 927263 | 19 | 8 | 15 |
| A5SS | 833 | ENSG00000185359 | HGS | chr17 | + | 81693869 | 81694050 | 81693869 | 81693962 | 81694814 | 81694853 | 2 | 2 | 6 |
| A5SS | 3908 | ENSG00000055609 | KMT2C | chr7 | - | 152163273 | 152163826 | 152163699 | 152163826 | 152162116 | 152163177 | 34 | 18 | 16 |
| A5SS | 5687 | ENSG00000115392 | FANCL | chr2 | - | 58198578 | 58198662 | 58198593 | 58198662 | 58165723 | 58165874 | 2 | 3 | 3 |

| | | | | | | | | | | | | | | | | | | | | |
|---|---|---|---|---|---|---|---|---|---|---|---|---|---|---|---|---|---|---|---|---|
| A5SS | 4170 | ENSG00000161618 | ALDH16A1 | chr19 | + | 49461618 | 49462036 | 49461618 | 49461800 | 49462569 | 49462755 | 24 | 6 | 12 | 18 | 2 | 2 | 0 | 1 | 2 | 0 |
| A5SS | 4234 | ENSG00000132781 | MUTYH | chr1 | - | 45338185 | 45338656 | 45338506 | 45338656 | 45334390 | 45334511 | 34 | 37 | 41 | 56 | 46 | 0 | 0 | 1 | 0 | 3 |
| A5SS | 4730 | ENSG00000110200 | ANAPC15 | chr11 | - | 72112649 | 72112780 | 72112733 | 72112780 | 72111411 | 72111496 | 45 | 22 | 21 | 29 | 34 | 2 | 0 | 1 | 0 | 0 |
| A5SS | 4918 | ENSG00000118961 | LDAH | chr2 | - | 20701569 | 20701652 | 20701574 | 20701652 | 20686906 | 20687094 | 10 | 19 | 15 | 21 | 31 | 0 | 0 | 0 | 2 | 0 |
| A5SS | 5224 | ENSG00000163644 | PPM1K | chr4 | - | 88278143 | 88278642 | 88278535 | 88278642 | 88277142 | 88277243 | 114 | 74 | 104 | 129 | 169 | 2 | 0 | 1 | 1 | 2 |
| A5SS | 5579 | ENSG00000134262 | AP4B1 | chr1 | - | 113899903 | 113900400 | 113900269 | 113900400 | 113898717 | 113898801 | 158 | 75 | 138 | 144 | 100 | 0 | 0 | 0 | 0 | 5 |
| A5SS | 6257 | ENSG00000164880 | INTS1 | chr7 | - | 1472242 | 1472386 | 1472272 | 1472386 | 1471570 | 1471641 | 0 | 0 | 0 | 2 | 0 | 42 | 16 | 44 | 50 | 14 |
| A5SS | 6514 | ENSG00000179818 | PCBP1-AS1 | chr2 | - | 70083506 | 70083687 | 70083537 | 70083687 | 70081369 | 70081490 | 1 | 1 | 3 | 7 | 6 | 1 | 0 | 1 | 0 | 0 |
| A5SS | 8081 | ENSG00000149577 | SIDT2 | chr11 | + | 117181415 | 117181537 | 117181415 | 117181514 | 117181806 | 117181971 | 147 | 66 | 44 | 131 | 116 | 1 | 2 | 2 | 0 | 0 |
| A5SS | 8156 | ENSG00000171204 | TMEM126B | chr11 | + | 85629131 | 85629262 | 85629131 | 85629203 | 85631686 | 85631808 | 14 | 8 | 8 | 14 | 13 | 0 | 0 | 0 | 1 | 0 |
| A5SS | 8268 | ENSG00000150433 | TMEM218 | chr11 | - | 125102636 | 125102809 | 125102729 | 125102809 | 125102131 | 125102317 | 16 | 15 | 11 | 11 | 8 | 1 | 0 | 0 | 0 | 0 |
| A5SS | 58 | ENSG00000182944 | EWSR1 | chr22 | + | 29288605 | 29288786 | 29288605 | 29288755 | 29290412 | 29290684 | 13 | 9 | 13 | 23 | 1 | 1 | 0 | 0 | 0 | 0 |
| A5SS | 426 | ENSG00000105135 | ILVBL | chr19 | - | 15116407 | 15118015 | 15117880 | 15118015 | 15116151 | 15116321 | 40 | 15 | 19 | 22 | 15 | 0 | 0 | 0 | 1 | 0 |
| A5SS | 780 | ENSG00000185624 | P4HB | chr17 | - | 81859178 | 81859387 | 81859180 | 81859387 | 81847242 | 81847347 | 0 | 0 | 2 | 1 | 4 | 3 | 0 | 2 | 0 | 1 |
| A5SS | 1236 | ENSG00000090861 | AARS1 | chr16 | - | 70253665 | 70253800 | 70253713 | 70253800 | 70253267 | 70253381 | 2 | 0 | 1 | 0 | 0 | 79 | 40 | 95 | 53 | 107 |
| A5SS | 1343 | ENSG00000102910 | LONP2 | chr16 | + | 48351580 | 48351798 | 48351580 | 48351685 | 48361704 | 48363122 | 58 | 50 | 55 | 86 | 74 | 0 | 0 | 0 | 0 | 4 |
| A5SS | 3655 | ENSG00000018510 | AGPS | chr2 | + | 177440964 | 177442235 | 177440964 | 177441040 | 177442406 | 177442486 | 14 | 22 | 24 | 12 | 20 | 0 | 1 | 0 | 0 | 0 |
| A5SS | 3885 | ENSG00000138386 | NAB1 | chr2 | + | 190655976 | 190656159 | 190655976 | 190656153 | 190659276 | 190659585 | 3 | 2 | 0 | 3 | 0 | 0 | 0 | 0 | 1 | 0 |
| A5SS | 4749 | ENSG00000160688 | FLAD1 | chr1 | + | 154990158 | 154990528 | 154990158 | 154990257 | 154992712 | 154992786 | 38 | 15 | 17 | 14 | 22 | 3 | 1 | 0 | 0 | 0 |
| A5SS | 4881 | ENSG00000163534 | FCRL1 | chr1 | - | 157802376 | 157802664 | 157802614 | 157802664 | 157801914 | 157802162 | 69 | 55 | 66 | 99 | 92 | 1 | 1 | 0 | 2 | 0 |
| A5SS | 4920 | ENSG00000118961 | LDAH | chr2 | - | 20801291 | 20801465 | 20801309 | 20801465 | 20790254 | 20790398 | 4 | 3 | 7 | 4 | 11 | 23 | 9 | 17 | 18 | 52 |
| A5SS | 4922 | ENSG00000187741 | FANCA | chr16 | - | 89739015 | 89739289 | 89739132 | 89739289 | 89738881 | 89738926 | 75 | 39 | 55 | 65 | 42 | 0 | 2 | 0 | 0 | 0 |
| A5SS | 4938 | ENSG00000177542 | SLC25A22 | chr11 | - | 794457 | 794513 | 794469 | 794513 | 793528 | 793619 | 15 | 3 | 12 | 24 | 4 | 0 | 0 | 0 | 2 | 0 |
| A5SS | 5250 | ENSG00000285756 | NA | chrX | - | 3828505 | 3829392 | 3829363 | 3829392 | 3818656 | 3818703 | 22 | 26 | 24 | 25 | 35 | 1 | 0 | 0 | 0 | 0 |
| A5SS | 5353 | ENSG00000064703 | DDX20 | chr1 | + | 111760705 | 111761125 | 111760705 | 111760848 | 111761225 | 111761284 | 61 | 38 | 65 | 64 | 87 | 0 | 5 | 3 | 0 | 0 |
| A5SS | 5751 | ENSG00000122203 | KIAA1191 | chr5 | - | 176359461 | 176359567 | 176359480 | 176359567 | 176348249 | 176348356 | 0 | 0 | 1 | 0 | 0 | 4 | 2 | 8 | 13 | 1 |
| A5SS | 6095 | ENSG00000135316 | SYNCRIP | chr6 | - | 85614980 | 85615347 | 85615018 | 85615347 | 85608334 | 85612920 | 93 | 52 | 91 | 67 | 128 | 0 | 0 | 1 | 4 | 0 |
| A5SS | 7103 | ENSG00000102393 | GLA | chrX | - | 101401232 | 101401809 | 101401667 | 101401809 | 101400665 | 101400757 | 18 | 26 | 30 | 33 | 22 | 1 | 0 | 0 | 0 | 1 |
| A5SS | 7569 | ENSG00000131089 | ARHGEF9 | chrX | - | 63754303 | 63755113 | 63755018 | 63755113 | 63724531 | 63724711 | 32 | 18 | 21 | 19 | 43 | 0 | 0 | 1 | 0 | 0 |
| A5SS | 7755 | ENSG00000165699 | TSC1 | chr9 | - | 132910570 | 132911568 | 132911452 | 132911568 | 132907300 | 132907370 | 44 | 41 | 44 | 64 | 37 | 2 | 0 | 0 | 0 | 0 |
| A5SS | 8259 | ENSG00000110330 | BIRC2 | chr11 | + | 102348597 | 102348764 | 102348597 | 102348719 | 102350345 | 102350565 | 17 | 20 | 14 | 13 | 41 | 0 | 0 | 1 | 0 | 0 |
| A5SS | 767 | ENSG00000073536 | NLE1 | chr17 | - | 35139819 | 35140066 | 35139848 | 35140066 | 35139234 | 35139314 | 1 | 0 | 8 | 0 | 2 | 6 | 6 | 7 | 5 | 10 |
| A5SS | 933 | ENSG00000154265 | ABCA5 | chr17 | - | 69273942 | 69274128 | 69273958 | 69274128 | 69271161 | 69271289 | 14 | 13 | 7 | 8 | 7 | 17 | 15 | 16 | 20 | 37 |
| A5SS | 1153 | ENSG00000121274 | TENT4B | chr16 | + | 50224890 | 50225011 | 50224890 | 50224994 | 50225238 | 50225285 | 0 | 0 | 0 | 1 | 2 | 30 | 20 | 20 | 14 | 67 |
| A5SS | 1181 | ENSG00000172775 | PSME3IP1 | chr16 | - | 57178501 | 57178852 | 57178805 | 57178852 | 57173727 | 57173869 | 112 | 92 | 126 | 115 | 121 | 1 | 3 | 0 | 0 | 2 |
| A5SS | 3228 | ENSG00000184014 | DENND5A | chr11 | - | 9141939 | 9142108 | 9141973 | 9142108 | 9140188 | 9140248 | 31 | 26 | 29 | 11 | 46 | 0 | 0 | 2 | 0 | 0 |
| A5SS | 3305 | ENSG00000173171 | MTX1 | chr1 | + | 155210547 | 155212219 | 155210547 | 155210627 | 155212384 | 155212567 | 58 | 29 | 40 | 48 | 45 | 1 | 0 | 1 | 0 | 0 |
| A5SS | 3561 | ENSG00000104381 | GDAP1 | chr8 | + | 74350394 | 74350578 | 74350394 | 74350530 | 74351273 | 74351466 | 15 | 6 | 4 | 16 | 15 | 0 | 0 | 0 | 0 | 1 |
| A5SS | 3820 | ENSG00000166971 | AKTIP | chr16 | - | 53502699 | 53503206 | 53503146 | 53503206 | 53500217 | 53500329 | 0 | 5 | 2 | 1 | 4 | 8 | 2 | 19 | 10 | 5 |
| A5SS | 3898 | ENSG00000183291 | SELENOF | chr1 | - | 86914027 | 86914424 | 86914314 | 86914424 | 86903280 | 86903448 | 145 | 86 | 155 | 156 | 174 | 0 | 2 | 0 | 1 | 2 |
| A5SS | 4050 | ENSG00000066279 | ASPM | chr1 | - | 197091906 | 197092056 | 197091930 | 197092056 | 197090849 | 197091041 | 13 | 4 | 17 | 6 | 6 | 1 | 0 | 0 | 0 | 0 |
| A5SS | 4165 | ENSG00000003393 | ALS2 | chr2 | - | 201746565 | 201746748 | 201746627 | 201746748 | 201744257 | 201744429 | 30 | 7 | 12 | 26 | 33 | 2 | 0 | 1 | 1 | 0 |
| A5SS | 5292 | ENSG00000224078 | SNHG14 | chr15 | + | 25099144 | 25102147 | 25099144 | 25099278 | 25103858 | 25103970 | 1 | 1 | 3 | 0 | 0 | 16 | 23 | 43 | 12 | 29 |
| A5SS | 6146 | ENSG00000137312 | FLOT1 | chr6 | - | 30741613 | 30741704 | 30741661 | 30741704 | 30741189 | 30741333 | 126 | 51 | 96 | 100 | 94 | 0 | 2 | 3 | 0 | 0 |
| A5SS | 6436 | ENSG00000119616 | FCF1 | chr14 | + | 74713484 | 74713620 | 74713484 | 74713507 | 74714871 | 74714943 | 23 | 11 | 15 | 24 | 29 | 1 | 1 | 0 | 1 | 0 |
| A5SS | 7964 | ENSG00000110711 | AIP | chr11 | + | 67490315 | 67490457 | 67490315 | 67490449 | 67490787 | 67491103 | 199 | 63 | 144 | 152 | 105 | 1 | 3 | 1 | 4 | 0 |
| A5SS | 8031 | ENSG00000177697 | CD151 | chr11 | + | 834529 | 834745 | 834529 | 834591 | 836062 | 836153 | 0 | 1 | 1 | 0 | 1 | 9 | 2 | 3 | 5 | 4 |
| A5SS | 8032 | ENSG00000177697 | CD151 | chr11 | + | 834529 | 834803 | 834529 | 834591 | 836062 | 836153 | 0 | 1 | 1 | 0 | 1 | 9 | 2 | 3 | 5 | 4 |
| A5SS | 8414 | ENSG00000180198 | RCC1 | chr1 | + | 28516724 | 28516867 | 28516724 | 28516853 | 28529857 | 28529939 | 33 | 8 | 22 | 34 | 53 | 2 | 0 | 0 | 0 | 1 |
| A5SS | 8429 | ENSG00000221817 | PPP3CB-AS1 | chr10 | + | 73496141 | 73496906 | 73496141 | 73496483 | 73499594 | 73499687 | 143 | 82 | 135 | 78 | 151 | 1 | 0 | 2 | 0 | 3 |
| A5SS | 116 | ENSG00000244625 | MIATNB | chr22 | + | 26726643 | 26726794 | 26726643 | 26726720 | 26738061 | 26738240 | 34 | 38 | 17 | 43 | 26 | 0 | 2 | 0 | 0 | 0 |
| A5SS | 772 | ENSG00000108799 | EZH1 | chr17 | - | 42727634 | 42727763 | 42727661 | 42727763 | 42724424 | 42724614 | 11 | 14 | 15 | 15 | 24 | 0 | 0 | 0 | 1 | 0 |
| A5SS | 1429 | ENSG00000198690 | FAN1 | chr15 | + | 30904511 | 30905897 | 30904511 | 30904590 | 30906370 | 30906504 | 60 | 24 | 18 | 46 | 54 | 0 | 1 | 0 | 0 | 0 |
| A5SS | 1866 | ENSG00000112679 | DUSP22 | chr6 | + | 311879 | 311962 | 311879 | 311914 | 335113 | 335163 | 94 | 49 | 100 | 132 | 101 | 0 | 3 | 0 | 0 | 0 |
| A5SS | 2354 | ENSG00000214655 | ZSWIM8 | chr10 | + | 73793587 | 73793719 | 73793587 | 73793698 | 73793864 | 73794044 | 29 | 20 | 25 | 39 | 26 | 2 | 0 | 0 | 0 | 0 |
| A5SS | 3394 | ENSG00000170581 | STAT2 | chr12 | - | 56349344 | 56349509 | 56349425 | 56349509 | 56348751 | 56348804 | 35 | 34 | 41 | 43 | 43 | 0 | 2 | 1 | 2 | 0 |
| A5SS | 3446 | ENSG00000144591 | GMPPA | chr2 | + | 219505457 | 219505761 | 219505457 | 219505555 | 219505979 | 219506072 | 34 | 14 | 19 | 24 | 19 | 0 | 0 | 0 | 1 | 0 |
| A5SS | 4344 | ENSG00000141552 | ANAPC11 | chr17 | + | 81894466 | 81894624 | 81894466 | 81894586 | 81899919 | 81900254 | 0 | 0 | 0 | 0 | 0 | 34 | 16 | 22 | 49 | 25 |
| A5SS | 4406 | ENSG00000143851 | PTPN7 | chr1 | - | 202148553 | 202148699 | 202148580 | 202148699 | 202147014 | 202148466 | 13 | 5 | 4 | 18 | 11 | 2 | 0 | 0 | 0 | 0 |
| A5SS | 5055 | ENSG00000004866 | ST7 | chr7 | + | 117209786 | 117209937 | 117209786 | 117209922 | 117219083 | 117219176 | 24 | 25 | 9 | 32 | 37 | 0 | 1 | 0 | 1 | 0 |
| A5SS | 5110 | ENSG00000126870 | DYNC2I1 | chr7 | + | 158902375 | 158905283 | 158902375 | 158902595 | 158905988 | 158906091 | 2 | 3 | 1 | 5 | 3 | 21 | 16 | 23 | 13 | 33 |
| A5SS | 5484 | ENSG00000125630 | POLR1B | chr2 | + | 112564365 | 112564499 | 112564365 | 112564404 | 112567966 | 112568137 | 55 | 24 | 49 | 37 | 36 | 0 | 0 | 1 | 1 | 0 |
| A5SS | 5697 | ENSG00000119912 | IDE | chr10 | - | 92468878 | 92470345 | 92470253 | 92470345 | 92465675 | 92465843 | 48 | 30 | 36 | 50 | 62 | 2 | 0 | 0 | 0 | 0 |
| A5SS | 6381 | ENSG00000196465 | MYL6B | chr12 | + | 56155418 | 56155592 | 56155418 | 56155493 | 56157467 | 56157545 | 36 | 8 | 43 | 41 | 21 | 0 | 0 | 0 | 0 | 1 |
| A5SS | 6404 | ENSG00000145191 | EIF2B5 | chr3 | + | 184140079 | 184140730 | 184140079 | 184140157 | 184141924 | 184142070 | 73 | 65 | 84 | 99 | 48 | 0 | 0 | 0 | 0 | 2 |
| A5SS | 7861 | ENSG00000142731 | PLK4 | chr4 | + | 127892364 | 127892514 | 127892364 | 127892398 | 127893284 | 127893418 | 17 | 22 | 16 | 20 | 25 | 0 | 1 | 0 | 0 | 0 |
| A5SS | 247 | ENSG00000099860 | GADD45B | chr19 | + | 2476528 | 2476630 | 2476528 | 2476628 | 2477028 | 2477251 | 70 | 26 | 57 | 66 | 72 | 0 | 2 | 0 | 0 | 0 |
| A5SS | 923 | ENSG00000273018 | FAM106A | chr17 | - | 18523650 | 18523877 | 18523776 | 18523877 | 18519481 | 18519674 | 2 | 0 | 3 | 3 | 3 | 0 | 4 | 1 | 8 | 2 |
| A5SS | 1706 | ENSG00000092108 | SCFD1 | chr14 | + | 30674983 | 30676855 | 30674983 | 30675065 | 30694772 | 30694869 | 47 | 73 | 33 | 47 | 17 | 109 | 77 | 80 | 98 | 147 |
| A5SS | 2829 | ENSG00000211456 | SACM1L | chr3 | + | 45689458 | 45689545 | 45689458 | 45689497 | 45703437 | 45703535 | 0 | 1 | 0 | 2 | 0 | 55 | 24 | 52 | 37 | 16 |
| A5SS | 3271 | ENSG00000198925 | ATG9A | chr2 | - | 219223868 | 219224022 | 219223927 | 219224022 | 219223584 | 219223764 | 62 | 34 | 74 | 62 | 41 | 1 | 0 | 0 | 1 | 3 |
| A5SS | 3621 | ENSG00000177728 | TMEM94 | chr17 | + | 75498911 | 75499118 | 75498911 | 75499082 | 75499261 | 75500090 | 0 | 0 | 0 | 0 | 0 | 37 | 11 | 17 | 26 | 5 |
| A5SS | 3689 | ENSG00000077782 | FGFR1 | chr8 | - | 38424316 | 38424699 | 38424508 | 38424699 | 38421796 | 38421941 | 2 | 2 | 2 | 0 | 0 | 7 | 5 | 6 | 2 | 0 |
| A5SS | 3952 | ENSG00000160051 | IQCC | chr1 | + | 32205670 | 32205963 | 32205670 | 32205723 | 32206153 | 32206297 | 2 | 0 | 1 | 3 | 0 | 1 | 0 | 1 | 0 | 0 |
| A5SS | 4516 | ENSG00000168610 | STAT3 | chr17 | - | 42323259 | 42323625 | 42323572 | 42323625 | 42322281 | 42322494 | 10 | 18 | 18 | 24 | 12 | 0 | 0 | 0 | 2 | 0 |
| A5SS | 4579 | ENSG00000177200 | CHD9 | chr16 | + | 53307680 | 53307953 | 53307680 | 53307905 | 53308685 | 53308854 | 70 | 34 | 39 | 41 | 79 | 13 | 8 | 6 | 5 | 7 |
| A5SS | 5005 | ENSG00000174371 | EXO1 | chr1 | + | 241853357 | 241853481 | 241853357 | 241853472 | 241857344 | 241857482 | 3 | 0 | 3 | 4 | 5 | 0 | 0 | 0 | 0 | 2 |
| A5SS | 5006 | ENSG00000196975 | ANXA4 | chr2 | + | 69806384 | 69806498 | 69806384 | 69806432 | 69807905 | 69807996 | 61 | 38 | 42 | 41 | 41 | 0 | 1 | 0 | 1 | 0 |
| A5SS | 5429 | ENSG00000068654 | POLR1A | chr2 | - | 86031329 | 86031635 | 86031512 | 86031635 | 86030195 | 86030396 | 43 | 44 | 59 | 71 | 48 | 1 | 0 | 0 | 1 | 0 |
| A5SS | 5766 | ENSG00000170325 | PRDM10 | chr11 | - | 129912084 | 129912225 | 129912123 | 129912225 | 129910475 | 129910656 | 7 | 1 | 2 | 6 | 0 | 0 | 0 | 0 | 2 | 0 |
| A5SS | 6182 | ENSG00000112237 | CCNC | chr6 | - | 99558496 | 99558548 | 99558500 | 99558548 | 99551839 | 99551895 | 59 | 39 | 59 | 64 | 106 | 0 | 1 | 1 | 0 | 3 |
| A5SS | 6374 | ENSG00000105854 | PON2 | chr7 | - | 95409900 | 95410101 | 95409953 | 95410101 | 95406986 | 95407068 | 64 | 46 | 18 | 35 | 63 | 2 | 1 | 0 | 0 | 0 |
| A5SS | 7490 | ENSG00000149084 | HSD17B12 | chr11 | + | 43680794 | 43681183 | 43680794 | 43680987 | 43750910 | 43750957 | 1 | 2 | 0 | 0 | 1 | 27 | 26 | 39 | 37 | 33 |
| A5SS | 7791 | ENSG00000130714 | POMT1 | chr9 | + | 131507367 | 131509022 | 131507367 | 131507514 | 131509742 | 131509808 | 19 | 12 | 26 | 18 | 10 | 0 | 0 | 1 | 0 | 0 |
| A5SS | 8436 | ENSG00000107959 | PITRM1 | chr10 | - | 3148167 | 3148291 | 3148170 | 3148291 | 3147986 | 3148063 | 0 | 0 | 0 | 0 | 0 | 14 | 15 | 20 | 20 | 14 |
| A5SS | 8520 | ENSG00000139620 | KANSL2 | chr12 | - | 48681381 | 48681641 | 48681570 | 48681641 | 48679654 | 48679833 | 117 | 85 | 106 | 172 | 171 | 2 | 2 | 0 | 0 | 0 |
| A5SS | 896 | ENSG00000178971 | CTC1 | chr17 | - | 8236057 | 8236342 | 8236082 | 8236342 | 8235830 | 8235959 | 38 | 32 | 44 | 62 | 34 | 1 | 1 | 0 | 0 | 0 |
| A5SS | 1453 | ENSG00000185658 | BRWD1 | chr21 | - | 39264890 | 39265019 | 39264985 | 39265019 | 39264459 | 39264685 | 277 | 254 | 257 | 302 | 301 | 6 | 2 | 1 | 1 | 1 |
| A5SS | 1713 | ENSG00000072042 | RDH11 | chr14 | - | 67692933 | 67693052 | 67692972 | 67693052 | 67691139 | 67691244 | 24 | 36 | 19 | 36 | 6 | 0 | 0 | 0 | 2 | 0 |
| A5SS | 1797 | ENSG00000150527 | MIA2 | chr14 | + | 39276933 | 39277065 | 39276933 | 39276955 | 39291018 | 39291111 | 54 | 25 | 30 | 42 | 43 | 0 | 1 | 0 | 0 | 0 |
| A5SS | 2109 | ENSG00000168268 | NT5DC2 | chr3 | - | 52528445 | 52528539 | 52528511 | 52528539 | 52528182 | 52528311 | 5 | 2 | 15 | 20 | 10 | 0 | 0 | 0 | 1 | 0 |
| A5SS | 2357 | ENSG00000038382 | TRIO | chr5 | + | 14487463 | 14488260 | 14487463 | 14487730 | 14492566 | 14492814 | 29 | 10 | 15 | 27 | 7 | 1 | 0 | 0 | 0 | 0 |
| A5SS | 2466 | ENSG00000198707 | CEP290 | chr12 | - | 88120113 | 88120276 | 88120117 | 88120276 | 88118642 | 88118743 | 14 | 11 | 8 | 11 | 46 | 0 | 0 | 0 | 0 | 3 |
| A5SS | 2533 | ENSG00000144959 | NCEH1 | chr3 | - | 172647861 | 172648114 | 172647885 | 172648114 | 172645622 | 172645692 | 0 | 1 | 1 | 0 | 0 | 16 | 10 | 20 | 10 | 36 |
| A5SS | 2595 | ENSG00000129187 | DCTD | chr4 | - | 182916672 | 182916857 | 182916722 | 182916857 | 182916418 | 182916539 | 12 | 4 | 8 | 20 | 7 | 0 | 0 | 0 | 1 | 0 |
| A5SS | 3195 | ENSG00000151576 | QTRT2 | chr3 | + | 114065236 | 114065457 | 114065236 | 114065278 | 114066227 | 114066283 | 150 | 115 | 169 | 149 | 354 | 0 | 5 | 0 | 1 | 0 |
| A5SS | 4121 | ENSG00000116337 | AMPD2 | chr1 | + | 109629326 | 109629490 | 109629326 | 109629486 | 109629795 | 109629916 | 49 | 17 | 37 | 32 | 29 | 0 | 2 | 0 | 0 | 0 |
| A5SS | 5150 | ENSG00000181163 | NPM1 | chr5 | + | 171392913 | 171392982 | 171392913 | 171392978 | 171400152 | 171400210 | 1 | 5 | 4 | 0 | 6 | 1295 | 919 | 1713 | 1527 | 2962 |
| A5SS | 5291 | ENSG00000224078 | SNHG14 | chr15 | + | 25099144 | 25100290 | 25099144 | 25099278 | 25100778 | 25100890 | 1 | 1 | 3 | 0 | 0 | 12 | 5 | 17 | 6 | 17 |
| A5SS | 5937 | ENSG00000124356 | STAMBP | chr2 | + | 73862202 | 73862945 | 73862202 | 73862304 | 73870142 | 73870352 | 50 | 44 | 28 | 38 | 88 | 0 | 0 | 0 | 0 | 3 |
| A5SS | 5938 | ENSG00000124356 | STAMBP | chr2 | + | 73862202 | 73867174 | 73862202 | 73862304 | 73870142 | 73870352 | 50 | 44 | 28 | 38 | 88 | 0 | 0 | 0 | 0 | 3 |
| A5SS | 6390 | ENSG00000183735 | TBK1 | chr12 | + | 64481841 | 64482021 | 64481841 | 64481956 | 64484302 | 64484499 | 211 | 112 | 104 | 182 | 170 | 2 | 0 | 1 | 3 | 6 |
| A5SS | 6967 | ENSG00000163389 | POGLUT1 | chr3 | + | 119468962 | 119469106 | 119468962 | 119469006 | 119469819 | 119469910 | 47 | 20 | 35 | 37 | 42 | 2 | 0 | 0 | 0 | 0 |
| A5SS | 7751 | ENSG00000165699 | TSC1 | chr9 | - | 132897422 | 132897610 | 132897427 | 132897610 | 132897183 | 132897345 | 46 | 34 | 47 | 54 | 76 | 1 | 0 | 0 | 1 | 2 |
| A5SS | 7752 | ENSG00000165699 | TSC1 | chr9 | - | 132901588 | 132901699 | 132901630 | 132901699 | 132897422 | 132897610 | 3 | 2 | 6 | 3 | 0 | 0 | 0 | 0 | 0 | 4 |
| A5SS | 7827 | ENSG00000083799 | CYLD | chr16 | + | 50742761 | 50742841 | 50742761 | 50742822 | 50744729 | 50744866 | 39 | 33 | 41 | 26 | 26 | 0 | 2 | 0 | 0 | 0 |
| A5SS | 8635 | ENSG00000084110 | HAL | chr12 | - | 95980797 | 95980863 | 95980814 | 95980863 | 95980555 | 95980721 | 12 | 15 | 17 | 13 | 19 | 1 | 0 | 0 | 0 | 0 |
| A5SS | 179 | ENSG00000239382 | ALKBH6 | chr19 | - | 36013020 | 36013422 | 36013343 | 36013422 | 36011403 | 36011464 | 39 | 29 | 44 | 32 | 66 | 0 | 0 | 0 | 1 | 0 |
| A5SS | 198 | ENSG00000126457 | PRMT1 | chr19 | + | 49680139 | 49680253 | 49680139 | 49680238 | 49680486 | 49680588 | 10 | 4 | 4 | 7 | 13 | 2 | 0 | 2 | 3 | 2 |
| A5SS | 535 | ENSG00000215440 | NPEPL1 | chr20 | + | 58713419 | 58714093 | 58713419 | 58713543 | 58714559 | 58714670 | 32 | 15 | 33 | 27 | 20 | 1 | 0 | 0 | 0 | 0 |
| A5SS | 715 | ENSG00000161944 | ASGR2 | chr17 | - | 7108756 | 7108888 | 7108771 | 7108888 | 7108461 | 7108557 | 0 | 1 | 0 | 0 | 0 | 17 | 9 | 2 | 13 | 4 |
| A5SS | 756 | ENSG00000182173 | TSEN54 | chr17 | + | 75521704 | 75523335 | 75521704 | 75522521 | 75523662 | 75523779 | 56 | 31 | 63 | 64 | 59 | 0 | 0 | 0 | 2 | 0 |
| A5SS | 818 | ENSG00000108298 | RPL19 | chr17 | + | 39202316 | 39202445 | 39202316 | 39202443 | 39202993 | 39203109 | 30 | 11 | 30 | 16 | 28 | 0 | 0 | 0 | 1 | 0 |
| A5SS | 1229 | ENSG00000214078 | CPNE1 | chr20 | - | 35630438 | 35631034 | 35630900 | 35631034 | 35627279 | 35627413 | 250 | 111 | 293 | 162 | 251 | 0 | 4 | 0 | 0 | 0 |
| A5SS | 1388 | ENSG00000167969 | ECI1 | chr16 | - | 2243317 | 2243439 | 2243368 | 2243439 | 2243045 | 2243224 | 39 | 13 | 21 | 27 | 16 | 0 | 1 | 0 | 0 | 0 |
| A5SS | 2637 | ENSG00000196209 | SIRPB2 | chr20 | - | 1479699 | 1480065 | 1479993 | 1480065 | 1477337 | 1477403 | 95 | 73 | 18 | 62 | 12 | 3 | 0 | 1 | 0 | 0 |
| A5SS | 2689 | ENSG00000164934 | DCAF13 | chr8 | + | 103427096 | 103430689 | 103427096 | 103427252 | 103432658 | 103432741 | 99 | 91 | 90 | 108 | 144 | 0 | 0 | 3 | 0 | 0 |
| A5SS | 2717 | ENSG00000005436 | GCFC2 | chr2 | - | 75702198 | 75702423 | 75702312 | 75702423 | 75701189 | 75701287 | 43 | 36 | 78 | 42 | 80 | 1 | 0 | 0 | 0 | 1 |
| A5SS | 3104 | ENSG00000165272 | AQP3 | chr9 | - | 33443178 | 33443458 | 33443320 | 33443458 | 33442851 | 33442970 | 7 | 1 | 8 | 1 | 8 | 50 | 18 | 70 | 25 | 33 |
| A5SS | 3137 | ENSG00000173889 | PHC3 | chr3 | - | 170171372 | 170171450 | 170171389 | 170171450 | 170145422 | 170145521 | 29 | 12 | 14 | 35 | 29 | 0 | 0 | 0 | 2 | 0 |
| A5SS | 4141 | ENSG00000154222 | CC2D1B | chr1 | - | 52355397 | 52355666 | 52355607 | 52355666 | 52354839 | 52354939 | 67 | 46 | 64 | 63 | 64 | 0 | 0 | 0 | 2 | 0 |
| A5SS | 4855 | ENSG00000137496 | IL18BP | chr11 | + | 72001404 | 72001552 | 72001404 | 72001424 | 72001783 | 72002804 | 57 | 19 | 33 | 30 | 13 | 0 | 0 | 0 | 1 | 0 |
| A5SS | 4932 | ENSG00000182362 | YBEY | chr21 | + | 46286869 | 46287123 | 46286869 | 46286988 | 46291333 | 46291462 | 311 | 157 | 105 | 277 | 189 | 1 | 0 | 2 | 3 | 0 |
| A5SS | 5100 | ENSG00000106105 | GARS1 | chr7 | + | 30601058 | 30601224 | 30601058 | 30601200 | 30603033 | 30603122 | 0 | 1 | 0 | 0 | 0 | 144 | 102 | 141 | 134 | 148 |
| A5SS | 5316 | ENSG00000136718 | IMP4 | chr2 | + | 130345566 | 130345933 | 130345566 | 130345699 | 130346017 | 130346112 | 75 | 39 | 94 | 44 | 65 | 0 | 0 | 0 | 0 | 2 |
| A5SS | 5406 | ENSG00000003509 | NDUFAF7 | chr2 | + | 37241577 | 37241791 | 37241577 | 37241637 | 37242634 | 37242693 | 23 | 24 | 23 | 41 | 62 | 2 | 0 | 2 | 0 | 0 |
| A5SS | 5446 | ENSG00000135912 | TTLL4 | chr2 | + | 218752762 | 218752973 | 218752762 | 218752847 | 218753114 | 218753185 | 42 | 29 | 35 | 48 | 56 | 0 | 1 | 0 | 0 | 0 |
| A5SS | 6734 | ENSG00000122783 | CYREN | chr7 | - | 135168744 | 135169060 | 135168930 | 135169060 | 135167731 | 135167807 | 139 | 108 | 117 | 136 | 131 | 0 | 0 | 0 | 2 | 2 |
| A5SS | 6739 | ENSG00000122783 | CYREN | chr7 | - | 135168785 | 135169060 | 135168930 | 135169060 | 135167731 | 135167807 | 146 | 111 | 123 | 144 | 125 | 0 | 0 | 0 | 2 | 2 |
| A5SS | 6828 | ENSG00000140543 | DET1 | chr15 | - | 88530622 | 88531715 | 88531608 | 88531715 | 88527598 | 88527786 | 36 | 31 | 52 | 32 | 29 | 0 | 0 | 0 | 1 | 0 |
| A5SS | 7137 | ENSG00000248323 | LUCAT1 | chr5 | - | 91313961 | 91314404 | 91314079 | 91314404 | 91313077 | 91313229 | 16 | 10 | 16 | 27 | 23 | 0 | 0 | 0 | 1 | 0 |
| A5SS | 7662 | ENSG00000132740 | IGHMBP2 | chr11 | + | 68906068 | 68906238 | 68906068 | 68906135 | 68908144 | 68908337 | 48 | 16 | 27 | 21 | 18 | 0 | 0 | 1 | 0 | 0 |
| A5SS | 7819 | ENSG00000188352 | FOCAD | chr9 | + | 20820940 | 20821071 | 20820940 | 20821061 | 20822988 | 20823115 | 35 | 17 | 34 | 48 | 37 | 0 | 0 | 2 | 0 | 0 |
| A5SS | 8048 | ENSG00000049449 | RCN1 | chr11 | + | 32103003 | 32103202 | 32103003 | 32103158 | 32103280 | 32103480 | 2 | 1 | 3 | 0 | 3 | 0 | 0 | 1 | 0 | 0 |
| A5SS | 8740 | ENSG00000111752 | PHC1 | chr12 | + | 8932562 | 8933350 | 8932562 | 8932831 | 8933864 | 8934012 | 31 | 12 | 36 | 37 | 21 | 1 | 0 | 0 | 0 | 0 |
| A5SS | 885 | ENSG00000215041 | NEURL4 | chr17 | - | 7324798 | 7324980 | 7324910 | 7324980 | 7324394 | 7324480 | 91 | 54 | 75 | 90 | 55 | 3 | 0 | 0 | 0 | 0 |
| A5SS | 1086 | ENSG00000125124 | BBS2 | chr16 | - | 56501944 | 56502456 | 56502316 | 56502456 | 56500853 | 56501025 | 84 | 93 | 83 | 110 | 49 | 0 | 1 | 2 | 1 | 0 |
| A5SS | 1204 | ENSG00000197912 | SPG7 | chr16 | + | 89540943 | 89542279 | 89540943 | 89541346 | 89544647 | 89544772 | 98 | 110 | 87 | 111 | 65 | 2 | 0 | 2 | 0 | 0 |
| A5SS | 1243 | ENSG00000090861 | AARS1 | chr16 | - | 70289420 | 70289555 | 70289529 | 70289555 | 70282619 | 70282784 | 28 | 8 | 30 | 27 | 25 | 0 | 0 | 1 | 0 | 0 |
| A5SS | 1669 | ENSG00000119682 | AREL1 | chr14 | - | 74675698 | 74675946 | 74675740 | 74675946 | 74674033 | 74674111 | 96 | 63 | 92 | 80 | 126 | 0 | 0 | 2 | 0 | 3 |
| A5SS | 1753 | ENSG00000099797 | TECR | chr19 | + | 14564785 | 14564992 | 14564785 | 14564858 | 14565065 | 14565123 | 77 | 33 | 86 | 69 | 63 | 0 | 0 | 0 | 1 | 1 |
| A5SS | 2077 | ENSG00000154153 | RETREG1 | chr5 | - | 16481008 | 16483472 | 16483345 | 16483472 | 16478849 | 16478987 | 48 | 29 | 81 | 49 | 134 | 0 | 0 | 0 | 0 | 4 |
| A5SS | 2276 | ENSG00000134644 | PUM1 | chr1 | - | 31059203 | 31059577 | 31059298 | 31059577 | 31055312 | 31055380 | 243 | 117 | 193 | 208 | 100 | 2 | 3 | 1 | 1 | 0 |
| A5SS | 2721 | ENSG00000155508 | CNOT8 | chr5 | + | 154858638 | 154858816 | 154858638 | 154858797 | 154863206 | 154863395 | 34 | 38 | 25 | 20 | 10 | 0 | 0 | 0 | 1 | 0 |
| A5SS | 3383 | ENSG00000114544 | SLC41A3 | chr3 | - | 126026334 | 126026479 | 126026392 | 126026479 | 126022785 | 126022932 | 17 | 6 | 24 | 25 | 24 | 0 | 0 | 1 | 0 | 0 |
| A5SS | 3544 | ENSG00000138780 | GSTCD | chr4 | + | 105717592 | 105718039 | 105717592 | 105717778 | 105719059 | 105719527 | 27 | 24 | 64 | 29 | 149 | 0 | 0 | 1 | 0 | 2 |
| A5SS | 3950 | ENSG00000156873 | PHKG2 | chr16 | + | 30751105 | 30751603 | 30751105 | 30751281 | 30753231 | 30753297 | 80 | 35 | 56 | 75 | 25 | 0 | 1 | 0 | 0 | 0 |
| A5SS | 4836 | ENSG00000048162 | NOP16 | chr5 | - | 176388234 | 176388598 | 176388432 | 176388598 | 176386839 | 176386909 | 20 | 30 | 33 | 33 | 15 | 0 | 0 | 0 | 1 | 0 |
| A5SS | 4880 | ENSG00000163534 | FCRL1 | chr1 | - | 157801914 | 157802664 | 157802376 | 157802664 | 157800057 | 157800085 | 50 | 29 | 48 | 33 | 20 | 0 | 0 | 0 | 1 | 0 |
| A5SS | 5160 | ENSG00000115464 | USP34 | chr2 | - | 61206759 | 61208977 | 61208898 | 61208977 | 61206016 | 61206124 | 278 | 218 | 250 | 242 | 592 | 7 | 0 | 0 | 0 | 2 |
| A5SS | 5217 | ENSG00000104907 | TRMT1 | chr19 | - | 13116145 | 13116340 | 13116186 | 13116340 | 13115996 | 13116052 | 72 | 70 | 77 | 88 | 89 | 1 | 0 | 0 | 2 | 2 |
| A5SS | 5347 | ENSG00000180488 | MIGA1 | chr1 | + | 77783237 | 77783351 | 77783237 | 77783267 | 77801330 | 77801508 | 68 | 45 | 68 | 57 | 34 | 0 | 0 | 0 | 1 | 1 |
| A5SS | 5440 | ENSG00000176978 | DPP7 | chr9 | - | 137114242 | 137114576 | 137114462 | 137114576 | 137113864 | 137114028 | 48 | 37 | 68 | 39 | 25 | 0 | 0 | 2 | 0 | 0 |
| A5SS | 5699 | ENSG00000119912 | IDE | chr10 | - | 92507574 | 92508205 | 92508112 | 92508205 | 92506441 | 92506522 | 141 | 88 | 117 | 138 | 148 | 0 | 2 | 0 | 1 | 0 |
| A5SS | 6745 | ENSG00000196305 | IARS1 | chr9 | - | 92253361 | 92256800 | 92256679 | 92256800 | 92251807 | 92251885 | 86 | 57 | 117 | 80 | 152 | 3 | 0 | 0 | 1 | 0 |
| A5SS | 7003 | ENSG00000144161 | ZC3H8 | chr2 | - | 112233257 | 112233371 | 112233259 | 112233371 | 112231837 | 112231945 | 23 | 20 | 25 | 35 | 28 | 0 | 0 | 0 | 2 | 0 |
| A5SS | 7405 | ENSG00000100023 | PPIL2 | chr22 | + | 21695393 | 21695496 | 21695393 | 21695492 | 21696884 | 21698220 | 18 | 9 | 4 | 5 | 5 | 1 | 0 | 0 | 0 | 0 |
| A5SS | 7594 | ENSG00000095303 | PTGS1 | chr9 | + | 122378774 | 122378910 | 122378774 | 122378851 | 122381370 | 122381552 | 128 | 119 | 20 | 180 | 188 | 0 | 1 | 1 | 0 | 0 |
| A5SS | 8343 | ENSG00000156650 | KAT6B | chr10 | + | 74975398 | 74977437 | 74975398 | 74976330 | 74979223 | 74979339 | 115 | 58 | 94 | 93 | 106 | 0 | 0 | 0 | 0 | 3 |
| A5SS | 182 | ENSG00000239382 | ALKBH6 | chr19 | - | 36013327 | 36013422 | 36013343 | 36013422 | 36013020 | 36013089 | 0 | 0 | 0 | 0 | 0 | 29 | 21 | 25 | 26 | 47 |
| A5SS | 344 | ENSG00000268027 | NA | chr19 | + | 41551179 | 41551494 | 41551179 | 41551302 | 41551752 | 41551828 | 37 | 14 | 38 | 35 | 23 | 1 | 0 | 0 | 0 | 0 |
| A5SS | 966 | ENSG00000108406 | DHX40 | chr17 | + | 59566626 | 59566794 | 59566626 | 59566709 | 59570517 | 59570663 | 142 | 103 | 142 | 142 | 164 | 0 | 0 | 0 | 1 | 4 |
| A5SS | 1097 | ENSG00000100336 | APOL4 | chr22 | - | 36195094 | 36195437 | 36195310 | 36195437 | 36189127 | 36191912 | 0 | 0 | 0 | 0 | 0 | 0 | 2 | 5 | 9 | 0 |
| A5SS | 1328 | ENSG00000102908 | NFAT5 | chr16 | + | 69647027 | 69647586 | 69647027 | 69647167 | 69653235 | 69653428 | 169 | 159 | 159 | 200 | 110 | 0 | 4 | 0 | 0 | 0 |
| A5SS | 1432 | ENSG00000198690 | FAN1 | chr15 | + | 30904511 | 30905897 | 30904511 | 30905353 | 30908117 | 30908258 | 45 | 45 | 59 | 39 | 70 | 0 | 0 | 0 | 1 | 0 |
| A5SS | 1518 | ENSG00000067369 | TP53BP1 | chr15 | - | 43420304 | 43420735 | 43420617 | 43420735 | 43416224 | 43416416 | 41 | 24 | 57 | 39 | 55 | 0 | 0 | 0 | 1 | 0 |
| A5SS | 2054 | ENSG00000131844 | MCCC2 | chr5 | + | 71634942 | 71635042 | 71634942 | 71634952 | 71635150 | 71635246 | 96 | 60 | 107 | 102 | 119 | 0 | 2 | 0 | 0 | 0 |
| A5SS | 3138 | ENSG00000173889 | PHC3 | chr3 | - | 170171372 | 170171450 | 170171389 | 170171450 | 170149085 | 170149244 | 57 | 39 | 42 | 71 | 35 | 0 | 1 | 0 | 3 | 0 |
| A5SS | 3482 | ENSG00000126107 | HECTD3 | chr1 | - | 45005793 | 45006116 | 45005996 | 45006116 | 45004599 | 45004806 | 70 | 36 | 62 | 75 | 70 | 0 | 0 | 0 | 2 | 0 |
| A5SS | 3931 | ENSG00000182134 | TDRKH | chr1 | - | 151779950 | 151780140 | 151780050 | 151780140 | 151776438 | 151776599 | 21 | 17 | 26 | 33 | 26 | 0 | 0 | 0 | 1 | 0 |
| A5SS | 3949 | ENSG00000156873 | PHKG2 | chr16 | + | 30751105 | 30751603 | 30751105 | 30751217 | 30753231 | 30753297 | 97 | 37 | 68 | 96 | 34 | 1 | 1 | 0 | 0 | 0 |
| A5SS | 5126 | ENSG00000112855 | HARS2 | chr5 | + | 140691452 | 140691756 | 140691452 | 140691515 | 140693590 | 140693665 | 55 | 31 | 65 | 39 | 24 | 0 | 0 | 0 | 1 | 0 |
| A5SS | 5333 | ENSG00000091527 | CDV3 | chr3 | + | 133574637 | 133574756 | 133574637 | 133574742 | 133575038 | 133575115 | 5 | 1 | 4 | 1 | 0 | 2 | 2 | 0 | 0 | 0 |
| A5SS | 5508 | ENSG00000085978 | ATG16L1 | chr2 | + | 233251672 | 233251942 | 233251672 | 233251940 | 233256101 | 233256195 | 77 | 30 | 62 | 69 | 25 | 1 | 0 | 0 | 0 | 2 |
| A5SS | 5885 | ENSG00000143436 | MRPL9 | chr1 | - | 151762104 | 151762155 | 151762112 | 151762155 | 151761961 | 151762025 | 19 | 15 | 32 | 21 | 14 | 0 | 0 | 0 | 1 | 0 |
| A5SS | 5908 | ENSG00000115307 | AUP1 | chr2 | - | 74528750 | 74529282 | 74529131 | 74529282 | 74527939 | 74528006 | 33 | 13 | 24 | 40 | 16 | 0 | 0 | 0 | 1 | 0 |
| A5SS | 6258 | ENSG00000108352 | RAPGEFL1 | chr17 | + | 40181298 | 40181486 | 40181298 | 40181373 | 40181615 | 40181694 | 2 | 3 | 2 | 3 | 0 | 0 | 0 | 1 | 1 | 0 |
| A5SS | 6407 | ENSG00000145191 | EIF2B5 | chr3 | + | 184140079 | 184140740 | 184140079 | 184140221 | 184141924 | 184142070 | 49 | 39 | 35 | 40 | 31 | 0 | 0 | 0 | 1 | 0 |
| A5SS | 6742 | ENSG00000170356 | OR2A20P | chr7 | - | 144254433 | 144254696 | 144254603 | 144254696 | 144253532 | 144253588 | 60 | 14 | 32 | 7 | 40 | 0 | 0 | 0 | 0 | 1 |
| A5SS | 7525 | ENSG00000099810 | MTAP | chr9 | + | 21815432 | 21815544 | 21815432 | 21815519 | 21816713 | 21816772 | 0 | 0 | 0 | 0 | 0 | 50 | 26 | 48 | 54 | 47 |
| A5SS | 908 | ENSG00000108439 | PNPO | chr17 | + | 47944615 | 47945612 | 47944615 | 47944715 | 47945860 | 47945989 | 43 | 16 | 32 | 44 | 35 | 1 | 0 | 0 | 0 | 0 |
| A5SS | 1185 | ENSG00000172775 | PSME3IP1 | chr16 | - | 57178709 | 57178852 | 57178805 | 57178852 | 57174168 | 57174286 | 46 | 37 | 50 | 67 | 58 | 0 | 1 | 0 | 0 | 0 |
| A5SS | 1186 | ENSG00000172775 | PSME3IP1 | chr16 | - | 57178501 | 57178852 | 57178709 | 57178852 | 57174619 | 57174688 | 77 | 77 | 112 | 107 | 111 | 1 | 0 | 0 | 1 | 0 |
| A5SS | 2023 | ENSG00000141279 | NPEPPS | chr17 | + | 47579389 | 47579511 | 47579389 | 47579455 | 47585499 | 47585745 | 71 | 46 | 56 | 77 | 229 | 0 | 1 | 0 | 2 | 0 |
| A5SS | 2167 | ENSG00000113758 | DBN1 | chr5 | - | 177467480 | 177467817 | 177467742 | 177467817 | 177467254 | 177467332 | 35 | 30 | 34 | 31 | 45 | 0 | 0 | 0 | 0 | 1 |
| A5SS | 2915 | ENSG00000010295 | IFFO1 | chr12 | - | 6548064 | 6548545 | 6548424 | 6548545 | 6541511 | 6541642 | 131 | 63 | 90 | 61 | 113 | 3 | 0 | 0 | 0 | 0 |
| A5SS | 2977 | ENSG00000163660 | CCNL1 | chr3 | - | 157156935 | 157157083 | 157157006 | 157157083 | 157153035 | 157153156 | 113 | 123 | 96 | 117 | 226 | 0 | 1 | 0 | 1 | 3 |
| A5SS | 3505 | ENSG00000214106 | PAXIP1-AS2 | chr7 | + | 154944026 | 154944620 | 154944026 | 154944398 | 154946359 | 154946493 | 77 | 54 | 74 | 76 | 115 | 0 | 0 | 1 | 0 | 1 |
| A5SS | 4117 | ENSG00000116337 | AMPD2 | chr1 | + | 109625661 | 109625805 | 109625661 | 109625792 | 109626159 | 109626228 | 0 | 0 | 0 | 0 | 0 | 67 | 25 | 44 | 73 | 2 |
| A5SS | 4394 | ENSG00000145390 | USP53 | chr4 | + | 119267319 | 119267482 | 119267319 | 119267435 | 119268267 | 119268420 | 64 | 58 | 88 | 82 | 126 | 0 | 2 | 0 | 0 | 0 |
| A5SS | 4514 | ENSG00000177225 | GATD1 | chr11 | - | 773427 | 773629 | 773521 | 773629 | 772426 | 772521 | 3 | 0 | 4 | 1 | 0 | 9 | 7 | 11 | 5 | 6 |
| A5SS | 4965 | ENSG00000131408 | NR1H2 | chr19 | + | 50378148 | 50378439 | 50378148 | 50378312 | 50378521 | 50378796 | 224 | 140 | 129 | 174 | 115 | 0 | 0 | 4 | 0 | 0 |
| A5SS | 5002 | ENSG00000163382 | NAXE | chr1 | + | 156592556 | 156593555 | 156592556 | 156592670 | 156593881 | 156594090 | 101 | 43 | 55 | 84 | 123 | 0 | 0 | 0 | 0 | 3 |
| A5SS | 5234 | ENSG00000115355 | CCDC88A | chr2 | - | 55299827 | 55299919 | 55299838 | 55299919 | 55296257 | 55296523 | 2 | 3 | 0 | 0 | 0 | 87 | 82 | 54 | 101 | 121 |
| A5SS | 5455 | ENSG00000039650 | PNKP | chr19 | - | 49861607 | 49861881 | 49861771 | 49861881 | 49861448 | 49861510 | 20 | 11 | 41 | 28 | 11 | 0 | 0 | 1 | 0 | 0 |
| A5SS | 5558 | ENSG00000112367 | FIG4 | chr6 | + | 109735149 | 109738453 | 109735149 | 109735298 | 109741443 | 109741544 | 140 | 80 | 98 | 96 | 207 | 0 | 0 | 0 | 0 | 5 |
| A5SS | 6451 | ENSG00000091106 | NLRC4 | chr2 | - | 32240941 | 32241125 | 32241032 | 32241125 | 32238131 | 32238302 | 1 | 1 | 1 | 1 | 0 | 62 | 37 | 27 | 36 | 30 |
| A5SS | 6678 | ENSG00000157191 | NECAP2 | chr1 | + | 16440723 | 16440853 | 16440723 | 16440775 | 16443631 | 16443732 | 270 | 139 | 228 | 200 | 186 | 1 | 1 | 0 | 4 | 0 |
| A5SS | 6737 | ENSG00000122783 | CYREN | chr7 | - | 135168748 | 135169060 | 135168930 | 135169060 | 135167731 | 135167807 | 191 | 151 | 165 | 167 | 166 | 0 | 0 | 0 | 2 | 2 |
| A5SS | 7287 | ENSG00000174197 | MGA | chr15 | + | 41736180 | 41736698 | 41736180 | 41736368 | 41748636 | 41748927 | 132 | 89 | 90 | 120 | 88 | 0 | 3 | 1 | 0 | 0 |
| A5SS | 7591 | ENSG00000196781 | TLE1 | chr9 | - | 81615981 | 81616134 | 81616020 | 81616134 | 81613376 | 81613521 | 125 | 39 | 69 | 90 | 71 | 1 | 2 | 0 | 0 | 1 |
| A5SS | 7891 | ENSG00000109911 | ELP4 | chr11 | + | 31594769 | 31594905 | 31594769 | 31594901 | 31603767 | 31603907 | 0 | 0 | 1 | 0 | 0 | 59 | 50 | 54 | 44 | 184 |
| A5SS | 7933 | ENSG00000162300 | ZFPL1 | chr11 | + | 65084690 | 65084800 | 65084690 | 65084782 | 65085114 | 65085226 | 43 | 29 | 52 | 47 | 55 | 0 | 0 | 0 | 2 | 0 |
| A5SS | 7982 | ENSG00000197798 | FAM118B | chr11 | + | 126229224 | 126229336 | 126229224 | 126229293 | 126234994 | 126235087 | 0 | 0 | 0 | 0 | 0 | 28 | 13 | 10 | 24 | 5 |
| A5SS | 8082 | ENSG00000149577 | SIDT2 | chr11 | + | 117189334 | 117190025 | 117189334 | 117189401 | 117190165 | 117190289 | 87 | 54 | 43 | 133 | 44 | 0 | 0 | 1 | 0 | 0 |
| A5SS | 8254 | ENSG00000100600 | LGMN | chr14 | - | 92718746 | 92718844 | 92718773 | 92718844 | 92717379 | 92717461 | 32 | 25 | 45 | 49 | 28 | 0 | 0 | 0 | 2 | 0 |
| A5SS | 8671 | ENSG00000102225 | CDK16 | chrX | + | 47224384 | 47224518 | 47224384 | 47224476 | 47224617 | 47224743 | 109 | 36 | 109 | 107 | 114 | 0 | 0 | 2 | 2 | 0 |
| A5SS | 386 | ENSG00000187994 | RINL | chr19 | - | 38876330 | 38876781 | 38876692 | 38876781 | 38873885 | 38873988 | 74 | 17 | 30 | 46 | 25 | 0 | 0 | 0 | 1 | 0 |
| A5SS | 693 | ENSG00000073605 | GSDMB | chr17 | - | 39906110 | 39906987 | 39906960 | 39906987 | 39905846 | 39905985 | 67 | 51 | 53 | 95 | 39 | 0 | 0 | 0 | 2 | 0 |

| | | | | | | | | | | | | | | | | | | | | | | | | | | | | | | | | | | | | | | | | | | | | | | |
|---|---|---|---|---|---|---|---|---|---|---|---|---|---|---|---|---|---|---|---|---|---|---|---|---|---|---|---|---|---|---|---|---|---|---|---|---|---|---|---|---|---|---|---|---|---|---|
| A5SS | 958 | ENSG00000170921 | TANC2 | chr17 | + | 63099174 | 63099357 | 63099174 | 63099246 | 63151269 | 63151380 | 40 | 14 | 15 | 15 | 19 | 1 | 0 | 0 | 0 | 0 | 15 | 21 | 7 | 25 | 22 | 0 | 0 | 0 | 0 | 0 | 260 | 149 | 1 | 1 | 0.958 | 1 | 1 | 1 | 1 | 1 | 1 | 1 | 1 | 1 | -0.008 |
| A5SS | 1022 | ENSG00000283103 | NA | chr19 | - | 58314670 | 58315036 | 58314805 | 58315036 | 58312969 | 58313113 | 63 | 32 | 73 | 65 | 88 | 3 | 0 | 1 | 1 | 0 | 25 | 58 | 69 | 79 | 10 | 0 | 0 | 1 | 3 | 0 | 284 | 149 | 1 | 1 | 0.917 | 1 | 0.975 | 0.972 | 1 | 1 | 1 | 0.973 | 0.933 | 1 | -0.008 |
| A5SS | 1184 | ENSG00000172775 | PSME3IP1 | chr16 | - | 57178501 | 57178852 | 57178805 | 57178852 | 57174168 | 57174286 | 68 | 51 | 86 | 90 | 85 | 0 | 1 | 0 | 0 | 0 | 47 | 113 | 66 | 75 | 30 | 0 | 0 | 0 | 0 | 0 | 298 | 149 | 1 | 1 | 1 | 0.962 | 1 | 1 | 1 | 1 | 1 | 1 | 1 | 1 | -0.008 |
| A5SS | 1281 | ENSG00000122390 | NAA60 | chr16 | + | 3443705 | 3443893 | 3443705 | 3443827 | 3448470 | 3448540 | 1 | 4 | 2 | 3 | 0 | 11 | 11 | 18 | 13 | 15 | 0 | 3 | 4 | 1 | 12 | 0 | 25 | 17 | 20 | 45 | 215 | 149 | 1 | 1 | 0.059 | 0.201 | 0.071 | 0.138 | 0 | NA | 0.077 | 0.14 | 0.033 | 0.156 | -0.008 |
| A5SS | 1735 | ENSG00000032219 | ARID4A | chr14 | + | 58366023 | 58366230 | 58366023 | 58366068 | 58366882 | 58367029 | 237 | 215 | 235 | 243 | 284 | 5 | 3 | 3 | 5 | 6 | 15 | 132 | 108 | 11 | 0 | 0 | 5 | 2 | 0 | 0 | 298 | 149 | 1 | 1 | 0.96 | 0.973 | 0.975 | 0.96 | 0.959 | 1 | 0.93 | 0.964 | 1 | NA | -0.008 |
| A5SS | 1933 | ENSG00000126214 | KLC1 | chr14 | + | 103679383 | 103679545 | 103679383 | 103679518 | 103693552 | 103693743 | 44 | 45 | 45 | 38 | 52 | 1 | 2 | 2 | 0 | 0 | 124 | 33 | 27 | 71 | 87 | 6 | 0 | 0 | 2 | 0 | 176 | 149 | 1 | 1 | 0.974 | 0.95 | 0.95 | 1 | 1 | 0.946 | 1 | 1 | 0.968 | 1 | -0.008 |
| A5SS | 2101 | ENSG00000108262 | GIT1 | chr17 | - | 29577135 | 29577247 | 29577183 | 29577247 | 29576862 | 29576996 | 43 | 30 | 50 | 70 | 52 | 0 | 1 | 0 | 0 | 0 | 83 | 126 | 45 | 91 | 122 | 0 | 0 | 0 | 0 | 0 | 197 | 149 | 1 | 1 | 1 | 0.958 | 1 | 1 | 1 | 1 | 1 | 1 | 1 | 1 | -0.008 |
| A5SS | 2280 | ENSG00000092020 | PPP2R3C | chr14 | - | 35108094 | 35108236 | 35108138 | 35108236 | 35107303 | 35107374 | 1 | 1 | 1 | 0 | 3 | 250 | 157 | 175 | 198 | 191 | 0 | 2 | 5 | 1 | 0 | 87 | 121 | 97 | 59 | 59 | 193 | 149 | 1 | 1 | 0.003 | 0.005 | 0.004 | 0 | 0.012 | 0 | 0.013 | 0.038 | 0.013 | 0 | -0.008 |
| A5SS | 2358 | ENSG00000140471 | LINS1 | chr15 | - | 100580443 | 100580945 | 100580800 | 100580945 | 100580262 | 100580352 | 246 | 184 | 218 | 190 | 372 | 1 | 0 | 0 | 2 | 2 | 47 | 59 | 125 | 64 | 9 | 0 | 0 | 0 | 0 | 0 | 298 | 149 | 1 | 1 | 0.992 | 1 | 1 | 0.979 | 0.989 | 1 | 1 | 1 | 1 | 1 | -0.008 |
| A5SS | 2966 | ENSG00000114867 | EIF4G1 | chr3 | + | 184317320 | 184317497 | 184317320 | 184317431 | 184317716 | 184317816 | 204 | 83 | 118 | 146 | 120 | 1 | 0 | 3 | 0 | 0 | 192 | 154 | 69 | 106 | 188 | 0 | 0 | 0 | 0 | 0 | 215 | 149 | 1 | 1 | 0.993 | 1 | 0.965 | 1 | 1 | 1 | 1 | 1 | 1 | 1 | -0.008 |
| A5SS | 3021 | ENSG00000156709 | AIFM1 | chrX | - | 130131666 | 130131799 | 130131674 | 130131799 | 130129969 | 130130166 | 0 | 0 | 0 | 0 | 0 | 38 | 34 | 55 | 31 | 46 | 0 | 0 | 1 | 0 | 0 | 92 | 53 | 24 | 58 | 54 | 157 | 149 | 1 | 1 | 0 | 0 | 0 | 0 | 0 | 0 | 0 | 0.038 | 0 | 0 | -0.008 |
| A5SS | 3359 | ENSG00000271723 | MROH7-TTC4 | chr1 | + | 54731485 | 54731700 | 54731485 | 54731549 | 54733628 | 54733710 | 85 | 82 | 101 | 99 | 194 | 0 | 0 | 2 | 0 | 0 | 60 | 87 | 54 | 52 | 91 | 0 | 0 | 0 | 0 | 0 | 298 | 149 | 1 | 1 | 1 | 1 | 0.962 | 1 | 1 | 1 | 1 | 1 | 1 | 1 | -0.008 |
| A5SS | 3749 | ENSG00000143493 | INTS7 | chr1 | - | 211952568 | 211952701 | 211952610 | 211952701 | 211946606 | 211946705 | 42 | 18 | 30 | 32 | 66 | 0 | 0 | 1 | 0 | 0 | 0 | 20 | 9 | 11 | 0 | 0 | 0 | 0 | 0 | 0 | 191 | 149 | 1 | 1 | 1 | 1 | 0.959 | 1 | 1 | NA | 1 | 1 | 1 | NA | -0.008 |
| A5SS | 3753 | ENSG00000182095 | TNRC18 | chr7 | - | 5314979 | 5315148 | 5314983 | 5315148 | 5313110 | 5313863 | 0 | 0 | 2 | 0 | 0 | 109 | 54 | 75 | 83 | 53 | 0 | 0 | 4 | 0 | 0 | 221 | 140 | 56 | 112 | 224 | 153 | 149 | 1 | 1 | 0 | 0 | 0.025 | 0 | 0 | 0 | 0 | 0.065 | 0 | 0 | -0.008 |
| A5SS | 4039 | ENSG00000106610 | STAG3L4 | chr7 | + | 67308576 | 67309139 | 67308576 | 67308652 | 67309563 | 67309715 | 74 | 42 | 65 | 47 | 66 | 0 | 0 | 0 | 1 | 0 | 140 | 76 | 90 | 97 | 193 | 0 | 0 | 0 | 0 | 0 | 298 | 149 | 1 | 1 | 1 | 1 | 1 | 0.959 | 1 | 1 | 1 | 1 | 1 | 1 | -0.008 |
| A5SS | 4055 | ENSG00000158773 | USF1 | chr1 | - | 161041734 | 161041846 | 161041833 | 161041846 | 161041323 | 161041411 | 144 | 78 | 138 | 113 | 69 | 0 | 2 | 0 | 0 | 0 | 467 | 240 | 214 | 291 | 387 | 0 | 0 | 0 | 0 | 0 | 248 | 149 | 1 | 1 | 1 | 0.959 | 1 | 1 | 1 | 1 | 1 | 1 | 1 | 1 | -0.008 |
| A5SS | 4214 | ENSG00000123064 | DDX54 | chr12 | - | 113180928 | 113181058 | 113180980 | 113181058 | 113179934 | 113180005 | 142 | 80 | 115 | 149 | 81 | 0 | 0 | 3 | 0 | 1 | 138 | 155 | 112 | 145 | 204 | 0 | 0 | 1 | 0 | 0 | 201 | 149 | 1 | 1 | 1 | 1 | 0.966 | 1 | 0.984 | 1 | 1 | 0.988 | 1 | 1 | -0.008 |
| A5SS | 4441 | ENSG00000158796 | DEDD | chr1 | - | 161124339 | 161124526 | 161124413 | 161124526 | 161124137 | 161124210 | 72 | 21 | 27 | 35 | 18 | 0 | 0 | 0 | 1 | 0 | 52 | 28 | 38 | 44 | 78 | 0 | 0 | 0 | 0 | 0 | 223 | 149 | 1 | 1 | 1 | 1 | 1 | 0.959 | 1 | 1 | 1 | 1 | 1 | 1 | -0.008 |
| A5SS | 5183 | ENSG00000177000 | MTHFR | chr1 | - | 11794728 | 11795348 | 11795097 | 11795348 | 11794357 | 11794538 | 48 | 30 | 32 | 48 | 41 | 0 | 0 | 0 | 1 | 0 | 240 | 72 | 30 | 71 | 102 | 0 | 0 | 0 | 0 | 0 | 298 | 149 | 1 | 1 | 1 | 1 | 1 | 0.96 | 1 | 1 | 1 | 1 | 1 | 1 | -0.008 |
| A5SS | 5442 | ENSG00000163848 | ZNF148 | chr3 | - | 125323308 | 125323444 | 125323387 | 125323444 | 125314936 | 125315053 | 121 | 68 | 91 | 142 | 99 | 0 | 0 | 2 | 1 | 3 | 15 | 61 | 95 | 32 | 59 | 0 | 0 | 3 | 0 | 0 | 228 | 149 | 1 | 1 | 1 | 1 | 0.967 | 0.989 | 0.956 | 1 | 1 | 0.954 | 1 | 1 | -0.008 |
| A5SS | 5526 | ENSG00000137947 | GTF2B | chr1 | - | 88887260 | 88887367 | 88887275 | 88887367 | 88863980 | 88864114 | 249 | 177 | 186 | 221 | 111 | 3 | 1 | 0 | 5 | 0 | 375 | 365 | 307 | 360 | 158 | 0 | 1 | 0 | 0 | 0 | 164 | 149 | 1 | 1 | 0.987 | 0.994 | 1 | 0.976 | 1 | 1 | 0.997 | 1 | 1 | 1 | -0.008 |
| A5SS | 5756 | ENSG00000123983 | ACSL3 | chr2 | + | 222916318 | 222918155 | 222916318 | 222916496 | 222919063 | 222919202 | 128 | 98 | 90 | 132 | 115 | 0 | 1 | 2 | 0 | 2 | 46 | 102 | 106 | 46 | 16 | 0 | 1 | 2 | 0 | 0 | 298 | 149 | 1 | 1 | 1 | 0.98 | 0.957 | 1 | 0.966 | 1 | 0.981 | 0.964 | 1 | 1 | -0.008 |
| A5SS | 5791 | ENSG00000160710 | ADAR | chr1 | - | 154590261 | 154590409 | 154590341 | 154590409 | 154589756 | 154589928 | 281 | 152 | 271 | 300 | 202 | 0 | 0 | 4 | 0 | 4 | 480 | 359 | 243 | 531 | 431 | 0 | 0 | 0 | 2 | 2 | 229 | 149 | 1 | 1 | 1 | 1 | 0.978 | 1 | 0.97 | 1 | 1 | 1 | 0.994 | 0.993 | -0.008 |
| A5SS | 5797 | ENSG00000079785 | DDX1 | chr2 | + | 15627053 | 15628839 | 15627053 | 15627145 | 15629601 | 15629697 | 122 | 91 | 112 | 107 | 185 | 1 | 0 | 0 | 3 | 0 | 15 | 40 | 73 | 57 | 5 | 0 | 0 | 1 | 0 | 0 | 298 | 149 | 1 | 1 | 0.984 | 1 | 1 | 0.947 | 1 | 1 | 1 | 0.973 | 1 | 1 | -0.008 |
| A5SS | 6306 | ENSG00000233614 | DDX11L10 | chr16 | + | 12293 | 12402 | 12293 | 12378 | 12901 | 14090 | 7 | 3 | 5 | 27 | 14 | 0 | 0 | 0 | 1 | 0 | 16 | 0 | 0 | 11 | 48 | 0 | 0 | 0 | 0 | 0 | 173 | 149 | 1 | 1 | 1 | 1 | 1 | 0.959 | 1 | 1 | NA | NA | 1 | 1 | -0.008 |
| A5SS | 6442 | ENSG00000134905 | CARS2 | chr13 | - | 110645966 | 110647239 | 110647100 | 110647239 | 110644384 | 110644483 | 127 | 76 | 90 | 101 | 62 | 1 | 0 | 1 | 0 | 0 | 110 | 79 | 57 | 104 | 157 | 0 | 0 | 0 | 0 | 0 | 298 | 149 | 1 | 1 | 0.984 | 1 | 0.978 | 1 | 1 | 1 | 1 | 1 | 1 | 1 | -0.008 |
| A5SS | 6667 | ENSG00000114503 | NCBP2 | chr3 | - | 196939250 | 196939432 | 196939392 | 196939432 | 196937509 | 196937631 | 335 | 282 | 321 | 328 | 424 | 3 | 0 | 1 | 0 | 4 | 112 | 212 | 213 | 167 | 51 | 0 | 0 | 0 | 0 | 0 | 291 | 149 | 1 | 1 | 0.983 | 1 | 0.994 | 1 | 0.982 | 1 | 1 | 1 | 1 | 1 | -0.008 |
| A5SS | 7220 | ENSG00000102309 | PIN4 | chrX | + | 72181764 | 72181925 | 72181764 | 72181828 | 72186460 | 72186534 | 0 | 0 | 0 | 0 | 0 | 55 | 62 | 46 | 28 | 44 | 0 | 0 | 0 | 4 | 0 | 66 | 38 | 15 | 56 | 30 | 246 | 149 | 1 | 1 | 0 | 0 | 0 | 0 | 0 | 0 | 0 | 0 | 0.041 | 0 | -0.008 |

[illegible]

| | | | | | | | | | | | | | | | | | | | | | | | | | | | | | | | | | | | | | | | | | | | | | | |
|---|---|---|---|---|---|---|---|---|---|---|---|---|---|---|---|---|---|---|---|---|---|---|---|---|---|---|---|---|---|---|---|---|---|---|---|---|---|---|---|---|---|---|---|---|---|---|
| A5SS | 680 | ENSG00000101019 | UQCC1 | chr20 | - | 35381917 | 35382025 | 35381983 | 35382025 | 35374183 | 35374256 | 119 | 113 | 92 | 94 | 107 | 0 | 0 | 1 | 0 | 0 | 98 | 107 | 111 | 149 | 106 | 0 | 0 | 0 | 0 | 0 | 215 | 149 | 1 | 1 | 1 | 1 | 0.985 | 1 | 1 | 1 | 1 | 1 | 1 | 1 | -0.003 |
| A5SS | 1041 | ENSG00000177731 | FLII | chr17 | - | 18245353 | 18245660 | 18245554 | 18245660 | 18245237 | 18245272 | 204 | 99 | 158 | 144 | 168 | 0 | 0 | 0 | 1 | 0 | 185 | 154 | 120 | 150 | 302 | 0 | 0 | 0 | 0 | 0 | 298 | 149 | 1 | 1 | 1 | 1 | 1 | 0.986 | 1 | 1 | 1 | 1 | 1 | 1 | -0.003 |
| A5SS | 1461 | ENSG00000137876 | RSL24D1 | chr15 | - | 55192710 | 55192833 | 55192719 | 55192833 | 55190974 | 55191047 | 0 | 1 | 1 | 0 | 0 | 439 | 229 | 396 | 380 | 569 | 0 | 0 | 6 | 0 | 0 | 147 | 213 | 251 | 112 | 50 | 158 | 149 | 1 | 1 | 0 | 0.004 | 0.002 | 0 | 0 | 0 | 0 | 0.022 | 0 | 0 | -0.003 |
| A5SS | 1463 | ENSG00000140479 | PCSK6 | chr15 | - | 101318184 | 101318422 | 101318318 | 101318422 | 101313375 | 101313505 | 0 | 0 | 0 | 2 | 0 | 14 | 16 | 6 | 32 | 25 | 0 | 0 | 0 | 2 | 11 | 33 | 8 | 6 | 28 | 481 | 283 | 149 | 1 | 1 | 0 | 0 | 0 | 0.032 | 0 | 0 | 0 | 0 | 0.036 | 0.012 | -0.003 |
| A5SS | 1702 | ENSG00000171469 | ZNF561 | chr19 | - | 9617044 | 9617171 | 9617048 | 9617171 | 9614020 | 9614103 | 90 | 96 | 77 | 62 | 124 | 2 | 0 | 2 | 2 | 2 | 70 | 68 | 86 | 56 | 16 | 0 | 6 | 0 | 0 | 0 | 153 | 149 | 1 | 1 | 0.978 | 1 | 0.974 | 0.968 | 0.984 | 1 | 0.917 | 1 | 1 | 1 | -0.003 |
| A5SS | 1733 | ENSG00000100938 | GMPR2 | chr14 | + | 24235966 | 24236140 | 24235966 | 24236041 | 24237070 | 24237152 | 226 | 127 | 200 | 242 | 200 | 0 | 1 | 0 | 0 | 0 | 285 | 254 | 166 | 224 | 279 | 0 | 0 | 0 | 0 | 0 | 248 | 149 | 1 | 1 | 1 | 0.987 | 1 | 1 | 1 | 1 | 1 | 1 | 1 | 1 | -0.003 |
| A5SS | 1867 | ENSG00000112679 | DUSP22 | chr6 | + | 311879 | 311962 | 311879 | 311914 | 345853 | 345928 | 114 | 72 | 76 | 140 | 100 | 0 | 0 | 1 | 0 | 0 | 364 | 141 | 94 | 139 | 325 | 0 | 0 | 0 | 0 | 0 | 197 | 149 | 1 | 1 | 1 | 1 | 0.983 | 1 | 1 | 1 | 1 | 1 | 1 | 1 | -0.003 |
| A5SS | 1871 | ENSG00000138081 | FBXO11 | chr2 | - | 47808328 | 47809266 | 47809157 | 47809266 | 47806997 | 47808247 | 240 | 247 | 210 | 273 | 492 | 2 | 0 | 0 | 0 | 0 | 39 | 84 | 165 | 79 | 10 | 0 | 0 | 0 | 0 | 0 | 298 | 149 | 1 | 1 | 0.984 | 1 | 1 | 1 | 1 | 1 | 1 | 1 | 1 | 1 | -0.003 |
| A5SS | 1987 | ENSG00000079616 | KIF22 | chr16 | + | 29796892 | 29797093 | 29796892 | 29797088 | 29798373 | 29798501 | 0 | 0 | 0 | 0 | 0 | 61 | 45 | 71 | 59 | 35 | 0 | 0 | 1 | 0 | 0 | 50 | 78 | 71 | 65 | 128 | 154 | 149 | 1 | 1 | 0 | 0 | 0 | 0 | 0 | 0 | 0 | 0.013 | 0 | 0 | -0.003 |
| A5SS | 2177 | ENSG00000133835 | HSD17B4 | chr5 | + | 119477416 | 119477501 | 119477416 | 119477492 | 119478833 | 119479021 | 214 | 141 | 182 | 211 | 249 | 0 | 1 | 1 | 0 | 0 | 118 | 146 | 95 | 140 | 36 | 0 | 0 | 0 | 0 | 0 | 158 | 149 | 1 | 1 | 1 | 0.993 | 0.994 | 1 | 1 | 1 | 1 | 1 | 1 | 1 | -0.003 |
| A5SS | 2377 | ENSG00000174720 | LARP7 | chr4 | + | 112644667 | 112644871 | 112644667 | 112644699 | 112646350 | 112646451 | 262 | 159 | 166 | 210 | 355 | 2 | 0 | 0 | 0 | 0 | 36 | 93 | 32 | 29 | 8 | 0 | 0 | 0 | 0 | 0 | 298 | 149 | 1 | 1 | 0.985 | 1 | 1 | 1 | 1 | 1 | 1 | 1 | 1 | 1 | -0.003 |
| A5SS | 2632 | ENSG00000064886 | CHI3L2 | chr1 | + | 111234906 | 111235057 | 111234906 | 111235000 | 111235638 | 111235763 | 153 | 36 | 205 | 75 | 260 | 1 | 0 | 1 | 0 | 0 | 83 | 249 | 67 | 70 | 213 | 0 | 0 | 0 | 0 | 0 | 206 | 149 | 1 | 1 | 0.991 | 1 | 0.993 | 1 | 1 | 1 | 1 | 1 | 1 | 1 | -0.003 |
| A5SS | 2751 | ENSG00000143669 | LYST | chr1 | - | 235766078 | 235766277 | 235766238 | 235766277 | 235762719 | 235762851 | 470 | 466 | 304 | 433 | 506 | 0 | 0 | 0 | 0 | 4 | 21 | 295 | 181 | 25 | 50 | 0 | 0 | 0 | 0 | 0 | 298 | 149 | 1 | 1 | 1 | 1 | 1 | 1 | 0.984 | 1 | 1 | 1 | 1 | 1 | -0.003 |
| A5SS | 3245 | ENSG00000137074 | APTX | chr9 | - | 32987543 | 32988129 | 32988082 | 32988129 | 32985970 | 32986030 | 61 | 43 | 33 | 51 | 115 | 0 | 0 | 0 | 0 | 1 | 59 | 70 | 75 | 42 | 18 | 0 | 0 | 0 | 0 | 0 | 298 | 149 | 1 | 1 | 1 | 1 | 1 | 1 | 0.983 | 1 | 1 | 1 | 1 | 1 | -0.003 |
| A5SS | 3291 | ENSG00000117054 | ACADM | chr1 | + | 75732852 | 75734871 | 75732852 | 75732922 | 75739979 | 75740110 | 169 | 108 | 151 | 195 | 248 | 0 | 0 | 1 | 0 | 0 | 11 | 96 | 62 | 48 | 9 | 0 | 0 | 0 | 0 | 0 | 298 | 149 | 1 | 1 | 1 | 1 | 0.987 | 1 | 1 | 1 | 1 | 1 | 1 | 1 | -0.003 |
| A5SS | 3299 | ENSG00000101457 | DNTTIP1 | chr20 | + | 45801998 | 45802057 | 45801998 | 45802055 | 45803332 | 45803378 | 208 | 127 | 174 | 145 | 203 | 0 | 0 | 2 | 0 | 2 | 395 | 189 | 162 | 236 | 458 | 0 | 0 | 0 | 0 | 4 | 151 | 149 | 1 | 1 | 1 | 1 | 0.988 | 1 | 0.99 | 1 | 1 | 1 | 1 | 0.991 | -0.003 |
| A5SS | 3348 | ENSG00000181555 | SETD2 | chr3 | - | 47080800 | 47081083 | 47080915 | 47081083 | 47067069 | 47067118 | 60 | 23 | 27 | 35 | 67 | 12 | 10 | 10 | 20 | 28 | 0 | 42 | 40 | 24 | 11 | 0 | 10 | 24 | 9 | 4 | 264 | 149 | 1 | 1 | 0.738 | 0.565 | 0.604 | 0.497 | 0.575 | NA | 0.703 | 0.485 | 0.601 | 0.608 | -0.003 |
| A5SS | 3651 | ENSG00000099622 | CIRBP | chr19 | + | 1271139 | 1271467 | 1271139 | 1271365 | 1271550 | 1271632 | 870 | 400 | 793 | 756 | 933 | 2 | 1 | 5 | 3 | 0 | 790 | 793 | 938 | 762 | 1522 | 0 | 1 | 1 | 4 | 0 | 251 | 149 | 1 | 1 | 0.996 | 0.996 | 0.989 | 0.993 | 1 | 1 | 0.998 | 0.998 | 0.991 | 1 | -0.003 |
| A5SS | 4112 | ENSG00000111331 | OAS3 | chr12 | + | 112950692 | 112950975 | 112950692 | 112950971 | 112961070 | 112961246 | 66 | 80 | 47 | 114 | 25 | 1 | 0 | 0 | 0 | 0 | 415 | 145 | 44 | 334 | 127 | 0 | 0 | 0 | 0 | 0 | 153 | 149 | 1 | 1 | 0.985 | 1 | 1 | 1 | 1 | 1 | 1 | 1 | 1 | 1 | -0.003 |
| A5SS | 4313 | ENSG00000127463 | EMC1 | chr1 | - | 19244738 | 19245030 | 19244905 | 19245030 | 19243949 | 19244015 | 4 | 7 | 0 | 3 | 7 | 80 | 43 | 58 | 40 | 44 | 7 | 3 | 5 | 9 | 9 | 133 | 85 | 32 | 50 | 157 | 298 | 149 | 1 | 1 | 0.024 | 0.075 | 0 | 0.036 | 0.074 | 0.026 | 0.017 | 0.072 | 0.083 | 0.028 | -0.003 |
| A5SS | 4410 | ENSG00000143575 | HAX1 | chr1 | + | 154273335 | 154273598 | 154273335 | 154273534 | 154273773 | 154273961 | 310 | 187 | 348 | 304 | 380 | 0 | 0 | 1 | 2 | 0 | 306 | 346 | 178 | 276 | 417 | 0 | 0 | 0 | 0 | 0 | 213 | 149 | 1 | 1 | 1 | 1 | 0.996 | 0.991 | 1 | 1 | 1 | 1 | 1 | 1 | -0.003 |
| A5SS | 4475 | ENSG00000177409 | SAMD9L | chr7 | - | 93146882 | 93147146 | 93147087 | 93147146 | 93145384 | 93145532 | 190 | 155 | 125 | 234 | 128 | 0 | 0 | 1 | 0 | 0 | 36 | 73 | 25 | 62 | 20 | 0 | 0 | 0 | 0 | 0 | 298 | 149 | 1 | 1 | 1 | 1 | 0.984 | 1 | 1 | 1 | 1 | 1 | 1 | 1 | -0.003 |
| A5SS | 4535 | ENSG00000116688 | MFN2 | chr1 | + | 11998769 | 11999095 | 11998769 | 11998878 | 12001400 | 12001554 | 100 | 100 | 83 | 147 | 119 | 0 | 0 | 0 | 1 | 0 | 160 | 148 | 113 | 139 | 635 | 0 | 0 | 0 | 0 | 0 | 298 | 149 | 1 | 1 | 1 | 1 | 1 | 0.987 | 1 | 1 | 1 | 1 | 1 | 1 | -0.003 |
| A5SS | 4929 | ENSG00000138398 | PPIG | chr2 | + | 169636091 | 169636228 | 169636091 | 169636162 | 169636412 | 169637523 | 257 | 291 | 225 | 229 | 406 | 0 | 3 | 0 | 0 | 0 | 43 | 76 | 112 | 27 | 5 | 0 | 0 | 0 | 0 | 0 | 215 | 149 | 1 | 1 | 1 | 0.985 | 1 | 1 | 1 | 1 | 1 | 1 | 1 | 1 | -0.003 |
| A5SS | 5137 | ENSG00000166986 | MARS1 | chr12 | + | 57489266 | 57489558 | 57489266 | 57489345 | 57489895 | 57489971 | 82 | 64 | 74 | 94 | 76 | 0 | 0 | 1 | 0 | 0 | 181 | 120 | 95 | 121 | 206 | 0 | 0 | 0 | 0 | 1 | 298 | 149 | 1 | 1 | 1 | 1 | 0.974 | 1 | 1 | 1 | 1 | 1 | 1 | 0.99 | -0.003 |
| A5SS | 5225 | ENSG00000121579 | NAA50 | chr3 | - | 113723958 | 113724095 | 113723980 | 113724095 | 113723421 | 113723541 | 153 | 83 | 112 | 89 | 165 | 0 | 0 | 0 | 1 | 0 | 24 | 61 | 126 | 58 | 37 | 0 | 0 | 0 | 0 | 0 | 171 | 149 | 1 | 1 | 1 | 1 | 1 | 0.987 | 1 | 1 | 1 | 1 | 1 | 1 | -0.003 |
| A5SS | 5249 | ENSG00000169621 | APLF | chr2 | + | 68513547 | 68513684 | 68513547 | 68513680 | 68526060 | 68526242 | 0 | 0 | 0 | 1 | 0 | 44 | 47 | 38 | 40 | 96 | 0 | 0 | 0 | 0 | 3 | 102 | 71 | 24 | 20 | 77 | 153 | 149 | 1 | 1 | 0 | 0 | 0 | 0.024 | 0 | 0 | 0 | 0 | 0 | 0.037 | -0.003 |
| A5SS | 5430 | ENSG00000115159 | GPD2 | chr2 | + | 156557388 | 156557582 | 156557388 | 156557551 | 156568824 | 156568959 | 86 | 87 | 53 | 115 | 87 | 1 | 0 | 0 | 0 | 0 | 81 | 110 | 90 | 101 | 121 | 0 | 0 | 0 | 0 | 0 | 180 | 149 | 1 | 1 | 0.986 | 1 | 1 | 1 | 1 | 1 | 1 | 1 | 1 | 1 | -0.003 |
| A5SS | 5497 | ENSG00000164032 | H2AZ1 | chr4 | - | 99948810 | 99948940 | 99948813 | 99948940 | 99948087 | 99948523 | 418 | 194 | 405 | 407 | 486 | 1 | 2 | 1 | 0 | 0 | 402 | 387 | 297 | 416 | 393 | 0 | 0 | 0 | 0 | 0 | 152 | 149 | 1 | 1 | 0.998 | 0.99 | 0.997 | 1 | 1 | 1 | 1 | 1 | 1 | 1 | -0.003 |
| A5SS | 5758 | ENSG00000123983 | ACSL3 | chr2 | + | 222927016 | 222928936 | 222927016 | 222927189 | 222930620 | 222930812 | 103 | 105 | 106 | 152 | 124 | 0 | 0 | 0 | 1 | 0 | 21 | 141 | 141 | 87 | 10 | 0 | 0 | 0 | 0 | 0 | 298 | 149 | 1 | 1 | 1 | 1 | 1 | 0.987 | 1 | 1 | 1 | 1 | 1 | 1 | -0.003 |
| A5SS | 5787 | ENSG00000115540 | MOB4 | chr2 | + | 197535529 | 197535630 | 197535529 | 197535605 | 197540110 | 197540153 | 186 | 92 | 137 | 153 | 300 | 0 | 0 | 2 | 0 | 0 | 10 | 68 | 81 | 40 | 34 | 0 | 0 | 0 | 0 | 0 | 174 | 149 | 1 | 1 | 1 | 1 | 0.983 | 1 | 1 | 1 | 1 | 1 | 1 | 1 | -0.003 |
| A5SS | 5798 | ENSG00000079785 | DDX1 | chr2 | + | 15628444 | 15628839 | 15628444 | 15628517 | 15629601 | 15629697 | 122 | 87 | 107 | 108 | 199 | 1 | 0 | 0 | 0 | 0 | 8 | 35 | 61 | 53 | 6 | 0 | 0 | 0 | 0 | 0 | 298 | 149 | 1 | 1 | 0.984 | 1 | 1 | 1 | 1 | 1 | 1 | 1 | 1 | 1 | -0.003 |
| A5SS | 5824 | ENSG00000180228 | PRKRA | chr2 | - | 178450241 | 178450411 | 178450365 | 178450411 | 178447455 | 178447586 | 123 | 64 | 120 | 129 | 199 | 0 | 0 | 0 | 1 | 0 | 45 | 55 | 53 | 71 | 12 | 0 | 0 | 0 | 0 | 0 | 273 | 149 | 1 | 1 | 1 | 1 | 1 | 0.986 | 1 | 1 | 1 | 1 | 1 | 1 | -0.003 |
| A5SS | 6039 | ENSG00000163939 | PBRM1 | chr3 | - | 52563282 | 52563493 | 52563363 | 52563493 | 52561766 | 52561968 | 206 | 162 | 200 | 227 | 383 | 0 | 0 | 1 | 1 | 0 | 53 | 135 | 113 | 96 | 9 | 0 | 0 | 0 | 0 | 0 | 230 | 149 | 1 | 1 | 1 | 1 | 0.992 | 0.993 | 1 | 1 | 1 | 1 | 1 | 1 | -0.003 |
| A5SS | 6070 | ENSG00000204410 | MSH5 | chr6 | + | 31761817 | 31761955 | 31761817 | 31761865 | 31762111 | 31762185 | 18 | 16 | 16 | 30 | 10 | 2 | 0 | 0 | 0 | 0 | 29 | 33 | 14 | 32 | 30 | 0 | 1 | 0 | 2 | 0 | 239 | 149 | 1 | 1 | 0.849 | 1 | 1 | 1 | 1 | 1 | 0.954 | 1 | 0.909 | 1 | -0.003 |
| A5SS | 6168 | ENSG00000135486 | HNRNPA1 | chr12 | + | 54282799 | 54282823 | 54282799 | 54282820 | 54283826 | 54283967 | 20 | 5 | 15 | 16 | 10 | 1 | 0 | 1 | 0 | 0 | 41 | 22 | 12 | 4 | 0 | 0 | 0 | 1 | 0 | 0 | 152 | 149 | 1 | 1 | 0.951 | 1 | 0.936 | 1 | 1 | 1 | 1 | 0.922 | 1 | NA | -0.003 |
| A5SS | 6419 | ENSG00000001631 | KRIT1 | chr7 | - | 92235402 | 92236542 | 92236412 | 92236542 | 92234807 | 92234923 | 117 | 111 | 71 | 84 | 155 | 0 | 3 | 0 | 0 | 0 | 19 | 20 | 53 | 33 | 17 | 0 | 0 | 1 | 0 | 0 | 298 | 149 | 1 | 1 | 1 | 0.949 | 1 | 1 | 1 | 1 | 1 | 0.964 | 1 | 1 | -0.003 |
| A5SS | 6591 | ENSG00000129103 | SUMF2 | chr7 | + | 56074585 | 56074736 | 56074585 | 56074639 | 56076833 | 56076889 | 194 | 124 | 255 | 204 | 386 | 0 | 0 | 0 | 2 | 0 | 194 | 273 | 89 | 110 | 286 | 0 | 0 | 0 | 0 | 0 | 246 | 149 | 1 | 1 | 1 | 1 | 1 | 0.984 | 1 | 1 | 1 | 1 | 1 | 1 | -0.003 |
| A5SS | 6664 | ENSG00000197150 | ABCB8 | chr7 | + | 151040497 | 151040644 | 151040497 | 151040634 | 151040827 | 151040922 | 0 | 2 | 2 | 0 | 0 | 7 | 3 | 11 | 7 | 1 | 0 | 3 | 0 | 0 | 0 | 0 | 10 | 0 | 1 | 0 | 159 | 149 | 1 | 1 | 0 | 0.385 | 0.146 | 0 | 0 | NA | 0.219 | NA | 0 | NA | -0.003 |
| A5SS | 7009 | ENSG00000244754 | N4BP2L2 | chr13 | - | 32536932 | 32537027 | 32537005 | 32537027 | 32522181 | 32522270 | 280 | 277 | 238 | 279 | 297 | 0 | 0 | 0 | 0 | 3 | 75 | 178 | 185 | 82 | 37 | 0 | 0 | 0 | 0 | 0 | 222 | 149 | 1 | 1 | 1 | 1 | 1 | 1 | 0.985 | 1 | 1 | 1 | 1 | 1 | -0.003 |
| A5SS | 7097 | ENSG00000124275 | MTRR | chr5 | + | 7873372 | 7873526 | 7873372 | 7873512 | 7875257 | 7875375 | 35 | 23 | 49 | 58 | 52 | 2 | 0 | 0 | 4 | 0 | 10 | 66 | 22 | 5 | 0 | 0 | 6 | 0 | 0 | 0 | 163 | 149 | 1 | 1 | 0.941 | 1 | 1 | 0.93 | 1 | 1 | 0.91 | 1 | 1 | NA | -0.003 |
| A5SS | 7341 | ENSG00000164961 | WASHC5 | chr8 | - | 125083712 | 125084022 | 125083834 | 125084022 | 125083112 | 125083258 | 153 | 78 | 136 | 146 | 148 | 0 | 0 | 1 | 0 | 0 | 55 | 73 | 63 | 125 | 14 | 0 | 0 | 0 | 0 | 0 | 271 | 149 | 1 | 1 | 1 | 1 | 0.987 | 1 | 1 | 1 | 1 | 1 | 1 | 1 | -0.003 |
| A5SS | 7448 | ENSG00000121022 | COPS5 | chr8 | - | 67047684 | 67047859 | 67047767 | 67047859 | 67045811 | 67045960 | 25 | 26 | 29 | 31 | 42 | 0 | 4 | 0 | 0 | 0 | 0 | 17 | 26 | 27 | 0 | 0 | 0 | 2 | 0 | 0 | 232 | 149 | 1 | 1 | 1 | 0.807 | 1 | 1 | 1 | NA | 1 | 0.893 | 1 | NA | -0.003 |
| A5SS | 7903 | ENSG00000136270 | TBRG4 | chr7 | - | 45102280 | 45102491 | 45102346 | 45102491 | 45101824 | 45102070 | 0 | 0 | 3 | 0 | 0 | 33 | 12 | 31 | 28 | 8 | 0 | 0 | 0 | 1 | 0 | 4 | 3 | 2 | 8 | 63 | 215 | 149 | 1 | 1 | 0 | 0 | 0.063 | 0 | 0 | 0 | 0 | 0 | 0.08 | 0 | -0.003 |
| A5SS | 7915 | ENSG00000110025 | SNX15 | chr11 | + | 65032167 | 65032271 | 65032167 | 65032203 | 65032430 | 65032551 | 4 | 0 | 1 | 0 | 0 | 59 | 35 | 36 | 39 | 24 | 0 | 0 | 0 | 3 | 3 | 55 | 68 | 28 | 32 | 104 | 217 | 149 | 1 | 1 | 0.044 | 0 | 0.019 | 0 | 0 | 0 | 0 | 0 | 0.06 | 0.019 | -0.003 |
| A5SS | 8035 | ENSG00000177697 | CD151 | chr11 | + | 837459 | 837618 | 837459 | 837612 | 837941 | 838028 | 36 | 23 | 21 | 62 | 5 | 0 | 0 | 0 | 1 | 0 | 28 | 29 | 14 | 28 | 200 | 0 | 0 | 0 | 0 | 0 | 155 | 149 | 1 | 1 | 1 | 1 | 1 | 0.983 | 1 | 1 | 1 | 1 | 1 | 1 | -0.003 |
| A5SS | 8185 | ENSG00000165916 | PSMC3 | chr11 | - | 47424045 | 47424183 | 47424108 | 47424183 | 47422829 | 47422973 | 186 | 102 | 156 | 181 | 99 | 1 | 1 | 0 | 0 | 2 | 334 | 286 | 180 | 262 | 362 | 0 | 1 | 0 | 1 | 6 | 212 | 149 | 1 | 1 | 0.992 | 0.986 | 1 | 1 | 0.972 | 1 | 0.995 | 1 | 0.995 | 0.977 | -0.003 |
| A5SS | 8225 | ENSG00000130830 | MPP1 | chrX | - | 154792141 | 154792285 | 154792192 | 154792285 | 154791768 | 154791847 | 114 | 111 | 78 | 205 | 101 | 0 | 1 | 0 | 1 | 0 | 409 | 249 | 157 | 215 | 1558 | 0 | 0 | 0 | 0 | 6 | 200 | 149 | 1 | 1 | 1 | 0.988 | 1 | 0.993 | 1 | 1 | 1 | 1 | 1 | 0.995 | -0.003 |
| A5SS | 8705 | ENSG00000132341 | RAN | chr12 | + | 130872583 | 130872629 | 130872583 | 130872617 | 130872835 | 130872920 | 338 | 217 | 323 | 361 | 253 | 0 | 0 | 0 | 0 | 3 | 464 | 357 | 260 | 387 | 489 | 0 | 0 | 0 | 0 | 0 | 161 | 149 | 1 | 1 | 1 | 1 | 1 | 1 | 0.987 | 1 | 1 | 1 | 1 | 1 | -0.003 |
| A5SS | 8722 | ENSG00000166012 | TAF1D | chr11 | - | 93731145 | 93731216 | 93731149 | 93731216 | 93730681 | 93730712 | 0 | 3 | 1 | 0 | 1 | 150 | 155 | 141 | 134 | 301 | 0 | 2 | 0 | 0 | 0 | 10 | 40 | 58 | 17 | 28 | 153 | 149 | 1 | 1 | 0 | 0.019 | 0.007 | 0 | 0.003 | 0 | 0.046 | 0 | 0 | 0 | -0.003 |
| A5SS | 237 | ENSG00000142541 | RPL13A | chr19 | + | 49489849 | 49490090 | 49489849 | 49489922 | 49490231 | 49490297 | 62 | 26 | 74 | 56 | 38 | 12021 | 4703 | 13109 | 9474 | 13052 | 78 | 35 | 41 | 45 | 127 | 5818 | 7716 | 5279 | 4104 | 9979 | 298 | 149 | 1 | 1 | 0.003 | 0.003 | 0.003 | 0.003 | 0.001 | 0.007 | 0.002 | 0.004 | 0.005 | 0.006 | -0.002 |
| A5SS | 468 | ENSG00000064601 | CTSA | chr20 | + | 45891568 | 45892027 | 45891568 | 45891762 | 45892272 | 45892323 | 229 | 99 | 112 | 176 | 140 | 0 | 0 | 0 | 1 | 0 | 350 | 496 | 132 | 337 | 2196 | 0 | 0 | 0 | 0 | 0 | 298 | 149 | 1 | 1 | 1 | 1 | 1 | 0.989 | 1 | 1 | 1 | 1 | 1 | 1 | -0.002 |
| A5SS | 779 | ENSG00000185624 | P4HB | chr17 | - | 81845817 | 81845991 | 81845870 | 81845991 | 81845560 | 81845742 | 1 | 3 | 1 | 4 | 0 | 1242 | 744 | 1021 | 1002 | 1415 | 20 | 1 | 3 | 20 | 7 | 2889 | 1624 | 1248 | 1918 | 3143 | 202 | 149 | 1 | 1 | 0.001 | 0.003 | 0.001 | 0.003 | 0 | 0.005 | 0 | 0.002 | 0.008 | 0.002 | -0.002 |
| A5SS | 849 | ENSG00000136490 | LIMD2 | chr17 | - | 63698798 | 63699069 | 63699027 | 63699069 | 63697815 | 63698711 | 230 | 72 | 223 | 160 | 48 | 0 | 0 | 1 | 0 | 0 | 100 | 172 | 110 | 90 | 241 | 0 | 0 | 0 | 0 | 0 | 298 | 149 | 1 | 1 | 1 | 1 | 0.991 | 1 | 1 | 1 | 1 | 1 | 1 | 1 | -0.002 |
| A5SS | 1020 | ENSG00000188895 | MSL1 | chr17 | + | 40133041 | 40133658 | 40133041 | 40133109 | 40133826 | 40133899 | 166 | 198 | 197 | 226 | 473 | 0 | 0 | 2 | 2 | 0 | 261 | 275 | 243 | 337 | 267 | 0 | 0 | 2 | 2 | 0 | 298 | 149 | 1 | 1 | 1 | 1 | 0.98 | 0.983 | 1 | 1 | 1 | 0.984 | 0.988 | 1 | -0.002 |
| A5SS | 1468 | ENSG00000166855 | CLPX | chr15 | - | 65158574 | 65158751 | 65158686 | 65158751 | 65157745 | 65157910 | 214 | 163 | 203 | 141 | 316 | 0 | 0 | 1 | 1 | 7 | 56 | 101 | 133 | 147 | 48 | 0 | 3 | 0 | 0 | 0 | 261 | 149 | 1 | 1 | 1 | 1 | 0.991 | 0.988 | 0.963 | 1 | 0.951 | 1 | 1 | 1 | -0.002 |
| A5SS | 1731 | ENSG00000100938 | GMPR2 | chr14 | + | 24233478 | 24233598 | 24233478 | 24233586 | 24235736 | 24235820 | 77 | 38 | 89 | 91 | 113 | 0 | 0 | 0 | 1 | 0 | 132 | 111 | 47 | 92 | 136 | 0 | 0 | 0 | 0 | 0 | 161 | 149 | 1 | 1 | 1 | 1 | 1 | 0.988 | 1 | 1 | 1 | 1 | 1 | 1 | -0.002 |
| A5SS | 1872 | ENSG00000138081 | FBXO11 | chr2 | - | 47810315 | 47813377 | 47813233 | 47813377 | 47809599 | 47809707 | 281 | 259 | 237 | 313 | 354 | 1 | 1 | 0 | 1 | 0 | 102 | 212 | 218 | 119 | 31 | 0 | 0 | 1 | 0 | 0 | 298 | 149 | 1 | 1 | 0.993 | 0.992 | 1 | 0.994 | 1 | 1 | 1 | 0.991 | 1 | 1 | -0.002 |
| A5SS | 1930 | ENSG00000126214 | KLC1 | chr14 | + | 103662701 | 103662927 | 103662701 | 103662921 | 103669510 | 103669598 | 112 | 84 | 114 | 145 | 88 | 0 | 0 | 1 | 0 | 1 | 263 | 117 | 128 | 171 | 157 | 0 | 0 | 0 | 2 | 0 | 155 | 149 | 1 | 1 | 1 | 1 | 0.991 | 1 | 0.988 | 1 | 1 | 1 | 0.988 | 1 | -0.002 |
| A5SS | 1972 | ENSG00000126777 | KTN1 | chr14 | + | 55671565 | 55671877 | 55671565 | 55671655 | 55672629 | 55672701 | 281 | 288 | 240 | 307 | 465 | 0 | 0 | 1 | 1 | 0 | 17 | 49 | 27 | 14 | 28 | 0 | 0 | 0 | 0 | 0 | 298 | 149 | 1 | 1 | 1 | 1 | 0.992 | 1 | 1 | 1 | 1 | 1 | 1 | 1 | -0.002 |
| A5SS | 1977 | ENSG00000139433 | GLTP | chr12 | - | 109857525 | 109857659 | 109857582 | 109857659 | 109855618 | 109855769 | 123 | 77 | 87 | 149 | 108 | 0 | 0 | 1 | 0 | 2 | 338 | 152 | 100 | 190 | 331 | 0 | 0 | 0 | 0 | 8 | 206 | 149 | 1 | 1 | 1 | 1 | 0.984 | 1 | 0.975 | 1 | 1 | 1 | 1 | 0.968 | -0.002 |
| A5SS | 2214 | ENSG00000170144 | HNRNPA3 | chr2 | + | 177215977 | 177216592 | 177215977 | 177216188 | 177216675 | 177216771 | 939 | 1025 | 818 | 938 | 944 | 2 | 1 | 0 | 0 | 2 | 878 | 842 | 1270 | 912 | 244 | 0 | 0 | 0 | 0 | 0 | 298 | 149 | 1 | 1 | 0.996 | 0.998 | 1 | 1 | 0.996 | 1 | 1 | 1 | 1 | 1 | -0.002 |
| A5SS | 2306 | ENSG00000127184 | COX7C | chr5 | + | 86619352 | 86619496 | 86619352 | 86619442 | 86620667 | 86620766 | 1557 | 725 | 1415 | 1229 | 1974 | 4 | 5 | 2 | 1 | 3 | 856 | 976 | 801 | 742 | 1061 | 0 | 4 | 0 | 0 | 0 | 203 | 149 | 1 | 1 | 0.997 | 0.991 | 0.998 | 0.999 | 0.998 | 1 | 0.994 | 1 | 1 | 1 | -0.002 |
| A5SS | 2546 | ENSG00000120708 | TGFBI | chr5 | + | 136046334 | 136046556 | 136046334 | 136046495 | 136046850 | 136047015 | 4 | 2 | 5 | 3 | 0 | 483 | 193 | 277 | 253 | 355 | 5 | 0 | 2 | 8 | 9 | 487 | 178 | 186 | 255 | 829 | 210 | 149 | 1 | 1 | 0.006 | 0.007 | 0.013 | 0.008 | 0 | 0.007 | 0 | 0.008 | 0.022 | 0.008 | -0.002 |
| A5SS | 2570 | ENSG00000203875 | SNHG5 | chr6 | - | 85677953 | 85678188 | 85678135 | 85678188 | 85677794 | 85677875 | 288 | 647 | 380 | 302 | 493 | 0 | 0 | 0 | 2 | 0 | 230 | 427 | 529 | 621 | 984 | 0 | 0 | 1 | 0 | 0 | 298 | 149 | 1 | 1 | 1 | 1 | 1 | 0.987 | 1 | 1 | 1 | 0.996 | 1 | 1 | -0.002 |
| A5SS | 2576 | ENSG00000203875 | SNHG5 | chr6 | - | 85678693 | 85678748 | 85678697 | 85678748 | 85678135 | 85678188 | 43 | 62 | 47 | 32 | 27 | 0 | 1 | 1 | 1 | 0 | 23 | 35 | 55 | 75 | 90 | 0 | 0 | 1 | 3 | 0 | 153 | 149 | 1 | 1 | 1 | 0.984 | 0.979 | 0.969 | 1 | 1 | 1 | 0.982 | 0.961 | 1 | -0.002 |
| A5SS | 2883 | ENSG00000004766 | VPS50 | chr7 | + | 93296741 | 93296959 | 93296741 | 93296836 | 93297144 | 93297243 | 2 | 1 | 0 | 0 | 2 | 32 | 45 | 39 | 50 | 77 | 0 | 0 | 1 | 0 | 0 | 0 | 9 | 9 | 5 | 10 | 272 | 149 | 1 | 1 | 0.033 | 0.012 | 0 | 0 | 0.014 | NA | 0 | 0.057 | 0 | 0 | -0.002 |
| A5SS | 2886 | ENSG00000221978 | CCNL2 | chr1 | - | 1392678 | 1392803 | 1392789 | 1392803 | 1390765 | 1390865 | 58 | 63 | 81 | 85 | 76 | 0 | 0 | 2 | 0 | 0 | 82 | 56 | 49 | 49 | 95 | 0 | 1 | 0 | 0 | 0 | 260 | 149 | 1 | 1 | 1 | 1 | 0.959 | 1 | 1 | 1 | 0.97 | 1 | 1 | 1 | -0.002 |
| A5SS | 2970 | ENSG00000114867 | EIF4G1 | chr3 | + | 184325850 | 184325957 | 184325850 | 184325951 | 184326526 | 184326629 | 0 | 1 | 0 | 0 | 1 | 278 | 179 | 241 | 287 | 181 | 6 | 0 | 0 | 2 | 0 | 533 | 311 | 233 | 275 | 618 | 155 | 149 | 1 | 1 | 0 | 0.005 | 0 | 0 | 0.005 | 0.011 | 0 | 0 | 0.007 | 0 | -0.002 |
| A5SS | 3308 | ENSG00000134352 | IL6ST | chr5 | - | 55969549 | 55969855 | 55969759 | 55969855 | 55968275 | 55968396 | 863 | 1019 | 1240 | 1053 | 1720 | 2 | 0 | 2 | 1 | 2 | 275 | 604 | 640 | 312 | 166 | 0 | 0 | 0 | 0 | 0 | 298 | 149 | 1 | 1 | 0.995 | 1 | 0.997 | 0.998 | 0.998 | 1 | 1 | 1 | 1 | 1 | -0.002 |
| A5SS | 3323 | ENSG00000145388 | METTL14 | chr4 | + | 118689369 | 118689457 | 118689369 | 118689404 | 118691531 | 118691612 | 289 | 237 | 211 | 277 | 314 | 0 | 0 | 0 | 0 | 2 | 51 | 177 | 179 | 94 | 24 | 0 | 0 | 0 | 0 | 0 | 202 | 149 | 1 | 1 | 1 | 1 | 1 | 1 | 0.991 | 1 | 1 | 1 | 1 | 1 | -0.002 |
| A5SS | 3326 | ENSG00000158864 | NDUFS2 | chr1 | + | 161209482 | 161209931 | 161209482 | 161209595 | 161210110 | 161210188 | 234 | 166 | 241 | 229 | 275 | 0 | 0 | 0 | 1 | 0 | 618 | 318 | 262 | 437 | 637 | 0 | 0 | 0 | 0 | 0 | 298 | 149 | 1 | 1 | 1 | 1 | 1 | 0.991 | 1 | 1 | 1 | 1 | 1 | 1 | -0.002 |
| A5SS | 3366 | ENSG00000164253 | WDR41 | chr5 | - | 77459049 | 77459124 | 77459061 | 77459124 | 77453816 | 77453928 | 0 | 1 | 1 | 0 | 0 | 144 | 46 | 60 | 92 | 157 | 0 | 3 | 0 | 0 | 0 | 98 | 59 | 43 | 41 | 28 | 161 | 149 | 1 | 1 | 0 | 0.02 | 0.015 | 0 | 0 | 0 | 0.045 | 0 | 0 | 0 | -0.002 |
| A5SS | 3372 | ENSG00000125347 | IRF1 | chr5 | - | 132484361 | 132484497 | 132484394 | 132484497 | 132483827 | 132484075 | 507 | 317 | 520 | 570 | 495 | 3 | 1 | 5 | 7 | 2 | 1879 | 412 | 736 | 1221 | 1153 | 8 | 4 | 0 | 14 | 4 | 182 | 149 | 1 | 1 | 0.993 | 0.996 | 0.988 | 0.985 | 0.995 | 0.995 | 0.988 | 1 | 0.986 | 0.996 | -0.002 |
| A5SS | 3414 | ENSG00000163719 | MTMR14 | chr3 | + | 9662266 | 9662375 | 9662266 | 9662285 | 9669431 | 9669492 | 118 | 73 | 132 | 141 | 147 | 0 | 0 | 0 | 1 | 0 | 461 | 189 | 179 | 269 | 570 | 0 | 0 | 0 | 0 | 0 | 239 | 149 | 1 | 1 | 1 | 1 | 1 | 0.989 | 1 | 1 | 1 | 1 | 1 | 1 | -0.002 |
| A5SS | 3603 | ENSG00000114861 | FOXP1 | chr3 | - | 70977642 | 70977719 | 70977646 | 70977719 | 70976940 | 70977042 | 411 | 352 | 546 | 520 | 391 | 4 | 0 | 1 | 2 | 0 | 263 | 371 | 505 | 357 | 183 | 0 | 2 | 0 | 0 | 0 | 153 | 149 | 1 | 1 | 0.99 | 1 | 0.998 | 0.996 | 1 | 1 | 0.994 | 1 | 1 | 1 | -0.002 |
| A5SS | 3684 | ENSG00000179335 | CLK3 | chr15 | + | 74625801 | 74625968 | 74625801 | 74625967 | 74627351 | 74627446 | 258 | 140 | 245 | 285 | 330 | 2 | 0 | 0 | 0 | 0 | 499 | 410 | 358 | 305 | 556 | 0 | 0 | 0 | 0 | 0 | 150 | 149 | 1 | 1 | 0.992 | 1 | 1 | 1 | 1 | 1 | 1 | 1 | 1 | 1 | -0.002 |
| A5SS | 3697 | ENSG00000182923 | CEP63 | chr3 | + | 134532777 | 134533030 | 134532777 | 134532900 | 134537154 | 134537268 | 7 | 6 | 4 | 5 | 4 | 66 | 62 | 62 | 77 | 83 | 0 | 10 | 6 | 1 | 0 | 29 | 50 | 63 | 31 | 0 | 279 | 149 | 1 | 1 | 0.054 | 0.049 | 0.033 | 0.034 | 0.025 | 0 | 0.097 | 0.048 | 0.017 | NA | -0.002 |
| A5SS | 3705 | ENSG00000185825 | BCAP31 | chrX | - | 153702934 | 153703058 | 153703039 | 153703058 | 153702006 | 153702107 | 696 | 395 | 553 | 704 | 994 | 0 | 1 | 0 | 1 | 4 | 933 | 757 | 423 | 606 | 2396 | 0 | 0 | 0 | 0 | 4 | 254 | 149 | 1 | 1 | 1 | 0.996 | 1 | 0.998 | 0.993 | 1 | 1 | 1 | 1 | 0.997 | -0.002 |
| A5SS | 3720 | ENSG00000163346 | PBXIP1 | chr1 | - | 154948132 | 154948366 | 154948225 | 154948366 | 154947981 | 154948005 | 159 | 75 | 156 | 140 | 52 | 0 | 0 | 1 | 0 | 0 | 28 | 164 | 71 | 50 | 111 | 0 | 0 | 0 | 0 | 0 | 242 | 149 | 1 | 1 | 1 | 1 | 0.99 | 1 | 1 | 1 | 1 | 1 | 1 | 1 | -0.002 |
| A5SS | 3899 | ENSG00000162894 | FCMR | chr1 | - | 206913758 | 206914094 | 206913861 | 206914094 | 206911729 | 206911952 | 601 | 335 | 701 | 545 | 923 | 1 | 0 | 1 | 1 | 0 | 248 | 546 | 205 | 164 | 396 | 0 | 0 | 0 | 0 | 0 | 252 | 149 | 1 | 1 | 0.997 | 1 | 0.998 | 0.997 | 1 | 1 | 1 | 1 | 1 | 1 | -0.002 |
| A5SS | 3901 | ENSG00000055609 | KMT2C | chr7 | - | 152144712 | 152145295 | 152145152 | 152145295 | 152139674 | 152139791 | 250 | 203 | 215 | 303 | 336 | 0 | 1 | 0 | 0 | 0 | 96 | 281 | 227 | 118 | 40 | 0 | 0 | 0 | 0 | 0 | 298 | 149 | 1 | 1 | 1 | 0.99 | 1 | 1 | 1 | 1 | 1 | 1 | 1 | 1 | -0.002 |
| A5SS | 3923 | ENSG00000095787 | WAC | chr10 | + | 28614566 | 28616362 | 28614566 | 28614685 | 28617656 | 28617784 | 590 | 425 | 520 | 621 | 817 | 2 | 5 | 0 | 2 | 0 | 246 | 498 | 679 | 384 | 79 | 1 | 2 | 3 | 0 | 0 | 298 | 149 | 1 | 1 | 0.993 | 0.977 | 1 | 0.994 | 1 | 0.992 | 0.992 | 0.991 | 1 | 1 | -0.002 |
| A5SS | 4028 | ENSG00000138095 | LRPPRC | chr2 | - | 43976142 | 43977052 | 43976993 | 43977052 | 43975090 | 43975217 | 253 | 175 | 244 | 210 | 338 | 0 | 1 | 0 | 0 | 2 | 49 | 136 | 130 | 81 | 57 | 0 | 0 | 1 | 0 | 0 | 298 | 149 | 1 | 1 | 1 | 0.989 | 1 | 1 | 0.988 | 1 | 1 | 0.985 | 1 | 1 | -0.002 |
| A5SS | 4339 | ENSG00000153283 | CD96 | chr3 | + | 111545045 | 111545402 | 111545045 | 111545328 | 111567522 | 111567647 | 941 | 732 | 1118 | 1045 | 1499 | 1 | 2 | 2 | 0 | 1 | 102 | 545 | 704 | 238 | 145 | 0 | 0 | 0 | 0 | 0 | 223 | 149 | 1 | 1 | 0.998 | 0.996 | 0.997 | 1 | 0.999 | 1 | 1 | 1 | 1 | 1 | -0.002 |
| A5SS | 4379 | ENSG00000172531 | PPP1CA | chr11 | - | 67398939 | 67399665 | 67399560 | 67399665 | 67398721 | 67398856 | 315 | 111 | 214 | 251 | 245 | 0 | 0 | 1 | 0 | 0 | 295 | 360 | 153 | 263 | 530 | 0 | 0 | 0 | 0 | 0 | 298 | 149 | 1 | 1 | 1 | 1 | 0.991 | 1 | 1 | 1 | 1 | 1 | 1 | 1 | -0.002 |
| A5SS | 4517 | ENSG00000168610 | STAT3 | chr17 | - | 42323259 | 42323354 | 42323286 | 42323354 | 42323003 | 42323143 | 523 | 356 | 411 | 535 | 657 | 2 | 0 | 6 | 1 | 1 | 1401 | 777 | 648 | 926 | 1157 | 1 | 4 | 0 | 0 | 8 | 176 | 149 | 1 | 1 | 0.996 | 1 | 0.983 | 0.998 | 0.998 | 0.999 | 0.994 | 1 | 1 | 0.992 | -0.002 |
| A5SS | 4532 | ENSG00000185885 | IFITM1 | chr11 | + | 314039 | 314392 | 314039 | 314356 | 314921 | 315272 | 6 | 3 | 1 | 5 | 0 | 1289 | 463 | 877 | 871 | 636 | 14 | 8 | 6 | 0 | 0 | 852 | 1173 | 897 | 1229 | 755 | 185 | 149 | 1 | 1 | 0.004 | 0.005 | 0.001 | 0.005 | 0 | 0.013 | 0.005 | 0.005 | 0 | 0 | -0.002 |
| A5SS | 4578 | ENSG00000177200 | CHD9 | chr16 | + | 53209481 | 53209813 | 53209481 | 53209541 | 53222643 | 53222755 | 193 | 248 | 168 | 249 | 219 | 0 | 0 | 0 | 1 | 0 | 24 | 141 | 81 | 41 | 41 | 0 | 0 | 0 | 0 | 0 | 298 | 149 | 1 | 1 | 1 | 1 | 1 | 0.992 | 1 | 1 | 1 | 1 | 1 | 1 | -0.002 |
| A5SS | 4603 | ENSG00000084733 | RAB10 | chr2 | + | 26127143 | 26127278 | 26127143 | 26127233 | 26127849 | 26127951 | 0 | 0 | 0 | 4 | 3 | 1158 | 1254 | 706 | 1030 | 1800 | 0 | 0 | 8 | 0 | 0 | 1760 | 946 | 527 | 1393 | 1299 | 194 | 149 | 1 | 1 | 0 | 0 | 0 | 0.003 | 0.001 | 0 | 0 | 0.012 | 0 | 0 | -0.002 |
| A5SS | 4627 | ENSG00000108523 | RNF167 | chr17 | + | 4940486 | 4940993 | 4940486 | 4940726 | 4941076 | 4941157 | 225 | 66 | 155 | 175 | 186 | 1 | 0 | 0 | 0 | 0 | 163 | 196 | 114 | 163 | 318 | 0 | 0 | 0 | 0 | 0 | 298 | 149 | 1 | 1 | 0.991 | 1 | 1 | 1 | 1 | 1 | 1 | 1 | 1 | 1 | -0.002 |
| A5SS | 5667 | ENSG00000095059 | DHPS | chr19 | - | 12679456 | 12679719 | 12679622 | 12679719 | 12677290 | 12677396 | 258 | 171 | 230 | 228 | 288 | 1 | 0 | 1 | 0 | 0 | 338 | 271 | 146 | 264 | 418 | 0 | 0 | 0 | 0 | 0 | 298 | 149 | 1 | 1 | 1 | 1 | 0.991 | 1 | 1 | 1 | 1 | 1 | 1 | 1 | -0.002 |
| A5SS | 5782 | ENSG00000064666 | CNN2 | chr19 | + | 1032558 | 1032696 | 1032558 | 1032669 | 1036129 | 1036246 | 847 | 502 | 1050 | 843 | 729 | 1 | 0 | 0 | 7 | 2 | 809 | 1074 | 751 | 966 | 1861 | 0 | 1 | 2 | 0 | 0 | 176 | 149 | 1 | 1 | 0.999 | 1 | 1 | 0.99 | 0.997 | 1 | 0.999 | 0.997 | 1 | 1 | -0.002 |
| A5SS | 5808 | ENSG00000130414 | NDUFA10 | chr2 | - | 240012331 | 240014860 | 240014840 | 240014860 | 240011616 | 240011696 | 375 | 253 | 319 | 260 | 787 | 0 | 1 | 0 | 0 | 0 | 487 | 389 | 283 | 269 | 358 | 0 | 0 | 0 | 0 | 0 | 298 | 149 | 1 | 1 | 1 | 0.992 | 1 | 1 | 1 | 1 | 1 | 1 | 1 | 1 | -0.002 |
| A5SS | 5811 | ENSG00000130414 | NDUFA10 | chr2 | - | 240014694 | 240014860 | 240014840 | 240014860 | 240011616 | 240011696 | 375 | 253 | 319 | 261 | 787 | 0 | 1 | 0 | 0 | 0 | 487 | 389 | 283 | 270 | 358 | 0 | 0 | 0 | 0 | 0 | 295 | 149 | 1 | 1 | 1 | 0.992 | 1 | 1 | 1 | 1 | 1 | 1 | 1 | 1 | -0.002 |
| A5SS | 6169 | ENSG00000135486 | HNRNPA1 | chr12 | + | 54282799 | 54282841 | 54282799 | 54282820 | 54283826 | 54283967 | 169 | 124 | 168 | 169 | 169 | 1 | 0 | 1 | 0 | 0 | 138 | 171 | 244 | 188 | 58 | 0 | 0 | 1 | 0 | 0 | 170 | 149 | 1 | 1 | 0.993 | 1 | 0.993 | 1 | 1 | 1 | 1 | 0.995 | 1 | 1 | -0.002 |
| A5SS | 6276 | ENSG00000089820 | ARHGAP4 | chrX | - | 153913217 | 153913600 | 153913408 | 153913600 | 153913023 | 153913051 | 248 | 109 | 241 | 238 | 137 | 1 | 0 | 0 | 0 | 0 | 193 | 199 | 166 | 224 | 406 | 0 | 0 | 0 | 0 | 0 | 298 | 149 | 1 | 1 | 0.992 | 1 | 1 | 1 | 1 | 1 | 1 | 1 | 1 | 1 | -0.002 |
| A5SS | 6756 | ENSG00000234741 | GAS5 | chr1 | - | 173865228 | 173865547 | 173865470 | 173865547 | 173864674 | 173864704 | 850 | 484 | 1023 | 780 | 1345 | 3 | 3 | 4 | 5 | 11 | 315 | 345 | 496 | 296 | 321 | 0 | 0 | 10 | 1 | 0 | 298 | 149 | 1 | 1 | 0.993 | 0.988 | 0.992 | 0.987 | 0.984 | 1 | 1 | 0.961 | 0.993 | 1 | -0.002 |
| A5SS | 6757 | ENSG00000234741 | GAS5 | chr1 | - | 173865228 | 173865547 | 173865509 | 173865547 | 173864674 | 173864704 | 877 | 513 | 1067 | 803 | 1418 | 2 | 1 | 1 | 0 | 0 | 312 | 370 | 520 | 318 | 335 | 0 | 0 | 0 | 0 | 0 | 298 | 149 | 1 | 1 | 0.995 | 0.996 | 0.998 | 1 | 1 | 1 | 1 | 1 | 1 | 1 | -0.002 |
| A5SS | 6780 | ENSG00000122545 | SEPTIN7 | chr7 | + | 35832797 | 35832900 | 35832797 | 35832893 | 35863551 | 35863658 | 715 | 532 | 475 | 696 | 857 | 0 | 0 | 2 | 0 | 3 | 55 | 287 | 121 | 38 | 27 | 0 | 0 | 0 | 0 | 0 | 156 | 149 | 1 | 1 | 1 | 1 | 0.996 | 1 | 0.996 | 1 | 1 | 1 | 1 | 1 | -0.002 |
| A5SS | 7007 | ENSG00000244754 | N4BP2L2 | chr13 | - | 32535768 | 32537027 | 32536932 | 32537027 | 32522181 | 32522270 | 797 | 794 | 722 | 788 | 1220 | 0 | 1 | 2 | 0 | 0 | 181 | 419 | 494 | 197 | 91 | 0 | 0 | 0 | 0 | 0 | 298 | 149 | 1 | 1 | 1 | 0.997 | 0.994 | 1 | 1 | 1 | 1 | 1 | 1 | 1 | -0.002 |
| A5SS | 7008 | ENSG00000244754 | N4BP2L2 | chr13 | - | 32535768 | 32537027 | 32537005 | 32537027 | 32522181 | 32522270 | 499 | 502 | 432 | 482 | 613 | 0 | 0 | 0 | 0 | 3 | 112 | 324 | 329 | 132 | 46 | 0 | 0 | 0 | 0 | 0 | 298 | 149 | 1 | 1 | 1 | 1 | 1 | 1 | 0.99 | 1 | 1 | 1 | 1 | 1 | -0.002 |
| A5SS | 7064 | ENSG00000135048 | CEMIP2 | chr9 | - | 71730704 | 71730914 | 71730738 | 71730914 | 71730047 | 71730253 | 305 | 124 | 179 | 233 | 200 | 0 | 1 | 0 | 4 | 0 | 94 | 360 | 518 | 208 | 39 | 0 | 1 | 7 | 0 | 0 | 183 | 149 | 1 | 1 | 1 | 0.99 | 1 | 0.979 | 1 | 1 | 0.997 | 0.984 | 1 | 1 | -0.002 |
| A5SS | 7087 | ENSG00000196924 | FLNA | chrX | - | 154365744 | 154366224 | 154366023 | 154366224 | 154365348 | 154365486 | 11 | 5 | 9 | 12 | 4 | 1882 | 700 | 982 | 1543 | 446 | 36 | 38 | 12 | 27 | 16 | 2582 | 1900 | 1144 | 1414 | 9433 | 298 | 149 | 1 | 1 | 0.003 | 0.004 | 0.005 | 0.004 | 0.004 | 0.007 | 0.01 | 0.005 | 0.009 | 0.001 | -0.002 |
| A5SS | 7092 | ENSG00000147140 | NONO | chrX | + | 71294226 | 71294528 | 71294226 | 71294348 | 71296564 | 71296660 | 903 | 651 | 901 | 891 | 1187 | 0 | 2 | 0 | 0 | 1 | 1404 | 922 | 721 | 832 | 1276 | 0 | 0 | 0 | 0 | 0 | 298 | 149 | 1 | 1 | 1 | 0.994 | 1 | 1 | 0.998 | 1 | 1 | 1 | 1 | 1 | -0.002 |
| A5SS | 7190 | ENSG00000147050 | KDM6A | chrX | + | 45082575 | 45082789 | 45082575 | 45082640 | 45083459 | 45083608 | 248 | 274 | 183 | 224 | 450 | 0 | 0 | 1 | 0 | 0 | 161 | 140 | 256 | 306 | 56 | 0 | 0 | 0 | 0 | 0 | 298 | 149 | 1 | 1 | 1 | 1 | 0.989 | 1 | 1 | 1 | 1 | 1 | 1 | 1 | -0.002 |
| A5SS | 7434 | ENSG00000160948 | VPS28 | chr8 | - | 144424717 | 144425772 | 144424945 | 144425772 | 144424214 | 144424268 | 239 | 115 | 177 | 177 | 161 | 1 | 0 | 0 | 0 | 0 | 111 | 259 | 146 | 164 | 309 | 0 | 0 | 0 | 0 | 0 | 298 | 149 | 1 | 1 | 0.992 | 1 | 1 | 1 | 1 | 1 | 1 | 1 | 1 | 1 | -0.002 |
| A5SS | 7435 | ENSG00000160948 | VPS28 | chr8 | - | 144424945 | 144425772 | 144425682 | 144425772 | 144424717 | 144424819 | 243 | 113 | 170 | 202 | 192 | 1 | 0 | 0 | 0 | 0 | 202 | 260 | 144 | 231 | 323 | 0 | 0 | 0 | 0 | 0 | 298 | 149 | 1 | 1 | 0.992 | 1 | 1 | 1 | 1 | 1 | 1 | 1 | 1 | 1 | -0.002 |
| A5SS | 7459 | ENSG00000104408 | EIF3E | chr8 | - | 108202982 | 108203117 | 108202989 | 108203117 | 108201742 | 108201923 | 319 | 271 | 440 | 343 | 868 | 3 | 0 | 0 | 0 | 2 | 25 | 128 | 91 | 41 | 16 | 0 | 0 | 0 | 0 | 0 | 156 | 149 | 1 | 1 | 0.99 | 1 | 1 | 1 | 0.998 | 1 | 1 | 1 | 1 | 1 | -0.002 |
| A5SS | 7495 | ENSG00000133794 | ARNTL | chr11 | + | 13378336 | 13381254 | 13378336 | 13378445 | 13385688 | 13385791 | 144 | 60 | 126 | 140 | 184 | 0 | 0 | 2 | 0 | 0 | 76 | 195 | 228 | 142 | 38 | 0 | 0 | 1 | 1 | 0 | 298 | 149 | 1 | 1 | 1 | 1 | 0.969 | 1 | 1 | 1 | 1 | 0.991 | 0.986 | 1 | -0.002 |
| A5SS | 7579 | ENSG00000113328 | CCNG1 | chr5 | + | 163439256 | 163439520 | 163439256 | 163439346 | 163441077 | 163441331 | 238 | 124 | 238 | 213 | 554 | 0 | 0 | 0 | 1 | 0 | 6 | 104 | 132 | 44 | 67 | 0 | 0 | 0 | 0 | 0 | 298 | 149 | 1 | 1 | 1 | 1 | 1 | 0.991 | 1 | 1 | 1 | 1 | 1 | 1 | -0.002 |
| A5SS | 7605 | ENSG00000119487 | MAPKAP1 | chr9 | - | 125672315 | 125672643 | 125672399 | 125672643 | 125669817 | 125669907 | 502 | 343 | 325 | 380 | 724 | 1 | 0 | 4 | 0 | 4 | 489 | 397 | 298 | 361 | 559 | 0 | 1 | 3 | 0 | 0 | 233 | 149 | 1 | 1 | 0.997 | 1 | 0.981 | 1 | 0.991 | 1 | 0.996 | 0.985 | 1 | 1 | -0.002 |
| A5SS | 7632 | ENSG00000100296 | THOC5 | chr22 | - | 29528936 | 29529239 | 29529161 | 29529239 | 29528425 | 29528466 | 2 | 0 | 0 | 0 | 1 | 91 | 35 | 79 | 57 | 58 | 0 | 0 | 0 | 4 | 0 | 127 | 56 | 37 | 60 | 134 | 298 | 149 | 1 | 1 | 0.011 | 0 | 0 | 0 | 0.009 | 0 | 0 | 0 | 0.032 | 0 | -0.002 |
| A5SS | 8068 | ENSG00000075239 | ACAT1 | chr11 | + | 108134220 | 108134316 | 108134220 | 108134296 | 108135141 | 108135242 | 96 | 70 | 97 | 94 | 72 | 1 | 0 | 0 | 0 | 0 | 52 | 118 | 104 | 123 | 19 | 0 | 0 | 0 | 0 | 0 | 169 | 149 | 1 | 1 | 0.988 | 1 | 1 | 1 | 1 | 1 | 1 | 1 | 1 | 1 | -0.002 |
| A5SS | 8558 | ENSG00000130119 | GNL3L | chrX | + | 54540134 | 54540242 | 54540134 | 54540238 | 54541272 | 54541389 | 119 | 105 | 112 | 108 | 92 | 1 | 0 | 0 | 0 | 0 | 350 | 123 | 98 | 211 | 260 | 0 | 0 | 0 | 0 | 0 | 153 | 149 | 1 | 1 | 0.991 | 1 | 1 | 1 | 1 | 1 | 1 | 1 | 1 | 1 | -0.002 |
| A5SS | 8626 | ENSG00000110719 | TCIRG1 | chr11 | + | 68047646 | 68047972 | 68047646 | 68047804 | 68048878 | 68048997 | 231 | 105 | 184 | 196 | 185 | 0 | 0 | 0 | 1 | 0 | 345 | 368 | 214 | 317 | 353 | 0 | 0 | 0 | 0 | 0 | 298 | 149 | 1 | 1 | 1 | 1 | 1 | 0.99 | 1 | 1 | 1 | 1 | 1 | 1 | -0.002 |
| A5SS | 8857 | ENSG00000111335 | OAS2 | chr12 | + | 112997519 | 112998410 | 112997519 | 112997755 | 113002931 | 113003102 | 498 | 503 | 407 | 613 | 747 | 0 | 0 | 2 | 0 | 0 | 2213 | 834 | 262 | 1444 | 335 | 0 | 0 | 0 | 0 | 0 | 298 | 149 | 1 | 1 | 1 | 1 | 0.99 | 1 | 1 | 1 | 1 | 1 | 1 | 1 | -0.002 |
| A5SS | 8885 | ENSG00000111300 | NAA25 | chr12 | - | 112039228 | 112040578 | 112040480 | 112040578 | 112033232 | 112033379 | 87 | 65 | 75 | 93 | 144 | 0 | 1 | 0 | 0 | 0 | 41 | 71 | 95 | 49 | 30 | 0 | 0 | 1 | 0 | 0 | 298 | 149 | 1 | 1 | 1 | 0.97 | 1 | 1 | 1 | 1 | 1 | 0.979 | 1 | 1 | -0.002 |
| A5SS | 110 | ENSG00000100242 | SUN2 | chr22 | - | 38740266 | 38741050 | 38740888 | 38741050 | 38739721 | 38739943 | 1027 | 400 | 675 | 768 | 363 | 0 | 1 | 0 | 0 | 0 | 690 | 845 | 546 | 552 | 1025 | 0 | 0 | 0 | 0 | 0 | 298 | 149 | 1 | 1 | 1 | 0.995 | 1 | 1 | 1 | 1 | 1 | 1 | 1 | 1 | -0.001 |
| A5SS | 136 | ENSG00000093072 | ADA2 | chr22 | - | 17189941 | 17191810 | 17191682 | 17191810 | 17188338 | 17188447 | 713 | 395 | 521 | 621 | 1019 | 1 | 1 | 0 | 4 | 2 | 1809 | 909 | 370 | 883 | 1498 | 0 | 0 | 2 | 5 | 0 | 298 | 149 | 1 | 1 | 0.997 | 0.995 | 1 | 0.987 | 0.996 | 1 | 1 | 0.989 | 0.989 | 1 | -0.001 |
| A5SS | 242 | ENSG00000142541 | RPL13A | chr19 | + | 49490474 | 49490600 | 49490474 | 49490576 | 49490778 | 49490864 | 68 | 34 | 74 | 46 | 26 | 12159 | 4971 | 13467 | 9814 | 15587 | 30 | 55 | 28 | 24 | 62 | 6724 | 7723 | 5307 | 4187 | 11129 | 173 | 149 | 1 | 1 | 0.005 | 0.006 | 0.005 | 0.004 | 0.001 | 0.004 | 0.006 | 0.005 | 0.005 | 0.005 | -0.001 |
| A5SS | 331 | ENSG00000142227 | EMP3 | chr19 | + | 48327520 | 48327664 | 48327520 | 48327561 | 48329351 | 48329492 | 284 | 113 | 155 | 206 | 104 | 0 | 0 | 0 | 0 | 1 | 497 | 277 | 159 | 328 | 587 | 0 | 0 | 0 | 2 | 0 | 252 | 149 | 1 | 1 | 1 | 1 | 1 | 1 | 0.984 | 1 | 1 | 1 | 0.99 | 1 | -0.001 |
| A5SS | 374 | ENSG00000105771 | SMG9 | chr19 | - | 43739672 | 43740218 | 43740106 | 43740218 | 43738121 | 43738217 | 0 | 0 | 2 | 2 | 1 | 25 | 24 | 30 | 48 | 27 | 0 | 0 | 2 | 0 | 0 | 47 | 28 | 12 | 25 | 52 | 298 | 149 | 1 | 1 | 0 | 0 | 0.032 | 0.02 | 0.018 | 0 | 0 | 0.077 | 0 | 0 | -0.001 |
| A5SS | 463 | ENSG00000101347 | SAMHD1 | chr20 | - | 36892578 | 36893066 | 36892762 | 36893066 | 36890903 | 36891807 | 373 | 275 | 286 | 293 | 575 | 0 | 0 | 1 | 0 | 0 | 290 | 208 | 102 | 178 | 144 | 0 | 0 | 0 | 0 | 0 | 298 | 149 | 1 | 1 | 1 | 1 | 0.993 | 1 | 1 | 1 | 1 | 1 | 1 | 1 | -0.001 |
| A5SS | 599 | ENSG00000166347 | CYB5A | chr18 | - | 74255740 | 74260944 | 74260914 | 74260944 | 74253296 | 74253665 | 44 | 26 | 64 | 73 | 45 | 0 | 1 | 0 | 0 | 0 | 25 | 56 | 14 | 13 | 41 | 0 | 2 | 0 | 0 | 0 | 298 | 149 | 1 | 1 | 1 | 0.929 | 1 | 1 | 1 | 1 | 0.933 | 1 | 1 | 1 | -0.001 |
| A5SS | 698 | ENSG00000169718 | DUS1L | chr17 | - | 82061556 | 82061721 | 82061617 | 82061721 | 82061208 | 82061353 | 1 | 2 | 2 | 2 | 0 | 66 | 15 | 57 | 53 | 34 | 0 | 4 | 2 | 3 | 0 | 36 | 77 | 19 | 44 | 93 | 210 | 149 | 1 | 1 | 0.011 | 0.086 | 0.024 | 0.026 | 0 | 0 | 0.036 | 0.069 | 0.046 | 0 | -0.001 |
| A5SS | 878 | ENSG00000007168 | PAFAH1B1 | chr17 | + | 2674059 | 2674288 | 2674059 | 2674096 | 2676504 | 2676606 | 562 | 339 | 477 | 486 | 637 | 0 | 0 | 0 | 1 | 0 | 354 | 576 | 546 | 583 | 298 | 0 | 0 | 0 | 0 | 0 | 298 | 149 | 1 | 1 | 1 | 1 | 1 | 0.996 | 1 | 1 | 1 | 1 | 1 | 1 | -0.001 |
| A5SS | 988 | ENSG00000168259 | DNAJC7 | chr17 | - | 41982245 | 41982401 | 41982254 | 41982401 | 41981854 | 41982007 | 0 | 0 | 0 | 0 | 5 | 331 | 276 | 286 | 345 | 356 | 2 | 2 | 5 | 0 | 0 | 870 | 353 | 523 | 665 | 392 | 158 | 149 | 1 | 1 | 0 | 0 | 0 | 0 | 0.013 | 0.002 | 0.005 | 0.009 | 0 | 0 | -0.001 |
| A5SS | 991 | ENSG00000168259 | DNAJC7 | chr17 | - | 41997114 | 41997239 | 41997136 | 41997239 | 41996310 | 41996424 | 155 | 72 | 99 | 97 | 176 | 0 | 0 | 2 | 0 | 0 | 165 | 101 | 136 | 181 | 120 | 3 | 0 | 0 | 0 | 0 | 171 | 149 | 1 | 1 | 1 | 1 | 0.977 | 1 | 1 | 0.98 | 1 | 1 | 1 | 1 | -0.001 |
| A5SS | 1159 | ENSG00000005844 | ITGAL | chr16 | + | 30483826 | 30483964 | 30483826 | 30483959 | 30484112 | 30484263 | 9 | 3 | 0 | 4 | 2 | 1129 | 513 | 595 | 790 | 650 | 4 | 6 | 6 | 2 | 9 | 1138 | 1152 | 568 | 802 | 1501 | 154 | 149 | 1 | 1 | 0.008 | 0.006 | 0 | 0.005 | 0.003 | 0.003 | 0.005 | 0.01 | 0.002 | 0.006 | -0.001 |
| A5SS | 1161 | ENSG00000005844 | ITGAL | chr16 | + | 30496096 | 30496294 | 30496096 | 30496201 | 30496435 | 30496566 | 1445 | 654 | 683 | 968 | 814 | 0 | 0 | 2 | 0 | 0 | 1236 | 1434 | 629 | 916 | 1795 | 0 | 0 | 0 | 0 | 0 | 242 | 149 | 1 | 1 | 1 | 1 | 0.995 | 1 | 1 | 1 | 1 | 1 | 1 | 1 | -0.001 |
| A5SS | 1162 | ENSG00000005844 | ITGAL | chr16 | + | 30496096 | 30496566 | 30496096 | 30496294 | 30499073 | 30499234 | 795 | 383 | 520 | 578 | 721 | 0 | 0 | 0 | 0 | 2 | 818 | 852 | 407 | 557 | 1106 | 0 | 0 | 0 | 0 | 0 | 298 | 149 | 1 | 1 | 1 | 1 | 1 | 1 | 0.994 | 1 | 1 | 1 | 1 | 1 | -0.001 |
| A5SS | 1191 | ENSG00000141002 | TCF25 | chr16 | + | 89887651 | 89887717 | 89887651 | 89887681 | 89892192 | 89892275 | 390 | 240 | 305 | 322 | 433 | 1 | 0 | 0 | 0 | 1 | 319 | 417 | 226 | 286 | 462 | 0 | 0 | 0 | 0 | 0 | 185 | 149 | 1 | 1 | 0.997 | 1 | 1 | 1 | 0.997 | 1 | 1 | 1 | 1 | 1 | -0.001 |
| A5SS | 1255 | ENSG00000157106 | SMG1 | chr16 | - | 18868522 | 18868719 | 18868632 | 18868719 | 18868189 | 18868354 | 241 | 170 | 258 | 241 | 333 | 1 | 0 | 0 | 1 | 0 | 96 | 256 | 211 | 44 | 0 | 0 | 0 | 1 | 0 | 0 | 259 | 149 | 1 | 1 | 0.993 | 1 | 1 | 0.993 | 1 | 1 | 1 | 0.992 | 1 | NA | -0.001 |
| A5SS | 1540 | ENSG00000137845 | ADAM10 | chr15 | - | 58633195 | 58633359 | 58633314 | 58633359 | 58627711 | 58627883 | 1917 | 1901 | 1290 | 1823 | 2033 | 0 | 1 | 2 | 0 | 1 | 503 | 1226 | 1339 | 484 | 283 | 0 | 0 | 0 | 0 | 0 | 268 | 149 | 1 | 1 | 1 | 0.999 | 0.997 | 1 | 0.999 | 1 | 1 | 1 | 1 | 1 | -0.001 |
| A5SS | 1665 | ENSG00000100949 | RABGGTA | chr14 | - | 24267865 | 24267958 | 24267869 | 24267958 | 24267659 | 24267776 | 1 | 0 | 0 | 0 | 0 | 59 | 26 | 44 | 48 | 60 | 0 | 2 | 0 | 0 | 0 | 57 | 82 | 26 | 46 | 60 | 153 | 149 | 1 | 1 | 0.016 | 0 | 0 | 0 | 0 | 0 | 0.023 | 0 | 0 | 0 | -0.001 |
| A5SS | 1910 | ENSG00000130779 | CLIP1 | chr12 | - | 122340752 | 122341697 | 122340980 | 122341697 | 122336631 | 122336748 | 277 | 290 | 184 | 305 | 267 | 0 | 0 | 0 | 1 | 2 | 39 | 237 | 351 | 132 | 25 | 0 | 1 | 2 | 0 | 0 | 298 | 149 | 1 | 1 | 1 | 1 | 1 | 0.993 | 0.985 | 1 | 0.992 | 0.989 | 1 | 1 | -0.001 |
| A5SS | 2164 | ENSG00000104805 | NUCB1 | chr19 | + | 48913473 | 48913564 | 48913473 | 48913532 | 48918725 | 48918784 | 820 | 579 | 593 | 762 | 764 | 1 | 2 | 0 | 0 | 0 | 2159 | 1539 | 931 | 1366 | 3714 | 0 | 1 | 0 | 0 | 0 | 181 | 149 | 1 | 1 | 0.999 | 0.996 | 1 | 1 | 1 | 1 | 0.999 | 1 | 1 | 1 | -0.001 |
| A5SS | 2285 | ENSG00000011243 | AKAP8L | chr19 | - | 15403474 | 15403715 | 15403657 | 15403715 | 15401149 | 15401603 | 334 | 243 | 394 | 422 | 413 | 0 | 0 | 1 | 0 | 0 | 596 | 454 | 323 | 359 | 672 | 0 | 0 | 0 | 0 | 0 | 298 | 149 | 1 | 1 | 1 | 1 | 0.995 | 1 | 1 | 1 | 1 | 1 | 1 | 1 | -0.001 |
| A5SS | 2331 | ENSG00000048140 | TSPAN17 | chr5 | + | 176656077 | 176656134 | 176656077 | 176656125 | 176656699 | 176656816 | 1 | 0 | 0 | 1 | 0 | 39 | 11 | 29 | 24 | 9 | 0 | 0 | 1 | 0 | 0 | 35 | 47 | 13 | 28 | 43 | 158 | 149 | 1 | 1 | 0.024 | 0 | 0 | 0.038 | 0 | 0 | 0 | 0.068 | 0 | 0 | -0.001 |
| A5SS | 2663 | ENSG00000131042 | LILRB2 | chr19 | - | 54276806 | 54277600 | 54277549 | 54277600 | 54276380 | 54276456 | 277 | 192 | 127 | 319 | 149 | 0 | 1 | 0 | 0 | 0 | 552 | 448 | 201 | 362 | 575 | 0 | 1 | 0 | 0 | 0 | 298 | 149 | 1 | 1 | 1 | 0.99 | 1 | 1 | 1 | 1 | 0.996 | 1 | 1 | 1 | -0.001 |
| A5SS | 2763 | ENSG00000012223 | LTF | chr3 | - | 46448862 | 46449017 | 46448868 | 46449017 | 46447307 | 46447398 | 10 | 2 | 5 | 157 | 5 | 0 | 0 | 0 | 1 | 0 | 152 | 139 | 54 | 3069 | 723 | 0 | 0 | 0 | 1 | 0 | 155 | 149 | 1 | 1 | 1 | 1 | 1 | 0.993 | 1 | 1 | 1 | 1 | 1 | 1 | -0.001 |
| A5SS | 2804 | ENSG00000013441 | CLK1 | chr2 | - | 200859679 | 200861466 | 200861237 | 200861466 | 200857972 | 200858089 | 1023 | 833 | 979 | 1154 | 1874 | 2 | 0 | 0 | 0 | 0 | 267 | 551 | 496 | 330 | 89 | 0 | 0 | 0 | 0 | 0 | 298 | 149 | 1 | 1 | 0.996 | 1 | 1 | 1 | 1 | 1 | 1 | 1 | 1 | 1 | -0.001 |
| A5SS | 3001 | ENSG00000116350 | SRSF4 | chr1 | - | 29159373 | 29160517 | 29160374 | 29160517 | 29154695 | 29154910 | 370 | 330 | 312 | 358 | 323 | 2 | 0 | 0 | 0 | 0 | 752 | 566 | 1005 | 776 | 678 | 0 | 0 | 2 | 0 | 0 | 298 | 149 | 1 | 1 | 0.989 | 1 | 1 | 1 | 1 | 1 | 1 | 0.996 | 1 | 1 | -0.001 |
| A5SS | 3011 | ENSG00000206503 | HLA-A | chr6 | + | 29944499 | 29945091 | 29944499 | 29944616 | 29945233 | 29945281 | 1345 | 410 | 1928 | 1146 | 2146 | 0 | 1 | 0 | 0 | 0 | 2229 | 995 | 2818 | 2493 | 2384 | 0 | 0 | 0 | 0 | 0 | 298 | 149 | 1 | 1 | 1 | 0.995 | 1 | 1 | 1 | 1 | 1 | 1 | 1 | 1 | -0.001 |
| A5SS | 3115 | ENSG00000109861 | CTSC | chr11 | - | 88312387 | 88312554 | 88312449 | 88312554 | 88309162 | 88309318 | 997 | 573 | 585 | 742 | 1285 | 3 | 0 | 3 | 0 | 4 | 1096 | 650 | 480 | 737 | 826 | 0 | 0 | 2 | 2 | 0 | 211 | 149 | 1 | 1 | 0.996 | 1 | 0.993 | 1 | 0.996 | 1 | 1 | 0.994 | 0.996 | 1 | -0.001 |
| A5SS | 3321 | ENSG00000113810 | SMC4 | chr3 | + | 160411919 | 160412084 | 160411919 | 160411977 | 160412325 | 160412453 | 181 | 215 | 158 | 175 | 367 | 0 | 0 | 0 | 2 | 4 | 60 | 122 | 99 | 26 | 36 | 0 | 0 | 2 | 0 | 0 | 256 | 149 | 1 | 1 | 1 | 1 | 1 | 0.981 | 0.982 | 1 | 1 | 0.966 | 1 | 1 | -0.001 |
| A5SS | 3796 | ENSG00000117335 | CD46 | chr1 | + | 207767012 | 207770362 | 207767012 | 207767195 | 207783291 | 207783330 | 594 | 481 | 613 | 641 | 847 | 1 | 0 | 0 | 0 | 0 | 141 | 353 | 329 | 265 | 71 | 0 | 0 | 0 | 0 | 0 | 298 | 149 | 1 | 1 | 0.997 | 1 | 1 | 1 | 1 | 1 | 1 | 1 | 1 | 1 | -0.001 |
| A5SS | 3838 | ENSG00000162434 | JAK1 | chr1 | - | 64841450 | 64841601 | 64841555 | 64841601 | 64841244 | 64841339 | 1232 | 769 | 992 | 1198 | 1262 | 0 | 1 | 0 | 1 | 0 | 739 | 770 | 945 | 963 | 344 | 0 | 0 | 0 | 0 | 0 | 254 | 149 | 1 | 1 | 1 | 0.998 | 1 | 0.999 | 1 | 1 | 1 | 1 | 1 | 1 | -0.001 |
| A5SS | 3853 | ENSG00000158517 | NCF1 | chr7 | + | 74782938 | 74783632 | 74782938 | 74783061 | 74785181 | 74785299 | 517 | 289 | 253 | 196 | 110 | 1 | 0 | 0 | 0 | 0 | 554 | 569 | 236 | 561 | 902 | 0 | 0 | 0 | 0 | 0 | 298 | 149 | 1 | 1 | 0.996 | 1 | 1 | 1 | 1 | 1 | 1 | 1 | 1 | 1 | -0.001 |
| A5SS | 3854 | ENSG00000158517 | NCF1 | chr7 | + | 74785181 | 74785299 | 74785181 | 74785261 | 74787983 | 74788088 | 644 | 382 | 356 | 317 | 185 | 0 | 2 | 0 | 0 | 0 | 716 | 726 | 263 | 615 | 992 | 0 | 0 | 0 | 0 | 0 | 187 | 149 | 1 | 1 | 1 | 0.993 | 1 | 1 | 1 | 1 | 1 | 1 | 1 | 1 | -0.001 |
| A5SS | 3906 | ENSG00000055609 | KMT2C | chr7 | - | 152158856 | 152159072 | 152158862 | 152159072 | 152157760 | 152157922 | 0 | 0 | 0 | 4 | 0 | 25 | 21 | 28 | 35 | 58 | 0 | 0 | 2 | 0 | 0 | 12 | 26 | 17 | 9 | 9 | 155 | 149 | 1 | 1 | 0 | 0 | 0 | 0.099 | 0 | 0 | 0 | 0.102 | 0 | 0 | -0.001 |
| A5SS | 3959 | ENSG00000163743 | RCHY1 | chr4 | - | 75491582 | 75491637 | 75491610 | 75491637 | 75490580 | 75490701 | 3 | 1 | 2 | 1 | 0 | 127 | 68 | 98 | 113 | 193 | 0 | 1 | 0 | 3 | 0 | 23 | 134 | 82 | 43 | 19 | 177 | 149 | 1 | 1 | 0.019 | 0.012 | 0.017 | 0.007 | 0 | 0 | 0.006 | 0 | 0.055 | 0 | -0.001 |
| A5SS | 4023 | ENSG00000138095 | LRPPRC | chr2 | - | 43894544 | 43899334 | 43899218 | 43899334 | 43889733 | 43889876 | 417 | 286 | 335 | 383 | 832 | 0 | 0 | 0 | 0 | 3 | 24 | 132 | 199 | 42 | 4 | 0 | 0 | 0 | 0 | 0 | 298 | 149 | 1 | 1 | 1 | 1 | 1 | 1 | 0.993 | 1 | 1 | 1 | 1 | 1 | -0.001 |
| A5SS | 4045 | ENSG00000142534 | RPS11 | chr19 | + | 49497193 | 49498046 | 49497193 | 49497595 | 49499511 | 49499689 | 1560 | 672 | 1641 | 1394 | 1328 | 0 | 0 | 1 | 0 | 1 | 1470 | 1368 | 933 | 1067 | 1915 | 0 | 0 | 0 | 0 | 0 | 298 | 149 | 1 | 1 | 1 | 1 | 0.999 | 1 | 0.998 | 1 | 1 | 1 | 1 | 1 | -0.001 |
| A5SS | 4273 | ENSG00000168159 | RNF187 | chr1 | + | 228488959 | 228489056 | 228488959 | 228489052 | 228493052 | 228493274 | 0 | 0 | 0 | 0 | 0 | 148 | 86 | 108 | 170 | 153 | 0 | 0 | 0 | 0 | 1 | 364 | 219 | 110 | 166 | 258 | 153 | 149 | 1 | 1 | 0 | 0 | 0 | 0 | 0 | 0 | 0 | 0 | 0 | 0.004 | -0.001 |
| A5SS | 4296 | ENSG00000071054 | MAP4K4 | chr2 | + | 101867211 | 101867318 | 101867211 | 101867309 | 101869621 | 101869797 | 2 | 0 | 1 | 0 | 0 | 223 | 159 | 198 | 274 | 305 | 0 | 0 | 5 | 0 | 0 | 363 | 260 | 245 | 324 | 235 | 158 | 149 | 1 | 1 | 0.008 | 0 | 0.005 | 0 | 0 | 0 | 0 | 0.019 | 0 | 0 | -0.001 |
| A5SS | 4444 | ENSG00000124126 | PREX1 | chr20 | - | 48632276 | 48632639 | 48632495 | 48632639 | 48630727 | 48630794 | 310 | 82 | 150 | 209 | 137 | 0 | 0 | 1 | 0 | 0 | 239 | 206 | 222 | 163 | 434 | 0 | 0 | 1 | 0 | 0 | 298 | 149 | 1 | 1 | 1 | 1 | 0.987 | 1 | 1 | 1 | 1 | 0.991 | 1 | 1 | -0.001 |
| A5SS | 4453 | ENSG00000164305 | CASP3 | chr4 | - | 184648464 | 184648631 | 184648502 | 184648631 | 184638400 | 184638468 | 25 | 12 | 18 | 9 | 1 | 2 | 1 | 0 | 1 | 0 | 16 | 18 | 4 | 3 | 4 | 0 | 1 | 1 | 0 | 0 | 187 | 149 | 1 | 1 | 0.909 | 0.905 | 1 | 0.878 | 1 | 1 | 0.935 | 0.761 | 1 | 1 | -0.001 |
| A5SS | 4538 | ENSG00000100612 | DHRS7 | chr14 | - | 60149352 | 60149568 | 60149449 | 60149568 | 60144805 | 60145013 | 1744 | 1008 | 872 | 1027 | 1130 | 0 | 3 | 0 | 1 | 0 | 2437 | 946 | 1209 | 1658 | 1594 | 0 | 0 | 0 | 0 | 0 | 246 | 149 | 1 | 1 | 1 | 0.995 | 1 | 0.998 | 1 | 1 | 1 | 1 | 1 | 1 | -0.001 |
| A5SS | 4655 | ENSG00000117616 | RSRP1 | chr1 | - | 25243549 | 25245301 | 25245149 | 25245301 | 25242248 | 25242705 | 878 | 697 | 708 | 807 | 1316 | 0 | 0 | 0 | 0 | 2 | 309 | 272 | 316 | 267 | 279 | 0 | 0 | 0 | 0 | 0 | 298 | 149 | 1 | 1 | 1 | 1 | 1 | 1 | 0.997 | 1 | 1 | 1 | 1 | 1 | -0.001 |
| A5SS | 4805 | ENSG00000159658 | EFCAB14 | chr1 | - | 46686783 | 46689691 | 46689586 | 46689691 | 46684490 | 46684602 | 559 | 402 | 469 | 454 | 1053 | 1 | 0 | 0 | 0 | 0 | 560 | 495 | 401 | 576 | 272 | 0 | 0 | 0 | 0 | 0 | 298 | 149 | 1 | 1 | 0.996 | 1 | 1 | 1 | 1 | 1 | 1 | 1 | 1 | 1 | -0.001 |
| A5SS | 4997 | ENSG00000009307 | CSDE1 | chr1 | - | 114749820 | 114750207 | 114749973 | 114750207 | 114739752 | 114739890 | 2174 | 1358 | 1937 | 2035 | 4685 | 2 | 0 | 0 | 2 | 0 | 317 | 1455 | 1521 | 979 | 321 | 0 | 0 | 1 | 0 | 0 | 298 | 149 | 1 | 1 | 0.998 | 1 | 1 | 0.998 | 1 | 1 | 1 | 0.999 | 1 | 1 | -0.001 |
| A5SS | 5103 | ENSG00000106105 | GARS1 | chr7 | + | 30631447 | 30631559 | 30631447 | 30631541 | 30632246 | 30632437 | 0 | 0 | 2 | 0 | 0 | 168 | 141 | 157 | 170 | 355 | 0 | 2 | 0 | 1 | 0 | 249 | 207 | 212 | 191 | 157 | 167 | 149 | 1 | 1 | 0 | 0 | 0.011 | 0 | 0 | 0 | 0.009 | 0 | 0.005 | 0 | -0.001 |
| A5SS | 5112 | ENSG00000118263 | KLF7 | chr2 | - | 207123773 | 207124404 | 207123975 | 207124404 | 207088457 | 207088581 | 107 | 93 | 141 | 146 | 130 | 0 | 0 | 0 | 1 | 0 | 121 | 146 | 228 | 206 | 118 | 0 | 0 | 1 | 0 | 0 | 298 | 149 | 1 | 1 | 1 | 1 | 1 | 0.986 | 1 | 1 | 1 | 0.991 | 1 | 1 | -0.001 |
| A5SS | 5135 | ENSG00000162191 | UBXN1 | chr11 | - | 62677517 | 62677832 | 62677780 | 62677832 | 62676812 | 62677005 | 195 | 103 | 226 | 184 | 210 | 0 | 1 | 0 | 0 | 0 | 373 | 272 | 179 | 249 | 390 | 0 | 0 | 0 | 2 | 0 | 298 | 149 | 1 | 1 | 1 | 0.981 | 1 | 1 | 1 | 1 | 1 | 1 | 0.984 | 1 | -0.001 |
| A5SS | 5274 | ENSG00000148303 | RPL7A | chr9 | + | 133348217 | 133348573 | 133348217 | 133348246 | 133348921 | 133349042 | 3 | 0 | 10 | 10 | 10 | 1828 | 905 | 2450 | 2023 | 3329 | 0 | 3 | 7 | 9 | 25 | 2079 | 1689 | 1517 | 1321 | 2698 | 298 | 149 | 1 | 1 | 0.001 | 0 | 0.002 | 0.002 | 0.001 | 0 | 0.001 | 0.002 | 0.003 | 0.005 | -0.001 |
| A5SS | 5606 | ENSG00000108883 | EFTUD2 | chr17 | - | 44894244 | 44894525 | 44894416 | 44894525 | 44886584 | 44886750 | 2 | 0 | 3 | 4 | 0 | 167 | 99 | 118 | 145 | 172 | 4 | 6 | 3 | 1 | 0 | 362 | 180 | 124 | 196 | 358 | 298 | 149 | 1 | 1 | 0.006 | 0 | 0.013 | 0.014 | 0 | 0.005 | 0.016 | 0.012 | 0.003 | 0 | -0.001 |
| A5SS | 5804 | ENSG00000115053 | NCL | chr2 | - | 231458265 | 231459125 | 231459000 | 231459125 | 231457642 | 231457800 | 1672 | 1455 | 1640 | 1850 | 2330 | 0 | 0 | 5 | 0 | 0 | 1349 | 1388 | 1554 | 1710 | 378 | 0 | 0 | 0 | 0 | 0 | 298 | 149 | 1 | 1 | 1 | 1 | 0.994 | 1 | 1 | 1 | 1 | 1 | 1 | 1 | -0.001 |
| A5SS | 5813 | ENSG00000130414 | NDUFA10 | chr2 | - | 240014738 | 240014860 | 240014840 | 240014860 | 240011616 | 240011696 | 430 | 276 | 411 | 326 | 880 | 0 | 1 | 0 | 0 | 0 | 591 | 453 | 316 | 324 | 423 | 0 | 0 | 0 | 0 | 0 | 251 | 149 | 1 | 1 | 1 | 0.994 | 1 | 1 | 1 | 1 | 1 | 1 | 1 | 1 | -0.001 |
| A5SS | 5814 | ENSG00000130414 | NDUFA10 | chr2 | - | 240014768 | 240014860 | 240014840 | 240014860 | 240011616 | 240011696 | 340 | 227 | 286 | 211 | 774 | 0 | 1 | 0 | 0 | 0 | 432 | 336 | 239 | 229 | 319 | 0 | 0 | 0 | 0 | 0 | 221 | 149 | 1 | 1 | 1 | 0.994 | 1 | 1 | 1 | 1 | 1 | 1 | 1 | 1 | -0.001 |
| A5SS | 5852 | ENSG00000169045 | HNRNPH1 | chr5 | - | 179617050 | 179617110 | 179617059 | 179617110 | 179616868 | 179616958 | 2633 | 2795 | 2533 | 2677 | 2719 | 27 | 11 | 10 | 15 | 16 | 918 | 1754 | 2005 | 1479 | 363 | 7 | 14 | 10 | 10 | 0 | 158 | 149 | 1 | 1 | 0.989 | 0.996 | 0.996 | 0.994 | 0.994 | 0.992 | 0.992 | 0.995 | 0.993 | 1 | -0.001 |
| A5SS | 5997 | ENSG00000137959 | IFI44L | chr1 | + | 78627905 | 78628393 | 78627905 | 78628018 | 78628950 | 78628999 | 343 | 991 | 251 | 580 | 565 | 0 | 0 | 0 | 1 | 0 | 1209 | 2004 | 803 | 2276 | 42 | 0 | 0 | 0 | 0 | 0 | 298 | 149 | 1 | 1 | 1 | 1 | 1 | 0.997 | 1 | 1 | 1 | 1 | 1 | 1 | -0.001 |
| A5SS | 5998 | ENSG00000137959 | IFI44L | chr1 | + | 78627905 | 78628999 | 78627905 | 78628393 | 78629719 | 78629915 | 167 | 477 | 122 | 397 | 145 | 0 | 3 | 0 | 1 | 0 | 394 | 843 | 371 | 901 | 7 | 0 | 4 | 0 | 2 | 0 | 298 | 149 | 1 | 1 | 1 | 0.988 | 1 | 0.995 | 1 | 1 | 0.991 | 1 | 0.996 | 1 | -0.001 |
| A5SS | 6153 | ENSG00000135486 | HNRNPA1 | chr12 | + | 54282572 | 54282665 | 54282572 | 54282662 | 54282799 | 54282874 | 2805 | 2712 | 3192 | 3023 | 3033 | 1 | 0 | 5 | 2 | 3 | 3272 | 2332 | 4030 | 3062 | 872 | 0 | 0 | 0 | 0 | 0 | 152 | 149 | 1 | 1 | 1 | 1 | 0.998 | 0.999 | 0.999 | 1 | 1 | 1 | 1 | 1 | -0.001 |
| A5SS | 6586 | ENSG00000141480 | ARRB2 | chr17 | + | 4716411 | 4716671 | 4716411 | 4716608 | 4717216 | 4717276 | 3 | 3 | 1 | 1 | 0 | 192 | 96 | 141 | 135 | 98 | 0 | 4 | 2 | 8 | 4 | 419 | 240 | 204 | 276 | 468 | 212 | 149 | 1 | 1 | 0.011 | 0.021 | 0.005 | 0.005 | 0 | 0 | 0.012 | 0.007 | 0.02 | 0.006 | -0.001 |
| A5SS | 6661 | ENSG00000122566 | HNRNPA2B1 | chr7 | - | 26191421 | 26192338 | 26192300 | 26192338 | 26190992 | 26191128 | 1621 | 1684 | 1319 | 1588 | 1073 | 1 | 3 | 0 | 0 | 0 | 480 | 823 | 1117 | 983 | 242 | 0 | 0 | 0 | 0 | 0 | 298 | 149 | 1 | 1 | 0.999 | 0.996 | 1 | 1 | 1 | 1 | 1 | 1 | 1 | 1 | -0.001 |
| A5SS | 6793 | ENSG00000107897 | ACBD5 | chr10 | - | 27210813 | 27211081 | 27210921 | 27211081 | 27208245 | 27208445 | 261 | 236 | 220 | 285 | 248 | 1 | 0 | 0 | 0 | 0 | 149 | 216 | 300 | 243 | 17 | 0 | 0 | 0 | 0 | 0 | 257 | 149 | 1 | 1 | 0.993 | 1 | 1 | 1 | 1 | 1 | 1 | 1 | 1 | 1 | -0.001 |
| A5SS | 6889 | ENSG00000125676 | THOC2 | chrX | - | 123706853 | 123706949 | 123706857 | 123706949 | 123703453 | 123703505 | 0 | 0 | 2 | 1 | 0 | 121 | 114 | 88 | 113 | 179 | 0 | 0 | 2 | 0 | 0 | 15 | 53 | 49 | 32 | 5 | 153 | 149 | 1 | 1 | 0 | 0 | 0.022 | 0.009 | 0 | 0 | 0 | 0.038 | 0 | 0 | -0.001 |
| A5SS | 6907 | ENSG00000139436 | GIT2 | chr12 | - | 109951166 | 109951316 | 109951211 | 109951316 | 109945259 | 109945349 | 39 | 19 | 22 | 25 | 18 | 0 | 0 | 0 | 0 | 2 | 22 | 19 | 28 | 33 | 24 | 0 | 0 | 3 | 0 | 0 | 194 | 149 | 1 | 1 | 1 | 1 | 1 | 1 | 0.874 | 1 | 1 | 0.878 | 1 | 1 | -0.001 |
| A5SS | 7086 | ENSG00000196924 | FLNA | chrX | - | 154350908 | 154351696 | 154351580 | 154351696 | 154350030 | 154350207 | 3387 | 1488 | 1719 | 2632 | 1075 | 0 | 0 | 5 | 0 | 0 | 4460 | 3022 | 1694 | 2418 | 14418 | 0 | 0 | 0 | 0 | 0 | 298 | 149 | 1 | 1 | 1 | 1 | 0.994 | 1 | 1 | 1 | 1 | 1 | 1 | 1 | -0.001 |
| A5SS | 7130 | ENSG00000197114 | ZGPAT | chr20 | + | 63733586 | 63733739 | 63733586 | 63733712 | 63734704 | 63734824 | 66 | 29 | 33 | 58 | 0 | 0 | 0 | 5 | 3 | 0 | 95 | 86 | 31 | 46 | 119 | 0 | 5 | 2 | 2 | 8 | 176 | 149 | 1 | 1 | 1 | 1 | 0.848 | 0.942 | NA | 1 | 0.936 | 0.929 | 0.951 | 0.926 | -0.001 |
| A5SS | 7171 | ENSG00000215301 | DDX3X | chrX | + | 41343215 | 41344128 | 41343215 | 41343351 | 41344238 | 41344399 | 793 | 529 | 703 | 924 | 984 | 1 | 0 | 0 | 2 | 0 | 117 | 419 | 557 | 393 | 53 | 0 | 0 | 0 | 0 | 0 | 298 | 149 | 1 | 1 | 0.997 | 1 | 1 | 0.996 | 1 | 1 | 1 | 1 | 1 | 1 | -0.001 |
| A5SS | 7245 | ENSG00000168300 | PCMTD1 | chr8 | - | 51860844 | 51861246 | 51860860 | 51861246 | 51833517 | 51833689 | 449 | 262 | 283 | 401 | 246 | 0 | 0 | 0 | 2 | 0 | 404 | 210 | 194 | 246 | 361 | 0 | 0 | 0 | 0 | 0 | 165 | 149 | 1 | 1 | 1 | 1 | 1 | 0.995 | 1 | 1 | 1 | 1 | 1 | 1 | -0.001 |
| A5SS | 7248 | ENSG00000087088 | BAX | chr19 | + | 48955547 | 48955833 | 48955547 | 48955599 | 48956197 | 48956333 | 98 | 46 | 68 | 117 | 50 | 1 | 0 | 0 | 0 | 0 | 165 | 121 | 58 | 138 | 282 | 0 | 0 | 0 | 1 | 0 | 298 | 149 | 1 | 1 | 0.98 | 1 | 1 | 1 | 1 | 1 | 1 | 1 | 0.986 | 1 | -0.001 |
| A5SS | 7280 | ENSG00000205581 | HMGN1 | chr21 | - | 39348421 | 39348577 | 39348544 | 39348577 | 39348291 | 39348339 | 256 | 225 | 327 | 329 | 183 | 0 | 0 | 1 | 0 | 0 | 390 | 254 | 261 | 415 | 322 | 0 | 0 | 0 | 0 | 0 | 272 | 149 | 1 | 1 | 1 | 1 | 0.994 | 1 | 1 | 1 | 1 | 1 | 1 | 1 | -0.001 |
| A5SS | 7462 | ENSG00000104408 | EIF3E | chr8 | - | 108241798 | 108241913 | 108241885 | 108241913 | 108239957 | 108240075 | 4600 | 2963 | 4658 | 3875 | 5149 | 2 | 3 | 4 | 1 | 3 | 631 | 1473 | 1432 | 495 | 640 | 0 | 0 | 0 | 0 | 0 | 236 | 149 | 1 | 1 | 0.999 | 0.998 | 0.999 | 1 | 0.999 | 1 | 1 | 1 | 1 | 1 | -0.001 |
| A5SS | 7470 | ENSG00000283189 | NA | chr3 | - | 49420210 | 49420342 | 49420214 | 49420342 | 49419709 | 49419788 | 10 | 3 | 4 | 7 | 5 | 0 | 0 | 0 | 1 | 0 | 10 | 13 | 22 | 18 | 27 | 0 | 0 | 3 | 0 | 0 | 153 | 149 | 1 | 1 | 1 | 1 | 1 | 0.872 | 1 | 1 | 1 | 0.877 | 1 | 1 | -0.001 |
| A5SS | 7586 | ENSG00000056586 | RC3H2 | chr9 | - | 122865348 | 122865657 | 122865623 | 122865657 | 122859916 | 122860131 | 224 | 174 | 235 | 264 | 502 | 0 | 0 | 0 | 0 | 1 | 39 | 218 | 177 | 54 | 132 | 0 | 0 | 0 | 0 | 0 | 298 | 149 | 1 | 1 | 1 | 1 | 1 | 1 | 0.996 | 1 | 1 | 1 | 1 | 1 | -0.001 |
| A5SS | 7781 | ENSG00000108424 | KPNB1 | chr17 | + | 47678307 | 47680134 | 47678307 | 47678413 | 47680507 | 47680669 | 816 | 702 | 813 | 970 | 1136 | 0 | 1 | 0 | 0 | 0 | 735 | 750 | 844 | 829 | 269 | 0 | 0 | 0 | 0 | 0 | 298 | 149 | 1 | 1 | 1 | 0.997 | 1 | 1 | 1 | 1 | 1 | 1 | 1 | 1 | -0.001 |
| A5SS | 7892 | ENSG00000152270 | PDE3B | chr11 | + | 14786436 | 14786685 | 14786436 | 14786532 | 14789105 | 14789242 | 525 | 525 | 978 | 673 | 1718 | 0 | 0 | 0 | 2 | 0 | 125 | 226 | 465 | 281 | 128 | 0 | 0 | 0 | 0 | 0 | 298 | 149 | 1 | 1 | 1 | 1 | 1 | 0.994 | 1 | 1 | 1 | 1 | 1 | 1 | -0.001 |
| A5SS | 8179 | ENSG00000089597 | GANAB | chr11 | - | 62639626 | 62639731 | 62639630 | 62639731 | 62639358 | 62639467 | 499 | 256 | 445 | 445 | 405 | 2 | 1 | 2 | 0 | 0 | 463 | 453 | 336 | 308 | 714 | 0 | 2 | 1 | 0 | 0 | 153 | 149 | 1 | 1 | 0.996 | 0.996 | 0.995 | 1 | 1 | 1 | 0.995 | 0.997 | 1 | 1 | -0.001 |
| A5SS | 8551 | ENSG00000213809 | KLRK1 | chr12 | - | 10379446 | 10379792 | 10379699 | 10379792 | 10378131 | 10378235 | 362 | 275 | 247 | 341 | 535 | 1 | 0 | 0 | 0 | 0 | 25 | 27 | 64 | 9 | 14 | 0 | 0 | 0 | 0 | 0 | 298 | 149 | 1 | 1 | 0.995 | 1 | 1 | 1 | 1 | 1 | 1 | 1 | 1 | 1 | -0.001 |
| A5SS | 8562 | ENSG00000087502 | ERGIC2 | chr12 | - | 29343119 | 29343282 | 29343162 | 29343282 | 29341733 | 29341816 | 209 | 134 | 172 | 163 | 260 | 0 | 0 | 0 | 0 | 1 | 24 | 52 | 32 | 14 | 1 | 0 | 0 | 0 | 0 | 0 | 192 | 149 | 1 | 1 | 1 | 1 | 1 | 1 | 0.995 | 1 | 1 | 1 | 1 | 1 | -0.001 |
| A5SS | 60 | ENSG00000182944 | EWSR1 | chr22 | + | 29288605 | 29291599 | 29288605 | 29288755 | 29292136 | 29292169 | 257 | 172 | 230 | 199 | 155 | 0 | 0 | 0 | 1 | 0 | 290 | 269 | 192 | 344 | 298 | 0 | 0 | 0 | 2 | 0 | 298 | 149 | 1 | 1 | 1 | 1 | 1 | 0.99 | 1 | 1 | 1 | 1 | 0.989 | 1 | 0 |
| A5SS | 239 | ENSG00000142541 | RPL13A | chr19 | + | 49489849 | 49490297 | 49489849 | 49489912 | 49490474 | 49490576 | 11939 | 4801 | 12468 | 9259 | 14143 | 2 | 0 | 0 | 1 | 0 | 6232 | 7518 | 5156 | 4217 | 10233 | 0 | 0 | 0 | 0 | 6 | 298 | 149 | 1 | 1 | 1 | 1 | 1 | 1 | 1 | 1 | 1 | 1 | 1 | 0.999 | 0 |
| A5SS | 240 | ENSG00000142541 | RPL13A | chr19 | + | 49489849 | 49490297 | 49489849 | 49489922 | 49490474 | 49490576 | 11938 | 4801 | 12471 | 9260 | 14136 | 1 | 0 | 4 | 0 | 0 | 6210 | 7513 | 5156 | 4208 | 10223 | 0 | 0 | 0 | 0 | 0 | 298 | 149 | 1 | 1 | 1 | 1 | 0.999 | 1 | 1 | 1 | 1 | 1 | 1 | 1 | 0 |
| A5SS | 280 | ENSG00000108107 | RPL28 | chr19 | + | 55386569 | 55386693 | 55386569 | 55386674 | 55387929 | 55388048 | 9597 | 4552 | 8040 | 8235 | 8016 | 0 | 0 | 0 | 1 | 0 | 7898 | 9109 | 6515 | 6430 | 10137 | 0 | 14 | 0 | 0 | 0 | 168 | 149 | 1 | 1 | 1 | 1 | 1 | 1 | 1 | 1 | 0.998 | 1 | 1 | 1 | 0 |
| A5SS | 313 | ENSG00000105640 | RPL18A | chr19 | + | 17861292 | 17861472 | 17861292 | 17861307 | 17862093 | 17862223 | 11431 | 4137 | 11963 | 8877 | 16985 | 1 | 0 | 2 | 0 | 2 | 6541 | 8020 | 4894 | 4173 | 9776 | 0 | 0 | 0 | 1 | 0 | 298 | 149 | 1 | 1 | 1 | 1 | 1 | 1 | 1 | 1 | 1 | 1 | 1 | 1 | 0 |
| A5SS | 330 | ENSG00000142227 | EMP3 | chr19 | + | 48327520 | 48327623 | 48327520 | 48327561 | 48329351 | 48329492 | 864 | 342 | 465 | 599 | 353 | 0 | 0 | 0 | 0 | 1 | 1229 | 895 | 455 | 1108 | 1533 | 0 | 0 | 0 | 2 | 0 | 211 | 149 | 1 | 1 | 1 | 1 | 1 | 1 | 0.996 | 1 | 1 | 1 | 0.997 | 1 | 0 |
| A5SS | 434 | ENSG00000142347 | MYO1F | chr19 | - | 8553138 | 8553437 | 8553349 | 8553437 | 8552032 | 8552164 | 1492 | 767 | 826 | 1120 | 789 | 1 | 0 | 0 | 0 | 0 | 3353 | 2174 | 1369 | 2405 | 3796 | 0 | 0 | 2 | 0 | 0 | 298 | 149 | 1 | 1 | 0.999 | 1 | 1 | 1 | 1 | 1 | 1 | 0.997 | 1 | 1 | 0 |
| A5SS | 464 | ENSG00000101347 | SAMHD1 | chr20 | - | 36935028 | 36935189 | 36935080 | 36935189 | 36930759 | 36930875 | 2263 | 1613 | 1533 | 1967 | 2266 | 0 | 0 | 0 | 0 | 0 | 3854 | 1424 | 839 | 2236 | 1363 | 0 | 0 | 0 | 1 | 0 | 201 | 149 | 1 | 1 | 1 | 1 | 1 | 1 | 1 | 1 | 1 | 1 | 0.999 | 1 | 0 |
| A5SS | 1043 | ENSG00000177731 | FLII | chr17 | - | 18253543 | 18253719 | 18253598 | 18253719 | 18253300 | 18253458 | 161 | 70 | 136 | 135 | 110 | 0 | 0 | 0 | 1 | 0 | 148 | 168 | 90 | 83 | 357 | 0 | 1 | 0 | 0 | 0 | 204 | 149 | 1 | 1 | 1 | 1 | 1 | 0.99 | 1 | 1 | 0.992 | 1 | 1 | 1 | 0 |
| A5SS | 1117 | ENSG00000125166 | GOT2 | chr16 | - | 58722118 | 58722278 | 58722149 | 58722278 | 58719195 | 58719255 | 0 | 1 | 1 | 0 | 1 | 96 | 67 | 111 | 77 | 105 | 0 | 0 | 1 | 1 | 0 | 43 | 88 | 53 | 57 | 86 | 180 | 149 | 1 | 1 | 0 | 0.012 | 0.007 | 0 | 0.008 | 0 | 0 | 0.015 | 0.014 | 0 | 0 |
| A5SS | 1221 | ENSG00000149925 | ALDOA | chr16 | + | 30067233 | 30067366 | 30067233 | 30067333 | 30067449 | 30067661 | 1436 | 659 | 1323 | 1413 | 849 | 0 | 0 | 0 | 2 | 0 | 2375 | 2272 | 1233 | 1806 | 4337 | 0 | 5 | 0 | 0 | 0 | 182 | 149 | 1 | 1 | 1 | 1 | 1 | 0.998 | 1 | 1 | 0.997 | 1 | 1 | 1 | 0 |
| A5SS | 1222 | ENSG00000149925 | ALDOA | chr16 | + | 30068817 | 30068990 | 30068817 | 30068978 | 30069305 | 30069389 | 4 | 2 | 1 | 0 | 1 | 1679 | 692 | 1179 | 1310 | 2137 | 4 | 6 | 0 | 0 | 4 | 2105 | 2093 | 1023 | 1842 | 4381 | 161 | 149 | 1 | 1 | 0.002 | 0.003 | 0.001 | 0 | 0 | 0.002 | 0.003 | 0 | 0 | 0.001 | 0 |
| A5SS | 1262 | ENSG00000102879 | CORO1A | chr16 | + | 30185208 | 30186720 | 30185208 | 30185407 | 30186815 | 30186945 | 1274 | 476 | 992 | 952 | 410 | 4 | 0 | 0 | 0 | 0 | 1237 | 1465 | 694 | 1223 | 1662 | 0 | 0 | 0 | 3 | 0 | 298 | 149 | 1 | 1 | 0.994 | 1 | 1 | 1 | 1 | 1 | 1 | 1 | 0.995 | 1 | 0 |
| A5SS | 1263 | ENSG00000160014 | CALM3 | chr19 | + | 46608196 | 46608340 | 46608196 | 46608327 | 46608481 | 46608981 | 1444 | 1076 | 1008 | 1634 | 1408 | 0 | 0 | 0 | 0 | 0 | 3654 | 1699 | 1267 | 1855 | 10669 | 0 | 0 | 0 | 0 | 12 | 162 | 149 | 1 | 1 | 1 | 1 | 1 | 1 | 1 | 1 | 1 | 1 | 1 | 0.999 | 0 |
| A5SS | 1469 | ENSG00000182718 | ANXA2 | chr15 | - | 60352378 | 60352476 | 60352382 | 60352476 | 60351723 | 60351819 | 0 | 0 | 0 | 0 | 0 | 1239 | 856 | 748 | 814 | 1546 | 0 | 2 | 0 | 0 | 0 | 2525 | 1179 | 737 | 1427 | 1999 | 153 | 149 | 1 | 1 | 0 | 0 | 0 | 0 | 0 | 0 | 0.002 | 0 | 0 | 0 | 0 |

| | | | | | | | | | |
|---|---|---|---|---|---|---|---|---|---|
| A5SS | 1483 | ENSG00000157823 | AP3S2 | chr15 | - | 89893876 | 89893994 | 89893880 | [illegible] |
| A5SS | 1699 | ENSG00000197249 | SERPINA1 | chr14 | - | 94380650 | 94381141 | 94380870 | [illegible] |
| A5SS | 1809 | ENSG00000100902 | PSMA6 | chr14 | + | 35312880 | 35313059 | 35312880 | [illegible] |
| A5SS | 2031 | ENSG00000130811 | EIF3G | chr19 | - | 10118861 | 10118956 | 10118867 | [illegible] |
| A5SS | 2135 | ENSG00000166888 | STAT6 | chr12 | - | 57107604 | 57107743 | 57107658 | [illegible] |
| A5SS | 2152 | ENSG00000113263 | ITK | chr5 | + | 157240061 | 157241720 | 157240061 | [illegible] |
| A5SS | 2228 | ENSG00000204628 | RACK1 | chr5 | - | 181241491 | 181242345 | 181242169 | [illegible] |
| A5SS | 2233 | ENSG00000204628 | RACK1 | chr5 | - | 181242169 | 181242345 | 181242285 | [illegible] |
| A5SS | 2234 | ENSG00000204628 | RACK1 | chr5 | - | 181242173 | 181242345 | 181242285 | [illegible] |
| A5SS | 2281 | ENSG00000092020 | PPP2R3C | chr14 | - | 35109818 | 35109931 | 35109847 | [illegible] |
| A5SS | 2401 | ENSG00000070756 | PABPC1 | chr8 | - | 100712651 | 100713181 | 100713086 | [illegible] |
| A5SS | 2405 | ENSG00000070756 | PABPC1 | chr8 | - | 100718086 | 100718280 | 100718230 | [illegible] |
| A5SS | 2406 | ENSG00000070756 | PABPC1 | chr8 | - | 100718182 | 100718280 | 100718230 | [illegible] |
| A5SS | 2561 | ENSG00000100650 | SRSF5 | chr14 | + | 69768797 | 69769251 | 69768797 | [illegible] |
| A5SS | 2610 | ENSG00000168397 | ATG4B | chr2 | + | 241651009 | 241651335 | 241651009 | [illegible] |
| A5SS | 2740 | ENSG00000168026 | TTC21A | chr3 | + | 39137194 | 39137387 | 39137194 | [illegible] |
| A5SS | 2803 | ENSG00000013441 | CLK1 | chr2 | - | 200856890 | 200857884 | 200857717 | [illegible] |
| A5SS | 2988 | ENSG00000163634 | THOC7 | chr3 | - | 63839655 | 63839773 | 63839682 | [illegible] |
| A5SS | 3007 | ENSG00000206503 | HLA-A | chr6 | + | 29942756 | 29943543 | 29942756 | [illegible] |
| A5SS | 3026 | ENSG00000198034 | RPS4X | chrX | - | 72273800 | 72275150 | 72275052 | [illegible] |
| A5SS | 3196 | ENSG00000168028 | RPSA | chr3 | + | 39408605 | 39408739 | 39408605 | [illegible] |
| A5SS | 3532 | ENSG00000163931 | TKT | chr3 | - | 53231349 | 53231550 | 53231356 | [illegible] |
| A5SS | 3706 | ENSG00000185825 | BCAP31 | chrX | - | 153724260 | 153724388 | 153724333 | [illegible] |
| A5SS | 3816 | ENSG00000175390 | EIF3F | chr11 | + | 7994425 | 7995118 | 7994425 | [illegible] |
| A5SS | 4044 | ENSG00000142534 | RPS11 | chr19 | + | 49497193 | 49497325 | 49497193 | [illegible] |
| A5SS | 4109 | ENSG00000145741 | BTF3 | chr5 | + | 73502487 | 73503117 | 73502487 | [illegible] |
| A5SS | 4149 | ENSG00000156482 | RPL30 | chr8 | - | 98044942 | 98045399 | 98045346 | [illegible] |
| A5SS | 4177 | ENSG00000137575 | SDCBP | chr8 | + | 58565018 | 58565084 | 58565018 | [illegible] |
| A5SS | 4366 | ENSG00000144381 | HSPD1 | chr2 | - | 197494656 | 197495376 | 197495293 | [illegible] |
| A5SS | 4432 | ENSG00000197989 | SNHG12 | chr1 | - | 28579532 | 28579981 | 28579911 | [illegible] |
| A5SS | 4631 | ENSG00000131165 | CHMP1A | chr16 | - | 89649350 | 89649497 | 89649377 | [illegible] |
| A5SS | 4951 | ENSG00000067225 | PKM | chr15 | - | 72208616 | 72208891 | 72208620 | [illegible] |
| A5SS | 5119 | ENSG00000142937 | RPS8 | chr1 | + | 44776674 | 44776774 | 44776674 | [illegible] |
| A5SS | 5147 | ENSG00000181163 | NPM1 | chr5 | + | 171390050 | 171391424 | 171390050 | [illegible] |
| A5SS | 5149 | ENSG00000181163 | NPM1 | chr5 | + | 171391304 | 171391799 | 171391304 | [illegible] |
| A5SS | 5153 | ENSG00000109332 | UBE2D3 | chr4 | - | 102801453 | 102802638 | 102802560 | [illegible] |
| A5SS | 5388 | ENSG00000204681 | GABBR1 | chr6 | - | 29630457 | 29630643 | 29630521 | [illegible] |
| A5SS | 5479 | ENSG00000204387 | SNHG32 | chr6 | + | 31835410 | 31835455 | 31835410 | [illegible] |
| A5SS | 5685 | ENSG00000122026 | RPL21 | chr13 | + | 27255241 | 27255354 | 27255241 | [illegible] |
| A5SS | 5781 | ENSG00000064666 | CNN2 | chr19 | + | 1032558 | 1032696 | 1032558 | [illegible] |
| A5SS | 5835 | ENSG00000145425 | RPS3A | chr4 | + | 151100974 | 151101162 | 151100974 | [illegible] |
| A5SS | 5837 | ENSG00000145425 | RPS3A | chr4 | + | 151102870 | 151103079 | 151102870 | [illegible] |
| A5SS | 6037 | ENSG00000114383 | TUSC2 | chr3 | - | 50327027 | 50327219 | 50327037 | [illegible] |
| A5SS | 6076 | ENSG00000204256 | BRD2 | chr6 | + | 32976561 | 32977570 | 32976561 | [illegible] |
| A5SS | 6082 | ENSG00000156508 | EEF1A1 | chr6 | - | 73518029 | 73518610 | 73518353 | [illegible] |
| A5SS | 6083 | ENSG00000156508 | EEF1A1 | chr6 | - | 73519336 | 73520056 | 73519882 | [illegible] |
| A5SS | 6084 | ENSG00000156508 | EEF1A1 | chr6 | - | 73519882 | 73520056 | 73519891 | [illegible] |
| A5SS | 6089 | ENSG00000169314 | C22orf15 | chr22 | + | 23764259 | 23764412 | 23764259 | [illegible] |
| A5SS | 6155 | ENSG00000135486 | HNRNPA1 | chr12 | + | 54282393 | 54282874 | 54282393 | [illegible] |
| A5SS | 6435 | ENSG00000119616 | FCF1 | chr14 | + | 74713484 | 74713552 | 74713484 | [illegible] |
| A5SS | 6587 | ENSG00000141480 | ARRB2 | chr17 | + | 4719282 | 4719420 | 4719282 | [illegible] |
| A5SS | 6603 | ENSG00000071462 | BUD23 | chr7 | + | 73683766 | 73684000 | 73683766 | [illegible] |
| A5SS | 6779 | ENSG00000166716 | ZNF592 | chr15 | + | 84790704 | 84790883 | 84790704 | [illegible] |
| A5SS | 7144 | ENSG00000152332 | UHMK1 | chr1 | + | 162499954 | 162501104 | 162499954 | [illegible] |
| A5SS | 7170 | ENSG00000215301 | DDX3X | chrX | + | 41342494 | 41342836 | 41342494 | [illegible] |
| A5SS | 7384 | ENSG00000070081 | NUCB2 | chr11 | + | 17286633 | 17286717 | 17286633 | [illegible] |
| A5SS | 7422 | ENSG00000104763 | ASAH1 | chr8 | - | 18067219 | 18067298 | 18067249 | [illegible] |
| A5SS | 7423 | ENSG00000104763 | ASAH1 | chr8 | - | 18071299 | 18071390 | 18071330 | [illegible] |
| A5SS | 7461 | ENSG00000104408 | EIF3E | chr8 | - | 108236160 | 108236203 | 108236162 | [illegible] |
| A5SS | 7463 | ENSG00000104408 | EIF3E | chr8 | - | 108241798 | 108241913 | 108241885 | [illegible] |
| A5SS | 7466 | ENSG00000008988 | RPS20 | chr8 | - | 56074253 | 56074510 | 56074380 | [illegible] |
| A5SS | 7615 | ENSG00000156042 | CFAP70 | chr10 | - | 73299600 | 73299665 | 73299604 | [illegible] |
| A5SS | 7625 | ENSG00000140988 | RPS2 | chr16 | - | 1962496 | 1962909 | 1962735 | [illegible] |
| A5SS | 7626 | ENSG00000140988 | RPS2 | chr16 | - | 1962496 | 1963256 | 1963148 | [illegible] |
| A5SS | 7628 | ENSG00000140988 | RPS2 | chr16 | - | 1963652 | 1964365 | 1964275 | [illegible] |
| A5SS | 7796 | ENSG00000137073 | UBAP2 | chr9 | - | 33996222 | 33998864 | 33998786 | [illegible] |
| A5SS | 7905 | ENSG00000254772 | EEF1G | chr11 | - | 62571539 | 62571682 | 62571623 | [illegible] |
| A5SS | 7963 | ENSG00000122218 | COPA | chr1 | - | 160332447 | 160333679 | 160333602 | [illegible] |
| A5SS | 8001 | ENSG00000149273 | RPS3 | chr11 | + | 75400693 | 75400824 | 75400693 | [illegible] |
| A5SS | 8116 | ENSG00000110321 | EIF4G2 | chr11 | - | 10799551 | 10800348 | 10800089 | [illegible] |
| A5SS | 8173 | ENSG00000084234 | APLP2 | chr11 | + | 130109428 | 130109617 | 130109428 | [illegible] |
| A5SS | 8346 | ENSG00000196531 | NACA | chr12 | - | 56714361 | 56714439 | 56714370 | [illegible] |
| A5SS | 8380 | ENSG00000105287 | PRKD2 | chr19 | - | 46678365 | 46678663 | 46678395 | [illegible] |
| A5SS | 8433 | ENSG00000107959 | PITRM1 | chr10 | - | 3147147 | 3147250 | 3147149 | [illegible] |
| A5SS | 8483 | ENSG00000118518 | RNF146 | chr6 | + | 127266882 | 127267095 | 127266882 | [illegible] |
| A5SS | 8550 | ENSG00000213809 | KLRK1 | chr12 | - | 10378553 | 10379792 | 10379699 | [illegible] |
| A5SS | 8651 | ENSG00000028203 | VEZT | chr12 | + | 95242163 | 95242314 | 95242163 | [illegible] |
| A5SS | 8664 | ENSG00000111144 | LTA4H | chr12 | - | 96009093 | 96013258 | 96013187 | [illegible] |
| A5SS | 53 | ENSG00000254413 | CHKB-CPT1B | chr22 | - | 50579726 | 50579830 | 50579790 | [illegible] |
| A5SS | 167 | ENSG00000104824 | HNRNPL | chr19 | - | 38840095 | 38840559 | 38840487 | [illegible] |
| A5SS | 351 | ENSG00000088053 | GP6 | chr19 | - | 55015678 | 55015733 | 55015682 | [illegible] |
| A5SS | 355 | ENSG00000090013 | BLVRB | chr19 | - | 40457498 | 40458244 | 40458154 | [illegible] |
| A5SS | 472 | ENSG00000064601 | CTSA | chr20 | + | 45892724 | 45894072 | 45892724 | [illegible] |
| A5SS | 731 | ENSG00000030582 | GRN | chr17 | + | 44351036 | 44351163 | 44351036 | [illegible] |
| A5SS | 881 | ENSG00000132471 | WBP2 | chr17 | - | 75855179 | 75855335 | 75855238 | [illegible] |
| A5SS | 930 | ENSG00000173821 | RNF213 | chr17 | + | 80381546 | 80383070 | 80381546 | [illegible] |
| A5SS | 1005 | ENSG00000126581 | BECN1 | chr17 | - | 42818786 | 42819609 | 42819547 | [illegible] |
| A5SS | 1014 | ENSG00000125970 | RALY | chr20 | + | 34072065 | 34072330 | 34072065 | [illegible] |
| A5SS | 1471 | ENSG00000182718 | ANXA2 | chr15 | - | 60360930 | 60361054 | 60360940 | [illegible] |
| A5SS | 1509 | ENSG00000102901 | CENPT | chr16 | - | 67829764 | 67830088 | 67829929 | [illegible] |
| A5SS | 1793 | ENSG00000213983 | AP1G2 | chr14 | - | 23567110 | 23567319 | 23567236 | [illegible] |
| A5SS | 1835 | ENSG00000103811 | CTSH | chr15 | - | 78926174 | 78927781 | 78927712 | [illegible] |
| A5SS | 1976 | ENSG00000118181 | RPS25 | chr11 | - | 119017924 | 119018053 | 119017957 | [illegible] |
| A5SS | 2086 | ENSG00000065357 | DGKA | chr12 | + | 55941997 | 55942083 | 55941997 | [illegible] |
| A5SS | 2184 | ENSG00000161960 | EIF4A1 | chr17 | + | 7576523 | 7576692 | 7576523 | [illegible] |
| A5SS | 2229 | ENSG00000204628 | RACK1 | chr5 | - | 181241491 | 181242345 | 181242173 | [illegible] |
| A5SS | 2461 | ENSG00000049656 | CLPTM1L | chr5 | - | 1331798 | 1331883 | 1331802 | [illegible] |
| A5SS | 2805 | ENSG00000188010 | MORN2 | chr2 | + | 38875990 | 38876211 | 38875990 | [illegible] |
| A5SS | 2833 | ENSG00000072756 | TRNT1 | chr3 | + | 3140509 | 3140648 | 3140509 | [illegible] |
| A5SS | 3182 | ENSG00000005812 | FBXL3 | chr13 | - | 77021512 | 77021861 | 77021656 | [illegible] |
| A5SS | 3272 | ENSG00000198925 | ATG9A | chr2 | - | 219228423 | 219228485 | 219228433 | [illegible] |
| A5SS | 3367 | ENSG00000080822 | CLDND1 | chr3 | - | 98521132 | 98521442 | 98521329 | [illegible] |
| A5SS | 3530 | ENSG00000111640 | GAPDH | chr12 | + | 6534809 | 6534991 | 6534809 | [illegible] |
| A5SS | 3639 | ENSG00000092010 | PSME1 | chr14 | + | 24138344 | 24138403 | 24138344 | [illegible] |
| A5SS | 3653 | ENSG00000018510 | AGPS | chr2 | + | 177440964 | 177441040 | 177440964 | [illegible] |
| A5SS | 3735 | ENSG00000026025 | VIM | chr10 | + | 17235845 | 17236379 | 17235845 | [illegible] |
| A5SS | 3839 | ENSG00000162434 | JAK1 | chr1 | - | 64845512 | 64846736 | 64846648 | [illegible] |
| A5SS | 4018 | ENSG00000131238 | PPT1 | chr1 | - | 40092044 | 40092507 | 40092397 | [illegible] |
| A5SS | 4151 | ENSG00000178467 | P4HTM | chr3 | + | 49002499 | 49006187 | 49002499 | [illegible] |
| A5SS | 4172 | ENSG00000165671 | NSD1 | chr5 | + | 177135086 | 177136030 | 177135086 | [illegible] |
| A5SS | 4301 | ENSG00000153187 | HNRNPU | chr1 | - | 244860334 | 244860474 | 244860357 | [illegible] |
| A5SS | 4373 | ENSG00000138688 | KIAA1109 | chr4 | + | 122209158 | 122209337 | 122209158 | [illegible] |
| A5SS | 4741 | ENSG00000082701 | GSK3B | chr3 | - | 119916039 | 119916174 | 119916043 | [illegible] |
| A5SS | 4891 | ENSG00000143418 | CERS2 | chr1 | - | 150967391 | 150967714 | 150967663 | [illegible] |
| A5SS | 5312 | ENSG00000088179 | PTPN4 | chr2 | + | 119915178 | 119916690 | 119915178 | [illegible] |
| A5SS | 5492 | ENSG00000198589 | LRBA | chr4 | - | 150798076 | 150798142 | 150798080 | [illegible] |
| A5SS | 5525 | ENSG00000137947 | GTF2B | chr1 | - | 88863980 | 88864114 | 88863985 | [illegible] |
| A5SS | 5739 | ENSG00000121741 | ZMYM2 | chr13 | + | 19993062 | 19993919 | 19993062 | [illegible] |
| A5SS | 5834 | ENSG00000145425 | RPS3A | chr4 | + | 151100974 | 151101085 | 151100974 | [illegible] |
| A5SS | 5853 | ENSG00000169045 | HNRNPH1 | chr5 | - | 179620891 | 179621397 | 179621241 | [illegible] |
| A5SS | 5873 | ENSG00000105372 | RPS19 | chr19 | + | 41860254 | 41860445 | 41860254 | [illegible] |
| A5SS | 6150 | ENSG00000135486 | HNRNPA1 | chr12 | + | 54281385 | 54281941 | 54281385 | [illegible] |
| A5SS | 6151 | ENSG00000135486 | HNRNPA1 | chr12 | + | 54281794 | 54281945 | 54281794 | [illegible] |
| A5SS | 6326 | ENSG00000126524 | SBDS | chr7 | - | 66994211 | 66994341 | 66994219 | [illegible] |
| A5SS | 6475 | ENSG00000142657 | PGD | chr1 | + | 10404160 | 10404279 | 10404160 | [illegible] |
| A5SS | 6543 | ENSG00000122557 | HERPUD2 | chr7 | - | 35694183 | 35695135 | 35694800 | [illegible] |
| A5SS | 6980 | ENSG00000072778 | ACADVL | chr17 | + | 7223812 | 7223875 | 7223812 | [illegible] |
| A5SS | 7158 | ENSG00000046651 | OFD1 | chrX | + | 13756577 | 13756771 | 13756577 | [illegible] |
| A5SS | 7160 | ENSG00000147133 | TAF1 | chrX | + | 71458240 | 71458366 | 71458240 | [illegible] |
| A5SS | 7307 | ENSG00000143321 | HDGF | chr1 | - | 156745007 | 156745373 | 156745296 | [illegible] |
| A5SS | 7627 | ENSG00000140988 | RPS2 | chr16 | - | 1962735 | 1963256 | 1963148 | [illegible] |
| A5SS | 7962 | ENSG00000122218 | COPA | chr1 | - | 160306353 | 160306493 | 160306454 | [illegible] |
| A5SS | 8021 | ENSG00000132274 | TRIM22 | chr11 | + | 5698314 | 5698545 | 5698314 | [illegible] |
| A5SS | 8026 | ENSG00000099783 | HNRNPM | chr19 | + | 8463496 | 8463504 | 8463496 | [illegible] |
| A5SS | 8150 | ENSG00000185187 | SIGIRR | chr11 | - | 409859 | 410027 | 409867 | [illegible] |
| A5SS | 8477 | ENSG00000150093 | ITGB1 | chr10 | - | 32932510 | 32932600 | 32932514 | [illegible] |
| A5SS | 8504 | ENSG00000099246 | RAB18 | chr10 | + | 27504285 | 27504448 | 27504285 | [illegible] |
| A5SS | 8676 | ENSG00000123384 | LRP1 | chr12 | + | 57201779 | 57201947 | 57201779 | [illegible] |
| A5SS | 8680 | ENSG00000089157 | RPLP0 | chr12 | - | 120198853 | 120199000 | 120198894 | [illegible] |
| A5SS | 8814 | ENSG00000123349 | PFDN5 | chr12 | + | 53295838 | 53297924 | 53295838 | [illegible] |
| A5SS | 8815 | ENSG00000123349 | PFDN5 | chr12 | + | 53296429 | 53297924 | 53296429 | [illegible] |
| A5SS | 8823 | ENSG00000111596 | CNOT2 | chr12 | + | 70344127 | 70344239 | 70344127 | [illegible] |
| A5SS | 8855 | ENSG00000188906 | LRRK2 | chr12 | + | 40364841 | 40365050 | 40364841 | [illegible] |
| A5SS | 8894 | ENSG00000067167 | TRAM1 | chr8 | - | 70597890 | 70598011 | 70597894 | [illegible] |
| A5SS | 8914 | ENSG00000111684 | LPCAT3 | chr12 | - | 6978340 | 6978507 | 6978420 | [illegible] |
| A5SS | 8954 | ENSG00000127022 | CANX | chr5 | + | 179705678 | 179705862 | 179705678 | [illegible] |
| A5SS | 146 | ENSG00000198131 | ZNF544 | chr19 | + | 58230393 | 58230497 | 58230393 | [illegible] |
| A5SS | 431 | ENSG00000105401 | CDC37 | chr19 | - | 10395227 | 10395543 | 10395434 | [illegible] |
| A5SS | 778 | ENSG00000185624 | P4HB | chr17 | - | 81845560 | 81845742 | 81845626 | [illegible] |
| A5SS | 787 | ENSG00000185624 | P4HB | chr17 | - | 81859178 | 81859387 | 81859180 | [illegible] |
| A5SS | 975 | ENSG00000073584 | SMARCE1 | chr17 | - | 40631591 | 40632367 | 40632194 | [illegible] |
| A5SS | 1110 | ENSG00000187555 | USP7 | chr16 | - | 8923214 | 8923413 | 8923353 | [illegible] |
| A5SS | 1175 | ENSG00000205250 | E2F4 | chr16 | + | 67192760 | 67192870 | 67192760 | [illegible] |
| A5SS | 1256 | ENSG00000089280 | FUS | chr16 | + | 31184938 | 31188357 | 31184938 | [illegible] |
| A5SS | 1418 | ENSG00000213614 | HEXA | chr15 | - | 72347685 | 72348134 | 72348030 | [illegible] |
| A5SS | 1715 | ENSG00000100568 | VTI1B | chr14 | - | 67656411 | 67656589 | 67656415 | [illegible] |
| A5SS | 1837 | ENSG00000103811 | CTSH | chr15 | - | 78932275 | 78932458 | 78932371 | [illegible] |
| A5SS | 2165 | ENSG00000113721 | PDGFRB | chr5 | - | 150134715 | 150135016 | 150134749 | [illegible] |
| A5SS | 2734 | ENSG00000129071 | MBD4 | chr3 | - | 129437719 | 129437950 | 129437807 | [illegible] |
| A5SS | 2972 | ENSG00000075785 | RAB7A | chr3 | + | 128806371 | 128806590 | 128806371 | [illegible] |
| A5SS | 3003 | ENSG00000186298 | PPP1CC | chr12 | - | 110722471 | 110722695 | 110722623 | [illegible] |
| A5SS | 3275 | ENSG00000167085 | PHB | chr17 | - | 49414739 | 49414858 | 49414834 | [illegible] |
| A5SS | 3403 | ENSG00000170581 | STAT2 | chr12 | - | 56354777 | 56355351 | 56355275 | [illegible] |
| A5SS | 3437 | ENSG00000055332 | EIF2AK2 | chr2 | - | 37109193 | 37109295 | 37109223 | [illegible] |
| A5SS | 3470 | ENSG00000117118 | SDHB | chr1 | - | 17028599 | 17028736 | 17028653 | [illegible] |
| A5SS | 3487 | ENSG00000071127 | WDR1 | chr4 | - | 10084440 | 10084530 | 10084442 | [illegible] |
| A5SS | 3559 | ENSG00000113194 | FAF2 | chr5 | + | 176493998 | 176494275 | 176493998 | [illegible] |
| A5SS | 3652 | ENSG00000099622 | CIRBP | chr19 | + | 1271139 | 1271467 | 1271139 | [illegible] |
| A5SS | 3716 | ENSG00000115524 | SF3B1 | chr2 | - | 197423807 | 197423974 | 197423893 | [illegible] |
| A5SS | 4543 | ENSG00000082014 | SMARCD3 | chr7 | - | 151242704 | 151242843 | 151242720 | [illegible] |
| A5SS | 4609 | ENSG00000204406 | MBD5 | chr2 | + | 148483109 | 148484135 | 148483109 | [illegible] |
| A5SS | 5470 | ENSG00000082153 | BZW1 | chr2 | + | 200819981 | 200820120 | 200819981 | [illegible] |
| A5SS | 5519 | ENSG00000223960 | CHROMR | chr2 | + | 178426365 | 178426703 | 178426365 | [illegible] |
| A5SS | 5594 | ENSG00000143390 | RFX5 | chr1 | - | 151346488 | 151346606 | 151346578 | [illegible] |
| A5SS | 6152 | ENSG00000135486 | HNRNPA1 | chr12 | + | 54282393 | 54282665 | 54282393 | [illegible] |
| A5SS | 6236 | ENSG00000168036 | CTNNB1 | chr3 | + | 41224525 | 41224759 | 41224525 | [illegible] |
| A5SS | 6465 | ENSG00000006530 | AGK | chr7 | + | 141615470 | 141615569 | 141615470 | [illegible] |
| A5SS | 6652 | ENSG00000123066 | MED13L | chr12 | - | 115980749 | 115982603 | 115982383 | [illegible] |
| A5SS | 6663 | ENSG00000122566 | HNRNPA2B1 | chr7 | - | 26196556 | 26197017 | 26196877 | [illegible] |
| A5SS | 6674 | ENSG00000105835 | NAMPT | chr7 | - | 106274945 | 106277179 | 106277083 | [illegible] |
| A5SS | 7191 | ENSG00000147050 | KDM6A | chrX | + | 45090722 | 45090864 | 45090722 | [illegible] |
| A5SS | 7202 | ENSG00000102119 | EMD | chrX | + | 154379484 | 154379566 | 154379484 | [illegible] |
| A5SS | 7246 | ENSG00000168300 | PCMTD1 | chr8 | - | 51860844 | 51861246 | 51860860 | [illegible] |
| A5SS | 7249 | ENSG00000145041 | DCAF1 | chr3 | - | 51466802 | 51466876 | 51466805 | [illegible] |
| A5SS | 7346 | ENSG00000034677 | RNF19A | chr8 | - | 100287500 | 100288267 | 100288206 | [illegible] |
| A5SS | 7426 | ENSG00000104763 | ASAH1 | chr8 | - | 18083940 | 18084120 | 18083980 | [illegible] |
| A5SS | 7496 | ENSG00000156738 | MS4A1 | chr11 | + | 60462184 | 60462533 | 60462184 | [illegible] |
| A5SS | 7837 | ENSG00000133943 | DGLUCY | chr14 | + | 91215404 | 91215793 | 91215404 | [illegible] |
| A5SS | 8053 | ENSG00000175550 | DRAP1 | chr11 | + | 65919947 | 65920041 | 65919947 | [illegible] |
| A5SS | 8172 | ENSG00000084234 | APLP2 | chr11 | + | 130069893 | 130070082 | 130069893 | [illegible] |
| A5SS | 8300 | ENSG00000198258 | UBL5 | chr19 | + | 9828326 | 9828393 | 9828326 | [illegible] |
| A5SS | 8314 | ENSG00000090621 | PABPC4 | chr1 | - | 39563613 | 39564542 | 39564470 | [illegible] |
| A5SS | 8738 | ENSG00000173442 | EHBP1L1 | chr11 | + | 65581538 | 65583765 | 65581538 | [illegible] |
| A5SS | 65 | ENSG00000196588 | MRTFA | chr22 | - | 40416975 | 40417046 | 40416985 | [illegible] |
| A5SS | 82 | ENSG00000100209 | HSCB | chr22 | + | 28743881 | 28743978 | 28743881 | [illegible] |
| A5SS | 111 | ENSG00000100242 | SUN2 | chr22 | - | 38740266 | 38741050 | 38741006 | [illegible] |
| A5SS | 304 | ENSG00000105193 | RPS16 | chr19 | - | 39433521 | 39433761 | 39433664 | [illegible] |
| A5SS | 415 | ENSG00000186010 | NDUFA13 | chr19 | + | 19527280 | 19527372 | 19527280 | [illegible] |
| A5SS | 418 | ENSG00000268790 | NA | chr19 | - | 16659954 | 16660000 | 16659958 | [illegible] |
| A5SS | 473 | ENSG00000064601 | CTSA | chr20 | + | 45894993 | 45895138 | 45894993 | [illegible] |
| A5SS | 1180 | ENSG00000103051 | COG4 | chr16 | - | 70519648 | 70519731 | 70519676 | [illegible] |
| A5SS | 1190 | ENSG00000260528 | FAM157C | chr16 | + | 90111371 | 90111485 | 90111371 | [illegible] |
| A5SS | 1368 | ENSG00000140694 | PARN | chr16 | - | 14627105 | 14628251 | 14628171 | [illegible] |
| A5SS | 2153 | ENSG00000113263 | ITK | chr5 | + | 157240061 | 157241720 | 157240061 | [illegible] |
| A5SS | 2394 | ENSG00000026297 | RNASET2 | chr6 | - | 166948565 | 166948625 | 166948569 | [illegible] |
| A5SS | 3094 | ENSG00000161204 | ABCF3 | chr3 | + | 184186506 | 184186654 | 184186506 | [illegible] |
| A5SS | 3114 | ENSG00000109861 | CTSC | chr11 | - | 88312387 | 88312554 | 88312426 | [illegible] |
| A5SS | 3527 | ENSG00000265241 | RBM8A | chr1 | - | 145926808 | 145927077 | 145927017 | [illegible] |
| A5SS | 3732 | ENSG00000026508 | CD44 | chr11 | + | 35189834 | 35190065 | 35189834 | [illegible] |

| | | | | | | | | | | | | | | | | | | | | | | |
|---|---|---|---|---|---|---|---|---|---|---|---|---|---|---|---|---|---|---|---|---|---|---|
| A5SS | 3859 | ENSG00000146007 | ZMAT2 | chr5 | + | 140703917 | 140704004 | 140703917 | 140703991 | 140704425 | 140704571 | 4 | 2 | 0 | [illegible] | [illegible] | [illegible] | [illegible] | [illegible] | [illegible] | [illegible] | [illegible] |
| A5SS | 4057 | ENSG00000158773 | USF1 | chr1 | - | 161043252 | 161043360 | 161043267 | 161043360 | 161042832 | 161042882 | 0 | 3 | 2 | [illegible] | [illegible] | [illegible] | [illegible] | [illegible] | [illegible] | [illegible] | [illegible] |
| A5SS | 4278 | ENSG00000135821 | GLUL | chr1 | - | 182386255 | 182386402 | 182386361 | 182386402 | 182385402 | 182385556 | 581 | 426 | 355 | [illegible] | [illegible] | [illegible] | [illegible] | [illegible] | [illegible] | [illegible] | [illegible] |
| A5SS | 5018 | ENSG00000112514 | CUTA | chr6 | - | 33417246 | 33417310 | 33417250 | 33417310 | 33417096 | 33417142 | 1 | 0 | 1 | [illegible] | [illegible] | [illegible] | [illegible] | [illegible] | [illegible] | [illegible] | [illegible] |
| A5SS | 5136 | ENSG00000162191 | UBXN1 | chr11 | - | 62677926 | 62678118 | 62677938 | 62678118 | 62677780 | 62677832 | 233 | 115 | 198 | 152 | 236 | 0 | 0 | 0 | 0 | 0 | 411 | 230 |
| A5SS | 5355 | ENSG00000084623 | EIF3I | chr1 | + | 32226170 | 32226320 | 32226170 | 32226283 | 32226402 | 32226530 | 299 | 160 | 287 | 285 | 280 | 0 | 0 | 0 | 0 | 0 | 398 | 316 |
| A5SS | 5529 | ENSG00000144895 | EIF2A | chr3 | + | 150567901 | 150568046 | 150567901 | 150567929 | 150568175 | 150568292 | 458 | 383 | 621 | 451 | 692 | 0 | 0 | 0 | 0 | 0 | 81 | 299 |
| A5SS | 5773 | ENSG00000184182 | UBE2F | chr2 | + | 238016565 | 238016633 | 238016565 | 238016630 | 238025341 | 238025412 | 327 | 190 | 195 | 296 | 249 | 0 | 0 | 0 | 0 | 0 | 389 | 243 |
| A5SS | 5870 | ENSG00000153094 | BCL2L11 | chr2 | + | 111123732 | 111124139 | 111123732 | 111123869 | 111164132 | 111168445 | 106 | 66 | 88 | 100 | 113 | 0 | 0 | 0 | 2 | 0 | 206 | 141 |
| A5SS | 6077 | ENSG00000204256 | BRD2 | chr6 | + | 32976561 | 32977570 | 32976561 | 32976936 | 32979827 | 32980132 | 100 | 95 | 96 | 149 | 69 | 0 | 0 | 0 | 0 | 0 | 558 | 135 |
| A5SS | 6526 | ENSG00000179818 | PCBP1-AS1 | chr2 | - | 70086054 | 70086273 | 70086123 | 70086273 | 70085815 | 70085903 | 10 | 8 | 21 | 6 | 0 | 0 | 0 | 1 | 1 | 0 | 11 | 10 |
| A5SS | 6642 | ENSG00000086730 | LAT2 | chr7 | + | 74216824 | 74216864 | 74216824 | 74216834 | 74219743 | 74219787 | 315 | 125 | 139 | 206 | 151 | 0 | 0 | 2 | 1 | 0 | 421 | 240 |
| A5SS | 6998 | ENSG00000005302 | MSL3 | chrX | + | 11759633 | 11759875 | 11759633 | 11759712 | 11760402 | 11760498 | 351 | 334 | 270 | 346 | 408 | 0 | 0 | 0 | 0 | 0 | 516 | 378 |
| A5SS | 7365 | ENSG00000035928 | RFC1 | chr4 | - | 39351347 | 39351476 | 39351431 | 39351476 | 39345400 | 39345476 | 126 | 119 | 142 | 153 | 128 | 0 | 0 | 0 | 2 | 0 | 31 | 168 |
| A5SS | 7409 | ENSG00000110077 | MS4A6A | chr11 | - | 60182977 | 60183331 | 60183135 | 60183331 | 60181580 | 60181741 | 1287 | 980 | 717 | 1011 | 2287 | 2 | 3 | 3 | 7 | 0 | 4392 | 1151 |
| A5SS | 7633 | ENSG00000100296 | THOC5 | chr22 | - | 29543414 | 29543542 | 29543428 | 29543542 | 29542858 | 29542956 | 0 | 0 | 0 | 0 | 2 | 61 | 58 | 75 | 78 | 102 | 0 | 0 |
| A5SS | 7680 | ENSG00000070061 | ELP1 | chr9 | - | 108881662 | 108881765 | 108881704 | 108881765 | 108880051 | 108880165 | 0 | 1 | 0 | 0 | 6 | 72 | 54 | 61 | 65 | 97 | 0 | 0 |
| A5SS | 8062 | ENSG00000101158 | NELFCD | chr20 | + | 58994109 | 58994239 | 58994109 | 58994191 | 58994641 | 58995133 | 118 | 75 | 111 | 137 | 99 | 1 | 0 | 1 | 0 | 0 | 156 | 139 |
| A5SS | 8384 | ENSG00000177239 | MAN1B1 | chr9 | + | 137086984 | 137087363 | 137086984 | 137087218 | 137088074 | 137088183 | 3 | 0 | 1 | 0 | 0 | 86 | 44 | 77 | 56 | 52 | 0 | 0 |
| A5SS | 8632 | ENSG00000179889 | PDXDC1 | chr16 | + | 15001775 | 15001887 | 15001775 | 15001856 | 15004186 | 15004333 | 0 | 0 | 1 | 2 | 0 | 88 | 55 | 105 | 76 | 66 | 0 | 0 |
| A5SS | 8882 | ENSG00000075415 | SLC25A3 | chr12 | + | 98593974 | 98595848 | 98593974 | 98594135 | 98597855 | 98598035 | 601 | 339 | 607 | 553 | 710 | 2 | 0 | 0 | 0 | 0 | 1016 | 725 |
| A5SS | 16 | ENSG00000184900 | SUMO3 | chr21 | - | 44808996 | 44809118 | 44809046 | 44809118 | 44805616 | 44807040 | 1 | 0 | 3 | 5 | 6 | 256 | 189 | 216 | 290 | 219 | 4 | 0 |
| A5SS | 100 | ENSG00000093000 | NUP50 | chr22 | + | 45170987 | 45171177 | 45170987 | 45171143 | 45171599 | 45171683 | 40 | 3 | 74 | 32 | 10 | 0 | 0 | 0 | 1 | 0 | 29 | 3 |
| A5SS | 228 | ENSG00000011600 | TYROBP | chr19 | - | 35907445 | 35907762 | 35907729 | 35907762 | 35907217 | 35907264 | 1018 | 343 | 430 | 615 | 412 | 0 | 0 | 0 | 1 | 0 | 1719 | 967 |
| A5SS | 298 | ENSG00000125755 | SYMPK | chr19 | - | 45827836 | 45829205 | 45828969 | 45829205 | 45827509 | 45827623 | 82 | 36 | 77 | 89 | 47 | 0 | 0 | 0 | 0 | 0 | 151 | 119 |
| A5SS | 359 | ENSG00000157540 | DYRK1A | chr21 | + | 37472683 | 37472880 | 37472683 | 37472829 | 37478207 | 37478300 | 426 | 281 | 419 | 484 | 532 | 0 | 0 | 0 | 0 | 3 | 244 | 458 |
| A5SS | 545 | ENSG00000101365 | IDH3B | chr20 | - | 2660456 | 2660829 | 2660696 | 2660829 | 2660262 | 2660365 | 142 | 76 | 161 | 203 | 292 | 0 | 0 | 0 | 0 | 0 | 284 | 204 |
| A5SS | 1130 | ENSG00000162066 | AMDHD2 | chr16 | + | 2528451 | 2528718 | 2528451 | 2528559 | 2528993 | 2529095 | 22 | 2 | 16 | 6 | 0 | 0 | 0 | 1 | 0 | 0 | 30 | 17 |
| A5SS | 1237 | ENSG00000090861 | AARS1 | chr16 | - | 70255719 | 70255836 | 70255727 | 70255836 | 70254620 | 70254734 | 0 | 0 | 0 | 1 | 0 | 46 | 31 | 35 | 46 | 48 | 0 | 0 |
| A5SS | 1839 | ENSG00000103811 | CTSH | chr15 | - | 78932582 | 78935082 | 78933730 | 78935082 | 78932371 | 78932458 | 14 | 14 | 15 | 31 | 12 | 1 | 0 | 0 | 0 | 0 | 53 | 32 |
| A5SS | 1937 | ENSG00000135506 | OS9 | chr12 | + | 57715759 | 57716193 | 57715759 | 57715970 | 57716411 | 57716512 | 193 | 148 | 192 | 266 | 152 | 0 | 0 | 0 | 2 | 0 | 760 | 448 |
| A5SS | 1986 | ENSG00000188994 | ZNF292 | chr6 | + | 87215902 | 87216057 | 87215902 | 87215962 | 87216298 | 87216377 | 274 | 172 | 171 | 232 | 356 | 0 | 0 | 0 | 0 | 0 | 285 | 157 |
| A5SS | 1998 | ENSG00000068796 | KIF2A | chr5 | + | 62361465 | 62362541 | 62361465 | 62361529 | 62363177 | 62363320 | 905 | 1190 | 633 | 1465 | 1710 | 0 | 0 | 0 | 0 | 0 | 550 | 728 |
| A5SS | 2598 | ENSG00000141562 | NARF | chr17 | + | 82483715 | 82484950 | 82483715 | 82483779 | 82485496 | 82485654 | 124 | 64 | 94 | 101 | 52 | 0 | 0 | 0 | 0 | 0 | 338 | 186 |
| A5SS | 2873 | ENSG00000172053 | QARS1 | chr3 | - | 49098352 | 49098480 | 49098372 | 49098480 | 49098191 | 49098258 | 141 | 90 | 126 | 151 | 131 | 0 | 0 | 0 | 0 | 0 | 148 | 126 |
| A5SS | 3072 | ENSG00000111642 | CHD4 | chr12 | - | 6577784 | 6578546 | 6578408 | 6578546 | 6573073 | 6573269 | 235 | 172 | 269 | 266 | 396 | 0 | 0 | 0 | 0 | 0 | 488 | 231 |
| A5SS | 3518 | ENSG00000089327 | FXYD5 | chr19 | + | 35166141 | 35166325 | 35166141 | 35166171 | 35169565 | 35169880 | 1154 | 620 | 1112 | 971 | 843 | 4 | 0 | 0 | 2 | 2 | 1612 | 1152 |
| A5SS | 3601 | ENSG00000125637 | PSD4 | chr2 | + | 113185364 | 113186255 | 113185364 | 113185440 | 113193047 | 113193128 | 23 | 76 | 57 | 109 | 33 | 0 | 0 | 0 | 0 | 0 | 183 | 110 |
| A5SS | 3644 | ENSG00000160789 | LMNA | chr1 | + | 156134402 | 156134528 | 156134402 | 156134523 | 156134804 | 156134975 | 37 | 57 | 44 | 79 | 28 | 0 | 0 | 0 | 0 | 0 | 230 | 74 |
| A5SS | 3795 | ENSG00000117335 | CD46 | chr1 | + | 207752067 | 207752309 | 207752067 | 207752097 | 207757013 | 207757202 | 301 | 160 | 243 | 305 | 119 | 2 | 0 | 1 | 0 | 0 | 63 | 186 |
| A5SS | 3872 | ENSG00000248932 | NA | chr3 | + | 139389875 | 139390429 | 139389875 | 139390247 | 139419556 | 139419680 | 6 | 0 | 3 | 0 | 0 | 1 | 2 | 1 | 0 | 2 | 8 | 0 |
| A5SS | 3890 | ENSG00000141556 | TBCD | chr17 | + | 82932657 | 82932855 | 82932657 | 82932735 | 82937270 | 82937360 | 1 | 1 | 5 | 3 | 6 | 22 | 7 | 15 | 15 | 11 | 0 | 2 |
| A5SS | 4025 | ENSG00000138095 | LRPPRC | chr2 | - | 43934187 | 43934296 | 43934189 | 43934296 | 43925892 | 43925961 | 1 | 0 | 0 | 4 | 0 | 279 | 206 | 279 | 258 | 339 | 0 | 0 |
| A5SS | 4123 | ENSG00000116337 | AMPD2 | chr1 | + | 109630232 | 109630439 | 109630232 | 109630406 | 109630682 | 109630793 | 2 | 0 | 0 | 0 | 2 | 103 | 67 | 88 | 86 | 30 | 0 | 1 |
| A5SS | 4157 | ENSG00000179632 | MAF1 | chr8 | + | 144106342 | 144106674 | 144106342 | 144106464 | 144106834 | 144106963 | 425 | 190 | 359 | 394 | 346 | 0 | 0 | 0 | 0 | 0 | 731 | 621 |
| A5SS | 4752 | ENSG00000089127 | OAS1 | chr12 | + | 112906961 | 112908824 | 112906961 | 112907219 | 112911050 | 112911235 | 128 | 109 | 77 | 140 | 90 | 2 | 0 | 0 | 0 | 0 | 494 | 252 |
| A5SS | 5109 | ENSG00000151092 | NGLY1 | chr3 | - | 25782892 | 25783414 | 25783259 | 25783414 | 25778573 | 25778688 | 2 | 0 | 0 | 0 | 0 | 55 | 30 | 59 | 71 | 88 | 0 | 0 |
| A5SS | 5215 | ENSG00000104907 | TRMT1 | chr19 | - | 13115996 | 13116431 | 13116145 | 13116431 | 13115625 | 13115768 | 81 | 72 | 73 | 98 | 95 | 0 | 0 | 0 | 0 | 0 | 162 | 69 |
| A5SS | 5362 | ENSG00000005483 | KMT2E | chr7 | + | 105105858 | 105106003 | 105105858 | 105105955 | 105106521 | 105106772 | 477 | 476 | 436 | 531 | 1110 | 1 | 1 | 1 | 2 | 1 | 61 | 375 |
| A5SS | 5936 | ENSG00000124356 | STAMBP | chr2 | + | 73849362 | 73849578 | 73849362 | 73849487 | 73850375 | 73850513 | 1 | 0 | 0 | 0 | 6 | 119 | 64 | 77 | 106 | 156 | 0 | 0 |
| A5SS | 5975 | ENSG00000120896 | SORBS3 | chr8 | + | 22561334 | 22561536 | 22561334 | 22561373 | 22561864 | 22561931 | 0 | 0 | 0 | 1 | 0 | 30 | 16 | 13 | 24 | 4 | 0 | 0 |
| A5SS | 6299 | ENSG00000214021 | TTLL3 | chr3 | + | 9817644 | 9820741 | 9817644 | 9817759 | 9826996 | 9827240 | 71 | 55 | 54 | 70 | 45 | 0 | 0 | 0 | 2 | 0 | 141 | 77 |
| A5SS | 6427 | ENSG00000130021 | PUDP | chrX | - | 7105619 | 7105838 | 7105748 | 7105838 | 7077219 | 7077449 | 125 | 126 | 142 | 108 | 140 | 0 | 0 | 1 | 0 | 0 | 289 | 126 |

[illegible]

| | | | | | | | | | | | | | | | | | | | | | | | | | | | | | | | | | | | | | | | | | | | | | | |
|---|---|---|---|---|---|---|---|---|---|---|---|---|---|---|---|---|---|---|---|---|---|---|---|---|---|---|---|---|---|---|---|---|---|---|---|---|---|---|---|---|---|---|---|---|---|---|
| A5SS | 6359 | ENSG00000088038 | CNOT3 | chr19 | + | 54143659 | 54144134 | 54143659 | 54143749 | 54144236 | 54144332 | 182 | 159 | 153 | 208 | 227 | 0 | 0 | 0 | 0 | 2 | 330 | 225 | 186 | 166 | 286 | 0 | 8 | 0 | 0 | 0 | 298 | 149 | 1 | 1 | 1 | 1 | 1 | 1 | 0.983 | 1 | 0.934 | 1 | 1 | 1 | 0.01 |
| A5SS | 6746 | ENSG00000196305 | IARS1 | chr9 | - | 92269877 | 92269983 | 92269884 | 92269983 | 92268173 | 92268300 | 0 | 4 | 2 | 3 | 0 | 161 | 93 | 144 | 124 | 141 | 0 | 0 | 3 | 0 | 0 | 71 | 98 | 108 | 114 | 32 | 156 | 149 | 1 | 1 | 0 | 0.039 | 0.013 | 0.023 | 0 | 0 | 0 | 0.026 | 0 | 0 | 0.01 |
| A5SS | 6886 | ENSG00000130734 | ATG4D | chr19 | + | 10546838 | 10547253 | 10546838 | 10547115 | 10548903 | 10549034 | 26 | 8 | 28 | 5 | 13 | 0 | 0 | 2 | 0 | 0 | 0 | 38 | 37 | 24 | 43 | 0 | 0 | 3 | 0 | 0 | 287 | 149 | 1 | 1 | 1 | 1 | 0.879 | 1 | 1 | NA | 1 | 0.865 | 1 | 1 | 0.01 |
| A5SS | 7425 | ENSG00000104763 | ASAH1 | chr8 | - | 18083916 | 18084073 | 18083980 | 18084073 | 18075540 | 18075587 | 1 | 0 | 1 | 3 | 0 | 114 | 47 | 79 | 59 | 24 | 0 | 0 | 0 | 0 | 0 | 13 | 73 | 89 | 95 | 85 | 213 | 149 | 1 | 1 | 0.006 | 0 | 0.009 | 0.034 | 0 | 0 | 0 | 0 | 0 | 0 | 0.01 |
| A5SS | 8279 | ENSG00000107537 | PHYH | chr10 | - | 13291824 | 13291912 | 13291830 | 13291912 | 13288359 | 13288541 | 0 | 0 | 0 | 1 | 0 | 28 | 23 | 30 | 19 | 43 | 0 | 0 | 0 | 0 | 0 | 43 | 23 | 29 | 11 | 6 | 155 | 149 | 1 | 1 | 0 | 0 | 0 | 0.048 | 0 | 0 | 0 | 0 | 0 | 0 | 0.01 |
| A5SS | 8455 | ENSG00000107862 | GBF1 | chr10 | + | 102362474 | 102363396 | 102362474 | 102362628 | 102363709 | 102363798 | 180 | 133 | 105 | 182 | 162 | 0 | 0 | 0 | 0 | 0 | 372 | 193 | 139 | 192 | 428 | 0 | 3 | 0 | 2 | 0 | 298 | 149 | 1 | 1 | 1 | 1 | 1 | 1 | 1 | 1 | 0.97 | 1 | 0.98 | 1 | 0.01 |
| A5SS | 8891 | ENSG00000110851 | PRDM4 | chr12 | - | 107751414 | 107752209 | 107751646 | 107752209 | 107746274 | 107746424 | 86 | 53 | 82 | 119 | 117 | 0 | 0 | 0 | 1 | 0 | 68 | 122 | 103 | 70 | 81 | 0 | 2 | 2 | 0 | 0 | 298 | 149 | 1 | 1 | 1 | 1 | 1 | 0.983 | 1 | 1 | 0.968 | 0.963 | 1 | 1 | 0.01 |
| A5SS | 31 | ENSG00000205930 | C21orf62-AS1 | chr21 | + | 32772160 | 32772320 | 32772160 | 32772308 | 32784796 | 32784919 | 2 | 1 | 10 | 7 | 1 | 3 | 12 | 9 | 9 | 26 | 0 | 1 | 1 | 12 | 0 | 0 | 8 | 1 | 11 | 22 | 161 | 149 | 1 | 1 | 0.382 | 0.072 | 0.507 | 0.419 | 0.034 | NA | 0.104 | 0.481 | 0.502 | 0 | 0.011 |
| A5SS | 175 | ENSG00000105618 | PRPF31 | chr19 | + | 54128072 | 54128377 | 54128072 | 54128200 | 54129056 | 54129370 | 84 | 37 | 83 | 84 | 60 | 1 | 0 | 0 | 0 | 0 | 153 | 92 | 50 | 93 | 213 | 0 | 0 | 0 | 4 | 0 | 298 | 149 | 1 | 1 | 0.977 | 1 | 1 | 1 | 1 | 1 | 1 | 1 | 0.921 | 1 | 0.011 |
| A5SS | 200 | ENSG00000196684 | HSH2D | chr19 | + | 16148723 | 16148875 | 16148723 | 16148800 | 16152551 | 16152641 | 78 | 29 | 59 | 71 | 11 | 0 | 0 | 0 | 0 | 0 | 97 | 81 | 87 | 127 | 130 | 0 | 0 | 0 | 5 | 0 | 224 | 149 | 1 | 1 | 1 | 1 | 1 | 1 | 1 | 1 | 1 | 1 | 0.944 | 1 | 0.011 |
| A5SS | 314 | ENSG00000129347 | KRI1 | chr19 | - | 10557551 | 10557895 | 10557768 | 10557895 | 10555284 | 10555349 | 82 | 62 | 129 | 97 | 60 | 0 | 0 | 0 | 0 | 0 | 77 | 87 | 36 | 72 | 158 | 0 | 0 | 0 | 2 | 0 | 298 | 149 | 1 | 1 | 1 | 1 | 1 | 1 | 1 | 1 | 1 | 1 | 0.947 | 1 | 0.011 |
| A5SS | 615 | ENSG00000124155 | PIGT | chr20 | + | 45420527 | 45421583 | 45420527 | 45420693 | 45424215 | 45424381 | 66 | 28 | 51 | 66 | 54 | 0 | 0 | 0 | 0 | 0 | 63 | 70 | 85 | 63 | 103 | 0 | 2 | 0 | 0 | 0 | 298 | 149 | 1 | 1 | 1 | 1 | 1 | 1 | 1 | 1 | 0.946 | 1 | 1 | 1 | 0.011 |
| A5SS | 967 | ENSG00000108406 | DHX40 | chr17 | + | 59566626 | 59566794 | 59566626 | 59566709 | 59573115 | 59573235 | 62 | 43 | 81 | 71 | 37 | 0 | 0 | 0 | 0 | 0 | 70 | 96 | 67 | 85 | 10 | 0 | 0 | 0 | 3 | 0 | 234 | 149 | 1 | 1 | 1 | 1 | 1 | 1 | 1 | 1 | 1 | 1 | 0.947 | 1 | 0.011 |
| A5SS | 1294 | ENSG00000118900 | UBN1 | chr16 | + | 4860663 | 4861102 | 4860663 | 4860953 | 4868832 | 4868903 | 191 | 105 | 129 | 134 | 110 | 1 | 0 | 0 | 0 | 0 | 254 | 212 | 192 | 273 | 423 | 0 | 0 | 3 | 5 | 0 | 298 | 149 | 1 | 1 | 0.99 | 1 | 1 | 1 | 1 | 1 | 1 | 0.97 | 0.965 | 1 | 0.011 |
| A5SS | 1659 | ENSG00000197102 | DYNC1H1 | chr14 | + | 102040597 | 102041734 | 102040597 | 102040673 | 102042012 | 102042124 | 81 | 59 | 87 | 100 | 75 | 0 | 0 | 0 | 0 | 0 | 84 | 110 | 66 | 102 | 68 | 0 | 1 | 0 | 2 | 0 | 298 | 149 | 1 | 1 | 1 | 1 | 1 | 1 | 1 | 1 | 0.982 | 1 | 0.962 | 1 | 0.011 |
| A5SS | 2268 | ENSG00000184584 | STING1 | chr5 | - | 139481158 | 139482302 | 139482209 | 139482302 | 139480789 | 139480898 | 87 | 41 | 88 | 65 | 35 | 1 | 0 | 0 | 0 | 0 | 149 | 100 | 66 | 72 | 160 | 0 | 0 | 0 | 3 | 0 | 298 | 149 | 1 | 1 | 0.978 | 1 | 1 | 1 | 1 | 1 | 1 | 1 | 0.923 | 1 | 0.011 |
| A5SS | 2636 | ENSG00000196209 | SIRPB2 | chr20 | - | 1479699 | 1480065 | 1479993 | 1480065 | 1473791 | 1473870 | 94 | 72 | 17 | 62 | 12 | 0 | 0 | 0 | 0 | 0 | 57 | 24 | 49 | 104 | 72 | 0 | 0 | 0 | 3 | 0 | 298 | 149 | 1 | 1 | 1 | 1 | 1 | 1 | 1 | 1 | 1 | 1 | 0.945 | 1 | 0.011 |
| A5SS | 2791 | ENSG00000156931 | VPS8 | chr3 | + | 184839697 | 184839758 | 184839697 | 184839752 | 184849070 | 184849195 | 0 | 0 | 1 | 0 | 0 | 42 | 29 | 17 | 42 | 44 | 0 | 0 | 0 | 0 | 0 | 18 | 80 | 22 | 34 | 10 | 155 | 149 | 1 | 1 | 0 | 0 | 0.054 | 0 | 0 | 0 | 0 | 0 | 0 | 0 | 0.011 |
| A5SS | 2837 | ENSG00000058056 | USP13 | chr3 | + | 179719939 | 179720039 | 179719939 | 179720034 | 179721401 | 179721589 | 0 | 0 | 1 | 0 | 0 | 17 | 7 | 16 | 17 | 25 | 0 | 0 | 0 | 0 | 0 | 10 | 10 | 14 | 6 | 36 | 154 | 149 | 1 | 1 | 0 | 0 | 0.057 | 0 | 0 | 0 | 0 | 0 | 0 | 0 | 0.011 |
| A5SS | 3986 | ENSG00000169398 | PTK2 | chr8 | - | 140675429 | 140675499 | 140675459 | 140675499 | 140674297 | 140674404 | 0 | 2 | 0 | 0 | 2 | 18 | 35 | 19 | 43 | 40 | 0 | 1 | 0 | 0 | 0 | 61 | 26 | 34 | 41 | 35 | 179 | 149 | 1 | 1 | 0 | 0.045 | 0 | 0 | 0.04 | 0 | 0.031 | 0 | 0 | 0 | 0.011 |
| A5SS | 4387 | ENSG00000145020 | AMT | chr3 | - | 49419709 | 49420342 | 49420210 | 49420342 | 49418970 | 49419405 | 26 | 12 | 22 | 27 | 16 | 0 | 0 | 0 | 0 | 0 | 31 | 29 | 43 | 34 | 68 | 0 | 0 | 0 | 0 | 2 | 298 | 149 | 1 | 1 | 1 | 1 | 1 | 1 | 1 | 1 | 1 | 1 | 1 | 0.944 | 0.011 |
| A5SS | 4905 | ENSG00000116984 | MTR | chr1 | + | 236886291 | 236889336 | 236886291 | 236886367 | 236891132 | 236891329 | 46 | 41 | 53 | 44 | 36 | 0 | 2 | 0 | 0 | 0 | 41 | 41 | 47 | 36 | 34 | 0 | 0 | 4 | 0 | 0 | 298 | 149 | 1 | 1 | 1 | 0.911 | 1 | 1 | 1 | 1 | 1 | 0.855 | 1 | 1 | 0.011 |
| A5SS | 5129 | ENSG00000112855 | HARS2 | chr5 | + | 140691794 | 140692329 | 140691794 | 140692017 | 140693590 | 140693665 | 16 | 8 | 11 | 22 | 35 | 0 | 0 | 0 | 0 | 2 | 43 | 42 | 28 | 16 | 2 | 0 | 4 | 0 | 0 | 0 | 298 | 149 | 1 | 1 | 1 | 1 | 1 | 1 | 0.897 | 1 | 0.84 | 1 | 1 | 1 | 0.011 |
| A5SS | 5265 | ENSG00000125875 | TBC1D20 | chr20 | - | 441587 | 441689 | 441619 | 441689 | 440247 | 440389 | 117 | 72 | 103 | 156 | 136 | 0 | 0 | 0 | 1 | 2 | 156 | 108 | 57 | 115 | 174 | 0 | 0 | 4 | 0 | 0 | 181 | 149 | 1 | 1 | 1 | 1 | 1 | 0.992 | 0.982 | 1 | 1 | 0.921 | 1 | 1 | 0.011 |
| A5SS | 5469 | ENSG00000082153 | BZW1 | chr2 | + | 200811545 | 200811946 | 200811545 | 200811727 | 200813207 | 200813281 | 7 | 2 | 5 | 6 | 7 | 20 | 7 | 3 | 5 | 2 | 0 | 2 | 8 | 6 | 0 | 5 | 11 | 11 | 0 | 0 | 298 | 149 | 1 | 1 | 0.149 | 0.125 | 0.455 | 0.375 | 0.636 | 0 | 0.083 | 0.267 | 1 | NA | 0.011 |
| A5SS | 5705 | ENSG00000178719 | GRINA | chr8 | + | 143991877 | 143992373 | 143991877 | 143992078 | 143992464 | 143992608 | 114 | 52 | 81 | 71 | 44 | 0 | 0 | 0 | 0 | 0 | 187 | 170 | 124 | 137 | 234 | 0 | 5 | 0 | 0 | 0 | 298 | 149 | 1 | 1 | 1 | 1 | 1 | 1 | 1 | 1 | 0.944 | 1 | 1 | 1 | 0.011 |
| A5SS | 7145 | ENSG00000152332 | UHMK1 | chr1 | + | 162499954 | 162501104 | 162499954 | 162500247 | 162512499 | 162512576 | 45 | 27 | 59 | 43 | 34 | 0 | 0 | 0 | 0 | 0 | 50 | 75 | 112 | 72 | 26 | 0 | 0 | 0 | 2 | 0 | 298 | 149 | 1 | 1 | 1 | 1 | 1 | 1 | 1 | 1 | 1 | 1 | 0.947 | 1 | 0.011 |
| A5SS | 7613 | ENSG00000168003 | SLC3A2 | chr11 | + | 62881892 | 62882999 | 62881892 | 62882066 | 62884456 | 62884525 | 177 | 103 | 115 | 162 | 209 | 0 | 0 | 0 | 1 | 0 | 223 | 155 | 137 | 251 | 331 | 0 | 2 | 2 | 2 | 0 | 298 | 149 | 1 | 1 | 1 | 1 | 1 | 0.988 | 1 | 1 | 0.975 | 0.972 | 0.984 | 1 | 0.011 |

[illegible]

| | | | | | | | | | | | | | | | | |
|---|---|---|---|---|---|---|---|---|---|---|---|---|---|---|---|---|
| A5SS | 2909 | ENSG00000175455 | CCDC14 | chr3 | - | 123955842 | 123956115 | 123956045 | 123956115 | 123948895 | 123949132 | 124 | 118 | 99 | 99 | 231 |
| A5SS | 3873 | ENSG00000164010 | ERMAP | chr1 | + | 42835037 | 42835165 | 42835037 | 42835154 | 42835731 | 42835760 | 0 | 0 | 0 | 0 | 4 |
| A5SS | 7289 | ENSG00000167904 | TMEM68 | chr8 | - | 55773226 | 55773378 | 55773268 | 55773378 | 55763935 | 55763980 | 0 | 1 | 0 | 1 | 0 |
| A5SS | 7667 | ENSG00000132740 | IGHMBP2 | chr11 | + | 68914822 | 68915023 | 68914822 | 68915017 | 68917735 | 68917883 | 20 | 26 | 18 | 18 | 14 |
| A5SS | 8242 | ENSG00000133895 | MEN1 | chr11 | - | 64809649 | 64810132 | 64809664 | 64810132 | 64807890 | 64808099 | 1 | 1 | 0 | 1 | 0 |
| A5SS | 7439 | ENSG00000185803 | SLC52A2 | chr8 | + | 144359187 | 144359521 | 144359187 | 144359423 | 144359622 | 144360493 | 1 | 0 | 0 | 2 | 0 |
| A5SS | 290 | ENSG00000198521 | ZNF43 | chr19 | - | 21845391 | 21845506 | 21845429 | 21845506 | 21819094 | 21819221 | 5 | 8 | 8 | 13 | 16 |
| A5SS | 1109 | ENSG00000086504 | MRPL28 | chr16 | - | 370484 | 370509 | 370498 | 370509 | 369930 | 370225 | 4 | 0 | 0 | 2 | 0 |
| A5SS | 1481 | ENSG00000182718 | ANXA2 | chr15 | - | 60396279 | 60396427 | 60396372 | 60396427 | 60390212 | 60390317 | 11 | 8 | 4 | 13 | 12 |
| A5SS | 1539 | ENSG00000168904 | LRRC28 | chr15 | + | 99255897 | 99256125 | 99255897 | 99255973 | 99287256 | 99287294 | 12 | 21 | 22 | 24 | 18 |
| A5SS | 6103 | ENSG00000135678 | CPM | chr12 | - | 68933141 | 68933202 | 68933159 | 68933202 | 68932677 | 68932840 | 5 | 0 | 1 | 5 | 0 |
| A5SS | 1172 | ENSG00000168701 | TMEM208 | chr16 | + | 67228796 | 67228881 | 67228796 | 67228836 | 67228975 | 67229278 | 36 | 29 | 45 | 35 | 12 |
| A5SS | 2179 | ENSG00000205309 | NT5M | chr17 | + | 17304363 | 17304481 | 17304363 | 17304477 | 17306542 | 17306643 | 0 | 0 | 0 | 1 | 0 |
| A5SS | 2781 | ENSG00000242086 | MUC20-OT1 | chr3 | + | 195708153 | 195708513 | 195708153 | 195708435 | 195708746 | 195708841 | 2 | 2 | 3 | 0 | 3 |
| A5SS | 3701 | ENSG00000182923 | CEP63 | chr3 | + | 134559149 | 134559429 | 134559149 | 134559305 | 134561376 | 134565031 | 13 | 16 | 3 | 6 | 27 |
| A5SS | 7578 | ENSG00000113328 | CCNG1 | chr5 | + | 163437594 | 163437804 | 163437594 | 163437750 | 163439256 | 163439520 | 42 | 7 | 36 | 25 | 32 |
| A5SS | 2353 | ENSG00000158290 | CUL4B | chrX | - | 120561132 | 120561453 | 120561371 | 120561453 | 120560082 | 120560625 | 0 | 0 | 0 | 1 | 0 |
| A5SS | 2562 | ENSG00000156876 | SASS6 | chr1 | - | 100107627 | 100108004 | 100107809 | 100108004 | 100107373 | 100107553 | 93 | 76 | 70 | 80 | 86 |
| A5SS | 3700 | ENSG00000182923 | CEP63 | chr3 | + | 134547334 | 134547472 | 134547334 | 134547468 | 134549061 | 134549176 | 30 | 41 | 27 | 56 | 54 |
| A5SS | 4232 | ENSG00000132781 | MUTYH | chr1 | - | 45331660 | 45332086 | 45332022 | 45332086 | 45331419 | 45331556 | 12 | 14 | 20 | 12 | 13 |
| A5SS | 6610 | ENSG00000133195 | SLC39A11 | chr17 | - | 72947527 | 72947875 | 72947730 | 72947875 | 72849633 | 72849804 | 4 | 2 | 2 | 6 | 2 |
| A5SS | 6901 | ENSG00000156970 | BUB1B | chr15 | + | 40170061 | 40170163 | 40170061 | 40170121 | 40170536 | 40170681 | 1 | 0 | 0 | 0 | 0 |
| A5SS | 4629 | ENSG00000171960 | PPIH | chr1 | + | 42659227 | 42659370 | 42659227 | 42659251 | 42660861 | 42660904 | 12 | 15 | 5 | 17 | 5 |
| A5SS | 6502 | ENSG00000006534 | ALDH3B1 | chr11 | + | 68015296 | 68018637 | 68015296 | 68015459 | 68018772 | 68018893 | 33 | 12 | 15 | 24 | 11 |
| A5SS | 515 | ENSG00000125885 | MCM8 | chr20 | + | 5950651 | 5951077 | 5950651 | 5951023 | 5952010 | 5952163 | 4 | 0 | 2 | 3 | 2 |
| A5SS | 2674 | ENSG00000102796 | DHRS12 | chr13 | - | 51773920 | 51774034 | 51773929 | 51774034 | 51769155 | 51769293 | 4 | 1 | 2 | 0 | 5 |
| A5SS | 3208 | ENSG00000163681 | SLMAP | chr3 | + | 57831382 | 57831536 | 57831382 | 57831530 | 57841298 | 57841371 | 8 | 1 | 5 | 6 | 8 |
| A5SS | 3427 | ENSG00000112294 | ALDH5A1 | chr6 | + | 24502522 | 24502763 | 24502522 | 24502606 | 24503262 | 24503433 | 2 | 1 | 2 | 2 | 1 |
| A5SS | 6948 | ENSG00000155966 | AFF2 | chrX | + | 148843381 | 148843433 | 148843381 | 148843415 | 148885888 | 148885985 | 0 | 0 | 0 | 3 | 0 |
| A5SS | 5557 | ENSG00000168385 | SEPTIN2 | chr2 | + | 241316421 | 241316555 | 241316421 | 241316540 | 241317498 | 241317599 | 1 | 1 | 3 | 0 | 0 |
| A5SS | 7817 | ENSG00000148396 | SEC16A | chr9 | - | 136463585 | 136463740 | 136463678 | 136463740 | 136461176 | 136461274 | 11 | 2 | 3 | 0 | 0 |
| A5SS | 2303 | ENSG00000118965 | WDR35 | chr2 | - | 19935466 | 19935603 | 19935470 | 19935603 | 19933400 | 19933511 | 0 | 0 | 3 | 1 | 0 |
| A5SS | 197 | ENSG00000214046 | SMIM7 | chr19 | - | 16647095 | 16647261 | 16647119 | 16647261 | 16631376 | 16631664 | 11 | 4 | 9 | 16 | 2 |
| A5SS | 1485 | ENSG00000185880 | TRIM69 | chr15 | + | 44756367 | 44758854 | 44756367 | 44756463 | 44759639 | 44759662 | 7 | 6 | 8 | 12 | 25 |
| A5SS | 1604 | ENSG00000127586 | CHTF18 | chr16 | + | 790524 | 792367 | 790524 | 792347 | 792438 | 792590 | 1 | 0 | 0 | 0 | 0 |
| A5SS | 8119 | ENSG00000149743 | TRPT1 | chr11 | - | 64224794 | 64224970 | 64224800 | 64224970 | 64224542 | 64224717 | 0 | 1 | 0 | 1 | 0 |
| A5SS | 8398 | ENSG00000148498 | PARD3 | chr10 | - | 34347955 | 34348115 | 34347964 | 34348115 | 34346197 | 34346449 | 1 | 0 | 0 | 2 | 0 |
| A5SS | 8574 | ENSG00000284770 | TBCE | chr1 | + | 235367412 | 235367552 | 235367412 | 235367504 | 235380018 | 235380149 | 1 | 0 | 0 | 4 | 0 |
| A5SS | 2836 | ENSG00000058056 | USP13 | chr3 | + | 179701007 | 179701179 | 179701007 | 179701129 | 179706933 | 179707076 | 0 | 1 | 0 | 8 | 0 |
| A5SS | 453 | ENSG00000174306 | ZHX3 | chr20 | - | 41268978 | 41269083 | 41268989 | 41269083 | 41202056 | 41205066 | 0 | 1 | 0 | 0 | 0 |
| A5SS | 611 | ENSG00000124155 | PIGT | chr20 | + | 45416516 | 45416694 | 45416516 | 45416654 | 45419294 | 45419395 | 30 | 11 | 21 | 44 | 35 |
| A5SS | 6719 | ENSG00000122786 | CALD1 | chr7 | + | 134932987 | 134934077 | 134932987 | 134933390 | 134941091 | 134941237 | 0 | 1 | 0 | 1 | 0 |
| A5SS | 8267 | ENSG00000150433 | TMEM218 | chr11 | - | 125102636 | 125102809 | 125102733 | 125102809 | 125101443 | 125101521 | 16 | 10 | 10 | 12 | 6 |
| A5SS | 92 | ENSG00000100395 | L3MBTL2 | chr22 | + | 41217122 | 41219536 | 41217122 | 41217202 | 41220733 | 41220868 | 55 | 20 | 45 | 64 | 48 |
| A5SS | 6045 | ENSG00000162585 | FAAP20 | chr1 | - | 2198753 | 2201225 | 2201049 | 2201225 | 2198006 | 2198142 | 4 | 1 | 0 | 1 | 0 |
| A5SS | 2342 | ENSG00000150756 | ATPSCKMT | chr5 | - | 10239041 | 10239356 | 10239066 | 10239356 | 10236477 | 10236615 | 5 | 0 | 1 | 2 | 0 |
| A5SS | 8538 | ENSG00000171792 | RHNO1 | chr12 | + | 2885282 | 2885534 | 2885282 | 2885492 | 2886977 | 2887111 | 4 | 2 | 14 | 3 | 4 |
| A5SS | 1066 | ENSG00000213918 | DNASE1 | chr16 | + | 3655848 | 3655966 | 3655848 | 3655937 | 3656101 | 3656185 | 0 | 0 | 2 | 0 | 0 |
| A5SS | 1563 | ENSG00000255571 | MIR9-3HG | chr15 | + | 89387816 | 89389482 | 89387816 | 89387881 | 89395027 | 89398484 | 17 | 7 | 12 | 6 | 7 |
| A5SS | 1752 | ENSG00000186648 | CARMIL3 | chr14 | + | 24064957 | 24065750 | 24064957 | 24065273 | 24066397 | 24066464 | 3 | 0 | 0 | 0 | 1 |
| A5SS | 5216 | ENSG00000104907 | TRMT1 | chr19 | - | 13116105 | 13116431 | 13116145 | 13116431 | 13115625 | 13115768 | 1 | 0 | 3 | 2 | 0 |
| A5SS | 7859 | ENSG00000142731 | PLK4 | chr4 | + | 127885707 | 127886728 | 127885707 | 127886626 | 127887395 | 127887496 | 16 | 9 | 19 | 18 | 53 |
| A5SS | 8212 | ENSG00000179532 | DNHD1 | chr11 | + | 6533680 | 6534173 | 6533680 | 6533806 | 6538382 | 6538510 | 13 | 7 | 5 | 4 | 8 |
| A5SS | 8216 | ENSG00000179532 | DNHD1 | chr11 | + | 6570008 | 6571423 | 6570008 | 6570396 | 6571635 | 6572027 | 1 | 1 | 4 | 8 | 4 |
| A5SS | 819 | ENSG00000108298 | RPL19 | chr17 | + | 39202316 | 39202448 | 39202316 | 39202443 | 39202999 | 39203109 | 30 | 11 | 32 | 16 | 28 |
| A5SS | 2102 | ENSG00000145604 | SKP2 | chr5 | + | 36163644 | 36164695 | 36163644 | 36163756 | 36166518 | 36166662 | 4 | 4 | 2 | 8 | 4 |
| A5SS | 4556 | ENSG00000115590 | IL1R2 | chr2 | + | 101991959 | 101992033 | 101991959 | 101992011 | 102008514 | 102008642 | 0 | 0 | 1 | 0 | 1 |
| A5SS | 5996 | ENSG00000067248 | DHX29 | chr5 | - | 55283115 | 55283811 | 55283202 | 55283811 | 55281371 | 55281515 | 7 | 1 | 5 | 7 | 14 |
| A5SS | 8297 | ENSG00000168283 | BMI1 | chr10 | + | 22326430 | 22326986 | 22326430 | 22326561 | 22327949 | 22328058 | 22 | 10 | 19 | 27 | 30 |
| A5SS | 8845 | ENSG00000125629 | INSIG2 | chr2 | + | 118096418 | 118096800 | 118096418 | 118096426 | 118103196 | 118103321 | 47 | 36 | 38 | 60 | 56 |
| A5SS | 8887 | ENSG00000111181 | SLC6A12 | chr12 | - | 209772 | 210043 | 209890 | 210043 | 204563 | 204698 | 26 | 6 | 8 | 3 | 3 |
| A5SS | 4701 | ENSG00000135913 | USP37 | chr2 | - | 218562672 | 218562813 | 218562717 | 218562813 | 218558497 | 218558677 | 8 | 4 | 3 | 8 | 3 |
| A5SS | 8496 | ENSG00000055950 | MRPL43 | chr10 | - | 100986686 | 100986975 | 100986748 | 100986975 | 100983754 | 100983817 | 16 | 4 | 8 | 1 | 4 |
| A5SS | 259 | ENSG00000102575 | ACP5 | chr19 | - | 11578479 | 11578657 | 11578584 | 11578657 | 11577592 | 11577679 | 0 | 0 | 1 | 7 | 0 |
| A5SS | 2942 | ENSG00000109065 | NAT9 | chr17 | - | 74775598 | 74775707 | 74775621 | 74775707 | 74773575 | 74773685 | 5 | 1 | 7 | 4 | 0 |
| A5SS | 5412 | ENSG00000205702 | CYP2D7 | chr22 | - | 42141511 | 42141675 | 42141624 | 42141675 | 42141151 | 42141339 | 1 | 1 | 3 | 3 | 0 |
| A5SS | 7511 | ENSG00000283239 | KBTBD11-OT1 | chr8 | + | 1869193 | 1869316 | 1869193 | 1869250 | 1876570 | 1876734 | 0 | 0 | 8 | 1 | 0 |
| A5SS | 3837 | ENSG00000158488 | CD1E | chr1 | + | 158355826 | 158356105 | 158355826 | 158356001 | 158356497 | 158356591 | 3 | 5 | 3 | 5 | 4 |
| A5SS | 6325 | ENSG00000106638 | TBL2 | chr7 | - | 73573937 | 73574122 | 73573960 | 73574122 | 73573319 | 73573471 | 24 | 9 | 20 | 12 | 12 |
| A5SS | 2062 | ENSG00000131504 | DIAPH1 | chr5 | - | 141527570 | 141527697 | 141527572 | 141527697 | 141526296 | 141526459 | 3 | 3 | 2 | 7 | 10 |
| A5SS | 3093 | ENSG00000138246 | DNAJC13 | chr3 | + | 132484362 | 132484493 | 132484362 | 132484488 | 132484587 | 132484672 | 0 | 0 | 0 | 1 | 1 |
| A5SS | 6384 | ENSG00000184307 | ZDHHC23 | chr3 | + | 113948685 | 113948963 | 113948685 | 113948734 | 113953699 | 113954410 | 13 | 17 | 18 | 17 | 34 |
| A5SS | 525 | ENSG00000244687 | UBE2V1 | chr20 | - | 50115484 | 50115954 | 50115698 | 50115954 | 50084128 | 50084254 | 10 | 7 | 9 | 7 | 13 |
| A5SS | 3597 | ENSG00000001460 | STPG1 | chr1 | - | 24383866 | 24384003 | 24383901 | 24384003 | 24379652 | 24379823 | 0 | 0 | 2 | 1 | 0 |
| A5SS | 4733 | ENSG00000242193 | CRYZL2P | chr1 | - | 178026071 | 178026255 | 178026154 | 178026255 | 178022439 | 178022668 | 1 | 0 | 0 | 0 | 0 |
| A5SS | 5728 | ENSG00000153253 | SCN3A | chr2 | - | 165140701 | 165140998 | 165140797 | 165140998 | 165139475 | 165139608 | 12 | 12 | 7 | 12 | 0 |
| A5SS | 7643 | ENSG00000136811 | ODF2 | chr9 | + | 128460941 | 128461067 | 128460941 | 128461010 | 128472912 | 128473042 | 20 | 7 | 4 | 12 | 1 |
| A5SS | 8772 | ENSG00000257221 | NA | chr12 | + | 108634330 | 108634629 | 108634330 | 108634422 | 108636394 | 108636455 | 8 | 3 | 15 | 6 | 0 |
| A5SS | 1046 | ENSG00000074370 | ATP2A3 | chr17 | - | 3928226 | 3928327 | 3928239 | 3928327 | 3927980 | 3928038 | 2 | 0 | 0 | 0 | 0 |
| A5SS | 2936 | ENSG00000009830 | POMT2 | chr14 | - | 77284881 | 77285041 | 77284949 | 77285041 | 77283796 | 77283873 | 0 | 0 | 1 | 1 | 1 |
| A5SS | 4029 | ENSG00000138095 | LRPPRC | chr2 | - | 43995773 | 43995962 | 43995798 | 43995962 | 43982237 | 43982434 | 0 | 2 | 4 | 3 | 0 |
| A5SS | 999 | ENSG00000188825 | LINC00910 | chr17 | - | 43388288 | 43388312 | 43388290 | 43388312 | 43381272 | 43381660 | 8 | 3 | 4 | 5 | 15 |
| A5SS | 4449 | ENSG00000117139 | KDM5B | chr1 | - | 202741366 | 202741722 | 202741545 | 202741722 | 202740673 | 202740812 | 64 | 54 | 70 | 74 | 27 |
| A5SS | 4606 | ENSG00000077152 | UBE2T | chr1 | - | 202335641 | 202335818 | 202335645 | 202335818 | 202334988 | 202335058 | 0 | 0 | 2 | 0 | 0 |
| A5SS | 5902 | ENSG00000115137 | DNAJC27 | chr2 | - | 24971415 | 24971616 | 24971555 | 24971616 | 24967210 | 24967293 | 9 | 15 | 11 | 15 | 9 |
| A5SS | 1598 | ENSG00000206149 | HERC2P9 | chr15 | + | 28655272 | 28655691 | 28655272 | 28655420 | 28655795 | 28656026 | 5 | 3 | 9 | 11 | 8 |
| A5SS | 5912 | ENSG00000168280 | KIF5C | chr2 | + | 148991009 | 148994538 | 148991009 | 148991198 | 148997263 | 148997340 | 9 | 19 | 16 | 17 | 20 |
| A5SS | 2498 | ENSG00000174652 | ZNF266 | chr19 | - | 9434064 | 9434235 | 9434074 | 9434235 | 9420064 | 9420218 | 0 | 2 | 2 | 2 | 7 |
| A5SS | 2501 | ENSG00000106692 | FKTN | chr9 | + | 105558137 | 105559109 | 105558137 | 105558165 | 105573654 | 105573746 | 0 | 0 | 1 | 1 | 0 |
| A5SS | 5413 | ENSG00000205702 | CYP2D7 | chr22 | - | 42141533 | 42141675 | 42141624 | 42141675 | 42141151 | 42141339 | 1 | 1 | 2 | 2 | 1 |
| A5SS | 8190 | ENSG00000186166 | CENATAC | chr11 | + | 119011220 | 119011311 | 119011220 | 119011283 | 119014993 | 119015083 | 4 | 2 | 2 | 5 | 1 |
| A5SS | 4681 | ENSG00000172915 | NBEA | chr13 | + | 35655578 | 35655755 | 35655578 | 35655749 | 35665084 | 35665186 | 0 | 0 | 1 | 0 | 0 |
| A5SS | 1364 | ENSG00000090905 | TNRC6A | chr16 | + | 24805013 | 24805733 | 24805013 | 24805151 | 24806205 | 24806283 | 68 | 43 | 44 | 61 | 55 |
| A5SS | 1576 | ENSG00000140474 | ULK3 | chr15 | - | 74842534 | 74842676 | 74842538 | 74842676 | 74842279 | 74842420 | 6 | 4 | 3 | 4 | 4 |
| A5SS | 4661 | ENSG00000133619 | KRBA1 | chr7 | + | 149730427 | 149730713 | 149730427 | 149730568 | 149731671 | 149731776 | 0 | 0 | 0 | 2 | 0 |
| A5SS | 4870 | ENSG00000160326 | SLC2A6 | chr9 | - | 133473414 | 133474088 | 133473979 | 133474088 | 133473104 | 133473250 | 5 | 4 | 5 | 9 | 3 |
| A5SS | 5513 | ENSG00000223960 | CHROMR | chr2 | + | 178426365 | 178426680 | 178426365 | 178426661 | 178430694 | 178430889 | 13 | 11 | 8 | 5 | 8 |
| A5SS | 6253 | ENSG00000137404 | NRM | chr6 | - | 30690024 | 30690243 | 30690046 | 30690243 | 30689275 | 30689452 | 0 | 0 | 1 | 0 | 1 |
| A5SS | 1430 | ENSG00000198690 | FAN1 | chr15 | + | 30904511 | 30905353 | 30904511 | 30904590 | 30908117 | 30908258 | 60 | 24 | 18 | 47 | 53 |
| A5SS | 1677 | ENSG00000100749 | VRK1 | chr14 | + | 96855223 | 96855356 | 96855223 | 96855350 | 96856527 | 96856586 | 5 | 1 | 1 | 10 | 5 |
| A5SS | 8133 | ENSG00000213619 | NDUFS3 | chr11 | + | 47579268 | 47579338 | 47579268 | 47579334 | 47580524 | 47580622 | 1 | 1 | 1 | 1 | 0 |
| A5SS | 565 | ENSG00000196756 | SNHG17 | chr20 | - | 38422086 | 38422241 | 38422095 | 38422241 | 38421005 | 38421098 | 8 | 2 | 5 | 3 | 1 |
| A5SS | 770 | ENSG00000108799 | EZH1 | chr17 | - | 42727634 | 42727763 | 42727661 | 42727763 | 42722794 | 42722915 | 29 | 21 | 23 | 29 | 36 |
| A5SS | 1123 | ENSG00000099364 | FBXL19 | chr16 | + | 30928466 | 30928630 | 30928466 | 30928628 | 30930072 | 30930584 | 1 | 0 | 0 | 0 | 2 |
| A5SS | 2391 | ENSG00000144659 | SLC25A38 | chr3 | + | 39390422 | 39390507 | 39390422 | 39390459 | 39391852 | 39392021 | 17 | 4 | 12 | 10 | 2 |
| A5SS | 5486 | ENSG00000244026 | FAM86DP | chr3 | - | 75434724 | 75435048 | 75434917 | 75435048 | 75432807 | 75432870 | 0 | 1 | 1 | 0 | 0 |
| A5SS | 1959 | ENSG00000228624 | HDAC2-AS2 | chr6 | + | 113991892 | 113992120 | 113991892 | 113992014 | 113993342 | 113993487 | 2 | 0 | 0 | 0 | 0 |
| A5SS | 2340 | ENSG00000101911 | PRPS2 | chrX | + | 12799206 | 12799399 | 12799206 | 12799390 | 12809233 | 12809332 | 11 | 2 | 10 | 2 | 15 |
| A5SS | 3006 | ENSG00000163755 | HPS3 | chr3 | + | 149129637 | 149130453 | 149129637 | 149129940 | 149140003 | 149140498 | 4 | 0 | 1 | 3 | 3 |
| A5SS | 4209 | ENSG00000048052 | HDAC9 | chr7 | + | 18495745 | 18496023 | 18495745 | 18495854 | 18496261 | 18496324 | 22 | 17 | 16 | 21 | 34 |
| A5SS | 2417 | ENSG00000073331 | ALPK1 | chr4 | + | 112302338 | 112302485 | 112302338 | 112302408 | 112315800 | 112315852 | 2 | 8 | 8 | 4 | 2 |
| A5SS | 4107 | ENSG00000256269 | HMBS | chr11 | + | 119092124 | 119092523 | 119092124 | 119092163 | 119092757 | 119092811 | 17 | 3 | 5 | 9 | 5 |
| A5SS | 8568 | ENSG00000257261 | NA | chr12 | + | 46383672 | 46383713 | 46383672 | 46383695 | 46387746 | 46387972 | 8 | 1 | 2 | 0 | 2 |
| A5SS | 1078 | ENSG00000102984 | ZNF821 | chr16 | - | 71867892 | 71868037 | 71867911 | 71868037 | 71859684 | 71860672 | 0 | 2 | 4 | 6 | 0 |
| A5SS | 6130 | ENSG00000132436 | FIGNL1 | chr7 | - | 50449544 | 50449723 | 50449555 | 50449723 | 50448180 | 50448289 | 4 | 0 | 1 | 10 | 0 |
| A5SS | 7649 | ENSG00000198000 | NOL8 | chr9 | - | 92325305 | 92325350 | 92325317 | 92325350 | 92324022 | 92324209 | 19 | 15 | 14 | 15 | 26 |
| A5SS | 8095 | ENSG00000086848 | ALG9 | chr11 | - | 111870227 | 111870370 | 111870231 | 111870370 | 111868601 | 111868736 | 2 | 0 | 1 | 5 | 5 |
| A5SS | 1134 | ENSG00000007038 | PRSS21 | chr16 | + | 2820954 | 2821109 | 2820954 | 2821067 | 2821365 | 2821719 | 0 | 2 | 3 | 1 | 6 |
| A5SS | 1387 | ENSG00000059145 | UNKL | chr16 | - | 1369933 | 1370357 | 1370129 | 1370357 | 1367655 | 1367858 | 3 | 0 | 0 | 2 | 0 |
| A5SS | 2434 | ENSG00000109654 | TRIM2 | chr4 | + | 153322647 | 153322828 | 153322647 | 153322816 | 153324077 | 153324148 | 2 | 0 | 0 | 0 | 0 |
| A5SS | 6350 | ENSG00000255036 | SUGT1P4-STRA6LP-CCDC180 | chr9 | + | 97323780 | 97323938 | 97323780 | 97323903 | 97325018 | 97325192 | 0 | 1 | 1 | 0 | 0 |
| A5SS | 7744 | ENSG00000172716 | SLFN11 | chr17 | - | 35367540 | 35367700 | 35367602 | 35367700 | 35366946 | 35367043 | 6 | 6 | 2 | 3 | 2 |
| A5SS | 3556 | ENSG00000048740 | CELF2 | chr10 | + | 11314138 | 11314270 | 11314138 | 11314258 | 11325835 | 11325979 | 5 | 3 | 4 | 2 | 0 |
| A5SS | 5294 | ENSG00000224078 | SNHG14 | chr15 | + | 25108997 | 25109103 | 25108997 | 25109099 | 25109197 | 25109403 | 37 | 8 | 25 | 20 | 60 |
| A5SS | 28 | ENSG00000241837 | ATP5PO | chr21 | - | 33915135 | 33915724 | 33915651 | 33915724 | 33914449 | 33914500 | 7 | 2 | 4 | 3 | 7 |
| A5SS | 1028 | ENSG00000161395 | PGAP3 | chr17 | - | 39673513 | 39674054 | 39673992 | 39674054 | 39671122 | 39672866 | 13 | 5 | 13 | 21 | 26 |
| A5SS | 3033 | ENSG00000159433 | STARD9 | chr15 | + | 42662793 | 42663917 | 42662793 | 42662891 | 42665252 | 42665330 | 5 | 17 | 7 | 16 | 15 |
| A5SS | 5106 | ENSG00000171132 | PRKCE | chr2 | + | 45701382 | 45701524 | 45701382 | 45701491 | 45840379 | 45840551 | 0 | 0 | 2 | 1 | 4 |
| A5SS | 1431 | ENSG00000198690 | FAN1 | chr15 | + | 30904511 | 30905897 | 30904511 | 30904590 | 30908117 | 30908258 | 72 | 38 | 38 | 63 | 66 |
| A5SS | 8742 | ENSG00000135452 | TSPAN31 | chr12 | + | 57745038 | 57745231 | 57745038 | 57745217 | 57745744 | 57745912 | 0 | 0 | 0 | 8 | 0 |
| A5SS | 8950 | ENSG00000121644 | DESI2 | chr1 | + | 244653125 | 244654025 | 244653125 | 244653355 | 244656295 | 244656521 | 7 | 1 | 6 | 5 | 2 |
| A5SS | 2493 | ENSG00000174652 | ZNF266 | chr19 | - | 9418761 | 9418895 | 9418765 | 9418895 | 9418504 | 9418631 | 39 | 13 | 8 | 19 | 14 |
| A5SS | 6112 | ENSG00000078269 | SYNJ2 | chr6 | + | 158059256 | 158059537 | 158059256 | 158059353 | 158062054 | 158062164 | 2 | 0 | 2 | 2 | 0 |
| A5SS | 2311 | ENSG00000013375 | PGM3 | chr6 | - | 83169062 | 83169323 | 83169241 | 83169323 | 83166129 | 83166466 | 32 | 30 | 29 | 37 | 36 |
| A5SS | 8374 | ENSG00000235823 | OLMALINC | chr10 | + | 100373567 | 100374083 | 100373567 | 100374012 | 100380982 | 100381109 | 1 | 0 | 0 | 2 | 0 |
| A5SS | 1439 | ENSG00000137871 | ZNF280D | chr15 | - | 56666678 | 56666986 | 56666799 | 56666986 | 56658423 | 56658486 | 65 | 62 | 63 | 50 | 63 |
| A5SS | 1472 | ENSG00000182718 | ANXA2 | chr15 | - | 60393012 | 60393440 | 60393037 | 60393440 | 60386027 | 60386086 | 0 | 3 | 5 | 6 | 6 |
| A5SS | 3596 | ENSG00000141577 | CEP131 | chr17 | - | 81198105 | 81198297 | 81198114 | 81198297 | 81197711 | 81197888 | 0 | 4 | 0 | 1 | 0 |
| A5SS | 6132 | ENSG00000132436 | FIGNL1 | chr7 | - | 50449555 | 50449723 | 50449562 | 50449723 | 50448180 | 50448289 | 0 | 0 | 2 | 1 | 4 |
| A5SS | 2739 | ENSG00000168026 | TTC21A | chr3 | + | 39118068 | 39118177 | 39118068 | 39118153 | 39119921 | 39120020 | 0 | 1 | 0 | 1 | 0 |
| A5SS | 5037 | ENSG00000172888 | ZNF621 | chr3 | + | 40524877 | 40525274 | 40524877 | 40525038 | 40525778 | 40525864 | 8 | 8 | 17 | 4 | 7 |
| A5SS | 5504 | ENSG00000239779 | WBP1 | chr2 | + | 74459437 | 74459562 | 74459437 | 74459552 | 74459642 | 74459745 | 18 | 18 | 26 | 20 | 15 |
| A5SS | 6175 | ENSG00000135486 | HNRNPA1 | chr12 | + | 54283078 | 54283234 | 54283078 | 54283225 | 54283826 | 54283967 | 4 | 0 | 1 | 0 | 0 |
| A5SS | 6792 | ENSG00000006757 | PNPLA4 | chrX | - | 7927670 | 7927739 | 7927685 | 7927739 | 7925939 | 7926132 | 4 | 1 | 10 | 3 | 12 |
| A5SS | 3186 | ENSG00000121578 | B4GALT4 | chr3 | - | 119240764 | 119240878 | 119240849 | 119240878 | 119236852 | 119237070 | 2 | 0 | 2 | 0 | 0 |
| A5SS | 3562 | ENSG00000104381 | GDAP1 | chr8 | + | 74351273 | 74351418 | 74351273 | 74351321 | 74360136 | 74360310 | 6 | 3 | 8 | 10 | 4 |
| A5SS | 4734 | ENSG00000232528 | NA | chr20 | - | 1778519 | 1778848 | 1778538 | 1778848 | 1774510 | 1774665 | 1 | 2 | 0 | 4 | 0 |
| A5SS | 961 | ENSG00000006282 | SPATA20 | chr17 | + | 50550997 | 50551222 | 50550997 | 50551190 | 50551510 | 50551679 | 1 | 0 | 2 | 0 | 0 |
| A5SS | 6114 | ENSG00000146054 | TRIM7 | chr5 | - | 181203476 | 181203640 | 181203544 | 181203640 | 181199850 | 181200081 | 2 | 0 | 0 | 3 | 0 |
| A5SS | 8614 | ENSG00000241370 | RPP21 | chr6 | + | 30345297 | 30345573 | 30345297 | 30345398 | 30346431 | 30346557 | 49 | 30 | 15 | 26 | 17 |
| A5SS | 1836 | ENSG00000103811 | CTSH | chr15 | - | 78931868 | 78932038 | 78931900 | 78932038 | 78931450 | 78931506 | 4 | 1 | 6 | 2 | 2 |
| A5SS | 8154 | ENSG00000221926 | TRIM16 | chr17 | - | 15682853 | 15682951 | 15682860 | 15682951 | 15680864 | 15680953 | 15 | 5 | 16 | 18 | 6 |
| A5SS | 8240 | ENSG00000133895 | MEN1 | chr11 | - | 64807890 | 64808099 | 64807995 | 64808099 | 64807551 | 64807680 | 46 | 19 | 53 | 50 | 24 |
| A5SS | 4666 | ENSG00000244005 | NFS1 | chr20 | - | 35699191 | 35699352 | 35699309 | 35699352 | 35698480 | 35698590 | 9 | 1 | 2 | 1 | 6 |
| A5SS | 6818 | ENSG00000149809 | TM7SF2 | chr11 | + | 65112810 | 65112894 | 65112810 | 65112865 | 65113219 | 65113414 | 0 | 1 | 0 | 0 | 0 |
| A5SS | 7835 | ENSG00000133943 | DGLUCY | chr14 | + | 91173288 | 91173439 | 91173288 | 91173403 | 91181185 | 91181389 | 12 | 3 | 9 | 9 | 0 |
| A5SS | 4702 | ENSG00000135913 | USP37 | chr2 | - | 218562672 | 218562813 | 218562717 | 218562813 | 218560819 | 218560883 | 28 | 23 | 33 | 54 | 21 |
| A5SS | 7113 | ENSG00000090316 | MAEA | chr4 | + | 1311978 | 1312238 | 1311978 | 1312161 | 1332756 | 1332865 | 0 | 2 | 2 | 1 | 0 |
| A5SS | 2807 | ENSG00000225697 | SLC26A6 | chr3 | - | 48631648 | 48631801 | 48631762 | 48631801 | 48631223 | 48631306 | 7 | 4 | 6 | 4 | 7 |
| A5SS | 5976 | ENSG00000144134 | RABL2A | chr2 | + | 113641352 | 113641480 | 113641352 | 113641450 | 113641780 | 113641864 | 3 | 1 | 1 | 1 | 0 |
| A5SS | 6056 | ENSG00000168000 | BSCL2 | chr11 | - | 62706202 | 62706344 | 62706258 | 62706344 | 62705300 | 62705617 | 3 | 0 | 1 | 0 | 0 |
| A5SS | 1825 | ENSG00000119723 | COQ6 | chr14 | + | 73950309 | 73950597 | 73950309 | 73950495 | 73953434 | 73953569 | 3 | 0 | 0 | 2 | 0 |
| A5SS | 3782 | ENSG00000144320 | LNPK | chr2 | - | 175995552 | 175995646 | 175995557 | 175995646 | 175994133 | 175994280 | 0 | 0 | 4 | 2 | 0 |
| A5SS | 7475 | ENSG00000139546 | TARBP2 | chr12 | + | 53503026 | 53503212 | 53503026 | 53503129 | 53503729 | 53503808 | 4 | 0 | 13 | 9 | 0 |
| A5SS | 2187 | ENSG00000179869 | ABCA13 | chr7 | + | 48507871 | 48508049 | 48507871 | 48507956 | 48511083 | 48511199 | 0 | 0 | 0 | 3 | 0 |
| A5SS | 7241 | ENSG00000104756 | KCTD9 | chr8 | - | 25446079 | 25446250 | 25446128 | 25446250 | 25440576 | 25440673 | 3 | 1 | 0 | 0 | 6 |
| A5SS | 5531 | ENSG00000124920 | MYRF | chr11 | + | 61783850 | 61784027 | 61783850 | 61783925 | 61784279 | 61784385 | 2 | 1 | 0 | 10 | 0 |
| A5SS | 6782 | ENSG00000156831 | NSMCE2 | chr8 | + | 125091843 | 125091958 | 125091843 | 125091900 | 125102050 | 125102146 | 7 | 7 | 6 | 8 | 4 |
| A5SS | 8830 | ENSG00000111247 | RAD51AP1 | chr12 | + | 4543762 | 4543911 | 4543762 | 4543904 | 4546308 | 4546418 | 2 | 0 | 0 | 0 | 0 |
| A5SS | 1523 | ENSG00000100321 | SYNGR1 | chr22 | + | 39374315 | 39374660 | 39374315 | 39374553 | 39376051 | 39376197 | 1 | 0 | 1 | 4 | 1 |
| A5SS | 949 | ENSG00000185813 | PCYT2 | chr17 | - | 81905669 | 81906177 | 81906099 | 81906177 | 81905381 | 81905447 | 18 | 8 | 17 | 18 | 12 |
| A5SS | 5971 | ENSG00000123600 | METTL8 | chr2 | - | 171331764 | 171331867 | 171331803 | 171331867 | 171326041 | 171326148 | 0 | 1 | 0 | 3 | 0 |
| A5SS | 7540 | ENSG00000163378 | EOGT | chr3 | - | 69012367 | 69012799 | 69012514 | 69012799 | 69011935 | 69011991 | 1 | 5 | 3 | 5 | 6 |
| A5SS | 8553 | ENSG00000211455 | STK38L | chr12 | + | 27302136 | 27302255 | 27302136 | 27302188 | 27308338 | 27308461 | 9 | 10 | 9 | 6 | 3 |
| A5SS | 2782 | ENSG00000242086 | MUC20-OT1 | chr3 | + | 195708428 | 195708555 | 195708428 | 195708435 | 195708746 | 195708841 | 5 | 4 | 6 | 2 | 3 |
| A5SS | 4936 | ENSG00000177542 | SLC25A22 | chr11 | - | 794452 | 794513 | 794457 | 794513 | 793528 | 793619 | 2 | 4 | 11 | 2 | 6 |
| A5SS | 2212 | ENSG00000146067 | FAM193B | chr5 | - | 177523158 | 177523265 | 177523162 | 177523265 | 177521973 | 177522071 | 7 | 5 | 9 | 4 | 7 |
| A5SS | 7833 | ENSG00000133943 | DGLUCY | chr14 | + | 91170002 | 91170201 | 91170002 | 91170186 | 91173288 | 91173439 | 26 | 14 | 31 | 37 | 18 |
| A5SS | 3353 | ENSG00000286061 | RCAN3AS | chr1 | - | 24502347 | 24502382 | 24502372 | 24502382 | 24499652 | 24499711 | 3 | 0 | 1 | 6 | 3 |
| A5SS | 8648 | ENSG00000238105 | GOLGA2P5 | chr12 | - | 100158678 | 100158873 | 100158797 | 100158873 | 100157996 | 100158077 | 9 | 1 | 5 | 2 | 4 |
| A5SS | 549 | ENSG00000101194 | SLC17A9 | chr20 | + | 62963583 | 62964022 | 62963583 | 62963680 | 62964227 | 62964315 | 0 | 1 | 2 | 2 | 0 |
| A5SS | 3888 | ENSG00000058272 | PPP1R12A | chr12 | - | 79786776 | 79788783 | 79788647 | 79788783 | 79781814 | 79781862 | 25 | 50 | 27 | 31 | 15 |
| A5SS | 4287 | ENSG00000181135 | ZNF707 | chr8 | + | 143691072 | 143691232 | 143691072 | 143691199 | 143691599 | 143691713 | 11 | 2 | 2 | 15 | 1 |
| A5SS | 7539 | ENSG00000104853 | CLPTM1 | chr19 | + | 44954590 | 44955085 | 44954590 | 44954701 | 44961962 | 44962075 | 4 | 7 | 10 | 6 | 0 |
| A5SS | 2363 | ENSG00000198826 | ARHGAP11A | chr15 | + | 32629594 | 32629762 | 32629594 | 32629675 | 32632978 | 32633108 | 5 | 12 | 25 | 20 | 16 |
| A5SS | 7081 | ENSG00000012048 | BRCA1 | chr17 | - | 43091434 | 43094860 | 43094743 | 43094860 | 43090943 | 43091032 | 8 | 13 | 6 | 26 | 17 |
| A5SS | 3953 | ENSG00000121766 | ZCCHC17 | chr1 | + | 31296981 | 31297213 | 31296981 | 31297075 | 31319108 | 31319166 | 2 | 4 | 3 | 3 | 0 |
| A5SS | 4678 | ENSG00000011021 | CLCN6 | chr1 | + | 11826155 | 11826224 | 11826155 | 11826214 | 11827114 | 11827221 | 0 | 0 | 0 | 0 | 1 |
| A5SS | 8638 | ENSG00000224429 | LINC00539 | chr13 | - | 21312150 | 21312365 | 21312278 | 21312365 | 21311041 | 21311118 | 3 | 1 | 4 | 0 | 0 |
| A5SS | 1365 | ENSG00000090905 | TNRC6A | chr16 | + | 24805013 | 24805733 | 24805013 | 24805151 | 24806573 | 24806784 | 5 | 6 | 7 | 1 | 4 |
| A5SS | 3501 | ENSG00000163449 | TMEM169 | chr2 | + | 216081918 | 216082232 | 216081918 | 216081979 | 216095837 | 216096234 | 0 | 2 | 1 | 2 | 0 |
| A5SS | 5972 | ENSG00000123600 | METTL8 | chr2 | - | 171331764 | 171331867 | 171331803 | 171331867 | 171330558 | 171330616 | 1 | 1 | 0 | 2 | 0 |
| A5SS | 7519 | ENSG00000283580 | NA | chr1 | + | 42775208 | 42775522 | 42775208 | 42775479 | 42789795 | 42789934 | 8 | 3 | 9 | 1 | 10 |
| A5SS | 3687 | ENSG00000077782 | FGFR1 | chr8 | - | 38419532 | 38419735 | 38419538 | 38419735 | 38418227 | 38418373 | 1 | 0 | 4 | 3 | 8 |
| A5SS | 4452 | ENSG00000141127 | PRPSAP2 | chr17 | + | 18856298 | 18856476 | 18856298 | 18856429 | 18865801 | 18865952 | 1 | 0 | 0 | 0 | 1 |
| A5SS | 8785 | ENSG00000186815 | TPCN1 | chr12 | + | 113221463 | 113221626 | 113221463 | 113221583 | 113226727 | 113226964 | 10 | 0 | 10 | 11 | 6 |
| A5SS | 5374 | ENSG00000169432 | SCN9A | chr2 | - | 166375696 | 166375944 | 166375734 | 166375944 | 166311498 | 166311806 | 3 | 2 | 0 | 2 | 0 |
| A5SS | 2090 | ENSG00000079215 | SLC1A3 | chr5 | + | 36629449 | 36630482 | 36629449 | 36629587 | 36671028 | 36671233 | 1 | 0 | 0 | 0 | 0 |
| A5SS | 5679 | ENSG00000102125 | TAZ | chrX | + | 154413206 | 154413358 | 154413206 | 154413339 | 154413481 | 154413567 | 1 | 5 | 2 | 3 | 5 |
| A5SS | 478 | ENSG00000000419 | DPM1 | chr20 | - | 50941104 | 50941209 | 50941128 | 50941209 | 50940864 | 50940955 | 3 | 7 | 1 | 0 | 0 |
| A5SS | 2270 | ENSG00000134644 | PUM1 | chr1 | - | 30950126 | 30950261 | 30950132 | 30950261 | 30941150 | 30941272 | 3 | 7 | 1 | 4 | 6 |
| A5SS | 2422 | ENSG00000167216 | KATNAL2 | chr18 | + | 46946056 | 46946546 | 46946056 | 46946275 | 46946853 | 46946923 | 2 | 4 | 7 | 9 | 8 |
| A5SS | 4558 | ENSG00000114491 | UMPS | chr3 | + | 124737567 | 124738443 | 124737567 | 124738239 | 124742151 | 124742266 | 12 | 6 | 11 | 13 | 23 |
| A5SS | 6928 | ENSG00000196152 | ZNF79 | chr9 | + | 127424373 | 127424910 | 127424373 | 127424803 | 127428551 | 127428660 | 9 | 8 | 4 | 4 | 5 |
| A5SS | 401 | ENSG00000261824 | LINC00662 | chr19 | - | 27746020 | 27746340 | 27746097 | 27746340 | 27732795 | 27732967 | 4 | 0 | 1 | 1 | 0 |
| A5SS | 1074 | ENSG00000172824 | CES4A | chr16 | + | 67000856 | 67001002 | 67000856 | 67000990 | 67001307 | 67001461 | 0 | 0 | 0 | 1 | 0 |
| A5SS | 3379 | ENSG00000095321 | CRAT | chr9 | - | 129108898 | 129109277 | 129109102 | 129109277 | 129107813 | 129108077 | 2 | 1 | 0 | 3 | 0 |
| A5SS | 6458 | ENSG00000068885 | IFT80 | chr3 | - | 160383436 | 160384646 | 160384405 | 160384646 | 160381502 | 160381718 | 30 | 34 | 37 | 55 | 91 |
| A5SS | 7776 | ENSG00000148356 | LRSAM1 | chr9 | + | 127497252 | 127497362 | 127497252 | 127497334 | 127501009 | 127501143 | 2 | 0 | 0 | 1 | 0 |

| | | | | | | | | | | | | | | |
|---|---|---|---|---|---|---|---|---|---|---|---|---|---|---|
| A5SS | 1632 | ENSG00000165795 | NDRG2 | chr14 | - | 21024029 | 21025061 | 21025028 | 21025061 | 21023240 | 21023321 | 3 | 1 | 1 |
| A5SS | 7205 | ENSG00000107815 | TWNK | chr10 | + | 100990868 | 100991054 | 100990868 | 100991010 | 100993189 | 100994401 | 5 | 3 | 2 |
| A5SS | 8241 | ENSG00000133895 | MEN1 | chr11 | - | 64807904 | 64808099 | 64807995 | 64808099 | 64807551 | 64807680 | 20 | 9 | 28 |
| A5SS | 1683 | ENSG00000006432 | MAP3K9 | chr14 | - | 70739897 | 70740164 | 70740041 | 70740164 | 70738244 | 70738398 | 4 | 3 | 2 |
| A5SS | 3213 | ENSG00000130270 | ATP8B3 | chr19 | - | 1811488 | 1811884 | 1811600 | 1811884 | 1810621 | 1810683 | 2 | 7 | 0 |
| A5SS | 5736 | ENSG00000079263 | SP140 | chr2 | + | 230244987 | 230245080 | 230244987 | 230245071 | 230255451 | 230255532 | 10 | 6 | 11 |
| A5SS | 8029 | ENSG00000177697 | CD151 | chr11 | + | 832951 | 833026 | 832951 | 833022 | 834529 | 834591 | 0 | 0 | 0 |
| A5SS | 8153 | ENSG00000166261 | ZNF202 | chr11 | - | 123740392 | 123740536 | 123740416 | 123740536 | 123740116 | 123740232 | 0 | 0 | 2 |
| A5SS | 6481 | ENSG00000010318 | PHF7 | chr3 | + | 52422759 | 52422881 | 52422759 | 52422836 | 52423090 | 52423640 | 4 | 2 | 0 |
| A5SS | 4619 | ENSG00000106462 | EZH2 | chr7 | - | 148816683 | 148818117 | 148817876 | 148818117 | 148815505 | 148815546 | 23 | 16 | 25 |
| A5SS | 5978 | ENSG00000072163 | LIMS2 | chr2 | - | 127640269 | 127640988 | 127640895 | 127640988 | 127640069 | 127640145 | 2 | 2 | 7 |
| A5SS | 6252 | ENSG00000137404 | NRM | chr6 | - | 30690024 | 30690243 | 30690046 | 30690243 | 30688046 | 30688942 | 0 | 0 | 1 |
| A5SS | 6768 | ENSG00000137501 | SYTL2 | chr11 | - | 85724031 | 85727967 | 85727895 | 85727967 | 85720857 | 85720959 | 29 | 11 | 18 |
| A5SS | 2188 | ENSG00000170074 | FAM153A | chr5 | - | 177714053 | 177714177 | 177714068 | 177714177 | 177713747 | 177713916 | 12 | 18 | 21 |
| A5SS | 6431 | ENSG00000105298 | CACTIN | chr19 | - | 3614385 | 3614589 | 3614396 | 3614589 | 3613463 | 3613586 | 0 | 2 | 1 |
| A5SS | 717 | ENSG00000132514 | CLEC10A | chr17 | - | 7075975 | 7076071 | 7075984 | 7076071 | 7075730 | 7075885 | 0 | 15 | 4 |
| A5SS | 1631 | ENSG00000165795 | NDRG2 | chr14 | - | 21023985 | 21024096 | 21024029 | 21024096 | 21023240 | 21023321 | 1 | 1 | 7 |
| A5SS | 2844 | ENSG00000157601 | MX1 | chr21 | + | 41426294 | 41426546 | 41426294 | 41426397 | 41427205 | 41427304 | 0 | 2 | 0 |
| A5SS | 5063 | ENSG00000183762 | KREMEN1 | chr22 | + | 29138623 | 29138833 | 29138623 | 29138782 | 29140281 | 29140366 | 0 | 0 | 0 |
| A5SS | 8656 | ENSG00000090006 | LTBP4 | chr19 | + | 40616888 | 40617042 | 40616888 | 40617020 | 40619346 | 40619493 | 1 | 0 | 0 |
| [illegible] | | | | | | | | | | | | | | |

| | | | | | | | | | | | | | | |
|---|---|---|---|---|---|---|---|---|---|---|---|---|---|---|
| A5SS | 380 | ENSG00000105176 | URI1 | chr19 | + | 29980158 | 29980364 | 29980158 | 29980250 | 29985222 | 29985301 | 8 | 5 | 5 |
| A5SS | 6098 | ENSG00000221838 | AP4M1 | chr7 | + | 100101879 | 100101989 | 100101879 | 100101968 | 100102674 | 100102781 | 0 | 2 | 1 |
| A5SS | 6167 | ENSG00000135486 | HNRNPA1 | chr12 | + | 54282572 | 54282665 | 54282572 | 54282662 | 54283826 | 54283967 | 2 | 0 | 0 |
| A5SS | 2916 | ENSG00000176261 | ZBTB8OS | chr1 | - | 32650401 | 32650525 | 32650432 | 32650525 | 32627507 | 32627544 | 0 | 1 | 2 |
| A5SS | 5327 | ENSG00000197037 | ZSCAN25 | chr7 | + | 99619560 | 99619997 | 99619560 | 99619993 | 99622548 | 99622640 | 3 | 1 | 4 |
| A5SS | 3411 | ENSG00000234663 | LINC01934 | chr2 | + | 181226470 | 181226539 | 181226470 | 181226535 | 181322660 | 181322730 | 5 | 1 | 0 |
| A5SS | 8540 | ENSG00000228705 | LINC00659 | chr20 | - | 62776455 | 62776921 | 62776805 | 62776921 | 62775314 | 62775520 | 0 | 1 | 7 |
| A5SS | 8282 | ENSG00000095066 | HOOK2 | chr19 | - | 12774560 | 12774741 | 12774634 | 12774741 | 12772993 | 12773044 | 2 | 3 | 0 |
| A5SS | 7669 | ENSG00000178445 | GLDC | chr9 | - | 6535946 | 6536236 | 6536063 | 6536236 | 6534707 | 6534788 | 0 | 0 | 0 |
| A5SS | 2770 | ENSG00000163702 | IL17RC | chr3 | + | 9923880 | 9924137 | 9923880 | 9924020 | 9924231 | 9924291 | 0 | 6 | 0 |
| A5SS | 3435 | ENSG00000154269 | ENPP3 | chr6 | + | 131674161 | 131674410 | 131674161 | 131674281 | 131675079 | 131675189 | 1 | 5 | 2 |
| A5SS | 8323 | ENSG00000157404 | KIT | chr4 | + | 54725856 | 54726050 | 54725856 | 54726038 | 54727217 | 54727324 | 2 | 4 | 3 |
| A5SS | 3994 | ENSG00000225746 | MEG8 | chr14 | + | 100944808 | 100944925 | 100944808 | 100944872 | 100947033 | 100947116 | 1 | 0 | 0 |
| A5SS | 6332 | ENSG00000164970 | FAM219A | chr9 | - | 34458203 | 34458552 | 34458236 | 34458552 | 34405864 | 34405964 | 8 | 4 | 2 |
| A5SS | 4748 | ENSG00000273136 | NBPF26 | chr1 | + | 120822103 | 120822267 | 120822103 | 120822156 | 120823308 | 120823360 | 7 | 6 | 9 |
| A5SS | 5050 | ENSG00000033867 | SLC4A7 | chr3 | - | 27484066 | 27484347 | 27484078 | 27484347 | 27452416 | 27452498 | 9 | 5 | 8 |
| A5SS | 7825 | ENSG00000074416 | MGLL | chr3 | - | 127822308 | 127823185 | 127823083 | 127823185 | 127821693 | 127821838 | 1 | 3 | 1 |
| A5SS | 8213 | ENSG00000179532 | DNHD1 | chr11 | + | 6533680 | 6534173 | 6533680 | 6533806 | 6538611 | 6538810 | 7 | 7 | 3 |
| A5SS | 2870 | ENSG00000171649 | ZIK1 | chr19 | + | 57584144 | 57584612 | 57584144 | 57584389 | 57584951 | 57584990 | 2 | 10 | 3 |
| A5SS | 2880 | ENSG00000172053 | QARS1 | chr3 | - | 49104356 | 49104471 | 49104372 | 49104471 | 49103862 | 49103972 | 10 | 0 | 2 |
| A5SS | 6483 | ENSG00000113303 | BTNL8 | chr5 | + | 180947511 | 180947755 | 180947511 | 180947625 | 180948354 | 180948375 | 1 | 0 | 3 |
| A5SS | 3157 | ENSG00000164211 | STARD4 | chr5 | - | 111512384 | 111512535 | 111512390 | 111512535 | 111507328 | 111507442 | 38 | 8 | 11 |
| A5SS | 400 | ENSG00000261824 | LINC00662 | chr19 | - | 27746020 | 27746340 | 27746080 | 27746340 | 27732795 | 27732967 | 4 | 0 | 1 |
| A5SS | 1961 | ENSG00000228624 | HDAC2-AS2 | chr6 | + | 113991892 | 113992120 | 113991892 | 113992018 | 113993342 | 113993487 | 2 | 0 | 0 |
| A5SS | 6212 | ENSG00000153406 | NMRAL1 | chr16 | - | 4471313 | 4471449 | 4471356 | 4471449 | 4469226 | 4469465 | 3 | 4 | 4 |
| A5SS | 361 | ENSG00000159917 | ZNF235 | chr19 | - | 44298807 | 44298903 | 44298819 | 44298903 | 44234754 | 44234869 | 0 | 1 | 1 |
| A5SS | 2543 | ENSG00000109762 | SNX25 | chr4 | + | 185332594 | 185332759 | 185332594 | 185332713 | 185339378 | 185339510 | 48 | 39 | 51 |
| A5SS | 3588 | ENSG00000116353 | MECR | chr1 | - | 29220833 | 29220921 | 29220837 | 29220921 | 29216587 | 29216812 | 0 | 0 | 0 |
| A5SS | 8497 | ENSG00000055950 | MRPL43 | chr10 | - | 100986686 | 100986975 | 100986748 | 100986975 | 100984033 | 100984074 | 14 | 4 | 8 |
| A5SS | 7564 | ENSG00000228340 | MIR646HG | chr20 | + | 60154406 | 60154506 | 60154406 | 60154501 | 60180834 | 60180913 | 4 | 4 | 0 |
| A5SS | 7992 | ENSG00000239523 | MYLK-AS1 | chr3 | + | 123585646 | 123585801 | 123585646 | 123585794 | 123629490 | 123629681 | 1 | 0 | 4 |
| A5SS | 3867 | ENSG00000251623 | FAM153B | chr5 | + | 176123996 | 176124120 | 176123996 | 176124105 | 176124262 | 176124432 | 8 | 3 | 8 |
| A5SS | 2460 | ENSG00000049656 | CLPTM1L | chr5 | - | 1331798 | 1331883 | 1331802 | 1331883 | 1325750 | 1325816 | 5 | 1 | 5 |
| A5SS | 2468 | ENSG00000267364 | NA | chr17 | + | 35323147 | 35323343 | 35323147 | 35323268 | 35324653 | 35324909 | 32 | 6 | 30 |
| A5SS | 3382 | ENSG00000095321 | CRAT | chr9 | - | 129109102 | 129109277 | 129109178 | 129109277 | 129108762 | 129108903 | 0 | 1 | 1 |
| A5SS | 5251 | ENSG00000285756 | NA | chrX | - | 3866921 | 3867806 | 3866949 | 3867806 | 3865042 | 3865142 | 0 | 12 | 0 |
| A5SS | 7689 | ENSG00000011454 | RABGAP1 | chr9 | + | 123065347 | 123065601 | 123065347 | 123065461 | 123073551 | 123073677 | 4 | 11 | 16 |
| A5SS | 6497 | ENSG00000142303 | ADAMTS10 | chr19 | - | 8595761 | 8595903 | 8595772 | 8595903 | 8592762 | 8592870 | 9 | 5 | 7 |
| A5SS | 8372 | ENSG00000180229 | HERC2P3 | chr15 | - | 20453619 | 20453762 | 20453705 | 20453762 | 20453350 | 20453502 | 3 | 15 | 6 |
| A5SS | 6808 | ENSG00000118513 | MYB | chr6 | + | 135200289 | 135200674 | 135200289 | 135200557 | 135201638 | 135201749 | 3 | 1 | 3 |
| A5SS | 8204 | ENSG00000054654 | SYNE2 | chr14 | + | 64216247 | 64216424 | 64216247 | 64216387 | 64219207 | 64219410 | 1 | 4 | 4 |
| A5SS | 7719 | ENSG00000135047 | CTSL | chr9 | + | 87727360 | 87727468 | 87727360 | 87727395 | 87727593 | 87727729 | 1 | 0 | 1 |
| A5SS | 6985 | ENSG00000147130 | ZMYM3 | chrX | - | 71246594 | 71246692 | 71246630 | 71246692 | 71246352 | 71246512 | 7 | 6 | 10 |
| A5SS | 5 | ENSG00000183570 | PCBP3 | chr21 | + | 45913950 | 45914094 | 45913950 | 45914025 | 45917587 | 45917629 | 0 | 0 | 0 |
| A5SS | 69 | ENSG00000185386 | MAPK11 | chr22 | - | 50267252 | 50267286 | 50267256 | 50267286 | 50267126 | 50267174 | 0 | 0 | 2 |
| A5SS | 70 | ENSG00000188130 | MAPK12 | chr22 | - | 50256787 | 50256964 | 50256934 | 50256964 | 50256598 | 50256646 | 0 | 0 | 2 |
| A5SS | 853 | ENSG00000159640 | ACE | chr17 | + | 63491208 | 63491381 | 63491208 | 63491368 | 63493435 | 63493659 | 1 | 1 | 2 |
| A5SS | 1252 | ENSG00000155666 | KDM8 | chr16 | + | 27218960 | 27219110 | 27218960 | 27219046 | 27220392 | 27220485 | 18 | 9 | 17 |
| A5SS | 1710 | ENSG00000075413 | MARK3 | chr14 | + | 103411644 | 103411765 | 103411644 | 103411749 | 103428386 | 103428440 | 1 | 0 | 0 |
| A5SS | 1862 | ENSG00000214900 | LINC01588 | chr14 | - | 50005363 | 50007520 | 50005593 | 50007520 | 49992777 | 49992873 | 4 | 2 | 2 |
| A5SS | 3107 | ENSG00000182853 | VMO1 | chr17 | - | 4785900 | 4786052 | 4785936 | 4786052 | 4785284 | 4785659 | 3 | 0 | 0 |
| A5SS | 3750 | ENSG00000167384 | ZNF180 | chr19 | - | 44500197 | 44500522 | 44500274 | 44500522 | 44497283 | 44497372 | 3 | 4 | 10 |
| A5SS | 3785 | ENSG00000156463 | SH3RF2 | chr5 | + | 146055980 | 146056231 | 146055980 | 146056213 | 146059865 | 146060224 | 0 | 0 | 0 |
| A5SS | 3982 | ENSG00000188004 | SNHG28 | chr1 | - | 159840710 | 159840866 | 159840768 | 159840866 | 159838364 | 159838451 | 5 | 0 | 0 |
| A5SS | 4188 | ENSG00000121988 | ZRANB3 | chr2 | - | 135227811 | 135228015 | 135227817 | 135228015 | 135219076 | 135219178 | 1 | 0 | 3 |
| A5SS | 4886 | ENSG00000182400 | TRAPPC6B | chr14 | - | 39159449 | 39159550 | 39159482 | 39159550 | 39151745 | 39151839 | 2 | 7 | 1 |
| A5SS | 5929 | ENSG00000124356 | STAMBP | chr2 | + | 73828966 | 73829070 | 73828966 | 73828996 | 73829404 | 73829510 | 1 | 1 | 0 |
| A5SS | 7377 | ENSG00000249859 | PVT1 | chr8 | + | 128049408 | 128049748 | 128049408 | 128049441 | 128070159 | 128070361 | 2 | 1 | 5 |
| A5SS | 7654 | ENSG00000137098 | SPAG8 | chr9 | - | 35811756 | 35812001 | 35811923 | 35812001 | 35810882 | 35811057 | 0 | 0 | 0 |
| A5SS | 7780 | ENSG00000175287 | PHYHD1 | chr9 | + | 128937756 | 128937944 | 128937756 | 128937778 | 128940368 | 128940497 | 0 | 2 | 3 |
| A5SS | 8721 | ENSG00000103356 | EARS2 | chr16 | - | 23553006 | 23553123 | 23553043 | 23553123 | 23552148 | 23552304 | 0 | 0 | 0 |
| A5SS | 8829 | ENSG00000111247 | RAD51AP1 | chr12 | + | 4543762 | 4543911 | 4543762 | 4543904 | 4545781 | 4545832 | 1 | 0 | 0 |
| A5SS | 8728 | ENSG00000139350 | NEDD1 | chr12 | + | 96907603 | 96907856 | 96907603 | 96907624 | 96919984 | 96920125 | 15 | 1 | 10 |
| A5SS | 8782 | ENSG00000127328 | RAB3IP | chr12 | + | 69800208 | 69800337 | 69800208 | 69800332 | 69812963 | 69813033 | 5 | 5 | 5 |
| A5SS | 6079 | ENSG00000092096 | SLC22A17 | chr14 | - | 23347459 | 23347731 | 23347513 | 23347731 | 23347100 | 23347212 | 1 | 0 | 3 |
| A5SS | 8470 | ENSG00000107862 | GBF1 | chr10 | + | 102376934 | 102377140 | 102376934 | 102377128 | 102379852 | 102379954 | 0 | 0 | 1 |
| A5SS | 8726 | ENSG00000139350 | NEDD1 | chr12 | + | 96907603 | 96907856 | 96907603 | 96907624 | 96909751 | 96909895 | 38 | 21 | 22 |
| A5SS | 8801 | ENSG00000130396 | AFDN | chr6 | + | 167965756 | 167966276 | 167965756 | 167966270 | 167969113 | 167969198 | 1 | 0 | 0 |
| A5SS | 6292 | ENSG00000109339 | MAPK10 | chr4 | - | 86194308 | 86194407 | 86194335 | 86194407 | 86159297 | 86159467 | 4 | 1 | 0 |
| A5SS | 5640 | ENSG00000132323 | ILKAP | chr2 | - | 238189852 | 238189972 | 238189862 | 238189972 | 238185180 | 238185287 | 3 | 1 | 9 |
| A5SS | 402 | ENSG00000261824 | LINC00662 | chr19 | - | 27746080 | 27746340 | 27746097 | 27746340 | 27732795 | 27732967 | 0 | 2 | 1 |
| A5SS | 8529 | ENSG00000155256 | ZFYVE27 | chr10 | + | 97738476 | 97738674 | 97738476 | 97738649 | 97749473 | 97749586 | 2 | 1 | 1 |
| A5SS | 1891 | ENSG00000105479 | ODAD1 | chr19 | - | 48320754 | 48320811 | 48320771 | 48320811 | 48320298 | 48320391 | 0 | 0 | 1 |
| A5SS | 2384 | ENSG00000103995 | CEP152 | chr15 | - | 48741600 | 48741704 | 48741630 | 48741704 | 48738150 | 48739288 | 6 | 10 | 7 |
| A5SS | 339 | ENSG00000160469 | BRSK1 | chr19 | + | 55287213 | 55287303 | 55287213 | 55287299 | 55289479 | 55289620 | 0 | 0 | 1 |
| A5SS | 3903 | ENSG00000055609 | KMT2C | chr7 | - | 152145152 | 152145295 | 152145164 | 152145295 | 152139674 | 152139791 | 1 | 1 | 2 |
| A5SS | 2271 | ENSG00000134644 | PUM1 | chr1 | - | 30950126 | 30950261 | 30950132 | 30950261 | 30941997 | 30942123 | 2 | 8 | 2 |
| A5SS | 5955 | ENSG00000110080 | ST3GAL4 | chr11 | + | 126406472 | 126406693 | 126406472 | 126406545 | 126406942 | 126407023 | 4 | 2 | 2 |
| A5SS | 6965 | ENSG00000100532 | CGRRF1 | chr14 | + | 54530048 | 54530763 | 54530048 | 54530405 | 54530902 | 54531050 | 15 | 3 | 6 |
| A5SS | 1446 | ENSG00000166794 | PPIB | chr15 | - | 64162036 | 64162154 | 64162040 | 64162154 | 64160103 | 64160193 | 1 | 0 | 1 |
| A5SS | 3587 | ENSG00000116353 | MECR | chr1 | - | 29220833 | 29220921 | 29220837 | 29220921 | 29216587 | 29216685 | 0 | 1 | 1 |
| A5SS | 2754 | ENSG00000163703 | CRELD1 | chr3 | + | 9933809 | 9933920 | 9933809 | 9933852 | 9934419 | 9934612 | 0 | 5 | 1 |
| A5SS | 2951 | ENSG00000157036 | EXOG | chr3 | + | 38497628 | 38497689 | 38497628 | 38497684 | 38501354 | 38501494 | 0 | 1 | 1 |
| A5SS | 6830 | ENSG00000140543 | DET1 | chr15 | - | 88531608 | 88531715 | 88531612 | 88531715 | 88527598 | 88527786 | 0 | 0 | 1 |
| A5SS | 6129 | ENSG00000132436 | FIGNL1 | chr7 | - | 50449499 | 50449723 | 50449562 | 50449723 | 50448180 | 50448289 | 7 | 0 | 4 |
| A5SS | 469 | ENSG00000064601 | CTSA | chr20 | + | 45892724 | 45892887 | 45892724 | 45892880 | 45893039 | 45893145 | 0 | 0 | 0 |
| A5SS | 2890 | ENSG00000237940 | LINC01238 | chr2 | + | 241971374 | 241971493 | 241971374 | 241971462 | 241971684 | 241971775 | 3 | 0 | 4 |
| A5SS | 3579 | ENSG00000169231 | THBS3 | chr1 | - | 155203085 | 155203137 | 155203092 | 155203137 | 155202811 | 155202960 | 6 | 5 | 3 |
| A5SS | 2070 | ENSG00000154134 | ROBO3 | chr11 | + | 124877518 | 124877752 | 124877518 | 124877658 | 124878297 | 124878436 | 3 | 0 | 2 |
| A5SS | 5330 | ENSG00000185480 | PARPBP | chr12 | + | 102153868 | 102153989 | 102153868 | 102153976 | 102154826 | 102155280 | 2 | 0 | 1 |
| A5SS | 1689 | ENSG00000092200 | RPGRIP1 | chr14 | + | 21321258 | 21321402 | 21321258 | 21321300 | 21321853 | 21322004 | 0 | 4 | 7 |
| A5SS | 5376 | ENSG00000137075 | RNF38 | chr9 | - | 36400935 | 36401198 | 36401097 | 36401198 | 36390466 | 36390616 | 5 | 1 | 6 |
| A5SS | 5922 | ENSG00000138380 | CARF | chr2 | + | 202913033 | 202913114 | 202913033 | 202913102 | 202917876 | 202918043 | 10 | 9 | 6 |
| A5SS | 8079 | ENSG00000087053 | MTMR2 | chr11 | - | 95866383 | 95866486 | 95866402 | 95866486 | 95865600 | 95865676 | 0 | 0 | 0 |
| A5SS | 144 | ENSG00000160336 | ZNF761 | chr19 | + | 53445000 | 53445219 | 53445000 | 53445123 | 53446226 | 53446337 | 0 | 2 | 4 |
| A5SS | 2278 | ENSG00000112992 | NNT | chr5 | + | 43647923 | 43648440 | 43647923 | 43648182 | 43649146 | 43649308 | 27 | 15 | 10 |
| A5SS | 3989 | ENSG00000110046 | ATG2A | chr11 | - | 64902509 | 64902680 | 64902515 | 64902680 | 64902259 | 64902386 | 10 | 0 | 5 |
| A5SS | 266 | ENSG00000196724 | ZNF418 | chr19 | - | 57935030 | 57935135 | 57935079 | 57935135 | 57933816 | 57933902 | 0 | 0 | 2 |
| A5SS | 2072 | ENSG00000154134 | ROBO3 | chr11 | + | 124877518 | 124878131 | 124877518 | 124877752 | 124878297 | 124878436 | 10 | 3 | 6 |
| A5SS | 3946 | ENSG00000169129 | AFAP1L2 | chr10 | - | 114297219 | 114297413 | 114297231 | 114297413 | 114296977 | 114297100 | 7 | 11 | 7 |
| A5SS | 8808 | ENSG00000215039 | CD27-AS1 | chr12 | - | 6450869 | 6450980 | 6450892 | 6450980 | 6448639 | 6448737 | 5 | 2 | 2 |
| A5SS | 3586 | ENSG00000116353 | MECR | chr1 | - | 29220716 | 29220921 | 29220837 | 29220921 | 29216587 | 29216685 | 0 | 5 | 1 |
| A5SS | 2764 | ENSG00000283646 | LINC02009 | chr3 | - | 46421372 | 46421492 | 46421421 | 46421492 | 46419516 | 46419663 | 0 | 0 | 0 |
| A5SS | 5731 | ENSG00000153253 | SCN3A | chr2 | - | 165140788 | 165140998 | 165140797 | 165140998 | 165139475 | 165139608 | 1 | 0 | 0 |
| A5SS | 7378 | ENSG00000249859 | PVT1 | chr8 | + | 128070159 | 128070361 | 128070159 | 128070272 | 128082752 | 128082848 | 2 | 2 | 3 |
| A5SS | 8881 | ENSG00000075089 | ACTR6 | chr12 | + | 100205675 | 100205757 | 100205675 | 100205744 | 100210072 | 100210208 | 3 | 1 | 1 |
| A5SS | 806 | ENSG00000141736 | ERBB2 | chr17 | + | 39706989 | 39707204 | 39706989 | 39707141 | 39708320 | 39708534 | 0 | 0 | 1 |
| A5SS | 6883 | ENSG00000130734 | ATG4D | chr19 | + | 10546838 | 10547082 | 10546838 | 10547075 | 10547188 | 10547253 | 1 | 0 | 1 |
| A5SS | 6564 | ENSG00000137731 | FXYD2 | chr11 | - | 117824161 | 117824744 | 117824653 | 117824744 | 117822678 | 117822717 | 0 | 0 | 2 |
| A5SS | 1065 | ENSG00000168488 | ATXN2L | chr16 | + | 28835932 | 28836485 | 28835932 | 28836122 | 28836727 | 28837237 | 7 | 4 | 8 |
| A5SS | 3174 | ENSG00000261040 | WFDC21P | chr17 | - | 60085054 | 60085203 | 60085112 | 60085203 | 60083565 | 60083766 | 0 | 5 | 0 |
| A5SS | 7927 | ENSG00000135424 | ITGA7 | chr12 | - | 55700565 | 55701154 | 55700898 | 55701154 | 55699869 | 55699989 | 1 | 0 | 1 |
| A5SS | 6331 | ENSG00000164970 | FAM219A | chr9 | - | 34405864 | 34405964 | 34405915 | 34405964 | 34402704 | 34402807 | 2 | 2 | 2 |
| A5SS | 4878 | ENSG00000163534 | FCRL1 | chr1 | - | 157797680 | 157797820 | 157797758 | 157797820 | 157797100 | 157797129 | 15 | 7 | 5 |
| A5SS | 6127 | ENSG00000132436 | FIGNL1 | chr7 | - | 50449499 | 50449723 | 50449544 | 50449723 | 50448180 | 50448289 | 7 | 0 | 4 |
| A5SS | 150 | ENSG00000131848 | ZSCAN5A | chr19 | - | 56314619 | 56314839 | 56314680 | 56314839 | 56313282 | 56313384 | 3 | 1 | 7 |
| A5SS | 3211 | ENSG00000284959 | NA | chr14 | - | 81053682 | 81053816 | 81053716 | 81053816 | 81052860 | 81053180 | 0 | 3 | 0 |
| A5SS | 1690 | ENSG00000092200 | RPGRIP1 | chr14 | + | 21321853 | 21322004 | 21321853 | 21321971 | 21324617 | 21325070 | 4 | 3 | 5 |
| A5SS | 8693 | ENSG00000116198 | CEP104 | chr1 | - | 3829132 | 3829373 | 3829265 | 3829373 | 3828884 | 3828971 | 6 | 2 | 4 |
| A5SS | 7813 | ENSG00000215126 | CBWD6 | chr9 | - | 41170278 | 41170423 | 41170283 | 41170423 | 41166783 | 41166830 | 1 | 0 | 0 |
| A5SS | 1991 | ENSG00000125459 | MSTO1 | chr1 | + | 155613656 | 155613766 | 155613656 | 155613733 | 155614058 | 155614935 | 3 | 5 | 3 |
| A5SS | 6425 | ENSG00000147894 | C9orf72 | chr9 | - | 27573708 | 27573866 | 27573786 | 27573866 | 27566676 | 27567164 | 13 | 3 | 3 |
| A5SS | 6128 | ENSG00000132436 | FIGNL1 | chr7 | - | 50449499 | 50449723 | 50449555 | 50449723 | 50448180 | 50448289 | 7 | 0 | 4 |
| A5SS | 4560 | ENSG00000215883 | CYB5RL | chr1 | - | 54190737 | 54190896 | 54190747 | 54190896 | 54187651 | 54187739 | 2 | 1 | 5 |
| A5SS | 358 | ENSG00000105146 | AURKC | chr19 | + | 57232541 | 57232685 | 57232541 | 57232680 | 57233459 | 57233608 | 0 | 0 | 1 |
| A5SS | 3293 | ENSG00000117054 | ACADM | chr1 | + | 75734790 | 75734875 | 75734790 | 75734869 | 75739979 | 75740110 | 1 | 1 | 0 |
| A5SS | 6126 | ENSG00000132436 | FIGNL1 | chr7 | - | 50449117 | 50449723 | 50449562 | 50449723 | 50448180 | 50448289 | 11 | 5 | 9 |
| A5SS | 6140 | ENSG00000251364 | NA | chr11 | - | 7461232 | 7461336 | 7461238 | 7461336 | 7450372 | 7450442 | 1 | 1 | 0 |
| A5SS | 5872 | ENSG00000148680 | HTR7 | chr10 | - | 90748833 | 90749594 | 90748838 | 90749594 | 90740822 | 90742528 | 1 | 1 | 0 |
| A5SS | 467 | ENSG00000064601 | CTSA | chr20 | + | 45891375 | 45891449 | 45891375 | 45891425 | 45891568 | 45891762 | 0 | 0 | 0 |
| A5SS | 507 | ENSG00000088836 | SLC4A11 | chr20 | - | 3238979 | 3239190 | 3239094 | 3239190 | 3237543 | 3237588 | 1 | 0 | 0 |
| A5SS | 604 | ENSG00000154040 | CABYR | chr18 | + | 24155700 | 24156999 | 24155700 | 24156042 | 24159471 | 24160069 | 0 | 0 | 0 |
| A5SS | 920 | ENSG00000244045 | TMEM199 | chr17 | + | 28357646 | 28357905 | 28357646 | 28357881 | 28359314 | 28359404 | 1 | 0 | 1 |
| A5SS | 1401 | ENSG00000161996 | WDR90 | chr16 | + | 660611 | 660870 | 660611 | 660714 | 661050 | 661172 | 1 | 0 | 0 |
| A5SS | 1415 | ENSG00000166145 | SPINT1 | chr15 | + | 40853710 | 40853929 | 40853710 | 40853881 | 40854396 | 40854522 | 2 | 0 | 1 |
| A5SS | 1427 | ENSG00000198690 | FAN1 | chr15 | + | 30903863 | 30903894 | 30903863 | 30903887 | 30904511 | 30904590 | 2 | 0 | 0 |
| A5SS | 2082 | ENSG00000183474 | GTF2H2C | chr5 | + | 69560222 | 69560403 | 69560222 | 69560242 | 69565099 | 69565188 | 4 | 2 | 2 |
| A5SS | 2389 | ENSG00000161912 | ADCY10P1 | chr6 | + | 41132293 | 41132525 | 41132293 | 41132513 | 41132874 | 41133000 | 0 | 0 | 0 |
| A5SS | 2855 | ENSG00000114346 | ECT2 | chr3 | + | 172752149 | 172752472 | 172752149 | 172752436 | 172754508 | 172754660 | 0 | 1 | 0 |
| A5SS | 2914 | ENSG00000173402 | DAG1 | chr3 | + | 49476848 | 49476905 | 49476848 | 49476900 | 49510418 | 49510819 | 0 | 1 | 0 |
| A5SS | 3286 | ENSG00000170667 | RASA4B | chr7 | - | 102487785 | 102488598 | 102488219 | 102488598 | 102485038 | 102485159 | 2 | 0 | 2 |
| A5SS | 3999 | ENSG00000137821 | LRRC49 | chr15 | + | 70893583 | 70893655 | 70893583 | 70893640 | 70895848 | 70895936 | 1 | 0 | 0 |
| A5SS | 4221 | ENSG00000147180 | ZNF711 | chrX | + | 85246929 | 85247188 | 85246929 | 85247183 | 85247546 | 85247651 | 1 | 0 | 0 |
| A5SS | 4397 | ENSG00000145979 | TBC1D7 | chr6 | - | 13307595 | 13307745 | 13307599 | 13307745 | 13305061 | 13305187 | 1 | 1 | 0 |
| A5SS | 4864 | ENSG00000149532 | CPSF7 | chr11 | - | 61429840 | 61429920 | 61429886 | 61429920 | 61429181 | 61429290 | 0 | 1 | 0 |
| A5SS | 5627 | ENSG00000070159 | PTPN3 | chr9 | - | 109383306 | 109383551 | 109383422 | 109383551 | 109382301 | 109382447 | 0 | 0 | 0 |
| A5SS | 5864 | ENSG00000118495 | PLAGL1 | chr6 | - | 144064450 | 144064598 | 144064467 | 144064598 | 143985134 | 143985174 | 0 | 0 | 0 |
| A5SS | 5923 | ENSG00000151164 | RAD9B | chr12 | + | 110502330 | 110502383 | 110502330 | 110502376 | 110503805 | 110503876 | 1 | 0 | 0 |
| A5SS | 6697 | ENSG00000127914 | AKAP9 | chr7 | + | 92062273 | 92065463 | 92062273 | 92062486 | 92066426 | 92066546 | 85 | 104 | 59 |
| A5SS | 7271 | ENSG00000160781 | PAQR6 | chr1 | - | 156247914 | 156248056 | 156247956 | 156248056 | 156246680 | 156246756 | 1 | 0 | 0 |
| A5SS | 7999 | ENSG00000235903 | CPB2-AS1 | chr13 | + | 46098397 | 46098810 | 46098397 | 46098781 | 46113183 | 46113507 | 1 | 0 | 0 |
| A5SS | 3156 | ENSG00000164211 | STARD4 | chr5 | - | 111512384 | 111512535 | 111512390 | 111512535 | 111506329 | 111506379 | 4 | 0 | 3 |
| A5SS | 1887 | ENSG00000006744 | ELAC2 | chr17 | - | 13002440 | 13002579 | 13002445 | 13002579 | 13000155 | 13000274 | 3 | 1 | 1 |
| A5SS | 1824 | ENSG00000119681 | LTBP2 | chr14 | - | 74502563 | 74502934 | 74502652 | 74502934 | 74501440 | 74501590 | 0 | 0 | 1 |
| A5SS | 8811 | ENSG00000135476 | ESPL1 | chr12 | + | 53268754 | 53268928 | 53268754 | 53268847 | 53269023 | 53270085 | 1 | 0 | 1 |
| A5SS | 2069 | ENSG00000154134 | ROBO3 | chr11 | + | 124877518 | 124877752 | 124877518 | 124877658 | 124877936 | 124878131 | 2 | 0 | 1 |
| A5SS | 2920 | ENSG00000176261 | ZBTB8OS | chr1 | - | 32650401 | 32650525 | 32650432 | 32650525 | 32633950 | 32634072 | 1 | 1 | 2 |
| A5SS | 78 | ENSG00000183765 | CHEK2 | chr22 | - | 28724564 | 28724923 | 28724885 | 28724923 | 28719394 | 28719485 | 1 | 1 | 1 |
| A5SS | 781 | ENSG00000185624 | P4HB | chr17 | - | 81859178 | 81859387 | 81859180 | 81859387 | 81855141 | 81855279 | 2 | 0 | 3 |
| A5SS | 6312 | ENSG00000078053 | AMPH | chr7 | - | 38427837 | 38429285 | 38429055 | 38429285 | 38426953 | 38426986 | 0 | 0 | 0 |
| A5SS | 6313 | ENSG00000078053 | AMPH | chr7 | - | 38428749 | 38429285 | 38429055 | 38429285 | 38426953 | 38426986 | 0 | 0 | 0 |
| A5SS | 2644 | ENSG00000163138 | PACRGL | chr4 | + | 20700457 | 20700747 | 20700457 | 20700611 | 20704465 | 20704533 | 4 | 0 | 2 |
| A5SS | 8368 | ENSG00000248538 | NA | chr8 | + | 9189045 | 9189597 | 9189045 | 9189586 | 9198923 | 9198991 | 0 | 0 | 0 |
| A5SS | 2938 | ENSG00000009830 | POMT2 | chr14 | - | 77306245 | 77306441 | 77306336 | 77306441 | 77304691 | 77304800 | 0 | 2 | 2 |
| A5SS | 4286 | ENSG00000181135 | ZNF707 | chr8 | + | 143691072 | 143691232 | 143691072 | 143691199 | 143691596 | 143691713 | 3 | 0 | 0 |
| A5SS | 8515 | ENSG00000165626 | BEND7 | chr10 | - | 13499738 | 13500080 | 13499777 | 13500080 | 13496765 | 13496888 | 0 | 0 | 0 |
| A5SS | 5730 | ENSG00000153253 | SCN3A | chr2 | - | 165140784 | 165140998 | 165140797 | 165140998 | 165139475 | 165139608 | 3 | 0 | 1 |
| A5SS | 8784 | ENSG00000177675 | CD163L1 | chr12 | - | 7433343 | 7433694 | 7433373 | 7433694 | 7432415 | 7432736 | 0 | 1 | 0 |
| A5SS | 2933 | ENSG00000009830 | POMT2 | chr14 | - | 77279683 | 77279928 | 77279822 | 77279928 | 77278728 | 77278869 | 1 | 0 | 2 |
| A5SS | 2645 | ENSG00000163138 | PACRGL | chr4 | + | 20700457 | 20700787 | 20700457 | 20700611 | 20704465 | 20704533 | 7 | 0 | 4 |
| A5SS | 7560 | ENSG00000228340 | MIR646HG | chr20 | + | 60096534 | 60097664 | 60096534 | 60097521 | 60100170 | 60100219 | 2 | 1 | 3 |
| A5SS | 32 | ENSG00000237484 | LINC01684 | chr21 | + | 24438252 | 24438443 | 24438252 | 24438433 | 24441168 | 24441353 | 2 | 5 | 0 |
| A5SS | 5161 | ENSG00000186281 | GPAT2 | chr2 | - | 96023105 | 96024000 | 96023922 | 96024000 | 96022957 | 96023023 | 0 | 0 | 2 |
| A5SS | 2671 | ENSG00000125375 | DMAC2L | chr14 | + | 50312323 | 50313060 | 50312323 | 50312389 | 50321482 | 50321594 | 3 | 0 | 5 |
| A5SS | 8479 | ENSG00000150093 | ITGB1 | chr10 | - | 32936241 | 32936423 | 32936349 | 32936423 | 32935491 | 32935558 | 1 | 6 | 3 |
| A5SS | 6014 | ENSG00000123219 | CENPK | chr5 | - | 65561462 | 65561564 | 65561496 | 65561564 | 65554796 | 65554946 | 32 | 18 | 46 |
| A5SS | 5874 | ENSG00000228203 | GRASLND | chr2 | - | 6916815 | 6917224 | 6916849 | 6917224 | 6915822 | 6915967 | 2 | 0 | 1 |
| A5SS | 5236 | ENSG00000178038 | ALS2CL | chr3 | - | 46681499 | 46681598 | 46681503 | 46681598 | 46681245 | 46681407 | 4 | 4 | 6 |
| A5SS | 11 | ENSG00000160207 | HSF2BP | chr21 | - | 43633262 | 43633421 | 43633271 | 43633421 | 43630321 | 43630454 | 1 | 0 | 0 |
| A5SS | 730 | ENSG00000030582 | GRN | chr17 | + | 44345301 | 44345605 | 44345301 | 44345334 | 44346088 | 44346228 | 2 | 0 | 0 |
| A5SS | 1962 | ENSG00000228624 | HDAC2-AS2 | chr6 | + | 113991892 | 113992187 | 113991892 | 113992018 | 113993342 | 113993487 | 0 | 1 | 0 |
| A5SS | 3670 | ENSG00000182389 | CACNB4 | chr2 | - | 151869176 | 151870611 | 151870530 | 151870611 | 151860710 | 151860820 | 3 | 5 | 4 |
| A5SS | 6916 | ENSG00000204632 | HLA-G | chr6 | + | 29829417 | 29829932 | 29829417 | 29829693 | 29830377 | 29830410 | 0 | 0 | 2 |
| A5SS | 7756 | ENSG00000165699 | TSC1 | chr9 | - | 132911409 | 132911568 | 132911452 | 132911568 | 132910570 | 132910692 | 3 | 2 | 5 |
| A5SS | 8753 | ENSG00000111077 | TNS2 | chr12 | + | 53058713 | 53058854 | 53058713 | 53058827 | 53059046 | 53060258 | 1 | 1 | 0 |
| A5SS | 6554 | ENSG00000152207 | CYSLTR2 | chr13 | + | 48691211 | 48691301 | 48691211 | 48691278 | 48693437 | 48693510 | 12 | 14 | 4 |
| A5SS | 8463 | ENSG00000107862 | GBF1 | chr10 | + | 102376559 | 102376788 | 102376559 | 102376784 | 102376934 | 102377140 | 1 | 4 | 3 |
| A5SS | 3879 | ENSG00000109184 | DCUN1D4 | chr4 | + | 51886567 | 51887244 | 51886567 | 51886638 | 51899269 | 51899378 | 19 | 11 | 17 |
| A5SS | 1577 | ENSG00000166839 | ANKDD1A | chr15 | + | 64917385 | 64917514 | 64917385 | 64917484 | 64921920 | 64922019 | 1 | 0 | 1 |
| A5SS | 6026 | ENSG00000114388 | NPRL2 | chr3 | - | 50348522 | 50348563 | 50348526 | 50348563 | 50347758 | 50347901 | 1 | 0 | 4 |
| A5SS | 2620 | ENSG00000020577 | SAMD4A | chr14 | + | 54784339 | 54784447 | 54784339 | 54784381 | 54784536 | 54784620 | 4 | 3 | 4 |
| A5SS | 4779 | ENSG00000140416 | TPM1 | chr15 | + | 63043705 | 63044152 | 63043705 | 63043831 | 63056984 | 63057118 | 1 | 1 | 1 |
| A5SS | 7653 | ENSG00000137098 | SPAG8 | chr9 | - | 35811181 | 35812001 | 35811923 | 35812001 | 35810882 | 35811057 | 2 | 1 | 7 |
| A5SS | 1189 | ENSG00000070915 | SLC12A3 | chr16 | + | 56892082 | 56892160 | 56892082 | 56892133 | 56892952 | 56893054 | 6 | 0 | 5 |
| A5SS | 8177 | ENSG00000118894 | EEF2KMT | chr16 | - | 5095374 | 5095514 | 5095451 | 5095514 | 5091793 | 5091895 | 1 | 0 | 3 |
| A5SS | 8444 | ENSG00000133678 | TMEM254 | chr10 | + | 80082144 | 80082217 | 80082144 | 80082204 | 80090379 | 80090431 | 2 | 2 | 0 |
| A5SS | 1676 | ENSG00000100749 | VRK1 | chr14 | + | 96855223 | 96855352 | 96855223 | 96855350 | 96856527 | 96856586 | 0 | 0 | 1 |
| A5SS | 7565 | ENSG00000228340 | MIR646HG | chr20 | + | 60378074 | 60378255 | 60378074 | 60378220 | 60379311 | 60380706 | 0 | 3 | 0 |
| A5SS | 560 | ENSG00000174365 | SNHG11 | chr20 | + | 38446687 | 38446967 | 38446687 | 38446948 | 38447465 | 38447623 | 0 | 1 | 0 |
| A5SS | 705 | ENSG00000131095 | GFAP | chr17 | - | 44910091 | 44910658 | 44910614 | 44910658 | 44908063 | 44908149 | 0 | 0 | 1 |
| A5SS | 983 | ENSG00000267248 | NA | chr17 | - | 60128955 | 60129090 | 60128992 | 60129090 | 60128540 | 60128659 | 2 | 0 | 3 |
| A5SS | 1392 | ENSG00000188039 | NWD1 | chr19 | + | 16744420 | 16745147 | 16744420 | 16744718 | 16749138 | 16750411 | 0 | 0 | 2 |
| A5SS | 2279 | ENSG00000113296 | THBS4 | chr5 | + | 79998331 | 79998427 | 79998331 | 79998375 | 80040076 | 80040280 | 1 | 0 | 0 |
| A5SS | 2336 | ENSG00000141084 | RANBP10 | chr16 | - | 67744264 | 67744455 | 67744287 | 67744455 | 67734857 | 67735042 | 0 | 2 | 1 |
| A5SS | 3119 | ENSG00000144395 | CCDC150 | chr2 | + | 196719496 | 196719666 | 196719496 | 196719584 | 196720173 | 196720234 | 0 | 0 | 1 |
| A5SS | 3419 | ENSG00000008128 | CDK11A | chr1 | - | 1709478 | 1709640 | 1709517 | 1709640 | 1709037 | 1709133 | 0 | 0 | 0 |
| A5SS | 3671 | ENSG00000182389 | CACNB4 | chr2 | - | 151869176 | 151872493 | 151872416 | 151872493 | 151860710 | 151860820 | 3 | 4 | 6 |
| A5SS | 3833 | ENSG00000118162 | KPTN | chr19 | - | 47483125 | 47483215 | 47483160 | 47483215 | 47481578 | 47481634 | 0 | 0 | 0 |
| A5SS | 4355 | ENSG00000169981 | ZNF35 | chr3 | + | 44648740 | 44648834 | 44648740 | 44648811 | 44650943 | 44651259 | 1 | 0 | 0 |

| | | | | | | | | | | | | | | | | | | | | | | | | | | | | | | | | | | | | | | | | | | | | | | |
|---|---|---|---|---|---|---|---|---|---|---|---|---|---|---|---|---|---|---|---|---|---|---|---|---|---|---|---|---|---|---|---|---|---|---|---|---|---|---|---|---|---|---|---|---|---|---|
| A5SS | 5572 | ENSG00000238009 | NA | chr1 | - | 120720 | 120932 | 120774 | 120932 | 112699 | 112804 | 0 | 0 | 0 | 1 | 0 | 0 | 0 | 0 | 0 | 0 | 0 | 0 | 0 | 0 | 0 | 0 | 0 | 0 | 0 | 3 | 203 | 149 | 1 | 1 | NA | NA | NA | 1 | NA | NA | NA | NA | NA | 0 | 1 |
| A5SS | 6290 | ENSG00000109339 | MAPK10 | chr4 | - | 86194308 | 86194407 | 86194335 | 86194407 | 86107222 | 86107352 | 2 | 0 | 0 | 0 | 0 | 0 | 0 | 0 | 0 | 0 | 0 | 0 | 0 | 0 | 0 | 0 | 0 | 0 | 0 | 5 | 176 | 149 | 1 | 1 | 1 | NA | NA | NA | NA | NA | NA | NA | NA | 0 | 1 |
| A5SS | 6434 | ENSG00000119616 | FCF1 | chr14 | + | 74713169 | 74713259 | 74713169 | 74713200 | 74714871 | 74714943 | 0 | 0 | 0 | 2 | 0 | 0 | 0 | 0 | 0 | 0 | 0 | 0 | 0 | 0 | 0 | 0 | 0 | 0 | 2 | 0 | 208 | 149 | 1 | 1 | NA | NA | NA | 1 | NA | NA | NA | NA | 0 | NA | 1 |
| A5SS | 6583 | ENSG00000115677 | HDLBP | chr2 | - | 241269200 | 241269295 | 241269205 | 241269295 | 241268953 | 241269105 | 0 | 0 | 1 | 0 | 2 | 0 | 0 | 0 | 0 | 0 | 0 | 0 | 0 | 0 | 0 | 0 | 0 | 0 | 2 | 0 | 154 | 149 | 1 | 1 | NA | NA | 1 | NA | | 1 | NA | NA | 0 | NA | 1 |
| A5SS | 6718 | ENSG00000122786 | CALD1 | chr7 | + | 134932987 | 134934077 | 134932987 | 134933390 | 134935687 | 134935765 | 0 | 1 | 0 | 0 | 0 | 0 | 0 | 0 | 0 | 0 | 0 | 0 | 0 | 0 | 0 | 0 | 0 | 0 | 3 | 0 | 298 | 149 | 1 | 1 | NA | 1 | NA | NA | NA | NA | NA | NA | 0 | NA | 1 |
| A5SS | 6930 | ENSG00000162669 | HFM1 | chr1 | - | 91379058 | 91379214 | 91379062 | 91379214 | 91378402 | 91378480 | 0 | 0 | 0 | 4 | 0 | 0 | 0 | 0 | 0 | 0 | 0 | 0 | 0 | 0 | 0 | 1 | 0 | 0 | 0 | 0 | 153 | 149 | 1 | 1 | NA | NA | NA | 1 | NA | NA | 0 | NA | NA | NA | 1 |
| A5SS | 7031 | ENSG00000184343 | SRPK3 | chrX | + | 153783973 | 153784024 | 153783973 | 153784021 | 153784120 | 153784213 | 0 | 0 | 0 | 7 | 0 | 0 | 0 | 0 | 0 | 0 | 0 | 0 | 0 | 0 | 0 | 0 | 0 | 0 | 5 | 0 | 152 | 149 | 1 | 1 | NA | NA | NA | 1 | NA | NA | NA | NA | 0 | NA | 1 |
| A5SS | 7178 | ENSG00000234665 | LERFS | chr9 | - | 62869076 | 62869304 | 62869189 | 62869304 | 62867696 | 62867876 | 1 | 0 | 0 | 2 | 0 | 0 | 0 | 0 | 0 | 0 | 0 | 0 | 0 | 0 | 0 | 0 | 0 | 0 | 2 | 0 | 262 | 149 | 1 | 1 | 1 | NA | NA | 1 | NA | NA | NA | NA | 0 | NA | 1 |
| A5SS | 7619 | ENSG00000116580 | GON4L | chr1 | - | 155857146 | 155857295 | 155857158 | 155857295 | 155826836 | 155827028 | 3 | 0 | 0 | 0 | 0 | 0 | 0 | 0 | 0 | 0 | 0 | 0 | 0 | 0 | 0 | 0 | 1 | 0 | 0 | 0 | 161 | 149 | 1 | 1 | 1 | NA | NA | NA | NA | NA | NA | 0 | NA | NA | 1 |
| A5SS | 8805 | ENSG00000111319 | SCNN1A | chr12 | - | 6362050 | 6362241 | 6362067 | 6362241 | 6355776 | 6355880 | 0 | 2 | 0 | 7 | 0 | 0 | 0 | 0 | 0 | 0 | 0 | 0 | 0 | 0 | 0 | 0 | 0 | 0 | 0 | 1 | 166 | 149 | 1 | 1 | NA | 1 | NA | 1 | NA | NA | NA | NA | NA | 0 | 1 |
| A5SS | 8933 | ENSG00000184497 | TMEM255B | chr13 | + | 113800826 | 113800912 | 113800826 | 113800865 | 113801652 | 113801812 | 0 | 0 | 0 | 1 | 0 | 0 | 0 | 0 | 0 | 0 | 0 | 0 | 0 | 0 | 0 | 0 | 2 | 0 | 0 | 9 | 196 | 149 | 1 | 1 | NA | NA | NA | 1 | NA | NA | NA | 0 | NA | 0 | 1 |

U12_A5SS_filtered

| Splice_type | ID | Gene_ID | Gene_Symbol | Chromosome | Strand | LongExonStart_0base | LongExonEnd | ShortES | ShortEE | FlankingES | FlankingEE | Inclusion_counts_HC SRR13214146 | SRR13214147 | SRR13214148 | SRR13214149 | SRR13214150 | Skipping_counts_HC SRR13214146 | SRR13214147 |
|---|---|---|---|---|---|---|---|---|---|---|---|---|---|---|---|---|---|---|
| A5SS | 5360 | ENSG00000084623 | EIF3I | chr1 | + | 32228726 | 32229024 | 32228726 | 32228816 | 32229134 | 32229208 | 17 | 11 | 18 | 19 | 12 | 136 | 82 |
| A5SS | 6633 | ENSG00000104133 | SPG11 | chr15 | - | 44596059 | 44596355 | 44596082 | 44596355 | 44595258 | 44595459 | 42 | 23 | 22 | 27 | 37 | 353 | 293 |
| A5SS | 1516 | ENSG00000140199 | SLC12A6 | chr15 | - | 34237389 | 34237550 | 34237418 | 34237550 | 34236707 | 34236815 | 6 | 7 | 10 | 12 | 10 | 273 | 231 |
| A5SS | 5356 | ENSG00000084623 | EIF3I | chr1 | + | 32224409 | 32226530 | 32224409 | 32224475 | 32228498 | 32228609 | 253 | 143 | 234 | 228 | 363 | 5 | 4 |
| A5SS | 8922 | ENSG00000117226 | GBP3 | chr1 | - | 89014404 | 89014656 | 89014546 | 89014656 | 89014082 | 89014279 | 28 | 34 | 47 | 42 | 55 | 61 | 75 |
| A5SS | 3077 | ENSG00000111642 | CHD4 | chr12 | - | 6587660 | 6587949 | 6587711 | 6587949 | 6587383 | 6587559 | 10 | 6 | 7 | 8 | 2 | 359 | 260 |
| A5SS | 4677 | ENSG00000011021 | CLCN6 | chr1 | + | 11826155 | 11826224 | 11826155 | 11826214 | 11827088 | 11827221 | 0 | 0 | 0 | 2 | 0 | 10 | 10 |
| A5SS | 5361 | ENSG00000084623 | EIF3I | chr1 | + | 32228726 | 32229208 | 32228726 | 32228816 | 32230926 | 32231019 | 161 | 113 | 180 | 196 | 136 | 0 | 0 |
| A5SS | 1648 | ENSG00000185024 | BRF1 | chr14 | - | 105246865 | 105247193 | 105247070 | 105247193 | 105241264 | 105241414 | 6 | 2 | 13 | 6 | 7 | 1 | 0 |
| A5SS | 5359 | ENSG00000084623 | EIF3I | chr1 | + | 32226170 | 32226530 | 32226170 | 32226320 | 32228498 | 32228609 | 236 | 145 | 231 | 223 | 356 | 1 | 2 |
| A5SS | 5592 | ENSG00000143390 | RFX5 | chr1 | - | 151346484 | 151346606 | 151346488 | 151346606 | 151346204 | 151346333 | 3 | 0 | 0 | 4 | 1 | 282 | 159 |
| A5SS | 1585 | ENSG00000137841 | PLCB2 | chr15 | - | 40296745 | 40296908 | 40296757 | 40296908 | 40296295 | 40296392 | 12 | 5 | 3 | 1 | 1 | 1 | 0 |
| A5SS | 573 | ENSG00000105576 | TNPO2 | chr19 | - | 12723768 | 12723932 | 12723772 | 12723932 | 12723248 | 12723365 | 3 | 1 | 3 | 8 | 20 | 1 | 0 |
| A5SS | 3076 | ENSG00000111642 | CHD4 | chr12 | - | 6583195 | 6583378 | 6583197 | 6583378 | 6583026 | 6583113 | 1 | 1 | 2 | 3 | 0 | 323 | 203 |
| A5SS | 1661 | ENSG00000174373 | RALGAPA1 | chr14 | - | 35688458 | 35690003 | 35689835 | 35690003 | 35686541 | 35686666 | 20 | 11 | 30 | 11 | 57 | 62 | 37 |
| A5SS | 3808 | ENSG00000163608 | NEPRO | chr3 | - | 113013948 | 113014032 | 113013952 | 113014032 | 113013271 | 113013403 | 2 | 2 | 3 | 6 | 11 | 21 | 22 |
| A5SS | 126 | ENSG00000100325 | ASCC2 | chr22 | - | 29834441 | 29834550 | 29834488 | 29834550 | 29825621 | 29825780 | 5 | 1 | 2 | 4 | 9 | 4 | 0 |
| A5SS | 8426 | ENSG00000148688 | RPP30 | chr10 | + | 90894774 | 90895483 | 90894774 | 90894891 | 90895879 | 90895917 | 95 | 50 | 105 | 69 | 100 | 6 | 12 |
| A5SS | 8086 | ENSG00000110274 | CEP164 | chr11 | + | 117408889 | 117409028 | 117408889 | 117409004 | 117409617 | 117409965 | 25 | 15 | 16 | 22 | 20 | 9 | 5 |
| A5SS | 3071 | ENSG00000111642 | CHD4 | chr12 | - | 6573070 | 6573269 | 6573073 | 6573269 | 6570868 | 6571032 | 13 | 2 | 8 | 16 | 22 | 232 | 109 |
| A5SS | 2903 | ENSG00000164068 | RNF123 | chr3 | + | 49689556 | 49689822 | 49689556 | 49689606 | 49691129 | 49691247 | 2 | 0 | 0 | 4 | 6 | 11 | 8 |
| A5SS | 3763 | ENSG00000088035 | ALG6 | chr1 | + | 63411145 | 63411331 | 63411145 | 63411155 | 63411681 | 63411806 | 30 | 25 | 36 | 41 | 92 | 0 | 0 |
| A5SS | 6964 | ENSG00000100532 | CGRRF1 | chr14 | + | 54530048 | 54530763 | 54530048 | 54530226 | 54530902 | 54531050 | 13 | 2 | 5 | 11 | 13 | 38 | 34 |
| A5SS | 3762 | ENSG00000088035 | ALG6 | chr1 | + | 63370770 | 63371287 | 63370770 | 63371059 | 63396512 | 63396597 | 11 | 7 | 13 | 21 | 21 | 98 | 32 |
| A5SS | 7111 | ENSG00000090316 | MAEA | chr4 | + | 1311978 | 1312238 | 1311978 | 1312161 | 1315396 | 1315600 | 2 | 5 | 8 | 0 | 0 | 40 | 22 |
| A5SS | 5240 | ENSG00000010244 | ZNF207 | chr17 | + | 32369598 | 32369782 | 32369598 | 32369745 | 32373398 | 32373457 | 190 | 138 | 201 | 268 | 182 | 3 | 2 |
| A5SS | 5344 | ENSG00000132676 | DAP3 | chr1 | + | 155689090 | 155689243 | 155689090 | 155689174 | 155709772 | 155709824 | 77 | 39 | 87 | 74 | 39 | 168 | 107 |
| A5SS | 8084 | ENSG00000110274 | CEP164 | chr11 | + | 117380613 | 117380714 | 117380613 | 117380705 | 117381700 | 117381868 | 3 | 3 | 6 | 3 | 1 | 39 | 23 |
| A5SS | 3971 | ENSG00000163479 | SSR2 | chr1 | - | 156018131 | 156018368 | 156018269 | 156018368 | 156014960 | 156015069 | 11 | 4 | 9 | 26 | 6 | 1147 | 420 |
| A5SS | 5358 | ENSG00000084623 | EIF3I | chr1 | + | 32226170 | 32226530 | 32226170 | 32226283 | 32228498 | 32228609 | 258 | 167 | 258 | 250 | 386 | 1 | 0 |
| A5SS | 874 | ENSG00000108587 | GOSR1 | chr17 | + | 30481142 | 30481263 | 30481142 | 30481257 | 30484213 | 30484301 | 46 | 33 | 43 | 43 | 41 | 77 | 68 |
| A5SS | 4816 | ENSG00000162909 | CAPN2 | chr1 | + | 223761580 | 223761739 | 223761580 | 223761617 | 223762185 | 223762251 | 37 | 27 | 34 | 10 | 12 | 557 | 300 |
| A5SS | 8428 | ENSG00000108094 | CUL2 | chr10 | - | 35090178 | 35090642 | 35090429 | 35090642 | 35071198 | 35071339 | 14 | 11 | 8 | 8 | 13 | 27 | 21 |
| A5SS | 2799 | ENSG00000068745 | IP6K2 | chr3 | - | 48694693 | 48695421 | 48695089 | 48695421 | 48694458 | 48694525 | 36 | 9 | 13 | 23 | 10 | 19 | 6 |
| A5SS | 8548 | ENSG00000079974 | RABL2B | chr22 | - | 50783038 | 50783286 | 50783044 | 50783286 | 50782187 | 50782351 | 3 | 4 | 7 | 8 | 14 | 1 | 0 |
| A5SS | 6341 | ENSG00000173715 | C11orf80 | chr11 | + | 66801030 | 66801161 | 66801030 | 66801102 | 66803994 | 66804194 | 1 | 0 | 3 | 1 | 2 | 15 | 2 |
| A5SS | 2344 | ENSG00000105875 | WDR91 | chr7 | - | 135197043 | 135198151 | 135197992 | 135198151 | 135196143 | 135196337 | 8 | 8 | 4 | 1 | 5 | 30 | 16 |
| A5SS | 5357 | ENSG00000084623 | EIF3I | chr1 | + | 32226170 | 32226320 | 32226170 | 32226283 | 32228498 | 32228609 | 23 | 25 | 29 | 34 | 37 | 1 | 0 |
| A5SS | 2624 | ENSG00000108591 | DRG2 | chr17 | + | 18093812 | 18093973 | 18093812 | 18093968 | 18098269 | 18098359 | 55 | 29 | 32 | 46 | 34 | 9 | 0 |
| A5SS | 8296 | ENSG00000143702 | CEP170 | chr1 | - | 243185778 | 243186422 | 243186258 | 243186422 | 243172696 | 243172846 | 157 | 158 | 113 | 130 | 216 | 24 | 24 |
| A5SS | 4787 | ENSG00000131778 | CHD1L | chr1 | + | 147270931 | 147271195 | 147270931 | 147271005 | 147272170 | 147272281 | 36 | 32 | 17 | 18 | 16 | 42 | 17 |
| A5SS | 7417 | ENSG00000174744 | BRMS1 | chr11 | - | 66341205 | 66341333 | 66341212 | 66341333 | 66340966 | 66341046 | 180 | 128 | 130 | 152 | 160 | 0 | 1 |
| A5SS | 6920 | ENSG00000147459 | DOCK5 | chr8 | + | 25368570 | 25368725 | 25368570 | 25368687 | 25369555 | 25369641 | 176 | 159 | 80 | 177 | 197 | 0 | 0 |
| A5SS | 6051 | ENSG00000105393 | BABAM1 | chr19 | + | 17267442 | 17267527 | 17267442 | 17267516 | 17268793 | 17269091 | 55 | 27 | 57 | 53 | 23 | 2 | 2 |
| A5SS | 3235 | ENSG00000197386 | HTT | chr4 | + | 3172907 | 3173331 | 3172907 | 3173131 | 3174720 | 3174799 | 5 | 2 | 6 | 7 | 0 | 64 | 38 |
| A5SS | 5272 | ENSG00000186141 | POLR3C | chr1 | + | 145826453 | 145826709 | 145826453 | 145826492 | 145826819 | 145827005 | 134 | 106 | 163 | 178 | 245 | 0 | 0 |
| A5SS | 759 | ENSG00000214087 | ARL16 | chr17 | - | 81682442 | 81682675 | 81682490 | 81682675 | 81682033 | 81682149 | 16 | 11 | 19 | 6 | 15 | 10 | 2 |
| A5SS | 2625 | ENSG00000108591 | DRG2 | chr17 | + | 18099016 | 18099093 | 18099016 | 18099077 | 18099632 | 18100435 | 4 | 4 | 2 | 6 | 5 | 40 | 17 |
| A5SS | 4103 | ENSG00000256269 | HMBS | chr11 | + | 119088634 | 119088848 | 119088634 | 119088707 | 119089081 | 119089131 | 0 | 0 | 1 | 2 | 8 | 29 | 18 |
| A5SS | 2906 | ENSG00000164068 | RNF123 | chr3 | + | 49712478 | 49712892 | 49712478 | 49712656 | 49713512 | 49713587 | 14 | 2 | 5 | 3 | 6 | 9 | 9 |
| A5SS | 4342 | ENSG00000111445 | RFC5 | chr12 | + | 118017689 | 118018053 | 118017689 | 118017815 | 118019071 | 118019136 | 9 | 14 | 14 | 8 | 29 | 2 | 1 |
| A5SS | 8812 | ENSG00000123349 | PFDN5 | chr12 | + | 53295838 | 53296275 | 53295838 | 53295941 | 53297849 | 53297924 | 1689 | 998 | 1549 | 1475 | 1772 | 3 | 13 |
| A5SS | 2700 | ENSG00000099949 | LZTR1 | chr22 | + | 20983026 | 20983122 | 20983026 | 20983089 | 20985840 | 20985897 | 5 | 0 | 2 | 2 | 0 | 36 | 11 |
| A5SS | 7252 | ENSG00000221914 | PPP2R2A | chr8 | + | 26293665 | 26293860 | 26293665 | 26293740 | 26338889 | 26338987 | 24 | 20 | 18 | 25 | 9 | 95 | 95 |
| A5SS | 720 | ENSG00000170004 | CHD3 | chr17 | + | 7909142 | 7909384 | 7909142 | 7909338 | 7910427 | 7910591 | 7 | 3 | 23 | 24 | 30 | 78 | 57 |
| A5SS | 4179 | ENSG00000139636 | LMBR1L | chr12 | - | 49106510 | 49106746 | 49106544 | 49106746 | 49105923 | 49105957 | 8 | 6 | 9 | 16 | 29 | 1 | 0 |
| A5SS | 8519 | ENSG00000139620 | KANSL2 | chr12 | - | 48660365 | 48660619 | 48660467 | 48660619 | 48654940 | 48655060 | 110 | 89 | 105 | 108 | 104 | 5 | 3 |
| A5SS | 7391 | ENSG00000136518 | ACTL6A | chr3 | + | 179562922 | 179563117 | 179562922 | 179562973 | 179569823 | 179569900 | 22 | 3 | 13 | 9 | 1 | 4 | 7 |
| A5SS | 6963 | ENSG00000100532 | CGRRF1 | chr14 | + | 54530048 | 54530405 | 54530048 | 54530226 | 54530902 | 54531050 | 7 | 2 | 5 | 9 | 9 | 38 | 34 |
| A5SS | 5396 | ENSG00000089060 | SLC8B1 | chr12 | - | 113332722 | 113332960 | 113332748 | 113332960 | 113321195 | 113321348 | 37 | 22 | 58 | 28 | 9 | 3 | 4 |
| A5SS | 3074 | ENSG00000111642 | CHD4 | chr12 | - | 6581037 | 6581173 | 6581043 | 6581173 | 6578845 | 6578917 | 36 | 19 | 18 | 24 | 13 | 320 | 254 |
| A5SS | 6593 | ENSG00000106477 | CEP41 | chr7 | - | 130404563 | 130404708 | 130404639 | 130404708 | 130402647 | 130402799 | 24 | 23 | 22 | 27 | 50 | 0 | 0 |
| A5SS | 7251 | ENSG00000221914 | PPP2R2A | chr8 | + | 26293665 | 26293860 | 26293665 | 26293740 | 26333485 | 26333570 | 19 | 16 | 17 | 22 | 9 | 12 | 0 |
| A5SS | 6367 | ENSG00000168014 | C2CD3 | chr11 | - | 74085521 | 74085886 | 74085617 | 74085886 | 74084880 | 74084970 | 10 | 3 | 21 | 13 | 10 | 66 | 73 |
| A5SS | 5587 | ENSG00000143390 | RFX5 | chr1 | - | 151346421 | 151346606 | 151346488 | 151346606 | 151346204 | 151346333 | 12 | 15 | 15 | 22 | 25 | 282 | 159 |
| A5SS | 3073 | ENSG00000111642 | CHD4 | chr12 | - | 6578337 | 6578546 | 6578408 | 6578546 | 6578028 | 6578117 | 15 | 17 | 5 | 7 | 4 | 1 | 1 |
| A5SS | 1282 | ENSG00000122390 | NAA60 | chr16 | + | 3484698 | 3485083 | 3484698 | 3485061 | 3485466 | 3486952 | 8 | 0 | 4 | 6 | 0 | 11 | 4 |
| A5SS | 227 | ENSG00000005007 | UPF1 | chr19 | + | 18852986 | 18853104 | 18852986 | 18853071 | 18853251 | 18853350 | 44 | 17 | 46 | 51 | 37 | 99 | 50 |
| A5SS | 4341 | ENSG00000111445 | RFC5 | chr12 | + | 118017689 | 118017852 | 118017689 | 118017815 | 118019071 | 118019136 | 1 | 6 | 1 | 5 | 15 | 2 | 1 |
| A5SS | 194 | ENSG00000178982 | EIF3K | chr19 | + | 38619195 | 38619327 | 38619195 | 38619317 | 38620336 | 38620435 | 1266 | 498 | 1045 | 932 | 508 | 24 | 19 |
| A5SS | 5127 | ENSG00000112855 | HARS2 | chr5 | + | 140691794 | 140692017 | 140691794 | 140691937 | 140693590 | 140693665 | 7 | 11 | 4 | 10 | 4 | 2 | 4 |
| A5SS | 3234 | ENSG00000197386 | HTT | chr4 | + | 3172907 | 3173218 | 3172907 | 3173131 | 3174720 | 3174799 | 4 | 4 | 4 | 9 | 6 | 64 | 38 |
| A5SS | 2697 | ENSG00000113504 | SLC12A7 | chr5 | - | 1075890 | 1076236 | 1076137 | 1076236 | 1075370 | 1075490 | 10 | 4 | 0 | 2 | 0 | 29 | 7 |
| A5SS | 8163 | ENSG00000159063 | ALG8 | chr11 | - | 78138709 | 78139624 | 78139493 | 78139624 | 78127357 | 78127436 | 4 | 4 | 2 | 2 | 10 | 13 | 9 |
| A5SS | 4178 | ENSG00000139636 | LMBR1L | chr12 | - | 49104718 | 49104885 | 49104745 | 49104885 | 49104447 | 49104551 | 0 | 3 | 2 | 1 | 2 | 24 | 28 |
| A5SS | 4786 | ENSG00000131778 | CHD1L | chr1 | + | 147270931 | 147271113 | 147270931 | 147271005 | 147272170 | 147272281 | 22 | 14 | 18 | 10 | 10 | 42 | 17 |
| A5SS | 3869 | ENSG00000095564 | BTAF1 | chr10 | + | 91959064 | 91959282 | 91959064 | 91959154 | 91959784 | 91959880 | 380 | 388 | 386 | 376 | 512 | 55 | 37 |
| A5SS | 8813 | ENSG00000123349 | PFDN5 | chr12 | + | 53296429 | 53296551 | 53296429 | 53296467 | 53297849 | 53297924 | 21 | 17 | 32 | 28 | 14 | 0 | 2 |
| A5SS | 5239 | ENSG00000152689 | RASGRP3 | chr2 | + | 33539093 | 33539323 | 33539093 | 33539210 | 33543511 | 33543627 | 1 | 3 | 6 | 10 | 2 | 13 | 3 |
| A5SS | 5128 | ENSG00000112855 | HARS2 | chr5 | + | 140691794 | 140692329 | 140691794 | 140691937 | 140693590 | 140693665 | 19 | 14 | 10 | 22 | 11 | 2 | 4 |
| A5SS | 4674 | ENSG00000124571 | XPO5 | chr6 | - | 43573397 | 43573601 | 43573479 | 43573601 | 43572505 | 43572578 | 0 | 1 | 2 | 0 | 1 | 71 | 65 |
| A5SS | 5590 | ENSG00000143390 | RFX5 | chr1 | - | 151346481 | 151346606 | 151346488 | 151346606 | 151346204 | 151346333 | 64 | 61 | 54 | 48 | 118 | 282 | 159 |
| A5SS | 7074 | ENSG00000215021 | PHB2 | chr12 | - | 6967170 | 6968021 | 6967891 | 6968021 | 6966464 | 6966500 | 640 | 313 | 630 | 535 | 978 | 0 | 2 |
| A5SS | 5544 | ENSG00000107521 | HPS1 | chr10 | - | 98433939 | 98434091 | 98433982 | 98434091 | 98431130 | 98431291 | 33 | 10 | 21 | 15 | 13 | 59 | 22 |
| A5SS | 3233 | ENSG00000197386 | HTT | chr4 | + | 3129923 | 3130047 | 3129923 | 3129999 | 3130304 | 3130423 | 199 | 134 | 165 | 199 | 155 | 3 | 5 |
| A5SS | 8334 | ENSG00000197771 | MCMBP | chr10 | - | 119843247 | 119843426 | 119843253 | 119843426 | 119842471 | 119842595 | 87 | 45 | 54 | 66 | 134 | 157 | 77 |
| A5SS | 3764 | ENSG00000088035 | ALG6 | chr1 | + | 63411145 | 63411331 | 63411145 | 63411155 | 63411925 | 63412061 | 47 | 28 | 59 | 60 | 130 | 10 | 0 |
| A5SS | 6635 | ENSG00000104133 | SPG11 | chr15 | - | 44620124 | 44620403 | 44620189 | 44620403 | 44615362 | 44615566 | 20 | 11 | 18 | 11 | 16 | 213 | 142 |
| A5SS | 7452 | ENSG00000164941 | INTS8 | chr8 | + | 94865505 | 94865700 | 94865505 | 94865690 | 94866157 | 94866191 | 5 | 2 | 6 | 8 | 7 | 177 | 93 |
| A5SS | 3678 | ENSG00000162946 | DISC1 | chr1 | + | 232008784 | 232009049 | 232008784 | 232008983 | 232026434 | 232026552 | 12 | 10 | 9 | 12 | 17 | 4 | 9 |

| | | | | | | | | | | | | | | | | | | | | | | | | | | | | | | | | | | | | | | | | | | | | | | |
|---|---|---|---|---|---|---|---|---|---|---|---|---|---|---|---|---|---|---|---|---|---|---|---|---|---|---|---|---|---|---|---|---|---|---|---|---|---|---|---|---|---|---|---|---|---|---|
| A5SS | 5583 | ENSG00000143390 | RFX5 | chr1 | - | 151346204 | 151346333 | 151346214 | 151346333 | 151345105 | 151345188 | 20 | 7 | 20 | 12 | 24 | 0 | 0 | 0 | 0 | 0 | 23 | 8 | 19 | 26 | 30 | 0 | 0 | 0 | 2 | 0 | 159 | 149 | 1 | 1 | 1 | 1 | 1 | 1 | 1 | 1 | 1 | 1 | 0.924 | 1 | 0.015 |
| A5SS | 5722 | ENSG00000153253 | SCN3A | chr2 | - | 165140650 | 165140998 | 165140701 | 165140998 | 165139475 | 165139608 | 2 | 0 | 0 | 0 | 1 | 1 | 6 | 1 | 6 | 0 | 0 | 6 | 6 | 0 | 0 | 0 | 0 | 0 | 0 | 0 | 200 | 149 | 1 | 1 | 0.598 | 0 | 0 | 0 | 1 | NA | 1 | 1 | NA | NA | -0.68 |
| A5SS | 5373 | ENSG00000169432 | SCN9A | chr2 | - | 166284452 | 166284824 | 166284485 | 166284824 | 166281678 | 166281808 | 0 | 1 | 2 | 0 | 0 | 6 | 1 | 9 | 5 | 1 | 0 | 0 | 1 | 0 | 0 | 5 | 0 | 0 | 8 | 0 | 182 | 149 | 1 | 1 | 0 | 0.45 | 0.154 | 0 | 0 | 0 | NA | 1 | 0 | NA | -0.213 |
| A5SS | 4343 | ENSG00000111445 | RFC5 | chr12 | + | 118017689 | 118018053 | 118017689 | 118017852 | 118019071 | 118019136 | 10 | 10 | 14 | 5 | 15 | 0 | 0 | 0 | 2 | 0 | 8 | 8 | 23 | 22 | 10 | 0 | 0 | 0 | 0 | 0 | 298 | 149 | 1 | 1 | 1 | 1 | 1 | 0.556 | 1 | 1 | 1 | 1 | 1 | 1 | -0.089 |
| A5SS | 7113 | ENSG00000090316 | MAEA | chr4 | + | 1311978 | 1312238 | 1311978 | 1312161 | 1332756 | 1332865 | 0 | 2 | 2 | 1 | 0 | 0 | 0 | 0 | 0 | 0 | 5 | 0 | 10 | 8 | 0 | 0 | 0 | 0 | 2 | 0 | 226 | 149 | 1 | 1 | NA | 1 | 1 | 1 | NA | 1 | NA | 1 | 0.725 | NA | 0.092 |
| A5SS | 4678 | ENSG00000011021 | CLCN6 | chr1 | + | 11826155 | 11826224 | 11826155 | 11826214 | 11827114 | 11827221 | 0 | 0 | 0 | 0 | 1 | 0 | 0 | 6 | 2 | 0 | 9 | 0 | 0 | 0 | 0 | 1 | 1 | 1 | 5 | 0 | 159 | 149 | 1 | 1 | NA | NA | 0 | 0 | 1 | 0.894 | 0 | 0 | 0 | NA | 0.11 |
| A5SS | 2062 | ENSG00000131504 | DIAPH1 | chr5 | - | 141527570 | 141527697 | 141527572 | 141527697 | 141526296 | 141526459 | 3 | 3 | 2 | 7 | 10 | 0 | 0 | 0 | 0 | 1 | 9 | 4 | 5 | 9 | 11 | 0 | 0 | 3 | 0 | 0 | 151 | 149 | 1 | 1 | 1 | 1 | 1 | 1 | 0.908 | 1 | 1 | 0.622 | 1 | 1 | 0.057 |
| A5SS | 5016 | ENSG00000124067 | SLC12A4 | chr16 | - | 67963436 | 67963559 | 67963464 | 67963559 | 67961574 | 67961706 | 0 | 0 | 0 | 0 | 0 | 3 | 2 | 2 | 1 | 5 | 0 | 0 | 1 | 0 | 0 | 0 | 0 | 5 | 3 | 18 | 177 | 149 | 1 | 1 | 0 | 0 | 0 | 0 | 0 | NA | NA | 0.144 | 0 | 0 | -0.048 |
| A5SS | 6098 | ENSG00000221838 | AP4M1 | chr7 | + | 100101879 | 100101989 | 100101879 | 100101968 | 100102674 | 100102781 | 0 | 2 | 1 | 1 | 0 | 6 | 0 | 5 | 5 | 2 | 0 | 0 | 0 | 0 | 0 | 8 | 1 | 2 | 0 | 0 | 170 | 149 | 1 | 1 | 0 | 1 | 0.149 | 0.149 | 0 | 0 | 0 | 0 | NA | NA | 0.26 |
| A5SS | 6099 | ENSG00000221838 | AP4M1 | chr7 | + | 100101879 | 100102393 | 100101879 | 100101968 | 100102674 | 100102781 | 3 | 1 | 0 | 0 | 1 | 6 | 0 | 5 | 5 | 2 | 0 | 1 | 0 | 0 | 0 | 8 | 1 | 2 | 0 | 0 | 298 | 149 | 1 | 1 | 0.2 | 1 | 0 | 0 | 0.2 | 0 | 0.333 | 0 | NA | NA | 0.169 |
| A5SS | 4306 | ENSG00000043514 | TRIT1 | chr1 | - | 39850118 | 39850261 | 39850122 | 39850261 | 39847985 | 39848091 | 1 | 0 | 2 | 0 | 0 | 2 | 2 | 0 | 0 | 0 | 0 | 0 | 1 | 0 | 10 | 0 | 0 | 0 | 0 | 0 | 153 | 149 | 1 | 1 | 0.327 | 0 | 1 | NA | NA | NA | NA | 1 | NA | 1 | -0.558 |
| A5SS | 6962 | ENSG00000100802 | C14orf93 | chr14 | - | 23010016 | 23010143 | 23010100 | 23010143 | 23000958 | 23001037 | 0 | 0 | 0 | 0 | 0 | 1 | 1 | 3 | 1 | 3 | 0 | 0 | 3 | 0 | 0 | 6 | 0 | 1 | 2 | 0 | 233 | 149 | 1 | 1 | 0 | 0 | 0 | 0 | 0 | 0 | NA | 0.657 | 0 | NA | -0.219 |
| A5SS | 5015 | ENSG00000124067 | SLC12A4 | chr16 | - | 67962243 | 67963559 | 67963464 | 67963559 | 67961574 | 67961706 | 4 | 0 | 4 | 1 | 1 | 3 | 2 | 2 | 1 | 5 | 0 | 1 | 4 | 3 | 0 | 0 | 0 | 5 | 3 | 18 | 298 | 149 | 1 | 1 | 0.4 | 0 | 0.5 | 0.333 | 0.091 | NA | 1 | 0.286 | 0.333 | 0 | -0.14 |
| A5SS | 5374 | ENSG00000169432 | SCN9A | chr2 | - | 166375696 | 166375944 | 166375734 | 166375944 | 166311498 | 166311806 | 3 | 2 | 0 | 2 | 0 | 0 | 0 | 0 | 0 | 0 | 0 | 9 | 2 | 3 | 4 | 0 | 0 | 0 | 2 | 0 | 187 | 149 | 1 | 1 | 1 | 1 | NA | 1 | NA | NA | 1 | 1 | 0.544 | 1 | 0.114 |
| A5SS | 4100 | ENSG00000136895 | GARNL3 | chr9 | + | 127243078 | 127243253 | 127243078 | 127243249 | 127264951 | 127265021 | 1 | 0 | 0 | 1 | 0 | 1 | 2 | 4 | 1 | 1 | 0 | 0 | 3 | 0 | 0 | 0 | 0 | 0 | 0 | 0 | 153 | 149 | 1 | 1 | 0.493 | 0 | 0 | 0.493 | 0 | NA | NA | 1 | NA | NA | -0.803 |
| A5SS | 5560 | ENSG00000112367 | FIG4 | chr6 | + | 109788797 | 109788956 | 109788797 | 109788913 | 109789593 | 109789677 | 4 | 2 | 2 | 0 | 3 | 1 | 0 | 1 | 3 | 0 | 14 | 0 | 2 | 0 | 0 | 0 | 3 | 0 | 0 | 0 | 192 | 149 | 1 | 1 | 0.756 | 1 | 0.608 | 0 | 1 | 1 | 0 | 1 | NA | NA | 0.006 |
| A5SS | 1524 | ENSG00000167077 | MEI1 | chr22 | + | 41758364 | 41758547 | 41758364 | 41758533 | 41763173 | 41763321 | 0 | 0 | 0 | 0 | 0 | 0 | 1 | 1 | 1 | 0 | 0 | 4 | 1 | 0 | 0 | 0 | 2 | 2 | 0 | 0 | 163 | 149 | 1 | 1 | NA | 0 | 0 | 0 | NA | NA | 0.646 | 0.314 | NA | NA | -0.48 |
| A5SS | 2627 | ENSG00000108591 | DRG2 | chr17 | + | 18099016 | 18099093 | 18099016 | 18099077 | 18100362 | 18100435 | 1 | 3 | 0 | 6 | 1 | 3 | 0 | 1 | 1 | 0 | 5 | 1 | 8 | 0 | 0 | 0 | 0 | 0 | 0 | 0 | 165 | 149 | 1 | 1 | 0.231 | 1 | 0 | 0.844 | 1 | 1 | 1 | 1 | NA | NA | -0.385 |
| A5SS | 4382 | ENSG00000111358 | GTF2H3 | chr12 | + | 123650993 | 123651156 | 123650993 | 123651056 | 123652523 | 123652561 | 2 | 1 | 3 | 0 | 0 | 0 | 0 | 1 | 0 | 3 | 0 | 0 | 5 | 1 | 4 | 0 | 0 | 2 | 0 | 0 | 249 | 149 | 1 | 1 | 1 | 1 | 0.642 | NA | 0 | NA | NA | 0.599 | 1 | 1 | -0.206 |
| A5SS | 8652 | ENSG00000028203 | VEZT | chr12 | + | 95258244 | 95258287 | 95258244 | 95258282 | 95262905 | 95263081 | 5 | 3 | 1 | 0 | 16 | 8 | 1 | 3 | 2 | 15 | 0 | 0 | 0 | 4 | 0 | 0 | 0 | 1 | 0 | 0 | 154 | 149 | 1 | 1 | 0.377 | 0.744 | 0.244 | 0 | 0.508 | NA | NA | 0 | 1 | NA | -0.125 |
| A5SS | 1537 | ENSG00000103852 | TTC23 | chr15 | - | 99249170 | 99249799 | 99249621 | 99249799 | 99228532 | 99228732 | 0 | 2 | 2 | 0 | 1 | 0 | 0 | 1 | 0 | 0 | 0 | 3 | 0 | 0 | 8 | 0 | 0 | 0 | 0 | 0 | 298 | 149 | 1 | 1 | NA | 1 | 0.5 | NA | 1 | NA | 1 | NA | NA | 1 | -0.167 |
| A5SS | 8651 | ENSG00000028203 | VEZT | chr12 | + | 95242163 | 95242314 | 95242163 | 95242264 | 95243940 | 95244009 | 0 | 0 | 1 | 0 | 0 | 1 | 0 | 0 | 0 | 0 | 7 | 0 | 0 | 0 | 0 | 0 | 0 | 2 | 0 | 0 | 199 | 149 | 1 | 1 | 0 | NA | 1 | NA | NA | 1 | NA | 0 | NA | NA | 0 |
| A5SS | 3598 | ENSG00000139842 | CUL4A | chr13 | + | 113209972 | 113210088 | 113209972 | 113210063 | 113218944 | 113219048 | 43 | 7 | 31 | 34 | 11 | 0 | 0 | 0 | 1 | 0 | 50 | 41 | 17 | 16 | 6 | 0 | 0 | 0 | 0 | 0 | 174 | 149 | 1 | 1 | 1 | 1 | 1 | 0.967 | 1 | 1 | 1 | 1 | 1 | 1 | -0.007 |
| A5SS | 7648 | ENSG00000198000 | NOL8 | chr9 | - | 92324905 | 92325350 | 92325317 | 92325350 | 92324022 | 92324209 | 3 | 2 | 1 | 6 | 8 | 3 | 0 | 2 | 0 | 2 | 0 | 11 | 2 | 5 | 0 | 0 | 6 | 0 | 0 | 0 | 298 | 149 | 1 | 1 | 0.333 | 1 | 0.2 | 1 | 0.667 | NA | 0.478 | 1 | 1 | NA | -0.186 |
| A5SS | 69 | ENSG00000185386 | MAPK11 | chr22 | - | 50267252 | 50267286 | 50267256 | 50267286 | 50267126 | 50267174 | 0 | 0 | 2 | 0 | 0 | 0 | 0 | 0 | 4 | 2 | 0 | 0 | 0 | 0 | 0 | 0 | 3 | 4 | 0 | 0 | 153 | 149 | 1 | 1 | NA | NA | 1 | 0 | 0 | NA | 0 | 0 | NA | NA | 0.333 |
| A5SS | 4371 | ENSG00000149527 | PLCH2 | chr1 | + | 2502111 | 2502910 | 2502111 | 2502409 | 2503921 | 2505532 | 2 | 1 | 1 | 4 | 0 | 3 | 0 | 3 | 0 | 0 | 0 | 2 | 4 | 0 | 0 | 0 | 0 | 0 | 0 | 0 | 298 | 149 | 1 | 1 | 0.25 | 1 | 0.143 | 1 | NA | NA | 1 | 1 | NA | NA | -0.402 |
| A5SS | 2846 | ENSG00000184470 | TXNRD2 | chr22 | - | 19877039 | 19877234 | 19877108 | 19877234 | 19875521 | 19875807 | 24 | 10 | 20 | 16 | 7 | 0 | 1 | 0 | 0 | 2 | 16 | 10 | 8 | 16 | 27 | 0 | 0 | 0 | 0 | 2 | 218 | 149 | 1 | 1 | 1 | 0.872 | 1 | 1 | 0.705 | 1 | 1 | 1 | 1 | 0.902 | -0.065 |
| A5SS | 4340 | ENSG00000111445 | RFC5 | chr12 | + | 118017689 | 118017852 | 118017689 | 118017815 | 118017994 | 118018053 | 1 | 6 | 1 | 3 | 15 | 2 | 0 | 0 | 1 | 0 | 4 | 11 | 3 | 0 | 8 | 0 | 0 | 0 | 0 | 0 | 186 | 149 | 1 | 1 | 0.286 | 1 | 1 | 0.706 | 1 | 1 | 1 | 1 | NA | 1 | -0.202 |
| A5SS | 5731 | ENSG00000153253 | SCN3A | chr2 | - | 165140788 | 165140998 | 165140797 | 165140998 | 165139475 | 165139608 | 1 | 0 | 0 | 2 | 0 | 3 | 0 | 3 | 0 | 0 | 0 | 0 | 0 | 0 | 0 | 0 | 7 | 0 | 0 | 0 | 158 | 149 | 1 | 1 | 0.239 | NA | 0 | 1 | NA | NA | 0 | NA | NA | NA | 0.413 |
| A5SS | 8333 | ENSG00000197771 | MCMBP | chr10 | - | 119843247 | 119843426 | 119843253 | 119843426 | 119840842 | 119840960 | 3 | 1 | 3 | 10 | 9 | 4 | 1 | 3 | 2 | 4 | 0 | 10 | 0 | 5 | 0 | 0 | 0 | 0 | 0 | 0 | 155 | 149 | 1 | 1 | 0.419 | 0.49 | 0.49 | 0.828 | 0.684 | NA | 1 | NA | 1 | NA | -0.418 |
| A5SS | 70 | ENSG00000188130 | MAPK12 | chr22 | - | 50256787 | 50256964 | 50256934 | 50256964 | 50256598 | 50256646 | 0 | 0 | 2 | 1 | 0 | 0 | 2 | 0 | 0 | 0 | 0 | 0 | 0 | 2 | 0 | 4 | 1 | 0 | 0 | 0 | 296 | 149 | 1 | 1 | NA | 0 | 1 | 1 | NA | 0 | 0 | NA | 1 | NA | 0.333 |
| A5SS | 5730 | ENSG00000153253 | SCN3A | chr2 | - | 165140784 | 165140998 | 165140797 | 165140998 | 165139475 | 165139608 | 3 | 0 | 1 | 1 | 0 | 3 | 0 | 3 | 0 | 0 | 0 | 0 | 0 | 0 | 0 | 0 | 7 | 0 | 0 | 0 | 162 | 149 | 1 | 1 | 0.479 | NA | 0.235 | 1 | NA | NA | 0 | NA | NA | NA | 0.571 |
| A5SS | 8159 | ENSG00000149262 | INTS4 | chr11 | - | 77987528 | 77987698 | 77987603 | 77987698 | 77981458 | 77981576 | 5 | 5 | 14 | 10 | 11 | 2 | 0 | 0 | 0 | 0 | 4 | 20 | 11 | 9 | 0 | 0 | 0 | 2 | 0 | 0 | 224 | 149 | 1 | 1 | 0.624 | 1 | 1 | 1 | 1 | 1 | 1 | 0.785 | 1 | NA | -0.021 |
| A5SS | 5725 | ENSG00000153253 | SCN3A | chr2 | - | 165140650 | 165140998 | 165140797 | 165140998 | 165139475 | 165139608 | 13 | 6 | 6 | 6 | 1 | 3 | 0 | 3 | 0 | 0 | 0 | 19 | 6 | 1 | 0 | 0 | 7 | 0 | 0 | 0 | 296 | 149 | 1 | 1 | 0.686 | 1 | 0.502 | 1 | 1 | NA | 0.577 | 1 | 1 | NA | -0.021 |
| A5SS | 6100 | ENSG00000221838 | AP4M1 | chr7 | + | 100101879 | 100102393 | 100101879 | 100101989 | 100102674 | 100102781 | 3 | 1 | 0 | 0 | 1 | 0 | 2 | 1 | 1 | 0 | 0 | 1 | 0 | 0 | 0 | 0 | 0 | 0 | 0 | 0 | 298 | 149 | 1 | 1 | 1 | 0.2 | 0 | 0 | 1 | NA | 1 | NA | NA | NA | -0.56 |
| A5SS | 7067 | ENSG00000102001 | CACNA1F | chrX | - | 49226936 | 49227127 | 49226969 | 49227127 | 49226609 | 49226702 | 2 | 0 | 0 | 1 | 0 | 0 | 1 | 0 | 0 | 0 | 0 | 1 | 0 | 1 | 0 | 0 | 0 | 0 | 0 | 0 | 182 | 149 | 1 | 1 | 1 | 0 | NA | 1 | NA | NA | 1 | NA | 1 | NA | -0.333 |
| A5SS | 5723 | ENSG00000153253 | SCN3A | chr2 | - | 165140650 | 165140998 | 165140784 | 165140998 | 165139475 | 165139608 | 12 | 6 | 5 | 5 | 1 | 2 | 0 | 0 | 0 | 0 | 0 | 19 | 6 | 1 | 0 | 0 | 0 | 0 | 0 | 0 | 283 | 149 | 1 | 1 | 0.76 | 1 | 1 | 1 | 1 | NA | 1 | 1 | 1 | NA | -0.048 |
| A5SS | 5840 | ENSG00000072195 | SPEG | chr2 | + | 219443107 | 219443440 | 219443107 | 219443150 | 219444024 | 219444059 | 2 | 0 | 3 | 6 | 5 | 3 | 0 | 3 | 0 | 2 | 0 | 0 | 0 | 4 | 2 | 0 | 0 | 0 | 0 | 0 | 298 | 149 | 1 | 1 | 0.25 | NA | 0.333 | 1 | 0.556 | NA | NA | NA | 1 | 1 | -0.465 |
| A5SS | 6290 | ENSG00000109339 | MAPK10 | chr4 | - | 86194308 | 86194407 | 86194335 | 86194407 | 86107222 | 86107352 | 2 | 0 | 0 | 0 | 0 | 0 | 0 | 0 | 0 | 0 | 0 | 0 | 0 | 0 | 0 | 0 | 0 | 0 | 5 | 0 | 176 | 149 | 1 | 1 | 1 | NA | NA | NA | NA | NA | NA | NA | NA | 0 | 1 |
| A5SS | 5724 | ENSG00000153253 | SCN3A | chr2 | - | 165140650 | 165140998 | 165140788 | 165140998 | 165139475 | 165139608 | 13 | 6 | 6 | 6 | 1 | 1 | 0 | 0 | 2 | 0 | 0 | 19 | 6 | 1 | 0 | 0 | 0 | 0 | 0 | 0 | 287 | 149 | 1 | 1 | 0.871 | 1 | 1 | 0.609 | 1 | NA | 1 | 1 | 1 | NA | -0.104 |
| A5SS | 6930 | ENSG00000162669 | HFM1 | chr1 | - | 91379058 | 91379214 | 91379062 | 91379214 | 91378402 | 91378480 | 0 | 0 | 0 | 4 | 0 | 0 | 0 | 0 | 0 | 0 | 0 | 0 | 0 | 0 | 0 | 0 | 1 | 0 | 0 | 0 | 153 | 149 | 1 | 1 | NA | NA | NA | 1 | NA | NA | 0 | NA | NA | NA | 1 |
| A5SS | 4107 | ENSG00000256269 | HMBS | chr11 | + | 119092124 | 119092523 | 119092124 | 119092163 | 119092757 | 119092811 | 17 | 3 | 5 | 9 | 5 | 2 | 0 | 2 | 0 | 1 | 1 | 16 | 7 | 5 | 0 | 0 | 22 | 0 | 1 | 0 | 298 | 149 | 1 | 1 | 0.81 | 1 | 0.556 | 1 | 0.714 | 1 | 0.267 | 1 | 0.714 | NA | 0.071 |
| A5SS | 1526 | ENSG00000103978 | TMEM87A | chr15 | - | 42272888 | 42273078 | 42272962 | 42273078 | 42272062 | 42272123 | 17 | 7 | 11 | 6 | 7 | 0 | 0 | 0 | 0 | 0 | 0 | 21 | 10 | 7 | 4 | 0 | 3 | 0 | 3 | 0 | 223 | 149 | 1 | 1 | 1 | 1 | 1 | 1 | 1 | NA | 0.824 | 1 | 0.609 | 1 | 0.142 |
| A5SS | 5841 | ENSG00000072195 | SPEG | chr2 | + | 219443107 | 219443440 | 219443107 | 219443150 | 219444029 | 219444059 | 2 | 0 | 3 | 6 | 5 | 0 | 0 | 2 | 1 | 0 | 0 | 0 | 0 | 4 | 2 | 0 | 0 | 0 | 0 | 0 | 298 | 149 | 1 | 1 | 1 | NA | 0.429 | 0.75 | 1 | NA | NA | NA | 1 | 1 | -0.205 |
| A5SS | 7031 | ENSG00000184343 | SRPK3 | chrX | + | 153783973 | 153784024 | 153783973 | 153784021 | 153784120 | 153784213 | 0 | 0 | 0 | 7 | 0 | 0 | 0 | 0 | 0 | 0 | 0 | 0 | 0 | 0 | 0 | 0 | 0 | 0 | 5 | 0 | 152 | 149 | 1 | 1 | NA | NA | NA | 1 | NA | NA | NA | NA | 0 | NA | 1 |
| A5SS | 5842 | ENSG00000072195 | SPEG | chr2 | + | 219443107 | 219443637 | 219443107 | 219443150 | 219444652 | 219444742 | 2 | 0 | 3 | 6 | 5 | 0 | 0 | 0 | 1 | 0 | 0 | 0 | 0 | 4 | 2 | 0 | 0 | 0 | 0 | 0 | 298 | 149 | 1 | 1 | 1 | NA | 1 | 0.75 | 1 | NA | NA | NA | 1 | 1 | -0.062 |
| A5SS | 1970 | ENSG00000164180 | TMEM161B | chr5 | - | 88268720 | 88268848 | 88268787 | 88268848 | 88228444 | 88228528 | 12 | 8 | 15 | 4 | 1 | 1 | 0 | 0 | 0 | 0 | 16 | 10 | 0 | 0 | 0 | 0 | 0 | 0 | 0 | 0 | 216 | 149 | 1 | 1 | 0.892 | 1 | 1 | 1 | 1 | 1 | 1 | NA | NA | NA | -0.022 |
| A5SS | 5543 | ENSG00000107521 | HPS1 | chr10 | - | 98433939 | 98434091 | 98433982 | 98434091 | 98430570 | 98430670 | 32 | 11 | 16 | 16 | 15 | 3 | 1 | 1 | 5 | 0 | 0 | 15 | 1 | 21 | 30 | 0 | 3 | 0 | 0 | 7 | 192 | 149 | 1 | 1 | 0.892 | 0.895 | 0.925 | 0.713 | 1 | NA | 0.795 | 1 | 1 | 0.769 | -0.006 |
| A5SS | 950 | ENSG00000185813 | PCYT2 | chr17 | - | 81907553 | 81907857 | 81907772 | 81907857 | 81906759 | 81906898 | 17 | 5 | 27 | 22 | 16 | 0 | 0 | 0 | 2 | 0 | 8 | 19 | 6 | 9 | 28 | 0 | 0 | 0 | 0 | 0 | 298 | 149 | 1 | 1 | 1 | 1 | 1 | 0.846 | 1 | 1 | 1 | 1 | 1 | 1 | -0.031 |
| A5SS | 6965 | ENSG00000100532 | CGRRF1 | chr14 | + | 54530048 | 54530763 | 54530048 | 54530405 | 54530902 | 54531050 | 15 | 3 | 6 | 6 | 14 | 0 | 0 | 0 | 0 | 0 | 0 | 17 | 10 | 0 | 0 | 0 | 1 | 0 | 2 | 0 | 298 | 149 | 1 | 1 | 1 | 1 | 1 | 1 | 1 | NA | 0.895 | 1 | 0 | NA | 0.368 |
| A5SS | 8427 | ENSG00000148688 | RPP30 | chr10 | + | 90900569 | 90900668 | 90900569 | 90900666 | 90902242 | 90902344 | 3 | 5 | 16 | 3 | 32 | 0 | 2 | 1 | 0 | 0 | 20 | 4 | 7 | 3 | 5 | 0 | 0 | 0 | 0 | 0 | 151 | 149 | 1 | 1 | 1 | 0.712 | 0.94 | 1 | 1 | 1 | 1 | 1 | 1 | 1 | -0.07 |
| A5SS | 5035 | ENSG00000116922 | C1orf109 | chr1 | - | 37689605 | 37689883 | 37689657 | 37689883 | 37686272 | 37686452 | 18 | 12 | 20 | 16 | 7 | 1 | 0 | 0 | 1 | 0 | 13 | 13 | 8 | 7 | 9 | 0 | 0 | 0 | 0 | 0 | 201 | 149 | 1 | 1 | 0.93 | 1 | 1 | 0.922 | 1 | 1 | 1 | 1 | 1 | 1 | -0.03 |
| A5SS | 6227 | ENSG00000223501 | VPS52 | chr6 | - | 33267864 | 33267997 | 33267870 | 33267997 | 33267681 | 33267739 | 92 | 76 | 87 | 99 | 176 | 0 | 0 | 0 | 2 | 0 | 91 | 86 | 57 | 63 | 209 | 0 | 0 | 0 | 0 | 0 | 155 | 149 | 1 | 1 | 1 | 1 | 1 | 0.979 | 1 | 1 | 1 | 1 | 1 | 1 | -0.004 |
| A5SS | 5728 | ENSG00000153253 | SCN3A | chr2 | - | 165140701 | 165140998 | 165140797 | 165140998 | 165139475 | 165139608 | 12 | 12 | 7 | 12 | 0 | 3 | 0 | 3 | 0 | 0 | 0 | 13 | 0 | 1 | 0 | 0 | 7 | 0 | 0 | 0 | 245 | 149 | 1 | 1 | 0.709 | 1 | 0.587 | 1 | NA | NA | 0.53 | NA | 1 | NA | 0.059 |
| A5SS | 5726 | ENSG00000153253 | SCN3A | chr2 | - | 165140701 | 165140998 | 165140784 | 165140998 | 165139475 | 165139608 | 11 | 12 | 6 | 11 | 0 | 2 | 0 | 0 | 0 | 0 | 0 | 13 | 0 | 1 | 0 | 0 | 0 | 0 | 0 | 0 | 232 | 149 | 1 | 1 | 0.779 | 1 | 1 | 1 | NA | NA | 1 | NA | 1 | NA | -0.055 |
| A5SS | 5727 | ENSG00000153253 | SCN3A | chr2 | - | 165140701 | 165140998 | 165140788 | 165140998 | 165139475 | 165139608 | 12 | 12 | 7 | 12 | 0 | 1 | 0 | 0 | 2 | 0 | 0 | 13 | 0 | 1 | 0 | 0 | 0 | 0 | 0 | 0 | 236 | 149 | 1 | 1 | 0.883 | 1 | 1 | 0.791 | NA | NA | 1 | NA | 1 | NA | -0.082 |
| A5SS | 5126 | ENSG00000112855 | HARS2 | chr5 | + | 140691452 | 140691756 | 140691452 | 140691515 | 140693590 | 140693665 | 55 | 31 | 65 | 39 | 24 | 0 | 0 | 0 | 1 | 0 | 13 | 20 | 38 | 29 | 78 | 0 | 0 | 0 | 0 | 0 | 298 | 149 | 1 | 1 | 1 | 1 | 1 | 0.951 | 1 | 1 | 1 | 1 | 1 | 1 | -0.01 |
| A5SS | 949 | ENSG00000185813 | PCYT2 | chr17 | - | 81905669 | 81906177 | 81906099 | 81906177 | 81905381 | 81905447 | 18 | 8 | 17 | 18 | 12 | 0 | 0 | 0 | 0 | 0 | 7 | 1 | 2 | 2 | 11 | 0 | 0 | 1 | 0 | 0 | 298 | 149 | 1 | 1 | 1 | 1 | 1 | 1 | 1 | 1 | 1 | 0.5 | 1 | 1 | 0.1 |
| A5SS | 8728 | ENSG00000139350 | NEDD1 | chr12 | + | 96907603 | 96907856 | 96907603 | 96907624 | 96919984 | 96920125 | 15 | 1 | 10 | 22 | 29 | 0 | 0 | 0 | 0 | 0 | 0 | 15 | 0 | 5 | 0 | 0 | 0 | 5 | 0 | 0 | 298 | 149 | 1 | 1 | 1 | 1 | 1 | 1 | 1 | NA | 1 | NA | 0.333 | NA | 0.334 |
| A5SS | 8300 | ENSG00000198258 | UBL5 | chr19 | + | 9828326 | 9828393 | 9828326 | 9828389 | 9828591 | 9828675 | 139 | 76 | 100 | 133 | 149 | 1 | 2 | 0 | 0 | 1 | 100 | 106 | 48 | 113 | 163 | 0 | 0 | 0 | 0 | 8 | 153 | 149 | 1 | 1 | 0.993 | 0.974 | 1 | 1 | 0.993 | 1 | 1 | 1 | 1 | 0.952 | 0.002 |
| A5SS | 7649 | ENSG00000198000 | NOL8 | chr9 | - | 92325305 | 92325350 | 92325317 | 92325350 | 92324022 | 92324209 | 19 | 15 | 14 | 15 | 26 | 3 | 0 | 2 | 0 | 2 | 0 | 16 | 2 | 0 | 0 | 0 | 6 | 0 | 0 | 0 | 161 | 149 | 1 | 1 | 0.854 | 1 | 0.866 | 1 | 0.923 | NA | 0.712 | 1 | NA | NA | 0.073 |
| A5SS | 8727 | ENSG00000139350 | NEDD1 | chr12 | + | 96907603 | 96907856 | 96907603 | 96907624 | 96917620 | 96917737 | 16 | 5 | 18 | 23 | 35 | 3 | 0 | 2 | 0 | 0 | 0 | 15 | 0 | 5 | 0 | 0 | 0 | 0 | 0 | 0 | 298 | 149 | 1 | 1 | 0.727 | 1 | 1 | 1 | 1 | NA | 1 | NA | 1 | NA | -0.055 |
| A5SS | 8294 | ENSG00000143702 | CEP170 | chr1 | - | 243142285 | 243142463 | 243142366 | 243142463 | 243139936 | 243140107 | 24 | 31 | 30 | 24 | 37 | 13 | 2 | 8 | 14 | 8 | 0 | 12 | 9 | 0 | 0 | 0 | 0 | 5 | 3 | 0 | 230 | 149 | 1 | 1 | 0.545 | 0.909 | 0.708 | 0.526 | 0.75 | NA | 1 | 0.538 | 0 | NA | 0.175 |
| A5SS | 7893 | ENSG00000100731 | PCNX1 | chr14 | + | 70995740 | 70995925 | 70995740 | 70995843 | 71009633 | 71009724 | 45 | 30 | 44 | 46 | 38 | 2 | 0 | 0 | 1 | 0 | 0 | 40 | 32 | 27 | 20 | 0 | 5 | 0 | 1 | 0 | 231 | 149 | 1 | 1 | 0.936 | 1 | 1 | 0.967 | 1 | NA | 0.838 | 1 | 0.946 | 1 | 0.035 |
| A5SS | 8491 | ENSG00000119977 | TCTN3 | chr10 | - | 95682650 | 95683629 | 95683100 | 95683629 | 95680471 | 95680609 | 26 | 18 | 33 | 28 | 41 | 2 | 3 | 3 | 0 | 3 | 0 | 19 | 20 | 10 | 4 | 0 | 0 | 3 | 0 | 0 | 298 | 149 | 1 | 1 | 0.867 | 0.75 | 0.846 | 1 | 0.872 | NA | 1 | 0.769 | 1 | 1 | -0.075 |
| A5SS | 8026 | ENSG00000099783 | HNRNPM | chr19 | + | 8463496 | 8463504 | 8463496 | 8463500 | 8463592 | 8463686 | 465 | 332 | 320 | 432 | 698 | 0 | 0 | 2 | 0 | 0 | 631 | 363 | 285 | 475 | 408 | 0 | 0 | 0 | 4 | 0 | 153 | 149 | 1 | 1 | 1 | 1 | 0.994 | 1 | 1 | 1 | 1 | 1 | 0.991 | 1 | 0.001 |
| A5SS | 7003 | ENSG00000144161 | ZC3H8 | chr2 | - | 112233257 | 112233371 | 112233259 | 112233371 | 112231837 | 112231945 | 23 | 20 | 25 | 35 | 28 | 0 | 0 | 0 | 2 | 0 | 0 | 6 | 18 | 13 | 0 | 0 | 0 | 0 | 0 | 0 | 151 | 149 | 1 | 1 | 1 | 1 | 1 | 0.945 | 1 | NA | 1 | 1 | 1 | NA | -0.011 |
| A5SS | 8295 | ENSG00000143702 | CEP170 | chr1 | - | 243142315 | 243142463 | 243142366 | 243142463 | 243139936 | 243140107 | 34 | 39 | 26 | 33 | 49 | 13 | 2 | 8 | 14 | 8 | 0 | 15 | 14 | 0 | 0 | 0 | 0 | 5 | 3 | 0 | 200 | 149 | 1 | 1 | 0.661 | 0.936 | 0.708 | 0.637 | 0.82 | NA | 1 | 0.676 | 0 | NA | 0.194 |
| A5SS | 8845 | ENSG00000125629 | INSIG2 | chr2 | + | 118096418 | 118096800 | 118096418 | 118096426 | 118103196 | 118103321 | 47 | 36 | 38 | 60 | 56 | 5 | 0 | 3 | 5 | 3 | 24 | 37 | 23 | 26 | 0 | 0 | 8 | 6 | 0 | 0 | 298 | 149 | 1 | 1 | 0.825 | 1 | 0.864 | 0.857 | 0.903 | 1 | 0.698 | 0.657 | 1 | NA | 0.051 |
| A5SS | 8726 | ENSG00000139350 | NEDD1 | chr12 | + | 96907603 | 96907856 | 96907603 | 96907624 | 96909751 | 96909895 | 38 | 21 | 22 | 49 | 59 | 3 | 1 | 3 | 1 | 0 | 0 | 23 | 1 | 8 | 0 | 2 | 1 | 1 | 0 | 0 | 298 | 149 | 1 | 1 | 0.864 | 0.913 | 0.786 | 0.961 | 1 | 0 | 0.92 | 0.333 | 1 | NA | 0.342 |
| A5SS | 8921 | ENSG00000117226 | GBP3 | chr1 | - | 89013184 | 89014279 | 89014082 | 89014279 | 89011746 | 89012027 | 98 | 101 | 84 | 80 | 66 | 0 | 0 | 6 | 11 | 0 | 0 | 88 | 57 | 43 | 26 | 0 | 0 | 2 | 0 | 0 | 298 | 149 | 1 | 1 | 1 | 1 | 0.875 | 0.784 | 1 | NA | 1 | 0.934 | 1 | 1 | -0.052 |
| A5SS | 6458 | ENSG00000068885 | IFT80 | chr3 | - | 160383436 | 160384646 | 160384405 | 160384646 | 160381502 | 160381718 | 30 | 34 | 37 | 55 | 91 | 0 | 0 | 0 | 0 | 0 | 8 | 16 | 9 | 9 | 0 | 0 | 0 | 4 | 0 | 0 | 298 | 149 | 1 | 1 | 1 | 1 | 1 | 1 | 1 | 1 | 1 | 1 | 0.529 | NA | 0.118 |
| A5SS | 2631 | ENSG00000145348 | TBCK | chr4 | - | 106251865 | 106252007 | 106251982 | 106252007 | 106250417 | 106250478 | 80 | 71 | 62 | 85 | 85 | 0 | 0 | 0 | 0 | 0 | 6 | 41 | 26 | 24 | 1 | 0 | 0 | 3 | 0 | 0 | 266 | 149 | 1 | 1 | 1 | 1 | 1 | 1 | 1 | 1 | 1 | 0.829 | 1 | 1 | 0.034 |
| A5SS | 5558 | ENSG00000112367 | FIG4 | chr6 | + | 109735149 | 109738453 | 109735149 | 109735298 | 109741443 | 109741544 | 140 | 80 | 98 | 96 | 207 | 0 | 0 | 0 | 0 | 5 | 86 | 49 | 23 | 85 | 35 | 0 | 0 | 0 | 0 | 0 | 298 | 149 | 1 | 1 | 1 | 1 | 1 | 1 | 0.954 | 1 | 1 | 1 | 1 | 1 | -0.009 |
| A5SS | 8752 | ENSG00000133858 | ZFC3H1 | chr12 | - | 71611057 | 71611887 | 71611785 | 71611887 | 71610694 | 71610757 | 93 | 67 | 63 | 64 | 180 | 0 | 1 | 0 | 1 | 3 | 0 | 63 | 13 | 1 | 4 | 0 | 0 | 1 | 0 | 0 | 298 | 149 | 1 | 1 | 1 | 0.971 | 1 | 0.97 | 0.968 | NA | 1 | 0.867 | 1 | 1 | 0.015 |
| A5SS | 7148 | ENSG00000104613 | INTS10 | chr8 | + | 19823872 | 19824044 | 19823872 | 19823981 | 19824802 | 19824972 | 204 | 132 | 215 | 215 | 586 | 2 | 3 | 0 | 0 | 2 | 175 | 165 | 171 | 218 | 66 | 0 | 0 | 4 | 0 | 0 | 212 | 149 | 1 | 1 | 0.986 | 0.969 | 1 | 1 | 0.995 | 1 | 1 | 0.968 | 1 | 1 | -0.004 |
| A5SS | 4235 | ENSG00000233355 | CHRM3-AS2 | chr1 | - | 239707533 | 239707760 | 239707614 | 239707760 | 239703891 | 239705715 | 93 | 128 | 226 | 120 | 104 | 0 | 0 | 1 | 1 | 0 | 22 | 22 | 47 | 21 | 0 | 0 | 1 | 0 | 0 | 0 | 230 | 149 | 1 | 1 | 1 | 1 | 0.993 | 0.987 | 1 | 1 | 0.934 | 1 | 1 | NA | 0.013 |
| A5SS | 4236 | ENSG00000233355 | CHRM3-AS2 | chr1 | - | 239707533 | 239707760 | 239707614 | 239707760 | 239706424 | 239706496 | 102 | 129 | 236 | 122 | 111 | 0 | 5 | 7 | 5 | 7 | 22 | 22 | 47 | 21 | 0 | 0 | 0 | 0 | 0 | 0 | 230 | 149 | 1 | 1 | 1 | 0.944 | 0.956 | 0.941 | 0.911 | 1 | 1 | 1 | 1 | NA | -0.05 |
| A5SS | 6667 | ENSG00000114503 | NCBP2 | chr3 | - | 196939250 | 196939432 | 196939392 | 196939432 | 196937509 | 196937631 | 335 | 282 | 321 | 328 | 424 | 3 | 0 | 1 | 0 | 4 | 112 | 212 | 213 | 167 | 51 | 0 | 0 | 0 | 0 | 0 | 291 | 149 | 1 | 1 | 0.983 | 1 | 0.994 | 1 | 0.982 | 1 | 1 | 1 | 1 | 1 | -0.008 |

**Supplementary_data_5S**

| Seq ID | Sample ID | Group | Cell type |
|---|---|---|---|
| s01.fq.gz | CTR_007_TTAGGC_L006_R1_001.fastq.gz | Healthy donor | Normal dense neutrophils |
| s02.fq.gz | CTR_150_ACTTGA_L007_R1_001.fastq.gz | Healthy donor | Normal dense neutrophils |
| s03.fq.gz | CTR_152_ATCACG_L007_R1_001.fastq.gz | Healthy donor | Normal dense neutrophils |
| s04.fq.gz | CTR_153_GGCTAC_L007_R1_001.fastq.gz | Healthy donor | Normal dense neutrophils |
| s05.fq.gz | CTR_154_ATCACG_L006_R1_001.fastq.gz | Healthy donor | Normal dense neutrophils |
| s06.fq.gz | CTR_161_CTTGTA_L008_R1_001.fastq.gz | Healthy donor | Normal dense neutrophils |
| s07.fq.gz | CTR_165_ACAGTG_L006_R1_001.fastq.gz | Healthy donor | Normal dense neutrophils |
| s08.fq.gz | CTR_22_ACTTGA_L008_R1_001.fastq.gz | Healthy donor | Normal dense neutrophils |
| s09.fq.gz | CTR_311_ACAGTG_L008_R1_001.fastq.gz | Healthy donor | Normal dense neutrophils |
| s10.fq.gz | CTR_997_CAGATC_L006_R1_001.fastq.gz | Healthy donor | Normal dense neutrophils |
| s11.fq.gz | LE_041_GCCAAT_L006_R1_001.fastq.gz | SLE patient | Normal dense neutrophils |
| s12.fq.gz | LE_145_LDG_CTTGTA_L007_R1_001.fastq.gz | SLE patient 145 | Low density neutrophils |
| s13.fq.gz | LE_145_PMN_GCCAAT_L008_R1_001.fastq.gz | SLE patient 145 | Normal dense neutrophils |
| s14.fq.gz | LE_147_LDG_TGACCA_L008_R1_001.fastq.gz | SLE patient 147 | Low density neutrophils |
| s15.fq.gz | LE_147_PMN_CGATGT_L006_R1_001.fastq.gz | SLE patient 147 | Normal dense neutrophils |
| s16.fq.gz | LE_150_LDG_GATCAG_L008_R1_001.fastq.gz | SLE patient 150 | Low density neutrophils |
| s17.fq.gz | LE_150_PMN_GGCTAC_L008_R1_001.fastq.gz | SLE patient 150 | Normal dense neutrophils |
| s18.fq.gz | LE_197_TTAGGC_L008_R1_001.fastq.gz | SLE patient | Normal dense neutrophils |
| s19.fq.gz | LE_237N_TAGCTT_L008_R1_001.fastq.gz | SLE patient | Normal dense neutrophils |
| s20.fq.gz | LE_245_CAGATC_L008_R1_001.fastq.gz | SLE patient | Normal dense neutrophils |
| s21.fq.gz | LE_263_GATCAG_L007_R1_001.fastq.gz | SLE patient | Normal dense neutrophils |
| s22.fq.gz | LE_277_TGACCA_L006_R1_001.fastq.gz | SLE patient | Normal dense neutrophils |
| s23.fq.gz | LE_278_CGATGT_L007_R1_001.fastq.gz | SLE patient | Normal dense neutrophils |
| s24.fq.gz | LE_278_TAGCTT_L007_R1_001.fastq.gz | SLE patient | Normal dense neutrophils |
| s25.fq.gz | LE_279_ATCACG_L008_R1_001.fastq.gz | SLE patient | Normal dense neutrophils |

| Demographics | | | | | | Complement | | |
|---|---|---|---|---|---|---|---|---|
| **Type** | **Sample number** | **Gender** | **Age** | **RACE** | **ETHNICITY** | **C3** | **C4** | **Seizure** |
| Healthy control | s1 (NDG) | F | 42 | | Not reported | | | |
| Healthy control | s3 (NDG) | M | 34 | | 3 (HISPANIC/LATINO) | | | |
| Healthy control | s4 (NDG) | M | 60 | | 1 (WHITE) | | | |
| Healthy control | s5 (NDG) | M | 57 | | 1 (WHITE) | | | |
| Healthy control | s6 (NDG) | F | 35 | | 1 (WHITE) | | | |
| Healthy control | s7 (NDG) | M | 34 | | 3 (HISPANIC/LATINO) | | | |
| Healthy control | s8 (NDG) | M | 49 | | 1 (WHITE) | | | |
| Healthy control | s10 (NDG) | F | 48 | | 1 (WHITE) | | | |
| SLE patient | s11 (NDG) | F | 30 | WHITE | LATINO OR HISPANIC | 91.4 | 25.4 | 0 |
| SLE patient | s12 (LDG) and s13 (NDG) | M | 25 | WHITE | NOT LATINO OR HISPANIC | 126.4 | 23.4 | 0 |
| SLE patient | s14 (LDG) and s15 (NDG) | F | 30 | AFRICAN | NOT LATINO OR HISPANIC | 93.6 | 18.3 | 0 |
| SLE patient | s16 (LDG) and s17 (NDG) | F | 56 | UNKNOWN | LATINO OR HISPANIC | 73.8 | 9.6 | 0 |
| SLE patient | s19 (NDG) | F | 34 | UNKNOWN | LATINO OR HISPANIC | 117.5 | 18 | 0 |
| SLE patient | s21 (NDG) | F | 60 | ASIAN | NOT LATINO OR HISPANIC | 112.8 | 28 | 0 |
| SLE patient | s22 (NDG) | F | 14 | WHITE | NOT LATINO OR HISPANIC | 103.2 | 20.6 | 0 |

| Psychosis | Organic_Brain_ Syndrome | Visual_Disturbance | Cranial_Nerve_ Disorder | Lupus_Headache | CVA | Vasculitis | Arthritis | Myositis |
|---|---|---|---|---|---|---|---|---|
| 0 | 0 | 0 | 0 | 0 | 0 | 0 | 0 | 0 |
| 0 | 0 | 0 | 0 | 0 | 0 | 0 | 0 | 0 |
| 0 | 0 | 0 | 0 | 0 | 0 | 0 | 0 | 0 |
| 0 | 0 | 0 | 0 | 0 | 0 | 0 | 4 | 0 |
| 0 | 0 | 0 | 0 | 0 | 0 | 0 | 0 | 0 |
| 0 | 0 | 0 | 0 | 0 | 0 | 0 | 0 | 0 |
| 0 | 0 | 0 | 0 | 0 | 0 | 0 | 0 | 0 |

| SLEDAI SCORE | | | | | | | | |
|---|---|---|---|---|---|---|---|---|
| Urinary_Casts | Hematuria | Proteinuria | Pyuria | Rash | Alopecia | Mucosal_Ulcers | Pleurisy | Pericarditis |
| 0 | 0 | 0 | 0 | 0 | 0 | 0 | 0 | 0 |
| 0 | 0 | 0 | 0 | 0 | 0 | 0 | 0 | 0 |
| 0 | 0 | 0 | 0 | 0 | 0 | 0 | 0 | 0 |
| 0 | 0 | 0 | 0 | 0 | 0 | 0 | 0 | 0 |
| 0 | 0 | 0 | 0 | 0 | 0 | 0 | 0 | 0 |
| 0 | 0 | 0 | 0 | 0 | 0 | 0 | 0 | 0 |
| 0 | 0 | 4 | 0 | 0 | 0 | 0 | 0 | 0 |

| Low_Complement | Increased_DNA_Binding | Fever | Thrombocytopenia | Leukopenia | TOTAL_SCORE |
|---|---|---|---|---|---|
| 0 | 2 | 0 | 0 | 0 | 2 |
| 0 | 0 | 0 | 0 | 0 | 0 |
| 0 | 2 | 0 | 0 | 0 | 2 |
| 2 | 2 | 0 | 0 | 0 | 8 |
| 0 | 0 | 0 | 0 | 0 | 0 |
| 0 | 0 | 0 | 0 | 0 | 0 |
| 0 | 0 | 0 | 0 | 0 | 4 |

| Autoantibodies | | | | | |
|---|---|---|---|---|---|
| ANA | Anti-ENA Screen | Anti-smith(SM) Ab | Anti-smRNP Ab | SS-A/Ro Ab, IgG (Anti-Sjogren's y. A) Ro | SS-B/La Ab, IgG (Anti-Sjogren's y. B) La |
| >12.0 (12/20/2012) | Neg | Neg | Neg | Neg | Neg |
| 9 [0.0-0.9] (03/24/2015) | >177 | 0.3 | 0.2 | > 8.0 | > 8.0 |
| > 12.0 [0.0-0.9] (03/18/2010) | >200 | >130 | >200 | Neg | Neg |
| > 12.0 [0.0-0.9] (03/22/2005) | >190 | 38 | 203 | 220 | 30 |
| 2.0 (8/7/2009) | 76(8/17/2004) | 67(8/17/2004) | 109(8/17/2004) | Neg | Neg |
| 4.7 (1/9/2014) | 106 (12/11/2014) | <0.2 (12/11/2014) | 1.1 (12/11/2014) | >8.0 (12/11/2014) | <0.2 (12/11/2014) |
| 0.2 | Neg | <0.2 | <0.2 | <0.2 | <0.2 |

| Rheumatoid Factor | Anti-CCP Ab | Anti-dsDNA | HCQ [mg/day] | Prednisone [mg/day] | AZA [mg/day] | MTX [mg/week] | MMF [mg/day] |
|---|---|---|---|---|---|---|---|
| <15 | | 103 | 400 | 5 | 100 | No | No |
| <15 | <20 | Neg | 400 | 30 | No | No | 3000 |
| <15 | N/A | 156 | 400 | 10 | No | No | 2000 |
| 45 | 55 | 206 | 400 | 5 | 100 | 15 | No |
| <15 | | Neg | 400 | 5 | No | No | No |
| <15 | | Neg | No | 10 | No | 20 | No |
| <15 | | <12.3 | No | 5 | No | No | 720 |

| Medication |
|---|
| **Other medications** |
| Coulmadin 15 mg p.o. daily. Recived cyclophosphamide infusion. Patient received cyclophosphamide for lupus nephritis date last cyclophosphamide infusio |
| Lisinopril 20 mg a day. |
| Cytoxan 1 gram/m2 IV, monthly pulse. Started9/2009, stopped, 3/2010. Usage past; Cytoxan 1 gram/m2 IV, NIH regimen, started, 4/2013, stopped, 4/2015. Usage past; synthroid 50 mcg PO, daily in AM, usage current; MVI PO, daily in AM, ssage current. In 2009, for CNS Lupus treated with pulse |
| Omeprazole 20 mg a day; amlodipine 5 mg once a day; pregabalin 20 mg once a day; used to take 5 mg once and 2.5 mg alternating once a day until Marcl |
| Ranitidine 75 milligrams twice a day; lisinopril 2.5 milligrams daily; xopenex p.r.n. for cough. |

n: 5/17/2023.

h 2015). She was never treated with cyclophosphamide.
